# Octanol-water partition coefficient measurements for the SAMPL6 Blind Prediction Challenge

**DOI:** 10.1101/757393

**Authors:** Mehtap Işık, Dorothy Levorse, David L. Mobley, Timothy Rhodes, John D. Chodera

## Abstract

Partition coefficients describe the equilibrium partitioning of a single, defined charge state of a solute between two liquid phases in contact, typically a neutral solute. Octanol-water partition coefficients (***K***_ow_), or their logarithms (log *P*), are frequently used as a measure of lipophilicity in drug discovery. The partition coefficient is a physicochemical property that captures the thermodynamics of relative solvation between aqueous and nonpolar phases, and therefore provides an excellent test for physics-based computational models that predict properties of pharmaceutical relevance such as protein-ligand binding affinities or hydration/solvation free energies. The SAMPL6 Part II Octanol-Water Partition Coefficient Prediction Challenge used a subset of kinase inhibitor fragment-like compounds from the SAMPL6 p*K*_a_ Prediction Challenge in a blind experimental benchmark. Following experimental data collection, the partition coefficient dataset was kept blinded until all predictions were collected from participating computational chemistry groups. A total of 91 submissions were received from 27 participating research groups. This paper presents the octanol-water log *P* dataset for this SAMPL6 Part II Partition Coefficient Challenge, which consisted of 11 compounds (six 4-aminoquinazolines, two benzimidazole, one pyrazolo[3,4-d]pyrimidine, one pyridine, one 2-oxoquinoline substructure containing compounds) with log *P* values in the range of 1.95–4.09. We describe the potentiometric log *P* measurement protocol used to collect this dataset using a Sirius T3, discuss the limitations of this experimental approach, and share suggestions for future log *P* data collection efforts for the evaluation of computational methods.

## 1 Introduction

The SAMPL (Statistical Assessment of the Modeling of Proteins and Ligands) Challenges [http://samplchallenges.github.io] are a series of blind prediction challenges for the computational chemistry community that aim to evaluate and advance computational tools for rational drug design [1]. These challenges focus the community on specific phenomena relevant to drug discovery—such as the contribution of force field inaccuracy to binding affinity prediction failures—and, using carefully-selected test systems, isolate these phenomena from other confounding factors. Through recurring community exercises involving blind prediction followed by data sharing and discussion, these challenges evaluate tools and methodologies prospectively, enforce data sharing to learn from failures, and generate high-quality datasets into the community as benchmark sets. As a result, SAMPL has driven progress in a number of areas over six previous rounds of challenge cycles [2–15].

To assess the accuracy of different computational methods, SAMPL has relied on the measurement of simple host-guest association affinities [6, 8, 11, 15–19] and other physical properties that isolate issues such as failing to capture relevant chemical effects, computationally-intensive conformational sampling, and force field accuracy. In SAMPL5, for example, a log *D* challenge was devised with the goal of isolating the accuracy of protein-ligand force fields from the difficulties of configurational sampling [20, 21]. In addition to being a useful surrogate for the accuracy of force fields in predicting binding free energies, partition or distribution coefficients are frequently used as a measure of lipophilicity in pharmacology [22], or as surrogates for solubility, permeability [23], and contributors to affinity [22, 24]. Lipophilicity is a critical physicochemical property that affects ADMET (absorption, distribution, metabolism, excretion, and toxicity) [22, 25, 26]. Since log *P* is utilized as a predictor for good drug-like properties in terms of pharmacokinetics and toxicity [25], accurate log *P* predictions of virtual molecules have high potential to benefit drug discovery and design.

Surprisingly, the cyclohexane-water log *D* challenge proved to be particularly problematic due to the necessity to account for protonation state effects to correctly compute the distribution coefficients, which assess the partitioning of all ionization states between phases [20]; failing to account for these protonation state effects led to modeling errors up to several log units [27]. As a result, the SAMPL6 Part II log *P* Prediction Challenge [28] aimed to further isolate the assessment of force field accuracy from the issues of conformational sampling and the modeling of ionization state equilibria by inviting participants to predict the partitioning of *neutral* drug-like molecules between aqueous and nonaqueous phases^1^. For maximum synergy with previous competitions, the challenge compound set was constructed to be a subset of kinase inhibitor fragment-like small molecules drawn from the SAMPL6 p*K*_a_ Challenge set [29], where the accuracy of participants to predict p*K*_a_ values was assessed. A blind challenge (the SAMPL6 Part II log *P* Blind Prediction Challenge) was run from November 1, 2018 to March 22, 2019 in which participants were given molecular structures and experimental details and asked to predict octanol-water partition coefficients before the data was unblinded on March 25, 2019. All primary and processed data was made available at https://github.com/MobleyLab/SAMPL6 immediately following the close of the competition.

### Partition coefficients and principles of their measurement

The partition coefficient describes the equilibrium partitioning of a molecule in a single, defined, charge state between two liquid phases in contact. Unless stated otherwise, in common usage partition coefficient (*P* or *P*^0^) refers to the partitioning of the neutral state of a molecule. In particular, the octanol-water partition coefficient of neutral species (frequently written as ***K_ow_*** or ***P***) is defined as

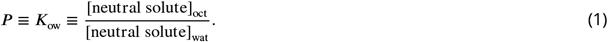

This quantity is often written in its log_10_ form, which we denote here as log *P*,

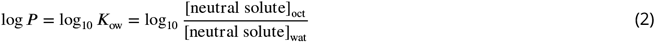

However, ionic species can also partition between phases [30–32]. The partition coefficients of ionic species is calculated using the same equation, *e.g. P*^+1^ refers to the partition equilibrium of +1 charge state of a molecule. Based on the experimental measurement method this value may be defined for a single tautomer or may involve multiple tautomers.

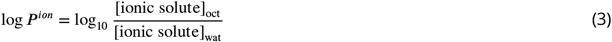

A closely related concept is that of the distribution coefficient (***D_ow_***, often written in log_10_ form as log *D*) which should not be confused with log *P*. log *D* is the logarithm of the sum of *all* species (both neutral and ionized) concentrations in the organic phase divided by the sum of neutral and ionic species concentrations in aqueous phase. Both octanol-water log *P* and log *D* values are frequently used as lipophilicity estimates [22]. However, while log *D* is pH-dependent, log *P* is independent of the pH of the aqueous phase. As log *P* is defined as the partition coefficient of neutral species, it would include all neutral tautomer populations if a compound can tautomerize.

The gold standard of partition coefficient measurement experimentation is the shake-flask method, according to the Organization for Economic Cooperation and Development(OECD)[33]. Methods developed as experimental refinements on the shake-flask method are high-throughput microscale shake flask [34, 35] and slow stirring methods [36]. Other direct methods for log *P* or log *D* determination include dialysis chamber-based methods [37], micellar electrokinetic capillary chromatography [38, 39], and counter-current chromatography [39]. An indirect experimental method that is widely used—despite being less reliable—is log *P* estimation based on reversed-phase high-performance liquid chromatography (HPLC) retention times [40–44]. The measurement principle for all of these methods is the measurement of log *D*—the equilibrium distribution coefficient for both neutral and ionized species—in a pH-dependent manner. As a result, in order to measure log *P* with these methods it is necessary to conduct the log *D* measurements at a pH where the analyte is completely un-ionized. At a pH where the analyte is at a neutral state, log *P* is *equal* to log *D*; however, accurately predicting or measuring the equilibrium ionization constant (p*K*_a_) of a substance is a prerequisite. Here in this study, however, we pursued an alternate approach for experimental determination of log *P*,which is potentiometric measurements.

### Potentiometric measurement of log *P* with the Sirius T3

The potentiometric log *P* measurement method determines log *P* values directly using potentiometric titrations in an immiscible biphasic system [45, 46]. The shift of apparent p*K*_a_ values when the aqueous phase is in contact with the octanol phase is used to estimate log *P* values. Experimental log *P* values presented in this study were collected using this potentiometric method, and they refer to the partition coefficient of the neutral species.

The potentiometric log *P* measurement method used by the Sirius T3 instrument (Pion) [46–51] is based on determination of the partition profile directly from acid-base titrations in a dual-phase water-partition solvent system consisting of two liquid phases in contact (Fig. 1). In this method, multiple potentiometric acid-base titrations are performed in the aqueous phase at various equilibrium volumetric ratios of octanol and water to observe the ionization and partitioning equilibrium behavior of the analyte. As the relative volume ratio of octanol to water changes, a shift in apparent p*K*_a_ (p_o_*K*_a_) is observed due to partitioning of neutral and ionic species—which have distinct octanol-water partitioning equilibria—into the octanol-rich phase. Equations describing this coupled partitioning and ionization equilibria are then solved to determine the log *P* of the neutral and ionic species. To use this method, aqueous p*K*_a_ value(s) must be known, and analytes must be fully water soluble at the highest concentration they reach during the titrations throughout the entire range of pH titration selected for the potentiometric log *P* measurement protocol. The largest pH range selected for titration can be pH 2–12 and the minimum range should include ±2 pH units around the p*K*_a_ and p_o_*K*_a_.

**Figure 1.**
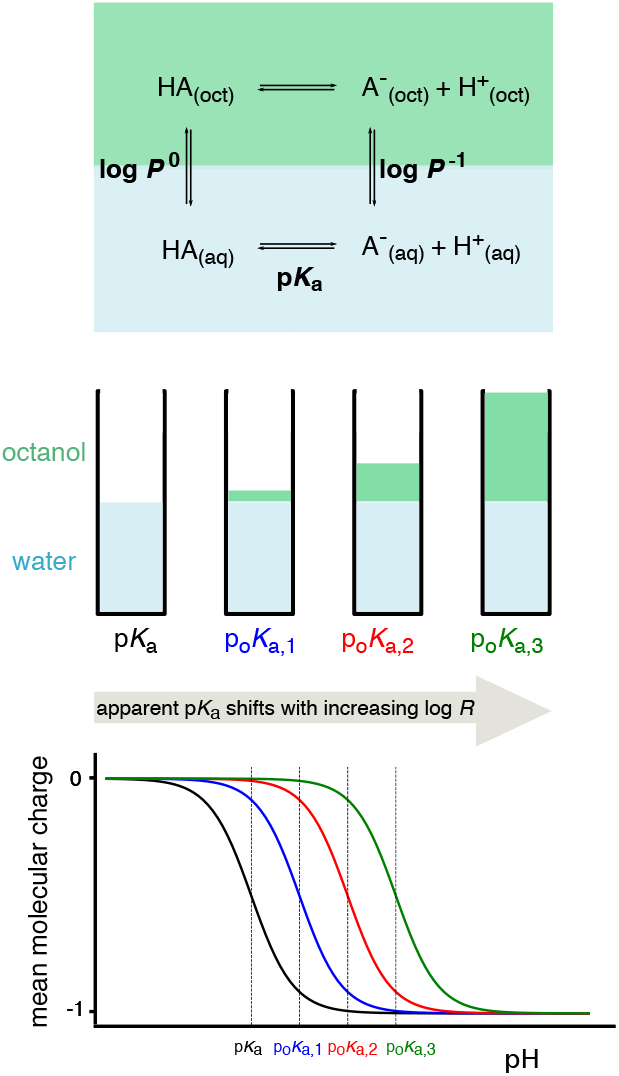
Potentiometric log *P* measurements are based on a model of ionization and partitioning equilibria [50]. Measurements of the p*K*_a_ and apparent p*K*_a_ (p_o_*K*_a_) at three octanol-water volumetric ratios (log *R*) are performed to estimate the partition coefficients of neutral and ionized species, log *P*^0^ and log *P*^-1^, respectively. An ionization and partitioning equilibria model, along with estimated potentiometric titration curves, are shown for a monoprotic acid in this figure.

When an ionizable substance is titrated in a two-phase system, the apparent p*K*_a_—here, denoted p_o_*K*_a_—observed in the titration shifts due to differential partitioning of neutral and ionized species into the nonaqueous phase. The p_o_*K*_a_ value is the apparent p*K*_a_ in the presence of partition solvent octanol. Its shift is dependent on the volumetric ratio of the water and octanol phases. The p_o_*K*_a_ value increases with increasing partition solvent volume for monoprotic acids and decreases with monoprotic bases. The shift in p_o_*K*_a_ is directly proportional to the log P of the compound and the ratio of octanol to water. For a monoprotic acid or base, the partition coefficient of neutral (*P*^0^) and ionic species (*P*^-1^, *P*^+1^) relates to p*K*_a_ and p_o_*K*_a_ as [50],

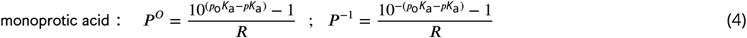

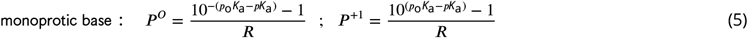

Here, ***R*** is the volume ratio of nonaqueous phase (***V***_nonaq_) to aqueous phase (***V***_aq_),

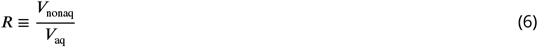

## 2 Methods

### 2.1 Compound selection and procurement

For the SAMPL6 Part II log *P* Challenge, we attempted to collect log *P* measurements for the entire set of 24 kinase inhibitor fragment-like compounds selected for the SAMPL6 p*K*_a_ Challenge [29, 52]. Details of compound selection criteria for the SAMPL6 p*K*_a_ set—driven in large part by cheminformatics filtering for experimental tractability and rapid, inexpensive compound procurement—can be found in the SAMPL6 p*K*_a_ experimental data collection paper [29]. Compounds with publicly available experimental log *P* measurements were excluded by checking the following sources: DrugBank [53], ChemSpider [54], NCI Open Database August 2006 release [55], Enhanced NCI Database Browser [56], and PubChem [57]. However, not all molecules selected for SAMPL6 were suitable for log *P* measurements using the Sirius T3, due to various reasons such as low solubility, apparent p*K*_a_ value shifting out of experimental range, or log *P* values out of experimental range limited by the sample vial. These limitations are explained in more detail in the Discussion section. Only 11 small molecules proved to be suitable for potentiometric log *P* measurements.

Molecule IDs assigned to these compounds for the SAMPL6 p*K*_a_ Challenge were preserved in the SAMPL6 Part II log *P* Challenge. A list of SAMPL6 log *P* Challenge small molecules, SMILES, and molecule IDs can be found in Table 1. Counterions, where present in solid formulations (see “Potentiometric log *P* measurements” section below), were included in SMILES for the sake of completeness, although no significant effect is expected from the presence of chloride counterions as experiments were conducted using KCl to maintain constant ionic strength. Procurement details for all compounds in the SAMPL6 log *P* Challenge compounds are presented in Table S1.

**Table 1.**
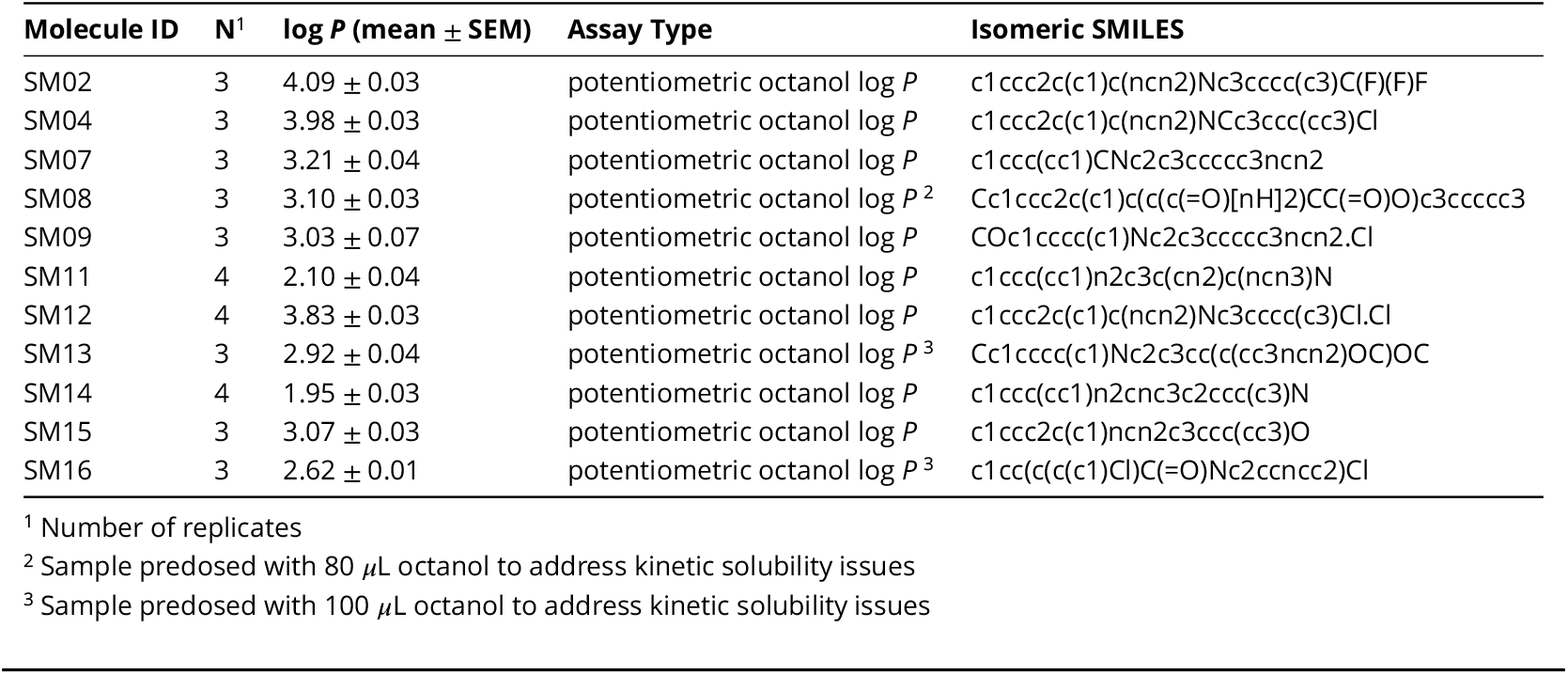
Experimental log *P* measurements for the SAMPL6 Part II log *P* Challenge. Potentiometric log *P* measurements were performed with the Sirius T3 in ISA water. Triplicate measurements were performed at 25.0 ± 0.5 and in the presence of 150 mM KCl to control ionic strength. log *P* values are reported as mean ± SEM of at least three independent replicates. log *P* values of independent replicate measurements are presented in Table S2. A computer readable form of this table can be found in the SI documents bundle (*logP_experimental_values.csv*).

### 2.2 Potentiometric log *P* measurements

Experimental octanol-water log *P* values of neutral species were collected using potentiometric log *P* (pH-metric log *P*) measurements [50] at 25.0±0.5 °C and constant ionic strength (0.15 M KCl). Aqueous p*K*_a_ values are required for log *P* determination with the Sirius T3, and were previously determined for all compounds in this set [29] using UV-metric p*K*_a_ measurements [58, 59] with the same instrument.

Three independent replicates were performed for each log *P* measurement using 1-octanol and water biphasic systems at 25.0°C, starting with solid material. General guidance of according to the instrument manual suggests optimal analyte mass should be in the range of 1–10 mg. “Sample weight” is the terminology used to describe analyte mass in Sirius T3 manuals, software, and reports. Due to solubility limitations of the SAMPL6 compounds, we tried to use analyte masses less than 3 mg. There was not much flexibility to adjust aqueous phase volume, since this is limited by the minimum volume required for the pH probe (1.4–1.5 mL) and the volume that must be spared for the octanol phase in the sample vial. Therefore, we adjusted analyte mass instead of aqueous phase volume when reducing sample concentration was necessary to achieve solubility.

For molecules with low solubility, target analyte mass was reduced, but not below a minimum of 1 mg. Samples were prepared by weighing 1–3 mg of analyte in solid powder form into Sirius T3 analysis vials using a Sartorius Analytical Balance (Model: ME235P) equipped with an antistatic ionizer. It was difficult to transfer powder compounds to achieve target masses in 1–3 mg range exactly. Instead, we opted to weigh out approximate target mass (±40% of the target mass was considered acceptable) and record the resulting sample mass. For instance, when aiming for 1 mg of compound, if 1.29 mg of compound was transferred to the balance, that was recorded as analyte mass and 1.29 mg was provided in to the Sirius T3 software for analysis. Reporting accurate analyte mass was important since analyte mass and purity are part of the Sirius T3 refinement model, although the analysis software doesn’t accept analyte purity as an input. Analyte purity (“sample concentration factor” according to Sirius T3) is estimated from the refinement model fit to experimental data given the reported analyte mass by the user. The remaining steps in sample preparation were performed by the automated titrator: addition of ionic-strength adjusted (ISA) water (typically 1.5 mL) and partition solvent (ISA water-saturated octanol), mixing, sonication,and titration with acid (0.5 M KCl) and base (0.5 M KOH) solutions targeting steps of 0.2 pH units. ISA water is 0.15 M KCl solution which was used to keep ionic strength constant during the experiment. ISA water was prepared by dissolving KCl salt in distilled water.

ISA water-saturated octanol was prepared by mixing 500 mL 1-octanol (Fisher Chemical, cat no A402-500, lot no 168525) with 26.3 mL ISA water (targeting 5% ISA water-octanol mixture by volume) and letting the mixture phases separate before attaching it to the automated titrator. Titrations were performed under argon flow on the liquid surface to minimize carbon dioxide absorption from the air.

In some cases, to help with kinetic solubility issues of the analytes, solid samples were predosed manually with 80–100 μL ISA water-saturated octanol prior to the addition of ISA water and partition solvent—these are noted in Table 1. Predosed volumes were provided to the analysis software as an input and were accounted for in the total octanol volume calculation. Whenever mean molecular charge *vs* pH plots showed experimental data points that deviated from the expected sigmoidal curve shape (oscillatory shape or steeper descent), we suspected solubility problems and attempted to prevent them by predosing octanol, which can only help the cases in which the solubility issue is a kinetic and not an equilibrium solubility issue. The only way to alleviate an equilibrium solubility issue entirely is to lower the analyte concentration by starting the experiment with a smaller analyte mass.

For each replicate log *P* measurement,three sequential automated acid-base titrations were performed in the same vial at three different volume ratios of octanol and water, using 0.5 M KOH and HCl solutions as titrants while monitoring pH with a pH electrode (Ag/AgCl double-junction reference electrode). Additional volumes of octanol were dispensed before each titration to achieve target octanol-water ratios. The sequence of three octanol-water ratios were determined using predicted log *R* profiles (apparent p*K*_a_ shift *vs* log_10_ of the volumetric ratios of partition solvents, as shown in Fig. 2C,D) or experimental log *R* profile if a previous iteration of the experiment is available during protocol optimization, with the goal of selecting three volumes that will maximize the |p*K*_a_ - p_o_*K*_a_| values between each titration. Experiments were designed so that maximum separation of p_o_*K*_a_ values can be achieved while the total liquid volume in the analysis vial did not exceed 3 mL by the end of the third titration.

**Figure 2.**
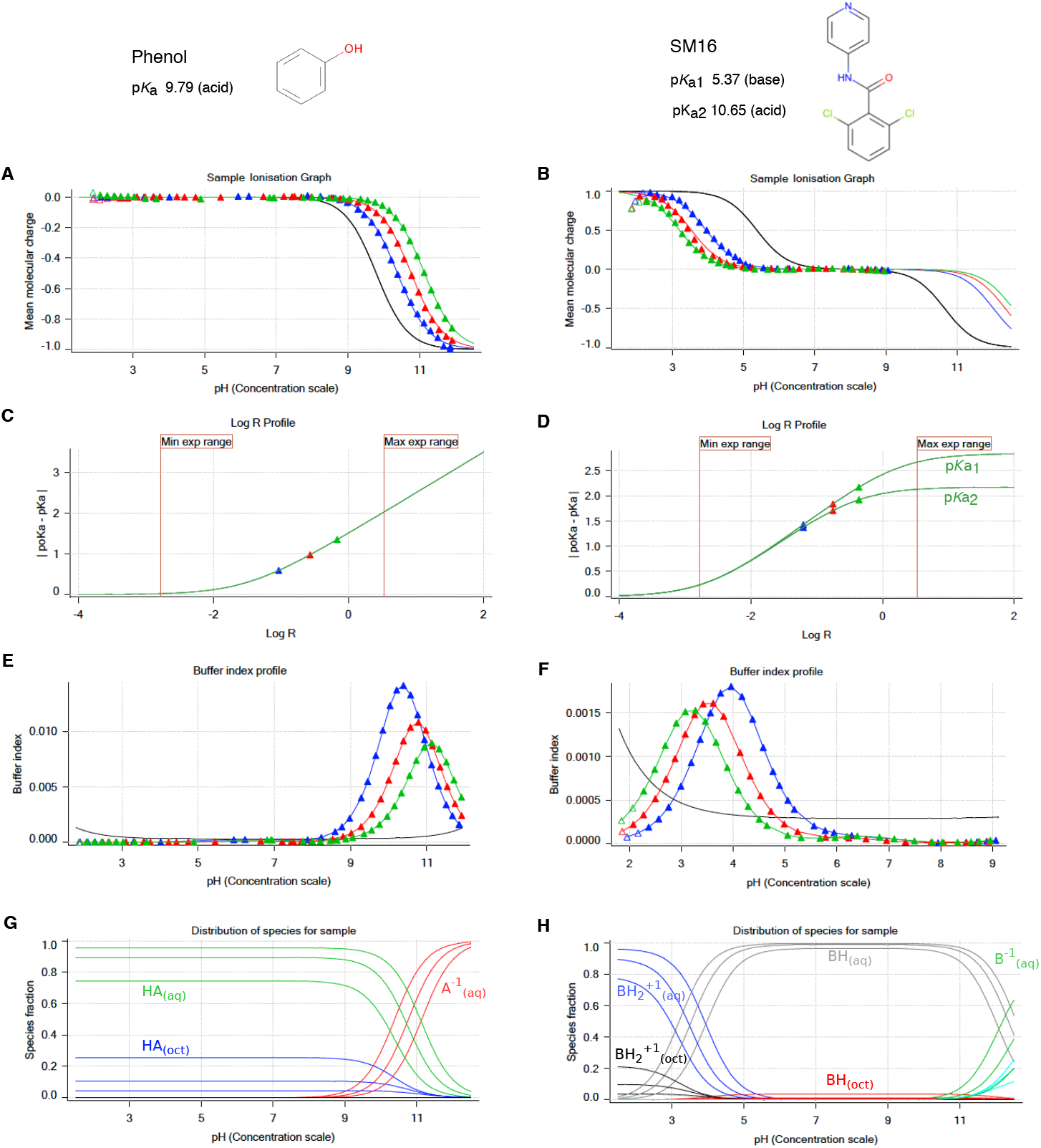
Illustrative potentiometric log *P* measurements of phenol (monoprotic, acid, log *P* 1.49) and SM16 (diprotic, amphoteric, log *P* 2.62) with the Sirius T3. Triangles represent experimental data points collected during the octanol-ISA water titrations and solid lines represent the ionization and partitioning model fit to the data. **A, B:** Computed mean molecular charge vs pH. Mean molecular charge is calculated based on experimental p*K*_a_ values and types (acid or base type) of the analyte. The black line is the model titration curve in aqueous media and based on the aqueous p*K*_a_. Blue, red, and green triangles represent three sequential titrations with increasing log *R* (increasing octanol) that show shifted p_o_*K*_a_ values. The inflection point of titration curves indicates the p*K*_a_ or p_o_*K*_a_, though these values are obtained by a global fit. For titration of acidic species, partitioning into the octanol phase increases the observed p_o_*K*_a_. In the titration of the basic p*K*_a_ of SM16, increasing log *R*causes a decrease in p_o_*K*_a_. The pH range of the experiment was determined such that only the titration of basic p*K*_a_ was captured (molecular charge between+1 and 0). **C, D:** log *R* profiles show a shift in p_o_*K*_a_ with respect to increasing relative octanol volume. These plots aid in the design of the experiment and selection of optimal octanol volumes that aim to maximize separation between p_o_*K*_a_ values for better model fit within experimental limitations (pH and analysis vial volume). **E, F:** Buffer index profiles show buffering capacity observed in three titrations with increasing log *R* (blue to green). The black line is the intrinsic buffering capacity of water. For an accurate potentiometric measurement, buffering capacity signal of the analyte must be above the buffering capacity of water. As octanol volume increases, the concentration of the analyte in aqueous phase, and thus buffering capacity, decreases. **G, H:** Predicted relative populations of ionization states in octanol and water phases as a function of pH, based on the equilibrium model fit to experimental data.

Two Sirius T3 software programs were used to execute measurement protocols (Sirius T3 Control v1.1.3.0) and analyze experiments (Sirius T3 Refine v1.1.3.0). The Sirius T3 Refine software has the capability of fitting partitioning and ionization equilibrium models to potentiometric data collected from a biphasic system to estimate log *P* values. The starting point for the model fit is simulated titration curves constructed using aqueous p*K*_a_ values (using prior p*K*_a_ measurements, here taken from [29]), predicted log *P* values, input analyte mass, and volumes of aqueous and organic phases dispensed to prepare the sample. Collected experimental measurements (pH vs dispensed volume of acid and base solutions) were used to refine the model parameters (log *P* of neutral species, log *P* of ionic species, analyte concentration factor, carbonate content, acidity error) to determine the log *P* values of neutral species and ions [48]. Potentiometric log *P* measurements have the potential to determine the partition coefficients of the ionic species (log *P*^1^) in addition to log *P* of the neutral species (log *P*^0^). It was, however, very challenging to design experiments to capture log *P* values of the ionic species due to volumetric limitations of the glass analysis vial and measurable pH range. Therefore, while optimizing experimental protocols, we prioritized the accuracy for only log *P* of the neutral species. Experimental protocols were optimized iteratively by adjusting octanol-water ratios, analyte concentration, and pH interval of the titration.

A partitioning and ionization equilibrium model [48] was fit to potentiometric measurements to estimate log *P* values of the neutral species and also the charged species, as implemented in Sirius T3 Refine Software. Experiments were optimized to determine log *P* of neutral species with good precision. log *P* estimates of charged species had high variance between replicate experiments performed in this study and were judged to be unreliable. Optimizing experiments further to be able to capture log *P* values of ionic species accurately would require larger log *R* values, which was limited by sample vial volume. Therefore, we decided not to pursue experimental data collection for ionic partition coefficients further.

### 2.3 Reporting uncertainty of log *P* measurements

Experimental uncertainties of log *P* measurements were reported as the standard error of the mean (SEM) of three or four replicates. The standard error of the mean (SEM) was estimated as

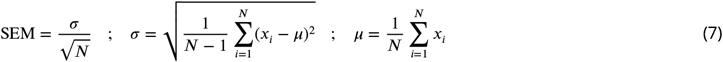

where ***σ*** denotes the unbiased sample estimator for the true standard deviation and ***μ*** denotes the sample mean. *x_i_* are observations and N is the number of observations.

The SEM calculated from independent replicate experiments as above was found to be larger than non-linear fit error reported by the Sirius T3 Refine Software from potentiometric log *P* model fit of a single experiment, thus leading us to believe that running replicate measurements and reporting mean and SEM of log *P* measurements better captured all sources of experimental uncertainty. We caution, however, that the statistical error estimated from three replicates is only determined to an order of magnitude [60].

### 2.4 Quality control of analytes

Purities of all SAMPL6 p*K*_a_ Challenge compounds—a subset of which formed the log *P* set used here—were determined by LC-MS and reported elsewhere [29]. The same lots of compounds were used for p*K*_a_ and log *P* measurements. LC-MS assessment showed that the 11 compounds reported in this study have a minimum of 96.5% purity and matching molecular weight to supplier reported values (Table S1).

When questions were raised about the accuracy of log *P* measurements for SM13 by a participant of SAMPL6 log *P* Challenge, we had additional quality control experiments performed to confirm the compound identity of SM13. LC-MS and NMR data were fully consistent with the structure of SM13 as originally provided (Figure S1, S2). High-Resolution Mass Spectrometry (HRMS)data was acquired using an Agilent 6560 Q-ToF by +ESI. NMR data were acquired for the sample dissolved in pyridine-*d*5. ^1^H, DQF-COSY, and ROESY spectra were acquired using a 600 MHz Bruker AVANCE III HD spectrometer equipped with a liquid nitrogen-cooled broadband Prodigy probe. Chemical shifts were assigned to validate the structure of SM13.

## 3 Results

In this study, we attempted to use the potentiometric log *P* measurement method of the Sirius T3 to measure log *P* values for 24 compounds of the SAMPL6 p*K*_a_ Challenge set. For 13 of the selected compounds, experimental constraints set by solubility, lipophilicity, p*K*_a_ properties of the analytes, and experiment analysis volume limitations of the Sirius T3 instrument resulted in an inability to achieve reliable log *P* measurements suitable for the blind challenge (Table S4). For example, SM24 has a basic p*K*_a_ of 2.60 and we could not optimize log *P* measurement protocol because in the presence of octanol phase apparent p*K*_a_ was shifting beyond the measurable pH range of the Sirius T3. On the other hand SM03 log *P* could not measured with potentiomentric method due to its low aqueous solubility. Only 11 of 24 compounds from the SAMPL6 p*K*_a_ Challenge set were found to be suitable for potentiometric log *P* measurements with the Sirius T3. The resulting challenge dataset presented here has a log *P* range of 1.95–4.09. Six of these represent the 4-amino quinazoline scaffold (SM02, SM04, SM07, SM09, SM12, SM13). There are two benzimidazoles (SM14, SM15), one pyrazolo[3,4-d] pyrimidine (SM11), one pyridine (SM16), and one 2-oxoquinoline (SM08) (Fig. 3). The mean and SEM of replicate log *P* measurements, SAMPL6 compound IDs (SMXX), and SMILES identifiers of these compounds are presented in Table 1. In all cases, the SEM of the log *P* measurements ranged between 0.01–0.07 log_10_ units.

**Figure 3.**
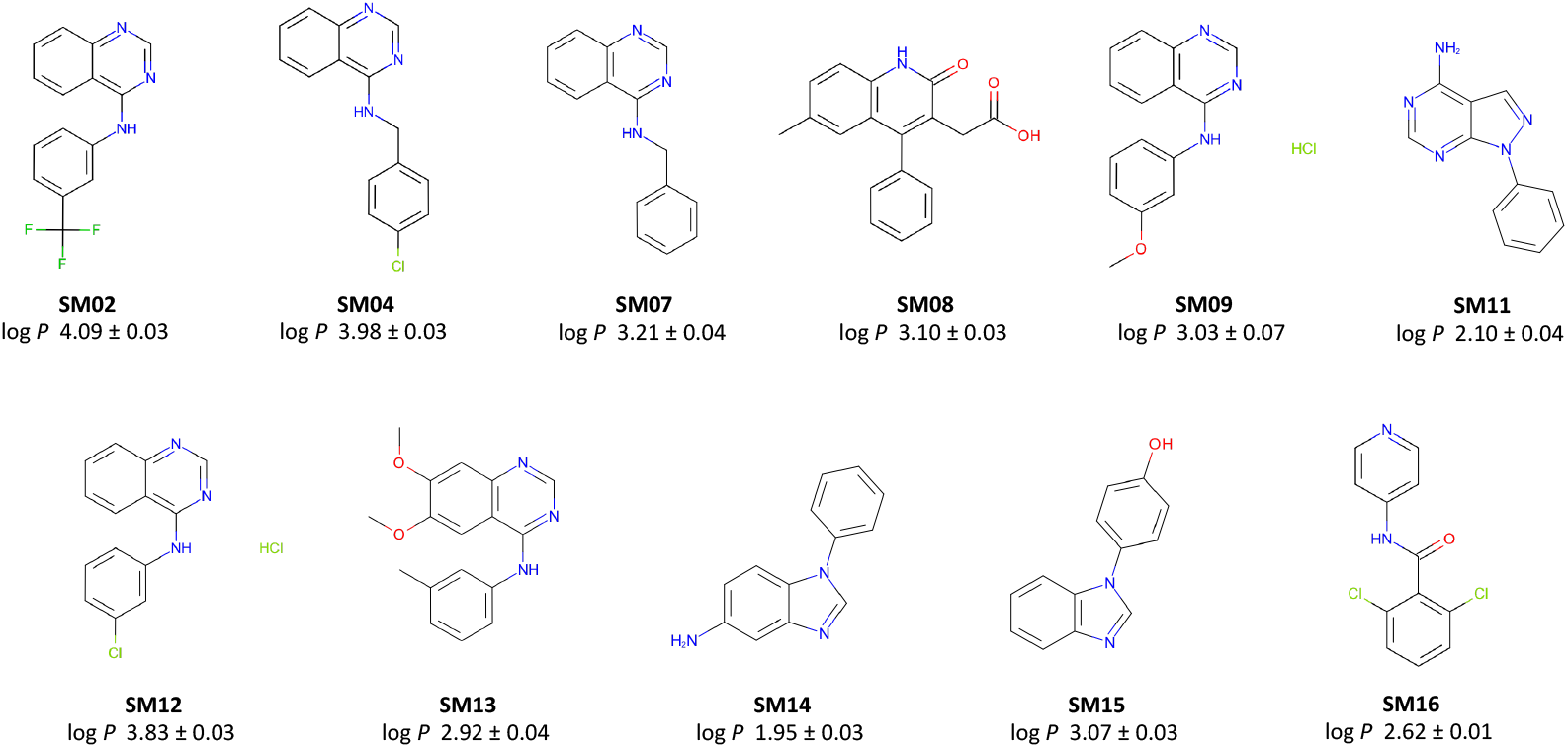
Molecules included in the SAMPL6 Part II log *P* Challenge set. Reliable experimental potentiometric log *P* measurements were collected for the 11 molecules depicted here. Reported uncertainties are expressed as the standard error of the mean (SEM) of replicate measurements. Molecules are depicted using OpenEye OEDepict Tool [61]. Canonical isomeric SMILES strings of all compounds are given in Table 1, and replicate log *P* measurements can be found in Table S2.

Results of independent replicate measurements are presented in Table S2. Preparation of each replicate sample started from weighing dry powder of the same analyte lot. The log *P* estimates from potentiometric titrations were evaluated using the partitioning and ionization equilibrium model as implemented in the Sirius T3 Refine software, which produces log *P* estimates for both neutral and ionic species. We observed that log *P* values of neutral species were highly reproducible, while variance of log *P* of ionized species between replicate experiments was high. It was also not possible to measure log *P* values of the ionized species reliably as doing so would require sampling higher log *R* values. Since it was probihitively difficult to optimize experimental protocols to capture partitioning of ionic species accurately, we optimized the experiments to prioritize accurate measurement of neutral species log *P* (log *P*^0^) and constructed the blind computational prediction challenge based on log *P*^0^ values.

## 4 Discussion

### 4.1 Dynamic range of log *P* measurements and solubility limitations

We attempted to measure the log *P* for all 24 SAMPL6 p*K*_a_ Challenge compounds, but the Sirius T3 potentiometric log *P* measurement method was able to provide reliable measurements for only a subset of 11 molecules which were included in the blind challenge. We only included molecules that yielded reliable, precise log *P* measurements in the computational blind challenge.

A number of factors restricted the ability to perform reliable log *P* measurements and led to elimination of some compounds from the initial set of 24: low water solubility within the pH range of the titration, the limited volume capacity of the glass sample vial which limits the maximum achievable octanol:water ratio, the octanol-dependent p_o_*K*_a_ values shifting outside the measurable pH range of 2–12 (especially high acidic p*K*_a_s and low basic p*K*_a_s). If an analyte does not suffer from the issues mentioned above, dynamic range of this log *P* measurement method is limited by smallest (related to dispensing accuracy and evaporation rate) and largest octanol volumes (related to analysis vial volume) that can be dispensed.

### 4.2 Optimizing experimental protocols for each compound

For the set of compounds in SAMPL6 Challenge, we observed that the Sirius T3 potentiometric log *P* measurement experiments were in practice very low throughput because of the necessary iterative protocol optimization for each compound. The parameters determining a potentiometric log *P* experiment are: mass of analyte, initial volume of ISA water, three target volumes of octanol for sequential titrations with increasing log *R*, and pH range of the pH titration. Factors that were considered in this optimization and limitations of choice are discussed below.

#### 4.2.1 Optimizing the sequence of octanol-water volumetric ratios and range of pH titration

To obtain reliable and precise log *P* estimates from experimental data, it is recommended to fit the ionization and partitioning equilibrium model to at least three potentiometric titrations with well separated p_o_*K*_a_ values (Figure 2A, B). log *P* values can also be estimated from two potentiometric titrations, but not as accurately. p_o_*K*_a_ values of sequential titrations need to be at least 0.3 p*K*_a_ units separated from one another and from the aqueous p*K*_a_. To achieve this, selecting an optimal set of octanol-water volumetric ratios is key.

It is logical to target the largest difference in octanol volumes, but the minimum volume of aqueous phase that provides enough depth for the pH probe (1.4 mL) and maximum analysis vial volume (3 mL) result in only 1.6 mL of available volume for the octanol phase, limiting the maximum octanol:water volume ratio ***R*** to ~ 1.1. Typically, one would pick octanol volumes for each of three sequential titrations that maximize the difference in p_o_*K*_a_ by maximizing the difference in log *R* values as much as possible considering the other experimental constraints. Simulated log *R* profiles based on predicted log *P* and experimental p*K*_a_ values provide guidance in the selection of octanol volumes (Fig. 2C, D). These plots show how much |p*K*_a_-p_o_*K*_a_| difference can be gained with respect to a change in log *R*, based on the titration and ionization propensity of each molecule, but they are only as useful as the accuracy of log *P* prediction. For that reason, potentiometric log *P* measurements needs to be optimized with an iterative process where the first experimental protocol is designed using predicted log *P* and experimental p*K*_a_ of the analyte. Based on the p_o_*K*_a_ shifts and quality of titration curves observed, a second experiment is designed to improve p_o_*K*_a_ shifts by adjusting the octanol volumes after consulting the log *R* profile and using the estimated log *P* from the previous experiment as a guide. Sometimes 3 or 4 iterations were necessary to reach an optimal protocol that results in a good fit between predicted and experimental titration curves and produces reproducible log *P* estimates. An example protocol optimization for SM02 guided by log *R* values is shown in Fig. 4.

**Figure 4.**
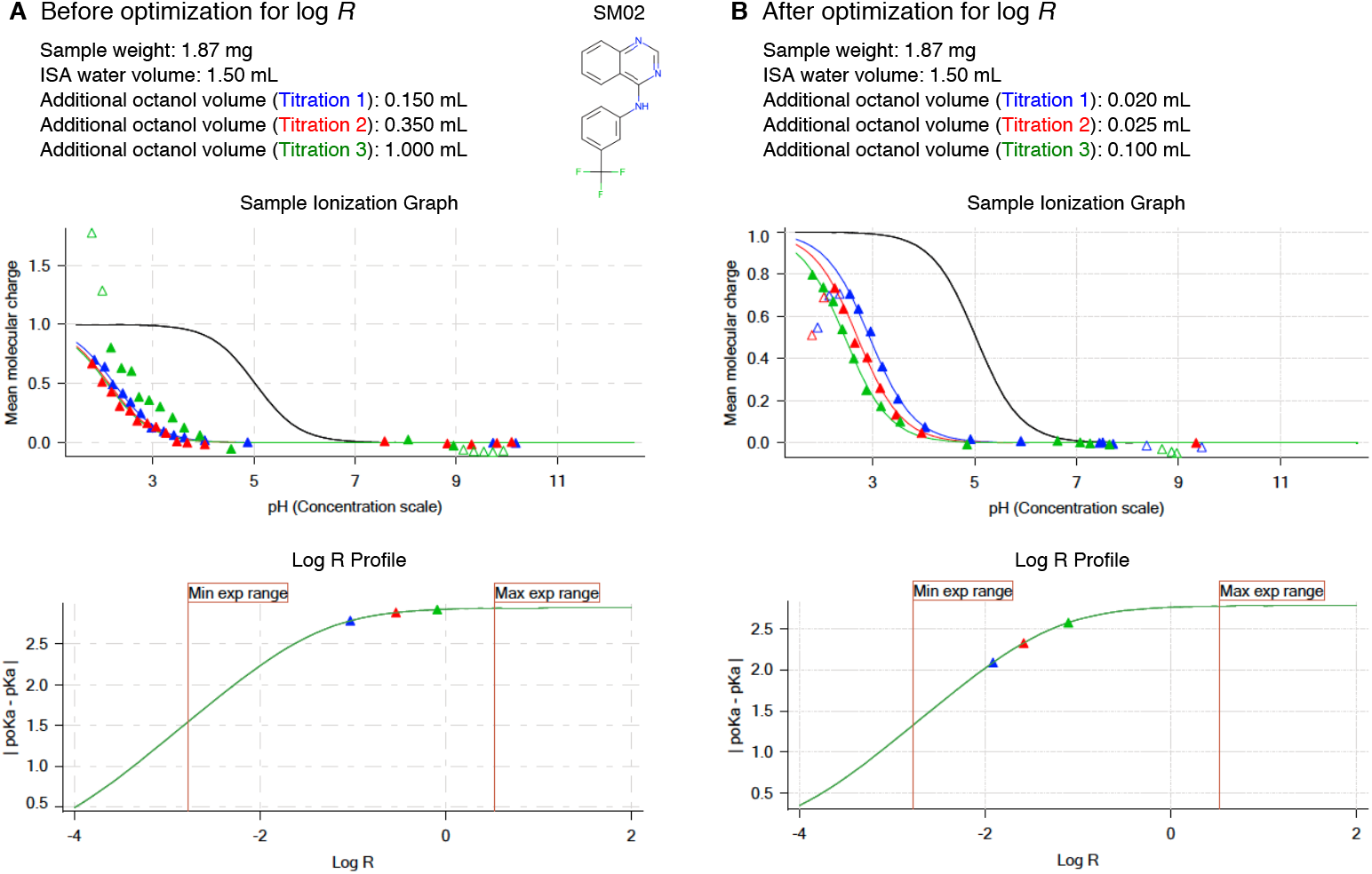
Potentiometric log *P* protocol optimization of SM02 based on log *R*. Experimental results of initial trial (**A**) and optimized protocol (**B**) are shown for SM02. log *R* profile before optimization (**A**lower panel) shows insufficient apparent *pK_a_* shift due to poor choice of octanol-water volume ratios. This experiment led to log *P* and log *P*^+1^ estimates of 4.32 and 1.38. For a good measurement, triangles that indicate |p*K*_a_-p_o_*K*_a_| of each titration in log *R* profiles must fall on the slope region of the log *R* profile instead of the plateau region. Adjusting log *R* by decreasing octanol volumes in each titration led to a better experiment with distinct titration curves and well separated p_o_*K*_a_values (**B**). log *P* and log *P*^+1^ were measuredas4.10and1.32withtheoptimizedprotocol. Once we achieved optimization of potentiometric log *P* protocol, triplicate measurements were collected using the same protocol.

While maximizing the difference in p_o_*K*_a_ values from each other and from the aqueous p*K*_a_ is desirable, sometimes it is necessary to reduce the octanol volume to limit the shift in p_o_*K*_a_ so that itremains within a measurable range. This would be necessary when the aqueous p*K*_a_ is a weak acid (p*K*_a_>9) or weak base (p*K*_a_<5),since the presence of the octanol phase causes p_o_*K*_a_ shifts towards higher and lower values, respectively, approaching the limit of the measurable pH range of the instrument. Measurable pH range is mainly limited by the acid and base strength of titration solutions against the increasing buffering capacity of water at pH values below 2 and above 12. It is also important to mention that even if the p_o_*K*_a_ value itself is within the stated measurement range of 2–12, if a large portion of the titration curve is beyond limits (i.e., saturation of fractional population on both sides of the p_o_*K*_a_), then the experimental titration curve may not be fit to the model titration curve exactly and p_o_*K*_a_ cannot be determined as precisely. When the dynamic part of the titration curve (p_o_*K*_a_ ± 2) shifts outside of the measureable pH range, it reduces the confidence in p_o_*K*_a_ estimates of the fit. Therefore, p_o_*K*_a_ values should ideally be at least ± 1 unit, and preferably ± 2 units away from the limits of pH measurement with this instrument, which can be extended to pH 1 and 13 at most. For this reason, it is easier to optimize log *P* experiments for monoprotic molecules which have acidic p*K*_a_s between 3–10 and basic p*K*_a_s between 4–11. Some molecules in the SAMPL6 set which were not suitable for potentiometric log *P* measurements because of this criteria were: SM01, SM17, SM18, SM19, and SM24 (Table S4).

#### 4.2.2 Sample preparation considerations and determination of appropriate starting concentration

Sample preparation starts with the weighing of solid powder material to analysis vials. How much analyte to use is another important decision that requires optimization. General guidance according to the Sirius T3 manual is to use 1–10 mg, and the aqueous phase volume is typically adjusted to the minimum volume (1.4–1.5 mL). The buffering capacity and compound solubility are the two factors that guide lower and higher limits of suitable analyte concentration. The Sirius T3 produces buffer index vs pH plots (Fig. 2E, D) which provide guidance on how much analyte is needed for sufficient potentiometric signal. To guide the first experiment, these plots can be simulated based on analyte mass, experimental p*K*_a_, predicted log *P*, and selected octanol volumes. In further iterations of experiments, the buffer index profiles of the previous experiment guides the decisions about how to optimize the protocol. On the other hand, aqueous solubility limits the maximum concentration of the analyte in the aqueous phase. Moreover, since the experimental methodology depends on measuring the p_o_*K*_a_ shift during pH titrations as species partition into the nonaqueous phase, the analyte must stay in solution over the titrated pH range for the entire experiment, as the presence of an insoluble phase represents another reservoir for compound partitioning that would invalidate the coupled ionization-and-partitioning model used to compute the log *P*. The pH titration range is adjusted to capture a sufficient region below and above the p_o_*K*_a_ to ensure ionization states with lower solubility are also visited (neutral and zwitterionic states).

For these compounds resembling fragments of kinase inhibitors–the compounds considered in the SAMPL6 p*K*_a_ Challenge [29] and this study–this solubility criterion turned out to be very challenging to meet. A large portion of compounds in the SAMPL6 p*K*_a_ Challenge set were found to be insufficiently soluble for potentiometric log *P* experiments at some region of the pH range that needs to be titrated during the experiment, more likely the pH region where the neutral population of analytes are prominent. These compounds for which potentiometric log *P* measurement could not be optimized due to solubility limitations are listed in Table S4. For other compounds, we had to try reducing the analyte sample quantity from 3 mg to 1 mg of compound to find the optimum balance between ensuring the compound remained fully soluble and ensuring sufficiently high buffering capacity signal. The rate of change of pH vs. volume of acid or base titrated in analyte solution must differ from the rate of pH change in just water. This quantity is expressed as a buffering index in buffer index profiles generated by Sirius T3 (Fig. 2E, F), where a black solid line describes the theoretical buffering capacity of water and colored triangles describe the experimental buffering capacity of the analyte. For high quality measurements, reaching at least 0.001 buffer index at the maximum point of the titration (at pH that equals p_o_*K*_a_) is recommended.

In our case, the exact solubility of compounds was not known prior to log *P* measurements. We had to evaluate precipitation issues based on the distortion of mean molecular charge vs pH profiles (Fig. 2E, D) from ideal shape by adjusting starting analyte masses until the distortions disappear. Distortions manifest as very steep drops or oscillations in relative ionization state populations with respect to pH. An example is shown in Fig 5A Sample Ionization Graph. The turbidity indicator of Sirius T3 can not be utilized for solubility detection during log *P* experiments since the presence of octanol causes turbidity in the aqueous phase due to vigorous stirring during titrations. Predosing 80-100 ***μ***L octanol before addition of ISA water, as well as sonication and stirring after titrant addition, were also helpful for overcoming kinetic solubility problems. An example protocol optimization for SM08 to overcome solubility problems is shown in Fig 5.

**Figure 5.**
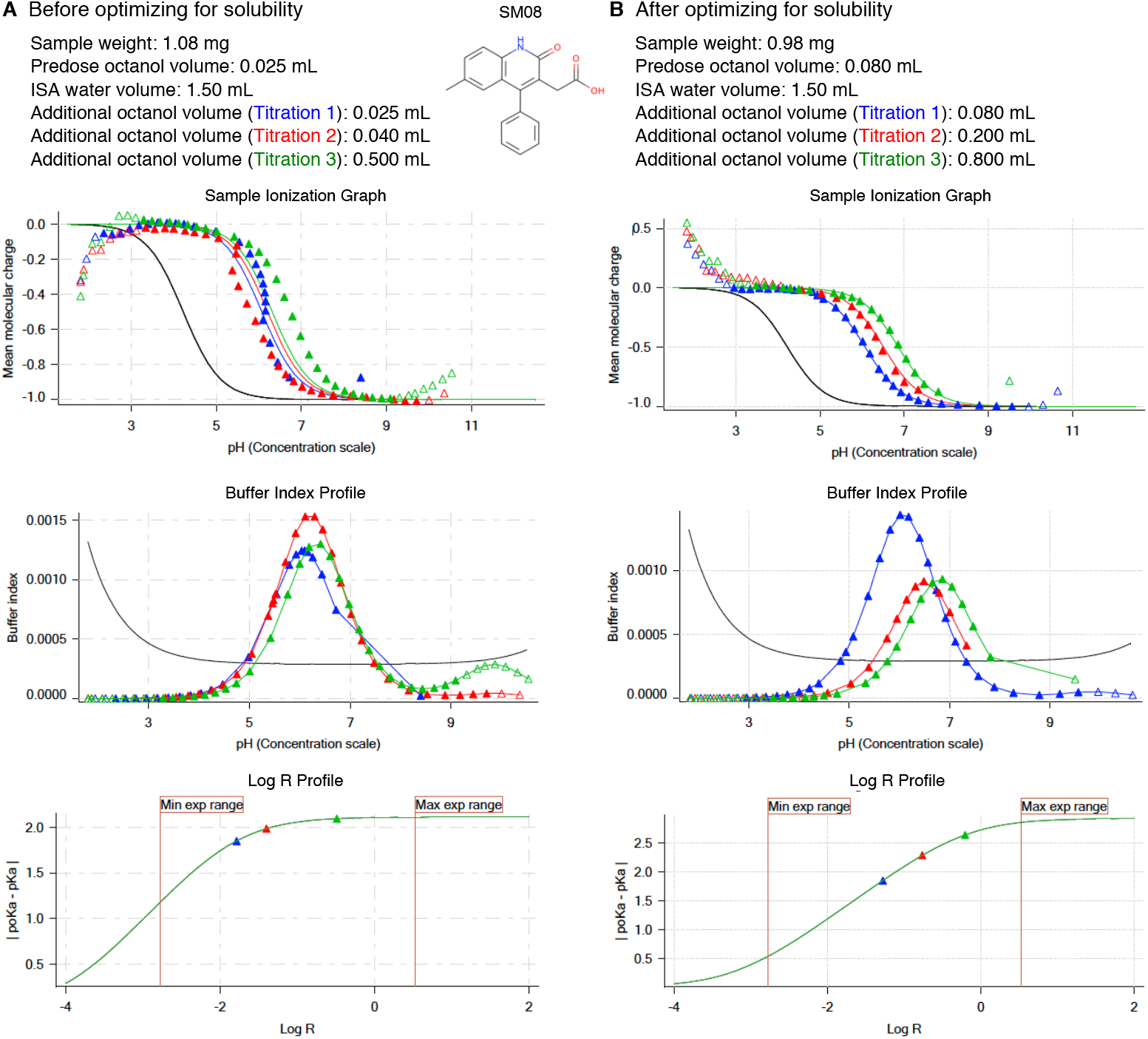
Potentiometric log *P* protocol optimization for SM08to alleviate aqueous solubility problems. Experimental results of initial protocol (**A**) and the optimized protocol (**B**) are shown for SM08. Both first (blue) and second (red) titrations in Sample Ionization Graph before optimization (**A** lower panel) show deviation from expected sigmoidal shape which is an indication ofan insoluble analyte. This experiment with solubility issuesled to log *P* and log*P*^+1^ estimates of 3.97 and 1.86. To eliminate precipitation, we could not lower analyte mass below 1 mg. Instead we were able to optimize the experimental protocol by increasing the predosed octanol volume and increasing additional octanol volumes added in each titration. Predosing octanol helps only with kinetic solubility issues. Larger octanol volumes can help to improve the experiment when thermo dynamic solubility is the limitation, by allowing larger amounts of analyte partitioning into the octanol phase and reducing the analyte concentration in aqueous phase. Additional octanol volumes were selected such that they would also improve log *R* profile of the measurement. With optimized protocol (**B**) we achieved sample ionization profiles without any precipitation effects. log *P* and log *P*^+1^ were measured as 3.16 and 0.23. Once we achieved optimization of potentiometric log *P* protocol, triplicate measurements were collected using the same protocol.

If possible, measuring solubility of compounds prior to potentiometric log *P* measurements can provide helpful information for more efficient log *P* measurement protocol optimization. However, since solubility is pH-dependent, the lowest solubility of the compound during the entire pH 2-12 range would be the information necessary to guide the experimental design. An experiment for a compound with 400 g/mol molecular weight using the minimum analysis made of 1 mg and 1.5 mL of aqueous phase corresponds to 1.67 mM. To be suitable for potentiometric log *P* measurements with the Sirius T3, at least 1.67 mM aqueous solubility is necessary throughout the pH range of the analysis.

One way to increase the dynamic range of potentiometric log *P* measurement with the Sirius T3 is to increase the range of log *R* that can be sampled by performing three different p_o_*K*_a_ measurements in three different analysis vials instead of three sequential titrations in one vial. But since log *R* is dependent on the cumulative octanol volume in sequential titrations, the advantage of the single titration approach is not significant. The single titration approach can only allow a small additional volume for octanol phase which would be used to dispense multiple acid and base stock solution volumes (~0.2 mL). We did not elect to investigate this design because we did not want to introduce another source of error: the variance in sample mass between measurements. Since the initial sample mass is an input parameter to the experimental model, using three different sample masses would introduce effects coming from the inaccuracy of the analytical balance to log *P* estimates.

Another way to prepare analyte samples for Sirius T3 measurements is to start from DMSO stock solutions instead of dry powder stocks. However, potentiometric measurements require 1-10 mg/mL analyte concentration in order to reach sufficient buffering capacity. The required concentration of the DMSO stock solution would be quite high, and sometimes impossible due to solubility limits in DMSO. Typical DMSO stock solution concentrations are 10 mM. For an analyte with 400 g/mol molecular weight, the concentration of 10 mM DMSO stock solution corresponds to 4 mg/mL. In order to achieve the minimum required 1 mg/mL analyte solution for the Sirius T3 experiment, the aqueous phase would have to consist of 25% DMSO which would cause significant cosolvent effects. On the other hand, achieving lower cosolvent presence, such as 2.5% DMSO, would require DMSO stock solutions of 100 mM at which concentration the analyte may not be soluble. Presence of cosolvent at even low amounts is undesirable due to the potential the effect on the log *P* measurements. Therefore, it is not recommended to perform these experiments starting from DMSO stock solutions.

### 4.3 Reliable determination of log *P* values of ionized species was not possible

Although it is possible to use Sirius T3 potentiometric log *P* measurements to determine the partition coefficients of ionic species as well, in practice, we were not able to achieve log *P*^1^ estimates with low variance between experiments. The partitioning of ionic species into the organic phase is typically much lower than that of the neutral species, and to capture this accurately by measuring sufficiently large p_o_*K*_a_ shifts, it would be necessary to use much larger octanol to water volumetric ratios ***R***. The Sirius T3 glass analyte vials can hold up to 3 mL, which limits the maximum achievable octanol to water volumetric ratio. Since at least 1.4 mL must be devoted to the aqueous phase for the pH probe, this leaves only 1.6 mL for the octanol phase, producing a maximum achievable ***R*** ~ 1.1. Another limitation was the measureable pH range. Since log *P* measurements rely on determining well-separated p_o_*K*_a_ values at different log *R* values to get a good model fit, the octanol to water volumetric ratio needs to be selected such that p_o_*K*_a_ values are well separated but not out of the measurable pH range (2-12).

To capture the partitioning of ionic species to the octanol layer reliably, experiments need to be set up with larger log *R* ratios which is problematic if this causes p_o_*K*_a_ to shift outside of the measureable pH range. Therefore, we designed the experiments to capture only the partition coefficient of the neutral species (log *P*^0^) accurately. The SAMPL6 log *P* Prediction Challenge was constructed only on prediction of neutral species.

The lack of reliable determination of partition coefficient values of the ionic species (log *P*^+1^ or log *P*^-1^) may be a source of systematic error in the estimate of log *P* of the neutral species (log *P*^0^). For hydrophobic compounds with negligible partitioning of the ionic species into the octanol-rich phase (log *P*^+1^, log *P*^−1^ ≤ 2), log *P*^0^ estimates would still be accurate even if ion partitioning is not captured well. For compounds that may have higher levels of ionic partitioning, to minimize the impact of inaccurate log *P*^+1^ or log *P*^-1^ experimental estimates on log *P*^0^ measurements, we used ACD/Labs predicted log *P*^+1^ and log *P*^-1^ values as the starting point for the refinement of the ionization and partitioning equilibrium model parameters (performed with Sirius T3 Refine Software).

### 4.4 Absence of zwitterions allowed accurate log *P* measurements of amphoteric molecules

Multiple publications point out discrepancies between log *P* values determined by the potentiometric method and the shake-flask experiments for zwitterionic compounds [62, 63]. There are multiprotic compounds in the SAMPL6 dataset (SM14, SM15, and SM16), but we believe these measurements were not affected by this problem because they are not zwitterionic. Zwitterionic molecules have a zwitterion as the dominant neutral state in the pH region between the two p*K*_a_s (a lower acidic p*K*_a_ and a higher basic p*K*_a_). SM14 has two basic p*K*_a_s and is not found as a zwitterion at any pH between 2-12. SM15 and SM16 are amphoteric compounds that possess both acidic and basic titratable groups, however, according to spectrophotometric p*K*_a_ measurements in the presence of cosolvent their acidic p*K*_a_ values are higher than their basic p*K*_a_ values. This means the major neutral form of these compound is the non-charged state, not a zwitterion. Spectrophotometric p*K*_a_ measurements with varying percentage of methanol as cosolvent were performed with the Sirius T3 and included in supplementary documents. Acidic or basic character of macroscopic p*K*_a_ values was assigned based on the slope of Yasuda-Shedlovsky plots.

In addition, quantum mechanics calculations [64] do not predict the presence of multiple tautomers of the neutral state at significant populations for any of the molecules in the SAMPL6 log *P* Challenge set. Possible tautomers, such as the zwitterionic state, are predicted to be much higher in energy and thus unlikely to play a significant role even if we considered a prediction error margin for quantum mechanics-based calculations. Therefore, we do not think our potentiometric log *P* measurements are influenced by presence of zwitterions or minor tautomeric forms.

### 4.5 Suggestions for future log *P* data collection

High quality datasets of experimental physicochemical property measurements are valuable for testing computational predictions. Benchmarking and evaluation efforts like the SAMPL challenges benefit from large experimental datasets with diverse chemical species. The quality of log *P* measurements collected with the Sirius T3 potentiometric method are satisfactory and comparable to gold standard shake flask measurements [45, 49, 51]. The Sirius T3 potentiometric log *P* method requires aqueous p*K*_a_s to be measured experimentally ahead of time. The ability to obtain log *P* measurements of neutral and charged species separately, instead of measuring pH dependent log *D*, is a unique advantage of the Sirius T3 approach compared to shake-flask or HPLC-based methods where ionization effects are involved with partitioning behaviour. However, due to previously discussed limitations and the necessity for extensive protocol optimization for each analyte, we are reluctant to suggest potentiometric log *P* measurements with the Sirius T3 as a general and high-throughput method for future log *P* data collection unless significant resources and work hours of a human expert can be dedicated to protocol optimization and data collection.

Informed selection of analytes can help improve the success of Sirius T3 experiments. For example, this approach is easier to apply to highly soluble compounds (more than 1 mg/ml solubulity in 0.15 M KCl through the entire range of pH titration range at room temperature) with p*K*_a_ values in the midrange (3<acidic p*K*_a_<10 and 4<basic p*K*_a_<11). There is no significant difference in difficulty between the measurements of monoprotic vs multiprotic compounds, as long as one of the p*K*_a_ values of the multiprotic compound is in the midrange. For determining the log *P* of neutral species, it is sufficient to collect potentiometric titration data between the neutral state and the +1 or −1 charged states by titrating the pH region that captures the relevant p_o_*K*_a_ values. It is not necessary to capture the titration of a second p*K*_a_ (Fig. 2B).

Our opinion is that log *D* measurements at a buffered pH can be much more easily obtained in a higher throughput fashion using miniaturized shake-flask measurements, such as those used in SAMPL5 log *D* Challenge experimental data collection [21]. To obtain log *P* values from experiments that were designed to measure log *D*s, it is necessary to measure the p*K*_a_ of compounds (such as with the Sirius T3) and conduct log *D* measurements using a buffered aqueous phase at a pH that will ensure that the analyte is completely in the neutral state. According to our experience, optimizing p*K*_a_ measurements with the Sirius T3 is significantly easier than optimizing log *P* measurements, especially if a spectrophotometric (UV-metric) p*K*_a_ method can be used instead of potentiometric, which is not an option for log *P* measurements.

## 5 Conclusion

This study reports the collection of experimental data for the SAMPL6 Part II log *P* Blind Prediction Challenge. In the physico-chemical property prediction challenge components of SAMPL6, we aimed to separately evaluate performance of computational methods for predicting ionization (p*K*_a_)and nonaqeuous partitioning(log *P*)of small molecules, collecting experimental data for these properties on the same set of compounds and fielding sequential, independent prediction challenges. While we attempted to measure octanol-water log *P* for all compounds in the SAMPL6 p*K*_a_ Challenge set—consisting of 24 compounds that resemble fragment so of kinase inhibitors— experimental limitations of the Sirius T3 potentiometric log *P* method meant that reliable log *P* measurement scould only be performed for 11 of these compounds. The resulting compound set had meaured log *P* values in the range of 1.95-4.09. This set included six molecules with 4-aminoquinazoline scaffolds, and two molecules with benzimidazole scaffolds. Although the chemical diversity and number of compounds was rather limited, blind high-quality log *P* datasets are rare, and still highly valuable for evaluating the performance of computational predictions. Therefore, the SAMPL6 Part II log *P* Blind Prediction Challenge was held between November 1, 2018 and March 22, 2019 using the log *P* measurements presented in this paper. This data set can be utilized as part of a benchmark set for the assessment of future log *P* predictions methods.

## 0.2 Abbreviations

SAMPL: Statistical Assessment of the Modeling of Proteins and Ligands
log *P*: log_10_ of the organic solvent-water partition coefficient (***K_ow_***, refers to partition of neutral species unless stated otherwise)
log *D*: log_10_ of organic solvent-water distribution coefficient (***D_ow_***)
log *R*: log_10_ of the volumetric ratios of partition solvents (octanol to water)
p*K*_a_: −log_10_ of the acid dissociation equilibrium constant
p_o_*K*_a_: −log_10_ apparent acid dissociation equilibrium constant in octanol-water biphasic system
ISA: Ionic-strength adjusted solution with 0.15 M KCl
SEM: Standard error of the mean
LC-MS: Liquid chromatography-mass spectrometry
NMR: Nuclear magnetic resonance spectroscopy
HRMS: High-resolution mass spectrometry
octanol: 1-octanol, also known as n-octanol

## 6 Code and data availability

All SAMPL6 log *P* Challenge instructions, submissions, experimental data and analysisare available at https://github.com/samplchallenges/SAMPL6/tree/master/physical_properties/logP

## 7 Overview of Supplementary Information

### Supplementary tables and figures appearing in the Supplementary Information document

- **Table S1:** Procurement details of SAMPL6 Part II log *P* Challenge compounds.
- **Table S2:** Replicate potentiometric log *P* measurements performed with Sirius T3 for octanol and ISA water biphasic system.
- **Table S3:** SMILES and InChI identifiers of SAMPL6 log *P* Challenge molecules.
- **Table S4:** Molecules from the SAMPL6 p*K*_a_ Challenge not included in the SAMPL6 log *P* Challenge.
- **Figure S1:** HRMS determination of SM13 molecular weight.
- **Figure S2:** NMR determination of SM13 structure.

### Additional supplementary files

*logP_experiment_reports.zip* file includes:

- Sirius T3 reports for all measurements in PDF and T3 formats: log *P* measurements of SAMPL6 log *P* Challenge set and cosolvent p*K*_a_ measurements of SM14, SM15, and SM16.
- Table 1 in CSV format (*logP_experimental_values.csv*)
- Table S2 in CSV format (*replicate-logP-measurements-table.csv*)
- Table S3 in CSV format (*chemical-identifiers-table.csv*)
- Table S4 in CSV format (*molecules_with_unsuccessful_logP_measurements.csv*)

## 8 Author Contributions

Conceptualization, MI, JDC, TR, DLM; Methodology, MI, DL; Software, MI; Formal Analysis, MI; Investigation, MI, DL; Resources, TR, DL; Data Curation, MI; Writing-Original Draft, MI, DL; Writing - Review and Editing, MI, DL, JDC, TR, DLM; Visualization, MI; Supervision, JDC, TR, DLM; Project Administration, MI; Funding Acquisition, DLM, JDC, TR, MI.

## 9 Acknowledgments

MI and JDC acknowledge support from the Sloan Kettering Institute. JDC acknowledges partial support from NIH grant P30 CA008748. MI, JDC, ASR, and DLM gratefully acknowledge support from NIH grant R01GM124270 supporting the SAMPL Blind Challenges. MI acknowledges support from a DorisJ. Hutchinson Fellowship. DLM appreciates financial support from the National Institutes of Health (1R01GM108889-01), the National Science Foundation (CHE 1352608). The authors are extremely grateful for the assistance and support from the MRL Preformulations and NMR Structure Elucidation groups for materials, expertise, and instrument time, without which this SAMPL challenge would not have been possible. The authors would like to thank Ryan Cohen from the NMR Structure Elucidation group for the NMR and LC-MS analysis of SM13. MI and DL are grateful to Pion/Sirius Analytical for their technical support in the planning and execution of this study. We are especially thankful to Karl Box (Sirius Analytical) for the guidance on optimization and interpretation of log *P* measurements with the Sirius T3. We thank Brad Sherborne (MRL; ORCID:0000-0002-0037-3427) for his valuable insights at the conception of the log *P* challenge and connecting us with TR and DL who were able to provide resources for experimental measurements. We acknowledge contributions from Caitlin Bannan who provided feedback on experimental data collection and structure of log *P* challenge from a computational chemist’s perspective. MI and JDC are grateful to OpenEye Scientific for providing a free academic software license for use in this work. The content is solely the responsibility of the authors and does not necessarily represent the official views of the National Institutes of Health. We thank anonymous reviewers for their input and constructive comments that improved this manuscript.

Research reported in this publication was supported by National Institute for General Medical Sciences of the National Institutes of Health under award number R01GM124270 and R01GM108889, and from the National Cancer Institute of the National Institutes of Health under P30CA008748.

## 10 Disclaimers

The content is solely the responsibility of the authors and does not necessarily represent the official views of the National Institutes of Health.

## Supplementary Information

**Table S1.** Procurement details of SAMPL6 Part II Octanol-Water Partition Coefficient Challenge compounds. ^1^Purities for these compounds were determined byLC-MS methods and reported elsewhere [29]

| **SAMPL6 Part II**<br>**Molecule ID** | **Supplier** | **Lot no** | **Cat no** | **Supplier**<br>**Reported**<br>**Purity** | **Purity**<br>**Detected**<br>**by LC-MS** | **CAS no** | **eMolecules ID** |
| --- | --- | --- | --- | --- | --- | --- | --- |
| SM02 | ChemDiv | CM02432403 | 3232-0333 |  | 97.62% |  | 1327907 |
| SM04 | ChemDiv |  | Z126957826 |  | 99.88% |  | 30719859 |
| SM07 | Enamine | 2017-0168841 | Z57161635 | 95% | 99.38% | 100818-54-0 | 1327878 |
| SM08 | Enamine | 2017-0168838 | Z57157353 | 95% | 99.24% | 65418_08_8 | 1367649 |
| SM09 | Enamine | 2017_0168839 | Z220564816 | 95% | 98.97% |  | 1865544 |
| SM11 | Maybridge | 142989 | RJC00689SC | 90% | 98.02% | 5334-30-5 | 719540 |
| SM12 | Maybridge | 265423 | DP00818SC |  | 98.13% |  | 1859493 |
| SM13 | Maybridge | 248841 | GK03474SC |  | 97.51% |  | 5828805 |
| SM14 | Enamine |  | Z57290870 |  | 96.48% |  | 31653344 |
| SM15 | Enamine |  | Z1318268952 |  | 98.70% |  | 37095168 |
| SM16 | VitaScreen |  | STK098832 |  | 97.73% |  | 1284691 |

**Table S2.**
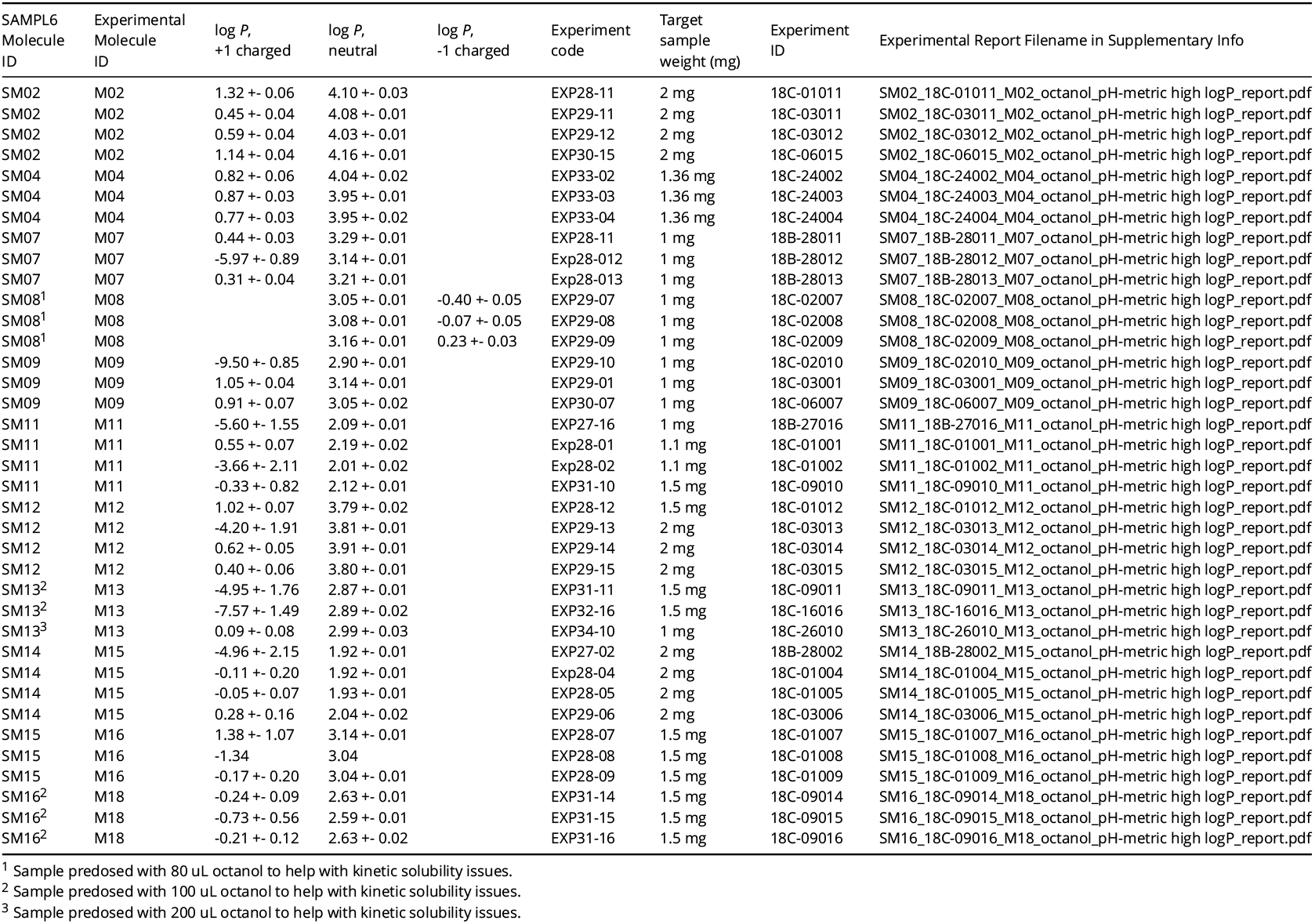
Replicate potentiometric log *P* measurements performed with Sirius T3 for octanol and ISA water biphasic system. Three or four independent replicate experiments were performed starting from powder samples. Measurements were performed at 25.0 ± 0.5°C and in the presence of approximately 150 mM KCl to adjust ionic strength. A partitioning and ionization equilibrium model was fit to potentiometric measurements to estimate log *P* values of the neutral species and also the charged species. Experiments were optimizied to be able to determine log *P* of neutral species with good precision. log *P* estimates of charged species have high variance between replicates and is unreliable.

**Table S3.**
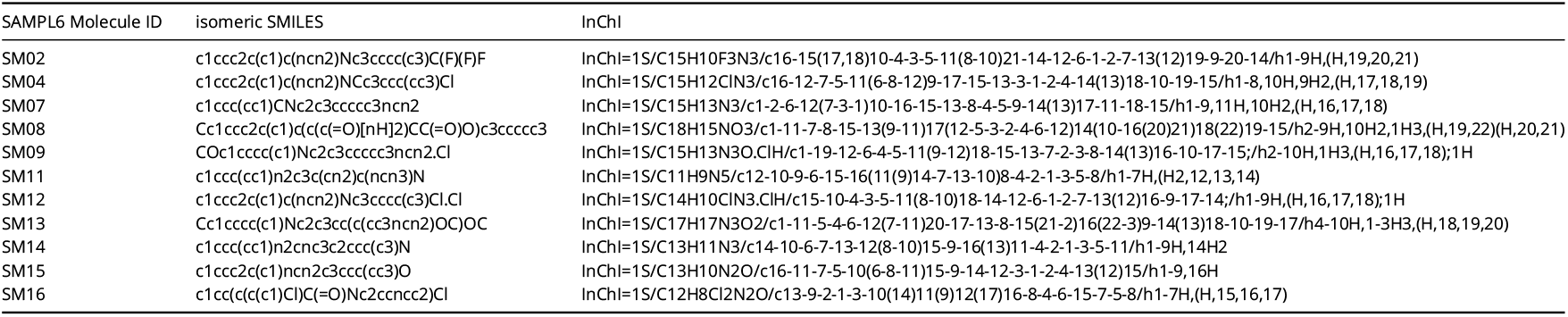
SMILES and InChı identifiers of SAMPL6 log *P* Challenge molecules.

**Table S4.**
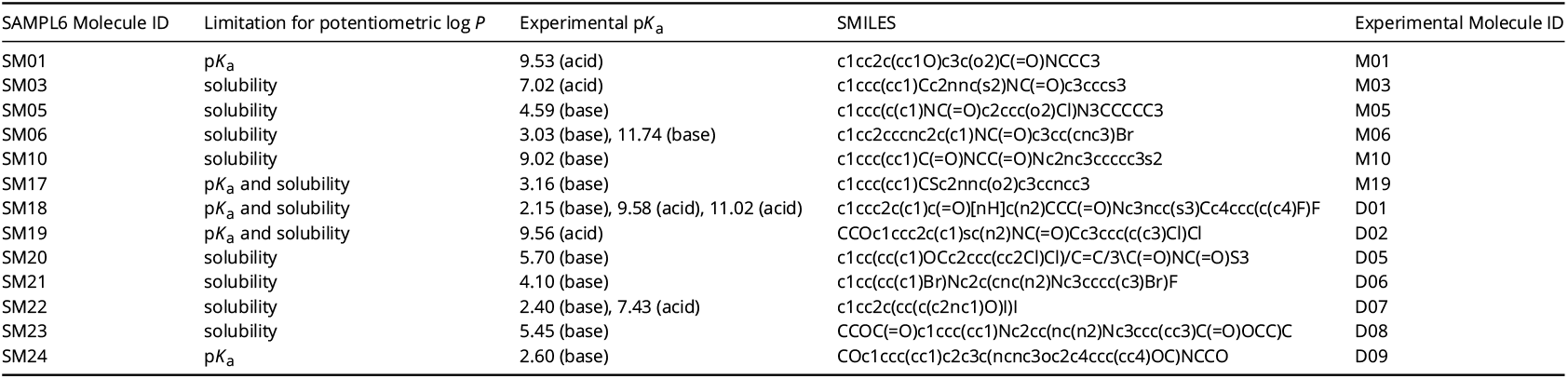
Molecules from SAMPL6 p*K*_a_ Challenge not included in SAMPL6 log *P* Challenge. These are molecules for which potentiometric log *P* experiments could not be optimized. Suspected reasons why good log *P* measurements could not be collected for these molecules are listed in the “Limitation for potentiometric log *P*” column. Limitation of p*K*_a_ value indicates that apparent p*K*_a_ shifts outside of measureable range in the presence of the octanol phase. Solubility limitation indicates that we could not find a potentiometric log *P* protocol that can avoid precipitation issues. Experimental p*K*_a_ values were originally reported elsewhere [29].

**Figure S1.**
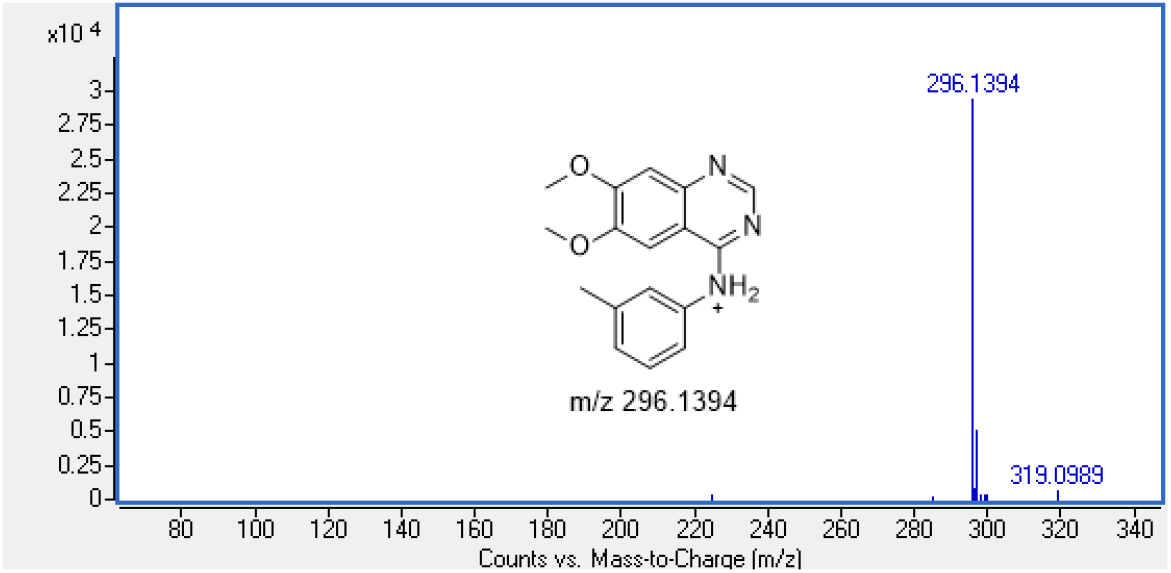
HRMS determination of SM13 molecular weight confirmed the supplier reported molecular weight.

**Figure S2.**
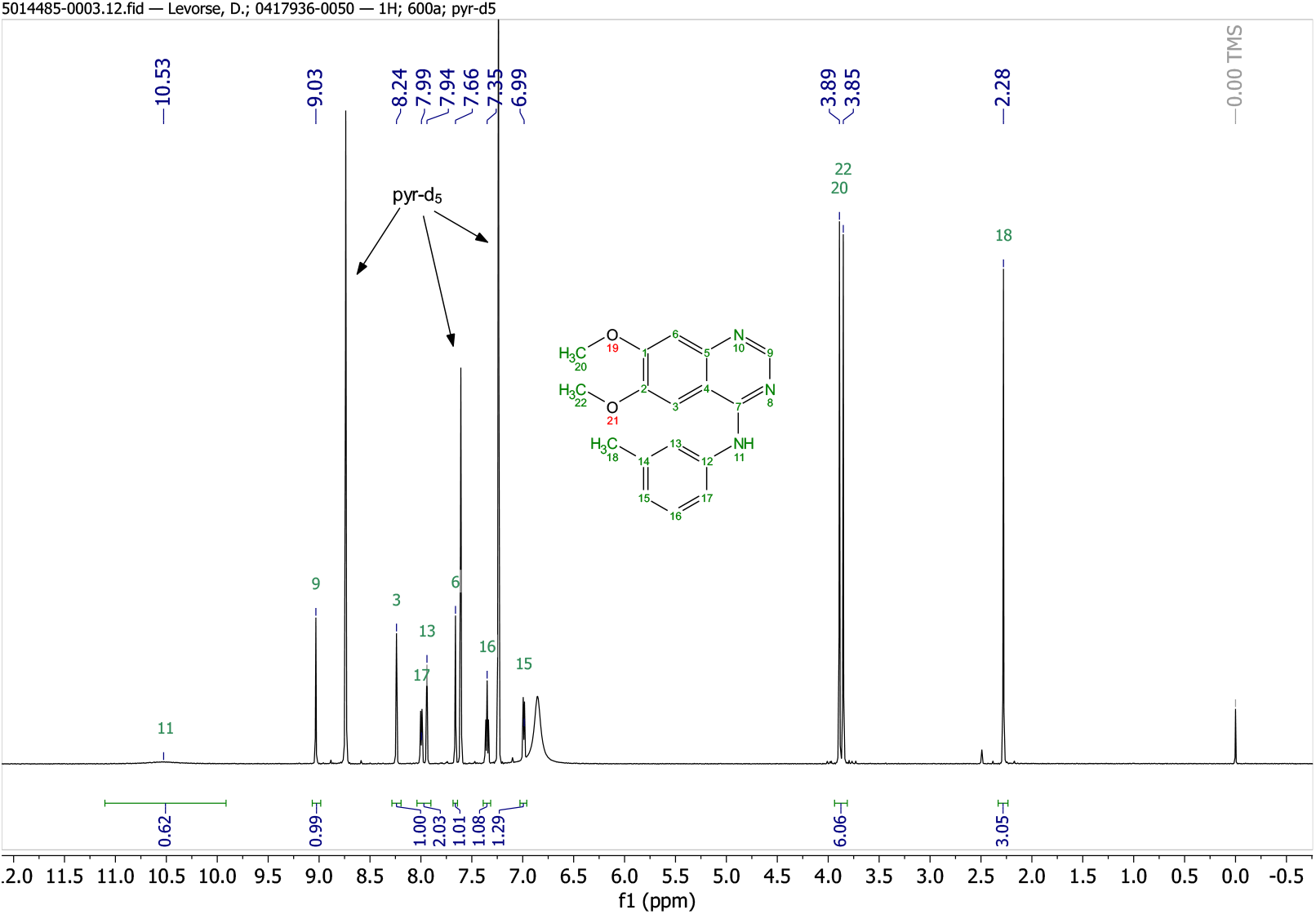
The ^1^H 1D NMR spectrum of SM13 confirms compound identity and structure.

## Footnotes

1 SAMPL6 was originally announced as featuring a log *D* prediction challenge, but there were difficulties in the collection of experimental data. The original plan was to measure log *P*^0^, log *P*^-1^, and log *P*^+1^ and calculate log *D* values at the experimental pH using these values. However, we were able to measure the partition coefficients of neutral species (log *P*^0^) much more reliably than ionic species with potentiometric log *P* method of Sirius T3, as elaborated further below.

