## Supplementary material for "Octanol-water partition coefficient measurements for the SAMPL6 Blind Prediction Challenge": SM02_18C-01011_M02_octanol_pH-metric high logP_report.pdf

| | | | |
|---|---|---|---|
| Sample name: | **M02_octanol** | Experiment start time: | **3/1/2018 2:44:22 PM** |
| Assay name: | **pH-metric high logP** | Analyst: | **Dorothy Levorse** |
| Assay ID: | **18C-01011** | Instrument ID: | **T312060** |
| Filename: | **C:\Sirius_T3\Mehtap\20180228_exp28_logP_T3-2\18C-01011_M02_octanol_pH-metric high logP.t3r** | | |

#### pH-metric Result

| | | |
|---|---|---|
| logP (XH +) | 1.32 | ±0.06 (n=50) |
| logP (neutral X) | 4.10 | ±0.03 (n=50) |

#### 18C-01011 Points 1 to 16

| | |
|---|---|
| M02_octanol concentration factor | 0.688 |
| Carbonate | 0.0278 mM |
| Acidity error | -0.23518 mM |

#### 18C-01011 Points 17 to 26

| | |
|---|---|
| M02_octanol concentration factor | 0.751 |
| Carbonate | 0.1291 mM |
| Acidity error | -0.36000 mM |

#### 18C-01011 Points 27 to 42

| | |
|---|---|
| M02_octanol concentration factor | 0.599 |
| Carbonate | 0.0511 mM |
| Acidity error | -0.26646 mM |

#### Warnings and errors

| | |
|---|---|
| Errors | None |
| Warnings | None |

#### Sample logD and percent species

| pH | M02_octanol logD | M02_octanol M02_octanolH | M02_octanol M02_octanol | M02_octanol M02_octanolH* | M02_octanol M02_octanol* | Comment |
|---|---|---|---|---|---|---|
| 1.000 | 1.34 | 4.34 % | 0.00 % | 90.61 % | 5.05 % | |
| 1.200 | 1.36 | 4.22 % | 0.00 % | 88.01 % | 7.77 % | Stomach pH |
| 2.000 | 1.51 | 2.99 % | 0.00 % | 62.31 % | 34.70 % | |
| 3.000 | 2.13 | 0.72 % | 0.01 % | 15.11 % | 84.16 % | |
| 4.000 | 3.03 | 0.08 % | 0.01 % | 1.76 % | 98.15 % | |
| 5.000 | 3.78 | 0.01 % | 0.01 % | 0.18 % | 99.80 % | |
| 6.000 | 4.05 | 0.00 % | 0.01 % | 0.02 % | 99.97 % | |
| 6.500 | 4.08 | 0.00 % | 0.01 % | 0.01 % | 99.99 % | |
| 7.000 | 4.09 | 0.00 % | 0.01 % | 0.00 % | 99.99 % | |
| 7.400 | 4.09 | 0.00 % | 0.01 % | 0.00 % | 99.99 % | Blood pH |
| 8.000 | 4.09 | 0.00 % | 0.01 % | 0.00 % | 99.99 % | |
| 9.000 | 4.10 | 0.00 % | 0.01 % | 0.00 % | 99.99 % | |
| 10.000 | 4.10 | 0.00 % | 0.01 % | 0.00 % | 99.99 % | |
| 11.000 | 4.10 | 0.00 % | 0.01 % | 0.00 % | 99.99 % | |
| 12.000 | 4.10 | 0.00 % | 0.01 % | 0.00 % | 99.99 % | |

Sample name: **M02_octanol** Experiment start time: **3/1/2018 2:44:22 PM**
Assay name: **pH-metric high logP** Analyst: **Dorothy Levorse**
Assay ID: **18C-01011** Instrument ID: **T312060**
Filename: **C:\Sirius_T3\Mehtap\20180228_exp28_logP_T3-2\18C-01011_M02_octanol_pH-metric high logP.t3r**

#### Graphs

Charge balance residuals

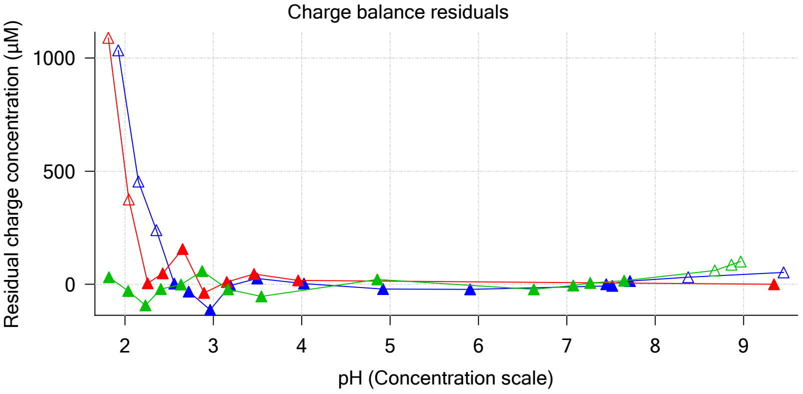


Ionic strength

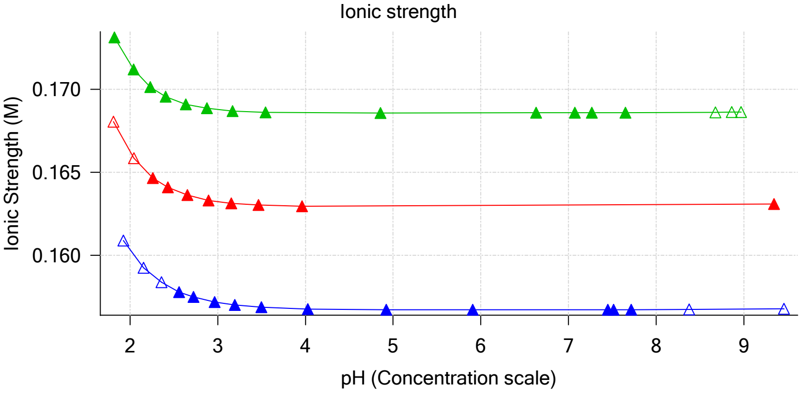


Buffer index profile

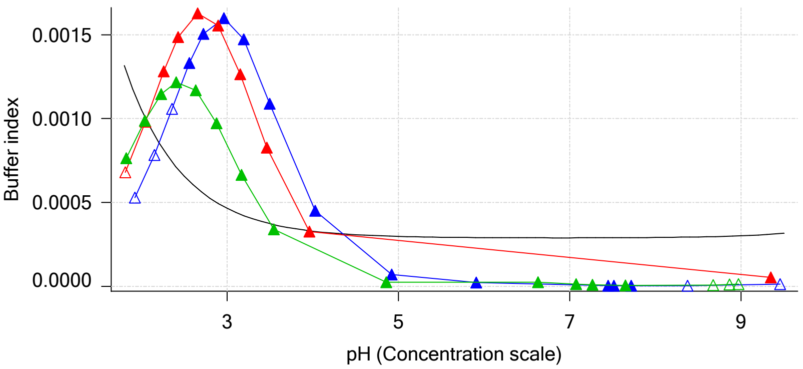


Sample M02_octanol Ionisation Graph

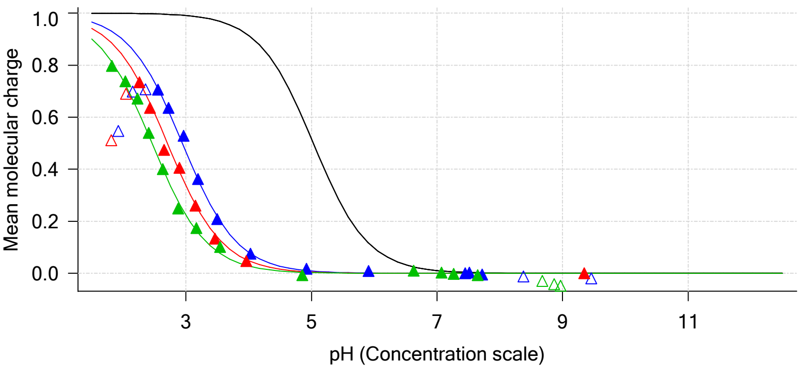


Distribution of species for sample M02_octanol

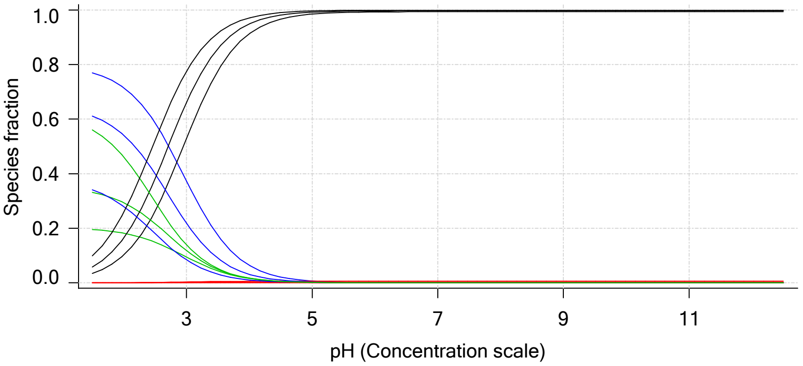


Lipophilicity Profile

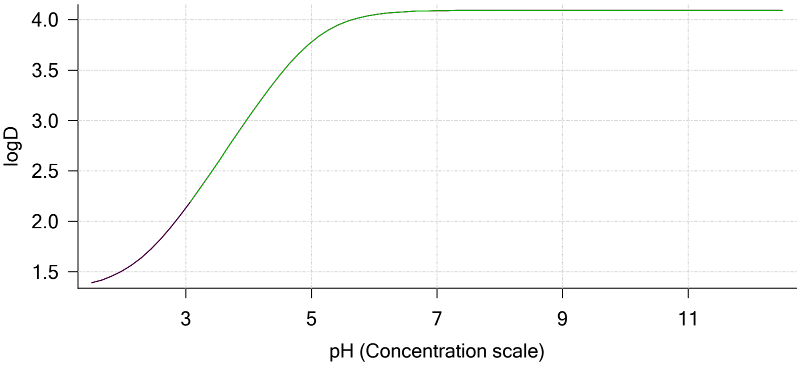


Log R Profile

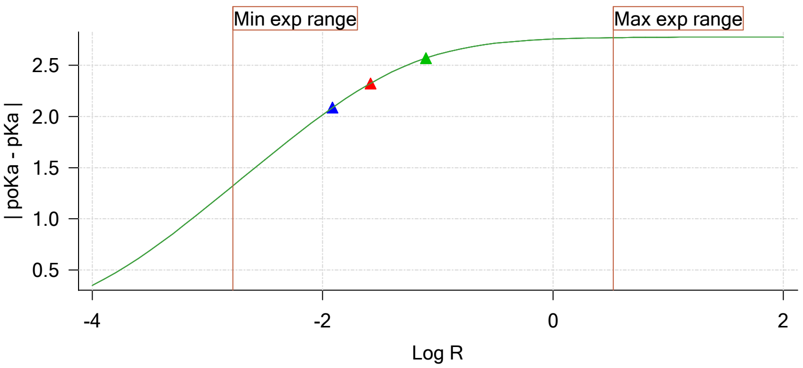


Sample Carbonate Ionisation Graph

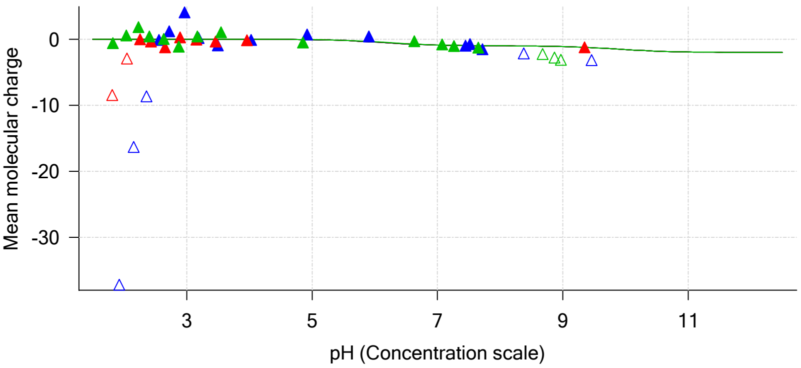

| | | | |
|---|---|---|---|
| Sample name: | **M02_octanol** | Experiment start time: | **3/1/2018 2:44:22 PM** |
| Assay name: | **pH-metric high logP** | Analyst: | **Dorothy Levorse** |
| Assay ID: | **18C-01011** | Instrument ID: | **T312060** |
| Filename: | **C:\Sirius_T3\Mehtap\20180228_exp28_logP_T3-2\18C-01011_M02_octanol_pH-metric high logP.t3r** | | |

#### Graphs (continued)

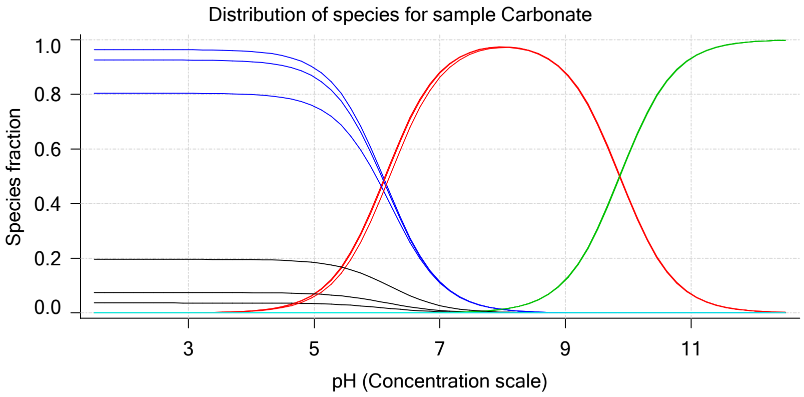

| | | | |
|---|---|---|---|
| Sample name: | **M02_octanol** | Experiment start time: | **3/1/2018 2:44:22 PM** |
| Assay name: | **pH-metric high logP** | Analyst: | **Dorothy Levorse** |
| Assay ID: | **18C-01011** | Instrument ID: | **T312060** |
| Filename: | **C:\Sirius_T3\Mehtap\20180228_exp28_logP_T3-2\18C-01011_M02_octanol_pH-metric high logP.t3r** | | |

# pH-metric high logP   Titration 1 of 3   18C-01011 Points 1 to 16

#### Overall results

| | |
|---|---|
| RMSD | 1.164 |
| Average ionic strength | 0.157 M |
| Average temperature | 24.9°C |
| Partition ratio | 0.0122 : 1 |
| Analyte concentration range | 3971.1 μM to 4077.5 μM |
| Total points considered | 11 of 16 |

#### Warnings and errors

| | |
|---|---|
| Errors | None |
| Warnings | Sample concentration factor out of range |

#### Four-Plus parameters

| | | | |
|---|---|---|---|
| Alpha | 0.130 | 3/1/2018 2:44:22 PM | C:\Sirius_T3\HCl18B27.t3r |
| S | 0.9970 | 3/1/2018 2:44:22 PM | C:\Sirius_T3\HCl18B27.t3r |
| jH | 0.8 | 3/1/2018 2:44:22 PM | C:\Sirius_T3\HCl18B27.t3r |
| jOH | -0.4 | 3/1/2018 2:44:22 PM | C:\Sirius_T3\HCl18B27.t3r |

#### Titrants

| | | | |
|---|---|---|---|
| 0.50 M HCl | 0.993513 | 3/1/2018 2:44:22 PM | C:\Sirius_T3\HCl18B27.t3r |
| 0.50 M KOH | 0.999845 | 3/1/2018 2:44:22 PM | C:\Sirius_T3\KOH18B27.t3r |

#### Sample

| | |
|---|---|
| M02_octanol concentration factor | 0.688 |
| Base pKa 1 | 5.03 |
| logP (XH +) | 1.60 |
| logP (neutral X) | 4.15 |

#### Sample graphs

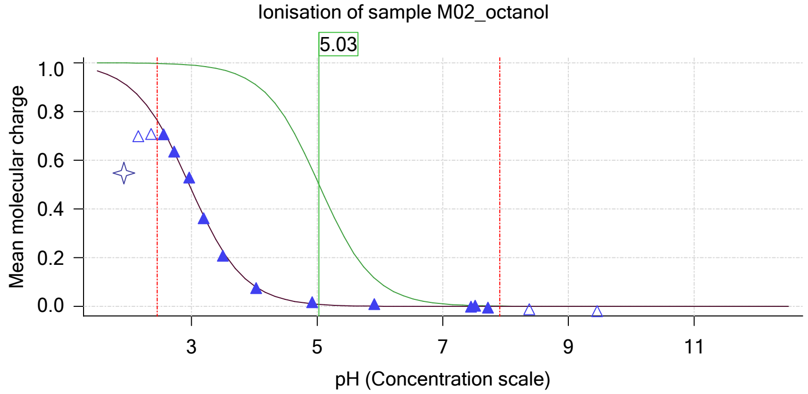


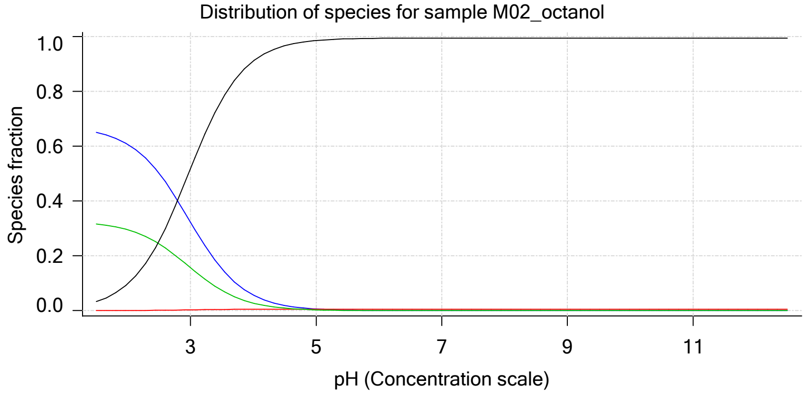

Sample name: **M02_octanol**
Assay name: **pH-metric high logP**
Assay ID: **18C-01011**
Filename: **C:\Sirius_T3\Mehtap\20180228_exp28_logP_T3-2\18C-01011_M02_octanol_pH-metric high logP.t3r**

Experiment start time: **3/1/2018 2:44:22 PM**
Analyst: **Dorothy Levorse**
Instrument ID: **T312060**

#### Sample graphs (continued)

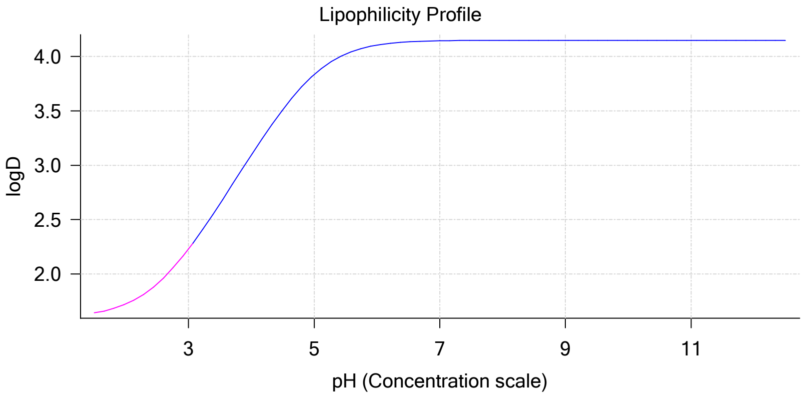


#### Sample logD and percent species

| pH | M02_octanol logD | M02_octanol M02_octanolH | M02_octanol M02_octanol | M02_octanol M02_octanolH* | M02_octanol M02_octanol* | Comment |
|---|---|---|---|---|---|---|
| 1.000 | 1.61 | 66.58 % | 0.01 % | 32.35 % | 1.07 % | |
| 1.200 | 1.62 | 66.16 % | 0.01 % | 32.15 % | 1.68 % | Stomach pH |
| 2.000 | 1.72 | 60.71 % | 0.06 % | 29.50 % | 9.74 % | |
| 3.000 | 2.23 | 32.27 % | 0.30 % | 15.68 % | 51.75 % | |
| 4.000 | 3.09 | 5.68 % | 0.53 % | 2.76 % | 91.04 % | |
| 5.000 | 3.83 | 0.61 % | 0.57 % | 0.30 % | 98.51 % | |
| 6.000 | 4.10 | 0.06 % | 0.58 % | 0.03 % | 99.33 % | |
| 6.500 | 4.13 | 0.02 % | 0.58 % | 0.01 % | 99.39 % | |
| 7.000 | 4.14 | 0.01 % | 0.58 % | 0.00 % | 99.41 % | |
| 7.400 | 4.15 | 0.00 % | 0.58 % | 0.00 % | 99.42 % | Blood pH |
| 8.000 | 4.15 | 0.00 % | 0.58 % | 0.00 % | 99.42 % | |
| 9.000 | 4.15 | 0.00 % | 0.58 % | 0.00 % | 99.42 % | |
| 10.000 | 4.15 | 0.00 % | 0.58 % | 0.00 % | 99.42 % | |
| 11.000 | 4.15 | 0.00 % | 0.58 % | 0.00 % | 99.42 % | |
| 12.000 | 4.15 | 0.00 % | 0.58 % | 0.00 % | 99.42 % | |

#### Carbonate and acidity

Carbonate 0.028 mM
Acidity error -0.235 mM

#### Other graphs

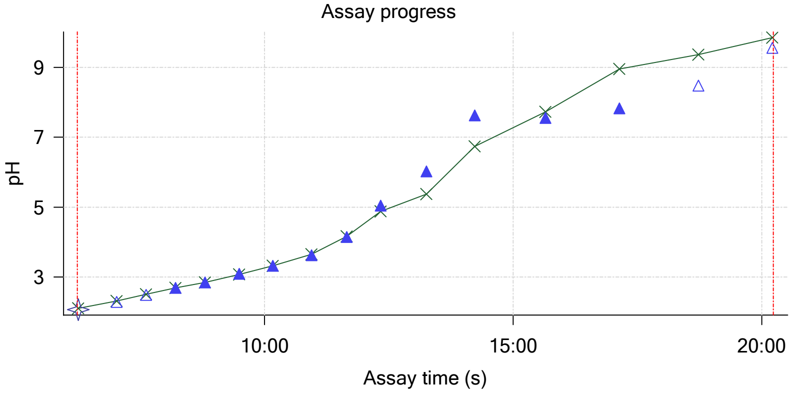


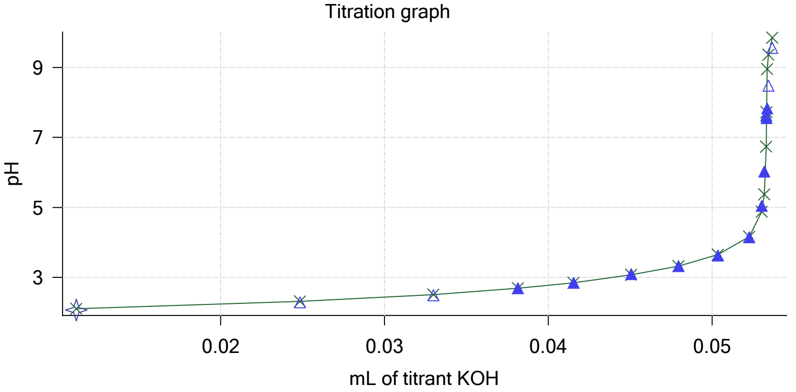

Sample name: **M02_octanol**
Assay name: **pH-metric high logP**
Assay ID: **18C-01011**
Filename: **C:\Sirius_T3\Mehtap\20180228_exp28_logP_T3-2\18C-01011_M02_octanol_pH-metric high logP.t3r**

Experiment start time: **3/1/2018 2:44:22 PM**
Analyst: **Dorothy Levorse**
Instrument ID: **T312060**

#### Other graphs (continued)

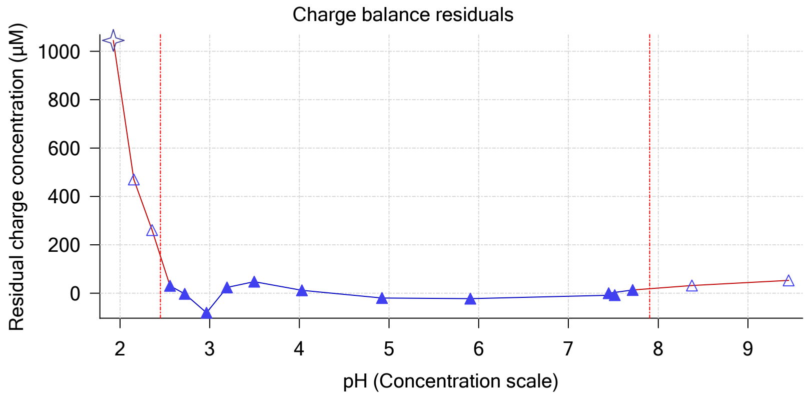


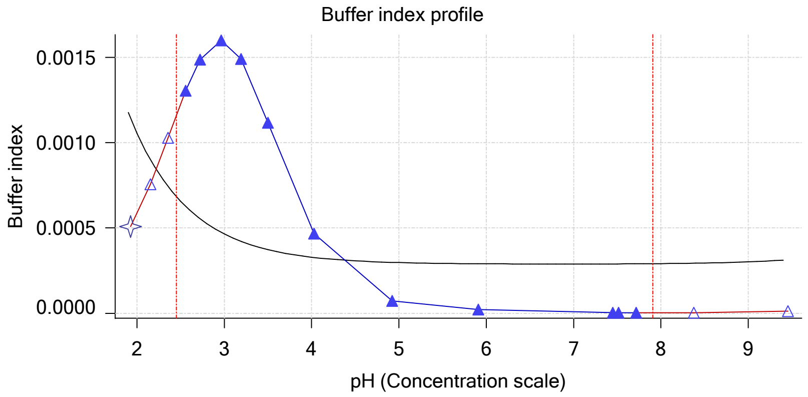


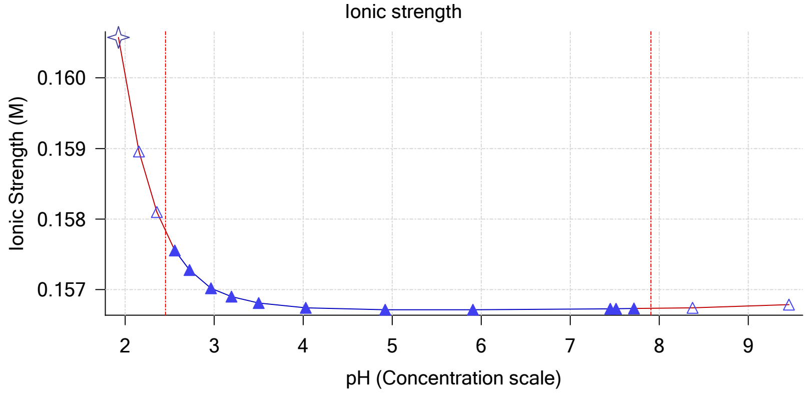


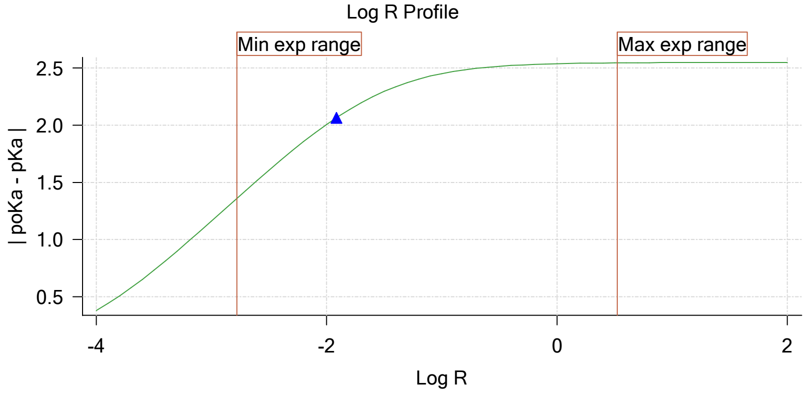


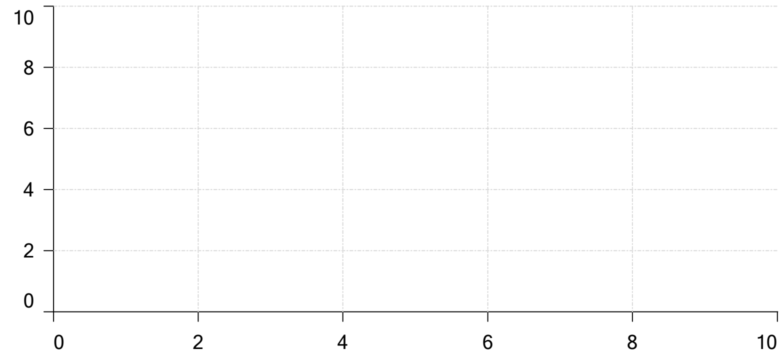


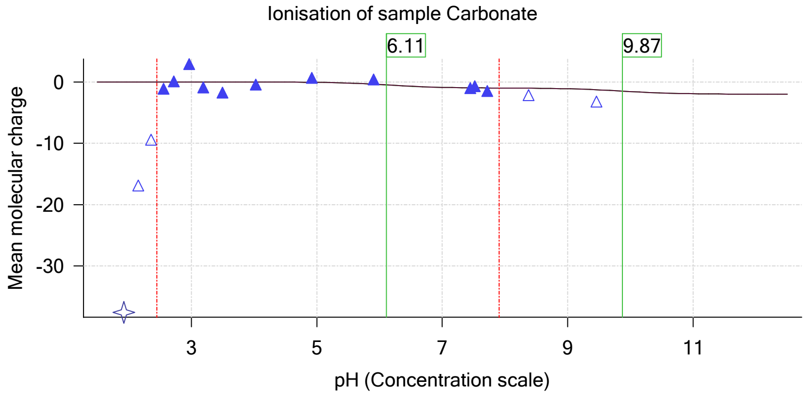

| | | | |
|---|---|---|---|
| Sample name: | **M02_octanol** | Experiment start time: | **3/1/2018 2:44:22 PM** |
| Assay name: | **pH-metric high logP** | Analyst: | **Dorothy Levorse** |
| Assay ID: | **18C-01011** | Instrument ID: | **T312060** |
| Filename: | **C:\Sirius_T3\Mehtap\20180228_exp28_logP_T3-2\18C-01011_M02_octanol_pH-metric high logP.t3r** | | |

### pH-metric high logP  Titration 2 of 3  18C-01011 Points 17 to 26

#### Overall results

| | |
|---|---|
| RMSD | 0.012 |
| Average ionic strength | 0.163 M |
| Average temperature | 25.0°C |
| Partition ratio | 0.0260 : 1 |
| Analyte concentration range | 3657.5 μM to 3779.3 μM |
| Total points considered | 8 of 10 |

#### Warnings and errors

| | |
|---|---|
| Errors | None |
| Warnings | None |

#### Four-Plus parameters

| | | | |
|---|---|---|---|
| Alpha | 0.130 | 3/1/2018 2:44:22 PM | C:\Sirius_T3\HCl18B27.t3r |
| S | 0.9970 | 3/1/2018 2:44:22 PM | C:\Sirius_T3\HCl18B27.t3r |
| jH | 0.8 | 3/1/2018 2:44:22 PM | C:\Sirius_T3\HCl18B27.t3r |
| jOH | -0.4 | 3/1/2018 2:44:22 PM | C:\Sirius_T3\HCl18B27.t3r |

#### Titrants

| | | | |
|---|---|---|---|
| 0.50 M HCl | 0.993513 | 3/1/2018 2:44:22 PM | C:\Sirius_T3\HCl18B27.t3r |
| 0.50 M KOH | 0.999845 | 3/1/2018 2:44:22 PM | C:\Sirius_T3\KOH18B27.t3r |

#### Sample

| | |
|---|---|
| M02_octanol concentration factor | 0.751 |
| Base pKa 1 | 5.03 |
| logP (XH +) | 1.60 |
| logP (neutral X) | 4.25 |

#### Sample graphs

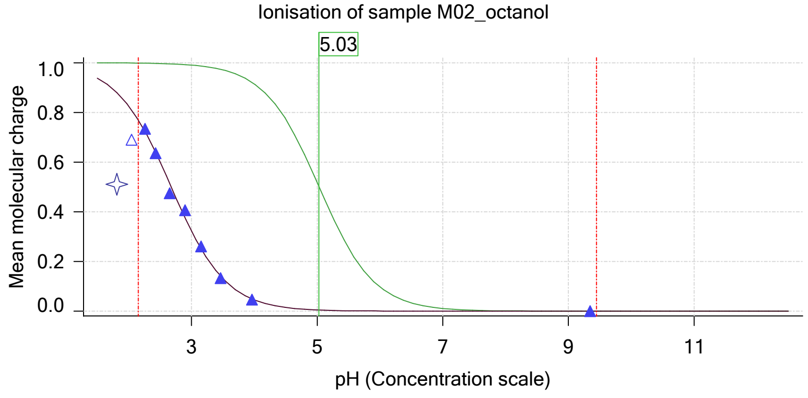


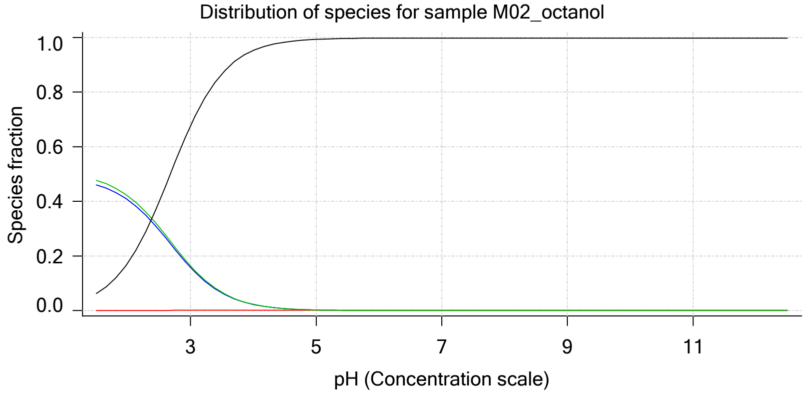

Sample name: **M02_octanol**
Assay name: **pH-metric high logP**
Assay ID: **18C-01011**
Filename: **C:\Sirius_T3\Mehtap\20180228_exp28_logP_T3-2\18C-01011_M02_octanol_pH-metric high logP.t3r**

Experiment start time: **3/1/2018 2:44:22 PM**
Analyst: **Dorothy Levorse**
Instrument ID: **T312060**

#### Sample graphs (continued)

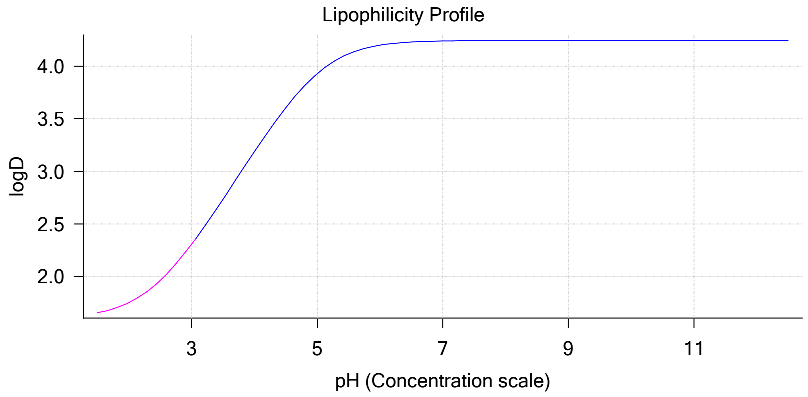


#### Sample logD and percent species

| pH | M02_octanol logD | M02_octanol M02_octanolH | M02_octanol M02_octanol | M02_octanol M02_octanolH* | M02_octanol M02_octanol* | Comment |
|---|---|---|---|---|---|---|
| 1.000 | 1.62 | 48.12 % | 0.00 % | 49.81 % | 2.06 % | |
| 1.200 | 1.63 | 47.55 % | 0.01 % | 49.22 % | 3.23 % | Stomach pH |
| 2.000 | 1.75 | 40.58 % | 0.04 % | 42.01 % | 17.38 % | |
| 3.000 | 2.31 | 15.81 % | 0.15 % | 16.36 % | 67.68 % | |
| 4.000 | 3.19 | 2.22 % | 0.21 % | 2.30 % | 95.26 % | |
| 5.000 | 3.93 | 0.23 % | 0.22 % | 0.24 % | 99.31 % | |
| 6.000 | 4.20 | 0.02 % | 0.22 % | 0.02 % | 99.74 % | |
| 6.500 | 4.23 | 0.01 % | 0.22 % | 0.01 % | 99.77 % | |
| 7.000 | 4.24 | 0.00 % | 0.22 % | 0.00 % | 99.78 % | |
| 7.400 | 4.24 | 0.00 % | 0.22 % | 0.00 % | 99.78 % | Blood pH |
| 8.000 | 4.25 | 0.00 % | 0.22 % | 0.00 % | 99.78 % | |
| 9.000 | 4.25 | 0.00 % | 0.22 % | 0.00 % | 99.78 % | |
| 10.000 | 4.25 | 0.00 % | 0.22 % | 0.00 % | 99.78 % | |
| 11.000 | 4.25 | 0.00 % | 0.22 % | 0.00 % | 99.78 % | |
| 12.000 | 4.25 | 0.00 % | 0.22 % | 0.00 % | 99.78 % | |

#### Carbonate and acidity

Carbonate 0.129 mM
Acidity error -0.360 mM

#### Other graphs

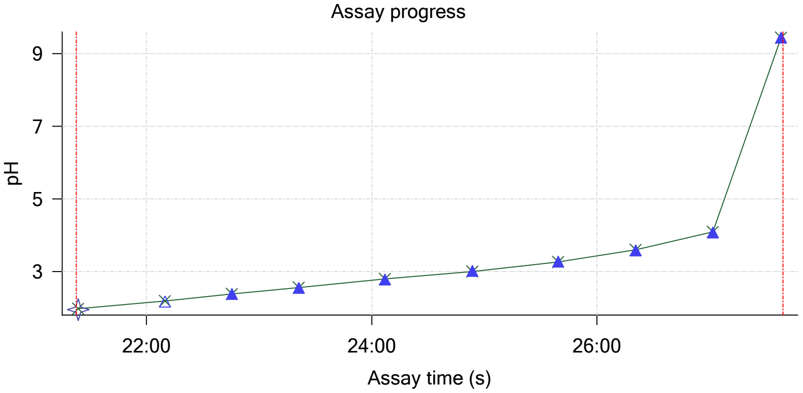


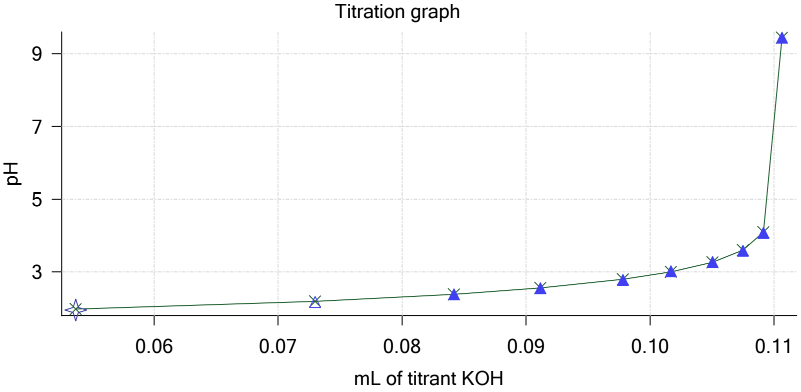

Sample name: **M02_octanol**
Assay name: **pH-metric high logP**
Assay ID: **18C-01011**
Filename: **C:\Sirius_T3\Mehtap\20180228_exp28_logP_T3-2\18C-01011_M02_octanol_pH-metric high logP.t3r**

Experiment start time: **3/1/2018 2:44:22 PM**
Analyst: **Dorothy Levorse**
Instrument ID: **T312060**

#### Other graphs (continued)

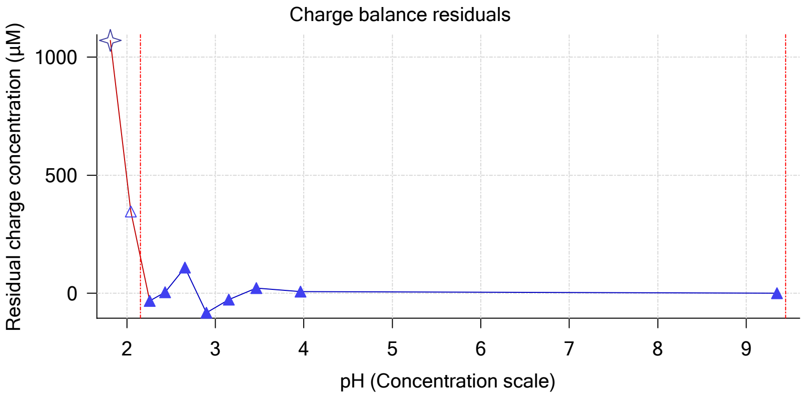


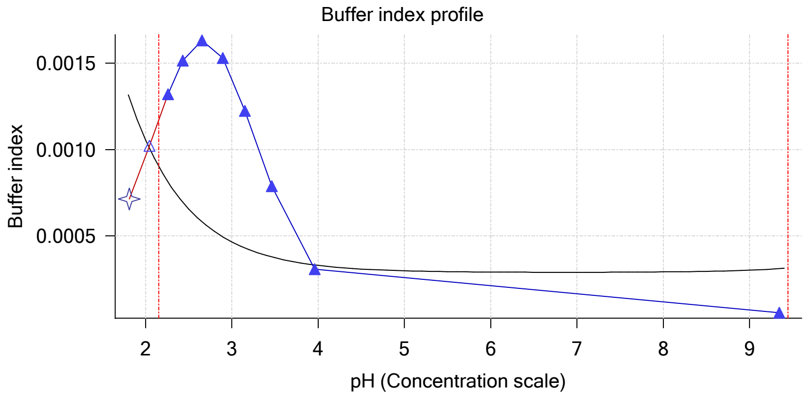


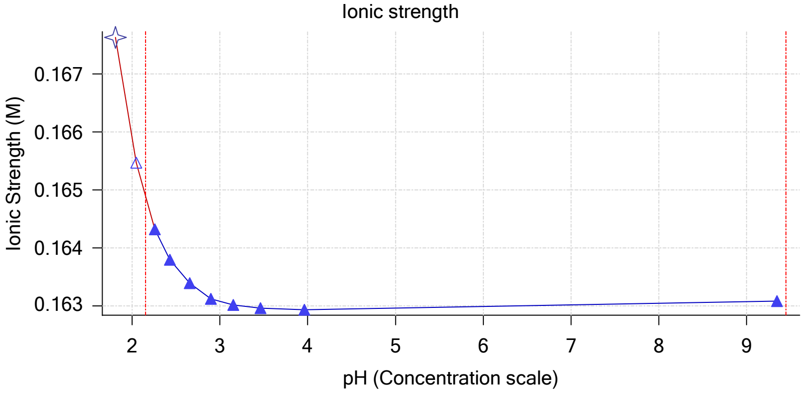


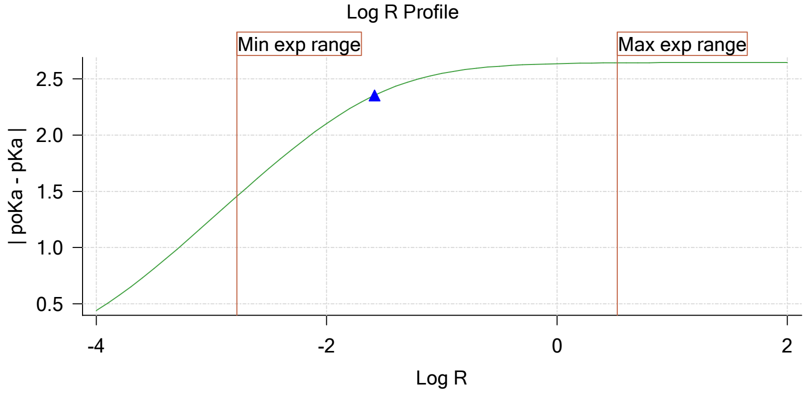


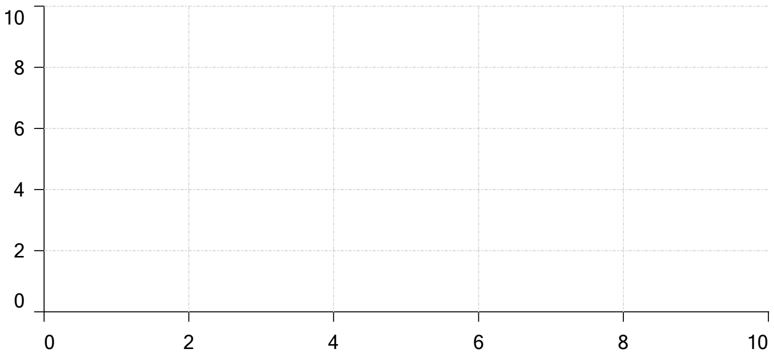


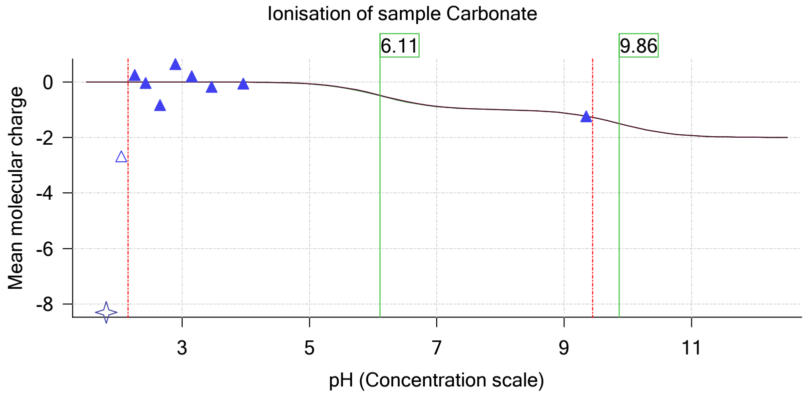

| | | | |
|---|---|---|---|
| Sample name: | **M02_octanol** | Experiment start time: | **3/1/2018 2:44:22 PM** |
| Assay name: | **pH-metric high logP** | Analyst: | **Dorothy Levorse** |
| Assay ID: | **18C-01011** | Instrument ID: | **T312060** |
| Filename: | **C:\Sirius_T3\Mehtap\20180228_exp28_logP_T3-2\18C-01011_M02_octanol_pH-metric high logP.t3r** | | |

### pH-metric high logP Titration 3 of 3 18C-01011 Points 27 to 42

#### Overall results

| | |
|---|---|
| RMSD | 0.734 |
| Average ionic strength | 0.169 M |
| Average temperature | 25.0°C |
| Partition ratio | 0.0789 : 1 |
| Analyte concentration range | 3250.6 µM to 3352.6 µM |
| Total points considered | 13 of 16 |

#### Warnings and errors

| | |
|---|---|
| Errors | None |
| Warnings | Sample concentration factor out of range |

#### Four-Plus parameters

| | | | |
|---|---|---|---|
| Alpha | 0.130 | 3/1/2018 2:44:22 PM | C:\Sirius_T3\HCl18B27.t3r |
| S | 0.9970 | 3/1/2018 2:44:22 PM | C:\Sirius_T3\HCl18B27.t3r |
| jH | 0.8 | 3/1/2018 2:44:22 PM | C:\Sirius_T3\HCl18B27.t3r |
| jOH | -0.4 | 3/1/2018 2:44:22 PM | C:\Sirius_T3\HCl18B27.t3r |

#### Titrants

| | | | |
|---|---|---|---|
| 0.50 M HCl | 0.993513 | 3/1/2018 2:44:22 PM | C:\Sirius_T3\HCl18B27.t3r |
| 0.50 M KOH | 0.999845 | 3/1/2018 2:44:22 PM | C:\Sirius_T3\KOH18B27.t3r |

#### Sample

| | |
|---|---|
| M02_octanol concentration factor | 0.599 |
| Base pKa 1 | 5.03 |
| logP (XH +) | 1.60 |
| logP (neutral X) | 4.27 |

#### Sample graphs

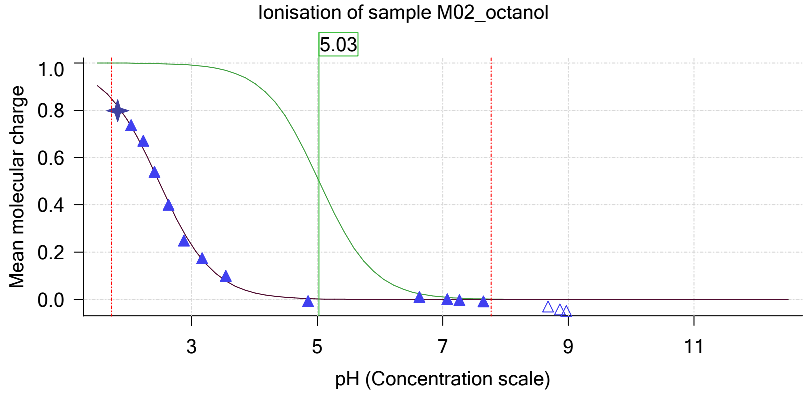


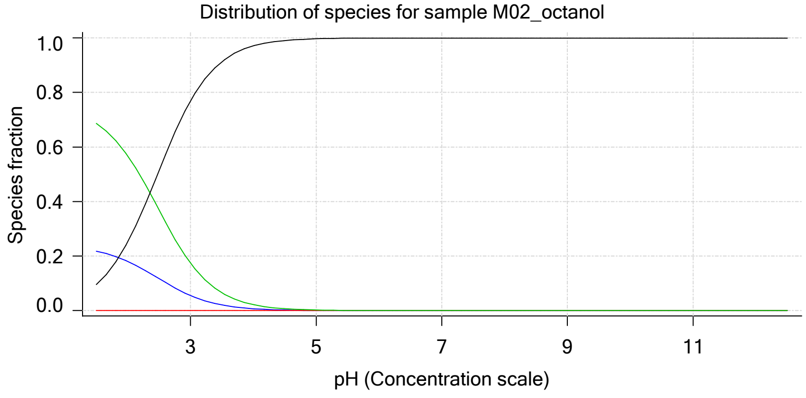

| | | | |
|---|---|---|---|
| Sample name: | **M02_octanol** | Experiment start time: | **3/1/2018 2:44:22 PM** |
| Assay name: | **pH-metric high logP** | Analyst: | **Dorothy Levorse** |
| Assay ID: | **18C-01011** | Instrument ID: | **T312060** |
| Filename: | **C:\Sirius_T3\Mehtap\20180228_exp28_logP_T3-2\18C-01011_M02_octanol_pH-metric high logP.t3r** | | |

#### Sample graphs (continued)

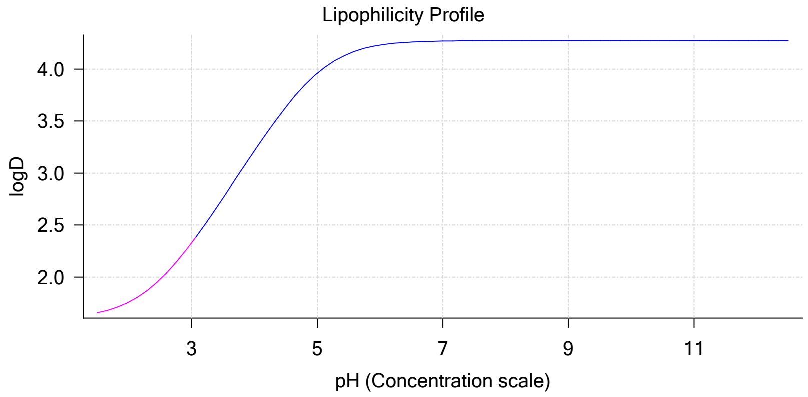


#### Sample logD and percent species

| pH | M02_octanol logD | M02_octanol M02_octanolH | M02_octanol M02_octanol | M02_octanol M02_octanolH* | M02_octanol M02_octanol* | Comment |
|---|---|---|---|---|---|---|
| 1.000 | 1.62 | 23.36 % | 0.00 % | 73.40 % | 3.24 % | |
| 1.200 | 1.63 | 22.92 % | 0.00 % | 72.04 % | 5.04 % | Stomach pH |
| 2.000 | 1.76 | 18.08 % | 0.02 % | 56.83 % | 25.06 % | |
| 3.000 | 2.33 | 5.55 % | 0.05 % | 17.45 % | 76.95 % | |
| 4.000 | 3.22 | 0.70 % | 0.07 % | 2.20 % | 97.03 % | |
| 5.000 | 3.96 | 0.07 % | 0.07 % | 0.23 % | 99.64 % | |
| 6.000 | 4.23 | 0.01 % | 0.07 % | 0.02 % | 99.90 % | |
| 6.500 | 4.26 | 0.00 % | 0.07 % | 0.01 % | 99.92 % | |
| 7.000 | 4.27 | 0.00 % | 0.07 % | 0.00 % | 99.93 % | |
| 7.400 | 4.27 | 0.00 % | 0.07 % | 0.00 % | 99.93 % | Blood pH |
| 8.000 | 4.27 | 0.00 % | 0.07 % | 0.00 % | 99.93 % | |
| 9.000 | 4.27 | 0.00 % | 0.07 % | 0.00 % | 99.93 % | |
| 10.000 | 4.27 | 0.00 % | 0.07 % | 0.00 % | 99.93 % | |
| 11.000 | 4.27 | 0.00 % | 0.07 % | 0.00 % | 99.93 % | |
| 12.000 | 4.27 | 0.00 % | 0.07 % | 0.00 % | 99.93 % | |

#### Carbonate and acidity

| | |
|---|---|
| Carbonate | 0.051 mM |
| Acidity error | -0.266 mM |

#### Other graphs

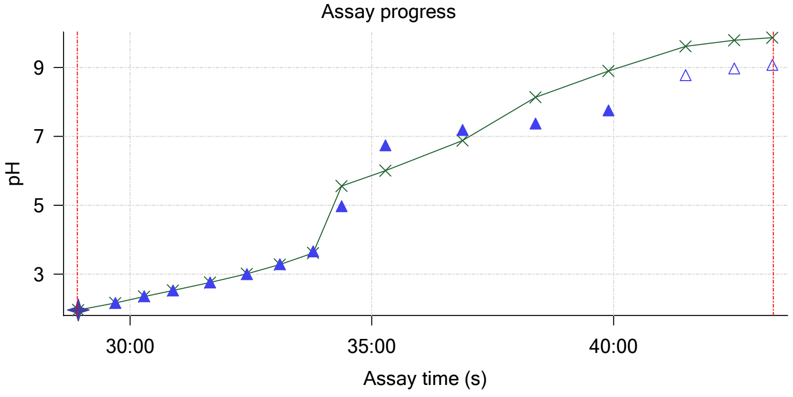


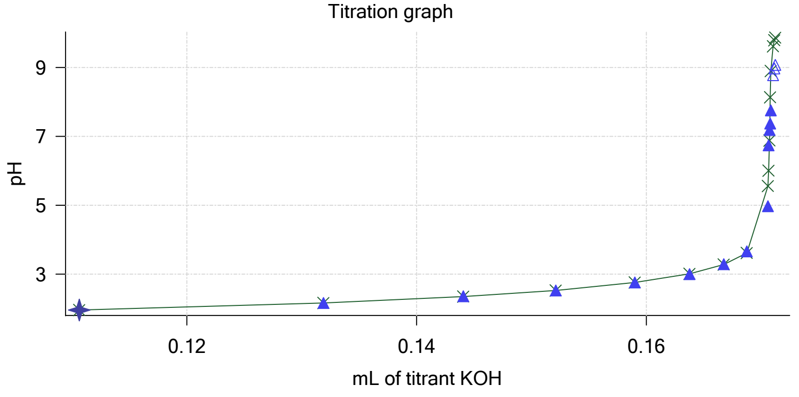

| | | | |
|---|---|---|---|
| Sample name: | **M02_octanol** | Experiment start time: | **3/1/2018 2:44:22 PM** |
| Assay name: | **pH-metric high logP** | Analyst: | **Dorothy Levorse** |
| Assay ID: | **18C-01011** | Instrument ID: | **T312060** |
| Filename: | **C:\Sirius_T3\Mehtap\20180228_exp28_logP_T3-2\18C-01011_M02_octanol_pH-metric high logP.t3r** | | |

#### Other graphs (continued)

Charge balance residuals

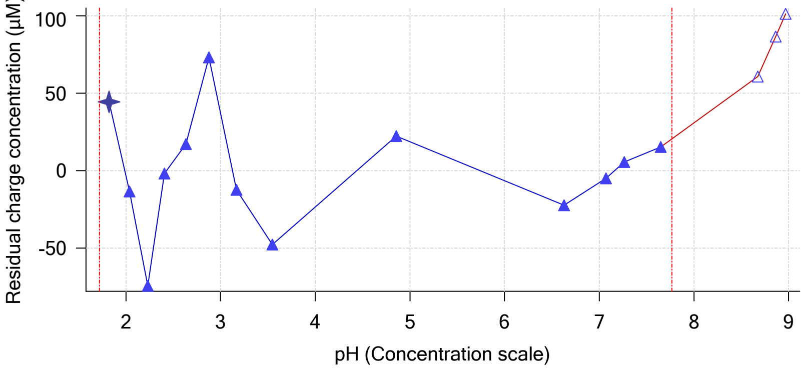


Buffer index profile

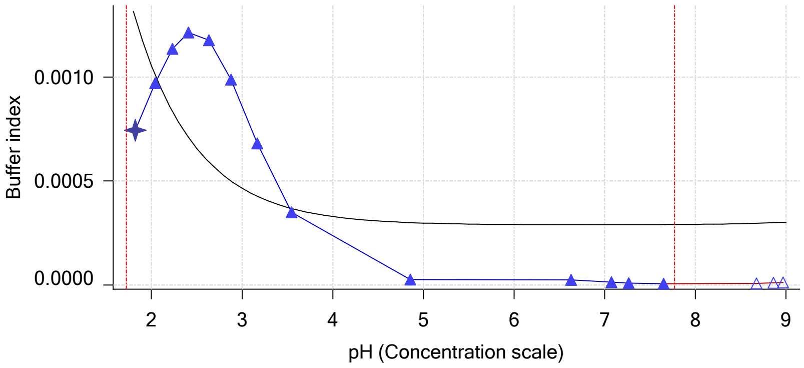


Ionic strength

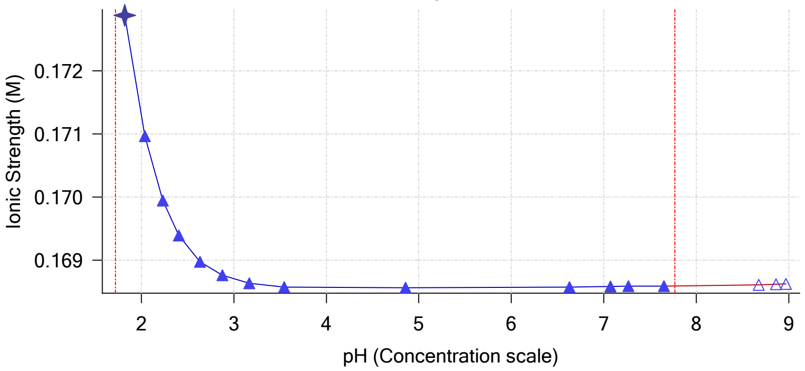


Log R Profile

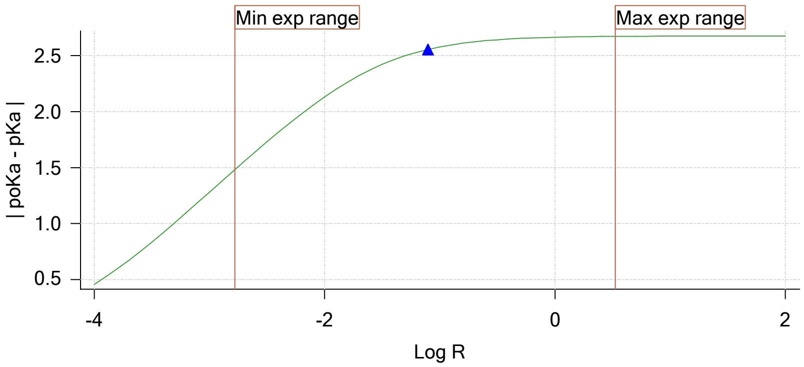


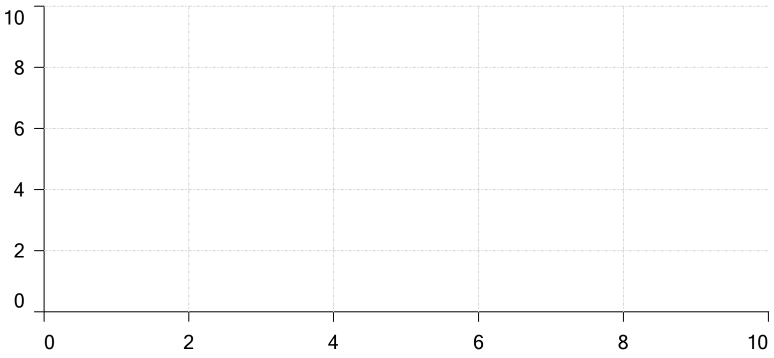


Ionisation of sample Carbonate

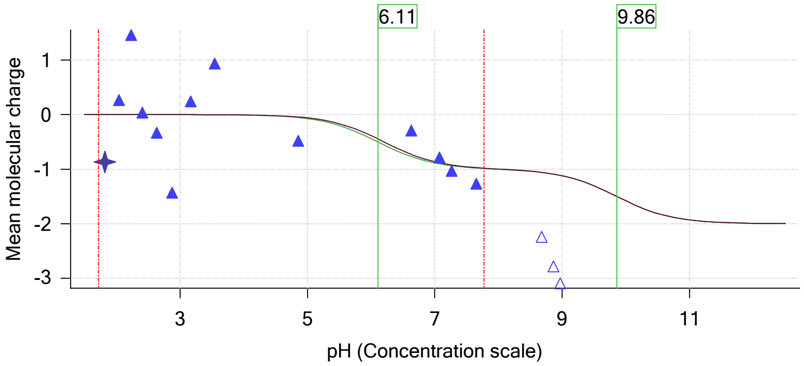

| | | | |
|---|---|---|---|
| Sample name: | **M02_octanol** | Experiment start time: | **3/1/2018 2:44:22 PM** |
| Assay name: | **pH-metric high logP** | Analyst: | **Dorothy Levorse** |
| Assay ID: | **18C-01011** | Instrument ID: | **T312060** |
| Filename: | **C:\Sirius_T3\Mehtap\20180228_exp28_logP_T3-2\18C-01011_M02_octanol_pH-metric high logP.t3r** | | |

#### Assay Model

| Settings | Value | Date/Time changed | Imported from |
|---|---|---|---|
| Sample name | M02_octanol | 12/6/2017 4:20:03 PM | User entered value |
| Sample by | Weight | | Default value |
| Sample weight | 0.001870 g | 2/28/2018 4:51:34 PM | User entered value |
| Formula weight | 289.26 g/mol | 12/6/2017 4:20:03 PM | User entered value |
| Solubility | Unknown | | Default value |
| Molecular weight | 289.26 | 12/6/2017 4:20:03 PM | User entered value |
| Individual pKa ionic environments | No | | Default value |
| Number of pKas | 1 | 12/6/2017 4:20:03 PM | User entered value |
| Sample is a | Base | 12/6/2017 4:20:03 PM | User entered value |
| pKa 1 | 5.03 | 12/6/2017 4:20:03 PM | User entered value |
| logp (XH +) | 1.60 | 12/6/2017 4:20:17 PM | User entered value |
| logP (neutral X) | 3.00 | 2/28/2018 2:50:25 PM | User entered value |

#### Events

| Time | Event | Water | Acid | Base | Octanol | pH | dpH/dt | pH R-squared | pH SD | dpH/dt time |
|---|---|---|---|---|---|---|---|---|---|---|
| 3:15.5 | Initial pH = 6.92 | | | | | | | | | |
| 6:15.3 | Data point 1 | 1.50000 mL | 0.05433 mL | 0.01119 mL | 0.01999 mL | 2.059 | 0.00260 | 0.18062 | 0.00030 | 10.0 s |
| 7:01.6 | Data point 2 | 1.50000 mL | 0.05433 mL | 0.02484 mL | 0.01999 mL | 2.282 | 0.00555 | 0.77597 | 0.00031 | 10.0 s |
| 7:37.2 | Data point 3 | 1.50000 mL | 0.05433 mL | 0.03300 mL | 0.01999 mL | 2.484 | 0.01434 | 0.77758 | 0.00080 | 10.0 s |
| 8:12.8 | Data point 4 | 1.50000 mL | 0.05433 mL | 0.03815 mL | 0.01999 mL | 2.683 | 0.00592 | 0.63815 | 0.00037 | 10.0 s |
| 8:48.3 | Data point 5 | 1.50000 mL | 0.05433 mL | 0.04153 mL | 0.01999 mL | 2.846 | -0.00121 | 0.24285 | 0.00012 | 10.5 s |
| 9:29.5 | Data point 6 | 1.50000 mL | 0.05433 mL | 0.04506 mL | 0.01999 mL | 3.087 | -0.00967 | 0.98095 | 0.00048 | 10.0 s |
| 10:10.2 | Data point 7 | 1.50000 mL | 0.05433 mL | 0.04795 mL | 0.01999 mL | 3.314 | -0.01605 | 0.70172 | 0.00095 | 16.0 s |
| 10:56.9 | Data point 8 | 1.50000 mL | 0.05433 mL | 0.05035 mL | 0.01999 mL | 3.619 | -0.01833 | 0.92570 | 0.00094 | 11.5 s |
| 11:39.2 | Data point 9 | 1.50000 mL | 0.05433 mL | 0.05226 mL | 0.01999 mL | 4.146 | -0.01442 | 0.66678 | 0.00087 | 15.5 s |
| 12:20.1 | Data point 10 | 1.50000 mL | 0.05433 mL | 0.05303 mL | 0.01999 mL | 5.037 | -0.01988 | 0.98479 | 0.00099 | 25.0 s |
| 13:15.6 | Data point 11 | 1.50000 mL | 0.05433 mL | 0.05318 mL | 0.01999 mL | 6.020 | -0.01461 | 0.67162 | 0.00088 | 27.5 s |
| 14:13.7 | Data point 12 | 1.50000 mL | 0.05433 mL | 0.05329 mL | 0.01999 mL | 7.623 | -0.08314 | 0.99820 | 0.00411 | Timed out at 59.5 s |
| 15:39.0 | Data point 13 | 1.50000 mL | 0.05433 mL | 0.05332 mL | 0.01999 mL | 7.556 | -0.01960 | 0.99189 | 0.00097 | 58.5 s |
| 17:08.1 | Data point 14 | 1.50000 mL | 0.05433 mL | 0.05336 mL | 0.01999 mL | 7.824 | -0.04269 | 0.97447 | 0.00214 | Timed out at 59.5 s |
| 18:43.7 | Data point 15 | 1.50000 mL | 0.05433 mL | 0.05343 mL | 0.01999 mL | 8.480 | -0.01896 | 0.89770 | 0.00099 | 53.5 s |
| 20:12.9 | Data point 16 | 1.50000 mL | 0.05433 mL | 0.05367 mL | 0.01999 mL | 9.557 | -0.01673 | 0.69489 | 0.00099 | 17.0 s |
| 21:23.5 | Data point 17 | 1.50000 mL | 0.11192 mL | 0.05367 mL | 0.04499 mL | 1.948 | 0.00696 | 0.77037 | 0.00039 | 10.0 s |
| 22:09.8 | Data point 18 | 1.50000 mL | 0.11192 mL | 0.07298 mL | 0.04499 mL | 2.177 | 0.01234 | 0.90932 | 0.00064 | 10.0 s |
| 22:45.5 | Data point 19 | 1.50000 mL | 0.11192 mL | 0.08420 mL | 0.04499 mL | 2.387 | -0.00583 | 0.35938 | 0.00048 | 10.0 s |
| 23:21.1 | Data point 20 | 1.50000 mL | 0.11192 mL | 0.09116 mL | 0.04499 mL | 2.556 | 0.00696 | 0.30662 | 0.00062 | 10.0 s |
| 24:07.1 | Data point 21 | 1.50000 mL | 0.11192 mL | 0.09781 mL | 0.04499 mL | 2.779 | 0.00229 | 0.34291 | 0.00019 | 10.5 s |
| 24:53.5 | Data point 22 | 1.50000 mL | 0.11192 mL | 0.10169 mL | 0.04499 mL | 3.019 | -0.00417 | 0.40847 | 0.00032 | 10.0 s |
| 25:39.4 | Data point 23 | 1.50000 mL | 0.11192 mL | 0.10503 mL | 0.04499 mL | 3.274 | -0.00304 | 0.48058 | 0.00022 | 10.5 s |
| 26:20.5 | Data point 24 | 1.50000 mL | 0.11192 mL | 0.10750 mL | 0.04499 mL | 3.583 | -0.01595 | 0.64072 | 0.00098 | 10.5 s |
| 27:01.7 | Data point 25 | 1.50000 mL | 0.11192 mL | 0.10915 mL | 0.04499 mL | 4.080 | -0.00893 | 0.27765 | 0.00084 | 11.0 s |
| 27:38.1 | Data point 26 | 1.50000 mL | 0.11192 mL | 0.11065 mL | 0.04499 mL | 9.447 | -0.01272 | 0.80417 | 0.00070 | 22.0 s |
| 28:55.6 | Data point 27 | 1.50000 mL | 0.17265 mL | 0.11065 mL | 0.14499 mL | 1.958 | -0.00711 | 0.87945 | 0.00037 | 10.0 s |
| 29:41.9 | Data point 28 | 1.50000 mL | 0.17265 mL | 0.13189 mL | 0.14499 mL | 2.170 | 0.00288 | 0.03386 | 0.00077 | 10.0 s |
| 30:17.6 | Data point 29 | 1.50000 mL | 0.17265 mL | 0.14407 mL | 0.14499 mL | 2.360 | 0.00992 | 0.49242 | 0.00070 | 10.0 s |
| 30:53.2 | Data point 30 | 1.50000 mL | 0.17265 mL | 0.15212 mL | 0.14499 mL | 2.533 | -0.00071 | 0.03478 | 0.00019 | 10.0 s |
| 31:39.2 | Data point 31 | 1.50000 mL | 0.17265 mL | 0.15898 mL | 0.14499 mL | 2.759 | -0.00280 | 0.03542 | 0.00073 | 10.0 s |
| 32:25.2 | Data point 32 | 1.50000 mL | 0.17265 mL | 0.16376 mL | 0.14499 mL | 2.999 | 0.00572 | 0.11987 | 0.00082 | 10.0 s |
| 33:05.9 | Data point 33 | 1.50000 mL | 0.17265 mL | 0.16672 mL | 0.14499 mL | 3.289 | -0.00045 | 0.04094 | 0.00011 | 10.5 s |
| 33:47.1 | Data point 34 | 1.50000 mL | 0.17265 mL | 0.16874 mL | 0.14499 mL | 3.666 | -0.00646 | 0.91671 | 0.00033 | 10.0 s |
| 34:22.5 | Data point 35 | 1.50000 mL | 0.17265 mL | 0.17058 mL | 0.14499 mL | 4.972 | -0.01889 | 0.93768 | 0.00096 | 29.0 s |
| 35:16.9 | Data point 36 | 1.50000 mL | 0.17265 mL | 0.17063 mL | 0.14499 mL | 6.740 | -0.05158 | 0.98694 | 0.00256 | Timed out at 59.5 s |
| 36:52.5 | Data point 37 | 1.50000 mL | 0.17265 mL | 0.17072 mL | 0.14499 mL | 7.181 | -0.03986 | 0.97488 | 0.00199 | Timed out at 59.5 s |

| | | | |
|---|---|---|---|
| Sample name: | **M02_octanol** | Experiment start time: | **3/1/2018 2:44:22 PM** |
| Assay name: | **pH-metric high logP** | Analyst: | **Dorothy Levorse** |
| Assay ID: | **18C-01011** | Instrument ID: | **T312060** |
| Filename: | **C:\Sirius_T3\Mehtap\20180228_exp28_logP_T3-2\18C-01011_M02_octanol_pH-metric high logP.t3r** | | |

#### Events (continued)

| Time | Event | Water | Acid | Base | Octanol | pH | dpH/dt | pH R-squared | pH SD | dpH/dt time |
|---|---|---|---|---|---|---|---|---|---|---|
| 38:23.1 | Data point 38 | 1.50000 mL | 0.17265 mL | 0.17077 mL | 0.14499 mL | 7.373 | -0.03900 | 0.95207 | 0.00197 | Timed out at 59.5 s |
| 39:53.6 | Data point 39 | 1.50000 mL | 0.17265 mL | 0.17081 mL | 0.14499 mL | 7.756 | -0.05606 | 0.98714 | 0.00279 | Timed out at 59.5 s |
| 41:29.3 | Data point 40 | 1.50000 mL | 0.17265 mL | 0.17103 mL | 0.14499 mL | 8.778 | -0.01761 | 0.76339 | 0.00100 | 24.5 s |
| 42:29.5 | Data point 41 | 1.50000 mL | 0.17265 mL | 0.17114 mL | 0.14499 mL | 8.966 | -0.01729 | 0.75749 | 0.00098 | 17.0 s |
| 43:17.1 | Data point 42 | 1.50000 mL | 0.17265 mL | 0.17121 mL | 0.14499 mL | 9.071 | -0.01579 | 0.68970 | 0.00094 | 23.0 s |
| 43:49.1 | Assay volumes | 1.50000 mL | 0.17265 mL | 0.17121 mL | 0.14499 mL | | | | | |

| | | | |
|---|---|---|---|
| Sample name: | **M02_octanol** | Experiment start time: | **3/1/2018 2:44:22 PM** |
| Assay name: | **pH-metric high logP** | Analyst: | **Dorothy Levorse** |
| Assay ID: | **18C-01011** | Instrument ID: | **T312060** |
| Filename: | **C:\Sirius_T3\Mehtap\20180228_exp28_logP_T3-2\18C-01011_M02_octanol_pH-metric high logP.t3r** | | |

#### Assay Settings

| Setting | Value | Original Value | Date/Time changed | Imported from |
|---|---|---|---|---|
| ***General Settings*** | | | | |
| Analyst name | Dorothy Levorse | | | |
| ***Standard Experiment Settings*** | | | | |
| Number of titrations | 3 | | | |
| Minimum pH | 2.000 | | | |
| Maximum pH | 9.000 | | | |
| pH step between points of | 0.200 | | | |
| Minimum titrant addition | 0.00002 mL | | | |
| Maximum titrant addition | 0.10000 mL | | | |
| Argon flow rate | 100% | | | |
| Start titration using | Cautious pH adjust | | | |
| ***Advanced General Settings*** | | | | |
| Detect turbidity using | None | | | |
| Collect turbidity sensor data | No | | | |
| Collect UV spectra | No | | | |
| Stir after titrant addition for | 5 seconds | | | |
| For titrant addition, stir at | 10% | | | |
| ***Titrant Pre-Dose*** | | | | |
| Titrant pre-dose | None | | | |
| ***Assay Medium*** | | | | |
| ISA water volume | 1.50 mL | | | |
| Water added | Automatic | | | |
| After water addition, stir for | 5 seconds | | | |
| At a speed of | 10% | | | |
| Partition solvent type | Octanol | | | |
| Partition volume | 0.020 mL | | | |
| Partition solvent added | Automatic | | | |
| After partition addition, stir for | 1 seconds | | | |
| ***Sample Sonication*** | | | | |
| Sonicate | No | | | |
| ***Sample Dissolution*** | | | | |
| Perform a dissolution stage | Yes | | | |
| Adjust and hold pH for dissolution | To start pH | | | |
| Stir to dissolve for | 120 seconds | | | |
| For dissolution, stir at | 10% | | | |
| ***Carbonate purge*** | | | | |
| Perform a carbonate purge | No | | | |
| ***Temperature Control*** | | | | |
| Wait for temperature | Yes | | | |
| Required start temperature | 25.0°C | | | |
| Acceptable deviation | 0.5°C | | | |
| Time to wait | 60 seconds | | | |
| Stir speed of | 50% | | | |
| ***Titration 1*** | | | | |
| Titrate from | Low to high pH | | | |
| Adjust to start pH | Yes | | | |
| After pH adjust stir for | 30 seconds | | | |
| Stir to allow partitioning for | 15 seconds | | | |
| Stirrer speed for partitioning | 50% | | | |
| ***Titration 2*** | | | | |
| Titrate from | Low to high pH | | | |
| Add additional water | 0.00 mL | | | |
| Additional partition solvent volume | 0.025 mL | | | |
| Additional partition solvent added | Automatic | | | |
| After pH adjust stir for | 30 seconds | | | |
| Stir to allow partitioning for | 15 seconds | | | |
| Stirrer speed for partitioning | 55% | | | |
| ***Titration 3*** | | | | |

| | | | |
|---|---|---|---|
| Sample name: | **M02_octanol** | Experiment start time: | **3/1/2018 2:44:22 PM** |
| Assay name: | **pH-metric high logP** | Analyst: | **Dorothy Levorse** |
| Assay ID: | **18C-01011** | Instrument ID: | **T312060** |
| Filename: | **C:\Sirius_T3\Mehtap\20180228_exp28_logP_T3-2\18C-01011_M02_octanol_pH-metric high logP.t3r** | | |

#### Assay Settings (continued)

| Setting | Value | Original Value | Date/Time changed | Imported from |
|---|---|---|---|---|
| Titrate from | Low to high pH | | | |
| Add additional water | 0.00 mL | | | |
| Additional partition solvent volume | 0.100 mL | | | |
| Additional partition solvent added | Automatic | | | |
| After pH adjust stir for | 30 seconds | | | |
| Stir to allow partitioning for | 15 seconds | | | |
| Stirrer speed for partitioning | 60% | | | |
| ***Data Point Stability*** | | | | |
| Stir during data point collection | No | | | |
| Delay before data point collection | 0 seconds | | | |
| Number of points to average | 20 points | | | |
| Time interval between points | 0.50 seconds | | | |
| Required maximum standard deviation | 0.00100 dpH/dt | | | |
| Stability timeout after | 60 seconds | | | |

#### Calibration Settings

| Setting | Value | Date/Time changed | Imported from |
|---|---|---|---|
| Four-Plus alpha | 0.130 | 3/1/2018 2:44:22 PM | C:\Sirius_T3\HCl18B27.t3r |
| Four-Plus S | 0.9970 | 3/1/2018 2:44:22 PM | C:\Sirius_T3\HCl18B27.t3r |
| Four-Plus jH | 0.8 | 3/1/2018 2:44:22 PM | C:\Sirius_T3\HCl18B27.t3r |
| Four-Plus jOH | -0.4 | 3/1/2018 2:44:22 PM | C:\Sirius_T3\HCl18B27.t3r |
| Base concentration factor | 1.000 | 3/1/2018 2:44:22 PM | C:\Sirius_T3\KOH18B27.t3r |
| Acid concentration factor | 0.994 | 3/1/2018 2:44:22 PM | C:\Sirius_T3\HCl18B27.t3r |

#### Instrument Settings

| Setting | Value | Batch Id | Install date |
|---|---|---|---|
| Instrument owner | Merck | | |
| Instrument ID | T312060 | | |
| Instrument type | T3 Simulator | | |
| Software version | 1.1.3.0 | | |
| Dispenser module | | T3DM1200361 | 3/31/2009 5:24:52 AM |
| Dispenser 0 | Water | | 3/31/2009 5:25:05 AM |
| Syringe volume | 2.5 mL | | |
| Firmware version | 1.2.1(r2) | | |
| Titrant | Water (0.15 M KCl) | 02-06-2018 | 2/27/2018 10:05:59 AM |
| Dispenser 2 | Acid | | 3/31/2009 5:25:11 AM |
| Syringe volume | 0.5 mL | | |
| Firmware version | 1.2.1(r2) | | |
| Titrant | Acid (0.5 M HCl) | 02-27-2018 | 2/27/2018 10:27:22 AM |
| Dispenser 1 | Base | | 3/31/2009 5:25:21 AM |
| Syringe volume | 0.5 mL | | |
| Firmware version | 1.2.1(r2) | | |
| Titrant | Base (0.5 M KOH) | 9/22/2017 | 2/27/2018 10:21:22 AM |
| Dispenser 5 | Cosolvent | | 3/31/2009 5:26:24 AM |
| Syringe volume | 2.5 mL | | |
| Firmware version | 1.2.1(r2) | | |
| Distribution valve 5 | Distribution Valve | | 3/31/2009 5:28:19 AM |
| Firmware version | 1.1.3 | | |
| Port A | Methanol (80%, 0.15 M KCl) | 09-26-17 | 2/7/2018 9:42:01 AM |
| Port B | Cyclohexane | 11-01-17 | 2/27/2018 10:37:57 AM |
| Dispenser 3 | Buffer | | 8/3/2010 5:05:16 AM |
| Syringe volume | 0.5 mL | | |
| Firmware version | 1.2.1(r2) | | |
| Titrant | Dodecane | 2018/01/31 | 2/28/2018 10:18:04 AM |
| Dispenser 6 | Octanol | | 10/22/2010 10:52:43 AM |
| Syringe volume | 0.5 mL | | |

Sample name: **M02_octanol**
Assay name: **pH-metric high logP**
Assay ID: **18C-01011**
Filename: **C:\Sirius_T3\Mehtap\20180228_exp28_logP_T3-2\18C-01011_M02_octanol_pH-metric high logP.t3r**

Experiment start time: **3/1/2018 2:44:22 PM**
Analyst: **Dorothy Levorse**
Instrument ID: **T312060**

#### Instrument Settings (continued)

| Setting | Value | Batch Id | Install date |
|---|---|---|---|
| Firmware version | 1.2.1(r2) | | |
| Titrant | Octanol | 01-31-2018 | 2/27/2018 9:59:35 AM |
| Titrator | | T3TM1200161 | 3/31/2009 5:24:17 AM |
| Horizontal axis firmware version | 1.17 AI1DI2DO2 Stepper 2 | | |
| Vertical axis firmware version | 1.17 AI1DI2DO2 Stepper 2 | | |
| Chassis I/O firmware version | 1.11 AI1DI0DO4 Norgren I/O | | |
| Probe I/O firmware version | 1.1.1 | | |
| Electrode | T3 Electrode | T3E0923 | 1/23/2018 2:01:00 PM |
| E0 calibration | +3.14 mV | | 3/1/2018 2:44:50 PM |
| Filling solution | 3M KCl | KCL097 | 2/27/2018 9:49:43 AM |
| Liquids | | | |
| Wash 1 | 50% IPA:50% Water | | 2/28/2018 10:23:32 AM |
| Wash 2 | 0.5% Trition X-100 in H20 | | 2/28/2018 10:23:34 AM |
| Buffer position 1 | pH7 Wash | | 2/28/2018 10:24:06 AM |
| Buffer position 2 | pH 7 | | 2/28/2018 10:24:08 AM |
| Storage position | | | 2/28/2018 10:21:14 AM |
| Wash water | 8.1e+003 mL | 02-27-2018 | 2/27/2018 9:54:39 AM |
| Waste | 7.3e+003 mL | | 11/28/2017 10:36:29 AM |
| Temperature controller | | | 8/5/2010 6:35:13 AM |
| Turbidity detector | | | 3/31/2009 5:24:45 AM |
| Spectrometer | | 074811 | 11/23/2010 11:22:28 AM |
| Dip probe | | 10196 | |
| Wavelength coefficient A0 | 183.333 | | |
| Wavelength coefficient A1 | 2.21568 | | |
| Wavelength coefficient A2 | -0.000289308 | | |
| Total lamp lit time | 112:08:55 | | 11/23/2010 11:22:28 AM |
| Calibrated on | 2/27/2018 10:40:38 AM | | |
| Integration time | 40 | | |
| Scans averaged | 10 | | |
| Autoloader | | T3AL1200345 | 11/10/2015 9:34:13 AM |
| Left-right axis firmware version | 1.17 AI1DI2DO2 Stepper 2 | | |
| Front-back axis firmware version | 1.17 AI1DI2DO2 Stepper 2 | | |
| Vertical axis firmware version | 1.17 AI1DI2DO2 Stepper 2 | | |
| Chassis I/O firmware version | 1.11 AI1DI0DO4 Norgren I/O | | |
| Configuration | | | |
| Alternate titration position | Titration position | | |
| Alternate reference position | Reference position | | |
| Maximum standard vial volume | 3.50 mL | | |
| Maximum alternate vial volume | 25.00 mL | | |
| Automatic action idle period | 5 minute(s) | | |
| Titrant tube volume | 1.3 mL | | |
| Syringe flush count | 3.50 | | |
| Flowing wash pump volume | 20.0 mL | | |
| Flowing wash stir duration | 5 s | | |
| Flowing wash stir speed | 30% | | |
| Solvent wash stir duration | 5 s | | |
| Solvent wash stir speed | 30% | | |
| Surfactant wash stir duration | 5 s | | |
| Surfactant wash stir speed | 30% | | |
| E0 calibration minimum number of points | 10 | | |
| E0 calibration maximum standard deviation | 0.01500 | | |
| E0 calibration timeout period | 60 s | | |
| E0 calibration stir duration | 5 s | | |
| E0 calibration preparation stir speed | 30% | | |
| E0 calibration buffer wash stir duration | 5 s | | |
| E0 calibration buffer wash stir speed | 30% | | |
| E0 calibration reading stir speed | 0% | | |
| Spectrometer calibration stir duration | 5 s | | |

| | | | |
|---|---|---|---|
| Sample name: | **M02_octanol** | Experiment start time: | **3/1/2018 2:44:22 PM** |
| Assay name: | **pH-metric high logP** | Analyst: | **Dorothy Levorse** |
| Assay ID: | **18C-01011** | Instrument ID: | **T312060** |
| Filename: | **C:\Sirius_T3\Mehtap\20180228_exp28_logP_T3-2\18C-01011_M02_octanol_pH-metric high logP.t3r** | | |

#### Instrument Settings (continued)

| Setting | Value | Batch Id | Install date |
|---|---|---|---|
| Spectrometer calibration stir speed | 30% | | |
| Spectrometer calibration wash pump volume | 20.0 mL | | |
| Spectrometer calibration wash stir duration | 5 s | | |
| Spectrometer calibration wash stir speed | 30% | | |
| Overhead dispense height | 10000 | | |

#### Refinement Settings

| Setting | Value | Default value |
|---|---|---|
| Turbidity detection method | None | None |
| Turbidity wavelength to assess | 500.0 nm | 500.0 nm |
| Turbidity maximum absorbance | 0.100 | 0.100 |
| Turbidity probe threshold | 50.00 | 50.00 |

#### Experiment Log

[2:37] Air gap created for Water (0.15 M KCl)
[2:38] Air gap created for Acid (0.5 M HCl)
[2:38] Air gap created for Base (0.5 M KOH)
[2:38] Air gap released for Water (0.15 M KCl)
[2:43] Titrator arm moved to Titration position
[2:43] Argon flow rate set to 100
[2:43] Titration 1 of 3
[2:43] Adding initial titrants
[2:43] Automatically add 1.50000 mL of water
[3:09] Dispensed 1.500000 mL of Water (0.15 M KCl)
[3:09] Stirrer speed set to 10
[3:14] Automatically add 0.02000 mL of Octanol
[3:14] Dispensed 0.019991 mL of Octanol
[3:15] Initial pH = 6.92
[3:15] Iterative adjust 6.92 -> 2.00
[3:15] pH 6.92 -> 2.00
[3:17] Air gap released for Acid (0.5 M HCl)
[3:18] Dispensed 0.054327 mL of Acid (0.5 M HCl)
[3:23] Holding pH 2.00
[5:23] Stirrer speed set to 0
[5:23] Stirrer speed set to 50
[5:23] Iterative adjust 1.90 -> 2.00
[5:23] pH 1.90 -> 2.00
[5:24] Air gap released for Base (0.5 M KOH)
[5:25] Dispensed 0.011195 mL of Base (0.5 M KOH)
[6:15] Stirrer speed set to 0
[6:25] Datapoint id 1 collected
[6:25] Stirrer speed set to 50
[6:30] pH 2.07 -> 2.27
[6:30] Using cautious pH adjust
[6:31] Dispensed 0.006867 mL of Base (0.5 M KOH)
[6:36] Stepping pH = 2.15
[6:36] Dispensed 0.005386 mL of Base (0.5 M KOH)
[6:41] Stepping pH = 2.24
[6:41] Dispensed 0.001388 mL of Base (0.5 M KOH)
[6:46] Stepping pH = 2.27
[7:02] Stirrer speed set to 0
[7:12] Datapoint id 2 collected
[7:12] Charge balance equation is out by 0.6%
[7:12] Stirrer speed set to 50
[7:17] pH 2.29 -> 2.49
[7:17] Using charge balance adjust
[7:17] Dispensed 0.008161 mL of Base (0.5 M KOH)

| | | | |
|---|---|---|---|
| Sample name: | **M02_octanol** | Experiment start time: | **3/1/2018 2:44:22 PM** |
| Assay name: | **pH-metric high logP** | Analyst: | **Dorothy Levorse** |
| Assay ID: | **18C-01011** | Instrument ID: | **T312060** |
| Filename: | **C:\Sirius_T3\Mehtap\20180228_exp28_logP_T3-2\18C-01011_M02_octanol_pH-metric high logP.t3r** | | |

#### Experiment Log (continued)

[7:37] Stirrer speed set to 0
[7:47] Datapoint id 3 collected
[7:47] Charge balance equation is out by -4.6%
[7:47] Stirrer speed set to 50
[7:52] pH 2.50 -> 2.70
[7:52] Using charge balance adjust
[7:53] Dispensed 0.005151 mL of Base (0.5 M KOH)
[8:13] Stirrer speed set to 0
[8:23] Datapoint id 4 collected
[8:23] Charge balance equation is out by -7.3%
[8:23] Stirrer speed set to 50
[8:28] pH 2.70 -> 2.90
[8:28] Using charge balance adjust
[8:28] Dispensed 0.003387 mL of Base (0.5 M KOH)
[8:48] Stirrer speed set to 0
[8:59] Datapoint id 5 collected
[8:59] Charge balance equation is out by -24.9%
[8:59] Stirrer speed set to 50
[9:04] pH 2.86 -> 3.06
[9:04] Using cautious pH adjust
[9:04] Dispensed 0.001246 mL of Base (0.5 M KOH)
[9:09] Stepping pH = 2.89
[9:09] Dispensed 0.002281 mL of Base (0.5 M KOH)
[9:14] Stepping pH = 3.07
[9:29] Stirrer speed set to 0
[9:39] Datapoint id 6 collected
[9:39] Charge balance equation is out by -41.5%
[9:39] Stirrer speed set to 50
[9:45] pH 3.10 -> 3.30
[9:45] Using cautious pH adjust
[9:45] Dispensed 0.000870 mL of Base (0.5 M KOH)
[9:50] Stepping pH = 3.12
[9:50] Dispensed 0.002023 mL of Base (0.5 M KOH)
[9:55] Stepping pH = 3.29
[10:10] Stirrer speed set to 0
[10:26] Datapoint id 7 collected
[10:26] Charge balance equation is out by -66.6%
[10:26] Stirrer speed set to 50
[10:31] pH 3.33 -> 3.53
[10:31] Using cautious pH adjust
[10:31] Dispensed 0.000729 mL of Base (0.5 M KOH)
[10:36] Stepping pH = 3.36
[10:37] Dispensed 0.001670 mL of Base (0.5 M KOH)
[10:42] Stepping pH = 3.58
[10:57] Stirrer speed set to 0
[11:08] Datapoint id 8 collected
[11:08] Charge balance equation is out by -63.8%
[11:08] Stirrer speed set to 50
[11:13] pH 3.65 -> 3.85
[11:13] Using cautious pH adjust
[11:14] Dispensed 0.000753 mL of Base (0.5 M KOH)
[11:19] Stepping pH = 3.70
[11:19] Dispensed 0.001152 mL of Base (0.5 M KOH)
[11:24] Stepping pH = 4.01
[11:39] Stirrer speed set to 0
[11:55] Datapoint id 9 collected
[11:55] Charge balance equation is out by -26.9%
[11:55] Stirrer speed set to 50
[12:00] pH 4.22 -> 4.42

| | | | |
|---|---|---|---|
| Sample name: | **M02_octanol** | Experiment start time: | **3/1/2018 2:44:22 PM** |
| Assay name: | **pH-metric high logP** | Analyst: | **Dorothy Levorse** |
| Assay ID: | **18C-01011** | Instrument ID: | **T312060** |
| Filename: | **C:\Sirius_T3\Mehtap\20180228_exp28_logP_T3-2\18C-01011_M02_octanol_pH-metric high logP.t3r** | | |

#### Experiment Log (continued)

[12:00] Using cautious pH adjust
[12:00] Dispensed 0.000776 mL of Base (0.5 M KOH)
[12:05] Stepping pH = 4.49
[12:20] Stirrer speed set to 0
[12:45] Datapoint id 10 collected
[12:45] Charge balance equation is out by 50.0%
[12:45] Stirrer speed set to 50
[12:50] pH 5.50 -> 5.70
[12:50] Using cautious pH adjust
[12:50] Dispensed 0.000118 mL of Base (0.5 M KOH)
[12:55] Stepping pH = 5.66
[12:55] Dispensed 0.000024 mL of Base (0.5 M KOH)
[13:00] Stepping pH = 5.88
[13:16] Stirrer speed set to 0
[13:43] Datapoint id 11 collected
[13:43] Charge balance equation is out by 40.8%
[13:43] Stirrer speed set to 50
[13:48] pH 6.03 -> 6.23
[13:48] Using cautious pH adjust
[13:48] Dispensed 0.000047 mL of Base (0.5 M KOH)
[13:53] Stepping pH = 6.09
[13:53] Dispensed 0.000071 mL of Base (0.5 M KOH)
[13:59] Stepping pH = 6.65
[14:14] Stirrer speed set to 0
[15:14] Datapoint id 12 collected
[15:14] Charge balance equation is out by -9.8%
[15:14] Stirrer speed set to 50
[15:19] pH 7.55 -> 7.75
[15:19] Using charge balance adjust
[15:19] Dispensed 0.000024 mL of Base (0.5 M KOH)
[15:39] Stirrer speed set to 0
[16:38] Datapoint id 13 collected
[16:38] Charge balance equation is out by -97.4%
[16:38] Stirrer speed set to 50
[16:43] pH 7.40 -> 7.60
[16:43] Using cautious pH adjust
[16:43] Dispensed 0.000024 mL of Base (0.5 M KOH)
[16:48] Stepping pH = 7.41
[16:48] Dispensed 0.000024 mL of Base (0.5 M KOH)
[16:53] Stepping pH = 7.62
[17:08] Stirrer speed set to 0
[18:08] Datapoint id 14 collected
[18:08] Charge balance equation is out by -248.2%
[18:08] Stirrer speed set to 50
[18:13] pH 7.67 -> 7.87
[18:13] Using cautious pH adjust
[18:13] Dispensed 0.000024 mL of Base (0.5 M KOH)
[18:18] Stepping pH = 7.62
[18:18] Dispensed 0.000024 mL of Base (0.5 M KOH)
[18:23] Stepping pH = 7.86
[18:23] Dispensed 0.000024 mL of Base (0.5 M KOH)
[18:29] Stepping pH = 8.40
[18:44] Stirrer speed set to 0
[19:37] Datapoint id 15 collected
[19:37] Charge balance equation is out by -741.2%
[19:37] Stirrer speed set to 50
[19:42] pH 8.40 -> 8.60
[19:42] Using cautious pH adjust
[19:42] Dispensed 0.000024 mL of Base (0.5 M KOH)

Sample name: **M02_octanol**
Assay name: **pH-metric high logP**
Assay ID: **18C-01011**
Filename: **C:\Sirius_T3\Mehtap\20180228_exp28_logP_T3-2\18C-01011_M02_octanol_pH-metric high logP.t3r**

Experiment start time: **3/1/2018 2:44:22 PM**
Analyst: **Dorothy Levorse**
Instrument ID: **T312060**

#### Experiment Log (continued)

[19:47] Stepping pH = 8.38
[19:48] Dispensed 0.000024 mL of Base (0.5 M KOH)
[19:53] Stepping pH = 8.35
[19:53] Dispensed 0.000188 mL of Base (0.5 M KOH)
[19:58] Stepping pH = 9.41
[20:13] Stirrer speed set to 0
[20:30] Datapoint id 16 collected
[20:30] Charge balance equation is out by -1,840.2%
[20:30] Titration 2 of 3
[20:30] Adding initial titrants
[20:30] Automatically add 0.02500 mL of Octanol
[20:31] Dispensed 0.025000 mL of Octanol
[20:31] Stirrer speed set to 10
[20:32] Stirrer speed set to 55
[20:32] Iterative adjust 9.57 -> 2.00
[20:32] pH 9.57 -> 2.00
[20:33] Dispensed 0.057596 mL of Acid (0.5 M HCl)
[21:23] Stirrer speed set to 0
[21:33] Datapoint id 17 collected
[21:33] Stirrer speed set to 55
[21:39] pH 1.96 -> 2.16
[21:39] Using cautious pH adjust
[21:39] Dispensed 0.009454 mL of Base (0.5 M KOH)
[21:44] Stepping pH = 2.04
[21:44] Dispensed 0.007502 mL of Base (0.5 M KOH)
[21:49] Stepping pH = 2.12
[21:50] Dispensed 0.002352 mL of Base (0.5 M KOH)
[21:55] Stepping pH = 2.16
[22:10] Stirrer speed set to 0
[22:20] Datapoint id 18 collected
[22:20] Charge balance equation is out by -2.0%
[22:20] Stirrer speed set to 55
[22:25] pH 2.19 -> 2.39
[22:25] Using charge balance adjust
[22:25] Dispensed 0.011218 mL of Base (0.5 M KOH)
[22:45] Stirrer speed set to 0
[22:55] Datapoint id 19 collected
[22:55] Charge balance equation is out by 1.0%
[22:55] Stirrer speed set to 55
[23:01] pH 2.40 -> 2.60
[23:01] Using charge balance adjust
[23:01] Dispensed 0.006961 mL of Base (0.5 M KOH)
[23:21] Stirrer speed set to 0
[23:31] Datapoint id 20 collected
[23:31] Charge balance equation is out by -20.4%
[23:31] Stirrer speed set to 55
[23:36] pH 2.57 -> 2.77
[23:36] Using cautious pH adjust
[23:36] Dispensed 0.002422 mL of Base (0.5 M KOH)
[23:41] Stepping pH = 2.62
[23:42] Dispensed 0.003316 mL of Base (0.5 M KOH)
[23:47] Stepping pH = 2.73
[23:47] Dispensed 0.000917 mL of Base (0.5 M KOH)
[23:52] Stepping pH = 2.76
[24:07] Stirrer speed set to 0
[24:18] Datapoint id 21 collected
[24:18] Charge balance equation is out by -37.8%
[24:18] Stirrer speed set to 55
[24:23] pH 2.79 -> 2.99

| | | | |
|---|---|---|---|
| Sample name: | **M02_octanol** | Experiment start time: | **3/1/2018 2:44:22 PM** |
| Assay name: | **pH-metric high logP** | Analyst: | **Dorothy Levorse** |
| Assay ID: | **18C-01011** | Instrument ID: | **T312060** |
| Filename: | **C:\Sirius_T3\Mehtap\20180228_exp28_logP_T3-2\18C-01011_M02_octanol_pH-metric high logP.t3r** | | |

#### Experiment Log (continued)

[24:23] Using cautious pH adjust
[24:23] Dispensed 0.001552 mL of Base (0.5 M KOH)
[24:28] Stepping pH = 2.84
[24:28] Dispensed 0.002187 mL of Base (0.5 M KOH)
[24:33] Stepping pH = 2.98
[24:33] Dispensed 0.000141 mL of Base (0.5 M KOH)
[24:38] Stepping pH = 3.00
[24:53] Stirrer speed set to 0
[25:03] Datapoint id 22 collected
[25:03] Charge balance equation is out by -25.1%
[25:03] Stirrer speed set to 55
[25:09] pH 3.03 -> 3.23
[25:09] Using cautious pH adjust
[25:09] Dispensed 0.001082 mL of Base (0.5 M KOH)
[25:14] Stepping pH = 3.07
[25:14] Dispensed 0.001952 mL of Base (0.5 M KOH)
[25:19] Stepping pH = 3.20
[25:19] Dispensed 0.000306 mL of Base (0.5 M KOH)
[25:24] Stepping pH = 3.25
[25:39] Stirrer speed set to 0
[25:50] Datapoint id 23 collected
[25:50] Charge balance equation is out by -53.6%
[25:50] Stirrer speed set to 55
[25:55] pH 3.29 -> 3.49
[25:55] Using cautious pH adjust
[25:55] Dispensed 0.000894 mL of Base (0.5 M KOH)
[26:00] Stepping pH = 3.33
[26:00] Dispensed 0.001576 mL of Base (0.5 M KOH)
[26:05] Stepping pH = 3.51
[26:20] Stirrer speed set to 0
[26:31] Datapoint id 24 collected
[26:31] Charge balance equation is out by -39.1%
[26:31] Stirrer speed set to 55
[26:36] pH 3.61 -> 3.81
[26:36] Using cautious pH adjust
[26:36] Dispensed 0.000847 mL of Base (0.5 M KOH)
[26:41] Stepping pH = 3.69
[26:41] Dispensed 0.000800 mL of Base (0.5 M KOH)
[26:46] Stepping pH = 3.90
[27:02] Stirrer speed set to 0
[27:13] Datapoint id 25 collected
[27:13] Charge balance equation is out by 2.9%
[27:13] Stirrer speed set to 55
[27:18] pH 4.16 -> 4.36
[27:18] Using charge balance adjust
[27:18] Dispensed 0.001505 mL of Base (0.5 M KOH)
[27:38] Stirrer speed set to 0
[28:00] Datapoint id 26 collected
[28:00] Charge balance equation is out by 2,543.0%
[28:00] Titration 3 of 3
[28:00] Adding initial titrants
[28:00] Automatically add 0.10000 mL of Octanol
[28:03] Dispensed 0.100000 mL of Octanol
[28:03] Stirrer speed set to 10
[28:04] Stirrer speed set to 60
[28:04] Iterative adjust 9.54 -> 2.00
[28:04] pH 9.54 -> 2.00
[28:05] Dispensed 0.060724 mL of Acid (0.5 M HCl)
[28:55] Stirrer speed set to 0

| | | | |
|---|---|---|---|
| Sample name: | **M02_octanol** | Experiment start time: | **3/1/2018 2:44:22 PM** |
| Assay name: | **pH-metric high logP** | Analyst: | **Dorothy Levorse** |
| Assay ID: | **18C-01011** | Instrument ID: | **T312060** |
| Filename: | **C:\Sirius_T3\Mehtap\20180228_exp28_logP_T3-2\18C-01011_M02_octanol_pH-metric high logP.t3r** | | |

#### Experiment Log (continued)

[29:06] Datapoint id 27 collected
[29:06] Stirrer speed set to 60
[29:11] pH 1.96 -> 2.16
[29:11] Using cautious pH adjust
[29:11] Dispensed 0.010042 mL of Base (0.5 M KOH)
[29:16] Stepping pH = 2.04
[29:16] Dispensed 0.008937 mL of Base (0.5 M KOH)
[29:22] Stepping pH = 2.13
[29:22] Dispensed 0.002258 mL of Base (0.5 M KOH)
[29:27] Stepping pH = 2.16
[29:42] Stirrer speed set to 0
[29:52] Datapoint id 28 collected
[29:52] Charge balance equation is out by -5.7%
[29:52] Stirrer speed set to 60
[29:57] pH 2.18 -> 2.38
[29:57] Using charge balance adjust
[29:57] Dispensed 0.012183 mL of Base (0.5 M KOH)
[30:18] Stirrer speed set to 0
[30:28] Datapoint id 29 collected
[30:28] Charge balance equation is out by -10.3%
[30:28] Stirrer speed set to 60
[30:33] pH 2.37 -> 2.57
[30:33] Using charge balance adjust
[30:33] Dispensed 0.008043 mL of Base (0.5 M KOH)
[30:53] Stirrer speed set to 0
[31:03] Datapoint id 30 collected
[31:03] Charge balance equation is out by -16.1%
[31:03] Stirrer speed set to 60
[31:08] pH 2.54 -> 2.74
[31:08] Using cautious pH adjust
[31:08] Dispensed 0.002775 mL of Base (0.5 M KOH)
[31:14] Stepping pH = 2.59
[31:14] Dispensed 0.003716 mL of Base (0.5 M KOH)
[31:19] Stepping pH = 2.72
[31:19] Dispensed 0.000376 mL of Base (0.5 M KOH)
[31:24] Stepping pH = 2.75
[31:39] Stirrer speed set to 0
[31:49] Datapoint id 31 collected
[31:49] Charge balance equation is out by -24.3%
[31:49] Stirrer speed set to 60
[31:54] pH 2.77 -> 2.97
[31:54] Using cautious pH adjust
[31:54] Dispensed 0.001787 mL of Base (0.5 M KOH)
[31:59] Stepping pH = 2.82
[32:00] Dispensed 0.002563 mL of Base (0.5 M KOH)
[32:05] Stepping pH = 2.94
[32:05] Dispensed 0.000423 mL of Base (0.5 M KOH)
[32:10] Stepping pH = 2.98
[32:25] Stirrer speed set to 0
[32:35] Datapoint id 32 collected
[32:35] Charge balance equation is out by -33.2%
[32:35] Stirrer speed set to 60
[32:40] pH 3.01 -> 3.21
[32:40] Using cautious pH adjust
[32:40] Dispensed 0.001246 mL of Base (0.5 M KOH)
[32:45] Stepping pH = 3.06
[32:46] Dispensed 0.001717 mL of Base (0.5 M KOH)
[32:51] Stepping pH = 3.23
[33:06] Stirrer speed set to 0

Sample name: **M02_octanol** Experiment start time: **3/1/2018 2:44:22 PM**
Assay name: **pH-metric high logP** Analyst: **Dorothy Levorse**
Assay ID: **18C-01011** Instrument ID: **T312060**
Filename: **C:\Sirius_T3\Mehtap\20180228_exp28_logP_T3-2\18C-01011_M02_octanol_pH-metric high logP.t3r**

#### Experiment Log (continued)

[33:16] Datapoint id 33 collected
[33:16] Charge balance equation is out by -18.3%
[33:16] Stirrer speed set to 60
[33:21] pH 3.31 -> 3.51
[33:21] Using cautious pH adjust
[33:22] Dispensed 0.001011 mL of Base (0.5 M KOH)
[33:27] Stepping pH = 3.38
[33:27] Dispensed 0.001011 mL of Base (0.5 M KOH)
[33:32] Stepping pH = 3.56
[33:47] Stirrer speed set to 0
[33:57] Datapoint id 34 collected
[33:57] Charge balance equation is out by 0.3%
[33:57] Stirrer speed set to 60
[34:02] pH 3.70 -> 3.90
[34:02] Using charge balance adjust
[34:02] Dispensed 0.001834 mL of Base (0.5 M KOH)
[34:22] Stirrer speed set to 0
[34:52] Datapoint id 35 collected
[34:52] Charge balance equation is out by 535.4%
[34:52] Stirrer speed set to 60
[34:57] pH 6.04 -> 6.24
[34:57] Using cautious pH adjust
[34:57] Dispensed 0.000047 mL of Base (0.5 M KOH)
[35:02] Stepping pH = 6.25
[35:17] Stirrer speed set to 0
[36:17] Datapoint id 36 collected
[36:17] Charge balance equation is out by 50.0%
[36:17] Stirrer speed set to 60
[36:22] pH 6.68 -> 6.88
[36:22] Using cautious pH adjust
[36:22] Dispensed 0.000024 mL of Base (0.5 M KOH)
[36:27] Stepping pH = 6.69
[36:27] Dispensed 0.000047 mL of Base (0.5 M KOH)
[36:32] Stepping pH = 6.81
[36:32] Dispensed 0.000024 mL of Base (0.5 M KOH)
[36:37] Stepping pH = 6.99
[36:52] Stirrer speed set to 0
[37:52] Datapoint id 37 collected
[37:52] Charge balance equation is out by -157.1%
[37:52] Stirrer speed set to 60
[37:58] pH 7.09 -> 7.29
[37:58] Using cautious pH adjust
[37:58] Dispensed 0.000024 mL of Base (0.5 M KOH)
[38:03] Stepping pH = 7.18
[38:03] Dispensed 0.000024 mL of Base (0.5 M KOH)
[38:08] Stepping pH = 7.33
[38:23] Stirrer speed set to 0
[39:23] Datapoint id 38 collected
[39:23] Charge balance equation is out by -88.9%
[39:23] Stirrer speed set to 60
[39:28] pH 7.33 -> 7.53
[39:28] Using cautious pH adjust
[39:28] Dispensed 0.000024 mL of Base (0.5 M KOH)
[39:33] Stepping pH = 7.45
[39:33] Dispensed 0.000024 mL of Base (0.5 M KOH)
[39:38] Stepping pH = 7.64
[39:54] Stirrer speed set to 0
[40:54] Datapoint id 39 collected
[40:54] Charge balance equation is out by -184.4%

| | | | |
|---|---|---|---|
| Sample name: | **M02_octanol** | Experiment start time: | **3/1/2018 2:44:22 PM** |
| Assay name: | **pH-metric high logP** | Analyst: | **Dorothy Levorse** |
| Assay ID: | **18C-01011** | Instrument ID: | **T312060** |
| Filename: | **C:\Sirius_T3\Mehtap\20180228_exp28_logP_T3-2\18C-01011_M02_octanol_pH-metric high logP.t3r** | | |

#### Experiment Log (continued)

[40:54] Stirrer speed set to 60
[40:59] pH 7.66 -> 7.86
[40:59] Using cautious pH adjust
[40:59] Dispensed 0.000024 mL of Base (0.5 M KOH)
[41:04] Stepping pH = 7.62
[41:04] Dispensed 0.000024 mL of Base (0.5 M KOH)
[41:09] Stepping pH = 7.58
[41:09] Dispensed 0.000165 mL of Base (0.5 M KOH)
[41:14] Stepping pH = 8.20
[41:29] Stirrer speed set to 0
[41:54] Datapoint id 40 collected
[41:54] Charge balance equation is out by -2,249.9%
[41:54] Stirrer speed set to 60
[41:59] pH 8.79 -> 8.99
[41:59] Using cautious pH adjust
[41:59] Dispensed 0.000024 mL of Base (0.5 M KOH)
[42:04] Stepping pH = 8.81
[42:04] Dispensed 0.000047 mL of Base (0.5 M KOH)
[42:09] Stepping pH = 8.88
[42:09] Dispensed 0.000047 mL of Base (0.5 M KOH)
[42:14] Stepping pH = 9.00
[42:29] Stirrer speed set to 0
[42:47] Datapoint id 41 collected
[42:47] Charge balance equation is out by -246.2%
[42:47] Stirrer speed set to 60
[42:52] pH 8.96 -> 9.05
[42:52] Using cautious pH adjust
[42:52] Dispensed 0.000024 mL of Base (0.5 M KOH)
[42:57] Stepping pH = 8.95
[42:57] Dispensed 0.000047 mL of Base (0.5 M KOH)
[43:02] Stepping pH = 9.06
[43:17] Stirrer speed set to 0
[43:40] Datapoint id 42 collected
[43:40] Charge balance equation is out by -272.1%
[43:40] Argon flow rate set to 0
[43:44] Titrator arm moved over Titration position
