## Supplementary material for "Octanol-water partition coefficient measurements for the SAMPL6 Blind Prediction Challenge": SM02_18C-03011_M02_octanol_pH-metric high logP_report.pdf

| | | | |
|---|---|---|---|
| Sample name: | **M02_octanol** | Experiment start time: | **3/3/2018 1:46:01 PM** |
| Assay name: | **pH-metric high logP** | Analyst: | **Dorothy Levorse** |
| Assay ID: | **18C-03011** | Instrument ID: | **T312060** |
| Filename: | **C:\Sirius_T3\Mehtap\20180302_exp29_logP_T3-2\18C-03011_M02_octanol_pH-metric high logP.t3r** | | |

#### pH-metric Result

| | | |
|---|---|---|
| logP (XH +) | 0.45 | ±0.04 (n=50) |
| logP (neutral X) | 4.08 | ±0.01 (n=50) |

#### 18C-03011 Points 1 to 20

| | |
|---|---|
| M02_octanol concentration factor | 0.967 |
| Carbonate | 0.1632 mM |
| Acidity error | -1.49856 mM |

#### 18C-03011 Points 21 to 48

| | |
|---|---|
| M02_octanol concentration factor | 1.157 |
| Carbonate | 0.1227 mM |
| Acidity error | -1.56956 mM |

#### 18C-03011 Points 49 to 70

| | |
|---|---|
| M02_octanol concentration factor | 1.597 |
| Carbonate | 0.1477 mM |
| Acidity error | -1.25180 mM |

#### Warnings and errors

| | |
|---|---|
| Errors | None |
| Warnings | None |

#### Sample logD and percent species

| pH | M02_octanol logD | M02_octanol M02_octanolH | M02_octanol M02_octanol | M02_octanol M02_octanolH* | M02_octanol M02_octanol* | Comment |
|---|---|---|---|---|---|---|
| 1.000 | 0.59 | 20.31 % | 0.00 % | 56.73 % | 22.95 % | |
| 1.200 | 0.66 | 17.91 % | 0.00 % | 50.02 % | 32.07 % | Stomach pH |
| 2.000 | 1.15 | 6.63 % | 0.01 % | 18.50 % | 74.87 % | |
| 3.000 | 2.06 | 0.86 % | 0.01 % | 2.39 % | 96.74 % | |
| 4.000 | 3.02 | 0.09 % | 0.01 % | 0.25 % | 99.66 % | |
| 5.000 | 3.77 | 0.01 % | 0.01 % | 0.02 % | 99.96 % | |
| 6.000 | 4.04 | 0.00 % | 0.01 % | 0.00 % | 99.99 % | |
| 6.500 | 4.07 | 0.00 % | 0.01 % | 0.00 % | 99.99 % | |
| 7.000 | 4.08 | 0.00 % | 0.01 % | 0.00 % | 99.99 % | |
| 7.400 | 4.08 | 0.00 % | 0.01 % | 0.00 % | 99.99 % | Blood pH |
| 8.000 | 4.08 | 0.00 % | 0.01 % | 0.00 % | 99.99 % | |
| 9.000 | 4.08 | 0.00 % | 0.01 % | 0.00 % | 99.99 % | |
| 10.000 | 4.08 | 0.00 % | 0.01 % | 0.00 % | 99.99 % | |
| 11.000 | 4.08 | 0.00 % | 0.01 % | 0.00 % | 99.99 % | |
| 12.000 | 4.08 | 0.00 % | 0.01 % | 0.00 % | 99.99 % | |

Sample name: **M02_octanol**
Assay name: **pH-metric high logP**
Assay ID: **18C-03011**
Filename: **C:\Sirius_T3\Mehtap\20180302_exp29_logP_T3-2\18C-03011_M02_octanol_pH-metric high logP.t3r**

Experiment start time: **3/3/2018 1:46:01 PM**
Analyst: **Dorothy Levorse**
Instrument ID: **T312060**

#### Graphs

Charge balance residuals

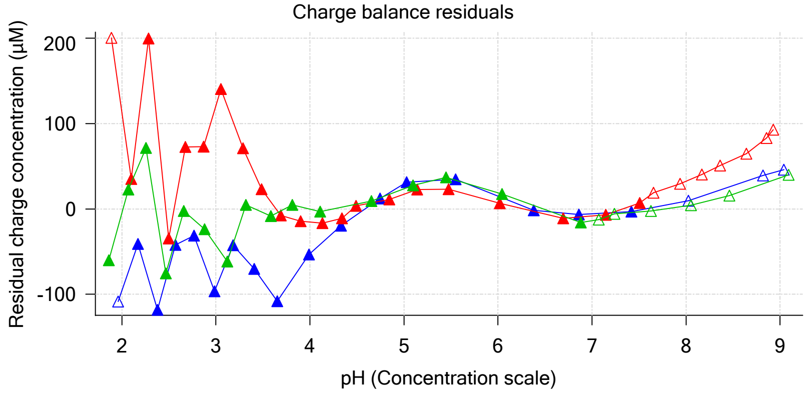


Ionic strength

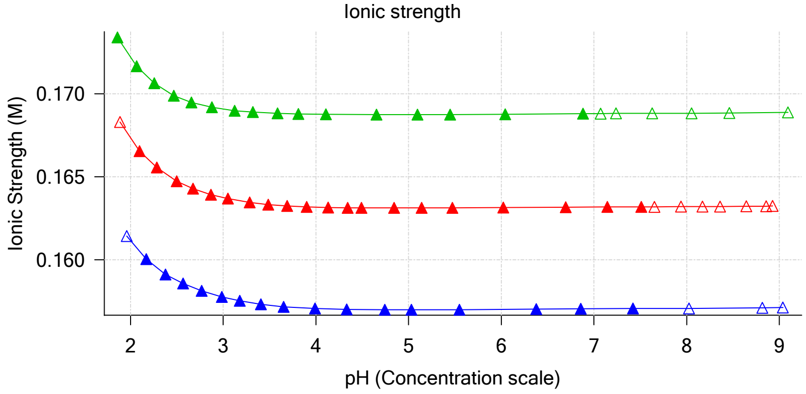


Buffer index profile

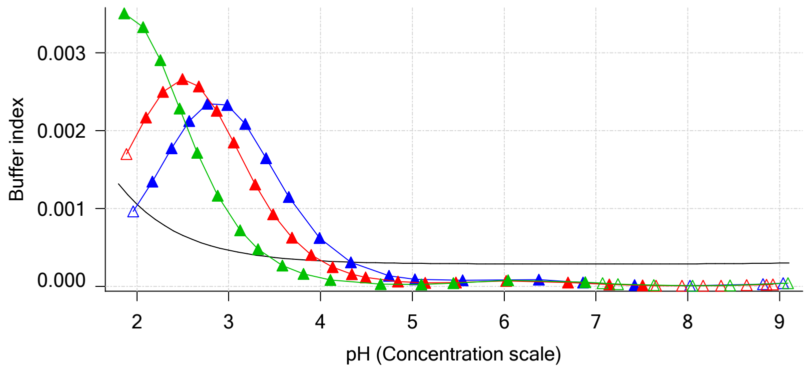


Sample M02_octanol Ionisation Graph

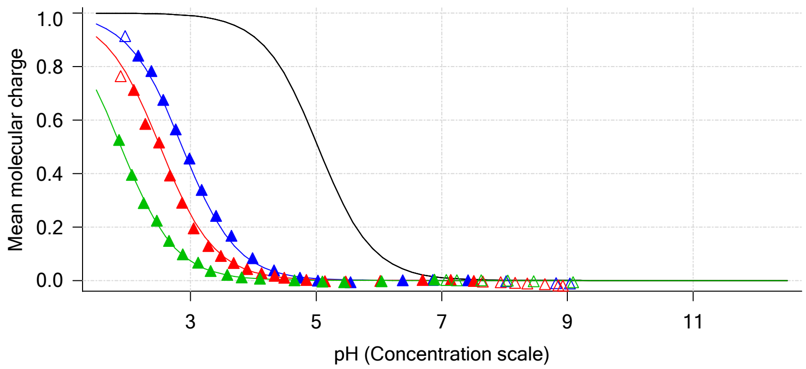


Distribution of species for sample M02_octanol

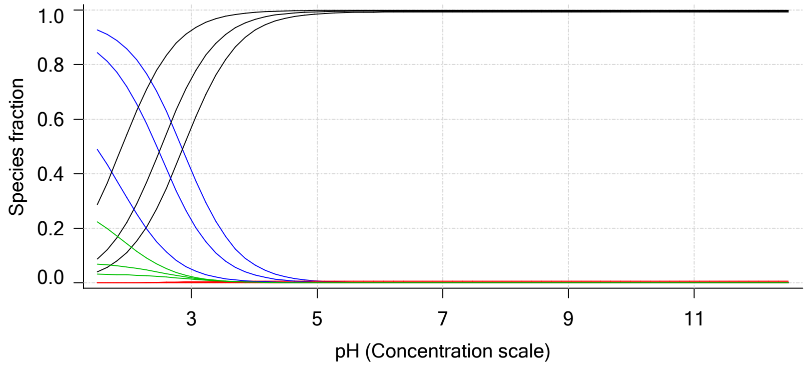


Lipophilicity Profile

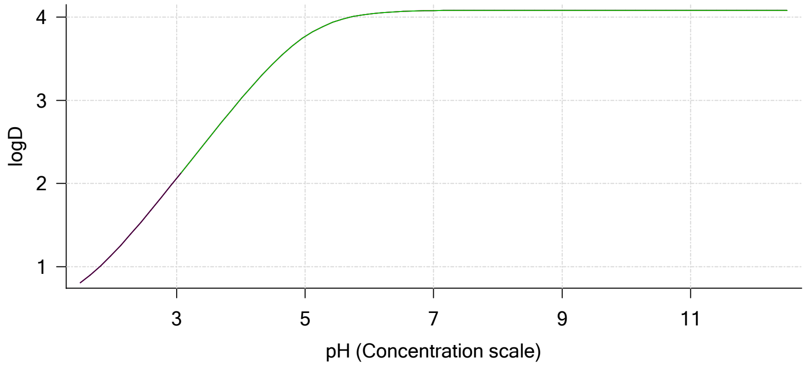


Log R Profile

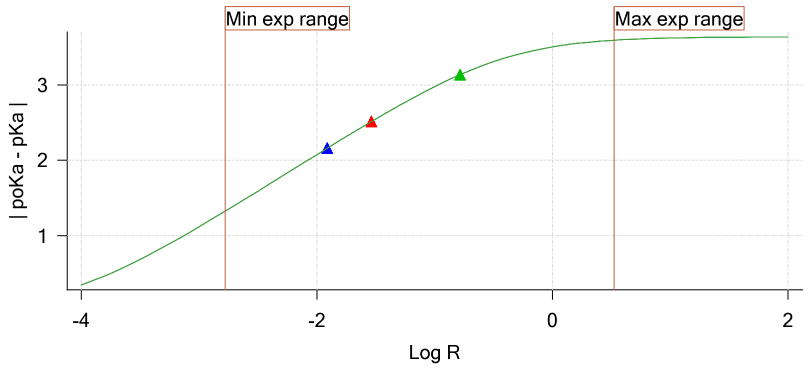


Sample Carbonate Ionisation Graph

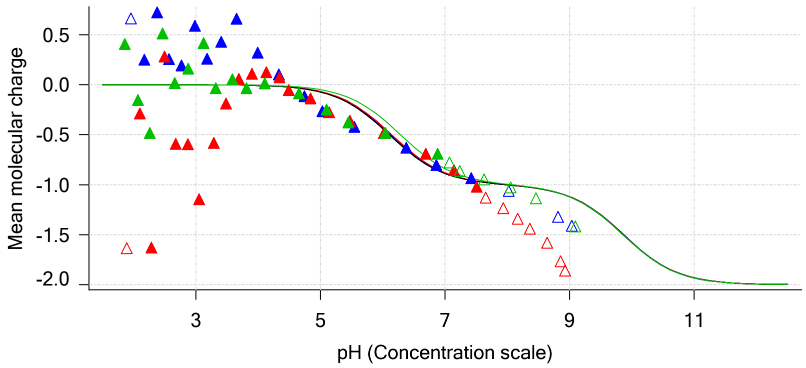

Sample name: **M02_octanol**
Assay name: **pH-metric high logP**
Assay ID: **18C-03011**
Filename: **C:\Sirius_T3\Mehtap\20180302_exp29_logP_T3-2\18C-03011_M02_octanol_pH-metric high logP.t3r**

Experiment start time: **3/3/2018 1:46:01 PM**
Analyst: **Dorothy Levorse**
Instrument ID: **T312060**

#### Graphs (continued)

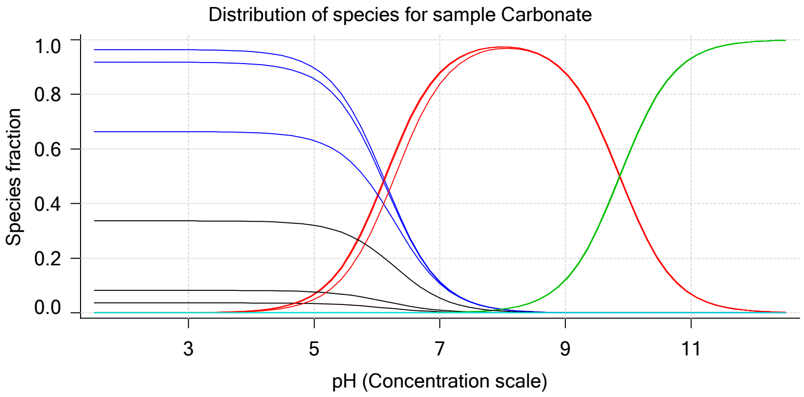

| | | | |
|---|---|---|---|
| Sample name: | **M02_octanol** | Experiment start time: | **3/3/2018 1:46:01 PM** |
| Assay name: | **pH-metric high logP** | Analyst: | **Dorothy Levorse** |
| Assay ID: | **18C-03011** | Instrument ID: | **T312060** |
| Filename: | **C:\Sirius_T3\Mehtap\20180302_exp29_logP_T3-2\18C-03011_M02_octanol_pH-metric high logP.t3r** | | |

### pH-metric high logP Titration 1 of 3 18C-03011 Points 1 to 20

#### Overall results

| | |
|---|---|
| RMSD | 0.135 |
| Average ionic strength | 0.157 M |
| Average temperature | 24.9°C |
| Partition ratio | 0.0123 : 1 |
| Analyte concentration range | 4180.2 µM to 4304.6 µM |
| Total points considered | 16 of 20 |

#### Warnings and errors

| | |
|---|---|
| Errors | None |
| Warnings | Excessive acidity error present |

#### Four-Plus parameters

| | | | |
|---|---|---|---|
| Alpha | 0.111 | 3/3/2018 1:46:01 PM | C:\Sirius_T3\HCl18C02.t3r |
| S | 0.9988 | 3/3/2018 1:46:01 PM | C:\Sirius_T3\HCl18C02.t3r |
| jH | 1.0 | 3/3/2018 1:46:01 PM | C:\Sirius_T3\HCl18C02.t3r |
| jOH | -0.8 | 3/3/2018 1:46:01 PM | C:\Sirius_T3\HCl18C02.t3r |

#### Titrants

| | | | |
|---|---|---|---|
| 0.50 M HCl | 0.999058 | 3/3/2018 1:46:01 PM | C:\Sirius_T3\HCl18C02.t3r |
| 0.50 M KOH | 0.999845 | 3/3/2018 1:46:01 PM | C:\Sirius_T3\KOH18B27.t3r |

#### Sample

| | |
|---|---|
| M02_octanol concentration factor | 0.967 |
| Base pKa 1 | 5.03 |
| logP (XH +) | 1.31 |
| logP (neutral X) | 4.13 |

#### Sample graphs

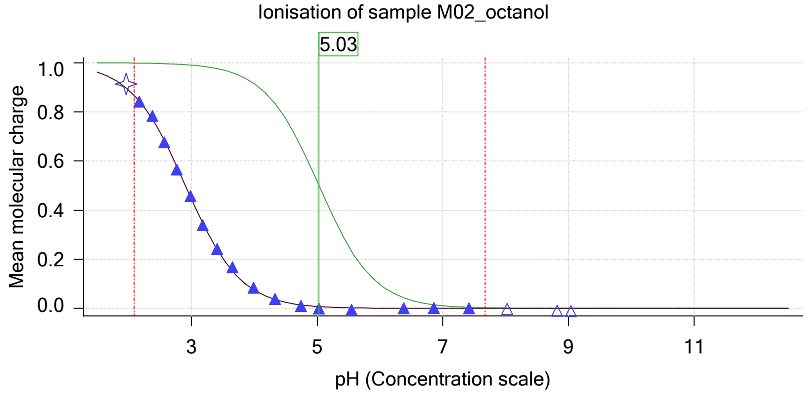


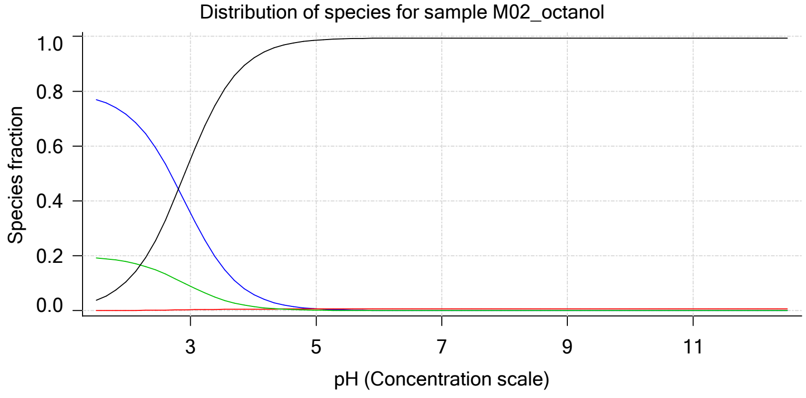

| | | | |
|---|---|---|---|
| Sample name: | **M02_octanol** | Experiment start time: | **3/3/2018 1:46:01 PM** |
| Assay name: | **pH-metric high logP** | Analyst: | **Dorothy Levorse** |
| Assay ID: | **18C-03011** | Instrument ID: | **T312060** |
| Filename: | **C:\Sirius_T3\Mehtap\20180302_exp29_logP_T3-2\18C-03011_M02_octanol_pH-metric high logP.t3r** | | |

#### Sample graphs (continued)

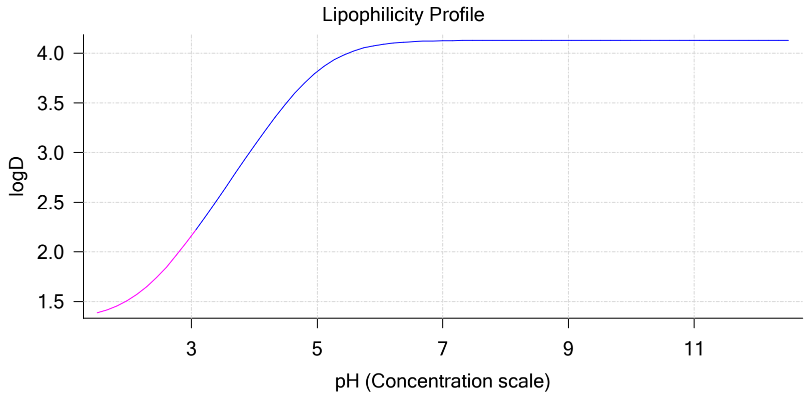


#### Sample logD and percent species

| pH | M02_octanol logD | M02_octanol M02_octanolH | M02_octanol M02_octanol | M02_octanol M02_octanolH* | M02_octanol M02_octanol* | Comment |
|---|---|---|---|---|---|---|
| 1.000 | 1.33 | 79.02 % | 0.01 % | 19.74 % | 1.22 % | |
| 1.200 | 1.35 | 78.46 % | 0.01 % | 19.60 % | 1.93 % | Stomach pH |
| 2.000 | 1.52 | 71.14 % | 0.07 % | 17.77 % | 11.02 % | |
| 3.000 | 2.16 | 35.60 % | 0.33 % | 8.90 % | 55.17 % | |
| 4.000 | 3.07 | 5.94 % | 0.55 % | 1.48 % | 92.02 % | |
| 5.000 | 3.82 | 0.64 % | 0.59 % | 0.16 % | 98.61 % | |
| 6.000 | 4.09 | 0.06 % | 0.60 % | 0.02 % | 99.32 % | |
| 6.500 | 4.12 | 0.02 % | 0.60 % | 0.01 % | 99.38 % | |
| 7.000 | 4.13 | 0.01 % | 0.60 % | 0.00 % | 99.39 % | |
| 7.400 | 4.13 | 0.00 % | 0.60 % | 0.00 % | 99.40 % | Blood pH |
| 8.000 | 4.13 | 0.00 % | 0.60 % | 0.00 % | 99.40 % | |
| 9.000 | 4.13 | 0.00 % | 0.60 % | 0.00 % | 99.40 % | |
| 10.000 | 4.13 | 0.00 % | 0.60 % | 0.00 % | 99.40 % | |
| 11.000 | 4.13 | 0.00 % | 0.60 % | 0.00 % | 99.40 % | |
| 12.000 | 4.13 | 0.00 % | 0.60 % | 0.00 % | 99.40 % | |

#### Carbonate and acidity

| | |
|---|---|
| Carbonate | 0.163 mM |
| Acidity error | -1.499 mM |

#### Other graphs

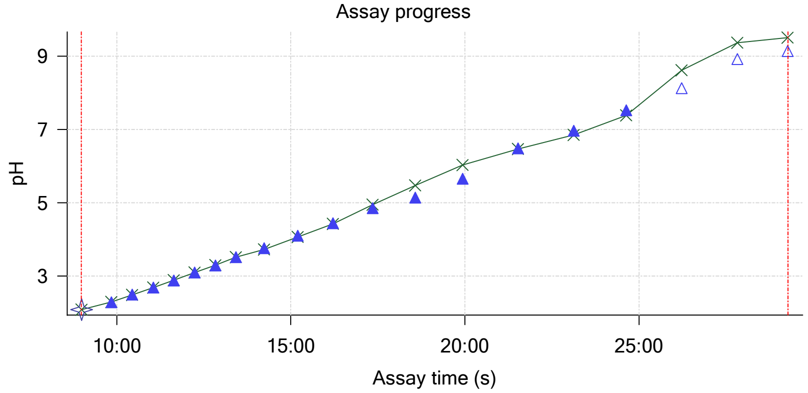


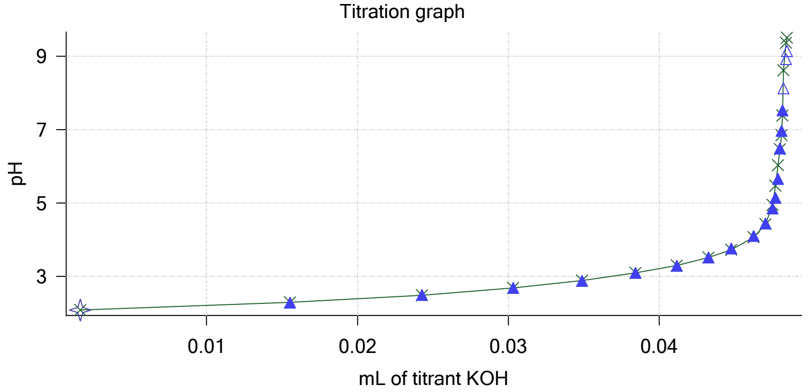

Sample name: **M02_octanol**
Assay name: **pH-metric high logP**
Assay ID: **18C-03011**
Filename: **C:\Sirius_T3\Mehtap\20180302_exp29_logP_T3-2\18C-03011_M02_octanol_pH-metric high logP.t3r**

Experiment start time: **3/3/2018 1:46:01 PM**
Analyst: **Dorothy Levorse**
Instrument ID: **T312060**

#### Other graphs (continued)

Charge balance residuals

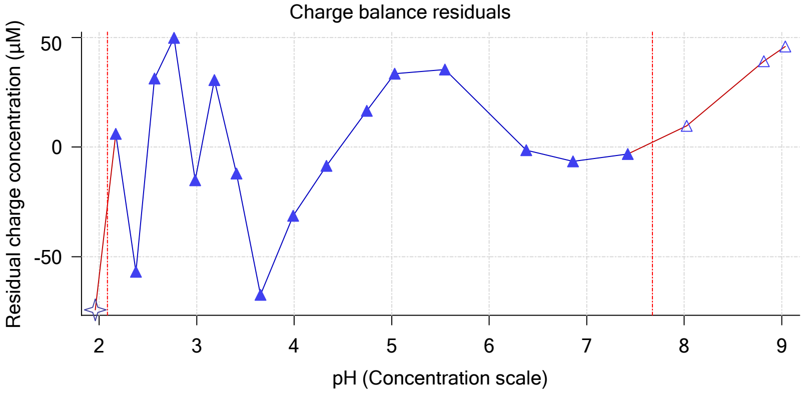


Buffer index profile

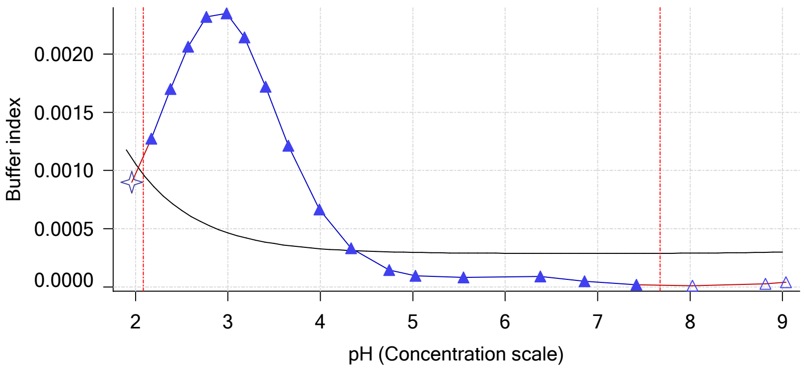


Ionic strength

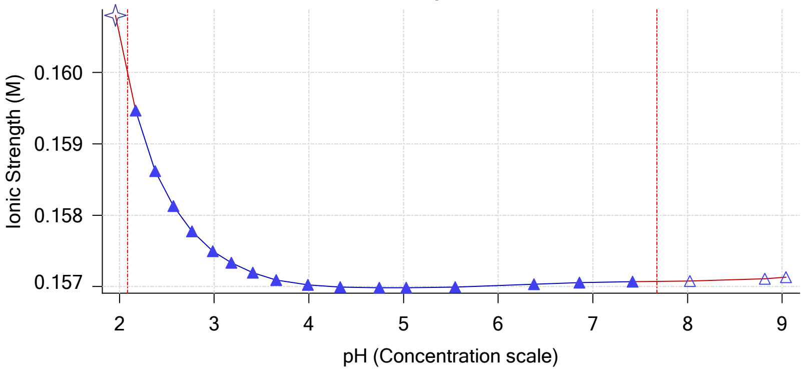


Log R Profile

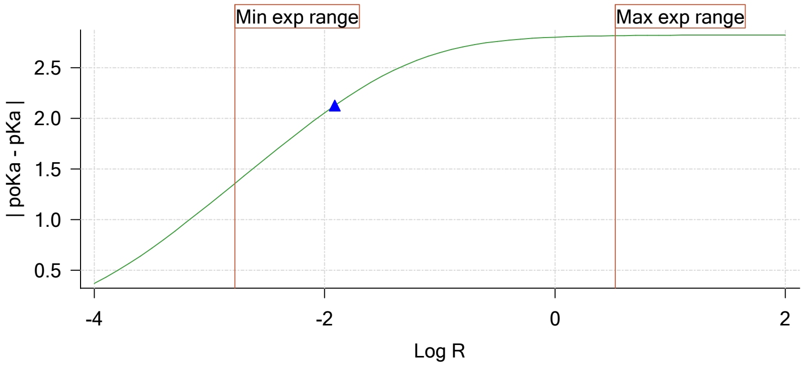


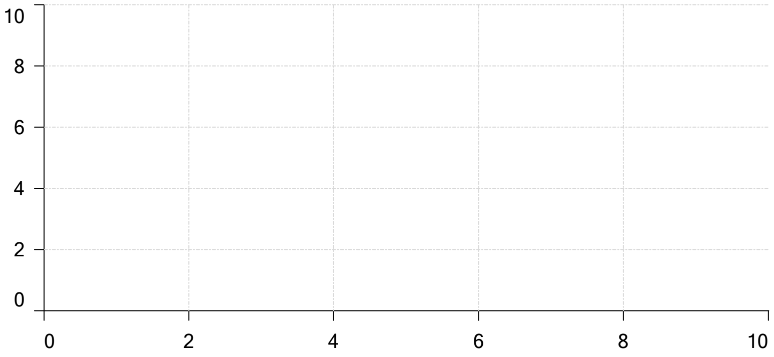


Ionisation of sample Carbonate

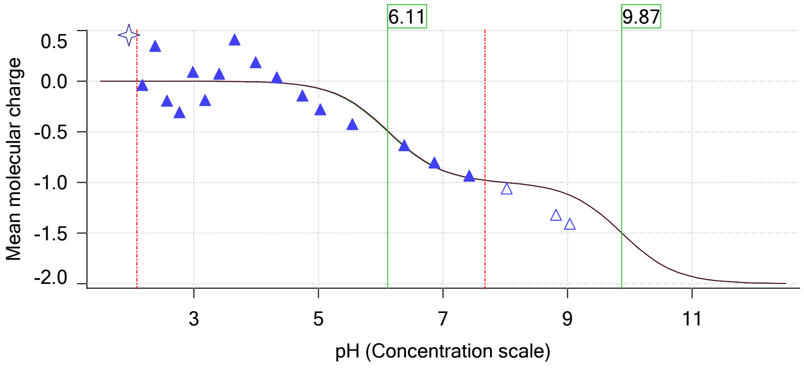

| | | | |
|---|---|---|---|
| Sample name: | **M02_octanol** | Experiment start time: | **3/3/2018 1:46:01 PM** |
| Assay name: | **pH-metric high logP** | Analyst: | **Dorothy Levorse** |
| Assay ID: | **18C-03011** | Instrument ID: | **T312060** |
| Filename: | **C:\Sirius_T3\Mehtap\20180302_exp29_logP_T3-2\18C-03011_M02_octanol_pH-metric high logP.t3r** | | |

### pH-metric high logP Titration 2 of 3 18C-03011 Points 21 to 48

#### Overall results

| | |
|---|---|
| RMSD | 0.183 |
| Average ionic strength | 0.163 M |
| Average temperature | 25.0°C |
| Partition ratio | 0.0290 : 1 |
| Analyte concentration range | 3843.8 µM to 3969.0 µM |
| Total points considered | 20 of 28 |

#### Warnings and errors

| | |
|---|---|
| Errors | None |
| Warnings | Excessive acidity error present |

#### Four-Plus parameters

| | | | |
|---|---|---|---|
| Alpha | 0.111 | 3/3/2018 1:46:01 PM | C:\Sirius_T3\HCl18C02.t3r |
| S | 0.9988 | 3/3/2018 1:46:01 PM | C:\Sirius_T3\HCl18C02.t3r |
| jH | 1.0 | 3/3/2018 1:46:01 PM | C:\Sirius_T3\HCl18C02.t3r |
| jOH | -0.8 | 3/3/2018 1:46:01 PM | C:\Sirius_T3\HCl18C02.t3r |

#### Titrants

| | | | |
|---|---|---|---|
| 0.50 M HCl | 0.999058 | 3/3/2018 1:46:01 PM | C:\Sirius_T3\HCl18C02.t3r |
| 0.50 M KOH | 0.999845 | 3/3/2018 1:46:01 PM | C:\Sirius_T3\KOH18B27.t3r |

#### Sample

| | |
|---|---|
| M02_octanol concentration factor | 1.157 |
| Base pKa 1 | 5.03 |
| logP (XH +) | 1.35 |
| logP (neutral X) | 4.30 |

#### Sample graphs

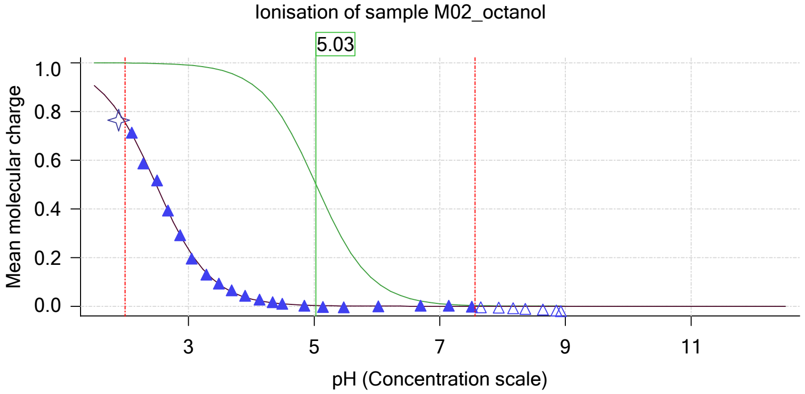


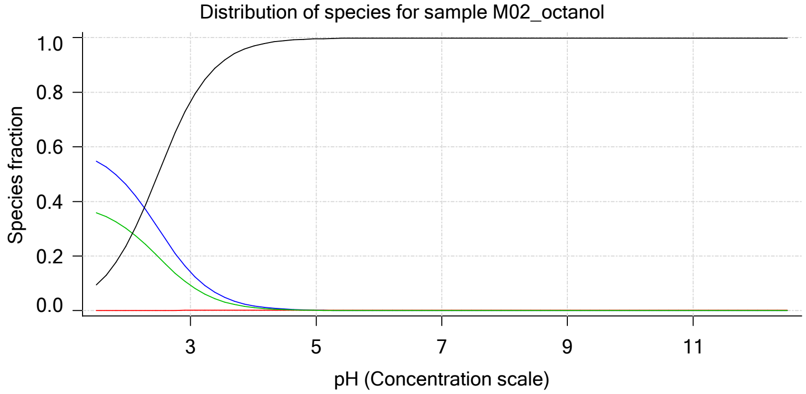

| | |
|---|---|
| Sample name: | **M02_octanol** |
| Assay name: | **pH-metric high logP** |
| Assay ID: | **18C-03011** |
| Filename: | **C:\Sirius_T3\Mehtap\20180302_exp29_logP_T3-2\18C-03011_M02_octanol_pH-metric high logP.t3r** |
| Experiment start time: | **3/3/2018 1:46:01 PM** |
| Analyst: | **Dorothy Levorse** |
| Instrument ID: | **T312060** |

#### Sample graphs (continued)

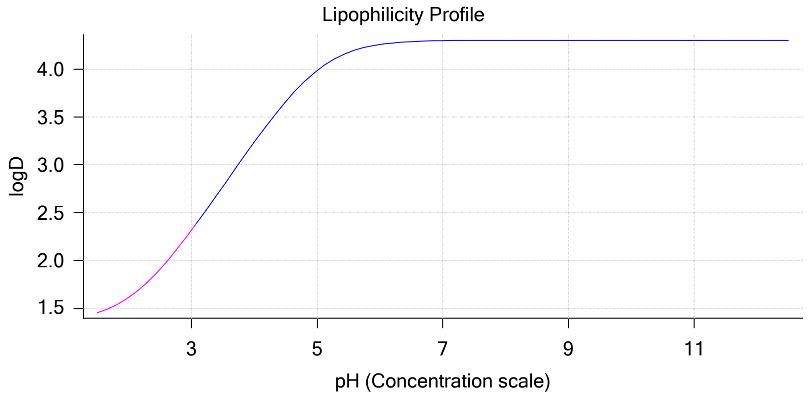


#### Sample logD and percent species

| pH | M02_octanol logD | M02_octanol M02_octanolH | M02_octanol M02_octanol | M02_octanol M02_octanolH* | M02_octanol M02_octanol* | Comment |
|---|---|---|---|---|---|---|
| 1.000 | 1.39 | 58.55 % | 0.01 % | 38.26 % | 3.18 % | |
| 1.200 | 1.41 | 57.48 % | 0.01 % | 37.56 % | 4.95 % | Stomach pH |
| 2.000 | 1.61 | 45.50 % | 0.04 % | 29.73 % | 24.72 % | |
| 3.000 | 2.32 | 14.10 % | 0.13 % | 9.21 % | 76.56 % | |
| 4.000 | 3.24 | 1.78 % | 0.17 % | 1.17 % | 96.88 % | |
| 5.000 | 3.99 | 0.18 % | 0.17 % | 0.12 % | 99.53 % | |
| 6.000 | 4.26 | 0.02 % | 0.17 % | 0.01 % | 99.80 % | |
| 6.500 | 4.29 | 0.01 % | 0.17 % | 0.00 % | 99.82 % | |
| 7.000 | 4.30 | 0.00 % | 0.17 % | 0.00 % | 99.83 % | |
| 7.400 | 4.30 | 0.00 % | 0.17 % | 0.00 % | 99.83 % | Blood pH |
| 8.000 | 4.30 | 0.00 % | 0.17 % | 0.00 % | 99.83 % | |
| 9.000 | 4.30 | 0.00 % | 0.17 % | 0.00 % | 99.83 % | |
| 10.000 | 4.30 | 0.00 % | 0.17 % | 0.00 % | 99.83 % | |
| 11.000 | 4.30 | 0.00 % | 0.17 % | 0.00 % | 99.83 % | |
| 12.000 | 4.30 | 0.00 % | 0.17 % | 0.00 % | 99.83 % | |

#### Carbonate and acidity

| | |
|---|---|
| Carbonate | 0.123 mM |
| Acidity error | -1.570 mM |

#### Other graphs

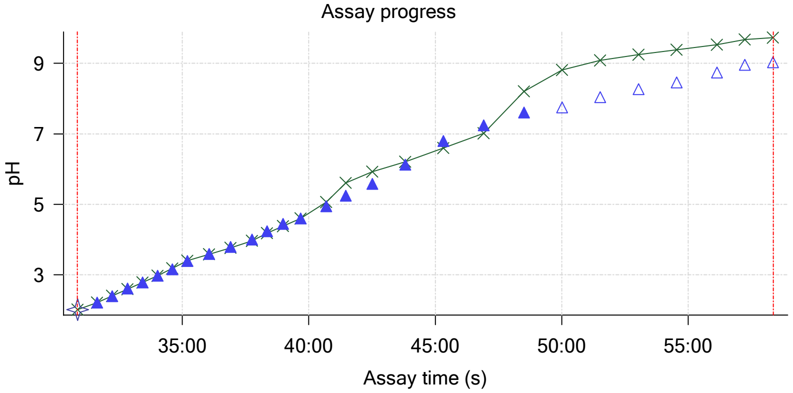


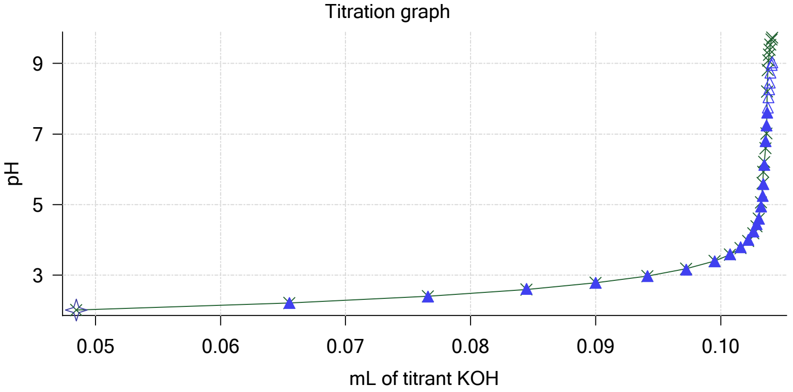

| | | | |
|---|---|---|---|
| Sample name: | **M02_octanol** | Experiment start time: | **3/3/2018 1:46:01 PM** |
| Assay name: | **pH-metric high logP** | Analyst: | **Dorothy Levorse** |
| Assay ID: | **18C-03011** | Instrument ID: | **T312060** |
| Filename: | **C:\Sirius_T3\Mehtap\20180302_exp29_logP_T3-2\18C-03011_M02_octanol_pH-metric high logP.t3r** | | |

#### Other graphs (continued)

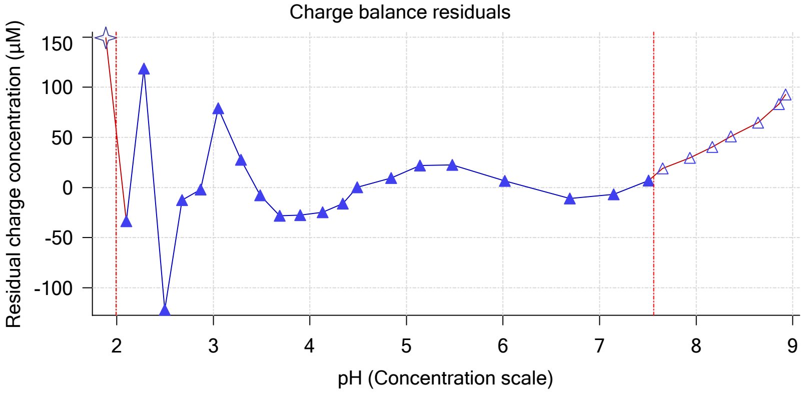


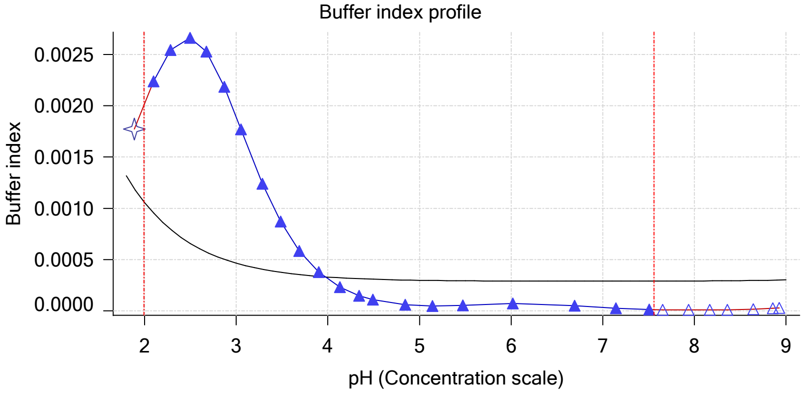


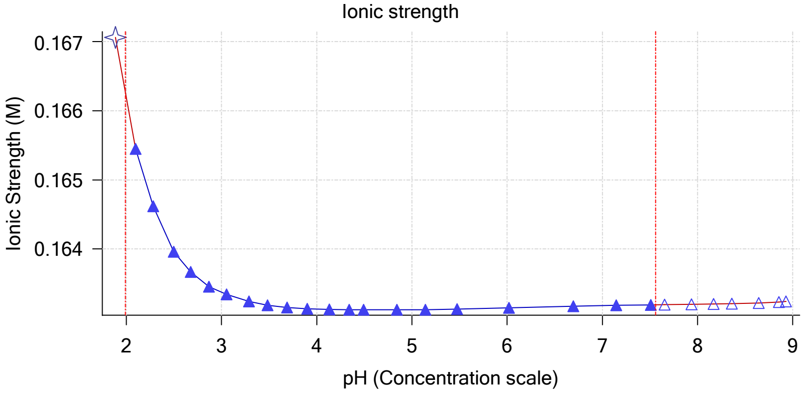


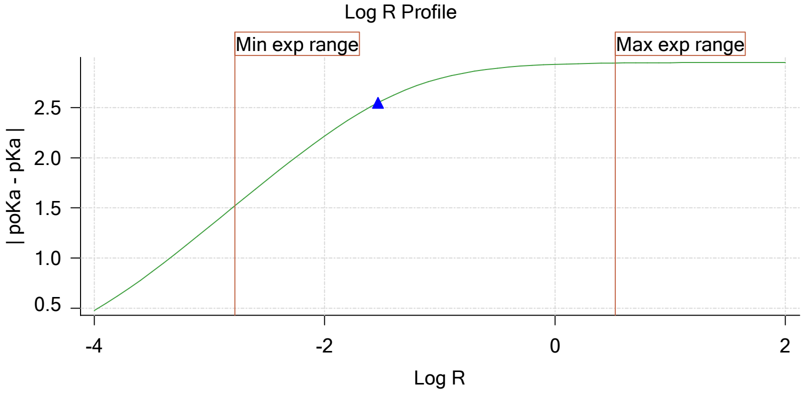


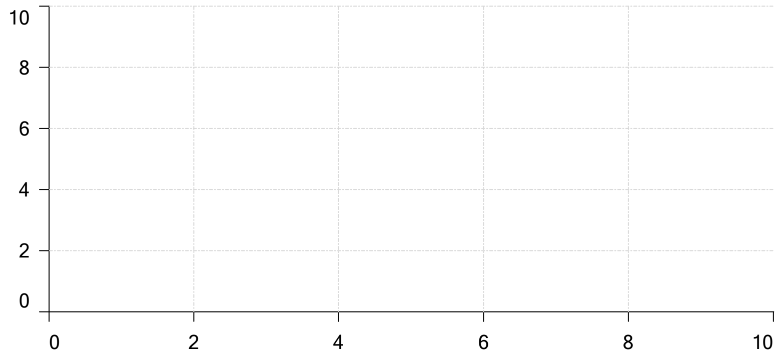

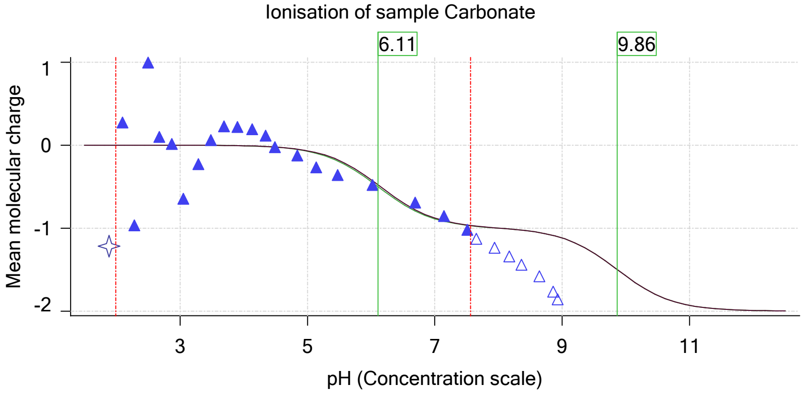

| | | | |
|---|---|---|---|
| Sample name: | **M02_octanol** | Experiment start time: | **3/3/2018 1:46:01 PM** |
| Assay name: | **pH-metric high logP** | Analyst: | **Dorothy Levorse** |
| Assay ID: | **18C-03011** | Instrument ID: | **T312060** |
| Filename: | **C:\Sirius_T3\Mehtap\20180302_exp29_logP_T3-2\18C-03011_M02_octanol_pH-metric high logP.t3r** | | |

### pH-metric high logP Titration 3 of 3 18C-03011 Points 49 to 70

#### Overall results

| | |
|---|---|
| RMSD | 0.256 |
| Average ionic strength | 0.169 M |
| Average temperature | 25.0°C |
| Partition ratio | 0.1643 : 1 |
| Analyte concentration range | 3175.3 µM to 3268.6 µM |
| Total points considered | 16 of 22 |

#### Warnings and errors

| | |
|---|---|
| Errors | None |
| Warnings | Sample concentration factor out of range |
| | Excessive acidity error present |

#### Four-Plus parameters

| | | | |
|---|---|---|---|
| Alpha | 0.111 | 3/3/2018 1:46:01 PM | C:\Sirius_T3\HCl18C02.t3r |
| S | 0.9988 | 3/3/2018 1:46:01 PM | C:\Sirius_T3\HCl18C02.t3r |
| jH | 1.0 | 3/3/2018 1:46:01 PM | C:\Sirius_T3\HCl18C02.t3r |
| jOH | -0.8 | 3/3/2018 1:46:01 PM | C:\Sirius_T3\HCl18C02.t3r |

#### Titrants

| | | | |
|---|---|---|---|
| 0.50 M HCl | 0.999058 | 3/3/2018 1:46:01 PM | C:\Sirius_T3\HCl18C02.t3r |
| 0.50 M KOH | 0.999845 | 3/3/2018 1:46:01 PM | C:\Sirius_T3\KOH18B27.t3r |

#### Sample

| | |
|---|---|
| M02_octanol concentration factor | 1.597 |
| Base pKa 1 | 5.03 |
| logP (XH +) | 1.32 |
| logP (neutral X) | 4.56 |

#### Sample graphs

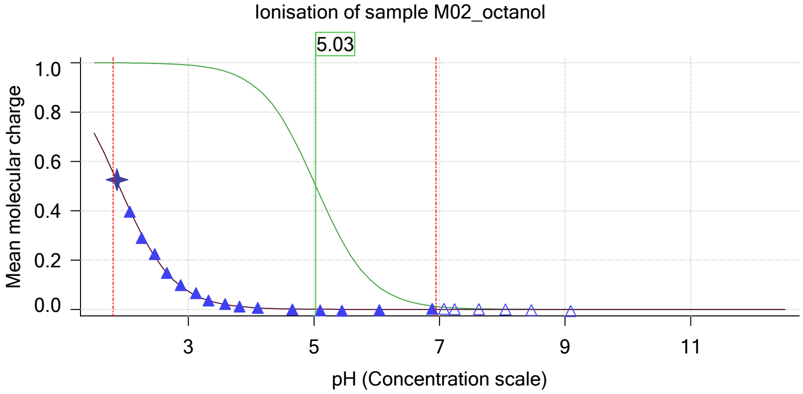


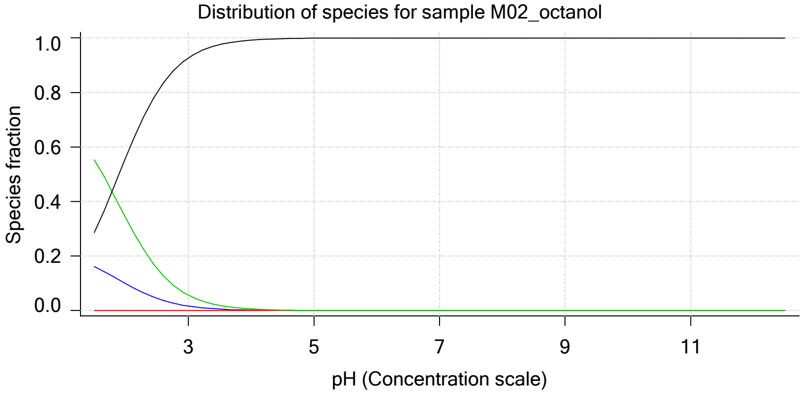

| | | | |
|---|---|---|---|
| Sample name: | **M02_octanol** | Experiment start time: | **3/3/2018 1:46:01 PM** |
| Assay name: | **pH-metric high logP** | Analyst: | **Dorothy Levorse** |
| Assay ID: | **18C-03011** | Instrument ID: | **T312060** |
| Filename: | **C:\Sirius_T3\Mehtap\20180302_exp29_logP_T3-2\18C-03011_M02_octanol_pH-metric high logP.t3r** | | |

#### Sample graphs (continued)

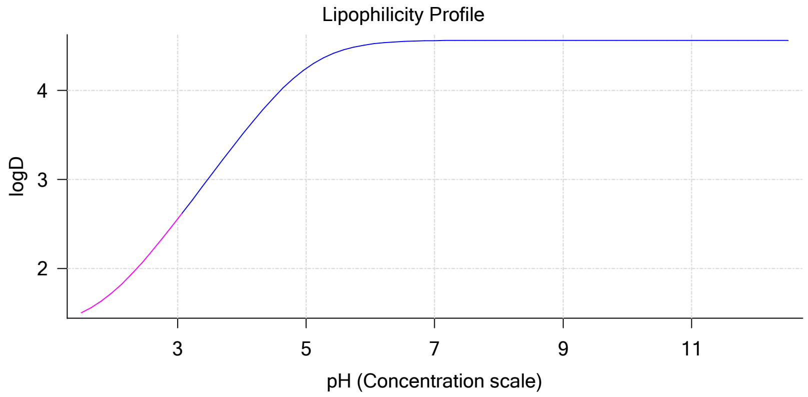


#### Sample logD and percent species

| pH | M02_octanol logD | M02_octanol M02_octanolH | M02_octanol M02_octanol | M02_octanol M02_octanolH* | M02_octanol M02_octanol* | Comment |
|---|---|---|---|---|---|---|
| 1.000 | 1.39 | 20.02 % | 0.00 % | 68.72 % | 11.26 % | |
| 1.200 | 1.42 | 18.78 % | 0.00 % | 64.47 % | 16.75 % | Stomach pH |
| 2.000 | 1.74 | 9.94 % | 0.01 % | 34.13 % | 55.92 % | |
| 3.000 | 2.56 | 1.65 % | 0.02 % | 5.66 % | 92.68 % | |
| 4.000 | 3.50 | 0.18 % | 0.02 % | 0.61 % | 99.20 % | |
| 5.000 | 4.25 | 0.02 % | 0.02 % | 0.06 % | 99.90 % | |
| 6.000 | 4.52 | 0.00 % | 0.02 % | 0.01 % | 99.98 % | |
| 6.500 | 4.55 | 0.00 % | 0.02 % | 0.00 % | 99.98 % | |
| 7.000 | 4.56 | 0.00 % | 0.02 % | 0.00 % | 99.98 % | |
| 7.400 | 4.56 | 0.00 % | 0.02 % | 0.00 % | 99.98 % | Blood pH |
| 8.000 | 4.56 | 0.00 % | 0.02 % | 0.00 % | 99.98 % | |
| 9.000 | 4.56 | 0.00 % | 0.02 % | 0.00 % | 99.98 % | |
| 10.000 | 4.56 | 0.00 % | 0.02 % | 0.00 % | 99.98 % | |
| 11.000 | 4.56 | 0.00 % | 0.02 % | 0.00 % | 99.98 % | |
| 12.000 | 4.56 | 0.00 % | 0.02 % | 0.00 % | 99.98 % | |

#### Carbonate and acidity

| | |
|---|---|
| Carbonate | 0.148 mM |
| Acidity error | -1.252 mM |

#### Other graphs

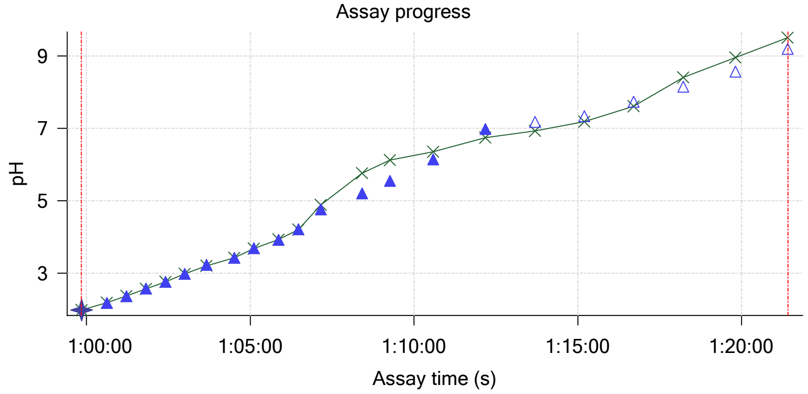


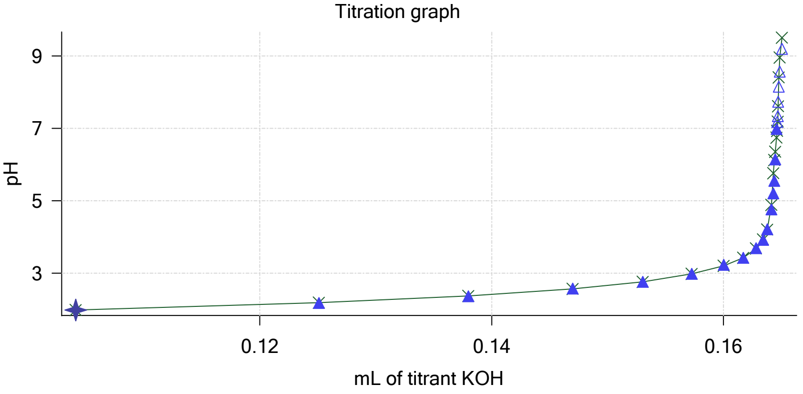

| | | | |
|---|---|---|---|
| Sample name: | **M02_octanol** | Experiment start time: | **3/3/2018 1:46:01 PM** |
| Assay name: | **pH-metric high logP** | Analyst: | **Dorothy Levorse** |
| Assay ID: | **18C-03011** | Instrument ID: | **T312060** |
| Filename: | **C:\Sirius_T3\Mehtap\20180302_exp29_logP_T3-2\18C-03011_M02_octanol_pH-metric high logP.t3r** | | |

#### Other graphs (continued)

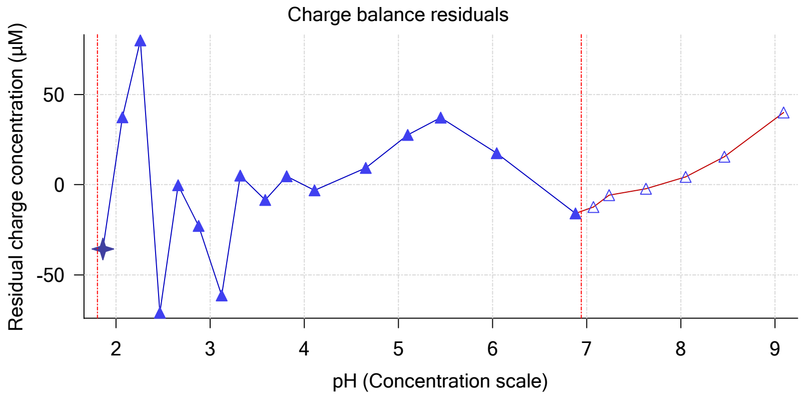


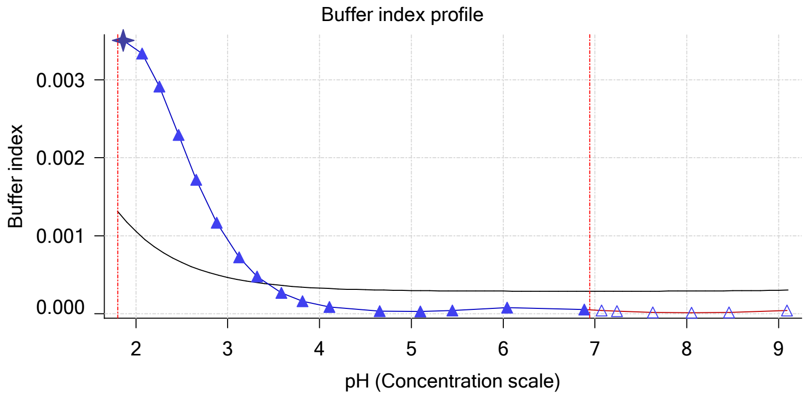


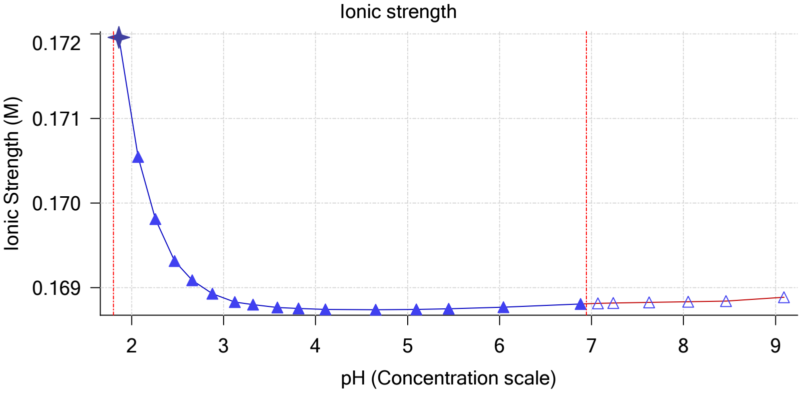


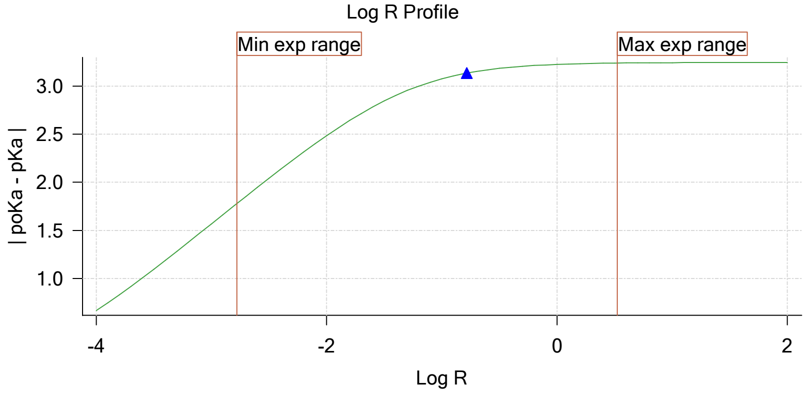


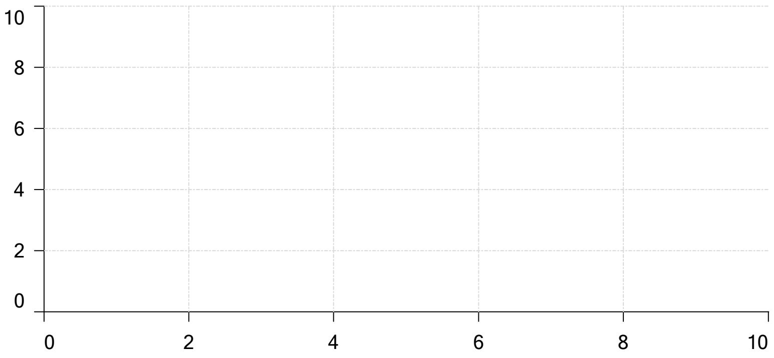

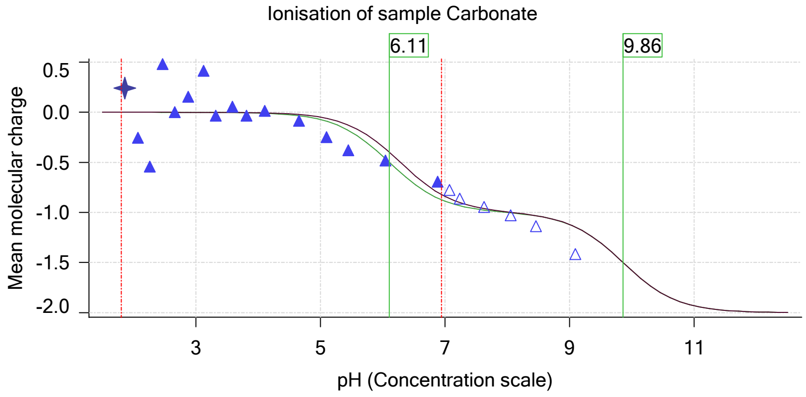

Sample name: **M02_octanol**
Assay name: **pH-metric high logP**
Assay ID: **18C-03011**
Filename: **C:\Sirius_T3\Mehtap\20180302_exp29_logP_T3-2\18C-03011_M02_octanol_pH-metric high logP.t3r**

Experiment start time: **3/3/2018 1:46:01 PM**
Analyst: **Dorothy Levorse**
Instrument ID: **T312060**

#### Assay Model

| Settings | Value | Date/Time changed | Imported from |
|---|---|---|---|
| Sample name | M02_octanol | 12/6/2017 4:20:03 PM | User entered value |
| Sample by | Weight | | Default value |
| Sample weight | 0.001960 g | 3/2/2018 5:09:57 PM | User entered value |
| Formula weight | 289.26 g/mol | 12/6/2017 4:20:03 PM | User entered value |
| Solubility | Unknown | | Default value |
| Molecular weight | 289.26 | 12/6/2017 4:20:03 PM | User entered value |
| Individual pKa ionic environments | No | | Default value |
| Number of pKas | 1 | 12/6/2017 4:20:03 PM | User entered value |
| Sample is a | Base | 12/6/2017 4:20:03 PM | User entered value |
| pKa 1 | 5.03 | 12/6/2017 4:20:03 PM | User entered value |
| logp (XH +) | 1.32 | 3/2/2018 3:38:13 PM | User entered value |
| logP (neutral X) | 4.10 | 3/2/2018 3:38:07 PM | User entered value |

#### Events

| Time | Event | Water | Acid | Base | Octanol | pH | dpH/dt | pH R-squared | pH SD | dpH/dt time |
|---|---|---|---|---|---|---|---|---|---|---|
| 6:00.1 | Initial pH = 8.70 | | | | | | | | | |
| 8:59.7 | Data point 1 | 1.50000 mL | 0.05252 mL | 0.00162 mL | 0.01999 mL | 2.078 | 0.01636 | 0.75364 | 0.00093 | 15.0 s |
| 9:50.7 | Data point 2 | 1.50000 mL | 0.05252 mL | 0.01552 mL | 0.01999 mL | 2.285 | -0.01511 | 0.81500 | 0.00083 | 10.5 s |
| 10:26.8 | Data point 3 | 1.50000 mL | 0.05252 mL | 0.02427 mL | 0.01999 mL | 2.490 | -0.00148 | 0.44552 | 0.00011 | 10.5 s |
| 11:02.9 | Data point 4 | 1.50000 mL | 0.05252 mL | 0.03032 mL | 0.01999 mL | 2.678 | -0.00348 | 0.65646 | 0.00021 | 10.0 s |
| 11:38.4 | Data point 5 | 1.50000 mL | 0.05252 mL | 0.03488 mL | 0.01999 mL | 2.876 | -0.01455 | 0.84506 | 0.00078 | 10.0 s |
| 12:13.9 | Data point 6 | 1.50000 mL | 0.05252 mL | 0.03843 mL | 0.01999 mL | 3.094 | -0.01527 | 0.85358 | 0.00082 | 10.5 s |
| 12:49.9 | Data point 7 | 1.50000 mL | 0.05252 mL | 0.04116 mL | 0.01999 mL | 3.288 | -0.01663 | 0.96111 | 0.00084 | 10.0 s |
| 13:25.4 | Data point 8 | 1.50000 mL | 0.05252 mL | 0.04327 mL | 0.01999 mL | 3.515 | -0.01850 | 0.86172 | 0.00098 | 23.0 s |
| 14:13.9 | Data point 9 | 1.50000 mL | 0.05252 mL | 0.04478 mL | 0.01999 mL | 3.760 | -0.01848 | 0.86901 | 0.00098 | 17.0 s |
| 15:11.8 | Data point 10 | 1.50000 mL | 0.05252 mL | 0.04626 mL | 0.01999 mL | 4.096 | -0.01860 | 0.90784 | 0.00096 | 20.0 s |
| 16:12.8 | Data point 11 | 1.50000 mL | 0.05252 mL | 0.04704 mL | 0.01999 mL | 4.437 | -0.01860 | 0.90771 | 0.00096 | 37.5 s |
| 17:20.9 | Data point 12 | 1.50000 mL | 0.05252 mL | 0.04751 mL | 0.01999 mL | 4.850 | -0.01809 | 0.87203 | 0.00096 | 43.0 s |
| 18:34.5 | Data point 13 | 1.50000 mL | 0.05252 mL | 0.04770 mL | 0.01999 mL | 5.134 | -0.01884 | 0.90212 | 0.00098 | 51.0 s |
| 19:56.1 | Data point 14 | 1.50000 mL | 0.05252 mL | 0.04786 mL | 0.01999 mL | 5.651 | -0.02196 | 0.93556 | 0.00112 | Timed out at 59.5 s |
| 21:31.8 | Data point 15 | 1.50000 mL | 0.05252 mL | 0.04800 mL | 0.01999 mL | 6.482 | -0.04707 | 0.98476 | 0.00234 | Timed out at 59.5 s |
| 23:07.4 | Data point 16 | 1.50000 mL | 0.05252 mL | 0.04809 mL | 0.01999 mL | 6.961 | -0.05553 | 0.98035 | 0.00277 | Timed out at 59.5 s |
| 24:38.0 | Data point 17 | 1.50000 mL | 0.05252 mL | 0.04817 mL | 0.01999 mL | 7.521 | -0.08676 | 0.99361 | 0.00430 | Timed out at 59.5 s |
| 26:13.5 | Data point 18 | 1.50000 mL | 0.05252 mL | 0.04824 mL | 0.01999 mL | 8.125 | -0.07978 | 0.98996 | 0.00396 | Timed out at 59.5 s |
| 27:49.3 | Data point 19 | 1.50000 mL | 0.05252 mL | 0.04840 mL | 0.01999 mL | 8.917 | -0.01848 | 0.88522 | 0.00097 | 56.0 s |
| 29:15.8 | Data point 20 | 1.50000 mL | 0.05252 mL | 0.04847 mL | 0.01999 mL | 9.138 | -0.01766 | 0.91576 | 0.00091 | 37.0 s |
| 30:51.7 | Data point 21 | 1.50000 mL | 0.10875 mL | 0.04847 mL | 0.05000 mL | 2.009 | -0.00919 | 0.60040 | 0.00059 | 10.0 s |
| 31:38.0 | Data point 22 | 1.50000 mL | 0.10875 mL | 0.06552 mL | 0.05000 mL | 2.215 | -0.00727 | 0.64690 | 0.00045 | 10.0 s |
| 32:13.7 | Data point 23 | 1.50000 mL | 0.10875 mL | 0.07660 mL | 0.05000 mL | 2.396 | -0.00146 | 0.31714 | 0.00013 | 10.5 s |
| 32:49.9 | Data point 24 | 1.50000 mL | 0.10875 mL | 0.08448 mL | 0.05000 mL | 2.609 | -0.00346 | 0.43326 | 0.00026 | 10.0 s |
| 33:25.5 | Data point 25 | 1.50000 mL | 0.10875 mL | 0.08998 mL | 0.05000 mL | 2.785 | -0.00991 | 0.29494 | 0.00090 | 10.0 s |
| 34:01.0 | Data point 26 | 1.50000 mL | 0.10875 mL | 0.09414 mL | 0.05000 mL | 2.978 | -0.00636 | 0.79094 | 0.00035 | 10.0 s |
| 34:36.5 | Data point 27 | 1.50000 mL | 0.10875 mL | 0.09722 mL | 0.05000 mL | 3.160 | -0.00652 | 0.78690 | 0.00036 | 10.0 s |
| 35:11.9 | Data point 28 | 1.50000 mL | 0.10875 mL | 0.09951 mL | 0.05000 mL | 3.395 | -0.01414 | 0.94922 | 0.00072 | 10.5 s |
| 36:03.3 | Data point 29 | 1.50000 mL | 0.10875 mL | 0.10073 mL | 0.05000 mL | 3.592 | -0.01172 | 0.70688 | 0.00069 | 10.0 s |
| 36:54.3 | Data point 30 | 1.50000 mL | 0.10875 mL | 0.10158 mL | 0.05000 mL | 3.795 | -0.01307 | 0.64357 | 0.00080 | 10.0 s |
| 37:45.1 | Data point 31 | 1.50000 mL | 0.10875 mL | 0.10219 mL | 0.05000 mL | 4.005 | -0.01735 | 0.75875 | 0.00098 | 10.5 s |
| 38:21.1 | Data point 32 | 1.50000 mL | 0.10875 mL | 0.10261 mL | 0.05000 mL | 4.238 | -0.01660 | 0.84161 | 0.00089 | 12.5 s |
| 38:59.1 | Data point 33 | 1.50000 mL | 0.10875 mL | 0.10287 mL | 0.05000 mL | 4.447 | -0.01934 | 0.93729 | 0.00099 | 16.0 s |
| 39:40.4 | Data point 34 | 1.50000 mL | 0.10875 mL | 0.10303 mL | 0.05000 mL | 4.596 | -0.01901 | 0.88676 | 0.00100 | 30.5 s |
| 40:41.5 | Data point 35 | 1.50000 mL | 0.10875 mL | 0.10322 mL | 0.05000 mL | 4.948 | -0.01691 | 0.77907 | 0.00095 | 16.0 s |

| | | | |
|---|---|---|---|
| Sample name: | **M02_octanol** | Experiment start time: | **3/3/2018 1:46:01 PM** |
| Assay name: | **pH-metric high logP** | Analyst: | **Dorothy Levorse** |
| Assay ID: | **18C-03011** | Instrument ID: | **T312060** |
| Filename: | **C:\Sirius_T3\Mehtap\20180302_exp29_logP_T3-2\18C-03011_M02_octanol_pH-metric high logP.t3r** | | |

#### Events (continued)

| Time | Event | Water | Acid | Base | Octanol | pH | dpH/dt | pH R-squared | pH SD | dpH/dt time |
|---|---|---|---|---|---|---|---|---|---|---|
| 41:28.0 | Data point 36 | 1.50000 mL | 0.10875 mL | 0.10334 mL | 0.05000 mL | 5.244 | -0.01809 | 0.82366 | 0.00098 | 32.5 s |
| 42:31.1 | Data point 37 | 1.50000 mL | 0.10875 mL | 0.10341 mL | 0.05000 mL | 5.578 | -0.01894 | 0.89362 | 0.00099 | 47.0 s |
| 43:48.7 | Data point 38 | 1.50000 mL | 0.10875 mL | 0.10348 mL | 0.05000 mL | 6.124 | -0.02064 | 0.93374 | 0.00106 | Timed out at 59.5 s |
| 45:19.2 | Data point 39 | 1.50000 mL | 0.10875 mL | 0.10358 mL | 0.05000 mL | 6.796 | -0.05867 | 0.98597 | 0.00292 | Timed out at 59.5 s |
| 46:54.9 | Data point 40 | 1.50000 mL | 0.10875 mL | 0.10365 mL | 0.05000 mL | 7.247 | -0.06483 | 0.98904 | 0.00322 | Timed out at 59.5 s |
| 48:30.7 | Data point 41 | 1.50000 mL | 0.10875 mL | 0.10372 mL | 0.05000 mL | 7.610 | -0.08577 | 0.99139 | 0.00426 | Timed out at 59.5 s |
| 50:01.2 | Data point 42 | 1.50000 mL | 0.10875 mL | 0.10376 mL | 0.05000 mL | 7.754 | -0.06230 | 0.98106 | 0.00311 | Timed out at 59.5 s |
| 51:31.7 | Data point 43 | 1.50000 mL | 0.10875 mL | 0.10381 mL | 0.05000 mL | 8.038 | -0.05883 | 0.98974 | 0.00292 | Timed out at 59.5 s |
| 53:02.2 | Data point 44 | 1.50000 mL | 0.10875 mL | 0.10386 mL | 0.05000 mL | 8.269 | -0.04245 | 0.97500 | 0.00212 | Timed out at 59.5 s |
| 54:32.7 | Data point 45 | 1.50000 mL | 0.10875 mL | 0.10390 mL | 0.05000 mL | 8.461 | -0.02925 | 0.94322 | 0.00149 | Timed out at 59.5 s |
| 56:08.3 | Data point 46 | 1.50000 mL | 0.10875 mL | 0.10398 mL | 0.05000 mL | 8.742 | -0.01524 | 0.61499 | 0.00096 | 30.5 s |
| 57:14.5 | Data point 47 | 1.50000 mL | 0.10875 mL | 0.10407 mL | 0.05000 mL | 8.956 | -0.01863 | 0.86242 | 0.00099 | 36.0 s |
| 58:21.1 | Data point 48 | 1.50000 mL | 0.10875 mL | 0.10412 mL | 0.05000 mL | 9.027 | -0.01667 | 0.74240 | 0.00096 | 26.0 s |
| 59:51.2 | Data point 49 | 1.50000 mL | 0.16893 mL | 0.10412 mL | 0.30000 mL | 1.983 | 0.00363 | 0.07579 | 0.00065 | 10.0 s |
| 1:00:37.5 | Data point 50 | 1.50000 mL | 0.16893 mL | 0.12509 mL | 0.30000 mL | 2.185 | -0.01327 | 0.52474 | 0.00091 | 10.0 s |
| 1:01:13.3 | Data point 51 | 1.50000 mL | 0.16893 mL | 0.13801 mL | 0.30000 mL | 2.371 | -0.01259 | 0.50607 | 0.00087 | 10.0 s |
| 1:01:48.9 | Data point 52 | 1.50000 mL | 0.16893 mL | 0.14699 mL | 0.30000 mL | 2.577 | -0.00872 | 0.34259 | 0.00074 | 10.0 s |
| 1:02:24.5 | Data point 53 | 1.50000 mL | 0.16893 mL | 0.15303 mL | 0.30000 mL | 2.768 | -0.00055 | 0.00118 | 0.00079 | 10.0 s |
| 1:02:59.9 | Data point 54 | 1.50000 mL | 0.16893 mL | 0.15727 mL | 0.30000 mL | 2.987 | -0.00399 | 0.04207 | 0.00096 | 14.5 s |
| 1:03:39.9 | Data point 55 | 1.50000 mL | 0.16893 mL | 0.16002 mL | 0.30000 mL | 3.230 | -0.00293 | 0.03061 | 0.00083 | 10.0 s |
| 1:04:30.9 | Data point 56 | 1.50000 mL | 0.16893 mL | 0.16169 mL | 0.30000 mL | 3.426 | 0.01036 | 0.28200 | 0.00096 | 10.0 s |
| 1:05:06.3 | Data point 57 | 1.50000 mL | 0.16893 mL | 0.16279 mL | 0.30000 mL | 3.692 | -0.01047 | 0.32693 | 0.00090 | 15.0 s |
| 1:05:51.9 | Data point 58 | 1.50000 mL | 0.16893 mL | 0.16340 mL | 0.30000 mL | 3.919 | -0.00471 | 0.52606 | 0.00032 | 10.5 s |
| 1:06:27.8 | Data point 59 | 1.50000 mL | 0.16893 mL | 0.16378 mL | 0.30000 mL | 4.213 | -0.01245 | 0.91405 | 0.00064 | 10.5 s |
| 1:07:09.1 | Data point 60 | 1.50000 mL | 0.16893 mL | 0.16414 mL | 0.30000 mL | 4.759 | -0.01882 | 0.91127 | 0.00097 | 45.0 s |
| 1:08:24.9 | Data point 61 | 1.50000 mL | 0.16893 mL | 0.16430 mL | 0.30000 mL | 5.201 | -0.01315 | 0.55571 | 0.00087 | 15.5 s |
| 1:09:16.0 | Data point 62 | 1.50000 mL | 0.16893 mL | 0.16439 mL | 0.30000 mL | 5.551 | -0.01841 | 0.93035 | 0.00094 | 48.5 s |
| 1:10:35.1 | Data point 63 | 1.50000 mL | 0.16893 mL | 0.16446 mL | 0.30000 mL | 6.146 | -0.03968 | 0.91076 | 0.00205 | Timed out at 59.5 s |
| 1:12:10.7 | Data point 64 | 1.50000 mL | 0.16893 mL | 0.16458 mL | 0.30000 mL | 6.984 | -0.09924 | 0.98030 | 0.00495 | Timed out at 59.5 s |
| 1:13:41.3 | Data point 65 | 1.50000 mL | 0.16893 mL | 0.16463 mL | 0.30000 mL | 7.173 | -0.07212 | 0.97043 | 0.00362 | Timed out at 59.5 s |
| 1:15:11.8 | Data point 66 | 1.50000 mL | 0.16893 mL | 0.16467 mL | 0.30000 mL | 7.339 | -0.06600 | 0.98427 | 0.00328 | Timed out at 59.5 s |
| 1:16:42.3 | Data point 67 | 1.50000 mL | 0.16893 mL | 0.16472 mL | 0.30000 mL | 7.729 | -0.07810 | 0.99269 | 0.00387 | Timed out at 59.5 s |
| 1:18:12.8 | Data point 68 | 1.50000 mL | 0.16893 mL | 0.16477 mL | 0.30000 mL | 8.153 | -0.05661 | 0.98395 | 0.00282 | Timed out at 59.5 s |
| 1:19:48.4 | Data point 69 | 1.50000 mL | 0.16893 mL | 0.16484 mL | 0.30000 mL | 8.561 | -0.03628 | 0.93717 | 0.00185 | Timed out at 59.5 s |
| 1:21:24.0 | Data point 70 | 1.50000 mL | 0.16893 mL | 0.16505 mL | 0.30000 mL | 9.190 | -0.01034 | 0.34587 | 0.00087 | 15.0 s |
| 1:21:48.1 | Assay volumes | 1.50000 mL | 0.16893 mL | 0.16505 mL | 0.30000 mL | | | | | |

| | | | |
|---|---|---|---|
| Sample name: | **M02_octanol** | Experiment start time: | **3/3/2018 1:46:01 PM** |
| Assay name: | **pH-metric high logP** | Analyst: | **Dorothy Levorse** |
| Assay ID: | **18C-03011** | Instrument ID: | **T312060** |
| Filename: | **C:\Sirius_T3\Mehtap\20180302_exp29_logP_T3-2\18C-03011_M02_octanol_pH-metric high logP.t3r** | | |

#### Assay Settings

| Setting | Value | Original Value | Date/Time changed | Imported from |
|---|---|---|---|---|
| ***General Settings*** | | | | |
| Analyst name | Dorothy Levorse | | | |
| ***Standard Experiment Settings*** | | | | |
| Number of titrations | 3 | | | |
| Minimum pH | 2.000 | | | |
| Maximum pH | 9.000 | | | |
| pH step between points of | 0.200 | | | |
| Minimum titrant addition | 0.00002 mL | | | |
| Maximum titrant addition | 0.10000 mL | | | |
| Argon flow rate | 100% | | | |
| Start titration using | Cautious pH adjust | | | |
| ***Advanced General Settings*** | | | | |
| Detect turbidity using | None | | | |
| Collect turbidity sensor data | No | | | |
| Collect UV spectra | No | | | |
| Stir after titrant addition for | 5 seconds | | | |
| For titrant addition, stir at | 10% | | | |
| ***Titrant Pre-Dose*** | | | | |
| Titrant pre-dose | None | | | |
| ***Assay Medium*** | | | | |
| ISA water volume | 1.50 mL | | | |
| Water added | Automatic | | | |
| Partition solvent type | Octanol | | | |
| Partition volume | 0.020 mL | | | |
| Partition solvent added | Automatic | | | |
| After partition addition, stir for | 1 seconds | | | |
| ***Sample Sonication*** | | | | |
| Sonicate | Yes | | | |
| Adjust pH for sonication | No | | | |
| Sonicate for | 120 seconds | | | |
| After sonication stir for | 5 seconds | | | |
| ***Sample Dissolution*** | | | | |
| Perform a dissolution stage | Yes | | | |
| Adjust and hold pH for dissolution | To start pH | | | |
| Stir to dissolve for | 120 seconds | | | |
| For dissolution, stir at | 10% | | | |
| ***Carbonate purge*** | | | | |
| Perform a carbonate purge | No | | | |
| ***Temperature Control*** | | | | |
| Wait for temperature | Yes | | | |
| Required start temperature | 25.0°C | | | |
| Acceptable deviation | 0.5°C | | | |
| Time to wait | 60 seconds | | | |
| Stir speed of | 50% | | | |
| ***Titration 1*** | | | | |
| Titrate from | Low to high pH | | | |
| Adjust to start pH | Yes | | | |
| After pH adjust stir for | 30 seconds | | | |
| Stir to allow partitioning for | 15 seconds | | | |
| Stirrer speed for partitioning | 50% | | | |
| ***Titration 2*** | | | | |
| Titrate from | Low to high pH | | | |
| Add additional water | 0.00 mL | | | |
| Additional partition solvent volume | 0.030 mL | | | |
| Additional partition solvent added | Automatic | | | |
| After pH adjust stir for | 30 seconds | | | |
| Stir to allow partitioning for | 15 seconds | | | |
| Stirrer speed for partitioning | 55% | | | |

| | | | |
|---|---|---|---|
| Sample name: | **M02_octanol** | Experiment start time: | **3/3/2018 1:46:01 PM** |
| Assay name: | **pH-metric high logP** | Analyst: | **Dorothy Levorse** |
| Assay ID: | **18C-03011** | Instrument ID: | **T312060** |
| Filename: | **C:\Sirius_T3\Mehtap\20180302_exp29_logP_T3-2\18C-03011_M02_octanol_pH-metric high logP.t3r** | | |

#### Assay Settings (continued)

| Setting | Value | Original Value | Date/Time changed | Imported from |
|---|---|---|---|---|
| ***Titration 3*** | | | | |
| Titrate from | Low to high pH | | | |
| Add additional water | 0.00 mL | | | |
| Additional partition solvent volume | 0.250 mL | | | |
| Additional partition solvent added | Automatic | | | |
| After pH adjust stir for | 30 seconds | | | |
| Stir to allow partitioning for | 15 seconds | | | |
| Stirrer speed for partitioning | 60% | | | |
| ***Data Point Stability*** | | | | |
| Stir during data point collection | No | | | |
| Delay before data point collection | 0 seconds | | | |
| Number of points to average | 20 points | | | |
| Time interval between points | 0.50 seconds | | | |
| Required maximum standard deviation | 0.00100 dpH/dt | | | |
| Stability timeout after | 60 seconds | | | |

#### Calibration Settings

| Setting | Value | Date/Time changed | Imported from |
|---|---|---|---|
| Four-Plus alpha | 0.111 | 3/3/2018 1:46:01 PM | C:\Sirius_T3\HCl18C02.t3r |
| Four-Plus S | 0.9988 | 3/3/2018 1:46:01 PM | C:\Sirius_T3\HCl18C02.t3r |
| Four-Plus jH | 1.0 | 3/3/2018 1:46:01 PM | C:\Sirius_T3\HCl18C02.t3r |
| Four-Plus jOH | -0.8 | 3/3/2018 1:46:01 PM | C:\Sirius_T3\HCl18C02.t3r |
| Base concentration factor | 1.000 | 3/3/2018 1:46:01 PM | C:\Sirius_T3\KOH18B27.t3r |
| Acid concentration factor | 0.999 | 3/3/2018 1:46:01 PM | C:\Sirius_T3\HCl18C02.t3r |

#### Instrument Settings

| Setting | Value | Batch Id | Install date |
|---|---|---|---|
| Instrument owner | Merck | | |
| Instrument ID | T312060 | | |
| Instrument type | T3 Simulator | | |
| Software version | 1.1.3.0 | | |
| Dispenser module | | T3DM1200361 | 3/31/2009 5:24:52 AM |
| Dispenser 0 | Water | | 3/31/2009 5:25:05 AM |
| Syringe volume | 2.5 mL | | |
| Firmware version | 1.2.1(r2) | | |
| Titrant | Water (0.15 M KCl) | 02-06-2018 | 2/27/2018 10:05:59 AM |
| Dispenser 2 | Acid | | 3/31/2009 5:25:11 AM |
| Syringe volume | 0.5 mL | | |
| Firmware version | 1.2.1(r2) | | |
| Titrant | Acid (0.5 M HCl) | 02-27-2018 | 2/27/2018 10:27:22 AM |
| Dispenser 1 | Base | | 3/31/2009 5:25:21 AM |
| Syringe volume | 0.5 mL | | |
| Firmware version | 1.2.1(r2) | | |
| Titrant | Base (0.5 M KOH) | 9/22/2017 | 2/27/2018 10:21:22 AM |
| Dispenser 5 | Cosolvent | | 3/31/2009 5:26:24 AM |
| Syringe volume | 2.5 mL | | |
| Firmware version | 1.2.1(r2) | | |
| Distribution valve 5 | Distribution Valve | | 3/31/2009 5:28:19 AM |
| Firmware version | 1.1.3 | | |
| Port A | Methanol (80%, 0.15 M KCl) | 09-26-17 | 2/7/2018 9:42:01 AM |
| Port B | Cyclohexane | 11-01-17 | 2/27/2018 10:37:57 AM |
| Dispenser 3 | Buffer | | 8/3/2010 5:05:16 AM |
| Syringe volume | 0.5 mL | | |
| Firmware version | 1.2.1(r2) | | |
| Titrant | Dodecane | 2018/01/31 | 2/28/2018 10:18:04 AM |
| Dispenser 6 | Octanol | | 10/22/2010 10:52:43 AM |

| | | | |
|---|---|---|---|
| Sample name: | **M02_octanol** | Experiment start time: | **3/3/2018 1:46:01 PM** |
| Assay name: | **pH-metric high logP** | Analyst: | **Dorothy Levorse** |
| Assay ID: | **18C-03011** | Instrument ID: | **T312060** |
| Filename: | **C:\Sirius_T3\Mehtap\20180302_exp29_logP_T3-2\18C-03011_M02_octanol_pH-metric high logP.t3r** | | |

#### Instrument Settings (continued)

| Setting | Value | Batch Id | Install date |
|---|---|---|---|
| Syringe volume | 0.5 mL | | |
| Firmware version | 1.2.1(r2) | | |
| Titrant | Octanol | 01-31-2018 | 2/27/2018 9:59:35 AM |
| Titrator | | T3TM1200161 | 3/31/2009 5:24:17 AM |
| Horizontal axis firmware version | 1.17 AI1DI2DO2 Stepper 2 | | |
| Vertical axis firmware version | 1.17 AI1DI2DO2 Stepper 2 | | |
| Chassis I/O firmware version | 1.11 AI1DI0DO4 Norgren I/O | | |
| Probe I/O firmware version | 1.1.1 | | |
| Electrode | T3 Electrode | T3E0923 | 1/23/2018 2:01:00 PM |
| E0 calibration | +6.59 mV | | 3/3/2018 1:46:29 PM |
| Filling solution | 3M KCl | KCL097 | 3/2/2018 9:43:24 AM |
| Liquids | | | |
| Wash 1 | 50% IPA:50% Water | | 3/2/2018 9:45:12 AM |
| Wash 2 | 0.5% Trition X-100 in H20 | | 3/2/2018 9:45:15 AM |
| Buffer position 1 | pH7 Wash | | 3/2/2018 9:45:18 AM |
| Buffer position 2 | pH 7 | | 3/2/2018 9:45:21 AM |
| Storage position | | | 3/2/2018 9:44:44 AM |
| Wash water | 6.7e+003 mL | 02-27-2018 | 2/27/2018 9:54:39 AM |
| Waste | 8.7e+003 mL | | 11/28/2017 10:36:29 AM |
| Temperature controller | | | 8/5/2010 6:35:13 AM |
| Turbidity detector | | | 3/31/2009 5:24:45 AM |
| Spectrometer | | 074811 | 11/23/2010 11:22:28 AM |
| Dip probe | | 10196 | |
| Wavelength coefficient A0 | 183.333 | | |
| Wavelength coefficient A1 | 2.21568 | | |
| Wavelength coefficient A2 | -0.000289308 | | |
| Total lamp lit time | 120:41:49 | | 11/23/2010 11:22:28 AM |
| Calibrated on | 2/27/2018 10:40:38 AM | | |
| Integration time | 40 | | |
| Scans averaged | 10 | | |
| Autoloader | | T3AL1200345 | 11/10/2015 9:34:13 AM |
| Left-right axis firmware version | 1.17 AI1DI2DO2 Stepper 2 | | |
| Front-back axis firmware version | 1.17 AI1DI2DO2 Stepper 2 | | |
| Vertical axis firmware version | 1.17 AI1DI2DO2 Stepper 2 | | |
| Chassis I/O firmware version | 1.11 AI1DI0DO4 Norgren I/O | | |
| Configuration | | | |
| Alternate titration position | Titration position | | |
| Alternate reference position | Reference position | | |
| Maximum standard vial volume | 3.50 mL | | |
| Maximum alternate vial volume | 25.00 mL | | |
| Automatic action idle period | 5 minute(s) | | |
| Titrant tube volume | 1.3 mL | | |
| Syringe flush count | 3.50 | | |
| Flowing wash pump volume | 20.0 mL | | |
| Flowing wash stir duration | 5 s | | |
| Flowing wash stir speed | 30% | | |
| Solvent wash stir duration | 5 s | | |
| Solvent wash stir speed | 30% | | |
| Surfactant wash stir duration | 5 s | | |
| Surfactant wash stir speed | 30% | | |
| E0 calibration minimum number of points | 10 | | |
| E0 calibration maximum standard deviation | 0.01500 | | |
| E0 calibration timeout period | 60 s | | |
| E0 calibration stir duration | 5 s | | |
| E0 calibration preparation stir speed | 30% | | |
| E0 calibration buffer wash stir duration | 5 s | | |
| E0 calibration buffer wash stir speed | 30% | | |
| E0 calibration reading stir speed | 0% | | |

| | | | |
|---|---|---|---|
| Sample name: | **M02_octanol** | Experiment start time: | **3/3/2018 1:46:01 PM** |
| Assay name: | **pH-metric high logP** | Analyst: | **Dorothy Levorse** |
| Assay ID: | **18C-03011** | Instrument ID: | **T312060** |
| Filename: | **C:\Sirius_T3\Mehtap\20180302_exp29_logP_T3-2\18C-03011_M02_octanol_pH-metric high logP.t3r** | | |

#### Instrument Settings (continued)

| Setting | Value | Batch Id | Install date |
|---|---|---|---|
| Spectrometer calibration stir duration | 5 s | | |
| Spectrometer calibration stir speed | 30% | | |
| Spectrometer calibration wash pump volume | 20.0 mL | | |
| Spectrometer calibration wash stir duration | 5 s | | |
| Spectrometer calibration wash stir speed | 30% | | |
| Overhead dispense height | 10000 | | |

#### Refinement Settings

| Setting | Value | Default value |
|---|---|---|
| Turbidity detection method | None | None |
| Turbidity wavelength to assess | 500.0 nm | 500.0 nm |
| Turbidity maximum absorbance | 0.100 | 0.100 |
| Turbidity probe threshold | 50.00 | 50.00 |

#### Experiment Log

[2:38] Air gap created for Water (0.15 M KCl)
[2:38] Air gap created for Acid (0.5 M HCl)
[2:38] Air gap created for Base (0.5 M KOH)
[2:39] Air gap released for Water (0.15 M KCl)
[2:42] Titrator arm moved over Titration position
[2:42] Titration 1 of 3
[2:42] Adding initial titrants
[2:42] Automatically add 1.50000 mL of water
[3:08] Dispensed 1.500000 mL of Water (0.15 M KCl)
[3:12] Titrator arm moved over Drain
[5:54] Titrator arm moved to Titration position
[5:54] Argon flow rate set to 100
[5:54] Stirrer speed set to 10
[5:59] Automatically add 0.02000 mL of Octanol
[5:59] Dispensed 0.019991 mL of Octanol
[6:00] Initial pH = 8.70
[6:00] Iterative adjust 8.70 -> 2.00
[6:00] pH 8.70 -> 2.00
[6:02] Air gap released for Acid (0.5 M HCl)
[6:03] Dispensed 0.052516 mL of Acid (0.5 M HCl)
[6:08] Holding pH 2.00
[8:08] Stirrer speed set to 0
[8:08] Stirrer speed set to 50
[8:08] Iterative adjust 1.98 -> 2.00
[8:08] pH 1.98 -> 2.00
[8:09] Air gap released for Base (0.5 M KOH)
[8:10] Dispensed 0.001623 mL of Base (0.5 M KOH)
[9:00] Stirrer speed set to 0
[9:15] Datapoint id 1 collected
[9:15] Stirrer speed set to 50
[9:20] pH 2.08 -> 2.28
[9:20] Using cautious pH adjust
[9:20] Dispensed 0.006656 mL of Base (0.5 M KOH)
[9:25] Stepping pH = 2.17
[9:26] Dispensed 0.005456 mL of Base (0.5 M KOH)
[9:31] Stepping pH = 2.25
[9:31] Dispensed 0.001787 mL of Base (0.5 M KOH)
[9:36] Stepping pH = 2.29
[9:51] Stirrer speed set to 0
[10:02] Datapoint id 2 collected
[10:02] Charge balance equation is out by -4.4%
[10:02] Stirrer speed set to 50

| | | | |
|---|---|---|---|
| Sample name: | **M02_octanol** | Experiment start time: | **3/3/2018 1:46:01 PM** |
| Assay name: | **pH-metric high logP** | Analyst: | **Dorothy Levorse** |
| Assay ID: | **18C-03011** | Instrument ID: | **T312060** |
| Filename: | **C:\Sirius_T3\Mehtap\20180302_exp29_logP_T3-2\18C-03011_M02_octanol_pH-metric high logP.t3r** | | |

#### Experiment Log (continued)

[10:07] pH 2.29 -> 2.49
[10:07] Using charge balance adjust
[10:07] Dispensed 0.008749 mL of Base (0.5 M KOH)
[10:27] Stirrer speed set to 0
[10:38] Datapoint id 3 collected
[10:38] Charge balance equation is out by -0.3%
[10:38] Stirrer speed set to 50
[10:43] pH 2.50 -> 2.70
[10:43] Using charge balance adjust
[10:43] Dispensed 0.006044 mL of Base (0.5 M KOH)
[11:03] Stirrer speed set to 0
[11:13] Datapoint id 4 collected
[11:13] Charge balance equation is out by -8.9%
[11:13] Stirrer speed set to 50
[11:18] pH 2.68 -> 2.88
[11:18] Using charge balance adjust
[11:19] Dispensed 0.004563 mL of Base (0.5 M KOH)
[11:39] Stirrer speed set to 0
[11:49] Datapoint id 5 collected
[11:49] Charge balance equation is out by -3.6%
[11:49] Stirrer speed set to 50
[11:54] pH 2.88 -> 3.08
[11:54] Using charge balance adjust
[11:54] Dispensed 0.003551 mL of Base (0.5 M KOH)
[12:14] Stirrer speed set to 0
[12:25] Datapoint id 6 collected
[12:25] Charge balance equation is out by 8.3%
[12:25] Stirrer speed set to 50
[12:30] pH 3.10 -> 3.30
[12:30] Using charge balance adjust
[12:30] Dispensed 0.002728 mL of Base (0.5 M KOH)
[12:50] Stirrer speed set to 0
[13:00] Datapoint id 7 collected
[13:00] Charge balance equation is out by -4.3%
[13:00] Stirrer speed set to 50
[13:05] pH 3.29 -> 3.49
[13:05] Using charge balance adjust
[13:06] Dispensed 0.002117 mL of Base (0.5 M KOH)
[13:26] Stirrer speed set to 0
[13:49] Datapoint id 8 collected
[13:49] Charge balance equation is out by 12.1%
[13:49] Stirrer speed set to 50
[13:54] pH 3.51 -> 3.71
[13:54] Using charge balance adjust
[13:54] Dispensed 0.001505 mL of Base (0.5 M KOH)
[14:14] Stirrer speed set to 0
[14:31] Datapoint id 9 collected
[14:31] Charge balance equation is out by 23.9%
[14:31] Stirrer speed set to 50
[14:36] pH 3.76 -> 3.96
[14:36] Using cautious pH adjust
[14:36] Dispensed 0.000494 mL of Base (0.5 M KOH)
[14:42] Stepping pH = 3.86
[14:42] Dispensed 0.000306 mL of Base (0.5 M KOH)
[14:47] Stepping pH = 3.94
[14:47] Dispensed 0.000047 mL of Base (0.5 M KOH)
[14:52] Stepping pH = 3.92
[14:52] Dispensed 0.000635 mL of Base (0.5 M KOH)
[14:57] Stepping pH = 4.16

| | | | |
|---|---|---|---|
| Sample name: | **M02_octanol** | Experiment start time: | **3/3/2018 1:46:01 PM** |
| Assay name: | **pH-metric high logP** | Analyst: | **Dorothy Levorse** |
| Assay ID: | **18C-03011** | Instrument ID: | **T312060** |
| Filename: | **C:\Sirius_T3\Mehtap\20180302_exp29_logP_T3-2\18C-03011_M02_octanol_pH-metric high logP.t3r** | | |

#### Experiment Log (continued)

[15:12] Stirrer speed set to 0
[15:32] Datapoint id 10 collected
[15:32] Charge balance equation is out by -51.7%
[15:32] Stirrer speed set to 50
[15:37] pH 4.11 -> 4.31
[15:37] Using cautious pH adjust
[15:37] Dispensed 0.000259 mL of Base (0.5 M KOH)
[15:42] Stepping pH = 4.20
[15:43] Dispensed 0.000188 mL of Base (0.5 M KOH)
[15:48] Stepping pH = 4.28
[15:48] Dispensed 0.000047 mL of Base (0.5 M KOH)
[15:53] Stepping pH = 4.28
[15:53] Dispensed 0.000282 mL of Base (0.5 M KOH)
[15:58] Stepping pH = 4.50
[16:13] Stirrer speed set to 0
[16:51] Datapoint id 11 collected
[16:51] Charge balance equation is out by -54.0%
[16:51] Stirrer speed set to 50
[16:56] pH 4.45 -> 4.65
[16:56] Using cautious pH adjust
[16:56] Dispensed 0.000118 mL of Base (0.5 M KOH)
[17:01] Stepping pH = 4.46
[17:01] Dispensed 0.000353 mL of Base (0.5 M KOH)
[17:06] Stepping pH = 4.97
[17:21] Stirrer speed set to 0
[18:04] Datapoint id 12 collected
[18:04] Charge balance equation is out by -97.0%
[18:04] Stirrer speed set to 50
[18:09] pH 4.93 -> 5.13
[18:09] Using cautious pH adjust
[18:09] Dispensed 0.000047 mL of Base (0.5 M KOH)
[18:15] Stepping pH = 4.94
[18:15] Dispensed 0.000141 mL of Base (0.5 M KOH)
[18:20] Stepping pH = 5.22
[18:35] Stirrer speed set to 0
[19:26] Datapoint id 13 collected
[19:26] Charge balance equation is out by -95.2%
[19:26] Stirrer speed set to 50
[19:31] pH 5.23 -> 5.43
[19:31] Using cautious pH adjust
[19:31] Dispensed 0.000024 mL of Base (0.5 M KOH)
[19:36] Stepping pH = 5.23
[19:36] Dispensed 0.000141 mL of Base (0.5 M KOH)
[19:41] Stepping pH = 5.64
[19:56] Stirrer speed set to 0
[20:56] Datapoint id 14 collected
[20:56] Charge balance equation is out by -204.5%
[20:56] Stirrer speed set to 50
[21:02] pH 5.71 -> 5.91
[21:02] Using cautious pH adjust
[21:02] Dispensed 0.000024 mL of Base (0.5 M KOH)
[21:07] Stepping pH = 5.75
[21:07] Dispensed 0.000047 mL of Base (0.5 M KOH)
[21:12] Stepping pH = 5.78
[21:12] Dispensed 0.000071 mL of Base (0.5 M KOH)
[21:17] Stepping pH = 6.41
[21:32] Stirrer speed set to 0
[22:32] Datapoint id 15 collected
[22:32] Charge balance equation is out by -213.0%

| | | | |
|---|---|---|---|
| Sample name: | **M02_octanol** | Experiment start time: | **3/3/2018 1:46:01 PM** |
| Assay name: | **pH-metric high logP** | Analyst: | **Dorothy Levorse** |
| Assay ID: | **18C-03011** | Instrument ID: | **T312060** |
| Filename: | **C:\Sirius_T3\Mehtap\20180302_exp29_logP_T3-2\18C-03011_M02_octanol_pH-metric high logP.t3r** | | |

#### Experiment Log (continued)

[22:32] Stirrer speed set to 50
[22:37] pH 6.50 -> 6.70
[22:37] Using cautious pH adjust
[22:37] Dispensed 0.000024 mL of Base (0.5 M KOH)
[22:42] Stepping pH = 6.50
[22:42] Dispensed 0.000047 mL of Base (0.5 M KOH)
[22:47] Stepping pH = 6.64
[22:48] Dispensed 0.000024 mL of Base (0.5 M KOH)
[22:53] Stepping pH = 6.88
[23:08] Stirrer speed set to 0
[24:08] Datapoint id 16 collected
[24:08] Charge balance equation is out by -164.9%
[24:08] Stirrer speed set to 50
[24:13] pH 6.94 -> 7.14
[24:13] Using cautious pH adjust
[24:13] Dispensed 0.000024 mL of Base (0.5 M KOH)
[24:18] Stepping pH = 6.91
[24:18] Dispensed 0.000047 mL of Base (0.5 M KOH)
[24:23] Stepping pH = 7.15
[24:38] Stirrer speed set to 0
[25:38] Datapoint id 17 collected
[25:38] Charge balance equation is out by -314.2%
[25:38] Stirrer speed set to 50
[25:43] pH 7.53 -> 7.73
[25:43] Using cautious pH adjust
[25:43] Dispensed 0.000024 mL of Base (0.5 M KOH)
[25:49] Stepping pH = 7.51
[25:49] Dispensed 0.000024 mL of Base (0.5 M KOH)
[25:54] Stepping pH = 7.55
[25:54] Dispensed 0.000024 mL of Base (0.5 M KOH)
[25:59] Stepping pH = 7.84
[26:14] Stirrer speed set to 0
[27:14] Datapoint id 18 collected
[27:14] Charge balance equation is out by -916.6%
[27:14] Stirrer speed set to 50
[27:19] pH 8.10 -> 8.30
[27:19] Using cautious pH adjust
[27:19] Dispensed 0.000024 mL of Base (0.5 M KOH)
[27:24] Stepping pH = 8.05
[27:24] Dispensed 0.000024 mL of Base (0.5 M KOH)
[27:29] Stepping pH = 8.01
[27:29] Dispensed 0.000118 mL of Base (0.5 M KOH)
[27:35] Stepping pH = 8.98
[27:50] Stirrer speed set to 0
[28:46] Datapoint id 19 collected
[28:46] Charge balance equation is out by -2,236.6%
[28:46] Stirrer speed set to 50
[28:51] pH 8.92 -> 9.05
[28:51] Using cautious pH adjust
[28:51] Dispensed 0.000024 mL of Base (0.5 M KOH)
[28:56] Stepping pH = 8.92
[28:56] Dispensed 0.000047 mL of Base (0.5 M KOH)
[29:01] Stepping pH = 9.11
[29:16] Stirrer speed set to 0
[29:53] Datapoint id 20 collected
[29:53] Charge balance equation is out by -259.1%
[29:53] Titration 2 of 3
[29:53] Adding initial titrants
[29:53] Automatically add 0.03000 mL of Octanol

| | | | |
|---|---|---|---|
| Sample name: | **M02_octanol** | Experiment start time: | **3/3/2018 1:46:01 PM** |
| Assay name: | **pH-metric high logP** | Analyst: | **Dorothy Levorse** |
| Assay ID: | **18C-03011** | Instrument ID: | **T312060** |
| Filename: | **C:\Sirius_T3\Mehtap\20180302_exp29_logP_T3-2\18C-03011_M02_octanol_pH-metric high logP.t3r** | | |

#### Experiment Log (continued)

[29:54] Dispensed 0.030009 mL of Octanol
[29:54] Stirrer speed set to 10
[29:55] Stirrer speed set to 55
[29:55] Iterative adjust 9.14 -> 2.00
[29:55] pH 9.14 -> 2.00
[29:57] Dispensed 0.054492 mL of Acid (0.5 M HCl)
[30:02] pH 2.02 -> 2.00
[30:02] Dispensed 0.001740 mL of Acid (0.5 M HCl)
[30:52] Stirrer speed set to 0
[31:02] Datapoint id 21 collected
[31:02] Stirrer speed set to 55
[31:07] pH 2.02 -> 2.22
[31:07] Using cautious pH adjust
[31:08] Dispensed 0.008396 mL of Base (0.5 M KOH)
[31:13] Stepping pH = 2.10
[31:13] Dispensed 0.007079 mL of Base (0.5 M KOH)
[31:18] Stepping pH = 2.19
[31:18] Dispensed 0.001576 mL of Base (0.5 M KOH)
[31:23] Stepping pH = 2.21
[31:38] Stirrer speed set to 0
[31:48] Datapoint id 22 collected
[31:48] Charge balance equation is out by -1.5%
[31:48] Stirrer speed set to 55
[31:54] pH 2.22 -> 2.42
[31:54] Using charge balance adjust
[31:54] Dispensed 0.011077 mL of Base (0.5 M KOH)
[32:14] Stirrer speed set to 0
[32:25] Datapoint id 23 collected
[32:25] Charge balance equation is out by -11.5%
[32:25] Stirrer speed set to 55
[32:30] pH 2.40 -> 2.60
[32:30] Using charge balance adjust
[32:30] Dispensed 0.007879 mL of Base (0.5 M KOH)
[32:50] Stirrer speed set to 0
[33:00] Datapoint id 24 collected
[33:00] Charge balance equation is out by 4.0%
[33:00] Stirrer speed set to 55
[33:05] pH 2.61 -> 2.81
[33:05] Using charge balance adjust
[33:06] Dispensed 0.005503 mL of Base (0.5 M KOH)
[33:26] Stirrer speed set to 0
[33:36] Datapoint id 25 collected
[33:36] Charge balance equation is out by -14.7%
[33:36] Stirrer speed set to 55
[33:41] pH 2.79 -> 2.99
[33:41] Using charge balance adjust
[33:41] Dispensed 0.004163 mL of Base (0.5 M KOH)
[34:01] Stirrer speed set to 0
[34:11] Datapoint id 26 collected
[34:11] Charge balance equation is out by -5.7%
[34:11] Stirrer speed set to 55
[34:16] pH 2.98 -> 3.18
[34:16] Using charge balance adjust
[34:17] Dispensed 0.003081 mL of Base (0.5 M KOH)
[34:37] Stirrer speed set to 0
[34:47] Datapoint id 27 collected
[34:47] Charge balance equation is out by -10.6%
[34:47] Stirrer speed set to 55
[34:52] pH 3.16 -> 3.36

| | | | |
|---|---|---|---|
| Sample name: | **M02_octanol** | Experiment start time: | **3/3/2018 1:46:01 PM** |
| Assay name: | **pH-metric high logP** | Analyst: | **Dorothy Levorse** |
| Assay ID: | **18C-03011** | Instrument ID: | **T312060** |
| Filename: | **C:\Sirius_T3\Mehtap\20180302_exp29_logP_T3-2\18C-03011_M02_octanol_pH-metric high logP.t3r** | | |

#### Experiment Log (continued)

[34:52] Using charge balance adjust
[34:52] Dispensed 0.002281 mL of Base (0.5 M KOH)
[35:12] Stirrer speed set to 0
[35:23] Datapoint id 28 collected
[35:23] Charge balance equation is out by 15.1%
[35:23] Stirrer speed set to 55
[35:28] pH 3.40 -> 3.60
[35:28] Using cautious pH adjust
[35:28] Dispensed 0.000753 mL of Base (0.5 M KOH)
[35:33] Stepping pH = 3.52
[35:33] Dispensed 0.000306 mL of Base (0.5 M KOH)
[35:38] Stepping pH = 3.58
[35:38] Dispensed 0.000047 mL of Base (0.5 M KOH)
[35:43] Stepping pH = 3.59
[35:43] Dispensed 0.000118 mL of Base (0.5 M KOH)
[35:49] Stepping pH = 3.61
[36:04] Stirrer speed set to 0
[36:14] Datapoint id 29 collected
[36:14] Charge balance equation is out by 17.7%
[36:14] Stirrer speed set to 55
[36:19] pH 3.60 -> 3.80
[36:19] Using cautious pH adjust
[36:19] Dispensed 0.000517 mL of Base (0.5 M KOH)
[36:24] Stepping pH = 3.73
[36:24] Dispensed 0.000188 mL of Base (0.5 M KOH)
[36:29] Stepping pH = 3.77
[36:29] Dispensed 0.000071 mL of Base (0.5 M KOH)
[36:34] Stepping pH = 3.78
[36:34] Dispensed 0.000071 mL of Base (0.5 M KOH)
[36:40] Stepping pH = 3.80
[36:55] Stirrer speed set to 0
[37:05] Datapoint id 30 collected
[37:05] Charge balance equation is out by 16.3%
[37:05] Stirrer speed set to 55
[37:10] pH 3.80 -> 4.00
[37:10] Using cautious pH adjust
[37:10] Dispensed 0.000329 mL of Base (0.5 M KOH)
[37:15] Stepping pH = 3.91
[37:15] Dispensed 0.000188 mL of Base (0.5 M KOH)
[37:20] Stepping pH = 3.99
[37:20] Dispensed 0.000024 mL of Base (0.5 M KOH)
[37:25] Stepping pH = 3.99
[37:25] Dispensed 0.000071 mL of Base (0.5 M KOH)
[37:30] Stepping pH = 4.02
[37:45] Stirrer speed set to 0
[37:56] Datapoint id 31 collected
[37:56] Charge balance equation is out by 9.2%
[37:56] Stirrer speed set to 55
[38:01] pH 4.02 -> 4.22
[38:01] Using charge balance adjust
[38:01] Dispensed 0.000423 mL of Base (0.5 M KOH)
[38:21] Stirrer speed set to 0
[38:34] Datapoint id 32 collected
[38:34] Charge balance equation is out by 10.4%
[38:34] Stirrer speed set to 55
[38:39] pH 4.26 -> 4.46
[38:39] Using charge balance adjust
[38:39] Dispensed 0.000259 mL of Base (0.5 M KOH)
[38:59] Stirrer speed set to 0

| | | | |
|---|---|---|---|
| Sample name: | **M02_octanol** | Experiment start time: | **3/3/2018 1:46:01 PM** |
| Assay name: | **pH-metric high logP** | Analyst: | **Dorothy Levorse** |
| Assay ID: | **18C-03011** | Instrument ID: | **T312060** |
| Filename: | **C:\Sirius_T3\Mehtap\20180302_exp29_logP_T3-2\18C-03011_M02_octanol_pH-metric high logP.t3r** | | |

#### Experiment Log (continued)

[39:15] Datapoint id 33 collected
[39:15] Charge balance equation is out by -4.0%
[39:15] Stirrer speed set to 55
[39:21] pH 4.47 -> 4.67
[39:21] Using charge balance adjust
[39:21] Dispensed 0.000165 mL of Base (0.5 M KOH)
[39:41] Stirrer speed set to 0
[40:11] Datapoint id 34 collected
[40:11] Charge balance equation is out by -37.4%
[40:11] Stirrer speed set to 55
[40:16] pH 4.64 -> 4.84
[40:16] Using cautious pH adjust
[40:17] Dispensed 0.000047 mL of Base (0.5 M KOH)
[40:22] Stepping pH = 4.65
[40:22] Dispensed 0.000141 mL of Base (0.5 M KOH)
[40:27] Stepping pH = 4.98
[40:42] Stirrer speed set to 0
[40:58] Datapoint id 35 collected
[40:58] Charge balance equation is out by -87.0%
[40:58] Stirrer speed set to 55
[41:03] pH 5.00 -> 5.20
[41:03] Using cautious pH adjust
[41:03] Dispensed 0.000024 mL of Base (0.5 M KOH)
[41:08] Stepping pH = 5.00
[41:08] Dispensed 0.000094 mL of Base (0.5 M KOH)
[41:13] Stepping pH = 5.27
[41:28] Stirrer speed set to 0
[42:01] Datapoint id 36 collected
[42:01] Charge balance equation is out by -98.9%
[42:01] Stirrer speed set to 55
[42:06] pH 5.33 -> 5.53
[42:06] Using cautious pH adjust
[42:06] Dispensed 0.000024 mL of Base (0.5 M KOH)
[42:11] Stepping pH = 5.34
[42:11] Dispensed 0.000047 mL of Base (0.5 M KOH)
[42:16] Stepping pH = 5.59
[42:31] Stirrer speed set to 0
[43:19] Datapoint id 37 collected
[43:19] Charge balance equation is out by -92.9%
[43:19] Stirrer speed set to 55
[43:24] pH 5.70 -> 5.90
[43:24] Using cautious pH adjust
[43:24] Dispensed 0.000024 mL of Base (0.5 M KOH)
[43:29] Stepping pH = 5.72
[43:29] Dispensed 0.000047 mL of Base (0.5 M KOH)
[43:34] Stepping pH = 6.12
[43:49] Stirrer speed set to 0
[44:49] Datapoint id 38 collected
[44:49] Charge balance equation is out by -87.7%
[44:49] Stirrer speed set to 55
[44:54] pH 6.16 -> 6.36
[44:54] Using cautious pH adjust
[44:54] Dispensed 0.000024 mL of Base (0.5 M KOH)
[44:59] Stepping pH = 6.16
[44:59] Dispensed 0.000071 mL of Base (0.5 M KOH)
[45:04] Stepping pH = 6.69
[45:20] Stirrer speed set to 0
[46:20] Datapoint id 39 collected
[46:20] Charge balance equation is out by -96.1%

Sample name: **M02_octanol**
Assay name: **pH-metric high logP**
Assay ID: **18C-03011**
Filename: **C:\Sirius_T3\Mehtap\20180302_exp29_logP_T3-2\18C-03011_M02_octanol_pH-metric high logP.t3r**

Experiment start time: **3/3/2018 1:46:01 PM**
Analyst: **Dorothy Levorse**
Instrument ID: **T312060**

#### Experiment Log (continued)

[46:20] Stirrer speed set to 55
[46:25] pH 6.83 -> 7.03
[46:25] Using cautious pH adjust
[46:25] Dispensed 0.000024 mL of Base (0.5 M KOH)
[46:30] Stepping pH = 6.85
[46:30] Dispensed 0.000024 mL of Base (0.5 M KOH)
[46:35] Stepping pH = 6.96
[46:35] Dispensed 0.000024 mL of Base (0.5 M KOH)
[46:40] Stepping pH = 7.20
[46:55] Stirrer speed set to 0
[47:55] Datapoint id 40 collected
[47:55] Charge balance equation is out by -198.9%
[47:55] Stirrer speed set to 55
[48:00] pH 7.31 -> 7.51
[48:00] Using cautious pH adjust
[48:00] Dispensed 0.000024 mL of Base (0.5 M KOH)
[48:05] Stepping pH = 7.34
[48:06] Dispensed 0.000024 mL of Base (0.5 M KOH)
[48:11] Stepping pH = 7.46
[48:11] Dispensed 0.000024 mL of Base (0.5 M KOH)
[48:16] Stepping pH = 7.68
[48:31] Stirrer speed set to 0
[49:31] Datapoint id 41 collected
[49:31] Charge balance equation is out by -452.4%
[49:31] Stirrer speed set to 55
[49:36] pH 7.54 -> 7.74
[49:36] Using cautious pH adjust
[49:36] Dispensed 0.000024 mL of Base (0.5 M KOH)
[49:41] Stepping pH = 7.55
[49:41] Dispensed 0.000024 mL of Base (0.5 M KOH)
[49:46] Stepping pH = 7.73
[50:02] Stirrer speed set to 0
[51:02] Datapoint id 42 collected
[51:02] Charge balance equation is out by -414.6%
[51:02] Stirrer speed set to 55
[51:07] pH 7.69 -> 7.89
[51:07] Using cautious pH adjust
[51:07] Dispensed 0.000024 mL of Base (0.5 M KOH)
[51:12] Stepping pH = 7.66
[51:12] Dispensed 0.000024 mL of Base (0.5 M KOH)
[51:17] Stepping pH = 7.90
[51:32] Stirrer speed set to 0
[52:32] Datapoint id 43 collected
[52:32] Charge balance equation is out by -498.4%
[52:32] Stirrer speed set to 55
[52:37] pH 8.03 -> 8.23
[52:37] Using cautious pH adjust
[52:37] Dispensed 0.000024 mL of Base (0.5 M KOH)
[52:42] Stepping pH = 8.05
[52:42] Dispensed 0.000024 mL of Base (0.5 M KOH)
[52:47] Stepping pH = 8.24
[53:02] Stirrer speed set to 0
[54:03] Datapoint id 44 collected
[54:03] Charge balance equation is out by -454.8%
[54:03] Stirrer speed set to 55
[54:08] pH 8.27 -> 8.47
[54:08] Using cautious pH adjust
[54:08] Dispensed 0.000024 mL of Base (0.5 M KOH)
[54:13] Stepping pH = 8.30

| | | | |
|---|---|---|---|
| Sample name: | **M02_octanol** | Experiment start time: | **3/3/2018 1:46:01 PM** |
| Assay name: | **pH-metric high logP** | Analyst: | **Dorothy Levorse** |
| Assay ID: | **18C-03011** | Instrument ID: | **T312060** |
| Filename: | **C:\Sirius_T3\Mehtap\20180302_exp29_logP_T3-2\18C-03011_M02_octanol_pH-metric high logP.t3r** | | |

#### Experiment Log (continued)

[54:13] Dispensed 0.000024 mL of Base (0.5 M KOH)
[54:18] Stepping pH = 8.46
[54:33] Stirrer speed set to 0
[55:33] Datapoint id 45 collected
[55:33] Charge balance equation is out by -298.4%
[55:33] Stirrer speed set to 55
[55:38] pH 8.50 -> 8.70
[55:38] Using cautious pH adjust
[55:38] Dispensed 0.000024 mL of Base (0.5 M KOH)
[55:43] Stepping pH = 8.53
[55:43] Dispensed 0.000024 mL of Base (0.5 M KOH)
[55:48] Stepping pH = 8.65
[55:48] Dispensed 0.000024 mL of Base (0.5 M KOH)
[55:54] Stepping pH = 8.76
[56:09] Stirrer speed set to 0
[56:39] Datapoint id 46 collected
[56:39] Charge balance equation is out by -284.7%
[56:39] Stirrer speed set to 55
[56:44] pH 8.74 -> 8.94
[56:44] Using cautious pH adjust
[56:44] Dispensed 0.000024 mL of Base (0.5 M KOH)
[56:49] Stepping pH = 8.74
[56:50] Dispensed 0.000047 mL of Base (0.5 M KOH)
[56:55] Stepping pH = 8.90
[56:55] Dispensed 0.000024 mL of Base (0.5 M KOH)
[57:00] Stepping pH = 8.96
[57:15] Stirrer speed set to 0
[57:51] Datapoint id 47 collected
[57:51] Charge balance equation is out by -204.3%
[57:51] Stirrer speed set to 55
[57:56] pH 8.98 -> 9.05
[57:56] Using cautious pH adjust
[57:56] Dispensed 0.000024 mL of Base (0.5 M KOH)
[58:01] Stepping pH = 8.98
[58:01] Dispensed 0.000024 mL of Base (0.5 M KOH)
[58:06] Stepping pH = 9.02
[58:21] Stirrer speed set to 0
[58:48] Datapoint id 48 collected
[58:48] Charge balance equation is out by -216.1%
[58:48] Titration 3 of 3
[58:48] Adding initial titrants
[58:48] Automatically add 0.25000 mL of Octanol
[58:53] Dispensed 0.250000 mL of Octanol
[58:53] Stirrer speed set to 10
[58:55] Stirrer speed set to 60
[58:55] Iterative adjust 9.03 -> 2.00
[58:55] pH 9.03 -> 2.00
[58:56] Dispensed 0.056044 mL of Acid (0.5 M HCl)
[59:01] pH 2.04 -> 2.00
[59:01] Dispensed 0.004139 mL of Acid (0.5 M HCl)
[59:52] Stirrer speed set to 0
[1:00:02] Datapoint id 49 collected
[1:00:02] Stirrer speed set to 60
[1:00:07] pH 1.99 -> 2.19
[1:00:07] Using cautious pH adjust
[1:00:07] Dispensed 0.009666 mL of Base (0.5 M KOH)
[1:00:12] Stepping pH = 2.07
[1:00:12] Dispensed 0.009055 mL of Base (0.5 M KOH)
[1:00:17] Stepping pH = 2.16

| | | | |
|---|---|---|---|
| Sample name: | **M02_octanol** | Experiment start time: | **3/3/2018 1:46:01 PM** |
| Assay name: | **pH-metric high logP** | Analyst: | **Dorothy Levorse** |
| Assay ID: | **18C-03011** | Instrument ID: | **T312060** |
| Filename: | **C:\Sirius_T3\Mehtap\20180302_exp29_logP_T3-2\18C-03011_M02_octanol_pH-metric high logP.t3r** | | |

#### Experiment Log (continued)

[1:00:18] Dispensed 0.002258 mL of Base (0.5 M KOH)
[1:00:23] Stepping pH = 2.19
[1:00:38] Stirrer speed set to 0
[1:00:48] Datapoint id 50 collected
[1:00:48] Charge balance equation is out by -8.6%
[1:00:48] Stirrer speed set to 60
[1:00:53] pH 2.19 -> 2.39
[1:00:53] Using charge balance adjust
[1:00:53] Dispensed 0.012912 mL of Base (0.5 M KOH)
[1:01:14] Stirrer speed set to 0
[1:01:24] Datapoint id 51 collected
[1:01:24] Charge balance equation is out by -8.4%
[1:01:24] Stirrer speed set to 60
[1:01:29] pH 2.37 -> 2.57
[1:01:29] Using charge balance adjust
[1:01:29] Dispensed 0.008984 mL of Base (0.5 M KOH)
[1:01:49] Stirrer speed set to 0
[1:01:59] Datapoint id 52 collected
[1:01:59] Charge balance equation is out by 2.2%
[1:01:59] Stirrer speed set to 60
[1:02:04] pH 2.58 -> 2.78
[1:02:04] Using charge balance adjust
[1:02:05] Dispensed 0.006044 mL of Base (0.5 M KOH)
[1:02:25] Stirrer speed set to 0
[1:02:35] Datapoint id 53 collected
[1:02:35] Charge balance equation is out by -7.6%
[1:02:35] Stirrer speed set to 60
[1:02:40] pH 2.77 -> 2.97
[1:02:40] Using charge balance adjust
[1:02:40] Dispensed 0.004233 mL of Base (0.5 M KOH)
[1:03:00] Stirrer speed set to 0
[1:03:15] Datapoint id 54 collected
[1:03:15] Charge balance equation is out by 7.6%
[1:03:15] Stirrer speed set to 60
[1:03:20] pH 2.99 -> 3.19
[1:03:20] Using charge balance adjust
[1:03:20] Dispensed 0.002752 mL of Base (0.5 M KOH)
[1:03:40] Stirrer speed set to 0
[1:03:50] Datapoint id 55 collected
[1:03:50] Charge balance equation is out by 19.2%
[1:03:50] Stirrer speed set to 60
[1:03:55] pH 3.23 -> 3.43
[1:03:55] Using cautious pH adjust
[1:03:55] Dispensed 0.000847 mL of Base (0.5 M KOH)
[1:04:01] Stepping pH = 3.33
[1:04:01] Dispensed 0.000541 mL of Base (0.5 M KOH)
[1:04:06] Stepping pH = 3.41
[1:04:06] Dispensed 0.000118 mL of Base (0.5 M KOH)
[1:04:11] Stepping pH = 3.42
[1:04:11] Dispensed 0.000165 mL of Base (0.5 M KOH)
[1:04:16] Stepping pH = 3.43
[1:04:31] Stirrer speed set to 0
[1:04:41] Datapoint id 56 collected
[1:04:41] Charge balance equation is out by 1.6%
[1:04:41] Stirrer speed set to 60
[1:04:46] pH 3.44 -> 3.64
[1:04:46] Using charge balance adjust
[1:04:46] Dispensed 0.001105 mL of Base (0.5 M KOH)
[1:05:07] Stirrer speed set to 0

| | | | |
|---|---|---|---|
| Sample name: | **M02_octanol** | Experiment start time: | **3/3/2018 1:46:01 PM** |
| Assay name: | **pH-metric high logP** | Analyst: | **Dorothy Levorse** |
| Assay ID: | **18C-03011** | Instrument ID: | **T312060** |
| Filename: | **C:\Sirius_T3\Mehtap\20180302_exp29_logP_T3-2\18C-03011_M02_octanol_pH-metric high logP.t3r** | | |

#### Experiment Log (continued)

[1:05:22] Datapoint id 57 collected
[1:05:22] Charge balance equation is out by 27.7%
[1:05:22] Stirrer speed set to 60
[1:05:27] pH 3.69 -> 3.89
[1:05:27] Using cautious pH adjust
[1:05:27] Dispensed 0.000306 mL of Base (0.5 M KOH)
[1:05:32] Stepping pH = 3.76
[1:05:32] Dispensed 0.000306 mL of Base (0.5 M KOH)
[1:05:37] Stepping pH = 3.92
[1:05:52] Stirrer speed set to 0
[1:06:03] Datapoint id 58 collected
[1:06:03] Charge balance equation is out by -0.4%
[1:06:03] Stirrer speed set to 60
[1:06:08] pH 3.93 -> 4.13
[1:06:08] Using charge balance adjust
[1:06:08] Dispensed 0.000376 mL of Base (0.5 M KOH)
[1:06:28] Stirrer speed set to 0
[1:06:39] Datapoint id 59 collected
[1:06:39] Charge balance equation is out by 41.3%
[1:06:39] Stirrer speed set to 60
[1:06:44] pH 4.23 -> 4.43
[1:06:44] Using cautious pH adjust
[1:06:44] Dispensed 0.000094 mL of Base (0.5 M KOH)
[1:06:49] Stepping pH = 4.24
[1:06:49] Dispensed 0.000259 mL of Base (0.5 M KOH)
[1:06:54] Stepping pH = 4.79
[1:07:09] Stirrer speed set to 0
[1:07:54] Datapoint id 60 collected
[1:07:54] Charge balance equation is out by -83.8%
[1:07:54] Stirrer speed set to 60
[1:08:00] pH 4.78 -> 4.98
[1:08:00] Using cautious pH adjust
[1:08:00] Dispensed 0.000024 mL of Base (0.5 M KOH)
[1:08:05] Stepping pH = 4.77
[1:08:05] Dispensed 0.000141 mL of Base (0.5 M KOH)
[1:08:10] Stepping pH = 5.22
[1:08:25] Stirrer speed set to 0
[1:08:41] Datapoint id 61 collected
[1:08:41] Charge balance equation is out by -208.6%
[1:08:41] Stirrer speed set to 60
[1:08:46] pH 5.30 -> 5.50
[1:08:46] Using cautious pH adjust
[1:08:46] Dispensed 0.000024 mL of Base (0.5 M KOH)
[1:08:51] Stepping pH = 5.31
[1:08:51] Dispensed 0.000047 mL of Base (0.5 M KOH)
[1:08:56] Stepping pH = 5.45
[1:08:56] Dispensed 0.000024 mL of Base (0.5 M KOH)
[1:09:01] Stepping pH = 5.56
[1:09:16] Stirrer speed set to 0
[1:10:05] Datapoint id 62 collected
[1:10:05] Charge balance equation is out by -166.1%
[1:10:05] Stirrer speed set to 60
[1:10:10] pH 5.72 -> 5.92
[1:10:10] Using cautious pH adjust
[1:10:10] Dispensed 0.000024 mL of Base (0.5 M KOH)
[1:10:15] Stepping pH = 5.75
[1:10:15] Dispensed 0.000047 mL of Base (0.5 M KOH)
[1:10:20] Stepping pH = 6.09
[1:10:35] Stirrer speed set to 0

#### Experiment Log (continued)

[1:11:35] Datapoint id 63 collected
[1:11:35] Charge balance equation is out by -74.3%
[1:11:35] Stirrer speed set to 60
[1:11:41] pH 6.22 -> 6.42
[1:11:41] Using cautious pH adjust
[1:11:41] Dispensed 0.000024 mL of Base (0.5 M KOH)
[1:11:46] Stepping pH = 6.24
[1:11:46] Dispensed 0.000047 mL of Base (0.5 M KOH)
[1:11:51] Stepping pH = 6.31
[1:11:51] Dispensed 0.000047 mL of Base (0.5 M KOH)
[1:11:56] Stepping pH = 6.87
[1:12:11] Stirrer speed set to 0
[1:13:11] Datapoint id 64 collected
[1:13:11] Charge balance equation is out by -176.8%
[1:13:11] Stirrer speed set to 60
[1:13:16] pH 7.12 -> 7.32
[1:13:16] Using cautious pH adjust
[1:13:16] Dispensed 0.000024 mL of Base (0.5 M KOH)
[1:13:21] Stepping pH = 7.20
[1:13:21] Dispensed 0.000024 mL of Base (0.5 M KOH)
[1:13:27] Stepping pH = 7.33
[1:13:42] Stirrer speed set to 0
[1:14:42] Datapoint id 65 collected
[1:14:42] Charge balance equation is out by -102.3%
[1:14:42] Stirrer speed set to 60
[1:14:47] pH 7.16 -> 7.36
[1:14:47] Using cautious pH adjust
[1:14:47] Dispensed 0.000024 mL of Base (0.5 M KOH)
[1:14:52] Stepping pH = 7.24
[1:14:52] Dispensed 0.000024 mL of Base (0.5 M KOH)
[1:14:57] Stepping pH = 7.41
[1:15:12] Stirrer speed set to 0
[1:16:12] Datapoint id 66 collected
[1:16:12] Charge balance equation is out by -115.3%
[1:16:12] Stirrer speed set to 60
[1:16:17] pH 7.58 -> 7.78
[1:16:17] Using cautious pH adjust
[1:16:17] Dispensed 0.000024 mL of Base (0.5 M KOH)
[1:16:22] Stepping pH = 7.73
[1:16:22] Dispensed 0.000024 mL of Base (0.5 M KOH)
[1:16:28] Stepping pH = 7.93
[1:16:43] Stirrer speed set to 0
[1:17:43] Datapoint id 67 collected
[1:17:43] Charge balance equation is out by -302.3%
[1:17:43] Stirrer speed set to 60
[1:17:48] pH 7.75 -> 7.95
[1:17:48] Using cautious pH adjust
[1:17:48] Dispensed 0.000024 mL of Base (0.5 M KOH)
[1:17:53] Stepping pH = 7.87
[1:17:53] Dispensed 0.000024 mL of Base (0.5 M KOH)
[1:17:58] Stepping pH = 8.15
[1:18:13] Stirrer speed set to 0
[1:19:13] Datapoint id 68 collected
[1:19:13] Charge balance equation is out by -373.8%
[1:19:13] Stirrer speed set to 60
[1:19:18] pH 8.24 -> 8.44
[1:19:18] Using cautious pH adjust
[1:19:18] Dispensed 0.000024 mL of Base (0.5 M KOH)
[1:19:23] Stepping pH = 8.26

| | | | |
|---|---|---|---|
| Sample name: | **M02_octanol** | Experiment start time: | **3/3/2018 1:46:01 PM** |
| Assay name: | **pH-metric high logP** | Analyst: | **Dorothy Levorse** |
| Assay ID: | **18C-03011** | Instrument ID: | **T312060** |
| Filename: | **C:\Sirius_T3\Mehtap\20180302_exp29_logP_T3-2\18C-03011_M02_octanol_pH-metric high logP.t3r** | | |

#### Experiment Log (continued)

[1:19:23] Dispensed 0.000024 mL of Base (0.5 M KOH)
[1:19:28] Stepping pH = 8.42
[1:19:29] Dispensed 0.000024 mL of Base (0.5 M KOH)
[1:19:34] Stepping pH = 8.58
[1:19:49] Stirrer speed set to 0
[1:20:49] Datapoint id 69 collected
[1:20:49] Charge balance equation is out by -452.3%
[1:20:49] Stirrer speed set to 60
[1:20:54] pH 8.61 -> 8.81
[1:20:54] Using cautious pH adjust
[1:20:54] Dispensed 0.000024 mL of Base (0.5 M KOH)
[1:20:59] Stepping pH = 8.61
[1:20:59] Dispensed 0.000024 mL of Base (0.5 M KOH)
[1:21:04] Stepping pH = 8.60
[1:21:04] Dispensed 0.000165 mL of Base (0.5 M KOH)
[1:21:09] Stepping pH = 9.23
[1:21:24] Stirrer speed set to 0
[1:21:39] Datapoint id 70 collected
[1:21:39] Charge balance equation is out by -904.7%
[1:21:39] Argon flow rate set to 0
[1:21:43] Titrator arm moved over Titration position
