## Supplementary material for "Octanol-water partition coefficient measurements for the SAMPL6 Blind Prediction Challenge": SM02_18C-03012_M02_octanol_pH-metric high logP_report.pdf

| | | | |
|---|---|---|---|
| Sample name: | **M02_octanol** | Experiment start time: | **3/3/2018 3:08:32 PM** |
| Assay name: | **pH-metric high logP** | Analyst: | **Dorothy Levorse** |
| Assay ID: | **18C-03012** | Instrument ID: | **T312060** |
| Filename: | **C:\Sirius_T3\Mehtap\20180302_exp29_logP_T3-2\18C-03012_M02_octanol_pH-metric high logP.t3r** | | |

#### pH-metric Result

| | | |
|---|---|---|
| logP (XH +) | 0.59 | ±0.04 (n=50) |
| logP (neutral X) | 4.03 | ±0.01 (n=50) |

#### 18C-03012 Points 1 to 26

| | |
|---|---|
| M02_octanol concentration factor | 0.955 |
| Carbonate | 0.2235 mM |
| Acidity error | -1.42826 mM |

#### 18C-03012 Points 27 to 50

| | |
|---|---|
| M02_octanol concentration factor | 0.892 |
| Carbonate | 0.1033 mM |
| Acidity error | -1.59633 mM |

#### 18C-03012 Points 51 to 75

| | |
|---|---|
| M02_octanol concentration factor | 1.218 |
| Carbonate | 0.1516 mM |
| Acidity error | -1.31894 mM |

#### Warnings and errors

| | |
|---|---|
| Errors | None |
| Warnings | None |

#### Sample logD and percent species

| pH | M02_octanol logD | M02_octanol M02_octanolH | M02_octanol M02_octanol | M02_octanol M02_octanolH* | M02_octanol M02_octanol* | Comment |
|---|---|---|---|---|---|---|
| 1.000 | 0.69 | 17.03 % | 0.00 % | 65.96 % | 17.01 % | |
| 1.200 | 0.74 | 15.49 % | 0.00 % | 59.99 % | 24.51 % | Stomach pH |
| 2.000 | 1.14 | 6.73 % | 0.01 % | 26.06 % | 67.20 % | |
| 3.000 | 2.01 | 0.95 % | 0.01 % | 3.70 % | 95.34 % | |
| 4.000 | 2.96 | 0.10 % | 0.01 % | 0.39 % | 99.51 % | |
| 5.000 | 3.71 | 0.01 % | 0.01 % | 0.04 % | 99.94 % | |
| 6.000 | 3.99 | 0.00 % | 0.01 % | 0.00 % | 99.99 % | |
| 6.500 | 4.01 | 0.00 % | 0.01 % | 0.00 % | 99.99 % | |
| 7.000 | 4.02 | 0.00 % | 0.01 % | 0.00 % | 99.99 % | |
| 7.400 | 4.03 | 0.00 % | 0.01 % | 0.00 % | 99.99 % | Blood pH |
| 8.000 | 4.03 | 0.00 % | 0.01 % | 0.00 % | 99.99 % | |
| 9.000 | 4.03 | 0.00 % | 0.01 % | 0.00 % | 99.99 % | |
| 10.000 | 4.03 | 0.00 % | 0.01 % | 0.00 % | 99.99 % | |
| 11.000 | 4.03 | 0.00 % | 0.01 % | 0.00 % | 99.99 % | |
| 12.000 | 4.03 | 0.00 % | 0.01 % | 0.00 % | 99.99 % | |

| | | | |
|---|---|---|---|
| Sample name: | **M02_octanol** | Experiment start time: | **3/3/2018 3:08:32 PM** |
| Assay name: | **pH-metric high logP** | Analyst: | **Dorothy Levorse** |
| Assay ID: | **18C-03012** | Instrument ID: | **T312060** |
| Filename: | **C:\Sirius_T3\Mehtap\20180302_exp29_logP_T3-2\18C-03012_M02_octanol_pH-metric high logP.t3r** | | |

#### Graphs

Charge balance residuals

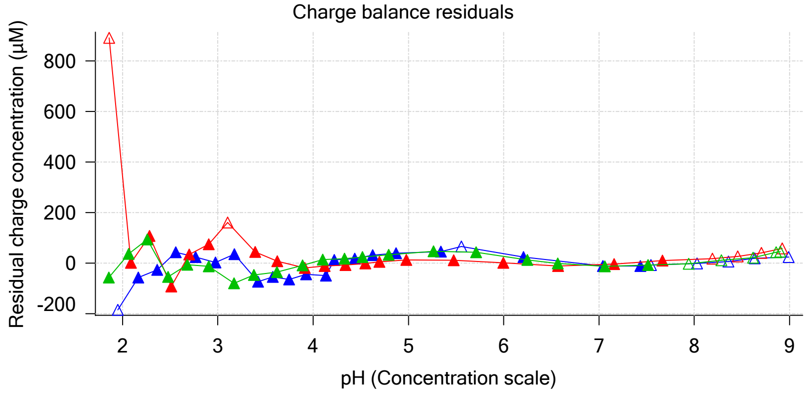

Ionic strength

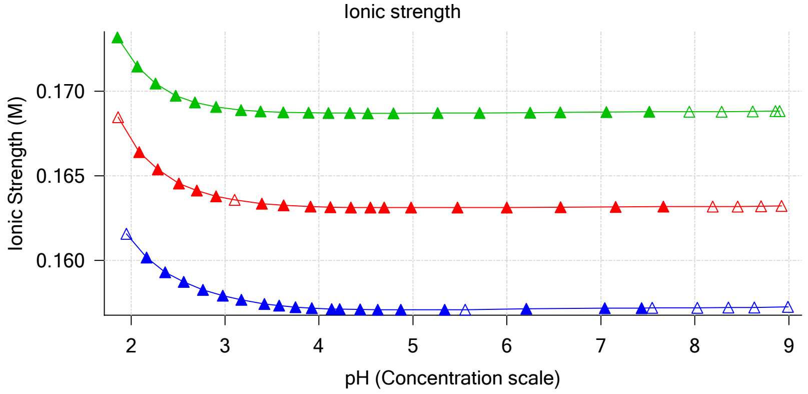

Buffer index profile

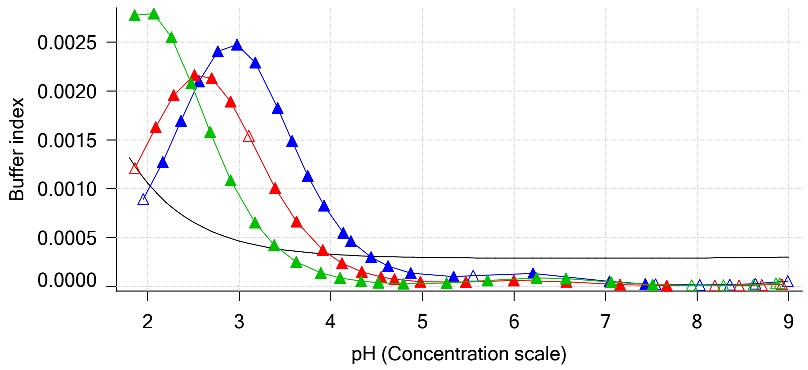

Sample M02_octanol Ionisation Graph

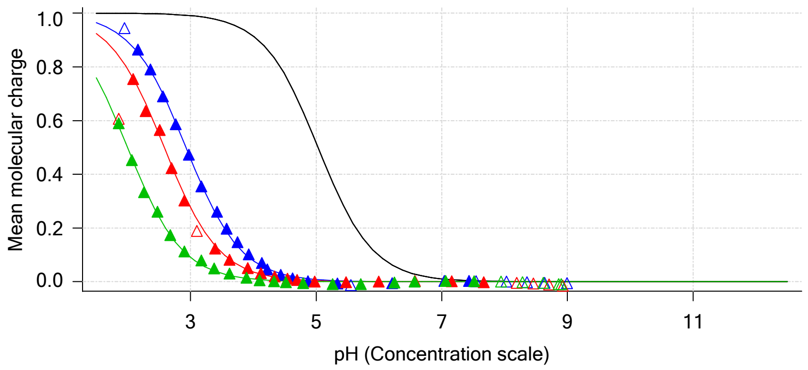

Distribution of species for sample M02_octanol

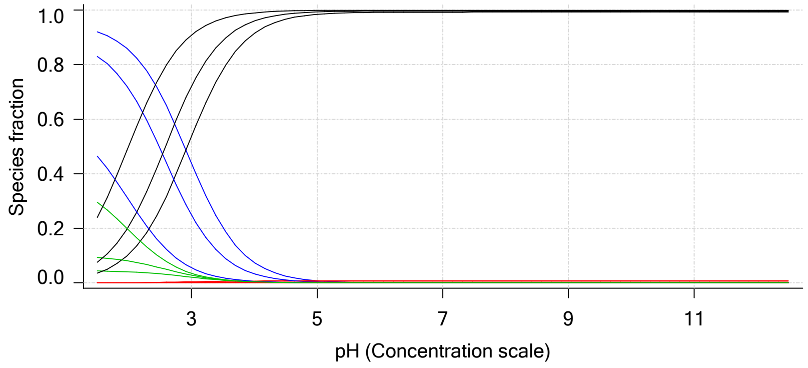

Lipophilicity Profile

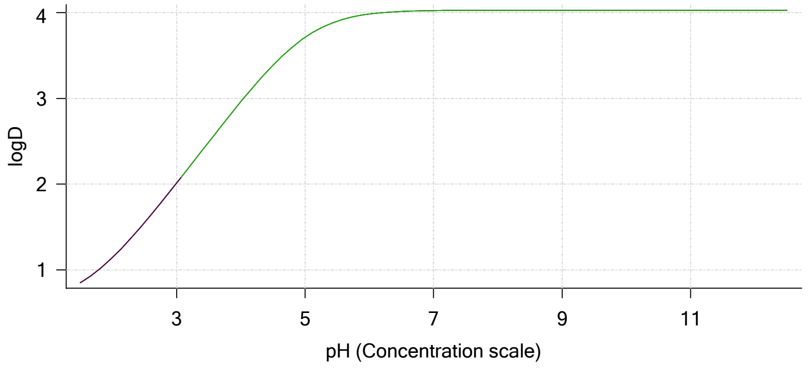

Log R Profile

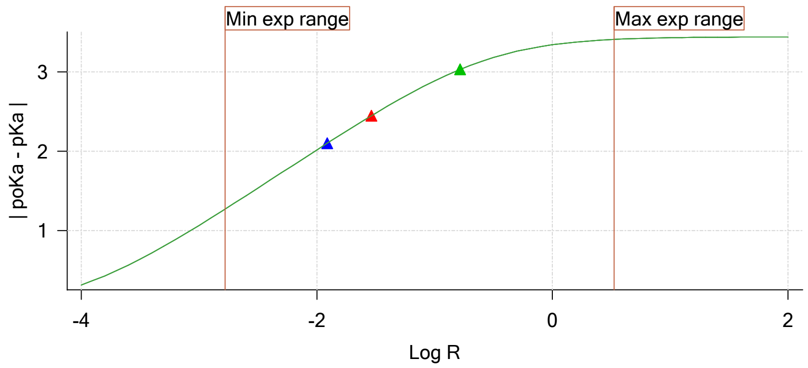

Sample Carbonate Ionisation Graph

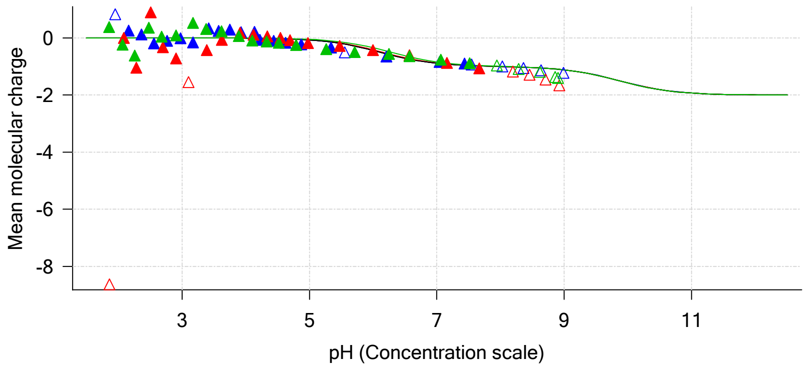

| | | | |
|---|---|---|---|
| Sample name: | **M02_octanol** | Experiment start time: | **3/3/2018 3:08:32 PM** |
| Assay name: | **pH-metric high logP** | Analyst: | **Dorothy Levorse** |
| Assay ID: | **18C-03012** | Instrument ID: | **T312060** |
| Filename: | **C:\Sirius_T3\Mehtap\20180302_exp29_logP_T3-2\18C-03012_M02_octanol_pH-metric high logP.t3r** | | |

#### Graphs (continued)

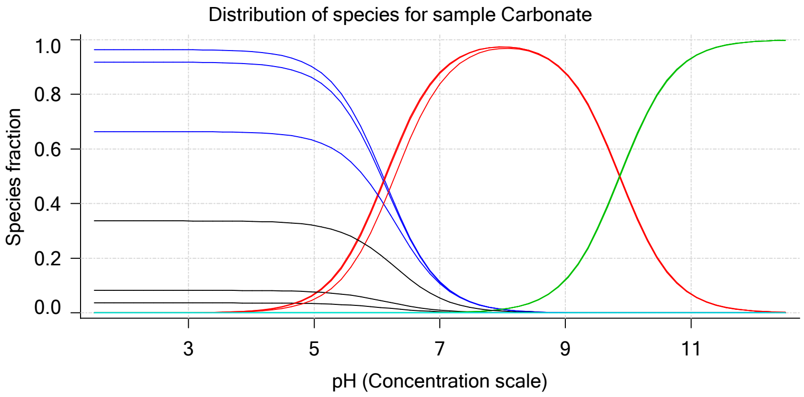

| | | | |
|---|---|---|---|
| Sample name: | **M02_octanol** | Experiment start time: | **3/3/2018 3:08:32 PM** |
| Assay name: | **pH-metric high logP** | Analyst: | **Dorothy Levorse** |
| Assay ID: | **18C-03012** | Instrument ID: | **T312060** |
| Filename: | **C:\Sirius_T3\Mehtap\20180302_exp29_logP_T3-2\18C-03012_M02_octanol_pH-metric high logP.t3r** | | |

### pH-metric high logP Titration 1 of 3 18C-03012 Points 1 to 26

#### Overall results

| | |
|---|---|
| RMSD | 0.169 |
| Average ionic strength | 0.158 M |
| Average temperature | 24.9°C |
| Partition ratio | 0.0123 : 1 |
| Analyte concentration range | 4430.5 μM to 4567.1 μM |
| Total points considered | 19 of 26 |

#### Warnings and errors

| | |
|---|---|
| Errors | None |
| Warnings | Excessive acidity error present |

#### Four-Plus parameters

| | | | |
|---|---|---|---|
| Alpha | 0.111 | 3/3/2018 3:08:32 PM | C:\Sirius_T3\HCl18C02.t3r |
| S | 0.9988 | 3/3/2018 3:08:32 PM | C:\Sirius_T3\HCl18C02.t3r |
| jH | 1.0 | 3/3/2018 3:08:32 PM | C:\Sirius_T3\HCl18C02.t3r |
| jOH | -0.8 | 3/3/2018 3:08:32 PM | C:\Sirius_T3\HCl18C02.t3r |

#### Titrants

| | | | |
|---|---|---|---|
| 0.50 M HCl | 0.999058 | 3/3/2018 3:08:32 PM | C:\Sirius_T3\HCl18C02.t3r |
| 0.50 M KOH | 0.999845 | 3/3/2018 3:08:32 PM | C:\Sirius_T3\KOH18B27.t3r |

#### Sample

| | |
|---|---|
| M02_octanol concentration factor | 0.955 |
| Base pKa 1 | 5.03 |
| logP (XH +) | 1.12 |
| logP (neutral X) | 4.07 |

#### Sample graphs

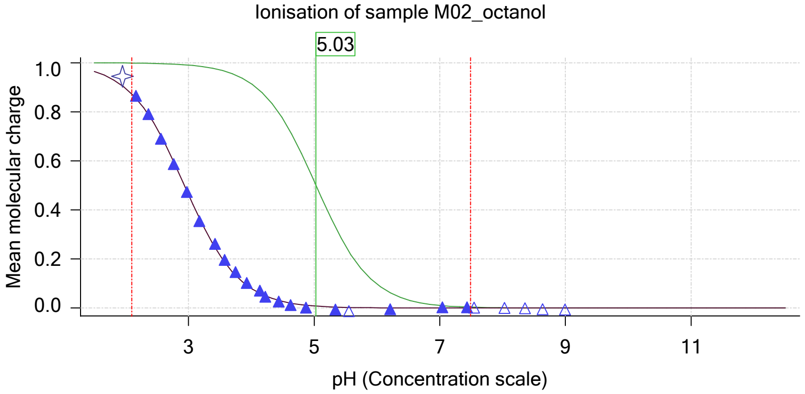

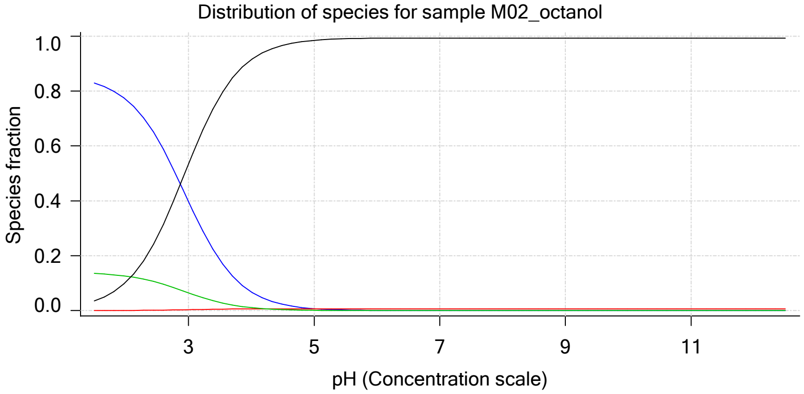

| | | | |
|---|---|---|---|
| Sample name: | **M02_octanol** | Experiment start time: | **3/3/2018 3:08:32 PM** |
| Assay name: | **pH-metric high logP** | Analyst: | **Dorothy Levorse** |
| Assay ID: | **18C-03012** | Instrument ID: | **T312060** |
| Filename: | **C:\Sirius_T3\Mehtap\20180302_exp29_logP_T3-2\18C-03012_M02_octanol_pH-metric high logP.t3r** | | |

#### Sample graphs (continued)

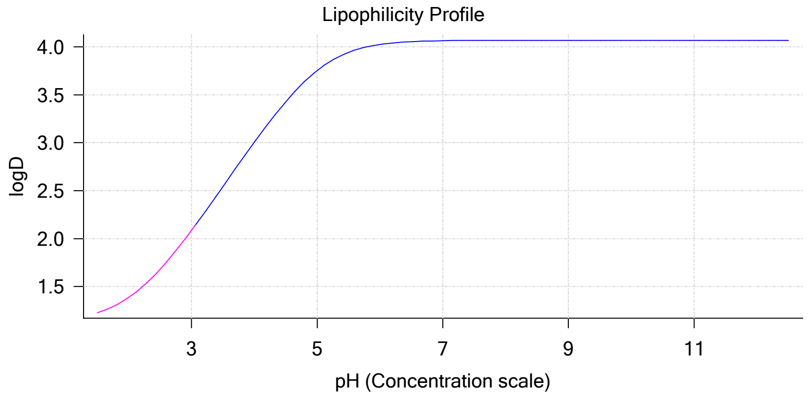

#### Sample logD and percent species

| pH | M02_octanol logD | M02_octanol M02_octanolH | M02_octanol M02_octanol | M02_octanol M02_octanolH* | M02_octanol M02_octanol* | Comment |
|---|---|---|---|---|---|---|
| 1.000 | 1.16 | 84.96 % | 0.01 % | 13.89 % | 1.14 % | |
| 1.200 | 1.18 | 84.39 % | 0.01 % | 13.80 % | 1.79 % | Stomach pH |
| 2.000 | 1.38 | 77.00 % | 0.07 % | 12.59 % | 10.33 % | |
| 3.000 | 2.08 | 39.77 % | 0.37 % | 6.50 % | 53.36 % | |
| 4.000 | 3.01 | 6.81 % | 0.64 % | 1.11 % | 91.44 % | |
| 5.000 | 3.75 | 0.73 % | 0.68 % | 0.12 % | 98.46 % | |
| 6.000 | 4.02 | 0.07 % | 0.69 % | 0.01 % | 99.22 % | |
| 6.500 | 4.05 | 0.02 % | 0.69 % | 0.00 % | 99.28 % | |
| 7.000 | 4.06 | 0.01 % | 0.69 % | 0.00 % | 99.30 % | |
| 7.400 | 4.07 | 0.00 % | 0.69 % | 0.00 % | 99.31 % | Blood pH |
| 8.000 | 4.07 | 0.00 % | 0.69 % | 0.00 % | 99.31 % | |
| 9.000 | 4.07 | 0.00 % | 0.69 % | 0.00 % | 99.31 % | |
| 10.000 | 4.07 | 0.00 % | 0.69 % | 0.00 % | 99.31 % | |
| 11.000 | 4.07 | 0.00 % | 0.69 % | 0.00 % | 99.31 % | |
| 12.000 | 4.07 | 0.00 % | 0.69 % | 0.00 % | 99.31 % | |

#### Carbonate and acidity

| | |
|---|---|
| Carbonate | 0.223 mM |
| Acidity error | -1.428 mM |

#### Other graphs

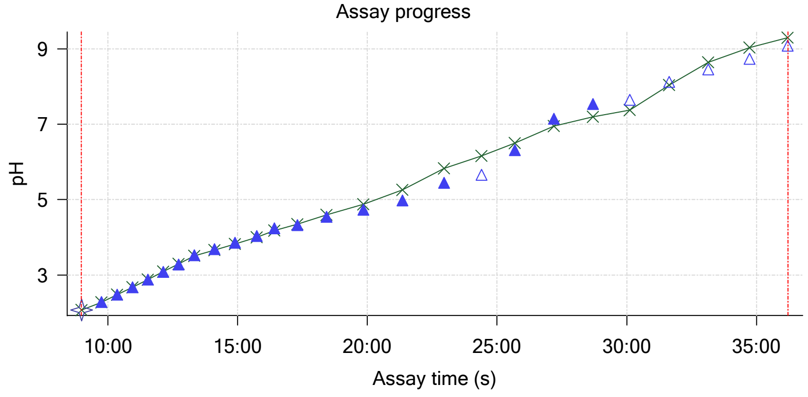

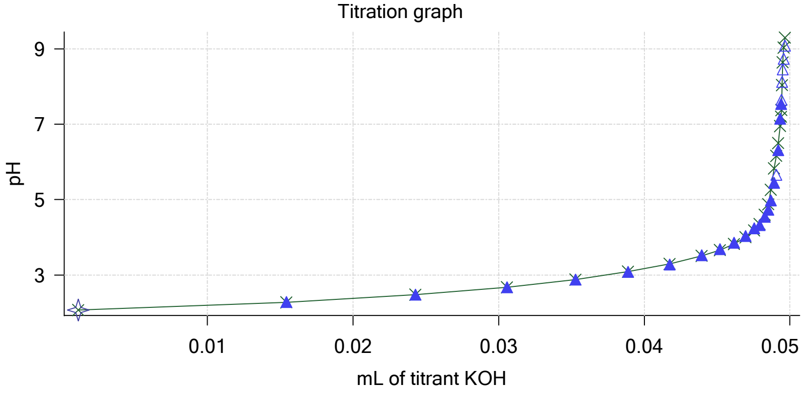

| | | | |
|---|---|---|---|
| Sample name: | **M02_octanol** | Experiment start time: | **3/3/2018 3:08:32 PM** |
| Assay name: | **pH-metric high logP** | Analyst: | **Dorothy Levorse** |
| Assay ID: | **18C-03012** | Instrument ID: | **T312060** |
| Filename: | **C:\Sirius_T3\Mehtap\20180302_exp29_logP_T3-2\18C-03012_M02_octanol_pH-metric high logP.t3r** | | |

#### Other graphs (continued)

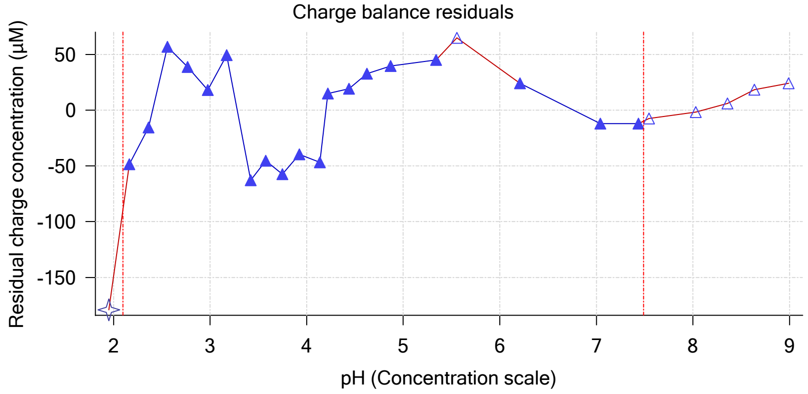

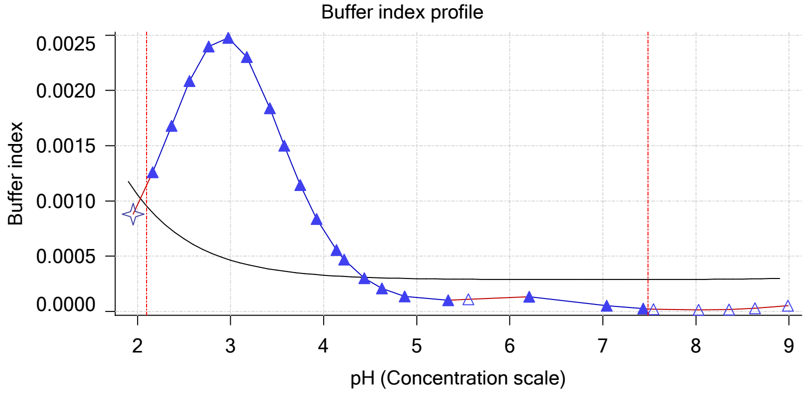

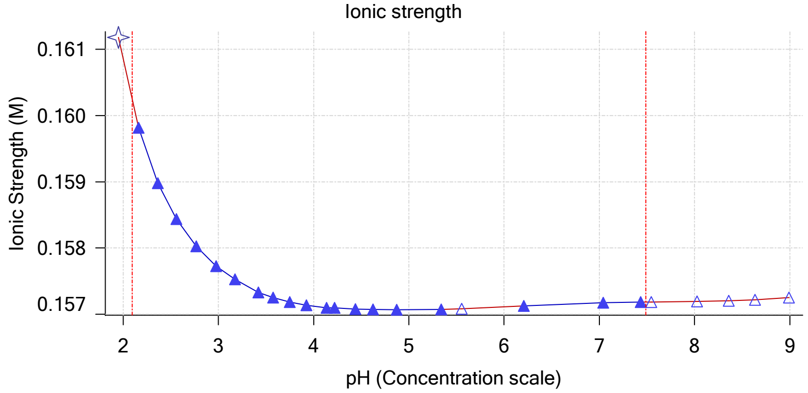

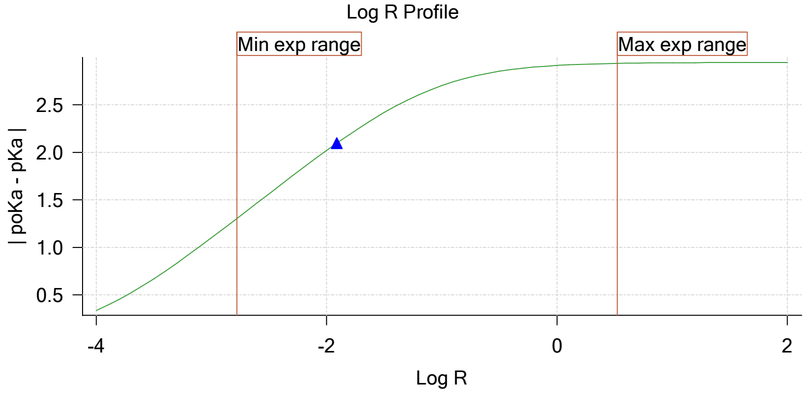

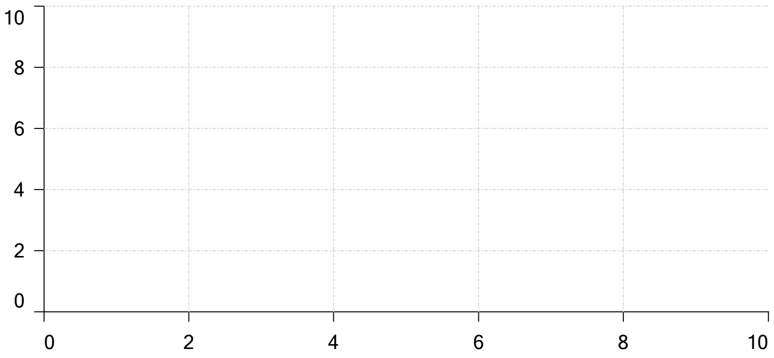

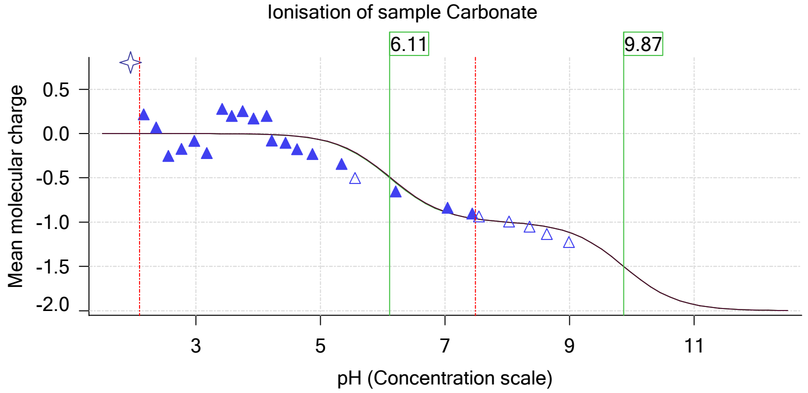

| | | | |
|---|---|---|---|
| Sample name: | **M02_octanol** | Experiment start time: | **3/3/2018 3:08:32 PM** |
| Assay name: | **pH-metric high logP** | Analyst: | **Dorothy Levorse** |
| Assay ID: | **18C-03012** | Instrument ID: | **T312060** |
| Filename: | **C:\Sirius_T3\Mehtap\20180302_exp29_logP_T3-2\18C-03012_M02_octanol_pH-metric high logP.t3r** | | |

### pH-metric high logP Titration 2 of 3 18C-03012 Points 27 to 50

#### Overall results

| | |
|---|---|
| RMSD | 0.235 |
| Average ionic strength | 0.163 M |
| Average temperature | 25.0°C |
| Partition ratio | 0.0290 : 1 |
| Analyte concentration range | 4080.4 µM to 4209.4 µM |
| Total points considered | 18 of 24 |

#### Warnings and errors

| | |
|---|---|
| Errors | None |
| Warnings | Excessive acidity error present |

#### Four-Plus parameters

| | | | |
|---|---|---|---|
| Alpha | 0.111 | 3/3/2018 3:08:32 PM | C:\Sirius_T3\HCl18C02.t3r |
| S | 0.9988 | 3/3/2018 3:08:32 PM | C:\Sirius_T3\HCl18C02.t3r |
| jH | 1.0 | 3/3/2018 3:08:32 PM | C:\Sirius_T3\HCl18C02.t3r |
| jOH | -0.8 | 3/3/2018 3:08:32 PM | C:\Sirius_T3\HCl18C02.t3r |

#### Titrants

| | | | |
|---|---|---|---|
| 0.50 M HCl | 0.999058 | 3/3/2018 3:08:32 PM | C:\Sirius_T3\HCl18C02.t3r |
| 0.50 M KOH | 0.999845 | 3/3/2018 3:08:32 PM | C:\Sirius_T3\KOH18B27.t3r |

#### Sample

| | |
|---|---|
| M02_octanol concentration factor | 0.892 |
| Base pKa 1 | 5.03 |
| logP (XH +) | 1.47 |
| logP (neutral X) | 4.27 |

#### Sample graphs

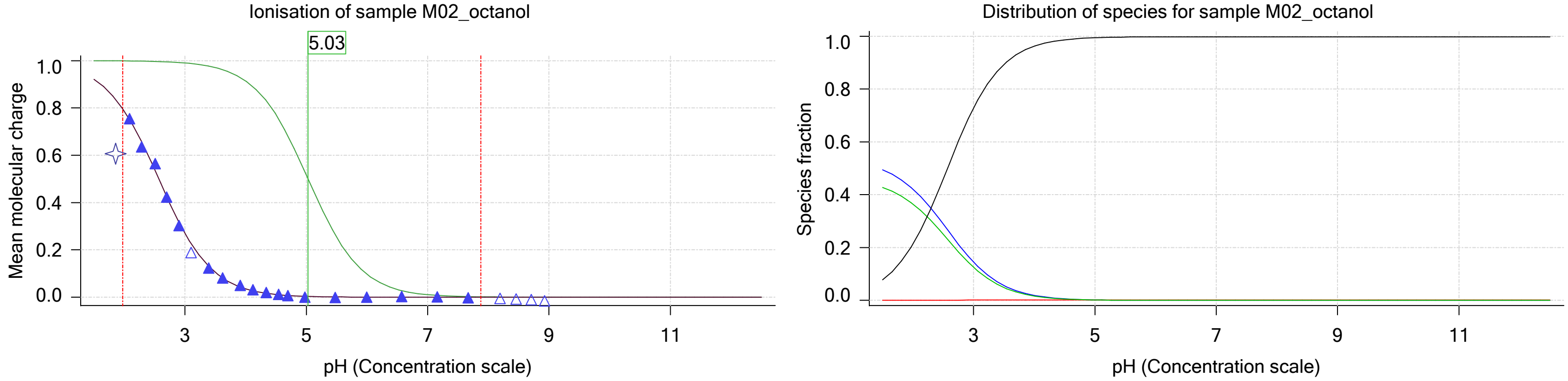

| | | | |
|---|---|---|---|
| Sample name: | **M02_octanol** | Experiment start time: | **3/3/2018 3:08:32 PM** |
| Assay name: | **pH-metric high logP** | Analyst: | **Dorothy Levorse** |
| Assay ID: | **18C-03012** | Instrument ID: | **T312060** |
| Filename: | **C:\Sirius_T3\Mehtap\20180302_exp29_logP_T3-2\18C-03012_M02_octanol_pH-metric high logP.t3r** | | |

#### Sample graphs (continued)

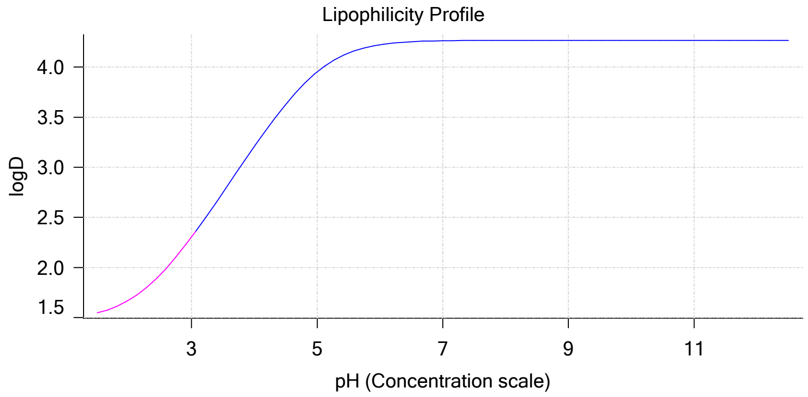

#### Sample logD and percent species

| pH | M02_octanol logD | M02_octanol M02_octanolH | M02_octanol M02_octanol | M02_octanol M02_octanolH* | M02_octanol M02_octanol* | Comment |
|---|---|---|---|---|---|---|
| 1.000 | 1.50 | 52.20 % | 0.00 % | 45.17 % | 2.62 % | |
| 1.200 | 1.51 | 51.41 % | 0.01 % | 44.49 % | 4.09 % | Stomach pH |
| 2.000 | 1.67 | 42.22 % | 0.04 % | 36.54 % | 21.21 % | |
| 3.000 | 2.30 | 14.50 % | 0.14 % | 12.55 % | 72.82 % | |
| 4.000 | 3.21 | 1.92 % | 0.18 % | 1.66 % | 96.25 % | |
| 5.000 | 3.95 | 0.20 % | 0.18 % | 0.17 % | 99.45 % | |
| 6.000 | 4.22 | 0.02 % | 0.19 % | 0.02 % | 99.78 % | |
| 6.500 | 4.25 | 0.01 % | 0.19 % | 0.01 % | 99.80 % | |
| 7.000 | 4.26 | 0.00 % | 0.19 % | 0.00 % | 99.81 % | |
| 7.400 | 4.27 | 0.00 % | 0.19 % | 0.00 % | 99.81 % | Blood pH |
| 8.000 | 4.27 | 0.00 % | 0.19 % | 0.00 % | 99.81 % | |
| 9.000 | 4.27 | 0.00 % | 0.19 % | 0.00 % | 99.81 % | |
| 10.000 | 4.27 | 0.00 % | 0.19 % | 0.00 % | 99.81 % | |
| 11.000 | 4.27 | 0.00 % | 0.19 % | 0.00 % | 99.81 % | |
| 12.000 | 4.27 | 0.00 % | 0.19 % | 0.00 % | 99.81 % | |

#### Carbonate and acidity

| | |
|---|---|
| Carbonate | 0.103 mM |
| Acidity error | -1.596 mM |

#### Other graphs

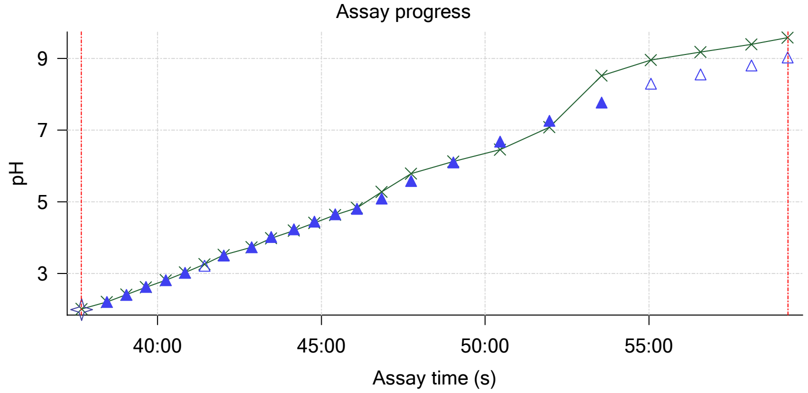

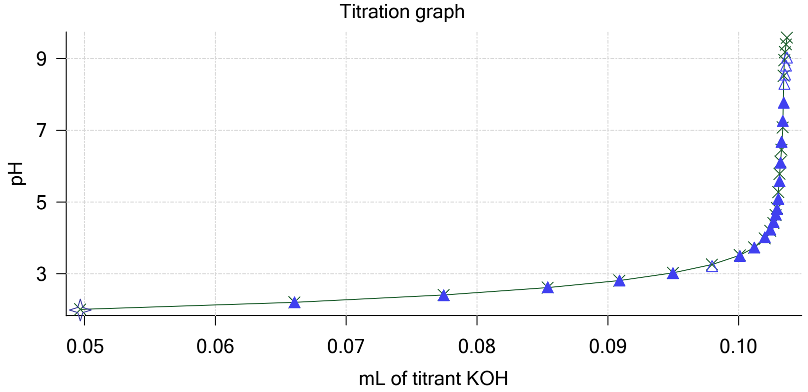

| | | | |
|---|---|---|---|
| Sample name: | **M02_octanol** | Experiment start time: | **3/3/2018 3:08:32 PM** |
| Assay name: | **pH-metric high logP** | Analyst: | **Dorothy Levorse** |
| Assay ID: | **18C-03012** | Instrument ID: | **T312060** |
| Filename: | **C:\Sirius_T3\Mehtap\20180302_exp29_logP_T3-2\18C-03012_M02_octanol_pH-metric high logP.t3r** | | |

#### Other graphs (continued)

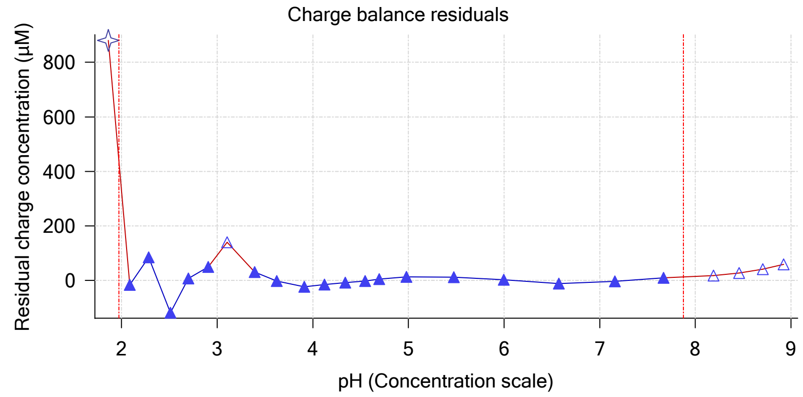

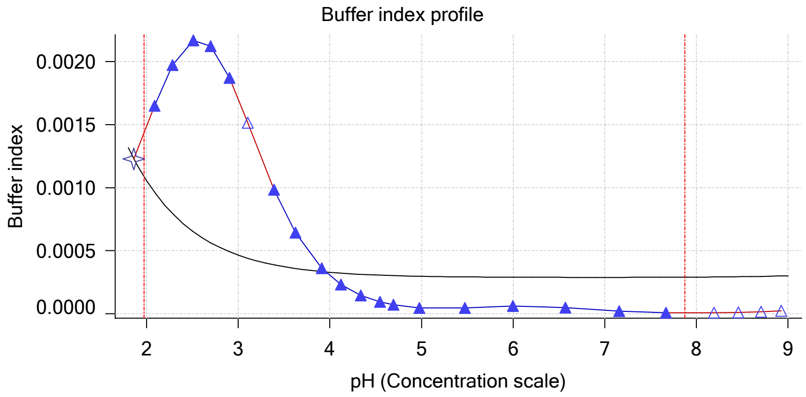

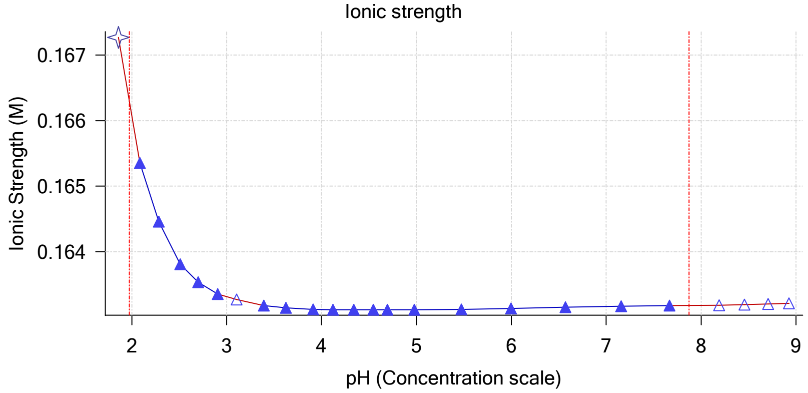

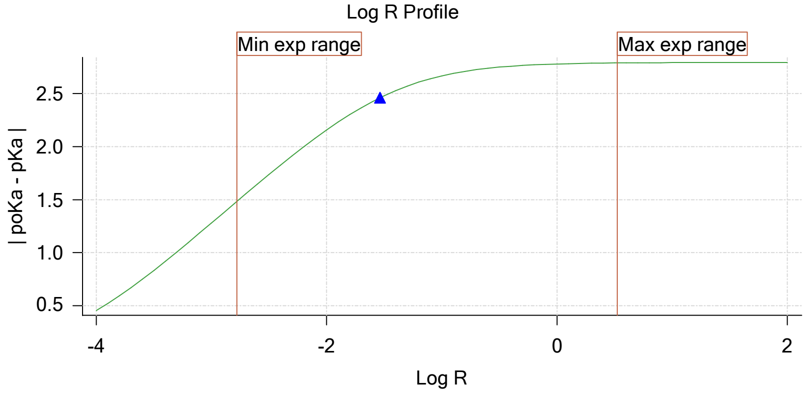

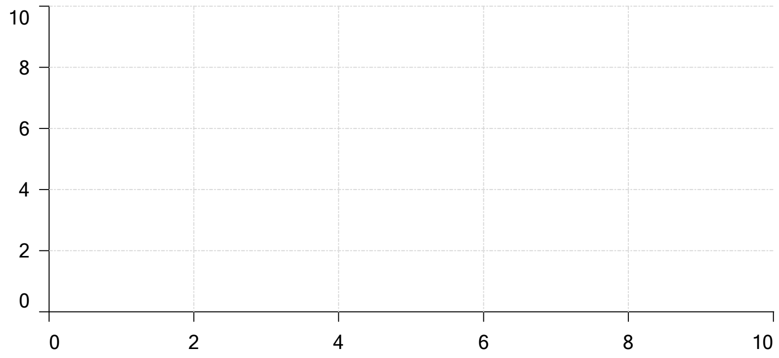

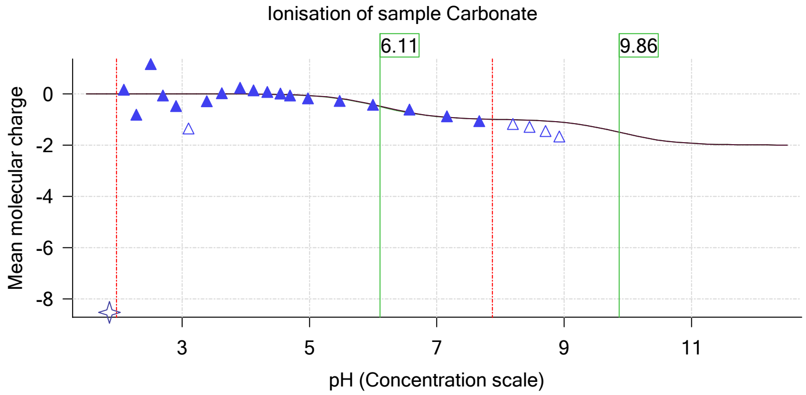

| | | | |
|---|---|---|---|
| Sample name: | **M02_octanol** | Experiment start time: | **3/3/2018 3:08:32 PM** |
| Assay name: | **pH-metric high logP** | Analyst: | **Dorothy Levorse** |
| Assay ID: | **18C-03012** | Instrument ID: | **T312060** |
| Filename: | **C:\Sirius_T3\Mehtap\20180302_exp29_logP_T3-2\18C-03012_M02_octanol_pH-metric high logP.t3r** | | |

# pH-metric high logP   Titration 3 of 3   18C-03012 Points 51 to 75

#### Overall results

| | |
|---|---|
| RMSD | 0.321 |
| Average ionic strength | 0.169 M |
| Average temperature | 25.0°C |
| Partition ratio | 0.1643 : 1 |
| Analyte concentration range | 3372.3 µM to 3470.6 µM |
| Total points considered | 20 of 25 |

#### Warnings and errors

| | |
|---|---|
| Errors | None |
| Warnings | Excessive acidity error present |

#### Four-Plus parameters

| | | | |
|---|---|---|---|
| Alpha | 0.111 | 3/3/2018 3:08:32 PM | C:\Sirius_T3\HCl18C02.t3r |
| S | 0.9988 | 3/3/2018 3:08:32 PM | C:\Sirius_T3\HCl18C02.t3r |
| jH | 1.0 | 3/3/2018 3:08:32 PM | C:\Sirius_T3\HCl18C02.t3r |
| jOH | -0.8 | 3/3/2018 3:08:32 PM | C:\Sirius_T3\HCl18C02.t3r |

#### Titrants

| | | | |
|---|---|---|---|
| 0.50 M HCl | 0.999058 | 3/3/2018 3:08:32 PM | C:\Sirius_T3\HCl18C02.t3r |
| 0.50 M KOH | 0.999845 | 3/3/2018 3:08:32 PM | C:\Sirius_T3\KOH18B27.t3r |

#### Sample

| | |
|---|---|
| M02_octanol concentration factor | 1.218 |
| Base pKa 1 | 5.03 |
| logP (XH +) | 1.33 |
| logP (neutral X) | 4.47 |

#### Sample graphs

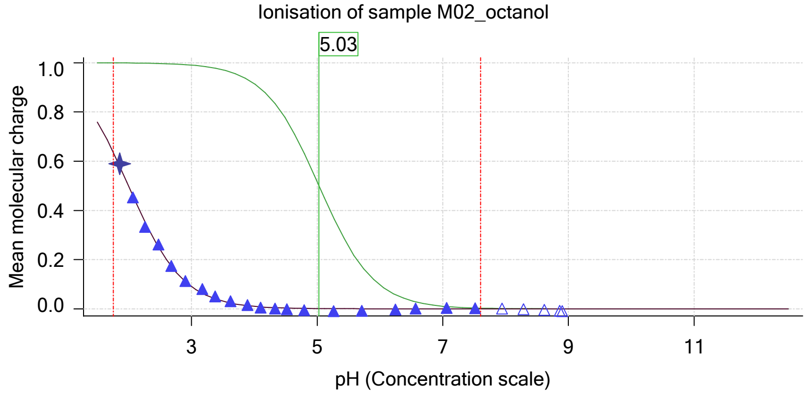

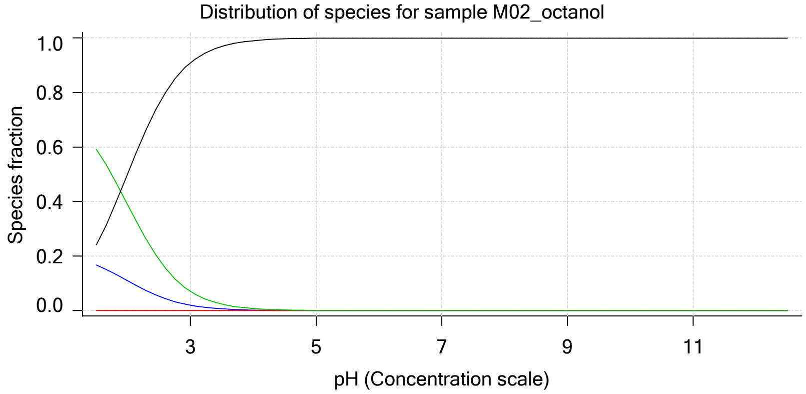

Sample name: **M02_octanol**
Assay name: **pH-metric high logP**
Assay ID: **18C-03012**
Filename: **C:\Sirius_T3\Mehtap\20180302_exp29_logP_T3-2\18C-03012_M02_octanol_pH-metric high logP.t3r**

Experiment start time: **3/3/2018 3:08:32 PM**
Analyst: **Dorothy Levorse**
Instrument ID: **T312060**

#### Sample graphs (continued)

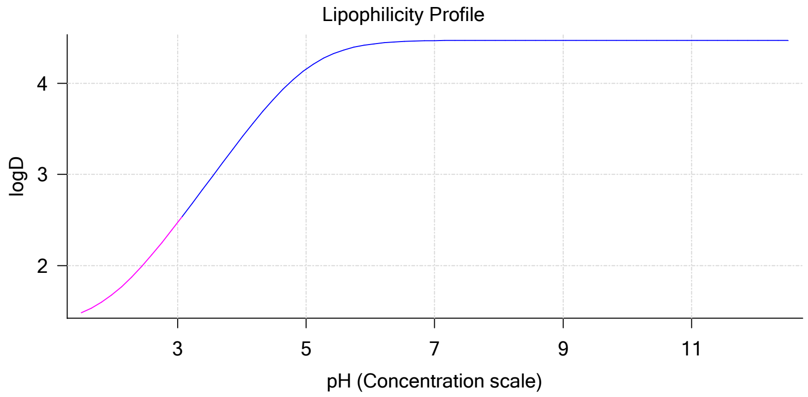

#### Sample logD and percent species

| pH | M02_octanol logD | M02_octanol M02_octanolH | M02_octanol M02_octanol | M02_octanol M02_octanolH* | M02_octanol M02_octanol* | Comment |
|---|---|---|---|---|---|---|
| 1.000 | 1.39 | 20.05 % | 0.00 % | 70.82 % | 9.14 % | |
| 1.200 | 1.41 | 19.03 % | 0.00 % | 67.22 % | 13.74 % | Stomach pH |
| 2.000 | 1.69 | 11.00 % | 0.01 % | 38.86 % | 50.13 % | |
| 3.000 | 2.47 | 2.00 % | 0.02 % | 7.05 % | 90.94 % | |
| 4.000 | 3.41 | 0.22 % | 0.02 % | 0.77 % | 99.00 % | |
| 5.000 | 4.16 | 0.02 % | 0.02 % | 0.08 % | 99.88 % | |
| 6.000 | 4.43 | 0.00 % | 0.02 % | 0.01 % | 99.97 % | |
| 6.500 | 4.46 | 0.00 % | 0.02 % | 0.00 % | 99.98 % | |
| 7.000 | 4.47 | 0.00 % | 0.02 % | 0.00 % | 99.98 % | |
| 7.400 | 4.47 | 0.00 % | 0.02 % | 0.00 % | 99.98 % | Blood pH |
| 8.000 | 4.47 | 0.00 % | 0.02 % | 0.00 % | 99.98 % | |
| 9.000 | 4.47 | 0.00 % | 0.02 % | 0.00 % | 99.98 % | |
| 10.000 | 4.47 | 0.00 % | 0.02 % | 0.00 % | 99.98 % | |
| 11.000 | 4.47 | 0.00 % | 0.02 % | 0.00 % | 99.98 % | |
| 12.000 | 4.47 | 0.00 % | 0.02 % | 0.00 % | 99.98 % | |

#### Carbonate and acidity

Carbonate 0.152 mM
Acidity error -1.319 mM

#### Other graphs

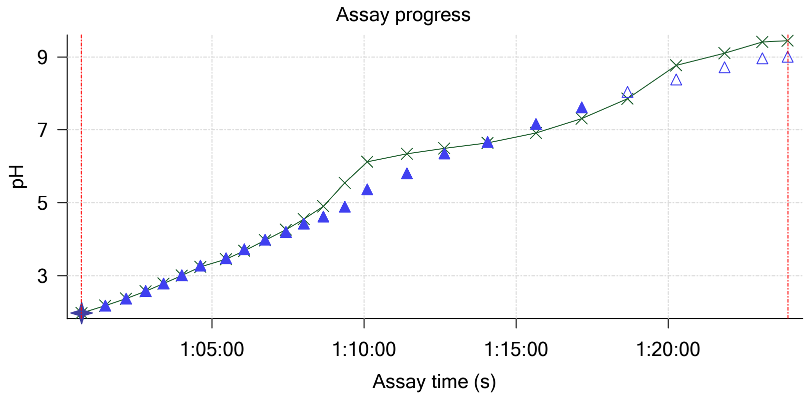

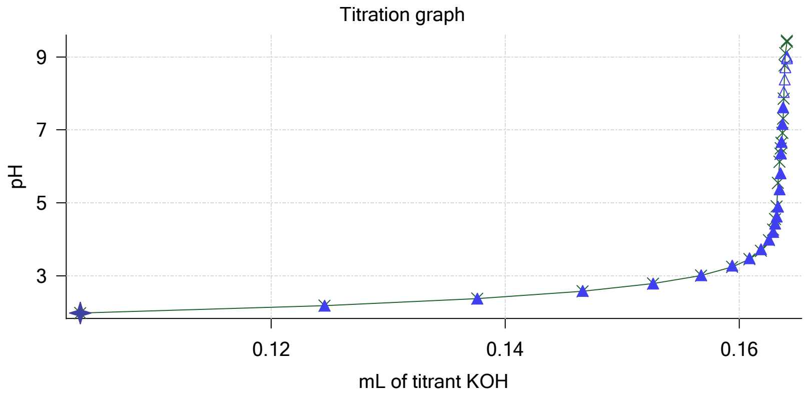

| | | | |
|---|---|---|---|
| Sample name: | **M02_octanol** | Experiment start time: | **3/3/2018 3:08:32 PM** |
| Assay name: | **pH-metric high logP** | Analyst: | **Dorothy Levorse** |
| Assay ID: | **18C-03012** | Instrument ID: | **T312060** |
| Filename: | **C:\Sirius_T3\Mehtap\20180302_exp29_logP_T3-2\18C-03012_M02_octanol_pH-metric high logP.t3r** | | |

#### Other graphs (continued)

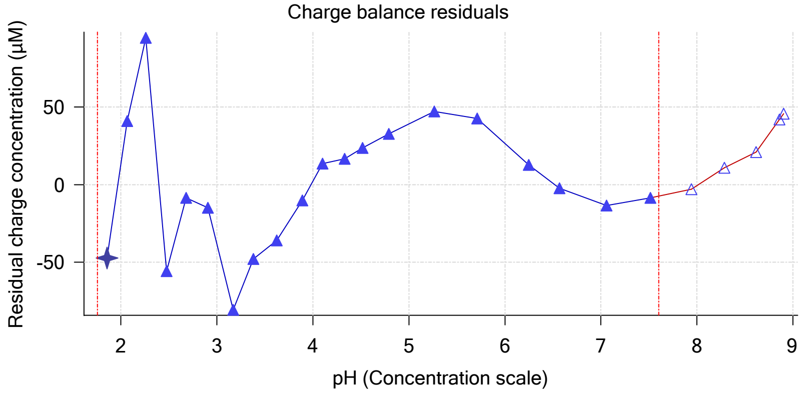

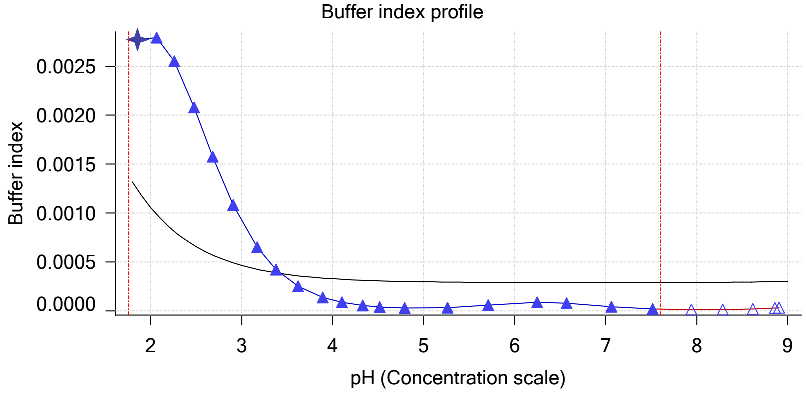

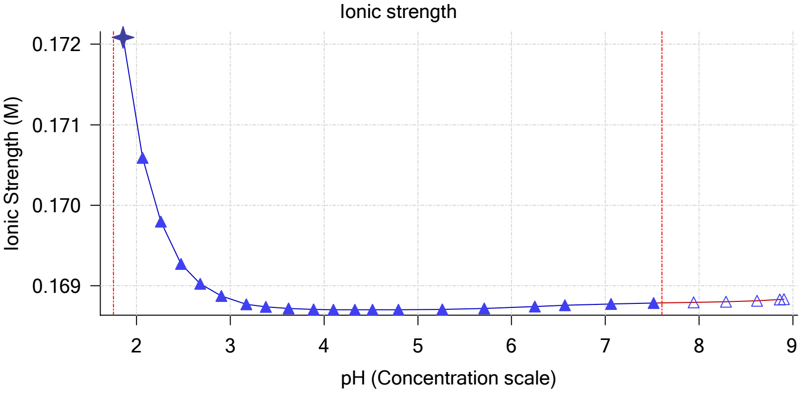

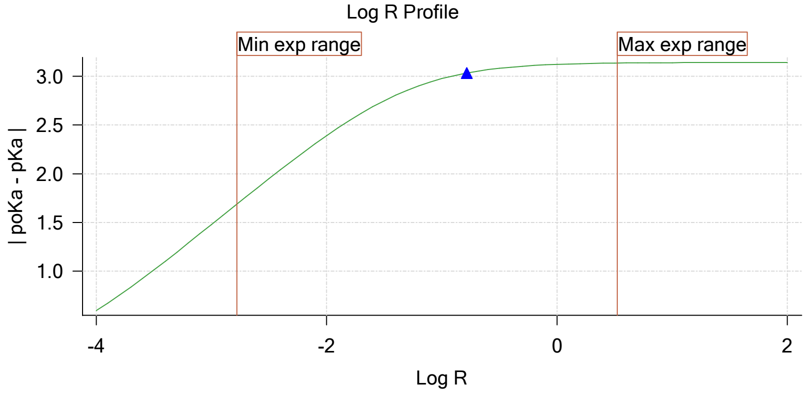

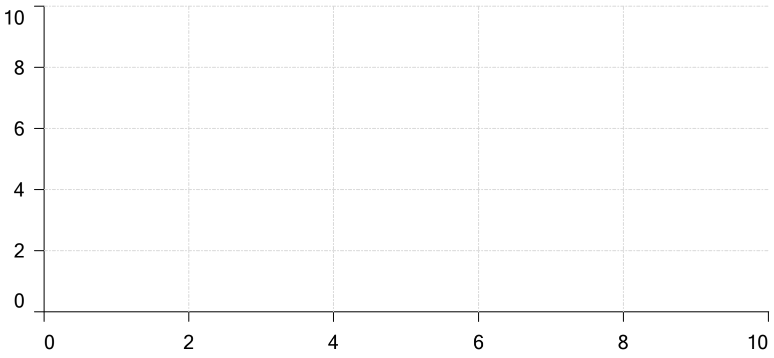

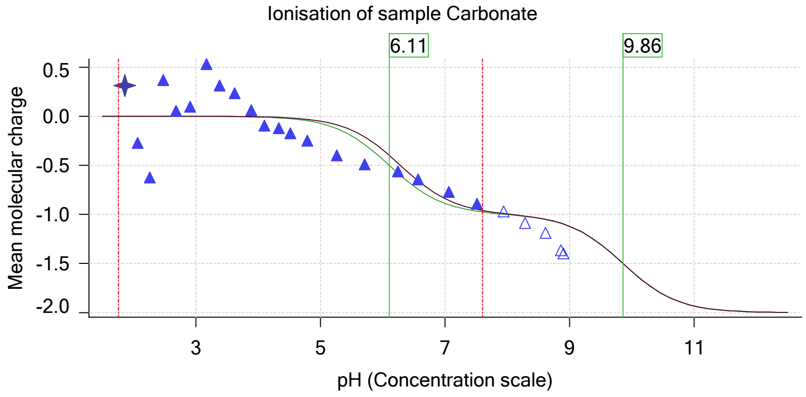

| | | | |
|---|---|---|---|
| Sample name: | **M02_octanol** | Experiment start time: | **3/3/2018 3:08:32 PM** |
| Assay name: | **pH-metric high logP** | Analyst: | **Dorothy Levorse** |
| Assay ID: | **18C-03012** | Instrument ID: | **T312060** |
| Filename: | **C:\Sirius_T3\Mehtap\20180302_exp29_logP_T3-2\18C-03012_M02_octanol_pH-metric high logP.t3r** | | |

#### Assay Model

| Settings | Value | Date/Time changed | Imported from |
|---|---|---|---|
| Sample name | M02_octanol | 12/6/2017 4:20:03 PM | User entered value |
| Sample by | Weight | | Default value |
| Sample weight | 0.002080 g | 3/2/2018 5:10:06 PM | User entered value |
| Formula weight | 289.26 g/mol | 12/6/2017 4:20:03 PM | User entered value |
| Solubility | Unknown | | Default value |
| Molecular weight | 289.26 | 12/6/2017 4:20:03 PM | User entered value |
| Individual pKa ionic environments | No | | Default value |
| Number of pKas | 1 | 12/6/2017 4:20:03 PM | User entered value |
| Sample is a | Base | 12/6/2017 4:20:03 PM | User entered value |
| pKa 1 | 5.03 | 12/6/2017 4:20:03 PM | User entered value |
| logp (XH +) | 1.32 | 3/2/2018 3:38:13 PM | User entered value |
| logP (neutral X) | 4.10 | 3/2/2018 3:38:07 PM | User entered value |

#### Events

| Time | Event | Water | Acid | Base | Octanol | pH | dpH/dt | pH R-squared | pH SD | dpH/dt time |
|---|---|---|---|---|---|---|---|---|---|---|
| 5:59.7 | Initial pH = 9.32 | | | | | | | | | |
| 8:59.2 | Data point 1 | 1.50000 mL | 0.05336 mL | 0.00113 mL | 0.01999 mL | 2.071 | 0.00065 | 0.01575 | 0.00025 | 10.0 s |
| 9:45.5 | Data point 2 | 1.50000 mL | 0.05336 mL | 0.01543 mL | 0.01999 mL | 2.279 | 0.00438 | 0.13165 | 0.00060 | 10.0 s |
| 10:21.2 | Data point 3 | 1.50000 mL | 0.05336 mL | 0.02429 mL | 0.01999 mL | 2.476 | -0.00354 | 0.41455 | 0.00027 | 10.5 s |
| 10:57.3 | Data point 4 | 1.50000 mL | 0.05336 mL | 0.03057 mL | 0.01999 mL | 2.669 | -0.00471 | 0.73726 | 0.00027 | 10.0 s |
| 11:32.8 | Data point 5 | 1.50000 mL | 0.05336 mL | 0.03528 mL | 0.01999 mL | 2.875 | -0.00270 | 0.03691 | 0.00069 | 10.0 s |
| 12:08.2 | Data point 6 | 1.50000 mL | 0.05336 mL | 0.03890 mL | 0.01999 mL | 3.083 | -0.01251 | 0.86143 | 0.00067 | 10.0 s |
| 12:43.7 | Data point 7 | 1.50000 mL | 0.05336 mL | 0.04175 mL | 0.01999 mL | 3.280 | -0.01215 | 0.94698 | 0.00062 | 10.5 s |
| 13:19.6 | Data point 8 | 1.50000 mL | 0.05336 mL | 0.04396 mL | 0.01999 mL | 3.527 | -0.01947 | 0.93803 | 0.00099 | 11.0 s |
| 14:06.4 | Data point 9 | 1.50000 mL | 0.05336 mL | 0.04520 mL | 0.01999 mL | 3.683 | -0.01786 | 0.92590 | 0.00092 | 11.5 s |
| 14:53.7 | Data point 10 | 1.50000 mL | 0.05336 mL | 0.04617 mL | 0.01999 mL | 3.855 | -0.01936 | 0.96090 | 0.00098 | 14.5 s |
| 15:44.0 | Data point 11 | 1.50000 mL | 0.05336 mL | 0.04697 mL | 0.01999 mL | 4.031 | -0.01779 | 0.90907 | 0.00092 | 15.5 s |
| 16:24.9 | Data point 12 | 1.50000 mL | 0.05336 mL | 0.04755 mL | 0.01999 mL | 4.242 | -0.01825 | 0.89567 | 0.00095 | 28.0 s |
| 17:18.3 | Data point 13 | 1.50000 mL | 0.05336 mL | 0.04793 mL | 0.01999 mL | 4.326 | -0.01954 | 0.97542 | 0.00098 | 36.5 s |
| 18:25.4 | Data point 14 | 1.50000 mL | 0.05336 mL | 0.04828 mL | 0.01999 mL | 4.543 | -0.01825 | 0.97150 | 0.00091 | 55.0 s |
| 19:50.9 | Data point 15 | 1.50000 mL | 0.05336 mL | 0.04852 mL | 0.01999 mL | 4.730 | -0.02010 | 0.91878 | 0.00103 | Timed out at 59.5 s |
| 21:21.5 | Data point 16 | 1.50000 mL | 0.05336 mL | 0.04871 mL | 0.01999 mL | 4.975 | -0.02384 | 0.95065 | 0.00121 | Timed out at 59.5 s |
| 22:57.2 | Data point 17 | 1.50000 mL | 0.05336 mL | 0.04892 mL | 0.01999 mL | 5.443 | -0.01935 | 0.93030 | 0.00099 | 56.0 s |
| 24:23.8 | Data point 18 | 1.50000 mL | 0.05336 mL | 0.04906 mL | 0.01999 mL | 5.658 | -0.01989 | 0.98952 | 0.00099 | 46.5 s |
| 25:40.9 | Data point 19 | 1.50000 mL | 0.05336 mL | 0.04920 mL | 0.01999 mL | 6.311 | -0.02340 | 0.95639 | 0.00118 | Timed out at 59.5 s |
| 27:11.5 | Data point 20 | 1.50000 mL | 0.05336 mL | 0.04934 mL | 0.01999 mL | 7.143 | -0.06220 | 0.99094 | 0.00309 | Timed out at 59.5 s |
| 28:41.9 | Data point 21 | 1.50000 mL | 0.05336 mL | 0.04939 mL | 0.01999 mL | 7.536 | -0.06100 | 0.98752 | 0.00303 | Timed out at 59.5 s |
| 30:07.3 | Data point 22 | 1.50000 mL | 0.05336 mL | 0.04941 mL | 0.01999 mL | 7.646 | -0.04673 | 0.98883 | 0.00232 | Timed out at 59.5 s |
| 31:37.8 | Data point 23 | 1.50000 mL | 0.05336 mL | 0.04946 mL | 0.01999 mL | 8.129 | -0.05480 | 0.98555 | 0.00273 | Timed out at 59.5 s |
| 33:08.3 | Data point 24 | 1.50000 mL | 0.05336 mL | 0.04951 mL | 0.01999 mL | 8.458 | -0.03117 | 0.96668 | 0.00157 | Timed out at 59.5 s |
| 34:43.9 | Data point 25 | 1.50000 mL | 0.05336 mL | 0.04958 mL | 0.01999 mL | 8.734 | -0.01915 | 0.91772 | 0.00099 | 57.5 s |
| 36:12.1 | Data point 26 | 1.50000 mL | 0.05336 mL | 0.04967 mL | 0.01999 mL | 9.090 | -0.01873 | 0.92804 | 0.00096 | 35.5 s |
| 37:41.3 | Data point 27 | 1.50000 mL | 0.10861 mL | 0.04967 mL | 0.05000 mL | 1.983 | -0.00848 | 0.30185 | 0.00076 | 10.0 s |
| 38:27.6 | Data point 28 | 1.50000 mL | 0.10861 mL | 0.06604 mL | 0.05000 mL | 2.203 | -0.01236 | 0.74273 | 0.00071 | 10.0 s |
| 39:03.3 | Data point 29 | 1.50000 mL | 0.10861 mL | 0.07745 mL | 0.05000 mL | 2.397 | -0.00234 | 0.07442 | 0.00042 | 10.0 s |
| 39:38.9 | Data point 30 | 1.50000 mL | 0.10861 mL | 0.08540 mL | 0.05000 mL | 2.621 | -0.00166 | 0.44144 | 0.00012 | 10.5 s |
| 40:15.0 | Data point 31 | 1.50000 mL | 0.10861 mL | 0.09087 mL | 0.05000 mL | 2.809 | -0.00504 | 0.92869 | 0.00026 | 10.0 s |
| 40:50.5 | Data point 32 | 1.50000 mL | 0.10861 mL | 0.09497 mL | 0.05000 mL | 3.014 | -0.00539 | 0.66005 | 0.00033 | 10.0 s |

| | | | |
|---|---|---|---|
| Sample name: | **M02_octanol** | Experiment start time: | **3/3/2018 3:08:32 PM** |
| Assay name: | **pH-metric high logP** | Analyst: | **Dorothy Levorse** |
| Assay ID: | **18C-03012** | Instrument ID: | **T312060** |
| Filename: | **C:\Sirius_T3\Mehtap\20180302_exp29_logP_T3-2\18C-03012_M02_octanol_pH-metric high logP.t3r** | | |

#### Events (continued)

| Time | Event | Water | Acid | Base | Octanol | pH | dpH/dt | pH R-squared | pH SD | dpH/dt time |
|---|---|---|---|---|---|---|---|---|---|---|
| 41:26.0 | Data point 33 | 1.50000 mL | 0.10861 mL | 0.09795 mL | 0.05000 mL | 3.211 | -0.00572 | 0.74071 | 0.00033 | 10.0 s |
| 42:01.6 | Data point 34 | 1.50000 mL | 0.10861 mL | 0.10009 mL | 0.05000 mL | 3.497 | -0.01360 | 0.48807 | 0.00096 | 10.0 s |
| 42:52.5 | Data point 35 | 1.50000 mL | 0.10861 mL | 0.10120 mL | 0.05000 mL | 3.730 | 0.00574 | 0.19061 | 0.00065 | 10.5 s |
| 43:28.4 | Data point 36 | 1.50000 mL | 0.10861 mL | 0.10200 mL | 0.05000 mL | 4.017 | -0.01503 | 0.86567 | 0.00080 | 11.0 s |
| 44:09.9 | Data point 37 | 1.50000 mL | 0.10861 mL | 0.10240 mL | 0.05000 mL | 4.227 | -0.01947 | 0.92718 | 0.00100 | 12.0 s |
| 44:47.3 | Data point 38 | 1.50000 mL | 0.10861 mL | 0.10266 mL | 0.05000 mL | 4.444 | -0.01821 | 0.92624 | 0.00093 | 13.0 s |
| 45:25.7 | Data point 39 | 1.50000 mL | 0.10861 mL | 0.10282 mL | 0.05000 mL | 4.652 | -0.01914 | 0.89982 | 0.00100 | 14.0 s |
| 46:05.1 | Data point 40 | 1.50000 mL | 0.10861 mL | 0.10292 mL | 0.05000 mL | 4.800 | -0.01859 | 0.91342 | 0.00096 | 15.0 s |
| 46:50.7 | Data point 41 | 1.50000 mL | 0.10861 mL | 0.10303 mL | 0.05000 mL | 5.083 | -0.01985 | 0.98349 | 0.00099 | 23.0 s |
| 47:44.2 | Data point 42 | 1.50000 mL | 0.10861 mL | 0.10313 mL | 0.05000 mL | 5.579 | -0.01759 | 0.91486 | 0.00091 | 42.0 s |
| 49:02.1 | Data point 43 | 1.50000 mL | 0.10861 mL | 0.10320 mL | 0.05000 mL | 6.101 | -0.01900 | 0.90215 | 0.00099 | 54.5 s |
| 50:27.2 | Data point 44 | 1.50000 mL | 0.10861 mL | 0.10327 mL | 0.05000 mL | 6.674 | -0.03446 | 0.99241 | 0.00171 | Timed out at 59.5 s |
| 51:57.7 | Data point 45 | 1.50000 mL | 0.10861 mL | 0.10336 mL | 0.05000 mL | 7.259 | -0.06329 | 0.99069 | 0.00314 | Timed out at 59.5 s |
| 53:33.3 | Data point 46 | 1.50000 mL | 0.10861 mL | 0.10343 mL | 0.05000 mL | 7.768 | -0.05820 | 0.98124 | 0.00290 | Timed out at 59.5 s |
| 55:03.8 | Data point 47 | 1.50000 mL | 0.10861 mL | 0.10348 mL | 0.05000 mL | 8.292 | -0.03946 | 0.97572 | 0.00197 | Timed out at 59.5 s |
| 56:34.3 | Data point 48 | 1.50000 mL | 0.10861 mL | 0.10353 mL | 0.05000 mL | 8.557 | -0.01940 | 0.93969 | 0.00099 | 57.5 s |
| 58:07.5 | Data point 49 | 1.50000 mL | 0.10861 mL | 0.10360 mL | 0.05000 mL | 8.805 | -0.00522 | 0.06822 | 0.00099 | 30.5 s |
| 59:13.7 | Data point 50 | 1.50000 mL | 0.10861 mL | 0.10369 mL | 0.05000 mL | 9.024 | -0.01835 | 0.91843 | 0.00094 | 25.0 s |
| 1:00:42.8 | Data point 51 | 1.50000 mL | 0.16823 mL | 0.10369 mL | 0.30000 mL | 1.978 | -0.00289 | 0.10876 | 0.00043 | 10.0 s |
| 1:01:29.1 | Data point 52 | 1.50000 mL | 0.16823 mL | 0.12458 mL | 0.30000 mL | 2.182 | -0.01606 | 0.64737 | 0.00099 | 15.5 s |
| 1:02:10.3 | Data point 53 | 1.50000 mL | 0.16823 mL | 0.13763 mL | 0.30000 mL | 2.373 | 0.00806 | 0.16357 | 0.00099 | 13.0 s |
| 1:02:49.0 | Data point 54 | 1.50000 mL | 0.16823 mL | 0.14661 mL | 0.30000 mL | 2.587 | -0.00022 | 0.00759 | 0.00012 | 10.0 s |
| 1:03:24.5 | Data point 55 | 1.50000 mL | 0.16823 mL | 0.15263 mL | 0.30000 mL | 2.790 | -0.00959 | 0.75034 | 0.00055 | 10.0 s |
| 1:04:00.1 | Data point 56 | 1.50000 mL | 0.16823 mL | 0.15675 mL | 0.30000 mL | 3.013 | 0.01134 | 0.31455 | 0.00100 | 11.5 s |
| 1:04:37.1 | Data point 57 | 1.50000 mL | 0.16823 mL | 0.15941 mL | 0.30000 mL | 3.278 | -0.01412 | 0.56497 | 0.00093 | 10.0 s |
| 1:05:27.9 | Data point 58 | 1.50000 mL | 0.16823 mL | 0.16087 mL | 0.30000 mL | 3.486 | 0.00232 | 0.15736 | 0.00029 | 10.5 s |
| 1:06:03.8 | Data point 59 | 1.50000 mL | 0.16823 mL | 0.16185 mL | 0.30000 mL | 3.728 | -0.00733 | 0.87321 | 0.00039 | 10.0 s |
| 1:06:44.4 | Data point 60 | 1.50000 mL | 0.16823 mL | 0.16254 mL | 0.30000 mL | 3.996 | -0.01008 | 0.94298 | 0.00051 | 10.5 s |
| 1:07:25.5 | Data point 61 | 1.50000 mL | 0.16823 mL | 0.16289 mL | 0.30000 mL | 4.206 | -0.01231 | 0.75314 | 0.00070 | 10.0 s |
| 1:08:00.9 | Data point 62 | 1.50000 mL | 0.16823 mL | 0.16308 mL | 0.30000 mL | 4.435 | 0.00196 | 0.01015 | 0.00096 | 13.5 s |
| 1:08:39.8 | Data point 63 | 1.50000 mL | 0.16823 mL | 0.16319 mL | 0.30000 mL | 4.623 | -0.01066 | 0.39352 | 0.00084 | 12.0 s |
| 1:09:22.3 | Data point 64 | 1.50000 mL | 0.16823 mL | 0.16331 mL | 0.30000 mL | 4.896 | -0.01695 | 0.88825 | 0.00089 | 13.5 s |
| 1:10:06.3 | Data point 65 | 1.50000 mL | 0.16823 mL | 0.16345 mL | 0.30000 mL | 5.367 | -0.01896 | 0.91841 | 0.00098 | 47.5 s |
| 1:11:24.4 | Data point 66 | 1.50000 mL | 0.16823 mL | 0.16352 mL | 0.30000 mL | 5.813 | 0.00169 | 0.01098 | 0.00080 | 43.0 s |
| 1:12:38.1 | Data point 67 | 1.50000 mL | 0.16823 mL | 0.16357 mL | 0.30000 mL | 6.350 | -0.02811 | 0.94564 | 0.00143 | Timed out at 59.5 s |
| 1:14:03.5 | Data point 68 | 1.50000 mL | 0.16823 mL | 0.16362 mL | 0.30000 mL | 6.671 | -0.04606 | 0.97751 | 0.00230 | Timed out at 59.5 s |
| 1:15:39.1 | Data point 69 | 1.50000 mL | 0.16823 mL | 0.16369 mL | 0.30000 mL | 7.163 | -0.06931 | 0.98791 | 0.00344 | Timed out at 59.5 s |
| 1:17:09.6 | Data point 70 | 1.50000 mL | 0.16823 mL | 0.16376 mL | 0.30000 mL | 7.616 | -0.08650 | 0.98058 | 0.00432 | Timed out at 59.5 s |
| 1:18:40.1 | Data point 71 | 1.50000 mL | 0.16823 mL | 0.16381 mL | 0.30000 mL | 8.042 | -0.06118 | 0.94917 | 0.00310 | Timed out at 59.5 s |
| 1:20:15.7 | Data point 72 | 1.50000 mL | 0.16823 mL | 0.16388 mL | 0.30000 mL | 8.385 | -0.04572 | 0.94304 | 0.00233 | Timed out at 59.5 s |
| 1:21:51.3 | Data point 73 | 1.50000 mL | 0.16823 mL | 0.16395 mL | 0.30000 mL | 8.716 | -0.01968 | 0.98797 | 0.00098 | 33.5 s |
| 1:23:05.7 | Data point 74 | 1.50000 mL | 0.16823 mL | 0.16406 mL | 0.30000 mL | 8.959 | -0.01414 | 0.95962 | 0.00071 | 24.5 s |
| 1:23:55.6 | Data point 75 | 1.50000 mL | 0.16823 mL | 0.16409 mL | 0.30000 mL | 9.002 | -0.01249 | 0.43204 | 0.00094 | 13.5 s |
| 1:24:18.2 | Assay volumes | 1.50000 mL | 0.16823 mL | 0.16409 mL | 0.30000 mL | | | | | |

| | | | |
|---|---|---|---|
| Sample name: | **M02_octanol** | Experiment start time: | **3/3/2018 3:08:32 PM** |
| Assay name: | **pH-metric high logP** | Analyst: | **Dorothy Levorse** |
| Assay ID: | **18C-03012** | Instrument ID: | **T312060** |
| Filename: | **C:\Sirius_T3\Mehtap\20180302_exp29_logP_T3-2\18C-03012_M02_octanol_pH-metric high logP.t3r** | | |

| | | | |
|---|---|---|---|
| Sample name: | **M02_octanol** | Experiment start time: | **3/3/2018 3:08:32 PM** |
| Assay name: | **pH-metric high logP** | Analyst: | **Dorothy Levorse** |
| Assay ID: | **18C-03012** | Instrument ID: | **T312060** |
| Filename: | **C:\Sirius_T3\Mehtap\20180302_exp29_logP_T3-2\18C-03012_M02_octanol_pH-metric high logP.t3r** | | |

#### Assay Settings (continued)

| Setting | Value | Original Value | Date/Time changed | Imported from |
|---|---|---|---|---|
| ***Titration 3*** | | | | |
| Titrate from | Low to high pH | | | |
| Add additional water | 0.00 mL | | | |
| Additional partition solvent volume | 0.250 mL | | | |
| Additional partition solvent added | Automatic | | | |
| After pH adjust stir for | 30 seconds | | | |
| Stir to allow partitioning for | 15 seconds | | | |
| Stirrer speed for partitioning | 60% | | | |
| ***Data Point Stability*** | | | | |
| Stir during data point collection | No | | | |
| Delay before data point collection | 0 seconds | | | |
| Number of points to average | 20 points | | | |
| Time interval between points | 0.50 seconds | | | |
| Required maximum standard deviation | 0.00100 dpH/dt | | | |
| Stability timeout after | 60 seconds | | | |

#### Calibration Settings

| Setting | Value | Date/Time changed | Imported from |
|---|---|---|---|
| Four-Plus alpha | 0.111 | 3/3/2018 3:08:32 PM | C:\Sirius_T3\HCl18C02.t3r |
| Four-Plus S | 0.9988 | 3/3/2018 3:08:32 PM | C:\Sirius_T3\HCl18C02.t3r |
| Four-Plus jH | 1.0 | 3/3/2018 3:08:32 PM | C:\Sirius_T3\HCl18C02.t3r |
| Four-Plus jOH | -0.8 | 3/3/2018 3:08:32 PM | C:\Sirius_T3\HCl18C02.t3r |
| Base concentration factor | 1.000 | 3/3/2018 3:08:32 PM | C:\Sirius_T3\KOH18B27.t3r |
| Acid concentration factor | 0.999 | 3/3/2018 3:08:32 PM | C:\Sirius_T3\HCl18C02.t3r |

#### Instrument Settings

| Setting | Value | Batch Id | Install date |
|---|---|---|---|
| Instrument owner | Merck | | |
| Instrument ID | T312060 | | |
| Instrument type | T3 Simulator | | |
| Software version | 1.1.3.0 | | |
| Dispenser module | | T3DM1200361 | 3/31/2009 5:24:52 AM |
| Dispenser 0 | Water | | 3/31/2009 5:25:05 AM |
| Syringe volume | 2.5 mL | | |
| Firmware version | 1.2.1(r2) | | |
| Titrant | Water (0.15 M KCl) | 02-06-2018 | 2/27/2018 10:05:59 AM |
| Dispenser 2 | Acid | | 3/31/2009 5:25:11 AM |
| Syringe volume | 0.5 mL | | |
| Firmware version | 1.2.1(r2) | | |
| Titrant | Acid (0.5 M HCl) | 02-27-2018 | 2/27/2018 10:27:22 AM |
| Dispenser 1 | Base | | 3/31/2009 5:25:21 AM |
| Syringe volume | 0.5 mL | | |
| Firmware version | 1.2.1(r2) | | |
| Titrant | Base (0.5 M KOH) | 9/22/2017 | 2/27/2018 10:21:22 AM |
| Dispenser 5 | Cosolvent | | 3/31/2009 5:26:24 AM |
| Syringe volume | 2.5 mL | | |
| Firmware version | 1.2.1(r2) | | |
| Distribution valve 5 | Distribution Valve | | 3/31/2009 5:28:19 AM |
| Firmware version | 1.1.3 | | |
| Port A | Methanol (80%, 0.15 M KCl) | 09-26-17 | 2/7/2018 9:42:01 AM |
| Port B | Cyclohexane | 11-01-17 | 2/27/2018 10:37:57 AM |
| Dispenser 3 | Buffer | | 8/3/2010 5:05:16 AM |
| Syringe volume | 0.5 mL | | |
| Firmware version | 1.2.1(r2) | | |
| Titrant | Dodecane | 2018/01/31 | 2/28/2018 10:18:04 AM |
| Dispenser 6 | Octanol | | 10/22/2010 10:52:43 AM |

| | | | |
|---|---|---|---|
| Sample name: | **M02_octanol** | Experiment start time: | **3/3/2018 3:08:32 PM** |
| Assay name: | **pH-metric high logP** | Analyst: | **Dorothy Levorse** |
| Assay ID: | **18C-03012** | Instrument ID: | **T312060** |
| Filename: | **C:\Sirius_T3\Mehtap\20180302_exp29_logP_T3-2\18C-03012_M02_octanol_pH-metric high logP.t3r** | | |

| | | | |
|---|---|---|---|
| Sample name: | **M02_octanol** | Experiment start time: | **3/3/2018 3:08:32 PM** |
| Assay name: | **pH-metric high logP** | Analyst: | **Dorothy Levorse** |
| Assay ID: | **18C-03012** | Instrument ID: | **T312060** |
| Filename: | **C:\Sirius_T3\Mehtap\20180302_exp29_logP_T3-2\18C-03012_M02_octanol_pH-metric high logP.t3r** | | |

#### Instrument Settings (continued)

| Setting | Value | Batch Id | Install date |
|---|---|---|---|
| Spectrometer calibration stir duration | 5 s | | |
| Spectrometer calibration stir speed | 30% | | |
| Spectrometer calibration wash pump volume | 20.0 mL | | |
| Spectrometer calibration wash stir duration | 5 s | | |
| Spectrometer calibration wash stir speed | 30% | | |
| Overhead dispense height | 10000 | | |

#### Refinement Settings

| Setting | Value | Default value |
|---|---|---|
| Turbidity detection method | None | None |
| Turbidity wavelength to assess | 500.0 nm | 500.0 nm |
| Turbidity maximum absorbance | 0.100 | 0.100 |
| Turbidity probe threshold | 50.00 | 50.00 |

#### Experiment Log

[2:37] Air gap created for Water (0.15 M KCl)
[2:38] Air gap created for Acid (0.5 M HCl)
[2:38] Air gap created for Base (0.5 M KOH)
[2:39] Air gap released for Water (0.15 M KCl)
[2:42] Titrator arm moved over Titration position
[2:42] Titration 1 of 3
[2:42] Adding initial titrants
[2:42] Automatically add 1.50000 mL of water
[3:08] Dispensed 1.500000 mL of Water (0.15 M KCl)
[3:12] Titrator arm moved over Drain
[5:53] Titrator arm moved to Titration position
[5:53] Argon flow rate set to 100
[5:53] Stirrer speed set to 10
[5:58] Automatically add 0.02000 mL of Octanol
[5:59] Dispensed 0.019991 mL of Octanol
[6:00] Initial pH = 9.32
[6:00] Iterative adjust 9.32 -> 2.00
[6:00] pH 9.32 -> 2.00
[6:02] Air gap released for Acid (0.5 M HCl)
[6:02] Dispensed 0.053363 mL of Acid (0.5 M HCl)
[6:08] Holding pH 2.00
[8:08] Stirrer speed set to 0
[8:08] Stirrer speed set to 50
[8:08] Iterative adjust 1.99 -> 2.00
[8:08] pH 1.99 -> 2.00
[8:08] Air gap released for Base (0.5 M KOH)
[8:09] Dispensed 0.001129 mL of Base (0.5 M KOH)
[8:59] Stirrer speed set to 0
[9:09] Datapoint id 1 collected
[9:09] Stirrer speed set to 50
[9:15] pH 2.08 -> 2.28
[9:15] Using cautious pH adjust
[9:15] Dispensed 0.006726 mL of Base (0.5 M KOH)
[9:20] Stepping pH = 2.16
[9:20] Dispensed 0.005738 mL of Base (0.5 M KOH)
[9:25] Stepping pH = 2.25
[9:25] Dispensed 0.001834 mL of Base (0.5 M KOH)
[9:31] Stepping pH = 2.28
[9:46] Stirrer speed set to 0
[9:56] Datapoint id 2 collected
[9:56] Charge balance equation is out by -6.2%
[9:56] Stirrer speed set to 50

| | | | |
|---|---|---|---|
| Sample name: | **M02_octanol** | Experiment start time: | **3/3/2018 3:08:32 PM** |
| Assay name: | **pH-metric high logP** | Analyst: | **Dorothy Levorse** |
| Assay ID: | **18C-03012** | Instrument ID: | **T312060** |
| Filename: | **C:\Sirius_T3\Mehtap\20180302_exp29_logP_T3-2\18C-03012_M02_octanol_pH-metric high logP.t3r** | | |

#### Experiment Log (continued)

[10:01] pH 2.29 -> 2.49
[10:01] Using charge balance adjust
[10:01] Dispensed 0.008866 mL of Base (0.5 M KOH)
[10:21] Stirrer speed set to 0
[10:32] Datapoint id 3 collected
[10:32] Charge balance equation is out by -5.7%
[10:32] Stirrer speed set to 50
[10:37] pH 2.48 -> 2.68
[10:37] Using charge balance adjust
[10:37] Dispensed 0.006279 mL of Base (0.5 M KOH)
[10:57] Stirrer speed set to 0
[11:07] Datapoint id 4 collected
[11:07] Charge balance equation is out by -6.6%
[11:07] Stirrer speed set to 50
[11:13] pH 2.68 -> 2.88
[11:13] Using charge balance adjust
[11:13] Dispensed 0.004704 mL of Base (0.5 M KOH)
[11:33] Stirrer speed set to 0
[11:43] Datapoint id 5 collected
[11:43] Charge balance equation is out by -0.6%
[11:43] Stirrer speed set to 50
[11:48] pH 2.88 -> 3.08
[11:48] Using charge balance adjust
[11:48] Dispensed 0.003622 mL of Base (0.5 M KOH)
[12:08] Stirrer speed set to 0
[12:18] Datapoint id 6 collected
[12:18] Charge balance equation is out by 0.1%
[12:18] Stirrer speed set to 50
[12:24] pH 3.09 -> 3.29
[12:24] Using charge balance adjust
[12:24] Dispensed 0.002846 mL of Base (0.5 M KOH)
[12:44] Stirrer speed set to 0
[12:54] Datapoint id 7 collected
[12:54] Charge balance equation is out by -4.2%
[12:54] Stirrer speed set to 50
[12:59] pH 3.28 -> 3.48
[12:59] Using charge balance adjust
[13:01] Dispensed 0.002211 mL of Base (0.5 M KOH)
[13:20] Stirrer speed set to 0
[13:31] Datapoint id 8 collected
[13:31] Charge balance equation is out by 21.6%
[13:31] Stirrer speed set to 50
[13:36] pH 3.53 -> 3.73
[13:36] Using cautious pH adjust
[13:36] Dispensed 0.000776 mL of Base (0.5 M KOH)
[13:41] Stepping pH = 3.67
[13:41] Dispensed 0.000282 mL of Base (0.5 M KOH)
[13:46] Stepping pH = 3.70
[13:46] Dispensed 0.000188 mL of Base (0.5 M KOH)
[13:51] Stepping pH = 3.72
[14:07] Stirrer speed set to 0
[14:18] Datapoint id 9 collected
[14:18] Charge balance equation is out by 19.9%
[14:18] Stirrer speed set to 50
[14:23] pH 3.69 -> 3.89
[14:23] Using cautious pH adjust
[14:23] Dispensed 0.000588 mL of Base (0.5 M KOH)
[14:28] Stepping pH = 3.82
[14:29] Dispensed 0.000235 mL of Base (0.5 M KOH)

Sample name: **M02_octanol**
Assay name: **pH-metric high logP**
Assay ID: **18C-03012**
Filename: **C:\Sirius_T3\Mehtap\20180302_exp29_logP_T3-2\18C-03012_M02_octanol_pH-metric high logP.t3r**
Experiment start time: **3/3/2018 3:08:32 PM**
Analyst: **Dorothy Levorse**
Instrument ID: **T312060**

#### Experiment Log (continued)

[14:34] Stepping pH = 3.86
[14:34] Dispensed 0.000141 mL of Base (0.5 M KOH)
[14:39] Stepping pH = 3.88
[14:54] Stirrer speed set to 0
[15:08] Datapoint id 10 collected
[15:08] Charge balance equation is out by 17.0%
[15:08] Stirrer speed set to 50
[15:14] pH 3.87 -> 4.07
[15:14] Using cautious pH adjust
[15:14] Dispensed 0.000423 mL of Base (0.5 M KOH)
[15:19] Stepping pH = 4.00
[15:19] Dispensed 0.000165 mL of Base (0.5 M KOH)
[15:24] Stepping pH = 4.02
[15:24] Dispensed 0.000212 mL of Base (0.5 M KOH)
[15:29] Stepping pH = 4.09
[15:44] Stirrer speed set to 0
[16:00] Datapoint id 11 collected
[16:00] Charge balance equation is out by 4.0%
[16:00] Stirrer speed set to 50
[16:05] pH 4.05 -> 4.25
[16:05] Using charge balance adjust
[16:05] Dispensed 0.000588 mL of Base (0.5 M KOH)
[16:25] Stirrer speed set to 0
[16:53] Datapoint id 12 collected
[16:53] Charge balance equation is out by -3.7%
[16:53] Stirrer speed set to 50
[16:58] pH 4.27 -> 4.47
[16:58] Using charge balance adjust
[16:58] Dispensed 0.000376 mL of Base (0.5 M KOH)
[17:19] Stirrer speed set to 0
[17:55] Datapoint id 13 collected
[17:55] Charge balance equation is out by -71.4%
[17:55] Stirrer speed set to 50
[18:00] pH 4.37 -> 4.57
[18:00] Using cautious pH adjust
[18:00] Dispensed 0.000141 mL of Base (0.5 M KOH)
[18:05] Stepping pH = 4.42
[18:05] Dispensed 0.000212 mL of Base (0.5 M KOH)
[18:11] Stepping pH = 4.67
[18:26] Stirrer speed set to 0
[19:21] Datapoint id 14 collected
[19:21] Charge balance equation is out by -17.2%
[19:21] Stirrer speed set to 50
[19:26] pH 4.58 -> 4.78
[19:26] Using cautious pH adjust
[19:26] Dispensed 0.000094 mL of Base (0.5 M KOH)
[19:31] Stepping pH = 4.63
[19:31] Dispensed 0.000141 mL of Base (0.5 M KOH)
[19:36] Stepping pH = 4.85
[19:51] Stirrer speed set to 0
[20:51] Datapoint id 15 collected
[20:51] Charge balance equation is out by -20.7%
[20:51] Stirrer speed set to 50
[20:56] pH 4.78 -> 4.98
[20:56] Using cautious pH adjust
[20:56] Dispensed 0.000071 mL of Base (0.5 M KOH)
[21:01] Stepping pH = 4.82
[21:01] Dispensed 0.000118 mL of Base (0.5 M KOH)
[21:07] Stepping pH = 5.08

| | | | |
|---|---|---|---|
| Sample name: | **M02_octanol** | Experiment start time: | **3/3/2018 3:08:32 PM** |
| Assay name: | **pH-metric high logP** | Analyst: | **Dorothy Levorse** |
| Assay ID: | **18C-03012** | Instrument ID: | **T312060** |
| Filename: | **C:\Sirius_T3\Mehtap\20180302_exp29_logP_T3-2\18C-03012_M02_octanol_pH-metric high logP.t3r** | | |

#### Experiment Log (continued)

[21:22] Stirrer speed set to 0
[22:22] Datapoint id 16 collected
[22:22] Charge balance equation is out by -35.6%
[22:22] Stirrer speed set to 50
[22:27] pH 5.00 -> 5.20
[22:27] Using cautious pH adjust
[22:27] Dispensed 0.000047 mL of Base (0.5 M KOH)
[22:32] Stepping pH = 5.11
[22:32] Dispensed 0.000024 mL of Base (0.5 M KOH)
[22:37] Stepping pH = 5.09
[22:37] Dispensed 0.000141 mL of Base (0.5 M KOH)
[22:42] Stepping pH = 5.51
[22:57] Stirrer speed set to 0
[23:53] Datapoint id 17 collected
[23:53] Charge balance equation is out by -140.4%
[23:53] Stirrer speed set to 50
[23:59] pH 5.46 -> 5.66
[23:59] Using cautious pH adjust
[23:59] Dispensed 0.000024 mL of Base (0.5 M KOH)
[24:04] Stepping pH = 5.45
[24:04] Dispensed 0.000118 mL of Base (0.5 M KOH)
[24:09] Stepping pH = 5.76
[24:24] Stirrer speed set to 0
[25:11] Datapoint id 18 collected
[25:11] Charge balance equation is out by -211.0%
[25:11] Stirrer speed set to 50
[25:16] pH 5.83 -> 6.03
[25:16] Using cautious pH adjust
[25:16] Dispensed 0.000024 mL of Base (0.5 M KOH)
[25:21] Stepping pH = 5.82
[25:21] Dispensed 0.000118 mL of Base (0.5 M KOH)
[25:26] Stepping pH = 6.32
[25:41] Stirrer speed set to 0
[26:41] Datapoint id 19 collected
[26:41] Charge balance equation is out by -217.8%
[26:41] Stirrer speed set to 50
[26:46] pH 6.25 -> 6.45
[26:46] Using cautious pH adjust
[26:46] Dispensed 0.000024 mL of Base (0.5 M KOH)
[26:51] Stepping pH = 6.23
[26:51] Dispensed 0.000118 mL of Base (0.5 M KOH)
[26:57] Stepping pH = 7.03
[27:12] Stirrer speed set to 0
[28:12] Datapoint id 20 collected
[28:12] Charge balance equation is out by -227.6%
[28:12] Stirrer speed set to 50
[28:17] pH 7.10 -> 7.30
[28:17] Using cautious pH adjust
[28:17] Dispensed 0.000024 mL of Base (0.5 M KOH)
[28:22] Stepping pH = 7.16
[28:22] Dispensed 0.000024 mL of Base (0.5 M KOH)
[28:27] Stepping pH = 7.53
[28:42] Stirrer speed set to 0
[29:42] Datapoint id 21 collected
[29:42] Charge balance equation is out by -180.3%
[29:42] Stirrer speed set to 50
[29:47] pH 7.45 -> 7.65
[29:47] Using cautious pH adjust
[29:47] Dispensed 0.000024 mL of Base (0.5 M KOH)

| | | | |
|---|---|---|---|
| Sample name: | **M02_octanol** | Experiment start time: | **3/3/2018 3:08:32 PM** |
| Assay name: | **pH-metric high logP** | Analyst: | **Dorothy Levorse** |
| Assay ID: | **18C-03012** | Instrument ID: | **T312060** |
| Filename: | **C:\Sirius_T3\Mehtap\20180302_exp29_logP_T3-2\18C-03012_M02_octanol_pH-metric high logP.t3r** | | |

#### Experiment Log (continued)

[29:52] Stepping pH = 7.65
[30:07] Stirrer speed set to 0
[31:08] Datapoint id 22 collected
[31:08] Charge balance equation is out by -150.3%
[31:08] Stirrer speed set to 50
[31:13] pH 7.60 -> 7.80
[31:13] Using cautious pH adjust
[31:13] Dispensed 0.000024 mL of Base (0.5 M KOH)
[31:18] Stepping pH = 7.75
[31:18] Dispensed 0.000024 mL of Base (0.5 M KOH)
[31:23] Stepping pH = 8.16
[31:38] Stirrer speed set to 0
[32:38] Datapoint id 23 collected
[32:38] Charge balance equation is out by -495.8%
[32:38] Stirrer speed set to 50
[32:43] pH 8.06 -> 8.26
[32:43] Using cautious pH adjust
[32:43] Dispensed 0.000024 mL of Base (0.5 M KOH)
[32:48] Stepping pH = 8.02
[32:48] Dispensed 0.000024 mL of Base (0.5 M KOH)
[32:53] Stepping pH = 8.38
[33:08] Stirrer speed set to 0
[34:09] Datapoint id 24 collected
[34:09] Charge balance equation is out by -485.1%
[34:09] Stirrer speed set to 50
[34:14] pH 8.42 -> 8.62
[34:14] Using cautious pH adjust
[34:14] Dispensed 0.000024 mL of Base (0.5 M KOH)
[34:19] Stepping pH = 8.39
[34:19] Dispensed 0.000024 mL of Base (0.5 M KOH)
[34:24] Stepping pH = 8.57
[34:24] Dispensed 0.000024 mL of Base (0.5 M KOH)
[34:29] Stepping pH = 8.82
[34:44] Stirrer speed set to 0
[35:42] Datapoint id 25 collected
[35:42] Charge balance equation is out by -501.8%
[35:42] Stirrer speed set to 50
[35:47] pH 8.72 -> 8.92
[35:47] Using cautious pH adjust
[35:47] Dispensed 0.000024 mL of Base (0.5 M KOH)
[35:52] Stepping pH = 8.69
[35:52] Dispensed 0.000071 mL of Base (0.5 M KOH)
[35:57] Stepping pH = 9.04
[36:12] Stirrer speed set to 0
[36:48] Datapoint id 26 collected
[36:48] Charge balance equation is out by -260.2%
[36:48] Titration 2 of 3
[36:48] Adding initial titrants
[36:48] Automatically add 0.03000 mL of Octanol
[36:49] Dispensed 0.030009 mL of Octanol
[36:49] Stirrer speed set to 10
[36:50] Stirrer speed set to 55
[36:50] Iterative adjust 9.10 -> 2.00
[36:50] pH 9.10 -> 2.00
[36:51] Dispensed 0.055245 mL of Acid (0.5 M HCl)
[37:41] Stirrer speed set to 0
[37:52] Datapoint id 27 collected
[37:52] Stirrer speed set to 55
[37:57] pH 1.99 -> 2.19

| | | | |
|---|---|---|---|
| Sample name: | **M02_octanol** | Experiment start time: | **3/3/2018 3:08:32 PM** |
| Assay name: | **pH-metric high logP** | Analyst: | **Dorothy Levorse** |
| Assay ID: | **18C-03012** | Instrument ID: | **T312060** |
| Filename: | **C:\Sirius_T3\Mehtap\20180302_exp29_logP_T3-2\18C-03012_M02_octanol_pH-metric high logP.t3r** | | |

#### Experiment Log (continued)

[37:57] Using cautious pH adjust
[37:57] Dispensed 0.008913 mL of Base (0.5 M KOH)
[38:02] Stepping pH = 2.08
[38:02] Dispensed 0.006279 mL of Base (0.5 M KOH)
[38:07] Stepping pH = 2.17
[38:08] Dispensed 0.001176 mL of Base (0.5 M KOH)
[38:13] Stepping pH = 2.19
[38:28] Stirrer speed set to 0
[38:38] Datapoint id 28 collected
[38:38] Charge balance equation is out by 8.2%
[38:38] Stirrer speed set to 55
[38:43] pH 2.21 -> 2.41
[38:43] Using charge balance adjust
[38:43] Dispensed 0.011406 mL of Base (0.5 M KOH)
[39:04] Stirrer speed set to 0
[39:14] Datapoint id 29 collected
[39:14] Charge balance equation is out by -5.6%
[39:14] Stirrer speed set to 55
[39:19] pH 2.40 -> 2.60
[39:19] Using charge balance adjust
[39:19] Dispensed 0.007949 mL of Base (0.5 M KOH)
[39:39] Stirrer speed set to 0
[39:50] Datapoint id 30 collected
[39:50] Charge balance equation is out by 8.8%
[39:50] Stirrer speed set to 55
[39:55] pH 2.63 -> 2.83
[39:55] Using charge balance adjust
[39:55] Dispensed 0.005480 mL of Base (0.5 M KOH)
[40:15] Stirrer speed set to 0
[40:25] Datapoint id 31 collected
[40:25] Charge balance equation is out by -9.5%
[40:25] Stirrer speed set to 55
[40:30] pH 2.82 -> 3.02
[40:30] Using charge balance adjust
[40:31] Dispensed 0.004092 mL of Base (0.5 M KOH)
[40:51] Stirrer speed set to 0
[41:01] Datapoint id 32 collected
[41:01] Charge balance equation is out by -0.9%
[41:01] Stirrer speed set to 55
[41:06] pH 3.02 -> 3.22
[41:06] Using charge balance adjust
[41:06] Dispensed 0.002987 mL of Base (0.5 M KOH)
[41:26] Stirrer speed set to 0
[41:36] Datapoint id 33 collected
[41:36] Charge balance equation is out by -4.3%
[41:36] Stirrer speed set to 55
[41:41] pH 3.22 -> 3.42
[41:41] Using charge balance adjust
[41:42] Dispensed 0.002140 mL of Base (0.5 M KOH)
[42:02] Stirrer speed set to 0
[42:12] Datapoint id 34 collected
[42:12] Charge balance equation is out by 40.8%
[42:12] Stirrer speed set to 55
[42:17] pH 3.51 -> 3.71
[42:17] Using cautious pH adjust
[42:17] Dispensed 0.000635 mL of Base (0.5 M KOH)
[42:22] Stepping pH = 3.63
[42:22] Dispensed 0.000259 mL of Base (0.5 M KOH)
[42:27] Stepping pH = 3.69

Sample name: **M02_octanol**
Assay name: **pH-metric high logP**
Assay ID: **18C-03012**
Filename: **C:\Sirius_T3\Mehtap\20180302_exp29_logP_T3-2\18C-03012_M02_octanol_pH-metric high logP.t3r**

Experiment start time: **3/3/2018 3:08:32 PM**
Analyst: **Dorothy Levorse**
Instrument ID: **T312060**

#### Experiment Log (continued)

[42:27] Dispensed 0.000047 mL of Base (0.5 M KOH)
[42:32] Stepping pH = 3.69
[42:32] Dispensed 0.000165 mL of Base (0.5 M KOH)
[42:38] Stepping pH = 3.74
[42:53] Stirrer speed set to 0
[43:03] Datapoint id 35 collected
[43:03] Charge balance equation is out by 12.2%
[43:03] Stirrer speed set to 55
[43:08] pH 3.74 -> 3.94
[43:08] Using charge balance adjust
[43:08] Dispensed 0.000800 mL of Base (0.5 M KOH)
[43:29] Stirrer speed set to 0
[43:40] Datapoint id 36 collected
[43:40] Charge balance equation is out by 38.4%
[43:40] Stirrer speed set to 55
[43:45] pH 4.03 -> 4.23
[43:45] Using cautious pH adjust
[43:45] Dispensed 0.000212 mL of Base (0.5 M KOH)
[43:50] Stepping pH = 4.11
[43:50] Dispensed 0.000188 mL of Base (0.5 M KOH)
[43:55] Stepping pH = 4.23
[44:10] Stirrer speed set to 0
[44:22] Datapoint id 37 collected
[44:22] Charge balance equation is out by 7.1%
[44:22] Stirrer speed set to 55
[44:27] pH 4.25 -> 4.45
[44:27] Using charge balance adjust
[44:27] Dispensed 0.000259 mL of Base (0.5 M KOH)
[44:48] Stirrer speed set to 0
[45:01] Datapoint id 38 collected
[45:01] Charge balance equation is out by -1.3%
[45:01] Stirrer speed set to 55
[45:06] pH 4.47 -> 4.67
[45:06] Using charge balance adjust
[45:06] Dispensed 0.000165 mL of Base (0.5 M KOH)
[45:26] Stirrer speed set to 0
[45:40] Datapoint id 39 collected
[45:40] Charge balance equation is out by -8.1%
[45:40] Stirrer speed set to 55
[45:45] pH 4.69 -> 4.89
[45:45] Using charge balance adjust
[45:45] Dispensed 0.000094 mL of Base (0.5 M KOH)
[46:05] Stirrer speed set to 0
[46:20] Datapoint id 40 collected
[46:20] Charge balance equation is out by -45.8%
[46:20] Stirrer speed set to 55
[46:25] pH 4.84 -> 5.04
[46:25] Using cautious pH adjust
[46:26] Dispensed 0.000047 mL of Base (0.5 M KOH)
[46:31] Stepping pH = 4.87
[46:31] Dispensed 0.000071 mL of Base (0.5 M KOH)
[46:36] Stepping pH = 5.07
[46:51] Stirrer speed set to 0
[47:14] Datapoint id 41 collected
[47:14] Charge balance equation is out by -57.4%
[47:14] Stirrer speed set to 55
[47:19] pH 5.15 -> 5.35
[47:19] Using cautious pH adjust
[47:19] Dispensed 0.000024 mL of Base (0.5 M KOH)

| | | | |
|---|---|---|---|
| Sample name: | **M02_octanol** | Experiment start time: | **3/3/2018 3:08:32 PM** |
| Assay name: | **pH-metric high logP** | Analyst: | **Dorothy Levorse** |
| Assay ID: | **18C-03012** | Instrument ID: | **T312060** |
| Filename: | **C:\Sirius_T3\Mehtap\20180302_exp29_logP_T3-2\18C-03012_M02_octanol_pH-metric high logP.t3r** | | |

#### Experiment Log (continued)

[47:24] Stepping pH = 5.16
[47:24] Dispensed 0.000071 mL of Base (0.5 M KOH)
[47:29] Stepping pH = 5.51
[47:44] Stirrer speed set to 0
[48:27] Datapoint id 42 collected
[48:27] Charge balance equation is out by -88.9%
[48:27] Stirrer speed set to 55
[48:32] pH 5.71 -> 5.91
[48:32] Using cautious pH adjust
[48:32] Dispensed 0.000024 mL of Base (0.5 M KOH)
[48:37] Stepping pH = 5.76
[48:37] Dispensed 0.000024 mL of Base (0.5 M KOH)
[48:42] Stepping pH = 5.83
[48:42] Dispensed 0.000024 mL of Base (0.5 M KOH)
[48:47] Stepping pH = 6.06
[49:02] Stirrer speed set to 0
[49:57] Datapoint id 43 collected
[49:57] Charge balance equation is out by -92.8%
[49:57] Stirrer speed set to 55
[50:02] pH 6.20 -> 6.40
[50:02] Using cautious pH adjust
[50:02] Dispensed 0.000024 mL of Base (0.5 M KOH)
[50:07] Stepping pH = 6.23
[50:07] Dispensed 0.000047 mL of Base (0.5 M KOH)
[50:12] Stepping pH = 6.63
[50:27] Stirrer speed set to 0
[51:27] Datapoint id 44 collected
[51:27] Charge balance equation is out by -69.3%
[51:27] Stirrer speed set to 55
[51:32] pH 6.63 -> 6.83
[51:32] Using cautious pH adjust
[51:33] Dispensed 0.000024 mL of Base (0.5 M KOH)
[51:38] Stepping pH = 6.63
[51:38] Dispensed 0.000071 mL of Base (0.5 M KOH)
[51:43] Stepping pH = 7.07
[51:58] Stirrer speed set to 0
[52:58] Datapoint id 45 collected
[52:58] Charge balance equation is out by -214.8%
[52:58] Stirrer speed set to 55
[53:03] pH 7.25 -> 7.45
[53:03] Using cautious pH adjust
[53:03] Dispensed 0.000024 mL of Base (0.5 M KOH)
[53:08] Stepping pH = 7.27
[53:08] Dispensed 0.000024 mL of Base (0.5 M KOH)
[53:13] Stepping pH = 7.40
[53:13] Dispensed 0.000024 mL of Base (0.5 M KOH)
[53:18] Stepping pH = 7.65
[53:33] Stirrer speed set to 0
[54:34] Datapoint id 46 collected
[54:34] Charge balance equation is out by -403.0%
[54:34] Stirrer speed set to 55
[54:39] pH 7.78 -> 7.98
[54:39] Using cautious pH adjust
[54:39] Dispensed 0.000024 mL of Base (0.5 M KOH)
[54:44] Stepping pH = 7.88
[54:44] Dispensed 0.000024 mL of Base (0.5 M KOH)
[54:49] Stepping pH = 8.12
[55:04] Stirrer speed set to 0
[56:04] Datapoint id 47 collected

Sample name: **M02_octanol**
Assay name: **pH-metric high logP**
Assay ID: **18C-03012**
Filename: **C:\Sirius_T3\Mehtap\20180302_exp29_logP_T3-2\18C-03012_M02_octanol_pH-metric high logP.t3r**

Experiment start time: **3/3/2018 3:08:32 PM**
Analyst: **Dorothy Levorse**
Instrument ID: **T312060**

#### Experiment Log (continued)

[56:04] Charge balance equation is out by -511.7%
[56:04] Stirrer speed set to 55
[56:09] pH 8.34 -> 8.54
[56:09] Using cautious pH adjust
[56:09] Dispensed 0.000024 mL of Base (0.5 M KOH)
[56:14] Stepping pH = 8.38
[56:14] Dispensed 0.000024 mL of Base (0.5 M KOH)
[56:19] Stepping pH = 8.53
[56:35] Stirrer speed set to 0
[57:32] Datapoint id 48 collected
[57:32] Charge balance equation is out by -253.0%
[57:32] Stirrer speed set to 55
[57:37] pH 8.57 -> 8.77
[57:37] Using cautious pH adjust
[57:37] Dispensed 0.000024 mL of Base (0.5 M KOH)
[57:42] Stepping pH = 8.59
[57:42] Dispensed 0.000024 mL of Base (0.5 M KOH)
[57:47] Stepping pH = 8.67
[57:48] Dispensed 0.000024 mL of Base (0.5 M KOH)
[57:53] Stepping pH = 8.77
[58:08] Stirrer speed set to 0
[58:38] Datapoint id 49 collected
[58:38] Charge balance equation is out by -278.9%
[58:38] Stirrer speed set to 55
[58:43] pH 8.82 -> 9.02
[58:43] Using cautious pH adjust
[58:43] Dispensed 0.000024 mL of Base (0.5 M KOH)
[58:49] Stepping pH = 8.83
[58:49] Dispensed 0.000047 mL of Base (0.5 M KOH)
[58:54] Stepping pH = 8.96
[58:54] Dispensed 0.000024 mL of Base (0.5 M KOH)
[58:59] Stepping pH = 9.04
[59:14] Stirrer speed set to 0
[59:39] Datapoint id 50 collected
[59:39] Charge balance equation is out by -176.9%
[59:39] Titration 3 of 3
[59:39] Adding initial titrants
[59:39] Automatically add 0.25000 mL of Octanol
[59:45] Dispensed 0.250000 mL of Octanol
[59:45] Stirrer speed set to 10
[59:46] Stirrer speed set to 60
[59:46] Iterative adjust 9.02 -> 2.00
[59:46] pH 9.02 -> 2.00
[59:47] Dispensed 0.056656 mL of Acid (0.5 M HCl)
[59:53] pH 2.03 -> 2.00
[59:53] Dispensed 0.002963 mL of Acid (0.5 M HCl)
[1:00:43] Stirrer speed set to 0
[1:00:53] Datapoint id 51 collected
[1:00:53] Stirrer speed set to 60
[1:00:58] pH 1.99 -> 2.19
[1:00:58] Using cautious pH adjust
[1:00:58] Dispensed 0.009784 mL of Base (0.5 M KOH)
[1:01:04] Stepping pH = 2.06
[1:01:04] Dispensed 0.008984 mL of Base (0.5 M KOH)
[1:01:09] Stepping pH = 2.16
[1:01:09] Dispensed 0.002117 mL of Base (0.5 M KOH)
[1:01:14] Stepping pH = 2.19
[1:01:29] Stirrer speed set to 0
[1:01:45] Datapoint id 52 collected

| | | | |
|---|---|---|---|
| Sample name: | **M02_octanol** | Experiment start time: | **3/3/2018 3:08:32 PM** |
| Assay name: | **pH-metric high logP** | Analyst: | **Dorothy Levorse** |
| Assay ID: | **18C-03012** | Instrument ID: | **T312060** |
| Filename: | **C:\Sirius_T3\Mehtap\20180302_exp29_logP_T3-2\18C-03012_M02_octanol_pH-metric high logP.t3r** | | |

#### Experiment Log (continued)

[1:01:45] Charge balance equation is out by -6.7%
[1:01:45] Stirrer speed set to 60
[1:01:50] pH 2.19 -> 2.39
[1:01:50] Using charge balance adjust
[1:01:50] Dispensed 0.013053 mL of Base (0.5 M KOH)
[1:02:11] Stirrer speed set to 0
[1:02:24] Datapoint id 53 collected
[1:02:24] Charge balance equation is out by -6.1%
[1:02:24] Stirrer speed set to 60
[1:02:29] pH 2.38 -> 2.58
[1:02:29] Using charge balance adjust
[1:02:29] Dispensed 0.008984 mL of Base (0.5 M KOH)
[1:02:49] Stirrer speed set to 0
[1:02:59] Datapoint id 54 collected
[1:02:59] Charge balance equation is out by 4.6%
[1:02:59] Stirrer speed set to 60
[1:03:04] pH 2.59 -> 2.79
[1:03:04] Using charge balance adjust
[1:03:05] Dispensed 0.006021 mL of Base (0.5 M KOH)
[1:03:25] Stirrer speed set to 0
[1:03:35] Datapoint id 55 collected
[1:03:35] Charge balance equation is out by -0.8%
[1:03:35] Stirrer speed set to 60
[1:03:40] pH 2.79 -> 2.99
[1:03:40] Using charge balance adjust
[1:03:40] Dispensed 0.004116 mL of Base (0.5 M KOH)
[1:04:00] Stirrer speed set to 0
[1:04:12] Datapoint id 56 collected
[1:04:12] Charge balance equation is out by 9.3%
[1:04:12] Stirrer speed set to 60
[1:04:17] pH 3.02 -> 3.22
[1:04:17] Using charge balance adjust
[1:04:17] Dispensed 0.002658 mL of Base (0.5 M KOH)
[1:04:37] Stirrer speed set to 0
[1:04:47] Datapoint id 57 collected
[1:04:47] Charge balance equation is out by 29.0%
[1:04:47] Stirrer speed set to 60
[1:04:52] pH 3.29 -> 3.49
[1:04:52] Using cautious pH adjust
[1:04:52] Dispensed 0.000776 mL of Base (0.5 M KOH)
[1:04:57] Stepping pH = 3.39
[1:04:58] Dispensed 0.000494 mL of Base (0.5 M KOH)
[1:05:03] Stepping pH = 3.47
[1:05:03] Dispensed 0.000094 mL of Base (0.5 M KOH)
[1:05:08] Stepping pH = 3.48
[1:05:08] Dispensed 0.000094 mL of Base (0.5 M KOH)
[1:05:13] Stepping pH = 3.49
[1:05:28] Stirrer speed set to 0
[1:05:39] Datapoint id 58 collected
[1:05:39] Charge balance equation is out by 4.0%
[1:05:39] Stirrer speed set to 60
[1:05:44] pH 3.50 -> 3.70
[1:05:44] Using charge balance adjust
[1:05:44] Dispensed 0.000988 mL of Base (0.5 M KOH)
[1:06:04] Stirrer speed set to 0
[1:06:14] Datapoint id 59 collected
[1:06:14] Charge balance equation is out by 16.0%
[1:06:14] Stirrer speed set to 60
[1:06:19] pH 3.74 -> 3.94

#### Experiment Log (continued)

[1:06:19] Using cautious pH adjust
[1:06:19] Dispensed 0.000282 mL of Base (0.5 M KOH)
[1:06:24] Stepping pH = 3.79
[1:06:24] Dispensed 0.000400 mL of Base (0.5 M KOH)
[1:06:29] Stepping pH = 3.99
[1:06:45] Stirrer speed set to 0
[1:06:55] Datapoint id 60 collected
[1:06:55] Charge balance equation is out by -17.6%
[1:06:55] Stirrer speed set to 60
[1:07:00] pH 4.01 -> 4.21
[1:07:00] Using cautious pH adjust
[1:07:00] Dispensed 0.000165 mL of Base (0.5 M KOH)
[1:07:05] Stepping pH = 4.07
[1:07:05] Dispensed 0.000188 mL of Base (0.5 M KOH)
[1:07:11] Stepping pH = 4.21
[1:07:26] Stirrer speed set to 0
[1:07:36] Datapoint id 61 collected
[1:07:36] Charge balance equation is out by -10.9%
[1:07:36] Stirrer speed set to 60
[1:07:41] pH 4.22 -> 4.42
[1:07:41] Using charge balance adjust
[1:07:41] Dispensed 0.000188 mL of Base (0.5 M KOH)
[1:08:01] Stirrer speed set to 0
[1:08:15] Datapoint id 62 collected
[1:08:15] Charge balance equation is out by 8.4%
[1:08:15] Stirrer speed set to 60
[1:08:20] pH 4.47 -> 4.67
[1:08:20] Using charge balance adjust
[1:08:20] Dispensed 0.000118 mL of Base (0.5 M KOH)
[1:08:40] Stirrer speed set to 0
[1:08:52] Datapoint id 63 collected
[1:08:52] Charge balance equation is out by -24.3%
[1:08:52] Stirrer speed set to 60
[1:08:57] pH 4.65 -> 4.85
[1:08:57] Using cautious pH adjust
[1:08:57] Dispensed 0.000047 mL of Base (0.5 M KOH)
[1:09:02] Stepping pH = 4.69
[1:09:02] Dispensed 0.000071 mL of Base (0.5 M KOH)
[1:09:07] Stepping pH = 4.89
[1:09:22] Stirrer speed set to 0
[1:09:36] Datapoint id 64 collected
[1:09:36] Charge balance equation is out by -31.4%
[1:09:36] Stirrer speed set to 60
[1:09:41] pH 4.94 -> 5.14
[1:09:41] Using cautious pH adjust
[1:09:41] Dispensed 0.000024 mL of Base (0.5 M KOH)
[1:09:46] Stepping pH = 4.94
[1:09:46] Dispensed 0.000118 mL of Base (0.5 M KOH)
[1:09:51] Stepping pH = 5.39
[1:10:07] Stirrer speed set to 0
[1:10:54] Datapoint id 65 collected
[1:10:54] Charge balance equation is out by -203.1%
[1:10:54] Stirrer speed set to 60
[1:10:59] pH 5.56 -> 5.76
[1:10:59] Using cautious pH adjust
[1:10:59] Dispensed 0.000024 mL of Base (0.5 M KOH)
[1:11:04] Stepping pH = 5.59
[1:11:04] Dispensed 0.000047 mL of Base (0.5 M KOH)
[1:11:10] Stepping pH = 5.78

| | | | |
|---|---|---|---|
| Sample name: | **M02_octanol** | Experiment start time: | **3/3/2018 3:08:32 PM** |
| Assay name: | **pH-metric high logP** | Analyst: | **Dorothy Levorse** |
| Assay ID: | **18C-03012** | Instrument ID: | **T312060** |
| Filename: | **C:\Sirius_T3\Mehtap\20180302_exp29_logP_T3-2\18C-03012_M02_octanol_pH-metric high logP.t3r** | | |

#### Experiment Log (continued)

[1:11:25] Stirrer speed set to 0
[1:12:08] Datapoint id 66 collected
[1:12:08] Charge balance equation is out by -75.9%
[1:12:08] Stirrer speed set to 60
[1:12:13] pH 6.06 -> 6.26
[1:12:13] Using cautious pH adjust
[1:12:13] Dispensed 0.000024 mL of Base (0.5 M KOH)
[1:12:18] Stepping pH = 6.18
[1:12:18] Dispensed 0.000024 mL of Base (0.5 M KOH)
[1:12:23] Stepping pH = 6.42
[1:12:38] Stirrer speed set to 0
[1:13:38] Datapoint id 67 collected
[1:13:38] Charge balance equation is out by 2.6%
[1:13:38] Stirrer speed set to 60
[1:13:43] pH 6.36 -> 6.56
[1:13:43] Using charge balance adjust
[1:13:44] Dispensed 0.000047 mL of Base (0.5 M KOH)
[1:14:04] Stirrer speed set to 0
[1:15:04] Datapoint id 68 collected
[1:15:04] Charge balance equation is out by 56.6%
[1:15:04] Stirrer speed set to 60
[1:15:09] pH 6.79 -> 6.99
[1:15:09] Using cautious pH adjust
[1:15:09] Dispensed 0.000024 mL of Base (0.5 M KOH)
[1:15:14] Stepping pH = 6.87
[1:15:14] Dispensed 0.000024 mL of Base (0.5 M KOH)
[1:15:19] Stepping pH = 6.91
[1:15:19] Dispensed 0.000024 mL of Base (0.5 M KOH)
[1:15:24] Stepping pH = 7.04
[1:15:39] Stirrer speed set to 0
[1:16:39] Datapoint id 69 collected
[1:16:39] Charge balance equation is out by -89.8%
[1:16:39] Stirrer speed set to 60
[1:16:44] pH 7.20 -> 7.40
[1:16:44] Using cautious pH adjust
[1:16:44] Dispensed 0.000024 mL of Base (0.5 M KOH)
[1:16:50] Stepping pH = 7.19
[1:16:50] Dispensed 0.000047 mL of Base (0.5 M KOH)
[1:16:55] Stepping pH = 7.44
[1:17:10] Stirrer speed set to 0
[1:18:10] Datapoint id 70 collected
[1:18:10] Charge balance equation is out by -272.1%
[1:18:10] Stirrer speed set to 60
[1:18:15] pH 7.69 -> 7.89
[1:18:15] Using cautious pH adjust
[1:18:15] Dispensed 0.000024 mL of Base (0.5 M KOH)
[1:18:20] Stepping pH = 7.81
[1:18:20] Dispensed 0.000024 mL of Base (0.5 M KOH)
[1:18:25] Stepping pH = 8.01
[1:18:40] Stirrer speed set to 0
[1:19:40] Datapoint id 71 collected
[1:19:40] Charge balance equation is out by -351.4%
[1:19:40] Stirrer speed set to 60
[1:19:45] pH 8.10 -> 8.30
[1:19:45] Using cautious pH adjust
[1:19:45] Dispensed 0.000024 mL of Base (0.5 M KOH)
[1:19:51] Stepping pH = 8.16
[1:19:51] Dispensed 0.000024 mL of Base (0.5 M KOH)
[1:19:56] Stepping pH = 8.26

Sample name: **M02_octanol**
Assay name: **pH-metric high logP**
Assay ID: **18C-03012**
Filename: **C:\Sirius_T3\Mehtap\20180302_exp29_logP_T3-2\18C-03012_M02_octanol_pH-metric high logP.t3r**
Experiment start time: **3/3/2018 3:08:32 PM**
Analyst: **Dorothy Levorse**
Instrument ID: **T312060**

#### Experiment Log (continued)

[1:19:56] Dispensed 0.000024 mL of Base (0.5 M KOH)
[1:20:01] Stepping pH = 8.39
[1:20:16] Stirrer speed set to 0
[1:21:16] Datapoint id 72 collected
[1:21:16] Charge balance equation is out by -565.1%
[1:21:16] Stirrer speed set to 60
[1:21:21] pH 8.46 -> 8.66
[1:21:21] Using cautious pH adjust
[1:21:21] Dispensed 0.000024 mL of Base (0.5 M KOH)
[1:21:26] Stepping pH = 8.52
[1:21:26] Dispensed 0.000024 mL of Base (0.5 M KOH)
[1:21:31] Stepping pH = 8.61
[1:21:31] Dispensed 0.000024 mL of Base (0.5 M KOH)
[1:21:36] Stepping pH = 8.71
[1:21:52] Stirrer speed set to 0
[1:22:25] Datapoint id 73 collected
[1:22:25] Charge balance equation is out by -283.1%
[1:22:25] Stirrer speed set to 60
[1:22:30] pH 8.76 -> 8.96
[1:22:30] Using cautious pH adjust
[1:22:30] Dispensed 0.000024 mL of Base (0.5 M KOH)
[1:22:35] Stepping pH = 8.80
[1:22:35] Dispensed 0.000024 mL of Base (0.5 M KOH)
[1:22:41] Stepping pH = 8.85
[1:22:41] Dispensed 0.000047 mL of Base (0.5 M KOH)
[1:22:46] Stepping pH = 8.91
[1:22:46] Dispensed 0.000024 mL of Base (0.5 M KOH)
[1:22:51] Stepping pH = 8.98
[1:23:06] Stirrer speed set to 0
[1:23:30] Datapoint id 74 collected
[1:23:30] Charge balance equation is out by -271.8%
[1:23:30] Stirrer speed set to 60
[1:23:36] pH 8.99 -> 9.05
[1:23:36] Using cautious pH adjust
[1:23:36] Dispensed 0.000024 mL of Base (0.5 M KOH)
[1:23:41] Stepping pH = 9.01
[1:23:56] Stirrer speed set to 0
[1:24:09] Datapoint id 75 collected
[1:24:09] Charge balance equation is out by -70.9%
[1:24:09] Argon flow rate set to 0
[1:24:13] Titrator arm moved over Titration position
