## Supplementary material for "Octanol-water partition coefficient measurements for the SAMPL6 Blind Prediction Challenge": SM02_18C-06015_M02_octanol_pH-metric high logP_report.pdf

| | | | |
|---|---|---|---|
| Sample name: | **M02_octanol** | Experiment start time: | **3/6/2018 5:41:59 PM** |
| Assay name: | **pH-metric high logP** | Analyst: | **Dorothy Levorse** |
| Assay ID: | **18C-06015** | Instrument ID: | **T312060** |
| Filename: | **C:\Sirius_T3\Mehtap\20180306_exp30_logP_T3-2\18C-06015_M02_octanol_pH-metric high logP.t3r** | | |

#### pH-metric Result

| | | |
|---|---|---|
| logP (XH +) | 1.14 | ±0.04 (n=50) |
| logP (neutral X) | 4.16 | ±0.01 (n=50) |

#### 18C-06015 Points 1 to 20

| | |
|---|---|
| M02_octanol concentration factor | 0.918 |
| Carbonate | 0.0000 mM |
| Acidity error | -0.42538 mM |

#### 18C-06015 Points 21 to 34

| | |
|---|---|
| M02_octanol concentration factor | 0.898 |
| Carbonate | 0.0001 mM |
| Acidity error | -0.16274 mM |

#### 18C-06015 Points 35 to 48

| | |
|---|---|
| M02_octanol concentration factor | 0.730 |
| Carbonate | 0.1987 mM |
| Acidity error | -0.19134 mM |

#### Warnings and errors

| | |
|---|---|
| Errors | None |
| Warnings | None |

#### Sample logD and percent species

| pH | M02_octanol logD | M02_octanol M02_octanolH | M02_octanol M02_octanol | M02_octanol M02_octanolH* | M02_octanol M02_octanol* | Comment |
|---|---|---|---|---|---|---|
| 1.000 | 1.18 | 6.22 % | 0.00 % | 85.42 % | 8.36 % | |
| 1.200 | 1.20 | 5.93 % | 0.00 % | 81.44 % | 12.64 % | Stomach pH |
| 2.000 | 1.43 | 3.55 % | 0.00 % | 48.74 % | 47.71 % | |
| 3.000 | 2.17 | 0.67 % | 0.01 % | 9.21 % | 90.12 % | |
| 4.000 | 3.09 | 0.07 % | 0.01 % | 1.01 % | 98.91 % | |
| 5.000 | 3.84 | 0.01 % | 0.01 % | 0.10 % | 99.88 % | |
| 6.000 | 4.11 | 0.00 % | 0.01 % | 0.01 % | 99.98 % | |
| 6.500 | 4.14 | 0.00 % | 0.01 % | 0.00 % | 99.99 % | |
| 7.000 | 4.15 | 0.00 % | 0.01 % | 0.00 % | 99.99 % | |
| 7.400 | 4.16 | 0.00 % | 0.01 % | 0.00 % | 99.99 % | Blood pH |
| 8.000 | 4.16 | 0.00 % | 0.01 % | 0.00 % | 99.99 % | |
| 9.000 | 4.16 | 0.00 % | 0.01 % | 0.00 % | 99.99 % | |
| 10.000 | 4.16 | 0.00 % | 0.01 % | 0.00 % | 99.99 % | |
| 11.000 | 4.16 | 0.00 % | 0.01 % | 0.00 % | 99.99 % | |
| 12.000 | 4.16 | 0.00 % | 0.01 % | 0.00 % | 99.99 % | |

| | | | |
|---|---|---|---|
| Sample name: | **M02_octanol** | Experiment start time: | **3/6/2018 5:41:59 PM** |
| Assay name: | **pH-metric high logP** | Analyst: | **Dorothy Levorse** |
| Assay ID: | **18C-06015** | Instrument ID: | **T312060** |
| Filename: | **C:\Sirius_T3\Mehtap\20180306_exp30_logP_T3-2\18C-06015_M02_octanol_pH-metric high logP.t3r** | | |

#### Graphs

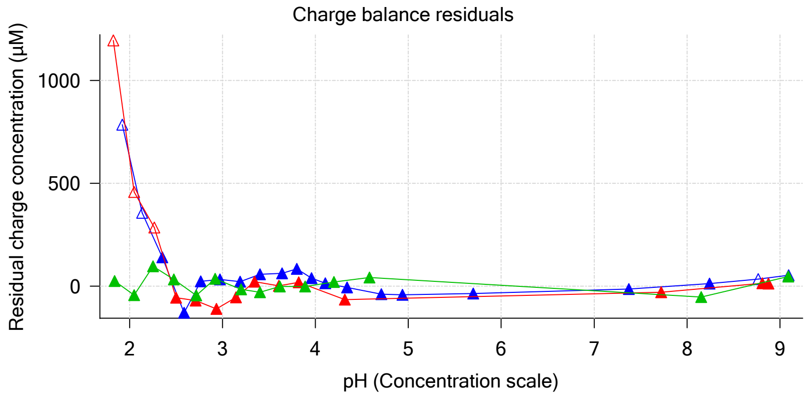


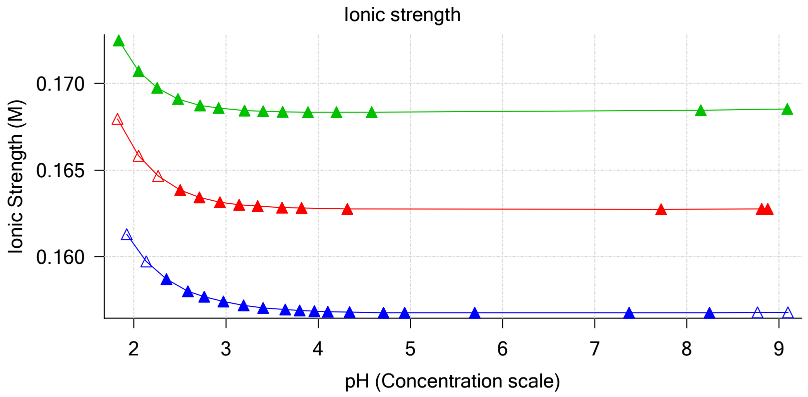


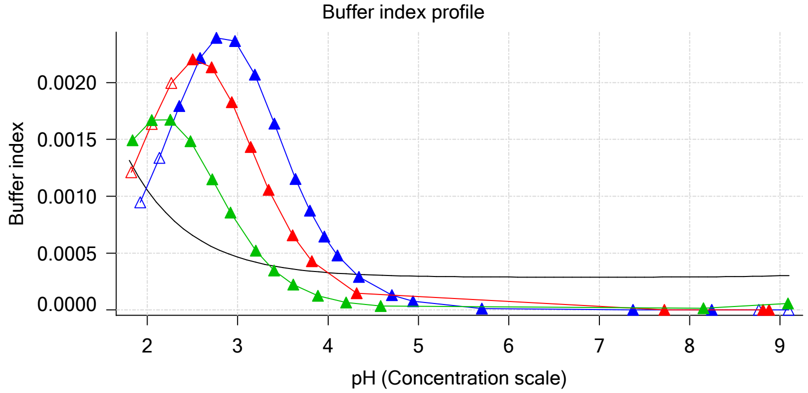


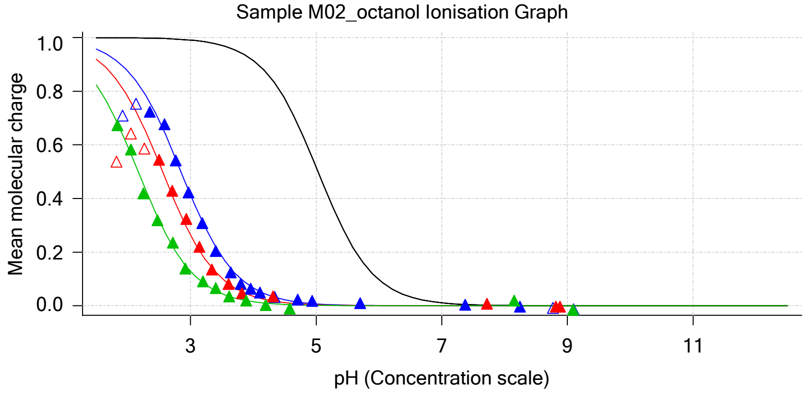


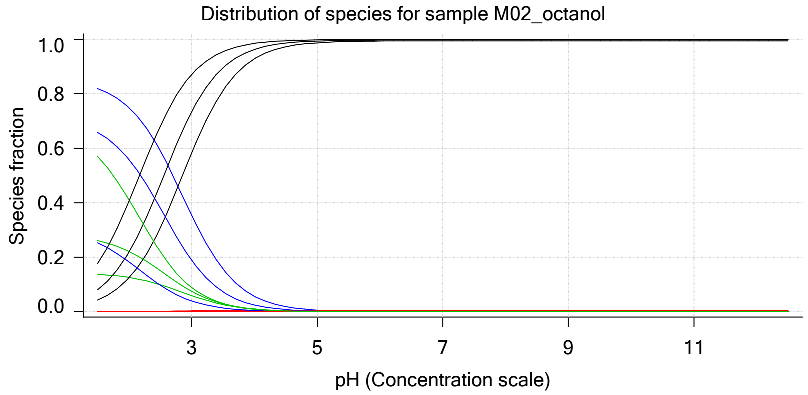


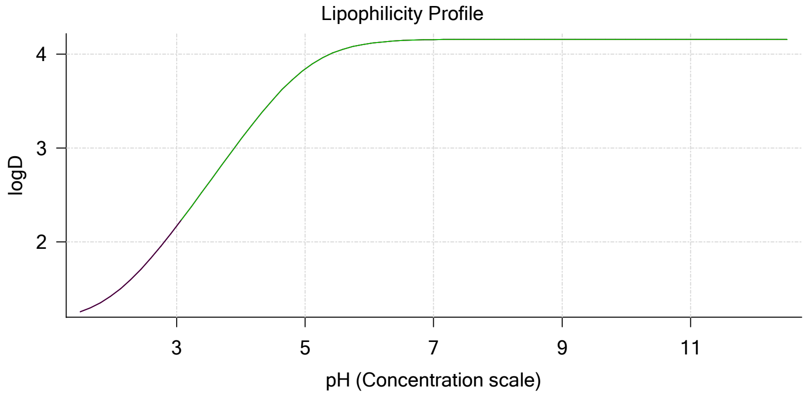


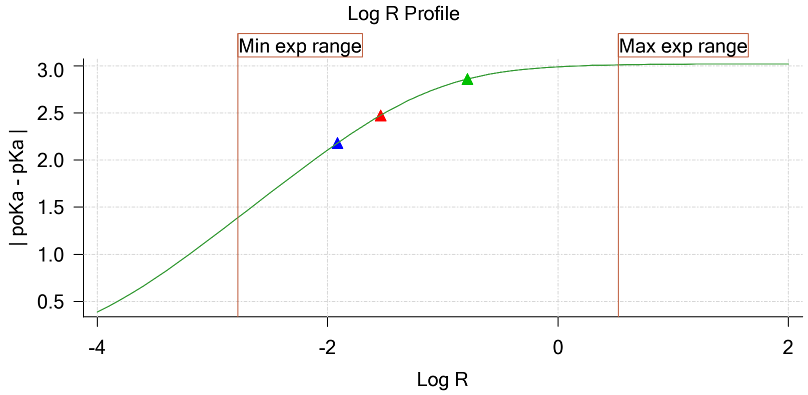


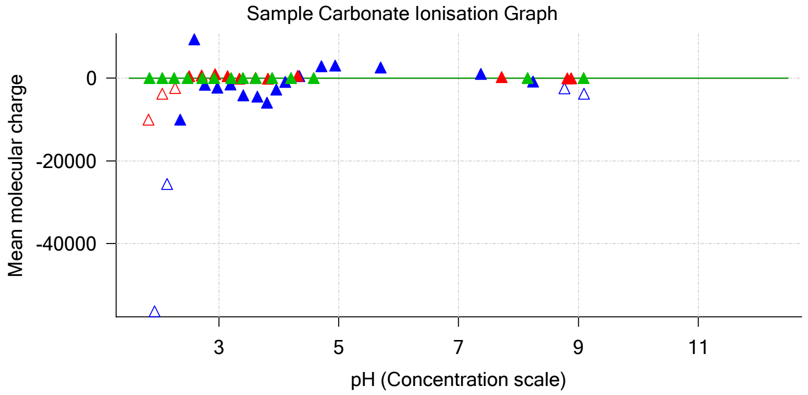

Sample name: **M02_octanol**
Assay name: **pH-metric high logP**
Assay ID: **18C-06015**
Filename: **C:\Sirius_T3\Mehtap\20180306_exp30_logP_T3-2\18C-06015_M02_octanol_pH-metric high logP.t3r**

Experiment start time: **3/6/2018 5:41:59 PM**
Analyst: **Dorothy Levorse**
Instrument ID: **T312060**

#### Graphs (continued)

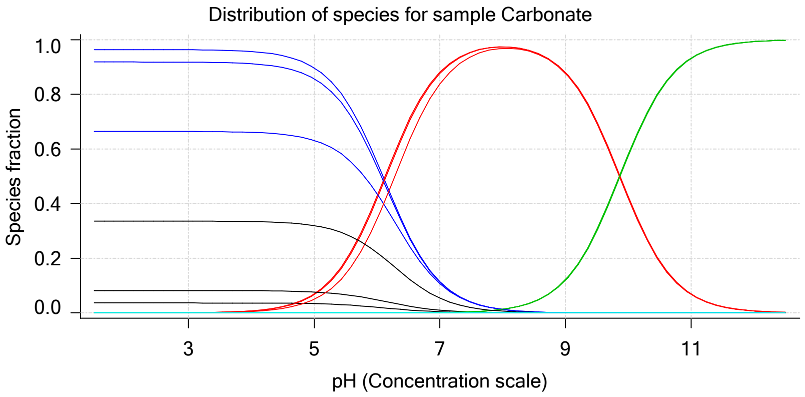

| | | | |
|---|---|---|---|
| Sample name: | **M02_octanol** | Experiment start time: | **3/6/2018 5:41:59 PM** |
| Assay name: | **pH-metric high logP** | Analyst: | **Dorothy Levorse** |
| Assay ID: | **18C-06015** | Instrument ID: | **T312060** |
| Filename: | **C:\Sirius_T3\Mehtap\20180306_exp30_logP_T3-2\18C-06015_M02_octanol_pH-metric high logP.t3r** | | |

### pH-metric high logP Titration 1 of 3 18C-06015 Points 1 to 20

#### Overall results

| | |
|---|---|
| RMSD | 2.836 |
| Average ionic strength | 0.157 M |
| Average temperature | 24.9°C |
| Partition ratio | 0.0122 : 1 |
| Analyte concentration range | 4481.9 µM to 4615.4 µM |
| Total points considered | 16 of 20 |

#### Warnings and errors

| | |
|---|---|
| Errors | None |
| Warnings | None |

#### Four-Plus parameters

| | | | |
|---|---|---|---|
| Alpha | 0.124 | 3/6/2018 5:41:59 PM | C:\Sirius_T3\18C-06006_Blank standardisation.t3r |
| S | 0.9973 | 3/6/2018 5:41:59 PM | C:\Sirius_T3\18C-06006_Blank standardisation.t3r |
| jH | 0.9 | 3/6/2018 5:41:59 PM | C:\Sirius_T3\18C-06006_Blank standardisation.t3r |
| jOH | -0.7 | 3/6/2018 5:41:59 PM | C:\Sirius_T3\18C-06006_Blank standardisation.t3r |

#### Titrants

| | | | |
|---|---|---|---|
| 0.50 M HCl | 0.989131 | 3/6/2018 5:41:59 PM | C:\Sirius_T3\18C-06006_Blank standardisation.t3r |
| 0.50 M KOH | 0.999845 | 3/6/2018 5:41:59 PM | C:\Sirius_T3\KOH18B27.t3r |

#### Sample

| | |
|---|---|
| M02_octanol concentration factor | 0.918 |
| Base pKa 1 | 5.03 |
| logP (XH +) | 1.32 |
| logP (neutral X) | 4.20 |

#### Sample graphs

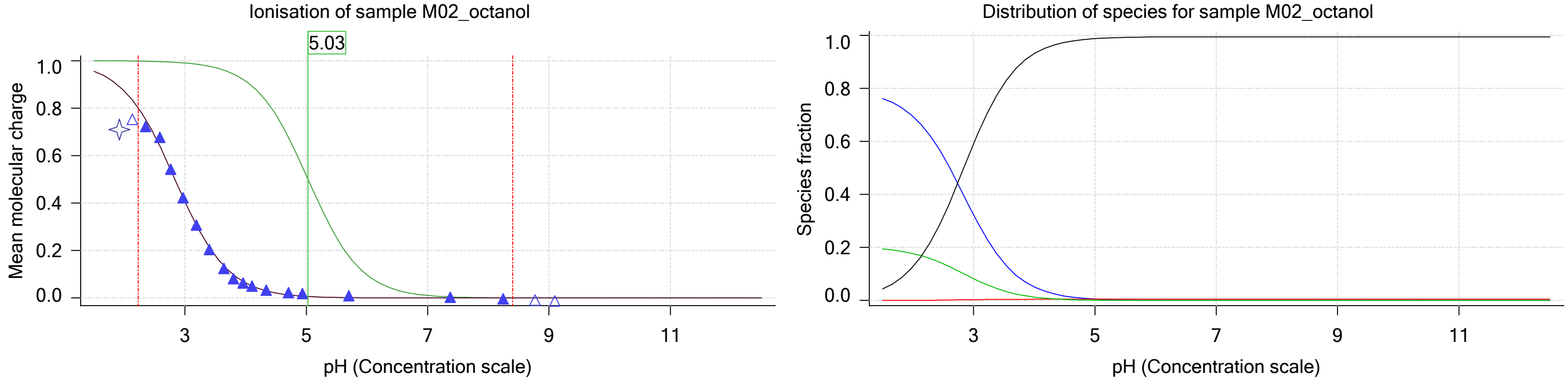

| | | | |
|---|---|---|---|
| Sample name: | **M02_octanol** | Experiment start time: | **3/6/2018 5:41:59 PM** |
| Assay name: | **pH-metric high logP** | Analyst: | **Dorothy Levorse** |
| Assay ID: | **18C-06015** | Instrument ID: | **T312060** |
| Filename: | **C:\Sirius_T3\Mehtap\20180306_exp30_logP_T3-2\18C-06015_M02_octanol_pH-metric high logP.t3r** | | |

#### Sample graphs (continued)

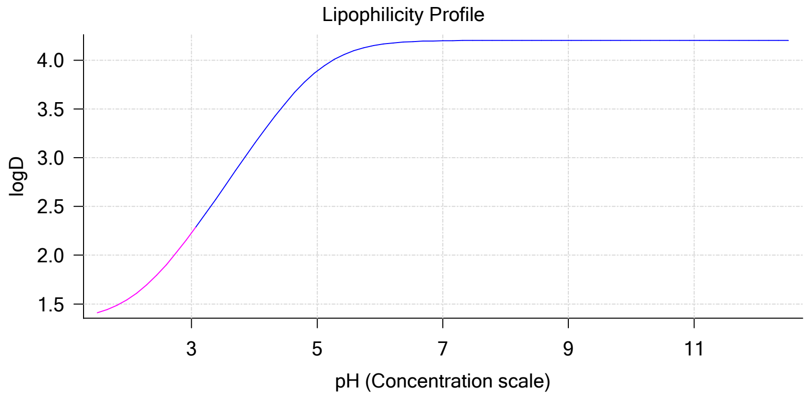


#### Sample logD and percent species

| pH | M02_octanol logD | M02_octanol M02_octanolH | M02_octanol M02_octanol | M02_octanol M02_octanolH* | M02_octanol M02_octanol* | Comment |
|---|---|---|---|---|---|---|
| 1.000 | 1.35 | 78.52 % | 0.01 % | 20.04 % | 1.43 % | |
| 1.200 | 1.37 | 77.87 % | 0.01 % | 19.87 % | 2.25 % | Stomach pH |
| 2.000 | 1.55 | 69.52 % | 0.06 % | 17.74 % | 12.68 % | |
| 3.000 | 2.23 | 32.38 % | 0.30 % | 8.26 % | 59.05 % | |
| 4.000 | 3.14 | 5.11 % | 0.48 % | 1.30 % | 93.11 % | |
| 5.000 | 3.89 | 0.54 % | 0.51 % | 0.14 % | 98.81 % | |
| 6.000 | 4.16 | 0.05 % | 0.51 % | 0.01 % | 99.42 % | |
| 6.500 | 4.19 | 0.02 % | 0.51 % | 0.00 % | 99.47 % | |
| 7.000 | 4.20 | 0.01 % | 0.51 % | 0.00 % | 99.48 % | |
| 7.400 | 4.20 | 0.00 % | 0.51 % | 0.00 % | 99.49 % | Blood pH |
| 8.000 | 4.20 | 0.00 % | 0.51 % | 0.00 % | 99.49 % | |
| 9.000 | 4.20 | 0.00 % | 0.51 % | 0.00 % | 99.49 % | |
| 10.000 | 4.20 | 0.00 % | 0.51 % | 0.00 % | 99.49 % | |
| 11.000 | 4.20 | 0.00 % | 0.51 % | 0.00 % | 99.49 % | |
| 12.000 | 4.20 | 0.00 % | 0.51 % | 0.00 % | 99.49 % | |

#### Carbonate and acidity

| | |
|---|---|
| Carbonate | 0.000 mM |
| Acidity error | -0.425 mM |

#### Other graphs

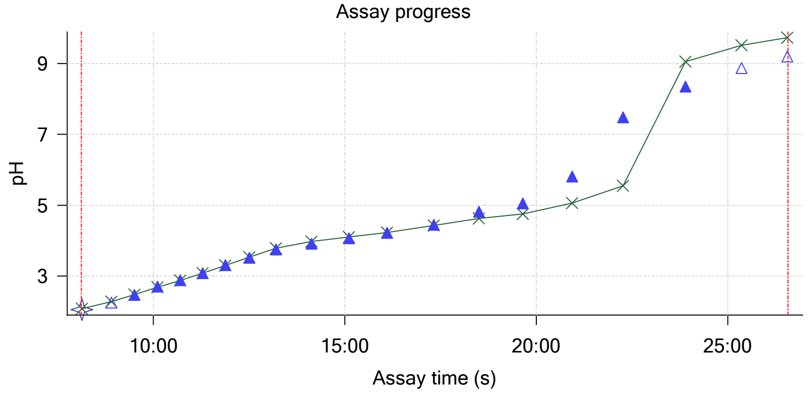


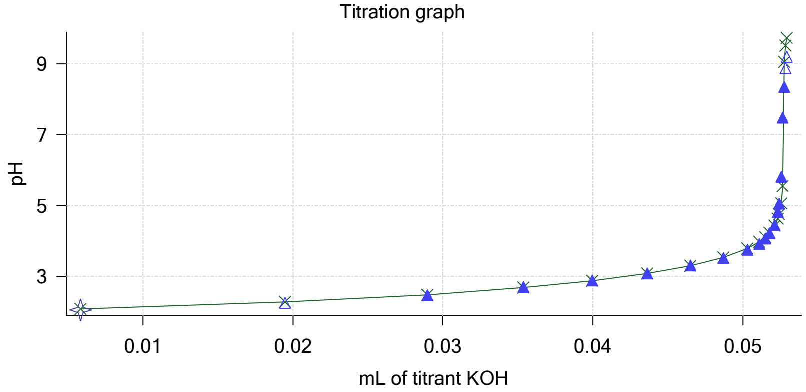

Sample name: **M02_octanol**
Assay name: **pH-metric high logP**
Assay ID: **18C-06015**
Filename: **C:\Sirius_T3\Mehtap\20180306_exp30_logP_T3-2\18C-06015_M02_octanol_pH-metric high logP.t3r**

Experiment start time: **3/6/2018 5:41:59 PM**
Analyst: **Dorothy Levorse**
Instrument ID: **T312060**

#### Other graphs (continued)

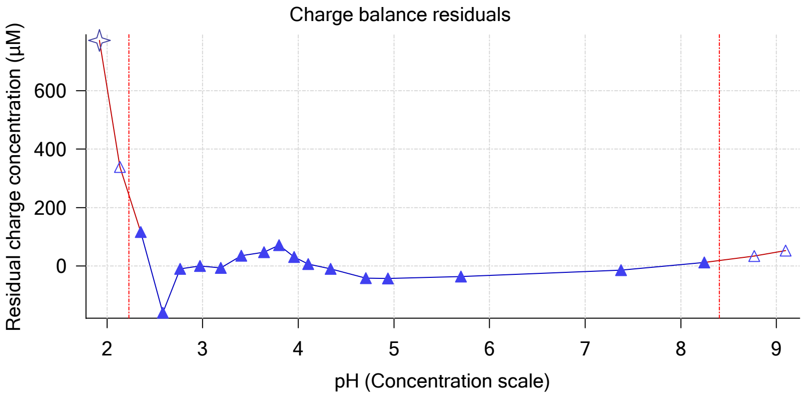


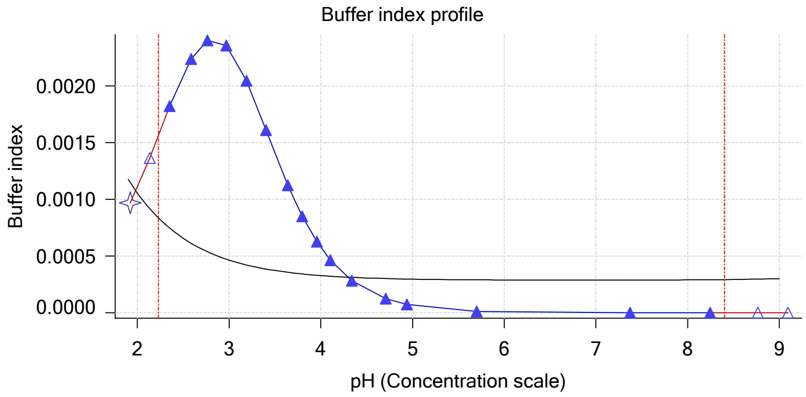


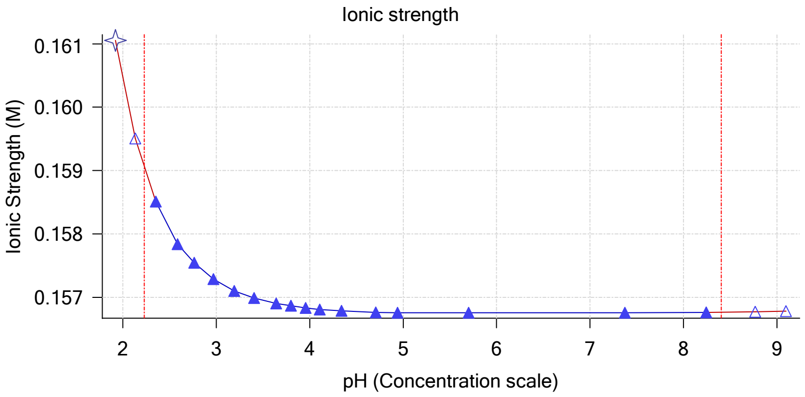


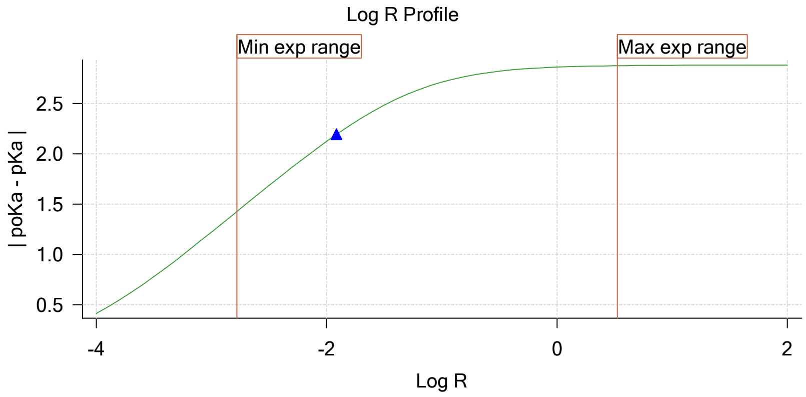


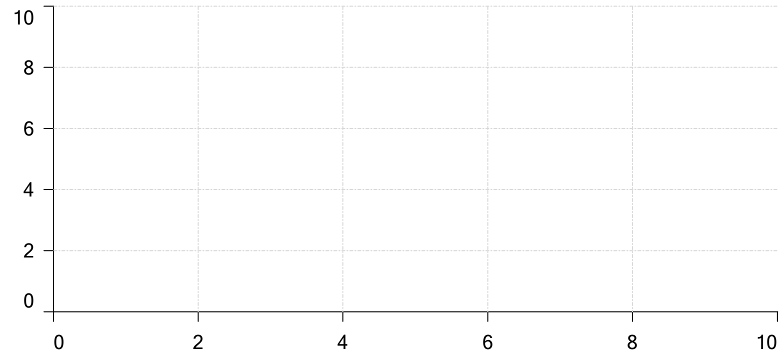


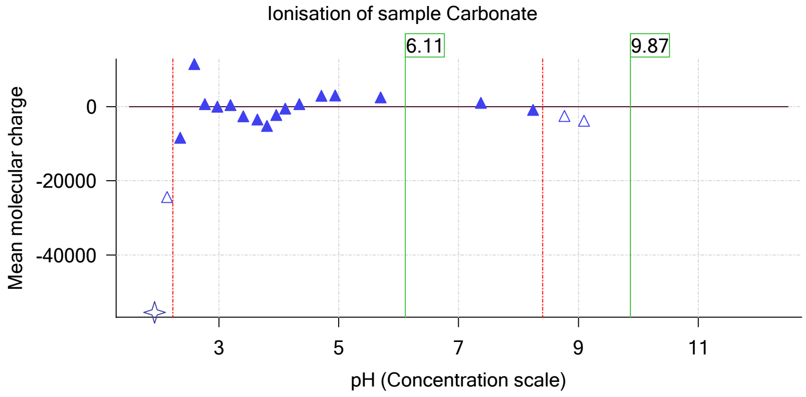

| | | | |
|---|---|---|---|
| Sample name: | **M02_octanol** | Experiment start time: | **3/6/2018 5:41:59 PM** |
| Assay name: | **pH-metric high logP** | Analyst: | **Dorothy Levorse** |
| Assay ID: | **18C-06015** | Instrument ID: | **T312060** |
| Filename: | **C:\Sirius_T3\Mehtap\20180306_exp30_logP_T3-2\18C-06015_M02_octanol_pH-metric high logP.t3r** | | |

# pH-metric high logP   Titration 2 of 3   18C-06015 Points 21 to 34

#### Overall results

| | |
|---|---|
| RMSD | 4.143 |
| Average ionic strength | 0.163 M |
| Average temperature | 25.0°C |
| Partition ratio | 0.0289 : 1 |
| Analyte concentration range | 4116.8 µM to 4254.1 µM |
| Total points considered | 11 of 14 |

#### Warnings and errors

| | |
|---|---|
| Errors | None |
| Warnings | None |

#### Four-Plus parameters

| | | | |
|---|---|---|---|
| Alpha | 0.124 | 3/6/2018 5:41:59 PM | C:\Sirius_T3\18C-06006_Blank standardisation.t3r |
| S | 0.9973 | 3/6/2018 5:41:59 PM | C:\Sirius_T3\18C-06006_Blank standardisation.t3r |
| jH | 0.9 | 3/6/2018 5:41:59 PM | C:\Sirius_T3\18C-06006_Blank standardisation.t3r |
| jOH | -0.7 | 3/6/2018 5:41:59 PM | C:\Sirius_T3\18C-06006_Blank standardisation.t3r |

#### Titrants

| | | | |
|---|---|---|---|
| 0.50 M HCl | 0.989131 | 3/6/2018 5:41:59 PM | C:\Sirius_T3\18C-06006_Blank standardisation.t3r |
| 0.50 M KOH | 0.999845 | 3/6/2018 5:41:59 PM | C:\Sirius_T3\KOH18B27.t3r |

#### Sample

| | |
|---|---|
| M02_octanol concentration factor | 0.898 |
| Base pKa 1 | 5.03 |
| logP (XH +) | 1.32 |
| logP (neutral X) | 4.19 |

#### Sample graphs

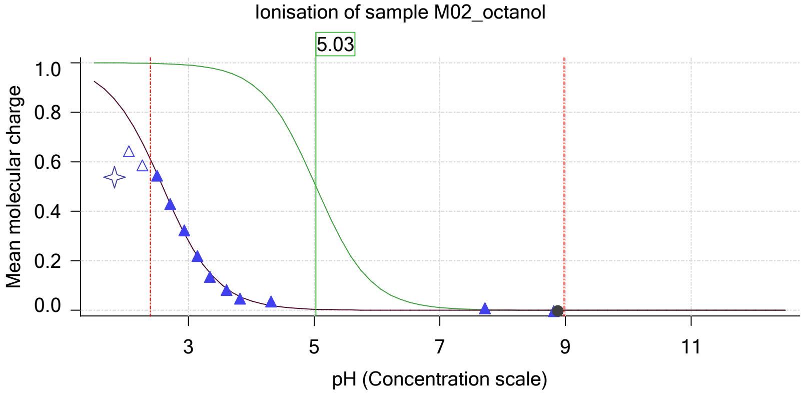


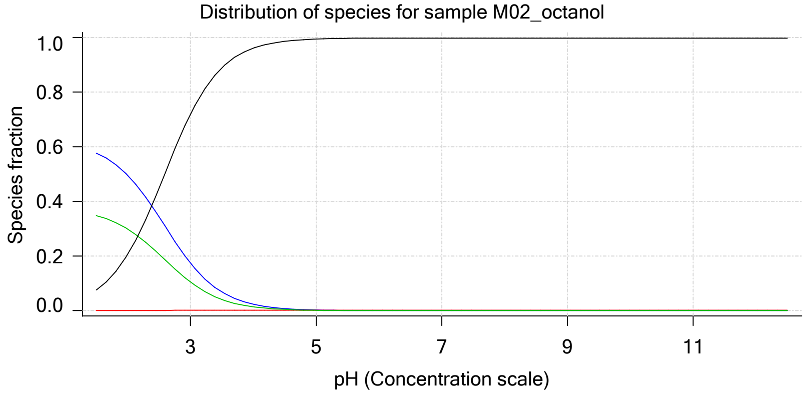

| | | | |
|---|---|---|---|
| Sample name: | **M02_octanol** | Experiment start time: | **3/6/2018 5:41:59 PM** |
| Assay name: | **pH-metric high logP** | Analyst: | **Dorothy Levorse** |
| Assay ID: | **18C-06015** | Instrument ID: | **T312060** |
| Filename: | **C:\Sirius_T3\Mehtap\20180306_exp30_logP_T3-2\18C-06015_M02_octanol_pH-metric high logP.t3r** | | |

#### Sample graphs (continued)

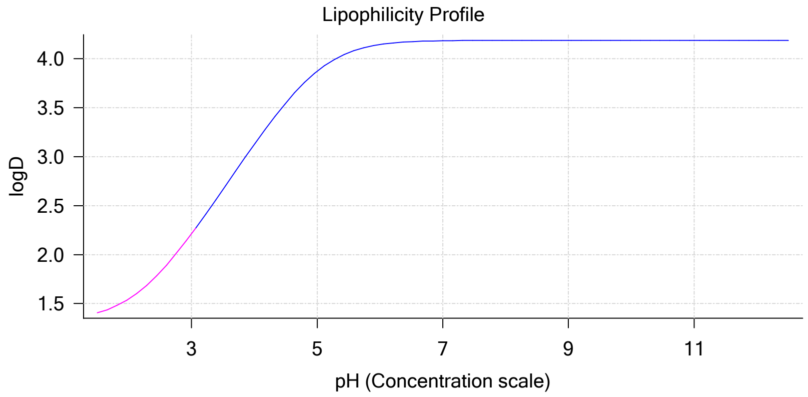


#### Sample logD and percent species

| pH | M02_octanol logD | M02_octanol M02_octanolH | M02_octanol M02_octanol | M02_octanol M02_octanolH* | M02_octanol M02_octanol* | Comment |
|---|---|---|---|---|---|---|
| 1.000 | 1.35 | 60.82 % | 0.01 % | 36.66 % | 2.52 % | |
| 1.200 | 1.36 | 59.93 % | 0.01 % | 36.13 % | 3.93 % | Stomach pH |
| 2.000 | 1.55 | 49.56 % | 0.05 % | 29.87 % | 20.52 % | |
| 3.000 | 2.21 | 17.38 % | 0.16 % | 10.48 % | 71.97 % | |
| 4.000 | 3.12 | 2.32 % | 0.22 % | 1.40 % | 96.06 % | |
| 5.000 | 3.87 | 0.24 % | 0.22 % | 0.14 % | 99.39 % | |
| 6.000 | 4.14 | 0.02 % | 0.22 % | 0.01 % | 99.74 % | |
| 6.500 | 4.17 | 0.01 % | 0.22 % | 0.00 % | 99.76 % | |
| 7.000 | 4.18 | 0.00 % | 0.22 % | 0.00 % | 99.77 % | |
| 7.400 | 4.19 | 0.00 % | 0.22 % | 0.00 % | 99.77 % | Blood pH |
| 8.000 | 4.19 | 0.00 % | 0.22 % | 0.00 % | 99.77 % | |
| 9.000 | 4.19 | 0.00 % | 0.22 % | 0.00 % | 99.78 % | |
| 10.000 | 4.19 | 0.00 % | 0.22 % | 0.00 % | 99.78 % | |
| 11.000 | 4.19 | 0.00 % | 0.22 % | 0.00 % | 99.78 % | |
| 12.000 | 4.19 | 0.00 % | 0.22 % | 0.00 % | 99.78 % | |

#### Carbonate and acidity

| | |
|---|---|
| Carbonate | 0.000 mM |
| Acidity error | -0.163 mM |

#### Other graphs

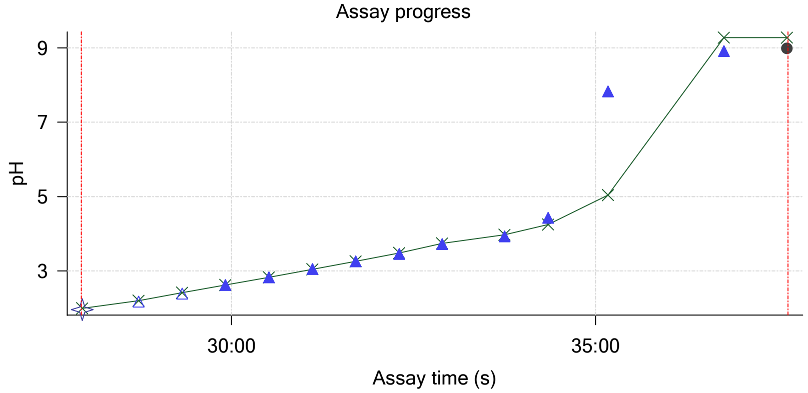


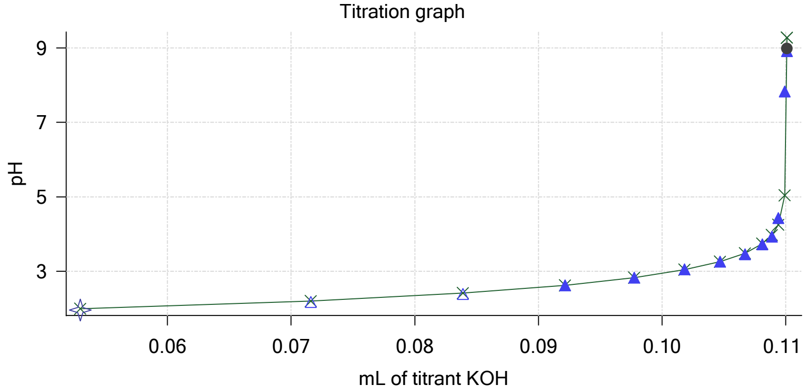

Sample name: **M02_octanol**
Assay name: **pH-metric high logP**
Assay ID: **18C-06015**
Filename: **C:\Sirius_T3\Mehtap\20180306_exp30_logP_T3-2\18C-06015_M02_octanol_pH-metric high logP.t3r**

Experiment start time: **3/6/2018 5:41:59 PM**
Analyst: **Dorothy Levorse**
Instrument ID: **T312060**

#### Other graphs (continued)

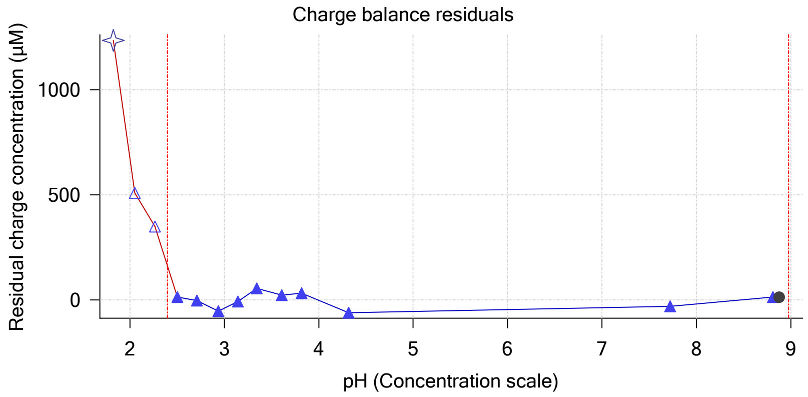


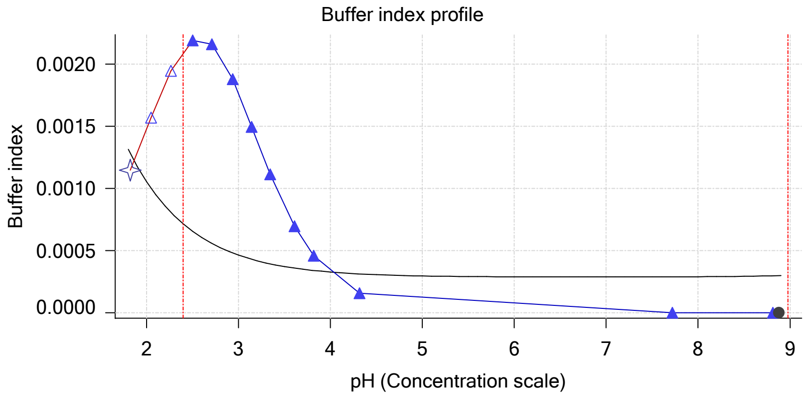


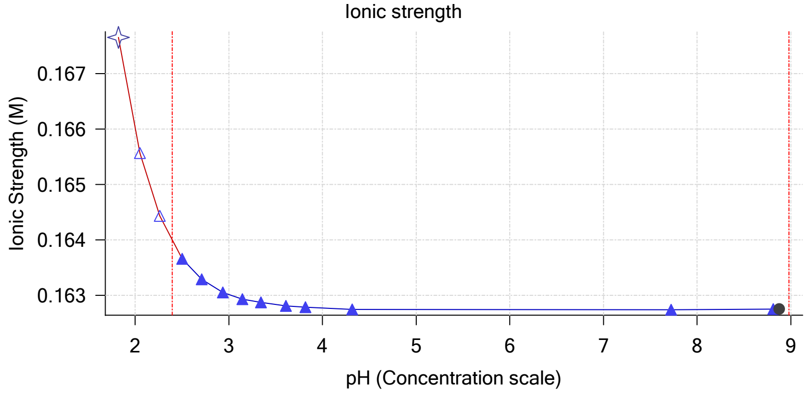


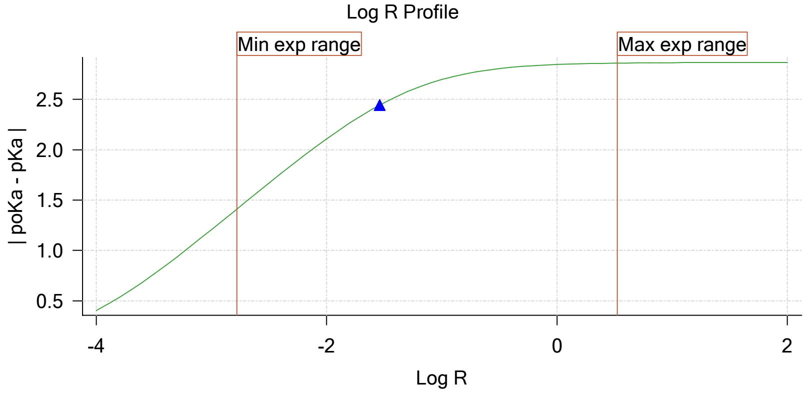


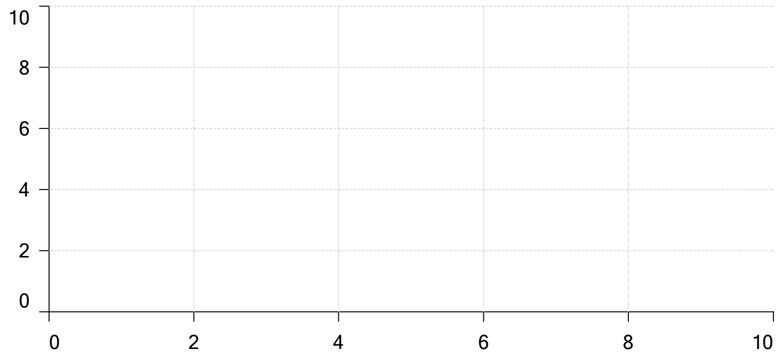


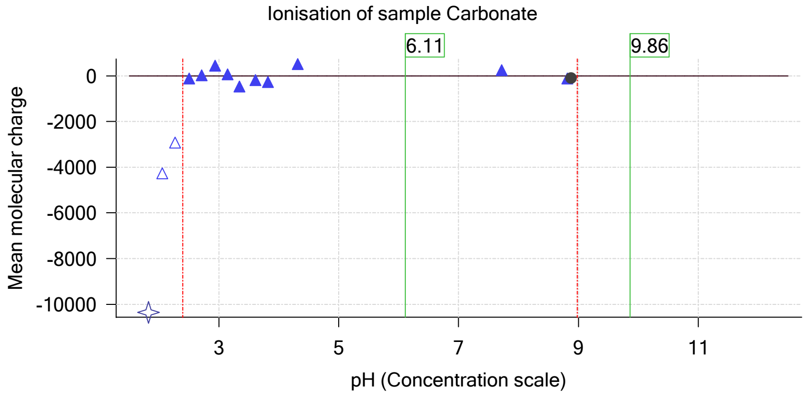

| | | | |
|---|---|---|---|
| Sample name: | **M02_octanol** | Experiment start time: | **3/6/2018 5:41:59 PM** |
| Assay name: | **pH-metric high logP** | Analyst: | **Dorothy Levorse** |
| Assay ID: | **18C-06015** | Instrument ID: | **T312060** |
| Filename: | **C:\Sirius_T3\Mehtap\20180306_exp30_logP_T3-2\18C-06015_M02_octanol_pH-metric high logP.t3r** | | |

### pH-metric high logP Titration 3 of 3 18C-06015 Points 35 to 48

#### Overall results

| | |
|---|---|
| RMSD | 0.729 |
| Average ionic strength | 0.169 M |
| Average temperature | 25.0°C |
| Partition ratio | 0.1637 : 1 |
| Analyte concentration range | 3405.3 μM to 3503.8 μM |
| Total points considered | 14 of 14 |

#### Warnings and errors

| | |
|---|---|
| Errors | None |
| Warnings | None |

#### Four-Plus parameters

| | | | |
|---|---|---|---|
| Alpha | 0.124 | 3/6/2018 5:41:59 PM | C:\Sirius_T3\18C-06006_Blank standardisation.t3r |
| S | 0.9973 | 3/6/2018 5:41:59 PM | C:\Sirius_T3\18C-06006_Blank standardisation.t3r |
| jH | 0.9 | 3/6/2018 5:41:59 PM | C:\Sirius_T3\18C-06006_Blank standardisation.t3r |
| jOH | -0.7 | 3/6/2018 5:41:59 PM | C:\Sirius_T3\18C-06006_Blank standardisation.t3r |

#### Titrants

| | | | |
|---|---|---|---|
| 0.50 M HCl | 0.989131 | 3/6/2018 5:41:59 PM | C:\Sirius_T3\18C-06006_Blank standardisation.t3r |
| 0.50 M KOH | 0.999845 | 3/6/2018 5:41:59 PM | C:\Sirius_T3\KOH18B27.t3r |

#### Sample

| | |
|---|---|
| M02_octanol concentration factor | 0.730 |
| Base pKa 1 | 5.03 |
| logP (XH +) | 1.32 |
| logP (neutral X) | 4.31 |

#### Sample graphs

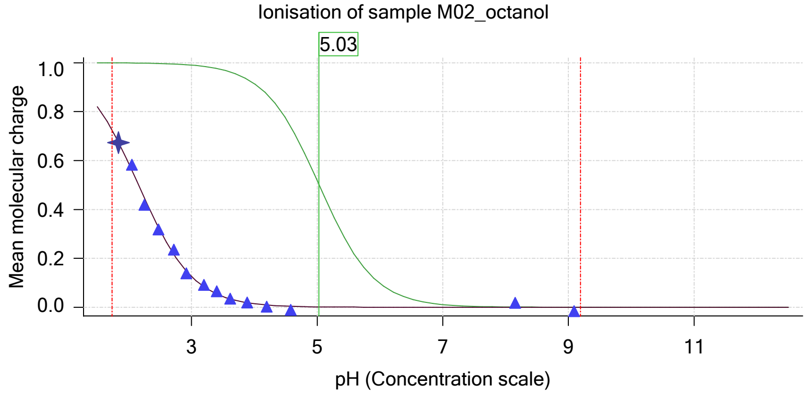


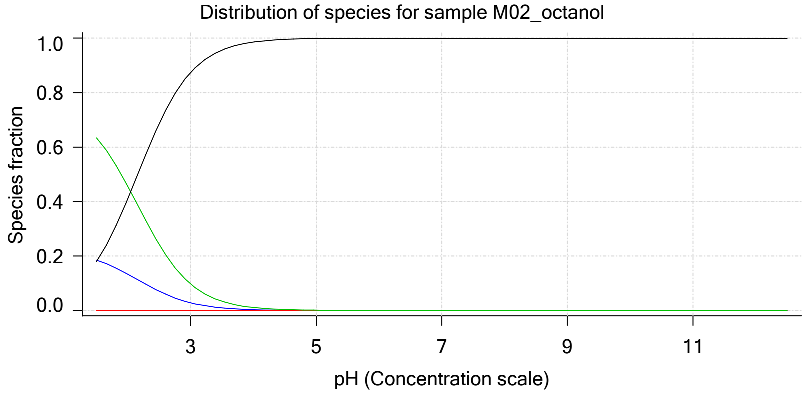

Sample name: **M02_octanol** Experiment start time: **3/6/2018 5:41:59 PM**
Assay name: **pH-metric high logP** Analyst: **Dorothy Levorse**
Assay ID: **18C-06015** Instrument ID: **T312060**
Filename: **C:\Sirius_T3\Mehtap\20180306_exp30_logP_T3-2\18C-06015_M02_octanol_pH-metric high logP.t3r**

#### Sample graphs (continued)

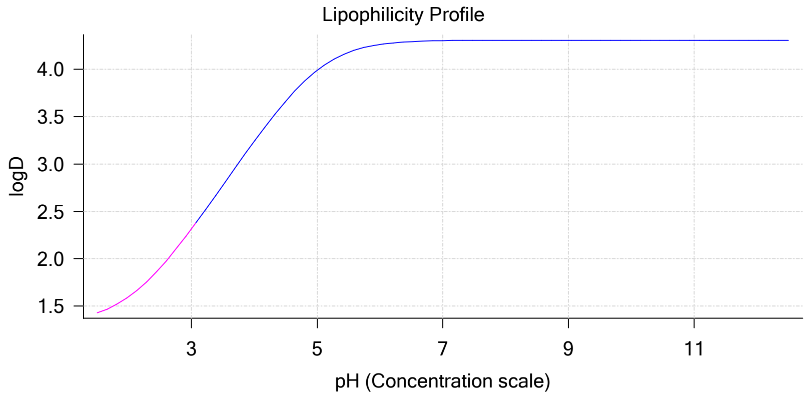


#### Sample logD and percent species

| pH | M02_octanol logD | M02_octanol M02_octanolH | M02_octanol M02_octanol | M02_octanol M02_octanolH* | M02_octanol M02_octanol* | Comment |
|---|---|---|---|---|---|---|
| 1.000 | 1.36 | 21.15 % | 0.00 % | 72.32 % | 6.53 % | |
| 1.200 | 1.38 | 20.37 % | 0.00 % | 69.66 % | 9.97 % | Stomach pH |
| 2.000 | 1.60 | 13.32 % | 0.01 % | 45.54 % | 41.14 % | |
| 3.000 | 2.32 | 2.83 % | 0.03 % | 9.68 % | 87.46 % | |
| 4.000 | 3.24 | 0.32 % | 0.03 % | 1.09 % | 98.56 % | |
| 5.000 | 3.99 | 0.03 % | 0.03 % | 0.11 % | 99.83 % | |
| 6.000 | 4.26 | 0.00 % | 0.03 % | 0.01 % | 99.96 % | |
| 6.500 | 4.29 | 0.00 % | 0.03 % | 0.00 % | 99.97 % | |
| 7.000 | 4.30 | 0.00 % | 0.03 % | 0.00 % | 99.97 % | |
| 7.400 | 4.30 | 0.00 % | 0.03 % | 0.00 % | 99.97 % | Blood pH |
| 8.000 | 4.31 | 0.00 % | 0.03 % | 0.00 % | 99.97 % | |
| 9.000 | 4.31 | 0.00 % | 0.03 % | 0.00 % | 99.97 % | |
| 10.000 | 4.31 | 0.00 % | 0.03 % | 0.00 % | 99.97 % | |
| 11.000 | 4.31 | 0.00 % | 0.03 % | 0.00 % | 99.97 % | |
| 12.000 | 4.31 | 0.00 % | 0.03 % | 0.00 % | 99.97 % | |

#### Carbonate and acidity

Carbonate 0.199 mM
Acidity error -0.191 mM

#### Other graphs

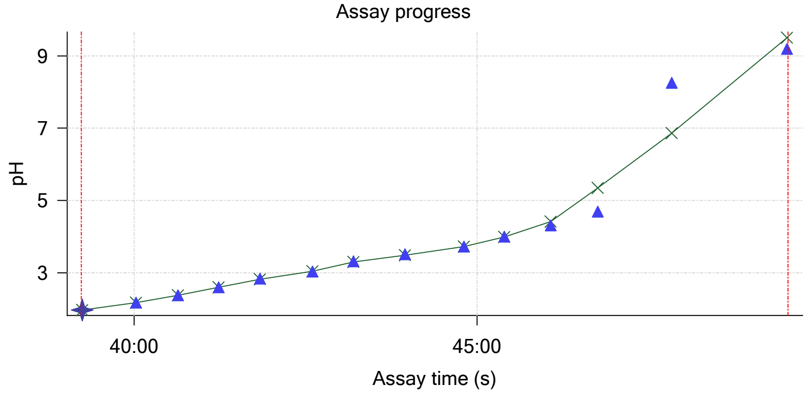


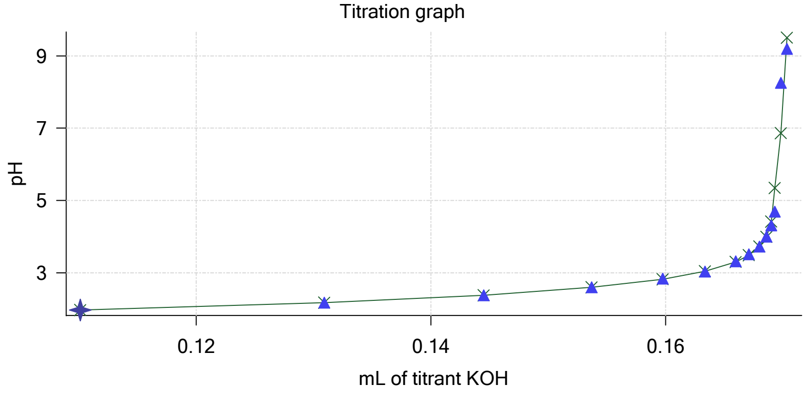

| | | | |
|---|---|---|---|
| Sample name: | **M02_octanol** | Experiment start time: | **3/6/2018 5:41:59 PM** |
| Assay name: | **pH-metric high logP** | Analyst: | **Dorothy Levorse** |
| Assay ID: | **18C-06015** | Instrument ID: | **T312060** |
| Filename: | **C:\Sirius_T3\Mehtap\20180306_exp30_logP_T3-2\18C-06015_M02_octanol_pH-metric high logP.t3r** | | |

#### Other graphs (continued)

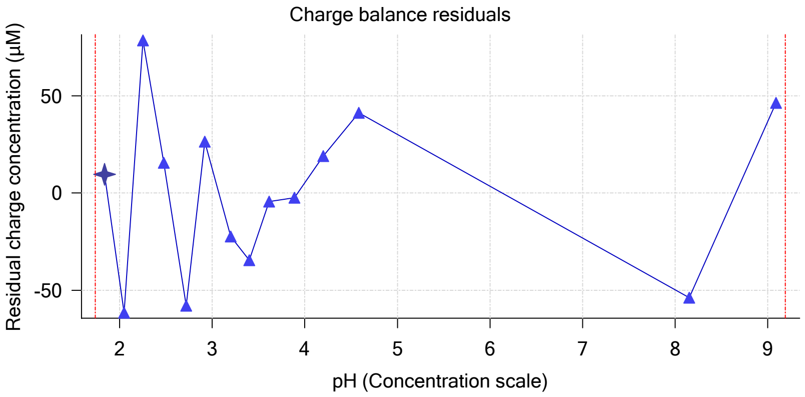


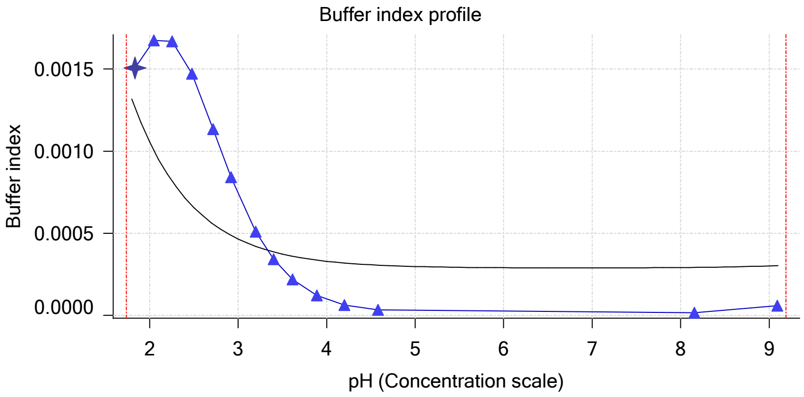


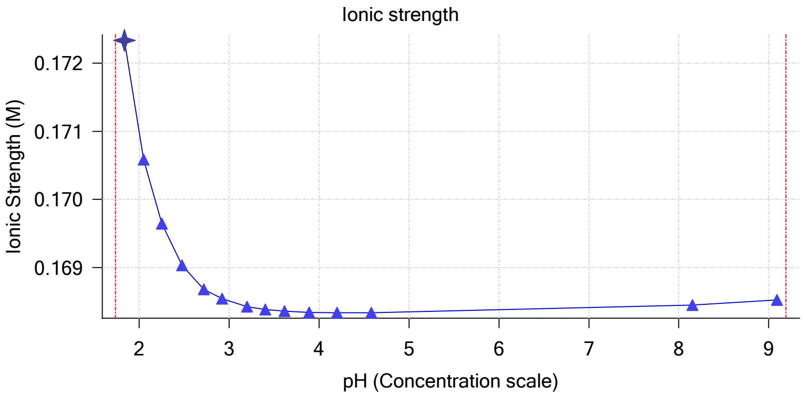


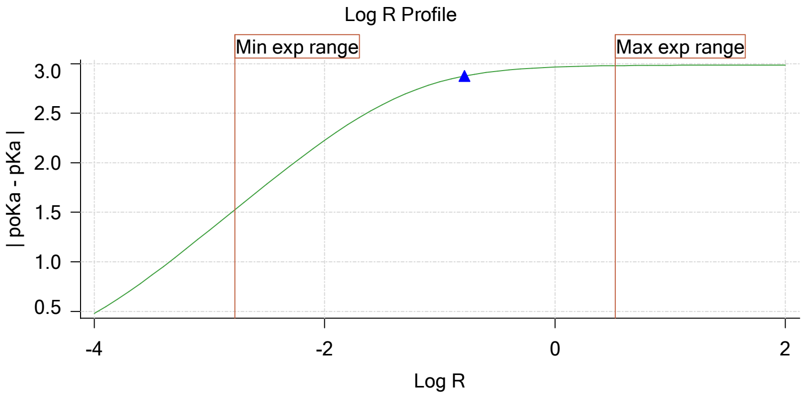


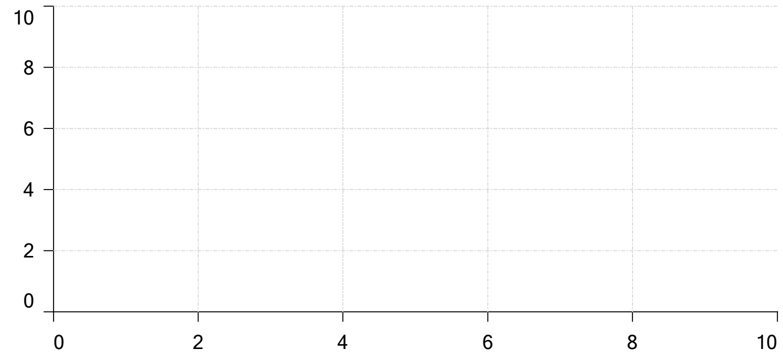

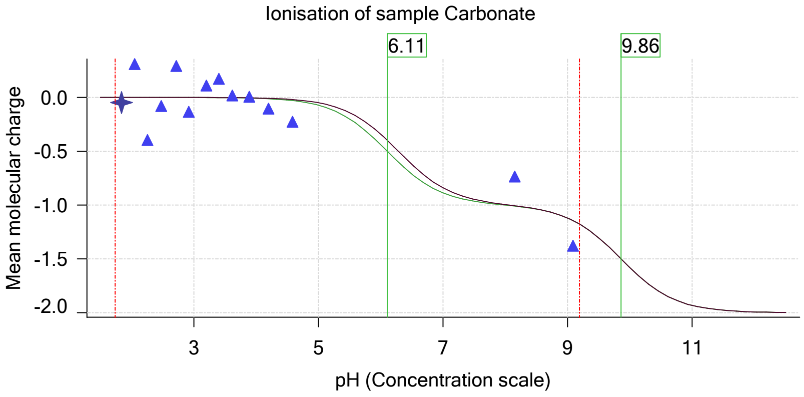

| | | | |
|---|---|---|---|
| Sample name: | **M02_octanol** | Experiment start time: | **3/6/2018 5:41:59 PM** |
| Assay name: | **pH-metric high logP** | Analyst: | **Dorothy Levorse** |
| Assay ID: | **18C-06015** | Instrument ID: | **T312060** |
| Filename: | **C:\Sirius_T3\Mehtap\20180306_exp30_logP_T3-2\18C-06015_M02_octanol_pH-metric high logP.t3r** | | |

#### Assay Model

| Settings | Value | Date/Time changed | Imported from |
|---|---|---|---|
| Sample name | M02_octanol | 12/6/2017 4:20:03 PM | User entered value |
| Sample by | Weight | | Default value |
| Sample weight | 0.002110 g | 3/6/2018 3:40:26 PM | User entered value |
| Formula weight | 289.26 g/mol | 12/6/2017 4:20:03 PM | User entered value |
| Solubility | Unknown | | Default value |
| Molecular weight | 289.26 | 12/6/2017 4:20:03 PM | User entered value |
| Individual pKa ionic environments | No | | Default value |
| Number of pKas | 1 | 12/6/2017 4:20:03 PM | User entered value |
| Sample is a | Base | 12/6/2017 4:20:03 PM | User entered value |
| pKa 1 | 5.03 | 12/6/2017 4:20:03 PM | User entered value |
| logp (XH +) | 1.32 | 3/2/2018 3:38:13 PM | User entered value |
| logP (neutral X) | 4.10 | 3/2/2018 3:38:07 PM | User entered value |

#### Events

| Time | Event | Water | Acid | Base | Octanol | pH | dpH/dt | pH R-squared | pH SD | dpH/dt time |
|---|---|---|---|---|---|---|---|---|---|---|
| 5:08.1 | Initial pH = 5.88 | | | | | | | | | |
| 8:07.7 | Data point 1 | 1.50000 mL | 0.05466 mL | 0.00583 mL | 0.01999 mL | 2.051 | 0.00164 | 0.09342 | 0.00026 | 10.0 s |
| 8:53.9 | Data point 2 | 1.50000 mL | 0.05466 mL | 0.01947 mL | 0.01999 mL | 2.258 | -0.00031 | 0.00286 | 0.00029 | 10.5 s |
| 9:30.0 | Data point 3 | 1.50000 mL | 0.05466 mL | 0.02897 mL | 0.01999 mL | 2.475 | -0.00173 | 0.32401 | 0.00015 | 10.5 s |
| 10:06.1 | Data point 4 | 1.50000 mL | 0.05466 mL | 0.03537 mL | 0.01999 mL | 2.705 | -0.01180 | 0.86985 | 0.00062 | 10.0 s |
| 10:41.7 | Data point 5 | 1.50000 mL | 0.05466 mL | 0.03996 mL | 0.01999 mL | 2.883 | -0.00877 | 0.93188 | 0.00045 | 10.0 s |
| 11:17.2 | Data point 6 | 1.50000 mL | 0.05466 mL | 0.04363 mL | 0.01999 mL | 3.088 | -0.01598 | 0.80563 | 0.00088 | 10.0 s |
| 11:52.7 | Data point 7 | 1.50000 mL | 0.05466 mL | 0.04652 mL | 0.01999 mL | 3.307 | -0.01840 | 0.83749 | 0.00099 | 12.0 s |
| 12:30.2 | Data point 8 | 1.50000 mL | 0.05466 mL | 0.04871 mL | 0.01999 mL | 3.520 | -0.01804 | 0.93551 | 0.00092 | 16.0 s |
| 13:11.8 | Data point 9 | 1.50000 mL | 0.05466 mL | 0.05031 mL | 0.01999 mL | 3.755 | -0.01934 | 0.98232 | 0.00096 | 20.0 s |
| 14:07.6 | Data point 10 | 1.50000 mL | 0.05466 mL | 0.05111 mL | 0.01999 mL | 3.913 | -0.01849 | 0.91937 | 0.00095 | 33.5 s |
| 15:06.5 | Data point 11 | 1.50000 mL | 0.05466 mL | 0.05151 mL | 0.01999 mL | 4.071 | -0.01887 | 0.88004 | 0.00099 | 34.0 s |
| 16:06.0 | Data point 12 | 1.50000 mL | 0.05466 mL | 0.05179 mL | 0.01999 mL | 4.218 | -0.01661 | 0.84054 | 0.00089 | 37.5 s |
| 17:19.3 | Data point 13 | 1.50000 mL | 0.05466 mL | 0.05212 mL | 0.01999 mL | 4.452 | -0.01927 | 0.92356 | 0.00099 | 40.0 s |
| 18:29.8 | Data point 14 | 1.50000 mL | 0.05466 mL | 0.05233 mL | 0.01999 mL | 4.819 | -0.01879 | 0.88303 | 0.00099 | 43.5 s |
| 19:38.8 | Data point 15 | 1.50000 mL | 0.05466 mL | 0.05242 mL | 0.01999 mL | 5.049 | -0.01780 | 0.88852 | 0.00093 | 46.5 s |
| 20:55.9 | Data point 16 | 1.50000 mL | 0.05466 mL | 0.05256 mL | 0.01999 mL | 5.809 | -0.01703 | 0.77107 | 0.00096 | 49.5 s |
| 22:16.0 | Data point 17 | 1.50000 mL | 0.05466 mL | 0.05266 mL | 0.01999 mL | 7.478 | -0.01823 | 0.89028 | 0.00095 | 57.0 s |
| 23:53.8 | Data point 18 | 1.50000 mL | 0.05466 mL | 0.05275 mL | 0.01999 mL | 8.346 | -0.01804 | 0.80184 | 0.00099 | 52.0 s |
| 25:21.6 | Data point 19 | 1.50000 mL | 0.05466 mL | 0.05285 mL | 0.01999 mL | 8.868 | -0.01663 | 0.74418 | 0.00095 | 36.0 s |
| 26:33.4 | Data point 20 | 1.50000 mL | 0.05466 mL | 0.05294 mL | 0.01999 mL | 9.195 | -0.01836 | 0.90519 | 0.00095 | 30.0 s |
| 27:57.2 | Data point 21 | 1.50000 mL | 0.11178 mL | 0.05294 mL | 0.05000 mL | 1.956 | -0.00466 | 0.28331 | 0.00043 | 10.0 s |
| 28:43.5 | Data point 22 | 1.50000 mL | 0.11178 mL | 0.07159 mL | 0.05000 mL | 2.177 | -0.01190 | 0.78202 | 0.00066 | 10.0 s |
| 29:19.2 | Data point 23 | 1.50000 mL | 0.11178 mL | 0.08389 mL | 0.05000 mL | 2.387 | -0.00289 | 0.10680 | 0.00044 | 10.0 s |
| 29:54.8 | Data point 24 | 1.50000 mL | 0.11178 mL | 0.09214 mL | 0.05000 mL | 2.622 | -0.00997 | 0.76942 | 0.00056 | 10.5 s |
| 30:30.9 | Data point 25 | 1.50000 mL | 0.11178 mL | 0.09774 mL | 0.05000 mL | 2.829 | -0.01166 | 0.83145 | 0.00063 | 10.5 s |
| 31:06.8 | Data point 26 | 1.50000 mL | 0.11178 mL | 0.10181 mL | 0.05000 mL | 3.053 | -0.01274 | 0.93127 | 0.00065 | 10.0 s |
| 31:42.3 | Data point 27 | 1.50000 mL | 0.11178 mL | 0.10468 mL | 0.05000 mL | 3.260 | -0.01427 | 0.92797 | 0.00073 | 10.5 s |
| 32:18.2 | Data point 28 | 1.50000 mL | 0.11178 mL | 0.10670 mL | 0.05000 mL | 3.459 | -0.00720 | 0.86970 | 0.00038 | 10.0 s |
| 32:53.6 | Data point 29 | 1.50000 mL | 0.11178 mL | 0.10811 mL | 0.05000 mL | 3.723 | -0.01606 | 0.92694 | 0.00082 | 10.5 s |
| 33:45.0 | Data point 30 | 1.50000 mL | 0.11178 mL | 0.10889 mL | 0.05000 mL | 3.933 | -0.01092 | 0.83346 | 0.00059 | 10.5 s |
| 34:20.9 | Data point 31 | 1.50000 mL | 0.11178 mL | 0.10941 mL | 0.05000 mL | 4.428 | -0.01931 | 0.91819 | 0.00100 | 18.5 s |
| 35:10.1 | Data point 32 | 1.50000 mL | 0.11178 mL | 0.10992 mL | 0.05000 mL | 7.825 | -0.06475 | 0.99712 | 0.00320 | Timed out at 59.5 s |
| 36:45.7 | Data point 33 | 1.50000 mL | 0.11178 mL | 0.11011 mL | 0.05000 mL | 8.913 | -0.01987 | 0.97328 | 0.00099 | 31.5 s |
| 37:37.5 | Data point 34 | 1.50000 mL | 0.11178 mL | 0.11011 mL | 0.05000 mL | 8.979 | -0.01914 | 0.92668 | 0.00098 | 33.0 s |
| 39:14.6 | Data point 35 | 1.50000 mL | 0.17180 mL | 0.11011 mL | 0.30000 mL | 1.969 | 0.00151 | 0.00822 | 0.00082 | 10.5 s |
| 40:01.6 | Data point 36 | 1.50000 mL | 0.17180 mL | 0.13090 mL | 0.30000 mL | 2.177 | -0.01405 | 0.55011 | 0.00094 | 11.0 s |
| 40:38.3 | Data point 37 | 1.50000 mL | 0.17180 mL | 0.14450 mL | 0.30000 mL | 2.376 | -0.01463 | 0.74877 | 0.00083 | 10.0 s |
| 41:13.9 | Data point 38 | 1.50000 mL | 0.17180 mL | 0.15369 mL | 0.30000 mL | 2.598 | -0.01418 | 0.73162 | 0.00082 | 10.5 s |
| 41:50.1 | Data point 39 | 1.50000 mL | 0.17180 mL | 0.15974 mL | 0.30000 mL | 2.837 | 0.01255 | 0.62249 | 0.00079 | 10.0 s |
| 42:35.9 | Data point 40 | 1.50000 mL | 0.17180 mL | 0.16336 mL | 0.30000 mL | 3.037 | 0.01290 | 0.63323 | 0.00080 | 10.5 s |

Sample name: **M02_octanol**
Assay name: **pH-metric high logP**
Assay ID: **18C-06015**
Filename: **C:\Sirius_T3\Mehtap\20180306_exp30_logP_T3-2\18C-06015_M02_octanol_pH-metric high logP.t3r**

Experiment start time: **3/6/2018 5:41:59 PM**
Analyst: **Dorothy Levorse**
Instrument ID: **T312060**

#### Events (continued)

| Time | Event | Water | Acid | Base | Octanol | pH | dpH/dt | pH R-squared | pH SD | dpH/dt time |
|---|---|---|---|---|---|---|---|---|---|---|
| 43:11.9 | Data point 41 | 1.50000 mL | 0.17180 mL | 0.16597 mL | 0.30000 mL | 3.316 | -0.01737 | 0.82670 | 0.00094 | 14.5 s |
| 43:57.2 | Data point 42 | 1.50000 mL | 0.17180 mL | 0.16710 mL | 0.30000 mL | 3.518 | -0.01034 | 0.83937 | 0.00056 | 10.5 s |
| 44:48.5 | Data point 43 | 1.50000 mL | 0.17180 mL | 0.16799 mL | 0.30000 mL | 3.731 | 0.01219 | 0.49410 | 0.00086 | 10.0 s |
| 45:23.9 | Data point 44 | 1.50000 mL | 0.17180 mL | 0.16858 mL | 0.30000 mL | 4.002 | -0.01494 | 0.80259 | 0.00082 | 10.0 s |
| 46:04.5 | Data point 45 | 1.50000 mL | 0.17180 mL | 0.16900 mL | 0.30000 mL | 4.312 | -0.01556 | 0.81871 | 0.00085 | 10.5 s |
| 46:45.7 | Data point 46 | 1.50000 mL | 0.17180 mL | 0.16928 mL | 0.30000 mL | 4.694 | -0.01928 | 0.95047 | 0.00098 | 29.0 s |
| 47:50.4 | Data point 47 | 1.50000 mL | 0.17180 mL | 0.16980 mL | 0.30000 mL | 8.255 | -0.04492 | 0.97778 | 0.00224 | Timed out at 59.5 s |
| 49:31.3 | Data point 48 | 1.50000 mL | 0.17180 mL | 0.17034 mL | 0.30000 mL | 9.190 | -0.01473 | 0.61216 | 0.00093 | 15.0 s |
| 49:55.4 | Assay volumes | 1.50000 mL | 0.17180 mL | 0.17034 mL | 0.30000 mL | | | | | |

| | | | |
|---|---|---|---|
| Sample name: | **M02_octanol** | Experiment start time: | **3/6/2018 5:41:59 PM** |
| Assay name: | **pH-metric high logP** | Analyst: | **Dorothy Levorse** |
| Assay ID: | **18C-06015** | Instrument ID: | **T312060** |
| Filename: | **C:\Sirius_T3\Mehtap\20180306_exp30_logP_T3-2\18C-06015_M02_octanol_pH-metric high logP.t3r** | | |

| | | | |
|---|---|---|---|
| Sample name: | **M02_octanol** | Experiment start time: | **3/6/2018 5:41:59 PM** |
| Assay name: | **pH-metric high logP** | Analyst: | **Dorothy Levorse** |
| Assay ID: | **18C-06015** | Instrument ID: | **T312060** |
| Filename: | **C:\Sirius_T3\Mehtap\20180306_exp30_logP_T3-2\18C-06015_M02_octanol_pH-metric high logP.t3r** | | |

#### Assay Settings (continued)

| Setting | Value | Original Value | Date/Time changed | Imported from |
|---|---|---|---|---|
| ***Titration 3*** | | | | |
| Titrate from | Low to high pH | | | |
| Add additional water | 0.00 mL | | | |
| Additional partition solvent volume | 0.250 mL | | | |
| Additional partition solvent added | Automatic | | | |
| After pH adjust stir for | 30 seconds | | | |
| Stir to allow partitioning for | 15 seconds | | | |
| Stirrer speed for partitioning | 60% | | | |
| ***Data Point Stability*** | | | | |
| Stir during data point collection | No | | | |
| Delay before data point collection | 0 seconds | | | |
| Number of points to average | 20 points | | | |
| Time interval between points | 0.50 seconds | | | |
| Required maximum standard deviation | 0.00100 dpH/dt | | | |
| Stability timeout after | 60 seconds | | | |

#### Calibration Settings

| Setting | Value | Date/Time changed | Imported from |
|---|---|---|---|
| Four-Plus alpha | 0.124 | 3/6/2018 5:41:59 PM | C:\Sirius_T3\18C-06006_Blank standardisation.t3r |
| Four-Plus S | 0.9973 | 3/6/2018 5:41:59 PM | C:\Sirius_T3\18C-06006_Blank standardisation.t3r |
| Four-Plus jH | 0.9 | 3/6/2018 5:41:59 PM | C:\Sirius_T3\18C-06006_Blank standardisation.t3r |
| Four-Plus jOH | -0.7 | 3/6/2018 5:41:59 PM | C:\Sirius_T3\18C-06006_Blank standardisation.t3r |
| Base concentration factor | 1.000 | 3/6/2018 5:41:59 PM | C:\Sirius_T3\KOH18B27.t3r |
| Acid concentration factor | 0.989 | 3/6/2018 5:41:59 PM | C:\Sirius_T3\18C-06006_Blank standardisation.t3r |

#### Instrument Settings

| Setting | Value | Batch Id | Install date |
|---|---|---|---|
| Instrument owner | Merck | | |
| Instrument ID | T312060 | | |
| Instrument type | T3 Simulator | | |
| Software version | 1.1.3.0 | | |
| Dispenser module | | T3DM1200361 | 3/31/2009 5:24:52 AM |
| Dispenser 0 | Water | | 3/31/2009 5:25:05 AM |
| Syringe volume | 2.5 mL | | |
| Firmware version | 1.2.1(r2) | | |
| Titrant | Water (0.15 M KCl) | 02-06-2018 | 2/27/2018 10:05:59 AM |
| Dispenser 2 | Acid | | 3/31/2009 5:25:11 AM |
| Syringe volume | 0.5 mL | | |
| Firmware version | 1.2.1(r2) | | |
| Titrant | Acid (0.5 M HCl) | 02-27-2018 | 2/27/2018 10:27:22 AM |
| Dispenser 1 | Base | | 3/31/2009 5:25:21 AM |
| Syringe volume | 0.5 mL | | |
| Firmware version | 1.2.1(r2) | | |
| Titrant | Base (0.5 M KOH) | 9/22/2017 | 2/27/2018 10:21:22 AM |
| Dispenser 5 | Cosolvent | | 3/31/2009 5:26:24 AM |
| Syringe volume | 2.5 mL | | |
| Firmware version | 1.2.1(r2) | | |
| Distribution valve 5 | Distribution Valve | | 3/31/2009 5:28:19 AM |
| Firmware version | 1.1.3 | | |
| Port A | Methanol (80%, 0.15 M KCl) | 02-08-2018 | 3/6/2018 9:28:59 AM |
| Port B | Cyclohexane | 11-01-17 | 2/27/2018 10:37:57 AM |
| Dispenser 3 | Buffer | | 8/3/2010 5:05:16 AM |
| Syringe volume | 0.5 mL | | |
| Firmware version | 1.2.1(r2) | | |
| Titrant | Dodecane | 2018/01/31 | 2/28/2018 10:18:04 AM |
| Dispenser 6 | Octanol | | 10/22/2010 10:52:43 AM |

| | | | |
|---|---|---|---|
| Sample name: | **M02_octanol** | Experiment start time: | **3/6/2018 5:41:59 PM** |
| Assay name: | **pH-metric high logP** | Analyst: | **Dorothy Levorse** |
| Assay ID: | **18C-06015** | Instrument ID: | **T312060** |
| Filename: | **C:\Sirius_T3\Mehtap\20180306_exp30_logP_T3-2\18C-06015_M02_octanol_pH-metric high logP.t3r** | | |

#### Instrument Settings (continued)

| Setting | Value | Batch Id | Install date |
|---|---|---|---|
| Syringe volume | 0.5 mL | | |
| Firmware version | 1.2.1(r2) | | |
| Titrant | Octanol | 01-31-2018 | 2/27/2018 9:59:35 AM |
| Titrator | | T3TM1200161 | 3/31/2009 5:24:17 AM |
| Horizontal axis firmware version | 1.17 AI1DI2DO2 Stepper 2 | | |
| Vertical axis firmware version | 1.17 AI1DI2DO2 Stepper 2 | | |
| Chassis I/O firmware version | 1.11 AI1DI0DO4 Norgren I/O | | |
| Probe I/O firmware version | 1.1.1 | | |
| Electrode | T3 Electrode | T3E0923 | 1/23/2018 2:01:00 PM |
| E0 calibration | +7.02 mV | | 3/6/2018 5:42:27 PM |
| Filling solution | 3M KCl | KCL097 | 3/6/2018 9:23:20 AM |
| Liquids | | | |
| Wash 1 | 50% IPA:50% Water | | 3/6/2018 9:24:32 AM |
| Wash 2 | 0.5% Trition X-100 in H20 | | 3/6/2018 9:24:35 AM |
| Buffer position 1 | pH7 Wash | | 3/6/2018 9:24:38 AM |
| Buffer position 2 | pH 7 | | 3/6/2018 9:24:40 AM |
| Storage position | | | 3/6/2018 9:24:07 AM |
| Wash water | 5.8e+003 mL | 02-27-2018 | 2/27/2018 9:54:39 AM |
| Waste | 9.8e+003 mL | | 11/28/2017 10:36:29 AM |
| Temperature controller | | | 8/5/2010 6:35:13 AM |
| Turbidity detector | | | 3/31/2009 5:24:45 AM |
| Spectrometer | | 074811 | 11/23/2010 11:22:28 AM |
| Dip probe | | 10196 | |
| Wavelength coefficient A0 | 183.333 | | |
| Wavelength coefficient A1 | 2.21568 | | |
| Wavelength coefficient A2 | -0.000289308 | | |
| Total lamp lit time | 123:01:40 | | 11/23/2010 11:22:28 AM |
| Calibrated on | 2/27/2018 10:40:38 AM | | |
| Integration time | 40 | | |
| Scans averaged | 10 | | |
| Autoloader | | T3AL1200345 | 11/10/2015 9:34:13 AM |
| Left-right axis firmware version | 1.17 AI1DI2DO2 Stepper 2 | | |
| Front-back axis firmware version | 1.17 AI1DI2DO2 Stepper 2 | | |
| Vertical axis firmware version | 1.17 AI1DI2DO2 Stepper 2 | | |
| Chassis I/O firmware version | 1.11 AI1DI0DO4 Norgren I/O | | |
| Configuration | | | |
| Alternate titration position | Titration position | | |
| Alternate reference position | Reference position | | |
| Maximum standard vial volume | 3.50 mL | | |
| Maximum alternate vial volume | 25.00 mL | | |
| Automatic action idle period | 5 minute(s) | | |
| Titrant tube volume | 1.3 mL | | |
| Syringe flush count | 3.50 | | |
| Flowing wash pump volume | 20.0 mL | | |
| Flowing wash stir duration | 5 s | | |
| Flowing wash stir speed | 30% | | |
| Solvent wash stir duration | 5 s | | |
| Solvent wash stir speed | 30% | | |
| Surfactant wash stir duration | 5 s | | |
| Surfactant wash stir speed | 30% | | |
| E0 calibration minimum number of points | 10 | | |
| E0 calibration maximum standard deviation | 0.01500 | | |
| E0 calibration timeout period | 60 s | | |
| E0 calibration stir duration | 5 s | | |
| E0 calibration preparation stir speed | 30% | | |
| E0 calibration buffer wash stir duration | 5 s | | |
| E0 calibration buffer wash stir speed | 30% | | |
| E0 calibration reading stir speed | 0% | | |

| | | | |
|---|---|---|---|
| Sample name: | **M02_octanol** | Experiment start time: | **3/6/2018 5:41:59 PM** |
| Assay name: | **pH-metric high logP** | Analyst: | **Dorothy Levorse** |
| Assay ID: | **18C-06015** | Instrument ID: | **T312060** |
| Filename: | **C:\Sirius_T3\Mehtap\20180306_exp30_logP_T3-2\18C-06015_M02_octanol_pH-metric high logP.t3r** | | |

#### Instrument Settings (continued)

| Setting | Value | Batch Id | Install date |
|---|---|---|---|
| Spectrometer calibration stir duration | 5 s | | |
| Spectrometer calibration stir speed | 30% | | |
| Spectrometer calibration wash pump volume | 20.0 mL | | |
| Spectrometer calibration wash stir duration | 5 s | | |
| Spectrometer calibration wash stir speed | 30% | | |
| Overhead dispense height | 10000 | | |

#### Refinement Settings

| Setting | Value | Default value |
|---|---|---|
| Turbidity detection method | None | None |
| Turbidity wavelength to assess | 500.0 nm | 500.0 nm |
| Turbidity maximum absorbance | 0.100 | 0.100 |
| Turbidity probe threshold | 50.00 | 50.00 |

#### Experiment Log

[49] Air gap released for Acid (0.5 M HCl)
[49] Air gap released for Base (0.5 M KOH)
[1:45] Air gap created for Water (0.15 M KCl)
[1:46] Air gap created for Acid (0.5 M HCl)
[1:46] Air gap created for Base (0.5 M KOH)
[1:46] Air gap released for Water (0.15 M KCl)
[1:50] Titrator arm moved over Titration position
[1:50] Titration 1 of 3
[1:50] Adding initial titrants
[1:50] Automatically add 1.50000 mL of water
[2:15] Dispensed 1.500000 mL of Water (0.15 M KCl)
[2:20] Titrator arm moved over Drain
[5:01] Titrator arm moved to Titration position
[5:01] Argon flow rate set to 100
[5:01] Stirrer speed set to 10
[5:06] Automatically add 0.02000 mL of Octanol
[5:07] Dispensed 0.019991 mL of Octanol
[5:08] Initial pH = 5.88
[5:08] Iterative adjust 5.88 -> 2.00
[5:08] pH 5.88 -> 2.00
[5:10] Air gap released for Acid (0.5 M HCl)
[5:10] Dispensed 0.054657 mL of Acid (0.5 M HCl)
[5:15] Holding pH 2.00
[7:15] Stirrer speed set to 0
[7:15] Stirrer speed set to 50
[7:15] Iterative adjust 1.95 -> 2.00
[7:15] pH 1.95 -> 2.00
[7:16] Air gap released for Base (0.5 M KOH)
[7:17] Dispensed 0.005833 mL of Base (0.5 M KOH)
[8:07] Stirrer speed set to 0
[8:17] Datapoint id 1 collected
[8:17] Stirrer speed set to 50
[8:22] pH 2.06 -> 2.26
[8:22] Using cautious pH adjust
[8:23] Dispensed 0.007197 mL of Base (0.5 M KOH)
[8:28] Stepping pH = 2.15
[8:28] Dispensed 0.005127 mL of Base (0.5 M KOH)
[8:33] Stepping pH = 2.23
[8:33] Dispensed 0.001317 mL of Base (0.5 M KOH)
[8:38] Stepping pH = 2.26
[8:54] Stirrer speed set to 0
[9:04] Datapoint id 2 collected

Sample name: **M02_octanol** Experiment start time: **3/6/2018 5:41:59 PM**
Assay name: **pH-metric high logP** Analyst: **Dorothy Levorse**
Assay ID: **18C-06015** Instrument ID: **T312060**
Filename: **C:\Sirius_T3\Mehtap\20180306_exp30_logP_T3-2\18C-06015_M02_octanol_pH-metric high logP.t3r**

#### Experiment Log (continued)

[9:04] Charge balance equation is out by 5.2%
[9:04] Stirrer speed set to 50
[9:09] pH 2.26 -> 2.46
[9:09] Using charge balance adjust
[9:09] Dispensed 0.009501 mL of Base (0.5 M KOH)
[9:30] Stirrer speed set to 0
[9:40] Datapoint id 3 collected
[9:40] Charge balance equation is out by 5.9%
[9:40] Stirrer speed set to 50
[9:45] pH 2.48 -> 2.68
[9:45] Using charge balance adjust
[9:46] Dispensed 0.006397 mL of Base (0.5 M KOH)
[10:06] Stirrer speed set to 0
[10:16] Datapoint id 4 collected
[10:16] Charge balance equation is out by 10.6%
[10:16] Stirrer speed set to 50
[10:21] pH 2.71 -> 2.91
[10:21] Using charge balance adjust
[10:21] Dispensed 0.004586 mL of Base (0.5 M KOH)
[10:41] Stirrer speed set to 0
[10:51] Datapoint id 5 collected
[10:51] Charge balance equation is out by -13.3%
[10:51] Stirrer speed set to 50
[10:56] pH 2.89 -> 3.09
[10:56] Using charge balance adjust
[10:57] Dispensed 0.003669 mL of Base (0.5 M KOH)
[11:17] Stirrer speed set to 0
[11:27] Datapoint id 6 collected
[11:27] Charge balance equation is out by -1.0%
[11:27] Stirrer speed set to 50
[11:32] pH 3.09 -> 3.29
[11:32] Using charge balance adjust
[11:32] Dispensed 0.002893 mL of Base (0.5 M KOH)
[11:52] Stirrer speed set to 0
[12:04] Datapoint id 7 collected
[12:04] Charge balance equation is out by 7.5%
[12:04] Stirrer speed set to 50
[12:09] pH 3.31 -> 3.51
[12:09] Using charge balance adjust
[12:10] Dispensed 0.002187 mL of Base (0.5 M KOH)
[12:30] Stirrer speed set to 0
[12:46] Datapoint id 8 collected
[12:46] Charge balance equation is out by 5.8%
[12:46] Stirrer speed set to 50
[12:51] pH 3.52 -> 3.72
[12:51] Using charge balance adjust
[12:51] Dispensed 0.001599 mL of Base (0.5 M KOH)
[13:11] Stirrer speed set to 0
[13:32] Datapoint id 9 collected
[13:32] Charge balance equation is out by 17.6%
[13:32] Stirrer speed set to 50
[13:37] pH 3.76 -> 3.96
[13:37] Using cautious pH adjust
[13:37] Dispensed 0.000541 mL of Base (0.5 M KOH)
[13:42] Stepping pH = 3.90
[13:42] Dispensed 0.000165 mL of Base (0.5 M KOH)
[13:47] Stepping pH = 3.93
[13:47] Dispensed 0.000094 mL of Base (0.5 M KOH)
[13:52] Stepping pH = 3.95

| | | | |
|---|---|---|---|
| Sample name: | **M02_octanol** | Experiment start time: | **3/6/2018 5:41:59 PM** |
| Assay name: | **pH-metric high logP** | Analyst: | **Dorothy Levorse** |
| Assay ID: | **18C-06015** | Instrument ID: | **T312060** |
| Filename: | **C:\Sirius_T3\Mehtap\20180306_exp30_logP_T3-2\18C-06015_M02_octanol_pH-metric high logP.t3r** | | |

#### Experiment Log (continued)

[14:07] Stirrer speed set to 0
[14:41] Datapoint id 10 collected
[14:41] Charge balance equation is out by 24.5%
[14:41] Stirrer speed set to 50
[14:46] pH 3.92 -> 4.12
[14:46] Using cautious pH adjust
[14:46] Dispensed 0.000400 mL of Base (0.5 M KOH)
[14:51] Stepping pH = 4.14
[15:06] Stirrer speed set to 0
[15:40] Datapoint id 11 collected
[15:40] Charge balance equation is out by 50.0%
[15:40] Stirrer speed set to 50
[15:45] pH 4.08 -> 4.28
[15:45] Using cautious pH adjust
[15:45] Dispensed 0.000282 mL of Base (0.5 M KOH)
[15:51] Stepping pH = 4.29
[16:06] Stirrer speed set to 0
[16:43] Datapoint id 12 collected
[16:43] Charge balance equation is out by 50.0%
[16:43] Stirrer speed set to 50
[16:48] pH 4.23 -> 4.43
[16:48] Using cautious pH adjust
[16:48] Dispensed 0.000212 mL of Base (0.5 M KOH)
[16:53] Stepping pH = 4.38
[16:54] Dispensed 0.000047 mL of Base (0.5 M KOH)
[16:59] Stepping pH = 4.39
[16:59] Dispensed 0.000071 mL of Base (0.5 M KOH)
[17:04] Stepping pH = 4.48
[17:19] Stirrer speed set to 0
[17:59] Datapoint id 13 collected
[17:59] Charge balance equation is out by 17.5%
[17:59] Stirrer speed set to 50
[18:04] pH 4.49 -> 4.69
[18:04] Using cautious pH adjust
[18:04] Dispensed 0.000118 mL of Base (0.5 M KOH)
[18:09] Stepping pH = 4.58
[18:09] Dispensed 0.000094 mL of Base (0.5 M KOH)
[18:14] Stepping pH = 4.81
[18:29] Stirrer speed set to 0
[19:13] Datapoint id 14 collected
[19:13] Charge balance equation is out by 11.3%
[19:13] Stirrer speed set to 50
[19:18] pH 4.89 -> 5.09
[19:18] Using charge balance adjust
[19:18] Dispensed 0.000094 mL of Base (0.5 M KOH)
[19:38] Stirrer speed set to 0
[20:25] Datapoint id 15 collected
[20:25] Charge balance equation is out by -19.7%
[20:25] Stirrer speed set to 50
[20:30] pH 5.10 -> 5.30
[20:30] Using cautious pH adjust
[20:30] Dispensed 0.000047 mL of Base (0.5 M KOH)
[20:35] Stepping pH = 5.11
[20:35] Dispensed 0.000094 mL of Base (0.5 M KOH)
[20:40] Stepping pH = 5.54
[20:56] Stirrer speed set to 0
[21:45] Datapoint id 16 collected
[21:45] Charge balance equation is out by -87.8%
[21:45] Stirrer speed set to 50

| | | | |
|---|---|---|---|
| Sample name: | **M02_octanol** | Experiment start time: | **3/6/2018 5:41:59 PM** |
| Assay name: | **pH-metric high logP** | Analyst: | **Dorothy Levorse** |
| Assay ID: | **18C-06015** | Instrument ID: | **T312060** |
| Filename: | **C:\Sirius_T3\Mehtap\20180306_exp30_logP_T3-2\18C-06015_M02_octanol_pH-metric high logP.t3r** | | |

#### Experiment Log (continued)

[21:50] pH 6.10 -> 6.30
[21:50] Using cautious pH adjust
[21:50] Dispensed 0.000024 mL of Base (0.5 M KOH)
[21:55] Stepping pH = 6.11
[21:55] Dispensed 0.000071 mL of Base (0.5 M KOH)
[22:01] Stepping pH = 6.60
[22:16] Stirrer speed set to 0
[23:13] Datapoint id 17 collected
[23:13] Charge balance equation is out by -96.1%
[23:13] Stirrer speed set to 50
[23:18] pH 7.61 -> 7.81
[23:18] Using cautious pH adjust
[23:18] Dispensed 0.000024 mL of Base (0.5 M KOH)
[23:23] Stepping pH = 7.64
[23:23] Dispensed 0.000024 mL of Base (0.5 M KOH)
[23:28] Stepping pH = 7.64
[23:28] Dispensed 0.000024 mL of Base (0.5 M KOH)
[23:33] Stepping pH = 7.71
[23:33] Dispensed 0.000024 mL of Base (0.5 M KOH)
[23:38] Stepping pH = 7.95
[23:53] Stirrer speed set to 0
[24:46] Datapoint id 18 collected
[24:46] Charge balance equation is out by -1,094.0%
[24:46] Stirrer speed set to 50
[24:51] pH 8.41 -> 8.61
[24:51] Using cautious pH adjust
[24:51] Dispensed 0.000024 mL of Base (0.5 M KOH)
[24:56] Stepping pH = 8.43
[24:56] Dispensed 0.000024 mL of Base (0.5 M KOH)
[25:01] Stepping pH = 8.43
[25:01] Dispensed 0.000047 mL of Base (0.5 M KOH)
[25:06] Stepping pH = 8.72
[25:21] Stirrer speed set to 0
[25:57] Datapoint id 19 collected
[25:57] Charge balance equation is out by -649.6%
[25:57] Stirrer speed set to 50
[26:02] pH 8.94 -> 9.05
[26:02] Using cautious pH adjust
[26:03] Dispensed 0.000024 mL of Base (0.5 M KOH)
[26:08] Stepping pH = 8.95
[26:08] Dispensed 0.000024 mL of Base (0.5 M KOH)
[26:13] Stepping pH = 8.97
[26:13] Dispensed 0.000047 mL of Base (0.5 M KOH)
[26:18] Stepping pH = 9.10
[26:33] Stirrer speed set to 0
[27:03] Datapoint id 20 collected
[27:03] Charge balance equation is out by -476.6%
[27:03] Titration 2 of 3
[27:03] Adding initial titrants
[27:03] Automatically add 0.03000 mL of Octanol
[27:04] Dispensed 0.030009 mL of Octanol
[27:04] Stirrer speed set to 10
[27:05] Stirrer speed set to 55
[27:05] Iterative adjust 9.21 -> 2.00
[27:05] pH 9.21 -> 2.00
[27:06] Dispensed 0.057126 mL of Acid (0.5 M HCl)
[27:57] Stirrer speed set to 0
[28:07] Datapoint id 21 collected
[28:07] Stirrer speed set to 55

| | | | |
|---|---|---|---|
| Sample name: | **M02_octanol** | Experiment start time: | **3/6/2018 5:41:59 PM** |
| Assay name: | **pH-metric high logP** | Analyst: | **Dorothy Levorse** |
| Assay ID: | **18C-06015** | Instrument ID: | **T312060** |
| Filename: | **C:\Sirius_T3\Mehtap\20180306_exp30_logP_T3-2\18C-06015_M02_octanol_pH-metric high logP.t3r** | | |

#### Experiment Log (continued)

[28:12] pH 1.97 -> 2.17
[28:12] Using cautious pH adjust
[28:12] Dispensed 0.009666 mL of Base (0.5 M KOH)
[28:17] Stepping pH = 2.05
[28:18] Dispensed 0.007361 mL of Base (0.5 M KOH)
[28:23] Stepping pH = 2.14
[28:23] Dispensed 0.001623 mL of Base (0.5 M KOH)
[28:28] Stepping pH = 2.17
[28:43] Stirrer speed set to 0
[28:53] Datapoint id 22 collected
[28:53] Charge balance equation is out by 3.5%
[28:53] Stirrer speed set to 55
[28:58] pH 2.18 -> 2.38
[28:58] Using charge balance adjust
[28:59] Dispensed 0.012300 mL of Base (0.5 M KOH)
[29:19] Stirrer speed set to 0
[29:29] Datapoint id 23 collected
[29:29] Charge balance equation is out by 2.4%
[29:29] Stirrer speed set to 55
[29:34] pH 2.40 -> 2.60
[29:34] Using charge balance adjust
[29:34] Dispensed 0.008255 mL of Base (0.5 M KOH)
[29:54] Stirrer speed set to 0
[30:05] Datapoint id 24 collected
[30:05] Charge balance equation is out by 12.9%
[30:05] Stirrer speed set to 55
[30:10] pH 2.63 -> 2.83
[30:10] Using charge balance adjust
[30:10] Dispensed 0.005597 mL of Base (0.5 M KOH)
[30:31] Stirrer speed set to 0
[30:41] Datapoint id 25 collected
[30:41] Charge balance equation is out by 0.3%
[30:41] Stirrer speed set to 55
[30:46] pH 2.83 -> 3.03
[30:46] Using charge balance adjust
[30:46] Dispensed 0.004069 mL of Base (0.5 M KOH)
[31:06] Stirrer speed set to 0
[31:16] Datapoint id 26 collected
[31:16] Charge balance equation is out by 8.9%
[31:16] Stirrer speed set to 55
[31:22] pH 3.06 -> 3.26
[31:22] Using charge balance adjust
[31:22] Dispensed 0.002869 mL of Base (0.5 M KOH)
[31:42] Stirrer speed set to 0
[31:52] Datapoint id 27 collected
[31:52] Charge balance equation is out by 0.5%
[31:52] Stirrer speed set to 55
[31:57] pH 3.26 -> 3.46
[31:57] Using charge balance adjust
[31:58] Dispensed 0.002023 mL of Base (0.5 M KOH)
[32:18] Stirrer speed set to 0
[32:28] Datapoint id 28 collected
[32:28] Charge balance equation is out by -2.6%
[32:28] Stirrer speed set to 55
[32:33] pH 3.47 -> 3.67
[32:33] Using charge balance adjust
[32:33] Dispensed 0.001411 mL of Base (0.5 M KOH)
[32:53] Stirrer speed set to 0
[33:04] Datapoint id 29 collected

| | | | |
|---|---|---|---|
| Sample name: | **M02_octanol** | Experiment start time: | **3/6/2018 5:41:59 PM** |
| Assay name: | **pH-metric high logP** | Analyst: | **Dorothy Levorse** |
| Assay ID: | **18C-06015** | Instrument ID: | **T312060** |
| Filename: | **C:\Sirius_T3\Mehtap\20180306_exp30_logP_T3-2\18C-06015_M02_octanol_pH-metric high logP.t3r** | | |

#### Experiment Log (continued)

[33:04] Charge balance equation is out by 27.7%
[33:04] Stirrer speed set to 55
[33:09] pH 3.73 -> 3.93
[33:09] Using cautious pH adjust
[33:09] Dispensed 0.000423 mL of Base (0.5 M KOH)
[33:14] Stepping pH = 3.84
[33:14] Dispensed 0.000235 mL of Base (0.5 M KOH)
[33:19] Stepping pH = 3.92
[33:19] Dispensed 0.000047 mL of Base (0.5 M KOH)
[33:24] Stepping pH = 3.92
[33:24] Dispensed 0.000071 mL of Base (0.5 M KOH)
[33:30] Stepping pH = 3.94
[33:45] Stirrer speed set to 0
[33:55] Datapoint id 30 collected
[33:55] Charge balance equation is out by 7.6%
[33:55] Stirrer speed set to 55
[34:00] pH 3.95 -> 4.15
[34:00] Using charge balance adjust
[34:00] Dispensed 0.000517 mL of Base (0.5 M KOH)
[34:21] Stirrer speed set to 0
[34:39] Datapoint id 31 collected
[34:39] Charge balance equation is out by 139.7%
[34:39] Stirrer speed set to 55
[34:44] pH 4.47 -> 4.67
[34:44] Using cautious pH adjust
[34:44] Dispensed 0.000094 mL of Base (0.5 M KOH)
[34:49] Stepping pH = 4.47
[34:50] Dispensed 0.000423 mL of Base (0.5 M KOH)
[34:55] Stepping pH = 7.49
[35:10] Stirrer speed set to 0
[36:10] Datapoint id 32 collected
[36:10] Charge balance equation is out by -201.6%
[36:10] Stirrer speed set to 55
[36:15] pH 8.18 -> 8.38
[36:15] Using cautious pH adjust
[36:15] Dispensed 0.000024 mL of Base (0.5 M KOH)
[36:20] Stepping pH = 8.14
[36:20] Dispensed 0.000024 mL of Base (0.5 M KOH)
[36:25] Stepping pH = 8.10
[36:25] Dispensed 0.000141 mL of Base (0.5 M KOH)
[36:30] Stepping pH = 8.92
[36:45] Stirrer speed set to 0
[37:17] Datapoint id 33 collected
[37:17] Charge balance equation is out by -2,089.2%
[37:17] Stirrer speed set to 55
[37:22] pH 9.03 -> 9.05
[37:22] Using cautious pH adjust
[37:37] Stirrer speed set to 0
[38:10] Datapoint id 34 collected
[38:10] Charge balance equation is out by 100.0%
[38:10] Titration 3 of 3
[38:10] Adding initial titrants
[38:10] Automatically add 0.25000 mL of Octanol
[38:16] Dispensed 0.250000 mL of Octanol
[38:16] Stirrer speed set to 10
[38:17] Stirrer speed set to 60
[38:17] Iterative adjust 9.00 -> 2.00
[38:17] pH 9.00 -> 2.00
[38:19] Dispensed 0.058725 mL of Acid (0.5 M HCl)

| | | | |
|---|---|---|---|
| Sample name: | **M02_octanol** | Experiment start time: | **3/6/2018 5:41:59 PM** |
| Assay name: | **pH-metric high logP** | Analyst: | **Dorothy Levorse** |
| Assay ID: | **18C-06015** | Instrument ID: | **T312060** |
| Filename: | **C:\Sirius_T3\Mehtap\20180306_exp30_logP_T3-2\18C-06015_M02_octanol_pH-metric high logP.t3r** | | |

#### Experiment Log (continued)

[38:24] pH 2.01 -> 2.00
[38:24] Dispensed 0.001294 mL of Acid (0.5 M HCl)
[39:14] Stirrer speed set to 0
[39:25] Datapoint id 35 collected
[39:25] Stirrer speed set to 60
[39:30] pH 1.98 -> 2.18
[39:30] Using cautious pH adjust
[39:30] Dispensed 0.010254 mL of Base (0.5 M KOH)
[39:35] Stepping pH = 2.06
[39:36] Dispensed 0.008043 mL of Base (0.5 M KOH)
[39:41] Stepping pH = 2.15
[39:41] Dispensed 0.002493 mL of Base (0.5 M KOH)
[39:46] Stepping pH = 2.18
[40:01] Stirrer speed set to 0
[40:12] Datapoint id 36 collected
[40:12] Charge balance equation is out by -1.3%
[40:12] Stirrer speed set to 60
[40:17] pH 2.18 -> 2.38
[40:17] Using charge balance adjust
[40:18] Dispensed 0.013594 mL of Base (0.5 M KOH)
[40:38] Stirrer speed set to 0
[40:48] Datapoint id 37 collected
[40:48] Charge balance equation is out by -1.2%
[40:48] Stirrer speed set to 60
[40:53] pH 2.38 -> 2.58
[40:53] Using charge balance adjust
[40:53] Dispensed 0.009196 mL of Base (0.5 M KOH)
[41:14] Stirrer speed set to 0
[41:24] Datapoint id 38 collected
[41:24] Charge balance equation is out by 9.4%
[41:24] Stirrer speed set to 60
[41:29] pH 2.60 -> 2.80
[41:29] Using charge balance adjust
[41:30] Dispensed 0.006044 mL of Base (0.5 M KOH)
[41:50] Stirrer speed set to 0
[42:00] Datapoint id 39 collected
[42:00] Charge balance equation is out by 16.8%
[42:00] Stirrer speed set to 60
[42:05] pH 2.85 -> 3.05
[42:05] Using cautious pH adjust
[42:05] Dispensed 0.001929 mL of Base (0.5 M KOH)
[42:10] Stepping pH = 2.94
[42:10] Dispensed 0.001435 mL of Base (0.5 M KOH)
[42:15] Stepping pH = 3.03
[42:15] Dispensed 0.000259 mL of Base (0.5 M KOH)
[42:20] Stepping pH = 3.04
[42:36] Stirrer speed set to 0
[42:46] Datapoint id 40 collected
[42:46] Charge balance equation is out by 5.9%
[42:46] Stirrer speed set to 60
[42:51] pH 3.05 -> 3.25
[42:51] Using charge balance adjust
[42:51] Dispensed 0.002611 mL of Base (0.5 M KOH)
[43:12] Stirrer speed set to 0
[43:26] Datapoint id 41 collected
[43:26] Charge balance equation is out by 35.1%
[43:26] Stirrer speed set to 60
[43:31] pH 3.32 -> 3.52
[43:31] Using cautious pH adjust

| | | | |
|---|---|---|---|
| Sample name: | **M02_octanol** | Experiment start time: | **3/6/2018 5:41:59 PM** |
| Assay name: | **pH-metric high logP** | Analyst: | **Dorothy Levorse** |
| Assay ID: | **18C-06015** | Instrument ID: | **T312060** |
| Filename: | **C:\Sirius_T3\Mehtap\20180306_exp30_logP_T3-2\18C-06015_M02_octanol_pH-metric high logP.t3r** | | |

#### Experiment Log (continued)

[43:31] Dispensed 0.000729 mL of Base (0.5 M KOH)
[43:36] Stepping pH = 3.43
[43:37] Dispensed 0.000400 mL of Base (0.5 M KOH)
[43:42] Stepping pH = 3.52
[43:57] Stirrer speed set to 0
[44:07] Datapoint id 42 collected
[44:07] Charge balance equation is out by 22.8%
[44:07] Stirrer speed set to 60
[44:12] pH 3.53 -> 3.73
[44:12] Using cautious pH adjust
[44:12] Dispensed 0.000470 mL of Base (0.5 M KOH)
[44:18] Stepping pH = 3.63
[44:18] Dispensed 0.000306 mL of Base (0.5 M KOH)
[44:23] Stepping pH = 3.71
[44:23] Dispensed 0.000047 mL of Base (0.5 M KOH)
[44:28] Stepping pH = 3.72
[44:28] Dispensed 0.000071 mL of Base (0.5 M KOH)
[44:33] Stepping pH = 3.73
[44:48] Stirrer speed set to 0
[44:58] Datapoint id 43 collected
[44:58] Charge balance equation is out by 5.5%
[44:58] Stirrer speed set to 60
[45:03] pH 3.75 -> 3.95
[45:03] Using charge balance adjust
[45:03] Dispensed 0.000588 mL of Base (0.5 M KOH)
[45:24] Stirrer speed set to 0
[45:34] Datapoint id 44 collected
[45:34] Charge balance equation is out by 26.0%
[45:34] Stirrer speed set to 60
[45:39] pH 4.03 -> 4.23
[45:39] Using cautious pH adjust
[45:39] Dispensed 0.000165 mL of Base (0.5 M KOH)
[45:44] Stepping pH = 4.07
[45:44] Dispensed 0.000259 mL of Base (0.5 M KOH)
[45:49] Stepping pH = 4.31
[46:04] Stirrer speed set to 0
[46:15] Datapoint id 45 collected
[46:15] Charge balance equation is out by -34.4%
[46:15] Stirrer speed set to 60
[46:20] pH 4.37 -> 4.57
[46:20] Using cautious pH adjust
[46:20] Dispensed 0.000071 mL of Base (0.5 M KOH)
[46:25] Stepping pH = 4.37
[46:25] Dispensed 0.000212 mL of Base (0.5 M KOH)
[46:30] Stepping pH = 4.68
[46:45] Stirrer speed set to 0
[47:14] Datapoint id 46 collected
[47:14] Charge balance equation is out by -97.4%
[47:14] Stirrer speed set to 60
[47:19] pH 4.84 -> 5.04
[47:19] Using cautious pH adjust
[47:19] Dispensed 0.000024 mL of Base (0.5 M KOH)
[47:25] Stepping pH = 4.84
[47:25] Dispensed 0.000141 mL of Base (0.5 M KOH)
[47:30] Stepping pH = 4.86
[47:30] Dispensed 0.000353 mL of Base (0.5 M KOH)
[47:35] Stepping pH = 8.18
[47:50] Stirrer speed set to 0
[48:50] Datapoint id 47 collected

| | | | |
|---|---|---|---|
| Sample name: | **M02_octanol** | Experiment start time: | **3/6/2018 5:41:59 PM** |
| Assay name: | **pH-metric high logP** | Analyst: | **Dorothy Levorse** |
| Assay ID: | **18C-06015** | Instrument ID: | **T312060** |
| Filename: | **C:\Sirius_T3\Mehtap\20180306_exp30_logP_T3-2\18C-06015_M02_octanol_pH-metric high logP.t3r** | | |

#### Experiment Log (continued)

[48:50] Charge balance equation is out by -866.5%
[48:50] Stirrer speed set to 60
[48:55] pH 8.41 -> 8.61
[48:55] Using cautious pH adjust
[48:55] Dispensed 0.000024 mL of Base (0.5 M KOH)
[49:00] Stepping pH = 8.38
[49:00] Dispensed 0.000047 mL of Base (0.5 M KOH)
[49:05] Stepping pH = 8.35
[49:06] Dispensed 0.000235 mL of Base (0.5 M KOH)
[49:11] Stepping pH = 8.46
[49:11] Dispensed 0.000235 mL of Base (0.5 M KOH)
[49:16] Stepping pH = 9.22
[49:31] Stirrer speed set to 0
[49:46] Datapoint id 48 collected
[49:46] Charge balance equation is out by -3,486.6%
[49:46] Argon flow rate set to 0
[49:50] Titrator arm moved over Titration position
[50:12] The autoloader failed to pick at location "Sample position"
