## Supplementary material for "Octanol-water partition coefficient measurements for the SAMPL6 Blind Prediction Challenge": SM04_18C-24002_M04_octanol_pH-metric high logP_report.pdf

| | | | |
|---|---|---|---|
| Sample name: | **M04_octanol** | Experiment start time: | **3/24/2018 1:34:06 AM** |
| Assay name: | **pH-metric high logP** | Analyst: | **Dorothy Levorse** |
| Assay ID: | **18C-24002** | Instrument ID: | **T312060** |
| Filename: | **C:\Sirius_T3\Mehtap\20180323_exp33_logP_T3-2\18C-24002_M04_octanol_pH-metric high logP.t3r** | | |

#### pH-metric Result

| | | |
|---|---|---|
| logP (XH +) | 0.82 | ±0.06 (n=50) |
| logP (neutral X) | 4.04 | ±0.02 (n=50) |
| RMSD | 0.362 | |

#### 18C-24002 Points 1 to 24

| | |
|---|---|
| M04_octanol concentration factor | 0.923 |
| Carbonate | 0.1572 mM |
| Acidity error | -0.23457 mM |

#### 18C-24002 Points 25 to 51

| | |
|---|---|
| M04_octanol concentration factor | 0.920 |
| Carbonate | 0.1065 mM |
| Acidity error | -0.29812 mM |

#### 18C-24002 Points 52 to 75

| | |
|---|---|
| M04_octanol concentration factor | 0.952 |
| Carbonate | 0.1039 mM |
| Acidity error | -0.10230 mM |

#### Warnings and errors

| | |
|---|---|
| Errors | None |
| Warnings | None |

#### Sample logD and percent species

| pH | M04_octanol logD | M04_octanol M04_octanolH | M04_octanol M04_octanol | M04_octanol M04_octanolH* | M04_octanol M04_octanol* | Comment |
|---|---|---|---|---|---|---|
| 1.000 | 0.83 | 12.93 % | 0.00 % | 85.56 % | 1.52 % | |
| 1.200 | 0.83 | 12.81 % | 0.00 % | 84.80 % | 2.38 % | Stomach pH |
| 2.000 | 0.89 | 11.37 % | 0.00 % | 75.28 % | 13.34 % | |
| 3.000 | 1.26 | 5.17 % | 0.01 % | 34.21 % | 60.62 % | |
| 4.000 | 2.09 | 0.80 % | 0.01 % | 5.30 % | 93.89 % | |
| 5.000 | 3.03 | 0.08 % | 0.01 % | 0.56 % | 99.35 % | |
| 6.000 | 3.75 | 0.01 % | 0.01 % | 0.06 % | 99.93 % | |
| 6.500 | 3.93 | 0.00 % | 0.01 % | 0.02 % | 99.97 % | |
| 7.000 | 4.00 | 0.00 % | 0.01 % | 0.01 % | 99.98 % | |
| 7.400 | 4.02 | 0.00 % | 0.01 % | 0.00 % | 99.99 % | Blood pH |
| 8.000 | 4.04 | 0.00 % | 0.01 % | 0.00 % | 99.99 % | |
| 9.000 | 4.04 | 0.00 % | 0.01 % | 0.00 % | 99.99 % | |
| 10.000 | 4.04 | 0.00 % | 0.01 % | 0.00 % | 99.99 % | |
| 11.000 | 4.04 | 0.00 % | 0.01 % | 0.00 % | 99.99 % | |
| 12.000 | 4.04 | 0.00 % | 0.01 % | 0.00 % | 99.99 % | |

| | | | |
|---|---|---|---|
| Sample name: | **M04_octanol** | Experiment start time: | **3/24/2018 1:34:06 AM** |
| Assay name: | **pH-metric high logP** | Analyst: | **Dorothy Levorse** |
| Assay ID: | **18C-24002** | Instrument ID: | **T312060** |
| Filename: | **C:\Sirius_T3\Mehtap\20180323_exp33_logP_T3-2\18C-24002_M04_octanol_pH-metric high logP.t3r** | | |

#### Graphs

Charge balance residuals

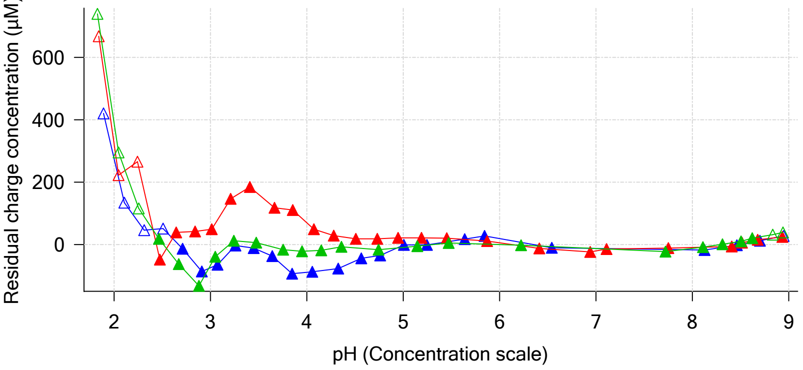


Ionic strength

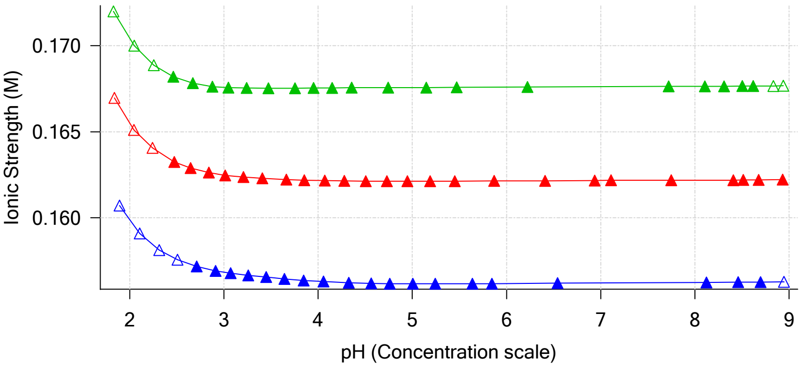


Buffer index profile

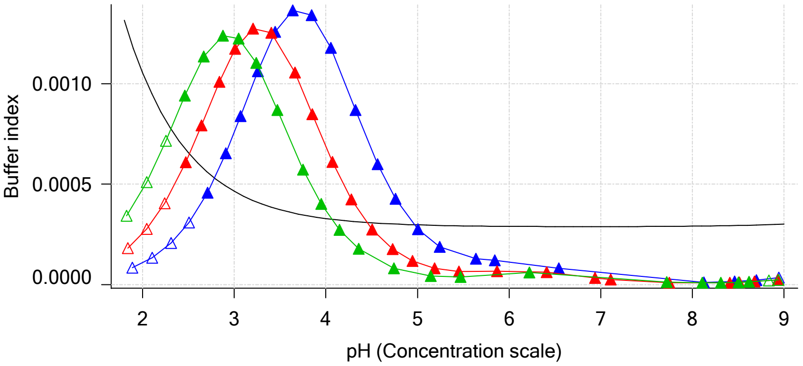


Sample M04_octanol Ionisation Graph

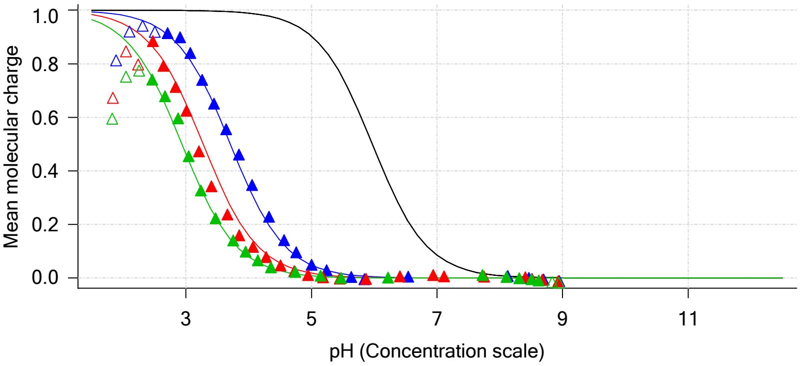


Distribution of species for sample M04_octanol

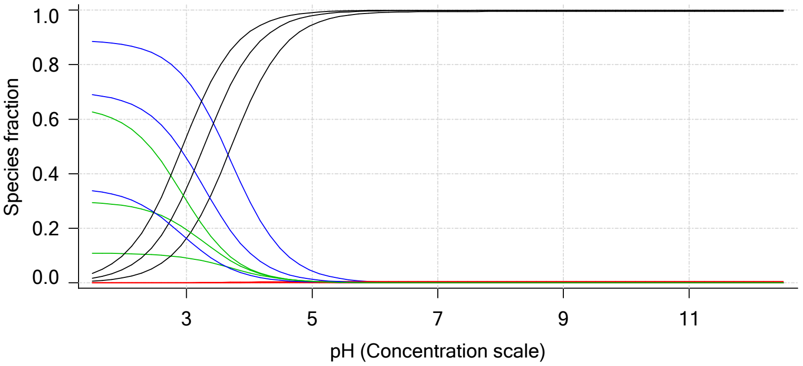


Lipophilicity Profile

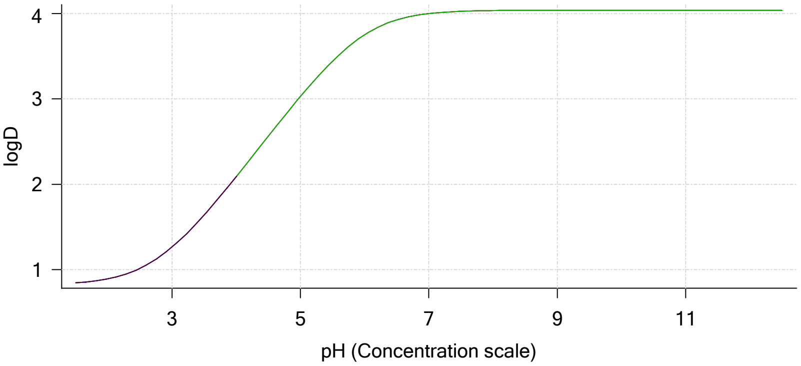


Log R Profile

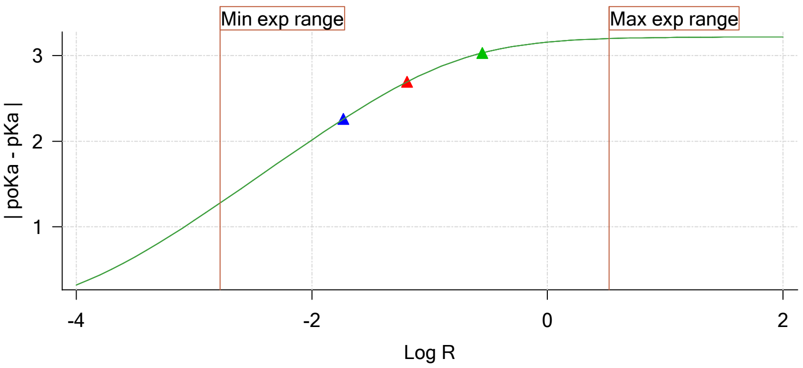


Sample Carbonate Ionisation Graph

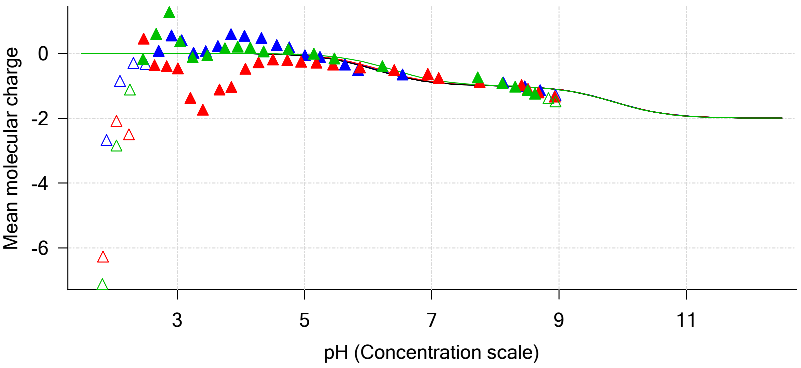

| | | | |
|---|---|---|---|
| Sample name: | **M04_octanol** | Experiment start time: | **3/24/2018 1:34:06 AM** |
| Assay name: | **pH-metric high logP** | Analyst: | **Dorothy Levorse** |
| Assay ID: | **18C-24002** | Instrument ID: | **T312060** |
| Filename: | **C:\Sirius_T3\Mehtap\20180323_exp33_logP_T3-2\18C-24002_M04_octanol_pH-metric high logP.t3r** | | |

#### Graphs (continued)

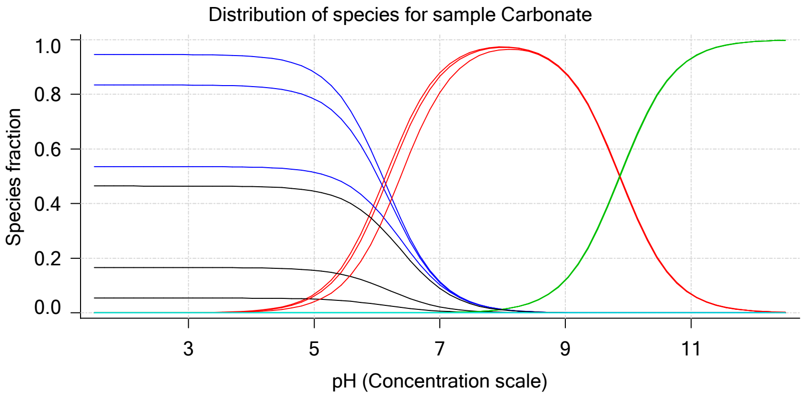

| | | | |
|---|---|---|---|
| Sample name: | **M04_octanol** | Experiment start time: | **3/24/2018 1:34:06 AM** |
| Assay name: | **pH-metric high logP** | Analyst: | **Dorothy Levorse** |
| Assay ID: | **18C-24002** | Instrument ID: | **T312060** |
| Filename: | **C:\Sirius_T3\Mehtap\20180323_exp33_logP_T3-2\18C-24002_M04_octanol_pH-metric high logP.t3r** | | |

### pH-metric high logP Titration 1 of 3 18C-24002 Points 1 to 24

#### Overall results

| | |
|---|---|
| RMSD | 0.282 |
| Average ionic strength | 0.156 M |
| Average temperature | 24.9°C |
| Partition ratio | 0.0186 : 1 |
| Analyte concentration range | 2528.6 μM to 2603.3 μM |
| Total points considered | 19 of 24 |

#### Warnings and errors

| | |
|---|---|
| Errors | None |
| Warnings | None |

#### Four-Plus parameters

| | | | |
|---|---|---|---|
| Alpha | 0.119 | 3/24/2018 1:34:06 AM | C:\Sirius_T3\HCl18C23.t3r |
| S | 0.9972 | 3/24/2018 1:34:06 AM | C:\Sirius_T3\HCl18C23.t3r |
| jH | 0.9 | 3/24/2018 1:34:06 AM | C:\Sirius_T3\HCl18C23.t3r |
| jOH | -0.3 | 3/24/2018 1:34:06 AM | C:\Sirius_T3\HCl18C23.t3r |

#### Titrants

| | | | |
|---|---|---|---|
| 0.50 M HCl | 0.997124 | 3/24/2018 1:34:06 AM | C:\Sirius_T3\HCl18C23.t3r |
| 0.50 M KOH | 1.003190 | 3/24/2018 1:34:06 AM | C:\Sirius_T3\KOH18C23.t3r |

#### Sample

| | |
|---|---|
| M04_octanol concentration factor | 0.923 |
| Base pKa 1 | 5.97 |
| logP (XH +) | 0.19 |
| logP (neutral X) | 3.95 |

#### Sample graphs

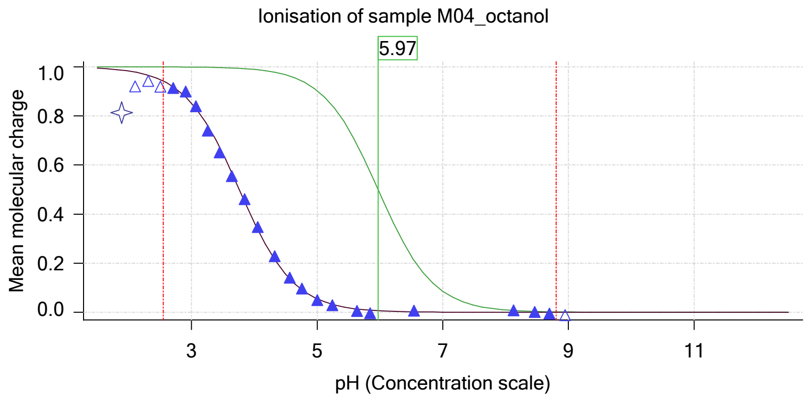


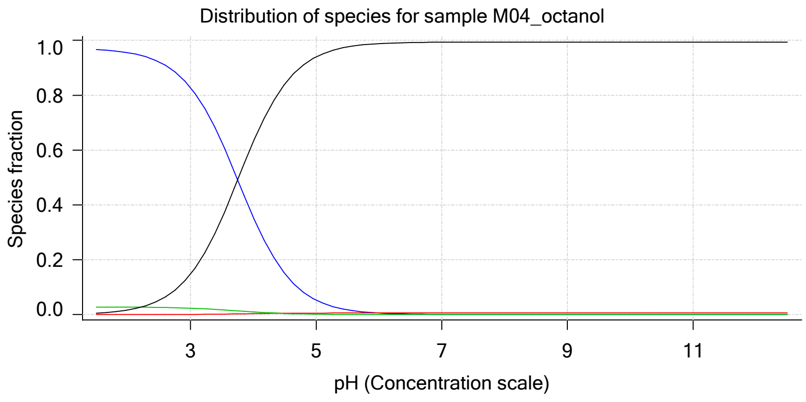

| | | | |
|---|---|---|---|
| Sample name: | **M04_octanol** | Experiment start time: | **3/24/2018 1:34:06 AM** |
| Assay name: | **pH-metric high logP** | Analyst: | **Dorothy Levorse** |
| Assay ID: | **18C-24002** | Instrument ID: | **T312060** |
| Filename: | **C:\Sirius_T3\Mehtap\20180323_exp33_logP_T3-2\18C-24002_M04_octanol_pH-metric high logP.t3r** | | |

#### Sample graphs (continued)

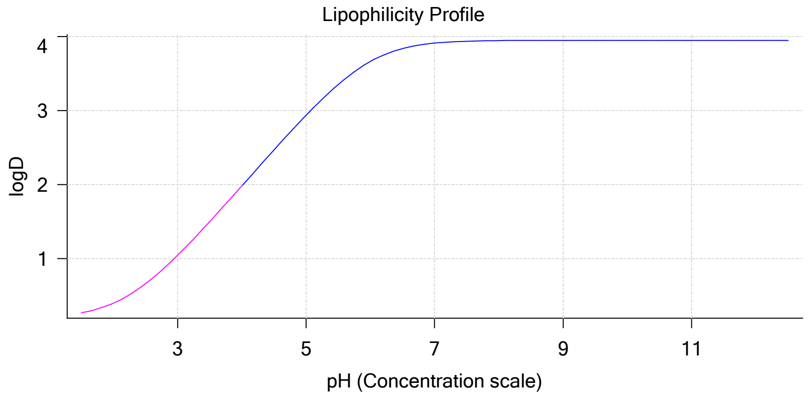


#### Sample logD and percent species

| pH | M04_octanol logD | M04_octanol M04_octanolH | M04_octanol M04_octanol | M04_octanol M04_octanolH* | M04_octanol M04_octanol* | Comment |
|---|---|---|---|---|---|---|
| 1.000 | 0.22 | 97.04 % | 0.00 % | 2.79 % | 0.17 % | |
| 1.200 | 0.23 | 96.94 % | 0.00 % | 2.79 % | 0.27 % | Stomach pH |
| 2.000 | 0.40 | 95.55 % | 0.01 % | 2.75 % | 1.70 % | |
| 3.000 | 1.05 | 82.83 % | 0.09 % | 2.38 % | 14.70 % | |
| 4.000 | 1.98 | 35.54 % | 0.38 % | 1.02 % | 63.06 % | |
| 5.000 | 2.94 | 5.30 % | 0.57 % | 0.15 % | 93.98 % | |
| 6.000 | 3.66 | 0.56 % | 0.60 % | 0.02 % | 98.83 % | |
| 6.500 | 3.84 | 0.18 % | 0.60 % | 0.01 % | 99.22 % | |
| 7.000 | 3.91 | 0.06 % | 0.60 % | 0.00 % | 99.34 % | |
| 7.400 | 3.93 | 0.02 % | 0.60 % | 0.00 % | 99.38 % | Blood pH |
| 8.000 | 3.95 | 0.01 % | 0.60 % | 0.00 % | 99.39 % | |
| 9.000 | 3.95 | 0.00 % | 0.60 % | 0.00 % | 99.40 % | |
| 10.000 | 3.95 | 0.00 % | 0.60 % | 0.00 % | 99.40 % | |
| 11.000 | 3.95 | 0.00 % | 0.60 % | 0.00 % | 99.40 % | |
| 12.000 | 3.95 | 0.00 % | 0.60 % | 0.00 % | 99.40 % | |

#### Carbonate and acidity

| | |
|---|---|
| Carbonate | 0.157 mM |
| Acidity error | -0.235 mM |

#### Other graphs

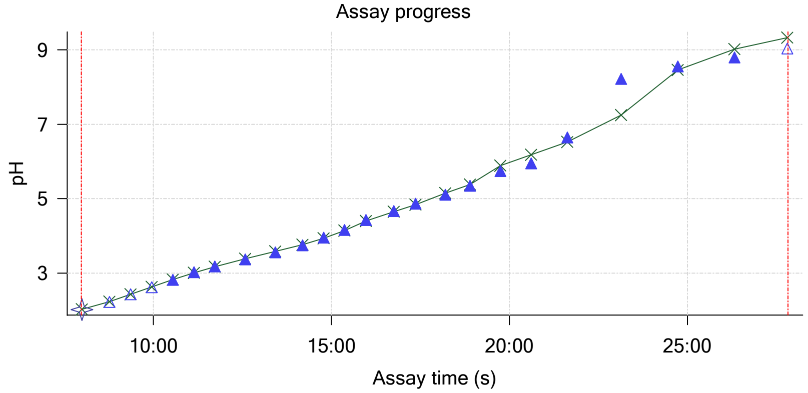


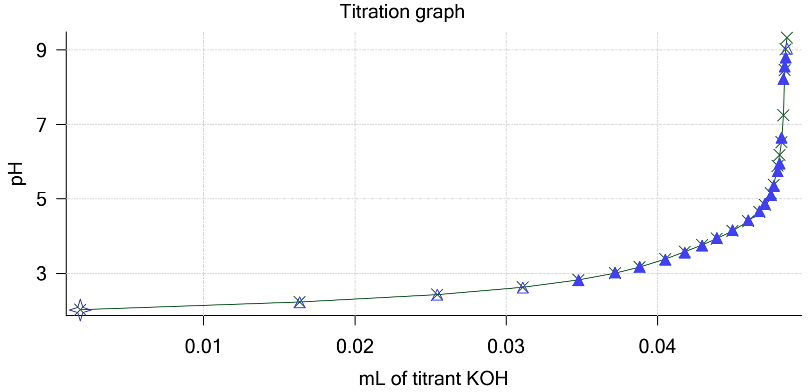

Sample name: **M04_octanol**
Assay name: **pH-metric high logP**
Assay ID: **18C-24002**
Filename: **C:\Sirius_T3\Mehtap\20180323_exp33_logP_T3-2\18C-24002_M04_octanol_pH-metric high logP.t3r**

Experiment start time: **3/24/2018 1:34:06 AM**
Analyst: **Dorothy Levorse**
Instrument ID: **T312060**

#### Other graphs (continued)

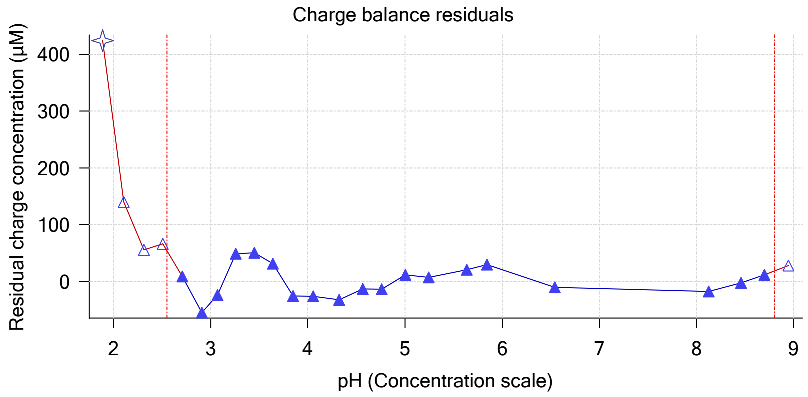


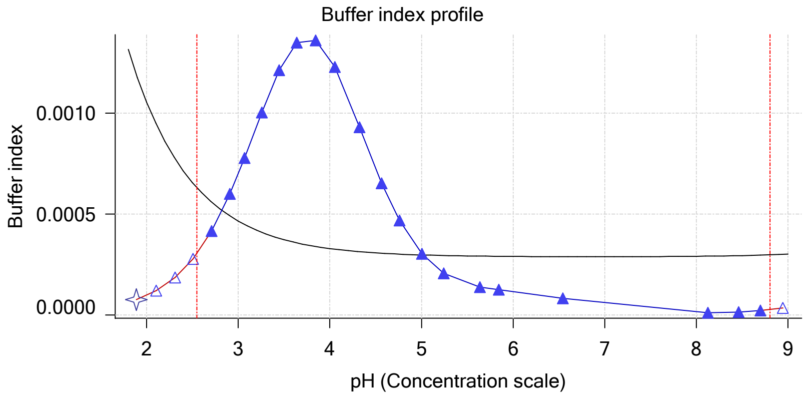


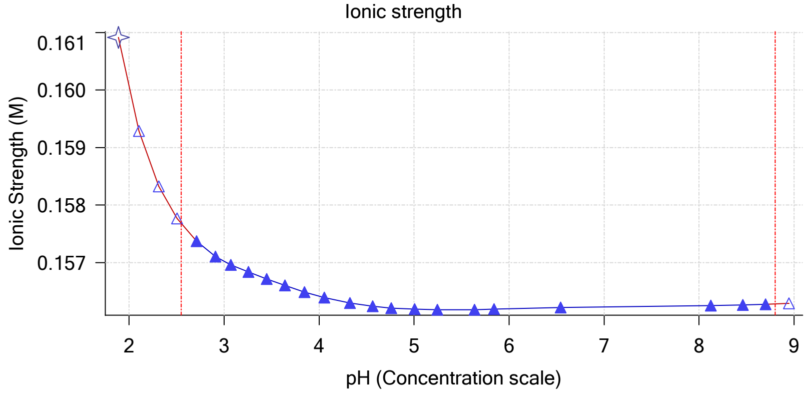


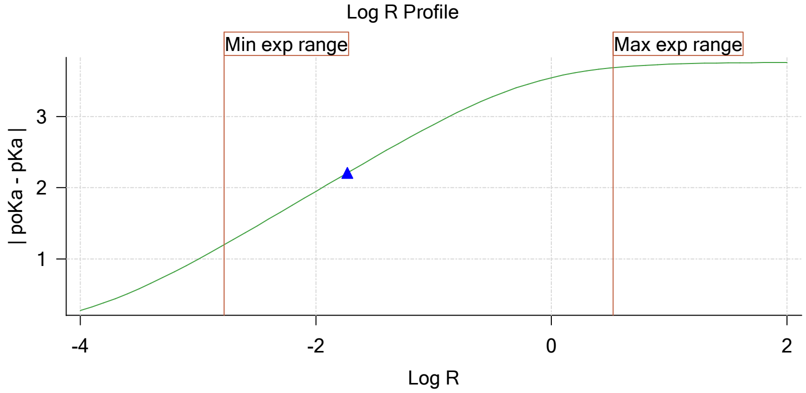


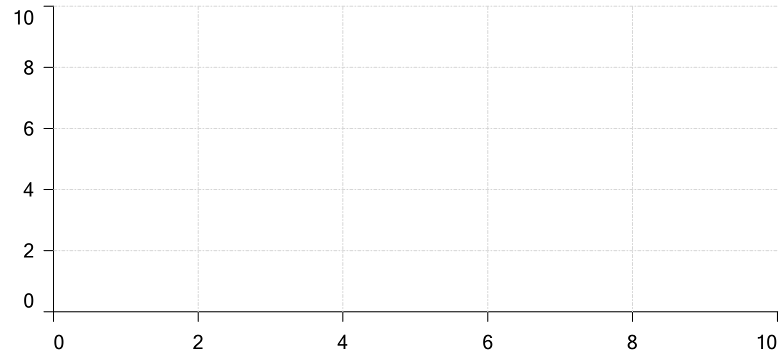

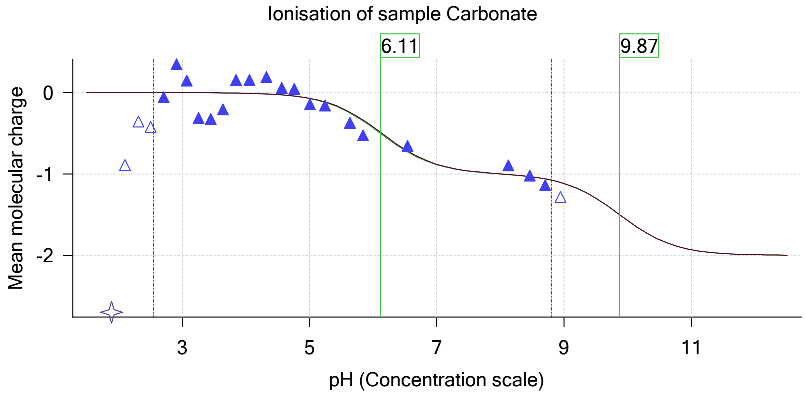

| | | | |
|---|---|---|---|
| Sample name: | **M04_octanol** | Experiment start time: | **3/24/2018 1:34:06 AM** |
| Assay name: | **pH-metric high logP** | Analyst: | **Dorothy Levorse** |
| Assay ID: | **18C-24002** | Instrument ID: | **T312060** |
| Filename: | **C:\Sirius_T3\Mehtap\20180323_exp33_logP_T3-2\18C-24002_M04_octanol_pH-metric high logP.t3r** | | |

### pH-metric high logP Titration 2 of 3 18C-24002 Points 25 to 51

#### Overall results

| | |
|---|---|
| RMSD | 0.334 |
| Average ionic strength | 0.162 M |
| Average temperature | 25.0°C |
| Partition ratio | 0.0644 : 1 |
| Analyte concentration range | 2267.2 µM to 2336.0 µM |
| Total points considered | 24 of 27 |

#### Warnings and errors

| | |
|---|---|
| Errors | None |
| Warnings | None |

#### Four-Plus parameters

| | | | |
|---|---|---|---|
| Alpha | 0.119 | 3/24/2018 1:34:06 AM | C:\Sirius_T3\HCl18C23.t3r |
| S | 0.9972 | 3/24/2018 1:34:06 AM | C:\Sirius_T3\HCl18C23.t3r |
| jH | 0.9 | 3/24/2018 1:34:06 AM | C:\Sirius_T3\HCl18C23.t3r |
| jOH | -0.3 | 3/24/2018 1:34:06 AM | C:\Sirius_T3\HCl18C23.t3r |

#### Titrants

| | | | |
|---|---|---|---|
| 0.50 M HCl | 0.997124 | 3/24/2018 1:34:06 AM | C:\Sirius_T3\HCl18C23.t3r |
| 0.50 M KOH | 1.003190 | 3/24/2018 1:34:06 AM | C:\Sirius_T3\KOH18C23.t3r |

#### Sample

| | |
|---|---|
| M04_octanol concentration factor | 0.920 |
| Base pKa 1 | 5.97 |
| logP (XH +) | 0.19 |
| logP (neutral X) | 4.01 |

#### Sample graphs

Ionisation of sample M04_octanol

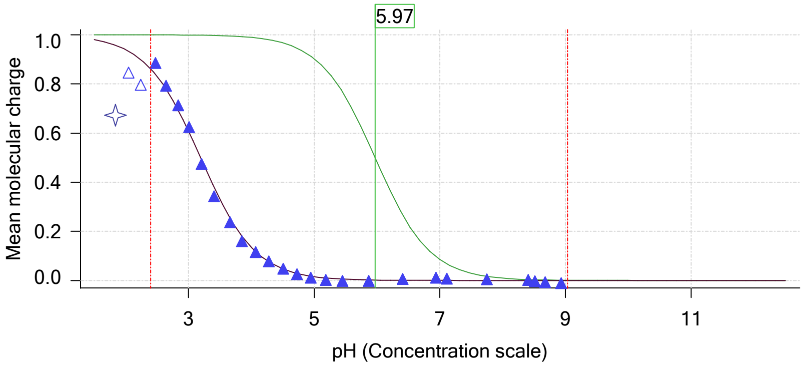


Distribution of species for sample M04_octanol

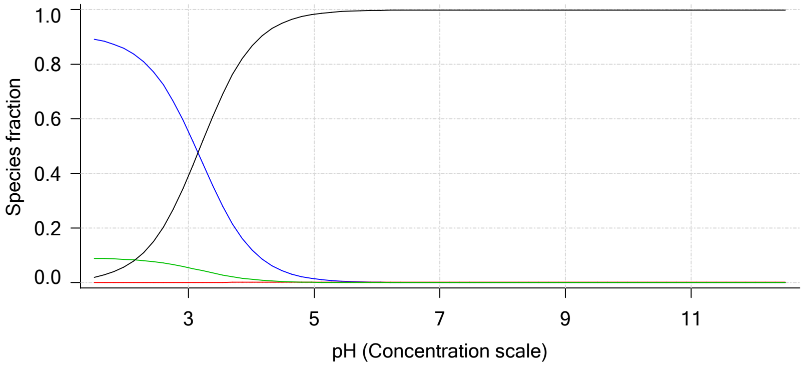

| | | | |
|---|---|---|---|
| Sample name: | **M04_octanol** | Experiment start time: | **3/24/2018 1:34:06 AM** |
| Assay name: | **pH-metric high logP** | Analyst: | **Dorothy Levorse** |
| Assay ID: | **18C-24002** | Instrument ID: | **T312060** |
| Filename: | **C:\Sirius_T3\Mehtap\20180323_exp33_logP_T3-2\18C-24002_M04_octanol_pH-metric high logP.t3r** | | |

#### Sample graphs (continued)

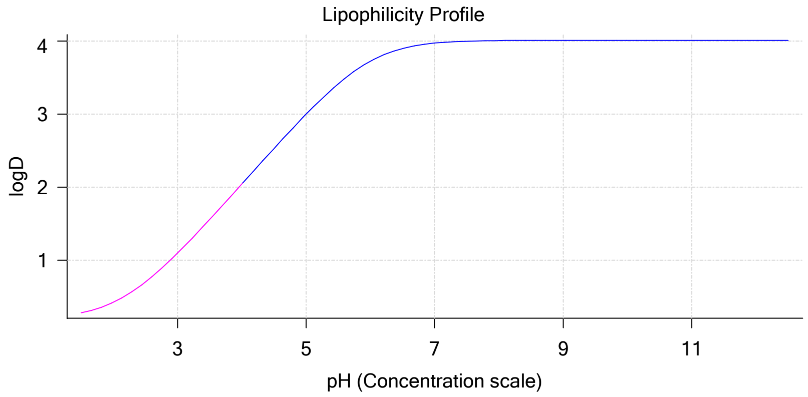


#### Sample logD and percent species

| pH | M04_octanol logD | M04_octanol M04_octanolH | M04_octanol M04_octanol | M04_octanol M04_octanolH* | M04_octanol M04_octanol* | Comment |
|---|---|---|---|---|---|---|
| 1.000 | 0.22 | 90.35 % | 0.00 % | 9.01 % | 0.64 % | |
| 1.200 | 0.24 | 90.02 % | 0.00 % | 8.97 % | 1.01 % | Stomach pH |
| 2.000 | 0.42 | 85.44 % | 0.01 % | 8.52 % | 6.03 % | |
| 3.000 | 1.10 | 55.36 % | 0.06 % | 5.52 % | 39.07 % | |
| 4.000 | 2.04 | 12.24 % | 0.13 % | 1.22 % | 86.40 % | |
| 5.000 | 3.00 | 1.39 % | 0.15 % | 0.14 % | 98.32 % | |
| 6.000 | 3.72 | 0.14 % | 0.15 % | 0.01 % | 99.69 % | |
| 6.500 | 3.90 | 0.04 % | 0.15 % | 0.00 % | 99.80 % | |
| 7.000 | 3.97 | 0.01 % | 0.15 % | 0.00 % | 99.83 % | |
| 7.400 | 3.99 | 0.01 % | 0.15 % | 0.00 % | 99.84 % | Blood pH |
| 8.000 | 4.01 | 0.00 % | 0.15 % | 0.00 % | 99.85 % | |
| 9.000 | 4.01 | 0.00 % | 0.15 % | 0.00 % | 99.85 % | |
| 10.000 | 4.01 | 0.00 % | 0.15 % | 0.00 % | 99.85 % | |
| 11.000 | 4.01 | 0.00 % | 0.15 % | 0.00 % | 99.85 % | |
| 12.000 | 4.01 | 0.00 % | 0.15 % | 0.00 % | 99.85 % | |

#### Carbonate and acidity

| | |
|---|---|
| Carbonate | 0.106 mM |
| Acidity error | -0.298 mM |

#### Other graphs

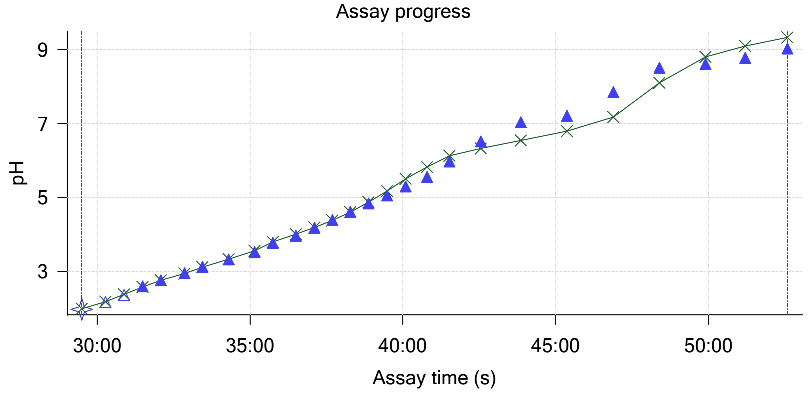


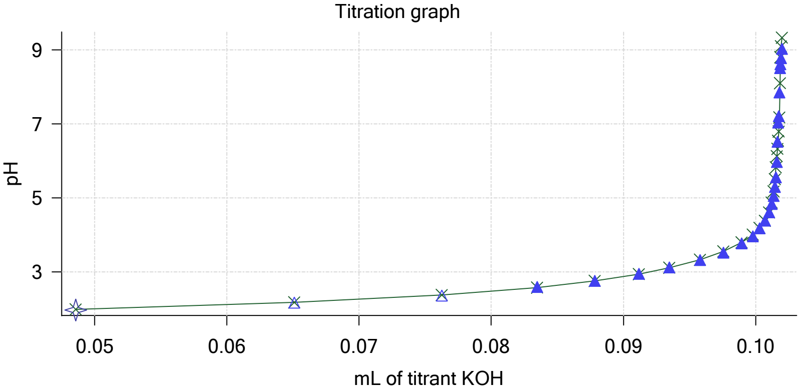

Sample name: **M04_octanol**
Assay name: **pH-metric high logP**
Assay ID: **18C-24002**
Filename: **C:\Sirius_T3\Mehtap\20180323_exp33_logP_T3-2\18C-24002_M04_octanol_pH-metric high logP.t3r**

Experiment start time: **3/24/2018 1:34:06 AM**
Analyst: **Dorothy Levorse**
Instrument ID: **T312060**

#### Other graphs (continued)

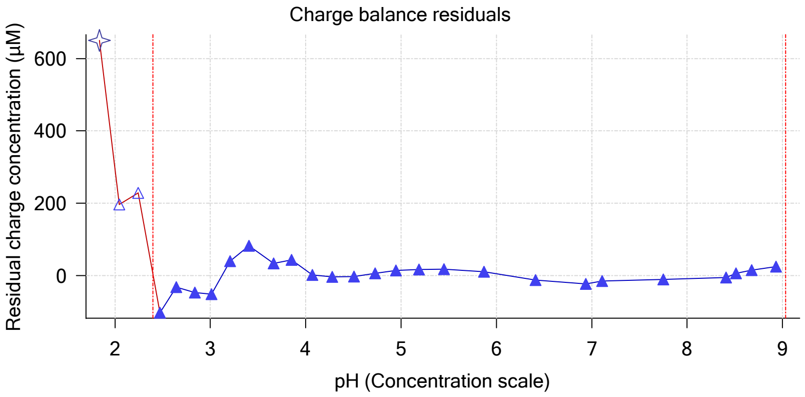


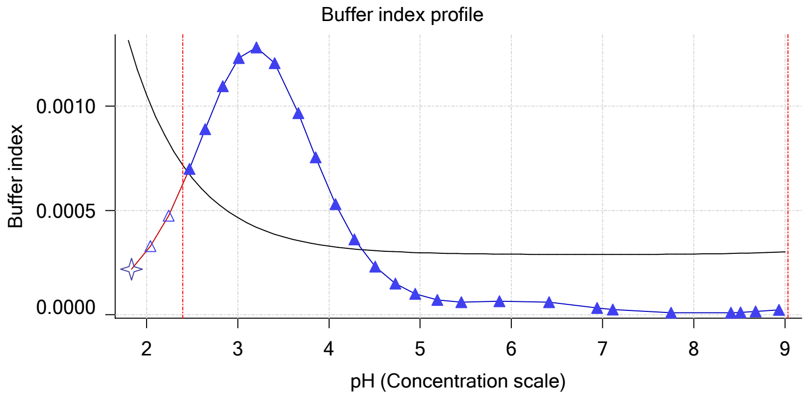


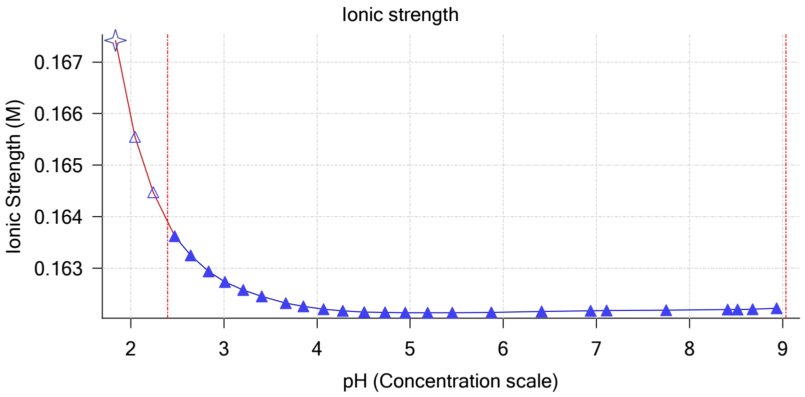


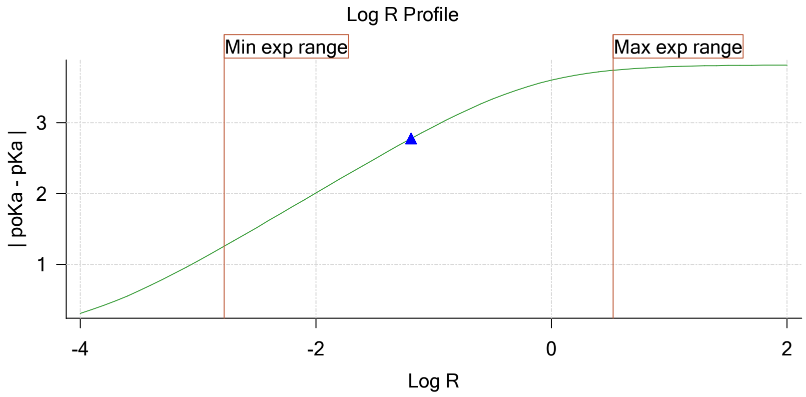


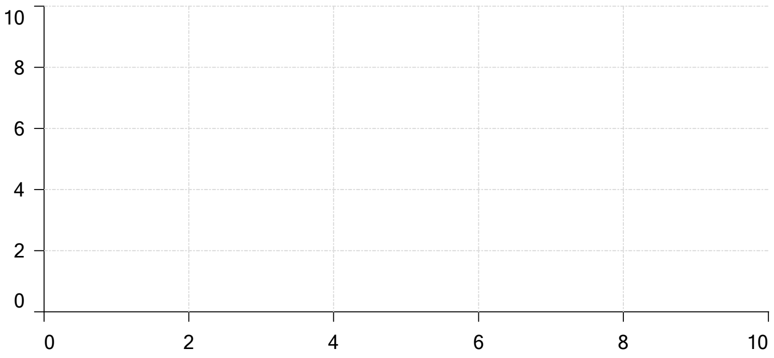

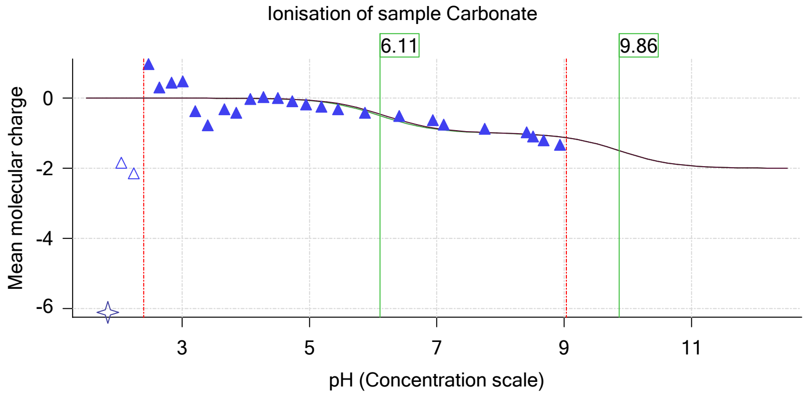

| | | | |
|---|---|---|---|
| Sample name: | **M04_octanol** | Experiment start time: | **3/24/2018 1:34:06 AM** |
| Assay name: | **pH-metric high logP** | Analyst: | **Dorothy Levorse** |
| Assay ID: | **18C-24002** | Instrument ID: | **T312060** |
| Filename: | **C:\Sirius_T3\Mehtap\20180323_exp33_logP_T3-2\18C-24002_M04_octanol_pH-metric high logP.t3r** | | |

### pH-metric high logP Titration 3 of 3 18C-24002 Points 52 to 75

#### Overall results

| | |
|---|---|
| RMSD | 0.452 |
| Average ionic strength | 0.168 M |
| Average temperature | 25.0°C |
| Partition ratio | 0.2805 : 1 |
| Analyte concentration range | 1766.3 µM to 1811.0 µM |
| Total points considered | 19 of 24 |

#### Warnings and errors

| | |
|---|---|
| Errors | None |
| Warnings | None |

#### Four-Plus parameters

| | | | |
|---|---|---|---|
| Alpha | 0.119 | 3/24/2018 1:34:06 AM | C:\Sirius_T3\HCl18C23.t3r |
| S | 0.9972 | 3/24/2018 1:34:06 AM | C:\Sirius_T3\HCl18C23.t3r |
| jH | 0.9 | 3/24/2018 1:34:06 AM | C:\Sirius_T3\HCl18C23.t3r |
| jOH | -0.3 | 3/24/2018 1:34:06 AM | C:\Sirius_T3\HCl18C23.t3r |

#### Titrants

| | | | |
|---|---|---|---|
| 0.50 M HCl | 0.997124 | 3/24/2018 1:34:06 AM | C:\Sirius_T3\HCl18C23.t3r |
| 0.50 M KOH | 1.003190 | 3/24/2018 1:34:06 AM | C:\Sirius_T3\KOH18C23.t3r |

#### Sample

| | |
|---|---|
| M04_octanol concentration factor | 0.952 |
| Base pKa 1 | 5.97 |
| logP (XH +) | 0.19 |
| logP (neutral X) | 3.70 |

#### Sample graphs

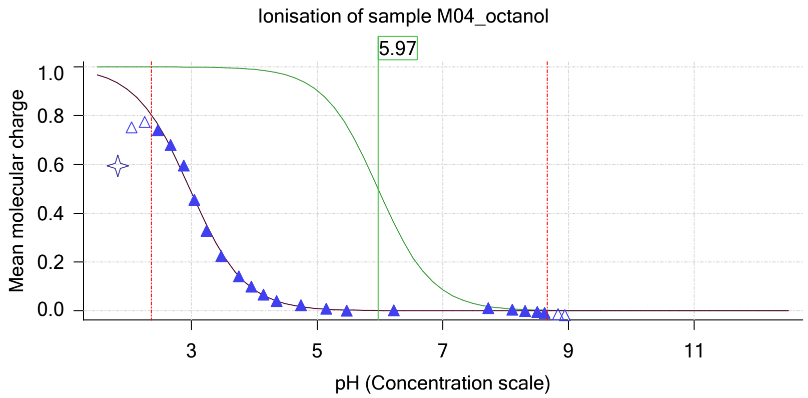


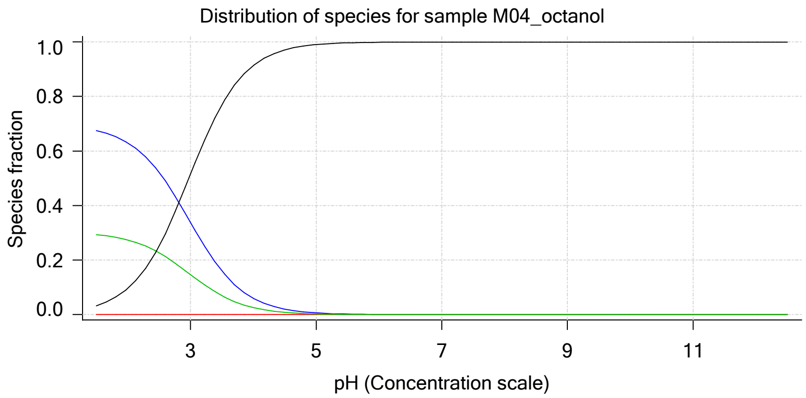

| | | | |
|---|---|---|---|
| Sample name: | **M04_octanol** | Experiment start time: | **3/24/2018 1:34:06 AM** |
| Assay name: | **pH-metric high logP** | Analyst: | **Dorothy Levorse** |
| Assay ID: | **18C-24002** | Instrument ID: | **T312060** |
| Filename: | **C:\Sirius_T3\Mehtap\20180323_exp33_logP_T3-2\18C-24002_M04_octanol_pH-metric high logP.t3r** | | |

#### Sample graphs (continued)

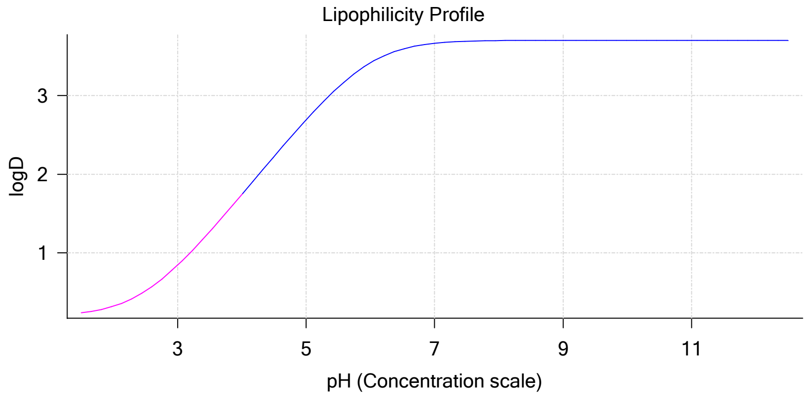


#### Sample logD and percent species

| pH | M04_octanol logD | M04_octanol M04_octanolH | M04_octanol M04_octanol | M04_octanol M04_octanolH* | M04_octanol M04_octanol* | Comment |
|---|---|---|---|---|---|---|
| 1.000 | 0.20 | 68.98 % | 0.00 % | 29.97 % | 1.05 % | |
| 1.200 | 0.21 | 68.56 % | 0.00 % | 29.78 % | 1.65 % | Stomach pH |
| 2.000 | 0.32 | 63.02 % | 0.01 % | 27.38 % | 9.60 % | |
| 3.000 | 0.84 | 33.81 % | 0.04 % | 14.69 % | 51.47 % | |
| 4.000 | 1.74 | 6.00 % | 0.06 % | 2.61 % | 91.33 % | |
| 5.000 | 2.69 | 0.65 % | 0.07 % | 0.28 % | 99.00 % | |
| 6.000 | 3.42 | 0.07 % | 0.07 % | 0.03 % | 99.84 % | |
| 6.500 | 3.59 | 0.02 % | 0.07 % | 0.01 % | 99.90 % | |
| 7.000 | 3.67 | 0.01 % | 0.07 % | 0.00 % | 99.92 % | |
| 7.400 | 3.69 | 0.00 % | 0.07 % | 0.00 % | 99.93 % | Blood pH |
| 8.000 | 3.70 | 0.00 % | 0.07 % | 0.00 % | 99.93 % | |
| 9.000 | 3.70 | 0.00 % | 0.07 % | 0.00 % | 99.93 % | |
| 10.000 | 3.70 | 0.00 % | 0.07 % | 0.00 % | 99.93 % | |
| 11.000 | 3.70 | 0.00 % | 0.07 % | 0.00 % | 99.93 % | |
| 12.000 | 3.70 | 0.00 % | 0.07 % | 0.00 % | 99.93 % | |

#### Carbonate and acidity

| | |
|---|---|
| Carbonate | 0.104 mM |
| Acidity error | -0.102 mM |

#### Other graphs

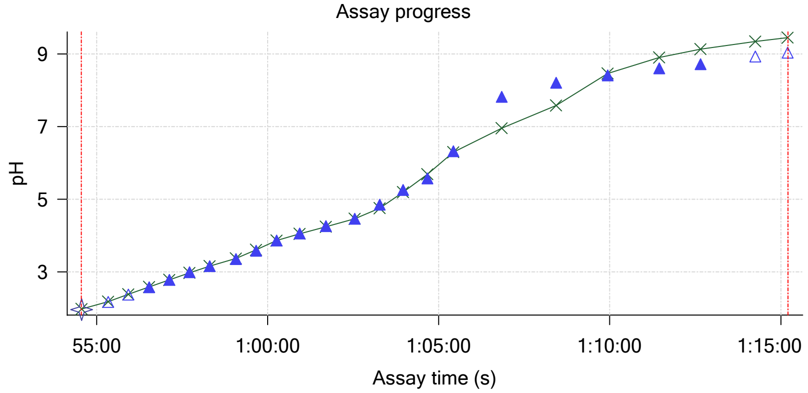


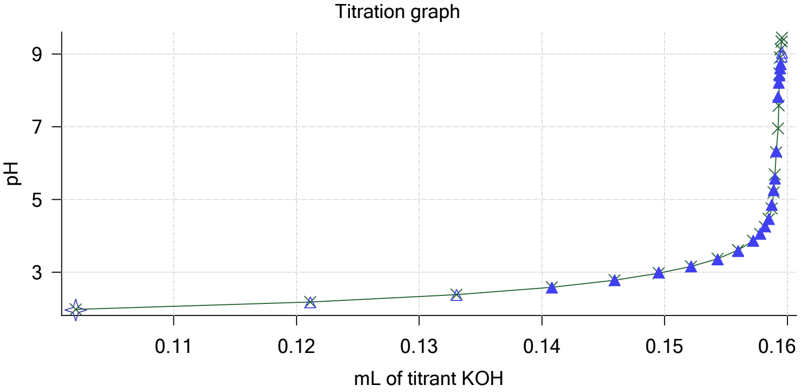

Sample name: **M04_octanol**
Assay name: **pH-metric high logP**
Assay ID: **18C-24002**
Filename: **C:\Sirius_T3\Mehtap\20180323_exp33_logP_T3-2\18C-24002_M04_octanol_pH-metric high logP.t3r**

Experiment start time: **3/24/2018 1:34:06 AM**
Analyst: **Dorothy Levorse**
Instrument ID: **T312060**

#### Other graphs (continued)

Charge balance residuals

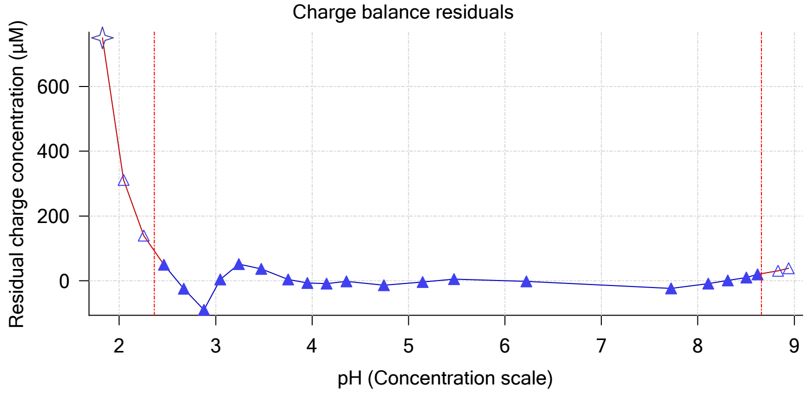


Buffer index profile

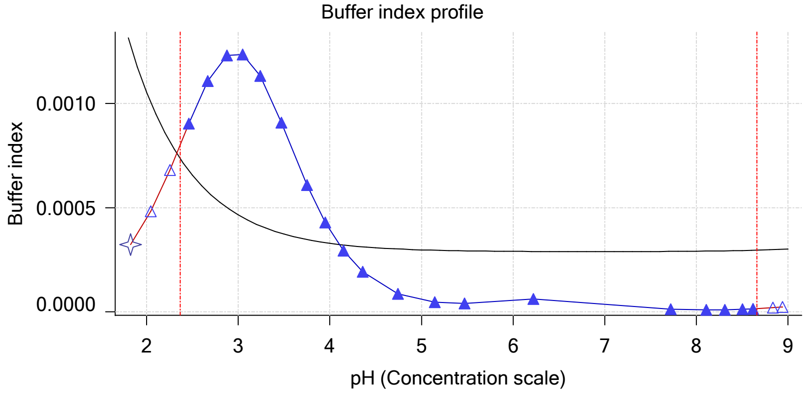


Ionic strength

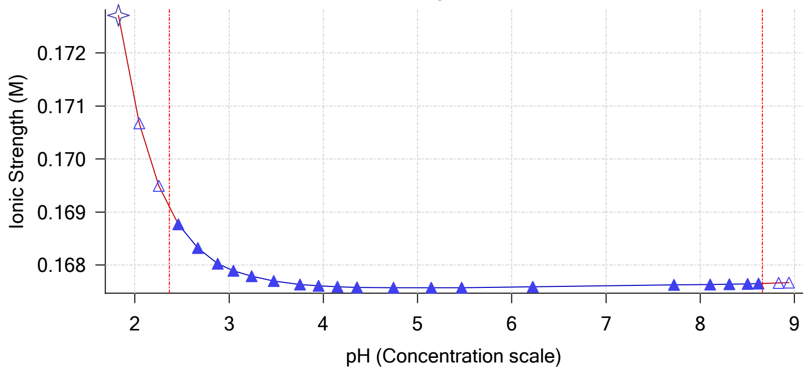


Log R Profile

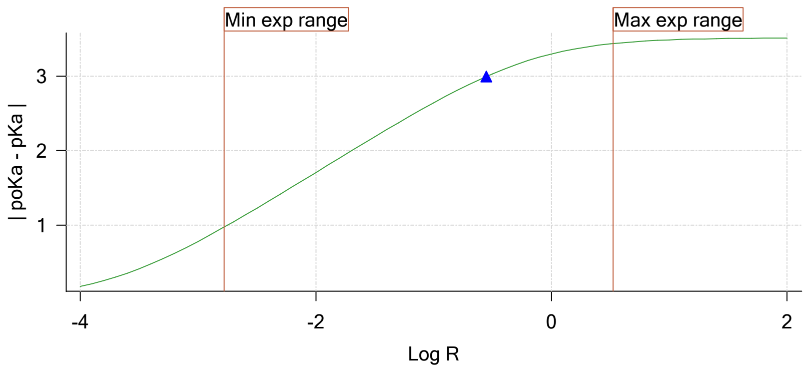


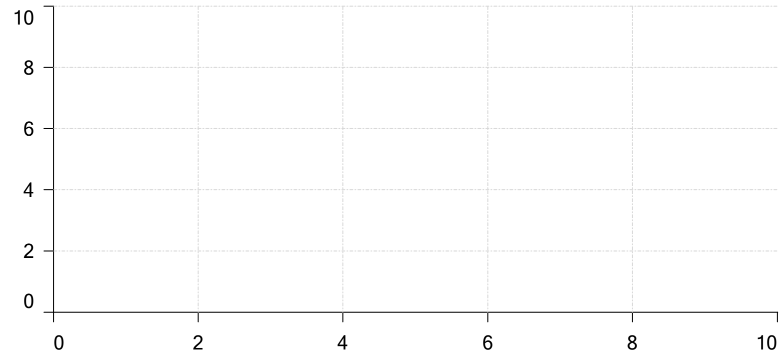


Ionisation of sample Carbonate

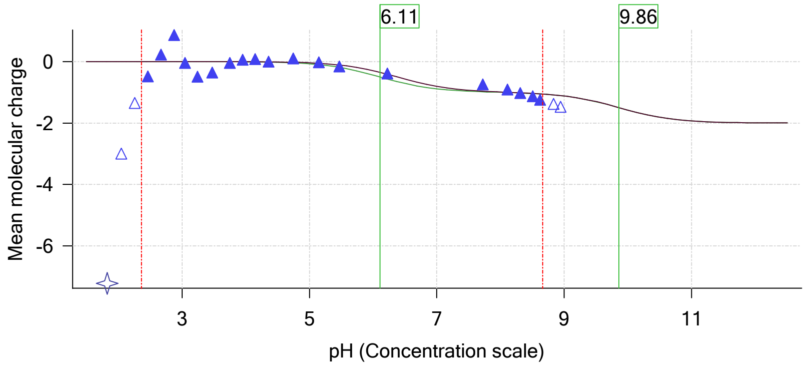

| | | | |
|---|---|---|---|
| Sample name: | **M04_octanol** | Experiment start time: | **3/24/2018 1:34:06 AM** |
| Assay name: | **pH-metric high logP** | Analyst: | **Dorothy Levorse** |
| Assay ID: | **18C-24002** | Instrument ID: | **T312060** |
| Filename: | **C:\Sirius_T3\Mehtap\20180323_exp33_logP_T3-2\18C-24002_M04_octanol_pH-metric high logP.t3r** | | |

#### Assay Model

| Settings | Value | Date/Time changed | Imported from |
|---|---|---|---|
| Sample name | M04_octanol | 3/9/2018 4:34:21 PM | User entered value |
| Sample by | Weight | | Default value |
| Sample weight | 0.001110 g | 3/23/2018 5:00:32 PM | User entered value |
| Formula weight | 269.73 g/mol | 3/9/2018 4:34:21 PM | User entered value |
| Solubility | Unknown | | Default value |
| Molecular weight | 269.73 | 3/9/2018 4:34:21 PM | User entered value |
| Individual pKa ionic environments | No | | Default value |
| Number of pKas | 1 | 3/9/2018 4:34:21 PM | User entered value |
| Sample is a | Base | 3/9/2018 4:34:21 PM | User entered value |
| pKa 1 | 5.97 | 3/9/2018 4:34:21 PM | User entered value |
| logp (XH +) | 0.19 | 3/9/2018 4:34:33 PM | User entered value |
| logP (neutral X) | 3.50 | 3/23/2018 2:32:46 PM | User entered value |

#### Events

| Time | Event | Water | Acid | Base | Octanol | pH | dpH/dt | pH R-squared | pH SD | dpH/dt time |
|---|---|---|---|---|---|---|---|---|---|---|
| 5:00.2 | Initial pH = 9.63 | | | | | | | | | |
| 7:59.7 | Data point 1 | 1.50000 mL | 0.04892 mL | 0.00183 mL | 0.03001 mL | 2.014 | -0.00160 | 0.32049 | 0.00014 | 10.5 s |
| 8:46.4 | Data point 2 | 1.50000 mL | 0.04892 mL | 0.01635 mL | 0.03001 mL | 2.224 | -0.00074 | 0.14269 | 0.00010 | 10.0 s |
| 9:21.9 | Data point 3 | 1.50000 mL | 0.04892 mL | 0.02545 mL | 0.03001 mL | 2.429 | 0.00290 | 0.05785 | 0.00060 | 10.0 s |
| 9:57.5 | Data point 4 | 1.50000 mL | 0.04892 mL | 0.03109 mL | 0.03001 mL | 2.620 | -0.00606 | 0.72611 | 0.00035 | 10.0 s |
| 10:33.1 | Data point 5 | 1.50000 mL | 0.04892 mL | 0.03478 mL | 0.03001 mL | 2.822 | -0.00318 | 0.34754 | 0.00027 | 10.0 s |
| 11:08.6 | Data point 6 | 1.50000 mL | 0.04892 mL | 0.03718 mL | 0.03001 mL | 3.022 | -0.00788 | 0.20158 | 0.00087 | 10.0 s |
| 11:44.0 | Data point 7 | 1.50000 mL | 0.04892 mL | 0.03883 mL | 0.03001 mL | 3.182 | -0.00246 | 0.53024 | 0.00017 | 10.0 s |
| 12:34.9 | Data point 8 | 1.50000 mL | 0.04892 mL | 0.04052 mL | 0.03001 mL | 3.368 | -0.01119 | 0.56806 | 0.00073 | 10.0 s |
| 13:25.9 | Data point 9 | 1.50000 mL | 0.04892 mL | 0.04182 mL | 0.03001 mL | 3.557 | -0.00577 | 0.09662 | 0.00092 | 10.0 s |
| 14:11.6 | Data point 10 | 1.50000 mL | 0.04892 mL | 0.04294 mL | 0.03001 mL | 3.749 | -0.00944 | 0.26941 | 0.00090 | 10.0 s |
| 14:47.0 | Data point 11 | 1.50000 mL | 0.04892 mL | 0.04393 mL | 0.03001 mL | 3.955 | -0.00902 | 0.80462 | 0.00050 | 10.0 s |
| 15:22.5 | Data point 12 | 1.50000 mL | 0.04892 mL | 0.04497 mL | 0.03001 mL | 4.163 | -0.00871 | 0.81314 | 0.00048 | 10.5 s |
| 15:58.4 | Data point 13 | 1.50000 mL | 0.04892 mL | 0.04600 mL | 0.03001 mL | 4.430 | -0.01386 | 0.79748 | 0.00077 | 11.0 s |
| 16:45.2 | Data point 14 | 1.50000 mL | 0.04892 mL | 0.04673 mL | 0.03001 mL | 4.671 | -0.00488 | 0.10408 | 0.00075 | 11.5 s |
| 17:22.1 | Data point 15 | 1.50000 mL | 0.04892 mL | 0.04711 mL | 0.03001 mL | 4.865 | -0.01667 | 0.86899 | 0.00088 | 14.5 s |
| 18:12.3 | Data point 16 | 1.50000 mL | 0.04892 mL | 0.04751 mL | 0.03001 mL | 5.110 | -0.01586 | 0.79314 | 0.00088 | 16.0 s |
| 18:53.6 | Data point 17 | 1.50000 mL | 0.04892 mL | 0.04770 mL | 0.03001 mL | 5.348 | -0.01986 | 0.98346 | 0.00099 | 20.5 s |
| 19:44.8 | Data point 18 | 1.50000 mL | 0.04892 mL | 0.04795 mL | 0.03001 mL | 5.739 | -0.01915 | 0.96564 | 0.00096 | 26.5 s |
| 20:36.6 | Data point 19 | 1.50000 mL | 0.04892 mL | 0.04807 mL | 0.03001 mL | 5.946 | -0.01903 | 0.92495 | 0.00098 | 30.5 s |
| 21:37.5 | Data point 20 | 1.50000 mL | 0.04892 mL | 0.04819 mL | 0.03001 mL | 6.643 | -0.02497 | 0.94842 | 0.00127 | Timed out at 59.5 s |
| 23:08.1 | Data point 21 | 1.50000 mL | 0.04892 mL | 0.04833 mL | 0.03001 mL | 8.221 | -0.07094 | 0.99161 | 0.00352 | Timed out at 59.5 s |
| 24:43.7 | Data point 22 | 1.50000 mL | 0.04892 mL | 0.04840 mL | 0.03001 mL | 8.555 | -0.03178 | 0.97425 | 0.00159 | Timed out at 59.5 s |
| 26:19.4 | Data point 23 | 1.50000 mL | 0.04892 mL | 0.04847 mL | 0.03001 mL | 8.794 | -0.01850 | 0.95661 | 0.00093 | 58.5 s |
| 27:48.5 | Data point 24 | 1.50000 mL | 0.04892 mL | 0.04857 mL | 0.03001 mL | 9.039 | -0.01975 | 0.95215 | 0.00100 | 41.5 s |
| 29:30.0 | Data point 25 | 1.50000 mL | 0.10310 mL | 0.04857 mL | 0.11002 mL | 1.964 | -0.00880 | 0.24648 | 0.00088 | 10.0 s |
| 30:16.3 | Data point 26 | 1.50000 mL | 0.10310 mL | 0.06510 mL | 0.11002 mL | 2.166 | -0.01564 | 0.64906 | 0.00096 | 11.0 s |
| 30:53.0 | Data point 27 | 1.50000 mL | 0.10310 mL | 0.07627 mL | 0.11002 mL | 2.359 | -0.00237 | 0.24657 | 0.00024 | 10.5 s |
| 31:29.0 | Data point 28 | 1.50000 mL | 0.10310 mL | 0.08347 mL | 0.11002 mL | 2.587 | 0.00566 | 0.08086 | 0.00098 | 10.5 s |
| 32:05.1 | Data point 29 | 1.50000 mL | 0.10310 mL | 0.08784 mL | 0.11002 mL | 2.757 | -0.00202 | 0.33078 | 0.00017 | 10.5 s |
| 32:51.5 | Data point 30 | 1.50000 mL | 0.10310 mL | 0.09118 mL | 0.11002 mL | 2.950 | -0.00478 | 0.66757 | 0.00029 | 10.0 s |
| 33:26.8 | Data point 31 | 1.50000 mL | 0.10310 mL | 0.09346 mL | 0.11002 mL | 3.123 | -0.00470 | 0.90243 | 0.00024 | 10.0 s |
| 34:17.8 | Data point 32 | 1.50000 mL | 0.10310 mL | 0.09579 mL | 0.11002 mL | 3.317 | -0.00344 | 0.59909 | 0.00022 | 10.5 s |
| 35:09.2 | Data point 33 | 1.50000 mL | 0.10310 mL | 0.09755 mL | 0.11002 mL | 3.515 | -0.00933 | 0.53854 | 0.00063 | 10.0 s |
| 35:44.5 | Data point 34 | 1.50000 mL | 0.10310 mL | 0.09894 mL | 0.11002 mL | 3.772 | -0.00653 | 0.77129 | 0.00037 | 10.0 s |
| 36:30.2 | Data point 35 | 1.50000 mL | 0.10310 mL | 0.09979 mL | 0.11002 mL | 3.961 | -0.00468 | 0.84923 | 0.00025 | 10.5 s |
| 37:06.1 | Data point 36 | 1.50000 mL | 0.10310 mL | 0.10031 mL | 0.11002 mL | 4.178 | -0.00915 | 0.61492 | 0.00058 | 10.0 s |
| 37:41.5 | Data point 37 | 1.50000 mL | 0.10310 mL | 0.10071 mL | 0.11002 mL | 4.385 | -0.00988 | 0.57323 | 0.00064 | 10.0 s |

| | | | |
|---|---|---|---|
| Sample name: | **M04_octanol** | Experiment start time: | **3/24/2018 1:34:06 AM** |
| Assay name: | **pH-metric high logP** | Analyst: | **Dorothy Levorse** |
| Assay ID: | **18C-24002** | Instrument ID: | **T312060** |
| Filename: | **C:\Sirius_T3\Mehtap\20180323_exp33_logP_T3-2\18C-24002_M04_octanol_pH-metric high logP.t3r** | | |

#### Events (continued)

| Time | Event | Water | Acid | Base | Octanol | pH | dpH/dt | pH R-squared | pH SD | dpH/dt time |
|---|---|---|---|---|---|---|---|---|---|---|
| 38:16.9 | Data point 38 | 1.50000 mL | 0.10310 mL | 0.10101 mL | 0.11002 mL | 4.612 | -0.00861 | 0.68608 | 0.00051 | 10.5 s |
| 38:52.8 | Data point 39 | 1.50000 mL | 0.10310 mL | 0.10122 mL | 0.11002 mL | 4.835 | -0.01226 | 0.66939 | 0.00074 | 10.5 s |
| 39:28.7 | Data point 40 | 1.50000 mL | 0.10310 mL | 0.10136 mL | 0.11002 mL | 5.051 | -0.01367 | 0.71800 | 0.00080 | 11.0 s |
| 40:05.1 | Data point 41 | 1.50000 mL | 0.10310 mL | 0.10146 mL | 0.11002 mL | 5.292 | -0.01155 | 0.59589 | 0.00074 | 12.0 s |
| 40:47.6 | Data point 42 | 1.50000 mL | 0.10310 mL | 0.10153 mL | 0.11002 mL | 5.555 | -0.01710 | 0.84028 | 0.00092 | 13.5 s |
| 41:31.6 | Data point 43 | 1.50000 mL | 0.10310 mL | 0.10160 mL | 0.11002 mL | 5.972 | -0.01840 | 0.92739 | 0.00094 | 31.0 s |
| 42:33.2 | Data point 44 | 1.50000 mL | 0.10310 mL | 0.10165 mL | 0.11002 mL | 6.512 | -0.01880 | 0.91936 | 0.00097 | 53.0 s |
| 43:51.7 | Data point 45 | 1.50000 mL | 0.10310 mL | 0.10169 mL | 0.11002 mL | 7.037 | -0.04193 | 0.98601 | 0.00209 | Timed out at 59.5 s |
| 45:22.1 | Data point 46 | 1.50000 mL | 0.10310 mL | 0.10174 mL | 0.11002 mL | 7.209 | -0.04636 | 0.98987 | 0.00230 | Timed out at 59.5 s |
| 46:52.7 | Data point 47 | 1.50000 mL | 0.10310 mL | 0.10179 mL | 0.11002 mL | 7.847 | -0.06759 | 0.99607 | 0.00335 | Timed out at 59.5 s |
| 48:23.2 | Data point 48 | 1.50000 mL | 0.10310 mL | 0.10183 mL | 0.11002 mL | 8.503 | -0.03401 | 0.99546 | 0.00168 | Timed out at 59.5 s |
| 49:53.7 | Data point 49 | 1.50000 mL | 0.10310 mL | 0.10188 mL | 0.11002 mL | 8.608 | -0.01529 | 0.60040 | 0.00097 | 47.5 s |
| 51:11.7 | Data point 50 | 1.50000 mL | 0.10310 mL | 0.10193 mL | 0.11002 mL | 8.772 | -0.01850 | 0.93390 | 0.00095 | 52.0 s |
| 52:34.3 | Data point 51 | 1.50000 mL | 0.10310 mL | 0.10200 mL | 0.11002 mL | 9.026 | -0.01279 | 0.96553 | 0.00064 | 36.5 s |
| 54:34.0 | Data point 52 | 1.50000 mL | 0.16030 mL | 0.10200 mL | 0.51002 mL | 1.954 | -0.00445 | 0.52317 | 0.00030 | 10.0 s |
| 55:20.3 | Data point 53 | 1.50000 mL | 0.16030 mL | 0.12112 mL | 0.51002 mL | 2.166 | 0.01490 | 0.84480 | 0.00080 | 10.0 s |
| 55:56.0 | Data point 54 | 1.50000 mL | 0.16030 mL | 0.13304 mL | 0.51002 mL | 2.372 | 0.00293 | 0.49233 | 0.00021 | 10.5 s |
| 56:32.0 | Data point 55 | 1.50000 mL | 0.16030 mL | 0.14080 mL | 0.51002 mL | 2.578 | 0.01404 | 0.59238 | 0.00090 | 10.0 s |
| 57:07.6 | Data point 56 | 1.50000 mL | 0.16030 mL | 0.14593 mL | 0.51002 mL | 2.782 | -0.00325 | 0.08785 | 0.00054 | 10.0 s |
| 57:43.1 | Data point 57 | 1.50000 mL | 0.16030 mL | 0.14953 mL | 0.51002 mL | 2.990 | -0.00261 | 0.10471 | 0.00040 | 10.0 s |
| 58:18.5 | Data point 58 | 1.50000 mL | 0.16030 mL | 0.15216 mL | 0.51002 mL | 3.158 | -0.00222 | 0.09026 | 0.00037 | 10.0 s |
| 59:04.3 | Data point 59 | 1.50000 mL | 0.16030 mL | 0.15433 mL | 0.51002 mL | 3.351 | -0.00880 | 0.28962 | 0.00081 | 10.0 s |
| 59:39.8 | Data point 60 | 1.50000 mL | 0.16030 mL | 0.15600 mL | 0.51002 mL | 3.582 | -0.01386 | 0.91128 | 0.00072 | 10.5 s |
| 1:00:15.8 | Data point 61 | 1.50000 mL | 0.16030 mL | 0.15722 mL | 0.51002 mL | 3.861 | -0.01252 | 0.59534 | 0.00080 | 10.0 s |
| 1:00:56.3 | Data point 62 | 1.50000 mL | 0.16030 mL | 0.15779 mL | 0.51002 mL | 4.058 | -0.00480 | 0.17045 | 0.00057 | 10.0 s |
| 1:01:42.1 | Data point 63 | 1.50000 mL | 0.16030 mL | 0.15818 mL | 0.51002 mL | 4.256 | -0.00431 | 0.26492 | 0.00041 | 10.0 s |
| 1:02:32.9 | Data point 64 | 1.50000 mL | 0.16030 mL | 0.15849 mL | 0.51002 mL | 4.464 | 0.00183 | 0.00838 | 0.00099 | 18.5 s |
| 1:03:16.7 | Data point 65 | 1.50000 mL | 0.16030 mL | 0.15873 mL | 0.51002 mL | 4.850 | -0.01054 | 0.48317 | 0.00075 | 10.5 s |
| 1:03:57.8 | Data point 66 | 1.50000 mL | 0.16030 mL | 0.15889 mL | 0.51002 mL | 5.251 | -0.01079 | 0.55670 | 0.00071 | 12.0 s |
| 1:04:40.3 | Data point 67 | 1.50000 mL | 0.16030 mL | 0.15898 mL | 0.51002 mL | 5.574 | -0.01750 | 0.78457 | 0.00098 | 15.0 s |
| 1:05:25.7 | Data point 68 | 1.50000 mL | 0.16030 mL | 0.15910 mL | 0.51002 mL | 6.324 | -0.01848 | 0.97545 | 0.00092 | 54.0 s |
| 1:06:50.3 | Data point 69 | 1.50000 mL | 0.16030 mL | 0.15924 mL | 0.51002 mL | 7.819 | -0.10642 | 0.99284 | 0.00528 | Timed out at 59.5 s |
| 1:08:25.9 | Data point 70 | 1.50000 mL | 0.16030 mL | 0.15931 mL | 0.51002 mL | 8.203 | -0.07202 | 0.98050 | 0.00359 | Timed out at 59.5 s |
| 1:09:56.4 | Data point 71 | 1.50000 mL | 0.16030 mL | 0.15936 mL | 0.51002 mL | 8.406 | -0.04547 | 0.97923 | 0.00227 | Timed out at 59.5 s |
| 1:11:26.8 | Data point 72 | 1.50000 mL | 0.16030 mL | 0.15941 mL | 0.51002 mL | 8.600 | -0.01814 | 0.81212 | 0.00099 | 42.0 s |
| 1:12:39.4 | Data point 73 | 1.50000 mL | 0.16030 mL | 0.15945 mL | 0.51002 mL | 8.714 | -0.02169 | 0.85709 | 0.00116 | Timed out at 59.5 s |
| 1:14:15.1 | Data point 74 | 1.50000 mL | 0.16030 mL | 0.15953 mL | 0.51002 mL | 8.928 | -0.01718 | 0.73352 | 0.00099 | 26.0 s |
| 1:15:11.8 | Data point 75 | 1.50000 mL | 0.16030 mL | 0.15957 mL | 0.51002 mL | 9.035 | -0.01441 | 0.58124 | 0.00093 | 19.5 s |
| 1:15:40.3 | Assay volumes | 1.50000 mL | 0.16030 mL | 0.15957 mL | 0.51002 mL | | | | | |

| | | | |
|---|---|---|---|
| Sample name: | **M04_octanol** | Experiment start time: | **3/24/2018 1:34:06 AM** |
| Assay name: | **pH-metric high logP** | Analyst: | **Dorothy Levorse** |
| Assay ID: | **18C-24002** | Instrument ID: | **T312060** |
| Filename: | **C:\Sirius_T3\Mehtap\20180323_exp33_logP_T3-2\18C-24002_M04_octanol_pH-metric high logP.t3r** | | |

| | | | |
|---|---|---|---|
| Sample name: | **M04_octanol** | Experiment start time: | **3/24/2018 1:34:06 AM** |
| Assay name: | **pH-metric high logP** | Analyst: | **Dorothy Levorse** |
| Assay ID: | **18C-24002** | Instrument ID: | **T312060** |
| Filename: | **C:\Sirius_T3\Mehtap\20180323_exp33_logP_T3-2\18C-24002_M04_octanol_pH-metric high logP.t3r** | | |

#### Assay Settings (continued)

| Setting | Value | Original Value | Date/Time changed | Imported from |
|---|---|---|---|---|
| ***Titration 3*** | | | | |
| Titrate from | Low to high pH | | | |
| Add additional water | 0.00 mL | | | |
| Additional partition solvent volume | 0.400 mL | | | |
| Additional partition solvent added | Automatic | | | |
| After pH adjust stir for | 30 seconds | | | |
| Stir to allow partitioning for | 15 seconds | | | |
| Stirrer speed for partitioning | 60% | | | |
| ***Data Point Stability*** | | | | |
| Stir during data point collection | No | | | |
| Delay before data point collection | 0 seconds | | | |
| Number of points to average | 20 points | | | |
| Time interval between points | 0.50 seconds | | | |
| Required maximum standard deviation | 0.00100 dpH/dt | | | |
| Stability timeout after | 60 seconds | | | |

#### Calibration Settings

| Setting | Value | Date/Time changed | Imported from |
|---|---|---|---|
| Four-Plus alpha | 0.119 | 3/24/2018 1:34:06 AM | C:\Sirius_T3\HCl18C23.t3r |
| Four-Plus S | 0.9972 | 3/24/2018 1:34:06 AM | C:\Sirius_T3\HCl18C23.t3r |
| Four-Plus jH | 0.9 | 3/24/2018 1:34:06 AM | C:\Sirius_T3\HCl18C23.t3r |
| Four-Plus jOH | -0.3 | 3/24/2018 1:34:06 AM | C:\Sirius_T3\HCl18C23.t3r |
| Base concentration factor | 1.003 | 3/24/2018 1:34:06 AM | C:\Sirius_T3\KOH18C23.t3r |
| Acid concentration factor | 0.997 | 3/24/2018 1:34:06 AM | C:\Sirius_T3\HCl18C23.t3r |

#### Instrument Settings

| Setting | Value | Batch Id | Install date |
|---|---|---|---|
| Instrument owner | Merck | | |
| Instrument ID | T312060 | | |
| Instrument type | T3 Simulator | | |
| Software version | 1.1.3.0 | | |
| Dispenser module | | T3DM1200361 | 3/31/2009 6:24:52 AM |
| Dispenser 0 | Water | | 3/31/2009 6:25:05 AM |
| Syringe volume | 2.5 mL | | |
| Firmware version | 1.2.1(r2) | | |
| Titrant | Water (0.15 M KCl) | 02-06-2018 | 3/16/2018 11:09:18 AM |
| Dispenser 2 | Acid | | 3/31/2009 6:25:11 AM |
| Syringe volume | 0.5 mL | | |
| Firmware version | 1.2.1(r2) | | |
| Titrant | Acid (0.5 M HCl) | 03-16-2018 | 3/16/2018 10:56:23 AM |
| Dispenser 1 | Base | | 3/31/2009 6:25:21 AM |
| Syringe volume | 0.5 mL | | |
| Firmware version | 1.2.1(r2) | | |
| Titrant | Base (0.5 M KOH) | 3/22/2018 | 3/23/2018 9:34:17 AM |
| Dispenser 5 | Cosolvent | | 3/31/2009 6:26:24 AM |
| Syringe volume | 2.5 mL | | |
| Firmware version | 1.2.1(r2) | | |
| Distribution valve 5 | Distribution Valve | | 3/31/2009 6:28:19 AM |
| Firmware version | 1.1.3 | | |
| Port A | Methanol (80%, 0.15 M KCl) | 02-08-2018 | 3/6/2018 10:28:59 AM |
| Port B | Cyclohexane | 11-01-17 | 2/27/2018 11:37:57 AM |
| Dispenser 3 | Buffer | | 8/3/2010 6:05:16 AM |
| Syringe volume | 0.5 mL | | |
| Firmware version | 1.2.1(r2) | | |
| Titrant | Dodecane | 2018/01/31 | 2/28/2018 11:18:04 AM |
| Dispenser 6 | Octanol | | 10/22/2010 11:52:43 AM |

| | | | |
|---|---|---|---|
| Sample name: | **M04_octanol** | Experiment start time: | **3/24/2018 1:34:06 AM** |
| Assay name: | **pH-metric high logP** | Analyst: | **Dorothy Levorse** |
| Assay ID: | **18C-24002** | Instrument ID: | **T312060** |
| Filename: | **C:\Sirius_T3\Mehtap\20180323_exp33_logP_T3-2\18C-24002_M04_octanol_pH-metric high logP.t3r** | | |

#### Instrument Settings (continued)

| Setting | Value | Batch Id | Install date |
|---|---|---|---|
| Syringe volume | 0.5 mL | | |
| Firmware version | 1.2.1(r2) | | |
| Titrant | Octanol | 01-31-2018 | 2/27/2018 10:59:35 AM |
| Titrator | | T3TM1200161 | 3/31/2009 6:24:17 AM |
| Horizontal axis firmware version | 1.17 AI1DI2DO2 Stepper 2 | | |
| Vertical axis firmware version | 1.17 AI1DI2DO2 Stepper 2 | | |
| Chassis I/O firmware version | 1.11 AI1DI0DO4 Norgren I/O | | |
| Probe I/O firmware version | 1.1.1 | | |
| Electrode | T3 Electrode | T3E0923 | 1/23/2018 3:01:00 PM |
| E0 calibration | +4.89 mV | | 3/24/2018 1:34:34 AM |
| Filling solution | 3M KCl | KCL097 | 3/23/2018 9:29:07 AM |
| Liquids | | | |
| Wash 1 | 50% IPA:50% Water | | 3/23/2018 9:29:12 AM |
| Wash 2 | 0.5% Trition X-100 in H20 | | 3/23/2018 9:29:15 AM |
| Buffer position 1 | pH7 Wash | | 3/23/2018 9:29:19 AM |
| Buffer position 2 | pH 7 | | 3/23/2018 9:29:21 AM |
| Storage position | | | 3/23/2018 9:30:23 AM |
| Wash water | 7.7e+003 mL | 03-12-2018 | 3/12/2018 9:25:04 AM |
| Waste | 2.5e+003 mL | | 3/12/2018 9:24:49 AM |
| Temperature controller | | | 8/5/2010 7:35:13 AM |
| Turbidity detector | | | 3/31/2009 6:24:45 AM |
| Spectrometer | | 074811 | 11/23/2010 12:22:28 PM |
| Dip probe | | 10196 | |
| Wavelength coefficient A0 | 183.333 | | |
| Wavelength coefficient A1 | 2.21568 | | |
| Wavelength coefficient A2 | -0.000289308 | | |
| Total lamp lit time | 162:53:01 | | 11/23/2010 12:22:28 PM |
| Calibrated on | 2/27/2018 11:40:38 AM | | |
| Integration time | 40 | | |
| Scans averaged | 10 | | |
| Autoloader | | T3AL1200345 | 11/10/2015 10:34:13 AM |
| Left-right axis firmware version | 1.17 AI1DI2DO2 Stepper 2 | | |
| Front-back axis firmware version | 1.17 AI1DI2DO2 Stepper 2 | | |
| Vertical axis firmware version | 1.17 AI1DI2DO2 Stepper 2 | | |
| Chassis I/O firmware version | 1.11 AI1DI0DO4 Norgren I/O | | |
| Configuration | | | |
| Alternate titration position | Titration position | | |
| Alternate reference position | Reference position | | |
| Maximum standard vial volume | 3.50 mL | | |
| Maximum alternate vial volume | 25.00 mL | | |
| Automatic action idle period | 5 minute(s) | | |
| Titrant tube volume | 1.3 mL | | |
| Syringe flush count | 3.50 | | |
| Flowing wash pump volume | 20.0 mL | | |
| Flowing wash stir duration | 5 s | | |
| Flowing wash stir speed | 30% | | |
| Solvent wash stir duration | 5 s | | |
| Solvent wash stir speed | 30% | | |
| Surfactant wash stir duration | 5 s | | |
| Surfactant wash stir speed | 30% | | |
| E0 calibration minimum number of points | 10 | | |
| E0 calibration maximum standard deviation | 0.01500 | | |
| E0 calibration timeout period | 60 s | | |
| E0 calibration stir duration | 5 s | | |
| E0 calibration preparation stir speed | 30% | | |
| E0 calibration buffer wash stir duration | 5 s | | |
| E0 calibration buffer wash stir speed | 30% | | |
| E0 calibration reading stir speed | 0% | | |

| | | | |
|---|---|---|---|
| Sample name: | **M04_octanol** | Experiment start time: | **3/24/2018 1:34:06 AM** |
| Assay name: | **pH-metric high logP** | Analyst: | **Dorothy Levorse** |
| Assay ID: | **18C-24002** | Instrument ID: | **T312060** |
| Filename: | **C:\Sirius_T3\Mehtap\20180323_exp33_logP_T3-2\18C-24002_M04_octanol_pH-metric high logP.t3r** | | |

#### Instrument Settings (continued)

| Setting | Value | Batch Id | Install date |
|---|---|---|---|
| Spectrometer calibration stir duration | 5 s | | |
| Spectrometer calibration stir speed | 30% | | |
| Spectrometer calibration wash pump volume | 20.0 mL | | |
| Spectrometer calibration wash stir duration | 5 s | | |
| Spectrometer calibration wash stir speed | 30% | | |
| Overhead dispense height | 10000 | | |

#### Refinement Settings

| Setting | Value | Default value |
|---|---|---|
| Turbidity detection method | None | None |
| Turbidity wavelength to assess | 500.0 nm | 500.0 nm |
| Turbidity maximum absorbance | 0.100 | 0.100 |
| Turbidity probe threshold | 50.00 | 50.00 |
