## Supplementary material for "Octanol-water partition coefficient measurements for the SAMPL6 Blind Prediction Challenge": SM04_18C-24003_M04_octanol_pH-metric high logP_report.pdf

| | | | |
|---|---|---|---|
| Sample name: | **M04_octanol** | Experiment start time: | **3/24/2018 2:50:31 AM** |
| Assay name: | **pH-metric high logP** | Analyst: | **Dorothy Levorse** |
| Assay ID: | **18C-24003** | Instrument ID: | **T312060** |
| Filename: | **C:\Sirius_T3\Mehtap\20180323_exp33_logP_T3-2\18C-24003_M04_octanol_pH-metric high logP.t3r** | | |

#### pH-metric Result

| | | |
|---|---|---|
| logP (XH +) | 0.87 | ±0.03 (n=50) |
| logP (neutral X) | 3.95 | ±0.01 (n=50) |
| RMSD | 0.342 | |

#### 18C-24003 Points 1 to 26

| | |
|---|---|
| M04_octanol concentration factor | 0.985 |
| Carbonate | 0.1126 mM |
| Acidity error | 0.07154 mM |

#### 18C-24003 Points 27 to 52

| | |
|---|---|
| M04_octanol concentration factor | 0.875 |
| Carbonate | 0.1064 mM |
| Acidity error | -0.19519 mM |

#### 18C-24003 Points 53 to 77

| | |
|---|---|
| M04_octanol concentration factor | 0.960 |
| Carbonate | 0.1188 mM |
| Acidity error | 0.08198 mM |

#### Warnings and errors

| | |
|---|---|
| Errors | None |
| Warnings | None |

#### Sample logD and percent species

| pH | M04_octanol logD | M04_octanol M04_octanolH | M04_octanol M04_octanol | M04_octanol M04_octanolH* | M04_octanol M04_octanol* | Comment |
|---|---|---|---|---|---|---|
| 1.000 | 0.88 | 11.71 % | 0.00 % | 87.17 % | 1.12 % | |
| 1.200 | 0.88 | 11.63 % | 0.00 % | 86.60 % | 1.77 % | Stomach pH |
| 2.000 | 0.92 | 10.63 % | 0.00 % | 79.16 % | 10.20 % | |
| 3.000 | 1.23 | 5.54 % | 0.01 % | 41.26 % | 53.19 % | |
| 4.000 | 2.01 | 0.96 % | 0.01 % | 7.13 % | 91.90 % | |
| 5.000 | 2.94 | 0.10 % | 0.01 % | 0.77 % | 99.12 % | |
| 6.000 | 3.67 | 0.01 % | 0.01 % | 0.08 % | 99.90 % | |
| 6.500 | 3.84 | 0.00 % | 0.01 % | 0.02 % | 99.96 % | |
| 7.000 | 3.91 | 0.00 % | 0.01 % | 0.01 % | 99.98 % | |
| 7.400 | 3.94 | 0.00 % | 0.01 % | 0.00 % | 99.99 % | Blood pH |
| 8.000 | 3.95 | 0.00 % | 0.01 % | 0.00 % | 99.99 % | |
| 9.000 | 3.95 | 0.00 % | 0.01 % | 0.00 % | 99.99 % | |
| 10.000 | 3.95 | 0.00 % | 0.01 % | 0.00 % | 99.99 % | |
| 11.000 | 3.95 | 0.00 % | 0.01 % | 0.00 % | 99.99 % | |
| 12.000 | 3.95 | 0.00 % | 0.01 % | 0.00 % | 99.99 % | |

| | | | |
|---|---|---|---|
| Sample name: | **M04_octanol** | Experiment start time: | **3/24/2018 2:50:31 AM** |
| Assay name: | **pH-metric high logP** | Analyst: | **Dorothy Levorse** |
| Assay ID: | **18C-24003** | Instrument ID: | **T312060** |
| Filename: | **C:\Sirius_T3\Mehtap\20180323_exp33_logP_T3-2\18C-24003_M04_octanol_pH-metric high logP.t3r** | | |

#### Graphs

Charge balance residuals

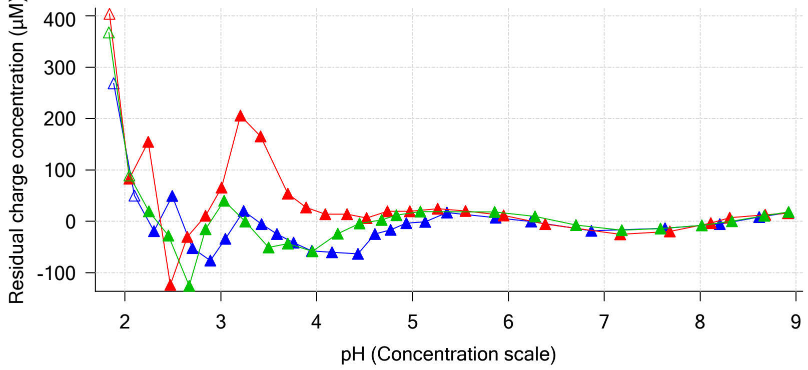


Ionic strength

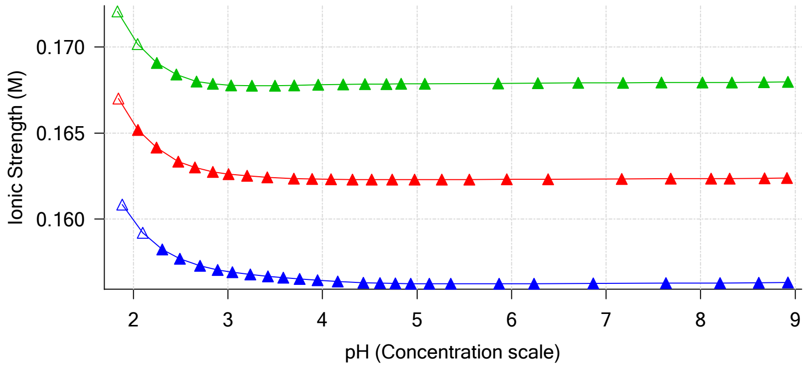


Buffer index profile

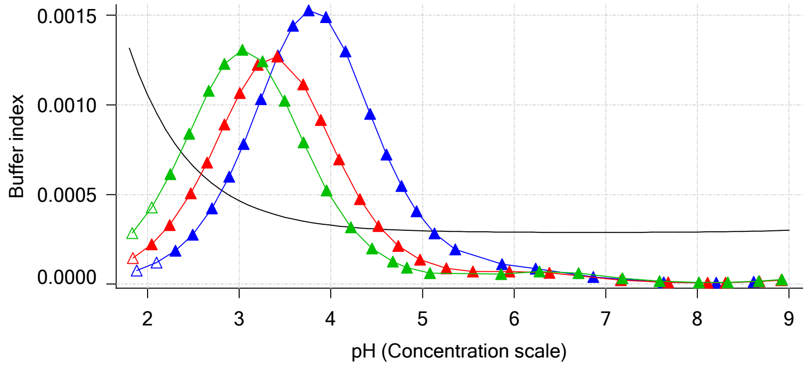


Sample M04_octanol Ionisation Graph

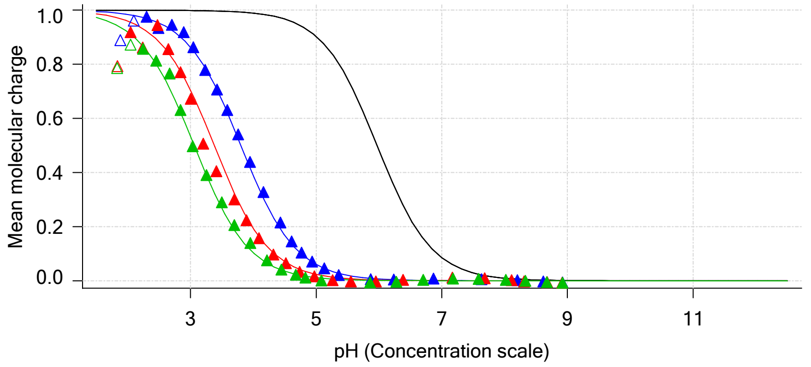


Distribution of species for sample M04_octanol

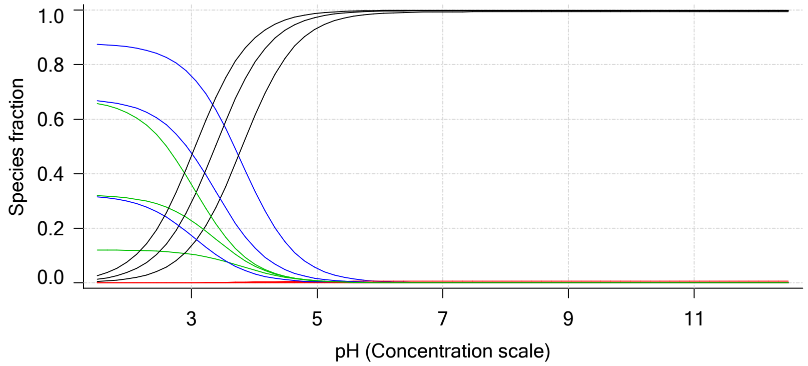


Lipophilicity Profile

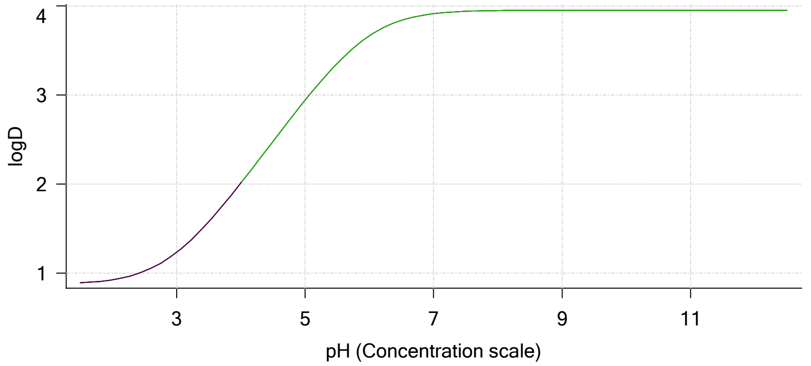


Log R Profile

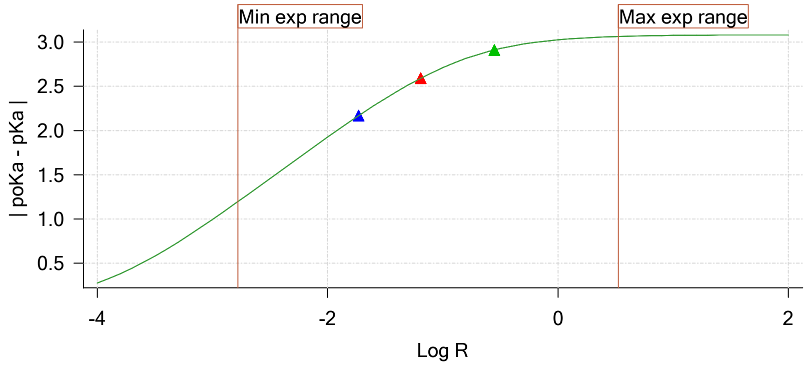


Sample Carbonate Ionisation Graph

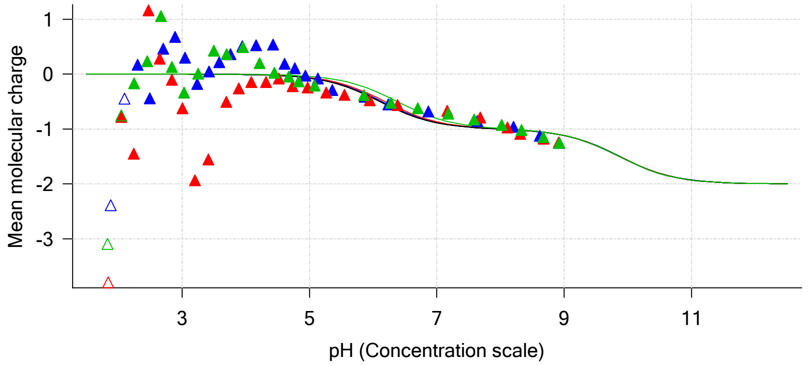

| | | | |
|---|---|---|---|
| Sample name: | **M04_octanol** | Experiment start time: | **3/24/2018 2:50:31 AM** |
| Assay name: | **pH-metric high logP** | Analyst: | **Dorothy Levorse** |
| Assay ID: | **18C-24003** | Instrument ID: | **T312060** |
| Filename: | **C:\Sirius_T3\Mehtap\20180323_exp33_logP_T3-2\18C-24003_M04_octanol_pH-metric high logP.t3r** | | |

#### Graphs (continued)

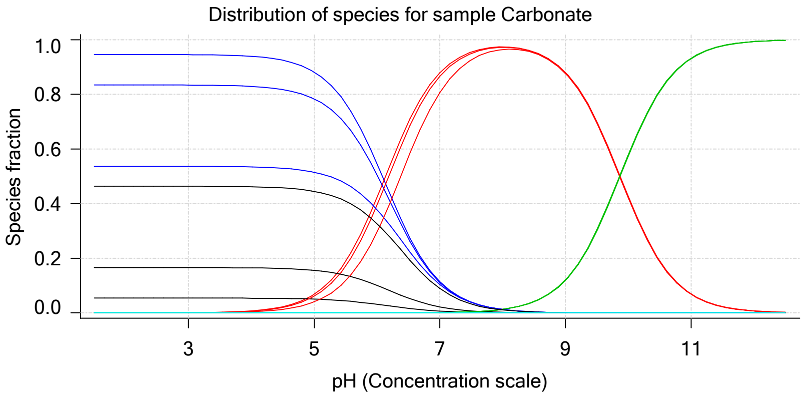

Sample name: **M04_octanol**
Assay name: **pH-metric high logP**
Assay ID: **18C-24003**
Filename: **C:\Sirius_T3\Mehtap\20180323_exp33_logP_T3-2\18C-24003_M04_octanol_pH-metric high logP.t3r**

Experiment start time: **3/24/2018 2:50:31 AM**
Analyst: **Dorothy Levorse**
Instrument ID: **T312060**

### pH-metric high logP Titration 1 of 3 18C-24003 Points 1 to 26

#### Overall results

| | |
|---|---|
| RMSD | 0.279 |
| Average ionic strength | 0.157 M |
| Average temperature | 24.9°C |
| Partition ratio | 0.0186 : 1 |
| Analyte concentration range | 2638.8 µM to 2719.9 µM |
| Total points considered | 24 of 26 |

#### Warnings and errors

| | |
|---|---|
| Errors | None |
| Warnings | None |

#### Four-Plus parameters

| | | | |
|---|---|---|---|
| Alpha | 0.119 | 3/24/2018 2:50:31 AM | C:\Sirius_T3\HCl18C23.t3r |
| S | 0.9972 | 3/24/2018 2:50:31 AM | C:\Sirius_T3\HCl18C23.t3r |
| jH | 0.9 | 3/24/2018 2:50:31 AM | C:\Sirius_T3\HCl18C23.t3r |
| jOH | -0.3 | 3/24/2018 2:50:31 AM | C:\Sirius_T3\HCl18C23.t3r |

#### Titrants

| | | | |
|---|---|---|---|
| 0.50 M HCl | 0.997124 | 3/24/2018 2:50:31 AM | C:\Sirius_T3\HCl18C23.t3r |
| 0.50 M KOH | 1.003190 | 3/24/2018 2:50:31 AM | C:\Sirius_T3\KOH18C23.t3r |

#### Sample

| | |
|---|---|
| M04_octanol concentration factor | 0.985 |
| Base pKa 1 | 5.97 |
| logP (XH +) | 0.19 |
| logP (neutral X) | 3.88 |

#### Sample graphs

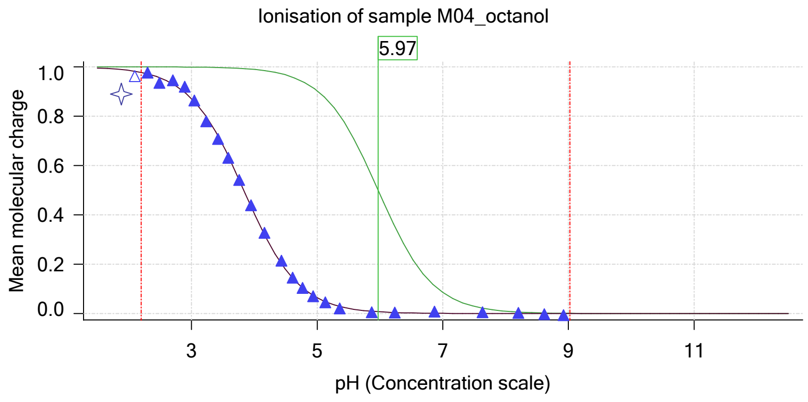


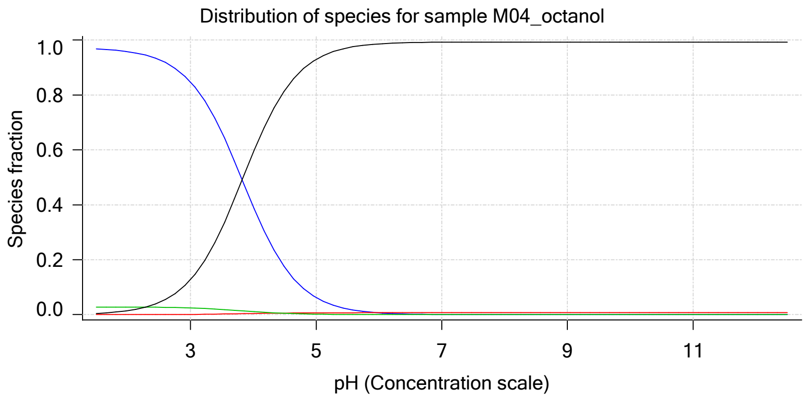

| | | | |
|---|---|---|---|
| Sample name: | **M04_octanol** | Experiment start time: | **3/24/2018 2:50:31 AM** |
| Assay name: | **pH-metric high logP** | Analyst: | **Dorothy Levorse** |
| Assay ID: | **18C-24003** | Instrument ID: | **T312060** |
| Filename: | **C:\Sirius_T3\Mehtap\20180323_exp33_logP_T3-2\18C-24003_M04_octanol_pH-metric high logP.t3r** | | |

#### Sample graphs (continued)

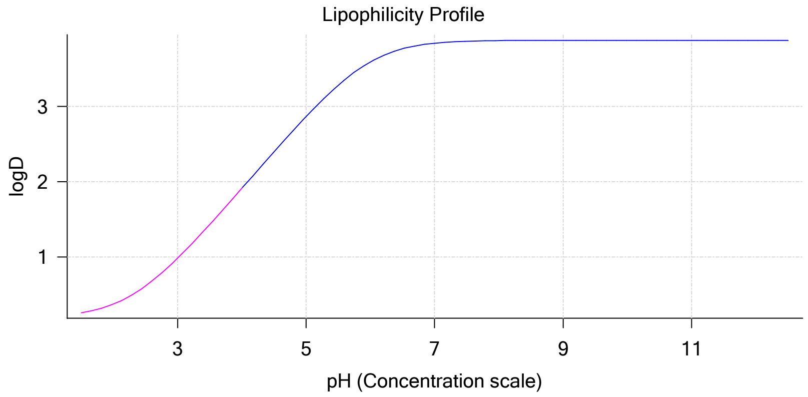


#### Sample logD and percent species

| pH | M04_octanol logD | M04_octanol M04_octanolH | M04_octanol M04_octanol | M04_octanol M04_octanolH* | M04_octanol M04_octanol* | Comment |
|---|---|---|---|---|---|---|
| 1.000 | 0.21 | 97.06 % | 0.00 % | 2.79 % | 0.15 % | |
| 1.200 | 0.22 | 96.98 % | 0.00 % | 2.79 % | 0.23 % | Stomach pH |
| 2.000 | 0.37 | 95.80 % | 0.01 % | 2.75 % | 1.44 % | |
| 3.000 | 0.98 | 84.73 % | 0.09 % | 2.44 % | 12.75 % | |
| 4.000 | 1.91 | 39.31 % | 0.42 % | 1.13 % | 59.14 % | |
| 5.000 | 2.87 | 6.18 % | 0.66 % | 0.18 % | 92.98 % | |
| 6.000 | 3.59 | 0.66 % | 0.70 % | 0.02 % | 98.62 % | |
| 6.500 | 3.77 | 0.21 % | 0.71 % | 0.01 % | 99.08 % | |
| 7.000 | 3.84 | 0.07 % | 0.71 % | 0.00 % | 99.23 % | |
| 7.400 | 3.86 | 0.03 % | 0.71 % | 0.00 % | 99.27 % | Blood pH |
| 8.000 | 3.87 | 0.01 % | 0.71 % | 0.00 % | 99.29 % | |
| 9.000 | 3.88 | 0.00 % | 0.71 % | 0.00 % | 99.29 % | |
| 10.000 | 3.88 | 0.00 % | 0.71 % | 0.00 % | 99.29 % | |
| 11.000 | 3.88 | 0.00 % | 0.71 % | 0.00 % | 99.29 % | |
| 12.000 | 3.88 | 0.00 % | 0.71 % | 0.00 % | 99.29 % | |

#### Carbonate and acidity

| | |
|---|---|
| Carbonate | 0.113 mM |
| Acidity error | 0.072 mM |

#### Other graphs

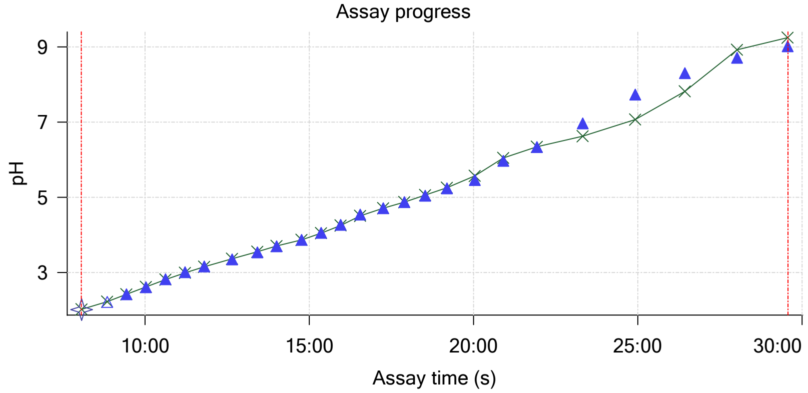


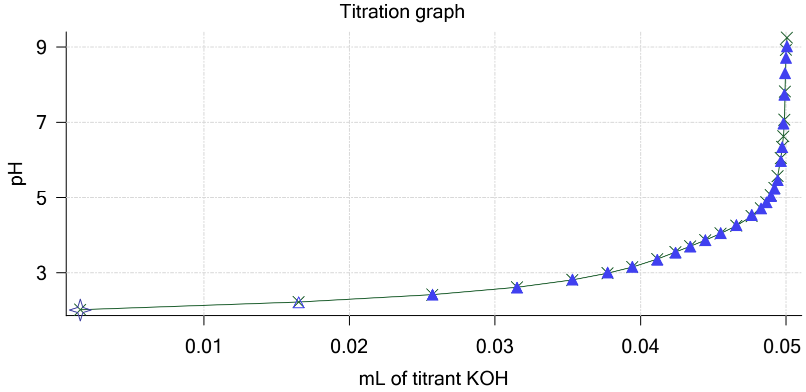

Sample name: **M04_octanol**
Assay name: **pH-metric high logP**
Assay ID: **18C-24003**
Filename: **C:\Sirius_T3\Mehtap\20180323_exp33_logP_T3-2\18C-24003_M04_octanol_pH-metric high logP.t3r**

Experiment start time: **3/24/2018 2:50:31 AM**
Analyst: **Dorothy Levorse**
Instrument ID: **T312060**

#### Other graphs (continued)

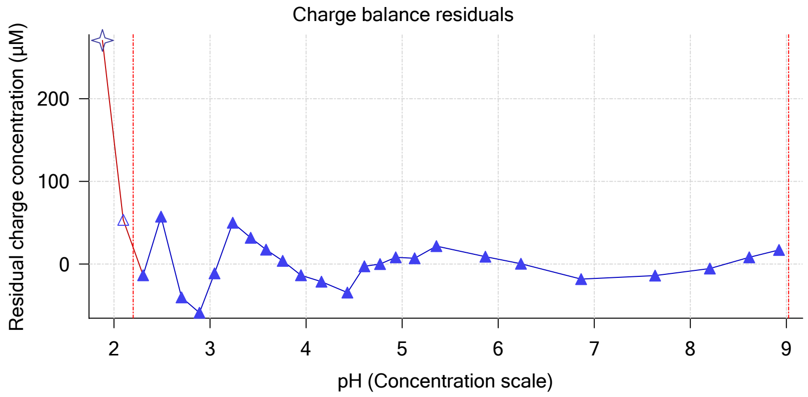


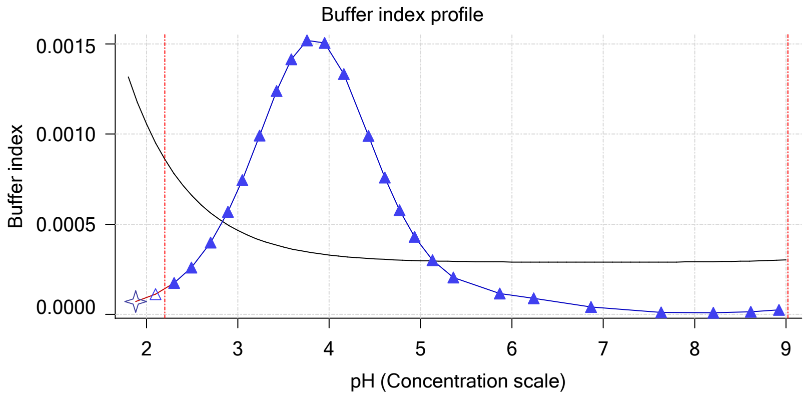


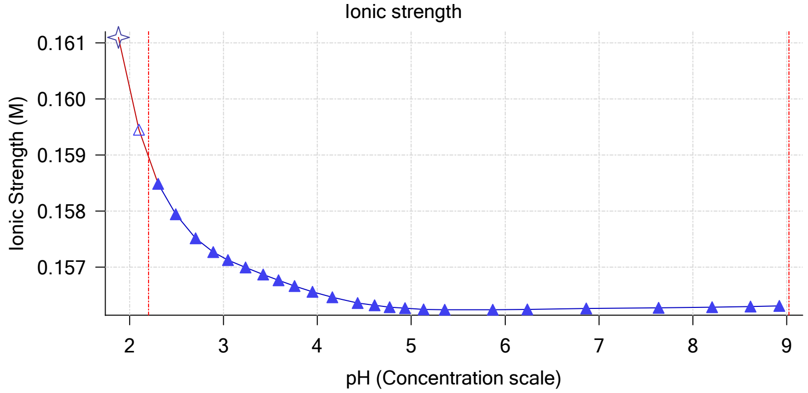


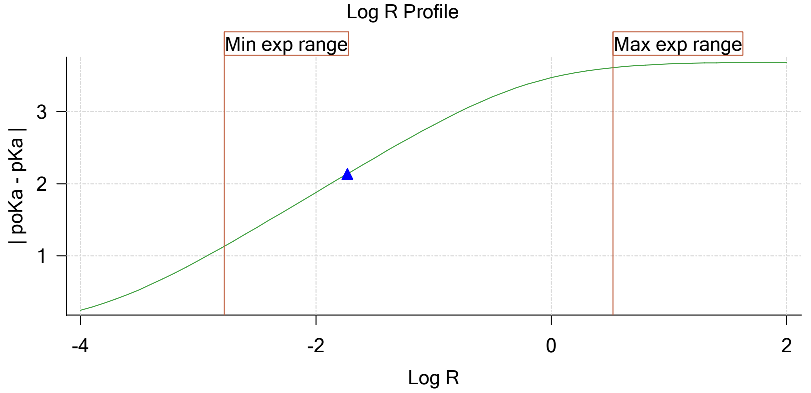


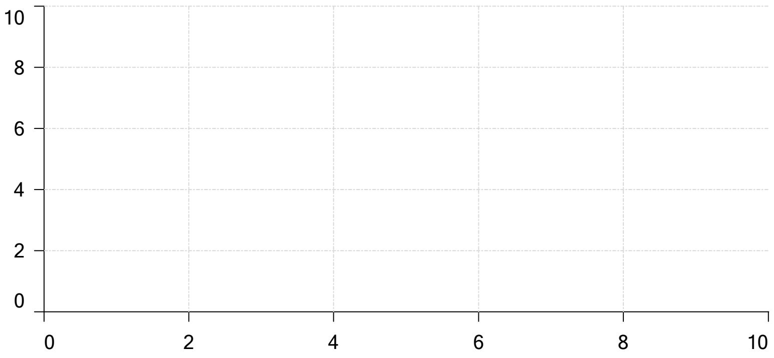


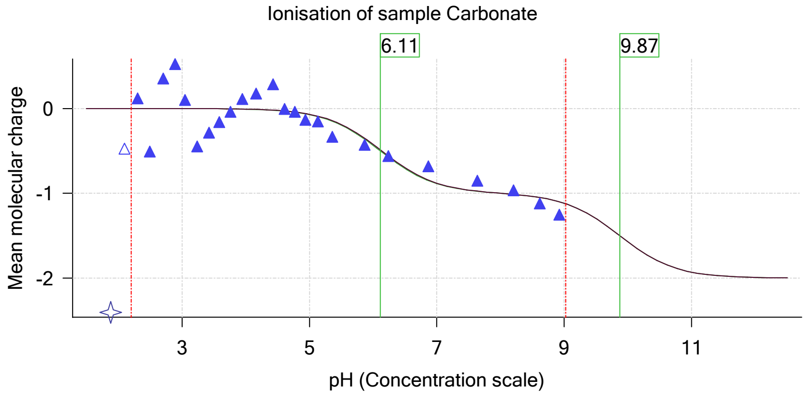

| | | | |
|---|---|---|---|
| Sample name: | **M04_octanol** | Experiment start time: | **3/24/2018 2:50:31 AM** |
| Assay name: | **pH-metric high logP** | Analyst: | **Dorothy Levorse** |
| Assay ID: | **18C-24003** | Instrument ID: | **T312060** |
| Filename: | **C:\Sirius_T3\Mehtap\20180323_exp33_logP_T3-2\18C-24003_M04_octanol_pH-metric high logP.t3r** | | |

# pH-metric high logP   Titration 2 of 3   18C-24003 Points 27 to 52

#### Overall results

| | |
|---|---|
| RMSD | 0.447 |
| Average ionic strength | 0.163 M |
| Average temperature | 25.0°C |
| Partition ratio | 0.0643 : 1 |
| Analyte concentration range | 2363.9 µM to 2436.4 µM |
| Total points considered | 25 of 26 |

#### Warnings and errors

| | |
|---|---|
| Errors | None |
| Warnings | None |

#### Four-Plus parameters

| | | | |
|---|---|---|---|
| Alpha | 0.119 | 3/24/2018 2:50:31 AM | C:\Sirius_T3\HCl18C23.t3r |
| S | 0.9972 | 3/24/2018 2:50:31 AM | C:\Sirius_T3\HCl18C23.t3r |
| jH | 0.9 | 3/24/2018 2:50:31 AM | C:\Sirius_T3\HCl18C23.t3r |
| jOH | -0.3 | 3/24/2018 2:50:31 AM | C:\Sirius_T3\HCl18C23.t3r |

#### Titrants

| | | | |
|---|---|---|---|
| 0.50 M HCl | 0.997124 | 3/24/2018 2:50:31 AM | C:\Sirius_T3\HCl18C23.t3r |
| 0.50 M KOH | 1.003190 | 3/24/2018 2:50:31 AM | C:\Sirius_T3\KOH18C23.t3r |

#### Sample

| | |
|---|---|
| M04_octanol concentration factor | 0.875 |
| Base pKa 1 | 5.97 |
| logP (XH +) | 0.19 |
| logP (neutral X) | 3.90 |

#### Sample graphs

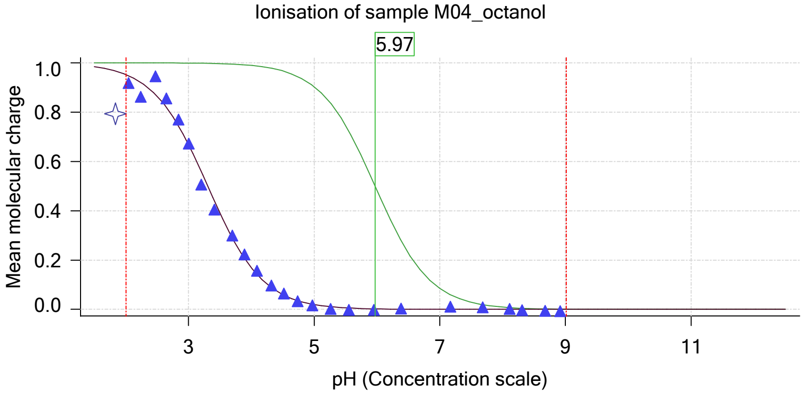


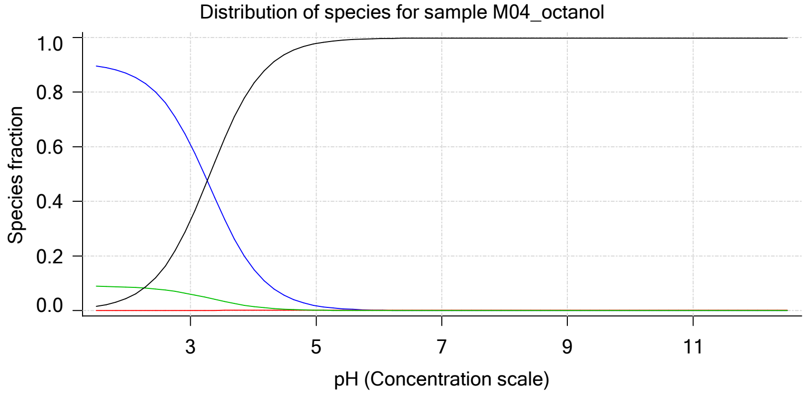

| | | | |
|---|---|---|---|
| Sample name: | **M04_octanol** | Experiment start time: | **3/24/2018 2:50:31 AM** |
| Assay name: | **pH-metric high logP** | Analyst: | **Dorothy Levorse** |
| Assay ID: | **18C-24003** | Instrument ID: | **T312060** |
| Filename: | **C:\Sirius_T3\Mehtap\20180323_exp33_logP_T3-2\18C-24003_M04_octanol_pH-metric high logP.t3r** | | |

#### Sample graphs (continued)

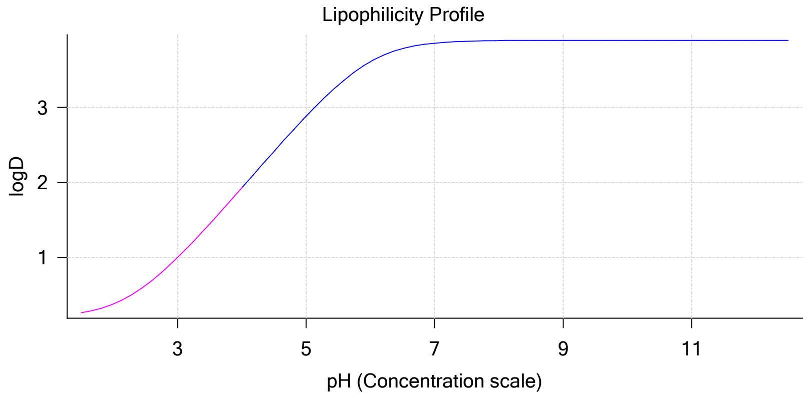


#### Sample logD and percent species

| pH | M04_octanol logD | M04_octanol M04_octanolH | M04_octanol M04_octanol | M04_octanol M04_octanolH* | M04_octanol M04_octanol* | Comment |
|---|---|---|---|---|---|---|
| 1.000 | 0.21 | 90.50 % | 0.00 % | 9.01 % | 0.49 % | |
| 1.200 | 0.23 | 90.24 % | 0.00 % | 8.99 % | 0.77 % | Stomach pH |
| 2.000 | 0.38 | 86.67 % | 0.01 % | 8.63 % | 4.69 % | |
| 3.000 | 1.00 | 60.90 % | 0.07 % | 6.07 % | 32.97 % | |
| 4.000 | 1.93 | 15.33 % | 0.16 % | 1.53 % | 82.98 % | |
| 5.000 | 2.88 | 1.81 % | 0.19 % | 0.18 % | 97.82 % | |
| 6.000 | 3.61 | 0.18 % | 0.20 % | 0.02 % | 99.60 % | |
| 6.500 | 3.78 | 0.06 % | 0.20 % | 0.01 % | 99.74 % | |
| 7.000 | 3.86 | 0.02 % | 0.20 % | 0.00 % | 99.78 % | |
| 7.400 | 3.88 | 0.01 % | 0.20 % | 0.00 % | 99.79 % | Blood pH |
| 8.000 | 3.89 | 0.00 % | 0.20 % | 0.00 % | 99.80 % | |
| 9.000 | 3.89 | 0.00 % | 0.20 % | 0.00 % | 99.80 % | |
| 10.000 | 3.90 | 0.00 % | 0.20 % | 0.00 % | 99.80 % | |
| 11.000 | 3.90 | 0.00 % | 0.20 % | 0.00 % | 99.80 % | |
| 12.000 | 3.90 | 0.00 % | 0.20 % | 0.00 % | 99.80 % | |

#### Carbonate and acidity

| | |
|---|---|
| Carbonate | 0.106 mM |
| Acidity error | -0.195 mM |

#### Other graphs

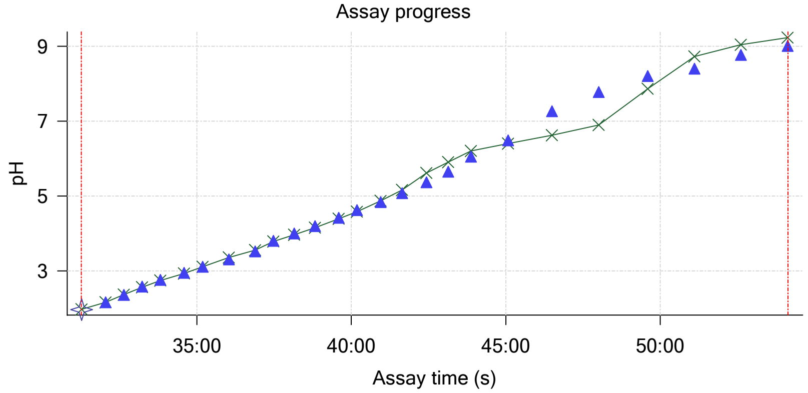


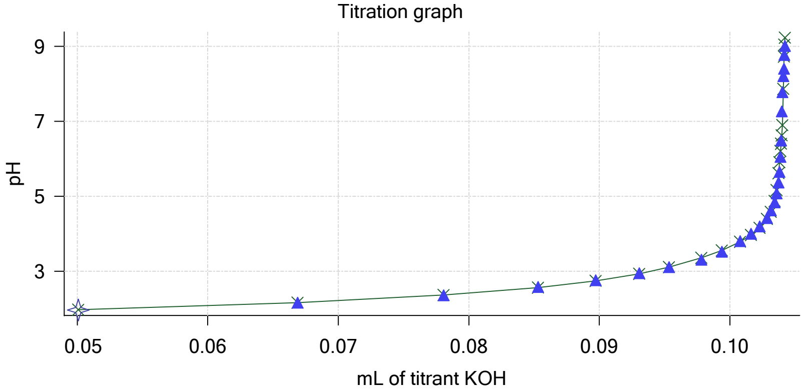

Sample name: **M04_octanol**
Assay name: **pH-metric high logP**
Assay ID: **18C-24003**
Filename: **C:\Sirius_T3\Mehtap\20180323_exp33_logP_T3-2\18C-24003_M04_octanol_pH-metric high logP.t3r**

Experiment start time: **3/24/2018 2:50:31 AM**
Analyst: **Dorothy Levorse**
Instrument ID: **T312060**

#### Other graphs (continued)

Charge balance residuals

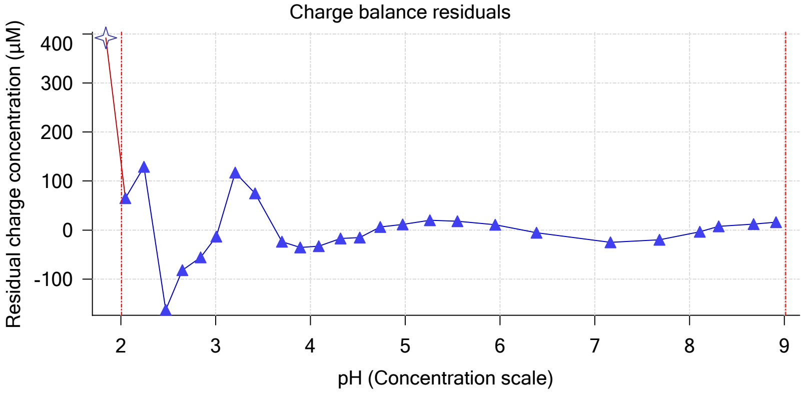


Buffer index profile

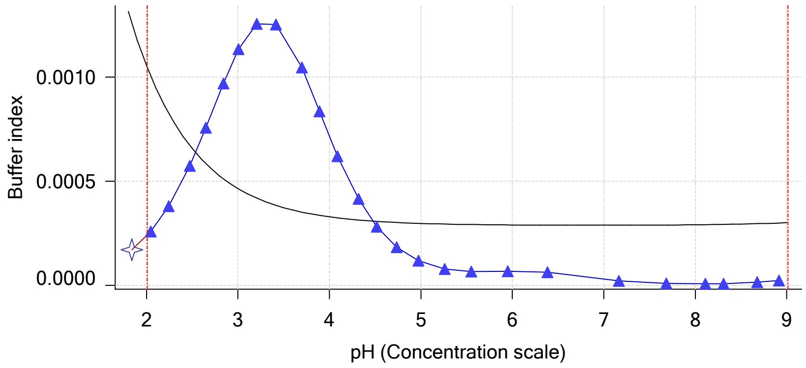


Ionic strength

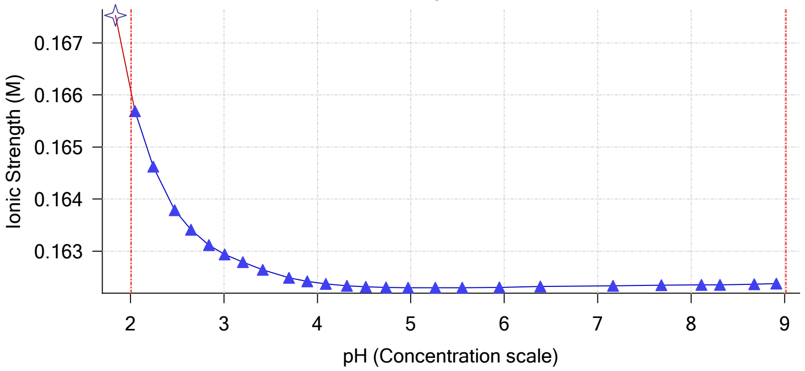


Log R Profile

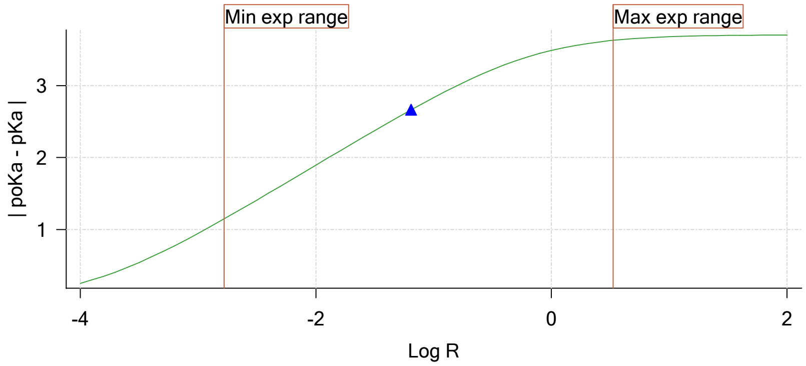


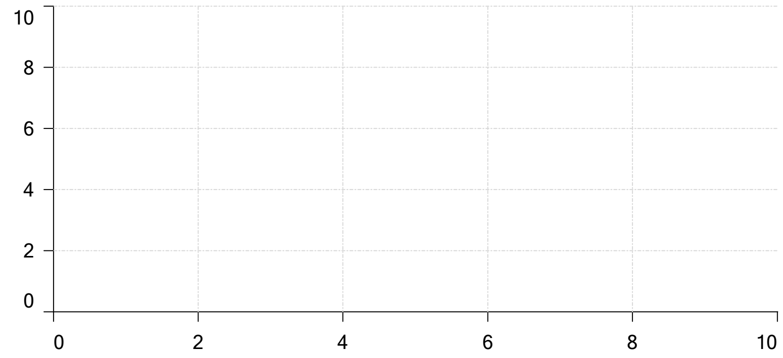

Ionisation of sample Carbonate

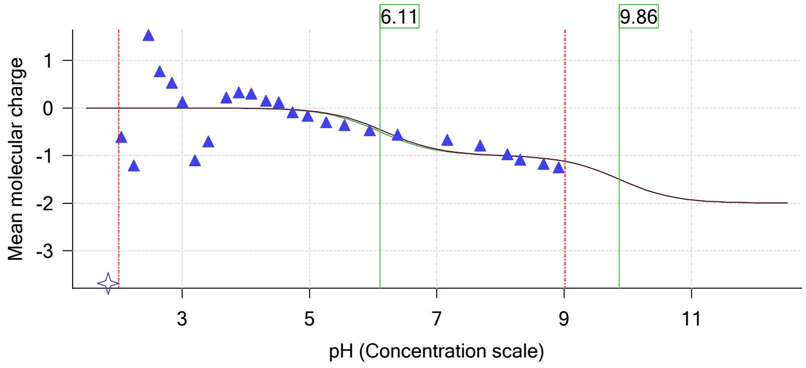

| | | | |
|---|---|---|---|
| Sample name: | **M04_octanol** | Experiment start time: | **3/24/2018 2:50:31 AM** |
| Assay name: | **pH-metric high logP** | Analyst: | **Dorothy Levorse** |
| Assay ID: | **18C-24003** | Instrument ID: | **T312060** |
| Filename: | **C:\Sirius_T3\Mehtap\20180323_exp33_logP_T3-2\18C-24003_M04_octanol_pH-metric high logP.t3r** | | |

### pH-metric high logP Titration 3 of 3 18C-24003 Points 53 to 77

#### Overall results

| | |
|---|---|
| RMSD | 0.260 |
| Average ionic strength | 0.168 M |
| Average temperature | 24.9°C |
| Partition ratio | 0.2796 : 1 |
| Analyte concentration range | 1840.5 μM to 1888.2 μM |
| Total points considered | 23 of 25 |

#### Warnings and errors

| | |
|---|---|
| Errors | None |
| Warnings | None |

#### Four-Plus parameters

| | | | |
|---|---|---|---|
| Alpha | 0.119 | 3/24/2018 2:50:31 AM | C:\Sirius_T3\HCl18C23.t3r |
| S | 0.9972 | 3/24/2018 2:50:31 AM | C:\Sirius_T3\HCl18C23.t3r |
| jH | 0.9 | 3/24/2018 2:50:31 AM | C:\Sirius_T3\HCl18C23.t3r |
| jOH | -0.3 | 3/24/2018 2:50:31 AM | C:\Sirius_T3\HCl18C23.t3r |

#### Titrants

| | | | |
|---|---|---|---|
| 0.50 M HCl | 0.997124 | 3/24/2018 2:50:31 AM | C:\Sirius_T3\HCl18C23.t3r |
| 0.50 M KOH | 1.003190 | 3/24/2018 2:50:31 AM | C:\Sirius_T3\KOH18C23.t3r |

#### Sample

| | |
|---|---|
| M04_octanol concentration factor | 0.960 |
| Base pKa 1 | 5.97 |
| logP (XH +) | 0.19 |
| logP (neutral X) | 3.59 |

#### Sample graphs

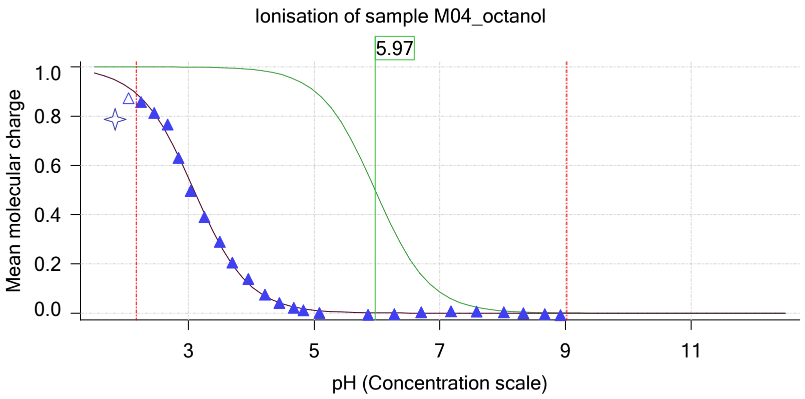


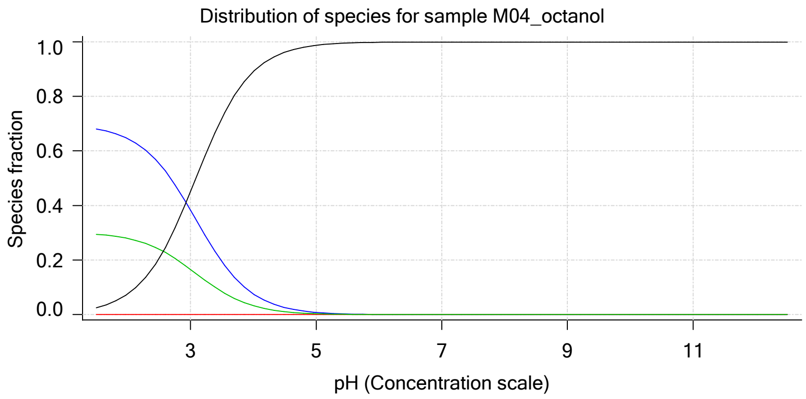

Sample name: **M04_octanol**
Assay name: **pH-metric high logP**
Assay ID: **18C-24003**
Filename: **C:\Sirius_T3\Mehtap\20180323_exp33_logP_T3-2\18C-24003_M04_octanol_pH-metric high logP.t3r**

Experiment start time: **3/24/2018 2:50:31 AM**
Analyst: **Dorothy Levorse**
Instrument ID: **T312060**

#### Sample graphs (continued)

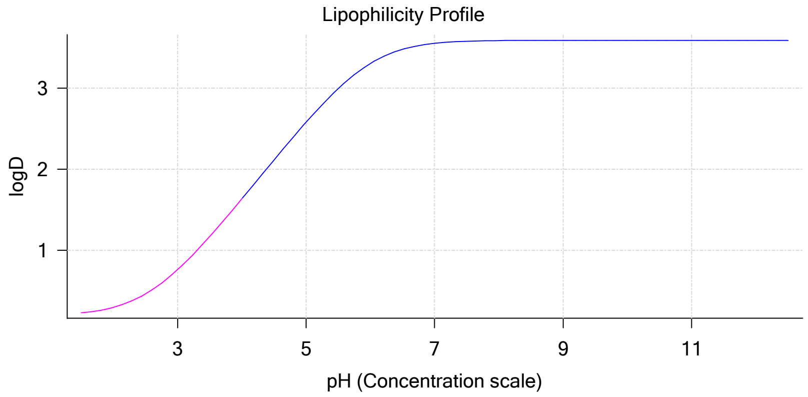


#### Sample logD and percent species

| pH | M04_octanol logD | M04_octanol M04_octanolH | M04_octanol M04_octanol | M04_octanol M04_octanolH* | M04_octanol M04_octanol* | Comment |
|---|---|---|---|---|---|---|
| 1.000 | 0.20 | 69.22 % | 0.00 % | 29.97 % | 0.81 % | |
| 1.200 | 0.21 | 68.89 % | 0.00 % | 29.83 % | 1.28 % | Stomach pH |
| 2.000 | 0.29 | 64.50 % | 0.01 % | 27.93 % | 7.57 % | |
| 3.000 | 0.76 | 38.35 % | 0.04 % | 16.61 % | 45.00 % | |
| 4.000 | 1.63 | 7.59 % | 0.08 % | 3.29 % | 89.04 % | |
| 5.000 | 2.58 | 0.84 % | 0.09 % | 0.36 % | 98.70 % | |
| 6.000 | 3.31 | 0.09 % | 0.09 % | 0.04 % | 99.79 % | |
| 6.500 | 3.48 | 0.03 % | 0.09 % | 0.01 % | 99.87 % | |
| 7.000 | 3.55 | 0.01 % | 0.09 % | 0.00 % | 99.90 % | |
| 7.400 | 3.58 | 0.00 % | 0.09 % | 0.00 % | 99.90 % | Blood pH |
| 8.000 | 3.59 | 0.00 % | 0.09 % | 0.00 % | 99.91 % | |
| 9.000 | 3.59 | 0.00 % | 0.09 % | 0.00 % | 99.91 % | |
| 10.000 | 3.59 | 0.00 % | 0.09 % | 0.00 % | 99.91 % | |
| 11.000 | 3.59 | 0.00 % | 0.09 % | 0.00 % | 99.91 % | |
| 12.000 | 3.59 | 0.00 % | 0.09 % | 0.00 % | 99.91 % | |

#### Carbonate and acidity

Carbonate 0.119 mM
Acidity error 0.082 mM

#### Other graphs

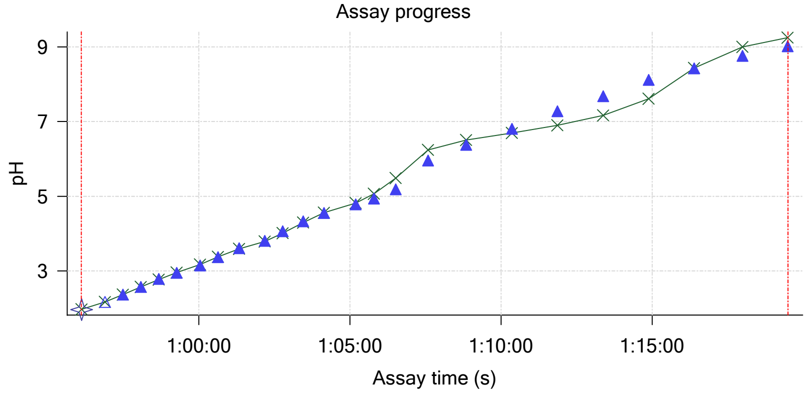


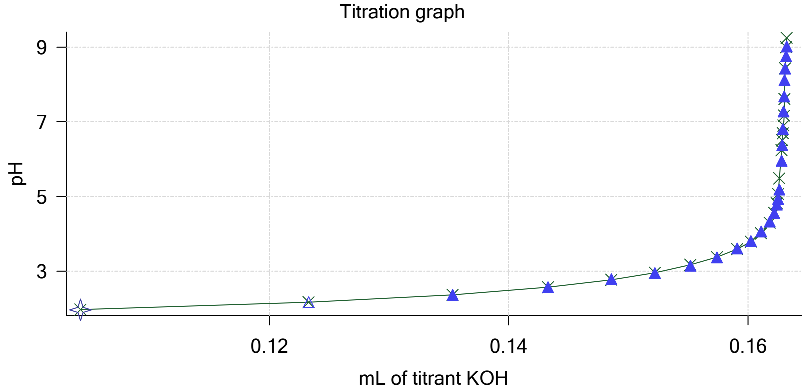

Sample name: **M04_octanol** Experiment start time: **3/24/2018 2:50:31 AM**
Assay name: **pH-metric high logP** Analyst: **Dorothy Levorse**
Assay ID: **18C-24003** Instrument ID: **T312060**
Filename: **C:\Sirius_T3\Mehtap\20180323_exp33_logP_T3-2\18C-24003_M04_octanol_pH-metric high logP.t3r**

#### Other graphs (continued)

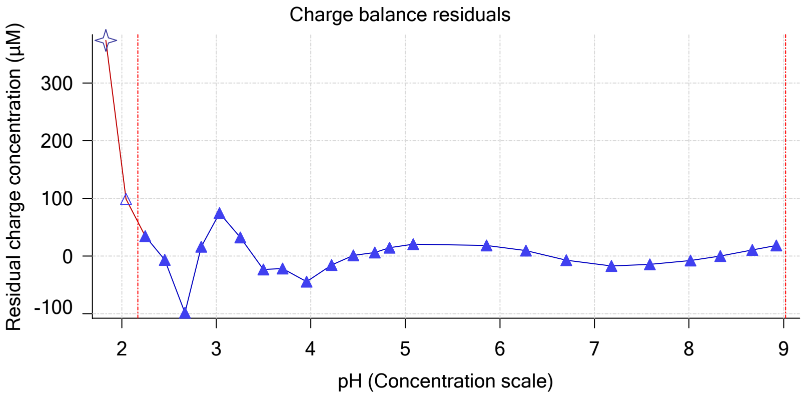


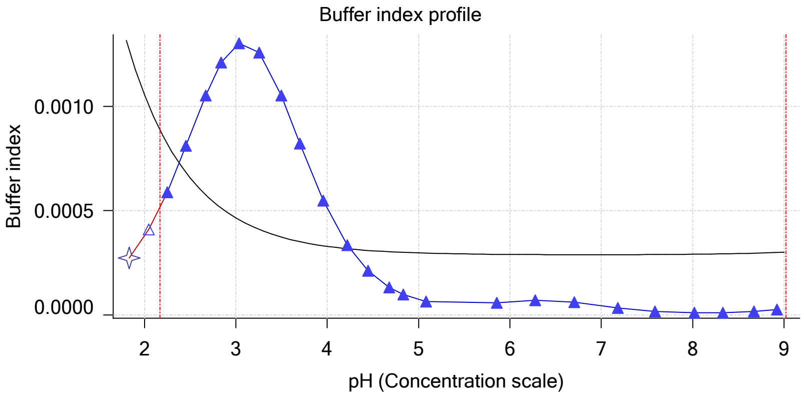


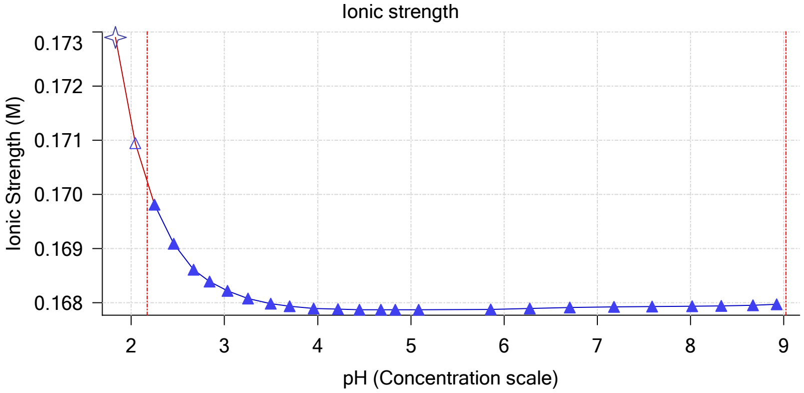


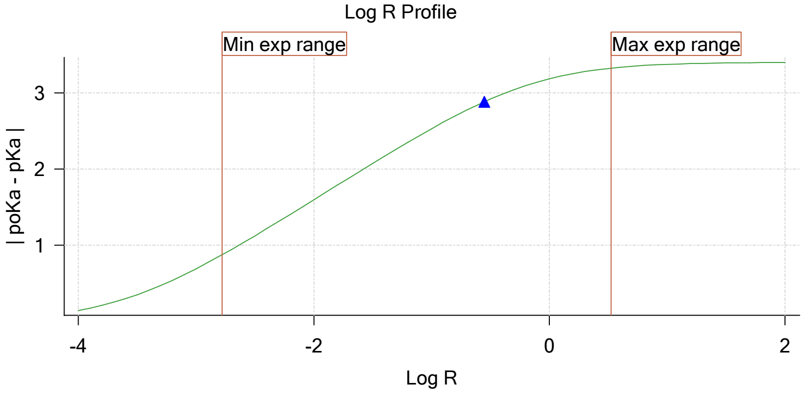


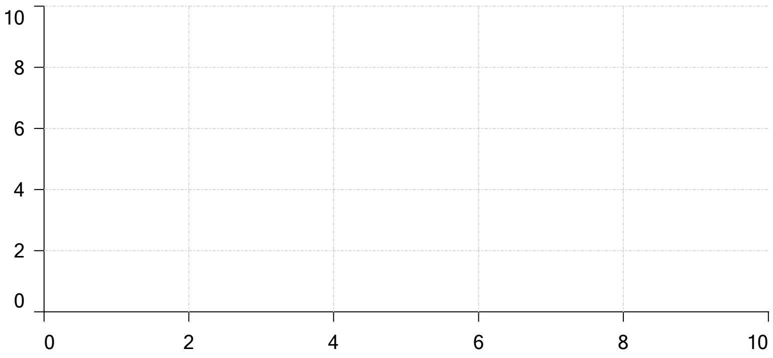


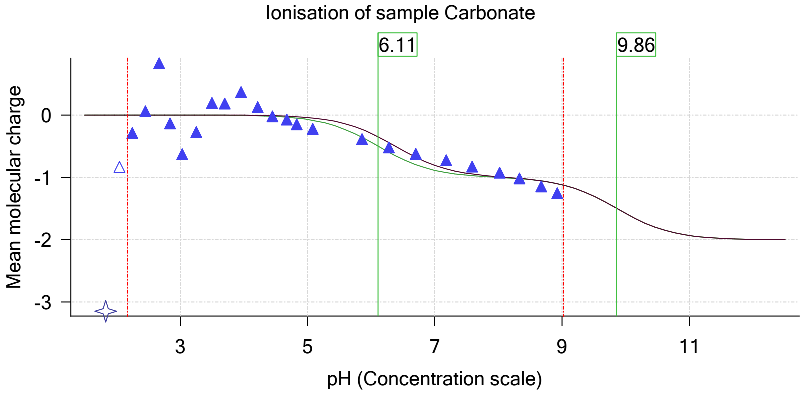

| | | | |
|---|---|---|---|
| Sample name: | **M04_octanol** | Experiment start time: | **3/24/2018 2:50:31 AM** |
| Assay name: | **pH-metric high logP** | Analyst: | **Dorothy Levorse** |
| Assay ID: | **18C-24003** | Instrument ID: | **T312060** |
| Filename: | **C:\Sirius_T3\Mehtap\20180323_exp33_logP_T3-2\18C-24003_M04_octanol_pH-metric high logP.t3r** | | |

#### Assay Model

| Settings | Value | Date/Time changed | Imported from |
|---|---|---|---|
| Sample name | M04_octanol | 3/9/2018 4:34:21 PM | User entered value |
| Sample by | Weight | | Default value |
| Sample weight | 0.001160 g | 3/23/2018 5:00:38 PM | User entered value |
| Formula weight | 269.73 g/mol | 3/9/2018 4:34:21 PM | User entered value |
| Solubility | Unknown | | Default value |
| Molecular weight | 269.73 | 3/9/2018 4:34:21 PM | User entered value |
| Individual pKa ionic environments | No | | Default value |
| Number of pKas | 1 | 3/9/2018 4:34:21 PM | User entered value |
| Sample is a | Base | 3/9/2018 4:34:21 PM | User entered value |
| pKa 1 | 5.97 | 3/9/2018 4:34:21 PM | User entered value |
| logp (XH +) | 0.19 | 3/9/2018 4:34:33 PM | User entered value |
| logP (neutral X) | 3.50 | 3/23/2018 2:32:46 PM | User entered value |

#### Events

| Time | Event | Water | Acid | Base | Octanol | pH | dpH/dt | pH R-squared | pH SD | dpH/dt time |
|---|---|---|---|---|---|---|---|---|---|---|
| 4:59.5 | Initial pH = 5.97 | | | | | | | | | |
| 8:04.2 | Data point 1 | 1.50000 mL | 0.04965 mL | 0.00151 mL | 0.03001 mL | 2.007 | 0.00427 | 0.35010 | 0.00036 | 10.0 s |
| 8:50.5 | Data point 2 | 1.50000 mL | 0.04965 mL | 0.01651 mL | 0.03001 mL | 2.218 | -0.00672 | 0.12678 | 0.00093 | 10.0 s |
| 9:26.1 | Data point 3 | 1.50000 mL | 0.04965 mL | 0.02571 mL | 0.03001 mL | 2.420 | 0.00425 | 0.38859 | 0.00034 | 10.0 s |
| 10:01.6 | Data point 4 | 1.50000 mL | 0.04965 mL | 0.03151 mL | 0.03001 mL | 2.606 | -0.00797 | 0.19529 | 0.00089 | 10.0 s |
| 10:37.2 | Data point 5 | 1.50000 mL | 0.04965 mL | 0.03532 mL | 0.03001 mL | 2.816 | -0.00186 | 0.44711 | 0.00014 | 10.0 s |
| 11:12.6 | Data point 6 | 1.50000 mL | 0.04965 mL | 0.03775 mL | 0.03001 mL | 3.003 | 0.00280 | 0.09562 | 0.00045 | 10.0 s |
| 11:48.1 | Data point 7 | 1.50000 mL | 0.04965 mL | 0.03944 mL | 0.03001 mL | 3.158 | 0.00375 | 0.14586 | 0.00048 | 10.0 s |
| 12:39.0 | Data point 8 | 1.50000 mL | 0.04965 mL | 0.04116 mL | 0.03001 mL | 3.347 | -0.00347 | 0.70036 | 0.00020 | 10.0 s |
| 13:24.7 | Data point 9 | 1.50000 mL | 0.04965 mL | 0.04240 mL | 0.03001 mL | 3.533 | -0.00887 | 0.63978 | 0.00055 | 10.0 s |
| 14:00.2 | Data point 10 | 1.50000 mL | 0.04965 mL | 0.04341 mL | 0.03001 mL | 3.695 | -0.00873 | 0.74697 | 0.00050 | 10.0 s |
| 14:46.0 | Data point 11 | 1.50000 mL | 0.04965 mL | 0.04445 mL | 0.03001 mL | 3.867 | -0.01180 | 0.86864 | 0.00063 | 10.0 s |
| 15:21.3 | Data point 12 | 1.50000 mL | 0.04965 mL | 0.04551 mL | 0.03001 mL | 4.056 | -0.01653 | 0.82606 | 0.00090 | 10.0 s |
| 15:56.8 | Data point 13 | 1.50000 mL | 0.04965 mL | 0.04659 mL | 0.03001 mL | 4.268 | -0.01706 | 0.91582 | 0.00088 | 10.5 s |
| 16:32.8 | Data point 14 | 1.50000 mL | 0.04965 mL | 0.04765 mL | 0.03001 mL | 4.537 | -0.01779 | 0.85452 | 0.00095 | 11.5 s |
| 17:14.9 | Data point 15 | 1.50000 mL | 0.04965 mL | 0.04828 mL | 0.03001 mL | 4.714 | -0.01769 | 0.95197 | 0.00089 | 13.0 s |
| 17:53.3 | Data point 16 | 1.50000 mL | 0.04965 mL | 0.04866 mL | 0.03001 mL | 4.876 | -0.01700 | 0.83581 | 0.00092 | 13.0 s |
| 18:31.7 | Data point 17 | 1.50000 mL | 0.04965 mL | 0.04897 mL | 0.03001 mL | 5.040 | -0.01747 | 0.89358 | 0.00091 | 14.0 s |
| 19:11.1 | Data point 18 | 1.50000 mL | 0.04965 mL | 0.04920 mL | 0.03001 mL | 5.236 | -0.01840 | 0.90934 | 0.00095 | 15.5 s |
| 20:02.3 | Data point 19 | 1.50000 mL | 0.04965 mL | 0.04944 mL | 0.03001 mL | 5.461 | -0.01934 | 0.94812 | 0.00098 | 21.5 s |
| 20:54.3 | Data point 20 | 1.50000 mL | 0.04965 mL | 0.04965 mL | 0.03001 mL | 5.969 | -0.01829 | 0.93298 | 0.00093 | 36.0 s |
| 21:55.7 | Data point 21 | 1.50000 mL | 0.04965 mL | 0.04974 mL | 0.03001 mL | 6.338 | -0.01998 | 0.98727 | 0.00099 | 53.0 s |
| 23:19.4 | Data point 22 | 1.50000 mL | 0.04965 mL | 0.04981 mL | 0.03001 mL | 6.965 | -0.05661 | 0.98918 | 0.00281 | Timed out at 59.5 s |
| 24:55.0 | Data point 23 | 1.50000 mL | 0.04965 mL | 0.04988 mL | 0.03001 mL | 7.732 | -0.10014 | 0.99537 | 0.00496 | Timed out at 59.5 s |
| 26:25.5 | Data point 24 | 1.50000 mL | 0.04965 mL | 0.04993 mL | 0.03001 mL | 8.300 | -0.05511 | 0.99507 | 0.00273 | Timed out at 59.5 s |
| 28:01.1 | Data point 25 | 1.50000 mL | 0.04965 mL | 0.05000 mL | 0.03001 mL | 8.709 | -0.01976 | 0.97372 | 0.00099 | 56.5 s |
| 29:33.2 | Data point 26 | 1.50000 mL | 0.04965 mL | 0.05007 mL | 0.03001 mL | 9.016 | -0.01909 | 0.93060 | 0.00098 | 43.0 s |
| 31:16.2 | Data point 27 | 1.50000 mL | 0.10503 mL | 0.05007 mL | 0.11002 mL | 1.966 | -0.00262 | 0.17546 | 0.00031 | 10.0 s |
| 32:02.5 | Data point 28 | 1.50000 mL | 0.10503 mL | 0.06689 mL | 0.11002 mL | 2.167 | -0.00002 | 0.00005 | 0.00013 | 10.0 s |
| 32:38.3 | Data point 29 | 1.50000 mL | 0.10503 mL | 0.07808 mL | 0.11002 mL | 2.360 | -0.00053 | 0.02080 | 0.00018 | 10.0 s |
| 33:13.7 | Data point 30 | 1.50000 mL | 0.10503 mL | 0.08530 mL | 0.11002 mL | 2.588 | 0.00069 | 0.01014 | 0.00034 | 10.0 s |
| 33:49.3 | Data point 31 | 1.50000 mL | 0.10503 mL | 0.08972 mL | 0.11002 mL | 2.762 | -0.00184 | 0.09470 | 0.00030 | 10.0 s |
| 34:35.2 | Data point 32 | 1.50000 mL | 0.10503 mL | 0.09306 mL | 0.11002 mL | 2.952 | -0.00238 | 0.48804 | 0.00017 | 10.5 s |
| 35:11.1 | Data point 33 | 1.50000 mL | 0.10503 mL | 0.09534 mL | 0.11002 mL | 3.117 | -0.00618 | 0.30787 | 0.00055 | 10.0 s |
| 36:02.2 | Data point 34 | 1.50000 mL | 0.10503 mL | 0.09781 mL | 0.11002 mL | 3.314 | -0.00705 | 0.70654 | 0.00041 | 10.0 s |
| 36:53.1 | Data point 35 | 1.50000 mL | 0.10503 mL | 0.09939 mL | 0.11002 mL | 3.524 | -0.00556 | 0.85447 | 0.00030 | 10.0 s |
| 37:28.6 | Data point 36 | 1.50000 mL | 0.10503 mL | 0.10080 mL | 0.11002 mL | 3.806 | -0.00526 | 0.75008 | 0.00030 | 10.0 s |
| 38:09.2 | Data point 37 | 1.50000 mL | 0.10503 mL | 0.10162 mL | 0.11002 mL | 3.999 | -0.01309 | 0.90328 | 0.00068 | 10.0 s |

| | | | |
|---|---|---|---|
| Sample name: | **M04_octanol** | Experiment start time: | **3/24/2018 2:50:31 AM** |
| Assay name: | **pH-metric high logP** | Analyst: | **Dorothy Levorse** |
| Assay ID: | **18C-24003** | Instrument ID: | **T312060** |
| Filename: | **C:\Sirius_T3\Mehtap\20180323_exp33_logP_T3-2\18C-24003_M04_octanol_pH-metric high logP.t3r** | | |

#### Events (continued)

| Time | Event | Water | Acid | Base | Octanol | pH | dpH/dt | pH R-squared | pH SD | dpH/dt time |
|---|---|---|---|---|---|---|---|---|---|---|
| 38:49.7 | Data point 38 | 1.50000 mL | 0.10503 mL | 0.10228 mL | 0.11002 mL | 4.196 | -0.00633 | 0.53369 | 0.00043 | 10.0 s |
| 39:35.4 | Data point 39 | 1.50000 mL | 0.10503 mL | 0.10285 mL | 0.11002 mL | 4.425 | -0.01381 | 0.73265 | 0.00080 | 10.0 s |
| 40:10.8 | Data point 40 | 1.50000 mL | 0.10503 mL | 0.10315 mL | 0.11002 mL | 4.625 | 0.00050 | 0.00129 | 0.00069 | 10.5 s |
| 40:57.0 | Data point 41 | 1.50000 mL | 0.10503 mL | 0.10343 mL | 0.11002 mL | 4.842 | -0.01579 | 0.77724 | 0.00088 | 11.0 s |
| 41:38.5 | Data point 42 | 1.50000 mL | 0.10503 mL | 0.10360 mL | 0.11002 mL | 5.078 | -0.00838 | 0.38707 | 0.00067 | 11.5 s |
| 42:25.7 | Data point 43 | 1.50000 mL | 0.10503 mL | 0.10374 mL | 0.11002 mL | 5.365 | -0.00881 | 0.23766 | 0.00089 | 12.0 s |
| 43:08.2 | Data point 44 | 1.50000 mL | 0.10503 mL | 0.10381 mL | 0.11002 mL | 5.655 | -0.01757 | 0.85957 | 0.00094 | 13.5 s |
| 43:52.2 | Data point 45 | 1.50000 mL | 0.10503 mL | 0.10388 mL | 0.11002 mL | 6.051 | -0.01851 | 0.93922 | 0.00094 | 41.5 s |
| 45:04.4 | Data point 46 | 1.50000 mL | 0.10503 mL | 0.10393 mL | 0.11002 mL | 6.487 | -0.02808 | 0.98839 | 0.00139 | Timed out at 59.5 s |
| 46:29.9 | Data point 47 | 1.50000 mL | 0.10503 mL | 0.10398 mL | 0.11002 mL | 7.265 | -0.09351 | 0.99809 | 0.00462 | Timed out at 59.5 s |
| 48:00.3 | Data point 48 | 1.50000 mL | 0.10503 mL | 0.10402 mL | 0.11002 mL | 7.780 | -0.09755 | 0.99711 | 0.00483 | Timed out at 59.5 s |
| 49:35.9 | Data point 49 | 1.50000 mL | 0.10503 mL | 0.10409 mL | 0.11002 mL | 8.206 | -0.05783 | 0.98965 | 0.00287 | Timed out at 59.5 s |
| 51:06.4 | Data point 50 | 1.50000 mL | 0.10503 mL | 0.10414 mL | 0.11002 mL | 8.404 | -0.03670 | 0.98260 | 0.00183 | Timed out at 59.5 s |
| 52:36.9 | Data point 51 | 1.50000 mL | 0.10503 mL | 0.10419 mL | 0.11002 mL | 8.772 | -0.02623 | 0.91998 | 0.00135 | Timed out at 59.5 s |
| 54:07.4 | Data point 52 | 1.50000 mL | 0.10503 mL | 0.10423 mL | 0.11002 mL | 9.008 | -0.01967 | 0.95537 | 0.00099 | 37.0 s |
| 56:07.6 | Data point 53 | 1.50000 mL | 0.16334 mL | 0.10423 mL | 0.51002 mL | 1.958 | -0.00498 | 0.69174 | 0.00030 | 10.0 s |
| 56:53.8 | Data point 54 | 1.50000 mL | 0.16334 mL | 0.12331 mL | 0.51002 mL | 2.165 | 0.01137 | 0.42530 | 0.00086 | 10.0 s |
| 57:29.6 | Data point 55 | 1.50000 mL | 0.16334 mL | 0.13533 mL | 0.51002 mL | 2.365 | -0.01105 | 0.37291 | 0.00089 | 10.0 s |
| 58:05.2 | Data point 56 | 1.50000 mL | 0.16334 mL | 0.14330 mL | 0.51002 mL | 2.569 | -0.00999 | 0.57469 | 0.00065 | 10.0 s |
| 58:40.8 | Data point 57 | 1.50000 mL | 0.16334 mL | 0.14859 mL | 0.51002 mL | 2.782 | -0.00724 | 0.84718 | 0.00039 | 10.0 s |
| 59:16.3 | Data point 58 | 1.50000 mL | 0.16334 mL | 0.15221 mL | 0.51002 mL | 2.951 | -0.00453 | 0.17701 | 0.00053 | 10.5 s |
| 1:00:02.6 | Data point 59 | 1.50000 mL | 0.16334 mL | 0.15520 mL | 0.51002 mL | 3.145 | -0.00569 | 0.12433 | 0.00080 | 10.0 s |
| 1:00:38.1 | Data point 60 | 1.50000 mL | 0.16334 mL | 0.15741 mL | 0.51002 mL | 3.364 | -0.01891 | 0.93813 | 0.00096 | 16.5 s |
| 1:01:20.1 | Data point 61 | 1.50000 mL | 0.16334 mL | 0.15910 mL | 0.51002 mL | 3.607 | -0.00838 | 0.90591 | 0.00043 | 10.0 s |
| 1:02:11.1 | Data point 62 | 1.50000 mL | 0.16334 mL | 0.16023 mL | 0.51002 mL | 3.809 | -0.01009 | 0.62944 | 0.00063 | 10.0 s |
| 1:02:46.4 | Data point 63 | 1.50000 mL | 0.16334 mL | 0.16110 mL | 0.51002 mL | 4.063 | -0.01406 | 0.72392 | 0.00082 | 10.0 s |
| 1:03:27.2 | Data point 64 | 1.50000 mL | 0.16334 mL | 0.16181 mL | 0.51002 mL | 4.326 | -0.01356 | 0.90136 | 0.00070 | 10.5 s |
| 1:04:08.2 | Data point 65 | 1.50000 mL | 0.16334 mL | 0.16218 mL | 0.51002 mL | 4.555 | -0.01747 | 0.81566 | 0.00096 | 37.5 s |
| 1:05:11.2 | Data point 66 | 1.50000 mL | 0.16334 mL | 0.16239 mL | 0.51002 mL | 4.783 | -0.01567 | 0.79185 | 0.00087 | 11.0 s |
| 1:05:47.6 | Data point 67 | 1.50000 mL | 0.16334 mL | 0.16251 mL | 0.51002 mL | 4.937 | -0.01579 | 0.62099 | 0.00099 | 12.5 s |
| 1:06:30.6 | Data point 68 | 1.50000 mL | 0.16334 mL | 0.16263 mL | 0.51002 mL | 5.187 | -0.01710 | 0.86918 | 0.00091 | 34.0 s |
| 1:07:35.2 | Data point 69 | 1.50000 mL | 0.16334 mL | 0.16279 mL | 0.51002 mL | 5.960 | -0.01912 | 0.94697 | 0.00097 | 45.0 s |
| 1:08:50.9 | Data point 70 | 1.50000 mL | 0.16334 mL | 0.16287 mL | 0.51002 mL | 6.378 | -0.03153 | 0.95619 | 0.00159 | Timed out at 59.5 s |
| 1:10:21.3 | Data point 71 | 1.50000 mL | 0.16334 mL | 0.16291 mL | 0.51002 mL | 6.803 | -0.06207 | 0.98528 | 0.00309 | Timed out at 59.5 s |
| 1:11:51.8 | Data point 72 | 1.50000 mL | 0.16334 mL | 0.16296 mL | 0.51002 mL | 7.279 | -0.09237 | 0.99346 | 0.00458 | Timed out at 59.5 s |
| 1:13:22.3 | Data point 73 | 1.50000 mL | 0.16334 mL | 0.16301 mL | 0.51002 mL | 7.684 | -0.09598 | 0.99125 | 0.00476 | Timed out at 59.5 s |
| 1:14:52.8 | Data point 74 | 1.50000 mL | 0.16334 mL | 0.16305 mL | 0.51002 mL | 8.114 | -0.08212 | 0.99372 | 0.00407 | Timed out at 59.5 s |
| 1:16:23.3 | Data point 75 | 1.50000 mL | 0.16334 mL | 0.16310 mL | 0.51002 mL | 8.426 | -0.04721 | 0.97560 | 0.00236 | Timed out at 59.5 s |
| 1:17:58.8 | Data point 76 | 1.50000 mL | 0.16334 mL | 0.16317 mL | 0.51002 mL | 8.764 | -0.01810 | 0.93699 | 0.00092 | 59.0 s |
| 1:19:28.4 | Data point 77 | 1.50000 mL | 0.16334 mL | 0.16324 mL | 0.51002 mL | 9.016 | -0.00920 | 0.71690 | 0.00054 | 30.0 s |
| 1:20:07.6 | Assay volumes | 1.50000 mL | 0.16334 mL | 0.16324 mL | 0.51002 mL | | | | | |

| | | | |
|---|---|---|---|
| Sample name: | **M04_octanol** | Experiment start time: | **3/24/2018 2:50:31 AM** |
| Assay name: | **pH-metric high logP** | Analyst: | **Dorothy Levorse** |
| Assay ID: | **18C-24003** | Instrument ID: | **T312060** |
| Filename: | **C:\Sirius_T3\Mehtap\20180323_exp33_logP_T3-2\18C-24003_M04_octanol_pH-metric high logP.t3r** | | |

| | | | |
|---|---|---|---|
| Sample name: | **M04_octanol** | Experiment start time: | **3/24/2018 2:50:31 AM** |
| Assay name: | **pH-metric high logP** | Analyst: | **Dorothy Levorse** |
| Assay ID: | **18C-24003** | Instrument ID: | **T312060** |
| Filename: | **C:\Sirius_T3\Mehtap\20180323_exp33_logP_T3-2\18C-24003_M04_octanol_pH-metric high logP.t3r** | | |

#### Assay Settings (continued)

| Setting | Value | Original Value | Date/Time changed | Imported from |
|---|---|---|---|---|
| ***Titration 3*** | | | | |
| Titrate from | Low to high pH | | | |
| Add additional water | 0.00 mL | | | |
| Additional partition solvent volume | 0.400 mL | | | |
| Additional partition solvent added | Automatic | | | |
| After pH adjust stir for | 30 seconds | | | |
| Stir to allow partitioning for | 15 seconds | | | |
| Stirrer speed for partitioning | 60% | | | |
| ***Data Point Stability*** | | | | |
| Stir during data point collection | No | | | |
| Delay before data point collection | 0 seconds | | | |
| Number of points to average | 20 points | | | |
| Time interval between points | 0.50 seconds | | | |
| Required maximum standard deviation | 0.00100 dpH/dt | | | |
| Stability timeout after | 60 seconds | | | |

#### Calibration Settings

| Setting | Value | Date/Time changed | Imported from |
|---|---|---|---|
| Four-Plus alpha | 0.119 | 3/24/2018 2:50:31 AM | C:\Sirius_T3\HCl18C23.t3r |
| Four-Plus S | 0.9972 | 3/24/2018 2:50:31 AM | C:\Sirius_T3\HCl18C23.t3r |
| Four-Plus jH | 0.9 | 3/24/2018 2:50:31 AM | C:\Sirius_T3\HCl18C23.t3r |
| Four-Plus jOH | -0.3 | 3/24/2018 2:50:31 AM | C:\Sirius_T3\HCl18C23.t3r |
| Base concentration factor | 1.003 | 3/24/2018 2:50:31 AM | C:\Sirius_T3\KOH18C23.t3r |
| Acid concentration factor | 0.997 | 3/24/2018 2:50:31 AM | C:\Sirius_T3\HCl18C23.t3r |

#### Instrument Settings

| Setting | Value | Batch Id | Install date |
|---|---|---|---|
| Instrument owner | Merck | | |
| Instrument ID | T312060 | | |
| Instrument type | T3 Simulator | | |
| Software version | 1.1.3.0 | | |
| Dispenser module | | T3DM1200361 | 3/31/2009 6:24:52 AM |
| Dispenser 0 | Water | | 3/31/2009 6:25:05 AM |
| Syringe volume | 2.5 mL | | |
| Firmware version | 1.2.1(r2) | | |
| Titrant | Water (0.15 M KCl) | 02-06-2018 | 3/16/2018 11:09:18 AM |
| Dispenser 2 | Acid | | 3/31/2009 6:25:11 AM |
| Syringe volume | 0.5 mL | | |
| Firmware version | 1.2.1(r2) | | |
| Titrant | Acid (0.5 M HCl) | 03-16-2018 | 3/16/2018 10:56:23 AM |
| Dispenser 1 | Base | | 3/31/2009 6:25:21 AM |
| Syringe volume | 0.5 mL | | |
| Firmware version | 1.2.1(r2) | | |
| Titrant | Base (0.5 M KOH) | 3/22/2018 | 3/23/2018 9:34:17 AM |
| Dispenser 5 | Cosolvent | | 3/31/2009 6:26:24 AM |
| Syringe volume | 2.5 mL | | |
| Firmware version | 1.2.1(r2) | | |
| Distribution valve 5 | Distribution Valve | | 3/31/2009 6:28:19 AM |
| Firmware version | 1.1.3 | | |
| Port A | Methanol (80%, 0.15 M KCl) | 02-08-2018 | 3/6/2018 10:28:59 AM |
| Port B | Cyclohexane | 11-01-17 | 2/27/2018 11:37:57 AM |
| Dispenser 3 | Buffer | | 8/3/2010 6:05:16 AM |
| Syringe volume | 0.5 mL | | |
| Firmware version | 1.2.1(r2) | | |
| Titrant | Dodecane | 2018/01/31 | 2/28/2018 11:18:04 AM |
| Dispenser 6 | Octanol | | 10/22/2010 11:52:43 AM |

| | | | |
|---|---|---|---|
| Sample name: | **M04_octanol** | Experiment start time: | **3/24/2018 2:50:31 AM** |
| Assay name: | **pH-metric high logP** | Analyst: | **Dorothy Levorse** |
| Assay ID: | **18C-24003** | Instrument ID: | **T312060** |
| Filename: | **C:\Sirius_T3\Mehtap\20180323_exp33_logP_T3-2\18C-24003_M04_octanol_pH-metric high logP.t3r** | | |

| | | | |
|---|---|---|---|
| Sample name: | **M04_octanol** | Experiment start time: | **3/24/2018 2:50:31 AM** |
| Assay name: | **pH-metric high logP** | Analyst: | **Dorothy Levorse** |
| Assay ID: | **18C-24003** | Instrument ID: | **T312060** |
| Filename: | **C:\Sirius_T3\Mehtap\20180323_exp33_logP_T3-2\18C-24003_M04_octanol_pH-metric high logP.t3r** | | |

#### Instrument Settings (continued)

| Setting | Value | Batch Id | Install date |
|---|---|---|---|
| Spectrometer calibration stir duration | 5 s | | |
| Spectrometer calibration stir speed | 30% | | |
| Spectrometer calibration wash pump volume | 20.0 mL | | |
| Spectrometer calibration wash stir duration | 5 s | | |
| Spectrometer calibration wash stir speed | 30% | | |
| Overhead dispense height | 10000 | | |

#### Refinement Settings

| Setting | Value | Default value |
|---|---|---|
| Turbidity detection method | None | None |
| Turbidity wavelength to assess | 500.0 nm | 500.0 nm |
| Turbidity maximum absorbance | 0.100 | 0.100 |
| Turbidity probe threshold | 50.00 | 50.00 |
