## Supplementary material for "Octanol-water partition coefficient measurements for the SAMPL6 Blind Prediction Challenge": SM04_18C-24004_M04_octanol_pH-metric high logP_report.pdf

| | | | |
|---|---|---|---|
| Sample name: | **M04_octanol** | Experiment start time: | **3/24/2018 4:11:23 AM** |
| Assay name: | **pH-metric high logP** | Analyst: | **Dorothy Levorse** |
| Assay ID: | **18C-24004** | Instrument ID: | **T312060** |
| Filename: | **C:\Sirius_T3\Mehtap\20180323_exp33_logP_T3-2\18C-24004_M04_octanol_pH-metric high logP.t3r** | | |

#### pH-metric Result

| | | |
|---|---|---|
| logP (XH +) | 0.77 | ±0.03 (n=50) |
| logP (neutral X) | 3.95 | ±0.02 (n=50) |
| RMSD | 0.422 | |

#### 18C-24004 Points 1 to 23

| | |
|---|---|
| M04_octanol concentration factor | 0.620 |
| Carbonate | 0.0570 mM |
| Acidity error | 0.07992 mM |

#### 18C-24004 Points 24 to 50

| | |
|---|---|
| M04_octanol concentration factor | 0.641 |
| Carbonate | 0.1306 mM |
| Acidity error | -0.26645 mM |

#### 18C-24004 Points 51 to 77

| | |
|---|---|
| M04_octanol concentration factor | 0.822 |
| Carbonate | 0.1176 mM |
| Acidity error | -0.01548 mM |

#### Warnings and errors

| | |
|---|---|
| Errors | None |
| Warnings | None |

#### Sample logD and percent species

| pH | M04_octanol logD | M04_octanol M04_octanolH | M04_octanol M04_octanol | M04_octanol M04_octanolH* | M04_octanol M04_octanol* | Comment |
|---|---|---|---|---|---|---|
| 1.000 | 0.78 | 14.18 % | 0.00 % | 84.45 % | 1.37 % | |
| 1.200 | 0.79 | 14.07 % | 0.00 % | 83.78 % | 2.15 % | Stomach pH |
| 2.000 | 0.84 | 12.63 % | 0.00 % | 75.19 % | 12.18 % | |
| 3.000 | 1.19 | 6.03 % | 0.01 % | 35.87 % | 58.10 % | |
| 4.000 | 2.01 | 0.97 % | 0.01 % | 5.76 % | 93.26 % | |
| 5.000 | 2.94 | 0.10 % | 0.01 % | 0.61 % | 99.27 % | |
| 6.000 | 3.67 | 0.01 % | 0.01 % | 0.06 % | 99.92 % | |
| 6.500 | 3.84 | 0.00 % | 0.01 % | 0.02 % | 99.97 % | |
| 7.000 | 3.92 | 0.00 % | 0.01 % | 0.01 % | 99.98 % | |
| 7.400 | 3.94 | 0.00 % | 0.01 % | 0.00 % | 99.99 % | Blood pH |
| 8.000 | 3.95 | 0.00 % | 0.01 % | 0.00 % | 99.99 % | |
| 9.000 | 3.95 | 0.00 % | 0.01 % | 0.00 % | 99.99 % | |
| 10.000 | 3.95 | 0.00 % | 0.01 % | 0.00 % | 99.99 % | |
| 11.000 | 3.95 | 0.00 % | 0.01 % | 0.00 % | 99.99 % | |
| 12.000 | 3.95 | 0.00 % | 0.01 % | 0.00 % | 99.99 % | |

| | |
|---|---|
| Sample name: | **M04_octanol** |
| Assay name: | **pH-metric high logP** |
| Assay ID: | **18C-24004** |
| Filename: | **C:\Sirius_T3\Mehtap\20180323_exp33_logP_T3-2\18C-24004_M04_octanol_pH-metric high logP.t3r** |
| Experiment start time: | **3/24/2018 4:11:23 AM** |
| Analyst: | **Dorothy Levorse** |
| Instrument ID: | **T312060** |

#### Graphs

Charge balance residuals

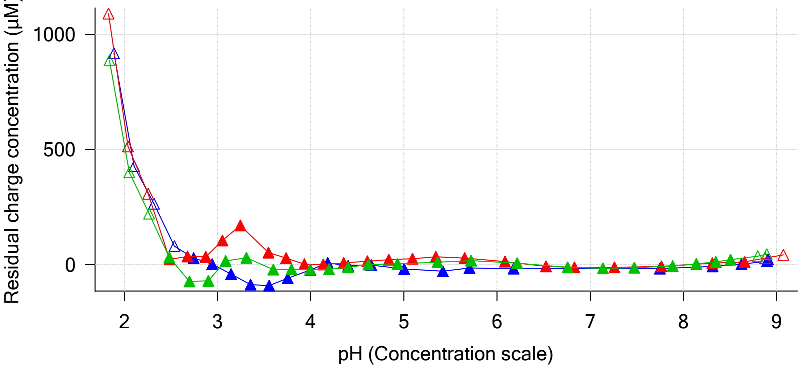


Ionic strength

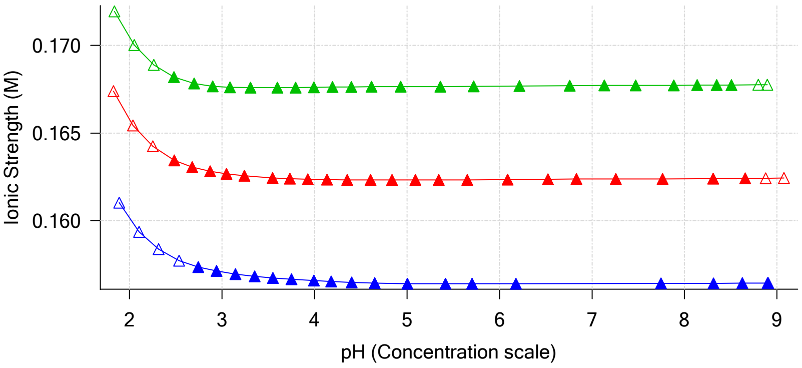


Buffer index profile

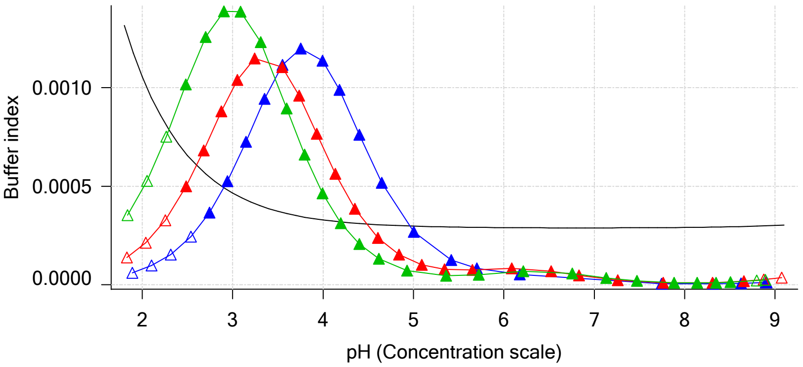


Sample M04_octanol Ionisation Graph

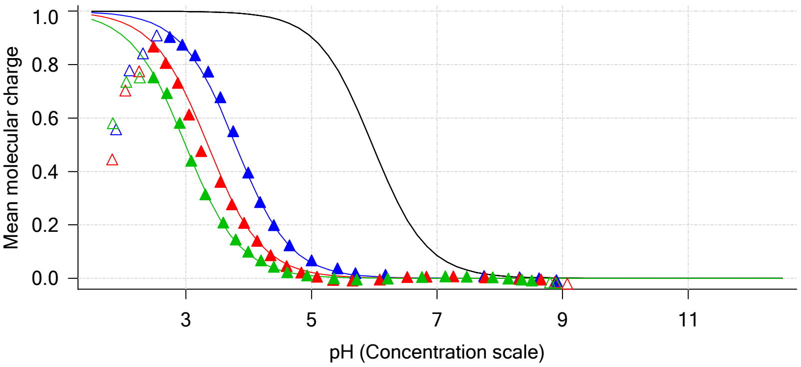


Distribution of species for sample M04_octanol

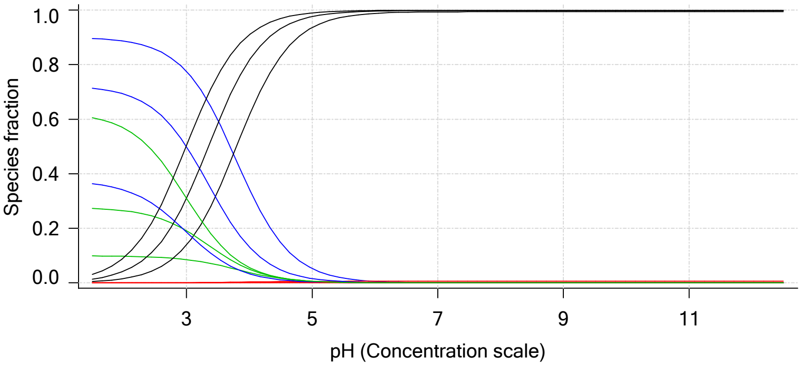


Lipophilicity Profile

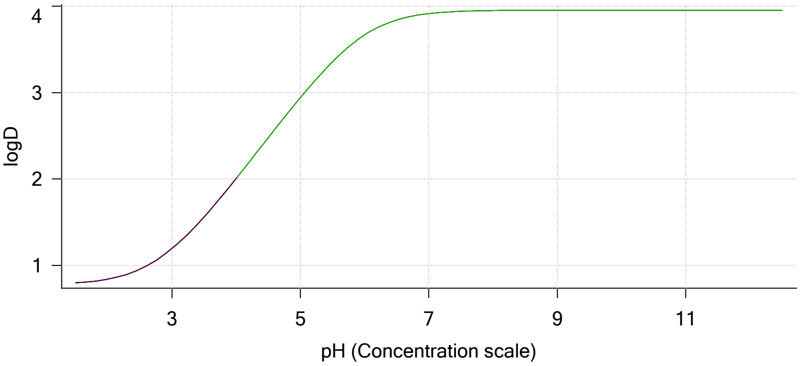


Log R Profile

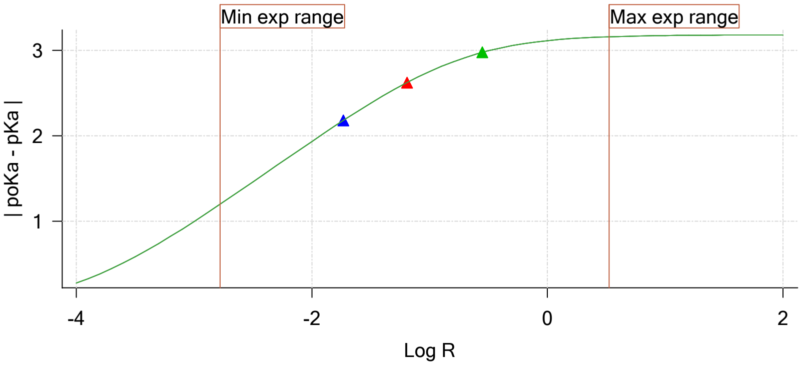


Sample Carbonate Ionisation Graph

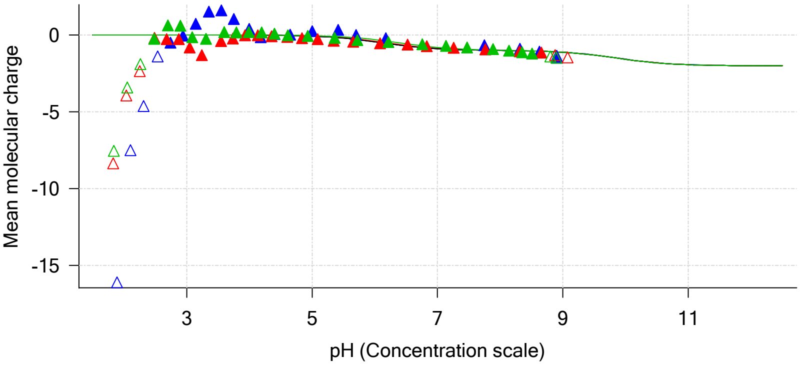

Sample name: **M04_octanol**
Assay name: **pH-metric high logP**
Assay ID: **18C-24004**
Filename: **C:\Sirius_T3\Mehtap\20180323_exp33_logP_T3-2\18C-24004_M04_octanol_pH-metric high logP.t3r**

Experiment start time: **3/24/2018 4:11:23 AM**
Analyst: **Dorothy Levorse**
Instrument ID: **T312060**

#### Graphs (continued)

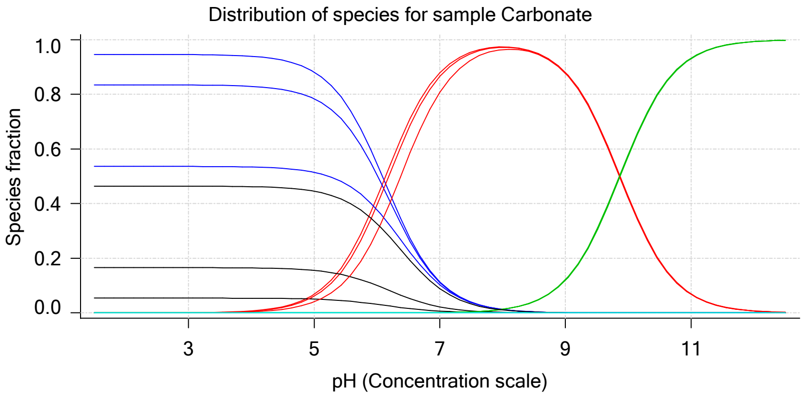

| | | | |
|---|---|---|---|
| Sample name: | **M04_octanol** | Experiment start time: | **3/24/2018 4:11:23 AM** |
| Assay name: | **pH-metric high logP** | Analyst: | **Dorothy Levorse** |
| Assay ID: | **18C-24004** | Instrument ID: | **T312060** |
| Filename: | **C:\Sirius_T3\Mehtap\20180323_exp33_logP_T3-2\18C-24004_M04_octanol_pH-metric high logP.t3r** | | |

### pH-metric high logP Titration 1 of 3 18C-24004 Points 1 to 23

#### Overall results

| | |
|---|---|
| RMSD | 0.635 |
| Average ionic strength | 0.157 M |
| Average temperature | 24.9°C |
| Partition ratio | 0.0185 : 1 |
| Analyte concentration range | 3292.8 μM to 3384.5 μM |
| Total points considered | 19 of 23 |

#### Warnings and errors

| | |
|---|---|
| Errors | None |
| Warnings | Sample concentration factor out of range |

#### Four-Plus parameters

| | | | |
|---|---|---|---|
| Alpha | 0.119 | 3/24/2018 4:11:23 AM | C:\Sirius_T3\HCl18C23.t3r |
| S | 0.9972 | 3/24/2018 4:11:23 AM | C:\Sirius_T3\HCl18C23.t3r |
| jH | 0.9 | 3/24/2018 4:11:23 AM | C:\Sirius_T3\HCl18C23.t3r |
| jOH | -0.3 | 3/24/2018 4:11:23 AM | C:\Sirius_T3\HCl18C23.t3r |

#### Titrants

| | | | |
|---|---|---|---|
| 0.50 M HCl | 0.997124 | 3/24/2018 4:11:23 AM | C:\Sirius_T3\HCl18C23.t3r |
| 0.50 M KOH | 1.003190 | 3/24/2018 4:11:23 AM | C:\Sirius_T3\KOH18C23.t3r |

#### Sample

| | |
|---|---|
| M04_octanol concentration factor | 0.620 |
| Base pKa 1 | 5.97 |
| logP (XH +) | 0.19 |
| logP (neutral X) | 3.88 |

#### Sample graphs

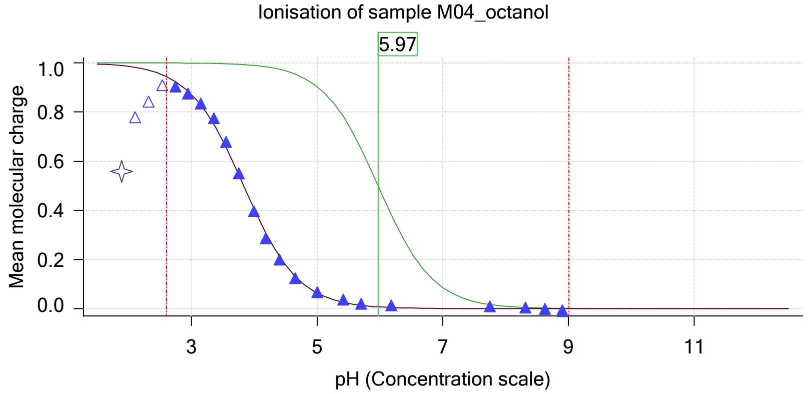


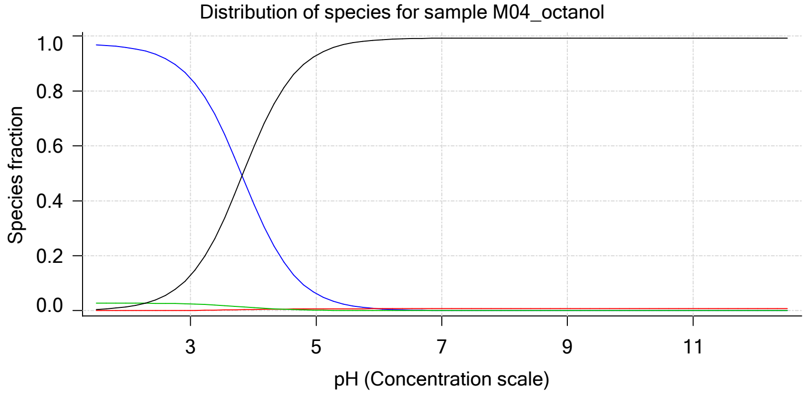

| | | | |
|---|---|---|---|
| Sample name: | **M04_octanol** | Experiment start time: | **3/24/2018 4:11:23 AM** |
| Assay name: | **pH-metric high logP** | Analyst: | **Dorothy Levorse** |
| Assay ID: | **18C-24004** | Instrument ID: | **T312060** |
| Filename: | **C:\Sirius_T3\Mehtap\20180323_exp33_logP_T3-2\18C-24004_M04_octanol_pH-metric high logP.t3r** | | |

#### Sample graphs (continued)

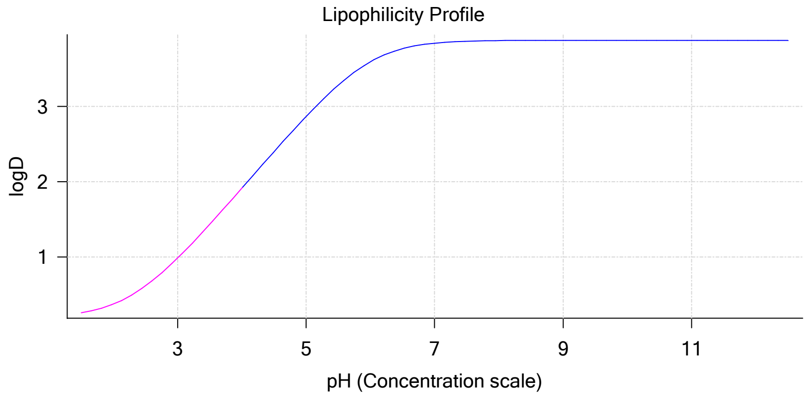


#### Sample logD and percent species

| pH | M04_octanol logD | M04_octanol M04_octanolH | M04_octanol M04_octanol | M04_octanol M04_octanolH* | M04_octanol M04_octanol* | Comment |
|---|---|---|---|---|---|---|
| 1.000 | 0.21 | 97.07 % | 0.00 % | 2.78 % | 0.15 % | |
| 1.200 | 0.22 | 96.99 % | 0.00 % | 2.78 % | 0.23 % | Stomach pH |
| 2.000 | 0.37 | 95.80 % | 0.01 % | 2.74 % | 1.44 % | |
| 3.000 | 0.99 | 84.72 % | 0.09 % | 2.43 % | 12.76 % | |
| 4.000 | 1.91 | 39.28 % | 0.42 % | 1.13 % | 59.18 % | |
| 5.000 | 2.87 | 6.17 % | 0.66 % | 0.18 % | 92.99 % | |
| 6.000 | 3.59 | 0.65 % | 0.70 % | 0.02 % | 98.63 % | |
| 6.500 | 3.77 | 0.21 % | 0.70 % | 0.01 % | 99.08 % | |
| 7.000 | 3.84 | 0.07 % | 0.71 % | 0.00 % | 99.23 % | |
| 7.400 | 3.87 | 0.03 % | 0.71 % | 0.00 % | 99.27 % | Blood pH |
| 8.000 | 3.88 | 0.01 % | 0.71 % | 0.00 % | 99.29 % | |
| 9.000 | 3.88 | 0.00 % | 0.71 % | 0.00 % | 99.29 % | |
| 10.000 | 3.88 | 0.00 % | 0.71 % | 0.00 % | 99.29 % | |
| 11.000 | 3.88 | 0.00 % | 0.71 % | 0.00 % | 99.29 % | |
| 12.000 | 3.88 | 0.00 % | 0.71 % | 0.00 % | 99.29 % | |

#### Carbonate and acidity

| | |
|---|---|
| Carbonate | 0.057 mM |
| Acidity error | 0.080 mM |

#### Other graphs

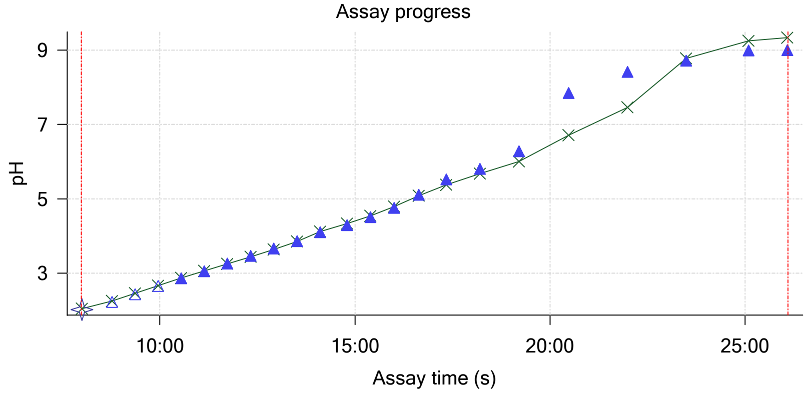


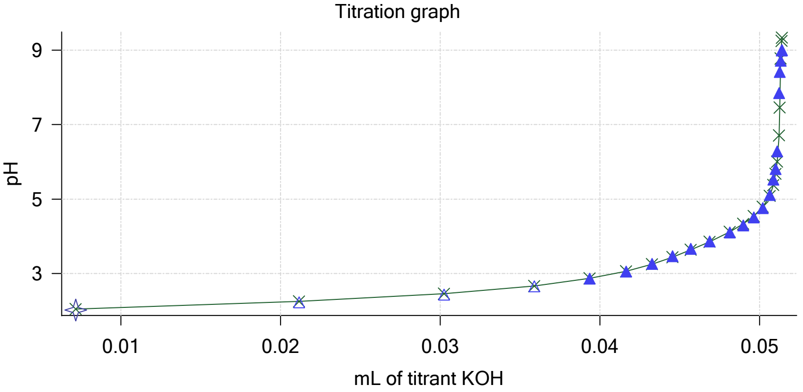

Sample name: **M04_octanol**
Assay name: **pH-metric high logP**
Assay ID: **18C-24004**
Filename: **C:\Sirius_T3\Mehtap\20180323_exp33_logP_T3-2\18C-24004_M04_octanol_pH-metric high logP.t3r**
Experiment start time: **3/24/2018 4:11:23 AM**
Analyst: **Dorothy Levorse**
Instrument ID: **T312060**

#### Other graphs (continued)

Charge balance residuals

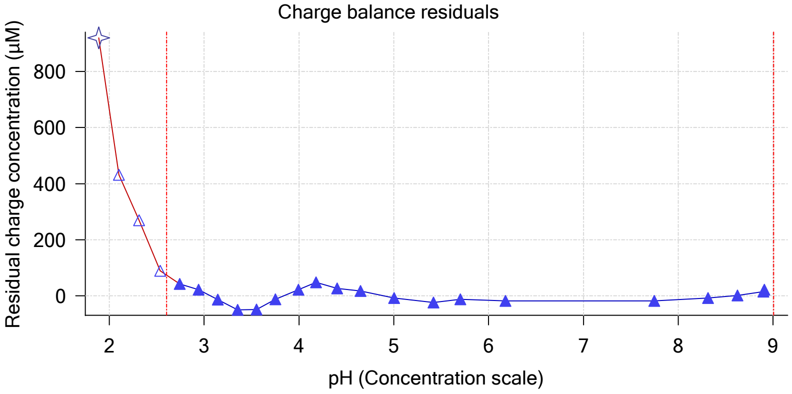


Buffer index profile

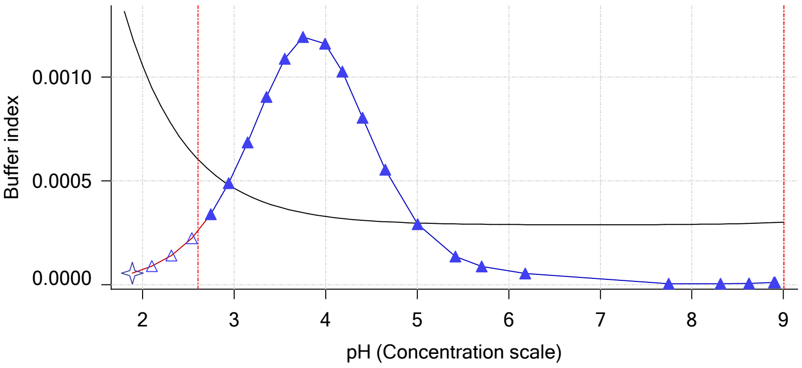


Ionic strength

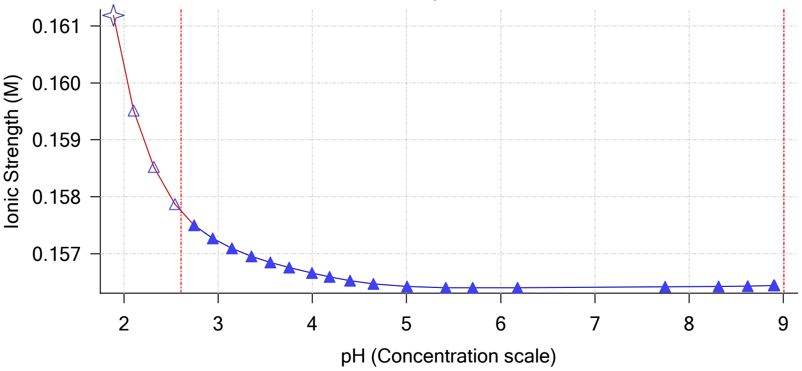


Log R Profile

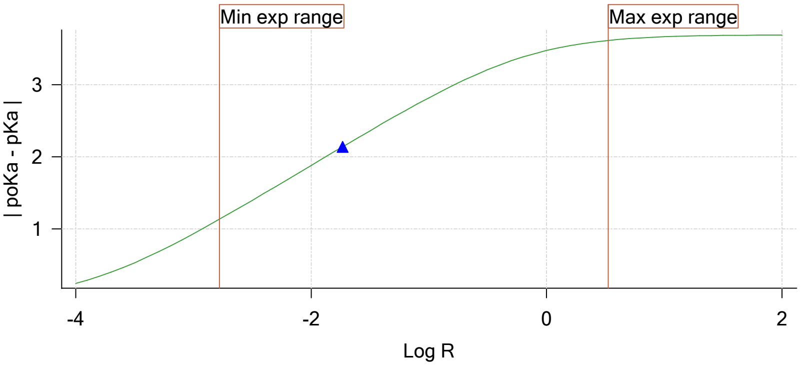


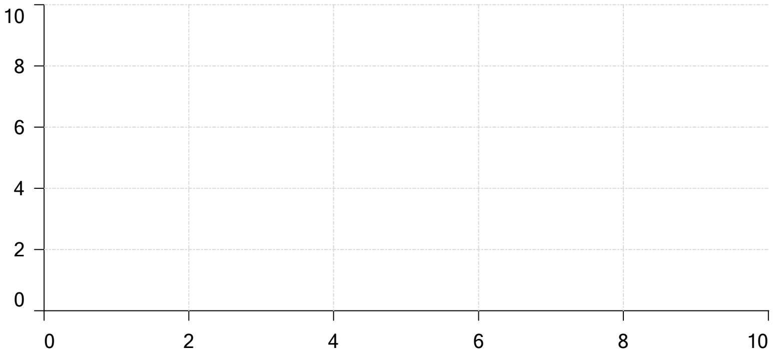


Ionisation of sample Carbonate

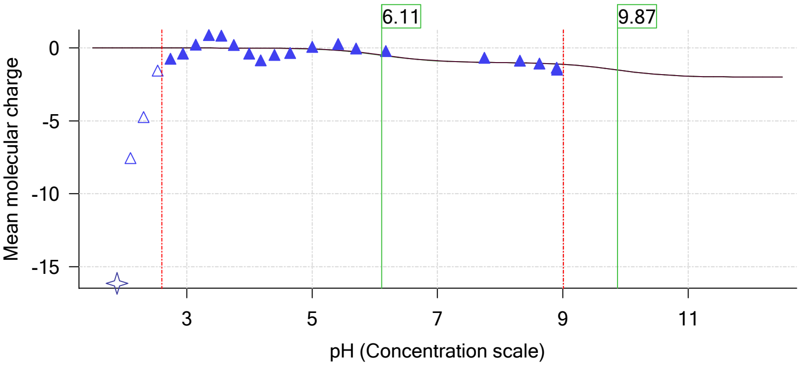

| | | | |
|---|---|---|---|
| Sample name: | **M04_octanol** | Experiment start time: | **3/24/2018 4:11:23 AM** |
| Assay name: | **pH-metric high logP** | Analyst: | **Dorothy Levorse** |
| Assay ID: | **18C-24004** | Instrument ID: | **T312060** |
| Filename: | **C:\Sirius_T3\Mehtap\20180323_exp33_logP_T3-2\18C-24004_M04_octanol_pH-metric high logP.t3r** | | |

# pH-metric high logP   Titration 2 of 3   18C-24004 Points 24 to 50

#### Overall results

| | |
|---|---|
| RMSD | 0.240 |
| Average ionic strength | 0.163 M |
| Average temperature | 25.0°C |
| Partition ratio | 0.0642 : 1 |
| Analyte concentration range | 2954.9 μM to 3043.2 μM |
| Total points considered | 22 of 27 |

#### Warnings and errors

| | |
|---|---|
| Errors | None |
| Warnings | Sample concentration factor out of range |

#### Four-Plus parameters

| | | | |
|---|---|---|---|
| Alpha | 0.119 | 3/24/2018 4:11:23 AM | C:\Sirius_T3\HCl18C23.t3r |
| S | 0.9972 | 3/24/2018 4:11:23 AM | C:\Sirius_T3\HCl18C23.t3r |
| jH | 0.9 | 3/24/2018 4:11:23 AM | C:\Sirius_T3\HCl18C23.t3r |
| jOH | -0.3 | 3/24/2018 4:11:23 AM | C:\Sirius_T3\HCl18C23.t3r |

#### Titrants

| | | | |
|---|---|---|---|
| 0.50 M HCl | 0.997124 | 3/24/2018 4:11:23 AM | C:\Sirius_T3\HCl18C23.t3r |
| 0.50 M KOH | 1.003190 | 3/24/2018 4:11:23 AM | C:\Sirius_T3\KOH18C23.t3r |

#### Sample

| | |
|---|---|
| M04_octanol concentration factor | 0.641 |
| Base pKa 1 | 5.97 |
| logP (XH +) | 0.19 |
| logP (neutral X) | 3.92 |

#### Sample graphs

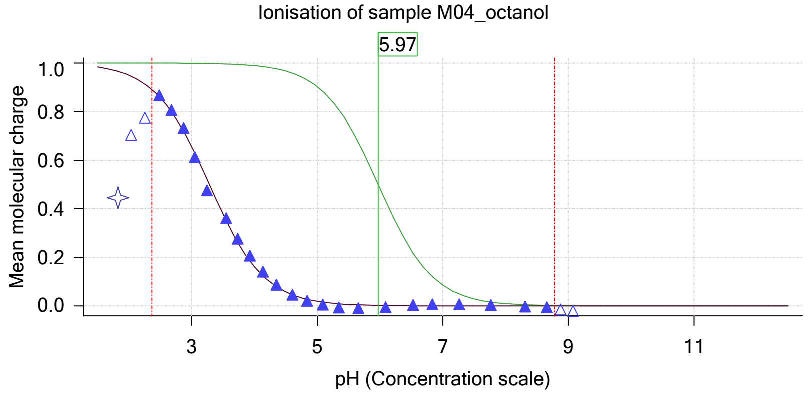


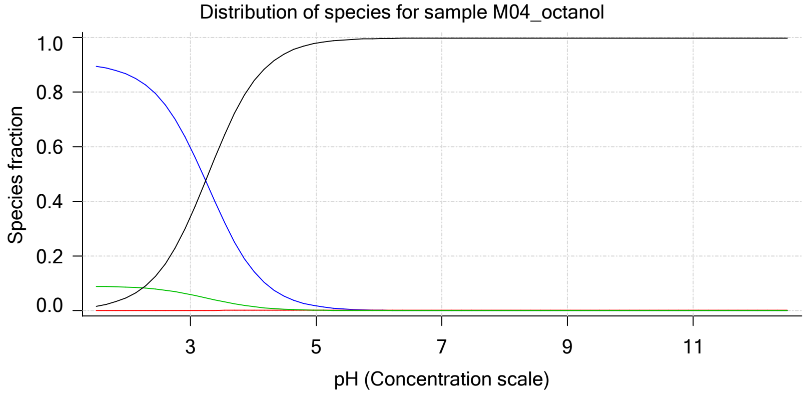

Sample name: **M04_octanol**
Assay name: **pH-metric high logP**
Assay ID: **18C-24004**
Filename: **C:\Sirius_T3\Mehtap\20180323_exp33_logP_T3-2\18C-24004_M04_octanol_pH-metric high logP.t3r**
Experiment start time: **3/24/2018 4:11:23 AM**
Analyst: **Dorothy Levorse**
Instrument ID: **T312060**

#### Sample graphs (continued)

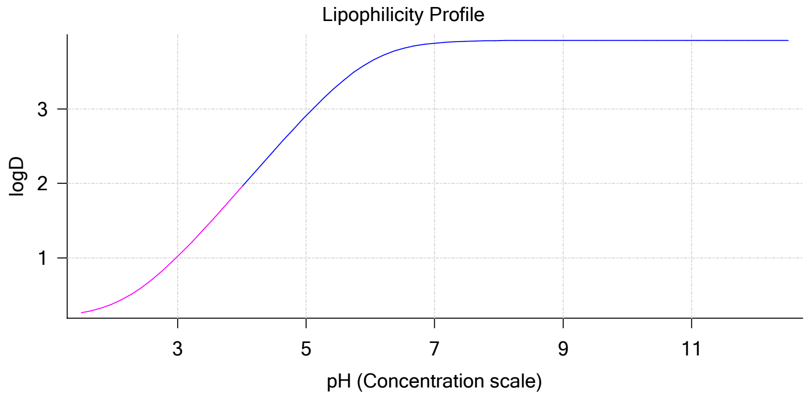


#### Sample logD and percent species

| pH | M04_octanol logD | M04_octanol M04_octanolH | M04_octanol M04_octanol | M04_octanol M04_octanolH* | M04_octanol M04_octanol* | Comment |
|---|---|---|---|---|---|---|
| 1.000 | 0.21 | 90.48 % | 0.00 % | 9.00 % | 0.52 % | |
| 1.200 | 0.23 | 90.21 % | 0.00 % | 8.97 % | 0.82 % | Stomach pH |
| 2.000 | 0.39 | 86.44 % | 0.01 % | 8.60 % | 4.96 % | |
| 3.000 | 1.02 | 59.74 % | 0.06 % | 5.94 % | 34.25 % | |
| 4.000 | 1.95 | 14.61 % | 0.16 % | 1.45 % | 83.78 % | |
| 5.000 | 2.91 | 1.71 % | 0.18 % | 0.17 % | 97.94 % | |
| 6.000 | 3.63 | 0.17 % | 0.19 % | 0.02 % | 99.62 % | |
| 6.500 | 3.81 | 0.06 % | 0.19 % | 0.01 % | 99.75 % | |
| 7.000 | 3.88 | 0.02 % | 0.19 % | 0.00 % | 99.79 % | |
| 7.400 | 3.90 | 0.01 % | 0.19 % | 0.00 % | 99.81 % | Blood pH |
| 8.000 | 3.92 | 0.00 % | 0.19 % | 0.00 % | 99.81 % | |
| 9.000 | 3.92 | 0.00 % | 0.19 % | 0.00 % | 99.81 % | |
| 10.000 | 3.92 | 0.00 % | 0.19 % | 0.00 % | 99.81 % | |
| 11.000 | 3.92 | 0.00 % | 0.19 % | 0.00 % | 99.81 % | |
| 12.000 | 3.92 | 0.00 % | 0.19 % | 0.00 % | 99.81 % | |

#### Carbonate and acidity

| | |
|---|---|
| Carbonate | 0.131 mM |
| Acidity error | -0.266 mM |

#### Other graphs

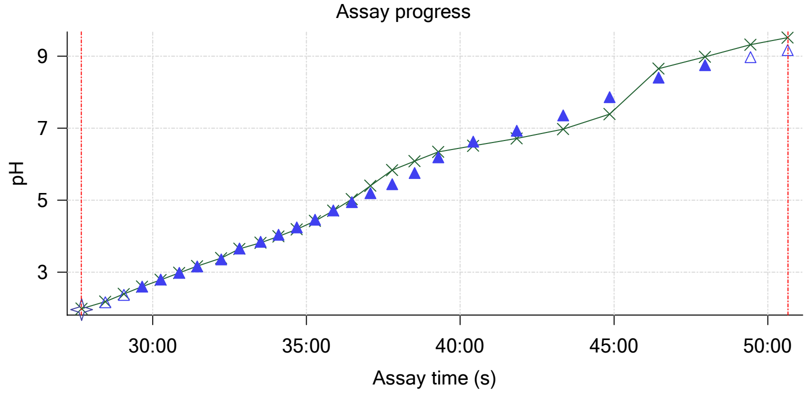


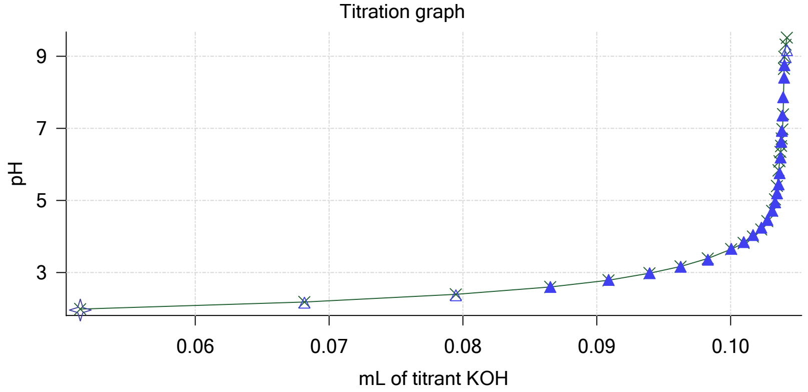

Sample name: **M04_octanol**
Assay name: **pH-metric high logP**
Assay ID: **18C-24004**
Filename: **C:\Sirius_T3\Mehtap\20180323_exp33_logP_T3-2\18C-24004_M04_octanol_pH-metric high logP.t3r**

Experiment start time: **3/24/2018 4:11:23 AM**
Analyst: **Dorothy Levorse**
Instrument ID: **T312060**

#### Other graphs (continued)

Charge balance residuals

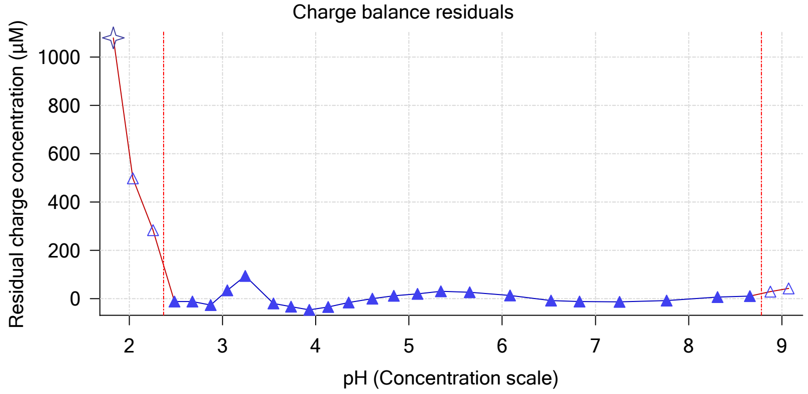


Buffer index profile

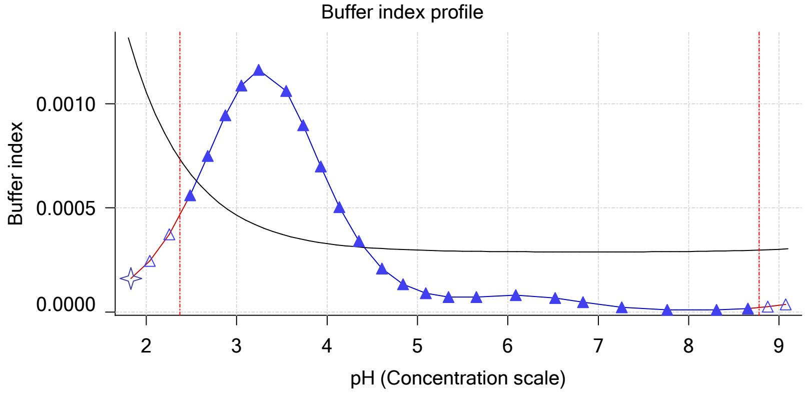


Ionic strength

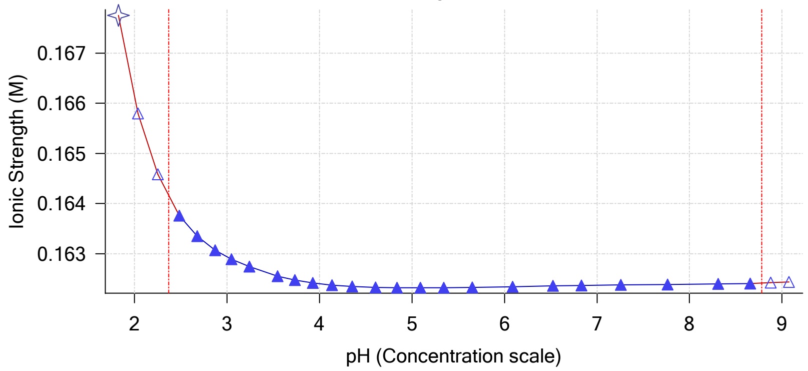


Log R Profile

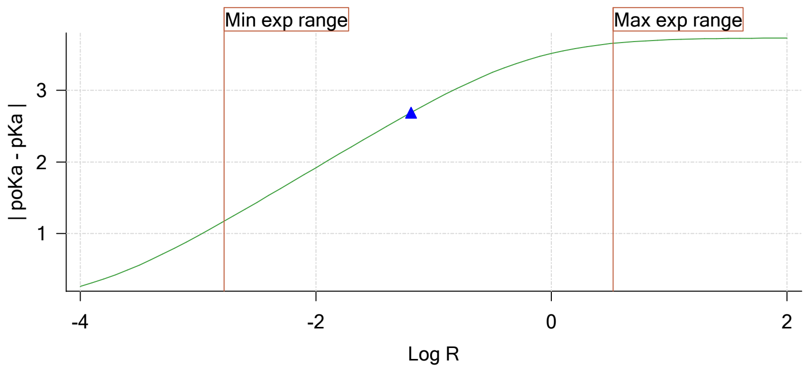


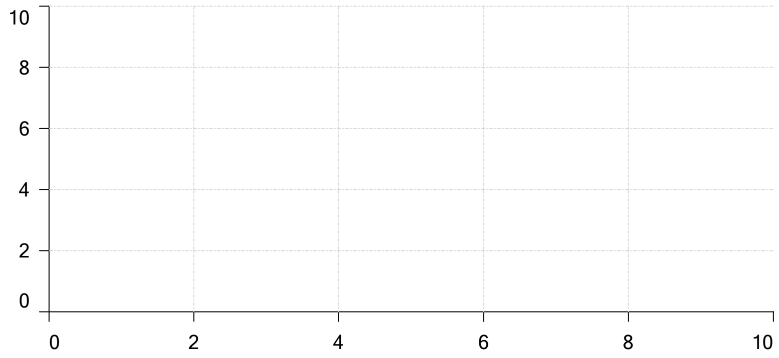

Ionisation of sample Carbonate

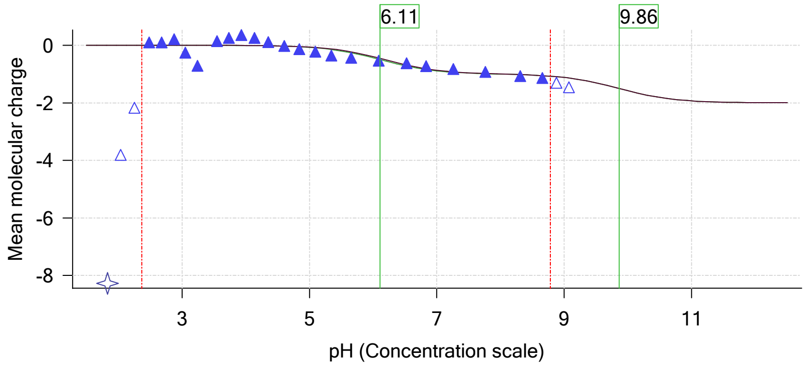

| | | | |
|---|---|---|---|
| Sample name: | **M04_octanol** | Experiment start time: | **3/24/2018 4:11:23 AM** |
| Assay name: | **pH-metric high logP** | Analyst: | **Dorothy Levorse** |
| Assay ID: | **18C-24004** | Instrument ID: | **T312060** |
| Filename: | **C:\Sirius_T3\Mehtap\20180323_exp33_logP_T3-2\18C-24004_M04_octanol_pH-metric high logP.t3r** | | |

# pH-metric high logP   Titration 3 of 3   18C-24004 Points 51 to 77

#### Overall results

| | |
|---|---|
| RMSD | 0.309 |
| Average ionic strength | 0.168 M |
| Average temperature | 25.0°C |
| Partition ratio | 0.2800 : 1 |
| Analyte concentration range | 2304.6 µM to 2362.2 µM |
| Total points considered | 22 of 27 |

#### Warnings and errors

| | |
|---|---|
| Errors | None |
| Warnings | None |

#### Four-Plus parameters

| | | | |
|---|---|---|---|
| Alpha | 0.119 | 3/24/2018 4:11:23 AM | C:\Sirius_T3\HCl18C23.t3r |
| S | 0.9972 | 3/24/2018 4:11:23 AM | C:\Sirius_T3\HCl18C23.t3r |
| jH | 0.9 | 3/24/2018 4:11:23 AM | C:\Sirius_T3\HCl18C23.t3r |
| jOH | -0.3 | 3/24/2018 4:11:23 AM | C:\Sirius_T3\HCl18C23.t3r |

#### Titrants

| | | | |
|---|---|---|---|
| 0.50 M HCl | 0.997124 | 3/24/2018 4:11:23 AM | C:\Sirius_T3\HCl18C23.t3r |
| 0.50 M KOH | 1.003190 | 3/24/2018 4:11:23 AM | C:\Sirius_T3\KOH18C23.t3r |

#### Sample

| | |
|---|---|
| M04_octanol concentration factor | 0.822 |
| Base pKa 1 | 5.97 |
| logP (XH +) | 0.19 |
| logP (neutral X) | 3.67 |

#### Sample graphs

Ionisation of sample M04_octanol

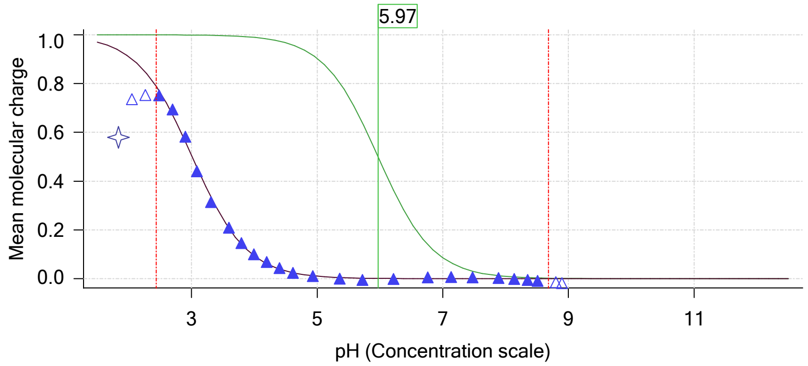


Distribution of species for sample M04_octanol

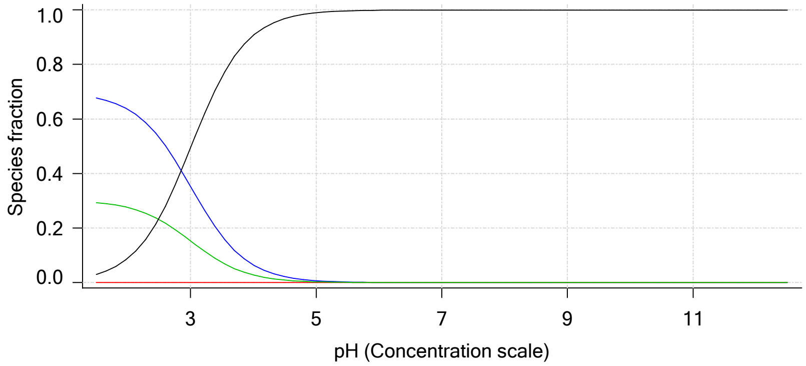

| | | | |
|---|---|---|---|
| Sample name: | **M04_octanol** | Experiment start time: | **3/24/2018 4:11:23 AM** |
| Assay name: | **pH-metric high logP** | Analyst: | **Dorothy Levorse** |
| Assay ID: | **18C-24004** | Instrument ID: | **T312060** |
| Filename: | **C:\Sirius_T3\Mehtap\20180323_exp33_logP_T3-2\18C-24004_M04_octanol_pH-metric high logP.t3r** | | |

#### Sample graphs (continued)

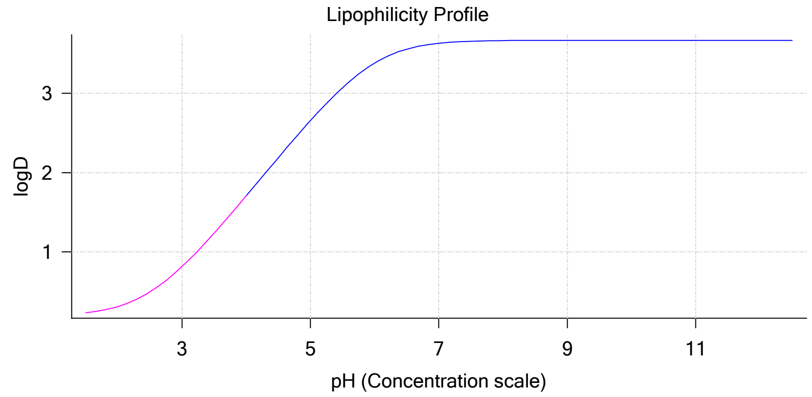


#### Sample logD and percent species

| pH | M04_octanol logD | M04_octanol M04_octanolH | M04_octanol M04_octanol | M04_octanol M04_octanolH* | M04_octanol M04_octanol* | Comment |
|---|---|---|---|---|---|---|
| 1.000 | 0.20 | 69.08 % | 0.00 % | 29.96 % | 0.97 % | |
| 1.200 | 0.21 | 68.69 % | 0.00 % | 29.79 % | 1.52 % | Stomach pH |
| 2.000 | 0.31 | 63.55 % | 0.01 % | 27.56 % | 8.88 % | |
| 3.000 | 0.81 | 35.31 % | 0.04 % | 15.31 % | 49.33 % | |
| 4.000 | 1.71 | 6.49 % | 0.07 % | 2.81 % | 90.63 % | |
| 5.000 | 2.66 | 0.71 % | 0.08 % | 0.31 % | 98.91 % | |
| 6.000 | 3.38 | 0.07 % | 0.08 % | 0.03 % | 99.82 % | |
| 6.500 | 3.56 | 0.02 % | 0.08 % | 0.01 % | 99.89 % | |
| 7.000 | 3.63 | 0.01 % | 0.08 % | 0.00 % | 99.91 % | |
| 7.400 | 3.65 | 0.00 % | 0.08 % | 0.00 % | 99.92 % | Blood pH |
| 8.000 | 3.66 | 0.00 % | 0.08 % | 0.00 % | 99.92 % | |
| 9.000 | 3.67 | 0.00 % | 0.08 % | 0.00 % | 99.92 % | |
| 10.000 | 3.67 | 0.00 % | 0.08 % | 0.00 % | 99.92 % | |
| 11.000 | 3.67 | 0.00 % | 0.08 % | 0.00 % | 99.92 % | |
| 12.000 | 3.67 | 0.00 % | 0.08 % | 0.00 % | 99.92 % | |

#### Carbonate and acidity

| | |
|---|---|
| Carbonate | 0.118 mM |
| Acidity error | -0.015 mM |

#### Other graphs

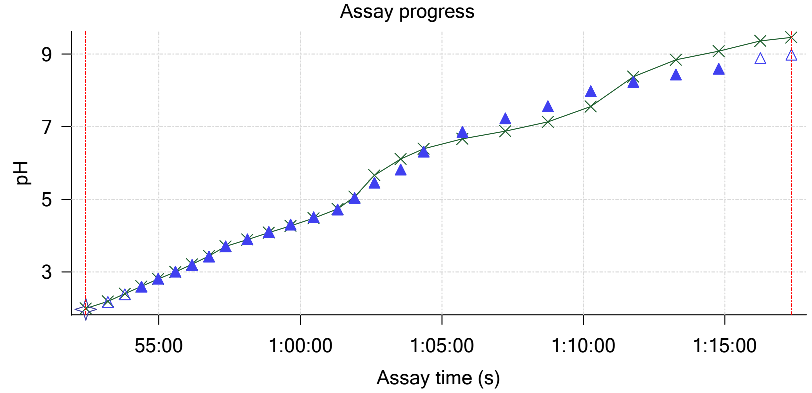


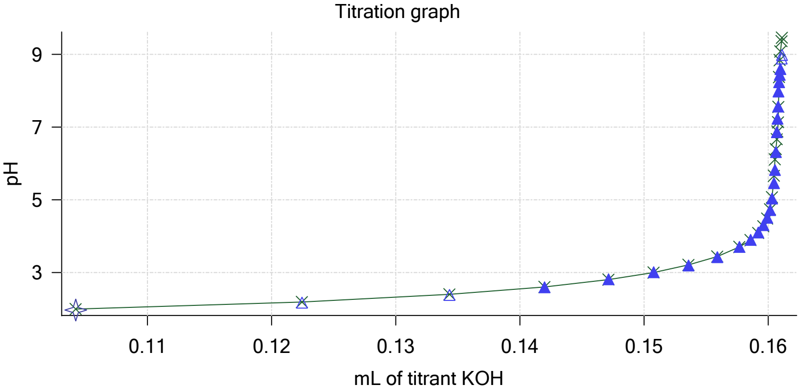

Sample name: **M04_octanol**
Assay name: **pH-metric high logP**
Assay ID: **18C-24004**
Filename: **C:\Sirius_T3\Mehtap\20180323_exp33_logP_T3-2\18C-24004_M04_octanol_pH-metric high logP.t3r**

Experiment start time: **3/24/2018 4:11:23 AM**
Analyst: **Dorothy Levorse**
Instrument ID: **T312060**

#### Other graphs (continued)

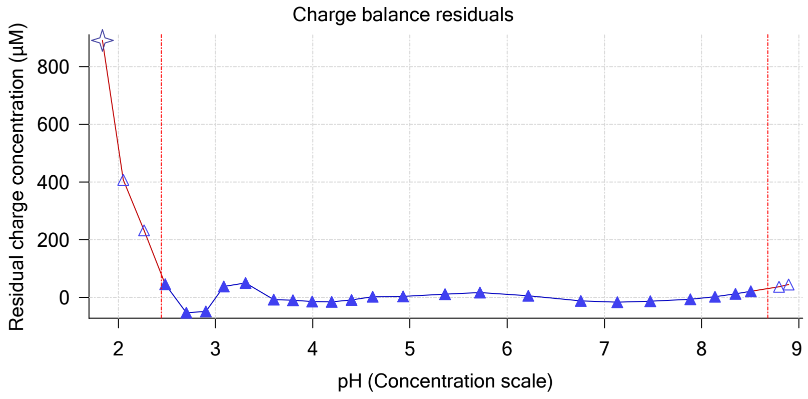


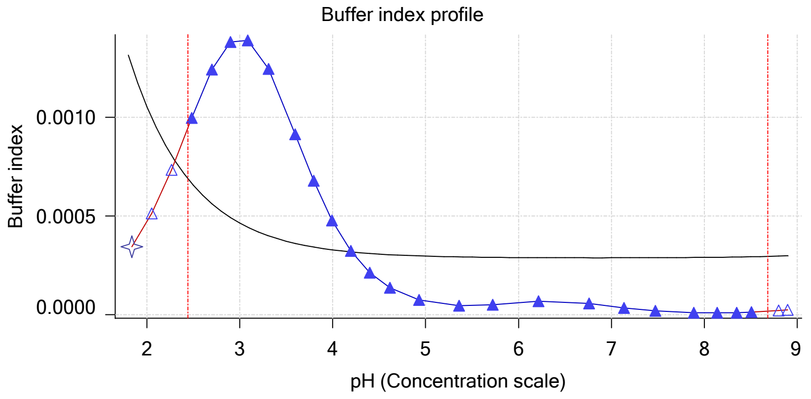


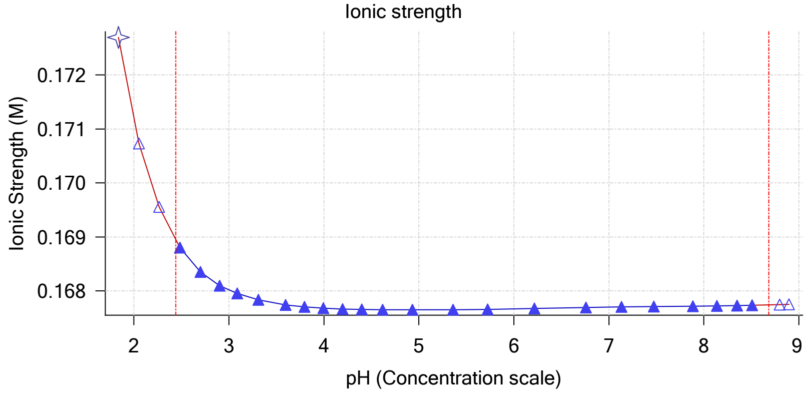


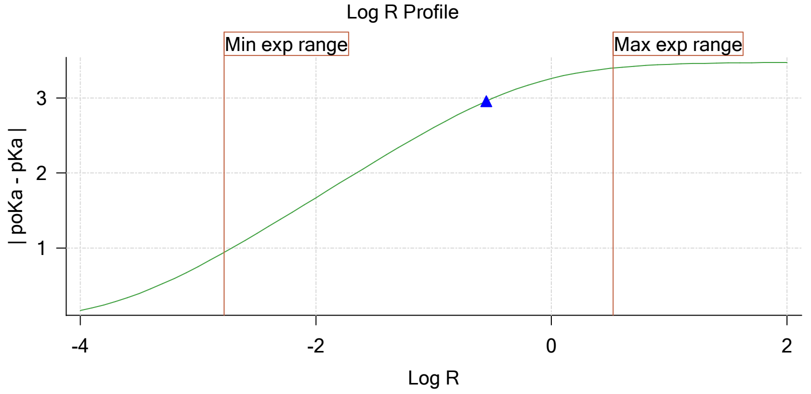


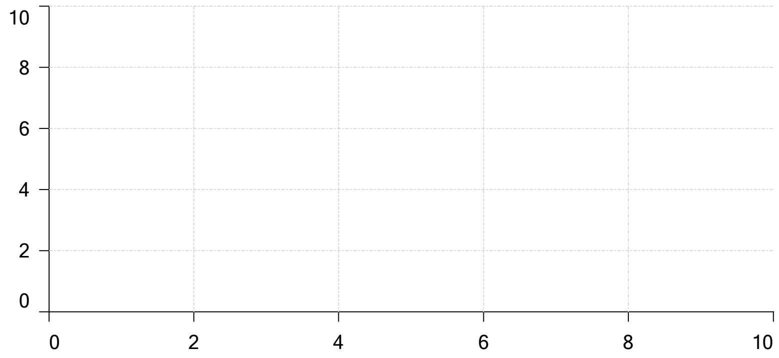


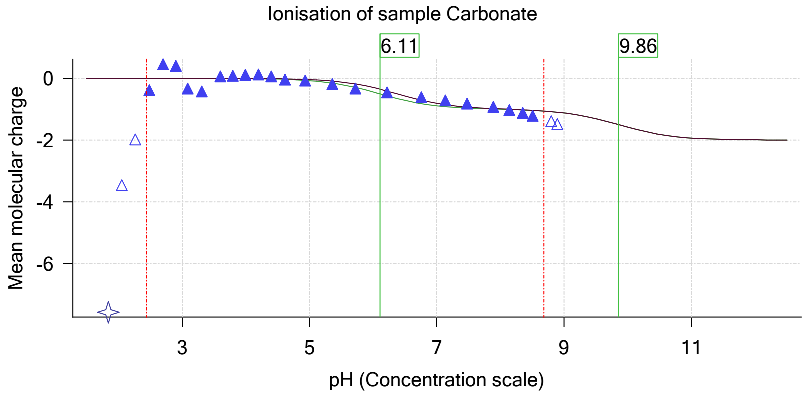

| | | | |
|---|---|---|---|
| Sample name: | **M04_octanol** | Experiment start time: | **3/24/2018 4:11:23 AM** |
| Assay name: | **pH-metric high logP** | Analyst: | **Dorothy Levorse** |
| Assay ID: | **18C-24004** | Instrument ID: | **T312060** |
| Filename: | **C:\Sirius_T3\Mehtap\20180323_exp33_logP_T3-2\18C-24004_M04_octanol_pH-metric high logP.t3r** | | |

#### Assay Model

| Settings | Value | Date/Time changed | Imported from |
|---|---|---|---|
| Sample name | M04_octanol | 3/9/2018 4:34:21 PM | User entered value |
| Sample by | Weight | | Default value |
| Sample weight | 0.001450 g | 3/23/2018 5:00:45 PM | User entered value |
| Formula weight | 269.73 g/mol | 3/9/2018 4:34:21 PM | User entered value |
| Solubility | Unknown | | Default value |
| Molecular weight | 269.73 | 3/9/2018 4:34:21 PM | User entered value |
| Individual pKa ionic environments | No | | Default value |
| Number of pKas | 1 | 3/9/2018 4:34:21 PM | User entered value |
| Sample is a | Base | 3/9/2018 4:34:21 PM | User entered value |
| pKa 1 | 5.97 | 3/9/2018 4:34:21 PM | User entered value |
| logp (XH +) | 0.19 | 3/9/2018 4:34:33 PM | User entered value |
| logP (neutral X) | 3.50 | 3/23/2018 2:32:46 PM | User entered value |

#### Events

| Time | Event | Water | Acid | Base | Octanol | pH | dpH/dt | pH R-squared | pH SD | dpH/dt time |
|---|---|---|---|---|---|---|---|---|---|---|
| 5:00.3 | Initial pH = 8.35 | | | | | | | | | |
| 7:60.0 | Data point 1 | 1.50000 mL | 0.05115 mL | 0.00717 mL | 0.03001 mL | 2.013 | -0.00513 | 0.25311 | 0.00050 | 10.0 s |
| 8:46.2 | Data point 2 | 1.50000 mL | 0.05115 mL | 0.02117 mL | 0.03001 mL | 2.222 | 0.00235 | 0.41042 | 0.00018 | 10.0 s |
| 9:21.8 | Data point 3 | 1.50000 mL | 0.05115 mL | 0.03024 mL | 0.03001 mL | 2.432 | -0.00386 | 0.04058 | 0.00095 | 10.0 s |
| 9:57.4 | Data point 4 | 1.50000 mL | 0.05115 mL | 0.03589 mL | 0.03001 mL | 2.652 | -0.00793 | 0.70535 | 0.00047 | 10.0 s |
| 10:32.9 | Data point 5 | 1.50000 mL | 0.05115 mL | 0.03937 mL | 0.03001 mL | 2.857 | -0.00289 | 0.42030 | 0.00022 | 10.0 s |
| 11:08.3 | Data point 6 | 1.50000 mL | 0.05115 mL | 0.04165 mL | 0.03001 mL | 3.054 | 0.00006 | 0.00064 | 0.00011 | 10.0 s |
| 11:43.9 | Data point 7 | 1.50000 mL | 0.05115 mL | 0.04327 mL | 0.03001 mL | 3.257 | -0.00042 | 0.04360 | 0.00010 | 10.5 s |
| 12:19.7 | Data point 8 | 1.50000 mL | 0.05115 mL | 0.04454 mL | 0.03001 mL | 3.464 | -0.00064 | 0.06303 | 0.00013 | 10.0 s |
| 12:55.2 | Data point 9 | 1.50000 mL | 0.05115 mL | 0.04570 mL | 0.03001 mL | 3.661 | -0.00672 | 0.71058 | 0.00039 | 10.5 s |
| 13:31.1 | Data point 10 | 1.50000 mL | 0.05115 mL | 0.04687 mL | 0.03001 mL | 3.861 | -0.00883 | 0.75826 | 0.00050 | 10.0 s |
| 14:06.6 | Data point 11 | 1.50000 mL | 0.05115 mL | 0.04814 mL | 0.03001 mL | 4.103 | -0.01310 | 0.89589 | 0.00068 | 10.5 s |
| 14:47.7 | Data point 12 | 1.50000 mL | 0.05115 mL | 0.04899 mL | 0.03001 mL | 4.289 | -0.00937 | 0.84367 | 0.00050 | 10.5 s |
| 15:23.5 | Data point 13 | 1.50000 mL | 0.05115 mL | 0.04965 mL | 0.03001 mL | 4.508 | -0.01463 | 0.74199 | 0.00084 | 11.0 s |
| 15:59.9 | Data point 14 | 1.50000 mL | 0.05115 mL | 0.05021 mL | 0.03001 mL | 4.755 | -0.01555 | 0.81188 | 0.00085 | 13.0 s |
| 16:38.2 | Data point 15 | 1.50000 mL | 0.05115 mL | 0.05063 mL | 0.03001 mL | 5.109 | -0.01643 | 0.85150 | 0.00088 | 17.0 s |
| 17:20.6 | Data point 16 | 1.50000 mL | 0.05115 mL | 0.05087 mL | 0.03001 mL | 5.521 | -0.01966 | 0.95659 | 0.00099 | 21.0 s |
| 18:12.1 | Data point 17 | 1.50000 mL | 0.05115 mL | 0.05101 mL | 0.03001 mL | 5.806 | -0.01764 | 0.91085 | 0.00091 | 29.5 s |
| 19:12.2 | Data point 18 | 1.50000 mL | 0.05115 mL | 0.05111 mL | 0.03001 mL | 6.280 | -0.01792 | 0.93360 | 0.00092 | 45.5 s |
| 20:28.4 | Data point 19 | 1.50000 mL | 0.05115 mL | 0.05122 mL | 0.03001 mL | 7.844 | -0.08873 | 0.99672 | 0.00439 | Timed out at 59.5 s |
| 21:58.9 | Data point 20 | 1.50000 mL | 0.05115 mL | 0.05127 mL | 0.03001 mL | 8.410 | -0.04014 | 0.97727 | 0.00201 | Timed out at 59.5 s |
| 23:29.4 | Data point 21 | 1.50000 mL | 0.05115 mL | 0.05132 mL | 0.03001 mL | 8.719 | -0.02194 | 0.97381 | 0.00110 | Timed out at 59.5 s |
| 25:05.0 | Data point 22 | 1.50000 mL | 0.05115 mL | 0.05139 mL | 0.03001 mL | 8.993 | -0.01950 | 0.95962 | 0.00098 | 34.5 s |
| 26:05.0 | Data point 23 | 1.50000 mL | 0.05115 mL | 0.05141 mL | 0.03001 mL | 9.001 | -0.01921 | 0.96673 | 0.00097 | 42.0 s |
| 27:41.8 | Data point 24 | 1.50000 mL | 0.10503 mL | 0.05141 mL | 0.11002 mL | 1.955 | -0.00341 | 0.14587 | 0.00044 | 10.0 s |
| 28:28.0 | Data point 25 | 1.50000 mL | 0.10503 mL | 0.06816 mL | 0.11002 mL | 2.160 | 0.00273 | 0.60068 | 0.00017 | 10.5 s |
| 29:04.3 | Data point 26 | 1.50000 mL | 0.10503 mL | 0.07947 mL | 0.11002 mL | 2.371 | 0.00238 | 0.28453 | 0.00022 | 10.0 s |
| 29:39.9 | Data point 27 | 1.50000 mL | 0.10503 mL | 0.08652 mL | 0.11002 mL | 2.601 | 0.00077 | 0.08864 | 0.00013 | 10.5 s |
| 30:15.9 | Data point 28 | 1.50000 mL | 0.10503 mL | 0.09087 mL | 0.11002 mL | 2.792 | -0.01433 | 0.72766 | 0.00083 | 10.5 s |
| 30:51.8 | Data point 29 | 1.50000 mL | 0.10503 mL | 0.09393 mL | 0.11002 mL | 2.985 | 0.01516 | 0.73138 | 0.00088 | 10.0 s |
| 31:27.3 | Data point 30 | 1.50000 mL | 0.10503 mL | 0.09626 mL | 0.11002 mL | 3.161 | -0.01511 | 0.64124 | 0.00093 | 11.0 s |
| 32:14.1 | Data point 31 | 1.50000 mL | 0.10503 mL | 0.09828 mL | 0.11002 mL | 3.353 | -0.00936 | 0.31606 | 0.00082 | 10.0 s |
| 32:49.5 | Data point 32 | 1.50000 mL | 0.10503 mL | 0.10005 mL | 0.11002 mL | 3.657 | -0.00287 | 0.78697 | 0.00016 | 10.5 s |
| 33:30.6 | Data point 33 | 1.50000 mL | 0.10503 mL | 0.10096 mL | 0.11002 mL | 3.844 | -0.00635 | 0.80051 | 0.00035 | 10.0 s |
| 34:06.0 | Data point 34 | 1.50000 mL | 0.10503 mL | 0.10167 mL | 0.11002 mL | 4.038 | -0.00491 | 0.70908 | 0.00029 | 10.0 s |
| 34:41.4 | Data point 35 | 1.50000 mL | 0.10503 mL | 0.10228 mL | 0.11002 mL | 4.243 | -0.00496 | 0.59949 | 0.00032 | 10.0 s |
| 35:16.8 | Data point 36 | 1.50000 mL | 0.10503 mL | 0.10275 mL | 0.11002 mL | 4.459 | -0.00596 | 0.57925 | 0.00039 | 10.5 s |
| 35:52.7 | Data point 37 | 1.50000 mL | 0.10503 mL | 0.10310 mL | 0.11002 mL | 4.712 | 0.00092 | 0.00227 | 0.00095 | 10.5 s |

| | | | |
|---|---|---|---|
| Sample name: | **M04_octanol** | Experiment start time: | **3/24/2018 4:11:23 AM** |
| Assay name: | **pH-metric high logP** | Analyst: | **Dorothy Levorse** |
| Assay ID: | **18C-24004** | Instrument ID: | **T312060** |
| Filename: | **C:\Sirius_T3\Mehtap\20180323_exp33_logP_T3-2\18C-24004_M04_octanol_pH-metric high logP.t3r** | | |

#### Events (continued)

| Time | Event | Water | Acid | Base | Octanol | pH | dpH/dt | pH R-squared | pH SD | dpH/dt time |
|---|---|---|---|---|---|---|---|---|---|---|
| 36:28.6 | Data point 38 | 1.50000 mL | 0.10503 mL | 0.10332 mL | 0.11002 mL | 4.945 | -0.00605 | 0.33921 | 0.00051 | 11.0 s |
| 37:05.0 | Data point 39 | 1.50000 mL | 0.10503 mL | 0.10346 mL | 0.11002 mL | 5.195 | -0.01462 | 0.70458 | 0.00086 | 12.0 s |
| 37:47.6 | Data point 40 | 1.50000 mL | 0.10503 mL | 0.10358 mL | 0.11002 mL | 5.448 | -0.00465 | 0.09101 | 0.00076 | 13.0 s |
| 38:31.1 | Data point 41 | 1.50000 mL | 0.10503 mL | 0.10365 mL | 0.11002 mL | 5.756 | -0.01450 | 0.74805 | 0.00083 | 15.5 s |
| 39:17.2 | Data point 42 | 1.50000 mL | 0.10503 mL | 0.10372 mL | 0.11002 mL | 6.188 | -0.01751 | 0.88071 | 0.00092 | 38.0 s |
| 40:25.8 | Data point 43 | 1.50000 mL | 0.10503 mL | 0.10376 mL | 0.11002 mL | 6.624 | -0.01828 | 0.88671 | 0.00096 | 59.5 s |
| 41:50.7 | Data point 44 | 1.50000 mL | 0.10503 mL | 0.10381 mL | 0.11002 mL | 6.931 | -0.02021 | 0.98765 | 0.00100 | Timed out at 59.5 s |
| 43:21.2 | Data point 45 | 1.50000 mL | 0.10503 mL | 0.10386 mL | 0.11002 mL | 7.358 | -0.05882 | 0.99495 | 0.00291 | Timed out at 59.5 s |
| 44:51.6 | Data point 46 | 1.50000 mL | 0.10503 mL | 0.10390 mL | 0.11002 mL | 7.860 | -0.06638 | 0.98979 | 0.00330 | Timed out at 59.5 s |
| 46:27.2 | Data point 47 | 1.50000 mL | 0.10503 mL | 0.10398 mL | 0.11002 mL | 8.405 | -0.03574 | 0.99078 | 0.00177 | Timed out at 59.5 s |
| 47:57.6 | Data point 48 | 1.50000 mL | 0.10503 mL | 0.10402 mL | 0.11002 mL | 8.751 | -0.01947 | 0.94555 | 0.00099 | 58.0 s |
| 49:26.2 | Data point 49 | 1.50000 mL | 0.10503 mL | 0.10412 mL | 0.11002 mL | 8.970 | -0.01588 | 0.97546 | 0.00079 | 42.0 s |
| 50:38.7 | Data point 50 | 1.50000 mL | 0.10503 mL | 0.10421 mL | 0.11002 mL | 9.167 | -0.01890 | 0.92626 | 0.00097 | 28.5 s |
| 52:25.2 | Data point 51 | 1.50000 mL | 0.16148 mL | 0.10421 mL | 0.51002 mL | 1.964 | -0.00413 | 0.66600 | 0.00025 | 10.5 s |
| 53:11.9 | Data point 52 | 1.50000 mL | 0.16148 mL | 0.12244 mL | 0.51002 mL | 2.173 | 0.00880 | 0.67097 | 0.00053 | 10.0 s |
| 53:47.6 | Data point 53 | 1.50000 mL | 0.16148 mL | 0.13434 mL | 0.51002 mL | 2.382 | 0.01052 | 0.53800 | 0.00071 | 10.0 s |
| 54:23.2 | Data point 54 | 1.50000 mL | 0.16148 mL | 0.14198 mL | 0.51002 mL | 2.597 | -0.00151 | 0.05969 | 0.00031 | 10.0 s |
| 54:58.7 | Data point 55 | 1.50000 mL | 0.16148 mL | 0.14713 mL | 0.51002 mL | 2.812 | -0.00306 | 0.69301 | 0.00018 | 10.5 s |
| 55:34.7 | Data point 56 | 1.50000 mL | 0.16148 mL | 0.15078 mL | 0.51002 mL | 3.013 | -0.00746 | 0.20419 | 0.00082 | 10.0 s |
| 56:10.2 | Data point 57 | 1.50000 mL | 0.16148 mL | 0.15360 mL | 0.51002 mL | 3.197 | 0.00123 | 0.02683 | 0.00037 | 10.5 s |
| 56:46.2 | Data point 58 | 1.50000 mL | 0.16148 mL | 0.15590 mL | 0.51002 mL | 3.420 | -0.00821 | 0.19314 | 0.00092 | 10.0 s |
| 57:21.7 | Data point 59 | 1.50000 mL | 0.16148 mL | 0.15769 mL | 0.51002 mL | 3.707 | 0.00459 | 0.14616 | 0.00059 | 10.0 s |
| 58:07.4 | Data point 60 | 1.50000 mL | 0.16148 mL | 0.15858 mL | 0.51002 mL | 3.903 | -0.00128 | 0.00501 | 0.00089 | 10.0 s |
| 58:53.1 | Data point 61 | 1.50000 mL | 0.16148 mL | 0.15920 mL | 0.51002 mL | 4.102 | -0.00123 | 0.00430 | 0.00093 | 10.0 s |
| 59:38.9 | Data point 62 | 1.50000 mL | 0.16148 mL | 0.15962 mL | 0.51002 mL | 4.303 | -0.01215 | 0.38768 | 0.00096 | 13.5 s |
| 1:00:28.1 | Data point 63 | 1.50000 mL | 0.16148 mL | 0.15993 mL | 0.51002 mL | 4.507 | -0.00113 | 0.00611 | 0.00072 | 10.0 s |
| 1:01:18.9 | Data point 64 | 1.50000 mL | 0.16148 mL | 0.16016 mL | 0.51002 mL | 4.723 | 0.00118 | 0.00902 | 0.00061 | 10.5 s |
| 1:01:54.8 | Data point 65 | 1.50000 mL | 0.16148 mL | 0.16033 mL | 0.51002 mL | 5.036 | -0.01870 | 0.92089 | 0.00096 | 11.5 s |
| 1:02:36.9 | Data point 66 | 1.50000 mL | 0.16148 mL | 0.16047 mL | 0.51002 mL | 5.464 | -0.01865 | 0.92187 | 0.00096 | 25.0 s |
| 1:03:32.5 | Data point 67 | 1.50000 mL | 0.16148 mL | 0.16056 mL | 0.51002 mL | 5.825 | -0.01533 | 0.67266 | 0.00092 | 18.5 s |
| 1:04:21.6 | Data point 68 | 1.50000 mL | 0.16148 mL | 0.16063 mL | 0.51002 mL | 6.318 | -0.01905 | 0.96692 | 0.00096 | 51.5 s |
| 1:05:43.7 | Data point 69 | 1.50000 mL | 0.16148 mL | 0.16070 mL | 0.51002 mL | 6.858 | -0.05249 | 0.96745 | 0.00264 | Timed out at 59.5 s |
| 1:07:14.2 | Data point 70 | 1.50000 mL | 0.16148 mL | 0.16075 mL | 0.51002 mL | 7.233 | -0.05610 | 0.97962 | 0.00280 | Timed out at 59.5 s |
| 1:08:44.7 | Data point 71 | 1.50000 mL | 0.16148 mL | 0.16079 mL | 0.51002 mL | 7.571 | -0.05662 | 0.99069 | 0.00281 | Timed out at 59.5 s |
| 1:10:15.2 | Data point 72 | 1.50000 mL | 0.16148 mL | 0.16084 mL | 0.51002 mL | 7.982 | -0.05547 | 0.99203 | 0.00275 | Timed out at 59.5 s |
| 1:11:45.7 | Data point 73 | 1.50000 mL | 0.16148 mL | 0.16089 mL | 0.51002 mL | 8.235 | -0.05197 | 0.96437 | 0.00261 | Timed out at 59.5 s |
| 1:13:16.2 | Data point 74 | 1.50000 mL | 0.16148 mL | 0.16094 mL | 0.51002 mL | 8.444 | -0.02913 | 0.98679 | 0.00145 | Timed out at 59.5 s |
| 1:14:46.7 | Data point 75 | 1.50000 mL | 0.16148 mL | 0.16098 mL | 0.51002 mL | 8.602 | -0.01977 | 0.98733 | 0.00098 | 47.0 s |
| 1:16:14.7 | Data point 76 | 1.50000 mL | 0.16148 mL | 0.16108 mL | 0.51002 mL | 8.892 | -0.01379 | 0.56644 | 0.00091 | 35.5 s |
| 1:17:20.7 | Data point 77 | 1.50000 mL | 0.16148 mL | 0.16112 mL | 0.51002 mL | 8.988 | -0.01977 | 0.96525 | 0.00099 | 28.5 s |
| 1:17:58.3 | Assay volumes | 1.50000 mL | 0.16148 mL | 0.16112 mL | 0.51002 mL | | | | | |

| | | | |
|---|---|---|---|
| Sample name: | **M04_octanol** | Experiment start time: | **3/24/2018 4:11:23 AM** |
| Assay name: | **pH-metric high logP** | Analyst: | **Dorothy Levorse** |
| Assay ID: | **18C-24004** | Instrument ID: | **T312060** |
| Filename: | **C:\Sirius_T3\Mehtap\20180323_exp33_logP_T3-2\18C-24004_M04_octanol_pH-metric high logP.t3r** | | |

| | | | |
|---|---|---|---|
| Sample name: | **M04_octanol** | Experiment start time: | **3/24/2018 4:11:23 AM** |
| Assay name: | **pH-metric high logP** | Analyst: | **Dorothy Levorse** |
| Assay ID: | **18C-24004** | Instrument ID: | **T312060** |
| Filename: | **C:\Sirius_T3\Mehtap\20180323_exp33_logP_T3-2\18C-24004_M04_octanol_pH-metric high logP.t3r** | | |

#### Assay Settings (continued)

| Setting | Value | Original Value | Date/Time changed | Imported from |
|---|---|---|---|---|
| ***Titration 3*** | | | | |
| Titrate from | Low to high pH | | | |
| Add additional water | 0.00 mL | | | |
| Additional partition solvent volume | 0.400 mL | | | |
| Additional partition solvent added | Automatic | | | |
| After pH adjust stir for | 30 seconds | | | |
| Stir to allow partitioning for | 15 seconds | | | |
| Stirrer speed for partitioning | 60% | | | |
| ***Data Point Stability*** | | | | |
| Stir during data point collection | No | | | |
| Delay before data point collection | 0 seconds | | | |
| Number of points to average | 20 points | | | |
| Time interval between points | 0.50 seconds | | | |
| Required maximum standard deviation | 0.00100 dpH/dt | | | |
| Stability timeout after | 60 seconds | | | |

#### Calibration Settings

| Setting | Value | Date/Time changed | Imported from |
|---|---|---|---|
| Four-Plus alpha | 0.119 | 3/24/2018 4:11:23 AM | C:\Sirius_T3\HCl18C23.t3r |
| Four-Plus S | 0.9972 | 3/24/2018 4:11:23 AM | C:\Sirius_T3\HCl18C23.t3r |
| Four-Plus jH | 0.9 | 3/24/2018 4:11:23 AM | C:\Sirius_T3\HCl18C23.t3r |
| Four-Plus jOH | -0.3 | 3/24/2018 4:11:23 AM | C:\Sirius_T3\HCl18C23.t3r |
| Base concentration factor | 1.003 | 3/24/2018 4:11:23 AM | C:\Sirius_T3\KOH18C23.t3r |
| Acid concentration factor | 0.997 | 3/24/2018 4:11:23 AM | C:\Sirius_T3\HCl18C23.t3r |

#### Instrument Settings

| Setting | Value | Batch Id | Install date |
|---|---|---|---|
| Instrument owner | Merck | | |
| Instrument ID | T312060 | | |
| Instrument type | T3 Simulator | | |
| Software version | 1.1.3.0 | | |
| Dispenser module | | T3DM1200361 | 3/31/2009 6:24:52 AM |
| Dispenser 0 | Water | | 3/31/2009 6:25:05 AM |
| Syringe volume | 2.5 mL | | |
| Firmware version | 1.2.1(r2) | | |
| Titrant | Water (0.15 M KCl) | 02-06-2018 | 3/16/2018 11:09:18 AM |
| Dispenser 2 | Acid | | 3/31/2009 6:25:11 AM |
| Syringe volume | 0.5 mL | | |
| Firmware version | 1.2.1(r2) | | |
| Titrant | Acid (0.5 M HCl) | 03-16-2018 | 3/16/2018 10:56:23 AM |
| Dispenser 1 | Base | | 3/31/2009 6:25:21 AM |
| Syringe volume | 0.5 mL | | |
| Firmware version | 1.2.1(r2) | | |
| Titrant | Base (0.5 M KOH) | 3/22/2018 | 3/23/2018 9:34:17 AM |
| Dispenser 5 | Cosolvent | | 3/31/2009 6:26:24 AM |
| Syringe volume | 2.5 mL | | |
| Firmware version | 1.2.1(r2) | | |
| Distribution valve 5 | Distribution Valve | | 3/31/2009 6:28:19 AM |
| Firmware version | 1.1.3 | | |
| Port A | Methanol (80%, 0.15 M KCl) | 02-08-2018 | 3/6/2018 10:28:59 AM |
| Port B | Cyclohexane | 11-01-17 | 2/27/2018 11:37:57 AM |
| Dispenser 3 | Buffer | | 8/3/2010 6:05:16 AM |
| Syringe volume | 0.5 mL | | |
| Firmware version | 1.2.1(r2) | | |
| Titrant | Dodecane | 2018/01/31 | 2/28/2018 11:18:04 AM |
| Dispenser 6 | Octanol | | 10/22/2010 11:52:43 AM |

| | | | |
|---|---|---|---|
| Sample name: | **M04_octanol** | Experiment start time: | **3/24/2018 4:11:23 AM** |
| Assay name: | **pH-metric high logP** | Analyst: | **Dorothy Levorse** |
| Assay ID: | **18C-24004** | Instrument ID: | **T312060** |
| Filename: | **C:\Sirius_T3\Mehtap\20180323_exp33_logP_T3-2\18C-24004_M04_octanol_pH-metric high logP.t3r** | | |

| | | | |
|---|---|---|---|
| Sample name: | **M04_octanol** | Experiment start time: | **3/24/2018 4:11:23 AM** |
| Assay name: | **pH-metric high logP** | Analyst: | **Dorothy Levorse** |
| Assay ID: | **18C-24004** | Instrument ID: | **T312060** |
| Filename: | **C:\Sirius_T3\Mehtap\20180323_exp33_logP_T3-2\18C-24004_M04_octanol_pH-metric high logP.t3r** | | |

#### Instrument Settings (continued)

| Setting | Value | Batch Id | Install date |
|---|---|---|---|
| Spectrometer calibration stir duration | 5 s | | |
| Spectrometer calibration stir speed | 30% | | |
| Spectrometer calibration wash pump volume | 20.0 mL | | |
| Spectrometer calibration wash stir duration | 5 s | | |
| Spectrometer calibration wash stir speed | 30% | | |
| Overhead dispense height | 10000 | | |

#### Refinement Settings

| Setting | Value | Default value |
|---|---|---|
| Turbidity detection method | None | None |
| Turbidity wavelength to assess | 500.0 nm | 500.0 nm |
| Turbidity maximum absorbance | 0.100 | 0.100 |
| Turbidity probe threshold | 50.00 | 50.00 |
