## Supplementary material for "Octanol-water partition coefficient measurements for the SAMPL6 Blind Prediction Challenge": SM07_18B-28011_M07_octanol_pH-metric high logP_report.pdf

| | | | |
|---|---|---|---|
| Sample name: | **M07_octanol** | Experiment start time: | **2/28/2018 4:26:20 PM** |
| Assay name: | **pH-metric high logP** | Analyst: | **Pion** |
| Assay ID: | **18B-28011** | Instrument ID: | **T312060** |
| Filename: | **C:\Sirius_T3\Mehtap\20180228_exp28_logP_T3-2\18B-28011_M07_octanol_pH-metric high logP.t3r** | | |

#### pH-metric Result

| | | |
|---|---|---|
| logP (XH +) | 0.44 | ±0.03 (n=50) |
| logP (neutral X) | 3.29 | ±0.01 (n=50) |

#### 18B-28011 Points 1 to 22

| | |
|---|---|
| M07_octanol concentration factor | 1.039 |
| Carbonate | 0.1202 mM |
| Acidity error | 0.21327 mM |

#### 18B-28011 Points 23 to 45

| | |
|---|---|
| M07_octanol concentration factor | 1.041 |
| Carbonate | 0.1876 mM |
| Acidity error | 0.19535 mM |

#### 18B-28011 Points 46 to 79

| | |
|---|---|
| M07_octanol concentration factor | 0.988 |
| Carbonate | 0.1406 mM |
| Acidity error | 2.18588 mM |

#### Warnings and errors

| | |
|---|---|
| Errors | None |
| Warnings | None |

#### Sample logD and percent species

| pH | M07_octanol logD | M07_octanol M07_octanolH | M07_octanol M07_octanol | M07_octanol M07_octanolH* | M07_octanol M07_octanol* | Comment |
|---|---|---|---|---|---|---|
| 1.000 | 0.45 | 26.38 % | 0.00 % | 73.19 % | 0.43 % | |
| 1.200 | 0.45 | 26.31 % | 0.00 % | 73.00 % | 0.69 % | Stomach pH |
| 2.000 | 0.47 | 25.38 % | 0.00 % | 70.44 % | 4.18 % | |
| 3.000 | 0.65 | 18.44 % | 0.02 % | 51.18 % | 30.36 % | |
| 4.000 | 1.28 | 4.94 % | 0.04 % | 13.70 % | 81.31 % | |
| 5.000 | 2.19 | 0.59 % | 0.05 % | 1.65 % | 97.71 % | |
| 6.000 | 2.95 | 0.06 % | 0.05 % | 0.17 % | 99.72 % | |
| 6.500 | 3.15 | 0.02 % | 0.05 % | 0.05 % | 99.88 % | |
| 7.000 | 3.24 | 0.01 % | 0.05 % | 0.02 % | 99.93 % | |
| 7.400 | 3.27 | 0.00 % | 0.05 % | 0.01 % | 99.94 % | Blood pH |
| 8.000 | 3.28 | 0.00 % | 0.05 % | 0.00 % | 99.95 % | |
| 9.000 | 3.29 | 0.00 % | 0.05 % | 0.00 % | 99.95 % | |
| 10.000 | 3.29 | 0.00 % | 0.05 % | 0.00 % | 99.95 % | |
| 11.000 | 3.29 | 0.00 % | 0.05 % | 0.00 % | 99.95 % | |
| 12.000 | 3.29 | 0.00 % | 0.05 % | 0.00 % | 99.95 % | |

| | | | |
|---|---|---|---|
| Sample name: | **M07_octanol** | Experiment start time: | **2/28/2018 4:26:20 PM** |
| Assay name: | **pH-metric high logP** | Analyst: | **Pion** |
| Assay ID: | **18B-28011** | Instrument ID: | **T312060** |
| Filename: | **C:\Sirius_T3\Mehtap\20180228_exp28_logP_T3-2\18B-28011_M07_octanol_pH-metric high logP.t3r** | | |

#### Graphs

Charge balance residuals

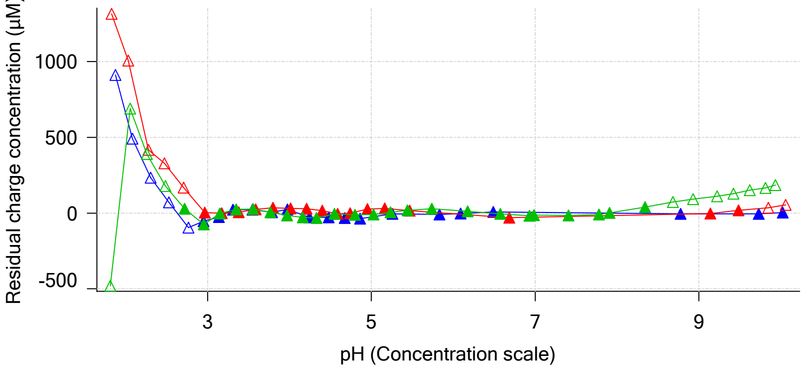


Ionic strength

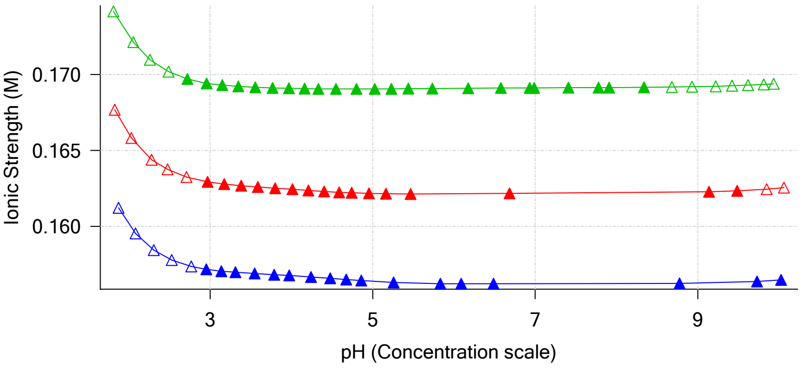


Buffer index profile

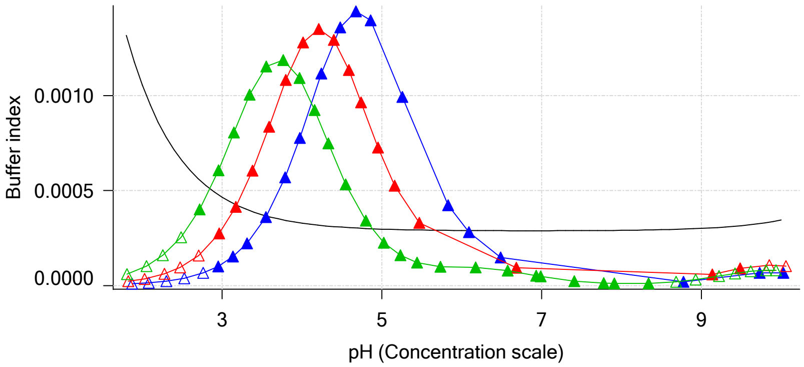


Sample M07_octanol Ionisation Graph

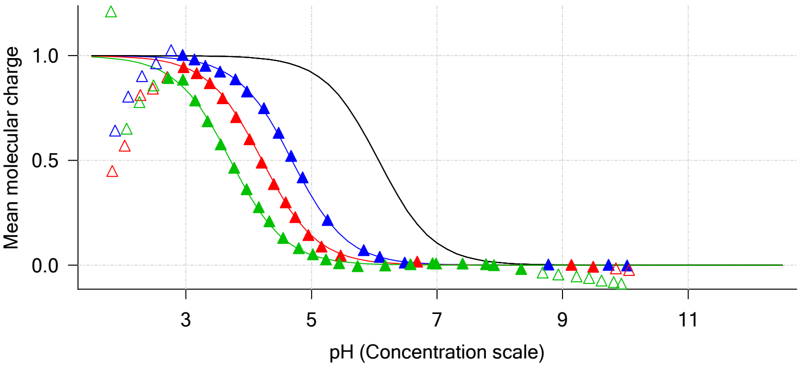


Distribution of species for sample M07_octanol

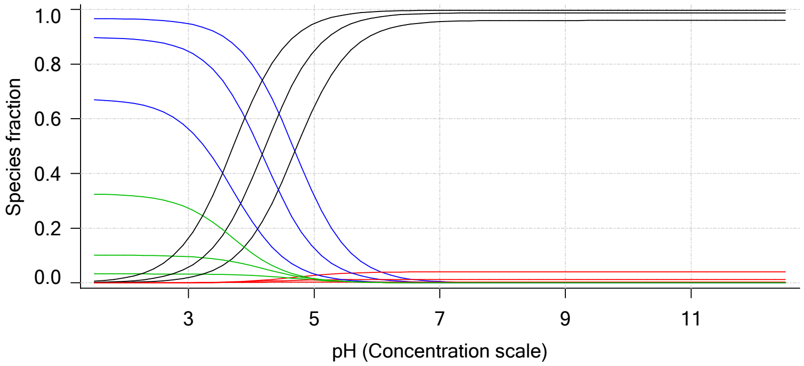


Lipophilicity Profile

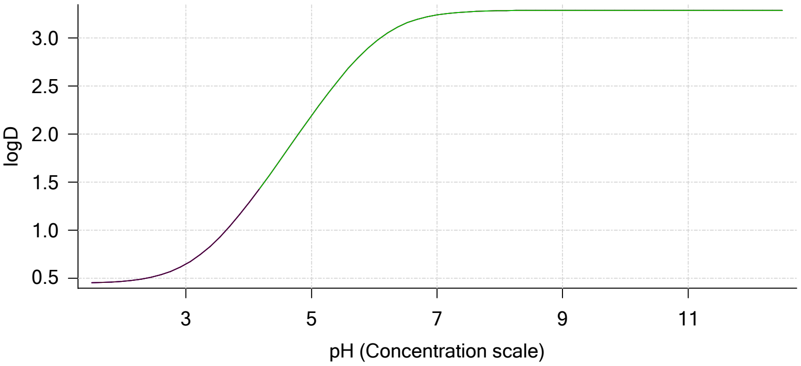


Log R Profile

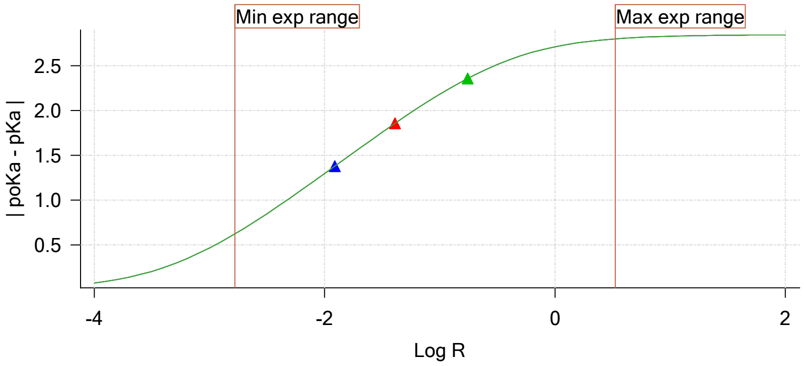


Sample Carbonate Ionisation Graph

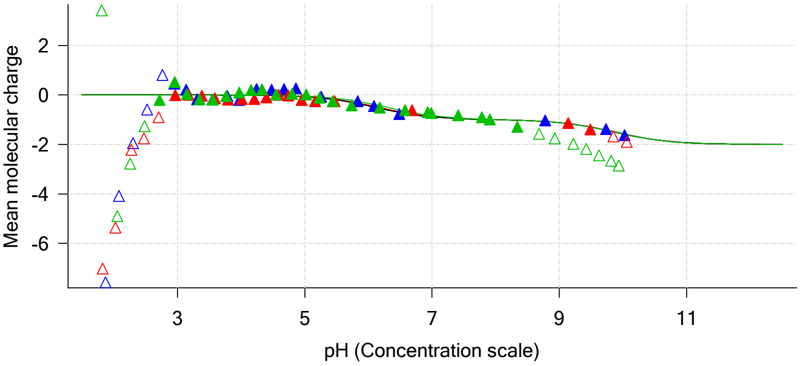

Sample name: **M07_octanol**
Assay name: **pH-metric high logP**
Assay ID: **18B-28011**
Filename: **C:\Sirius_T3\Mehtap\20180228_exp28_logP_T3-2\18B-28011_M07_octanol_pH-metric high logP.t3r**

Experiment start time: **2/28/2018 4:26:20 PM**
Analyst: **Pion**
Instrument ID: **T312060**

#### Graphs (continued)

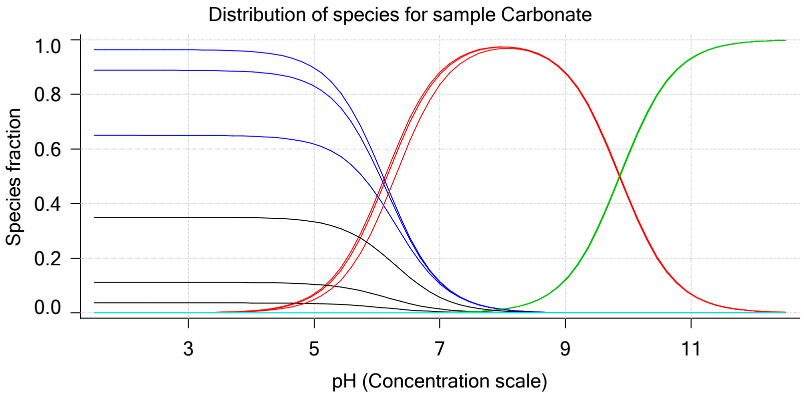

| | | | |
|---|---|---|---|
| Sample name: | **M07_octanol** | Experiment start time: | **2/28/2018 4:26:20 PM** |
| Assay name: | **pH-metric high logP** | Analyst: | **Pion** |
| Assay ID: | **18B-28011** | Instrument ID: | **T312060** |
| Filename: | **C:\Sirius_T3\Mehtap\20180228_exp28_logP_T3-2\18B-28011_M07_octanol_pH-metric high logP.t3r** | | |

# pH-metric high logP   Titration 1 of 3   18B-28011 Points 1 to 22

#### Overall results

| | |
|---|---|
| RMSD | 0.037 |
| Average ionic strength | 0.157 M |
| Average temperature | 25.0°C |
| Partition ratio | 0.0123 : 1 |
| Analyte concentration range | 2362.2 µM to 2434.4 µM |
| Total points considered | 17 of 22 |

#### Warnings and errors

| | |
|---|---|
| Errors | None |
| Warnings | None |

#### Four-Plus parameters

| | | | |
|---|---|---|---|
| Alpha | 0.130 | 2/28/2018 4:26:20 PM | C:\Sirius_T3\HCl18B27.t3r |
| S | 0.9970 | 2/28/2018 4:26:20 PM | C:\Sirius_T3\HCl18B27.t3r |
| jH | 0.8 | 2/28/2018 4:26:20 PM | C:\Sirius_T3\HCl18B27.t3r |
| jOH | -0.4 | 2/28/2018 4:26:20 PM | C:\Sirius_T3\HCl18B27.t3r |

#### Titrants

| | | | |
|---|---|---|---|
| 0.50 M HCl | 0.993513 | 2/28/2018 4:26:20 PM | C:\Sirius_T3\HCl18B27.t3r |
| 0.50 M KOH | 0.999845 | 2/28/2018 4:26:20 PM | C:\Sirius_T3\KOH18B27.t3r |

#### Sample

| | |
|---|---|
| M07_octanol concentration factor | 1.039 |
| Base pKa 1 | 6.07 |
| logP (XH +) | 0.50 |
| logP (neutral X) | 3.27 |

#### Sample graphs

Ionisation of sample M07_octanol

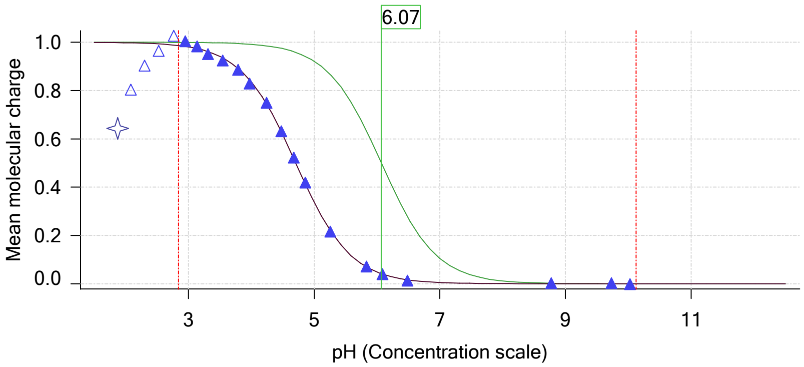


Distribution of species for sample M07_octanol

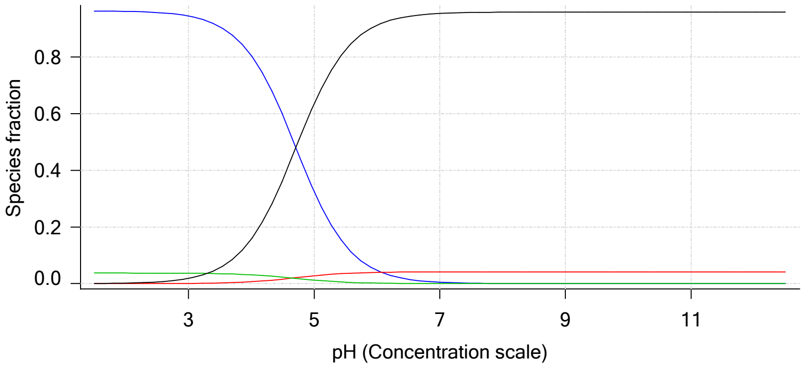

| | | | |
|---|---|---|---|
| Sample name: | **M07_octanol** | Experiment start time: | **2/28/2018 4:26:20 PM** |
| Assay name: | **pH-metric high logP** | Analyst: | **Pion** |
| Assay ID: | **18B-28011** | Instrument ID: | **T312060** |
| Filename: | **C:\Sirius_T3\Mehtap\20180228_exp28_logP_T3-2\18B-28011_M07_octanol_pH-metric high logP.t3r** | | |

#### Sample graphs (continued)

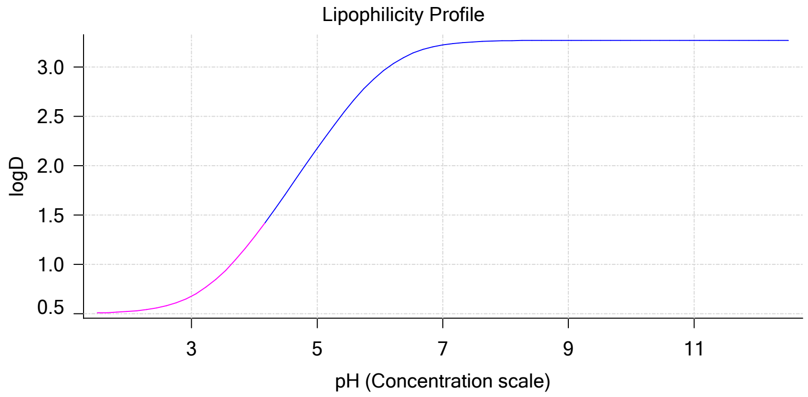


#### Sample logD and percent species

| pH | M07_octanol logD | M07_octanol M07_octanolH | M07_octanol M07_octanol | M07_octanol M07_octanolH* | M07_octanol M07_octanol* | Comment |
|---|---|---|---|---|---|---|
| 1.000 | 0.50 | 96.24 % | 0.00 % | 3.74 % | 0.02 % | |
| 1.200 | 0.50 | 96.23 % | 0.00 % | 3.74 % | 0.03 % | Stomach pH |
| 2.000 | 0.52 | 96.07 % | 0.01 % | 3.73 % | 0.19 % | |
| 3.000 | 0.68 | 94.41 % | 0.08 % | 3.67 % | 1.84 % | |
| 4.000 | 1.28 | 80.48 % | 0.69 % | 3.13 % | 15.71 % | |
| 5.000 | 2.17 | 32.51 % | 2.77 % | 1.26 % | 63.46 % | |
| 6.000 | 2.93 | 4.67 % | 3.98 % | 0.18 % | 91.17 % | |
| 6.500 | 3.13 | 1.53 % | 4.11 % | 0.06 % | 94.30 % | |
| 7.000 | 3.22 | 0.49 % | 4.16 % | 0.02 % | 95.33 % | |
| 7.400 | 3.25 | 0.20 % | 4.17 % | 0.01 % | 95.63 % | Blood pH |
| 8.000 | 3.27 | 0.05 % | 4.18 % | 0.00 % | 95.77 % | |
| 9.000 | 3.27 | 0.00 % | 4.18 % | 0.00 % | 95.82 % | |
| 10.000 | 3.27 | 0.00 % | 4.18 % | 0.00 % | 95.82 % | |
| 11.000 | 3.27 | 0.00 % | 4.18 % | 0.00 % | 95.82 % | |
| 12.000 | 3.27 | 0.00 % | 4.18 % | 0.00 % | 95.82 % | |

#### Carbonate and acidity

| | |
|---|---|
| Carbonate | 0.120 mM |
| Acidity error | 0.213 mM |

#### Other graphs

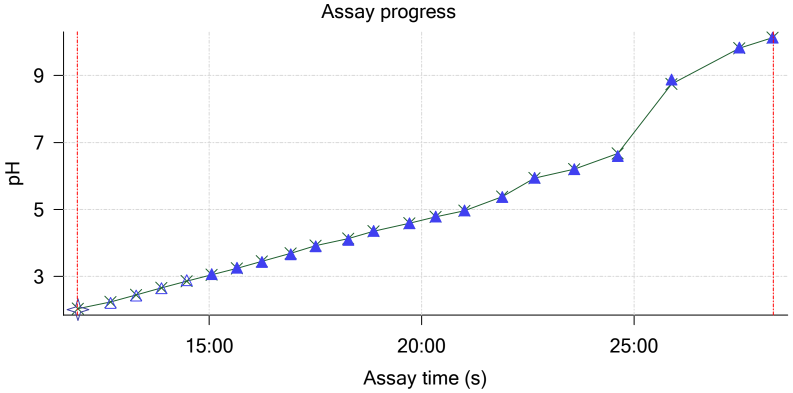


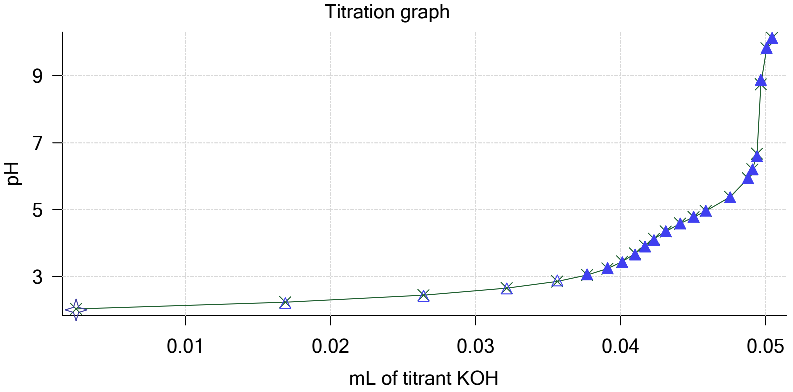

| | | | |
|---|---|---|---|
| Sample name: | **M07_octanol** | Experiment start time: | **2/28/2018 4:26:20 PM** |
| Assay name: | **pH-metric high logP** | Analyst: | **Pion** |
| Assay ID: | **18B-28011** | Instrument ID: | **T312060** |
| Filename: | **C:\Sirius_T3\Mehtap\20180228_exp28_logP_T3-2\18B-28011_M07_octanol_pH-metric high logP.t3r** | | |

#### Other graphs (continued)

Charge balance residuals

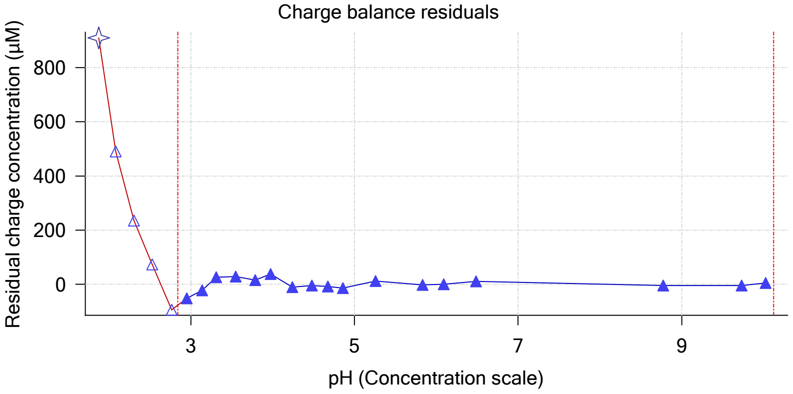


Buffer index profile

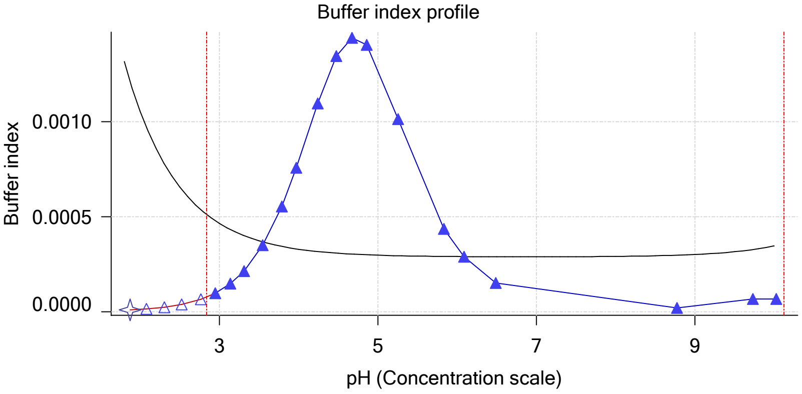


Ionic strength

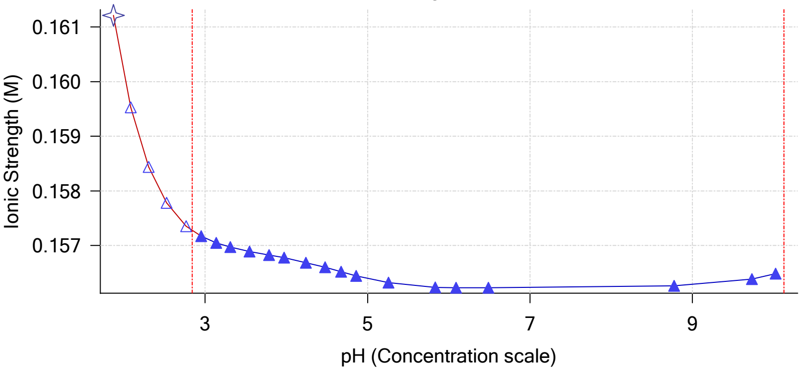


Log R Profile

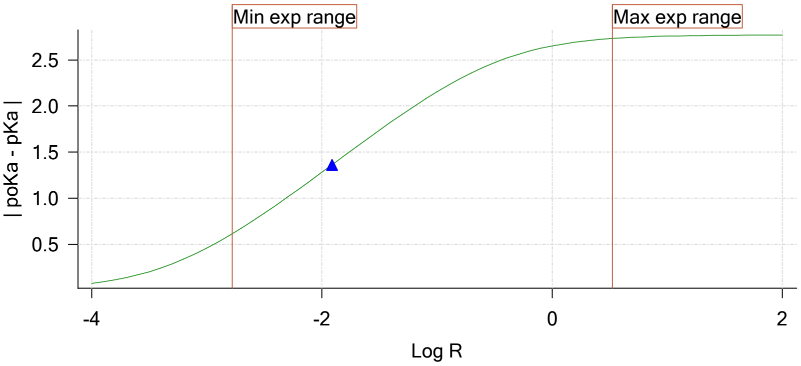


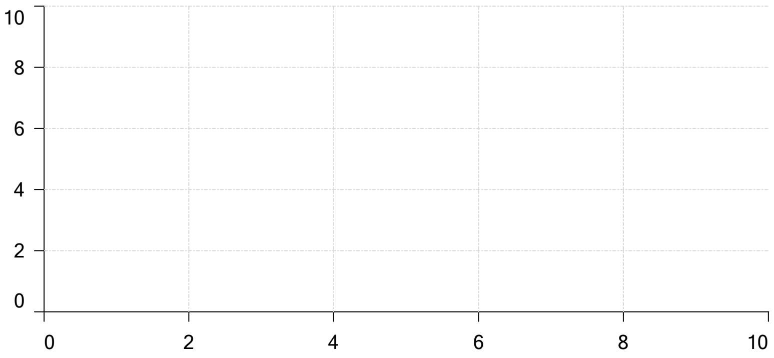


Ionisation of sample Carbonate

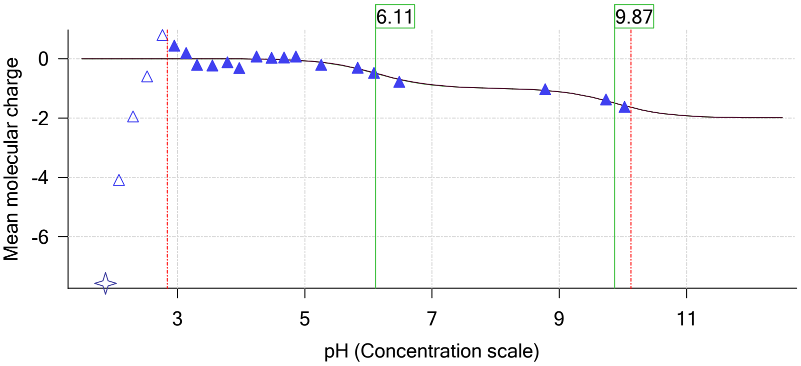

| | | | |
|---|---|---|---|
| Sample name: | **M07_octanol** | Experiment start time: | **2/28/2018 4:26:20 PM** |
| Assay name: | **pH-metric high logP** | Analyst: | **Pion** |
| Assay ID: | **18B-28011** | Instrument ID: | **T312060** |
| Filename: | **C:\Sirius_T3\Mehtap\20180228_exp28_logP_T3-2\18B-28011_M07_octanol_pH-metric high logP.t3r** | | |

### pH-metric high logP Titration 2 of 3 18B-28011 Points 23 to 45

#### Overall results

| | |
|---|---|
| RMSD | 0.084 |
| Average ionic strength | 0.162 M |
| Average temperature | 25.0°C |
| Partition ratio | 0.0408 : 1 |
| Analyte concentration range | 2150.3 µM to 2218.8 µM |
| Total points considered | 16 of 23 |

#### Warnings and errors

| | |
|---|---|
| Errors | None |
| Warnings | None |

#### Four-Plus parameters

| | | | |
|---|---|---|---|
| Alpha | 0.130 | 2/28/2018 4:26:20 PM | C:\Sirius_T3\HCl18B27.t3r |
| S | 0.9970 | 2/28/2018 4:26:20 PM | C:\Sirius_T3\HCl18B27.t3r |
| jH | 0.8 | 2/28/2018 4:26:20 PM | C:\Sirius_T3\HCl18B27.t3r |
| jOH | -0.4 | 2/28/2018 4:26:20 PM | C:\Sirius_T3\HCl18B27.t3r |

#### Titrants

| | | | |
|---|---|---|---|
| 0.50 M HCl | 0.993513 | 2/28/2018 4:26:20 PM | C:\Sirius_T3\HCl18B27.t3r |
| 0.50 M KOH | 0.999845 | 2/28/2018 4:26:20 PM | C:\Sirius_T3\KOH18B27.t3r |

#### Sample

| | |
|---|---|
| M07_octanol concentration factor | 1.041 |
| Base pKa 1 | 6.07 |
| logP (XH +) | 0.50 |
| logP (neutral X) | 3.31 |

#### Sample graphs

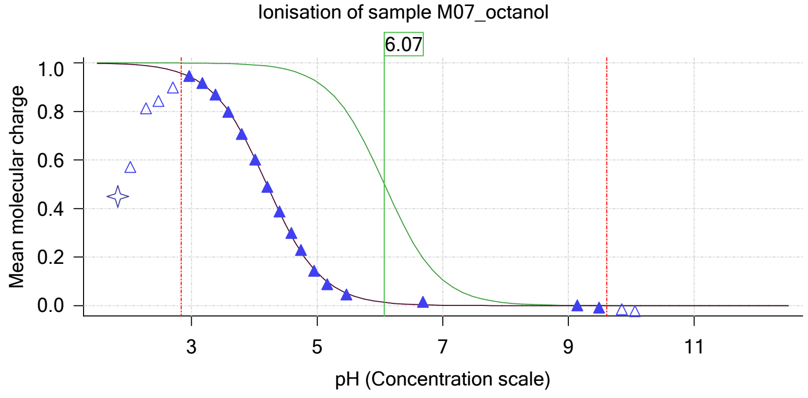


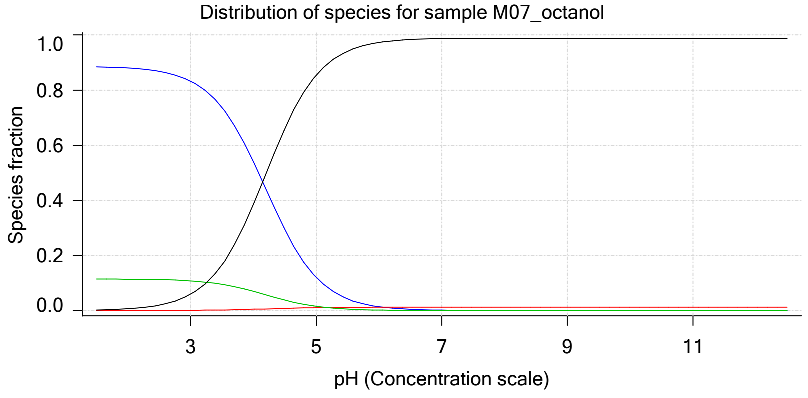

| | | | |
|---|---|---|---|
| Sample name: | **M07_octanol** | Experiment start time: | **2/28/2018 4:26:20 PM** |
| Assay name: | **pH-metric high logP** | Analyst: | **Pion** |
| Assay ID: | **18B-28011** | Instrument ID: | **T312060** |
| Filename: | **C:\Sirius_T3\Mehtap\20180228_exp28_logP_T3-2\18B-28011_M07_octanol_pH-metric high logP.t3r** | | |

#### Sample graphs (continued)

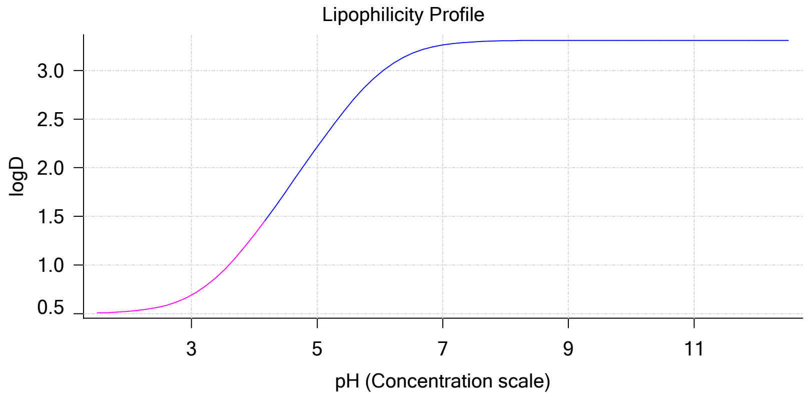


#### Sample logD and percent species

| pH | M07_octanol logD | M07_octanol M07_octanolH | M07_octanol M07_octanol | M07_octanol M07_octanolH* | M07_octanol M07_octanol* | Comment |
|---|---|---|---|---|---|---|
| 1.000 | 0.50 | 88.51 % | 0.00 % | 11.43 % | 0.06 % | |
| 1.200 | 0.50 | 88.48 % | 0.00 % | 11.42 % | 0.10 % | Stomach pH |
| 2.000 | 0.52 | 88.01 % | 0.01 % | 11.36 % | 0.63 % | |
| 3.000 | 0.69 | 83.26 % | 0.07 % | 10.75 % | 5.92 % | |
| 4.000 | 1.31 | 54.08 % | 0.46 % | 6.98 % | 38.48 % | |
| 5.000 | 2.21 | 12.01 % | 1.02 % | 1.55 % | 85.42 % | |
| 6.000 | 2.97 | 1.37 % | 1.16 % | 0.18 % | 97.29 % | |
| 6.500 | 3.17 | 0.44 % | 1.18 % | 0.06 % | 98.33 % | |
| 7.000 | 3.26 | 0.14 % | 1.18 % | 0.02 % | 98.66 % | |
| 7.400 | 3.29 | 0.06 % | 1.18 % | 0.01 % | 98.76 % | Blood pH |
| 8.000 | 3.31 | 0.01 % | 1.18 % | 0.00 % | 98.80 % | |
| 9.000 | 3.31 | 0.00 % | 1.18 % | 0.00 % | 98.82 % | |
| 10.000 | 3.31 | 0.00 % | 1.18 % | 0.00 % | 98.82 % | |
| 11.000 | 3.31 | 0.00 % | 1.18 % | 0.00 % | 98.82 % | |
| 12.000 | 3.31 | 0.00 % | 1.18 % | 0.00 % | 98.82 % | |

#### Carbonate and acidity

| | |
|---|---|
| Carbonate | 0.188 mM |
| Acidity error | 0.195 mM |

#### Other graphs

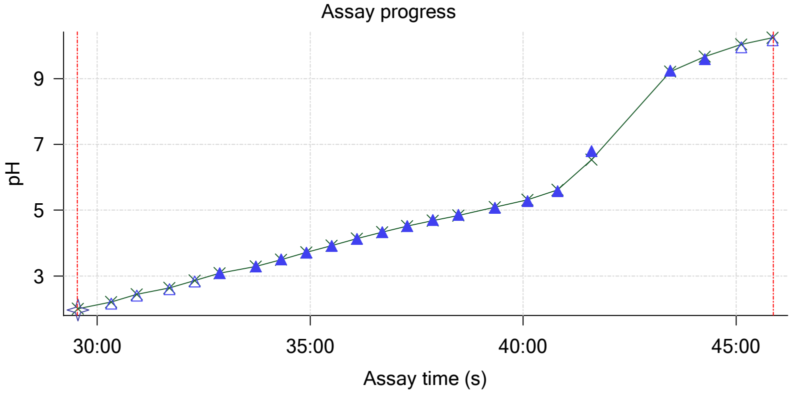


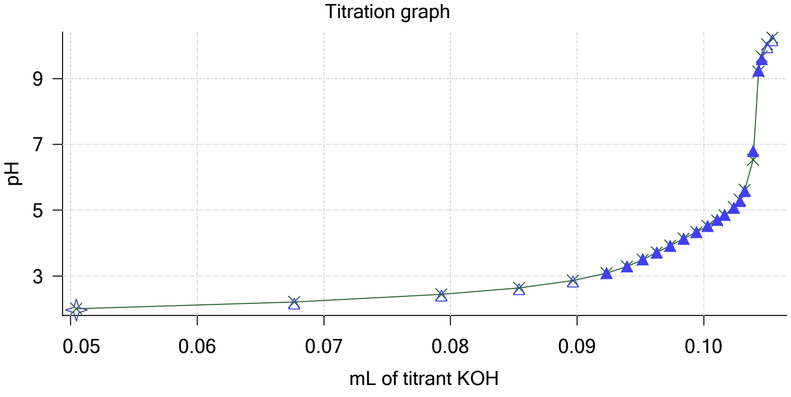

| | | | |
|---|---|---|---|
| Sample name: | **M07_octanol** | Experiment start time: | **2/28/2018 4:26:20 PM** |
| Assay name: | **pH-metric high logP** | Analyst: | **Pion** |
| Assay ID: | **18B-28011** | Instrument ID: | **T312060** |
| Filename: | **C:\Sirius_T3\Mehtap\20180228_exp28_logP_T3-2\18B-28011_M07_octanol_pH-metric high logP.t3r** | | |

#### Other graphs (continued)

Charge balance residuals

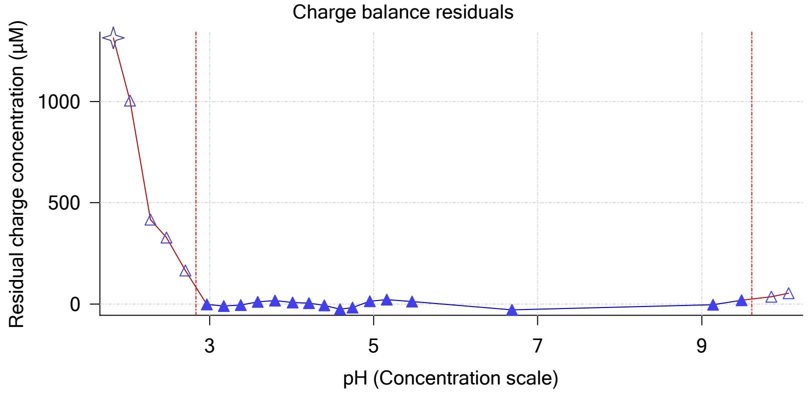


Buffer index profile

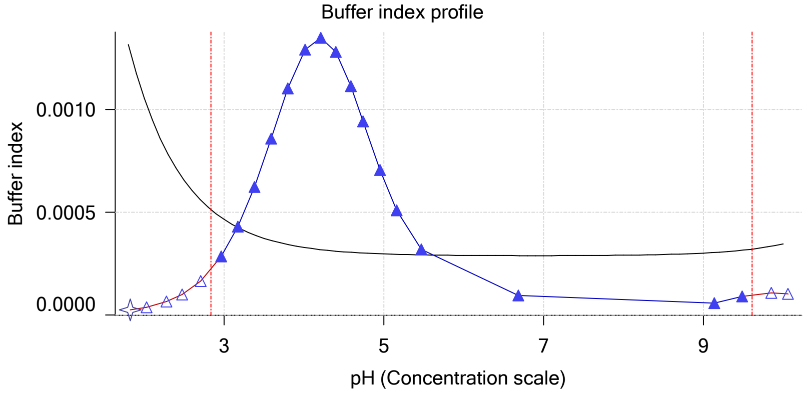


Ionic strength

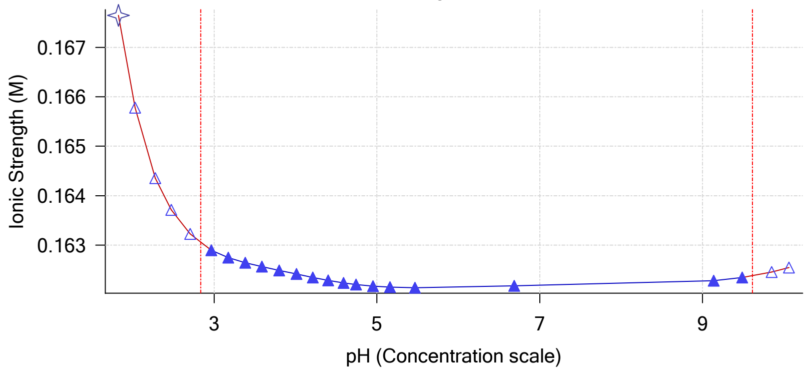


Log R Profile

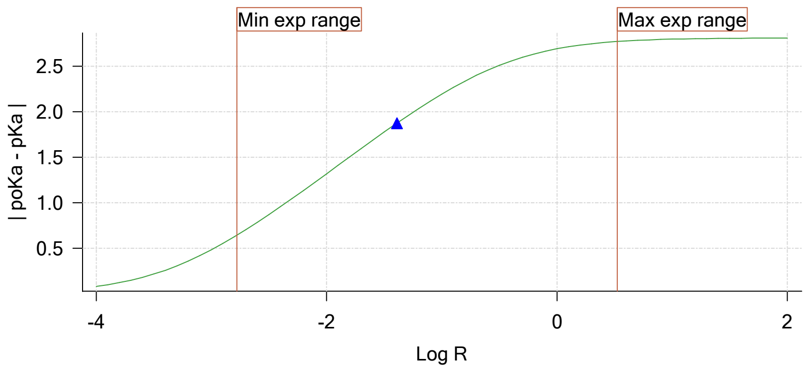


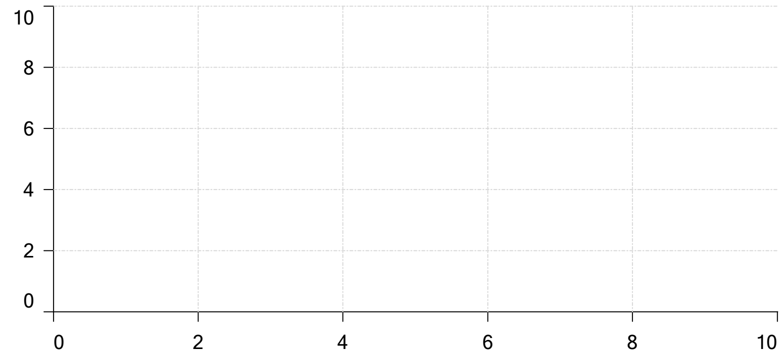


Ionisation of sample Carbonate

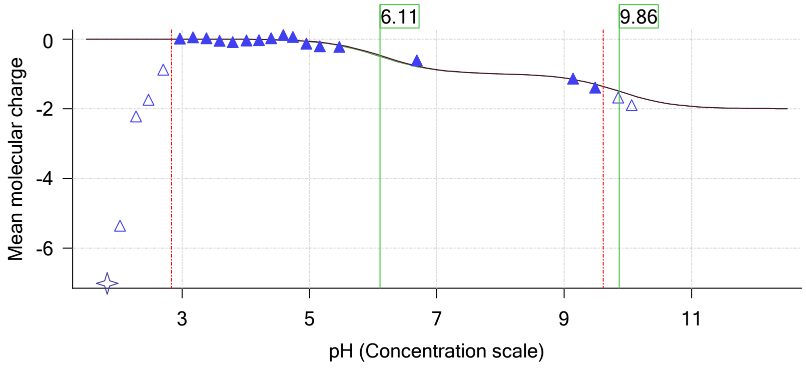

| | |
|---|---|
| Sample name: | **M07_octanol** |
| Assay name: | **pH-metric high logP** |
| Assay ID: | **18B-28011** |
| Filename: | **C:\Sirius_T3\Mehtap\20180228_exp28_logP_T3-2\18B-28011_M07_octanol_pH-metric high logP.t3r** |
| Experiment start time: | **2/28/2018 4:26:20 PM** |
| Analyst: | **Pion** |
| Instrument ID: | **T312060** |

# pH-metric high logP   Titration 3 of 3   18B-28011 Points 46 to 79

#### Overall results

| | |
|---|---|
| RMSD | 0.443 |
| Average ionic strength | 0.169 M |
| Average temperature | 25.0°C |
| Partition ratio | 0.1746 : 1 |
| Analyte concentration range | 1775.6 μM to 1831.9 μM |
| Total points considered | 23 of 34 |

#### Warnings and errors

| | |
|---|---|
| Errors | None |
| Warnings | Excessive acidity error present |

#### Four-Plus parameters

| | | | |
|---|---|---|---|
| Alpha | 0.130 | 2/28/2018 4:26:20 PM | C:\Sirius_T3\HCl18B27.t3r |
| S | 0.9970 | 2/28/2018 4:26:20 PM | C:\Sirius_T3\HCl18B27.t3r |
| jH | 0.8 | 2/28/2018 4:26:20 PM | C:\Sirius_T3\HCl18B27.t3r |
| jOH | -0.4 | 2/28/2018 4:26:20 PM | C:\Sirius_T3\HCl18B27.t3r |

#### Titrants

| | | | |
|---|---|---|---|
| 0.50 M HCl | 0.993513 | 2/28/2018 4:26:20 PM | C:\Sirius_T3\HCl18B27.t3r |
| 0.50 M KOH | 0.999845 | 2/28/2018 4:26:20 PM | C:\Sirius_T3\KOH18B27.t3r |

#### Sample

| | |
|---|---|
| M07_octanol concentration factor | 0.988 |
| Base pKa 1 | 6.07 |
| logP (XH +) | 0.50 |
| logP (neutral X) | 3.30 |

#### Sample graphs

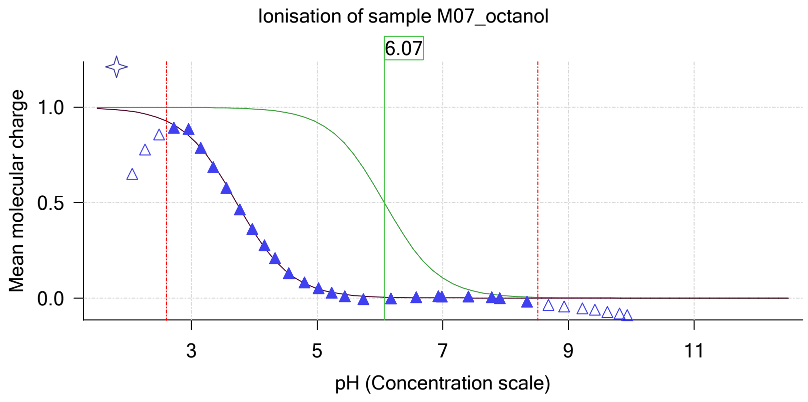


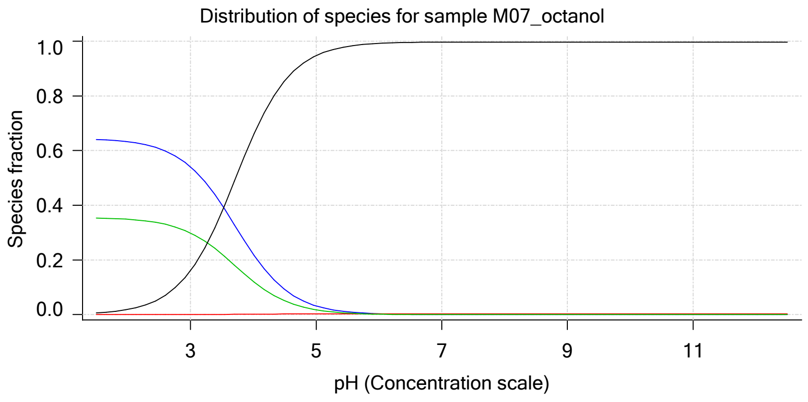

| | | | |
|---|---|---|---|
| Sample name: | **M07_octanol** | Experiment start time: | **2/28/2018 4:26:20 PM** |
| Assay name: | **pH-metric high logP** | Analyst: | **Pion** |
| Assay ID: | **18B-28011** | Instrument ID: | **T312060** |
| Filename: | **C:\Sirius_T3\Mehtap\20180228_exp28_logP_T3-2\18B-28011_M07_octanol_pH-metric high logP.t3r** | | |

#### Sample graphs (continued)

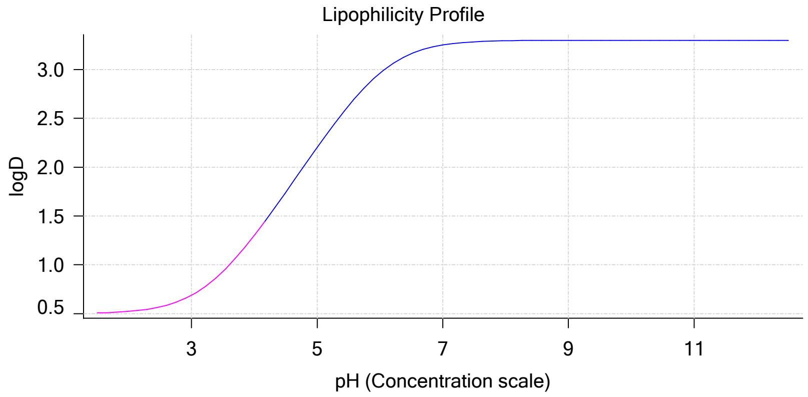


#### Sample logD and percent species

| pH | M07_octanol logD | M07_octanol M07_octanolH | M07_octanol M07_octanol | M07_octanol M07_octanolH* | M07_octanol M07_octanol* | Comment |
|---|---|---|---|---|---|---|
| 1.000 | 0.50 | 64.31 % | 0.00 % | 35.50 % | 0.19 % | |
| 1.200 | 0.50 | 64.24 % | 0.00 % | 35.46 % | 0.30 % | Stomach pH |
| 2.000 | 0.52 | 63.22 % | 0.01 % | 34.90 % | 1.88 % | |
| 3.000 | 0.69 | 54.07 % | 0.05 % | 29.85 % | 16.04 % | |
| 4.000 | 1.30 | 22.09 % | 0.19 % | 12.19 % | 65.53 % | |
| 5.000 | 2.20 | 3.20 % | 0.27 % | 1.76 % | 94.77 % | |
| 6.000 | 2.96 | 0.33 % | 0.28 % | 0.18 % | 99.20 % | |
| 6.500 | 3.16 | 0.11 % | 0.29 % | 0.06 % | 99.55 % | |
| 7.000 | 3.25 | 0.03 % | 0.29 % | 0.02 % | 99.66 % | |
| 7.400 | 3.28 | 0.01 % | 0.29 % | 0.01 % | 99.69 % | Blood pH |
| 8.000 | 3.30 | 0.00 % | 0.29 % | 0.00 % | 99.71 % | |
| 9.000 | 3.30 | 0.00 % | 0.29 % | 0.00 % | 99.71 % | |
| 10.000 | 3.30 | 0.00 % | 0.29 % | 0.00 % | 99.71 % | |
| 11.000 | 3.30 | 0.00 % | 0.29 % | 0.00 % | 99.71 % | |
| 12.000 | 3.30 | 0.00 % | 0.29 % | 0.00 % | 99.71 % | |

#### Carbonate and acidity

| | |
|---|---|
| Carbonate | 0.141 mM |
| Acidity error | 2.186 mM |

#### Other graphs

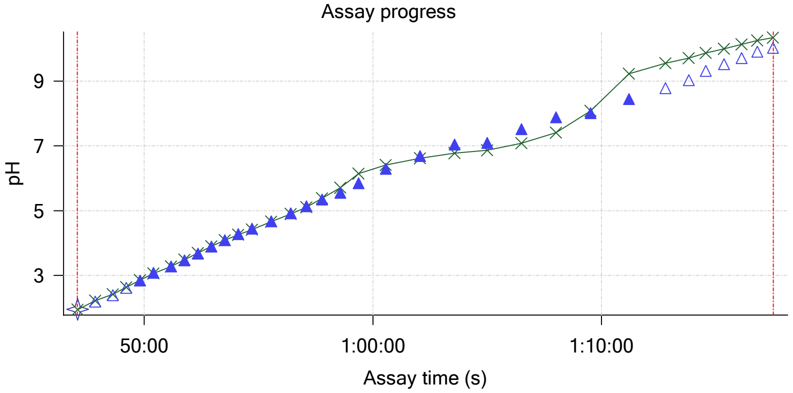


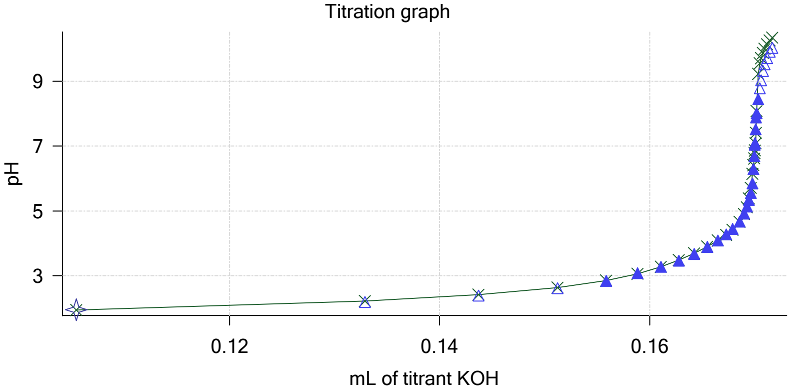

| | | | |
|---|---|---|---|
| Sample name: | **M07_octanol** | Experiment start time: | **2/28/2018 4:26:20 PM** |
| Assay name: | **pH-metric high logP** | Analyst: | **Pion** |
| Assay ID: | **18B-28011** | Instrument ID: | **T312060** |
| Filename: | **C:\Sirius_T3\Mehtap\20180228_exp28_logP_T3-2\18B-28011_M07_octanol_pH-metric high logP.t3r** | | |

#### Other graphs (continued)

Charge balance residuals

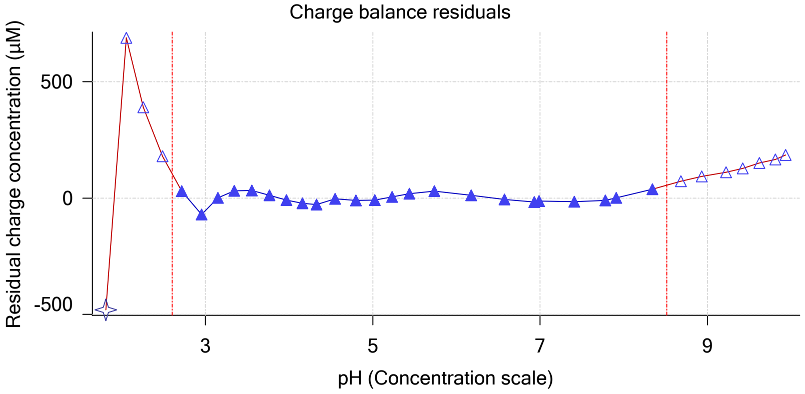


Buffer index profile

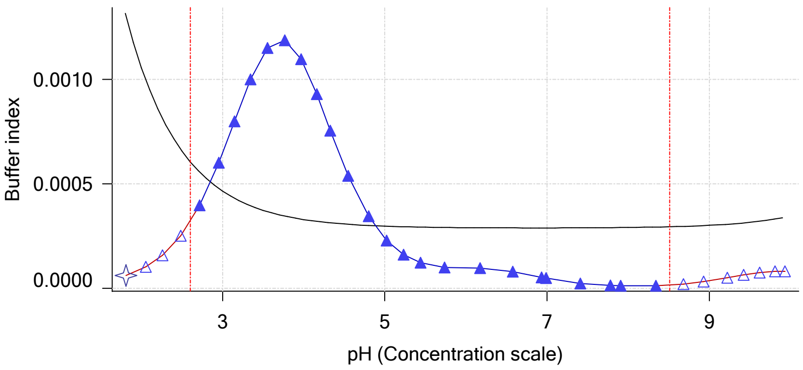


Ionic strength

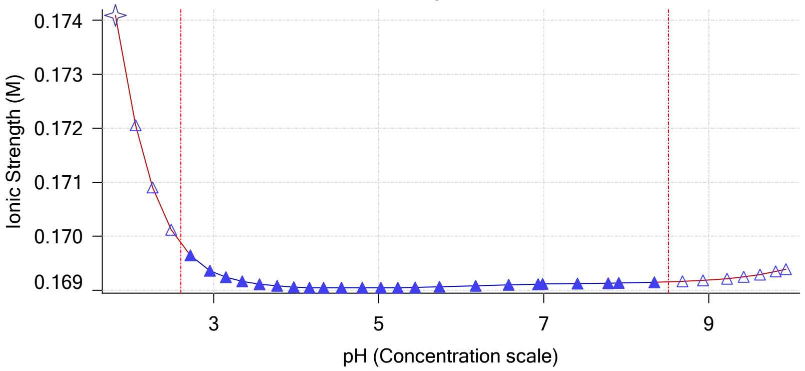


Log R Profile

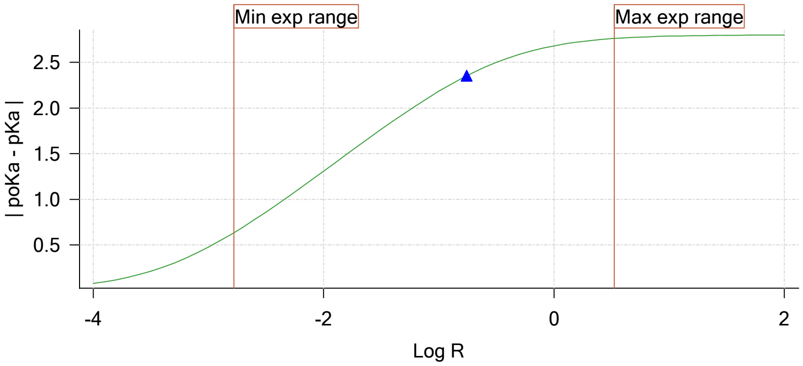


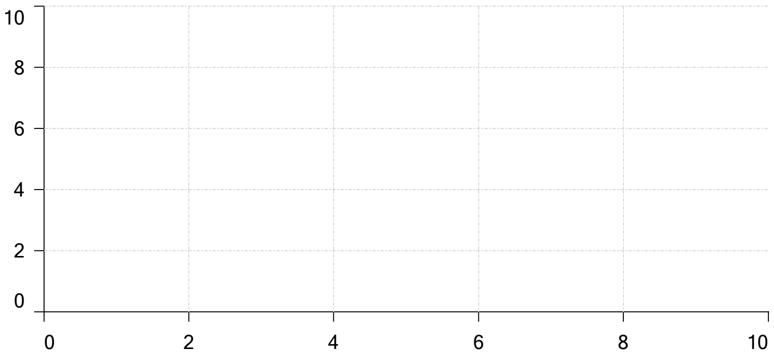


Ionisation of sample Carbonate

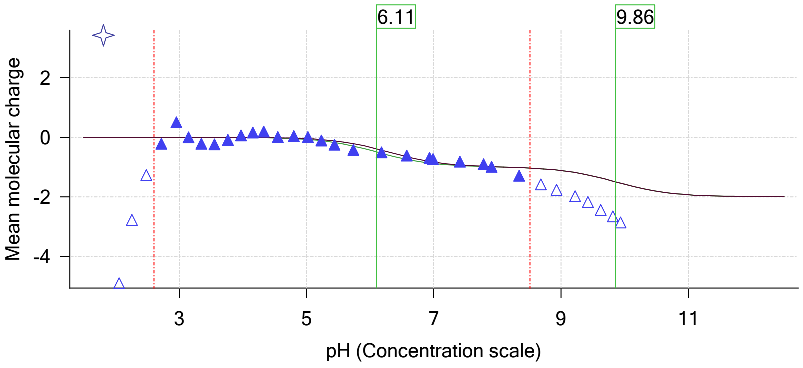

| | | | |
|---|---|---|---|
| Sample name: | **M07_octanol** | Experiment start time: | **2/28/2018 4:26:20 PM** |
| Assay name: | **pH-metric high logP** | Analyst: | **Pion** |
| Assay ID: | **18B-28011** | Instrument ID: | **T312060** |
| Filename: | **C:\Sirius_T3\Mehtap\20180228_exp28_logP_T3-2\18B-28011_M07_octanol_pH-metric high logP.t3r** | | |

#### Assay Model

| Settings | Value | Date/Time changed | Imported from |
|---|---|---|---|
| Sample name | M07_octanol | 2/27/2018 4:29:24 PM | User entered value |
| Sample by | Weight | | Default value |
| Sample weight | 0.000900 g | 2/28/2018 4:24:06 PM | User entered value |
| Formula weight | 235.28 g/mol | 2/27/2018 4:29:24 PM | User entered value |
| Solubility | Unknown | | Default value |
| Molecular weight | 235.28 | 2/27/2018 4:29:24 PM | User entered value |
| Individual pKa ionic environments | No | | Default value |
| Number of pKas | 1 | 2/27/2018 4:29:24 PM | User entered value |
| Sample is a | Base | 2/27/2018 4:29:24 PM | User entered value |
| pKa 1 | 6.07 | 2/27/2018 4:29:24 PM | User entered value |
| logp (XH +) | 0.50 | 2/28/2018 1:33:04 PM | User entered value |
| logP (neutral X) | 3.44 | 2/28/2018 1:33:10 PM | User entered value |

#### Events

| Time | Event | Water | Acid | Base | Octanol | pH | dpH/dt | pH R-squared | pH SD | dpH/dt time |
|---|---|---|---|---|---|---|---|---|---|---|
| 8:55.6 | Initial pH = 6.32 | | | | | | | | | |
| 11:55.1 | Data point 1 | 1.50000 mL | 0.04887 mL | 0.00245 mL | 0.01999 mL | 2.007 | -0.00759 | 0.43163 | 0.00057 | 10.0 s |
| 12:41.2 | Data point 2 | 1.50000 mL | 0.04887 mL | 0.01689 mL | 0.01999 mL | 2.212 | -0.00650 | 0.83933 | 0.00035 | 10.5 s |
| 13:17.4 | Data point 3 | 1.50000 mL | 0.04887 mL | 0.02641 mL | 0.01999 mL | 2.431 | -0.00986 | 0.77807 | 0.00055 | 10.0 s |
| 13:52.9 | Data point 4 | 1.50000 mL | 0.04887 mL | 0.03215 mL | 0.01999 mL | 2.651 | -0.00451 | 0.66680 | 0.00027 | 10.0 s |
| 14:28.4 | Data point 5 | 1.50000 mL | 0.04887 mL | 0.03563 mL | 0.01999 mL | 2.888 | -0.00773 | 0.36187 | 0.00063 | 10.0 s |
| 15:03.8 | Data point 6 | 1.50000 mL | 0.04887 mL | 0.03770 mL | 0.01999 mL | 3.072 | -0.00302 | 0.21499 | 0.00032 | 10.0 s |
| 15:39.3 | Data point 7 | 1.50000 mL | 0.04887 mL | 0.03911 mL | 0.01999 mL | 3.259 | -0.00406 | 0.30171 | 0.00036 | 10.0 s |
| 16:14.6 | Data point 8 | 1.50000 mL | 0.04887 mL | 0.04012 mL | 0.01999 mL | 3.431 | -0.00336 | 0.75731 | 0.00019 | 10.0 s |
| 16:55.2 | Data point 9 | 1.50000 mL | 0.04887 mL | 0.04099 mL | 0.01999 mL | 3.667 | -0.00529 | 0.80865 | 0.00029 | 10.0 s |
| 17:30.5 | Data point 10 | 1.50000 mL | 0.04887 mL | 0.04167 mL | 0.01999 mL | 3.906 | -0.01649 | 0.89884 | 0.00086 | 10.5 s |
| 18:16.6 | Data point 11 | 1.50000 mL | 0.04887 mL | 0.04231 mL | 0.01999 mL | 4.090 | -0.00742 | 0.86888 | 0.00039 | 10.0 s |
| 18:52.0 | Data point 12 | 1.50000 mL | 0.04887 mL | 0.04311 mL | 0.01999 mL | 4.359 | -0.01526 | 0.79653 | 0.00084 | 10.0 s |
| 19:43.0 | Data point 13 | 1.50000 mL | 0.04887 mL | 0.04412 mL | 0.01999 mL | 4.593 | -0.01458 | 0.78492 | 0.00081 | 11.5 s |
| 20:19.9 | Data point 14 | 1.50000 mL | 0.04887 mL | 0.04504 mL | 0.01999 mL | 4.792 | -0.01834 | 0.83155 | 0.00099 | 15.0 s |
| 21:00.3 | Data point 15 | 1.50000 mL | 0.04887 mL | 0.04588 mL | 0.01999 mL | 4.975 | -0.01816 | 0.89144 | 0.00095 | 12.5 s |
| 21:53.9 | Data point 16 | 1.50000 mL | 0.04887 mL | 0.04755 mL | 0.01999 mL | 5.371 | -0.01774 | 0.89163 | 0.00093 | 15.0 s |
| 22:39.5 | Data point 17 | 1.50000 mL | 0.04887 mL | 0.04880 mL | 0.01999 mL | 5.944 | -0.01937 | 0.91808 | 0.00100 | 25.5 s |
| 23:35.6 | Data point 18 | 1.50000 mL | 0.04887 mL | 0.04911 mL | 0.01999 mL | 6.201 | -0.01860 | 0.87008 | 0.00098 | 25.5 s |
| 24:36.8 | Data point 19 | 1.50000 mL | 0.04887 mL | 0.04941 mL | 0.01999 mL | 6.598 | -0.01882 | 0.95235 | 0.00095 | 40.0 s |
| 25:52.5 | Data point 20 | 1.50000 mL | 0.04887 mL | 0.04967 mL | 0.01999 mL | 8.879 | -0.02286 | 0.13576 | 0.00307 | Timed out at 59.5 s |
| 27:28.2 | Data point 21 | 1.50000 mL | 0.04887 mL | 0.05007 mL | 0.01999 mL | 9.833 | -0.01571 | 0.67713 | 0.00094 | 16.5 s |
| 28:15.3 | Data point 22 | 1.50000 mL | 0.04887 mL | 0.05045 mL | 0.01999 mL | 10.125 | -0.01833 | 0.90348 | 0.00095 | 18.5 s |
| 29:33.1 | Data point 23 | 1.50000 mL | 0.10350 mL | 0.05045 mL | 0.06999 mL | 1.961 | -0.00524 | 0.41258 | 0.00040 | 10.5 s |
| 30:19.8 | Data point 24 | 1.50000 mL | 0.10350 mL | 0.06766 mL | 0.06999 mL | 2.160 | -0.01414 | 0.83789 | 0.00076 | 10.5 s |
| 30:56.0 | Data point 25 | 1.50000 mL | 0.10350 mL | 0.07928 mL | 0.06999 mL | 2.405 | 0.00314 | 0.09186 | 0.00051 | 10.0 s |
| 31:42.0 | Data point 26 | 1.50000 mL | 0.10350 mL | 0.08542 mL | 0.06999 mL | 2.598 | -0.00349 | 0.57578 | 0.00023 | 10.0 s |
| 32:17.4 | Data point 27 | 1.50000 mL | 0.10350 mL | 0.08968 mL | 0.06999 mL | 2.829 | -0.00230 | 0.56610 | 0.00015 | 10.0 s |
| 32:52.8 | Data point 28 | 1.50000 mL | 0.10350 mL | 0.09231 mL | 0.06999 mL | 3.088 | -0.00226 | 0.48854 | 0.00016 | 10.0 s |
| 33:43.8 | Data point 29 | 1.50000 mL | 0.10350 mL | 0.09393 mL | 0.06999 mL | 3.295 | 0.00205 | 0.02071 | 0.00070 | 10.0 s |
| 34:19.2 | Data point 30 | 1.50000 mL | 0.10350 mL | 0.09518 mL | 0.06999 mL | 3.501 | -0.00570 | 0.60687 | 0.00036 | 10.0 s |
| 34:54.6 | Data point 31 | 1.50000 mL | 0.10350 mL | 0.09628 mL | 0.06999 mL | 3.707 | -0.00228 | 0.44952 | 0.00017 | 10.5 s |
| 35:30.6 | Data point 32 | 1.50000 mL | 0.10350 mL | 0.09734 mL | 0.06999 mL | 3.917 | -0.00812 | 0.27411 | 0.00077 | 10.0 s |
| 36:06.0 | Data point 33 | 1.50000 mL | 0.10350 mL | 0.09840 mL | 0.06999 mL | 4.130 | -0.00941 | 0.30499 | 0.00084 | 10.0 s |
| 36:41.4 | Data point 34 | 1.50000 mL | 0.10350 mL | 0.09941 mL | 0.06999 mL | 4.327 | 0.00167 | 0.00821 | 0.00091 | 10.0 s |
| 37:16.9 | Data point 35 | 1.50000 mL | 0.10350 mL | 0.10031 mL | 0.06999 mL | 4.517 | -0.00558 | 0.51690 | 0.00038 | 10.5 s |
| 37:52.8 | Data point 36 | 1.50000 mL | 0.10350 mL | 0.10106 mL | 0.06999 mL | 4.705 | -0.00463 | 0.40740 | 0.00036 | 10.5 s |
| 38:28.8 | Data point 37 | 1.50000 mL | 0.10350 mL | 0.10165 mL | 0.06999 mL | 4.857 | -0.00336 | 0.10816 | 0.00050 | 10.5 s |
| 39:20.2 | Data point 38 | 1.50000 mL | 0.10350 mL | 0.10237 mL | 0.06999 mL | 5.067 | -0.00648 | 0.11307 | 0.00095 | 10.5 s |
| 40:06.4 | Data point 39 | 1.50000 mL | 0.10350 mL | 0.10285 mL | 0.06999 mL | 5.275 | 0.00498 | 0.07222 | 0.00091 | 11.5 s |

| | | | |
|---|---|---|---|
| Sample name: | **M07_octanol** | Experiment start time: | **2/28/2018 4:26:20 PM** |
| Assay name: | **pH-metric high logP** | Analyst: | **Pion** |
| Assay ID: | **18B-28011** | Instrument ID: | **T312060** |
| Filename: | **C:\Sirius_T3\Mehtap\20180228_exp28_logP_T3-2\18B-28011_M07_octanol_pH-metric high logP.t3r** | | |

#### Events (continued)

| Time | Event | Water | Acid | Base | Octanol | pH | dpH/dt | pH R-squared | pH SD | dpH/dt time |
|---|---|---|---|---|---|---|---|---|---|---|
| 40:48.5 | Data point 40 | 1.50000 mL | 0.10350 mL | 0.10325 mL | 0.06999 mL | 5.581 | -0.01086 | 0.46682 | 0.00078 | 12.0 s |
| 41:36.2 | Data point 41 | 1.50000 mL | 0.10350 mL | 0.10388 mL | 0.06999 mL | 6.795 | -0.04882 | 0.99621 | 0.00241 | Timed out at 59.5 s |
| 43:27.1 | Data point 42 | 1.50000 mL | 0.10350 mL | 0.10433 mL | 0.06999 mL | 9.241 | -0.00847 | 0.26017 | 0.00082 | 18.5 s |
| 44:16.0 | Data point 43 | 1.50000 mL | 0.10350 mL | 0.10459 mL | 0.06999 mL | 9.587 | -0.01617 | 0.74261 | 0.00093 | 21.0 s |
| 45:07.4 | Data point 44 | 1.50000 mL | 0.10350 mL | 0.10501 mL | 0.06999 mL | 9.949 | -0.01841 | 0.93681 | 0.00094 | 13.5 s |
| 45:51.4 | Data point 45 | 1.50000 mL | 0.10350 mL | 0.10541 mL | 0.06999 mL | 10.160 | -0.00819 | 0.56417 | 0.00054 | 10.0 s |
| 47:05.4 | Data point 46 | 1.50000 mL | 0.16268 mL | 0.10541 mL | 0.31999 mL | 1.946 | -0.00597 | 0.52848 | 0.00041 | 10.0 s |
| 47:51.7 | Data point 47 | 1.50000 mL | 0.16268 mL | 0.13290 mL | 0.31999 mL | 2.185 | 0.00614 | 0.38583 | 0.00049 | 10.0 s |
| 48:37.7 | Data point 48 | 1.50000 mL | 0.16268 mL | 0.14372 mL | 0.31999 mL | 2.386 | -0.00203 | 0.41833 | 0.00016 | 10.0 s |
| 49:13.2 | Data point 49 | 1.50000 mL | 0.16268 mL | 0.15122 mL | 0.31999 mL | 2.611 | 0.01311 | 0.84025 | 0.00071 | 10.0 s |
| 49:48.7 | Data point 50 | 1.50000 mL | 0.16268 mL | 0.15588 mL | 0.31999 mL | 2.842 | 0.00403 | 0.34986 | 0.00034 | 10.0 s |
| 50:24.2 | Data point 51 | 1.50000 mL | 0.16268 mL | 0.15887 mL | 0.31999 mL | 3.077 | -0.00378 | 0.75203 | 0.00022 | 10.0 s |
| 51:10.1 | Data point 52 | 1.50000 mL | 0.16268 mL | 0.16108 mL | 0.31999 mL | 3.269 | 0.00384 | 0.15940 | 0.00048 | 10.0 s |
| 51:45.5 | Data point 53 | 1.50000 mL | 0.16268 mL | 0.16279 mL | 0.31999 mL | 3.466 | -0.00301 | 0.05747 | 0.00062 | 10.0 s |
| 52:20.9 | Data point 54 | 1.50000 mL | 0.16268 mL | 0.16425 mL | 0.31999 mL | 3.673 | 0.00201 | 0.08022 | 0.00035 | 10.0 s |
| 52:56.4 | Data point 55 | 1.50000 mL | 0.16268 mL | 0.16550 mL | 0.31999 mL | 3.886 | -0.00573 | 0.82166 | 0.00031 | 10.0 s |
| 53:31.7 | Data point 56 | 1.50000 mL | 0.16268 mL | 0.16651 mL | 0.31999 mL | 4.087 | -0.00755 | 0.87150 | 0.00040 | 10.0 s |
| 54:07.0 | Data point 57 | 1.50000 mL | 0.16268 mL | 0.16731 mL | 0.31999 mL | 4.277 | -0.00644 | 0.13717 | 0.00086 | 10.0 s |
| 54:42.4 | Data point 58 | 1.50000 mL | 0.16268 mL | 0.16790 mL | 0.31999 mL | 4.446 | -0.01161 | 0.50856 | 0.00080 | 10.0 s |
| 55:33.2 | Data point 59 | 1.50000 mL | 0.16268 mL | 0.16858 mL | 0.31999 mL | 4.665 | -0.00119 | 0.00535 | 0.00080 | 10.5 s |
| 56:24.4 | Data point 60 | 1.50000 mL | 0.16268 mL | 0.16900 mL | 0.31999 mL | 4.916 | -0.00586 | 0.09627 | 0.00093 | 10.5 s |
| 57:05.5 | Data point 61 | 1.50000 mL | 0.16268 mL | 0.16926 mL | 0.31999 mL | 5.139 | 0.00326 | 0.04495 | 0.00076 | 11.0 s |
| 57:47.1 | Data point 62 | 1.50000 mL | 0.16268 mL | 0.16947 mL | 0.31999 mL | 5.345 | -0.00065 | 0.00209 | 0.00070 | 11.5 s |
| 58:34.2 | Data point 63 | 1.50000 mL | 0.16268 mL | 0.16964 mL | 0.31999 mL | 5.553 | 0.00764 | 0.15851 | 0.00095 | 12.5 s |
| 59:22.4 | Data point 64 | 1.50000 mL | 0.16268 mL | 0.16980 mL | 0.31999 mL | 5.848 | -0.01710 | 0.92021 | 0.00088 | 35.5 s |
| 1:00:33.7 | Data point 65 | 1.50000 mL | 0.16268 mL | 0.16990 mL | 0.31999 mL | 6.288 | -0.01881 | 0.94810 | 0.00095 | 59.5 s |
| 1:02:03.8 | Data point 66 | 1.50000 mL | 0.16268 mL | 0.16997 mL | 0.31999 mL | 6.686 | -0.05982 | 0.98248 | 0.00298 | Timed out at 59.5 s |
| 1:03:34.3 | Data point 67 | 1.50000 mL | 0.16268 mL | 0.17001 mL | 0.31999 mL | 7.040 | -0.07343 | 0.98170 | 0.00366 | Timed out at 59.5 s |
| 1:04:59.6 | Data point 68 | 1.50000 mL | 0.16268 mL | 0.17004 mL | 0.31999 mL | 7.096 | -0.04310 | 0.98785 | 0.00214 | Timed out at 59.5 s |
| 1:06:30.2 | Data point 69 | 1.50000 mL | 0.16268 mL | 0.17008 mL | 0.31999 mL | 7.517 | -0.07608 | 0.99474 | 0.00377 | Timed out at 59.5 s |
| 1:08:00.6 | Data point 70 | 1.50000 mL | 0.16268 mL | 0.17013 mL | 0.31999 mL | 7.886 | -0.06164 | 0.99333 | 0.00306 | Timed out at 59.5 s |
| 1:09:31.1 | Data point 71 | 1.50000 mL | 0.16268 mL | 0.17018 mL | 0.31999 mL | 8.015 | -0.04517 | 0.98471 | 0.00225 | Timed out at 59.5 s |
| 1:11:11.9 | Data point 72 | 1.50000 mL | 0.16268 mL | 0.17034 mL | 0.31999 mL | 8.445 | -0.02344 | 0.95432 | 0.00119 | Timed out at 59.5 s |
| 1:12:47.5 | Data point 73 | 1.50000 mL | 0.16268 mL | 0.17051 mL | 0.31999 mL | 8.787 | -0.01486 | 0.61766 | 0.00093 | 20.5 s |
| 1:13:48.8 | Data point 74 | 1.50000 mL | 0.16268 mL | 0.17063 mL | 0.31999 mL | 9.033 | -0.01484 | 0.60375 | 0.00094 | 13.5 s |
| 1:14:33.0 | Data point 75 | 1.50000 mL | 0.16268 mL | 0.17079 mL | 0.31999 mL | 9.324 | -0.01384 | 0.49067 | 0.00098 | 17.5 s |
| 1:15:21.1 | Data point 76 | 1.50000 mL | 0.16268 mL | 0.17096 mL | 0.31999 mL | 9.524 | -0.01653 | 0.72433 | 0.00096 | 10.5 s |
| 1:16:07.3 | Data point 77 | 1.50000 mL | 0.16268 mL | 0.17119 mL | 0.31999 mL | 9.720 | -0.01049 | 0.45814 | 0.00077 | 10.5 s |
| 1:16:48.4 | Data point 78 | 1.50000 mL | 0.16268 mL | 0.17145 mL | 0.31999 mL | 9.913 | -0.00202 | 0.02625 | 0.00062 | 10.5 s |
| 1:17:29.5 | Data point 79 | 1.50000 mL | 0.16268 mL | 0.17168 mL | 0.31999 mL | 10.034 | -0.01309 | 0.45959 | 0.00095 | 13.0 s |
| 1:17:51.5 | Assay volumes | 1.50000 mL | 0.16268 mL | 0.17168 mL | 0.31999 mL | | | | | |

| | | | |
|---|---|---|---|
| Sample name: | **M07_octanol** | Experiment start time: | **2/28/2018 4:26:20 PM** |
| Assay name: | **pH-metric high logP** | Analyst: | **Pion** |
| Assay ID: | **18B-28011** | Instrument ID: | **T312060** |
| Filename: | **C:\Sirius_T3\Mehtap\20180228_exp28_logP_T3-2\18B-28011_M07_octanol_pH-metric high logP.t3r** | | |

#### Assay Settings

| Setting | Value | Original Value | Date/Time changed | Imported from |
|---|---|---|---|---|
| ***General Settings*** | | | | |
| Analyst name | Pion | | | |
| ***Standard Experiment Settings*** | | | | |
| Number of titrations | 3 | | | |
| Minimum pH | 2.000 | | | |
| Maximum pH | 10.000 | | | |
| pH step between points of | 0.200 | | | |
| Minimum titrant addition | 0.00002 mL | | | |
| Maximum titrant addition | 0.10000 mL | | | |
| Argon flow rate | 100% | | | |
| Start titration using | Cautious pH adjust | | | |
| ***Advanced General Settings*** | | | | |
| Detect turbidity using | None | | | |
| Collect turbidity sensor data | No | | | |
| Collect UV spectra | No | | | |
| Stir after titrant addition for | 5 seconds | | | |
| For titrant addition, stir at | 10% | | | |
| ***Titrant Pre-Dose*** | | | | |
| Titrant pre-dose | None | | | |
| ***Assay Medium*** | | | | |
| ISA water volume | 1.50 mL | | | |
| Water added | Automatic | | | |
| Partition solvent type | Octanol | | | |
| Partition volume | 0.020 mL | | | |
| Partition solvent added | Automatic | | | |
| After partition addition, stir for | 1 seconds | | | |
| ***Sample Sonication*** | | | | |
| Sonicate | Yes | | | |
| Adjust pH for sonication | No | | | |
| Sonicate for | 300 seconds | | | |
| After sonication stir for | 5 seconds | | | |
| ***Sample Dissolution*** | | | | |
| Perform a dissolution stage | Yes | | | |
| Adjust and hold pH for dissolution | To start pH | | | |
| Stir to dissolve for | 120 seconds | | | |
| For dissolution, stir at | 10% | | | |
| ***Carbonate purge*** | | | | |
| Perform a carbonate purge | No | | | |
| ***Temperature Control*** | | | | |
| Wait for temperature | Yes | | | |
| Required start temperature | 25.0°C | | | |
| Acceptable deviation | 0.5°C | | | |
| Time to wait | 60 seconds | | | |
| Stir speed of | 50% | | | |
| ***Titration 1*** | | | | |
| Titrate from | Low to high pH | | | |
| Adjust to start pH | Yes | | | |
| After pH adjust stir for | 30 seconds | | | |
| Stir to allow partitioning for | 15 seconds | | | |
| Stirrer speed for partitioning | 50% | | | |
| ***Titration 2*** | | | | |
| Titrate from | Low to high pH | | | |
| Add additional water | 0.00 mL | | | |
| Additional partition solvent volume | 0.050 mL | | | |
| Additional partition solvent added | Automatic | | | |
| After pH adjust stir for | 30 seconds | | | |
| Stir to allow partitioning for | 15 seconds | | | |
| Stirrer speed for partitioning | 55% | | | |

| | | | |
|---|---|---|---|
| Sample name: | **M07_octanol** | Experiment start time: | **2/28/2018 4:26:20 PM** |
| Assay name: | **pH-metric high logP** | Analyst: | **Pion** |
| Assay ID: | **18B-28011** | Instrument ID: | **T312060** |
| Filename: | **C:\Sirius_T3\Mehtap\20180228_exp28_logP_T3-2\18B-28011_M07_octanol_pH-metric high logP.t3r** | | |

#### Assay Settings (continued)

| Setting | Value | Original Value | Date/Time changed | Imported from |
|---|---|---|---|---|
| ***Titration 3*** | | | | |
| Titrate from | Low to high pH | | | |
| Add additional water | 0.00 mL | | | |
| Additional partition solvent volume | 0.250 mL | | | |
| Additional partition solvent added | Automatic | | | |
| After pH adjust stir for | 30 seconds | | | |
| Stir to allow partitioning for | 15 seconds | | | |
| Stirrer speed for partitioning | 60% | | | |
| ***Data Point Stability*** | | | | |
| Stir during data point collection | No | | | |
| Delay before data point collection | 0 seconds | | | |
| Number of points to average | 20 points | | | |
| Time interval between points | 0.50 seconds | | | |
| Required maximum standard deviation | 0.00100 dpH/dt | | | |
| Stability timeout after | 60 seconds | | | |

#### Calibration Settings

| Setting | Value | Date/Time changed | Imported from |
|---|---|---|---|
| Four-Plus alpha | 0.130 | 2/28/2018 4:26:20 PM | C:\Sirius_T3\HCl18B27.t3r |
| Four-Plus S | 0.9970 | 2/28/2018 4:26:20 PM | C:\Sirius_T3\HCl18B27.t3r |
| Four-Plus jH | 0.8 | 2/28/2018 4:26:20 PM | C:\Sirius_T3\HCl18B27.t3r |
| Four-Plus jOH | -0.4 | 2/28/2018 4:26:20 PM | C:\Sirius_T3\HCl18B27.t3r |
| Base concentration factor | 1.000 | 2/28/2018 4:26:20 PM | C:\Sirius_T3\KOH18B27.t3r |
| Acid concentration factor | 0.994 | 2/28/2018 4:26:20 PM | C:\Sirius_T3\HCl18B27.t3r |

#### Instrument Settings

| Setting | Value | Batch Id | Install date |
|---|---|---|---|
| Instrument owner | Merck | | |
| Instrument ID | T312060 | | |
| Instrument type | T3 Simulator | | |
| Software version | 1.1.3.0 | | |
| Dispenser module | | T3DM1200361 | 3/31/2009 5:24:52 AM |
| Dispenser 0 | Water | | 3/31/2009 5:25:05 AM |
| Syringe volume | 2.5 mL | | |
| Firmware version | 1.2.1(r2) | | |
| Titrant | Water (0.15 M KCl) | 02-06-2018 | 2/27/2018 10:05:59 AM |
| Dispenser 2 | Acid | | 3/31/2009 5:25:11 AM |
| Syringe volume | 0.5 mL | | |
| Firmware version | 1.2.1(r2) | | |
| Titrant | Acid (0.5 M HCl) | 02-27-2018 | 2/27/2018 10:27:22 AM |
| Dispenser 1 | Base | | 3/31/2009 5:25:21 AM |
| Syringe volume | 0.5 mL | | |
| Firmware version | 1.2.1(r2) | | |
| Titrant | Base (0.5 M KOH) | 9/22/2017 | 2/27/2018 10:21:22 AM |
| Dispenser 5 | Cosolvent | | 3/31/2009 5:26:24 AM |
| Syringe volume | 2.5 mL | | |
| Firmware version | 1.2.1(r2) | | |
| Distribution valve 5 | Distribution Valve | | 3/31/2009 5:28:19 AM |
| Firmware version | 1.1.3 | | |
| Port A | Methanol (80%, 0.15 M KCl) | 09-26-17 | 2/7/2018 9:42:01 AM |
| Port B | Cyclohexane | 11-01-17 | 2/27/2018 10:37:57 AM |
| Dispenser 3 | Buffer | | 8/3/2010 5:05:16 AM |
| Syringe volume | 0.5 mL | | |
| Firmware version | 1.2.1(r2) | | |
| Titrant | Dodecane | 2018/01/31 | 2/28/2018 10:18:04 AM |
| Dispenser 6 | Octanol | | 10/22/2010 10:52:43 AM |

| | | | |
|---|---|---|---|
| Sample name: | **M07_octanol** | Experiment start time: | **2/28/2018 4:26:20 PM** |
| Assay name: | **pH-metric high logP** | Analyst: | **Pion** |
| Assay ID: | **18B-28011** | Instrument ID: | **T312060** |
| Filename: | **C:\Sirius_T3\Mehtap\20180228_exp28_logP_T3-2\18B-28011_M07_octanol_pH-metric high logP.t3r** | | |

#### Instrument Settings (continued)

| Setting | Value | Batch Id | Install date |
|---|---|---|---|
| Syringe volume | 0.5 mL | | |
| Firmware version | 1.2.1(r2) | | |
| Titrant | Octanol | 01-31-2018 | 2/27/2018 9:59:35 AM |
| Titrator | | T3TM1200161 | 3/31/2009 5:24:17 AM |
| Horizontal axis firmware version | 1.17 AI1DI2DO2 Stepper 2 | | |
| Vertical axis firmware version | 1.17 AI1DI2DO2 Stepper 2 | | |
| Chassis I/O firmware version | 1.11 AI1DI0DO4 Norgren I/O | | |
| Probe I/O firmware version | 1.1.1 | | |
| Electrode | T3 Electrode | T3E0923 | 1/23/2018 2:01:00 PM |
| E0 calibration | +3.96 mV | | 2/28/2018 4:27:04 PM |
| Filling solution | 3M KCl | KCL097 | 2/27/2018 9:49:43 AM |
| Liquids | | | |
| Wash 1 | 50% IPA:50% Water | | 2/28/2018 10:23:32 AM |
| Wash 2 | 0.5% Trition X-100 in H20 | | 2/28/2018 10:23:34 AM |
| Buffer position 1 | pH7 Wash | | 2/28/2018 10:24:06 AM |
| Buffer position 2 | pH 7 | | 2/28/2018 10:24:08 AM |
| Storage position | | | 2/28/2018 10:21:14 AM |
| Wash water | 8.8e+003 mL | 02-27-2018 | 2/27/2018 9:54:39 AM |
| Waste | 6.7e+003 mL | | 11/28/2017 10:36:29 AM |
| Temperature controller | | | 8/5/2010 6:35:13 AM |
| Turbidity detector | | | 3/31/2009 5:24:45 AM |
| Spectrometer | | 074811 | 11/23/2010 11:22:28 AM |
| Dip probe | | 10196 | |
| Wavelength coefficient A0 | 183.333 | | |
| Wavelength coefficient A1 | 2.21568 | | |
| Wavelength coefficient A2 | -0.000289308 | | |
| Total lamp lit time | 112:08:55 | | 11/23/2010 11:22:28 AM |
| Calibrated on | 2/27/2018 10:40:38 AM | | |
| Integration time | 40 | | |
| Scans averaged | 10 | | |
| Autoloader | | T3AL1200345 | 11/10/2015 9:34:13 AM |
| Left-right axis firmware version | 1.17 AI1DI2DO2 Stepper 2 | | |
| Front-back axis firmware version | 1.17 AI1DI2DO2 Stepper 2 | | |
| Vertical axis firmware version | 1.17 AI1DI2DO2 Stepper 2 | | |
| Chassis I/O firmware version | 1.11 AI1DI0DO4 Norgren I/O | | |
| Configuration | | | |
| Alternate titration position | Titration position | | |
| Alternate reference position | Reference position | | |
| Maximum standard vial volume | 3.50 mL | | |
| Maximum alternate vial volume | 25.00 mL | | |
| Automatic action idle period | 5 minute(s) | | |
| Titrant tube volume | 1.3 mL | | |
| Syringe flush count | 3.50 | | |
| Flowing wash pump volume | 20.0 mL | | |
| Flowing wash stir duration | 5 s | | |
| Flowing wash stir speed | 30% | | |
| Solvent wash stir duration | 5 s | | |
| Solvent wash stir speed | 30% | | |
| Surfactant wash stir duration | 5 s | | |
| Surfactant wash stir speed | 30% | | |
| E0 calibration minimum number of points | 10 | | |
| E0 calibration maximum standard deviation | 0.01500 | | |
| E0 calibration timeout period | 60 s | | |
| E0 calibration stir duration | 5 s | | |
| E0 calibration preparation stir speed | 30% | | |
| E0 calibration buffer wash stir duration | 5 s | | |
| E0 calibration buffer wash stir speed | 30% | | |
| E0 calibration reading stir speed | 0% | | |

| | | | |
|---|---|---|---|
| Sample name: | **M07_octanol** | Experiment start time: | **2/28/2018 4:26:20 PM** |
| Assay name: | **pH-metric high logP** | Analyst: | **Pion** |
| Assay ID: | **18B-28011** | Instrument ID: | **T312060** |
| Filename: | **C:\Sirius_T3\Mehtap\20180228_exp28_logP_T3-2\18B-28011_M07_octanol_pH-metric high logP.t3r** | | |

#### Instrument Settings (continued)

| Setting | Value | Batch Id | Install date |
|---|---|---|---|
| Spectrometer calibration stir duration | 5 s | | |
| Spectrometer calibration stir speed | 30% | | |
| Spectrometer calibration wash pump volume | 20.0 mL | | |
| Spectrometer calibration wash stir duration | 5 s | | |
| Spectrometer calibration wash stir speed | 30% | | |
| Overhead dispense height | 10000 | | |

#### Refinement Settings

| Setting | Value | Default value |
|---|---|---|
| Turbidity detection method | None | None |
| Turbidity wavelength to assess | 500.0 nm | 500.0 nm |
| Turbidity maximum absorbance | 0.100 | 0.100 |
| Turbidity probe threshold | 50.00 | 50.00 |

#### Experiment Log

[1:57] Air gap released for Acid (0.5 M HCl)
[1:57] Air gap released for Base (0.5 M KOH)
[2:33] Air gap created for Water (0.15 M KCl)
[2:33] Air gap created for Acid (0.5 M HCl)
[2:33] Air gap created for Base (0.5 M KOH)
[2:34] Air gap released for Water (0.15 M KCl)
[2:38] Titrator arm moved over Titration position
[2:38] Titration 1 of 3
[2:38] Adding initial titrants
[2:38] Automatically add 1.50000 mL of water
[3:03] Dispensed 1.500000 mL of Water (0.15 M KCl)
[3:07] Titrator arm moved over Drain
[8:49] Titrator arm moved to Titration position
[8:49] Argon flow rate set to 100
[8:49] Stirrer speed set to 10
[8:54] Automatically add 0.02000 mL of Octanol
[8:54] Dispensed 0.019991 mL of Octanol
[8:55] Initial pH = 6.32
[8:55] Iterative adjust 6.32 -> 2.00
[8:55] pH 6.32 -> 2.00
[8:57] Air gap released for Acid (0.5 M HCl)
[8:58] Dispensed 0.048871 mL of Acid (0.5 M HCl)
[9:03] Holding pH 2.00
[11:03] Stirrer speed set to 0
[11:03] Stirrer speed set to 50
[11:03] Iterative adjust 1.98 -> 2.00
[11:03] pH 1.98 -> 2.00
[11:04] Air gap released for Base (0.5 M KOH)
[11:04] Dispensed 0.002446 mL of Base (0.5 M KOH)
[11:55] Stirrer speed set to 0
[12:05] Datapoint id 1 collected
[12:05] Stirrer speed set to 50
[12:10] pH 2.02 -> 2.22
[12:10] Using cautious pH adjust
[12:10] Dispensed 0.007643 mL of Base (0.5 M KOH)
[12:15] Stepping pH = 2.11
[12:16] Dispensed 0.005339 mL of Base (0.5 M KOH)
[12:21] Stepping pH = 2.19
[12:21] Dispensed 0.001458 mL of Base (0.5 M KOH)
[12:26] Stepping pH = 2.22
[12:41] Stirrer speed set to 0
[12:52] Datapoint id 2 collected

| | | | |
|---|---|---|---|
| Sample name: | **M07_octanol** | Experiment start time: | **2/28/2018 4:26:20 PM** |
| Assay name: | **pH-metric high logP** | Analyst: | **Pion** |
| Assay ID: | **18B-28011** | Instrument ID: | **T312060** |
| Filename: | **C:\Sirius_T3\Mehtap\20180228_exp28_logP_T3-2\18B-28011_M07_octanol_pH-metric high logP.t3r** | | |

#### Experiment Log (continued)

[12:52] Charge balance equation is out by 5.5%
[12:52] Stirrer speed set to 50
[12:57] pH 2.22 -> 2.42
[12:57] Using charge balance adjust
[12:57] Dispensed 0.009525 mL of Base (0.5 M KOH)
[13:17] Stirrer speed set to 0
[13:27] Datapoint id 3 collected
[13:27] Charge balance equation is out by 6.0%
[13:27] Stirrer speed set to 50
[13:32] pH 2.44 -> 2.64
[13:32] Using charge balance adjust
[13:32] Dispensed 0.005738 mL of Base (0.5 M KOH)
[13:53] Stirrer speed set to 0
[14:03] Datapoint id 4 collected
[14:03] Charge balance equation is out by 5.9%
[14:03] Stirrer speed set to 50
[14:08] pH 2.66 -> 2.86
[14:08] Using charge balance adjust
[14:08] Dispensed 0.003481 mL of Base (0.5 M KOH)
[14:28] Stirrer speed set to 0
[14:38] Datapoint id 5 collected
[14:38] Charge balance equation is out by 14.5%
[14:38] Stirrer speed set to 50
[14:43] pH 2.89 -> 3.09
[14:43] Using charge balance adjust
[14:43] Dispensed 0.002070 mL of Base (0.5 M KOH)
[15:04] Stirrer speed set to 0
[15:14] Datapoint id 6 collected
[15:14] Charge balance equation is out by -10.8%
[15:14] Stirrer speed set to 50
[15:19] pH 3.08 -> 3.28
[15:19] Using charge balance adjust
[15:19] Dispensed 0.001411 mL of Base (0.5 M KOH)
[15:39] Stirrer speed set to 0
[15:49] Datapoint id 7 collected
[15:49] Charge balance equation is out by -9.7%
[15:49] Stirrer speed set to 50
[15:54] pH 3.26 -> 3.46
[15:54] Using charge balance adjust
[15:54] Dispensed 0.001011 mL of Base (0.5 M KOH)
[16:14] Stirrer speed set to 0
[16:24] Datapoint id 8 collected
[16:24] Charge balance equation is out by -16.8%
[16:24] Stirrer speed set to 50
[16:29] pH 3.44 -> 3.64
[16:29] Using cautious pH adjust
[16:30] Dispensed 0.000400 mL of Base (0.5 M KOH)
[16:35] Stepping pH = 3.50
[16:35] Dispensed 0.000470 mL of Base (0.5 M KOH)
[16:40] Stepping pH = 3.66
[16:55] Stirrer speed set to 0
[17:05] Datapoint id 9 collected
[17:05] Charge balance equation is out by -8.3%
[17:05] Stirrer speed set to 50
[17:10] pH 3.67 -> 3.87
[17:10] Using charge balance adjust
[17:10] Dispensed 0.000682 mL of Base (0.5 M KOH)
[17:30] Stirrer speed set to 0
[17:41] Datapoint id 10 collected

| | | | |
|---|---|---|---|
| Sample name: | **M07_octanol** | Experiment start time: | **2/28/2018 4:26:20 PM** |
| Assay name: | **pH-metric high logP** | Analyst: | **Pion** |
| Assay ID: | **18B-28011** | Instrument ID: | **T312060** |
| Filename: | **C:\Sirius_T3\Mehtap\20180228_exp28_logP_T3-2\18B-28011_M07_octanol_pH-metric high logP.t3r** | | |

#### Experiment Log (continued)

[17:41] Charge balance equation is out by 15.6%
[17:41] Stirrer speed set to 50
[17:46] pH 3.91 -> 4.11
[17:46] Using cautious pH adjust
[17:46] Dispensed 0.000353 mL of Base (0.5 M KOH)
[17:51] Stepping pH = 4.02
[17:51] Dispensed 0.000235 mL of Base (0.5 M KOH)
[17:56] Stepping pH = 4.10
[17:56] Dispensed 0.000047 mL of Base (0.5 M KOH)
[18:01] Stepping pH = 4.11
[18:16] Stirrer speed set to 0
[18:26] Datapoint id 11 collected
[18:26] Charge balance equation is out by 11.4%
[18:26] Stirrer speed set to 50
[18:31] pH 4.10 -> 4.30
[18:31] Using charge balance adjust
[18:32] Dispensed 0.000800 mL of Base (0.5 M KOH)
[18:52] Stirrer speed set to 0
[19:02] Datapoint id 12 collected
[19:02] Charge balance equation is out by 29.2%
[19:02] Stirrer speed set to 50
[19:07] pH 4.36 -> 4.56
[19:07] Using cautious pH adjust
[19:07] Dispensed 0.000447 mL of Base (0.5 M KOH)
[19:12] Stepping pH = 4.48
[19:12] Dispensed 0.000259 mL of Base (0.5 M KOH)
[19:17] Stepping pH = 4.55
[19:17] Dispensed 0.000047 mL of Base (0.5 M KOH)
[19:22] Stepping pH = 4.55
[19:22] Dispensed 0.000259 mL of Base (0.5 M KOH)
[19:28] Stepping pH = 4.62
[19:43] Stirrer speed set to 0
[19:54] Datapoint id 13 collected
[19:54] Charge balance equation is out by -12.3%
[19:54] Stirrer speed set to 50
[19:59] pH 4.61 -> 4.81
[19:59] Using charge balance adjust
[19:59] Dispensed 0.000917 mL of Base (0.5 M KOH)
[20:20] Stirrer speed set to 0
[20:35] Datapoint id 14 collected
[20:35] Charge balance equation is out by -8.1%
[20:35] Stirrer speed set to 50
[20:40] pH 4.81 -> 5.01
[20:40] Using charge balance adjust
[20:40] Dispensed 0.000847 mL of Base (0.5 M KOH)
[21:00] Stirrer speed set to 0
[21:13] Datapoint id 15 collected
[21:13] Charge balance equation is out by -18.0%
[21:13] Stirrer speed set to 50
[21:18] pH 5.00 -> 5.20
[21:18] Using cautious pH adjust
[21:18] Dispensed 0.000353 mL of Base (0.5 M KOH)
[21:23] Stepping pH = 5.07
[21:23] Dispensed 0.000376 mL of Base (0.5 M KOH)
[21:28] Stepping pH = 5.17
[21:28] Dispensed 0.000071 mL of Base (0.5 M KOH)
[21:33] Stepping pH = 5.14
[21:33] Dispensed 0.000870 mL of Base (0.5 M KOH)
[21:39] Stepping pH = 5.45

| | | | |
|---|---|---|---|
| Sample name: | **M07_octanol** | Experiment start time: | **2/28/2018 4:26:20 PM** |
| Assay name: | **pH-metric high logP** | Analyst: | **Pion** |
| Assay ID: | **18B-28011** | Instrument ID: | **T312060** |
| Filename: | **C:\Sirius_T3\Mehtap\20180228_exp28_logP_T3-2\18B-28011_M07_octanol_pH-metric high logP.t3r** | | |

#### Experiment Log (continued)

[21:54] Stirrer speed set to 0
[22:09] Datapoint id 16 collected
[22:09] Charge balance equation is out by -134.1%
[22:09] Stirrer speed set to 50
[22:14] pH 5.41 -> 5.61
[22:14] Using cautious pH adjust
[22:14] Dispensed 0.000212 mL of Base (0.5 M KOH)
[22:19] Stepping pH = 5.41
[22:19] Dispensed 0.001035 mL of Base (0.5 M KOH)
[22:24] Stepping pH = 6.09
[22:39] Stirrer speed set to 0
[23:05] Datapoint id 17 collected
[23:05] Charge balance equation is out by -201.4%
[23:05] Stirrer speed set to 50
[23:10] pH 5.99 -> 6.19
[23:10] Using cautious pH adjust
[23:10] Dispensed 0.000071 mL of Base (0.5 M KOH)
[23:15] Stepping pH = 5.99
[23:15] Dispensed 0.000235 mL of Base (0.5 M KOH)
[23:20] Stepping pH = 6.23
[23:35] Stirrer speed set to 0
[24:01] Datapoint id 18 collected
[24:01] Charge balance equation is out by -98.3%
[24:01] Stirrer speed set to 50
[24:06] pH 6.26 -> 6.46
[24:06] Using cautious pH adjust
[24:06] Dispensed 0.000047 mL of Base (0.5 M KOH)
[24:11] Stepping pH = 6.26
[24:11] Dispensed 0.000165 mL of Base (0.5 M KOH)
[24:16] Stepping pH = 6.37
[24:16] Dispensed 0.000094 mL of Base (0.5 M KOH)
[24:21] Stepping pH = 6.59
[24:37] Stirrer speed set to 0
[25:17] Datapoint id 19 collected
[25:17] Charge balance equation is out by -178.7%
[25:17] Stirrer speed set to 50
[25:22] pH 6.71 -> 6.91
[25:22] Using cautious pH adjust
[25:22] Dispensed 0.000024 mL of Base (0.5 M KOH)
[25:27] Stepping pH = 6.72
[25:27] Dispensed 0.000071 mL of Base (0.5 M KOH)
[25:32] Stepping pH = 6.75
[25:32] Dispensed 0.000165 mL of Base (0.5 M KOH)
[25:37] Stepping pH = 8.75
[25:52] Stirrer speed set to 0
[26:52] Datapoint id 20 collected
[26:52] Charge balance equation is out by -408.8%
[26:52] Stirrer speed set to 50
[26:57] pH 8.97 -> 9.17
[26:57] Using cautious pH adjust
[26:57] Dispensed 0.000024 mL of Base (0.5 M KOH)
[27:03] Stepping pH = 8.97
[27:03] Dispensed 0.000094 mL of Base (0.5 M KOH)
[27:08] Stepping pH = 8.98
[27:08] Dispensed 0.000282 mL of Base (0.5 M KOH)
[27:13] Stepping pH = 9.83
[27:28] Stirrer speed set to 0
[27:45] Datapoint id 21 collected
[27:45] Charge balance equation is out by -942.4%

| | | | |
|---|---|---|---|
| Sample name: | **M07_octanol** | Experiment start time: | **2/28/2018 4:26:20 PM** |
| Assay name: | **pH-metric high logP** | Analyst: | **Pion** |
| Assay ID: | **18B-28011** | Instrument ID: | **T312060** |
| Filename: | **C:\Sirius_T3\Mehtap\20180228_exp28_logP_T3-2\18B-28011_M07_octanol_pH-metric high logP.t3r** | | |

#### Experiment Log (continued)

[27:45] Stirrer speed set to 50
[27:50] pH 9.85 -> 10.05
[27:50] Using cautious pH adjust
[27:50] Dispensed 0.000118 mL of Base (0.5 M KOH)
[27:55] Stepping pH = 9.87
[27:55] Dispensed 0.000259 mL of Base (0.5 M KOH)
[28:00] Stepping pH = 10.13
[28:15] Stirrer speed set to 0
[28:34] Datapoint id 22 collected
[28:34] Charge balance equation is out by -73.7%
[28:34] Titration 2 of 3
[28:34] Adding initial titrants
[28:34] Automatically add 0.05000 mL of Octanol
[28:35] Dispensed 0.050000 mL of Octanol
[28:35] Stirrer speed set to 10
[28:36] Stirrer speed set to 55
[28:36] Iterative adjust 10.14 -> 2.00
[28:36] pH 10.14 -> 2.00
[28:37] Dispensed 0.052658 mL of Acid (0.5 M HCl)
[28:42] pH 2.02 -> 2.00
[28:43] Dispensed 0.001976 mL of Acid (0.5 M HCl)
[29:33] Stirrer speed set to 0
[29:43] Datapoint id 23 collected
[29:43] Stirrer speed set to 55
[29:48] pH 1.97 -> 2.17
[29:48] Using cautious pH adjust
[29:49] Dispensed 0.009196 mL of Base (0.5 M KOH)
[29:54] Stepping pH = 2.06
[29:54] Dispensed 0.006515 mL of Base (0.5 M KOH)
[29:59] Stepping pH = 2.14
[29:59] Dispensed 0.001505 mL of Base (0.5 M KOH)
[30:04] Stepping pH = 2.16
[30:20] Stirrer speed set to 0
[30:30] Datapoint id 24 collected
[30:30] Charge balance equation is out by 6.5%
[30:30] Stirrer speed set to 55
[30:35] pH 2.16 -> 2.36
[30:35] Using charge balance adjust
[30:36] Dispensed 0.011618 mL of Base (0.5 M KOH)
[30:56] Stirrer speed set to 0
[31:06] Datapoint id 25 collected
[31:06] Charge balance equation is out by 20.5%
[31:06] Stirrer speed set to 55
[31:11] pH 2.41 -> 2.61
[31:11] Using cautious pH adjust
[31:11] Dispensed 0.003293 mL of Base (0.5 M KOH)
[31:16] Stepping pH = 2.51
[31:16] Dispensed 0.002023 mL of Base (0.5 M KOH)
[31:21] Stepping pH = 2.58
[31:21] Dispensed 0.000823 mL of Base (0.5 M KOH)
[31:27] Stepping pH = 2.60
[31:42] Stirrer speed set to 0
[31:52] Datapoint id 26 collected
[31:52] Charge balance equation is out by 6.8%
[31:52] Stirrer speed set to 55
[31:57] pH 2.61 -> 2.81
[31:57] Using charge balance adjust
[31:57] Dispensed 0.004257 mL of Base (0.5 M KOH)
[32:17] Stirrer speed set to 0

Sample name: **M07_octanol**
Assay name: **pH-metric high logP**
Assay ID: **18B-28011**
Filename: **C:\Sirius_T3\Mehtap\20180228_exp28_logP_T3-2\18B-28011_M07_octanol_pH-metric high logP.t3r**

Experiment start time: **2/28/2018 4:26:20 PM**
Analyst: **Pion**
Instrument ID: **T312060**

#### Experiment Log (continued)

[32:27] Datapoint id 27 collected
[32:27] Charge balance equation is out by 11.6%
[32:27] Stirrer speed set to 55
[32:32] pH 2.83 -> 3.03
[32:32] Using charge balance adjust
[32:32] Dispensed 0.002634 mL of Base (0.5 M KOH)
[32:53] Stirrer speed set to 0
[33:03] Datapoint id 28 collected
[33:03] Charge balance equation is out by 26.9%
[33:03] Stirrer speed set to 55
[33:08] pH 3.09 -> 3.29
[33:08] Using cautious pH adjust
[33:08] Dispensed 0.000823 mL of Base (0.5 M KOH)
[33:13] Stepping pH = 3.19
[33:13] Dispensed 0.000588 mL of Base (0.5 M KOH)
[33:18] Stepping pH = 3.27
[33:18] Dispensed 0.000118 mL of Base (0.5 M KOH)
[33:23] Stepping pH = 3.28
[33:23] Dispensed 0.000094 mL of Base (0.5 M KOH)
[33:28] Stepping pH = 3.30
[33:44] Stirrer speed set to 0
[33:54] Datapoint id 29 collected
[33:54] Charge balance equation is out by 1.7%
[33:54] Stirrer speed set to 55
[33:59] pH 3.30 -> 3.50
[33:59] Using charge balance adjust
[33:59] Dispensed 0.001246 mL of Base (0.5 M KOH)
[34:19] Stirrer speed set to 0
[34:29] Datapoint id 30 collected
[34:29] Charge balance equation is out by -0.4%
[34:29] Stirrer speed set to 55
[34:34] pH 3.51 -> 3.71
[34:34] Using charge balance adjust
[34:34] Dispensed 0.001105 mL of Base (0.5 M KOH)
[34:54] Stirrer speed set to 0
[35:05] Datapoint id 31 collected
[35:05] Charge balance equation is out by 1.1%
[35:05] Stirrer speed set to 55
[35:10] pH 3.71 -> 3.91
[35:10] Using charge balance adjust
[35:10] Dispensed 0.001058 mL of Base (0.5 M KOH)
[35:30] Stirrer speed set to 0
[35:40] Datapoint id 32 collected
[35:40] Charge balance equation is out by 2.5%
[35:40] Stirrer speed set to 55
[35:45] pH 3.92 -> 4.12
[35:45] Using charge balance adjust
[35:46] Dispensed 0.001058 mL of Base (0.5 M KOH)
[36:06] Stirrer speed set to 0
[36:16] Datapoint id 33 collected
[36:16] Charge balance equation is out by 3.6%
[36:16] Stirrer speed set to 55
[36:21] pH 4.13 -> 4.33
[36:21] Using charge balance adjust
[36:21] Dispensed 0.001011 mL of Base (0.5 M KOH)
[36:41] Stirrer speed set to 0
[36:51] Datapoint id 34 collected
[36:51] Charge balance equation is out by -3.3%
[36:51] Stirrer speed set to 55

| | | | |
|---|---|---|---|
| Sample name: | **M07_octanol** | Experiment start time: | **2/28/2018 4:26:20 PM** |
| Assay name: | **pH-metric high logP** | Analyst: | **Pion** |
| Assay ID: | **18B-28011** | Instrument ID: | **T312060** |
| Filename: | **C:\Sirius_T3\Mehtap\20180228_exp28_logP_T3-2\18B-28011_M07_octanol_pH-metric high logP.t3r** | | |

#### Experiment Log (continued)

[36:56] pH 4.34 -> 4.54
[36:56] Using charge balance adjust
[36:56] Dispensed 0.000894 mL of Base (0.5 M KOH)
[37:17] Stirrer speed set to 0
[37:27] Datapoint id 35 collected
[37:27] Charge balance equation is out by -9.8%
[37:27] Stirrer speed set to 55
[37:32] pH 4.53 -> 4.73
[37:32] Using charge balance adjust
[37:32] Dispensed 0.000753 mL of Base (0.5 M KOH)
[37:53] Stirrer speed set to 0
[38:03] Datapoint id 36 collected
[38:03] Charge balance equation is out by -11.7%
[38:03] Stirrer speed set to 55
[38:08] pH 4.72 -> 4.92
[38:08] Using charge balance adjust
[38:08] Dispensed 0.000588 mL of Base (0.5 M KOH)
[38:29] Stirrer speed set to 0
[38:39] Datapoint id 37 collected
[38:39] Charge balance equation is out by -30.8%
[38:39] Stirrer speed set to 55
[38:44] pH 4.87 -> 5.07
[38:44] Using cautious pH adjust
[38:44] Dispensed 0.000235 mL of Base (0.5 M KOH)
[38:49] Stepping pH = 4.93
[38:49] Dispensed 0.000282 mL of Base (0.5 M KOH)
[38:54] Stepping pH = 5.01
[38:55] Dispensed 0.000141 mL of Base (0.5 M KOH)
[39:00] Stepping pH = 5.05
[39:00] Dispensed 0.000071 mL of Base (0.5 M KOH)
[39:05] Stepping pH = 5.07
[39:20] Stirrer speed set to 0
[39:30] Datapoint id 38 collected
[39:30] Charge balance equation is out by -55.1%
[39:30] Stirrer speed set to 55
[39:36] pH 5.09 -> 5.29
[39:36] Using cautious pH adjust
[39:36] Dispensed 0.000165 mL of Base (0.5 M KOH)
[39:41] Stepping pH = 5.13
[39:41] Dispensed 0.000282 mL of Base (0.5 M KOH)
[39:46] Stepping pH = 5.28
[39:46] Dispensed 0.000024 mL of Base (0.5 M KOH)
[39:51] Stepping pH = 5.29
[40:06] Stirrer speed set to 0
[40:18] Datapoint id 39 collected
[40:18] Charge balance equation is out by -46.4%
[40:18] Stirrer speed set to 55
[40:23] pH 5.30 -> 5.50
[40:23] Using cautious pH adjust
[40:23] Dispensed 0.000118 mL of Base (0.5 M KOH)
[40:28] Stepping pH = 5.32
[40:28] Dispensed 0.000282 mL of Base (0.5 M KOH)
[40:33] Stepping pH = 5.58
[40:48] Stirrer speed set to 0
[41:00] Datapoint id 40 collected
[41:00] Charge balance equation is out by -83.4%
[41:00] Stirrer speed set to 55
[41:05] pH 5.62 -> 5.82
[41:05] Using cautious pH adjust

Sample name: **M07_octanol**
Assay name: **pH-metric high logP**
Assay ID: **18B-28011**
Filename: **C:\Sirius_T3\Mehtap\20180228_exp28_logP_T3-2\18B-28011_M07_octanol_pH-metric high logP.t3r**
Experiment start time: **2/28/2018 4:26:20 PM**
Analyst: **Pion**
Instrument ID: **T312060**

#### Experiment Log (continued)

[41:05] Dispensed 0.000071 mL of Base (0.5 M KOH)
[41:11] Stepping pH = 5.63
[41:11] Dispensed 0.000188 mL of Base (0.5 M KOH)
[41:16] Stepping pH = 5.66
[41:16] Dispensed 0.000376 mL of Base (0.5 M KOH)
[41:21] Stepping pH = 6.69
[41:36] Stirrer speed set to 0
[42:36] Datapoint id 41 collected
[42:36] Charge balance equation is out by -392.1%
[42:36] Stirrer speed set to 55
[42:41] pH 6.97 -> 7.17
[42:41] Using cautious pH adjust
[42:41] Dispensed 0.000024 mL of Base (0.5 M KOH)
[42:46] Stepping pH = 7.02
[42:46] Dispensed 0.000024 mL of Base (0.5 M KOH)
[42:51] Stepping pH = 7.05
[42:51] Dispensed 0.000024 mL of Base (0.5 M KOH)
[42:56] Stepping pH = 7.08
[42:56] Dispensed 0.000047 mL of Base (0.5 M KOH)
[43:02] Stepping pH = 7.10
[43:02] Dispensed 0.000047 mL of Base (0.5 M KOH)
[43:07] Stepping pH = 7.10
[43:07] Dispensed 0.000282 mL of Base (0.5 M KOH)
[43:12] Stepping pH = 9.27
[43:27] Stirrer speed set to 0
[43:45] Datapoint id 42 collected
[43:45] Charge balance equation is out by -1,782.6%
[43:45] Stirrer speed set to 55
[43:50] pH 9.30 -> 9.50
[43:50] Using cautious pH adjust
[43:50] Dispensed 0.000047 mL of Base (0.5 M KOH)
[43:56] Stepping pH = 9.30
[43:56] Dispensed 0.000212 mL of Base (0.5 M KOH)
[44:01] Stepping pH = 9.60
[44:16] Stirrer speed set to 0
[44:37] Datapoint id 43 collected
[44:37] Charge balance equation is out by -200.1%
[44:37] Stirrer speed set to 55
[44:42] pH 9.58 -> 9.78
[44:42] Using cautious pH adjust
[44:42] Dispensed 0.000071 mL of Base (0.5 M KOH)
[44:47] Stepping pH = 9.58
[44:47] Dispensed 0.000353 mL of Base (0.5 M KOH)
[44:52] Stepping pH = 9.97
[45:07] Stirrer speed set to 0
[45:21] Datapoint id 44 collected
[45:21] Charge balance equation is out by -201.1%
[45:21] Stirrer speed set to 55
[45:26] pH 9.95 -> 10.05
[45:26] Using cautious pH adjust
[45:26] Dispensed 0.000071 mL of Base (0.5 M KOH)
[45:31] Stepping pH = 9.94
[45:31] Dispensed 0.000329 mL of Base (0.5 M KOH)
[45:36] Stepping pH = 10.17
[45:51] Stirrer speed set to 0
[46:01] Datapoint id 45 collected
[46:01] Charge balance equation is out by -210.8%
[46:01] Titration 3 of 3
[46:01] Adding initial titrants

| | | | |
|---|---|---|---|
| Sample name: | **M07_octanol** | Experiment start time: | **2/28/2018 4:26:20 PM** |
| Assay name: | **pH-metric high logP** | Analyst: | **Pion** |
| Assay ID: | **18B-28011** | Instrument ID: | **T312060** |
| Filename: | **C:\Sirius_T3\Mehtap\20180228_exp28_logP_T3-2\18B-28011_M07_octanol_pH-metric high logP.t3r** | | |

#### Experiment Log (continued)

[46:01] Automatically add 0.25000 mL of Octanol
[46:07] Dispensed 0.250000 mL of Octanol
[46:07] Stirrer speed set to 10
[46:08] Stirrer speed set to 60
[46:08] Iterative adjust 10.17 -> 2.00
[46:08] pH 10.17 -> 2.00
[46:10] Dispensed 0.055715 mL of Acid (0.5 M HCl)
[46:15] pH 2.03 -> 2.00
[46:15] Dispensed 0.003457 mL of Acid (0.5 M HCl)
[47:05] Stirrer speed set to 0
[47:15] Datapoint id 46 collected
[47:15] Stirrer speed set to 60
[47:20] pH 1.96 -> 2.16
[47:20] Using cautious pH adjust
[47:21] Dispensed 0.010113 mL of Base (0.5 M KOH)
[47:26] Stepping pH = 2.02
[47:26] Dispensed 0.010865 mL of Base (0.5 M KOH)
[47:31] Stepping pH = 2.09
[47:31] Dispensed 0.006515 mL of Base (0.5 M KOH)
[47:36] Stepping pH = 2.18
[47:51] Stirrer speed set to 0
[48:01] Datapoint id 47 collected
[48:01] Charge balance equation is out by -35.9%
[48:01] Stirrer speed set to 60
[48:06] pH 2.19 -> 2.39
[48:06] Using cautious pH adjust
[48:07] Dispensed 0.005903 mL of Base (0.5 M KOH)
[48:12] Stepping pH = 2.29
[48:12] Dispensed 0.003481 mL of Base (0.5 M KOH)
[48:17] Stepping pH = 2.36
[48:17] Dispensed 0.001435 mL of Base (0.5 M KOH)
[48:22] Stepping pH = 2.39
[48:37] Stirrer speed set to 0
[48:47] Datapoint id 48 collected
[48:47] Charge balance equation is out by 8.3%
[48:47] Stirrer speed set to 60
[48:52] pH 2.39 -> 2.59
[48:52] Using charge balance adjust
[48:53] Dispensed 0.007502 mL of Base (0.5 M KOH)
[49:13] Stirrer speed set to 0
[49:23] Datapoint id 49 collected
[49:23] Charge balance equation is out by 9.3%
[49:23] Stirrer speed set to 60
[49:28] pH 2.62 -> 2.82
[49:28] Using charge balance adjust
[49:28] Dispensed 0.004657 mL of Base (0.5 M KOH)
[49:48] Stirrer speed set to 0
[49:59] Datapoint id 50 collected
[49:59] Charge balance equation is out by 13.0%
[49:59] Stirrer speed set to 60
[50:04] pH 2.85 -> 3.05
[50:04] Using charge balance adjust
[50:04] Dispensed 0.002987 mL of Base (0.5 M KOH)
[50:24] Stirrer speed set to 0
[50:34] Datapoint id 51 collected
[50:34] Charge balance equation is out by 15.0%
[50:34] Stirrer speed set to 60
[50:39] pH 3.08 -> 3.28
[50:39] Using cautious pH adjust

| | | | |
|---|---|---|---|
| Sample name: | **M07_octanol** | Experiment start time: | **2/28/2018 4:26:20 PM** |
| Assay name: | **pH-metric high logP** | Analyst: | **Pion** |
| Assay ID: | **18B-28011** | Instrument ID: | **T312060** |
| Filename: | **C:\Sirius_T3\Mehtap\20180228_exp28_logP_T3-2\18B-28011_M07_octanol_pH-metric high logP.t3r** | | |

#### Experiment Log (continued)

[50:39] Dispensed 0.001058 mL of Base (0.5 M KOH)
[50:44] Stepping pH = 3.17
[50:44] Dispensed 0.000823 mL of Base (0.5 M KOH)
[50:50] Stepping pH = 3.24
[50:50] Dispensed 0.000329 mL of Base (0.5 M KOH)
[50:55] Stepping pH = 3.27
[51:10] Stirrer speed set to 0
[51:20] Datapoint id 52 collected
[51:20] Charge balance equation is out by -4.5%
[51:20] Stirrer speed set to 60
[51:25] pH 3.28 -> 3.48
[51:25] Using charge balance adjust
[51:25] Dispensed 0.001717 mL of Base (0.5 M KOH)
[51:45] Stirrer speed set to 0
[51:55] Datapoint id 53 collected
[51:55] Charge balance equation is out by -5.2%
[51:55] Stirrer speed set to 60
[52:00] pH 3.47 -> 3.67
[52:00] Using charge balance adjust
[52:00] Dispensed 0.001458 mL of Base (0.5 M KOH)
[52:21] Stirrer speed set to 0
[52:31] Datapoint id 54 collected
[52:31] Charge balance equation is out by -0.7%
[52:31] Stirrer speed set to 60
[52:36] pH 3.68 -> 3.88
[52:36] Using charge balance adjust
[52:36] Dispensed 0.001246 mL of Base (0.5 M KOH)
[52:56] Stirrer speed set to 0
[53:06] Datapoint id 55 collected
[53:06] Charge balance equation is out by 3.3%
[53:06] Stirrer speed set to 60
[53:11] pH 3.89 -> 4.09
[53:11] Using charge balance adjust
[53:11] Dispensed 0.001011 mL of Base (0.5 M KOH)
[53:31] Stirrer speed set to 0
[53:41] Datapoint id 56 collected
[53:41] Charge balance equation is out by -1.8%
[53:41] Stirrer speed set to 60
[53:47] pH 4.10 -> 4.30
[53:47] Using charge balance adjust
[53:47] Dispensed 0.000800 mL of Base (0.5 M KOH)
[54:07] Stirrer speed set to 0
[54:17] Datapoint id 57 collected
[54:17] Charge balance equation is out by -9.4%
[54:17] Stirrer speed set to 60
[54:22] pH 4.29 -> 4.49
[54:22] Using charge balance adjust
[54:22] Dispensed 0.000588 mL of Base (0.5 M KOH)
[54:42] Stirrer speed set to 0
[54:52] Datapoint id 58 collected
[54:52] Charge balance equation is out by -20.0%
[54:52] Stirrer speed set to 60
[54:57] pH 4.45 -> 4.65
[54:57] Using cautious pH adjust
[54:57] Dispensed 0.000235 mL of Base (0.5 M KOH)
[55:02] Stepping pH = 4.52
[55:02] Dispensed 0.000235 mL of Base (0.5 M KOH)
[55:08] Stepping pH = 4.61
[55:08] Dispensed 0.000094 mL of Base (0.5 M KOH)

Sample name: **M07_octanol**
Assay name: **pH-metric high logP**
Assay ID: **18B-28011**
Filename: **C:\Sirius_T3\Mehtap\20180228_exp28_logP_T3-2\18B-28011_M07_octanol_pH-metric high logP.t3r**
Experiment start time: **2/28/2018 4:26:20 PM**
Analyst: **Pion**
Instrument ID: **T312060**

#### Experiment Log (continued)

[55:13] Stepping pH = 4.62
[55:13] Dispensed 0.000118 mL of Base (0.5 M KOH)
[55:18] Stepping pH = 4.67
[55:33] Stirrer speed set to 0
[55:43] Datapoint id 59 collected
[55:43] Charge balance equation is out by -49.9%
[55:43] Stirrer speed set to 60
[55:49] pH 4.68 -> 4.88
[55:49] Using cautious pH adjust
[55:49] Dispensed 0.000141 mL of Base (0.5 M KOH)
[55:54] Stepping pH = 4.75
[55:54] Dispensed 0.000165 mL of Base (0.5 M KOH)
[55:59] Stepping pH = 4.86
[55:59] Dispensed 0.000024 mL of Base (0.5 M KOH)
[56:04] Stepping pH = 4.86
[56:04] Dispensed 0.000094 mL of Base (0.5 M KOH)
[56:09] Stepping pH = 4.92
[56:24] Stirrer speed set to 0
[56:35] Datapoint id 60 collected
[56:35] Charge balance equation is out by -46.6%
[56:35] Stirrer speed set to 60
[56:40] pH 4.93 -> 5.13
[56:40] Using cautious pH adjust
[56:40] Dispensed 0.000094 mL of Base (0.5 M KOH)
[56:45] Stepping pH = 4.97
[56:45] Dispensed 0.000165 mL of Base (0.5 M KOH)
[56:50] Stepping pH = 5.15
[57:05] Stirrer speed set to 0
[57:16] Datapoint id 61 collected
[57:16] Charge balance equation is out by -38.7%
[57:16] Stirrer speed set to 60
[57:21] pH 5.16 -> 5.36
[57:21] Using cautious pH adjust
[57:21] Dispensed 0.000047 mL of Base (0.5 M KOH)
[57:27] Stepping pH = 5.17
[57:27] Dispensed 0.000165 mL of Base (0.5 M KOH)
[57:32] Stepping pH = 5.35
[57:47] Stirrer speed set to 0
[57:58] Datapoint id 62 collected
[57:58] Charge balance equation is out by -87.8%
[57:58] Stirrer speed set to 60
[58:03] pH 5.38 -> 5.58
[58:03] Using cautious pH adjust
[58:03] Dispensed 0.000047 mL of Base (0.5 M KOH)
[58:09] Stepping pH = 5.41
[58:09] Dispensed 0.000094 mL of Base (0.5 M KOH)
[58:14] Stepping pH = 5.54
[58:14] Dispensed 0.000024 mL of Base (0.5 M KOH)
[58:19] Stepping pH = 5.57
[58:34] Stirrer speed set to 0
[58:47] Datapoint id 63 collected
[58:47] Charge balance equation is out by -84.7%
[58:47] Stirrer speed set to 60
[58:52] pH 5.63 -> 5.83
[58:52] Using cautious pH adjust
[58:52] Dispensed 0.000024 mL of Base (0.5 M KOH)
[58:57] Stepping pH = 5.64
[58:57] Dispensed 0.000094 mL of Base (0.5 M KOH)
[59:02] Stepping pH = 5.74

Sample name: **M07_octanol**
Assay name: **pH-metric high logP**
Assay ID: **18B-28011**
Filename: **C:\Sirius_T3\Mehtap\20180228_exp28_logP_T3-2\18B-28011_M07_octanol_pH-metric high logP.t3r**

Experiment start time: **2/28/2018 4:26:20 PM**
Analyst: **Pion**
Instrument ID: **T312060**

#### Experiment Log (continued)

[59:02] Dispensed 0.000047 mL of Base (0.5 M KOH)
[59:07] Stepping pH = 5.86
[59:22] Stirrer speed set to 0
[59:58] Datapoint id 64 collected
[59:58] Charge balance equation is out by -167.1%
[59:58] Stirrer speed set to 60
[1:00:03] pH 5.99 -> 6.19
[1:00:03] Using cautious pH adjust
[1:00:03] Dispensed 0.000024 mL of Base (0.5 M KOH)
[1:00:08] Stepping pH = 6.02
[1:00:08] Dispensed 0.000047 mL of Base (0.5 M KOH)
[1:00:13] Stepping pH = 6.15
[1:00:13] Dispensed 0.000024 mL of Base (0.5 M KOH)
[1:00:18] Stepping pH = 6.30
[1:00:33] Stirrer speed set to 0
[1:01:33] Datapoint id 65 collected
[1:01:33] Charge balance equation is out by -96.8%
[1:01:33] Stirrer speed set to 60
[1:01:38] pH 6.35 -> 6.55
[1:01:38] Using cautious pH adjust
[1:01:38] Dispensed 0.000024 mL of Base (0.5 M KOH)
[1:01:43] Stepping pH = 6.40
[1:01:43] Dispensed 0.000047 mL of Base (0.5 M KOH)
[1:01:48] Stepping pH = 6.63
[1:02:04] Stirrer speed set to 0
[1:03:04] Datapoint id 66 collected
[1:03:04] Charge balance equation is out by -27.0%
[1:03:04] Stirrer speed set to 60
[1:03:09] pH 6.73 -> 6.93
[1:03:09] Using cautious pH adjust
[1:03:09] Dispensed 0.000024 mL of Base (0.5 M KOH)
[1:03:14] Stepping pH = 6.79
[1:03:14] Dispensed 0.000024 mL of Base (0.5 M KOH)
[1:03:19] Stepping pH = 7.01
[1:03:34] Stirrer speed set to 0
[1:04:34] Datapoint id 67 collected
[1:04:34] Charge balance equation is out by -13.9%
[1:04:34] Stirrer speed set to 60
[1:04:39] pH 7.04 -> 7.24
[1:04:39] Using charge balance adjust
[1:04:39] Dispensed 0.000024 mL of Base (0.5 M KOH)
[1:04:59] Stirrer speed set to 0
[1:05:59] Datapoint id 68 collected
[1:05:59] Charge balance equation is out by -70.5%
[1:05:59] Stirrer speed set to 60
[1:06:05] pH 7.21 -> 7.41
[1:06:05] Using cautious pH adjust
[1:06:05] Dispensed 0.000024 mL of Base (0.5 M KOH)
[1:06:10] Stepping pH = 7.21
[1:06:10] Dispensed 0.000024 mL of Base (0.5 M KOH)
[1:06:15] Stepping pH = 7.44
[1:06:30] Stirrer speed set to 0
[1:07:30] Datapoint id 69 collected
[1:07:30] Charge balance equation is out by -155.1%
[1:07:30] Stirrer speed set to 60
[1:07:35] pH 7.73 -> 7.93
[1:07:35] Using cautious pH adjust
[1:07:35] Dispensed 0.000024 mL of Base (0.5 M KOH)
[1:07:40] Stepping pH = 7.83

| | | | |
|---|---|---|---|
| Sample name: | **M07_octanol** | Experiment start time: | **2/28/2018 4:26:20 PM** |
| Assay name: | **pH-metric high logP** | Analyst: | **Pion** |
| Assay ID: | **18B-28011** | Instrument ID: | **T312060** |
| Filename: | **C:\Sirius_T3\Mehtap\20180228_exp28_logP_T3-2\18B-28011_M07_octanol_pH-metric high logP.t3r** | | |

#### Experiment Log (continued)

[1:07:40] Dispensed 0.000024 mL of Base (0.5 M KOH)
[1:07:45] Stepping pH = 8.04
[1:08:00] Stirrer speed set to 0
[1:09:00] Datapoint id 70 collected
[1:09:00] Charge balance equation is out by -346.1%
[1:09:00] Stirrer speed set to 60
[1:09:06] pH 7.76 -> 7.96
[1:09:06] Using cautious pH adjust
[1:09:06] Dispensed 0.000024 mL of Base (0.5 M KOH)
[1:09:11] Stepping pH = 7.79
[1:09:11] Dispensed 0.000024 mL of Base (0.5 M KOH)
[1:09:16] Stepping pH = 8.01
[1:09:31] Stirrer speed set to 0
[1:10:31] Datapoint id 71 collected
[1:10:31] Charge balance equation is out by -354.4%
[1:10:31] Stirrer speed set to 60
[1:10:36] pH 8.11 -> 8.31
[1:10:36] Using cautious pH adjust
[1:10:36] Dispensed 0.000024 mL of Base (0.5 M KOH)
[1:10:41] Stepping pH = 8.17
[1:10:41] Dispensed 0.000024 mL of Base (0.5 M KOH)
[1:10:46] Stepping pH = 8.19
[1:10:46] Dispensed 0.000024 mL of Base (0.5 M KOH)
[1:10:51] Stepping pH = 8.17
[1:10:51] Dispensed 0.000094 mL of Base (0.5 M KOH)
[1:10:57] Stepping pH = 8.47
[1:11:12] Stirrer speed set to 0
[1:12:12] Datapoint id 72 collected
[1:12:12] Charge balance equation is out by -1,435.6%
[1:12:12] Stirrer speed set to 60
[1:12:17] pH 8.43 -> 8.63
[1:12:17] Using cautious pH adjust
[1:12:17] Dispensed 0.000024 mL of Base (0.5 M KOH)
[1:12:22] Stepping pH = 8.42
[1:12:22] Dispensed 0.000047 mL of Base (0.5 M KOH)
[1:12:27] Stepping pH = 8.45
[1:12:27] Dispensed 0.000094 mL of Base (0.5 M KOH)
[1:12:32] Stepping pH = 8.81
[1:12:47] Stirrer speed set to 0
[1:13:08] Datapoint id 73 collected
[1:13:08] Charge balance equation is out by -948.3%
[1:13:08] Stirrer speed set to 60
[1:13:13] pH 8.83 -> 9.03
[1:13:13] Using cautious pH adjust
[1:13:13] Dispensed 0.000024 mL of Base (0.5 M KOH)
[1:13:18] Stepping pH = 8.84
[1:13:18] Dispensed 0.000047 mL of Base (0.5 M KOH)
[1:13:23] Stepping pH = 8.94
[1:13:23] Dispensed 0.000024 mL of Base (0.5 M KOH)
[1:13:28] Stepping pH = 9.00
[1:13:28] Dispensed 0.000024 mL of Base (0.5 M KOH)
[1:13:33] Stepping pH = 9.04
[1:13:49] Stirrer speed set to 0
[1:14:02] Datapoint id 74 collected
[1:14:02] Charge balance equation is out by -262.5%
[1:14:02] Stirrer speed set to 60
[1:14:07] pH 9.05 -> 9.25
[1:14:07] Using cautious pH adjust
[1:14:07] Dispensed 0.000024 mL of Base (0.5 M KOH)

Sample name: **M07_octanol**
Assay name: **pH-metric high logP**
Assay ID: **18B-28011**
Filename: **C:\Sirius_T3\Mehtap\20180228_exp28_logP_T3-2\18B-28011_M07_octanol_pH-metric high logP.t3r**
Experiment start time: **2/28/2018 4:26:20 PM**
Analyst: **Pion**
Instrument ID: **T312060**

#### Experiment Log (continued)

[1:14:13] Stepping pH = 9.04
[1:14:13] Dispensed 0.000141 mL of Base (0.5 M KOH)
[1:14:18] Stepping pH = 9.33
[1:14:33] Stirrer speed set to 0
[1:14:50] Datapoint id 75 collected
[1:14:50] Charge balance equation is out by -206.3%
[1:14:50] Stirrer speed set to 60
[1:14:55] pH 9.34 -> 9.54
[1:14:55] Using cautious pH adjust
[1:14:56] Dispensed 0.000047 mL of Base (0.5 M KOH)
[1:15:01] Stepping pH = 9.37
[1:15:01] Dispensed 0.000118 mL of Base (0.5 M KOH)
[1:15:06] Stepping pH = 9.53
[1:15:21] Stirrer speed set to 0
[1:15:31] Datapoint id 76 collected
[1:15:31] Charge balance equation is out by -78.6%
[1:15:31] Stirrer speed set to 60
[1:15:37] pH 9.54 -> 9.74
[1:15:37] Using cautious pH adjust
[1:15:37] Dispensed 0.000071 mL of Base (0.5 M KOH)
[1:15:42] Stepping pH = 9.57
[1:15:42] Dispensed 0.000141 mL of Base (0.5 M KOH)
[1:15:47] Stepping pH = 9.72
[1:15:47] Dispensed 0.000024 mL of Base (0.5 M KOH)
[1:15:52] Stepping pH = 9.73
[1:16:07] Stirrer speed set to 0
[1:16:18] Datapoint id 77 collected
[1:16:18] Charge balance equation is out by -62.5%
[1:16:18] Stirrer speed set to 60
[1:16:23] pH 9.73 -> 9.93
[1:16:23] Using cautious pH adjust
[1:16:23] Dispensed 0.000094 mL of Base (0.5 M KOH)
[1:16:28] Stepping pH = 9.78
[1:16:28] Dispensed 0.000165 mL of Base (0.5 M KOH)
[1:16:33] Stepping pH = 9.93
[1:16:48] Stirrer speed set to 0
[1:16:59] Datapoint id 78 collected
[1:16:59] Charge balance equation is out by -29.9%
[1:16:59] Stirrer speed set to 60
[1:17:04] pH 9.92 -> 10.05
[1:17:04] Using cautious pH adjust
[1:17:04] Dispensed 0.000094 mL of Base (0.5 M KOH)
[1:17:09] Stepping pH = 9.95
[1:17:09] Dispensed 0.000141 mL of Base (0.5 M KOH)
[1:17:14] Stepping pH = 10.04
[1:17:29] Stirrer speed set to 0
[1:17:42] Datapoint id 79 collected
[1:17:42] Charge balance equation is out by -36.7%
[1:17:42] Argon flow rate set to 0
[1:17:46] Titrator arm moved over Titration position
