## Supplementary material for "Octanol-water partition coefficient measurements for the SAMPL6 Blind Prediction Challenge": SM07_18B-28012_M07_octanol_pH-metric high logP_report.pdf

| | | | |
|---|---|---|---|
| Sample name: | **M07_octanol** | Experiment start time: | **2/28/2018 5:44:56 PM** |
| Assay name: | **pH-metric high logP** | Analyst: | **Pion** |
| Assay ID: | **18B-28012** | Instrument ID: | **T312060** |
| Filename: | **C:\Sirius_T3\Mehtap\20180228_exp28_logP_T3-2\18B-28012_M07_octanol_pH-metric high logP.t3r** | | |

#### pH-metric Result

| | | |
|---|---|---|
| logP (XH +) | -5.97 | ±0.89 (n=49) |
| logP (neutral X) | 3.14 | ±0.01 (n=49) |

#### 18B-28012 Points 1 to 30

| | |
|---|---|
| M07_octanol concentration factor | 0.885 |
| Carbonate | 0.0000 mM |
| Acidity error | 0.24941 mM |

#### 18B-28012 Points 31 to 62

| | |
|---|---|
| M07_octanol concentration factor | 0.786 |
| Carbonate | 0.0623 mM |
| Acidity error | 0.05292 mM |

#### 18B-28012 Points 63 to 95

| | |
|---|---|
| M07_octanol concentration factor | 0.842 |
| Carbonate | 0.0998 mM |
| Acidity error | 0.05919 mM |

#### Warnings and errors

| | |
|---|---|
| Errors | None |
| Warnings | One or more logP values out of range |

#### Sample logD and percent species

| pH | M07_octanol logD | M07_octanol M07_octanolH | M07_octanol M07_octanol | M07_octanol M07_octanolH* | M07_octanol M07_octanol* | Comment |
|---|---|---|---|---|---|---|
| 1.000 | -1.93 | 98.83 % | 0.00 % | 0.00 % | 1.16 % | |
| 1.200 | -1.73 | 98.17 % | 0.00 % | 0.00 % | 1.83 % | Stomach pH |
| 2.000 | -0.93 | 89.45 % | 0.01 % | 0.00 % | 10.54 % | |
| 3.000 | 0.07 | 45.89 % | 0.04 % | 0.00 % | 54.07 % | |
| 4.000 | 1.07 | 7.82 % | 0.07 % | 0.00 % | 92.12 % | |
| 5.000 | 2.04 | 0.84 % | 0.07 % | 0.00 % | 99.09 % | |
| 6.000 | 2.80 | 0.08 % | 0.07 % | 0.00 % | 99.84 % | |
| 6.500 | 3.00 | 0.03 % | 0.07 % | 0.00 % | 99.90 % | |
| 7.000 | 3.09 | 0.01 % | 0.07 % | 0.00 % | 99.92 % | |
| 7.400 | 3.12 | 0.00 % | 0.07 % | 0.00 % | 99.92 % | Blood pH |
| 8.000 | 3.14 | 0.00 % | 0.07 % | 0.00 % | 99.93 % | |
| 9.000 | 3.14 | 0.00 % | 0.07 % | 0.00 % | 99.93 % | |
| 10.000 | 3.14 | 0.00 % | 0.07 % | 0.00 % | 99.93 % | |
| 11.000 | 3.14 | 0.00 % | 0.07 % | 0.00 % | 99.93 % | |
| 12.000 | 3.14 | 0.00 % | 0.07 % | 0.00 % | 99.93 % | |

| | | | |
|---|---|---|---|
| Sample name: | **M07_octanol** | Experiment start time: | **2/28/2018 5:44:56 PM** |
| Assay name: | **pH-metric high logP** | Analyst: | **Pion** |
| Assay ID: | **18B-28012** | Instrument ID: | **T312060** |
| Filename: | **C:\Sirius_T3\Mehtap\20180228_exp28_logP_T3-2\18B-28012_M07_octanol_pH-metric high logP.t3r** | | |

#### Graphs

Charge balance residuals

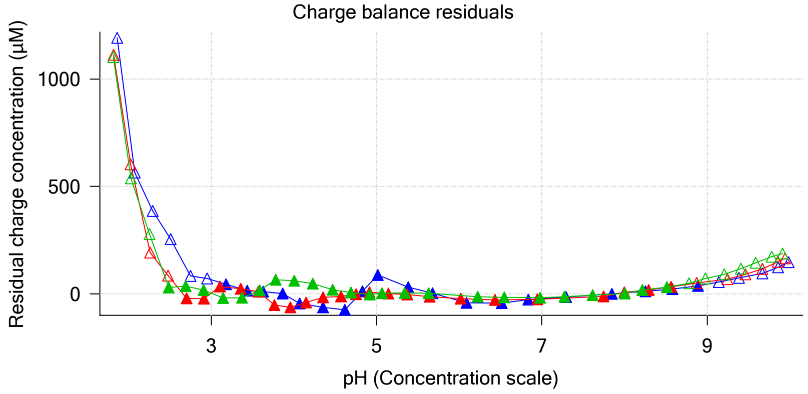


Ionic strength

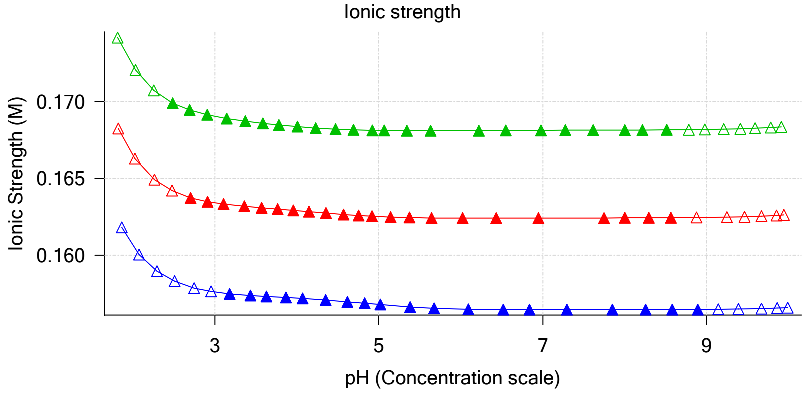


Buffer index profile

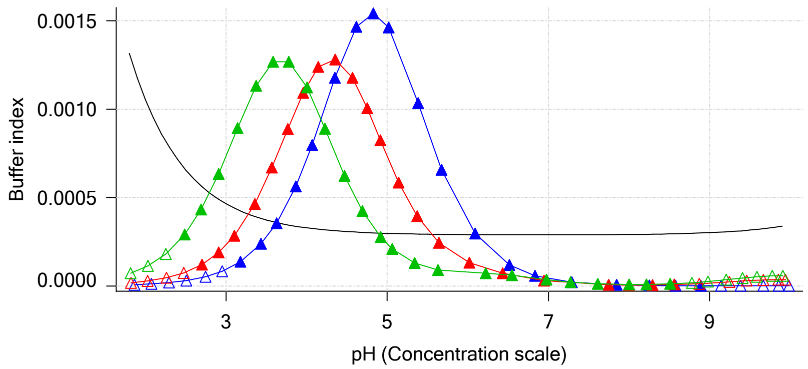


Sample M07_octanol Ionisation Graph

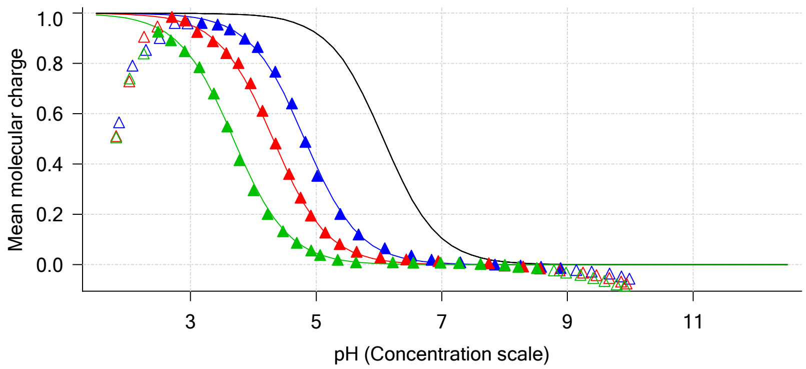


Distribution of species for sample M07_octanol

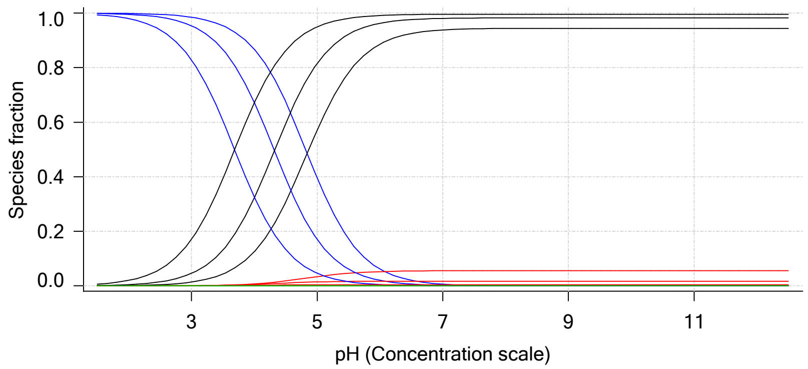


Lipophilicity Profile

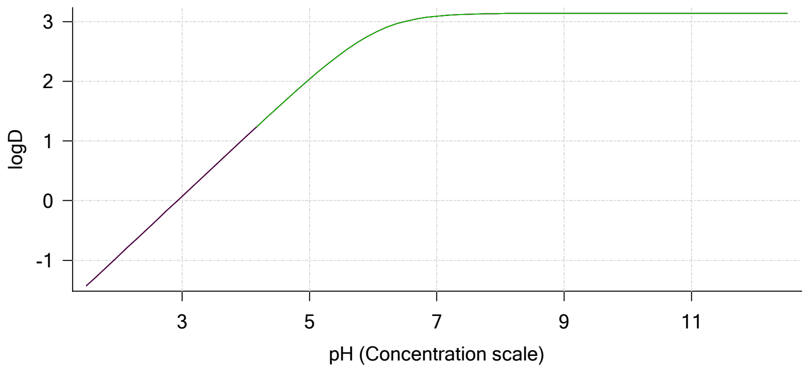


Log R Profile

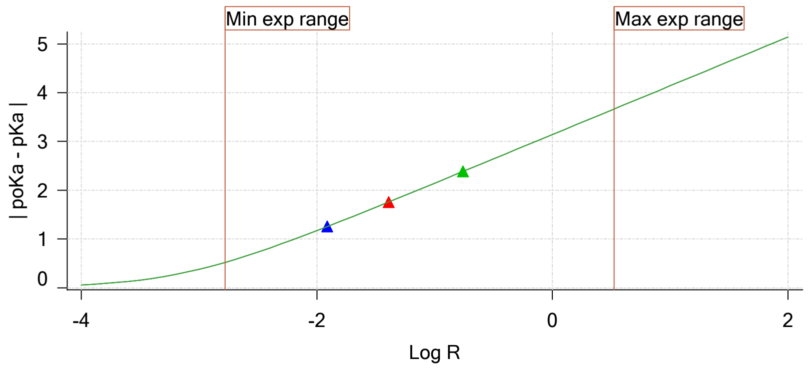


Sample Carbonate Ionisation Graph

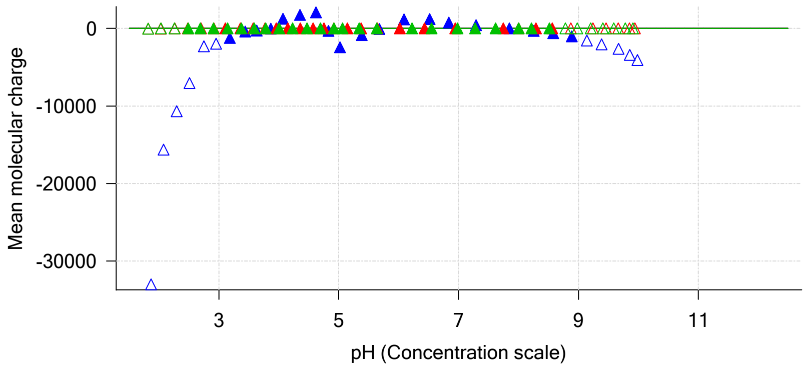

| | | | |
|---|---|---|---|
| Sample name: | **M07_octanol** | Experiment start time: | **2/28/2018 5:44:56 PM** |
| Assay name: | **pH-metric high logP** | Analyst: | **Pion** |
| Assay ID: | **18B-28012** | Instrument ID: | **T312060** |
| Filename: | **C:\Sirius_T3\Mehtap\20180228_exp28_logP_T3-2\18B-28012_M07_octanol_pH-metric high logP.t3r** | | |

#### Graphs (continued)

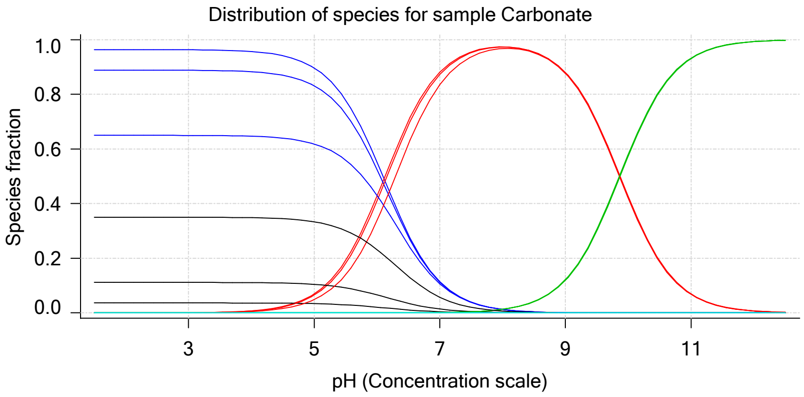

| | |
|---|---|
| Sample name: | **M07_octanol** |
| Assay name: | **pH-metric high logP** |
| Assay ID: | **18B-28012** |
| Filename: | **C:\Sirius_T3\Mehtap\20180228_exp28_logP_T3-2\18B-28012_M07_octanol_pH-metric high logP.t3r** |
| Experiment start time: | **2/28/2018 5:44:56 PM** |
| Analyst: | **Pion** |
| Instrument ID: | **T312060** |

### pH-metric high logP Titration 1 of 3 18B-28012 Points 1 to 30

#### Overall results

| | |
|---|---|
| RMSD | 0.538 |
| Average ionic strength | 0.157 M |
| Average temperature | 25.0°C |
| Partition ratio | 0.0122 : 1 |
| Analyte concentration range | 2983.5 µM to 3075.2 µM |
| Total points considered | 19 of 30 |

#### Warnings and errors

| | |
|---|---|
| Errors | None |
| Warnings | None |

#### Four-Plus parameters

| | | | |
|---|---|---|---|
| Alpha | 0.130 | 2/28/2018 5:44:56 PM | C:\Sirius_T3\HCl18B27.t3r |
| S | 0.9970 | 2/28/2018 5:44:56 PM | C:\Sirius_T3\HCl18B27.t3r |
| jH | 0.8 | 2/28/2018 5:44:56 PM | C:\Sirius_T3\HCl18B27.t3r |
| jOH | -0.4 | 2/28/2018 5:44:56 PM | C:\Sirius_T3\HCl18B27.t3r |

#### Titrants

| | | | |
|---|---|---|---|
| 0.50 M HCl | 0.993513 | 2/28/2018 5:44:56 PM | C:\Sirius_T3\HCl18B27.t3r |
| 0.50 M KOH | 0.999845 | 2/28/2018 5:44:56 PM | C:\Sirius_T3\KOH18B27.t3r |

#### Sample

| | |
|---|---|
| M07_octanol concentration factor | 0.885 |
| Base pKa 1 | 6.07 |
| logP (XH +) | 0.46 |
| logP (neutral X) | 3.15 |

#### Sample graphs

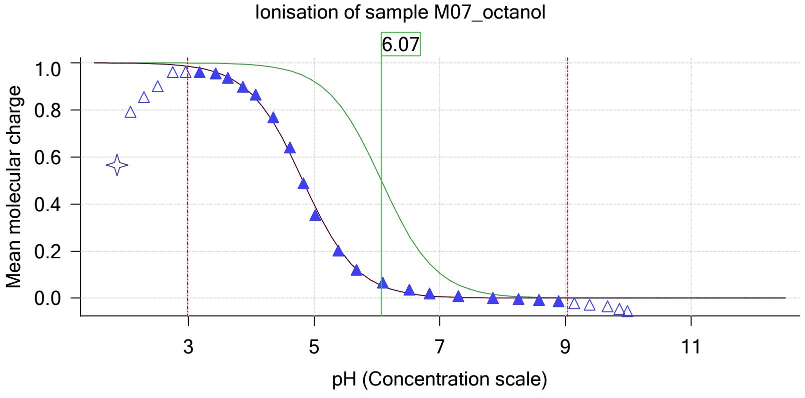


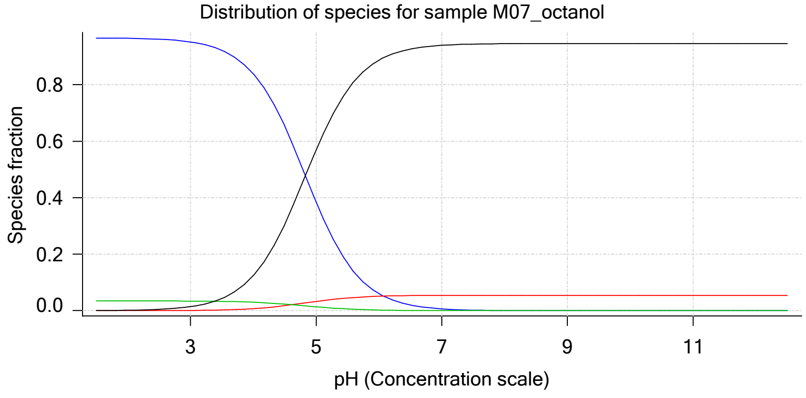

| | | | |
|---|---|---|---|
| Sample name: | **M07_octanol** | Experiment start time: | **2/28/2018 5:44:56 PM** |
| Assay name: | **pH-metric high logP** | Analyst: | **Pion** |
| Assay ID: | **18B-28012** | Instrument ID: | **T312060** |
| Filename: | **C:\Sirius_T3\Mehtap\20180228_exp28_logP_T3-2\18B-28012_M07_octanol_pH-metric high logP.t3r** | | |

#### Sample graphs (continued)

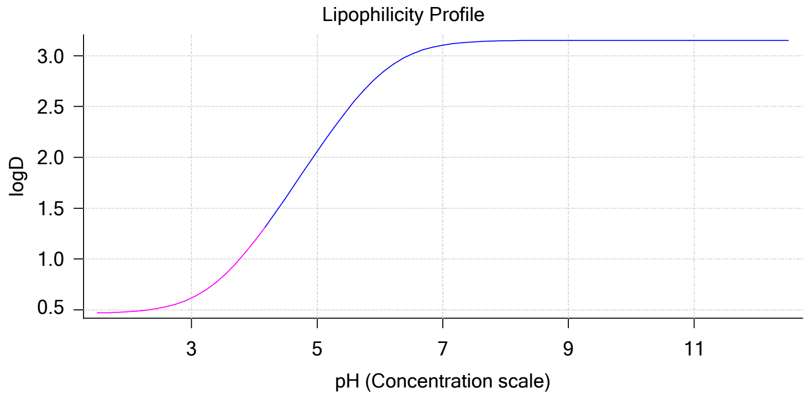


#### Sample logD and percent species

| pH | M07_octanol logD | M07_octanol M07_octanolH | M07_octanol M07_octanol | M07_octanol M07_octanolH* | M07_octanol M07_octanol* | Comment |
|---|---|---|---|---|---|---|
| 1.000 | 0.46 | 96.55 % | 0.00 % | 3.43 % | 0.01 % | |
| 1.200 | 0.47 | 96.55 % | 0.00 % | 3.43 % | 0.02 % | Stomach pH |
| 2.000 | 0.48 | 96.42 % | 0.01 % | 3.43 % | 0.14 % | |
| 3.000 | 0.61 | 95.13 % | 0.08 % | 3.38 % | 1.41 % | |
| 4.000 | 1.17 | 83.88 % | 0.71 % | 2.98 % | 12.42 % | |
| 5.000 | 2.06 | 38.44 % | 3.27 % | 1.37 % | 56.92 % | |
| 6.000 | 2.82 | 5.99 % | 5.10 % | 0.21 % | 88.70 % | |
| 6.500 | 3.02 | 1.98 % | 5.32 % | 0.07 % | 92.63 % | |
| 7.000 | 3.10 | 0.63 % | 5.40 % | 0.02 % | 93.94 % | |
| 7.400 | 3.13 | 0.25 % | 5.42 % | 0.01 % | 94.32 % | Blood pH |
| 8.000 | 3.15 | 0.06 % | 5.43 % | 0.00 % | 94.50 % | |
| 9.000 | 3.15 | 0.01 % | 5.43 % | 0.00 % | 94.56 % | |
| 10.000 | 3.15 | 0.00 % | 5.44 % | 0.00 % | 94.56 % | |
| 11.000 | 3.15 | 0.00 % | 5.44 % | 0.00 % | 94.56 % | |
| 12.000 | 3.15 | 0.00 % | 5.44 % | 0.00 % | 94.56 % | |

#### Carbonate and acidity

| | |
|---|---|
| Carbonate | 0.000 mM |
| Acidity error | 0.249 mM |

#### Other graphs

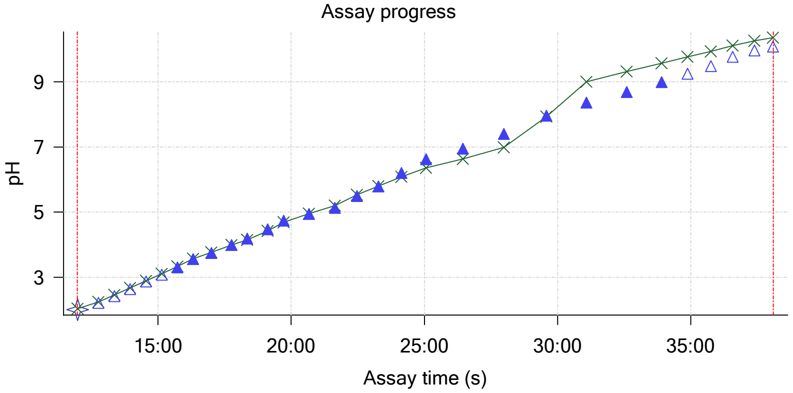


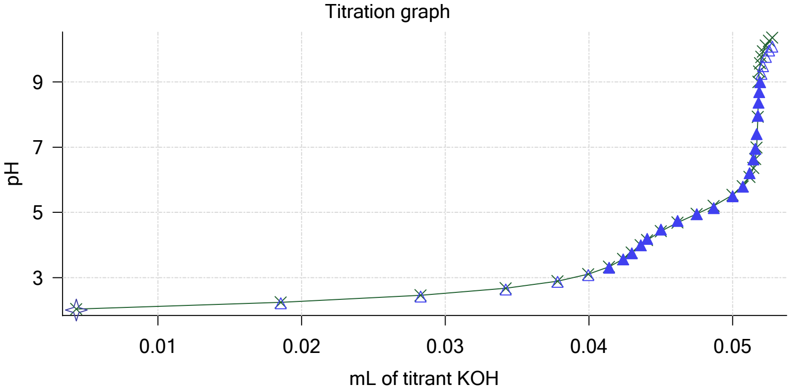

Sample name: **M07_octanol**
Assay name: **pH-metric high logP**
Assay ID: **18B-28012**
Filename: **C:\Sirius_T3\Mehtap\20180228_exp28_logP_T3-2\18B-28012_M07_octanol_pH-metric high logP.t3r**

Experiment start time: **2/28/2018 5:44:56 PM**
Analyst: **Pion**
Instrument ID: **T312060**

#### Other graphs (continued)

Charge balance residuals

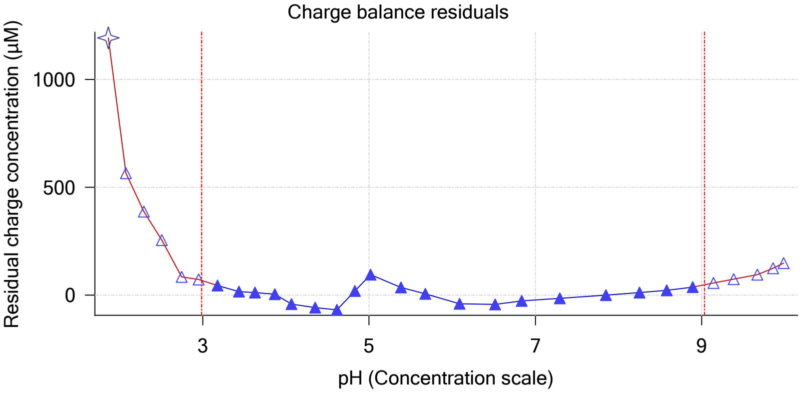


Buffer index profile

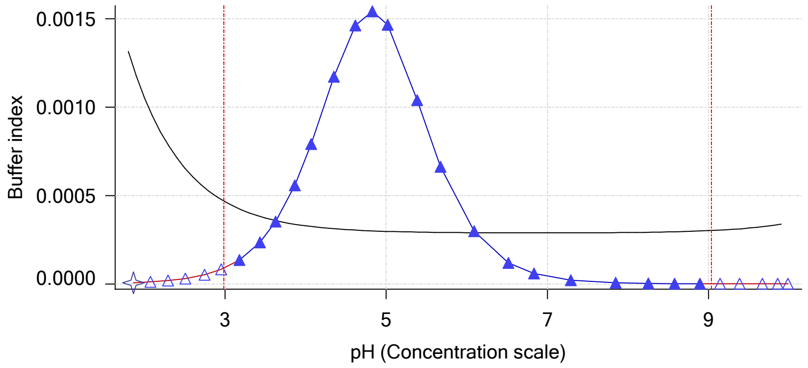


Ionic strength

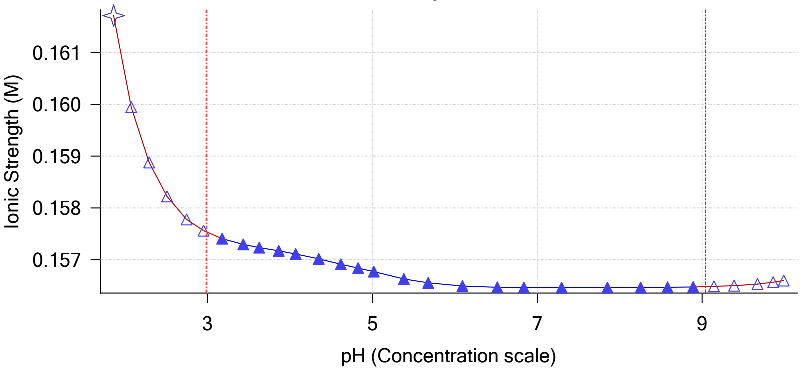


Log R Profile

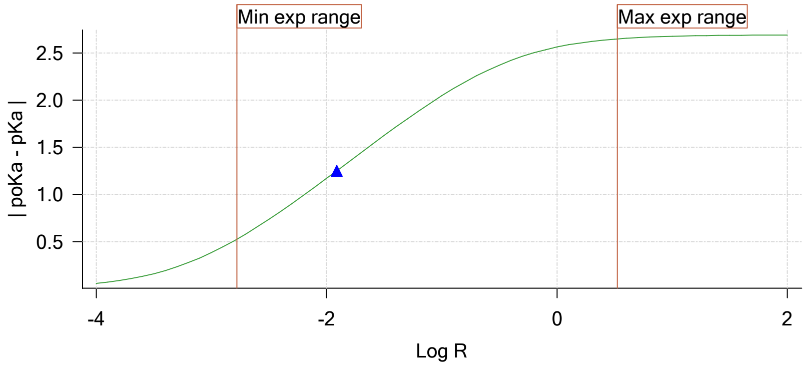


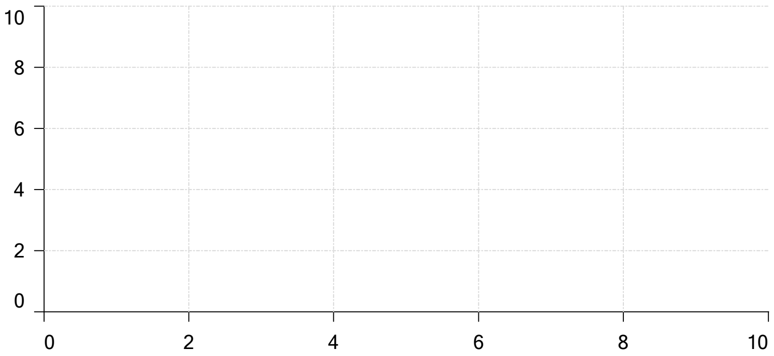


Ionisation of sample Carbonate

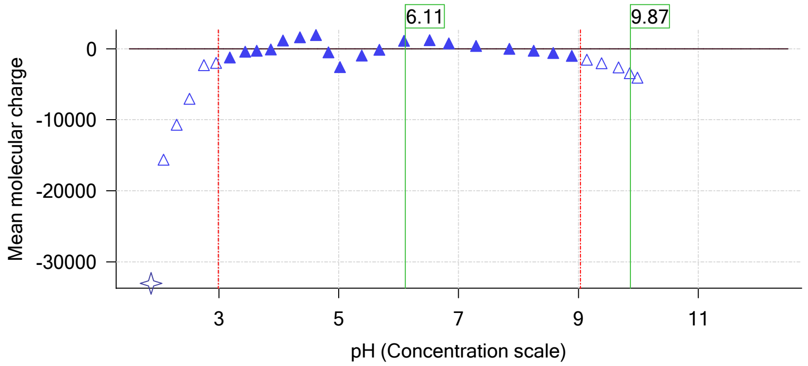

| | | | |
|---|---|---|---|
| Sample name: | **M07_octanol** | Experiment start time: | **2/28/2018 5:44:56 PM** |
| Assay name: | **pH-metric high logP** | Analyst: | **Pion** |
| Assay ID: | **18B-28012** | Instrument ID: | **T312060** |
| Filename: | **C:\Sirius_T3\Mehtap\20180228_exp28_logP_T3-2\18B-28012_M07_octanol_pH-metric high logP.t3r** | | |

# pH-metric high logP   Titration 2 of 3   18B-28012 Points 31 to 62

#### Overall results

| | |
|---|---|
| RMSD | 0.578 |
| Average ionic strength | 0.163 M |
| Average temperature | 25.0°C |
| Partition ratio | 0.0406 : 1 |
| Analyte concentration range | 2713.2 µM to 2800.4 µM |
| Total points considered | 22 of 32 |

#### Warnings and errors

| | |
|---|---|
| Errors | None |
| Warnings | None |

#### Four-Plus parameters

| | | | |
|---|---|---|---|
| Alpha | 0.130 | 2/28/2018 5:44:56 PM | C:\Sirius_T3\HCl18B27.t3r |
| S | 0.9970 | 2/28/2018 5:44:56 PM | C:\Sirius_T3\HCl18B27.t3r |
| jH | 0.8 | 2/28/2018 5:44:56 PM | C:\Sirius_T3\HCl18B27.t3r |
| jOH | -0.4 | 2/28/2018 5:44:56 PM | C:\Sirius_T3\HCl18B27.t3r |

#### Titrants

| | | | |
|---|---|---|---|
| 0.50 M HCl | 0.993513 | 2/28/2018 5:44:56 PM | C:\Sirius_T3\HCl18B27.t3r |
| 0.50 M KOH | 0.999845 | 2/28/2018 5:44:56 PM | C:\Sirius_T3\KOH18B27.t3r |

#### Sample

| | |
|---|---|
| M07_octanol concentration factor | 0.786 |
| Base pKa 1 | 6.07 |
| logP (XH +) | 0.52 |
| logP (neutral X) | 3.18 |

#### Sample graphs

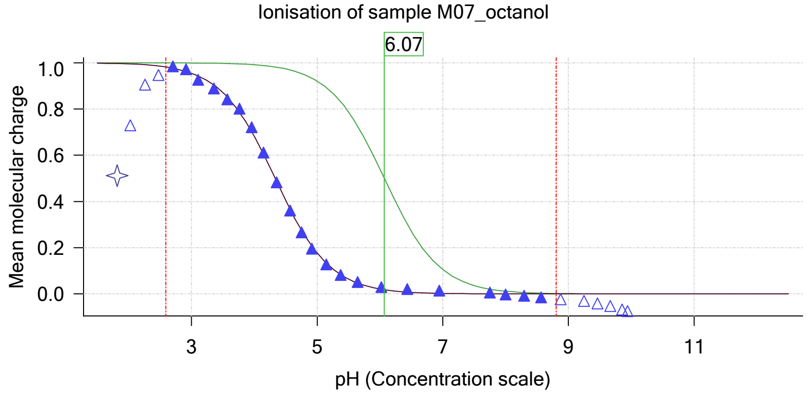


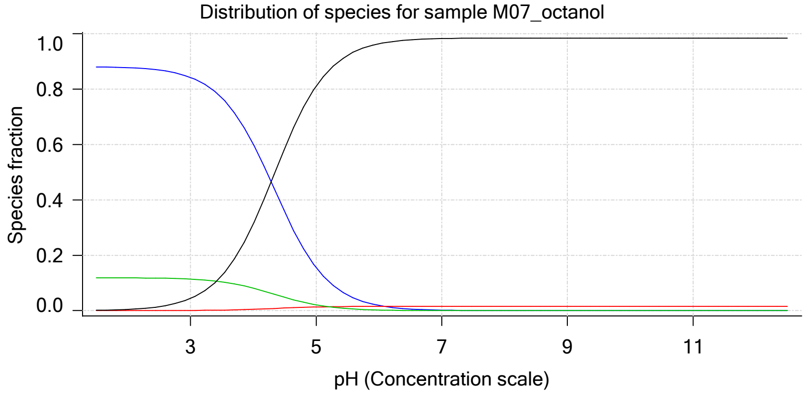

| | | | |
|---|---|---|---|
| Sample name: | **M07_octanol** | Experiment start time: | **2/28/2018 5:44:56 PM** |
| Assay name: | **pH-metric high logP** | Analyst: | **Pion** |
| Assay ID: | **18B-28012** | Instrument ID: | **T312060** |
| Filename: | **C:\Sirius_T3\Mehtap\20180228_exp28_logP_T3-2\18B-28012_M07_octanol_pH-metric high logP.t3r** | | |

#### Sample graphs (continued)

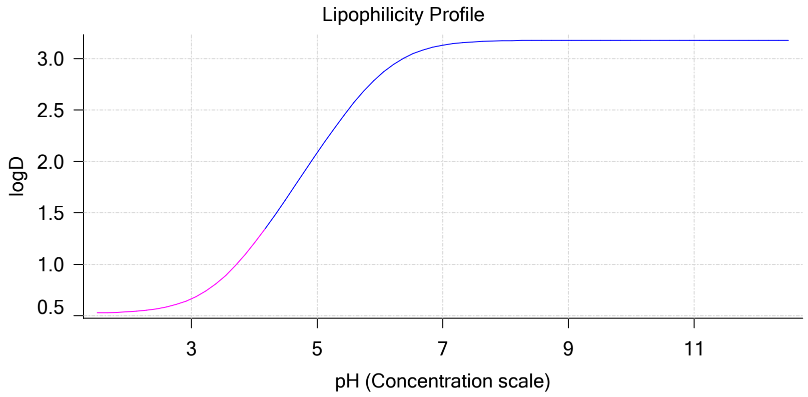


#### Sample logD and percent species

| pH | M07_octanol logD | M07_octanol M07_octanolH | M07_octanol M07_octanol | M07_octanol M07_octanolH* | M07_octanol M07_octanol* | Comment |
|---|---|---|---|---|---|---|
| 1.000 | 0.52 | 88.06 % | 0.00 % | 11.89 % | 0.05 % | |
| 1.200 | 0.52 | 88.04 % | 0.00 % | 11.89 % | 0.07 % | Stomach pH |
| 2.000 | 0.54 | 87.69 % | 0.01 % | 11.84 % | 0.46 % | |
| 3.000 | 0.66 | 84.18 % | 0.07 % | 11.37 % | 4.38 % | |
| 4.000 | 1.20 | 60.09 % | 0.51 % | 8.11 % | 31.29 % | |
| 5.000 | 2.08 | 15.56 % | 1.32 % | 2.10 % | 81.02 % | |
| 6.000 | 2.84 | 1.85 % | 1.57 % | 0.25 % | 96.33 % | |
| 6.500 | 3.04 | 0.59 % | 1.60 % | 0.08 % | 97.73 % | |
| 7.000 | 3.13 | 0.19 % | 1.60 % | 0.03 % | 98.18 % | |
| 7.400 | 3.16 | 0.08 % | 1.61 % | 0.01 % | 98.31 % | Blood pH |
| 8.000 | 3.17 | 0.02 % | 1.61 % | 0.00 % | 98.37 % | |
| 9.000 | 3.18 | 0.00 % | 1.61 % | 0.00 % | 98.39 % | |
| 10.000 | 3.18 | 0.00 % | 1.61 % | 0.00 % | 98.39 % | |
| 11.000 | 3.18 | 0.00 % | 1.61 % | 0.00 % | 98.39 % | |
| 12.000 | 3.18 | 0.00 % | 1.61 % | 0.00 % | 98.39 % | |

#### Carbonate and acidity

| | |
|---|---|
| Carbonate | 0.062 mM |
| Acidity error | 0.053 mM |

#### Other graphs

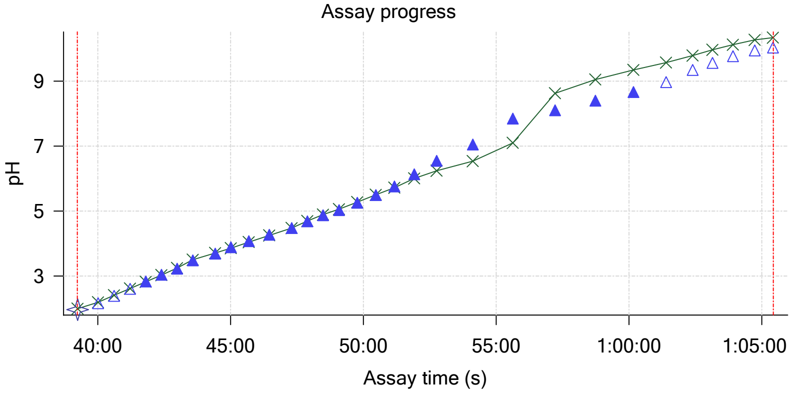


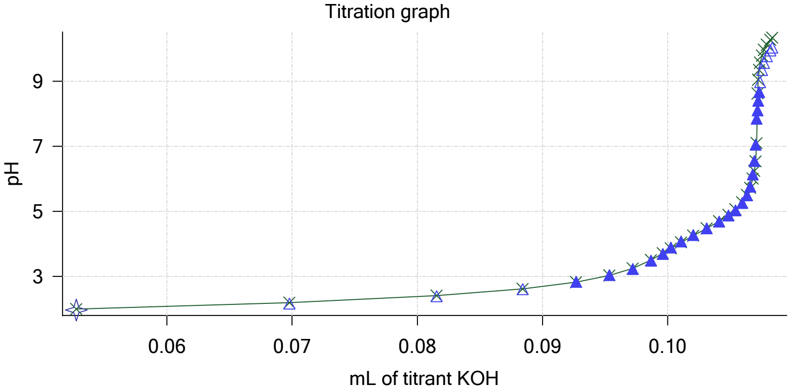

| | | | |
|---|---|---|---|
| Sample name: | **M07_octanol** | Experiment start time: | **2/28/2018 5:44:56 PM** |
| Assay name: | **pH-metric high logP** | Analyst: | **Pion** |
| Assay ID: | **18B-28012** | Instrument ID: | **T312060** |
| Filename: | **C:\Sirius_T3\Mehtap\20180228_exp28_logP_T3-2\18B-28012_M07_octanol_pH-metric high logP.t3r** | | |

#### Other graphs (continued)

Charge balance residuals

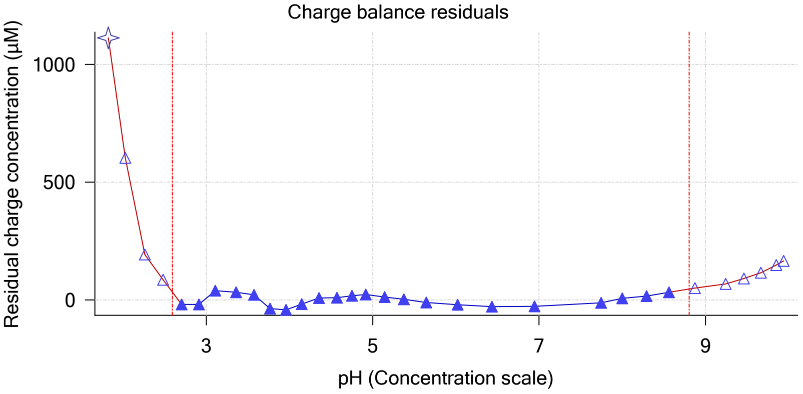


Buffer index profile

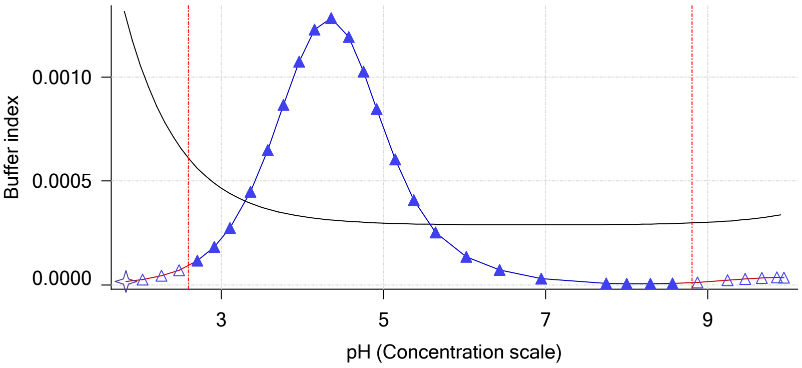


Ionic strength

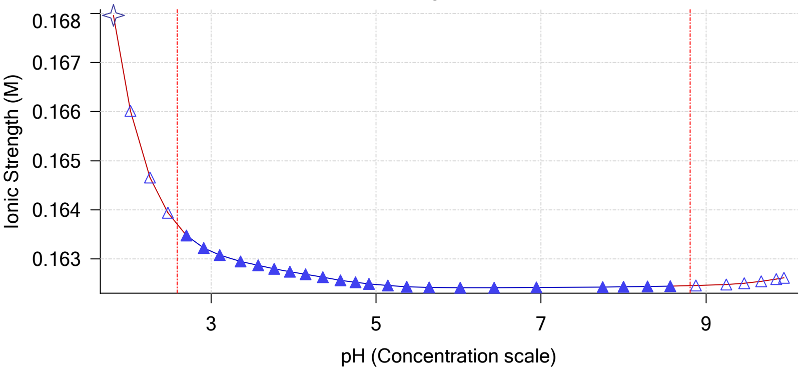


Log R Profile

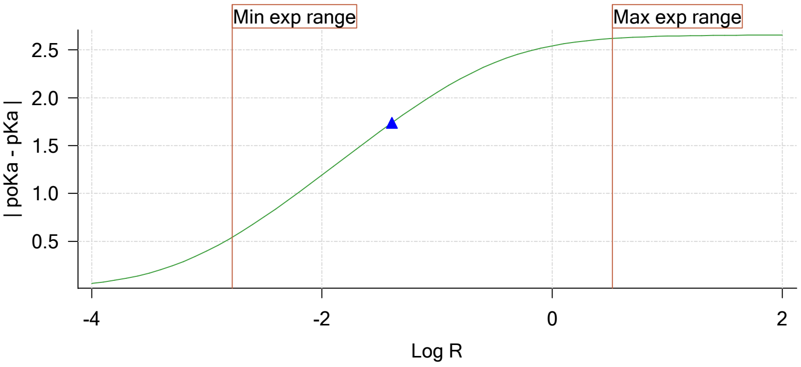


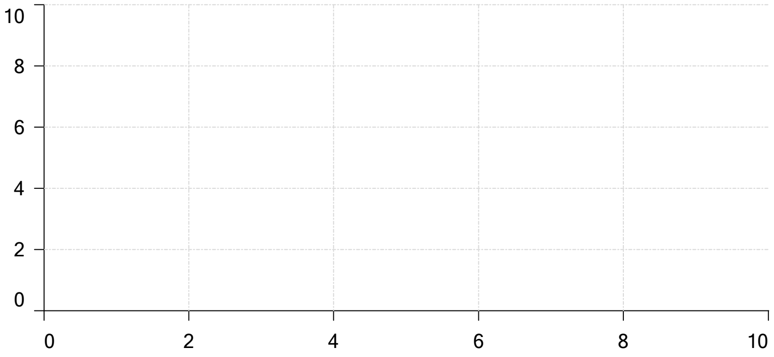


Ionisation of sample Carbonate

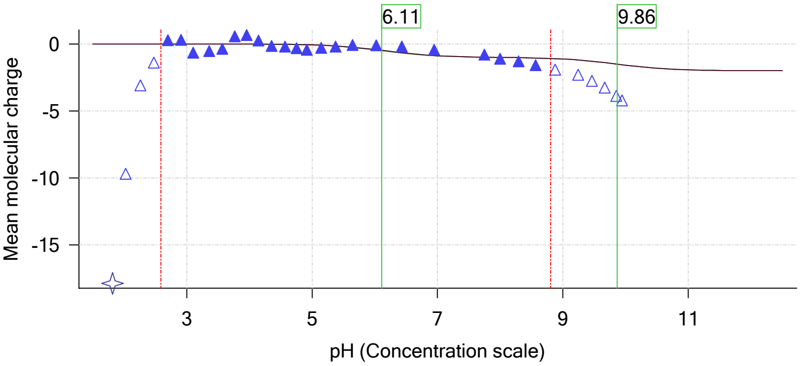

| | | | |
|---|---|---|---|
| Sample name: | **M07_octanol** | Experiment start time: | **2/28/2018 5:44:56 PM** |
| Assay name: | **pH-metric high logP** | Analyst: | **Pion** |
| Assay ID: | **18B-28012** | Instrument ID: | **T312060** |
| Filename: | **C:\Sirius_T3\Mehtap\20180228_exp28_logP_T3-2\18B-28012_M07_octanol_pH-metric high logP.t3r** | | |

### pH-metric high logP Titration 3 of 3 18B-28012 Points 63 to 95

#### Overall results

| | |
|---|---|
| RMSD | 0.432 |
| Average ionic strength | 0.168 M |
| Average temperature | 25.0°C |
| Partition ratio | 0.1744 : 1 |
| Analyte concentration range | 2246.9 µM to 2311.5 µM |
| Total points considered | 23 of 33 |

#### Warnings and errors

| | |
|---|---|
| Errors | None |
| Warnings | None |

#### Four-Plus parameters

| | | | |
|---|---|---|---|
| Alpha | 0.130 | 2/28/2018 5:44:56 PM | C:\Sirius_T3\HCl18B27.t3r |
| S | 0.9970 | 2/28/2018 5:44:56 PM | C:\Sirius_T3\HCl18B27.t3r |
| jH | 0.8 | 2/28/2018 5:44:56 PM | C:\Sirius_T3\HCl18B27.t3r |
| jOH | -0.4 | 2/28/2018 5:44:56 PM | C:\Sirius_T3\HCl18B27.t3r |

#### Titrants

| | | | |
|---|---|---|---|
| 0.50 M HCl | 0.993513 | 2/28/2018 5:44:56 PM | C:\Sirius_T3\HCl18B27.t3r |
| 0.50 M KOH | 0.999845 | 2/28/2018 5:44:56 PM | C:\Sirius_T3\KOH18B27.t3r |

#### Sample

| | |
|---|---|
| M07_octanol concentration factor | 0.842 |
| Base pKa 1 | 6.07 |
| logP (XH +) | 0.57 |
| logP (neutral X) | 3.38 |

#### Sample graphs

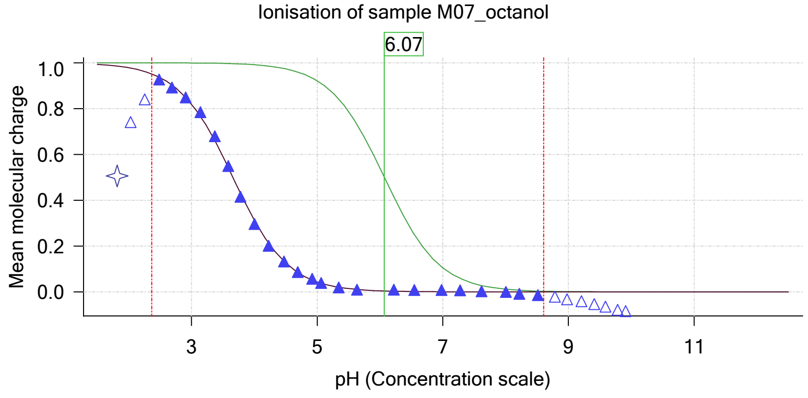


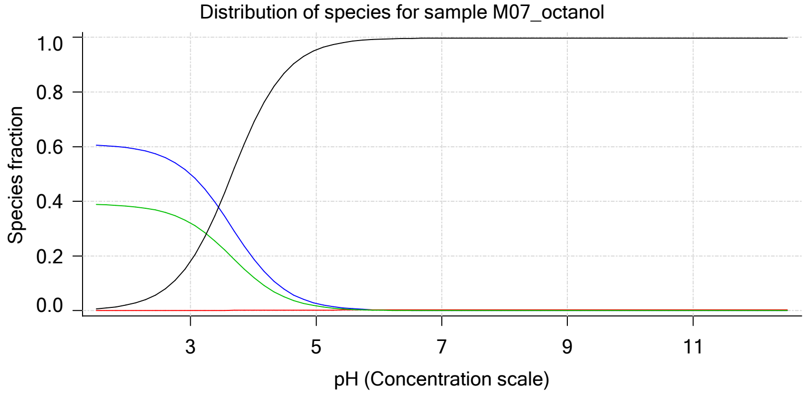

| | | | |
|---|---|---|---|
| Sample name: | **M07_octanol** | Experiment start time: | **2/28/2018 5:44:56 PM** |
| Assay name: | **pH-metric high logP** | Analyst: | **Pion** |
| Assay ID: | **18B-28012** | Instrument ID: | **T312060** |
| Filename: | **C:\Sirius_T3\Mehtap\20180228_exp28_logP_T3-2\18B-28012_M07_octanol_pH-metric high logP.t3r** | | |

#### Sample graphs (continued)

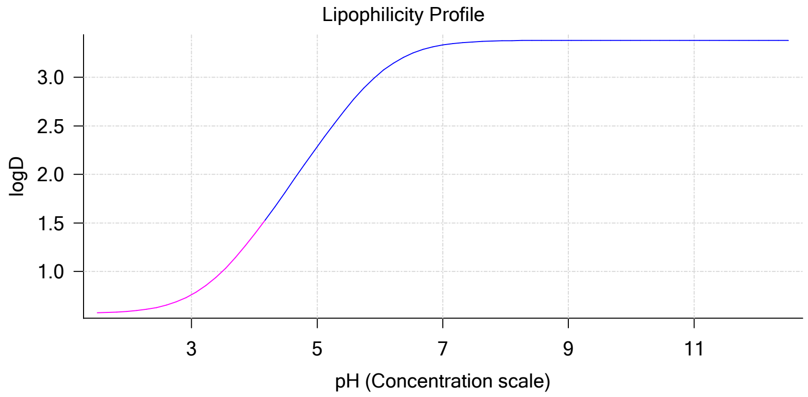


#### Sample logD and percent species

| pH | M07_octanol logD | M07_octanol M07_octanolH | M07_octanol M07_octanol | M07_octanol M07_octanolH* | M07_octanol M07_octanol* | Comment |
|---|---|---|---|---|---|---|
| 1.000 | 0.57 | 60.78 % | 0.00 % | 39.00 % | 0.22 % | |
| 1.200 | 0.57 | 60.70 % | 0.00 % | 38.95 % | 0.34 % | Stomach pH |
| 2.000 | 0.59 | 59.61 % | 0.01 % | 38.25 % | 2.14 % | |
| 3.000 | 0.76 | 49.98 % | 0.04 % | 32.07 % | 17.91 % | |
| 4.000 | 1.38 | 19.11 % | 0.16 % | 12.26 % | 68.47 % | |
| 5.000 | 2.28 | 2.66 % | 0.23 % | 1.71 % | 95.40 % | |
| 6.000 | 3.05 | 0.28 % | 0.24 % | 0.18 % | 99.31 % | |
| 6.500 | 3.25 | 0.09 % | 0.24 % | 0.06 % | 99.62 % | |
| 7.000 | 3.33 | 0.03 % | 0.24 % | 0.02 % | 99.72 % | |
| 7.400 | 3.36 | 0.01 % | 0.24 % | 0.01 % | 99.74 % | Blood pH |
| 8.000 | 3.38 | 0.00 % | 0.24 % | 0.00 % | 99.76 % | |
| 9.000 | 3.38 | 0.00 % | 0.24 % | 0.00 % | 99.76 % | |
| 10.000 | 3.38 | 0.00 % | 0.24 % | 0.00 % | 99.76 % | |
| 11.000 | 3.38 | 0.00 % | 0.24 % | 0.00 % | 99.76 % | |
| 12.000 | 3.38 | 0.00 % | 0.24 % | 0.00 % | 99.76 % | |

#### Carbonate and acidity

| | |
|---|---|
| Carbonate | 0.100 mM |
| Acidity error | 0.059 mM |

#### Other graphs

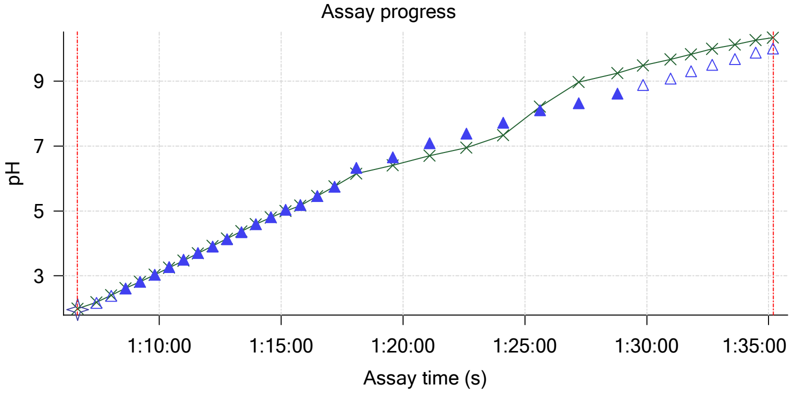


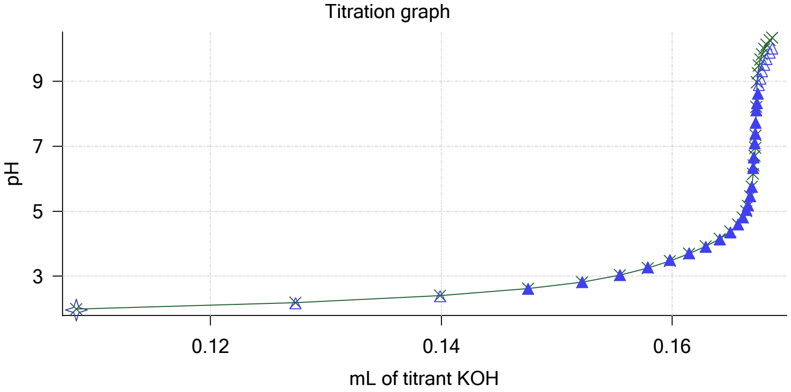

| | | | |
|---|---|---|---|
| Sample name: | **M07_octanol** | Experiment start time: | **2/28/2018 5:44:56 PM** |
| Assay name: | **pH-metric high logP** | Analyst: | **Pion** |
| Assay ID: | **18B-28012** | Instrument ID: | **T312060** |
| Filename: | **C:\Sirius_T3\Mehtap\20180228_exp28_logP_T3-2\18B-28012_M07_octanol_pH-metric high logP.t3r** | | |

#### Other graphs (continued)

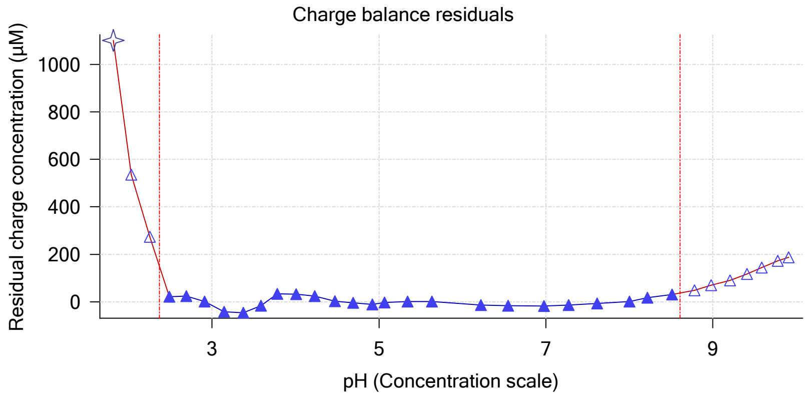


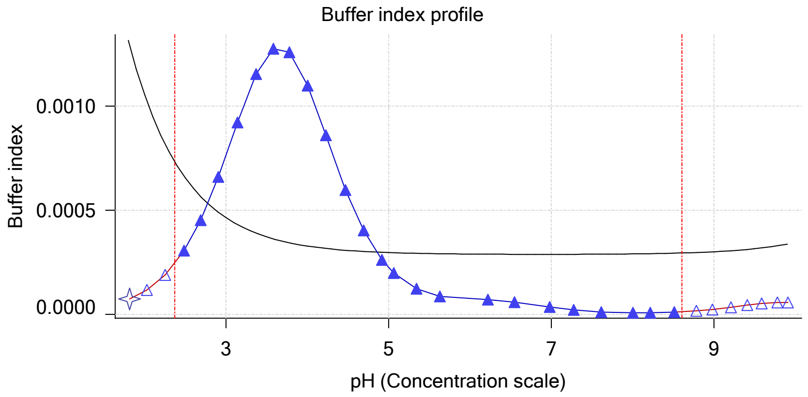


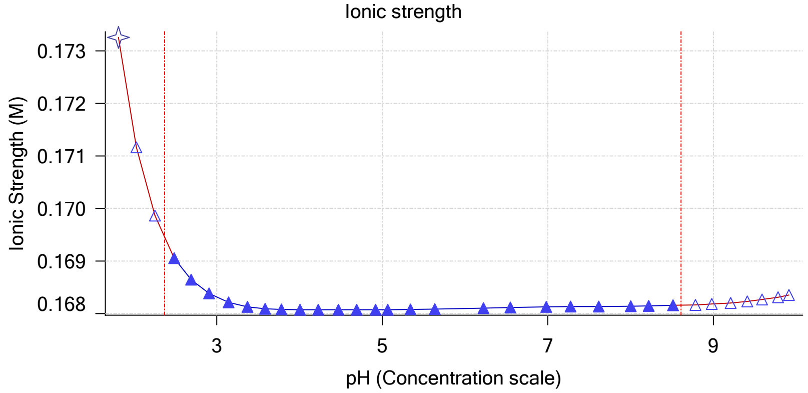


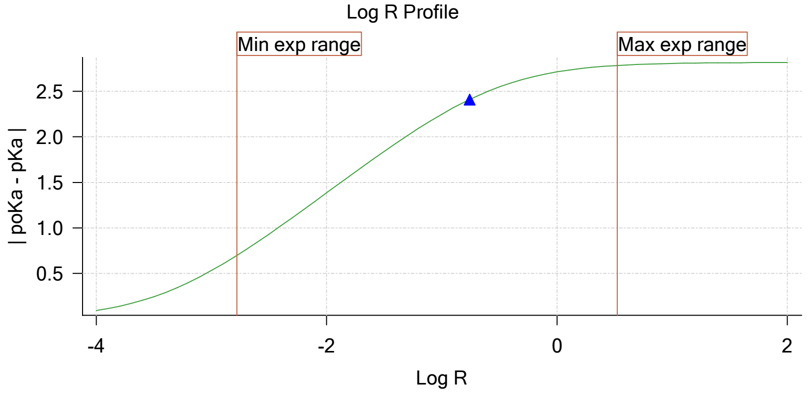


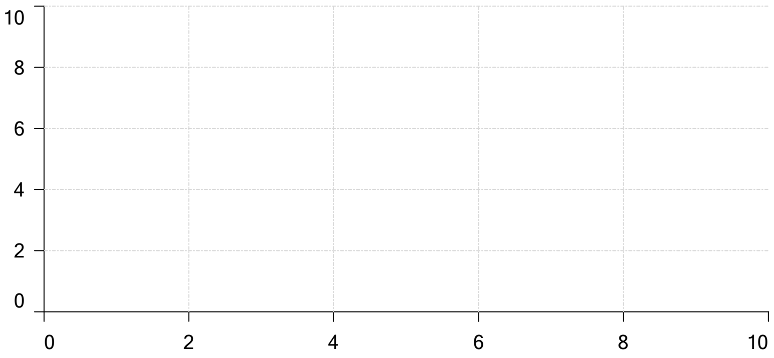

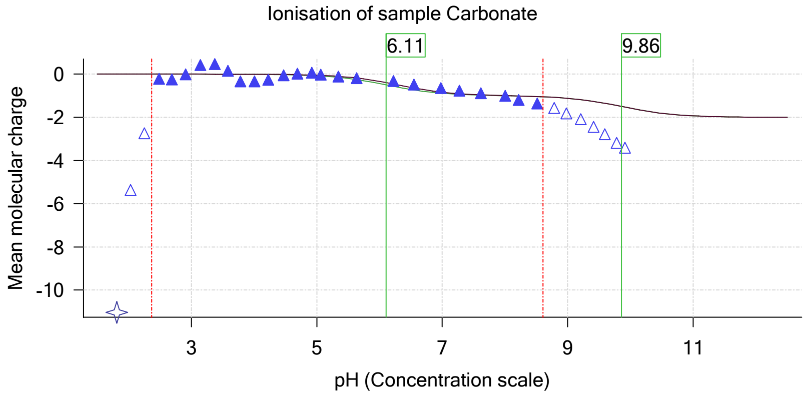

| | | | |
|---|---|---|---|
| Sample name: | **M07_octanol** | Experiment start time: | **2/28/2018 5:44:56 PM** |
| Assay name: | **pH-metric high logP** | Analyst: | **Pion** |
| Assay ID: | **18B-28012** | Instrument ID: | **T312060** |
| Filename: | **C:\Sirius_T3\Mehtap\20180228_exp28_logP_T3-2\18B-28012_M07_octanol_pH-metric high logP.t3r** | | |

#### Assay Model

| Settings | Value | Date/Time changed | Imported from |
|---|---|---|---|
| Sample name | M07_octanol | 2/27/2018 4:29:24 PM | User entered value |
| Sample by | Weight | | Default value |
| Sample weight | 0.001140 g | 2/28/2018 4:24:13 PM | User entered value |
| Formula weight | 235.28 g/mol | 2/27/2018 4:29:24 PM | User entered value |
| Solubility | Unknown | | Default value |
| Molecular weight | 235.28 | 2/27/2018 4:29:24 PM | User entered value |
| Individual pKa ionic environments | No | | Default value |
| Number of pKas | 1 | 2/27/2018 4:29:24 PM | User entered value |
| Sample is a | Base | 2/27/2018 4:29:24 PM | User entered value |
| pKa 1 | 6.07 | 2/27/2018 4:29:24 PM | User entered value |
| logp (XH +) | 0.50 | 2/28/2018 1:33:04 PM | User entered value |
| logP (neutral X) | 3.44 | 2/28/2018 1:33:10 PM | User entered value |

#### Events

| Time | Event | Water | Acid | Base | Octanol | pH | dpH/dt | pH R-squared | pH SD | dpH/dt time |
|---|---|---|---|---|---|---|---|---|---|---|
| 8:59.8 | Initial pH = 8.94 | | | | | | | | | |
| 11:59.4 | Data point 1 | 1.50000 mL | 0.05127 mL | 0.00433 mL | 0.01999 mL | 1.999 | -0.00679 | 0.88781 | 0.00036 | 10.5 s |
| 12:46.0 | Data point 2 | 1.50000 mL | 0.05127 mL | 0.01856 mL | 0.01999 mL | 2.206 | -0.00488 | 0.10282 | 0.00075 | 10.0 s |
| 13:21.7 | Data point 3 | 1.50000 mL | 0.05127 mL | 0.02829 mL | 0.01999 mL | 2.420 | -0.00179 | 0.10235 | 0.00028 | 10.0 s |
| 13:57.3 | Data point 4 | 1.50000 mL | 0.05127 mL | 0.03420 mL | 0.01999 mL | 2.634 | 0.00674 | 0.36990 | 0.00055 | 10.5 s |
| 14:33.3 | Data point 5 | 1.50000 mL | 0.05127 mL | 0.03782 mL | 0.01999 mL | 2.871 | -0.00703 | 0.23966 | 0.00071 | 10.0 s |
| 15:08.8 | Data point 6 | 1.50000 mL | 0.05127 mL | 0.03996 mL | 0.01999 mL | 3.075 | -0.00875 | 0.22861 | 0.00090 | 10.0 s |
| 15:44.1 | Data point 7 | 1.50000 mL | 0.05127 mL | 0.04139 mL | 0.01999 mL | 3.301 | -0.00343 | 0.34132 | 0.00029 | 10.0 s |
| 16:19.5 | Data point 8 | 1.50000 mL | 0.05127 mL | 0.04238 mL | 0.01999 mL | 3.554 | -0.00525 | 0.72590 | 0.00030 | 10.0 s |
| 17:00.2 | Data point 9 | 1.50000 mL | 0.05127 mL | 0.04297 mL | 0.01999 mL | 3.748 | -0.00822 | 0.25539 | 0.00080 | 10.0 s |
| 17:45.9 | Data point 10 | 1.50000 mL | 0.05127 mL | 0.04360 mL | 0.01999 mL | 3.985 | 0.00207 | 0.02036 | 0.00072 | 10.0 s |
| 18:21.3 | Data point 11 | 1.50000 mL | 0.05127 mL | 0.04405 mL | 0.01999 mL | 4.187 | -0.00914 | 0.64694 | 0.00056 | 10.0 s |
| 19:07.1 | Data point 12 | 1.50000 mL | 0.05127 mL | 0.04501 mL | 0.01999 mL | 4.469 | -0.01050 | 0.85079 | 0.00056 | 10.5 s |
| 19:43.0 | Data point 13 | 1.50000 mL | 0.05127 mL | 0.04617 mL | 0.01999 mL | 4.732 | -0.00875 | 0.19263 | 0.00099 | 11.0 s |
| 20:40.1 | Data point 14 | 1.50000 mL | 0.05127 mL | 0.04751 mL | 0.01999 mL | 4.944 | -0.00471 | 0.06280 | 0.00093 | 12.5 s |
| 21:38.6 | Data point 15 | 1.50000 mL | 0.05127 mL | 0.04868 mL | 0.01999 mL | 5.133 | -0.01539 | 0.58584 | 0.00099 | 13.5 s |
| 22:27.9 | Data point 16 | 1.50000 mL | 0.05127 mL | 0.05000 mL | 0.01999 mL | 5.497 | -0.01785 | 0.90166 | 0.00093 | 18.0 s |
| 23:16.5 | Data point 17 | 1.50000 mL | 0.05127 mL | 0.05071 mL | 0.01999 mL | 5.789 | -0.01719 | 0.91631 | 0.00089 | 21.0 s |
| 24:08.1 | Data point 18 | 1.50000 mL | 0.05127 mL | 0.05118 mL | 0.01999 mL | 6.202 | -0.01867 | 0.95635 | 0.00094 | 25.0 s |
| 25:03.7 | Data point 19 | 1.50000 mL | 0.05127 mL | 0.05143 mL | 0.01999 mL | 6.626 | -0.01776 | 0.83266 | 0.00096 | 47.5 s |
| 26:26.9 | Data point 20 | 1.50000 mL | 0.05127 mL | 0.05158 mL | 0.01999 mL | 6.945 | -0.01819 | 0.87488 | 0.00096 | 56.5 s |
| 27:59.1 | Data point 21 | 1.50000 mL | 0.05127 mL | 0.05167 mL | 0.01999 mL | 7.402 | -0.04008 | 0.97402 | 0.00201 | Timed out at 59.5 s |
| 29:34.7 | Data point 22 | 1.50000 mL | 0.05127 mL | 0.05174 mL | 0.01999 mL | 7.954 | -0.04746 | 0.99119 | 0.00236 | Timed out at 59.5 s |
| 31:05.2 | Data point 23 | 1.50000 mL | 0.05127 mL | 0.05179 mL | 0.01999 mL | 8.360 | -0.02983 | 0.96268 | 0.00150 | Timed out at 59.5 s |
| 32:35.7 | Data point 24 | 1.50000 mL | 0.05127 mL | 0.05183 mL | 0.01999 mL | 8.683 | -0.01860 | 0.94531 | 0.00095 | 43.0 s |
| 33:54.3 | Data point 25 | 1.50000 mL | 0.05127 mL | 0.05191 mL | 0.01999 mL | 8.994 | -0.01985 | 0.97488 | 0.00099 | 28.0 s |
| 34:52.9 | Data point 26 | 1.50000 mL | 0.05127 mL | 0.05200 mL | 0.01999 mL | 9.245 | -0.01948 | 0.97749 | 0.00097 | 22.0 s |
| 35:45.4 | Data point 27 | 1.50000 mL | 0.05127 mL | 0.05212 mL | 0.01999 mL | 9.487 | -0.01908 | 0.98282 | 0.00095 | 18.0 s |
| 36:34.0 | Data point 28 | 1.50000 mL | 0.05127 mL | 0.05230 mL | 0.01999 mL | 9.769 | -0.01911 | 0.95487 | 0.00097 | 13.5 s |
| 37:23.1 | Data point 29 | 1.50000 mL | 0.05127 mL | 0.05254 mL | 0.01999 mL | 9.959 | -0.01930 | 0.93400 | 0.00099 | 11.0 s |
| 38:04.7 | Data point 30 | 1.50000 mL | 0.05127 mL | 0.05275 mL | 0.01999 mL | 10.086 | -0.01735 | 0.90522 | 0.00090 | 10.5 s |
| 39:14.5 | Data point 31 | 1.50000 mL | 0.10745 mL | 0.05275 mL | 0.06999 mL | 1.956 | -0.00499 | 0.33277 | 0.00043 | 10.0 s |
| 40:00.8 | Data point 32 | 1.50000 mL | 0.10745 mL | 0.06978 mL | 0.06999 mL | 2.157 | -0.00137 | 0.09377 | 0.00022 | 10.0 s |
| 40:36.5 | Data point 33 | 1.50000 mL | 0.10745 mL | 0.08154 mL | 0.06999 mL | 2.389 | 0.01008 | 0.68241 | 0.00060 | 10.5 s |
| 41:12.6 | Data point 34 | 1.50000 mL | 0.10745 mL | 0.08843 mL | 0.06999 mL | 2.603 | -0.00003 | 0.00007 | 0.00018 | 10.0 s |
| 41:48.1 | Data point 35 | 1.50000 mL | 0.10745 mL | 0.09269 mL | 0.06999 mL | 2.825 | -0.00269 | 0.22934 | 0.00028 | 10.0 s |
| 42:23.5 | Data point 36 | 1.50000 mL | 0.10745 mL | 0.09537 mL | 0.06999 mL | 3.033 | -0.00407 | 0.55814 | 0.00027 | 10.0 s |
| 42:58.9 | Data point 37 | 1.50000 mL | 0.10745 mL | 0.09722 mL | 0.06999 mL | 3.227 | -0.00358 | 0.59751 | 0.00023 | 10.0 s |

| | | | |
|---|---|---|---|
| Sample name: | **M07_octanol** | Experiment start time: | **2/28/2018 5:44:56 PM** |
| Assay name: | **pH-metric high logP** | Analyst: | **Pion** |
| Assay ID: | **18B-28012** | Instrument ID: | **T312060** |
| Filename: | **C:\Sirius_T3\Mehtap\20180228_exp28_logP_T3-2\18B-28012_M07_octanol_pH-metric high logP.t3r** | | |

#### Events (continued)

| Time | Event | Water | Acid | Base | Octanol | pH | dpH/dt | pH R-squared | pH SD | dpH/dt time |
|---|---|---|---|---|---|---|---|---|---|---|
| 43:34.4 | Data point 38 | 1.50000 mL | 0.10745 mL | 0.09868 mL | 0.06999 mL | 3.478 | -0.00403 | 0.43861 | 0.00030 | 10.0 s |
| 44:25.3 | Data point 39 | 1.50000 mL | 0.10745 mL | 0.09962 mL | 0.06999 mL | 3.690 | -0.00498 | 0.42701 | 0.00038 | 10.0 s |
| 45:00.7 | Data point 40 | 1.50000 mL | 0.10745 mL | 0.10026 mL | 0.06999 mL | 3.885 | -0.00655 | 0.31183 | 0.00058 | 10.0 s |
| 45:41.4 | Data point 41 | 1.50000 mL | 0.10745 mL | 0.10108 mL | 0.06999 mL | 4.076 | -0.00350 | 0.03966 | 0.00087 | 10.0 s |
| 46:27.2 | Data point 42 | 1.50000 mL | 0.10745 mL | 0.10205 mL | 0.06999 mL | 4.264 | -0.00534 | 0.73313 | 0.00031 | 10.0 s |
| 47:18.1 | Data point 43 | 1.50000 mL | 0.10745 mL | 0.10313 mL | 0.06999 mL | 4.472 | -0.00695 | 0.55723 | 0.00046 | 10.0 s |
| 47:53.5 | Data point 44 | 1.50000 mL | 0.10745 mL | 0.10412 mL | 0.06999 mL | 4.687 | 0.00135 | 0.00555 | 0.00090 | 10.0 s |
| 48:28.9 | Data point 45 | 1.50000 mL | 0.10745 mL | 0.10487 mL | 0.06999 mL | 4.868 | -0.00089 | 0.00956 | 0.00045 | 10.5 s |
| 49:04.9 | Data point 46 | 1.50000 mL | 0.10745 mL | 0.10543 mL | 0.06999 mL | 5.032 | 0.00810 | 0.39796 | 0.00063 | 11.0 s |
| 49:46.4 | Data point 47 | 1.50000 mL | 0.10745 mL | 0.10597 mL | 0.06999 mL | 5.258 | 0.00679 | 0.13828 | 0.00090 | 11.0 s |
| 50:28.0 | Data point 48 | 1.50000 mL | 0.10745 mL | 0.10635 mL | 0.06999 mL | 5.487 | 0.00018 | 0.00010 | 0.00089 | 11.5 s |
| 51:10.0 | Data point 49 | 1.50000 mL | 0.10745 mL | 0.10661 mL | 0.06999 mL | 5.756 | -0.01673 | 0.85338 | 0.00089 | 14.5 s |
| 51:55.1 | Data point 50 | 1.50000 mL | 0.10745 mL | 0.10682 mL | 0.06999 mL | 6.132 | -0.01877 | 0.87402 | 0.00099 | 19.5 s |
| 52:45.2 | Data point 51 | 1.50000 mL | 0.10745 mL | 0.10694 mL | 0.06999 mL | 6.543 | -0.01943 | 0.92774 | 0.00100 | 51.5 s |
| 54:07.2 | Data point 52 | 1.50000 mL | 0.10745 mL | 0.10703 mL | 0.06999 mL | 7.054 | -0.04976 | 0.98613 | 0.00247 | Timed out at 59.5 s |
| 55:37.7 | Data point 53 | 1.50000 mL | 0.10745 mL | 0.10713 mL | 0.06999 mL | 7.854 | -0.06673 | 0.98909 | 0.00331 | Timed out at 59.5 s |
| 57:13.3 | Data point 54 | 1.50000 mL | 0.10745 mL | 0.10720 mL | 0.06999 mL | 8.106 | -0.04215 | 0.99160 | 0.00209 | Timed out at 59.5 s |
| 58:43.8 | Data point 55 | 1.50000 mL | 0.10745 mL | 0.10724 mL | 0.06999 mL | 8.397 | -0.02007 | 0.98697 | 0.00100 | 50.5 s |
| 1:00:10.1 | Data point 56 | 1.50000 mL | 0.10745 mL | 0.10731 mL | 0.06999 mL | 8.669 | -0.01679 | 0.97495 | 0.00084 | 38.0 s |
| 1:01:23.7 | Data point 57 | 1.50000 mL | 0.10745 mL | 0.10741 mL | 0.06999 mL | 8.978 | -0.01933 | 0.92055 | 0.00099 | 29.5 s |
| 1:02:23.8 | Data point 58 | 1.50000 mL | 0.10745 mL | 0.10755 mL | 0.06999 mL | 9.347 | -0.01767 | 0.84096 | 0.00095 | 14.5 s |
| 1:03:08.8 | Data point 59 | 1.50000 mL | 0.10745 mL | 0.10771 mL | 0.06999 mL | 9.564 | -0.01365 | 0.77349 | 0.00077 | 15.5 s |
| 1:03:54.9 | Data point 60 | 1.50000 mL | 0.10745 mL | 0.10793 mL | 0.06999 mL | 9.768 | -0.01979 | 0.95642 | 0.00100 | 13.5 s |
| 1:04:44.1 | Data point 61 | 1.50000 mL | 0.10745 mL | 0.10821 mL | 0.06999 mL | 9.953 | -0.01717 | 0.91485 | 0.00089 | 10.5 s |
| 1:05:25.1 | Data point 62 | 1.50000 mL | 0.10745 mL | 0.10837 mL | 0.06999 mL | 10.042 | -0.01543 | 0.86770 | 0.00082 | 10.0 s |
| 1:06:39.2 | Data point 63 | 1.50000 mL | 0.16776 mL | 0.10837 mL | 0.31999 mL | 1.951 | -0.00886 | 0.78453 | 0.00049 | 10.0 s |
| 1:07:25.4 | Data point 64 | 1.50000 mL | 0.16776 mL | 0.12737 mL | 0.31999 mL | 2.161 | 0.01497 | 0.60547 | 0.00095 | 10.5 s |
| 1:08:01.6 | Data point 65 | 1.50000 mL | 0.16776 mL | 0.13989 mL | 0.31999 mL | 2.381 | 0.00407 | 0.78143 | 0.00023 | 10.0 s |
| 1:08:37.2 | Data point 66 | 1.50000 mL | 0.16776 mL | 0.14751 mL | 0.31999 mL | 2.612 | 0.00821 | 0.51467 | 0.00056 | 10.0 s |
| 1:09:12.7 | Data point 67 | 1.50000 mL | 0.16776 mL | 0.15221 mL | 0.31999 mL | 2.814 | -0.00211 | 0.44340 | 0.00016 | 10.5 s |
| 1:09:48.8 | Data point 68 | 1.50000 mL | 0.16776 mL | 0.15548 mL | 0.31999 mL | 3.031 | -0.00824 | 0.29769 | 0.00075 | 10.0 s |
| 1:10:24.3 | Data point 69 | 1.50000 mL | 0.16776 mL | 0.15788 mL | 0.31999 mL | 3.265 | -0.00300 | 0.73677 | 0.00017 | 10.0 s |
| 1:10:59.7 | Data point 70 | 1.50000 mL | 0.16776 mL | 0.15981 mL | 0.31999 mL | 3.493 | -0.00479 | 0.74608 | 0.00027 | 10.0 s |
| 1:11:35.2 | Data point 71 | 1.50000 mL | 0.16776 mL | 0.16148 mL | 0.31999 mL | 3.705 | -0.00484 | 0.90505 | 0.00025 | 10.5 s |
| 1:12:11.2 | Data point 72 | 1.50000 mL | 0.16776 mL | 0.16291 mL | 0.31999 mL | 3.900 | -0.00806 | 0.48548 | 0.00057 | 10.5 s |
| 1:12:47.1 | Data point 73 | 1.50000 mL | 0.16776 mL | 0.16414 mL | 0.31999 mL | 4.124 | -0.01087 | 0.62887 | 0.00068 | 10.0 s |
| 1:13:22.5 | Data point 74 | 1.50000 mL | 0.16776 mL | 0.16505 mL | 0.31999 mL | 4.346 | -0.01000 | 0.57912 | 0.00065 | 10.0 s |
| 1:13:58.0 | Data point 75 | 1.50000 mL | 0.16776 mL | 0.16571 mL | 0.31999 mL | 4.587 | -0.00486 | 0.06935 | 0.00091 | 11.0 s |
| 1:14:34.4 | Data point 76 | 1.50000 mL | 0.16776 mL | 0.16613 mL | 0.31999 mL | 4.808 | 0.00762 | 0.26942 | 0.00072 | 11.0 s |
| 1:15:10.8 | Data point 77 | 1.50000 mL | 0.16776 mL | 0.16642 mL | 0.31999 mL | 5.034 | -0.01627 | 0.71284 | 0.00095 | 11.0 s |
| 1:15:47.2 | Data point 78 | 1.50000 mL | 0.16776 mL | 0.16658 mL | 0.31999 mL | 5.179 | 0.01179 | 0.55572 | 0.00078 | 11.5 s |
| 1:16:29.3 | Data point 79 | 1.50000 mL | 0.16776 mL | 0.16677 mL | 0.31999 mL | 5.456 | -0.00292 | 0.02167 | 0.00098 | 11.5 s |
| 1:17:11.3 | Data point 80 | 1.50000 mL | 0.16776 mL | 0.16689 mL | 0.31999 mL | 5.746 | -0.01772 | 0.96453 | 0.00089 | 23.5 s |
| 1:18:05.3 | Data point 81 | 1.50000 mL | 0.16776 mL | 0.16700 mL | 0.31999 mL | 6.334 | -0.01888 | 0.94530 | 0.00096 | 59.0 s |
| 1:19:35.0 | Data point 82 | 1.50000 mL | 0.16776 mL | 0.16707 mL | 0.31999 mL | 6.658 | -0.04000 | 0.97540 | 0.00200 | Timed out at 59.5 s |
| 1:21:05.5 | Data point 83 | 1.50000 mL | 0.16776 mL | 0.16714 mL | 0.31999 mL | 7.088 | -0.06280 | 0.99503 | 0.00311 | Timed out at 59.5 s |
| 1:22:36.0 | Data point 84 | 1.50000 mL | 0.16776 mL | 0.16719 mL | 0.31999 mL | 7.382 | -0.07007 | 0.99379 | 0.00347 | Timed out at 59.5 s |
| 1:24:06.5 | Data point 85 | 1.50000 mL | 0.16776 mL | 0.16724 mL | 0.31999 mL | 7.722 | -0.06487 | 0.99756 | 0.00321 | Timed out at 59.5 s |
| 1:25:37.0 | Data point 86 | 1.50000 mL | 0.16776 mL | 0.16729 mL | 0.31999 mL | 8.108 | -0.04105 | 0.99278 | 0.00203 | Timed out at 59.5 s |

Sample name: **M07_octanol**
Assay name: **pH-metric high logP**
Assay ID: **18B-28012**
Filename: **C:\Sirius_T3\Mehtap\20180228_exp28_logP_T3-2\18B-28012_M07_octanol_pH-metric high logP.t3r**
Experiment start time: **2/28/2018 5:44:56 PM**
Analyst: **Pion**
Instrument ID: **T312060**

#### Events (continued)

| Time | Event | Water | Acid | Base | Octanol | pH | dpH/dt | pH R-squared | pH SD | dpH/dt time |
|---|---|---|---|---|---|---|---|---|---|---|
| 1:27:12.6 | Data point 87 | 1.50000 mL | 0.16776 mL | 0.16736 mL | 0.31999 mL | 8.322 | -0.02438 | 0.97344 | 0.00122 | Timed out at 59.5 s |
| 1:28:48.2 | Data point 88 | 1.50000 mL | 0.16776 mL | 0.16743 mL | 0.31999 mL | 8.617 | -0.01662 | 0.76643 | 0.00094 | 22.0 s |
| 1:29:51.0 | Data point 89 | 1.50000 mL | 0.16776 mL | 0.16752 mL | 0.31999 mL | 8.885 | -0.01802 | 0.91131 | 0.00093 | 27.0 s |
| 1:30:58.8 | Data point 90 | 1.50000 mL | 0.16776 mL | 0.16764 mL | 0.31999 mL | 9.081 | -0.01665 | 0.80009 | 0.00092 | 14.5 s |
| 1:31:49.0 | Data point 91 | 1.50000 mL | 0.16776 mL | 0.16778 mL | 0.31999 mL | 9.309 | -0.01346 | 0.47170 | 0.00097 | 11.5 s |
| 1:32:41.3 | Data point 92 | 1.50000 mL | 0.16776 mL | 0.16797 mL | 0.31999 mL | 9.511 | -0.01581 | 0.68192 | 0.00095 | 15.0 s |
| 1:33:37.1 | Data point 93 | 1.50000 mL | 0.16776 mL | 0.16818 mL | 0.31999 mL | 9.689 | -0.00057 | 0.00339 | 0.00048 | 10.5 s |
| 1:34:28.5 | Data point 94 | 1.50000 mL | 0.16776 mL | 0.16846 mL | 0.31999 mL | 9.881 | -0.01835 | 0.83849 | 0.00099 | 11.5 s |
| 1:35:10.5 | Data point 95 | 1.50000 mL | 0.16776 mL | 0.16867 mL | 0.31999 mL | 10.010 | 0.00024 | 0.00025 | 0.00075 | 10.0 s |
| 1:35:29.7 | Assay volumes | 1.50000 mL | 0.16776 mL | 0.16867 mL | 0.31999 mL | | | | | |

| | | | |
|---|---|---|---|
| Sample name: | **M07_octanol** | Experiment start time: | **2/28/2018 5:44:56 PM** |
| Assay name: | **pH-metric high logP** | Analyst: | **Pion** |
| Assay ID: | **18B-28012** | Instrument ID: | **T312060** |
| Filename: | **C:\Sirius_T3\Mehtap\20180228_exp28_logP_T3-2\18B-28012_M07_octanol_pH-metric high logP.t3r** | | |

| | | | |
|---|---|---|---|
| Sample name: | **M07_octanol** | Experiment start time: | **2/28/2018 5:44:56 PM** |
| Assay name: | **pH-metric high logP** | Analyst: | **Pion** |
| Assay ID: | **18B-28012** | Instrument ID: | **T312060** |
| Filename: | **C:\Sirius_T3\Mehtap\20180228_exp28_logP_T3-2\18B-28012_M07_octanol_pH-metric high logP.t3r** | | |

#### Assay Settings (continued)

| Setting | Value | Original Value | Date/Time changed | Imported from |
|---|---|---|---|---|
| ***Titration 3*** | | | | |
| Titrate from | Low to high pH | | | |
| Add additional water | 0.00 mL | | | |
| Additional partition solvent volume | 0.250 mL | | | |
| Additional partition solvent added | Automatic | | | |
| After pH adjust stir for | 30 seconds | | | |
| Stir to allow partitioning for | 15 seconds | | | |
| Stirrer speed for partitioning | 60% | | | |
| ***Data Point Stability*** | | | | |
| Stir during data point collection | No | | | |
| Delay before data point collection | 0 seconds | | | |
| Number of points to average | 20 points | | | |
| Time interval between points | 0.50 seconds | | | |
| Required maximum standard deviation | 0.00100 dpH/dt | | | |
| Stability timeout after | 60 seconds | | | |

#### Calibration Settings

| Setting | Value | Date/Time changed | Imported from |
|---|---|---|---|
| Four-Plus alpha | 0.130 | 2/28/2018 5:44:56 PM | C:\Sirius_T3\HCl18B27.t3r |
| Four-Plus S | 0.9970 | 2/28/2018 5:44:56 PM | C:\Sirius_T3\HCl18B27.t3r |
| Four-Plus jH | 0.8 | 2/28/2018 5:44:56 PM | C:\Sirius_T3\HCl18B27.t3r |
| Four-Plus jOH | -0.4 | 2/28/2018 5:44:56 PM | C:\Sirius_T3\HCl18B27.t3r |
| Base concentration factor | 1.000 | 2/28/2018 5:44:56 PM | C:\Sirius_T3\KOH18B27.t3r |
| Acid concentration factor | 0.994 | 2/28/2018 5:44:56 PM | C:\Sirius_T3\HCl18B27.t3r |

#### Instrument Settings

| Setting | Value | Batch Id | Install date |
|---|---|---|---|
| Instrument owner | Merck | | |
| Instrument ID | T312060 | | |
| Instrument type | T3 Simulator | | |
| Software version | 1.1.3.0 | | |
| Dispenser module | | T3DM1200361 | 3/31/2009 5:24:52 AM |
| Dispenser 0 | Water | | 3/31/2009 5:25:05 AM |
| Syringe volume | 2.5 mL | | |
| Firmware version | 1.2.1(r2) | | |
| Titrant | Water (0.15 M KCl) | 02-06-2018 | 2/27/2018 10:05:59 AM |
| Dispenser 2 | Acid | | 3/31/2009 5:25:11 AM |
| Syringe volume | 0.5 mL | | |
| Firmware version | 1.2.1(r2) | | |
| Titrant | Acid (0.5 M HCl) | 02-27-2018 | 2/27/2018 10:27:22 AM |
| Dispenser 1 | Base | | 3/31/2009 5:25:21 AM |
| Syringe volume | 0.5 mL | | |
| Firmware version | 1.2.1(r2) | | |
| Titrant | Base (0.5 M KOH) | 9/22/2017 | 2/27/2018 10:21:22 AM |
| Dispenser 5 | Cosolvent | | 3/31/2009 5:26:24 AM |
| Syringe volume | 2.5 mL | | |
| Firmware version | 1.2.1(r2) | | |
| Distribution valve 5 | Distribution Valve | | 3/31/2009 5:28:19 AM |
| Firmware version | 1.1.3 | | |
| Port A | Methanol (80%, 0.15 M KCl) | 09-26-17 | 2/7/2018 9:42:01 AM |
| Port B | Cyclohexane | 11-01-17 | 2/27/2018 10:37:57 AM |
| Dispenser 3 | Buffer | | 8/3/2010 5:05:16 AM |
| Syringe volume | 0.5 mL | | |
| Firmware version | 1.2.1(r2) | | |
| Titrant | Dodecane | 2018/01/31 | 2/28/2018 10:18:04 AM |
| Dispenser 6 | Octanol | | 10/22/2010 10:52:43 AM |

| | | | |
|---|---|---|---|
| Sample name: | **M07_octanol** | Experiment start time: | **2/28/2018 5:44:56 PM** |
| Assay name: | **pH-metric high logP** | Analyst: | **Pion** |
| Assay ID: | **18B-28012** | Instrument ID: | **T312060** |
| Filename: | **C:\Sirius_T3\Mehtap\20180228_exp28_logP_T3-2\18B-28012_M07_octanol_pH-metric high logP.t3r** | | |

| | | | |
|---|---|---|---|
| Sample name: | **M07_octanol** | Experiment start time: | **2/28/2018 5:44:56 PM** |
| Assay name: | **pH-metric high logP** | Analyst: | **Pion** |
| Assay ID: | **18B-28012** | Instrument ID: | **T312060** |
| Filename: | **C:\Sirius_T3\Mehtap\20180228_exp28_logP_T3-2\18B-28012_M07_octanol_pH-metric high logP.t3r** | | |

#### Instrument Settings (continued)

| Setting | Value | Batch Id | Install date |
|---|---|---|---|
| Spectrometer calibration stir duration | 5 s | | |
| Spectrometer calibration stir speed | 30% | | |
| Spectrometer calibration wash pump volume | 20.0 mL | | |
| Spectrometer calibration wash stir duration | 5 s | | |
| Spectrometer calibration wash stir speed | 30% | | |
| Overhead dispense height | 10000 | | |

#### Refinement Settings

| Setting | Value | Default value |
|---|---|---|
| Turbidity detection method | None | None |
| Turbidity wavelength to assess | 500.0 nm | 500.0 nm |
| Turbidity maximum absorbance | 0.100 | 0.100 |
| Turbidity probe threshold | 50.00 | 50.00 |

#### Experiment Log

[2:37] Air gap created for Water (0.15 M KCl)
[2:38] Air gap created for Acid (0.5 M HCl)
[2:38] Air gap created for Base (0.5 M KOH)
[2:39] Air gap released for Water (0.15 M KCl)
[2:42] Titrator arm moved over Titration position
[2:42] Titration 1 of 3
[2:42] Adding initial titrants
[2:42] Automatically add 1.50000 mL of water
[3:08] Dispensed 1.500000 mL of Water (0.15 M KCl)
[3:12] Titrator arm moved over Drain
[8:53] Titrator arm moved to Titration position
[8:53] Argon flow rate set to 100
[8:53] Stirrer speed set to 10
[8:58] Automatically add 0.02000 mL of Octanol
[8:59] Dispensed 0.019991 mL of Octanol
[9:00] Initial pH = 8.94
[9:00] Iterative adjust 8.94 -> 2.00
[9:00] pH 8.94 -> 2.00
[9:02] Air gap released for Acid (0.5 M HCl)
[9:02] Dispensed 0.051270 mL of Acid (0.5 M HCl)
[9:08] Holding pH 2.00
[11:08] Stirrer speed set to 0
[11:08] Stirrer speed set to 50
[11:08] Iterative adjust 1.96 -> 2.00
[11:08] pH 1.96 -> 2.00
[11:08] Air gap released for Base (0.5 M KOH)
[11:09] Dispensed 0.004327 mL of Base (0.5 M KOH)
[12:00] Stirrer speed set to 0
[12:10] Datapoint id 1 collected
[12:10] Stirrer speed set to 50
[12:15] pH 2.01 -> 2.21
[12:15] Using cautious pH adjust
[12:15] Dispensed 0.007808 mL of Base (0.5 M KOH)
[12:21] Stepping pH = 2.10
[12:21] Dispensed 0.005550 mL of Base (0.5 M KOH)
[12:26] Stepping pH = 2.19
[12:26] Dispensed 0.000870 mL of Base (0.5 M KOH)
[12:31] Stepping pH = 2.21
[12:46] Stirrer speed set to 0
[12:56] Datapoint id 2 collected
[12:56] Charge balance equation is out by 8.8%
[12:56] Stirrer speed set to 50

| | | | |
|---|---|---|---|
| Sample name: | **M07_octanol** | Experiment start time: | **2/28/2018 5:44:56 PM** |
| Assay name: | **pH-metric high logP** | Analyst: | **Pion** |
| Assay ID: | **18B-28012** | Instrument ID: | **T312060** |
| Filename: | **C:\Sirius_T3\Mehtap\20180228_exp28_logP_T3-2\18B-28012_M07_octanol_pH-metric high logP.t3r** | | |

#### Experiment Log (continued)

[13:01] pH 2.21 -> 2.41
[13:01] Using charge balance adjust
[13:02] Dispensed 0.009737 mL of Base (0.5 M KOH)
[13:22] Stirrer speed set to 0
[13:32] Datapoint id 3 collected
[13:32] Charge balance equation is out by 4.0%
[13:32] Stirrer speed set to 50
[13:37] pH 2.43 -> 2.63
[13:37] Using charge balance adjust
[13:37] Dispensed 0.005903 mL of Base (0.5 M KOH)
[13:57] Stirrer speed set to 0
[14:08] Datapoint id 4 collected
[14:08] Charge balance equation is out by 3.4%
[14:08] Stirrer speed set to 50
[14:13] pH 2.64 -> 2.84
[14:13] Using charge balance adjust
[14:13] Dispensed 0.003622 mL of Base (0.5 M KOH)
[14:33] Stirrer speed set to 0
[14:43] Datapoint id 5 collected
[14:43] Charge balance equation is out by 13.9%
[14:43] Stirrer speed set to 50
[14:49] pH 2.88 -> 3.08
[14:49] Using charge balance adjust
[14:49] Dispensed 0.002140 mL of Base (0.5 M KOH)
[15:09] Stirrer speed set to 0
[15:19] Datapoint id 6 collected
[15:19] Charge balance equation is out by -3.1%
[15:19] Stirrer speed set to 50
[15:24] pH 3.08 -> 3.28
[15:24] Using charge balance adjust
[15:24] Dispensed 0.001435 mL of Base (0.5 M KOH)
[15:44] Stirrer speed set to 0
[15:54] Datapoint id 7 collected
[15:54] Charge balance equation is out by 8.5%
[15:54] Stirrer speed set to 50
[15:59] pH 3.31 -> 3.51
[15:59] Using charge balance adjust
[16:00] Dispensed 0.000988 mL of Base (0.5 M KOH)
[16:20] Stirrer speed set to 0
[16:30] Datapoint id 8 collected
[16:30] Charge balance equation is out by 22.7%
[16:30] Stirrer speed set to 50
[16:35] pH 3.56 -> 3.76
[16:35] Using cautious pH adjust
[16:35] Dispensed 0.000400 mL of Base (0.5 M KOH)
[16:40] Stepping pH = 3.68
[16:40] Dispensed 0.000188 mL of Base (0.5 M KOH)
[16:45] Stepping pH = 3.75
[17:00] Stirrer speed set to 0
[17:10] Datapoint id 9 collected
[17:10] Charge balance equation is out by 25.4%
[17:10] Stirrer speed set to 50
[17:15] pH 3.76 -> 3.96
[17:15] Using cautious pH adjust
[17:16] Dispensed 0.000400 mL of Base (0.5 M KOH)
[17:21] Stepping pH = 3.92
[17:21] Dispensed 0.000071 mL of Base (0.5 M KOH)
[17:26] Stepping pH = 3.93
[17:26] Dispensed 0.000165 mL of Base (0.5 M KOH)

Sample name: **M07_octanol**
Assay name: **pH-metric high logP**
Assay ID: **18B-28012**
Filename: **C:\Sirius_T3\Mehtap\20180228_exp28_logP_T3-2\18B-28012_M07_octanol_pH-metric high logP.t3r**
Experiment start time: **2/28/2018 5:44:56 PM**
Analyst: **Pion**
Instrument ID: **T312060**

#### Experiment Log (continued)

[17:31] Stepping pH = 3.99
[17:46] Stirrer speed set to 0
[17:56] Datapoint id 10 collected
[17:56] Charge balance equation is out by 18.0%
[17:56] Stirrer speed set to 50
[18:01] pH 4.00 -> 4.20
[18:01] Using cautious pH adjust
[18:01] Dispensed 0.000447 mL of Base (0.5 M KOH)
[18:06] Stepping pH = 4.20
[18:21] Stirrer speed set to 0
[18:32] Datapoint id 11 collected
[18:32] Charge balance equation is out by 50.0%
[18:32] Stirrer speed set to 50
[18:37] pH 4.20 -> 4.40
[18:37] Using cautious pH adjust
[18:37] Dispensed 0.000517 mL of Base (0.5 M KOH)
[18:42] Stepping pH = 4.38
[18:42] Dispensed 0.000071 mL of Base (0.5 M KOH)
[18:47] Stepping pH = 4.37
[18:47] Dispensed 0.000376 mL of Base (0.5 M KOH)
[18:52] Stepping pH = 4.48
[19:07] Stirrer speed set to 0
[19:18] Datapoint id 12 collected
[19:18] Charge balance equation is out by 7.7%
[19:18] Stirrer speed set to 50
[19:23] pH 4.48 -> 4.68
[19:23] Using charge balance adjust
[19:23] Dispensed 0.001152 mL of Base (0.5 M KOH)
[19:43] Stirrer speed set to 0
[19:54] Datapoint id 13 collected
[19:54] Charge balance equation is out by 27.1%
[19:54] Stirrer speed set to 50
[19:59] pH 4.74 -> 4.94
[19:59] Using cautious pH adjust
[19:59] Dispensed 0.000541 mL of Base (0.5 M KOH)
[20:04] Stepping pH = 4.85
[20:05] Dispensed 0.000353 mL of Base (0.5 M KOH)
[20:10] Stepping pH = 4.90
[20:10] Dispensed 0.000212 mL of Base (0.5 M KOH)
[20:15] Stepping pH = 4.93
[20:15] Dispensed 0.000071 mL of Base (0.5 M KOH)
[20:20] Stepping pH = 4.93
[20:20] Dispensed 0.000165 mL of Base (0.5 M KOH)
[20:25] Stepping pH = 4.98
[20:40] Stirrer speed set to 0
[20:53] Datapoint id 14 collected
[20:53] Charge balance equation is out by -21.6%
[20:53] Stirrer speed set to 50
[20:58] pH 4.96 -> 5.16
[20:58] Using cautious pH adjust
[20:58] Dispensed 0.000470 mL of Base (0.5 M KOH)
[21:03] Stepping pH = 5.07
[21:03] Dispensed 0.000282 mL of Base (0.5 M KOH)
[21:08] Stepping pH = 5.12
[21:08] Dispensed 0.000188 mL of Base (0.5 M KOH)
[21:13] Stepping pH = 5.14
[21:13] Dispensed 0.000118 mL of Base (0.5 M KOH)
[21:19] Stepping pH = 5.15
[21:19] Dispensed 0.000118 mL of Base (0.5 M KOH)

#### Experiment Log (continued)

[21:24] Stepping pH = 5.18
[21:39] Stirrer speed set to 0
[21:52] Datapoint id 15 collected
[21:52] Charge balance equation is out by -25.4%
[21:52] Stirrer speed set to 50
[21:57] pH 5.17 -> 5.37
[21:57] Using cautious pH adjust
[21:58] Dispensed 0.000376 mL of Base (0.5 M KOH)
[22:03] Stepping pH = 5.30
[22:03] Dispensed 0.000141 mL of Base (0.5 M KOH)
[22:08] Stepping pH = 5.29
[22:08] Dispensed 0.000800 mL of Base (0.5 M KOH)
[22:13] Stepping pH = 5.59
[22:28] Stirrer speed set to 0
[22:46] Datapoint id 16 collected
[22:46] Charge balance equation is out by -79.4%
[22:46] Stirrer speed set to 50
[22:51] pH 5.54 -> 5.74
[22:51] Using cautious pH adjust
[22:51] Dispensed 0.000212 mL of Base (0.5 M KOH)
[22:56] Stepping pH = 5.57
[22:56] Dispensed 0.000494 mL of Base (0.5 M KOH)
[23:02] Stepping pH = 5.89
[23:17] Stirrer speed set to 0
[23:38] Datapoint id 17 collected
[23:38] Charge balance equation is out by -72.2%
[23:38] Stirrer speed set to 50
[23:43] pH 5.84 -> 6.04
[23:43] Using cautious pH adjust
[23:43] Dispensed 0.000118 mL of Base (0.5 M KOH)
[23:48] Stepping pH = 5.85
[23:48] Dispensed 0.000353 mL of Base (0.5 M KOH)
[23:53] Stepping pH = 6.29
[24:08] Stirrer speed set to 0
[24:33] Datapoint id 18 collected
[24:33] Charge balance equation is out by -89.2%
[24:33] Stirrer speed set to 50
[24:38] pH 6.26 -> 6.46
[24:38] Using cautious pH adjust
[24:38] Dispensed 0.000071 mL of Base (0.5 M KOH)
[24:44] Stepping pH = 6.26
[24:44] Dispensed 0.000188 mL of Base (0.5 M KOH)
[24:49] Stepping pH = 6.68
[25:04] Stirrer speed set to 0
[25:51] Datapoint id 19 collected
[25:51] Charge balance equation is out by -96.9%
[25:51] Stirrer speed set to 50
[25:57] pH 6.69 -> 6.89
[25:57] Using cautious pH adjust
[25:57] Dispensed 0.000024 mL of Base (0.5 M KOH)
[26:02] Stepping pH = 6.70
[26:02] Dispensed 0.000094 mL of Base (0.5 M KOH)
[26:07] Stepping pH = 6.87
[26:07] Dispensed 0.000024 mL of Base (0.5 M KOH)
[26:12] Stepping pH = 6.96
[26:27] Stirrer speed set to 0
[27:24] Datapoint id 20 collected
[27:24] Charge balance equation is out by -124.8%
[27:24] Stirrer speed set to 50

Sample name: **M07_octanol**
Assay name: **pH-metric high logP**
Assay ID: **18B-28012**
Filename: **C:\Sirius_T3\Mehtap\20180228_exp28_logP_T3-2\18B-28012_M07_octanol_pH-metric high logP.t3r**
Experiment start time: **2/28/2018 5:44:56 PM**
Analyst: **Pion**
Instrument ID: **T312060**

#### Experiment Log (continued)

[27:29] pH 7.02 -> 7.22
[27:29] Using cautious pH adjust
[27:29] Dispensed 0.000024 mL of Base (0.5 M KOH)
[27:34] Stepping pH = 7.04
[27:34] Dispensed 0.000047 mL of Base (0.5 M KOH)
[27:39] Stepping pH = 7.19
[27:39] Dispensed 0.000024 mL of Base (0.5 M KOH)
[27:44] Stepping pH = 7.38
[27:59] Stirrer speed set to 0
[28:59] Datapoint id 21 collected
[28:59] Charge balance equation is out by -161.1%
[28:59] Stirrer speed set to 50
[29:04] pH 7.46 -> 7.66
[29:04] Using cautious pH adjust
[29:04] Dispensed 0.000024 mL of Base (0.5 M KOH)
[29:09] Stepping pH = 7.49
[29:10] Dispensed 0.000024 mL of Base (0.5 M KOH)
[29:15] Stepping pH = 7.60
[29:15] Dispensed 0.000024 mL of Base (0.5 M KOH)
[29:20] Stepping pH = 7.84
[29:35] Stirrer speed set to 0
[30:35] Datapoint id 22 collected
[30:35] Charge balance equation is out by -351.4%
[30:35] Stirrer speed set to 50
[30:40] pH 7.98 -> 8.18
[30:40] Using cautious pH adjust
[30:40] Dispensed 0.000024 mL of Base (0.5 M KOH)
[30:45] Stepping pH = 8.02
[30:45] Dispensed 0.000024 mL of Base (0.5 M KOH)
[30:50] Stepping pH = 8.26
[31:05] Stirrer speed set to 0
[32:05] Datapoint id 23 collected
[32:05] Charge balance equation is out by -398.6%
[32:05] Stirrer speed set to 50
[32:10] pH 8.42 -> 8.62
[32:10] Using cautious pH adjust
[32:10] Dispensed 0.000024 mL of Base (0.5 M KOH)
[32:16] Stepping pH = 8.45
[32:16] Dispensed 0.000024 mL of Base (0.5 M KOH)
[32:21] Stepping pH = 8.64
[32:36] Stirrer speed set to 0
[33:19] Datapoint id 24 collected
[33:19] Charge balance equation is out by -209.9%
[33:19] Stirrer speed set to 50
[33:24] pH 8.71 -> 8.91
[33:24] Using cautious pH adjust
[33:24] Dispensed 0.000024 mL of Base (0.5 M KOH)
[33:29] Stepping pH = 8.73
[33:29] Dispensed 0.000024 mL of Base (0.5 M KOH)
[33:34] Stepping pH = 8.84
[33:34] Dispensed 0.000024 mL of Base (0.5 M KOH)
[33:39] Stepping pH = 8.98
[33:55] Stirrer speed set to 0
[34:23] Datapoint id 25 collected
[34:23] Charge balance equation is out by -216.2%
[34:23] Stirrer speed set to 50
[34:28] pH 9.01 -> 9.21
[34:28] Using cautious pH adjust
[34:28] Dispensed 0.000024 mL of Base (0.5 M KOH)

| | | | |
|---|---|---|---|
| Sample name: | **M07_octanol** | Experiment start time: | **2/28/2018 5:44:56 PM** |
| Assay name: | **pH-metric high logP** | Analyst: | **Pion** |
| Assay ID: | **18B-28012** | Instrument ID: | **T312060** |
| Filename: | **C:\Sirius_T3\Mehtap\20180228_exp28_logP_T3-2\18B-28012_M07_octanol_pH-metric high logP.t3r** | | |

#### Experiment Log (continued)

[34:33] Stepping pH = 9.01
[34:33] Dispensed 0.000071 mL of Base (0.5 M KOH)
[34:38] Stepping pH = 9.23
[34:53] Stirrer speed set to 0
[35:15] Datapoint id 26 collected
[35:15] Charge balance equation is out by -96.5%
[35:15] Stirrer speed set to 50
[35:20] pH 9.25 -> 9.45
[35:20] Using cautious pH adjust
[35:20] Dispensed 0.000024 mL of Base (0.5 M KOH)
[35:25] Stepping pH = 9.25
[35:25] Dispensed 0.000094 mL of Base (0.5 M KOH)
[35:30] Stepping pH = 9.48
[35:46] Stirrer speed set to 0
[36:04] Datapoint id 27 collected
[36:04] Charge balance equation is out by -96.5%
[36:04] Stirrer speed set to 50
[36:09] pH 9.49 -> 9.69
[36:09] Using cautious pH adjust
[36:09] Dispensed 0.000047 mL of Base (0.5 M KOH)
[36:14] Stepping pH = 9.51
[36:14] Dispensed 0.000141 mL of Base (0.5 M KOH)
[36:19] Stepping pH = 9.77
[36:34] Stirrer speed set to 0
[36:48] Datapoint id 28 collected
[36:48] Charge balance equation is out by -78.5%
[36:48] Stirrer speed set to 50
[36:53] pH 9.77 -> 9.97
[36:53] Using cautious pH adjust
[36:53] Dispensed 0.000094 mL of Base (0.5 M KOH)
[36:58] Stepping pH = 9.84
[36:58] Dispensed 0.000118 mL of Base (0.5 M KOH)
[37:03] Stepping pH = 9.96
[37:03] Dispensed 0.000024 mL of Base (0.5 M KOH)
[37:08] Stepping pH = 9.97
[37:23] Stirrer speed set to 0
[37:34] Datapoint id 29 collected
[37:34] Charge balance equation is out by -18.4%
[37:34] Stirrer speed set to 50
[37:39] pH 9.96 -> 10.05
[37:39] Using cautious pH adjust
[37:39] Dispensed 0.000047 mL of Base (0.5 M KOH)
[37:45] Stepping pH = 9.96
[37:45] Dispensed 0.000165 mL of Base (0.5 M KOH)
[37:50] Stepping pH = 10.10
[38:05] Stirrer speed set to 0
[38:15] Datapoint id 30 collected
[38:15] Charge balance equation is out by -90.3%
[38:15] Titration 2 of 3
[38:15] Adding initial titrants
[38:15] Automatically add 0.05000 mL of Octanol
[38:17] Dispensed 0.050000 mL of Octanol
[38:17] Stirrer speed set to 10
[38:18] Stirrer speed set to 55
[38:18] Iterative adjust 10.09 -> 2.00
[38:18] pH 10.09 -> 2.00
[38:19] Dispensed 0.054774 mL of Acid (0.5 M HCl)
[38:24] pH 2.01 -> 2.00
[38:24] Dispensed 0.001411 mL of Acid (0.5 M HCl)

| | | | |
|---|---|---|---|
| Sample name: | **M07_octanol** | Experiment start time: | **2/28/2018 5:44:56 PM** |
| Assay name: | **pH-metric high logP** | Analyst: | **Pion** |
| Assay ID: | **18B-28012** | Instrument ID: | **T312060** |
| Filename: | **C:\Sirius_T3\Mehtap\20180228_exp28_logP_T3-2\18B-28012_M07_octanol_pH-metric high logP.t3r** | | |

#### Experiment Log (continued)

[39:15] Stirrer speed set to 0
[39:25] Datapoint id 31 collected
[39:25] Stirrer speed set to 55
[39:30] pH 1.96 -> 2.16
[39:30] Using cautious pH adjust
[39:30] Dispensed 0.009337 mL of Base (0.5 M KOH)
[39:35] Stepping pH = 2.06
[39:35] Dispensed 0.005809 mL of Base (0.5 M KOH)
[39:41] Stepping pH = 2.13
[39:41] Dispensed 0.001881 mL of Base (0.5 M KOH)
[39:46] Stepping pH = 2.16
[40:01] Stirrer speed set to 0
[40:11] Datapoint id 32 collected
[40:11] Charge balance equation is out by 8.8%
[40:11] Stirrer speed set to 55
[40:16] pH 2.16 -> 2.36
[40:16] Using charge balance adjust
[40:16] Dispensed 0.011759 mL of Base (0.5 M KOH)
[40:37] Stirrer speed set to 0
[40:47] Datapoint id 33 collected
[40:47] Charge balance equation is out by 14.2%
[40:47] Stirrer speed set to 55
[40:52] pH 2.39 -> 2.59
[40:52] Using charge balance adjust
[40:53] Dispensed 0.006891 mL of Base (0.5 M KOH)
[41:13] Stirrer speed set to 0
[41:23] Datapoint id 34 collected
[41:23] Charge balance equation is out by 4.7%
[41:23] Stirrer speed set to 55
[41:28] pH 2.61 -> 2.81
[41:28] Using charge balance adjust
[41:28] Dispensed 0.004257 mL of Base (0.5 M KOH)
[41:48] Stirrer speed set to 0
[41:58] Datapoint id 35 collected
[41:58] Charge balance equation is out by 7.3%
[41:58] Stirrer speed set to 55
[42:03] pH 2.83 -> 3.03
[42:03] Using charge balance adjust
[42:03] Dispensed 0.002681 mL of Base (0.5 M KOH)
[42:24] Stirrer speed set to 0
[42:34] Datapoint id 36 collected
[42:34] Charge balance equation is out by 0.1%
[42:34] Stirrer speed set to 55
[42:39] pH 3.04 -> 3.24
[42:39] Using charge balance adjust
[42:39] Dispensed 0.001858 mL of Base (0.5 M KOH)
[42:59] Stirrer speed set to 0
[43:09] Datapoint id 37 collected
[43:09] Charge balance equation is out by -6.5%
[43:09] Stirrer speed set to 55
[43:14] pH 3.23 -> 3.43
[43:14] Using charge balance adjust
[43:14] Dispensed 0.001458 mL of Base (0.5 M KOH)
[43:35] Stirrer speed set to 0
[43:45] Datapoint id 38 collected
[43:45] Charge balance equation is out by 21.9%
[43:45] Stirrer speed set to 55
[43:50] pH 3.49 -> 3.69
[43:50] Using cautious pH adjust

| | | | |
|---|---|---|---|
| Sample name: | **M07_octanol** | Experiment start time: | **2/28/2018 5:44:56 PM** |
| Assay name: | **pH-metric high logP** | Analyst: | **Pion** |
| Assay ID: | **18B-28012** | Instrument ID: | **T312060** |
| Filename: | **C:\Sirius_T3\Mehtap\20180228_exp28_logP_T3-2\18B-28012_M07_octanol_pH-metric high logP.t3r** | | |

#### Experiment Log (continued)

[43:50] Dispensed 0.000635 mL of Base (0.5 M KOH)
[43:55] Stepping pH = 3.64
[43:55] Dispensed 0.000165 mL of Base (0.5 M KOH)
[44:00] Stepping pH = 3.67
[44:00] Dispensed 0.000047 mL of Base (0.5 M KOH)
[44:05] Stepping pH = 3.68
[44:05] Dispensed 0.000094 mL of Base (0.5 M KOH)
[44:10] Stepping pH = 3.70
[44:25] Stirrer speed set to 0
[44:36] Datapoint id 39 collected
[44:36] Charge balance equation is out by 24.8%
[44:36] Stirrer speed set to 55
[44:41] pH 3.70 -> 3.90
[44:41] Using cautious pH adjust
[44:41] Dispensed 0.000635 mL of Base (0.5 M KOH)
[44:46] Stepping pH = 3.89
[45:01] Stirrer speed set to 0
[45:11] Datapoint id 40 collected
[45:11] Charge balance equation is out by 50.0%
[45:11] Stirrer speed set to 55
[45:16] pH 3.89 -> 4.09
[45:16] Using cautious pH adjust
[45:16] Dispensed 0.000635 mL of Base (0.5 M KOH)
[45:21] Stepping pH = 4.04
[45:21] Dispensed 0.000188 mL of Base (0.5 M KOH)
[45:26] Stepping pH = 4.08
[45:42] Stirrer speed set to 0
[45:52] Datapoint id 41 collected
[45:52] Charge balance equation is out by 35.8%
[45:52] Stirrer speed set to 55
[45:57] pH 4.08 -> 4.28
[45:57] Using cautious pH adjust
[45:57] Dispensed 0.000635 mL of Base (0.5 M KOH)
[46:02] Stepping pH = 4.22
[46:02] Dispensed 0.000235 mL of Base (0.5 M KOH)
[46:07] Stepping pH = 4.26
[46:07] Dispensed 0.000094 mL of Base (0.5 M KOH)
[46:12] Stepping pH = 4.27
[46:27] Stirrer speed set to 0
[46:37] Datapoint id 42 collected
[46:37] Charge balance equation is out by 24.0%
[46:37] Stirrer speed set to 55
[46:42] pH 4.27 -> 4.47
[46:42] Using cautious pH adjust
[46:43] Dispensed 0.000588 mL of Base (0.5 M KOH)
[46:48] Stepping pH = 4.40
[46:48] Dispensed 0.000282 mL of Base (0.5 M KOH)
[46:53] Stepping pH = 4.45
[46:53] Dispensed 0.000094 mL of Base (0.5 M KOH)
[46:58] Stepping pH = 4.46
[46:58] Dispensed 0.000118 mL of Base (0.5 M KOH)
[47:03] Stepping pH = 4.48
[47:18] Stirrer speed set to 0
[47:28] Datapoint id 43 collected
[47:28] Charge balance equation is out by 8.5%
[47:28] Stirrer speed set to 55
[47:33] pH 4.48 -> 4.68
[47:33] Using charge balance adjust
[47:33] Dispensed 0.000988 mL of Base (0.5 M KOH)

| | | | |
|---|---|---|---|
| Sample name: | **M07_octanol** | Experiment start time: | **2/28/2018 5:44:56 PM** |
| Assay name: | **pH-metric high logP** | Analyst: | **Pion** |
| Assay ID: | **18B-28012** | Instrument ID: | **T312060** |
| Filename: | **C:\Sirius_T3\Mehtap\20180228_exp28_logP_T3-2\18B-28012_M07_octanol_pH-metric high logP.t3r** | | |

#### Experiment Log (continued)

[47:54] Stirrer speed set to 0
[48:04] Datapoint id 44 collected
[48:04] Charge balance equation is out by 3.3%
[48:04] Stirrer speed set to 55
[48:09] pH 4.70 -> 4.90
[48:09] Using charge balance adjust
[48:09] Dispensed 0.000753 mL of Base (0.5 M KOH)
[48:29] Stirrer speed set to 0
[48:40] Datapoint id 45 collected
[48:40] Charge balance equation is out by -14.9%
[48:40] Stirrer speed set to 55
[48:45] pH 4.88 -> 5.08
[48:45] Using charge balance adjust
[48:45] Dispensed 0.000564 mL of Base (0.5 M KOH)
[49:05] Stirrer speed set to 0
[49:16] Datapoint id 46 collected
[49:16] Charge balance equation is out by -25.1%
[49:16] Stirrer speed set to 55
[49:21] pH 5.05 -> 5.25
[49:21] Using cautious pH adjust
[49:21] Dispensed 0.000212 mL of Base (0.5 M KOH)
[49:26] Stepping pH = 5.10
[49:26] Dispensed 0.000329 mL of Base (0.5 M KOH)
[49:31] Stepping pH = 5.27
[49:47] Stirrer speed set to 0
[49:58] Datapoint id 47 collected
[49:58] Charge balance equation is out by -26.5%
[49:58] Stirrer speed set to 55
[50:03] pH 5.28 -> 5.48
[50:03] Using cautious pH adjust
[50:03] Dispensed 0.000141 mL of Base (0.5 M KOH)
[50:08] Stepping pH = 5.32
[50:08] Dispensed 0.000235 mL of Base (0.5 M KOH)
[50:13] Stepping pH = 5.50
[50:28] Stirrer speed set to 0
[50:40] Datapoint id 48 collected
[50:40] Charge balance equation is out by -34.5%
[50:40] Stirrer speed set to 55
[50:45] pH 5.52 -> 5.72
[50:45] Using cautious pH adjust
[50:45] Dispensed 0.000094 mL of Base (0.5 M KOH)
[50:50] Stepping pH = 5.55
[50:50] Dispensed 0.000165 mL of Base (0.5 M KOH)
[50:55] Stepping pH = 5.77
[51:10] Stirrer speed set to 0
[51:25] Datapoint id 49 collected
[51:25] Charge balance equation is out by -41.4%
[51:25] Stirrer speed set to 55
[51:30] pH 5.79 -> 5.99
[51:30] Using cautious pH adjust
[51:30] Dispensed 0.000071 mL of Base (0.5 M KOH)
[51:35] Stepping pH = 5.82
[51:35] Dispensed 0.000141 mL of Base (0.5 M KOH)
[51:40] Stepping pH = 6.15
[51:55] Stirrer speed set to 0
[52:15] Datapoint id 50 collected
[52:15] Charge balance equation is out by -59.1%
[52:15] Stirrer speed set to 55
[52:20] pH 6.18 -> 6.38

Sample name: **M07_octanol**
Assay name: **pH-metric high logP**
Assay ID: **18B-28012**
Filename: **C:\Sirius_T3\Mehtap\20180228_exp28_logP_T3-2\18B-28012_M07_octanol_pH-metric high logP.t3r**

Experiment start time: **2/28/2018 5:44:56 PM**
Analyst: **Pion**
Instrument ID: **T312060**

#### Experiment Log (continued)

[52:20] Using cautious pH adjust
[52:20] Dispensed 0.000047 mL of Base (0.5 M KOH)
[52:25] Stepping pH = 6.22
[52:25] Dispensed 0.000071 mL of Base (0.5 M KOH)
[52:30] Stepping pH = 6.55
[52:45] Stirrer speed set to 0
[53:37] Datapoint id 51 collected
[53:37] Charge balance equation is out by -52.2%
[53:37] Stirrer speed set to 55
[53:42] pH 6.63 -> 6.83
[53:42] Using cautious pH adjust
[53:42] Dispensed 0.000024 mL of Base (0.5 M KOH)
[53:47] Stepping pH = 6.64
[53:47] Dispensed 0.000071 mL of Base (0.5 M KOH)
[53:52] Stepping pH = 7.11
[54:07] Stirrer speed set to 0
[55:07] Datapoint id 52 collected
[55:07] Charge balance equation is out by -86.4%
[55:07] Stirrer speed set to 55
[55:13] pH 7.04 -> 7.24
[55:13] Using cautious pH adjust
[55:13] Dispensed 0.000024 mL of Base (0.5 M KOH)
[55:18] Stepping pH = 7.01
[55:18] Dispensed 0.000071 mL of Base (0.5 M KOH)
[55:23] Stepping pH = 7.76
[55:38] Stirrer speed set to 0
[56:38] Datapoint id 53 collected
[56:38] Charge balance equation is out by -279.5%
[56:38] Stirrer speed set to 55
[56:43] pH 7.97 -> 8.17
[56:43] Using cautious pH adjust
[56:43] Dispensed 0.000024 mL of Base (0.5 M KOH)
[56:48] Stepping pH = 8.01
[56:48] Dispensed 0.000024 mL of Base (0.5 M KOH)
[56:53] Stepping pH = 8.15
[56:53] Dispensed 0.000024 mL of Base (0.5 M KOH)
[56:58] Stepping pH = 8.29
[57:14] Stirrer speed set to 0
[58:14] Datapoint id 54 collected
[58:14] Charge balance equation is out by -699.4%
[58:14] Stirrer speed set to 55
[58:19] pH 8.07 -> 8.27
[58:19] Using cautious pH adjust
[58:19] Dispensed 0.000024 mL of Base (0.5 M KOH)
[58:24] Stepping pH = 8.04
[58:24] Dispensed 0.000024 mL of Base (0.5 M KOH)
[58:29] Stepping pH = 8.33
[58:44] Stirrer speed set to 0
[59:35] Datapoint id 55 collected
[59:35] Charge balance equation is out by -428.0%
[59:35] Stirrer speed set to 55
[59:40] pH 8.41 -> 8.61
[59:40] Using cautious pH adjust
[59:40] Dispensed 0.000024 mL of Base (0.5 M KOH)
[59:45] Stepping pH = 8.45
[59:45] Dispensed 0.000024 mL of Base (0.5 M KOH)
[59:50] Stepping pH = 8.56
[59:50] Dispensed 0.000024 mL of Base (0.5 M KOH)
[59:55] Stepping pH = 8.69

Sample name: **M07_octanol** Experiment start time: **2/28/2018 5:44:56 PM**
Assay name: **pH-metric high logP** Analyst: **Pion**
Assay ID: **18B-28012** Instrument ID: **T312060**
Filename: **C:\Sirius_T3\Mehtap\20180228_exp28_logP_T3-2\18B-28012_M07_octanol_pH-metric high logP.t3r**

#### Experiment Log (continued)

[1:00:10] Stirrer speed set to 0
[1:00:48] Datapoint id 56 collected
[1:00:48] Charge balance equation is out by -357.1%
[1:00:48] Stirrer speed set to 55
[1:00:53] pH 8.73 -> 8.93
[1:00:53] Using cautious pH adjust
[1:00:53] Dispensed 0.000024 mL of Base (0.5 M KOH)
[1:00:59] Stepping pH = 8.73
[1:00:59] Dispensed 0.000047 mL of Base (0.5 M KOH)
[1:01:04] Stepping pH = 8.91
[1:01:04] Dispensed 0.000024 mL of Base (0.5 M KOH)
[1:01:09] Stepping pH = 8.99
[1:01:24] Stirrer speed set to 0
[1:01:53] Datapoint id 57 collected
[1:01:53] Charge balance equation is out by -204.7%
[1:01:53] Stirrer speed set to 55
[1:01:59] pH 9.00 -> 9.20
[1:01:59] Using cautious pH adjust
[1:01:59] Dispensed 0.000024 mL of Base (0.5 M KOH)
[1:02:04] Stepping pH = 8.99
[1:02:04] Dispensed 0.000118 mL of Base (0.5 M KOH)
[1:02:09] Stepping pH = 9.35
[1:02:24] Stirrer speed set to 0
[1:02:38] Datapoint id 58 collected
[1:02:38] Charge balance equation is out by -205.9%
[1:02:38] Stirrer speed set to 55
[1:02:44] pH 9.36 -> 9.56
[1:02:44] Using cautious pH adjust
[1:02:44] Dispensed 0.000047 mL of Base (0.5 M KOH)
[1:02:49] Stepping pH = 9.37
[1:02:49] Dispensed 0.000118 mL of Base (0.5 M KOH)
[1:02:54] Stepping pH = 9.58
[1:03:09] Stirrer speed set to 0
[1:03:25] Datapoint id 59 collected
[1:03:25] Charge balance equation is out by -88.1%
[1:03:25] Stirrer speed set to 55
[1:03:30] pH 9.57 -> 9.77
[1:03:30] Using cautious pH adjust
[1:03:30] Dispensed 0.000071 mL of Base (0.5 M KOH)
[1:03:35] Stepping pH = 9.61
[1:03:35] Dispensed 0.000141 mL of Base (0.5 M KOH)
[1:03:40] Stepping pH = 9.77
[1:03:55] Stirrer speed set to 0
[1:04:09] Datapoint id 60 collected
[1:04:09] Charge balance equation is out by -44.9%
[1:04:09] Stirrer speed set to 55
[1:04:14] pH 9.77 -> 9.97
[1:04:14] Using cautious pH adjust
[1:04:14] Dispensed 0.000094 mL of Base (0.5 M KOH)
[1:04:19] Stepping pH = 9.83
[1:04:19] Dispensed 0.000141 mL of Base (0.5 M KOH)
[1:04:24] Stepping pH = 9.93
[1:04:24] Dispensed 0.000047 mL of Base (0.5 M KOH)
[1:04:29] Stepping pH = 9.96
[1:04:44] Stirrer speed set to 0
[1:04:55] Datapoint id 61 collected
[1:04:55] Charge balance equation is out by -40.3%
[1:04:55] Stirrer speed set to 55
[1:05:00] pH 9.95 -> 10.05

| | | | |
|---|---|---|---|
| Sample name: | **M07_octanol** | Experiment start time: | **2/28/2018 5:44:56 PM** |
| Assay name: | **pH-metric high logP** | Analyst: | **Pion** |
| Assay ID: | **18B-28012** | Instrument ID: | **T312060** |
| Filename: | **C:\Sirius_T3\Mehtap\20180228_exp28_logP_T3-2\18B-28012_M07_octanol_pH-metric high logP.t3r** | | |

#### Experiment Log (continued)

[1:05:00] Using cautious pH adjust
[1:05:00] Dispensed 0.000071 mL of Base (0.5 M KOH)
[1:05:05] Stepping pH = 9.98
[1:05:05] Dispensed 0.000094 mL of Base (0.5 M KOH)
[1:05:10] Stepping pH = 10.04
[1:05:25] Stirrer speed set to 0
[1:05:35] Datapoint id 62 collected
[1:05:35] Charge balance equation is out by -24.0%
[1:05:35] Titration 3 of 3
[1:05:35] Adding initial titrants
[1:05:35] Automatically add 0.25000 mL of Octanol
[1:05:41] Dispensed 0.250000 mL of Octanol
[1:05:41] Stirrer speed set to 10
[1:05:42] Stirrer speed set to 60
[1:05:42] Iterative adjust 10.05 -> 2.00
[1:05:42] pH 10.05 -> 2.00
[1:05:44] Dispensed 0.057714 mL of Acid (0.5 M HCl)
[1:05:49] pH 2.02 -> 2.00
[1:05:49] Dispensed 0.002587 mL of Acid (0.5 M HCl)
[1:06:39] Stirrer speed set to 0
[1:06:49] Datapoint id 63 collected
[1:06:49] Stirrer speed set to 60
[1:06:54] pH 1.96 -> 2.16
[1:06:54] Using cautious pH adjust
[1:06:55] Dispensed 0.010113 mL of Base (0.5 M KOH)
[1:07:00] Stepping pH = 2.05
[1:07:00] Dispensed 0.007620 mL of Base (0.5 M KOH)
[1:07:05] Stepping pH = 2.14
[1:07:05] Dispensed 0.001270 mL of Base (0.5 M KOH)
[1:07:10] Stepping pH = 2.16
[1:07:26] Stirrer speed set to 0
[1:07:36] Datapoint id 64 collected
[1:07:36] Charge balance equation is out by 6.0%
[1:07:36] Stirrer speed set to 60
[1:07:41] pH 2.17 -> 2.37
[1:07:41] Using charge balance adjust
[1:07:42] Dispensed 0.012512 mL of Base (0.5 M KOH)
[1:08:02] Stirrer speed set to 0
[1:08:12] Datapoint id 65 collected
[1:08:12] Charge balance equation is out by 7.3%
[1:08:12] Stirrer speed set to 60
[1:08:17] pH 2.39 -> 2.59
[1:08:17] Using charge balance adjust
[1:08:17] Dispensed 0.007620 mL of Base (0.5 M KOH)
[1:08:37] Stirrer speed set to 0
[1:08:47] Datapoint id 66 collected
[1:08:47] Charge balance equation is out by 11.9%
[1:08:47] Stirrer speed set to 60
[1:08:52] pH 2.62 -> 2.82
[1:08:52] Using charge balance adjust
[1:08:53] Dispensed 0.004704 mL of Base (0.5 M KOH)
[1:09:13] Stirrer speed set to 0
[1:09:23] Datapoint id 67 collected
[1:09:23] Charge balance equation is out by -2.4%
[1:09:23] Stirrer speed set to 60
[1:09:29] pH 2.82 -> 3.02
[1:09:29] Using charge balance adjust
[1:09:29] Dispensed 0.003269 mL of Base (0.5 M KOH)
[1:09:49] Stirrer speed set to 0

| | | | |
|---|---|---|---|
| Sample name: | **M07_octanol** | Experiment start time: | **2/28/2018 5:44:56 PM** |
| Assay name: | **pH-metric high logP** | Analyst: | **Pion** |
| Assay ID: | **18B-28012** | Instrument ID: | **T312060** |
| Filename: | **C:\Sirius_T3\Mehtap\20180228_exp28_logP_T3-2\18B-28012_M07_octanol_pH-metric high logP.t3r** | | |

#### Experiment Log (continued)

[1:09:59] Datapoint id 68 collected
[1:09:59] Charge balance equation is out by 6.0%
[1:09:59] Stirrer speed set to 60
[1:10:04] pH 3.04 -> 3.24
[1:10:04] Using charge balance adjust
[1:10:04] Dispensed 0.002399 mL of Base (0.5 M KOH)
[1:10:24] Stirrer speed set to 0
[1:10:34] Datapoint id 69 collected
[1:10:34] Charge balance equation is out by 13.8%
[1:10:34] Stirrer speed set to 60
[1:10:40] pH 3.27 -> 3.47
[1:10:40] Using charge balance adjust
[1:10:40] Dispensed 0.001929 mL of Base (0.5 M KOH)
[1:11:00] Stirrer speed set to 0
[1:11:10] Datapoint id 70 collected
[1:11:10] Charge balance equation is out by 10.7%
[1:11:10] Stirrer speed set to 60
[1:11:15] pH 3.50 -> 3.70
[1:11:15] Using charge balance adjust
[1:11:15] Dispensed 0.001670 mL of Base (0.5 M KOH)
[1:11:35] Stirrer speed set to 0
[1:11:46] Datapoint id 71 collected
[1:11:46] Charge balance equation is out by 1.9%
[1:11:46] Stirrer speed set to 60
[1:11:51] pH 3.71 -> 3.91
[1:11:51] Using charge balance adjust
[1:11:51] Dispensed 0.001435 mL of Base (0.5 M KOH)
[1:12:11] Stirrer speed set to 0
[1:12:22] Datapoint id 72 collected
[1:12:22] Charge balance equation is out by -6.8%
[1:12:22] Stirrer speed set to 60
[1:12:27] pH 3.91 -> 4.11
[1:12:27] Using charge balance adjust
[1:12:27] Dispensed 0.001223 mL of Base (0.5 M KOH)
[1:12:47] Stirrer speed set to 0
[1:12:57] Datapoint id 73 collected
[1:12:57] Charge balance equation is out by 8.5%
[1:12:57] Stirrer speed set to 60
[1:13:02] pH 4.13 -> 4.33
[1:13:02] Using charge balance adjust
[1:13:03] Dispensed 0.000917 mL of Base (0.5 M KOH)
[1:13:23] Stirrer speed set to 0
[1:13:33] Datapoint id 74 collected
[1:13:33] Charge balance equation is out by 8.2%
[1:13:33] Stirrer speed set to 60
[1:13:38] pH 4.36 -> 4.56
[1:13:38] Using charge balance adjust
[1:13:38] Dispensed 0.000659 mL of Base (0.5 M KOH)
[1:13:58] Stirrer speed set to 0
[1:14:09] Datapoint id 75 collected
[1:14:09] Charge balance equation is out by 14.5%
[1:14:09] Stirrer speed set to 60
[1:14:14] pH 4.60 -> 4.80
[1:14:14] Using charge balance adjust
[1:14:14] Dispensed 0.000423 mL of Base (0.5 M KOH)
[1:14:35] Stirrer speed set to 0
[1:14:46] Datapoint id 76 collected
[1:14:46] Charge balance equation is out by 1.5%
[1:14:46] Stirrer speed set to 60

| | | | |
|---|---|---|---|
| Sample name: | **M07_octanol** | Experiment start time: | **2/28/2018 5:44:56 PM** |
| Assay name: | **pH-metric high logP** | Analyst: | **Pion** |
| Assay ID: | **18B-28012** | Instrument ID: | **T312060** |
| Filename: | **C:\Sirius_T3\Mehtap\20180228_exp28_logP_T3-2\18B-28012_M07_octanol_pH-metric high logP.t3r** | | |

#### Experiment Log (continued)

[1:14:51] pH 4.82 -> 5.02
[1:14:51] Using charge balance adjust
[1:14:51] Dispensed 0.000282 mL of Base (0.5 M KOH)
[1:15:11] Stirrer speed set to 0
[1:15:22] Datapoint id 77 collected
[1:15:22] Charge balance equation is out by 6.9%
[1:15:22] Stirrer speed set to 60
[1:15:27] pH 5.04 -> 5.24
[1:15:27] Using charge balance adjust
[1:15:27] Dispensed 0.000165 mL of Base (0.5 M KOH)
[1:15:47] Stirrer speed set to 0
[1:15:59] Datapoint id 78 collected
[1:15:59] Charge balance equation is out by -32.4%
[1:15:59] Stirrer speed set to 60
[1:16:04] pH 5.20 -> 5.40
[1:16:04] Using cautious pH adjust
[1:16:04] Dispensed 0.000071 mL of Base (0.5 M KOH)
[1:16:09] Stepping pH = 5.23
[1:16:09] Dispensed 0.000118 mL of Base (0.5 M KOH)
[1:16:14] Stepping pH = 5.46
[1:16:29] Stirrer speed set to 0
[1:16:41] Datapoint id 79 collected
[1:16:41] Charge balance equation is out by -41.0%
[1:16:41] Stirrer speed set to 60
[1:16:46] pH 5.50 -> 5.70
[1:16:46] Using cautious pH adjust
[1:16:46] Dispensed 0.000047 mL of Base (0.5 M KOH)
[1:16:51] Stepping pH = 5.54
[1:16:51] Dispensed 0.000071 mL of Base (0.5 M KOH)
[1:16:56] Stepping pH = 5.75
[1:17:11] Stirrer speed set to 0
[1:17:35] Datapoint id 80 collected
[1:17:35] Charge balance equation is out by -44.5%
[1:17:35] Stirrer speed set to 60
[1:17:40] pH 5.82 -> 6.02
[1:17:40] Using cautious pH adjust
[1:17:40] Dispensed 0.000024 mL of Base (0.5 M KOH)
[1:17:45] Stepping pH = 5.83
[1:17:45] Dispensed 0.000094 mL of Base (0.5 M KOH)
[1:17:50] Stepping pH = 6.40
[1:18:05] Stirrer speed set to 0
[1:19:05] Datapoint id 81 collected
[1:19:05] Charge balance equation is out by -88.8%
[1:19:05] Stirrer speed set to 60
[1:19:10] pH 6.36 -> 6.56
[1:19:10] Using cautious pH adjust
[1:19:10] Dispensed 0.000024 mL of Base (0.5 M KOH)
[1:19:15] Stepping pH = 6.40
[1:19:15] Dispensed 0.000047 mL of Base (0.5 M KOH)
[1:19:20] Stepping pH = 6.66
[1:19:35] Stirrer speed set to 0
[1:20:35] Datapoint id 82 collected
[1:20:35] Charge balance equation is out by -45.8%
[1:20:35] Stirrer speed set to 60
[1:20:40] pH 6.67 -> 6.87
[1:20:40] Using cautious pH adjust
[1:20:40] Dispensed 0.000024 mL of Base (0.5 M KOH)
[1:20:45] Stepping pH = 6.70
[1:20:45] Dispensed 0.000047 mL of Base (0.5 M KOH)

Sample name: **M07_octanol**
Assay name: **pH-metric high logP**
Assay ID: **18B-28012**
Filename: **C:\Sirius_T3\Mehtap\20180228_exp28_logP_T3-2\18B-28012_M07_octanol_pH-metric high logP.t3r**
Experiment start time: **2/28/2018 5:44:56 PM**
Analyst: **Pion**
Instrument ID: **T312060**

#### Experiment Log (continued)

[1:20:51] Stepping pH = 7.15
[1:21:06] Stirrer speed set to 0
[1:22:06] Datapoint id 83 collected
[1:22:06] Charge balance equation is out by -73.9%
[1:22:06] Stirrer speed set to 60
[1:22:11] pH 7.13 -> 7.33
[1:22:11] Using cautious pH adjust
[1:22:11] Dispensed 0.000024 mL of Base (0.5 M KOH)
[1:22:16] Stepping pH = 7.19
[1:22:16] Dispensed 0.000024 mL of Base (0.5 M KOH)
[1:22:21] Stepping pH = 7.37
[1:22:36] Stirrer speed set to 0
[1:23:36] Datapoint id 84 collected
[1:23:36] Charge balance equation is out by -87.9%
[1:23:36] Stirrer speed set to 60
[1:23:41] pH 7.36 -> 7.56
[1:23:41] Using cautious pH adjust
[1:23:41] Dispensed 0.000024 mL of Base (0.5 M KOH)
[1:23:46] Stepping pH = 7.41
[1:23:46] Dispensed 0.000024 mL of Base (0.5 M KOH)
[1:23:52] Stepping pH = 7.71
[1:24:07] Stirrer speed set to 0
[1:25:07] Datapoint id 85 collected
[1:25:07] Charge balance equation is out by -173.4%
[1:25:07] Stirrer speed set to 60
[1:25:12] pH 7.90 -> 8.10
[1:25:12] Using cautious pH adjust
[1:25:12] Dispensed 0.000024 mL of Base (0.5 M KOH)
[1:25:17] Stepping pH = 7.97
[1:25:17] Dispensed 0.000024 mL of Base (0.5 M KOH)
[1:25:22] Stepping pH = 8.16
[1:25:37] Stirrer speed set to 0
[1:26:37] Datapoint id 86 collected
[1:26:37] Charge balance equation is out by -375.0%
[1:26:37] Stirrer speed set to 60
[1:26:42] pH 8.13 -> 8.33
[1:26:42] Using cautious pH adjust
[1:26:42] Dispensed 0.000024 mL of Base (0.5 M KOH)
[1:26:47] Stepping pH = 8.15
[1:26:47] Dispensed 0.000024 mL of Base (0.5 M KOH)
[1:26:53] Stepping pH = 8.23
[1:26:53] Dispensed 0.000024 mL of Base (0.5 M KOH)
[1:26:58] Stepping pH = 8.34
[1:27:13] Stirrer speed set to 0
[1:28:13] Datapoint id 87 collected
[1:28:13] Charge balance equation is out by -530.1%
[1:28:13] Stirrer speed set to 60
[1:28:18] pH 8.40 -> 8.60
[1:28:18] Using cautious pH adjust
[1:28:18] Dispensed 0.000024 mL of Base (0.5 M KOH)
[1:28:23] Stepping pH = 8.46
[1:28:23] Dispensed 0.000024 mL of Base (0.5 M KOH)
[1:28:28] Stepping pH = 8.56
[1:28:28] Dispensed 0.000024 mL of Base (0.5 M KOH)
[1:28:33] Stepping pH = 8.61
[1:28:48] Stirrer speed set to 0
[1:29:10] Datapoint id 88 collected
[1:29:10] Charge balance equation is out by -322.0%
[1:29:10] Stirrer speed set to 60

| | | | |
|---|---|---|---|
| Sample name: | **M07_octanol** | Experiment start time: | **2/28/2018 5:44:56 PM** |
| Assay name: | **pH-metric high logP** | Analyst: | **Pion** |
| Assay ID: | **18B-28012** | Instrument ID: | **T312060** |
| Filename: | **C:\Sirius_T3\Mehtap\20180228_exp28_logP_T3-2\18B-28012_M07_octanol_pH-metric high logP.t3r** | | |

#### Experiment Log (continued)

[1:29:16] pH 8.63 -> 8.83
[1:29:16] Using cautious pH adjust
[1:29:16] Dispensed 0.000024 mL of Base (0.5 M KOH)
[1:29:21] Stepping pH = 8.66
[1:29:21] Dispensed 0.000024 mL of Base (0.5 M KOH)
[1:29:26] Stepping pH = 8.73
[1:29:26] Dispensed 0.000024 mL of Base (0.5 M KOH)
[1:29:31] Stepping pH = 8.80
[1:29:31] Dispensed 0.000024 mL of Base (0.5 M KOH)
[1:29:36] Stepping pH = 8.89
[1:29:51] Stirrer speed set to 0
[1:30:18] Datapoint id 89 collected
[1:30:18] Charge balance equation is out by -324.1%
[1:30:18] Stirrer speed set to 60
[1:30:23] pH 8.89 -> 9.09
[1:30:23] Using cautious pH adjust
[1:30:23] Dispensed 0.000024 mL of Base (0.5 M KOH)
[1:30:28] Stepping pH = 8.90
[1:30:29] Dispensed 0.000047 mL of Base (0.5 M KOH)
[1:30:34] Stepping pH = 9.01
[1:30:34] Dispensed 0.000024 mL of Base (0.5 M KOH)
[1:30:39] Stepping pH = 9.04
[1:30:39] Dispensed 0.000024 mL of Base (0.5 M KOH)
[1:30:44] Stepping pH = 9.09
[1:30:59] Stirrer speed set to 0
[1:31:14] Datapoint id 90 collected
[1:31:14] Charge balance equation is out by -257.9%
[1:31:14] Stirrer speed set to 60
[1:31:19] pH 9.11 -> 9.31
[1:31:19] Using cautious pH adjust
[1:31:19] Dispensed 0.000024 mL of Base (0.5 M KOH)
[1:31:24] Stepping pH = 9.12
[1:31:24] Dispensed 0.000094 mL of Base (0.5 M KOH)
[1:31:29] Stepping pH = 9.30
[1:31:29] Dispensed 0.000024 mL of Base (0.5 M KOH)
[1:31:34] Stepping pH = 9.32
[1:31:49] Stirrer speed set to 0
[1:32:01] Datapoint id 91 collected
[1:32:01] Charge balance equation is out by -124.8%
[1:32:01] Stirrer speed set to 60
[1:32:06] pH 9.31 -> 9.51
[1:32:06] Using cautious pH adjust
[1:32:06] Dispensed 0.000047 mL of Base (0.5 M KOH)
[1:32:11] Stepping pH = 9.35
[1:32:11] Dispensed 0.000094 mL of Base (0.5 M KOH)
[1:32:16] Stepping pH = 9.49
[1:32:16] Dispensed 0.000024 mL of Base (0.5 M KOH)
[1:32:21] Stepping pH = 9.50
[1:32:21] Dispensed 0.000024 mL of Base (0.5 M KOH)
[1:32:26] Stepping pH = 9.52
[1:32:41] Stirrer speed set to 0
[1:32:57] Datapoint id 92 collected
[1:32:57] Charge balance equation is out by -93.9%
[1:32:57] Stirrer speed set to 60
[1:33:02] pH 9.50 -> 9.70
[1:33:02] Using cautious pH adjust
[1:33:02] Dispensed 0.000071 mL of Base (0.5 M KOH)
[1:33:07] Stepping pH = 9.58
[1:33:07] Dispensed 0.000071 mL of Base (0.5 M KOH)

| | | | |
|---|---|---|---|
| Sample name: | **M07_octanol** | Experiment start time: | **2/28/2018 5:44:56 PM** |
| Assay name: | **pH-metric high logP** | Analyst: | **Pion** |
| Assay ID: | **18B-28012** | Instrument ID: | **T312060** |
| Filename: | **C:\Sirius_T3\Mehtap\20180228_exp28_logP_T3-2\18B-28012_M07_octanol_pH-metric high logP.t3r** | | |

#### Experiment Log (continued)

[1:33:12] Stepping pH = 9.65
[1:33:12] Dispensed 0.000047 mL of Base (0.5 M KOH)
[1:33:17] Stepping pH = 9.68
[1:33:17] Dispensed 0.000024 mL of Base (0.5 M KOH)
[1:33:22] Stepping pH = 9.70
[1:33:37] Stirrer speed set to 0
[1:33:48] Datapoint id 93 collected
[1:33:48] Charge balance equation is out by -50.3%
[1:33:48] Stirrer speed set to 60
[1:33:53] pH 9.69 -> 9.89
[1:33:53] Using cautious pH adjust
[1:33:53] Dispensed 0.000094 mL of Base (0.5 M KOH)
[1:33:58] Stepping pH = 9.76
[1:33:58] Dispensed 0.000118 mL of Base (0.5 M KOH)
[1:34:03] Stepping pH = 9.85
[1:34:03] Dispensed 0.000047 mL of Base (0.5 M KOH)
[1:34:08] Stepping pH = 9.87
[1:34:08] Dispensed 0.000024 mL of Base (0.5 M KOH)
[1:34:14] Stepping pH = 9.89
[1:34:29] Stirrer speed set to 0
[1:34:40] Datapoint id 94 collected
[1:34:40] Charge balance equation is out by -41.2%
[1:34:40] Stirrer speed set to 60
[1:34:45] pH 9.89 -> 10.05
[1:34:45] Using cautious pH adjust
[1:34:45] Dispensed 0.000094 mL of Base (0.5 M KOH)
[1:34:50] Stepping pH = 9.94
[1:34:51] Dispensed 0.000118 mL of Base (0.5 M KOH)
[1:34:56] Stepping pH = 10.02
[1:35:11] Stirrer speed set to 0
[1:35:21] Datapoint id 95 collected
[1:35:21] Charge balance equation is out by -9.4%
[1:35:21] Argon flow rate set to 0
[1:35:25] Titrator arm moved over Titration position
