## Supplementary material for "Octanol-water partition coefficient measurements for the SAMPL6 Blind Prediction Challenge": SM07_18B-28013_M07_octanol_pH-metric high logP_report.pdf

| | | | |
|---|---|---|---|
| Sample name: | **M07_octanol** | Experiment start time: | **2/28/2018 7:21:13 PM** |
| Assay name: | **pH-metric high logP** | Analyst: | **Pion** |
| Assay ID: | **18B-28013** | Instrument ID: | **T312060** |
| Filename: | **C:\Sirius_T3\Mehtap\20180228_exp28_logP_T3-2\18B-28013_M07_octanol_pH-metric high logP.t3r** | | |

#### pH-metric Result

| | | |
|---|---|---|
| logP (XH +) | 0.31 | ±0.04 (n=50) |
| logP (neutral X) | 3.21 | ±0.01 (n=50) |

#### 18B-28013 Points 1 to 31

| | |
|---|---|
| M07_octanol concentration factor | 1.061 |
| Carbonate | 0.0000 mM |
| Acidity error | 0.26960 mM |

#### 18B-28013 Points 32 to 65

| | |
|---|---|
| M07_octanol concentration factor | 0.967 |
| Carbonate | 0.0076 mM |
| Acidity error | 0.04628 mM |

#### 18B-28013 Points 66 to 95

| | |
|---|---|
| M07_octanol concentration factor | 0.957 |
| Carbonate | 0.0568 mM |
| Acidity error | 0.17006 mM |

#### Warnings and errors

| | |
|---|---|
| Errors | None |
| Warnings | None |

#### Sample logD and percent species

| pH | M07_octanol logD | M07_octanol M07_octanolH | M07_octanol M07_octanol | M07_octanol M07_octanolH* | M07_octanol M07_octanol* | Comment |
|---|---|---|---|---|---|---|
| 1.000 | 0.31 | 32.68 % | 0.00 % | 66.87 % | 0.45 % | |
| 1.200 | 0.32 | 32.60 % | 0.00 % | 66.69 % | 0.71 % | Stomach pH |
| 2.000 | 0.34 | 31.42 % | 0.00 % | 64.28 % | 4.29 % | |
| 3.000 | 0.53 | 22.66 % | 0.02 % | 46.35 % | 30.97 % | |
| 4.000 | 1.19 | 5.98 % | 0.05 % | 12.23 % | 81.74 % | |
| 5.000 | 2.11 | 0.72 % | 0.06 % | 1.46 % | 97.76 % | |
| 6.000 | 2.87 | 0.07 % | 0.06 % | 0.15 % | 99.72 % | |
| 6.500 | 3.07 | 0.02 % | 0.06 % | 0.05 % | 99.87 % | |
| 7.000 | 3.16 | 0.01 % | 0.06 % | 0.01 % | 99.92 % | |
| 7.400 | 3.19 | 0.00 % | 0.06 % | 0.01 % | 99.93 % | Blood pH |
| 8.000 | 3.20 | 0.00 % | 0.06 % | 0.00 % | 99.94 % | |
| 9.000 | 3.21 | 0.00 % | 0.06 % | 0.00 % | 99.94 % | |
| 10.000 | 3.21 | 0.00 % | 0.06 % | 0.00 % | 99.94 % | |
| 11.000 | 3.21 | 0.00 % | 0.06 % | 0.00 % | 99.94 % | |
| 12.000 | 3.21 | 0.00 % | 0.06 % | 0.00 % | 99.94 % | |

Sample name: **M07_octanol**
Assay name: **pH-metric high logP**
Assay ID: **18B-28013**
Filename: **C:\Sirius_T3\Mehtap\20180228_exp28_logP_T3-2\18B-28013_M07_octanol_pH-metric high logP.t3r**

Experiment start time: **2/28/2018 7:21:13 PM**
Analyst: **Pion**
Instrument ID: **T312060**

#### Graphs

Charge balance residuals

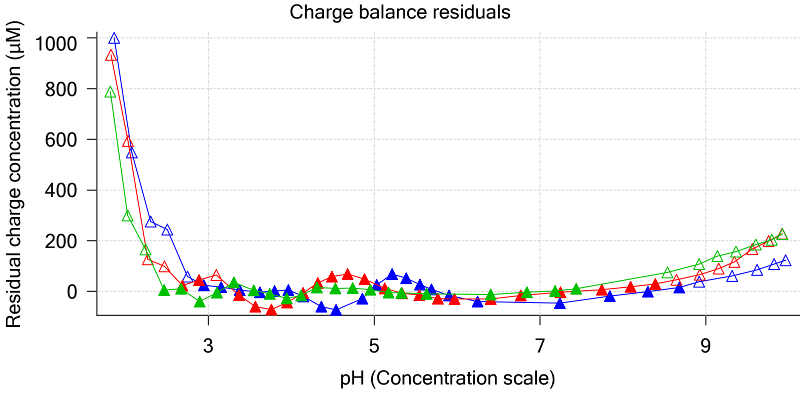


Ionic strength

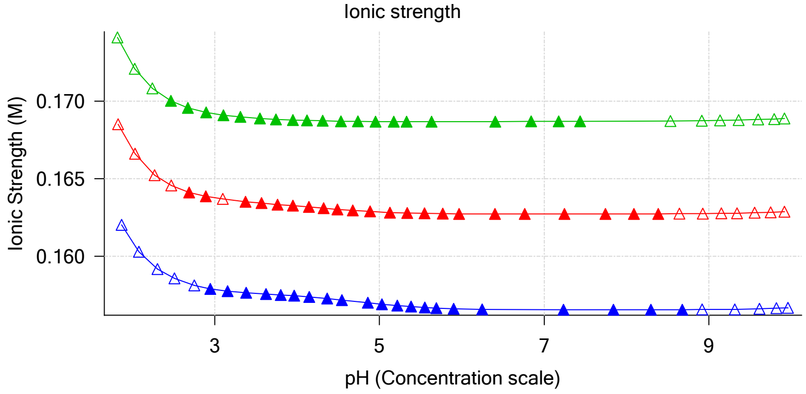


Buffer index profile

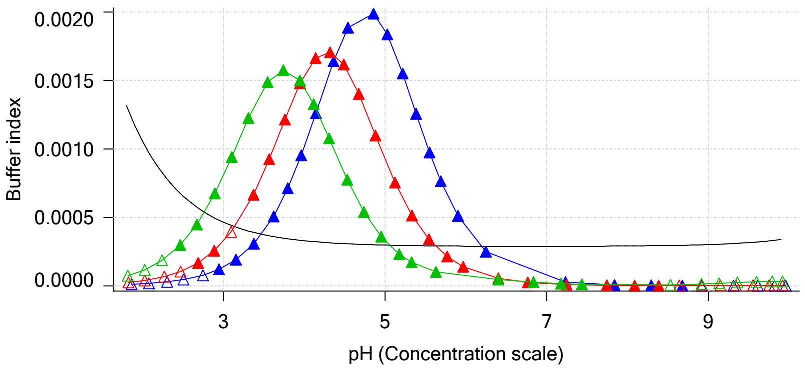


Sample M07_octanol Ionisation Graph

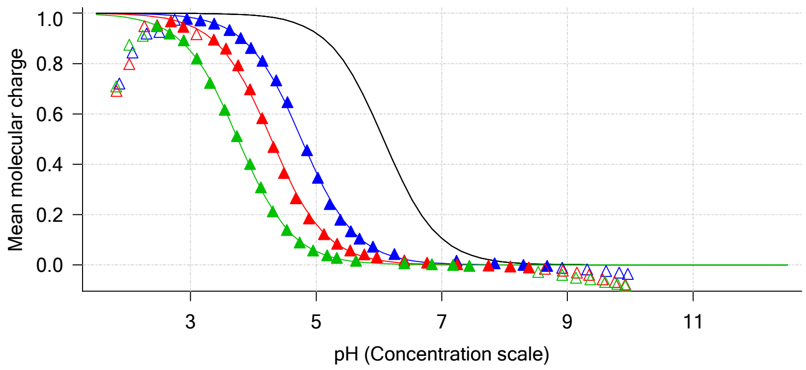


Distribution of species for sample M07_octanol

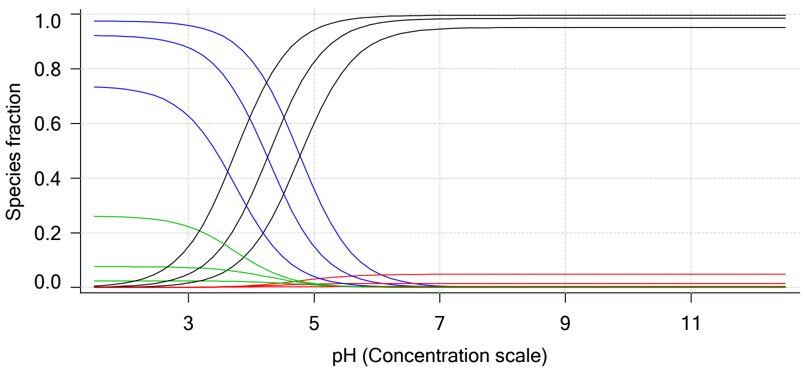


Lipophilicity Profile

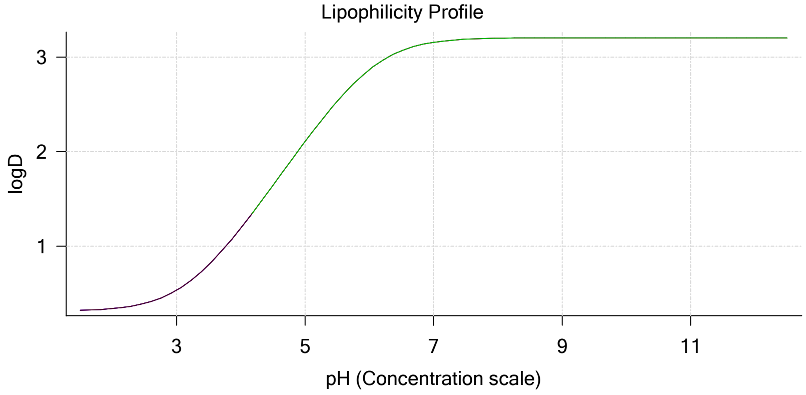


Log R Profile

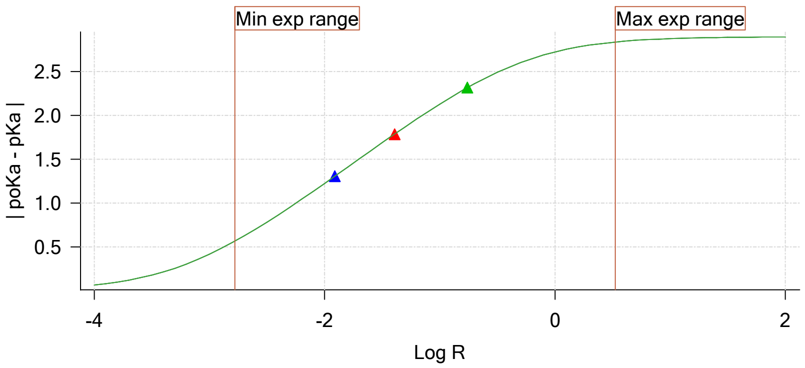


Sample Carbonate Ionisation Graph

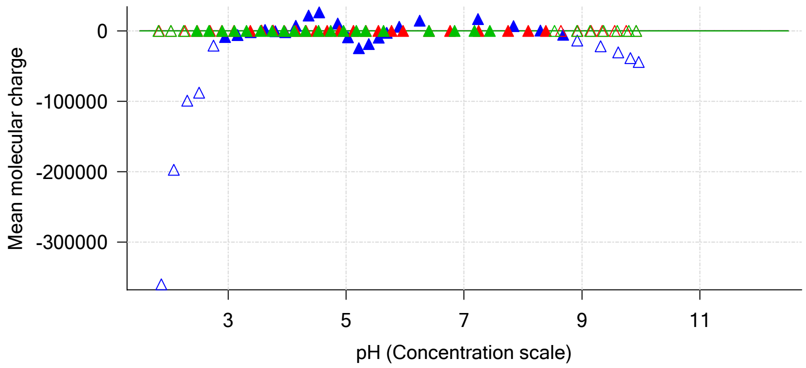

| | | | |
|---|---|---|---|
| Sample name: | **M07_octanol** | Experiment start time: | **2/28/2018 7:21:13 PM** |
| Assay name: | **pH-metric high logP** | Analyst: | **Pion** |
| Assay ID: | **18B-28013** | Instrument ID: | **T312060** |
| Filename: | **C:\Sirius_T3\Mehtap\20180228_exp28_logP_T3-2\18B-28013_M07_octanol_pH-metric high logP.t3r** | | |

#### Graphs (continued)

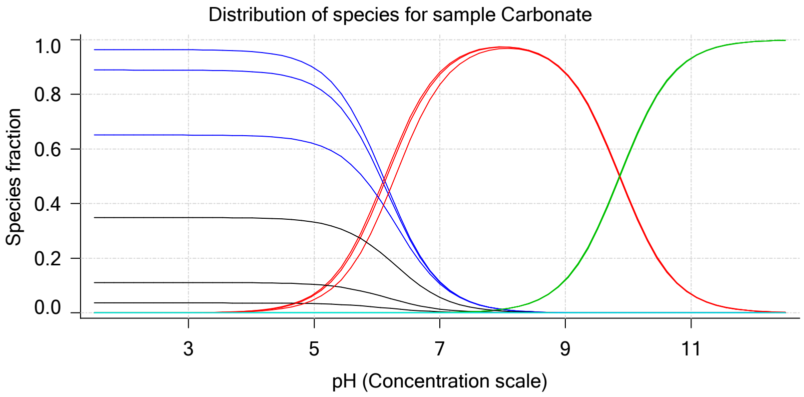

| | | | |
|---|---|---|---|
| Sample name: | **M07_octanol** | Experiment start time: | **2/28/2018 7:21:13 PM** |
| Assay name: | **pH-metric high logP** | Analyst: | **Pion** |
| Assay ID: | **18B-28013** | Instrument ID: | **T312060** |
| Filename: | **C:\Sirius_T3\Mehtap\20180228_exp28_logP_T3-2\18B-28013_M07_octanol_pH-metric high logP.t3r** | | |

### pH-metric high logP Titration 1 of 3 18B-28013 Points 1 to 31

#### Overall results

| | |
|---|---|
| RMSD | 0.587 |
| Average ionic strength | 0.157 M |
| Average temperature | 25.0°C |
| Partition ratio | 0.0122 : 1 |
| Analyte concentration range | 3242.7 µM to 3348.4 µM |
| Total points considered | 21 of 31 |

#### Warnings and errors

| | |
|---|---|
| Errors | None |
| Warnings | None |

#### Four-Plus parameters

| | | | |
|---|---|---|---|
| Alpha | 0.130 | 2/28/2018 7:21:12 PM | C:\Sirius_T3\HCl18B27.t3r |
| S | 0.9970 | 2/28/2018 7:21:12 PM | C:\Sirius_T3\HCl18B27.t3r |
| jH | 0.8 | 2/28/2018 7:21:12 PM | C:\Sirius_T3\HCl18B27.t3r |
| jOH | -0.4 | 2/28/2018 7:21:12 PM | C:\Sirius_T3\HCl18B27.t3r |

#### Titrants

| | | | |
|---|---|---|---|
| 0.50 M HCl | 0.993513 | 2/28/2018 7:21:12 PM | C:\Sirius_T3\HCl18B27.t3r |
| 0.50 M KOH | 0.999845 | 2/28/2018 7:21:13 PM | C:\Sirius_T3\KOH18B27.t3r |

#### Sample

| | |
|---|---|
| M07_octanol concentration factor | 1.061 |
| Base pKa 1 | 6.07 |
| logP (XH +) | 0.43 |
| logP (neutral X) | 3.21 |

#### Sample graphs

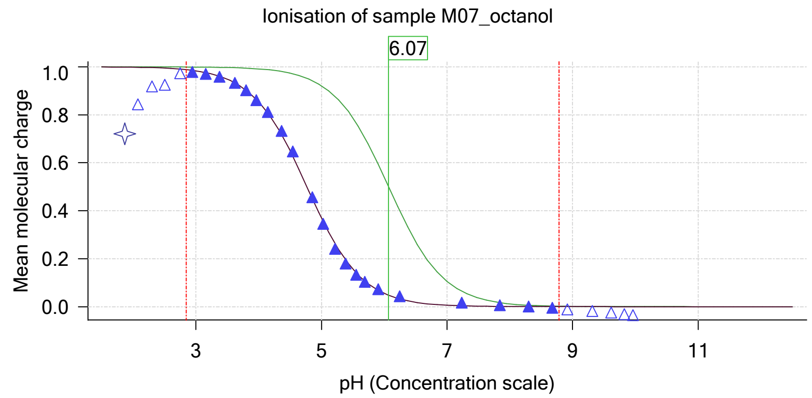


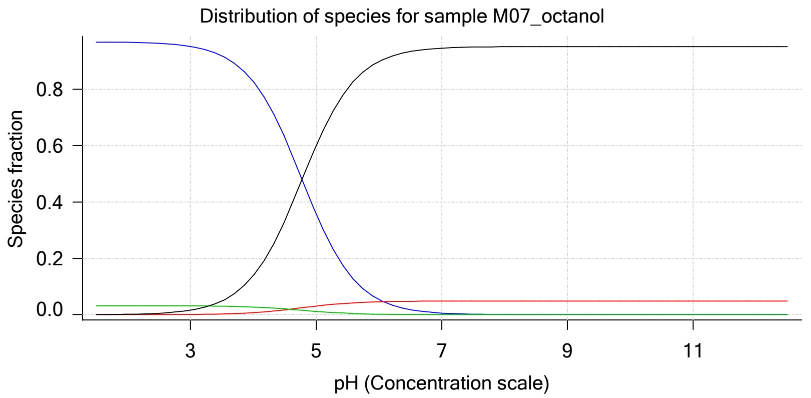

| | | | |
|---|---|---|---|
| Sample name: | **M07_octanol** | Experiment start time: | **2/28/2018 7:21:13 PM** |
| Assay name: | **pH-metric high logP** | Analyst: | **Pion** |
| Assay ID: | **18B-28013** | Instrument ID: | **T312060** |
| Filename: | **C:\Sirius_T3\Mehtap\20180228_exp28_logP_T3-2\18B-28013_M07_octanol_pH-metric high logP.t3r** | | |

#### Sample graphs (continued)

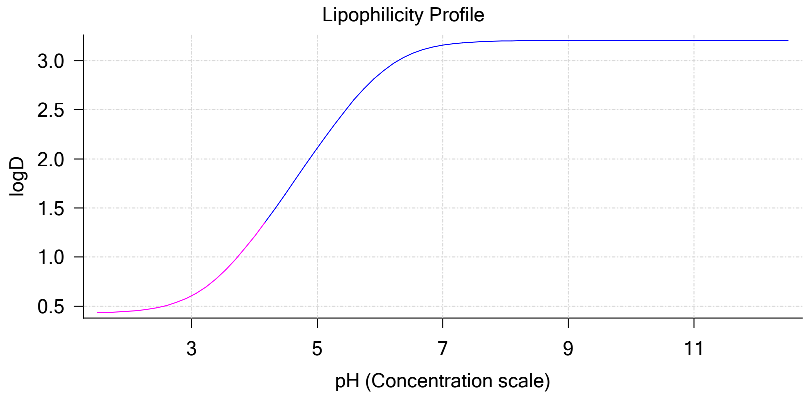


#### Sample logD and percent species

| pH | M07_octanol logD | M07_octanol M07_octanolH | M07_octanol M07_octanol | M07_octanol M07_octanolH* | M07_octanol M07_octanol* | Comment |
|---|---|---|---|---|---|---|
| 1.000 | 0.43 | 96.82 % | 0.00 % | 3.16 % | 0.02 % | |
| 1.200 | 0.43 | 96.81 % | 0.00 % | 3.16 % | 0.03 % | Stomach pH |
| 2.000 | 0.45 | 96.67 % | 0.01 % | 3.16 % | 0.16 % | |
| 3.000 | 0.61 | 95.22 % | 0.08 % | 3.11 % | 1.59 % | |
| 4.000 | 1.21 | 82.74 % | 0.70 % | 2.70 % | 13.85 % | |
| 5.000 | 2.11 | 35.82 % | 3.05 % | 1.17 % | 59.96 % | |
| 6.000 | 2.87 | 5.37 % | 4.57 % | 0.18 % | 89.88 % | |
| 6.500 | 3.07 | 1.77 % | 4.75 % | 0.06 % | 93.43 % | |
| 7.000 | 3.16 | 0.57 % | 4.81 % | 0.02 % | 94.61 % | |
| 7.400 | 3.19 | 0.23 % | 4.83 % | 0.01 % | 94.94 % | Blood pH |
| 8.000 | 3.20 | 0.06 % | 4.84 % | 0.00 % | 95.11 % | |
| 9.000 | 3.21 | 0.01 % | 4.84 % | 0.00 % | 95.16 % | |
| 10.000 | 3.21 | 0.00 % | 4.84 % | 0.00 % | 95.16 % | |
| 11.000 | 3.21 | 0.00 % | 4.84 % | 0.00 % | 95.16 % | |
| 12.000 | 3.21 | 0.00 % | 4.84 % | 0.00 % | 95.16 % | |

#### Carbonate and acidity

| | |
|---|---|
| Carbonate | 0.000 mM |
| Acidity error | 0.270 mM |

#### Other graphs

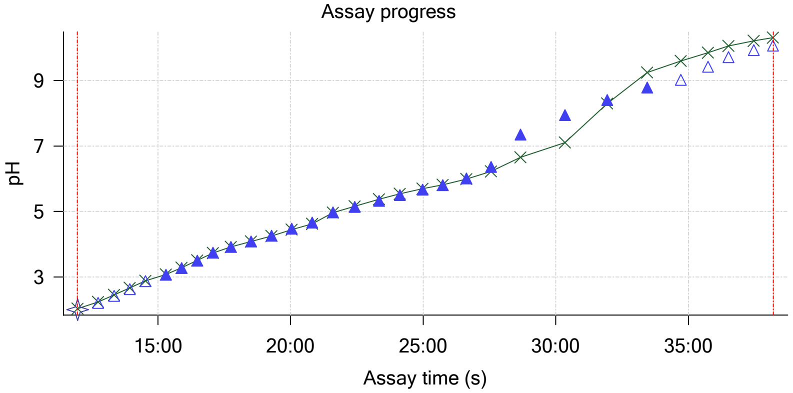


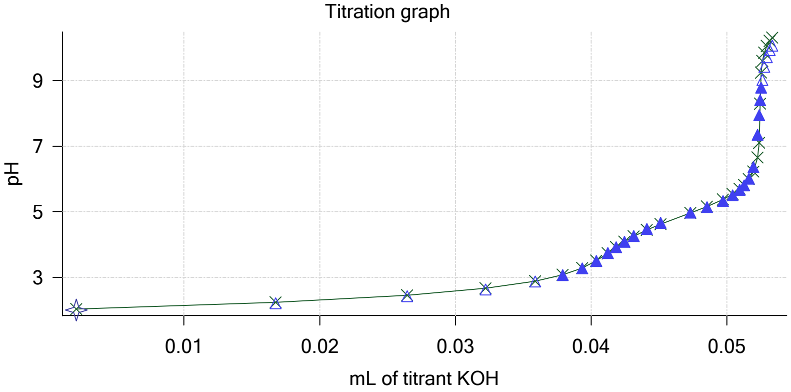

Sample name: **M07_octanol**
Assay name: **pH-metric high logP**
Assay ID: **18B-28013**
Filename: **C:\Sirius_T3\Mehtap\20180228_exp28_logP_T3-2\18B-28013_M07_octanol_pH-metric high logP.t3r**

Experiment start time: **2/28/2018 7:21:13 PM**
Analyst: **Pion**
Instrument ID: **T312060**

#### Other graphs (continued)

Charge balance residuals

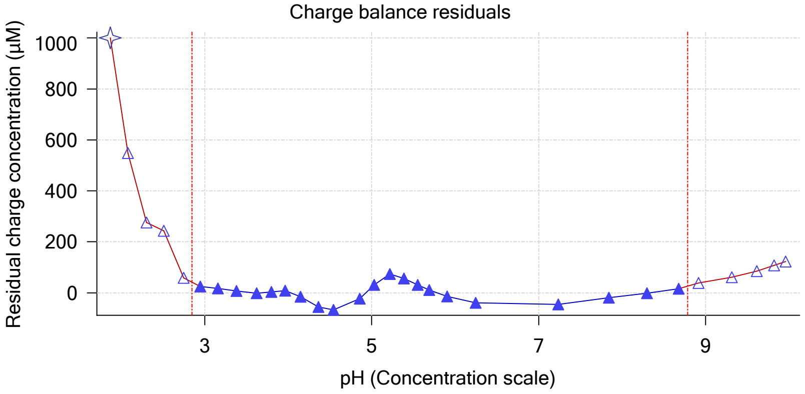


Buffer index profile

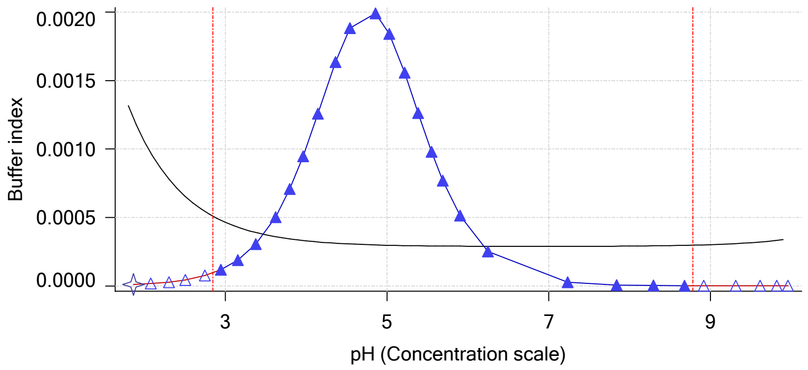


Ionic strength

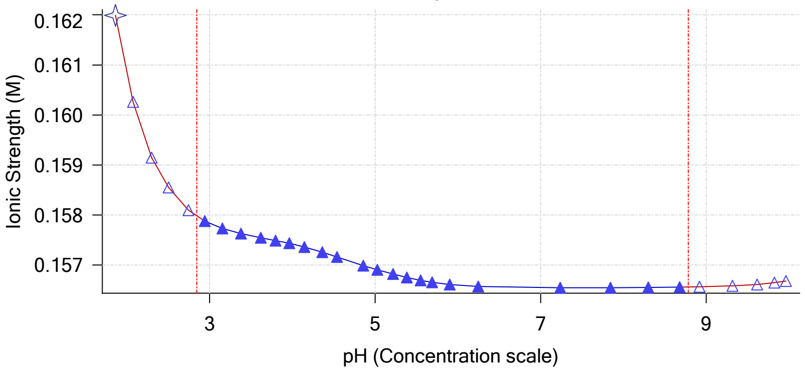


Log R Profile

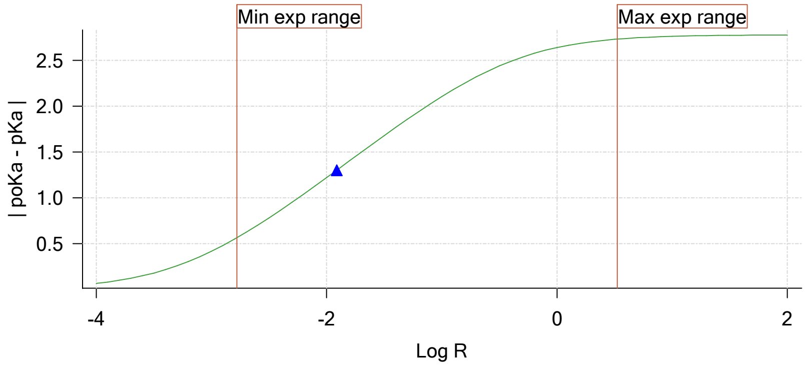


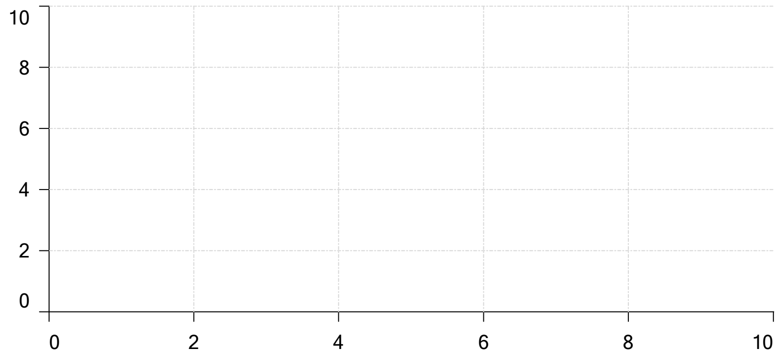


Ionisation of sample Carbonate

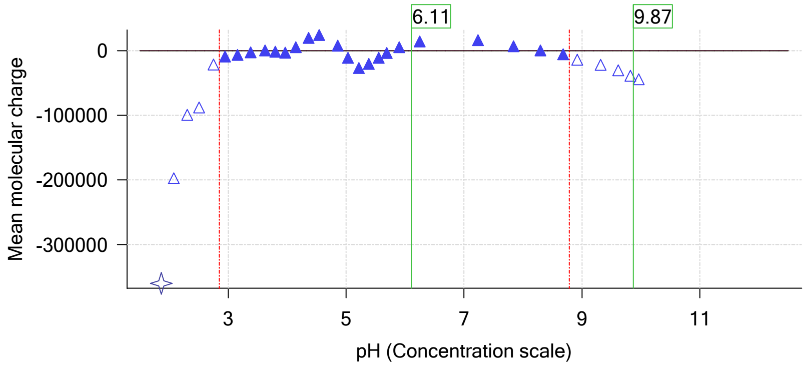

| | | | |
|---|---|---|---|
| Sample name: | **M07_octanol** | Experiment start time: | **2/28/2018 7:21:13 PM** |
| Assay name: | **pH-metric high logP** | Analyst: | **Pion** |
| Assay ID: | **18B-28013** | Instrument ID: | **T312060** |
| Filename: | **C:\Sirius_T3\Mehtap\20180228_exp28_logP_T3-2\18B-28013_M07_octanol_pH-metric high logP.t3r** | | |

### pH-metric high logP Titration 2 of 3 18B-28013 Points 32 to 65

#### Overall results

| | |
|---|---|
| RMSD | 0.919 |
| Average ionic strength | 0.163 M |
| Average temperature | 25.0°C |
| Partition ratio | 0.0405 : 1 |
| Analyte concentration range | 2940.2 µM to 3039.0 µM |
| Total points considered | 22 of 34 |

#### Warnings and errors

| | |
|---|---|
| Errors | None |
| Warnings | None |

#### Four-Plus parameters

| | | | |
|---|---|---|---|
| Alpha | 0.130 | 2/28/2018 7:21:12 PM | C:\Sirius_T3\HCl18B27.t3r |
| S | 0.9970 | 2/28/2018 7:21:12 PM | C:\Sirius_T3\HCl18B27.t3r |
| jH | 0.8 | 2/28/2018 7:21:12 PM | C:\Sirius_T3\HCl18B27.t3r |
| jOH | -0.4 | 2/28/2018 7:21:12 PM | C:\Sirius_T3\HCl18B27.t3r |

#### Titrants

| | | | |
|---|---|---|---|
| 0.50 M HCl | 0.993513 | 2/28/2018 7:21:12 PM | C:\Sirius_T3\HCl18B27.t3r |
| 0.50 M KOH | 0.999845 | 2/28/2018 7:21:13 PM | C:\Sirius_T3\KOH18B27.t3r |

#### Sample

| | |
|---|---|
| M07_octanol concentration factor | 0.967 |
| Base pKa 1 | 6.07 |
| logP (XH +) | 0.52 |
| logP (neutral X) | 3.23 |

#### Sample graphs

Ionisation of sample M07_octanol

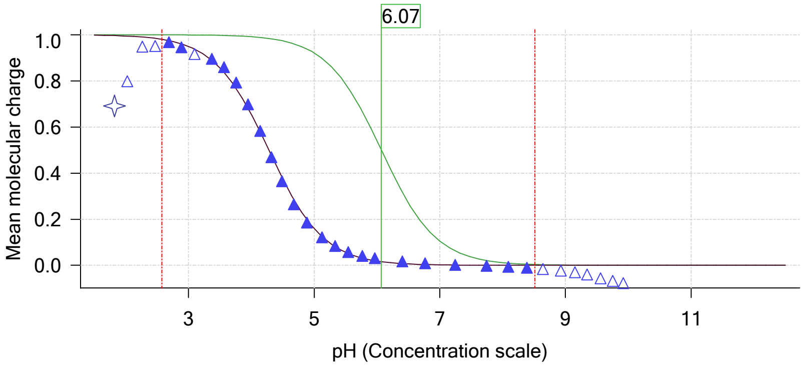


Distribution of species for sample M07_octanol

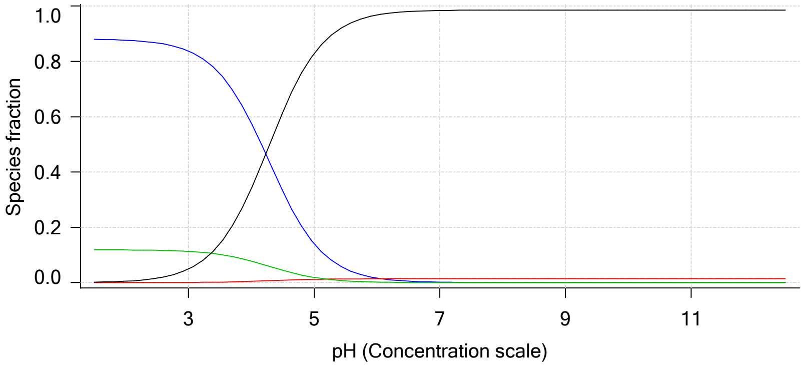

| | | | |
|---|---|---|---|
| Sample name: | **M07_octanol** | Experiment start time: | **2/28/2018 7:21:13 PM** |
| Assay name: | **pH-metric high logP** | Analyst: | **Pion** |
| Assay ID: | **18B-28013** | Instrument ID: | **T312060** |
| Filename: | **C:\Sirius_T3\Mehtap\20180228_exp28_logP_T3-2\18B-28013_M07_octanol_pH-metric high logP.t3r** | | |

#### Sample graphs (continued)

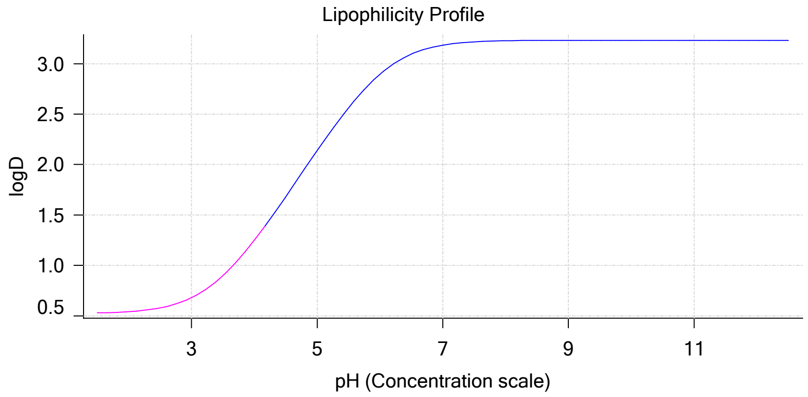


#### Sample logD and percent species

| pH | M07_octanol logD | M07_octanol M07_octanolH | M07_octanol M07_octanol | M07_octanol M07_octanolH* | M07_octanol M07_octanol* | Comment |
|---|---|---|---|---|---|---|
| 1.000 | 0.52 | 88.06 % | 0.00 % | 11.88 % | 0.05 % | |
| 1.200 | 0.53 | 88.04 % | 0.00 % | 11.88 % | 0.08 % | Stomach pH |
| 2.000 | 0.54 | 87.65 % | 0.01 % | 11.83 % | 0.52 % | |
| 3.000 | 0.68 | 83.70 % | 0.07 % | 11.30 % | 4.93 % | |
| 4.000 | 1.25 | 57.73 % | 0.49 % | 7.79 % | 33.99 % | |
| 5.000 | 2.14 | 14.07 % | 1.20 % | 1.90 % | 82.84 % | |
| 6.000 | 2.90 | 1.64 % | 1.40 % | 0.22 % | 96.74 % | |
| 6.500 | 3.10 | 0.53 % | 1.42 % | 0.07 % | 97.99 % | |
| 7.000 | 3.18 | 0.17 % | 1.42 % | 0.02 % | 98.39 % | |
| 7.400 | 3.21 | 0.07 % | 1.42 % | 0.01 % | 98.50 % | Blood pH |
| 8.000 | 3.23 | 0.02 % | 1.42 % | 0.00 % | 98.56 % | |
| 9.000 | 3.23 | 0.00 % | 1.42 % | 0.00 % | 98.57 % | |
| 10.000 | 3.23 | 0.00 % | 1.42 % | 0.00 % | 98.57 % | |
| 11.000 | 3.23 | 0.00 % | 1.42 % | 0.00 % | 98.58 % | |
| 12.000 | 3.23 | 0.00 % | 1.42 % | 0.00 % | 98.58 % | |

#### Carbonate and acidity

| | |
|---|---|
| Carbonate | 0.008 mM |
| Acidity error | 0.046 mM |

#### Other graphs

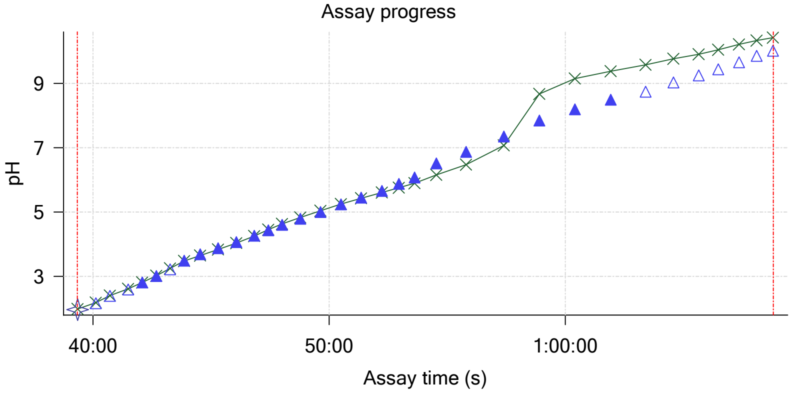


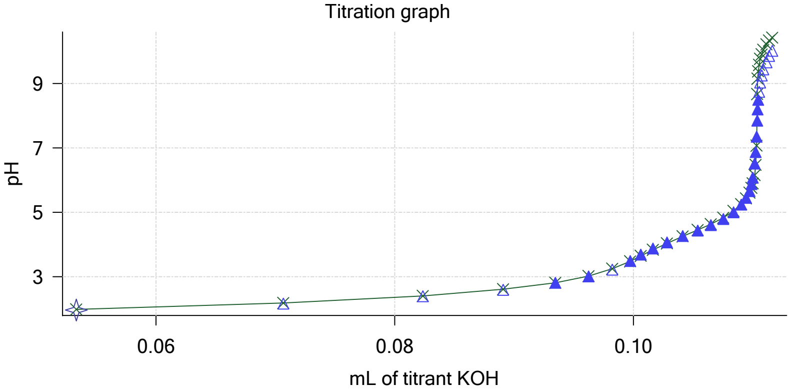

Sample name: **M07_octanol**
Assay name: **pH-metric high logP**
Assay ID: **18B-28013**
Filename: **C:\Sirius_T3\Mehtap\20180228_exp28_logP_T3-2\18B-28013_M07_octanol_pH-metric high logP.t3r**

Experiment start time: **2/28/2018 7:21:13 PM**
Analyst: **Pion**
Instrument ID: **T312060**

#### Other graphs (continued)

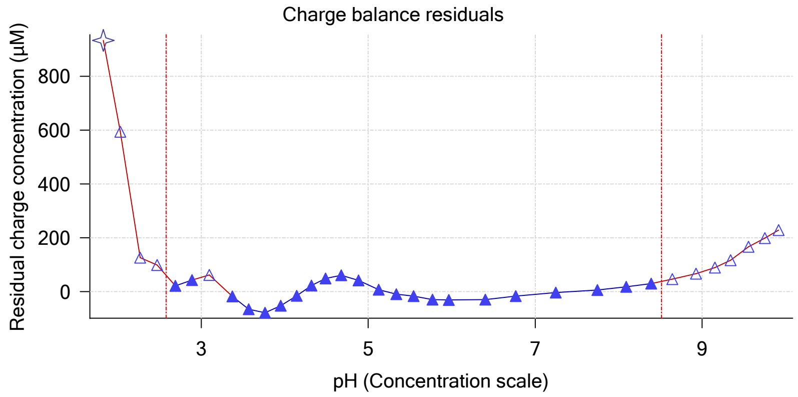


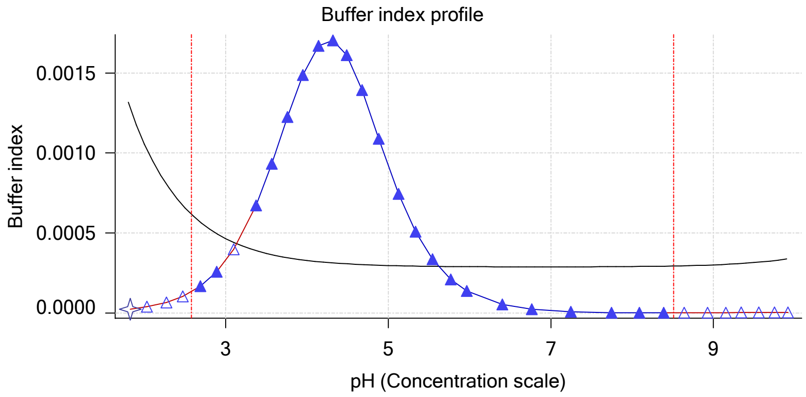


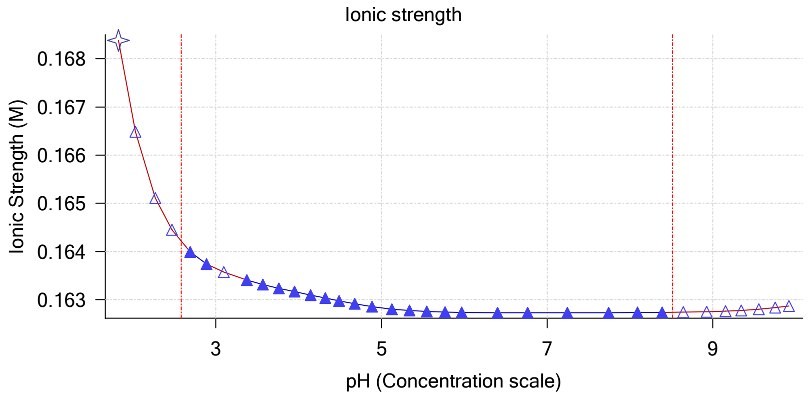


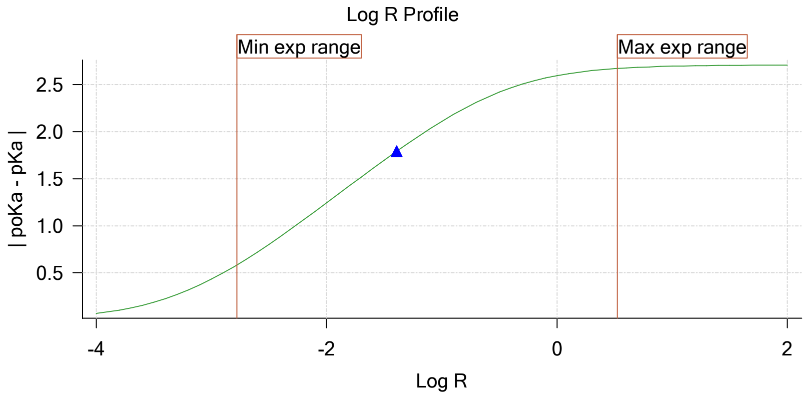


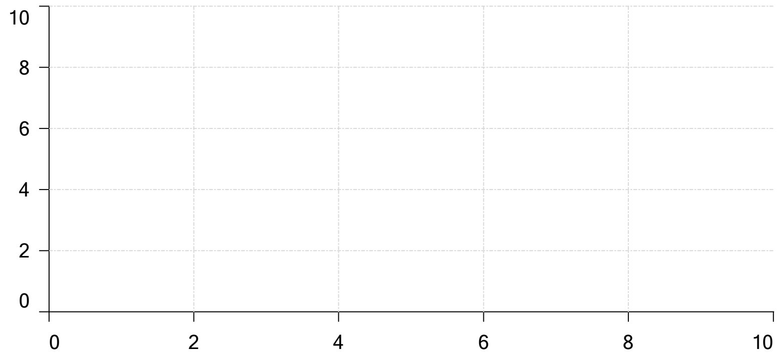


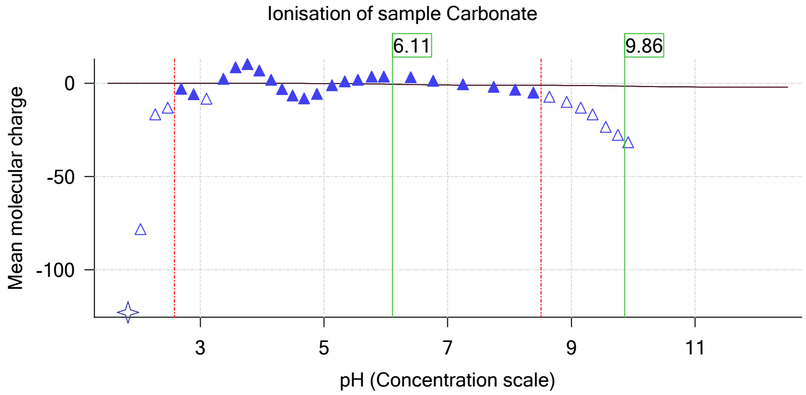

| | | | |
|---|---|---|---|
| Sample name: | **M07_octanol** | Experiment start time: | **2/28/2018 7:21:13 PM** |
| Assay name: | **pH-metric high logP** | Analyst: | **Pion** |
| Assay ID: | **18B-28013** | Instrument ID: | **T312060** |
| Filename: | **C:\Sirius_T3\Mehtap\20180228_exp28_logP_T3-2\18B-28013_M07_octanol_pH-metric high logP.t3r** | | |

# pH-metric high logP   Titration 3 of 3   18B-28013 Points 66 to 95

#### Overall results

| | |
|---|---|
| RMSD | 0.230 |
| Average ionic strength | 0.169 M |
| Average temperature | 25.0°C |
| Partition ratio | 0.1734 : 1 |
| Analyte concentration range | 2430.3 μM to 2503.3 μM |
| Total points considered | 20 of 30 |

#### Warnings and errors

| | |
|---|---|
| Errors | None |
| Warnings | None |

#### Four-Plus parameters

| | | | |
|---|---|---|---|
| Alpha | 0.130 | 2/28/2018 7:21:12 PM | C:\Sirius_T3\HCl18B27.t3r |
| S | 0.9970 | 2/28/2018 7:21:12 PM | C:\Sirius_T3\HCl18B27.t3r |
| jH | 0.8 | 2/28/2018 7:21:12 PM | C:\Sirius_T3\HCl18B27.t3r |
| jOH | -0.4 | 2/28/2018 7:21:12 PM | C:\Sirius_T3\HCl18B27.t3r |

#### Titrants

| | | | |
|---|---|---|---|
| 0.50 M HCl | 0.993513 | 2/28/2018 7:21:12 PM | C:\Sirius_T3\HCl18B27.t3r |
| 0.50 M KOH | 0.999845 | 2/28/2018 7:21:13 PM | C:\Sirius_T3\KOH18B27.t3r |

#### Sample

| | |
|---|---|
| M07_octanol concentration factor | 0.957 |
| Base pKa 1 | 6.07 |
| logP (XH +) | 0.49 |
| logP (neutral X) | 3.26 |

#### Sample graphs

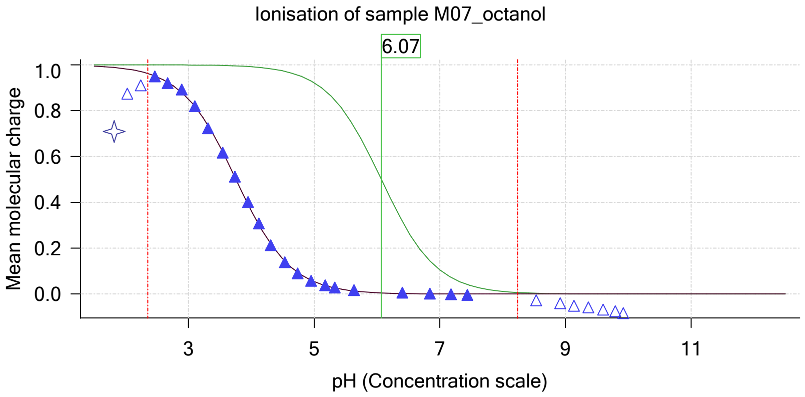


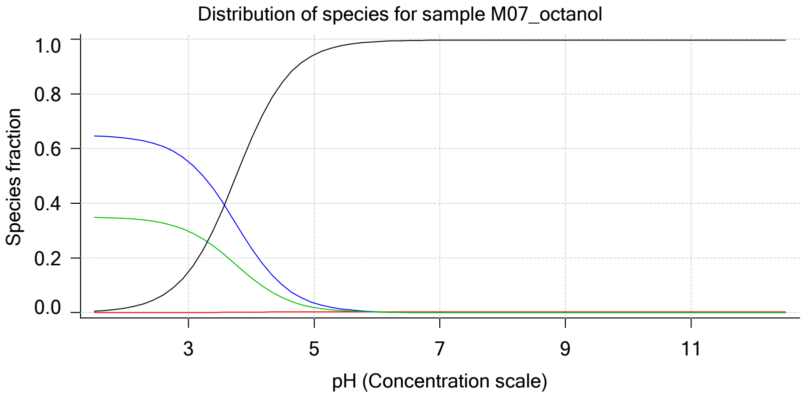

| | | | |
|---|---|---|---|
| Sample name: | **M07_octanol** | Experiment start time: | **2/28/2018 7:21:13 PM** |
| Assay name: | **pH-metric high logP** | Analyst: | **Pion** |
| Assay ID: | **18B-28013** | Instrument ID: | **T312060** |
| Filename: | **C:\Sirius_T3\Mehtap\20180228_exp28_logP_T3-2\18B-28013_M07_octanol_pH-metric high logP.t3r** | | |

#### Sample graphs (continued)

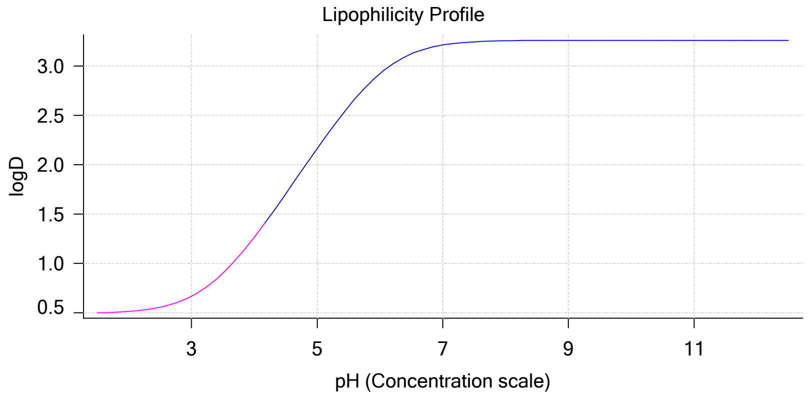


#### Sample logD and percent species

| pH | M07_octanol logD | M07_octanol M07_octanolH | M07_octanol M07_octanol | M07_octanol M07_octanolH* | M07_octanol M07_octanol* | Comment |
|---|---|---|---|---|---|---|
| 1.000 | 0.49 | 64.86 % | 0.00 % | 34.97 % | 0.17 % | |
| 1.200 | 0.50 | 64.79 % | 0.00 % | 34.93 % | 0.28 % | Stomach pH |
| 2.000 | 0.51 | 63.85 % | 0.01 % | 34.43 % | 1.72 % | |
| 3.000 | 0.67 | 55.29 % | 0.05 % | 29.81 % | 14.85 % | |
| 4.000 | 1.27 | 23.62 % | 0.20 % | 12.73 % | 63.45 % | |
| 5.000 | 2.16 | 3.51 % | 0.30 % | 1.89 % | 94.30 % | |
| 6.000 | 2.92 | 0.37 % | 0.31 % | 0.20 % | 99.12 % | |
| 6.500 | 3.12 | 0.12 % | 0.32 % | 0.06 % | 99.50 % | |
| 7.000 | 3.21 | 0.04 % | 0.32 % | 0.02 % | 99.63 % | |
| 7.400 | 3.24 | 0.01 % | 0.32 % | 0.01 % | 99.66 % | Blood pH |
| 8.000 | 3.25 | 0.00 % | 0.32 % | 0.00 % | 99.68 % | |
| 9.000 | 3.26 | 0.00 % | 0.32 % | 0.00 % | 99.68 % | |
| 10.000 | 3.26 | 0.00 % | 0.32 % | 0.00 % | 99.68 % | |
| 11.000 | 3.26 | 0.00 % | 0.32 % | 0.00 % | 99.68 % | |
| 12.000 | 3.26 | 0.00 % | 0.32 % | 0.00 % | 99.68 % | |

#### Carbonate and acidity

| | |
|---|---|
| Carbonate | 0.057 mM |
| Acidity error | 0.170 mM |

#### Other graphs

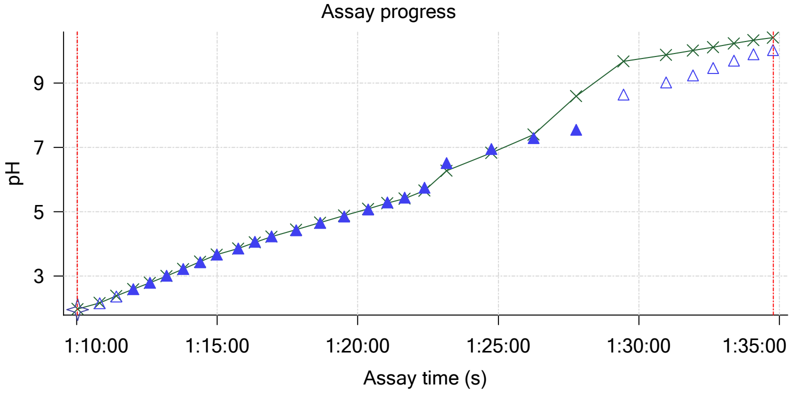


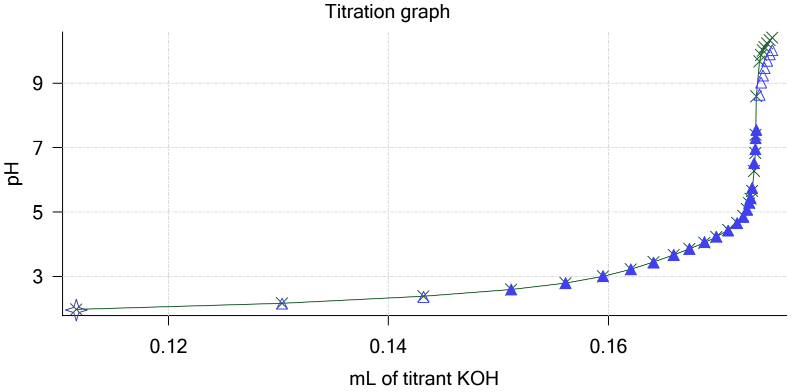

Sample name: **M07_octanol**
Assay name: **pH-metric high logP**
Assay ID: **18B-28013**
Filename: **C:\Sirius_T3\Mehtap\20180228_exp28_logP_T3-2\18B-28013_M07_octanol_pH-metric high logP.t3r**

Experiment start time: **2/28/2018 7:21:13 PM**
Analyst: **Pion**
Instrument ID: **T312060**

#### Other graphs (continued)

Charge balance residuals

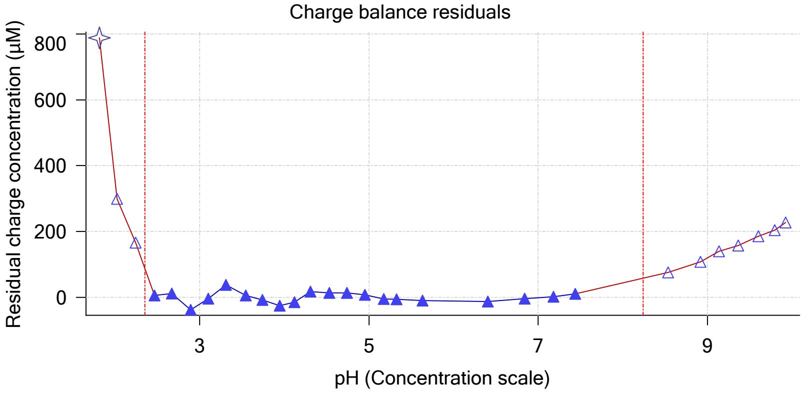


Buffer index profile

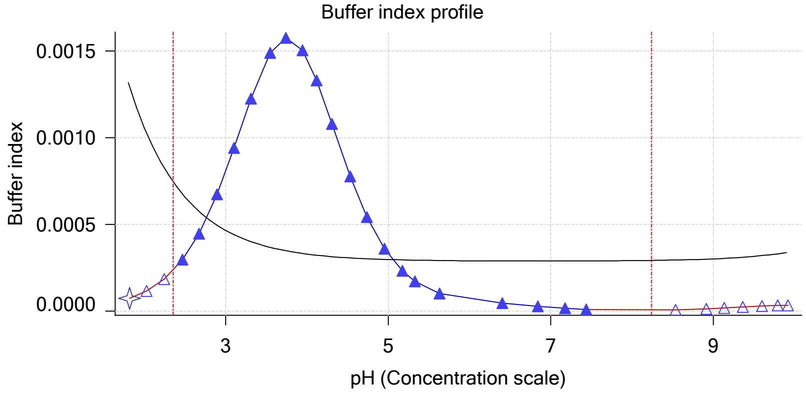


Ionic strength

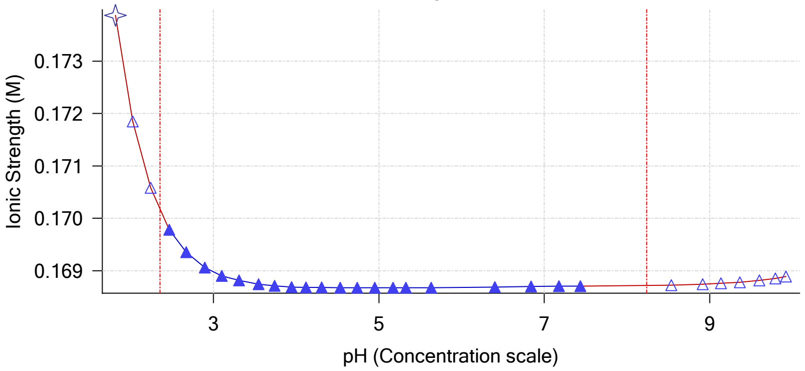


Log R Profile

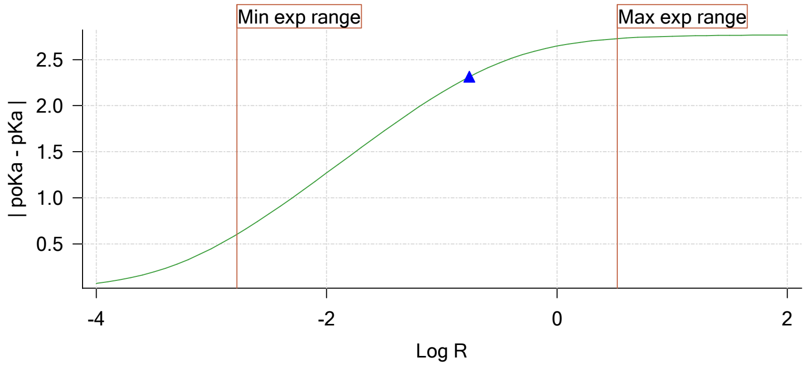


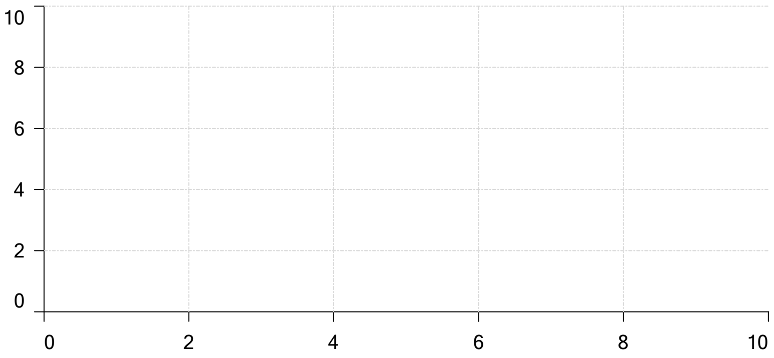


Ionisation of sample Carbonate

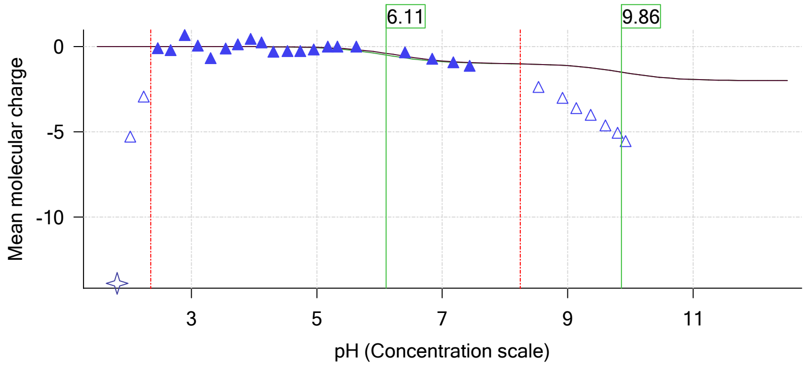

| | | | |
|---|---|---|---|
| Sample name: | **M07_octanol** | Experiment start time: | **2/28/2018 7:21:13 PM** |
| Assay name: | **pH-metric high logP** | Analyst: | **Pion** |
| Assay ID: | **18B-28013** | Instrument ID: | **T312060** |
| Filename: | **C:\Sirius_T3\Mehtap\20180228_exp28_logP_T3-2\18B-28013_M07_octanol_pH-metric high logP.t3r** | | |

#### Assay Model

| Settings | Value | Date/Time changed | Imported from |
|---|---|---|---|
| Sample name | M07_octanol | 2/27/2018 4:29:24 PM | User entered value |
| Sample by | Weight | | Default value |
| Sample weight | 0.001240 g | 2/28/2018 4:24:19 PM | User entered value |
| Formula weight | 235.28 g/mol | 2/27/2018 4:29:24 PM | User entered value |
| Solubility | Unknown | | Default value |
| Molecular weight | 235.28 | 2/27/2018 4:29:24 PM | User entered value |
| Individual pKa ionic environments | No | | Default value |
| Number of pKas | 1 | 2/27/2018 4:29:24 PM | User entered value |
| Sample is a | Base | 2/27/2018 4:29:24 PM | User entered value |
| pKa 1 | 6.07 | 2/27/2018 4:29:24 PM | User entered value |
| logp (XH +) | 0.50 | 2/28/2018 1:33:04 PM | User entered value |
| logP (neutral X) | 3.44 | 2/28/2018 1:33:10 PM | User entered value |

#### Events

| Time | Event | Water | Acid | Base | Octanol | pH | dpH/dt | pH R-squared | pH SD | dpH/dt time |
|---|---|---|---|---|---|---|---|---|---|---|
| 8:59.2 | Initial pH = 6.72 | | | | | | | | | |
| 11:58.8 | Data point 1 | 1.50000 mL | 0.05191 mL | 0.00207 mL | 0.01999 mL | 2.001 | 0.00132 | 0.04152 | 0.00032 | 10.0 s |
| 12:44.9 | Data point 2 | 1.50000 mL | 0.05191 mL | 0.01675 mL | 0.01999 mL | 2.208 | -0.01261 | 0.85529 | 0.00067 | 10.5 s |
| 13:21.1 | Data point 3 | 1.50000 mL | 0.05191 mL | 0.02643 mL | 0.01999 mL | 2.430 | -0.01070 | 0.78000 | 0.00060 | 10.0 s |
| 13:56.6 | Data point 4 | 1.50000 mL | 0.05191 mL | 0.03222 mL | 0.01999 mL | 2.632 | -0.00370 | 0.50717 | 0.00026 | 10.0 s |
| 14:32.2 | Data point 5 | 1.50000 mL | 0.05191 mL | 0.03587 mL | 0.01999 mL | 2.871 | -0.00475 | 0.47439 | 0.00034 | 10.0 s |
| 15:18.0 | Data point 6 | 1.50000 mL | 0.05191 mL | 0.03789 mL | 0.01999 mL | 3.068 | -0.00916 | 0.24962 | 0.00091 | 10.0 s |
| 15:53.4 | Data point 7 | 1.50000 mL | 0.05191 mL | 0.03935 mL | 0.01999 mL | 3.277 | -0.00524 | 0.87796 | 0.00028 | 10.5 s |
| 16:29.4 | Data point 8 | 1.50000 mL | 0.05191 mL | 0.04038 mL | 0.01999 mL | 3.499 | -0.00471 | 0.65043 | 0.00029 | 10.0 s |
| 17:04.8 | Data point 9 | 1.50000 mL | 0.05191 mL | 0.04123 mL | 0.01999 mL | 3.741 | -0.00484 | 0.50306 | 0.00034 | 10.0 s |
| 17:45.4 | Data point 10 | 1.50000 mL | 0.05191 mL | 0.04184 mL | 0.01999 mL | 3.917 | -0.00416 | 0.61481 | 0.00026 | 10.0 s |
| 18:31.2 | Data point 11 | 1.50000 mL | 0.05191 mL | 0.04245 mL | 0.01999 mL | 4.082 | -0.00448 | 0.70035 | 0.00026 | 10.0 s |
| 19:16.9 | Data point 12 | 1.50000 mL | 0.05191 mL | 0.04313 mL | 0.01999 mL | 4.264 | -0.01077 | 0.67467 | 0.00065 | 10.0 s |
| 20:02.6 | Data point 13 | 1.50000 mL | 0.05191 mL | 0.04410 mL | 0.01999 mL | 4.480 | -0.01584 | 0.85617 | 0.00085 | 10.5 s |
| 20:48.8 | Data point 14 | 1.50000 mL | 0.05191 mL | 0.04511 mL | 0.01999 mL | 4.658 | -0.01680 | 0.77140 | 0.00095 | 11.5 s |
| 21:36.2 | Data point 15 | 1.50000 mL | 0.05191 mL | 0.04730 mL | 0.01999 mL | 4.972 | -0.01791 | 0.88784 | 0.00094 | 13.5 s |
| 22:25.5 | Data point 16 | 1.50000 mL | 0.05191 mL | 0.04854 mL | 0.01999 mL | 5.144 | -0.01931 | 0.92268 | 0.00099 | 14.0 s |
| 23:20.4 | Data point 17 | 1.50000 mL | 0.05191 mL | 0.04972 mL | 0.01999 mL | 5.332 | -0.01736 | 0.89270 | 0.00091 | 16.5 s |
| 24:07.7 | Data point 18 | 1.50000 mL | 0.05191 mL | 0.05042 mL | 0.01999 mL | 5.498 | -0.01945 | 0.93126 | 0.00100 | 20.5 s |
| 24:58.7 | Data point 19 | 1.50000 mL | 0.05191 mL | 0.05094 mL | 0.01999 mL | 5.664 | -0.01909 | 0.92243 | 0.00098 | 20.5 s |
| 25:44.7 | Data point 20 | 1.50000 mL | 0.05191 mL | 0.05127 mL | 0.01999 mL | 5.804 | -0.01742 | 0.80585 | 0.00096 | 23.0 s |
| 26:38.3 | Data point 21 | 1.50000 mL | 0.05191 mL | 0.05162 mL | 0.01999 mL | 6.014 | -0.01944 | 0.96457 | 0.00098 | 24.5 s |
| 27:33.3 | Data point 22 | 1.50000 mL | 0.05191 mL | 0.05195 mL | 0.01999 mL | 6.358 | -0.01926 | 0.90710 | 0.00100 | 36.0 s |
| 28:39.9 | Data point 23 | 1.50000 mL | 0.05191 mL | 0.05226 mL | 0.01999 mL | 7.345 | -0.04577 | 0.99604 | 0.00226 | Timed out at 59.5 s |
| 30:20.7 | Data point 24 | 1.50000 mL | 0.05191 mL | 0.05238 mL | 0.01999 mL | 7.948 | -0.04715 | 0.98406 | 0.00235 | Timed out at 59.5 s |
| 31:56.4 | Data point 25 | 1.50000 mL | 0.05191 mL | 0.05245 mL | 0.01999 mL | 8.403 | -0.03639 | 0.99199 | 0.00180 | Timed out at 59.5 s |
| 33:26.9 | Data point 26 | 1.50000 mL | 0.05191 mL | 0.05252 mL | 0.01999 mL | 8.783 | -0.01951 | 0.93124 | 0.00100 | 45.5 s |
| 34:42.9 | Data point 27 | 1.50000 mL | 0.05191 mL | 0.05261 mL | 0.01999 mL | 9.021 | -0.01783 | 0.95357 | 0.00090 | 31.0 s |
| 35:44.5 | Data point 28 | 1.50000 mL | 0.05191 mL | 0.05275 mL | 0.01999 mL | 9.417 | -0.01646 | 0.77635 | 0.00092 | 15.5 s |
| 36:30.7 | Data point 29 | 1.50000 mL | 0.05191 mL | 0.05294 mL | 0.01999 mL | 9.717 | -0.01888 | 0.92426 | 0.00097 | 21.5 s |
| 37:28.0 | Data point 30 | 1.50000 mL | 0.05191 mL | 0.05315 mL | 0.01999 mL | 9.923 | -0.01891 | 0.96558 | 0.00095 | 12.5 s |
| 38:11.1 | Data point 31 | 1.50000 mL | 0.05191 mL | 0.05334 mL | 0.01999 mL | 10.061 | -0.01803 | 0.97803 | 0.00090 | 10.5 s |
| 39:21.0 | Data point 32 | 1.50000 mL | 0.11084 mL | 0.05334 mL | 0.06999 mL | 1.960 | -0.00708 | 0.60704 | 0.00045 | 10.0 s |
| 40:07.3 | Data point 33 | 1.50000 mL | 0.11084 mL | 0.07067 mL | 0.06999 mL | 2.160 | -0.01155 | 0.76739 | 0.00065 | 10.0 s |
| 40:43.0 | Data point 34 | 1.50000 mL | 0.11084 mL | 0.08236 mL | 0.06999 mL | 2.395 | -0.00810 | 0.72844 | 0.00047 | 10.5 s |
| 41:29.5 | Data point 35 | 1.50000 mL | 0.11084 mL | 0.08909 mL | 0.06999 mL | 2.596 | -0.00385 | 0.58762 | 0.00025 | 10.0 s |
| 42:05.0 | Data point 36 | 1.50000 mL | 0.11084 mL | 0.09344 mL | 0.06999 mL | 2.811 | -0.00364 | 0.09675 | 0.00058 | 10.0 s |
| 42:40.5 | Data point 37 | 1.50000 mL | 0.11084 mL | 0.09624 mL | 0.06999 mL | 3.010 | -0.00760 | 0.20188 | 0.00084 | 10.0 s |

| | | | |
|---|---|---|---|
| Sample name: | **M07_octanol** | Experiment start time: | **2/28/2018 7:21:13 PM** |
| Assay name: | **pH-metric high logP** | Analyst: | **Pion** |
| Assay ID: | **18B-28013** | Instrument ID: | **T312060** |
| Filename: | **C:\Sirius_T3\Mehtap\20180228_exp28_logP_T3-2\18B-28013_M07_octanol_pH-metric high logP.t3r** | | |

#### Events (continued)

| Time | Event | Water | Acid | Base | Octanol | pH | dpH/dt | pH R-squared | pH SD | dpH/dt time |
|---|---|---|---|---|---|---|---|---|---|---|
| 43:16.0 | Data point 38 | 1.50000 mL | 0.11084 mL | 0.09821 mL | 0.06999 mL | 3.218 | -0.00906 | 0.28704 | 0.00084 | 10.0 s |
| 43:51.4 | Data point 39 | 1.50000 mL | 0.11084 mL | 0.09972 mL | 0.06999 mL | 3.492 | 0.00211 | 0.05894 | 0.00043 | 10.5 s |
| 44:32.6 | Data point 40 | 1.50000 mL | 0.11084 mL | 0.10061 mL | 0.06999 mL | 3.685 | -0.00509 | 0.57053 | 0.00033 | 10.0 s |
| 45:18.3 | Data point 41 | 1.50000 mL | 0.11084 mL | 0.10162 mL | 0.06999 mL | 3.880 | -0.00602 | 0.45174 | 0.00044 | 10.0 s |
| 46:04.1 | Data point 42 | 1.50000 mL | 0.11084 mL | 0.10280 mL | 0.06999 mL | 4.068 | 0.00238 | 0.05728 | 0.00049 | 10.0 s |
| 46:49.8 | Data point 43 | 1.50000 mL | 0.11084 mL | 0.10412 mL | 0.06999 mL | 4.260 | -0.01117 | 0.50124 | 0.00078 | 10.0 s |
| 47:25.3 | Data point 44 | 1.50000 mL | 0.11084 mL | 0.10536 mL | 0.06999 mL | 4.437 | -0.00931 | 0.43556 | 0.00070 | 10.0 s |
| 48:00.7 | Data point 45 | 1.50000 mL | 0.11084 mL | 0.10647 mL | 0.06999 mL | 4.605 | -0.00382 | 0.51934 | 0.00026 | 10.5 s |
| 48:47.0 | Data point 46 | 1.50000 mL | 0.11084 mL | 0.10753 mL | 0.06999 mL | 4.793 | 0.00774 | 0.33101 | 0.00067 | 10.5 s |
| 49:38.5 | Data point 47 | 1.50000 mL | 0.11084 mL | 0.10837 mL | 0.06999 mL | 4.999 | 0.00150 | 0.03062 | 0.00042 | 10.5 s |
| 50:29.8 | Data point 48 | 1.50000 mL | 0.11084 mL | 0.10903 mL | 0.06999 mL | 5.240 | -0.00100 | 0.00811 | 0.00055 | 11.0 s |
| 51:21.7 | Data point 49 | 1.50000 mL | 0.11084 mL | 0.10943 mL | 0.06999 mL | 5.450 | -0.00888 | 0.38198 | 0.00071 | 11.5 s |
| 52:14.0 | Data point 50 | 1.50000 mL | 0.11084 mL | 0.10971 mL | 0.06999 mL | 5.658 | -0.01542 | 0.74055 | 0.00089 | 13.0 s |
| 52:57.5 | Data point 51 | 1.50000 mL | 0.11084 mL | 0.10988 mL | 0.06999 mL | 5.881 | -0.01855 | 0.89943 | 0.00097 | 14.5 s |
| 53:37.4 | Data point 52 | 1.50000 mL | 0.11084 mL | 0.11000 mL | 0.06999 mL | 6.079 | -0.01880 | 0.89624 | 0.00098 | 24.5 s |
| 54:32.4 | Data point 53 | 1.50000 mL | 0.11084 mL | 0.11014 mL | 0.06999 mL | 6.513 | -0.01832 | 0.91525 | 0.00095 | 45.5 s |
| 55:48.5 | Data point 54 | 1.50000 mL | 0.11084 mL | 0.11023 mL | 0.06999 mL | 6.875 | -0.02824 | 0.99129 | 0.00140 | Timed out at 59.5 s |
| 57:24.1 | Data point 55 | 1.50000 mL | 0.11084 mL | 0.11030 mL | 0.06999 mL | 7.353 | -0.05669 | 0.99082 | 0.00281 | Timed out at 59.5 s |
| 58:54.6 | Data point 56 | 1.50000 mL | 0.11084 mL | 0.11035 mL | 0.06999 mL | 7.851 | -0.05247 | 0.99574 | 0.00260 | Timed out at 59.5 s |
| 1:00:25.1 | Data point 57 | 1.50000 mL | 0.11084 mL | 0.11039 mL | 0.06999 mL | 8.195 | -0.03651 | 0.95479 | 0.00184 | Timed out at 59.5 s |
| 1:01:55.6 | Data point 58 | 1.50000 mL | 0.11084 mL | 0.11044 mL | 0.06999 mL | 8.493 | -0.01930 | 0.96296 | 0.00097 | 53.0 s |
| 1:03:24.4 | Data point 59 | 1.50000 mL | 0.11084 mL | 0.11051 mL | 0.06999 mL | 8.746 | -0.01430 | 0.96004 | 0.00072 | 35.0 s |
| 1:04:35.2 | Data point 60 | 1.50000 mL | 0.11084 mL | 0.11061 mL | 0.06999 mL | 9.030 | -0.01839 | 0.90118 | 0.00096 | 29.0 s |
| 1:05:39.8 | Data point 61 | 1.50000 mL | 0.11084 mL | 0.11072 mL | 0.06999 mL | 9.255 | -0.01963 | 0.96157 | 0.00099 | 13.5 s |
| 1:06:29.0 | Data point 62 | 1.50000 mL | 0.11084 mL | 0.11087 mL | 0.06999 mL | 9.445 | -0.01933 | 0.93954 | 0.00098 | 17.0 s |
| 1:07:21.8 | Data point 63 | 1.50000 mL | 0.11084 mL | 0.11112 mL | 0.06999 mL | 9.658 | -0.01953 | 0.96611 | 0.00098 | 14.0 s |
| 1:08:06.4 | Data point 64 | 1.50000 mL | 0.11084 mL | 0.11136 mL | 0.06999 mL | 9.854 | -0.01512 | 0.84020 | 0.00082 | 11.0 s |
| 1:08:47.9 | Data point 65 | 1.50000 mL | 0.11084 mL | 0.11162 mL | 0.06999 mL | 10.018 | -0.01750 | 0.92297 | 0.00090 | 10.5 s |
| 1:10:02.6 | Data point 66 | 1.50000 mL | 0.17368 mL | 0.11162 mL | 0.31999 mL | 1.953 | -0.00769 | 0.84527 | 0.00041 | 10.5 s |
| 1:10:49.3 | Data point 67 | 1.50000 mL | 0.17368 mL | 0.13034 mL | 0.31999 mL | 2.156 | -0.00310 | 0.19456 | 0.00035 | 10.0 s |
| 1:11:25.0 | Data point 68 | 1.50000 mL | 0.17368 mL | 0.14318 mL | 0.31999 mL | 2.369 | -0.00632 | 0.23843 | 0.00064 | 10.5 s |
| 1:12:01.1 | Data point 69 | 1.50000 mL | 0.17368 mL | 0.15115 mL | 0.31999 mL | 2.591 | 0.00518 | 0.21450 | 0.00055 | 10.0 s |
| 1:12:36.7 | Data point 70 | 1.50000 mL | 0.17368 mL | 0.15612 mL | 0.31999 mL | 2.794 | -0.00339 | 0.46040 | 0.00025 | 10.0 s |
| 1:13:12.2 | Data point 71 | 1.50000 mL | 0.17368 mL | 0.15953 mL | 0.31999 mL | 3.017 | -0.00260 | 0.03972 | 0.00065 | 10.0 s |
| 1:13:47.7 | Data point 72 | 1.50000 mL | 0.17368 mL | 0.16204 mL | 0.31999 mL | 3.224 | -0.01399 | 0.69435 | 0.00083 | 10.5 s |
| 1:14:23.6 | Data point 73 | 1.50000 mL | 0.17368 mL | 0.16411 mL | 0.31999 mL | 3.429 | -0.01099 | 0.70364 | 0.00065 | 10.0 s |
| 1:14:59.0 | Data point 74 | 1.50000 mL | 0.17368 mL | 0.16595 mL | 0.31999 mL | 3.666 | -0.00580 | 0.58863 | 0.00037 | 10.0 s |
| 1:15:44.8 | Data point 75 | 1.50000 mL | 0.17368 mL | 0.16738 mL | 0.31999 mL | 3.860 | 0.00294 | 0.10993 | 0.00044 | 10.0 s |
| 1:16:20.3 | Data point 76 | 1.50000 mL | 0.17368 mL | 0.16874 mL | 0.31999 mL | 4.065 | -0.01443 | 0.61135 | 0.00091 | 10.5 s |
| 1:16:56.2 | Data point 77 | 1.50000 mL | 0.17368 mL | 0.16983 mL | 0.31999 mL | 4.237 | -0.01769 | 0.77303 | 0.00099 | 16.5 s |
| 1:17:48.5 | Data point 78 | 1.50000 mL | 0.17368 mL | 0.17088 mL | 0.31999 mL | 4.426 | -0.01278 | 0.47370 | 0.00092 | 10.0 s |
| 1:18:39.5 | Data point 79 | 1.50000 mL | 0.17368 mL | 0.17171 mL | 0.31999 mL | 4.649 | -0.00486 | 0.31109 | 0.00043 | 10.5 s |
| 1:19:31.0 | Data point 80 | 1.50000 mL | 0.17368 mL | 0.17225 mL | 0.31999 mL | 4.855 | -0.00462 | 0.06310 | 0.00091 | 10.5 s |
| 1:20:22.3 | Data point 81 | 1.50000 mL | 0.17368 mL | 0.17260 mL | 0.31999 mL | 5.067 | -0.00056 | 0.00122 | 0.00080 | 10.5 s |
| 1:21:03.3 | Data point 82 | 1.50000 mL | 0.17368 mL | 0.17281 mL | 0.31999 mL | 5.289 | -0.01845 | 0.83141 | 0.00100 | 11.5 s |
| 1:21:40.3 | Data point 83 | 1.50000 mL | 0.17368 mL | 0.17293 mL | 0.31999 mL | 5.442 | -0.01376 | 0.70225 | 0.00081 | 11.5 s |
| 1:22:22.3 | Data point 84 | 1.50000 mL | 0.17368 mL | 0.17307 mL | 0.31999 mL | 5.745 | -0.01592 | 0.71364 | 0.00093 | 16.5 s |
| 1:23:09.4 | Data point 85 | 1.50000 mL | 0.17368 mL | 0.17326 mL | 0.31999 mL | 6.515 | -0.03653 | 0.99353 | 0.00181 | Timed out at 59.5 s |
| 1:24:45.1 | Data point 86 | 1.50000 mL | 0.17368 mL | 0.17335 mL | 0.31999 mL | 6.950 | -0.06050 | 0.97491 | 0.00303 | Timed out at 59.5 s |
| 1:26:15.6 | Data point 87 | 1.50000 mL | 0.17368 mL | 0.17340 mL | 0.31999 mL | 7.287 | -0.05907 | 0.99436 | 0.00292 | Timed out at 59.5 s |

Sample name: **M07_octanol**
Assay name: **pH-metric high logP**
Assay ID: **18B-28013**
Filename: **C:\Sirius_T3\Mehtap\20180228_exp28_logP_T3-2\18B-28013_M07_octanol_pH-metric high logP.t3r**

Experiment start time: **2/28/2018 7:21:13 PM**
Analyst: **Pion**
Instrument ID: **T312060**

#### Events (continued)

| Time | Event | Water | Acid | Base | Octanol | pH | dpH/dt | pH R-squared | pH SD | dpH/dt time |
|---|---|---|---|---|---|---|---|---|---|---|
| 1:27:46.0 | Data point 88 | 1.50000 mL | 0.17368 mL | 0.17345 mL | 0.31999 mL | 7.543 | -0.06077 | 0.98579 | 0.00302 | Timed out at 59.5 s |
| 1:29:26.9 | Data point 89 | 1.50000 mL | 0.17368 mL | 0.17373 mL | 0.31999 mL | 8.640 | -0.01815 | 0.88900 | 0.00095 | 55.0 s |
| 1:30:57.7 | Data point 90 | 1.50000 mL | 0.17368 mL | 0.17389 mL | 0.31999 mL | 9.020 | -0.01693 | 0.78905 | 0.00094 | 16.5 s |
| 1:31:55.0 | Data point 91 | 1.50000 mL | 0.17368 mL | 0.17406 mL | 0.31999 mL | 9.240 | -0.01696 | 0.79810 | 0.00094 | 13.0 s |
| 1:32:38.5 | Data point 92 | 1.50000 mL | 0.17368 mL | 0.17420 mL | 0.31999 mL | 9.465 | -0.01825 | 0.92964 | 0.00094 | 13.5 s |
| 1:33:22.6 | Data point 93 | 1.50000 mL | 0.17368 mL | 0.17444 mL | 0.31999 mL | 9.701 | -0.00732 | 0.21315 | 0.00078 | 11.5 s |
| 1:34:04.7 | Data point 94 | 1.50000 mL | 0.17368 mL | 0.17467 mL | 0.31999 mL | 9.894 | -0.01667 | 0.74381 | 0.00095 | 10.5 s |
| 1:34:45.7 | Data point 95 | 1.50000 mL | 0.17368 mL | 0.17491 mL | 0.31999 mL | 10.021 | -0.01410 | 0.74350 | 0.00081 | 10.0 s |
| 1:35:04.9 | Assay volumes | 1.50000 mL | 0.17368 mL | 0.17491 mL | 0.31999 mL | | | | | |

| | | | |
|---|---|---|---|
| Sample name: | **M07_octanol** | Experiment start time: | **2/28/2018 7:21:13 PM** |
| Assay name: | **pH-metric high logP** | Analyst: | **Pion** |
| Assay ID: | **18B-28013** | Instrument ID: | **T312060** |
| Filename: | **C:\Sirius_T3\Mehtap\20180228_exp28_logP_T3-2\18B-28013_M07_octanol_pH-metric high logP.t3r** | | |

| | | | |
|---|---|---|---|
| Sample name: | **M07_octanol** | Experiment start time: | **2/28/2018 7:21:13 PM** |
| Assay name: | **pH-metric high logP** | Analyst: | **Pion** |
| Assay ID: | **18B-28013** | Instrument ID: | **T312060** |
| Filename: | **C:\Sirius_T3\Mehtap\20180228_exp28_logP_T3-2\18B-28013_M07_octanol_pH-metric high logP.t3r** | | |

#### Assay Settings (continued)

| Setting | Value | Original Value | Date/Time changed | Imported from |
|---|---|---|---|---|
| ***Titration 3*** | | | | |
| Titrate from | Low to high pH | | | |
| Add additional water | 0.00 mL | | | |
| Additional partition solvent volume | 0.250 mL | | | |
| Additional partition solvent added | Automatic | | | |
| After pH adjust stir for | 30 seconds | | | |
| Stir to allow partitioning for | 15 seconds | | | |
| Stirrer speed for partitioning | 60% | | | |
| ***Data Point Stability*** | | | | |
| Stir during data point collection | No | | | |
| Delay before data point collection | 0 seconds | | | |
| Number of points to average | 20 points | | | |
| Time interval between points | 0.50 seconds | | | |
| Required maximum standard deviation | 0.00100 dpH/dt | | | |
| Stability timeout after | 60 seconds | | | |

#### Calibration Settings

| Setting | Value | Date/Time changed | Imported from |
|---|---|---|---|
| Four-Plus alpha | 0.130 | 2/28/2018 7:21:12 PM | C:\Sirius_T3\HCl18B27.t3r |
| Four-Plus S | 0.9970 | 2/28/2018 7:21:12 PM | C:\Sirius_T3\HCl18B27.t3r |
| Four-Plus jH | 0.8 | 2/28/2018 7:21:12 PM | C:\Sirius_T3\HCl18B27.t3r |
| Four-Plus jOH | -0.4 | 2/28/2018 7:21:12 PM | C:\Sirius_T3\HCl18B27.t3r |
| Base concentration factor | 1.000 | 2/28/2018 7:21:13 PM | C:\Sirius_T3\KOH18B27.t3r |
| Acid concentration factor | 0.994 | 2/28/2018 7:21:12 PM | C:\Sirius_T3\HCl18B27.t3r |

#### Instrument Settings

| Setting | Value | Batch Id | Install date |
|---|---|---|---|
| Instrument owner | Merck | | |
| Instrument ID | T312060 | | |
| Instrument type | T3 Simulator | | |
| Software version | 1.1.3.0 | | |
| Dispenser module | | T3DM1200361 | 3/31/2009 5:24:52 AM |
| Dispenser 0 | Water | | 3/31/2009 5:25:05 AM |
| Syringe volume | 2.5 mL | | |
| Firmware version | 1.2.1(r2) | | |
| Titrant | Water (0.15 M KCl) | 02-06-2018 | 2/27/2018 10:05:59 AM |
| Dispenser 2 | Acid | | 3/31/2009 5:25:11 AM |
| Syringe volume | 0.5 mL | | |
| Firmware version | 1.2.1(r2) | | |
| Titrant | Acid (0.5 M HCl) | 02-27-2018 | 2/27/2018 10:27:22 AM |
| Dispenser 1 | Base | | 3/31/2009 5:25:21 AM |
| Syringe volume | 0.5 mL | | |
| Firmware version | 1.2.1(r2) | | |
| Titrant | Base (0.5 M KOH) | 9/22/2017 | 2/27/2018 10:21:22 AM |
| Dispenser 5 | Cosolvent | | 3/31/2009 5:26:24 AM |
| Syringe volume | 2.5 mL | | |
| Firmware version | 1.2.1(r2) | | |
| Distribution valve 5 | Distribution Valve | | 3/31/2009 5:28:19 AM |
| Firmware version | 1.1.3 | | |
| Port A | Methanol (80%, 0.15 M KCl) | 09-26-17 | 2/7/2018 9:42:01 AM |
| Port B | Cyclohexane | 11-01-17 | 2/27/2018 10:37:57 AM |
| Dispenser 3 | Buffer | | 8/3/2010 5:05:16 AM |
| Syringe volume | 0.5 mL | | |
| Firmware version | 1.2.1(r2) | | |
| Titrant | Dodecane | 2018/01/31 | 2/28/2018 10:18:04 AM |
| Dispenser 6 | Octanol | | 10/22/2010 10:52:43 AM |

| | | | |
|---|---|---|---|
| Sample name: | **M07_octanol** | Experiment start time: | **2/28/2018 7:21:13 PM** |
| Assay name: | **pH-metric high logP** | Analyst: | **Pion** |
| Assay ID: | **18B-28013** | Instrument ID: | **T312060** |
| Filename: | **C:\Sirius_T3\Mehtap\20180228_exp28_logP_T3-2\18B-28013_M07_octanol_pH-metric high logP.t3r** | | |

Sample name: **M07_octanol**
Assay name: **pH-metric high logP**
Assay ID: **18B-28013**
Filename: **C:\Sirius_T3\Mehtap\20180228_exp28_logP_T3-2\18B-28013_M07_octanol_pH-metric high logP.t3r**
Experiment start time: **2/28/2018 7:21:13 PM**
Analyst: **Pion**
Instrument ID: **T312060**

#### Instrument Settings (continued)

| Setting | Value | Batch Id | Install date |
|---|---|---|---|
| Spectrometer calibration stir duration | 5 s | | |
| Spectrometer calibration stir speed | 30% | | |
| Spectrometer calibration wash pump volume | 20.0 mL | | |
| Spectrometer calibration wash stir duration | 5 s | | |
| Spectrometer calibration wash stir speed | 30% | | |
| Overhead dispense height | 10000 | | |

#### Refinement Settings

| Setting | Value | Default value |
|---|---|---|
| Turbidity detection method | None | None |
| Turbidity wavelength to assess | 500.0 nm | 500.0 nm |
| Turbidity maximum absorbance | 0.100 | 0.100 |
| Turbidity probe threshold | 50.00 | 50.00 |

#### Experiment Log

[2:37] Air gap created for Water (0.15 M KCl)
[2:37] Air gap created for Acid (0.5 M HCl)
[2:38] Air gap created for Base (0.5 M KOH)
[2:38] Air gap released for Water (0.15 M KCl)
[2:42] Titrator arm moved over Titration position
[2:42] Titration 1 of 3
[2:42] Adding initial titrants
[2:42] Automatically add 1.50000 mL of water
[3:07] Dispensed 1.500000 mL of Water (0.15 M KCl)
[3:11] Titrator arm moved over Drain
[8:52] Titrator arm moved to Titration position
[8:52] Argon flow rate set to 100
[8:52] Stirrer speed set to 10
[8:57] Automatically add 0.02000 mL of Octanol
[8:58] Dispensed 0.019991 mL of Octanol
[8:59] Initial pH = 6.72
[8:59] Iterative adjust 6.72 -> 2.00
[8:59] pH 6.72 -> 2.00
[9:01] Air gap released for Acid (0.5 M HCl)
[9:01] Dispensed 0.051905 mL of Acid (0.5 M HCl)
[9:06] Holding pH 2.00
[11:07] Stirrer speed set to 0
[11:07] Stirrer speed set to 50
[11:07] Iterative adjust 1.98 -> 2.00
[11:07] pH 1.98 -> 2.00
[11:07] Air gap released for Base (0.5 M KOH)
[11:08] Dispensed 0.002070 mL of Base (0.5 M KOH)
[11:58] Stirrer speed set to 0
[12:08] Datapoint id 1 collected
[12:08] Stirrer speed set to 50
[12:13] pH 2.01 -> 2.21
[12:13] Using cautious pH adjust
[12:14] Dispensed 0.007761 mL of Base (0.5 M KOH)
[12:19] Stepping pH = 2.10
[12:19] Dispensed 0.005644 mL of Base (0.5 M KOH)
[12:24] Stepping pH = 2.19
[12:24] Dispensed 0.001270 mL of Base (0.5 M KOH)
[12:29] Stepping pH = 2.21
[12:44] Stirrer speed set to 0
[12:55] Datapoint id 2 collected
[12:55] Charge balance equation is out by 5.6%
[12:55] Stirrer speed set to 50

| | | | |
|---|---|---|---|
| Sample name: | **M07_octanol** | Experiment start time: | **2/28/2018 7:21:13 PM** |
| Assay name: | **pH-metric high logP** | Analyst: | **Pion** |
| Assay ID: | **18B-28013** | Instrument ID: | **T312060** |
| Filename: | **C:\Sirius_T3\Mehtap\20180228_exp28_logP_T3-2\18B-28013_M07_octanol_pH-metric high logP.t3r** | | |

#### Experiment Log (continued)

[13:00] pH 2.21 -> 2.41
[13:00] Using charge balance adjust
[13:00] Dispensed 0.009690 mL of Base (0.5 M KOH)
[13:21] Stirrer speed set to 0
[13:31] Datapoint id 3 collected
[13:31] Charge balance equation is out by 8.2%
[13:31] Stirrer speed set to 50
[13:36] pH 2.44 -> 2.64
[13:36] Using charge balance adjust
[13:36] Dispensed 0.005786 mL of Base (0.5 M KOH)
[13:56] Stirrer speed set to 0
[14:06] Datapoint id 4 collected
[14:06] Charge balance equation is out by -2.6%
[14:06] Stirrer speed set to 50
[14:11] pH 2.64 -> 2.84
[14:11] Using charge balance adjust
[14:12] Dispensed 0.003645 mL of Base (0.5 M KOH)
[14:32] Stirrer speed set to 0
[14:42] Datapoint id 5 collected
[14:42] Charge balance equation is out by 15.5%
[14:42] Stirrer speed set to 50
[14:47] pH 2.88 -> 3.08
[14:47] Using cautious pH adjust
[14:47] Dispensed 0.001082 mL of Base (0.5 M KOH)
[14:52] Stepping pH = 2.97
[14:52] Dispensed 0.000729 mL of Base (0.5 M KOH)
[14:57] Stepping pH = 3.05
[14:57] Dispensed 0.000212 mL of Base (0.5 M KOH)
[15:02] Stepping pH = 3.07
[15:18] Stirrer speed set to 0
[15:28] Datapoint id 6 collected
[15:28] Charge balance equation is out by 6.5%
[15:28] Stirrer speed set to 50
[15:33] pH 3.07 -> 3.27
[15:33] Using charge balance adjust
[15:33] Dispensed 0.001458 mL of Base (0.5 M KOH)
[15:53] Stirrer speed set to 0
[16:04] Datapoint id 7 collected
[16:04] Charge balance equation is out by 1.7%
[16:04] Stirrer speed set to 50
[16:09] pH 3.28 -> 3.48
[16:09] Using charge balance adjust
[16:09] Dispensed 0.001035 mL of Base (0.5 M KOH)
[16:29] Stirrer speed set to 0
[16:39] Datapoint id 8 collected
[16:39] Charge balance equation is out by 8.8%
[16:39] Stirrer speed set to 50
[16:44] pH 3.50 -> 3.70
[16:44] Using charge balance adjust
[16:44] Dispensed 0.000847 mL of Base (0.5 M KOH)
[17:04] Stirrer speed set to 0
[17:14] Datapoint id 9 collected
[17:14] Charge balance equation is out by 18.0%
[17:14] Stirrer speed set to 50
[17:19] pH 3.74 -> 3.94
[17:19] Using cautious pH adjust
[17:20] Dispensed 0.000423 mL of Base (0.5 M KOH)
[17:25] Stepping pH = 3.87
[17:25] Dispensed 0.000188 mL of Base (0.5 M KOH)

Sample name: **M07_octanol**
Assay name: **pH-metric high logP**
Assay ID: **18B-28013**
Filename: **C:\Sirius_T3\Mehtap\20180228_exp28_logP_T3-2\18B-28013_M07_octanol_pH-metric high logP.t3r**

Experiment start time: **2/28/2018 7:21:13 PM**
Analyst: **Pion**
Instrument ID: **T312060**

#### Experiment Log (continued)

[17:30] Stepping pH = 3.94
[17:45] Stirrer speed set to 0
[17:55] Datapoint id 10 collected
[17:55] Charge balance equation is out by 27.2%
[17:55] Stirrer speed set to 50
[18:00] pH 3.92 -> 4.12
[18:00] Using cautious pH adjust
[18:00] Dispensed 0.000470 mL of Base (0.5 M KOH)
[18:05] Stepping pH = 4.08
[18:05] Dispensed 0.000094 mL of Base (0.5 M KOH)
[18:10] Stepping pH = 4.11
[18:11] Dispensed 0.000047 mL of Base (0.5 M KOH)
[18:16] Stepping pH = 4.12
[18:31] Stirrer speed set to 0
[18:41] Datapoint id 11 collected
[18:41] Charge balance equation is out by 32.3%
[18:41] Stirrer speed set to 50
[18:46] pH 4.09 -> 4.29
[18:46] Using cautious pH adjust
[18:46] Dispensed 0.000517 mL of Base (0.5 M KOH)
[18:51] Stepping pH = 4.26
[18:51] Dispensed 0.000094 mL of Base (0.5 M KOH)
[18:56] Stepping pH = 4.27
[18:56] Dispensed 0.000071 mL of Base (0.5 M KOH)
[19:01] Stepping pH = 4.28
[19:16] Stirrer speed set to 0
[19:26] Datapoint id 12 collected
[19:26] Charge balance equation is out by 33.6%
[19:26] Stirrer speed set to 50
[19:32] pH 4.27 -> 4.47
[19:32] Using cautious pH adjust
[19:32] Dispensed 0.000588 mL of Base (0.5 M KOH)
[19:37] Stepping pH = 4.44
[19:37] Dispensed 0.000094 mL of Base (0.5 M KOH)
[19:42] Stepping pH = 4.44
[19:42] Dispensed 0.000282 mL of Base (0.5 M KOH)
[19:47] Stepping pH = 4.52
[20:02] Stirrer speed set to 0
[20:13] Datapoint id 13 collected
[20:13] Charge balance equation is out by 16.4%
[20:13] Stirrer speed set to 50
[20:18] pH 4.48 -> 4.68
[20:18] Using cautious pH adjust
[20:18] Dispensed 0.000635 mL of Base (0.5 M KOH)
[20:23] Stepping pH = 4.63
[20:23] Dispensed 0.000188 mL of Base (0.5 M KOH)
[20:28] Stepping pH = 4.65
[20:28] Dispensed 0.000188 mL of Base (0.5 M KOH)
[20:33] Stepping pH = 4.68
[20:48] Stirrer speed set to 0
[21:00] Datapoint id 14 collected
[21:00] Charge balance equation is out by 20.8%
[21:00] Stirrer speed set to 50
[21:05] pH 4.68 -> 4.88
[21:05] Using cautious pH adjust
[21:05] Dispensed 0.000611 mL of Base (0.5 M KOH)
[21:10] Stepping pH = 4.81
[21:10] Dispensed 0.000235 mL of Base (0.5 M KOH)
[21:15] Stepping pH = 4.80

| | | | |
|---|---|---|---|
| Sample name: | **M07_octanol** | Experiment start time: | **2/28/2018 7:21:13 PM** |
| Assay name: | **pH-metric high logP** | Analyst: | **Pion** |
| Assay ID: | **18B-28013** | Instrument ID: | **T312060** |
| Filename: | **C:\Sirius_T3\Mehtap\20180228_exp28_logP_T3-2\18B-28013_M07_octanol_pH-metric high logP.t3r** | | |

#### Experiment Log (continued)

[21:16] Dispensed 0.001341 mL of Base (0.5 M KOH)
[21:21] Stepping pH = 5.06
[21:36] Stirrer speed set to 0
[21:49] Datapoint id 15 collected
[21:49] Charge balance equation is out by -79.3%
[21:49] Stirrer speed set to 50
[21:54] pH 5.00 -> 5.20
[21:54] Using cautious pH adjust
[21:55] Dispensed 0.000494 mL of Base (0.5 M KOH)
[22:00] Stepping pH = 5.09
[22:00] Dispensed 0.000353 mL of Base (0.5 M KOH)
[22:05] Stepping pH = 5.13
[22:05] Dispensed 0.000400 mL of Base (0.5 M KOH)
[22:10] Stepping pH = 5.20
[22:25] Stirrer speed set to 0
[22:39] Datapoint id 16 collected
[22:39] Charge balance equation is out by -26.8%
[22:39] Stirrer speed set to 50
[22:44] pH 5.18 -> 5.38
[22:44] Using cautious pH adjust
[22:44] Dispensed 0.000400 mL of Base (0.5 M KOH)
[22:49] Stepping pH = 5.27
[22:49] Dispensed 0.000306 mL of Base (0.5 M KOH)
[22:55] Stepping pH = 5.32
[22:55] Dispensed 0.000235 mL of Base (0.5 M KOH)
[23:00] Stepping pH = 5.34
[23:00] Dispensed 0.000235 mL of Base (0.5 M KOH)
[23:05] Stepping pH = 5.39
[23:20] Stirrer speed set to 0
[23:37] Datapoint id 17 collected
[23:37] Charge balance equation is out by -49.3%
[23:37] Stirrer speed set to 50
[23:42] pH 5.37 -> 5.57
[23:42] Using cautious pH adjust
[23:42] Dispensed 0.000306 mL of Base (0.5 M KOH)
[23:47] Stepping pH = 5.42
[23:47] Dispensed 0.000400 mL of Base (0.5 M KOH)
[23:52] Stepping pH = 5.57
[24:07] Stirrer speed set to 0
[24:28] Datapoint id 18 collected
[24:28] Charge balance equation is out by -17.6%
[24:28] Stirrer speed set to 50
[24:33] pH 5.54 -> 5.74
[24:33] Using cautious pH adjust
[24:33] Dispensed 0.000235 mL of Base (0.5 M KOH)
[24:38] Stepping pH = 5.60
[24:38] Dispensed 0.000282 mL of Base (0.5 M KOH)
[24:43] Stepping pH = 5.75
[24:58] Stirrer speed set to 0
[25:19] Datapoint id 19 collected
[25:19] Charge balance equation is out by -11.9%
[25:19] Stirrer speed set to 50
[25:24] pH 5.71 -> 5.91
[25:24] Using charge balance adjust
[25:24] Dispensed 0.000329 mL of Base (0.5 M KOH)
[25:44] Stirrer speed set to 0
[26:07] Datapoint id 20 collected
[26:07] Charge balance equation is out by -52.3%
[26:07] Stirrer speed set to 50

Sample name: **M07_octanol**
Assay name: **pH-metric high logP**
Assay ID: **18B-28013**
Filename: **C:\Sirius_T3\Mehtap\20180228_exp28_logP_T3-2\18B-28013_M07_octanol_pH-metric high logP.t3r**

Experiment start time: **2/28/2018 7:21:13 PM**
Analyst: **Pion**
Instrument ID: **T312060**

#### Experiment Log (continued)

[26:12] pH 5.85 -> 6.05
[26:12] Using cautious pH adjust
[26:12] Dispensed 0.000141 mL of Base (0.5 M KOH)
[26:18] Stepping pH = 5.89
[26:18] Dispensed 0.000212 mL of Base (0.5 M KOH)
[26:23] Stepping pH = 6.08
[26:38] Stirrer speed set to 0
[27:02] Datapoint id 21 collected
[27:02] Charge balance equation is out by -33.5%
[27:02] Stirrer speed set to 50
[27:07] pH 6.07 -> 6.27
[27:07] Using cautious pH adjust
[27:08] Dispensed 0.000094 mL of Base (0.5 M KOH)
[27:13] Stepping pH = 6.09
[27:13] Dispensed 0.000235 mL of Base (0.5 M KOH)
[27:18] Stepping pH = 6.43
[27:33] Stirrer speed set to 0
[28:09] Datapoint id 22 collected
[28:09] Charge balance equation is out by -82.8%
[28:09] Stirrer speed set to 50
[28:14] pH 6.40 -> 6.60
[28:14] Using cautious pH adjust
[28:14] Dispensed 0.000047 mL of Base (0.5 M KOH)
[28:19] Stepping pH = 6.39
[28:19] Dispensed 0.000259 mL of Base (0.5 M KOH)
[28:24] Stepping pH = 7.32
[28:39] Stirrer speed set to 0
[29:40] Datapoint id 23 collected
[29:40] Charge balance equation is out by -208.7%
[29:40] Stirrer speed set to 50
[29:45] pH 7.40 -> 7.60
[29:45] Using cautious pH adjust
[29:45] Dispensed 0.000024 mL of Base (0.5 M KOH)
[29:50] Stepping pH = 7.44
[29:50] Dispensed 0.000024 mL of Base (0.5 M KOH)
[29:55] Stepping pH = 7.46
[29:55] Dispensed 0.000024 mL of Base (0.5 M KOH)
[30:00] Stepping pH = 7.49
[30:00] Dispensed 0.000047 mL of Base (0.5 M KOH)
[30:05] Stepping pH = 7.77
[30:20] Stirrer speed set to 0
[31:20] Datapoint id 24 collected
[31:20] Charge balance equation is out by -679.3%
[31:20] Stirrer speed set to 50
[31:25] pH 8.02 -> 8.22
[31:25] Using cautious pH adjust
[31:25] Dispensed 0.000024 mL of Base (0.5 M KOH)
[31:31] Stepping pH = 8.06
[31:31] Dispensed 0.000024 mL of Base (0.5 M KOH)
[31:36] Stepping pH = 8.12
[31:36] Dispensed 0.000024 mL of Base (0.5 M KOH)
[31:41] Stepping pH = 8.37
[31:56] Stirrer speed set to 0
[32:56] Datapoint id 25 collected
[32:56] Charge balance equation is out by -627.8%
[32:56] Stirrer speed set to 50
[33:01] pH 8.46 -> 8.66
[33:01] Using cautious pH adjust
[33:01] Dispensed 0.000024 mL of Base (0.5 M KOH)

Sample name: **M07_octanol**
Assay name: **pH-metric high logP**
Assay ID: **18B-28013**
Filename: **C:\Sirius_T3\Mehtap\20180228_exp28_logP_T3-2\18B-28013_M07_octanol_pH-metric high logP.t3r**
Experiment start time: **2/28/2018 7:21:13 PM**
Analyst: **Pion**
Instrument ID: **T312060**

#### Experiment Log (continued)

[33:06] Stepping pH = 8.44
[33:06] Dispensed 0.000047 mL of Base (0.5 M KOH)
[33:11] Stepping pH = 8.80
[33:26] Stirrer speed set to 0
[34:12] Datapoint id 26 collected
[34:12] Charge balance equation is out by -310.7%
[34:12] Stirrer speed set to 50
[34:17] pH 8.74 -> 8.94
[34:17] Using cautious pH adjust
[34:17] Dispensed 0.000024 mL of Base (0.5 M KOH)
[34:22] Stepping pH = 8.71
[34:22] Dispensed 0.000071 mL of Base (0.5 M KOH)
[34:27] Stepping pH = 9.00
[34:43] Stirrer speed set to 0
[35:14] Datapoint id 27 collected
[35:14] Charge balance equation is out by -261.3%
[35:14] Stirrer speed set to 50
[35:19] pH 9.03 -> 9.23
[35:19] Using cautious pH adjust
[35:19] Dispensed 0.000024 mL of Base (0.5 M KOH)
[35:24] Stepping pH = 9.03
[35:24] Dispensed 0.000118 mL of Base (0.5 M KOH)
[35:29] Stepping pH = 9.41
[35:44] Stirrer speed set to 0
[36:00] Datapoint id 28 collected
[36:00] Charge balance equation is out by -201.0%
[36:00] Stirrer speed set to 50
[36:05] pH 9.42 -> 9.62
[36:05] Using cautious pH adjust
[36:05] Dispensed 0.000047 mL of Base (0.5 M KOH)
[36:10] Stepping pH = 9.44
[36:10] Dispensed 0.000141 mL of Base (0.5 M KOH)
[36:15] Stepping pH = 9.72
[36:30] Stirrer speed set to 0
[36:52] Datapoint id 29 collected
[36:52] Charge balance equation is out by -86.2%
[36:52] Stirrer speed set to 50
[36:57] pH 9.73 -> 9.93
[36:57] Using cautious pH adjust
[36:57] Dispensed 0.000094 mL of Base (0.5 M KOH)
[37:02] Stepping pH = 9.79
[37:02] Dispensed 0.000094 mL of Base (0.5 M KOH)
[37:07] Stepping pH = 9.92
[37:07] Dispensed 0.000024 mL of Base (0.5 M KOH)
[37:12] Stepping pH = 9.93
[37:28] Stirrer speed set to 0
[37:40] Datapoint id 30 collected
[37:40] Charge balance equation is out by -22.0%
[37:40] Stirrer speed set to 50
[37:45] pH 9.92 -> 10.05
[37:45] Using cautious pH adjust
[37:45] Dispensed 0.000071 mL of Base (0.5 M KOH)
[37:50] Stepping pH = 9.95
[37:50] Dispensed 0.000118 mL of Base (0.5 M KOH)
[37:56] Stepping pH = 10.06
[38:11] Stirrer speed set to 0
[38:21] Datapoint id 31 collected
[38:21] Charge balance equation is out by -30.0%
[38:21] Titration 2 of 3

| | | | |
|---|---|---|---|
| Sample name: | **M07_octanol** | Experiment start time: | **2/28/2018 7:21:13 PM** |
| Assay name: | **pH-metric high logP** | Analyst: | **Pion** |
| Assay ID: | **18B-28013** | Instrument ID: | **T312060** |
| Filename: | **C:\Sirius_T3\Mehtap\20180228_exp28_logP_T3-2\18B-28013_M07_octanol_pH-metric high logP.t3r** | | |

#### Experiment Log (continued)

[38:21] Adding initial titrants
[38:21] Automatically add 0.05000 mL of Octanol
[38:23] Dispensed 0.050000 mL of Octanol
[38:23] Stirrer speed set to 10
[38:24] Stirrer speed set to 55
[38:24] Iterative adjust 10.07 -> 2.00
[38:24] pH 10.07 -> 2.00
[38:25] Dispensed 0.055621 mL of Acid (0.5 M HCl)
[38:30] pH 2.03 -> 2.00
[38:30] Dispensed 0.003316 mL of Acid (0.5 M HCl)
[39:21] Stirrer speed set to 0
[39:31] Datapoint id 32 collected
[39:31] Stirrer speed set to 55
[39:36] pH 1.96 -> 2.16
[39:36] Using cautious pH adjust
[39:36] Dispensed 0.009313 mL of Base (0.5 M KOH)
[39:41] Stepping pH = 2.06
[39:41] Dispensed 0.005997 mL of Base (0.5 M KOH)
[39:47] Stepping pH = 2.13
[39:47] Dispensed 0.002023 mL of Base (0.5 M KOH)
[39:52] Stepping pH = 2.16
[40:07] Stirrer speed set to 0
[40:17] Datapoint id 33 collected
[40:17] Charge balance equation is out by 7.0%
[40:17] Stirrer speed set to 55
[40:22] pH 2.16 -> 2.36
[40:22] Using charge balance adjust
[40:22] Dispensed 0.011689 mL of Base (0.5 M KOH)
[40:43] Stirrer speed set to 0
[40:53] Datapoint id 34 collected
[40:53] Charge balance equation is out by 15.3%
[40:53] Stirrer speed set to 55
[40:58] pH 2.40 -> 2.60
[40:58] Using cautious pH adjust
[40:58] Dispensed 0.003387 mL of Base (0.5 M KOH)
[41:03] Stepping pH = 2.49
[41:04] Dispensed 0.002446 mL of Base (0.5 M KOH)
[41:09] Stepping pH = 2.57
[41:09] Dispensed 0.000894 mL of Base (0.5 M KOH)
[41:14] Stepping pH = 2.60
[41:29] Stirrer speed set to 0
[41:39] Datapoint id 35 collected
[41:39] Charge balance equation is out by 0.8%
[41:39] Stirrer speed set to 55
[41:44] pH 2.60 -> 2.80
[41:44] Using charge balance adjust
[41:44] Dispensed 0.004351 mL of Base (0.5 M KOH)
[42:05] Stirrer speed set to 0
[42:15] Datapoint id 36 collected
[42:15] Charge balance equation is out by 4.1%
[42:15] Stirrer speed set to 55
[42:20] pH 2.82 -> 3.02
[42:20] Using charge balance adjust
[42:20] Dispensed 0.002799 mL of Base (0.5 M KOH)
[42:40] Stirrer speed set to 0
[42:50] Datapoint id 37 collected
[42:50] Charge balance equation is out by -3.5%
[42:50] Stirrer speed set to 55
[42:55] pH 3.02 -> 3.22

| | | | |
|---|---|---|---|
| Sample name: | **M07_octanol** | Experiment start time: | **2/28/2018 7:21:13 PM** |
| Assay name: | **pH-metric high logP** | Analyst: | **Pion** |
| Assay ID: | **18B-28013** | Instrument ID: | **T312060** |
| Filename: | **C:\Sirius_T3\Mehtap\20180228_exp28_logP_T3-2\18B-28013_M07_octanol_pH-metric high logP.t3r** | | |

#### Experiment Log (continued)

[42:55] Using charge balance adjust
[42:55] Dispensed 0.001976 mL of Base (0.5 M KOH)
[43:16] Stirrer speed set to 0
[43:26] Datapoint id 38 collected
[43:26] Charge balance equation is out by 1.2%
[43:26] Stirrer speed set to 55
[43:31] pH 3.22 -> 3.42
[43:31] Using charge balance adjust
[43:31] Dispensed 0.001505 mL of Base (0.5 M KOH)
[43:51] Stirrer speed set to 0
[44:02] Datapoint id 39 collected
[44:02] Charge balance equation is out by 33.9%
[44:02] Stirrer speed set to 55
[44:07] pH 3.50 -> 3.70
[44:07] Using cautious pH adjust
[44:07] Dispensed 0.000659 mL of Base (0.5 M KOH)
[44:12] Stepping pH = 3.64
[44:12] Dispensed 0.000235 mL of Base (0.5 M KOH)
[44:17] Stepping pH = 3.69
[44:32] Stirrer speed set to 0
[44:42] Datapoint id 40 collected
[44:42] Charge balance equation is out by 32.5%
[44:42] Stirrer speed set to 55
[44:47] pH 3.69 -> 3.89
[44:47] Using cautious pH adjust
[44:47] Dispensed 0.000659 mL of Base (0.5 M KOH)
[44:52] Stepping pH = 3.82
[44:52] Dispensed 0.000259 mL of Base (0.5 M KOH)
[44:58] Stepping pH = 3.87
[44:58] Dispensed 0.000094 mL of Base (0.5 M KOH)
[45:03] Stepping pH = 3.89
[45:18] Stirrer speed set to 0
[45:28] Datapoint id 41 collected
[45:28] Charge balance equation is out by 22.8%
[45:28] Stirrer speed set to 55
[45:33] pH 3.88 -> 4.08
[45:33] Using cautious pH adjust
[45:33] Dispensed 0.000682 mL of Base (0.5 M KOH)
[45:38] Stepping pH = 4.00
[45:38] Dispensed 0.000353 mL of Base (0.5 M KOH)
[45:43] Stepping pH = 4.06
[45:43] Dispensed 0.000141 mL of Base (0.5 M KOH)
[45:49] Stepping pH = 4.08
[46:04] Stirrer speed set to 0
[46:14] Datapoint id 42 collected
[46:14] Charge balance equation is out by 15.0%
[46:14] Stirrer speed set to 55
[46:19] pH 4.07 -> 4.27
[46:19] Using cautious pH adjust
[46:19] Dispensed 0.000682 mL of Base (0.5 M KOH)
[46:24] Stepping pH = 4.18
[46:24] Dispensed 0.000423 mL of Base (0.5 M KOH)
[46:29] Stepping pH = 4.24
[46:29] Dispensed 0.000212 mL of Base (0.5 M KOH)
[46:34] Stepping pH = 4.27
[46:49] Stirrer speed set to 0
[46:59] Datapoint id 43 collected
[46:59] Charge balance equation is out by 3.8%
[46:59] Stirrer speed set to 55

| | | | |
|---|---|---|---|
| Sample name: | **M07_octanol** | Experiment start time: | **2/28/2018 7:21:13 PM** |
| Assay name: | **pH-metric high logP** | Analyst: | **Pion** |
| Assay ID: | **18B-28013** | Instrument ID: | **T312060** |
| Filename: | **C:\Sirius_T3\Mehtap\20180228_exp28_logP_T3-2\18B-28013_M07_octanol_pH-metric high logP.t3r** | | |

#### Experiment Log (continued)

[47:04] pH 4.27 -> 4.47
[47:04] Using charge balance adjust
[47:05] Dispensed 0.001246 mL of Base (0.5 M KOH)
[47:25] Stirrer speed set to 0
[47:35] Datapoint id 44 collected
[47:35] Charge balance equation is out by -14.4%
[47:35] Stirrer speed set to 55
[47:40] pH 4.44 -> 4.64
[47:40] Using charge balance adjust
[47:40] Dispensed 0.001105 mL of Base (0.5 M KOH)
[48:00] Stirrer speed set to 0
[48:11] Datapoint id 45 collected
[48:11] Charge balance equation is out by -17.5%
[48:11] Stirrer speed set to 55
[48:16] pH 4.61 -> 4.81
[48:16] Using cautious pH adjust
[48:16] Dispensed 0.000447 mL of Base (0.5 M KOH)
[48:21] Stepping pH = 4.69
[48:21] Dispensed 0.000447 mL of Base (0.5 M KOH)
[48:26] Stepping pH = 4.77
[48:26] Dispensed 0.000165 mL of Base (0.5 M KOH)
[48:31] Stepping pH = 4.80
[48:47] Stirrer speed set to 0
[48:57] Datapoint id 46 collected
[48:57] Charge balance equation is out by -16.8%
[48:57] Stirrer speed set to 55
[49:02] pH 4.81 -> 5.01
[49:02] Using cautious pH adjust
[49:02] Dispensed 0.000353 mL of Base (0.5 M KOH)
[49:07] Stepping pH = 4.89
[49:08] Dispensed 0.000306 mL of Base (0.5 M KOH)
[49:13] Stepping pH = 4.97
[49:13] Dispensed 0.000094 mL of Base (0.5 M KOH)
[49:18] Stepping pH = 4.99
[49:18] Dispensed 0.000094 mL of Base (0.5 M KOH)
[49:23] Stepping pH = 5.01
[49:38] Stirrer speed set to 0
[49:49] Datapoint id 47 collected
[49:49] Charge balance equation is out by -19.3%
[49:49] Stirrer speed set to 55
[49:54] pH 5.01 -> 5.21
[49:54] Using cautious pH adjust
[49:54] Dispensed 0.000235 mL of Base (0.5 M KOH)
[49:59] Stepping pH = 5.09
[49:59] Dispensed 0.000235 mL of Base (0.5 M KOH)
[50:04] Stepping pH = 5.19
[50:04] Dispensed 0.000047 mL of Base (0.5 M KOH)
[50:09] Stepping pH = 5.19
[50:09] Dispensed 0.000141 mL of Base (0.5 M KOH)
[50:14] Stepping pH = 5.25
[50:29] Stirrer speed set to 0
[50:40] Datapoint id 48 collected
[50:40] Charge balance equation is out by -35.3%
[50:40] Stirrer speed set to 55
[50:46] pH 5.26 -> 5.46
[50:46] Using cautious pH adjust
[50:46] Dispensed 0.000165 mL of Base (0.5 M KOH)
[50:51] Stepping pH = 5.33
[50:51] Dispensed 0.000165 mL of Base (0.5 M KOH)

Sample name: **M07_octanol** Experiment start time: **2/28/2018 7:21:13 PM**
Assay name: **pH-metric high logP** Analyst: **Pion**
Assay ID: **18B-28013** Instrument ID: **T312060**
Filename: **C:\Sirius_T3\Mehtap\20180228_exp28_logP_T3-2\18B-28013_M07_octanol_pH-metric high logP.t3r**

#### Experiment Log (continued)

[50:56] Stepping pH = 5.44
[50:56] Dispensed 0.000024 mL of Base (0.5 M KOH)
[51:01] Stepping pH = 5.44
[51:01] Dispensed 0.000047 mL of Base (0.5 M KOH)
[51:06] Stepping pH = 5.46
[51:21] Stirrer speed set to 0
[51:33] Datapoint id 49 collected
[51:33] Charge balance equation is out by -26.8%
[51:33] Stirrer speed set to 55
[51:38] pH 5.48 -> 5.68
[51:38] Using cautious pH adjust
[51:38] Dispensed 0.000118 mL of Base (0.5 M KOH)
[51:43] Stepping pH = 5.54
[51:43] Dispensed 0.000118 mL of Base (0.5 M KOH)
[51:48] Stepping pH = 5.64
[51:48] Dispensed 0.000024 mL of Base (0.5 M KOH)
[51:53] Stepping pH = 5.65
[51:53] Dispensed 0.000024 mL of Base (0.5 M KOH)
[51:58] Stepping pH = 5.67
[52:14] Stirrer speed set to 0
[52:27] Datapoint id 50 collected
[52:27] Charge balance equation is out by -33.5%
[52:27] Stirrer speed set to 55
[52:32] pH 5.69 -> 5.89
[52:32] Using cautious pH adjust
[52:32] Dispensed 0.000071 mL of Base (0.5 M KOH)
[52:37] Stepping pH = 5.75
[52:37] Dispensed 0.000094 mL of Base (0.5 M KOH)
[52:42] Stepping pH = 5.89
[52:57] Stirrer speed set to 0
[53:12] Datapoint id 51 collected
[53:12] Charge balance equation is out by -9.3%
[53:12] Stirrer speed set to 55
[53:17] pH 5.91 -> 6.11
[53:17] Using charge balance adjust
[53:17] Dispensed 0.000118 mL of Base (0.5 M KOH)
[53:37] Stirrer speed set to 0
[54:02] Datapoint id 52 collected
[54:02] Charge balance equation is out by -17.4%
[54:02] Stirrer speed set to 55
[54:07] pH 6.13 -> 6.33
[54:07] Using cautious pH adjust
[54:07] Dispensed 0.000047 mL of Base (0.5 M KOH)
[54:12] Stepping pH = 6.17
[54:12] Dispensed 0.000094 mL of Base (0.5 M KOH)
[54:17] Stepping pH = 6.53
[54:32] Stirrer speed set to 0
[55:18] Datapoint id 53 collected
[55:18] Charge balance equation is out by -53.9%
[55:18] Stirrer speed set to 55
[55:23] pH 6.56 -> 6.76
[55:23] Using cautious pH adjust
[55:23] Dispensed 0.000024 mL of Base (0.5 M KOH)
[55:28] Stepping pH = 6.57
[55:28] Dispensed 0.000071 mL of Base (0.5 M KOH)
[55:33] Stepping pH = 6.85
[55:48] Stirrer speed set to 0
[56:48] Datapoint id 54 collected
[56:48] Charge balance equation is out by -91.3%

Sample name: **M07_octanol** Experiment start time: **2/28/2018 7:21:13 PM**
Assay name: **pH-metric high logP** Analyst: **Pion**
Assay ID: **18B-28013** Instrument ID: **T312060**
Filename: **C:\Sirius_T3\Mehtap\20180228_exp28_logP_T3-2\18B-28013_M07_octanol_pH-metric high logP.t3r**

#### Experiment Log (continued)

[56:48] Stirrer speed set to 55
[56:53] pH 7.00 -> 7.20
[56:53] Using cautious pH adjust
[56:53] Dispensed 0.000024 mL of Base (0.5 M KOH)
[56:58] Stepping pH = 7.03
[56:58] Dispensed 0.000024 mL of Base (0.5 M KOH)
[57:03] Stepping pH = 7.13
[57:03] Dispensed 0.000024 mL of Base (0.5 M KOH)
[57:09] Stepping pH = 7.33
[57:24] Stirrer speed set to 0
[58:24] Datapoint id 55 collected
[58:24] Charge balance equation is out by -155.9%
[58:24] Stirrer speed set to 55
[58:29] pH 7.44 -> 7.64
[58:29] Using cautious pH adjust
[58:29] Dispensed 0.000024 mL of Base (0.5 M KOH)
[58:34] Stepping pH = 7.50
[58:34] Dispensed 0.000024 mL of Base (0.5 M KOH)
[58:39] Stepping pH = 7.75
[58:54] Stirrer speed set to 0
[59:54] Datapoint id 56 collected
[59:54] Charge balance equation is out by -253.5%
[59:54] Stirrer speed set to 55
[59:59] pH 7.87 -> 8.07
[59:59] Using cautious pH adjust
[59:59] Dispensed 0.000024 mL of Base (0.5 M KOH)
[1:00:04] Stepping pH = 7.93
[1:00:04] Dispensed 0.000024 mL of Base (0.5 M KOH)
[1:00:10] Stepping pH = 8.14
[1:00:25] Stirrer speed set to 0
[1:01:25] Datapoint id 57 collected
[1:01:25] Charge balance equation is out by -437.3%
[1:01:25] Stirrer speed set to 55
[1:01:30] pH 8.27 -> 8.47
[1:01:30] Using cautious pH adjust
[1:01:30] Dispensed 0.000024 mL of Base (0.5 M KOH)
[1:01:35] Stepping pH = 8.31
[1:01:35] Dispensed 0.000024 mL of Base (0.5 M KOH)
[1:01:40] Stepping pH = 8.47
[1:01:55] Stirrer speed set to 0
[1:02:48] Datapoint id 58 collected
[1:02:48] Charge balance equation is out by -280.7%
[1:02:48] Stirrer speed set to 55
[1:02:53] pH 8.50 -> 8.70
[1:02:53] Using cautious pH adjust
[1:02:53] Dispensed 0.000024 mL of Base (0.5 M KOH)
[1:02:59] Stepping pH = 8.50
[1:02:59] Dispensed 0.000024 mL of Base (0.5 M KOH)
[1:03:04] Stepping pH = 8.60
[1:03:04] Dispensed 0.000024 mL of Base (0.5 M KOH)
[1:03:09] Stepping pH = 8.73
[1:03:24] Stirrer speed set to 0
[1:03:59] Datapoint id 59 collected
[1:03:59] Charge balance equation is out by -300.8%
[1:03:59] Stirrer speed set to 55
[1:04:04] pH 8.78 -> 8.98
[1:04:04] Using cautious pH adjust
[1:04:04] Dispensed 0.000024 mL of Base (0.5 M KOH)
[1:04:09] Stepping pH = 8.79

Sample name: **M07_octanol** Experiment start time: **2/28/2018 7:21:13 PM**
Assay name: **pH-metric high logP** Analyst: **Pion**
Assay ID: **18B-28013** Instrument ID: **T312060**
Filename: **C:\Sirius_T3\Mehtap\20180228_exp28_logP_T3-2\18B-28013_M07_octanol_pH-metric high logP.t3r**

#### Experiment Log (continued)

[1:04:09] Dispensed 0.000047 mL of Base (0.5 M KOH)
[1:04:14] Stepping pH = 8.94
[1:04:15] Dispensed 0.000024 mL of Base (0.5 M KOH)
[1:04:20] Stepping pH = 9.02
[1:04:35] Stirrer speed set to 0
[1:05:04] Datapoint id 60 collected
[1:05:04] Charge balance equation is out by -188.4%
[1:05:04] Stirrer speed set to 55
[1:05:09] pH 9.05 -> 9.25
[1:05:09] Using cautious pH adjust
[1:05:09] Dispensed 0.000024 mL of Base (0.5 M KOH)
[1:05:14] Stepping pH = 9.05
[1:05:14] Dispensed 0.000071 mL of Base (0.5 M KOH)
[1:05:19] Stepping pH = 9.21
[1:05:19] Dispensed 0.000024 mL of Base (0.5 M KOH)
[1:05:24] Stepping pH = 9.25
[1:05:39] Stirrer speed set to 0
[1:05:53] Datapoint id 61 collected
[1:05:53] Charge balance equation is out by -138.2%
[1:05:53] Stirrer speed set to 55
[1:05:58] pH 9.25 -> 9.45
[1:05:58] Using cautious pH adjust
[1:05:58] Dispensed 0.000047 mL of Base (0.5 M KOH)
[1:06:03] Stepping pH = 9.29
[1:06:03] Dispensed 0.000071 mL of Base (0.5 M KOH)
[1:06:08] Stepping pH = 9.42
[1:06:08] Dispensed 0.000024 mL of Base (0.5 M KOH)
[1:06:13] Stepping pH = 9.45
[1:06:29] Stirrer speed set to 0
[1:06:46] Datapoint id 62 collected
[1:06:46] Charge balance equation is out by -77.2%
[1:06:46] Stirrer speed set to 55
[1:06:51] pH 9.45 -> 9.65
[1:06:51] Using cautious pH adjust
[1:06:51] Dispensed 0.000047 mL of Base (0.5 M KOH)
[1:06:56] Stepping pH = 9.48
[1:06:56] Dispensed 0.000118 mL of Base (0.5 M KOH)
[1:07:01] Stepping pH = 9.56
[1:07:01] Dispensed 0.000094 mL of Base (0.5 M KOH)
[1:07:06] Stepping pH = 9.66
[1:07:21] Stirrer speed set to 0
[1:07:35] Datapoint id 63 collected
[1:07:35] Charge balance equation is out by -136.0%
[1:07:35] Stirrer speed set to 55
[1:07:41] pH 9.66 -> 9.86
[1:07:41] Using cautious pH adjust
[1:07:41] Dispensed 0.000071 mL of Base (0.5 M KOH)
[1:07:46] Stepping pH = 9.69
[1:07:46] Dispensed 0.000165 mL of Base (0.5 M KOH)
[1:07:51] Stepping pH = 9.86
[1:08:06] Stirrer speed set to 0
[1:08:17] Datapoint id 64 collected
[1:08:17] Charge balance equation is out by -55.4%
[1:08:17] Stirrer speed set to 55
[1:08:22] pH 9.86 -> 10.05
[1:08:22] Using cautious pH adjust
[1:08:22] Dispensed 0.000118 mL of Base (0.5 M KOH)
[1:08:27] Stepping pH = 9.92
[1:08:27] Dispensed 0.000141 mL of Base (0.5 M KOH)

Sample name: **M07_octanol**
Assay name: **pH-metric high logP**
Assay ID: **18B-28013**
Filename: **C:\Sirius_T3\Mehtap\20180228_exp28_logP_T3-2\18B-28013_M07_octanol_pH-metric high logP.t3r**

Experiment start time: **2/28/2018 7:21:13 PM**
Analyst: **Pion**
Instrument ID: **T312060**

#### Experiment Log (continued)

[1:08:32] Stepping pH = 10.02
[1:08:48] Stirrer speed set to 0
[1:08:58] Datapoint id 65 collected
[1:08:58] Charge balance equation is out by -13.4%
[1:08:58] Titration 3 of 3
[1:08:58] Adding initial titrants
[1:08:58] Automatically add 0.25000 mL of Octanol
[1:09:04] Dispensed 0.250000 mL of Octanol
[1:09:04] Stirrer speed set to 10
[1:09:05] Stirrer speed set to 60
[1:09:05] Iterative adjust 10.02 -> 2.00
[1:09:05] pH 10.02 -> 2.00
[1:09:07] Dispensed 0.058702 mL of Acid (0.5 M HCl)
[1:09:12] pH 2.04 -> 2.00
[1:09:12] Dispensed 0.004139 mL of Acid (0.5 M HCl)
[1:10:02] Stirrer speed set to 0
[1:10:13] Datapoint id 66 collected
[1:10:13] Stirrer speed set to 60
[1:10:18] pH 1.96 -> 2.16
[1:10:18] Using cautious pH adjust
[1:10:18] Dispensed 0.010183 mL of Base (0.5 M KOH)
[1:10:23] Stepping pH = 2.05
[1:10:23] Dispensed 0.006632 mL of Base (0.5 M KOH)
[1:10:29] Stepping pH = 2.13
[1:10:29] Dispensed 0.001905 mL of Base (0.5 M KOH)
[1:10:34] Stepping pH = 2.16
[1:10:49] Stirrer speed set to 0
[1:10:59] Datapoint id 67 collected
[1:10:59] Charge balance equation is out by 8.1%
[1:10:59] Stirrer speed set to 60
[1:11:04] pH 2.16 -> 2.36
[1:11:04] Using charge balance adjust
[1:11:04] Dispensed 0.012841 mL of Base (0.5 M KOH)
[1:11:25] Stirrer speed set to 0
[1:11:35] Datapoint id 68 collected
[1:11:35] Charge balance equation is out by 5.6%
[1:11:35] Stirrer speed set to 60
[1:11:40] pH 2.37 -> 2.57
[1:11:40] Using charge balance adjust
[1:11:40] Dispensed 0.007973 mL of Base (0.5 M KOH)
[1:12:01] Stirrer speed set to 0
[1:12:11] Datapoint id 69 collected
[1:12:11] Charge balance equation is out by 9.9%
[1:12:11] Stirrer speed set to 60
[1:12:16] pH 2.60 -> 2.80
[1:12:16] Using charge balance adjust
[1:12:16] Dispensed 0.004962 mL of Base (0.5 M KOH)
[1:12:36] Stirrer speed set to 0
[1:12:46] Datapoint id 70 collected
[1:12:46] Charge balance equation is out by -2.1%
[1:12:46] Stirrer speed set to 60
[1:12:51] pH 2.80 -> 3.00
[1:12:51] Using charge balance adjust
[1:12:52] Dispensed 0.003410 mL of Base (0.5 M KOH)
[1:13:12] Stirrer speed set to 0
[1:13:22] Datapoint id 71 collected
[1:13:22] Charge balance equation is out by 7.6%
[1:13:22] Stirrer speed set to 60
[1:13:27] pH 3.02 -> 3.22

Sample name: **M07_octanol** Experiment start time: **2/28/2018 7:21:13 PM**
Assay name: **pH-metric high logP** Analyst: **Pion**
Assay ID: **18B-28013** Instrument ID: **T312060**
Filename: **C:\Sirius_T3\Mehtap\20180228_exp28_logP_T3-2\18B-28013_M07_octanol_pH-metric high logP.t3r**

#### Experiment Log (continued)

[1:13:27] Using charge balance adjust
[1:13:27] Dispensed 0.002516 mL of Base (0.5 M KOH)
[1:13:47] Stirrer speed set to 0
[1:13:58] Datapoint id 72 collected
[1:13:58] Charge balance equation is out by 0.5%
[1:13:58] Stirrer speed set to 60
[1:14:03] pH 3.23 -> 3.43
[1:14:03] Using charge balance adjust
[1:14:03] Dispensed 0.002070 mL of Base (0.5 M KOH)
[1:14:23] Stirrer speed set to 0
[1:14:33] Datapoint id 73 collected
[1:14:33] Charge balance equation is out by 1.7%
[1:14:33] Stirrer speed set to 60
[1:14:38] pH 3.43 -> 3.63
[1:14:38] Using charge balance adjust
[1:14:38] Dispensed 0.001834 mL of Base (0.5 M KOH)
[1:14:59] Stirrer speed set to 0
[1:15:09] Datapoint id 74 collected
[1:15:09] Charge balance equation is out by 15.9%
[1:15:09] Stirrer speed set to 60
[1:15:14] pH 3.67 -> 3.87
[1:15:14] Using cautious pH adjust
[1:15:14] Dispensed 0.000800 mL of Base (0.5 M KOH)
[1:15:19] Stepping pH = 3.78
[1:15:19] Dispensed 0.000494 mL of Base (0.5 M KOH)
[1:15:24] Stepping pH = 3.85
[1:15:24] Dispensed 0.000141 mL of Base (0.5 M KOH)
[1:15:29] Stepping pH = 3.87
[1:15:44] Stirrer speed set to 0
[1:15:54] Datapoint id 75 collected
[1:15:54] Charge balance equation is out by 10.4%
[1:15:54] Stirrer speed set to 60
[1:16:00] pH 3.87 -> 4.07
[1:16:00] Using charge balance adjust
[1:16:00] Dispensed 0.001364 mL of Base (0.5 M KOH)
[1:16:20] Stirrer speed set to 0
[1:16:30] Datapoint id 76 collected
[1:16:30] Charge balance equation is out by -1.5%
[1:16:30] Stirrer speed set to 60
[1:16:35] pH 4.07 -> 4.27
[1:16:35] Using charge balance adjust
[1:16:36] Dispensed 0.001082 mL of Base (0.5 M KOH)
[1:16:56] Stirrer speed set to 0
[1:17:12] Datapoint id 77 collected
[1:17:12] Charge balance equation is out by -15.8%
[1:17:12] Stirrer speed set to 60
[1:17:17] pH 4.24 -> 4.44
[1:17:17] Using cautious pH adjust
[1:17:18] Dispensed 0.000423 mL of Base (0.5 M KOH)
[1:17:23] Stepping pH = 4.31
[1:17:23] Dispensed 0.000423 mL of Base (0.5 M KOH)
[1:17:28] Stepping pH = 4.39
[1:17:28] Dispensed 0.000212 mL of Base (0.5 M KOH)
[1:17:33] Stepping pH = 4.43
[1:17:48] Stirrer speed set to 0
[1:17:58] Datapoint id 78 collected
[1:17:58] Charge balance equation is out by -24.6%
[1:17:58] Stirrer speed set to 60
[1:18:03] pH 4.44 -> 4.64

| | | | |
|---|---|---|---|
| Sample name: | **M07_octanol** | Experiment start time: | **2/28/2018 7:21:13 PM** |
| Assay name: | **pH-metric high logP** | Analyst: | **Pion** |
| Assay ID: | **18B-28013** | Instrument ID: | **T312060** |
| Filename: | **C:\Sirius_T3\Mehtap\20180228_exp28_logP_T3-2\18B-28013_M07_octanol_pH-metric high logP.t3r** | | |

#### Experiment Log (continued)

[1:18:03] Using cautious pH adjust
[1:18:03] Dispensed 0.000306 mL of Base (0.5 M KOH)
[1:18:08] Stepping pH = 4.51
[1:18:09] Dispensed 0.000306 mL of Base (0.5 M KOH)
[1:18:14] Stepping pH = 4.60
[1:18:14] Dispensed 0.000094 mL of Base (0.5 M KOH)
[1:18:19] Stepping pH = 4.61
[1:18:19] Dispensed 0.000118 mL of Base (0.5 M KOH)
[1:18:24] Stepping pH = 4.65
[1:18:39] Stirrer speed set to 0
[1:18:50] Datapoint id 79 collected
[1:18:50] Charge balance equation is out by -33.9%
[1:18:50] Stirrer speed set to 60
[1:18:55] pH 4.66 -> 4.86
[1:18:55] Using cautious pH adjust
[1:18:55] Dispensed 0.000212 mL of Base (0.5 M KOH)
[1:19:00] Stepping pH = 4.74
[1:19:00] Dispensed 0.000188 mL of Base (0.5 M KOH)
[1:19:05] Stepping pH = 4.82
[1:19:05] Dispensed 0.000071 mL of Base (0.5 M KOH)
[1:19:10] Stepping pH = 4.84
[1:19:10] Dispensed 0.000071 mL of Base (0.5 M KOH)
[1:19:15] Stepping pH = 4.86
[1:19:31] Stirrer speed set to 0
[1:19:41] Datapoint id 80 collected
[1:19:41] Charge balance equation is out by -26.0%
[1:19:41] Stirrer speed set to 60
[1:19:46] pH 4.87 -> 5.07
[1:19:46] Using cautious pH adjust
[1:19:46] Dispensed 0.000141 mL of Base (0.5 M KOH)
[1:19:51] Stepping pH = 4.95
[1:19:51] Dispensed 0.000118 mL of Base (0.5 M KOH)
[1:19:56] Stepping pH = 5.02
[1:19:56] Dispensed 0.000047 mL of Base (0.5 M KOH)
[1:20:02] Stepping pH = 5.05
[1:20:02] Dispensed 0.000047 mL of Base (0.5 M KOH)
[1:20:07] Stepping pH = 5.08
[1:20:22] Stirrer speed set to 0
[1:20:32] Datapoint id 81 collected
[1:20:32] Charge balance equation is out by -32.6%
[1:20:32] Stirrer speed set to 60
[1:20:37] pH 5.09 -> 5.29
[1:20:37] Using cautious pH adjust
[1:20:38] Dispensed 0.000094 mL of Base (0.5 M KOH)
[1:20:43] Stepping pH = 5.15
[1:20:43] Dispensed 0.000118 mL of Base (0.5 M KOH)
[1:20:48] Stepping pH = 5.29
[1:21:03] Stirrer speed set to 0
[1:21:14] Datapoint id 82 collected
[1:21:14] Charge balance equation is out by -12.9%
[1:21:14] Stirrer speed set to 60
[1:21:20] pH 5.30 -> 5.50
[1:21:20] Using charge balance adjust
[1:21:20] Dispensed 0.000118 mL of Base (0.5 M KOH)
[1:21:40] Stirrer speed set to 0
[1:21:51] Datapoint id 83 collected
[1:21:51] Charge balance equation is out by -30.6%
[1:21:51] Stirrer speed set to 60
[1:21:56] pH 5.46 -> 5.66

| | | | |
|---|---|---|---|
| Sample name: | **M07_octanol** | Experiment start time: | **2/28/2018 7:21:13 PM** |
| Assay name: | **pH-metric high logP** | Analyst: | **Pion** |
| Assay ID: | **18B-28013** | Instrument ID: | **T312060** |
| Filename: | **C:\Sirius_T3\Mehtap\20180228_exp28_logP_T3-2\18B-28013_M07_octanol_pH-metric high logP.t3r** | | |

#### Experiment Log (continued)

[1:21:56] Using cautious pH adjust
[1:21:57] Dispensed 0.000047 mL of Base (0.5 M KOH)
[1:22:02] Stepping pH = 5.50
[1:22:02] Dispensed 0.000094 mL of Base (0.5 M KOH)
[1:22:07] Stepping pH = 5.75
[1:22:22] Stirrer speed set to 0
[1:22:38] Datapoint id 84 collected
[1:22:38] Charge balance equation is out by -42.0%
[1:22:38] Stirrer speed set to 60
[1:22:43] pH 5.77 -> 5.97
[1:22:43] Using cautious pH adjust
[1:22:44] Dispensed 0.000024 mL of Base (0.5 M KOH)
[1:22:49] Stepping pH = 5.77
[1:22:49] Dispensed 0.000165 mL of Base (0.5 M KOH)
[1:22:54] Stepping pH = 6.64
[1:23:09] Stirrer speed set to 0
[1:24:09] Datapoint id 85 collected
[1:24:09] Charge balance equation is out by -201.0%
[1:24:09] Stirrer speed set to 60
[1:24:14] pH 6.65 -> 6.85
[1:24:14] Using cautious pH adjust
[1:24:14] Dispensed 0.000024 mL of Base (0.5 M KOH)
[1:24:19] Stepping pH = 6.68
[1:24:19] Dispensed 0.000047 mL of Base (0.5 M KOH)
[1:24:24] Stepping pH = 6.82
[1:24:24] Dispensed 0.000024 mL of Base (0.5 M KOH)
[1:24:30] Stepping pH = 6.97
[1:24:45] Stirrer speed set to 0
[1:25:45] Datapoint id 86 collected
[1:25:45] Charge balance equation is out by -106.3%
[1:25:45] Stirrer speed set to 60
[1:25:50] pH 7.10 -> 7.30
[1:25:50] Using cautious pH adjust
[1:25:50] Dispensed 0.000024 mL of Base (0.5 M KOH)
[1:25:55] Stepping pH = 7.14
[1:25:55] Dispensed 0.000024 mL of Base (0.5 M KOH)
[1:26:00] Stepping pH = 7.29
[1:26:15] Stirrer speed set to 0
[1:27:15] Datapoint id 87 collected
[1:27:15] Charge balance equation is out by -76.1%
[1:27:15] Stirrer speed set to 60
[1:27:20] pH 7.31 -> 7.51
[1:27:20] Using cautious pH adjust
[1:27:20] Dispensed 0.000024 mL of Base (0.5 M KOH)
[1:27:25] Stepping pH = 7.39
[1:27:25] Dispensed 0.000024 mL of Base (0.5 M KOH)
[1:27:31] Stepping pH = 7.60
[1:27:46] Stirrer speed set to 0
[1:28:46] Datapoint id 88 collected
[1:28:46] Charge balance equation is out by -151.3%
[1:28:46] Stirrer speed set to 60
[1:28:51] pH 7.53 -> 7.73
[1:28:51] Using cautious pH adjust
[1:28:51] Dispensed 0.000024 mL of Base (0.5 M KOH)
[1:28:56] Stepping pH = 7.52
[1:28:56] Dispensed 0.000024 mL of Base (0.5 M KOH)
[1:29:01] Stepping pH = 7.53
[1:29:01] Dispensed 0.000094 mL of Base (0.5 M KOH)
[1:29:06] Stepping pH = 7.58

Sample name: **M07_octanol**
Assay name: **pH-metric high logP**
Assay ID: **18B-28013**
Filename: **C:\Sirius_T3\Mehtap\20180228_exp28_logP_T3-2\18B-28013_M07_octanol_pH-metric high logP.t3r**

Experiment start time: **2/28/2018 7:21:13 PM**
Analyst: **Pion**
Instrument ID: **T312060**

#### Experiment Log (continued)

[1:29:06] Dispensed 0.000141 mL of Base (0.5 M KOH)
[1:29:11] Stepping pH = 8.69
[1:29:26] Stirrer speed set to 0
[1:30:22] Datapoint id 89 collected
[1:30:22] Charge balance equation is out by -2,279.8%
[1:30:22] Stirrer speed set to 60
[1:30:27] pH 8.69 -> 8.89
[1:30:27] Using cautious pH adjust
[1:30:27] Dispensed 0.000024 mL of Base (0.5 M KOH)
[1:30:32] Stepping pH = 8.69
[1:30:32] Dispensed 0.000047 mL of Base (0.5 M KOH)
[1:30:37] Stepping pH = 8.71
[1:30:37] Dispensed 0.000094 mL of Base (0.5 M KOH)
[1:30:42] Stepping pH = 9.03
[1:30:57] Stirrer speed set to 0
[1:31:14] Datapoint id 90 collected
[1:31:14] Charge balance equation is out by -541.2%
[1:31:14] Stirrer speed set to 60
[1:31:19] pH 9.03 -> 9.23
[1:31:19] Using cautious pH adjust
[1:31:19] Dispensed 0.000024 mL of Base (0.5 M KOH)
[1:31:24] Stepping pH = 9.03
[1:31:24] Dispensed 0.000071 mL of Base (0.5 M KOH)
[1:31:29] Stepping pH = 9.14
[1:31:29] Dispensed 0.000047 mL of Base (0.5 M KOH)
[1:31:34] Stepping pH = 9.21
[1:31:34] Dispensed 0.000024 mL of Base (0.5 M KOH)
[1:31:39] Stepping pH = 9.24
[1:31:55] Stirrer speed set to 0
[1:32:08] Datapoint id 91 collected
[1:32:08] Charge balance equation is out by -236.4%
[1:32:08] Stirrer speed set to 60
[1:32:13] pH 9.26 -> 9.46
[1:32:13] Using cautious pH adjust
[1:32:13] Dispensed 0.000047 mL of Base (0.5 M KOH)
[1:32:18] Stepping pH = 9.29
[1:32:18] Dispensed 0.000094 mL of Base (0.5 M KOH)
[1:32:23] Stepping pH = 9.46
[1:32:38] Stirrer speed set to 0
[1:32:52] Datapoint id 92 collected
[1:32:52] Charge balance equation is out by -62.5%
[1:32:52] Stirrer speed set to 60
[1:32:57] pH 9.48 -> 9.68
[1:32:57] Using cautious pH adjust
[1:32:57] Dispensed 0.000071 mL of Base (0.5 M KOH)
[1:33:02] Stepping pH = 9.50
[1:33:02] Dispensed 0.000165 mL of Base (0.5 M KOH)
[1:33:07] Stepping pH = 9.70
[1:33:22] Stirrer speed set to 0
[1:33:34] Datapoint id 93 collected
[1:33:34] Charge balance equation is out by -82.3%
[1:33:34] Stirrer speed set to 60
[1:33:39] pH 9.71 -> 9.91
[1:33:39] Using cautious pH adjust
[1:33:39] Dispensed 0.000094 mL of Base (0.5 M KOH)
[1:33:44] Stepping pH = 9.76
[1:33:44] Dispensed 0.000141 mL of Base (0.5 M KOH)
[1:33:49] Stepping pH = 9.90
[1:34:04] Stirrer speed set to 0

| | | | |
|---|---|---|---|
| Sample name: | **M07_octanol** | Experiment start time: | **2/28/2018 7:21:13 PM** |
| Assay name: | **pH-metric high logP** | Analyst: | **Pion** |
| Assay ID: | **18B-28013** | Instrument ID: | **T312060** |
| Filename: | **C:\Sirius_T3\Mehtap\20180228_exp28_logP_T3-2\18B-28013_M07_octanol_pH-metric high logP.t3r** | | |

#### Experiment Log (continued)

[1:34:15] Datapoint id 94 collected
[1:34:15] Charge balance equation is out by -20.8%
[1:34:15] Stirrer speed set to 60
[1:34:20] pH 9.90 -> 10.05
[1:34:20] Using cautious pH adjust
[1:34:20] Dispensed 0.000094 mL of Base (0.5 M KOH)
[1:34:25] Stepping pH = 9.94
[1:34:25] Dispensed 0.000141 mL of Base (0.5 M KOH)
[1:34:30] Stepping pH = 10.03
[1:34:45] Stirrer speed set to 0
[1:34:55] Datapoint id 95 collected
[1:34:55] Charge balance equation is out by -25.1%
[1:34:55] Argon flow rate set to 0
[1:34:59] Titrator arm moved over Titration position
