## Supplementary material for "Octanol-water partition coefficient measurements for the SAMPL6 Blind Prediction Challenge": SM08_18C-02007_M08_octanol_pH-metric high logP_report.pdf

| | | | |
|---|---|---|---|
| Sample name: | **M08_octanol** | Experiment start time: | **3/2/2018 5:10:52 PM** |
| Assay name: | **pH-metric high logP** | Analyst: | **Pion** |
| Assay ID: | **18C-02007** | Instrument ID: | **T312060** |
| Filename: | **C:\Sirius_T3\Mehtap\20180302_exp29_logP_T3-2\18C-02007_M08_octanol_pH-metric high logP.t3r** | | |

#### pH-metric Result

| | | |
|---|---|---|
| logP (neutral XH) | 3.05 | ±0.01 (n=50) |
| logP (X -) | -0.40 | ±0.05 (n=50) |

#### 18C-02007 Points 2 to 38

| | |
|---|---|
| M08_octanol concentration factor | 1.137 |
| Carbonate | 0.0754 mM |
| Acidity error | -1.80478 mM |

#### 18C-02007 Points 39 to 77

| | |
|---|---|
| M08_octanol concentration factor | 0.855 |
| Carbonate | 0.3938 mM |
| Acidity error | -1.60863 mM |

#### 18C-02007 Points 78 to 114

| | |
|---|---|
| M08_octanol concentration factor | 0.850 |
| Carbonate | 0.3937 mM |
| Acidity error | -1.48346 mM |

#### Warnings and errors

| | |
|---|---|
| Errors | None |
| Warnings | None |

#### Sample logD and percent species

| pH | M08_octanol logD | M08_octanol M08_octanolH | M08_octanol M08_octanol | M08_octanol M08_octanolH* | M08_octanol M08_octanol* | Comment |
|---|---|---|---|---|---|---|
| 1.000 | 3.05 | 0.09 % | 0.00 % | 99.91 % | 0.00 % | |
| 1.200 | 3.05 | 0.09 % | 0.00 % | 99.91 % | 0.00 % | Stomach pH |
| 2.000 | 3.05 | 0.09 % | 0.00 % | 99.91 % | 0.00 % | |
| 3.000 | 3.03 | 0.09 % | 0.01 % | 99.90 % | 0.00 % | |
| 4.000 | 2.85 | 0.09 % | 0.05 % | 99.84 % | 0.02 % | |
| 5.000 | 2.21 | 0.09 % | 0.53 % | 99.18 % | 0.21 % | |
| 6.000 | 1.28 | 0.08 % | 4.94 % | 93.03 % | 1.95 % | |
| 6.500 | 0.80 | 0.07 % | 13.61 % | 80.97 % | 5.36 % | |
| 7.000 | 0.36 | 0.05 % | 30.52 % | 57.42 % | 12.01 % | |
| 7.400 | 0.06 | 0.03 % | 46.65 % | 34.95 % | 18.36 % | Blood pH |
| 8.000 | -0.24 | 0.01 % | 63.21 % | 11.90 % | 24.88 % | |
| 9.000 | -0.38 | 0.00 % | 70.80 % | 1.33 % | 27.87 % | |
| 10.000 | -0.40 | 0.00 % | 71.66 % | 0.13 % | 28.21 % | |
| 11.000 | -0.40 | 0.00 % | 71.74 % | 0.01 % | 28.24 % | |
| 12.000 | -0.40 | 0.00 % | 71.75 % | 0.00 % | 28.25 % | |

| | | | |
|---|---|---|---|
| Sample name: | **M08_octanol** | Experiment start time: | **3/2/2018 5:10:52 PM** |
| Assay name: | **pH-metric high logP** | Analyst: | **Pion** |
| Assay ID: | **18C-02007** | Instrument ID: | **T312060** |
| Filename: | **C:\Sirius_T3\Mehtap\20180302_exp29_logP_T3-2\18C-02007_M08_octanol_pH-metric high logP.t3r** | | |

#### Graphs

Charge balance residuals

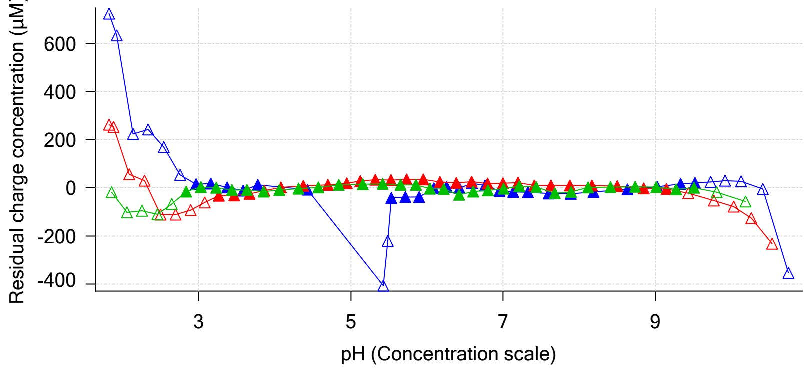


Ionic strength

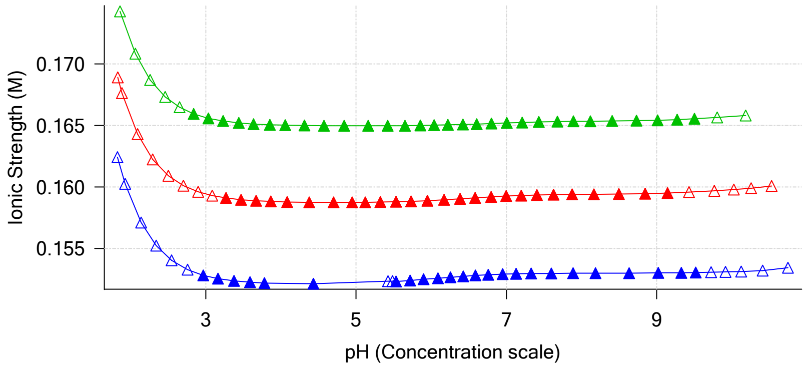


Buffer index profile

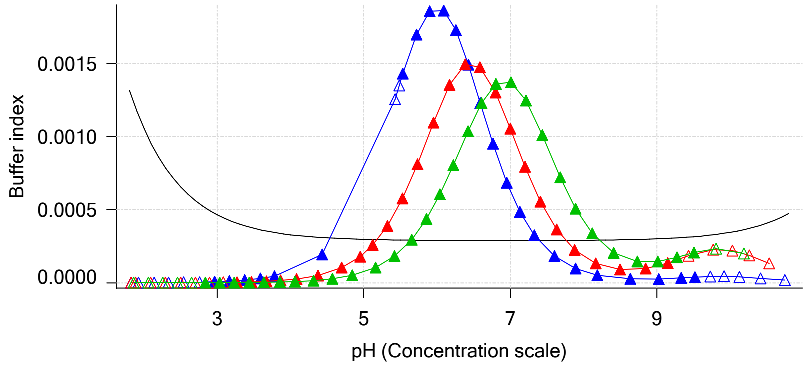


Sample M08_octanol Ionisation Graph

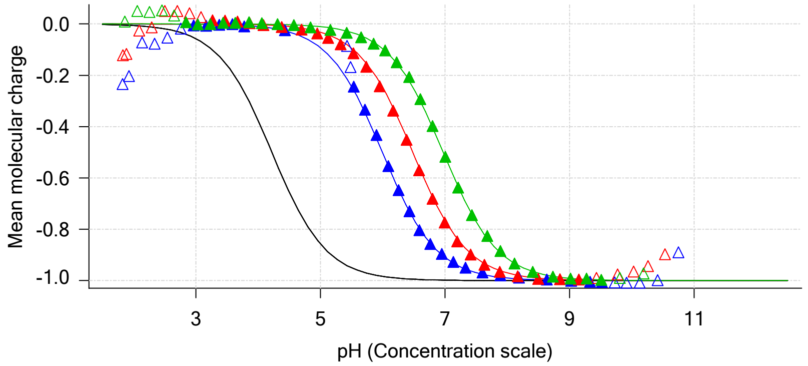


Distribution of species for sample M08_octanol

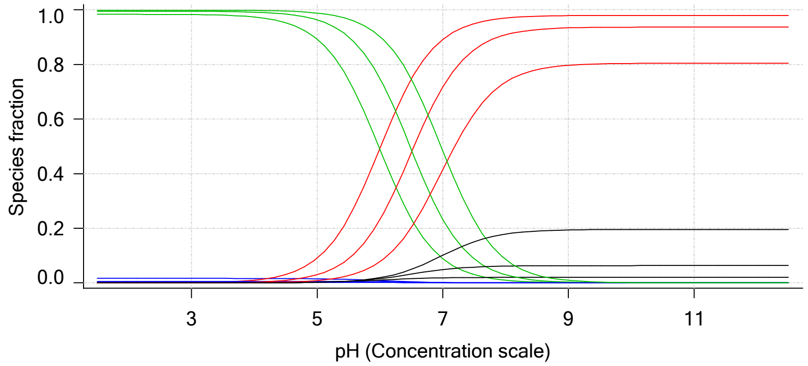


Lipophilicity Profile

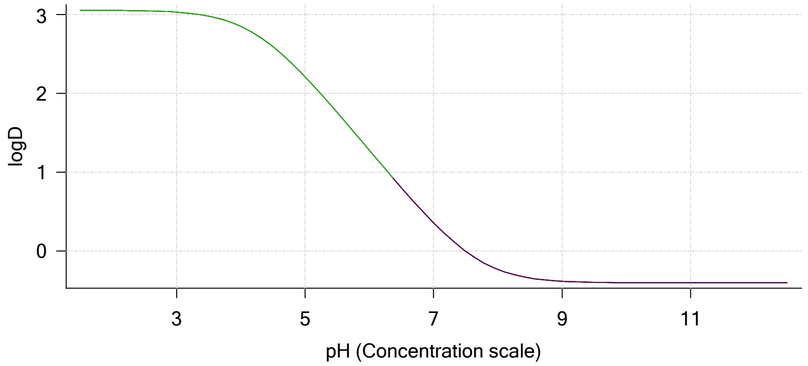


Log R Profile

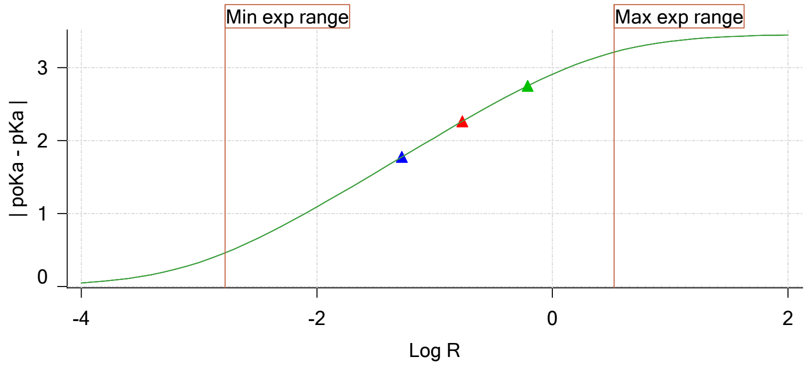


Sample Carbonate Ionisation Graph

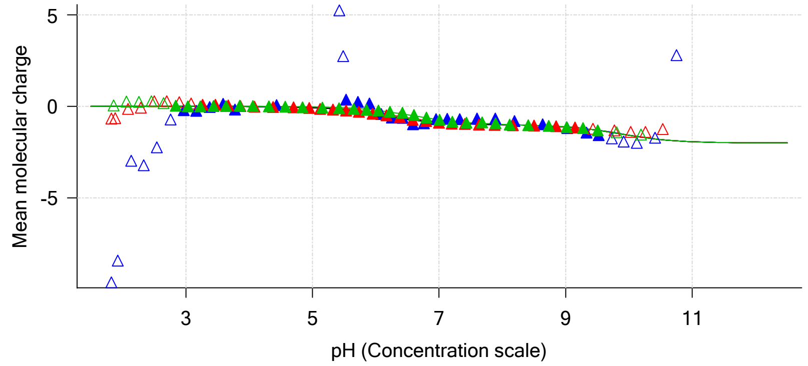

| | | | |
|---|---|---|---|
| Sample name: | **M08_octanol** | Experiment start time: | **3/2/2018 5:10:52 PM** |
| Assay name: | **pH-metric high logP** | Analyst: | **Pion** |
| Assay ID: | **18C-02007** | Instrument ID: | **T312060** |
| Filename: | **C:\Sirius_T3\Mehtap\20180302_exp29_logP_T3-2\18C-02007_M08_octanol_pH-metric high logP.t3r** | | |

#### Graphs (continued)

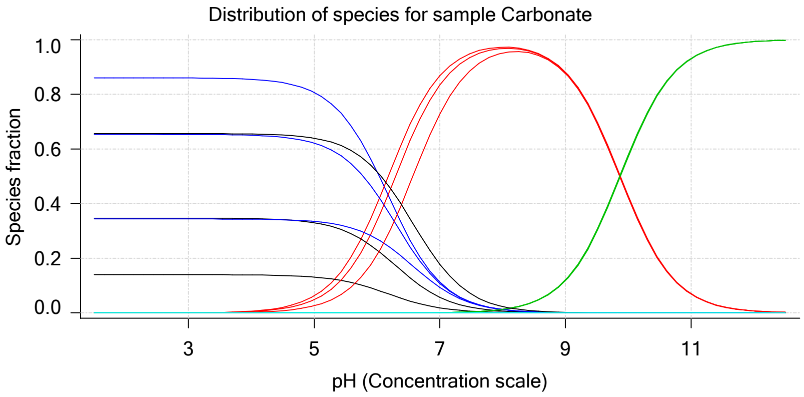

| | | | |
|---|---|---|---|
| Sample name: | **M08_octanol** | Experiment start time: | **3/2/2018 5:10:52 PM** |
| Assay name: | **pH-metric high logP** | Analyst: | **Pion** |
| Assay ID: | **18C-02007** | Instrument ID: | **T312060** |
| Filename: | **C:\Sirius_T3\Mehtap\20180302_exp29_logP_T3-2\18C-02007_M08_octanol_pH-metric high logP.t3r** | | |

### pH-metric high logP Titration 1 of 3 18C-02007 Points 2 to 38

#### Overall results

| | |
|---|---|
| RMSD | 0.092 |
| Average ionic strength | 0.153 M |
| Average temperature | 25.0°C |
| Partition ratio | 0.0526 : 1 |
| Analyte concentration range | 2593.3 µM to 2686.2 µM |
| Total points considered | 24 of 37 |

#### Warnings and errors

| | |
|---|---|
| Errors | None |
| Warnings | One or more logP values out of range<br>Excessive acidity error present |

#### Four-Plus parameters

| | | | |
|---|---|---|---|
| Alpha | 0.111 | 3/2/2018 5:10:52 PM | C:\Sirius_T3\HCl18C02.t3r |
| S | 0.9988 | 3/2/2018 5:10:52 PM | C:\Sirius_T3\HCl18C02.t3r |
| jH | 1.0 | 3/2/2018 5:10:52 PM | C:\Sirius_T3\HCl18C02.t3r |
| jOH | -0.8 | 3/2/2018 5:10:52 PM | C:\Sirius_T3\HCl18C02.t3r |

#### Titrants

| | | | |
|---|---|---|---|
| 0.50 M HCl | 0.999058 | 3/2/2018 5:10:52 PM | C:\Sirius_T3\HCl18C02.t3r |
| 0.50 M KOH | 0.999845 | 3/2/2018 5:10:52 PM | C:\Sirius_T3\KOH18B27.t3r |

#### Sample

| | |
|---|---|
| M08_octanol concentration factor | 1.137 |
| Acid pKa 1 | 4.22 |
| logP (neutral XH) | 3.06 |
| logP (X -) | -5.22 |

#### Sample graphs

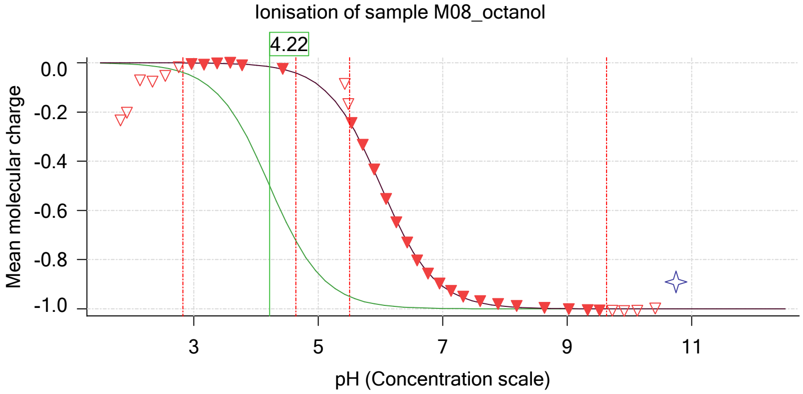


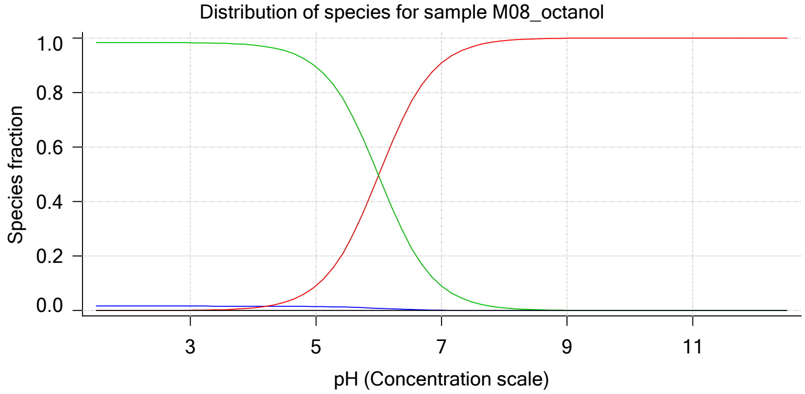

Sample name: **M08_octanol**
Assay name: **pH-metric high logP**
Assay ID: **18C-02007**
Filename: **C:\Sirius_T3\Mehtap\20180302_exp29_logP_T3-2\18C-02007_M08_octanol_pH-metric high logP.t3r**

Experiment start time: **3/2/2018 5:10:52 PM**
Analyst: **Pion**
Instrument ID: **T312060**

#### Sample graphs (continued)

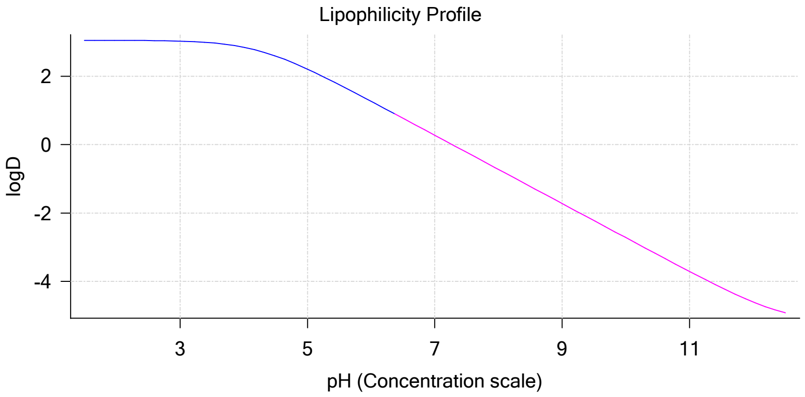


#### Sample logD and percent species

| pH | M08_octanol logD | M08_octanol M08_octanolH | M08_octanol M08_octanol | M08_octanol M08_octanolH* | M08_octanol M08_octanol* | Comment |
|---|---|---|---|---|---|---|
| 1.000 | 3.05 | 1.65 % | 0.00 % | 98.35 % | 0.00 % | |
| 1.200 | 3.05 | 1.65 % | 0.00 % | 98.35 % | 0.00 % | Stomach pH |
| 2.000 | 3.05 | 1.65 % | 0.01 % | 98.34 % | 0.00 % | |
| 3.000 | 3.03 | 1.65 % | 0.10 % | 98.26 % | 0.00 % | |
| 4.000 | 2.85 | 1.63 % | 0.98 % | 97.39 % | 0.00 % | |
| 5.000 | 2.21 | 1.50 % | 9.03 % | 89.48 % | 0.00 % | |
| 6.000 | 1.27 | 0.83 % | 49.80 % | 49.37 % | 0.00 % | |
| 6.500 | 0.77 | 0.40 % | 75.83 % | 23.77 % | 0.00 % | |
| 7.000 | 0.27 | 0.15 % | 90.84 % | 9.00 % | 0.00 % | |
| 7.400 | -0.13 | 0.06 % | 96.14 % | 3.79 % | 0.00 % | Blood pH |
| 8.000 | -0.72 | 0.02 % | 99.00 % | 0.98 % | 0.00 % | |
| 9.000 | -1.72 | 0.00 % | 99.90 % | 0.10 % | 0.00 % | |
| 10.000 | -2.72 | 0.00 % | 99.99 % | 0.01 % | 0.00 % | |
| 11.000 | -3.71 | 0.00 % | 100.00 % | 0.00 % | 0.00 % | |
| 12.000 | -4.60 | 0.00 % | 100.00 % | 0.00 % | 0.00 % | |

#### Carbonate and acidity

Carbonate 0.075 mM
Acidity error -1.805 mM

#### Other graphs

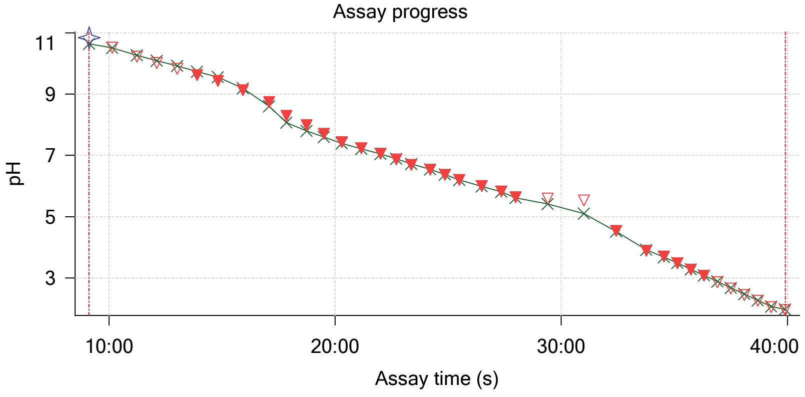


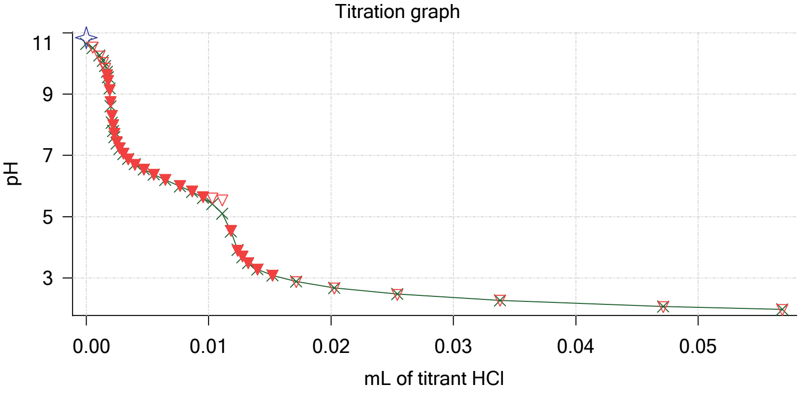

Sample name: **M08_octanol**
Assay name: **pH-metric high logP**
Assay ID: **18C-02007**
Filename: **C:\Sirius_T3\Mehtap\20180302_exp29_logP_T3-2\18C-02007_M08_octanol_pH-metric high logP.t3r**

Experiment start time: **3/2/2018 5:10:52 PM**
Analyst: **Pion**
Instrument ID: **T312060**

#### Other graphs (continued)

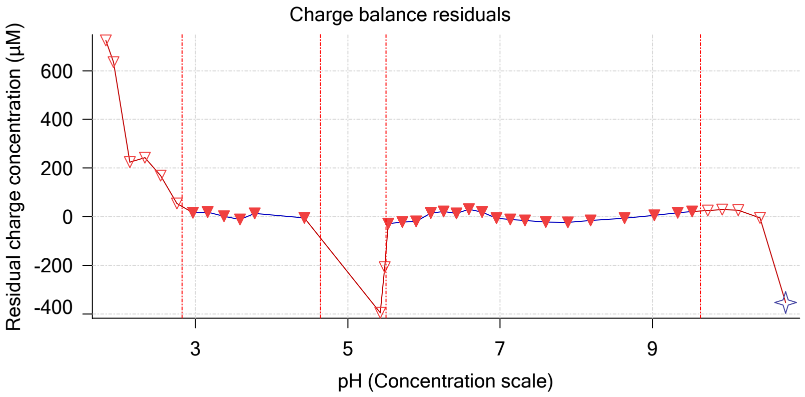


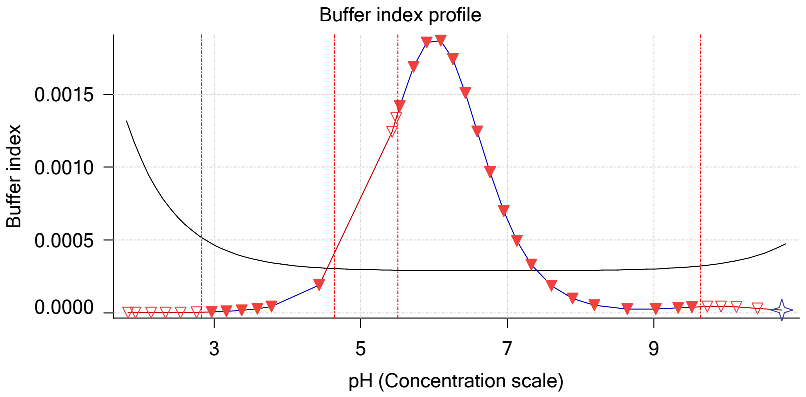


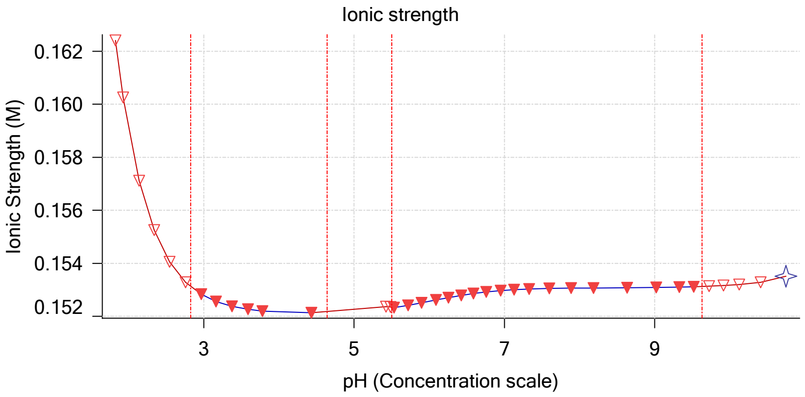


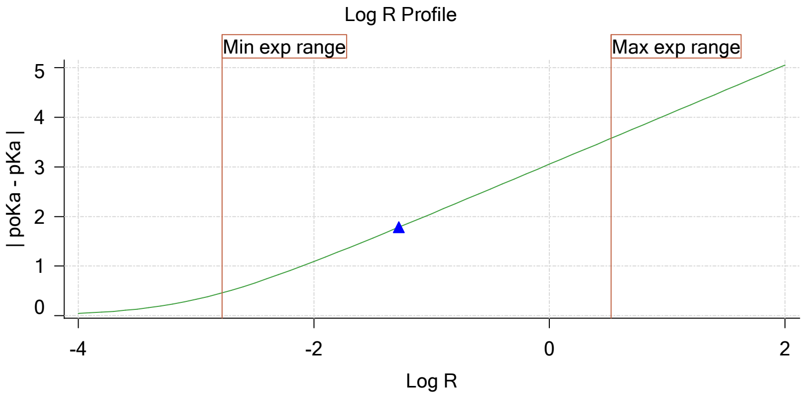


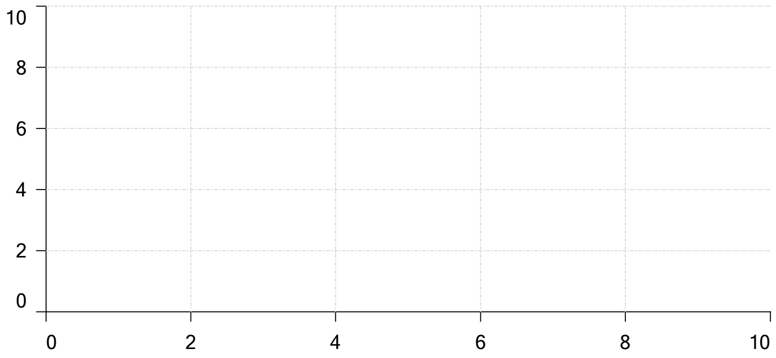


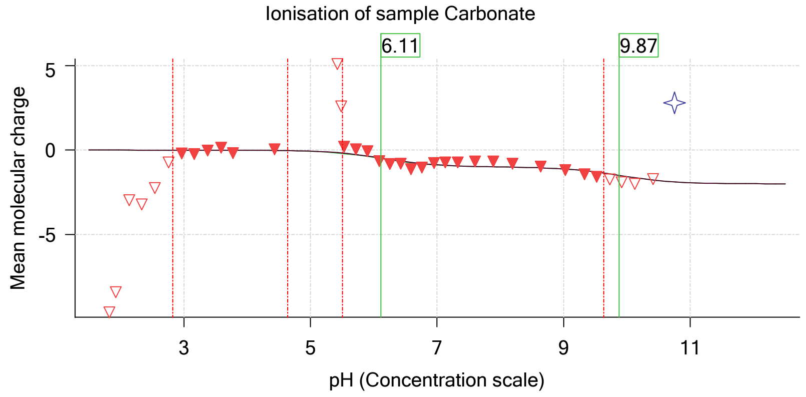

Sample name: **M08_octanol**
Assay name: **pH-metric high logP**
Assay ID: **18C-02007**
Filename: **C:\Sirius_T3\Mehtap\20180302_exp29_logP_T3-2\18C-02007_M08_octanol_pH-metric high logP.t3r**
Experiment start time: **3/2/2018 5:10:52 PM**
Analyst: **Pion**
Instrument ID: **T312060**

### pH-metric high logP Titration 2 of 3 18C-02007 Points 39 to 77

#### Overall results

| | |
|---|---|
| RMSD | 0.038 |
| Average ionic strength | 0.159 M |
| Average temperature | 25.0°C |
| Partition ratio | 0.1719 : 1 |
| Analyte concentration range | 2174.7 µM to 2243.9 µM |
| Total points considered | 26 of 39 |

#### Warnings and errors

| | |
|---|---|
| Errors | None |
| Warnings | One or more logP values out of range<br>Excessive acidity error present |

#### Four-Plus parameters

| | | | |
|---|---|---|---|
| Alpha | 0.111 | 3/2/2018 5:10:52 PM | C:\Sirius_T3\HCl18C02.t3r |
| S | 0.9988 | 3/2/2018 5:10:52 PM | C:\Sirius_T3\HCl18C02.t3r |
| jH | 1.0 | 3/2/2018 5:10:52 PM | C:\Sirius_T3\HCl18C02.t3r |
| jOH | -0.8 | 3/2/2018 5:10:52 PM | C:\Sirius_T3\HCl18C02.t3r |

#### Titrants

| | | | |
|---|---|---|---|
| 0.50 M HCl | 0.999058 | 3/2/2018 5:10:52 PM | C:\Sirius_T3\HCl18C02.t3r |
| 0.50 M KOH | 0.999845 | 3/2/2018 5:10:52 PM | C:\Sirius_T3\KOH18B27.t3r |

#### Sample

| | |
|---|---|
| M08_octanol concentration factor | 0.855 |
| Acid pKa 1 | 4.22 |
| logP (neutral XH) | 3.00 |
| logP (X -) | -5.22 |

#### Sample graphs

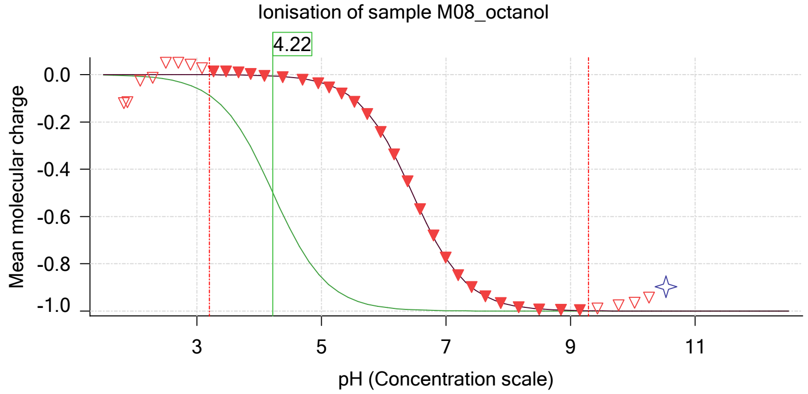


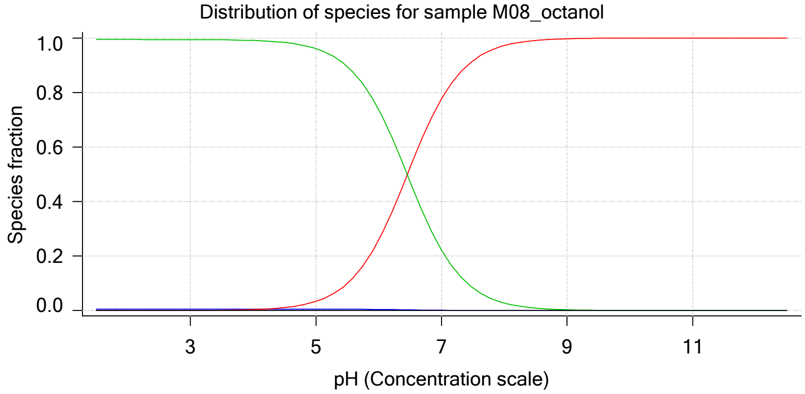

Sample name: **M08_octanol**
Assay name: **pH-metric high logP**
Assay ID: **18C-02007**
Filename: **C:\Sirius_T3\Mehtap\20180302_exp29_logP_T3-2\18C-02007_M08_octanol_pH-metric high logP.t3r**

Experiment start time: **3/2/2018 5:10:52 PM**
Analyst: **Pion**
Instrument ID: **T312060**

#### Sample graphs (continued)

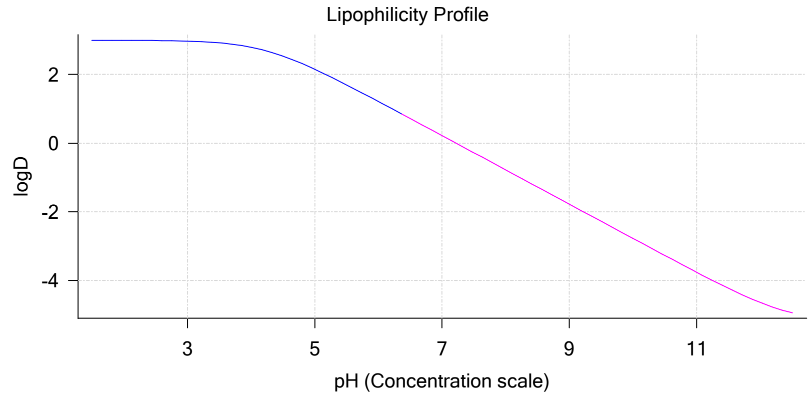


#### Sample logD and percent species

| pH | M08_octanol logD | M08_octanol M08_octanolH | M08_octanol M08_octanol | M08_octanol M08_octanolH* | M08_octanol M08_octanol* | Comment |
|---|---|---|---|---|---|---|
| 1.000 | 3.00 | 0.58 % | 0.00 % | 99.42 % | 0.00 % | |
| 1.200 | 3.00 | 0.58 % | 0.00 % | 99.42 % | 0.00 % | Stomach pH |
| 2.000 | 3.00 | 0.58 % | 0.00 % | 99.42 % | 0.00 % | |
| 3.000 | 2.97 | 0.58 % | 0.03 % | 99.39 % | 0.00 % | |
| 4.000 | 2.79 | 0.58 % | 0.35 % | 99.07 % | 0.00 % | |
| 5.000 | 2.15 | 0.56 % | 3.37 % | 96.07 % | 0.00 % | |
| 6.000 | 1.21 | 0.43 % | 25.88 % | 73.69 % | 0.00 % | |
| 6.500 | 0.72 | 0.28 % | 52.47 % | 47.25 % | 0.00 % | |
| 7.000 | 0.22 | 0.13 % | 77.74 % | 22.14 % | 0.00 % | |
| 7.400 | -0.18 | 0.06 % | 89.76 % | 10.18 % | 0.00 % | Blood pH |
| 8.000 | -0.78 | 0.02 % | 97.22 % | 2.77 % | 0.00 % | |
| 9.000 | -1.78 | 0.00 % | 99.71 % | 0.28 % | 0.00 % | |
| 10.000 | -2.78 | 0.00 % | 99.97 % | 0.03 % | 0.00 % | |
| 11.000 | -3.77 | 0.00 % | 100.00 % | 0.00 % | 0.00 % | |
| 12.000 | -4.65 | 0.00 % | 100.00 % | 0.00 % | 0.00 % | |

#### Carbonate and acidity

| | |
|---|---|
| Carbonate | 0.394 mM |
| Acidity error | -1.609 mM |

#### Other graphs

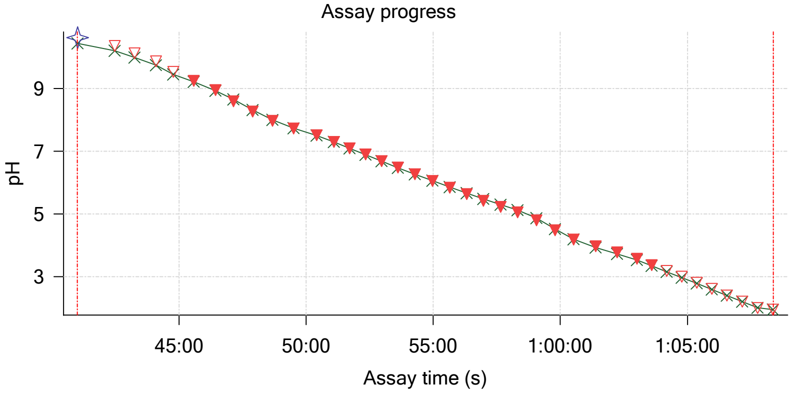


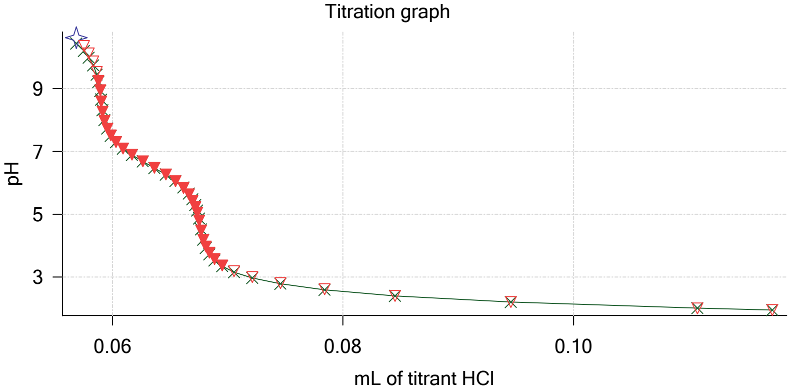

| | | | |
|---|---|---|---|
| Sample name: | **M08_octanol** | Experiment start time: | **3/2/2018 5:10:52 PM** |
| Assay name: | **pH-metric high logP** | Analyst: | **Pion** |
| Assay ID: | **18C-02007** | Instrument ID: | **T312060** |
| Filename: | **C:\Sirius_T3\Mehtap\20180302_exp29_logP_T3-2\18C-02007_M08_octanol_pH-metric high logP.t3r** | | |

#### Other graphs (continued)

Charge balance residuals

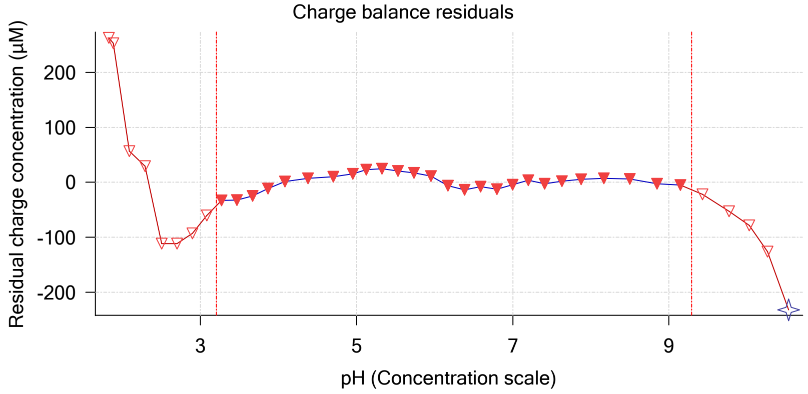


Buffer index profile

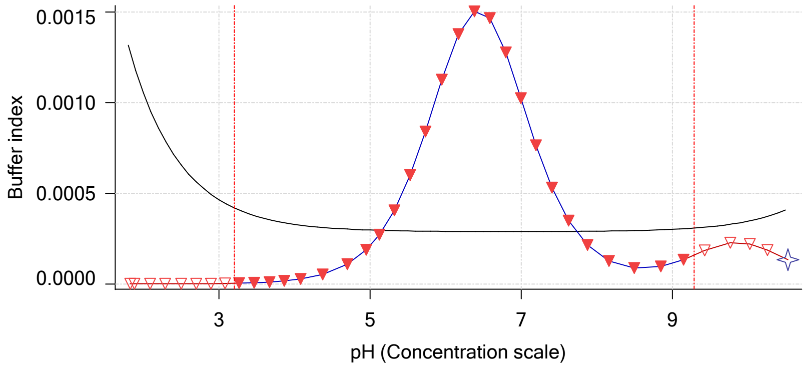


Ionic strength

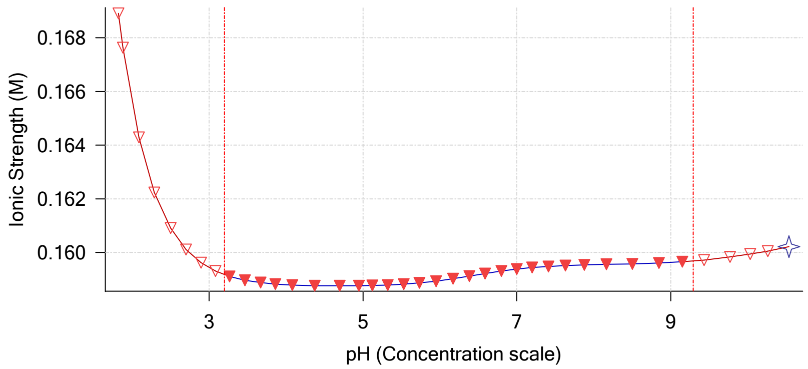


Log R Profile

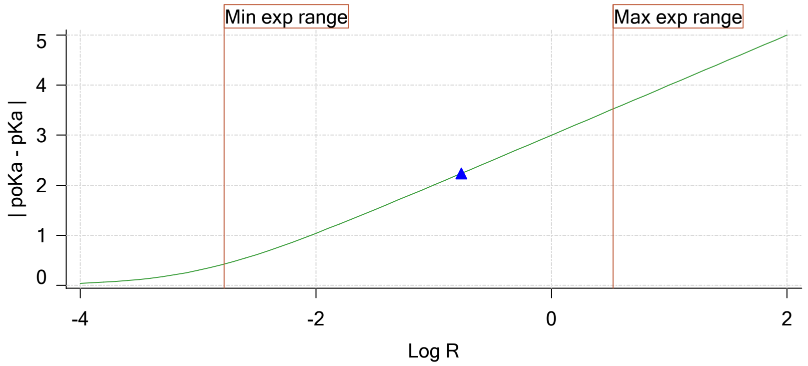


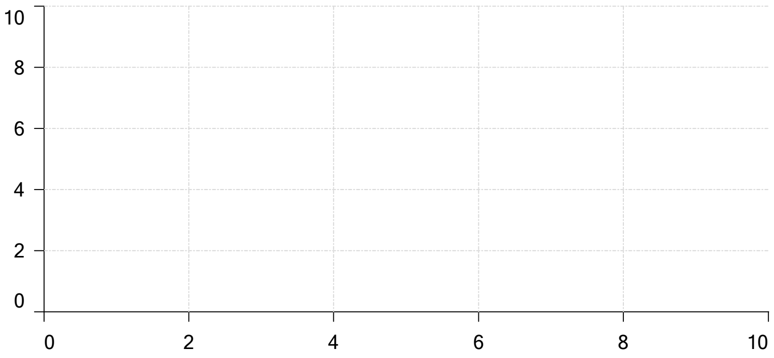


Ionisation of sample Carbonate

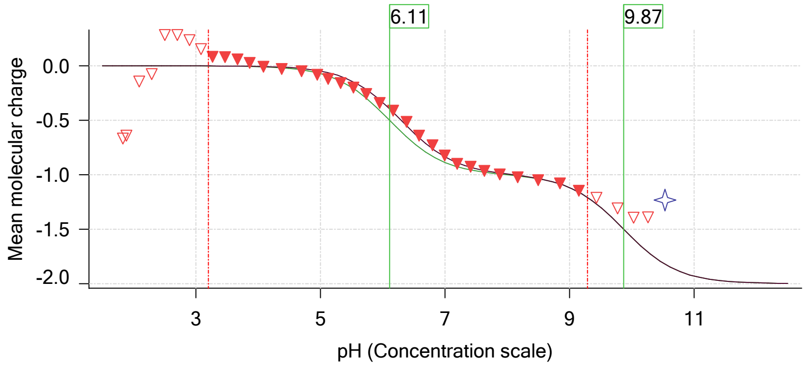

| | | | |
|---|---|---|---|
| Sample name: | **M08_octanol** | Experiment start time: | **3/2/2018 5:10:52 PM** |
| Assay name: | **pH-metric high logP** | Analyst: | **Pion** |
| Assay ID: | **18C-02007** | Instrument ID: | **T312060** |
| Filename: | **C:\Sirius_T3\Mehtap\20180302_exp29_logP_T3-2\18C-02007_M08_octanol_pH-metric high logP.t3r** | | |

### pH-metric high logP Titration 3 of 3 18C-02007 Points 78 to 114

#### Overall results

| | |
|---|---|
| RMSD | 0.046 |
| Average ionic strength | 0.165 M |
| Average temperature | 25.0°C |
| Partition ratio | 0.6178 : 1 |
| Analyte concentration range | 1479.7 μM to 1511.5 μM |
| Total points considered | 30 of 37 |

#### Warnings and errors

| | |
|---|---|
| Errors | None |
| Warnings | One or more logP values out of range<br>Excessive acidity error present |

#### Four-Plus parameters

| | | | |
|---|---|---|---|
| Alpha | 0.111 | 3/2/2018 5:10:52 PM | C:\Sirius_T3\HCl18C02.t3r |
| S | 0.9988 | 3/2/2018 5:10:52 PM | C:\Sirius_T3\HCl18C02.t3r |
| jH | 1.0 | 3/2/2018 5:10:52 PM | C:\Sirius_T3\HCl18C02.t3r |
| jOH | -0.8 | 3/2/2018 5:10:52 PM | C:\Sirius_T3\HCl18C02.t3r |

#### Titrants

| | | | |
|---|---|---|---|
| 0.50 M HCl | 0.999058 | 3/2/2018 5:10:52 PM | C:\Sirius_T3\HCl18C02.t3r |
| 0.50 M KOH | 0.999845 | 3/2/2018 5:10:52 PM | C:\Sirius_T3\KOH18B27.t3r |

#### Sample

| | |
|---|---|
| M08_octanol concentration factor | 0.850 |
| Acid pKa 1 | 4.22 |
| logP (neutral XH) | 2.97 |
| logP (X -) | -6.01 |

#### Sample graphs

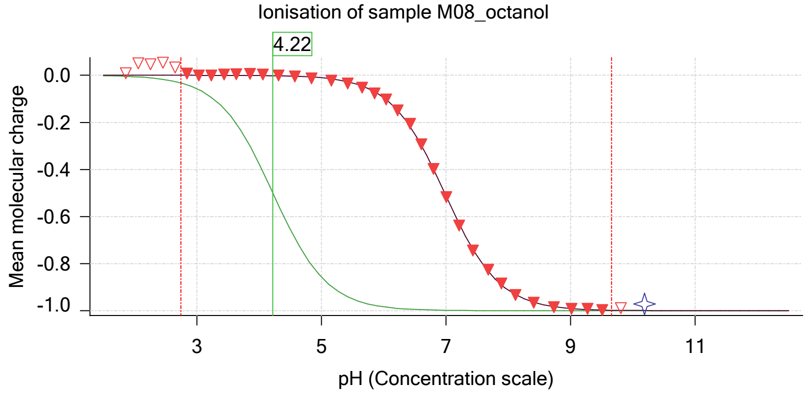


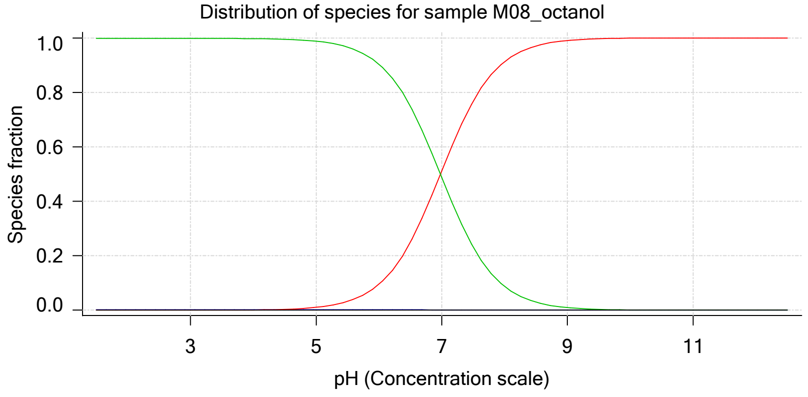

| | | | |
|---|---|---|---|
| Sample name: | **M08_octanol** | Experiment start time: | **3/2/2018 5:10:52 PM** |
| Assay name: | **pH-metric high logP** | Analyst: | **Pion** |
| Assay ID: | **18C-02007** | Instrument ID: | **T312060** |
| Filename: | **C:\Sirius_T3\Mehtap\20180302_exp29_logP_T3-2\18C-02007_M08_octanol_pH-metric high logP.t3r** | | |

#### Sample graphs (continued)

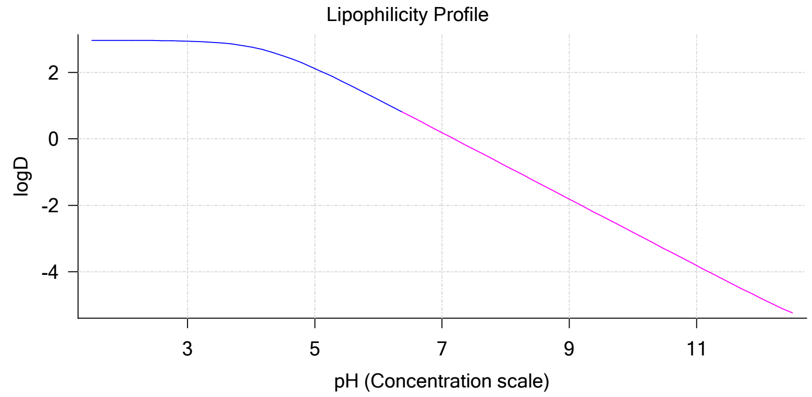


#### Sample logD and percent species

| pH | M08_octanol logD | M08_octanol M08_octanolH | M08_octanol M08_octanol | M08_octanol M08_octanolH* | M08_octanol M08_octanol* | Comment |
|---|---|---|---|---|---|---|
| 1.000 | 2.97 | 0.17 % | 0.00 % | 99.83 % | 0.00 % | |
| 1.200 | 2.97 | 0.17 % | 0.00 % | 99.83 % | 0.00 % | Stomach pH |
| 2.000 | 2.96 | 0.17 % | 0.00 % | 99.82 % | 0.00 % | |
| 3.000 | 2.94 | 0.17 % | 0.01 % | 99.82 % | 0.00 % | |
| 4.000 | 2.76 | 0.17 % | 0.10 % | 99.72 % | 0.00 % | |
| 5.000 | 2.12 | 0.17 % | 1.04 % | 98.79 % | 0.00 % | |
| 6.000 | 1.18 | 0.16 % | 9.50 % | 90.34 % | 0.00 % | |
| 6.500 | 0.68 | 0.13 % | 24.93 % | 74.94 % | 0.00 % | |
| 7.000 | 0.19 | 0.09 % | 51.23 % | 48.69 % | 0.00 % | |
| 7.400 | -0.21 | 0.05 % | 72.51 % | 27.44 % | 0.00 % | Blood pH |
| 8.000 | -0.81 | 0.02 % | 91.31 % | 8.68 % | 0.00 % | |
| 9.000 | -1.81 | 0.00 % | 99.06 % | 0.94 % | 0.00 % | |
| 10.000 | -2.81 | 0.00 % | 99.90 % | 0.09 % | 0.00 % | |
| 11.000 | -3.81 | 0.00 % | 99.99 % | 0.01 % | 0.00 % | |
| 12.000 | -4.79 | 0.00 % | 100.00 % | 0.00 % | 0.00 % | |

#### Carbonate and acidity

| | |
|---|---|
| Carbonate | 0.394 mM |
| Acidity error | -1.483 mM |

#### Other graphs

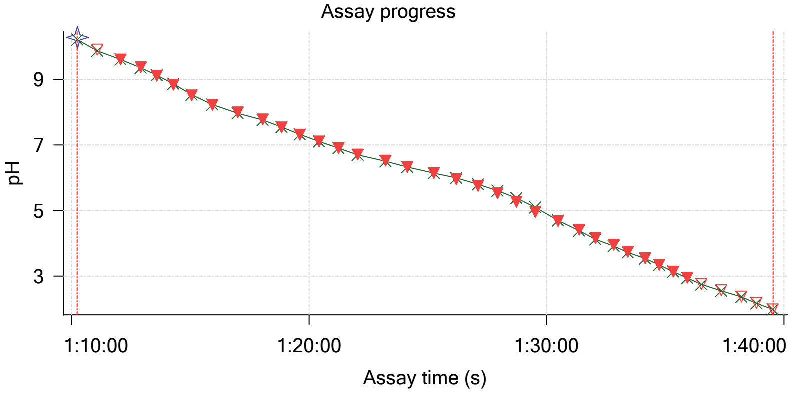


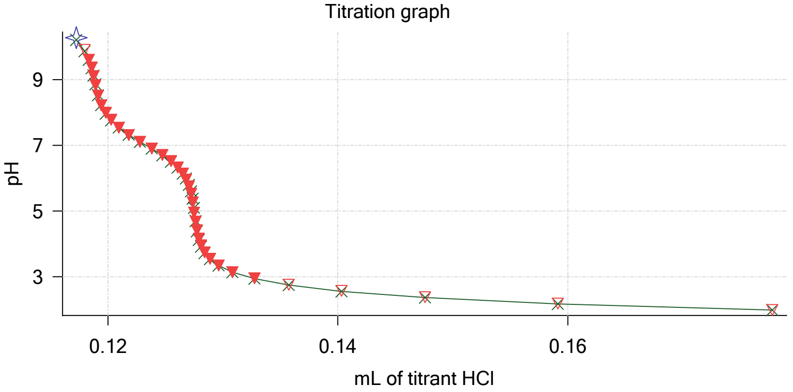

Sample name: **M08_octanol**
Assay name: **pH-metric high logP**
Assay ID: **18C-02007**
Filename: **C:\Sirius_T3\Mehtap\20180302_exp29_logP_T3-2\18C-02007_M08_octanol_pH-metric high logP.t3r**

Experiment start time: **3/2/2018 5:10:52 PM**
Analyst: **Pion**
Instrument ID: **T312060**

#### Other graphs (continued)

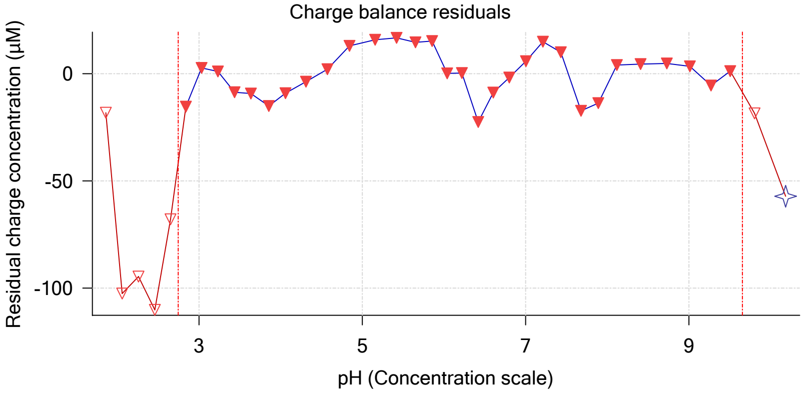


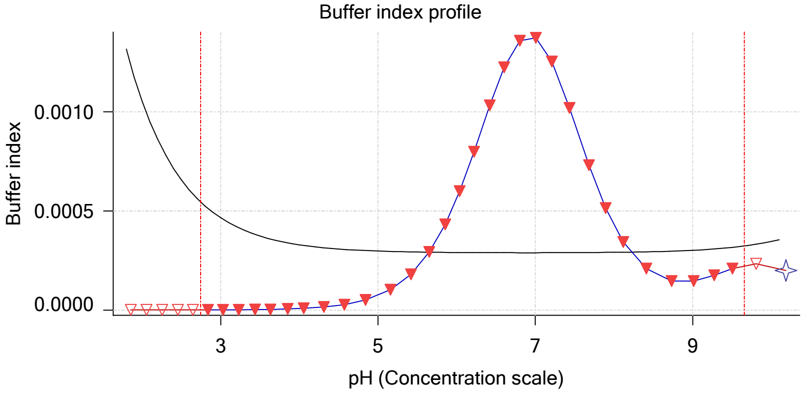


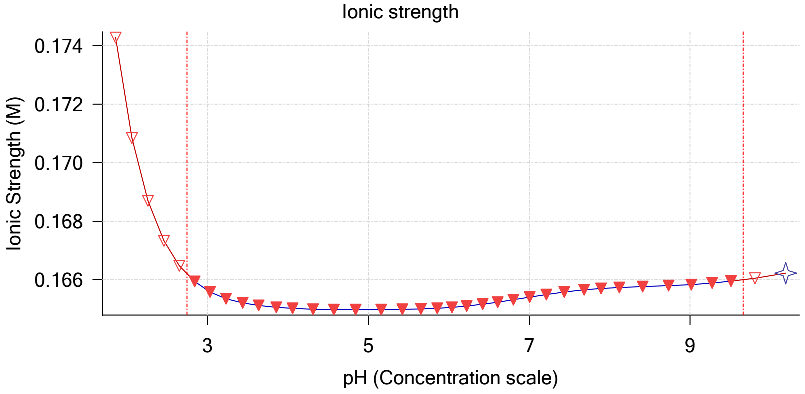


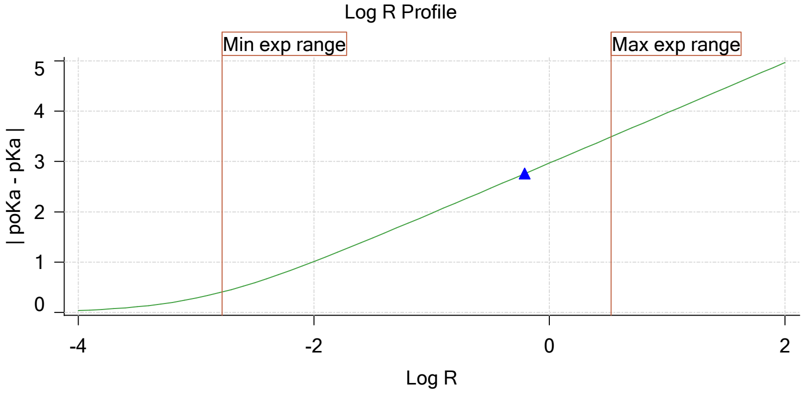


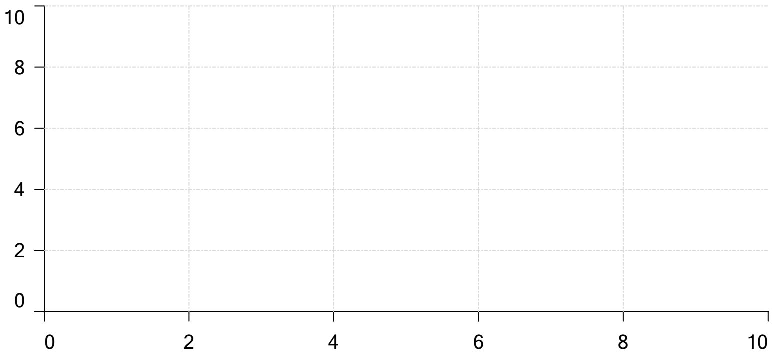


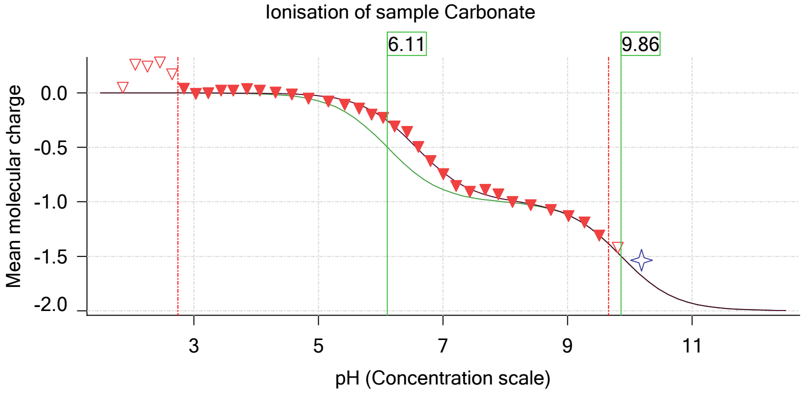

| | | | |
|---|---|---|---|
| Sample name: | **M08_octanol** | Experiment start time: | **3/2/2018 5:10:52 PM** |
| Assay name: | **pH-metric high logP** | Analyst: | **Pion** |
| Assay ID: | **18C-02007** | Instrument ID: | **T312060** |
| Filename: | **C:\Sirius_T3\Mehtap\20180302_exp29_logP_T3-2\18C-02007_M08_octanol_pH-metric high logP.t3r** | | |

#### Assay Model

| Settings | Value | Date/Time changed | Imported from |
|---|---|---|---|
| Sample name | M08_octanol | 2/27/2018 4:33:51 PM | User entered value |
| Sample by | Weight | | Default value |
| Sample weight | 0.001250 g | 3/2/2018 5:08:08 PM | User entered value |
| Formula weight | 293.32 g/mol | 2/27/2018 4:33:51 PM | User entered value |
| Solubility | Unknown | | Default value |
| Molecular weight | 293.32 | 2/27/2018 4:33:51 PM | User entered value |
| Individual pKa ionic environments | No | | Default value |
| Number of pKas | 1 | 2/27/2018 4:33:51 PM | User entered value |
| Sample is a | Acid | 2/27/2018 4:33:51 PM | User entered value |
| pKa 1 | 4.22 | 2/27/2018 4:33:51 PM | User entered value |
| logP (neutral XH) | 2.98 | 3/2/2018 3:22:58 PM | User entered value |
| logP (X -) | -5.22 | 3/2/2018 3:23:03 PM | User entered value |

#### Events

| Time | Event | Water | Acid | Base | Octanol | pH | dpH/dt | pH R-squared | pH SD | dpH time |
|---|---|---|---|---|---|---|---|---|---|---|
| 6:14.2 | Manual volume addition | | | | 0.08000 mL | | | | | |
| 6:15.3 | Initial pH = 5.74 | | | | | | | | | |
| 9:07.3 | Data point 2 | 1.50000 mL | 0.00000 mL | 0.00647 mL | 0.08000 mL | 10.845 | -0.01910 | 0.96118 | 0.00097 | 34.5 s |
| 10:08.5 | Data point 3 | 1.50000 mL | 0.00052 mL | 0.00647 mL | 0.08000 mL | 10.511 | 0.01702 | 0.82617 | 0.00093 | 35.0 s |
| 11:14.1 | Data point 4 | 1.50000 mL | 0.00108 mL | 0.00647 mL | 0.08000 mL | 10.223 | -0.01634 | 0.68740 | 0.00097 | 27.0 s |
| 12:06.6 | Data point 5 | 1.50000 mL | 0.00136 mL | 0.00647 mL | 0.08000 mL | 10.016 | -0.01334 | 0.51589 | 0.00092 | 28.5 s |
| 13:00.5 | Data point 6 | 1.50000 mL | 0.00155 mL | 0.00647 mL | 0.08000 mL | 9.825 | -0.01728 | 0.80734 | 0.00095 | 28.0 s |
| 13:53.9 | Data point 7 | 1.50000 mL | 0.00169 mL | 0.00647 mL | 0.08000 mL | 9.619 | -0.01617 | 0.77696 | 0.00091 | 29.5 s |
| 14:48.8 | Data point 8 | 1.50000 mL | 0.00179 mL | 0.00647 mL | 0.08000 mL | 9.430 | -0.01709 | 0.89746 | 0.00089 | 30.5 s |
| 15:54.8 | Data point 9 | 1.50000 mL | 0.00190 mL | 0.00647 mL | 0.08000 mL | 9.124 | -0.01954 | 0.95895 | 0.00100 | 34.0 s |
| 17:04.8 | Data point 10 | 1.50000 mL | 0.00200 mL | 0.00647 mL | 0.08000 mL | 8.735 | 0.01424 | 0.54643 | 0.00095 | 10.5 s |
| 17:50.7 | Data point 11 | 1.50000 mL | 0.00209 mL | 0.00647 mL | 0.08000 mL | 8.289 | 0.01419 | 0.56162 | 0.00094 | 17.5 s |
| 18:43.8 | Data point 12 | 1.50000 mL | 0.00219 mL | 0.00647 mL | 0.08000 mL | 7.989 | 0.01768 | 0.82202 | 0.00096 | 16.0 s |
| 19:30.3 | Data point 13 | 1.50000 mL | 0.00230 mL | 0.00647 mL | 0.08000 mL | 7.701 | 0.01538 | 0.74821 | 0.00089 | 17.0 s |
| 20:17.7 | Data point 14 | 1.50000 mL | 0.00249 mL | 0.00647 mL | 0.08000 mL | 7.430 | 0.01705 | 0.74628 | 0.00097 | 16.0 s |
| 21:09.3 | Data point 15 | 1.50000 mL | 0.00273 mL | 0.00647 mL | 0.08000 mL | 7.233 | 0.01759 | 0.83369 | 0.00095 | 16.0 s |
| 22:00.8 | Data point 16 | 1.50000 mL | 0.00303 mL | 0.00647 mL | 0.08000 mL | 7.053 | 0.01636 | 0.81366 | 0.00091 | 15.5 s |
| 22:41.9 | Data point 17 | 1.50000 mL | 0.00343 mL | 0.00647 mL | 0.08000 mL | 6.864 | 0.01785 | 0.82800 | 0.00097 | 15.0 s |
| 23:22.4 | Data point 18 | 1.50000 mL | 0.00397 mL | 0.00647 mL | 0.08000 mL | 6.692 | 0.01764 | 0.77148 | 0.00099 | 14.0 s |
| 24:12.1 | Data point 19 | 1.50000 mL | 0.00470 mL | 0.00647 mL | 0.08000 mL | 6.530 | 0.01783 | 0.78830 | 0.00099 | 13.5 s |
| 24:51.2 | Data point 20 | 1.50000 mL | 0.00553 mL | 0.00647 mL | 0.08000 mL | 6.359 | 0.01727 | 0.78473 | 0.00097 | 13.0 s |

| | | | |
|---|---|---|---|
| Sample name: | **M08_octanol** | Experiment start time: | **3/2/2018 5:10:52 PM** |
| Assay name: | **pH-metric high logP** | Analyst: | **Pion** |
| Assay ID: | **18C-02007** | Instrument ID: | **T312060** |
| Filename: | **C:\Sirius_T3\Mehtap\20180302_exp29_logP_T3-2\18C-02007_M08_octanol_pH-metric high logP.t3r** | | |

#### Events (continued)

| Time | Event | Water | Acid | Base | Octanol | pH | dpH/dt | pH R-squared | pH SD | dpH/dt time |
|---|---|---|---|---|---|---|---|---|---|---|
| 25:29.8 | Data point 21 | 1.50000 mL | 0.00647 mL | 0.00647 mL | 0.08000 mL | 6.192 | 0.01558 | 0.76055 | 0.00089 | 13.0 s |
| 26:28.7 | Data point 22 | 1.50000 mL | 0.00767 mL | 0.00647 mL | 0.08000 mL | 6.001 | 0.01489 | 0.64323 | 0.00092 | 11.0 s |
| 27:20.5 | Data point 23 | 1.50000 mL | 0.00865 mL | 0.00647 mL | 0.08000 mL | 5.821 | 0.01662 | 0.89411 | 0.00087 | 13.0 s |
| 27:58.9 | Data point 24 | 1.50000 mL | 0.00955 mL | 0.00647 mL | 0.08000 mL | 5.637 | 0.05313 | 0.99810 | 0.00262 | Timed out at 59.0 s |
| 29:24.2 | Data point 25 | 1.50000 mL | 0.01030 mL | 0.00647 mL | 0.08000 mL | 5.588 | 0.13136 | 0.99843 | 0.00649 | Timed out at 59.5 s |
| 31:00.0 | Data point 26 | 1.50000 mL | 0.01110 mL | 0.00647 mL | 0.08000 mL | 5.531 | 0.09941 | 0.99750 | 0.00492 | Timed out at 59.5 s |
| 32:25.5 | Data point 27 | 1.50000 mL | 0.01183 mL | 0.00647 mL | 0.08000 mL | 4.537 | 0.01986 | 0.98034 | 0.00099 | 49.0 s |
| 33:45.2 | Data point 28 | 1.50000 mL | 0.01237 mL | 0.00647 mL | 0.08000 mL | 3.885 | 0.01681 | 0.92569 | 0.00086 | 11.5 s |
| 34:32.4 | Data point 29 | 1.50000 mL | 0.01275 mL | 0.00647 mL | 0.08000 mL | 3.693 | 0.00140 | 0.02989 | 0.00040 | 10.0 s |
| 35:08.0 | Data point 30 | 1.50000 mL | 0.01322 mL | 0.00647 mL | 0.08000 mL | 3.482 | 0.01095 | 0.91600 | 0.00057 | 10.0 s |
| 35:43.5 | Data point 31 | 1.50000 mL | 0.01399 mL | 0.00647 mL | 0.08000 mL | 3.270 | 0.00378 | 0.79588 | 0.00021 | 10.0 s |
| 36:19.0 | Data point 32 | 1.50000 mL | 0.01522 mL | 0.00647 mL | 0.08000 mL | 3.073 | -0.00000 | 0.00000 | 0.00009 | 10.0 s |
| 36:54.5 | Data point 33 | 1.50000 mL | 0.01714 mL | 0.00647 mL | 0.08000 mL | 2.866 | -0.00657 | 0.26783 | 0.00063 | 10.0 s |
| 37:30.0 | Data point 34 | 1.50000 mL | 0.02027 mL | 0.00647 mL | 0.08000 mL | 2.651 | -0.00505 | 0.84244 | 0.00027 | 10.0 s |
| 38:05.6 | Data point 35 | 1.50000 mL | 0.02542 mL | 0.00647 mL | 0.08000 mL | 2.448 | -0.00911 | 0.70428 | 0.00054 | 10.0 s |
| 38:41.2 | Data point 36 | 1.50000 mL | 0.03380 mL | 0.00647 mL | 0.08000 mL | 2.253 | -0.00829 | 0.95487 | 0.00042 | 10.5 s |
| 39:17.3 | Data point 37 | 1.50000 mL | 0.04713 mL | 0.00647 mL | 0.08000 mL | 2.045 | -0.00957 | 0.94345 | 0.00049 | 10.5 s |
| 39:53.4 | Data point 38 | 1.50000 mL | 0.05687 mL | 0.00647 mL | 0.08000 mL | 1.947 | -0.01349 | 0.79563 | 0.00075 | 10.0 s |
| 41:01.1 | Data point 39 | 1.50000 mL | 0.05687 mL | 0.06235 mL | 0.28000 mL | 10.631 | -0.01759 | 0.84525 | 0.00094 | 46.5 s |
| 42:28.5 | Data point 40 | 1.50000 mL | 0.05753 mL | 0.06235 mL | 0.28000 mL | 10.359 | -0.01582 | 0.80255 | 0.00087 | 21.0 s |
| 43:14.9 | Data point 41 | 1.50000 mL | 0.05795 mL | 0.06235 mL | 0.28000 mL | 10.128 | -0.01731 | 0.77064 | 0.00098 | 20.5 s |
| 44:06.0 | Data point 42 | 1.50000 mL | 0.05833 mL | 0.06235 mL | 0.28000 mL | 9.868 | -0.01482 | 0.86960 | 0.00079 | 10.0 s |
| 44:46.6 | Data point 43 | 1.50000 mL | 0.05863 mL | 0.06235 mL | 0.28000 mL | 9.530 | 0.01648 | 0.70393 | 0.00097 | 18.0 s |
| 45:35.3 | Data point 44 | 1.50000 mL | 0.05880 mL | 0.06235 mL | 0.28000 mL | 9.248 | 0.00953 | 0.92650 | 0.00049 | 10.0 s |
| 46:26.1 | Data point 45 | 1.50000 mL | 0.05894 mL | 0.06235 mL | 0.28000 mL | 8.947 | 0.01716 | 0.79689 | 0.00095 | 12.0 s |
| 47:08.7 | Data point 46 | 1.50000 mL | 0.05903 mL | 0.06235 mL | 0.28000 mL | 8.599 | 0.01796 | 0.79610 | 0.00099 | 15.0 s |
| 47:54.2 | Data point 47 | 1.50000 mL | 0.05915 mL | 0.06235 mL | 0.28000 mL | 8.266 | 0.01550 | 0.73917 | 0.00089 | 16.0 s |
| 48:40.8 | Data point 48 | 1.50000 mL | 0.05931 mL | 0.06235 mL | 0.28000 mL | 7.978 | 0.01748 | 0.85525 | 0.00093 | 14.5 s |
| 49:30.9 | Data point 49 | 1.50000 mL | 0.05955 mL | 0.06235 mL | 0.28000 mL | 7.732 | 0.01742 | 0.81873 | 0.00095 | 13.5 s |
| 50:25.3 | Data point 50 | 1.50000 mL | 0.05988 mL | 0.06235 mL | 0.28000 mL | 7.511 | 0.01772 | 0.84280 | 0.00095 | 15.0 s |
| 51:05.7 | Data point 51 | 1.50000 mL | 0.06030 mL | 0.06235 mL | 0.28000 mL | 7.300 | 0.01794 | 0.80992 | 0.00098 | 12.0 s |
| 51:43.1 | Data point 52 | 1.50000 mL | 0.06091 mL | 0.06235 mL | 0.28000 mL | 7.099 | 0.01591 | 0.80910 | 0.00087 | 12.0 s |
| 52:20.5 | Data point 53 | 1.50000 mL | 0.06169 mL | 0.06235 mL | 0.28000 mL | 6.900 | 0.01723 | 0.80221 | 0.00095 | 12.5 s |
| 52:58.5 | Data point 54 | 1.50000 mL | 0.06263 mL | 0.06235 mL | 0.28000 mL | 6.688 | 0.01881 | 0.86957 | 0.00100 | 12.5 s |
| 53:36.4 | Data point 55 | 1.50000 mL | 0.06364 mL | 0.06235 mL | 0.28000 mL | 6.485 | 0.01949 | 0.96513 | 0.00098 | 14.5 s |
| 54:16.5 | Data point 56 | 1.50000 mL | 0.06463 mL | 0.06235 mL | 0.28000 mL | 6.272 | 0.01746 | 0.81817 | 0.00095 | 16.0 s |
| 54:58.0 | Data point 57 | 1.50000 mL | 0.06548 mL | 0.06235 mL | 0.28000 mL | 6.054 | 0.01509 | 0.72773 | 0.00087 | 15.5 s |
| 55:38.9 | Data point 58 | 1.50000 mL | 0.06616 mL | 0.06235 mL | 0.28000 mL | 5.836 | 0.01598 | 0.67878 | 0.00096 | 14.5 s |
| 56:18.8 | Data point 59 | 1.50000 mL | 0.06663 mL | 0.06235 mL | 0.28000 mL | 5.633 | 0.01711 | 0.75074 | 0.00098 | 14.5 s |
| 56:58.7 | Data point 60 | 1.50000 mL | 0.06696 mL | 0.06235 mL | 0.28000 mL | 5.426 | 0.01163 | 0.43580 | 0.00087 | 15.0 s |
| 57:39.1 | Data point 61 | 1.50000 mL | 0.06719 mL | 0.06235 mL | 0.28000 mL | 5.227 | 0.01748 | 0.74548 | 0.00100 | 14.5 s |
| 58:19.1 | Data point 62 | 1.50000 mL | 0.06736 mL | 0.06235 mL | 0.28000 mL | 5.053 | 0.01418 | 0.59508 | 0.00091 | 14.0 s |
| 59:03.7 | Data point 63 | 1.50000 mL | 0.06752 mL | 0.06235 mL | 0.28000 mL | 4.804 | 0.01416 | 0.51436 | 0.00098 | 13.0 s |
| 59:47.2 | Data point 64 | 1.50000 mL | 0.06769 mL | 0.06235 mL | 0.28000 mL | 4.481 | -0.01418 | 0.54917 | 0.00095 | 13.5 s |
| 1:00:31.3 | Data point 65 | 1.50000 mL | 0.06787 mL | 0.06235 mL | 0.28000 mL | 4.188 | 0.00796 | 0.27623 | 0.00075 | 11.5 s |
| 1:01:23.6 | Data point 66 | 1.50000 mL | 0.06811 mL | 0.06235 mL | 0.28000 mL | 3.970 | 0.00985 | 0.32707 | 0.00085 | 10.0 s |
| 1:02:14.3 | Data point 67 | 1.50000 mL | 0.06841 mL | 0.06235 mL | 0.28000 mL | 3.775 | 0.00616 | 0.11203 | 0.00091 | 10.5 s |
| 1:03:00.6 | Data point 68 | 1.50000 mL | 0.06886 mL | 0.06235 mL | 0.28000 mL | 3.573 | -0.00222 | 0.09420 | 0.00036 | 10.0 s |
| 1:03:36.0 | Data point 69 | 1.50000 mL | 0.06952 mL | 0.06235 mL | 0.28000 mL | 3.375 | -0.00763 | 0.55206 | 0.00051 | 10.0 s |
| 1:04:11.4 | Data point 70 | 1.50000 mL | 0.07056 mL | 0.06235 mL | 0.28000 mL | 3.190 | -0.00162 | 0.01741 | 0.00061 | 10.0 s |
| 1:04:46.9 | Data point 71 | 1.50000 mL | 0.07213 mL | 0.06235 mL | 0.28000 mL | 3.003 | -0.00478 | 0.16896 | 0.00058 | 10.0 s |
| 1:05:22.5 | Data point 72 | 1.50000 mL | 0.07458 mL | 0.06235 mL | 0.28000 mL | 2.809 | -0.01075 | 0.85851 | 0.00057 | 10.0 s |
| 1:05:57.9 | Data point 73 | 1.50000 mL | 0.07841 mL | 0.06235 mL | 0.28000 mL | 2.612 | -0.00808 | 0.55458 | 0.00054 | 10.0 s |
| 1:06:33.4 | Data point 74 | 1.50000 mL | 0.08452 mL | 0.06235 mL | 0.28000 mL | 2.403 | -0.01513 | 0.90224 | 0.00079 | 10.5 s |

| | | | |
|---|---|---|---|
| Sample name: | **M08_octanol** | Experiment start time: | **3/2/2018 5:10:52 PM** |
| Assay name: | **pH-metric high logP** | Analyst: | **Pion** |
| Assay ID: | **18C-02007** | Instrument ID: | **T312060** |
| Filename: | **C:\Sirius_T3\Mehtap\20180302_exp29_logP_T3-2\18C-02007_M08_octanol_pH-metric high logP.t3r** | | |

#### Events (continued)

| Time | Event | Water | Acid | Base | Octanol | pH | dpH/dt | pH R-squared | pH SD | dpH/dt time |
|---|---|---|---|---|---|---|---|---|---|---|
| 1:07:09.6 | Data point 75 | 1.50000 mL | 0.09457 mL | 0.06235 mL | 0.28000 mL | 2.205 | -0.01575 | 0.80047 | 0.00087 | 10.5 s |
| 1:07:46.0 | Data point 76 | 1.50000 mL | 0.11075 mL | 0.06235 mL | 0.28000 mL | 2.005 | -0.01140 | 0.94279 | 0.00058 | 10.0 s |
| 1:08:21.6 | Data point 77 | 1.50000 mL | 0.11724 mL | 0.06235 mL | 0.28000 mL | 1.949 | -0.01059 | 0.59925 | 0.00068 | 10.5 s |
| 1:10:15.2 | Data point 78 | 1.50000 mL | 0.11724 mL | 0.12227 mL | 1.08000 mL | 10.284 | -0.01559 | 0.68044 | 0.00093 | 19.0 s |
| 1:11:04.8 | Data point 79 | 1.50000 mL | 0.11797 mL | 0.12227 mL | 1.08000 mL | 9.905 | -0.01823 | 0.85938 | 0.00097 | 29.0 s |
| 1:12:04.4 | Data point 80 | 1.50000 mL | 0.11832 mL | 0.12227 mL | 1.08000 mL | 9.606 | 0.00322 | 0.20225 | 0.00035 | 10.0 s |
| 1:12:55.2 | Data point 81 | 1.50000 mL | 0.11858 mL | 0.12227 mL | 1.08000 mL | 9.372 | 0.01018 | 0.33175 | 0.00087 | 10.0 s |
| 1:13:35.8 | Data point 82 | 1.50000 mL | 0.11874 mL | 0.12227 mL | 1.08000 mL | 9.113 | 0.01675 | 0.72883 | 0.00097 | 11.5 s |
| 1:14:17.8 | Data point 83 | 1.50000 mL | 0.11891 mL | 0.12227 mL | 1.08000 mL | 8.833 | 0.01720 | 0.72480 | 0.00100 | 15.0 s |
| 1:15:03.3 | Data point 84 | 1.50000 mL | 0.11912 mL | 0.12227 mL | 1.08000 mL | 8.510 | 0.01867 | 0.92540 | 0.00096 | 22.0 s |
| 1:15:55.9 | Data point 85 | 1.50000 mL | 0.11940 mL | 0.12227 mL | 1.08000 mL | 8.218 | 0.01492 | 0.63364 | 0.00092 | 23.5 s |
| 1:17:00.2 | Data point 86 | 1.50000 mL | 0.11980 mL | 0.12227 mL | 1.08000 mL | 7.993 | 0.01343 | 0.47459 | 0.00096 | 22.0 s |
| 1:18:03.1 | Data point 87 | 1.50000 mL | 0.12027 mL | 0.12227 mL | 1.08000 mL | 7.782 | 0.00779 | 0.17186 | 0.00093 | 22.5 s |
| 1:18:51.0 | Data point 88 | 1.50000 mL | 0.12093 mL | 0.12227 mL | 1.08000 mL | 7.536 | 0.01513 | 0.63465 | 0.00094 | 20.5 s |
| 1:19:36.9 | Data point 89 | 1.50000 mL | 0.12180 mL | 0.12227 mL | 1.08000 mL | 7.314 | 0.01545 | 0.64305 | 0.00095 | 23.0 s |
| 1:20:25.4 | Data point 90 | 1.50000 mL | 0.12279 mL | 0.12227 mL | 1.08000 mL | 7.108 | 0.01486 | 0.61616 | 0.00093 | 24.0 s |
| 1:21:14.9 | Data point 91 | 1.50000 mL | 0.12380 mL | 0.12227 mL | 1.08000 mL | 6.904 | 0.01585 | 0.65380 | 0.00097 | 23.0 s |
| 1:22:03.3 | Data point 92 | 1.50000 mL | 0.12472 mL | 0.12227 mL | 1.08000 mL | 6.708 | 0.01765 | 0.76772 | 0.00100 | 29.0 s |
| 1:23:13.3 | Data point 93 | 1.50000 mL | 0.12549 mL | 0.12227 mL | 1.08000 mL | 6.525 | 0.01641 | 0.69561 | 0.00097 | 29.5 s |
| 1:24:08.2 | Data point 94 | 1.50000 mL | 0.12606 mL | 0.12227 mL | 1.08000 mL | 6.325 | 0.01752 | 0.80208 | 0.00097 | 31.5 s |
| 1:25:15.4 | Data point 95 | 1.50000 mL | 0.12651 mL | 0.12227 mL | 1.08000 mL | 6.143 | 0.01142 | 0.37473 | 0.00092 | 31.5 s |
| 1:26:12.4 | Data point 96 | 1.50000 mL | 0.12679 mL | 0.12227 mL | 1.08000 mL | 5.958 | 0.01161 | 0.45352 | 0.00085 | 19.0 s |
| 1:27:07.1 | Data point 97 | 1.50000 mL | 0.12705 mL | 0.12227 mL | 1.08000 mL | 5.756 | 0.01128 | 0.44370 | 0.00084 | 18.5 s |
| 1:27:56.1 | Data point 98 | 1.50000 mL | 0.12723 mL | 0.12227 mL | 1.08000 mL | 5.523 | 0.00514 | 0.07395 | 0.00093 | 17.5 s |
| 1:28:44.2 | Data point 99 | 1.50000 mL | 0.12737 mL | 0.12227 mL | 1.08000 mL | 5.263 | 0.00888 | 0.39531 | 0.00070 | 17.0 s |
| 1:29:31.7 | Data point 100 | 1.50000 mL | 0.12749 mL | 0.12227 mL | 1.08000 mL | 4.946 | 0.01013 | 0.28860 | 0.00093 | 22.0 s |
| 1:30:29.4 | Data point 101 | 1.50000 mL | 0.12761 mL | 0.12227 mL | 1.08000 mL | 4.676 | 0.01394 | 0.55015 | 0.00093 | 17.5 s |
| 1:31:22.6 | Data point 102 | 1.50000 mL | 0.12773 mL | 0.12227 mL | 1.08000 mL | 4.416 | 0.01242 | 0.40861 | 0.00096 | 11.0 s |
| 1:32:04.1 | Data point 103 | 1.50000 mL | 0.12789 mL | 0.12227 mL | 1.08000 mL | 4.164 | 0.00814 | 0.19383 | 0.00091 | 10.0 s |
| 1:32:49.9 | Data point 104 | 1.50000 mL | 0.12810 mL | 0.12227 mL | 1.08000 mL | 3.959 | 0.00234 | 0.03684 | 0.00060 | 10.0 s |
| 1:33:25.3 | Data point 105 | 1.50000 mL | 0.12841 mL | 0.12227 mL | 1.08000 mL | 3.741 | -0.01363 | 0.76827 | 0.00077 | 18.5 s |
| 1:34:09.1 | Data point 106 | 1.50000 mL | 0.12888 mL | 0.12227 mL | 1.08000 mL | 3.543 | -0.01018 | 0.83349 | 0.00055 | 10.0 s |
| 1:34:44.5 | Data point 107 | 1.50000 mL | 0.12963 mL | 0.12227 mL | 1.08000 mL | 3.337 | -0.01423 | 0.90918 | 0.00074 | 10.0 s |
| 1:35:20.0 | Data point 108 | 1.50000 mL | 0.13083 mL | 0.12227 mL | 1.08000 mL | 3.139 | -0.01823 | 0.90719 | 0.00095 | 10.0 s |
| 1:35:55.6 | Data point 109 | 1.50000 mL | 0.13274 mL | 0.12227 mL | 1.08000 mL | 2.947 | -0.01750 | 0.91940 | 0.00090 | 10.0 s |
| 1:36:31.1 | Data point 110 | 1.50000 mL | 0.13572 mL | 0.12227 mL | 1.08000 mL | 2.759 | -0.01737 | 0.77482 | 0.00097 | 24.5 s |
| 1:37:21.2 | Data point 111 | 1.50000 mL | 0.14033 mL | 0.12227 mL | 1.08000 mL | 2.568 | -0.01758 | 0.81168 | 0.00096 | 25.5 s |
| 1:38:12.4 | Data point 112 | 1.50000 mL | 0.14758 mL | 0.12227 mL | 1.08000 mL | 2.370 | -0.01653 | 0.73042 | 0.00096 | 12.0 s |
| 1:38:50.1 | Data point 113 | 1.50000 mL | 0.15915 mL | 0.12227 mL | 1.08000 mL | 2.176 | -0.01588 | 0.71562 | 0.00093 | 15.5 s |
| 1:39:31.5 | Data point 114 | 1.50000 mL | 0.17780 mL | 0.12227 mL | 1.08000 mL | 1.980 | -0.01565 | 0.88128 | 0.00082 | 11.5 s |
| 1:39:52.1 | Assay volumes | 1.50000 mL | 0.17780 mL | 0.12227 mL | 1.08000 mL | | | | | |

| | | | |
|---|---|---|---|
| Sample name: | **M08_octanol** | Experiment start time: | **3/2/2018 5:10:52 PM** |
| Assay name: | **pH-metric high logP** | Analyst: | **Pion** |
| Assay ID: | **18C-02007** | Instrument ID: | **T312060** |
| Filename: | **C:\Sirius_T3\Mehtap\20180302_exp29_logP_T3-2\18C-02007_M08_octanol_pH-metric high logP.t3r** | | |

#### Assay Settings

| Setting | Value | Original Value | Date/Time changed | Imported from |
|---|---|---|---|---|
| ***General Settings*** | | | | |
| Analyst name | Pion | | | |
| ***Standard Experiment Settings*** | | | | |
| Number of titrations | 3 | | | |
| Minimum pH | 2.000 | | | |
| Maximum pH | 10.000 | | | |
| pH step between points of | 0.200 | | | |
| Minimum titrant addition | 0.00002 mL | | | |
| Maximum titrant addition | 0.10000 mL | | | |
| Argon flow rate | 100% | | | |
| Start titration using | Cautious pH adjust | | | |
| ***Advanced General Settings*** | | | | |
| Detect turbidity using | None | | | |
| Collect turbidity sensor data | No | | | |
| Collect UV spectra | No | | | |
| Stir after titrant addition for | 5 seconds | | | |
| For titrant addition, stir at | 10% | | | |
| ***Titrant Pre-Dose*** | | | | |
| Titrant pre-dose | None | | | |
| ***Assay Medium*** | | | | |
| ISA water volume | 1.50 mL | | | |
| Water added | Automatic | | | |
| Partition solvent type | Octanol | | | |
| Partition volume | 0.080 mL | | | |
| Partition solvent added | Manual in advance | | | |
| After partition addition, stir for | 1 seconds | | | |
| ***Sample Sonication*** | | | | |
| Sonicate | Yes | | | |
| Adjust pH for sonication | No | | | |
| Sonicate for | 120 seconds | | | |
| After sonication stir for | 5 seconds | | | |
| ***Sample Dissolution*** | | | | |
| Perform a dissolution stage | Yes | | | |
| Adjust and hold pH for dissolution | To start pH | | | |
| Stir to dissolve for | 120 seconds | | | |
| For dissolution, stir at | 10% | | | |
| ***Carbonate purge*** | | | | |
| Perform a carbonate purge | No | | | |
| ***Temperature Control*** | | | | |
| Wait for temperature | Yes | | | |
| Required start temperature | 25.0°C | | | |
| Acceptable deviation | 0.5°C | | | |
| Time to wait | 60 seconds | | | |
| Stir speed of | 50% | | | |
| ***Titration 1*** | | | | |
| Titrate from | High to low pH | | | |
| Adjust to start pH | Yes | | | |
| After pH adjust stir for | 30 seconds | | | |
| Stir to allow partitioning for | 15 seconds | | | |
| Stirrer speed for partitioning | 50% | | | |
| ***Titration 2*** | | | | |
| Titrate from | High to low pH | | | |
| Add additional water | 0.00 mL | | | |
| Additional partition solvent volume | 0.200 mL | | | |
| Additional partition solvent added | Automatic | | | |
| After pH adjust stir for | 30 seconds | | | |
| Stir to allow partitioning for | 15 seconds | | | |
| Stirrer speed for partitioning | 55% | | | |

| | | | |
|---|---|---|---|
| Sample name: | **M08_octanol** | Experiment start time: | **3/2/2018 5:10:52 PM** |
| Assay name: | **pH-metric high logP** | Analyst: | **Pion** |
| Assay ID: | **18C-02007** | Instrument ID: | **T312060** |
| Filename: | **C:\Sirius_T3\Mehtap\20180302_exp29_logP_T3-2\18C-02007_M08_octanol_pH-metric high logP.t3r** | | |

#### Assay Settings (continued)

| Setting | Value | Original Value | Date/Time changed | Imported from |
|---|---|---|---|---|
| ***Titration 3*** | | | | |
| Titrate from | High to low pH | | | |
| Add additional water | 0.00 mL | | | |
| Additional partition solvent volume | 0.800 mL | | | |
| Additional partition solvent added | Automatic | | | |
| After pH adjust stir for | 30 seconds | | | |
| Stir to allow partitioning for | 15 seconds | | | |
| Stirrer speed for partitioning | 60% | | | |
| ***Data Point Stability*** | | | | |
| Stir during data point collection | No | | | |
| Delay before data point collection | 0 seconds | | | |
| Number of points to average | 20 points | | | |
| Time interval between points | 0.50 seconds | | | |
| Required maximum standard deviation | 0.00100 dpH/dt | | | |
| Stability timeout after | 60 seconds | | | |

#### Calibration Settings

| Setting | Value | Date/Time changed | Imported from |
|---|---|---|---|
| Four-Plus alpha | 0.111 | 3/2/2018 5:10:52 PM | C:\Sirius_T3\HCl18C02.t3r |
| Four-Plus S | 0.9988 | 3/2/2018 5:10:52 PM | C:\Sirius_T3\HCl18C02.t3r |
| Four-Plus jH | 1.0 | 3/2/2018 5:10:52 PM | C:\Sirius_T3\HCl18C02.t3r |
| Four-Plus jOH | -0.8 | 3/2/2018 5:10:52 PM | C:\Sirius_T3\HCl18C02.t3r |
| Base concentration factor | 1.000 | 3/2/2018 5:10:52 PM | C:\Sirius_T3\KOH18B27.t3r |
| Acid concentration factor | 0.999 | 3/2/2018 5:10:52 PM | C:\Sirius_T3\HCl18C02.t3r |

#### Instrument Settings

| Setting | Value | Batch Id | Install date |
|---|---|---|---|
| Instrument owner | Merck | | |
| Instrument ID | T312060 | | |
| Instrument type | T3 Simulator | | |
| Software version | 1.1.3.0 | | |
| Dispenser module | | T3DM1200361 | 3/31/2009 5:24:52 AM |
| Dispenser 0 | Water | | 3/31/2009 5:25:05 AM |
| Syringe volume | 2.5 mL | | |
| Firmware version | 1.2.1(r2) | | |
| Titrant | Water (0.15 M KCl) | 02-06-2018 | 2/27/2018 10:05:59 AM |
| Dispenser 2 | Acid | | 3/31/2009 5:25:11 AM |
| Syringe volume | 0.5 mL | | |
| Firmware version | 1.2.1(r2) | | |
| Titrant | Acid (0.5 M HCl) | 02-27-2018 | 2/27/2018 10:27:22 AM |
| Dispenser 1 | Base | | 3/31/2009 5:25:21 AM |
| Syringe volume | 0.5 mL | | |
| Firmware version | 1.2.1(r2) | | |
| Titrant | Base (0.5 M KOH) | 9/22/2017 | 2/27/2018 10:21:22 AM |
| Dispenser 5 | Cosolvent | | 3/31/2009 5:26:24 AM |
| Syringe volume | 2.5 mL | | |
| Firmware version | 1.2.1(r2) | | |
| Distribution valve 5 | Distribution Valve | | 3/31/2009 5:28:19 AM |
| Firmware version | 1.1.3 | | |
| Port A | Methanol (80%, 0.15 M KCl) | 09-26-17 | 2/7/2018 9:42:01 AM |
| Port B | Cyclohexane | 11-01-17 | 2/27/2018 10:37:57 AM |
| Dispenser 3 | Buffer | | 8/3/2010 5:05:16 AM |
| Syringe volume | 0.5 mL | | |
| Firmware version | 1.2.1(r2) | | |
| Titrant | Dodecane | 2018/01/31 | 2/28/2018 10:18:04 AM |
| Dispenser 6 | Octanol | | 10/22/2010 10:52:43 AM |

Sample name: **M08_octanol**
Assay name: **pH-metric high logP**
Assay ID: **18C-02007**
Filename: **C:\Sirius_T3\Mehtap\20180302_exp29_logP_T3-2\18C-02007_M08_octanol_pH-metric high logP.t3r**

Experiment start time: **3/2/2018 5:10:52 PM**
Analyst: **Pion**
Instrument ID: **T312060**

#### Instrument Settings (continued)

| Setting | Value | Batch Id | Install date |
|---|---|---|---|
| Syringe volume | 0.5 mL | | |
| Firmware version | 1.2.1(r2) | | |
| Titrant | Octanol | 01-31-2018 | 2/27/2018 9:59:35 AM |
| Titrator | | T3TM1200161 | 3/31/2009 5:24:17 AM |
| Horizontal axis firmware version | 1.17 AI1DI2DO2 Stepper 2 | | |
| Vertical axis firmware version | 1.17 AI1DI2DO2 Stepper 2 | | |
| Chassis I/O firmware version | 1.11 AI1DI0DO4 Norgren I/O | | |
| Probe I/O firmware version | 1.1.1 | | |
| Electrode | T3 Electrode | T3E0923 | 1/23/2018 2:01:00 PM |
| E0 calibration | +3.92 mV | | 3/2/2018 5:11:36 PM |
| Filling solution | 3M KCl | KCL097 | 3/2/2018 9:43:24 AM |
| Liquids | | | |
| Wash 1 | 50% IPA:50% Water | | 3/2/2018 9:45:12 AM |
| Wash 2 | 0.5% Trition X-100 in H20 | | 3/2/2018 9:45:15 AM |
| Buffer position 1 | pH7 Wash | | 3/2/2018 9:45:18 AM |
| Buffer position 2 | pH 7 | | 3/2/2018 9:45:21 AM |
| Storage position | | | 3/2/2018 9:44:44 AM |
| Wash water | 7.4e+003 mL | 02-27-2018 | 2/27/2018 9:54:39 AM |
| Waste | 8.1e+003 mL | | 11/28/2017 10:36:29 AM |
| Temperature controller | | | 8/5/2010 6:35:13 AM |
| Turbidity detector | | | 3/31/2009 5:24:45 AM |
| Spectrometer | | 074811 | 11/23/2010 11:22:28 AM |
| Dip probe | | 10196 | |
| Wavelength coefficient A0 | 183.333 | | |
| Wavelength coefficient A1 | 2.21568 | | |
| Wavelength coefficient A2 | -0.000289308 | | |
| Total lamp lit time | 120:41:49 | | 11/23/2010 11:22:28 AM |
| Calibrated on | 2/27/2018 10:40:38 AM | | |
| Integration time | 40 | | |
| Scans averaged | 10 | | |
| Autoloader | | T3AL1200345 | 11/10/2015 9:34:13 AM |
| Left-right axis firmware version | 1.17 AI1DI2DO2 Stepper 2 | | |
| Front-back axis firmware version | 1.17 AI1DI2DO2 Stepper 2 | | |
| Vertical axis firmware version | 1.17 AI1DI2DO2 Stepper 2 | | |
| Chassis I/O firmware version | 1.11 AI1DI0DO4 Norgren I/O | | |
| Configuration | | | |
| Alternate titration position | Titration position | | |
| Alternate reference position | Reference position | | |
| Maximum standard vial volume | 3.50 mL | | |
| Maximum alternate vial volume | 25.00 mL | | |
| Automatic action idle period | 5 minute(s) | | |
| Titrant tube volume | 1.3 mL | | |
| Syringe flush count | 3.50 | | |
| Flowing wash pump volume | 20.0 mL | | |
| Flowing wash stir duration | 5 s | | |
| Flowing wash stir speed | 30% | | |
| Solvent wash stir duration | 5 s | | |
| Solvent wash stir speed | 30% | | |
| Surfactant wash stir duration | 5 s | | |
| Surfactant wash stir speed | 30% | | |
| E0 calibration minimum number of points | 10 | | |
| E0 calibration maximum standard deviation | 0.01500 | | |
| E0 calibration timeout period | 60 s | | |
| E0 calibration stir duration | 5 s | | |
| E0 calibration preparation stir speed | 30% | | |
| E0 calibration buffer wash stir duration | 5 s | | |
| E0 calibration buffer wash stir speed | 30% | | |
| E0 calibration reading stir speed | 0% | | |

| | | | |
|---|---|---|---|
| Sample name: | **M08_octanol** | Experiment start time: | **3/2/2018 5:10:52 PM** |
| Assay name: | **pH-metric high logP** | Analyst: | **Pion** |
| Assay ID: | **18C-02007** | Instrument ID: | **T312060** |
| Filename: | **C:\Sirius_T3\Mehtap\20180302_exp29_logP_T3-2\18C-02007_M08_octanol_pH-metric high logP.t3r** | | |

#### Instrument Settings (continued)

| Setting | Value | Batch Id | Install date |
|---|---|---|---|
| Spectrometer calibration stir duration | 5 s | | |
| Spectrometer calibration stir speed | 30% | | |
| Spectrometer calibration wash pump volume | 20.0 mL | | |
| Spectrometer calibration wash stir duration | 5 s | | |
| Spectrometer calibration wash stir speed | 30% | | |
| Overhead dispense height | 10000 | | |

#### Refinement Settings

| Setting | Value | Default value |
|---|---|---|
| Turbidity detection method | None | None |
| Turbidity wavelength to assess | 500.0 nm | 500.0 nm |
| Turbidity maximum absorbance | 0.100 | 0.100 |
| Turbidity probe threshold | 50.00 | 50.00 |

#### Experiment Log

[1:59] Air gap released for Acid (0.5 M HCl)
[2:54] Air gap created for Water (0.15 M KCl)
[2:54] Air gap created for Acid (0.5 M HCl)
[2:55] Air gap created for Base (0.5 M KOH)
[2:55] Air gap released for Water (0.15 M KCl)
[2:59] Titrator arm moved over Titration position
[2:59] Titration 1 of 3
[2:59] Adding initial titrants
[2:59] Automatically add 1.50000 mL of water
[3:24] Dispensed 1.500000 mL of Water (0.15 M KCl)
[3:28] Titrator arm moved over Drain
[6:09] Titrator arm moved to Titration position
[6:09] Argon flow rate set to 100
[6:09] Stirrer speed set to 10
[6:16] Initial pH = 5.74
[6:16] Iterative adjust 5.74 -> 10.00
[6:16] pH 5.74 -> 10.00
[6:16] Air gap released for Base (0.5 M KOH)
[6:17] Dispensed 0.006468 mL of Base (0.5 M KOH)
[6:22] Holding pH 10.00
[8:22] Stirrer speed set to 0
[8:22] Stirrer speed set to 50
[8:22] Iterative adjust 11.25 -> 10.00
[9:07] Stirrer speed set to 0
[9:42] Datapoint id 2 collected
[9:42] Stirrer speed set to 50
[9:47] pH 10.83 -> 10.63
[9:47] Using cautious pH adjust
[9:48] Air gap released for Acid (0.5 M HCl)
[9:49] Dispensed 0.000517 mL of Acid (0.5 M HCl)
[9:54] Stepping pH = 10.62
[10:09] Stirrer speed set to 0
[10:44] Datapoint id 3 collected
[10:44] Charge balance equation is out by 50.1%
[10:44] Stirrer speed set to 50
[10:49] pH 10.50 -> 10.30
[10:49] Using cautious pH adjust
[10:49] Dispensed 0.000259 mL of Acid (0.5 M HCl)
[10:54] Stepping pH = 10.44
[10:54] Dispensed 0.000306 mL of Acid (0.5 M HCl)
[10:59] Stepping pH = 10.31
[11:14] Stirrer speed set to 0

#### Experiment Log (continued)

[11:41] Datapoint id 4 collected
[11:41] Charge balance equation is out by -9.4%
[11:41] Stirrer speed set to 50
[11:46] pH 10.22 -> 10.02
[11:46] Using charge balance adjust
[11:47] Dispensed 0.000282 mL of Acid (0.5 M HCl)
[12:07] Stirrer speed set to 0
[12:35] Datapoint id 5 collected
[12:35] Charge balance equation is out by 4.0%
[12:35] Stirrer speed set to 50
[12:40] pH 10.00 -> 9.80
[12:40] Using charge balance adjust
[12:40] Dispensed 0.000188 mL of Acid (0.5 M HCl)
[13:01] Stirrer speed set to 0
[13:29] Datapoint id 6 collected
[13:29] Charge balance equation is out by -10.5%
[13:29] Stirrer speed set to 50
[13:34] pH 9.81 -> 9.61
[13:34] Using charge balance adjust
[13:34] Dispensed 0.000141 mL of Acid (0.5 M HCl)
[13:54] Stirrer speed set to 0
[14:24] Datapoint id 7 collected
[14:24] Charge balance equation is out by -4.1%
[14:24] Stirrer speed set to 50
[14:29] pH 9.60 -> 9.40
[14:29] Using charge balance adjust
[14:29] Dispensed 0.000094 mL of Acid (0.5 M HCl)
[14:49] Stirrer speed set to 0
[15:20] Datapoint id 8 collected
[15:20] Charge balance equation is out by -16.0%
[15:20] Stirrer speed set to 50
[15:25] pH 9.40 -> 9.20
[15:25] Using cautious pH adjust
[15:25] Dispensed 0.000024 mL of Acid (0.5 M HCl)
[15:30] Stepping pH = 9.38
[15:30] Dispensed 0.000071 mL of Acid (0.5 M HCl)
[15:35] Stepping pH = 9.22
[15:35] Dispensed 0.000024 mL of Acid (0.5 M HCl)
[15:40] Stepping pH = 9.17
[15:55] Stirrer speed set to 0
[16:29] Datapoint id 9 collected
[16:29] Charge balance equation is out by -121.4%
[16:29] Stirrer speed set to 50
[16:34] pH 9.07 -> 8.87
[16:34] Using cautious pH adjust
[16:34] Dispensed 0.000024 mL of Acid (0.5 M HCl)
[16:40] Stepping pH = 9.05
[16:40] Dispensed 0.000047 mL of Acid (0.5 M HCl)
[16:45] Stepping pH = 8.91
[16:45] Dispensed 0.000024 mL of Acid (0.5 M HCl)
[16:50] Stepping pH = 8.80
[17:05] Stirrer speed set to 0
[17:15] Datapoint id 10 collected
[17:15] Charge balance equation is out by -150.8%
[17:15] Stirrer speed set to 50
[17:20] pH 8.72 -> 8.52
[17:20] Using cautious pH adjust
[17:21] Dispensed 0.000024 mL of Acid (0.5 M HCl)
[17:26] Stepping pH = 8.71

| | | | |
|---|---|---|---|
| Sample name: | **M08_octanol** | Experiment start time: | **3/2/2018 5:10:52 PM** |
| Assay name: | **pH-metric high logP** | Analyst: | **Pion** |
| Assay ID: | **18C-02007** | Instrument ID: | **T312060** |
| Filename: | **C:\Sirius_T3\Mehtap\20180302_exp29_logP_T3-2\18C-02007_M08_octanol_pH-metric high logP.t3r** | | |

#### Experiment Log (continued)

[17:26] Dispensed 0.000024 mL of Acid (0.5 M HCl)
[17:31] Stepping pH = 8.66
[17:31] Dispensed 0.000047 mL of Acid (0.5 M HCl)
[17:36] Stepping pH = 8.37
[17:51] Stirrer speed set to 0
[18:08] Datapoint id 11 collected
[18:08] Charge balance equation is out by -335.3%
[18:08] Stirrer speed set to 50
[18:13] pH 8.27 -> 8.07
[18:13] Using cautious pH adjust
[18:14] Dispensed 0.000024 mL of Acid (0.5 M HCl)
[18:19] Stepping pH = 8.26
[18:19] Dispensed 0.000047 mL of Acid (0.5 M HCl)
[18:24] Stepping pH = 8.12
[18:24] Dispensed 0.000024 mL of Acid (0.5 M HCl)
[18:29] Stepping pH = 8.04
[18:44] Stirrer speed set to 0
[19:00] Datapoint id 12 collected
[19:00] Charge balance equation is out by -155.8%
[19:00] Stirrer speed set to 50
[19:05] pH 7.96 -> 7.76
[19:05] Using cautious pH adjust
[19:05] Dispensed 0.000024 mL of Acid (0.5 M HCl)
[19:10] Stepping pH = 7.95
[19:10] Dispensed 0.000094 mL of Acid (0.5 M HCl)
[19:15] Stepping pH = 7.75
[19:31] Stirrer speed set to 0
[19:48] Datapoint id 13 collected
[19:48] Charge balance equation is out by -93.4%
[19:48] Stirrer speed set to 50
[19:53] pH 7.69 -> 7.49
[19:53] Using cautious pH adjust
[19:53] Dispensed 0.000047 mL of Acid (0.5 M HCl)
[19:58] Stepping pH = 7.66
[19:58] Dispensed 0.000141 mL of Acid (0.5 M HCl)
[20:03] Stepping pH = 7.44
[20:18] Stirrer speed set to 0
[20:34] Datapoint id 14 collected
[20:34] Charge balance equation is out by -74.1%
[20:34] Stirrer speed set to 50
[20:39] pH 7.42 -> 7.22
[20:39] Using cautious pH adjust
[20:39] Dispensed 0.000094 mL of Acid (0.5 M HCl)
[20:44] Stepping pH = 7.35
[20:44] Dispensed 0.000118 mL of Acid (0.5 M HCl)
[20:49] Stepping pH = 7.23
[20:49] Dispensed 0.000024 mL of Acid (0.5 M HCl)
[20:54] Stepping pH = 7.22
[21:10] Stirrer speed set to 0
[21:26] Datapoint id 15 collected
[21:26] Charge balance equation is out by -18.8%
[21:26] Stirrer speed set to 50
[21:31] pH 7.22 -> 7.02
[21:31] Using cautious pH adjust
[21:31] Dispensed 0.000141 mL of Acid (0.5 M HCl)
[21:36] Stepping pH = 7.12
[21:36] Dispensed 0.000094 mL of Acid (0.5 M HCl)
[21:41] Stepping pH = 7.07
[21:41] Dispensed 0.000071 mL of Acid (0.5 M HCl)

Sample name: **M08_octanol** Experiment start time: **3/2/2018 5:10:52 PM**
Assay name: **pH-metric high logP** Analyst: **Pion**
Assay ID: **18C-02007** Instrument ID: **T312060**
Filename: **C:\Sirius_T3\Mehtap\20180302_exp29_logP_T3-2\18C-02007_M08_octanol_pH-metric high logP.t3r**

#### Experiment Log (continued)

[21:46] Stepping pH = 7.03
[22:01] Stirrer speed set to 0
[22:17] Datapoint id 16 collected
[22:17] Charge balance equation is out by -9.3%
[22:17] Stirrer speed set to 50
[22:22] pH 7.05 -> 6.85
[22:22] Using charge balance adjust
[22:22] Dispensed 0.000400 mL of Acid (0.5 M HCl)
[22:42] Stirrer speed set to 0
[22:57] Datapoint id 17 collected
[22:57] Charge balance equation is out by -6.8%
[22:57] Stirrer speed set to 50
[23:02] pH 6.86 -> 6.66
[23:02] Using charge balance adjust
[23:02] Dispensed 0.000541 mL of Acid (0.5 M HCl)
[23:23] Stirrer speed set to 0
[23:37] Datapoint id 18 collected
[23:37] Charge balance equation is out by -17.6%
[23:37] Stirrer speed set to 50
[23:42] pH 6.69 -> 6.49
[23:42] Using cautious pH adjust
[23:42] Dispensed 0.000353 mL of Acid (0.5 M HCl)
[23:47] Stepping pH = 6.57
[23:47] Dispensed 0.000188 mL of Acid (0.5 M HCl)
[23:52] Stepping pH = 6.54
[23:52] Dispensed 0.000188 mL of Acid (0.5 M HCl)
[23:57] Stepping pH = 6.50
[24:12] Stirrer speed set to 0
[24:26] Datapoint id 19 collected
[24:26] Charge balance equation is out by -4.7%
[24:26] Stirrer speed set to 50
[24:31] pH 6.53 -> 6.33
[24:31] Using charge balance adjust
[24:31] Dispensed 0.000823 mL of Acid (0.5 M HCl)
[24:51] Stirrer speed set to 0
[25:05] Datapoint id 20 collected
[25:05] Charge balance equation is out by -13.9%
[25:05] Stirrer speed set to 50
[25:10] pH 6.36 -> 6.16
[25:10] Using charge balance adjust
[25:10] Dispensed 0.000941 mL of Acid (0.5 M HCl)
[25:30] Stirrer speed set to 0
[25:43] Datapoint id 21 collected
[25:43] Charge balance equation is out by -17.0%
[25:43] Stirrer speed set to 50
[25:48] pH 6.19 -> 5.99
[25:48] Using cautious pH adjust
[25:48] Dispensed 0.000494 mL of Acid (0.5 M HCl)
[25:53] Stepping pH = 6.09
[25:53] Dispensed 0.000306 mL of Acid (0.5 M HCl)
[25:59] Stepping pH = 6.04
[25:59] Dispensed 0.000188 mL of Acid (0.5 M HCl)
[26:04] Stepping pH = 6.01
[26:04] Dispensed 0.000118 mL of Acid (0.5 M HCl)
[26:09] Stepping pH = 6.00
[26:09] Dispensed 0.000094 mL of Acid (0.5 M HCl)
[26:14] Stepping pH = 5.99
[26:29] Stirrer speed set to 0
[26:40] Datapoint id 22 collected

| | | | |
|---|---|---|---|
| Sample name: | **M08_octanol** | Experiment start time: | **3/2/2018 5:10:52 PM** |
| Assay name: | **pH-metric high logP** | Analyst: | **Pion** |
| Assay ID: | **18C-02007** | Instrument ID: | **T312060** |
| Filename: | **C:\Sirius_T3\Mehtap\20180302_exp29_logP_T3-2\18C-02007_M08_octanol_pH-metric high logP.t3r** | | |

#### Experiment Log (continued)

[26:40] Charge balance equation is out by -22.6%
[26:40] Stirrer speed set to 50
[26:45] pH 6.00 -> 5.80
[26:45] Using cautious pH adjust
[26:45] Dispensed 0.000494 mL of Acid (0.5 M HCl)
[26:50] Stepping pH = 5.88
[26:50] Dispensed 0.000235 mL of Acid (0.5 M HCl)
[26:55] Stepping pH = 5.85
[26:55] Dispensed 0.000188 mL of Acid (0.5 M HCl)
[27:00] Stepping pH = 5.82
[27:01] Dispensed 0.000071 mL of Acid (0.5 M HCl)
[27:06] Stepping pH = 5.81
[27:21] Stirrer speed set to 0
[27:34] Datapoint id 23 collected
[27:34] Charge balance equation is out by -0.9%
[27:34] Stirrer speed set to 50
[27:39] pH 5.82 -> 5.62
[27:39] Using charge balance adjust
[27:39] Dispensed 0.000894 mL of Acid (0.5 M HCl)
[27:59] Stirrer speed set to 0
[28:59] Datapoint id 24 collected
[28:59] Charge balance equation is out by -7.3%
[28:59] Stirrer speed set to 50
[29:04] pH 5.65 -> 5.45
[29:04] Using charge balance adjust
[29:04] Dispensed 0.000753 mL of Acid (0.5 M HCl)
[29:24] Stirrer speed set to 0
[30:24] Datapoint id 25 collected
[30:24] Charge balance equation is out by -68.1%
[30:24] Stirrer speed set to 50
[30:30] pH 5.66 -> 5.46
[30:30] Using cautious pH adjust
[30:30] Dispensed 0.000376 mL of Acid (0.5 M HCl)
[30:35] Stepping pH = 5.52
[30:35] Dispensed 0.000118 mL of Acid (0.5 M HCl)
[30:40] Stepping pH = 5.51
[30:40] Dispensed 0.000306 mL of Acid (0.5 M HCl)
[30:45] Stepping pH = 5.31
[31:00] Stirrer speed set to 0
[32:00] Datapoint id 26 collected
[32:00] Charge balance equation is out by -6.1%
[32:00] Stirrer speed set to 50
[32:05] pH 5.62 -> 5.42
[32:05] Using charge balance adjust
[32:06] Dispensed 0.000729 mL of Acid (0.5 M HCl)
[32:26] Stirrer speed set to 0
[33:15] Datapoint id 27 collected
[33:15] Charge balance equation is out by 443.4%
[33:15] Stirrer speed set to 50
[33:20] pH 4.60 -> 4.40
[33:20] Using cautious pH adjust
[33:20] Dispensed 0.000094 mL of Acid (0.5 M HCl)
[33:25] Stepping pH = 4.61
[33:25] Dispensed 0.000447 mL of Acid (0.5 M HCl)
[33:30] Stepping pH = 3.88
[33:45] Stirrer speed set to 0
[33:57] Datapoint id 28 collected
[33:57] Charge balance equation is out by -208.0%
[33:57] Stirrer speed set to 50

| | | | |
|---|---|---|---|
| Sample name: | **M08_octanol** | Experiment start time: | **3/2/2018 5:10:52 PM** |
| Assay name: | **pH-metric high logP** | Analyst: | **Pion** |
| Assay ID: | **18C-02007** | Instrument ID: | **T312060** |
| Filename: | **C:\Sirius_T3\Mehtap\20180302_exp29_logP_T3-2\18C-02007_M08_octanol_pH-metric high logP.t3r** | | |

#### Experiment Log (continued)

[34:02] pH 3.89 -> 3.69
[34:02] Using cautious pH adjust
[34:02] Dispensed 0.000165 mL of Acid (0.5 M HCl)
[34:07] Stepping pH = 3.81
[34:07] Dispensed 0.000188 mL of Acid (0.5 M HCl)
[34:12] Stepping pH = 3.70
[34:12] Dispensed 0.000024 mL of Acid (0.5 M HCl)
[34:18] Stepping pH = 3.70
[34:33] Stirrer speed set to 0
[34:43] Datapoint id 29 collected
[34:43] Charge balance equation is out by -13.2%
[34:43] Stirrer speed set to 50
[34:48] pH 3.69 -> 3.49
[34:48] Using charge balance adjust
[34:48] Dispensed 0.000470 mL of Acid (0.5 M HCl)
[35:08] Stirrer speed set to 0
[35:18] Datapoint id 30 collected
[35:18] Charge balance equation is out by 6.0%
[35:18] Stirrer speed set to 50
[35:23] pH 3.48 -> 3.28
[35:23] Using charge balance adjust
[35:24] Dispensed 0.000776 mL of Acid (0.5 M HCl)
[35:44] Stirrer speed set to 0
[35:54] Datapoint id 31 collected
[35:54] Charge balance equation is out by 3.3%
[35:54] Stirrer speed set to 50
[35:59] pH 3.28 -> 3.08
[35:59] Using charge balance adjust
[35:59] Dispensed 0.001223 mL of Acid (0.5 M HCl)
[36:19] Stirrer speed set to 0
[36:29] Datapoint id 32 collected
[36:29] Charge balance equation is out by 2.2%
[36:29] Stirrer speed set to 50
[36:34] pH 3.08 -> 2.88
[36:34] Using charge balance adjust
[36:34] Dispensed 0.001929 mL of Acid (0.5 M HCl)
[36:55] Stirrer speed set to 0
[37:05] Datapoint id 33 collected
[37:05] Charge balance equation is out by 6.6%
[37:05] Stirrer speed set to 50
[37:10] pH 2.87 -> 2.67
[37:10] Using charge balance adjust
[37:10] Dispensed 0.003128 mL of Acid (0.5 M HCl)
[37:30] Stirrer speed set to 0
[37:40] Datapoint id 34 collected
[37:40] Charge balance equation is out by 9.6%
[37:40] Stirrer speed set to 50
[37:45] pH 2.66 -> 2.46
[37:45] Using charge balance adjust
[37:46] Dispensed 0.005151 mL of Acid (0.5 M HCl)
[38:06] Stirrer speed set to 0
[38:16] Datapoint id 35 collected
[38:16] Charge balance equation is out by 4.2%
[38:16] Stirrer speed set to 50
[38:21] pH 2.45 -> 2.25
[38:21] Using charge balance adjust
[38:21] Dispensed 0.008373 mL of Acid (0.5 M HCl)
[38:41] Stirrer speed set to 0
[38:52] Datapoint id 36 collected

Sample name: **M08_octanol**
Assay name: **pH-metric high logP**
Assay ID: **18C-02007**
Filename: **C:\Sirius_T3\Mehtap\20180302_exp29_logP_T3-2\18C-02007_M08_octanol_pH-metric high logP.t3r**
Experiment start time: **3/2/2018 5:10:52 PM**
Analyst: **Pion**
Instrument ID: **T312060**

#### Experiment Log (continued)

[38:52] Charge balance equation is out by -0.7%
[38:52] Stirrer speed set to 50
[38:57] pH 2.26 -> 2.06
[38:57] Using charge balance adjust
[38:57] Dispensed 0.013335 mL of Acid (0.5 M HCl)
[39:18] Stirrer speed set to 0
[39:28] Datapoint id 37 collected
[39:28] Charge balance equation is out by 6.9%
[39:28] Stirrer speed set to 50
[39:33] pH 2.05 -> 1.95
[39:33] Using charge balance adjust
[39:33] Dispensed 0.009737 mL of Acid (0.5 M HCl)
[39:54] Stirrer speed set to 0
[40:04] Datapoint id 38 collected
[40:04] Charge balance equation is out by -48.9%
[40:04] Titration 2 of 3
[40:04] Adding initial titrants
[40:04] Automatically add 0.20000 mL of Octanol
[40:08] Dispensed 0.200000 mL of Octanol
[40:08] Stirrer speed set to 10
[40:09] Stirrer speed set to 55
[40:09] Iterative adjust 1.95 -> 10.00
[40:09] pH 1.95 -> 10.00
[40:11] Dispensed 0.055880 mL of Base (0.5 M KOH)
[41:01] Stirrer speed set to 0
[41:48] Datapoint id 39 collected
[41:48] Stirrer speed set to 55
[41:53] pH 10.59 -> 10.39
[41:53] Using cautious pH adjust
[41:53] Dispensed 0.000329 mL of Acid (0.5 M HCl)
[41:58] Stepping pH = 10.50
[41:58] Dispensed 0.000259 mL of Acid (0.5 M HCl)
[42:03] Stepping pH = 10.41
[42:03] Dispensed 0.000047 mL of Acid (0.5 M HCl)
[42:08] Stepping pH = 10.40
[42:08] Dispensed 0.000024 mL of Acid (0.5 M HCl)
[42:14] Stepping pH = 10.39
[42:29] Stirrer speed set to 0
[42:50] Datapoint id 40 collected
[42:50] Charge balance equation is out by -0.8%
[42:50] Stirrer speed set to 55
[42:55] pH 10.36 -> 10.16
[42:55] Using charge balance adjust
[42:55] Dispensed 0.000423 mL of Acid (0.5 M HCl)
[43:15] Stirrer speed set to 0
[43:36] Datapoint id 41 collected
[43:36] Charge balance equation is out by 17.8%
[43:36] Stirrer speed set to 55
[43:41] pH 10.13 -> 9.93
[43:41] Using cautious pH adjust
[43:41] Dispensed 0.000118 mL of Acid (0.5 M HCl)
[43:46] Stepping pH = 10.09
[43:46] Dispensed 0.000259 mL of Acid (0.5 M HCl)
[43:51] Stepping pH = 9.88
[44:06] Stirrer speed set to 0
[44:16] Datapoint id 42 collected
[44:16] Charge balance equation is out by -46.0%
[44:16] Stirrer speed set to 55
[44:21] pH 9.86 -> 9.66

#### Experiment Log (continued)

[44:21] Using cautious pH adjust
[44:21] Dispensed 0.000071 mL of Acid (0.5 M HCl)
[44:26] Stepping pH = 9.85
[44:27] Dispensed 0.000235 mL of Acid (0.5 M HCl)
[44:32] Stepping pH = 9.55
[44:47] Stirrer speed set to 0
[45:05] Datapoint id 43 collected
[45:05] Charge balance equation is out by -94.9%
[45:05] Stirrer speed set to 55
[45:10] pH 9.51 -> 9.31
[45:10] Using cautious pH adjust
[45:10] Dispensed 0.000047 mL of Acid (0.5 M HCl)
[45:15] Stepping pH = 9.50
[45:15] Dispensed 0.000118 mL of Acid (0.5 M HCl)
[45:20] Stepping pH = 9.30
[45:36] Stirrer speed set to 0
[45:46] Datapoint id 44 collected
[45:46] Charge balance equation is out by -88.8%
[45:46] Stirrer speed set to 55
[45:51] pH 9.23 -> 9.03
[45:51] Using cautious pH adjust
[45:51] Dispensed 0.000024 mL of Acid (0.5 M HCl)
[45:56] Stepping pH = 9.22
[45:56] Dispensed 0.000071 mL of Acid (0.5 M HCl)
[46:01] Stepping pH = 9.10
[46:01] Dispensed 0.000024 mL of Acid (0.5 M HCl)
[46:06] Stepping pH = 9.06
[46:06] Dispensed 0.000024 mL of Acid (0.5 M HCl)
[46:11] Stepping pH = 9.02
[46:26] Stirrer speed set to 0
[46:38] Datapoint id 45 collected
[46:38] Charge balance equation is out by -183.9%
[46:38] Stirrer speed set to 55
[46:43] pH 8.90 -> 8.70
[46:43] Using cautious pH adjust
[46:44] Dispensed 0.000024 mL of Acid (0.5 M HCl)
[46:49] Stepping pH = 8.89
[46:49] Dispensed 0.000071 mL of Acid (0.5 M HCl)
[46:54] Stepping pH = 8.63
[47:09] Stirrer speed set to 0
[47:24] Datapoint id 46 collected
[47:24] Charge balance equation is out by -94.4%
[47:24] Stirrer speed set to 55
[47:29] pH 8.54 -> 8.34
[47:29] Using cautious pH adjust
[47:29] Dispensed 0.000024 mL of Acid (0.5 M HCl)
[47:34] Stepping pH = 8.53
[47:34] Dispensed 0.000094 mL of Acid (0.5 M HCl)
[47:39] Stepping pH = 8.30
[47:54] Stirrer speed set to 0
[48:10] Datapoint id 47 collected
[48:10] Charge balance equation is out by -96.0%
[48:10] Stirrer speed set to 55
[48:16] pH 8.23 -> 8.03
[48:16] Using cautious pH adjust
[48:16] Dispensed 0.000047 mL of Acid (0.5 M HCl)
[48:21] Stepping pH = 8.20
[48:21] Dispensed 0.000118 mL of Acid (0.5 M HCl)
[48:26] Stepping pH = 8.01

#### Experiment Log (continued)

[48:41] Stirrer speed set to 0
[48:55] Datapoint id 48 collected
[48:55] Charge balance equation is out by -69.2%
[48:55] Stirrer speed set to 55
[49:01] pH 7.95 -> 7.75
[49:01] Using cautious pH adjust
[49:01] Dispensed 0.000094 mL of Acid (0.5 M HCl)
[49:06] Stepping pH = 7.87
[49:06] Dispensed 0.000094 mL of Acid (0.5 M HCl)
[49:11] Stepping pH = 7.78
[49:11] Dispensed 0.000047 mL of Acid (0.5 M HCl)
[49:16] Stepping pH = 7.75
[49:31] Stirrer speed set to 0
[49:45] Datapoint id 49 collected
[49:45] Charge balance equation is out by -25.6%
[49:45] Stirrer speed set to 55
[49:50] pH 7.72 -> 7.52
[49:50] Using cautious pH adjust
[49:50] Dispensed 0.000141 mL of Acid (0.5 M HCl)
[49:55] Stepping pH = 7.62
[49:55] Dispensed 0.000118 mL of Acid (0.5 M HCl)
[50:00] Stepping pH = 7.55
[50:00] Dispensed 0.000047 mL of Acid (0.5 M HCl)
[50:05] Stepping pH = 7.53
[50:05] Dispensed 0.000024 mL of Acid (0.5 M HCl)
[50:10] Stepping pH = 7.52
[50:25] Stirrer speed set to 0
[50:41] Datapoint id 50 collected
[50:41] Charge balance equation is out by -10.2%
[50:41] Stirrer speed set to 55
[50:46] pH 7.50 -> 7.30
[50:46] Using charge balance adjust
[50:46] Dispensed 0.000423 mL of Acid (0.5 M HCl)
[51:06] Stirrer speed set to 0
[51:18] Datapoint id 51 collected
[51:18] Charge balance equation is out by -1.5%
[51:18] Stirrer speed set to 55
[51:23] pH 7.29 -> 7.09
[51:23] Using charge balance adjust
[51:23] Dispensed 0.000611 mL of Acid (0.5 M HCl)
[51:43] Stirrer speed set to 0
[51:55] Datapoint id 52 collected
[51:55] Charge balance equation is out by -3.4%
[51:55] Stirrer speed set to 55
[52:00] pH 7.09 -> 6.89
[52:00] Using charge balance adjust
[52:01] Dispensed 0.000776 mL of Acid (0.5 M HCl)
[52:21] Stirrer speed set to 0
[52:33] Datapoint id 53 collected
[52:33] Charge balance equation is out by -3.7%
[52:33] Stirrer speed set to 55
[52:38] pH 6.89 -> 6.69
[52:38] Using charge balance adjust
[52:38] Dispensed 0.000941 mL of Acid (0.5 M HCl)
[52:59] Stirrer speed set to 0
[53:11] Datapoint id 54 collected
[53:11] Charge balance equation is out by 3.2%
[53:11] Stirrer speed set to 55
[53:16] pH 6.69 -> 6.49

Sample name: **M08_octanol** Experiment start time: **3/2/2018 5:10:52 PM**
Assay name: **pH-metric high logP** Analyst: **Pion**
Assay ID: **18C-02007** Instrument ID: **T312060**
Filename: **C:\Sirius_T3\Mehtap\20180302_exp29_logP_T3-2\18C-02007_M08_octanol_pH-metric high logP.t3r**

#### Experiment Log (continued)

[53:16] Using charge balance adjust
[53:16] Dispensed 0.001011 mL of Acid (0.5 M HCl)
[53:37] Stirrer speed set to 0
[53:51] Datapoint id 55 collected
[53:51] Charge balance equation is out by 1.7%
[53:51] Stirrer speed set to 55
[53:56] pH 6.48 -> 6.28
[53:56] Using charge balance adjust
[53:56] Dispensed 0.000988 mL of Acid (0.5 M HCl)
[54:17] Stirrer speed set to 0
[54:33] Datapoint id 56 collected
[54:33] Charge balance equation is out by 3.6%
[54:33] Stirrer speed set to 55
[54:38] pH 6.26 -> 6.06
[54:38] Using charge balance adjust
[54:38] Dispensed 0.000847 mL of Acid (0.5 M HCl)
[54:58] Stirrer speed set to 0
[55:14] Datapoint id 57 collected
[55:14] Charge balance equation is out by 4.4%
[55:14] Stirrer speed set to 55
[55:19] pH 6.04 -> 5.84
[55:19] Using charge balance adjust
[55:19] Dispensed 0.000682 mL of Acid (0.5 M HCl)
[55:39] Stirrer speed set to 0
[55:54] Datapoint id 58 collected
[55:54] Charge balance equation is out by 4.2%
[55:54] Stirrer speed set to 55
[55:59] pH 5.82 -> 5.62
[55:59] Using charge balance adjust
[55:59] Dispensed 0.000470 mL of Acid (0.5 M HCl)
[56:19] Stirrer speed set to 0
[56:34] Datapoint id 59 collected
[56:34] Charge balance equation is out by -5.4%
[56:34] Stirrer speed set to 55
[56:39] pH 5.61 -> 5.41
[56:39] Using charge balance adjust
[56:39] Dispensed 0.000329 mL of Acid (0.5 M HCl)
[56:59] Stirrer speed set to 0
[57:14] Datapoint id 60 collected
[57:14] Charge balance equation is out by -8.3%
[57:14] Stirrer speed set to 55
[57:19] pH 5.41 -> 5.21
[57:19] Using charge balance adjust
[57:19] Dispensed 0.000235 mL of Acid (0.5 M HCl)
[57:39] Stirrer speed set to 0
[57:54] Datapoint id 61 collected
[57:54] Charge balance equation is out by -7.7%
[57:54] Stirrer speed set to 55
[57:59] pH 5.20 -> 5.00
[57:59] Using charge balance adjust
[57:59] Dispensed 0.000165 mL of Acid (0.5 M HCl)
[58:19] Stirrer speed set to 0
[58:33] Datapoint id 62 collected
[58:33] Charge balance equation is out by -25.6%
[58:33] Stirrer speed set to 55
[58:38] pH 5.02 -> 4.82
[58:38] Using cautious pH adjust
[58:38] Dispensed 0.000071 mL of Acid (0.5 M HCl)
[58:44] Stepping pH = 4.97

Sample name: **M08_octanol**
Assay name: **pH-metric high logP**
Assay ID: **18C-02007**
Filename: **C:\Sirius_T3\Mehtap\20180302_exp29_logP_T3-2\18C-02007_M08_octanol_pH-metric high logP.t3r**
Experiment start time: **3/2/2018 5:10:52 PM**
Analyst: **Pion**
Instrument ID: **T312060**

#### Experiment Log (continued)

[58:44] Dispensed 0.000094 mL of Acid (0.5 M HCl)
[58:49] Stepping pH = 4.80
[59:04] Stirrer speed set to 0
[59:17] Datapoint id 63 collected
[59:17] Charge balance equation is out by -23.5%
[59:17] Stirrer speed set to 55
[59:22] pH 4.77 -> 4.57
[59:22] Using cautious pH adjust
[59:22] Dispensed 0.000047 mL of Acid (0.5 M HCl)
[59:27] Stepping pH = 4.74
[59:27] Dispensed 0.000118 mL of Acid (0.5 M HCl)
[59:32] Stepping pH = 4.49
[59:47] Stirrer speed set to 0
[1:00:01] Datapoint id 64 collected
[1:00:01] Charge balance equation is out by -69.6%
[1:00:01] Stirrer speed set to 55
[1:00:06] pH 4.47 -> 4.27
[1:00:06] Using cautious pH adjust
[1:00:06] Dispensed 0.000047 mL of Acid (0.5 M HCl)
[1:00:11] Stepping pH = 4.45
[1:00:11] Dispensed 0.000141 mL of Acid (0.5 M HCl)
[1:00:16] Stepping pH = 4.20
[1:00:31] Stirrer speed set to 0
[1:00:43] Datapoint id 65 collected
[1:00:43] Charge balance equation is out by -85.9%
[1:00:43] Stirrer speed set to 55
[1:00:48] pH 4.19 -> 3.99
[1:00:48] Using cautious pH adjust
[1:00:48] Dispensed 0.000094 mL of Acid (0.5 M HCl)
[1:00:53] Stepping pH = 4.10
[1:00:53] Dispensed 0.000071 mL of Acid (0.5 M HCl)
[1:00:58] Stepping pH = 4.03
[1:00:58] Dispensed 0.000047 mL of Acid (0.5 M HCl)
[1:01:04] Stepping pH = 4.00
[1:01:04] Dispensed 0.000024 mL of Acid (0.5 M HCl)
[1:01:09] Stepping pH = 3.98
[1:01:24] Stirrer speed set to 0
[1:01:34] Datapoint id 66 collected
[1:01:34] Charge balance equation is out by -32.8%
[1:01:34] Stirrer speed set to 55
[1:01:39] pH 3.97 -> 3.77
[1:01:39] Using cautious pH adjust
[1:01:39] Dispensed 0.000141 mL of Acid (0.5 M HCl)
[1:01:44] Stepping pH = 3.87
[1:01:44] Dispensed 0.000094 mL of Acid (0.5 M HCl)
[1:01:49] Stepping pH = 3.81
[1:01:49] Dispensed 0.000047 mL of Acid (0.5 M HCl)
[1:01:54] Stepping pH = 3.78
[1:01:54] Dispensed 0.000024 mL of Acid (0.5 M HCl)
[1:01:59] Stepping pH = 3.78
[1:02:14] Stirrer speed set to 0
[1:02:25] Datapoint id 67 collected
[1:02:25] Charge balance equation is out by -15.5%
[1:02:25] Stirrer speed set to 55
[1:02:30] pH 3.77 -> 3.57
[1:02:30] Using cautious pH adjust
[1:02:30] Dispensed 0.000212 mL of Acid (0.5 M HCl)
[1:02:35] Stepping pH = 3.66
[1:02:35] Dispensed 0.000141 mL of Acid (0.5 M HCl)

Sample name: **M08_octanol** Experiment start time: **3/2/2018 5:10:52 PM**
Assay name: **pH-metric high logP** Analyst: **Pion**
Assay ID: **18C-02007** Instrument ID: **T312060**
Filename: **C:\Sirius_T3\Mehtap\20180302_exp29_logP_T3-2\18C-02007_M08_octanol_pH-metric high logP.t3r**

#### Experiment Log (continued)

[1:02:40] Stepping pH = 3.61
[1:02:41] Dispensed 0.000094 mL of Acid (0.5 M HCl)
[1:02:46] Stepping pH = 3.58
[1:03:01] Stirrer speed set to 0
[1:03:11] Datapoint id 68 collected
[1:03:11] Charge balance equation is out by -6.3%
[1:03:11] Stirrer speed set to 55
[1:03:16] pH 3.57 -> 3.37
[1:03:16] Using charge balance adjust
[1:03:16] Dispensed 0.000659 mL of Acid (0.5 M HCl)
[1:03:36] Stirrer speed set to 0
[1:03:46] Datapoint id 69 collected
[1:03:46] Charge balance equation is out by -2.7%
[1:03:46] Stirrer speed set to 55
[1:03:51] pH 3.37 -> 3.17
[1:03:51] Using charge balance adjust
[1:03:51] Dispensed 0.001035 mL of Acid (0.5 M HCl)
[1:04:12] Stirrer speed set to 0
[1:04:22] Datapoint id 70 collected
[1:04:22] Charge balance equation is out by -8.0%
[1:04:22] Stirrer speed set to 55
[1:04:27] pH 3.19 -> 2.99
[1:04:27] Using charge balance adjust
[1:04:27] Dispensed 0.001576 mL of Acid (0.5 M HCl)
[1:04:47] Stirrer speed set to 0
[1:04:57] Datapoint id 71 collected
[1:04:57] Charge balance equation is out by -5.6%
[1:04:57] Stirrer speed set to 55
[1:05:02] pH 3.01 -> 2.81
[1:05:02] Using charge balance adjust
[1:05:02] Dispensed 0.002446 mL of Acid (0.5 M HCl)
[1:05:23] Stirrer speed set to 0
[1:05:33] Datapoint id 72 collected
[1:05:33] Charge balance equation is out by -1.4%
[1:05:33] Stirrer speed set to 55
[1:05:38] pH 2.81 -> 2.61
[1:05:38] Using charge balance adjust
[1:05:38] Dispensed 0.003833 mL of Acid (0.5 M HCl)
[1:05:58] Stirrer speed set to 0
[1:06:08] Datapoint id 73 collected
[1:06:08] Charge balance equation is out by -0.1%
[1:06:08] Stirrer speed set to 55
[1:06:13] pH 2.61 -> 2.41
[1:06:13] Using charge balance adjust
[1:06:13] Dispensed 0.006115 mL of Acid (0.5 M HCl)
[1:06:34] Stirrer speed set to 0
[1:06:44] Datapoint id 74 collected
[1:06:44] Charge balance equation is out by 5.7%
[1:06:44] Stirrer speed set to 55
[1:06:49] pH 2.41 -> 2.21
[1:06:49] Using charge balance adjust
[1:06:50] Dispensed 0.010042 mL of Acid (0.5 M HCl)
[1:07:10] Stirrer speed set to 0
[1:07:20] Datapoint id 75 collected
[1:07:20] Charge balance equation is out by -0.1%
[1:07:20] Stirrer speed set to 55
[1:07:25] pH 2.21 -> 2.01
[1:07:25] Using charge balance adjust
[1:07:26] Dispensed 0.016181 mL of Acid (0.5 M HCl)

| | | | |
|---|---|---|---|
| Sample name: | **M08_octanol** | Experiment start time: | **3/2/2018 5:10:52 PM** |
| Assay name: | **pH-metric high logP** | Analyst: | **Pion** |
| Assay ID: | **18C-02007** | Instrument ID: | **T312060** |
| Filename: | **C:\Sirius_T3\Mehtap\20180302_exp29_logP_T3-2\18C-02007_M08_octanol_pH-metric high logP.t3r** | | |

#### Experiment Log (continued)

[1:07:46] Stirrer speed set to 0
[1:07:56] Datapoint id 76 collected
[1:07:56] Charge balance equation is out by 1.9%
[1:07:56] Stirrer speed set to 55
[1:08:01] pH 2.01 -> 1.95
[1:08:01] Using charge balance adjust
[1:08:02] Dispensed 0.006491 mL of Acid (0.5 M HCl)
[1:08:22] Stirrer speed set to 0
[1:08:32] Datapoint id 77 collected
[1:08:32] Charge balance equation is out by -70.0%
[1:08:32] Titration 3 of 3
[1:08:32] Adding initial titrants
[1:08:32] Automatically add 0.80000 mL of Octanol
[1:09:23] Dispensed 0.800000 mL of Octanol
[1:09:23] Stirrer speed set to 10
[1:09:24] Stirrer speed set to 60
[1:09:24] Iterative adjust 1.95 -> 10.00
[1:09:24] pH 1.95 -> 10.00
[1:09:25] Dispensed 0.059925 mL of Base (0.5 M KOH)
[1:10:15] Stirrer speed set to 0
[1:10:34] Datapoint id 78 collected
[1:10:34] Stirrer speed set to 60
[1:10:40] pH 10.30 -> 10.10
[1:10:40] Using cautious pH adjust
[1:10:40] Dispensed 0.000188 mL of Acid (0.5 M HCl)
[1:10:45] Stepping pH = 10.28
[1:10:45] Dispensed 0.000541 mL of Acid (0.5 M HCl)
[1:10:50] Stepping pH = 9.94
[1:11:05] Stirrer speed set to 0
[1:11:34] Datapoint id 79 collected
[1:11:34] Charge balance equation is out by -81.9%
[1:11:34] Stirrer speed set to 60
[1:11:39] pH 9.90 -> 9.70
[1:11:39] Using cautious pH adjust
[1:11:39] Dispensed 0.000094 mL of Acid (0.5 M HCl)
[1:11:44] Stepping pH = 9.90
[1:11:44] Dispensed 0.000259 mL of Acid (0.5 M HCl)
[1:11:50] Stepping pH = 9.63
[1:12:05] Stirrer speed set to 0
[1:12:15] Datapoint id 80 collected
[1:12:15] Charge balance equation is out by -91.9%
[1:12:15] Stirrer speed set to 60
[1:12:20] pH 9.60 -> 9.40
[1:12:20] Using cautious pH adjust
[1:12:20] Dispensed 0.000047 mL of Acid (0.5 M HCl)
[1:12:25] Stepping pH = 9.60
[1:12:25] Dispensed 0.000165 mL of Acid (0.5 M HCl)
[1:12:30] Stepping pH = 9.43
[1:12:30] Dispensed 0.000024 mL of Acid (0.5 M HCl)
[1:12:35] Stepping pH = 9.42
[1:12:35] Dispensed 0.000024 mL of Acid (0.5 M HCl)
[1:12:40] Stepping pH = 9.40
[1:12:55] Stirrer speed set to 0
[1:13:05] Datapoint id 81 collected
[1:13:05] Charge balance equation is out by -132.7%
[1:13:05] Stirrer speed set to 60
[1:13:11] pH 9.37 -> 9.17
[1:13:11] Using cautious pH adjust
[1:13:11] Dispensed 0.000047 mL of Acid (0.5 M HCl)

| | | | |
|---|---|---|---|
| Sample name: | **M08_octanol** | Experiment start time: | **3/2/2018 5:10:52 PM** |
| Assay name: | **pH-metric high logP** | Analyst: | **Pion** |
| Assay ID: | **18C-02007** | Instrument ID: | **T312060** |
| Filename: | **C:\Sirius_T3\Mehtap\20180302_exp29_logP_T3-2\18C-02007_M08_octanol_pH-metric high logP.t3r** | | |

#### Experiment Log (continued)

[1:13:16] Stepping pH = 9.36
[1:13:16] Dispensed 0.000118 mL of Acid (0.5 M HCl)
[1:13:21] Stepping pH = 9.14
[1:13:36] Stirrer speed set to 0
[1:13:47] Datapoint id 82 collected
[1:13:47] Charge balance equation is out by -88.9%
[1:13:47] Stirrer speed set to 60
[1:13:53] pH 9.10 -> 8.90
[1:13:53] Using cautious pH adjust
[1:13:53] Dispensed 0.000047 mL of Acid (0.5 M HCl)
[1:13:58] Stepping pH = 9.08
[1:13:58] Dispensed 0.000118 mL of Acid (0.5 M HCl)
[1:14:03] Stepping pH = 8.86
[1:14:18] Stirrer speed set to 0
[1:14:33] Datapoint id 83 collected
[1:14:33] Charge balance equation is out by -89.5%
[1:14:33] Stirrer speed set to 60
[1:14:38] pH 8.81 -> 8.61
[1:14:38] Using cautious pH adjust
[1:14:38] Dispensed 0.000047 mL of Acid (0.5 M HCl)
[1:14:43] Stepping pH = 8.80
[1:14:43] Dispensed 0.000165 mL of Acid (0.5 M HCl)
[1:14:48] Stepping pH = 8.51
[1:15:04] Stirrer speed set to 0
[1:15:26] Datapoint id 84 collected
[1:15:26] Charge balance equation is out by -94.1%
[1:15:26] Stirrer speed set to 60
[1:15:31] pH 8.48 -> 8.28
[1:15:31] Using cautious pH adjust
[1:15:31] Dispensed 0.000094 mL of Acid (0.5 M HCl)
[1:15:36] Stepping pH = 8.44
[1:15:36] Dispensed 0.000188 mL of Acid (0.5 M HCl)
[1:15:41] Stepping pH = 8.22
[1:15:56] Stirrer speed set to 0
[1:16:20] Datapoint id 85 collected
[1:16:20] Charge balance equation is out by -44.0%
[1:16:20] Stirrer speed set to 60
[1:16:25] pH 8.19 -> 7.99
[1:16:25] Using cautious pH adjust
[1:16:25] Dispensed 0.000165 mL of Acid (0.5 M HCl)
[1:16:30] Stepping pH = 8.11
[1:16:30] Dispensed 0.000165 mL of Acid (0.5 M HCl)
[1:16:35] Stepping pH = 8.01
[1:16:35] Dispensed 0.000024 mL of Acid (0.5 M HCl)
[1:16:40] Stepping pH = 8.00
[1:16:40] Dispensed 0.000047 mL of Acid (0.5 M HCl)
[1:16:45] Stepping pH = 7.99
[1:17:00] Stirrer speed set to 0
[1:17:23] Datapoint id 86 collected
[1:17:23] Charge balance equation is out by -19.6%
[1:17:23] Stirrer speed set to 60
[1:17:28] pH 7.97 -> 7.77
[1:17:28] Using cautious pH adjust
[1:17:28] Dispensed 0.000235 mL of Acid (0.5 M HCl)
[1:17:33] Stepping pH = 7.85
[1:17:33] Dispensed 0.000141 mL of Acid (0.5 M HCl)
[1:17:38] Stepping pH = 7.79
[1:17:38] Dispensed 0.000047 mL of Acid (0.5 M HCl)
[1:17:43] Stepping pH = 7.79

| | | | |
|---|---|---|---|
| Sample name: | **M08_octanol** | Experiment start time: | **3/2/2018 5:10:52 PM** |
| Assay name: | **pH-metric high logP** | Analyst: | **Pion** |
| Assay ID: | **18C-02007** | Instrument ID: | **T312060** |
| Filename: | **C:\Sirius_T3\Mehtap\20180302_exp29_logP_T3-2\18C-02007_M08_octanol_pH-metric high logP.t3r** | | |

#### Experiment Log (continued)

[1:17:43] Dispensed 0.000047 mL of Acid (0.5 M HCl)
[1:17:48] Stepping pH = 7.78
[1:18:03] Stirrer speed set to 0
[1:18:26] Datapoint id 87 collected
[1:18:26] Charge balance equation is out by 3.4%
[1:18:26] Stirrer speed set to 60
[1:18:31] pH 7.76 -> 7.56
[1:18:31] Using charge balance adjust
[1:18:31] Dispensed 0.000659 mL of Acid (0.5 M HCl)
[1:18:51] Stirrer speed set to 0
[1:19:12] Datapoint id 88 collected
[1:19:12] Charge balance equation is out by 13.2%
[1:19:12] Stirrer speed set to 60
[1:19:17] pH 7.52 -> 7.32
[1:19:17] Using charge balance adjust
[1:19:17] Dispensed 0.000870 mL of Acid (0.5 M HCl)
[1:19:37] Stirrer speed set to 0
[1:20:00] Datapoint id 89 collected
[1:20:00] Charge balance equation is out by 2.1%
[1:20:00] Stirrer speed set to 60
[1:20:05] pH 7.30 -> 7.10
[1:20:05] Using charge balance adjust
[1:20:05] Dispensed 0.000988 mL of Acid (0.5 M HCl)
[1:20:26] Stirrer speed set to 0
[1:20:50] Datapoint id 90 collected
[1:20:50] Charge balance equation is out by -5.8%
[1:20:50] Stirrer speed set to 60
[1:20:55] pH 7.09 -> 6.89
[1:20:55] Using charge balance adjust
[1:20:55] Dispensed 0.001011 mL of Acid (0.5 M HCl)
[1:21:15] Stirrer speed set to 0
[1:21:38] Datapoint id 91 collected
[1:21:38] Charge balance equation is out by -9.0%
[1:21:38] Stirrer speed set to 60
[1:21:43] pH 6.88 -> 6.68
[1:21:43] Using charge balance adjust
[1:21:43] Dispensed 0.000917 mL of Acid (0.5 M HCl)
[1:22:04] Stirrer speed set to 0
[1:22:33] Datapoint id 92 collected
[1:22:33] Charge balance equation is out by -16.1%
[1:22:33] Stirrer speed set to 60
[1:22:38] pH 6.67 -> 6.47
[1:22:38] Using cautious pH adjust
[1:22:38] Dispensed 0.000376 mL of Acid (0.5 M HCl)
[1:22:43] Stepping pH = 6.57
[1:22:43] Dispensed 0.000259 mL of Acid (0.5 M HCl)
[1:22:48] Stepping pH = 6.50
[1:22:48] Dispensed 0.000094 mL of Acid (0.5 M HCl)
[1:22:53] Stepping pH = 6.48
[1:22:53] Dispensed 0.000047 mL of Acid (0.5 M HCl)
[1:22:58] Stepping pH = 6.47
[1:23:13] Stirrer speed set to 0
[1:23:43] Datapoint id 93 collected
[1:23:43] Charge balance equation is out by -7.8%
[1:23:43] Stirrer speed set to 60
[1:23:48] pH 6.47 -> 6.27
[1:23:48] Using charge balance adjust
[1:23:48] Dispensed 0.000564 mL of Acid (0.5 M HCl)
[1:24:08] Stirrer speed set to 0

| | | | |
|---|---|---|---|
| Sample name: | **M08_octanol** | Experiment start time: | **3/2/2018 5:10:52 PM** |
| Assay name: | **pH-metric high logP** | Analyst: | **Pion** |
| Assay ID: | **18C-02007** | Instrument ID: | **T312060** |
| Filename: | **C:\Sirius_T3\Mehtap\20180302_exp29_logP_T3-2\18C-02007_M08_octanol_pH-metric high logP.t3r** | | |

#### Experiment Log (continued)

[1:24:40] Datapoint id 94 collected
[1:24:40] Charge balance equation is out by -25.7%
[1:24:40] Stirrer speed set to 60
[1:24:45] pH 6.26 -> 6.06
[1:24:45] Using cautious pH adjust
[1:24:45] Dispensed 0.000212 mL of Acid (0.5 M HCl)
[1:24:50] Stepping pH = 6.18
[1:24:50] Dispensed 0.000165 mL of Acid (0.5 M HCl)
[1:24:55] Stepping pH = 6.10
[1:24:55] Dispensed 0.000071 mL of Acid (0.5 M HCl)
[1:25:01] Stepping pH = 6.07
[1:25:16] Stirrer speed set to 0
[1:25:47] Datapoint id 95 collected
[1:25:47] Charge balance equation is out by -6.6%
[1:25:47] Stirrer speed set to 60
[1:25:52] pH 6.08 -> 5.88
[1:25:52] Using charge balance adjust
[1:25:52] Dispensed 0.000282 mL of Acid (0.5 M HCl)
[1:26:13] Stirrer speed set to 0
[1:26:32] Datapoint id 96 collected
[1:26:32] Charge balance equation is out by -37.9%
[1:26:32] Stirrer speed set to 60
[1:26:37] pH 5.89 -> 5.69
[1:26:37] Using cautious pH adjust
[1:26:37] Dispensed 0.000094 mL of Acid (0.5 M HCl)
[1:26:42] Stepping pH = 5.84
[1:26:42] Dispensed 0.000141 mL of Acid (0.5 M HCl)
[1:26:47] Stepping pH = 5.71
[1:26:47] Dispensed 0.000024 mL of Acid (0.5 M HCl)
[1:26:52] Stepping pH = 5.70
[1:27:07] Stirrer speed set to 0
[1:27:26] Datapoint id 97 collected
[1:27:26] Charge balance equation is out by -33.9%
[1:27:26] Stirrer speed set to 60
[1:27:31] pH 5.69 -> 5.49
[1:27:31] Using cautious pH adjust
[1:27:31] Dispensed 0.000071 mL of Acid (0.5 M HCl)
[1:27:36] Stepping pH = 5.64
[1:27:36] Dispensed 0.000118 mL of Acid (0.5 M HCl)
[1:27:41] Stepping pH = 5.45
[1:27:56] Stirrer speed set to 0
[1:28:14] Datapoint id 98 collected
[1:28:14] Charge balance equation is out by -30.0%
[1:28:14] Stirrer speed set to 60
[1:28:19] pH 5.45 -> 5.25
[1:28:19] Using cautious pH adjust
[1:28:19] Dispensed 0.000047 mL of Acid (0.5 M HCl)
[1:28:24] Stepping pH = 5.42
[1:28:24] Dispensed 0.000094 mL of Acid (0.5 M HCl)
[1:28:29] Stepping pH = 5.22
[1:28:44] Stirrer speed set to 0
[1:29:01] Datapoint id 99 collected
[1:29:01] Charge balance equation is out by -55.8%
[1:29:01] Stirrer speed set to 60
[1:29:07] pH 5.20 -> 5.00
[1:29:07] Using cautious pH adjust
[1:29:07] Dispensed 0.000024 mL of Acid (0.5 M HCl)
[1:29:12] Stepping pH = 5.19
[1:29:12] Dispensed 0.000094 mL of Acid (0.5 M HCl)

Sample name: **M08_octanol**
Assay name: **pH-metric high logP**
Assay ID: **18C-02007**
Filename: **C:\Sirius_T3\Mehtap\20180302_exp29_logP_T3-2\18C-02007_M08_octanol_pH-metric high logP.t3r**
Experiment start time: **3/2/2018 5:10:52 PM**
Analyst: **Pion**
Instrument ID: **T312060**

#### Experiment Log (continued)

[1:29:17] Stepping pH = 4.93
[1:29:32] Stirrer speed set to 0
[1:29:54] Datapoint id 100 collected
[1:29:54] Charge balance equation is out by -94.2%
[1:29:54] Stirrer speed set to 60
[1:29:59] pH 4.89 -> 4.69
[1:29:59] Using cautious pH adjust
[1:29:59] Dispensed 0.000024 mL of Acid (0.5 M HCl)
[1:30:04] Stepping pH = 4.89
[1:30:04] Dispensed 0.000071 mL of Acid (0.5 M HCl)
[1:30:09] Stepping pH = 4.71
[1:30:09] Dispensed 0.000024 mL of Acid (0.5 M HCl)
[1:30:14] Stepping pH = 4.65
[1:30:30] Stirrer speed set to 0
[1:30:47] Datapoint id 101 collected
[1:30:47] Charge balance equation is out by -132.2%
[1:30:47] Stirrer speed set to 60
[1:30:52] pH 4.63 -> 4.43
[1:30:52] Using cautious pH adjust
[1:30:52] Dispensed 0.000047 mL of Acid (0.5 M HCl)
[1:30:57] Stepping pH = 4.56
[1:30:57] Dispensed 0.000047 mL of Acid (0.5 M HCl)
[1:31:03] Stepping pH = 4.46
[1:31:03] Dispensed 0.000024 mL of Acid (0.5 M HCl)
[1:31:08] Stepping pH = 4.42
[1:31:23] Stirrer speed set to 0
[1:31:34] Datapoint id 102 collected
[1:31:34] Charge balance equation is out by -38.2%
[1:31:34] Stirrer speed set to 60
[1:31:39] pH 4.41 -> 4.21
[1:31:39] Using cautious pH adjust
[1:31:39] Dispensed 0.000047 mL of Acid (0.5 M HCl)
[1:31:44] Stepping pH = 4.38
[1:31:44] Dispensed 0.000118 mL of Acid (0.5 M HCl)
[1:31:49] Stepping pH = 4.16
[1:32:04] Stirrer speed set to 0
[1:32:14] Datapoint id 103 collected
[1:32:14] Charge balance equation is out by -54.7%
[1:32:14] Stirrer speed set to 60
[1:32:19] pH 4.15 -> 3.95
[1:32:19] Using cautious pH adjust
[1:32:20] Dispensed 0.000094 mL of Acid (0.5 M HCl)
[1:32:25] Stepping pH = 4.07
[1:32:25] Dispensed 0.000094 mL of Acid (0.5 M HCl)
[1:32:30] Stepping pH = 3.98
[1:32:30] Dispensed 0.000024 mL of Acid (0.5 M HCl)
[1:32:35] Stepping pH = 3.96
[1:32:50] Stirrer speed set to 0
[1:33:00] Datapoint id 104 collected
[1:33:00] Charge balance equation is out by -11.2%
[1:33:00] Stirrer speed set to 60
[1:33:05] pH 3.95 -> 3.75
[1:33:05] Using charge balance adjust
[1:33:05] Dispensed 0.000306 mL of Acid (0.5 M HCl)
[1:33:25] Stirrer speed set to 0
[1:33:44] Datapoint id 105 collected
[1:33:44] Charge balance equation is out by 5.1%
[1:33:44] Stirrer speed set to 60
[1:33:49] pH 3.74 -> 3.54

#### Experiment Log (continued)

[1:33:49] Using charge balance adjust
[1:33:49] Dispensed 0.000470 mL of Acid (0.5 M HCl)
[1:34:09] Stirrer speed set to 0
[1:34:19] Datapoint id 106 collected
[1:34:19] Charge balance equation is out by -1.6%
[1:34:19] Stirrer speed set to 60
[1:34:24] pH 3.54 -> 3.34
[1:34:24] Using charge balance adjust
[1:34:25] Dispensed 0.000753 mL of Acid (0.5 M HCl)
[1:34:45] Stirrer speed set to 0
[1:34:55] Datapoint id 107 collected
[1:34:55] Charge balance equation is out by 2.6%
[1:34:55] Stirrer speed set to 60
[1:35:00] pH 3.34 -> 3.14
[1:35:00] Using charge balance adjust
[1:35:00] Dispensed 0.001199 mL of Acid (0.5 M HCl)
[1:35:20] Stirrer speed set to 0
[1:35:30] Datapoint id 108 collected
[1:35:30] Charge balance equation is out by 2.0%
[1:35:30] Stirrer speed set to 60
[1:35:35] pH 3.14 -> 2.94
[1:35:35] Using charge balance adjust
[1:35:36] Dispensed 0.001905 mL of Acid (0.5 M HCl)
[1:35:56] Stirrer speed set to 0
[1:36:06] Datapoint id 109 collected
[1:36:06] Charge balance equation is out by -1.1%
[1:36:06] Stirrer speed set to 60
[1:36:11] pH 2.95 -> 2.75
[1:36:11] Using charge balance adjust
[1:36:11] Dispensed 0.002987 mL of Acid (0.5 M HCl)
[1:36:31] Stirrer speed set to 0
[1:36:56] Datapoint id 110 collected
[1:36:56] Charge balance equation is out by -5.0%
[1:36:56] Stirrer speed set to 60
[1:37:01] pH 2.76 -> 2.56
[1:37:01] Using charge balance adjust
[1:37:01] Dispensed 0.004610 mL of Acid (0.5 M HCl)
[1:37:21] Stirrer speed set to 0
[1:37:47] Datapoint id 111 collected
[1:37:47] Charge balance equation is out by -2.3%
[1:37:47] Stirrer speed set to 60
[1:37:52] pH 2.57 -> 2.37
[1:37:52] Using charge balance adjust
[1:37:52] Dispensed 0.007244 mL of Acid (0.5 M HCl)
[1:38:13] Stirrer speed set to 0
[1:38:25] Datapoint id 112 collected
[1:38:25] Charge balance equation is out by 1.1%
[1:38:25] Stirrer speed set to 60
[1:38:30] pH 2.38 -> 2.18
[1:38:30] Using charge balance adjust
[1:38:30] Dispensed 0.011571 mL of Acid (0.5 M HCl)
[1:38:50] Stirrer speed set to 0
[1:39:06] Datapoint id 113 collected
[1:39:06] Charge balance equation is out by -0.0%
[1:39:06] Stirrer speed set to 60
[1:39:11] pH 2.18 -> 1.98
[1:39:11] Using charge balance adjust
[1:39:11] Dispensed 0.018650 mL of Acid (0.5 M HCl)
[1:39:32] Stirrer speed set to 0

| | | | |
|---|---|---|---|
| Sample name: | **M08_octanol** | Experiment start time: | **3/2/2018 5:10:52 PM** |
| Assay name: | **pH-metric high logP** | Analyst: | **Pion** |
| Assay ID: | **18C-02007** | Instrument ID: | **T312060** |
| Filename: | **C:\Sirius_T3\Mehtap\20180302_exp29_logP_T3-2\18C-02007_M08_octanol_pH-metric high logP.t3r** | | |

#### Experiment Log (continued)

[1:39:43] Datapoint id 114 collected
[1:39:43] Charge balance equation is out by 0.2%
[1:39:43] Argon flow rate set to 0
[1:39:47] Titrator arm moved over Titration position
