## Supplementary material for "Octanol-water partition coefficient measurements for the SAMPL6 Blind Prediction Challenge": SM08_18C-02008_M08_octanol_pH-metric high logP_report.pdf

| | | | |
|---|---|---|---|
| Sample name: | **M08_octanol** | Experiment start time: | **3/2/2018 6:51:30 PM** |
| Assay name: | **pH-metric high logP** | Analyst: | **Pion** |
| Assay ID: | **18C-02008** | Instrument ID: | **T312060** |
| Filename: | **C:\Sirius_T3\Mehtap\20180302_exp29_logP_T3-2\18C-02008_M08_octanol_pH-metric high logP.t3r** | | |

#### pH-metric Result

| | | |
|---|---|---|
| logP (neutral XH) | 3.08 | ±0.01 (n=50) |
| logP (X -) | -0.07 | ±0.05 (n=50) |

#### 18C-02008 Points 2 to 33

| | |
|---|---|
| M08_octanol concentration factor | 1.008 |
| Carbonate | 0.0000 mM |
| Acidity error | -0.21439 mM |

#### 18C-02008 Points 34 to 67

| | |
|---|---|
| M08_octanol concentration factor | 1.106 |
| Carbonate | 0.0000 mM |
| Acidity error | -0.33976 mM |

#### 18C-02008 Points 68 to 106

| | |
|---|---|
| M08_octanol concentration factor | 0.984 |
| Carbonate | 0.2479 mM |
| Acidity error | -0.08487 mM |

#### Warnings and errors

| | |
|---|---|
| Errors | None |
| Warnings | None |

#### Sample logD and percent species

| pH | M08_octanol logD | M08_octanol M08_octanolH | M08_octanol M08_octanol | M08_octanol M08_octanolH* | M08_octanol M08_octanol* | Comment |
|---|---|---|---|---|---|---|
| 1.000 | 3.08 | 0.08 % | 0.00 % | 99.92 % | 0.00 % | |
| 1.200 | 3.08 | 0.08 % | 0.00 % | 99.92 % | 0.00 % | Stomach pH |
| 2.000 | 3.08 | 0.08 % | 0.00 % | 99.92 % | 0.00 % | |
| 3.000 | 3.06 | 0.08 % | 0.00 % | 99.91 % | 0.00 % | |
| 4.000 | 2.88 | 0.08 % | 0.05 % | 99.82 % | 0.04 % | |
| 5.000 | 2.24 | 0.08 % | 0.50 % | 99.00 % | 0.42 % | |
| 6.000 | 1.31 | 0.08 % | 4.58 % | 91.44 % | 3.91 % | |
| 6.500 | 0.85 | 0.06 % | 12.23 % | 77.26 % | 10.45 % | |
| 7.000 | 0.45 | 0.04 % | 25.94 % | 51.84 % | 22.18 % | |
| 7.400 | 0.22 | 0.02 % | 37.72 % | 30.01 % | 32.25 % | Blood pH |
| 8.000 | 0.02 | 0.01 % | 48.66 % | 9.72 % | 41.61 % | |
| 9.000 | -0.06 | 0.00 % | 53.33 % | 1.07 % | 45.60 % | |
| 10.000 | -0.07 | 0.00 % | 53.85 % | 0.11 % | 46.04 % | |
| 11.000 | -0.07 | 0.00 % | 53.90 % | 0.01 % | 46.09 % | |
| 12.000 | -0.07 | 0.00 % | 53.91 % | 0.00 % | 46.09 % | |

Sample name: **M08_octanol**
Assay name: **pH-metric high logP**
Assay ID: **18C-02008**
Filename: **C:\Sirius_T3\Mehtap\20180302_exp29_logP_T3-2\18C-02008_M08_octanol_pH-metric high logP.t3r**

Experiment start time: **3/2/2018 6:51:30 PM**
Analyst: **Pion**
Instrument ID: **T312060**

#### Graphs

Charge balance residuals

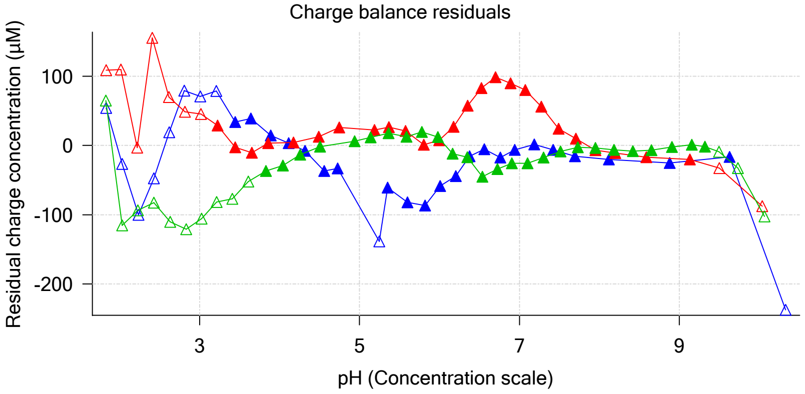


Ionic strength

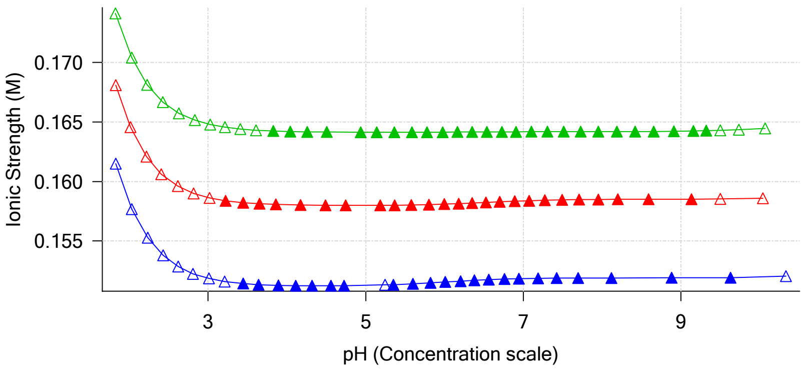


Buffer index profile

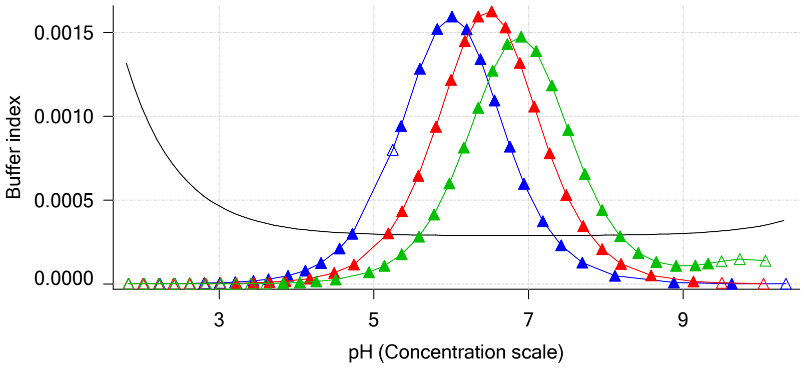


Sample M08_octanol Ionisation Graph

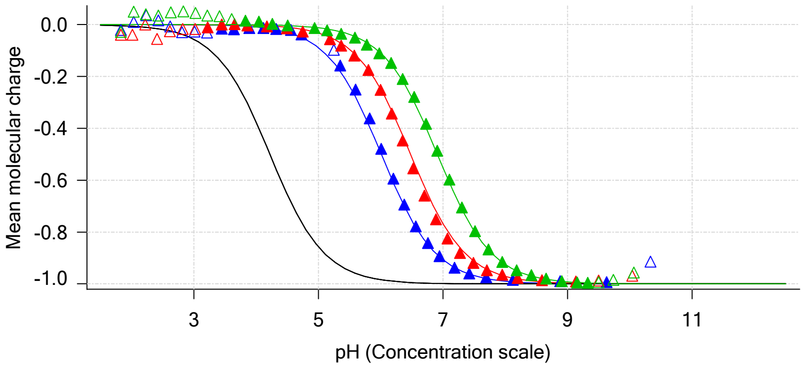


Distribution of species for sample M08_octanol

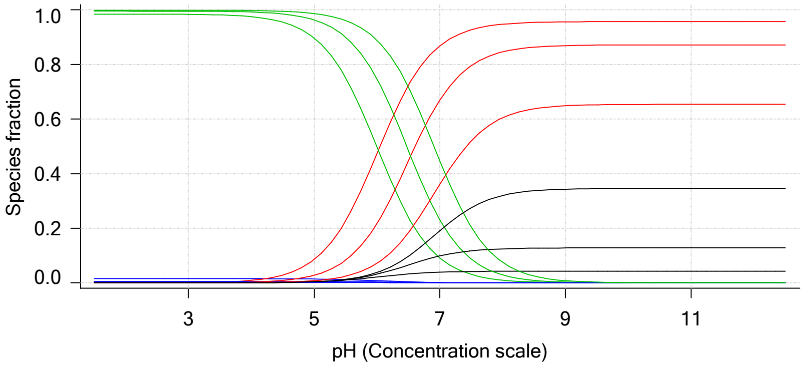


Lipophilicity Profile

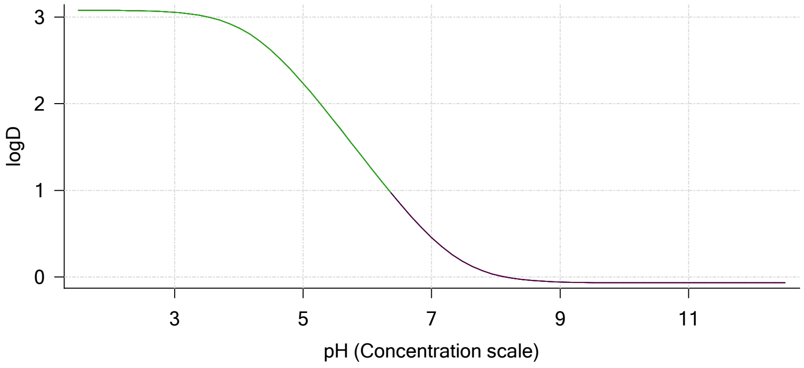


Log R Profile

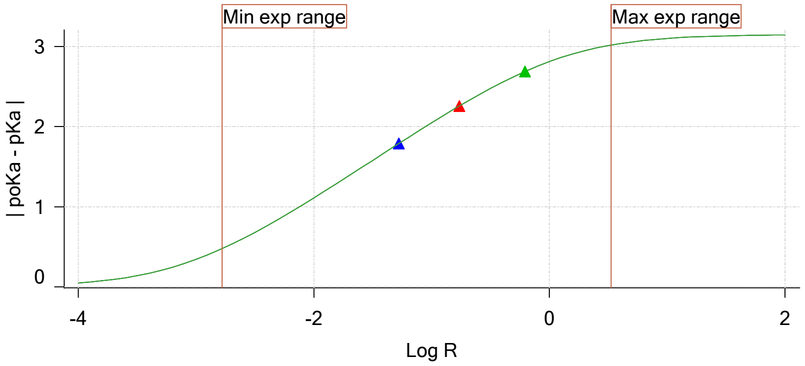


Sample Carbonate Ionisation Graph

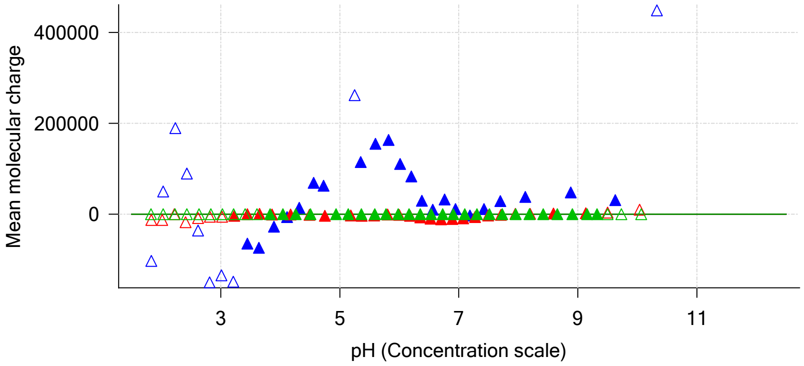

Sample name: **M08_octanol**
Assay name: **pH-metric high logP**
Assay ID: **18C-02008**
Filename: **C:\Sirius_T3\Mehtap\20180302_exp29_logP_T3-2\18C-02008_M08_octanol_pH-metric high logP.t3r**

Experiment start time: **3/2/2018 6:51:30 PM**
Analyst: **Pion**
Instrument ID: **T312060**

#### Graphs (continued)

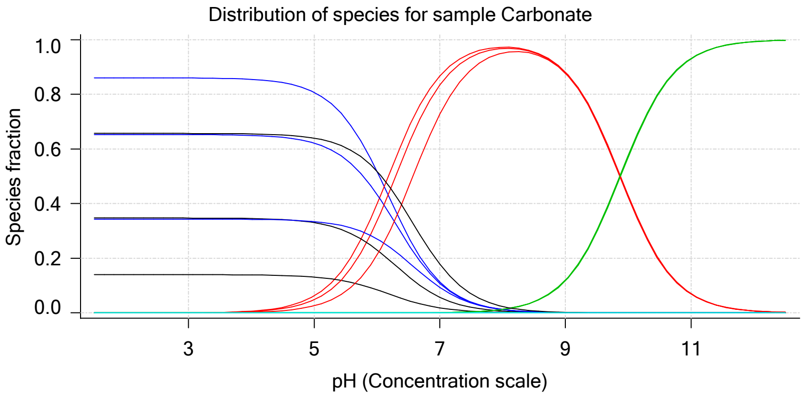

| | | | |
|---|---|---|---|
| Sample name: | **M08_octanol** | Experiment start time: | **3/2/2018 6:51:30 PM** |
| Assay name: | **pH-metric high logP** | Analyst: | **Pion** |
| Assay ID: | **18C-02008** | Instrument ID: | **T312060** |
| Filename: | **C:\Sirius_T3\Mehtap\20180302_exp29_logP_T3-2\18C-02008_M08_octanol_pH-metric high logP.t3r** | | |

### pH-metric high logP Titration 1 of 3 18C-02008 Points 2 to 33

#### Overall results

| | |
|---|---|
| RMSD | 0.161 |
| Average ionic strength | 0.152 M |
| Average temperature | 24.9°C |
| Partition ratio | 0.0526 : 1 |
| Analyte concentration range | 2531.4 µM to 2619.0 µM |
| Total points considered | 22 of 32 |

#### Warnings and errors

| | |
|---|---|
| Errors | None |
| Warnings | One or more logP values out of range |

#### Four-Plus parameters

| | | | |
|---|---|---|---|
| Alpha | 0.111 | 3/2/2018 6:51:30 PM | C:\Sirius_T3\HCl18C02.t3r |
| S | 0.9988 | 3/2/2018 6:51:30 PM | C:\Sirius_T3\HCl18C02.t3r |
| jH | 1.0 | 3/2/2018 6:51:30 PM | C:\Sirius_T3\HCl18C02.t3r |
| jOH | -0.8 | 3/2/2018 6:51:30 PM | C:\Sirius_T3\HCl18C02.t3r |

#### Titrants

| | | | |
|---|---|---|---|
| 0.50 M HCl | 0.999058 | 3/2/2018 6:51:30 PM | C:\Sirius_T3\HCl18C02.t3r |
| 0.50 M KOH | 0.999845 | 3/2/2018 6:51:30 PM | C:\Sirius_T3\KOH18B27.t3r |

#### Sample

| | |
|---|---|
| M08_octanol concentration factor | 1.008 |
| Acid pKa 1 | 4.22 |
| logP (neutral XH) | 3.10 |
| logP (X -) | -5.13 |

#### Sample graphs

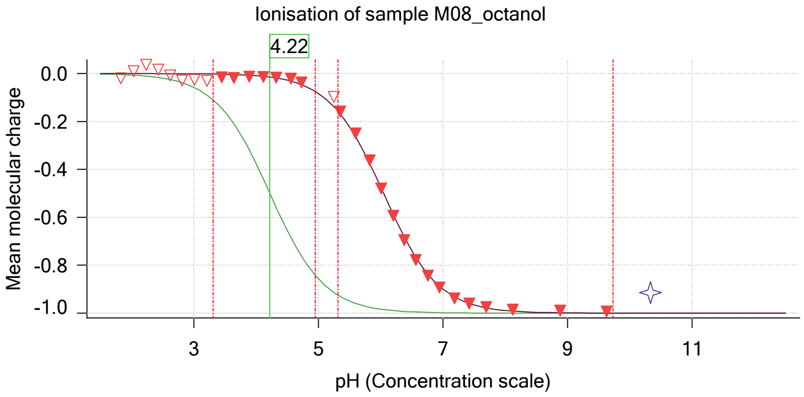


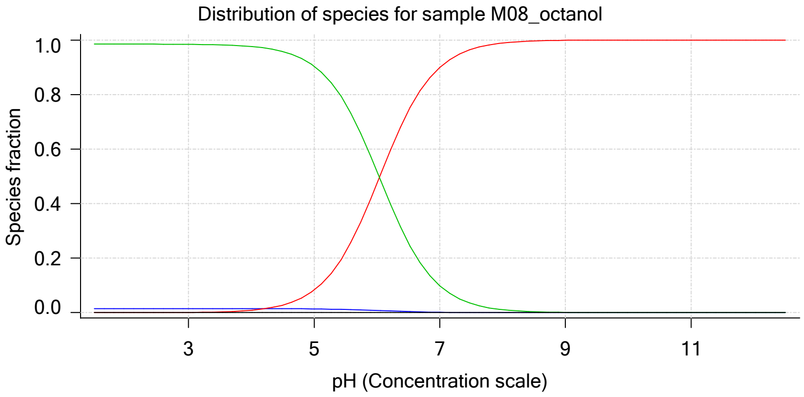

| | | | |
|---|---|---|---|
| Sample name: | **M08_octanol** | Experiment start time: | **3/2/2018 6:51:30 PM** |
| Assay name: | **pH-metric high logP** | Analyst: | **Pion** |
| Assay ID: | **18C-02008** | Instrument ID: | **T312060** |
| Filename: | **C:\Sirius_T3\Mehtap\20180302_exp29_logP_T3-2\18C-02008_M08_octanol_pH-metric high logP.t3r** | | |

#### Sample graphs (continued)

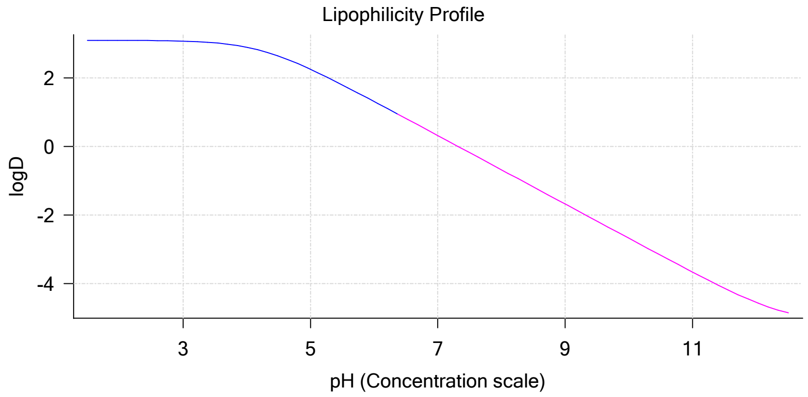


#### Sample logD and percent species

| pH | M08_octanol logD | M08_octanol M08_octanolH | M08_octanol M08_octanol | M08_octanol M08_octanolH* | M08_octanol M08_octanol* | Comment |
|---|---|---|---|---|---|---|
| 1.000 | 3.10 | 1.49 % | 0.00 % | 98.50 % | 0.00 % | |
| 1.200 | 3.10 | 1.49 % | 0.00 % | 98.50 % | 0.00 % | Stomach pH |
| 2.000 | 3.10 | 1.49 % | 0.01 % | 98.50 % | 0.00 % | |
| 3.000 | 3.07 | 1.49 % | 0.09 % | 98.42 % | 0.00 % | |
| 4.000 | 2.89 | 1.48 % | 0.89 % | 97.63 % | 0.00 % | |
| 5.000 | 2.25 | 1.37 % | 8.26 % | 90.37 % | 0.00 % | |
| 6.000 | 1.31 | 0.79 % | 47.38 % | 51.83 % | 0.00 % | |
| 6.500 | 0.82 | 0.39 % | 74.01 % | 25.60 % | 0.00 % | |
| 7.000 | 0.32 | 0.15 % | 90.01 % | 9.84 % | 0.00 % | |
| 7.400 | -0.08 | 0.06 % | 95.77 % | 4.17 % | 0.00 % | Blood pH |
| 8.000 | -0.68 | 0.02 % | 98.90 % | 1.08 % | 0.00 % | |
| 9.000 | -1.68 | 0.00 % | 99.89 % | 0.11 % | 0.00 % | |
| 10.000 | -2.68 | 0.00 % | 99.99 % | 0.01 % | 0.00 % | |
| 11.000 | -3.67 | 0.00 % | 100.00 % | 0.00 % | 0.00 % | |
| 12.000 | -4.55 | 0.00 % | 100.00 % | 0.00 % | 0.00 % | |

#### Carbonate and acidity

| | |
|---|---|
| Carbonate | 0.000 mM |
| Acidity error | -0.214 mM |

#### Other graphs

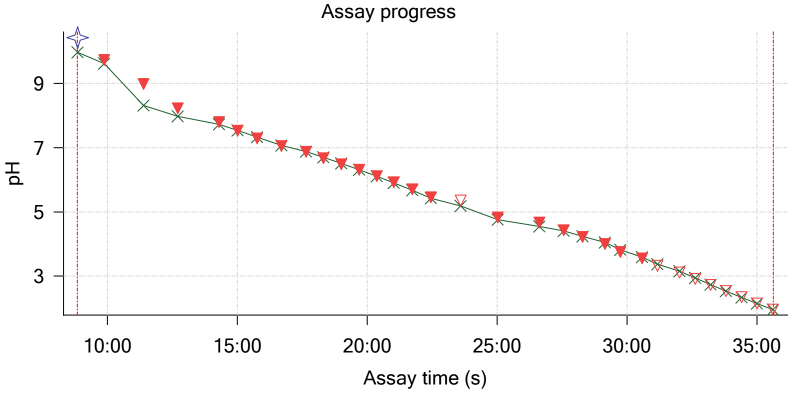


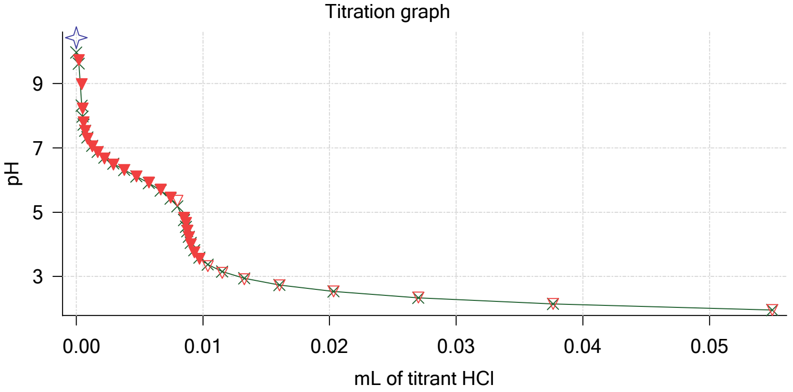

| | | | |
|---|---|---|---|
| Sample name: | **M08_octanol** | Experiment start time: | **3/2/2018 6:51:30 PM** |
| Assay name: | **pH-metric high logP** | Analyst: | **Pion** |
| Assay ID: | **18C-02008** | Instrument ID: | **T312060** |
| Filename: | **C:\Sirius_T3\Mehtap\20180302_exp29_logP_T3-2\18C-02008_M08_octanol_pH-metric high logP.t3r** | | |

#### Other graphs (continued)

Charge balance residuals

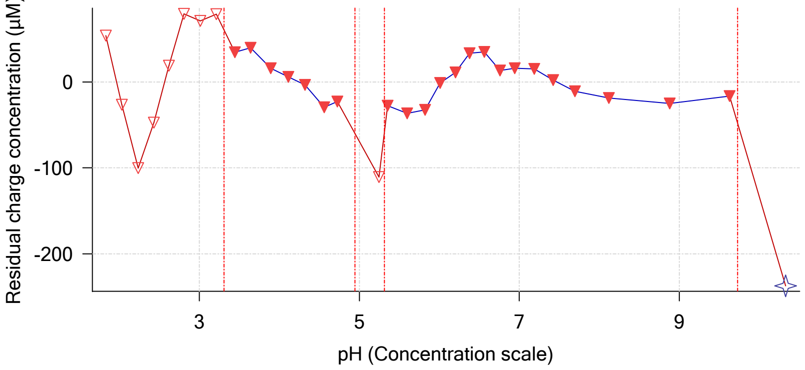


Buffer index profile

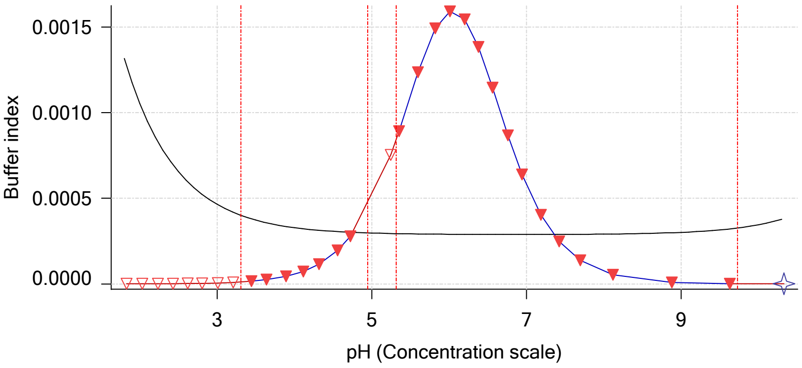


Ionic strength

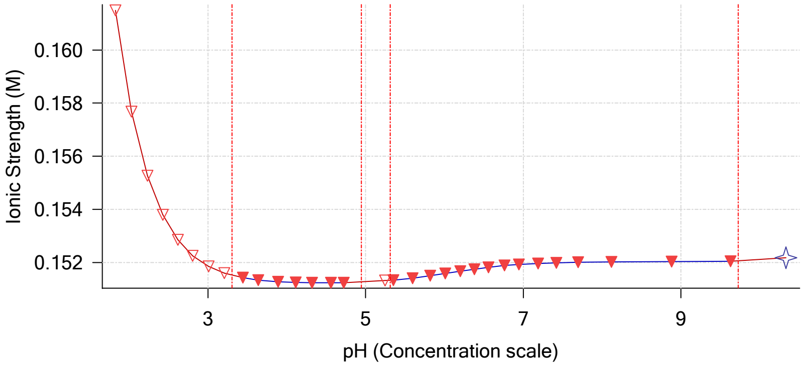


Log R Profile

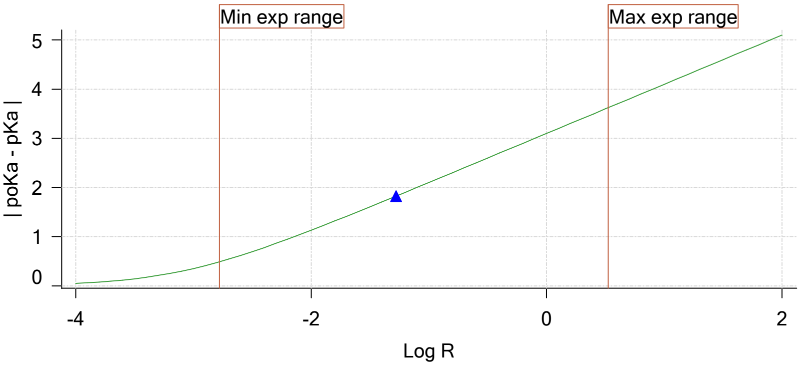


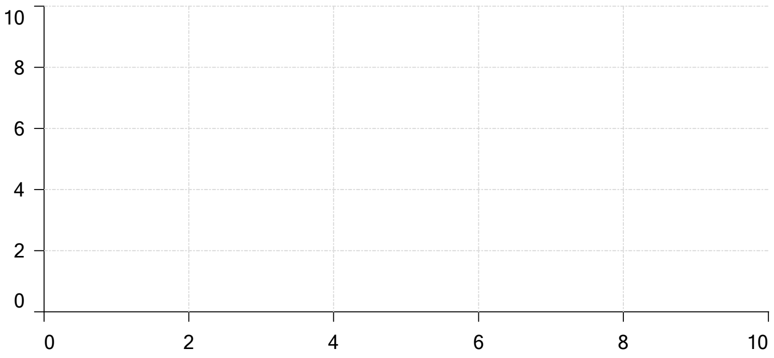


Ionisation of sample Carbonate

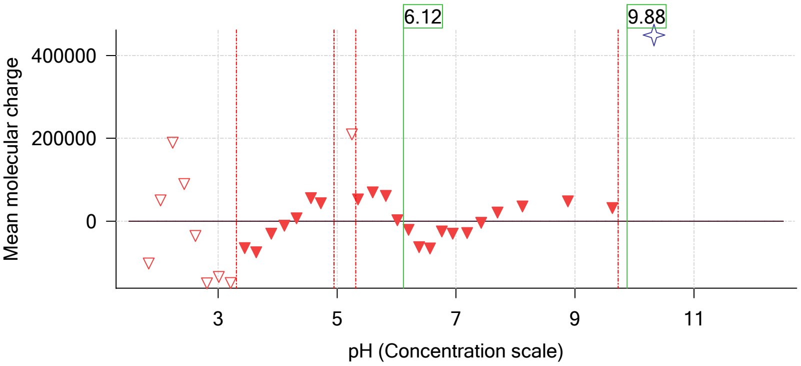

| | | | |
|---|---|---|---|
| Sample name: | **M08_octanol** | Experiment start time: | **3/2/2018 6:51:30 PM** |
| Assay name: | **pH-metric high logP** | Analyst: | **Pion** |
| Assay ID: | **18C-02008** | Instrument ID: | **T312060** |
| Filename: | **C:\Sirius_T3\Mehtap\20180302_exp29_logP_T3-2\18C-02008_M08_octanol_pH-metric high logP.t3r** | | |

# pH-metric high logP   Titration 2 of 3   18C-02008 Points 34 to 67

#### Overall results

| | |
|---|---|
| RMSD | 0.101 |
| Average ionic strength | 0.158 M |
| Average temperature | 25.0°C |
| Partition ratio | 0.1721 : 1 |
| Analyte concentration range | 2125.1 µM to 2191.1 µM |
| Total points considered | 25 of 34 |

#### Warnings and errors

| | |
|---|---|
| Errors | None |
| Warnings | One or more logP values out of range |

#### Four-Plus parameters

| | | | |
|---|---|---|---|
| Alpha | 0.111 | 3/2/2018 6:51:30 PM | C:\Sirius_T3\HCl18C02.t3r |
| S | 0.9988 | 3/2/2018 6:51:30 PM | C:\Sirius_T3\HCl18C02.t3r |
| jH | 1.0 | 3/2/2018 6:51:30 PM | C:\Sirius_T3\HCl18C02.t3r |
| jOH | -0.8 | 3/2/2018 6:51:30 PM | C:\Sirius_T3\HCl18C02.t3r |

#### Titrants

| | | | |
|---|---|---|---|
| 0.50 M HCl | 0.999058 | 3/2/2018 6:51:30 PM | C:\Sirius_T3\HCl18C02.t3r |
| 0.50 M KOH | 0.999845 | 3/2/2018 6:51:30 PM | C:\Sirius_T3\KOH18B27.t3r |

#### Sample

| | |
|---|---|
| M08_octanol concentration factor | 1.106 |
| Acid pKa 1 | 4.22 |
| logP (neutral XH) | 2.98 |
| logP (X -) | -4.72 |

#### Sample graphs

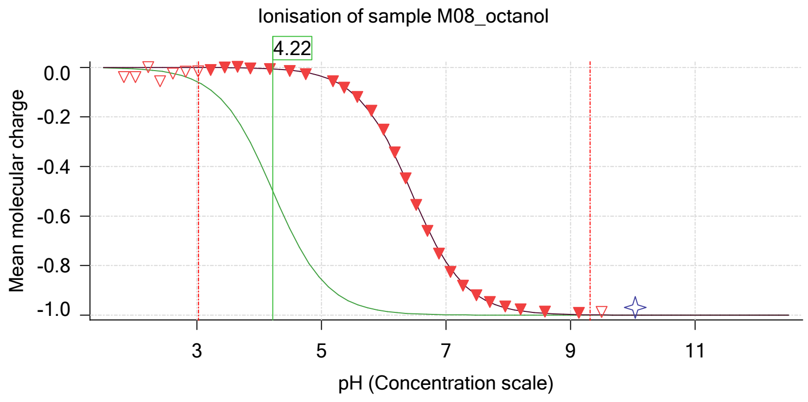


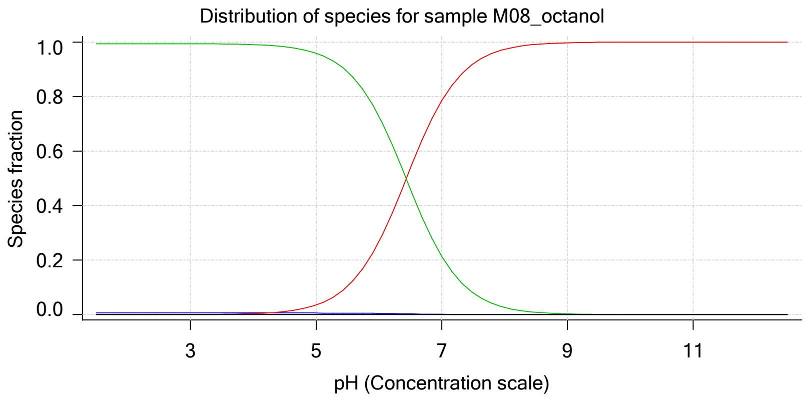

| | | | |
|---|---|---|---|
| Sample name: | **M08_octanol** | Experiment start time: | **3/2/2018 6:51:30 PM** |
| Assay name: | **pH-metric high logP** | Analyst: | **Pion** |
| Assay ID: | **18C-02008** | Instrument ID: | **T312060** |
| Filename: | **C:\Sirius_T3\Mehtap\20180302_exp29_logP_T3-2\18C-02008_M08_octanol_pH-metric high logP.t3r** | | |

#### Sample graphs (continued)

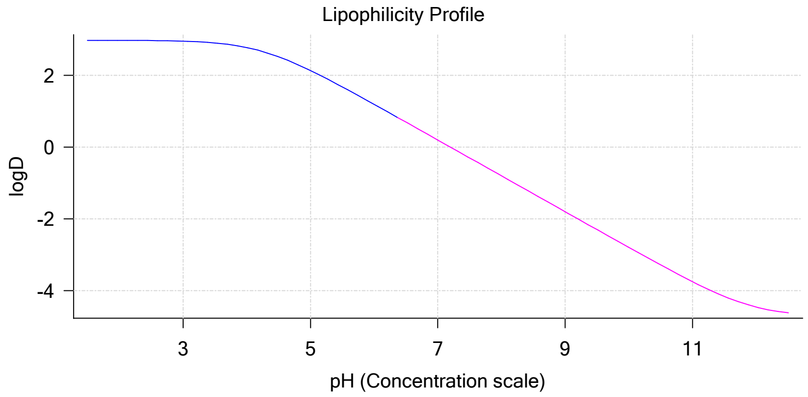


#### Sample logD and percent species

| pH | M08_octanol logD | M08_octanol M08_octanolH | M08_octanol M08_octanol | M08_octanol M08_octanolH* | M08_octanol M08_octanol* | Comment |
|---|---|---|---|---|---|---|
| 1.000 | 2.98 | 0.61 % | 0.00 % | 99.39 % | 0.00 % | |
| 1.200 | 2.98 | 0.61 % | 0.00 % | 99.39 % | 0.00 % | Stomach pH |
| 2.000 | 2.98 | 0.61 % | 0.00 % | 99.39 % | 0.00 % | |
| 3.000 | 2.95 | 0.61 % | 0.04 % | 99.36 % | 0.00 % | |
| 4.000 | 2.77 | 0.61 % | 0.36 % | 99.03 % | 0.00 % | |
| 5.000 | 2.13 | 0.59 % | 3.53 % | 95.88 % | 0.00 % | |
| 6.000 | 1.19 | 0.44 % | 26.80 % | 72.76 % | 0.00 % | |
| 6.500 | 0.70 | 0.28 % | 53.65 % | 46.06 % | 0.00 % | |
| 7.000 | 0.20 | 0.13 % | 78.54 % | 21.32 % | 0.00 % | |
| 7.400 | -0.20 | 0.06 % | 90.19 % | 9.75 % | 0.00 % | Blood pH |
| 8.000 | -0.80 | 0.02 % | 97.34 % | 2.64 % | 0.00 % | |
| 9.000 | -1.80 | 0.00 % | 99.73 % | 0.27 % | 0.00 % | |
| 10.000 | -2.80 | 0.00 % | 99.97 % | 0.03 % | 0.00 % | |
| 11.000 | -3.75 | 0.00 % | 100.00 % | 0.00 % | 0.00 % | |
| 12.000 | -4.46 | 0.00 % | 100.00 % | 0.00 % | 0.00 % | |

#### Carbonate and acidity

| | |
|---|---|
| Carbonate | 0.000 mM |
| Acidity error | -0.340 mM |

#### Other graphs

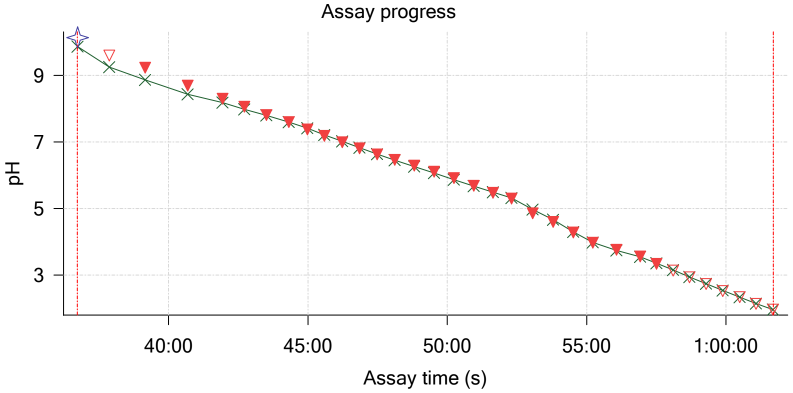


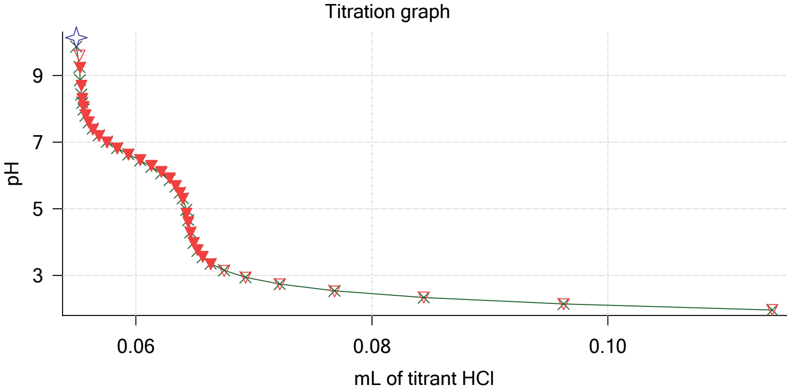

| | | | |
|---|---|---|---|
| Sample name: | **M08_octanol** | Experiment start time: | **3/2/2018 6:51:30 PM** |
| Assay name: | **pH-metric high logP** | Analyst: | **Pion** |
| Assay ID: | **18C-02008** | Instrument ID: | **T312060** |
| Filename: | **C:\Sirius_T3\Mehtap\20180302_exp29_logP_T3-2\18C-02008_M08_octanol_pH-metric high logP.t3r** | | |

#### Other graphs (continued)

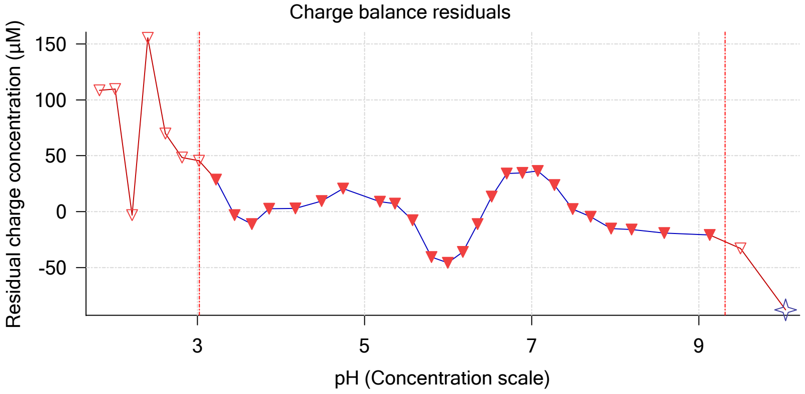


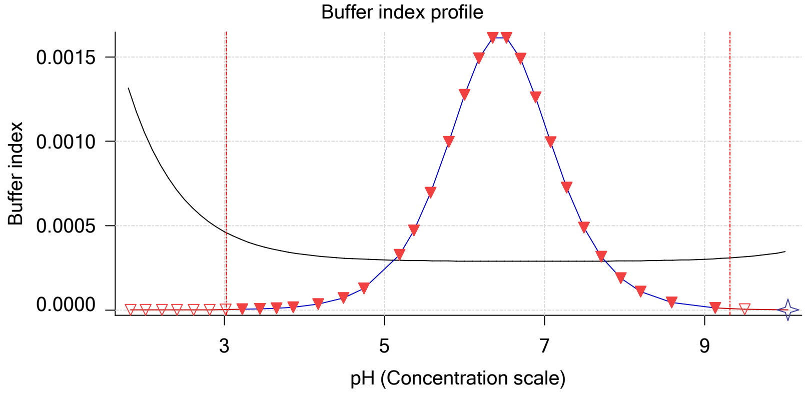


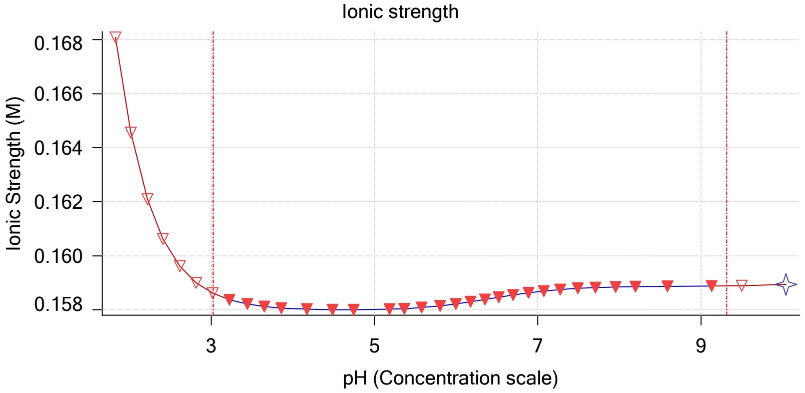


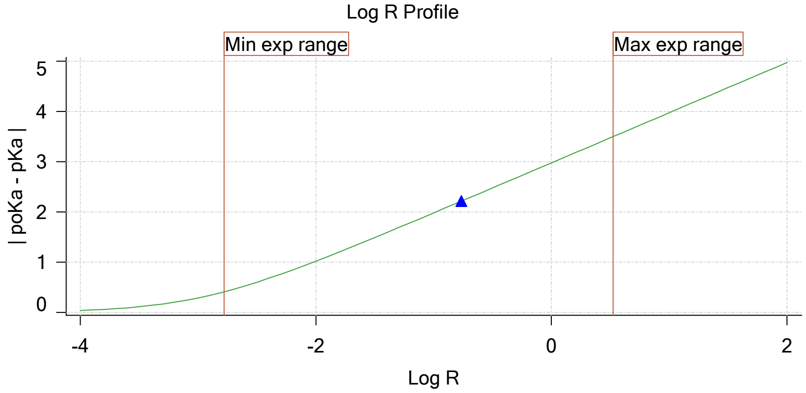


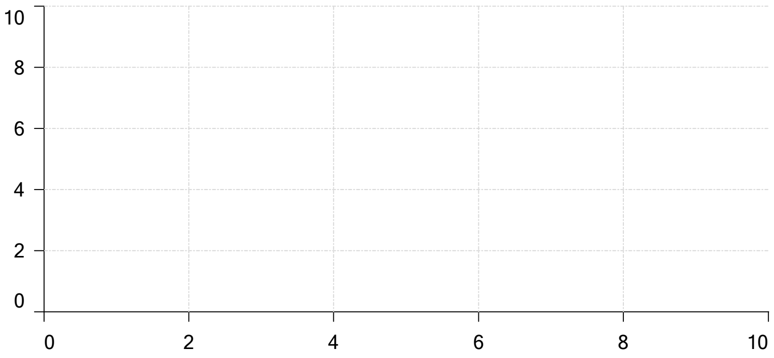

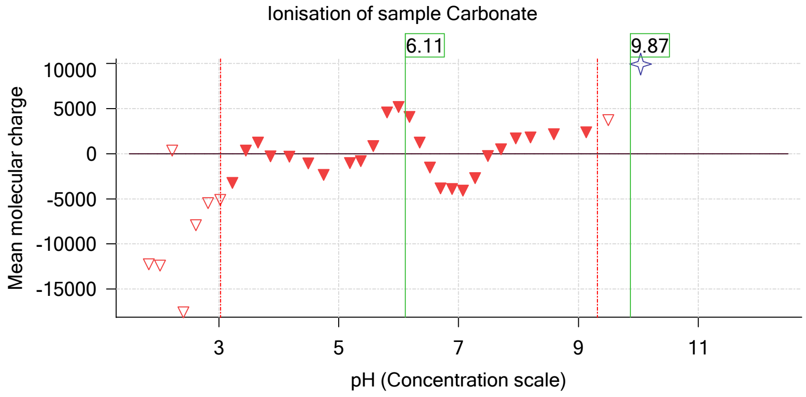

| | | | |
|---|---|---|---|
| Sample name: | **M08_octanol** | Experiment start time: | **3/2/2018 6:51:30 PM** |
| Assay name: | **pH-metric high logP** | Analyst: | **Pion** |
| Assay ID: | **18C-02008** | Instrument ID: | **T312060** |
| Filename: | **C:\Sirius_T3\Mehtap\20180302_exp29_logP_T3-2\18C-02008_M08_octanol_pH-metric high logP.t3r** | | |

### pH-metric high logP Titration 3 of 3 18C-02008 Points 68 to 106

#### Overall results

| | |
|---|---|
| RMSD | 0.055 |
| Average ionic strength | 0.165 M |
| Average temperature | 25.0°C |
| Partition ratio | 0.6191 : 1 |
| Analyte concentration range | 1443.8 µM to 1476.3 µM |
| Total points considered | 26 of 39 |

#### Warnings and errors

| | |
|---|---|
| Errors | None |
| Warnings | One or more logP values out of range |

#### Four-Plus parameters

| | | | |
|---|---|---|---|
| Alpha | 0.111 | 3/2/2018 6:51:30 PM | C:\Sirius_T3\HCl18C02.t3r |
| S | 0.9988 | 3/2/2018 6:51:30 PM | C:\Sirius_T3\HCl18C02.t3r |
| jH | 1.0 | 3/2/2018 6:51:30 PM | C:\Sirius_T3\HCl18C02.t3r |
| jOH | -0.8 | 3/2/2018 6:51:30 PM | C:\Sirius_T3\HCl18C02.t3r |

#### Titrants

| | | | |
|---|---|---|---|
| 0.50 M HCl | 0.999058 | 3/2/2018 6:51:30 PM | C:\Sirius_T3\HCl18C02.t3r |
| 0.50 M KOH | 0.999845 | 3/2/2018 6:51:30 PM | C:\Sirius_T3\KOH18B27.t3r |

#### Sample

| | |
|---|---|
| M08_octanol concentration factor | 0.984 |
| Acid pKa 1 | 4.22 |
| logP (neutral XH) | 2.91 |
| logP (X -) | -5.22 |

#### Sample graphs

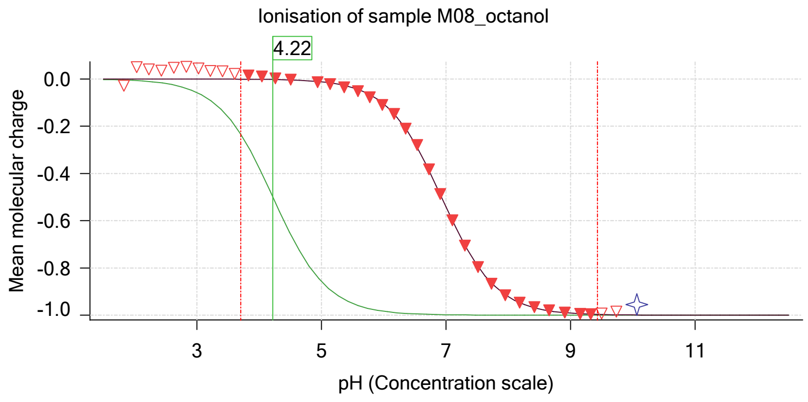


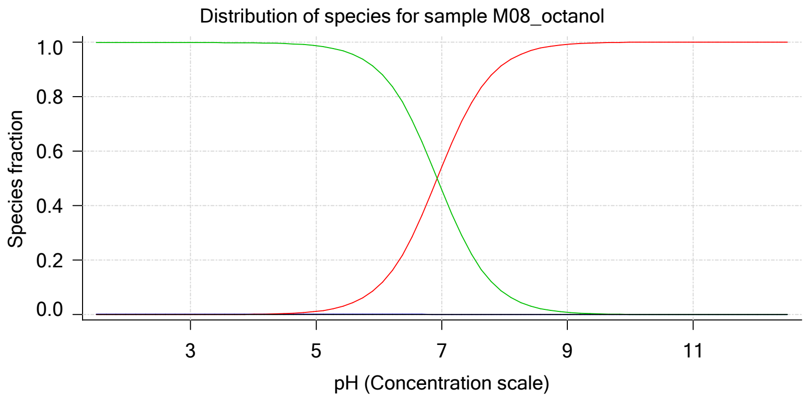

| | | | |
|---|---|---|---|
| Sample name: | **M08_octanol** | Experiment start time: | **3/2/2018 6:51:30 PM** |
| Assay name: | **pH-metric high logP** | Analyst: | **Pion** |
| Assay ID: | **18C-02008** | Instrument ID: | **T312060** |
| Filename: | **C:\Sirius_T3\Mehtap\20180302_exp29_logP_T3-2\18C-02008_M08_octanol_pH-metric high logP.t3r** | | |

#### Sample graphs (continued)

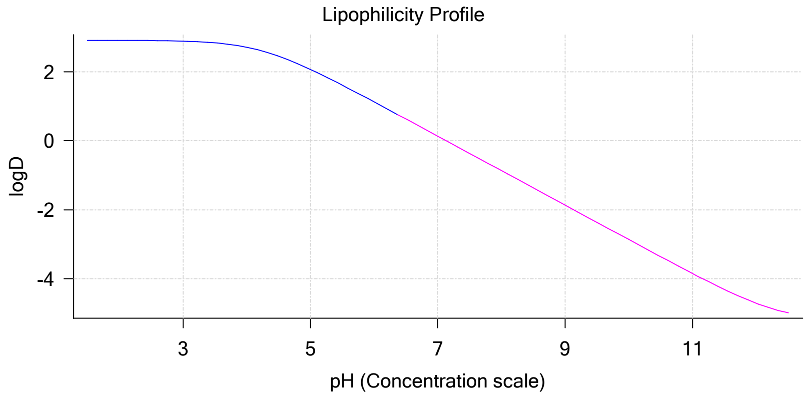


#### Sample logD and percent species

| pH | M08_octanol logD | M08_octanol M08_octanolH | M08_octanol M08_octanol | M08_octanol M08_octanolH* | M08_octanol M08_octanol* | Comment |
|---|---|---|---|---|---|---|
| 1.000 | 2.91 | 0.20 % | 0.00 % | 99.80 % | 0.00 % | |
| 1.200 | 2.91 | 0.20 % | 0.00 % | 99.80 % | 0.00 % | Stomach pH |
| 2.000 | 2.91 | 0.20 % | 0.00 % | 99.80 % | 0.00 % | |
| 3.000 | 2.89 | 0.20 % | 0.01 % | 99.79 % | 0.00 % | |
| 4.000 | 2.71 | 0.20 % | 0.12 % | 99.69 % | 0.00 % | |
| 5.000 | 2.07 | 0.19 % | 1.17 % | 98.64 % | 0.00 % | |
| 6.000 | 1.13 | 0.18 % | 10.57 % | 89.25 % | 0.00 % | |
| 6.500 | 0.63 | 0.14 % | 27.22 % | 72.64 % | 0.00 % | |
| 7.000 | 0.13 | 0.09 % | 54.18 % | 45.73 % | 0.00 % | |
| 7.400 | -0.27 | 0.05 % | 74.81 % | 25.14 % | 0.00 % | Blood pH |
| 8.000 | -0.87 | 0.02 % | 92.20 % | 7.78 % | 0.00 % | |
| 9.000 | -1.87 | 0.00 % | 99.16 % | 0.84 % | 0.00 % | |
| 10.000 | -2.86 | 0.00 % | 99.92 % | 0.08 % | 0.00 % | |
| 11.000 | -3.85 | 0.00 % | 99.99 % | 0.01 % | 0.00 % | |
| 12.000 | -4.71 | 0.00 % | 100.00 % | 0.00 % | 0.00 % | |

#### Carbonate and acidity

| | |
|---|---|
| Carbonate | 0.248 mM |
| Acidity error | -0.085 mM |

#### Other graphs

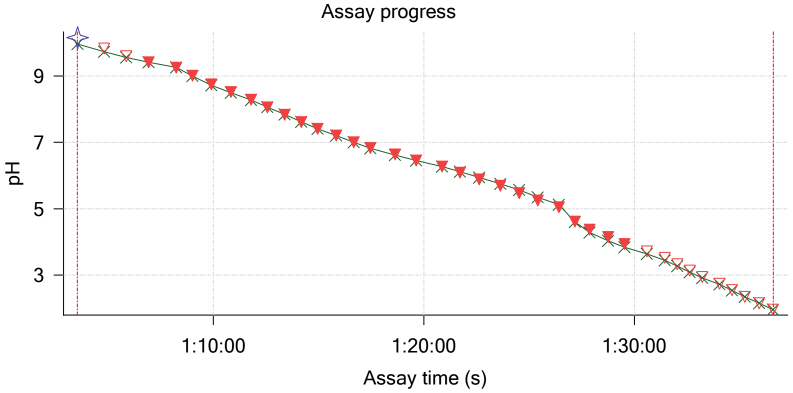


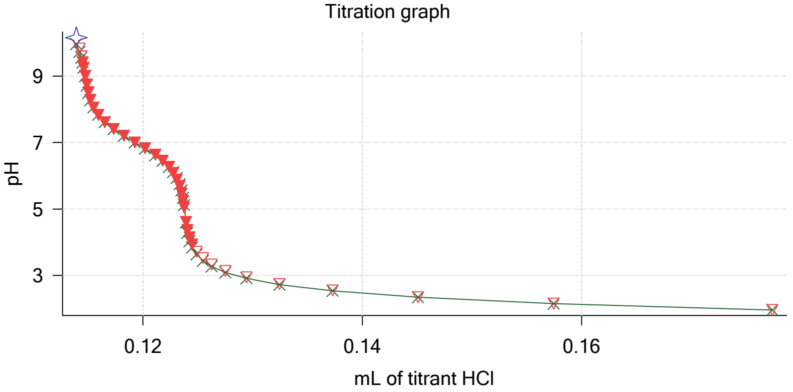

Sample name: **M08_octanol**
Assay name: **pH-metric high logP**
Assay ID: **18C-02008**
Filename: **C:\Sirius_T3\Mehtap\20180302_exp29_logP_T3-2\18C-02008_M08_octanol_pH-metric high logP.t3r**
Experiment start time: **3/2/2018 6:51:30 PM**
Analyst: **Pion**
Instrument ID: **T312060**

#### Other graphs (continued)

Charge balance residuals

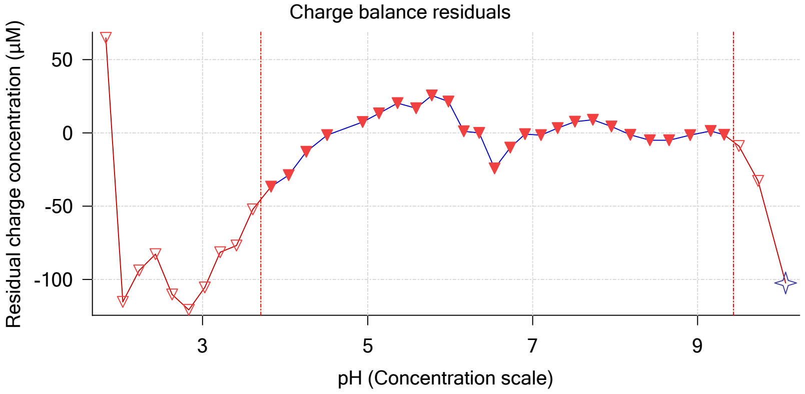


Buffer index profile

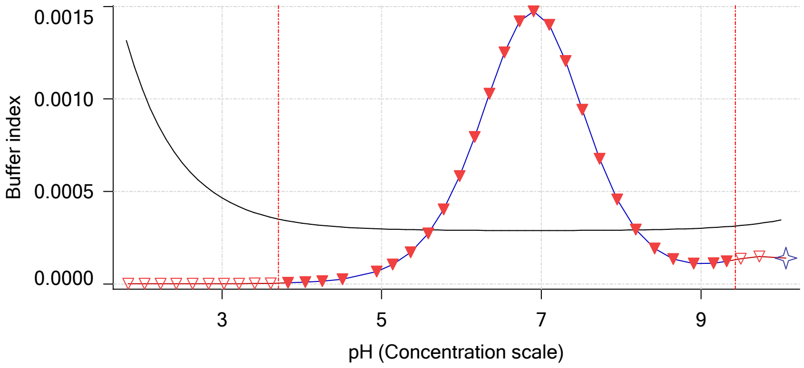


Ionic strength

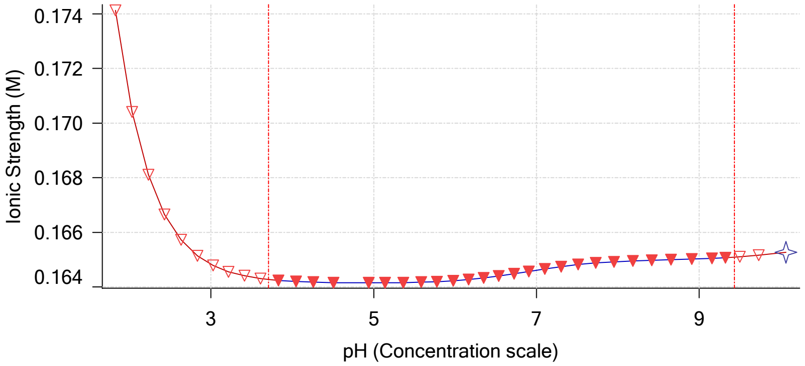


Log R Profile

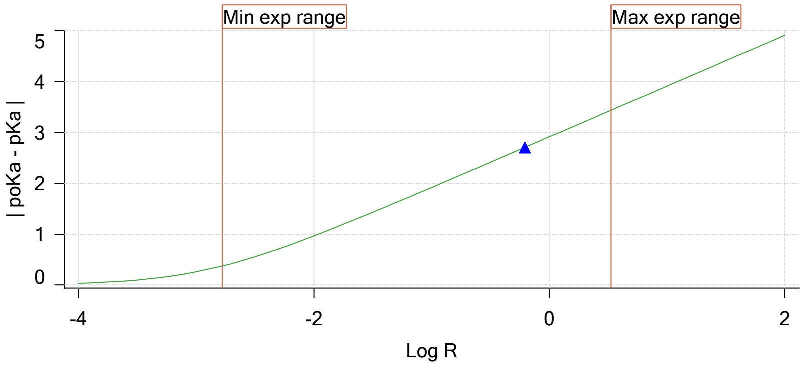


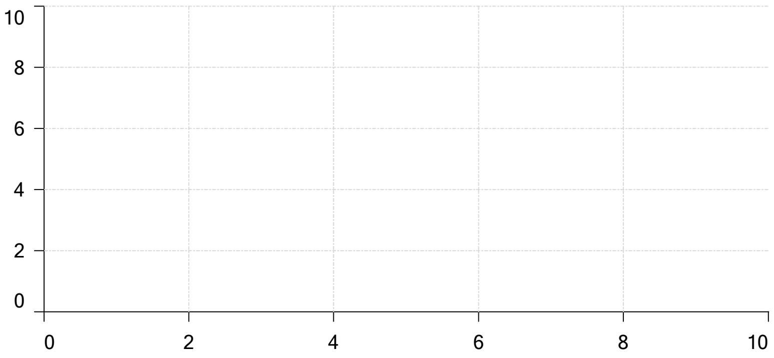


Ionisation of sample Carbonate

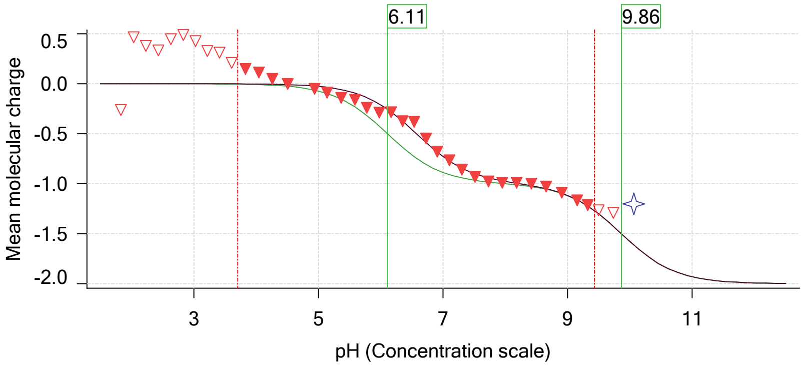

| | | | |
|---|---|---|---|
| Sample name: | **M08_octanol** | Experiment start time: | **3/2/2018 6:51:30 PM** |
| Assay name: | **pH-metric high logP** | Analyst: | **Pion** |
| Assay ID: | **18C-02008** | Instrument ID: | **T312060** |
| Filename: | **C:\Sirius_T3\Mehtap\20180302_exp29_logP_T3-2\18C-02008_M08_octanol_pH-metric high logP.t3r** | | |

#### Assay Model

| Settings | Value | Date/Time changed | Imported from |
|---|---|---|---|
| Sample name | M08_octanol | 2/27/2018 4:33:51 PM | User entered value |
| Sample by | Weight | | Default value |
| Sample weight | 0.001220 g | 3/2/2018 5:08:15 PM | User entered value |
| Formula weight | 293.32 g/mol | 2/27/2018 4:33:51 PM | User entered value |
| Solubility | Unknown | | Default value |
| Molecular weight | 293.32 | 2/27/2018 4:33:51 PM | User entered value |
| Individual pKa ionic environments | No | | Default value |
| Number of pKas | 1 | 2/27/2018 4:33:51 PM | User entered value |
| Sample is a | Acid | 2/27/2018 4:33:51 PM | User entered value |
| pKa 1 | 4.22 | 2/27/2018 4:33:51 PM | User entered value |
| logP (neutral XH) | 2.98 | 3/2/2018 3:22:58 PM | User entered value |
| logP (X -) | -5.22 | 3/2/2018 3:23:03 PM | User entered value |

#### Events

| Time | Event | Water | Acid | Base | Octanol | pH | dpH/dt | pH R-squared | pH SD | dpH time |
|---|---|---|---|---|---|---|---|---|---|---|
| 5:58.6 | Manual volume addition | | | | 0.08000 mL | | | | | |
| 5:59.7 | Initial pH = 5.21 | | | | | | | | | |
| 8:51.7 | Data point 2 | 1.50000 mL | 0.00000 mL | 0.00811 mL | 0.08000 mL | 10.428 | 0.01958 | 0.93900 | 0.00100 | 34.0 s |
| 9:52.4 | Data point 3 | 1.50000 mL | 0.00021 mL | 0.00811 mL | 0.08000 mL | 9.727 | 0.01586 | 0.62733 | 0.00099 | 55.5 s |
| 11:23.7 | Data point 4 | 1.50000 mL | 0.00042 mL | 0.00811 mL | 0.08000 mL | 8.981 | -0.01653 | 0.82499 | 0.00090 | 43.0 s |
| 12:42.4 | Data point 5 | 1.50000 mL | 0.00049 mL | 0.00811 mL | 0.08000 mL | 8.220 | -0.04412 | 0.98170 | 0.00220 | Time out at |
| 14:17.9 | Data point 6 | 1.50000 mL | 0.00059 mL | 0.00811 mL | 0.08000 mL | 7.798 | 0.01512 | 0.58186 | 0.00098 | 12.5 s |
| 15:00.9 | Data point 7 | 1.50000 mL | 0.00071 mL | 0.00811 mL | 0.08000 mL | 7.523 | 0.01618 | 0.67470 | 0.00097 | 14.5 s |
| 15:46.0 | Data point 8 | 1.50000 mL | 0.00089 mL | 0.00811 mL | 0.08000 mL | 7.289 | 0.01805 | 0.80919 | 0.00099 | 15.0 s |
| 16:41.8 | Data point 9 | 1.50000 mL | 0.00127 mL | 0.00811 mL | 0.08000 mL | 7.046 | 0.01754 | 0.75358 | 0.00100 | 16.0 s |
| 17:38.7 | Data point 10 | 1.50000 mL | 0.00169 mL | 0.00811 mL | 0.08000 mL | 6.863 | 0.01849 | 0.84615 | 0.00099 | 14.5 s |
| 18:18.6 | Data point 11 | 1.50000 mL | 0.00223 mL | 0.00811 mL | 0.08000 mL | 6.664 | 0.01638 | 0.78161 | 0.00092 | 16.0 s |
| 18:59.9 | Data point 12 | 1.50000 mL | 0.00294 mL | 0.00811 mL | 0.08000 mL | 6.481 | 0.01849 | 0.86308 | 0.00098 | 16.5 s |
| 19:41.9 | Data point 13 | 1.50000 mL | 0.00379 mL | 0.00811 mL | 0.08000 mL | 6.307 | 0.01848 | 0.90886 | 0.00096 | 15.0 s |
| 20:22.3 | Data point 14 | 1.50000 mL | 0.00475 mL | 0.00811 mL | 0.08000 mL | 6.113 | 0.01882 | 0.90673 | 0.00098 | 14.0 s |
| 21:01.8 | Data point 15 | 1.50000 mL | 0.00574 mL | 0.00811 mL | 0.08000 mL | 5.924 | 0.01677 | 0.83278 | 0.00091 | 17.0 s |
| 21:44.2 | Data point 16 | 1.50000 mL | 0.00668 mL | 0.00811 mL | 0.08000 mL | 5.702 | 0.01845 | 0.92173 | 0.00095 | 17.5 s |
| 22:27.2 | Data point 17 | 1.50000 mL | 0.00746 mL | 0.00811 mL | 0.08000 mL | 5.455 | 0.01862 | 0.94817 | 0.00094 | 28.0 s |
| 23:36.2 | Data point 18 | 1.50000 mL | 0.00797 mL | 0.00811 mL | 0.08000 mL | 5.350 | 0.09983 | 0.99423 | 0.00495 | Time out at |
| 25:01.6 | Data point 19 | 1.50000 mL | 0.00851 mL | 0.00811 mL | 0.08000 mL | 4.830 | 0.04926 | 0.98780 | 0.00245 | Time out at |

| | | | |
|---|---|---|---|
| Sample name: | **M08_octanol** | Experiment start time: | **3/2/2018 6:51:30 PM** |
| Assay name: | **pH-metric high logP** | Analyst: | **Pion** |
| Assay ID: | **18C-02008** | Instrument ID: | **T312060** |
| Filename: | **C:\Sirius_T3\Mehtap\20180302_exp29_logP_T3-2\18C-02008_M08_octanol_pH-metric high logP.t3r** | | |

#### Events (continued)

| Time | Event | Water | Acid | Base | Octanol | pH | dpH/dt | pH R-squared | pH SD | dpH/dt time |
|---|---|---|---|---|---|---|---|---|---|---|
| 26:37.2 | Data point 20 | 1.50000 mL | 0.00868 mL | 0.00811 mL | 0.08000 mL | 4.665 | 0.01816 | 0.93657 | 0.00093 | 31.5 s |
| 27:34.0 | Data point 21 | 1.50000 mL | 0.00877 mL | 0.00811 mL | 0.08000 mL | 4.425 | 0.01775 | 0.90251 | 0.00092 | 12.5 s |
| 28:17.1 | Data point 22 | 1.50000 mL | 0.00889 mL | 0.00811 mL | 0.08000 mL | 4.219 | 0.01953 | 0.93075 | 0.00100 | 11.0 s |
| 29:08.8 | Data point 23 | 1.50000 mL | 0.00905 mL | 0.00811 mL | 0.08000 mL | 3.997 | 0.00018 | 0.00008 | 0.00098 | 10.0 s |
| 29:44.2 | Data point 24 | 1.50000 mL | 0.00931 mL | 0.00811 mL | 0.08000 mL | 3.747 | 0.00337 | 0.08202 | 0.00058 | 10.0 s |
| 30:35.1 | Data point 25 | 1.50000 mL | 0.00974 mL | 0.00811 mL | 0.08000 mL | 3.552 | 0.00143 | 0.03293 | 0.00039 | 10.0 s |
| 31:10.5 | Data point 26 | 1.50000 mL | 0.01040 mL | 0.00811 mL | 0.08000 mL | 3.317 | 0.00181 | 0.11529 | 0.00026 | 10.0 s |
| 32:01.5 | Data point 27 | 1.50000 mL | 0.01152 mL | 0.00811 mL | 0.08000 mL | 3.118 | -0.00447 | 0.58442 | 0.00029 | 10.5 s |
| 32:37.5 | Data point 28 | 1.50000 mL | 0.01326 mL | 0.00811 mL | 0.08000 mL | 2.917 | -0.00516 | 0.25417 | 0.00051 | 10.0 s |
| 33:13.0 | Data point 29 | 1.50000 mL | 0.01604 mL | 0.00811 mL | 0.08000 mL | 2.730 | -0.01202 | 0.42201 | 0.00091 | 10.0 s |
| 33:48.5 | Data point 30 | 1.50000 mL | 0.02032 mL | 0.00811 mL | 0.08000 mL | 2.540 | -0.00759 | 0.75110 | 0.00043 | 10.5 s |
| 34:24.6 | Data point 31 | 1.50000 mL | 0.02702 mL | 0.00811 mL | 0.08000 mL | 2.348 | -0.00583 | 0.78723 | 0.00032 | 10.0 s |
| 35:00.3 | Data point 32 | 1.50000 mL | 0.03765 mL | 0.00811 mL | 0.08000 mL | 2.149 | -0.01355 | 0.91748 | 0.00070 | 10.5 s |
| 35:36.6 | Data point 33 | 1.50000 mL | 0.05496 mL | 0.00811 mL | 0.08000 mL | 1.953 | -0.01411 | 0.85512 | 0.00075 | 10.5 s |
| 36:44.7 | Data point 34 | 1.50000 mL | 0.05496 mL | 0.06333 mL | 0.28000 mL | 10.137 | -0.00357 | 0.03271 | 0.00098 | 37.5 s |
| 37:52.8 | Data point 35 | 1.50000 mL | 0.05522 mL | 0.06333 mL | 0.28000 mL | 9.598 | 0.01355 | 0.57771 | 0.00088 | 51.5 s |
| 39:09.8 | Data point 36 | 1.50000 mL | 0.05529 mL | 0.06333 mL | 0.28000 mL | 9.232 | 0.01873 | 0.90967 | 0.00097 | 55.5 s |
| 40:41.0 | Data point 37 | 1.50000 mL | 0.05539 mL | 0.06333 mL | 0.28000 mL | 8.688 | -0.00364 | 0.04343 | 0.00086 | 40.0 s |
| 41:56.7 | Data point 38 | 1.50000 mL | 0.05548 mL | 0.06333 mL | 0.28000 mL | 8.299 | 0.01075 | 0.31366 | 0.00095 | 10.5 s |
| 42:43.0 | Data point 39 | 1.50000 mL | 0.05560 mL | 0.06333 mL | 0.28000 mL | 8.053 | 0.01326 | 0.61367 | 0.00084 | 12.0 s |
| 43:30.7 | Data point 40 | 1.50000 mL | 0.05576 mL | 0.06333 mL | 0.28000 mL | 7.808 | 0.01660 | 0.85424 | 0.00089 | 12.5 s |
| 44:18.9 | Data point 41 | 1.50000 mL | 0.05602 mL | 0.06333 mL | 0.28000 mL | 7.592 | 0.01619 | 0.81503 | 0.00089 | 14.5 s |
| 44:58.8 | Data point 42 | 1.50000 mL | 0.05637 mL | 0.06333 mL | 0.28000 mL | 7.376 | 0.01819 | 0.83297 | 0.00098 | 11.5 s |
| 45:35.7 | Data point 43 | 1.50000 mL | 0.05689 mL | 0.06333 mL | 0.28000 mL | 7.176 | 0.01797 | 0.88644 | 0.00094 | 12.5 s |
| 46:13.6 | Data point 44 | 1.50000 mL | 0.05757 mL | 0.06333 mL | 0.28000 mL | 6.991 | 0.01928 | 0.96006 | 0.00097 | 12.0 s |
| 46:51.1 | Data point 45 | 1.50000 mL | 0.05842 mL | 0.06333 mL | 0.28000 mL | 6.803 | 0.01836 | 0.93846 | 0.00094 | 12.5 s |
| 47:29.0 | Data point 46 | 1.50000 mL | 0.05938 mL | 0.06333 mL | 0.28000 mL | 6.627 | 0.01804 | 0.92322 | 0.00093 | 12.5 s |
| 48:06.9 | Data point 47 | 1.50000 mL | 0.06037 mL | 0.06333 mL | 0.28000 mL | 6.455 | 0.01858 | 0.93024 | 0.00095 | 16.5 s |
| 48:48.9 | Data point 48 | 1.50000 mL | 0.06134 mL | 0.06333 mL | 0.28000 mL | 6.281 | 0.01664 | 0.78265 | 0.00093 | 17.5 s |
| 49:31.9 | Data point 49 | 1.50000 mL | 0.06218 mL | 0.06333 mL | 0.28000 mL | 6.101 | 0.01692 | 0.74989 | 0.00096 | 17.0 s |
| 50:14.3 | Data point 50 | 1.50000 mL | 0.06289 mL | 0.06333 mL | 0.28000 mL | 5.907 | 0.01653 | 0.76306 | 0.00093 | 17.5 s |
| 50:57.3 | Data point 51 | 1.50000 mL | 0.06341 mL | 0.06333 mL | 0.28000 mL | 5.681 | 0.01220 | 0.53640 | 0.00082 | 15.5 s |
| 51:38.2 | Data point 52 | 1.50000 mL | 0.06376 mL | 0.06333 mL | 0.28000 mL | 5.471 | 0.01147 | 0.51173 | 0.00079 | 15.0 s |
| 52:18.6 | Data point 53 | 1.50000 mL | 0.06399 mL | 0.06333 mL | 0.28000 mL | 5.290 | 0.01455 | 0.56277 | 0.00096 | 14.5 s |
| 53:03.6 | Data point 54 | 1.50000 mL | 0.06430 mL | 0.06333 mL | 0.28000 mL | 4.851 | 0.01470 | 0.53131 | 0.00100 | 14.0 s |
| 53:48.2 | Data point 55 | 1.50000 mL | 0.06446 mL | 0.06333 mL | 0.28000 mL | 4.594 | 0.01132 | 0.40356 | 0.00088 | 12.5 s |
| 54:31.4 | Data point 56 | 1.50000 mL | 0.06465 mL | 0.06333 mL | 0.28000 mL | 4.279 | 0.00590 | 0.14884 | 0.00076 | 11.5 s |
| 55:13.4 | Data point 57 | 1.50000 mL | 0.06491 mL | 0.06333 mL | 0.28000 mL | 3.965 | 0.00967 | 0.40120 | 0.00075 | 10.0 s |
| 56:04.3 | Data point 58 | 1.50000 mL | 0.06524 mL | 0.06333 mL | 0.28000 mL | 3.759 | 0.00813 | 0.49244 | 0.00057 | 10.0 s |
| 56:55.2 | Data point 59 | 1.50000 mL | 0.06566 mL | 0.06333 mL | 0.28000 mL | 3.552 | -0.00215 | 0.16107 | 0.00027 | 10.0 s |
| 57:30.6 | Data point 60 | 1.50000 mL | 0.06635 mL | 0.06333 mL | 0.28000 mL | 3.331 | -0.00414 | 0.34668 | 0.00035 | 10.0 s |
| 58:06.0 | Data point 61 | 1.50000 mL | 0.06747 mL | 0.06333 mL | 0.28000 mL | 3.126 | -0.00603 | 0.61652 | 0.00038 | 10.0 s |
| 58:41.5 | Data point 62 | 1.50000 mL | 0.06931 mL | 0.06333 mL | 0.28000 mL | 2.927 | -0.00487 | 0.09075 | 0.00080 | 10.0 s |
| 59:17.0 | Data point 63 | 1.50000 mL | 0.07220 mL | 0.06333 mL | 0.28000 mL | 2.726 | 0.00082 | 0.04244 | 0.00020 | 10.5 s |
| 59:53.1 | Data point 64 | 1.50000 mL | 0.07683 mL | 0.06333 mL | 0.28000 mL | 2.519 | -0.00715 | 0.90269 | 0.00037 | 10.5 s |
| 1:00:29.1 | Data point 65 | 1.50000 mL | 0.08441 mL | 0.06333 mL | 0.28000 mL | 2.334 | -0.01263 | 0.94471 | 0.00064 | 10.0 s |
| 1:01:04.8 | Data point 66 | 1.50000 mL | 0.09624 mL | 0.06333 mL | 0.28000 mL | 2.133 | -0.01155 | 0.84750 | 0.00062 | 10.0 s |
| 1:01:40.7 | Data point 67 | 1.50000 mL | 0.11392 mL | 0.06333 mL | 0.28000 mL | 1.953 | -0.00652 | 0.61375 | 0.00041 | 10.5 s |
| 1:03:34.1 | Data point 68 | 1.50000 mL | 0.11392 mL | 0.12345 mL | 1.08000 mL | 10.164 | -0.01244 | 0.38157 | 0.00099 | 45.0 s |
| 1:04:49.8 | Data point 69 | 1.50000 mL | 0.11423 mL | 0.12345 mL | 1.08000 mL | 9.832 | -0.00830 | 0.23375 | 0.00085 | 37.5 s |
| 1:05:52.7 | Data point 70 | 1.50000 mL | 0.11439 mL | 0.12345 mL | 1.08000 mL | 9.598 | -0.01486 | 0.60594 | 0.00094 | 38.0 s |
| 1:06:56.1 | Data point 71 | 1.50000 mL | 0.11451 mL | 0.12345 mL | 1.08000 mL | 9.421 | -0.00966 | 0.27141 | 0.00092 | 53.0 s |
| 1:08:14.6 | Data point 72 | 1.50000 mL | 0.11461 mL | 0.12345 mL | 1.08000 mL | 9.256 | -0.00823 | 0.36404 | 0.00067 | 11.0 s |
| 1:09:01.3 | Data point 73 | 1.50000 mL | 0.11475 mL | 0.12345 mL | 1.08000 mL | 9.009 | 0.01389 | 0.51931 | 0.00095 | 18.0 s |
| 1:09:55.0 | Data point 74 | 1.50000 mL | 0.11489 mL | 0.12345 mL | 1.08000 mL | 8.754 | 0.00577 | 0.08155 | 0.00100 | 20.0 s |
| 1:10:50.8 | Data point 75 | 1.50000 mL | 0.11503 mL | 0.12345 mL | 1.08000 mL | 8.520 | 0.01828 | 0.95610 | 0.00092 | 21.5 s |
| 1:11:48.1 | Data point 76 | 1.50000 mL | 0.11522 mL | 0.12345 mL | 1.08000 mL | 8.286 | 0.01567 | 0.71329 | 0.00092 | 20.5 s |

| | | | |
|---|---|---|---|
| Sample name: | **M08_octanol** | Experiment start time: | **3/2/2018 6:51:30 PM** |
| Assay name: | **pH-metric high logP** | Analyst: | **Pion** |
| Assay ID: | **18C-02008** | Instrument ID: | **T312060** |
| Filename: | **C:\Sirius_T3\Mehtap\20180302_exp29_logP_T3-2\18C-02008_M08_octanol_pH-metric high logP.t3r** | | |

#### Events (continued)

| Time | Event | Water | Acid | Base | Octanol | pH | dpH/dt | pH R-squared | pH SD | dpH/dt time |
|---|---|---|---|---|---|---|---|---|---|---|
| 1:12:34.1 | Data point 77 | 1.50000 mL | 0.11550 mL | 0.12345 mL | 1.08000 mL | 8.054 | 0.00873 | 0.25062 | 0.00086 | 24.0 s |
| 1:13:23.5 | Data point 78 | 1.50000 mL | 0.11592 mL | 0.12345 mL | 1.08000 mL | 7.830 | 0.01471 | 0.68425 | 0.00088 | 22.0 s |
| 1:14:11.0 | Data point 79 | 1.50000 mL | 0.11653 mL | 0.12345 mL | 1.08000 mL | 7.615 | 0.01082 | 0.31604 | 0.00095 | 21.0 s |
| 1:14:57.5 | Data point 80 | 1.50000 mL | 0.11733 mL | 0.12345 mL | 1.08000 mL | 7.406 | 0.01311 | 0.67015 | 0.00079 | 27.5 s |
| 1:15:50.4 | Data point 81 | 1.50000 mL | 0.11827 mL | 0.12345 mL | 1.08000 mL | 7.204 | 0.01583 | 0.70594 | 0.00093 | 24.5 s |
| 1:16:40.4 | Data point 82 | 1.50000 mL | 0.11926 mL | 0.12345 mL | 1.08000 mL | 7.008 | 0.01015 | 0.26007 | 0.00098 | 22.0 s |
| 1:17:27.9 | Data point 83 | 1.50000 mL | 0.12020 mL | 0.12345 mL | 1.08000 mL | 6.829 | 0.00824 | 0.26586 | 0.00079 | 29.0 s |
| 1:18:37.8 | Data point 84 | 1.50000 mL | 0.12114 mL | 0.12345 mL | 1.08000 mL | 6.639 | 0.01737 | 0.74638 | 0.00099 | 35.0 s |
| 1:19:38.3 | Data point 85 | 1.50000 mL | 0.12180 mL | 0.12345 mL | 1.08000 mL | 6.453 | 0.01561 | 0.66863 | 0.00094 | 33.0 s |
| 1:20:52.2 | Data point 86 | 1.50000 mL | 0.12239 mL | 0.12345 mL | 1.08000 mL | 6.268 | 0.01016 | 0.29466 | 0.00092 | 25.5 s |
| 1:21:43.1 | Data point 87 | 1.50000 mL | 0.12277 mL | 0.12345 mL | 1.08000 mL | 6.082 | 0.01424 | 0.61244 | 0.00090 | 24.0 s |
| 1:22:37.7 | Data point 88 | 1.50000 mL | 0.12310 mL | 0.12345 mL | 1.08000 mL | 5.882 | 0.00792 | 0.16581 | 0.00096 | 19.0 s |
| 1:23:37.7 | Data point 89 | 1.50000 mL | 0.12335 mL | 0.12345 mL | 1.08000 mL | 5.690 | 0.00115 | 0.00389 | 0.00091 | 23.5 s |
| 1:24:31.7 | Data point 90 | 1.50000 mL | 0.12352 mL | 0.12345 mL | 1.08000 mL | 5.467 | 0.01785 | 0.83193 | 0.00097 | 22.5 s |
| 1:25:24.8 | Data point 91 | 1.50000 mL | 0.12366 mL | 0.12345 mL | 1.08000 mL | 5.243 | 0.01854 | 0.86754 | 0.00098 | 29.5 s |
| 1:26:24.8 | Data point 92 | 1.50000 mL | 0.12375 mL | 0.12345 mL | 1.08000 mL | 5.042 | 0.00911 | 0.28586 | 0.00084 | 14.5 s |
| 1:27:09.9 | Data point 93 | 1.50000 mL | 0.12392 mL | 0.12345 mL | 1.08000 mL | 4.612 | -0.01412 | 0.49010 | 0.00100 | 12.0 s |
| 1:27:52.4 | Data point 94 | 1.50000 mL | 0.12406 mL | 0.12345 mL | 1.08000 mL | 4.363 | 0.01967 | 0.96830 | 0.00099 | 17.0 s |
| 1:28:45.1 | Data point 95 | 1.50000 mL | 0.12425 mL | 0.12345 mL | 1.08000 mL | 4.149 | 0.00177 | 0.01471 | 0.00072 | 10.5 s |
| 1:29:31.3 | Data point 96 | 1.50000 mL | 0.12448 mL | 0.12345 mL | 1.08000 mL | 3.934 | -0.00929 | 0.21345 | 0.00099 | 23.5 s |
| 1:30:35.6 | Data point 97 | 1.50000 mL | 0.12488 mL | 0.12345 mL | 1.08000 mL | 3.715 | -0.00629 | 0.73094 | 0.00036 | 10.0 s |
| 1:31:26.6 | Data point 98 | 1.50000 mL | 0.12547 mL | 0.12345 mL | 1.08000 mL | 3.517 | -0.01016 | 0.69269 | 0.00060 | 10.0 s |
| 1:32:02.1 | Data point 99 | 1.50000 mL | 0.12627 mL | 0.12345 mL | 1.08000 mL | 3.321 | -0.01475 | 0.90060 | 0.00077 | 10.0 s |
| 1:32:37.6 | Data point 100 | 1.50000 mL | 0.12752 mL | 0.12345 mL | 1.08000 mL | 3.134 | -0.00934 | 0.64893 | 0.00057 | 10.5 s |
| 1:33:13.4 | Data point 101 | 1.50000 mL | 0.12945 mL | 0.12345 mL | 1.08000 mL | 2.940 | 0.00025 | 0.00017 | 0.00095 | 23.5 s |
| 1:34:02.5 | Data point 102 | 1.50000 mL | 0.13245 mL | 0.12345 mL | 1.08000 mL | 2.741 | -0.01740 | 0.87852 | 0.00092 | 10.0 s |
| 1:34:38.0 | Data point 103 | 1.50000 mL | 0.13730 mL | 0.12345 mL | 1.08000 mL | 2.539 | -0.01964 | 0.96661 | 0.00099 | 10.5 s |
| 1:35:14.2 | Data point 104 | 1.50000 mL | 0.14508 mL | 0.12345 mL | 1.08000 mL | 2.343 | 0.00480 | 0.07839 | 0.00085 | 15.0 s |
| 1:35:54.9 | Data point 105 | 1.50000 mL | 0.15746 mL | 0.12345 mL | 1.08000 mL | 2.149 | -0.01890 | 0.90387 | 0.00098 | 13.5 s |
| 1:36:34.4 | Data point 106 | 1.50000 mL | 0.17738 mL | 0.12345 mL | 1.08000 mL | 1.950 | -0.01112 | 0.54218 | 0.00075 | 20.5 s |
| 1:37:04.0 | Assay volumes | 1.50000 mL | 0.17738 mL | 0.12345 mL | 1.08000 mL | | | | | |

| | | | |
|---|---|---|---|
| Sample name: | **M08_octanol** | Experiment start time: | **3/2/2018 6:51:30 PM** |
| Assay name: | **pH-metric high logP** | Analyst: | **Pion** |
| Assay ID: | **18C-02008** | Instrument ID: | **T312060** |
| Filename: | **C:\Sirius_T3\Mehtap\20180302_exp29_logP_T3-2\18C-02008_M08_octanol_pH-metric high logP.t3r** | | |

#### Assay Settings

| Setting | Value | Original Value | Date/Time changed | Imported from |
|---|---|---|---|---|
| ***General Settings*** | | | | |
| Analyst name | Pion | | | |
| ***Standard Experiment Settings*** | | | | |
| Number of titrations | 3 | | | |
| Minimum pH | 2.000 | | | |
| Maximum pH | 10.000 | | | |
| pH step between points of | 0.200 | | | |
| Minimum titrant addition | 0.00002 mL | | | |
| Maximum titrant addition | 0.10000 mL | | | |
| Argon flow rate | 100% | | | |
| Start titration using | Cautious pH adjust | | | |
| ***Advanced General Settings*** | | | | |
| Detect turbidity using | None | | | |
| Collect turbidity sensor data | No | | | |
| Collect UV spectra | No | | | |
| Stir after titrant addition for | 5 seconds | | | |
| For titrant addition, stir at | 10% | | | |
| ***Titrant Pre-Dose*** | | | | |
| Titrant pre-dose | None | | | |
| ***Assay Medium*** | | | | |
| ISA water volume | 1.50 mL | | | |
| Water added | Automatic | | | |
| Partition solvent type | Octanol | | | |
| Partition volume | 0.080 mL | | | |
| Partition solvent added | Manual in advance | | | |
| After partition addition, stir for | 1 seconds | | | |
| ***Sample Sonication*** | | | | |
| Sonicate | Yes | | | |
| Adjust pH for sonication | No | | | |
| Sonicate for | 120 seconds | | | |
| After sonication stir for | 5 seconds | | | |
| ***Sample Dissolution*** | | | | |
| Perform a dissolution stage | Yes | | | |
| Adjust and hold pH for dissolution | To start pH | | | |
| Stir to dissolve for | 120 seconds | | | |
| For dissolution, stir at | 10% | | | |
| ***Carbonate purge*** | | | | |
| Perform a carbonate purge | No | | | |
| ***Temperature Control*** | | | | |
| Wait for temperature | Yes | | | |
| Required start temperature | 25.0°C | | | |
| Acceptable deviation | 0.5°C | | | |
| Time to wait | 60 seconds | | | |
| Stir speed of | 50% | | | |
| ***Titration 1*** | | | | |
| Titrate from | High to low pH | | | |
| Adjust to start pH | Yes | | | |
| After pH adjust stir for | 30 seconds | | | |
| Stir to allow partitioning for | 15 seconds | | | |
| Stirrer speed for partitioning | 50% | | | |
| ***Titration 2*** | | | | |
| Titrate from | High to low pH | | | |
| Add additional water | 0.00 mL | | | |
| Additional partition solvent volume | 0.200 mL | | | |
| Additional partition solvent added | Automatic | | | |
| After pH adjust stir for | 30 seconds | | | |
| Stir to allow partitioning for | 15 seconds | | | |
| Stirrer speed for partitioning | 55% | | | |

| | | | |
|---|---|---|---|
| Sample name: | **M08_octanol** | Experiment start time: | **3/2/2018 6:51:30 PM** |
| Assay name: | **pH-metric high logP** | Analyst: | **Pion** |
| Assay ID: | **18C-02008** | Instrument ID: | **T312060** |
| Filename: | **C:\Sirius_T3\Mehtap\20180302_exp29_logP_T3-2\18C-02008_M08_octanol_pH-metric high logP.t3r** | | |

#### Assay Settings (continued)

| Setting | Value | Original Value | Date/Time changed | Imported from |
|---|---|---|---|---|
| ***Titration 3*** | | | | |
| Titrate from | High to low pH | | | |
| Add additional water | 0.00 mL | | | |
| Additional partition solvent volume | 0.800 mL | | | |
| Additional partition solvent added | Automatic | | | |
| After pH adjust stir for | 30 seconds | | | |
| Stir to allow partitioning for | 15 seconds | | | |
| Stirrer speed for partitioning | 60% | | | |
| ***Data Point Stability*** | | | | |
| Stir during data point collection | No | | | |
| Delay before data point collection | 0 seconds | | | |
| Number of points to average | 20 points | | | |
| Time interval between points | 0.50 seconds | | | |
| Required maximum standard deviation | 0.00100 dpH/dt | | | |
| Stability timeout after | 60 seconds | | | |

#### Calibration Settings

| Setting | Value | Date/Time changed | Imported from |
|---|---|---|---|
| Four-Plus alpha | 0.111 | 3/2/2018 6:51:30 PM | C:\Sirius_T3\HCl18C02.t3r |
| Four-Plus S | 0.9988 | 3/2/2018 6:51:30 PM | C:\Sirius_T3\HCl18C02.t3r |
| Four-Plus jH | 1.0 | 3/2/2018 6:51:30 PM | C:\Sirius_T3\HCl18C02.t3r |
| Four-Plus jOH | -0.8 | 3/2/2018 6:51:30 PM | C:\Sirius_T3\HCl18C02.t3r |
| Base concentration factor | 1.000 | 3/2/2018 6:51:30 PM | C:\Sirius_T3\KOH18B27.t3r |
| Acid concentration factor | 0.999 | 3/2/2018 6:51:30 PM | C:\Sirius_T3\HCl18C02.t3r |

#### Instrument Settings

| Setting | Value | Batch Id | Install date |
|---|---|---|---|
| Instrument owner | Merck | | |
| Instrument ID | T312060 | | |
| Instrument type | T3 Simulator | | |
| Software version | 1.1.3.0 | | |
| Dispenser module | | T3DM1200361 | 3/31/2009 5:24:52 AM |
| Dispenser 0 | Water | | 3/31/2009 5:25:05 AM |
| Syringe volume | 2.5 mL | | |
| Firmware version | 1.2.1(r2) | | |
| Titrant | Water (0.15 M KCl) | 02-06-2018 | 2/27/2018 10:05:59 AM |
| Dispenser 2 | Acid | | 3/31/2009 5:25:11 AM |
| Syringe volume | 0.5 mL | | |
| Firmware version | 1.2.1(r2) | | |
| Titrant | Acid (0.5 M HCl) | 02-27-2018 | 2/27/2018 10:27:22 AM |
| Dispenser 1 | Base | | 3/31/2009 5:25:21 AM |
| Syringe volume | 0.5 mL | | |
| Firmware version | 1.2.1(r2) | | |
| Titrant | Base (0.5 M KOH) | 9/22/2017 | 2/27/2018 10:21:22 AM |
| Dispenser 5 | Cosolvent | | 3/31/2009 5:26:24 AM |
| Syringe volume | 2.5 mL | | |
| Firmware version | 1.2.1(r2) | | |
| Distribution valve 5 | Distribution Valve | | 3/31/2009 5:28:19 AM |
| Firmware version | 1.1.3 | | |
| Port A | Methanol (80%, 0.15 M KCl) | 09-26-17 | 2/7/2018 9:42:01 AM |
| Port B | Cyclohexane | 11-01-17 | 2/27/2018 10:37:57 AM |
| Dispenser 3 | Buffer | | 8/3/2010 5:05:16 AM |
| Syringe volume | 0.5 mL | | |
| Firmware version | 1.2.1(r2) | | |
| Titrant | Dodecane | 2018/01/31 | 2/28/2018 10:18:04 AM |
| Dispenser 6 | Octanol | | 10/22/2010 10:52:43 AM |

| | | | |
|---|---|---|---|
| Sample name: | **M08_octanol** | Experiment start time: | **3/2/2018 6:51:30 PM** |
| Assay name: | **pH-metric high logP** | Analyst: | **Pion** |
| Assay ID: | **18C-02008** | Instrument ID: | **T312060** |
| Filename: | **C:\Sirius_T3\Mehtap\20180302_exp29_logP_T3-2\18C-02008_M08_octanol_pH-metric high logP.t3r** | | |

#### Instrument Settings (continued)

| Setting | Value | Batch Id | Install date |
|---|---|---|---|
| Syringe volume | 0.5 mL | | |
| Firmware version | 1.2.1(r2) | | |
| Titrant | Octanol | 01-31-2018 | 2/27/2018 9:59:35 AM |
| Titrator | | T3TM1200161 | 3/31/2009 5:24:17 AM |
| Horizontal axis firmware version | 1.17 AI1DI2DO2 Stepper 2 | | |
| Vertical axis firmware version | 1.17 AI1DI2DO2 Stepper 2 | | |
| Chassis I/O firmware version | 1.11 AI1DI0DO4 Norgren I/O | | |
| Probe I/O firmware version | 1.1.1 | | |
| Electrode | T3 Electrode | T3E0923 | 1/23/2018 2:01:00 PM |
| E0 calibration | +2.88 mV | | 3/2/2018 6:51:58 PM |
| Filling solution | 3M KCl | KCL097 | 3/2/2018 9:43:24 AM |
| Liquids | | | |
| Wash 1 | 50% IPA:50% Water | | 3/2/2018 9:45:12 AM |
| Wash 2 | 0.5% Trition X-100 in H20 | | 3/2/2018 9:45:15 AM |
| Buffer position 1 | pH7 Wash | | 3/2/2018 9:45:18 AM |
| Buffer position 2 | pH 7 | | 3/2/2018 9:45:21 AM |
| Storage position | | | 3/2/2018 9:44:44 AM |
| Wash water | 7.3e+003 mL | 02-27-2018 | 2/27/2018 9:54:39 AM |
| Waste | 8.1e+003 mL | | 11/28/2017 10:36:29 AM |
| Temperature controller | | | 8/5/2010 6:35:13 AM |
| Turbidity detector | | | 3/31/2009 5:24:45 AM |
| Spectrometer | | 074811 | 11/23/2010 11:22:28 AM |
| Dip probe | | 10196 | |
| Wavelength coefficient A0 | 183.333 | | |
| Wavelength coefficient A1 | 2.21568 | | |
| Wavelength coefficient A2 | -0.000289308 | | |
| Total lamp lit time | 120:41:49 | | 11/23/2010 11:22:28 AM |
| Calibrated on | 2/27/2018 10:40:38 AM | | |
| Integration time | 40 | | |
| Scans averaged | 10 | | |
| Autoloader | | T3AL1200345 | 11/10/2015 9:34:13 AM |
| Left-right axis firmware version | 1.17 AI1DI2DO2 Stepper 2 | | |
| Front-back axis firmware version | 1.17 AI1DI2DO2 Stepper 2 | | |
| Vertical axis firmware version | 1.17 AI1DI2DO2 Stepper 2 | | |
| Chassis I/O firmware version | 1.11 AI1DI0DO4 Norgren I/O | | |
| Configuration | | | |
| Alternate titration position | Titration position | | |
| Alternate reference position | Reference position | | |
| Maximum standard vial volume | 3.50 mL | | |
| Maximum alternate vial volume | 25.00 mL | | |
| Automatic action idle period | 5 minute(s) | | |
| Titrant tube volume | 1.3 mL | | |
| Syringe flush count | 3.50 | | |
| Flowing wash pump volume | 20.0 mL | | |
| Flowing wash stir duration | 5 s | | |
| Flowing wash stir speed | 30% | | |
| Solvent wash stir duration | 5 s | | |
| Solvent wash stir speed | 30% | | |
| Surfactant wash stir duration | 5 s | | |
| Surfactant wash stir speed | 30% | | |
| E0 calibration minimum number of points | 10 | | |
| E0 calibration maximum standard deviation | 0.01500 | | |
| E0 calibration timeout period | 60 s | | |
| E0 calibration stir duration | 5 s | | |
| E0 calibration preparation stir speed | 30% | | |
| E0 calibration buffer wash stir duration | 5 s | | |
| E0 calibration buffer wash stir speed | 30% | | |
| E0 calibration reading stir speed | 0% | | |

| | | | |
|---|---|---|---|
| Sample name: | **M08_octanol** | Experiment start time: | **3/2/2018 6:51:30 PM** |
| Assay name: | **pH-metric high logP** | Analyst: | **Pion** |
| Assay ID: | **18C-02008** | Instrument ID: | **T312060** |
| Filename: | **C:\Sirius_T3\Mehtap\20180302_exp29_logP_T3-2\18C-02008_M08_octanol_pH-metric high logP.t3r** | | |

#### Instrument Settings (continued)

| Setting | Value | Batch Id | Install date |
|---|---|---|---|
| Spectrometer calibration stir duration | 5 s | | |
| Spectrometer calibration stir speed | 30% | | |
| Spectrometer calibration wash pump volume | 20.0 mL | | |
| Spectrometer calibration wash stir duration | 5 s | | |
| Spectrometer calibration wash stir speed | 30% | | |
| Overhead dispense height | 10000 | | |

#### Refinement Settings

| Setting | Value | Default value |
|---|---|---|
| Turbidity detection method | None | None |
| Turbidity wavelength to assess | 500.0 nm | 500.0 nm |
| Turbidity maximum absorbance | 0.100 | 0.100 |
| Turbidity probe threshold | 50.00 | 50.00 |

#### Experiment Log

[2:38] Air gap created for Water (0.15 M KCl)
[2:38] Air gap created for Acid (0.5 M HCl)
[2:39] Air gap created for Base (0.5 M KOH)
[2:39] Air gap released for Water (0.15 M KCl)
[2:43] Titrator arm moved over Titration position
[2:43] Titration 1 of 3
[2:43] Adding initial titrants
[2:43] Automatically add 1.50000 mL of water
[3:08] Dispensed 1.500000 mL of Water (0.15 M KCl)
[3:12] Titrator arm moved over Drain
[5:54] Titrator arm moved to Titration position
[5:54] Argon flow rate set to 100
[5:54] Stirrer speed set to 10
[6:00] Initial pH = 5.21
[6:00] Iterative adjust 5.21 -> 10.00
[6:00] pH 5.21 -> 10.00
[6:01] Air gap released for Base (0.5 M KOH)
[6:02] Dispensed 0.008114 mL of Base (0.5 M KOH)
[6:07] Holding pH 10.00
[8:07] Stirrer speed set to 0
[8:07] Stirrer speed set to 50
[8:07] Iterative adjust 11.12 -> 10.00
[8:52] Stirrer speed set to 0
[9:26] Datapoint id 2 collected
[9:26] Stirrer speed set to 50
[9:31] pH 10.39 -> 10.19
[9:31] Using cautious pH adjust
[9:32] Air gap released for Acid (0.5 M HCl)
[9:33] Dispensed 0.000212 mL of Acid (0.5 M HCl)
[9:38] Stepping pH = 10.07
[9:53] Stirrer speed set to 0
[10:48] Datapoint id 3 collected
[10:48] Charge balance equation is out by 50.1%
[10:48] Stirrer speed set to 50
[10:53] pH 9.64 -> 9.44
[10:53] Using cautious pH adjust
[10:54] Dispensed 0.000047 mL of Acid (0.5 M HCl)
[10:59] Stepping pH = 9.60
[10:59] Dispensed 0.000071 mL of Acid (0.5 M HCl)
[11:04] Stepping pH = 9.55
[11:04] Dispensed 0.000094 mL of Acid (0.5 M HCl)
[11:09] Stepping pH = 9.35

#### Experiment Log (continued)

[11:24] Stirrer speed set to 0
[12:07] Datapoint id 4 collected
[12:07] Charge balance equation is out by -137.2%
[12:07] Stirrer speed set to 50
[12:12] pH 8.90 -> 8.70
[12:12] Using cautious pH adjust
[12:12] Dispensed 0.000024 mL of Acid (0.5 M HCl)
[12:17] Stepping pH = 8.82
[12:17] Dispensed 0.000024 mL of Acid (0.5 M HCl)
[12:22] Stepping pH = 8.74
[12:22] Dispensed 0.000024 mL of Acid (0.5 M HCl)
[12:28] Stepping pH = 8.61
[12:43] Stirrer speed set to 0
[13:43] Datapoint id 5 collected
[13:43] Charge balance equation is out by -121.9%
[13:43] Stirrer speed set to 50
[13:48] pH 8.13 -> 7.93
[13:48] Using cautious pH adjust
[13:48] Dispensed 0.000024 mL of Acid (0.5 M HCl)
[13:53] Stepping pH = 8.08
[13:53] Dispensed 0.000047 mL of Acid (0.5 M HCl)
[13:58] Stepping pH = 8.00
[13:58] Dispensed 0.000024 mL of Acid (0.5 M HCl)
[14:03] Stepping pH = 7.90
[14:18] Stirrer speed set to 0
[14:31] Datapoint id 6 collected
[14:31] Charge balance equation is out by -91.6%
[14:31] Stirrer speed set to 50
[14:36] pH 7.77 -> 7.57
[14:36] Using cautious pH adjust
[14:36] Dispensed 0.000047 mL of Acid (0.5 M HCl)
[14:41] Stepping pH = 7.72
[14:41] Dispensed 0.000071 mL of Acid (0.5 M HCl)
[14:46] Stepping pH = 7.57
[15:01] Stirrer speed set to 0
[15:16] Datapoint id 7 collected
[15:16] Charge balance equation is out by -35.0%
[15:16] Stirrer speed set to 50
[15:21] pH 7.51 -> 7.31
[15:21] Using cautious pH adjust
[15:21] Dispensed 0.000071 mL of Acid (0.5 M HCl)
[15:26] Stepping pH = 7.44
[15:26] Dispensed 0.000118 mL of Acid (0.5 M HCl)
[15:31] Stepping pH = 7.28
[15:46] Stirrer speed set to 0
[16:01] Datapoint id 8 collected
[16:01] Charge balance equation is out by -17.3%
[16:01] Stirrer speed set to 50
[16:06] pH 7.29 -> 7.09
[16:06] Using cautious pH adjust
[16:06] Dispensed 0.000118 mL of Acid (0.5 M HCl)
[16:12] Stepping pH = 7.20
[16:12] Dispensed 0.000118 mL of Acid (0.5 M HCl)
[16:17] Stepping pH = 7.10
[16:17] Dispensed 0.000024 mL of Acid (0.5 M HCl)
[16:22] Stepping pH = 7.10
[16:22] Dispensed 0.000118 mL of Acid (0.5 M HCl)
[16:27] Stepping pH = 7.02
[16:42] Stirrer speed set to 0

| | | | |
|---|---|---|---|
| Sample name: | **M08_octanol** | Experiment start time: | **3/2/2018 6:51:30 PM** |
| Assay name: | **pH-metric high logP** | Analyst: | **Pion** |
| Assay ID: | **18C-02008** | Instrument ID: | **T312060** |
| Filename: | **C:\Sirius_T3\Mehtap\20180302_exp29_logP_T3-2\18C-02008_M08_octanol_pH-metric high logP.t3r** | | |

#### Experiment Log (continued)

[16:58] Datapoint id 9 collected
[16:58] Charge balance equation is out by -48.6%
[16:58] Stirrer speed set to 50
[17:03] pH 7.04 -> 6.84
[17:03] Using cautious pH adjust
[17:03] Dispensed 0.000188 mL of Acid (0.5 M HCl)
[17:08] Stepping pH = 6.92
[17:09] Dispensed 0.000118 mL of Acid (0.5 M HCl)
[17:14] Stepping pH = 6.88
[17:14] Dispensed 0.000071 mL of Acid (0.5 M HCl)
[17:19] Stepping pH = 6.86
[17:19] Dispensed 0.000047 mL of Acid (0.5 M HCl)
[17:24] Stepping pH = 6.85
[17:39] Stirrer speed set to 0
[17:54] Datapoint id 10 collected
[17:54] Charge balance equation is out by -11.0%
[17:54] Stirrer speed set to 50
[17:59] pH 6.86 -> 6.66
[17:59] Using charge balance adjust
[17:59] Dispensed 0.000541 mL of Acid (0.5 M HCl)
[18:19] Stirrer speed set to 0
[18:35] Datapoint id 11 collected
[18:35] Charge balance equation is out by -2.2%
[18:35] Stirrer speed set to 50
[18:40] pH 6.66 -> 6.46
[18:40] Using charge balance adjust
[18:40] Dispensed 0.000706 mL of Acid (0.5 M HCl)
[19:00] Stirrer speed set to 0
[19:17] Datapoint id 12 collected
[19:17] Charge balance equation is out by -10.0%
[19:17] Stirrer speed set to 50
[19:22] pH 6.48 -> 6.28
[19:22] Using charge balance adjust
[19:22] Dispensed 0.000847 mL of Acid (0.5 M HCl)
[19:42] Stirrer speed set to 0
[19:57] Datapoint id 13 collected
[19:57] Charge balance equation is out by -11.9%
[19:57] Stirrer speed set to 50
[20:02] pH 6.31 -> 6.11
[20:02] Using charge balance adjust
[20:02] Dispensed 0.000964 mL of Acid (0.5 M HCl)
[20:23] Stirrer speed set to 0
[20:37] Datapoint id 14 collected
[20:37] Charge balance equation is out by -3.1%
[20:37] Stirrer speed set to 50
[20:42] pH 6.12 -> 5.92
[20:42] Using charge balance adjust
[20:42] Dispensed 0.000988 mL of Acid (0.5 M HCl)
[21:02] Stirrer speed set to 0
[21:19] Datapoint id 15 collected
[21:19] Charge balance equation is out by -3.5%
[21:19] Stirrer speed set to 50
[21:24] pH 5.93 -> 5.73
[21:24] Using charge balance adjust
[21:24] Dispensed 0.000941 mL of Acid (0.5 M HCl)
[21:45] Stirrer speed set to 0
[22:02] Datapoint id 16 collected
[22:02] Charge balance equation is out by 12.1%
[22:02] Stirrer speed set to 50

| | | | |
|---|---|---|---|
| Sample name: | **M08_octanol** | Experiment start time: | **3/2/2018 6:51:30 PM** |
| Assay name: | **pH-metric high logP** | Analyst: | **Pion** |
| Assay ID: | **18C-02008** | Instrument ID: | **T312060** |
| Filename: | **C:\Sirius_T3\Mehtap\20180302_exp29_logP_T3-2\18C-02008_M08_octanol_pH-metric high logP.t3r** | | |

#### Experiment Log (continued)

[22:07] pH 5.71 -> 5.51
[22:07] Using charge balance adjust
[22:07] Dispensed 0.000776 mL of Acid (0.5 M HCl)
[22:28] Stirrer speed set to 0
[22:56] Datapoint id 17 collected
[22:56] Charge balance equation is out by 25.0%
[22:56] Stirrer speed set to 50
[23:01] pH 5.46 -> 5.26
[23:01] Using cautious pH adjust
[23:01] Dispensed 0.000282 mL of Acid (0.5 M HCl)
[23:06] Stepping pH = 5.35
[23:06] Dispensed 0.000141 mL of Acid (0.5 M HCl)
[23:11] Stepping pH = 5.28
[23:11] Dispensed 0.000024 mL of Acid (0.5 M HCl)
[23:16] Stepping pH = 5.28
[23:16] Dispensed 0.000071 mL of Acid (0.5 M HCl)
[23:21] Stepping pH = 5.24
[23:36] Stirrer speed set to 0
[24:36] Datapoint id 18 collected
[24:36] Charge balance equation is out by 10.7%
[24:36] Stirrer speed set to 50
[24:42] pH 5.41 -> 5.21
[24:42] Using charge balance adjust
[24:42] Dispensed 0.000541 mL of Acid (0.5 M HCl)
[25:02] Stirrer speed set to 0
[26:02] Datapoint id 19 collected
[26:02] Charge balance equation is out by 187.7%
[26:02] Stirrer speed set to 50
[26:07] pH 4.89 -> 4.69
[26:07] Using cautious pH adjust
[26:07] Dispensed 0.000118 mL of Acid (0.5 M HCl)
[26:12] Stepping pH = 4.73
[26:12] Dispensed 0.000024 mL of Acid (0.5 M HCl)
[26:17] Stepping pH = 4.70
[26:17] Dispensed 0.000024 mL of Acid (0.5 M HCl)
[26:22] Stepping pH = 4.65
[26:38] Stirrer speed set to 0
[27:09] Datapoint id 20 collected
[27:09] Charge balance equation is out by 31.0%
[27:09] Stirrer speed set to 50
[27:14] pH 4.68 -> 4.48
[27:14] Using cautious pH adjust
[27:14] Dispensed 0.000094 mL of Acid (0.5 M HCl)
[27:19] Stepping pH = 4.44
[27:34] Stirrer speed set to 0
[27:47] Datapoint id 21 collected
[27:47] Charge balance equation is out by 50.0%
[27:47] Stirrer speed set to 50
[27:52] pH 4.43 -> 4.23
[27:52] Using cautious pH adjust
[27:52] Dispensed 0.000071 mL of Acid (0.5 M HCl)
[27:57] Stepping pH = 4.32
[27:57] Dispensed 0.000047 mL of Acid (0.5 M HCl)
[28:02] Stepping pH = 4.23
[28:17] Stirrer speed set to 0
[28:28] Datapoint id 22 collected
[28:28] Charge balance equation is out by 19.5%
[28:28] Stirrer speed set to 50
[28:34] pH 4.22 -> 4.02

| | | | |
|---|---|---|---|
| Sample name: | **M08_octanol** | Experiment start time: | **3/2/2018 6:51:30 PM** |
| Assay name: | **pH-metric high logP** | Analyst: | **Pion** |
| Assay ID: | **18C-02008** | Instrument ID: | **T312060** |
| Filename: | **C:\Sirius_T3\Mehtap\20180302_exp29_logP_T3-2\18C-02008_M08_octanol_pH-metric high logP.t3r** | | |

#### Experiment Log (continued)

[28:34] Using cautious pH adjust
[28:34] Dispensed 0.000094 mL of Acid (0.5 M HCl)
[28:39] Stepping pH = 4.08
[28:39] Dispensed 0.000024 mL of Acid (0.5 M HCl)
[28:44] Stepping pH = 4.05
[28:44] Dispensed 0.000024 mL of Acid (0.5 M HCl)
[28:49] Stepping pH = 4.04
[28:49] Dispensed 0.000024 mL of Acid (0.5 M HCl)
[28:54] Stepping pH = 4.01
[29:09] Stirrer speed set to 0
[29:19] Datapoint id 23 collected
[29:19] Charge balance equation is out by 5.7%
[29:19] Stirrer speed set to 50
[29:24] pH 4.00 -> 3.80
[29:24] Using charge balance adjust
[29:24] Dispensed 0.000259 mL of Acid (0.5 M HCl)
[29:45] Stirrer speed set to 0
[29:55] Datapoint id 24 collected
[29:55] Charge balance equation is out by 26.5%
[29:55] Stirrer speed set to 50
[30:00] pH 3.75 -> 3.55
[30:00] Using cautious pH adjust
[30:00] Dispensed 0.000212 mL of Acid (0.5 M HCl)
[30:05] Stepping pH = 3.64
[30:05] Dispensed 0.000141 mL of Acid (0.5 M HCl)
[30:10] Stepping pH = 3.57
[30:10] Dispensed 0.000047 mL of Acid (0.5 M HCl)
[30:15] Stepping pH = 3.56
[30:15] Dispensed 0.000024 mL of Acid (0.5 M HCl)
[30:20] Stepping pH = 3.56
[30:35] Stirrer speed set to 0
[30:45] Datapoint id 25 collected
[30:45] Charge balance equation is out by 1.3%
[30:45] Stirrer speed set to 50
[30:51] pH 3.55 -> 3.35
[30:51] Using charge balance adjust
[30:51] Dispensed 0.000659 mL of Acid (0.5 M HCl)
[31:11] Stirrer speed set to 0
[31:21] Datapoint id 26 collected
[31:21] Charge balance equation is out by 18.1%
[31:21] Stirrer speed set to 50
[31:26] pH 3.32 -> 3.12
[31:26] Using cautious pH adjust
[31:26] Dispensed 0.000541 mL of Acid (0.5 M HCl)
[31:31] Stepping pH = 3.21
[31:31] Dispensed 0.000353 mL of Acid (0.5 M HCl)
[31:36] Stepping pH = 3.15
[31:36] Dispensed 0.000141 mL of Acid (0.5 M HCl)
[31:42] Stepping pH = 3.13
[31:42] Dispensed 0.000094 mL of Acid (0.5 M HCl)
[31:47] Stepping pH = 3.12
[32:02] Stirrer speed set to 0
[32:12] Datapoint id 27 collected
[32:12] Charge balance equation is out by -5.1%
[32:12] Stirrer speed set to 50
[32:17] pH 3.12 -> 2.92
[32:17] Using charge balance adjust
[32:18] Dispensed 0.001740 mL of Acid (0.5 M HCl)
[32:38] Stirrer speed set to 0

| | | | |
|---|---|---|---|
| Sample name: | **M08_octanol** | Experiment start time: | **3/2/2018 6:51:30 PM** |
| Assay name: | **pH-metric high logP** | Analyst: | **Pion** |
| Assay ID: | **18C-02008** | Instrument ID: | **T312060** |
| Filename: | **C:\Sirius_T3\Mehtap\20180302_exp29_logP_T3-2\18C-02008_M08_octanol_pH-metric high logP.t3r** | | |

#### Experiment Log (continued)

[32:48] Datapoint id 28 collected
[32:48] Charge balance equation is out by 2.3%
[32:48] Stirrer speed set to 50
[32:53] pH 2.92 -> 2.72
[32:53] Using charge balance adjust
[32:53] Dispensed 0.002775 mL of Acid (0.5 M HCl)
[33:13] Stirrer speed set to 0
[33:23] Datapoint id 29 collected
[33:23] Charge balance equation is out by -4.1%
[33:23] Stirrer speed set to 50
[33:28] pH 2.73 -> 2.53
[33:28] Using charge balance adjust
[33:29] Dispensed 0.004280 mL of Acid (0.5 M HCl)
[33:49] Stirrer speed set to 0
[33:59] Datapoint id 30 collected
[33:59] Charge balance equation is out by -2.8%
[33:59] Stirrer speed set to 50
[34:04] pH 2.54 -> 2.34
[34:04] Using charge balance adjust
[34:05] Dispensed 0.006703 mL of Acid (0.5 M HCl)
[34:25] Stirrer speed set to 0
[34:35] Datapoint id 31 collected
[34:35] Charge balance equation is out by -1.5%
[34:35] Stirrer speed set to 50
[34:40] pH 2.35 -> 2.15
[34:40] Using charge balance adjust
[34:40] Dispensed 0.010630 mL of Acid (0.5 M HCl)
[35:01] Stirrer speed set to 0
[35:11] Datapoint id 32 collected
[35:11] Charge balance equation is out by 1.5%
[35:11] Stirrer speed set to 50
[35:16] pH 2.15 -> 1.95
[35:16] Using charge balance adjust
[35:17] Dispensed 0.017310 mL of Acid (0.5 M HCl)
[35:37] Stirrer speed set to 0
[35:47] Datapoint id 33 collected
[35:47] Charge balance equation is out by -0.1%
[35:47] Titration 2 of 3
[35:47] Adding initial titrants
[35:47] Automatically add 0.20000 mL of Octanol
[35:52] Dispensed 0.200000 mL of Octanol
[35:52] Stirrer speed set to 10
[35:53] Stirrer speed set to 55
[35:53] Iterative adjust 1.96 -> 10.00
[35:53] pH 1.96 -> 10.00
[35:55] Dispensed 0.055221 mL of Base (0.5 M KOH)
[36:45] Stirrer speed set to 0
[37:23] Datapoint id 34 collected
[37:23] Stirrer speed set to 55
[37:28] pH 10.05 -> 9.85
[37:28] Using cautious pH adjust
[37:28] Dispensed 0.000118 mL of Acid (0.5 M HCl)
[37:33] Stepping pH = 9.99
[37:33] Dispensed 0.000141 mL of Acid (0.5 M HCl)
[37:38] Stepping pH = 9.81
[37:53] Stirrer speed set to 0
[38:45] Datapoint id 35 collected
[38:45] Charge balance equation is out by -10.4%
[38:45] Stirrer speed set to 55

#### Experiment Log (continued)

[38:50] pH 9.48 -> 9.28
[38:50] Using charge balance adjust
[38:50] Dispensed 0.000071 mL of Acid (0.5 M HCl)
[39:10] Stirrer speed set to 0
[40:06] Datapoint id 36 collected
[40:06] Charge balance equation is out by 26.4%
[40:06] Stirrer speed set to 55
[40:11] pH 9.11 -> 8.91
[40:11] Using cautious pH adjust
[40:11] Dispensed 0.000024 mL of Acid (0.5 M HCl)
[40:16] Stepping pH = 9.08
[40:16] Dispensed 0.000047 mL of Acid (0.5 M HCl)
[40:21] Stepping pH = 8.95
[40:21] Dispensed 0.000024 mL of Acid (0.5 M HCl)
[40:26] Stepping pH = 8.86
[40:41] Stirrer speed set to 0
[41:21] Datapoint id 37 collected
[41:21] Charge balance equation is out by -100.8%
[41:21] Stirrer speed set to 55
[41:27] pH 8.60 -> 8.40
[41:27] Using cautious pH adjust
[41:27] Dispensed 0.000024 mL of Acid (0.5 M HCl)
[41:32] Stepping pH = 8.56
[41:32] Dispensed 0.000047 mL of Acid (0.5 M HCl)
[41:37] Stepping pH = 8.42
[41:37] Dispensed 0.000024 mL of Acid (0.5 M HCl)
[41:42] Stepping pH = 8.36
[41:57] Stirrer speed set to 0
[42:08] Datapoint id 38 collected
[42:08] Charge balance equation is out by -78.4%
[42:08] Stirrer speed set to 55
[42:13] pH 8.28 -> 8.08
[42:13] Using cautious pH adjust
[42:13] Dispensed 0.000047 mL of Acid (0.5 M HCl)
[42:18] Stepping pH = 8.21
[42:18] Dispensed 0.000047 mL of Acid (0.5 M HCl)
[42:23] Stepping pH = 8.11
[42:23] Dispensed 0.000024 mL of Acid (0.5 M HCl)
[42:28] Stepping pH = 8.08
[42:43] Stirrer speed set to 0
[42:55] Datapoint id 39 collected
[42:55] Charge balance equation is out by -33.8%
[42:55] Stirrer speed set to 55
[43:00] pH 8.04 -> 7.84
[43:00] Using cautious pH adjust
[43:00] Dispensed 0.000071 mL of Acid (0.5 M HCl)
[43:06] Stepping pH = 7.94
[43:06] Dispensed 0.000047 mL of Acid (0.5 M HCl)
[43:11] Stepping pH = 7.89
[43:11] Dispensed 0.000047 mL of Acid (0.5 M HCl)
[43:16] Stepping pH = 7.83
[43:31] Stirrer speed set to 0
[43:44] Datapoint id 40 collected
[43:44] Charge balance equation is out by -18.6%
[43:44] Stirrer speed set to 55
[43:49] pH 7.80 -> 7.60
[43:49] Using cautious pH adjust
[43:49] Dispensed 0.000118 mL of Acid (0.5 M HCl)
[43:54] Stepping pH = 7.70

| | | | |
|---|---|---|---|
| Sample name: | **M08_octanol** | Experiment start time: | **3/2/2018 6:51:30 PM** |
| Assay name: | **pH-metric high logP** | Analyst: | **Pion** |
| Assay ID: | **18C-02008** | Instrument ID: | **T312060** |
| Filename: | **C:\Sirius_T3\Mehtap\20180302_exp29_logP_T3-2\18C-02008_M08_octanol_pH-metric high logP.t3r** | | |

#### Experiment Log (continued)

[43:54] Dispensed 0.000094 mL of Acid (0.5 M HCl)
[43:59] Stepping pH = 7.63
[43:59] Dispensed 0.000047 mL of Acid (0.5 M HCl)
[44:04] Stepping pH = 7.60
[44:19] Stirrer speed set to 0
[44:34] Datapoint id 41 collected
[44:34] Charge balance equation is out by -7.6%
[44:34] Stirrer speed set to 55
[44:39] pH 7.59 -> 7.39
[44:39] Using charge balance adjust
[44:39] Dispensed 0.000353 mL of Acid (0.5 M HCl)
[44:59] Stirrer speed set to 0
[45:11] Datapoint id 42 collected
[45:11] Charge balance equation is out by 4.7%
[45:11] Stirrer speed set to 55
[45:16] pH 7.37 -> 7.17
[45:16] Using charge balance adjust
[45:16] Dispensed 0.000517 mL of Acid (0.5 M HCl)
[45:36] Stirrer speed set to 0
[45:49] Datapoint id 43 collected
[45:49] Charge balance equation is out by -1.7%
[45:49] Stirrer speed set to 55
[45:54] pH 7.17 -> 6.97
[45:54] Using charge balance adjust
[45:54] Dispensed 0.000682 mL of Acid (0.5 M HCl)
[46:14] Stirrer speed set to 0
[46:26] Datapoint id 44 collected
[46:26] Charge balance equation is out by -8.7%
[46:26] Stirrer speed set to 55
[46:31] pH 6.99 -> 6.79
[46:31] Using charge balance adjust
[46:31] Dispensed 0.000847 mL of Acid (0.5 M HCl)
[46:51] Stirrer speed set to 0
[47:04] Datapoint id 45 collected
[47:04] Charge balance equation is out by -5.3%
[47:04] Stirrer speed set to 55
[47:09] pH 6.80 -> 6.60
[47:09] Using charge balance adjust
[47:09] Dispensed 0.000964 mL of Acid (0.5 M HCl)
[47:29] Stirrer speed set to 0
[47:42] Datapoint id 46 collected
[47:42] Charge balance equation is out by -11.4%
[47:42] Stirrer speed set to 55
[47:47] pH 6.63 -> 6.43
[47:47] Using charge balance adjust
[47:47] Dispensed 0.000988 mL of Acid (0.5 M HCl)
[48:07] Stirrer speed set to 0
[48:24] Datapoint id 47 collected
[48:24] Charge balance equation is out by -13.9%
[48:24] Stirrer speed set to 55
[48:29] pH 6.45 -> 6.25
[48:29] Using charge balance adjust
[48:29] Dispensed 0.000964 mL of Acid (0.5 M HCl)
[48:49] Stirrer speed set to 0
[49:07] Datapoint id 48 collected
[49:07] Charge balance equation is out by -14.7%
[49:07] Stirrer speed set to 55
[49:12] pH 6.28 -> 6.08
[49:12] Using charge balance adjust

| | | | |
|---|---|---|---|
| Sample name: | **M08_octanol** | Experiment start time: | **3/2/2018 6:51:30 PM** |
| Assay name: | **pH-metric high logP** | Analyst: | **Pion** |
| Assay ID: | **18C-02008** | Instrument ID: | **T312060** |
| Filename: | **C:\Sirius_T3\Mehtap\20180302_exp29_logP_T3-2\18C-02008_M08_octanol_pH-metric high logP.t3r** | | |

#### Experiment Log (continued)

[49:12] Dispensed 0.000847 mL of Acid (0.5 M HCl)
[49:32] Stirrer speed set to 0
[49:49] Datapoint id 49 collected
[49:49] Charge balance equation is out by -13.2%
[49:49] Stirrer speed set to 55
[49:54] pH 6.09 -> 5.89
[49:54] Using charge balance adjust
[49:54] Dispensed 0.000706 mL of Acid (0.5 M HCl)
[50:15] Stirrer speed set to 0
[50:32] Datapoint id 50 collected
[50:32] Charge balance equation is out by -7.5%
[50:32] Stirrer speed set to 55
[50:37] pH 5.90 -> 5.70
[50:37] Using charge balance adjust
[50:37] Dispensed 0.000517 mL of Acid (0.5 M HCl)
[50:58] Stirrer speed set to 0
[51:13] Datapoint id 51 collected
[51:13] Charge balance equation is out by 7.5%
[51:13] Stirrer speed set to 55
[51:18] pH 5.67 -> 5.47
[51:18] Using charge balance adjust
[51:18] Dispensed 0.000353 mL of Acid (0.5 M HCl)
[51:38] Stirrer speed set to 0
[51:54] Datapoint id 52 collected
[51:54] Charge balance equation is out by -2.1%
[51:54] Stirrer speed set to 55
[51:59] pH 5.46 -> 5.26
[51:59] Using charge balance adjust
[51:59] Dispensed 0.000235 mL of Acid (0.5 M HCl)
[52:19] Stirrer speed set to 0
[52:33] Datapoint id 53 collected
[52:33] Charge balance equation is out by -17.6%
[52:33] Stirrer speed set to 55
[52:39] pH 5.27 -> 5.07
[52:39] Using cautious pH adjust
[52:39] Dispensed 0.000094 mL of Acid (0.5 M HCl)
[52:44] Stepping pH = 5.25
[52:44] Dispensed 0.000212 mL of Acid (0.5 M HCl)
[52:49] Stepping pH = 4.83
[53:04] Stirrer speed set to 0
[53:18] Datapoint id 54 collected
[53:18] Charge balance equation is out by -65.9%
[53:18] Stirrer speed set to 55
[53:23] pH 4.84 -> 4.64
[53:23] Using cautious pH adjust
[53:23] Dispensed 0.000047 mL of Acid (0.5 M HCl)
[53:28] Stepping pH = 4.81
[53:28] Dispensed 0.000118 mL of Acid (0.5 M HCl)
[53:33] Stepping pH = 4.59
[53:48] Stirrer speed set to 0
[54:01] Datapoint id 55 collected
[54:01] Charge balance equation is out by -62.4%
[54:01] Stirrer speed set to 55
[54:06] pH 4.59 -> 4.39
[54:06] Using cautious pH adjust
[54:06] Dispensed 0.000047 mL of Acid (0.5 M HCl)
[54:11] Stepping pH = 4.59
[54:12] Dispensed 0.000141 mL of Acid (0.5 M HCl)
[54:17] Stepping pH = 4.27

| | | | |
|---|---|---|---|
| Sample name: | **M08_octanol** | Experiment start time: | **3/2/2018 6:51:30 PM** |
| Assay name: | **pH-metric high logP** | Analyst: | **Pion** |
| Assay ID: | **18C-02008** | Instrument ID: | **T312060** |
| Filename: | **C:\Sirius_T3\Mehtap\20180302_exp29_logP_T3-2\18C-02008_M08_octanol_pH-metric high logP.t3r** | | |

#### Experiment Log (continued)

[54:32] Stirrer speed set to 0
[54:43] Datapoint id 56 collected
[54:43] Charge balance equation is out by -100.0%
[54:43] Stirrer speed set to 55
[54:48] pH 4.28 -> 4.08
[54:48] Using cautious pH adjust
[54:48] Dispensed 0.000071 mL of Acid (0.5 M HCl)
[54:54] Stepping pH = 4.25
[54:54] Dispensed 0.000188 mL of Acid (0.5 M HCl)
[54:59] Stepping pH = 3.97
[55:14] Stirrer speed set to 0
[55:24] Datapoint id 57 collected
[55:24] Charge balance equation is out by -70.8%
[55:24] Stirrer speed set to 55
[55:29] pH 3.96 -> 3.76
[55:29] Using cautious pH adjust
[55:29] Dispensed 0.000141 mL of Acid (0.5 M HCl)
[55:34] Stepping pH = 3.87
[55:34] Dispensed 0.000118 mL of Acid (0.5 M HCl)
[55:39] Stepping pH = 3.79
[55:39] Dispensed 0.000047 mL of Acid (0.5 M HCl)
[55:44] Stepping pH = 3.77
[55:44] Dispensed 0.000024 mL of Acid (0.5 M HCl)
[55:49] Stepping pH = 3.76
[56:05] Stirrer speed set to 0
[56:15] Datapoint id 58 collected
[56:15] Charge balance equation is out by -19.0%
[56:15] Stirrer speed set to 55
[56:20] pH 3.76 -> 3.56
[56:20] Using cautious pH adjust
[56:20] Dispensed 0.000212 mL of Acid (0.5 M HCl)
[56:25] Stepping pH = 3.63
[56:25] Dispensed 0.000118 mL of Acid (0.5 M HCl)
[56:30] Stepping pH = 3.59
[56:30] Dispensed 0.000047 mL of Acid (0.5 M HCl)
[56:35] Stepping pH = 3.57
[56:35] Dispensed 0.000047 mL of Acid (0.5 M HCl)
[56:40] Stepping pH = 3.56
[56:55] Stirrer speed set to 0
[57:06] Datapoint id 59 collected
[57:06] Charge balance equation is out by 3.2%
[57:06] Stirrer speed set to 55
[57:11] pH 3.55 -> 3.35
[57:11] Using charge balance adjust
[57:11] Dispensed 0.000682 mL of Acid (0.5 M HCl)
[57:31] Stirrer speed set to 0
[57:41] Datapoint id 60 collected
[57:41] Charge balance equation is out by 10.1%
[57:41] Stirrer speed set to 55
[57:46] pH 3.33 -> 3.13
[57:46] Using charge balance adjust
[57:46] Dispensed 0.001129 mL of Acid (0.5 M HCl)
[58:06] Stirrer speed set to 0
[58:16] Datapoint id 61 collected
[58:16] Charge balance equation is out by 3.9%
[58:16] Stirrer speed set to 55
[58:21] pH 3.13 -> 2.93
[58:21] Using charge balance adjust
[58:22] Dispensed 0.001834 mL of Acid (0.5 M HCl)

| | | | |
|---|---|---|---|
| Sample name: | **M08_octanol** | Experiment start time: | **3/2/2018 6:51:30 PM** |
| Assay name: | **pH-metric high logP** | Analyst: | **Pion** |
| Assay ID: | **18C-02008** | Instrument ID: | **T312060** |
| Filename: | **C:\Sirius_T3\Mehtap\20180302_exp29_logP_T3-2\18C-02008_M08_octanol_pH-metric high logP.t3r** | | |

#### Experiment Log (continued)

[58:42] Stirrer speed set to 0
[58:52] Datapoint id 62 collected
[58:52] Charge balance equation is out by 1.2%
[58:52] Stirrer speed set to 55
[58:57] pH 2.93 -> 2.73
[58:57] Using charge balance adjust
[58:57] Dispensed 0.002893 mL of Acid (0.5 M HCl)
[59:17] Stirrer speed set to 0
[59:28] Datapoint id 63 collected
[59:28] Charge balance equation is out by 3.2%
[59:28] Stirrer speed set to 55
[59:33] pH 2.73 -> 2.53
[59:33] Using charge balance adjust
[59:33] Dispensed 0.004633 mL of Acid (0.5 M HCl)
[59:53] Stirrer speed set to 0
[1:00:04] Datapoint id 64 collected
[1:00:04] Charge balance equation is out by 5.3%
[1:00:04] Stirrer speed set to 55
[1:00:09] pH 2.52 -> 2.32
[1:00:09] Using charge balance adjust
[1:00:09] Dispensed 0.007573 mL of Acid (0.5 M HCl)
[1:00:29] Stirrer speed set to 0
[1:00:39] Datapoint id 65 collected
[1:00:39] Charge balance equation is out by -5.8%
[1:00:39] Stirrer speed set to 55
[1:00:45] pH 2.34 -> 2.14
[1:00:45] Using charge balance adjust
[1:00:45] Dispensed 0.011830 mL of Acid (0.5 M HCl)
[1:01:05] Stirrer speed set to 0
[1:01:15] Datapoint id 66 collected
[1:01:15] Charge balance equation is out by 1.7%
[1:01:15] Stirrer speed set to 55
[1:01:20] pH 2.14 -> 1.95
[1:01:20] Using charge balance adjust
[1:01:21] Dispensed 0.017686 mL of Acid (0.5 M HCl)
[1:01:41] Stirrer speed set to 0
[1:01:52] Datapoint id 67 collected
[1:01:52] Charge balance equation is out by -8.3%
[1:01:52] Titration 3 of 3
[1:01:52] Adding initial titrants
[1:01:52] Automatically add 0.80000 mL of Octanol
[1:02:42] Dispensed 0.800000 mL of Octanol
[1:02:42] Stirrer speed set to 10
[1:02:43] Stirrer speed set to 60
[1:02:43] Iterative adjust 1.94 -> 10.00
[1:02:43] pH 1.94 -> 10.00
[1:02:44] Dispensed 0.060113 mL of Base (0.5 M KOH)
[1:03:34] Stirrer speed set to 0
[1:04:20] Datapoint id 68 collected
[1:04:20] Stirrer speed set to 60
[1:04:25] pH 10.13 -> 9.93
[1:04:25] Using cautious pH adjust
[1:04:25] Dispensed 0.000141 mL of Acid (0.5 M HCl)
[1:04:30] Stepping pH = 10.07
[1:04:30] Dispensed 0.000165 mL of Acid (0.5 M HCl)
[1:04:35] Stepping pH = 9.92
[1:04:50] Stirrer speed set to 0
[1:05:28] Datapoint id 69 collected
[1:05:28] Charge balance equation is out by -4.4%

#### Experiment Log (continued)

[1:05:28] Stirrer speed set to 60
[1:05:33] pH 9.82 -> 9.62
[1:05:33] Using charge balance adjust
[1:05:33] Dispensed 0.000165 mL of Acid (0.5 M HCl)
[1:05:53] Stirrer speed set to 0
[1:06:31] Datapoint id 70 collected
[1:06:31] Charge balance equation is out by 12.1%
[1:06:31] Stirrer speed set to 60
[1:06:36] pH 9.60 -> 9.40
[1:06:36] Using charge balance adjust
[1:06:36] Dispensed 0.000118 mL of Acid (0.5 M HCl)
[1:06:56] Stirrer speed set to 0
[1:07:50] Datapoint id 71 collected
[1:07:50] Charge balance equation is out by -11.8%
[1:07:50] Stirrer speed set to 60
[1:07:55] pH 9.39 -> 9.19
[1:07:55] Using charge balance adjust
[1:07:55] Dispensed 0.000094 mL of Acid (0.5 M HCl)
[1:08:15] Stirrer speed set to 0
[1:08:26] Datapoint id 72 collected
[1:08:26] Charge balance equation is out by -31.9%
[1:08:26] Stirrer speed set to 60
[1:08:31] pH 9.25 -> 9.05
[1:08:31] Using cautious pH adjust
[1:08:31] Dispensed 0.000047 mL of Acid (0.5 M HCl)
[1:08:36] Stepping pH = 9.20
[1:08:36] Dispensed 0.000071 mL of Acid (0.5 M HCl)
[1:08:41] Stepping pH = 9.08
[1:08:41] Dispensed 0.000024 mL of Acid (0.5 M HCl)
[1:08:47] Stepping pH = 9.04
[1:09:02] Stirrer speed set to 0
[1:09:20] Datapoint id 73 collected
[1:09:20] Charge balance equation is out by -49.6%
[1:09:20] Stirrer speed set to 60
[1:09:25] pH 9.00 -> 8.80
[1:09:25] Using cautious pH adjust
[1:09:25] Dispensed 0.000047 mL of Acid (0.5 M HCl)
[1:09:30] Stepping pH = 8.94
[1:09:30] Dispensed 0.000047 mL of Acid (0.5 M HCl)
[1:09:35] Stepping pH = 8.86
[1:09:35] Dispensed 0.000047 mL of Acid (0.5 M HCl)
[1:09:40] Stepping pH = 8.78
[1:09:55] Stirrer speed set to 0
[1:10:15] Datapoint id 74 collected
[1:10:15] Charge balance equation is out by -58.1%
[1:10:15] Stirrer speed set to 60
[1:10:20] pH 8.72 -> 8.52
[1:10:20] Using cautious pH adjust
[1:10:21] Dispensed 0.000071 mL of Acid (0.5 M HCl)
[1:10:26] Stepping pH = 8.63
[1:10:26] Dispensed 0.000047 mL of Acid (0.5 M HCl)
[1:10:31] Stepping pH = 8.56
[1:10:31] Dispensed 0.000024 mL of Acid (0.5 M HCl)
[1:10:36] Stepping pH = 8.52
[1:10:51] Stirrer speed set to 0
[1:11:13] Datapoint id 75 collected
[1:11:13] Charge balance equation is out by -17.8%
[1:11:13] Stirrer speed set to 60
[1:11:18] pH 8.50 -> 8.30

| | | | |
|---|---|---|---|
| Sample name: | **M08_octanol** | Experiment start time: | **3/2/2018 6:51:30 PM** |
| Assay name: | **pH-metric high logP** | Analyst: | **Pion** |
| Assay ID: | **18C-02008** | Instrument ID: | **T312060** |
| Filename: | **C:\Sirius_T3\Mehtap\20180302_exp29_logP_T3-2\18C-02008_M08_octanol_pH-metric high logP.t3r** | | |

#### Experiment Log (continued)

[1:11:18] Using cautious pH adjust
[1:11:18] Dispensed 0.000094 mL of Acid (0.5 M HCl)
[1:11:23] Stepping pH = 8.39
[1:11:23] Dispensed 0.000071 mL of Acid (0.5 M HCl)
[1:11:28] Stepping pH = 8.31
[1:11:28] Dispensed 0.000024 mL of Acid (0.5 M HCl)
[1:11:33] Stepping pH = 8.29
[1:11:48] Stirrer speed set to 0
[1:12:09] Datapoint id 76 collected
[1:12:09] Charge balance equation is out by 5.6%
[1:12:09] Stirrer speed set to 60
[1:12:14] pH 8.27 -> 8.07
[1:12:14] Using charge balance adjust
[1:12:14] Dispensed 0.000282 mL of Acid (0.5 M HCl)
[1:12:34] Stirrer speed set to 0
[1:12:58] Datapoint id 77 collected
[1:12:58] Charge balance equation is out by 6.1%
[1:12:58] Stirrer speed set to 60
[1:13:04] pH 8.03 -> 7.83
[1:13:04] Using charge balance adjust
[1:13:04] Dispensed 0.000423 mL of Acid (0.5 M HCl)
[1:13:24] Stirrer speed set to 0
[1:13:46] Datapoint id 78 collected
[1:13:46] Charge balance equation is out by -1.0%
[1:13:46] Stirrer speed set to 60
[1:13:51] pH 7.81 -> 7.61
[1:13:51] Using charge balance adjust
[1:13:51] Dispensed 0.000611 mL of Acid (0.5 M HCl)
[1:14:11] Stirrer speed set to 0
[1:14:32] Datapoint id 79 collected
[1:14:32] Charge balance equation is out by -0.6%
[1:14:32] Stirrer speed set to 60
[1:14:37] pH 7.59 -> 7.39
[1:14:37] Using charge balance adjust
[1:14:38] Dispensed 0.000800 mL of Acid (0.5 M HCl)
[1:14:58] Stirrer speed set to 0
[1:15:25] Datapoint id 80 collected
[1:15:25] Charge balance equation is out by -5.5%
[1:15:25] Stirrer speed set to 60
[1:15:30] pH 7.38 -> 7.18
[1:15:30] Using charge balance adjust
[1:15:31] Dispensed 0.000941 mL of Acid (0.5 M HCl)
[1:15:51] Stirrer speed set to 0
[1:16:15] Datapoint id 81 collected
[1:16:15] Charge balance equation is out by -10.4%
[1:16:15] Stirrer speed set to 60
[1:16:20] pH 7.18 -> 6.98
[1:16:20] Using charge balance adjust
[1:16:21] Dispensed 0.000988 mL of Acid (0.5 M HCl)
[1:16:41] Stirrer speed set to 0
[1:17:03] Datapoint id 82 collected
[1:17:03] Charge balance equation is out by -12.8%
[1:17:03] Stirrer speed set to 60
[1:17:08] pH 6.99 -> 6.79
[1:17:08] Using charge balance adjust
[1:17:08] Dispensed 0.000941 mL of Acid (0.5 M HCl)
[1:17:28] Stirrer speed set to 0
[1:17:57] Datapoint id 83 collected
[1:17:57] Charge balance equation is out by -21.6%

Sample name: **M08_octanol** Experiment start time: **3/2/2018 6:51:30 PM**
Assay name: **pH-metric high logP** Analyst: **Pion**
Assay ID: **18C-02008** Instrument ID: **T312060**
Filename: **C:\Sirius_T3\Mehtap\20180302_exp29_logP_T3-2\18C-02008_M08_octanol_pH-metric high logP.t3r**

#### Experiment Log (continued)

[1:17:57] Stirrer speed set to 60
[1:18:02] pH 6.79 -> 6.59
[1:18:02] Using cautious pH adjust
[1:18:02] Dispensed 0.000423 mL of Acid (0.5 M HCl)
[1:18:08] Stepping pH = 6.70
[1:18:08] Dispensed 0.000329 mL of Acid (0.5 M HCl)
[1:18:13] Stepping pH = 6.62
[1:18:13] Dispensed 0.000094 mL of Acid (0.5 M HCl)
[1:18:18] Stepping pH = 6.61
[1:18:18] Dispensed 0.000094 mL of Acid (0.5 M HCl)
[1:18:23] Stepping pH = 6.59
[1:18:38] Stirrer speed set to 0
[1:19:13] Datapoint id 84 collected
[1:19:13] Charge balance equation is out by -11.6%
[1:19:13] Stirrer speed set to 60
[1:19:18] pH 6.59 -> 6.39
[1:19:18] Using charge balance adjust
[1:19:18] Dispensed 0.000659 mL of Acid (0.5 M HCl)
[1:19:39] Stirrer speed set to 0
[1:20:12] Datapoint id 85 collected
[1:20:12] Charge balance equation is out by -31.3%
[1:20:12] Stirrer speed set to 60
[1:20:17] pH 6.40 -> 6.20
[1:20:17] Using cautious pH adjust
[1:20:17] Dispensed 0.000259 mL of Acid (0.5 M HCl)
[1:20:22] Stepping pH = 6.31
[1:20:22] Dispensed 0.000212 mL of Acid (0.5 M HCl)
[1:20:27] Stepping pH = 6.23
[1:20:27] Dispensed 0.000071 mL of Acid (0.5 M HCl)
[1:20:32] Stepping pH = 6.21
[1:20:32] Dispensed 0.000047 mL of Acid (0.5 M HCl)
[1:20:37] Stepping pH = 6.20
[1:20:53] Stirrer speed set to 0
[1:21:18] Datapoint id 86 collected
[1:21:18] Charge balance equation is out by -13.3%
[1:21:18] Stirrer speed set to 60
[1:21:23] pH 6.22 -> 6.02
[1:21:23] Using charge balance adjust
[1:21:23] Dispensed 0.000376 mL of Acid (0.5 M HCl)
[1:21:43] Stirrer speed set to 0
[1:22:08] Datapoint id 87 collected
[1:22:08] Charge balance equation is out by -33.3%
[1:22:08] Stirrer speed set to 60
[1:22:13] pH 6.01 -> 5.81
[1:22:13] Using cautious pH adjust
[1:22:13] Dispensed 0.000118 mL of Acid (0.5 M HCl)
[1:22:18] Stepping pH = 5.97
[1:22:18] Dispensed 0.000212 mL of Acid (0.5 M HCl)
[1:22:23] Stepping pH = 5.81
[1:22:38] Stirrer speed set to 0
[1:22:57] Datapoint id 88 collected
[1:22:57] Charge balance equation is out by -32.4%
[1:22:57] Stirrer speed set to 60
[1:23:02] pH 5.82 -> 5.62
[1:23:02] Using cautious pH adjust
[1:23:02] Dispensed 0.000094 mL of Acid (0.5 M HCl)
[1:23:07] Stepping pH = 5.76
[1:23:08] Dispensed 0.000118 mL of Acid (0.5 M HCl)
[1:23:13] Stepping pH = 5.65

| | | | |
|---|---|---|---|
| Sample name: | **M08_octanol** | Experiment start time: | **3/2/2018 6:51:30 PM** |
| Assay name: | **pH-metric high logP** | Analyst: | **Pion** |
| Assay ID: | **18C-02008** | Instrument ID: | **T312060** |
| Filename: | **C:\Sirius_T3\Mehtap\20180302_exp29_logP_T3-2\18C-02008_M08_octanol_pH-metric high logP.t3r** | | |

#### Experiment Log (continued)

[1:23:13] Dispensed 0.000024 mL of Acid (0.5 M HCl)
[1:23:18] Stepping pH = 5.63
[1:23:18] Dispensed 0.000024 mL of Acid (0.5 M HCl)
[1:23:23] Stepping pH = 5.62
[1:23:38] Stirrer speed set to 0
[1:24:02] Datapoint id 89 collected
[1:24:02] Charge balance equation is out by -35.8%
[1:24:02] Stirrer speed set to 60
[1:24:07] pH 5.62 -> 5.42
[1:24:07] Using cautious pH adjust
[1:24:07] Dispensed 0.000047 mL of Acid (0.5 M HCl)
[1:24:12] Stepping pH = 5.59
[1:24:12] Dispensed 0.000118 mL of Acid (0.5 M HCl)
[1:24:17] Stepping pH = 5.42
[1:24:32] Stirrer speed set to 0
[1:24:55] Datapoint id 90 collected
[1:24:55] Charge balance equation is out by -48.7%
[1:24:55] Stirrer speed set to 60
[1:25:00] pH 5.41 -> 5.21
[1:25:00] Using cautious pH adjust
[1:25:00] Dispensed 0.000047 mL of Acid (0.5 M HCl)
[1:25:05] Stepping pH = 5.39
[1:25:05] Dispensed 0.000094 mL of Acid (0.5 M HCl)
[1:25:10] Stepping pH = 5.20
[1:25:25] Stirrer speed set to 0
[1:25:55] Datapoint id 91 collected
[1:25:55] Charge balance equation is out by -72.9%
[1:25:55] Stirrer speed set to 60
[1:26:00] pH 5.20 -> 5.00
[1:26:00] Using cautious pH adjust
[1:26:00] Dispensed 0.000024 mL of Acid (0.5 M HCl)
[1:26:05] Stepping pH = 5.17
[1:26:05] Dispensed 0.000071 mL of Acid (0.5 M HCl)
[1:26:10] Stepping pH = 5.00
[1:26:25] Stirrer speed set to 0
[1:26:40] Datapoint id 92 collected
[1:26:40] Charge balance equation is out by -79.7%
[1:26:40] Stirrer speed set to 60
[1:26:45] pH 4.97 -> 4.77
[1:26:45] Using cautious pH adjust
[1:26:45] Dispensed 0.000024 mL of Acid (0.5 M HCl)
[1:26:50] Stepping pH = 4.97
[1:26:50] Dispensed 0.000141 mL of Acid (0.5 M HCl)
[1:26:55] Stepping pH = 4.59
[1:27:10] Stirrer speed set to 0
[1:27:22] Datapoint id 93 collected
[1:27:22] Charge balance equation is out by -206.8%
[1:27:22] Stirrer speed set to 60
[1:27:27] pH 4.58 -> 4.38
[1:27:27] Using cautious pH adjust
[1:27:27] Dispensed 0.000047 mL of Acid (0.5 M HCl)
[1:27:32] Stepping pH = 4.55
[1:27:33] Dispensed 0.000094 mL of Acid (0.5 M HCl)
[1:27:38] Stepping pH = 4.36
[1:27:53] Stirrer speed set to 0
[1:28:10] Datapoint id 94 collected
[1:28:10] Charge balance equation is out by -82.7%
[1:28:10] Stirrer speed set to 60
[1:28:15] pH 4.35 -> 4.15

| | | | |
|---|---|---|---|
| Sample name: | **M08_octanol** | Experiment start time: | **3/2/2018 6:51:30 PM** |
| Assay name: | **pH-metric high logP** | Analyst: | **Pion** |
| Assay ID: | **18C-02008** | Instrument ID: | **T312060** |
| Filename: | **C:\Sirius_T3\Mehtap\20180302_exp29_logP_T3-2\18C-02008_M08_octanol_pH-metric high logP.t3r** | | |

#### Experiment Log (continued)

[1:28:15] Using cautious pH adjust
[1:28:15] Dispensed 0.000071 mL of Acid (0.5 M HCl)
[1:28:20] Stepping pH = 4.29
[1:28:20] Dispensed 0.000094 mL of Acid (0.5 M HCl)
[1:28:25] Stepping pH = 4.17
[1:28:25] Dispensed 0.000024 mL of Acid (0.5 M HCl)
[1:28:30] Stepping pH = 4.15
[1:28:45] Stirrer speed set to 0
[1:28:56] Datapoint id 95 collected
[1:28:56] Charge balance equation is out by -35.8%
[1:28:56] Stirrer speed set to 60
[1:29:01] pH 4.13 -> 3.93
[1:29:01] Using cautious pH adjust
[1:29:01] Dispensed 0.000094 mL of Acid (0.5 M HCl)
[1:29:06] Stepping pH = 4.05
[1:29:06] Dispensed 0.000094 mL of Acid (0.5 M HCl)
[1:29:11] Stepping pH = 3.97
[1:29:11] Dispensed 0.000047 mL of Acid (0.5 M HCl)
[1:29:17] Stepping pH = 3.94
[1:29:32] Stirrer speed set to 0
[1:29:55] Datapoint id 96 collected
[1:29:55] Charge balance equation is out by -17.3%
[1:29:55] Stirrer speed set to 60
[1:30:00] pH 3.92 -> 3.72
[1:30:00] Using cautious pH adjust
[1:30:00] Dispensed 0.000165 mL of Acid (0.5 M HCl)
[1:30:05] Stepping pH = 3.83
[1:30:05] Dispensed 0.000141 mL of Acid (0.5 M HCl)
[1:30:11] Stepping pH = 3.75
[1:30:11] Dispensed 0.000047 mL of Acid (0.5 M HCl)
[1:30:16] Stepping pH = 3.74
[1:30:16] Dispensed 0.000047 mL of Acid (0.5 M HCl)
[1:30:21] Stepping pH = 3.72
[1:30:36] Stirrer speed set to 0
[1:30:46] Datapoint id 97 collected
[1:30:46] Charge balance equation is out by -21.4%
[1:30:46] Stirrer speed set to 60
[1:30:51] pH 3.72 -> 3.52
[1:30:51] Using cautious pH adjust
[1:30:51] Dispensed 0.000259 mL of Acid (0.5 M HCl)
[1:30:56] Stepping pH = 3.61
[1:30:56] Dispensed 0.000188 mL of Acid (0.5 M HCl)
[1:31:01] Stepping pH = 3.55
[1:31:02] Dispensed 0.000094 mL of Acid (0.5 M HCl)
[1:31:07] Stepping pH = 3.53
[1:31:07] Dispensed 0.000047 mL of Acid (0.5 M HCl)
[1:31:12] Stepping pH = 3.52
[1:31:27] Stirrer speed set to 0
[1:31:37] Datapoint id 98 collected
[1:31:37] Charge balance equation is out by -13.3%
[1:31:37] Stirrer speed set to 60
[1:31:42] pH 3.52 -> 3.32
[1:31:42] Using charge balance adjust
[1:31:42] Dispensed 0.000800 mL of Acid (0.5 M HCl)
[1:32:02] Stirrer speed set to 0
[1:32:12] Datapoint id 99 collected
[1:32:12] Charge balance equation is out by -0.9%
[1:32:12] Stirrer speed set to 60
[1:32:18] pH 3.32 -> 3.12

Sample name: **M08_octanol**
Assay name: **pH-metric high logP**
Assay ID: **18C-02008**
Filename: **C:\Sirius_T3\Mehtap\20180302_exp29_logP_T3-2\18C-02008_M08_octanol_pH-metric high logP.t3r**
Experiment start time: **3/2/2018 6:51:30 PM**
Analyst: **Pion**
Instrument ID: **T312060**

#### Experiment Log (continued)

[1:32:18] Using charge balance adjust
[1:32:18] Dispensed 0.001246 mL of Acid (0.5 M HCl)
[1:32:38] Stirrer speed set to 0
[1:32:48] Datapoint id 100 collected
[1:32:48] Charge balance equation is out by -4.4%
[1:32:48] Stirrer speed set to 60
[1:32:53] pH 3.14 -> 2.94
[1:32:53] Using charge balance adjust
[1:32:54] Dispensed 0.001929 mL of Acid (0.5 M HCl)
[1:33:14] Stirrer speed set to 0
[1:33:37] Datapoint id 101 collected
[1:33:37] Charge balance equation is out by -0.5%
[1:33:37] Stirrer speed set to 60
[1:33:42] pH 2.94 -> 2.74
[1:33:42] Using charge balance adjust
[1:33:43] Dispensed 0.003010 mL of Acid (0.5 M HCl)
[1:34:03] Stirrer speed set to 0
[1:34:13] Datapoint id 102 collected
[1:34:13] Charge balance equation is out by 1.9%
[1:34:13] Stirrer speed set to 60
[1:34:18] pH 2.74 -> 2.54
[1:34:18] Using charge balance adjust
[1:34:18] Dispensed 0.004845 mL of Acid (0.5 M HCl)
[1:34:38] Stirrer speed set to 0
[1:34:49] Datapoint id 103 collected
[1:34:49] Charge balance equation is out by 1.5%
[1:34:49] Stirrer speed set to 60
[1:34:54] pH 2.54 -> 2.34
[1:34:54] Using charge balance adjust
[1:34:54] Dispensed 0.007785 mL of Acid (0.5 M HCl)
[1:35:15] Stirrer speed set to 0
[1:35:30] Datapoint id 104 collected
[1:35:30] Charge balance equation is out by -0.7%
[1:35:30] Stirrer speed set to 60
[1:35:35] pH 2.35 -> 2.15
[1:35:35] Using charge balance adjust
[1:35:35] Dispensed 0.012371 mL of Acid (0.5 M HCl)
[1:35:55] Stirrer speed set to 0
[1:36:09] Datapoint id 105 collected
[1:36:09] Charge balance equation is out by -0.3%
[1:36:09] Stirrer speed set to 60
[1:36:14] pH 2.15 -> 1.95
[1:36:14] Using charge balance adjust
[1:36:14] Dispensed 0.019920 mL of Acid (0.5 M HCl)
[1:36:35] Stirrer speed set to 0
[1:36:55] Datapoint id 106 collected
[1:36:55] Charge balance equation is out by 1.4%
[1:36:55] Argon flow rate set to 0
[1:36:59] Titrator arm moved over Titration position
