## Supplementary material for "Octanol-water partition coefficient measurements for the SAMPL6 Blind Prediction Challenge": SM08_18C-02009_M08_octanol_pH-metric high logP_report.pdf

| | | | |
|---|---|---|---|
| Sample name: | **M08_octanol** | Experiment start time: | **3/2/2018 8:29:22 PM** |
| Assay name: | **pH-metric high logP** | Analyst: | **Pion** |
| Assay ID: | **18C-02009** | Instrument ID: | **T312060** |
| Filename: | **C:\Sirius_T3\Mehtap\20180302_exp29_logP_T3-2\18C-02009_M08_octanol_pH-metric high logP.t3r** | | |

#### pH-metric Result

| | | |
|---|---|---|
| logP (neutral XH) | 3.16 | ±0.01 (n=50) |
| logP (X -) | 0.23 | ±0.03 (n=50) |

#### 18C-02009 Points 2 to 38

| | |
|---|---|
| M08_octanol concentration factor | 1.095 |
| Carbonate | 0.0863 mM |
| Acidity error | -0.76941 mM |

#### 18C-02009 Points 39 to 63

| | |
|---|---|
| M08_octanol concentration factor | 0.769 |
| Carbonate | 0.0001 mM |
| Acidity error | -0.70585 mM |

#### 18C-02009 Points 64 to 92

| | |
|---|---|
| M08_octanol concentration factor | 0.703 |
| Carbonate | 0.2882 mM |
| Acidity error | -0.43720 mM |

#### Warnings and errors

| | |
|---|---|
| Errors | None |
| Warnings | None |

#### Sample logD and percent species

| pH | M08_octanol logD | M08_octanol M08_octanolH | M08_octanol M08_octanol | M08_octanol M08_octanolH* | M08_octanol M08_octanol* | Comment |
|---|---|---|---|---|---|---|
| 1.000 | 3.16 | 0.07 % | 0.00 % | 99.93 % | 0.00 % | |
| 1.200 | 3.16 | 0.07 % | 0.00 % | 99.93 % | 0.00 % | Stomach pH |
| 2.000 | 3.16 | 0.07 % | 0.00 % | 99.93 % | 0.00 % | |
| 3.000 | 3.13 | 0.07 % | 0.00 % | 99.92 % | 0.01 % | |
| 4.000 | 2.95 | 0.07 % | 0.04 % | 99.82 % | 0.07 % | |
| 5.000 | 2.32 | 0.07 % | 0.41 % | 98.81 % | 0.71 % | |
| 6.000 | 1.40 | 0.06 % | 3.75 % | 89.77 % | 6.42 % | |
| 6.500 | 0.97 | 0.05 % | 9.73 % | 73.58 % | 16.64 % | |
| 7.000 | 0.61 | 0.03 % | 19.59 % | 46.86 % | 33.52 % | |
| 7.400 | 0.43 | 0.02 % | 27.29 % | 25.99 % | 46.69 % | Blood pH |
| 8.000 | 0.29 | 0.01 % | 33.90 % | 8.11 % | 57.99 % | |
| 9.000 | 0.24 | 0.00 % | 36.57 % | 0.87 % | 62.56 % | |
| 10.000 | 0.23 | 0.00 % | 36.86 % | 0.09 % | 63.05 % | |
| 11.000 | 0.23 | 0.00 % | 36.89 % | 0.01 % | 63.10 % | |
| 12.000 | 0.23 | 0.00 % | 36.89 % | 0.00 % | 63.11 % | |

Sample name: **M08_octanol**
Assay name: **pH-metric high logP**
Assay ID: **18C-02009**
Filename: **C:\Sirius_T3\Mehtap\20180302_exp29_logP_T3-2\18C-02009_M08_octanol_pH-metric high logP.t3r**

Experiment start time: **3/2/2018 8:29:22 PM**
Analyst: **Pion**
Instrument ID: **T312060**

#### Graphs

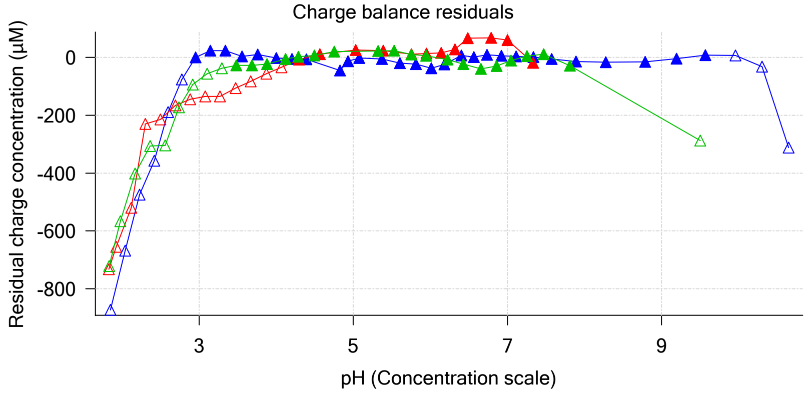


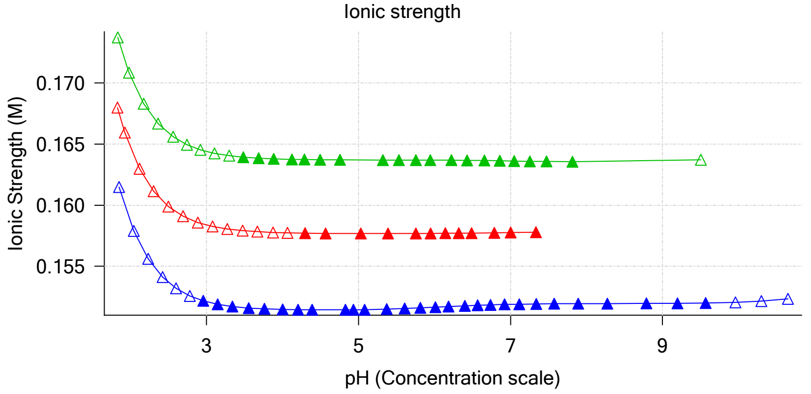


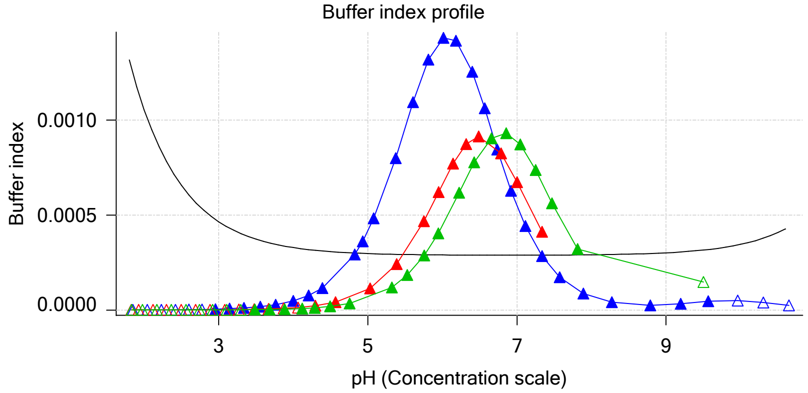


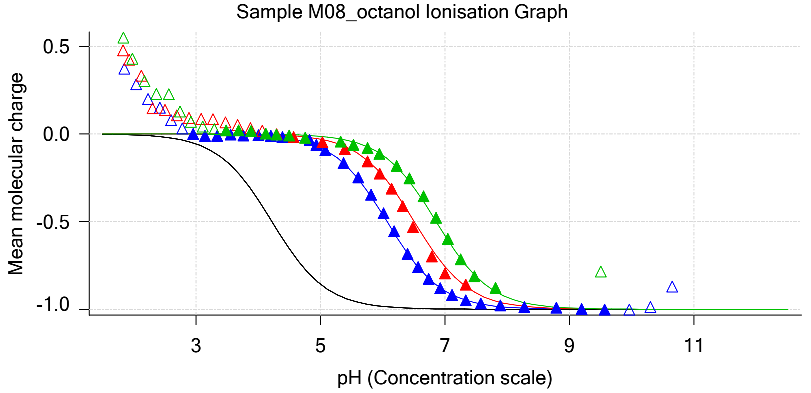


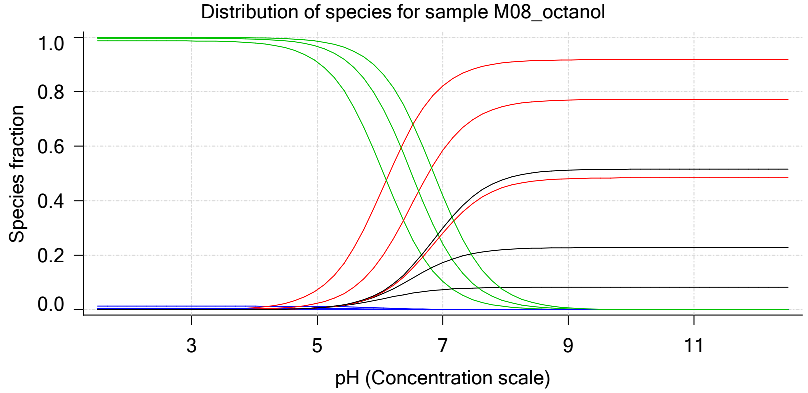


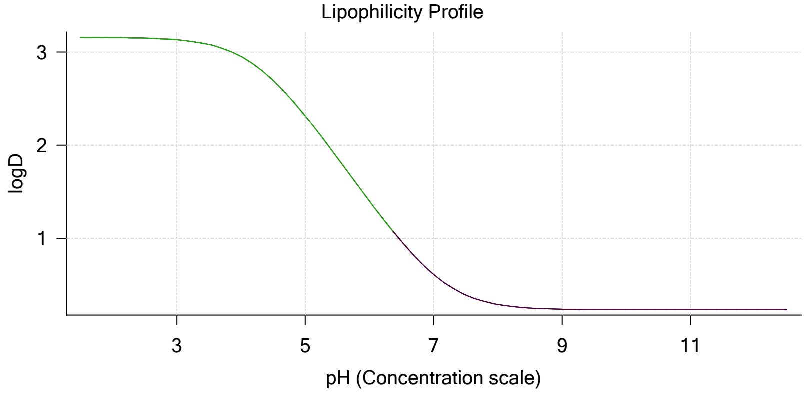


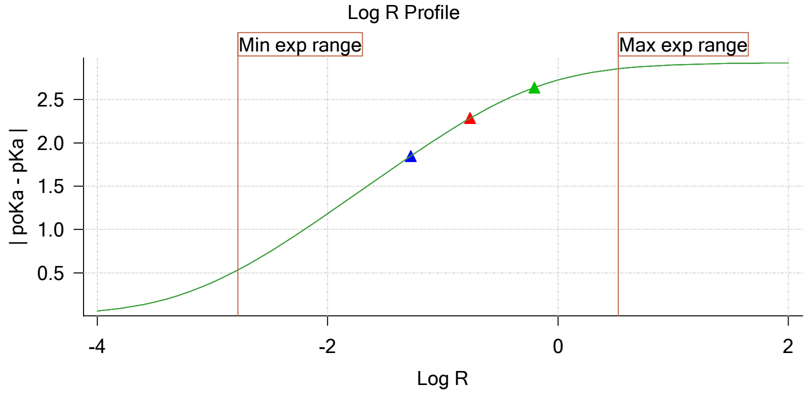


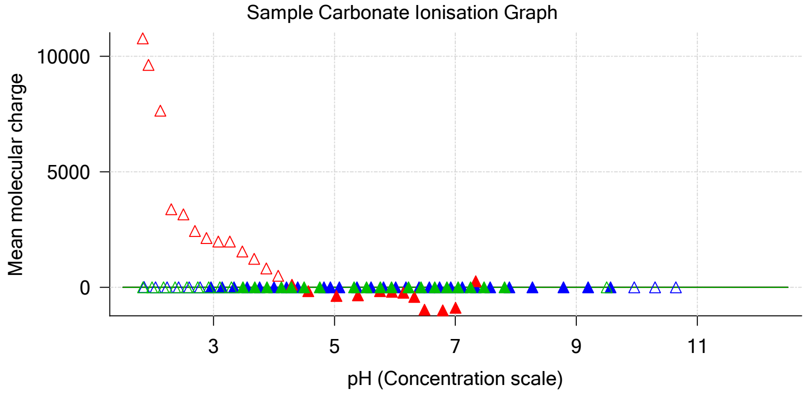

| | | | |
|---|---|---|---|
| Sample name: | **M08_octanol** | Experiment start time: | **3/2/2018 8:29:22 PM** |
| Assay name: | **pH-metric high logP** | Analyst: | **Pion** |
| Assay ID: | **18C-02009** | Instrument ID: | **T312060** |
| Filename: | **C:\Sirius_T3\Mehtap\20180302_exp29_logP_T3-2\18C-02009_M08_octanol_pH-metric high logP.t3r** | | |

#### Graphs (continued)

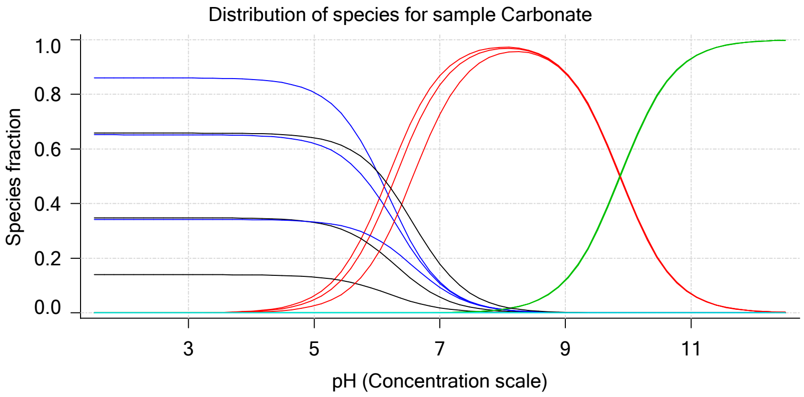

Sample name: **M08_octanol** | Experiment start time: **3/2/2018 8:29:22 PM**
Assay name: **pH-metric high logP** | Analyst: **Pion**
Assay ID: **18C-02009** | Instrument ID: **T312060**
Filename: **C:\Sirius_T3\Mehtap\20180302_exp29_logP_T3-2\18C-02009_M08_octanol_pH-metric high logP.t3r**

# pH-metric high logP   Titration 1 of 3   18C-02009 Points 2 to 38

#### Overall results

| | |
|---|---|
| RMSD | 0.094 |
| Average ionic strength | 0.152 M |
| Average temperature | 25.0°C |
| Partition ratio | 0.0526 : 1 |
| Analyte concentration range | 2033.3 µM to 2105.6 µM |
| Total points considered | 28 of 37 |

#### Warnings and errors

| | |
|---|---|
| Errors | None |
| Warnings | One or more logP values out of range |

#### Four-Plus parameters

| | | | |
|---|---|---|---|
| Alpha | 0.111 | 3/2/2018 8:29:22 PM | C:\Sirius_T3\HCl18C02.t3r |
| S | 0.9988 | 3/2/2018 8:29:22 PM | C:\Sirius_T3\HCl18C02.t3r |
| jH | 1.0 | 3/2/2018 8:29:22 PM | C:\Sirius_T3\HCl18C02.t3r |
| jOH | -0.8 | 3/2/2018 8:29:22 PM | C:\Sirius_T3\HCl18C02.t3r |

#### Titrants

| | | | |
|---|---|---|---|
| 0.50 M HCl | 0.999058 | 3/2/2018 8:29:22 PM | C:\Sirius_T3\HCl18C02.t3r |
| 0.50 M KOH | 0.999845 | 3/2/2018 8:29:22 PM | C:\Sirius_T3\KOH18B27.t3r |

#### Sample

| | |
|---|---|
| M08_octanol concentration factor | 1.095 |
| Acid pKa 1 | 4.22 |
| logP (neutral XH) | 3.13 |
| logP (X -) | -5.22 |

#### Sample graphs

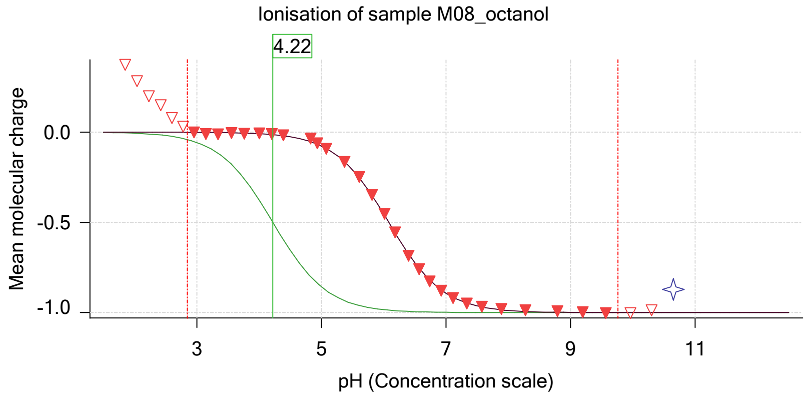


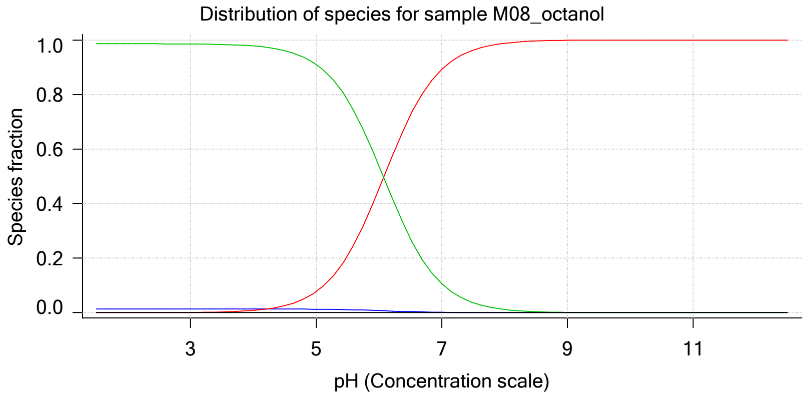

#### Sample graphs (continued)

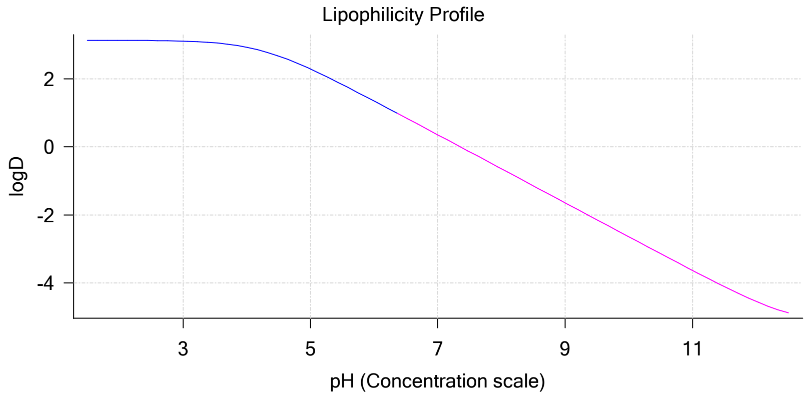


#### Sample logD and percent species

| pH | M08_octanol logD | M08_octanol M08_octanolH | M08_octanol M08_octanol | M08_octanol M08_octanolH* | M08_octanol M08_octanol* | Comment |
|---|---|---|---|---|---|---|
| 1.000 | 3.13 | 1.38 % | 0.00 % | 98.62 % | 0.00 % | |
| 1.200 | 3.13 | 1.38 % | 0.00 % | 98.62 % | 0.00 % | Stomach pH |
| 2.000 | 3.13 | 1.38 % | 0.01 % | 98.61 % | 0.00 % | |
| 3.000 | 3.11 | 1.38 % | 0.08 % | 98.54 % | 0.00 % | |
| 4.000 | 2.93 | 1.37 % | 0.82 % | 97.81 % | 0.00 % | |
| 5.000 | 2.29 | 1.27 % | 7.67 % | 91.06 % | 0.00 % | |
| 6.000 | 1.35 | 0.75 % | 45.38 % | 53.86 % | 0.00 % | |
| 6.500 | 0.85 | 0.38 % | 72.43 % | 27.19 % | 0.00 % | |
| 7.000 | 0.35 | 0.15 % | 89.26 % | 10.59 % | 0.00 % | |
| 7.400 | -0.05 | 0.06 % | 95.43 % | 4.51 % | 0.00 % | Blood pH |
| 8.000 | -0.65 | 0.02 % | 98.81 % | 1.17 % | 0.00 % | |
| 9.000 | -1.65 | 0.00 % | 99.88 % | 0.12 % | 0.00 % | |
| 10.000 | -2.65 | 0.00 % | 99.99 % | 0.01 % | 0.00 % | |
| 11.000 | -3.64 | 0.00 % | 100.00 % | 0.00 % | 0.00 % | |
| 12.000 | -4.54 | 0.00 % | 100.00 % | 0.00 % | 0.00 % | |

#### Carbonate and acidity

Carbonate 0.086 mM
Acidity error -0.769 mM

#### Other graphs

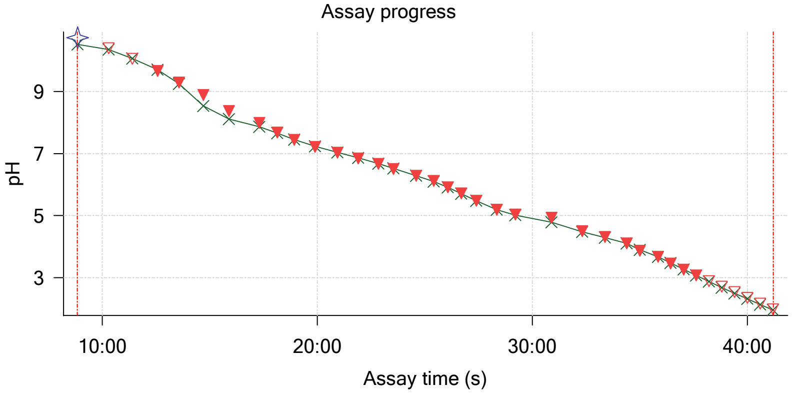


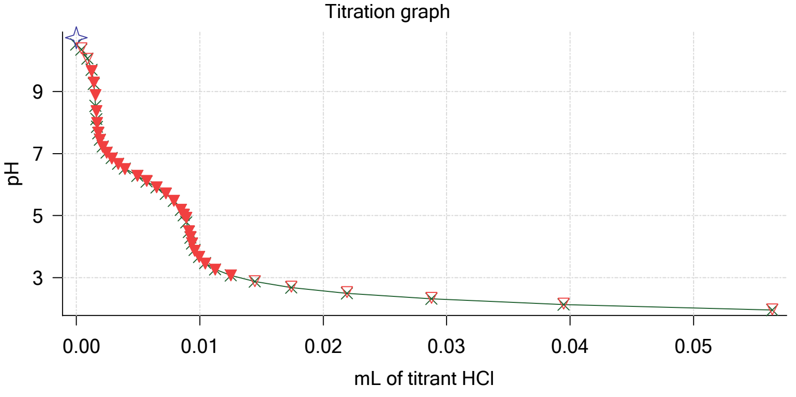

| | | | |
|---|---|---|---|
| Sample name: | **M08_octanol** | Experiment start time: | **3/2/2018 8:29:22 PM** |
| Assay name: | **pH-metric high logP** | Analyst: | **Pion** |
| Assay ID: | **18C-02009** | Instrument ID: | **T312060** |
| Filename: | **C:\Sirius_T3\Mehtap\20180302_exp29_logP_T3-2\18C-02009_M08_octanol_pH-metric high logP.t3r** | | |

#### Other graphs (continued)

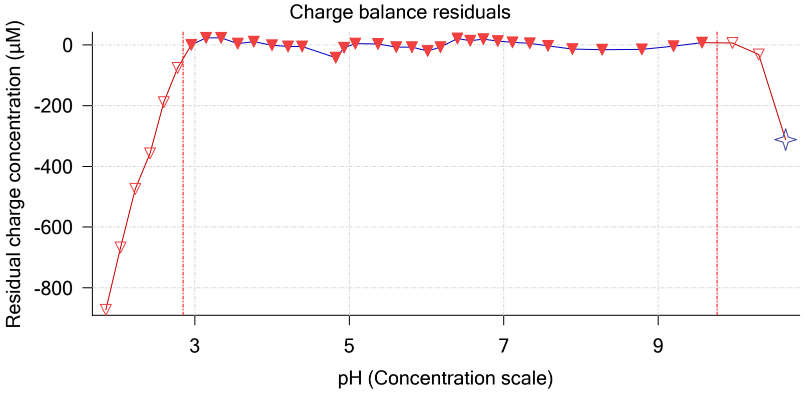


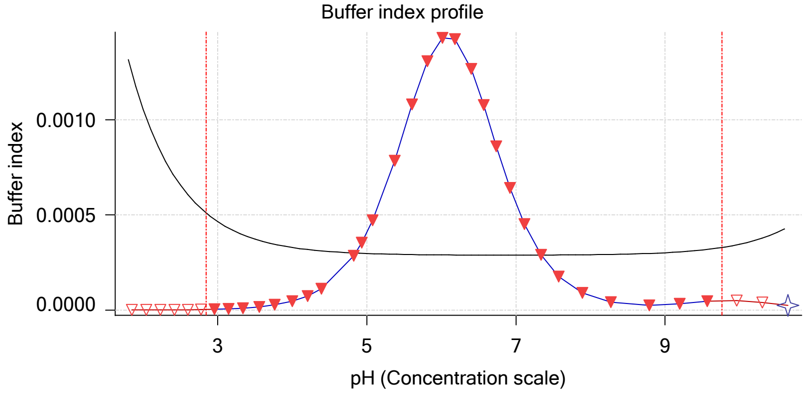


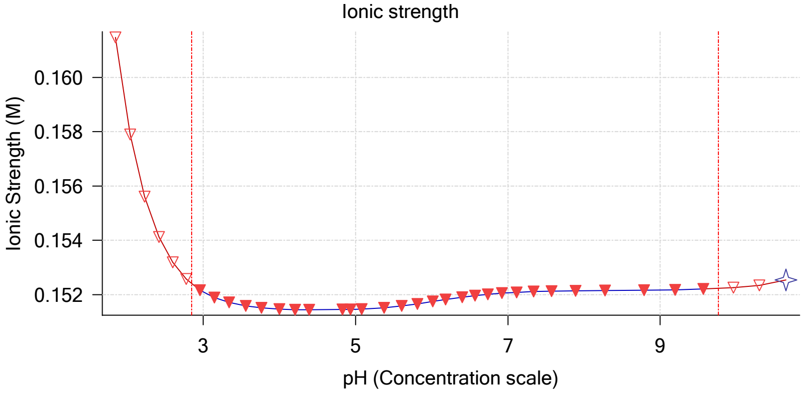


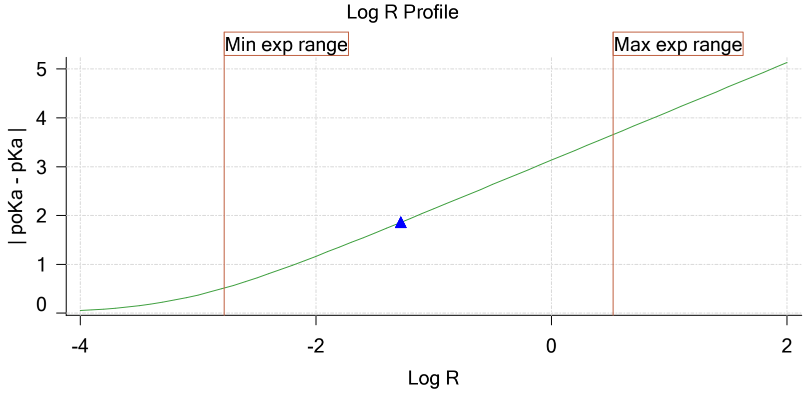


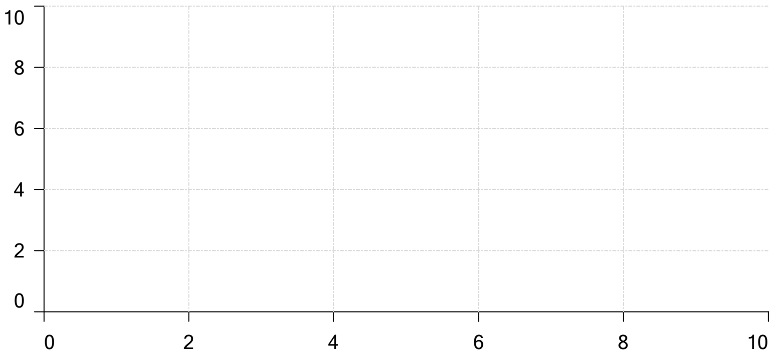


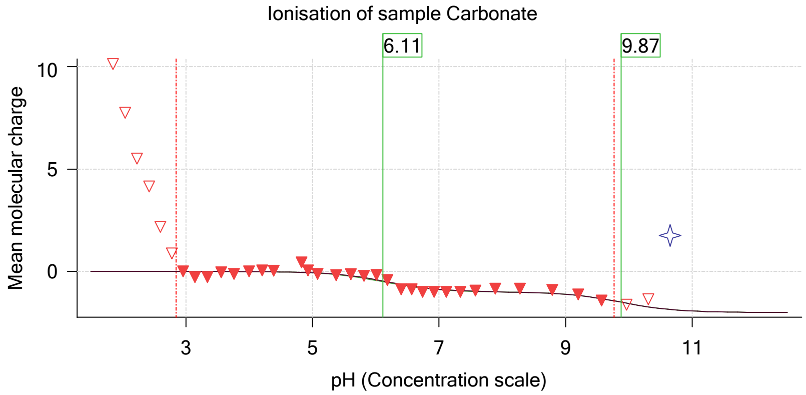

| | | | |
|---|---|---|---|
| Sample name: | **M08_octanol** | Experiment start time: | **3/2/2018 8:29:22 PM** |
| Assay name: | **pH-metric high logP** | Analyst: | **Pion** |
| Assay ID: | **18C-02009** | Instrument ID: | **T312060** |
| Filename: | **C:\Sirius_T3\Mehtap\20180302_exp29_logP_T3-2\18C-02009_M08_octanol_pH-metric high logP.t3r** | | |

### pH-metric high logP Titration 2 of 3 18C-02009 Points 39 to 63

#### Overall results

| | |
|---|---|
| RMSD | 0.061 |
| Average ionic strength | 0.158 M |
| Average temperature | 25.0°C |
| Partition ratio | 0.1728 : 1 |
| Analyte concentration range | 1712.1 µM to 1763.3 µM |
| Total points considered | 12 of 25 |

#### Warnings and errors

| | |
|---|---|
| Errors | None |
| Warnings | One or more logP values out of range |

#### Four-Plus parameters

| | | | |
|---|---|---|---|
| Alpha | 0.111 | 3/2/2018 8:29:22 PM | C:\Sirius_T3\HCl18C02.t3r |
| S | 0.9988 | 3/2/2018 8:29:22 PM | C:\Sirius_T3\HCl18C02.t3r |
| jH | 1.0 | 3/2/2018 8:29:22 PM | C:\Sirius_T3\HCl18C02.t3r |
| jOH | -0.8 | 3/2/2018 8:29:22 PM | C:\Sirius_T3\HCl18C02.t3r |

#### Titrants

| | | | |
|---|---|---|---|
| 0.50 M HCl | 0.999058 | 3/2/2018 8:29:22 PM | C:\Sirius_T3\HCl18C02.t3r |
| 0.50 M KOH | 0.999845 | 3/2/2018 8:29:22 PM | C:\Sirius_T3\KOH18B27.t3r |

#### Sample

| | |
|---|---|
| M08_octanol concentration factor | 0.769 |
| Acid pKa 1 | 4.22 |
| logP (neutral XH) | 3.00 |
| logP (X -) | -5.22 |

#### Sample graphs

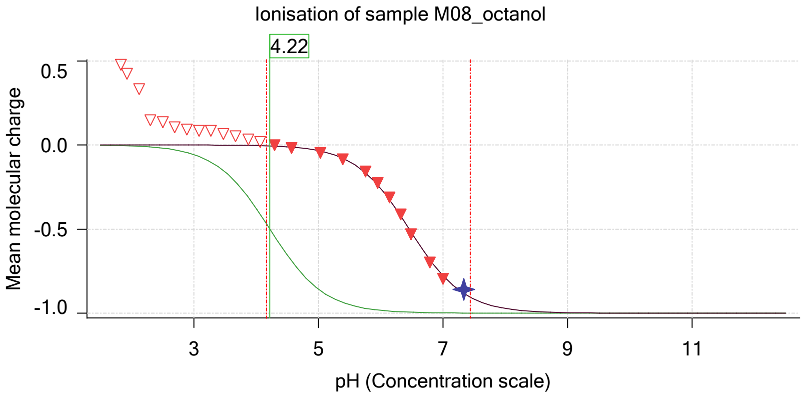


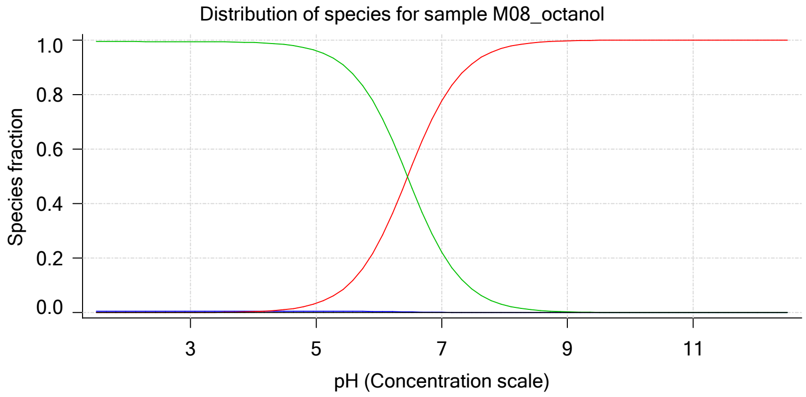

| | |
|---|---|
| Sample name: | **M08_octanol** |
| Assay name: | **pH-metric high logP** |
| Assay ID: | **18C-02009** |
| Filename: | **C:\Sirius_T3\Mehtap\20180302_exp29_logP_T3-2\18C-02009_M08_octanol_pH-metric high logP.t3r** |
| Experiment start time: | **3/2/2018 8:29:22 PM** |
| Analyst: | **Pion** |
| Instrument ID: | **T312060** |

#### Sample graphs (continued)

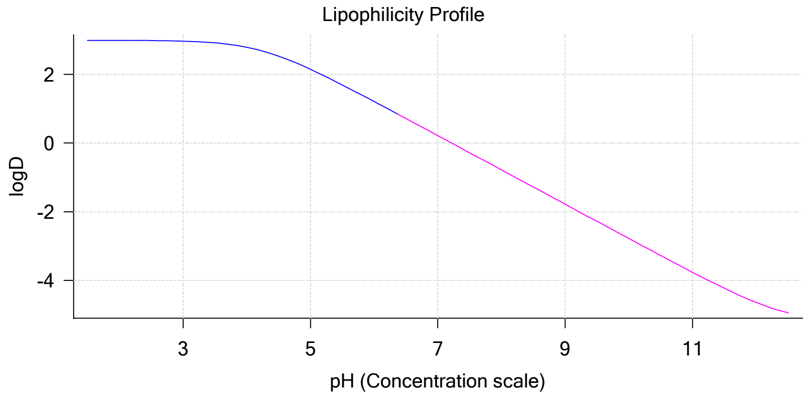


#### Sample logD and percent species

| pH | M08_octanol logD | M08_octanol M08_octanolH | M08_octanol M08_octanol | M08_octanol M08_octanolH* | M08_octanol M08_octanol* | Comment |
|---|---|---|---|---|---|---|
| 1.000 | 3.00 | 0.58 % | 0.00 % | 99.42 % | 0.00 % | |
| 1.200 | 3.00 | 0.58 % | 0.00 % | 99.42 % | 0.00 % | Stomach pH |
| 2.000 | 2.99 | 0.58 % | 0.00 % | 99.42 % | 0.00 % | |
| 3.000 | 2.97 | 0.58 % | 0.03 % | 99.39 % | 0.00 % | |
| 4.000 | 2.79 | 0.58 % | 0.35 % | 99.07 % | 0.00 % | |
| 5.000 | 2.15 | 0.56 % | 3.38 % | 96.06 % | 0.00 % | |
| 6.000 | 1.21 | 0.43 % | 25.90 % | 73.67 % | 0.00 % | |
| 6.500 | 0.71 | 0.28 % | 52.50 % | 47.23 % | 0.00 % | |
| 7.000 | 0.22 | 0.13 % | 77.75 % | 22.12 % | 0.00 % | |
| 7.400 | -0.18 | 0.06 % | 89.77 % | 10.17 % | 0.00 % | Blood pH |
| 8.000 | -0.78 | 0.02 % | 97.22 % | 2.77 % | 0.00 % | |
| 9.000 | -1.78 | 0.00 % | 99.71 % | 0.28 % | 0.00 % | |
| 10.000 | -2.78 | 0.00 % | 99.97 % | 0.03 % | 0.00 % | |
| 11.000 | -3.77 | 0.00 % | 100.00 % | 0.00 % | 0.00 % | |
| 12.000 | -4.65 | 0.00 % | 100.00 % | 0.00 % | 0.00 % | |

#### Carbonate and acidity

| | |
|---|---|
| Carbonate | 0.000 mM |
| Acidity error | -0.706 mM |

#### Other graphs

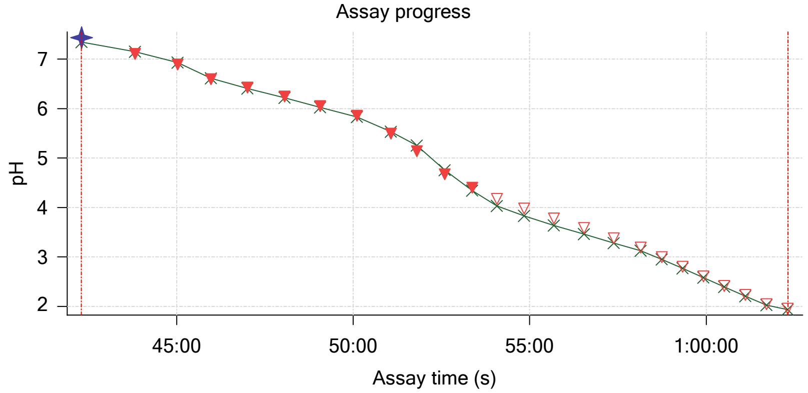


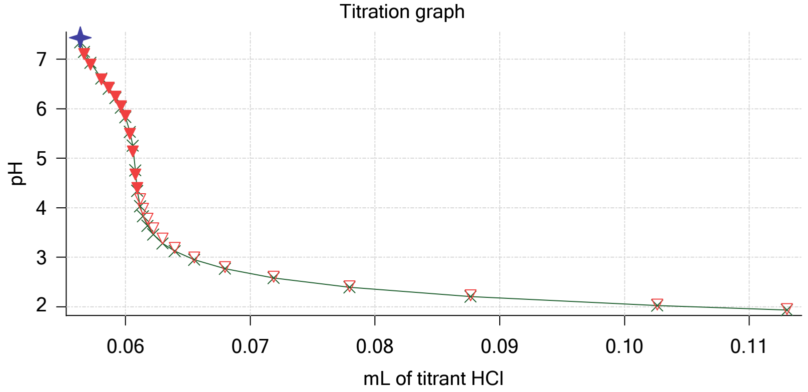

Sample name: **M08_octanol**
Assay name: **pH-metric high logP**
Assay ID: **18C-02009**
Filename: **C:\Sirius_T3\Mehtap\20180302_exp29_logP_T3-2\18C-02009_M08_octanol_pH-metric high logP.t3r**

Experiment start time: **3/2/2018 8:29:22 PM**
Analyst: **Pion**
Instrument ID: **T312060**

#### Other graphs (continued)

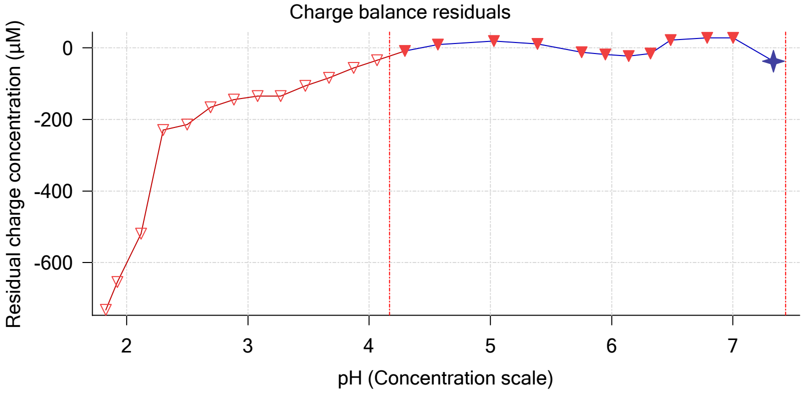


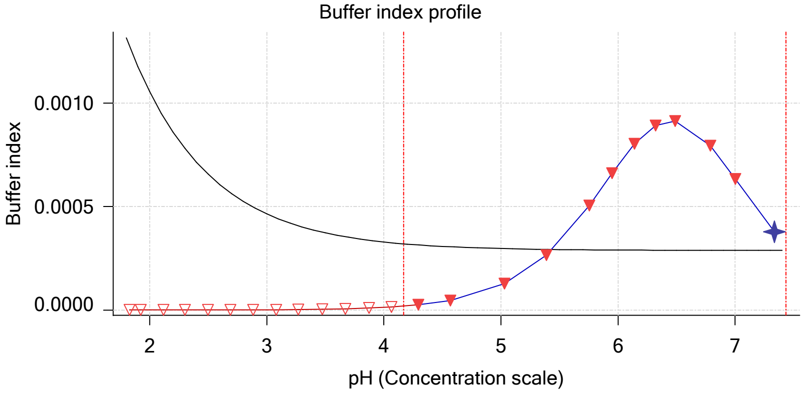


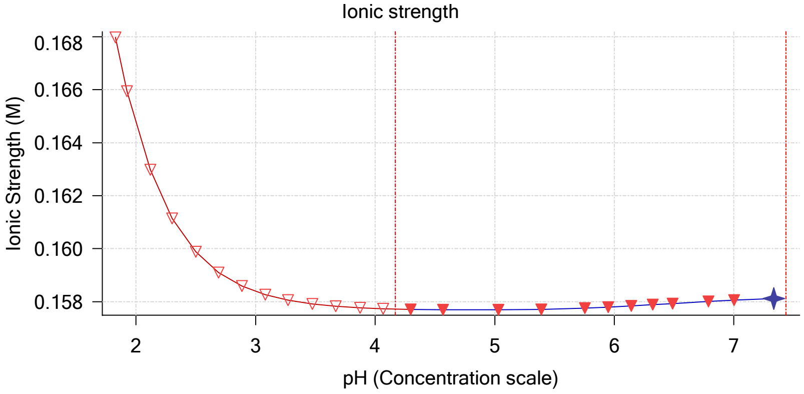


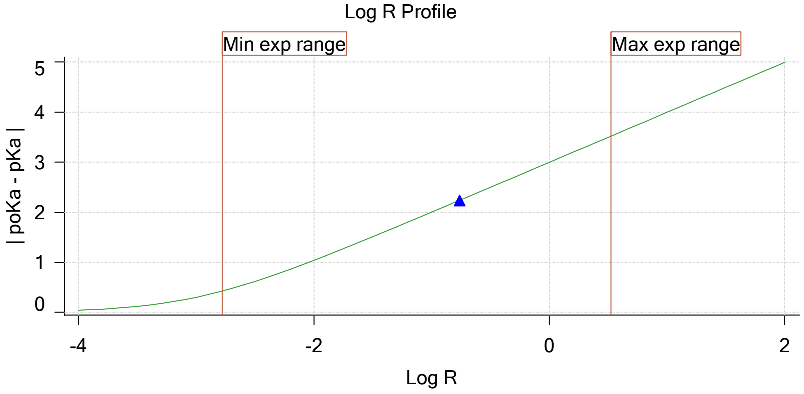


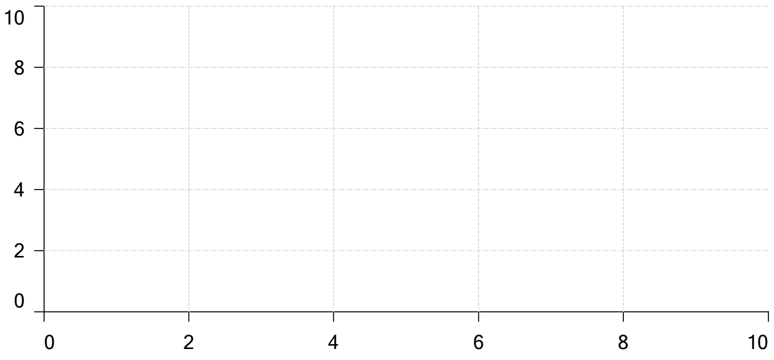


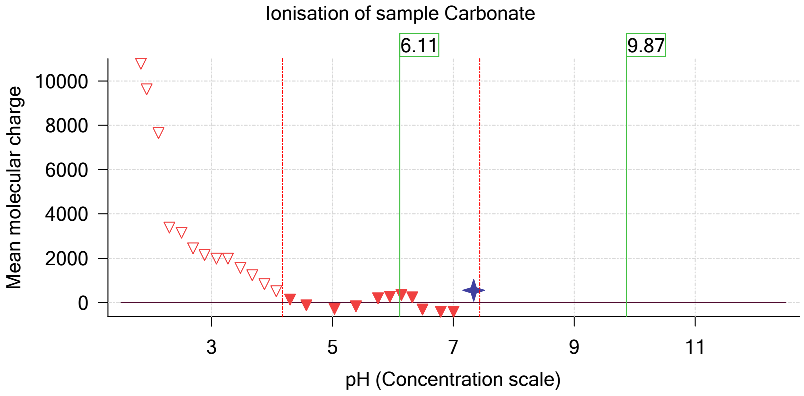

| | | | |
|---|---|---|---|
| Sample name: | **M08_octanol** | Experiment start time: | **3/2/2018 8:29:22 PM** |
| Assay name: | **pH-metric high logP** | Analyst: | **Pion** |
| Assay ID: | **18C-02009** | Instrument ID: | **T312060** |
| Filename: | **C:\Sirius_T3\Mehtap\20180302_exp29_logP_T3-2\18C-02009_M08_octanol_pH-metric high logP.t3r** | | |

### pH-metric high logP Titration 3 of 3 18C-02009 Points 64 to 92

#### Overall results

| | |
|---|---|
| RMSD | 0.092 |
| Average ionic strength | 0.164 M |
| Average temperature | 25.0°C |
| Partition ratio | 0.6226 : 1 |
| Analyte concentration range | 1164.0 µM to 1189.2 µM |
| Total points considered | 19 of 29 |

#### Warnings and errors

| | |
|---|---|
| Errors | None |
| Warnings | One or more logP values out of range |

#### Four-Plus parameters

| | | | |
|---|---|---|---|
| Alpha | 0.111 | 3/2/2018 8:29:22 PM | C:\Sirius_T3\HCl18C02.t3r |
| S | 0.9988 | 3/2/2018 8:29:22 PM | C:\Sirius_T3\HCl18C02.t3r |
| jH | 1.0 | 3/2/2018 8:29:22 PM | C:\Sirius_T3\HCl18C02.t3r |
| jOH | -0.8 | 3/2/2018 8:29:22 PM | C:\Sirius_T3\HCl18C02.t3r |

#### Titrants

| | | | |
|---|---|---|---|
| 0.50 M HCl | 0.999058 | 3/2/2018 8:29:22 PM | C:\Sirius_T3\HCl18C02.t3r |
| 0.50 M KOH | 0.999845 | 3/2/2018 8:29:22 PM | C:\Sirius_T3\KOH18B27.t3r |

#### Sample

| | |
|---|---|
| M08_octanol concentration factor | 0.703 |
| Acid pKa 1 | 4.22 |
| logP (neutral XH) | 2.87 |
| logP (X -) | -5.22 |

#### Sample graphs

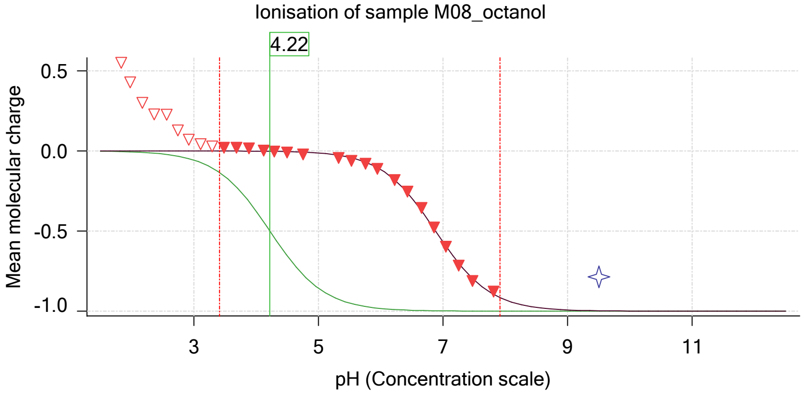


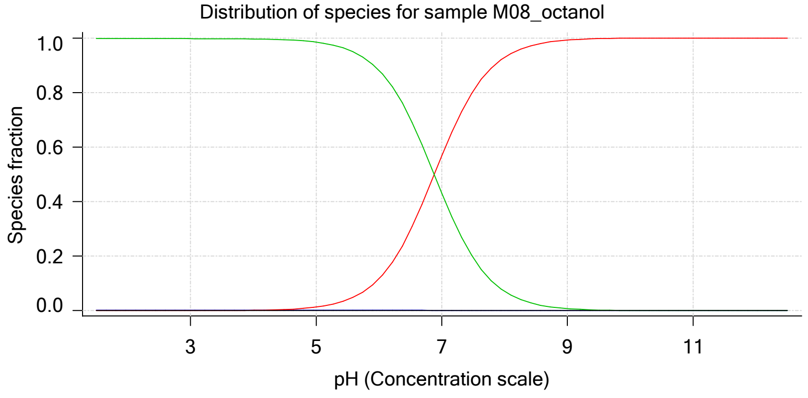

Sample name: **M08_octanol** Experiment start time: **3/2/2018 8:29:22 PM**
Assay name: **pH-metric high logP** Analyst: **Pion**
Assay ID: **18C-02009** Instrument ID: **T312060**
Filename: **C:\Sirius_T3\Mehtap\20180302_exp29_logP_T3-2\18C-02009_M08_octanol_pH-metric high logP.t3r**

#### Sample graphs (continued)

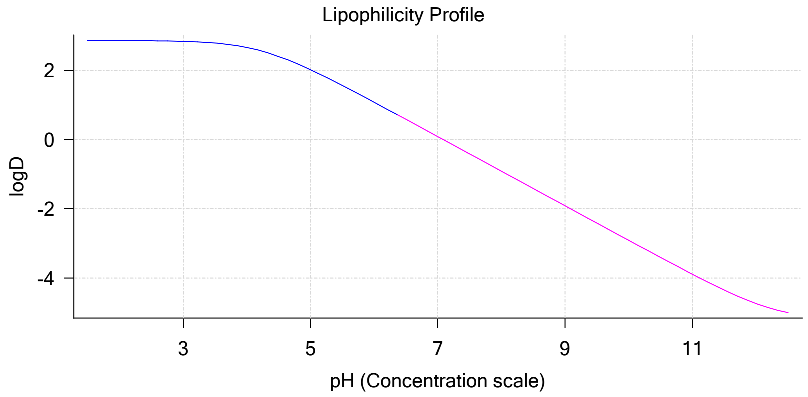


#### Sample logD and percent species

| pH | M08_octanol logD | M08_octanol M08_octanolH | M08_octanol M08_octanol | M08_octanol M08_octanolH* | M08_octanol M08_octanol* | Comment |
|---|---|---|---|---|---|---|
| 1.000 | 2.86 | 0.22 % | 0.00 % | 99.78 % | 0.00 % | |
| 1.200 | 2.86 | 0.22 % | 0.00 % | 99.78 % | 0.00 % | Stomach pH |
| 2.000 | 2.86 | 0.22 % | 0.00 % | 99.78 % | 0.00 % | |
| 3.000 | 2.84 | 0.22 % | 0.01 % | 99.77 % | 0.00 % | |
| 4.000 | 2.66 | 0.22 % | 0.13 % | 99.65 % | 0.00 % | |
| 5.000 | 2.02 | 0.22 % | 1.30 % | 98.48 % | 0.00 % | |
| 6.000 | 1.08 | 0.19 % | 11.64 % | 88.17 % | 0.00 % | |
| 6.500 | 0.58 | 0.15 % | 29.40 % | 70.44 % | 0.00 % | |
| 7.000 | 0.08 | 0.09 % | 56.84 % | 43.06 % | 0.00 % | |
| 7.400 | -0.32 | 0.05 % | 76.79 % | 23.16 % | 0.00 % | Blood pH |
| 8.000 | -0.91 | 0.02 % | 92.94 % | 7.04 % | 0.00 % | |
| 9.000 | -1.91 | 0.00 % | 99.25 % | 0.75 % | 0.00 % | |
| 10.000 | -2.91 | 0.00 % | 99.92 % | 0.08 % | 0.00 % | |
| 11.000 | -3.89 | 0.00 % | 99.99 % | 0.01 % | 0.00 % | |
| 12.000 | -4.74 | 0.00 % | 100.00 % | 0.00 % | 0.00 % | |

#### Carbonate and acidity

| | |
|---|---|
| Carbonate | 0.288 mM |
| Acidity error | -0.437 mM |

#### Other graphs

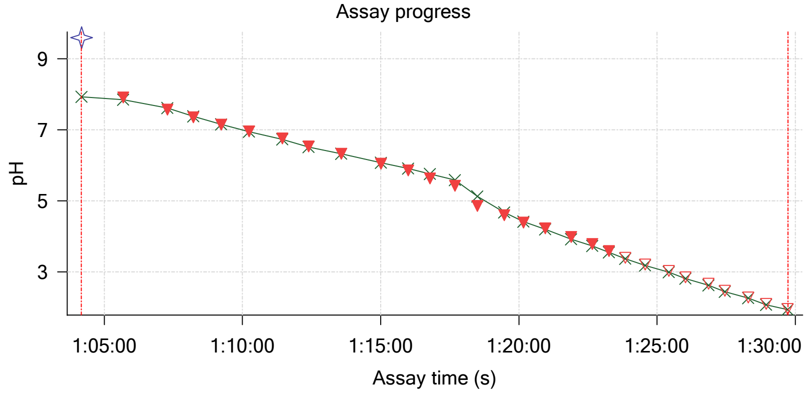


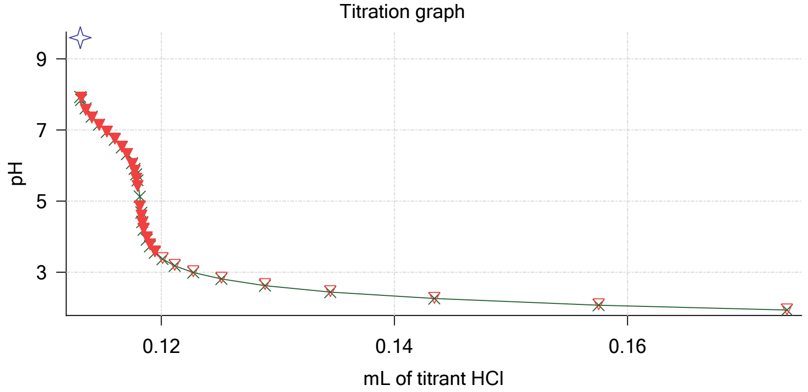

| | | | |
|---|---|---|---|
| Sample name: | **M08_octanol** | Experiment start time: | **3/2/2018 8:29:22 PM** |
| Assay name: | **pH-metric high logP** | Analyst: | **Pion** |
| Assay ID: | **18C-02009** | Instrument ID: | **T312060** |
| Filename: | **C:\Sirius_T3\Mehtap\20180302_exp29_logP_T3-2\18C-02009_M08_octanol_pH-metric high logP.t3r** | | |

#### Other graphs (continued)

Charge balance residuals

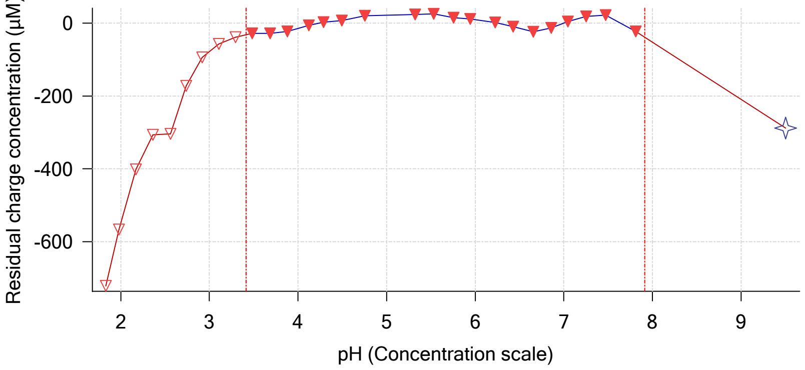


Buffer index profile

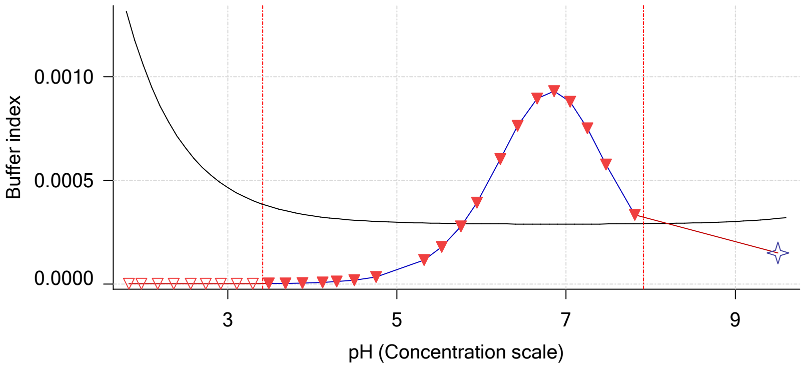


Ionic strength

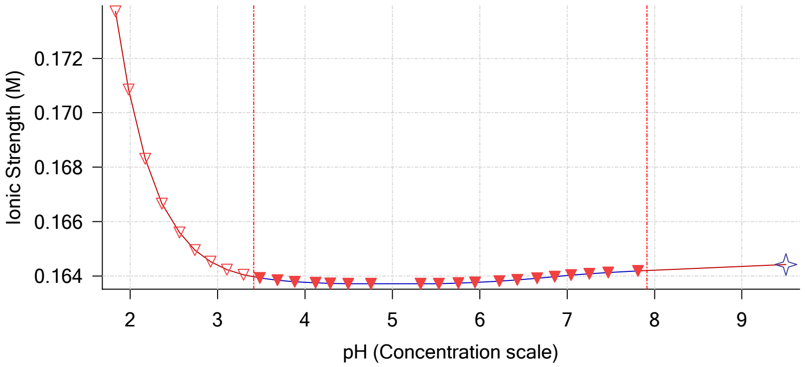


Log R Profile

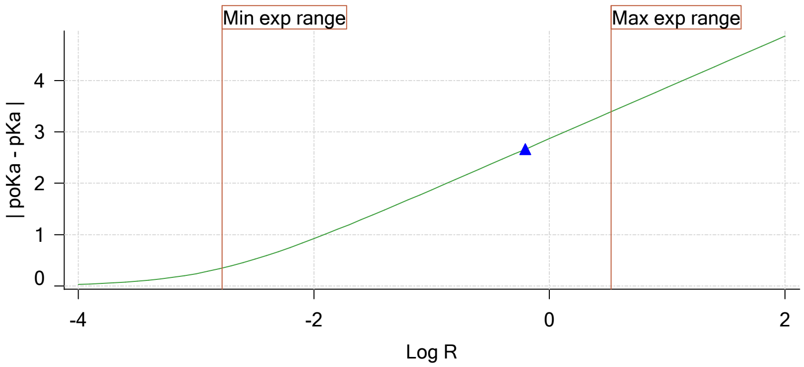


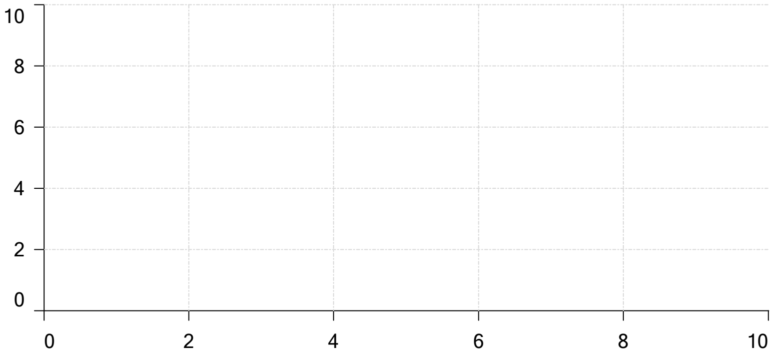


Ionisation of sample Carbonate

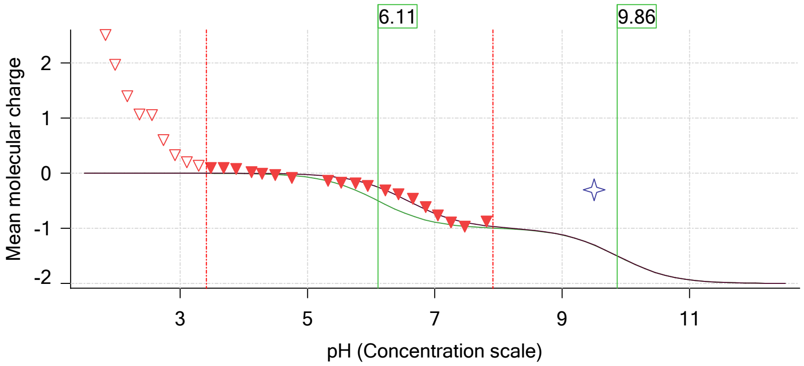

| | | | |
|---|---|---|---|
| Sample name: | **M08_octanol** | Experiment start time: | **3/2/2018 8:29:22 PM** |
| Assay name: | **pH-metric high logP** | Analyst: | **Pion** |
| Assay ID: | **18C-02009** | Instrument ID: | **T312060** |
| Filename: | **C:\Sirius_T3\Mehtap\20180302_exp29_logP_T3-2\18C-02009_M08_octanol_pH-metric high logP.t3r** | | |

#### Assay Model

| Settings | Value | Date/Time changed | Imported from |
|---|---|---|---|
| Sample name | M08_octanol | 2/27/2018 4:33:51 PM | User entered value |
| Sample by | Weight | | Default value |
| Sample weight | 0.000980 g | 3/2/2018 5:08:22 PM | User entered value |
| Formula weight | 293.32 g/mol | 2/27/2018 4:33:51 PM | User entered value |
| Solubility | Unknown | | Default value |
| Molecular weight | 293.32 | 2/27/2018 4:33:51 PM | User entered value |
| Individual pKa ionic environments | No | | Default value |
| Number of pKas | 1 | 2/27/2018 4:33:51 PM | User entered value |
| Sample is a | Acid | 2/27/2018 4:33:51 PM | User entered value |
| pKa 1 | 4.22 | 2/27/2018 4:33:51 PM | User entered value |
| logP (neutral XH) | 2.98 | 3/2/2018 3:22:58 PM | User entered value |
| logP (X -) | -5.22 | 3/2/2018 3:23:03 PM | User entered value |

#### Events

| Time | Event | Water | Acid | Base | Octanol | pH | dpH/dt | pH R-squared | pH SD | dpH time |
|---|---|---|---|---|---|---|---|---|---|---|
| 5:58.3 | Manual volume addition | | | | 0.08000 mL | | | | | |
| 5:59.4 | Initial pH = 5.13 | | | | | | | | | |
| 8:51.4 | Data point 2 | 1.50000 mL | 0.00000 mL | 0.00680 mL | 0.08000 mL | 10.746 | -0.11246 | 0.99456 | 0.00557 | Time out at |
| 10:18.0 | Data point 3 | 1.50000 mL | 0.00042 mL | 0.00680 mL | 0.08000 mL | 10.400 | 0.01868 | 0.91894 | 0.00096 | 35.0 s |
| 11:23.6 | Data point 4 | 1.50000 mL | 0.00092 mL | 0.00680 mL | 0.08000 mL | 10.058 | 0.01821 | 0.90743 | 0.00094 | 40.5 s |
| 12:34.8 | Data point 5 | 1.50000 mL | 0.00125 mL | 0.00680 mL | 0.08000 mL | 9.666 | -0.01373 | 0.55829 | 0.00091 | 28.5 s |
| 13:33.8 | Data point 6 | 1.50000 mL | 0.00143 mL | 0.00680 mL | 0.08000 mL | 9.296 | -0.01475 | 0.60041 | 0.00094 | 32.5 s |
| 14:42.1 | Data point 7 | 1.50000 mL | 0.00155 mL | 0.00680 mL | 0.08000 mL | 8.888 | -0.01820 | 0.87464 | 0.00096 | 36.0 s |
| 15:53.7 | Data point 8 | 1.50000 mL | 0.00162 mL | 0.00680 mL | 0.08000 mL | 8.376 | -0.01780 | 0.85957 | 0.00095 | 49.0 s |
| 17:18.5 | Data point 9 | 1.50000 mL | 0.00169 mL | 0.00680 mL | 0.08000 mL | 7.991 | 0.00108 | 0.00290 | 0.00099 | 19.5 s |
| 18:08.5 | Data point 10 | 1.50000 mL | 0.00179 mL | 0.00680 mL | 0.08000 mL | 7.676 | 0.01522 | 0.69377 | 0.00090 | 17.0 s |
| 18:56.0 | Data point 11 | 1.50000 mL | 0.00193 mL | 0.00680 mL | 0.08000 mL | 7.439 | 0.01692 | 0.86099 | 0.00090 | 17.0 s |
| 19:53.8 | Data point 12 | 1.50000 mL | 0.00216 mL | 0.00680 mL | 0.08000 mL | 7.216 | 0.01704 | 0.82585 | 0.00093 | 21.0 s |
| 20:55.8 | Data point 13 | 1.50000 mL | 0.00247 mL | 0.00680 mL | 0.08000 mL | 7.024 | 0.01724 | 0.88730 | 0.00090 | 17.0 s |
| 21:53.7 | Data point 14 | 1.50000 mL | 0.00287 mL | 0.00680 mL | 0.08000 mL | 6.840 | 0.01697 | 0.84626 | 0.00091 | 16.0 s |
| 22:50.5 | Data point 15 | 1.50000 mL | 0.00339 mL | 0.00680 mL | 0.08000 mL | 6.669 | 0.01841 | 0.84076 | 0.00099 | 16.5 s |
| 23:32.5 | Data point 16 | 1.50000 mL | 0.00395 mL | 0.00680 mL | 0.08000 mL | 6.504 | 0.01741 | 0.89494 | 0.00091 | 17.0 s |
| 24:35.6 | Data point 17 | 1.50000 mL | 0.00494 mL | 0.00680 mL | 0.08000 mL | 6.285 | 0.01783 | 0.92060 | 0.00092 | 13.5 s |
| 25:24.8 | Data point 18 | 1.50000 mL | 0.00571 mL | 0.00680 mL | 0.08000 mL | 6.116 | 0.01844 | 0.92964 | 0.00095 | 13.5 s |
| 26:03.8 | Data point 19 | 1.50000 mL | 0.00651 mL | 0.00680 mL | 0.08000 mL | 5.916 | 0.01893 | 0.91741 | 0.00098 | 12.5 s |
| 26:41.7 | Data point 20 | 1.50000 mL | 0.00727 mL | 0.00680 mL | 0.08000 mL | 5.710 | 0.01949 | 0.97281 | 0.00098 | 16.0 s |

| | | | |
|---|---|---|---|
| Sample name: | **M08_octanol** | Experiment start time: | **3/2/2018 8:29:22 PM** |
| Assay name: | **pH-metric high logP** | Analyst: | **Pion** |
| Assay ID: | **18C-02009** | Instrument ID: | **T312060** |
| Filename: | **C:\Sirius_T3\Mehtap\20180302_exp29_logP_T3-2\18C-02009_M08_octanol_pH-metric high logP.t3r** | | |

#### Events (continued)

| Time | Event | Water | Acid | Base | Octanol | pH | dpH/dt | pH R-squared | pH SD | dpH/dt time |
|---|---|---|---|---|---|---|---|---|---|---|
| 27:23.2 | Data point 21 | 1.50000 mL | 0.00790 mL | 0.00680 mL | 0.08000 mL | 5.477 | 0.01911 | 0.96079 | 0.00096 | 17.0 s |
| 28:21.0 | Data point 22 | 1.50000 mL | 0.00847 mL | 0.00680 mL | 0.08000 mL | 5.183 | 0.01836 | 0.88513 | 0.00096 | 21.0 s |
| 29:12.6 | Data point 23 | 1.50000 mL | 0.00870 mL | 0.00680 mL | 0.08000 mL | 5.036 | 0.05948 | 0.99622 | 0.00294 | Timed out at 59.5 s |
| 30:53.3 | Data point 24 | 1.50000 mL | 0.00891 mL | 0.00680 mL | 0.08000 mL | 4.931 | 0.05785 | 0.99230 | 0.00287 | Timed out at 59.5 s |
| 32:18.7 | Data point 25 | 1.50000 mL | 0.00913 mL | 0.00680 mL | 0.08000 mL | 4.496 | 0.01948 | 0.93912 | 0.00099 | 33.0 s |
| 33:22.3 | Data point 26 | 1.50000 mL | 0.00924 mL | 0.00680 mL | 0.08000 mL | 4.310 | 0.01919 | 0.92386 | 0.00099 | 25.5 s |
| 34:23.5 | Data point 27 | 1.50000 mL | 0.00938 mL | 0.00680 mL | 0.08000 mL | 4.106 | 0.01620 | 0.92889 | 0.00083 | 10.5 s |
| 34:59.4 | Data point 28 | 1.50000 mL | 0.00960 mL | 0.00680 mL | 0.08000 mL | 3.868 | 0.01215 | 0.93122 | 0.00062 | 10.0 s |
| 35:50.2 | Data point 29 | 1.50000 mL | 0.00995 mL | 0.00680 mL | 0.08000 mL | 3.663 | 0.01206 | 0.86827 | 0.00064 | 10.0 s |
| 36:25.7 | Data point 30 | 1.50000 mL | 0.01044 mL | 0.00680 mL | 0.08000 mL | 3.447 | 0.00079 | 0.06082 | 0.00016 | 10.5 s |
| 37:01.6 | Data point 31 | 1.50000 mL | 0.01124 mL | 0.00680 mL | 0.08000 mL | 3.252 | 0.00291 | 0.66841 | 0.00018 | 10.0 s |
| 37:37.1 | Data point 32 | 1.50000 mL | 0.01251 mL | 0.00680 mL | 0.08000 mL | 3.063 | 0.00379 | 0.17037 | 0.00045 | 10.0 s |
| 38:12.6 | Data point 33 | 1.50000 mL | 0.01446 mL | 0.00680 mL | 0.08000 mL | 2.886 | -0.00068 | 0.10900 | 0.00010 | 10.0 s |
| 38:48.2 | Data point 34 | 1.50000 mL | 0.01743 mL | 0.00680 mL | 0.08000 mL | 2.708 | -0.00988 | 0.27293 | 0.00093 | 10.0 s |
| 39:23.8 | Data point 35 | 1.50000 mL | 0.02192 mL | 0.00680 mL | 0.08000 mL | 2.530 | -0.00487 | 0.81957 | 0.00027 | 10.0 s |
| 39:59.3 | Data point 36 | 1.50000 mL | 0.02876 mL | 0.00680 mL | 0.08000 mL | 2.343 | -0.01064 | 0.51106 | 0.00073 | 10.0 s |
| 40:35.0 | Data point 37 | 1.50000 mL | 0.03949 mL | 0.00680 mL | 0.08000 mL | 2.157 | -0.01066 | 0.81291 | 0.00058 | 10.0 s |
| 41:10.9 | Data point 38 | 1.50000 mL | 0.05637 mL | 0.00680 mL | 0.08000 mL | 1.971 | -0.01287 | 0.96844 | 0.00065 | 10.5 s |
| 42:18.9 | Data point 39 | 1.50000 mL | 0.05637 mL | 0.05847 mL | 0.28000 mL | 7.437 | -0.14549 | 0.99322 | 0.00721 | Timed out at 59.5 s |
| 43:49.4 | Data point 40 | 1.50000 mL | 0.05670 mL | 0.05847 mL | 0.28000 mL | 7.103 | -0.01286 | 0.42347 | 0.00098 | 36.5 s |
| 45:01.7 | Data point 41 | 1.50000 mL | 0.05720 mL | 0.05847 mL | 0.28000 mL | 6.891 | 0.01894 | 0.94814 | 0.00096 | 21.0 s |
| 45:58.4 | Data point 42 | 1.50000 mL | 0.05807 mL | 0.05847 mL | 0.28000 mL | 6.590 | 0.01908 | 0.89638 | 0.00100 | 26.5 s |
| 47:00.7 | Data point 43 | 1.50000 mL | 0.05868 mL | 0.05847 mL | 0.28000 mL | 6.423 | 0.01918 | 0.91840 | 0.00099 | 27.0 s |
| 48:03.4 | Data point 44 | 1.50000 mL | 0.05920 mL | 0.05847 mL | 0.28000 mL | 6.245 | 0.01858 | 0.85197 | 0.00099 | 25.0 s |
| 49:04.1 | Data point 45 | 1.50000 mL | 0.05964 mL | 0.05847 mL | 0.28000 mL | 6.052 | 0.01835 | 0.93095 | 0.00094 | 26.5 s |
| 50:06.3 | Data point 46 | 1.50000 mL | 0.06000 mL | 0.05847 mL | 0.28000 mL | 5.858 | 0.01996 | 0.97913 | 0.00100 | 22.0 s |
| 51:04.1 | Data point 47 | 1.50000 mL | 0.06037 mL | 0.05847 mL | 0.28000 mL | 5.493 | 0.01776 | 0.89690 | 0.00093 | 19.0 s |
| 51:48.5 | Data point 48 | 1.50000 mL | 0.06058 mL | 0.05847 mL | 0.28000 mL | 5.135 | 0.01954 | 0.96361 | 0.00098 | 16.5 s |
| 52:35.6 | Data point 49 | 1.50000 mL | 0.06079 mL | 0.05847 mL | 0.28000 mL | 4.673 | 0.01944 | 0.94099 | 0.00099 | 16.0 s |
| 53:22.2 | Data point 50 | 1.50000 mL | 0.06096 mL | 0.05847 mL | 0.28000 mL | 4.401 | 0.01686 | 0.77173 | 0.00095 | 11.5 s |
| 54:04.2 | Data point 51 | 1.50000 mL | 0.06117 mL | 0.05847 mL | 0.28000 mL | 4.172 | 0.01841 | 0.83644 | 0.00099 | 10.5 s |
| 54:50.4 | Data point 52 | 1.50000 mL | 0.06141 mL | 0.05847 mL | 0.28000 mL | 3.979 | 0.00270 | 0.11201 | 0.00040 | 10.0 s |
| 55:41.2 | Data point 53 | 1.50000 mL | 0.06176 mL | 0.05847 mL | 0.28000 mL | 3.777 | 0.00199 | 0.23344 | 0.00020 | 10.0 s |
| 56:32.1 | Data point 54 | 1.50000 mL | 0.06223 mL | 0.05847 mL | 0.28000 mL | 3.582 | 0.00442 | 0.54450 | 0.00030 | 10.0 s |
| 57:23.0 | Data point 55 | 1.50000 mL | 0.06298 mL | 0.05847 mL | 0.28000 mL | 3.377 | -0.00183 | 0.13137 | 0.00025 | 10.0 s |
| 58:08.8 | Data point 56 | 1.50000 mL | 0.06395 mL | 0.05847 mL | 0.28000 mL | 3.187 | -0.00575 | 0.75760 | 0.00033 | 10.0 s |
| 58:44.2 | Data point 57 | 1.50000 mL | 0.06552 mL | 0.05847 mL | 0.28000 mL | 2.993 | -0.00768 | 0.67458 | 0.00046 | 10.0 s |
| 59:19.7 | Data point 58 | 1.50000 mL | 0.06799 mL | 0.05847 mL | 0.28000 mL | 2.800 | -0.00920 | 0.23870 | 0.00093 | 10.0 s |
| 59:55.3 | Data point 59 | 1.50000 mL | 0.07187 mL | 0.05847 mL | 0.28000 mL | 2.609 | -0.00893 | 0.93688 | 0.00046 | 10.0 s |
| 1:00:30.9 | Data point 60 | 1.50000 mL | 0.07796 mL | 0.05847 mL | 0.28000 mL | 2.414 | -0.01402 | 0.81271 | 0.00077 | 10.0 s |
| 1:01:06.6 | Data point 61 | 1.50000 mL | 0.08765 mL | 0.05847 mL | 0.28000 mL | 2.234 | -0.01252 | 0.94599 | 0.00064 | 10.0 s |
| 1:01:42.4 | Data point 62 | 1.50000 mL | 0.10263 mL | 0.05847 mL | 0.28000 mL | 2.045 | -0.01296 | 0.81107 | 0.00071 | 10.0 s |
| 1:02:18.1 | Data point 63 | 1.50000 mL | 0.11303 mL | 0.05847 mL | 0.28000 mL | 1.952 | -0.01017 | 0.74429 | 0.00058 | 10.0 s |
| 1:04:11.0 | Data point 64 | 1.50000 mL | 0.11303 mL | 0.11660 mL | 1.08000 mL | 9.602 | -0.07572 | 0.97633 | 0.00379 | Timed out at 59.5 s |
| 1:05:41.5 | Data point 65 | 1.50000 mL | 0.11312 mL | 0.11660 mL | 1.08000 mL | 7.914 | -0.01009 | 0.17302 | 0.00120 | Timed out at 59.5 s |
| 1:07:17.2 | Data point 66 | 1.50000 mL | 0.11350 mL | 0.11660 mL | 1.08000 mL | 7.574 | 0.00521 | 0.07080 | 0.00097 | 20.5 s |
| 1:08:13.4 | Data point 67 | 1.50000 mL | 0.11402 mL | 0.11660 mL | 1.08000 mL | 7.354 | 0.00770 | 0.15435 | 0.00097 | 25.0 s |
| 1:09:14.2 | Data point 68 | 1.50000 mL | 0.11465 mL | 0.11660 mL | 1.08000 mL | 7.149 | 0.01456 | 0.60986 | 0.00092 | 24.5 s |
| 1:10:14.4 | Data point 69 | 1.50000 mL | 0.11531 mL | 0.11660 mL | 1.08000 mL | 6.960 | 0.01422 | 0.79037 | 0.00079 | 31.5 s |
| 1:11:26.9 | Data point 70 | 1.50000 mL | 0.11599 mL | 0.11660 mL | 1.08000 mL | 6.759 | 0.01621 | 0.64775 | 0.00100 | 31.5 s |
| 1:12:23.9 | Data point 71 | 1.50000 mL | 0.11660 mL | 0.11660 mL | 1.08000 mL | 6.531 | 0.01843 | 0.87072 | 0.00098 | 45.0 s |
| 1:13:34.3 | Data point 72 | 1.50000 mL | 0.11705 mL | 0.11660 mL | 1.08000 mL | 6.327 | 0.01609 | 0.74484 | 0.00092 | 40.5 s |

| | | | |
|---|---|---|---|
| Sample name: | **M08_octanol** | Experiment start time: | **3/2/2018 8:29:22 PM** |
| Assay name: | **pH-metric high logP** | Analyst: | **Pion** |
| Assay ID: | **18C-02009** | Instrument ID: | **T312060** |
| Filename: | **C:\Sirius_T3\Mehtap\20180302_exp29_logP_T3-2\18C-02009_M08_octanol_pH-metric high logP.t3r** | | |

#### Events (continued)

| Time | Event | Water | Acid | Base | Octanol | pH | dpH/dt | pH R-squared | pH SD | dpH/dt time |
|---|---|---|---|---|---|---|---|---|---|---|
| 1:15:00.9 | Data point 73 | 1.50000 mL | 0.11750 mL | 0.11660 mL | 1.08000 mL | 6.048 | 0.00782 | 0.18941 | 0.00089 | 23.5 s |
| 1:16:00.0 | Data point 74 | 1.50000 mL | 0.11771 mL | 0.11660 mL | 1.08000 mL | 5.859 | 0.00920 | 0.23435 | 0.00094 | 21.5 s |
| 1:16:46.9 | Data point 75 | 1.50000 mL | 0.11785 mL | 0.11660 mL | 1.08000 mL | 5.634 | 0.01897 | 0.91284 | 0.00098 | 24.0 s |
| 1:17:41.5 | Data point 76 | 1.50000 mL | 0.11797 mL | 0.11660 mL | 1.08000 mL | 5.426 | 0.01127 | 0.33431 | 0.00097 | 18.0 s |
| 1:18:30.1 | Data point 77 | 1.50000 mL | 0.11816 mL | 0.11660 mL | 1.08000 mL | 4.860 | 0.01580 | 0.66582 | 0.00096 | 22.5 s |
| 1:19:28.2 | Data point 78 | 1.50000 mL | 0.11827 mL | 0.11660 mL | 1.08000 mL | 4.601 | -0.00526 | 0.07244 | 0.00097 | 11.5 s |
| 1:20:10.2 | Data point 79 | 1.50000 mL | 0.11837 mL | 0.11660 mL | 1.08000 mL | 4.394 | 0.01090 | 0.29750 | 0.00099 | 21.5 s |
| 1:20:57.2 | Data point 80 | 1.50000 mL | 0.11848 mL | 0.11660 mL | 1.08000 mL | 4.228 | 0.00338 | 0.03554 | 0.00089 | 10.5 s |
| 1:21:53.7 | Data point 81 | 1.50000 mL | 0.11874 mL | 0.11660 mL | 1.08000 mL | 3.988 | -0.01055 | 0.71176 | 0.00062 | 10.0 s |
| 1:22:39.4 | Data point 82 | 1.50000 mL | 0.11903 mL | 0.11660 mL | 1.08000 mL | 3.791 | -0.01849 | 0.93115 | 0.00095 | 10.5 s |
| 1:23:15.3 | Data point 83 | 1.50000 mL | 0.11945 mL | 0.11660 mL | 1.08000 mL | 3.591 | -0.01497 | 0.62245 | 0.00094 | 10.0 s |
| 1:23:50.8 | Data point 84 | 1.50000 mL | 0.12011 mL | 0.11660 mL | 1.08000 mL | 3.403 | 0.01050 | 0.27436 | 0.00099 | 18.0 s |
| 1:24:34.2 | Data point 85 | 1.50000 mL | 0.12114 mL | 0.11660 mL | 1.08000 mL | 3.213 | 0.00072 | 0.00159 | 0.00089 | 25.5 s |
| 1:25:25.3 | Data point 86 | 1.50000 mL | 0.12274 mL | 0.11660 mL | 1.08000 mL | 3.027 | -0.00532 | 0.13136 | 0.00073 | 10.5 s |
| 1:26:01.3 | Data point 87 | 1.50000 mL | 0.12517 mL | 0.11660 mL | 1.08000 mL | 2.849 | -0.00610 | 0.09602 | 0.00097 | 25.0 s |
| 1:26:51.9 | Data point 88 | 1.50000 mL | 0.12888 mL | 0.11660 mL | 1.08000 mL | 2.672 | -0.01765 | 0.96186 | 0.00089 | 10.0 s |
| 1:27:27.4 | Data point 89 | 1.50000 mL | 0.13450 mL | 0.11660 mL | 1.08000 mL | 2.475 | -0.00467 | 0.05593 | 0.00097 | 24.5 s |
| 1:28:17.7 | Data point 90 | 1.50000 mL | 0.14344 mL | 0.11660 mL | 1.08000 mL | 2.286 | -0.01613 | 0.73740 | 0.00093 | 13.5 s |
| 1:28:56.9 | Data point 91 | 1.50000 mL | 0.15753 mL | 0.11660 mL | 1.08000 mL | 2.098 | 0.00368 | 0.03491 | 0.00097 | 20.5 s |
| 1:29:43.3 | Data point 92 | 1.50000 mL | 0.17368 mL | 0.11660 mL | 1.08000 mL | 1.955 | 0.00356 | 0.03439 | 0.00095 | 19.5 s |
| 1:30:11.8 | Assay volumes | 1.50000 mL | 0.17368 mL | 0.11660 mL | 1.08000 mL | | | | | |

| | | | |
|---|---|---|---|
| Sample name: | **M08_octanol** | Experiment start time: | **3/2/2018 8:29:22 PM** |
| Assay name: | **pH-metric high logP** | Analyst: | **Pion** |
| Assay ID: | **18C-02009** | Instrument ID: | **T312060** |
| Filename: | **C:\Sirius_T3\Mehtap\20180302_exp29_logP_T3-2\18C-02009_M08_octanol_pH-metric high logP.t3r** | | |

| | | | |
|---|---|---|---|
| Sample name: | **M08_octanol** | Experiment start time: | **3/2/2018 8:29:22 PM** |
| Assay name: | **pH-metric high logP** | Analyst: | **Pion** |
| Assay ID: | **18C-02009** | Instrument ID: | **T312060** |
| Filename: | **C:\Sirius_T3\Mehtap\20180302_exp29_logP_T3-2\18C-02009_M08_octanol_pH-metric high logP.t3r** | | |

#### Assay Settings (continued)

| Setting | Value | Original Value | Date/Time changed | Imported from |
|---|---|---|---|---|
| ***Titration 3*** | | | | |
| Titrate from | High to low pH | | | |
| Add additional water | 0.00 mL | | | |
| Additional partition solvent volume | 0.800 mL | | | |
| Additional partition solvent added | Automatic | | | |
| After pH adjust stir for | 30 seconds | | | |
| Stir to allow partitioning for | 15 seconds | | | |
| Stirrer speed for partitioning | 60% | | | |
| ***Data Point Stability*** | | | | |
| Stir during data point collection | No | | | |
| Delay before data point collection | 0 seconds | | | |
| Number of points to average | 20 points | | | |
| Time interval between points | 0.50 seconds | | | |
| Required maximum standard deviation | 0.00100 dpH/dt | | | |
| Stability timeout after | 60 seconds | | | |

#### Calibration Settings

| Setting | Value | Date/Time changed | Imported from |
|---|---|---|---|
| Four-Plus alpha | 0.111 | 3/2/2018 8:29:22 PM | C:\Sirius_T3\HCl18C02.t3r |
| Four-Plus S | 0.9988 | 3/2/2018 8:29:22 PM | C:\Sirius_T3\HCl18C02.t3r |
| Four-Plus jH | 1.0 | 3/2/2018 8:29:22 PM | C:\Sirius_T3\HCl18C02.t3r |
| Four-Plus jOH | -0.8 | 3/2/2018 8:29:22 PM | C:\Sirius_T3\HCl18C02.t3r |
| Base concentration factor | 1.000 | 3/2/2018 8:29:22 PM | C:\Sirius_T3\KOH18B27.t3r |
| Acid concentration factor | 0.999 | 3/2/2018 8:29:22 PM | C:\Sirius_T3\HCl18C02.t3r |

#### Instrument Settings

| Setting | Value | Batch Id | Install date |
|---|---|---|---|
| Instrument owner | Merck | | |
| Instrument ID | T312060 | | |
| Instrument type | T3 Simulator | | |
| Software version | 1.1.3.0 | | |
| Dispenser module | | T3DM1200361 | 3/31/2009 5:24:52 AM |
| Dispenser 0 | Water | | 3/31/2009 5:25:05 AM |
| Syringe volume | 2.5 mL | | |
| Firmware version | 1.2.1(r2) | | |
| Titrant | Water (0.15 M KCl) | 02-06-2018 | 2/27/2018 10:05:59 AM |
| Dispenser 2 | Acid | | 3/31/2009 5:25:11 AM |
| Syringe volume | 0.5 mL | | |
| Firmware version | 1.2.1(r2) | | |
| Titrant | Acid (0.5 M HCl) | 02-27-2018 | 2/27/2018 10:27:22 AM |
| Dispenser 1 | Base | | 3/31/2009 5:25:21 AM |
| Syringe volume | 0.5 mL | | |
| Firmware version | 1.2.1(r2) | | |
| Titrant | Base (0.5 M KOH) | 9/22/2017 | 2/27/2018 10:21:22 AM |
| Dispenser 5 | Cosolvent | | 3/31/2009 5:26:24 AM |
| Syringe volume | 2.5 mL | | |
| Firmware version | 1.2.1(r2) | | |
| Distribution valve 5 | Distribution Valve | | 3/31/2009 5:28:19 AM |
| Firmware version | 1.1.3 | | |
| Port A | Methanol (80%, 0.15 M KCl) | 09-26-17 | 2/7/2018 9:42:01 AM |
| Port B | Cyclohexane | 11-01-17 | 2/27/2018 10:37:57 AM |
| Dispenser 3 | Buffer | | 8/3/2010 5:05:16 AM |
| Syringe volume | 0.5 mL | | |
| Firmware version | 1.2.1(r2) | | |
| Titrant | Dodecane | 2018/01/31 | 2/28/2018 10:18:04 AM |
| Dispenser 6 | Octanol | | 10/22/2010 10:52:43 AM |

| | | | |
|---|---|---|---|
| Sample name: | **M08_octanol** | Experiment start time: | **3/2/2018 8:29:22 PM** |
| Assay name: | **pH-metric high logP** | Analyst: | **Pion** |
| Assay ID: | **18C-02009** | Instrument ID: | **T312060** |
| Filename: | **C:\Sirius_T3\Mehtap\20180302_exp29_logP_T3-2\18C-02009_M08_octanol_pH-metric high logP.t3r** | | |

| | | | |
|---|---|---|---|
| Sample name: | **M08_octanol** | Experiment start time: | **3/2/2018 8:29:22 PM** |
| Assay name: | **pH-metric high logP** | Analyst: | **Pion** |
| Assay ID: | **18C-02009** | Instrument ID: | **T312060** |
| Filename: | **C:\Sirius_T3\Mehtap\20180302_exp29_logP_T3-2\18C-02009_M08_octanol_pH-metric high logP.t3r** | | |

#### Instrument Settings (continued)

| Setting | Value | Batch Id | Install date |
|---|---|---|---|
| Spectrometer calibration stir duration | 5 s | | |
| Spectrometer calibration stir speed | 30% | | |
| Spectrometer calibration wash pump volume | 20.0 mL | | |
| Spectrometer calibration wash stir duration | 5 s | | |
| Spectrometer calibration wash stir speed | 30% | | |
| Overhead dispense height | 10000 | | |

#### Refinement Settings

| Setting | Value | Default value |
|---|---|---|
| Turbidity detection method | None | None |
| Turbidity wavelength to assess | 500.0 nm | 500.0 nm |
| Turbidity maximum absorbance | 0.100 | 0.100 |
| Turbidity probe threshold | 50.00 | 50.00 |

#### Experiment Log

[2:37] Air gap created for Water (0.15 M KCl)
[2:38] Air gap created for Acid (0.5 M HCl)
[2:38] Air gap created for Base (0.5 M KOH)
[2:38] Air gap released for Water (0.15 M KCl)
[2:42] Titrator arm moved over Titration position
[2:42] Titration 1 of 3
[2:42] Adding initial titrants
[2:42] Automatically add 1.50000 mL of water
[3:07] Dispensed 1.500000 mL of Water (0.15 M KCl)
[3:12] Titrator arm moved over Drain
[5:53] Titrator arm moved to Titration position
[5:53] Argon flow rate set to 100
[5:53] Stirrer speed set to 10
[5:59] Initial pH = 5.13
[5:59] Iterative adjust 5.13 -> 10.00
[5:59] pH 5.13 -> 10.00
[6:00] Air gap released for Base (0.5 M KOH)
[6:01] Dispensed 0.006797 mL of Base (0.5 M KOH)
[6:06] Holding pH 10.00
[8:06] Stirrer speed set to 0
[8:06] Stirrer speed set to 50
[8:06] Iterative adjust 11.17 -> 10.00
[8:51] Stirrer speed set to 0
[9:51] Datapoint id 2 collected
[9:51] Stirrer speed set to 50
[9:57] pH 10.75 -> 10.55
[9:57] Using cautious pH adjust
[9:57] Air gap released for Acid (0.5 M HCl)
[9:58] Dispensed 0.000423 mL of Acid (0.5 M HCl)
[10:03] Stepping pH = 10.52
[10:18] Stirrer speed set to 0
[10:53] Datapoint id 3 collected
[10:53] Charge balance equation is out by 49.8%
[10:53] Stirrer speed set to 50
[10:58] pH 10.39 -> 10.19
[10:58] Using cautious pH adjust
[10:58] Dispensed 0.000188 mL of Acid (0.5 M HCl)
[11:03] Stepping pH = 10.34
[11:04] Dispensed 0.000306 mL of Acid (0.5 M HCl)
[11:09] Stepping pH = 10.16
[11:24] Stirrer speed set to 0
[12:04] Datapoint id 4 collected

| | | | |
|---|---|---|---|
| Sample name: | **M08_octanol** | Experiment start time: | **3/2/2018 8:29:22 PM** |
| Assay name: | **pH-metric high logP** | Analyst: | **Pion** |
| Assay ID: | **18C-02009** | Instrument ID: | **T312060** |
| Filename: | **C:\Sirius_T3\Mehtap\20180302_exp29_logP_T3-2\18C-02009_M08_octanol_pH-metric high logP.t3r** | | |

#### Experiment Log (continued)

[12:04] Charge balance equation is out by -24.4%
[12:04] Stirrer speed set to 50
[12:09] pH 10.05 -> 9.85
[12:09] Using cautious pH adjust
[12:09] Dispensed 0.000094 mL of Acid (0.5 M HCl)
[12:15] Stepping pH = 10.02
[12:15] Dispensed 0.000235 mL of Acid (0.5 M HCl)
[12:20] Stepping pH = 9.77
[12:35] Stirrer speed set to 0
[13:03] Datapoint id 5 collected
[13:03] Charge balance equation is out by -70.9%
[13:03] Stirrer speed set to 50
[13:09] pH 9.66 -> 9.46
[13:09] Using cautious pH adjust
[13:09] Dispensed 0.000047 mL of Acid (0.5 M HCl)
[13:14] Stepping pH = 9.64
[13:14] Dispensed 0.000141 mL of Acid (0.5 M HCl)
[13:19] Stepping pH = 9.39
[13:34] Stirrer speed set to 0
[14:07] Datapoint id 6 collected
[14:07] Charge balance equation is out by -86.9%
[14:07] Stirrer speed set to 50
[14:12] pH 9.28 -> 9.08
[14:12] Using cautious pH adjust
[14:12] Dispensed 0.000024 mL of Acid (0.5 M HCl)
[14:17] Stepping pH = 9.26
[14:17] Dispensed 0.000071 mL of Acid (0.5 M HCl)
[14:22] Stepping pH = 9.10
[14:22] Dispensed 0.000024 mL of Acid (0.5 M HCl)
[14:27] Stepping pH = 9.03
[14:42] Stirrer speed set to 0
[15:18] Datapoint id 7 collected
[15:18] Charge balance equation is out by -126.0%
[15:18] Stirrer speed set to 50
[15:23] pH 8.82 -> 8.62
[15:23] Using cautious pH adjust
[15:23] Dispensed 0.000024 mL of Acid (0.5 M HCl)
[15:28] Stepping pH = 8.78
[15:28] Dispensed 0.000024 mL of Acid (0.5 M HCl)
[15:34] Stepping pH = 8.69
[15:34] Dispensed 0.000024 mL of Acid (0.5 M HCl)
[15:39] Stepping pH = 8.55
[15:54] Stirrer speed set to 0
[16:43] Datapoint id 8 collected
[16:43] Charge balance equation is out by -159.4%
[16:43] Stirrer speed set to 50
[16:48] pH 8.28 -> 8.08
[16:48] Using cautious pH adjust
[16:48] Dispensed 0.000024 mL of Acid (0.5 M HCl)
[16:53] Stepping pH = 8.24
[16:53] Dispensed 0.000024 mL of Acid (0.5 M HCl)
[16:58] Stepping pH = 8.17
[16:58] Dispensed 0.000024 mL of Acid (0.5 M HCl)
[17:03] Stepping pH = 8.08
[17:19] Stirrer speed set to 0
[17:38] Datapoint id 9 collected
[17:38] Charge balance equation is out by -148.3%
[17:38] Stirrer speed set to 50
[17:43] pH 7.95 -> 7.75

| | | | |
|---|---|---|---|
| Sample name: | **M08_octanol** | Experiment start time: | **3/2/2018 8:29:22 PM** |
| Assay name: | **pH-metric high logP** | Analyst: | **Pion** |
| Assay ID: | **18C-02009** | Instrument ID: | **T312060** |
| Filename: | **C:\Sirius_T3\Mehtap\20180302_exp29_logP_T3-2\18C-02009_M08_octanol_pH-metric high logP.t3r** | | |

#### Experiment Log (continued)

[17:43] Using cautious pH adjust
[17:43] Dispensed 0.000024 mL of Acid (0.5 M HCl)
[17:48] Stepping pH = 7.92
[17:48] Dispensed 0.000071 mL of Acid (0.5 M HCl)
[17:53] Stepping pH = 7.72
[18:09] Stirrer speed set to 0
[18:26] Datapoint id 10 collected
[18:26] Charge balance equation is out by -62.6%
[18:26] Stirrer speed set to 50
[18:31] pH 7.65 -> 7.45
[18:31] Using cautious pH adjust
[18:31] Dispensed 0.000047 mL of Acid (0.5 M HCl)
[18:36] Stepping pH = 7.62
[18:36] Dispensed 0.000094 mL of Acid (0.5 M HCl)
[18:41] Stepping pH = 7.44
[18:56] Stirrer speed set to 0
[19:13] Datapoint id 11 collected
[19:13] Charge balance equation is out by -49.6%
[19:13] Stirrer speed set to 50
[19:18] pH 7.42 -> 7.22
[19:18] Using cautious pH adjust
[19:18] Dispensed 0.000071 mL of Acid (0.5 M HCl)
[19:23] Stepping pH = 7.36
[19:23] Dispensed 0.000094 mL of Acid (0.5 M HCl)
[19:29] Stepping pH = 7.25
[19:29] Dispensed 0.000024 mL of Acid (0.5 M HCl)
[19:34] Stepping pH = 7.24
[19:34] Dispensed 0.000047 mL of Acid (0.5 M HCl)
[19:39] Stepping pH = 7.21
[19:54] Stirrer speed set to 0
[20:15] Datapoint id 12 collected
[20:15] Charge balance equation is out by -60.4%
[20:15] Stirrer speed set to 50
[20:20] pH 7.21 -> 7.01
[20:20] Using cautious pH adjust
[20:20] Dispensed 0.000118 mL of Acid (0.5 M HCl)
[20:25] Stepping pH = 7.11
[20:25] Dispensed 0.000094 mL of Acid (0.5 M HCl)
[20:30] Stepping pH = 7.04
[20:31] Dispensed 0.000047 mL of Acid (0.5 M HCl)
[20:36] Stepping pH = 7.03
[20:36] Dispensed 0.000047 mL of Acid (0.5 M HCl)
[20:41] Stepping pH = 7.01
[20:56] Stirrer speed set to 0
[21:13] Datapoint id 13 collected
[21:13] Charge balance equation is out by -27.4%
[21:13] Stirrer speed set to 50
[21:18] pH 7.02 -> 6.82
[21:18] Using cautious pH adjust
[21:18] Dispensed 0.000165 mL of Acid (0.5 M HCl)
[21:23] Stepping pH = 6.90
[21:23] Dispensed 0.000094 mL of Acid (0.5 M HCl)
[21:28] Stepping pH = 6.86
[21:28] Dispensed 0.000071 mL of Acid (0.5 M HCl)
[21:34] Stepping pH = 6.84
[21:34] Dispensed 0.000071 mL of Acid (0.5 M HCl)
[21:39] Stepping pH = 6.82
[21:54] Stirrer speed set to 0
[22:10] Datapoint id 14 collected

Sample name: **M08_octanol** Experiment start time: **3/2/2018 8:29:22 PM**
Assay name: **pH-metric high logP** Analyst: **Pion**
Assay ID: **18C-02009** Instrument ID: **T312060**
Filename: **C:\Sirius_T3\Mehtap\20180302_exp29_logP_T3-2\18C-02009_M08_octanol_pH-metric high logP.t3r**

#### Experiment Log (continued)

[22:10] Charge balance equation is out by -25.3%
[22:10] Stirrer speed set to 50
[22:15] pH 6.84 -> 6.64
[22:15] Using cautious pH adjust
[22:15] Dispensed 0.000235 mL of Acid (0.5 M HCl)
[22:20] Stepping pH = 6.72
[22:20] Dispensed 0.000118 mL of Acid (0.5 M HCl)
[22:25] Stepping pH = 6.68
[22:25] Dispensed 0.000094 mL of Acid (0.5 M HCl)
[22:30] Stepping pH = 6.66
[22:30] Dispensed 0.000071 mL of Acid (0.5 M HCl)
[22:36] Stepping pH = 6.65
[22:51] Stirrer speed set to 0
[23:07] Datapoint id 15 collected
[23:07] Charge balance equation is out by -11.8%
[23:07] Stirrer speed set to 50
[23:12] pH 6.67 -> 6.47
[23:12] Using charge balance adjust
[23:12] Dispensed 0.000564 mL of Acid (0.5 M HCl)
[23:33] Stirrer speed set to 0
[23:50] Datapoint id 16 collected
[23:50] Charge balance equation is out by -18.0%
[23:50] Stirrer speed set to 50
[23:55] pH 6.50 -> 6.30
[23:55] Using cautious pH adjust
[23:55] Dispensed 0.000329 mL of Acid (0.5 M HCl)
[24:00] Stepping pH = 6.39
[24:00] Dispensed 0.000212 mL of Acid (0.5 M HCl)
[24:05] Stepping pH = 6.35
[24:05] Dispensed 0.000165 mL of Acid (0.5 M HCl)
[24:10] Stepping pH = 6.32
[24:10] Dispensed 0.000071 mL of Acid (0.5 M HCl)
[24:15] Stepping pH = 6.32
[24:15] Dispensed 0.000212 mL of Acid (0.5 M HCl)
[24:21] Stepping pH = 6.26
[24:36] Stirrer speed set to 0
[24:49] Datapoint id 17 collected
[24:49] Charge balance equation is out by -45.8%
[24:49] Stirrer speed set to 50
[24:54] pH 6.29 -> 6.09
[24:54] Using cautious pH adjust
[24:54] Dispensed 0.000400 mL of Acid (0.5 M HCl)
[25:00] Stepping pH = 6.17
[25:00] Dispensed 0.000212 mL of Acid (0.5 M HCl)
[25:05] Stepping pH = 6.13
[25:05] Dispensed 0.000165 mL of Acid (0.5 M HCl)
[25:10] Stepping pH = 6.10
[25:25] Stirrer speed set to 0
[25:38] Datapoint id 18 collected
[25:38] Charge balance equation is out by 3.6%
[25:38] Stirrer speed set to 50
[25:44] pH 6.12 -> 5.92
[25:44] Using charge balance adjust
[25:44] Dispensed 0.000800 mL of Acid (0.5 M HCl)
[26:04] Stirrer speed set to 0
[26:16] Datapoint id 19 collected
[26:16] Charge balance equation is out by 1.0%
[26:16] Stirrer speed set to 50
[26:22] pH 5.92 -> 5.72

| | | | |
|---|---|---|---|
| Sample name: | **M08_octanol** | Experiment start time: | **3/2/2018 8:29:22 PM** |
| Assay name: | **pH-metric high logP** | Analyst: | **Pion** |
| Assay ID: | **18C-02009** | Instrument ID: | **T312060** |
| Filename: | **C:\Sirius_T3\Mehtap\20180302_exp29_logP_T3-2\18C-02009_M08_octanol_pH-metric high logP.t3r** | | |

#### Experiment Log (continued)

[26:22] Using charge balance adjust
[26:22] Dispensed 0.000753 mL of Acid (0.5 M HCl)
[26:42] Stirrer speed set to 0
[26:58] Datapoint id 20 collected
[26:58] Charge balance equation is out by 4.1%
[26:58] Stirrer speed set to 50
[27:03] pH 5.71 -> 5.51
[27:03] Using charge balance adjust
[27:03] Dispensed 0.000635 mL of Acid (0.5 M HCl)
[27:23] Stirrer speed set to 0
[27:40] Datapoint id 21 collected
[27:40] Charge balance equation is out by 17.0%
[27:40] Stirrer speed set to 50
[27:45] pH 5.48 -> 5.28
[27:45] Using cautious pH adjust
[27:45] Dispensed 0.000235 mL of Acid (0.5 M HCl)
[27:51] Stepping pH = 5.37
[27:51] Dispensed 0.000141 mL of Acid (0.5 M HCl)
[27:56] Stepping pH = 5.29
[27:56] Dispensed 0.000024 mL of Acid (0.5 M HCl)
[28:01] Stepping pH = 5.30
[28:01] Dispensed 0.000165 mL of Acid (0.5 M HCl)
[28:06] Stepping pH = 5.18
[28:21] Stirrer speed set to 0
[28:42] Datapoint id 22 collected
[28:42] Charge balance equation is out by -22.3%
[28:42] Stirrer speed set to 50
[28:47] pH 5.18 -> 4.98
[28:47] Using cautious pH adjust
[28:47] Dispensed 0.000141 mL of Acid (0.5 M HCl)
[28:52] Stepping pH = 5.08
[28:52] Dispensed 0.000094 mL of Acid (0.5 M HCl)
[28:58] Stepping pH = 4.98
[29:13] Stirrer speed set to 0
[30:13] Datapoint id 23 collected
[30:13] Charge balance equation is out by 20.1%
[30:13] Stirrer speed set to 50
[30:18] pH 5.05 -> 4.85
[30:18] Using cautious pH adjust
[30:18] Dispensed 0.000118 mL of Acid (0.5 M HCl)
[30:23] Stepping pH = 4.93
[30:23] Dispensed 0.000047 mL of Acid (0.5 M HCl)
[30:28] Stepping pH = 4.88
[30:28] Dispensed 0.000024 mL of Acid (0.5 M HCl)
[30:33] Stepping pH = 4.87
[30:33] Dispensed 0.000024 mL of Acid (0.5 M HCl)
[30:38] Stepping pH = 4.83
[30:53] Stirrer speed set to 0
[31:53] Datapoint id 24 collected
[31:53] Charge balance equation is out by 5.3%
[31:53] Stirrer speed set to 50
[31:59] pH 4.99 -> 4.79
[31:59] Using charge balance adjust
[31:59] Dispensed 0.000212 mL of Acid (0.5 M HCl)
[32:19] Stirrer speed set to 0
[32:52] Datapoint id 25 collected
[32:52] Charge balance equation is out by 145.5%
[32:52] Stirrer speed set to 50
[32:57] pH 4.51 -> 4.31

Sample name: **M08_octanol** Experiment start time: **3/2/2018 8:29:22 PM**
Assay name: **pH-metric high logP** Analyst: **Pion**
Assay ID: **18C-02009** Instrument ID: **T312060**
Filename: **C:\Sirius_T3\Mehtap\20180302_exp29_logP_T3-2\18C-02009_M08_octanol_pH-metric high logP.t3r**

#### Experiment Log (continued)

[32:57] Using cautious pH adjust
[32:57] Dispensed 0.000071 mL of Acid (0.5 M HCl)
[33:02] Stepping pH = 4.40
[33:02] Dispensed 0.000047 mL of Acid (0.5 M HCl)
[33:07] Stepping pH = 4.32
[33:22] Stirrer speed set to 0
[33:48] Datapoint id 26 collected
[33:48] Charge balance equation is out by 19.0%
[33:48] Stirrer speed set to 50
[33:53] pH 4.31 -> 4.11
[33:53] Using cautious pH adjust
[33:53] Dispensed 0.000071 mL of Acid (0.5 M HCl)
[33:58] Stepping pH = 4.21
[33:58] Dispensed 0.000047 mL of Acid (0.5 M HCl)
[34:03] Stepping pH = 4.14
[34:03] Dispensed 0.000024 mL of Acid (0.5 M HCl)
[34:08] Stepping pH = 4.12
[34:24] Stirrer speed set to 0
[34:34] Datapoint id 27 collected
[34:34] Charge balance equation is out by 1.1%
[34:34] Stirrer speed set to 50
[34:39] pH 4.11 -> 3.91
[34:39] Using charge balance adjust
[34:39] Dispensed 0.000212 mL of Acid (0.5 M HCl)
[34:59] Stirrer speed set to 0
[35:09] Datapoint id 28 collected
[35:10] Charge balance equation is out by 21.2%
[35:10] Stirrer speed set to 50
[35:15] pH 3.87 -> 3.67
[35:15] Using cautious pH adjust
[35:15] Dispensed 0.000165 mL of Acid (0.5 M HCl)
[35:20] Stepping pH = 3.76
[35:20] Dispensed 0.000094 mL of Acid (0.5 M HCl)
[35:25] Stepping pH = 3.71
[35:25] Dispensed 0.000047 mL of Acid (0.5 M HCl)
[35:30] Stepping pH = 3.69
[35:30] Dispensed 0.000047 mL of Acid (0.5 M HCl)
[35:35] Stepping pH = 3.67
[35:50] Stirrer speed set to 0
[36:00] Datapoint id 29 collected
[36:00] Charge balance equation is out by -14.5%
[36:00] Stirrer speed set to 50
[36:05] pH 3.67 -> 3.47
[36:05] Using charge balance adjust
[36:06] Dispensed 0.000494 mL of Acid (0.5 M HCl)
[36:26] Stirrer speed set to 0
[36:36] Datapoint id 30 collected
[36:36] Charge balance equation is out by 10.8%
[36:36] Stirrer speed set to 50
[36:41] pH 3.45 -> 3.25
[36:41] Using charge balance adjust
[36:41] Dispensed 0.000800 mL of Acid (0.5 M HCl)
[37:02] Stirrer speed set to 0
[37:12] Datapoint id 31 collected
[37:12] Charge balance equation is out by 0.6%
[37:12] Stirrer speed set to 50
[37:17] pH 3.26 -> 3.06
[37:17] Using charge balance adjust
[37:17] Dispensed 0.001270 mL of Acid (0.5 M HCl)

#### Experiment Log (continued)

[37:37] Stirrer speed set to 0
[37:47] Datapoint id 32 collected
[37:47] Charge balance equation is out by -2.6%
[37:47] Stirrer speed set to 50
[37:52] pH 3.07 -> 2.87
[37:52] Using charge balance adjust
[37:52] Dispensed 0.001952 mL of Acid (0.5 M HCl)
[38:13] Stirrer speed set to 0
[38:23] Datapoint id 33 collected
[38:23] Charge balance equation is out by -7.9%
[38:23] Stirrer speed set to 50
[38:28] pH 2.89 -> 2.69
[38:28] Using charge balance adjust
[38:28] Dispensed 0.002963 mL of Acid (0.5 M HCl)
[38:48] Stirrer speed set to 0
[38:58] Datapoint id 34 collected
[38:58] Charge balance equation is out by -8.0%
[38:58] Stirrer speed set to 50
[39:03] pH 2.71 -> 2.51
[39:03] Using charge balance adjust
[39:04] Dispensed 0.004492 mL of Acid (0.5 M HCl)
[39:24] Stirrer speed set to 0
[39:34] Datapoint id 35 collected
[39:34] Charge balance equation is out by -8.4%
[39:34] Stirrer speed set to 50
[39:39] pH 2.54 -> 2.34
[39:39] Using charge balance adjust
[39:39] Dispensed 0.006844 mL of Acid (0.5 M HCl)
[39:59] Stirrer speed set to 0
[40:09] Datapoint id 36 collected
[40:09] Charge balance equation is out by -3.3%
[40:09] Stirrer speed set to 50
[40:15] pH 2.35 -> 2.15
[40:15] Using charge balance adjust
[40:15] Dispensed 0.010724 mL of Acid (0.5 M HCl)
[40:35] Stirrer speed set to 0
[40:45] Datapoint id 37 collected
[40:45] Charge balance equation is out by -4.4%
[40:45] Stirrer speed set to 50
[40:50] pH 2.16 -> 1.96
[40:50] Using charge balance adjust
[40:51] Dispensed 0.016886 mL of Acid (0.5 M HCl)
[41:11] Stirrer speed set to 0
[41:22] Datapoint id 38 collected
[41:22] Charge balance equation is out by -3.8%
[41:22] Titration 2 of 3
[41:22] Adding initial titrants
[41:22] Automatically add 0.20000 mL of Octanol
[41:26] Dispensed 0.200000 mL of Octanol
[41:26] Stirrer speed set to 10
[41:27] Stirrer speed set to 55
[41:27] Iterative adjust 1.97 -> 10.00
[41:27] pH 1.97 -> 10.00
[41:29] Dispensed 0.051670 mL of Base (0.5 M KOH)
[42:19] Stirrer speed set to 0
[43:19] Datapoint id 39 collected
[43:19] Stirrer speed set to 55
[43:24] pH 7.36 -> 7.16
[43:24] Using cautious pH adjust

| | | | |
|---|---|---|---|
| Sample name: | **M08_octanol** | Experiment start time: | **3/2/2018 8:29:22 PM** |
| Assay name: | **pH-metric high logP** | Analyst: | **Pion** |
| Assay ID: | **18C-02009** | Instrument ID: | **T312060** |
| Filename: | **C:\Sirius_T3\Mehtap\20180302_exp29_logP_T3-2\18C-02009_M08_octanol_pH-metric high logP.t3r** | | |

#### Experiment Log (continued)

[43:24] Dispensed 0.000212 mL of Acid (0.5 M HCl)
[43:29] Stepping pH = 7.24
[43:29] Dispensed 0.000118 mL of Acid (0.5 M HCl)
[43:34] Stepping pH = 7.17
[43:49] Stirrer speed set to 0
[44:26] Datapoint id 40 collected
[44:26] Charge balance equation is out by 21.6%
[44:26] Stirrer speed set to 55
[44:31] pH 7.10 -> 6.90
[44:31] Using cautious pH adjust
[44:31] Dispensed 0.000306 mL of Acid (0.5 M HCl)
[44:36] Stepping pH = 6.94
[44:36] Dispensed 0.000094 mL of Acid (0.5 M HCl)
[44:41] Stepping pH = 6.93
[44:42] Dispensed 0.000094 mL of Acid (0.5 M HCl)
[44:47] Stepping pH = 6.89
[45:02] Stirrer speed set to 0
[45:23] Datapoint id 41 collected
[45:23] Charge balance equation is out by 21.8%
[45:23] Stirrer speed set to 55
[45:28] pH 6.89 -> 6.69
[45:28] Using cautious pH adjust
[45:28] Dispensed 0.000376 mL of Acid (0.5 M HCl)
[45:33] Stepping pH = 6.72
[45:33] Dispensed 0.000047 mL of Acid (0.5 M HCl)
[45:38] Stepping pH = 6.74
[45:38] Dispensed 0.000447 mL of Acid (0.5 M HCl)
[45:43] Stepping pH = 6.56
[45:58] Stirrer speed set to 0
[46:25] Datapoint id 42 collected
[46:25] Charge balance equation is out by -19.0%
[46:25] Stirrer speed set to 55
[46:30] pH 6.59 -> 6.39
[46:30] Using cautious pH adjust
[46:30] Dispensed 0.000400 mL of Acid (0.5 M HCl)
[46:35] Stepping pH = 6.44
[46:35] Dispensed 0.000141 mL of Acid (0.5 M HCl)
[46:40] Stepping pH = 6.41
[46:41] Dispensed 0.000071 mL of Acid (0.5 M HCl)
[46:46] Stepping pH = 6.39
[47:01] Stirrer speed set to 0
[47:28] Datapoint id 43 collected
[47:28] Charge balance equation is out by 24.6%
[47:28] Stirrer speed set to 55
[47:33] pH 6.41 -> 6.21
[47:33] Using cautious pH adjust
[47:33] Dispensed 0.000376 mL of Acid (0.5 M HCl)
[47:38] Stepping pH = 6.25
[47:38] Dispensed 0.000071 mL of Acid (0.5 M HCl)
[47:43] Stepping pH = 6.24
[47:43] Dispensed 0.000071 mL of Acid (0.5 M HCl)
[47:48] Stepping pH = 6.21
[48:03] Stirrer speed set to 0
[48:29] Datapoint id 44 collected
[48:29] Charge balance equation is out by 30.2%
[48:29] Stirrer speed set to 55
[48:34] pH 6.24 -> 6.04
[48:34] Using cautious pH adjust
[48:34] Dispensed 0.000329 mL of Acid (0.5 M HCl)

| | | | |
|---|---|---|---|
| Sample name: | **M08_octanol** | Experiment start time: | **3/2/2018 8:29:22 PM** |
| Assay name: | **pH-metric high logP** | Analyst: | **Pion** |
| Assay ID: | **18C-02009** | Instrument ID: | **T312060** |
| Filename: | **C:\Sirius_T3\Mehtap\20180302_exp29_logP_T3-2\18C-02009_M08_octanol_pH-metric high logP.t3r** | | |

#### Experiment Log (continued)

[48:39] Stepping pH = 6.06
[48:39] Dispensed 0.000047 mL of Acid (0.5 M HCl)
[48:44] Stepping pH = 6.06
[48:44] Dispensed 0.000071 mL of Acid (0.5 M HCl)
[48:49] Stepping pH = 6.02
[49:04] Stirrer speed set to 0
[49:31] Datapoint id 45 collected
[49:31] Charge balance equation is out by 32.8%
[49:31] Stirrer speed set to 55
[49:36] pH 6.04 -> 5.84
[49:36] Using cautious pH adjust
[49:36] Dispensed 0.000259 mL of Acid (0.5 M HCl)
[49:41] Stepping pH = 5.87
[49:41] Dispensed 0.000024 mL of Acid (0.5 M HCl)
[49:46] Stepping pH = 5.86
[49:46] Dispensed 0.000071 mL of Acid (0.5 M HCl)
[49:51] Stepping pH = 5.82
[50:06] Stirrer speed set to 0
[50:28] Datapoint id 46 collected
[50:28] Charge balance equation is out by 30.4%
[50:28] Stirrer speed set to 55
[50:34] pH 5.85 -> 5.65
[50:34] Using cautious pH adjust
[50:34] Dispensed 0.000188 mL of Acid (0.5 M HCl)
[50:39] Stepping pH = 5.67
[50:39] Dispensed 0.000024 mL of Acid (0.5 M HCl)
[50:44] Stepping pH = 5.68
[50:44] Dispensed 0.000165 mL of Acid (0.5 M HCl)
[50:49] Stepping pH = 5.46
[51:04] Stirrer speed set to 0
[51:23] Datapoint id 47 collected
[51:23] Charge balance equation is out by 5.3%
[51:23] Stirrer speed set to 55
[51:28] pH 5.48 -> 5.28
[51:28] Using charge balance adjust
[51:28] Dispensed 0.000212 mL of Acid (0.5 M HCl)
[51:49] Stirrer speed set to 0
[52:05] Datapoint id 48 collected
[52:05] Charge balance equation is out by 71.8%
[52:05] Stirrer speed set to 55
[52:10] pH 5.12 -> 4.92
[52:10] Using cautious pH adjust
[52:10] Dispensed 0.000047 mL of Acid (0.5 M HCl)
[52:15] Stepping pH = 5.10
[52:15] Dispensed 0.000165 mL of Acid (0.5 M HCl)
[52:21] Stepping pH = 4.66
[52:36] Stirrer speed set to 0
[52:52] Datapoint id 49 collected
[52:52] Charge balance equation is out by -86.2%
[52:52] Stirrer speed set to 55
[52:57] pH 4.66 -> 4.46
[52:57] Using cautious pH adjust
[52:57] Dispensed 0.000047 mL of Acid (0.5 M HCl)
[53:02] Stepping pH = 4.65
[53:02] Dispensed 0.000118 mL of Acid (0.5 M HCl)
[53:07] Stepping pH = 4.41
[53:22] Stirrer speed set to 0
[53:34] Datapoint id 50 collected
[53:34] Charge balance equation is out by -89.3%

| | | | |
|---|---|---|---|
| Sample name: | **M08_octanol** | Experiment start time: | **3/2/2018 8:29:22 PM** |
| Assay name: | **pH-metric high logP** | Analyst: | **Pion** |
| Assay ID: | **18C-02009** | Instrument ID: | **T312060** |
| Filename: | **C:\Sirius_T3\Mehtap\20180302_exp29_logP_T3-2\18C-02009_M08_octanol_pH-metric high logP.t3r** | | |

#### Experiment Log (continued)

[53:34] Stirrer speed set to 55
[53:39] pH 4.40 -> 4.20
[53:39] Using cautious pH adjust
[53:39] Dispensed 0.000047 mL of Acid (0.5 M HCl)
[53:44] Stepping pH = 4.39
[53:44] Dispensed 0.000165 mL of Acid (0.5 M HCl)
[53:49] Stepping pH = 4.17
[54:04] Stirrer speed set to 0
[54:15] Datapoint id 51 collected
[54:15] Charge balance equation is out by -92.5%
[54:15] Stirrer speed set to 55
[54:20] pH 4.17 -> 3.97
[54:20] Using cautious pH adjust
[54:20] Dispensed 0.000094 mL of Acid (0.5 M HCl)
[54:25] Stepping pH = 4.12
[54:25] Dispensed 0.000118 mL of Acid (0.5 M HCl)
[54:30] Stepping pH = 3.99
[54:30] Dispensed 0.000024 mL of Acid (0.5 M HCl)
[54:35] Stepping pH = 3.98
[54:50] Stirrer speed set to 0
[55:00] Datapoint id 52 collected
[55:00] Charge balance equation is out by -30.4%
[55:00] Stirrer speed set to 55
[55:06] pH 3.98 -> 3.78
[55:06] Using cautious pH adjust
[55:06] Dispensed 0.000141 mL of Acid (0.5 M HCl)
[55:11] Stepping pH = 3.89
[55:11] Dispensed 0.000118 mL of Acid (0.5 M HCl)
[55:16] Stepping pH = 3.82
[55:16] Dispensed 0.000047 mL of Acid (0.5 M HCl)
[55:21] Stepping pH = 3.80
[55:21] Dispensed 0.000047 mL of Acid (0.5 M HCl)
[55:26] Stepping pH = 3.78
[55:41] Stirrer speed set to 0
[55:51] Datapoint id 53 collected
[55:51] Charge balance equation is out by -27.9%
[55:51] Stirrer speed set to 55
[55:56] pH 3.78 -> 3.58
[55:56] Using cautious pH adjust
[55:56] Dispensed 0.000212 mL of Acid (0.5 M HCl)
[56:02] Stepping pH = 3.67
[56:02] Dispensed 0.000141 mL of Acid (0.5 M HCl)
[56:07] Stepping pH = 3.62
[56:07] Dispensed 0.000094 mL of Acid (0.5 M HCl)
[56:12] Stepping pH = 3.59
[56:12] Dispensed 0.000024 mL of Acid (0.5 M HCl)
[56:17] Stepping pH = 3.59
[56:32] Stirrer speed set to 0
[56:42] Datapoint id 54 collected
[56:42] Charge balance equation is out by -15.7%
[56:42] Stirrer speed set to 55
[56:47] pH 3.59 -> 3.39
[56:47] Using cautious pH adjust
[56:47] Dispensed 0.000306 mL of Acid (0.5 M HCl)
[56:52] Stepping pH = 3.50
[56:53] Dispensed 0.000282 mL of Acid (0.5 M HCl)
[56:58] Stepping pH = 3.41
[56:58] Dispensed 0.000071 mL of Acid (0.5 M HCl)
[57:03] Stepping pH = 3.40

Sample name: **M08_octanol**
Assay name: **pH-metric high logP**
Assay ID: **18C-02009**
Filename: **C:\Sirius_T3\Mehtap\20180302_exp29_logP_T3-2\18C-02009_M08_octanol_pH-metric high logP.t3r**
Experiment start time: **3/2/2018 8:29:22 PM**
Analyst: **Pion**
Instrument ID: **T312060**

#### Experiment Log (continued)

[57:03] Dispensed 0.000094 mL of Acid (0.5 M HCl)
[57:08] Stepping pH = 3.38
[57:23] Stirrer speed set to 0
[57:33] Datapoint id 55 collected
[57:33] Charge balance equation is out by -22.9%
[57:33] Stirrer speed set to 55
[57:38] pH 3.38 -> 3.18
[57:38] Using cautious pH adjust
[57:38] Dispensed 0.000517 mL of Acid (0.5 M HCl)
[57:43] Stepping pH = 3.26
[57:43] Dispensed 0.000282 mL of Acid (0.5 M HCl)
[57:49] Stepping pH = 3.21
[57:49] Dispensed 0.000165 mL of Acid (0.5 M HCl)
[57:54] Stepping pH = 3.19
[58:09] Stirrer speed set to 0
[58:19] Datapoint id 56 collected
[58:19] Charge balance equation is out by 5.3%
[58:19] Stirrer speed set to 55
[58:24] pH 3.19 -> 2.99
[58:24] Using charge balance adjust
[58:24] Dispensed 0.001576 mL of Acid (0.5 M HCl)
[58:44] Stirrer speed set to 0
[58:54] Datapoint id 57 collected
[58:54] Charge balance equation is out by -0.7%
[58:54] Stirrer speed set to 55
[58:59] pH 3.00 -> 2.80
[58:59] Using charge balance adjust
[59:00] Dispensed 0.002469 mL of Acid (0.5 M HCl)
[59:20] Stirrer speed set to 0
[59:30] Datapoint id 58 collected
[59:30] Charge balance equation is out by -1.6%
[59:30] Stirrer speed set to 55
[59:35] pH 2.81 -> 2.61
[59:35] Using charge balance adjust
[59:35] Dispensed 0.003881 mL of Acid (0.5 M HCl)
[59:55] Stirrer speed set to 0
[1:00:05] Datapoint id 59 collected
[1:00:05] Charge balance equation is out by -2.2%
[1:00:05] Stirrer speed set to 55
[1:00:10] pH 2.61 -> 2.41
[1:00:10] Using charge balance adjust
[1:00:11] Dispensed 0.006091 mL of Acid (0.5 M HCl)
[1:00:31] Stirrer speed set to 0
[1:00:41] Datapoint id 60 collected
[1:00:41] Charge balance equation is out by -0.3%
[1:00:41] Stirrer speed set to 55
[1:00:46] pH 2.42 -> 2.22
[1:00:46] Using charge balance adjust
[1:00:46] Dispensed 0.009690 mL of Acid (0.5 M HCl)
[1:01:07] Stirrer speed set to 0
[1:01:17] Datapoint id 61 collected
[1:01:17] Charge balance equation is out by -8.4%
[1:01:17] Stirrer speed set to 55
[1:01:22] pH 2.24 -> 2.04
[1:01:22] Using charge balance adjust
[1:01:22] Dispensed 0.014981 mL of Acid (0.5 M HCl)
[1:01:42] Stirrer speed set to 0
[1:01:52] Datapoint id 62 collected
[1:01:52] Charge balance equation is out by -3.3%

Sample name: **M08_octanol** | Experiment start time: **3/2/2018 8:29:22 PM**
Assay name: **pH-metric high logP** | Analyst: **Pion**
Assay ID: **18C-02009** | Instrument ID: **T312060**
Filename: **C:\Sirius_T3\Mehtap\20180302_exp29_logP_T3-2\18C-02009_M08_octanol_pH-metric high logP.t3r**

#### Experiment Log (continued)

[1:01:52] Stirrer speed set to 55
[1:01:58] pH 2.05 -> 1.95
[1:01:58] Using charge balance adjust
[1:01:58] Dispensed 0.010395 mL of Acid (0.5 M HCl)
[1:02:18] Stirrer speed set to 0
[1:02:28] Datapoint id 63 collected
[1:02:28] Charge balance equation is out by -51.4%
[1:02:28] Titration 3 of 3
[1:02:28] Adding initial titrants
[1:02:28] Automatically add 0.80000 mL of Octanol
[1:03:18] Dispensed 0.800000 mL of Octanol
[1:03:18] Stirrer speed set to 10
[1:03:19] Stirrer speed set to 60
[1:03:19] Iterative adjust 1.94 -> 10.00
[1:03:19] pH 1.94 -> 10.00
[1:03:21] Dispensed 0.058137 mL of Base (0.5 M KOH)
[1:04:11] Stirrer speed set to 0
[1:05:11] Datapoint id 64 collected
[1:05:11] Stirrer speed set to 60
[1:05:16] pH 8.61 -> 8.41
[1:05:16] Using cautious pH adjust
[1:05:16] Dispensed 0.000071 mL of Acid (0.5 M HCl)
[1:05:21] Stepping pH = 8.48
[1:05:21] Dispensed 0.000024 mL of Acid (0.5 M HCl)
[1:05:26] Stepping pH = 8.35
[1:05:42] Stirrer speed set to 0
[1:06:42] Datapoint id 65 collected
[1:06:42] Charge balance equation is out by 28.7%
[1:06:42] Stirrer speed set to 60
[1:06:47] pH 7.80 -> 7.60
[1:06:47] Using cautious pH adjust
[1:06:47] Dispensed 0.000259 mL of Acid (0.5 M HCl)
[1:06:52] Stepping pH = 7.65
[1:06:52] Dispensed 0.000071 mL of Acid (0.5 M HCl)
[1:06:57] Stepping pH = 7.62
[1:06:57] Dispensed 0.000047 mL of Acid (0.5 M HCl)
[1:07:02] Stepping pH = 7.60
[1:07:17] Stirrer speed set to 0
[1:07:38] Datapoint id 66 collected
[1:07:38] Charge balance equation is out by 24.0%
[1:07:38] Stirrer speed set to 60
[1:07:43] pH 7.54 -> 7.34
[1:07:43] Using cautious pH adjust
[1:07:43] Dispensed 0.000329 mL of Acid (0.5 M HCl)
[1:07:48] Stepping pH = 7.40
[1:07:48] Dispensed 0.000141 mL of Acid (0.5 M HCl)
[1:07:53] Stepping pH = 7.36
[1:07:53] Dispensed 0.000047 mL of Acid (0.5 M HCl)
[1:07:58] Stepping pH = 7.35
[1:08:13] Stirrer speed set to 0
[1:08:39] Datapoint id 67 collected
[1:08:39] Charge balance equation is out by 23.3%
[1:08:39] Stirrer speed set to 60
[1:08:44] pH 7.32 -> 7.12
[1:08:44] Using cautious pH adjust
[1:08:44] Dispensed 0.000400 mL of Acid (0.5 M HCl)
[1:08:49] Stepping pH = 7.18
[1:08:49] Dispensed 0.000165 mL of Acid (0.5 M HCl)
[1:08:54] Stepping pH = 7.14

#### Experiment Log (continued)

[1:08:54] Dispensed 0.000071 mL of Acid (0.5 M HCl)
[1:08:59] Stepping pH = 7.12
[1:09:14] Stirrer speed set to 0
[1:09:39] Datapoint id 68 collected
[1:09:39] Charge balance equation is out by 21.8%
[1:09:39] Stirrer speed set to 60
[1:09:44] pH 7.11 -> 6.91
[1:09:44] Using cautious pH adjust
[1:09:44] Dispensed 0.000400 mL of Acid (0.5 M HCl)
[1:09:49] Stepping pH = 6.98
[1:09:49] Dispensed 0.000165 mL of Acid (0.5 M HCl)
[1:09:54] Stepping pH = 6.94
[1:09:54] Dispensed 0.000094 mL of Acid (0.5 M HCl)
[1:09:59] Stepping pH = 6.92
[1:10:15] Stirrer speed set to 0
[1:10:46] Datapoint id 69 collected
[1:10:46] Charge balance equation is out by 16.7%
[1:10:46] Stirrer speed set to 60
[1:10:51] pH 6.92 -> 6.72
[1:10:51] Using cautious pH adjust
[1:10:51] Dispensed 0.000376 mL of Acid (0.5 M HCl)
[1:10:56] Stepping pH = 6.79
[1:10:56] Dispensed 0.000165 mL of Acid (0.5 M HCl)
[1:11:02] Stepping pH = 6.74
[1:11:02] Dispensed 0.000071 mL of Acid (0.5 M HCl)
[1:11:07] Stepping pH = 6.73
[1:11:07] Dispensed 0.000071 mL of Acid (0.5 M HCl)
[1:11:12] Stepping pH = 6.71
[1:11:27] Stirrer speed set to 0
[1:11:59] Datapoint id 70 collected
[1:11:59] Charge balance equation is out by 9.2%
[1:11:59] Stirrer speed set to 60
[1:12:04] pH 6.71 -> 6.51
[1:12:04] Using charge balance adjust
[1:12:04] Dispensed 0.000611 mL of Acid (0.5 M HCl)
[1:12:24] Stirrer speed set to 0
[1:13:09] Datapoint id 71 collected
[1:13:09] Charge balance equation is out by -12.8%
[1:13:09] Stirrer speed set to 60
[1:13:14] pH 6.46 -> 6.26
[1:13:14] Using charge balance adjust
[1:13:14] Dispensed 0.000447 mL of Acid (0.5 M HCl)
[1:13:34] Stirrer speed set to 0
[1:14:15] Datapoint id 72 collected
[1:14:15] Charge balance equation is out by -34.1%
[1:14:15] Stirrer speed set to 60
[1:14:20] pH 6.25 -> 6.05
[1:14:20] Using cautious pH adjust
[1:14:20] Dispensed 0.000165 mL of Acid (0.5 M HCl)
[1:14:25] Stepping pH = 6.17
[1:14:25] Dispensed 0.000141 mL of Acid (0.5 M HCl)
[1:14:30] Stepping pH = 6.07
[1:14:30] Dispensed 0.000024 mL of Acid (0.5 M HCl)
[1:14:36] Stepping pH = 6.06
[1:14:36] Dispensed 0.000024 mL of Acid (0.5 M HCl)
[1:14:41] Stepping pH = 6.06
[1:14:41] Dispensed 0.000094 mL of Acid (0.5 M HCl)
[1:14:46] Stepping pH = 5.98
[1:15:01] Stirrer speed set to 0

| | | | |
|---|---|---|---|
| Sample name: | **M08_octanol** | Experiment start time: | **3/2/2018 8:29:22 PM** |
| Assay name: | **pH-metric high logP** | Analyst: | **Pion** |
| Assay ID: | **18C-02009** | Instrument ID: | **T312060** |
| Filename: | **C:\Sirius_T3\Mehtap\20180302_exp29_logP_T3-2\18C-02009_M08_octanol_pH-metric high logP.t3r** | | |

#### Experiment Log (continued)

[1:15:24] Datapoint id 73 collected
[1:15:24] Charge balance equation is out by -45.5%
[1:15:24] Stirrer speed set to 60
[1:15:30] pH 5.98 -> 5.78
[1:15:30] Using cautious pH adjust
[1:15:30] Dispensed 0.000094 mL of Acid (0.5 M HCl)
[1:15:35] Stepping pH = 5.91
[1:15:35] Dispensed 0.000094 mL of Acid (0.5 M HCl)
[1:15:40] Stepping pH = 5.80
[1:15:40] Dispensed 0.000024 mL of Acid (0.5 M HCl)
[1:15:45] Stepping pH = 5.78
[1:16:00] Stirrer speed set to 0
[1:16:22] Datapoint id 74 collected
[1:16:22] Charge balance equation is out by -10.2%
[1:16:22] Stirrer speed set to 60
[1:16:27] pH 5.79 -> 5.59
[1:16:27] Using charge balance adjust
[1:16:27] Dispensed 0.000141 mL of Acid (0.5 M HCl)
[1:16:47] Stirrer speed set to 0
[1:17:11] Datapoint id 75 collected
[1:17:11] Charge balance equation is out by -22.3%
[1:17:11] Stirrer speed set to 60
[1:17:16] pH 5.58 -> 5.38
[1:17:16] Using cautious pH adjust
[1:17:16] Dispensed 0.000047 mL of Acid (0.5 M HCl)
[1:17:21] Stepping pH = 5.54
[1:17:21] Dispensed 0.000071 mL of Acid (0.5 M HCl)
[1:17:26] Stepping pH = 5.39
[1:17:42] Stirrer speed set to 0
[1:18:00] Datapoint id 76 collected
[1:18:00] Charge balance equation is out by -38.4%
[1:18:00] Stirrer speed set to 60
[1:18:05] pH 5.37 -> 5.17
[1:18:05] Using cautious pH adjust
[1:18:05] Dispensed 0.000024 mL of Acid (0.5 M HCl)
[1:18:10] Stepping pH = 5.38
[1:18:10] Dispensed 0.000165 mL of Acid (0.5 M HCl)
[1:18:15] Stepping pH = 4.87
[1:18:30] Stirrer speed set to 0
[1:18:53] Datapoint id 77 collected
[1:18:53] Charge balance equation is out by -202.4%
[1:18:53] Stirrer speed set to 60
[1:18:58] pH 4.81 -> 4.61
[1:18:58] Using cautious pH adjust
[1:18:58] Dispensed 0.000024 mL of Acid (0.5 M HCl)
[1:19:03] Stepping pH = 4.80
[1:19:03] Dispensed 0.000071 mL of Acid (0.5 M HCl)
[1:19:08] Stepping pH = 4.64
[1:19:08] Dispensed 0.000024 mL of Acid (0.5 M HCl)
[1:19:13] Stepping pH = 4.58
[1:19:28] Stirrer speed set to 0
[1:19:40] Datapoint id 78 collected
[1:19:40] Charge balance equation is out by -130.6%
[1:19:40] Stirrer speed set to 60
[1:19:45] pH 4.59 -> 4.39
[1:19:45] Using cautious pH adjust
[1:19:45] Dispensed 0.000047 mL of Acid (0.5 M HCl)
[1:19:50] Stepping pH = 4.52
[1:19:50] Dispensed 0.000047 mL of Acid (0.5 M HCl)

| | | | |
|---|---|---|---|
| Sample name: | **M08_octanol** | Experiment start time: | **3/2/2018 8:29:22 PM** |
| Assay name: | **pH-metric high logP** | Analyst: | **Pion** |
| Assay ID: | **18C-02009** | Instrument ID: | **T312060** |
| Filename: | **C:\Sirius_T3\Mehtap\20180302_exp29_logP_T3-2\18C-02009_M08_octanol_pH-metric high logP.t3r** | | |

#### Experiment Log (continued)

[1:19:55] Stepping pH = 4.40
[1:20:10] Stirrer speed set to 0
[1:20:32] Datapoint id 79 collected
[1:20:32] Charge balance equation is out by -2.3%
[1:20:32] Stirrer speed set to 60
[1:20:37] pH 4.39 -> 4.19
[1:20:37] Using charge balance adjust
[1:20:37] Dispensed 0.000118 mL of Acid (0.5 M HCl)
[1:20:57] Stirrer speed set to 0
[1:21:08] Datapoint id 80 collected
[1:21:08] Charge balance equation is out by -19.7%
[1:21:08] Stirrer speed set to 60
[1:21:13] pH 4.22 -> 4.02
[1:21:13] Using cautious pH adjust
[1:21:13] Dispensed 0.000071 mL of Acid (0.5 M HCl)
[1:21:18] Stepping pH = 4.16
[1:21:18] Dispensed 0.000094 mL of Acid (0.5 M HCl)
[1:21:23] Stepping pH = 4.05
[1:21:23] Dispensed 0.000024 mL of Acid (0.5 M HCl)
[1:21:28] Stepping pH = 4.03
[1:21:28] Dispensed 0.000024 mL of Acid (0.5 M HCl)
[1:21:33] Stepping pH = 4.03
[1:21:34] Dispensed 0.000047 mL of Acid (0.5 M HCl)
[1:21:39] Stepping pH = 3.99
[1:21:54] Stirrer speed set to 0
[1:22:04] Datapoint id 81 collected
[1:22:04] Charge balance equation is out by -78.1%
[1:22:04] Stirrer speed set to 60
[1:22:09] pH 3.99 -> 3.79
[1:22:09] Using cautious pH adjust
[1:22:09] Dispensed 0.000141 mL of Acid (0.5 M HCl)
[1:22:14] Stepping pH = 3.88
[1:22:14] Dispensed 0.000094 mL of Acid (0.5 M HCl)
[1:22:19] Stepping pH = 3.82
[1:22:19] Dispensed 0.000047 mL of Acid (0.5 M HCl)
[1:22:24] Stepping pH = 3.80
[1:22:39] Stirrer speed set to 0
[1:22:50] Datapoint id 82 collected
[1:22:50] Charge balance equation is out by 0.9%
[1:22:50] Stirrer speed set to 60
[1:22:55] pH 3.80 -> 3.60
[1:22:55] Using charge balance adjust
[1:22:55] Dispensed 0.000423 mL of Acid (0.5 M HCl)
[1:23:15] Stirrer speed set to 0
[1:23:25] Datapoint id 83 collected
[1:23:25] Charge balance equation is out by 2.4%
[1:23:25] Stirrer speed set to 60
[1:23:31] pH 3.59 -> 3.39
[1:23:31] Using charge balance adjust
[1:23:31] Dispensed 0.000659 mL of Acid (0.5 M HCl)
[1:23:51] Stirrer speed set to 0
[1:24:09] Datapoint id 84 collected
[1:24:09] Charge balance equation is out by -5.5%
[1:24:09] Stirrer speed set to 60
[1:24:14] pH 3.41 -> 3.21
[1:24:14] Using charge balance adjust
[1:24:14] Dispensed 0.001035 mL of Acid (0.5 M HCl)
[1:24:34] Stirrer speed set to 0
[1:25:00] Datapoint id 85 collected

| | | | |
|---|---|---|---|
| Sample name: | **M08_octanol** | Experiment start time: | **3/2/2018 8:29:22 PM** |
| Assay name: | **pH-metric high logP** | Analyst: | **Pion** |
| Assay ID: | **18C-02009** | Instrument ID: | **T312060** |
| Filename: | **C:\Sirius_T3\Mehtap\20180302_exp29_logP_T3-2\18C-02009_M08_octanol_pH-metric high logP.t3r** | | |

#### Experiment Log (continued)

[1:25:00] Charge balance equation is out by -3.2%
[1:25:00] Stirrer speed set to 60
[1:25:05] pH 3.22 -> 3.02
[1:25:05] Using charge balance adjust
[1:25:05] Dispensed 0.001599 mL of Acid (0.5 M HCl)
[1:25:25] Stirrer speed set to 0
[1:25:36] Datapoint id 86 collected
[1:25:36] Charge balance equation is out by -5.0%
[1:25:36] Stirrer speed set to 60
[1:25:41] pH 3.03 -> 2.83
[1:25:41] Using charge balance adjust
[1:25:41] Dispensed 0.002422 mL of Acid (0.5 M HCl)
[1:26:01] Stirrer speed set to 0
[1:26:26] Datapoint id 87 collected
[1:26:26] Charge balance equation is out by -7.8%
[1:26:26] Stirrer speed set to 60
[1:26:32] pH 2.85 -> 2.65
[1:26:32] Using charge balance adjust
[1:26:32] Dispensed 0.003716 mL of Acid (0.5 M HCl)
[1:26:52] Stirrer speed set to 0
[1:27:02] Datapoint id 88 collected
[1:27:02] Charge balance equation is out by -9.4%
[1:27:02] Stirrer speed set to 60
[1:27:07] pH 2.68 -> 2.48
[1:27:07] Using charge balance adjust
[1:27:07] Dispensed 0.005621 mL of Acid (0.5 M HCl)
[1:27:27] Stirrer speed set to 0
[1:27:52] Datapoint id 89 collected
[1:27:52] Charge balance equation is out by 0.3%
[1:27:52] Stirrer speed set to 60
[1:27:57] pH 2.48 -> 2.28
[1:27:57] Using charge balance adjust
[1:27:58] Dispensed 0.008937 mL of Acid (0.5 M HCl)
[1:28:18] Stirrer speed set to 0
[1:28:31] Datapoint id 90 collected
[1:28:31] Charge balance equation is out by -2.8%
[1:28:31] Stirrer speed set to 60
[1:28:36] pH 2.29 -> 2.09
[1:28:36] Using charge balance adjust
[1:28:37] Dispensed 0.014087 mL of Acid (0.5 M HCl)
[1:28:57] Stirrer speed set to 0
[1:29:18] Datapoint id 91 collected
[1:29:18] Charge balance equation is out by -3.5%
[1:29:18] Stirrer speed set to 60
[1:29:23] pH 2.10 -> 1.95
[1:29:23] Using charge balance adjust
[1:29:23] Dispensed 0.016157 mL of Acid (0.5 M HCl)
[1:29:43] Stirrer speed set to 0
[1:30:03] Datapoint id 92 collected
[1:30:03] Charge balance equation is out by -25.8%
[1:30:03] Argon flow rate set to 0
[1:30:07] Titrator arm moved over Titration position
