## Supplementary material for "Octanol-water partition coefficient measurements for the SAMPL6 Blind Prediction Challenge": SM09_18C-02010_M09_octanol_pH-metric high logP_report.pdf

| | | | |
|---|---|---|---|
| Sample name: | **M09_octanol** | Experiment start time: | **3/2/2018 10:00:20 PM** |
| Assay name: | **pH-metric high logP** | Analyst: | **Pion** |
| Assay ID: | **18C-02010** | Instrument ID: | **T312060** |
| Filename: | **C:\Sirius_T3\Mehtap\20180302_exp29_logP_T3-2\18C-02010_M09_octanol_pH-metric high logP.t3r** | | |

#### pH-metric Result

| | | |
|---|---|---|
| logP (XH +) | -9.50 | ±0.85 (n=47) |
| logP (neutral X) | 2.90 | ±0.02 (n=47) |

#### 18C-02010 Points 1 to 22

| | |
|---|---|
| M09_octanol concentration factor | 1.064 |
| Carbonate | 0.0587 mM |
| Acidity error | -0.83134 mM |

#### 18C-02010 Points 23 to 44

| | |
|---|---|
| M09_octanol concentration factor | 0.800 |
| Carbonate | 0.0538 mM |
| Acidity error | -0.43636 mM |

#### 18C-02010 Points 45 to 64

| | |
|---|---|
| M09_octanol concentration factor | 1.103 |
| Carbonate | 0.1700 mM |
| Acidity error | -0.97715 mM |

#### Warnings and errors

| | |
|---|---|
| Errors | None |
| Warnings | One or more logP values out of range |

#### Sample logD and percent species

| pH | M09_octanol logD | M09_octanol M09_octanolH | M09_octanol M09_octanol | M09_octanol M09_octanolH* | M09_octanol M09_octanol* | Comment |
|---|---|---|---|---|---|---|
| 1.000 | -1.47 | 96.71 % | 0.00 % | 0.00 % | 3.29 % | |
| 1.200 | -1.27 | 94.88 % | 0.01 % | 0.00 % | 5.11 % | Stomach pH |
| 2.000 | -0.47 | 74.60 % | 0.03 % | 0.00 % | 25.37 % | |
| 3.000 | 0.53 | 22.70 % | 0.10 % | 0.00 % | 77.20 % | |
| 4.000 | 1.51 | 2.85 % | 0.12 % | 0.00 % | 97.03 % | |
| 5.000 | 2.38 | 0.29 % | 0.12 % | 0.00 % | 99.58 % | |
| 6.000 | 2.81 | 0.03 % | 0.13 % | 0.00 % | 99.85 % | |
| 6.500 | 2.87 | 0.01 % | 0.13 % | 0.00 % | 99.87 % | |
| 7.000 | 2.89 | 0.00 % | 0.13 % | 0.00 % | 99.87 % | |
| 7.400 | 2.90 | 0.00 % | 0.13 % | 0.00 % | 99.87 % | Blood pH |
| 8.000 | 2.90 | 0.00 % | 0.13 % | 0.00 % | 99.87 % | |
| 9.000 | 2.90 | 0.00 % | 0.13 % | 0.00 % | 99.87 % | |
| 10.000 | 2.90 | 0.00 % | 0.13 % | 0.00 % | 99.87 % | |
| 11.000 | 2.90 | 0.00 % | 0.13 % | 0.00 % | 99.87 % | |
| 12.000 | 2.90 | 0.00 % | 0.13 % | 0.00 % | 99.87 % | |

Sample name: **M09_octanol**
Assay name: **pH-metric high logP**
Assay ID: **18C-02010**
Filename: **C:\Sirius_T3\Mehtap\20180302_exp29_logP_T3-2\18C-02010_M09_octanol_pH-metric high logP.t3r**

Experiment start time: **3/2/2018 10:00:20 PM**
Analyst: **Pion**
Instrument ID: **T312060**

#### Graphs

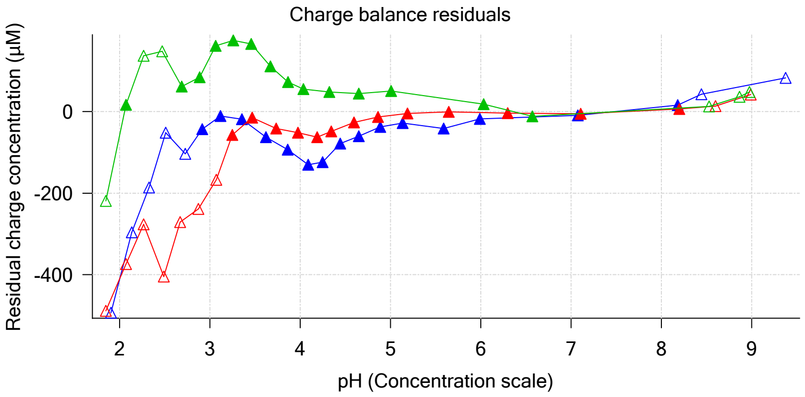


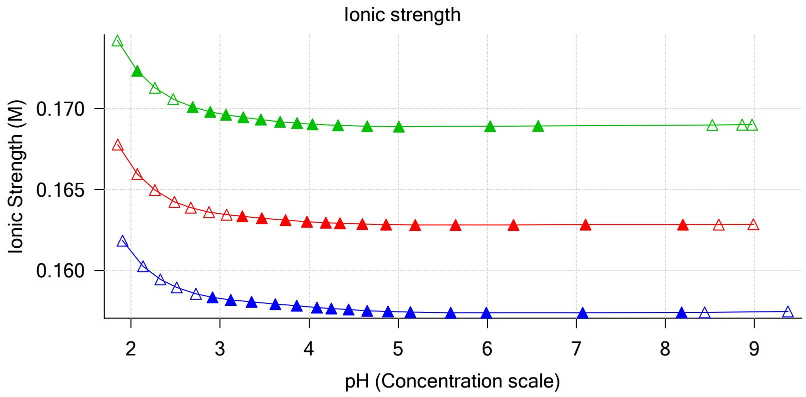


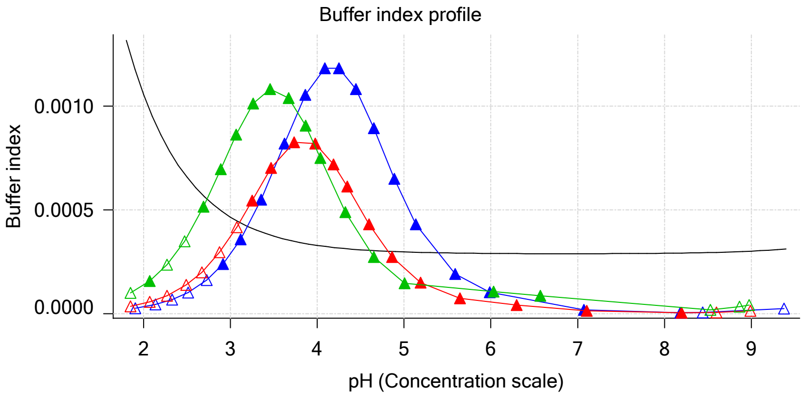


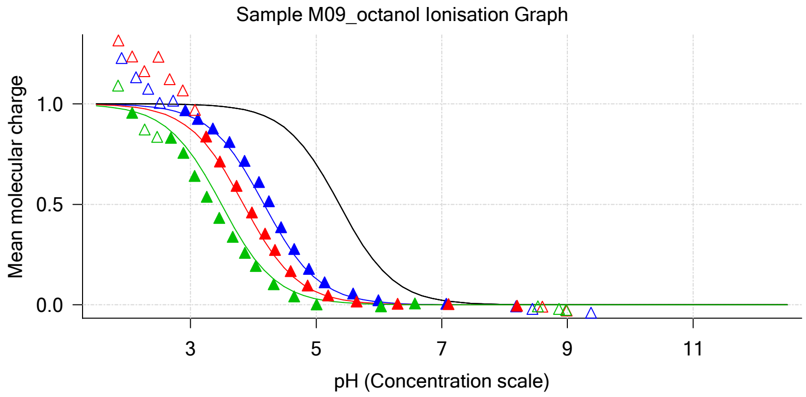


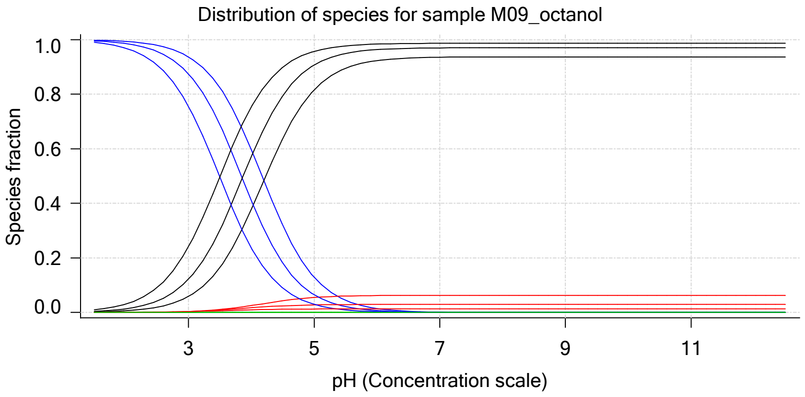


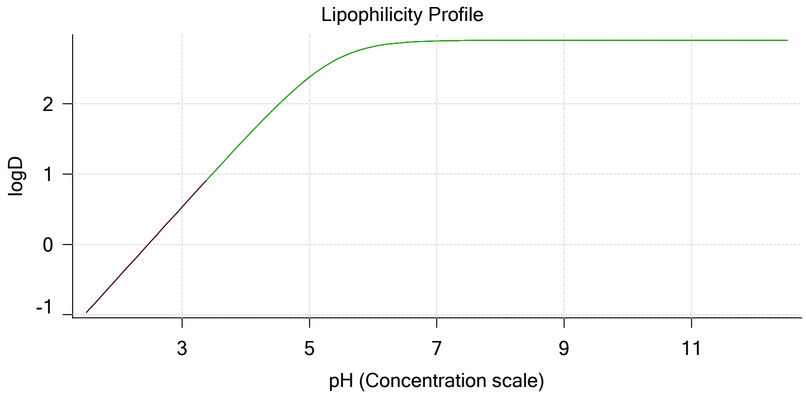


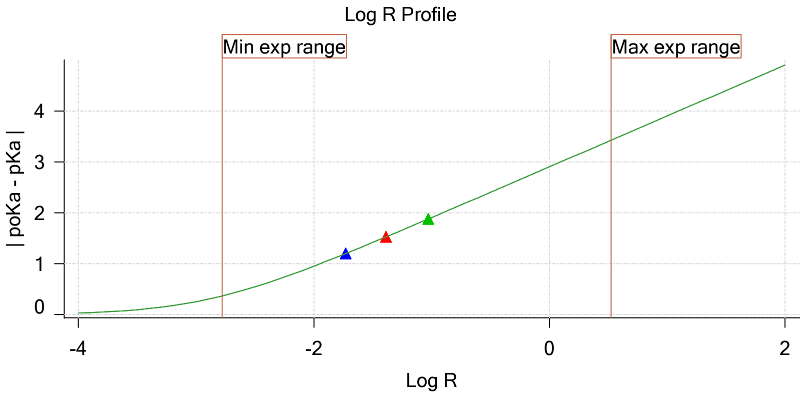


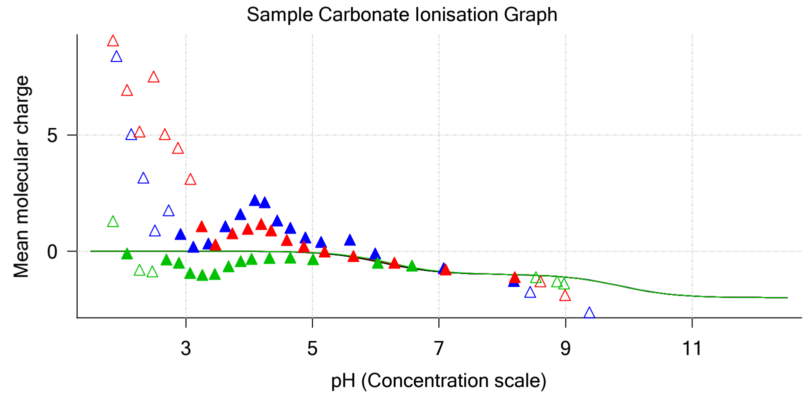

Sample name: **M09_octanol** Experiment start time: **3/2/2018 10:00:20 PM**
Assay name: **pH-metric high logP** Analyst: **Pion**
Assay ID: **18C-02010** Instrument ID: **T312060**
Filename: **C:\Sirius_T3\Mehtap\20180302_exp29_logP_T3-2\18C-02010_M09_octanol_pH-metric high logP.t3r**

#### Graphs (continued)

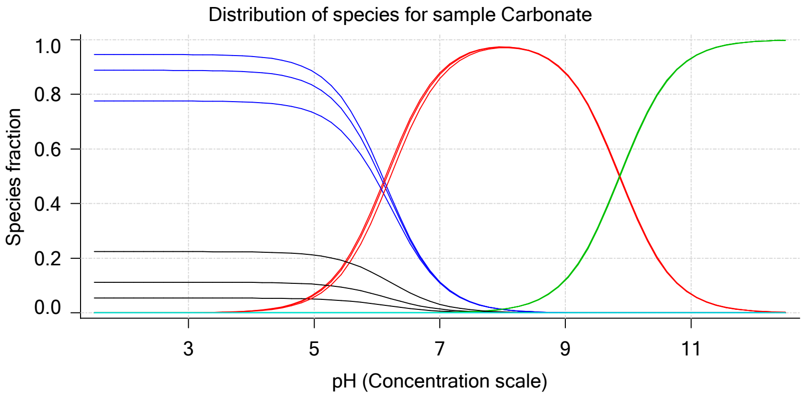

Sample name: **M09_octanol**
Assay name: **pH-metric high logP**
Assay ID: **18C-02010**
Filename: **C:\Sirius_T3\Mehtap\20180302_exp29_logP_T3-2\18C-02010_M09_octanol_pH-metric high logP.t3r**
Experiment start time: **3/2/2018 10:00:20 PM**
Analyst: **Pion**
Instrument ID: **T312060**

### pH-metric high logP Titration 1 of 3 18C-02010 Points 1 to 22

#### Overall results

| | |
|---|---|
| RMSD | 0.413 |
| Average ionic strength | 0.158 M |
| Average temperature | 24.9°C |
| Partition ratio | 0.0186 : 1 |
| Analyte concentration range | 1902.7 µM to 1959.6 µM |
| Total points considered | 15 of 22 |

#### Warnings and errors

| | |
|---|---|
| Errors | None |
| Warnings | None |

#### Four-Plus parameters

| | | | |
|---|---|---|---|
| Alpha | 0.111 | 3/2/2018 10:00:20 PM | C:\Sirius_T3\HCl18C02.t3r |
| S | 0.9988 | 3/2/2018 10:00:20 PM | C:\Sirius_T3\HCl18C02.t3r |
| jH | 1.0 | 3/2/2018 10:00:20 PM | C:\Sirius_T3\HCl18C02.t3r |
| jOH | -0.8 | 3/2/2018 10:00:20 PM | C:\Sirius_T3\HCl18C02.t3r |

#### Titrants

| | | | |
|---|---|---|---|
| 0.50 M HCl | 0.999058 | 3/2/2018 10:00:20 PM | C:\Sirius_T3\HCl18C02.t3r |
| 0.50 M KOH | 0.999845 | 3/2/2018 10:00:20 PM | C:\Sirius_T3\KOH18B27.t3r |

#### Sample

| | |
|---|---|
| M09_octanol concentration factor | 1.064 |
| M09_octanol stoichiometry | 1.000 |
| Chloride stoichiometry | 1.000 |
| Base pKa 1 | 5.37 |
| logP (XH +) | 0.48 |
| logP (neutral X) | 2.84 |

#### Sample graphs

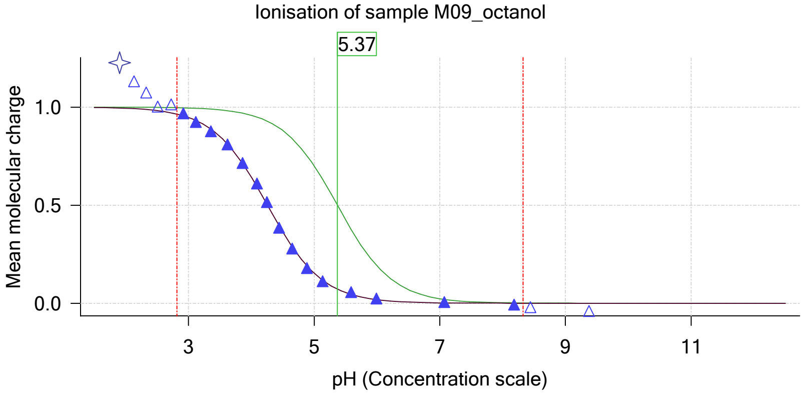


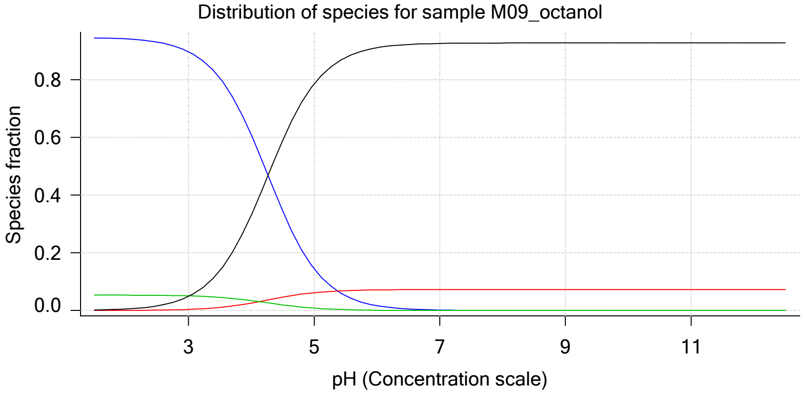

| | | | |
|---|---|---|---|
| Sample name: | **M09_octanol** | Experiment start time: | **3/2/2018 10:00:20 PM** |
| Assay name: | **pH-metric high logP** | Analyst: | **Pion** |
| Assay ID: | **18C-02010** | Instrument ID: | **T312060** |
| Filename: | **C:\Sirius_T3\Mehtap\20180302_exp29_logP_T3-2\18C-02010_M09_octanol_pH-metric high logP.t3r** | | |

#### Sample graphs (continued)

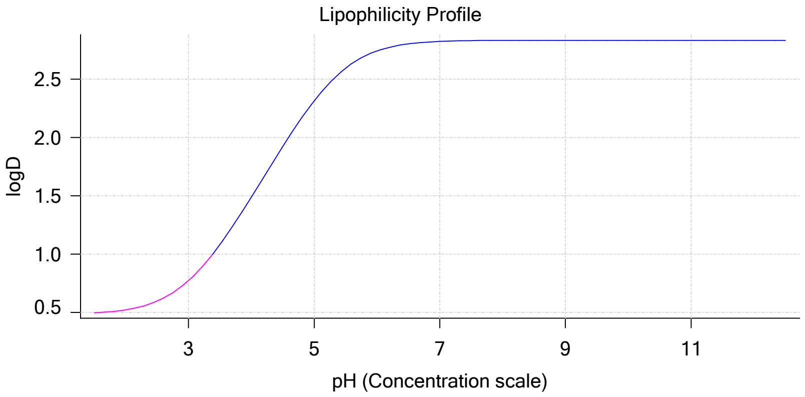


#### Sample logD and percent species

| pH | M09_octanol logD | M09_octanol M09_octanolH | M09_octanol M09_octanol | M09_octanol M09_octanolH* | M09_octanol M09_octanol* | Comment |
|---|---|---|---|---|---|---|
| 1.000 | 0.49 | 94.58 % | 0.00 % | 5.36 % | 0.05 % | |
| 1.200 | 0.49 | 94.55 % | 0.01 % | 5.36 % | 0.08 % | Stomach pH |
| 2.000 | 0.52 | 94.11 % | 0.04 % | 5.34 % | 0.51 % | |
| 3.000 | 0.77 | 89.67 % | 0.38 % | 5.08 % | 4.86 % | |
| 4.000 | 1.49 | 60.91 % | 2.60 % | 3.45 % | 33.04 % | |
| 5.000 | 2.32 | 14.48 % | 6.17 % | 0.82 % | 78.53 % | |
| 6.000 | 2.74 | 1.68 % | 7.16 % | 0.10 % | 91.07 % | |
| 6.500 | 2.80 | 0.54 % | 7.25 % | 0.03 % | 92.18 % | |
| 7.000 | 2.83 | 0.17 % | 7.28 % | 0.01 % | 92.54 % | |
| 7.400 | 2.83 | 0.07 % | 7.28 % | 0.00 % | 92.64 % | Blood pH |
| 8.000 | 2.83 | 0.02 % | 7.29 % | 0.00 % | 92.69 % | |
| 9.000 | 2.83 | 0.00 % | 7.29 % | 0.00 % | 92.71 % | |
| 10.000 | 2.84 | 0.00 % | 7.29 % | 0.00 % | 92.71 % | |
| 11.000 | 2.84 | 0.00 % | 7.29 % | 0.00 % | 92.71 % | |
| 12.000 | 2.84 | 0.00 % | 7.29 % | 0.00 % | 92.71 % | |

#### Carbonate and acidity

| | |
|---|---|
| Carbonate | 0.059 mM |
| Acidity error | -0.831 mM |

#### Other graphs

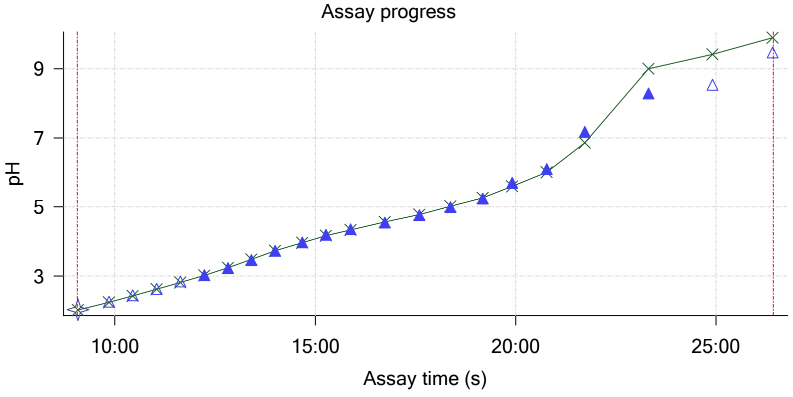


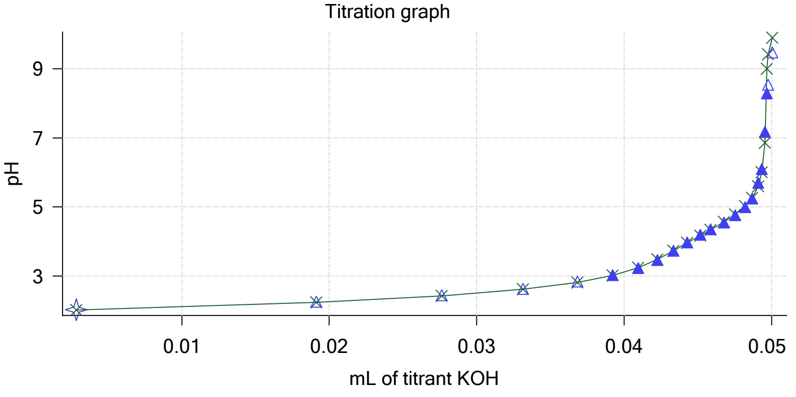

| | | | |
|---|---|---|---|
| Sample name: | **M09_octanol** | Experiment start time: | **3/2/2018 10:00:20 PM** |
| Assay name: | **pH-metric high logP** | Analyst: | **Pion** |
| Assay ID: | **18C-02010** | Instrument ID: | **T312060** |
| Filename: | **C:\Sirius_T3\Mehtap\20180302_exp29_logP_T3-2\18C-02010_M09_octanol_pH-metric high logP.t3r** | | |

#### Other graphs (continued)

Charge balance residuals

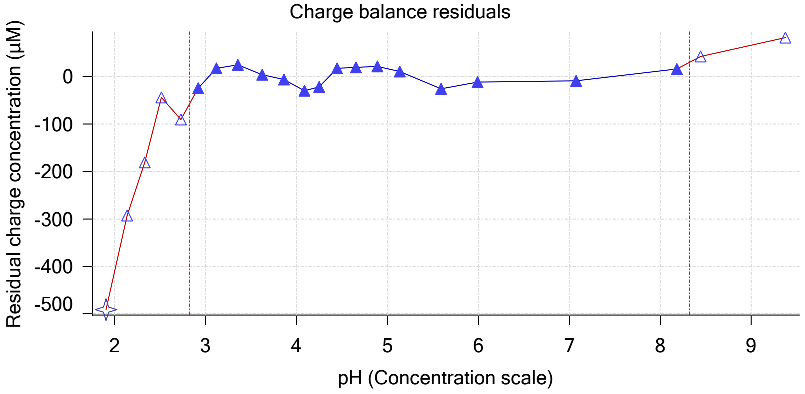


Buffer index profile

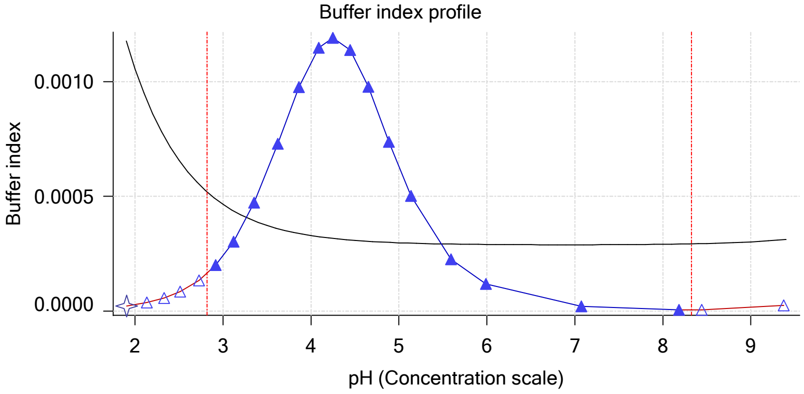


Ionic strength

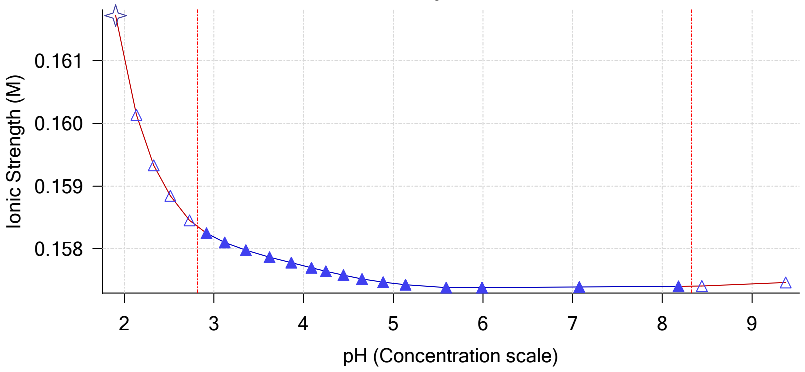


Log R Profile

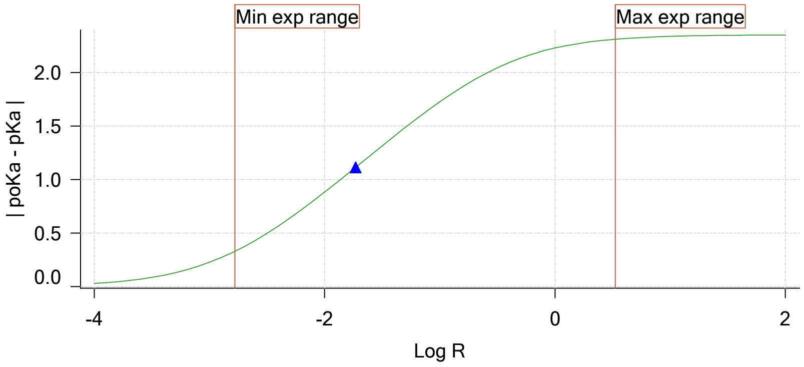


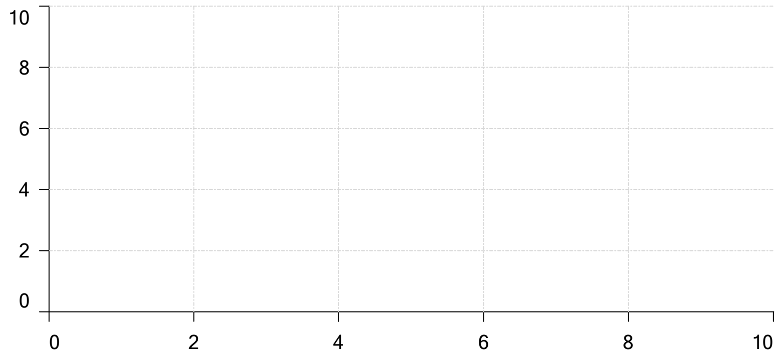

Ionisation of sample Carbonate

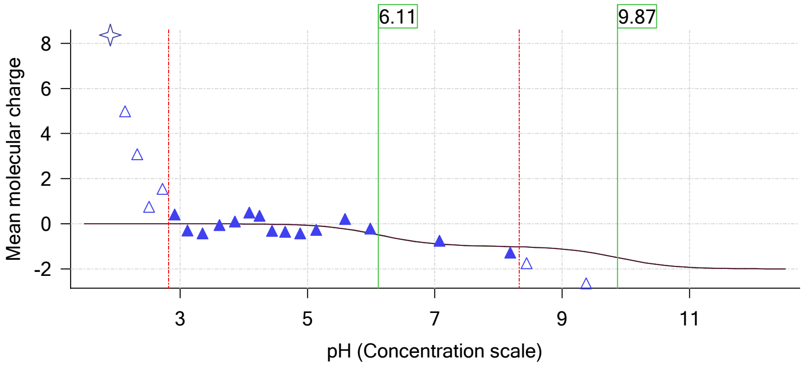

| | | | |
|---|---|---|---|
| Sample name: | **M09_octanol** | Experiment start time: | **3/2/2018 10:00:20 PM** |
| Assay name: | **pH-metric high logP** | Analyst: | **Pion** |
| Assay ID: | **18C-02010** | Instrument ID: | **T312060** |
| Filename: | **C:\Sirius_T3\Mehtap\20180302_exp29_logP_T3-2\18C-02010_M09_octanol_pH-metric high logP.t3r** | | |

### pH-metric high logP Titration 2 of 3 18C-02010 Points 23 to 44

#### Overall results

| | |
|---|---|
| RMSD | 0.195 |
| Average ionic strength | 0.163 M |
| Average temperature | 25.0°C |
| Partition ratio | 0.0409 : 1 |
| Analyte concentration range | 1744.4 μM to 1798.6 μM |
| Total points considered | 13 of 22 |

#### Warnings and errors

| | |
|---|---|
| Errors | None |
| Warnings | None |

#### Four-Plus parameters

| | | | |
|---|---|---|---|
| Alpha | 0.111 | 3/2/2018 10:00:20 PM | C:\Sirius_T3\HCl18C02.t3r |
| S | 0.9988 | 3/2/2018 10:00:20 PM | C:\Sirius_T3\HCl18C02.t3r |
| jH | 1.0 | 3/2/2018 10:00:20 PM | C:\Sirius_T3\HCl18C02.t3r |
| jOH | -0.8 | 3/2/2018 10:00:20 PM | C:\Sirius_T3\HCl18C02.t3r |

#### Titrants

| | | | |
|---|---|---|---|
| 0.50 M HCl | 0.999058 | 3/2/2018 10:00:20 PM | C:\Sirius_T3\HCl18C02.t3r |
| 0.50 M KOH | 0.999845 | 3/2/2018 10:00:20 PM | C:\Sirius_T3\KOH18B27.t3r |

#### Sample

| | |
|---|---|
| M09_octanol concentration factor | 0.800 |
| M09_octanol stoichiometry | 1.000 |
| Chloride stoichiometry | 1.000 |
| Base pKa 1 | 5.37 |
| logP (XH +) | 0.84 |
| logP (neutral X) | 2.95 |

#### Sample graphs

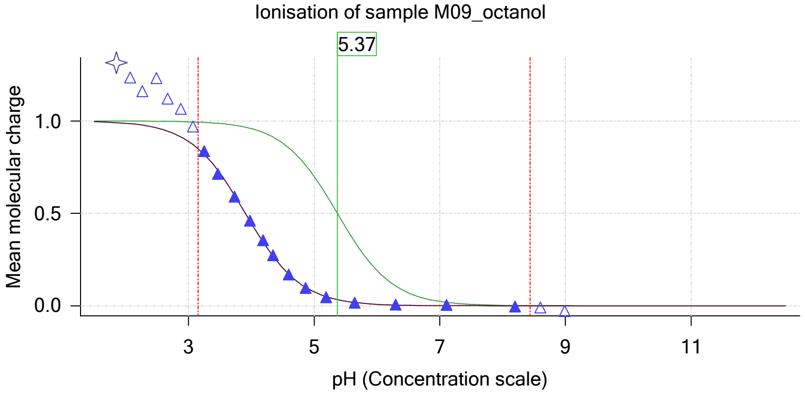


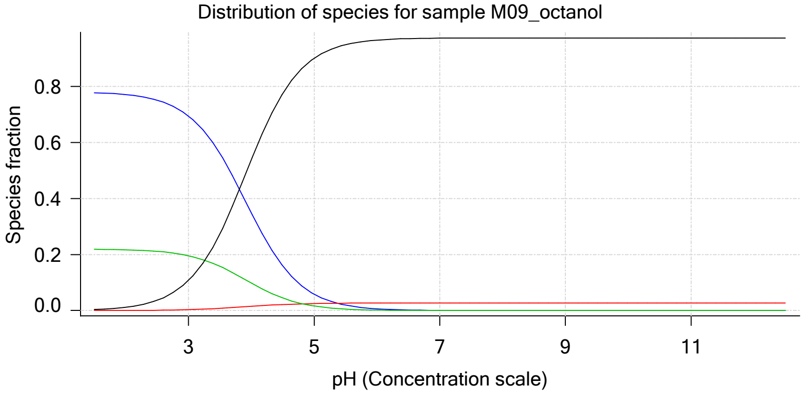

Sample name: **M09_octanol**
Assay name: **pH-metric high logP**
Assay ID: **18C-02010**
Filename: **C:\Sirius_T3\Mehtap\20180302_exp29_logP_T3-2\18C-02010_M09_octanol_pH-metric high logP.t3r**
Experiment start time: **3/2/2018 10:00:20 PM**
Analyst: **Pion**
Instrument ID: **T312060**

#### Sample graphs (continued)

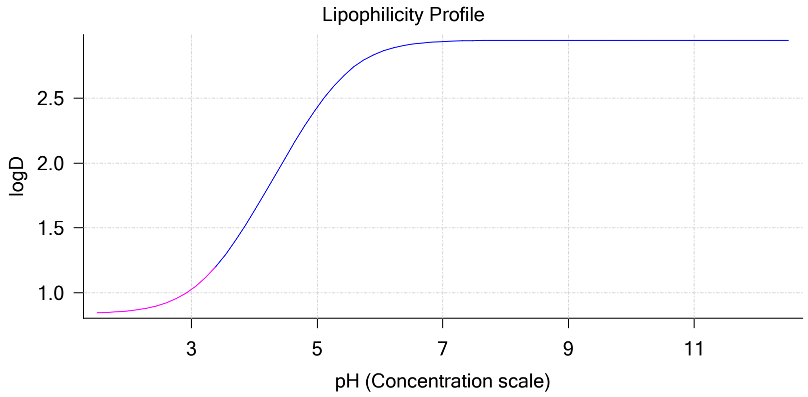


#### Sample logD and percent species

| pH | M09_octanol logD | M09_octanol M09_octanolH | M09_octanol M09_octanol | M09_octanol M09_octanolH* | M09_octanol M09_octanol* | Comment |
|---|---|---|---|---|---|---|
| 1.000 | 0.84 | 77.94 % | 0.00 % | 21.93 % | 0.12 % | |
| 1.200 | 0.84 | 77.89 % | 0.01 % | 21.92 % | 0.19 % | Stomach pH |
| 2.000 | 0.86 | 77.09 % | 0.03 % | 21.69 % | 1.19 % | |
| 3.000 | 1.03 | 69.45 % | 0.30 % | 19.54 % | 10.72 % | |
| 4.000 | 1.63 | 34.88 % | 1.49 % | 9.81 % | 53.82 % | |
| 5.000 | 2.43 | 5.83 % | 2.49 % | 1.64 % | 90.04 % | |
| 6.000 | 2.86 | 0.63 % | 2.67 % | 0.18 % | 96.53 % | |
| 6.500 | 2.92 | 0.20 % | 2.68 % | 0.06 % | 97.06 % | |
| 7.000 | 2.94 | 0.06 % | 2.69 % | 0.02 % | 97.23 % | |
| 7.400 | 2.94 | 0.03 % | 2.69 % | 0.01 % | 97.28 % | Blood pH |
| 8.000 | 2.95 | 0.01 % | 2.69 % | 0.00 % | 97.30 % | |
| 9.000 | 2.95 | 0.00 % | 2.69 % | 0.00 % | 97.31 % | |
| 10.000 | 2.95 | 0.00 % | 2.69 % | 0.00 % | 97.31 % | |
| 11.000 | 2.95 | 0.00 % | 2.69 % | 0.00 % | 97.31 % | |
| 12.000 | 2.95 | 0.00 % | 2.69 % | 0.00 % | 97.31 % | |

#### Carbonate and acidity

| | |
|---|---|
| Carbonate | 0.054 mM |
| Acidity error | -0.436 mM |

#### Other graphs

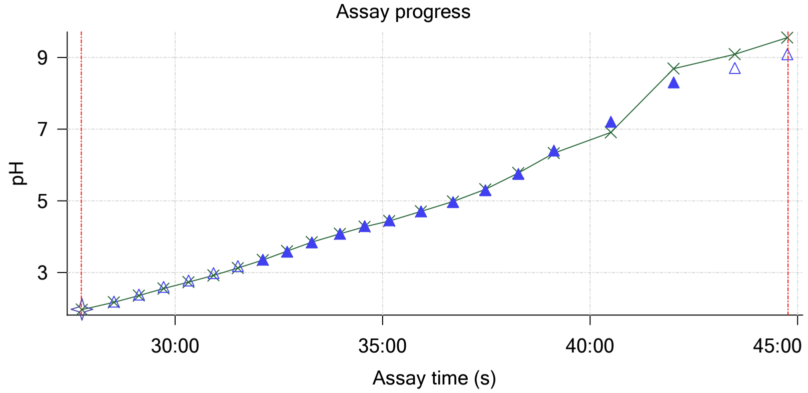


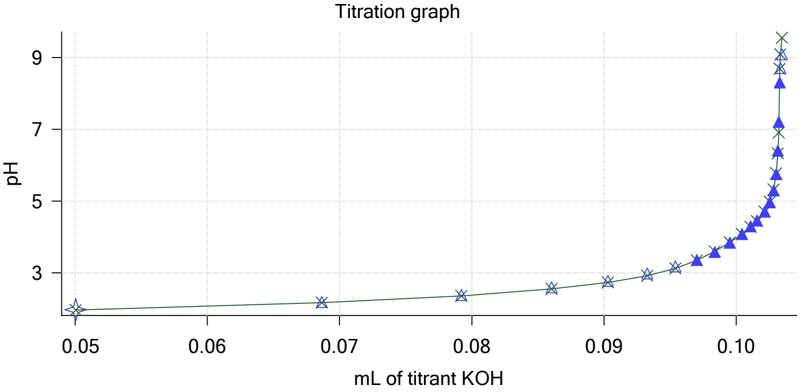

| | | | |
|---|---|---|---|
| Sample name: | **M09_octanol** | Experiment start time: | **3/2/2018 10:00:20 PM** |
| Assay name: | **pH-metric high logP** | Analyst: | **Pion** |
| Assay ID: | **18C-02010** | Instrument ID: | **T312060** |
| Filename: | **C:\Sirius_T3\Mehtap\20180302_exp29_logP_T3-2\18C-02010_M09_octanol_pH-metric high logP.t3r** | | |

#### Other graphs (continued)

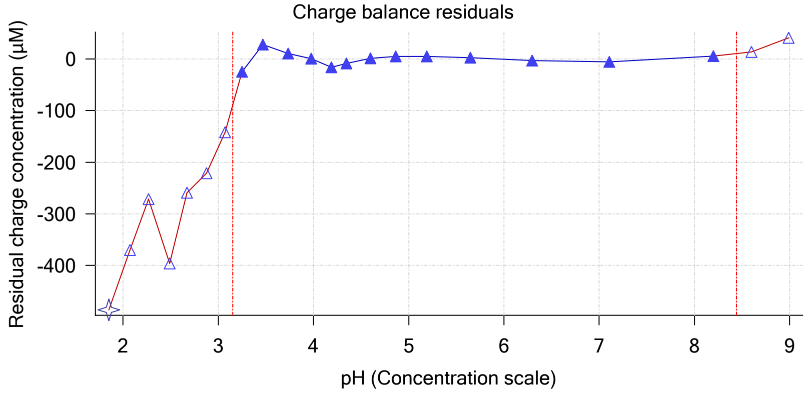


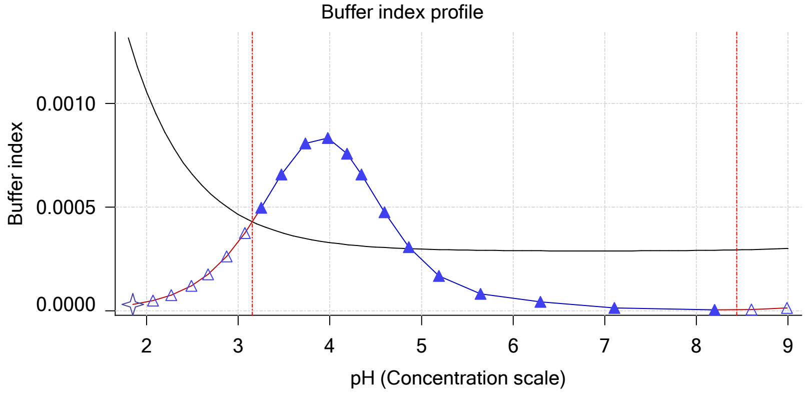


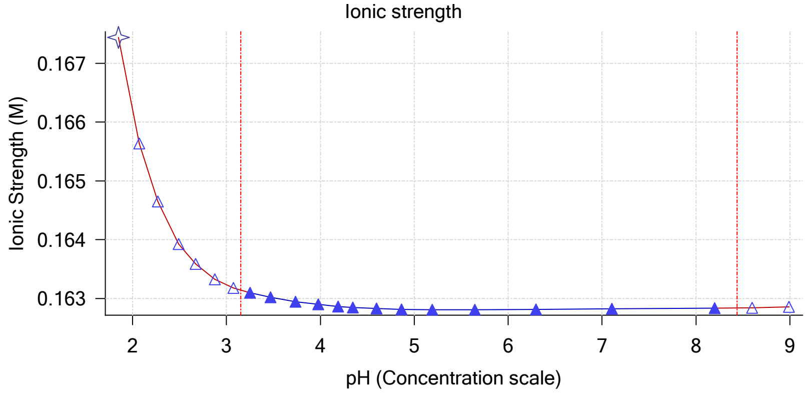


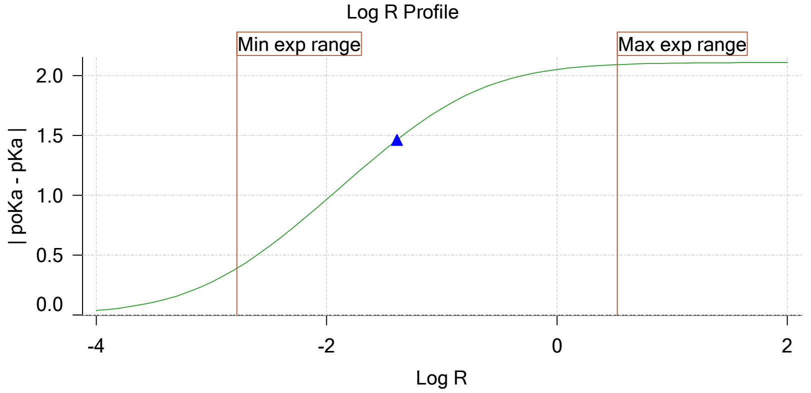


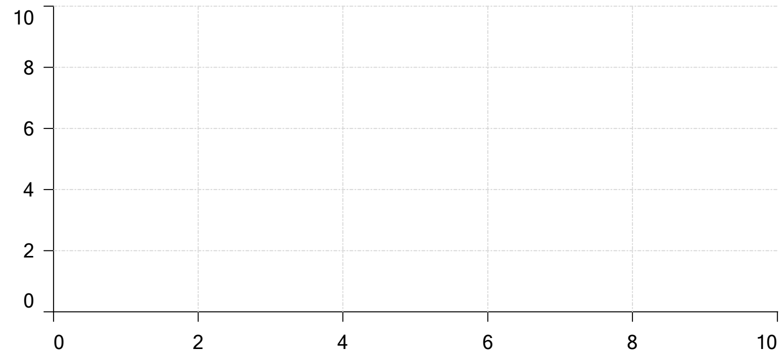

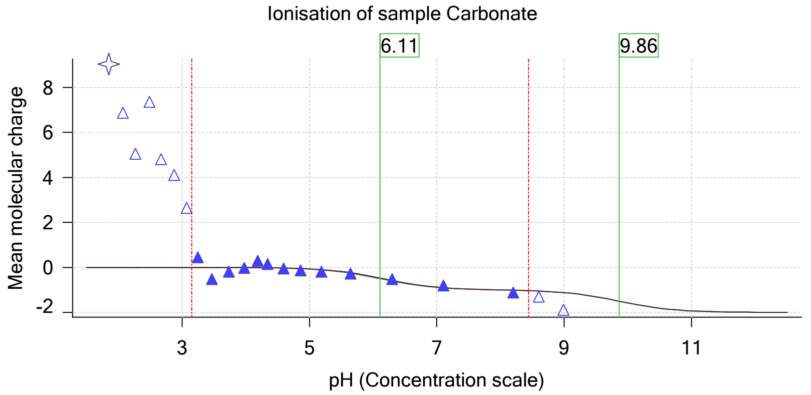

| | | | |
|---|---|---|---|
| Sample name: | **M09_octanol** | Experiment start time: | **3/2/2018 10:00:20 PM** |
| Assay name: | **pH-metric high logP** | Analyst: | **Pion** |
| Assay ID: | **18C-02010** | Instrument ID: | **T312060** |
| Filename: | **C:\Sirius_T3\Mehtap\20180302_exp29_logP_T3-2\18C-02010_M09_octanol_pH-metric high logP.t3r** | | |

### pH-metric high logP Titration 3 of 3 18C-02010 Points 45 to 64

#### Overall results

| | |
|---|---|
| RMSD | 0.092 |
| Average ionic strength | 0.169 M |
| Average temperature | 25.0°C |
| Partition ratio | 0.0935 : 1 |
| Analyte concentration range | 1555.0 μM to 1601.8 μM |
| Total points considered | 14 of 20 |

#### Warnings and errors

| | |
|---|---|
| Errors | None |
| Warnings | None |

#### Four-Plus parameters

| | | | |
|---|---|---|---|
| Alpha | 0.111 | 3/2/2018 10:00:20 PM | C:\Sirius_T3\HCl18C02.t3r |
| S | 0.9988 | 3/2/2018 10:00:20 PM | C:\Sirius_T3\HCl18C02.t3r |
| jH | 1.0 | 3/2/2018 10:00:20 PM | C:\Sirius_T3\HCl18C02.t3r |
| jOH | -0.8 | 3/2/2018 10:00:20 PM | C:\Sirius_T3\HCl18C02.t3r |

#### Titrants

| | | | |
|---|---|---|---|
| 0.50 M HCl | 0.999058 | 3/2/2018 10:00:20 PM | C:\Sirius_T3\HCl18C02.t3r |
| 0.50 M KOH | 0.999845 | 3/2/2018 10:00:20 PM | C:\Sirius_T3\KOH18B27.t3r |

#### Sample

| | |
|---|---|
| M09_octanol concentration factor | 1.103 |
| M09_octanol stoichiometry | 1.000 |
| Chloride stoichiometry | 1.000 |
| Base pKa 1 | 5.37 |
| logP (XH +) | 0.88 |
| logP (neutral X) | 3.27 |

#### Sample graphs

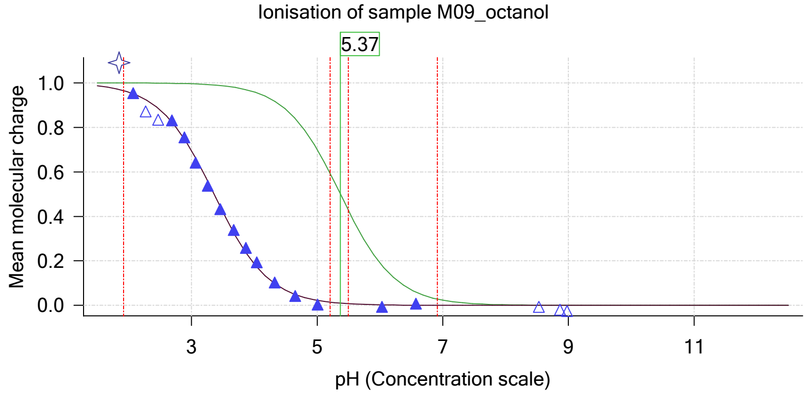


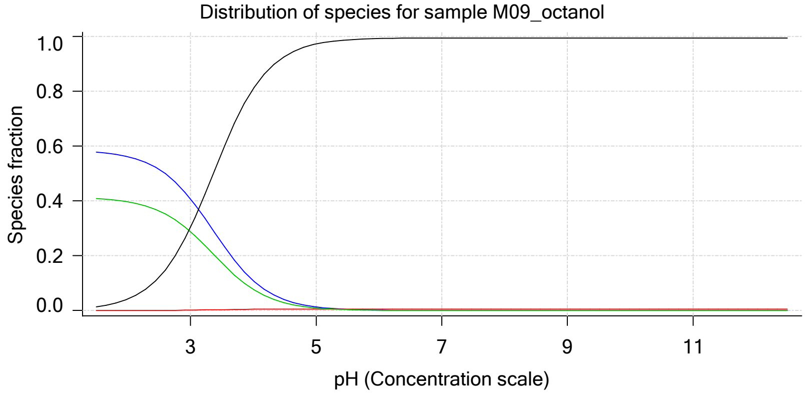

Sample name: **M09_octanol** Experiment start time: **3/2/2018 10:00:20 PM**
Assay name: **pH-metric high logP** Analyst: **Pion**
Assay ID: **18C-02010** Instrument ID: **T312060**
Filename: **C:\Sirius_T3\Mehtap\20180302_exp29_logP_T3-2\18C-02010_M09_octanol_pH-metric high logP.t3r**

#### Sample graphs (continued)

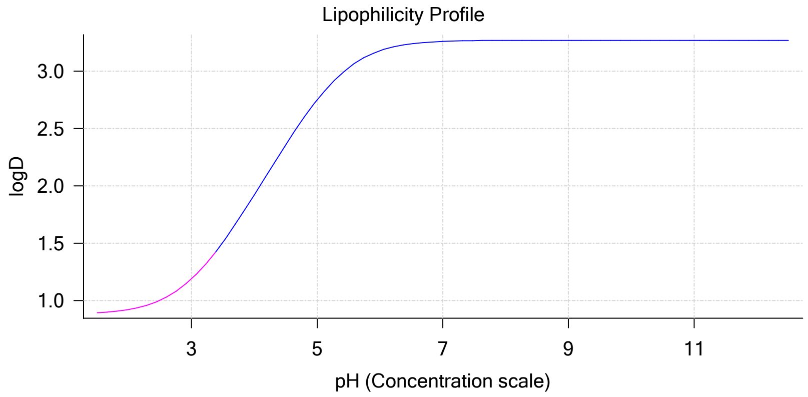


#### Sample logD and percent species

| pH | M09_octanol logD | M09_octanol M09_octanolH | M09_octanol M09_octanol | M09_octanol M09_octanolH* | M09_octanol M09_octanol* | Comment |
|---|---|---|---|---|---|---|
| 1.000 | 0.88 | 58.36 % | 0.00 % | 41.21 % | 0.43 % | |
| 1.200 | 0.89 | 58.21 % | 0.00 % | 41.10 % | 0.69 % | Stomach pH |
| 2.000 | 0.92 | 56.15 % | 0.02 % | 39.65 % | 4.17 % | |
| 3.000 | 1.19 | 40.76 % | 0.17 % | 28.78 % | 30.28 % | |
| 4.000 | 1.92 | 10.89 % | 0.46 % | 7.69 % | 80.95 % | |
| 5.000 | 2.75 | 1.31 % | 0.56 % | 0.92 % | 97.21 % | |
| 6.000 | 3.18 | 0.13 % | 0.57 % | 0.09 % | 99.20 % | |
| 6.500 | 3.24 | 0.04 % | 0.57 % | 0.03 % | 99.36 % | |
| 7.000 | 3.26 | 0.01 % | 0.57 % | 0.01 % | 99.41 % | |
| 7.400 | 3.27 | 0.01 % | 0.57 % | 0.00 % | 99.42 % | Blood pH |
| 8.000 | 3.27 | 0.00 % | 0.57 % | 0.00 % | 99.43 % | |
| 9.000 | 3.27 | 0.00 % | 0.57 % | 0.00 % | 99.43 % | |
| 10.000 | 3.27 | 0.00 % | 0.57 % | 0.00 % | 99.43 % | |
| 11.000 | 3.27 | 0.00 % | 0.57 % | 0.00 % | 99.43 % | |
| 12.000 | 3.27 | 0.00 % | 0.57 % | 0.00 % | 99.43 % | |

#### Carbonate and acidity

| | |
|---|---|
| Carbonate | 0.170 mM |
| Acidity error | -0.977 mM |

#### Other graphs

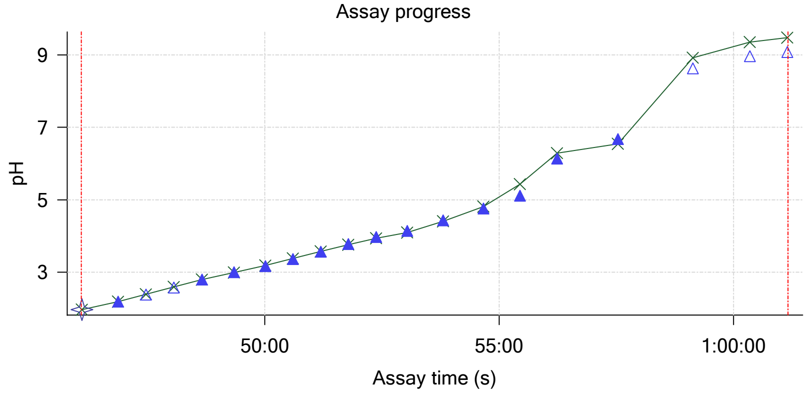


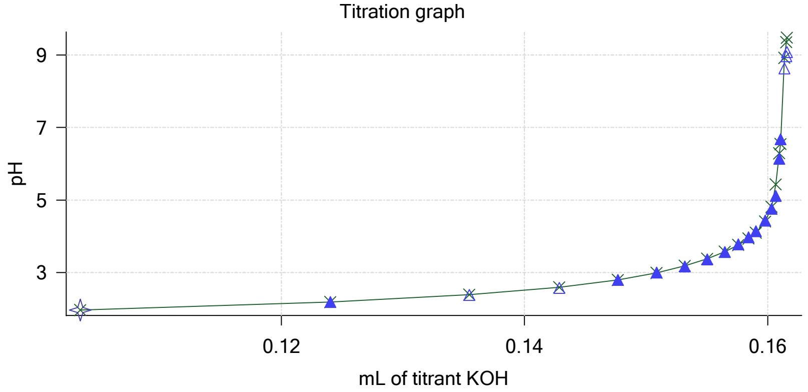

Sample name: **M09_octanol** Experiment start time: **3/2/2018 10:00:20 PM**
Assay name: **pH-metric high logP** Analyst: **Pion**
Assay ID: **18C-02010** Instrument ID: **T312060**
Filename: **C:\Sirius_T3\Mehtap\20180302_exp29_logP_T3-2\18C-02010_M09_octanol_pH-metric high logP.t3r**

#### Other graphs (continued)

Charge balance residuals

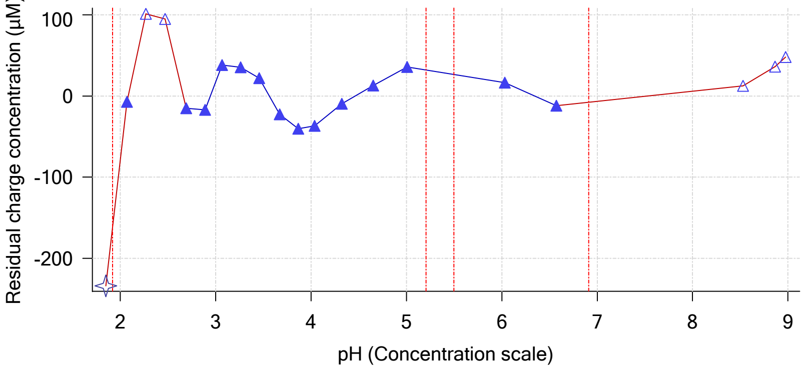


Buffer index profile

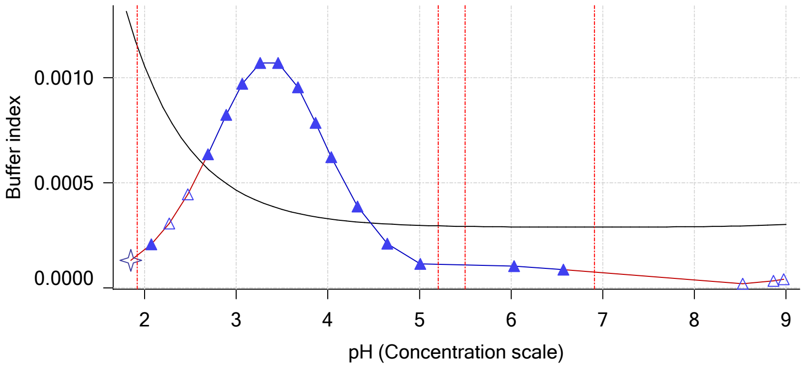


Ionic strength

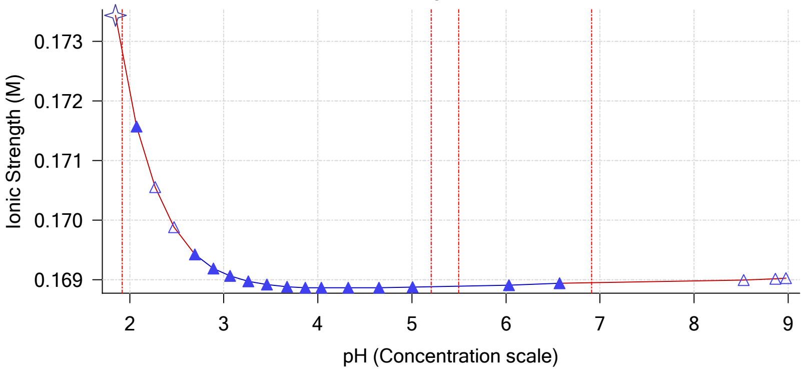


Log R Profile

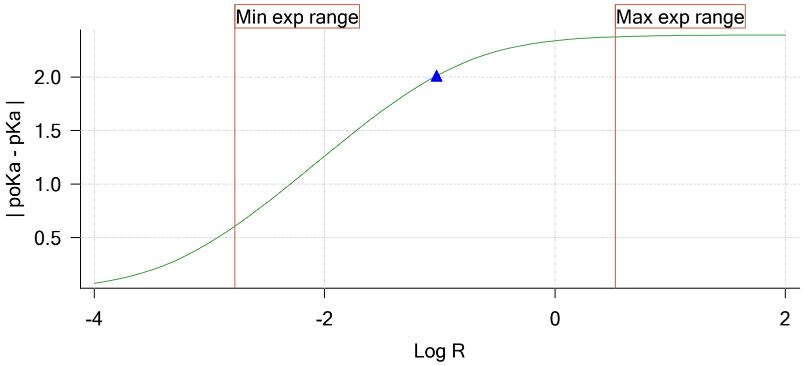


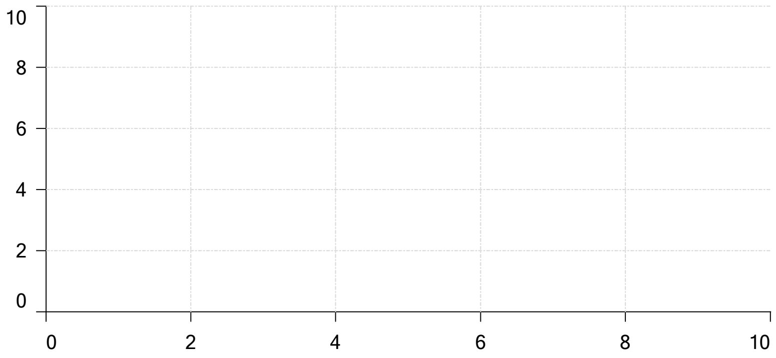


Ionisation of sample Carbonate

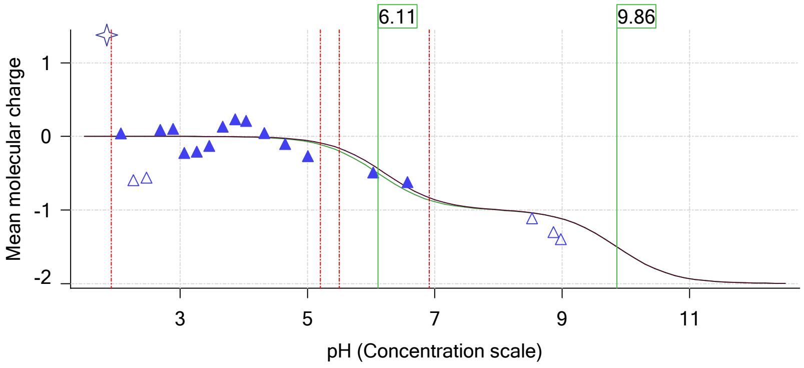

| | | | |
|---|---|---|---|
| Sample name: | **M09_octanol** | Experiment start time: | **3/2/2018 10:00:20 PM** |
| Assay name: | **pH-metric high logP** | Analyst: | **Pion** |
| Assay ID: | **18C-02010** | Instrument ID: | **T312060** |
| Filename: | **C:\Sirius_T3\Mehtap\20180302_exp29_logP_T3-2\18C-02010_M09_octanol_pH-metric high logP.t3r** | | |

#### Assay Model

| Settings | Value | Date/Time changed | Imported from |
|---|---|---|---|
| Sample name | M09_octanol | 2/27/2018 4:56:17 PM | User entered value |
| Sample by | Weight | | Default value |
| Sample weight | 0.000890 g | 3/2/2018 5:08:29 PM | User entered value |
| Formula weight | 287.74 g/mol | 2/27/2018 4:45:45 PM | User entered value |
| Solubility | Unknown | | Default value |
| Molecular weight | 251.28 | 2/27/2018 4:45:45 PM | User entered value |
| Individual pKa ionic environments | No | | Default value |
| Number of pKas | 1 | 2/27/2018 4:45:45 PM | User entered value |
| Sample is a | Base | 2/27/2018 4:45:45 PM | User entered value |
| pKa 1 | 5.37 | 2/27/2018 4:45:45 PM | User entered value |
| logp (XH +) | 0.76 | 3/2/2018 3:27:23 PM | User entered value |
| logP (neutral X) | 3.27 | 3/2/2018 3:27:17 PM | User entered value |
| Stoichiometry | 1.00000 | | Default value |
| Aprotic counterion name | Chloride | | From standards.xml file |
| Stoichiometry | 1.00 | | From standards.xml file |
| Charge per counterion | -1 | | From standards.xml file |

#### Events

| Time | Event | Water | Acid | Base | Octanol | pH | dpH/dt | pH R-squared | pH SD | dpH/dt time |
|---|---|---|---|---|---|---|---|---|---|---|
| 6:00.1 | Initial pH = 4.31 | | | | | | | | | |
| 9:04.7 | Data point 1 | 1.50000 mL | 0.04553 mL | 0.00285 mL | 0.03001 mL | 2.026 | -0.01138 | 0.33547 | 0.00097 | 10.0 s |
| 9:51.0 | Data point 2 | 1.50000 mL | 0.04553 mL | 0.01912 mL | 0.03001 mL | 2.251 | -0.00371 | 0.21517 | 0.00040 | 10.0 s |
| 10:26.6 | Data point 3 | 1.50000 mL | 0.04553 mL | 0.02763 mL | 0.03001 mL | 2.442 | -0.00300 | 0.28283 | 0.00028 | 10.5 s |
| 11:02.7 | Data point 4 | 1.50000 mL | 0.04553 mL | 0.03314 mL | 0.03001 mL | 2.623 | -0.00261 | 0.52506 | 0.00018 | 10.0 s |
| 11:38.3 | Data point 5 | 1.50000 mL | 0.04553 mL | 0.03683 mL | 0.03001 mL | 2.836 | -0.00461 | 0.55068 | 0.00031 | 10.0 s |
| 12:13.7 | Data point 6 | 1.50000 mL | 0.04553 mL | 0.03923 mL | 0.03001 mL | 3.025 | -0.00570 | 0.58263 | 0.00037 | 10.0 s |
| 12:49.2 | Data point 7 | 1.50000 mL | 0.04553 mL | 0.04095 mL | 0.03001 mL | 3.227 | -0.00639 | 0.77318 | 0.00036 | 10.0 s |
| 13:24.6 | Data point 8 | 1.50000 mL | 0.04553 mL | 0.04226 mL | 0.03001 mL | 3.462 | -0.00426 | 0.73568 | 0.00025 | 10.0 s |
| 14:00.0 | Data point 9 | 1.50000 mL | 0.04553 mL | 0.04334 mL | 0.03001 mL | 3.728 | -0.00671 | 0.78883 | 0.00037 | 10.0 s |
| 14:40.6 | Data point 10 | 1.50000 mL | 0.04553 mL | 0.04428 mL | 0.03001 mL | 3.968 | -0.01073 | 0.80668 | 0.00059 | 10.0 s |
| 15:16.1 | Data point 11 | 1.50000 mL | 0.04553 mL | 0.04516 mL | 0.03001 mL | 4.194 | -0.01846 | 0.90114 | 0.00096 | 11.5 s |
| 15:53.0 | Data point 12 | 1.50000 mL | 0.04553 mL | 0.04586 mL | 0.03001 mL | 4.354 | -0.01589 | 0.68146 | 0.00095 | 10.5 s |
| 16:44.4 | Data point 13 | 1.50000 mL | 0.04553 mL | 0.04678 mL | 0.03001 mL | 4.549 | -0.01839 | 0.87702 | 0.00097 | 10.5 s |
| 17:35.9 | Data point 14 | 1.50000 mL | 0.04553 mL | 0.04753 mL | 0.03001 mL | 4.756 | -0.01781 | 0.86166 | 0.00095 | 11.0 s |
| 18:22.6 | Data point 15 | 1.50000 mL | 0.04553 mL | 0.04821 mL | 0.03001 mL | 4.992 | -0.01766 | 0.92547 | 0.00091 | 12.5 s |
| 19:10.8 | Data point 16 | 1.50000 mL | 0.04553 mL | 0.04868 mL | 0.03001 mL | 5.240 | -0.01809 | 0.83242 | 0.00098 | 13.5 s |
| 19:54.9 | Data point 17 | 1.50000 mL | 0.04553 mL | 0.04908 mL | 0.03001 mL | 5.694 | -0.01649 | 0.87889 | 0.00087 | 16.0 s |
| 20:46.7 | Data point 18 | 1.50000 mL | 0.04553 mL | 0.04934 mL | 0.03001 mL | 6.092 | -0.01930 | 0.92959 | 0.00099 | 21.0 s |
| 21:43.4 | Data point 19 | 1.50000 mL | 0.04553 mL | 0.04955 mL | 0.03001 mL | 7.176 | -0.04769 | 0.99644 | 0.00236 | Timed out at 59.5 s |
| 23:19.0 | Data point 20 | 1.50000 mL | 0.04553 mL | 0.04967 mL | 0.03001 mL | 8.283 | -0.03268 | 0.96974 | 0.00164 | Timed out at 59.5 s |
| 24:54.6 | Data point 21 | 1.50000 mL | 0.04553 mL | 0.04976 mL | 0.03001 mL | 8.541 | -0.01854 | 0.89905 | 0.00097 | 54.5 s |
| 26:24.8 | Data point 22 | 1.50000 mL | 0.04553 mL | 0.05005 mL | 0.03001 mL | 9.474 | -0.01874 | 0.97230 | 0.00094 | 21.5 s |
| 27:45.4 | Data point 23 | 1.50000 mL | 0.09967 mL | 0.05005 mL | 0.07001 mL | 1.973 | -0.00556 | 0.20263 | 0.00061 | 10.0 s |
| 28:31.7 | Data point 24 | 1.50000 mL | 0.09967 mL | 0.06865 mL | 0.07001 mL | 2.188 | -0.00175 | 0.27969 | 0.00016 | 10.5 s |
| 29:07.8 | Data point 25 | 1.50000 mL | 0.09967 mL | 0.07921 mL | 0.07001 mL | 2.381 | -0.00137 | 0.05773 | 0.00028 | 10.0 s |
| 29:43.5 | Data point 26 | 1.50000 mL | 0.09967 mL | 0.08603 mL | 0.07001 mL | 2.600 | -0.00181 | 0.18269 | 0.00021 | 10.5 s |
| 30:19.5 | Data point 27 | 1.50000 mL | 0.09967 mL | 0.09029 mL | 0.07001 mL | 2.780 | -0.00959 | 0.84691 | 0.00052 | 10.5 s |
| 30:55.5 | Data point 28 | 1.50000 mL | 0.09967 mL | 0.09327 mL | 0.07001 mL | 2.984 | -0.00603 | 0.12688 | 0.00083 | 10.0 s |
| 31:31.0 | Data point 29 | 1.50000 mL | 0.09967 mL | 0.09539 mL | 0.07001 mL | 3.180 | -0.00463 | 0.68835 | 0.00028 | 10.5 s |
| 32:07.0 | Data point 30 | 1.50000 mL | 0.09967 mL | 0.09701 mL | 0.07001 mL | 3.357 | -0.00337 | 0.46236 | 0.00024 | 10.0 s |
| 32:42.4 | Data point 31 | 1.50000 mL | 0.09967 mL | 0.09838 mL | 0.07001 mL | 3.576 | -0.01194 | 0.46846 | 0.00086 | 10.0 s |
| 33:17.9 | Data point 32 | 1.50000 mL | 0.09967 mL | 0.09951 mL | 0.07001 mL | 3.840 | -0.01046 | 0.33603 | 0.00089 | 10.0 s |
| 33:58.5 | Data point 33 | 1.50000 mL | 0.09967 mL | 0.10042 mL | 0.07001 mL | 4.082 | -0.00936 | 0.85406 | 0.00050 | 10.5 s |
| 34:34.4 | Data point 34 | 1.50000 mL | 0.09967 mL | 0.10108 mL | 0.07001 mL | 4.294 | -0.00407 | 0.09219 | 0.00066 | 10.0 s |

| | | | |
|---|---|---|---|
| Sample name: | **M09_octanol** | Experiment start time: | **3/2/2018 10:00:20 PM** |
| Assay name: | **pH-metric high logP** | Analyst: | **Pion** |
| Assay ID: | **18C-02010** | Instrument ID: | **T312060** |
| Filename: | **C:\Sirius_T3\Mehtap\20180302_exp29_logP_T3-2\18C-02010_M09_octanol_pH-metric high logP.t3r** | | |

#### Events (continued)

| Time | Event | Water | Acid | Base | Octanol | pH | dpH/dt | pH R-squared | pH SD | dpH/dt time |
|---|---|---|---|---|---|---|---|---|---|---|
| 35:09.9 | Data point 35 | 1.50000 mL | 0.09967 mL | 0.10155 mL | 0.07001 mL | 4.450 | -0.01452 | 0.77733 | 0.00081 | 10.0 s |
| 35:55.6 | Data point 36 | 1.50000 mL | 0.09967 mL | 0.10214 mL | 0.07001 mL | 4.702 | -0.01411 | 0.84297 | 0.00076 | 10.5 s |
| 36:41.9 | Data point 37 | 1.50000 mL | 0.09967 mL | 0.10254 mL | 0.07001 mL | 4.968 | -0.01431 | 0.68972 | 0.00085 | 11.0 s |
| 37:28.6 | Data point 38 | 1.50000 mL | 0.09967 mL | 0.10282 mL | 0.07001 mL | 5.294 | -0.01678 | 0.70413 | 0.00099 | 12.0 s |
| 38:16.4 | Data point 39 | 1.50000 mL | 0.09967 mL | 0.10301 mL | 0.07001 mL | 5.750 | -0.01457 | 0.79249 | 0.00081 | 15.5 s |
| 39:07.6 | Data point 40 | 1.50000 mL | 0.09967 mL | 0.10313 mL | 0.07001 mL | 6.399 | -0.01829 | 0.88102 | 0.00096 | 47.0 s |
| 40:30.3 | Data point 41 | 1.50000 mL | 0.09967 mL | 0.10320 mL | 0.07001 mL | 7.207 | -0.05114 | 0.99091 | 0.00254 | Timed out at 59.5 s |
| 42:00.8 | Data point 42 | 1.50000 mL | 0.09967 mL | 0.10327 mL | 0.07001 mL | 8.299 | -0.01875 | 0.88424 | 0.00099 | 58.0 s |
| 43:29.5 | Data point 43 | 1.50000 mL | 0.09967 mL | 0.10332 mL | 0.07001 mL | 8.699 | -0.01983 | 0.98147 | 0.00099 | 35.0 s |
| 44:45.3 | Data point 44 | 1.50000 mL | 0.09967 mL | 0.10346 mL | 0.07001 mL | 9.089 | -0.01827 | 0.94147 | 0.00093 | 20.5 s |
| 46:06.3 | Data point 45 | 1.50000 mL | 0.15753 mL | 0.10346 mL | 0.17001 mL | 1.972 | -0.01076 | 0.86425 | 0.00057 | 10.0 s |
| 46:52.6 | Data point 46 | 1.50000 mL | 0.15753 mL | 0.12404 mL | 0.17001 mL | 2.189 | -0.00758 | 0.73014 | 0.00044 | 10.0 s |
| 47:28.3 | Data point 47 | 1.50000 mL | 0.15753 mL | 0.13547 mL | 0.17001 mL | 2.382 | -0.00182 | 0.17529 | 0.00021 | 10.0 s |
| 48:03.9 | Data point 48 | 1.50000 mL | 0.15753 mL | 0.14285 mL | 0.17001 mL | 2.583 | 0.00231 | 0.53001 | 0.00016 | 10.5 s |
| 48:39.9 | Data point 49 | 1.50000 mL | 0.15753 mL | 0.14767 mL | 0.17001 mL | 2.800 | 0.01463 | 0.61912 | 0.00092 | 15.5 s |
| 49:20.9 | Data point 50 | 1.50000 mL | 0.15753 mL | 0.15087 mL | 0.17001 mL | 2.998 | 0.01486 | 0.59518 | 0.00095 | 14.5 s |
| 50:00.9 | Data point 51 | 1.50000 mL | 0.15753 mL | 0.15320 mL | 0.17001 mL | 3.174 | -0.00236 | 0.06490 | 0.00046 | 10.0 s |
| 50:36.4 | Data point 52 | 1.50000 mL | 0.15753 mL | 0.15503 mL | 0.17001 mL | 3.369 | -0.00368 | 0.52645 | 0.00025 | 10.0 s |
| 51:11.8 | Data point 53 | 1.50000 mL | 0.15753 mL | 0.15647 mL | 0.17001 mL | 3.564 | -0.00334 | 0.04831 | 0.00075 | 10.0 s |
| 51:47.3 | Data point 54 | 1.50000 mL | 0.15753 mL | 0.15760 mL | 0.17001 mL | 3.779 | -0.00287 | 0.06293 | 0.00056 | 10.0 s |
| 52:22.7 | Data point 55 | 1.50000 mL | 0.15753 mL | 0.15842 mL | 0.17001 mL | 3.972 | 0.01222 | 0.41308 | 0.00094 | 14.5 s |
| 53:02.6 | Data point 56 | 1.50000 mL | 0.15753 mL | 0.15903 mL | 0.17001 mL | 4.143 | -0.00657 | 0.60274 | 0.00042 | 10.0 s |
| 53:48.4 | Data point 57 | 1.50000 mL | 0.15753 mL | 0.15981 mL | 0.17001 mL | 4.429 | 0.01474 | 0.60321 | 0.00094 | 16.0 s |
| 54:40.1 | Data point 58 | 1.50000 mL | 0.15753 mL | 0.16033 mL | 0.17001 mL | 4.755 | 0.01365 | 0.52664 | 0.00093 | 11.0 s |
| 55:26.8 | Data point 59 | 1.50000 mL | 0.15753 mL | 0.16065 mL | 0.17001 mL | 5.112 | -0.00952 | 0.31084 | 0.00084 | 11.5 s |
| 56:14.0 | Data point 60 | 1.50000 mL | 0.15753 mL | 0.16096 mL | 0.17001 mL | 6.134 | -0.01828 | 0.87737 | 0.00096 | 42.5 s |
| 57:32.1 | Data point 61 | 1.50000 mL | 0.15753 mL | 0.16105 mL | 0.17001 mL | 6.674 | -0.03582 | 0.95893 | 0.00181 | Timed out at 59.5 s |
| 59:07.8 | Data point 62 | 1.50000 mL | 0.15753 mL | 0.16138 mL | 0.17001 mL | 8.628 | -0.01937 | 0.92742 | 0.00099 | 32.0 s |
| 1:00:20.6 | Data point 63 | 1.50000 mL | 0.15753 mL | 0.16152 mL | 0.17001 mL | 8.963 | -0.00684 | 0.12695 | 0.00095 | 17.5 s |
| 1:01:08.6 | Data point 64 | 1.50000 mL | 0.15753 mL | 0.16159 mL | 0.17001 mL | 9.075 | -0.01927 | 0.95912 | 0.00097 | 15.0 s |
| 1:01:32.7 | Assay volumes | 1.50000 mL | 0.15753 mL | 0.16159 mL | 0.17001 mL | | | | | |

| | | | |
|---|---|---|---|
| Sample name: | **M09_octanol** | Experiment start time: | **3/2/2018 10:00:20 PM** |
| Assay name: | **pH-metric high logP** | Analyst: | **Pion** |
| Assay ID: | **18C-02010** | Instrument ID: | **T312060** |
| Filename: | **C:\Sirius_T3\Mehtap\20180302_exp29_logP_T3-2\18C-02010_M09_octanol_pH-metric high logP.t3r** | | |

| | | | |
|---|---|---|---|
| Sample name: | **M09_octanol** | Experiment start time: | **3/2/2018 10:00:20 PM** |
| Assay name: | **pH-metric high logP** | Analyst: | **Pion** |
| Assay ID: | **18C-02010** | Instrument ID: | **T312060** |
| Filename: | **C:\Sirius_T3\Mehtap\20180302_exp29_logP_T3-2\18C-02010_M09_octanol_pH-metric high logP.t3r** | | |

#### Assay Settings (continued)

| Setting | Value | Original Value | Date/Time changed | Imported from |
|---|---|---|---|---|
| ***Titration 3*** | | | | |
| Titrate from | Low to high pH | | | |
| Add additional water | 0.00 mL | | | |
| Additional partition solvent volume | 0.100 mL | | | |
| Additional partition solvent added | Automatic | | | |
| After pH adjust stir for | 30 seconds | | | |
| Stir to allow partitioning for | 15 seconds | | | |
| Stirrer speed for partitioning | 60% | | | |
| ***Data Point Stability*** | | | | |
| Stir during data point collection | No | | | |
| Delay before data point collection | 0 seconds | | | |
| Number of points to average | 20 points | | | |
| Time interval between points | 0.50 seconds | | | |
| Required maximum standard deviation | 0.00100 dpH/dt | | | |
| Stability timeout after | 60 seconds | | | |

#### Calibration Settings

| Setting | Value | Date/Time changed | Imported from |
|---|---|---|---|
| Four-Plus alpha | 0.111 | 3/2/2018 10:00:20 PM | C:\Sirius_T3\HCl18C02.t3r |
| Four-Plus S | 0.9988 | 3/2/2018 10:00:20 PM | C:\Sirius_T3\HCl18C02.t3r |
| Four-Plus jH | 1.0 | 3/2/2018 10:00:20 PM | C:\Sirius_T3\HCl18C02.t3r |
| Four-Plus jOH | -0.8 | 3/2/2018 10:00:20 PM | C:\Sirius_T3\HCl18C02.t3r |
| Base concentration factor | 1.000 | 3/2/2018 10:00:20 PM | C:\Sirius_T3\KOH18B27.t3r |
| Acid concentration factor | 0.999 | 3/2/2018 10:00:20 PM | C:\Sirius_T3\HCl18C02.t3r |

#### Instrument Settings

| Setting | Value | Batch Id | Install date |
|---|---|---|---|
| Instrument owner | Merck | | |
| Instrument ID | T312060 | | |
| Instrument type | T3 Simulator | | |
| Software version | 1.1.3.0 | | |
| Dispenser module | | T3DM1200361 | 3/31/2009 5:24:52 AM |
| Dispenser 0 | Water | | 3/31/2009 5:25:05 AM |
| Syringe volume | 2.5 mL | | |
| Firmware version | 1.2.1(r2) | | |
| Titrant | Water (0.15 M KCl) | 02-06-2018 | 2/27/2018 10:05:59 AM |
| Dispenser 2 | Acid | | 3/31/2009 5:25:11 AM |
| Syringe volume | 0.5 mL | | |
| Firmware version | 1.2.1(r2) | | |
| Titrant | Acid (0.5 M HCl) | 02-27-2018 | 2/27/2018 10:27:22 AM |
| Dispenser 1 | Base | | 3/31/2009 5:25:21 AM |
| Syringe volume | 0.5 mL | | |
| Firmware version | 1.2.1(r2) | | |
| Titrant | Base (0.5 M KOH) | 9/22/2017 | 2/27/2018 10:21:22 AM |
| Dispenser 5 | Cosolvent | | 3/31/2009 5:26:24 AM |
| Syringe volume | 2.5 mL | | |
| Firmware version | 1.2.1(r2) | | |
| Distribution valve 5 | Distribution Valve | | 3/31/2009 5:28:19 AM |
| Firmware version | 1.1.3 | | |
| Port A | Methanol (80%, 0.15 M KCl) | 09-26-17 | 2/7/2018 9:42:01 AM |
| Port B | Cyclohexane | 11-01-17 | 2/27/2018 10:37:57 AM |
| Dispenser 3 | Buffer | | 8/3/2010 5:05:16 AM |
| Syringe volume | 0.5 mL | | |
| Firmware version | 1.2.1(r2) | | |
| Titrant | Dodecane | 2018/01/31 | 2/28/2018 10:18:04 AM |
| Dispenser 6 | Octanol | | 10/22/2010 10:52:43 AM |

| | | | |
|---|---|---|---|
| Sample name: | **M09_octanol** | Experiment start time: | **3/2/2018 10:00:20 PM** |
| Assay name: | **pH-metric high logP** | Analyst: | **Pion** |
| Assay ID: | **18C-02010** | Instrument ID: | **T312060** |
| Filename: | **C:\Sirius_T3\Mehtap\20180302_exp29_logP_T3-2\18C-02010_M09_octanol_pH-metric high logP.t3r** | | |

| | | | |
|---|---|---|---|
| Sample name: | **M09_octanol** | Experiment start time: | **3/2/2018 10:00:20 PM** |
| Assay name: | **pH-metric high logP** | Analyst: | **Pion** |
| Assay ID: | **18C-02010** | Instrument ID: | **T312060** |
| Filename: | **C:\Sirius_T3\Mehtap\20180302_exp29_logP_T3-2\18C-02010_M09_octanol_pH-metric high logP.t3r** | | |

#### Instrument Settings (continued)

| Setting | Value | Batch Id | Install date |
|---|---|---|---|
| Spectrometer calibration stir duration | 5 s | | |
| Spectrometer calibration stir speed | 30% | | |
| Spectrometer calibration wash pump volume | 20.0 mL | | |
| Spectrometer calibration wash stir duration | 5 s | | |
| Spectrometer calibration wash stir speed | 30% | | |
| Overhead dispense height | 10000 | | |

#### Refinement Settings

| Setting | Value | Default value |
|---|---|---|
| Turbidity detection method | None | None |
| Turbidity wavelength to assess | 500.0 nm | 500.0 nm |
| Turbidity maximum absorbance | 0.100 | 0.100 |
| Turbidity probe threshold | 50.00 | 50.00 |

#### Experiment Log

[2:37] Air gap created for Water (0.15 M KCl)
[2:38] Air gap created for Acid (0.5 M HCl)
[2:38] Air gap created for Base (0.5 M KOH)
[2:39] Air gap released for Water (0.15 M KCl)
[2:42] Titrator arm moved over Titration position
[2:42] Titration 1 of 3
[2:42] Adding initial titrants
[2:42] Automatically add 1.50000 mL of water
[3:08] Dispensed 1.500000 mL of Water (0.15 M KCl)
[3:12] Titrator arm moved over Drain
[5:53] Titrator arm moved to Titration position
[5:53] Argon flow rate set to 100
[5:53] Stirrer speed set to 10
[5:58] Automatically add 0.03000 mL of Octanol
[5:59] Dispensed 0.030009 mL of Octanol
[6:00] Initial pH = 4.31
[6:00] Iterative adjust 4.31 -> 2.00
[6:00] pH 4.31 -> 2.00
[6:02] Air gap released for Acid (0.5 M HCl)
[6:03] Dispensed 0.043815 mL of Acid (0.5 M HCl)
[6:08] pH 2.02 -> 2.00
[6:08] Dispensed 0.001717 mL of Acid (0.5 M HCl)
[6:13] Holding pH 2.00
[8:13] Stirrer speed set to 0
[8:13] Stirrer speed set to 50
[8:13] Iterative adjust 1.97 -> 2.00
[8:13] pH 1.97 -> 2.00
[8:14] Air gap released for Base (0.5 M KOH)
[8:15] Dispensed 0.002846 mL of Base (0.5 M KOH)
[9:05] Stirrer speed set to 0
[9:15] Datapoint id 1 collected
[9:15] Stirrer speed set to 50
[9:20] pH 2.03 -> 2.23
[9:20] Using cautious pH adjust
[9:20] Dispensed 0.007173 mL of Base (0.5 M KOH)
[9:25] Stepping pH = 2.10
[9:26] Dispensed 0.007832 mL of Base (0.5 M KOH)
[9:31] Stepping pH = 2.21
[9:31] Dispensed 0.001270 mL of Base (0.5 M KOH)
[9:36] Stepping pH = 2.24
[9:51] Stirrer speed set to 0
[10:01] Datapoint id 2 collected

Sample name: **M09_octanol**
Assay name: **pH-metric high logP**
Assay ID: **18C-02010**
Filename: **C:\Sirius_T3\Mehtap\20180302_exp29_logP_T3-2\18C-02010_M09_octanol_pH-metric high logP.t3r**

Experiment start time: **3/2/2018 10:00:20 PM**
Analyst: **Pion**
Instrument ID: **T312060**

#### Experiment Log (continued)

[10:01] Charge balance equation is out by -13.4%
[10:01] Stirrer speed set to 50
[10:06] pH 2.26 -> 2.46
[10:06] Using charge balance adjust
[10:07] Dispensed 0.008514 mL of Base (0.5 M KOH)
[10:27] Stirrer speed set to 0
[10:37] Datapoint id 3 collected
[10:37] Charge balance equation is out by -7.5%
[10:37] Stirrer speed set to 50
[10:42] pH 2.45 -> 2.65
[10:42] Using charge balance adjust
[10:43] Dispensed 0.005503 mL of Base (0.5 M KOH)
[11:03] Stirrer speed set to 0
[11:13] Datapoint id 4 collected
[11:13] Charge balance equation is out by -12.9%
[11:13] Stirrer speed set to 50
[11:18] pH 2.63 -> 2.83
[11:18] Using charge balance adjust
[11:18] Dispensed 0.003692 mL of Base (0.5 M KOH)
[11:38] Stirrer speed set to 0
[11:48] Datapoint id 5 collected
[11:48] Charge balance equation is out by 3.3%
[11:48] Stirrer speed set to 50
[11:54] pH 2.84 -> 3.04
[11:54] Using charge balance adjust
[11:54] Dispensed 0.002399 mL of Base (0.5 M KOH)
[12:14] Stirrer speed set to 0
[12:24] Datapoint id 6 collected
[12:24] Charge balance equation is out by -8.2%
[12:24] Stirrer speed set to 50
[12:29] pH 3.03 -> 3.23
[12:29] Using charge balance adjust
[12:29] Dispensed 0.001717 mL of Base (0.5 M KOH)
[12:49] Stirrer speed set to 0
[12:59] Datapoint id 7 collected
[12:59] Charge balance equation is out by -2.1%
[12:59] Stirrer speed set to 50
[13:04] pH 3.23 -> 3.43
[13:04] Using charge balance adjust
[13:05] Dispensed 0.001317 mL of Base (0.5 M KOH)
[13:25] Stirrer speed set to 0
[13:35] Datapoint id 8 collected
[13:35] Charge balance equation is out by 15.0%
[13:35] Stirrer speed set to 50
[13:40] pH 3.47 -> 3.67
[13:40] Using charge balance adjust
[13:40] Dispensed 0.001082 mL of Base (0.5 M KOH)
[14:00] Stirrer speed set to 0
[14:10] Datapoint id 9 collected
[14:10] Charge balance equation is out by 31.3%
[14:10] Stirrer speed set to 50
[14:15] pH 3.73 -> 3.93
[14:15] Using cautious pH adjust
[14:15] Dispensed 0.000494 mL of Base (0.5 M KOH)
[14:21] Stepping pH = 3.81
[14:21] Dispensed 0.000447 mL of Base (0.5 M KOH)
[14:26] Stepping pH = 3.94
[14:41] Stirrer speed set to 0
[14:51] Datapoint id 10 collected

| | | | |
|---|---|---|---|
| Sample name: | **M09_octanol** | Experiment start time: | **3/2/2018 10:00:20 PM** |
| Assay name: | **pH-metric high logP** | Analyst: | **Pion** |
| Assay ID: | **18C-02010** | Instrument ID: | **T312060** |
| Filename: | **C:\Sirius_T3\Mehtap\20180302_exp29_logP_T3-2\18C-02010_M09_octanol_pH-metric high logP.t3r** | | |

#### Experiment Log (continued)

[14:51] Charge balance equation is out by 3.1%
[14:51] Stirrer speed set to 50
[14:56] pH 3.97 -> 4.17
[14:56] Using charge balance adjust
[14:56] Dispensed 0.000870 mL of Base (0.5 M KOH)
[15:16] Stirrer speed set to 0
[15:28] Datapoint id 11 collected
[15:28] Charge balance equation is out by 10.9%
[15:28] Stirrer speed set to 50
[15:33] pH 4.20 -> 4.40
[15:33] Using charge balance adjust
[15:33] Dispensed 0.000706 mL of Base (0.5 M KOH)
[15:53] Stirrer speed set to 0
[16:04] Datapoint id 12 collected
[16:04] Charge balance equation is out by -22.6%
[16:04] Stirrer speed set to 50
[16:09] pH 4.36 -> 4.56
[16:09] Using cautious pH adjust
[16:09] Dispensed 0.000306 mL of Base (0.5 M KOH)
[16:14] Stepping pH = 4.41
[16:14] Dispensed 0.000447 mL of Base (0.5 M KOH)
[16:19] Stepping pH = 4.52
[16:19] Dispensed 0.000118 mL of Base (0.5 M KOH)
[16:24] Stepping pH = 4.55
[16:24] Dispensed 0.000047 mL of Base (0.5 M KOH)
[16:29] Stepping pH = 4.56
[16:45] Stirrer speed set to 0
[16:55] Datapoint id 13 collected
[16:55] Charge balance equation is out by -53.1%
[16:55] Stirrer speed set to 50
[17:00] pH 4.56 -> 4.76
[17:00] Using cautious pH adjust
[17:00] Dispensed 0.000235 mL of Base (0.5 M KOH)
[17:06] Stepping pH = 4.60
[17:06] Dispensed 0.000329 mL of Base (0.5 M KOH)
[17:11] Stepping pH = 4.70
[17:11] Dispensed 0.000165 mL of Base (0.5 M KOH)
[17:16] Stepping pH = 4.75
[17:16] Dispensed 0.000024 mL of Base (0.5 M KOH)
[17:21] Stepping pH = 4.76
[17:36] Stirrer speed set to 0
[17:47] Datapoint id 14 collected
[17:47] Charge balance equation is out by -67.5%
[17:47] Stirrer speed set to 50
[17:52] pH 4.77 -> 4.97
[17:52] Using cautious pH adjust
[17:52] Dispensed 0.000165 mL of Base (0.5 M KOH)
[17:57] Stepping pH = 4.79
[17:57] Dispensed 0.000400 mL of Base (0.5 M KOH)
[18:03] Stepping pH = 4.92
[18:03] Dispensed 0.000118 mL of Base (0.5 M KOH)
[18:08] Stepping pH = 4.98
[18:23] Stirrer speed set to 0
[18:35] Datapoint id 15 collected
[18:35] Charge balance equation is out by -106.8%
[18:35] Stirrer speed set to 50
[18:40] pH 5.01 -> 5.21
[18:40] Using cautious pH adjust
[18:41] Dispensed 0.000094 mL of Base (0.5 M KOH)

Sample name: **M09_octanol** Experiment start time: **3/2/2018 10:00:20 PM**
Assay name: **pH-metric high logP** Analyst: **Pion**
Assay ID: **18C-02010** Instrument ID: **T312060**
Filename: **C:\Sirius_T3\Mehtap\20180302_exp29_logP_T3-2\18C-02010_M09_octanol_pH-metric high logP.t3r**

#### Experiment Log (continued)

[18:46] Stepping pH = 5.01
[18:46] Dispensed 0.000306 mL of Base (0.5 M KOH)
[18:51] Stepping pH = 5.16
[18:51] Dispensed 0.000071 mL of Base (0.5 M KOH)
[18:56] Stepping pH = 5.22
[19:11] Stirrer speed set to 0
[19:25] Datapoint id 16 collected
[19:25] Charge balance equation is out by -128.8%
[19:25] Stirrer speed set to 50
[19:30] pH 5.26 -> 5.46
[19:30] Using cautious pH adjust
[19:30] Dispensed 0.000071 mL of Base (0.5 M KOH)
[19:35] Stepping pH = 5.26
[19:35] Dispensed 0.000329 mL of Base (0.5 M KOH)
[19:40] Stepping pH = 5.52
[19:55] Stirrer speed set to 0
[20:11] Datapoint id 17 collected
[20:11] Charge balance equation is out by -200.5%
[20:11] Stirrer speed set to 50
[20:16] pH 5.72 -> 5.92
[20:16] Using cautious pH adjust
[20:16] Dispensed 0.000047 mL of Base (0.5 M KOH)
[20:21] Stepping pH = 5.73
[20:22] Dispensed 0.000094 mL of Base (0.5 M KOH)
[20:27] Stepping pH = 5.79
[20:27] Dispensed 0.000118 mL of Base (0.5 M KOH)
[20:32] Stepping pH = 5.94
[20:47] Stirrer speed set to 0
[21:08] Datapoint id 18 collected
[21:08] Charge balance equation is out by -265.1%
[21:08] Stirrer speed set to 50
[21:13] pH 6.11 -> 6.31
[21:13] Using cautious pH adjust
[21:13] Dispensed 0.000024 mL of Base (0.5 M KOH)
[21:18] Stepping pH = 6.11
[21:18] Dispensed 0.000141 mL of Base (0.5 M KOH)
[21:23] Stepping pH = 6.26
[21:23] Dispensed 0.000047 mL of Base (0.5 M KOH)
[21:29] Stepping pH = 6.65
[21:44] Stirrer speed set to 0
[22:44] Datapoint id 19 collected
[22:44] Charge balance equation is out by -268.6%
[22:44] Stirrer speed set to 50
[22:49] pH 7.06 -> 7.26
[22:49] Using cautious pH adjust
[22:49] Dispensed 0.000024 mL of Base (0.5 M KOH)
[22:54] Stepping pH = 7.04
[22:54] Dispensed 0.000047 mL of Base (0.5 M KOH)
[22:59] Stepping pH = 7.13
[22:59] Dispensed 0.000047 mL of Base (0.5 M KOH)
[23:04] Stepping pH = 7.50
[23:19] Stirrer speed set to 0
[24:19] Datapoint id 20 collected
[24:19] Charge balance equation is out by -526.7%
[24:19] Stirrer speed set to 50
[24:24] pH 8.19 -> 8.39
[24:24] Using cautious pH adjust
[24:24] Dispensed 0.000024 mL of Base (0.5 M KOH)
[24:29] Stepping pH = 8.16

| | | | |
|---|---|---|---|
| Sample name: | **M09_octanol** | Experiment start time: | **3/2/2018 10:00:20 PM** |
| Assay name: | **pH-metric high logP** | Analyst: | **Pion** |
| Assay ID: | **18C-02010** | Instrument ID: | **T312060** |
| Filename: | **C:\Sirius_T3\Mehtap\20180302_exp29_logP_T3-2\18C-02010_M09_octanol_pH-metric high logP.t3r** | | |

#### Experiment Log (continued)

[24:29] Dispensed 0.000024 mL of Base (0.5 M KOH)
[24:35] Stepping pH = 8.20
[24:35] Dispensed 0.000047 mL of Base (0.5 M KOH)
[24:40] Stepping pH = 8.49
[24:55] Stirrer speed set to 0
[25:49] Datapoint id 21 collected
[25:49] Charge balance equation is out by -997.9%
[25:49] Stirrer speed set to 50
[25:55] pH 8.47 -> 8.67
[25:55] Using cautious pH adjust
[25:55] Dispensed 0.000024 mL of Base (0.5 M KOH)
[26:00] Stepping pH = 8.43
[26:00] Dispensed 0.000047 mL of Base (0.5 M KOH)
[26:05] Stepping pH = 8.40
[26:05] Dispensed 0.000212 mL of Base (0.5 M KOH)
[26:10] Stepping pH = 9.23
[26:25] Stirrer speed set to 0
[26:47] Datapoint id 22 collected
[26:47] Charge balance equation is out by -1,935.7%
[26:47] Titration 2 of 3
[26:47] Adding initial titrants
[26:47] Automatically add 0.04000 mL of Octanol
[26:48] Dispensed 0.040005 mL of Octanol
[26:48] Stirrer speed set to 10
[26:49] Stirrer speed set to 55
[26:49] Iterative adjust 9.48 -> 2.00
[26:49] pH 9.48 -> 2.00
[26:50] Dispensed 0.048871 mL of Acid (0.5 M HCl)
[26:55] pH 2.05 -> 2.00
[26:55] Dispensed 0.005268 mL of Acid (0.5 M HCl)
[27:46] Stirrer speed set to 0
[27:56] Datapoint id 23 collected
[27:56] Stirrer speed set to 55
[28:01] pH 1.98 -> 2.18
[28:01] Using cautious pH adjust
[28:01] Dispensed 0.008702 mL of Base (0.5 M KOH)
[28:06] Stepping pH = 2.05
[28:06] Dispensed 0.008302 mL of Base (0.5 M KOH)
[28:12] Stepping pH = 2.15
[28:12] Dispensed 0.001599 mL of Base (0.5 M KOH)
[28:17] Stepping pH = 2.18
[28:32] Stirrer speed set to 0
[28:42] Datapoint id 24 collected
[28:42] Charge balance equation is out by -6.9%
[28:42] Stirrer speed set to 55
[28:47] pH 2.19 -> 2.39
[28:47] Using charge balance adjust
[28:48] Dispensed 0.010560 mL of Base (0.5 M KOH)
[29:08] Stirrer speed set to 0
[29:18] Datapoint id 25 collected
[29:18] Charge balance equation is out by -5.9%
[29:18] Stirrer speed set to 55
[29:23] pH 2.39 -> 2.59
[29:23] Using charge balance adjust
[29:23] Dispensed 0.006820 mL of Base (0.5 M KOH)
[29:44] Stirrer speed set to 0
[29:54] Datapoint id 26 collected
[29:54] Charge balance equation is out by 6.5%
[29:54] Stirrer speed set to 55

Sample name: **M09_octanol** Experiment start time: **3/2/2018 10:00:20 PM**
Assay name: **pH-metric high logP** Analyst: **Pion**
Assay ID: **18C-02010** Instrument ID: **T312060**
Filename: **C:\Sirius_T3\Mehtap\20180302_exp29_logP_T3-2\18C-02010_M09_octanol_pH-metric high logP.t3r**

#### Experiment Log (continued)

[29:59] pH 2.61 -> 2.81
[29:59] Using charge balance adjust
[30:00] Dispensed 0.004257 mL of Base (0.5 M KOH)
[30:20] Stirrer speed set to 0
[30:30] Datapoint id 27 collected
[30:30] Charge balance equation is out by -12.7%
[30:30] Stirrer speed set to 55
[30:35] pH 2.78 -> 2.98
[30:35] Using charge balance adjust
[30:36] Dispensed 0.002987 mL of Base (0.5 M KOH)
[30:56] Stirrer speed set to 0
[31:06] Datapoint id 28 collected
[31:06] Charge balance equation is out by 0.0%
[31:06] Stirrer speed set to 55
[31:11] pH 2.99 -> 3.19
[31:11] Using charge balance adjust
[31:11] Dispensed 0.002117 mL of Base (0.5 M KOH)
[31:31] Stirrer speed set to 0
[31:42] Datapoint id 29 collected
[31:42] Charge balance equation is out by -3.1%
[31:42] Stirrer speed set to 55
[31:47] pH 3.18 -> 3.38
[31:47] Using charge balance adjust
[31:47] Dispensed 0.001623 mL of Base (0.5 M KOH)
[32:07] Stirrer speed set to 0
[32:17] Datapoint id 30 collected
[32:17] Charge balance equation is out by -13.1%
[32:17] Stirrer speed set to 55
[32:22] pH 3.36 -> 3.56
[32:22] Using charge balance adjust
[32:22] Dispensed 0.001364 mL of Base (0.5 M KOH)
[32:43] Stirrer speed set to 0
[32:53] Datapoint id 31 collected
[32:53] Charge balance equation is out by 7.5%
[32:53] Stirrer speed set to 55
[32:58] pH 3.58 -> 3.78
[32:58] Using charge balance adjust
[32:58] Dispensed 0.001129 mL of Base (0.5 M KOH)
[33:18] Stirrer speed set to 0
[33:28] Datapoint id 32 collected
[33:28] Charge balance equation is out by 30.1%
[33:28] Stirrer speed set to 55
[33:33] pH 3.84 -> 4.04
[33:33] Using cautious pH adjust
[33:33] Dispensed 0.000447 mL of Base (0.5 M KOH)
[33:38] Stepping pH = 3.91
[33:38] Dispensed 0.000470 mL of Base (0.5 M KOH)
[33:44] Stepping pH = 4.03
[33:59] Stirrer speed set to 0
[34:09] Datapoint id 33 collected
[34:09] Charge balance equation is out by -3.9%
[34:09] Stirrer speed set to 55
[34:14] pH 4.08 -> 4.28
[34:14] Using charge balance adjust
[34:14] Dispensed 0.000659 mL of Base (0.5 M KOH)
[34:35] Stirrer speed set to 0
[34:45] Datapoint id 34 collected
[34:45] Charge balance equation is out by 5.3%
[34:45] Stirrer speed set to 55

Sample name: **M09_octanol**
Assay name: **pH-metric high logP**
Assay ID: **18C-02010**
Filename: **C:\Sirius_T3\Mehtap\20180302_exp29_logP_T3-2\18C-02010_M09_octanol_pH-metric high logP.t3r**
Experiment start time: **3/2/2018 10:00:20 PM**
Analyst: **Pion**
Instrument ID: **T312060**

#### Experiment Log (continued)

[34:50] pH 4.30 -> 4.50
[34:50] Using charge balance adjust
[34:50] Dispensed 0.000470 mL of Base (0.5 M KOH)
[35:10] Stirrer speed set to 0
[35:20] Datapoint id 35 collected
[35:20] Charge balance equation is out by -24.5%
[35:20] Stirrer speed set to 55
[35:25] pH 4.46 -> 4.66
[35:25] Using cautious pH adjust
[35:25] Dispensed 0.000188 mL of Base (0.5 M KOH)
[35:30] Stepping pH = 4.50
[35:30] Dispensed 0.000306 mL of Base (0.5 M KOH)
[35:36] Stepping pH = 4.61
[35:36] Dispensed 0.000094 mL of Base (0.5 M KOH)
[35:41] Stepping pH = 4.67
[35:56] Stirrer speed set to 0
[36:06] Datapoint id 36 collected
[36:06] Charge balance equation is out by -61.6%
[36:06] Stirrer speed set to 55
[36:11] pH 4.71 -> 4.91
[36:11] Using cautious pH adjust
[36:12] Dispensed 0.000118 mL of Base (0.5 M KOH)
[36:17] Stepping pH = 4.74
[36:17] Dispensed 0.000235 mL of Base (0.5 M KOH)
[36:22] Stepping pH = 4.88
[36:22] Dispensed 0.000047 mL of Base (0.5 M KOH)
[36:27] Stepping pH = 4.93
[36:42] Stirrer speed set to 0
[36:53] Datapoint id 37 collected
[36:53] Charge balance equation is out by -76.9%
[36:53] Stirrer speed set to 55
[36:58] pH 4.98 -> 5.18
[36:58] Using cautious pH adjust
[36:58] Dispensed 0.000071 mL of Base (0.5 M KOH)
[37:03] Stepping pH = 5.00
[37:03] Dispensed 0.000188 mL of Base (0.5 M KOH)
[37:09] Stepping pH = 5.17
[37:09] Dispensed 0.000024 mL of Base (0.5 M KOH)
[37:14] Stepping pH = 5.23
[37:29] Stirrer speed set to 0
[37:41] Datapoint id 38 collected
[37:41] Charge balance equation is out by -100.4%
[37:41] Stirrer speed set to 55
[37:46] pH 5.31 -> 5.51
[37:46] Using cautious pH adjust
[37:46] Dispensed 0.000047 mL of Base (0.5 M KOH)
[37:51] Stepping pH = 5.32
[37:51] Dispensed 0.000118 mL of Base (0.5 M KOH)
[37:56] Stepping pH = 5.49
[37:56] Dispensed 0.000024 mL of Base (0.5 M KOH)
[38:01] Stepping pH = 5.62
[38:17] Stirrer speed set to 0
[38:32] Datapoint id 39 collected
[38:32] Charge balance equation is out by -114.7%
[38:32] Stirrer speed set to 55
[38:37] pH 5.77 -> 5.97
[38:37] Using cautious pH adjust
[38:37] Dispensed 0.000024 mL of Base (0.5 M KOH)
[38:42] Stepping pH = 5.78

Sample name: **M09_octanol**
Assay name: **pH-metric high logP**
Assay ID: **18C-02010**
Filename: **C:\Sirius_T3\Mehtap\20180302_exp29_logP_T3-2\18C-02010_M09_octanol_pH-metric high logP.t3r**

Experiment start time: **3/2/2018 10:00:20 PM**
Analyst: **Pion**
Instrument ID: **T312060**

#### Experiment Log (continued)

[38:42] Dispensed 0.000071 mL of Base (0.5 M KOH)
[38:48] Stepping pH = 5.95
[38:48] Dispensed 0.000024 mL of Base (0.5 M KOH)
[38:53] Stepping pH = 6.16
[39:08] Stirrer speed set to 0
[39:55] Datapoint id 40 collected
[39:55] Charge balance equation is out by -131.1%
[39:55] Stirrer speed set to 55
[40:00] pH 6.40 -> 6.60
[40:00] Using cautious pH adjust
[40:00] Dispensed 0.000024 mL of Base (0.5 M KOH)
[40:05] Stepping pH = 6.45
[40:05] Dispensed 0.000024 mL of Base (0.5 M KOH)
[40:10] Stepping pH = 6.54
[40:10] Dispensed 0.000024 mL of Base (0.5 M KOH)
[40:15] Stepping pH = 6.77
[40:31] Stirrer speed set to 0
[41:31] Datapoint id 41 collected
[41:31] Charge balance equation is out by -75.8%
[41:31] Stirrer speed set to 55
[41:36] pH 7.09 -> 7.29
[41:36] Using cautious pH adjust
[41:36] Dispensed 0.000024 mL of Base (0.5 M KOH)
[41:41] Stepping pH = 7.03
[41:41] Dispensed 0.000047 mL of Base (0.5 M KOH)
[41:46] Stepping pH = 7.49
[42:01] Stirrer speed set to 0
[42:59] Datapoint id 42 collected
[42:59] Charge balance equation is out by -364.7%
[42:59] Stirrer speed set to 55
[43:04] pH 8.28 -> 8.48
[43:04] Using cautious pH adjust
[43:04] Dispensed 0.000024 mL of Base (0.5 M KOH)
[43:09] Stepping pH = 8.37
[43:09] Dispensed 0.000024 mL of Base (0.5 M KOH)
[43:15] Stepping pH = 8.52
[43:30] Stirrer speed set to 0
[44:05] Datapoint id 43 collected
[44:05] Charge balance equation is out by -286.1%
[44:05] Stirrer speed set to 55
[44:10] pH 8.68 -> 8.88
[44:10] Using cautious pH adjust
[44:10] Dispensed 0.000024 mL of Base (0.5 M KOH)
[44:15] Stepping pH = 8.68
[44:15] Dispensed 0.000047 mL of Base (0.5 M KOH)
[44:20] Stepping pH = 8.76
[44:20] Dispensed 0.000047 mL of Base (0.5 M KOH)
[44:25] Stepping pH = 8.85
[44:25] Dispensed 0.000024 mL of Base (0.5 M KOH)
[44:30] Stepping pH = 8.98
[44:46] Stirrer speed set to 0
[45:06] Datapoint id 44 collected
[45:06] Charge balance equation is out by -572.8%
[45:06] Titration 3 of 3
[45:06] Adding initial titrants
[45:06] Automatically add 0.10000 mL of Octanol
[45:08] Dispensed 0.100000 mL of Octanol
[45:08] Stirrer speed set to 10
[45:10] Stirrer speed set to 60

| | | | |
|---|---|---|---|
| Sample name: | **M09_octanol** | Experiment start time: | **3/2/2018 10:00:20 PM** |
| Assay name: | **pH-metric high logP** | Analyst: | **Pion** |
| Assay ID: | **18C-02010** | Instrument ID: | **T312060** |
| Filename: | **C:\Sirius_T3\Mehtap\20180302_exp29_logP_T3-2\18C-02010_M09_octanol_pH-metric high logP.t3r** | | |

#### Experiment Log (continued)

[45:10] Iterative adjust 9.09 -> 2.00
[45:10] pH 9.09 -> 2.00
[45:11] Dispensed 0.051529 mL of Acid (0.5 M HCl)
[45:16] pH 2.06 -> 2.00
[45:16] Dispensed 0.006326 mL of Acid (0.5 M HCl)
[46:07] Stirrer speed set to 0
[46:17] Datapoint id 45 collected
[46:17] Stirrer speed set to 60
[46:22] pH 1.97 -> 2.17
[46:22] Using cautious pH adjust
[46:22] Dispensed 0.009431 mL of Base (0.5 M KOH)
[46:27] Stepping pH = 2.04
[46:27] Dispensed 0.009525 mL of Base (0.5 M KOH)
[46:32] Stepping pH = 2.15
[46:33] Dispensed 0.001623 mL of Base (0.5 M KOH)
[46:38] Stepping pH = 2.18
[46:53] Stirrer speed set to 0
[47:03] Datapoint id 46 collected
[47:03] Charge balance equation is out by -9.2%
[47:03] Stirrer speed set to 60
[47:08] pH 2.19 -> 2.39
[47:08] Using charge balance adjust
[47:08] Dispensed 0.011430 mL of Base (0.5 M KOH)
[47:28] Stirrer speed set to 0
[47:38] Datapoint id 47 collected
[47:38] Charge balance equation is out by -4.3%
[47:38] Stirrer speed set to 60
[47:44] pH 2.39 -> 2.59
[47:44] Using charge balance adjust
[47:44] Dispensed 0.007385 mL of Base (0.5 M KOH)
[48:04] Stirrer speed set to 0
[48:15] Datapoint id 48 collected
[48:15] Charge balance equation is out by -1.9%
[48:15] Stirrer speed set to 60
[48:20] pH 2.59 -> 2.79
[48:20] Using charge balance adjust
[48:20] Dispensed 0.004821 mL of Base (0.5 M KOH)
[48:40] Stirrer speed set to 0
[48:56] Datapoint id 49 collected
[48:56] Charge balance equation is out by 5.6%
[48:56] Stirrer speed set to 60
[49:01] pH 2.81 -> 3.01
[49:01] Using charge balance adjust
[49:01] Dispensed 0.003198 mL of Base (0.5 M KOH)
[49:21] Stirrer speed set to 0
[49:36] Datapoint id 50 collected
[49:36] Charge balance equation is out by -4.8%
[49:36] Stirrer speed set to 60
[49:41] pH 3.00 -> 3.20
[49:41] Using charge balance adjust
[49:41] Dispensed 0.002328 mL of Base (0.5 M KOH)
[50:01] Stirrer speed set to 0
[50:11] Datapoint id 51 collected
[50:11] Charge balance equation is out by -14.6%
[50:11] Stirrer speed set to 60
[50:16] pH 3.18 -> 3.38
[50:16] Using charge balance adjust
[50:16] Dispensed 0.001834 mL of Base (0.5 M KOH)
[50:37] Stirrer speed set to 0

| | | | |
|---|---|---|---|
| Sample name: | **M09_octanol** | Experiment start time: | **3/2/2018 10:00:20 PM** |
| Assay name: | **pH-metric high logP** | Analyst: | **Pion** |
| Assay ID: | **18C-02010** | Instrument ID: | **T312060** |
| Filename: | **C:\Sirius_T3\Mehtap\20180302_exp29_logP_T3-2\18C-02010_M09_octanol_pH-metric high logP.t3r** | | |

#### Experiment Log (continued)

[50:47] Datapoint id 52 collected
[50:47] Charge balance equation is out by -5.0%
[50:47] Stirrer speed set to 60
[50:52] pH 3.37 -> 3.57
[50:52] Using charge balance adjust
[50:52] Dispensed 0.001435 mL of Base (0.5 M KOH)
[51:12] Stirrer speed set to 0
[51:22] Datapoint id 53 collected
[51:22] Charge balance equation is out by -4.5%
[51:22] Stirrer speed set to 60
[51:27] pH 3.57 -> 3.77
[51:27] Using charge balance adjust
[51:27] Dispensed 0.001129 mL of Base (0.5 M KOH)
[51:47] Stirrer speed set to 0
[51:57] Datapoint id 54 collected
[51:57] Charge balance equation is out by 5.2%
[51:57] Stirrer speed set to 60
[52:03] pH 3.78 -> 3.98
[52:03] Using charge balance adjust
[52:03] Dispensed 0.000823 mL of Base (0.5 M KOH)
[52:23] Stirrer speed set to 0
[52:37] Datapoint id 55 collected
[52:37] Charge balance equation is out by -5.8%
[52:37] Stirrer speed set to 60
[52:43] pH 3.98 -> 4.18
[52:43] Using charge balance adjust
[52:43] Dispensed 0.000611 mL of Base (0.5 M KOH)
[53:03] Stirrer speed set to 0
[53:13] Datapoint id 56 collected
[53:13] Charge balance equation is out by -17.3%
[53:13] Stirrer speed set to 60
[53:18] pH 4.15 -> 4.35
[53:18] Using cautious pH adjust
[53:18] Dispensed 0.000235 mL of Base (0.5 M KOH)
[53:23] Stepping pH = 4.19
[53:23] Dispensed 0.000400 mL of Base (0.5 M KOH)
[53:28] Stepping pH = 4.30
[53:28] Dispensed 0.000141 mL of Base (0.5 M KOH)
[53:33] Stepping pH = 4.38
[53:49] Stirrer speed set to 0
[54:05] Datapoint id 57 collected
[54:05] Charge balance equation is out by -71.6%
[54:05] Stirrer speed set to 60
[54:10] pH 4.43 -> 4.63
[54:10] Using cautious pH adjust
[54:10] Dispensed 0.000118 mL of Base (0.5 M KOH)
[54:15] Stepping pH = 4.44
[54:15] Dispensed 0.000353 mL of Base (0.5 M KOH)
[54:20] Stepping pH = 4.61
[54:20] Dispensed 0.000047 mL of Base (0.5 M KOH)
[54:25] Stepping pH = 4.70
[54:40] Stirrer speed set to 0
[54:51] Datapoint id 58 collected
[54:51] Charge balance equation is out by -107.6%
[54:51] Stirrer speed set to 60
[54:56] pH 4.76 -> 4.96
[54:56] Using cautious pH adjust
[54:56] Dispensed 0.000071 mL of Base (0.5 M KOH)
[55:02] Stepping pH = 4.77

| | | | |
|---|---|---|---|
| Sample name: | **M09_octanol** | Experiment start time: | **3/2/2018 10:00:20 PM** |
| Assay name: | **pH-metric high logP** | Analyst: | **Pion** |
| Assay ID: | **18C-02010** | Instrument ID: | **T312060** |
| Filename: | **C:\Sirius_T3\Mehtap\20180302_exp29_logP_T3-2\18C-02010_M09_octanol_pH-metric high logP.t3r** | | |

#### Experiment Log (continued)

[55:02] Dispensed 0.000188 mL of Base (0.5 M KOH)
[55:07] Stepping pH = 4.90
[55:07] Dispensed 0.000071 mL of Base (0.5 M KOH)
[55:12] Stepping pH = 5.01
[55:27] Stirrer speed set to 0
[55:39] Datapoint id 59 collected
[55:39] Charge balance equation is out by -151.8%
[55:39] Stirrer speed set to 60
[55:44] pH 5.13 -> 5.33
[55:44] Using cautious pH adjust
[55:44] Dispensed 0.000024 mL of Base (0.5 M KOH)
[55:49] Stepping pH = 5.14
[55:49] Dispensed 0.000094 mL of Base (0.5 M KOH)
[55:54] Stepping pH = 5.17
[55:54] Dispensed 0.000188 mL of Base (0.5 M KOH)
[55:59] Stepping pH = 5.59
[56:14] Stirrer speed set to 0
[56:57] Datapoint id 60 collected
[56:57] Charge balance equation is out by -376.9%
[56:57] Stirrer speed set to 60
[57:02] pH 6.13 -> 6.33
[57:02] Using cautious pH adjust
[57:02] Dispensed 0.000024 mL of Base (0.5 M KOH)
[57:07] Stepping pH = 6.17
[57:07] Dispensed 0.000047 mL of Base (0.5 M KOH)
[57:12] Stepping pH = 6.29
[57:12] Dispensed 0.000024 mL of Base (0.5 M KOH)
[57:17] Stepping pH = 6.46
[57:32] Stirrer speed set to 0
[58:32] Datapoint id 61 collected
[58:32] Charge balance equation is out by -74.4%
[58:32] Stirrer speed set to 60
[58:37] pH 6.61 -> 6.81
[58:37] Using cautious pH adjust
[58:37] Dispensed 0.000024 mL of Base (0.5 M KOH)
[58:43] Stepping pH = 6.59
[58:43] Dispensed 0.000094 mL of Base (0.5 M KOH)
[58:48] Stepping pH = 6.63
[58:48] Dispensed 0.000212 mL of Base (0.5 M KOH)
[58:53] Stepping pH = 7.68
[59:08] Stirrer speed set to 0
[59:40] Datapoint id 62 collected
[59:40] Charge balance equation is out by -805.8%
[59:40] Stirrer speed set to 60
[59:45] pH 8.62 -> 8.82
[59:45] Using cautious pH adjust
[59:45] Dispensed 0.000024 mL of Base (0.5 M KOH)
[59:50] Stepping pH = 8.65
[59:50] Dispensed 0.000024 mL of Base (0.5 M KOH)
[59:55] Stepping pH = 8.68
[59:55] Dispensed 0.000047 mL of Base (0.5 M KOH)
[1:00:01] Stepping pH = 8.75
[1:00:01] Dispensed 0.000047 mL of Base (0.5 M KOH)
[1:00:06] Stepping pH = 8.87
[1:00:21] Stirrer speed set to 0
[1:00:38] Datapoint id 63 collected
[1:00:38] Charge balance equation is out by -507.2%
[1:00:38] Stirrer speed set to 60
[1:00:43] pH 8.96 -> 9.05

| | | | |
|---|---|---|---|
| Sample name: | **M09_octanol** | Experiment start time: | **3/2/2018 10:00:20 PM** |
| Assay name: | **pH-metric high logP** | Analyst: | **Pion** |
| Assay ID: | **18C-02010** | Instrument ID: | **T312060** |
| Filename: | **C:\Sirius_T3\Mehtap\20180302_exp29_logP_T3-2\18C-02010_M09_octanol_pH-metric high logP.t3r** | | |

#### Experiment Log (continued)

[1:00:43] Using cautious pH adjust
[1:00:43] Dispensed 0.000024 mL of Base (0.5 M KOH)
[1:00:49] Stepping pH = 8.96
[1:00:49] Dispensed 0.000047 mL of Base (0.5 M KOH)
[1:00:54] Stepping pH = 9.00
[1:01:09] Stirrer speed set to 0
[1:01:24] Datapoint id 64 collected
[1:01:24] Charge balance equation is out by -267.3%
[1:01:24] Argon flow rate set to 0
[1:01:28] Titrator arm moved over Titration position
