## Supplementary material for "Octanol-water partition coefficient measurements for the SAMPL6 Blind Prediction Challenge": SM09_18C-03001_M09_octanol_pH-metric high logP_report.pdf

| | | | |
|---|---|---|---|
| Sample name: | **M09_octanol** | Experiment start time: | **3/3/2018 12:18:26 AM** |
| Assay name: | **pH-metric high logP** | Analyst: | **Pion** |
| Assay ID: | **18C-03001** | Instrument ID: | **T312060** |
| Filename: | **C:\Sirius_T3\Mehtap\20180302_exp29_logP_T3-2\18C-03001_M09_octanol_pH-metric high logP.t3r** | | |

#### pH-metric Result

| | | |
|---|---|---|
| logP (XH +) | 1.05 | ±0.04 (n=50) |
| logP (neutral X) | 3.14 | ±0.01 (n=50) |

#### 18C-03001 Points 1 to 25

| | |
|---|---|
| M09_octanol concentration factor | 1.208 |
| Carbonate | 0.0513 mM |
| Acidity error | -0.65776 mM |

#### 18C-03001 Points 26 to 53

| | |
|---|---|
| M09_octanol concentration factor | 1.218 |
| Carbonate | 0.1056 mM |
| Acidity error | -0.76829 mM |

#### 18C-03001 Points 54 to 80

| | |
|---|---|
| M09_octanol concentration factor | 1.072 |
| Carbonate | 0.0742 mM |
| Acidity error | -0.32205 mM |

#### Warnings and errors

| | |
|---|---|
| Errors | None |
| Warnings | None |

#### Sample logD and percent species

| pH | M09_octanol logD | M09_octanol M09_octanolH | M09_octanol M09_octanol | M09_octanol M09_octanolH* | M09_octanol M09_octanol* | Comment |
|---|---|---|---|---|---|---|
| 1.000 | 1.05 | 8.12 % | 0.00 % | 91.40 % | 0.48 % | |
| 1.200 | 1.05 | 8.10 % | 0.00 % | 91.14 % | 0.76 % | Stomach pH |
| 2.000 | 1.07 | 7.78 % | 0.00 % | 87.58 % | 4.63 % | |
| 3.000 | 1.23 | 5.49 % | 0.02 % | 61.80 % | 32.69 % | |
| 4.000 | 1.83 | 1.39 % | 0.06 % | 15.67 % | 82.88 % | |
| 5.000 | 2.63 | 0.16 % | 0.07 % | 1.85 % | 97.91 % | |
| 6.000 | 3.05 | 0.02 % | 0.07 % | 0.19 % | 99.72 % | |
| 6.500 | 3.11 | 0.01 % | 0.07 % | 0.06 % | 99.86 % | |
| 7.000 | 3.13 | 0.00 % | 0.07 % | 0.02 % | 99.91 % | |
| 7.400 | 3.14 | 0.00 % | 0.07 % | 0.01 % | 99.92 % | Blood pH |
| 8.000 | 3.14 | 0.00 % | 0.07 % | 0.00 % | 99.93 % | |
| 9.000 | 3.14 | 0.00 % | 0.07 % | 0.00 % | 99.93 % | |
| 10.000 | 3.14 | 0.00 % | 0.07 % | 0.00 % | 99.93 % | |
| 11.000 | 3.14 | 0.00 % | 0.07 % | 0.00 % | 99.93 % | |
| 12.000 | 3.14 | 0.00 % | 0.07 % | 0.00 % | 99.93 % | |

| | | | |
|---|---|---|---|
| Sample name: | **M09_octanol** | Experiment start time: | **3/3/2018 12:18:26 AM** |
| Assay name: | **pH-metric high logP** | Analyst: | **Pion** |
| Assay ID: | **18C-03001** | Instrument ID: | **T312060** |
| Filename: | **C:\Sirius_T3\Mehtap\20180302_exp29_logP_T3-2\18C-03001_M09_octanol_pH-metric high logP.t3r** | | |

#### Graphs

Charge balance residuals

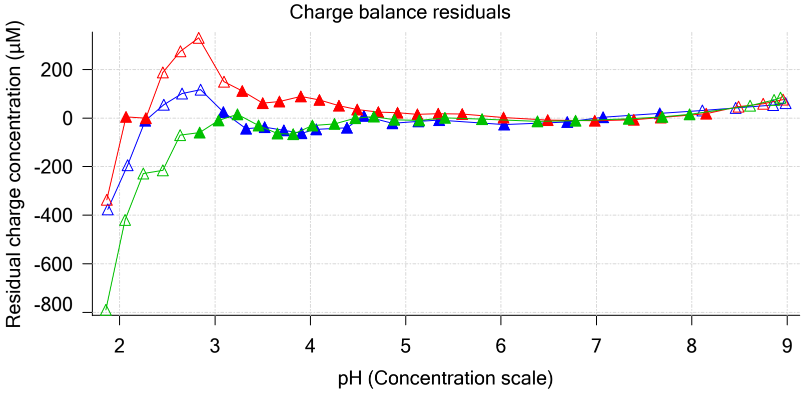


Ionic strength

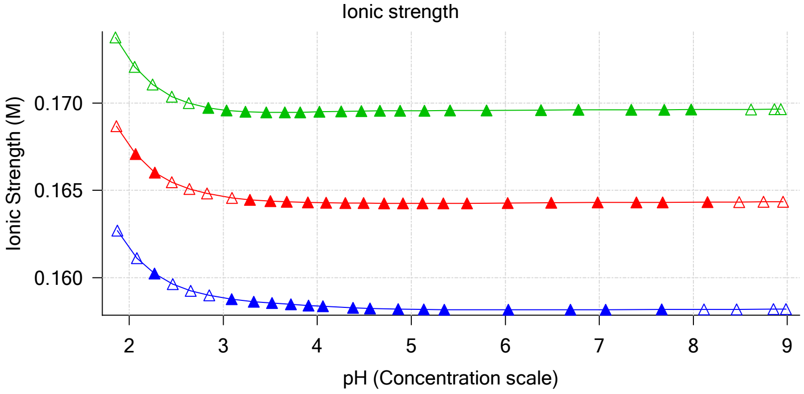


Buffer index profile

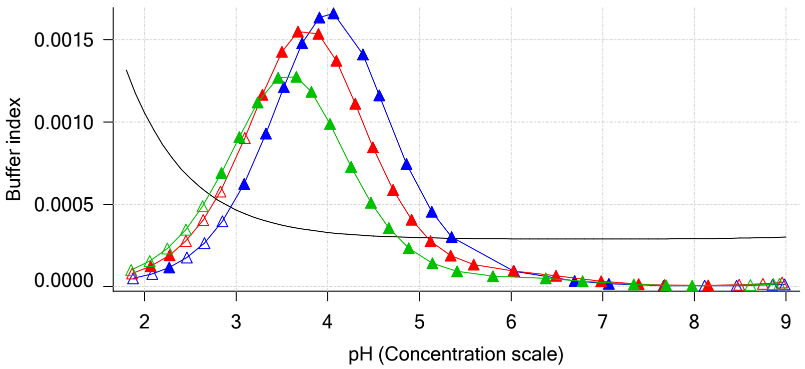


Sample M09_octanol Ionisation Graph

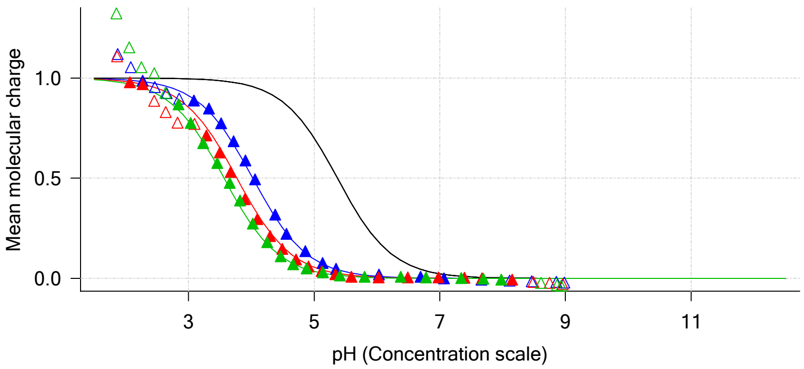


Distribution of species for sample M09_octanol

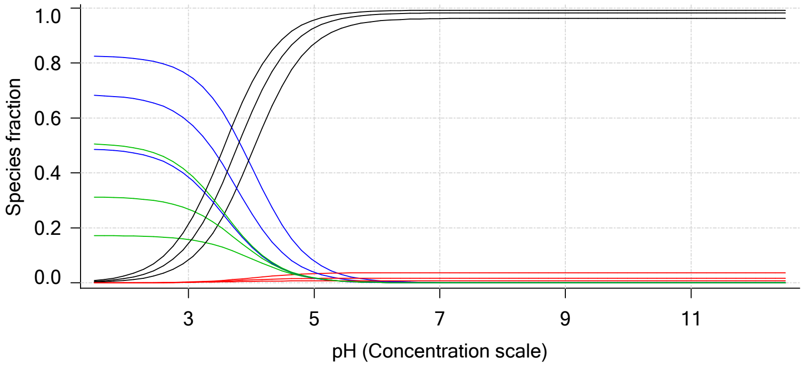


Lipophilicity Profile

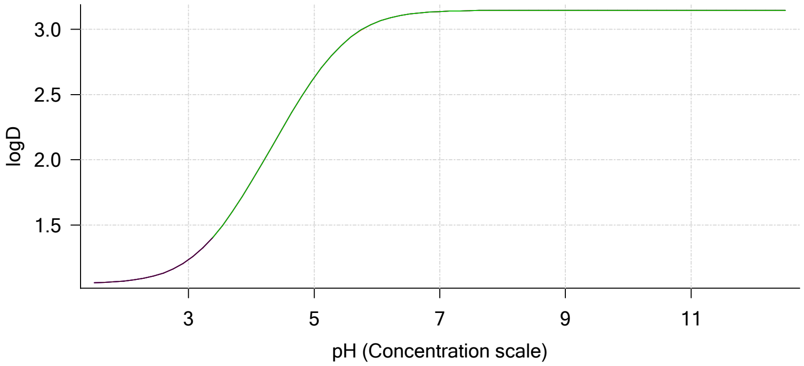


Log R Profile

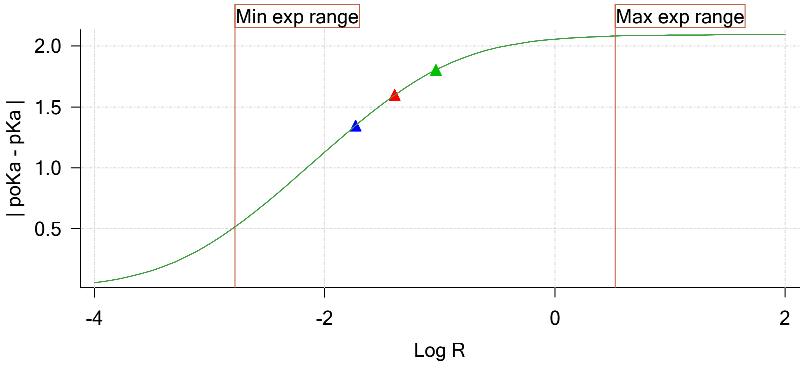


Sample Carbonate Ionisation Graph

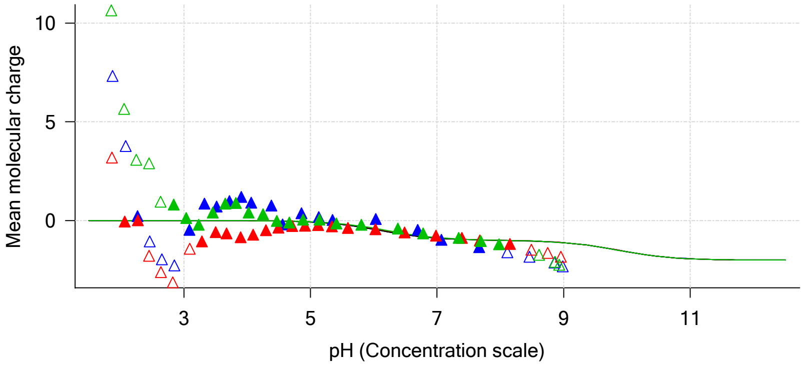

| | | | |
|---|---|---|---|
| Sample name: | **M09_octanol** | Experiment start time: | **3/3/2018 12:18:26 AM** |
| Assay name: | **pH-metric high logP** | Analyst: | **Pion** |
| Assay ID: | **18C-03001** | Instrument ID: | **T312060** |
| Filename: | **C:\Sirius_T3\Mehtap\20180302_exp29_logP_T3-2\18C-03001_M09_octanol_pH-metric high logP.t3r** | | |

#### Graphs (continued)

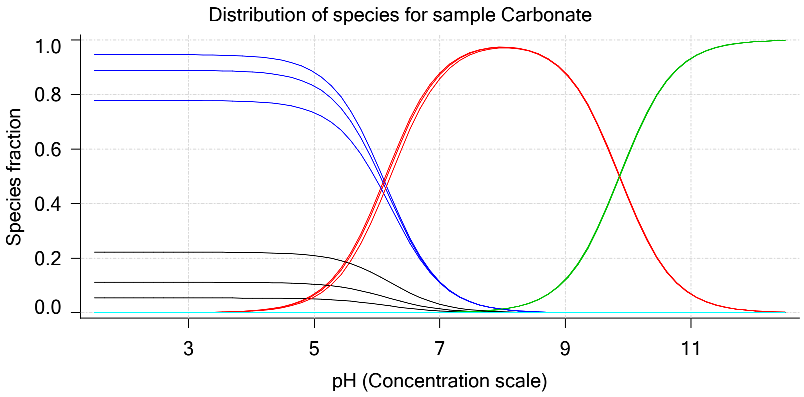

| | | | |
|---|---|---|---|
| Sample name: | **M09_octanol** | Experiment start time: | **3/3/2018 12:18:26 AM** |
| Assay name: | **pH-metric high logP** | Analyst: | **Pion** |
| Assay ID: | **18C-03001** | Instrument ID: | **T312060** |
| Filename: | **C:\Sirius_T3\Mehtap\20180302_exp29_logP_T3-2\18C-03001_M09_octanol_pH-metric high logP.t3r** | | |

### pH-metric high logP Titration 1 of 3 18C-03001 Points 1 to 25

#### Overall results

| | |
|---|---|
| RMSD | 0.656 |
| Average ionic strength | 0.158 M |
| Average temperature | 25.0°C |
| Partition ratio | 0.0185 : 1 |
| Analyte concentration range | 2340.8 µM to 2417.9 µM |
| Total points considered | 16 of 25 |

#### Warnings and errors

| | |
|---|---|
| Errors | None |
| Warnings | None |

#### Four-Plus parameters

| | | | |
|---|---|---|---|
| Alpha | 0.111 | 3/3/2018 12:18:25 AM | C:\Sirius_T3\HCl18C02.t3r |
| S | 0.9988 | 3/3/2018 12:18:25 AM | C:\Sirius_T3\HCl18C02.t3r |
| jH | 1.0 | 3/3/2018 12:18:25 AM | C:\Sirius_T3\HCl18C02.t3r |
| jOH | -0.8 | 3/3/2018 12:18:25 AM | C:\Sirius_T3\HCl18C02.t3r |

#### Titrants

| | | | |
|---|---|---|---|
| 0.50 M HCl | 0.999058 | 3/3/2018 12:18:25 AM | C:\Sirius_T3\HCl18C02.t3r |
| 0.50 M KOH | 0.999845 | 3/3/2018 12:18:26 AM | C:\Sirius_T3\KOH18B27.t3r |

#### Sample

| | |
|---|---|
| M09_octanol concentration factor | 1.208 |
| M09_octanol stoichiometry | 1.000 |
| Chloride stoichiometry | 1.000 |
| Base pKa 1 | 5.37 |
| logP (XH +) | 0.87 |
| logP (neutral X) | 3.09 |

#### Sample graphs

Ionisation of sample M09_octanol

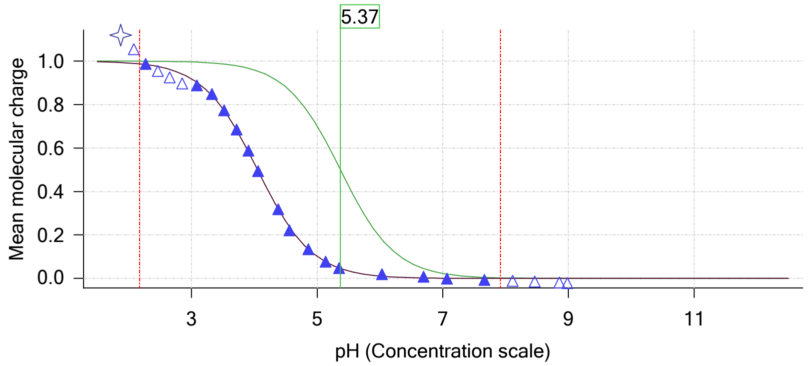


Distribution of species for sample M09_octanol

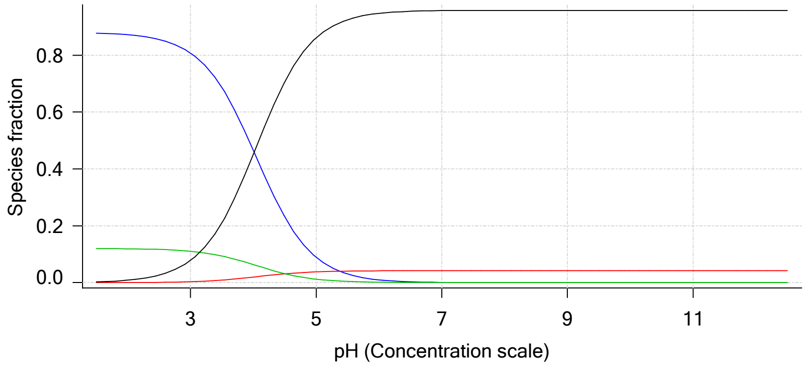

Sample name: **M09_octanol**
Assay name: **pH-metric high logP**
Assay ID: **18C-03001**
Filename: **C:\Sirius_T3\Mehtap\20180302_exp29_logP_T3-2\18C-03001_M09_octanol_pH-metric high logP.t3r**

Experiment start time: **3/3/2018 12:18:26 AM**
Analyst: **Pion**
Instrument ID: **T312060**

#### Sample graphs (continued)

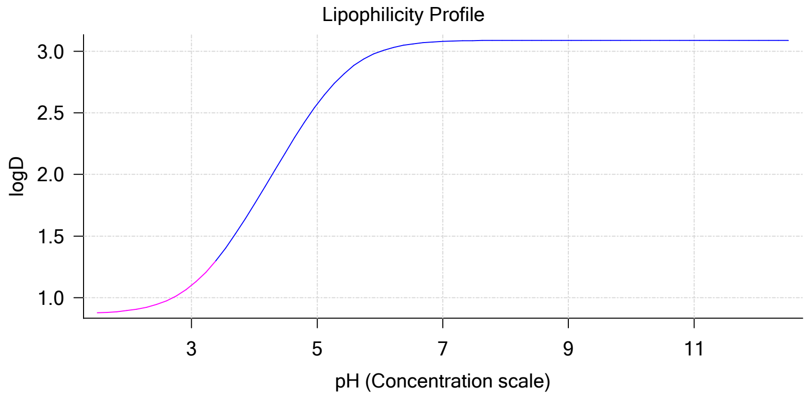


#### Sample logD and percent species

| pH | M09_octanol logD | M09_octanol M09_octanolH | M09_octanol M09_octanol | M09_octanol M09_octanolH* | M09_octanol M09_octanol* | Comment |
|---|---|---|---|---|---|---|
| 1.000 | 0.87 | 87.91 % | 0.00 % | 12.00 % | 0.09 % | |
| 1.200 | 0.87 | 87.86 % | 0.01 % | 12.00 % | 0.13 % | Stomach pH |
| 2.000 | 0.90 | 87.21 % | 0.04 % | 11.91 % | 0.84 % | |
| 3.000 | 1.10 | 80.80 % | 0.34 % | 11.03 % | 7.83 % | |
| 4.000 | 1.76 | 46.56 % | 1.99 % | 6.36 % | 45.10 % | |
| 5.000 | 2.57 | 8.89 % | 3.79 % | 1.21 % | 86.10 % | |
| 6.000 | 3.00 | 0.98 % | 4.17 % | 0.13 % | 94.72 % | |
| 6.500 | 3.06 | 0.31 % | 4.20 % | 0.04 % | 95.44 % | |
| 7.000 | 3.08 | 0.10 % | 4.21 % | 0.01 % | 95.67 % | |
| 7.400 | 3.08 | 0.04 % | 4.22 % | 0.01 % | 95.74 % | Blood pH |
| 8.000 | 3.09 | 0.01 % | 4.22 % | 0.00 % | 95.77 % | |
| 9.000 | 3.09 | 0.00 % | 4.22 % | 0.00 % | 95.78 % | |
| 10.000 | 3.09 | 0.00 % | 4.22 % | 0.00 % | 95.78 % | |
| 11.000 | 3.09 | 0.00 % | 4.22 % | 0.00 % | 95.78 % | |
| 12.000 | 3.09 | 0.00 % | 4.22 % | 0.00 % | 95.78 % | |

#### Carbonate and acidity

| | |
|---|---|
| Carbonate | 0.051 mM |
| Acidity error | -0.658 mM |

#### Other graphs

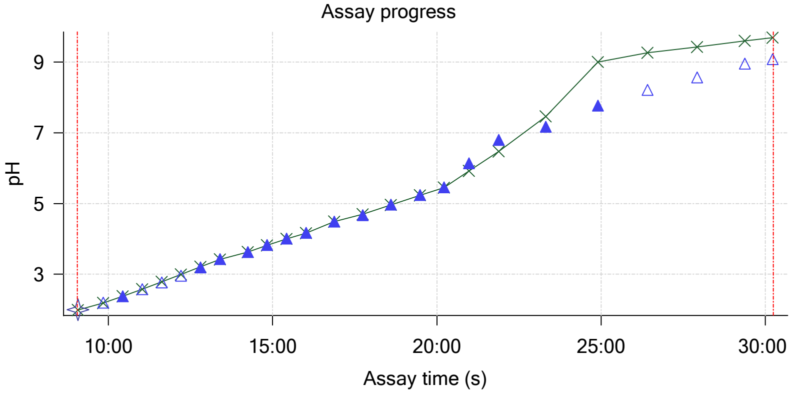


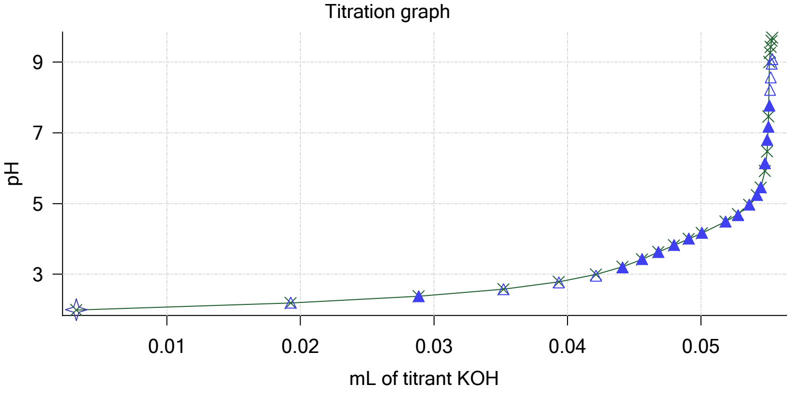

| | | | |
|---|---|---|---|
| Sample name: | **M09_octanol** | Experiment start time: | **3/3/2018 12:18:26 AM** |
| Assay name: | **pH-metric high logP** | Analyst: | **Pion** |
| Assay ID: | **18C-03001** | Instrument ID: | **T312060** |
| Filename: | **C:\Sirius_T3\Mehtap\20180302_exp29_logP_T3-2\18C-03001_M09_octanol_pH-metric high logP.t3r** | | |

#### Other graphs (continued)

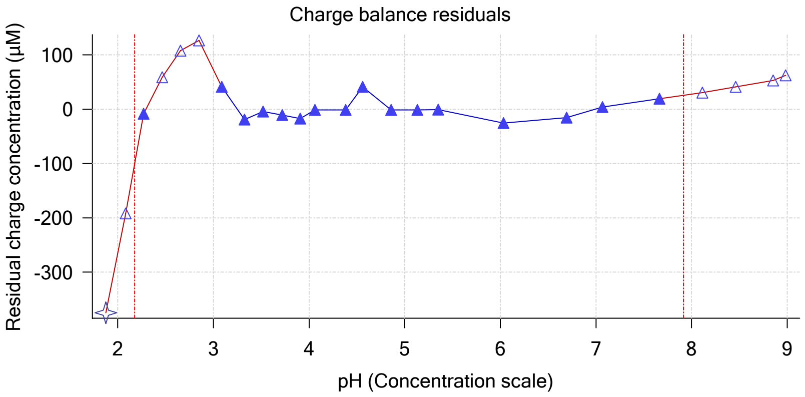


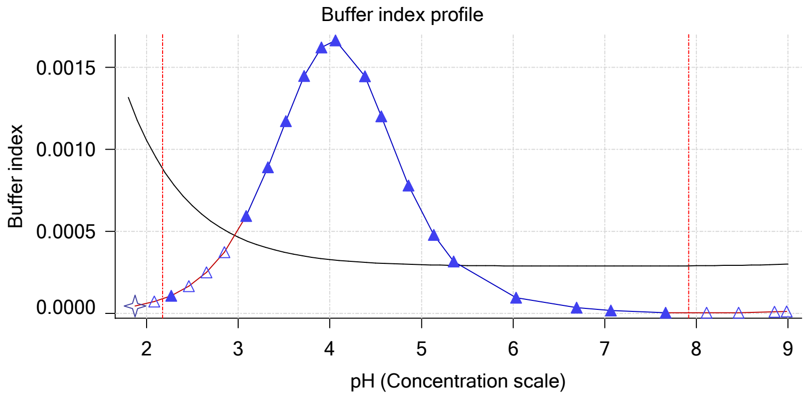


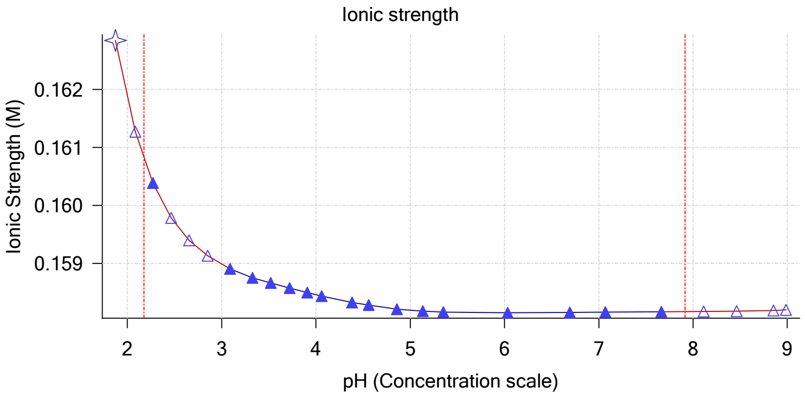


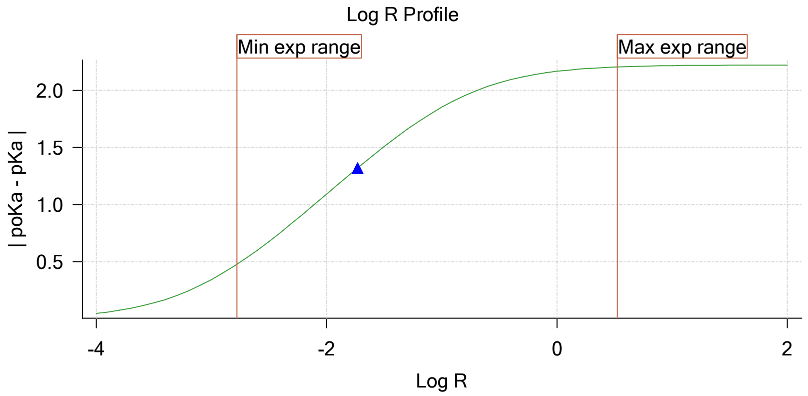


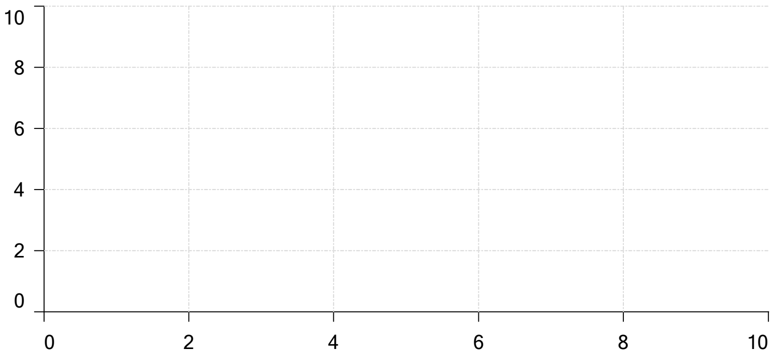


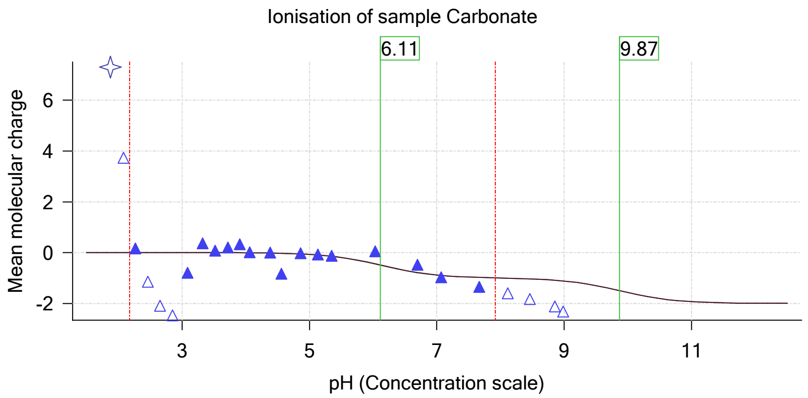

| | | | |
|---|---|---|---|
| Sample name: | **M09_octanol** | Experiment start time: | **3/3/2018 12:18:26 AM** |
| Assay name: | **pH-metric high logP** | Analyst: | **Pion** |
| Assay ID: | **18C-03001** | Instrument ID: | **T312060** |
| Filename: | **C:\Sirius_T3\Mehtap\20180302_exp29_logP_T3-2\18C-03001_M09_octanol_pH-metric high logP.t3r** | | |

# pH-metric high logP   Titration 2 of 3   18C-03001 Points 26 to 53

#### Overall results

| | |
|---|---|
| RMSD | 0.344 |
| Average ionic strength | 0.165 M |
| Average temperature | 25.0°C |
| Partition ratio | 0.0407 : 1 |
| Analyte concentration range | 2140.2 µM to 2210.0 µM |
| Total points considered | 20 of 28 |

#### Warnings and errors

| | |
|---|---|
| Errors | None |
| Warnings | None |

#### Four-Plus parameters

| | | | |
|---|---|---|---|
| Alpha | 0.111 | 3/3/2018 12:18:25 AM | C:\Sirius_T3\HCl18C02.t3r |
| S | 0.9988 | 3/3/2018 12:18:25 AM | C:\Sirius_T3\HCl18C02.t3r |
| jH | 1.0 | 3/3/2018 12:18:25 AM | C:\Sirius_T3\HCl18C02.t3r |
| jOH | -0.8 | 3/3/2018 12:18:25 AM | C:\Sirius_T3\HCl18C02.t3r |

#### Titrants

| | | | |
|---|---|---|---|
| 0.50 M HCl | 0.999058 | 3/3/2018 12:18:25 AM | C:\Sirius_T3\HCl18C02.t3r |
| 0.50 M KOH | 0.999845 | 3/3/2018 12:18:26 AM | C:\Sirius_T3\KOH18B27.t3r |

#### Sample

| | |
|---|---|
| M09_octanol concentration factor | 1.218 |
| M09_octanol stoichiometry | 1.000 |
| Chloride stoichiometry | 1.000 |
| Base pKa 1 | 5.37 |
| logP (XH +) | 0.73 |
| logP (neutral X) | 3.12 |

#### Sample graphs

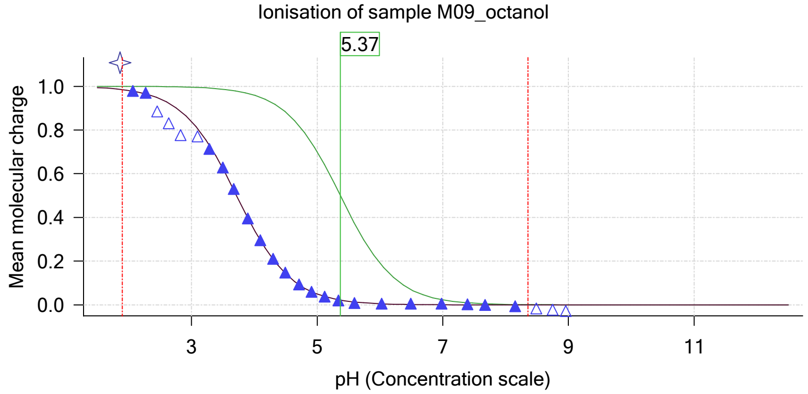


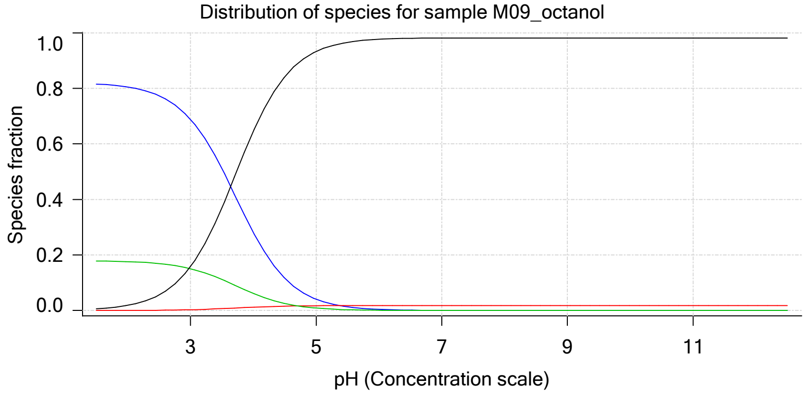

| | | | |
|---|---|---|---|
| Sample name: | **M09_octanol** | Experiment start time: | **3/3/2018 12:18:26 AM** |
| Assay name: | **pH-metric high logP** | Analyst: | **Pion** |
| Assay ID: | **18C-03001** | Instrument ID: | **T312060** |
| Filename: | **C:\Sirius_T3\Mehtap\20180302_exp29_logP_T3-2\18C-03001_M09_octanol_pH-metric high logP.t3r** | | |

#### Sample graphs (continued)

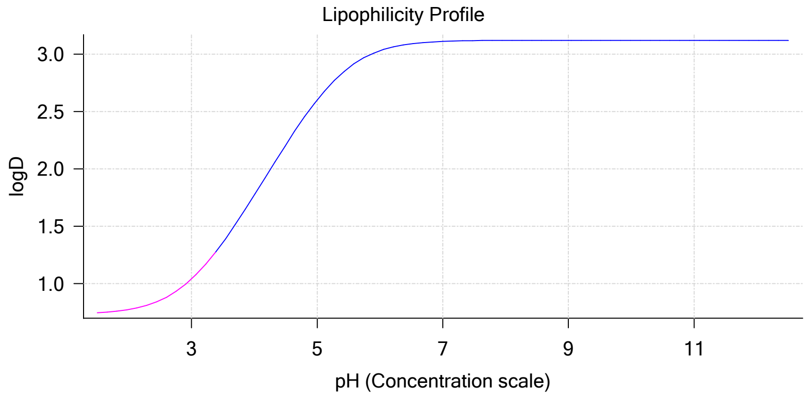


#### Sample logD and percent species

| pH | M09_octanol logD | M09_octanol M09_octanolH | M09_octanol M09_octanol | M09_octanol M09_octanolH* | M09_octanol M09_octanol* | Comment |
|---|---|---|---|---|---|---|
| 1.000 | 0.74 | 81.90 % | 0.00 % | 17.91 % | 0.19 % | |
| 1.200 | 0.74 | 81.80 % | 0.01 % | 17.89 % | 0.30 % | Stomach pH |
| 2.000 | 0.77 | 80.51 % | 0.03 % | 17.61 % | 1.85 % | |
| 3.000 | 1.04 | 68.86 % | 0.29 % | 15.06 % | 15.78 % | |
| 4.000 | 1.77 | 28.14 % | 1.20 % | 6.16 % | 64.50 % | |
| 5.000 | 2.60 | 4.07 % | 1.74 % | 0.89 % | 93.30 % | |
| 6.000 | 3.03 | 0.43 % | 1.82 % | 0.09 % | 97.66 % | |
| 6.500 | 3.09 | 0.14 % | 1.82 % | 0.03 % | 98.01 % | |
| 7.000 | 3.11 | 0.04 % | 1.83 % | 0.01 % | 98.12 % | |
| 7.400 | 3.12 | 0.02 % | 1.83 % | 0.00 % | 98.15 % | Blood pH |
| 8.000 | 3.12 | 0.00 % | 1.83 % | 0.00 % | 98.17 % | |
| 9.000 | 3.12 | 0.00 % | 1.83 % | 0.00 % | 98.17 % | |
| 10.000 | 3.12 | 0.00 % | 1.83 % | 0.00 % | 98.17 % | |
| 11.000 | 3.12 | 0.00 % | 1.83 % | 0.00 % | 98.17 % | |
| 12.000 | 3.12 | 0.00 % | 1.83 % | 0.00 % | 98.17 % | |

#### Carbonate and acidity

| | |
|---|---|
| Carbonate | 0.106 mM |
| Acidity error | -0.768 mM |

#### Other graphs

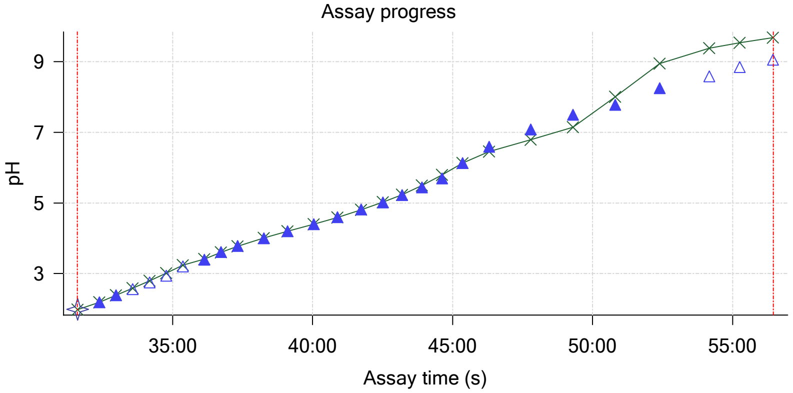


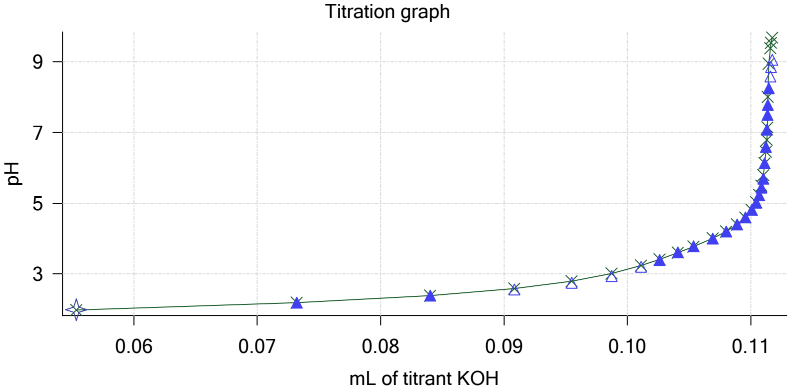

| | | | |
|---|---|---|---|
| Sample name: | **M09_octanol** | Experiment start time: | **3/3/2018 12:18:26 AM** |
| Assay name: | **pH-metric high logP** | Analyst: | **Pion** |
| Assay ID: | **18C-03001** | Instrument ID: | **T312060** |
| Filename: | **C:\Sirius_T3\Mehtap\20180302_exp29_logP_T3-2\18C-03001_M09_octanol_pH-metric high logP.t3r** | | |

#### Other graphs (continued)

Charge balance residuals

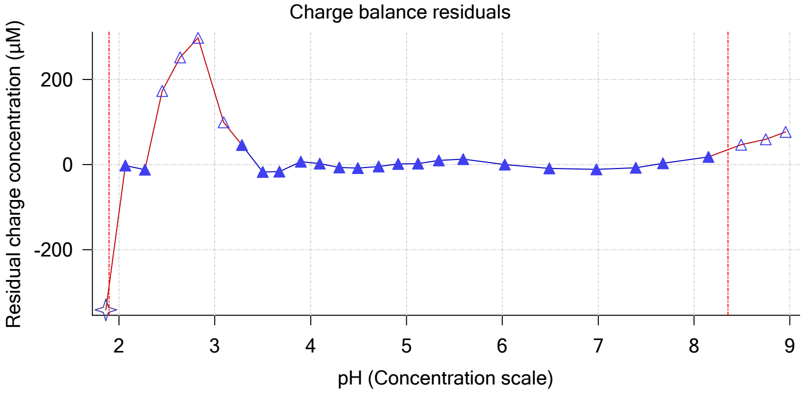


Buffer index profile

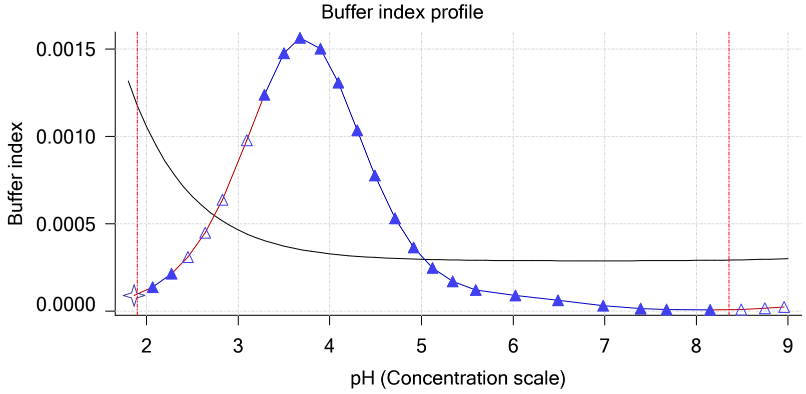


Ionic strength

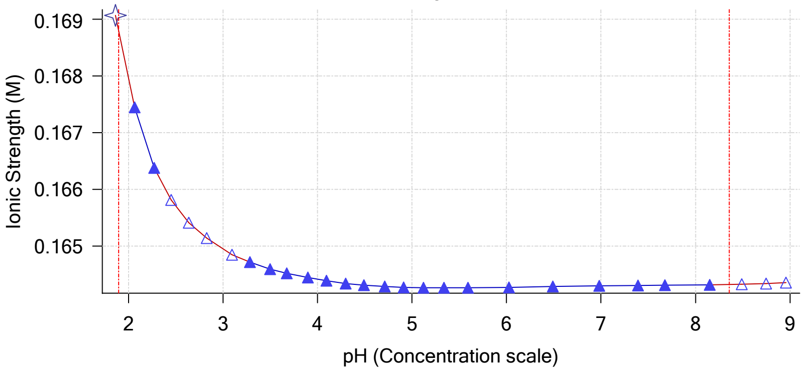


Log R Profile

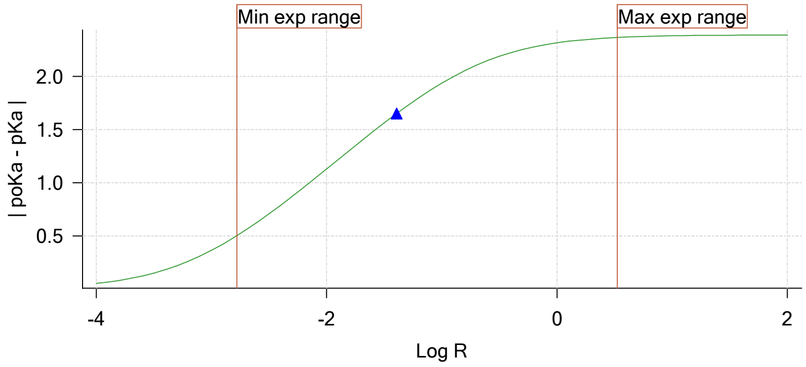


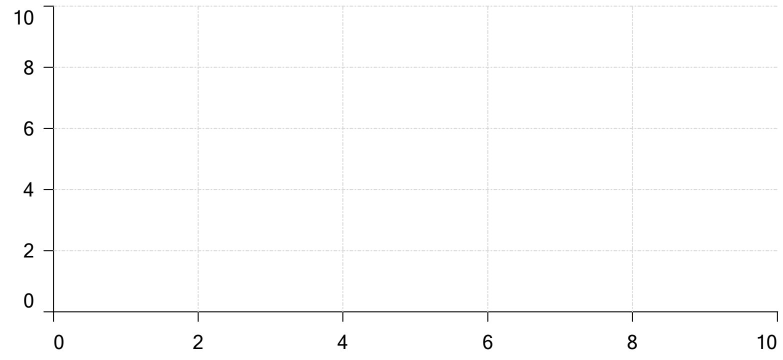


Ionisation of sample Carbonate

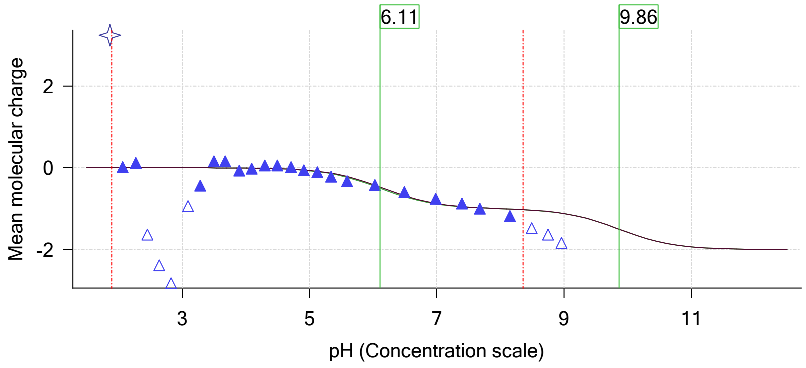

| | | | |
|---|---|---|---|
| Sample name: | **M09_octanol** | Experiment start time: | **3/3/2018 12:18:26 AM** |
| Assay name: | **pH-metric high logP** | Analyst: | **Pion** |
| Assay ID: | **18C-03001** | Instrument ID: | **T312060** |
| Filename: | **C:\Sirius_T3\Mehtap\20180302_exp29_logP_T3-2\18C-03001_M09_octanol_pH-metric high logP.t3r** | | |

### pH-metric high logP Titration 3 of 3 18C-03001 Points 54 to 80

#### Overall results

| | |
|---|---|
| RMSD | 0.436 |
| Average ionic strength | 0.170 M |
| Average temperature | 25.0°C |
| Partition ratio | 0.0924 : 1 |
| Analyte concentration range | 1903.5 μM to 1963.3 μM |
| Total points considered | 19 of 27 |

#### Warnings and errors

| | |
|---|---|
| Errors | None |
| Warnings | None |

#### Four-Plus parameters

| | | | |
|---|---|---|---|
| Alpha | 0.111 | 3/3/2018 12:18:25 AM | C:\Sirius_T3\HCl18C02.t3r |
| S | 0.9988 | 3/3/2018 12:18:25 AM | C:\Sirius_T3\HCl18C02.t3r |
| jH | 1.0 | 3/3/2018 12:18:25 AM | C:\Sirius_T3\HCl18C02.t3r |
| jOH | -0.8 | 3/3/2018 12:18:25 AM | C:\Sirius_T3\HCl18C02.t3r |

#### Titrants

| | | | |
|---|---|---|---|
| 0.50 M HCl | 0.999058 | 3/3/2018 12:18:25 AM | C:\Sirius_T3\HCl18C02.t3r |
| 0.50 M KOH | 0.999845 | 3/3/2018 12:18:26 AM | C:\Sirius_T3\KOH18B27.t3r |

#### Sample

| | |
|---|---|
| M09_octanol concentration factor | 1.072 |
| M09_octanol stoichiometry | 1.000 |
| Chloride stoichiometry | 1.000 |
| Base pKa 1 | 5.37 |
| logP (XH +) | 0.75 |
| logP (neutral X) | 2.98 |

#### Sample graphs

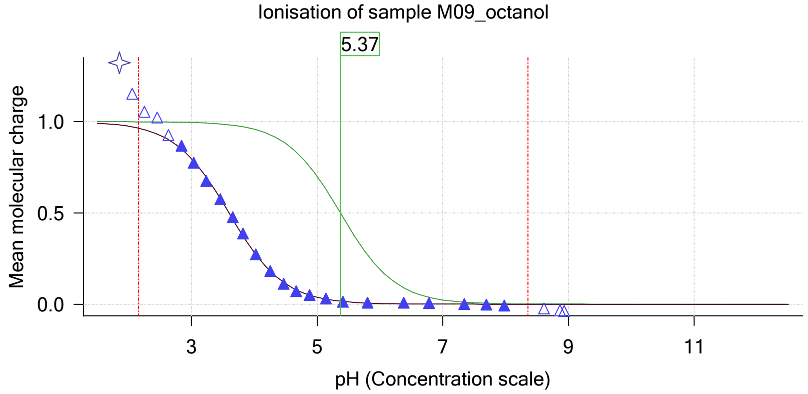


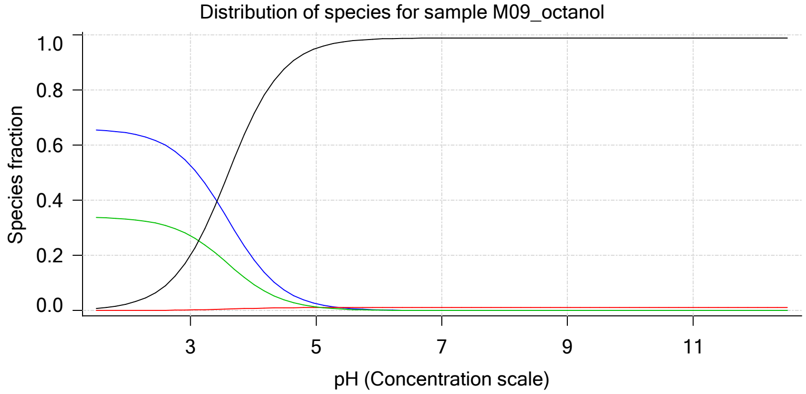

Sample name: **M09_octanol**
Assay name: **pH-metric high logP**
Assay ID: **18C-03001**
Filename: **C:\Sirius_T3\Mehtap\20180302_exp29_logP_T3-2\18C-03001_M09_octanol_pH-metric high logP.t3r**

Experiment start time: **3/3/2018 12:18:26 AM**
Analyst: **Pion**
Instrument ID: **T312060**

#### Sample graphs (continued)

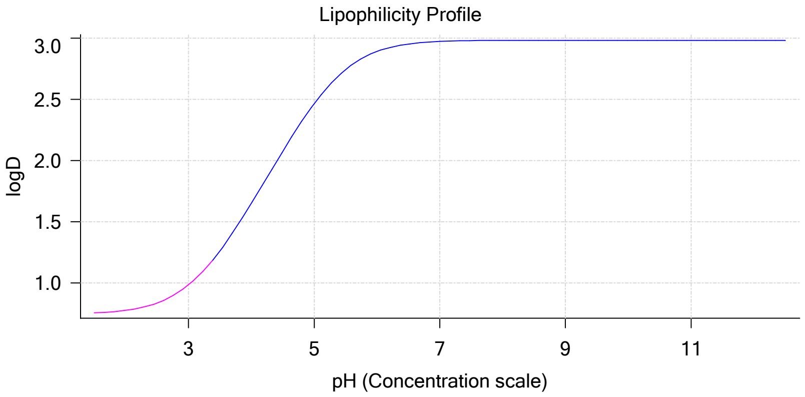


#### Sample logD and percent species

| pH | M09_octanol logD | M09_octanol M09_octanolH | M09_octanol M09_octanol | M09_octanol M09_octanolH* | M09_octanol M09_octanol* | Comment |
|---|---|---|---|---|---|---|
| 1.000 | 0.75 | 65.86 % | 0.00 % | 33.89 % | 0.25 % | |
| 1.200 | 0.75 | 65.76 % | 0.00 % | 33.84 % | 0.40 % | Stomach pH |
| 2.000 | 0.78 | 64.40 % | 0.03 % | 33.13 % | 2.44 % | |
| 3.000 | 0.98 | 52.69 % | 0.22 % | 27.11 % | 19.97 % | |
| 4.000 | 1.65 | 18.70 % | 0.80 % | 9.62 % | 70.88 % | |
| 5.000 | 2.46 | 2.51 % | 1.07 % | 1.29 % | 95.13 % | |
| 6.000 | 2.89 | 0.26 % | 1.11 % | 0.13 % | 98.50 % | |
| 6.500 | 2.95 | 0.08 % | 1.11 % | 0.04 % | 98.76 % | |
| 7.000 | 2.97 | 0.03 % | 1.11 % | 0.01 % | 98.85 % | |
| 7.400 | 2.98 | 0.01 % | 1.11 % | 0.01 % | 98.87 % | Blood pH |
| 8.000 | 2.98 | 0.00 % | 1.11 % | 0.00 % | 98.88 % | |
| 9.000 | 2.98 | 0.00 % | 1.11 % | 0.00 % | 98.89 % | |
| 10.000 | 2.98 | 0.00 % | 1.11 % | 0.00 % | 98.89 % | |
| 11.000 | 2.98 | 0.00 % | 1.11 % | 0.00 % | 98.89 % | |
| 12.000 | 2.98 | 0.00 % | 1.11 % | 0.00 % | 98.89 % | |

#### Carbonate and acidity

| | |
|---|---|
| Carbonate | 0.074 mM |
| Acidity error | -0.322 mM |

#### Other graphs

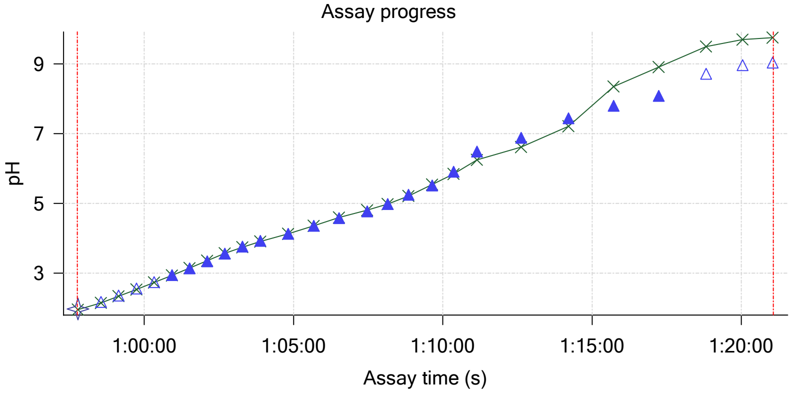


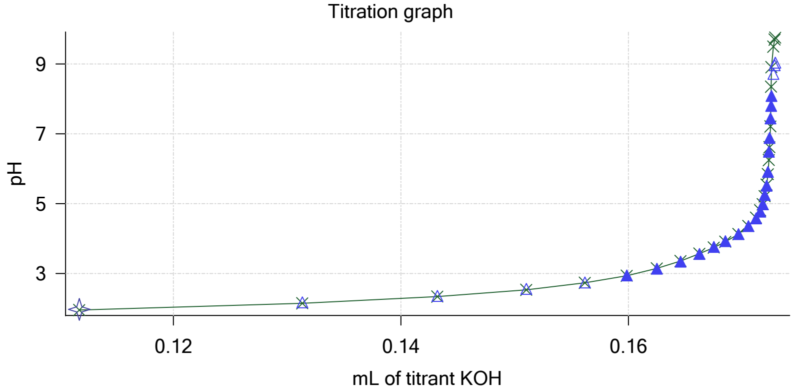

| | | | |
|---|---|---|---|
| Sample name: | **M09_octanol** | Experiment start time: | **3/3/2018 12:18:26 AM** |
| Assay name: | **pH-metric high logP** | Analyst: | **Pion** |
| Assay ID: | **18C-03001** | Instrument ID: | **T312060** |
| Filename: | **C:\Sirius_T3\Mehtap\20180302_exp29_logP_T3-2\18C-03001_M09_octanol_pH-metric high logP.t3r** | | |

#### Other graphs (continued)

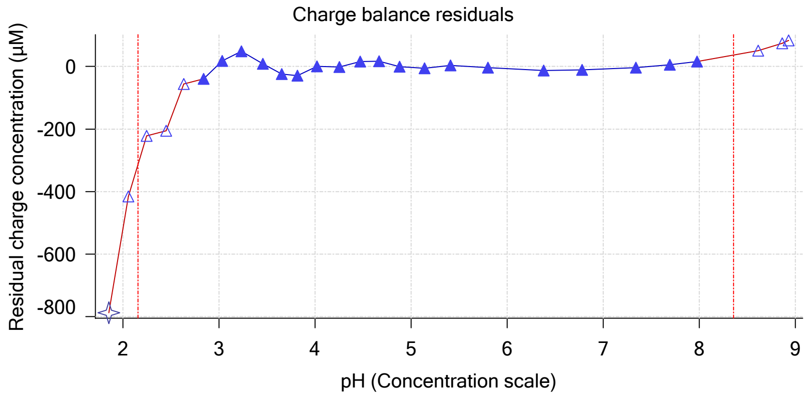


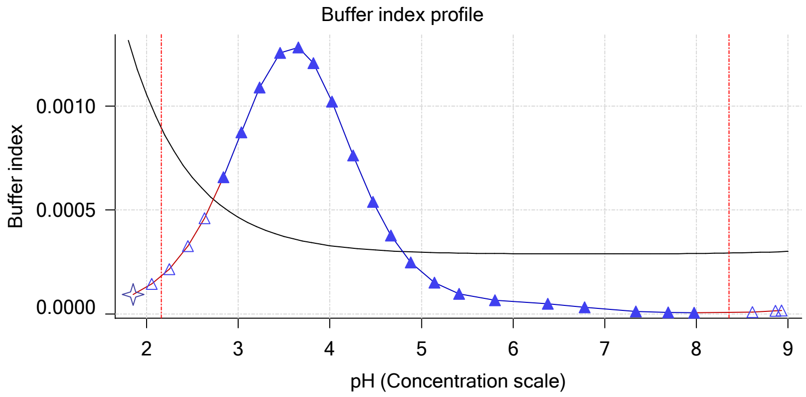


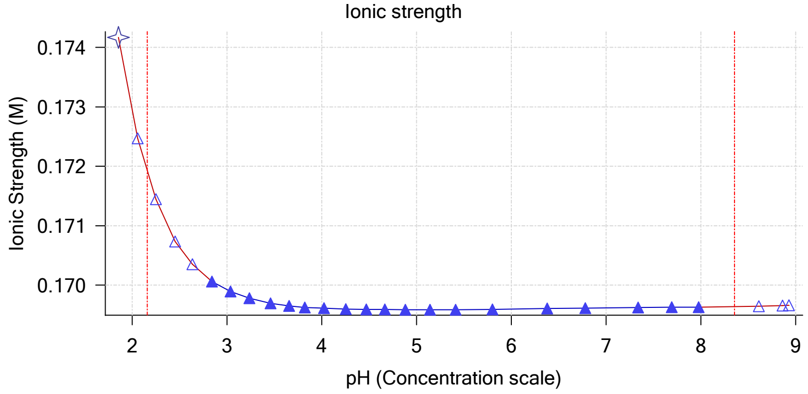


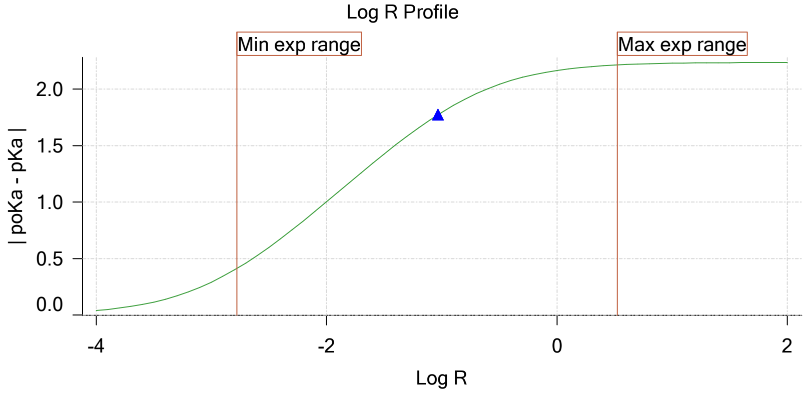


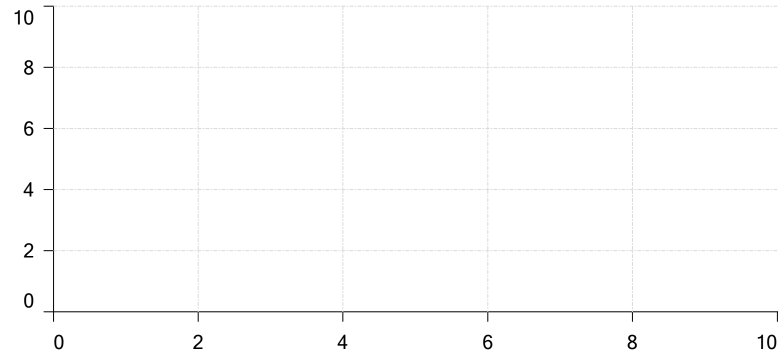

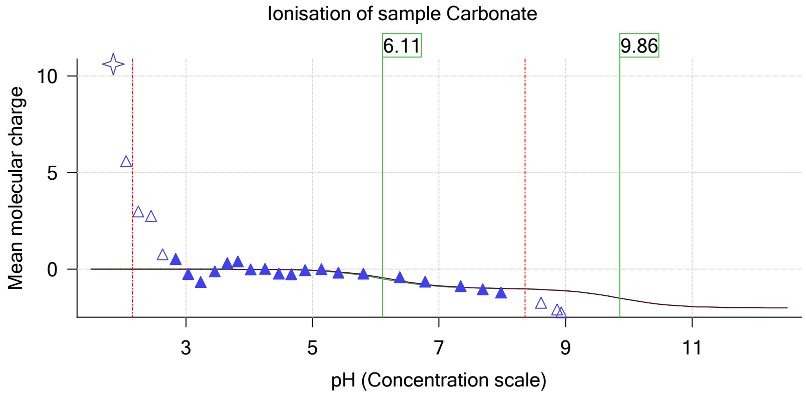

| | | | |
|---|---|---|---|
| Sample name: | **M09_octanol** | Experiment start time: | **3/3/2018 12:18:26 AM** |
| Assay name: | **pH-metric high logP** | Analyst: | **Pion** |
| Assay ID: | **18C-03001** | Instrument ID: | **T312060** |
| Filename: | **C:\Sirius_T3\Mehtap\20180302_exp29_logP_T3-2\18C-03001_M09_octanol_pH-metric high logP.t3r** | | |

#### Assay Model

| Settings | Value | Date/Time changed | Imported from |
|---|---|---|---|
| Sample name | M09_octanol | 2/27/2018 4:56:17 PM | User entered value |
| Sample by | Weight | | Default value |
| Sample weight | 0.001100 g | 3/2/2018 5:08:41 PM | User entered value |
| Formula weight | 287.74 g/mol | 2/27/2018 4:45:45 PM | User entered value |
| Solubility | Unknown | | Default value |
| Molecular weight | 251.28 | 2/27/2018 4:45:45 PM | User entered value |
| Individual pKa ionic environments | No | | Default value |
| Number of pKas | 1 | 2/27/2018 4:45:45 PM | User entered value |
| Sample is a | Base | 2/27/2018 4:45:45 PM | User entered value |
| pKa 1 | 5.37 | 2/27/2018 4:45:45 PM | User entered value |
| logp (XH +) | 0.76 | 3/2/2018 3:27:23 PM | User entered value |
| logP (neutral X) | 3.27 | 3/2/2018 3:27:17 PM | User entered value |
| Stoichiometry | 1.00000 | | Default value |
| Aprotic counterion name | Chloride | | From standards.xml file |
| Stoichiometry | 1.00 | | From standards.xml file |
| Charge per counterion | -1 | | From standards.xml file |

#### Events

| Time | Event | Water | Acid | Base | Octanol | pH | dpH/dt | pH R-squared | pH SD | dpH/dt time |
|---|---|---|---|---|---|---|---|---|---|---|
| 5:59.7 | Initial pH = 4.70 | | | | | | | | | |
| 9:04.4 | Data point 1 | 1.50000 mL | 0.04781 mL | 0.00322 mL | 0.03001 mL | 1.997 | 0.00021 | 0.00128 | 0.00028 | 10.0 s |
| 9:50.6 | Data point 2 | 1.50000 mL | 0.04781 mL | 0.01929 mL | 0.03001 mL | 2.199 | -0.00807 | 0.15986 | 0.00100 | 10.0 s |
| 10:26.3 | Data point 3 | 1.50000 mL | 0.04781 mL | 0.02886 mL | 0.03001 mL | 2.383 | -0.00750 | 0.76284 | 0.00042 | 10.0 s |
| 11:01.9 | Data point 4 | 1.50000 mL | 0.04781 mL | 0.03521 mL | 0.03001 mL | 2.574 | -0.00115 | 0.21590 | 0.00012 | 10.0 s |
| 11:37.4 | Data point 5 | 1.50000 mL | 0.04781 mL | 0.03935 mL | 0.03001 mL | 2.764 | -0.00311 | 0.44869 | 0.00023 | 10.0 s |
| 12:12.9 | Data point 6 | 1.50000 mL | 0.04781 mL | 0.04214 mL | 0.03001 mL | 2.959 | 0.00347 | 0.05811 | 0.00071 | 10.0 s |
| 12:48.4 | Data point 7 | 1.50000 mL | 0.04781 mL | 0.04412 mL | 0.03001 mL | 3.196 | -0.00193 | 0.10328 | 0.00030 | 10.0 s |
| 13:23.8 | Data point 8 | 1.50000 mL | 0.04781 mL | 0.04558 mL | 0.03001 mL | 3.433 | -0.00576 | 0.33769 | 0.00049 | 10.0 s |
| 14:14.7 | Data point 9 | 1.50000 mL | 0.04781 mL | 0.04680 mL | 0.03001 mL | 3.626 | 0.00424 | 0.07343 | 0.00077 | 10.0 s |
| 14:50.2 | Data point 10 | 1.50000 mL | 0.04781 mL | 0.04798 mL | 0.03001 mL | 3.827 | -0.00743 | 0.22433 | 0.00078 | 10.0 s |
| 15:25.6 | Data point 11 | 1.50000 mL | 0.04781 mL | 0.04908 mL | 0.03001 mL | 4.012 | -0.00611 | 0.18146 | 0.00071 | 10.0 s |
| 16:01.0 | Data point 12 | 1.50000 mL | 0.04781 mL | 0.05007 mL | 0.03001 mL | 4.168 | 0.00689 | 0.18082 | 0.00080 | 10.5 s |
| 16:52.5 | Data point 13 | 1.50000 mL | 0.04781 mL | 0.05183 mL | 0.03001 mL | 4.487 | 0.00440 | 0.05974 | 0.00089 | 11.0 s |
| 17:44.4 | Data point 14 | 1.50000 mL | 0.04781 mL | 0.05278 mL | 0.03001 mL | 4.667 | -0.01155 | 0.70656 | 0.00068 | 11.0 s |
| 18:36.3 | Data point 15 | 1.50000 mL | 0.04781 mL | 0.05362 mL | 0.03001 mL | 4.964 | -0.01372 | 0.57218 | 0.00090 | 12.0 s |
| 19:29.2 | Data point 16 | 1.50000 mL | 0.04781 mL | 0.05419 mL | 0.03001 mL | 5.238 | -0.01399 | 0.63621 | 0.00087 | 13.5 s |
| 20:13.3 | Data point 17 | 1.50000 mL | 0.04781 mL | 0.05447 mL | 0.03001 mL | 5.454 | -0.01800 | 0.81818 | 0.00098 | 14.5 s |
| 20:58.4 | Data point 18 | 1.50000 mL | 0.04781 mL | 0.05480 mL | 0.03001 mL | 6.136 | -0.01947 | 0.94607 | 0.00099 | 24.0 s |
| 21:53.1 | Data point 19 | 1.50000 mL | 0.04781 mL | 0.05496 mL | 0.03001 mL | 6.796 | -0.01955 | 0.98614 | 0.00097 | 55.0 s |
| 23:18.9 | Data point 20 | 1.50000 mL | 0.04781 mL | 0.05506 mL | 0.03001 mL | 7.170 | -0.02220 | 0.96314 | 0.00112 | Timed out at 59.5 s |
| 24:54.4 | Data point 21 | 1.50000 mL | 0.04781 mL | 0.05513 mL | 0.03001 mL | 7.765 | -0.05086 | 0.98182 | 0.00253 | Timed out at 59.5 s |
| 26:24.9 | Data point 22 | 1.50000 mL | 0.04781 mL | 0.05517 mL | 0.03001 mL | 8.213 | -0.03736 | 0.98947 | 0.00186 | Timed out at 59.5 s |
| 27:55.4 | Data point 23 | 1.50000 mL | 0.04781 mL | 0.05522 mL | 0.03001 mL | 8.561 | -0.01843 | 0.94257 | 0.00094 | 51.0 s |
| 29:22.2 | Data point 24 | 1.50000 mL | 0.04781 mL | 0.05529 mL | 0.03001 mL | 8.953 | -0.01480 | 0.58788 | 0.00095 | 20.5 s |
| 30:13.4 | Data point 25 | 1.50000 mL | 0.04781 mL | 0.05534 mL | 0.03001 mL | 9.083 | -0.01975 | 0.98237 | 0.00098 | 24.0 s |
| 31:36.5 | Data point 26 | 1.50000 mL | 0.10442 mL | 0.05534 mL | 0.07001 mL | 1.986 | -0.00842 | 0.45051 | 0.00062 | 10.0 s |
| 32:22.8 | Data point 27 | 1.50000 mL | 0.10442 mL | 0.07321 mL | 0.07001 mL | 2.184 | 0.00425 | 0.08074 | 0.00074 | 10.0 s |
| 32:58.4 | Data point 28 | 1.50000 mL | 0.10442 mL | 0.08401 mL | 0.07001 mL | 2.386 | -0.00574 | 0.55446 | 0.00038 | 10.5 s |
| 33:34.5 | Data point 29 | 1.50000 mL | 0.10442 mL | 0.09083 mL | 0.07001 mL | 2.563 | -0.00261 | 0.29606 | 0.00024 | 10.5 s |
| 34:10.6 | Data point 30 | 1.50000 mL | 0.10442 mL | 0.09548 mL | 0.07001 mL | 2.748 | -0.00851 | 0.22823 | 0.00088 | 10.0 s |
| 34:46.1 | Data point 31 | 1.50000 mL | 0.10442 mL | 0.09873 mL | 0.07001 mL | 2.936 | -0.00490 | 0.63259 | 0.00030 | 10.0 s |
| 35:21.6 | Data point 32 | 1.50000 mL | 0.10442 mL | 0.10111 mL | 0.07001 mL | 3.202 | 0.00315 | 0.37649 | 0.00025 | 10.5 s |
| 36:07.9 | Data point 33 | 1.50000 mL | 0.10442 mL | 0.10261 mL | 0.07001 mL | 3.392 | -0.00411 | 0.37428 | 0.00033 | 10.0 s |

| | | | |
|---|---|---|---|
| Sample name: | **M09_octanol** | Experiment start time: | **3/3/2018 12:18:26 AM** |
| Assay name: | **pH-metric high logP** | Analyst: | **Pion** |
| Assay ID: | **18C-03001** | Instrument ID: | **T312060** |
| Filename: | **C:\Sirius_T3\Mehtap\20180302_exp29_logP_T3-2\18C-03001_M09_octanol_pH-metric high logP.t3r** | | |

#### Events (continued)

| Time | Event | Water | Acid | Base | Octanol | pH | dpH/dt | pH R-squared | pH SD | dpH/dt time |
|---|---|---|---|---|---|---|---|---|---|---|
| 36:43.4 | Data point 34 | 1.50000 mL | 0.10442 mL | 0.10409 mL | 0.07001 mL | 3.607 | -0.00351 | 0.55022 | 0.00023 | 10.0 s |
| 37:18.9 | Data point 35 | 1.50000 mL | 0.10442 mL | 0.10536 mL | 0.07001 mL | 3.781 | -0.00697 | 0.45953 | 0.00051 | 10.0 s |
| 38:15.1 | Data point 36 | 1.50000 mL | 0.10442 mL | 0.10691 mL | 0.07001 mL | 4.005 | 0.00441 | 0.11123 | 0.00065 | 10.5 s |
| 39:06.5 | Data point 37 | 1.50000 mL | 0.10442 mL | 0.10800 mL | 0.07001 mL | 4.199 | -0.00237 | 0.07863 | 0.00042 | 10.0 s |
| 40:02.4 | Data point 38 | 1.50000 mL | 0.10442 mL | 0.10889 mL | 0.07001 mL | 4.404 | -0.00334 | 0.17918 | 0.00039 | 10.0 s |
| 40:53.4 | Data point 39 | 1.50000 mL | 0.10442 mL | 0.10955 mL | 0.07001 mL | 4.598 | -0.01051 | 0.50833 | 0.00073 | 10.0 s |
| 41:44.3 | Data point 40 | 1.50000 mL | 0.10442 mL | 0.11009 mL | 0.07001 mL | 4.816 | -0.00546 | 0.23998 | 0.00055 | 10.5 s |
| 42:30.4 | Data point 41 | 1.50000 mL | 0.10442 mL | 0.11044 mL | 0.07001 mL | 5.018 | -0.00850 | 0.19854 | 0.00094 | 11.0 s |
| 43:12.0 | Data point 42 | 1.50000 mL | 0.10442 mL | 0.11068 mL | 0.07001 mL | 5.225 | -0.01559 | 0.61445 | 0.00098 | 11.5 s |
| 43:54.1 | Data point 43 | 1.50000 mL | 0.10442 mL | 0.11087 mL | 0.07001 mL | 5.444 | -0.01372 | 0.82414 | 0.00075 | 12.5 s |
| 44:37.2 | Data point 44 | 1.50000 mL | 0.10442 mL | 0.11101 mL | 0.07001 mL | 5.696 | -0.01715 | 0.77305 | 0.00096 | 13.5 s |
| 45:21.1 | Data point 45 | 1.50000 mL | 0.10442 mL | 0.11112 mL | 0.07001 mL | 6.130 | -0.01954 | 0.96881 | 0.00098 | 26.5 s |
| 46:18.2 | Data point 46 | 1.50000 mL | 0.10442 mL | 0.11122 mL | 0.07001 mL | 6.592 | -0.01874 | 0.90315 | 0.00097 | 53.5 s |
| 47:47.4 | Data point 47 | 1.50000 mL | 0.10442 mL | 0.11129 mL | 0.07001 mL | 7.084 | -0.04097 | 0.96909 | 0.00205 | Timed out at 59.5 s |
| 49:17.9 | Data point 48 | 1.50000 mL | 0.10442 mL | 0.11134 mL | 0.07001 mL | 7.495 | -0.05100 | 0.98620 | 0.00253 | Timed out at 59.5 s |
| 50:48.4 | Data point 49 | 1.50000 mL | 0.10442 mL | 0.11138 mL | 0.07001 mL | 7.776 | -0.04776 | 0.98456 | 0.00238 | Timed out at 59.5 s |
| 52:24.0 | Data point 50 | 1.50000 mL | 0.10442 mL | 0.11145 mL | 0.07001 mL | 8.250 | -0.03292 | 0.98852 | 0.00163 | Timed out at 59.5 s |
| 54:10.0 | Data point 51 | 1.50000 mL | 0.10442 mL | 0.11157 mL | 0.07001 mL | 8.590 | -0.00702 | 0.18474 | 0.00081 | 30.0 s |
| 55:15.7 | Data point 52 | 1.50000 mL | 0.10442 mL | 0.11164 mL | 0.07001 mL | 8.848 | -0.01818 | 0.90325 | 0.00095 | 35.0 s |
| 56:26.4 | Data point 53 | 1.50000 mL | 0.10442 mL | 0.11174 mL | 0.07001 mL | 9.055 | -0.01929 | 0.94782 | 0.00098 | 19.5 s |
| 57:46.5 | Data point 54 | 1.50000 mL | 0.16536 mL | 0.11174 mL | 0.17001 mL | 1.977 | -0.01088 | 0.90338 | 0.00057 | 10.0 s |
| 58:32.8 | Data point 55 | 1.50000 mL | 0.16536 mL | 0.13133 mL | 0.17001 mL | 2.175 | 0.01443 | 0.67056 | 0.00087 | 10.0 s |
| 59:08.5 | Data point 56 | 1.50000 mL | 0.16536 mL | 0.14320 mL | 0.17001 mL | 2.362 | -0.00235 | 0.19580 | 0.00026 | 10.5 s |
| 59:44.6 | Data point 57 | 1.50000 mL | 0.16536 mL | 0.15104 mL | 0.17001 mL | 2.563 | 0.00080 | 0.04198 | 0.00019 | 10.0 s |
| 1:00:20.1 | Data point 58 | 1.50000 mL | 0.16536 mL | 0.15618 mL | 0.17001 mL | 2.744 | -0.00463 | 0.54415 | 0.00031 | 10.0 s |
| 1:00:55.6 | Data point 59 | 1.50000 mL | 0.16536 mL | 0.15985 mL | 0.17001 mL | 2.949 | 0.00206 | 0.02453 | 0.00065 | 10.0 s |
| 1:01:31.1 | Data point 60 | 1.50000 mL | 0.16536 mL | 0.16251 mL | 0.17001 mL | 3.143 | -0.00224 | 0.06525 | 0.00043 | 10.0 s |
| 1:02:06.6 | Data point 61 | 1.50000 mL | 0.16536 mL | 0.16458 mL | 0.17001 mL | 3.342 | -0.00396 | 0.72084 | 0.00023 | 10.0 s |
| 1:02:42.1 | Data point 62 | 1.50000 mL | 0.16536 mL | 0.16625 mL | 0.17001 mL | 3.564 | 0.00587 | 0.14501 | 0.00076 | 10.0 s |
| 1:03:17.5 | Data point 63 | 1.50000 mL | 0.16536 mL | 0.16752 mL | 0.17001 mL | 3.761 | 0.00890 | 0.28722 | 0.00082 | 10.5 s |
| 1:03:53.5 | Data point 64 | 1.50000 mL | 0.16536 mL | 0.16851 mL | 0.17001 mL | 3.925 | -0.00097 | 0.01530 | 0.00039 | 10.0 s |
| 1:04:49.6 | Data point 65 | 1.50000 mL | 0.16536 mL | 0.16966 mL | 0.17001 mL | 4.128 | -0.00722 | 0.41676 | 0.00055 | 10.0 s |
| 1:05:40.5 | Data point 66 | 1.50000 mL | 0.16536 mL | 0.17056 mL | 0.17001 mL | 4.359 | -0.00877 | 0.21278 | 0.00094 | 10.0 s |
| 1:06:31.4 | Data point 67 | 1.50000 mL | 0.16536 mL | 0.17121 mL | 0.17001 mL | 4.575 | 0.01310 | 0.51315 | 0.00090 | 16.0 s |
| 1:07:28.2 | Data point 68 | 1.50000 mL | 0.16536 mL | 0.17159 mL | 0.17001 mL | 4.771 | -0.01121 | 0.73701 | 0.00065 | 10.5 s |
| 1:08:09.3 | Data point 69 | 1.50000 mL | 0.16536 mL | 0.17180 mL | 0.17001 mL | 4.986 | -0.00731 | 0.13298 | 0.00099 | 11.0 s |
| 1:08:50.9 | Data point 70 | 1.50000 mL | 0.16536 mL | 0.17199 mL | 0.17001 mL | 5.245 | -0.00725 | 0.21815 | 0.00077 | 12.0 s |
| 1:09:38.5 | Data point 71 | 1.50000 mL | 0.16536 mL | 0.17215 mL | 0.17001 mL | 5.515 | -0.01092 | 0.47619 | 0.00078 | 13.0 s |
| 1:10:22.0 | Data point 72 | 1.50000 mL | 0.16536 mL | 0.17225 mL | 0.17001 mL | 5.905 | -0.01340 | 0.44094 | 0.00100 | 16.5 s |
| 1:11:09.0 | Data point 73 | 1.50000 mL | 0.16536 mL | 0.17234 mL | 0.17001 mL | 6.483 | -0.01687 | 0.87993 | 0.00089 | 57.5 s |
| 1:12:37.2 | Data point 74 | 1.50000 mL | 0.16536 mL | 0.17241 mL | 0.17001 mL | 6.882 | -0.04276 | 0.96454 | 0.00215 | Timed out at 59.5 s |
| 1:14:12.9 | Data point 75 | 1.50000 mL | 0.16536 mL | 0.17248 mL | 0.17001 mL | 7.441 | -0.07234 | 0.98643 | 0.00360 | Timed out at 59.5 s |
| 1:15:43.3 | Data point 76 | 1.50000 mL | 0.16536 mL | 0.17253 mL | 0.17001 mL | 7.793 | -0.06080 | 0.98083 | 0.00303 | Timed out at 59.5 s |
| 1:17:13.8 | Data point 77 | 1.50000 mL | 0.16536 mL | 0.17258 mL | 0.17001 mL | 8.077 | -0.04863 | 0.96800 | 0.00244 | Timed out at 59.5 s |
| 1:18:49.5 | Data point 78 | 1.50000 mL | 0.16536 mL | 0.17274 mL | 0.17001 mL | 8.712 | -0.01736 | 0.96206 | 0.00087 | 32.5 s |
| 1:20:02.8 | Data point 79 | 1.50000 mL | 0.16536 mL | 0.17286 mL | 0.17001 mL | 8.963 | -0.01640 | 0.89684 | 0.00086 | 30.0 s |
| 1:21:03.3 | Data point 80 | 1.50000 mL | 0.16536 mL | 0.17291 mL | 0.17001 mL | 9.029 | -0.00786 | 0.17453 | 0.00093 | 13.5 s |
| 1:21:25.8 | Assay volumes | 1.50000 mL | 0.16536 mL | 0.17291 mL | 0.17001 mL | | | | | |

| | | | |
|---|---|---|---|
| Sample name: | **M09_octanol** | Experiment start time: | **3/3/2018 12:18:26 AM** |
| Assay name: | **pH-metric high logP** | Analyst: | **Pion** |
| Assay ID: | **18C-03001** | Instrument ID: | **T312060** |
| Filename: | **C:\Sirius_T3\Mehtap\20180302_exp29_logP_T3-2\18C-03001_M09_octanol_pH-metric high logP.t3r** | | |

| | | | |
|---|---|---|---|
| Sample name: | **M09_octanol** | Experiment start time: | **3/3/2018 12:18:26 AM** |
| Assay name: | **pH-metric high logP** | Analyst: | **Pion** |
| Assay ID: | **18C-03001** | Instrument ID: | **T312060** |
| Filename: | **C:\Sirius_T3\Mehtap\20180302_exp29_logP_T3-2\18C-03001_M09_octanol_pH-metric high logP.t3r** | | |

#### Assay Settings (continued)

| Setting | Value | Original Value | Date/Time changed | Imported from |
|---|---|---|---|---|
| ***Titration 3*** | | | | |
| Titrate from | Low to high pH | | | |
| Add additional water | 0.00 mL | | | |
| Additional partition solvent volume | 0.100 mL | | | |
| Additional partition solvent added | Automatic | | | |
| After pH adjust stir for | 30 seconds | | | |
| Stir to allow partitioning for | 15 seconds | | | |
| Stirrer speed for partitioning | 60% | | | |
| ***Data Point Stability*** | | | | |
| Stir during data point collection | No | | | |
| Delay before data point collection | 0 seconds | | | |
| Number of points to average | 20 points | | | |
| Time interval between points | 0.50 seconds | | | |
| Required maximum standard deviation | 0.00100 dpH/dt | | | |
| Stability timeout after | 60 seconds | | | |

#### Calibration Settings

| Setting | Value | Date/Time changed | Imported from |
|---|---|---|---|
| Four-Plus alpha | 0.111 | 3/3/2018 12:18:25 AM | C:\Sirius_T3\HCl18C02.t3r |
| Four-Plus S | 0.9988 | 3/3/2018 12:18:25 AM | C:\Sirius_T3\HCl18C02.t3r |
| Four-Plus jH | 1.0 | 3/3/2018 12:18:25 AM | C:\Sirius_T3\HCl18C02.t3r |
| Four-Plus jOH | -0.8 | 3/3/2018 12:18:25 AM | C:\Sirius_T3\HCl18C02.t3r |
| Base concentration factor | 1.000 | 3/3/2018 12:18:26 AM | C:\Sirius_T3\KOH18B27.t3r |
| Acid concentration factor | 0.999 | 3/3/2018 12:18:25 AM | C:\Sirius_T3\HCl18C02.t3r |

#### Instrument Settings

| Setting | Value | Batch Id | Install date |
|---|---|---|---|
| Instrument owner | Merck | | |
| Instrument ID | T312060 | | |
| Instrument type | T3 Simulator | | |
| Software version | 1.1.3.0 | | |
| Dispenser module | | T3DM1200361 | 3/31/2009 5:24:52 AM |
| Dispenser 0 | Water | | 3/31/2009 5:25:05 AM |
| Syringe volume | 2.5 mL | | |
| Firmware version | 1.2.1(r2) | | |
| Titrant | Water (0.15 M KCl) | 02-06-2018 | 2/27/2018 10:05:59 AM |
| Dispenser 2 | Acid | | 3/31/2009 5:25:11 AM |
| Syringe volume | 0.5 mL | | |
| Firmware version | 1.2.1(r2) | | |
| Titrant | Acid (0.5 M HCl) | 02-27-2018 | 2/27/2018 10:27:22 AM |
| Dispenser 1 | Base | | 3/31/2009 5:25:21 AM |
| Syringe volume | 0.5 mL | | |
| Firmware version | 1.2.1(r2) | | |
| Titrant | Base (0.5 M KOH) | 9/22/2017 | 2/27/2018 10:21:22 AM |
| Dispenser 5 | Cosolvent | | 3/31/2009 5:26:24 AM |
| Syringe volume | 2.5 mL | | |
| Firmware version | 1.2.1(r2) | | |
| Distribution valve 5 | Distribution Valve | | 3/31/2009 5:28:19 AM |
| Firmware version | 1.1.3 | | |
| Port A | Methanol (80%, 0.15 M KCl) | 09-26-17 | 2/7/2018 9:42:01 AM |
| Port B | Cyclohexane | 11-01-17 | 2/27/2018 10:37:57 AM |
| Dispenser 3 | Buffer | | 8/3/2010 5:05:16 AM |
| Syringe volume | 0.5 mL | | |
| Firmware version | 1.2.1(r2) | | |
| Titrant | Dodecane | 2018/01/31 | 2/28/2018 10:18:04 AM |
| Dispenser 6 | Octanol | | 10/22/2010 10:52:43 AM |

| | | | |
|---|---|---|---|
| Sample name: | **M09_octanol** | Experiment start time: | **3/3/2018 12:18:26 AM** |
| Assay name: | **pH-metric high logP** | Analyst: | **Pion** |
| Assay ID: | **18C-03001** | Instrument ID: | **T312060** |
| Filename: | **C:\Sirius_T3\Mehtap\20180302_exp29_logP_T3-2\18C-03001_M09_octanol_pH-metric high logP.t3r** | | |

| | | | |
|---|---|---|---|
| Sample name: | **M09_octanol** | Experiment start time: | **3/3/2018 12:18:26 AM** |
| Assay name: | **pH-metric high logP** | Analyst: | **Pion** |
| Assay ID: | **18C-03001** | Instrument ID: | **T312060** |
| Filename: | **C:\Sirius_T3\Mehtap\20180302_exp29_logP_T3-2\18C-03001_M09_octanol_pH-metric high logP.t3r** | | |

#### Instrument Settings (continued)

| Setting | Value | Batch Id | Install date |
|---|---|---|---|
| Spectrometer calibration stir duration | 5 s | | |
| Spectrometer calibration stir speed | 30% | | |
| Spectrometer calibration wash pump volume | 20.0 mL | | |
| Spectrometer calibration wash stir duration | 5 s | | |
| Spectrometer calibration wash stir speed | 30% | | |
| Overhead dispense height | 10000 | | |

#### Refinement Settings

| Setting | Value | Default value |
|---|---|---|
| Turbidity detection method | None | None |
| Turbidity wavelength to assess | 500.0 nm | 500.0 nm |
| Turbidity maximum absorbance | 0.100 | 0.100 |
| Turbidity probe threshold | 50.00 | 50.00 |

#### Experiment Log

[2:37] Air gap created for Water (0.15 M KCl)
[2:37] Air gap created for Acid (0.5 M HCl)
[2:38] Air gap created for Base (0.5 M KOH)
[2:38] Air gap released for Water (0.15 M KCl)
[2:42] Titrator arm moved over Titration position
[2:42] Titration 1 of 3
[2:42] Adding initial titrants
[2:42] Automatically add 1.50000 mL of water
[3:07] Dispensed 1.500000 mL of Water (0.15 M KCl)
[3:11] Titrator arm moved over Drain
[5:52] Titrator arm moved to Titration position
[5:52] Argon flow rate set to 100
[5:52] Stirrer speed set to 10
[5:57] Automatically add 0.03000 mL of Octanol
[5:58] Dispensed 0.030009 mL of Octanol
[5:59] Initial pH = 4.70
[5:59] Iterative adjust 4.70 -> 2.00
[5:59] pH 4.70 -> 2.00
[6:01] Air gap released for Acid (0.5 M HCl)
[6:02] Dispensed 0.046096 mL of Acid (0.5 M HCl)
[6:07] pH 2.02 -> 2.00
[6:07] Dispensed 0.001717 mL of Acid (0.5 M HCl)
[6:12] Holding pH 2.00
[8:12] Stirrer speed set to 0
[8:12] Stirrer speed set to 50
[8:12] Iterative adjust 1.97 -> 2.00
[8:12] pH 1.97 -> 2.00
[8:13] Air gap released for Base (0.5 M KOH)
[8:14] Dispensed 0.003222 mL of Base (0.5 M KOH)
[9:04] Stirrer speed set to 0
[9:14] Datapoint id 1 collected
[9:14] Stirrer speed set to 50
[9:19] pH 2.00 -> 2.20
[9:19] Using cautious pH adjust
[9:19] Dispensed 0.007690 mL of Base (0.5 M KOH)
[9:24] Stepping pH = 2.08
[9:25] Dispensed 0.006867 mL of Base (0.5 M KOH)
[9:30] Stepping pH = 2.18
[9:30] Dispensed 0.001505 mL of Base (0.5 M KOH)
[9:35] Stepping pH = 2.20
[9:50] Stirrer speed set to 0
[10:00] Datapoint id 2 collected

Sample name: **M09_octanol** Experiment start time: **3/3/2018 12:18:26 AM**
Assay name: **pH-metric high logP** Analyst: **Pion**
Assay ID: **18C-03001** Instrument ID: **T312060**
Filename: **C:\Sirius_T3\Mehtap\20180302_exp29_logP_T3-2\18C-03001_M09_octanol_pH-metric high logP.t3r**

#### Experiment Log (continued)

[10:00] Charge balance equation is out by -4.5%
[10:00] Stirrer speed set to 50
[10:05] pH 2.21 -> 2.41
[10:05] Using charge balance adjust
[10:06] Dispensed 0.009572 mL of Base (0.5 M KOH)
[10:26] Stirrer speed set to 0
[10:36] Datapoint id 3 collected
[10:36] Charge balance equation is out by -12.3%
[10:36] Stirrer speed set to 50
[10:41] pH 2.39 -> 2.59
[10:41] Using charge balance adjust
[10:41] Dispensed 0.006350 mL of Base (0.5 M KOH)
[11:01] Stirrer speed set to 0
[11:11] Datapoint id 4 collected
[11:11] Charge balance equation is out by -7.1%
[11:11] Stirrer speed set to 50
[11:17] pH 2.58 -> 2.78
[11:17] Using charge balance adjust
[11:17] Dispensed 0.004139 mL of Base (0.5 M KOH)
[11:37] Stirrer speed set to 0
[11:47] Datapoint id 5 collected
[11:47] Charge balance equation is out by -8.6%
[11:47] Stirrer speed set to 50
[11:52] pH 2.77 -> 2.97
[11:52] Using charge balance adjust
[11:52] Dispensed 0.002799 mL of Base (0.5 M KOH)
[12:12] Stirrer speed set to 0
[12:22] Datapoint id 6 collected
[12:22] Charge balance equation is out by -6.9%
[12:22] Stirrer speed set to 50
[12:28] pH 2.97 -> 3.17
[12:28] Using charge balance adjust
[12:28] Dispensed 0.001976 mL of Base (0.5 M KOH)
[12:48] Stirrer speed set to 0
[12:58] Datapoint id 7 collected
[12:58] Charge balance equation is out by 13.6%
[12:58] Stirrer speed set to 50
[13:03] pH 3.20 -> 3.40
[13:03] Using charge balance adjust
[13:03] Dispensed 0.001458 mL of Base (0.5 M KOH)
[13:23] Stirrer speed set to 0
[13:33] Datapoint id 8 collected
[13:33] Charge balance equation is out by 15.2%
[13:33] Stirrer speed set to 50
[13:38] pH 3.44 -> 3.64
[13:38] Using cautious pH adjust
[13:39] Dispensed 0.000611 mL of Base (0.5 M KOH)
[13:44] Stepping pH = 3.54
[13:44] Dispensed 0.000423 mL of Base (0.5 M KOH)
[13:49] Stepping pH = 3.61
[13:49] Dispensed 0.000118 mL of Base (0.5 M KOH)
[13:54] Stepping pH = 3.63
[13:54] Dispensed 0.000071 mL of Base (0.5 M KOH)
[13:59] Stepping pH = 3.64
[14:14] Stirrer speed set to 0
[14:24] Datapoint id 9 collected
[14:24] Charge balance equation is out by 1.1%
[14:24] Stirrer speed set to 50
[14:29] pH 3.63 -> 3.83

| | | | |
|---|---|---|---|
| Sample name: | **M09_octanol** | Experiment start time: | **3/3/2018 12:18:26 AM** |
| Assay name: | **pH-metric high logP** | Analyst: | **Pion** |
| Assay ID: | **18C-03001** | Instrument ID: | **T312060** |
| Filename: | **C:\Sirius_T3\Mehtap\20180302_exp29_logP_T3-2\18C-03001_M09_octanol_pH-metric high logP.t3r** | | |

#### Experiment Log (continued)

[14:29] Using charge balance adjust
[14:29] Dispensed 0.001176 mL of Base (0.5 M KOH)
[14:50] Stirrer speed set to 0
[15:00] Datapoint id 10 collected
[15:00] Charge balance equation is out by -3.3%
[15:00] Stirrer speed set to 50
[15:05] pH 3.83 -> 4.03
[15:05] Using charge balance adjust
[15:05] Dispensed 0.001105 mL of Base (0.5 M KOH)
[15:25] Stirrer speed set to 0
[15:35] Datapoint id 11 collected
[15:35] Charge balance equation is out by -10.7%
[15:35] Stirrer speed set to 50
[15:40] pH 4.02 -> 4.22
[15:40] Using charge balance adjust
[15:40] Dispensed 0.000988 mL of Base (0.5 M KOH)
[16:01] Stirrer speed set to 0
[16:11] Datapoint id 12 collected
[16:11] Charge balance equation is out by -25.2%
[16:11] Stirrer speed set to 50
[16:16] pH 4.18 -> 4.38
[16:16] Using cautious pH adjust
[16:16] Dispensed 0.000447 mL of Base (0.5 M KOH)
[16:21] Stepping pH = 4.25
[16:21] Dispensed 0.000470 mL of Base (0.5 M KOH)
[16:27] Stepping pH = 4.35
[16:27] Dispensed 0.000118 mL of Base (0.5 M KOH)
[16:32] Stepping pH = 4.34
[16:32] Dispensed 0.000729 mL of Base (0.5 M KOH)
[16:37] Stepping pH = 4.51
[16:52] Stirrer speed set to 0
[17:03] Datapoint id 13 collected
[17:03] Charge balance equation is out by -100.8%
[17:03] Stirrer speed set to 50
[17:08] pH 4.50 -> 4.70
[17:08] Using cautious pH adjust
[17:08] Dispensed 0.000306 mL of Base (0.5 M KOH)
[17:13] Stepping pH = 4.56
[17:13] Dispensed 0.000376 mL of Base (0.5 M KOH)
[17:18] Stepping pH = 4.65
[17:19] Dispensed 0.000141 mL of Base (0.5 M KOH)
[17:24] Stepping pH = 4.67
[17:24] Dispensed 0.000118 mL of Base (0.5 M KOH)
[17:29] Stepping pH = 4.70
[17:44] Stirrer speed set to 0
[17:55] Datapoint id 14 collected
[17:55] Charge balance equation is out by -55.5%
[17:55] Stirrer speed set to 50
[18:00] pH 4.68 -> 4.88
[18:00] Using cautious pH adjust
[18:00] Dispensed 0.000212 mL of Base (0.5 M KOH)
[18:05] Stepping pH = 4.74
[18:05] Dispensed 0.000306 mL of Base (0.5 M KOH)
[18:10] Stepping pH = 4.86
[18:11] Dispensed 0.000047 mL of Base (0.5 M KOH)
[18:16] Stepping pH = 4.86
[18:16] Dispensed 0.000282 mL of Base (0.5 M KOH)
[18:21] Stepping pH = 4.99
[18:36] Stirrer speed set to 0

Sample name: **M09_octanol** Experiment start time: **3/3/2018 12:18:26 AM**
Assay name: **pH-metric high logP** Analyst: **Pion**
Assay ID: **18C-03001** Instrument ID: **T312060**
Filename: **C:\Sirius_T3\Mehtap\20180302_exp29_logP_T3-2\18C-03001_M09_octanol_pH-metric high logP.t3r**

#### Experiment Log (continued)

[18:48] Datapoint id 15 collected
[18:48] Charge balance equation is out by -89.5%
[18:48] Stirrer speed set to 50
[18:53] pH 4.99 -> 5.19
[18:53] Using cautious pH adjust
[18:53] Dispensed 0.000141 mL of Base (0.5 M KOH)
[18:58] Stepping pH = 5.02
[18:58] Dispensed 0.000235 mL of Base (0.5 M KOH)
[19:03] Stepping pH = 5.15
[19:03] Dispensed 0.000047 mL of Base (0.5 M KOH)
[19:08] Stepping pH = 5.16
[19:09] Dispensed 0.000141 mL of Base (0.5 M KOH)
[19:14] Stepping pH = 5.27
[19:29] Stirrer speed set to 0
[19:42] Datapoint id 16 collected
[19:42] Charge balance equation is out by -110.5%
[19:42] Stirrer speed set to 50
[19:47] pH 5.26 -> 5.46
[19:47] Using cautious pH adjust
[19:47] Dispensed 0.000071 mL of Base (0.5 M KOH)
[19:53] Stepping pH = 5.28
[19:53] Dispensed 0.000212 mL of Base (0.5 M KOH)
[19:58] Stepping pH = 5.48
[20:13] Stirrer speed set to 0
[20:27] Datapoint id 17 collected
[20:27] Charge balance equation is out by -85.7%
[20:27] Stirrer speed set to 50
[20:32] pH 5.48 -> 5.68
[20:32] Using cautious pH adjust
[20:33] Dispensed 0.000047 mL of Base (0.5 M KOH)
[20:38] Stepping pH = 5.48
[20:38] Dispensed 0.000282 mL of Base (0.5 M KOH)
[20:43] Stepping pH = 6.08
[20:58] Stirrer speed set to 0
[21:22] Datapoint id 18 collected
[21:22] Charge balance equation is out by -201.5%
[21:22] Stirrer speed set to 50
[21:27] pH 6.16 -> 6.36
[21:27] Using cautious pH adjust
[21:27] Dispensed 0.000024 mL of Base (0.5 M KOH)
[21:32] Stepping pH = 6.16
[21:32] Dispensed 0.000141 mL of Base (0.5 M KOH)
[21:38] Stepping pH = 6.76
[21:53] Stirrer speed set to 0
[22:48] Datapoint id 19 collected
[22:48] Charge balance equation is out by -210.1%
[22:48] Stirrer speed set to 50
[22:53] pH 6.82 -> 7.02
[22:53] Using cautious pH adjust
[22:53] Dispensed 0.000024 mL of Base (0.5 M KOH)
[22:58] Stepping pH = 6.82
[22:58] Dispensed 0.000071 mL of Base (0.5 M KOH)
[23:03] Stepping pH = 7.05
[23:18] Stirrer speed set to 0
[24:18] Datapoint id 20 collected
[24:18] Charge balance equation is out by -228.7%
[24:18] Stirrer speed set to 50
[24:23] pH 7.20 -> 7.40
[24:23] Using cautious pH adjust

Sample name: **M09_octanol**
Assay name: **pH-metric high logP**
Assay ID: **18C-03001**
Filename: **C:\Sirius_T3\Mehtap\20180302_exp29_logP_T3-2\18C-03001_M09_octanol_pH-metric high logP.t3r**

Experiment start time: **3/3/2018 12:18:26 AM**
Analyst: **Pion**
Instrument ID: **T312060**

#### Experiment Log (continued)

[24:24] Dispensed 0.000024 mL of Base (0.5 M KOH)
[24:29] Stepping pH = 7.22
[24:29] Dispensed 0.000024 mL of Base (0.5 M KOH)
[24:34] Stepping pH = 7.28
[24:34] Dispensed 0.000024 mL of Base (0.5 M KOH)
[24:39] Stepping pH = 7.54
[24:54] Stirrer speed set to 0
[25:54] Datapoint id 21 collected
[25:54] Charge balance equation is out by -355.4%
[25:54] Stirrer speed set to 50
[25:59] pH 7.96 -> 8.16
[25:59] Using cautious pH adjust
[25:59] Dispensed 0.000024 mL of Base (0.5 M KOH)
[26:04] Stepping pH = 7.99
[26:04] Dispensed 0.000024 mL of Base (0.5 M KOH)
[26:09] Stepping pH = 8.15
[26:24] Stirrer speed set to 0
[27:24] Datapoint id 22 collected
[27:24] Charge balance equation is out by -510.2%
[27:24] Stirrer speed set to 50
[27:30] pH 8.24 -> 8.44
[27:30] Using cautious pH adjust
[27:30] Dispensed 0.000024 mL of Base (0.5 M KOH)
[27:35] Stepping pH = 8.26
[27:35] Dispensed 0.000024 mL of Base (0.5 M KOH)
[27:40] Stepping pH = 8.48
[27:55] Stirrer speed set to 0
[28:46] Datapoint id 23 collected
[28:46] Charge balance equation is out by -341.8%
[28:46] Stirrer speed set to 50
[28:51] pH 8.60 -> 8.80
[28:51] Using cautious pH adjust
[28:51] Dispensed 0.000024 mL of Base (0.5 M KOH)
[28:56] Stepping pH = 8.61
[28:56] Dispensed 0.000024 mL of Base (0.5 M KOH)
[29:02] Stepping pH = 8.77
[29:02] Dispensed 0.000024 mL of Base (0.5 M KOH)
[29:07] Stepping pH = 8.94
[29:22] Stirrer speed set to 0
[29:42] Datapoint id 24 collected
[29:42] Charge balance equation is out by -267.1%
[29:42] Stirrer speed set to 50
[29:47] pH 8.96 -> 9.05
[29:47] Using cautious pH adjust
[29:47] Dispensed 0.000024 mL of Base (0.5 M KOH)
[29:53] Stepping pH = 8.97
[29:53] Dispensed 0.000024 mL of Base (0.5 M KOH)
[29:58] Stepping pH = 9.06
[30:13] Stirrer speed set to 0
[30:37] Datapoint id 25 collected
[30:37] Charge balance equation is out by -171.7%
[30:37] Titration 2 of 3
[30:37] Adding initial titrants
[30:37] Automatically add 0.04000 mL of Octanol
[30:38] Dispensed 0.040005 mL of Octanol
[30:38] Stirrer speed set to 10
[30:39] Stirrer speed set to 55
[30:39] Iterative adjust 9.09 -> 2.00
[30:39] pH 9.09 -> 2.00

| | | | |
|---|---|---|---|
| Sample name: | **M09_octanol** | Experiment start time: | **3/3/2018 12:18:26 AM** |
| Assay name: | **pH-metric high logP** | Analyst: | **Pion** |
| Assay ID: | **18C-03001** | Instrument ID: | **T312060** |
| Filename: | **C:\Sirius_T3\Mehtap\20180302_exp29_logP_T3-2\18C-03001_M09_octanol_pH-metric high logP.t3r** | | |

#### Experiment Log (continued)

[30:40] Dispensed 0.050376 mL of Acid (0.5 M HCl)
[30:46] pH 2.06 -> 2.00
[30:46] Dispensed 0.006232 mL of Acid (0.5 M HCl)
[31:36] Stirrer speed set to 0
[31:46] Datapoint id 26 collected
[31:46] Stirrer speed set to 55
[31:51] pH 1.99 -> 2.19
[31:51] Using cautious pH adjust
[31:51] Dispensed 0.008561 mL of Base (0.5 M KOH)
[31:57] Stepping pH = 2.08
[31:57] Dispensed 0.006515 mL of Base (0.5 M KOH)
[32:02] Stepping pH = 2.15
[32:02] Dispensed 0.002799 mL of Base (0.5 M KOH)
[32:07] Stepping pH = 2.18
[32:22] Stirrer speed set to 0
[32:32] Datapoint id 27 collected
[32:32] Charge balance equation is out by -4.4%
[32:32] Stirrer speed set to 55
[32:37] pH 2.19 -> 2.39
[32:37] Using charge balance adjust
[32:38] Dispensed 0.010795 mL of Base (0.5 M KOH)
[32:58] Stirrer speed set to 0
[33:08] Datapoint id 28 collected
[33:08] Charge balance equation is out by -0.6%
[33:08] Stirrer speed set to 55
[33:14] pH 2.39 -> 2.59
[33:14] Using charge balance adjust
[33:14] Dispensed 0.006820 mL of Base (0.5 M KOH)
[33:34] Stirrer speed set to 0
[33:45] Datapoint id 29 collected
[33:45] Charge balance equation is out by -14.0%
[33:45] Stirrer speed set to 55
[33:50] pH 2.57 -> 2.77
[33:50] Using charge balance adjust
[33:50] Dispensed 0.004657 mL of Base (0.5 M KOH)
[34:10] Stirrer speed set to 0
[34:20] Datapoint id 30 collected
[34:20] Charge balance equation is out by -11.3%
[34:20] Stirrer speed set to 55
[34:25] pH 2.76 -> 2.96
[34:25] Using charge balance adjust
[34:25] Dispensed 0.003246 mL of Base (0.5 M KOH)
[34:46] Stirrer speed set to 0
[34:56] Datapoint id 31 collected
[34:56] Charge balance equation is out by -10.5%
[34:56] Stirrer speed set to 55
[35:01] pH 2.94 -> 3.14
[35:01] Using charge balance adjust
[35:01] Dispensed 0.002375 mL of Base (0.5 M KOH)
[35:21] Stirrer speed set to 0
[35:32] Datapoint id 32 collected
[35:32] Charge balance equation is out by 29.9%
[35:32] Stirrer speed set to 55
[35:37] pH 3.21 -> 3.41
[35:37] Using cautious pH adjust
[35:37] Dispensed 0.000870 mL of Base (0.5 M KOH)
[35:42] Stepping pH = 3.32
[35:42] Dispensed 0.000517 mL of Base (0.5 M KOH)
[35:47] Stepping pH = 3.39

| | | | |
|---|---|---|---|
| Sample name: | **M09_octanol** | Experiment start time: | **3/3/2018 12:18:26 AM** |
| Assay name: | **pH-metric high logP** | Analyst: | **Pion** |
| Assay ID: | **18C-03001** | Instrument ID: | **T312060** |
| Filename: | **C:\Sirius_T3\Mehtap\20180302_exp29_logP_T3-2\18C-03001_M09_octanol_pH-metric high logP.t3r** | | |

#### Experiment Log (continued)

[35:47] Dispensed 0.000118 mL of Base (0.5 M KOH)
[35:52] Stepping pH = 3.40
[36:07] Stirrer speed set to 0
[36:18] Datapoint id 33 collected
[36:18] Charge balance equation is out by 13.6%
[36:18] Stirrer speed set to 55
[36:23] pH 3.40 -> 3.60
[36:23] Using charge balance adjust
[36:23] Dispensed 0.001482 mL of Base (0.5 M KOH)
[36:43] Stirrer speed set to 0
[36:53] Datapoint id 34 collected
[36:53] Charge balance equation is out by 4.3%
[36:53] Stirrer speed set to 55
[36:58] pH 3.61 -> 3.81
[36:58] Using charge balance adjust
[36:58] Dispensed 0.001270 mL of Base (0.5 M KOH)
[37:18] Stirrer speed set to 0
[37:28] Datapoint id 35 collected
[37:28] Charge balance equation is out by -15.5%
[37:28] Stirrer speed set to 55
[37:34] pH 3.78 -> 3.98
[37:34] Using cautious pH adjust
[37:34] Dispensed 0.000541 mL of Base (0.5 M KOH)
[37:39] Stepping pH = 3.86
[37:39] Dispensed 0.000541 mL of Base (0.5 M KOH)
[37:44] Stepping pH = 3.95
[37:44] Dispensed 0.000188 mL of Base (0.5 M KOH)
[37:49] Stepping pH = 3.97
[37:49] Dispensed 0.000071 mL of Base (0.5 M KOH)
[37:54] Stepping pH = 3.97
[37:54] Dispensed 0.000212 mL of Base (0.5 M KOH)
[38:00] Stepping pH = 4.01
[38:15] Stirrer speed set to 0
[38:25] Datapoint id 36 collected
[38:25] Charge balance equation is out by -41.9%
[38:25] Stirrer speed set to 55
[38:30] pH 4.01 -> 4.21
[38:30] Using cautious pH adjust
[38:30] Dispensed 0.000423 mL of Base (0.5 M KOH)
[38:35] Stepping pH = 4.09
[38:35] Dispensed 0.000400 mL of Base (0.5 M KOH)
[38:41] Stepping pH = 4.17
[38:41] Dispensed 0.000188 mL of Base (0.5 M KOH)
[38:46] Stepping pH = 4.20
[38:46] Dispensed 0.000071 mL of Base (0.5 M KOH)
[38:51] Stepping pH = 4.21
[39:06] Stirrer speed set to 0
[39:16] Datapoint id 37 collected
[39:16] Charge balance equation is out by -27.1%
[39:16] Stirrer speed set to 55
[39:21] pH 4.21 -> 4.41
[39:21] Using cautious pH adjust
[39:21] Dispensed 0.000329 mL of Base (0.5 M KOH)
[39:26] Stepping pH = 4.28
[39:26] Dispensed 0.000306 mL of Base (0.5 M KOH)
[39:31] Stepping pH = 4.36
[39:31] Dispensed 0.000141 mL of Base (0.5 M KOH)
[39:37] Stepping pH = 4.39
[39:37] Dispensed 0.000047 mL of Base (0.5 M KOH)

| | | | |
|---|---|---|---|
| Sample name: | **M09_octanol** | Experiment start time: | **3/3/2018 12:18:26 AM** |
| Assay name: | **pH-metric high logP** | Analyst: | **Pion** |
| Assay ID: | **18C-03001** | Instrument ID: | **T312060** |
| Filename: | **C:\Sirius_T3\Mehtap\20180302_exp29_logP_T3-2\18C-03001_M09_octanol_pH-metric high logP.t3r** | | |

#### Experiment Log (continued)

[39:42] Stepping pH = 4.40
[39:42] Dispensed 0.000071 mL of Base (0.5 M KOH)
[39:47] Stepping pH = 4.41
[40:02] Stirrer speed set to 0
[40:12] Datapoint id 38 collected
[40:12] Charge balance equation is out by -35.9%
[40:12] Stirrer speed set to 55
[40:17] pH 4.41 -> 4.61
[40:17] Using cautious pH adjust
[40:17] Dispensed 0.000235 mL of Base (0.5 M KOH)
[40:22] Stepping pH = 4.48
[40:22] Dispensed 0.000259 mL of Base (0.5 M KOH)
[40:27] Stepping pH = 4.57
[40:28] Dispensed 0.000094 mL of Base (0.5 M KOH)
[40:33] Stepping pH = 4.59
[40:33] Dispensed 0.000071 mL of Base (0.5 M KOH)
[40:38] Stepping pH = 4.60
[40:53] Stirrer speed set to 0
[41:03] Datapoint id 39 collected
[41:03] Charge balance equation is out by -42.4%
[41:03] Stirrer speed set to 55
[41:08] pH 4.61 -> 4.81
[41:08] Using cautious pH adjust
[41:08] Dispensed 0.000165 mL of Base (0.5 M KOH)
[41:13] Stepping pH = 4.66
[41:13] Dispensed 0.000212 mL of Base (0.5 M KOH)
[41:18] Stepping pH = 4.76
[41:18] Dispensed 0.000071 mL of Base (0.5 M KOH)
[41:24] Stepping pH = 4.78
[41:24] Dispensed 0.000094 mL of Base (0.5 M KOH)
[41:29] Stepping pH = 4.82
[41:44] Stirrer speed set to 0
[41:54] Datapoint id 40 collected
[41:54] Charge balance equation is out by -63.5%
[41:54] Stirrer speed set to 55
[41:59] pH 4.83 -> 5.03
[41:59] Using cautious pH adjust
[41:59] Dispensed 0.000094 mL of Base (0.5 M KOH)
[42:05] Stepping pH = 4.86
[42:05] Dispensed 0.000235 mL of Base (0.5 M KOH)
[42:10] Stepping pH = 5.02
[42:10] Dispensed 0.000024 mL of Base (0.5 M KOH)
[42:15] Stepping pH = 5.03
[42:30] Stirrer speed set to 0
[42:41] Datapoint id 41 collected
[42:41] Charge balance equation is out by -73.8%
[42:41] Stirrer speed set to 55
[42:46] pH 5.04 -> 5.24
[42:46] Using cautious pH adjust
[42:46] Dispensed 0.000071 mL of Base (0.5 M KOH)
[42:51] Stepping pH = 5.06
[42:51] Dispensed 0.000165 mL of Base (0.5 M KOH)
[42:56] Stepping pH = 5.23
[43:12] Stirrer speed set to 0
[43:23] Datapoint id 42 collected
[43:23] Charge balance equation is out by -73.2%
[43:23] Stirrer speed set to 55
[43:28] pH 5.25 -> 5.45
[43:28] Using cautious pH adjust

| | | | |
|---|---|---|---|
| Sample name: | **M09_octanol** | Experiment start time: | **3/3/2018 12:18:26 AM** |
| Assay name: | **pH-metric high logP** | Analyst: | **Pion** |
| Assay ID: | **18C-03001** | Instrument ID: | **T312060** |
| Filename: | **C:\Sirius_T3\Mehtap\20180302_exp29_logP_T3-2\18C-03001_M09_octanol_pH-metric high logP.t3r** | | |

#### Experiment Log (continued)

[43:28] Dispensed 0.000047 mL of Base (0.5 M KOH)
[43:33] Stepping pH = 5.25
[43:33] Dispensed 0.000141 mL of Base (0.5 M KOH)
[43:39] Stepping pH = 5.44
[43:54] Stirrer speed set to 0
[44:06] Datapoint id 43 collected
[44:06] Charge balance equation is out by -97.6%
[44:06] Stirrer speed set to 55
[44:11] pH 5.47 -> 5.67
[44:11] Using cautious pH adjust
[44:11] Dispensed 0.000047 mL of Base (0.5 M KOH)
[44:16] Stepping pH = 5.48
[44:17] Dispensed 0.000094 mL of Base (0.5 M KOH)
[44:22] Stepping pH = 5.69
[44:37] Stirrer speed set to 0
[44:50] Datapoint id 44 collected
[44:50] Charge balance equation is out by -89.8%
[44:50] Stirrer speed set to 55
[44:55] pH 5.73 -> 5.93
[44:55] Using cautious pH adjust
[44:55] Dispensed 0.000024 mL of Base (0.5 M KOH)
[45:00] Stepping pH = 5.74
[45:00] Dispensed 0.000094 mL of Base (0.5 M KOH)
[45:06] Stepping pH = 6.11
[45:21] Stirrer speed set to 0
[45:47] Datapoint id 45 collected
[45:47] Charge balance equation is out by -93.6%
[45:47] Stirrer speed set to 55
[45:52] pH 6.19 -> 6.39
[45:52] Using cautious pH adjust
[45:52] Dispensed 0.000024 mL of Base (0.5 M KOH)
[45:57] Stepping pH = 6.20
[45:57] Dispensed 0.000071 mL of Base (0.5 M KOH)
[46:03] Stepping pH = 6.56
[46:18] Stirrer speed set to 0
[47:11] Datapoint id 46 collected
[47:11] Charge balance equation is out by -89.9%
[47:11] Stirrer speed set to 55
[47:16] pH 6.72 -> 6.92
[47:16] Using cautious pH adjust
[47:16] Dispensed 0.000024 mL of Base (0.5 M KOH)
[47:22] Stepping pH = 6.76
[47:22] Dispensed 0.000024 mL of Base (0.5 M KOH)
[47:27] Stepping pH = 6.87
[47:27] Dispensed 0.000024 mL of Base (0.5 M KOH)
[47:32] Stepping pH = 7.07
[47:47] Stirrer speed set to 0
[48:47] Datapoint id 47 collected
[48:47] Charge balance equation is out by -117.3%
[48:47] Stirrer speed set to 55
[48:52] pH 7.14 -> 7.34
[48:52] Using cautious pH adjust
[48:52] Dispensed 0.000024 mL of Base (0.5 M KOH)
[48:57] Stepping pH = 7.19
[48:57] Dispensed 0.000024 mL of Base (0.5 M KOH)
[49:02] Stepping pH = 7.37
[49:17] Stirrer speed set to 0
[50:17] Datapoint id 48 collected
[50:17] Charge balance equation is out by -157.7%

| | | | |
|---|---|---|---|
| Sample name: | **M09_octanol** | Experiment start time: | **3/3/2018 12:18:26 AM** |
| Assay name: | **pH-metric high logP** | Analyst: | **Pion** |
| Assay ID: | **18C-03001** | Instrument ID: | **T312060** |
| Filename: | **C:\Sirius_T3\Mehtap\20180302_exp29_logP_T3-2\18C-03001_M09_octanol_pH-metric high logP.t3r** | | |

#### Experiment Log (continued)

[50:17] Stirrer speed set to 55
[50:23] pH 7.40 -> 7.60
[50:23] Using cautious pH adjust
[50:23] Dispensed 0.000024 mL of Base (0.5 M KOH)
[50:28] Stepping pH = 7.40
[50:28] Dispensed 0.000024 mL of Base (0.5 M KOH)
[50:33] Stepping pH = 7.65
[50:48] Stirrer speed set to 0
[51:48] Datapoint id 49 collected
[51:48] Charge balance equation is out by -351.7%
[51:48] Stirrer speed set to 55
[51:53] pH 7.81 -> 8.01
[51:53] Using cautious pH adjust
[51:53] Dispensed 0.000024 mL of Base (0.5 M KOH)
[51:58] Stepping pH = 7.82
[51:58] Dispensed 0.000024 mL of Base (0.5 M KOH)
[52:03] Stepping pH = 7.91
[52:03] Dispensed 0.000024 mL of Base (0.5 M KOH)
[52:08] Stepping pH = 8.19
[52:24] Stirrer speed set to 0
[53:24] Datapoint id 50 collected
[53:24] Charge balance equation is out by -782.1%
[53:24] Stirrer speed set to 55
[53:29] pH 8.25 -> 8.45
[53:29] Using cautious pH adjust
[53:29] Dispensed 0.000024 mL of Base (0.5 M KOH)
[53:34] Stepping pH = 8.27
[53:34] Dispensed 0.000024 mL of Base (0.5 M KOH)
[53:39] Stepping pH = 8.31
[53:39] Dispensed 0.000024 mL of Base (0.5 M KOH)
[53:44] Stepping pH = 8.38
[53:44] Dispensed 0.000024 mL of Base (0.5 M KOH)
[53:49] Stepping pH = 8.42
[53:49] Dispensed 0.000024 mL of Base (0.5 M KOH)
[53:54] Stepping pH = 8.58
[54:10] Stirrer speed set to 0
[54:40] Datapoint id 51 collected
[54:40] Charge balance equation is out by -941.0%
[54:40] Stirrer speed set to 55
[54:45] pH 8.62 -> 8.82
[54:45] Using cautious pH adjust
[54:45] Dispensed 0.000024 mL of Base (0.5 M KOH)
[54:50] Stepping pH = 8.65
[54:50] Dispensed 0.000024 mL of Base (0.5 M KOH)
[54:55] Stepping pH = 8.73
[54:55] Dispensed 0.000024 mL of Base (0.5 M KOH)
[55:00] Stepping pH = 8.85
[55:15] Stirrer speed set to 0
[55:50] Datapoint id 52 collected
[55:50] Charge balance equation is out by -223.8%
[55:50] Stirrer speed set to 55
[55:55] pH 8.86 -> 9.05
[55:55] Using cautious pH adjust
[55:55] Dispensed 0.000024 mL of Base (0.5 M KOH)
[56:01] Stepping pH = 8.86
[56:01] Dispensed 0.000047 mL of Base (0.5 M KOH)
[56:06] Stepping pH = 8.99
[56:06] Dispensed 0.000024 mL of Base (0.5 M KOH)
[56:11] Stepping pH = 9.06

Sample name: **M09_octanol** Experiment start time: **3/3/2018 12:18:26 AM**
Assay name: **pH-metric high logP** Analyst: **Pion**
Assay ID: **18C-03001** Instrument ID: **T312060**
Filename: **C:\Sirius_T3\Mehtap\20180302_exp29_logP_T3-2\18C-03001_M09_octanol_pH-metric high logP.t3r**

#### Experiment Log (continued)

[56:26] Stirrer speed set to 0
[56:45] Datapoint id 53 collected
[56:45] Charge balance equation is out by -177.4%
[56:45] Titration 3 of 3
[56:45] Adding initial titrants
[56:45] Automatically add 0.10000 mL of Octanol
[56:48] Dispensed 0.100000 mL of Octanol
[56:48] Stirrer speed set to 10
[56:49] Stirrer speed set to 60
[56:49] Iterative adjust 9.06 -> 2.00
[56:49] pH 9.06 -> 2.00
[56:50] Dispensed 0.053293 mL of Acid (0.5 M HCl)
[56:55] pH 2.07 -> 2.00
[56:56] Dispensed 0.007643 mL of Acid (0.5 M HCl)
[57:46] Stirrer speed set to 0
[57:56] Datapoint id 54 collected
[57:56] Stirrer speed set to 60
[58:01] pH 1.98 -> 2.18
[58:01] Using cautious pH adjust
[58:01] Dispensed 0.009431 mL of Base (0.5 M KOH)
[58:07] Stepping pH = 2.06
[58:07] Dispensed 0.007902 mL of Base (0.5 M KOH)
[58:12] Stepping pH = 2.15
[58:12] Dispensed 0.002258 mL of Base (0.5 M KOH)
[58:17] Stepping pH = 2.17
[58:32] Stirrer speed set to 0
[58:42] Datapoint id 55 collected
[58:42] Charge balance equation is out by -3.9%
[58:42] Stirrer speed set to 60
[58:47] pH 2.18 -> 2.38
[58:47] Using charge balance adjust
[58:48] Dispensed 0.011877 mL of Base (0.5 M KOH)
[59:08] Stirrer speed set to 0
[59:19] Datapoint id 56 collected
[59:19] Charge balance equation is out by -8.4%
[59:19] Stirrer speed set to 60
[59:24] pH 2.37 -> 2.57
[59:24] Using charge balance adjust
[59:24] Dispensed 0.007832 mL of Base (0.5 M KOH)
[59:44] Stirrer speed set to 0
[59:54] Datapoint id 57 collected
[59:54] Charge balance equation is out by -1.8%
[59:54] Stirrer speed set to 60
[59:59] pH 2.57 -> 2.77
[59:59] Using charge balance adjust
[59:59] Dispensed 0.005151 mL of Base (0.5 M KOH)
[1:00:20] Stirrer speed set to 0
[1:00:30] Datapoint id 58 collected
[1:00:30] Charge balance equation is out by -13.0%
[1:00:30] Stirrer speed set to 60
[1:00:35] pH 2.75 -> 2.95
[1:00:35] Using charge balance adjust
[1:00:35] Dispensed 0.003669 mL of Base (0.5 M KOH)
[1:00:55] Stirrer speed set to 0
[1:01:05] Datapoint id 59 collected
[1:01:05] Charge balance equation is out by -1.4%
[1:01:05] Stirrer speed set to 60
[1:01:10] pH 2.96 -> 3.16
[1:01:10] Using charge balance adjust

Sample name: **M09_octanol** Experiment start time: **3/3/2018 12:18:26 AM**
Assay name: **pH-metric high logP** Analyst: **Pion**
Assay ID: **18C-03001** Instrument ID: **T312060**
Filename: **C:\Sirius_T3\Mehtap\20180302_exp29_logP_T3-2\18C-03001_M09_octanol_pH-metric high logP.t3r**

#### Experiment Log (continued)

[1:01:10] Dispensed 0.002658 mL of Base (0.5 M KOH)
[1:01:31] Stirrer speed set to 0
[1:01:41] Datapoint id 60 collected
[1:01:41] Charge balance equation is out by -7.0%
[1:01:41] Stirrer speed set to 60
[1:01:46] pH 3.15 -> 3.35
[1:01:46] Using charge balance adjust
[1:01:46] Dispensed 0.002070 mL of Base (0.5 M KOH)
[1:02:06] Stirrer speed set to 0
[1:02:16] Datapoint id 61 collected
[1:02:16] Charge balance equation is out by -2.9%
[1:02:16] Stirrer speed set to 60
[1:02:21] pH 3.35 -> 3.55
[1:02:21] Using charge balance adjust
[1:02:21] Dispensed 0.001670 mL of Base (0.5 M KOH)
[1:02:42] Stirrer speed set to 0
[1:02:52] Datapoint id 62 collected
[1:02:52] Charge balance equation is out by 8.9%
[1:02:52] Stirrer speed set to 60
[1:02:57] pH 3.57 -> 3.77
[1:02:57] Using charge balance adjust
[1:02:57] Dispensed 0.001270 mL of Base (0.5 M KOH)
[1:03:17] Stirrer speed set to 0
[1:03:28] Datapoint id 63 collected
[1:03:28] Charge balance equation is out by -5.5%
[1:03:28] Stirrer speed set to 60
[1:03:33] pH 3.77 -> 3.97
[1:03:33] Using charge balance adjust
[1:03:33] Dispensed 0.000988 mL of Base (0.5 M KOH)
[1:03:53] Stirrer speed set to 0
[1:04:03] Datapoint id 64 collected
[1:04:03] Charge balance equation is out by -21.8%
[1:04:03] Stirrer speed set to 60
[1:04:08] pH 3.93 -> 4.13
[1:04:08] Using cautious pH adjust
[1:04:08] Dispensed 0.000376 mL of Base (0.5 M KOH)
[1:04:13] Stepping pH = 4.00
[1:04:13] Dispensed 0.000423 mL of Base (0.5 M KOH)
[1:04:19] Stepping pH = 4.08
[1:04:19] Dispensed 0.000188 mL of Base (0.5 M KOH)
[1:04:24] Stepping pH = 4.11
[1:04:24] Dispensed 0.000071 mL of Base (0.5 M KOH)
[1:04:29] Stepping pH = 4.12
[1:04:29] Dispensed 0.000094 mL of Base (0.5 M KOH)
[1:04:34] Stepping pH = 4.13
[1:04:49] Stirrer speed set to 0
[1:04:59] Datapoint id 65 collected
[1:04:59] Charge balance equation is out by -52.8%
[1:04:59] Stirrer speed set to 60
[1:05:04] pH 4.14 -> 4.34
[1:05:04] Using cautious pH adjust
[1:05:04] Dispensed 0.000259 mL of Base (0.5 M KOH)
[1:05:09] Stepping pH = 4.20
[1:05:09] Dispensed 0.000329 mL of Base (0.5 M KOH)
[1:05:15] Stepping pH = 4.28
[1:05:15] Dispensed 0.000141 mL of Base (0.5 M KOH)
[1:05:20] Stepping pH = 4.30
[1:05:20] Dispensed 0.000165 mL of Base (0.5 M KOH)
[1:05:25] Stepping pH = 4.36

#### Experiment Log (continued)

[1:05:40] Stirrer speed set to 0
[1:05:50] Datapoint id 66 collected
[1:05:50] Charge balance equation is out by -68.3%
[1:05:50] Stirrer speed set to 60
[1:05:55] pH 4.36 -> 4.56
[1:05:55] Using cautious pH adjust
[1:05:55] Dispensed 0.000165 mL of Base (0.5 M KOH)
[1:06:00] Stepping pH = 4.40
[1:06:00] Dispensed 0.000353 mL of Base (0.5 M KOH)
[1:06:06] Stepping pH = 4.54
[1:06:06] Dispensed 0.000047 mL of Base (0.5 M KOH)
[1:06:11] Stepping pH = 4.54
[1:06:11] Dispensed 0.000094 mL of Base (0.5 M KOH)
[1:06:16] Stepping pH = 4.57
[1:06:31] Stirrer speed set to 0
[1:06:47] Datapoint id 67 collected
[1:06:47] Charge balance equation is out by -89.4%
[1:06:47] Stirrer speed set to 60
[1:06:52] pH 4.59 -> 4.79
[1:06:52] Using cautious pH adjust
[1:06:52] Dispensed 0.000118 mL of Base (0.5 M KOH)
[1:06:57] Stepping pH = 4.63
[1:06:57] Dispensed 0.000188 mL of Base (0.5 M KOH)
[1:07:02] Stepping pH = 4.76
[1:07:02] Dispensed 0.000024 mL of Base (0.5 M KOH)
[1:07:07] Stepping pH = 4.76
[1:07:08] Dispensed 0.000047 mL of Base (0.5 M KOH)
[1:07:13] Stepping pH = 4.78
[1:07:28] Stirrer speed set to 0
[1:07:38] Datapoint id 68 collected
[1:07:38] Charge balance equation is out by -71.7%
[1:07:38] Stirrer speed set to 60
[1:07:43] pH 4.79 -> 4.99
[1:07:43] Using cautious pH adjust
[1:07:43] Dispensed 0.000071 mL of Base (0.5 M KOH)
[1:07:49] Stepping pH = 4.82
[1:07:49] Dispensed 0.000141 mL of Base (0.5 M KOH)
[1:07:54] Stepping pH = 4.99
[1:08:09] Stirrer speed set to 0
[1:08:20] Datapoint id 69 collected
[1:08:20] Charge balance equation is out by -48.7%
[1:08:20] Stirrer speed set to 60
[1:08:25] pH 5.01 -> 5.21
[1:08:25] Using cautious pH adjust
[1:08:25] Dispensed 0.000047 mL of Base (0.5 M KOH)
[1:08:30] Stepping pH = 5.01
[1:08:30] Dispensed 0.000141 mL of Base (0.5 M KOH)
[1:08:35] Stepping pH = 5.24
[1:08:50] Stirrer speed set to 0
[1:09:02] Datapoint id 70 collected
[1:09:02] Charge balance equation is out by -94.7%
[1:09:02] Stirrer speed set to 60
[1:09:07] pH 5.27 -> 5.47
[1:09:07] Using cautious pH adjust
[1:09:07] Dispensed 0.000024 mL of Base (0.5 M KOH)
[1:09:13] Stepping pH = 5.27
[1:09:13] Dispensed 0.000094 mL of Base (0.5 M KOH)
[1:09:18] Stepping pH = 5.39
[1:09:18] Dispensed 0.000047 mL of Base (0.5 M KOH)

| | | | |
|---|---|---|---|
| Sample name: | **M09_octanol** | Experiment start time: | **3/3/2018 12:18:26 AM** |
| Assay name: | **pH-metric high logP** | Analyst: | **Pion** |
| Assay ID: | **18C-03001** | Instrument ID: | **T312060** |
| Filename: | **C:\Sirius_T3\Mehtap\20180302_exp29_logP_T3-2\18C-03001_M09_octanol_pH-metric high logP.t3r** | | |

#### Experiment Log (continued)

[1:09:23] Stepping pH = 5.50
[1:09:38] Stirrer speed set to 0
[1:09:51] Datapoint id 71 collected
[1:09:51] Charge balance equation is out by -168.0%
[1:09:51] Stirrer speed set to 60
[1:09:56] pH 5.56 -> 5.76
[1:09:56] Using cautious pH adjust
[1:09:56] Dispensed 0.000024 mL of Base (0.5 M KOH)
[1:10:01] Stepping pH = 5.58
[1:10:01] Dispensed 0.000071 mL of Base (0.5 M KOH)
[1:10:06] Stepping pH = 5.86
[1:10:22] Stirrer speed set to 0
[1:10:38] Datapoint id 72 collected
[1:10:38] Charge balance equation is out by -88.7%
[1:10:38] Stirrer speed set to 60
[1:10:43] pH 5.96 -> 6.16
[1:10:43] Using cautious pH adjust
[1:10:43] Dispensed 0.000024 mL of Base (0.5 M KOH)
[1:10:48] Stepping pH = 5.98
[1:10:48] Dispensed 0.000071 mL of Base (0.5 M KOH)
[1:10:53] Stepping pH = 6.55
[1:11:09] Stirrer speed set to 0
[1:12:06] Datapoint id 73 collected
[1:12:06] Charge balance equation is out by -84.8%
[1:12:06] Stirrer speed set to 60
[1:12:11] pH 6.50 -> 6.70
[1:12:11] Using cautious pH adjust
[1:12:11] Dispensed 0.000024 mL of Base (0.5 M KOH)
[1:12:17] Stepping pH = 6.52
[1:12:17] Dispensed 0.000047 mL of Base (0.5 M KOH)
[1:12:22] Stepping pH = 6.85
[1:12:37] Stirrer speed set to 0
[1:13:37] Datapoint id 74 collected
[1:13:37] Charge balance equation is out by -88.4%
[1:13:37] Stirrer speed set to 60
[1:13:42] pH 6.98 -> 7.18
[1:13:42] Using cautious pH adjust
[1:13:42] Dispensed 0.000024 mL of Base (0.5 M KOH)
[1:13:47] Stepping pH = 7.04
[1:13:47] Dispensed 0.000024 mL of Base (0.5 M KOH)
[1:13:52] Stepping pH = 7.14
[1:13:52] Dispensed 0.000024 mL of Base (0.5 M KOH)
[1:13:57] Stepping pH = 7.42
[1:14:12] Stirrer speed set to 0
[1:15:12] Datapoint id 75 collected
[1:15:12] Charge balance equation is out by -161.4%
[1:15:12] Stirrer speed set to 60
[1:15:18] pH 7.51 -> 7.71
[1:15:18] Using cautious pH adjust
[1:15:18] Dispensed 0.000024 mL of Base (0.5 M KOH)
[1:15:23] Stepping pH = 7.56
[1:15:23] Dispensed 0.000024 mL of Base (0.5 M KOH)
[1:15:28] Stepping pH = 7.71
[1:15:43] Stirrer speed set to 0
[1:16:43] Datapoint id 76 collected
[1:16:43] Charge balance equation is out by -306.0%
[1:16:43] Stirrer speed set to 60
[1:16:48] pH 7.94 -> 8.14
[1:16:48] Using cautious pH adjust

Sample name: **M09_octanol** Experiment start time: **3/3/2018 12:18:26 AM**
Assay name: **pH-metric high logP** Analyst: **Pion**
Assay ID: **18C-03001** Instrument ID: **T312060**
Filename: **C:\Sirius_T3\Mehtap\20180302_exp29_logP_T3-2\18C-03001_M09_octanol_pH-metric high logP.t3r**

#### Experiment Log (continued)

[1:16:48] Dispensed 0.000024 mL of Base (0.5 M KOH)
[1:16:53] Stepping pH = 8.00
[1:16:53] Dispensed 0.000024 mL of Base (0.5 M KOH)
[1:16:58] Stepping pH = 8.19
[1:17:13] Stirrer speed set to 0
[1:18:13] Datapoint id 77 collected
[1:18:13] Charge balance equation is out by -415.5%
[1:18:13] Stirrer speed set to 60
[1:18:18] pH 8.04 -> 8.24
[1:18:18] Using cautious pH adjust
[1:18:19] Dispensed 0.000024 mL of Base (0.5 M KOH)
[1:18:24] Stepping pH = 8.02
[1:18:24] Dispensed 0.000024 mL of Base (0.5 M KOH)
[1:18:29] Stepping pH = 8.01
[1:18:29] Dispensed 0.000118 mL of Base (0.5 M KOH)
[1:18:34] Stepping pH = 8.74
[1:18:49] Stirrer speed set to 0
[1:19:22] Datapoint id 78 collected
[1:19:22] Charge balance equation is out by -1,836.6%
[1:19:22] Stirrer speed set to 60
[1:19:27] pH 8.74 -> 8.94
[1:19:27] Using cautious pH adjust
[1:19:27] Dispensed 0.000024 mL of Base (0.5 M KOH)
[1:19:32] Stepping pH = 8.74
[1:19:32] Dispensed 0.000047 mL of Base (0.5 M KOH)
[1:19:37] Stepping pH = 8.85
[1:19:37] Dispensed 0.000024 mL of Base (0.5 M KOH)
[1:19:42] Stepping pH = 8.90
[1:19:42] Dispensed 0.000024 mL of Base (0.5 M KOH)
[1:19:47] Stepping pH = 8.96
[1:20:02] Stirrer speed set to 0
[1:20:32] Datapoint id 79 collected
[1:20:32] Charge balance equation is out by -287.6%
[1:20:32] Stirrer speed set to 60
[1:20:37] pH 8.98 -> 9.05
[1:20:37] Using cautious pH adjust
[1:20:38] Dispensed 0.000024 mL of Base (0.5 M KOH)
[1:20:43] Stepping pH = 8.99
[1:20:43] Dispensed 0.000024 mL of Base (0.5 M KOH)
[1:20:48] Stepping pH = 9.02
[1:21:03] Stirrer speed set to 0
[1:21:16] Datapoint id 80 collected
[1:21:16] Charge balance equation is out by -194.9%
[1:21:16] Argon flow rate set to 0
[1:21:20] Titrator arm moved over Titration position
