## Supplementary material for "Octanol-water partition coefficient measurements for the SAMPL6 Blind Prediction Challenge": SM09_18C-06007_M09_octanol_pH-metric high logP_report.pdf

| | | | |
|---|---|---|---|
| Sample name: | **M09_octanol** | Experiment start time: | **3/6/2018 3:40:58 PM** |
| Assay name: | **pH-metric high logP** | Analyst: | **Pion** |
| Assay ID: | **18C-06007** | Instrument ID: | **T312060** |
| Filename: | **C:\Sirius_T3\Mehtap\20180306_exp30_logP_T3-2\18C-06007_M09_octanol_pH-metric high logP.t3r** | | |

#### pH-metric Result

| | | |
|---|---|---|
| logP (XH +) | 0.91 | ±0.07 (n=50) |
| logP (neutral X) | 3.05 | ±0.02 (n=50) |

#### 18C-06007 Points 1 to 23

| | |
|---|---|
| M09_octanol concentration factor | 1.046 |
| Carbonate | 0.0318 mM |
| Acidity error | 0.09711 mM |

#### 18C-06007 Points 24 to 50

| | |
|---|---|
| M09_octanol concentration factor | 0.952 |
| Carbonate | 0.0262 mM |
| Acidity error | 0.24086 mM |

#### 18C-06007 Points 51 to 74

| | |
|---|---|
| M09_octanol concentration factor | 0.937 |
| Carbonate | 0.0860 mM |
| Acidity error | 0.30528 mM |

#### Warnings and errors

| | |
|---|---|
| Errors | None |
| Warnings | None |

#### Sample logD and percent species

| pH | M09_octanol logD | M09_octanol M09_octanolH | M09_octanol M09_octanol | M09_octanol M09_octanolH* | M09_octanol M09_octanol* | Comment |
|---|---|---|---|---|---|---|
| 1.000 | 0.92 | 10.79 % | 0.00 % | 88.69 % | 0.52 % | |
| 1.200 | 0.92 | 10.76 % | 0.00 % | 88.42 % | 0.82 % | Stomach pH |
| 2.000 | 0.94 | 10.31 % | 0.00 % | 84.72 % | 4.96 % | |
| 3.000 | 1.11 | 7.13 % | 0.03 % | 58.56 % | 34.29 % | |
| 4.000 | 1.73 | 1.74 % | 0.07 % | 14.32 % | 83.86 % | |
| 5.000 | 2.54 | 0.20 % | 0.09 % | 1.67 % | 98.03 % | |
| 6.000 | 2.96 | 0.02 % | 0.09 % | 0.17 % | 99.72 % | |
| 6.500 | 3.02 | 0.01 % | 0.09 % | 0.05 % | 99.85 % | |
| 7.000 | 3.04 | 0.00 % | 0.09 % | 0.02 % | 99.89 % | |
| 7.400 | 3.05 | 0.00 % | 0.09 % | 0.01 % | 99.90 % | Blood pH |
| 8.000 | 3.05 | 0.00 % | 0.09 % | 0.00 % | 99.91 % | |
| 9.000 | 3.05 | 0.00 % | 0.09 % | 0.00 % | 99.91 % | |
| 10.000 | 3.05 | 0.00 % | 0.09 % | 0.00 % | 99.91 % | |
| 11.000 | 3.05 | 0.00 % | 0.09 % | 0.00 % | 99.91 % | |
| 12.000 | 3.05 | 0.00 % | 0.09 % | 0.00 % | 99.91 % | |

Sample name: **M09_octanol**
Assay name: **pH-metric high logP**
Assay ID: **18C-06007**
Filename: **C:\Sirius_T3\Mehtap\20180306_exp30_logP_T3-2\18C-06007_M09_octanol_pH-metric high logP.t3r**

Experiment start time: **3/6/2018 3:40:58 PM**
Analyst: **Pion**
Instrument ID: **T312060**

#### Graphs

Charge balance residuals

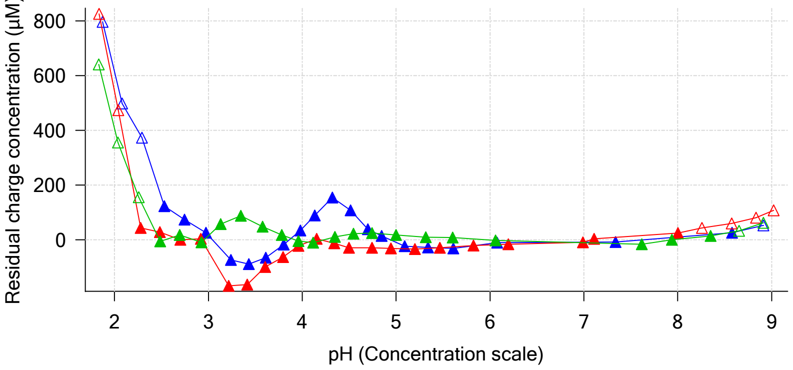


Ionic strength

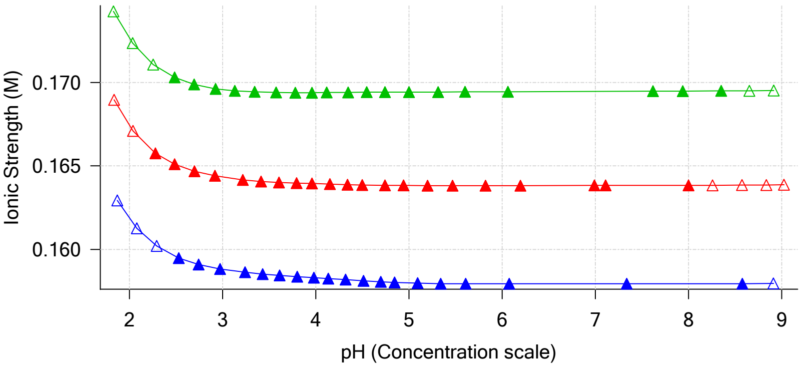


Buffer index profile

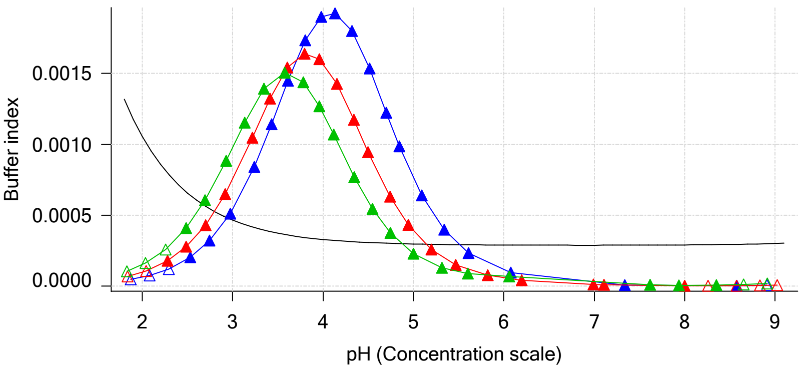


Sample M09_octanol Ionisation Graph

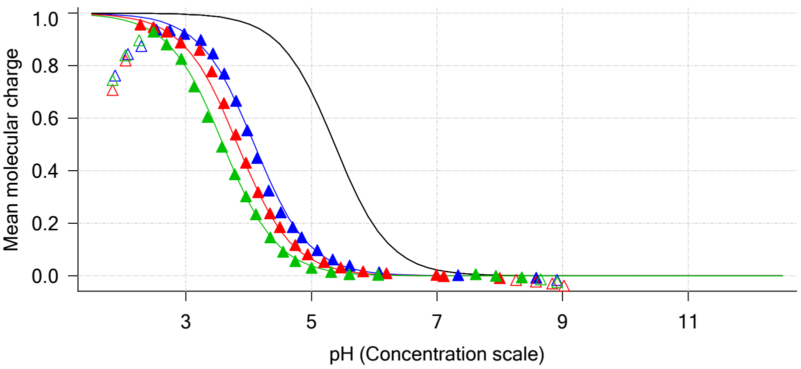


Distribution of species for sample M09_octanol

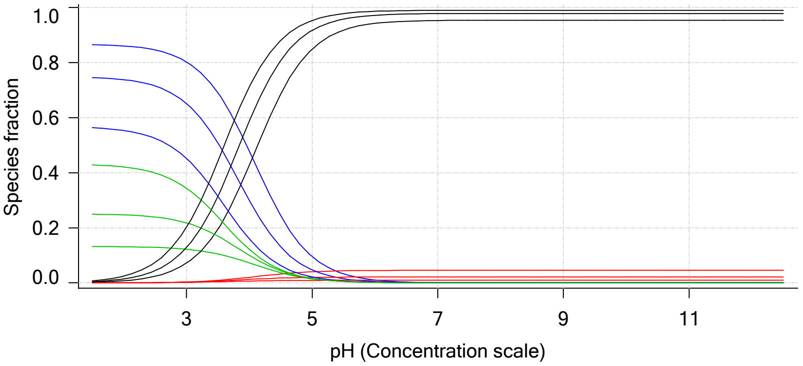


Lipophilicity Profile

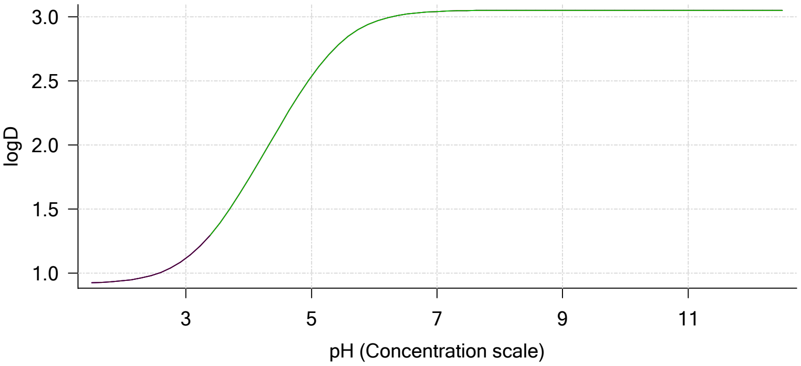


Log R Profile

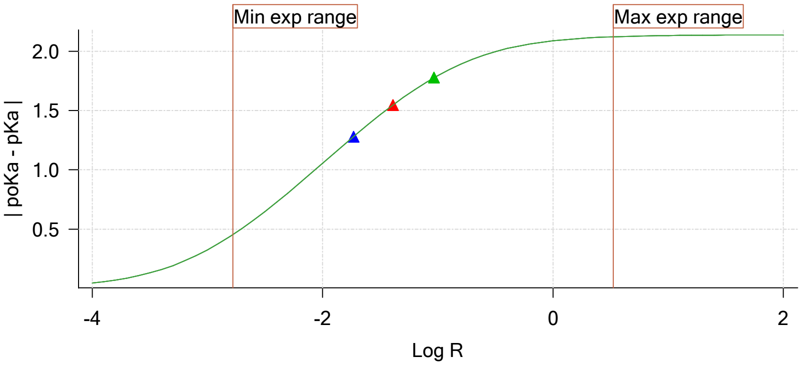


Sample Carbonate Ionisation Graph

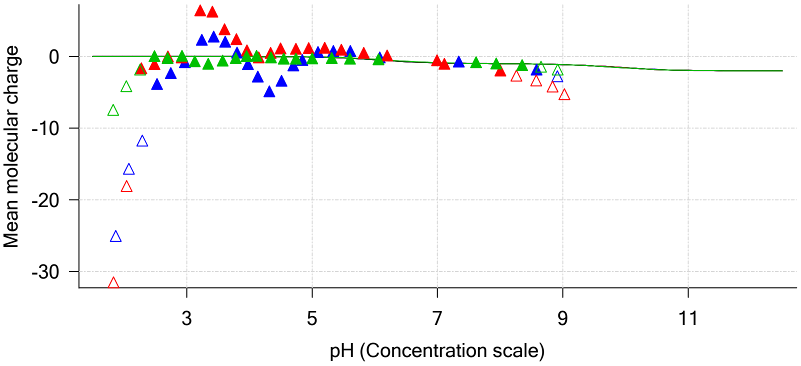

| | | | |
|---|---|---|---|
| Sample name: | **M09_octanol** | Experiment start time: | **3/6/2018 3:40:58 PM** |
| Assay name: | **pH-metric high logP** | Analyst: | **Pion** |
| Assay ID: | **18C-06007** | Instrument ID: | **T312060** |
| Filename: | **C:\Sirius_T3\Mehtap\20180306_exp30_logP_T3-2\18C-06007_M09_octanol_pH-metric high logP.t3r** | | |

#### Graphs (continued)

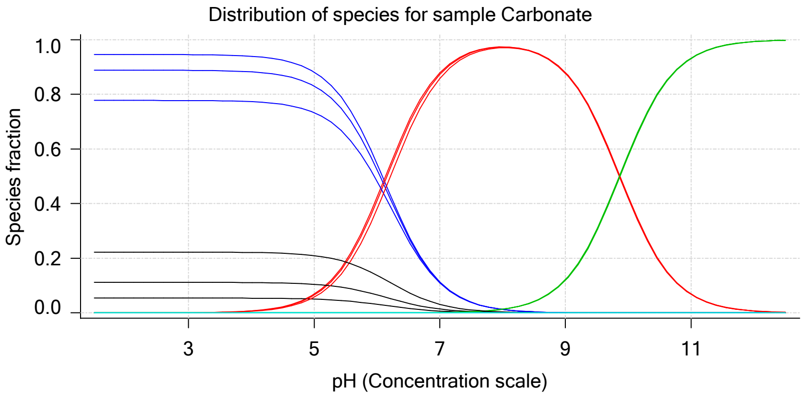

| | | | |
|---|---|---|---|
| Sample name: | **M09_octanol** | Experiment start time: | **3/6/2018 3:40:58 PM** |
| Assay name: | **pH-metric high logP** | Analyst: | **Pion** |
| Assay ID: | **18C-06007** | Instrument ID: | **T312060** |
| Filename: | **C:\Sirius_T3\Mehtap\20180306_exp30_logP_T3-2\18C-06007_M09_octanol_pH-metric high logP.t3r** | | |

### pH-metric high logP Titration 1 of 3 18C-06007 Points 1 to 23

#### Overall results

| | |
|---|---|
| RMSD | 0.369 |
| Average ionic strength | 0.158 M |
| Average temperature | 24.9°C |
| Partition ratio | 0.0185 : 1 |
| Analyte concentration range | 3130.9 µM to 3230.2 µM |
| Total points considered | 19 of 23 |

#### Warnings and errors

| | |
|---|---|
| Errors | None |
| Warnings | None |

#### Four-Plus parameters

| | | | |
|---|---|---|---|
| Alpha | 0.124 | 3/6/2018 3:40:58 PM | C:\Sirius_T3\18C-06006_Blank standardisation.t3r |
| S | 0.9973 | 3/6/2018 3:40:58 PM | C:\Sirius_T3\18C-06006_Blank standardisation.t3r |
| jH | 0.9 | 3/6/2018 3:40:58 PM | C:\Sirius_T3\18C-06006_Blank standardisation.t3r |
| jOH | -0.7 | 3/6/2018 3:40:58 PM | C:\Sirius_T3\18C-06006_Blank standardisation.t3r |

#### Titrants

| | | | |
|---|---|---|---|
| 0.50 M HCl | 0.989131 | 3/6/2018 3:40:58 PM | C:\Sirius_T3\18C-06006_Blank standardisation.t3r |
| 0.50 M KOH | 0.999845 | 3/6/2018 3:40:58 PM | C:\Sirius_T3\KOH18B27.t3r |

#### Sample

| | |
|---|---|
| M09_octanol concentration factor | 1.046 |
| M09_octanol stoichiometry | 1.000 |
| Chloride stoichiometry | 1.000 |
| Base pKa 1 | 5.37 |
| logP (XH +) | 0.75 |
| logP (neutral X) | 3.05 |

#### Sample graphs

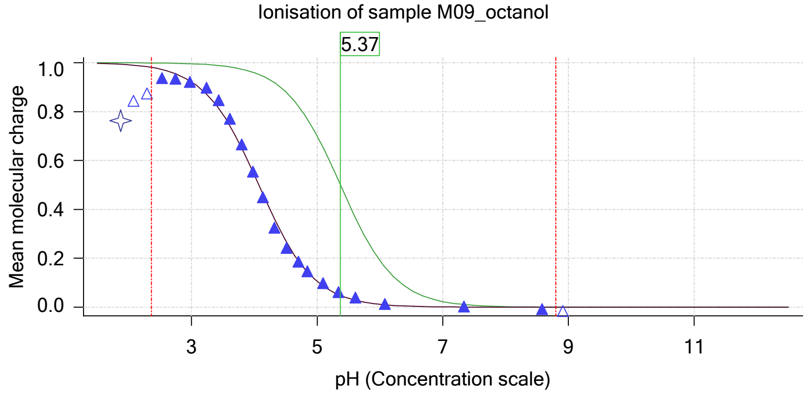


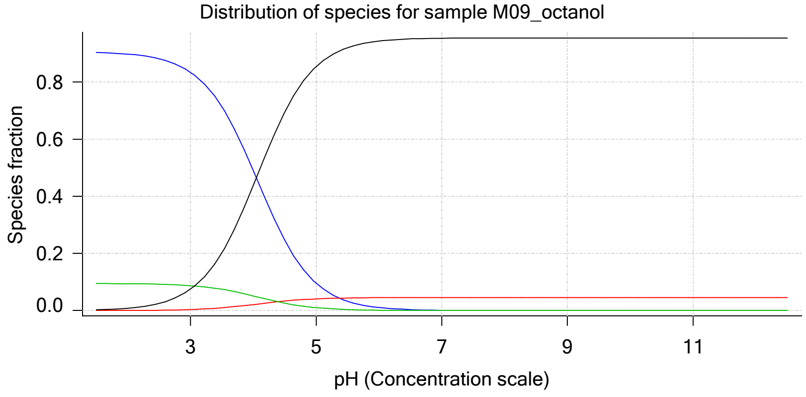

| | | | |
|---|---|---|---|
| Sample name: | **M09_octanol** | Experiment start time: | **3/6/2018 3:40:58 PM** |
| Assay name: | **pH-metric high logP** | Analyst: | **Pion** |
| Assay ID: | **18C-06007** | Instrument ID: | **T312060** |
| Filename: | **C:\Sirius_T3\Mehtap\20180306_exp30_logP_T3-2\18C-06007_M09_octanol_pH-metric high logP.t3r** | | |

#### Sample graphs (continued)

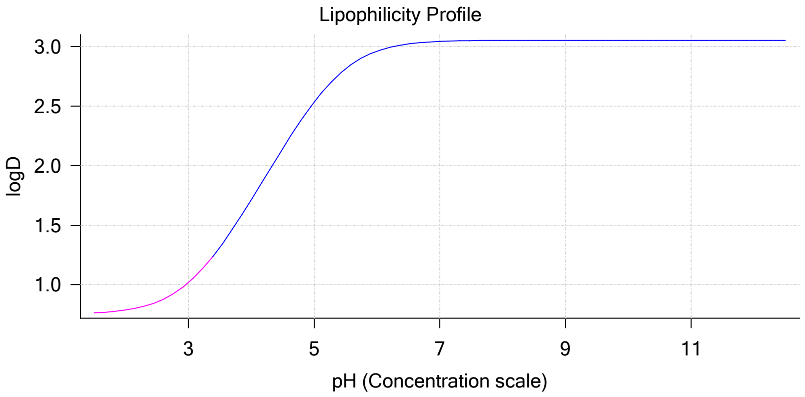


#### Sample logD and percent species

| pH | M09_octanol logD | M09_octanol M09_octanolH | M09_octanol M09_octanol | M09_octanol M09_octanolH* | M09_octanol M09_octanol* | Comment |
|---|---|---|---|---|---|---|
| 1.000 | 0.75 | 90.46 % | 0.00 % | 9.46 % | 0.08 % | |
| 1.200 | 0.76 | 90.41 % | 0.01 % | 9.45 % | 0.13 % | Stomach pH |
| 2.000 | 0.79 | 89.77 % | 0.04 % | 9.39 % | 0.80 % | |
| 3.000 | 1.02 | 83.46 % | 0.36 % | 8.72 % | 7.46 % | |
| 4.000 | 1.71 | 48.98 % | 2.09 % | 5.12 % | 43.81 % | |
| 5.000 | 2.53 | 9.55 % | 4.07 % | 1.00 % | 85.38 % | |
| 6.000 | 2.96 | 1.05 % | 4.50 % | 0.11 % | 94.34 % | |
| 6.500 | 3.02 | 0.34 % | 4.54 % | 0.04 % | 95.09 % | |
| 7.000 | 3.04 | 0.11 % | 4.55 % | 0.01 % | 95.34 % | |
| 7.400 | 3.05 | 0.04 % | 4.55 % | 0.00 % | 95.40 % | Blood pH |
| 8.000 | 3.05 | 0.01 % | 4.55 % | 0.00 % | 95.44 % | |
| 9.000 | 3.05 | 0.00 % | 4.55 % | 0.00 % | 95.45 % | |
| 10.000 | 3.05 | 0.00 % | 4.55 % | 0.00 % | 95.45 % | |
| 11.000 | 3.05 | 0.00 % | 4.55 % | 0.00 % | 95.45 % | |
| 12.000 | 3.05 | 0.00 % | 4.55 % | 0.00 % | 95.45 % | |

#### Carbonate and acidity

| | |
|---|---|
| Carbonate | 0.032 mM |
| Acidity error | 0.097 mM |

#### Other graphs

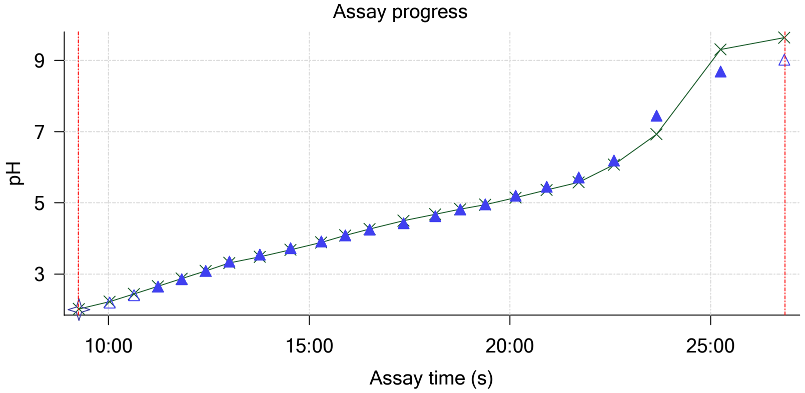


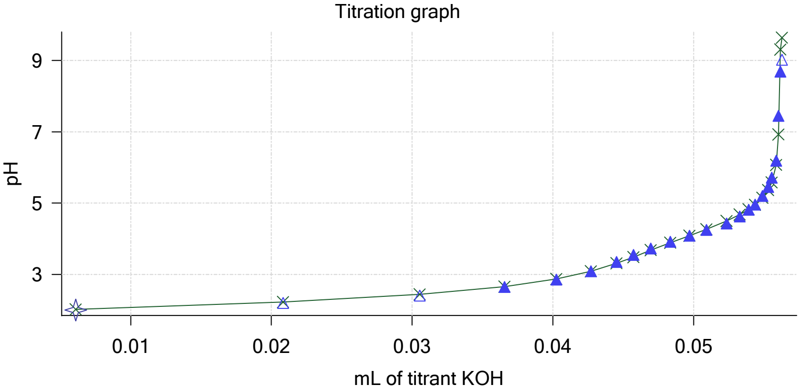

Sample name: **M09_octanol**
Assay name: **pH-metric high logP**
Assay ID: **18C-06007**
Filename: **C:\Sirius_T3\Mehtap\20180306_exp30_logP_T3-2\18C-06007_M09_octanol_pH-metric high logP.t3r**

Experiment start time: **3/6/2018 3:40:58 PM**
Analyst: **Pion**
Instrument ID: **T312060**

#### Other graphs (continued)

Charge balance residuals

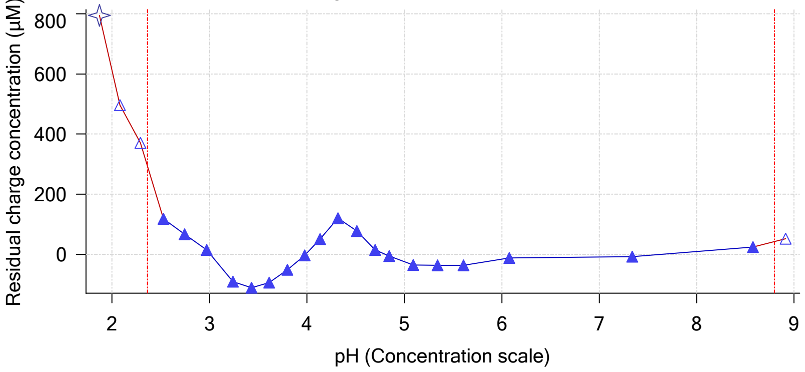


Buffer index profile

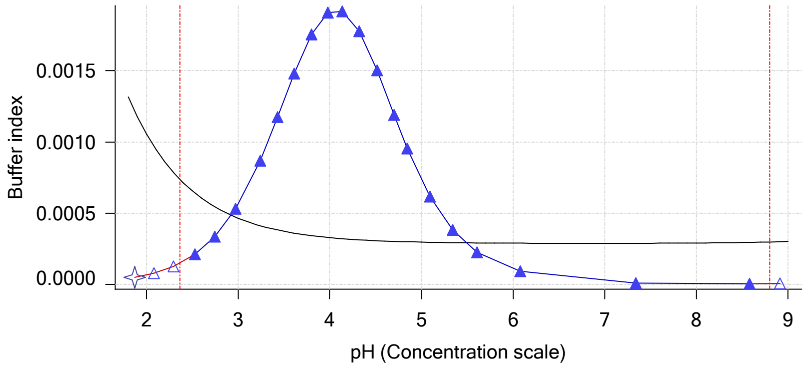


Ionic strength

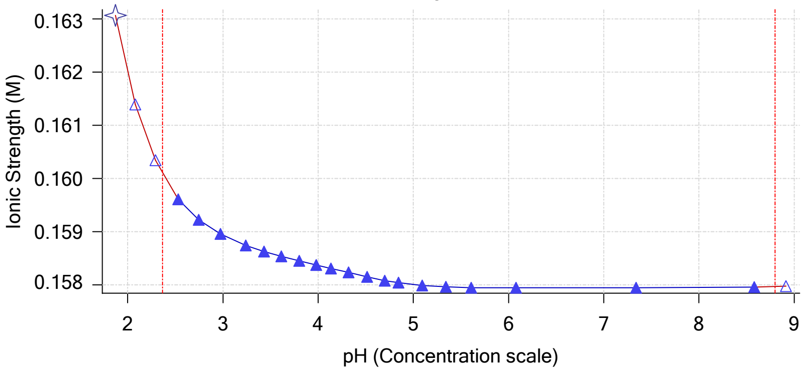


Log R Profile

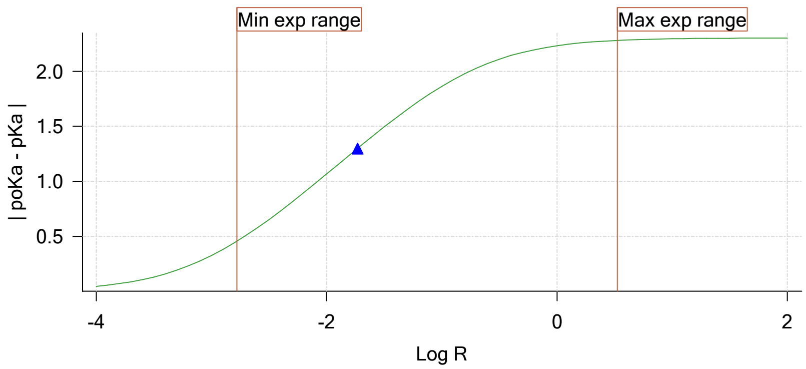


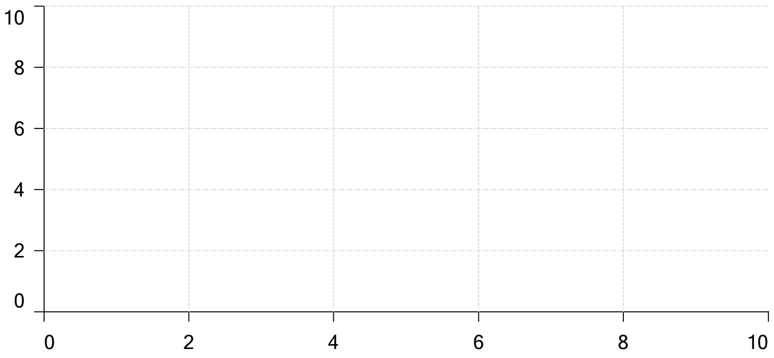


Ionisation of sample Carbonate

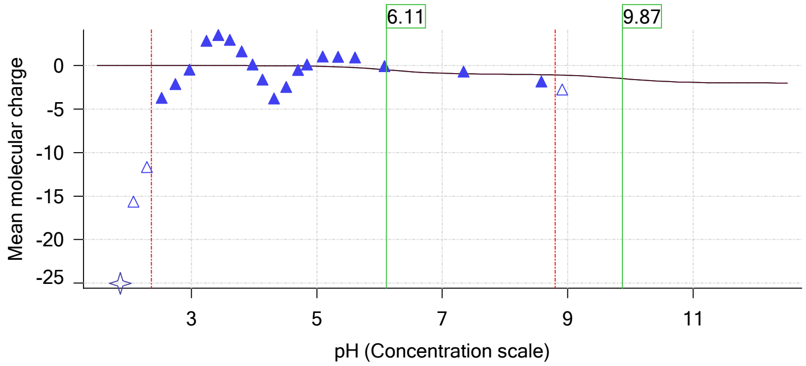

| | | | |
|---|---|---|---|
| Sample name: | **M09_octanol** | Experiment start time: | **3/6/2018 3:40:58 PM** |
| Assay name: | **pH-metric high logP** | Analyst: | **Pion** |
| Assay ID: | **18C-06007** | Instrument ID: | **T312060** |
| Filename: | **C:\Sirius_T3\Mehtap\20180306_exp30_logP_T3-2\18C-06007_M09_octanol_pH-metric high logP.t3r** | | |

# pH-metric high logP   Titration 2 of 3   18C-06007 Points 24 to 50

#### Overall results

| | |
|---|---|
| RMSD | 0.910 |
| Average ionic strength | 0.164 M |
| Average temperature | 25.0°C |
| Partition ratio | 0.0407 : 1 |
| Analyte concentration range | 2862.7 µM to 2955.4 µM |
| Total points considered | 21 of 27 |

#### Warnings and errors

| | |
|---|---|
| Errors | None |
| Warnings | None |

#### Four-Plus parameters

| | | | |
|---|---|---|---|
| Alpha | 0.124 | 3/6/2018 3:40:58 PM | C:\Sirius_T3\18C-06006_Blank standardisation.t3r |
| S | 0.9973 | 3/6/2018 3:40:58 PM | C:\Sirius_T3\18C-06006_Blank standardisation.t3r |
| jH | 0.9 | 3/6/2018 3:40:58 PM | C:\Sirius_T3\18C-06006_Blank standardisation.t3r |
| jOH | -0.7 | 3/6/2018 3:40:58 PM | C:\Sirius_T3\18C-06006_Blank standardisation.t3r |

#### Titrants

| | | | |
|---|---|---|---|
| 0.50 M HCl | 0.989131 | 3/6/2018 3:40:58 PM | C:\Sirius_T3\18C-06006_Blank standardisation.t3r |
| 0.50 M KOH | 0.999845 | 3/6/2018 3:40:58 PM | C:\Sirius_T3\KOH18B27.t3r |

#### Sample

| | |
|---|---|
| M09_octanol concentration factor | 0.952 |
| M09_octanol stoichiometry | 1.000 |
| Chloride stoichiometry | 1.000 |
| Base pKa 1 | 5.37 |
| logP (XH +) | 0.76 |
| logP (neutral X) | 2.97 |

#### Sample graphs

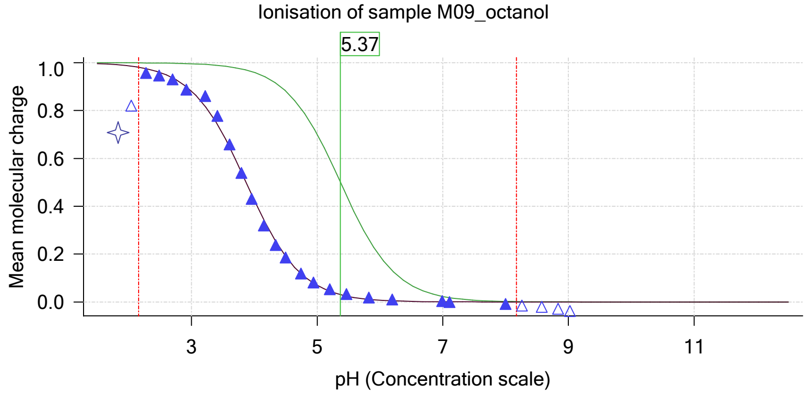


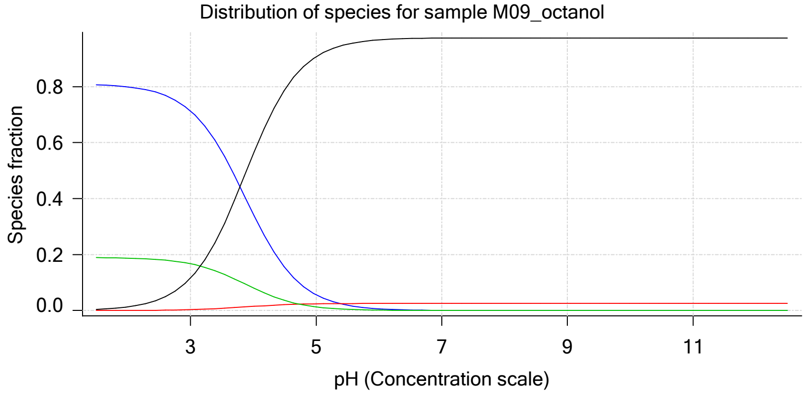

| | | | |
|---|---|---|---|
| Sample name: | **M09_octanol** | Experiment start time: | **3/6/2018 3:40:58 PM** |
| Assay name: | **pH-metric high logP** | Analyst: | **Pion** |
| Assay ID: | **18C-06007** | Instrument ID: | **T312060** |
| Filename: | **C:\Sirius_T3\Mehtap\20180306_exp30_logP_T3-2\18C-06007_M09_octanol_pH-metric high logP.t3r** | | |

#### Sample graphs (continued)

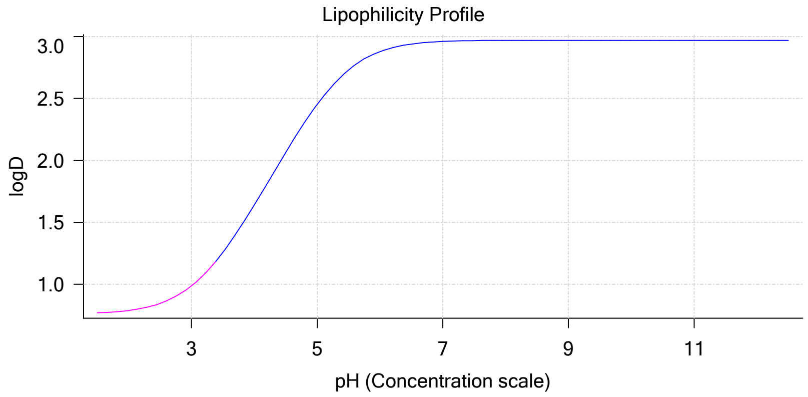


#### Sample logD and percent species

| pH | M09_octanol logD | M09_octanol M09_octanolH | M09_octanol M09_octanol | M09_octanol M09_octanolH* | M09_octanol M09_octanol* | Comment |
|---|---|---|---|---|---|---|
| 1.000 | 0.76 | 80.91 % | 0.00 % | 18.96 % | 0.13 % | |
| 1.200 | 0.76 | 80.84 % | 0.01 % | 18.94 % | 0.21 % | Stomach pH |
| 2.000 | 0.79 | 79.94 % | 0.03 % | 18.73 % | 1.30 % | |
| 3.000 | 0.99 | 71.38 % | 0.30 % | 16.73 % | 11.59 % | |
| 4.000 | 1.64 | 34.48 % | 1.47 % | 8.08 % | 55.97 % | |
| 5.000 | 2.45 | 5.59 % | 2.38 % | 1.31 % | 90.72 % | |
| 6.000 | 2.88 | 0.60 % | 2.54 % | 0.14 % | 96.72 % | |
| 6.500 | 2.94 | 0.19 % | 2.55 % | 0.04 % | 97.21 % | |
| 7.000 | 2.96 | 0.06 % | 2.56 % | 0.01 % | 97.37 % | |
| 7.400 | 2.97 | 0.02 % | 2.56 % | 0.01 % | 97.41 % | Blood pH |
| 8.000 | 2.97 | 0.01 % | 2.56 % | 0.00 % | 97.43 % | |
| 9.000 | 2.97 | 0.00 % | 2.56 % | 0.00 % | 97.44 % | |
| 10.000 | 2.97 | 0.00 % | 2.56 % | 0.00 % | 97.44 % | |
| 11.000 | 2.97 | 0.00 % | 2.56 % | 0.00 % | 97.44 % | |
| 12.000 | 2.97 | 0.00 % | 2.56 % | 0.00 % | 97.44 % | |

#### Carbonate and acidity

| | |
|---|---|
| Carbonate | 0.026 mM |
| Acidity error | 0.241 mM |

#### Other graphs

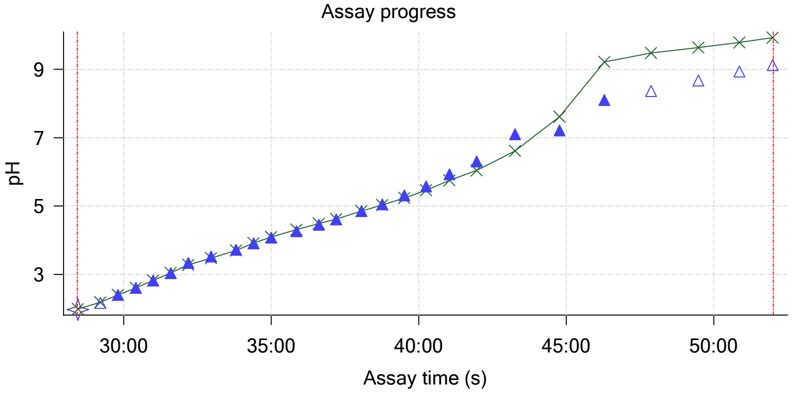


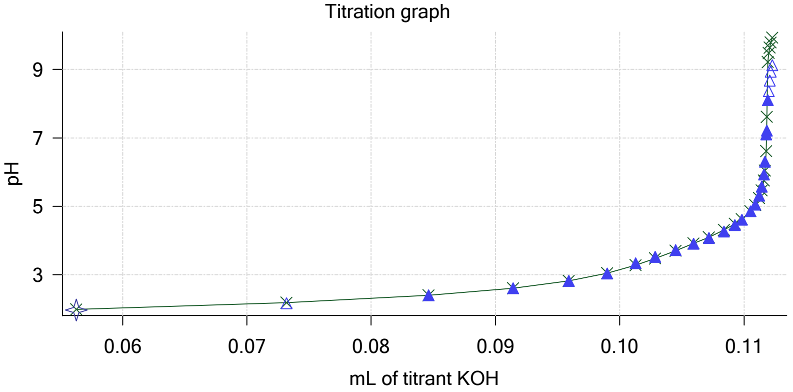

| | | | |
|---|---|---|---|
| Sample name: | **M09_octanol** | Experiment start time: | **3/6/2018 3:40:58 PM** |
| Assay name: | **pH-metric high logP** | Analyst: | **Pion** |
| Assay ID: | **18C-06007** | Instrument ID: | **T312060** |
| Filename: | **C:\Sirius_T3\Mehtap\20180306_exp30_logP_T3-2\18C-06007_M09_octanol_pH-metric high logP.t3r** | | |

#### Other graphs (continued)

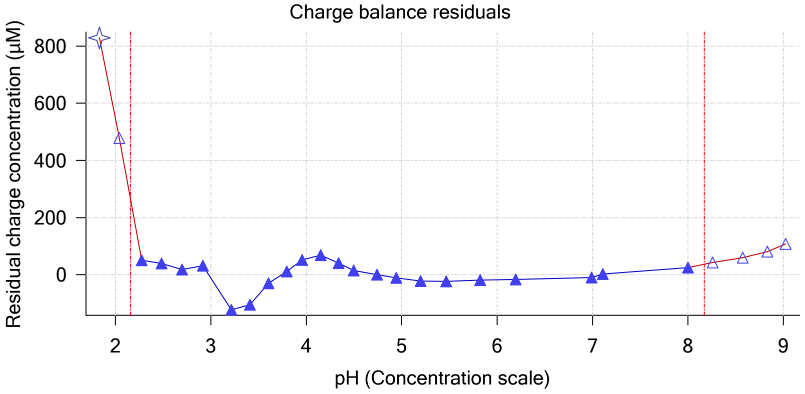


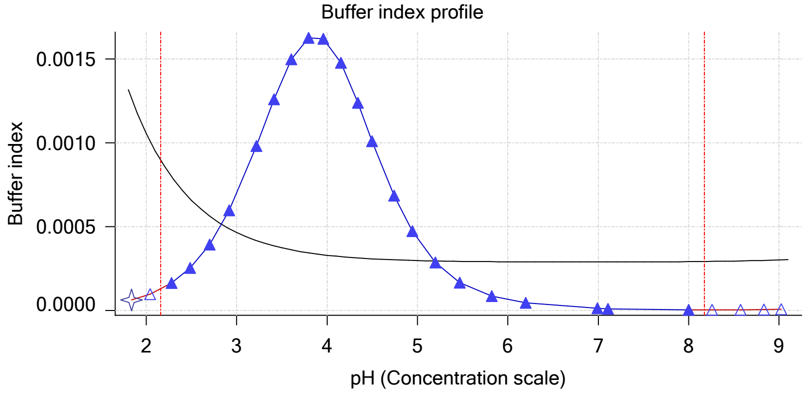


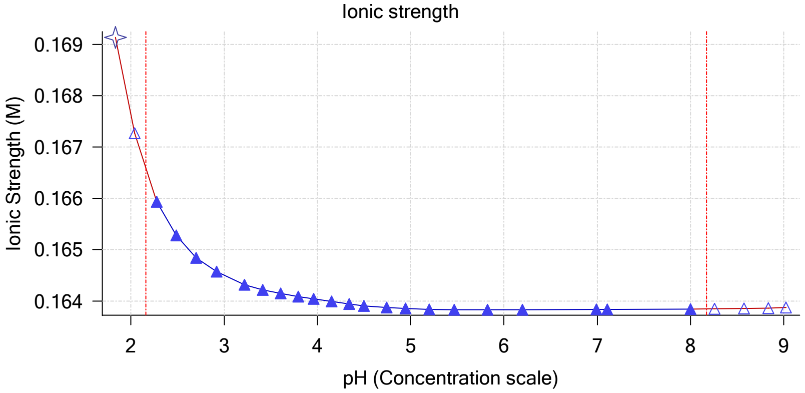


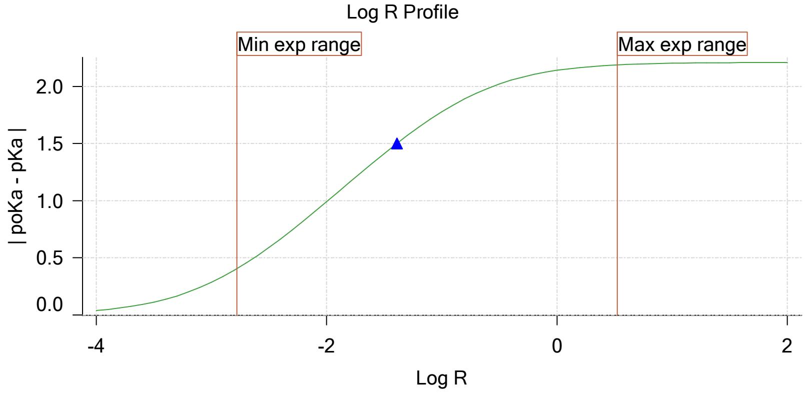


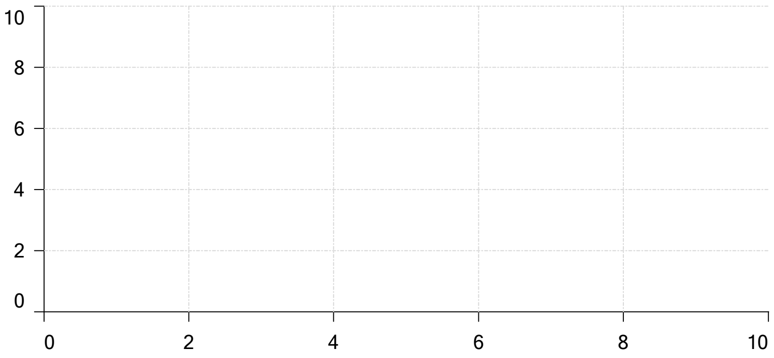

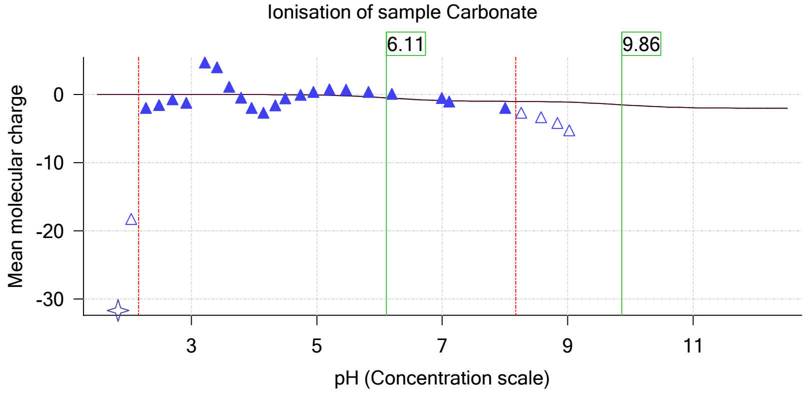

| | | | |
|---|---|---|---|
| Sample name: | **M09_octanol** | Experiment start time: | **3/6/2018 3:40:58 PM** |
| Assay name: | **pH-metric high logP** | Analyst: | **Pion** |
| Assay ID: | **18C-06007** | Instrument ID: | **T312060** |
| Filename: | **C:\Sirius_T3\Mehtap\20180306_exp30_logP_T3-2\18C-06007_M09_octanol_pH-metric high logP.t3r** | | |

### pH-metric high logP Titration 3 of 3 18C-06007 Points 51 to 74

#### Overall results

| | |
|---|---|
| RMSD | 0.405 |
| Average ionic strength | 0.170 M |
| Average temperature | 25.0°C |
| Partition ratio | 0.0926 : 1 |
| Analyte concentration range | 2547.3 µM to 2626.2 µM |
| Total points considered | 19 of 24 |

#### Warnings and errors

| | |
|---|---|
| Errors | None |
| Warnings | None |

#### Four-Plus parameters

| | | | |
|---|---|---|---|
| Alpha | 0.124 | 3/6/2018 3:40:58 PM | C:\Sirius_T3\18C-06006_Blank standardisation.t3r |
| S | 0.9973 | 3/6/2018 3:40:58 PM | C:\Sirius_T3\18C-06006_Blank standardisation.t3r |
| jH | 0.9 | 3/6/2018 3:40:58 PM | C:\Sirius_T3\18C-06006_Blank standardisation.t3r |
| jOH | -0.7 | 3/6/2018 3:40:58 PM | C:\Sirius_T3\18C-06006_Blank standardisation.t3r |

#### Titrants

| | | | |
|---|---|---|---|
| 0.50 M HCl | 0.989131 | 3/6/2018 3:40:58 PM | C:\Sirius_T3\18C-06006_Blank standardisation.t3r |
| 0.50 M KOH | 0.999845 | 3/6/2018 3:40:58 PM | C:\Sirius_T3\KOH18B27.t3r |

#### Sample

| | |
|---|---|
| M09_octanol concentration factor | 0.937 |
| M09_octanol stoichiometry | 1.000 |
| Chloride stoichiometry | 1.000 |
| Base pKa 1 | 5.37 |
| logP (XH +) | 0.76 |
| logP (neutral X) | 3.01 |

#### Sample graphs

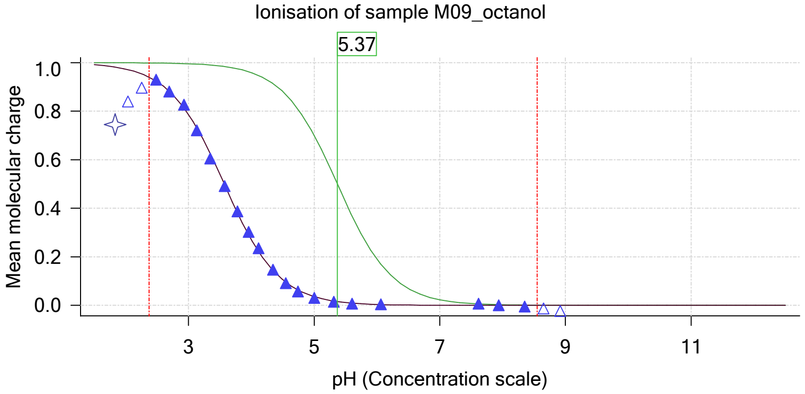


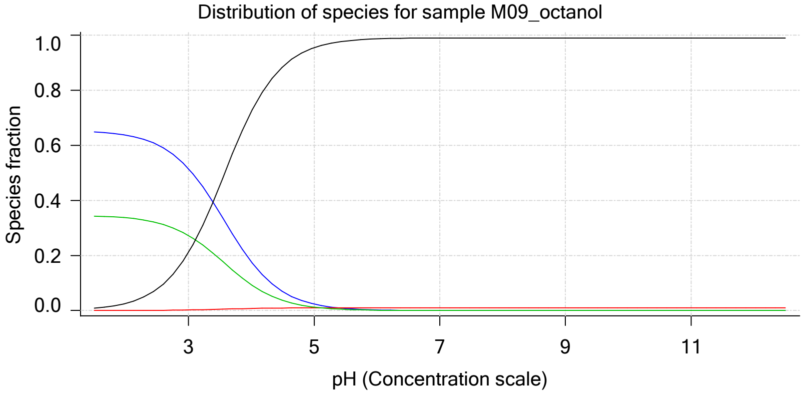

| | | | |
|---|---|---|---|
| Sample name: | **M09_octanol** | Experiment start time: | **3/6/2018 3:40:58 PM** |
| Assay name: | **pH-metric high logP** | Analyst: | **Pion** |
| Assay ID: | **18C-06007** | Instrument ID: | **T312060** |
| Filename: | **C:\Sirius_T3\Mehtap\20180306_exp30_logP_T3-2\18C-06007_M09_octanol_pH-metric high logP.t3r** | | |

#### Sample graphs (continued)

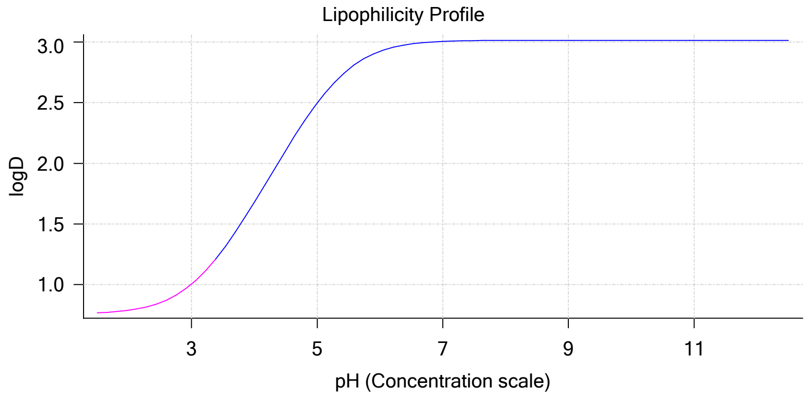


#### Sample logD and percent species

| pH | M09_octanol logD | M09_octanol M09_octanolH | M09_octanol M09_octanol | M09_octanol M09_octanolH* | M09_octanol M09_octanol* | Comment |
|---|---|---|---|---|---|---|
| 1.000 | 0.76 | 65.23 % | 0.00 % | 34.50 % | 0.27 % | |
| 1.200 | 0.76 | 65.13 % | 0.00 % | 34.45 % | 0.42 % | Stomach pH |
| 2.000 | 0.79 | 63.69 % | 0.03 % | 33.69 % | 2.60 % | |
| 3.000 | 1.00 | 51.51 % | 0.22 % | 27.25 % | 21.02 % | |
| 4.000 | 1.68 | 17.69 % | 0.75 % | 9.36 % | 72.19 % | |
| 5.000 | 2.50 | 2.34 % | 1.00 % | 1.24 % | 95.43 % | |
| 6.000 | 2.92 | 0.24 % | 1.03 % | 0.13 % | 98.60 % | |
| 6.500 | 2.98 | 0.08 % | 1.03 % | 0.04 % | 98.85 % | |
| 7.000 | 3.00 | 0.02 % | 1.03 % | 0.01 % | 98.93 % | |
| 7.400 | 3.01 | 0.01 % | 1.03 % | 0.01 % | 98.95 % | Blood pH |
| 8.000 | 3.01 | 0.00 % | 1.03 % | 0.00 % | 98.96 % | |
| 9.000 | 3.01 | 0.00 % | 1.03 % | 0.00 % | 98.97 % | |
| 10.000 | 3.01 | 0.00 % | 1.03 % | 0.00 % | 98.97 % | |
| 11.000 | 3.01 | 0.00 % | 1.03 % | 0.00 % | 98.97 % | |
| 12.000 | 3.01 | 0.00 % | 1.03 % | 0.00 % | 98.97 % | |

#### Carbonate and acidity

| | |
|---|---|
| Carbonate | 0.086 mM |
| Acidity error | 0.305 mM |

#### Other graphs

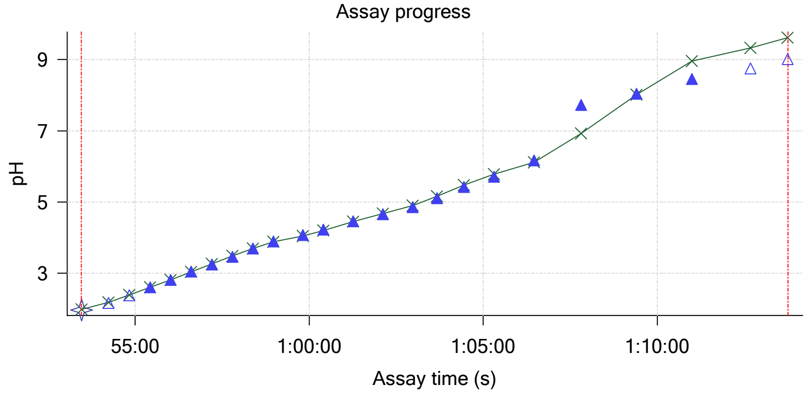


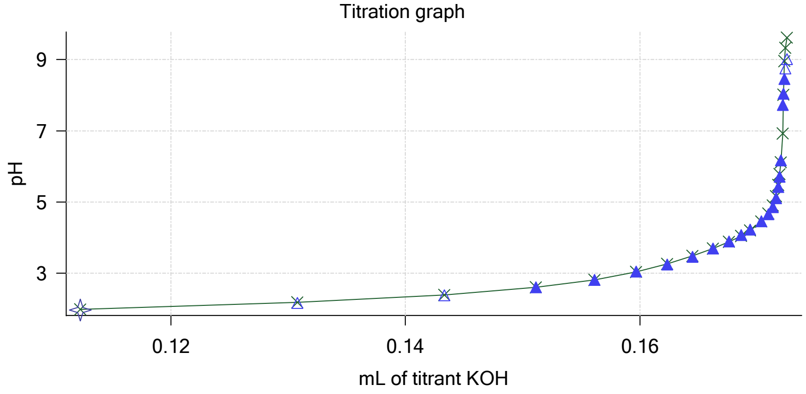

| | | | |
|---|---|---|---|
| Sample name: | **M09_octanol** | Experiment start time: | **3/6/2018 3:40:58 PM** |
| Assay name: | **pH-metric high logP** | Analyst: | **Pion** |
| Assay ID: | **18C-06007** | Instrument ID: | **T312060** |
| Filename: | **C:\Sirius_T3\Mehtap\20180306_exp30_logP_T3-2\18C-06007_M09_octanol_pH-metric high logP.t3r** | | |

#### Other graphs (continued)

Charge balance residuals

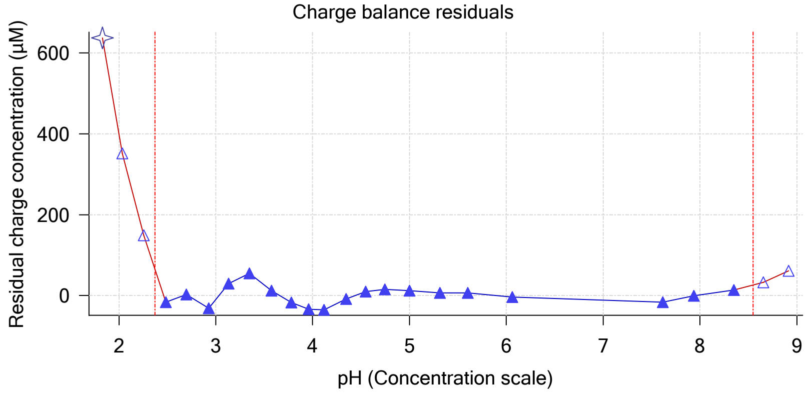


Buffer index profile

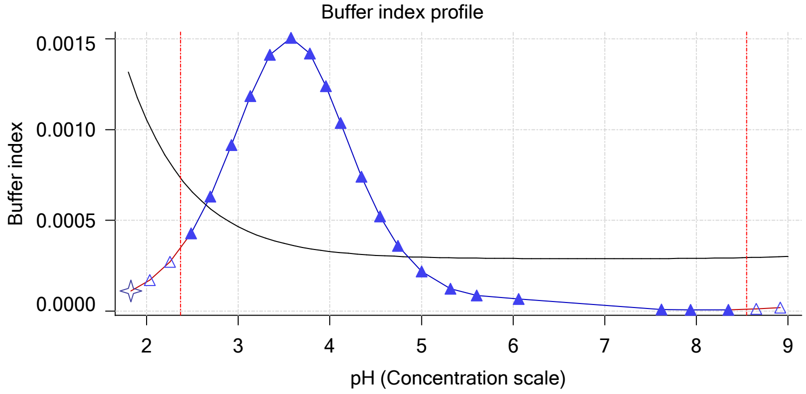


Ionic strength

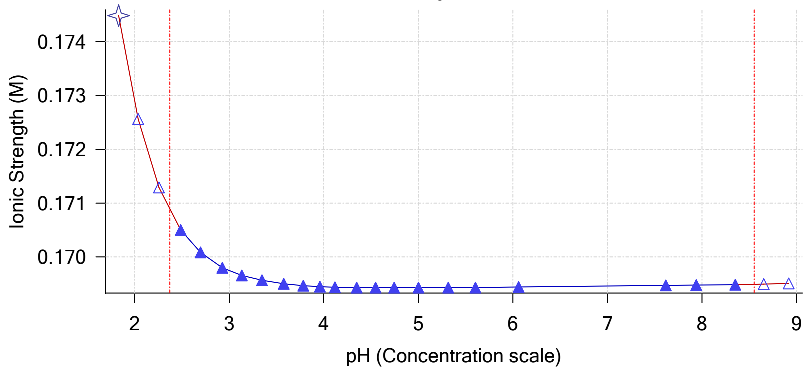


Log R Profile

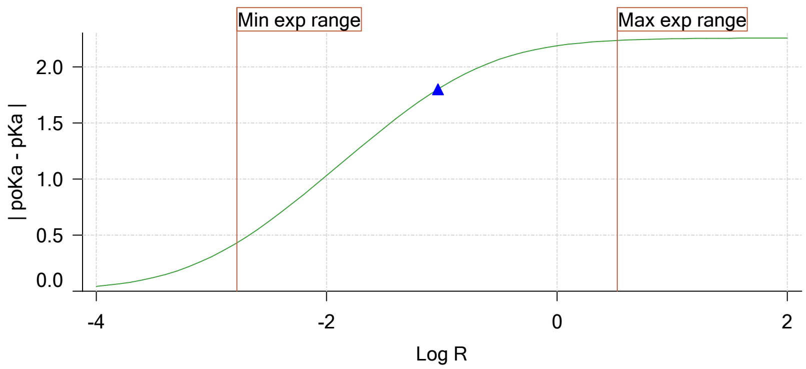


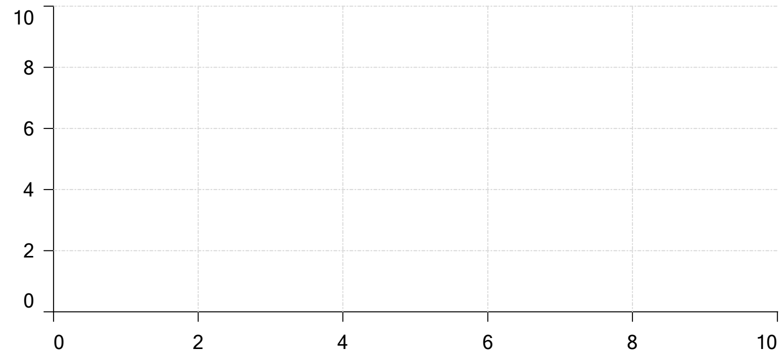


Ionisation of sample Carbonate

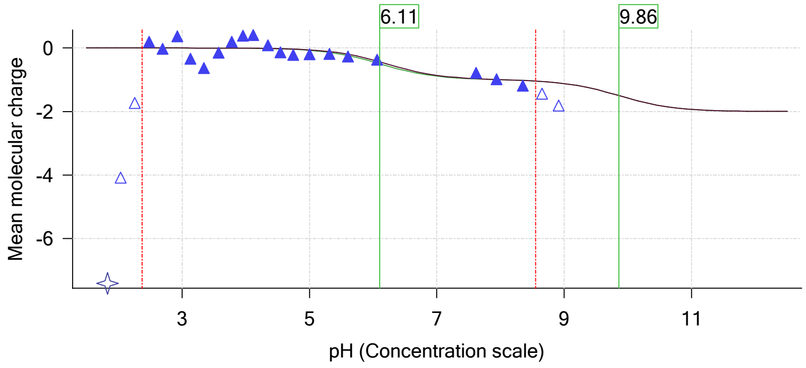

| | | | |
|---|---|---|---|
| Sample name: | **M09_octanol** | Experiment start time: | **3/6/2018 3:40:58 PM** |
| Assay name: | **pH-metric high logP** | Analyst: | **Pion** |
| Assay ID: | **18C-06007** | Instrument ID: | **T312060** |
| Filename: | **C:\Sirius_T3\Mehtap\20180306_exp30_logP_T3-2\18C-06007_M09_octanol_pH-metric high logP.t3r** | | |

#### Assay Model

| Settings | Value | Date/Time changed | Imported from |
|---|---|---|---|
| Sample name | M09_octanol | 2/27/2018 4:56:17 PM | User entered value |
| Sample by | Weight | | Default value |
| Sample weight | 0.001470 g | 3/6/2018 3:39:12 PM | User entered value |
| Formula weight | 287.74 g/mol | 2/27/2018 4:45:45 PM | User entered value |
| Solubility | Unknown | | Default value |
| Molecular weight | 251.28 | 2/27/2018 4:45:45 PM | User entered value |
| Individual pKa ionic environments | No | | Default value |
| Number of pKas | 1 | 2/27/2018 4:45:45 PM | User entered value |
| Sample is a | Base | 2/27/2018 4:45:45 PM | User entered value |
| pKa 1 | 5.37 | 2/27/2018 4:45:45 PM | User entered value |
| logp (XH +) | 0.76 | 3/2/2018 3:27:23 PM | User entered value |
| logP (neutral X) | 3.27 | 3/2/2018 3:27:17 PM | User entered value |
| Stoichiometry | 1.00000 | | Default value |
| Aprotic counterion name | Chloride | | From standards.xml file |
| Stoichiometry | 1.00 | | From standards.xml file |
| Charge per counterion | -1 | | From standards.xml file |

#### Events

| Time | Event | Water | Acid | Base | Octanol | pH | dpH/dt | pH R-squared | pH SD | dpH/dt time |
|---|---|---|---|---|---|---|---|---|---|---|
| 6:16.5 | Initial pH = 3.92 | | | | | | | | | |
| 9:16.0 | Data point 1 | 1.50000 mL | 0.04544 mL | 0.00609 mL | 0.03001 mL | 2.003 | -0.00765 | 0.85441 | 0.00041 | 10.0 s |
| 10:02.2 | Data point 2 | 1.50000 mL | 0.04544 mL | 0.02084 mL | 0.03001 mL | 2.206 | -0.00581 | 0.82838 | 0.00032 | 10.5 s |
| 10:38.4 | Data point 3 | 1.50000 mL | 0.04544 mL | 0.03053 mL | 0.03001 mL | 2.415 | -0.00308 | 0.68547 | 0.00018 | 10.5 s |
| 11:14.3 | Data point 4 | 1.50000 mL | 0.04544 mL | 0.03657 mL | 0.03001 mL | 2.649 | -0.00951 | 0.26483 | 0.00091 | 10.0 s |
| 11:49.8 | Data point 5 | 1.50000 mL | 0.04544 mL | 0.04024 mL | 0.03001 mL | 2.863 | -0.01634 | 0.89001 | 0.00086 | 10.0 s |
| 12:25.3 | Data point 6 | 1.50000 mL | 0.04544 mL | 0.04271 mL | 0.03001 mL | 3.089 | -0.00759 | 0.82120 | 0.00041 | 10.0 s |
| 13:00.8 | Data point 7 | 1.50000 mL | 0.04544 mL | 0.04452 mL | 0.03001 mL | 3.357 | -0.00824 | 0.86493 | 0.00044 | 10.0 s |
| 13:46.4 | Data point 8 | 1.50000 mL | 0.04544 mL | 0.04572 mL | 0.03001 mL | 3.546 | -0.00997 | 0.85301 | 0.00053 | 10.0 s |
| 14:32.2 | Data point 9 | 1.50000 mL | 0.04544 mL | 0.04694 mL | 0.03001 mL | 3.727 | -0.00887 | 0.87089 | 0.00047 | 10.5 s |
| 15:18.4 | Data point 10 | 1.50000 mL | 0.04544 mL | 0.04833 mL | 0.03001 mL | 3.916 | -0.01261 | 0.86654 | 0.00067 | 10.0 s |
| 15:54.0 | Data point 11 | 1.50000 mL | 0.04544 mL | 0.04969 mL | 0.03001 mL | 4.093 | -0.01852 | 0.93335 | 0.00095 | 11.0 s |
| 16:30.4 | Data point 12 | 1.50000 mL | 0.04544 mL | 0.05092 mL | 0.03001 mL | 4.247 | -0.01783 | 0.96763 | 0.00090 | 10.0 s |
| 17:21.2 | Data point 13 | 1.50000 mL | 0.04544 mL | 0.05233 mL | 0.03001 mL | 4.433 | -0.01718 | 0.93747 | 0.00088 | 11.5 s |
| 18:08.5 | Data point 14 | 1.50000 mL | 0.04544 mL | 0.05327 mL | 0.03001 mL | 4.627 | -0.01904 | 0.93322 | 0.00097 | 12.0 s |
| 18:45.9 | Data point 15 | 1.50000 mL | 0.04544 mL | 0.05390 mL | 0.03001 mL | 4.812 | -0.01725 | 0.84669 | 0.00093 | 12.0 s |
| 19:23.3 | Data point 16 | 1.50000 mL | 0.04544 mL | 0.05435 mL | 0.03001 mL | 4.956 | -0.01847 | 0.93434 | 0.00094 | 15.0 s |
| 20:08.9 | Data point 17 | 1.50000 mL | 0.04544 mL | 0.05489 mL | 0.03001 mL | 5.203 | -0.01881 | 0.92505 | 0.00097 | 16.0 s |
| 20:55.1 | Data point 18 | 1.50000 mL | 0.04544 mL | 0.05529 mL | 0.03001 mL | 5.450 | -0.01772 | 0.88375 | 0.00093 | 17.5 s |
| 21:43.1 | Data point 19 | 1.50000 mL | 0.04544 mL | 0.05555 mL | 0.03001 mL | 5.717 | -0.01934 | 0.94249 | 0.00098 | 17.0 s |
| 22:35.7 | Data point 20 | 1.50000 mL | 0.04544 mL | 0.05586 mL | 0.03001 mL | 6.186 | -0.01984 | 0.97180 | 0.00099 | 33.0 s |
| 23:39.3 | Data point 21 | 1.50000 mL | 0.04544 mL | 0.05602 mL | 0.03001 mL | 7.443 | -0.10565 | 0.99712 | 0.00522 | Timed out at 59.5 s |
| 25:15.0 | Data point 22 | 1.50000 mL | 0.04544 mL | 0.05616 mL | 0.03001 mL | 8.681 | -0.01903 | 0.92118 | 0.00098 | 59.5 s |
| 26:50.1 | Data point 23 | 1.50000 mL | 0.04544 mL | 0.05628 mL | 0.03001 mL | 9.012 | -0.01984 | 0.96660 | 0.00100 | 37.5 s |
| 28:26.7 | Data point 24 | 1.50000 mL | 0.10228 mL | 0.05628 mL | 0.07001 mL | 1.967 | -0.00938 | 0.82269 | 0.00051 | 10.0 s |
| 29:12.7 | Data point 25 | 1.50000 mL | 0.10228 mL | 0.07319 mL | 0.07001 mL | 2.167 | -0.00239 | 0.33662 | 0.00020 | 10.0 s |
| 29:48.4 | Data point 26 | 1.50000 mL | 0.10228 mL | 0.08462 mL | 0.07001 mL | 2.401 | -0.00220 | 0.08165 | 0.00038 | 10.5 s |
| 30:24.5 | Data point 27 | 1.50000 mL | 0.10228 mL | 0.09142 mL | 0.07001 mL | 2.605 | -0.00054 | 0.07623 | 0.00010 | 10.0 s |
| 31:00.1 | Data point 28 | 1.50000 mL | 0.10228 mL | 0.09591 mL | 0.07001 mL | 2.819 | -0.00633 | 0.20637 | 0.00069 | 10.0 s |
| 31:35.6 | Data point 29 | 1.50000 mL | 0.10228 mL | 0.09899 mL | 0.07001 mL | 3.036 | -0.01322 | 0.81377 | 0.00072 | 10.5 s |
| 32:11.6 | Data point 30 | 1.50000 mL | 0.10228 mL | 0.10129 mL | 0.07001 mL | 3.333 | -0.00473 | 0.38455 | 0.00038 | 10.0 s |
| 32:57.3 | Data point 31 | 1.50000 mL | 0.10228 mL | 0.10285 mL | 0.07001 mL | 3.527 | -0.00844 | 0.34234 | 0.00071 | 10.0 s |
| 33:48.3 | Data point 32 | 1.50000 mL | 0.10228 mL | 0.10449 mL | 0.07001 mL | 3.721 | -0.01104 | 0.52367 | 0.00075 | 10.5 s |
| 34:24.3 | Data point 33 | 1.50000 mL | 0.10228 mL | 0.10595 mL | 0.07001 mL | 3.910 | -0.00623 | 0.66641 | 0.00038 | 10.0 s |
| 34:59.8 | Data point 34 | 1.50000 mL | 0.10228 mL | 0.10717 mL | 0.07001 mL | 4.073 | -0.00589 | 0.76350 | 0.00033 | 10.5 s |
| 35:51.2 | Data point 35 | 1.50000 mL | 0.10228 mL | 0.10840 mL | 0.07001 mL | 4.265 | -0.01218 | 0.79615 | 0.00067 | 10.0 s |

| | | | |
|---|---|---|---|
| Sample name: | **M09_octanol** | Experiment start time: | **3/6/2018 3:40:58 PM** |
| Assay name: | **pH-metric high logP** | Analyst: | **Pion** |
| Assay ID: | **18C-06007** | Instrument ID: | **T312060** |
| Filename: | **C:\Sirius_T3\Mehtap\20180306_exp30_logP_T3-2\18C-06007_M09_octanol_pH-metric high logP.t3r** | | |

#### Events (continued)

| Time | Event | Water | Acid | Base | Octanol | pH | dpH/dt | pH R-squared | pH SD | dpH/dt time |
|---|---|---|---|---|---|---|---|---|---|---|
| 36:36.9 | Data point 36 | 1.50000 mL | 0.10228 mL | 0.10927 mL | 0.07001 mL | 4.453 | -0.01574 | 0.77656 | 0.00088 | 10.0 s |
| 37:12.3 | Data point 37 | 1.50000 mL | 0.10228 mL | 0.10983 mL | 0.07001 mL | 4.611 | -0.01114 | 0.70830 | 0.00065 | 10.5 s |
| 38:03.7 | Data point 38 | 1.50000 mL | 0.10228 mL | 0.11054 mL | 0.07001 mL | 4.855 | -0.01503 | 0.83195 | 0.00081 | 11.5 s |
| 38:45.7 | Data point 39 | 1.50000 mL | 0.10228 mL | 0.11091 mL | 0.07001 mL | 5.055 | -0.00135 | 0.00984 | 0.00067 | 14.5 s |
| 39:30.9 | Data point 40 | 1.50000 mL | 0.10228 mL | 0.11122 mL | 0.07001 mL | 5.310 | -0.01935 | 0.93748 | 0.00099 | 13.5 s |
| 40:15.0 | Data point 41 | 1.50000 mL | 0.10228 mL | 0.11143 mL | 0.07001 mL | 5.577 | -0.01835 | 0.93150 | 0.00094 | 17.0 s |
| 41:02.4 | Data point 42 | 1.50000 mL | 0.10228 mL | 0.11160 mL | 0.07001 mL | 5.932 | -0.01854 | 0.95517 | 0.00094 | 25.0 s |
| 41:58.0 | Data point 43 | 1.50000 mL | 0.10228 mL | 0.11169 mL | 0.07001 mL | 6.304 | -0.01938 | 0.96132 | 0.00098 | 47.5 s |
| 43:16.1 | Data point 44 | 1.50000 mL | 0.10228 mL | 0.11178 mL | 0.07001 mL | 7.096 | -0.08353 | 0.99184 | 0.00414 | Timed out at 59.5 s |
| 44:46.6 | Data point 45 | 1.50000 mL | 0.10228 mL | 0.11183 mL | 0.07001 mL | 7.214 | -0.05929 | 0.99332 | 0.00294 | Timed out at 59.5 s |
| 46:17.1 | Data point 46 | 1.50000 mL | 0.10228 mL | 0.11192 mL | 0.07001 mL | 8.104 | -0.05533 | 0.99034 | 0.00275 | Timed out at 59.5 s |
| 47:52.7 | Data point 47 | 1.50000 mL | 0.10228 mL | 0.11199 mL | 0.07001 mL | 8.361 | -0.02231 | 0.99013 | 0.00111 | Timed out at 59.5 s |
| 49:28.4 | Data point 48 | 1.50000 mL | 0.10228 mL | 0.11207 mL | 0.07001 mL | 8.675 | -0.01974 | 0.96662 | 0.00099 | 43.0 s |
| 50:52.3 | Data point 49 | 1.50000 mL | 0.10228 mL | 0.11216 mL | 0.07001 mL | 8.935 | -0.01905 | 0.95683 | 0.00096 | 32.0 s |
| 51:60.0 | Data point 50 | 1.50000 mL | 0.10228 mL | 0.11228 mL | 0.07001 mL | 9.123 | -0.01924 | 0.93622 | 0.00098 | 27.5 s |
| 53:28.1 | Data point 51 | 1.50000 mL | 0.16298 mL | 0.11228 mL | 0.17001 mL | 1.963 | -0.00291 | 0.11888 | 0.00042 | 10.0 s |
| 54:14.4 | Data point 52 | 1.50000 mL | 0.16298 mL | 0.13079 mL | 0.17001 mL | 2.162 | -0.00496 | 0.56826 | 0.00032 | 10.0 s |
| 54:50.1 | Data point 53 | 1.50000 mL | 0.16298 mL | 0.14332 mL | 0.17001 mL | 2.380 | -0.01003 | 0.32764 | 0.00087 | 10.0 s |
| 55:25.7 | Data point 54 | 1.50000 mL | 0.16298 mL | 0.15113 mL | 0.17001 mL | 2.605 | -0.00142 | 0.12787 | 0.00020 | 10.0 s |
| 56:01.3 | Data point 55 | 1.50000 mL | 0.16298 mL | 0.15614 mL | 0.17001 mL | 2.814 | -0.00875 | 0.26635 | 0.00084 | 10.0 s |
| 56:36.8 | Data point 56 | 1.50000 mL | 0.16298 mL | 0.15969 mL | 0.17001 mL | 3.044 | -0.00319 | 0.61153 | 0.00020 | 10.0 s |
| 57:12.3 | Data point 57 | 1.50000 mL | 0.16298 mL | 0.16232 mL | 0.17001 mL | 3.248 | -0.01113 | 0.63073 | 0.00069 | 10.0 s |
| 57:47.8 | Data point 58 | 1.50000 mL | 0.16298 mL | 0.16449 mL | 0.17001 mL | 3.462 | -0.00502 | 0.86633 | 0.00027 | 10.0 s |
| 58:23.2 | Data point 59 | 1.50000 mL | 0.16298 mL | 0.16625 mL | 0.17001 mL | 3.691 | -0.01227 | 0.89636 | 0.00064 | 10.0 s |
| 58:58.6 | Data point 60 | 1.50000 mL | 0.16298 mL | 0.16761 mL | 0.17001 mL | 3.895 | -0.01553 | 0.75391 | 0.00088 | 25.0 s |
| 59:49.0 | Data point 61 | 1.50000 mL | 0.16298 mL | 0.16863 mL | 0.17001 mL | 4.072 | -0.01667 | 0.79855 | 0.00092 | 10.0 s |
| 1:00:24.5 | Data point 62 | 1.50000 mL | 0.16298 mL | 0.16940 mL | 0.17001 mL | 4.230 | -0.00448 | 0.66946 | 0.00027 | 10.5 s |
| 1:01:15.9 | Data point 63 | 1.50000 mL | 0.16298 mL | 0.17037 mL | 0.17001 mL | 4.459 | -0.00209 | 0.01107 | 0.00098 | 10.5 s |
| 1:02:07.3 | Data point 64 | 1.50000 mL | 0.16298 mL | 0.17096 mL | 0.17001 mL | 4.659 | -0.01788 | 0.81862 | 0.00098 | 10.5 s |
| 1:02:58.6 | Data point 65 | 1.50000 mL | 0.16298 mL | 0.17133 mL | 0.17001 mL | 4.856 | -0.01893 | 0.90105 | 0.00099 | 11.0 s |
| 1:03:40.2 | Data point 66 | 1.50000 mL | 0.16298 mL | 0.17161 mL | 0.17001 mL | 5.111 | -0.01888 | 0.92060 | 0.00097 | 16.0 s |
| 1:04:26.8 | Data point 67 | 1.50000 mL | 0.16298 mL | 0.17180 mL | 0.17001 mL | 5.425 | -0.01926 | 0.91497 | 0.00099 | 21.0 s |
| 1:05:18.4 | Data point 68 | 1.50000 mL | 0.16298 mL | 0.17192 mL | 0.17001 mL | 5.713 | -0.01897 | 0.89914 | 0.00099 | 38.5 s |
| 1:06:27.4 | Data point 69 | 1.50000 mL | 0.16298 mL | 0.17201 mL | 0.17001 mL | 6.169 | -0.01908 | 0.94473 | 0.00097 | 50.5 s |
| 1:07:48.5 | Data point 70 | 1.50000 mL | 0.16298 mL | 0.17218 mL | 0.17001 mL | 7.721 | -0.10780 | 0.99169 | 0.00535 | Timed out at 59.5 s |
| 1:09:24.1 | Data point 71 | 1.50000 mL | 0.16298 mL | 0.17225 mL | 0.17001 mL | 8.041 | -0.05145 | 0.99163 | 0.00255 | Timed out at 59.5 s |
| 1:10:59.7 | Data point 72 | 1.50000 mL | 0.16298 mL | 0.17232 mL | 0.17001 mL | 8.452 | -0.02336 | 0.95422 | 0.00118 | Timed out at 59.5 s |
| 1:12:40.5 | Data point 73 | 1.50000 mL | 0.16298 mL | 0.17241 mL | 0.17001 mL | 8.754 | -0.01965 | 0.97546 | 0.00098 | 28.0 s |
| 1:13:44.2 | Data point 74 | 1.50000 mL | 0.16298 mL | 0.17255 mL | 0.17001 mL | 9.015 | -0.01623 | 0.82712 | 0.00088 | 23.0 s |
| 1:14:16.2 | Assay volumes | 1.50000 mL | 0.16298 mL | 0.17255 mL | 0.17001 mL | | | | | |

| | | | |
|---|---|---|---|
| Sample name: | **M09_octanol** | Experiment start time: | **3/6/2018 3:40:58 PM** |
| Assay name: | **pH-metric high logP** | Analyst: | **Pion** |
| Assay ID: | **18C-06007** | Instrument ID: | **T312060** |
| Filename: | **C:\Sirius_T3\Mehtap\20180306_exp30_logP_T3-2\18C-06007_M09_octanol_pH-metric high logP.t3r** | | |

| | | | |
|---|---|---|---|
| Sample name: | **M09_octanol** | Experiment start time: | **3/6/2018 3:40:58 PM** |
| Assay name: | **pH-metric high logP** | Analyst: | **Pion** |
| Assay ID: | **18C-06007** | Instrument ID: | **T312060** |
| Filename: | **C:\Sirius_T3\Mehtap\20180306_exp30_logP_T3-2\18C-06007_M09_octanol_pH-metric high logP.t3r** | | |

#### Assay Settings (continued)

| Setting | Value | Original Value | Date/Time changed | Imported from |
|---|---|---|---|---|
| ***Titration 3*** | | | | |
| Titrate from | Low to high pH | | | |
| Add additional water | 0.00 mL | | | |
| Additional partition solvent volume | 0.100 mL | | | |
| Additional partition solvent added | Automatic | | | |
| After pH adjust stir for | 30 seconds | | | |
| Stir to allow partitioning for | 15 seconds | | | |
| Stirrer speed for partitioning | 60% | | | |
| ***Data Point Stability*** | | | | |
| Stir during data point collection | No | | | |
| Delay before data point collection | 0 seconds | | | |
| Number of points to average | 20 points | | | |
| Time interval between points | 0.50 seconds | | | |
| Required maximum standard deviation | 0.00100 dpH/dt | | | |
| Stability timeout after | 60 seconds | | | |

#### Calibration Settings

| Setting | Value | Date/Time changed | Imported from |
|---|---|---|---|
| Four-Plus alpha | 0.124 | 3/6/2018 3:40:58 PM | C:\Sirius_T3\18C-06006_Blank standardisation.t3r |
| Four-Plus S | 0.9973 | 3/6/2018 3:40:58 PM | C:\Sirius_T3\18C-06006_Blank standardisation.t3r |
| Four-Plus jH | 0.9 | 3/6/2018 3:40:58 PM | C:\Sirius_T3\18C-06006_Blank standardisation.t3r |
| Four-Plus jOH | -0.7 | 3/6/2018 3:40:58 PM | C:\Sirius_T3\18C-06006_Blank standardisation.t3r |
| Base concentration factor | 1.000 | 3/6/2018 3:40:58 PM | C:\Sirius_T3\KOH18B27.t3r |
| Acid concentration factor | 0.989 | 3/6/2018 3:40:58 PM | C:\Sirius_T3\18C-06006_Blank standardisation.t3r |

#### Instrument Settings

| Setting | Value | Batch Id | Install date |
|---|---|---|---|
| Instrument owner | Merck | | |
| Instrument ID | T312060 | | |
| Instrument type | T3 Simulator | | |
| Software version | 1.1.3.0 | | |
| Dispenser module | | T3DM1200361 | 3/31/2009 5:24:52 AM |
| Dispenser 0 | Water | | 3/31/2009 5:25:05 AM |
| Syringe volume | 2.5 mL | | |
| Firmware version | 1.2.1(r2) | | |
| Titrant | Water (0.15 M KCl) | 02-06-2018 | 2/27/2018 10:05:59 AM |
| Dispenser 2 | Acid | | 3/31/2009 5:25:11 AM |
| Syringe volume | 0.5 mL | | |
| Firmware version | 1.2.1(r2) | | |
| Titrant | Acid (0.5 M HCl) | 02-27-2018 | 2/27/2018 10:27:22 AM |
| Dispenser 1 | Base | | 3/31/2009 5:25:21 AM |
| Syringe volume | 0.5 mL | | |
| Firmware version | 1.2.1(r2) | | |
| Titrant | Base (0.5 M KOH) | 9/22/2017 | 2/27/2018 10:21:22 AM |
| Dispenser 5 | Cosolvent | | 3/31/2009 5:26:24 AM |
| Syringe volume | 2.5 mL | | |
| Firmware version | 1.2.1(r2) | | |
| Distribution valve 5 | Distribution Valve | | 3/31/2009 5:28:19 AM |
| Firmware version | 1.1.3 | | |
| Port A | Methanol (80%, 0.15 M KCl) | 02-08-2018 | 3/6/2018 9:28:59 AM |
| Port B | Cyclohexane | 11-01-17 | 2/27/2018 10:37:57 AM |
| Dispenser 3 | Buffer | | 8/3/2010 5:05:16 AM |
| Syringe volume | 0.5 mL | | |
| Firmware version | 1.2.1(r2) | | |
| Titrant | Dodecane | 2018/01/31 | 2/28/2018 10:18:04 AM |
| Dispenser 6 | Octanol | | 10/22/2010 10:52:43 AM |

| | | | |
|---|---|---|---|
| Sample name: | **M09_octanol** | Experiment start time: | **3/6/2018 3:40:58 PM** |
| Assay name: | **pH-metric high logP** | Analyst: | **Pion** |
| Assay ID: | **18C-06007** | Instrument ID: | **T312060** |
| Filename: | **C:\Sirius_T3\Mehtap\20180306_exp30_logP_T3-2\18C-06007_M09_octanol_pH-metric high logP.t3r** | | |

| | | | |
|---|---|---|---|
| Sample name: | **M09_octanol** | Experiment start time: | **3/6/2018 3:40:58 PM** |
| Assay name: | **pH-metric high logP** | Analyst: | **Pion** |
| Assay ID: | **18C-06007** | Instrument ID: | **T312060** |
| Filename: | **C:\Sirius_T3\Mehtap\20180306_exp30_logP_T3-2\18C-06007_M09_octanol_pH-metric high logP.t3r** | | |

#### Instrument Settings (continued)

| Setting | Value | Batch Id | Install date |
|---|---|---|---|
| Spectrometer calibration stir duration | 5 s | | |
| Spectrometer calibration stir speed | 30% | | |
| Spectrometer calibration wash pump volume | 20.0 mL | | |
| Spectrometer calibration wash stir duration | 5 s | | |
| Spectrometer calibration wash stir speed | 30% | | |
| Overhead dispense height | 10000 | | |

#### Refinement Settings

| Setting | Value | Default value |
|---|---|---|
| Turbidity detection method | None | None |
| Turbidity wavelength to assess | 500.0 nm | 500.0 nm |
| Turbidity maximum absorbance | 0.100 | 0.100 |
| Turbidity probe threshold | 50.00 | 50.00 |

#### Experiment Log

[1:59] Air gap released for Acid (0.5 M HCl)
[2:54] Air gap created for Water (0.15 M KCl)
[2:54] Air gap created for Acid (0.5 M HCl)
[2:55] Air gap created for Base (0.5 M KOH)
[2:55] Air gap released for Water (0.15 M KCl)
[2:59] Titrator arm moved over Titration position
[2:59] Titration 1 of 3
[2:59] Adding initial titrants
[2:59] Automatically add 1.50000 mL of water
[3:24] Dispensed 1.500000 mL of Water (0.15 M KCl)
[3:28] Titrator arm moved over Drain
[6:10] Titrator arm moved to Titration position
[6:10] Argon flow rate set to 100
[6:10] Stirrer speed set to 10
[6:15] Automatically add 0.03000 mL of Octanol
[6:16] Dispensed 0.030009 mL of Octanol
[6:17] Initial pH = 3.92
[6:17] Iterative adjust 3.92 -> 2.00
[6:17] pH 3.92 -> 2.00
[6:18] Air gap released for Acid (0.5 M HCl)
[6:19] Dispensed 0.045437 mL of Acid (0.5 M HCl)
[6:24] Holding pH 2.00
[8:24] Stirrer speed set to 0
[8:24] Stirrer speed set to 50
[8:24] Iterative adjust 1.94 -> 2.00
[8:24] pH 1.94 -> 2.00
[8:25] Air gap released for Base (0.5 M KOH)
[8:26] Dispensed 0.006091 mL of Base (0.5 M KOH)
[9:16] Stirrer speed set to 0
[9:26] Datapoint id 1 collected
[9:26] Stirrer speed set to 50
[9:31] pH 2.01 -> 2.21
[9:31] Using cautious pH adjust
[9:32] Dispensed 0.007714 mL of Base (0.5 M KOH)
[9:37] Stepping pH = 2.10
[9:37] Dispensed 0.006185 mL of Base (0.5 M KOH)
[9:42] Stepping pH = 2.19
[9:42] Dispensed 0.000847 mL of Base (0.5 M KOH)
[9:47] Stepping pH = 2.21
[10:03] Stirrer speed set to 0
[10:13] Datapoint id 2 collected
[10:13] Charge balance equation is out by 4.3%

Sample name: **M09_octanol** Experiment start time: **3/6/2018 3:40:58 PM**
Assay name: **pH-metric high logP** Analyst: **Pion**
Assay ID: **18C-06007** Instrument ID: **T312060**
Filename: **C:\Sirius_T3\Mehtap\20180306_exp30_logP_T3-2\18C-06007_M09_octanol_pH-metric high logP.t3r**

#### Experiment Log (continued)

[10:13] Stirrer speed set to 50
[10:18] pH 2.21 -> 2.41
[10:18] Using charge balance adjust
[10:19] Dispensed 0.009690 mL of Base (0.5 M KOH)
[10:39] Stirrer speed set to 0
[10:49] Datapoint id 3 collected
[10:49] Charge balance equation is out by 1.3%
[10:49] Stirrer speed set to 50
[10:54] pH 2.42 -> 2.62
[10:54] Using charge balance adjust
[10:54] Dispensed 0.006044 mL of Base (0.5 M KOH)
[11:15] Stirrer speed set to 0
[11:25] Datapoint id 4 collected
[11:25] Charge balance equation is out by 13.5%
[11:25] Stirrer speed set to 50
[11:30] pH 2.66 -> 2.86
[11:30] Using charge balance adjust
[11:30] Dispensed 0.003669 mL of Base (0.5 M KOH)
[11:50] Stirrer speed set to 0
[12:00] Datapoint id 5 collected
[12:00] Charge balance equation is out by 3.7%
[12:00] Stirrer speed set to 50
[12:05] pH 2.87 -> 3.07
[12:05] Using charge balance adjust
[12:05] Dispensed 0.002469 mL of Base (0.5 M KOH)
[12:26] Stirrer speed set to 0
[12:36] Datapoint id 6 collected
[12:36] Charge balance equation is out by 10.9%
[12:36] Stirrer speed set to 50
[12:41] pH 3.10 -> 3.30
[12:41] Using charge balance adjust
[12:41] Dispensed 0.001811 mL of Base (0.5 M KOH)
[13:01] Stirrer speed set to 0
[13:11] Datapoint id 7 collected
[13:11] Charge balance equation is out by 30.3%
[13:11] Stirrer speed set to 50
[13:16] pH 3.36 -> 3.56
[13:16] Using cautious pH adjust
[13:16] Dispensed 0.000753 mL of Base (0.5 M KOH)
[13:21] Stepping pH = 3.49
[13:21] Dispensed 0.000329 mL of Base (0.5 M KOH)
[13:27] Stepping pH = 3.54
[13:27] Dispensed 0.000118 mL of Base (0.5 M KOH)
[13:32] Stepping pH = 3.55
[13:47] Stirrer speed set to 0
[13:57] Datapoint id 8 collected
[13:57] Charge balance equation is out by 19.6%
[13:57] Stirrer speed set to 50
[14:02] pH 3.55 -> 3.75
[14:02] Using cautious pH adjust
[14:02] Dispensed 0.000729 mL of Base (0.5 M KOH)
[14:07] Stepping pH = 3.68
[14:07] Dispensed 0.000329 mL of Base (0.5 M KOH)
[14:12] Stepping pH = 3.72
[14:12] Dispensed 0.000165 mL of Base (0.5 M KOH)
[14:17] Stepping pH = 3.74
[14:33] Stirrer speed set to 0
[14:43] Datapoint id 9 collected
[14:43] Charge balance equation is out by 16.9%

Sample name: **M09_octanol** Experiment start time: **3/6/2018 3:40:58 PM**
Assay name: **pH-metric high logP** Analyst: **Pion**
Assay ID: **18C-06007** Instrument ID: **T312060**
Filename: **C:\Sirius_T3\Mehtap\20180306_exp30_logP_T3-2\18C-06007_M09_octanol_pH-metric high logP.t3r**

#### Experiment Log (continued)

[14:43] Stirrer speed set to 50
[14:48] pH 3.73 -> 3.93
[14:48] Using cautious pH adjust
[14:48] Dispensed 0.000706 mL of Base (0.5 M KOH)
[14:53] Stepping pH = 3.84
[14:53] Dispensed 0.000423 mL of Base (0.5 M KOH)
[14:58] Stepping pH = 3.89
[14:59] Dispensed 0.000259 mL of Base (0.5 M KOH)
[15:04] Stepping pH = 3.92
[15:19] Stirrer speed set to 0
[15:29] Datapoint id 10 collected
[15:29] Charge balance equation is out by 2.0%
[15:29] Stirrer speed set to 50
[15:34] pH 3.92 -> 4.12
[15:34] Using charge balance adjust
[15:34] Dispensed 0.001364 mL of Base (0.5 M KOH)
[15:54] Stirrer speed set to 0
[16:05] Datapoint id 11 collected
[16:05] Charge balance equation is out by -13.7%
[16:05] Stirrer speed set to 50
[16:10] pH 4.10 -> 4.30
[16:10] Using charge balance adjust
[16:11] Dispensed 0.001223 mL of Base (0.5 M KOH)
[16:31] Stirrer speed set to 0
[16:41] Datapoint id 12 collected
[16:41] Charge balance equation is out by -26.8%
[16:41] Stirrer speed set to 50
[16:46] pH 4.25 -> 4.45
[16:46] Using cautious pH adjust
[16:46] Dispensed 0.000541 mL of Base (0.5 M KOH)
[16:51] Stepping pH = 4.33
[16:51] Dispensed 0.000541 mL of Base (0.5 M KOH)
[16:56] Stepping pH = 4.42
[16:56] Dispensed 0.000188 mL of Base (0.5 M KOH)
[17:01] Stepping pH = 4.43
[17:01] Dispensed 0.000141 mL of Base (0.5 M KOH)
[17:06] Stepping pH = 4.45
[17:22] Stirrer speed set to 0
[17:33] Datapoint id 13 collected
[17:33] Charge balance equation is out by -31.1%
[17:33] Stirrer speed set to 50
[17:38] pH 4.44 -> 4.64
[17:38] Using cautious pH adjust
[17:38] Dispensed 0.000423 mL of Base (0.5 M KOH)
[17:43] Stepping pH = 4.55
[17:43] Dispensed 0.000259 mL of Base (0.5 M KOH)
[17:49] Stepping pH = 4.59
[17:49] Dispensed 0.000259 mL of Base (0.5 M KOH)
[17:54] Stepping pH = 4.64
[18:09] Stirrer speed set to 0
[18:21] Datapoint id 14 collected
[18:21] Charge balance equation is out by -9.3%
[18:21] Stirrer speed set to 50
[18:26] pH 4.64 -> 4.84
[18:26] Using charge balance adjust
[18:26] Dispensed 0.000635 mL of Base (0.5 M KOH)
[18:46] Stirrer speed set to 0
[18:58] Datapoint id 15 collected
[18:58] Charge balance equation is out by -14.0%

Sample name: **M09_octanol** Experiment start time: **3/6/2018 3:40:58 PM**
Assay name: **pH-metric high logP** Analyst: **Pion**
Assay ID: **18C-06007** Instrument ID: **T312060**
Filename: **C:\Sirius_T3\Mehtap\20180306_exp30_logP_T3-2\18C-06007_M09_octanol_pH-metric high logP.t3r**

#### Experiment Log (continued)

[18:58] Stirrer speed set to 50
[19:03] pH 4.83 -> 5.03
[19:03] Using charge balance adjust
[19:03] Dispensed 0.000447 mL of Base (0.5 M KOH)
[19:24] Stirrer speed set to 0
[19:39] Datapoint id 16 collected
[19:39] Charge balance equation is out by -38.2%
[19:39] Stirrer speed set to 50
[19:44] pH 4.98 -> 5.18
[19:44] Using cautious pH adjust
[19:44] Dispensed 0.000165 mL of Base (0.5 M KOH)
[19:49] Stepping pH = 5.01
[19:49] Dispensed 0.000376 mL of Base (0.5 M KOH)
[19:54] Stepping pH = 5.25
[20:09] Stirrer speed set to 0
[20:25] Datapoint id 17 collected
[20:25] Charge balance equation is out by -62.0%
[20:25] Stirrer speed set to 50
[20:30] pH 5.23 -> 5.43
[20:30] Using cautious pH adjust
[20:30] Dispensed 0.000118 mL of Base (0.5 M KOH)
[20:35] Stepping pH = 5.25
[20:35] Dispensed 0.000282 mL of Base (0.5 M KOH)
[20:40] Stepping pH = 5.49
[20:55] Stirrer speed set to 0
[21:13] Datapoint id 18 collected
[21:13] Charge balance equation is out by -85.2%
[21:13] Stirrer speed set to 50
[21:18] pH 5.49 -> 5.69
[21:18] Using cautious pH adjust
[21:18] Dispensed 0.000071 mL of Base (0.5 M KOH)
[21:23] Stepping pH = 5.50
[21:23] Dispensed 0.000188 mL of Base (0.5 M KOH)
[21:28] Stepping pH = 5.74
[21:43] Stirrer speed set to 0
[22:00] Datapoint id 19 collected
[22:00] Charge balance equation is out by -94.3%
[22:00] Stirrer speed set to 50
[22:06] pH 5.76 -> 5.96
[22:06] Using cautious pH adjust
[22:06] Dispensed 0.000047 mL of Base (0.5 M KOH)
[22:11] Stepping pH = 5.77
[22:11] Dispensed 0.000141 mL of Base (0.5 M KOH)
[22:16] Stepping pH = 5.85
[22:16] Dispensed 0.000118 mL of Base (0.5 M KOH)
[22:21] Stepping pH = 6.19
[22:36] Stirrer speed set to 0
[23:09] Datapoint id 20 collected
[23:09] Charge balance equation is out by -210.5%
[23:09] Stirrer speed set to 50
[23:14] pH 6.23 -> 6.43
[23:14] Using cautious pH adjust
[23:14] Dispensed 0.000024 mL of Base (0.5 M KOH)
[23:19] Stepping pH = 6.23
[23:19] Dispensed 0.000141 mL of Base (0.5 M KOH)
[23:25] Stepping pH = 7.24
[23:40] Stirrer speed set to 0
[24:40] Datapoint id 21 collected
[24:40] Charge balance equation is out by -211.2%

| | | | |
|---|---|---|---|
| Sample name: | **M09_octanol** | Experiment start time: | **3/6/2018 3:40:58 PM** |
| Assay name: | **pH-metric high logP** | Analyst: | **Pion** |
| Assay ID: | **18C-06007** | Instrument ID: | **T312060** |
| Filename: | **C:\Sirius_T3\Mehtap\20180306_exp30_logP_T3-2\18C-06007_M09_octanol_pH-metric high logP.t3r** | | |

#### Experiment Log (continued)

[24:40] Stirrer speed set to 50
[24:45] pH 7.39 -> 7.59
[24:45] Using cautious pH adjust
[24:45] Dispensed 0.000024 mL of Base (0.5 M KOH)
[24:50] Stepping pH = 7.36
[24:50] Dispensed 0.000024 mL of Base (0.5 M KOH)
[24:55] Stepping pH = 7.36
[24:55] Dispensed 0.000094 mL of Base (0.5 M KOH)
[25:00] Stepping pH = 8.58
[25:15] Stirrer speed set to 0
[26:15] Datapoint id 22 collected
[26:15] Charge balance equation is out by -1,276.8%
[26:15] Stirrer speed set to 50
[26:20] pH 8.72 -> 8.92
[26:20] Using cautious pH adjust
[26:20] Dispensed 0.000024 mL of Base (0.5 M KOH)
[26:25] Stepping pH = 8.71
[26:25] Dispensed 0.000071 mL of Base (0.5 M KOH)
[26:30] Stepping pH = 8.89
[26:30] Dispensed 0.000024 mL of Base (0.5 M KOH)
[26:35] Stepping pH = 9.02
[26:50] Stirrer speed set to 0
[27:28] Datapoint id 23 collected
[27:28] Charge balance equation is out by -331.4%
[27:28] Titration 2 of 3
[27:28] Adding initial titrants
[27:28] Automatically add 0.04000 mL of Octanol
[27:29] Dispensed 0.040005 mL of Octanol
[27:29] Stirrer speed set to 10
[27:30] Stirrer speed set to 55
[27:30] Iterative adjust 9.02 -> 2.00
[27:30] pH 9.02 -> 2.00
[27:31] Dispensed 0.054351 mL of Acid (0.5 M HCl)
[27:36] pH 2.02 -> 2.00
[27:37] Dispensed 0.002493 mL of Acid (0.5 M HCl)
[28:27] Stirrer speed set to 0
[28:37] Datapoint id 24 collected
[28:37] Stirrer speed set to 55
[28:42] pH 1.97 -> 2.17
[28:42] Using cautious pH adjust
[28:42] Dispensed 0.009149 mL of Base (0.5 M KOH)
[28:47] Stepping pH = 2.07
[28:48] Dispensed 0.005691 mL of Base (0.5 M KOH)
[28:53] Stepping pH = 2.14
[28:53] Dispensed 0.002070 mL of Base (0.5 M KOH)
[28:58] Stepping pH = 2.17
[29:13] Stirrer speed set to 0
[29:23] Datapoint id 25 collected
[29:23] Charge balance equation is out by 7.6%
[29:23] Stirrer speed set to 55
[29:28] pH 2.17 -> 2.37
[29:28] Using charge balance adjust
[29:29] Dispensed 0.011430 mL of Base (0.5 M KOH)
[29:49] Stirrer speed set to 0
[29:59] Datapoint id 26 collected
[29:59] Charge balance equation is out by 13.7%
[29:59] Stirrer speed set to 55
[30:04] pH 2.41 -> 2.61
[30:04] Using charge balance adjust

| | | | |
|---|---|---|---|
| Sample name: | **M09_octanol** | Experiment start time: | **3/6/2018 3:40:58 PM** |
| Assay name: | **pH-metric high logP** | Analyst: | **Pion** |
| Assay ID: | **18C-06007** | Instrument ID: | **T312060** |
| Filename: | **C:\Sirius_T3\Mehtap\20180306_exp30_logP_T3-2\18C-06007_M09_octanol_pH-metric high logP.t3r** | | |

#### Experiment Log (continued)

[30:05] Dispensed 0.006797 mL of Base (0.5 M KOH)
[30:25] Stirrer speed set to 0
[30:35] Datapoint id 27 collected
[30:35] Charge balance equation is out by -1.0%
[30:35] Stirrer speed set to 55
[30:40] pH 2.61 -> 2.81
[30:40] Using charge balance adjust
[30:40] Dispensed 0.004492 mL of Base (0.5 M KOH)
[31:00] Stirrer speed set to 0
[31:10] Datapoint id 28 collected
[31:10] Charge balance equation is out by 4.7%
[31:10] Stirrer speed set to 55
[31:16] pH 2.82 -> 3.02
[31:16] Using charge balance adjust
[31:16] Dispensed 0.003081 mL of Base (0.5 M KOH)
[31:36] Stirrer speed set to 0
[31:46] Datapoint id 29 collected
[31:46] Charge balance equation is out by 6.2%
[31:46] Stirrer speed set to 55
[31:52] pH 3.04 -> 3.24
[31:52] Using charge balance adjust
[31:52] Dispensed 0.002305 mL of Base (0.5 M KOH)
[32:12] Stirrer speed set to 0
[32:22] Datapoint id 30 collected
[32:22] Charge balance equation is out by 46.1%
[32:22] Stirrer speed set to 55
[32:27] pH 3.34 -> 3.54
[32:27] Using cautious pH adjust
[32:27] Dispensed 0.000917 mL of Base (0.5 M KOH)
[32:32] Stepping pH = 3.46
[32:32] Dispensed 0.000447 mL of Base (0.5 M KOH)
[32:37] Stepping pH = 3.51
[32:37] Dispensed 0.000188 mL of Base (0.5 M KOH)
[32:43] Stepping pH = 3.53
[32:58] Stirrer speed set to 0
[33:08] Datapoint id 31 collected
[33:08] Charge balance equation is out by 15.3%
[33:08] Stirrer speed set to 55
[33:13] pH 3.53 -> 3.73
[33:13] Using cautious pH adjust
[33:13] Dispensed 0.000823 mL of Base (0.5 M KOH)
[33:18] Stepping pH = 3.64
[33:18] Dispensed 0.000494 mL of Base (0.5 M KOH)
[33:23] Stepping pH = 3.70
[33:23] Dispensed 0.000212 mL of Base (0.5 M KOH)
[33:28] Stepping pH = 3.72
[33:28] Dispensed 0.000118 mL of Base (0.5 M KOH)
[33:34] Stepping pH = 3.73
[33:49] Stirrer speed set to 0
[33:59] Datapoint id 32 collected
[33:59] Charge balance equation is out by -0.4%
[33:59] Stirrer speed set to 55
[34:04] pH 3.72 -> 3.92
[34:04] Using charge balance adjust
[34:04] Dispensed 0.001458 mL of Base (0.5 M KOH)
[34:25] Stirrer speed set to 0
[34:35] Datapoint id 33 collected
[34:35] Charge balance equation is out by -7.5%
[34:35] Stirrer speed set to 55

| | | | |
|---|---|---|---|
| Sample name: | **M09_octanol** | Experiment start time: | **3/6/2018 3:40:58 PM** |
| Assay name: | **pH-metric high logP** | Analyst: | **Pion** |
| Assay ID: | **18C-06007** | Instrument ID: | **T312060** |
| Filename: | **C:\Sirius_T3\Mehtap\20180306_exp30_logP_T3-2\18C-06007_M09_octanol_pH-metric high logP.t3r** | | |

#### Experiment Log (continued)

[34:40] pH 3.91 -> 4.11
[34:40] Using charge balance adjust
[34:40] Dispensed 0.001223 mL of Base (0.5 M KOH)
[35:00] Stirrer speed set to 0
[35:11] Datapoint id 34 collected
[35:11] Charge balance equation is out by -20.6%
[35:11] Stirrer speed set to 55
[35:16] pH 4.08 -> 4.28
[35:16] Using cautious pH adjust
[35:16] Dispensed 0.000517 mL of Base (0.5 M KOH)
[35:21] Stepping pH = 4.16
[35:21] Dispensed 0.000470 mL of Base (0.5 M KOH)
[35:26] Stepping pH = 4.24
[35:26] Dispensed 0.000165 mL of Base (0.5 M KOH)
[35:31] Stepping pH = 4.27
[35:31] Dispensed 0.000071 mL of Base (0.5 M KOH)
[35:36] Stepping pH = 4.27
[35:51] Stirrer speed set to 0
[36:02] Datapoint id 35 collected
[36:02] Charge balance equation is out by -18.3%
[36:02] Stirrer speed set to 55
[36:07] pH 4.27 -> 4.47
[36:07] Using cautious pH adjust
[36:07] Dispensed 0.000376 mL of Base (0.5 M KOH)
[36:12] Stepping pH = 4.35
[36:12] Dispensed 0.000329 mL of Base (0.5 M KOH)
[36:17] Stepping pH = 4.42
[36:17] Dispensed 0.000165 mL of Base (0.5 M KOH)
[36:22] Stepping pH = 4.46
[36:37] Stirrer speed set to 0
[36:47] Datapoint id 36 collected
[36:47] Charge balance equation is out by -13.4%
[36:47] Stirrer speed set to 55
[36:52] pH 4.46 -> 4.66
[36:52] Using charge balance adjust
[36:52] Dispensed 0.000564 mL of Base (0.5 M KOH)
[37:13] Stirrer speed set to 0
[37:23] Datapoint id 37 collected
[37:23] Charge balance equation is out by -25.1%
[37:23] Stirrer speed set to 55
[37:28] pH 4.62 -> 4.82
[37:28] Using cautious pH adjust
[37:28] Dispensed 0.000212 mL of Base (0.5 M KOH)
[37:33] Stepping pH = 4.67
[37:34] Dispensed 0.000306 mL of Base (0.5 M KOH)
[37:39] Stepping pH = 4.79
[37:39] Dispensed 0.000047 mL of Base (0.5 M KOH)
[37:44] Stepping pH = 4.80
[37:44] Dispensed 0.000141 mL of Base (0.5 M KOH)
[37:49] Stepping pH = 4.86
[38:04] Stirrer speed set to 0
[38:16] Datapoint id 38 collected
[38:16] Charge balance equation is out by -66.8%
[38:16] Stirrer speed set to 55
[38:21] pH 4.87 -> 5.07
[38:21] Using cautious pH adjust
[38:21] Dispensed 0.000118 mL of Base (0.5 M KOH)
[38:26] Stepping pH = 4.90
[38:26] Dispensed 0.000259 mL of Base (0.5 M KOH)

| | | | |
|---|---|---|---|
| Sample name: | **M09_octanol** | Experiment start time: | **3/6/2018 3:40:58 PM** |
| Assay name: | **pH-metric high logP** | Analyst: | **Pion** |
| Assay ID: | **18C-06007** | Instrument ID: | **T312060** |
| Filename: | **C:\Sirius_T3\Mehtap\20180306_exp30_logP_T3-2\18C-06007_M09_octanol_pH-metric high logP.t3r** | | |

#### Experiment Log (continued)

[38:31] Stepping pH = 5.07
[38:46] Stirrer speed set to 0
[39:01] Datapoint id 39 collected
[39:01] Charge balance equation is out by -50.1%
[39:01] Stirrer speed set to 55
[39:06] pH 5.08 -> 5.28
[39:06] Using cautious pH adjust
[39:06] Dispensed 0.000094 mL of Base (0.5 M KOH)
[39:11] Stepping pH = 5.10
[39:11] Dispensed 0.000212 mL of Base (0.5 M KOH)
[39:16] Stepping pH = 5.32
[39:31] Stirrer speed set to 0
[39:45] Datapoint id 40 collected
[39:45] Charge balance equation is out by -73.8%
[39:45] Stirrer speed set to 55
[39:50] pH 5.34 -> 5.54
[39:50] Using cautious pH adjust
[39:50] Dispensed 0.000047 mL of Base (0.5 M KOH)
[39:55] Stepping pH = 5.34
[39:55] Dispensed 0.000165 mL of Base (0.5 M KOH)
[40:00] Stepping pH = 5.59
[40:15] Stirrer speed set to 0
[40:32] Datapoint id 41 collected
[40:32] Charge balance equation is out by -98.5%
[40:32] Stirrer speed set to 55
[40:37] pH 5.61 -> 5.81
[40:37] Using cautious pH adjust
[40:37] Dispensed 0.000047 mL of Base (0.5 M KOH)
[40:43] Stepping pH = 5.62
[40:43] Dispensed 0.000118 mL of Base (0.5 M KOH)
[40:48] Stepping pH = 5.97
[41:03] Stirrer speed set to 0
[41:28] Datapoint id 42 collected
[41:28] Charge balance equation is out by -92.4%
[41:28] Stirrer speed set to 55
[41:33] pH 5.99 -> 6.19
[41:33] Using cautious pH adjust
[41:33] Dispensed 0.000024 mL of Base (0.5 M KOH)
[41:38] Stepping pH = 6.00
[41:38] Dispensed 0.000071 mL of Base (0.5 M KOH)
[41:43] Stepping pH = 6.30
[41:58] Stirrer speed set to 0
[42:46] Datapoint id 43 collected
[42:46] Charge balance equation is out by -90.5%
[42:46] Stirrer speed set to 55
[42:51] pH 6.33 -> 6.53
[42:51] Using cautious pH adjust
[42:51] Dispensed 0.000024 mL of Base (0.5 M KOH)
[42:56] Stepping pH = 6.35
[42:56] Dispensed 0.000071 mL of Base (0.5 M KOH)
[43:01] Stepping pH = 6.95
[43:16] Stirrer speed set to 0
[44:16] Datapoint id 44 collected
[44:16] Charge balance equation is out by -77.9%
[44:16] Stirrer speed set to 55
[44:22] pH 7.16 -> 7.36
[44:22] Using cautious pH adjust
[44:22] Dispensed 0.000024 mL of Base (0.5 M KOH)
[44:27] Stepping pH = 7.21

Sample name: **M09_octanol**
Assay name: **pH-metric high logP**
Assay ID: **18C-06007**
Filename: **C:\Sirius_T3\Mehtap\20180306_exp30_logP_T3-2\18C-06007_M09_octanol_pH-metric high logP.t3r**

Experiment start time: **3/6/2018 3:40:58 PM**
Analyst: **Pion**
Instrument ID: **T312060**

#### Experiment Log (continued)

[44:27] Dispensed 0.000024 mL of Base (0.5 M KOH)
[44:32] Stepping pH = 7.36
[44:47] Stirrer speed set to 0
[45:47] Datapoint id 45 collected
[45:47] Charge balance equation is out by -160.1%
[45:47] Stirrer speed set to 55
[45:52] pH 7.07 -> 7.27
[45:52] Using cautious pH adjust
[45:52] Dispensed 0.000024 mL of Base (0.5 M KOH)
[45:57] Stepping pH = 7.02
[45:57] Dispensed 0.000071 mL of Base (0.5 M KOH)
[46:02] Stepping pH = 7.98
[46:17] Stirrer speed set to 0
[47:17] Datapoint id 46 collected
[47:17] Charge balance equation is out by -347.1%
[47:17] Stirrer speed set to 55
[47:23] pH 8.04 -> 8.24
[47:23] Using cautious pH adjust
[47:23] Dispensed 0.000024 mL of Base (0.5 M KOH)
[47:28] Stepping pH = 8.03
[47:28] Dispensed 0.000024 mL of Base (0.5 M KOH)
[47:33] Stepping pH = 8.16
[47:33] Dispensed 0.000024 mL of Base (0.5 M KOH)
[47:38] Stepping pH = 8.35
[47:53] Stirrer speed set to 0
[48:53] Datapoint id 47 collected
[48:53] Charge balance equation is out by -699.5%
[48:53] Stirrer speed set to 55
[48:58] pH 8.43 -> 8.63
[48:58] Using cautious pH adjust
[48:58] Dispensed 0.000024 mL of Base (0.5 M KOH)
[49:03] Stepping pH = 8.46
[49:03] Dispensed 0.000024 mL of Base (0.5 M KOH)
[49:08] Stepping pH = 8.57
[49:09] Dispensed 0.000024 mL of Base (0.5 M KOH)
[49:14] Stepping pH = 8.69
[49:29] Stirrer speed set to 0
[50:12] Datapoint id 48 collected
[50:12] Charge balance equation is out by -340.0%
[50:12] Stirrer speed set to 55
[50:17] pH 8.69 -> 8.89
[50:17] Using cautious pH adjust
[50:17] Dispensed 0.000024 mL of Base (0.5 M KOH)
[50:22] Stepping pH = 8.69
[50:22] Dispensed 0.000024 mL of Base (0.5 M KOH)
[50:27] Stepping pH = 8.77
[50:27] Dispensed 0.000024 mL of Base (0.5 M KOH)
[50:32] Stepping pH = 8.86
[50:32] Dispensed 0.000024 mL of Base (0.5 M KOH)
[50:38] Stepping pH = 8.93
[50:53] Stirrer speed set to 0
[51:25] Datapoint id 49 collected
[51:25] Charge balance equation is out by -371.7%
[51:25] Stirrer speed set to 55
[51:30] pH 8.95 -> 9.05
[51:30] Using cautious pH adjust
[51:30] Dispensed 0.000024 mL of Base (0.5 M KOH)
[51:35] Stepping pH = 8.95
[51:35] Dispensed 0.000024 mL of Base (0.5 M KOH)

Sample name: **M09_octanol**
Assay name: **pH-metric high logP**
Assay ID: **18C-06007**
Filename: **C:\Sirius_T3\Mehtap\20180306_exp30_logP_T3-2\18C-06007_M09_octanol_pH-metric high logP.t3r**

Experiment start time: **3/6/2018 3:40:58 PM**
Analyst: **Pion**
Instrument ID: **T312060**

#### Experiment Log (continued)

[51:40] Stepping pH = 8.96
[51:40] Dispensed 0.000071 mL of Base (0.5 M KOH)
[51:45] Stepping pH = 9.13
[52:00] Stirrer speed set to 0
[52:28] Datapoint id 50 collected
[52:28] Charge balance equation is out by -530.3%
[52:28] Titration 3 of 3
[52:28] Adding initial titrants
[52:28] Automatically add 0.10000 mL of Octanol
[52:30] Dispensed 0.100000 mL of Octanol
[52:30] Stirrer speed set to 10
[52:31] Stirrer speed set to 60
[52:31] Iterative adjust 9.13 -> 2.00
[52:31] pH 9.13 -> 2.00
[52:33] Dispensed 0.057314 mL of Acid (0.5 M HCl)
[52:38] pH 2.03 -> 2.00
[52:38] Dispensed 0.003387 mL of Acid (0.5 M HCl)
[53:28] Stirrer speed set to 0
[53:39] Datapoint id 51 collected
[53:39] Stirrer speed set to 60
[53:44] pH 1.97 -> 2.17
[53:44] Using cautious pH adjust
[53:44] Dispensed 0.009901 mL of Base (0.5 M KOH)
[53:49] Stepping pH = 2.06
[53:49] Dispensed 0.006914 mL of Base (0.5 M KOH)
[53:54] Stepping pH = 2.14
[53:55] Dispensed 0.001693 mL of Base (0.5 M KOH)
[54:00] Stepping pH = 2.17
[54:15] Stirrer speed set to 0
[54:25] Datapoint id 52 collected
[54:25] Charge balance equation is out by 6.5%
[54:25] Stirrer speed set to 60
[54:30] pH 2.17 -> 2.37
[54:30] Using charge balance adjust
[54:30] Dispensed 0.012535 mL of Base (0.5 M KOH)
[54:50] Stirrer speed set to 0
[55:00] Datapoint id 53 collected
[55:00] Charge balance equation is out by 6.7%
[55:00] Stirrer speed set to 60
[55:06] pH 2.38 -> 2.58
[55:06] Using charge balance adjust
[55:06] Dispensed 0.007808 mL of Base (0.5 M KOH)
[55:26] Stirrer speed set to 0
[55:36] Datapoint id 54 collected
[55:36] Charge balance equation is out by 10.7%
[55:36] Stirrer speed set to 60
[55:41] pH 2.61 -> 2.81
[55:41] Using charge balance adjust
[55:41] Dispensed 0.005009 mL of Base (0.5 M KOH)
[56:02] Stirrer speed set to 0
[56:12] Datapoint id 55 collected
[56:12] Charge balance equation is out by 2.5%
[56:12] Stirrer speed set to 60
[56:17] pH 2.82 -> 3.02
[56:17] Using charge balance adjust
[56:17] Dispensed 0.003551 mL of Base (0.5 M KOH)
[56:37] Stirrer speed set to 0
[56:47] Datapoint id 56 collected
[56:47] Charge balance equation is out by 13.2%

| | | | |
|---|---|---|---|
| Sample name: | **M09_octanol** | Experiment start time: | **3/6/2018 3:40:58 PM** |
| Assay name: | **pH-metric high logP** | Analyst: | **Pion** |
| Assay ID: | **18C-06007** | Instrument ID: | **T312060** |
| Filename: | **C:\Sirius_T3\Mehtap\20180306_exp30_logP_T3-2\18C-06007_M09_octanol_pH-metric high logP.t3r** | | |

#### Experiment Log (continued)

[56:47] Stirrer speed set to 60
[56:52] pH 3.05 -> 3.25
[56:52] Using charge balance adjust
[56:52] Dispensed 0.002634 mL of Base (0.5 M KOH)
[57:13] Stirrer speed set to 0
[57:23] Datapoint id 57 collected
[57:23] Charge balance equation is out by 1.0%
[57:23] Stirrer speed set to 60
[57:28] pH 3.25 -> 3.45
[57:28] Using charge balance adjust
[57:28] Dispensed 0.002164 mL of Base (0.5 M KOH)
[57:48] Stirrer speed set to 0
[57:58] Datapoint id 58 collected
[57:58] Charge balance equation is out by 5.5%
[57:58] Stirrer speed set to 60
[58:03] pH 3.46 -> 3.66
[58:03] Using charge balance adjust
[58:03] Dispensed 0.001764 mL of Base (0.5 M KOH)
[58:24] Stirrer speed set to 0
[58:34] Datapoint id 59 collected
[58:34] Charge balance equation is out by 13.2%
[58:34] Stirrer speed set to 60
[58:39] pH 3.69 -> 3.89
[58:39] Using charge balance adjust
[58:39] Dispensed 0.001364 mL of Base (0.5 M KOH)
[58:59] Stirrer speed set to 0
[59:24] Datapoint id 60 collected
[59:24] Charge balance equation is out by 0.3%
[59:24] Stirrer speed set to 60
[59:29] pH 3.90 -> 4.10
[59:29] Using charge balance adjust
[59:29] Dispensed 0.001011 mL of Base (0.5 M KOH)
[59:49] Stirrer speed set to 0
[59:59] Datapoint id 61 collected
[59:59] Charge balance equation is out by -14.0%
[59:59] Stirrer speed set to 60
[1:00:05] pH 4.08 -> 4.28
[1:00:05] Using charge balance adjust
[1:00:05] Dispensed 0.000776 mL of Base (0.5 M KOH)
[1:00:25] Stirrer speed set to 0
[1:00:35] Datapoint id 62 collected
[1:00:35] Charge balance equation is out by -23.3%
[1:00:35] Stirrer speed set to 60
[1:00:40] pH 4.24 -> 4.44
[1:00:40] Using cautious pH adjust
[1:00:41] Dispensed 0.000282 mL of Base (0.5 M KOH)
[1:00:46] Stepping pH = 4.28
[1:00:46] Dispensed 0.000447 mL of Base (0.5 M KOH)
[1:00:51] Stepping pH = 4.41
[1:00:51] Dispensed 0.000071 mL of Base (0.5 M KOH)
[1:00:56] Stepping pH = 4.42
[1:00:56] Dispensed 0.000165 mL of Base (0.5 M KOH)
[1:01:01] Stepping pH = 4.46
[1:01:16] Stirrer speed set to 0
[1:01:27] Datapoint id 63 collected
[1:01:27] Charge balance equation is out by -64.5%
[1:01:27] Stirrer speed set to 60
[1:01:32] pH 4.46 -> 4.66
[1:01:32] Using cautious pH adjust

| | | | |
|---|---|---|---|
| Sample name: | **M09_octanol** | Experiment start time: | **3/6/2018 3:40:58 PM** |
| Assay name: | **pH-metric high logP** | Analyst: | **Pion** |
| Assay ID: | **18C-06007** | Instrument ID: | **T312060** |
| Filename: | **C:\Sirius_T3\Mehtap\20180306_exp30_logP_T3-2\18C-06007_M09_octanol_pH-metric high logP.t3r** | | |

#### Experiment Log (continued)

[1:01:32] Dispensed 0.000188 mL of Base (0.5 M KOH)
[1:01:37] Stepping pH = 4.52
[1:01:37] Dispensed 0.000235 mL of Base (0.5 M KOH)
[1:01:42] Stepping pH = 4.62
[1:01:42] Dispensed 0.000071 mL of Base (0.5 M KOH)
[1:01:47] Stepping pH = 4.63
[1:01:47] Dispensed 0.000094 mL of Base (0.5 M KOH)
[1:01:53] Stepping pH = 4.66
[1:02:08] Stirrer speed set to 0
[1:02:18] Datapoint id 64 collected
[1:02:18] Charge balance equation is out by -59.0%
[1:02:18] Stirrer speed set to 60
[1:02:23] pH 4.67 -> 4.87
[1:02:23] Using cautious pH adjust
[1:02:23] Dispensed 0.000118 mL of Base (0.5 M KOH)
[1:02:28] Stepping pH = 4.71
[1:02:28] Dispensed 0.000188 mL of Base (0.5 M KOH)
[1:02:34] Stepping pH = 4.84
[1:02:34] Dispensed 0.000024 mL of Base (0.5 M KOH)
[1:02:39] Stepping pH = 4.85
[1:02:39] Dispensed 0.000047 mL of Base (0.5 M KOH)
[1:02:44] Stepping pH = 4.86
[1:02:59] Stirrer speed set to 0
[1:03:10] Datapoint id 65 collected
[1:03:10] Charge balance equation is out by -59.7%
[1:03:10] Stirrer speed set to 60
[1:03:15] pH 4.87 -> 5.07
[1:03:15] Using cautious pH adjust
[1:03:15] Dispensed 0.000071 mL of Base (0.5 M KOH)
[1:03:20] Stepping pH = 4.88
[1:03:20] Dispensed 0.000212 mL of Base (0.5 M KOH)
[1:03:25] Stepping pH = 5.12
[1:03:41] Stirrer speed set to 0
[1:03:57] Datapoint id 66 collected
[1:03:57] Charge balance equation is out by -86.7%
[1:03:57] Stirrer speed set to 60
[1:04:02] pH 5.14 -> 5.34
[1:04:02] Using cautious pH adjust
[1:04:02] Dispensed 0.000047 mL of Base (0.5 M KOH)
[1:04:07] Stepping pH = 5.15
[1:04:07] Dispensed 0.000141 mL of Base (0.5 M KOH)
[1:04:12] Stepping pH = 5.43
[1:04:27] Stirrer speed set to 0
[1:04:48] Datapoint id 67 collected
[1:04:48] Charge balance equation is out by -88.8%
[1:04:48] Stirrer speed set to 60
[1:04:53] pH 5.48 -> 5.68
[1:04:53] Using cautious pH adjust
[1:04:53] Dispensed 0.000024 mL of Base (0.5 M KOH)
[1:04:59] Stepping pH = 5.49
[1:04:59] Dispensed 0.000094 mL of Base (0.5 M KOH)
[1:05:04] Stepping pH = 5.73
[1:05:19] Stirrer speed set to 0
[1:05:57] Datapoint id 68 collected
[1:05:57] Charge balance equation is out by -92.7%
[1:05:57] Stirrer speed set to 60
[1:06:02] pH 5.81 -> 6.01
[1:06:02] Using cautious pH adjust
[1:06:02] Dispensed 0.000024 mL of Base (0.5 M KOH)

| | | | |
|---|---|---|---|
| Sample name: | **M09_octanol** | Experiment start time: | **3/6/2018 3:40:58 PM** |
| Assay name: | **pH-metric high logP** | Analyst: | **Pion** |
| Assay ID: | **18C-06007** | Instrument ID: | **T312060** |
| Filename: | **C:\Sirius_T3\Mehtap\20180306_exp30_logP_T3-2\18C-06007_M09_octanol_pH-metric high logP.t3r** | | |

#### Experiment Log (continued)

[1:06:07] Stepping pH = 5.82
[1:06:08] Dispensed 0.000071 mL of Base (0.5 M KOH)
[1:06:13] Stepping pH = 6.20
[1:06:28] Stirrer speed set to 0
[1:07:18] Datapoint id 69 collected
[1:07:18] Charge balance equation is out by -91.3%
[1:07:18] Stirrer speed set to 60
[1:07:23] pH 6.15 -> 6.35
[1:07:23] Using cautious pH adjust
[1:07:23] Dispensed 0.000024 mL of Base (0.5 M KOH)
[1:07:29] Stepping pH = 6.11
[1:07:29] Dispensed 0.000141 mL of Base (0.5 M KOH)
[1:07:34] Stepping pH = 7.70
[1:07:49] Stirrer speed set to 0
[1:08:49] Datapoint id 70 collected
[1:08:49] Charge balance equation is out by -243.1%
[1:08:49] Stirrer speed set to 60
[1:08:54] pH 7.85 -> 8.05
[1:08:54] Using cautious pH adjust
[1:08:54] Dispensed 0.000024 mL of Base (0.5 M KOH)
[1:08:59] Stepping pH = 7.89
[1:08:59] Dispensed 0.000024 mL of Base (0.5 M KOH)
[1:09:04] Stepping pH = 7.95
[1:09:04] Dispensed 0.000024 mL of Base (0.5 M KOH)
[1:09:09] Stepping pH = 8.07
[1:09:24] Stirrer speed set to 0
[1:10:24] Datapoint id 71 collected
[1:10:24] Charge balance equation is out by -683.0%
[1:10:24] Stirrer speed set to 60
[1:10:30] pH 8.14 -> 8.34
[1:10:30] Using cautious pH adjust
[1:10:30] Dispensed 0.000024 mL of Base (0.5 M KOH)
[1:10:35] Stepping pH = 8.19
[1:10:35] Dispensed 0.000024 mL of Base (0.5 M KOH)
[1:10:40] Stepping pH = 8.32
[1:10:40] Dispensed 0.000024 mL of Base (0.5 M KOH)
[1:10:45] Stepping pH = 8.48
[1:11:00] Stirrer speed set to 0
[1:12:00] Datapoint id 72 collected
[1:12:00] Charge balance equation is out by -550.4%
[1:12:00] Stirrer speed set to 60
[1:12:05] pH 8.50 -> 8.70
[1:12:05] Using cautious pH adjust
[1:12:05] Dispensed 0.000024 mL of Base (0.5 M KOH)
[1:12:10] Stepping pH = 8.51
[1:12:10] Dispensed 0.000024 mL of Base (0.5 M KOH)
[1:12:15] Stepping pH = 8.60
[1:12:15] Dispensed 0.000024 mL of Base (0.5 M KOH)
[1:12:21] Stepping pH = 8.68
[1:12:21] Dispensed 0.000024 mL of Base (0.5 M KOH)
[1:12:26] Stepping pH = 8.77
[1:12:41] Stirrer speed set to 0
[1:13:09] Datapoint id 73 collected
[1:13:09] Charge balance equation is out by -398.4%
[1:13:09] Stirrer speed set to 60
[1:13:14] pH 8.76 -> 8.96
[1:13:14] Using cautious pH adjust
[1:13:14] Dispensed 0.000024 mL of Base (0.5 M KOH)
[1:13:19] Stepping pH = 8.75

Sample name: **M09_octanol**
Assay name: **pH-metric high logP**
Assay ID: **18C-06007**
Filename: **C:\Sirius_T3\Mehtap\20180306_exp30_logP_T3-2\18C-06007_M09_octanol_pH-metric high logP.t3r**

Experiment start time: **3/6/2018 3:40:58 PM**
Analyst: **Pion**
Instrument ID: **T312060**

#### Experiment Log (continued)

[1:13:19] Dispensed 0.000071 mL of Base (0.5 M KOH)
[1:13:24] Stepping pH = 8.87
[1:13:24] Dispensed 0.000047 mL of Base (0.5 M KOH)
[1:13:29] Stepping pH = 9.01
[1:13:45] Stirrer speed set to 0
[1:14:08] Datapoint id 74 collected
[1:14:08] Charge balance equation is out by -368.3%
[1:14:08] Argon flow rate set to 0
[1:14:11] Titrator arm moved over Titration position
