## Supplementary material for "Octanol-water partition coefficient measurements for the SAMPL6 Blind Prediction Challenge": SM11_18B-27016_M11_octanol_pH-metric high logP_report.pdf

| | | | |
|---|---|---|---|
| Sample name: | **M11_octanol** | Experiment start time: | **2/27/2018 10:54:30 PM** |
| Assay name: | **pH-metric high logP** | Analyst: | **Pion** |
| Assay ID: | **18B-27016** | Instrument ID: | **T312060** |
| Filename: | **C:\Sirius_T3\Mehtap\20180227_exp27_logP_T3-2\18B-27016_M11_octanol_pH-metric high logP.t3r** | | |

#### pH-metric Result

| | | |
|---|---|---|
| logP (XH +) | -5.60 | ±1.55 (n=48) |
| logP (neutral X) | 2.09 | ±0.01 (n=48) |

#### 18B-27016 Points 1 to 20

| | |
|---|---|
| M11_octanol concentration factor | 0.990 |
| Carbonate | 0.0000 mM |
| Acidity error | 0.67760 mM |

#### 18B-27016 Points 21 to 40

| | |
|---|---|
| M11_octanol concentration factor | 0.815 |
| Carbonate | 0.0822 mM |
| Acidity error | 0.32858 mM |

#### 18B-27016 Points 41 to 62

| | |
|---|---|
| M11_octanol concentration factor | 0.703 |
| Carbonate | 0.1416 mM |
| Acidity error | 0.69165 mM |

#### Warnings and errors

| | |
|---|---|
| Errors | None |
| Warnings | One or more logP values out of range |

#### Sample logD and percent species

| pH | M11_octanol logD | M11_octanol M11_octanolH | M11_octanol M11_octanol | M11_octanol M11_octanolH* | M11_octanol M11_octanol* | Comment |
|---|---|---|---|---|---|---|
| 1.000 | -0.80 | 86.28 % | 0.11 % | 0.00 % | 13.61 % | |
| 1.200 | -0.60 | 79.87 % | 0.16 % | 0.00 % | 19.97 % | Stomach pH |
| 2.000 | 0.19 | 38.61 % | 0.50 % | 0.00 % | 60.90 % | |
| 3.000 | 1.15 | 5.92 % | 0.76 % | 0.00 % | 93.32 % | |
| 4.000 | 1.84 | 0.62 % | 0.81 % | 0.00 % | 98.57 % | |
| 5.000 | 2.06 | 0.06 % | 0.81 % | 0.00 % | 99.13 % | |
| 6.000 | 2.08 | 0.01 % | 0.81 % | 0.00 % | 99.18 % | |
| 6.500 | 2.09 | 0.00 % | 0.81 % | 0.00 % | 99.19 % | |
| 7.000 | 2.09 | 0.00 % | 0.81 % | 0.00 % | 99.19 % | |
| 7.400 | 2.09 | 0.00 % | 0.81 % | 0.00 % | 99.19 % | Blood pH |
| 8.000 | 2.09 | 0.00 % | 0.81 % | 0.00 % | 99.19 % | |
| 9.000 | 2.09 | 0.00 % | 0.81 % | 0.00 % | 99.19 % | |
| 10.000 | 2.09 | 0.00 % | 0.81 % | 0.00 % | 99.19 % | |
| 11.000 | 2.09 | 0.00 % | 0.81 % | 0.00 % | 99.19 % | |
| 12.000 | 2.09 | 0.00 % | 0.81 % | 0.00 % | 99.19 % | |

Sample name: **M11_octanol**
Assay name: **pH-metric high logP**
Assay ID: **18B-27016**
Filename: **C:\Sirius_T3\Mehtap\20180227_exp27_logP_T3-2\18B-27016_M11_octanol_pH-metric high logP.t3r**

Experiment start time: **2/27/2018 10:54:30 PM**
Analyst: **Pion**
Instrument ID: **T312060**

#### Graphs

Charge balance residuals

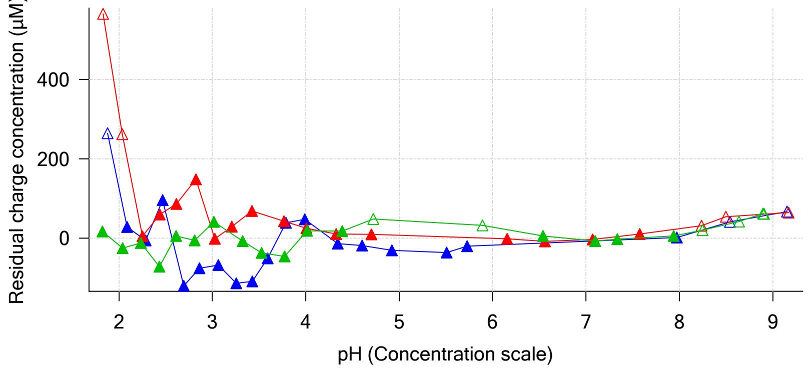


Ionic strength

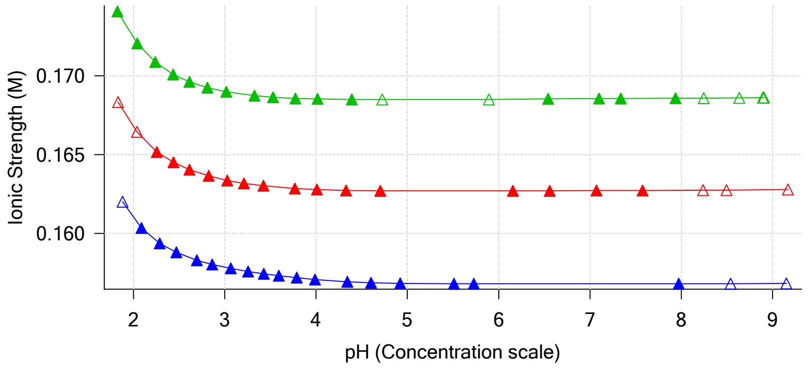


Buffer index profile

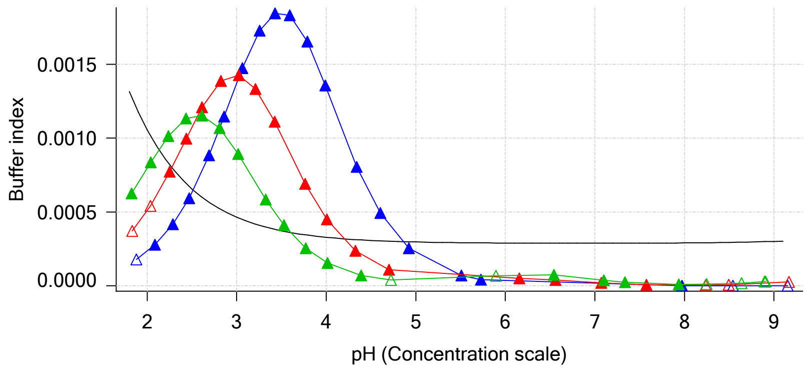


Sample M11_octanol Ionisation Graph

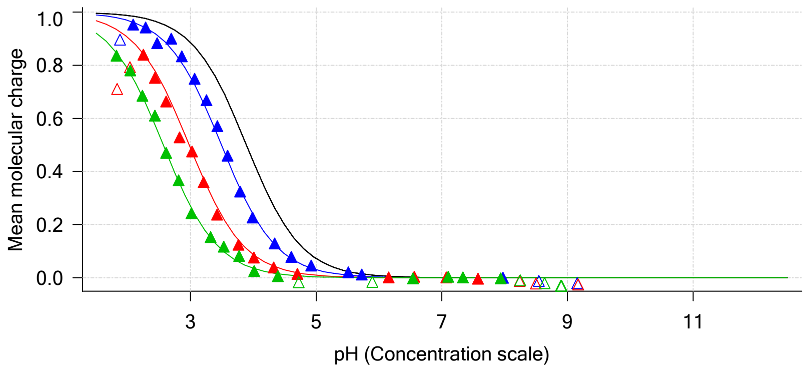


Distribution of species for sample M11_octanol

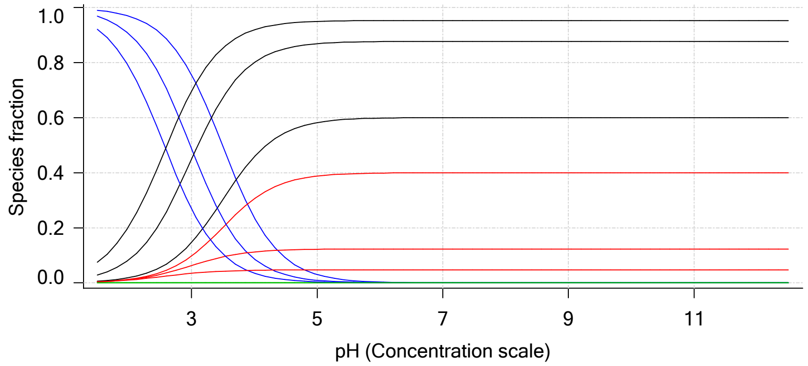


Lipophilicity Profile

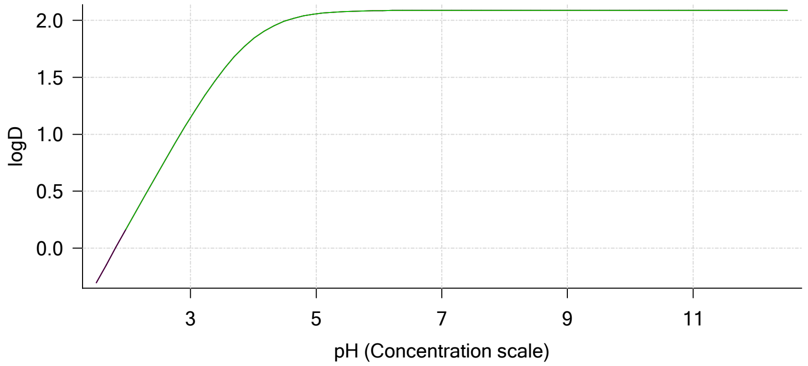


Log R Profile

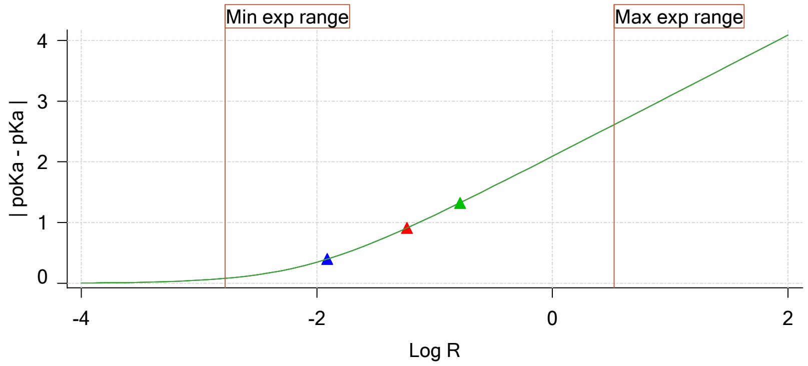


Sample Carbonate Ionisation Graph

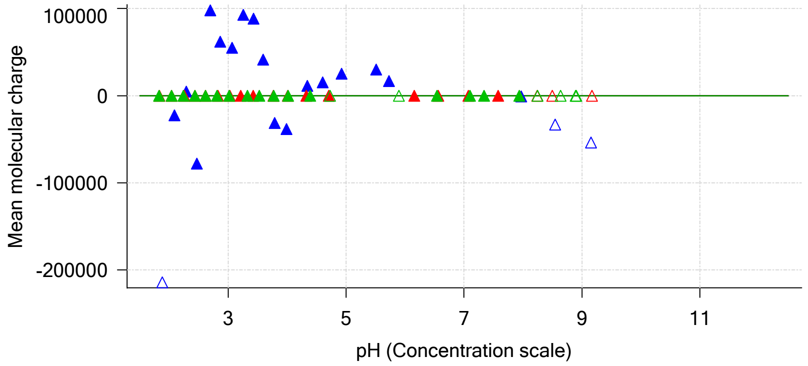

| | | | |
|---|---|---|---|
| Sample name: | **M11_octanol** | Experiment start time: | **2/27/2018 10:54:30 PM** |
| Assay name: | **pH-metric high logP** | Analyst: | **Pion** |
| Assay ID: | **18B-27016** | Instrument ID: | **T312060** |
| Filename: | **C:\Sirius_T3\Mehtap\20180227_exp27_logP_T3-2\18B-27016_M11_octanol_pH-metric high logP.t3r** | | |

#### Graphs (continued)

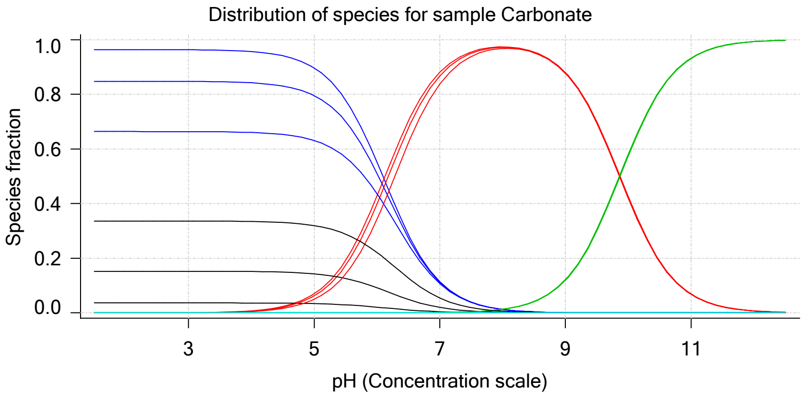

| | | | |
|---|---|---|---|
| Sample name: | **M11_octanol** | Experiment start time: | **2/27/2018 10:54:30 PM** |
| Assay name: | **pH-metric high logP** | Analyst: | **Pion** |
| Assay ID: | **18B-27016** | Instrument ID: | **T312060** |
| Filename: | **C:\Sirius_T3\Mehtap\20180227_exp27_logP_T3-2\18B-27016_M11_octanol_pH-metric high logP.t3r** | | |

### pH-metric high logP Titration 1 of 3 18B-27016 Points 1 to 20

#### Overall results

| | |
|---|---|
| RMSD | 0.163 |
| Average ionic strength | 0.158 M |
| Average temperature | 25.0°C |
| Partition ratio | 0.0123 : 1 |
| Analyte concentration range | 3203.9 μM to 3307.2 μM |
| Total points considered | 17 of 20 |

#### Warnings and errors

| | |
|---|---|
| Errors | None |
| Warnings | None |

#### Four-Plus parameters

| | | | |
|---|---|---|---|
| Alpha | 0.130 | 2/27/2018 10:54:30 PM | C:\Sirius_T3\HCl18B27.t3r |
| S | 0.9970 | 2/27/2018 10:54:30 PM | C:\Sirius_T3\HCl18B27.t3r |
| jH | 0.8 | 2/27/2018 10:54:30 PM | C:\Sirius_T3\HCl18B27.t3r |
| jOH | -0.4 | 2/27/2018 10:54:30 PM | C:\Sirius_T3\HCl18B27.t3r |

#### Titrants

| | | | |
|---|---|---|---|
| 0.50 M HCl | 0.993513 | 2/27/2018 10:54:30 PM | C:\Sirius_T3\HCl18B27.t3r |
| 0.50 M KOH | 0.999845 | 2/27/2018 10:54:30 PM | C:\Sirius_T3\KOH18B27.t3r |

#### Sample

| | |
|---|---|
| M11_octanol concentration factor | 0.990 |
| Base pKa 1 | 3.89 |
| logP (XH +) | -0.58 |
| logP (neutral X) | 2.04 |

#### Sample graphs

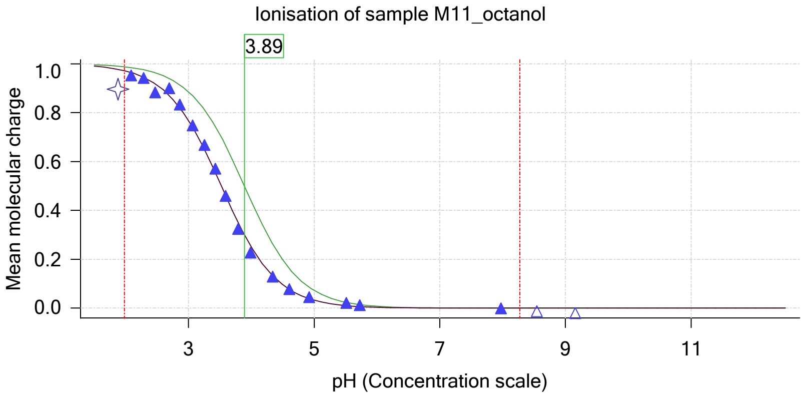


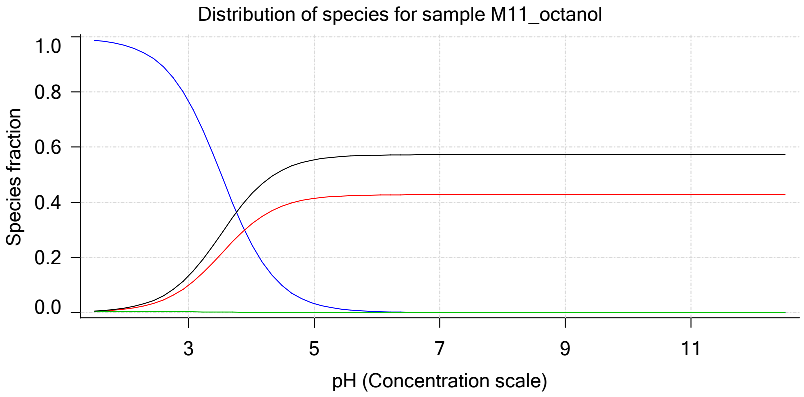

| | | | |
|---|---|---|---|
| Sample name: | **M11_octanol** | Experiment start time: | **2/27/2018 10:54:30 PM** |
| Assay name: | **pH-metric high logP** | Analyst: | **Pion** |
| Assay ID: | **18B-27016** | Instrument ID: | **T312060** |

Filename: **C:\Sirius_T3\Mehtap\20180227_exp27_logP_T3-2\18B-27016_M11_octanol_pH-metric high logP.t3r**

#### Sample graphs (continued)

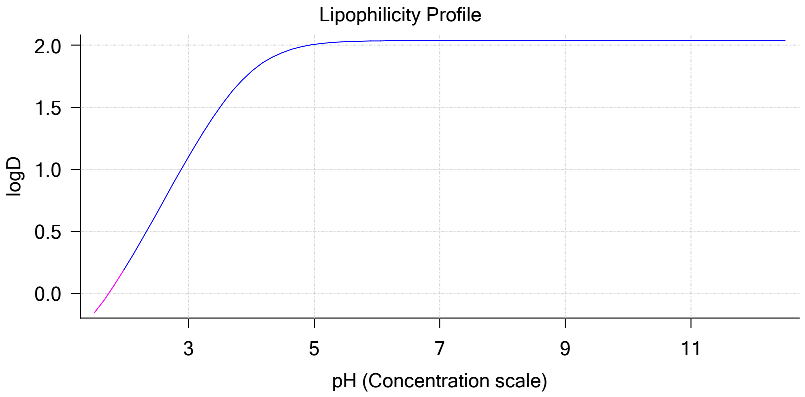


#### Sample logD and percent species

| pH | M11_octanol logD | M11_octanol M11_octanolH | M11_octanol M11_octanol | M11_octanol M11_octanolH* | M11_octanol M11_octanol* | Comment |
|---|---|---|---|---|---|---|
| 1.000 | -0.40 | 99.38 % | 0.13 % | 0.32 % | 0.17 % | |
| 1.200 | -0.31 | 99.21 % | 0.20 % | 0.32 % | 0.27 % | Stomach pH |
| 2.000 | 0.22 | 96.77 % | 1.25 % | 0.31 % | 1.67 % | |
| 3.000 | 1.10 | 76.66 % | 9.88 % | 0.25 % | 13.22 % | |
| 4.000 | 1.79 | 24.90 % | 32.08 % | 0.08 % | 42.94 % | |
| 5.000 | 2.01 | 3.21 % | 41.38 % | 0.01 % | 55.40 % | |
| 6.000 | 2.03 | 0.33 % | 42.61 % | 0.00 % | 57.05 % | |
| 6.500 | 2.04 | 0.10 % | 42.71 % | 0.00 % | 57.18 % | |
| 7.000 | 2.04 | 0.03 % | 42.74 % | 0.00 % | 57.22 % | |
| 7.400 | 2.04 | 0.01 % | 42.75 % | 0.00 % | 57.24 % | Blood pH |
| 8.000 | 2.04 | 0.00 % | 42.76 % | 0.00 % | 57.24 % | |
| 9.000 | 2.04 | 0.00 % | 42.76 % | 0.00 % | 57.24 % | |
| 10.000 | 2.04 | 0.00 % | 42.76 % | 0.00 % | 57.24 % | |
| 11.000 | 2.04 | 0.00 % | 42.76 % | 0.00 % | 57.24 % | |
| 12.000 | 2.04 | 0.00 % | 42.76 % | 0.00 % | 57.24 % | |

#### Carbonate and acidity

| | |
|---|---|
| Carbonate | 0.000 mM |
| Acidity error | 0.678 mM |

#### Other graphs

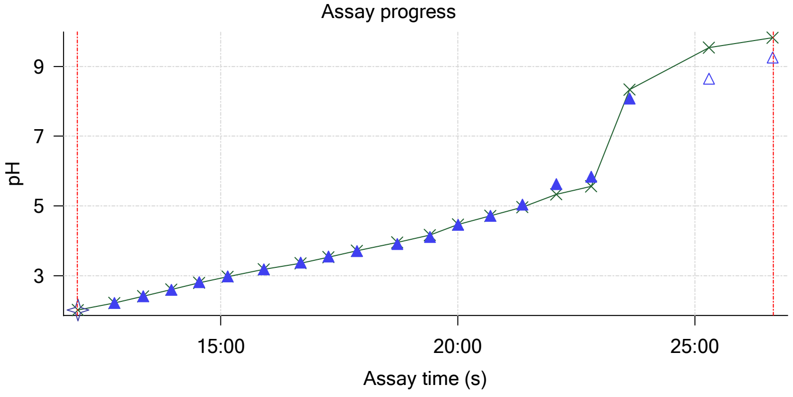


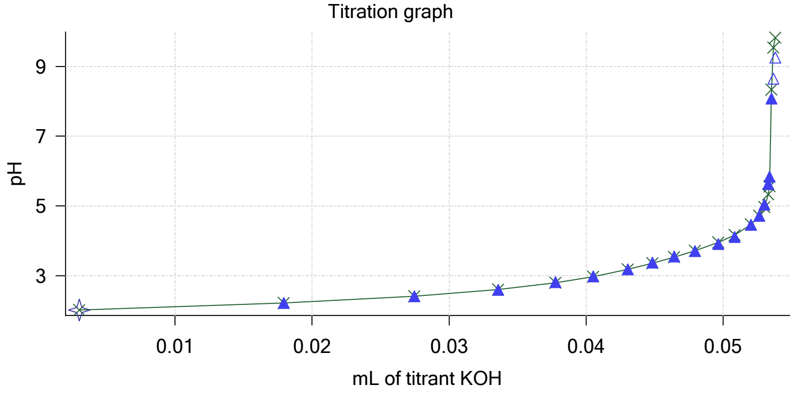

| | | | |
|---|---|---|---|
| Sample name: | **M11_octanol** | Experiment start time: | **2/27/2018 10:54:30 PM** |
| Assay name: | **pH-metric high logP** | Analyst: | **Pion** |
| Assay ID: | **18B-27016** | Instrument ID: | **T312060** |
| Filename: | **C:\Sirius_T3\Mehtap\20180227_exp27_logP_T3-2\18B-27016_M11_octanol_pH-metric high logP.t3r** | | |

#### Other graphs (continued)

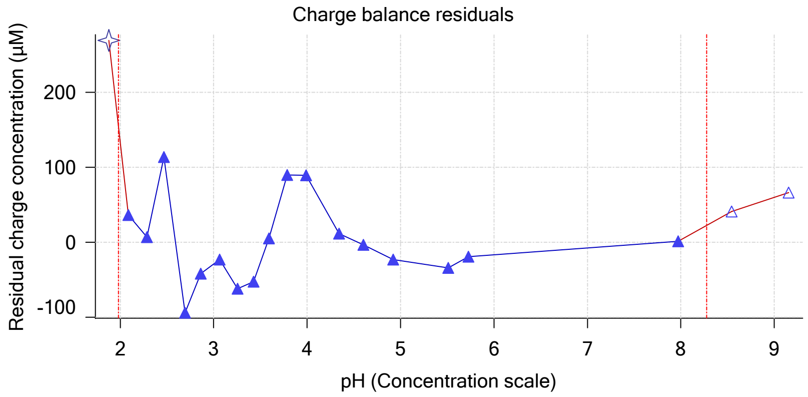


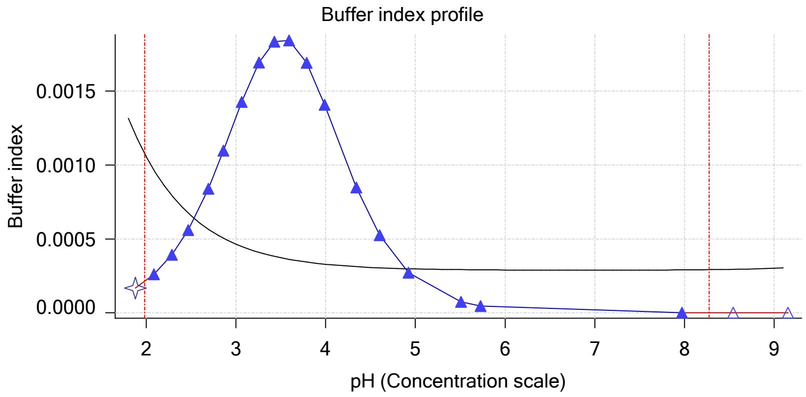


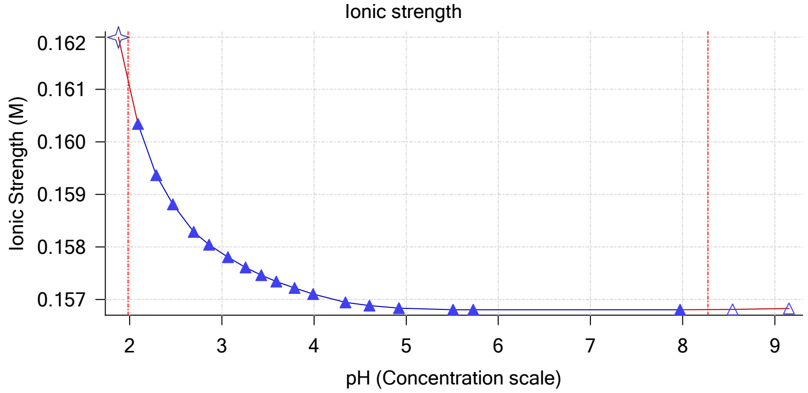


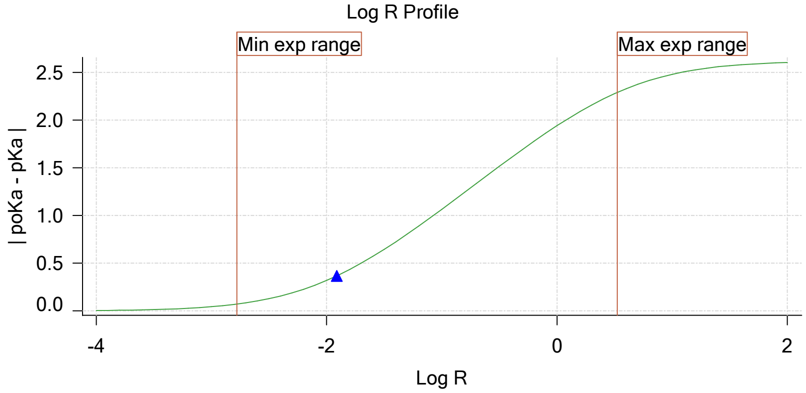


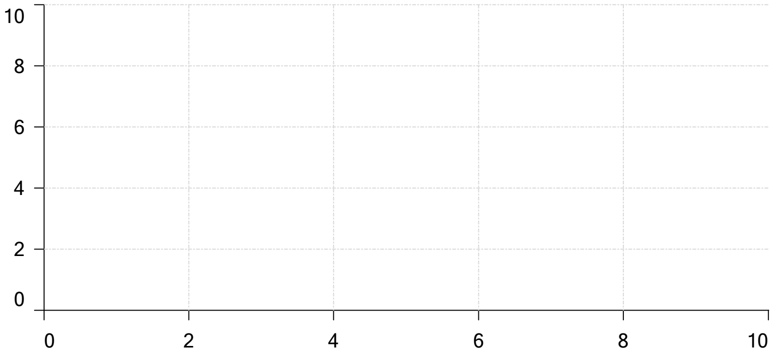

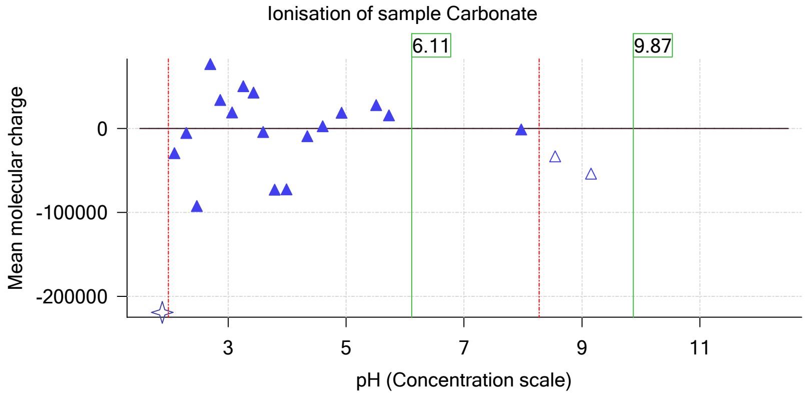

| | | | |
|---|---|---|---|
| Sample name: | **M11_octanol** | Experiment start time: | **2/27/2018 10:54:30 PM** |
| Assay name: | **pH-metric high logP** | Analyst: | **Pion** |
| Assay ID: | **18B-27016** | Instrument ID: | **T312060** |
| Filename: | **C:\Sirius_T3\Mehtap\20180227_exp27_logP_T3-2\18B-27016_M11_octanol_pH-metric high logP.t3r** | | |

### pH-metric high logP Titration 2 of 3 18B-27016 Points 21 to 40

#### Overall results

| | |
|---|---|
| RMSD | 0.275 |
| Average ionic strength | 0.163 M |
| Average temperature | 25.0°C |
| Partition ratio | 0.0582 : 1 |
| Analyte concentration range | 2864.7 μM to 2955.2 μM |
| Total points considered | 15 of 20 |

#### Warnings and errors

| | |
|---|---|
| Errors | None |
| Warnings | None |

#### Four-Plus parameters

| | | | |
|---|---|---|---|
| Alpha | 0.130 | 2/27/2018 10:54:30 PM | C:\Sirius_T3\HCl18B27.t3r |
| S | 0.9970 | 2/27/2018 10:54:30 PM | C:\Sirius_T3\HCl18B27.t3r |
| jH | 0.8 | 2/27/2018 10:54:30 PM | C:\Sirius_T3\HCl18B27.t3r |
| jOH | -0.4 | 2/27/2018 10:54:30 PM | C:\Sirius_T3\HCl18B27.t3r |

#### Titrants

| | | | |
|---|---|---|---|
| 0.50 M HCl | 0.993513 | 2/27/2018 10:54:30 PM | C:\Sirius_T3\HCl18B27.t3r |
| 0.50 M KOH | 0.999845 | 2/27/2018 10:54:30 PM | C:\Sirius_T3\KOH18B27.t3r |

#### Sample

| | |
|---|---|
| M11_octanol concentration factor | 0.815 |
| Base pKa 1 | 3.89 |
| logP (XH +) | -0.58 |
| logP (neutral X) | 2.15 |

#### Sample graphs

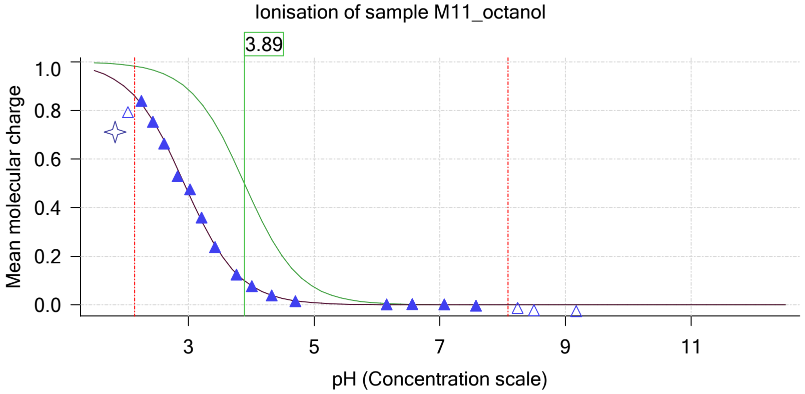


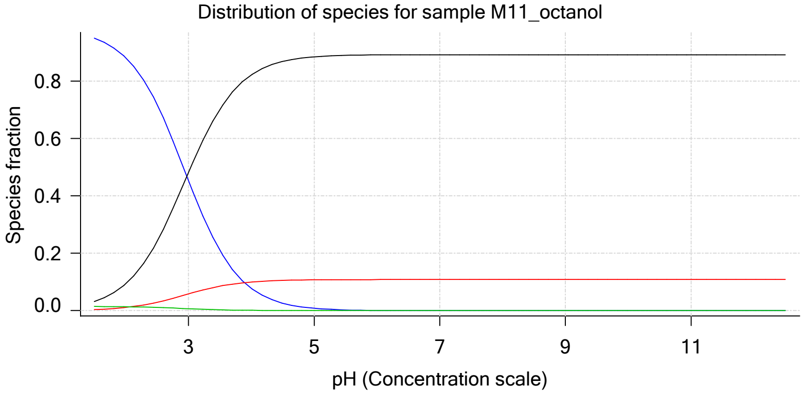

| | | | |
|---|---|---|---|
| Sample name: | **M11_octanol** | Experiment start time: | **2/27/2018 10:54:30 PM** |
| Assay name: | **pH-metric high logP** | Analyst: | **Pion** |
| Assay ID: | **18B-27016** | Instrument ID: | **T312060** |
| Filename: | **C:\Sirius_T3\Mehtap\20180227_exp27_logP_T3-2\18B-27016_M11_octanol_pH-metric high logP.t3r** | | |

#### Sample graphs (continued)

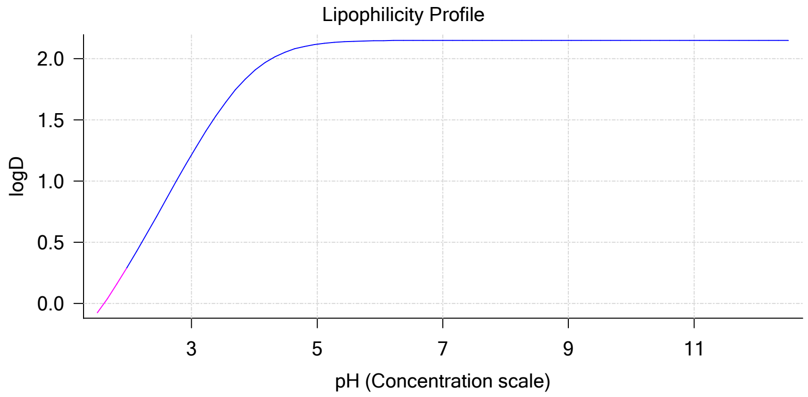


#### Sample logD and percent species

| pH | M11_octanol logD | M11_octanol M11_octanolH | M11_octanol M11_octanol | M11_octanol M11_octanolH* | M11_octanol M11_octanol* | Comment |
|---|---|---|---|---|---|---|
| 1.000 | -0.35 | 97.35 % | 0.13 % | 1.49 % | 1.03 % | |
| 1.200 | -0.26 | 96.70 % | 0.20 % | 1.48 % | 1.63 % | Stomach pH |
| 2.000 | 0.31 | 88.17 % | 1.14 % | 1.35 % | 9.35 % | |
| 3.000 | 1.21 | 45.36 % | 5.84 % | 0.69 % | 48.10 % | |
| 4.000 | 1.90 | 7.75 % | 9.98 % | 0.12 % | 82.15 % | |
| 5.000 | 2.12 | 0.83 % | 10.74 % | 0.01 % | 88.41 % | |
| 6.000 | 2.15 | 0.08 % | 10.82 % | 0.00 % | 89.09 % | |
| 6.500 | 2.15 | 0.03 % | 10.83 % | 0.00 % | 89.14 % | |
| 7.000 | 2.15 | 0.01 % | 10.83 % | 0.00 % | 89.16 % | |
| 7.400 | 2.15 | 0.00 % | 10.83 % | 0.00 % | 89.16 % | Blood pH |
| 8.000 | 2.15 | 0.00 % | 10.83 % | 0.00 % | 89.17 % | |
| 9.000 | 2.15 | 0.00 % | 10.83 % | 0.00 % | 89.17 % | |
| 10.000 | 2.15 | 0.00 % | 10.83 % | 0.00 % | 89.17 % | |
| 11.000 | 2.15 | 0.00 % | 10.83 % | 0.00 % | 89.17 % | |
| 12.000 | 2.15 | 0.00 % | 10.83 % | 0.00 % | 89.17 % | |

#### Carbonate and acidity

| | |
|---|---|
| Carbonate | 0.082 mM |
| Acidity error | 0.329 mM |

#### Other graphs

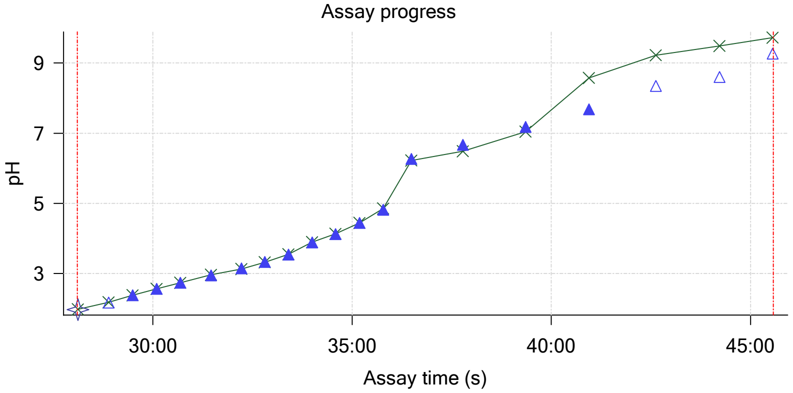


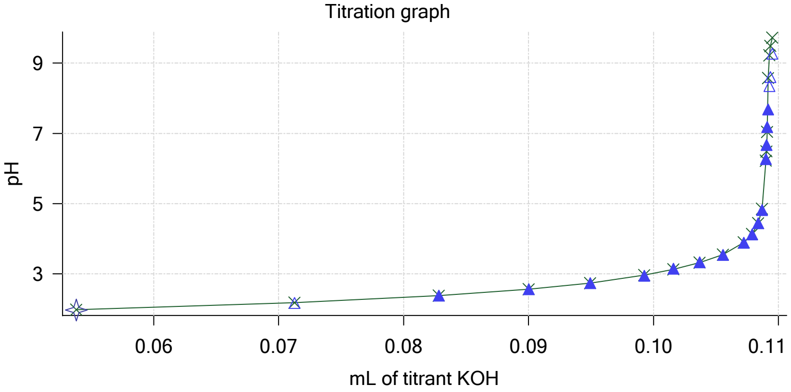

| | | | |
|---|---|---|---|
| Sample name: | **M11_octanol** | Experiment start time: | **2/27/2018 10:54:30 PM** |
| Assay name: | **pH-metric high logP** | Analyst: | **Pion** |
| Assay ID: | **18B-27016** | Instrument ID: | **T312060** |
| Filename: | **C:\Sirius_T3\Mehtap\20180227_exp27_logP_T3-2\18B-27016_M11_octanol_pH-metric high logP.t3r** | | |

#### Other graphs (continued)

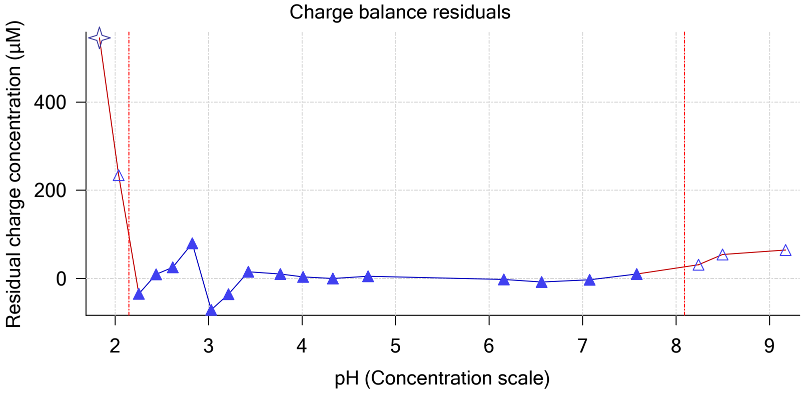


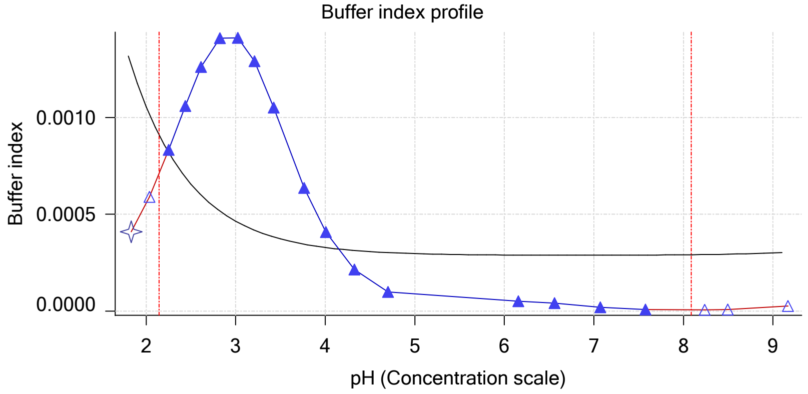


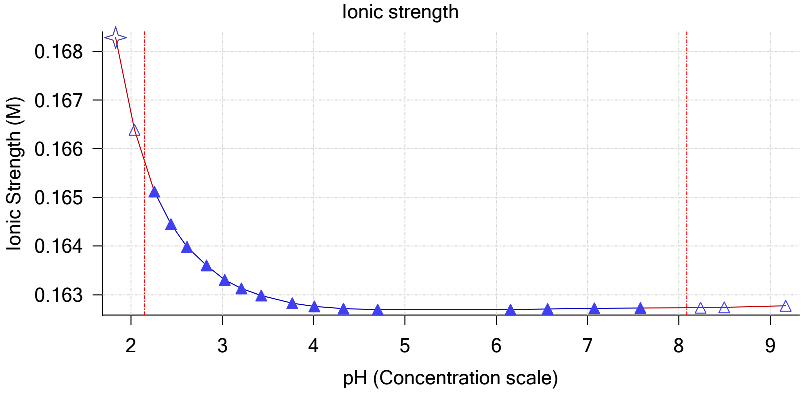


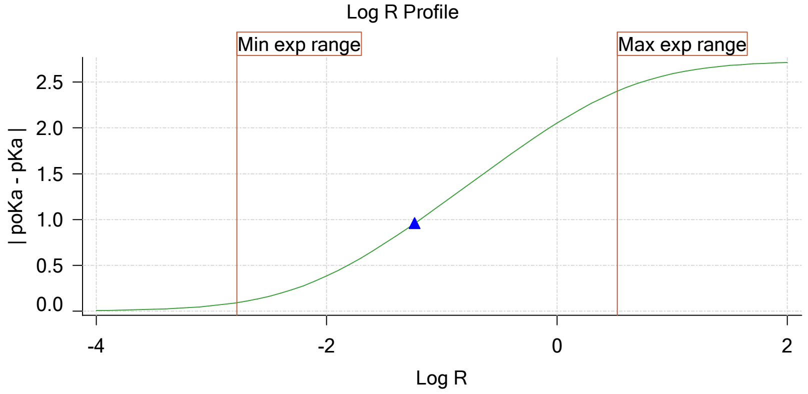


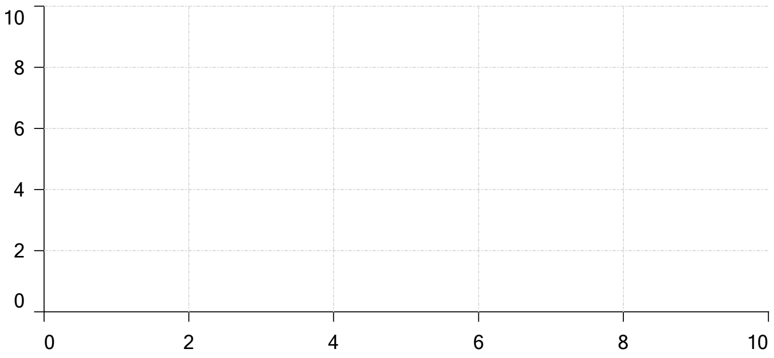

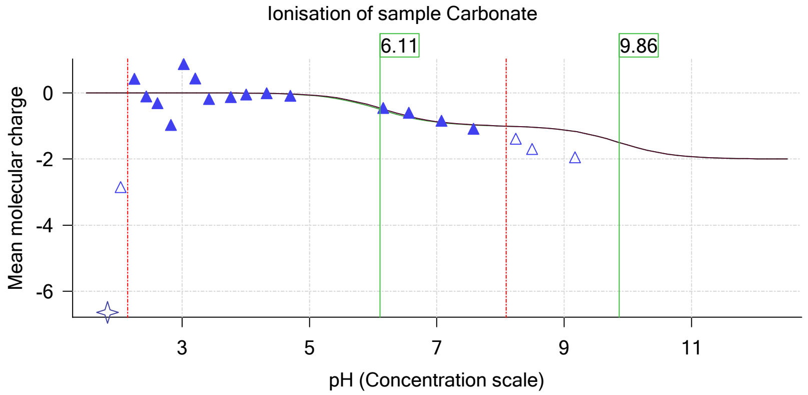

Sample name: **M11_octanol** | Experiment start time: **2/27/2018 10:54:30 PM**
Assay name: **pH-metric high logP** | Analyst: **Pion**
Assay ID: **18B-27016** | Instrument ID: **T312060**
Filename: **C:\Sirius_T3\Mehtap\20180227_exp27_logP_T3-2\18B-27016_M11_octanol_pH-metric high logP.t3r**

# pH-metric high logP   Titration 3 of 3   18B-27016 Points 41 to 62

#### Overall results

| | |
|---|---|
| RMSD | 0.122 |
| Average ionic strength | 0.169 M |
| Average temperature | 25.0°C |
| Partition ratio | 0.1640 : 1 |
| Analyte concentration range | 2436.6 µM to 2507.4 µM |
| Total points considered | 16 of 22 |

#### Warnings and errors

| | |
|---|---|
| Errors | None |
| Warnings | None |

#### Four-Plus parameters

| | | | |
|---|---|---|---|
| Alpha | 0.130 | 2/27/2018 10:54:30 PM | C:\Sirius_T3\HCl18B27.t3r |
| S | 0.9970 | 2/27/2018 10:54:30 PM | C:\Sirius_T3\HCl18B27.t3r |
| jH | 0.8 | 2/27/2018 10:54:30 PM | C:\Sirius_T3\HCl18B27.t3r |
| jOH | -0.4 | 2/27/2018 10:54:30 PM | C:\Sirius_T3\HCl18B27.t3r |

#### Titrants

| | | | |
|---|---|---|---|
| 0.50 M HCl | 0.993513 | 2/27/2018 10:54:30 PM | C:\Sirius_T3\HCl18B27.t3r |
| 0.50 M KOH | 0.999845 | 2/27/2018 10:54:30 PM | C:\Sirius_T3\KOH18B27.t3r |

#### Sample

| | |
|---|---|
| M11_octanol concentration factor | 0.703 |
| Base pKa 1 | 3.89 |
| logP (XH +) | -0.58 |
| logP (neutral X) | 2.09 |

#### Sample graphs

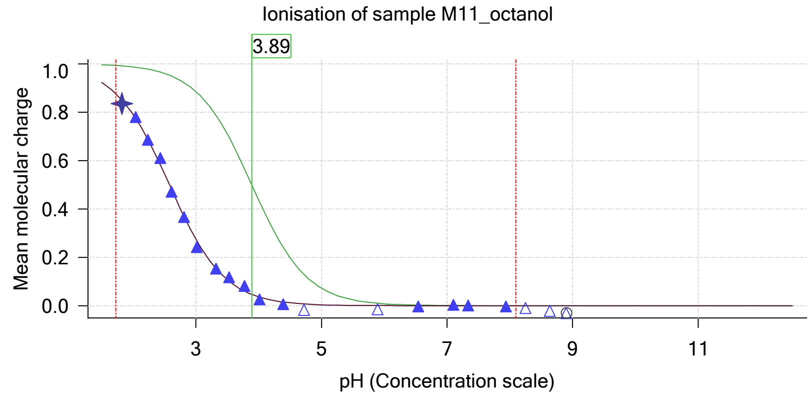


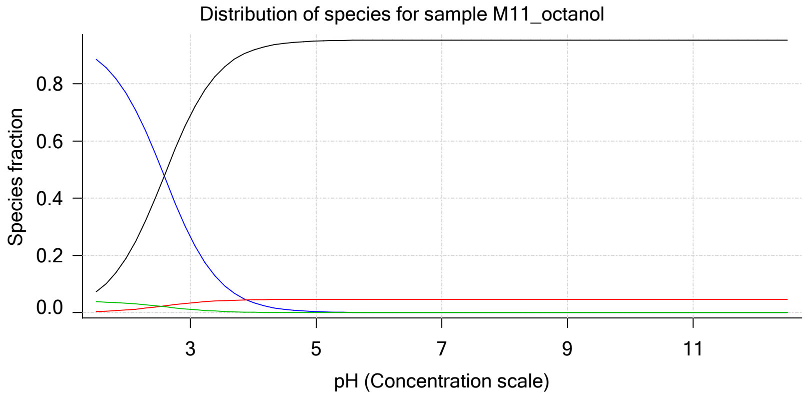

| | | | |
|---|---|---|---|
| Sample name: | **M11_octanol** | Experiment start time: | **2/27/2018 10:54:30 PM** |
| Assay name: | **pH-metric high logP** | Analyst: | **Pion** |
| Assay ID: | **18B-27016** | Instrument ID: | **T312060** |
| Filename: | **C:\Sirius_T3\Mehtap\20180227_exp27_logP_T3-2\18B-27016_M11_octanol_pH-metric high logP.t3r** | | |

#### Sample graphs (continued)

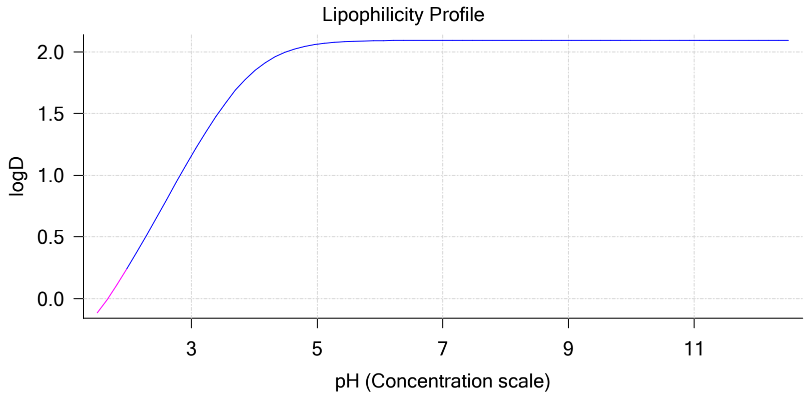


#### Sample logD and percent species

| pH | M11_octanol logD | M11_octanol M11_octanolH | M11_octanol M11_octanol | M11_octanol M11_octanolH* | M11_octanol M11_octanol* | Comment |
|---|---|---|---|---|---|---|
| 1.000 | -0.37 | 93.40 % | 0.12 % | 4.03 % | 2.45 % | |
| 1.200 | -0.29 | 92.02 % | 0.19 % | 3.97 % | 3.82 % | Stomach pH |
| 2.000 | 0.26 | 75.86 % | 0.98 % | 3.27 % | 19.89 % | |
| 3.000 | 1.16 | 26.36 % | 3.40 % | 1.14 % | 69.10 % | |
| 4.000 | 1.84 | 3.50 % | 4.51 % | 0.15 % | 91.83 % | |
| 5.000 | 2.06 | 0.36 % | 4.67 % | 0.02 % | 94.96 % | |
| 6.000 | 2.09 | 0.04 % | 4.68 % | 0.00 % | 95.28 % | |
| 6.500 | 2.09 | 0.01 % | 4.68 % | 0.00 % | 95.30 % | |
| 7.000 | 2.09 | 0.00 % | 4.68 % | 0.00 % | 95.31 % | |
| 7.400 | 2.09 | 0.00 % | 4.68 % | 0.00 % | 95.31 % | Blood pH |
| 8.000 | 2.09 | 0.00 % | 4.68 % | 0.00 % | 95.32 % | |
| 9.000 | 2.09 | 0.00 % | 4.68 % | 0.00 % | 95.32 % | |
| 10.000 | 2.09 | 0.00 % | 4.68 % | 0.00 % | 95.32 % | |
| 11.000 | 2.09 | 0.00 % | 4.68 % | 0.00 % | 95.32 % | |
| 12.000 | 2.09 | 0.00 % | 4.68 % | 0.00 % | 95.32 % | |

#### Carbonate and acidity

| | |
|---|---|
| Carbonate | 0.142 mM |
| Acidity error | 0.692 mM |

#### Other graphs

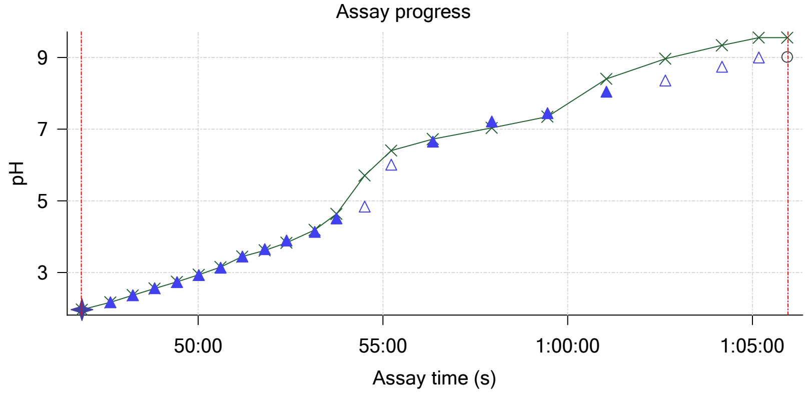


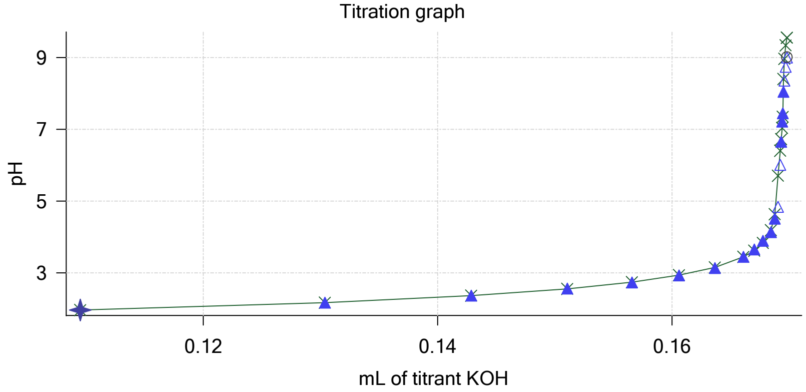

Sample name: **M11_octanol**
Assay name: **pH-metric high logP**
Assay ID: **18B-27016**
Filename: **C:\Sirius_T3\Mehtap\20180227_exp27_logP_T3-2\18B-27016_M11_octanol_pH-metric high logP.t3r**

Experiment start time: **2/27/2018 10:54:30 PM**
Analyst: **Pion**
Instrument ID: **T312060**

#### Other graphs (continued)

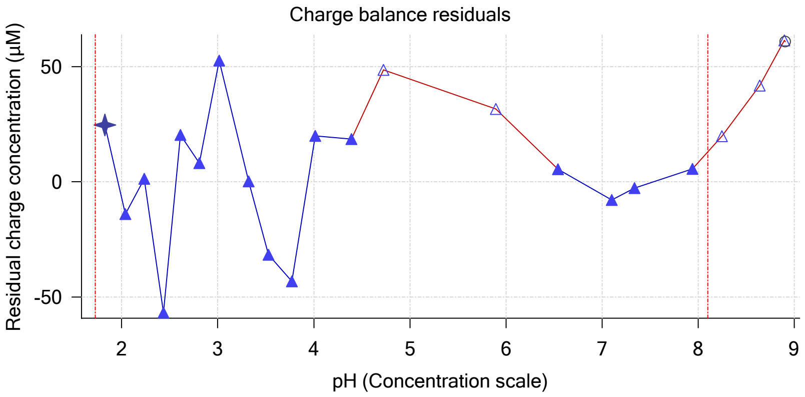


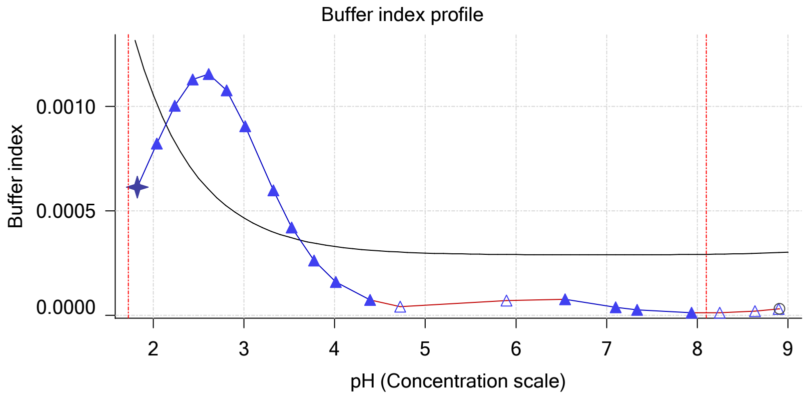


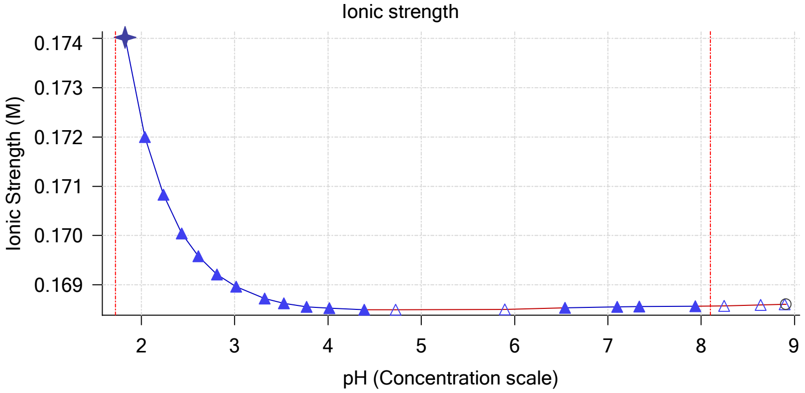


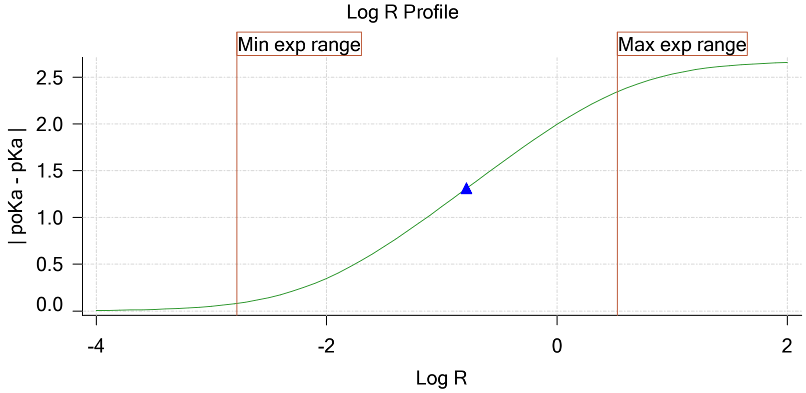


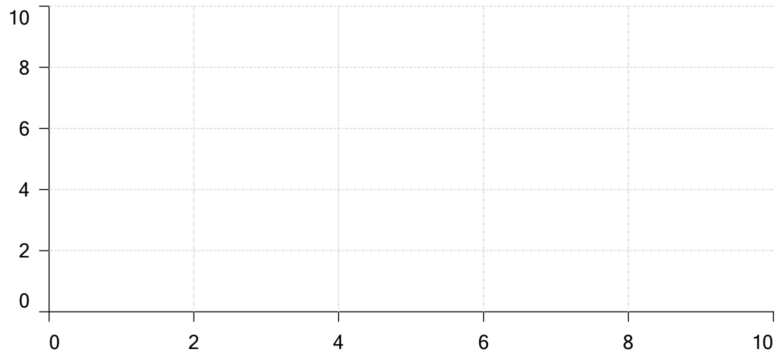


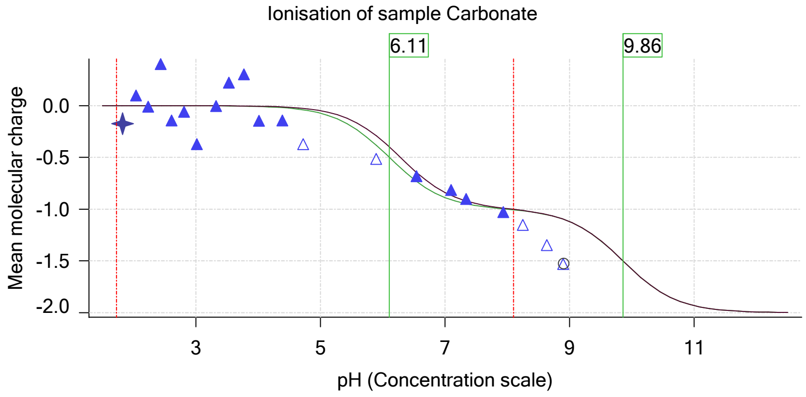

| | | | |
|---|---|---|---|
| Sample name: | **M11_octanol** | Experiment start time: | **2/27/2018 10:54:30 PM** |
| Assay name: | **pH-metric high logP** | Analyst: | **Pion** |
| Assay ID: | **18B-27016** | Instrument ID: | **T312060** |
| Filename: | **C:\Sirius_T3\Mehtap\20180227_exp27_logP_T3-2\18B-27016_M11_octanol_pH-metric high logP.t3r** | | |

#### Assay Model

| Settings | Value | Date/Time changed | Imported from |
|---|---|---|---|
| Sample name | M11_octanol | 2/27/2018 4:54:30 PM | User entered value |
| Sample by | Weight | | Default value |
| Sample weight | 0.001100 g | 2/27/2018 6:40:56 PM | User entered value |
| Formula weight | 211.22 g/mol | 2/27/2018 4:54:30 PM | User entered value |
| Solubility | Unknown | | Default value |
| Molecular weight | 211.22 | 2/27/2018 4:54:30 PM | User entered value |
| Individual pKa ionic environments | No | | Default value |
| Number of pKas | 1 | 2/27/2018 4:54:30 PM | User entered value |
| Sample is a | Base | 2/27/2018 4:54:30 PM | User entered value |
| pKa 1 | 3.89 | 2/27/2018 4:54:30 PM | User entered value |
| logp (XH +) | -0.58 | 2/27/2018 4:54:54 PM | User entered value |
| logP (neutral X) | 1.24 | 2/27/2018 4:54:49 PM | User entered value |

#### Events

| Time | Event | Water | Acid | Base | Octanol | pH | dpH/dt | pH R-squared | pH SD | dpH/dt time |
|---|---|---|---|---|---|---|---|---|---|---|
| 8:60.0 | Initial pH = 6.35 | | | | | | | | | |
| 11:59.5 | Data point 1 | 1.50000 mL | 0.05167 mL | 0.00301 mL | 0.01999 mL | 2.012 | -0.01321 | 0.59861 | 0.00084 | 10.0 s |
| 12:45.7 | Data point 2 | 1.50000 mL | 0.05167 mL | 0.01794 mL | 0.01999 mL | 2.217 | -0.01201 | 0.89280 | 0.00063 | 10.5 s |
| 13:21.8 | Data point 3 | 1.50000 mL | 0.05167 mL | 0.02747 mL | 0.01999 mL | 2.415 | -0.00642 | 0.73636 | 0.00037 | 10.0 s |
| 13:57.4 | Data point 4 | 1.50000 mL | 0.05167 mL | 0.03358 mL | 0.01999 mL | 2.593 | -0.00697 | 0.59547 | 0.00045 | 10.0 s |
| 14:32.9 | Data point 5 | 1.50000 mL | 0.05167 mL | 0.03777 mL | 0.01999 mL | 2.816 | -0.00524 | 0.75021 | 0.00030 | 10.5 s |
| 15:08.9 | Data point 6 | 1.50000 mL | 0.05167 mL | 0.04052 mL | 0.01999 mL | 2.984 | -0.01107 | 0.79804 | 0.00061 | 10.0 s |
| 15:54.8 | Data point 7 | 1.50000 mL | 0.05167 mL | 0.04304 mL | 0.01999 mL | 3.186 | -0.00468 | 0.88101 | 0.00025 | 10.5 s |
| 16:41.1 | Data point 8 | 1.50000 mL | 0.05167 mL | 0.04485 mL | 0.01999 mL | 3.377 | -0.00932 | 0.44692 | 0.00069 | 10.0 s |
| 17:16.6 | Data point 9 | 1.50000 mL | 0.05167 mL | 0.04643 mL | 0.01999 mL | 3.547 | -0.00399 | 0.79959 | 0.00022 | 10.5 s |
| 17:52.6 | Data point 10 | 1.50000 mL | 0.05167 mL | 0.04795 mL | 0.01999 mL | 3.710 | -0.00939 | 0.88441 | 0.00049 | 10.0 s |
| 18:43.6 | Data point 11 | 1.50000 mL | 0.05167 mL | 0.04965 mL | 0.01999 mL | 3.906 | 0.00461 | 0.20532 | 0.00050 | 10.5 s |
| 19:24.7 | Data point 12 | 1.50000 mL | 0.05167 mL | 0.05085 mL | 0.01999 mL | 4.108 | -0.01572 | 0.89585 | 0.00082 | 10.0 s |
| 20:00.3 | Data point 13 | 1.50000 mL | 0.05167 mL | 0.05205 mL | 0.01999 mL | 4.458 | -0.01283 | 0.78922 | 0.00071 | 10.5 s |
| 20:41.3 | Data point 14 | 1.50000 mL | 0.05167 mL | 0.05263 mL | 0.01999 mL | 4.719 | -0.01182 | 0.87243 | 0.00063 | 10.0 s |
| 21:21.9 | Data point 15 | 1.50000 mL | 0.05167 mL | 0.05301 mL | 0.01999 mL | 5.038 | -0.01739 | 0.94532 | 0.00088 | 12.5 s |
| 22:05.0 | Data point 16 | 1.50000 mL | 0.05167 mL | 0.05329 mL | 0.01999 mL | 5.624 | -0.01784 | 0.80310 | 0.00098 | 18.5 s |
| 22:48.9 | Data point 17 | 1.50000 mL | 0.05167 mL | 0.05339 mL | 0.01999 mL | 5.840 | -0.01669 | 0.76560 | 0.00094 | 18.0 s |
| 23:37.4 | Data point 18 | 1.50000 mL | 0.05167 mL | 0.05353 mL | 0.01999 mL | 8.078 | -0.05197 | 0.99587 | 0.00257 | Timed out at 59.5 s |
| 25:18.2 | Data point 19 | 1.50000 mL | 0.05167 mL | 0.05367 mL | 0.01999 mL | 8.647 | -0.01738 | 0.86596 | 0.00092 | 44.5 s |
| 26:38.3 | Data point 20 | 1.50000 mL | 0.05167 mL | 0.05381 mL | 0.01999 mL | 9.256 | -0.01881 | 0.92735 | 0.00096 | 29.0 s |
| 28:07.4 | Data point 21 | 1.50000 mL | 0.10842 mL | 0.05381 mL | 0.10000 mL | 1.967 | -0.00520 | 0.28864 | 0.00048 | 10.0 s |
| 28:53.7 | Data point 22 | 1.50000 mL | 0.10842 mL | 0.07126 mL | 0.10000 mL | 2.168 | -0.01928 | 0.91140 | 0.00100 | 10.5 s |
| 29:29.9 | Data point 23 | 1.50000 mL | 0.10842 mL | 0.08283 mL | 0.10000 mL | 2.381 | 0.00042 | 0.05097 | 0.00009 | 10.5 s |
| 30:06.0 | Data point 24 | 1.50000 mL | 0.10842 mL | 0.09000 mL | 0.10000 mL | 2.562 | -0.00366 | 0.61217 | 0.00023 | 10.0 s |
| 30:41.6 | Data point 25 | 1.50000 mL | 0.10842 mL | 0.09492 mL | 0.10000 mL | 2.738 | -0.01006 | 0.31818 | 0.00088 | 10.0 s |
| 31:27.5 | Data point 26 | 1.50000 mL | 0.10842 mL | 0.09925 mL | 0.10000 mL | 2.947 | -0.00503 | 0.77202 | 0.00028 | 10.0 s |
| 32:13.3 | Data point 27 | 1.50000 mL | 0.10842 mL | 0.10160 mL | 0.10000 mL | 3.147 | -0.00762 | 0.61374 | 0.00048 | 10.0 s |
| 32:48.8 | Data point 28 | 1.50000 mL | 0.10842 mL | 0.10369 mL | 0.10000 mL | 3.331 | -0.01378 | 0.87687 | 0.00073 | 10.0 s |
| 33:24.3 | Data point 29 | 1.50000 mL | 0.10842 mL | 0.10555 mL | 0.10000 mL | 3.544 | -0.00880 | 0.29642 | 0.00080 | 10.0 s |
| 33:59.7 | Data point 30 | 1.50000 mL | 0.10842 mL | 0.10722 mL | 0.10000 mL | 3.884 | -0.00929 | 0.26981 | 0.00088 | 10.0 s |
| 34:35.2 | Data point 31 | 1.50000 mL | 0.10842 mL | 0.10788 mL | 0.10000 mL | 4.125 | -0.00452 | 0.80460 | 0.00025 | 10.5 s |
| 35:11.1 | Data point 32 | 1.50000 mL | 0.10842 mL | 0.10837 mL | 0.10000 mL | 4.444 | -0.01177 | 0.87839 | 0.00062 | 10.5 s |
| 35:46.9 | Data point 33 | 1.50000 mL | 0.10842 mL | 0.10868 mL | 0.10000 mL | 4.818 | -0.01577 | 0.78042 | 0.00088 | 12.0 s |
| 36:29.4 | Data point 34 | 1.50000 mL | 0.10842 mL | 0.10898 mL | 0.10000 mL | 6.267 | -0.01927 | 0.97409 | 0.00096 | 52.0 s |
| 37:46.9 | Data point 35 | 1.50000 mL | 0.10842 mL | 0.10903 mL | 0.10000 mL | 6.671 | -0.01898 | 0.96189 | 0.00096 | 58.5 s |
| 39:21.1 | Data point 36 | 1.50000 mL | 0.10842 mL | 0.10910 mL | 0.10000 mL | 7.182 | -0.05514 | 0.98470 | 0.00274 | Timed out at 59.5 s |
| 40:56.7 | Data point 37 | 1.50000 mL | 0.10842 mL | 0.10917 mL | 0.10000 mL | 7.684 | -0.05926 | 0.99557 | 0.00293 | Timed out at 59.5 s |

| | | | |
|---|---|---|---|
| Sample name: | **M11_octanol** | Experiment start time: | **2/27/2018 10:54:30 PM** |
| Assay name: | **pH-metric high logP** | Analyst: | **Pion** |
| Assay ID: | **18B-27016** | Instrument ID: | **T312060** |
| Filename: | **C:\Sirius_T3\Mehtap\20180227_exp27_logP_T3-2\18B-27016_M11_octanol_pH-metric high logP.t3r** | | |

#### Events (continued)

| Time | Event | Water | Acid | Base | Octanol | pH | dpH/dt | pH R-squared | pH SD | dpH/dt time |
|---|---|---|---|---|---|---|---|---|---|---|
| 42:37.4 | Data point 38 | 1.50000 mL | 0.10842 mL | 0.10927 mL | 0.10000 mL | 8.344 | -0.02444 | 0.97413 | 0.00122 | Timed out at 59.5 s |
| 44:13.0 | Data point 39 | 1.50000 mL | 0.10842 mL | 0.10936 mL | 0.10000 mL | 8.601 | -0.01930 | 0.94203 | 0.00098 | 44.5 s |
| 45:33.1 | Data point 40 | 1.50000 mL | 0.10842 mL | 0.10950 mL | 0.10000 mL | 9.271 | -0.01442 | 0.69862 | 0.00085 | 15.0 s |
| 46:50.9 | Data point 41 | 1.50000 mL | 0.16747 mL | 0.10950 mL | 0.30000 mL | 1.960 | -0.00611 | 0.71181 | 0.00036 | 10.0 s |
| 47:37.4 | Data point 42 | 1.50000 mL | 0.16747 mL | 0.13039 mL | 0.30000 mL | 2.171 | -0.01056 | 0.60425 | 0.00067 | 10.5 s |
| 48:13.6 | Data point 43 | 1.50000 mL | 0.16747 mL | 0.14287 mL | 0.30000 mL | 2.364 | -0.00724 | 0.54028 | 0.00049 | 10.0 s |
| 48:49.2 | Data point 44 | 1.50000 mL | 0.16747 mL | 0.15106 mL | 0.30000 mL | 2.559 | -0.00372 | 0.68600 | 0.00022 | 10.5 s |
| 49:25.4 | Data point 45 | 1.50000 mL | 0.16747 mL | 0.15656 mL | 0.30000 mL | 2.736 | -0.00396 | 0.44516 | 0.00029 | 10.0 s |
| 50:00.9 | Data point 46 | 1.50000 mL | 0.16747 mL | 0.16058 mL | 0.30000 mL | 2.932 | -0.00663 | 0.33641 | 0.00056 | 10.0 s |
| 50:36.4 | Data point 47 | 1.50000 mL | 0.16747 mL | 0.16364 mL | 0.30000 mL | 3.137 | -0.01016 | 0.30029 | 0.00092 | 10.0 s |
| 51:11.9 | Data point 48 | 1.50000 mL | 0.16747 mL | 0.16609 mL | 0.30000 mL | 3.444 | -0.00956 | 0.44143 | 0.00071 | 10.5 s |
| 51:47.9 | Data point 49 | 1.50000 mL | 0.16747 mL | 0.16700 mL | 0.30000 mL | 3.647 | -0.00982 | 0.27260 | 0.00093 | 10.0 s |
| 52:23.4 | Data point 50 | 1.50000 mL | 0.16747 mL | 0.16773 mL | 0.30000 mL | 3.891 | -0.00230 | 0.16102 | 0.00028 | 10.0 s |
| 53:09.1 | Data point 51 | 1.50000 mL | 0.16747 mL | 0.16842 mL | 0.30000 mL | 4.133 | -0.00752 | 0.36746 | 0.00061 | 10.0 s |
| 53:44.5 | Data point 52 | 1.50000 mL | 0.16747 mL | 0.16877 mL | 0.30000 mL | 4.507 | -0.01597 | 0.71656 | 0.00093 | 10.0 s |
| 54:30.2 | Data point 53 | 1.50000 mL | 0.16747 mL | 0.16903 mL | 0.30000 mL | 4.840 | -0.00857 | 0.27004 | 0.00082 | 12.5 s |
| 55:13.3 | Data point 54 | 1.50000 mL | 0.16747 mL | 0.16921 mL | 0.30000 mL | 6.006 | -0.01875 | 0.96219 | 0.00094 | 37.5 s |
| 56:21.2 | Data point 55 | 1.50000 mL | 0.16747 mL | 0.16931 mL | 0.30000 mL | 6.652 | -0.05803 | 0.99479 | 0.00287 | Timed out at 59.5 s |
| 57:56.8 | Data point 56 | 1.50000 mL | 0.16747 mL | 0.16938 mL | 0.30000 mL | 7.209 | -0.06064 | 0.98448 | 0.00302 | Timed out at 59.5 s |
| 59:27.4 | Data point 57 | 1.50000 mL | 0.16747 mL | 0.16943 mL | 0.30000 mL | 7.445 | -0.06310 | 0.99629 | 0.00312 | Timed out at 59.5 s |
| 1:01:03.0 | Data point 58 | 1.50000 mL | 0.16747 mL | 0.16950 mL | 0.30000 mL | 8.044 | -0.04412 | 0.95409 | 0.00223 | Timed out at 59.5 s |
| 1:02:38.8 | Data point 59 | 1.50000 mL | 0.16747 mL | 0.16957 mL | 0.30000 mL | 8.352 | -0.01985 | 0.97791 | 0.00099 | 51.0 s |
| 1:04:10.8 | Data point 60 | 1.50000 mL | 0.16747 mL | 0.16969 mL | 0.30000 mL | 8.741 | -0.01671 | 0.79899 | 0.00092 | 24.0 s |
| 1:05:10.6 | Data point 61 | 1.50000 mL | 0.16747 mL | 0.16980 mL | 0.30000 mL | 8.999 | -0.01827 | 0.88409 | 0.00096 | 26.0 s |
| 1:05:56.8 | Data point 62 | 1.50000 mL | 0.16747 mL | 0.16980 mL | 0.30000 mL | 9.010 | -0.01788 | 0.90579 | 0.00093 | 19.0 s |
| 1:06:25.0 | Assay volumes | 1.50000 mL | 0.16747 mL | 0.16980 mL | 0.30000 mL | | | | | |

| | | | |
|---|---|---|---|
| Sample name: | **M11_octanol** | Experiment start time: | **2/27/2018 10:54:30 PM** |
| Assay name: | **pH-metric high logP** | Analyst: | **Pion** |
| Assay ID: | **18B-27016** | Instrument ID: | **T312060** |
| Filename: | **C:\Sirius_T3\Mehtap\20180227_exp27_logP_T3-2\18B-27016_M11_octanol_pH-metric high logP.t3r** | | |

| | | | |
|---|---|---|---|
| Sample name: | **M11_octanol** | Experiment start time: | **2/27/2018 10:54:30 PM** |
| Assay name: | **pH-metric high logP** | Analyst: | **Pion** |
| Assay ID: | **18B-27016** | Instrument ID: | **T312060** |
| Filename: | **C:\Sirius_T3\Mehtap\20180227_exp27_logP_T3-2\18B-27016_M11_octanol_pH-metric high logP.t3r** | | |

#### Assay Settings (continued)

| Setting | Value | Original Value | Date/Time changed | Imported from |
|---|---|---|---|---|
| ***Titration 3*** | | | | |
| Titrate from | Low to high pH | | | |
| Add additional water | 0.00 mL | | | |
| Additional partition solvent volume | 0.200 mL | | | |
| Additional partition solvent added | Automatic | | | |
| After pH adjust stir for | 30 seconds | | | |
| Stir to allow partitioning for | 15 seconds | | | |
| Stirrer speed for partitioning | 60% | | | |
| ***Data Point Stability*** | | | | |
| Stir during data point collection | No | | | |
| Delay before data point collection | 0 seconds | | | |
| Number of points to average | 20 points | | | |
| Time interval between points | 0.50 seconds | | | |
| Required maximum standard deviation | 0.00100 dpH/dt | | | |
| Stability timeout after | 60 seconds | | | |

#### Calibration Settings

| Setting | Value | Date/Time changed | Imported from |
|---|---|---|---|
| Four-Plus alpha | 0.130 | 2/27/2018 10:54:30 PM | C:\Sirius_T3\HCl18B27.t3r |
| Four-Plus S | 0.9970 | 2/27/2018 10:54:30 PM | C:\Sirius_T3\HCl18B27.t3r |
| Four-Plus jH | 0.8 | 2/27/2018 10:54:30 PM | C:\Sirius_T3\HCl18B27.t3r |
| Four-Plus jOH | -0.4 | 2/27/2018 10:54:30 PM | C:\Sirius_T3\HCl18B27.t3r |
| Base concentration factor | 1.000 | 2/27/2018 10:54:30 PM | C:\Sirius_T3\KOH18B27.t3r |
| Acid concentration factor | 0.994 | 2/27/2018 10:54:30 PM | C:\Sirius_T3\HCl18B27.t3r |

#### Instrument Settings

| Setting | Value | Batch Id | Install date |
|---|---|---|---|
| Instrument owner | Merck | | |
| Instrument ID | T312060 | | |
| Instrument type | T3 Simulator | | |
| Software version | 1.1.3.0 | | |
| Dispenser module | | T3DM1200361 | 3/31/2009 5:24:52 AM |
| Dispenser 0 | Water | | 3/31/2009 5:25:05 AM |
| Syringe volume | 2.5 mL | | |
| Firmware version | 1.2.1(r2) | | |
| Titrant | Water (0.15 M KCl) | 02-06-2018 | 2/27/2018 10:05:59 AM |
| Dispenser 2 | Acid | | 3/31/2009 5:25:11 AM |
| Syringe volume | 0.5 mL | | |
| Firmware version | 1.2.1(r2) | | |
| Titrant | Acid (0.5 M HCl) | 02-27-2018 | 2/27/2018 10:27:22 AM |
| Dispenser 1 | Base | | 3/31/2009 5:25:21 AM |
| Syringe volume | 0.5 mL | | |
| Firmware version | 1.2.1(r2) | | |
| Titrant | Base (0.5 M KOH) | 9/22/2017 | 2/27/2018 10:21:22 AM |
| Dispenser 5 | Cosolvent | | 3/31/2009 5:26:24 AM |
| Syringe volume | 2.5 mL | | |
| Firmware version | 1.2.1(r2) | | |
| Distribution valve 5 | Distribution Valve | | 3/31/2009 5:28:19 AM |
| Firmware version | 1.1.3 | | |
| Port A | Methanol (80%, 0.15 M KCl) | 09-26-17 | 2/7/2018 9:42:01 AM |
| Port B | Cyclohexane | 11-01-17 | 2/27/2018 10:37:57 AM |
| Dispenser 3 | Buffer | | 8/3/2010 5:05:16 AM |
| Syringe volume | 0.5 mL | | |
| Firmware version | 1.2.1(r2) | | |
| Titrant | Dodecane | 2018/01/31 | 1/31/2018 12:26:26 PM |
| Dispenser 6 | Octanol | | 10/22/2010 10:52:43 AM |

| | | | |
|---|---|---|---|
| Sample name: | **M11_octanol** | Experiment start time: | **2/27/2018 10:54:30 PM** |
| Assay name: | **pH-metric high logP** | Analyst: | **Pion** |
| Assay ID: | **18B-27016** | Instrument ID: | **T312060** |
| Filename: | **C:\Sirius_T3\Mehtap\20180227_exp27_logP_T3-2\18B-27016_M11_octanol_pH-metric high logP.t3r** | | |

| | | | |
|---|---|---|---|
| Sample name: | **M11_octanol** | Experiment start time: | **2/27/2018 10:54:30 PM** |
| Assay name: | **pH-metric high logP** | Analyst: | **Pion** |
| Assay ID: | **18B-27016** | Instrument ID: | **T312060** |
| Filename: | **C:\Sirius_T3\Mehtap\20180227_exp27_logP_T3-2\18B-27016_M11_octanol_pH-metric high logP.t3r** | | |

#### Instrument Settings (continued)

| Setting | Value | Batch Id | Install date |
|---|---|---|---|
| Spectrometer calibration stir duration | 5 s | | |
| Spectrometer calibration stir speed | 30% | | |
| Spectrometer calibration wash pump volume | 20.0 mL | | |
| Spectrometer calibration wash stir duration | 5 s | | |
| Spectrometer calibration wash stir speed | 30% | | |
| Overhead dispense height | 10000 | | |

#### Refinement Settings

| Setting | Value | Default value |
|---|---|---|
| Turbidity detection method | None | None |
| Turbidity wavelength to assess | 500.0 nm | 500.0 nm |
| Turbidity maximum absorbance | 0.100 | 0.100 |
| Turbidity probe threshold | 50.00 | 50.00 |

#### Experiment Log

[2:37] Air gap created for Water (0.15 M KCl)
[2:38] Air gap created for Acid (0.5 M HCl)
[2:38] Air gap created for Base (0.5 M KOH)
[2:39] Air gap released for Water (0.15 M KCl)
[2:42] Titrator arm moved over Titration position
[2:42] Titration 1 of 3
[2:42] Adding initial titrants
[2:42] Automatically add 1.50000 mL of water
[3:08] Dispensed 1.500000 mL of Water (0.15 M KCl)
[3:12] Titrator arm moved over Drain
[8:53] Titrator arm moved to Titration position
[8:53] Argon flow rate set to 100
[8:53] Stirrer speed set to 10
[8:58] Automatically add 0.02000 mL of Octanol
[8:59] Dispensed 0.019991 mL of Octanol
[9:00] Initial pH = 6.35
[9:00] Iterative adjust 6.35 -> 2.00
[9:00] pH 6.35 -> 2.00
[9:02] Air gap released for Acid (0.5 M HCl)
[9:02] Dispensed 0.051670 mL of Acid (0.5 M HCl)
[9:08] Holding pH 2.00
[11:08] Stirrer speed set to 0
[11:08] Stirrer speed set to 50
[11:08] Iterative adjust 1.97 -> 2.00
[11:08] pH 1.97 -> 2.00
[11:08] Air gap released for Base (0.5 M KOH)
[11:09] Dispensed 0.003010 mL of Base (0.5 M KOH)
[11:59] Stirrer speed set to 0
[12:10] Datapoint id 1 collected
[12:10] Stirrer speed set to 50
[12:15] pH 2.02 -> 2.22
[12:15] Using cautious pH adjust
[12:15] Dispensed 0.007573 mL of Base (0.5 M KOH)
[12:20] Stepping pH = 2.11
[12:20] Dispensed 0.005950 mL of Base (0.5 M KOH)
[12:25] Stepping pH = 2.20
[12:25] Dispensed 0.001411 mL of Base (0.5 M KOH)
[12:31] Stepping pH = 2.22
[12:46] Stirrer speed set to 0
[12:56] Datapoint id 2 collected
[12:56] Charge balance equation is out by 1.4%
[12:56] Stirrer speed set to 50

| | | | |
|---|---|---|---|
| Sample name: | **M11_octanol** | Experiment start time: | **2/27/2018 10:54:30 PM** |
| Assay name: | **pH-metric high logP** | Analyst: | **Pion** |
| Assay ID: | **18B-27016** | Instrument ID: | **T312060** |
| Filename: | **C:\Sirius_T3\Mehtap\20180227_exp27_logP_T3-2\18B-27016_M11_octanol_pH-metric high logP.t3r** | | |

#### Experiment Log (continued)

[13:01] pH 2.22 -> 2.42
[13:01] Using charge balance adjust
[13:02] Dispensed 0.009525 mL of Base (0.5 M KOH)
[13:22] Stirrer speed set to 0
[13:32] Datapoint id 3 collected
[13:32] Charge balance equation is out by -4.9%
[13:32] Stirrer speed set to 50
[13:37] pH 2.42 -> 2.62
[13:37] Using charge balance adjust
[13:37] Dispensed 0.006115 mL of Base (0.5 M KOH)
[13:57] Stirrer speed set to 0
[14:07] Datapoint id 4 collected
[14:07] Charge balance equation is out by -14.8%
[14:07] Stirrer speed set to 50
[14:12] pH 2.60 -> 2.80
[14:12] Using charge balance adjust
[14:13] Dispensed 0.004186 mL of Base (0.5 M KOH)
[14:33] Stirrer speed set to 0
[14:43] Datapoint id 5 collected
[14:43] Charge balance equation is out by 7.6%
[14:43] Stirrer speed set to 50
[14:48] pH 2.82 -> 3.02
[14:48] Using charge balance adjust
[14:49] Dispensed 0.002752 mL of Base (0.5 M KOH)
[15:09] Stirrer speed set to 0
[15:19] Datapoint id 6 collected
[15:19] Charge balance equation is out by -19.4%
[15:19] Stirrer speed set to 50
[15:24] pH 2.99 -> 3.19
[15:24] Using cautious pH adjust
[15:24] Dispensed 0.001058 mL of Base (0.5 M KOH)
[15:29] Stepping pH = 3.06
[15:29] Dispensed 0.001152 mL of Base (0.5 M KOH)
[15:34] Stepping pH = 3.16
[15:35] Dispensed 0.000306 mL of Base (0.5 M KOH)
[15:40] Stepping pH = 3.19
[15:55] Stirrer speed set to 0
[16:05] Datapoint id 7 collected
[16:05] Charge balance equation is out by -18.8%
[16:05] Stirrer speed set to 50
[16:10] pH 3.19 -> 3.39
[16:10] Using cautious pH adjust
[16:10] Dispensed 0.000870 mL of Base (0.5 M KOH)
[16:16] Stepping pH = 3.28
[16:16] Dispensed 0.000706 mL of Base (0.5 M KOH)
[16:21] Stepping pH = 3.36
[16:21] Dispensed 0.000235 mL of Base (0.5 M KOH)
[16:26] Stepping pH = 3.38
[16:41] Stirrer speed set to 0
[16:51] Datapoint id 8 collected
[16:51] Charge balance equation is out by -2.8%
[16:51] Stirrer speed set to 50
[16:56] pH 3.38 -> 3.58
[16:56] Using charge balance adjust
[16:56] Dispensed 0.001576 mL of Base (0.5 M KOH)
[17:17] Stirrer speed set to 0
[17:27] Datapoint id 9 collected
[17:27] Charge balance equation is out by -14.9%
[17:27] Stirrer speed set to 50

| | | | |
|---|---|---|---|
| Sample name: | **M11_octanol** | Experiment start time: | **2/27/2018 10:54:30 PM** |
| Assay name: | **pH-metric high logP** | Analyst: | **Pion** |
| Assay ID: | **18B-27016** | Instrument ID: | **T312060** |
| Filename: | **C:\Sirius_T3\Mehtap\20180227_exp27_logP_T3-2\18B-27016_M11_octanol_pH-metric high logP.t3r** | | |

#### Experiment Log (continued)

[17:32] pH 3.55 -> 3.75
[17:32] Using charge balance adjust
[17:32] Dispensed 0.001529 mL of Base (0.5 M KOH)
[17:53] Stirrer speed set to 0
[18:03] Datapoint id 10 collected
[18:03] Charge balance equation is out by -21.5%
[18:03] Stirrer speed set to 50
[18:08] pH 3.71 -> 3.91
[18:08] Using cautious pH adjust
[18:08] Dispensed 0.000729 mL of Base (0.5 M KOH)
[18:13] Stepping pH = 3.79
[18:13] Dispensed 0.000729 mL of Base (0.5 M KOH)
[18:18] Stepping pH = 3.89
[18:18] Dispensed 0.000118 mL of Base (0.5 M KOH)
[18:23] Stepping pH = 3.90
[18:23] Dispensed 0.000118 mL of Base (0.5 M KOH)
[18:28] Stepping pH = 3.91
[18:44] Stirrer speed set to 0
[18:54] Datapoint id 11 collected
[18:54] Charge balance equation is out by -16.0%
[18:54] Stirrer speed set to 50
[18:59] pH 3.91 -> 4.11
[18:59] Using cautious pH adjust
[18:59] Dispensed 0.000682 mL of Base (0.5 M KOH)
[19:04] Stepping pH = 4.01
[19:04] Dispensed 0.000517 mL of Base (0.5 M KOH)
[19:10] Stepping pH = 4.11
[19:25] Stirrer speed set to 0
[19:35] Datapoint id 12 collected
[19:35] Charge balance equation is out by 12.1%
[19:35] Stirrer speed set to 50
[19:40] pH 4.12 -> 4.32
[19:40] Using charge balance adjust
[19:40] Dispensed 0.001199 mL of Base (0.5 M KOH)
[20:00] Stirrer speed set to 0
[20:11] Datapoint id 13 collected
[20:11] Charge balance equation is out by 71.5%
[20:11] Stirrer speed set to 50
[20:16] pH 4.46 -> 4.66
[20:16] Using cautious pH adjust
[20:16] Dispensed 0.000400 mL of Base (0.5 M KOH)
[20:21] Stepping pH = 4.58
[20:21] Dispensed 0.000188 mL of Base (0.5 M KOH)
[20:26] Stepping pH = 4.68
[20:41] Stirrer speed set to 0
[20:51] Datapoint id 14 collected
[20:51] Charge balance equation is out by 26.5%
[20:51] Stirrer speed set to 50
[20:56] pH 4.73 -> 4.93
[20:56] Using cautious pH adjust
[20:56] Dispensed 0.000259 mL of Base (0.5 M KOH)
[21:02] Stepping pH = 4.86
[21:02] Dispensed 0.000118 mL of Base (0.5 M KOH)
[21:07] Stepping pH = 4.98
[21:22] Stirrer speed set to 0
[21:34] Datapoint id 15 collected
[21:34] Charge balance equation is out by 28.5%
[21:34] Stirrer speed set to 50
[21:39] pH 5.06 -> 5.26

Sample name: **M11_octanol**
Assay name: **pH-metric high logP**
Assay ID: **18B-27016**
Filename: **C:\Sirius_T3\Mehtap\20180227_exp27_logP_T3-2\18B-27016_M11_octanol_pH-metric high logP.t3r**
Experiment start time: **2/27/2018 10:54:30 PM**
Analyst: **Pion**
Instrument ID: **T312060**

#### Experiment Log (continued)

[21:39] Using cautious pH adjust
[21:40] Dispensed 0.000141 mL of Base (0.5 M KOH)
[21:45] Stepping pH = 5.13
[21:45] Dispensed 0.000141 mL of Base (0.5 M KOH)
[21:50] Stepping pH = 5.52
[22:05] Stirrer speed set to 0
[22:23] Datapoint id 16 collected
[22:23] Charge balance equation is out by -5.6%
[22:23] Stirrer speed set to 50
[22:29] pH 5.70 -> 5.90
[22:29] Using charge balance adjust
[22:29] Dispensed 0.000094 mL of Base (0.5 M KOH)
[22:49] Stirrer speed set to 0
[23:07] Datapoint id 17 collected
[23:07] Charge balance equation is out by -30.7%
[23:07] Stirrer speed set to 50
[23:12] pH 5.93 -> 6.13
[23:12] Using cautious pH adjust
[23:12] Dispensed 0.000047 mL of Base (0.5 M KOH)
[23:17] Stepping pH = 5.95
[23:17] Dispensed 0.000094 mL of Base (0.5 M KOH)
[23:22] Stepping pH = 7.00
[23:37] Stirrer speed set to 0
[24:37] Datapoint id 18 collected
[24:37] Charge balance equation is out by -81.0%
[24:37] Stirrer speed set to 50
[24:42] pH 8.28 -> 8.48
[24:42] Using cautious pH adjust
[24:42] Dispensed 0.000024 mL of Base (0.5 M KOH)
[24:48] Stepping pH = 8.30
[24:48] Dispensed 0.000024 mL of Base (0.5 M KOH)
[24:53] Stepping pH = 8.31
[24:53] Dispensed 0.000047 mL of Base (0.5 M KOH)
[24:58] Stepping pH = 8.37
[24:58] Dispensed 0.000047 mL of Base (0.5 M KOH)
[25:03] Stepping pH = 8.51
[25:18] Stirrer speed set to 0
[26:03] Datapoint id 19 collected
[26:03] Charge balance equation is out by -1,164.1%
[26:03] Stirrer speed set to 50
[26:08] pH 8.73 -> 8.93
[26:08] Using cautious pH adjust
[26:08] Dispensed 0.000024 mL of Base (0.5 M KOH)
[26:13] Stepping pH = 8.74
[26:13] Dispensed 0.000024 mL of Base (0.5 M KOH)
[26:18] Stepping pH = 8.75
[26:18] Dispensed 0.000094 mL of Base (0.5 M KOH)
[26:23] Stepping pH = 9.13
[26:38] Stirrer speed set to 0
[27:07] Datapoint id 20 collected
[27:07] Charge balance equation is out by -543.2%
[27:07] Titration 2 of 3
[27:07] Adding initial titrants
[27:07] Automatically add 0.08000 mL of Octanol
[27:09] Dispensed 0.080009 mL of Octanol
[27:09] Stirrer speed set to 10
[27:10] Stirrer speed set to 55
[27:10] Iterative adjust 9.27 -> 2.00
[27:10] pH 9.27 -> 2.00

Sample name: **M11_octanol** Experiment start time: **2/27/2018 10:54:30 PM**
Assay name: **pH-metric high logP** Analyst: **Pion**
Assay ID: **18B-27016** Instrument ID: **T312060**
Filename: **C:\Sirius_T3\Mehtap\20180227_exp27_logP_T3-2\18B-27016_M11_octanol_pH-metric high logP.t3r**

#### Experiment Log (continued)

[27:12] Dispensed 0.054798 mL of Acid (0.5 M HCl)
[27:17] pH 2.02 -> 2.00
[27:17] Dispensed 0.001952 mL of Acid (0.5 M HCl)
[28:07] Stirrer speed set to 0
[28:17] Datapoint id 21 collected
[28:17] Stirrer speed set to 55
[28:22] pH 1.97 -> 2.17
[28:22] Using cautious pH adjust
[28:23] Dispensed 0.009102 mL of Base (0.5 M KOH)
[28:28] Stepping pH = 2.07
[28:28] Dispensed 0.005833 mL of Base (0.5 M KOH)
[28:33] Stepping pH = 2.14
[28:33] Dispensed 0.002516 mL of Base (0.5 M KOH)
[28:39] Stepping pH = 2.17
[28:54] Stirrer speed set to 0
[29:04] Datapoint id 22 collected
[29:04] Charge balance equation is out by 4.1%
[29:04] Stirrer speed set to 55
[29:09] pH 2.17 -> 2.37
[29:09] Using charge balance adjust
[29:10] Dispensed 0.011571 mL of Base (0.5 M KOH)
[29:30] Stirrer speed set to 0
[29:40] Datapoint id 23 collected
[29:40] Charge balance equation is out by 4.1%
[29:40] Stirrer speed set to 55
[29:45] pH 2.39 -> 2.59
[29:45] Using charge balance adjust
[29:46] Dispensed 0.007173 mL of Base (0.5 M KOH)
[30:06] Stirrer speed set to 0
[30:16] Datapoint id 24 collected
[30:16] Charge balance equation is out by -13.1%
[30:16] Stirrer speed set to 55
[30:21] pH 2.57 -> 2.77
[30:21] Using charge balance adjust
[30:21] Dispensed 0.004915 mL of Base (0.5 M KOH)
[30:42] Stirrer speed set to 0
[30:52] Datapoint id 25 collected
[30:52] Charge balance equation is out by -15.1%
[30:52] Stirrer speed set to 55
[30:57] pH 2.75 -> 2.95
[30:57] Using cautious pH adjust
[30:57] Dispensed 0.001764 mL of Base (0.5 M KOH)
[31:02] Stepping pH = 2.81
[31:02] Dispensed 0.002093 mL of Base (0.5 M KOH)
[31:07] Stepping pH = 2.92
[31:07] Dispensed 0.000470 mL of Base (0.5 M KOH)
[31:12] Stepping pH = 2.95
[31:27] Stirrer speed set to 0
[31:37] Datapoint id 26 collected
[31:37] Charge balance equation is out by -22.6%
[31:37] Stirrer speed set to 55
[31:43] pH 2.95 -> 3.15
[31:43] Using cautious pH adjust
[31:43] Dispensed 0.001294 mL of Base (0.5 M KOH)
[31:48] Stepping pH = 3.06
[31:48] Dispensed 0.000753 mL of Base (0.5 M KOH)
[31:53] Stepping pH = 3.12
[31:53] Dispensed 0.000306 mL of Base (0.5 M KOH)
[31:58] Stepping pH = 3.15

Sample name: **M11_octanol** Experiment start time: **2/27/2018 10:54:30 PM**
Assay name: **pH-metric high logP** Analyst: **Pion**
Assay ID: **18B-27016** Instrument ID: **T312060**
Filename: **C:\Sirius_T3\Mehtap\20180227_exp27_logP_T3-2\18B-27016_M11_octanol_pH-metric high logP.t3r**

#### Experiment Log (continued)

[32:13] Stirrer speed set to 0
[32:23] Datapoint id 27 collected
[32:23] Charge balance equation is out by 9.1%
[32:23] Stirrer speed set to 55
[32:28] pH 3.15 -> 3.35
[32:28] Using charge balance adjust
[32:29] Dispensed 0.002093 mL of Base (0.5 M KOH)
[32:49] Stirrer speed set to 0
[32:59] Datapoint id 28 collected
[32:59] Charge balance equation is out by -11.4%
[32:59] Stirrer speed set to 55
[33:04] pH 3.34 -> 3.54
[33:04] Using charge balance adjust
[33:04] Dispensed 0.001858 mL of Base (0.5 M KOH)
[33:24] Stirrer speed set to 0
[33:34] Datapoint id 29 collected
[33:34] Charge balance equation is out by 3.5%
[33:34] Stirrer speed set to 55
[33:39] pH 3.55 -> 3.75
[33:39] Using charge balance adjust
[33:39] Dispensed 0.001670 mL of Base (0.5 M KOH)
[34:00] Stirrer speed set to 0
[34:10] Datapoint id 30 collected
[34:10] Charge balance equation is out by 67.8%
[34:10] Stirrer speed set to 55
[34:15] pH 3.89 -> 4.09
[34:15] Using cautious pH adjust
[34:15] Dispensed 0.000659 mL of Base (0.5 M KOH)
[34:20] Stepping pH = 4.11
[34:35] Stirrer speed set to 0
[34:46] Datapoint id 31 collected
[34:46] Charge balance equation is out by 50.0%
[34:46] Stirrer speed set to 55
[34:51] pH 4.14 -> 4.34
[34:51] Using cautious pH adjust
[34:51] Dispensed 0.000494 mL of Base (0.5 M KOH)
[34:56] Stepping pH = 4.42
[35:11] Stirrer speed set to 0
[35:22] Datapoint id 32 collected
[35:22] Charge balance equation is out by 50.0%
[35:22] Stirrer speed set to 55
[35:27] pH 4.45 -> 4.65
[35:27] Using cautious pH adjust
[35:27] Dispensed 0.000306 mL of Base (0.5 M KOH)
[35:32] Stepping pH = 4.69
[35:47] Stirrer speed set to 0
[35:59] Datapoint id 33 collected
[35:59] Charge balance equation is out by 50.0%
[35:59] Stirrer speed set to 55
[36:04] pH 4.85 -> 5.05
[36:04] Using cautious pH adjust
[36:04] Dispensed 0.000141 mL of Base (0.5 M KOH)
[36:09] Stepping pH = 4.91
[36:09] Dispensed 0.000165 mL of Base (0.5 M KOH)
[36:14] Stepping pH = 5.80
[36:29] Stirrer speed set to 0
[37:21] Datapoint id 34 collected
[37:21] Charge balance equation is out by -5.2%
[37:21] Stirrer speed set to 55

| | | | |
|---|---|---|---|
| Sample name: | **M11_octanol** | Experiment start time: | **2/27/2018 10:54:30 PM** |
| Assay name: | **pH-metric high logP** | Analyst: | **Pion** |
| Assay ID: | **18B-27016** | Instrument ID: | **T312060** |
| Filename: | **C:\Sirius_T3\Mehtap\20180227_exp27_logP_T3-2\18B-27016_M11_octanol_pH-metric high logP.t3r** | | |

#### Experiment Log (continued)

[37:27] pH 6.56 -> 6.76
[37:27] Using charge balance adjust
[37:27] Dispensed 0.000047 mL of Base (0.5 M KOH)
[37:47] Stirrer speed set to 0
[38:45] Datapoint id 35 collected
[38:45] Charge balance equation is out by -43.3%
[38:45] Stirrer speed set to 55
[38:51] pH 6.84 -> 7.04
[38:51] Using cautious pH adjust
[38:51] Dispensed 0.000024 mL of Base (0.5 M KOH)
[38:56] Stepping pH = 6.89
[38:56] Dispensed 0.000024 mL of Base (0.5 M KOH)
[39:01] Stepping pH = 6.95
[39:01] Dispensed 0.000024 mL of Base (0.5 M KOH)
[39:06] Stepping pH = 7.16
[39:21] Stirrer speed set to 0
[40:21] Datapoint id 36 collected
[40:21] Charge balance equation is out by -119.4%
[40:21] Stirrer speed set to 55
[40:26] pH 7.34 -> 7.54
[40:26] Using cautious pH adjust
[40:26] Dispensed 0.000024 mL of Base (0.5 M KOH)
[40:31] Stepping pH = 7.43
[40:31] Dispensed 0.000024 mL of Base (0.5 M KOH)
[40:36] Stepping pH = 7.48
[40:36] Dispensed 0.000024 mL of Base (0.5 M KOH)
[40:42] Stepping pH = 7.57
[40:57] Stirrer speed set to 0
[41:57] Datapoint id 37 collected
[41:57] Charge balance equation is out by -407.6%
[41:57] Stirrer speed set to 55
[42:02] pH 7.81 -> 8.01
[42:02] Using cautious pH adjust
[42:02] Dispensed 0.000024 mL of Base (0.5 M KOH)
[42:07] Stepping pH = 7.83
[42:07] Dispensed 0.000024 mL of Base (0.5 M KOH)
[42:12] Stepping pH = 7.87
[42:12] Dispensed 0.000024 mL of Base (0.5 M KOH)
[42:17] Stepping pH = 7.91
[42:17] Dispensed 0.000024 mL of Base (0.5 M KOH)
[42:22] Stepping pH = 8.04
[42:37] Stirrer speed set to 0
[43:37] Datapoint id 38 collected
[43:37] Charge balance equation is out by -1,031.4%
[43:37] Stirrer speed set to 55
[43:42] pH 8.38 -> 8.58
[43:42] Using cautious pH adjust
[43:42] Dispensed 0.000024 mL of Base (0.5 M KOH)
[43:48] Stepping pH = 8.40
[43:48] Dispensed 0.000024 mL of Base (0.5 M KOH)
[43:53] Stepping pH = 8.43
[43:53] Dispensed 0.000047 mL of Base (0.5 M KOH)
[43:58] Stepping pH = 8.58
[44:13] Stirrer speed set to 0
[44:58] Datapoint id 39 collected
[44:58] Charge balance equation is out by -519.7%
[44:58] Stirrer speed set to 55
[45:03] pH 8.70 -> 8.90
[45:03] Using cautious pH adjust

| | | | |
|---|---|---|---|
| Sample name: | **M11_octanol** | Experiment start time: | **2/27/2018 10:54:30 PM** |
| Assay name: | **pH-metric high logP** | Analyst: | **Pion** |
| Assay ID: | **18B-27016** | Instrument ID: | **T312060** |
| Filename: | **C:\Sirius_T3\Mehtap\20180227_exp27_logP_T3-2\18B-27016_M11_octanol_pH-metric high logP.t3r** | | |

#### Experiment Log (continued)

[45:03] Dispensed 0.000024 mL of Base (0.5 M KOH)
[45:08] Stepping pH = 8.70
[45:08] Dispensed 0.000024 mL of Base (0.5 M KOH)
[45:13] Stepping pH = 8.71
[45:13] Dispensed 0.000094 mL of Base (0.5 M KOH)
[45:18] Stepping pH = 9.07
[45:33] Stirrer speed set to 0
[45:48] Datapoint id 40 collected
[45:48] Charge balance equation is out by -554.5%
[45:48] Titration 3 of 3
[45:48] Adding initial titrants
[45:48] Automatically add 0.20000 mL of Octanol
[45:53] Dispensed 0.200000 mL of Octanol
[45:53] Stirrer speed set to 10
[45:54] Stirrer speed set to 60
[45:54] Iterative adjust 9.28 -> 2.00
[45:54] pH 9.28 -> 2.00
[45:55] Dispensed 0.057761 mL of Acid (0.5 M HCl)
[46:00] pH 2.01 -> 2.00
[46:01] Dispensed 0.001294 mL of Acid (0.5 M HCl)
[46:51] Stirrer speed set to 0
[47:01] Datapoint id 41 collected
[47:01] Stirrer speed set to 60
[47:06] pH 1.97 -> 2.17
[47:06] Using cautious pH adjust
[47:06] Dispensed 0.009901 mL of Base (0.5 M KOH)
[47:11] Stepping pH = 2.05
[47:12] Dispensed 0.007926 mL of Base (0.5 M KOH)
[47:17] Stepping pH = 2.13
[47:17] Dispensed 0.003057 mL of Base (0.5 M KOH)
[47:22] Stepping pH = 2.17
[47:37] Stirrer speed set to 0
[47:48] Datapoint id 42 collected
[47:48] Charge balance equation is out by -5.4%
[47:48] Stirrer speed set to 60
[47:53] pH 2.17 -> 2.37
[47:53] Using charge balance adjust
[47:53] Dispensed 0.012488 mL of Base (0.5 M KOH)
[48:14] Stirrer speed set to 0
[48:24] Datapoint id 43 collected
[48:24] Charge balance equation is out by -4.2%
[48:24] Stirrer speed set to 60
[48:29] pH 2.37 -> 2.57
[48:29] Using charge balance adjust
[48:29] Dispensed 0.008184 mL of Base (0.5 M KOH)
[48:49] Stirrer speed set to 0
[49:00] Datapoint id 44 collected
[49:00] Charge balance equation is out by -4.0%
[49:00] Stirrer speed set to 60
[49:05] pH 2.57 -> 2.77
[49:05] Using charge balance adjust
[49:05] Dispensed 0.005503 mL of Base (0.5 M KOH)
[49:25] Stirrer speed set to 0
[49:35] Datapoint id 45 collected
[49:35] Charge balance equation is out by -14.9%
[49:35] Stirrer speed set to 60
[49:40] pH 2.74 -> 2.94
[49:40] Using charge balance adjust
[49:41] Dispensed 0.004022 mL of Base (0.5 M KOH)

Sample name: **M11_octanol**
Assay name: **pH-metric high logP**
Assay ID: **18B-27016**
Filename: **C:\Sirius_T3\Mehtap\20180227_exp27_logP_T3-2\18B-27016_M11_octanol_pH-metric high logP.t3r**

Experiment start time: **2/27/2018 10:54:30 PM**
Analyst: **Pion**
Instrument ID: **T312060**

#### Experiment Log (continued)

[50:01] Stirrer speed set to 0
[50:11] Datapoint id 46 collected
[50:11] Charge balance equation is out by -5.7%
[50:11] Stirrer speed set to 60
[50:16] pH 2.94 -> 3.14
[50:16] Using charge balance adjust
[50:16] Dispensed 0.003057 mL of Base (0.5 M KOH)
[50:36] Stirrer speed set to 0
[50:46] Datapoint id 47 collected
[50:46] Charge balance equation is out by 0.9%
[50:46] Stirrer speed set to 60
[50:52] pH 3.14 -> 3.34
[50:52] Using charge balance adjust
[50:52] Dispensed 0.002446 mL of Base (0.5 M KOH)
[51:12] Stirrer speed set to 0
[51:22] Datapoint id 48 collected
[51:22] Charge balance equation is out by 52.1%
[51:22] Stirrer speed set to 60
[51:27] pH 3.45 -> 3.65
[51:27] Using cautious pH adjust
[51:28] Dispensed 0.000917 mL of Base (0.5 M KOH)
[51:33] Stepping pH = 3.65
[51:48] Stirrer speed set to 0
[51:58] Datapoint id 49 collected
[51:58] Charge balance equation is out by 50.0%
[51:58] Stirrer speed set to 60
[52:03] pH 3.65 -> 3.85
[52:03] Using cautious pH adjust
[52:03] Dispensed 0.000729 mL of Base (0.5 M KOH)
[52:08] Stepping pH = 3.89
[52:23] Stirrer speed set to 0
[52:33] Datapoint id 50 collected
[52:33] Charge balance equation is out by 50.0%
[52:33] Stirrer speed set to 60
[52:38] pH 3.90 -> 4.10
[52:38] Using cautious pH adjust
[52:39] Dispensed 0.000517 mL of Base (0.5 M KOH)
[52:44] Stepping pH = 4.06
[52:44] Dispensed 0.000094 mL of Base (0.5 M KOH)
[52:49] Stepping pH = 4.08
[52:49] Dispensed 0.000071 mL of Base (0.5 M KOH)
[52:54] Stepping pH = 4.12
[53:09] Stirrer speed set to 0
[53:19] Datapoint id 51 collected
[53:19] Charge balance equation is out by 33.8%
[53:19] Stirrer speed set to 60
[53:24] pH 4.14 -> 4.34
[53:24] Using cautious pH adjust
[53:24] Dispensed 0.000353 mL of Base (0.5 M KOH)
[53:29] Stepping pH = 4.47
[53:44] Stirrer speed set to 0
[53:54] Datapoint id 52 collected
[53:54] Charge balance equation is out by 50.0%
[53:54] Stirrer speed set to 60
[54:00] pH 4.52 -> 4.72
[54:00] Using cautious pH adjust
[54:00] Dispensed 0.000165 mL of Base (0.5 M KOH)
[54:05] Stepping pH = 4.67
[54:05] Dispensed 0.000047 mL of Base (0.5 M KOH)

| | | | |
|---|---|---|---|
| Sample name: | **M11_octanol** | Experiment start time: | **2/27/2018 10:54:30 PM** |
| Assay name: | **pH-metric high logP** | Analyst: | **Pion** |
| Assay ID: | **18B-27016** | Instrument ID: | **T312060** |
| Filename: | **C:\Sirius_T3\Mehtap\20180227_exp27_logP_T3-2\18B-27016_M11_octanol_pH-metric high logP.t3r** | | |

#### Experiment Log (continued)

[54:10] Stepping pH = 4.69
[54:10] Dispensed 0.000047 mL of Base (0.5 M KOH)
[54:15] Stepping pH = 4.78
[54:30] Stirrer speed set to 0
[54:43] Datapoint id 53 collected
[54:43] Charge balance equation is out by 22.1%
[54:43] Stirrer speed set to 60
[54:48] pH 4.89 -> 5.09
[54:48] Using cautious pH adjust
[54:48] Dispensed 0.000071 mL of Base (0.5 M KOH)
[54:53] Stepping pH = 4.94
[54:53] Dispensed 0.000118 mL of Base (0.5 M KOH)
[54:58] Stepping pH = 6.12
[55:13] Stirrer speed set to 0
[55:51] Datapoint id 54 collected
[55:51] Charge balance equation is out by -21.1%
[55:51] Stirrer speed set to 60
[55:56] pH 6.13 -> 6.33
[55:56] Using cautious pH adjust
[55:56] Dispensed 0.000024 mL of Base (0.5 M KOH)
[56:01] Stepping pH = 6.16
[56:01] Dispensed 0.000071 mL of Base (0.5 M KOH)
[56:06] Stepping pH = 6.51
[56:21] Stirrer speed set to 0
[57:21] Datapoint id 55 collected
[57:21] Charge balance equation is out by -82.3%
[57:21] Stirrer speed set to 60
[57:26] pH 6.85 -> 7.05
[57:26] Using cautious pH adjust
[57:26] Dispensed 0.000024 mL of Base (0.5 M KOH)
[57:31] Stepping pH = 6.94
[57:31] Dispensed 0.000024 mL of Base (0.5 M KOH)
[57:37] Stepping pH = 7.01
[57:37] Dispensed 0.000024 mL of Base (0.5 M KOH)
[57:42] Stepping pH = 7.10
[57:57] Stirrer speed set to 0
[58:57] Datapoint id 56 collected
[58:57] Charge balance equation is out by -92.6%
[58:57] Stirrer speed set to 60
[59:02] pH 7.25 -> 7.45
[59:02] Using cautious pH adjust
[59:02] Dispensed 0.000024 mL of Base (0.5 M KOH)
[59:07] Stepping pH = 7.33
[59:07] Dispensed 0.000024 mL of Base (0.5 M KOH)
[59:12] Stepping pH = 7.49
[59:27] Stirrer speed set to 0
[1:00:27] Datapoint id 57 collected
[1:00:27] Charge balance equation is out by -136.9%
[1:00:27] Stirrer speed set to 60
[1:00:32] pH 7.57 -> 7.77
[1:00:32] Using cautious pH adjust
[1:00:32] Dispensed 0.000024 mL of Base (0.5 M KOH)
[1:00:38] Stepping pH = 7.65
[1:00:38] Dispensed 0.000024 mL of Base (0.5 M KOH)
[1:00:43] Stepping pH = 7.74
[1:00:43] Dispensed 0.000024 mL of Base (0.5 M KOH)
[1:00:48] Stepping pH = 7.91
[1:01:03] Stirrer speed set to 0
[1:02:03] Datapoint id 58 collected

| | | | |
|---|---|---|---|
| Sample name: | **M11_octanol** | Experiment start time: | **2/27/2018 10:54:30 PM** |
| Assay name: | **pH-metric high logP** | Analyst: | **Pion** |
| Assay ID: | **18B-27016** | Instrument ID: | **T312060** |
| Filename: | **C:\Sirius_T3\Mehtap\20180227_exp27_logP_T3-2\18B-27016_M11_octanol_pH-metric high logP.t3r** | | |

#### Experiment Log (continued)

[1:02:03] Charge balance equation is out by -478.6%
[1:02:03] Stirrer speed set to 60
[1:02:08] pH 8.11 -> 8.31
[1:02:08] Using cautious pH adjust
[1:02:08] Dispensed 0.000024 mL of Base (0.5 M KOH)
[1:02:13] Stepping pH = 8.12
[1:02:13] Dispensed 0.000024 mL of Base (0.5 M KOH)
[1:02:18] Stepping pH = 8.20
[1:02:19] Dispensed 0.000024 mL of Base (0.5 M KOH)
[1:02:24] Stepping pH = 8.30
[1:02:39] Stirrer speed set to 0
[1:03:30] Datapoint id 59 collected
[1:03:30] Charge balance equation is out by -552.5%
[1:03:30] Stirrer speed set to 60
[1:03:35] pH 8.41 -> 8.61
[1:03:35] Using cautious pH adjust
[1:03:35] Dispensed 0.000024 mL of Base (0.5 M KOH)
[1:03:40] Stepping pH = 8.43
[1:03:40] Dispensed 0.000024 mL of Base (0.5 M KOH)
[1:03:45] Stepping pH = 8.46
[1:03:45] Dispensed 0.000047 mL of Base (0.5 M KOH)
[1:03:50] Stepping pH = 8.58
[1:03:51] Dispensed 0.000024 mL of Base (0.5 M KOH)
[1:03:56] Stepping pH = 8.71
[1:04:11] Stirrer speed set to 0
[1:04:35] Datapoint id 60 collected
[1:04:35] Charge balance equation is out by -623.7%
[1:04:35] Stirrer speed set to 60
[1:04:40] pH 8.77 -> 8.97
[1:04:40] Using cautious pH adjust
[1:04:40] Dispensed 0.000024 mL of Base (0.5 M KOH)
[1:04:45] Stepping pH = 8.78
[1:04:45] Dispensed 0.000047 mL of Base (0.5 M KOH)
[1:04:50] Stepping pH = 8.84
[1:04:50] Dispensed 0.000047 mL of Base (0.5 M KOH)
[1:04:55] Stepping pH = 8.97
[1:05:11] Stirrer speed set to 0
[1:05:37] Datapoint id 61 collected
[1:05:37] Charge balance equation is out by -302.6%
[1:05:37] Stirrer speed set to 60
[1:05:42] pH 9.03 -> 9.05
[1:05:42] Using cautious pH adjust
[1:05:57] Stirrer speed set to 0
[1:06:16] Datapoint id 62 collected
[1:06:16] Charge balance equation is out by 100.0%
[1:06:16] Argon flow rate set to 0
[1:06:20] Titrator arm moved over Titration position
