## Supplementary material for "Octanol-water partition coefficient measurements for the SAMPL6 Blind Prediction Challenge": SM11_18C-01001_M11_octanol_pH-metric high logP_report.pdf

| | | | |
|---|---|---|---|
| Sample name: | **M11_octanol** | Experiment start time: | **3/1/2018 12:11:22 AM** |
| Assay name: | **pH-metric high logP** | Analyst: | **Pion** |
| Assay ID: | **18C-01001** | Instrument ID: | **T312060** |
| Filename: | **C:\Sirius_T3\Mehtap\20180228_exp28_logP_T3-2\18C-01001_M11_octanol_pH-metric high logP.t3r** | | |

#### pH-metric Result

| | | |
|---|---|---|
| logP (XH +) | 0.55 | ±0.07 (n=50) |
| logP (neutral X) | 2.19 | ±0.02 (n=50) |

#### 18C-01001 Points 1 to 23

| | |
|---|---|
| M11_octanol concentration factor | 1.023 |
| Carbonate | 0.0000 mM |
| Acidity error | 0.03008 mM |

#### 18C-01001 Points 24 to 48

| | |
|---|---|
| M11_octanol concentration factor | 0.912 |
| Carbonate | 0.0957 mM |
| Acidity error | -0.30629 mM |

#### 18C-01001 Points 49 to 69

| | |
|---|---|
| M11_octanol concentration factor | 0.792 |
| Carbonate | 0.0733 mM |
| Acidity error | -0.81928 mM |

#### Warnings and errors

| | |
|---|---|
| Errors | None |
| Warnings | None |

#### Sample logD and percent species

| pH | M11_octanol logD | M11_octanol M11_octanolH | M11_octanol M11_octanol | M11_octanol M11_octanolH* | M11_octanol M11_octanol* | Comment |
|---|---|---|---|---|---|---|
| 1.000 | 0.57 | 21.08 % | 0.03 % | 74.70 % | 4.20 % | |
| 1.200 | 0.59 | 20.57 % | 0.04 % | 72.90 % | 6.49 % | Stomach pH |
| 2.000 | 0.74 | 15.27 % | 0.20 % | 54.12 % | 30.41 % | |
| 3.000 | 1.32 | 4.07 % | 0.52 % | 14.42 % | 80.99 % | |
| 4.000 | 1.95 | 0.49 % | 0.63 % | 1.73 % | 97.15 % | |
| 5.000 | 2.16 | 0.05 % | 0.64 % | 0.18 % | 99.13 % | |
| 6.000 | 2.19 | 0.00 % | 0.64 % | 0.02 % | 99.33 % | |
| 6.500 | 2.19 | 0.00 % | 0.64 % | 0.01 % | 99.35 % | |
| 7.000 | 2.19 | 0.00 % | 0.64 % | 0.00 % | 99.35 % | |
| 7.400 | 2.19 | 0.00 % | 0.64 % | 0.00 % | 99.36 % | Blood pH |
| 8.000 | 2.19 | 0.00 % | 0.64 % | 0.00 % | 99.36 % | |
| 9.000 | 2.19 | 0.00 % | 0.64 % | 0.00 % | 99.36 % | |
| 10.000 | 2.19 | 0.00 % | 0.64 % | 0.00 % | 99.36 % | |
| 11.000 | 2.19 | 0.00 % | 0.64 % | 0.00 % | 99.36 % | |
| 12.000 | 2.19 | 0.00 % | 0.64 % | 0.00 % | 99.36 % | |

| | | | |
|---|---|---|---|
| Sample name: | **M11_octanol** | Experiment start time: | **3/1/2018 12:11:22 AM** |
| Assay name: | **pH-metric high logP** | Analyst: | **Pion** |
| Assay ID: | **18C-01001** | Instrument ID: | **T312060** |
| Filename: | **C:\Sirius_T3\Mehtap\20180228_exp28_logP_T3-2\18C-01001_M11_octanol_pH-metric high logP.t3r** | | |

#### Graphs

Charge balance residuals

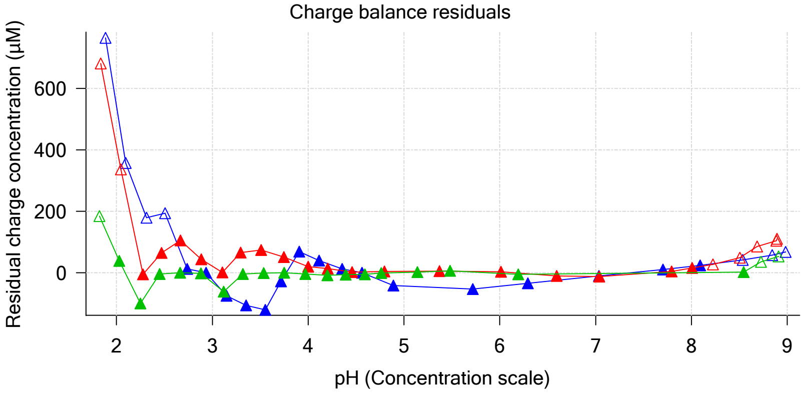


Ionic strength

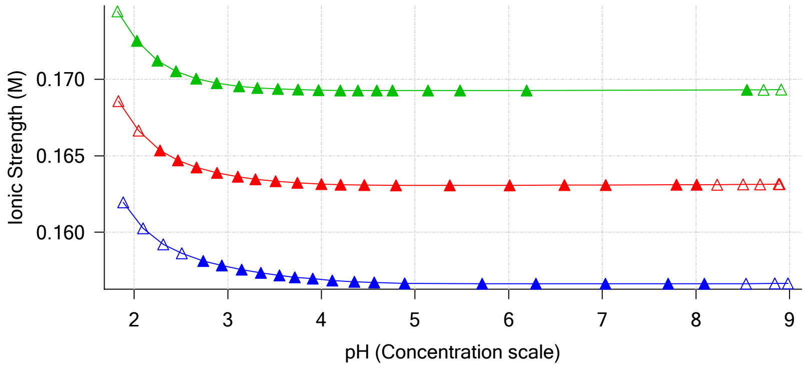


Buffer index profile

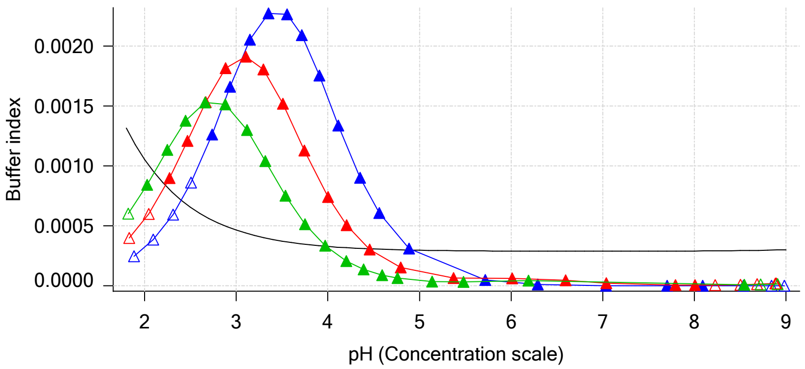


Sample M11_octanol Ionisation Graph

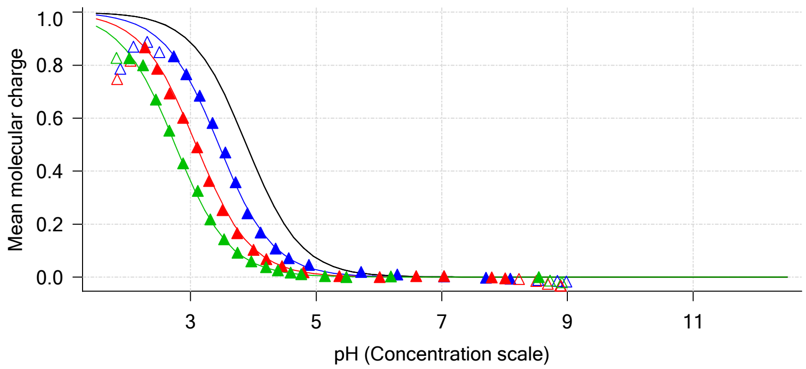


Distribution of species for sample M11_octanol

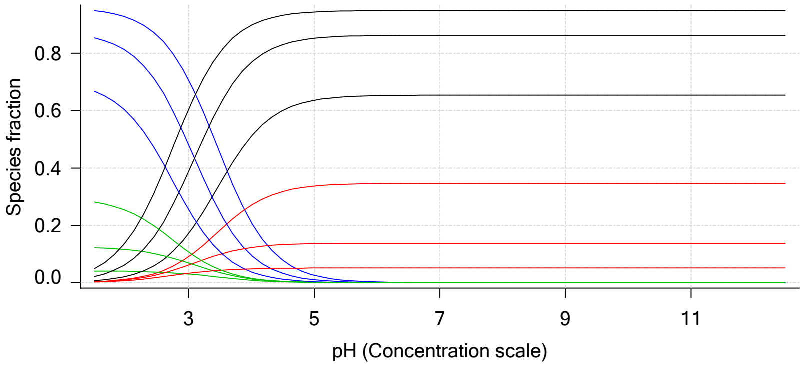


Lipophilicity Profile

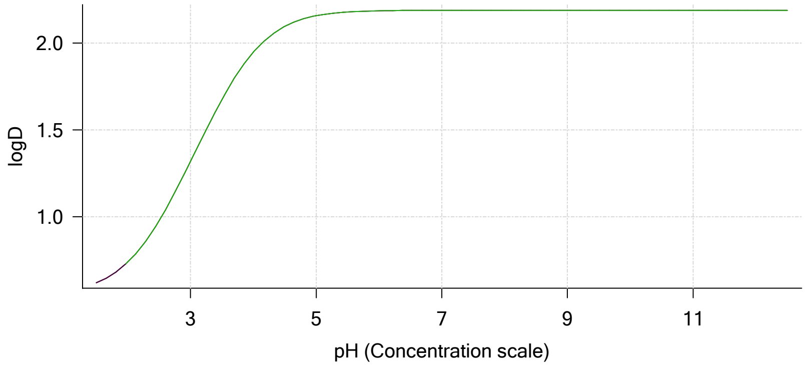


Log R Profile

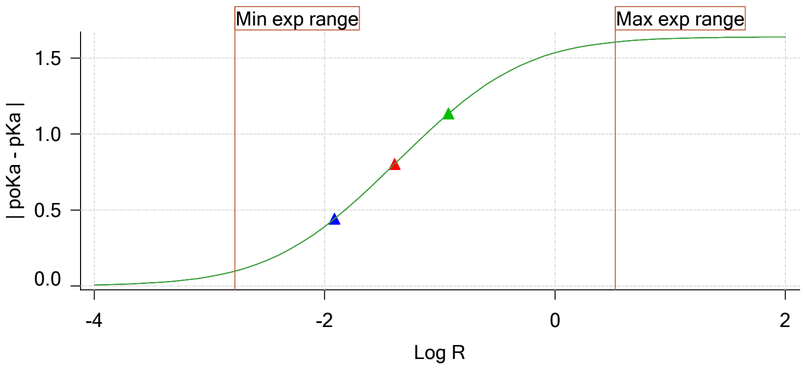


Sample Carbonate Ionisation Graph

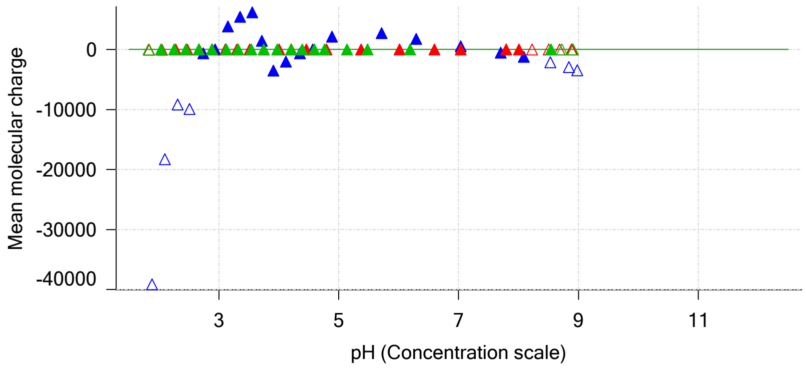

| | | | |
|---|---|---|---|
| Sample name: | **M11_octanol** | Experiment start time: | **3/1/2018 12:11:22 AM** |
| Assay name: | **pH-metric high logP** | Analyst: | **Pion** |
| Assay ID: | **18C-01001** | Instrument ID: | **T312060** |
| Filename: | **C:\Sirius_T3\Mehtap\20180228_exp28_logP_T3-2\18C-01001_M11_octanol_pH-metric high logP.t3r** | | |

#### Graphs (continued)

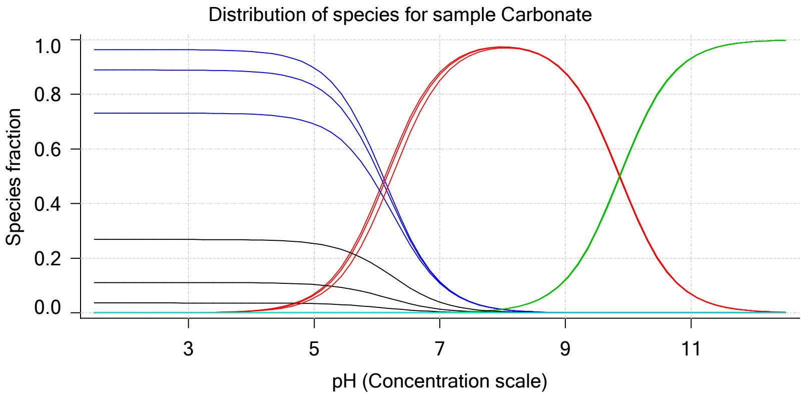

| | | | |
|---|---|---|---|
| Sample name: | **M11_octanol** | Experiment start time: | **3/1/2018 12:11:22 AM** |
| Assay name: | **pH-metric high logP** | Analyst: | **Pion** |
| Assay ID: | **18C-01001** | Instrument ID: | **T312060** |
| Filename: | **C:\Sirius_T3\Mehtap\20180228_exp28_logP_T3-2\18C-01001_M11_octanol_pH-metric high logP.t3r** | | |

# pH-metric high logP   Titration 1 of 3   18C-01001 Points 1 to 23

#### Overall results

| | |
|---|---|
| RMSD | 1.827 |
| Average ionic strength | 0.157 M |
| Average temperature | 25.0°C |
| Partition ratio | 0.0122 : 1 |
| Analyte concentration range | 3838.3 μM to 3962.6 μM |
| Total points considered | 16 of 23 |

#### Warnings and errors

| | |
|---|---|
| Errors | None |
| Warnings | One or more logP values out of range |

#### Four-Plus parameters

| | | | |
|---|---|---|---|
| Alpha | 0.130 | 3/1/2018 12:11:22 AM | C:\Sirius_T3\HCl18B27.t3r |
| S | 0.9970 | 3/1/2018 12:11:22 AM | C:\Sirius_T3\HCl18B27.t3r |
| jH | 0.8 | 3/1/2018 12:11:22 AM | C:\Sirius_T3\HCl18B27.t3r |
| jOH | -0.4 | 3/1/2018 12:11:22 AM | C:\Sirius_T3\HCl18B27.t3r |

#### Titrants

| | | | |
|---|---|---|---|
| 0.50 M HCl | 0.993513 | 3/1/2018 12:11:22 AM | C:\Sirius_T3\HCl18B27.t3r |
| 0.50 M KOH | 0.999845 | 3/1/2018 12:11:22 AM | C:\Sirius_T3\KOH18B27.t3r |

#### Sample

| | |
|---|---|
| M11_octanol concentration factor | 1.023 |
| Base pKa 1 | 3.89 |
| logP (XH +) | -8.26 |
| logP (neutral X) | 2.13 |

#### Sample graphs

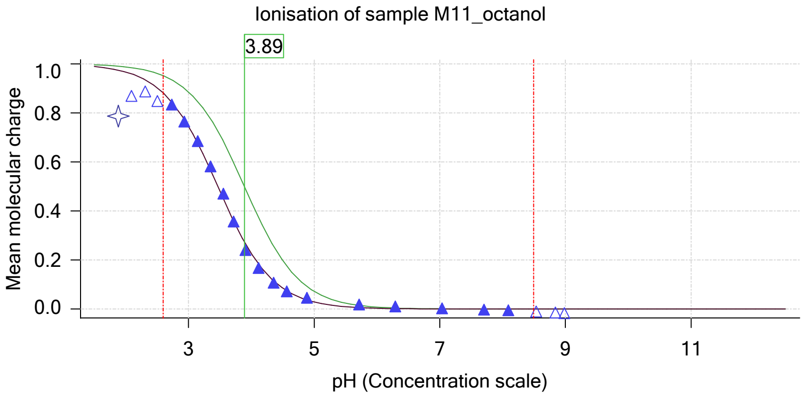


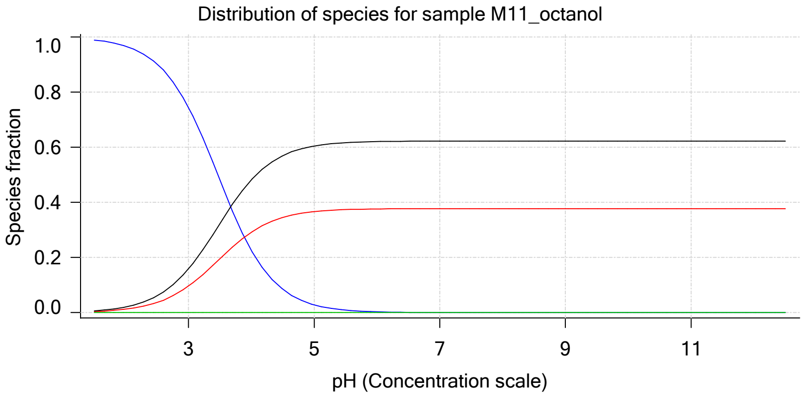

Sample name: **M11_octanol**
Assay name: **pH-metric high logP**
Assay ID: **18C-01001**
Filename: **C:\Sirius_T3\Mehtap\20180228_exp28_logP_T3-2\18C-01001_M11_octanol_pH-metric high logP.t3r**
Experiment start time: **3/1/2018 12:11:22 AM**
Analyst: **Pion**
Instrument ID: **T312060**

#### Sample graphs (continued)

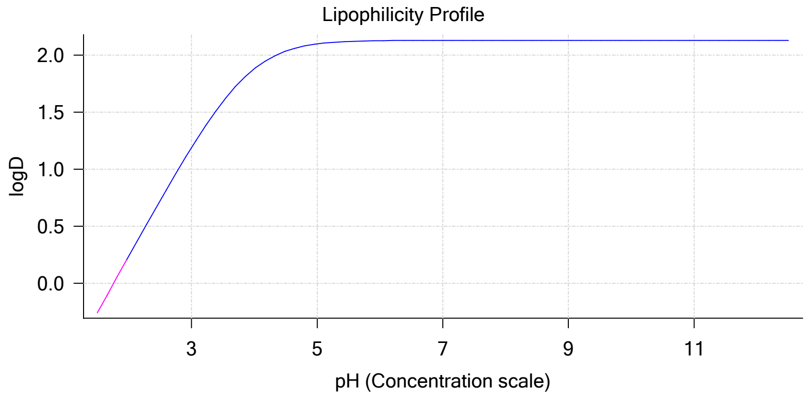


#### Sample logD and percent species

| pH | M11_octanol logD | M11_octanol M11_octanolH | M11_octanol M11_octanol | M11_octanol M11_octanolH* | M11_octanol M11_octanol* | Comment |
|---|---|---|---|---|---|---|
| 1.000 | -0.76 | 99.66 % | 0.13 % | 0.00 % | 0.21 % | |
| 1.200 | -0.56 | 99.46 % | 0.20 % | 0.00 % | 0.34 % | Stomach pH |
| 2.000 | 0.24 | 96.70 % | 1.25 % | 0.00 % | 2.06 % | |
| 3.000 | 1.19 | 74.55 % | 9.60 % | 0.00 % | 15.84 % | |
| 4.000 | 1.88 | 22.66 % | 29.19 % | 0.00 % | 48.16 % | |
| 5.000 | 2.10 | 2.85 % | 36.66 % | 0.00 % | 60.49 % | |
| 6.000 | 2.13 | 0.29 % | 37.63 % | 0.00 % | 62.08 % | |
| 6.500 | 2.13 | 0.09 % | 37.70 % | 0.00 % | 62.20 % | |
| 7.000 | 2.13 | 0.03 % | 37.73 % | 0.00 % | 62.24 % | |
| 7.400 | 2.13 | 0.01 % | 37.73 % | 0.00 % | 62.25 % | Blood pH |
| 8.000 | 2.13 | 0.00 % | 37.74 % | 0.00 % | 62.26 % | |
| 9.000 | 2.13 | 0.00 % | 37.74 % | 0.00 % | 62.26 % | |
| 10.000 | 2.13 | 0.00 % | 37.74 % | 0.00 % | 62.26 % | |
| 11.000 | 2.13 | 0.00 % | 37.74 % | 0.00 % | 62.26 % | |
| 12.000 | 2.13 | 0.00 % | 37.74 % | 0.00 % | 62.26 % | |

#### Carbonate and acidity

| | |
|---|---|
| Carbonate | 0.000 mM |
| Acidity error | 0.030 mM |

#### Other graphs

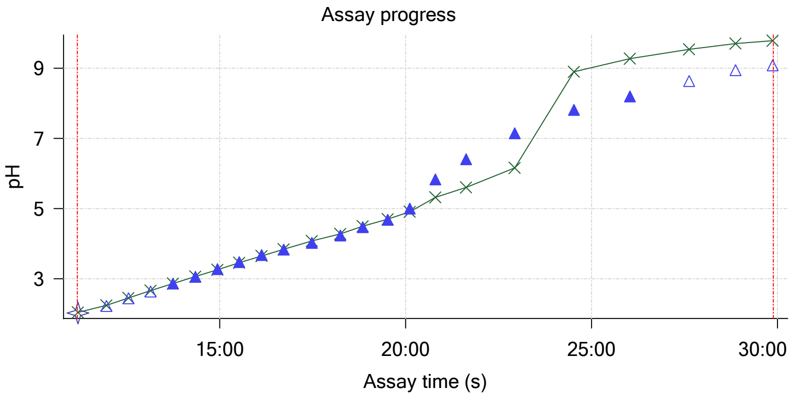


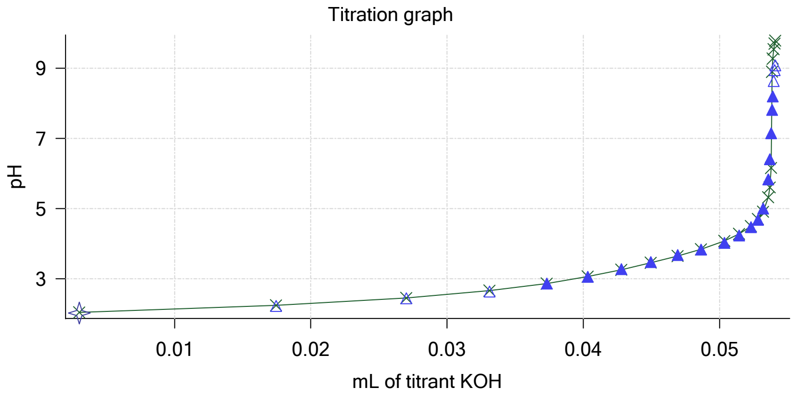

Sample name: **M11_octanol**
Assay name: **pH-metric high logP**
Assay ID: **18C-01001**
Filename: **C:\Sirius_T3\Mehtap\20180228_exp28_logP_T3-2\18C-01001_M11_octanol_pH-metric high logP.t3r**

Experiment start time: **3/1/2018 12:11:22 AM**
Analyst: **Pion**
Instrument ID: **T312060**

#### Other graphs (continued)

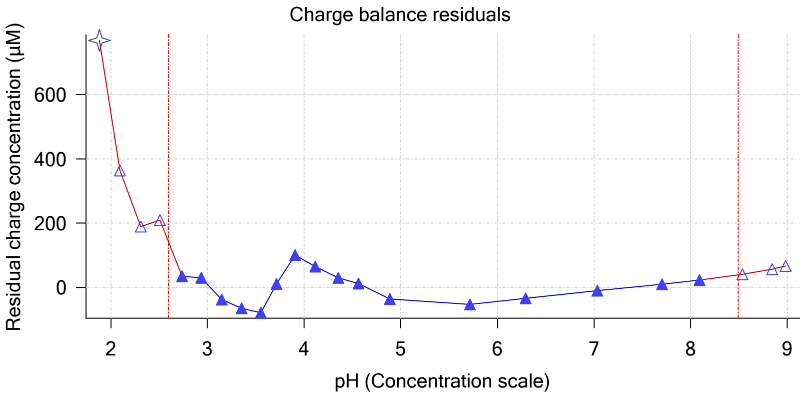


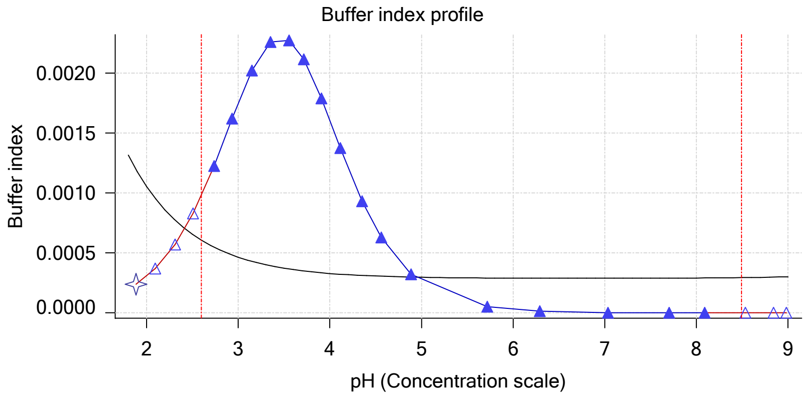


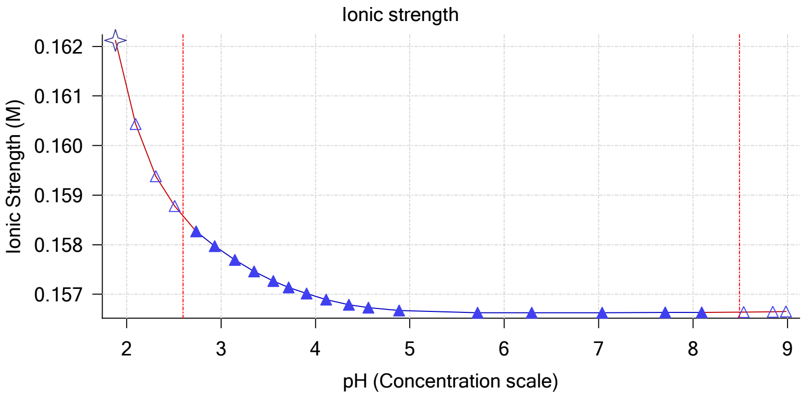


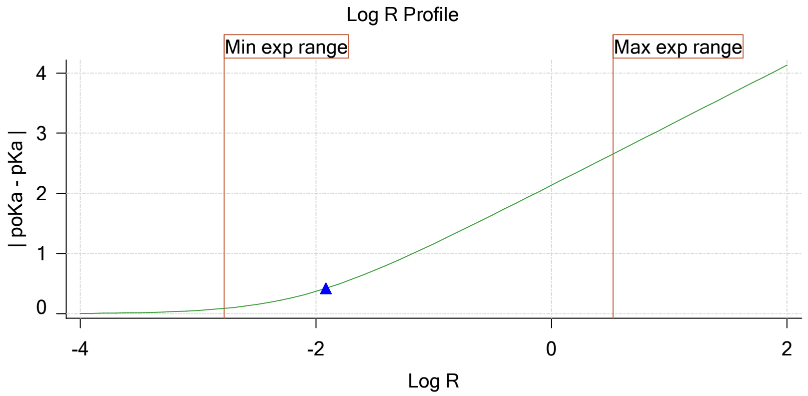


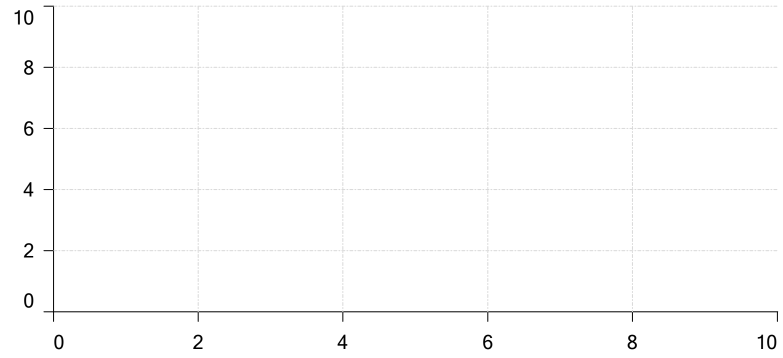


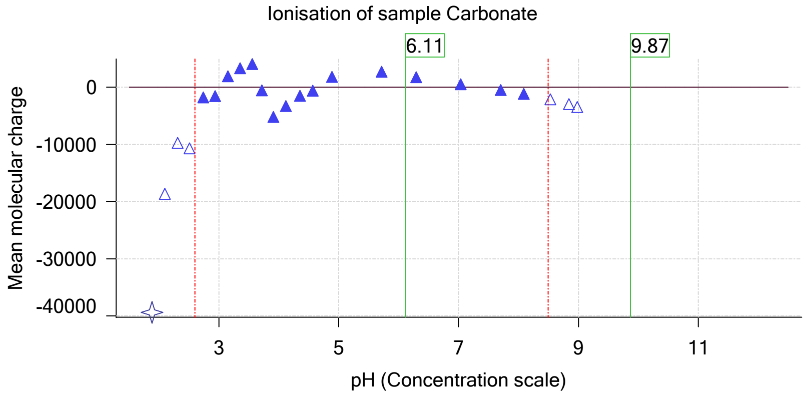

| | | | |
|---|---|---|---|
| Sample name: | **M11_octanol** | Experiment start time: | **3/1/2018 12:11:22 AM** |
| Assay name: | **pH-metric high logP** | Analyst: | **Pion** |
| Assay ID: | **18C-01001** | Instrument ID: | **T312060** |
| Filename: | **C:\Sirius_T3\Mehtap\20180228_exp28_logP_T3-2\18C-01001_M11_octanol_pH-metric high logP.t3r** | | |

### pH-metric high logP Titration 2 of 3 18C-01001 Points 24 to 48

#### Overall results

| | |
|---|---|
| RMSD | 0.387 |
| Average ionic strength | 0.164 M |
| Average temperature | 25.0°C |
| Partition ratio | 0.0404 : 1 |
| Analyte concentration range | 3480.1 µM to 3596.9 µM |
| Total points considered | 18 of 25 |

#### Warnings and errors

| | |
|---|---|
| Errors | None |
| Warnings | One or more logP values out of range |

#### Four-Plus parameters

| | | | |
|---|---|---|---|
| Alpha | 0.130 | 3/1/2018 12:11:22 AM | C:\Sirius_T3\HCl18B27.t3r |
| S | 0.9970 | 3/1/2018 12:11:22 AM | C:\Sirius_T3\HCl18B27.t3r |
| jH | 0.8 | 3/1/2018 12:11:22 AM | C:\Sirius_T3\HCl18B27.t3r |
| jOH | -0.4 | 3/1/2018 12:11:22 AM | C:\Sirius_T3\HCl18B27.t3r |

#### Titrants

| | | | |
|---|---|---|---|
| 0.50 M HCl | 0.993513 | 3/1/2018 12:11:22 AM | C:\Sirius_T3\HCl18B27.t3r |
| 0.50 M KOH | 0.999845 | 3/1/2018 12:11:22 AM | C:\Sirius_T3\KOH18B27.t3r |

#### Sample

| | |
|---|---|
| M11_octanol concentration factor | 0.912 |
| Base pKa 1 | 3.89 |
| logP (XH +) | -4.46 |
| logP (neutral X) | 2.16 |

#### Sample graphs

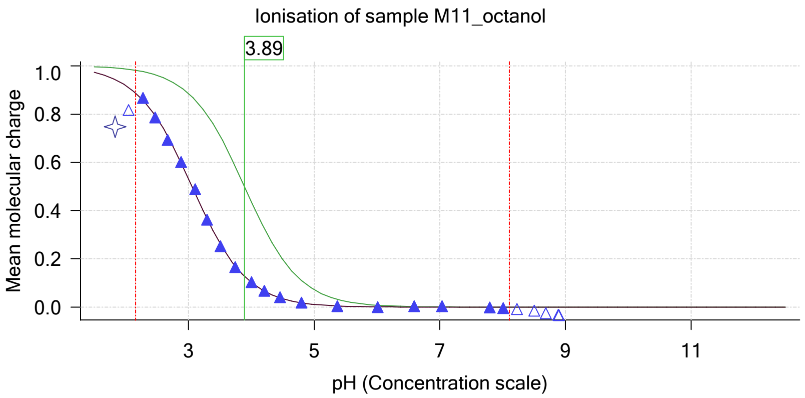


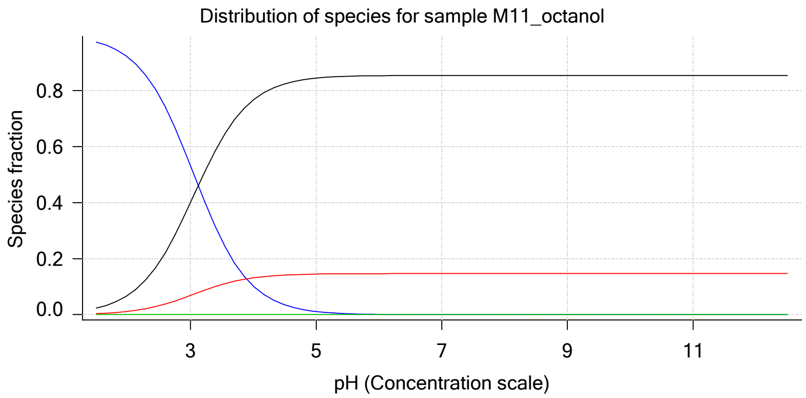

| | | | |
|---|---|---|---|
| Sample name: | **M11_octanol** | Experiment start time: | **3/1/2018 12:11:22 AM** |
| Assay name: | **pH-metric high logP** | Analyst: | **Pion** |
| Assay ID: | **18C-01001** | Instrument ID: | **T312060** |
| Filename: | **C:\Sirius_T3\Mehtap\20180228_exp28_logP_T3-2\18C-01001_M11_octanol_pH-metric high logP.t3r** | | |

#### Sample graphs (continued)

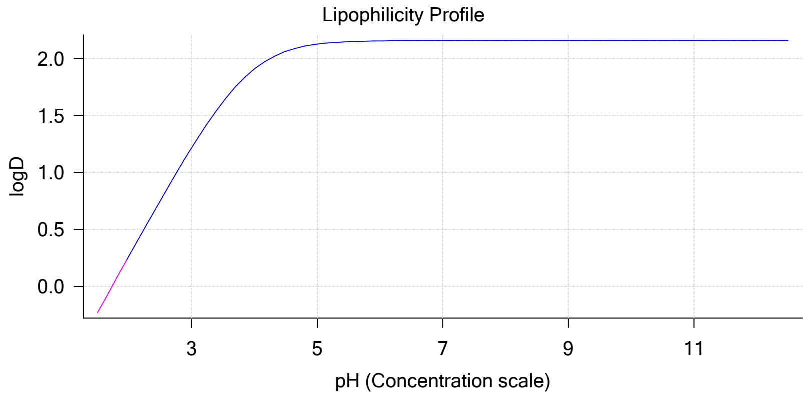


#### Sample logD and percent species

| pH | M11_octanol logD | M11_octanol M11_octanolH | M11_octanol M11_octanol | M11_octanol M11_octanolH* | M11_octanol M11_octanol* | Comment |
|---|---|---|---|---|---|---|
| 1.000 | -0.73 | 99.13 % | 0.13 % | 0.00 % | 0.74 % | |
| 1.200 | -0.53 | 98.62 % | 0.20 % | 0.00 % | 1.17 % | Stomach pH |
| 2.000 | 0.26 | 91.91 % | 1.18 % | 0.00 % | 6.90 % | |
| 3.000 | 1.22 | 53.20 % | 6.85 % | 0.00 % | 39.94 % | |
| 4.000 | 1.91 | 10.21 % | 13.15 % | 0.00 % | 76.64 % | |
| 5.000 | 2.13 | 1.12 % | 14.48 % | 0.00 % | 84.40 % | |
| 6.000 | 2.16 | 0.11 % | 14.63 % | 0.00 % | 85.26 % | |
| 6.500 | 2.16 | 0.04 % | 14.64 % | 0.00 % | 85.32 % | |
| 7.000 | 2.16 | 0.01 % | 14.64 % | 0.00 % | 85.34 % | |
| 7.400 | 2.16 | 0.00 % | 14.64 % | 0.00 % | 85.35 % | Blood pH |
| 8.000 | 2.16 | 0.00 % | 14.65 % | 0.00 % | 85.35 % | |
| 9.000 | 2.16 | 0.00 % | 14.65 % | 0.00 % | 85.35 % | |
| 10.000 | 2.16 | 0.00 % | 14.65 % | 0.00 % | 85.35 % | |
| 11.000 | 2.16 | 0.00 % | 14.65 % | 0.00 % | 85.35 % | |
| 12.000 | 2.16 | 0.00 % | 14.65 % | 0.00 % | 85.35 % | |

#### Carbonate and acidity

| | | |
|---|---|---|
| 💎 | Carbonate | 0.096 mM |
| 💎 | Acidity error | -0.306 mM |

#### Other graphs

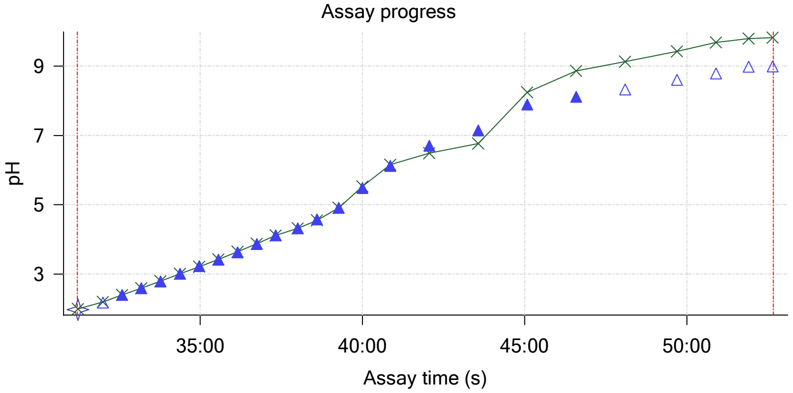


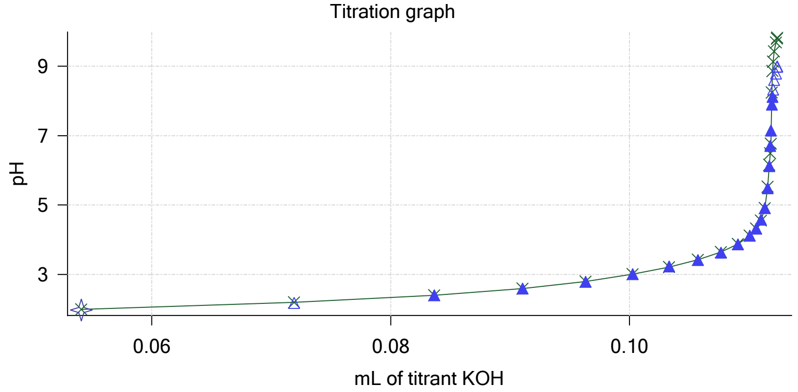

| | | | |
|---|---|---|---|
| Sample name: | **M11_octanol** | Experiment start time: | **3/1/2018 12:11:22 AM** |
| Assay name: | **pH-metric high logP** | Analyst: | **Pion** |
| Assay ID: | **18C-01001** | Instrument ID: | **T312060** |
| Filename: | **C:\Sirius_T3\Mehtap\20180228_exp28_logP_T3-2\18C-01001_M11_octanol_pH-metric high logP.t3r** | | |

#### Other graphs (continued)

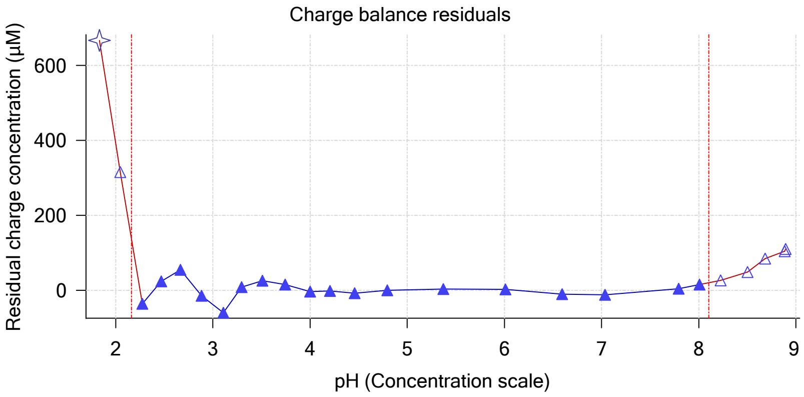


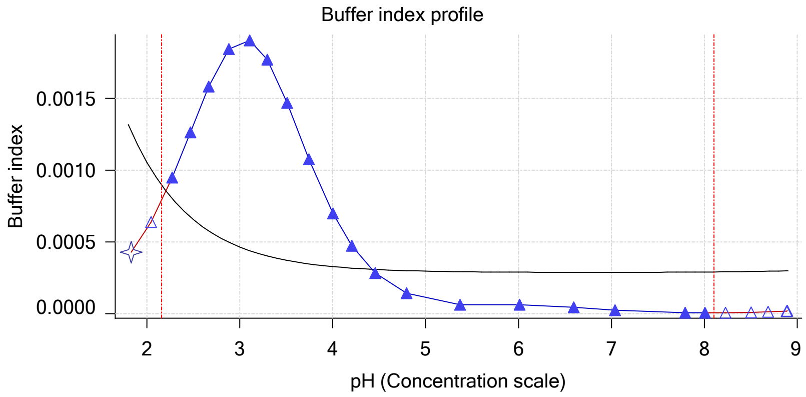


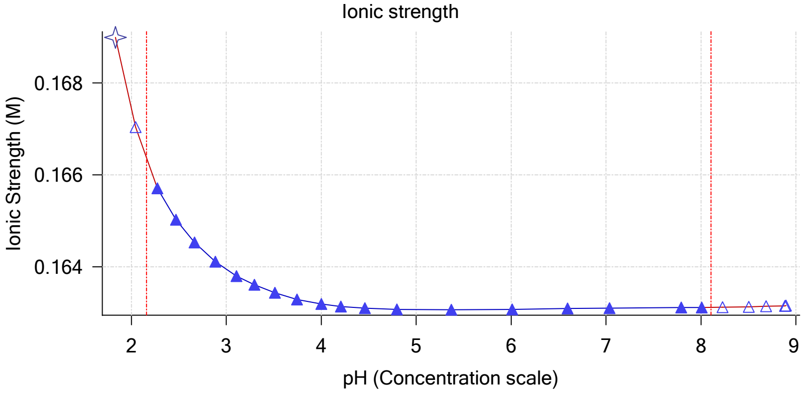


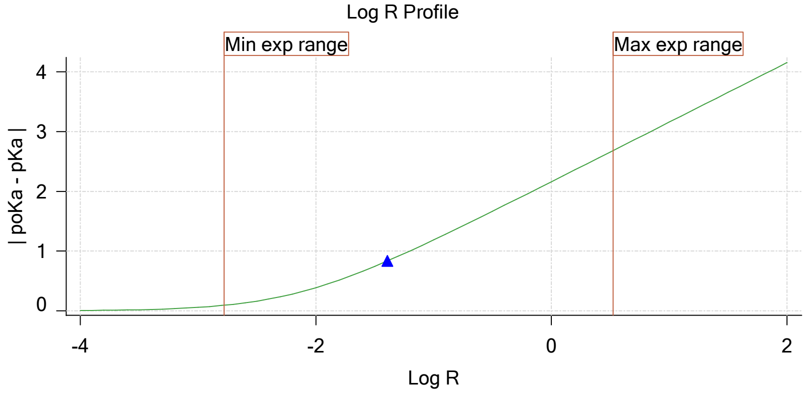


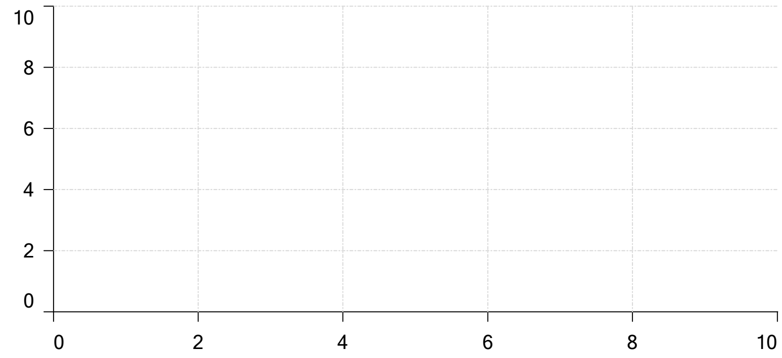


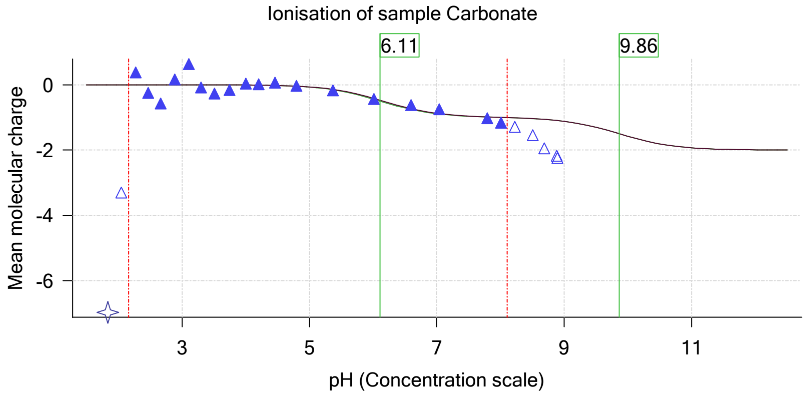

| | | | |
|---|---|---|---|
| Sample name: | **M11_octanol** | Experiment start time: | **3/1/2018 12:11:22 AM** |
| Assay name: | **pH-metric high logP** | Analyst: | **Pion** |
| Assay ID: | **18C-01001** | Instrument ID: | **T312060** |
| Filename: | **C:\Sirius_T3\Mehtap\20180228_exp28_logP_T3-2\18C-01001_M11_octanol_pH-metric high logP.t3r** | | |

### pH-metric high logP Titration 3 of 3 18C-01001 Points 49 to 69

#### Overall results

| | |
|---|---|
| RMSD | 0.050 |
| Average ionic strength | 0.170 M |
| Average temperature | 25.0°C |
| Partition ratio | 0.1189 : 1 |
| Analyte concentration range | 3013.9 μM to 3107.7 μM |
| Total points considered | 18 of 21 |

#### Warnings and errors

| | |
|---|---|
| Errors | None |
| Warnings | One or more logP values out of range |

#### Four-Plus parameters

| | | | |
|---|---|---|---|
| Alpha | 0.130 | 3/1/2018 12:11:22 AM | C:\Sirius_T3\HCl18B27.t3r |
| S | 0.9970 | 3/1/2018 12:11:22 AM | C:\Sirius_T3\HCl18B27.t3r |
| jH | 0.8 | 3/1/2018 12:11:22 AM | C:\Sirius_T3\HCl18B27.t3r |
| jOH | -0.4 | 3/1/2018 12:11:22 AM | C:\Sirius_T3\HCl18B27.t3r |

#### Titrants

| | | | |
|---|---|---|---|
| 0.50 M HCl | 0.993513 | 3/1/2018 12:11:22 AM | C:\Sirius_T3\HCl18B27.t3r |
| 0.50 M KOH | 0.999845 | 3/1/2018 12:11:22 AM | C:\Sirius_T3\KOH18B27.t3r |

#### Sample

| | |
|---|---|
| M11_octanol concentration factor | 0.792 |
| Base pKa 1 | 3.89 |
| logP (XH +) | -5.10 |
| logP (neutral X) | 2.01 |

#### Sample graphs

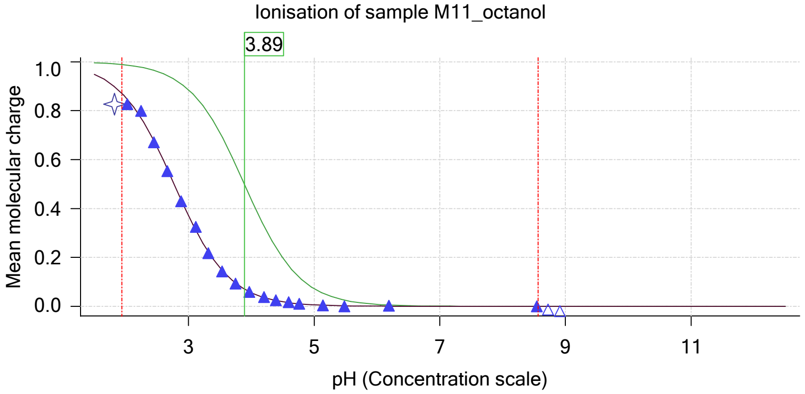


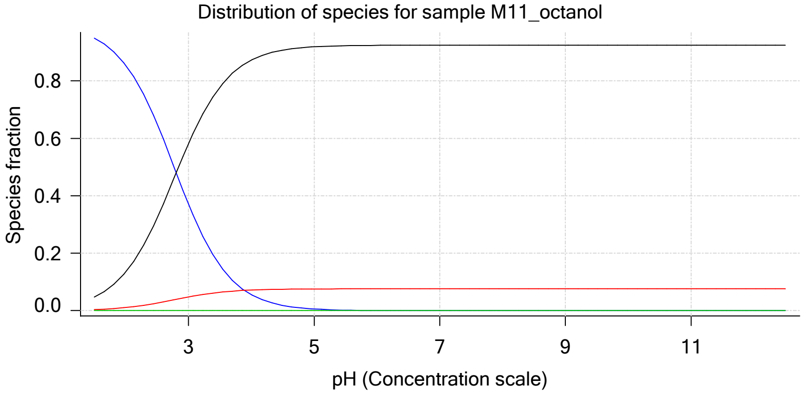

| | | | |
|---|---|---|---|
| Sample name: | **M11_octanol** | Experiment start time: | **3/1/2018 12:11:22 AM** |
| Assay name: | **pH-metric high logP** | Analyst: | **Pion** |
| Assay ID: | **18C-01001** | Instrument ID: | **T312060** |
| Filename: | **C:\Sirius_T3\Mehtap\20180228_exp28_logP_T3-2\18C-01001_M11_octanol_pH-metric high logP.t3r** | | |

#### Sample graphs (continued)

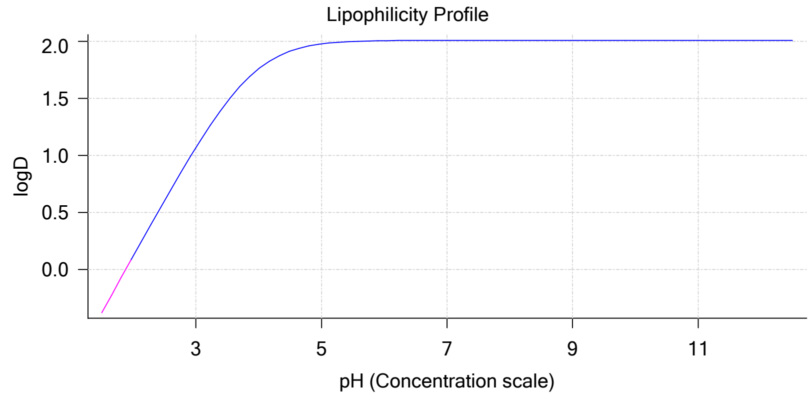


#### Sample logD and percent species

| pH | M11_octanol logD | M11_octanol M11_octanolH | M11_octanol M11_octanol | M11_octanol M11_octanolH* | M11_octanol M11_octanol* | Comment |
|---|---|---|---|---|---|---|
| 1.000 | -0.88 | 98.33 % | 0.13 % | 0.00 % | 1.54 % | |
| 1.200 | -0.68 | 97.38 % | 0.20 % | 0.00 % | 2.42 % | Stomach pH |
| 2.000 | 0.11 | 85.51 % | 1.10 % | 0.00 % | 13.39 % | |
| 3.000 | 1.07 | 37.11 % | 4.78 % | 0.00 % | 58.11 % | |
| 4.000 | 1.76 | 5.57 % | 7.18 % | 0.00 % | 87.25 % | |
| 5.000 | 1.98 | 0.59 % | 7.56 % | 0.00 % | 91.86 % | |
| 6.000 | 2.01 | 0.06 % | 7.60 % | 0.00 % | 92.34 % | |
| 6.500 | 2.01 | 0.02 % | 7.60 % | 0.00 % | 92.38 % | |
| 7.000 | 2.01 | 0.01 % | 7.60 % | 0.00 % | 92.39 % | |
| 7.400 | 2.01 | 0.00 % | 7.60 % | 0.00 % | 92.40 % | Blood pH |
| 8.000 | 2.01 | 0.00 % | 7.60 % | 0.00 % | 92.40 % | |
| 9.000 | 2.01 | 0.00 % | 7.60 % | 0.00 % | 92.40 % | |
| 10.000 | 2.01 | 0.00 % | 7.60 % | 0.00 % | 92.40 % | |
| 11.000 | 2.01 | 0.00 % | 7.60 % | 0.00 % | 92.40 % | |
| 12.000 | 2.01 | 0.00 % | 7.60 % | 0.00 % | 92.40 % | |

#### Carbonate and acidity

| | |
|---|---|
| Carbonate | 0.073 mM |
| Acidity error | -0.819 mM |

#### Other graphs

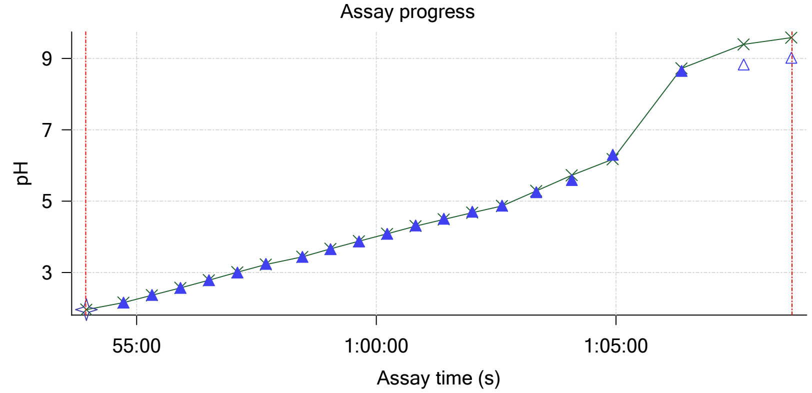


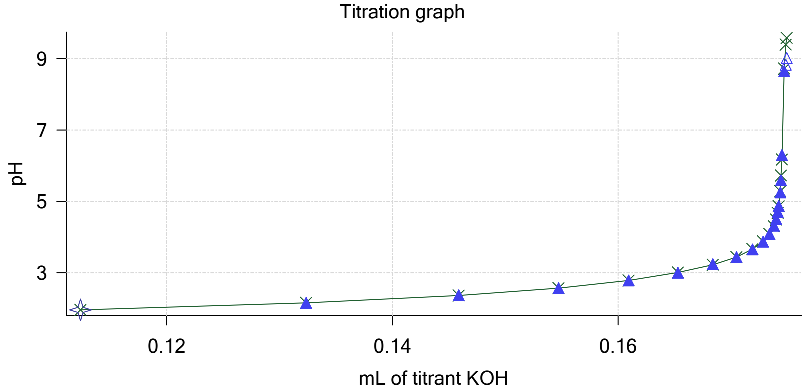

Sample name: **M11_octanol** | Experiment start time: **3/1/2018 12:11:22 AM**
Assay name: **pH-metric high logP** | Analyst: **Pion**
Assay ID: **18C-01001** | Instrument ID: **T312060**
Filename: **C:\Sirius_T3\Mehtap\20180228_exp28_logP_T3-2\18C-01001_M11_octanol_pH-metric high logP.t3r**

#### Other graphs (continued)

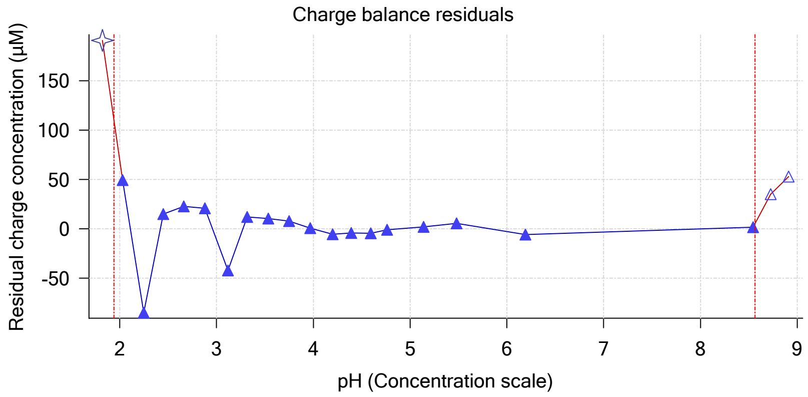


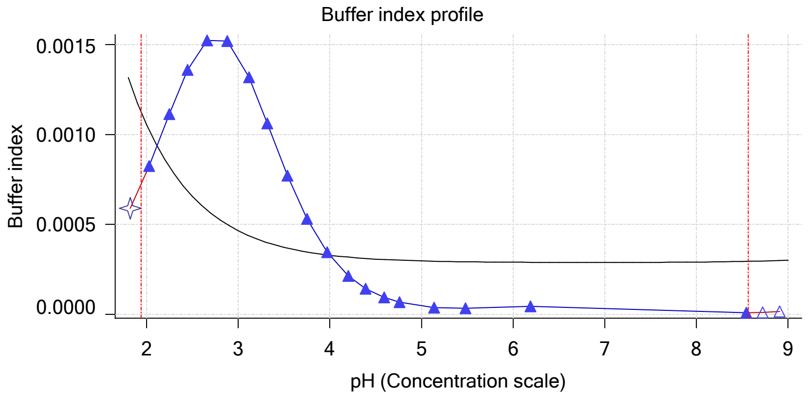


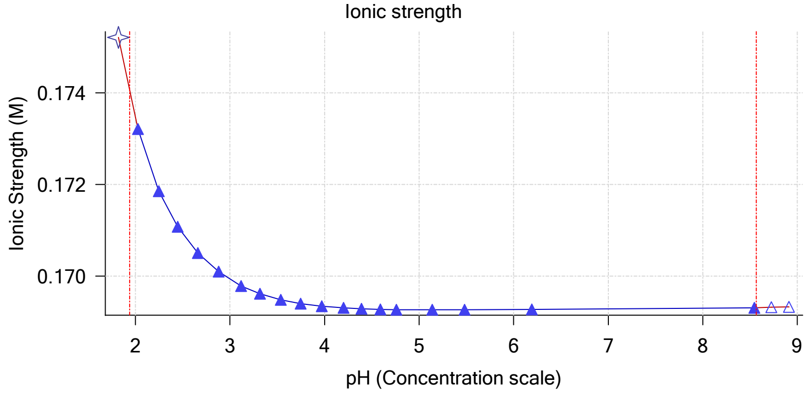


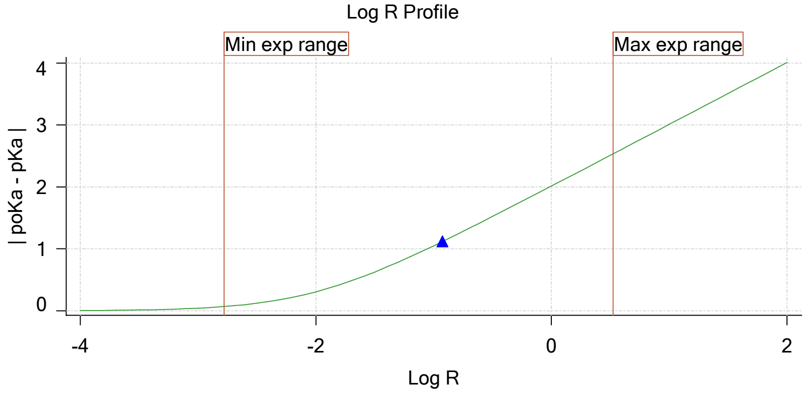


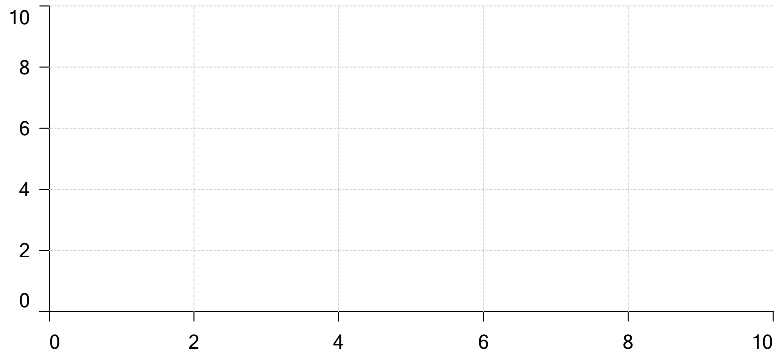

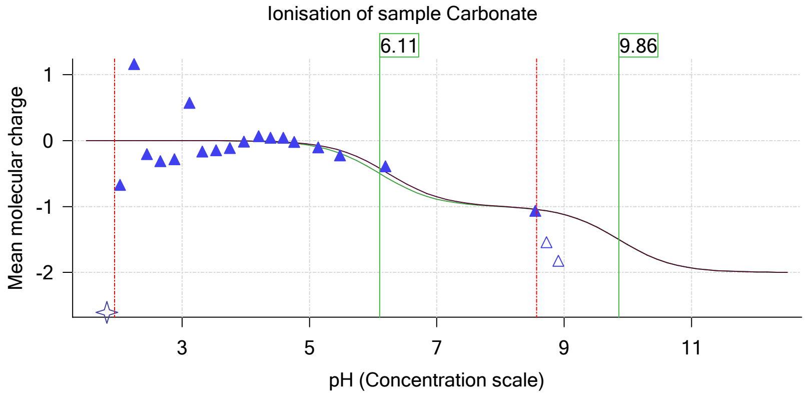

| | | | |
|---|---|---|---|
| Sample name: | **M11_octanol** | Experiment start time: | **3/1/2018 12:11:22 AM** |
| Assay name: | **pH-metric high logP** | Analyst: | **Pion** |
| Assay ID: | **18C-01001** | Instrument ID: | **T312060** |
| Filename: | **C:\Sirius_T3\Mehtap\20180228_exp28_logP_T3-2\18C-01001_M11_octanol_pH-metric high logP.t3r** | | |

#### Assay Model

| Settings | Value | Date/Time changed | Imported from |
|---|---|---|---|
| Sample name | M11_octanol | 2/27/2018 4:54:30 PM | User entered value |
| Sample by | Weight | | Default value |
| Sample weight | 0.001320 g | 2/28/2018 4:24:52 PM | User entered value |
| Formula weight | 211.22 g/mol | 2/27/2018 4:54:30 PM | User entered value |
| Solubility | Unknown | | Default value |
| Molecular weight | 211.22 | 2/27/2018 4:54:30 PM | User entered value |
| Individual pKa ionic environments | No | | Default value |
| Number of pKas | 1 | 2/27/2018 4:54:30 PM | User entered value |
| Sample is a | Base | 2/27/2018 4:54:30 PM | User entered value |
| pKa 1 | 3.89 | 2/27/2018 4:54:30 PM | User entered value |
| logp (XH +) | -5.60 | 2/28/2018 1:53:21 PM | User entered value |
| logP (neutral X) | 2.09 | 2/28/2018 1:53:11 PM | User entered value |

#### Events

| Time | Event | Water | Acid | Base | Octanol | pH | dpH/dt | pH R-squared | pH SD | dpH/dt time |
|---|---|---|---|---|---|---|---|---|---|---|
| 8:11.4 | Initial pH = 9.64 | | | | | | | | | |
| 11:11.0 | Data point 1 | 1.50000 mL | 0.05407 mL | 0.00301 mL | 0.01999 mL | 2.018 | -0.00907 | 0.79017 | 0.00050 | 10.0 s |
| 11:57.2 | Data point 2 | 1.50000 mL | 0.05407 mL | 0.01745 mL | 0.01999 mL | 2.225 | -0.00501 | 0.21160 | 0.00054 | 10.0 s |
| 12:32.9 | Data point 3 | 1.50000 mL | 0.05407 mL | 0.02702 mL | 0.01999 mL | 2.438 | -0.00560 | 0.18844 | 0.00064 | 10.0 s |
| 13:08.5 | Data point 4 | 1.50000 mL | 0.05407 mL | 0.03311 mL | 0.01999 mL | 2.633 | -0.00265 | 0.48730 | 0.00019 | 10.5 s |
| 13:44.5 | Data point 5 | 1.50000 mL | 0.05407 mL | 0.03732 mL | 0.01999 mL | 2.860 | -0.01106 | 0.66289 | 0.00067 | 10.5 s |
| 14:20.5 | Data point 6 | 1.50000 mL | 0.05407 mL | 0.04033 mL | 0.01999 mL | 3.057 | -0.01232 | 0.47963 | 0.00088 | 10.0 s |
| 14:56.0 | Data point 7 | 1.50000 mL | 0.05407 mL | 0.04280 mL | 0.01999 mL | 3.270 | -0.00677 | 0.19054 | 0.00077 | 10.0 s |
| 15:31.5 | Data point 8 | 1.50000 mL | 0.05407 mL | 0.04497 mL | 0.01999 mL | 3.472 | -0.00166 | 0.35149 | 0.00014 | 10.5 s |
| 16:07.5 | Data point 9 | 1.50000 mL | 0.05407 mL | 0.04692 mL | 0.01999 mL | 3.673 | -0.00781 | 0.35544 | 0.00065 | 10.0 s |
| 16:42.9 | Data point 10 | 1.50000 mL | 0.05407 mL | 0.04864 mL | 0.01999 mL | 3.835 | -0.01388 | 0.51424 | 0.00096 | 10.0 s |
| 17:28.7 | Data point 11 | 1.50000 mL | 0.05407 mL | 0.05035 mL | 0.01999 mL | 4.027 | -0.01025 | 0.38200 | 0.00082 | 10.0 s |
| 18:14.5 | Data point 12 | 1.50000 mL | 0.05407 mL | 0.05143 mL | 0.01999 mL | 4.232 | -0.00787 | 0.92490 | 0.00040 | 10.5 s |
| 18:50.5 | Data point 13 | 1.50000 mL | 0.05407 mL | 0.05230 mL | 0.01999 mL | 4.470 | -0.00601 | 0.45130 | 0.00044 | 10.0 s |
| 19:31.0 | Data point 14 | 1.50000 mL | 0.05407 mL | 0.05282 mL | 0.01999 mL | 4.679 | -0.00976 | 0.57077 | 0.00064 | 10.0 s |
| 20:06.4 | Data point 15 | 1.50000 mL | 0.05407 mL | 0.05320 mL | 0.01999 mL | 5.003 | -0.01866 | 0.86396 | 0.00099 | 11.0 s |
| 20:48.0 | Data point 16 | 1.50000 mL | 0.05407 mL | 0.05357 mL | 0.01999 mL | 5.829 | -0.01903 | 0.91633 | 0.00098 | 19.0 s |
| 21:37.5 | Data point 17 | 1.50000 mL | 0.05407 mL | 0.05369 mL | 0.01999 mL | 6.403 | -0.02005 | 0.98528 | 0.00100 | 42.5 s |
| 22:55.8 | Data point 18 | 1.50000 mL | 0.05407 mL | 0.05379 mL | 0.01999 mL | 7.144 | -0.04872 | 0.99706 | 0.00241 | Timed out at 59.5 s |
| 24:31.4 | Data point 19 | 1.50000 mL | 0.05407 mL | 0.05386 mL | 0.01999 mL | 7.810 | -0.04860 | 0.99788 | 0.00240 | Timed out at 59.5 s |
| 26:01.9 | Data point 20 | 1.50000 mL | 0.05407 mL | 0.05390 mL | 0.01999 mL | 8.196 | -0.02409 | 0.99122 | 0.00119 | Timed out at 59.5 s |
| 27:37.5 | Data point 21 | 1.50000 mL | 0.05407 mL | 0.05397 mL | 0.01999 mL | 8.638 | -0.02003 | 0.99205 | 0.00099 | 39.0 s |
| 28:52.2 | Data point 22 | 1.50000 mL | 0.05407 mL | 0.05405 mL | 0.01999 mL | 8.945 | -0.02002 | 0.98473 | 0.00100 | 29.5 s |
| 29:52.2 | Data point 23 | 1.50000 mL | 0.05407 mL | 0.05409 mL | 0.01999 mL | 9.084 | -0.01992 | 0.97892 | 0.00099 | 22.5 s |
| 31:14.1 | Data point 24 | 1.50000 mL | 0.11336 mL | 0.05409 mL | 0.06999 mL | 1.969 | -0.01287 | 0.76123 | 0.00073 | 10.0 s |
| 32:00.4 | Data point 25 | 1.50000 mL | 0.11336 mL | 0.07192 mL | 0.06999 mL | 2.176 | -0.00432 | 0.09399 | 0.00070 | 10.0 s |
| 32:36.1 | Data point 26 | 1.50000 mL | 0.11336 mL | 0.08365 mL | 0.06999 mL | 2.402 | 0.00539 | 0.49157 | 0.00038 | 10.0 s |
| 33:11.7 | Data point 27 | 1.50000 mL | 0.11336 mL | 0.09104 mL | 0.06999 mL | 2.594 | -0.00764 | 0.16492 | 0.00093 | 10.0 s |
| 33:47.2 | Data point 28 | 1.50000 mL | 0.11336 mL | 0.09631 mL | 0.06999 mL | 2.790 | -0.00104 | 0.07057 | 0.00019 | 10.0 s |
| 34:22.7 | Data point 29 | 1.50000 mL | 0.11336 mL | 0.10026 mL | 0.06999 mL | 3.007 | -0.00433 | 0.31139 | 0.00038 | 10.0 s |
| 34:58.2 | Data point 30 | 1.50000 mL | 0.11336 mL | 0.10332 mL | 0.06999 mL | 3.227 | -0.00212 | 0.02826 | 0.00062 | 10.0 s |
| 35:33.8 | Data point 31 | 1.50000 mL | 0.11336 mL | 0.10572 mL | 0.06999 mL | 3.417 | 0.00173 | 0.09499 | 0.00028 | 10.0 s |
| 36:09.3 | Data point 32 | 1.50000 mL | 0.11336 mL | 0.10764 mL | 0.06999 mL | 3.630 | -0.00346 | 0.60333 | 0.00022 | 10.0 s |
| 36:44.7 | Data point 33 | 1.50000 mL | 0.11336 mL | 0.10908 mL | 0.06999 mL | 3.864 | -0.00645 | 0.48290 | 0.00046 | 10.0 s |
| 37:20.1 | Data point 34 | 1.50000 mL | 0.11336 mL | 0.11007 mL | 0.06999 mL | 4.119 | -0.00379 | 0.54985 | 0.00025 | 10.0 s |
| 38:00.7 | Data point 35 | 1.50000 mL | 0.11336 mL | 0.11061 mL | 0.06999 mL | 4.322 | -0.00366 | 0.30464 | 0.00033 | 10.0 s |
| 38:36.1 | Data point 36 | 1.50000 mL | 0.11336 mL | 0.11101 mL | 0.06999 mL | 4.575 | -0.01448 | 0.78873 | 0.00080 | 10.0 s |
| 39:16.6 | Data point 37 | 1.50000 mL | 0.11336 mL | 0.11134 mL | 0.06999 mL | 4.912 | -0.01854 | 0.87629 | 0.00098 | 13.0 s |

| | | | |
|---|---|---|---|
| Sample name: | **M11_octanol** | Experiment start time: | **3/1/2018 12:11:22 AM** |
| Assay name: | **pH-metric high logP** | Analyst: | **Pion** |
| Assay ID: | **18C-01001** | Instrument ID: | **T312060** |
| Filename: | **C:\Sirius_T3\Mehtap\20180228_exp28_logP_T3-2\18C-01001_M11_octanol_pH-metric high logP.t3r** | | |

#### Events (continued)

| Time | Event | Water | Acid | Base | Octanol | pH | dpH/dt | pH R-squared | pH SD | dpH/dt time |
|---|---|---|---|---|---|---|---|---|---|---|
| 40:00.2 | Data point 38 | 1.50000 mL | 0.11336 mL | 0.11157 mL | 0.06999 mL | 5.485 | -0.01247 | 0.47642 | 0.00089 | 16.0 s |
| 40:51.8 | Data point 39 | 1.50000 mL | 0.11336 mL | 0.11171 mL | 0.06999 mL | 6.124 | -0.01978 | 0.97231 | 0.00099 | 36.5 s |
| 42:04.0 | Data point 40 | 1.50000 mL | 0.11336 mL | 0.11178 mL | 0.06999 mL | 6.704 | -0.02890 | 0.99478 | 0.00143 | Timed out at 59.5 s |
| 43:34.5 | Data point 41 | 1.50000 mL | 0.11336 mL | 0.11183 mL | 0.06999 mL | 7.146 | -0.04871 | 0.99489 | 0.00241 | Timed out at 59.5 s |
| 45:05.1 | Data point 42 | 1.50000 mL | 0.11336 mL | 0.11192 mL | 0.06999 mL | 7.900 | -0.05502 | 0.99571 | 0.00272 | Timed out at 59.5 s |
| 46:35.6 | Data point 43 | 1.50000 mL | 0.11336 mL | 0.11197 mL | 0.06999 mL | 8.113 | -0.02606 | 0.99186 | 0.00129 | Timed out at 59.5 s |
| 48:06.1 | Data point 44 | 1.50000 mL | 0.11336 mL | 0.11202 mL | 0.06999 mL | 8.331 | -0.01989 | 0.98982 | 0.00099 | 55.0 s |
| 49:41.9 | Data point 45 | 1.50000 mL | 0.11336 mL | 0.11211 mL | 0.06999 mL | 8.608 | -0.01909 | 0.97284 | 0.00096 | 31.0 s |
| 50:53.8 | Data point 46 | 1.50000 mL | 0.11336 mL | 0.11225 mL | 0.06999 mL | 8.789 | -0.01955 | 0.99182 | 0.00097 | 25.0 s |
| 51:54.4 | Data point 47 | 1.50000 mL | 0.11336 mL | 0.11235 mL | 0.06999 mL | 8.987 | -0.01750 | 0.90247 | 0.00091 | 19.0 s |
| 52:38.8 | Data point 48 | 1.50000 mL | 0.11336 mL | 0.11237 mL | 0.06999 mL | 8.997 | -0.01930 | 0.96484 | 0.00097 | 16.5 s |
| 53:57.2 | Data point 49 | 1.50000 mL | 0.17855 mL | 0.11237 mL | 0.21999 mL | 1.958 | -0.00753 | 0.72606 | 0.00044 | 10.0 s |
| 54:43.6 | Data point 50 | 1.50000 mL | 0.17855 mL | 0.13236 mL | 0.21999 mL | 2.160 | -0.00028 | 0.00344 | 0.00023 | 10.0 s |
| 55:19.4 | Data point 51 | 1.50000 mL | 0.17855 mL | 0.14588 mL | 0.21999 mL | 2.375 | 0.00030 | 0.00453 | 0.00022 | 10.0 s |
| 55:55.0 | Data point 52 | 1.50000 mL | 0.17855 mL | 0.15473 mL | 0.21999 mL | 2.574 | -0.00751 | 0.16614 | 0.00091 | 10.0 s |
| 56:30.6 | Data point 53 | 1.50000 mL | 0.17855 mL | 0.16091 mL | 0.21999 mL | 2.786 | -0.00360 | 0.47623 | 0.00026 | 10.0 s |
| 57:06.1 | Data point 54 | 1.50000 mL | 0.17855 mL | 0.16529 mL | 0.21999 mL | 3.003 | 0.00336 | 0.12158 | 0.00048 | 10.0 s |
| 57:41.6 | Data point 55 | 1.50000 mL | 0.17855 mL | 0.16837 mL | 0.21999 mL | 3.241 | -0.01332 | 0.90714 | 0.00069 | 10.0 s |
| 58:27.4 | Data point 56 | 1.50000 mL | 0.17855 mL | 0.17046 mL | 0.21999 mL | 3.438 | -0.00073 | 0.01099 | 0.00034 | 10.0 s |
| 59:02.8 | Data point 57 | 1.50000 mL | 0.17855 mL | 0.17190 mL | 0.21999 mL | 3.656 | -0.00412 | 0.83143 | 0.00022 | 10.0 s |
| 59:38.3 | Data point 58 | 1.50000 mL | 0.17855 mL | 0.17281 mL | 0.21999 mL | 3.868 | -0.00343 | 0.75983 | 0.00019 | 10.0 s |
| 1:00:13.7 | Data point 59 | 1.50000 mL | 0.17855 mL | 0.17340 mL | 0.21999 mL | 4.088 | -0.00449 | 0.80203 | 0.00025 | 10.0 s |
| 1:00:49.1 | Data point 60 | 1.50000 mL | 0.17855 mL | 0.17378 mL | 0.21999 mL | 4.318 | -0.00344 | 0.13702 | 0.00046 | 10.0 s |
| 1:01:24.5 | Data point 61 | 1.50000 mL | 0.17855 mL | 0.17399 mL | 0.21999 mL | 4.508 | -0.01637 | 0.76668 | 0.00092 | 10.0 s |
| 1:01:59.9 | Data point 62 | 1.50000 mL | 0.17855 mL | 0.17413 mL | 0.21999 mL | 4.709 | -0.01145 | 0.54777 | 0.00076 | 12.0 s |
| 1:02:37.3 | Data point 63 | 1.50000 mL | 0.17855 mL | 0.17422 mL | 0.21999 mL | 4.876 | -0.00617 | 0.20813 | 0.00067 | 12.5 s |
| 1:03:20.3 | Data point 64 | 1.50000 mL | 0.17855 mL | 0.17434 mL | 0.21999 mL | 5.253 | -0.01491 | 0.67066 | 0.00090 | 14.0 s |
| 1:04:04.7 | Data point 65 | 1.50000 mL | 0.17855 mL | 0.17441 mL | 0.21999 mL | 5.593 | -0.01977 | 0.97774 | 0.00099 | 20.5 s |
| 1:04:55.7 | Data point 66 | 1.50000 mL | 0.17855 mL | 0.17448 mL | 0.21999 mL | 6.302 | -0.01947 | 0.98219 | 0.00097 | 55.5 s |
| 1:06:21.7 | Data point 67 | 1.50000 mL | 0.17855 mL | 0.17469 mL | 0.21999 mL | 8.648 | -0.01798 | 0.88440 | 0.00094 | 37.0 s |
| 1:07:39.6 | Data point 68 | 1.50000 mL | 0.17855 mL | 0.17483 mL | 0.21999 mL | 8.828 | -0.01767 | 0.82826 | 0.00096 | 24.0 s |
| 1:08:39.2 | Data point 69 | 1.50000 mL | 0.17855 mL | 0.17493 mL | 0.21999 mL | 9.013 | -0.01774 | 0.89128 | 0.00093 | 16.0 s |
| 1:09:04.2 | Assay volumes | 1.50000 mL | 0.17855 mL | 0.17493 mL | 0.21999 mL | | | | | |

| | | | |
|---|---|---|---|
| Sample name: | **M11_octanol** | Experiment start time: | **3/1/2018 12:11:22 AM** |
| Assay name: | **pH-metric high logP** | Analyst: | **Pion** |
| Assay ID: | **18C-01001** | Instrument ID: | **T312060** |
| Filename: | **C:\Sirius_T3\Mehtap\20180228_exp28_logP_T3-2\18C-01001_M11_octanol_pH-metric high logP.t3r** | | |

| | | | |
|---|---|---|---|
| Sample name: | **M11_octanol** | Experiment start time: | **3/1/2018 12:11:22 AM** |
| Assay name: | **pH-metric high logP** | Analyst: | **Pion** |
| Assay ID: | **18C-01001** | Instrument ID: | **T312060** |
| Filename: | **C:\Sirius_T3\Mehtap\20180228_exp28_logP_T3-2\18C-01001_M11_octanol_pH-metric high logP.t3r** | | |

#### Assay Settings (continued)

| Setting | Value | Original Value | Date/Time changed | Imported from |
|---|---|---|---|---|
| ***Titration 3*** | | | | |
| Titrate from | Low to high pH | | | |
| Add additional water | 0.00 mL | | | |
| Additional partition solvent volume | 0.150 mL | | | |
| Additional partition solvent added | Automatic | | | |
| After pH adjust stir for | 30 seconds | | | |
| Stir to allow partitioning for | 15 seconds | | | |
| Stirrer speed for partitioning | 60% | | | |
| ***Data Point Stability*** | | | | |
| Stir during data point collection | No | | | |
| Delay before data point collection | 0 seconds | | | |
| Number of points to average | 20 points | | | |
| Time interval between points | 0.50 seconds | | | |
| Required maximum standard deviation | 0.00100 dpH/dt | | | |
| Stability timeout after | 60 seconds | | | |

#### Calibration Settings

| Setting | Value | Date/Time changed | Imported from |
|---|---|---|---|
| Four-Plus alpha | 0.130 | 3/1/2018 12:11:22 AM | C:\Sirius_T3\HCl18B27.t3r |
| Four-Plus S | 0.9970 | 3/1/2018 12:11:22 AM | C:\Sirius_T3\HCl18B27.t3r |
| Four-Plus jH | 0.8 | 3/1/2018 12:11:22 AM | C:\Sirius_T3\HCl18B27.t3r |
| Four-Plus jOH | -0.4 | 3/1/2018 12:11:22 AM | C:\Sirius_T3\HCl18B27.t3r |
| Base concentration factor | 1.000 | 3/1/2018 12:11:22 AM | C:\Sirius_T3\KOH18B27.t3r |
| Acid concentration factor | 0.994 | 3/1/2018 12:11:22 AM | C:\Sirius_T3\HCl18B27.t3r |

#### Instrument Settings

| Setting | Value | Batch Id | Install date |
|---|---|---|---|
| Instrument owner | Merck | | |
| Instrument ID | T312060 | | |
| Instrument type | T3 Simulator | | |
| Software version | 1.1.3.0 | | |
| Dispenser module | | T3DM1200361 | 3/31/2009 5:24:52 AM |
| Dispenser 0 | Water | | 3/31/2009 5:25:05 AM |
| Syringe volume | 2.5 mL | | |
| Firmware version | 1.2.1(r2) | | |
| Titrant | Water (0.15 M KCl) | 02-06-2018 | 2/27/2018 10:05:59 AM |
| Dispenser 2 | Acid | | 3/31/2009 5:25:11 AM |
| Syringe volume | 0.5 mL | | |
| Firmware version | 1.2.1(r2) | | |
| Titrant | Acid (0.5 M HCl) | 02-27-2018 | 2/27/2018 10:27:22 AM |
| Dispenser 1 | Base | | 3/31/2009 5:25:21 AM |
| Syringe volume | 0.5 mL | | |
| Firmware version | 1.2.1(r2) | | |
| Titrant | Base (0.5 M KOH) | 9/22/2017 | 2/27/2018 10:21:22 AM |
| Dispenser 5 | Cosolvent | | 3/31/2009 5:26:24 AM |
| Syringe volume | 2.5 mL | | |
| Firmware version | 1.2.1(r2) | | |
| Distribution valve 5 | Distribution Valve | | 3/31/2009 5:28:19 AM |
| Firmware version | 1.1.3 | | |
| Port A | Methanol (80%, 0.15 M KCl) | 09-26-17 | 2/7/2018 9:42:01 AM |
| Port B | Cyclohexane | 11-01-17 | 2/27/2018 10:37:57 AM |
| Dispenser 3 | Buffer | | 8/3/2010 5:05:16 AM |
| Syringe volume | 0.5 mL | | |
| Firmware version | 1.2.1(r2) | | |
| Titrant | Dodecane | 2018/01/31 | 2/28/2018 10:18:04 AM |
| Dispenser 6 | Octanol | | 10/22/2010 10:52:43 AM |

| | | | |
|---|---|---|---|
| Sample name: | **M11_octanol** | Experiment start time: | **3/1/2018 12:11:22 AM** |
| Assay name: | **pH-metric high logP** | Analyst: | **Pion** |
| Assay ID: | **18C-01001** | Instrument ID: | **T312060** |
| Filename: | **C:\Sirius_T3\Mehtap\20180228_exp28_logP_T3-2\18C-01001_M11_octanol_pH-metric high logP.t3r** | | |

Sample name: **M11_octanol**
Assay name: **pH-metric high logP**
Assay ID: **18C-01001**
Filename: **C:\Sirius_T3\Mehtap\20180228_exp28_logP_T3-2\18C-01001_M11_octanol_pH-metric high logP.t3r**
Experiment start time: **3/1/2018 12:11:22 AM**
Analyst: **Pion**
Instrument ID: **T312060**

#### Instrument Settings (continued)

| Setting | Value | Batch Id | Install date |
|---|---|---|---|
| Spectrometer calibration stir duration | 5 s | | |
| Spectrometer calibration stir speed | 30% | | |
| Spectrometer calibration wash pump volume | 20.0 mL | | |
| Spectrometer calibration wash stir duration | 5 s | | |
| Spectrometer calibration wash stir speed | 30% | | |
| Overhead dispense height | 10000 | | |

#### Refinement Settings

| Setting | Value | Default value |
|---|---|---|
| Turbidity detection method | None | None |
| Turbidity wavelength to assess | 500.0 nm | 500.0 nm |
| Turbidity maximum absorbance | 0.100 | 0.100 |
| Turbidity probe threshold | 50.00 | 50.00 |

#### Experiment Log

[1:49] Air gap created for Water (0.15 M KCl)
[1:49] Air gap created for Acid (0.5 M HCl)
[1:50] Air gap created for Base (0.5 M KOH)
[1:50] Air gap released for Water (0.15 M KCl)
[1:54] Titrator arm moved over Titration position
[1:54] Titration 1 of 3
[1:54] Adding initial titrants
[1:54] Automatically add 1.50000 mL of water
[2:19] Dispensed 1.500000 mL of Water (0.15 M KCl)
[2:23] Titrator arm moved over Drain
[8:05] Titrator arm moved to Titration position
[8:05] Argon flow rate set to 100
[8:05] Stirrer speed set to 10
[8:10] Automatically add 0.02000 mL of Octanol
[8:10] Dispensed 0.019991 mL of Octanol
[8:11] Initial pH = 9.64
[8:11] Iterative adjust 9.64 -> 2.00
[8:11] pH 9.64 -> 2.00
[8:13] Air gap released for Acid (0.5 M HCl)
[8:14] Dispensed 0.054069 mL of Acid (0.5 M HCl)
[8:19] Holding pH 2.00
[10:19] Stirrer speed set to 0
[10:19] Stirrer speed set to 50
[10:19] Iterative adjust 1.97 -> 2.00
[10:19] pH 1.97 -> 2.00
[10:20] Air gap released for Base (0.5 M KOH)
[10:21] Dispensed 0.003010 mL of Base (0.5 M KOH)
[11:11] Stirrer speed set to 0
[11:21] Datapoint id 1 collected
[11:21] Stirrer speed set to 50
[11:26] pH 2.02 -> 2.22
[11:26] Using cautious pH adjust
[11:26] Dispensed 0.007643 mL of Base (0.5 M KOH)
[11:32] Stepping pH = 2.11
[11:32] Dispensed 0.005833 mL of Base (0.5 M KOH)
[11:37] Stepping pH = 2.20
[11:37] Dispensed 0.000964 mL of Base (0.5 M KOH)
[11:42] Stepping pH = 2.22
[11:57] Stirrer speed set to 0
[12:07] Datapoint id 2 collected
[12:07] Charge balance equation is out by 5.6%
[12:07] Stirrer speed set to 50

Sample name: **M11_octanol**
Assay name: **pH-metric high logP**
Assay ID: **18C-01001**
Filename: **C:\Sirius_T3\Mehtap\20180228_exp28_logP_T3-2\18C-01001_M11_octanol_pH-metric high logP.t3r**

Experiment start time: **3/1/2018 12:11:22 AM**
Analyst: **Pion**
Instrument ID: **T312060**

#### Experiment Log (continued)

[12:12] pH 2.23 -> 2.43
[12:12] Using charge balance adjust
[12:13] Dispensed 0.009572 mL of Base (0.5 M KOH)
[12:33] Stirrer speed set to 0
[12:43] Datapoint id 3 collected
[12:43] Charge balance equation is out by 3.5%
[12:43] Stirrer speed set to 50
[12:48] pH 2.44 -> 2.64
[12:48] Using charge balance adjust
[12:48] Dispensed 0.006091 mL of Base (0.5 M KOH)
[13:09] Stirrer speed set to 0
[13:19] Datapoint id 4 collected
[13:19] Charge balance equation is out by -5.3%
[13:19] Stirrer speed set to 50
[13:24] pH 2.64 -> 2.84
[13:24] Using charge balance adjust
[13:24] Dispensed 0.004210 mL of Base (0.5 M KOH)
[13:45] Stirrer speed set to 0
[13:55] Datapoint id 5 collected
[13:55] Charge balance equation is out by 10.7%
[13:55] Stirrer speed set to 50
[14:00] pH 2.87 -> 3.07
[14:00] Using charge balance adjust
[14:00] Dispensed 0.003010 mL of Base (0.5 M KOH)
[14:21] Stirrer speed set to 0
[14:31] Datapoint id 6 collected
[14:31] Charge balance equation is out by -4.5%
[14:31] Stirrer speed set to 50
[14:36] pH 3.06 -> 3.26
[14:36] Using charge balance adjust
[14:36] Dispensed 0.002469 mL of Base (0.5 M KOH)
[14:56] Stirrer speed set to 0
[15:06] Datapoint id 7 collected
[15:06] Charge balance equation is out by 4.2%
[15:06] Stirrer speed set to 50
[15:11] pH 3.27 -> 3.47
[15:11] Using charge balance adjust
[15:11] Dispensed 0.002164 mL of Base (0.5 M KOH)
[15:32] Stirrer speed set to 0
[15:42] Datapoint id 8 collected
[15:42] Charge balance equation is out by 0.1%
[15:42] Stirrer speed set to 50
[15:47] pH 3.48 -> 3.68
[15:47] Using charge balance adjust
[15:47] Dispensed 0.001952 mL of Base (0.5 M KOH)
[16:07] Stirrer speed set to 0
[16:18] Datapoint id 9 collected
[16:18] Charge balance equation is out by -1.5%
[16:18] Stirrer speed set to 50
[16:23] pH 3.67 -> 3.87
[16:23] Using charge balance adjust
[16:23] Dispensed 0.001717 mL of Base (0.5 M KOH)
[16:43] Stirrer speed set to 0
[16:53] Datapoint id 10 collected
[16:53] Charge balance equation is out by -16.9%
[16:53] Stirrer speed set to 50
[16:58] pH 3.84 -> 4.04
[16:58] Using cautious pH adjust
[16:58] Dispensed 0.000729 mL of Base (0.5 M KOH)

| | | | |
|---|---|---|---|
| Sample name: | **M11_octanol** | Experiment start time: | **3/1/2018 12:11:22 AM** |
| Assay name: | **pH-metric high logP** | Analyst: | **Pion** |
| Assay ID: | **18C-01001** | Instrument ID: | **T312060** |
| Filename: | **C:\Sirius_T3\Mehtap\20180228_exp28_logP_T3-2\18C-01001_M11_octanol_pH-metric high logP.t3r** | | |

#### Experiment Log (continued)

[17:03] Stepping pH = 3.91
[17:03] Dispensed 0.000823 mL of Base (0.5 M KOH)
[17:08] Stepping pH = 4.01
[17:09] Dispensed 0.000165 mL of Base (0.5 M KOH)
[17:14] Stepping pH = 4.04
[17:29] Stirrer speed set to 0
[17:39] Datapoint id 11 collected
[17:39] Charge balance equation is out by -17.2%
[17:39] Stirrer speed set to 50
[17:44] pH 4.03 -> 4.23
[17:44] Using cautious pH adjust
[17:44] Dispensed 0.000588 mL of Base (0.5 M KOH)
[17:49] Stepping pH = 4.12
[17:49] Dispensed 0.000447 mL of Base (0.5 M KOH)
[17:54] Stepping pH = 4.22
[17:54] Dispensed 0.000047 mL of Base (0.5 M KOH)
[17:59] Stepping pH = 4.24
[18:14] Stirrer speed set to 0
[18:25] Datapoint id 12 collected
[18:25] Charge balance equation is out by 7.9%
[18:25] Stirrer speed set to 50
[18:30] pH 4.24 -> 4.44
[18:30] Using charge balance adjust
[18:30] Dispensed 0.000870 mL of Base (0.5 M KOH)
[18:50] Stirrer speed set to 0
[19:01] Datapoint id 13 collected
[19:01] Charge balance equation is out by 17.3%
[19:01] Stirrer speed set to 50
[19:06] pH 4.47 -> 4.67
[19:06] Using cautious pH adjust
[19:06] Dispensed 0.000282 mL of Base (0.5 M KOH)
[19:11] Stepping pH = 4.56
[19:11] Dispensed 0.000235 mL of Base (0.5 M KOH)
[19:16] Stepping pH = 4.67
[19:31] Stirrer speed set to 0
[19:41] Datapoint id 14 collected
[19:41] Charge balance equation is out by 9.3%
[19:41] Stirrer speed set to 50
[19:46] pH 4.69 -> 4.89
[19:46] Using charge balance adjust
[19:46] Dispensed 0.000376 mL of Base (0.5 M KOH)
[20:06] Stirrer speed set to 0
[20:17] Datapoint id 15 collected
[20:17] Charge balance equation is out by 58.7%
[20:17] Stirrer speed set to 50
[20:23] pH 5.01 -> 5.21
[20:23] Using cautious pH adjust
[20:23] Dispensed 0.000094 mL of Base (0.5 M KOH)
[20:28] Stepping pH = 5.02
[20:28] Dispensed 0.000282 mL of Base (0.5 M KOH)
[20:33] Stepping pH = 5.74
[20:48] Stirrer speed set to 0
[21:07] Datapoint id 16 collected
[21:07] Charge balance equation is out by -91.3%
[21:07] Stirrer speed set to 50
[21:12] pH 5.85 -> 6.05
[21:12] Using cautious pH adjust
[21:12] Dispensed 0.000024 mL of Base (0.5 M KOH)
[21:17] Stepping pH = 5.87

| | | | |
|---|---|---|---|
| Sample name: | **M11_octanol** | Experiment start time: | **3/1/2018 12:11:22 AM** |
| Assay name: | **pH-metric high logP** | Analyst: | **Pion** |
| Assay ID: | **18C-01001** | Instrument ID: | **T312060** |
| Filename: | **C:\Sirius_T3\Mehtap\20180228_exp28_logP_T3-2\18C-01001_M11_octanol_pH-metric high logP.t3r** | | |

#### Experiment Log (continued)

[21:17] Dispensed 0.000094 mL of Base (0.5 M KOH)
[21:22] Stepping pH = 6.25
[21:38] Stirrer speed set to 0
[22:20] Datapoint id 17 collected
[22:20] Charge balance equation is out by -85.8%
[22:20] Stirrer speed set to 50
[22:25] pH 6.40 -> 6.60
[22:25] Using cautious pH adjust
[22:25] Dispensed 0.000024 mL of Base (0.5 M KOH)
[22:30] Stepping pH = 6.43
[22:30] Dispensed 0.000047 mL of Base (0.5 M KOH)
[22:36] Stepping pH = 6.55
[22:36] Dispensed 0.000024 mL of Base (0.5 M KOH)
[22:41] Stepping pH = 6.90
[22:56] Stirrer speed set to 0
[23:56] Datapoint id 18 collected
[23:56] Charge balance equation is out by -113.3%
[23:56] Stirrer speed set to 50
[24:01] pH 7.16 -> 7.36
[24:01] Using cautious pH adjust
[24:01] Dispensed 0.000024 mL of Base (0.5 M KOH)
[24:06] Stepping pH = 7.21
[24:06] Dispensed 0.000024 mL of Base (0.5 M KOH)
[24:11] Stepping pH = 7.34
[24:11] Dispensed 0.000024 mL of Base (0.5 M KOH)
[24:16] Stepping pH = 7.64
[24:31] Stirrer speed set to 0
[25:31] Datapoint id 19 collected
[25:31] Charge balance equation is out by -330.3%
[25:31] Stirrer speed set to 50
[25:37] pH 7.74 -> 7.94
[25:37] Using cautious pH adjust
[25:37] Dispensed 0.000024 mL of Base (0.5 M KOH)
[25:42] Stepping pH = 7.80
[25:42] Dispensed 0.000024 mL of Base (0.5 M KOH)
[25:47] Stepping pH = 8.00
[26:02] Stirrer speed set to 0
[27:02] Datapoint id 20 collected
[27:02] Charge balance equation is out by -530.4%
[27:02] Stirrer speed set to 50
[27:07] pH 8.20 -> 8.40
[27:07] Using cautious pH adjust
[27:07] Dispensed 0.000024 mL of Base (0.5 M KOH)
[27:12] Stepping pH = 8.24
[27:12] Dispensed 0.000024 mL of Base (0.5 M KOH)
[27:17] Stepping pH = 8.38
[27:17] Dispensed 0.000024 mL of Base (0.5 M KOH)
[27:22] Stepping pH = 8.62
[27:38] Stirrer speed set to 0
[28:17] Datapoint id 21 collected
[28:17] Charge balance equation is out by -617.9%
[28:17] Stirrer speed set to 50
[28:22] pH 8.66 -> 8.86
[28:22] Using cautious pH adjust
[28:22] Dispensed 0.000024 mL of Base (0.5 M KOH)
[28:27] Stepping pH = 8.68
[28:27] Dispensed 0.000024 mL of Base (0.5 M KOH)
[28:32] Stepping pH = 8.76
[28:32] Dispensed 0.000024 mL of Base (0.5 M KOH)

Sample name: **M11_octanol** Experiment start time: **3/1/2018 12:11:22 AM**
Assay name: **pH-metric high logP** Analyst: **Pion**
Assay ID: **18C-01001** Instrument ID: **T312060**
Filename: **C:\Sirius_T3\Mehtap\20180228_exp28_logP_T3-2\18C-01001_M11_octanol_pH-metric high logP.t3r**

#### Experiment Log (continued)

[28:37] Stepping pH = 8.92
[28:52] Stirrer speed set to 0
[29:22] Datapoint id 22 collected
[29:22] Charge balance equation is out by -250.7%
[29:22] Stirrer speed set to 50
[29:27] pH 8.95 -> 9.05
[29:27] Using cautious pH adjust
[29:27] Dispensed 0.000024 mL of Base (0.5 M KOH)
[29:32] Stepping pH = 8.95
[29:32] Dispensed 0.000024 mL of Base (0.5 M KOH)
[29:37] Stepping pH = 9.06
[29:52] Stirrer speed set to 0
[30:15] Datapoint id 23 collected
[30:15] Charge balance equation is out by -155.3%
[30:15] Titration 2 of 3
[30:15] Adding initial titrants
[30:15] Automatically add 0.05000 mL of Octanol
[30:16] Dispensed 0.050000 mL of Octanol
[30:16] Stirrer speed set to 10
[30:17] Stirrer speed set to 55
[30:17] Iterative adjust 9.09 -> 2.00
[30:17] pH 9.09 -> 2.00
[30:19] Dispensed 0.056421 mL of Acid (0.5 M HCl)
[30:24] pH 2.03 -> 2.00
[30:24] Dispensed 0.002869 mL of Acid (0.5 M HCl)
[31:14] Stirrer speed set to 0
[31:24] Datapoint id 24 collected
[31:24] Stirrer speed set to 55
[31:29] pH 1.97 -> 2.17
[31:29] Using cautious pH adjust
[31:30] Dispensed 0.009313 mL of Base (0.5 M KOH)
[31:35] Stepping pH = 2.07
[31:35] Dispensed 0.006303 mL of Base (0.5 M KOH)
[31:40] Stepping pH = 2.14
[31:40] Dispensed 0.002211 mL of Base (0.5 M KOH)
[31:45] Stepping pH = 2.17
[32:00] Stirrer speed set to 0
[32:10] Datapoint id 25 collected
[32:10] Charge balance equation is out by 4.3%
[32:10] Stirrer speed set to 55
[32:16] pH 2.18 -> 2.38
[32:16] Using charge balance adjust
[32:16] Dispensed 0.011736 mL of Base (0.5 M KOH)
[32:36] Stirrer speed set to 0
[32:46] Datapoint id 26 collected
[32:46] Charge balance equation is out by 10.1%
[32:46] Stirrer speed set to 55
[32:51] pH 2.41 -> 2.61
[32:51] Using charge balance adjust
[32:52] Dispensed 0.007385 mL of Base (0.5 M KOH)
[33:12] Stirrer speed set to 0
[33:22] Datapoint id 27 collected
[33:22] Charge balance equation is out by -6.9%
[33:22] Stirrer speed set to 55
[33:27] pH 2.60 -> 2.80
[33:27] Using charge balance adjust
[33:27] Dispensed 0.005268 mL of Base (0.5 M KOH)
[33:47] Stirrer speed set to 0
[33:57] Datapoint id 28 collected

Sample name: **M11_octanol** Experiment start time: **3/1/2018 12:11:22 AM**
Assay name: **pH-metric high logP** Analyst: **Pion**
Assay ID: **18C-01001** Instrument ID: **T312060**
Filename: **C:\Sirius_T3\Mehtap\20180228_exp28_logP_T3-2\18C-01001_M11_octanol_pH-metric high logP.t3r**

#### Experiment Log (continued)

[33:57] Charge balance equation is out by -4.7%
[33:57] Stirrer speed set to 55
[34:02] pH 2.80 -> 3.00
[34:02] Using charge balance adjust
[34:03] Dispensed 0.003951 mL of Base (0.5 M KOH)
[34:23] Stirrer speed set to 0
[34:33] Datapoint id 29 collected
[34:33] Charge balance equation is out by 5.7%
[34:33] Stirrer speed set to 55
[34:38] pH 3.01 -> 3.21
[34:38] Using charge balance adjust
[34:38] Dispensed 0.003057 mL of Base (0.5 M KOH)
[34:58] Stirrer speed set to 0
[35:08] Datapoint id 30 collected
[35:08] Charge balance equation is out by 7.6%
[35:08] Stirrer speed set to 55
[35:13] pH 3.23 -> 3.43
[35:13] Using charge balance adjust
[35:14] Dispensed 0.002399 mL of Base (0.5 M KOH)
[35:34] Stirrer speed set to 0
[35:44] Datapoint id 31 collected
[35:44] Charge balance equation is out by -7.7%
[35:44] Stirrer speed set to 55
[35:49] pH 3.42 -> 3.62
[35:49] Using charge balance adjust
[35:49] Dispensed 0.001929 mL of Base (0.5 M KOH)
[36:09] Stirrer speed set to 0
[36:19] Datapoint id 32 collected
[36:19] Charge balance equation is out by 4.5%
[36:19] Stirrer speed set to 55
[36:24] pH 3.63 -> 3.83
[36:24] Using charge balance adjust
[36:25] Dispensed 0.001435 mL of Base (0.5 M KOH)
[36:45] Stirrer speed set to 0
[36:55] Datapoint id 33 collected
[36:55] Charge balance equation is out by 14.8%
[36:55] Stirrer speed set to 55
[37:00] pH 3.87 -> 4.07
[37:00] Using charge balance adjust
[37:00] Dispensed 0.000988 mL of Base (0.5 M KOH)
[37:20] Stirrer speed set to 0
[37:30] Datapoint id 34 collected
[37:30] Charge balance equation is out by 25.2%
[37:30] Stirrer speed set to 55
[37:35] pH 4.13 -> 4.33
[37:35] Using cautious pH adjust
[37:35] Dispensed 0.000306 mL of Base (0.5 M KOH)
[37:40] Stepping pH = 4.21
[37:41] Dispensed 0.000235 mL of Base (0.5 M KOH)
[37:46] Stepping pH = 4.32
[38:01] Stirrer speed set to 0
[38:11] Datapoint id 35 collected
[38:11] Charge balance equation is out by 9.9%
[38:11] Stirrer speed set to 55
[38:16] pH 4.33 -> 4.53
[38:16] Using charge balance adjust
[38:16] Dispensed 0.000400 mL of Base (0.5 M KOH)
[38:36] Stirrer speed set to 0
[38:46] Datapoint id 36 collected

Sample name: **M11_octanol** Experiment start time: **3/1/2018 12:11:22 AM**
Assay name: **pH-metric high logP** Analyst: **Pion**
Assay ID: **18C-01001** Instrument ID: **T312060**
Filename: **C:\Sirius_T3\Mehtap\20180228_exp28_logP_T3-2\18C-01001_M11_octanol_pH-metric high logP.t3r**

#### Experiment Log (continued)

[38:46] Charge balance equation is out by 23.7%
[38:46] Stirrer speed set to 55
[38:51] pH 4.58 -> 4.78
[38:51] Using cautious pH adjust
[38:51] Dispensed 0.000118 mL of Base (0.5 M KOH)
[38:56] Stepping pH = 4.62
[38:56] Dispensed 0.000212 mL of Base (0.5 M KOH)
[39:02] Stepping pH = 4.90
[39:17] Stirrer speed set to 0
[39:30] Datapoint id 37 collected
[39:30] Charge balance equation is out by -32.8%
[39:30] Stirrer speed set to 55
[39:35] pH 4.93 -> 5.13
[39:35] Using cautious pH adjust
[39:35] Dispensed 0.000071 mL of Base (0.5 M KOH)
[39:40] Stepping pH = 4.94
[39:40] Dispensed 0.000165 mL of Base (0.5 M KOH)
[39:45] Stepping pH = 5.45
[40:00] Stirrer speed set to 0
[40:16] Datapoint id 38 collected
[40:16] Charge balance equation is out by -85.1%
[40:16] Stirrer speed set to 55
[40:21] pH 5.53 -> 5.73
[40:21] Using cautious pH adjust
[40:21] Dispensed 0.000024 mL of Base (0.5 M KOH)
[40:26] Stepping pH = 5.56
[40:27] Dispensed 0.000047 mL of Base (0.5 M KOH)
[40:32] Stepping pH = 5.61
[40:32] Dispensed 0.000071 mL of Base (0.5 M KOH)
[40:37] Stepping pH = 6.14
[40:52] Stirrer speed set to 0
[41:28] Datapoint id 39 collected
[41:28] Charge balance equation is out by -212.4%
[41:28] Stirrer speed set to 55
[41:34] pH 6.23 -> 6.43
[41:34] Using cautious pH adjust
[41:34] Dispensed 0.000024 mL of Base (0.5 M KOH)
[41:39] Stepping pH = 6.29
[41:39] Dispensed 0.000024 mL of Base (0.5 M KOH)
[41:44] Stepping pH = 6.37
[41:44] Dispensed 0.000024 mL of Base (0.5 M KOH)
[41:49] Stepping pH = 6.60
[42:04] Stirrer speed set to 0
[43:04] Datapoint id 40 collected
[43:04] Charge balance equation is out by -59.8%
[43:04] Stirrer speed set to 55
[43:09] pH 6.71 -> 6.91
[43:09] Using cautious pH adjust
[43:09] Dispensed 0.000024 mL of Base (0.5 M KOH)
[43:14] Stepping pH = 6.85
[43:14] Dispensed 0.000024 mL of Base (0.5 M KOH)
[43:19] Stepping pH = 7.05
[43:35] Stirrer speed set to 0
[44:35] Datapoint id 41 collected
[44:35] Charge balance equation is out by -35.1%
[44:35] Stirrer speed set to 55
[44:40] pH 6.87 -> 7.07
[44:40] Using cautious pH adjust
[44:40] Dispensed 0.000024 mL of Base (0.5 M KOH)

| | | | |
|---|---|---|---|
| Sample name: | **M11_octanol** | Experiment start time: | **3/1/2018 12:11:22 AM** |
| Assay name: | **pH-metric high logP** | Analyst: | **Pion** |
| Assay ID: | **18C-01001** | Instrument ID: | **T312060** |
| Filename: | **C:\Sirius_T3\Mehtap\20180228_exp28_logP_T3-2\18C-01001_M11_octanol_pH-metric high logP.t3r** | | |

#### Experiment Log (continued)

[44:45] Stepping pH = 6.82
[44:45] Dispensed 0.000071 mL of Base (0.5 M KOH)
[44:50] Stepping pH = 7.44
[45:05] Stirrer speed set to 0
[46:05] Datapoint id 42 collected
[46:05] Charge balance equation is out by -316.2%
[46:05] Stirrer speed set to 55
[46:10] pH 7.84 -> 8.04
[46:10] Using cautious pH adjust
[46:10] Dispensed 0.000024 mL of Base (0.5 M KOH)
[46:15] Stepping pH = 7.89
[46:15] Dispensed 0.000024 mL of Base (0.5 M KOH)
[46:20] Stepping pH = 8.09
[46:36] Stirrer speed set to 0
[47:36] Datapoint id 43 collected
[47:36] Charge balance equation is out by -492.9%
[47:36] Stirrer speed set to 55
[47:41] pH 8.08 -> 8.28
[47:41] Using cautious pH adjust
[47:41] Dispensed 0.000024 mL of Base (0.5 M KOH)
[47:46] Stepping pH = 8.16
[47:46] Dispensed 0.000024 mL of Base (0.5 M KOH)
[47:51] Stepping pH = 8.30
[48:06] Stirrer speed set to 0
[49:01] Datapoint id 44 collected
[49:01] Charge balance equation is out by -412.6%
[49:01] Stirrer speed set to 55
[49:06] pH 8.32 -> 8.52
[49:06] Using cautious pH adjust
[49:06] Dispensed 0.000024 mL of Base (0.5 M KOH)
[49:11] Stepping pH = 8.36
[49:11] Dispensed 0.000024 mL of Base (0.5 M KOH)
[49:17] Stepping pH = 8.42
[49:17] Dispensed 0.000024 mL of Base (0.5 M KOH)
[49:22] Stepping pH = 8.50
[49:22] Dispensed 0.000024 mL of Base (0.5 M KOH)
[49:27] Stepping pH = 8.61
[49:42] Stirrer speed set to 0
[50:13] Datapoint id 45 collected
[50:13] Charge balance equation is out by -623.6%
[50:13] Stirrer speed set to 55
[50:18] pH 8.59 -> 8.79
[50:18] Using cautious pH adjust
[50:18] Dispensed 0.000024 mL of Base (0.5 M KOH)
[50:23] Stepping pH = 8.61
[50:23] Dispensed 0.000024 mL of Base (0.5 M KOH)
[50:28] Stepping pH = 8.66
[50:28] Dispensed 0.000047 mL of Base (0.5 M KOH)
[50:34] Stepping pH = 8.69
[50:34] Dispensed 0.000047 mL of Base (0.5 M KOH)
[50:39] Stepping pH = 8.79
[50:54] Stirrer speed set to 0
[51:19] Datapoint id 46 collected
[51:19] Charge balance equation is out by -667.4%
[51:19] Stirrer speed set to 55
[51:24] pH 8.78 -> 8.98
[51:24] Using cautious pH adjust
[51:24] Dispensed 0.000024 mL of Base (0.5 M KOH)
[51:29] Stepping pH = 8.79

| | | | |
|---|---|---|---|
| Sample name: | **M11_octanol** | Experiment start time: | **3/1/2018 12:11:22 AM** |
| Assay name: | **pH-metric high logP** | Analyst: | **Pion** |
| Assay ID: | **18C-01001** | Instrument ID: | **T312060** |
| Filename: | **C:\Sirius_T3\Mehtap\20180228_exp28_logP_T3-2\18C-01001_M11_octanol_pH-metric high logP.t3r** | | |

#### Experiment Log (continued)

[51:29] Dispensed 0.000047 mL of Base (0.5 M KOH)
[51:34] Stepping pH = 8.93
[51:34] Dispensed 0.000024 mL of Base (0.5 M KOH)
[51:39] Stepping pH = 9.00
[51:54] Stirrer speed set to 0
[52:14] Datapoint id 47 collected
[52:14] Charge balance equation is out by -185.9%
[52:14] Stirrer speed set to 55
[52:19] pH 8.99 -> 9.05
[52:19] Using cautious pH adjust
[52:19] Dispensed 0.000024 mL of Base (0.5 M KOH)
[52:24] Stepping pH = 9.00
[52:39] Stirrer speed set to 0
[52:55] Datapoint id 48 collected
[52:55] Charge balance equation is out by -80.3%
[52:55] Titration 3 of 3
[52:55] Adding initial titrants
[52:55] Automatically add 0.15000 mL of Octanol
[52:59] Dispensed 0.150000 mL of Octanol
[52:59] Stirrer speed set to 10
[53:00] Stirrer speed set to 60
[53:00] Iterative adjust 9.00 -> 2.00
[53:00] pH 9.00 -> 2.00
[53:02] Dispensed 0.058678 mL of Acid (0.5 M HCl)
[53:07] pH 2.06 -> 2.00
[53:07] Dispensed 0.006515 mL of Acid (0.5 M HCl)
[53:57] Stirrer speed set to 0
[54:07] Datapoint id 49 collected
[54:07] Stirrer speed set to 60
[54:12] pH 1.96 -> 2.16
[54:12] Using cautious pH adjust
[54:13] Dispensed 0.010466 mL of Base (0.5 M KOH)
[54:18] Stepping pH = 2.05
[54:18] Dispensed 0.007667 mL of Base (0.5 M KOH)
[54:23] Stepping pH = 2.13
[54:23] Dispensed 0.001858 mL of Base (0.5 M KOH)
[54:28] Stepping pH = 2.16
[54:44] Stirrer speed set to 0
[54:54] Datapoint id 50 collected
[54:54] Charge balance equation is out by 4.5%
[54:54] Stirrer speed set to 60
[54:59] pH 2.16 -> 2.36
[54:59] Using charge balance adjust
[54:59] Dispensed 0.013523 mL of Base (0.5 M KOH)
[55:19] Stirrer speed set to 0
[55:29] Datapoint id 51 collected
[55:29] Charge balance equation is out by 5.4%
[55:29] Stirrer speed set to 60
[55:35] pH 2.38 -> 2.58
[55:35] Using charge balance adjust
[55:35] Dispensed 0.008843 mL of Base (0.5 M KOH)
[55:55] Stirrer speed set to 0
[56:05] Datapoint id 52 collected
[56:05] Charge balance equation is out by -2.8%
[56:05] Stirrer speed set to 60
[56:10] pH 2.58 -> 2.78
[56:10] Using charge balance adjust
[56:10] Dispensed 0.006185 mL of Base (0.5 M KOH)
[56:31] Stirrer speed set to 0

| | | | |
|---|---|---|---|
| Sample name: | **M11_octanol** | Experiment start time: | **3/1/2018 12:11:22 AM** |
| Assay name: | **pH-metric high logP** | Analyst: | **Pion** |
| Assay ID: | **18C-01001** | Instrument ID: | **T312060** |
| Filename: | **C:\Sirius_T3\Mehtap\20180228_exp28_logP_T3-2\18C-01001_M11_octanol_pH-metric high logP.t3r** | | |

#### Experiment Log (continued)

[56:41] Datapoint id 53 collected
[56:41] Charge balance equation is out by 2.9%
[56:41] Stirrer speed set to 60
[56:46] pH 2.79 -> 2.99
[56:46] Using charge balance adjust
[56:46] Dispensed 0.004374 mL of Base (0.5 M KOH)
[57:06] Stirrer speed set to 0
[57:16] Datapoint id 54 collected
[57:16] Charge balance equation is out by 5.5%
[57:16] Stirrer speed set to 60
[57:21] pH 3.01 -> 3.21
[57:21] Using charge balance adjust
[57:21] Dispensed 0.003081 mL of Base (0.5 M KOH)
[57:42] Stirrer speed set to 0
[57:52] Datapoint id 55 collected
[57:52] Charge balance equation is out by 17.6%
[57:52] Stirrer speed set to 60
[57:57] pH 3.25 -> 3.45
[57:57] Using cautious pH adjust
[57:57] Dispensed 0.001035 mL of Base (0.5 M KOH)
[58:02] Stepping pH = 3.33
[58:02] Dispensed 0.000823 mL of Base (0.5 M KOH)
[58:07] Stepping pH = 3.42
[58:07] Dispensed 0.000235 mL of Base (0.5 M KOH)
[58:12] Stepping pH = 3.44
[58:27] Stirrer speed set to 0
[58:37] Datapoint id 56 collected
[58:37] Charge balance equation is out by -2.1%
[58:37] Stirrer speed set to 60
[58:42] pH 3.44 -> 3.64
[58:42] Using charge balance adjust
[58:43] Dispensed 0.001435 mL of Base (0.5 M KOH)
[59:03] Stirrer speed set to 0
[59:13] Datapoint id 57 collected
[59:13] Charge balance equation is out by 5.6%
[59:13] Stirrer speed set to 60
[59:18] pH 3.66 -> 3.86
[59:18] Using charge balance adjust
[59:18] Dispensed 0.000917 mL of Base (0.5 M KOH)
[59:38] Stirrer speed set to 0
[59:48] Datapoint id 58 collected
[59:48] Charge balance equation is out by 2.8%
[59:48] Stirrer speed set to 60
[59:53] pH 3.88 -> 4.08
[59:53] Using charge balance adjust
[59:53] Dispensed 0.000588 mL of Base (0.5 M KOH)
[1:00:14] Stirrer speed set to 0
[1:00:24] Datapoint id 59 collected
[1:00:24] Charge balance equation is out by 5.7%
[1:00:24] Stirrer speed set to 60
[1:00:29] pH 4.09 -> 4.29
[1:00:29] Using charge balance adjust
[1:00:29] Dispensed 0.000376 mL of Base (0.5 M KOH)
[1:00:49] Stirrer speed set to 0
[1:00:59] Datapoint id 60 collected
[1:00:59] Charge balance equation is out by 11.7%
[1:00:59] Stirrer speed set to 60
[1:01:04] pH 4.33 -> 4.53
[1:01:04] Using charge balance adjust

| | | | |
|---|---|---|---|
| Sample name: | **M11_octanol** | Experiment start time: | **3/1/2018 12:11:22 AM** |
| Assay name: | **pH-metric high logP** | Analyst: | **Pion** |
| Assay ID: | **18C-01001** | Instrument ID: | **T312060** |
| Filename: | **C:\Sirius_T3\Mehtap\20180228_exp28_logP_T3-2\18C-01001_M11_octanol_pH-metric high logP.t3r** | | |

#### Experiment Log (continued)

[1:01:04] Dispensed 0.000212 mL of Base (0.5 M KOH)
[1:01:24] Stirrer speed set to 0
[1:01:35] Datapoint id 61 collected
[1:01:35] Charge balance equation is out by -11.9%
[1:01:35] Stirrer speed set to 60
[1:01:40] pH 4.52 -> 4.72
[1:01:40] Using charge balance adjust
[1:01:40] Dispensed 0.000141 mL of Base (0.5 M KOH)
[1:02:00] Stirrer speed set to 0
[1:02:12] Datapoint id 62 collected
[1:02:12] Charge balance equation is out by -7.5%
[1:02:12] Stirrer speed set to 60
[1:02:17] pH 4.73 -> 4.93
[1:02:17] Using charge balance adjust
[1:02:17] Dispensed 0.000094 mL of Base (0.5 M KOH)
[1:02:37] Stirrer speed set to 0
[1:02:50] Datapoint id 63 collected
[1:02:50] Charge balance equation is out by -28.9%
[1:02:50] Stirrer speed set to 60
[1:02:55] pH 4.92 -> 5.12
[1:02:55] Using cautious pH adjust
[1:02:55] Dispensed 0.000024 mL of Base (0.5 M KOH)
[1:03:00] Stepping pH = 4.94
[1:03:00] Dispensed 0.000094 mL of Base (0.5 M KOH)
[1:03:05] Stepping pH = 5.24
[1:03:20] Stirrer speed set to 0
[1:03:34] Datapoint id 64 collected
[1:03:34] Charge balance equation is out by -82.8%
[1:03:34] Stirrer speed set to 60
[1:03:39] pH 5.35 -> 5.55
[1:03:39] Using cautious pH adjust
[1:03:39] Dispensed 0.000024 mL of Base (0.5 M KOH)
[1:03:44] Stepping pH = 5.38
[1:03:45] Dispensed 0.000047 mL of Base (0.5 M KOH)
[1:03:50] Stepping pH = 5.57
[1:04:05] Stirrer speed set to 0
[1:04:25] Datapoint id 65 collected
[1:04:25] Charge balance equation is out by -49.2%
[1:04:25] Stirrer speed set to 60
[1:04:30] pH 5.71 -> 5.91
[1:04:30] Using cautious pH adjust
[1:04:30] Dispensed 0.000024 mL of Base (0.5 M KOH)
[1:04:35] Stepping pH = 5.75
[1:04:36] Dispensed 0.000047 mL of Base (0.5 M KOH)
[1:04:41] Stepping pH = 6.27
[1:04:56] Stirrer speed set to 0
[1:05:51] Datapoint id 66 collected
[1:05:51] Charge balance equation is out by -57.7%
[1:05:51] Stirrer speed set to 60
[1:05:56] pH 6.32 -> 6.52
[1:05:56] Using cautious pH adjust
[1:05:56] Dispensed 0.000024 mL of Base (0.5 M KOH)
[1:06:01] Stepping pH = 6.20
[1:06:02] Dispensed 0.000188 mL of Base (0.5 M KOH)
[1:06:07] Stepping pH = 8.64
[1:06:22] Stirrer speed set to 0
[1:06:59] Datapoint id 67 collected
[1:06:59] Charge balance equation is out by -355.4%
[1:06:59] Stirrer speed set to 60

Sample name: **M11_octanol** | Experiment start time: **3/1/2018 12:11:22 AM**
Assay name: **pH-metric high logP** | Analyst: **Pion**
Assay ID: **18C-01001** | Instrument ID: **T312060**
Filename: **C:\Sirius_T3\Mehtap\20180228_exp28_logP_T3-2\18C-01001_M11_octanol_pH-metric high logP.t3r**

#### Experiment Log (continued)

[1:07:04] pH 8.65 -> 8.85
[1:07:04] Using cautious pH adjust
[1:07:04] Dispensed 0.000024 mL of Base (0.5 M KOH)
[1:07:09] Stepping pH = 8.61
[1:07:09] Dispensed 0.000071 mL of Base (0.5 M KOH)
[1:07:14] Stepping pH = 8.77
[1:07:14] Dispensed 0.000024 mL of Base (0.5 M KOH)
[1:07:19] Stepping pH = 8.81
[1:07:19] Dispensed 0.000024 mL of Base (0.5 M KOH)
[1:07:24] Stepping pH = 8.85
[1:07:40] Stirrer speed set to 0
[1:08:04] Datapoint id 68 collected
[1:08:04] Charge balance equation is out by -499.1%
[1:08:04] Stirrer speed set to 60
[1:08:09] pH 8.85 -> 9.05
[1:08:09] Using cautious pH adjust
[1:08:09] Dispensed 0.000024 mL of Base (0.5 M KOH)
[1:08:14] Stepping pH = 8.87
[1:08:14] Dispensed 0.000047 mL of Base (0.5 M KOH)
[1:08:19] Stepping pH = 8.98
[1:08:19] Dispensed 0.000024 mL of Base (0.5 M KOH)
[1:08:24] Stepping pH = 9.02
[1:08:39] Stirrer speed set to 0
[1:08:55] Datapoint id 69 collected
[1:08:55] Charge balance equation is out by -147.2%
[1:08:55] Argon flow rate set to 0
[1:08:59] Titrator arm moved over Titration position
