## Supplementary material for "Octanol-water partition coefficient measurements for the SAMPL6 Blind Prediction Challenge": SM11_18C-01002_M11_octanol_pH-metric high logP_report.pdf

| | | | |
|---|---|---|---|
| Sample name: | **M11_octanol** | Experiment start time: | **3/1/2018 1:21:08 AM** |
| Assay name: | **pH-metric high logP** | Analyst: | **Pion** |
| Assay ID: | **18C-01002** | Instrument ID: | **T312060** |
| Filename: | **C:\Sirius_T3\Mehtap\20180228_exp28_logP_T3-2\18C-01002_M11_octanol_pH-metric high logP.t3r** | | |

### pH-metric Result

| | | |
|---|---|---|
| logP (XH +) | -3.66 | ±2.11 (n=50) |
| logP (neutral X) | 2.01 | ±0.02 (n=50) |

### 18C-01002 Points 1 to 28

| | |
|---|---|
| M11_octanol concentration factor | 0.940 |
| Carbonate | 0.0379 mM |
| Acidity error | 0.00456 mM |

### 18C-01002 Points 29 to 49

| | |
|---|---|
| M11_octanol concentration factor | 0.777 |
| Carbonate | 0.1154 mM |
| Acidity error | -0.46703 mM |

### 18C-01002 Points 50 to 75

| | |
|---|---|
| M11_octanol concentration factor | 0.713 |
| Carbonate | 0.1025 mM |
| Acidity error | -1.33190 mM |

### Warnings and errors

| | |
|---|---|
| Errors | None |
| Warnings | One or more logP values out of range |

### Sample logD and percent species

| pH | M11_octanol logD | M11_octanol M11_octanolH | M11_octanol M11_octanol | M11_octanol M11_octanolH* | M11_octanol M11_octanol* | Comment |
|---|---|---|---|---|---|---|
| 1.000 | -0.88 | 88.32 % | 0.11 % | 0.02 % | 11.55 % | |
| 1.200 | -0.68 | 82.68 % | 0.17 % | 0.02 % | 17.13 % | Stomach pH |
| 2.000 | 0.11 | 43.10 % | 0.56 % | 0.01 % | 56.34 % | |
| 3.000 | 1.06 | 7.04 % | 0.91 % | 0.00 % | 92.05 % | |
| 4.000 | 1.76 | 0.75 % | 0.97 % | 0.00 % | 98.28 % | |
| 5.000 | 1.97 | 0.08 % | 0.98 % | 0.00 % | 98.95 % | |
| 6.000 | 2.00 | 0.01 % | 0.98 % | 0.00 % | 99.02 % | |
| 6.500 | 2.01 | 0.00 % | 0.98 % | 0.00 % | 99.02 % | |
| 7.000 | 2.01 | 0.00 % | 0.98 % | 0.00 % | 99.02 % | |
| 7.400 | 2.01 | 0.00 % | 0.98 % | 0.00 % | 99.02 % | Blood pH |
| 8.000 | 2.01 | 0.00 % | 0.98 % | 0.00 % | 99.02 % | |
| 9.000 | 2.01 | 0.00 % | 0.98 % | 0.00 % | 99.02 % | |
| 10.000 | 2.01 | 0.00 % | 0.98 % | 0.00 % | 99.02 % | |
| 11.000 | 2.01 | 0.00 % | 0.98 % | 0.00 % | 99.02 % | |
| 12.000 | 2.01 | 0.00 % | 0.98 % | 0.00 % | 99.02 % | |

Sample name: **M11_octanol**
Assay name: **pH-metric high logP**
Assay ID: **18C-01002**
Filename: **C:\Sirius_T3\Mehtap\20180228_exp28_logP_T3-2\18C-01002_M11_octanol_pH-metric high logP.t3r**

Experiment start time: **3/1/2018 1:21:08 AM**
Analyst: **Pion**
Instrument ID: **T312060**

### Graphs

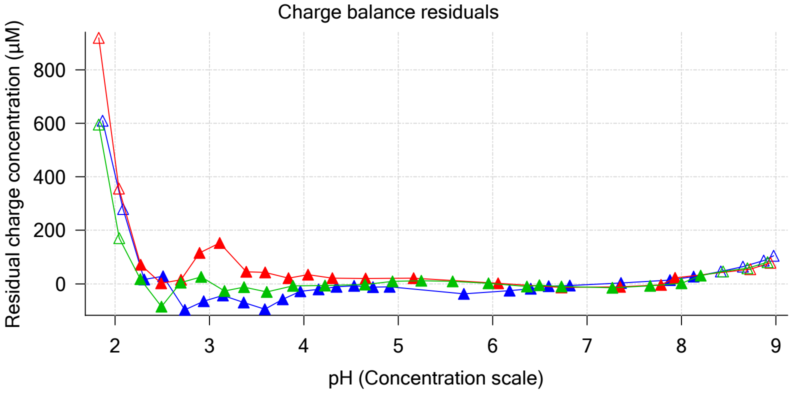


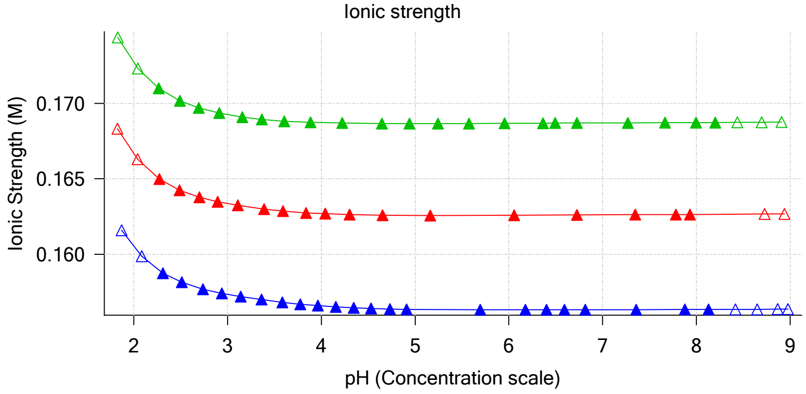


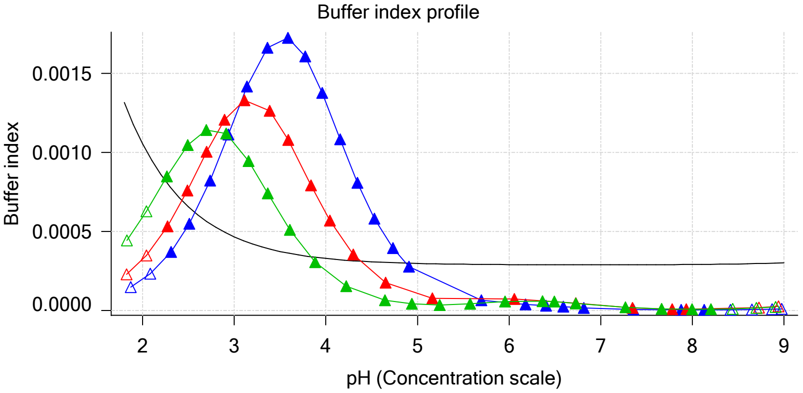


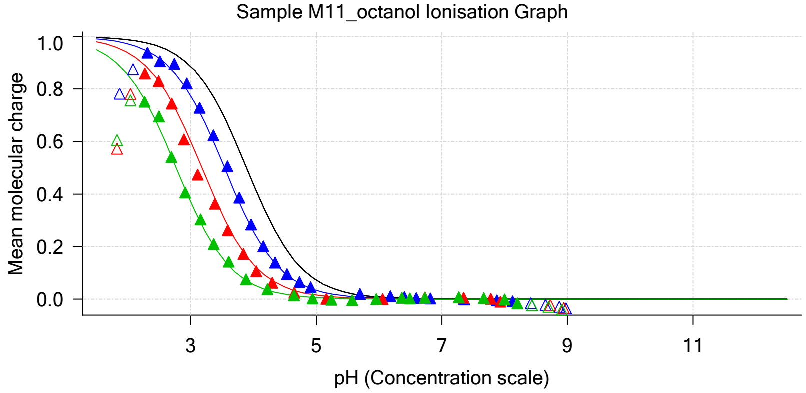


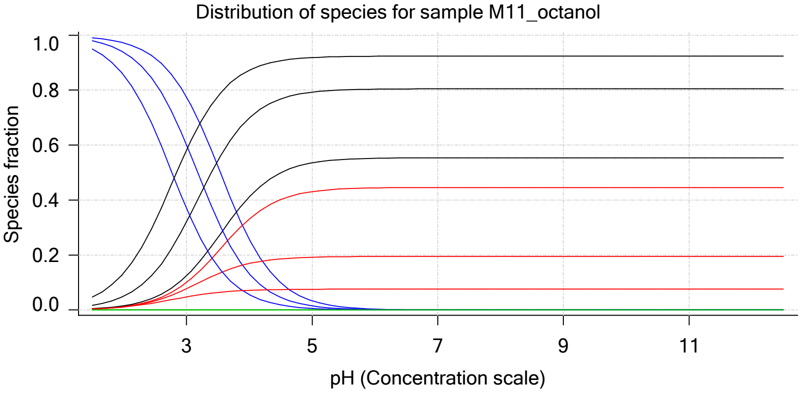


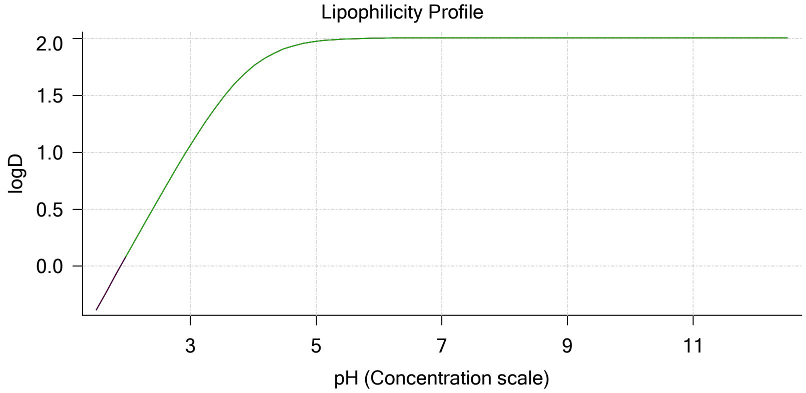


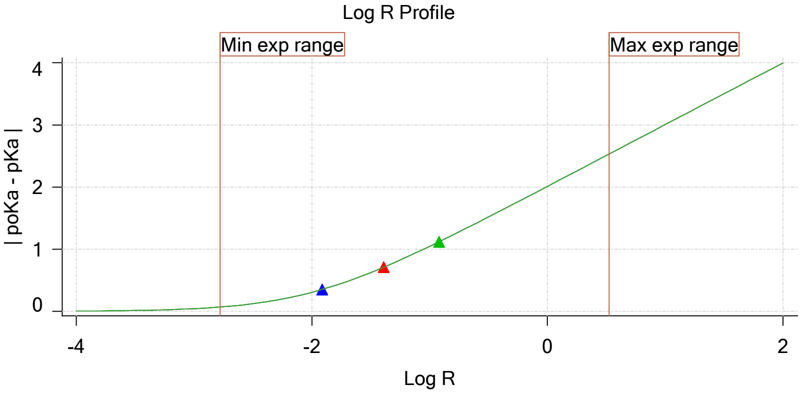


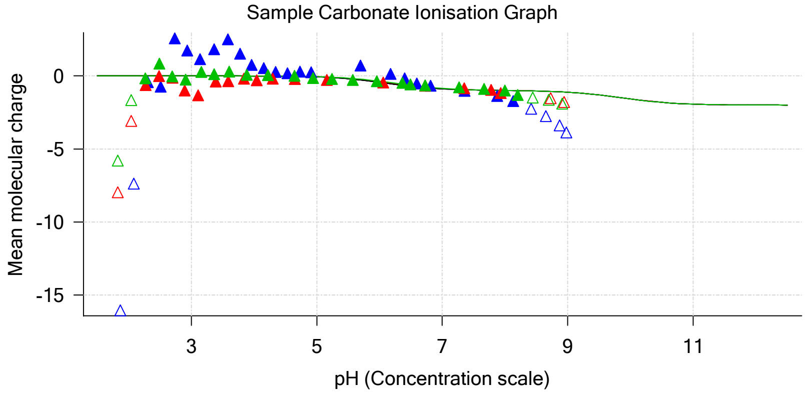

| | | | |
|---|---|---|---|
| Sample name: | **M11_octanol** | Experiment start time: | **3/1/2018 1:21:08 AM** |
| Assay name: | **pH-metric high logP** | Analyst: | **Pion** |
| Assay ID: | **18C-01002** | Instrument ID: | **T312060** |
| Filename: | **C:\Sirius_T3\Mehtap\20180228_exp28_logP_T3-2\18C-01002_M11_octanol_pH-metric high logP.t3r** | | |

### Graphs (continued)

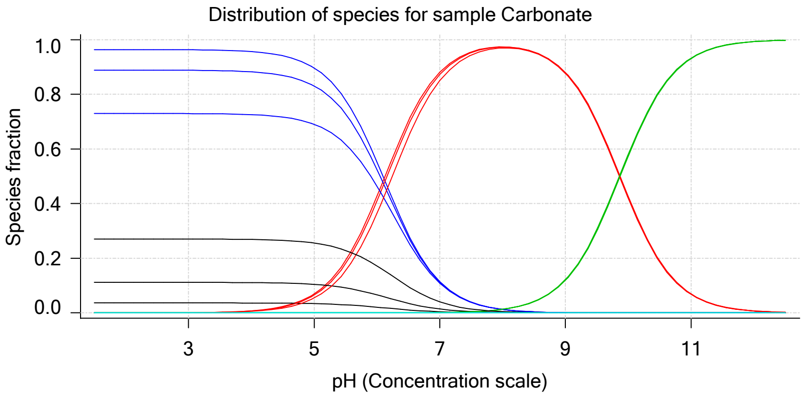

| | | | |
|---|---|---|---|
| Sample name: | **M11_octanol** | Experiment start time: | **3/1/2018 1:21:08 AM** |
| Assay name: | **pH-metric high logP** | Analyst: | **Pion** |
| Assay ID: | **18C-01002** | Instrument ID: | **T312060** |
| Filename: | **C:\Sirius_T3\Mehtap\20180228_exp28_logP_T3-2\18C-01002_M11_octanol_pH-metric high logP.t3r** | | |

# pH-metric high logP   Titration 1 of 3   18C-01002 Points 1 to 28

### Overall results

| | |
|---|---|
| RMSD | 0.953 |
| Average ionic strength | 0.157 M |
| Average temperature | 24.9°C |
| Partition ratio | 0.0122 : 1 |
| Analyte concentration range | 3149.9 µM to 3249.9 µM |
| Total points considered | 22 of 28 |

### Warnings and errors

| | |
|---|---|
| Errors | None |
| Warnings | One or more logP values out of range |

### Four-Plus parameters

| | | | |
|---|---|---|---|
| Alpha | 0.130 | 3/1/2018 1:21:08 AM | C:\Sirius_T3\HCl18B27.t3r |
| S | 0.9970 | 3/1/2018 1:21:08 AM | C:\Sirius_T3\HCl18B27.t3r |
| jH | 0.8 | 3/1/2018 1:21:08 AM | C:\Sirius_T3\HCl18B27.t3r |
| jOH | -0.4 | 3/1/2018 1:21:08 AM | C:\Sirius_T3\HCl18B27.t3r |

### Titrants

| | | | |
|---|---|---|---|
| 0.50 M HCl | 0.993513 | 3/1/2018 1:21:08 AM | C:\Sirius_T3\HCl18B27.t3r |
| 0.50 M KOH | 0.999845 | 3/1/2018 1:21:08 AM | C:\Sirius_T3\KOH18B27.t3r |

### Sample

| | |
|---|---|
| M11_octanol concentration factor | 0.940 |
| Base pKa 1 | 3.89 |
| logP (XH +) | -5.60 |
| logP (neutral X) | 1.94 |

### Sample graphs

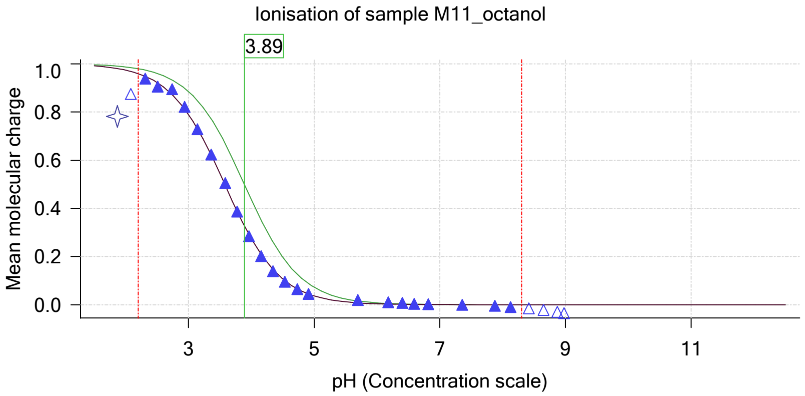


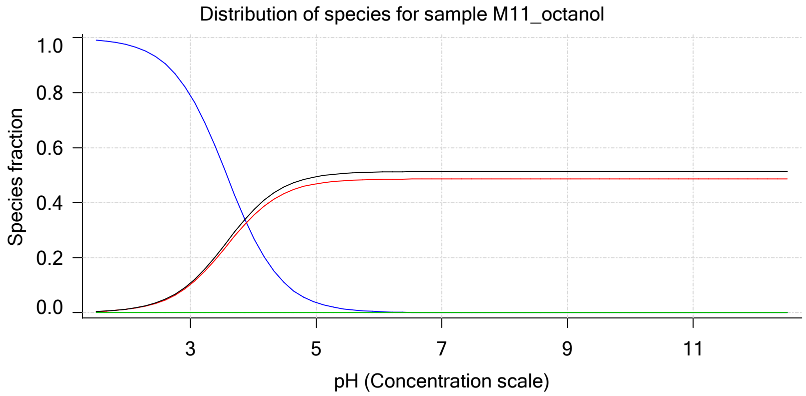

| | | | |
|---|---|---|---|
| Sample name: | **M11_octanol** | Experiment start time: | **3/1/2018 1:21:08 AM** |
| Assay name: | **pH-metric high logP** | Analyst: | **Pion** |
| Assay ID: | **18C-01002** | Instrument ID: | **T312060** |
| Filename: | **C:\Sirius_T3\Mehtap\20180228_exp28_logP_T3-2\18C-01002_M11_octanol_pH-metric high logP.t3r** | | |

### Sample graphs (continued)

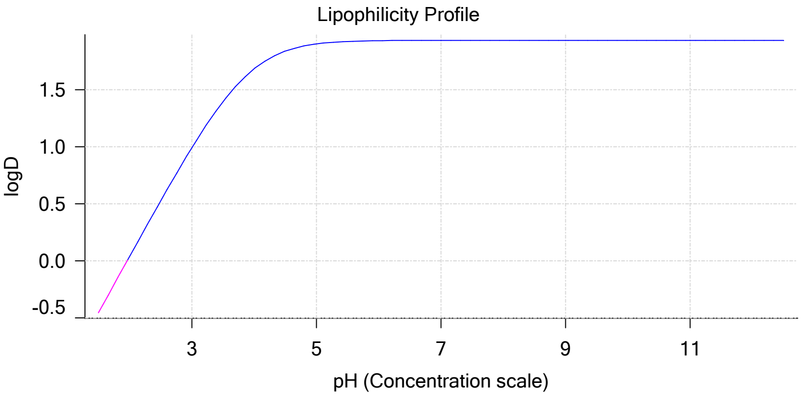


### Sample logD and percent species

| pH | M11_octanol logD | M11_octanol M11_octanolH | M11_octanol M11_octanol | M11_octanol M11_octanolH* | M11_octanol M11_octanol* | Comment |
|---|---|---|---|---|---|---|
| 1.000 | -0.96 | 99.74 % | 0.13 % | 0.00 % | 0.14 % | |
| 1.200 | -0.76 | 99.58 % | 0.20 % | 0.00 % | 0.21 % | Stomach pH |
| 2.000 | 0.04 | 97.42 % | 1.26 % | 0.00 % | 1.32 % | |
| 3.000 | 0.99 | 79.07 % | 10.19 % | 0.00 % | 10.74 % | |
| 4.000 | 1.69 | 27.42 % | 35.32 % | 0.00 % | 37.26 % | |
| 5.000 | 1.90 | 3.64 % | 46.90 % | 0.00 % | 49.46 % | |
| 6.000 | 1.93 | 0.38 % | 48.48 % | 0.00 % | 51.14 % | |
| 6.500 | 1.93 | 0.12 % | 48.61 % | 0.00 % | 51.27 % | |
| 7.000 | 1.93 | 0.04 % | 48.65 % | 0.00 % | 51.31 % | |
| 7.400 | 1.93 | 0.02 % | 48.66 % | 0.00 % | 51.33 % | Blood pH |
| 8.000 | 1.94 | 0.00 % | 48.67 % | 0.00 % | 51.33 % | |
| 9.000 | 1.94 | 0.00 % | 48.67 % | 0.00 % | 51.33 % | |
| 10.000 | 1.94 | 0.00 % | 48.67 % | 0.00 % | 51.33 % | |
| 11.000 | 1.94 | 0.00 % | 48.67 % | 0.00 % | 51.33 % | |
| 12.000 | 1.94 | 0.00 % | 48.67 % | 0.00 % | 51.33 % | |

### Carbonate and acidity

| | |
|---|---|
| Carbonate | 0.038 mM |
| Acidity error | 0.005 mM |

### Other graphs

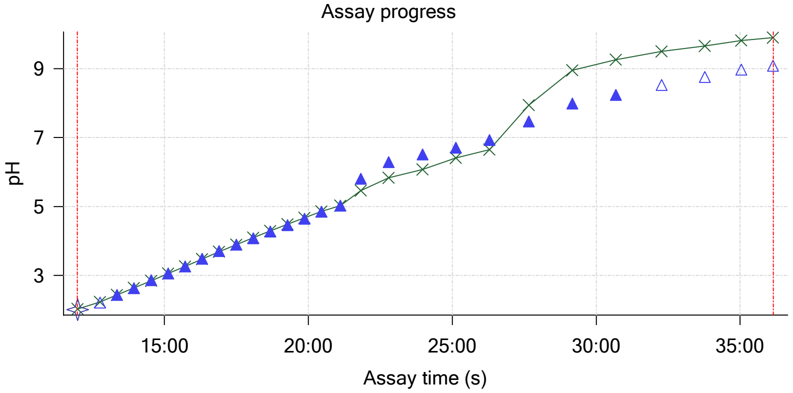


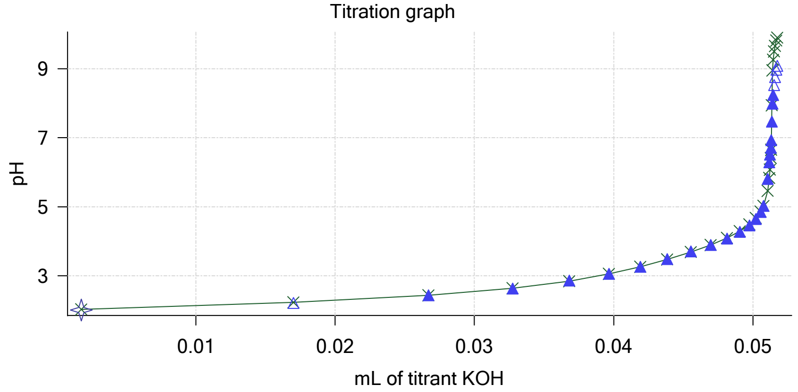

Sample name: **M11_octanol**
Assay name: **pH-metric high logP**
Assay ID: **18C-01002**
Filename: **C:\Sirius_T3\Mehtap\20180228_exp28_logP_T3-2\18C-01002_M11_octanol_pH-metric high logP.t3r**

Experiment start time: **3/1/2018 1:21:08 AM**
Analyst: **Pion**
Instrument ID: **T312060**

### Other graphs (continued)

Charge balance residuals

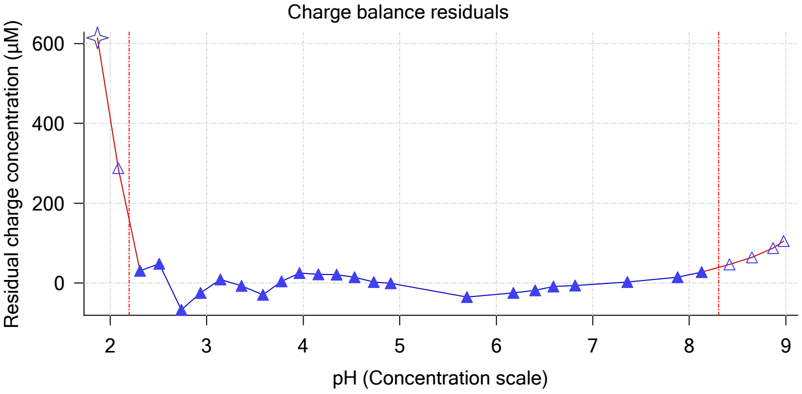


Buffer index profile

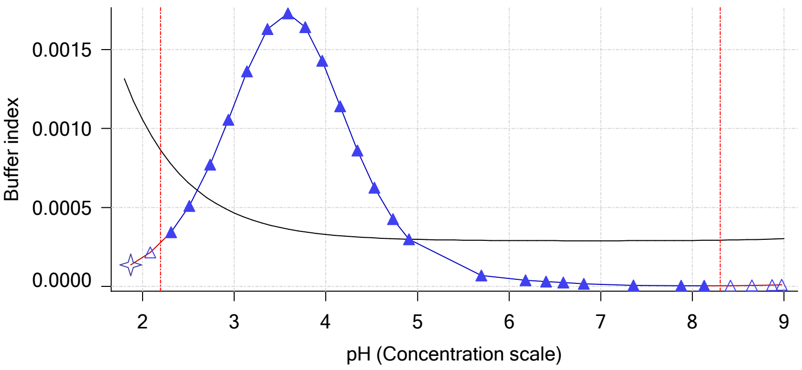


Ionic strength

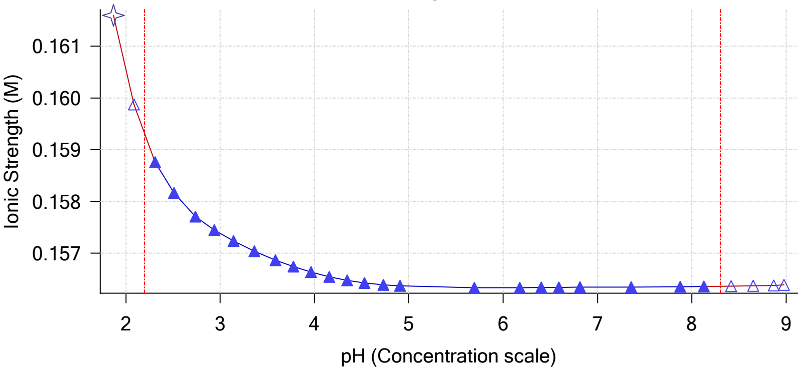


Log R Profile

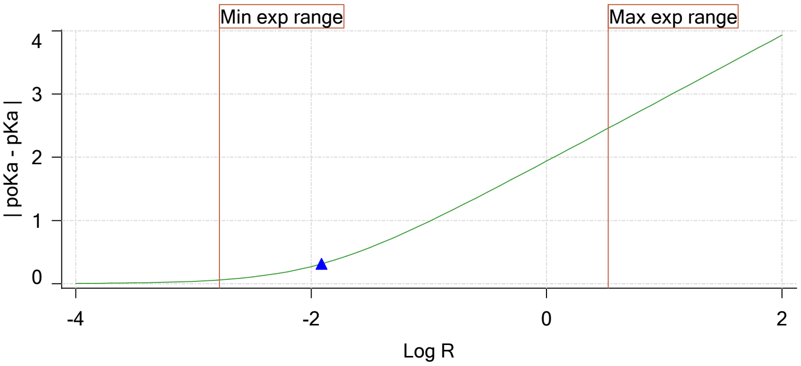


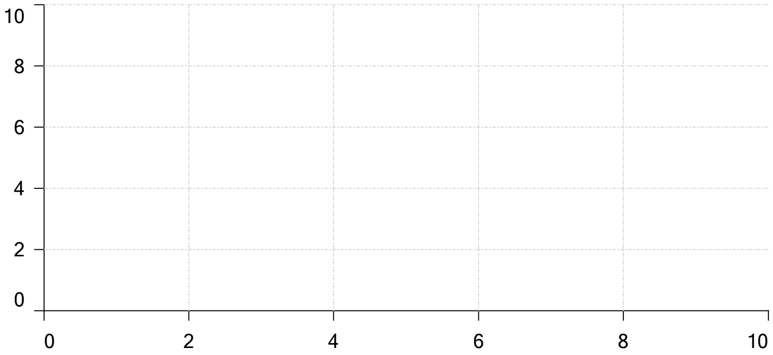


Ionisation of sample Carbonate

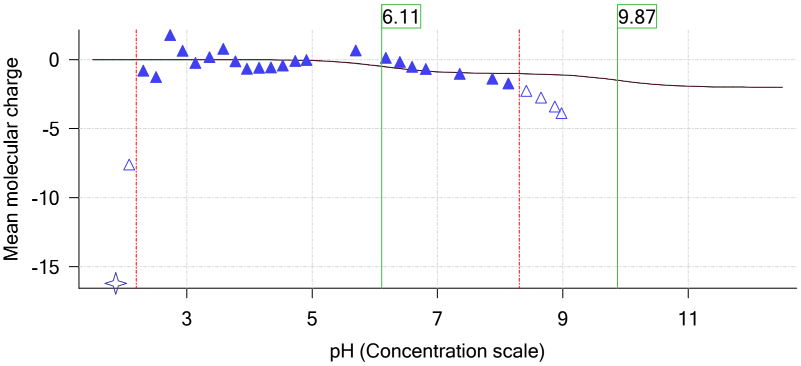

| | | | |
|---|---|---|---|
| Sample name: | **M11_octanol** | Experiment start time: | **3/1/2018 1:21:08 AM** |
| Assay name: | **pH-metric high logP** | Analyst: | **Pion** |
| Assay ID: | **18C-01002** | Instrument ID: | **T312060** |
| Filename: | **C:\Sirius_T3\Mehtap\20180228_exp28_logP_T3-2\18C-01002_M11_octanol_pH-metric high logP.t3r** | | |

# pH-metric high logP   Titration 2 of 3   18C-01002 Points 29 to 49

### Overall results

| | |
|---|---|
| RMSD | 0.536 |
| Average ionic strength | 0.163 M |
| Average temperature | 25.0°C |
| Partition ratio | 0.0407 : 1 |
| Analyte concentration range | 2865.3 µM to 2956.0 µM |
| Total points considered | 17 of 21 |

### Warnings and errors

| | |
|---|---|
| Errors | None |
| Warnings | One or more logP values out of range |

### Four-Plus parameters

| | | | |
|---|---|---|---|
| Alpha | 0.130 | 3/1/2018 1:21:08 AM | C:\Sirius_T3\HCl18B27.t3r |
| S | 0.9970 | 3/1/2018 1:21:08 AM | C:\Sirius_T3\HCl18B27.t3r |
| jH | 0.8 | 3/1/2018 1:21:08 AM | C:\Sirius_T3\HCl18B27.t3r |
| jOH | -0.4 | 3/1/2018 1:21:08 AM | C:\Sirius_T3\HCl18B27.t3r |

### Titrants

| | | | |
|---|---|---|---|
| 0.50 M HCl | 0.993513 | 3/1/2018 1:21:08 AM | C:\Sirius_T3\HCl18B27.t3r |
| 0.50 M KOH | 0.999845 | 3/1/2018 1:21:08 AM | C:\Sirius_T3\KOH18B27.t3r |

### Sample

| | |
|---|---|
| M11_octanol concentration factor | 0.777 |
| Base pKa 1 | 3.89 |
| logP (XH +) | -5.10 |
| logP (neutral X) | 2.08 |

### Sample graphs

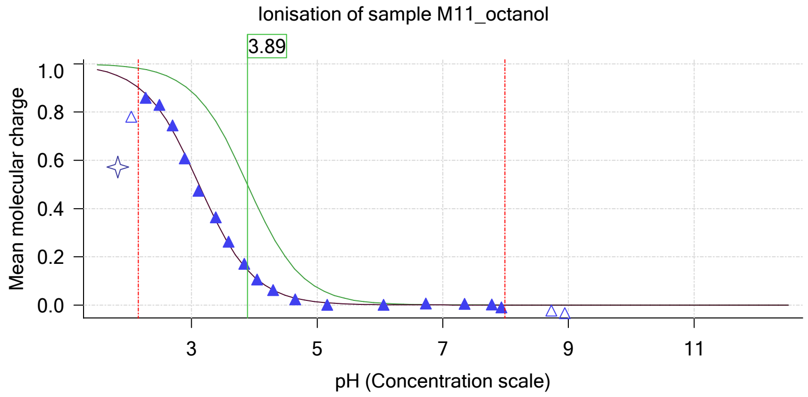


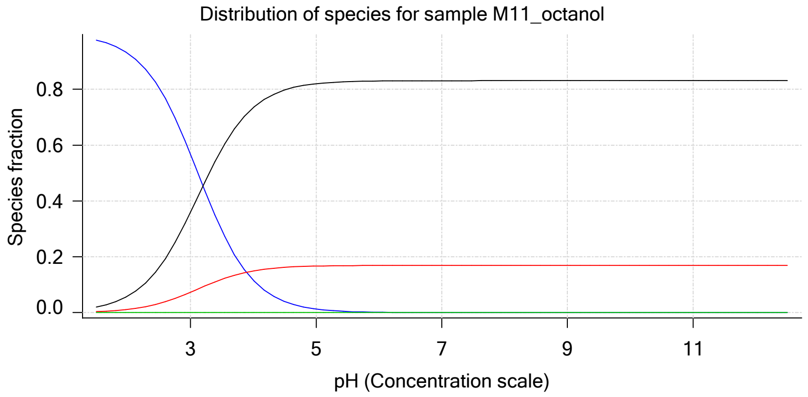

| | | | |
|---|---|---|---|
| Sample name: | **M11_octanol** | Experiment start time: | **3/1/2018 1:21:08 AM** |
| Assay name: | **pH-metric high logP** | Analyst: | **Pion** |
| Assay ID: | **18C-01002** | Instrument ID: | **T312060** |
| Filename: | **C:\Sirius_T3\Mehtap\20180228_exp28_logP_T3-2\18C-01002_M11_octanol_pH-metric high logP.t3r** | | |

### Sample graphs (continued)

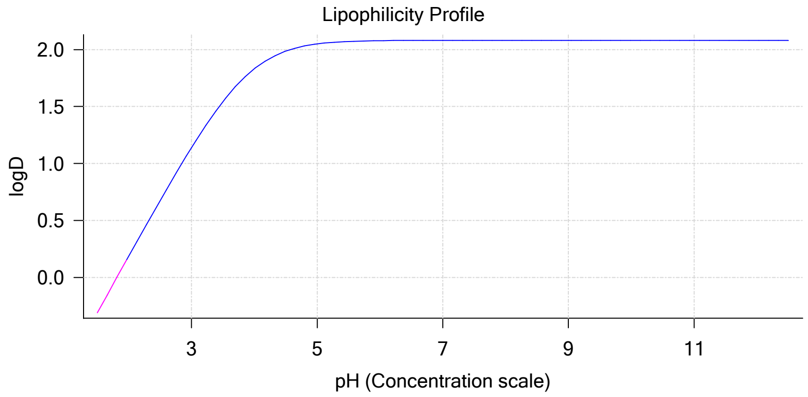


### Sample logD and percent species

| pH | M11_octanol logD | M11_octanol M11_octanolH | M11_octanol M11_octanol | M11_octanol M11_octanolH* | M11_octanol M11_octanol* | Comment |
|---|---|---|---|---|---|---|
| 1.000 | -0.81 | 99.24 % | 0.13 % | 0.00 % | 0.63 % | |
| 1.200 | -0.61 | 98.81 % | 0.20 % | 0.00 % | 0.99 % | Stomach pH |
| 2.000 | 0.19 | 92.92 % | 1.20 % | 0.00 % | 5.88 % | |
| 3.000 | 1.14 | 56.77 % | 7.31 % | 0.00 % | 35.91 % | |
| 4.000 | 1.83 | 11.61 % | 14.96 % | 0.00 % | 73.44 % | |
| 5.000 | 2.05 | 1.30 % | 16.70 % | 0.00 % | 82.00 % | |
| 6.000 | 2.08 | 0.13 % | 16.90 % | 0.00 % | 82.97 % | |
| 6.500 | 2.08 | 0.04 % | 16.91 % | 0.00 % | 83.05 % | |
| 7.000 | 2.08 | 0.01 % | 16.92 % | 0.00 % | 83.07 % | |
| 7.400 | 2.08 | 0.01 % | 16.92 % | 0.00 % | 83.08 % | Blood pH |
| 8.000 | 2.08 | 0.00 % | 16.92 % | 0.00 % | 83.08 % | |
| 9.000 | 2.08 | 0.00 % | 16.92 % | 0.00 % | 83.08 % | |
| 10.000 | 2.08 | 0.00 % | 16.92 % | 0.00 % | 83.08 % | |
| 11.000 | 2.08 | 0.00 % | 16.92 % | 0.00 % | 83.08 % | |
| 12.000 | 2.08 | 0.00 % | 16.92 % | 0.00 % | 83.08 % | |

### Carbonate and acidity

| | |
|---|---|
| Carbonate | 0.115 mM |
| Acidity error | -0.467 mM |

### Other graphs

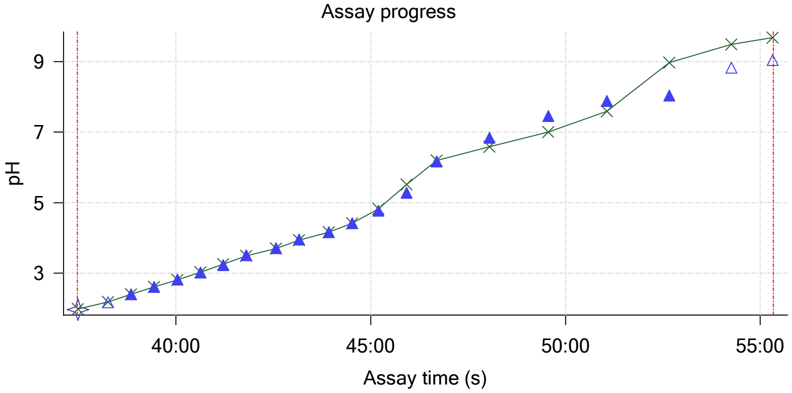


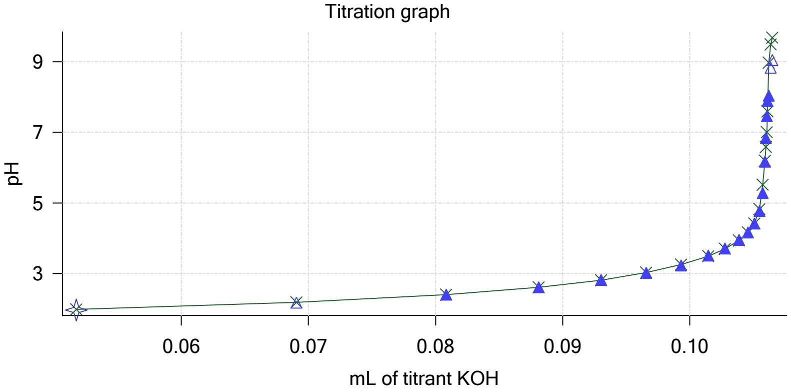

Sample name: **M11_octanol**
Assay name: **pH-metric high logP**
Assay ID: **18C-01002**
Filename: **C:\Sirius_T3\Mehtap\20180228_exp28_logP_T3-2\18C-01002_M11_octanol_pH-metric high logP.t3r**

Experiment start time: **3/1/2018 1:21:08 AM**
Analyst: **Pion**
Instrument ID: **T312060**

### Other graphs (continued)

Charge balance residuals

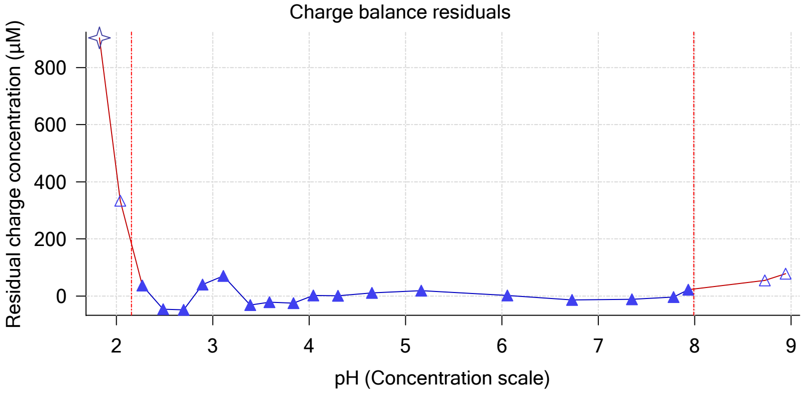


Buffer index profile

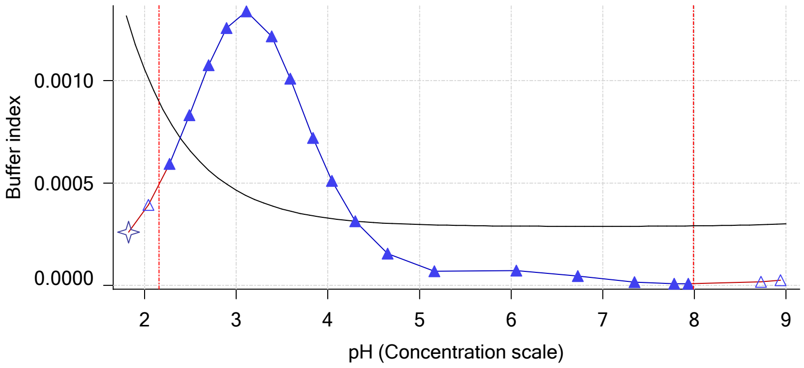


Ionic strength

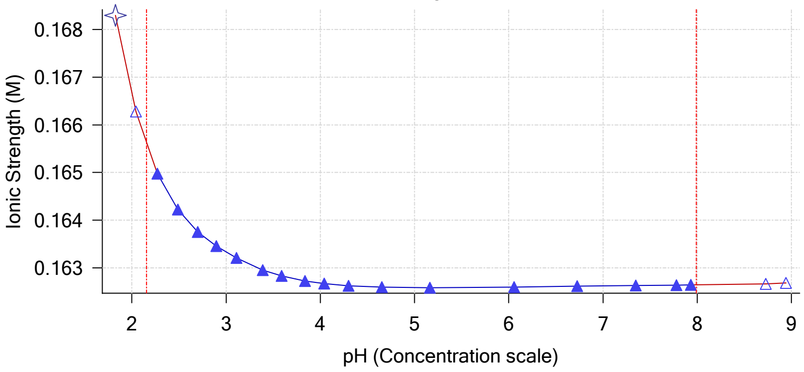


Log R Profile

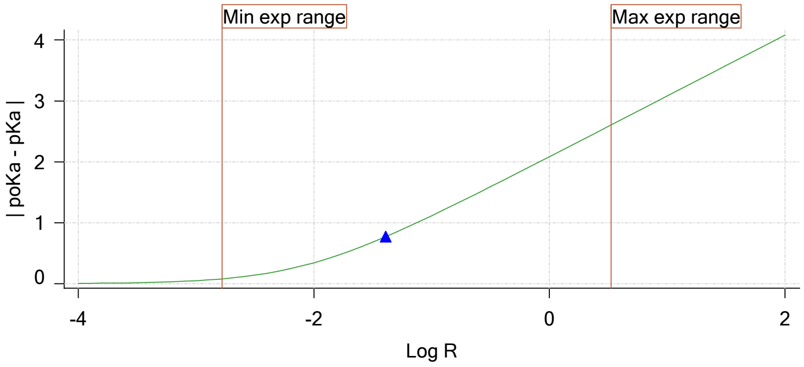


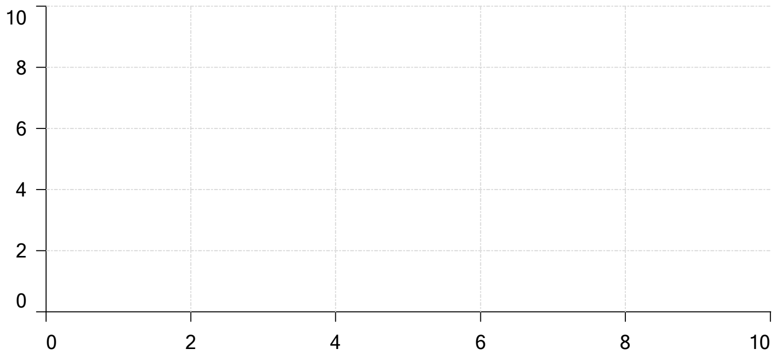

Ionisation of sample Carbonate

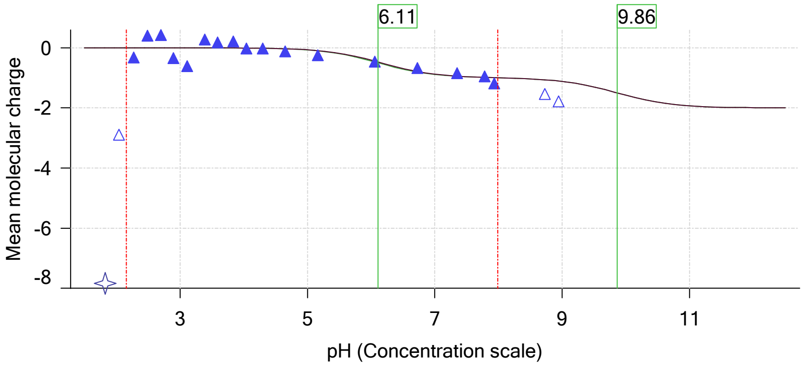

| | | | |
|---|---|---|---|
| Sample name: | **M11_octanol** | Experiment start time: | **3/1/2018 1:21:08 AM** |
| Assay name: | **pH-metric high logP** | Analyst: | **Pion** |
| Assay ID: | **18C-01002** | Instrument ID: | **T312060** |
| Filename: | **C:\Sirius_T3\Mehtap\20180228_exp28_logP_T3-2\18C-01002_M11_octanol_pH-metric high logP.t3r** | | |

# pH-metric high logP   Titration 3 of 3   18C-01002 Points 50 to 75

### Overall results

| | |
|---|---|
| RMSD | 0.550 |
| Average ionic strength | 0.169 M |
| Average temperature | 25.0°C |
| Partition ratio | 0.1199 : 1 |
| Analyte concentration range | 2489.4 µM to 2561.6 µM |
| Total points considered | 21 of 26 |

### Warnings and errors

| | |
|---|---|
| Errors | None |
| Warnings | One or more logP values out of range |
| | Excessive acidity error present |

### Four-Plus parameters

| | | | |
|---|---|---|---|
| Alpha | 0.130 | 3/1/2018 1:21:08 AM | C:\Sirius_T3\HCl18B27.t3r |
| S | 0.9970 | 3/1/2018 1:21:08 AM | C:\Sirius_T3\HCl18B27.t3r |
| jH | 0.8 | 3/1/2018 1:21:08 AM | C:\Sirius_T3\HCl18B27.t3r |
| jOH | -0.4 | 3/1/2018 1:21:08 AM | C:\Sirius_T3\HCl18B27.t3r |

### Titrants

| | | | |
|---|---|---|---|
| 0.50 M HCl | 0.993513 | 3/1/2018 1:21:08 AM | C:\Sirius_T3\HCl18B27.t3r |
| 0.50 M KOH | 0.999845 | 3/1/2018 1:21:08 AM | C:\Sirius_T3\KOH18B27.t3r |

### Sample

| | |
|---|---|
| M11_octanol concentration factor | 0.713 |
| Base pKa 1 | 3.89 |
| logP (XH +) | -5.10 |
| logP (neutral X) | 1.99 |

### Sample graphs

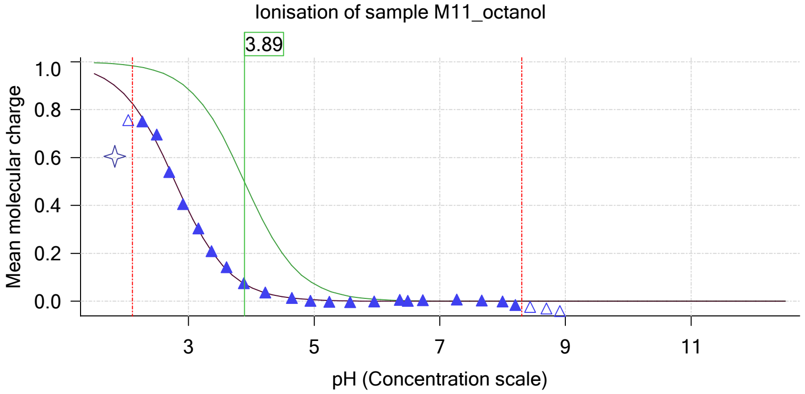


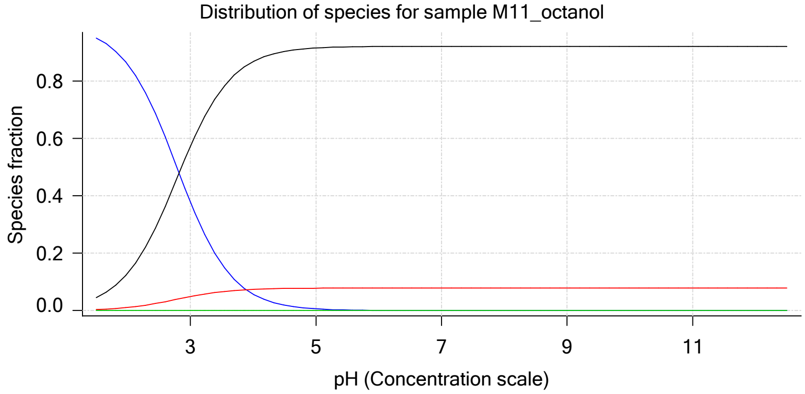

| | | | |
|---|---|---|---|
| Sample name: | **M11_octanol** | Experiment start time: | **3/1/2018 1:21:08 AM** |
| Assay name: | **pH-metric high logP** | Analyst: | **Pion** |
| Assay ID: | **18C-01002** | Instrument ID: | **T312060** |
| Filename: | **C:\Sirius_T3\Mehtap\20180228_exp28_logP_T3-2\18C-01002_M11_octanol_pH-metric high logP.t3r** | | |

### Sample graphs (continued)

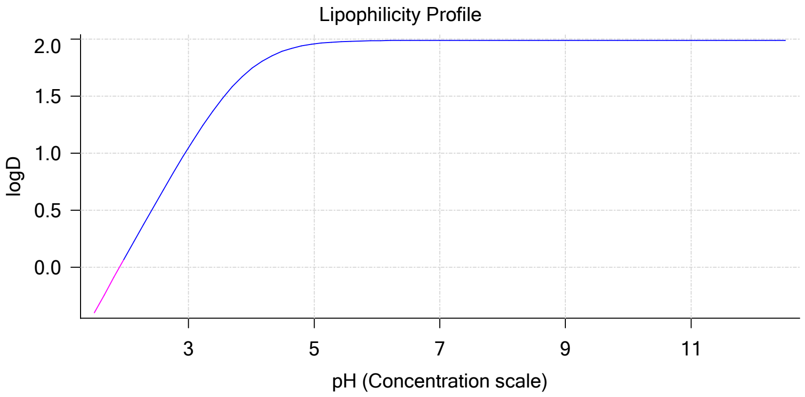


### Sample logD and percent species

| pH | M11_octanol logD | M11_octanol M11_octanolH | M11_octanol M11_octanol | M11_octanol M11_octanolH* | M11_octanol M11_octanol* | Comment |
|---|---|---|---|---|---|---|
| 1.000 | -0.90 | 98.39 % | 0.13 % | 0.00 % | 1.49 % | |
| 1.200 | -0.70 | 97.47 % | 0.20 % | 0.00 % | 2.33 % | Stomach pH |
| 2.000 | 0.09 | 85.92 % | 1.11 % | 0.00 % | 12.97 % | |
| 3.000 | 1.05 | 37.90 % | 4.88 % | 0.00 % | 57.21 % | |
| 4.000 | 1.74 | 5.75 % | 7.41 % | 0.00 % | 86.84 % | |
| 5.000 | 1.96 | 0.61 % | 7.82 % | 0.00 % | 91.58 % | |
| 6.000 | 1.99 | 0.06 % | 7.86 % | 0.00 % | 92.08 % | |
| 6.500 | 1.99 | 0.02 % | 7.86 % | 0.00 % | 92.12 % | |
| 7.000 | 1.99 | 0.01 % | 7.86 % | 0.00 % | 92.13 % | |
| 7.400 | 1.99 | 0.00 % | 7.86 % | 0.00 % | 92.13 % | Blood pH |
| 8.000 | 1.99 | 0.00 % | 7.86 % | 0.00 % | 92.14 % | |
| 9.000 | 1.99 | 0.00 % | 7.86 % | 0.00 % | 92.14 % | |
| 10.000 | 1.99 | 0.00 % | 7.86 % | 0.00 % | 92.14 % | |
| 11.000 | 1.99 | 0.00 % | 7.86 % | 0.00 % | 92.14 % | |
| 12.000 | 1.99 | 0.00 % | 7.86 % | 0.00 % | 92.14 % | |

### Carbonate and acidity

| | |
|---|---|
| Carbonate | 0.103 mM |
| Acidity error | -1.332 mM |

### Other graphs

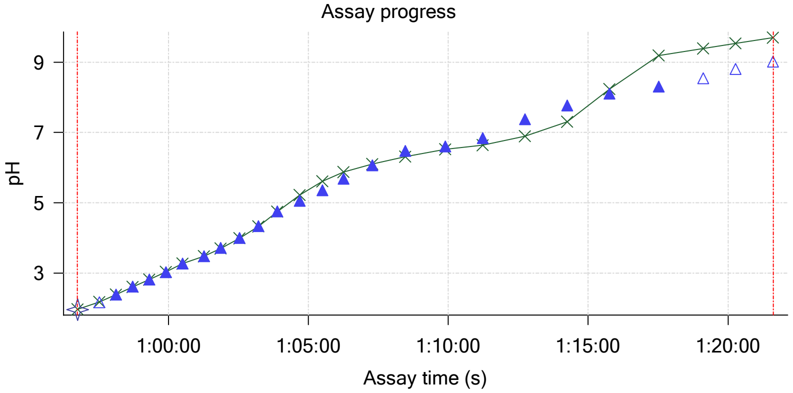


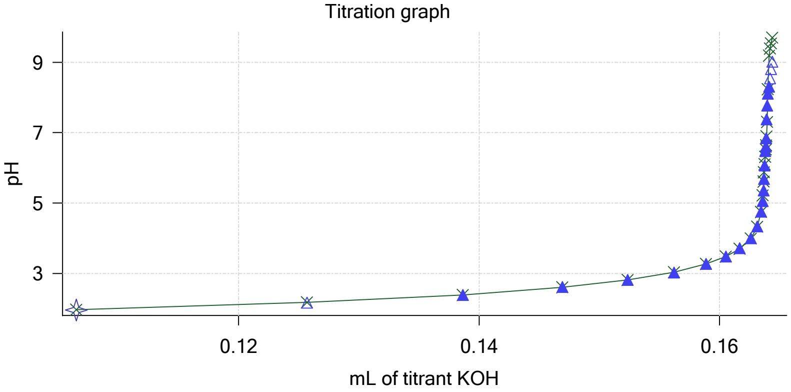

Sample name: **M11_octanol**
Assay name: **pH-metric high logP**
Assay ID: **18C-01002**
Filename: **C:\Sirius_T3\Mehtap\20180228_exp28_logP_T3-2\18C-01002_M11_octanol_pH-metric high logP.t3r**

Experiment start time: **3/1/2018 1:21:08 AM**
Analyst: **Pion**
Instrument ID: **T312060**

### Other graphs (continued)

Charge balance residuals

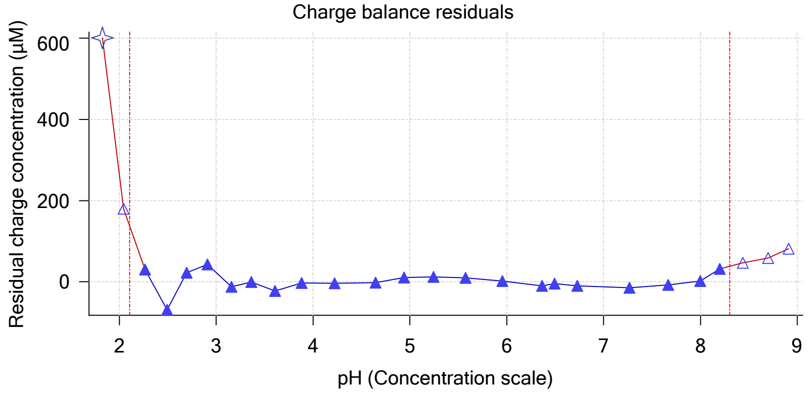


Buffer index profile

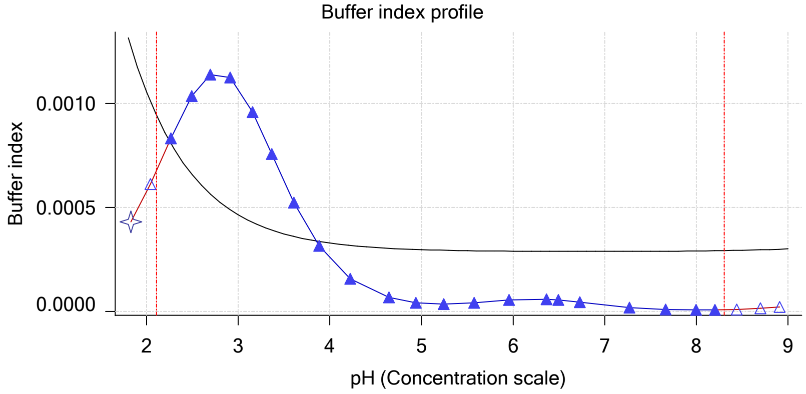


Ionic strength

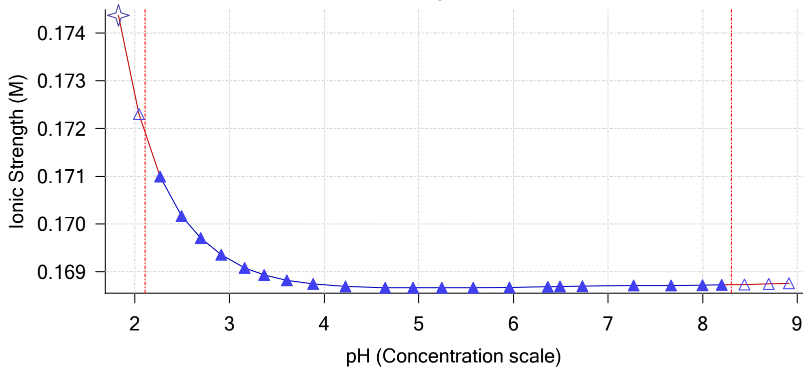


Log R Profile

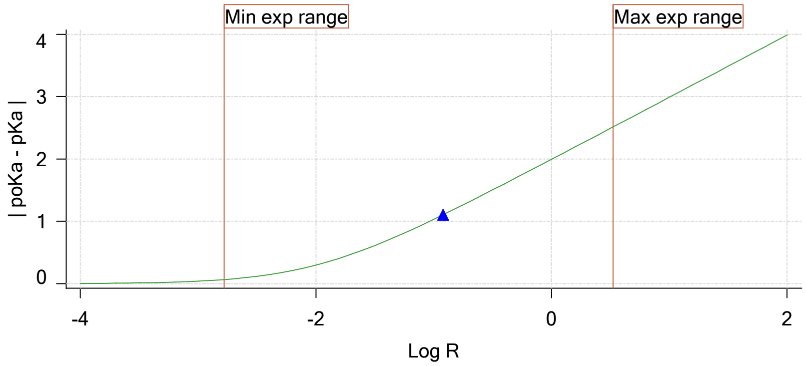


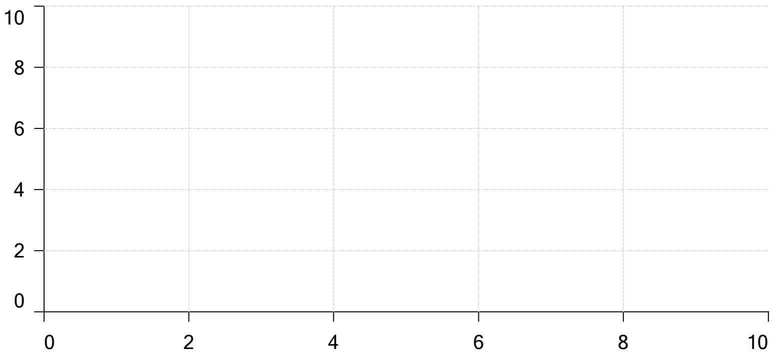


Ionisation of sample Carbonate

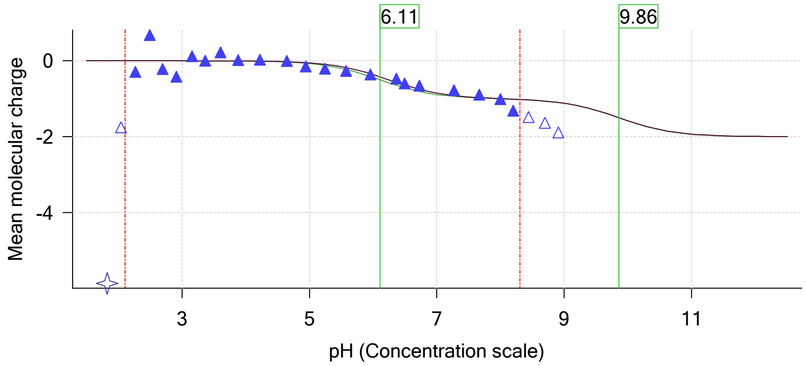

| | | | |
|---|---|---|---|
| Sample name: | **M11_octanol** | Experiment start time: | **3/1/2018 1:21:08 AM** |
| Assay name: | **pH-metric high logP** | Analyst: | **Pion** |
| Assay ID: | **18C-01002** | Instrument ID: | **T312060** |
| Filename: | **C:\Sirius_T3\Mehtap\20180228_exp28_logP_T3-2\18C-01002_M11_octanol_pH-metric high logP.t3r** | | |

### Assay Model

| Settings | Value | Date/Time changed | Imported from |
|---|---|---|---|
| Sample name | M11_octanol | 2/27/2018 4:54:30 PM | User entered value |
| Sample by | Weight | | Default value |
| Sample weight | 0.001080 g | 2/28/2018 4:24:59 PM | User entered value |
| Formula weight | 211.22 g/mol | 2/27/2018 4:54:30 PM | User entered value |
| Solubility | Unknown | | Default value |
| Molecular weight | 211.22 | 2/27/2018 4:54:30 PM | User entered value |
| Individual pKa ionic environments | No | | Default value |
| Number of pKas | 1 | 2/27/2018 4:54:30 PM | User entered value |
| Sample is a | Base | 2/27/2018 4:54:30 PM | User entered value |
| pKa 1 | 3.89 | 2/27/2018 4:54:30 PM | User entered value |
| logp (XH +) | -5.60 | 2/28/2018 1:53:21 PM | User entered value |
| logP (neutral X) | 2.09 | 2/28/2018 1:53:11 PM | User entered value |

### Events

| Time | Event | Water | Acid | Base | Octanol | pH | dpH/dt | pH R-squared | pH SD | dpH/dt time |
|---|---|---|---|---|---|---|---|---|---|---|
| 8:59.7 | Initial pH = 8.06 | | | | | | | | | |
| 11:59.3 | Data point 1 | 1.50000 mL | 0.05153 mL | 0.00179 mL | 0.01999 mL | 2.005 | -0.00768 | 0.87550 | 0.00040 | 10.0 s |
| 12:45.4 | Data point 2 | 1.50000 mL | 0.05153 mL | 0.01700 mL | 0.01999 mL | 2.215 | -0.00104 | 0.14554 | 0.00013 | 10.0 s |
| 13:21.1 | Data point 3 | 1.50000 mL | 0.05153 mL | 0.02672 mL | 0.01999 mL | 2.436 | -0.00042 | 0.01137 | 0.00020 | 10.0 s |
| 13:56.6 | Data point 4 | 1.50000 mL | 0.05153 mL | 0.03274 mL | 0.01999 mL | 2.635 | -0.00726 | 0.18423 | 0.00084 | 10.0 s |
| 14:32.2 | Data point 5 | 1.50000 mL | 0.05153 mL | 0.03681 mL | 0.01999 mL | 2.861 | -0.00222 | 0.27923 | 0.00021 | 10.0 s |
| 15:07.7 | Data point 6 | 1.50000 mL | 0.05153 mL | 0.03965 mL | 0.01999 mL | 3.060 | -0.00837 | 0.77297 | 0.00047 | 10.0 s |
| 15:43.1 | Data point 7 | 1.50000 mL | 0.05153 mL | 0.04191 mL | 0.01999 mL | 3.262 | -0.00454 | 0.61543 | 0.00029 | 10.0 s |
| 16:18.6 | Data point 8 | 1.50000 mL | 0.05153 mL | 0.04384 mL | 0.01999 mL | 3.482 | -0.00445 | 0.55554 | 0.00029 | 10.0 s |
| 16:54.1 | Data point 9 | 1.50000 mL | 0.05153 mL | 0.04553 mL | 0.01999 mL | 3.705 | -0.01127 | 0.51278 | 0.00078 | 10.0 s |
| 17:29.6 | Data point 10 | 1.50000 mL | 0.05153 mL | 0.04697 mL | 0.01999 mL | 3.894 | -0.00094 | 0.05467 | 0.00020 | 10.5 s |
| 18:05.4 | Data point 11 | 1.50000 mL | 0.05153 mL | 0.04814 mL | 0.01999 mL | 4.080 | -0.00544 | 0.86041 | 0.00029 | 10.0 s |
| 18:40.8 | Data point 12 | 1.50000 mL | 0.05153 mL | 0.04906 mL | 0.01999 mL | 4.274 | -0.01020 | 0.91768 | 0.00053 | 10.5 s |
| 19:16.8 | Data point 13 | 1.50000 mL | 0.05153 mL | 0.04974 mL | 0.01999 mL | 4.463 | -0.01138 | 0.78057 | 0.00064 | 10.0 s |
| 19:52.2 | Data point 14 | 1.50000 mL | 0.05153 mL | 0.05021 mL | 0.01999 mL | 4.648 | -0.01383 | 0.82707 | 0.00075 | 10.0 s |
| 20:27.6 | Data point 15 | 1.50000 mL | 0.05153 mL | 0.05054 mL | 0.01999 mL | 4.848 | -0.01852 | 0.92972 | 0.00095 | 13.5 s |
| 21:06.5 | Data point 16 | 1.50000 mL | 0.05153 mL | 0.05075 mL | 0.01999 mL | 5.023 | -0.01863 | 0.95790 | 0.00094 | 12.0 s |
| 21:49.0 | Data point 17 | 1.50000 mL | 0.05153 mL | 0.05106 mL | 0.01999 mL | 5.808 | -0.01925 | 0.92072 | 0.00099 | 22.5 s |
| 22:47.2 | Data point 18 | 1.50000 mL | 0.05153 mL | 0.05118 mL | 0.01999 mL | 6.289 | -0.01833 | 0.92758 | 0.00094 | 40.0 s |
| 23:57.7 | Data point 19 | 1.50000 mL | 0.05153 mL | 0.05122 mL | 0.01999 mL | 6.513 | -0.01738 | 0.88907 | 0.00091 | 44.5 s |
| 25:07.7 | Data point 20 | 1.50000 mL | 0.05153 mL | 0.05127 mL | 0.01999 mL | 6.702 | -0.01872 | 0.87409 | 0.00099 | 44.5 s |
| 26:17.6 | Data point 21 | 1.50000 mL | 0.05153 mL | 0.05129 mL | 0.01999 mL | 6.926 | -0.01864 | 0.99231 | 0.00092 | 51.5 s |
| 27:39.8 | Data point 22 | 1.50000 mL | 0.05153 mL | 0.05134 mL | 0.01999 mL | 7.465 | -0.05629 | 0.99724 | 0.00278 | Timed out at 59.5 s |
| 29:10.2 | Data point 23 | 1.50000 mL | 0.05153 mL | 0.05139 mL | 0.01999 mL | 7.984 | -0.04354 | 0.97478 | 0.00218 | Timed out at 59.5 s |
| 30:40.7 | Data point 24 | 1.50000 mL | 0.05153 mL | 0.05143 mL | 0.01999 mL | 8.234 | -0.03056 | 0.98451 | 0.00152 | Timed out at 59.5 s |
| 32:16.3 | Data point 25 | 1.50000 mL | 0.05153 mL | 0.05151 mL | 0.01999 mL | 8.520 | -0.01890 | 0.92260 | 0.00097 | 54.5 s |
| 33:46.5 | Data point 26 | 1.50000 mL | 0.05153 mL | 0.05158 mL | 0.01999 mL | 8.753 | -0.01846 | 0.87251 | 0.00098 | 40.5 s |
| 35:02.6 | Data point 27 | 1.50000 mL | 0.05153 mL | 0.05167 mL | 0.01999 mL | 8.972 | -0.01838 | 0.89623 | 0.00096 | 35.0 s |
| 36:08.2 | Data point 28 | 1.50000 mL | 0.05153 mL | 0.05174 mL | 0.01999 mL | 9.079 | -0.01907 | 0.95761 | 0.00096 | 21.5 s |
| 37:29.0 | Data point 29 | 1.50000 mL | 0.10802 mL | 0.05174 mL | 0.06999 mL | 1.962 | -0.01000 | 0.87704 | 0.00053 | 10.0 s |
| 38:15.3 | Data point 30 | 1.50000 mL | 0.10802 mL | 0.06905 mL | 0.06999 mL | 2.172 | -0.01070 | 0.38504 | 0.00085 | 10.0 s |
| 38:51.0 | Data point 31 | 1.50000 mL | 0.10802 mL | 0.08083 mL | 0.06999 mL | 2.398 | 0.00033 | 0.00320 | 0.00029 | 10.0 s |
| 39:26.6 | Data point 32 | 1.50000 mL | 0.10802 mL | 0.08812 mL | 0.06999 mL | 2.613 | -0.00001 | 0.00001 | 0.00013 | 10.0 s |
| 40:02.1 | Data point 33 | 1.50000 mL | 0.10802 mL | 0.09302 mL | 0.06999 mL | 2.822 | -0.00176 | 0.22522 | 0.00018 | 10.0 s |
| 40:37.6 | Data point 34 | 1.50000 mL | 0.10802 mL | 0.09657 mL | 0.06999 mL | 3.017 | -0.00360 | 0.20627 | 0.00039 | 10.0 s |
| 41:13.0 | Data point 35 | 1.50000 mL | 0.10802 mL | 0.09932 mL | 0.06999 mL | 3.231 | -0.00250 | 0.58779 | 0.00016 | 10.0 s |
| 41:48.5 | Data point 36 | 1.50000 mL | 0.10802 mL | 0.10146 mL | 0.06999 mL | 3.509 | -0.00677 | 0.28436 | 0.00063 | 10.0 s |
| 42:34.2 | Data point 37 | 1.50000 mL | 0.10802 mL | 0.10278 mL | 0.06999 mL | 3.709 | -0.00300 | 0.43965 | 0.00022 | 10.0 s |

| Sample name: | M11_octanol | Experiment start time: | 3/1/2018 1:21:08 AM |
|---|---|---|---|
| Assay name: | pH-metric high logP | Analyst: | Pion |
| Assay ID: | 18C-01002 | Instrument ID: | T312060 |
| Filename: | C:\Sirius_T3\Mehtap\20180228_exp28_logP_T3-2\18C-01002_M11_octanol_pH-metric high logP.t3r | | |

### Events (continued)

| Time | Event | Water | Acid | Base | Octanol | pH | dpH/dt | pH R-squared | pH SD | dpH/dt time |
|---|---|---|---|---|---|---|---|---|---|---|
| 43:09.6 | Data point 38 | 1.50000 mL | 0.10802 mL | 0.10388 mL | 0.06999 mL | 3.957 | -0.00406 | 0.29082 | 0.00037 | 10.0 s |
| 43:55.3 | Data point 39 | 1.50000 mL | 0.10802 mL | 0.10459 mL | 0.06999 mL | 4.161 | -0.00477 | 0.72196 | 0.00028 | 10.5 s |
| 44:31.3 | Data point 40 | 1.50000 mL | 0.10802 mL | 0.10508 mL | 0.06999 mL | 4.417 | -0.01592 | 0.90741 | 0.00083 | 10.0 s |
| 45:11.8 | Data point 41 | 1.50000 mL | 0.10802 mL | 0.10548 mL | 0.06999 mL | 4.769 | -0.01738 | 0.82458 | 0.00095 | 13.0 s |
| 45:55.4 | Data point 42 | 1.50000 mL | 0.10802 mL | 0.10574 mL | 0.06999 mL | 5.276 | -0.01305 | 0.55308 | 0.00087 | 15.5 s |
| 46:41.4 | Data point 43 | 1.50000 mL | 0.10802 mL | 0.10590 mL | 0.06999 mL | 6.170 | -0.01853 | 0.94972 | 0.00094 | 46.0 s |
| 48:03.1 | Data point 44 | 1.50000 mL | 0.10802 mL | 0.10600 mL | 0.06999 mL | 6.837 | -0.04872 | 0.98691 | 0.00242 | Timed out at 59.5 s |
| 49:33.5 | Data point 45 | 1.50000 mL | 0.10802 mL | 0.10607 mL | 0.06999 mL | 7.455 | -0.05872 | 0.98494 | 0.00292 | Timed out at 59.5 s |
| 51:04.0 | Data point 46 | 1.50000 mL | 0.10802 mL | 0.10612 mL | 0.06999 mL | 7.887 | -0.06009 | 0.99005 | 0.00298 | Timed out at 59.5 s |
| 52:39.7 | Data point 47 | 1.50000 mL | 0.10802 mL | 0.10621 mL | 0.06999 mL | 8.038 | -0.04607 | 0.98857 | 0.00229 | Timed out at 59.5 s |
| 54:15.4 | Data point 48 | 1.50000 mL | 0.10802 mL | 0.10637 mL | 0.06999 mL | 8.831 | -0.01813 | 0.88499 | 0.00095 | 28.0 s |
| 55:19.1 | Data point 49 | 1.50000 mL | 0.10802 mL | 0.10649 mL | 0.06999 mL | 9.044 | -0.01850 | 0.93380 | 0.00095 | 24.5 s |
| 56:45.5 | Data point 50 | 1.50000 mL | 0.16959 mL | 0.10649 mL | 0.21999 mL | 1.963 | -0.00309 | 0.05617 | 0.00064 | 10.0 s |
| 57:31.7 | Data point 51 | 1.50000 mL | 0.16959 mL | 0.12568 mL | 0.21999 mL | 2.174 | -0.01262 | 0.82319 | 0.00069 | 10.0 s |
| 58:07.5 | Data point 52 | 1.50000 mL | 0.16959 mL | 0.13864 mL | 0.21999 mL | 2.394 | -0.00228 | 0.23781 | 0.00023 | 10.0 s |
| 58:43.1 | Data point 53 | 1.50000 mL | 0.16959 mL | 0.14692 mL | 0.21999 mL | 2.618 | 0.00798 | 0.26467 | 0.00077 | 10.0 s |
| 59:18.7 | Data point 54 | 1.50000 mL | 0.16959 mL | 0.15235 mL | 0.21999 mL | 2.818 | -0.00238 | 0.60816 | 0.00015 | 10.5 s |
| 59:54.8 | Data point 55 | 1.50000 mL | 0.16959 mL | 0.15621 mL | 0.21999 mL | 3.034 | -0.00228 | 0.56070 | 0.00015 | 10.0 s |
| 1:00:30.3 | Data point 56 | 1.50000 mL | 0.16959 mL | 0.15889 mL | 0.21999 mL | 3.279 | 0.00108 | 0.00354 | 0.00090 | 10.0 s |
| 1:01:16.1 | Data point 57 | 1.50000 mL | 0.16959 mL | 0.16054 mL | 0.21999 mL | 3.485 | -0.00480 | 0.78494 | 0.00027 | 10.0 s |
| 1:01:51.6 | Data point 58 | 1.50000 mL | 0.16959 mL | 0.16169 mL | 0.21999 mL | 3.726 | -0.00042 | 0.00046 | 0.00096 | 10.0 s |
| 1:02:32.2 | Data point 59 | 1.50000 mL | 0.16959 mL | 0.16261 mL | 0.21999 mL | 4.002 | 0.00063 | 0.00106 | 0.00095 | 10.0 s |
| 1:03:12.8 | Data point 60 | 1.50000 mL | 0.16959 mL | 0.16315 mL | 0.21999 mL | 4.342 | 0.00232 | 0.02472 | 0.00073 | 10.0 s |
| 1:03:53.3 | Data point 61 | 1.50000 mL | 0.16959 mL | 0.16345 mL | 0.21999 mL | 4.762 | 0.00143 | 0.00712 | 0.00084 | 12.0 s |
| 1:04:41.0 | Data point 62 | 1.50000 mL | 0.16959 mL | 0.16359 mL | 0.21999 mL | 5.055 | 0.00471 | 0.07278 | 0.00086 | 13.5 s |
| 1:05:30.2 | Data point 63 | 1.50000 mL | 0.16959 mL | 0.16366 mL | 0.21999 mL | 5.358 | -0.00404 | 0.04516 | 0.00094 | 14.5 s |
| 1:06:15.2 | Data point 64 | 1.50000 mL | 0.16959 mL | 0.16371 mL | 0.21999 mL | 5.687 | -0.01776 | 0.89415 | 0.00093 | 36.5 s |
| 1:07:17.1 | Data point 65 | 1.50000 mL | 0.16959 mL | 0.16376 mL | 0.21999 mL | 6.068 | -0.01777 | 0.97728 | 0.00089 | 40.5 s |
| 1:08:28.0 | Data point 66 | 1.50000 mL | 0.16959 mL | 0.16381 mL | 0.21999 mL | 6.474 | -0.02868 | 0.97106 | 0.00144 | Timed out at 59.5 s |
| 1:09:53.3 | Data point 67 | 1.50000 mL | 0.16959 mL | 0.16385 mL | 0.21999 mL | 6.603 | -0.01652 | 0.86246 | 0.00088 | 55.0 s |
| 1:11:13.8 | Data point 68 | 1.50000 mL | 0.16959 mL | 0.16388 mL | 0.21999 mL | 6.837 | -0.03943 | 0.95981 | 0.00199 | Timed out at 59.5 s |
| 1:12:44.3 | Data point 69 | 1.50000 mL | 0.16959 mL | 0.16392 mL | 0.21999 mL | 7.376 | -0.07353 | 0.99451 | 0.00364 | Timed out at 59.5 s |
| 1:14:14.8 | Data point 70 | 1.50000 mL | 0.16959 mL | 0.16397 mL | 0.21999 mL | 7.773 | -0.06843 | 0.99215 | 0.00339 | Timed out at 59.5 s |
| 1:15:45.3 | Data point 71 | 1.50000 mL | 0.16959 mL | 0.16402 mL | 0.21999 mL | 8.104 | -0.04184 | 0.98451 | 0.00208 | Timed out at 59.5 s |
| 1:17:31.3 | Data point 72 | 1.50000 mL | 0.16959 mL | 0.16414 mL | 0.21999 mL | 8.306 | -0.03456 | 0.94740 | 0.00175 | Timed out at 59.5 s |
| 1:19:06.9 | Data point 73 | 1.50000 mL | 0.16959 mL | 0.16420 mL | 0.21999 mL | 8.544 | -0.01489 | 0.84815 | 0.00080 | 33.5 s |
| 1:20:16.2 | Data point 74 | 1.50000 mL | 0.16959 mL | 0.16428 mL | 0.21999 mL | 8.801 | -0.01803 | 0.81436 | 0.00099 | 44.0 s |
| 1:21:36.0 | Data point 75 | 1.50000 mL | 0.16959 mL | 0.16439 mL | 0.21999 mL | 9.012 | -0.01523 | 0.58113 | 0.00099 | 13.0 s |
| 1:21:58.0 | Assay volumes | 1.50000 mL | 0.16959 mL | 0.16439 mL | 0.21999 mL | | | | | |

| | | | |
|---|---|---|---|
| Sample name: | **M11_octanol** | Experiment start time: | **3/1/2018 1:21:08 AM** |
| Assay name: | **pH-metric high logP** | Analyst: | **Pion** |
| Assay ID: | **18C-01002** | Instrument ID: | **T312060** |
| Filename: | **C:\Sirius_T3\Mehtap\20180228_exp28_logP_T3-2\18C-01002_M11_octanol_pH-metric high logP.t3r** | | |

| | | | |
|---|---|---|---|
| Sample name: | **M11_octanol** | Experiment start time: | **3/1/2018 1:21:08 AM** |
| Assay name: | **pH-metric high logP** | Analyst: | **Pion** |
| Assay ID: | **18C-01002** | Instrument ID: | **T312060** |
| Filename: | **C:\Sirius_T3\Mehtap\20180228_exp28_logP_T3-2\18C-01002_M11_octanol_pH-metric high logP.t3r** | | |

### Assay Settings (continued)

| Setting | Value | Original Value | Date/Time changed | Imported from |
|---|---|---|---|---|
| ***Titration 3*** | | | | |
| Titrate from | Low to high pH | | | |
| Add additional water | 0.00 mL | | | |
| Additional partition solvent volume | 0.150 mL | | | |
| Additional partition solvent added | Automatic | | | |
| After pH adjust stir for | 30 seconds | | | |
| Stir to allow partitioning for | 15 seconds | | | |
| Stirrer speed for partitioning | 60% | | | |
| ***Data Point Stability*** | | | | |
| Stir during data point collection | No | | | |
| Delay before data point collection | 0 seconds | | | |
| Number of points to average | 20 points | | | |
| Time interval between points | 0.50 seconds | | | |
| Required maximum standard deviation | 0.00100 dpH/dt | | | |
| Stability timeout after | 60 seconds | | | |

### Calibration Settings

| Setting | Value | Date/Time changed | Imported from |
|---|---|---|---|
| Four-Plus alpha | 0.130 | 3/1/2018 1:21:08 AM | C:\Sirius_T3\HCl18B27.t3r |
| Four-Plus S | 0.9970 | 3/1/2018 1:21:08 AM | C:\Sirius_T3\HCl18B27.t3r |
| Four-Plus jH | 0.8 | 3/1/2018 1:21:08 AM | C:\Sirius_T3\HCl18B27.t3r |
| Four-Plus jOH | -0.4 | 3/1/2018 1:21:08 AM | C:\Sirius_T3\HCl18B27.t3r |
| Base concentration factor | 1.000 | 3/1/2018 1:21:08 AM | C:\Sirius_T3\KOH18B27.t3r |
| Acid concentration factor | 0.994 | 3/1/2018 1:21:08 AM | C:\Sirius_T3\HCl18B27.t3r |

### Instrument Settings

| Setting | Value | Batch Id | Install date |
|---|---|---|---|
| Instrument owner | Merck | | |
| Instrument ID | T312060 | | |
| Instrument type | T3 Simulator | | |
| Software version | 1.1.3.0 | | |
| Dispenser module | | T3DM1200361 | 3/31/2009 5:24:52 AM |
| Dispenser 0 | Water | | 3/31/2009 5:25:05 AM |
| Syringe volume | 2.5 mL | | |
| Firmware version | 1.2.1(r2) | | |
| Titrant | Water (0.15 M KCl) | 02-06-2018 | 2/27/2018 10:05:59 AM |
| Dispenser 2 | Acid | | 3/31/2009 5:25:11 AM |
| Syringe volume | 0.5 mL | | |
| Firmware version | 1.2.1(r2) | | |
| Titrant | Acid (0.5 M HCl) | 02-27-2018 | 2/27/2018 10:27:22 AM |
| Dispenser 1 | Base | | 3/31/2009 5:25:21 AM |
| Syringe volume | 0.5 mL | | |
| Firmware version | 1.2.1(r2) | | |
| Titrant | Base (0.5 M KOH) | 9/22/2017 | 2/27/2018 10:21:22 AM |
| Dispenser 5 | Cosolvent | | 3/31/2009 5:26:24 AM |
| Syringe volume | 2.5 mL | | |
| Firmware version | 1.2.1(r2) | | |
| Distribution valve 5 | Distribution Valve | | 3/31/2009 5:28:19 AM |
| Firmware version | 1.1.3 | | |
| Port A | Methanol (80%, 0.15 M KCl) | 09-26-17 | 2/7/2018 9:42:01 AM |
| Port B | Cyclohexane | 11-01-17 | 2/27/2018 10:37:57 AM |
| Dispenser 3 | Buffer | | 8/3/2010 5:05:16 AM |
| Syringe volume | 0.5 mL | | |
| Firmware version | 1.2.1(r2) | | |
| Titrant | Dodecane | 2018/01/31 | 2/28/2018 10:18:04 AM |
| Dispenser 6 | Octanol | | 10/22/2010 10:52:43 AM |

| | | | |
|---|---|---|---|
| Sample name: | **M11_octanol** | Experiment start time: | **3/1/2018 1:21:08 AM** |
| Assay name: | **pH-metric high logP** | Analyst: | **Pion** |
| Assay ID: | **18C-01002** | Instrument ID: | **T312060** |
| Filename: | **C:\Sirius_T3\Mehtap\20180228_exp28_logP_T3-2\18C-01002_M11_octanol_pH-metric high logP.t3r** | | |

| | | | |
|---|---|---|---|
| Sample name: | **M11_octanol** | Experiment start time: | **3/1/2018 1:21:08 AM** |
| Assay name: | **pH-metric high logP** | Analyst: | **Pion** |
| Assay ID: | **18C-01002** | Instrument ID: | **T312060** |
| Filename: | **C:\Sirius_T3\Mehtap\20180228_exp28_logP_T3-2\18C-01002_M11_octanol_pH-metric high logP.t3r** | | |

### Instrument Settings (continued)

| Setting | Value | Batch Id | Install date |
|---|---|---|---|
| Spectrometer calibration stir duration | 5 s | | |
| Spectrometer calibration stir speed | 30% | | |
| Spectrometer calibration wash pump volume | 20.0 mL | | |
| Spectrometer calibration wash stir duration | 5 s | | |
| Spectrometer calibration wash stir speed | 30% | | |
| Overhead dispense height | 10000 | | |

### Refinement Settings

| Setting | Value | Default value |
|---|---|---|
| Turbidity detection method | None | None |
| Turbidity wavelength to assess | 500.0 nm | 500.0 nm |
| Turbidity maximum absorbance | 0.100 | 0.100 |
| Turbidity probe threshold | 50.00 | 50.00 |

### Experiment Log

[2:37] Air gap created for Water (0.15 M KCl)
[2:37] Air gap created for Acid (0.5 M HCl)
[2:38] Air gap created for Base (0.5 M KOH)
[2:38] Air gap released for Water (0.15 M KCl)
[2:42] Titrator arm moved over Titration position
[2:42] Titration 1 of 3
[2:42] Adding initial titrants
[2:42] Automatically add 1.50000 mL of water
[3:07] Dispensed 1.500000 mL of Water (0.15 M KCl)
[3:11] Titrator arm moved over Drain
[8:53] Titrator arm moved to Titration position
[8:53] Argon flow rate set to 100
[8:53] Stirrer speed set to 10
[8:58] Automatically add 0.02000 mL of Octanol
[8:58] Dispensed 0.019991 mL of Octanol
[8:59] Initial pH = 8.06
[8:59] Iterative adjust 8.06 -> 2.00
[8:59] pH 8.06 -> 2.00
[9:01] Air gap released for Acid (0.5 M HCl)
[9:02] Dispensed 0.051529 mL of Acid (0.5 M HCl)
[9:07] Holding pH 2.00
[11:07] Stirrer speed set to 0
[11:07] Stirrer speed set to 50
[11:07] Iterative adjust 1.98 -> 2.00
[11:07] pH 1.98 -> 2.00
[11:08] Air gap released for Base (0.5 M KOH)
[11:09] Dispensed 0.001787 mL of Base (0.5 M KOH)
[11:59] Stirrer speed set to 0
[12:09] Datapoint id 1 collected
[12:09] Stirrer speed set to 50
[12:14] pH 2.01 -> 2.21
[12:14] Using cautious pH adjust
[12:14] Dispensed 0.007855 mL of Base (0.5 M KOH)
[12:19] Stepping pH = 2.09
[12:20] Dispensed 0.006232 mL of Base (0.5 M KOH)
[12:25] Stepping pH = 2.19
[12:25] Dispensed 0.001129 mL of Base (0.5 M KOH)
[12:30] Stepping pH = 2.21
[12:45] Stirrer speed set to 0
[12:55] Datapoint id 2 collected
[12:55] Charge balance equation is out by 3.1%
[12:55] Stirrer speed set to 50

Sample name: **M11_octanol** Experiment start time: **3/1/2018 1:21:08 AM**
Assay name: **pH-metric high logP** Analyst: **Pion**
Assay ID: **18C-01002** Instrument ID: **T312060**
Filename: **C:\Sirius_T3\Mehtap\20180228_exp28_logP_T3-2\18C-01002_M11_octanol_pH-metric high logP.t3r**

### Experiment Log (continued)

[13:00] pH 2.22 -> 2.42
[13:00] Using charge balance adjust
[13:01] Dispensed 0.009713 mL of Base (0.5 M KOH)
[13:21] Stirrer speed set to 0
[13:31] Datapoint id 3 collected
[13:31] Charge balance equation is out by 7.5%
[13:31] Stirrer speed set to 50
[13:36] pH 2.44 -> 2.64
[13:36] Using charge balance adjust
[13:36] Dispensed 0.006021 mL of Base (0.5 M KOH)
[13:56] Stirrer speed set to 0
[14:06] Datapoint id 4 collected
[14:06] Charge balance equation is out by -3.5%
[14:06] Stirrer speed set to 50
[14:11] pH 2.64 -> 2.84
[14:11] Using charge balance adjust
[14:12] Dispensed 0.004069 mL of Base (0.5 M KOH)
[14:32] Stirrer speed set to 0
[14:42] Datapoint id 5 collected
[14:42] Charge balance equation is out by 10.1%
[14:42] Stirrer speed set to 50
[14:47] pH 2.86 -> 3.06
[14:47] Using charge balance adjust
[14:47] Dispensed 0.002846 mL of Base (0.5 M KOH)
[15:07] Stirrer speed set to 0
[15:17] Datapoint id 6 collected
[15:17] Charge balance equation is out by -2.6%
[15:17] Stirrer speed set to 50
[15:22] pH 3.07 -> 3.27
[15:22] Using charge balance adjust
[15:23] Dispensed 0.002258 mL of Base (0.5 M KOH)
[15:43] Stirrer speed set to 0
[15:53] Datapoint id 7 collected
[15:53] Charge balance equation is out by -2.0%
[15:53] Stirrer speed set to 50
[15:58] pH 3.27 -> 3.47
[15:58] Using charge balance adjust
[15:58] Dispensed 0.001929 mL of Base (0.5 M KOH)
[16:18] Stirrer speed set to 0
[16:28] Datapoint id 8 collected
[16:28] Charge balance equation is out by 8.1%
[16:28] Stirrer speed set to 50
[16:33] pH 3.49 -> 3.69
[16:33] Using charge balance adjust
[16:34] Dispensed 0.001693 mL of Base (0.5 M KOH)
[16:54] Stirrer speed set to 0
[17:04] Datapoint id 9 collected
[17:04] Charge balance equation is out by 8.4%
[17:04] Stirrer speed set to 50
[17:09] pH 3.71 -> 3.91
[17:09] Using charge balance adjust
[17:09] Dispensed 0.001435 mL of Base (0.5 M KOH)
[17:29] Stirrer speed set to 0
[17:40] Datapoint id 10 collected
[17:40] Charge balance equation is out by -7.0%
[17:40] Stirrer speed set to 50
[17:45] pH 3.90 -> 4.10
[17:45] Using charge balance adjust
[17:45] Dispensed 0.001176 mL of Base (0.5 M KOH)

| | | | |
|---|---|---|---|
| Sample name: | **M11_octanol** | Experiment start time: | **3/1/2018 1:21:08 AM** |
| Assay name: | **pH-metric high logP** | Analyst: | **Pion** |
| Assay ID: | **18C-01002** | Instrument ID: | **T312060** |
| Filename: | **C:\Sirius_T3\Mehtap\20180228_exp28_logP_T3-2\18C-01002_M11_octanol_pH-metric high logP.t3r** | | |

### Experiment Log (continued)

[18:05] Stirrer speed set to 0
[18:15] Datapoint id 11 collected
[18:15] Charge balance equation is out by -8.8%
[18:15] Stirrer speed set to 50
[18:20] pH 4.09 -> 4.29
[18:20] Using charge balance adjust
[18:20] Dispensed 0.000917 mL of Base (0.5 M KOH)
[18:41] Stirrer speed set to 0
[18:51] Datapoint id 12 collected
[18:51] Charge balance equation is out by -6.3%
[18:51] Stirrer speed set to 50
[18:56] pH 4.28 -> 4.48
[18:56] Using charge balance adjust
[18:56] Dispensed 0.000682 mL of Base (0.5 M KOH)
[19:16] Stirrer speed set to 0
[19:26] Datapoint id 13 collected
[19:26] Charge balance equation is out by -8.5%
[19:26] Stirrer speed set to 50
[19:32] pH 4.47 -> 4.67
[19:32] Using charge balance adjust
[19:32] Dispensed 0.000470 mL of Base (0.5 M KOH)
[19:52] Stirrer speed set to 0
[20:02] Datapoint id 14 collected
[20:02] Charge balance equation is out by -10.3%
[20:02] Stirrer speed set to 50
[20:07] pH 4.65 -> 4.85
[20:07] Using charge balance adjust
[20:07] Dispensed 0.000329 mL of Base (0.5 M KOH)
[20:27] Stirrer speed set to 0
[20:41] Datapoint id 15 collected
[20:41] Charge balance equation is out by -2.0%
[20:41] Stirrer speed set to 50
[20:46] pH 4.86 -> 5.06
[20:46] Using charge balance adjust
[20:46] Dispensed 0.000212 mL of Base (0.5 M KOH)
[21:06] Stirrer speed set to 0
[21:18] Datapoint id 16 collected
[21:18] Charge balance equation is out by -17.2%
[21:18] Stirrer speed set to 50
[21:23] pH 5.03 -> 5.23
[21:23] Using cautious pH adjust
[21:23] Dispensed 0.000071 mL of Base (0.5 M KOH)
[21:28] Stepping pH = 5.04
[21:29] Dispensed 0.000235 mL of Base (0.5 M KOH)
[21:34] Stepping pH = 5.67
[21:49] Stirrer speed set to 0
[22:11] Datapoint id 17 collected
[22:11] Charge balance equation is out by -93.1%
[22:11] Stirrer speed set to 50
[22:16] pH 5.82 -> 6.02
[22:16] Using cautious pH adjust
[22:16] Dispensed 0.000024 mL of Base (0.5 M KOH)
[22:21] Stepping pH = 5.84
[22:22] Dispensed 0.000071 mL of Base (0.5 M KOH)
[22:27] Stepping pH = 5.97
[22:27] Dispensed 0.000024 mL of Base (0.5 M KOH)
[22:32] Stepping pH = 6.20
[22:47] Stirrer speed set to 0
[23:27] Datapoint id 18 collected

Sample name: **M11_octanol** Experiment start time: **3/1/2018 1:21:08 AM**
Assay name: **pH-metric high logP** Analyst: **Pion**
Assay ID: **18C-01002** Instrument ID: **T312060**
Filename: **C:\Sirius_T3\Mehtap\20180228_exp28_logP_T3-2\18C-01002_M11_octanol_pH-metric high logP.t3r**

### Experiment Log (continued)

[23:27] Charge balance equation is out by -130.2%
[23:27] Stirrer speed set to 50
[23:32] pH 6.32 -> 6.52
[23:32] Using cautious pH adjust
[23:32] Dispensed 0.000024 mL of Base (0.5 M KOH)
[23:37] Stepping pH = 6.38
[23:37] Dispensed 0.000024 mL of Base (0.5 M KOH)
[23:42] Stepping pH = 6.54
[23:57] Stirrer speed set to 0
[24:42] Datapoint id 19 collected
[24:42] Charge balance equation is out by -11.0%
[24:42] Stirrer speed set to 50
[24:47] pH 6.49 -> 6.69
[24:47] Using charge balance adjust
[24:47] Dispensed 0.000047 mL of Base (0.5 M KOH)
[25:07] Stirrer speed set to 0
[25:52] Datapoint id 20 collected
[25:52] Charge balance equation is out by 7.8%
[25:52] Stirrer speed set to 50
[25:57] pH 6.69 -> 6.89
[25:57] Using charge balance adjust
[25:57] Dispensed 0.000024 mL of Base (0.5 M KOH)
[26:17] Stirrer speed set to 0
[27:09] Datapoint id 21 collected
[27:09] Charge balance equation is out by 15.7%
[27:09] Stirrer speed set to 50
[27:14] pH 6.96 -> 7.16
[27:14] Using cautious pH adjust
[27:14] Dispensed 0.000024 mL of Base (0.5 M KOH)
[27:19] Stepping pH = 7.08
[27:19] Dispensed 0.000024 mL of Base (0.5 M KOH)
[27:24] Stepping pH = 7.29
[27:40] Stirrer speed set to 0
[28:40] Datapoint id 22 collected
[28:40] Charge balance equation is out by -110.3%
[28:40] Stirrer speed set to 50
[28:45] pH 7.43 -> 7.63
[28:45] Using cautious pH adjust
[28:45] Dispensed 0.000024 mL of Base (0.5 M KOH)
[28:50] Stepping pH = 7.47
[28:50] Dispensed 0.000024 mL of Base (0.5 M KOH)
[28:55] Stepping pH = 7.66
[29:10] Stirrer speed set to 0
[30:10] Datapoint id 23 collected
[30:10] Charge balance equation is out by -359.3%
[30:10] Stirrer speed set to 50
[30:15] pH 7.88 -> 8.08
[30:15] Using cautious pH adjust
[30:15] Dispensed 0.000024 mL of Base (0.5 M KOH)
[30:20] Stepping pH = 7.89
[30:20] Dispensed 0.000024 mL of Base (0.5 M KOH)
[30:25] Stepping pH = 8.10
[30:40] Stirrer speed set to 0
[31:40] Datapoint id 24 collected
[31:40] Charge balance equation is out by -547.1%
[31:40] Stirrer speed set to 50
[31:46] pH 8.15 -> 8.35
[31:46] Using cautious pH adjust
[31:46] Dispensed 0.000024 mL of Base (0.5 M KOH)

Sample name: **M11_octanol**
Assay name: **pH-metric high logP**
Assay ID: **18C-01002**
Filename: **C:\Sirius_T3\Mehtap\20180228_exp28_logP_T3-2\18C-01002_M11_octanol_pH-metric high logP.t3r**

Experiment start time: **3/1/2018 1:21:08 AM**
Analyst: **Pion**
Instrument ID: **T312060**

### Experiment Log (continued)

[31:51] Stepping pH = 8.14
[31:51] Dispensed 0.000024 mL of Base (0.5 M KOH)
[31:56] Stepping pH = 8.26
[31:56] Dispensed 0.000024 mL of Base (0.5 M KOH)
[32:01] Stepping pH = 8.46
[32:16] Stirrer speed set to 0
[33:11] Datapoint id 25 collected
[33:11] Charge balance equation is out by -675.1%
[33:11] Stirrer speed set to 50
[33:16] pH 8.52 -> 8.72
[33:16] Using cautious pH adjust
[33:16] Dispensed 0.000024 mL of Base (0.5 M KOH)
[33:21] Stepping pH = 8.57
[33:21] Dispensed 0.000024 mL of Base (0.5 M KOH)
[33:26] Stepping pH = 8.67
[33:26] Dispensed 0.000024 mL of Base (0.5 M KOH)
[33:31] Stepping pH = 8.76
[33:46] Stirrer speed set to 0
[34:27] Datapoint id 26 collected
[34:27] Charge balance equation is out by -301.4%
[34:27] Stirrer speed set to 50
[34:32] pH 8.73 -> 8.93
[34:32] Using cautious pH adjust
[34:32] Dispensed 0.000024 mL of Base (0.5 M KOH)
[34:37] Stepping pH = 8.74
[34:37] Dispensed 0.000024 mL of Base (0.5 M KOH)
[34:42] Stepping pH = 8.81
[34:42] Dispensed 0.000047 mL of Base (0.5 M KOH)
[34:47] Stepping pH = 8.96
[35:02] Stirrer speed set to 0
[35:37] Datapoint id 27 collected
[35:37] Charge balance equation is out by -310.7%
[35:37] Stirrer speed set to 50
[35:43] pH 8.95 -> 9.05
[35:43] Using cautious pH adjust
[35:43] Dispensed 0.000024 mL of Base (0.5 M KOH)
[35:48] Stepping pH = 8.95
[35:48] Dispensed 0.000047 mL of Base (0.5 M KOH)
[35:53] Stepping pH = 9.03
[36:08] Stirrer speed set to 0
[36:29] Datapoint id 28 collected
[36:29] Charge balance equation is out by -289.1%
[36:29] Titration 2 of 3
[36:29] Adding initial titrants
[36:29] Automatically add 0.05000 mL of Octanol
[36:31] Dispensed 0.050000 mL of Octanol
[36:31] Stirrer speed set to 10
[36:32] Stirrer speed set to 55
[36:32] Iterative adjust 9.08 -> 2.00
[36:32] pH 9.08 -> 2.00
[36:33] Dispensed 0.054139 mL of Acid (0.5 M HCl)
[36:38] pH 2.02 -> 2.00
[36:38] Dispensed 0.002352 mL of Acid (0.5 M HCl)
[37:29] Stirrer speed set to 0
[37:39] Datapoint id 29 collected
[37:39] Stirrer speed set to 55
[37:44] pH 1.97 -> 2.17
[37:44] Using cautious pH adjust
[37:44] Dispensed 0.009313 mL of Base (0.5 M KOH)

Sample name: **M11_octanol**
Assay name: **pH-metric high logP**
Assay ID: **18C-01002**
Filename: **C:\Sirius_T3\Mehtap\20180228_exp28_logP_T3-2\18C-01002_M11_octanol_pH-metric high logP.t3r**

Experiment start time: **3/1/2018 1:21:08 AM**
Analyst: **Pion**
Instrument ID: **T312060**

### Experiment Log (continued)

[37:49] Stepping pH = 2.06
[37:50] Dispensed 0.006538 mL of Base (0.5 M KOH)
[37:55] Stepping pH = 2.15
[37:55] Dispensed 0.001458 mL of Base (0.5 M KOH)
[38:00] Stepping pH = 2.17
[38:15] Stirrer speed set to 0
[38:25] Datapoint id 30 collected
[38:25] Charge balance equation is out by 7.0%
[38:25] Stirrer speed set to 55
[38:30] pH 2.17 -> 2.37
[38:30] Using charge balance adjust
[38:30] Dispensed 0.011783 mL of Base (0.5 M KOH)
[38:51] Stirrer speed set to 0
[39:01] Datapoint id 31 collected
[39:01] Charge balance equation is out by 12.3%
[39:01] Stirrer speed set to 55
[39:06] pH 2.40 -> 2.60
[39:06] Using charge balance adjust
[39:06] Dispensed 0.007291 mL of Base (0.5 M KOH)
[39:26] Stirrer speed set to 0
[39:36] Datapoint id 32 collected
[39:36] Charge balance equation is out by 5.2%
[39:36] Stirrer speed set to 55
[39:41] pH 2.62 -> 2.82
[39:41] Using charge balance adjust
[39:42] Dispensed 0.004892 mL of Base (0.5 M KOH)
[40:02] Stirrer speed set to 0
[40:12] Datapoint id 33 collected
[40:12] Charge balance equation is out by 2.4%
[40:12] Stirrer speed set to 55
[40:17] pH 2.83 -> 3.03
[40:17] Using charge balance adjust
[40:17] Dispensed 0.003551 mL of Base (0.5 M KOH)
[40:37] Stirrer speed set to 0
[40:47] Datapoint id 34 collected
[40:47] Charge balance equation is out by -4.4%
[40:47] Stirrer speed set to 55
[40:52] pH 3.02 -> 3.22
[40:52] Using charge balance adjust
[40:52] Dispensed 0.002752 mL of Base (0.5 M KOH)
[41:13] Stirrer speed set to 0
[41:23] Datapoint id 35 collected
[41:23] Charge balance equation is out by 5.0%
[41:23] Stirrer speed set to 55
[41:28] pH 3.24 -> 3.44
[41:28] Using charge balance adjust
[41:28] Dispensed 0.002140 mL of Base (0.5 M KOH)
[41:48] Stirrer speed set to 0
[41:58] Datapoint id 36 collected
[41:58] Charge balance equation is out by 37.0%
[41:58] Stirrer speed set to 55
[42:03] pH 3.51 -> 3.71
[42:03] Using cautious pH adjust
[42:03] Dispensed 0.000753 mL of Base (0.5 M KOH)
[42:08] Stepping pH = 3.61
[42:09] Dispensed 0.000517 mL of Base (0.5 M KOH)
[42:14] Stepping pH = 3.70
[42:14] Dispensed 0.000047 mL of Base (0.5 M KOH)
[42:19] Stepping pH = 3.71

Sample name: **M11_octanol**
Assay name: **pH-metric high logP**
Assay ID: **18C-01002**
Filename: **C:\Sirius_T3\Mehtap\20180228_exp28_logP_T3-2\18C-01002_M11_octanol_pH-metric high logP.t3r**

Experiment start time: **3/1/2018 1:21:08 AM**
Analyst: **Pion**
Instrument ID: **T312060**

### Experiment Log (continued)

[42:34] Stirrer speed set to 0
[42:44] Datapoint id 37 collected
[42:44] Charge balance equation is out by 11.6%
[42:44] Stirrer speed set to 55
[42:49] pH 3.71 -> 3.91
[42:49] Using charge balance adjust
[42:49] Dispensed 0.001105 mL of Base (0.5 M KOH)
[43:09] Stirrer speed set to 0
[43:19] Datapoint id 38 collected
[43:19] Charge balance equation is out by 21.3%
[43:19] Stirrer speed set to 55
[43:24] pH 3.96 -> 4.16
[43:24] Using cautious pH adjust
[43:24] Dispensed 0.000353 mL of Base (0.5 M KOH)
[43:30] Stepping pH = 4.04
[43:30] Dispensed 0.000306 mL of Base (0.5 M KOH)
[43:35] Stepping pH = 4.14
[43:35] Dispensed 0.000047 mL of Base (0.5 M KOH)
[43:40] Stepping pH = 4.16
[43:55] Stirrer speed set to 0
[44:06] Datapoint id 39 collected
[44:06] Charge balance equation is out by -1.4%
[44:06] Stirrer speed set to 55
[44:11] pH 4.17 -> 4.37
[44:11] Using charge balance adjust
[44:11] Dispensed 0.000494 mL of Base (0.5 M KOH)
[44:31] Stirrer speed set to 0
[44:41] Datapoint id 40 collected
[44:41] Charge balance equation is out by 26.0%
[44:41] Stirrer speed set to 55
[44:46] pH 4.42 -> 4.62
[44:46] Using cautious pH adjust
[44:46] Dispensed 0.000141 mL of Base (0.5 M KOH)
[44:51] Stepping pH = 4.46
[44:51] Dispensed 0.000259 mL of Base (0.5 M KOH)
[44:56] Stepping pH = 4.72
[45:12] Stirrer speed set to 0
[45:25] Datapoint id 41 collected
[45:25] Charge balance equation is out by -38.6%
[45:25] Stirrer speed set to 55
[45:30] pH 4.78 -> 4.98
[45:30] Using cautious pH adjust
[45:30] Dispensed 0.000071 mL of Base (0.5 M KOH)
[45:35] Stepping pH = 4.79
[45:35] Dispensed 0.000188 mL of Base (0.5 M KOH)
[45:40] Stepping pH = 5.22
[45:55] Stirrer speed set to 0
[46:11] Datapoint id 42 collected
[46:11] Charge balance equation is out by -92.6%
[46:11] Stirrer speed set to 55
[46:16] pH 5.29 -> 5.49
[46:16] Using cautious pH adjust
[46:16] Dispensed 0.000024 mL of Base (0.5 M KOH)
[46:21] Stepping pH = 5.29
[46:21] Dispensed 0.000141 mL of Base (0.5 M KOH)
[46:26] Stepping pH = 5.90
[46:41] Stirrer speed set to 0
[47:27] Datapoint id 43 collected
[47:27] Charge balance equation is out by -206.7%

| | | | |
|---|---|---|---|
| Sample name: | **M11_octanol** | Experiment start time: | **3/1/2018 1:21:08 AM** |
| Assay name: | **pH-metric high logP** | Analyst: | **Pion** |
| Assay ID: | **18C-01002** | Instrument ID: | **T312060** |
| Filename: | **C:\Sirius_T3\Mehtap\20180228_exp28_logP_T3-2\18C-01002_M11_octanol_pH-metric high logP.t3r** | | |

### Experiment Log (continued)

[47:27] Stirrer speed set to 55
[47:32] pH 6.12 -> 6.32
[47:32] Using cautious pH adjust
[47:32] Dispensed 0.000024 mL of Base (0.5 M KOH)
[47:37] Stepping pH = 6.14
[47:37] Dispensed 0.000047 mL of Base (0.5 M KOH)
[47:43] Stepping pH = 6.30
[47:43] Dispensed 0.000024 mL of Base (0.5 M KOH)
[47:48] Stepping pH = 6.62
[48:03] Stirrer speed set to 0
[49:03] Datapoint id 44 collected
[49:03] Charge balance equation is out by -118.6%
[49:03] Stirrer speed set to 55
[49:08] pH 6.74 -> 6.94
[49:08] Using cautious pH adjust
[49:08] Dispensed 0.000024 mL of Base (0.5 M KOH)
[49:13] Stepping pH = 6.76
[49:13] Dispensed 0.000047 mL of Base (0.5 M KOH)
[49:18] Stepping pH = 7.01
[49:33] Stirrer speed set to 0
[50:33] Datapoint id 45 collected
[50:33] Charge balance equation is out by -103.4%
[50:33] Stirrer speed set to 55
[50:38] pH 7.33 -> 7.53
[50:38] Using cautious pH adjust
[50:38] Dispensed 0.000024 mL of Base (0.5 M KOH)
[50:43] Stepping pH = 7.29
[50:43] Dispensed 0.000024 mL of Base (0.5 M KOH)
[50:49] Stepping pH = 7.56
[51:04] Stirrer speed set to 0
[52:04] Datapoint id 46 collected
[52:04] Charge balance equation is out by -392.4%
[52:04] Stirrer speed set to 55
[52:09] pH 7.78 -> 7.98
[52:09] Using cautious pH adjust
[52:09] Dispensed 0.000024 mL of Base (0.5 M KOH)
[52:14] Stepping pH = 7.73
[52:14] Dispensed 0.000024 mL of Base (0.5 M KOH)
[52:19] Stepping pH = 7.78
[52:19] Dispensed 0.000047 mL of Base (0.5 M KOH)
[52:24] Stepping pH = 8.11
[52:39] Stirrer speed set to 0
[53:39] Datapoint id 47 collected
[53:39] Charge balance equation is out by -1,044.0%
[53:39] Stirrer speed set to 55
[53:45] pH 7.92 -> 8.12
[53:45] Using cautious pH adjust
[53:45] Dispensed 0.000024 mL of Base (0.5 M KOH)
[53:50] Stepping pH = 7.88
[53:50] Dispensed 0.000024 mL of Base (0.5 M KOH)
[53:55] Stepping pH = 7.86
[53:55] Dispensed 0.000118 mL of Base (0.5 M KOH)
[54:00] Stepping pH = 8.83
[54:15] Stirrer speed set to 0
[54:43] Datapoint id 48 collected
[54:43] Charge balance equation is out by -2,120.7%
[54:43] Stirrer speed set to 55
[54:48] pH 8.82 -> 9.02
[54:48] Using cautious pH adjust

| | | | |
|---|---|---|---|
| Sample name: | **M11_octanol** | Experiment start time: | **3/1/2018 1:21:08 AM** |
| Assay name: | **pH-metric high logP** | Analyst: | **Pion** |
| Assay ID: | **18C-01002** | Instrument ID: | **T312060** |
| Filename: | **C:\Sirius_T3\Mehtap\20180228_exp28_logP_T3-2\18C-01002_M11_octanol_pH-metric high logP.t3r** | | |

### Experiment Log (continued)

[54:48] Dispensed 0.000024 mL of Base (0.5 M KOH)
[54:53] Stepping pH = 8.83
[54:53] Dispensed 0.000047 mL of Base (0.5 M KOH)
[54:58] Stepping pH = 8.90
[54:59] Dispensed 0.000047 mL of Base (0.5 M KOH)
[55:04] Stepping pH = 9.03
[55:19] Stirrer speed set to 0
[55:43] Datapoint id 49 collected
[55:43] Charge balance equation is out by -286.1%
[55:43] Titration 3 of 3
[55:43] Adding initial titrants
[55:43] Automatically add 0.15000 mL of Octanol
[55:47] Dispensed 0.150000 mL of Octanol
[55:47] Stirrer speed set to 10
[55:48] Stirrer speed set to 60
[55:48] Iterative adjust 9.05 -> 2.00
[55:48] pH 9.05 -> 2.00
[55:50] Dispensed 0.056397 mL of Acid (0.5 M HCl)
[55:55] pH 2.05 -> 2.00
[55:55] Dispensed 0.005174 mL of Acid (0.5 M HCl)
[56:45] Stirrer speed set to 0
[56:55] Datapoint id 50 collected
[56:55] Stirrer speed set to 60
[57:00] pH 1.97 -> 2.17
[57:00] Using cautious pH adjust
[57:01] Dispensed 0.010113 mL of Base (0.5 M KOH)
[57:06] Stepping pH = 2.05
[57:06] Dispensed 0.007949 mL of Base (0.5 M KOH)
[57:11] Stepping pH = 2.15
[57:11] Dispensed 0.001129 mL of Base (0.5 M KOH)
[57:16] Stepping pH = 2.17
[57:31] Stirrer speed set to 0
[57:41] Datapoint id 51 collected
[57:41] Charge balance equation is out by 5.1%
[57:41] Stirrer speed set to 60
[57:47] pH 2.17 -> 2.37
[57:47] Using charge balance adjust
[57:47] Dispensed 0.012959 mL of Base (0.5 M KOH)
[58:07] Stirrer speed set to 0
[58:17] Datapoint id 52 collected
[58:17] Charge balance equation is out by 9.6%
[58:17] Stirrer speed set to 60
[58:22] pH 2.40 -> 2.60
[58:22] Using charge balance adjust
[58:23] Dispensed 0.008278 mL of Base (0.5 M KOH)
[58:43] Stirrer speed set to 0
[58:53] Datapoint id 53 collected
[58:53] Charge balance equation is out by 10.7%
[58:53] Stirrer speed set to 60
[58:58] pH 2.62 -> 2.82
[58:58] Using charge balance adjust
[58:58] Dispensed 0.005433 mL of Base (0.5 M KOH)
[59:18] Stirrer speed set to 0
[59:29] Datapoint id 54 collected
[59:29] Charge balance equation is out by -3.5%
[59:29] Stirrer speed set to 60
[59:34] pH 2.82 -> 3.02
[59:34] Using charge balance adjust
[59:34] Dispensed 0.003857 mL of Base (0.5 M KOH)

| | | | |
|---|---|---|---|
| Sample name: | **M11_octanol** | Experiment start time: | **3/1/2018 1:21:08 AM** |
| Assay name: | **pH-metric high logP** | Analyst: | **Pion** |
| Assay ID: | **18C-01002** | Instrument ID: | **T312060** |
| Filename: | **C:\Sirius_T3\Mehtap\20180228_exp28_logP_T3-2\18C-01002_M11_octanol_pH-metric high logP.t3r** | | |

### Experiment Log (continued)

[59:55] Stirrer speed set to 0
[1:00:05] Datapoint id 55 collected
[1:00:05] Charge balance equation is out by 4.7%
[1:00:05] Stirrer speed set to 60
[1:00:10] pH 3.04 -> 3.24
[1:00:10] Using charge balance adjust
[1:00:10] Dispensed 0.002681 mL of Base (0.5 M KOH)
[1:00:30] Stirrer speed set to 0
[1:00:40] Datapoint id 56 collected
[1:00:40] Charge balance equation is out by 20.3%
[1:00:40] Stirrer speed set to 60
[1:00:45] pH 3.28 -> 3.48
[1:00:45] Using cautious pH adjust
[1:00:45] Dispensed 0.000870 mL of Base (0.5 M KOH)
[1:00:50] Stepping pH = 3.37
[1:00:50] Dispensed 0.000706 mL of Base (0.5 M KOH)
[1:00:56] Stepping pH = 3.47
[1:00:56] Dispensed 0.000071 mL of Base (0.5 M KOH)
[1:01:01] Stepping pH = 3.49
[1:01:16] Stirrer speed set to 0
[1:01:26] Datapoint id 57 collected
[1:01:26] Charge balance equation is out by 5.0%
[1:01:26] Stirrer speed set to 60
[1:01:31] pH 3.49 -> 3.69
[1:01:31] Using charge balance adjust
[1:01:31] Dispensed 0.001152 mL of Base (0.5 M KOH)
[1:01:51] Stirrer speed set to 0
[1:02:01] Datapoint id 58 collected
[1:02:01] Charge balance equation is out by 16.7%
[1:02:01] Stirrer speed set to 60
[1:02:06] pH 3.73 -> 3.93
[1:02:06] Using cautious pH adjust
[1:02:07] Dispensed 0.000353 mL of Base (0.5 M KOH)
[1:02:12] Stepping pH = 3.78
[1:02:12] Dispensed 0.000564 mL of Base (0.5 M KOH)
[1:02:17] Stepping pH = 3.98
[1:02:32] Stirrer speed set to 0
[1:02:42] Datapoint id 59 collected
[1:02:42] Charge balance equation is out by -28.5%
[1:02:42] Stirrer speed set to 60
[1:02:47] pH 4.01 -> 4.21
[1:02:47] Using cautious pH adjust
[1:02:47] Dispensed 0.000188 mL of Base (0.5 M KOH)
[1:02:52] Stepping pH = 4.05
[1:02:52] Dispensed 0.000353 mL of Base (0.5 M KOH)
[1:02:57] Stepping pH = 4.32
[1:03:12] Stirrer speed set to 0
[1:03:22] Datapoint id 60 collected
[1:03:22] Charge balance equation is out by -40.2%
[1:03:22] Stirrer speed set to 60
[1:03:28] pH 4.36 -> 4.56
[1:03:28] Using cautious pH adjust
[1:03:28] Dispensed 0.000094 mL of Base (0.5 M KOH)
[1:03:33] Stepping pH = 4.38
[1:03:33] Dispensed 0.000212 mL of Base (0.5 M KOH)
[1:03:38] Stepping pH = 4.73
[1:03:53] Stirrer speed set to 0
[1:04:05] Datapoint id 61 collected
[1:04:05] Charge balance equation is out by -65.1%

### Experiment Log (continued)

[1:04:05] Stirrer speed set to 60
[1:04:10] pH 4.80 -> 5.00
[1:04:10] Using cautious pH adjust
[1:04:10] Dispensed 0.000047 mL of Base (0.5 M KOH)
[1:04:15] Stepping pH = 4.83
[1:04:15] Dispensed 0.000071 mL of Base (0.5 M KOH)
[1:04:20] Stepping pH = 4.94
[1:04:21] Dispensed 0.000024 mL of Base (0.5 M KOH)
[1:04:26] Stepping pH = 5.04
[1:04:41] Stirrer speed set to 0
[1:04:54] Datapoint id 62 collected
[1:04:54] Charge balance equation is out by -98.4%
[1:04:54] Stirrer speed set to 60
[1:04:59] pH 5.08 -> 5.28
[1:04:59] Using cautious pH adjust
[1:04:59] Dispensed 0.000024 mL of Base (0.5 M KOH)
[1:05:04] Stepping pH = 5.13
[1:05:05] Dispensed 0.000024 mL of Base (0.5 M KOH)
[1:05:10] Stepping pH = 5.22
[1:05:10] Dispensed 0.000024 mL of Base (0.5 M KOH)
[1:05:15] Stepping pH = 5.32
[1:05:30] Stirrer speed set to 0
[1:05:44] Datapoint id 63 collected
[1:05:44] Charge balance equation is out by -64.2%
[1:05:44] Stirrer speed set to 60
[1:05:50] pH 5.43 -> 5.63
[1:05:50] Using cautious pH adjust
[1:05:50] Dispensed 0.000024 mL of Base (0.5 M KOH)
[1:05:55] Stepping pH = 5.51
[1:05:55] Dispensed 0.000024 mL of Base (0.5 M KOH)
[1:06:00] Stepping pH = 5.65
[1:06:15] Stirrer speed set to 0
[1:06:51] Datapoint id 64 collected
[1:06:51] Charge balance equation is out by -14.9%
[1:06:51] Stirrer speed set to 60
[1:06:57] pH 5.73 -> 5.93
[1:06:57] Using charge balance adjust
[1:06:57] Dispensed 0.000047 mL of Base (0.5 M KOH)
[1:07:17] Stirrer speed set to 0
[1:07:57] Datapoint id 65 collected
[1:07:57] Charge balance equation is out by 69.9%
[1:07:57] Stirrer speed set to 60
[1:08:02] pH 6.06 -> 6.26
[1:08:02] Using cautious pH adjust
[1:08:02] Dispensed 0.000024 mL of Base (0.5 M KOH)
[1:08:08] Stepping pH = 6.22
[1:08:08] Dispensed 0.000024 mL of Base (0.5 M KOH)
[1:08:13] Stepping pH = 6.45
[1:08:28] Stirrer speed set to 0
[1:09:28] Datapoint id 66 collected
[1:09:28] Charge balance equation is out by 4.3%
[1:09:28] Stirrer speed set to 60
[1:09:33] pH 6.45 -> 6.65
[1:09:33] Using charge balance adjust
[1:09:33] Dispensed 0.000047 mL of Base (0.5 M KOH)
[1:09:53] Stirrer speed set to 0
[1:10:48] Datapoint id 67 collected
[1:10:48] Charge balance equation is out by -24.7%
[1:10:48] Stirrer speed set to 60

| | | | |
|---|---|---|---|
| Sample name: | **M11_octanol** | Experiment start time: | **3/1/2018 1:21:08 AM** |
| Assay name: | **pH-metric high logP** | Analyst: | **Pion** |
| Assay ID: | **18C-01002** | Instrument ID: | **T312060** |
| Filename: | **C:\Sirius_T3\Mehtap\20180228_exp28_logP_T3-2\18C-01002_M11_octanol_pH-metric high logP.t3r** | | |

### Experiment Log (continued)

[1:10:53] pH 6.58 -> 6.78
[1:10:53] Using cautious pH adjust
[1:10:53] Dispensed 0.000024 mL of Base (0.5 M KOH)
[1:10:58] Stepping pH = 6.77
[1:11:13] Stirrer speed set to 0
[1:12:13] Datapoint id 68 collected
[1:12:13] Charge balance equation is out by 47.7%
[1:12:13] Stirrer speed set to 60
[1:12:19] pH 6.87 -> 7.07
[1:12:19] Using cautious pH adjust
[1:12:19] Dispensed 0.000024 mL of Base (0.5 M KOH)
[1:12:24] Stepping pH = 7.05
[1:12:24] Dispensed 0.000024 mL of Base (0.5 M KOH)
[1:12:29] Stepping pH = 7.33
[1:12:44] Stirrer speed set to 0
[1:13:44] Datapoint id 69 collected
[1:13:44] Charge balance equation is out by -44.9%
[1:13:44] Stirrer speed set to 60
[1:13:49] pH 7.36 -> 7.56
[1:13:49] Using cautious pH adjust
[1:13:49] Dispensed 0.000024 mL of Base (0.5 M KOH)
[1:13:54] Stepping pH = 7.45
[1:13:54] Dispensed 0.000024 mL of Base (0.5 M KOH)
[1:13:59] Stepping pH = 7.73
[1:14:14] Stirrer speed set to 0
[1:15:14] Datapoint id 70 collected
[1:15:14] Charge balance equation is out by -212.1%
[1:15:14] Stirrer speed set to 60
[1:15:20] pH 7.77 -> 7.97
[1:15:20] Using cautious pH adjust
[1:15:20] Dispensed 0.000024 mL of Base (0.5 M KOH)
[1:15:25] Stepping pH = 7.85
[1:15:25] Dispensed 0.000024 mL of Base (0.5 M KOH)
[1:15:30] Stepping pH = 8.04
[1:15:45] Stirrer speed set to 0
[1:16:45] Datapoint id 71 collected
[1:16:45] Charge balance equation is out by -401.0%
[1:16:45] Stirrer speed set to 60
[1:16:50] pH 8.05 -> 8.25
[1:16:50] Using cautious pH adjust
[1:16:50] Dispensed 0.000024 mL of Base (0.5 M KOH)
[1:16:55] Stepping pH = 8.15
[1:16:55] Dispensed 0.000024 mL of Base (0.5 M KOH)
[1:17:00] Stepping pH = 8.21
[1:17:00] Dispensed 0.000024 mL of Base (0.5 M KOH)
[1:17:06] Stepping pH = 8.19
[1:17:06] Dispensed 0.000024 mL of Base (0.5 M KOH)
[1:17:11] Stepping pH = 8.20
[1:17:11] Dispensed 0.000024 mL of Base (0.5 M KOH)
[1:17:16] Stepping pH = 8.32
[1:17:31] Stirrer speed set to 0
[1:18:31] Datapoint id 72 collected
[1:18:31] Charge balance equation is out by -1,251.5%
[1:18:31] Stirrer speed set to 60
[1:18:36] pH 8.28 -> 8.48
[1:18:36] Using cautious pH adjust
[1:18:36] Dispensed 0.000024 mL of Base (0.5 M KOH)
[1:18:41] Stepping pH = 8.27
[1:18:41] Dispensed 0.000024 mL of Base (0.5 M KOH)

Sample name: **M11_octanol** Experiment start time: **3/1/2018 1:21:08 AM**
Assay name: **pH-metric high logP** Analyst: **Pion**
Assay ID: **18C-01002** Instrument ID: **T312060**
Filename: **C:\Sirius_T3\Mehtap\20180228_exp28_logP_T3-2\18C-01002_M11_octanol_pH-metric high logP.t3r**

### Experiment Log (continued)

[1:18:46] Stepping pH = 8.39
[1:18:46] Dispensed 0.000024 mL of Base (0.5 M KOH)
[1:18:51] Stepping pH = 8.53
[1:19:07] Stirrer speed set to 0
[1:19:40] Datapoint id 73 collected
[1:19:40] Charge balance equation is out by -521.2%
[1:19:40] Stirrer speed set to 60
[1:19:45] pH 8.59 -> 8.79
[1:19:45] Using cautious pH adjust
[1:19:45] Dispensed 0.000024 mL of Base (0.5 M KOH)
[1:19:51] Stepping pH = 8.65
[1:19:51] Dispensed 0.000024 mL of Base (0.5 M KOH)
[1:19:56] Stepping pH = 8.73
[1:19:56] Dispensed 0.000024 mL of Base (0.5 M KOH)
[1:20:01] Stepping pH = 8.81
[1:20:16] Stirrer speed set to 0
[1:21:00] Datapoint id 74 collected
[1:21:00] Charge balance equation is out by -196.5%
[1:21:00] Stirrer speed set to 60
[1:21:05] pH 8.79 -> 8.99
[1:21:05] Using cautious pH adjust
[1:21:05] Dispensed 0.000024 mL of Base (0.5 M KOH)
[1:21:10] Stepping pH = 8.78
[1:21:10] Dispensed 0.000071 mL of Base (0.5 M KOH)
[1:21:15] Stepping pH = 8.93
[1:21:15] Dispensed 0.000024 mL of Base (0.5 M KOH)
[1:21:21] Stepping pH = 9.00
[1:21:36] Stirrer speed set to 0
[1:21:49] Datapoint id 75 collected
[1:21:49] Charge balance equation is out by -327.4%
[1:21:49] Argon flow rate set to 0
[1:21:53] Titrator arm moved over Titration position
