## Supplementary material for "Octanol-water partition coefficient measurements for the SAMPL6 Blind Prediction Challenge": SM11_18C-09010_M11_octanol_pH-metric high logP_report.pdf

| | | | |
|---|---|---|---|
| Sample name: | **M11_octanol** | Experiment start time: | **3/9/2018 4:01:24 PM** |
| Assay name: | **pH-metric high logP** | Analyst: | **Pion** |
| Assay ID: | **18C-09010** | Instrument ID: | **T312060** |
| Filename: | **C:\Sirius_T3\Mehtap\20180309_exp31_logP_T3-2\18C-09010_M11_octanol_pH-metric high logP.t3r** | | |

### pH-metric Result

| | | |
|---|---|---|
| logP (XH +) | -0.33 | ±0.82 (n=50) |
| logP (neutral X) | 2.12 | ±0.01 (n=50) |
| RMSD | 0.537 | |

### 18C-09010 Points 1 to 24

| | |
|---|---|
| M11_octanol concentration factor | 0.962 |
| Carbonate | 0.0600 mM |
| Acidity error | -0.46392 mM |

### 18C-09010 Points 25 to 45

| | |
|---|---|
| M11_octanol concentration factor | 0.930 |
| Carbonate | 0.0890 mM |
| Acidity error | -0.75734 mM |

### 18C-09010 Points 46 to 69

| | |
|---|---|
| M11_octanol concentration factor | 0.907 |
| Carbonate | 0.1290 mM |
| Acidity error | -0.81079 mM |

### Warnings and errors

| | |
|---|---|
| Errors | None |
| Warnings | None |

### Sample logD and percent species

| pH | M11_octanol logD | M11_octanol M11_octanolH | M11_octanol M11_octanol | M11_octanol M11_octanolH* | M11_octanol M11_octanol* | Comment |
|---|---|---|---|---|---|---|
| 1.000 | -0.20 | 61.06 % | 0.08 % | 28.50 % | 10.36 % | |
| 1.200 | -0.13 | 57.55 % | 0.12 % | 26.86 % | 15.48 % | Stomach pH |
| 2.000 | 0.33 | 31.48 % | 0.41 % | 14.69 % | 53.42 % | |
| 3.000 | 1.19 | 5.39 % | 0.69 % | 2.51 % | 91.40 % | |
| 4.000 | 1.87 | 0.58 % | 0.75 % | 0.27 % | 98.40 % | |
| 5.000 | 2.09 | 0.06 % | 0.75 % | 0.03 % | 99.16 % | |
| 6.000 | 2.12 | 0.01 % | 0.75 % | 0.00 % | 99.24 % | |
| 6.500 | 2.12 | 0.00 % | 0.75 % | 0.00 % | 99.24 % | |
| 7.000 | 2.12 | 0.00 % | 0.75 % | 0.00 % | 99.25 % | |
| 7.400 | 2.12 | 0.00 % | 0.75 % | 0.00 % | 99.25 % | Blood pH |
| 8.000 | 2.12 | 0.00 % | 0.75 % | 0.00 % | 99.25 % | |
| 9.000 | 2.12 | 0.00 % | 0.75 % | 0.00 % | 99.25 % | |
| 10.000 | 2.12 | 0.00 % | 0.75 % | 0.00 % | 99.25 % | |
| 11.000 | 2.12 | 0.00 % | 0.75 % | 0.00 % | 99.25 % | |
| 12.000 | 2.12 | 0.00 % | 0.75 % | 0.00 % | 99.25 % | |

Sample name: **M11_octanol** Experiment start time: **3/9/2018 4:01:24 PM**
Assay name: **pH-metric high logP** Analyst: **Pion**
Assay ID: **18C-09010** Instrument ID: **T312060**
Filename: **C:\Sirius_T3\Mehtap\20180309_exp31_logP_T3-2\18C-09010_M11_octanol_pH-metric high logP.t3r**

### Graphs

Charge balance residuals

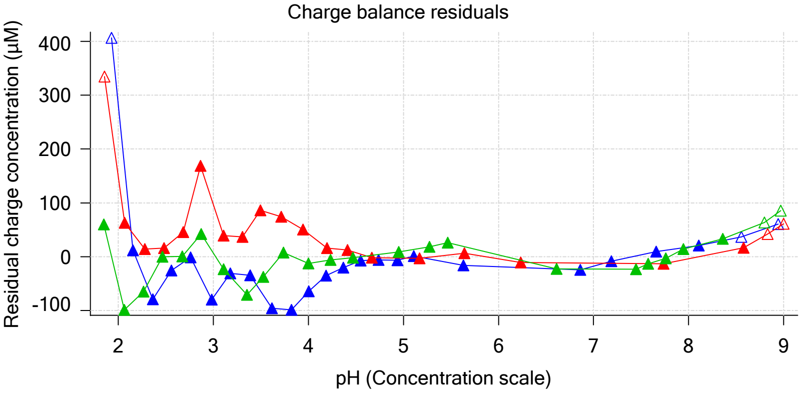


Ionic strength

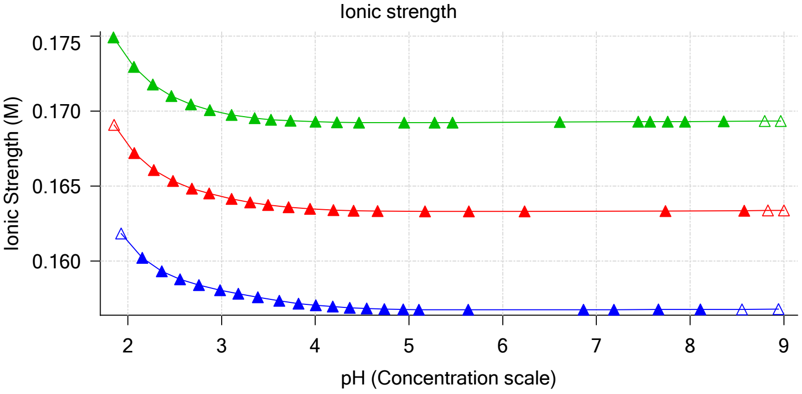


Buffer index profile

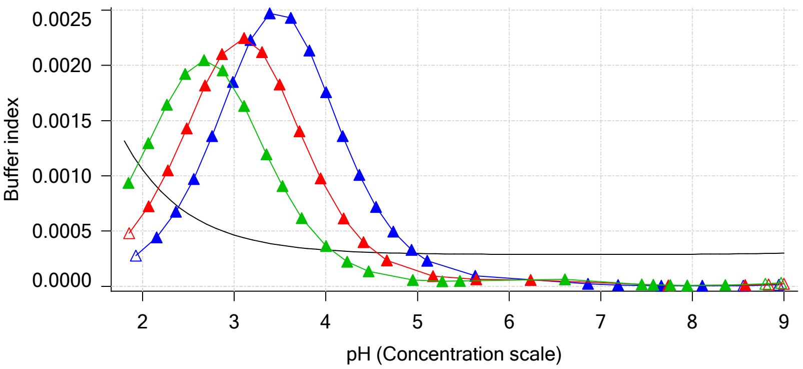


Sample M11_octanol Ionisation Graph

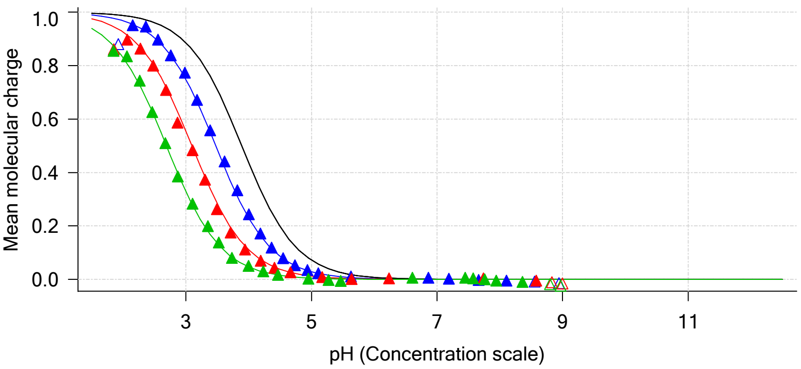


Distribution of species for sample M11_octanol

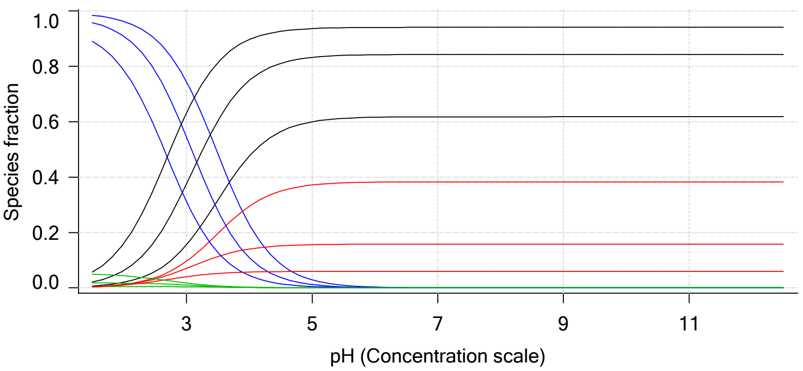


Lipophilicity Profile

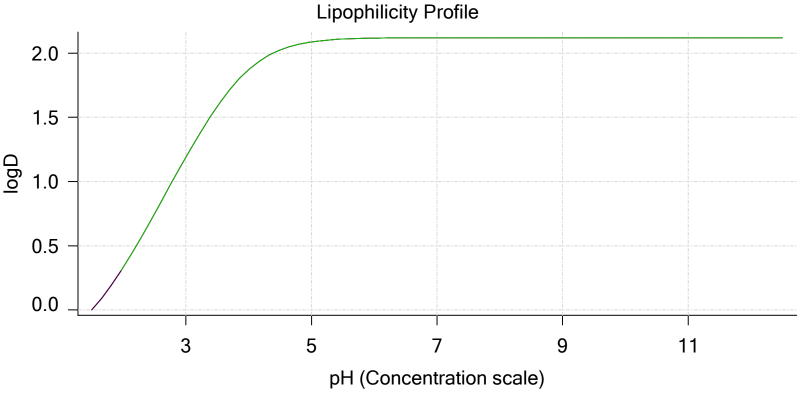


Log R Profile

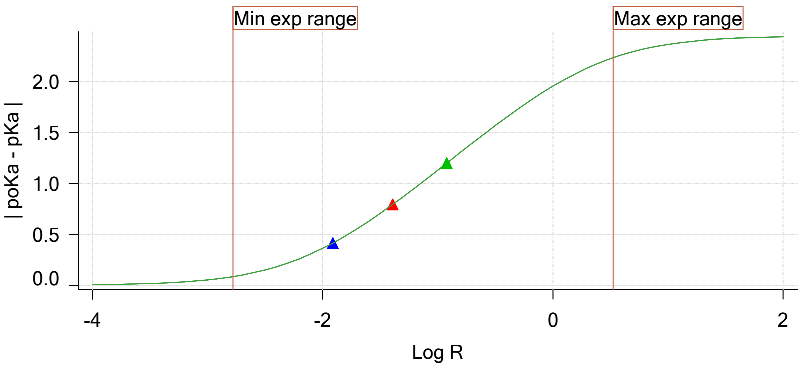


Sample Carbonate Ionisation Graph

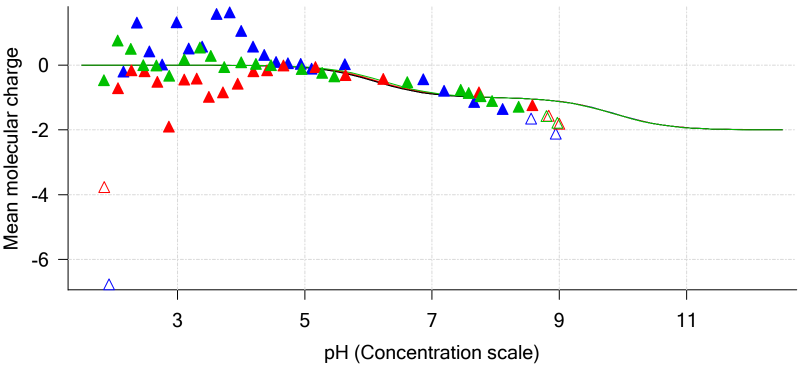

| | | | |
|---|---|---|---|
| Sample name: | **M11_octanol** | Experiment start time: | **3/9/2018 4:01:24 PM** |
| Assay name: | **pH-metric high logP** | Analyst: | **Pion** |
| Assay ID: | **18C-09010** | Instrument ID: | **T312060** |
| Filename: | **C:\Sirius_T3\Mehtap\20180309_exp31_logP_T3-2\18C-09010_M11_octanol_pH-metric high logP.t3r** | | |

### Graphs (continued)

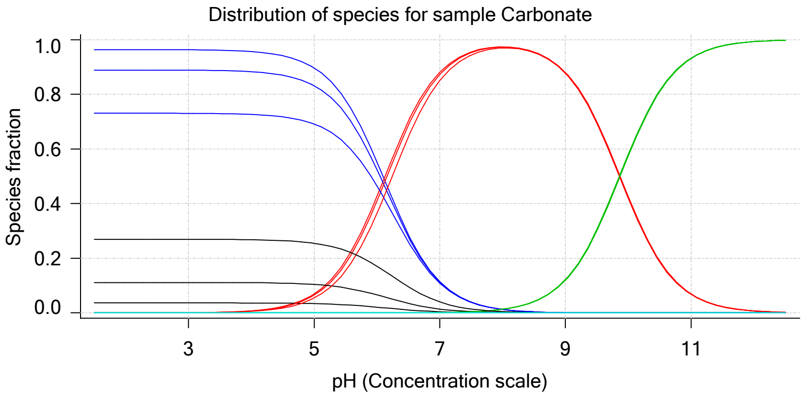

| | | | |
|---|---|---|---|
| Sample name: | **M11_octanol** | Experiment start time: | **3/9/2018 4:01:24 PM** |
| Assay name: | **pH-metric high logP** | Analyst: | **Pion** |
| Assay ID: | **18C-09010** | Instrument ID: | **T312060** |
| Filename: | **C:\Sirius_T3\Mehtap\20180309_exp31_logP_T3-2\18C-09010_M11_octanol_pH-metric high logP.t3r** | | |

# pH-metric high logP   Titration 1 of 3   18C-09010 Points 1 to 24

### Overall results

| | |
|---|---|
| RMSD | 0.632 |
| Average ionic strength | 0.157 M |
| Average temperature | 24.9°C |
| Partition ratio | 0.0123 : 1 |
| Analyte concentration range | 4427.7 µM to 4566.6 µM |
| Total points considered | 21 of 24 |

### Warnings and errors

| | |
|---|---|
| Errors | None |
| Warnings | None |

### Four-Plus parameters

| | | | |
|---|---|---|---|
| Alpha | 0.102 | 3/9/2018 4:01:24 PM | C:\Sirius_T3\HCl18C09.t3r |
| S | 0.9967 | 3/9/2018 4:01:24 PM | C:\Sirius_T3\HCl18C09.t3r |
| jH | 1.2 | 3/9/2018 4:01:24 PM | C:\Sirius_T3\HCl18C09.t3r |
| jOH | 0.0 | 3/9/2018 4:01:24 PM | C:\Sirius_T3\HCl18C09.t3r |

### Titrants

| | | | |
|---|---|---|---|
| 0.50 M HCl | 0.999843 | 3/9/2018 4:01:24 PM | C:\Sirius_T3\HCl18C09.t3r |
| 0.50 M KOH | 0.999845 | 3/9/2018 4:01:24 PM | C:\Sirius_T3\KOH18B27.t3r |

### Sample

| | |
|---|---|
| M11_octanol concentration factor | 0.962 |
| Base pKa 1 | 3.89 |
| logP (XH +) | 0.52 |
| logP (neutral X) | 2.10 |

### Sample graphs

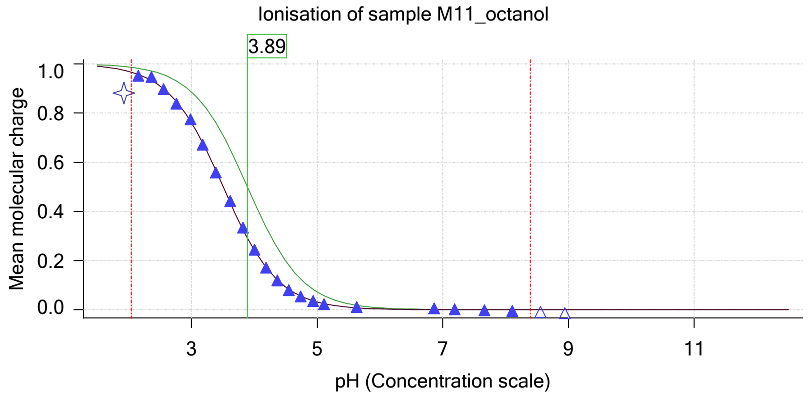


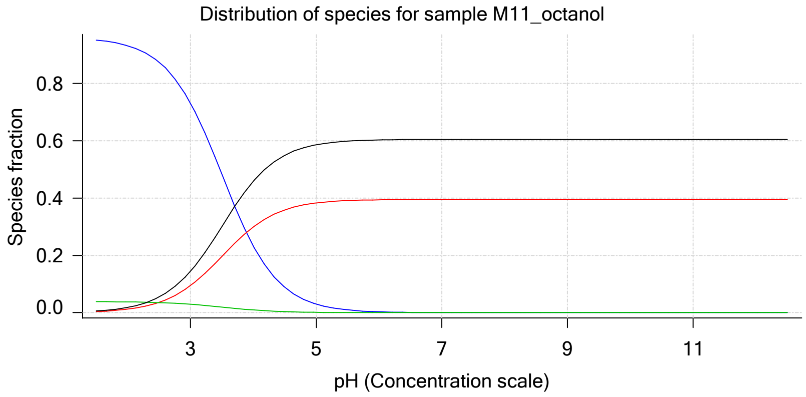

| | | | |
|---|---|---|---|
| Sample name: | **M11_octanol** | Experiment start time: | **3/9/2018 4:01:24 PM** |
| Assay name: | **pH-metric high logP** | Analyst: | **Pion** |
| Assay ID: | **18C-09010** | Instrument ID: | **T312060** |
| Filename: | **C:\Sirius_T3\Mehtap\20180309_exp31_logP_T3-2\18C-09010_M11_octanol_pH-metric high logP.t3r** | | |

### Sample graphs (continued)

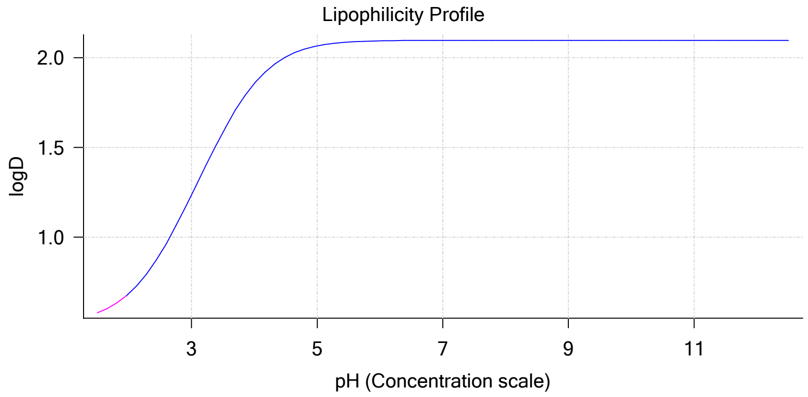


### Sample logD and percent species

| pH | M11_octanol logD | M11_octanol M11_octanolH | M11_octanol M11_octanol | M11_octanol M11_octanolH* | M11_octanol M11_octanol* | Comment |
|---|---|---|---|---|---|---|
| 1.000 | 0.54 | 95.82 % | 0.12 % | 3.86 % | 0.19 % | |
| 1.200 | 0.55 | 95.65 % | 0.20 % | 3.86 % | 0.30 % | Stomach pH |
| 2.000 | 0.68 | 93.20 % | 1.20 % | 3.76 % | 1.84 % | |
| 3.000 | 1.23 | 73.19 % | 9.43 % | 2.95 % | 14.43 % | |
| 4.000 | 1.86 | 23.25 % | 29.95 % | 0.94 % | 45.86 % | |
| 5.000 | 2.07 | 2.97 % | 38.29 % | 0.12 % | 58.62 % | |
| 6.000 | 2.09 | 0.31 % | 39.39 % | 0.01 % | 60.30 % | |
| 6.500 | 2.10 | 0.10 % | 39.47 % | 0.00 % | 60.43 % | |
| 7.000 | 2.10 | 0.03 % | 39.50 % | 0.00 % | 60.47 % | |
| 7.400 | 2.10 | 0.01 % | 39.51 % | 0.00 % | 60.48 % | Blood pH |
| 8.000 | 2.10 | 0.00 % | 39.51 % | 0.00 % | 60.49 % | |
| 9.000 | 2.10 | 0.00 % | 39.51 % | 0.00 % | 60.49 % | |
| 10.000 | 2.10 | 0.00 % | 39.51 % | 0.00 % | 60.49 % | |
| 11.000 | 2.10 | 0.00 % | 39.51 % | 0.00 % | 60.49 % | |
| 12.000 | 2.10 | 0.00 % | 39.51 % | 0.00 % | 60.49 % | |

### Carbonate and acidity

| | | |
|---|---|---|
| 💎 | Carbonate | 0.060 mM |
| 💎 | Acidity error | -0.464 mM |

### Other graphs

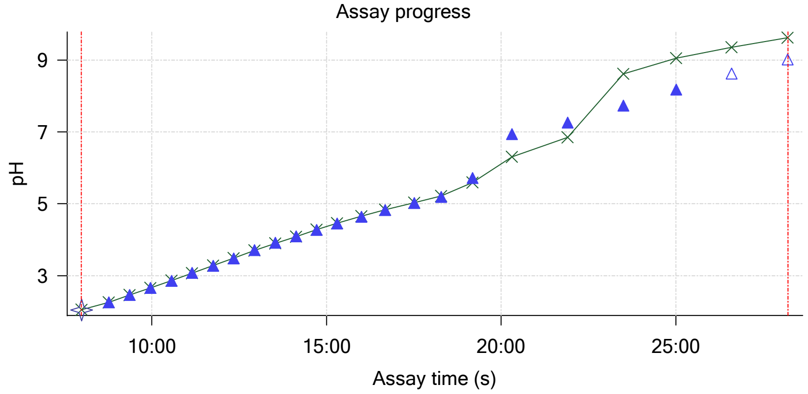


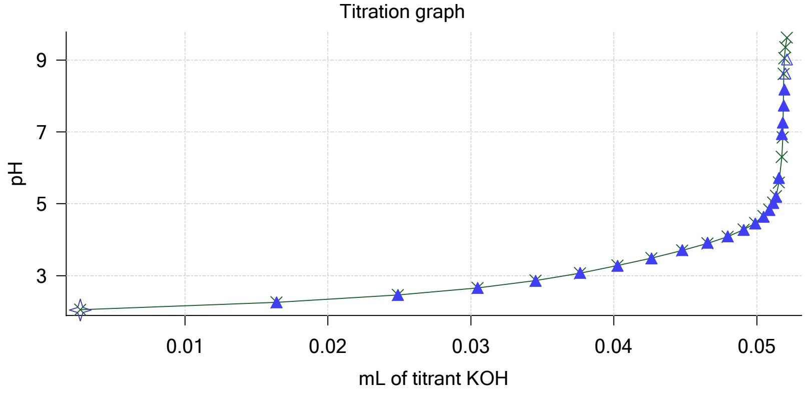

Sample name: **M11_octanol**
Assay name: **pH-metric high logP**
Assay ID: **18C-09010**
Filename: **C:\Sirius_T3\Mehtap\20180309_exp31_logP_T3-2\18C-09010_M11_octanol_pH-metric high logP.t3r**

Experiment start time: **3/9/2018 4:01:24 PM**
Analyst: **Pion**
Instrument ID: **T312060**

### Other graphs (continued)

Charge balance residuals

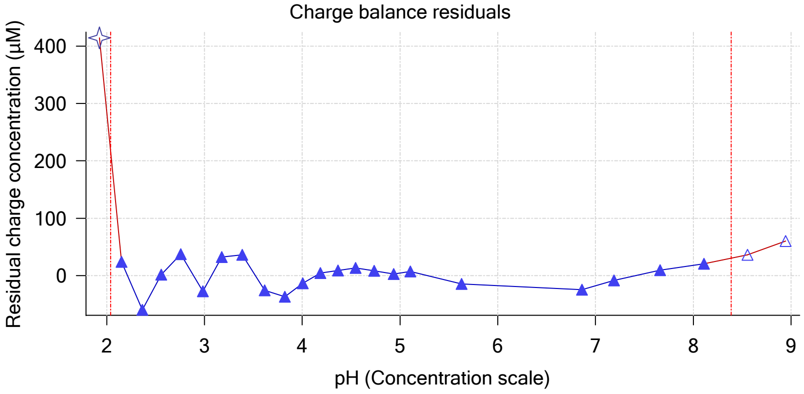


Buffer index profile

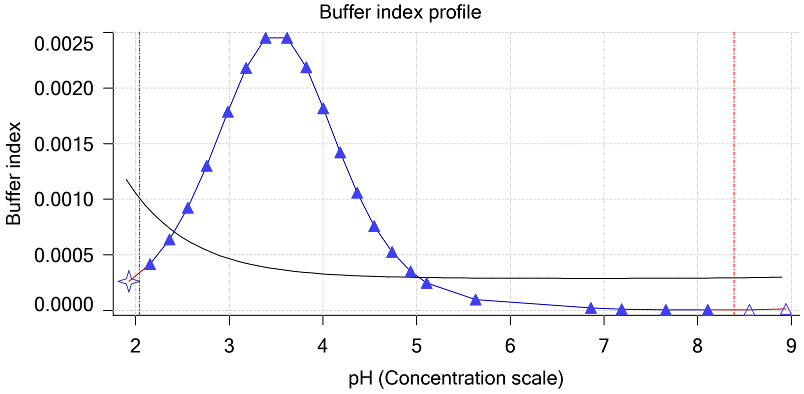


Ionic strength

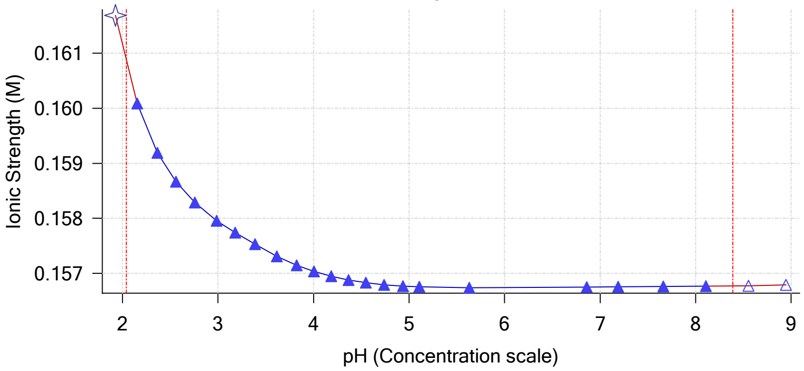


Log R Profile

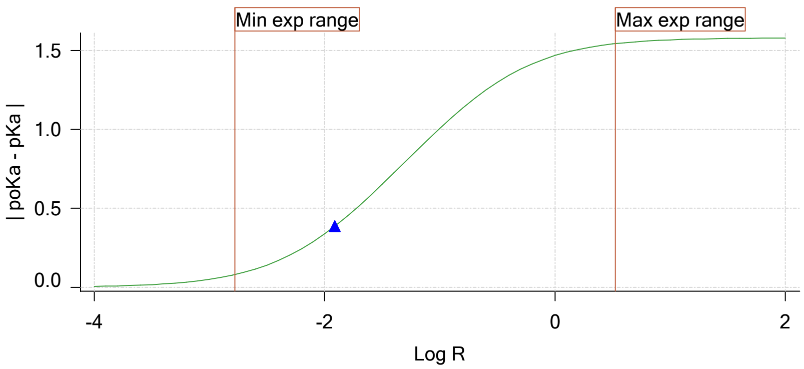


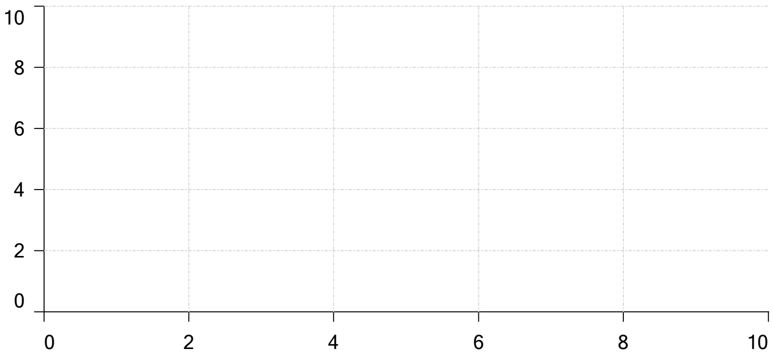

Ionisation of sample Carbonate

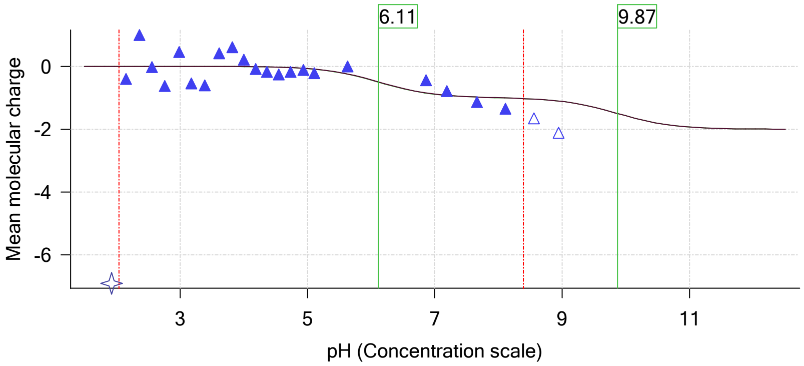

| | | | |
|---|---|---|---|
| Sample name: | **M11_octanol** | Experiment start time: | **3/9/2018 4:01:24 PM** |
| Assay name: | **pH-metric high logP** | Analyst: | **Pion** |
| Assay ID: | **18C-09010** | Instrument ID: | **T312060** |
| Filename: | **C:\Sirius_T3\Mehtap\20180309_exp31_logP_T3-2\18C-09010_M11_octanol_pH-metric high logP.t3r** | | |

# pH-metric high logP   Titration 2 of 3   18C-09010 Points 25 to 45

### Overall results

| | |
|---|---|
| RMSD | 0.373 |
| Average ionic strength | 0.164 M |
| Average temperature | 25.0°C |
| Partition ratio | 0.0405 : 1 |
| Analyte concentration range | 4011.7 µM to 4147.6 µM |
| Total points considered | 18 of 21 |

### Warnings and errors

| | |
|---|---|
| Errors | None |
| Warnings | None |

### Four-Plus parameters

| | | | |
|---|---|---|---|
| Alpha | 0.102 | 3/9/2018 4:01:24 PM | C:\Sirius_T3\HCl18C09.t3r |
| S | 0.9967 | 3/9/2018 4:01:24 PM | C:\Sirius_T3\HCl18C09.t3r |
| jH | 1.2 | 3/9/2018 4:01:24 PM | C:\Sirius_T3\HCl18C09.t3r |
| jOH | 0.0 | 3/9/2018 4:01:24 PM | C:\Sirius_T3\HCl18C09.t3r |

### Titrants

| | | | |
|---|---|---|---|
| 0.50 M HCl | 0.999843 | 3/9/2018 4:01:24 PM | C:\Sirius_T3\HCl18C09.t3r |
| 0.50 M KOH | 0.999845 | 3/9/2018 4:01:24 PM | C:\Sirius_T3\KOH18B27.t3r |

### Sample

| | |
|---|---|
| M11_octanol concentration factor | 0.930 |
| Base pKa 1 | 3.89 |
| logP (XH +) | 0.55 |
| logP (neutral X) | 2.22 |

### Sample graphs

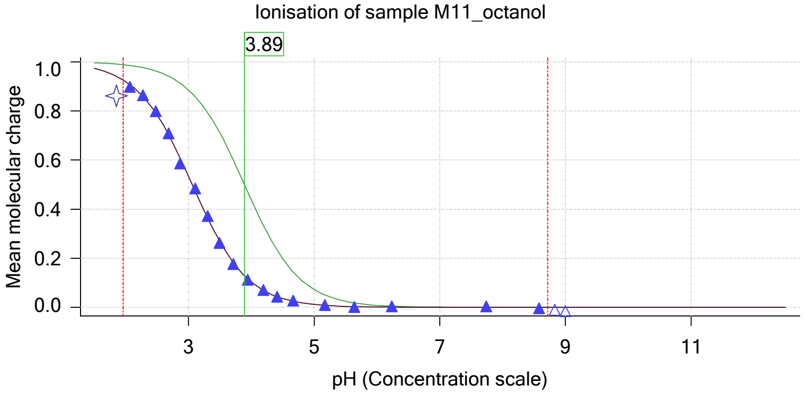


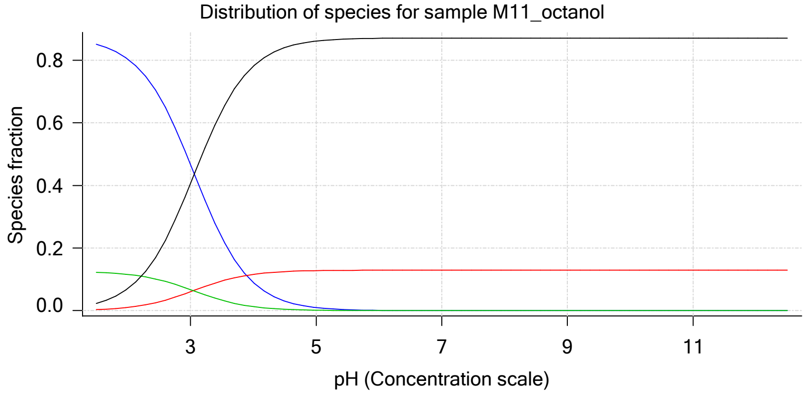

Sample name: **M11_octanol**
Assay name: **pH-metric high logP**
Assay ID: **18C-09010**
Filename: **C:\Sirius_T3\Mehtap\20180309_exp31_logP_T3-2\18C-09010_M11_octanol_pH-metric high logP.t3r**

Experiment start time: **3/9/2018 4:01:24 PM**
Analyst: **Pion**
Instrument ID: **T312060**

### Sample graphs (continued)

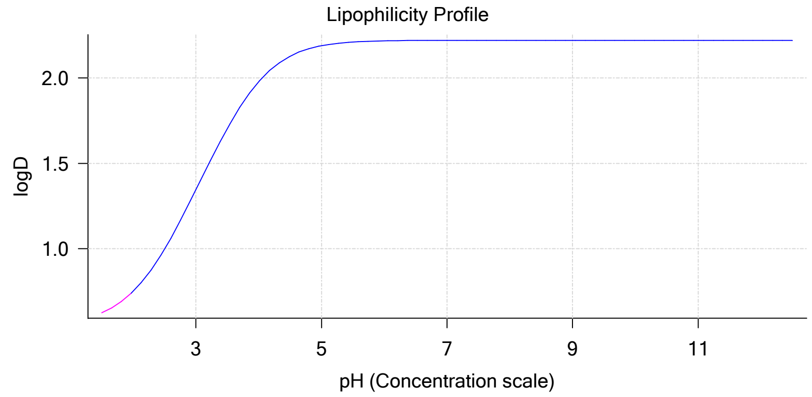


### Sample logD and percent species

| pH | M11_octanol logD | M11_octanol M11_octanolH | M11_octanol M11_octanol | M11_octanol M11_octanolH* | M11_octanol M11_octanol* | Comment |
|---|---|---|---|---|---|---|
| 1.000 | 0.57 | 86.67 % | 0.11 % | 12.47 % | 0.75 % | |
| 1.200 | 0.59 | 86.23 % | 0.18 % | 12.40 % | 1.19 % | Stomach pH |
| 2.000 | 0.75 | 80.41 % | 1.04 % | 11.57 % | 6.98 % | |
| 3.000 | 1.34 | 46.71 % | 6.02 % | 6.72 % | 40.55 % | |
| 4.000 | 1.98 | 9.00 % | 11.59 % | 1.29 % | 78.12 % | |
| 5.000 | 2.19 | 0.99 % | 12.77 % | 0.14 % | 86.09 % | |
| 6.000 | 2.22 | 0.10 % | 12.91 % | 0.01 % | 86.98 % | |
| 6.500 | 2.22 | 0.03 % | 12.92 % | 0.00 % | 87.05 % | |
| 7.000 | 2.22 | 0.01 % | 12.92 % | 0.00 % | 87.07 % | |
| 7.400 | 2.22 | 0.00 % | 12.92 % | 0.00 % | 87.08 % | Blood pH |
| 8.000 | 2.22 | 0.00 % | 12.92 % | 0.00 % | 87.08 % | |
| 9.000 | 2.22 | 0.00 % | 12.92 % | 0.00 % | 87.08 % | |
| 10.000 | 2.22 | 0.00 % | 12.92 % | 0.00 % | 87.08 % | |
| 11.000 | 2.22 | 0.00 % | 12.92 % | 0.00 % | 87.08 % | |
| 12.000 | 2.22 | 0.00 % | 12.92 % | 0.00 % | 87.08 % | |

### Carbonate and acidity

| | |
|---|---|
| Carbonate | 0.089 mM |
| Acidity error | -0.757 mM |

### Other graphs

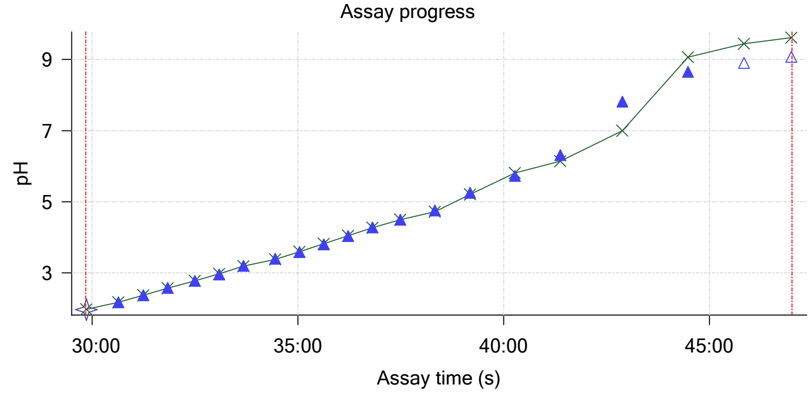


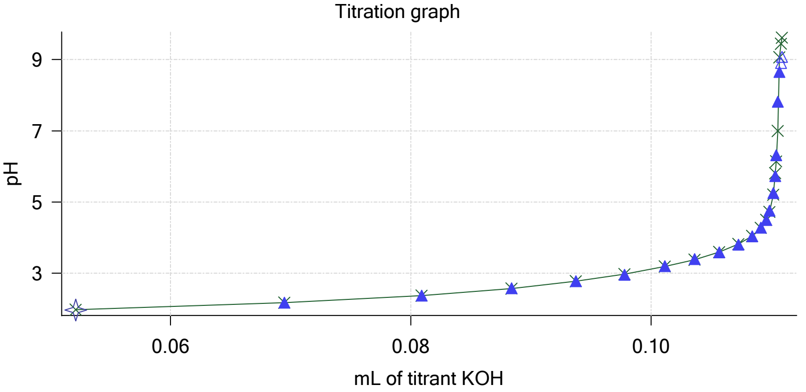

| | | | |
|---|---|---|---|
| Sample name: | **M11_octanol** | Experiment start time: | **3/9/2018 4:01:24 PM** |
| Assay name: | **pH-metric high logP** | Analyst: | **Pion** |
| Assay ID: | **18C-09010** | Instrument ID: | **T312060** |
| Filename: | **C:\Sirius_T3\Mehtap\20180309_exp31_logP_T3-2\18C-09010_M11_octanol_pH-metric high logP.t3r** | | |

### Other graphs (continued)

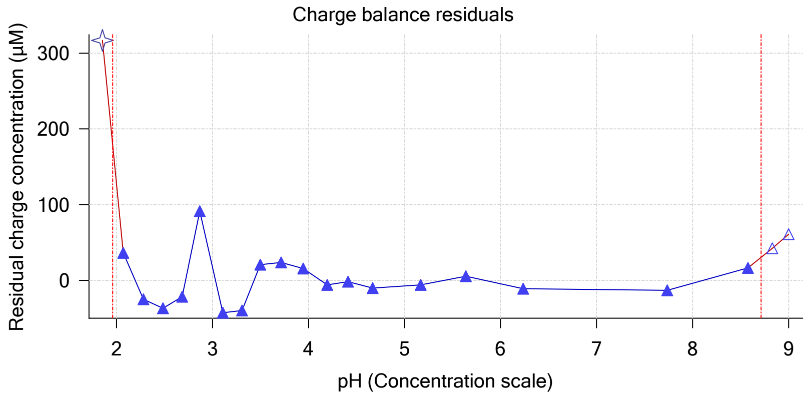


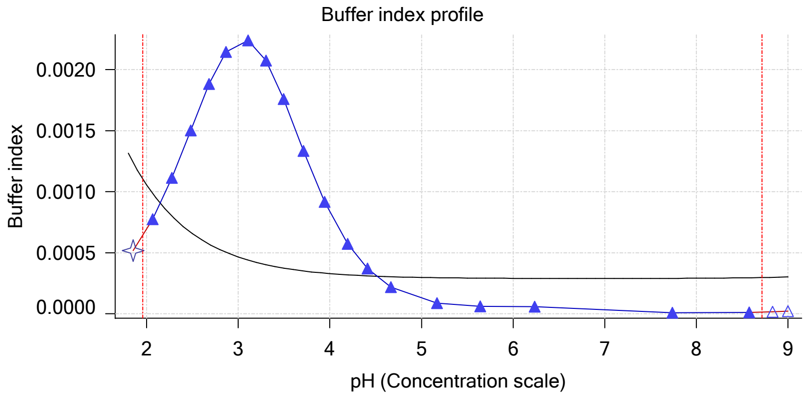


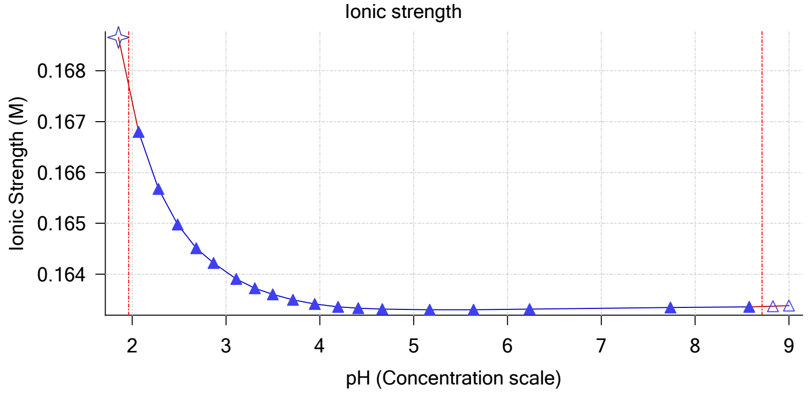


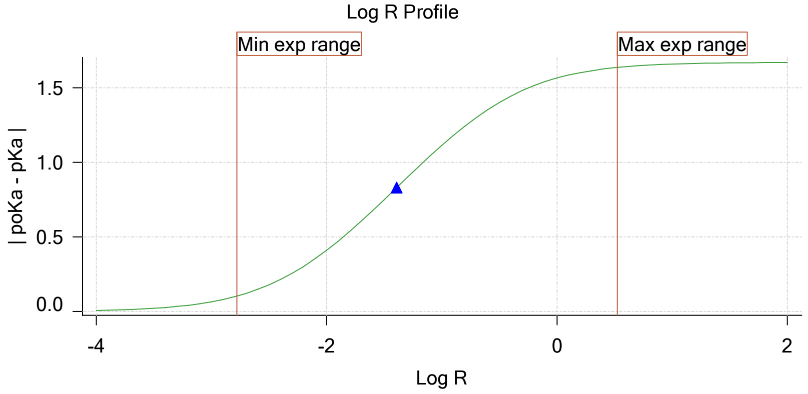


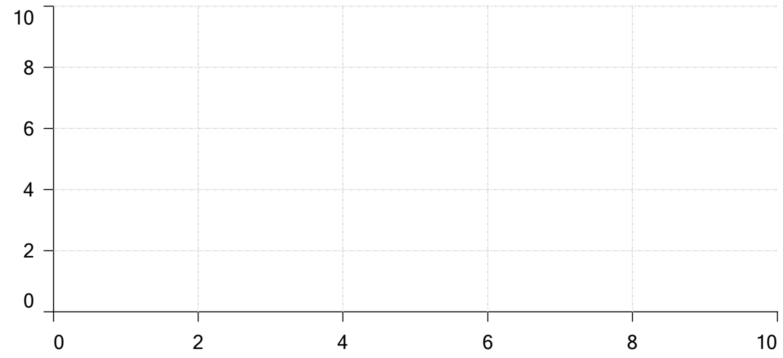


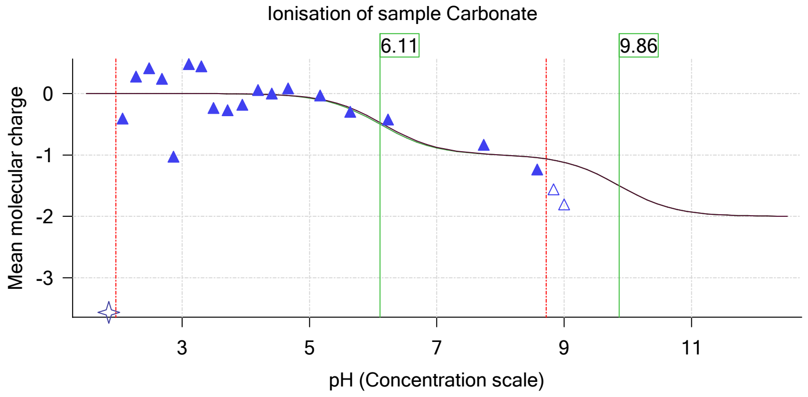

| | | | |
|---|---|---|---|
| Sample name: | **M11_octanol** | Experiment start time: | **3/9/2018 4:01:24 PM** |
| Assay name: | **pH-metric high logP** | Analyst: | **Pion** |
| Assay ID: | **18C-09010** | Instrument ID: | **T312060** |
| Filename: | **C:\Sirius_T3\Mehtap\20180309_exp31_logP_T3-2\18C-09010_M11_octanol_pH-metric high logP.t3r** | | |

# pH-metric high logP   Titration 3 of 3   18C-09010 Points 46 to 69

### Overall results

| | |
|---|---|
| RMSD | 0.548 |
| Average ionic strength | 0.170 M |
| Average temperature | 25.0°C |
| Partition ratio | 0.1193 : 1 |
| Analyte concentration range | 3477.2 µM to 3586.0 µM |
| Total points considered | 22 of 24 |

### Warnings and errors

| | |
|---|---|
| Errors | None |
| Warnings | None |

### Four-Plus parameters

| | | | |
|---|---|---|---|
| Alpha | 0.102 | 3/9/2018 4:01:24 PM | C:\Sirius_T3\HCl18C09.t3r |
| S | 0.9967 | 3/9/2018 4:01:24 PM | C:\Sirius_T3\HCl18C09.t3r |
| jH | 1.2 | 3/9/2018 4:01:24 PM | C:\Sirius_T3\HCl18C09.t3r |
| jOH | 0.0 | 3/9/2018 4:01:24 PM | C:\Sirius_T3\HCl18C09.t3r |

### Titrants

| | | | |
|---|---|---|---|
| 0.50 M HCl | 0.999843 | 3/9/2018 4:01:24 PM | C:\Sirius_T3\HCl18C09.t3r |
| 0.50 M KOH | 0.999845 | 3/9/2018 4:01:24 PM | C:\Sirius_T3\KOH18B27.t3r |

### Sample

| | |
|---|---|
| M11_octanol concentration factor | 0.907 |
| Base pKa 1 | 3.89 |
| logP (XH +) | 0.55 |
| logP (neutral X) | 2.25 |

### Sample graphs

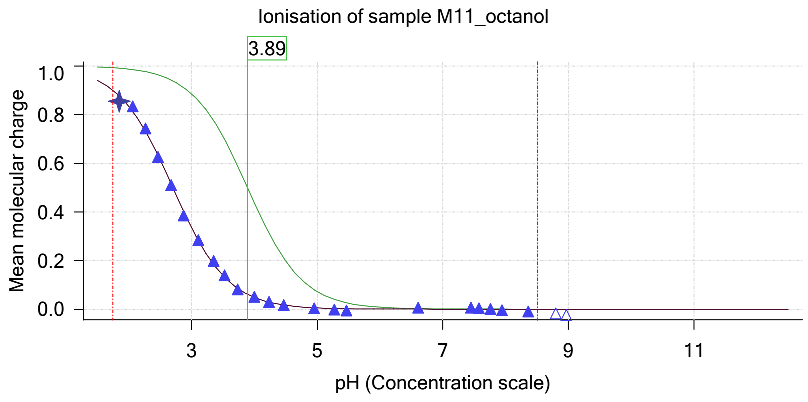


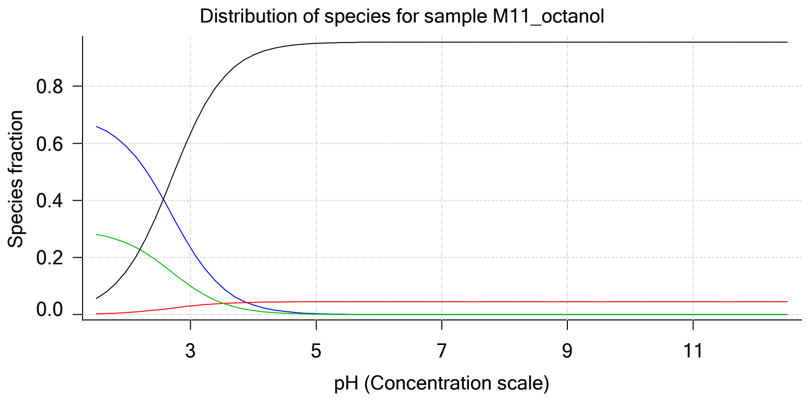

| | | | |
|---|---|---|---|
| Sample name: | **M11_octanol** | Experiment start time: | **3/9/2018 4:01:24 PM** |
| Assay name: | **pH-metric high logP** | Analyst: | **Pion** |
| Assay ID: | **18C-09010** | Instrument ID: | **T312060** |
| Filename: | **C:\Sirius_T3\Mehtap\20180309_exp31_logP_T3-2\18C-09010_M11_octanol_pH-metric high logP.t3r** | | |

### Sample graphs (continued)

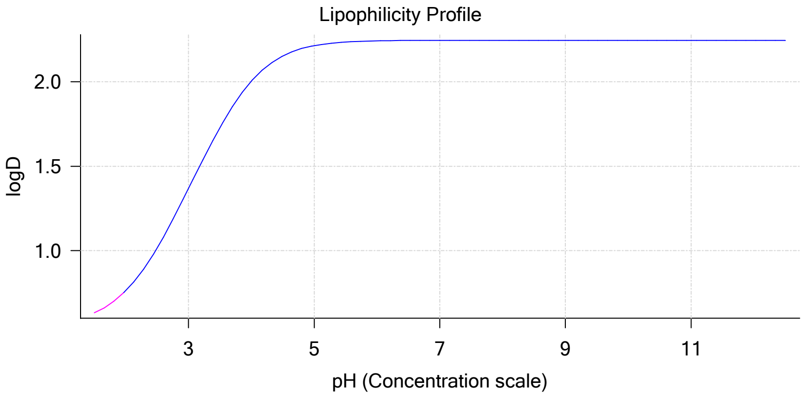


### Sample logD and percent species

| pH | M11_octanol logD | M11_octanol M11_octanolH | M11_octanol M11_octanol | M11_octanol M11_octanolH* | M11_octanol M11_octanol* | Comment |
|---|---|---|---|---|---|---|
| 1.000 | 0.58 | 68.74 % | 0.09 % | 29.31 % | 1.86 % | |
| 1.200 | 0.59 | 67.97 % | 0.14 % | 28.98 % | 2.92 % | Stomach pH |
| 2.000 | 0.76 | 58.48 % | 0.75 % | 24.93 % | 15.84 % | |
| 3.000 | 1.37 | 23.45 % | 3.02 % | 10.00 % | 63.53 % | |
| 4.000 | 2.00 | 3.36 % | 4.32 % | 1.43 % | 90.89 % | |
| 5.000 | 2.21 | 0.35 % | 4.52 % | 0.15 % | 94.98 % | |
| 6.000 | 2.24 | 0.04 % | 4.54 % | 0.02 % | 95.41 % | |
| 6.500 | 2.25 | 0.01 % | 4.54 % | 0.00 % | 95.44 % | |
| 7.000 | 2.25 | 0.00 % | 4.54 % | 0.00 % | 95.46 % | |
| 7.400 | 2.25 | 0.00 % | 4.54 % | 0.00 % | 95.46 % | Blood pH |
| 8.000 | 2.25 | 0.00 % | 4.54 % | 0.00 % | 95.46 % | |
| 9.000 | 2.25 | 0.00 % | 4.54 % | 0.00 % | 95.46 % | |
| 10.000 | 2.25 | 0.00 % | 4.54 % | 0.00 % | 95.46 % | |
| 11.000 | 2.25 | 0.00 % | 4.54 % | 0.00 % | 95.46 % | |
| 12.000 | 2.25 | 0.00 % | 4.54 % | 0.00 % | 95.46 % | |

### Carbonate and acidity

| | | |
|---|---|---|
| | Carbonate | 0.129 mM |
| | Acidity error | -0.811 mM |

### Other graphs

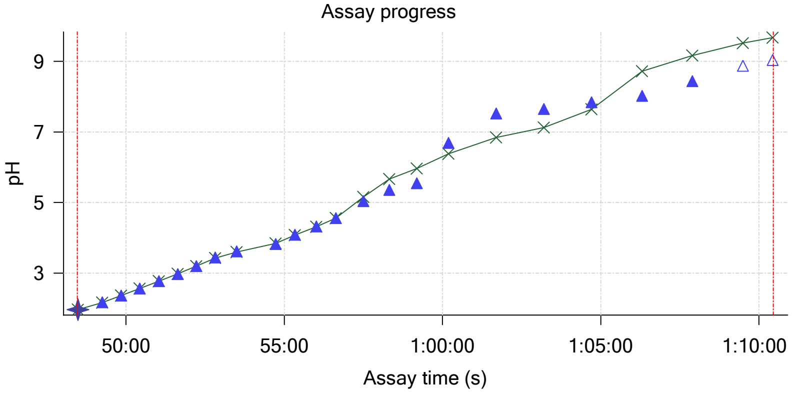


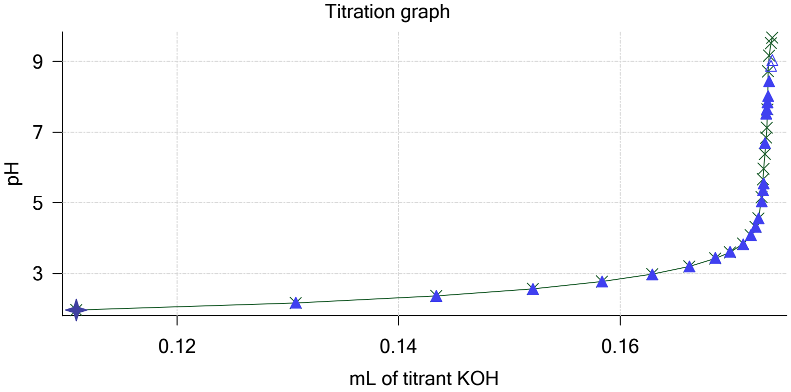

Sample name: **M11_octanol**
Assay name: **pH-metric high logP**
Assay ID: **18C-09010**
Filename: **C:\Sirius_T3\Mehtap\20180309_exp31_logP_T3-2\18C-09010_M11_octanol_pH-metric high logP.t3r**

Experiment start time: **3/9/2018 4:01:24 PM**
Analyst: **Pion**
Instrument ID: **T312060**

### Other graphs (continued)

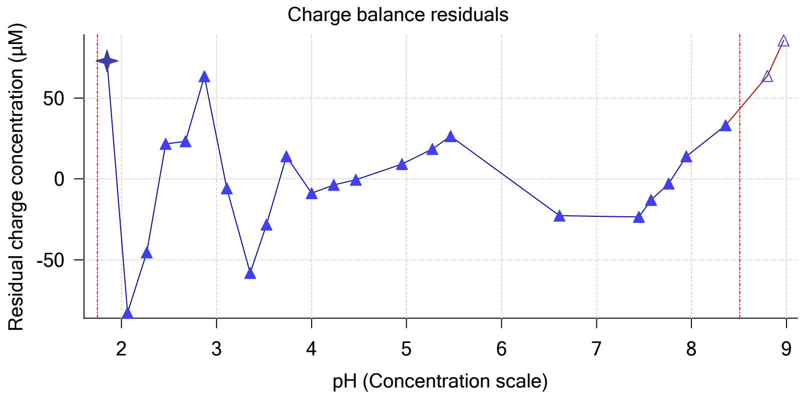


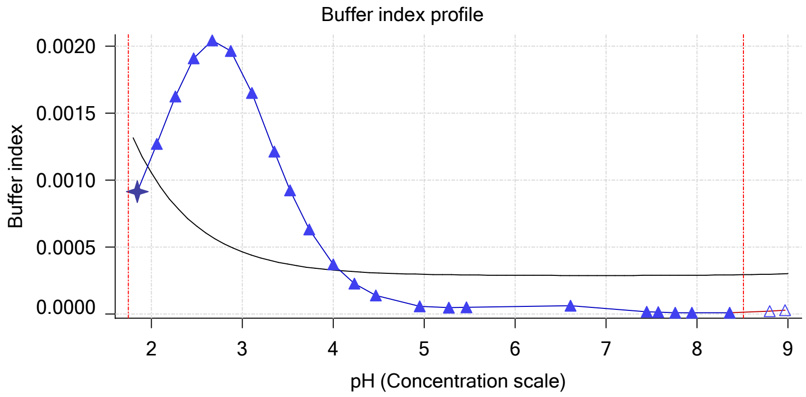


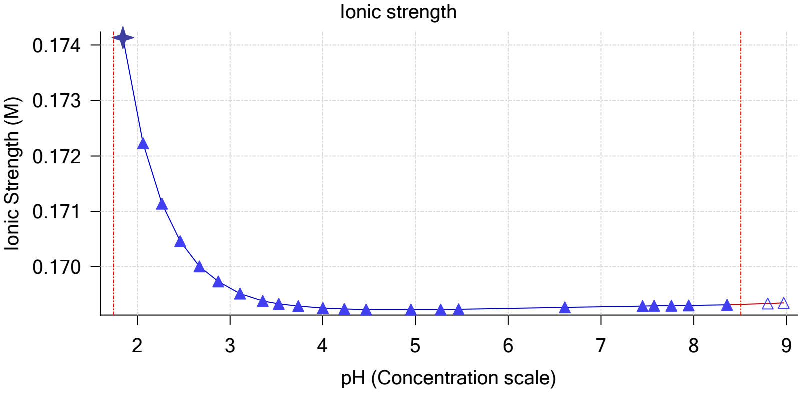


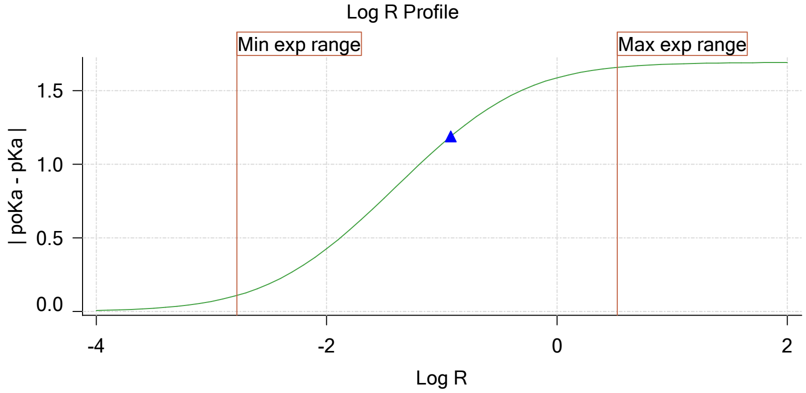


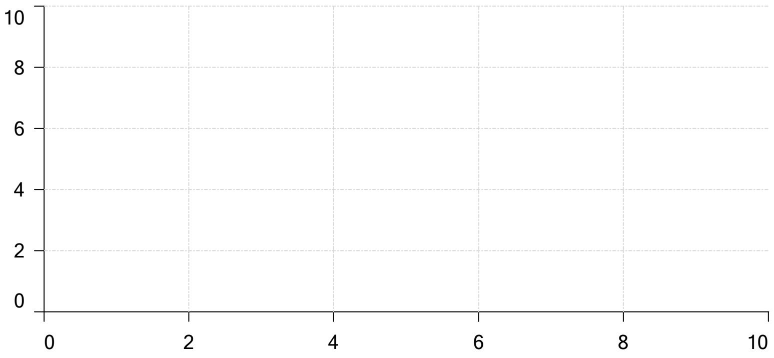


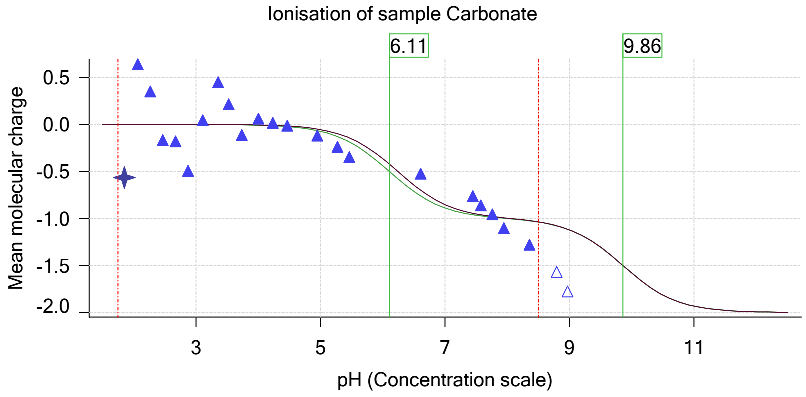

| | | | |
|---|---|---|---|
| Sample name: | **M11_octanol** | Experiment start time: | **3/9/2018 4:01:24 PM** |
| Assay name: | **pH-metric high logP** | Analyst: | **Pion** |
| Assay ID: | **18C-09010** | Instrument ID: | **T312060** |
| Filename: | **C:\Sirius_T3\Mehtap\20180309_exp31_logP_T3-2\18C-09010_M11_octanol_pH-metric high logP.t3r** | | |

### Assay Model

| Settings | Value | Date/Time changed | Imported from |
|---|---|---|---|
| Sample name | M11_octanol | 2/27/2018 5:54:30 PM | User entered value |
| Sample by | Weight | | Default value |
| Sample weight | 0.001520 g | 3/9/2018 2:22:09 PM | User entered value |
| Formula weight | 211.22 g/mol | 2/27/2018 5:54:30 PM | User entered value |
| Solubility | Unknown | | Default value |
| Molecular weight | 211.22 | 2/27/2018 5:54:30 PM | User entered value |
| Individual pKa ionic environments | No | | Default value |
| Number of pKas | 1 | 2/27/2018 5:54:30 PM | User entered value |
| Sample is a | Base | 2/27/2018 5:54:30 PM | User entered value |
| pKa 1 | 3.89 | 2/27/2018 5:54:30 PM | User entered value |
| logp (XH +) | 0.55 | 3/2/2018 4:29:35 PM | User entered value |
| logP (neutral X) | 2.19 | 3/2/2018 4:29:30 PM | User entered value |

### Events

| Time | Event | Water | Acid | Base | Octanol | pH | dpH/dt | pH R-squared | pH SD | dpH/dt time |
|---|---|---|---|---|---|---|---|---|---|---|
| 5:00.2 | Initial pH = 9.10 | | | | | | | | | |
| 7:59.8 | Data point 1 | 1.50000 mL | 0.05315 mL | 0.00268 mL | 0.01999 mL | 2.037 | -0.00511 | 0.30733 | 0.00046 | 10.0 s |
| 8:46.1 | Data point 2 | 1.50000 mL | 0.05315 mL | 0.01642 mL | 0.01999 mL | 2.256 | 0.00288 | 0.69530 | 0.00017 | 10.5 s |
| 9:22.2 | Data point 3 | 1.50000 mL | 0.05315 mL | 0.02491 mL | 0.01999 mL | 2.463 | 0.00374 | 0.32148 | 0.00033 | 10.0 s |
| 9:57.8 | Data point 4 | 1.50000 mL | 0.05315 mL | 0.03048 mL | 0.01999 mL | 2.655 | 0.00078 | 0.10371 | 0.00012 | 10.5 s |
| 10:33.9 | Data point 5 | 1.50000 mL | 0.05315 mL | 0.03452 mL | 0.01999 mL | 2.854 | -0.00337 | 0.19592 | 0.00038 | 10.0 s |
| 11:09.3 | Data point 6 | 1.50000 mL | 0.05315 mL | 0.03765 mL | 0.01999 mL | 3.078 | 0.00146 | 0.01563 | 0.00058 | 10.5 s |
| 11:45.3 | Data point 7 | 1.50000 mL | 0.05315 mL | 0.04026 mL | 0.01999 mL | 3.271 | -0.00477 | 0.53594 | 0.00032 | 10.0 s |
| 12:20.8 | Data point 8 | 1.50000 mL | 0.05315 mL | 0.04264 mL | 0.01999 mL | 3.480 | -0.00721 | 0.85954 | 0.00038 | 10.0 s |
| 12:56.3 | Data point 9 | 1.50000 mL | 0.05315 mL | 0.04478 mL | 0.01999 mL | 3.707 | -0.01214 | 0.77245 | 0.00068 | 10.5 s |
| 13:32.3 | Data point 10 | 1.50000 mL | 0.05315 mL | 0.04657 mL | 0.01999 mL | 3.912 | -0.01020 | 0.53326 | 0.00069 | 10.0 s |
| 14:07.7 | Data point 11 | 1.50000 mL | 0.05315 mL | 0.04798 mL | 0.01999 mL | 4.094 | -0.00754 | 0.73914 | 0.00043 | 10.0 s |
| 14:43.1 | Data point 12 | 1.50000 mL | 0.05315 mL | 0.04908 mL | 0.01999 mL | 4.274 | -0.01378 | 0.67663 | 0.00083 | 10.0 s |
| 15:18.5 | Data point 13 | 1.50000 mL | 0.05315 mL | 0.04988 mL | 0.01999 mL | 4.454 | -0.01794 | 0.90958 | 0.00093 | 16.5 s |
| 16:00.3 | Data point 14 | 1.50000 mL | 0.05315 mL | 0.05047 mL | 0.01999 mL | 4.635 | -0.01742 | 0.86808 | 0.00092 | 15.0 s |
| 16:40.7 | Data point 15 | 1.50000 mL | 0.05315 mL | 0.05087 mL | 0.01999 mL | 4.824 | -0.01787 | 0.84622 | 0.00096 | 24.5 s |
| 17:30.7 | Data point 16 | 1.50000 mL | 0.05315 mL | 0.05115 mL | 0.01999 mL | 5.023 | -0.01827 | 0.90373 | 0.00095 | 21.0 s |
| 18:17.2 | Data point 17 | 1.50000 mL | 0.05315 mL | 0.05134 mL | 0.01999 mL | 5.193 | -0.01878 | 0.91929 | 0.00097 | 23.0 s |
| 19:10.8 | Data point 18 | 1.50000 mL | 0.05315 mL | 0.05155 mL | 0.01999 mL | 5.715 | -0.01901 | 0.96541 | 0.00096 | 37.0 s |
| 20:18.6 | Data point 19 | 1.50000 mL | 0.05315 mL | 0.05174 mL | 0.01999 mL | 6.941 | -0.08865 | 0.99621 | 0.00438 | Timed out at 59.5 s |
| 21:54.2 | Data point 20 | 1.50000 mL | 0.05315 mL | 0.05181 mL | 0.01999 mL | 7.267 | -0.07065 | 0.99539 | 0.00350 | Timed out at 59.5 s |
| 23:29.8 | Data point 21 | 1.50000 mL | 0.05315 mL | 0.05188 mL | 0.01999 mL | 7.738 | -0.08731 | 0.99760 | 0.00432 | Timed out at 59.5 s |
| 25:00.3 | Data point 22 | 1.50000 mL | 0.05315 mL | 0.05193 mL | 0.01999 mL | 8.184 | -0.04618 | 0.99543 | 0.00229 | Timed out at 59.5 s |
| 26:36.0 | Data point 23 | 1.50000 mL | 0.05315 mL | 0.05200 mL | 0.01999 mL | 8.628 | -0.02509 | 0.99264 | 0.00124 | Timed out at 59.5 s |
| 28:11.7 | Data point 24 | 1.50000 mL | 0.05315 mL | 0.05212 mL | 0.01999 mL | 9.017 | -0.01906 | 0.93147 | 0.00097 | 40.5 s |
| 29:51.6 | Data point 25 | 1.50000 mL | 0.11291 mL | 0.05212 mL | 0.06999 mL | 1.966 | 0.00082 | 0.01226 | 0.00037 | 10.5 s |
| 30:38.4 | Data point 26 | 1.50000 mL | 0.11291 mL | 0.06950 mL | 0.06999 mL | 2.173 | -0.00810 | 0.69034 | 0.00048 | 10.5 s |
| 31:14.6 | Data point 27 | 1.50000 mL | 0.11291 mL | 0.08090 mL | 0.06999 mL | 2.379 | 0.00330 | 0.80719 | 0.00018 | 10.0 s |
| 31:50.1 | Data point 28 | 1.50000 mL | 0.11291 mL | 0.08841 mL | 0.06999 mL | 2.580 | -0.01357 | 0.47607 | 0.00097 | 14.0 s |
| 32:29.7 | Data point 29 | 1.50000 mL | 0.11291 mL | 0.09374 mL | 0.06999 mL | 2.779 | -0.00359 | 0.57678 | 0.00023 | 10.0 s |
| 33:05.2 | Data point 30 | 1.50000 mL | 0.11291 mL | 0.09781 mL | 0.06999 mL | 2.960 | -0.00538 | 0.43414 | 0.00040 | 10.0 s |
| 33:40.7 | Data point 31 | 1.50000 mL | 0.11291 mL | 0.10115 mL | 0.06999 mL | 3.200 | -0.01650 | 0.90031 | 0.00086 | 10.5 s |
| 34:27.1 | Data point 32 | 1.50000 mL | 0.11291 mL | 0.10362 mL | 0.06999 mL | 3.397 | -0.00830 | 0.30538 | 0.00074 | 10.0 s |
| 35:02.5 | Data point 33 | 1.50000 mL | 0.11291 mL | 0.10569 mL | 0.06999 mL | 3.587 | -0.01081 | 0.29933 | 0.00098 | 10.0 s |
| 35:38.0 | Data point 34 | 1.50000 mL | 0.11291 mL | 0.10729 mL | 0.06999 mL | 3.805 | -0.00516 | 0.79057 | 0.00029 | 10.0 s |
| 36:13.4 | Data point 35 | 1.50000 mL | 0.11291 mL | 0.10842 mL | 0.06999 mL | 4.034 | -0.00755 | 0.89227 | 0.00039 | 10.0 s |

| | | | |
|---|---|---|---|
| Sample name: | **M11_octanol** | Experiment start time: | **3/9/2018 4:01:24 PM** |
| Assay name: | **pH-metric high logP** | Analyst: | **Pion** |
| Assay ID: | **18C-09010** | Instrument ID: | **T312060** |
| Filename: | **C:\Sirius_T3\Mehtap\20180309_exp31_logP_T3-2\18C-09010_M11_octanol_pH-metric high logP.t3r** | | |

### Events (continued)

| Time | Event | Water | Acid | Base | Octanol | pH | dpH/dt | pH R-squared | pH SD | dpH/dt time |
|---|---|---|---|---|---|---|---|---|---|---|
| 36:48.8 | Data point 36 | 1.50000 mL | 0.11291 mL | 0.10915 mL | 0.06999 mL | 4.284 | -0.01862 | 0.91237 | 0.00096 | 10.0 s |
| 37:29.3 | Data point 37 | 1.50000 mL | 0.11291 mL | 0.10960 mL | 0.06999 mL | 4.498 | -0.01521 | 0.66710 | 0.00092 | 25.0 s |
| 38:19.8 | Data point 38 | 1.50000 mL | 0.11291 mL | 0.10988 mL | 0.06999 mL | 4.754 | -0.01754 | 0.82634 | 0.00095 | 20.5 s |
| 39:10.7 | Data point 39 | 1.50000 mL | 0.11291 mL | 0.11018 mL | 0.06999 mL | 5.255 | -0.01812 | 0.84415 | 0.00097 | 30.0 s |
| 40:16.4 | Data point 40 | 1.50000 mL | 0.11291 mL | 0.11035 mL | 0.06999 mL | 5.724 | -0.01903 | 0.93383 | 0.00097 | 35.5 s |
| 41:22.4 | Data point 41 | 1.50000 mL | 0.11291 mL | 0.11042 mL | 0.06999 mL | 6.316 | -0.03790 | 0.98438 | 0.00188 | Timed out at 59.5 s |
| 42:53.0 | Data point 42 | 1.50000 mL | 0.11291 mL | 0.11056 mL | 0.06999 mL | 7.814 | -0.06409 | 0.99645 | 0.00317 | Timed out at 59.5 s |
| 44:28.6 | Data point 43 | 1.50000 mL | 0.11291 mL | 0.11070 mL | 0.06999 mL | 8.652 | -0.02006 | 0.98249 | 0.00100 | 46.0 s |
| 45:50.3 | Data point 44 | 1.50000 mL | 0.11291 mL | 0.11082 mL | 0.06999 mL | 8.905 | -0.01828 | 0.84297 | 0.00098 | 38.5 s |
| 46:59.3 | Data point 45 | 1.50000 mL | 0.11291 mL | 0.11091 mL | 0.06999 mL | 9.073 | -0.01775 | 0.86902 | 0.00094 | 27.5 s |
| 48:28.6 | Data point 46 | 1.50000 mL | 0.17582 mL | 0.11091 mL | 0.21999 mL | 1.960 | -0.00614 | 0.61144 | 0.00039 | 10.0 s |
| 49:14.9 | Data point 47 | 1.50000 mL | 0.17582 mL | 0.13074 mL | 0.21999 mL | 2.169 | 0.01032 | 0.30311 | 0.00093 | 10.0 s |
| 49:50.6 | Data point 48 | 1.50000 mL | 0.17582 mL | 0.14341 mL | 0.21999 mL | 2.367 | -0.00532 | 0.40508 | 0.00041 | 10.0 s |
| 50:26.1 | Data point 49 | 1.50000 mL | 0.17582 mL | 0.15214 mL | 0.21999 mL | 2.563 | -0.00517 | 0.08291 | 0.00089 | 10.5 s |
| 51:02.2 | Data point 50 | 1.50000 mL | 0.17582 mL | 0.15837 mL | 0.21999 mL | 2.768 | -0.00887 | 0.21333 | 0.00095 | 10.0 s |
| 51:37.8 | Data point 51 | 1.50000 mL | 0.17582 mL | 0.16289 mL | 0.21999 mL | 2.967 | 0.00295 | 0.02140 | 0.00100 | 10.0 s |
| 52:13.3 | Data point 52 | 1.50000 mL | 0.17582 mL | 0.16625 mL | 0.21999 mL | 3.200 | -0.00340 | 0.17172 | 0.00041 | 10.0 s |
| 52:48.8 | Data point 53 | 1.50000 mL | 0.17582 mL | 0.16860 mL | 0.21999 mL | 3.446 | -0.01379 | 0.46424 | 0.00100 | 10.0 s |
| 53:29.5 | Data point 54 | 1.50000 mL | 0.17582 mL | 0.16992 mL | 0.21999 mL | 3.617 | 0.01450 | 0.85818 | 0.00077 | 33.5 s |
| 54:43.8 | Data point 55 | 1.50000 mL | 0.17582 mL | 0.17110 mL | 0.21999 mL | 3.827 | -0.00092 | 0.01708 | 0.00035 | 10.5 s |
| 55:19.9 | Data point 56 | 1.50000 mL | 0.17582 mL | 0.17180 mL | 0.21999 mL | 4.090 | 0.00886 | 0.25159 | 0.00087 | 10.5 s |
| 56:00.9 | Data point 57 | 1.50000 mL | 0.17582 mL | 0.17222 mL | 0.21999 mL | 4.321 | -0.01922 | 0.95151 | 0.00097 | 11.5 s |
| 56:37.7 | Data point 58 | 1.50000 mL | 0.17582 mL | 0.17248 mL | 0.21999 mL | 4.556 | -0.01818 | 0.96232 | 0.00092 | 21.5 s |
| 57:29.8 | Data point 59 | 1.50000 mL | 0.17582 mL | 0.17277 mL | 0.21999 mL | 5.036 | -0.00086 | 0.00356 | 0.00071 | 14.0 s |
| 58:19.3 | Data point 60 | 1.50000 mL | 0.17582 mL | 0.17288 mL | 0.21999 mL | 5.357 | -0.01461 | 0.57206 | 0.00095 | 16.0 s |
| 59:10.9 | Data point 61 | 1.50000 mL | 0.17582 mL | 0.17295 mL | 0.21999 mL | 5.549 | -0.01615 | 0.98210 | 0.00080 | 29.5 s |
| 1:00:11.1 | Data point 62 | 1.50000 mL | 0.17582 mL | 0.17307 mL | 0.21999 mL | 6.690 | -0.09840 | 0.99748 | 0.00486 | Timed out at 59.5 s |
| 1:01:41.6 | Data point 63 | 1.50000 mL | 0.17582 mL | 0.17319 mL | 0.21999 mL | 7.526 | -0.12365 | 0.99119 | 0.00614 | Timed out at 59.5 s |
| 1:03:12.0 | Data point 64 | 1.50000 mL | 0.17582 mL | 0.17324 mL | 0.21999 mL | 7.651 | -0.08733 | 0.99550 | 0.00433 | Timed out at 59.5 s |
| 1:04:42.5 | Data point 65 | 1.50000 mL | 0.17582 mL | 0.17328 mL | 0.21999 mL | 7.837 | -0.13623 | 0.43667 | 0.01018 | Timed out at 59.5 s |
| 1:06:18.2 | Data point 66 | 1.50000 mL | 0.17582 mL | 0.17335 mL | 0.21999 mL | 8.021 | -0.06575 | 0.99623 | 0.00325 | Timed out at 59.5 s |
| 1:07:53.8 | Data point 67 | 1.50000 mL | 0.17582 mL | 0.17345 mL | 0.21999 mL | 8.434 | -0.11867 | 0.41261 | 0.00912 | Timed out at 59.5 s |
| 1:09:29.4 | Data point 68 | 1.50000 mL | 0.17582 mL | 0.17361 mL | 0.21999 mL | 8.870 | -0.01721 | 0.79761 | 0.00095 | 21.0 s |
| 1:10:26.0 | Data point 69 | 1.50000 mL | 0.17582 mL | 0.17373 mL | 0.21999 mL | 9.041 | -0.01672 | 0.74463 | 0.00096 | 15.5 s |
| 1:10:50.6 | Assay volumes | 1.50000 mL | 0.17582 mL | 0.17373 mL | 0.21999 mL | | | | | |

| | | | |
|---|---|---|---|
| Sample name: | **M11_octanol** | Experiment start time: | **3/9/2018 4:01:24 PM** |
| Assay name: | **pH-metric high logP** | Analyst: | **Pion** |
| Assay ID: | **18C-09010** | Instrument ID: | **T312060** |
| Filename: | **C:\Sirius_T3\Mehtap\20180309_exp31_logP_T3-2\18C-09010_M11_octanol_pH-metric high logP.t3r** | | |

| | | | |
|---|---|---|---|
| Sample name: | **M11_octanol** | Experiment start time: | **3/9/2018 4:01:24 PM** |
| Assay name: | **pH-metric high logP** | Analyst: | **Pion** |
| Assay ID: | **18C-09010** | Instrument ID: | **T312060** |
| Filename: | **C:\Sirius_T3\Mehtap\20180309_exp31_logP_T3-2\18C-09010_M11_octanol_pH-metric high logP.t3r** | | |

### Assay Settings (continued)

| Setting | Value | Original Value | Date/Time changed | Imported from |
|---|---|---|---|---|
| ***Titration 3*** | | | | |
| Titrate from | Low to high pH | | | |
| Add additional water | 0.00 mL | | | |
| Additional partition solvent volume | 0.150 mL | | | |
| Additional partition solvent added | Automatic | | | |
| After pH adjust stir for | 30 seconds | | | |
| Stir to allow partitioning for | 15 seconds | | | |
| Stirrer speed for partitioning | 60% | | | |
| ***Data Point Stability*** | | | | |
| Stir during data point collection | No | | | |
| Delay before data point collection | 0 seconds | | | |
| Number of points to average | 20 points | | | |
| Time interval between points | 0.50 seconds | | | |
| Required maximum standard deviation | 0.00100 dpH/dt | | | |
| Stability timeout after | 60 seconds | | | |

### Calibration Settings

| Setting | Value | Date/Time changed | Imported from |
|---|---|---|---|
| Four-Plus alpha | 0.102 | 3/9/2018 4:01:24 PM | C:\Sirius_T3\HCl18C09.t3r |
| Four-Plus S | 0.9967 | 3/9/2018 4:01:24 PM | C:\Sirius_T3\HCl18C09.t3r |
| Four-Plus jH | 1.2 | 3/9/2018 4:01:24 PM | C:\Sirius_T3\HCl18C09.t3r |
| Four-Plus jOH | 0.0 | 3/9/2018 4:01:24 PM | C:\Sirius_T3\HCl18C09.t3r |
| Base concentration factor | 1.000 | 3/9/2018 4:01:24 PM | C:\Sirius_T3\KOH18B27.t3r |
| Acid concentration factor | 1.000 | 3/9/2018 4:01:24 PM | C:\Sirius_T3\HCl18C09.t3r |

### Instrument Settings

| Setting | Value | Batch Id | Install date |
|---|---|---|---|
| Instrument owner | Merck | | |
| Instrument ID | T312060 | | |
| Instrument type | T3 Simulator | | |
| Software version | 1.1.3.0 | | |
| Dispenser module | | T3DM1200361 | 3/31/2009 6:24:52 AM |
| Dispenser 0 | Water | | 3/31/2009 6:25:05 AM |
| Syringe volume | 2.5 mL | | |
| Firmware version | 1.2.1(r2) | | |
| Titrant | Water (0.15 M KCl) | 02-06-2018 | 2/27/2018 11:05:59 AM |
| Dispenser 2 | Acid | | 3/31/2009 6:25:11 AM |
| Syringe volume | 0.5 mL | | |
| Firmware version | 1.2.1(r2) | | |
| Titrant | Acid (0.5 M HCl) | 02-27-2018 | 2/27/2018 11:27:22 AM |
| Dispenser 1 | Base | | 3/31/2009 6:25:21 AM |
| Syringe volume | 0.5 mL | | |
| Firmware version | 1.2.1(r2) | | |
| Titrant | Base (0.5 M KOH) | 9/22/2017 | 2/27/2018 11:21:22 AM |
| Dispenser 5 | Cosolvent | | 3/31/2009 6:26:24 AM |
| Syringe volume | 2.5 mL | | |
| Firmware version | 1.2.1(r2) | | |
| Distribution valve 5 | Distribution Valve | | 3/31/2009 6:28:19 AM |
| Firmware version | 1.1.3 | | |
| Port A | Methanol (80%, 0.15 M KCl) | 02-08-2018 | 3/6/2018 10:28:59 AM |
| Port B | Cyclohexane | 11-01-17 | 2/27/2018 11:37:57 AM |
| Dispenser 3 | Buffer | | 8/3/2010 6:05:16 AM |
| Syringe volume | 0.5 mL | | |
| Firmware version | 1.2.1(r2) | | |
| Titrant | Dodecane | 2018/01/31 | 2/28/2018 11:18:04 AM |
| Dispenser 6 | Octanol | | 10/22/2010 11:52:43 AM |

| | | | |
|---|---|---|---|
| Sample name: | **M11_octanol** | Experiment start time: | **3/9/2018 4:01:24 PM** |
| Assay name: | **pH-metric high logP** | Analyst: | **Pion** |
| Assay ID: | **18C-09010** | Instrument ID: | **T312060** |
| Filename: | **C:\Sirius_T3\Mehtap\20180309_exp31_logP_T3-2\18C-09010_M11_octanol_pH-metric high logP.t3r** | | |

| | | | |
|---|---|---|---|
| Sample name: | **M11_octanol** | Experiment start time: | **3/9/2018 4:01:24 PM** |
| Assay name: | **pH-metric high logP** | Analyst: | **Pion** |
| Assay ID: | **18C-09010** | Instrument ID: | **T312060** |
| Filename: | **C:\Sirius_T3\Mehtap\20180309_exp31_logP_T3-2\18C-09010_M11_octanol_pH-metric high logP.t3r** | | |

### Instrument Settings (continued)

| Setting | Value | Batch Id | Install date |
|---|---|---|---|
| Spectrometer calibration stir duration | 5 s | | |
| Spectrometer calibration stir speed | 30% | | |
| Spectrometer calibration wash pump volume | 20.0 mL | | |
| Spectrometer calibration wash stir duration | 5 s | | |
| Spectrometer calibration wash stir speed | 30% | | |
| Overhead dispense height | 10000 | | |

### Refinement Settings

| Setting | Value | Default value |
|---|---|---|
| Turbidity detection method | None | None |
| Turbidity wavelength to assess | 500.0 nm | 500.0 nm |
| Turbidity maximum absorbance | 0.100 | 0.100 |
| Turbidity probe threshold | 50.00 | 50.00 |
