## Supplementary material for "Octanol-water partition coefficient measurements for the SAMPL6 Blind Prediction Challenge": SM12_18C-01012_M12_octanol_pH-metric high logP_report.pdf

| | | | |
|---|---|---|---|
| Sample name: | **M12_octanol** | Experiment start time: | **3/1/2018 3:28:52 PM** |
| Assay name: | **pH-metric high logP** | Analyst: | **Pion** |
| Assay ID: | **18C-01012** | Instrument ID: | **T312060** |
| Filename: | **C:\Sirius_T3\Mehtap\20180228_exp28_logP_T3-2\18C-01012_M12_octanol_pH-metric high logP.t3r** | | |

#### pH-metric Result

| | | |
|---|---|---|
| logP (XH +) | 1.02 | ±0.07 (n=50) |
| logP (neutral X) | 3.79 | ±0.02 (n=50) |

#### 18C-01012 Points 1 to 21

| | |
|---|---|
| M12_octanol concentration factor | 0.790 |
| Carbonate | 0.0885 mM |
| Acidity error | -0.75786 mM |

#### 18C-01012 Points 22 to 45

| | |
|---|---|
| M12_octanol concentration factor | 0.666 |
| Carbonate | 0.1221 mM |
| Acidity error | -0.39695 mM |

#### 18C-01012 Points 46 to 87

| | |
|---|---|
| M12_octanol concentration factor | 0.461 |
| Carbonate | 1.9011 mM |
| Acidity error | -0.34659 mM |

#### Warnings and errors

| | |
|---|---|
| Errors | None |
| Warnings | Excessive carbonate concentration present |

#### Sample logD and percent species

| pH | M12_octanol logD | M12_octanol M12_octanolH | M12_octanol M12_octanol | M12_octanol M12_octanolH* | M12_octanol M12_octanol* | Comment |
|---|---|---|---|---|---|---|
| 1.000 | 1.04 | 8.40 % | 0.00 % | 88.90 % | 2.71 % | |
| 1.200 | 1.05 | 8.26 % | 0.00 % | 87.51 % | 4.23 % | Stomach pH |
| 2.000 | 1.14 | 6.75 % | 0.00 % | 71.47 % | 21.78 % | |
| 3.000 | 1.63 | 2.28 % | 0.01 % | 24.14 % | 73.56 % | |
| 4.000 | 2.50 | 0.30 % | 0.02 % | 3.17 % | 96.52 % | |
| 5.000 | 3.33 | 0.03 % | 0.02 % | 0.33 % | 99.63 % | |
| 6.000 | 3.71 | 0.00 % | 0.02 % | 0.03 % | 99.95 % | |
| 6.500 | 3.76 | 0.00 % | 0.02 % | 0.01 % | 99.97 % | |
| 7.000 | 3.78 | 0.00 % | 0.02 % | 0.00 % | 99.98 % | |
| 7.400 | 3.79 | 0.00 % | 0.02 % | 0.00 % | 99.98 % | Blood pH |
| 8.000 | 3.79 | 0.00 % | 0.02 % | 0.00 % | 99.98 % | |
| 9.000 | 3.79 | 0.00 % | 0.02 % | 0.00 % | 99.98 % | |
| 10.000 | 3.79 | 0.00 % | 0.02 % | 0.00 % | 99.98 % | |
| 11.000 | 3.79 | 0.00 % | 0.02 % | 0.00 % | 99.98 % | |
| 12.000 | 3.79 | 0.00 % | 0.02 % | 0.00 % | 99.98 % | |

### pH-metric Result

| | | | |
|---|---|---|---|
| Sample name: | **M12_octanol** | Experiment start time: | **3/1/2018 3:28:52 PM** |
| Assay name: | **pH-metric high logP** | Analyst: | **Pion** |
| Assay ID: | **18C-01012** | Instrument ID: | **T312060** |
| Filename: | **C:\Sirius_T3\Mehtap\20180228_exp28_logP_T3-2\18C-01012_M12_octanol_pH-metric high logP.t3r** | | |

#### Graphs

Charge balance residuals

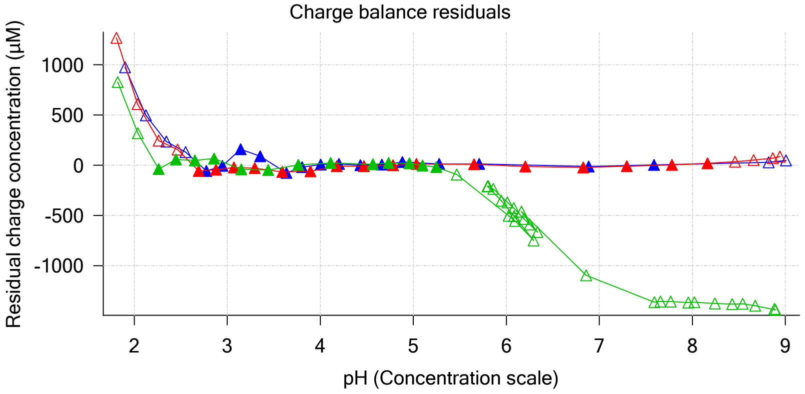


Ionic strength

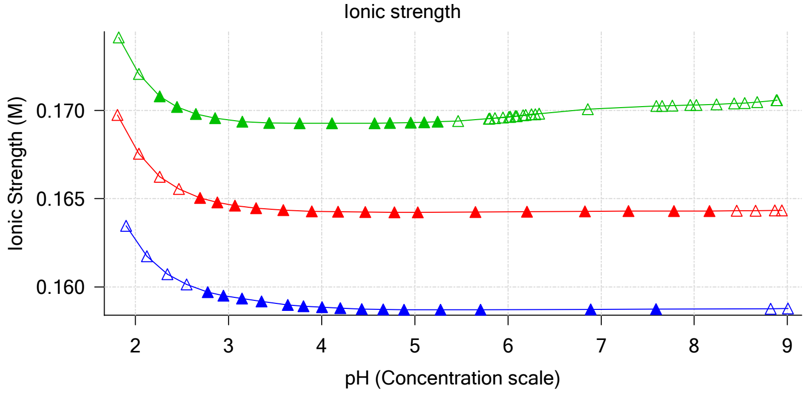


Buffer index profile

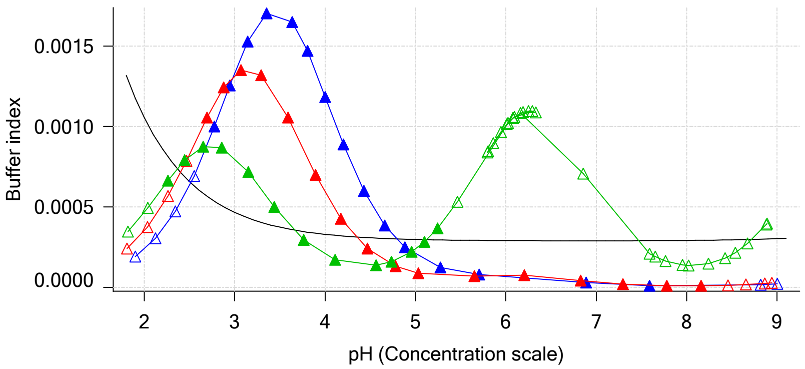


Sample M12_octanol Ionisation Graph

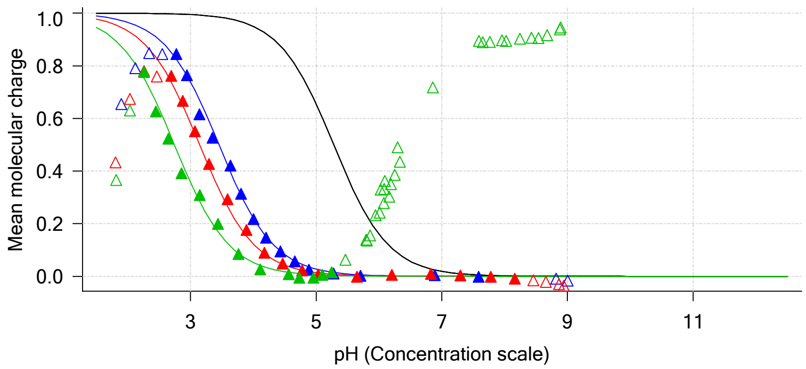


Distribution of species for sample M12_octanol

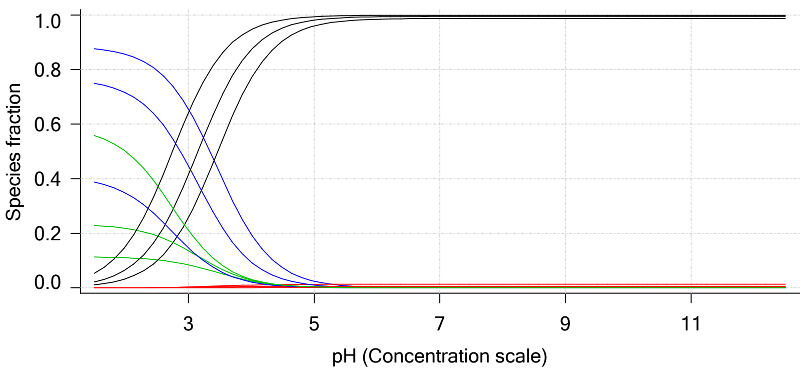


Lipophilicity Profile

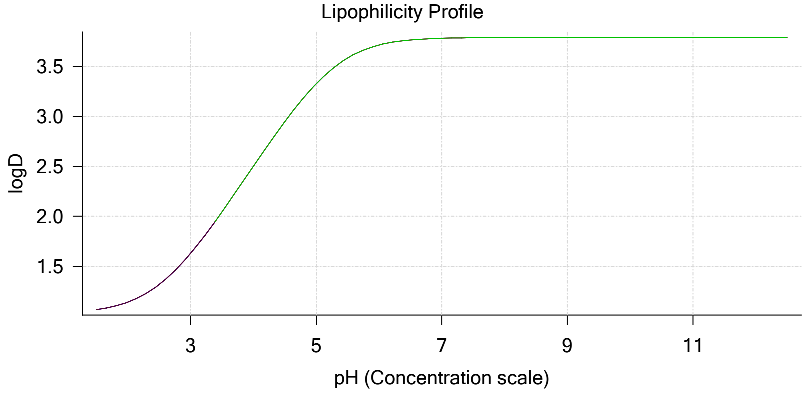


Log R Profile

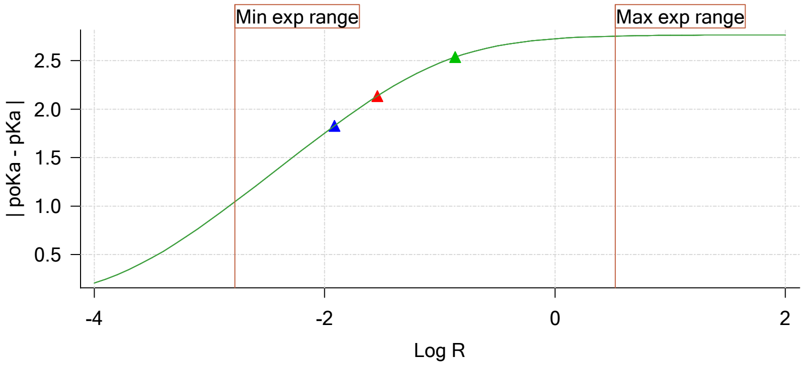


Sample Carbonate Ionisation Graph

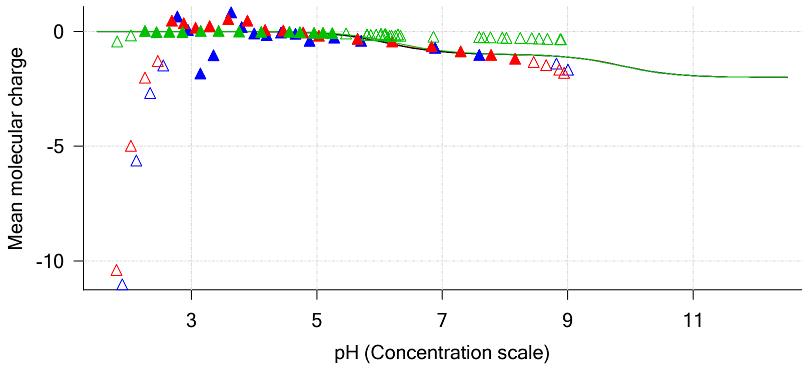

Sample name: **M12_octanol**
Assay name: **pH-metric high logP**
Assay ID: **18C-01012**
Filename: **C:\Sirius_T3\Mehtap\20180228_exp28_logP_T3-2\18C-01012_M12_octanol_pH-metric high logP.t3r**

Experiment start time: **3/1/2018 3:28:52 PM**
Analyst: **Pion**
Instrument ID: **T312060**

#### Graphs (continued)

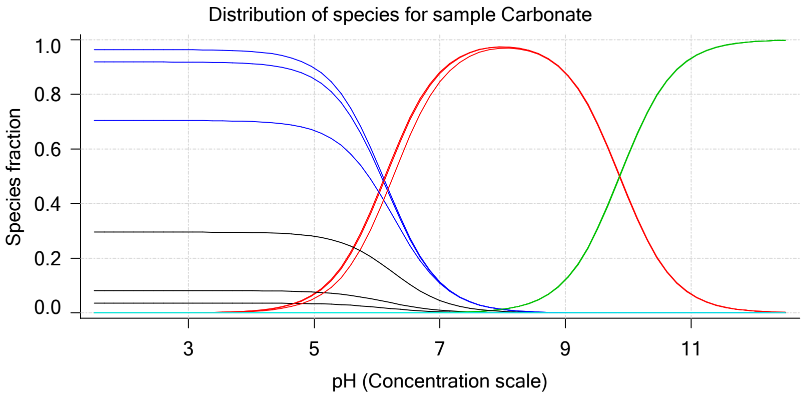

| | | | |
|---|---|---|---|
| Sample name: | **M12_octanol** | Experiment start time: | **3/1/2018 3:28:52 PM** |
| Assay name: | **pH-metric high logP** | Analyst: | **Pion** |
| Assay ID: | **18C-01012** | Instrument ID: | **T312060** |
| Filename: | **C:\Sirius_T3\Mehtap\20180228_exp28_logP_T3-2\18C-01012_M12_octanol_pH-metric high logP.t3r** | | |

# pH-metric high logP   Titration 1 of 3   18C-01012 Points 1 to 21

#### Overall results

| | |
|---|---|
| RMSD | 0.175 |
| Average ionic strength | 0.159 M |
| Average temperature | 25.0°C |
| Partition ratio | 0.0122 : 1 |
| Analyte concentration range | 3733.6 µM to 3842.7 µM |
| Total points considered | 15 of 21 |

#### Warnings and errors

| | |
|---|---|
| Errors | None |
| Warnings | None |

#### Four-Plus parameters

| | | | |
|---|---|---|---|
| Alpha | 0.130 | 3/1/2018 3:28:52 PM | C:\Sirius_T3\HCl18B27.t3r |
| S | 0.9970 | 3/1/2018 3:28:52 PM | C:\Sirius_T3\HCl18B27.t3r |
| jH | 0.8 | 3/1/2018 3:28:52 PM | C:\Sirius_T3\HCl18B27.t3r |
| jOH | -0.4 | 3/1/2018 3:28:52 PM | C:\Sirius_T3\HCl18B27.t3r |

#### Titrants

| | | | |
|---|---|---|---|
| 0.50 M HCl | 0.993513 | 3/1/2018 3:28:52 PM | C:\Sirius_T3\HCl18B27.t3r |
| 0.50 M KOH | 0.999845 | 3/1/2018 3:28:52 PM | C:\Sirius_T3\KOH18B27.t3r |

#### Sample

| | |
|---|---|
| M12_octanol concentration factor | 0.790 |
| M12_octanol stoichiometry | 1.000 |
| Chloride stoichiometry | 1.000 |
| Base pKa 1 | 5.28 |
| logP (XH +) | 1.53 |
| logP (neutral X) | 3.90 |

#### Sample graphs

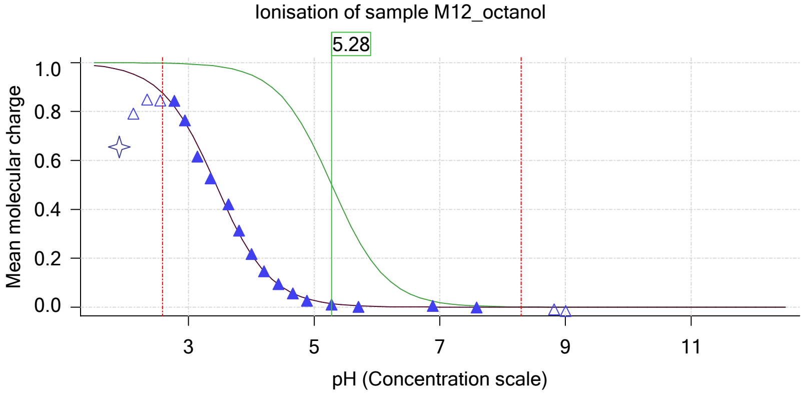


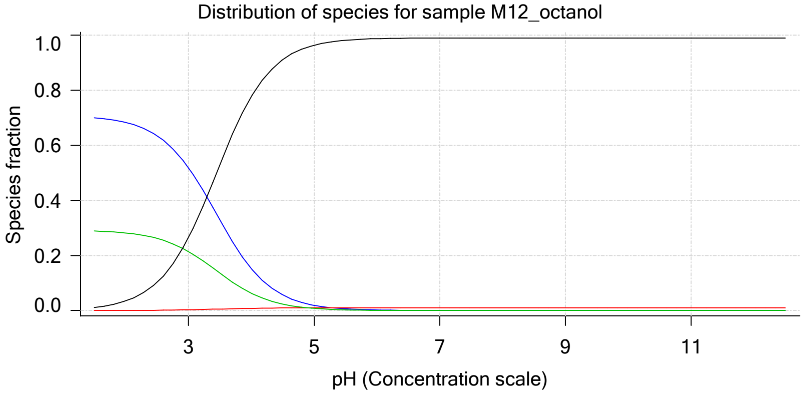

Sample name: **M12_octanol** Experiment start time: **3/1/2018 3:28:52 PM**
Assay name: **pH-metric high logP** Analyst: **Pion**
Assay ID: **18C-01012** Instrument ID: **T312060**
Filename: **C:\Sirius_T3\Mehtap\20180228_exp28_logP_T3-2\18C-01012_M12_octanol_pH-metric high logP.t3r**

#### Sample graphs (continued)

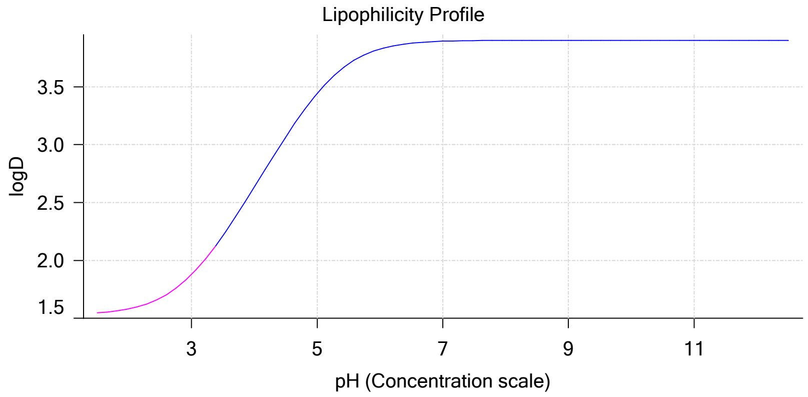


#### Sample logD and percent species

| pH | M12_octanol logD | M12_octanol M12_octanolH | M12_octanol M12_octanol | M12_octanol M12_octanolH* | M12_octanol M12_octanol* | Comment |
|---|---|---|---|---|---|---|
| 1.000 | 1.54 | 70.54 % | 0.00 % | 29.09 % | 0.36 % | |
| 1.200 | 1.54 | 70.39 % | 0.01 % | 29.03 % | 0.57 % | Stomach pH |
| 2.000 | 1.58 | 68.31 % | 0.04 % | 28.17 % | 3.48 % | |
| 3.000 | 1.88 | 51.88 % | 0.27 % | 21.40 % | 26.45 % | |
| 4.000 | 2.63 | 15.23 % | 0.80 % | 6.28 % | 77.68 % | |
| 5.000 | 3.44 | 1.89 % | 0.99 % | 0.78 % | 96.34 % | |
| 6.000 | 3.83 | 0.19 % | 1.02 % | 0.08 % | 98.71 % | |
| 6.500 | 3.88 | 0.06 % | 1.02 % | 0.03 % | 98.90 % | |
| 7.000 | 3.89 | 0.02 % | 1.02 % | 0.01 % | 98.95 % | |
| 7.400 | 3.90 | 0.01 % | 1.02 % | 0.00 % | 98.97 % | Blood pH |
| 8.000 | 3.90 | 0.00 % | 1.02 % | 0.00 % | 98.98 % | |
| 9.000 | 3.90 | 0.00 % | 1.02 % | 0.00 % | 98.98 % | |
| 10.000 | 3.90 | 0.00 % | 1.02 % | 0.00 % | 98.98 % | |
| 11.000 | 3.90 | 0.00 % | 1.02 % | 0.00 % | 98.98 % | |
| 12.000 | 3.90 | 0.00 % | 1.02 % | 0.00 % | 98.98 % | |

#### Carbonate and acidity

Carbonate 0.088 mM
Acidity error -0.758 mM

#### Other graphs

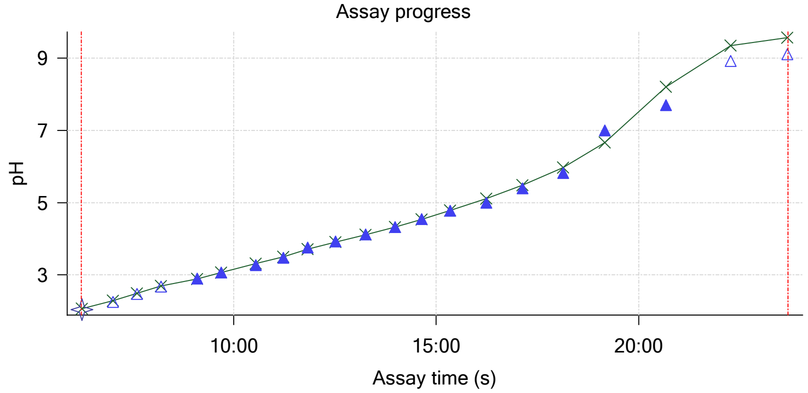


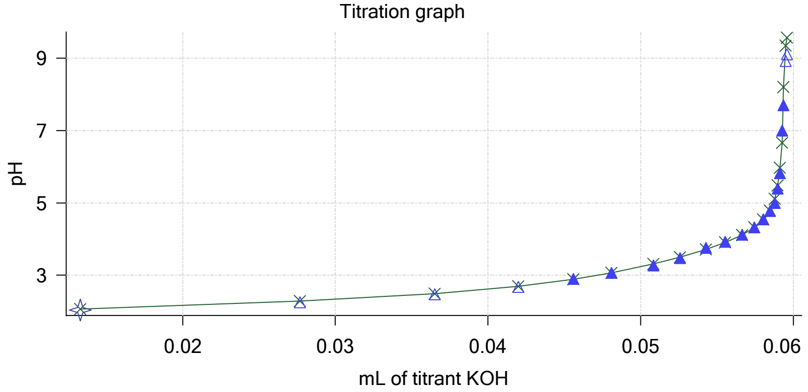

Sample name: **M12_octanol**
Assay name: **pH-metric high logP**
Assay ID: **18C-01012**
Filename: **C:\Sirius_T3\Mehtap\20180228_exp28_logP_T3-2\18C-01012_M12_octanol_pH-metric high logP.t3r**

Experiment start time: **3/1/2018 3:28:52 PM**
Analyst: **Pion**
Instrument ID: **T312060**

#### Other graphs (continued)

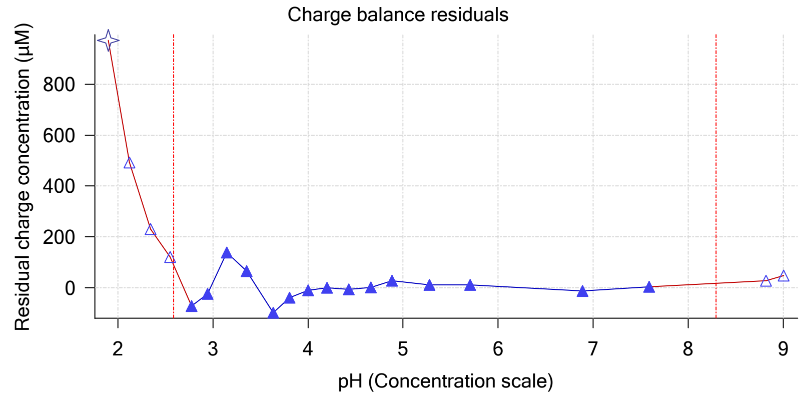


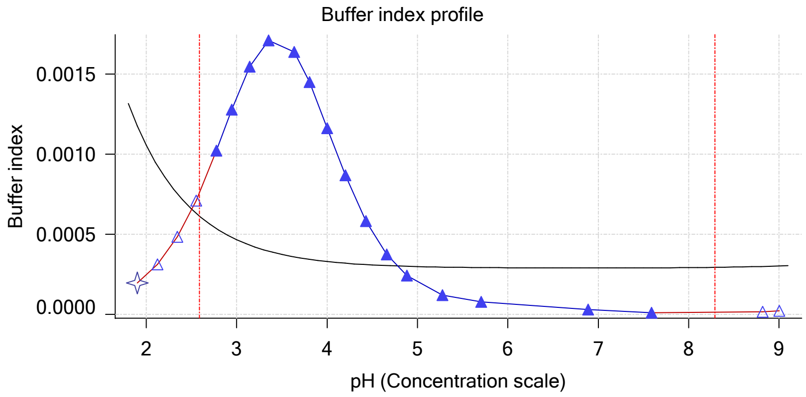


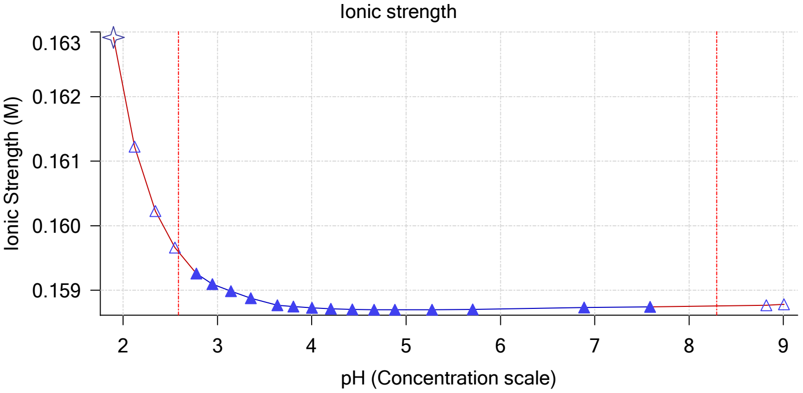


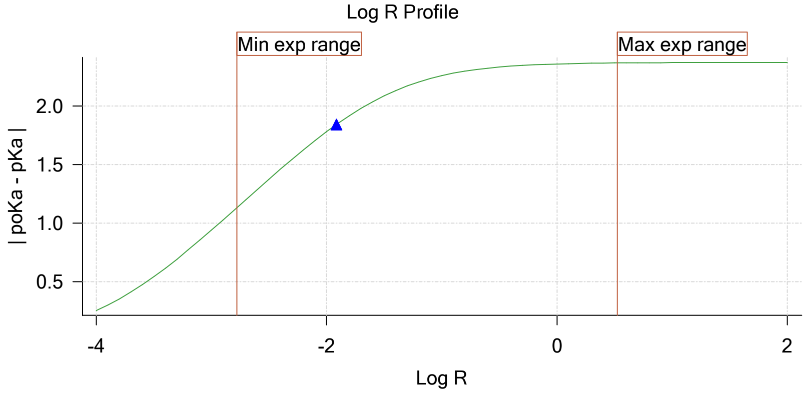


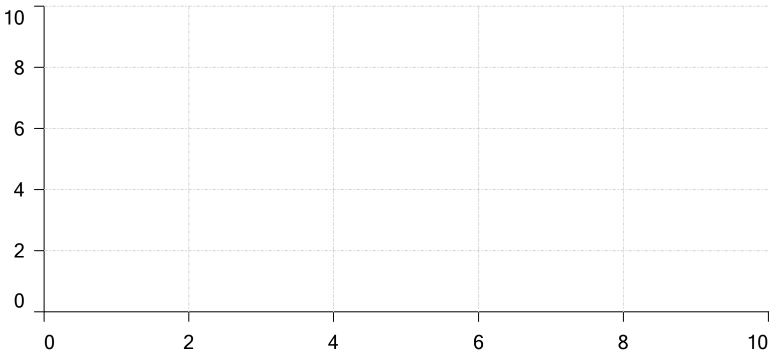

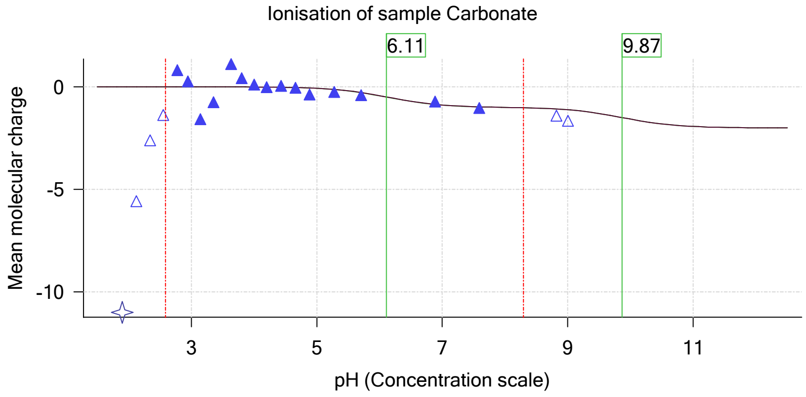

| | | | |
|---|---|---|---|
| Sample name: | **M12_octanol** | Experiment start time: | **3/1/2018 3:28:52 PM** |
| Assay name: | **pH-metric high logP** | Analyst: | **Pion** |
| Assay ID: | **18C-01012** | Instrument ID: | **T312060** |
| Filename: | **C:\Sirius_T3\Mehtap\20180228_exp28_logP_T3-2\18C-01012_M12_octanol_pH-metric high logP.t3r** | | |

### pH-metric high logP Titration 2 of 3 18C-01012 Points 22 to 45

#### Overall results

| | |
|---|---|
| RMSD | 0.444 |
| Average ionic strength | 0.164 M |
| Average temperature | 25.0°C |
| Partition ratio | 0.0288 : 1 |
| Analyte concentration range | 3433.0 μM to 3545.2 μM |
| Total points considered | 16 of 24 |

#### Warnings and errors

| | |
|---|---|
| Errors | None |
| Warnings | Sample concentration factor out of range |

#### Four-Plus parameters

| | | | |
|---|---|---|---|
| Alpha | 0.130 | 3/1/2018 3:28:52 PM | C:\Sirius_T3\HCl18B27.t3r |
| S | 0.9970 | 3/1/2018 3:28:52 PM | C:\Sirius_T3\HCl18B27.t3r |
| jH | 0.8 | 3/1/2018 3:28:52 PM | C:\Sirius_T3\HCl18B27.t3r |
| jOH | -0.4 | 3/1/2018 3:28:52 PM | C:\Sirius_T3\HCl18B27.t3r |

#### Titrants

| | | | |
|---|---|---|---|
| 0.50 M HCl | 0.993513 | 3/1/2018 3:28:52 PM | C:\Sirius_T3\HCl18B27.t3r |
| 0.50 M KOH | 0.999845 | 3/1/2018 3:28:52 PM | C:\Sirius_T3\KOH18B27.t3r |

#### Sample

| | |
|---|---|
| M12_octanol concentration factor | 0.666 |
| M12_octanol stoichiometry | 1.000 |
| Chloride stoichiometry | 1.000 |
| Base pKa 1 | 5.28 |
| logP (XH +) | 1.53 |
| logP (neutral X) | 3.93 |

#### Sample graphs

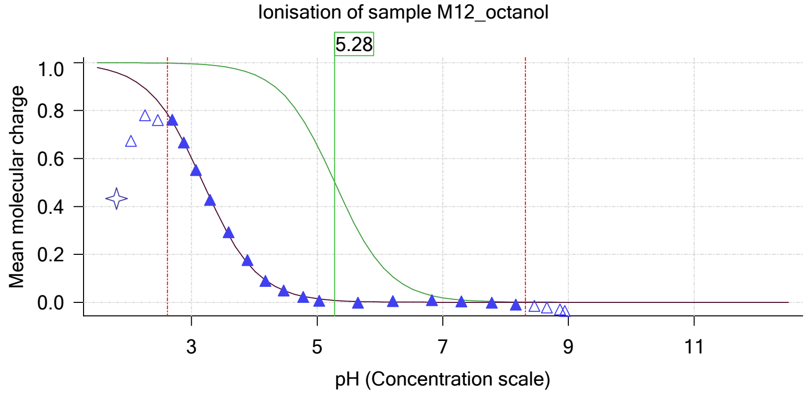


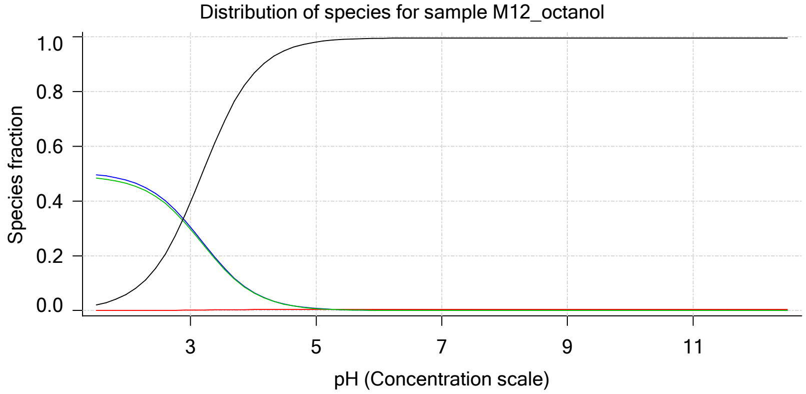

| | | | |
|---|---|---|---|
| Sample name: | **M12_octanol** | Experiment start time: | **3/1/2018 3:28:52 PM** |
| Assay name: | **pH-metric high logP** | Analyst: | **Pion** |
| Assay ID: | **18C-01012** | Instrument ID: | **T312060** |
| Filename: | **C:\Sirius_T3\Mehtap\20180228_exp28_logP_T3-2\18C-01012_M12_octanol_pH-metric high logP.t3r** | | |

#### Sample graphs (continued)

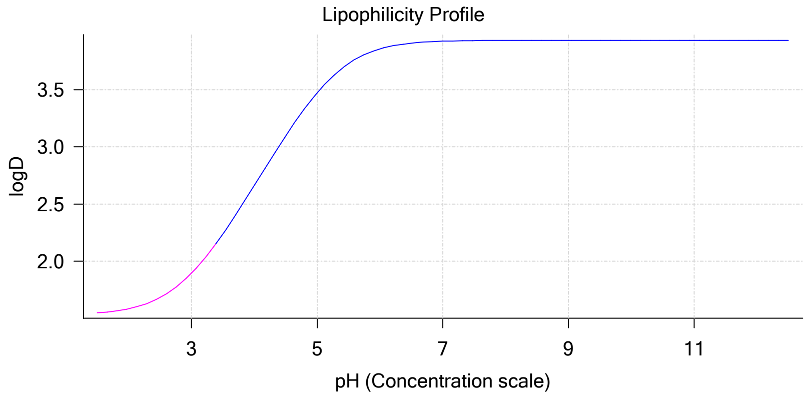


#### Sample logD and percent species

| pH | M12_octanol logD | M12_octanol M12_octanolH | M12_octanol M12_octanol | M12_octanol M12_octanolH* | M12_octanol M12_octanol* | Comment |
|---|---|---|---|---|---|---|
| 1.000 | 1.54 | 50.30 % | 0.00 % | 49.05 % | 0.65 % | |
| 1.200 | 1.54 | 50.10 % | 0.00 % | 48.86 % | 1.03 % | Stomach pH |
| 2.000 | 1.58 | 47.50 % | 0.02 % | 46.32 % | 6.15 % | |
| 3.000 | 1.89 | 30.53 % | 0.16 % | 29.77 % | 39.54 % | |
| 4.000 | 2.66 | 6.68 % | 0.35 % | 6.51 % | 86.46 % | |
| 5.000 | 3.47 | 0.76 % | 0.40 % | 0.74 % | 98.11 % | |
| 6.000 | 3.86 | 0.08 % | 0.40 % | 0.07 % | 99.45 % | |
| 6.500 | 3.91 | 0.02 % | 0.40 % | 0.02 % | 99.55 % | |
| 7.000 | 3.92 | 0.01 % | 0.40 % | 0.01 % | 99.58 % | |
| 7.400 | 3.93 | 0.00 % | 0.40 % | 0.00 % | 99.59 % | Blood pH |
| 8.000 | 3.93 | 0.00 % | 0.40 % | 0.00 % | 99.59 % | |
| 9.000 | 3.93 | 0.00 % | 0.40 % | 0.00 % | 99.60 % | |
| 10.000 | 3.93 | 0.00 % | 0.40 % | 0.00 % | 99.60 % | |
| 11.000 | 3.93 | 0.00 % | 0.40 % | 0.00 % | 99.60 % | |
| 12.000 | 3.93 | 0.00 % | 0.40 % | 0.00 % | 99.60 % | |

#### Carbonate and acidity

| | |
|---|---|
| Carbonate | 0.122 mM |
| Acidity error | -0.397 mM |

#### Other graphs

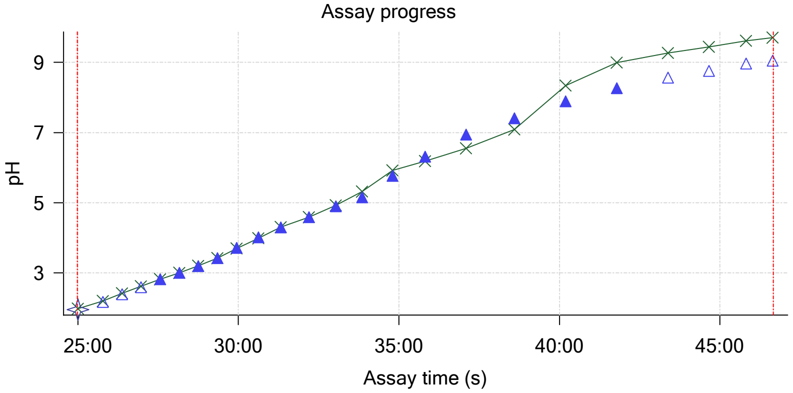


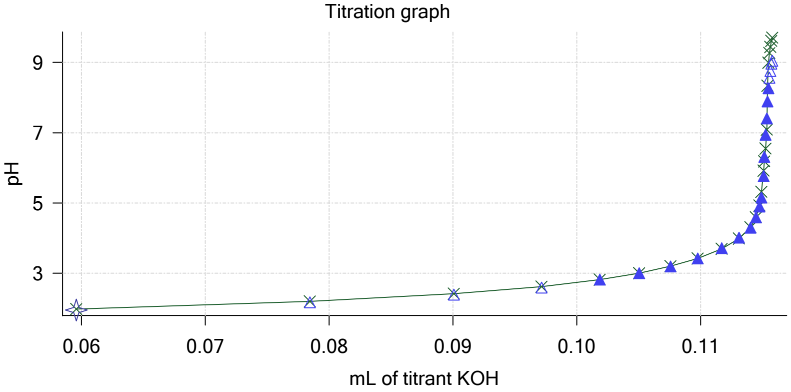

| | | | |
|---|---|---|---|
| Sample name: | **M12_octanol** | Experiment start time: | **3/1/2018 3:28:52 PM** |
| Assay name: | **pH-metric high logP** | Analyst: | **Pion** |
| Assay ID: | **18C-01012** | Instrument ID: | **T312060** |
| Filename: | **C:\Sirius_T3\Mehtap\20180228_exp28_logP_T3-2\18C-01012_M12_octanol_pH-metric high logP.t3r** | | |

#### Other graphs (continued)

Charge balance residuals

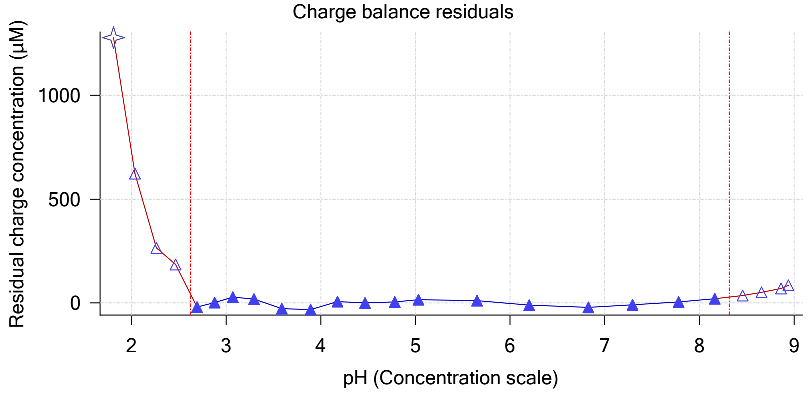


Buffer index profile

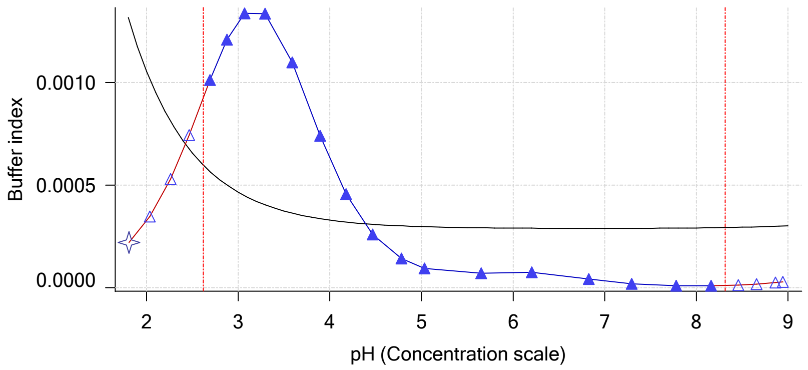


Ionic strength

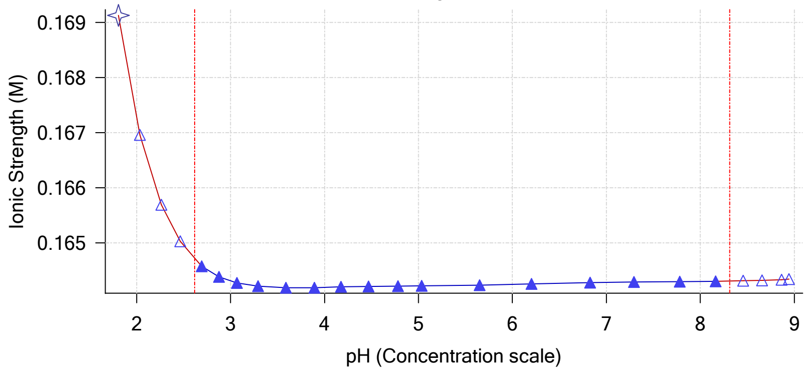


Log R Profile

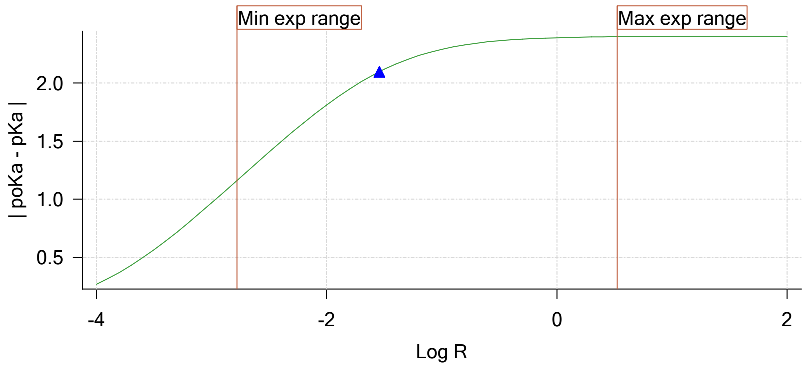


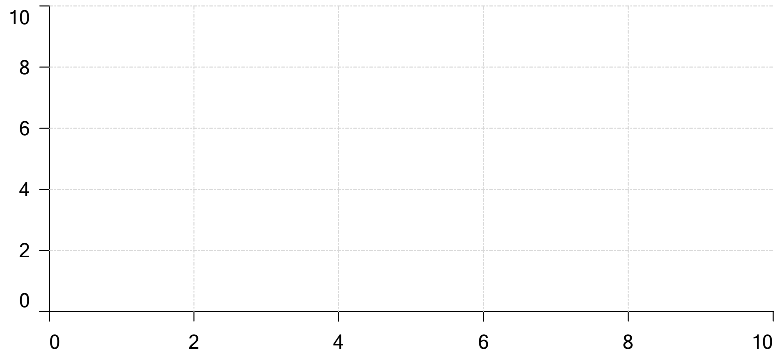


Ionisation of sample Carbonate

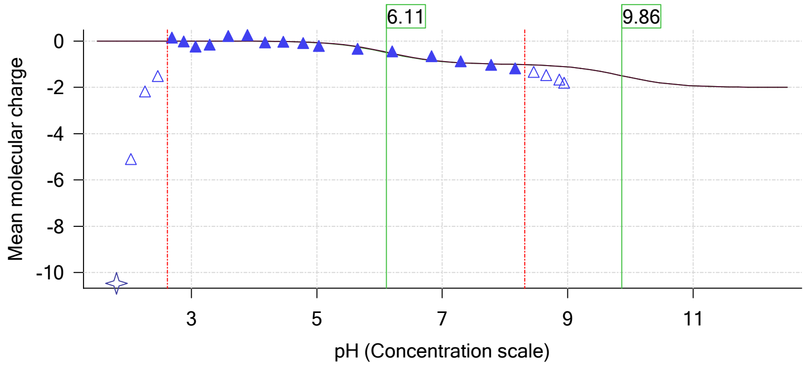

| | | | |
|---|---|---|---|
| Sample name: | **M12_octanol** | Experiment start time: | **3/1/2018 3:28:52 PM** |
| Assay name: | **pH-metric high logP** | Analyst: | **Pion** |
| Assay ID: | **18C-01012** | Instrument ID: | **T312060** |
| Filename: | **C:\Sirius_T3\Mehtap\20180228_exp28_logP_T3-2\18C-01012_M12_octanol_pH-metric high logP.t3r** | | |

### pH-metric high logP Titration 3 of 3 18C-01012 Points 46 to 87

#### Overall results

| | |
|---|---|
| RMSD | 0.047 |
| Average ionic strength | 0.169 M |
| Average temperature | 25.0°C |
| Partition ratio | 0.1359 : 1 |
| Analyte concentration range | 2911.0 μM to 2994.8 μM |
| Total points considered | 13 of 42 |

#### Warnings and errors

| | |
|---|---|
| Errors | None |
| Warnings | Sample concentration factor out of range |
| | Excessive carbonate concentration present |

#### Four-Plus parameters

| | | | |
|---|---|---|---|
| Alpha | 0.130 | 3/1/2018 3:28:52 PM | C:\Sirius_T3\HCl18B27.t3r |
| S | 0.9970 | 3/1/2018 3:28:52 PM | C:\Sirius_T3\HCl18B27.t3r |
| jH | 0.8 | 3/1/2018 3:28:52 PM | C:\Sirius_T3\HCl18B27.t3r |
| jOH | -0.4 | 3/1/2018 3:28:52 PM | C:\Sirius_T3\HCl18B27.t3r |

#### Titrants

| | | | |
|---|---|---|---|
| 0.50 M HCl | 0.993513 | 3/1/2018 3:28:52 PM | C:\Sirius_T3\HCl18B27.t3r |
| 0.50 M KOH | 0.999845 | 3/1/2018 3:28:52 PM | C:\Sirius_T3\KOH18B27.t3r |

#### Sample

| | |
|---|---|
| M12_octanol concentration factor | 0.461 |
| M12_octanol stoichiometry | 1.000 |
| Chloride stoichiometry | 1.000 |
| Base pKa 1 | 5.28 |
| logP (XH +) | 1.53 |
| logP (neutral X) | 4.17 |

#### Sample graphs

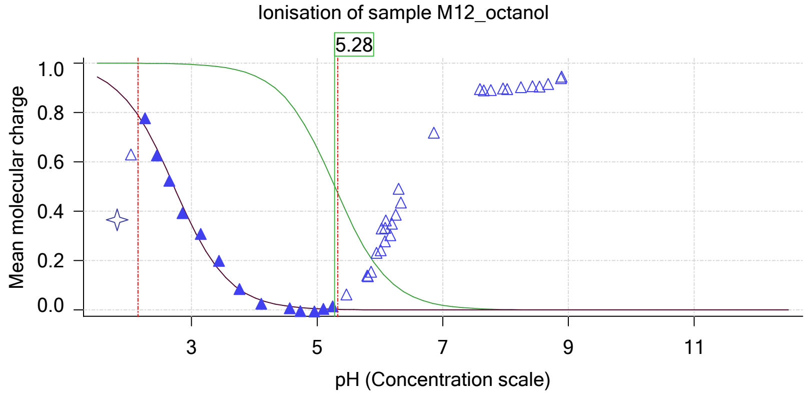


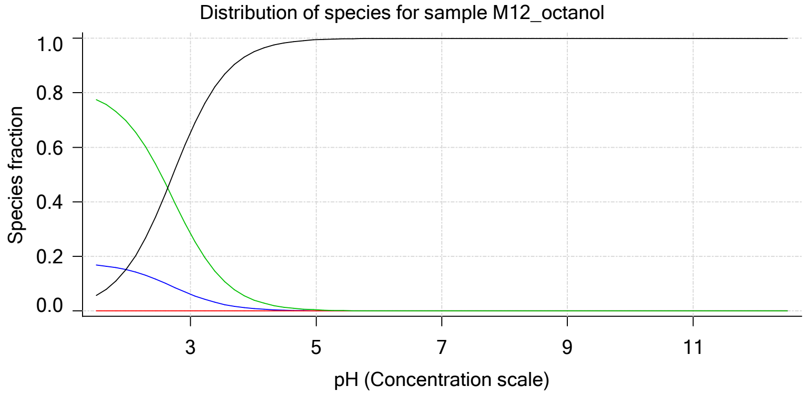

| | | | |
|---|---|---|---|
| Sample name: | **M12_octanol** | Experiment start time: | **3/1/2018 3:28:52 PM** |
| Assay name: | **pH-metric high logP** | Analyst: | **Pion** |
| Assay ID: | **18C-01012** | Instrument ID: | **T312060** |
| Filename: | **C:\Sirius_T3\Mehtap\20180228_exp28_logP_T3-2\18C-01012_M12_octanol_pH-metric high logP.t3r** | | |

#### Sample graphs (continued)

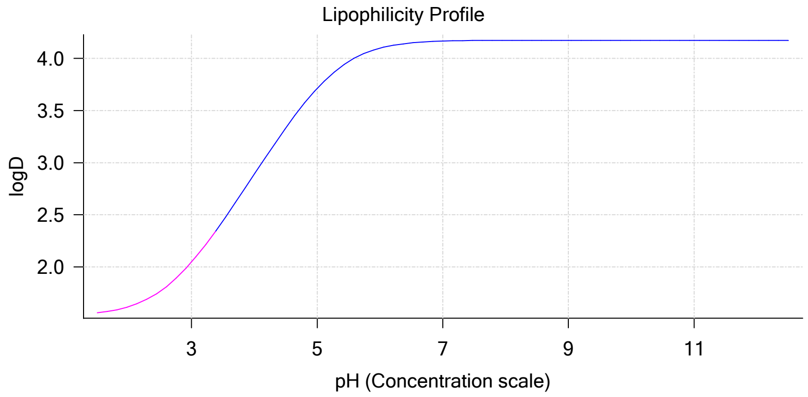


#### Sample logD and percent species

| pH | M12_octanol logD | M12_octanol M12_octanolH | M12_octanol M12_octanol | M12_octanol M12_octanolH* | M12_octanol M12_octanol* | Comment |
|---|---|---|---|---|---|---|
| 1.000 | 1.54 | 17.51 % | 0.00 % | 80.63 % | 1.86 % | |
| 1.200 | 1.55 | 17.32 % | 0.00 % | 79.76 % | 2.92 % | Stomach pH |
| 2.000 | 1.62 | 15.00 % | 0.01 % | 69.06 % | 15.93 % | |
| 3.000 | 2.05 | 6.16 % | 0.03 % | 28.37 % | 65.44 % | |
| 4.000 | 2.89 | 0.89 % | 0.05 % | 4.12 % | 94.94 % | |
| 5.000 | 3.71 | 0.09 % | 0.05 % | 0.43 % | 99.43 % | |
| 6.000 | 4.10 | 0.01 % | 0.05 % | 0.04 % | 99.90 % | |
| 6.500 | 4.15 | 0.00 % | 0.05 % | 0.01 % | 99.93 % | |
| 7.000 | 4.16 | 0.00 % | 0.05 % | 0.00 % | 99.95 % | |
| 7.400 | 4.17 | 0.00 % | 0.05 % | 0.00 % | 99.95 % | Blood pH |
| 8.000 | 4.17 | 0.00 % | 0.05 % | 0.00 % | 99.95 % | |
| 9.000 | 4.17 | 0.00 % | 0.05 % | 0.00 % | 99.95 % | |
| 10.000 | 4.17 | 0.00 % | 0.05 % | 0.00 % | 99.95 % | |
| 11.000 | 4.17 | 0.00 % | 0.05 % | 0.00 % | 99.95 % | |
| 12.000 | 4.17 | 0.00 % | 0.05 % | 0.00 % | 99.95 % | |

#### Carbonate and acidity

| | |
|---|---|
| Carbonate | 1.901 mM |
| Acidity error | -0.347 mM |

#### Other graphs

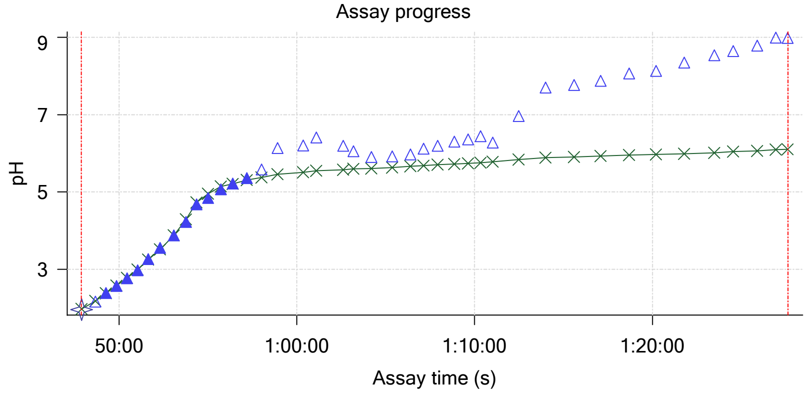


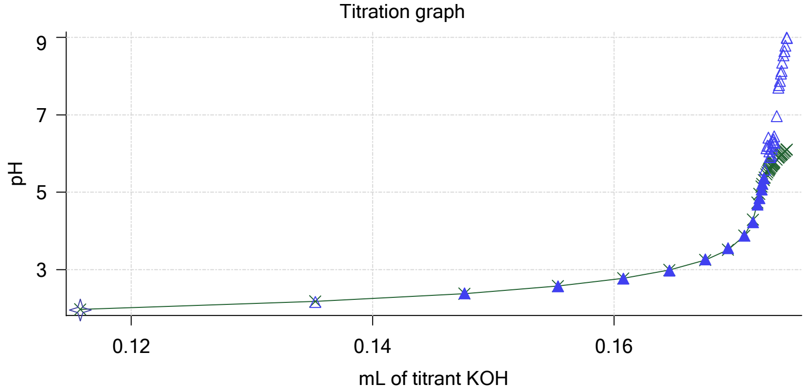

Sample name: **M12_octanol**
Assay name: **pH-metric high logP**
Assay ID: **18C-01012**
Filename: **C:\Sirius_T3\Mehtap\20180228_exp28_logP_T3-2\18C-01012_M12_octanol_pH-metric high logP.t3r**

Experiment start time: **3/1/2018 3:28:52 PM**
Analyst: **Pion**
Instrument ID: **T312060**

#### Other graphs (continued)

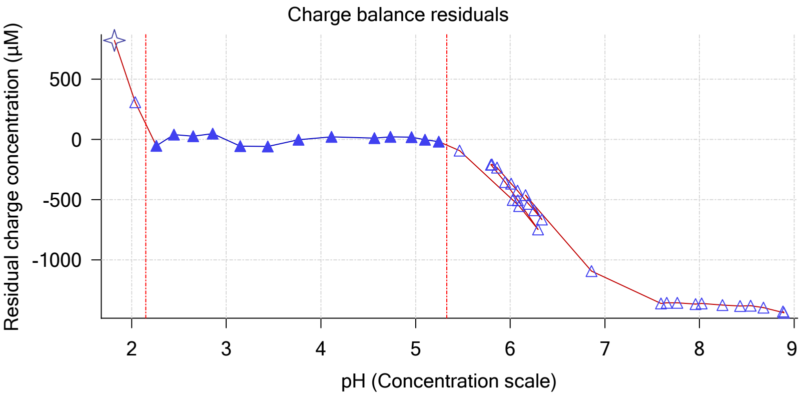


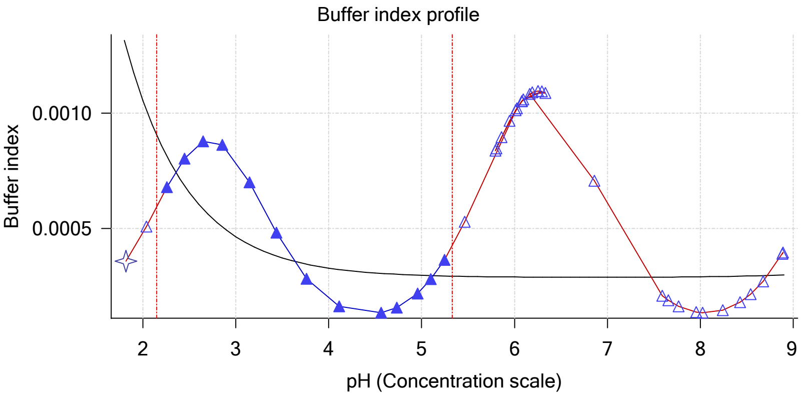


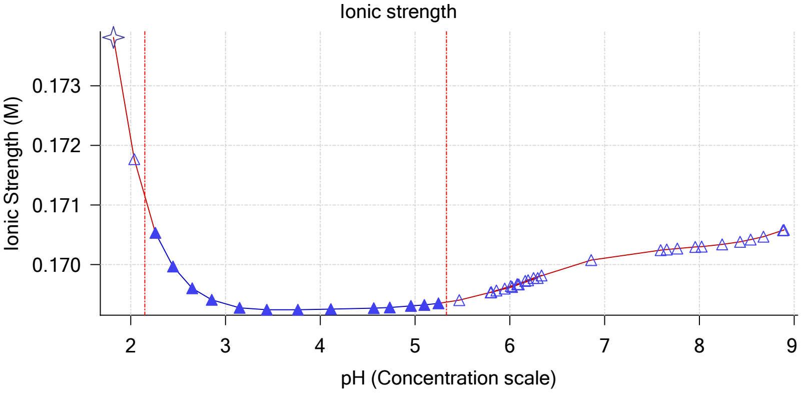


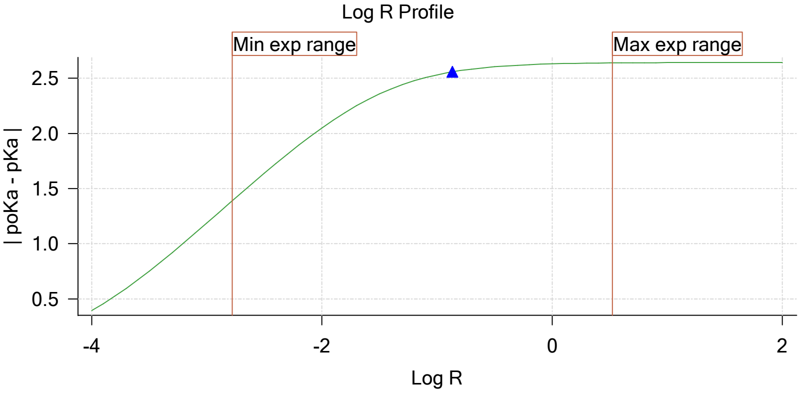


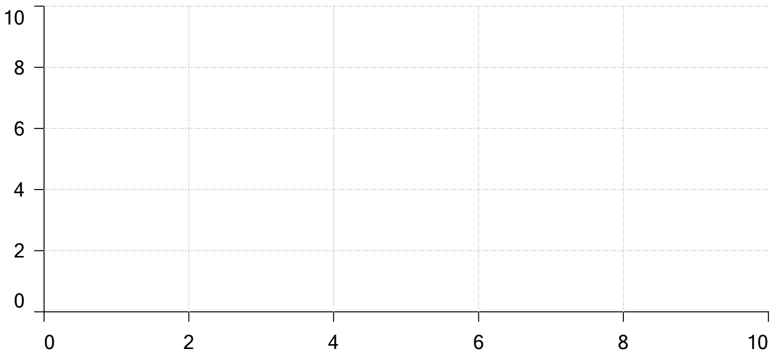

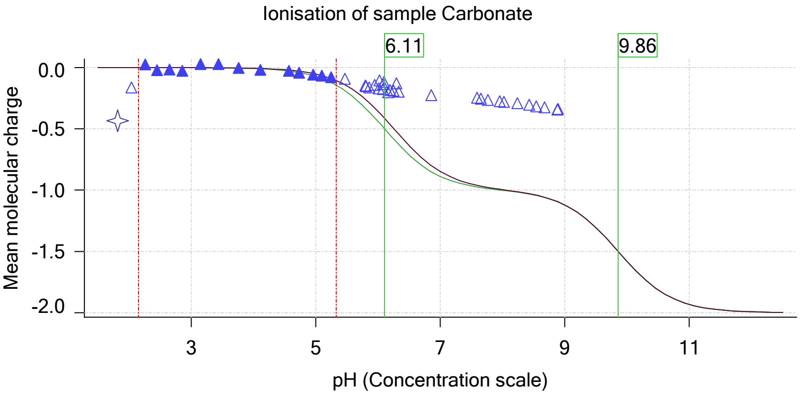

| | | | |
|---|---|---|---|
| Sample name: | **M12_octanol** | Experiment start time: | **3/1/2018 3:28:52 PM** |
| Assay name: | **pH-metric high logP** | Analyst: | **Pion** |
| Assay ID: | **18C-01012** | Instrument ID: | **T312060** |
| Filename: | **C:\Sirius_T3\Mehtap\20180228_exp28_logP_T3-2\18C-01012_M12_octanol_pH-metric high logP.t3r** | | |

#### Assay Model

| Settings | Value | Date/Time changed | Imported from |
|---|---|---|---|
| Sample name | M12_octanol | 2/28/2018 2:58:36 PM | User entered value |
| Sample by | Weight | | Default value |
| Sample weight | 0.001780 g | 2/28/2018 4:51:40 PM | User entered value |
| Formula weight | 292.16 g/mol | 2/28/2018 2:58:36 PM | User entered value |
| Solubility | Unknown | | Default value |
| Molecular weight | 255.70 | 2/28/2018 2:58:36 PM | User entered value |
| Individual pKa ionic environments | No | | Default value |
| Number of pKas | 1 | 2/28/2018 2:58:36 PM | User entered value |
| Sample is a | Base | 2/28/2018 2:58:36 PM | User entered value |
| pKa 1 | 5.28 | 2/28/2018 2:58:36 PM | User entered value |
| logp (XH +) | 1.53 | 2/28/2018 2:59:16 PM | User entered value |
| logP (neutral X) | 3.50 | 2/28/2018 2:59:06 PM | User entered value |
| Stoichiometry | 1.00000 | | Default value |
| Aprotic counterion name | Chloride | | From standards.xml file |
| Stoichiometry | 1.00 | | From standards.xml file |
| Charge per counterion | -1 | | From standards.xml file |

#### Events

| Time | Event | Water | Acid | Base | Octanol | pH | dpH/dt | pH R-squared | pH SD | dpH/dt time |
|---|---|---|---|---|---|---|---|---|---|---|
| 3:15.3 | Initial pH = 4.96 | | | | | | | | | |
| 6:15.1 | Data point 1 | 1.50000 mL | 0.05221 mL | 0.01329 mL | 0.01999 mL | 2.033 | -0.00115 | 0.13540 | 0.00015 | 10.0 s |
| 7:01.4 | Data point 2 | 1.50000 mL | 0.05221 mL | 0.02768 mL | 0.01999 mL | 2.252 | 0.00111 | 0.19272 | 0.00012 | 10.0 s |
| 7:37.0 | Data point 3 | 1.50000 mL | 0.05221 mL | 0.03652 mL | 0.01999 mL | 2.470 | -0.00312 | 0.03211 | 0.00086 | 10.0 s |
| 8:12.5 | Data point 4 | 1.50000 mL | 0.05221 mL | 0.04198 mL | 0.01999 mL | 2.675 | -0.00299 | 0.02511 | 0.00093 | 28.0 s |
| 9:06.1 | Data point 5 | 1.50000 mL | 0.05221 mL | 0.04560 mL | 0.01999 mL | 2.899 | 0.00724 | 0.28037 | 0.00067 | 10.0 s |
| 9:41.6 | Data point 6 | 1.50000 mL | 0.05221 mL | 0.04809 mL | 0.01999 mL | 3.067 | -0.00758 | 0.65584 | 0.00046 | 10.0 s |
| 10:32.6 | Data point 7 | 1.50000 mL | 0.05221 mL | 0.05085 mL | 0.01999 mL | 3.266 | -0.00574 | 0.79317 | 0.00032 | 10.5 s |
| 11:13.7 | Data point 8 | 1.50000 mL | 0.05221 mL | 0.05259 mL | 0.01999 mL | 3.474 | -0.01919 | 0.98927 | 0.00095 | 10.5 s |
| 11:49.7 | Data point 9 | 1.50000 mL | 0.05221 mL | 0.05428 mL | 0.01999 mL | 3.755 | -0.01633 | 0.83230 | 0.00088 | 10.5 s |
| 12:30.9 | Data point 10 | 1.50000 mL | 0.05221 mL | 0.05555 mL | 0.01999 mL | 3.923 | -0.01859 | 0.95788 | 0.00094 | 14.0 s |
| 13:15.5 | Data point 11 | 1.50000 mL | 0.05221 mL | 0.05666 mL | 0.01999 mL | 4.120 | -0.01675 | 0.71331 | 0.00098 | 13.0 s |
| 13:59.1 | Data point 12 | 1.50000 mL | 0.05221 mL | 0.05746 mL | 0.01999 mL | 4.319 | -0.01672 | 0.90369 | 0.00087 | 14.0 s |
| 14:38.5 | Data point 13 | 1.50000 mL | 0.05221 mL | 0.05804 mL | 0.01999 mL | 4.547 | -0.01987 | 0.96798 | 0.00100 | 17.0 s |
| 15:21.0 | Data point 14 | 1.50000 mL | 0.05221 mL | 0.05847 mL | 0.01999 mL | 4.777 | -0.01763 | 0.88444 | 0.00093 | 23.0 s |
| 16:14.5 | Data point 15 | 1.50000 mL | 0.05221 mL | 0.05880 mL | 0.01999 mL | 4.998 | -0.01818 | 0.93024 | 0.00093 | 28.0 s |
| 17:08.0 | Data point 16 | 1.50000 mL | 0.05221 mL | 0.05898 mL | 0.01999 mL | 5.391 | -0.01937 | 0.94705 | 0.00098 | 29.5 s |
| 18:08.0 | Data point 17 | 1.50000 mL | 0.05221 mL | 0.05913 mL | 0.01999 mL | 5.819 | -0.01888 | 0.96763 | 0.00095 | 31.5 s |
| 19:10.1 | Data point 18 | 1.50000 mL | 0.05221 mL | 0.05927 mL | 0.01999 mL | 6.996 | -0.07112 | 0.99199 | 0.00353 | Timed out at 59.5 s |
| 20:40.6 | Data point 19 | 1.50000 mL | 0.05221 mL | 0.05936 mL | 0.01999 mL | 7.695 | -0.08201 | 0.99679 | 0.00406 | Timed out at 59.5 s |
| 22:16.3 | Data point 20 | 1.50000 mL | 0.05221 mL | 0.05950 mL | 0.01999 mL | 8.922 | -0.00844 | 0.17826 | 0.00099 | 53.5 s |
| 23:40.4 | Data point 21 | 1.50000 mL | 0.05221 mL | 0.05960 mL | 0.01999 mL | 9.107 | -0.02002 | 0.98651 | 0.00100 | 26.0 s |
| 25:00.1 | Data point 22 | 1.50000 mL | 0.10891 mL | 0.05960 mL | 0.05000 mL | 1.945 | -0.00363 | 0.51184 | 0.00025 | 10.5 s |
| 25:46.8 | Data point 23 | 1.50000 mL | 0.10891 mL | 0.07846 mL | 0.05000 mL | 2.167 | 0.00370 | 0.60390 | 0.00024 | 10.0 s |
| 26:22.5 | Data point 24 | 1.50000 mL | 0.10891 mL | 0.09008 mL | 0.05000 mL | 2.390 | -0.00263 | 0.03650 | 0.00068 | 10.0 s |
| 26:58.0 | Data point 25 | 1.50000 mL | 0.10891 mL | 0.09715 mL | 0.05000 mL | 2.592 | 0.00780 | 0.23852 | 0.00079 | 10.5 s |
| 27:34.1 | Data point 26 | 1.50000 mL | 0.10891 mL | 0.10186 mL | 0.05000 mL | 2.817 | 0.00513 | 0.20428 | 0.00056 | 10.0 s |
| 28:09.6 | Data point 27 | 1.50000 mL | 0.10891 mL | 0.10503 mL | 0.05000 mL | 3.001 | -0.00273 | 0.39758 | 0.00021 | 10.0 s |
| 28:45.1 | Data point 28 | 1.50000 mL | 0.10891 mL | 0.10757 mL | 0.05000 mL | 3.191 | -0.00242 | 0.40166 | 0.00019 | 10.5 s |
| 29:21.2 | Data point 29 | 1.50000 mL | 0.10891 mL | 0.10976 mL | 0.05000 mL | 3.415 | -0.00337 | 0.31261 | 0.00030 | 10.0 s |
| 29:56.7 | Data point 30 | 1.50000 mL | 0.10891 mL | 0.11171 mL | 0.05000 mL | 3.709 | -0.00838 | 0.53036 | 0.00057 | 10.0 s |
| 30:37.3 | Data point 31 | 1.50000 mL | 0.10891 mL | 0.11310 mL | 0.05000 mL | 4.012 | -0.01267 | 0.50941 | 0.00088 | 11.5 s |
| 31:19.4 | Data point 32 | 1.50000 mL | 0.10891 mL | 0.11402 mL | 0.05000 mL | 4.293 | -0.01573 | 0.65616 | 0.00096 | 27.0 s |
| 32:11.9 | Data point 33 | 1.50000 mL | 0.10891 mL | 0.11446 mL | 0.05000 mL | 4.584 | -0.01886 | 0.90646 | 0.00098 | 25.0 s |
| 33:02.3 | Data point 34 | 1.50000 mL | 0.10891 mL | 0.11475 mL | 0.05000 mL | 4.896 | -0.01935 | 0.95587 | 0.00098 | 18.5 s |

| | | | |
|---|---|---|---|
| Sample name: | **M12_octanol** | Experiment start time: | **3/1/2018 3:28:52 PM** |
| Assay name: | **pH-metric high logP** | Analyst: | **Pion** |
| Assay ID: | **18C-01012** | Instrument ID: | **T312060** |
| Filename: | **C:\Sirius_T3\Mehtap\20180228_exp28_logP_T3-2\18C-01012_M12_octanol_pH-metric high logP.t3r** | | |

#### Events (continued)

| Time | Event | Water | Acid | Base | Octanol | pH | dpH/dt | pH R-squared | pH SD | dpH/dt time |
|---|---|---|---|---|---|---|---|---|---|---|
| 33:51.3 | Data point 35 | 1.50000 mL | 0.10891 mL | 0.11491 mL | 0.05000 mL | 5.148 | -0.01470 | 0.60540 | 0.00093 | 26.5 s |
| 34:48.4 | Data point 36 | 1.50000 mL | 0.10891 mL | 0.11508 mL | 0.05000 mL | 5.764 | -0.01973 | 0.94931 | 0.00100 | 30.0 s |
| 35:49.0 | Data point 37 | 1.50000 mL | 0.10891 mL | 0.11515 mL | 0.05000 mL | 6.314 | -0.01346 | 0.55470 | 0.00089 | 46.0 s |
| 37:05.6 | Data point 38 | 1.50000 mL | 0.10891 mL | 0.11524 mL | 0.05000 mL | 6.936 | -0.04576 | 0.98945 | 0.00227 | Timed out at 59.5 s |
| 38:36.1 | Data point 39 | 1.50000 mL | 0.10891 mL | 0.11533 mL | 0.05000 mL | 7.403 | -0.05091 | 0.98174 | 0.00254 | Timed out at 59.5 s |
| 40:11.7 | Data point 40 | 1.50000 mL | 0.10891 mL | 0.11540 mL | 0.05000 mL | 7.887 | -0.07516 | 0.99257 | 0.00373 | Timed out at 59.5 s |
| 41:47.3 | Data point 41 | 1.50000 mL | 0.10891 mL | 0.11547 mL | 0.05000 mL | 8.267 | -0.03226 | 0.96652 | 0.00162 | Timed out at 59.5 s |
| 43:22.8 | Data point 42 | 1.50000 mL | 0.10891 mL | 0.11555 mL | 0.05000 mL | 8.561 | -0.01882 | 0.90027 | 0.00098 | 41.0 s |
| 44:39.5 | Data point 43 | 1.50000 mL | 0.10891 mL | 0.11562 mL | 0.05000 mL | 8.761 | -0.02011 | 0.98771 | 0.00100 | 33.5 s |
| 45:48.7 | Data point 44 | 1.50000 mL | 0.10891 mL | 0.11571 mL | 0.05000 mL | 8.967 | -0.01575 | 0.73120 | 0.00091 | 19.5 s |
| 46:38.8 | Data point 45 | 1.50000 mL | 0.10891 mL | 0.11578 mL | 0.05000 mL | 9.045 | -0.01982 | 0.97953 | 0.00099 | 17.0 s |
| 47:53.5 | Data point 46 | 1.50000 mL | 0.16860 mL | 0.11578 mL | 0.25000 mL | 1.956 | -0.00773 | 0.91840 | 0.00040 | 10.5 s |
| 48:40.3 | Data point 47 | 1.50000 mL | 0.16860 mL | 0.13525 mL | 0.25000 mL | 2.168 | 0.00144 | 0.01980 | 0.00051 | 10.0 s |
| 49:16.0 | Data point 48 | 1.50000 mL | 0.16860 mL | 0.14760 mL | 0.25000 mL | 2.388 | 0.01561 | 0.79013 | 0.00087 | 10.0 s |
| 49:51.7 | Data point 49 | 1.50000 mL | 0.16860 mL | 0.15534 mL | 0.25000 mL | 2.573 | -0.01149 | 0.57763 | 0.00075 | 10.0 s |
| 50:27.2 | Data point 50 | 1.50000 mL | 0.16860 mL | 0.16075 mL | 0.25000 mL | 2.774 | 0.00396 | 0.06451 | 0.00077 | 10.0 s |
| 51:02.7 | Data point 51 | 1.50000 mL | 0.16860 mL | 0.16458 mL | 0.25000 mL | 2.979 | -0.00515 | 0.16315 | 0.00063 | 10.0 s |
| 51:38.2 | Data point 52 | 1.50000 mL | 0.16860 mL | 0.16755 mL | 0.25000 mL | 3.270 | 0.00396 | 0.26107 | 0.00038 | 10.0 s |
| 52:18.8 | Data point 53 | 1.50000 mL | 0.16860 mL | 0.16943 mL | 0.25000 mL | 3.558 | 0.01629 | 0.64793 | 0.00100 | 15.5 s |
| 53:05.0 | Data point 54 | 1.50000 mL | 0.16860 mL | 0.17079 mL | 0.25000 mL | 3.882 | 0.01336 | 0.49641 | 0.00094 | 10.5 s |
| 53:46.1 | Data point 55 | 1.50000 mL | 0.16860 mL | 0.17150 mL | 0.25000 mL | 4.230 | -0.01083 | 0.45645 | 0.00079 | 10.5 s |
| 54:22.0 | Data point 56 | 1.50000 mL | 0.16860 mL | 0.17187 mL | 0.25000 mL | 4.682 | -0.00156 | 0.00649 | 0.00095 | 13.0 s |
| 55:00.4 | Data point 57 | 1.50000 mL | 0.16860 mL | 0.17204 mL | 0.25000 mL | 4.849 | -0.00797 | 0.20959 | 0.00086 | 13.0 s |
| 55:43.9 | Data point 58 | 1.50000 mL | 0.16860 mL | 0.17220 mL | 0.25000 mL | 5.072 | -0.00963 | 0.30906 | 0.00086 | 14.5 s |
| 56:23.8 | Data point 59 | 1.50000 mL | 0.16860 mL | 0.17227 mL | 0.25000 mL | 5.214 | 0.01328 | 0.44826 | 0.00098 | 17.5 s |
| 57:11.9 | Data point 60 | 1.50000 mL | 0.16860 mL | 0.17239 mL | 0.25000 mL | 5.360 | -0.00926 | 0.25608 | 0.00090 | 13.5 s |
| 58:01.0 | Data point 61 | 1.50000 mL | 0.16860 mL | 0.17248 mL | 0.25000 mL | 5.579 | 0.01451 | 0.60856 | 0.00092 | 24.0 s |
| 58:55.6 | Data point 62 | 1.50000 mL | 0.16860 mL | 0.17260 mL | 0.25000 mL | 6.137 | -0.01945 | 0.97898 | 0.00097 | 55.5 s |
| 1:00:21.7 | Data point 63 | 1.50000 mL | 0.16860 mL | 0.17269 mL | 0.25000 mL | 6.206 | 0.00784 | 0.16179 | 0.00096 | 13.5 s |
| 1:01:05.7 | Data point 64 | 1.50000 mL | 0.16860 mL | 0.17277 mL | 0.25000 mL | 6.406 | -0.02142 | 0.96918 | 0.00107 | Timed out at 59.5 s |
| 1:02:36.2 | Data point 65 | 1.50000 mL | 0.16860 mL | 0.17281 mL | 0.25000 mL | 6.193 | 0.00801 | 0.28675 | 0.00074 | 10.0 s |
| 1:03:11.6 | Data point 66 | 1.50000 mL | 0.16860 mL | 0.17286 mL | 0.25000 mL | 6.057 | 0.00973 | 0.24043 | 0.00098 | 35.0 s |
| 1:04:12.0 | Data point 67 | 1.50000 mL | 0.16860 mL | 0.17288 mL | 0.25000 mL | 5.908 | 0.01936 | 0.92962 | 0.00099 | 39.0 s |
| 1:05:21.4 | Data point 68 | 1.50000 mL | 0.16860 mL | 0.17293 mL | 0.25000 mL | 5.918 | 0.01461 | 0.56025 | 0.00096 | 37.0 s |
| 1:06:23.9 | Data point 69 | 1.50000 mL | 0.16860 mL | 0.17300 mL | 0.25000 mL | 5.972 | -0.01320 | 0.59468 | 0.00085 | 13.0 s |
| 1:07:07.4 | Data point 70 | 1.50000 mL | 0.16860 mL | 0.17305 mL | 0.25000 mL | 6.123 | -0.01667 | 0.70517 | 0.00098 | 17.5 s |
| 1:07:55.5 | Data point 71 | 1.50000 mL | 0.16860 mL | 0.17310 mL | 0.25000 mL | 6.190 | 0.01867 | 0.87098 | 0.00099 | 30.5 s |
| 1:08:51.4 | Data point 72 | 1.50000 mL | 0.16860 mL | 0.17314 mL | 0.25000 mL | 6.303 | -0.00843 | 0.60418 | 0.00054 | 16.0 s |
| 1:09:38.0 | Data point 73 | 1.50000 mL | 0.16860 mL | 0.17319 mL | 0.25000 mL | 6.363 | -0.01074 | 0.33367 | 0.00092 | 17.0 s |
| 1:10:20.4 | Data point 74 | 1.50000 mL | 0.16860 mL | 0.17324 mL | 0.25000 mL | 6.444 | -0.01718 | 0.73214 | 0.00099 | 15.0 s |
| 1:11:00.8 | Data point 75 | 1.50000 mL | 0.16860 mL | 0.17328 mL | 0.25000 mL | 6.273 | 0.01956 | 0.96509 | 0.00098 | 57.5 s |
| 1:12:28.9 | Data point 76 | 1.50000 mL | 0.16860 mL | 0.17345 mL | 0.25000 mL | 6.967 | -0.09221 | 0.98961 | 0.00458 | Timed out at 59.5 s |
| 1:13:59.4 | Data point 77 | 1.50000 mL | 0.16860 mL | 0.17359 mL | 0.25000 mL | 7.698 | -0.10796 | 0.99667 | 0.00535 | Timed out at 59.5 s |
| 1:15:35.0 | Data point 78 | 1.50000 mL | 0.16860 mL | 0.17366 mL | 0.25000 mL | 7.761 | -0.05693 | 0.98901 | 0.00283 | Timed out at 59.5 s |
| 1:17:05.5 | Data point 79 | 1.50000 mL | 0.16860 mL | 0.17373 mL | 0.25000 mL | 7.873 | -0.04996 | 0.97617 | 0.00250 | Timed out at 59.5 s |
| 1:18:41.2 | Data point 80 | 1.50000 mL | 0.16860 mL | 0.17380 mL | 0.25000 mL | 8.062 | -0.05264 | 0.96602 | 0.00265 | Timed out at 59.5 s |

| | | | |
|---|---|---|---|
| Sample name: | **M12_octanol** | Experiment start time: | **3/1/2018 3:28:52 PM** |
| Assay name: | **pH-metric high logP** | Analyst: | **Pion** |
| Assay ID: | **18C-01012** | Instrument ID: | **T312060** |
| Filename: | **C:\Sirius_T3\Mehtap\20180228_exp28_logP_T3-2\18C-01012_M12_octanol_pH-metric high logP.t3r** | | |

#### Events (continued)

| Time | Event | Water | Acid | Base | Octanol | pH | dpH/dt | pH R-squared | pH SD | dpH/dt time |
|---|---|---|---|---|---|---|---|---|---|---|
| 1:20:11.7 | Data point 81 | 1.50000 mL | 0.16860 mL | 0.17385 mL | 0.25000 mL | 8.128 | -0.04768 | 0.99488 | 0.00236 | Timed out at 59.5 s |
| 1:21:47.4 | Data point 82 | 1.50000 mL | 0.16860 mL | 0.17392 mL | 0.25000 mL | 8.346 | -0.02426 | 0.96191 | 0.00122 | Timed out at 59.5 s |
| 1:23:28.3 | Data point 83 | 1.50000 mL | 0.16860 mL | 0.17401 mL | 0.25000 mL | 8.532 | -0.01603 | 0.65257 | 0.00098 | 28.0 s |
| 1:24:32.0 | Data point 84 | 1.50000 mL | 0.16860 mL | 0.17411 mL | 0.25000 mL | 8.645 | -0.01972 | 0.96650 | 0.00099 | 46.0 s |
| 1:25:53.8 | Data point 85 | 1.50000 mL | 0.16860 mL | 0.17418 mL | 0.25000 mL | 8.779 | -0.01801 | 0.80706 | 0.00099 | 22.0 s |
| 1:26:56.6 | Data point 86 | 1.50000 mL | 0.16860 mL | 0.17429 mL | 0.25000 mL | 8.994 | -0.01105 | 0.30301 | 0.00099 | 14.5 s |
| 1:27:36.5 | Data point 87 | 1.50000 mL | 0.16860 mL | 0.17432 mL | 0.25000 mL | 8.985 | -0.01268 | 0.48445 | 0.00090 | 22.5 s |
| 1:28:08.1 | Assay volumes | 1.50000 mL | 0.16860 mL | 0.17432 mL | 0.25000 mL | | | | | |

| | | | |
|---|---|---|---|
| Sample name: | **M12_octanol** | Experiment start time: | **3/1/2018 3:28:52 PM** |
| Assay name: | **pH-metric high logP** | Analyst: | **Pion** |
| Assay ID: | **18C-01012** | Instrument ID: | **T312060** |
| Filename: | **C:\Sirius_T3\Mehtap\20180228_exp28_logP_T3-2\18C-01012_M12_octanol_pH-metric high logP.t3r** | | |

| | | | |
|---|---|---|---|
| Sample name: | **M12_octanol** | Experiment start time: | **3/1/2018 3:28:52 PM** |
| Assay name: | **pH-metric high logP** | Analyst: | **Pion** |
| Assay ID: | **18C-01012** | Instrument ID: | **T312060** |
| Filename: | **C:\Sirius_T3\Mehtap\20180228_exp28_logP_T3-2\18C-01012_M12_octanol_pH-metric high logP.t3r** | | |

#### Assay Settings (continued)

| Setting | Value | Original Value | Date/Time changed | Imported from |
|---|---|---|---|---|
| Titrate from | Low to high pH | | | |
| Add additional water | 0.00 mL | | | |
| Additional partition solvent volume | 0.200 mL | | | |
| Additional partition solvent added | Automatic | | | |
| After pH adjust stir for | 30 seconds | | | |
| Stir to allow partitioning for | 15 seconds | | | |
| Stirrer speed for partitioning | 60% | | | |
| ***Data Point Stability*** | | | | |
| Stir during data point collection | No | | | |
| Delay before data point collection | 0 seconds | | | |
| Number of points to average | 20 points | | | |
| Time interval between points | 0.50 seconds | | | |
| Required maximum standard deviation | 0.00100 dpH/dt | | | |
| Stability timeout after | 60 seconds | | | |

#### Calibration Settings

| Setting | Value | Date/Time changed | Imported from |
|---|---|---|---|
| Four-Plus alpha | 0.130 | 3/1/2018 3:28:52 PM | C:\Sirius_T3\HCl18B27.t3r |
| Four-Plus S | 0.9970 | 3/1/2018 3:28:52 PM | C:\Sirius_T3\HCl18B27.t3r |
| Four-Plus jH | 0.8 | 3/1/2018 3:28:52 PM | C:\Sirius_T3\HCl18B27.t3r |
| Four-Plus jOH | -0.4 | 3/1/2018 3:28:52 PM | C:\Sirius_T3\HCl18B27.t3r |
| Base concentration factor | 1.000 | 3/1/2018 3:28:52 PM | C:\Sirius_T3\KOH18B27.t3r |
| Acid concentration factor | 0.994 | 3/1/2018 3:28:52 PM | C:\Sirius_T3\HCl18B27.t3r |

#### Instrument Settings

| Setting | Value | Batch Id | Install date |
|---|---|---|---|
| Instrument owner | Merck | | |
| Instrument ID | T312060 | | |
| Instrument type | T3 Simulator | | |
| Software version | 1.1.3.0 | | |
| Dispenser module | | T3DM1200361 | 3/31/2009 5:24:52 AM |
| Dispenser 0 | Water | | 3/31/2009 5:25:05 AM |
| Syringe volume | 2.5 mL | | |
| Firmware version | 1.2.1(r2) | | |
| Titrant | Water (0.15 M KCl) | 02-06-2018 | 2/27/2018 10:05:59 AM |
| Dispenser 2 | Acid | | 3/31/2009 5:25:11 AM |
| Syringe volume | 0.5 mL | | |
| Firmware version | 1.2.1(r2) | | |
| Titrant | Acid (0.5 M HCl) | 02-27-2018 | 2/27/2018 10:27:22 AM |
| Dispenser 1 | Base | | 3/31/2009 5:25:21 AM |
| Syringe volume | 0.5 mL | | |
| Firmware version | 1.2.1(r2) | | |
| Titrant | Base (0.5 M KOH) | 9/22/2017 | 2/27/2018 10:21:22 AM |
| Dispenser 5 | Cosolvent | | 3/31/2009 5:26:24 AM |
| Syringe volume | 2.5 mL | | |
| Firmware version | 1.2.1(r2) | | |
| Distribution valve 5 | Distribution Valve | | 3/31/2009 5:28:19 AM |
| Firmware version | 1.1.3 | | |
| Port A | Methanol (80%, 0.15 M KCl) | 09-26-17 | 2/7/2018 9:42:01 AM |
| Port B | Cyclohexane | 11-01-17 | 2/27/2018 10:37:57 AM |
| Dispenser 3 | Buffer | | 8/3/2010 5:05:16 AM |
| Syringe volume | 0.5 mL | | |
| Firmware version | 1.2.1(r2) | | |
| Titrant | Dodecane | 2018/01/31 | 2/28/2018 10:18:04 AM |
| Dispenser 6 | Octanol | | 10/22/2010 10:52:43 AM |
| Syringe volume | 0.5 mL | | |

| | | | |
|---|---|---|---|
| Sample name: | **M12_octanol** | Experiment start time: | **3/1/2018 3:28:52 PM** |
| Assay name: | **pH-metric high logP** | Analyst: | **Pion** |
| Assay ID: | **18C-01012** | Instrument ID: | **T312060** |
| Filename: | **C:\Sirius_T3\Mehtap\20180228_exp28_logP_T3-2\18C-01012_M12_octanol_pH-metric high logP.t3r** | | |

| | | | |
|---|---|---|---|
| Sample name: | **M12_octanol** | Experiment start time: | **3/1/2018 3:28:52 PM** |
| Assay name: | **pH-metric high logP** | Analyst: | **Pion** |
| Assay ID: | **18C-01012** | Instrument ID: | **T312060** |
| Filename: | **C:\Sirius_T3\Mehtap\20180228_exp28_logP_T3-2\18C-01012_M12_octanol_pH-metric high logP.t3r** | | |

#### Instrument Settings (continued)

| Setting | Value | Batch Id | Install date |
|---|---|---|---|
| Spectrometer calibration stir speed | 30% | | |
| Spectrometer calibration wash pump volume | 20.0 mL | | |
| Spectrometer calibration wash stir duration | 5 s | | |
| Spectrometer calibration wash stir speed | 30% | | |
| Overhead dispense height | 10000 | | |

#### Refinement Settings

| Setting | Value | Default value |
|---|---|---|
| Turbidity detection method | None | None |
| Turbidity wavelength to assess | 500.0 nm | 500.0 nm |
| Turbidity maximum absorbance | 0.100 | 0.100 |
| Turbidity probe threshold | 50.00 | 50.00 |

#### Experiment Log

[2:38] Air gap created for Water (0.15 M KCl)
[2:38] Air gap created for Acid (0.5 M HCl)
[2:38] Air gap created for Base (0.5 M KOH)
[2:39] Air gap released for Water (0.15 M KCl)
[2:44] Titrator arm moved to Titration position
[2:44] Argon flow rate set to 100
[2:44] Titration 1 of 3
[2:44] Adding initial titrants
[2:44] Automatically add 1.50000 mL of water
[3:09] Dispensed 1.500000 mL of Water (0.15 M KCl)
[3:09] Stirrer speed set to 10
[3:14] Automatically add 0.02000 mL of Octanol
[3:15] Dispensed 0.019991 mL of Octanol
[3:16] Initial pH = 4.96
[3:16] Iterative adjust 4.96 -> 2.00
[3:16] pH 4.96 -> 2.00
[3:18] Air gap released for Acid (0.5 M HCl)
[3:18] Dispensed 0.052211 mL of Acid (0.5 M HCl)
[3:23] Holding pH 2.00
[5:23] Stirrer speed set to 0
[5:23] Stirrer speed set to 50
[5:23] Iterative adjust 1.89 -> 2.00
[5:23] pH 1.89 -> 2.00
[5:24] Air gap released for Base (0.5 M KOH)
[5:25] Dispensed 0.013288 mL of Base (0.5 M KOH)
[6:15] Stirrer speed set to 0
[6:26] Datapoint id 1 collected
[6:26] Stirrer speed set to 50
[6:31] pH 2.04 -> 2.24
[6:31] Using cautious pH adjust
[6:31] Dispensed 0.007314 mL of Base (0.5 M KOH)
[6:36] Stepping pH = 2.12
[6:36] Dispensed 0.005974 mL of Base (0.5 M KOH)
[6:41] Stepping pH = 2.22
[6:42] Dispensed 0.001105 mL of Base (0.5 M KOH)
[6:47] Stepping pH = 2.24
[7:02] Stirrer speed set to 0
[7:12] Datapoint id 2 collected
[7:12] Charge balance equation is out by 1.6%
[7:12] Stirrer speed set to 50
[7:17] pH 2.26 -> 2.46
[7:17] Using charge balance adjust
[7:17] Dispensed 0.008843 mL of Base (0.5 M KOH)

| | | | |
|---|---|---|---|
| Sample name: | **M12_octanol** | Experiment start time: | **3/1/2018 3:28:52 PM** |
| Assay name: | **pH-metric high logP** | Analyst: | **Pion** |
| Assay ID: | **18C-01012** | Instrument ID: | **T312060** |
| Filename: | **C:\Sirius_T3\Mehtap\20180228_exp28_logP_T3-2\18C-01012_M12_octanol_pH-metric high logP.t3r** | | |

#### Experiment Log (continued)

[7:37] Stirrer speed set to 0
[7:47] Datapoint id 3 collected
[7:47] Charge balance equation is out by 4.9%
[7:47] Stirrer speed set to 50
[7:53] pH 2.48 -> 2.68
[7:53] Using charge balance adjust
[7:53] Dispensed 0.005456 mL of Base (0.5 M KOH)
[8:13] Stirrer speed set to 0
[8:41] Datapoint id 4 collected
[8:41] Charge balance equation is out by -2.2%
[8:41] Stirrer speed set to 50
[8:46] pH 2.68 -> 2.88
[8:46] Using charge balance adjust
[8:46] Dispensed 0.003622 mL of Base (0.5 M KOH)
[9:07] Stirrer speed set to 0
[9:17] Datapoint id 5 collected
[9:17] Charge balance equation is out by 9.1%
[9:17] Stirrer speed set to 50
[9:22] pH 2.90 -> 3.10
[9:22] Using charge balance adjust
[9:22] Dispensed 0.002493 mL of Base (0.5 M KOH)
[9:42] Stirrer speed set to 0
[9:52] Datapoint id 6 collected
[9:52] Charge balance equation is out by -16.5%
[9:52] Stirrer speed set to 50
[9:57] pH 3.07 -> 3.27
[9:57] Using cautious pH adjust
[9:57] Dispensed 0.001011 mL of Base (0.5 M KOH)
[10:02] Stepping pH = 3.12
[10:02] Dispensed 0.001317 mL of Base (0.5 M KOH)
[10:08] Stepping pH = 3.24
[10:08] Dispensed 0.000212 mL of Base (0.5 M KOH)
[10:13] Stepping pH = 3.25
[10:13] Dispensed 0.000212 mL of Base (0.5 M KOH)
[10:18] Stepping pH = 3.27
[10:33] Stirrer speed set to 0
[10:44] Datapoint id 7 collected
[10:44] Charge balance equation is out by -36.5%
[10:44] Stirrer speed set to 50
[10:49] pH 3.27 -> 3.47
[10:49] Using cautious pH adjust
[10:49] Dispensed 0.000894 mL of Base (0.5 M KOH)
[10:54] Stepping pH = 3.35
[10:54] Dispensed 0.000847 mL of Base (0.5 M KOH)
[10:59] Stepping pH = 3.46
[11:14] Stirrer speed set to 0
[11:25] Datapoint id 8 collected
[11:25] Charge balance equation is out by 2.1%
[11:25] Stirrer speed set to 50
[11:30] pH 3.48 -> 3.68
[11:30] Using charge balance adjust
[11:30] Dispensed 0.001693 mL of Base (0.5 M KOH)
[11:50] Stirrer speed set to 0
[12:01] Datapoint id 9 collected
[12:01] Charge balance equation is out by 39.1%
[12:01] Stirrer speed set to 50
[12:06] pH 3.73 -> 3.93
[12:06] Using cautious pH adjust
[12:06] Dispensed 0.000823 mL of Base (0.5 M KOH)

Sample name: **M12_octanol**
Assay name: **pH-metric high logP**
Assay ID: **18C-01012**
Filename: **C:\Sirius_T3\Mehtap\20180228_exp28_logP_T3-2\18C-01012_M12_octanol_pH-metric high logP.t3r**
Experiment start time: **3/1/2018 3:28:52 PM**
Analyst: **Pion**
Instrument ID: **T312060**

#### Experiment Log (continued)

[12:11] Stepping pH = 3.85
[12:11] Dispensed 0.000447 mL of Base (0.5 M KOH)
[12:16] Stepping pH = 3.94
[12:31] Stirrer speed set to 0
[12:45] Datapoint id 10 collected
[12:45] Charge balance equation is out by 22.7%
[12:45] Stirrer speed set to 50
[12:50] pH 3.90 -> 4.10
[12:50] Using cautious pH adjust
[12:51] Dispensed 0.000800 mL of Base (0.5 M KOH)
[12:56] Stepping pH = 4.04
[12:56] Dispensed 0.000306 mL of Base (0.5 M KOH)
[13:01] Stepping pH = 4.12
[13:16] Stirrer speed set to 0
[13:29] Datapoint id 11 collected
[13:29] Charge balance equation is out by 30.8%
[13:29] Stirrer speed set to 50
[13:34] pH 4.13 -> 4.33
[13:34] Using cautious pH adjust
[13:34] Dispensed 0.000706 mL of Base (0.5 M KOH)
[13:39] Stepping pH = 4.30
[13:39] Dispensed 0.000094 mL of Base (0.5 M KOH)
[13:44] Stepping pH = 4.35
[14:00] Stirrer speed set to 0
[14:14] Datapoint id 12 collected
[14:14] Charge balance equation is out by 42.4%
[14:14] Stirrer speed set to 50
[14:19] pH 4.31 -> 4.51
[14:19] Using cautious pH adjust
[14:19] Dispensed 0.000588 mL of Base (0.5 M KOH)
[14:24] Stepping pH = 4.55
[14:39] Stirrer speed set to 0
[14:56] Datapoint id 13 collected
[14:56] Charge balance equation is out by 50.0%
[14:56] Stirrer speed set to 50
[15:01] pH 4.55 -> 4.75
[15:01] Using cautious pH adjust
[15:01] Dispensed 0.000423 mL of Base (0.5 M KOH)
[15:06] Stepping pH = 4.83
[15:21] Stirrer speed set to 0
[15:44] Datapoint id 14 collected
[15:44] Charge balance equation is out by 50.0%
[15:44] Stirrer speed set to 50
[15:50] pH 4.82 -> 5.02
[15:50] Using cautious pH adjust
[15:50] Dispensed 0.000259 mL of Base (0.5 M KOH)
[15:55] Stepping pH = 4.97
[15:55] Dispensed 0.000071 mL of Base (0.5 M KOH)
[16:00] Stepping pH = 5.06
[16:15] Stirrer speed set to 0
[16:43] Datapoint id 15 collected
[16:43] Charge balance equation is out by 38.3%
[16:43] Stirrer speed set to 50
[16:48] pH 5.02 -> 5.22
[16:48] Using cautious pH adjust
[16:48] Dispensed 0.000188 mL of Base (0.5 M KOH)
[16:53] Stepping pH = 5.44
[17:08] Stirrer speed set to 0
[17:38] Datapoint id 16 collected

Sample name: **M12_octanol**
Assay name: **pH-metric high logP**
Assay ID: **18C-01012**
Filename: **C:\Sirius_T3\Mehtap\20180228_exp28_logP_T3-2\18C-01012_M12_octanol_pH-metric high logP.t3r**

Experiment start time: **3/1/2018 3:28:52 PM**
Analyst: **Pion**
Instrument ID: **T312060**

#### Experiment Log (continued)

[17:38] Charge balance equation is out by 50.0%
[17:38] Stirrer speed set to 50
[17:43] pH 5.43 -> 5.63
[17:43] Using cautious pH adjust
[17:43] Dispensed 0.000094 mL of Base (0.5 M KOH)
[17:48] Stepping pH = 5.54
[17:48] Dispensed 0.000047 mL of Base (0.5 M KOH)
[17:53] Stepping pH = 5.77
[18:08] Stirrer speed set to 0
[18:40] Datapoint id 17 collected
[18:40] Charge balance equation is out by 18.4%
[18:40] Stirrer speed set to 50
[18:45] pH 5.84 -> 6.04
[18:45] Using cautious pH adjust
[18:45] Dispensed 0.000047 mL of Base (0.5 M KOH)
[18:50] Stepping pH = 5.88
[18:50] Dispensed 0.000094 mL of Base (0.5 M KOH)
[18:55] Stepping pH = 6.68
[19:11] Stirrer speed set to 0
[20:11] Datapoint id 18 collected
[20:11] Charge balance equation is out by -44.4%
[20:11] Stirrer speed set to 50
[20:16] pH 6.88 -> 7.08
[20:16] Using cautious pH adjust
[20:16] Dispensed 0.000024 mL of Base (0.5 M KOH)
[20:21] Stepping pH = 6.87
[20:21] Dispensed 0.000071 mL of Base (0.5 M KOH)
[20:26] Stepping pH = 7.48
[20:41] Stirrer speed set to 0
[21:41] Datapoint id 19 collected
[21:41] Charge balance equation is out by -232.0%
[21:41] Stirrer speed set to 50
[21:46] pH 7.52 -> 7.72
[21:46] Using cautious pH adjust
[21:46] Dispensed 0.000024 mL of Base (0.5 M KOH)
[21:51] Stepping pH = 7.42
[21:51] Dispensed 0.000024 mL of Base (0.5 M KOH)
[21:56] Stepping pH = 7.45
[21:57] Dispensed 0.000094 mL of Base (0.5 M KOH)
[22:02] Stepping pH = 8.76
[22:17] Stirrer speed set to 0
[23:10] Datapoint id 20 collected
[23:10] Charge balance equation is out by -1,588.8%
[23:10] Stirrer speed set to 50
[23:15] pH 8.90 -> 9.05
[23:15] Using cautious pH adjust
[23:15] Dispensed 0.000024 mL of Base (0.5 M KOH)
[23:21] Stepping pH = 8.90
[23:21] Dispensed 0.000071 mL of Base (0.5 M KOH)
[23:26] Stepping pH = 9.05
[23:41] Stirrer speed set to 0
[24:07] Datapoint id 21 collected
[24:07] Charge balance equation is out by -237.3%
[24:07] Titration 2 of 3
[24:07] Adding initial titrants
[24:07] Automatically add 0.03000 mL of Octanol
[24:08] Dispensed 0.030009 mL of Octanol
[24:08] Stirrer speed set to 10
[24:09] Stirrer speed set to 55

| | | | |
|---|---|---|---|
| Sample name: | **M12_octanol** | Experiment start time: | **3/1/2018 3:28:52 PM** |
| Assay name: | **pH-metric high logP** | Analyst: | **Pion** |
| Assay ID: | **18C-01012** | Instrument ID: | **T312060** |
| Filename: | **C:\Sirius_T3\Mehtap\20180228_exp28_logP_T3-2\18C-01012_M12_octanol_pH-metric high logP.t3r** | | |

#### Experiment Log (continued)

[24:09] Iterative adjust 9.11 -> 2.00
[24:09] pH 9.11 -> 2.00
[24:10] Dispensed 0.056703 mL of Acid (0.5 M HCl)
[25:01] Stirrer speed set to 0
[25:11] Datapoint id 22 collected
[25:11] Stirrer speed set to 55
[25:16] pH 1.96 -> 2.16
[25:16] Using cautious pH adjust
[25:16] Dispensed 0.009572 mL of Base (0.5 M KOH)
[25:22] Stepping pH = 2.04
[25:22] Dispensed 0.007832 mL of Base (0.5 M KOH)
[25:27] Stepping pH = 2.13
[25:27] Dispensed 0.001458 mL of Base (0.5 M KOH)
[25:32] Stepping pH = 2.16
[25:47] Stirrer speed set to 0
[25:57] Datapoint id 23 collected
[25:57] Charge balance equation is out by 1.5%
[25:57] Stirrer speed set to 55
[26:02] pH 2.17 -> 2.37
[26:02] Using charge balance adjust
[26:03] Dispensed 0.011618 mL of Base (0.5 M KOH)
[26:23] Stirrer speed set to 0
[26:33] Datapoint id 24 collected
[26:33] Charge balance equation is out by 8.1%
[26:33] Stirrer speed set to 55
[26:38] pH 2.40 -> 2.60
[26:38] Using charge balance adjust
[26:38] Dispensed 0.007079 mL of Base (0.5 M KOH)
[26:58] Stirrer speed set to 0
[27:09] Datapoint id 25 collected
[27:09] Charge balance equation is out by -3.0%
[27:09] Stirrer speed set to 55
[27:14] pH 2.60 -> 2.80
[27:14] Using charge balance adjust
[27:14] Dispensed 0.004704 mL of Base (0.5 M KOH)
[27:35] Stirrer speed set to 0
[27:45] Datapoint id 26 collected
[27:45] Charge balance equation is out by 9.3%
[27:45] Stirrer speed set to 55
[27:50] pH 2.82 -> 3.02
[27:50] Using charge balance adjust
[27:50] Dispensed 0.003175 mL of Base (0.5 M KOH)
[28:10] Stirrer speed set to 0
[28:20] Datapoint id 27 collected
[28:20] Charge balance equation is out by -11.4%
[28:20] Stirrer speed set to 55
[28:25] pH 3.00 -> 3.20
[28:25] Using charge balance adjust
[28:25] Dispensed 0.002540 mL of Base (0.5 M KOH)
[28:46] Stirrer speed set to 0
[28:56] Datapoint id 28 collected
[28:56] Charge balance equation is out by -6.4%
[28:56] Stirrer speed set to 55
[29:01] pH 3.19 -> 3.39
[29:01] Using charge balance adjust
[29:01] Dispensed 0.002187 mL of Base (0.5 M KOH)
[29:22] Stirrer speed set to 0
[29:32] Datapoint id 29 collected
[29:32] Charge balance equation is out by 13.7%

Sample name: **M12_octanol**
Assay name: **pH-metric high logP**
Assay ID: **18C-01012**
Filename: **C:\Sirius_T3\Mehtap\20180228_exp28_logP_T3-2\18C-01012_M12_octanol_pH-metric high logP.t3r**
Experiment start time: **3/1/2018 3:28:52 PM**
Analyst: **Pion**
Instrument ID: **T312060**

#### Experiment Log (continued)

[29:32] Stirrer speed set to 55
[29:37] pH 3.42 -> 3.62
[29:37] Using charge balance adjust
[29:37] Dispensed 0.001952 mL of Base (0.5 M KOH)
[29:57] Stirrer speed set to 0
[30:07] Datapoint id 30 collected
[30:07] Charge balance equation is out by 46.6%
[30:07] Stirrer speed set to 55
[30:12] pH 3.71 -> 3.91
[30:12] Using cautious pH adjust
[30:12] Dispensed 0.000870 mL of Base (0.5 M KOH)
[30:17] Stepping pH = 3.82
[30:18] Dispensed 0.000517 mL of Base (0.5 M KOH)
[30:23] Stepping pH = 3.94
[30:38] Stirrer speed set to 0
[30:49] Datapoint id 31 collected
[30:49] Charge balance equation is out by 20.1%
[30:49] Stirrer speed set to 55
[30:54] pH 4.01 -> 4.21
[30:54] Using cautious pH adjust
[30:55] Dispensed 0.000659 mL of Base (0.5 M KOH)
[31:00] Stepping pH = 4.14
[31:00] Dispensed 0.000259 mL of Base (0.5 M KOH)
[31:05] Stepping pH = 4.27
[31:20] Stirrer speed set to 0
[31:47] Datapoint id 32 collected
[31:47] Charge balance equation is out by 30.0%
[31:47] Stirrer speed set to 55
[31:52] pH 4.31 -> 4.51
[31:52] Using cautious pH adjust
[31:52] Dispensed 0.000447 mL of Base (0.5 M KOH)
[31:57] Stepping pH = 4.55
[32:12] Stirrer speed set to 0
[32:37] Datapoint id 33 collected
[32:37] Charge balance equation is out by 50.0%
[32:37] Stirrer speed set to 55
[32:42] pH 4.60 -> 4.80
[32:42] Using cautious pH adjust
[32:43] Dispensed 0.000282 mL of Base (0.5 M KOH)
[32:48] Stepping pH = 4.90
[33:03] Stirrer speed set to 0
[33:21] Datapoint id 34 collected
[33:21] Charge balance equation is out by 50.0%
[33:21] Stirrer speed set to 55
[33:26] pH 4.92 -> 5.12
[33:26] Using cautious pH adjust
[33:26] Dispensed 0.000141 mL of Base (0.5 M KOH)
[33:32] Stepping pH = 5.08
[33:32] Dispensed 0.000024 mL of Base (0.5 M KOH)
[33:37] Stepping pH = 5.15
[33:52] Stirrer speed set to 0
[34:18] Datapoint id 35 collected
[34:18] Charge balance equation is out by 40.7%
[34:18] Stirrer speed set to 55
[34:23] pH 5.16 -> 5.36
[34:23] Using cautious pH adjust
[34:23] Dispensed 0.000094 mL of Base (0.5 M KOH)
[34:29] Stepping pH = 5.26
[34:29] Dispensed 0.000071 mL of Base (0.5 M KOH)

| | | | |
|---|---|---|---|
| Sample name: | **M12_octanol** | Experiment start time: | **3/1/2018 3:28:52 PM** |
| Assay name: | **pH-metric high logP** | Analyst: | **Pion** |
| Assay ID: | **18C-01012** | Instrument ID: | **T312060** |
| Filename: | **C:\Sirius_T3\Mehtap\20180228_exp28_logP_T3-2\18C-01012_M12_octanol_pH-metric high logP.t3r** | | |

#### Experiment Log (continued)

[34:34] Stepping pH = 5.69
[34:49] Stirrer speed set to 0
[35:19] Datapoint id 36 collected
[35:19] Charge balance equation is out by 17.2%
[35:19] Stirrer speed set to 55
[35:24] pH 5.79 -> 5.99
[35:24] Using cautious pH adjust
[35:24] Dispensed 0.000047 mL of Base (0.5 M KOH)
[35:29] Stepping pH = 5.93
[35:29] Dispensed 0.000024 mL of Base (0.5 M KOH)
[35:34] Stepping pH = 6.19
[35:49] Stirrer speed set to 0
[36:36] Datapoint id 37 collected
[36:36] Charge balance equation is out by 23.2%
[36:36] Stirrer speed set to 55
[36:41] pH 6.24 -> 6.44
[36:41] Using cautious pH adjust
[36:41] Dispensed 0.000024 mL of Base (0.5 M KOH)
[36:46] Stepping pH = 6.26
[36:46] Dispensed 0.000071 mL of Base (0.5 M KOH)
[36:51] Stepping pH = 6.71
[37:06] Stirrer speed set to 0
[38:06] Datapoint id 38 collected
[38:06] Charge balance equation is out by -67.6%
[38:06] Stirrer speed set to 55
[38:11] pH 6.80 -> 7.00
[38:11] Using cautious pH adjust
[38:11] Dispensed 0.000024 mL of Base (0.5 M KOH)
[38:16] Stepping pH = 6.78
[38:16] Dispensed 0.000071 mL of Base (0.5 M KOH)
[38:21] Stepping pH = 7.04
[38:37] Stirrer speed set to 0
[39:37] Datapoint id 39 collected
[39:37] Charge balance equation is out by -246.2%
[39:37] Stirrer speed set to 55
[39:42] pH 7.29 -> 7.49
[39:42] Using cautious pH adjust
[39:42] Dispensed 0.000024 mL of Base (0.5 M KOH)
[39:47] Stepping pH = 7.29
[39:47] Dispensed 0.000024 mL of Base (0.5 M KOH)
[39:52] Stepping pH = 7.41
[39:52] Dispensed 0.000024 mL of Base (0.5 M KOH)
[39:57] Stepping pH = 7.70
[40:12] Stirrer speed set to 0
[41:12] Datapoint id 40 collected
[41:12] Charge balance equation is out by -384.4%
[41:12] Stirrer speed set to 55
[41:17] pH 7.79 -> 7.99
[41:17] Using cautious pH adjust
[41:17] Dispensed 0.000024 mL of Base (0.5 M KOH)
[41:22] Stepping pH = 7.81
[41:22] Dispensed 0.000024 mL of Base (0.5 M KOH)
[41:28] Stepping pH = 7.95
[41:28] Dispensed 0.000024 mL of Base (0.5 M KOH)
[41:33] Stepping pH = 8.17
[41:48] Stirrer speed set to 0
[42:48] Datapoint id 41 collected
[42:48] Charge balance equation is out by -765.5%
[42:48] Stirrer speed set to 55

| | | | |
|---|---|---|---|
| Sample name: | **M12_octanol** | Experiment start time: | **3/1/2018 3:28:52 PM** |
| Assay name: | **pH-metric high logP** | Analyst: | **Pion** |
| Assay ID: | **18C-01012** | Instrument ID: | **T312060** |
| Filename: | **C:\Sirius_T3\Mehtap\20180228_exp28_logP_T3-2\18C-01012_M12_octanol_pH-metric high logP.t3r** | | |

#### Experiment Log (continued)

[42:53] pH 8.18 -> 8.38
[42:53] Using cautious pH adjust
[42:53] Dispensed 0.000024 mL of Base (0.5 M KOH)
[42:58] Stepping pH = 8.17
[42:58] Dispensed 0.000024 mL of Base (0.5 M KOH)
[43:03] Stepping pH = 8.32
[43:03] Dispensed 0.000024 mL of Base (0.5 M KOH)
[43:08] Stepping pH = 8.51
[43:23] Stirrer speed set to 0
[44:04] Datapoint id 42 collected
[44:04] Charge balance equation is out by -611.1%
[44:04] Stirrer speed set to 55
[44:09] pH 8.54 -> 8.74
[44:09] Using cautious pH adjust
[44:10] Dispensed 0.000024 mL of Base (0.5 M KOH)
[44:15] Stepping pH = 8.56
[44:15] Dispensed 0.000024 mL of Base (0.5 M KOH)
[44:20] Stepping pH = 8.64
[44:20] Dispensed 0.000024 mL of Base (0.5 M KOH)
[44:25] Stepping pH = 8.75
[44:40] Stirrer speed set to 0
[45:14] Datapoint id 43 collected
[45:14] Charge balance equation is out by -275.0%
[45:14] Stirrer speed set to 55
[45:19] pH 8.74 -> 8.94
[45:19] Using cautious pH adjust
[45:19] Dispensed 0.000024 mL of Base (0.5 M KOH)
[45:24] Stepping pH = 8.74
[45:24] Dispensed 0.000047 mL of Base (0.5 M KOH)
[45:29] Stepping pH = 8.86
[45:29] Dispensed 0.000024 mL of Base (0.5 M KOH)
[45:34] Stepping pH = 8.97
[45:49] Stirrer speed set to 0
[46:09] Datapoint id 44 collected
[46:09] Charge balance equation is out by -212.1%
[46:09] Stirrer speed set to 55
[46:14] pH 8.94 -> 9.05
[46:14] Using cautious pH adjust
[46:14] Dispensed 0.000024 mL of Base (0.5 M KOH)
[46:19] Stepping pH = 8.94
[46:19] Dispensed 0.000047 mL of Base (0.5 M KOH)
[46:24] Stepping pH = 9.01
[46:39] Stirrer speed set to 0
[46:56] Datapoint id 45 collected
[46:56] Charge balance equation is out by -261.0%
[46:56] Titration 3 of 3
[46:56] Adding initial titrants
[46:56] Automatically add 0.20000 mL of Octanol
[47:01] Dispensed 0.200000 mL of Octanol
[47:01] Stirrer speed set to 10
[47:02] Stirrer speed set to 60
[47:02] Iterative adjust 9.05 -> 2.00
[47:02] pH 9.05 -> 2.00
[47:04] Dispensed 0.059690 mL of Acid (0.5 M HCl)
[47:54] Stirrer speed set to 0
[48:04] Datapoint id 46 collected
[48:04] Stirrer speed set to 60
[48:10] pH 1.96 -> 2.16
[48:10] Using cautious pH adjust

Sample name: **M12_octanol**
Assay name: **pH-metric high logP**
Assay ID: **18C-01012**
Filename: **C:\Sirius_T3\Mehtap\20180228_exp28_logP_T3-2\18C-01012_M12_octanol_pH-metric high logP.t3r**
Experiment start time: **3/1/2018 3:28:52 PM**
Analyst: **Pion**
Instrument ID: **T312060**

#### Experiment Log (continued)

[48:10] Dispensed 0.010113 mL of Base (0.5 M KOH)
[48:15] Stepping pH = 2.05
[48:15] Dispensed 0.007361 mL of Base (0.5 M KOH)
[48:20] Stepping pH = 2.13
[48:21] Dispensed 0.001999 mL of Base (0.5 M KOH)
[48:26] Stepping pH = 2.16
[48:41] Stirrer speed set to 0
[48:51] Datapoint id 47 collected
[48:51] Charge balance equation is out by 3.8%
[48:51] Stirrer speed set to 60
[48:56] pH 2.18 -> 2.38
[48:56] Using charge balance adjust
[48:56] Dispensed 0.012347 mL of Base (0.5 M KOH)
[49:16] Stirrer speed set to 0
[49:27] Datapoint id 48 collected
[49:27] Charge balance equation is out by 4.8%
[49:27] Stirrer speed set to 60
[49:32] pH 2.39 -> 2.59
[49:32] Using charge balance adjust
[49:32] Dispensed 0.007738 mL of Base (0.5 M KOH)
[49:52] Stirrer speed set to 0
[50:02] Datapoint id 49 collected
[50:02] Charge balance equation is out by -10.9%
[50:02] Stirrer speed set to 60
[50:07] pH 2.58 -> 2.78
[50:07] Using charge balance adjust
[50:07] Dispensed 0.005409 mL of Base (0.5 M KOH)
[50:28] Stirrer speed set to 0
[50:38] Datapoint id 50 collected
[50:38] Charge balance equation is out by -1.8%
[50:38] Stirrer speed set to 60
[50:43] pH 2.78 -> 2.98
[50:43] Using charge balance adjust
[50:43] Dispensed 0.003833 mL of Base (0.5 M KOH)
[51:03] Stirrer speed set to 0
[51:13] Datapoint id 51 collected
[51:13] Charge balance equation is out by -0.7%
[51:13] Stirrer speed set to 60
[51:18] pH 2.99 -> 3.19
[51:18] Using charge balance adjust
[51:18] Dispensed 0.002963 mL of Base (0.5 M KOH)
[51:39] Stirrer speed set to 0
[51:49] Datapoint id 52 collected
[51:49] Charge balance equation is out by 41.2%
[51:49] Stirrer speed set to 60
[51:54] pH 3.28 -> 3.48
[51:54] Using cautious pH adjust
[51:54] Dispensed 0.001152 mL of Base (0.5 M KOH)
[51:59] Stepping pH = 3.38
[51:59] Dispensed 0.000729 mL of Base (0.5 M KOH)
[52:04] Stepping pH = 3.51
[52:19] Stirrer speed set to 0
[52:35] Datapoint id 53 collected
[52:35] Charge balance equation is out by 18.3%
[52:35] Stirrer speed set to 60
[52:40] pH 3.56 -> 3.76
[52:40] Using cautious pH adjust
[52:40] Dispensed 0.000917 mL of Base (0.5 M KOH)
[52:45] Stepping pH = 3.68

| | | | |
|---|---|---|---|
| Sample name: | **M12_octanol** | Experiment start time: | **3/1/2018 3:28:52 PM** |
| Assay name: | **pH-metric high logP** | Analyst: | **Pion** |
| Assay ID: | **18C-01012** | Instrument ID: | **T312060** |
| Filename: | **C:\Sirius_T3\Mehtap\20180228_exp28_logP_T3-2\18C-01012_M12_octanol_pH-metric high logP.t3r** | | |

#### Experiment Log (continued)

[52:45] Dispensed 0.000447 mL of Base (0.5 M KOH)
[52:50] Stepping pH = 3.81
[53:05] Stirrer speed set to 0
[53:16] Datapoint id 54 collected
[53:16] Charge balance equation is out by 25.4%
[53:16] Stirrer speed set to 60
[53:21] pH 3.89 -> 4.09
[53:21] Using cautious pH adjust
[53:21] Dispensed 0.000659 mL of Base (0.5 M KOH)
[53:26] Stepping pH = 4.07
[53:26] Dispensed 0.000047 mL of Base (0.5 M KOH)
[53:31] Stepping pH = 4.21
[53:47] Stirrer speed set to 0
[53:57] Datapoint id 55 collected
[53:57] Charge balance equation is out by 45.6%
[53:57] Stirrer speed set to 60
[54:02] pH 4.23 -> 4.43
[54:02] Using cautious pH adjust
[54:02] Dispensed 0.000376 mL of Base (0.5 M KOH)
[54:07] Stepping pH = 4.66
[54:22] Stirrer speed set to 0
[54:35] Datapoint id 56 collected
[54:35] Charge balance equation is out by 50.0%
[54:35] Stirrer speed set to 60
[54:41] pH 4.69 -> 4.89
[54:41] Using cautious pH adjust
[54:41] Dispensed 0.000165 mL of Base (0.5 M KOH)
[54:46] Stepping pH = 4.90
[55:01] Stirrer speed set to 0
[55:14] Datapoint id 57 collected
[55:14] Charge balance equation is out by 50.0%
[55:14] Stirrer speed set to 60
[55:19] pH 4.82 -> 5.02
[55:19] Using cautious pH adjust
[55:19] Dispensed 0.000118 mL of Base (0.5 M KOH)
[55:24] Stepping pH = 4.96
[55:24] Dispensed 0.000047 mL of Base (0.5 M KOH)
[55:29] Stepping pH = 5.07
[55:44] Stirrer speed set to 0
[55:59] Datapoint id 58 collected
[55:59] Charge balance equation is out by 35.4%
[55:59] Stirrer speed set to 60
[56:04] pH 5.04 -> 5.24
[56:04] Using cautious pH adjust
[56:04] Dispensed 0.000071 mL of Base (0.5 M KOH)
[56:09] Stepping pH = 5.24
[56:24] Stirrer speed set to 0
[56:42] Datapoint id 59 collected
[56:42] Charge balance equation is out by 50.0%
[56:42] Stirrer speed set to 60
[56:47] pH 5.18 -> 5.38
[56:47] Using cautious pH adjust
[56:47] Dispensed 0.000071 mL of Base (0.5 M KOH)
[56:52] Stepping pH = 5.28
[56:52] Dispensed 0.000047 mL of Base (0.5 M KOH)
[56:57] Stepping pH = 5.44
[57:12] Stirrer speed set to 0
[57:26] Datapoint id 60 collected
[57:26] Charge balance equation is out by 17.2%

| | | | |
|---|---|---|---|
| Sample name: | **M12_octanol** | Experiment start time: | **3/1/2018 3:28:52 PM** |
| Assay name: | **pH-metric high logP** | Analyst: | **Pion** |
| Assay ID: | **18C-01012** | Instrument ID: | **T312060** |
| Filename: | **C:\Sirius_T3\Mehtap\20180228_exp28_logP_T3-2\18C-01012_M12_octanol_pH-metric high logP.t3r** | | |

#### Experiment Log (continued)

[57:26] Stirrer speed set to 60
[57:31] pH 5.44 -> 5.64
[57:31] Using cautious pH adjust
[57:31] Dispensed 0.000047 mL of Base (0.5 M KOH)
[57:36] Stepping pH = 5.52
[57:36] Dispensed 0.000024 mL of Base (0.5 M KOH)
[57:41] Stepping pH = 5.57
[57:41] Dispensed 0.000024 mL of Base (0.5 M KOH)
[57:46] Stepping pH = 5.67
[58:01] Stirrer speed set to 0
[58:26] Datapoint id 61 collected
[58:26] Charge balance equation is out by -31.4%
[58:26] Stirrer speed set to 60
[58:31] pH 5.57 -> 5.77
[58:31] Using cautious pH adjust
[58:31] Dispensed 0.000024 mL of Base (0.5 M KOH)
[58:36] Stepping pH = 5.60
[58:36] Dispensed 0.000094 mL of Base (0.5 M KOH)
[58:41] Stepping pH = 6.29
[58:56] Stirrer speed set to 0
[59:52] Datapoint id 62 collected
[59:52] Charge balance equation is out by -71.8%
[59:52] Stirrer speed set to 60
[59:57] pH 5.94 -> 6.14
[59:57] Using cautious pH adjust
[59:57] Dispensed 0.000024 mL of Base (0.5 M KOH)
[1:00:02] Stepping pH = 5.95
[1:00:02] Dispensed 0.000071 mL of Base (0.5 M KOH)
[1:00:07] Stepping pH = 6.30
[1:00:22] Stirrer speed set to 0
[1:00:36] Datapoint id 63 collected
[1:00:36] Charge balance equation is out by -89.9%
[1:00:36] Stirrer speed set to 60
[1:00:41] pH 6.12 -> 6.32
[1:00:41] Using cautious pH adjust
[1:00:41] Dispensed 0.000024 mL of Base (0.5 M KOH)
[1:00:46] Stepping pH = 6.16
[1:00:46] Dispensed 0.000047 mL of Base (0.5 M KOH)
[1:00:51] Stepping pH = 6.48
[1:01:06] Stirrer speed set to 0
[1:02:06] Datapoint id 64 collected
[1:02:06] Charge balance equation is out by -38.2%
[1:02:06] Stirrer speed set to 60
[1:02:11] pH 6.11 -> 6.31
[1:02:11] Using cautious pH adjust
[1:02:11] Dispensed 0.000024 mL of Base (0.5 M KOH)
[1:02:16] Stepping pH = 6.18
[1:02:16] Dispensed 0.000024 mL of Base (0.5 M KOH)
[1:02:22] Stepping pH = 6.34
[1:02:37] Stirrer speed set to 0
[1:02:47] Datapoint id 65 collected
[1:02:47] Charge balance equation is out by -6.6%
[1:02:47] Stirrer speed set to 60
[1:02:52] pH 6.05 -> 6.25
[1:02:52] Using charge balance adjust
[1:02:52] Dispensed 0.000047 mL of Base (0.5 M KOH)
[1:03:12] Stirrer speed set to 0
[1:03:47] Datapoint id 66 collected
[1:03:47] Charge balance equation is out by -98.3%

| | | | |
|---|---|---|---|
| Sample name: | **M12_octanol** | Experiment start time: | **3/1/2018 3:28:52 PM** |
| Assay name: | **pH-metric high logP** | Analyst: | **Pion** |
| Assay ID: | **18C-01012** | Instrument ID: | **T312060** |
| Filename: | **C:\Sirius_T3\Mehtap\20180228_exp28_logP_T3-2\18C-01012_M12_octanol_pH-metric high logP.t3r** | | |

#### Experiment Log (continued)

[1:03:47] Stirrer speed set to 60
[1:03:52] pH 5.78 -> 5.98
[1:03:52] Using cautious pH adjust
[1:03:52] Dispensed 0.000024 mL of Base (0.5 M KOH)
[1:03:57] Stepping pH = 5.98
[1:04:12] Stirrer speed set to 0
[1:04:52] Datapoint id 67 collected
[1:04:52] Charge balance equation is out by 50.0%
[1:04:52] Stirrer speed set to 60
[1:04:57] pH 5.76 -> 5.96
[1:04:57] Using cautious pH adjust
[1:04:57] Dispensed 0.000024 mL of Base (0.5 M KOH)
[1:05:02] Stepping pH = 5.85
[1:05:02] Dispensed 0.000024 mL of Base (0.5 M KOH)
[1:05:07] Stepping pH = 6.07
[1:05:22] Stirrer speed set to 0
[1:05:59] Datapoint id 68 collected
[1:05:59] Charge balance equation is out by 7.8%
[1:05:59] Stirrer speed set to 60
[1:06:04] pH 5.71 -> 5.91
[1:06:04] Using charge balance adjust
[1:06:04] Dispensed 0.000071 mL of Base (0.5 M KOH)
[1:06:24] Stirrer speed set to 0
[1:06:37] Datapoint id 69 collected
[1:06:37] Charge balance equation is out by 33.1%
[1:06:37] Stirrer speed set to 60
[1:06:42] pH 5.84 -> 6.04
[1:06:42] Using cautious pH adjust
[1:06:43] Dispensed 0.000024 mL of Base (0.5 M KOH)
[1:06:48] Stepping pH = 5.96
[1:06:48] Dispensed 0.000024 mL of Base (0.5 M KOH)
[1:06:53] Stepping pH = 6.21
[1:07:08] Stirrer speed set to 0
[1:07:25] Datapoint id 70 collected
[1:07:25] Charge balance equation is out by 15.2%
[1:07:25] Stirrer speed set to 60
[1:07:30] pH 5.94 -> 6.14
[1:07:30] Using cautious pH adjust
[1:07:31] Dispensed 0.000024 mL of Base (0.5 M KOH)
[1:07:36] Stepping pH = 6.12
[1:07:36] Dispensed 0.000024 mL of Base (0.5 M KOH)
[1:07:41] Stepping pH = 6.28
[1:07:56] Stirrer speed set to 0
[1:08:27] Datapoint id 71 collected
[1:08:27] Charge balance equation is out by 14.3%
[1:08:27] Stirrer speed set to 60
[1:08:32] pH 6.04 -> 6.24
[1:08:32] Using charge balance adjust
[1:08:32] Dispensed 0.000047 mL of Base (0.5 M KOH)
[1:08:52] Stirrer speed set to 0
[1:09:08] Datapoint id 72 collected
[1:09:08] Charge balance equation is out by 29.3%
[1:09:08] Stirrer speed set to 60
[1:09:13] pH 6.21 -> 6.41
[1:09:13] Using cautious pH adjust
[1:09:13] Dispensed 0.000024 mL of Base (0.5 M KOH)
[1:09:18] Stepping pH = 6.32
[1:09:18] Dispensed 0.000024 mL of Base (0.5 M KOH)
[1:09:23] Stepping pH = 6.47

Sample name: **M12_octanol**
Assay name: **pH-metric high logP**
Assay ID: **18C-01012**
Filename: **C:\Sirius_T3\Mehtap\20180228_exp28_logP_T3-2\18C-01012_M12_octanol_pH-metric high logP.t3r**
Experiment start time: **3/1/2018 3:28:52 PM**
Analyst: **Pion**
Instrument ID: **T312060**

#### Experiment Log (continued)

[1:09:38] Stirrer speed set to 0
[1:09:55] Datapoint id 73 collected
[1:09:55] Charge balance equation is out by 11.9%
[1:09:55] Stirrer speed set to 60
[1:10:01] pH 6.23 -> 6.43
[1:10:01] Using charge balance adjust
[1:10:01] Dispensed 0.000047 mL of Base (0.5 M KOH)
[1:10:21] Stirrer speed set to 0
[1:10:36] Datapoint id 74 collected
[1:10:36] Charge balance equation is out by 6.5%
[1:10:36] Stirrer speed set to 60
[1:10:41] pH 6.39 -> 6.59
[1:10:41] Using charge balance adjust
[1:10:41] Dispensed 0.000047 mL of Base (0.5 M KOH)
[1:11:01] Stirrer speed set to 0
[1:11:59] Datapoint id 75 collected
[1:11:59] Charge balance equation is out by -41.6%
[1:11:59] Stirrer speed set to 60
[1:12:04] pH 5.94 -> 6.14
[1:12:04] Using cautious pH adjust
[1:12:04] Dispensed 0.000024 mL of Base (0.5 M KOH)
[1:12:09] Stepping pH = 5.92
[1:12:09] Dispensed 0.000141 mL of Base (0.5 M KOH)
[1:12:14] Stepping pH = 7.15
[1:12:29] Stirrer speed set to 0
[1:13:29] Datapoint id 76 collected
[1:13:29] Charge balance equation is out by -219.5%
[1:13:29] Stirrer speed set to 60
[1:13:34] pH 6.70 -> 6.90
[1:13:34] Using cautious pH adjust
[1:13:34] Dispensed 0.000024 mL of Base (0.5 M KOH)
[1:13:40] Stepping pH = 6.66
[1:13:40] Dispensed 0.000118 mL of Base (0.5 M KOH)
[1:13:45] Stepping pH = 7.66
[1:14:00] Stirrer speed set to 0
[1:15:00] Datapoint id 77 collected
[1:15:00] Charge balance equation is out by -255.5%
[1:15:00] Stirrer speed set to 60
[1:15:05] pH 7.53 -> 7.73
[1:15:05] Using cautious pH adjust
[1:15:05] Dispensed 0.000024 mL of Base (0.5 M KOH)
[1:15:10] Stepping pH = 7.55
[1:15:10] Dispensed 0.000024 mL of Base (0.5 M KOH)
[1:15:15] Stepping pH = 7.66
[1:15:15] Dispensed 0.000024 mL of Base (0.5 M KOH)
[1:15:20] Stepping pH = 7.84
[1:15:35] Stirrer speed set to 0
[1:16:35] Datapoint id 78 collected
[1:16:35] Charge balance equation is out by -467.5%
[1:16:35] Stirrer speed set to 60
[1:16:41] pH 7.53 -> 7.73
[1:16:41] Using cautious pH adjust
[1:16:41] Dispensed 0.000024 mL of Base (0.5 M KOH)
[1:16:46] Stepping pH = 7.43
[1:16:46] Dispensed 0.000047 mL of Base (0.5 M KOH)
[1:16:51] Stepping pH = 7.78
[1:17:06] Stirrer speed set to 0
[1:18:06] Datapoint id 79 collected
[1:18:06] Charge balance equation is out by -479.6%

| | | | |
|---|---|---|---|
| Sample name: | **M12_octanol** | Experiment start time: | **3/1/2018 3:28:52 PM** |
| Assay name: | **pH-metric high logP** | Analyst: | **Pion** |
| Assay ID: | **18C-01012** | Instrument ID: | **T312060** |
| Filename: | **C:\Sirius_T3\Mehtap\20180228_exp28_logP_T3-2\18C-01012_M12_octanol_pH-metric high logP.t3r** | | |

#### Experiment Log (continued)

[1:18:06] Stirrer speed set to 60
[1:18:11] pH 7.85 -> 8.05
[1:18:11] Using cautious pH adjust
[1:18:11] Dispensed 0.000024 mL of Base (0.5 M KOH)
[1:18:16] Stepping pH = 7.94
[1:18:16] Dispensed 0.000024 mL of Base (0.5 M KOH)
[1:18:21] Stepping pH = 8.03
[1:18:21] Dispensed 0.000024 mL of Base (0.5 M KOH)
[1:18:27] Stepping pH = 8.07
[1:18:42] Stirrer speed set to 0
[1:19:42] Datapoint id 80 collected
[1:19:42] Charge balance equation is out by -636.5%
[1:19:42] Stirrer speed set to 60
[1:19:47] pH 7.94 -> 8.14
[1:19:47] Using cautious pH adjust
[1:19:47] Dispensed 0.000024 mL of Base (0.5 M KOH)
[1:19:52] Stepping pH = 8.03
[1:19:52] Dispensed 0.000024 mL of Base (0.5 M KOH)
[1:19:57] Stepping pH = 8.17
[1:20:12] Stirrer speed set to 0
[1:21:12] Datapoint id 81 collected
[1:21:12] Charge balance equation is out by -388.4%
[1:21:12] Stirrer speed set to 60
[1:21:17] pH 8.07 -> 8.27
[1:21:17] Using cautious pH adjust
[1:21:17] Dispensed 0.000024 mL of Base (0.5 M KOH)
[1:21:22] Stepping pH = 8.13
[1:21:22] Dispensed 0.000024 mL of Base (0.5 M KOH)
[1:21:28] Stepping pH = 8.23
[1:21:28] Dispensed 0.000024 mL of Base (0.5 M KOH)
[1:21:33] Stepping pH = 8.37
[1:21:48] Stirrer speed set to 0
[1:22:48] Datapoint id 82 collected
[1:22:48] Charge balance equation is out by -583.2%
[1:22:48] Stirrer speed set to 60
[1:22:53] pH 8.38 -> 8.58
[1:22:53] Using cautious pH adjust
[1:22:53] Dispensed 0.000024 mL of Base (0.5 M KOH)
[1:22:58] Stepping pH = 8.43
[1:22:58] Dispensed 0.000024 mL of Base (0.5 M KOH)
[1:23:03] Stepping pH = 8.46
[1:23:03] Dispensed 0.000024 mL of Base (0.5 M KOH)
[1:23:08] Stepping pH = 8.54
[1:23:09] Dispensed 0.000024 mL of Base (0.5 M KOH)
[1:23:14] Stepping pH = 8.63
[1:23:29] Stirrer speed set to 0
[1:23:57] Datapoint id 83 collected
[1:23:57] Charge balance equation is out by -489.1%
[1:23:57] Stirrer speed set to 60
[1:24:02] pH 8.48 -> 8.68
[1:24:02] Using cautious pH adjust
[1:24:02] Dispensed 0.000024 mL of Base (0.5 M KOH)
[1:24:07] Stepping pH = 8.46
[1:24:07] Dispensed 0.000047 mL of Base (0.5 M KOH)
[1:24:12] Stepping pH = 8.59
[1:24:12] Dispensed 0.000024 mL of Base (0.5 M KOH)
[1:24:17] Stepping pH = 8.68
[1:24:32] Stirrer speed set to 0
[1:25:19] Datapoint id 84 collected

| | | | |
|---|---|---|---|
| Sample name: | **M12_octanol** | Experiment start time: | **3/1/2018 3:28:52 PM** |
| Assay name: | **pH-metric high logP** | Analyst: | **Pion** |
| Assay ID: | **18C-01012** | Instrument ID: | **T312060** |
| Filename: | **C:\Sirius_T3\Mehtap\20180228_exp28_logP_T3-2\18C-01012_M12_octanol_pH-metric high logP.t3r** | | |

#### Experiment Log (continued)

[1:25:19] Charge balance equation is out by -472.3%
[1:25:19] Stirrer speed set to 60
[1:25:24] pH 8.61 -> 8.81
[1:25:24] Using cautious pH adjust
[1:25:24] Dispensed 0.000024 mL of Base (0.5 M KOH)
[1:25:29] Stepping pH = 8.65
[1:25:29] Dispensed 0.000024 mL of Base (0.5 M KOH)
[1:25:34] Stepping pH = 8.72
[1:25:34] Dispensed 0.000024 mL of Base (0.5 M KOH)
[1:25:39] Stepping pH = 8.80
[1:25:54] Stirrer speed set to 0
[1:26:16] Datapoint id 85 collected
[1:26:16] Charge balance equation is out by -185.7%
[1:26:16] Stirrer speed set to 60
[1:26:21] pH 8.77 -> 8.97
[1:26:21] Using cautious pH adjust
[1:26:21] Dispensed 0.000024 mL of Base (0.5 M KOH)
[1:26:27] Stepping pH = 8.79
[1:26:27] Dispensed 0.000047 mL of Base (0.5 M KOH)
[1:26:32] Stepping pH = 8.90
[1:26:32] Dispensed 0.000024 mL of Base (0.5 M KOH)
[1:26:37] Stepping pH = 8.95
[1:26:37] Dispensed 0.000024 mL of Base (0.5 M KOH)
[1:26:42] Stepping pH = 9.00
[1:26:57] Stirrer speed set to 0
[1:27:12] Datapoint id 86 collected
[1:27:12] Charge balance equation is out by -250.7%
[1:27:12] Stirrer speed set to 60
[1:27:17] pH 8.99 -> 9.05
[1:27:17] Using cautious pH adjust
[1:27:17] Dispensed 0.000024 mL of Base (0.5 M KOH)
[1:27:22] Stepping pH = 9.00
[1:27:37] Stirrer speed set to 0
[1:27:59] Datapoint id 87 collected
[1:27:59] Charge balance equation is out by -76.9%
[1:27:59] Argon flow rate set to 0
[1:28:03] Titrator arm moved over Titration position
