## Supplementary material for "Octanol-water partition coefficient measurements for the SAMPL6 Blind Prediction Challenge": SM12_18C-03013_M12_octanol_pH-metric high logP_report.pdf

| | | | |
|---|---|---|---|
| Sample name: | **M12_octanol** | Experiment start time: | **3/3/2018 4:33:34 PM** |
| Assay name: | **pH-metric high logP** | Analyst: | **Pion** |
| Assay ID: | **18C-03013** | Instrument ID: | **T312060** |
| Filename: | **C:\Sirius_T3\Mehtap\20180302_exp29_logP_T3-2\18C-03013_M12_octanol_pH-metric high logP.t3r** | | |

### pH-metric Result

| | | |
|---|---|---|
| logP (XH +) | -4.20 | ±1.91 (n=49) |
| logP (neutral X) | 3.81 | ±0.01 (n=49) |

#### 18C-03013 Points 1 to 30

| | |
|---|---|
| M12_octanol concentration factor | 0.900 |
| Carbonate | 0.0742 mM |
| Acidity error | -0.41736 mM |

#### 18C-03013 Points 31 to 56

| | |
|---|---|
| M12_octanol concentration factor | 0.766 |
| Carbonate | 0.1690 mM |
| Acidity error | 0.13861 mM |

#### 18C-03013 Points 57 to 80

| | |
|---|---|
| M12_octanol concentration factor | 1.090 |
| Carbonate | 0.1802 mM |
| Acidity error | -0.94352 mM |

### Warnings and errors

| | |
|---|---|
| Errors | None |
| Warnings | One or more logP values out of range |

### Sample logD and percent species

| pH | M12_octanol logD | M12_octanol M12_octanolH | M12_octanol M12_octanol | M12_octanol M12_octanolH* | M12_octanol M12_octanol* | Comment |
|---|---|---|---|---|---|---|
| 1.000 | -0.47 | 74.87 % | 0.00 % | 0.00 % | 25.12 % | |
| 1.200 | -0.27 | 65.28 % | 0.01 % | 0.00 % | 34.71 % | Stomach pH |
| 2.000 | 0.53 | 22.96 % | 0.01 % | 0.00 % | 77.03 % | |
| 3.000 | 1.52 | 2.89 % | 0.02 % | 0.00 % | 97.09 % | |
| 4.000 | 2.50 | 0.30 % | 0.02 % | 0.00 % | 99.69 % | |
| 5.000 | 3.34 | 0.03 % | 0.02 % | 0.00 % | 99.95 % | |
| 6.000 | 3.73 | 0.00 % | 0.02 % | 0.00 % | 99.98 % | |
| 6.500 | 3.78 | 0.00 % | 0.02 % | 0.00 % | 99.98 % | |
| 7.000 | 3.80 | 0.00 % | 0.02 % | 0.00 % | 99.98 % | |
| 7.400 | 3.80 | 0.00 % | 0.02 % | 0.00 % | 99.98 % | Blood pH |
| 8.000 | 3.80 | 0.00 % | 0.02 % | 0.00 % | 99.98 % | |
| 9.000 | 3.81 | 0.00 % | 0.02 % | 0.00 % | 99.98 % | |
| 10.000 | 3.81 | 0.00 % | 0.02 % | 0.00 % | 99.98 % | |
| 11.000 | 3.81 | 0.00 % | 0.02 % | 0.00 % | 99.98 % | |
| 12.000 | 3.81 | 0.00 % | 0.02 % | 0.00 % | 99.98 % | |

| | | | |
|---|---|---|---|
| Sample name: | **M12_octanol** | Experiment start time: | **3/3/2018 4:33:34 PM** |
| Assay name: | **pH-metric high logP** | Analyst: | **Pion** |
| Assay ID: | **18C-03013** | Instrument ID: | **T312060** |
| Filename: | **C:\Sirius_T3\Mehtap\20180302_exp29_logP_T3-2\18C-03013_M12_octanol_pH-metric high logP.t3r** | | |

### Graphs

Charge balance residuals

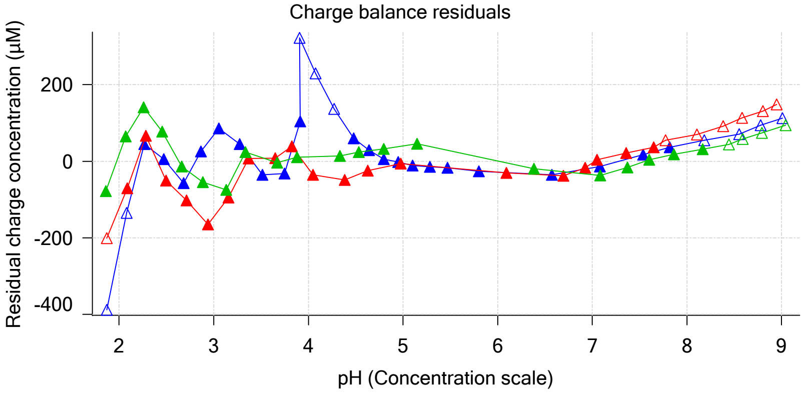


Ionic strength

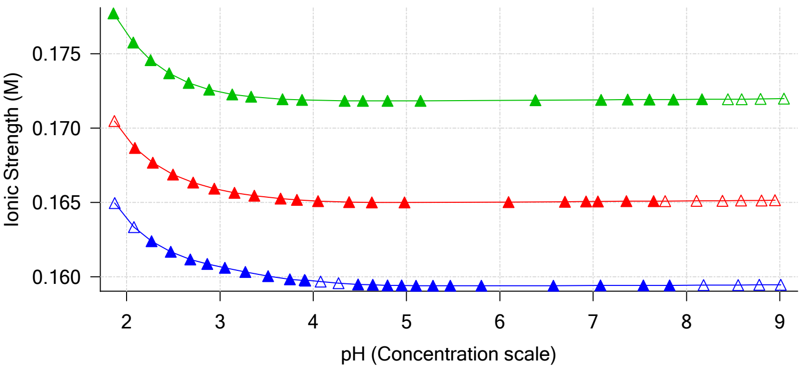


Buffer index profile

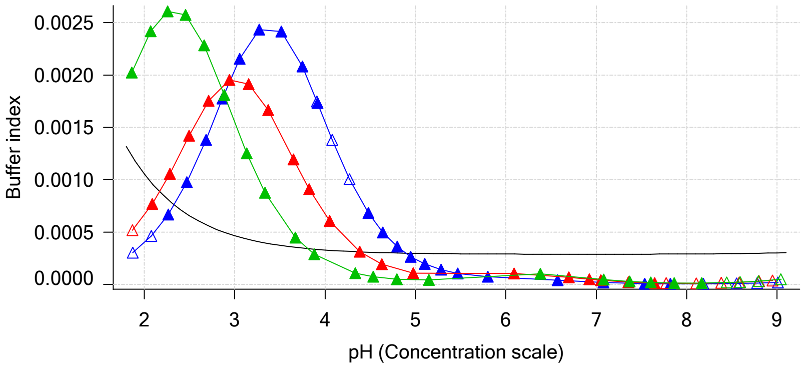


Sample M12_octanol Ionisation Graph

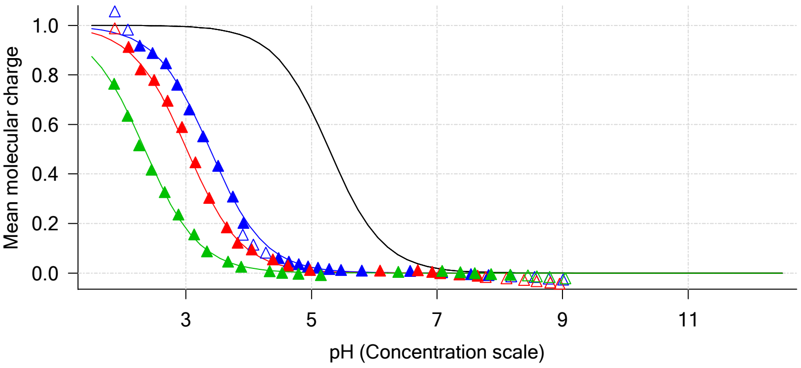


Distribution of species for sample M12_octanol

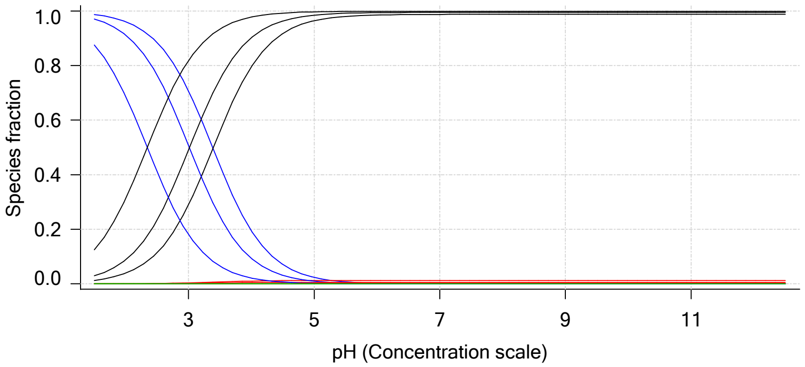


Lipophilicity Profile

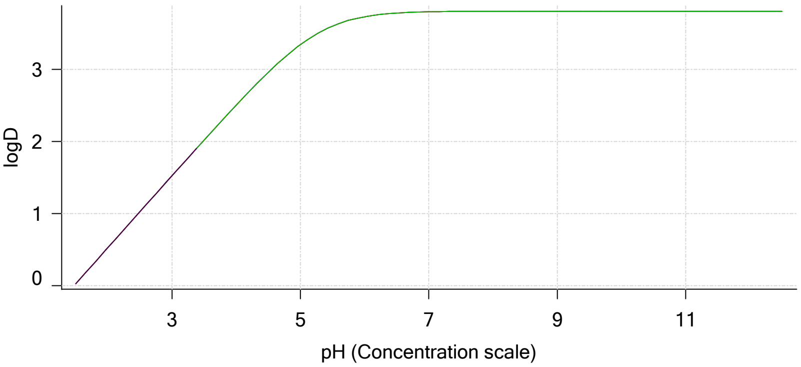


Log R Profile

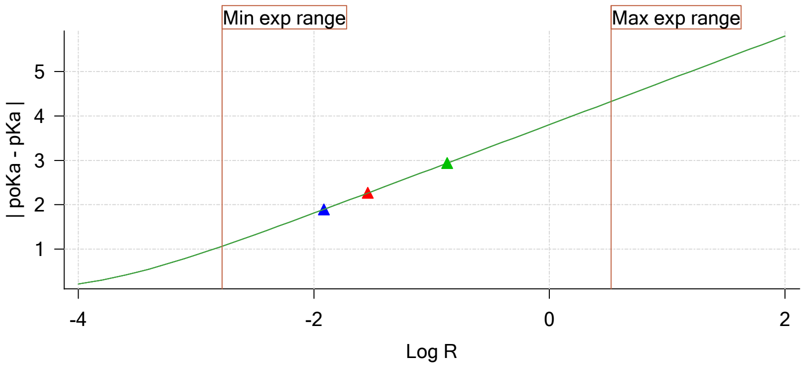


Sample Carbonate Ionisation Graph

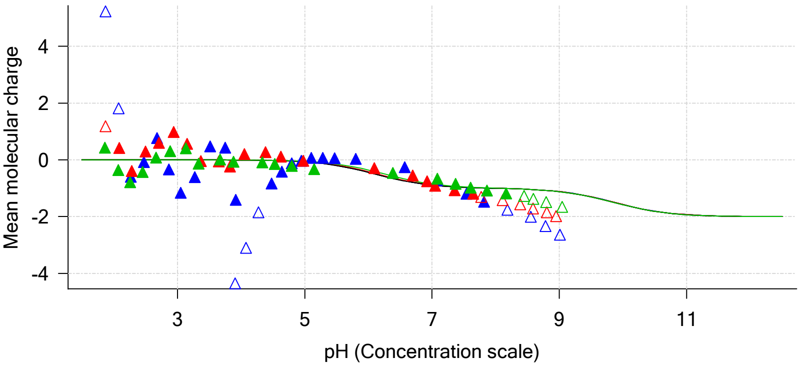

| | | | |
|---|---|---|---|
| Sample name: | **M12_octanol** | Experiment start time: | **3/3/2018 4:33:34 PM** |
| Assay name: | **pH-metric high logP** | Analyst: | **Pion** |
| Assay ID: | **18C-03013** | Instrument ID: | **T312060** |
| Filename: | **C:\Sirius_T3\Mehtap\20180302_exp29_logP_T3-2\18C-03013_M12_octanol_pH-metric high logP.t3r** | | |

### Graphs (continued)

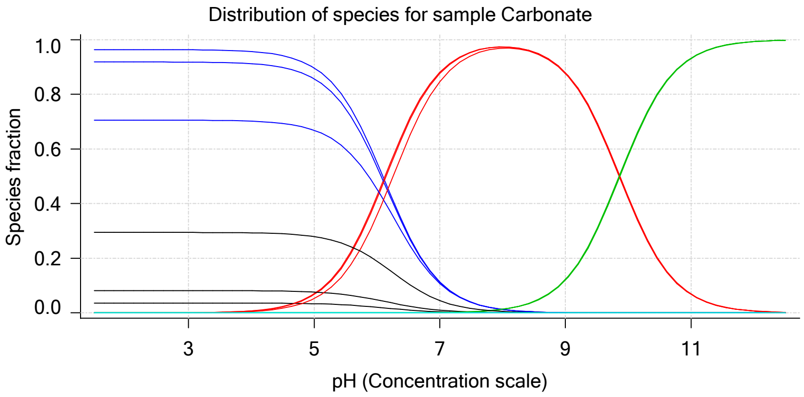

| | | | |
|---|---|---|---|
| Sample name: | **M12_octanol** | Experiment start time: | **3/3/2018 4:33:34 PM** |
| Assay name: | **pH-metric high logP** | Analyst: | **Pion** |
| Assay ID: | **18C-03013** | Instrument ID: | **T312060** |
| Filename: | **C:\Sirius_T3\Mehtap\20180302_exp29_logP_T3-2\18C-03013_M12_octanol_pH-metric high logP.t3r** | | |

# pH-metric high logP   Titration 1 of 3   18C-03013 Points 1 to 30

### Overall results

| | |
|---|---|
| RMSD | 1.132 |
| Average ionic strength | 0.160 M |
| Average temperature | 24.9°C |
| Partition ratio | 0.0122 : 1 |
| Analyte concentration range | 4689.3 µM to 4859.4 µM |
| Total points considered | 21 of 30 |

### Warnings and errors

| | |
|---|---|
| Errors | None |
| Warnings | None |

### Four-Plus parameters

| | | | |
|---|---|---|---|
| Alpha | 0.111 | 3/3/2018 4:33:34 PM | C:\Sirius_T3\HCl18C02.t3r |
| S | 0.9988 | 3/3/2018 4:33:34 PM | C:\Sirius_T3\HCl18C02.t3r |
| jH | 1.0 | 3/3/2018 4:33:34 PM | C:\Sirius_T3\HCl18C02.t3r |
| jOH | -0.8 | 3/3/2018 4:33:34 PM | C:\Sirius_T3\HCl18C02.t3r |

### Titrants

| | | | |
|---|---|---|---|
| 0.50 M HCl | 0.999058 | 3/3/2018 4:33:34 PM | C:\Sirius_T3\HCl18C02.t3r |
| 0.50 M KOH | 0.999845 | 3/3/2018 4:33:34 PM | C:\Sirius_T3\KOH18B27.t3r |

### Sample

| | |
|---|---|
| M12_octanol concentration factor | 0.900 |
| M12_octanol stoichiometry | 1.000 |
| Chloride stoichiometry | 1.000 |
| Base pKa 1 | 5.28 |
| logP (XH +) | 1.02 |
| logP (neutral X) | 3.87 |

### Sample graphs

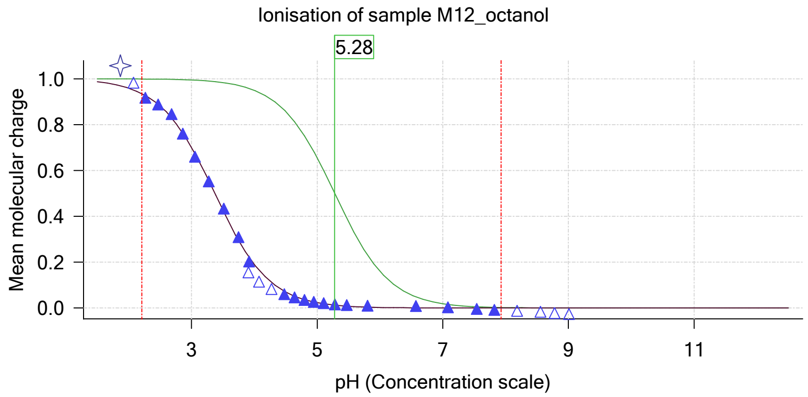


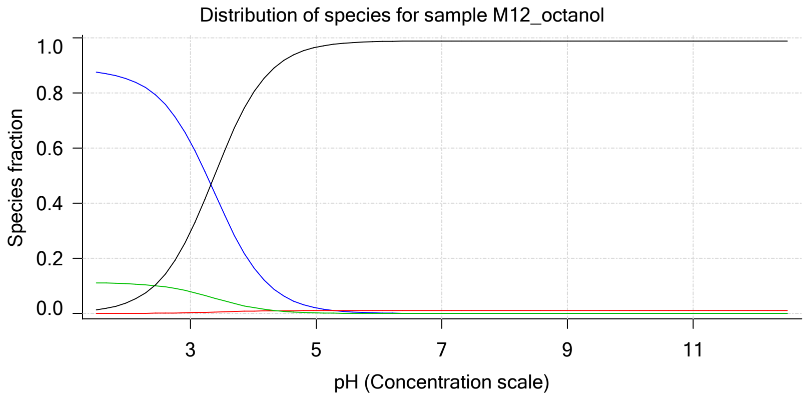

Sample name: **M12_octanol**
Assay name: **pH-metric high logP**
Assay ID: **18C-03013**
Filename: **C:\Sirius_T3\Mehtap\20180302_exp29_logP_T3-2\18C-03013_M12_octanol_pH-metric high logP.t3r**
Experiment start time: **3/3/2018 4:33:34 PM**
Analyst: **Pion**
Instrument ID: **T312060**

### Sample graphs (continued)

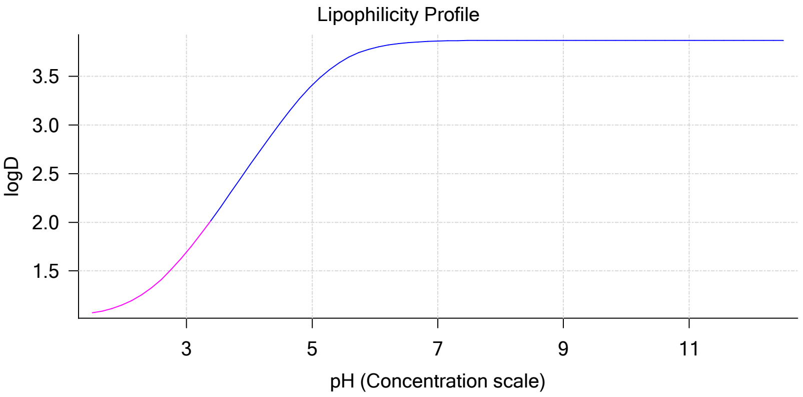


### Sample logD and percent species

| pH | M12_octanol logD | M12_octanol M12_octanolH | M12_octanol M12_octanol | M12_octanol M12_octanolH* | M12_octanol M12_octanol* | Comment |
|---|---|---|---|---|---|---|
| 1.000 | 1.04 | 88.32 % | 0.00 % | 11.25 % | 0.42 % | |
| 1.200 | 1.04 | 88.11 % | 0.01 % | 11.23 % | 0.66 % | Stomach pH |
| 2.000 | 1.16 | 85.08 % | 0.04 % | 10.84 % | 4.03 % | |
| 3.000 | 1.69 | 62.24 % | 0.33 % | 7.93 % | 29.50 % | |
| 4.000 | 2.58 | 16.89 % | 0.89 % | 2.15 % | 80.07 % | |
| 5.000 | 3.41 | 2.04 % | 1.07 % | 0.26 % | 96.63 % | |
| 6.000 | 3.80 | 0.21 % | 1.09 % | 0.03 % | 98.67 % | |
| 6.500 | 3.85 | 0.07 % | 1.09 % | 0.01 % | 98.83 % | |
| 7.000 | 3.86 | 0.02 % | 1.09 % | 0.00 % | 98.88 % | |
| 7.400 | 3.87 | 0.01 % | 1.09 % | 0.00 % | 98.90 % | Blood pH |
| 8.000 | 3.87 | 0.00 % | 1.09 % | 0.00 % | 98.90 % | |
| 9.000 | 3.87 | 0.00 % | 1.09 % | 0.00 % | 98.90 % | |
| 10.000 | 3.87 | 0.00 % | 1.09 % | 0.00 % | 98.91 % | |
| 11.000 | 3.87 | 0.00 % | 1.09 % | 0.00 % | 98.91 % | |
| 12.000 | 3.87 | 0.00 % | 1.09 % | 0.00 % | 98.91 % | |

### Carbonate and acidity

Carbonate 0.074 mM
Acidity error -0.417 mM

### Other graphs

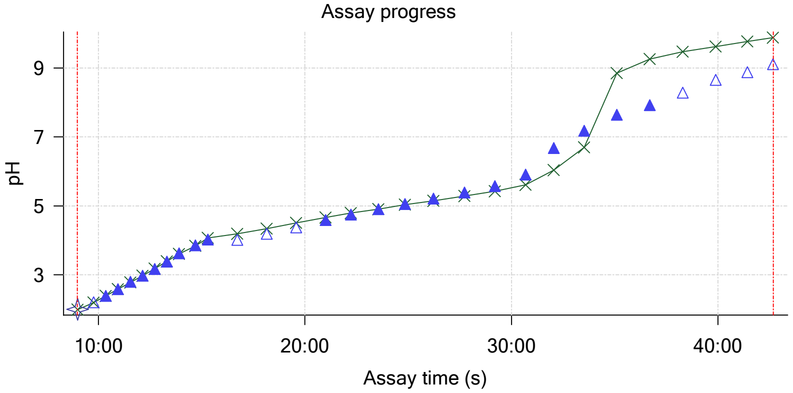


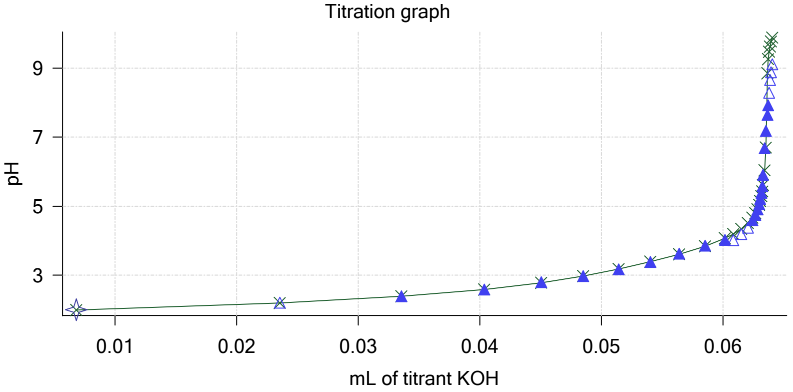

| | | | |
|---|---|---|---|
| Sample name: | **M12_octanol** | Experiment start time: | **3/3/2018 4:33:34 PM** |
| Assay name: | **pH-metric high logP** | Analyst: | **Pion** |
| Assay ID: | **18C-03013** | Instrument ID: | **T312060** |
| Filename: | **C:\Sirius_T3\Mehtap\20180302_exp29_logP_T3-2\18C-03013_M12_octanol_pH-metric high logP.t3r** | | |

### Other graphs (continued)

Charge balance residuals

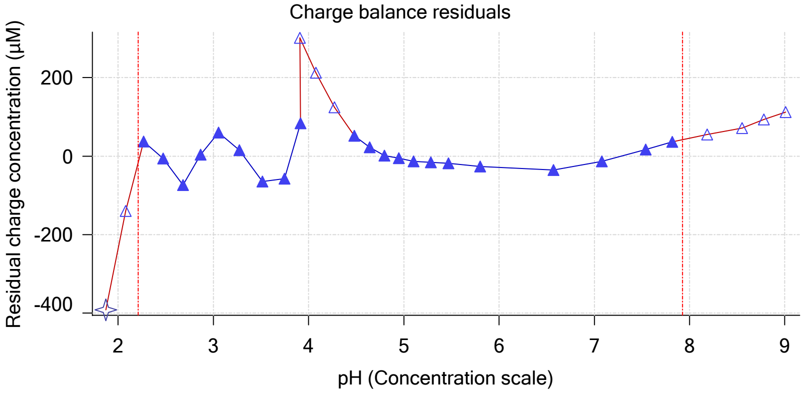


Buffer index profile

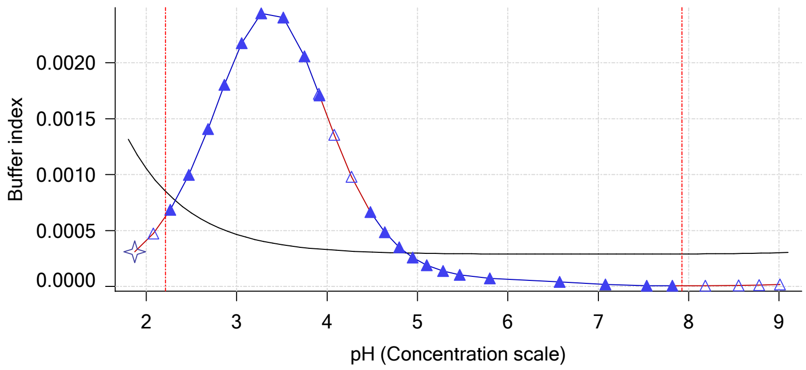


Ionic strength

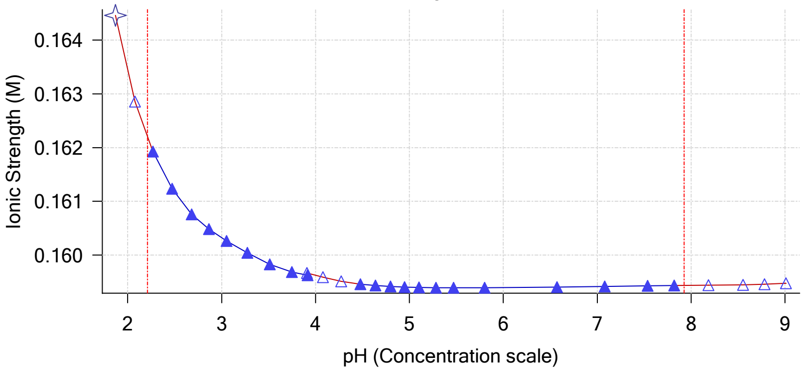


Log R Profile

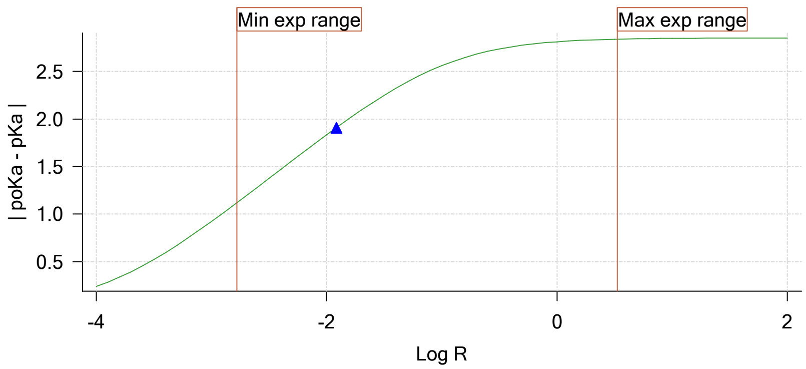


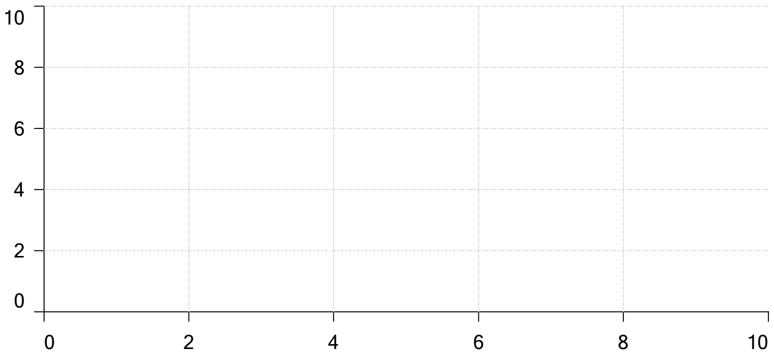


Ionisation of sample Carbonate

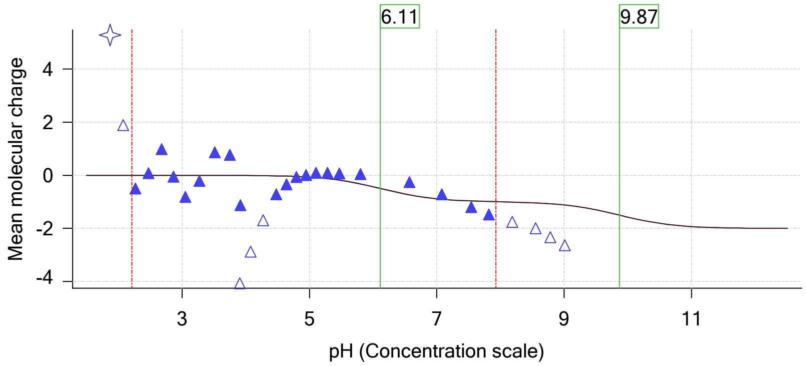

| | | | |
|---|---|---|---|
| Sample name: | **M12_octanol** | Experiment start time: | **3/3/2018 4:33:34 PM** |
| Assay name: | **pH-metric high logP** | Analyst: | **Pion** |
| Assay ID: | **18C-03013** | Instrument ID: | **T312060** |
| Filename: | **C:\Sirius_T3\Mehtap\20180302_exp29_logP_T3-2\18C-03013_M12_octanol_pH-metric high logP.t3r** | | |

# pH-metric high logP   Titration 2 of 3   18C-03013 Points 31 to 56

### Overall results

| | |
|---|---|
| RMSD | 0.588 |
| Average ionic strength | 0.165 M |
| Average temperature | 25.0°C |
| Partition ratio | 0.0288 : 1 |
| Analyte concentration range | 4305.5 µM to 4450.5 µM |
| Total points considered | 19 of 26 |

### Warnings and errors

| | |
|---|---|
| Errors | None |
| Warnings | None |

### Four-Plus parameters

| | | | |
|---|---|---|---|
| Alpha | 0.111 | 3/3/2018 4:33:34 PM | C:\Sirius_T3\HCl18C02.t3r |
| S | 0.9988 | 3/3/2018 4:33:34 PM | C:\Sirius_T3\HCl18C02.t3r |
| jH | 1.0 | 3/3/2018 4:33:34 PM | C:\Sirius_T3\HCl18C02.t3r |
| jOH | -0.8 | 3/3/2018 4:33:34 PM | C:\Sirius_T3\HCl18C02.t3r |

### Titrants

| | | | |
|---|---|---|---|
| 0.50 M HCl | 0.999058 | 3/3/2018 4:33:34 PM | C:\Sirius_T3\HCl18C02.t3r |
| 0.50 M KOH | 0.999845 | 3/3/2018 4:33:34 PM | C:\Sirius_T3\KOH18B27.t3r |

### Sample

| | |
|---|---|
| M12_octanol concentration factor | 0.766 |
| M12_octanol stoichiometry | 1.000 |
| Chloride stoichiometry | 1.000 |
| Base pKa 1 | 5.28 |
| logP (XH +) | 1.02 |
| logP (neutral X) | 3.88 |

### Sample graphs

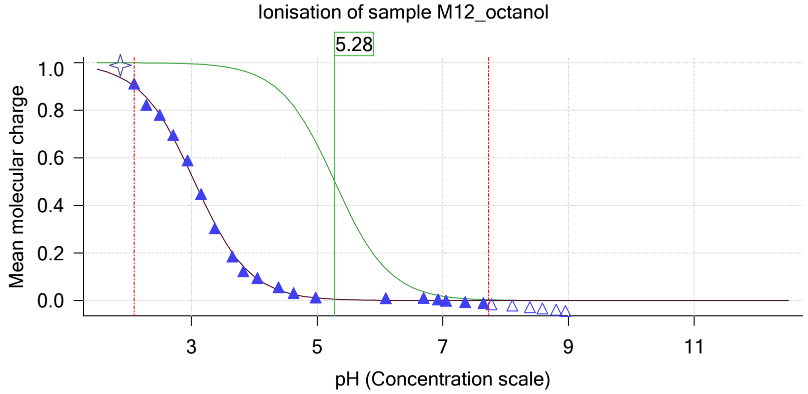


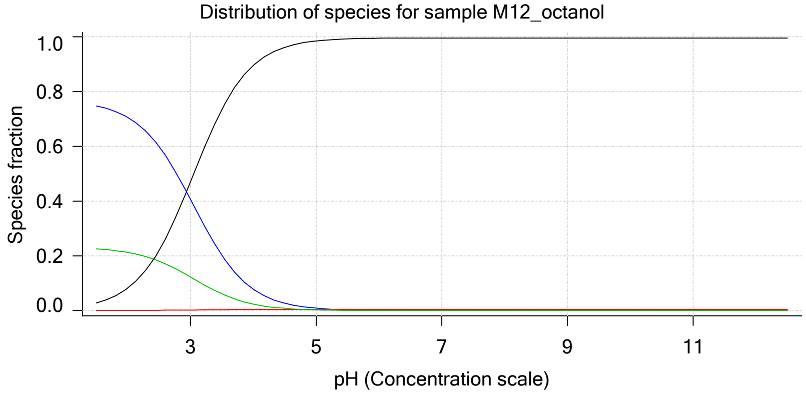

| | | | |
|---|---|---|---|
| Sample name: | **M12_octanol** | Experiment start time: | **3/3/2018 4:33:34 PM** |
| Assay name: | **pH-metric high logP** | Analyst: | **Pion** |
| Assay ID: | **18C-03013** | Instrument ID: | **T312060** |
| Filename: | **C:\Sirius_T3\Mehtap\20180302_exp29_logP_T3-2\18C-03013_M12_octanol_pH-metric high logP.t3r** | | |

### Sample graphs (continued)

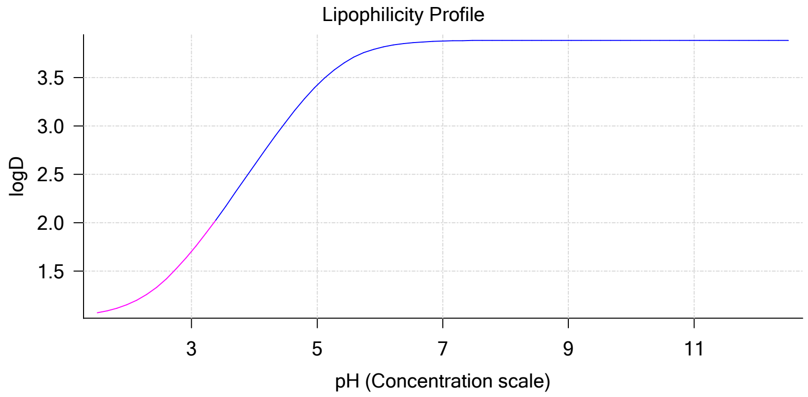


### Sample logD and percent species

| pH | M12_octanol logD | M12_octanol M12_octanolH | M12_octanol M12_octanol | M12_octanol M12_octanolH* | M12_octanol M12_octanol* | Comment |
|---|---|---|---|---|---|---|
| 1.000 | 1.04 | 76.18 % | 0.00 % | 22.94 % | 0.88 % | |
| 1.200 | 1.05 | 75.78 % | 0.01 % | 22.82 % | 1.39 % | Stomach pH |
| 2.000 | 1.16 | 70.56 % | 0.04 % | 21.25 % | 8.16 % | |
| 3.000 | 1.70 | 40.60 % | 0.21 % | 12.23 % | 46.96 % | |
| 4.000 | 2.59 | 7.74 % | 0.41 % | 2.33 % | 89.52 % | |
| 5.000 | 3.42 | 0.85 % | 0.45 % | 0.26 % | 98.45 % | |
| 6.000 | 3.81 | 0.09 % | 0.45 % | 0.03 % | 99.44 % | |
| 6.500 | 3.86 | 0.03 % | 0.45 % | 0.01 % | 99.51 % | |
| 7.000 | 3.88 | 0.01 % | 0.45 % | 0.00 % | 99.54 % | |
| 7.400 | 3.88 | 0.00 % | 0.45 % | 0.00 % | 99.54 % | Blood pH |
| 8.000 | 3.88 | 0.00 % | 0.45 % | 0.00 % | 99.55 % | |
| 9.000 | 3.88 | 0.00 % | 0.45 % | 0.00 % | 99.55 % | |
| 10.000 | 3.88 | 0.00 % | 0.45 % | 0.00 % | 99.55 % | |
| 11.000 | 3.88 | 0.00 % | 0.45 % | 0.00 % | 99.55 % | |
| 12.000 | 3.88 | 0.00 % | 0.45 % | 0.00 % | 99.55 % | |

### Carbonate and acidity

| | |
|---|---|
| Carbonate | 0.169 mM |
| Acidity error | 0.139 mM |

### Other graphs

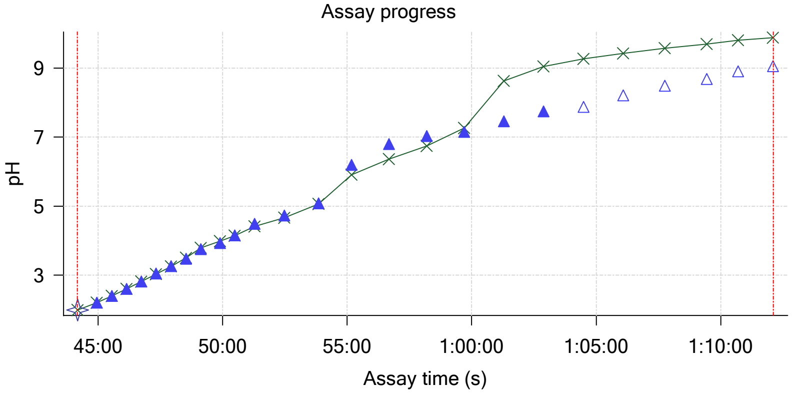


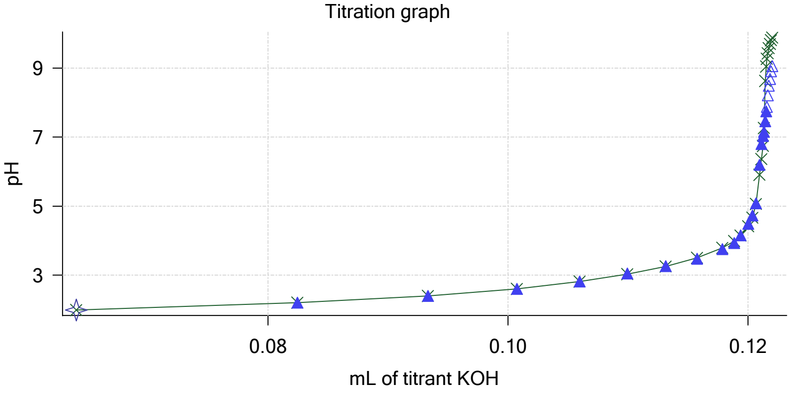

Sample name: **M12_octanol**
Assay name: **pH-metric high logP**
Assay ID: **18C-03013**
Filename: **C:\Sirius_T3\Mehtap\20180302_exp29_logP_T3-2\18C-03013_M12_octanol_pH-metric high logP.t3r**

Experiment start time: **3/3/2018 4:33:34 PM**
Analyst: **Pion**
Instrument ID: **T312060**

### Other graphs (continued)

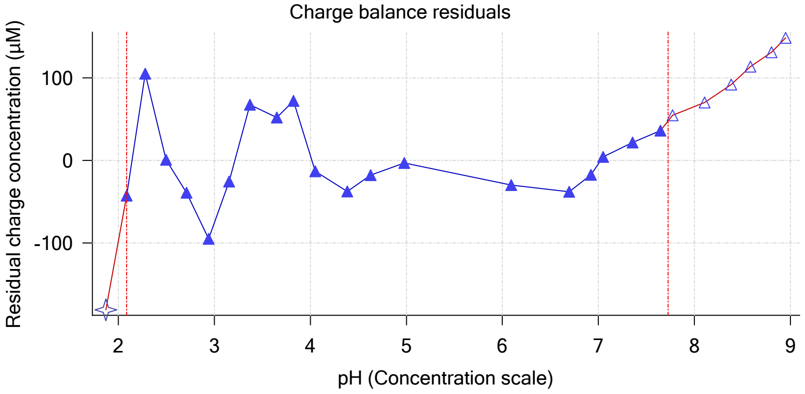


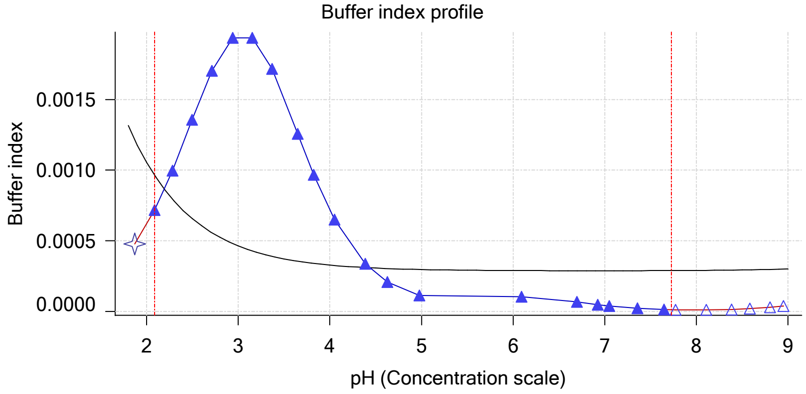


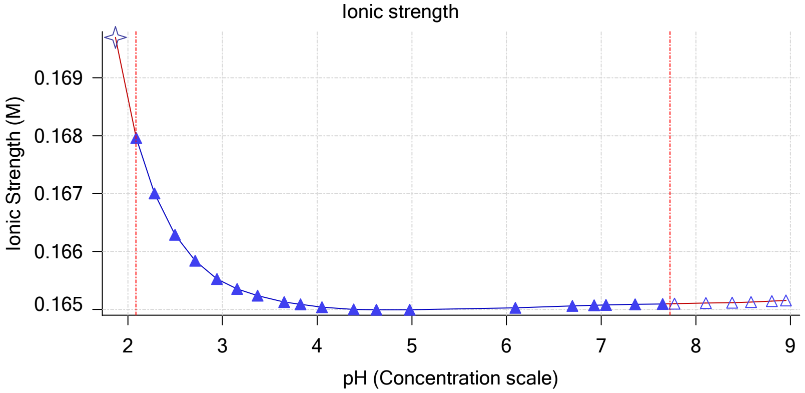


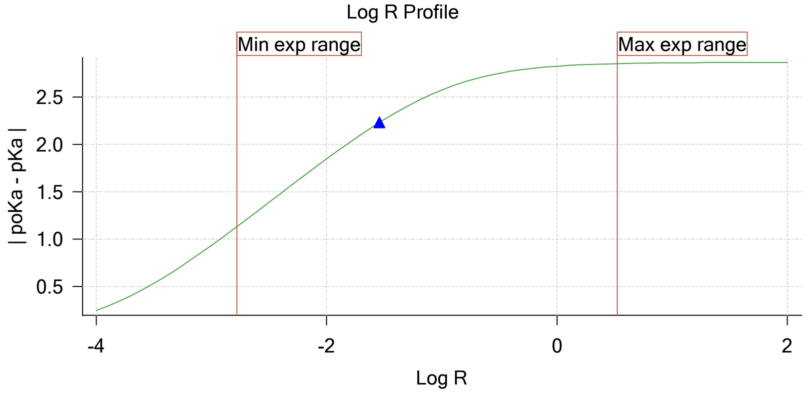


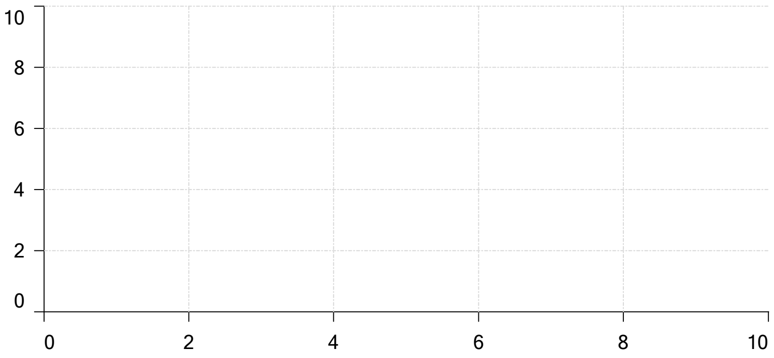


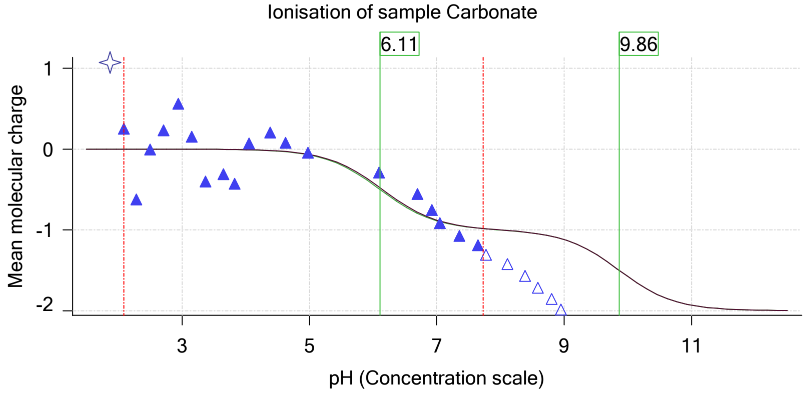

| | | | |
|---|---|---|---|
| Sample name: | **M12_octanol** | Experiment start time: | **3/3/2018 4:33:34 PM** |
| Assay name: | **pH-metric high logP** | Analyst: | **Pion** |
| Assay ID: | **18C-03013** | Instrument ID: | **T312060** |
| Filename: | **C:\Sirius_T3\Mehtap\20180302_exp29_logP_T3-2\18C-03013_M12_octanol_pH-metric high logP.t3r** | | |

# pH-metric high logP   Titration 3 of 3   18C-03013 Points 57 to 80

### Overall results

| | |
|---|---|
| RMSD | 0.527 |
| Average ionic strength | 0.172 M |
| Average temperature | 25.0°C |
| Partition ratio | 0.1351 : 1 |
| Analyte concentration range | 3639.4 µM to 3752.4 µM |
| Total points considered | 20 of 24 |

### Warnings and errors

| | |
|---|---|
| Errors | None |
| Warnings | None |

### Four-Plus parameters

| | | | |
|---|---|---|---|
| Alpha | 0.111 | 3/3/2018 4:33:34 PM | C:\Sirius_T3\HCl18C02.t3r |
| S | 0.9988 | 3/3/2018 4:33:34 PM | C:\Sirius_T3\HCl18C02.t3r |
| jH | 1.0 | 3/3/2018 4:33:34 PM | C:\Sirius_T3\HCl18C02.t3r |
| jOH | -0.8 | 3/3/2018 4:33:34 PM | C:\Sirius_T3\HCl18C02.t3r |

### Titrants

| | | | |
|---|---|---|---|
| 0.50 M HCl | 0.999058 | 3/3/2018 4:33:34 PM | C:\Sirius_T3\HCl18C02.t3r |
| 0.50 M KOH | 0.999845 | 3/3/2018 4:33:34 PM | C:\Sirius_T3\KOH18B27.t3r |

### Sample

| | |
|---|---|
| M12_octanol concentration factor | 1.090 |
| M12_octanol stoichiometry | 1.000 |
| Chloride stoichiometry | 1.000 |
| Base pKa 1 | 5.28 |
| logP (XH +) | 1.02 |
| logP (neutral X) | 4.20 |

### Sample graphs

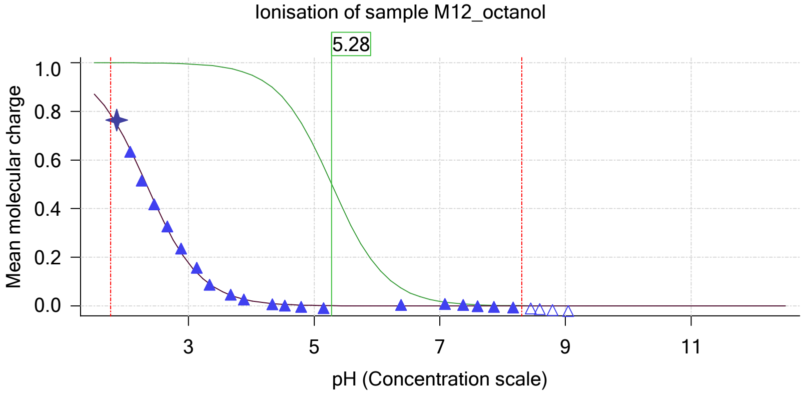


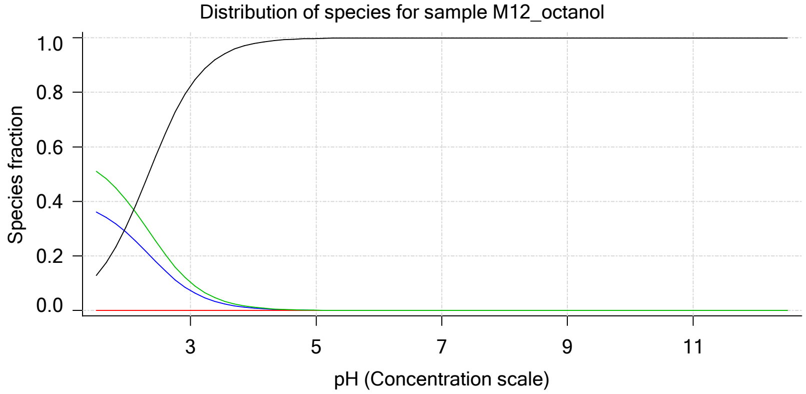

Sample name: **M12_octanol** Experiment start time: **3/3/2018 4:33:34 PM**
Assay name: **pH-metric high logP** Analyst: **Pion**
Assay ID: **18C-03013** Instrument ID: **T312060**
Filename: **C:\Sirius_T3\Mehtap\20180302_exp29_logP_T3-2\18C-03013_M12_octanol_pH-metric high logP.t3r**

### Sample graphs (continued)

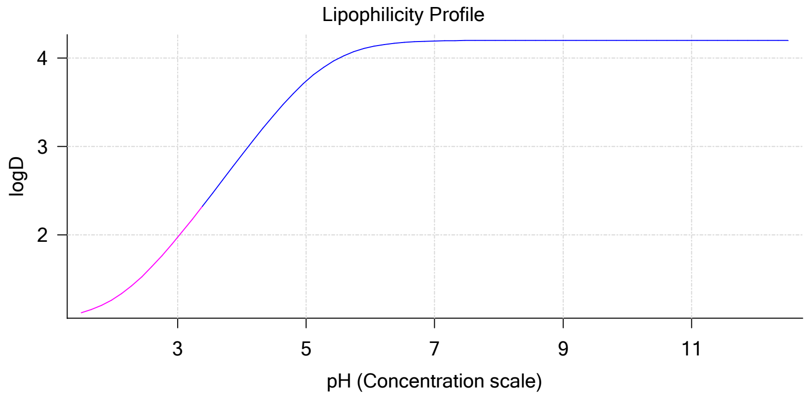


### Sample logD and percent species

| pH | M12_octanol logD | M12_octanol M12_octanolH | M12_octanol M12_octanol | M12_octanol M12_octanolH* | M12_octanol M12_octanol* | Comment |
|---|---|---|---|---|---|---|
| 1.000 | 1.05 | 39.56 % | 0.00 % | 55.97 % | 4.47 % | |
| 1.200 | 1.07 | 38.55 % | 0.00 % | 54.54 % | 6.90 % | Stomach pH |
| 2.000 | 1.27 | 28.21 % | 0.01 % | 39.91 % | 31.87 % | |
| 3.000 | 1.97 | 7.29 % | 0.04 % | 10.31 % | 82.36 % | |
| 4.000 | 2.91 | 0.87 % | 0.05 % | 1.23 % | 97.86 % | |
| 5.000 | 3.74 | 0.09 % | 0.05 % | 0.12 % | 99.74 % | |
| 6.000 | 4.13 | 0.01 % | 0.05 % | 0.01 % | 99.93 % | |
| 6.500 | 4.18 | 0.00 % | 0.05 % | 0.00 % | 99.95 % | |
| 7.000 | 4.19 | 0.00 % | 0.05 % | 0.00 % | 99.95 % | |
| 7.400 | 4.20 | 0.00 % | 0.05 % | 0.00 % | 99.95 % | Blood pH |
| 8.000 | 4.20 | 0.00 % | 0.05 % | 0.00 % | 99.95 % | |
| 9.000 | 4.20 | 0.00 % | 0.05 % | 0.00 % | 99.95 % | |
| 10.000 | 4.20 | 0.00 % | 0.05 % | 0.00 % | 99.95 % | |
| 11.000 | 4.20 | 0.00 % | 0.05 % | 0.00 % | 99.95 % | |
| 12.000 | 4.20 | 0.00 % | 0.05 % | 0.00 % | 99.95 % | |

### Carbonate and acidity

Carbonate 0.180 mM
Acidity error -0.944 mM

### Other graphs

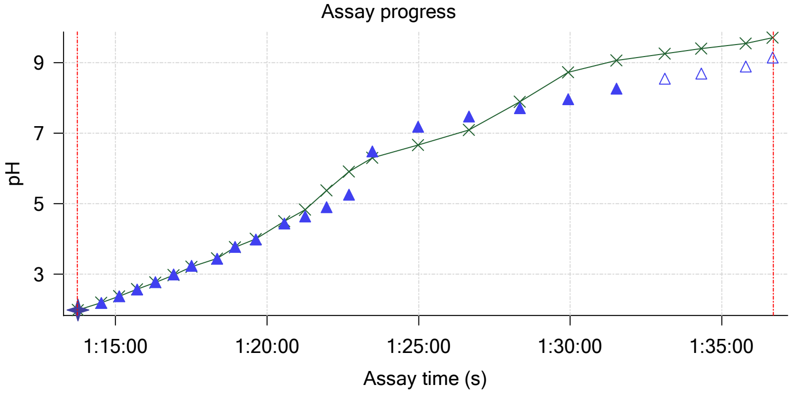


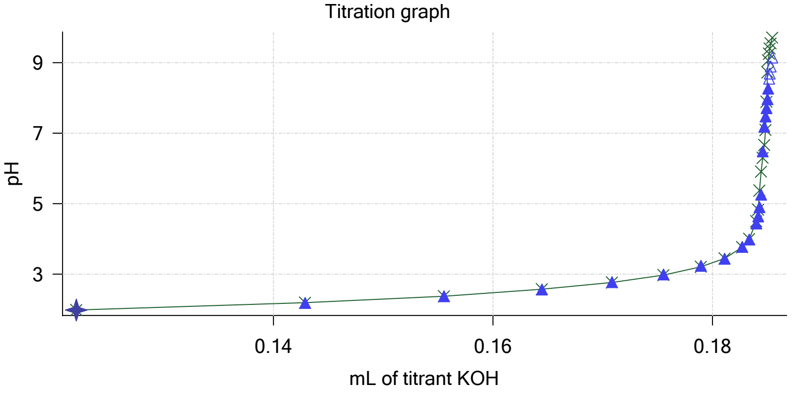

| | | | |
|---|---|---|---|
| Sample name: | **M12_octanol** | Experiment start time: | **3/3/2018 4:33:34 PM** |
| Assay name: | **pH-metric high logP** | Analyst: | **Pion** |
| Assay ID: | **18C-03013** | Instrument ID: | **T312060** |
| Filename: | **C:\Sirius_T3\Mehtap\20180302_exp29_logP_T3-2\18C-03013_M12_octanol_pH-metric high logP.t3r** | | |

### Other graphs (continued)

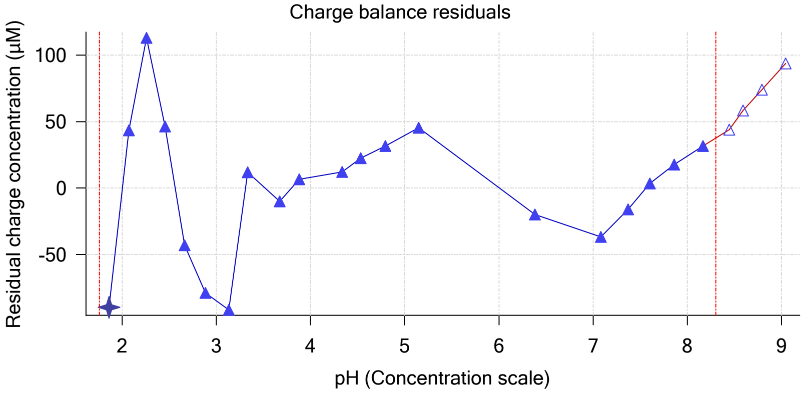


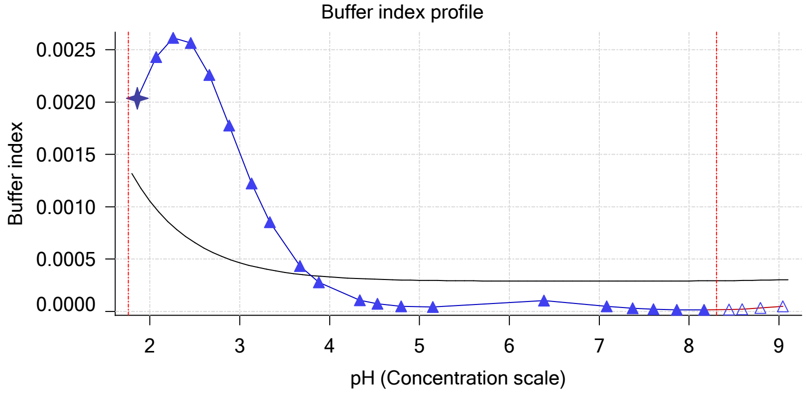


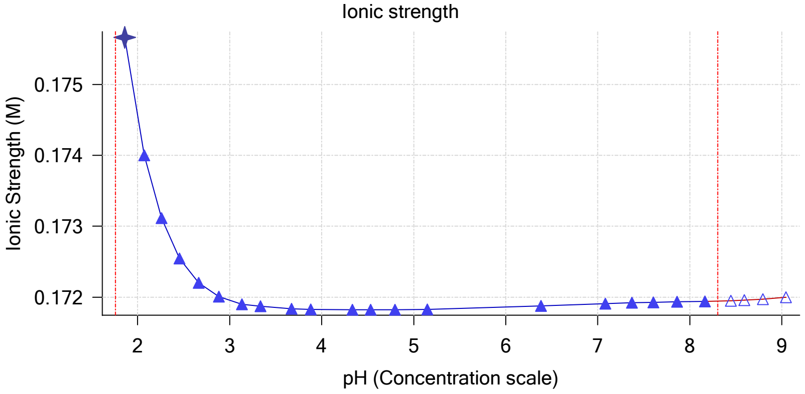


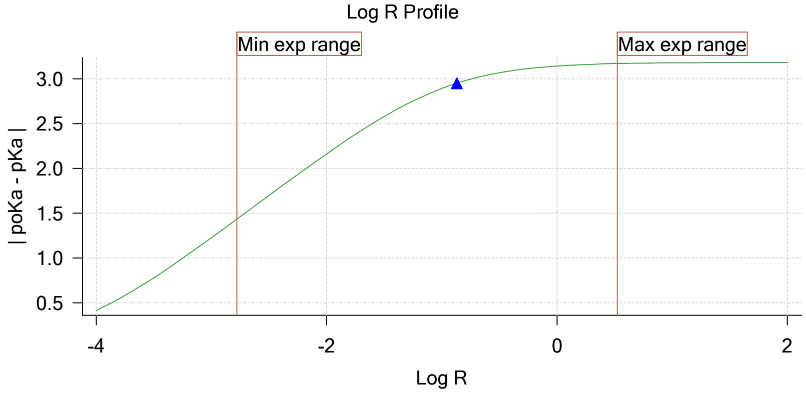


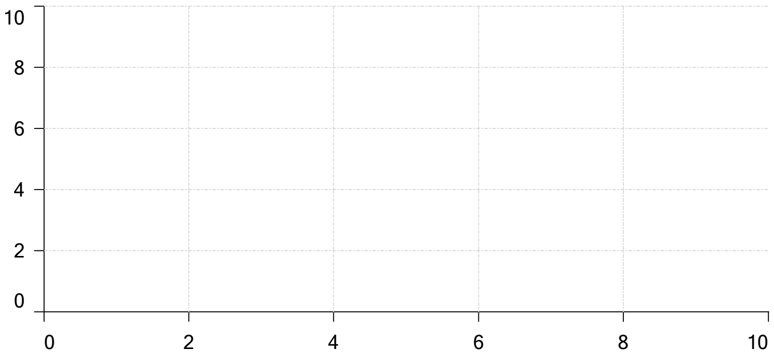

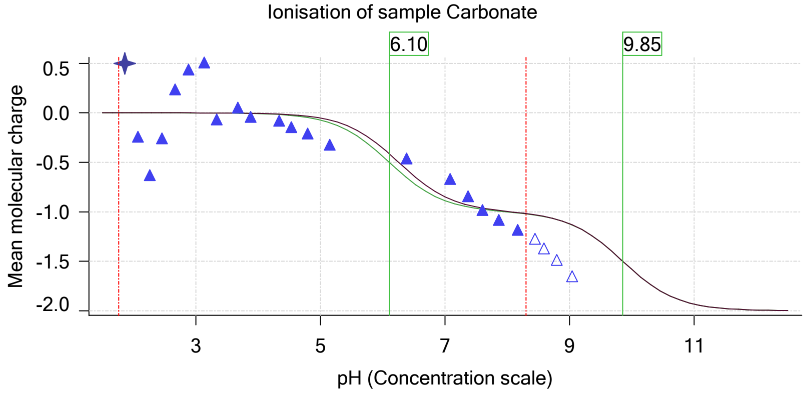

| | | | |
|---|---|---|---|
| Sample name: | **M12_octanol** | Experiment start time: | **3/3/2018 4:33:34 PM** |
| Assay name: | **pH-metric high logP** | Analyst: | **Pion** |
| Assay ID: | **18C-03013** | Instrument ID: | **T312060** |
| Filename: | **C:\Sirius_T3\Mehtap\20180302_exp29_logP_T3-2\18C-03013_M12_octanol_pH-metric high logP.t3r** | | |

### Assay Model

| Settings | Value | Date/Time changed | Imported from |
|---|---|---|---|
| Sample name | M12_octanol | 2/28/2018 2:58:36 PM | User entered value |
| Sample by | Weight | | Default value |
| Sample weight | 0.002240 g | 3/2/2018 5:10:19 PM | User entered value |
| Formula weight | 292.16 g/mol | 2/28/2018 2:58:36 PM | User entered value |
| Solubility | Unknown | | Default value |
| Molecular weight | 255.70 | 2/28/2018 2:58:36 PM | User entered value |
| Individual pKa ionic environments | No | | Default value |
| Number of pKas | 1 | 2/28/2018 2:58:36 PM | User entered value |
| Sample is a | Base | 2/28/2018 2:58:36 PM | User entered value |
| pKa 1 | 5.28 | 2/28/2018 2:58:36 PM | User entered value |
| logp (XH +) | 1.02 | 3/2/2018 3:44:28 PM | User entered value |
| logP (neutral X) | 3.79 | 3/2/2018 3:44:35 PM | User entered value |
| Stoichiometry | 1.00000 | | Default value |
| Aprotic counterion name | Chloride | | From standards.xml file |
| Stoichiometry | 1.00 | | From standards.xml file |
| Charge per counterion | -1 | | From standards.xml file |

### Events

| Time | Event | Water | Acid | Base | Octanol | pH | dpH/dt | pH R-squared | pH SD | dpH/dt time |
|---|---|---|---|---|---|---|---|---|---|---|
| 5:59.7 | Initial pH = 4.07 | | | | | | | | | |
| 8:59.3 | Data point 1 | 1.50000 mL | 0.05094 mL | 0.00682 mL | 0.01999 mL | 1.993 | 0.00166 | 0.09032 | 0.00027 | 10.0 s |
| 9:45.6 | Data point 2 | 1.50000 mL | 0.05094 mL | 0.02354 mL | 0.01999 mL | 2.195 | -0.01016 | 0.27478 | 0.00096 | 10.0 s |
| 10:21.2 | Data point 3 | 1.50000 mL | 0.05094 mL | 0.03356 mL | 0.01999 mL | 2.380 | -0.00393 | 0.49609 | 0.00028 | 10.0 s |
| 10:56.8 | Data point 4 | 1.50000 mL | 0.05094 mL | 0.04036 mL | 0.01999 mL | 2.583 | -0.01061 | 0.30160 | 0.00095 | 10.0 s |
| 11:32.3 | Data point 5 | 1.50000 mL | 0.05094 mL | 0.04504 mL | 0.01999 mL | 2.791 | -0.00615 | 0.53625 | 0.00041 | 10.0 s |
| 12:07.9 | Data point 6 | 1.50000 mL | 0.05094 mL | 0.04849 mL | 0.01999 mL | 2.973 | -0.01184 | 0.89998 | 0.00062 | 10.0 s |
| 12:43.3 | Data point 7 | 1.50000 mL | 0.05094 mL | 0.05141 mL | 0.01999 mL | 3.161 | -0.01641 | 0.89430 | 0.00086 | 10.0 s |
| 13:18.8 | Data point 8 | 1.50000 mL | 0.05094 mL | 0.05402 mL | 0.01999 mL | 3.380 | -0.01816 | 0.97797 | 0.00091 | 10.0 s |
| 13:54.4 | Data point 9 | 1.50000 mL | 0.05094 mL | 0.05640 mL | 0.01999 mL | 3.621 | -0.01999 | 0.98251 | 0.00100 | 11.0 s |
| 14:41.2 | Data point 10 | 1.50000 mL | 0.05094 mL | 0.05851 mL | 0.01999 mL | 3.854 | -0.01872 | 0.92596 | 0.00096 | 11.5 s |
| 15:18.1 | Data point 11 | 1.50000 mL | 0.05094 mL | 0.06016 mL | 0.01999 mL | 4.021 | -0.07914 | 0.99504 | 0.00392 | Timed out at 59.5 s |
| 16:43.6 | Data point 12 | 1.50000 mL | 0.05094 mL | 0.06082 mL | 0.01999 mL | 4.014 | -0.06809 | 0.99320 | 0.00337 | Timed out at 59.5 s |
| 18:09.0 | Data point 13 | 1.50000 mL | 0.05094 mL | 0.06150 mL | 0.01999 mL | 4.183 | -0.04322 | 0.98848 | 0.00215 | Timed out at 59.5 s |
| 19:34.3 | Data point 14 | 1.50000 mL | 0.05094 mL | 0.06204 mL | 0.01999 mL | 4.377 | -0.02931 | 0.97915 | 0.00146 | Timed out at 59.5 s |
| 20:59.7 | Data point 15 | 1.50000 mL | 0.05094 mL | 0.06242 mL | 0.01999 mL | 4.584 | -0.01998 | 0.99064 | 0.00099 | 48.5 s |
| 22:13.6 | Data point 16 | 1.50000 mL | 0.05094 mL | 0.06265 mL | 0.01999 mL | 4.745 | -0.01777 | 0.91711 | 0.00092 | 54.0 s |
| 23:33.1 | Data point 17 | 1.50000 mL | 0.05094 mL | 0.06282 mL | 0.01999 mL | 4.902 | -0.01909 | 0.98788 | 0.00095 | 46.5 s |
| 24:50.2 | Data point 18 | 1.50000 mL | 0.05094 mL | 0.06296 mL | 0.01999 mL | 5.053 | -0.01967 | 0.95681 | 0.00099 | 57.5 s |
| 26:13.2 | Data point 19 | 1.50000 mL | 0.05094 mL | 0.06305 mL | 0.01999 mL | 5.207 | -0.02130 | 0.95071 | 0.00108 | Timed out at 59.5 s |
| 27:43.7 | Data point 20 | 1.50000 mL | 0.05094 mL | 0.06315 mL | 0.01999 mL | 5.387 | -0.01948 | 0.92987 | 0.00100 | 57.5 s |
| 29:11.7 | Data point 21 | 1.50000 mL | 0.05094 mL | 0.06322 mL | 0.01999 mL | 5.573 | -0.01912 | 0.95211 | 0.00097 | 59.0 s |
| 30:41.3 | Data point 22 | 1.50000 mL | 0.05094 mL | 0.06329 mL | 0.01999 mL | 5.905 | -0.01889 | 0.90762 | 0.00098 | 51.5 s |
| 32:03.4 | Data point 23 | 1.50000 mL | 0.05094 mL | 0.06341 mL | 0.01999 mL | 6.675 | -0.01975 | 0.98665 | 0.00098 | 57.0 s |
| 33:31.0 | Data point 24 | 1.50000 mL | 0.05094 mL | 0.06352 mL | 0.01999 mL | 7.182 | -0.02923 | 0.95987 | 0.00147 | Timed out at 59.5 s |
| 35:06.7 | Data point 25 | 1.50000 mL | 0.05094 mL | 0.06364 mL | 0.01999 mL | 7.640 | -0.04416 | 0.98507 | 0.00220 | Timed out at 59.5 s |
| 36:42.3 | Data point 26 | 1.50000 mL | 0.05094 mL | 0.06371 mL | 0.01999 mL | 7.920 | -0.03822 | 0.98044 | 0.00191 | Timed out at 59.5 s |
| 38:17.9 | Data point 27 | 1.50000 mL | 0.05094 mL | 0.06378 mL | 0.01999 mL | 8.284 | -0.02815 | 0.97144 | 0.00141 | Timed out at 59.5 s |

| | | | |
|---|---|---|---|
| Sample name: | **M12_octanol** | Experiment start time: | **3/3/2018 4:33:34 PM** |
| Assay name: | **pH-metric high logP** | Analyst: | **Pion** |
| Assay ID: | **18C-03013** | Instrument ID: | **T312060** |
| Filename: | **C:\Sirius_T3\Mehtap\20180302_exp29_logP_T3-2\18C-03013_M12_octanol_pH-metric high logP.t3r** | | |

### Events (continued)

| Time | Event | Water | Acid | Base | Octanol | pH | dpH/dt | pH R-squared | pH SD | dpH/dt time |
|---|---|---|---|---|---|---|---|---|---|---|
| 39:53.6 | Data point 28 | 1.50000 mL | 0.05094 mL | 0.06385 mL | 0.01999 mL | 8.653 | -0.01851 | 0.94276 | 0.00094 | 56.5 s |
| 41:25.8 | Data point 29 | 1.50000 mL | 0.05094 mL | 0.06395 mL | 0.01999 mL | 8.880 | -0.01941 | 0.93233 | 0.00099 | 43.5 s |
| 42:40.0 | Data point 30 | 1.50000 mL | 0.05094 mL | 0.06404 mL | 0.01999 mL | 9.109 | -0.01865 | 0.94873 | 0.00095 | 37.5 s |
| 44:11.2 | Data point 31 | 1.50000 mL | 0.10868 mL | 0.06404 mL | 0.05000 mL | 1.992 | -0.01464 | 0.98078 | 0.00073 | 10.0 s |
| 44:57.5 | Data point 32 | 1.50000 mL | 0.10868 mL | 0.08246 mL | 0.05000 mL | 2.204 | -0.00623 | 0.39555 | 0.00049 | 10.0 s |
| 45:33.2 | Data point 33 | 1.50000 mL | 0.10868 mL | 0.09334 mL | 0.05000 mL | 2.395 | 0.00024 | 0.00175 | 0.00029 | 10.0 s |
| 46:08.8 | Data point 34 | 1.50000 mL | 0.10868 mL | 0.10075 mL | 0.05000 mL | 2.608 | 0.00510 | 0.08694 | 0.00085 | 10.0 s |
| 46:44.4 | Data point 35 | 1.50000 mL | 0.10868 mL | 0.10597 mL | 0.05000 mL | 2.821 | -0.00756 | 0.57280 | 0.00049 | 10.0 s |
| 47:19.9 | Data point 36 | 1.50000 mL | 0.10868 mL | 0.10995 mL | 0.05000 mL | 3.049 | -0.00595 | 0.80766 | 0.00033 | 10.5 s |
| 47:56.0 | Data point 37 | 1.50000 mL | 0.10868 mL | 0.11315 mL | 0.05000 mL | 3.262 | -0.00544 | 0.73730 | 0.00031 | 10.5 s |
| 48:32.0 | Data point 38 | 1.50000 mL | 0.10868 mL | 0.11578 mL | 0.05000 mL | 3.477 | -0.00488 | 0.14223 | 0.00064 | 10.0 s |
| 49:07.5 | Data point 39 | 1.50000 mL | 0.10868 mL | 0.11787 mL | 0.05000 mL | 3.757 | -0.00502 | 0.11123 | 0.00074 | 10.5 s |
| 49:53.8 | Data point 40 | 1.50000 mL | 0.10868 mL | 0.11886 mL | 0.05000 mL | 3.930 | -0.01303 | 0.75605 | 0.00074 | 10.5 s |
| 50:29.7 | Data point 41 | 1.50000 mL | 0.10868 mL | 0.11940 mL | 0.05000 mL | 4.156 | -0.01560 | 0.85505 | 0.00083 | 12.0 s |
| 51:17.4 | Data point 42 | 1.50000 mL | 0.10868 mL | 0.12004 mL | 0.05000 mL | 4.490 | -0.01911 | 0.93383 | 0.00098 | 46.0 s |
| 52:28.8 | Data point 43 | 1.50000 mL | 0.10868 mL | 0.12039 mL | 0.05000 mL | 4.732 | -0.01804 | 0.90169 | 0.00094 | 52.0 s |
| 53:51.5 | Data point 44 | 1.50000 mL | 0.10868 mL | 0.12067 mL | 0.05000 mL | 5.082 | -0.01877 | 0.92983 | 0.00096 | 48.5 s |
| 55:10.7 | Data point 45 | 1.50000 mL | 0.10868 mL | 0.12098 mL | 0.05000 mL | 6.196 | -0.03857 | 0.98173 | 0.00192 | Timed out at 59.5 s |
| 56:41.2 | Data point 46 | 1.50000 mL | 0.10868 mL | 0.12114 mL | 0.05000 mL | 6.798 | -0.04966 | 0.98696 | 0.00247 | Timed out at 59.5 s |
| 58:11.7 | Data point 47 | 1.50000 mL | 0.10868 mL | 0.12126 mL | 0.05000 mL | 7.026 | -0.03315 | 0.95815 | 0.00167 | Timed out at 59.5 s |
| 59:42.2 | Data point 48 | 1.50000 mL | 0.10868 mL | 0.12136 mL | 0.05000 mL | 7.152 | -0.03446 | 0.98537 | 0.00171 | Timed out at 59.5 s |
| 1:01:17.8 | Data point 49 | 1.50000 mL | 0.10868 mL | 0.12145 mL | 0.05000 mL | 7.458 | -0.05118 | 0.98692 | 0.00254 | Timed out at 59.5 s |
| 1:02:53.4 | Data point 50 | 1.50000 mL | 0.10868 mL | 0.12152 mL | 0.05000 mL | 7.749 | -0.09026 | 0.98237 | 0.00450 | Timed out at 59.5 s |
| 1:04:29.0 | Data point 51 | 1.50000 mL | 0.10868 mL | 0.12159 mL | 0.05000 mL | 7.874 | -0.07009 | 0.98935 | 0.00348 | Timed out at 59.5 s |
| 1:06:04.6 | Data point 52 | 1.50000 mL | 0.10868 mL | 0.12166 mL | 0.05000 mL | 8.208 | -0.05709 | 0.99176 | 0.00283 | Timed out at 59.5 s |
| 1:07:45.4 | Data point 53 | 1.50000 mL | 0.10868 mL | 0.12175 mL | 0.05000 mL | 8.485 | -0.02993 | 0.97232 | 0.00150 | Timed out at 59.5 s |
| 1:09:26.2 | Data point 54 | 1.50000 mL | 0.10868 mL | 0.12185 mL | 0.05000 mL | 8.684 | -0.01638 | 0.69410 | 0.00097 | 39.5 s |
| 1:10:41.4 | Data point 55 | 1.50000 mL | 0.10868 mL | 0.12194 mL | 0.05000 mL | 8.903 | -0.01932 | 0.91951 | 0.00099 | 53.5 s |
| 1:12:05.4 | Data point 56 | 1.50000 mL | 0.10868 mL | 0.12204 mL | 0.05000 mL | 9.050 | -0.01983 | 0.98693 | 0.00099 | 37.0 s |
| 1:13:45.4 | Data point 57 | 1.50000 mL | 0.17119 mL | 0.12204 mL | 0.25000 mL | 1.982 | -0.00868 | 0.51882 | 0.00059 | 10.0 s |
| 1:14:31.7 | Data point 58 | 1.50000 mL | 0.17119 mL | 0.14287 mL | 0.25000 mL | 2.187 | -0.00313 | 0.29423 | 0.00028 | 10.0 s |
| 1:15:07.5 | Data point 59 | 1.50000 mL | 0.17119 mL | 0.15553 mL | 0.25000 mL | 2.372 | 0.00043 | 0.02077 | 0.00015 | 10.0 s |
| 1:15:43.1 | Data point 60 | 1.50000 mL | 0.17119 mL | 0.16446 mL | 0.25000 mL | 2.567 | -0.00212 | 0.35123 | 0.00018 | 10.5 s |
| 1:16:19.2 | Data point 61 | 1.50000 mL | 0.17119 mL | 0.17084 mL | 0.25000 mL | 2.773 | 0.00623 | 0.11536 | 0.00091 | 10.0 s |
| 1:16:54.7 | Data point 62 | 1.50000 mL | 0.17119 mL | 0.17552 mL | 0.25000 mL | 2.992 | -0.00796 | 0.85989 | 0.00042 | 10.0 s |
| 1:17:30.2 | Data point 63 | 1.50000 mL | 0.17119 mL | 0.17893 mL | 0.25000 mL | 3.240 | -0.00517 | 0.42305 | 0.00039 | 10.0 s |
| 1:18:21.2 | Data point 64 | 1.50000 mL | 0.17119 mL | 0.18109 mL | 0.25000 mL | 3.441 | 0.00207 | 0.09593 | 0.00033 | 10.0 s |
| 1:18:56.8 | Data point 65 | 1.50000 mL | 0.17119 mL | 0.18271 mL | 0.25000 mL | 3.778 | -0.00557 | 0.68049 | 0.00033 | 10.5 s |
| 1:19:37.9 | Data point 66 | 1.50000 mL | 0.17119 mL | 0.18335 mL | 0.25000 mL | 3.987 | -0.00947 | 0.83986 | 0.00051 | 10.0 s |
| 1:20:33.9 | Data point 67 | 1.50000 mL | 0.17119 mL | 0.18398 mL | 0.25000 mL | 4.440 | -0.01073 | 0.32598 | 0.00093 | 11.0 s |
| 1:21:15.4 | Data point 68 | 1.50000 mL | 0.17119 mL | 0.18415 mL | 0.25000 mL | 4.637 | -0.01956 | 0.95178 | 0.00099 | 17.0 s |
| 1:21:57.9 | Data point 69 | 1.50000 mL | 0.17119 mL | 0.18429 mL | 0.25000 mL | 4.901 | -0.01572 | 0.83949 | 0.00085 | 13.5 s |
| 1:22:41.9 | Data point 70 | 1.50000 mL | 0.17119 mL | 0.18443 mL | 0.25000 mL | 5.253 | -0.01459 | 0.61037 | 0.00092 | 15.5 s |
| 1:23:28.0 | Data point 71 | 1.50000 mL | 0.17119 mL | 0.18457 mL | 0.25000 mL | 6.486 | -0.02584 | 0.99076 | 0.00128 | Timed out at 59.5 s |
| 1:24:58.6 | Data point 72 | 1.50000 mL | 0.17119 mL | 0.18471 mL | 0.25000 mL | 7.185 | -0.06008 | 0.98340 | 0.00299 | Timed out at 59.5 s |

| | | | |
|---|---|---|---|
| Sample name: | **M12_octanol** | Experiment start time: | **3/3/2018 4:33:34 PM** |
| Assay name: | **pH-metric high logP** | Analyst: | **Pion** |
| Assay ID: | **18C-03013** | Instrument ID: | **T312060** |
| Filename: | **C:\Sirius_T3\Mehtap\20180302_exp29_logP_T3-2\18C-03013_M12_octanol_pH-metric high logP.t3r** | | |

### Events (continued)

| Time | Event | Water | Acid | Base | Octanol | pH | dpH/dt | pH R-squared | pH SD | dpH/dt time |
|---|---|---|---|---|---|---|---|---|---|---|
| 1:26:39.3 | Data point 73 | 1.50000 mL | 0.17119 mL | 0.18483 mL | 0.25000 mL | 7.473 | -0.04422 | 0.98321 | 0.00220 | Timed out at 59.5 s |
| 1:28:20.1 | Data point 74 | 1.50000 mL | 0.17119 mL | 0.18493 mL | 0.25000 mL | 7.705 | -0.05489 | 0.98906 | 0.00273 | Timed out at 59.5 s |
| 1:29:55.7 | Data point 75 | 1.50000 mL | 0.17119 mL | 0.18499 mL | 0.25000 mL | 7.963 | -0.05454 | 0.98638 | 0.00271 | Timed out at 59.5 s |
| 1:31:31.4 | Data point 76 | 1.50000 mL | 0.17119 mL | 0.18507 mL | 0.25000 mL | 8.267 | -0.04359 | 0.97797 | 0.00218 | Timed out at 59.5 s |
| 1:33:06.9 | Data point 77 | 1.50000 mL | 0.17119 mL | 0.18514 mL | 0.25000 mL | 8.545 | -0.01535 | 0.64494 | 0.00094 | 37.0 s |
| 1:34:19.6 | Data point 78 | 1.50000 mL | 0.17119 mL | 0.18521 mL | 0.25000 mL | 8.691 | -0.01787 | 0.92709 | 0.00092 | 57.5 s |
| 1:35:47.7 | Data point 79 | 1.50000 mL | 0.17119 mL | 0.18530 mL | 0.25000 mL | 8.893 | -0.01116 | 0.33380 | 0.00095 | 17.5 s |
| 1:36:40.9 | Data point 80 | 1.50000 mL | 0.17119 mL | 0.18544 mL | 0.25000 mL | 9.143 | -0.01321 | 0.57777 | 0.00086 | 13.0 s |
| 1:37:02.9 | Assay volumes | 1.50000 mL | 0.17119 mL | 0.18544 mL | 0.25000 mL | | | | | |

| | | | |
|---|---|---|---|
| Sample name: | **M12_octanol** | Experiment start time: | **3/3/2018 4:33:34 PM** |
| Assay name: | **pH-metric high logP** | Analyst: | **Pion** |
| Assay ID: | **18C-03013** | Instrument ID: | **T312060** |
| Filename: | **C:\Sirius_T3\Mehtap\20180302_exp29_logP_T3-2\18C-03013_M12_octanol_pH-metric high logP.t3r** | | |

| | | | |
|---|---|---|---|
| Sample name: | **M12_octanol** | Experiment start time: | **3/3/2018 4:33:34 PM** |
| Assay name: | **pH-metric high logP** | Analyst: | **Pion** |
| Assay ID: | **18C-03013** | Instrument ID: | **T312060** |
| Filename: | **C:\Sirius_T3\Mehtap\20180302_exp29_logP_T3-2\18C-03013_M12_octanol_pH-metric high logP.t3r** | | |

### Assay Settings (continued)

| Setting | Value | Original Value | Date/Time changed | Imported from |
|---|---|---|---|---|
| ***Titration 3*** | | | | |
| Titrate from | Low to high pH | | | |
| Add additional water | 0.00 mL | | | |
| Additional partition solvent volume | 0.200 mL | | | |
| Additional partition solvent added | Automatic | | | |
| After pH adjust stir for | 30 seconds | | | |
| Stir to allow partitioning for | 15 seconds | | | |
| Stirrer speed for partitioning | 60% | | | |
| ***Data Point Stability*** | | | | |
| Stir during data point collection | No | | | |
| Delay before data point collection | 0 seconds | | | |
| Number of points to average | 20 points | | | |
| Time interval between points | 0.50 seconds | | | |
| Required maximum standard deviation | 0.00100 dpH/dt | | | |
| Stability timeout after | 60 seconds | | | |

### Calibration Settings

| Setting | Value | Date/Time changed | Imported from |
|---|---|---|---|
| Four-Plus alpha | 0.111 | 3/3/2018 4:33:34 PM | C:\Sirius_T3\HCl18C02.t3r |
| Four-Plus S | 0.9988 | 3/3/2018 4:33:34 PM | C:\Sirius_T3\HCl18C02.t3r |
| Four-Plus jH | 1.0 | 3/3/2018 4:33:34 PM | C:\Sirius_T3\HCl18C02.t3r |
| Four-Plus jOH | -0.8 | 3/3/2018 4:33:34 PM | C:\Sirius_T3\HCl18C02.t3r |
| Base concentration factor | 1.000 | 3/3/2018 4:33:34 PM | C:\Sirius_T3\KOH18B27.t3r |
| Acid concentration factor | 0.999 | 3/3/2018 4:33:34 PM | C:\Sirius_T3\HCl18C02.t3r |

### Instrument Settings

| Setting | Value | Batch Id | Install date |
|---|---|---|---|
| Instrument owner | Merck | | |
| Instrument ID | T312060 | | |
| Instrument type | T3 Simulator | | |
| Software version | 1.1.3.0 | | |
| Dispenser module | | T3DM1200361 | 3/31/2009 5:24:52 AM |
| Dispenser 0 | Water | | 3/31/2009 5:25:05 AM |
| Syringe volume | 2.5 mL | | |
| Firmware version | 1.2.1(r2) | | |
| Titrant | Water (0.15 M KCl) | 02-06-2018 | 2/27/2018 10:05:59 AM |
| Dispenser 2 | Acid | | 3/31/2009 5:25:11 AM |
| Syringe volume | 0.5 mL | | |
| Firmware version | 1.2.1(r2) | | |
| Titrant | Acid (0.5 M HCl) | 02-27-2018 | 2/27/2018 10:27:22 AM |
| Dispenser 1 | Base | | 3/31/2009 5:25:21 AM |
| Syringe volume | 0.5 mL | | |
| Firmware version | 1.2.1(r2) | | |
| Titrant | Base (0.5 M KOH) | 9/22/2017 | 2/27/2018 10:21:22 AM |
| Dispenser 5 | Cosolvent | | 3/31/2009 5:26:24 AM |
| Syringe volume | 2.5 mL | | |
| Firmware version | 1.2.1(r2) | | |
| Distribution valve 5 | Distribution Valve | | 3/31/2009 5:28:19 AM |
| Firmware version | 1.1.3 | | |
| Port A | Methanol (80%, 0.15 M KCl) | 09-26-17 | 2/7/2018 9:42:01 AM |
| Port B | Cyclohexane | 11-01-17 | 2/27/2018 10:37:57 AM |
| Dispenser 3 | Buffer | | 8/3/2010 5:05:16 AM |
| Syringe volume | 0.5 mL | | |
| Firmware version | 1.2.1(r2) | | |
| Titrant | Dodecane | 2018/01/31 | 2/28/2018 10:18:04 AM |
| Dispenser 6 | Octanol | | 10/22/2010 10:52:43 AM |

| | | | |
|---|---|---|---|
| Sample name: | **M12_octanol** | Experiment start time: | **3/3/2018 4:33:34 PM** |
| Assay name: | **pH-metric high logP** | Analyst: | **Pion** |
| Assay ID: | **18C-03013** | Instrument ID: | **T312060** |
| Filename: | **C:\Sirius_T3\Mehtap\20180302_exp29_logP_T3-2\18C-03013_M12_octanol_pH-metric high logP.t3r** | | |

| | | | |
|---|---|---|---|
| Sample name: | **M12_octanol** | Experiment start time: | **3/3/2018 4:33:34 PM** |
| Assay name: | **pH-metric high logP** | Analyst: | **Pion** |
| Assay ID: | **18C-03013** | Instrument ID: | **T312060** |
| Filename: | **C:\Sirius_T3\Mehtap\20180302_exp29_logP_T3-2\18C-03013_M12_octanol_pH-metric high logP.t3r** | | |

### Instrument Settings (continued)

| Setting | Value | Batch Id | Install date |
|---|---|---|---|
| Spectrometer calibration stir duration | 5 s | | |
| Spectrometer calibration stir speed | 30% | | |
| Spectrometer calibration wash pump volume | 20.0 mL | | |
| Spectrometer calibration wash stir duration | 5 s | | |
| Spectrometer calibration wash stir speed | 30% | | |
| Overhead dispense height | 10000 | | |

### Refinement Settings

| Setting | Value | Default value |
|---|---|---|
| Turbidity detection method | None | None |
| Turbidity wavelength to assess | 500.0 nm | 500.0 nm |
| Turbidity maximum absorbance | 0.100 | 0.100 |
| Turbidity probe threshold | 50.00 | 50.00 |

### Experiment Log

[2:37] Air gap created for Water (0.15 M KCl)
[2:38] Air gap created for Acid (0.5 M HCl)
[2:38] Air gap created for Base (0.5 M KOH)
[2:38] Air gap released for Water (0.15 M KCl)
[2:42] Titrator arm moved over Titration position
[2:42] Titration 1 of 3
[2:42] Adding initial titrants
[2:42] Automatically add 1.50000 mL of water
[3:07] Dispensed 1.500000 mL of Water (0.15 M KCl)
[3:12] Titrator arm moved over Drain
[5:53] Titrator arm moved to Titration position
[5:53] Argon flow rate set to 100
[5:53] Stirrer speed set to 10
[5:58] Automatically add 0.02000 mL of Octanol
[5:59] Dispensed 0.019991 mL of Octanol
[6:00] Initial pH = 4.07
[6:00] Iterative adjust 4.07 -> 2.00
[6:00] pH 4.07 -> 2.00
[6:01] Air gap released for Acid (0.5 M HCl)
[6:02] Dispensed 0.050941 mL of Acid (0.5 M HCl)
[6:07] Holding pH 2.00
[8:07] Stirrer speed set to 0
[8:07] Stirrer speed set to 50
[8:07] Iterative adjust 1.94 -> 2.00
[8:07] pH 1.94 -> 2.00
[8:08] Air gap released for Base (0.5 M KOH)
[8:09] Dispensed 0.006820 mL of Base (0.5 M KOH)
[8:59] Stirrer speed set to 0
[9:09] Datapoint id 1 collected
[9:09] Stirrer speed set to 50
[9:14] pH 2.00 -> 2.20
[9:14] Using cautious pH adjust
[9:15] Dispensed 0.007902 mL of Base (0.5 M KOH)
[9:20] Stepping pH = 2.08
[9:20] Dispensed 0.006985 mL of Base (0.5 M KOH)
[9:25] Stepping pH = 2.17
[9:25] Dispensed 0.001834 mL of Base (0.5 M KOH)
[9:30] Stepping pH = 2.20
[9:45] Stirrer speed set to 0
[9:55] Datapoint id 2 collected
[9:55] Charge balance equation is out by -5.8%
[9:55] Stirrer speed set to 50

Sample name: **M12_octanol**
Assay name: **pH-metric high logP**
Assay ID: **18C-03013**
Filename: **C:\Sirius_T3\Mehtap\20180302_exp29_logP_T3-2\18C-03013_M12_octanol_pH-metric high logP.t3r**
Experiment start time: **3/3/2018 4:33:34 PM**
Analyst: **Pion**
Instrument ID: **T312060**

### Experiment Log (continued)

[10:01] pH 2.20 -> 2.40
[10:01] Using charge balance adjust
[10:01] Dispensed 0.010019 mL of Base (0.5 M KOH)
[10:21] Stirrer speed set to 0
[10:31] Datapoint id 3 collected
[10:31] Charge balance equation is out by -11.0%
[10:31] Stirrer speed set to 50
[10:36] pH 2.39 -> 2.59
[10:36] Using charge balance adjust
[10:36] Dispensed 0.006797 mL of Base (0.5 M KOH)
[10:57] Stirrer speed set to 0
[11:07] Datapoint id 4 collected
[11:07] Charge balance equation is out by -2.1%
[11:07] Stirrer speed set to 50
[11:12] pH 2.59 -> 2.79
[11:12] Using charge balance adjust
[11:12] Dispensed 0.004680 mL of Base (0.5 M KOH)
[11:32] Stirrer speed set to 0
[11:42] Datapoint id 5 collected
[11:42] Charge balance equation is out by 0.9%
[11:42] Stirrer speed set to 50
[11:47] pH 2.80 -> 3.00
[11:47] Using charge balance adjust
[11:48] Dispensed 0.003457 mL of Base (0.5 M KOH)
[12:08] Stirrer speed set to 0
[12:18] Datapoint id 6 collected
[12:18] Charge balance equation is out by -12.7%
[12:18] Stirrer speed set to 50
[12:23] pH 2.98 -> 3.18
[12:23] Using charge balance adjust
[12:23] Dispensed 0.002916 mL of Base (0.5 M KOH)
[12:43] Stirrer speed set to 0
[12:53] Datapoint id 7 collected
[12:53] Charge balance equation is out by -8.1%
[12:53] Stirrer speed set to 50
[12:58] pH 3.17 -> 3.37
[12:58] Using charge balance adjust
[12:58] Dispensed 0.002611 mL of Base (0.5 M KOH)
[13:19] Stirrer speed set to 0
[13:29] Datapoint id 8 collected
[13:29] Charge balance equation is out by 7.3%
[13:29] Stirrer speed set to 50
[13:34] pH 3.38 -> 3.58
[13:34] Using charge balance adjust
[13:34] Dispensed 0.002375 mL of Base (0.5 M KOH)
[13:54] Stirrer speed set to 0
[14:05] Datapoint id 9 collected
[14:05] Charge balance equation is out by 18.9%
[14:05] Stirrer speed set to 50
[14:10] pH 3.62 -> 3.82
[14:10] Using cautious pH adjust
[14:10] Dispensed 0.001035 mL of Base (0.5 M KOH)
[14:16] Stepping pH = 3.77
[14:16] Dispensed 0.000329 mL of Base (0.5 M KOH)
[14:21] Stepping pH = 3.77
[14:21] Dispensed 0.000753 mL of Base (0.5 M KOH)
[14:26] Stepping pH = 3.89
[14:41] Stirrer speed set to 0
[14:53] Datapoint id 10 collected

### Experiment Log (continued)

[14:53] Charge balance equation is out by -3.0%
[14:53] Stirrer speed set to 50
[14:58] pH 3.86 -> 4.06
[14:58] Using charge balance adjust
[14:58] Dispensed 0.001646 mL of Base (0.5 M KOH)
[15:18] Stirrer speed set to 0
[16:18] Datapoint id 11 collected
[16:18] Charge balance equation is out by -20.9%
[16:18] Stirrer speed set to 50
[16:23] pH 4.02 -> 4.22
[16:23] Using cautious pH adjust
[16:23] Dispensed 0.000659 mL of Base (0.5 M KOH)
[16:28] Stepping pH = 4.23
[16:43] Stirrer speed set to 0
[17:43] Datapoint id 12 collected
[17:43] Charge balance equation is out by 50.0%
[17:43] Stirrer speed set to 50
[17:49] pH 4.01 -> 4.21
[17:49] Using cautious pH adjust
[17:49] Dispensed 0.000682 mL of Base (0.5 M KOH)
[17:54] Stepping pH = 4.38
[18:09] Stirrer speed set to 0
[19:09] Datapoint id 13 collected
[19:09] Charge balance equation is out by 50.0%
[19:09] Stirrer speed set to 50
[19:14] pH 4.18 -> 4.38
[19:14] Using cautious pH adjust
[19:14] Dispensed 0.000541 mL of Base (0.5 M KOH)
[19:19] Stepping pH = 4.58
[19:34] Stirrer speed set to 0
[20:34] Datapoint id 14 collected
[20:34] Charge balance equation is out by 50.0%
[20:34] Stirrer speed set to 50
[20:39] pH 4.38 -> 4.58
[20:39] Using cautious pH adjust
[20:39] Dispensed 0.000376 mL of Base (0.5 M KOH)
[20:44] Stepping pH = 4.77
[21:00] Stirrer speed set to 0
[21:48] Datapoint id 15 collected
[21:48] Charge balance equation is out by 50.0%
[21:48] Stirrer speed set to 50
[21:53] pH 4.61 -> 4.81
[21:53] Using cautious pH adjust
[21:53] Dispensed 0.000235 mL of Base (0.5 M KOH)
[21:58] Stepping pH = 4.90
[22:14] Stirrer speed set to 0
[23:08] Datapoint id 16 collected
[23:08] Charge balance equation is out by 50.0%
[23:08] Stirrer speed set to 50
[23:13] pH 4.77 -> 4.97
[23:13] Using cautious pH adjust
[23:13] Dispensed 0.000165 mL of Base (0.5 M KOH)
[23:18] Stepping pH = 5.01
[23:33] Stirrer speed set to 0
[24:20] Datapoint id 17 collected
[24:20] Charge balance equation is out by 50.0%
[24:20] Stirrer speed set to 50
[24:25] pH 4.93 -> 5.13
[24:25] Using cautious pH adjust

| | | | |
|---|---|---|---|
| Sample name: | **M12_octanol** | Experiment start time: | **3/3/2018 4:33:34 PM** |
| Assay name: | **pH-metric high logP** | Analyst: | **Pion** |
| Assay ID: | **18C-03013** | Instrument ID: | **T312060** |
| Filename: | **C:\Sirius_T3\Mehtap\20180302_exp29_logP_T3-2\18C-03013_M12_octanol_pH-metric high logP.t3r** | | |

### Experiment Log (continued)

[24:25] Dispensed 0.000118 mL of Base (0.5 M KOH)
[24:30] Stepping pH = 5.12
[24:30] Dispensed 0.000024 mL of Base (0.5 M KOH)
[24:35] Stepping pH = 5.14
[24:50] Stirrer speed set to 0
[25:48] Datapoint id 18 collected
[25:48] Charge balance equation is out by 42.0%
[25:48] Stirrer speed set to 50
[25:53] pH 5.09 -> 5.29
[25:53] Using cautious pH adjust
[25:53] Dispensed 0.000094 mL of Base (0.5 M KOH)
[25:58] Stepping pH = 5.30
[26:13] Stirrer speed set to 0
[27:13] Datapoint id 19 collected
[27:13] Charge balance equation is out by 50.0%
[27:13] Stirrer speed set to 50
[27:18] pH 5.22 -> 5.42
[27:18] Using cautious pH adjust
[27:18] Dispensed 0.000071 mL of Base (0.5 M KOH)
[27:23] Stepping pH = 5.35
[27:23] Dispensed 0.000024 mL of Base (0.5 M KOH)
[27:28] Stepping pH = 5.44
[27:44] Stirrer speed set to 0
[28:41] Datapoint id 20 collected
[28:41] Charge balance equation is out by 30.4%
[28:41] Stirrer speed set to 50
[28:46] pH 5.39 -> 5.59
[28:46] Using cautious pH adjust
[28:46] Dispensed 0.000047 mL of Base (0.5 M KOH)
[28:51] Stepping pH = 5.50
[28:51] Dispensed 0.000024 mL of Base (0.5 M KOH)
[28:56] Stepping pH = 5.60
[29:12] Stirrer speed set to 0
[30:11] Datapoint id 21 collected
[30:11] Charge balance equation is out by 20.0%
[30:11] Stirrer speed set to 50
[30:16] pH 5.58 -> 5.78
[30:16] Using cautious pH adjust
[30:16] Dispensed 0.000047 mL of Base (0.5 M KOH)
[30:21] Stepping pH = 5.73
[30:21] Dispensed 0.000024 mL of Base (0.5 M KOH)
[30:26] Stepping pH = 5.90
[30:41] Stirrer speed set to 0
[31:33] Datapoint id 22 collected
[31:33] Charge balance equation is out by 25.8%
[31:33] Stirrer speed set to 50
[31:38] pH 5.90 -> 6.10
[31:38] Using cautious pH adjust
[31:38] Dispensed 0.000024 mL of Base (0.5 M KOH)
[31:43] Stepping pH = 5.91
[31:43] Dispensed 0.000094 mL of Base (0.5 M KOH)
[31:48] Stepping pH = 6.63
[32:03] Stirrer speed set to 0
[33:00] Datapoint id 23 collected
[33:00] Charge balance equation is out by -90.2%
[33:00] Stirrer speed set to 50
[33:06] pH 6.63 -> 6.83
[33:06] Using cautious pH adjust
[33:06] Dispensed 0.000024 mL of Base (0.5 M KOH)

| | | | |
|---|---|---|---|
| Sample name: | **M12_octanol** | Experiment start time: | **3/3/2018 4:33:34 PM** |
| Assay name: | **pH-metric high logP** | Analyst: | **Pion** |
| Assay ID: | **18C-03013** | Instrument ID: | **T312060** |
| Filename: | **C:\Sirius_T3\Mehtap\20180302_exp29_logP_T3-2\18C-03013_M12_octanol_pH-metric high logP.t3r** | | |

### Experiment Log (continued)

[33:11] Stepping pH = 6.61
[33:11] Dispensed 0.000094 mL of Base (0.5 M KOH)
[33:16] Stepping pH = 7.08
[33:31] Stirrer speed set to 0
[34:31] Datapoint id 24 collected
[34:31] Charge balance equation is out by -229.9%
[34:31] Stirrer speed set to 50
[34:36] pH 7.12 -> 7.32
[34:36] Using cautious pH adjust
[34:36] Dispensed 0.000024 mL of Base (0.5 M KOH)
[34:41] Stepping pH = 7.09
[34:41] Dispensed 0.000047 mL of Base (0.5 M KOH)
[34:46] Stepping pH = 7.18
[34:46] Dispensed 0.000047 mL of Base (0.5 M KOH)
[34:51] Stepping pH = 7.57
[35:07] Stirrer speed set to 0
[36:07] Datapoint id 25 collected
[36:07] Charge balance equation is out by -576.8%
[36:07] Stirrer speed set to 50
[36:12] pH 7.58 -> 7.78
[36:12] Using cautious pH adjust
[36:12] Dispensed 0.000024 mL of Base (0.5 M KOH)
[36:17] Stepping pH = 7.57
[36:17] Dispensed 0.000024 mL of Base (0.5 M KOH)
[36:22] Stepping pH = 7.63
[36:22] Dispensed 0.000024 mL of Base (0.5 M KOH)
[36:27] Stepping pH = 7.82
[36:42] Stirrer speed set to 0
[37:42] Datapoint id 26 collected
[37:42] Charge balance equation is out by -808.4%
[37:42] Stirrer speed set to 50
[37:47] pH 7.85 -> 8.05
[37:47] Using cautious pH adjust
[37:47] Dispensed 0.000024 mL of Base (0.5 M KOH)
[37:52] Stepping pH = 7.82
[37:52] Dispensed 0.000024 mL of Base (0.5 M KOH)
[37:58] Stepping pH = 7.89
[37:58] Dispensed 0.000024 mL of Base (0.5 M KOH)
[38:03] Stepping pH = 8.14
[38:18] Stirrer speed set to 0
[39:18] Datapoint id 27 collected
[39:18] Charge balance equation is out by -902.2%
[39:18] Stirrer speed set to 50
[39:23] pH 8.29 -> 8.49
[39:23] Using cautious pH adjust
[39:23] Dispensed 0.000024 mL of Base (0.5 M KOH)
[39:28] Stepping pH = 8.33
[39:28] Dispensed 0.000024 mL of Base (0.5 M KOH)
[39:33] Stepping pH = 8.43
[39:33] Dispensed 0.000024 mL of Base (0.5 M KOH)
[39:38] Stepping pH = 8.64
[39:53] Stirrer speed set to 0
[40:50] Datapoint id 28 collected
[40:50] Charge balance equation is out by -511.3%
[40:50] Stirrer speed set to 50
[40:55] pH 8.63 -> 8.83
[40:55] Using cautious pH adjust
[40:55] Dispensed 0.000024 mL of Base (0.5 M KOH)
[41:00] Stepping pH = 8.59

Sample name: **M12_octanol** Experiment start time: **3/3/2018 4:33:34 PM**
Assay name: **pH-metric high logP** Analyst: **Pion**
Assay ID: **18C-03013** Instrument ID: **T312060**
Filename: **C:\Sirius_T3\Mehtap\20180302_exp29_logP_T3-2\18C-03013_M12_octanol_pH-metric high logP.t3r**

### Experiment Log (continued)

[41:00] Dispensed 0.000047 mL of Base (0.5 M KOH)
[41:05] Stepping pH = 8.78
[41:05] Dispensed 0.000024 mL of Base (0.5 M KOH)
[41:11] Stepping pH = 8.93
[41:26] Stirrer speed set to 0
[42:09] Datapoint id 29 collected
[42:09] Charge balance equation is out by -393.5%
[42:09] Stirrer speed set to 50
[42:14] pH 8.86 -> 9.05
[42:14] Using cautious pH adjust
[42:14] Dispensed 0.000024 mL of Base (0.5 M KOH)
[42:20] Stepping pH = 8.86
[42:20] Dispensed 0.000071 mL of Base (0.5 M KOH)
[42:25] Stepping pH = 9.12
[42:40] Stirrer speed set to 0
[43:17] Datapoint id 30 collected
[43:17] Charge balance equation is out by -226.3%
[43:17] Titration 2 of 3
[43:17] Adding initial titrants
[43:17] Automatically add 0.03000 mL of Octanol
[43:18] Dispensed 0.030009 mL of Octanol
[43:18] Stirrer speed set to 10
[43:19] Stirrer speed set to 55
[43:19] Iterative adjust 9.11 -> 2.00
[43:19] pH 9.11 -> 2.00
[43:21] Dispensed 0.057738 mL of Acid (0.5 M HCl)
[44:11] Stirrer speed set to 0
[44:21] Datapoint id 31 collected
[44:21] Stirrer speed set to 55
[44:26] pH 2.00 -> 2.20
[44:26] Using cautious pH adjust
[44:27] Dispensed 0.008537 mL of Base (0.5 M KOH)
[44:32] Stepping pH = 2.08
[44:32] Dispensed 0.007855 mL of Base (0.5 M KOH)
[44:37] Stepping pH = 2.17
[44:37] Dispensed 0.002023 mL of Base (0.5 M KOH)
[44:42] Stepping pH = 2.21
[44:57] Stirrer speed set to 0
[45:07] Datapoint id 32 collected
[45:07] Charge balance equation is out by -7.9%
[45:07] Stirrer speed set to 55
[45:13] pH 2.21 -> 2.41
[45:13] Using charge balance adjust
[45:13] Dispensed 0.010889 mL of Base (0.5 M KOH)
[45:33] Stirrer speed set to 0
[45:43] Datapoint id 33 collected
[45:43] Charge balance equation is out by -6.8%
[45:43] Stirrer speed set to 55
[45:48] pH 2.40 -> 2.60
[45:48] Using charge balance adjust
[45:48] Dispensed 0.007408 mL of Base (0.5 M KOH)
[46:09] Stirrer speed set to 0
[46:19] Datapoint id 34 collected
[46:19] Charge balance equation is out by 2.4%
[46:19] Stirrer speed set to 55
[46:24] pH 2.61 -> 2.81
[46:24] Using charge balance adjust
[46:24] Dispensed 0.005221 mL of Base (0.5 M KOH)
[46:44] Stirrer speed set to 0

| | | | |
|---|---|---|---|
| Sample name: | **M12_octanol** | Experiment start time: | **3/3/2018 4:33:34 PM** |
| Assay name: | **pH-metric high logP** | Analyst: | **Pion** |
| Assay ID: | **18C-03013** | Instrument ID: | **T312060** |
| Filename: | **C:\Sirius_T3\Mehtap\20180302_exp29_logP_T3-2\18C-03013_M12_octanol_pH-metric high logP.t3r** | | |

### Experiment Log (continued)

[46:54] Datapoint id 35 collected
[46:54] Charge balance equation is out by 3.1%
[46:54] Stirrer speed set to 55
[46:59] pH 2.83 -> 3.03
[46:59] Using charge balance adjust
[47:00] Dispensed 0.003975 mL of Base (0.5 M KOH)
[47:20] Stirrer speed set to 0
[47:30] Datapoint id 36 collected
[47:30] Charge balance equation is out by 11.3%
[47:30] Stirrer speed set to 55
[47:35] pH 3.05 -> 3.25
[47:35] Using charge balance adjust
[47:36] Dispensed 0.003198 mL of Base (0.5 M KOH)
[47:56] Stirrer speed set to 0
[48:06] Datapoint id 37 collected
[48:06] Charge balance equation is out by 5.8%
[48:06] Stirrer speed set to 55
[48:12] pH 3.26 -> 3.46
[48:12] Using charge balance adjust
[48:12] Dispensed 0.002634 mL of Base (0.5 M KOH)
[48:32] Stirrer speed set to 0
[48:42] Datapoint id 38 collected
[48:42] Charge balance equation is out by 8.2%
[48:42] Stirrer speed set to 55
[48:47] pH 3.48 -> 3.68
[48:47] Using charge balance adjust
[48:47] Dispensed 0.002093 mL of Base (0.5 M KOH)
[49:07] Stirrer speed set to 0
[49:18] Datapoint id 39 collected
[49:18] Charge balance equation is out by 39.2%
[49:18] Stirrer speed set to 55
[49:23] pH 3.74 -> 3.94
[49:23] Using cautious pH adjust
[49:23] Dispensed 0.000729 mL of Base (0.5 M KOH)
[49:28] Stepping pH = 3.89
[49:28] Dispensed 0.000188 mL of Base (0.5 M KOH)
[49:33] Stepping pH = 3.93
[49:33] Dispensed 0.000071 mL of Base (0.5 M KOH)
[49:39] Stepping pH = 3.94
[49:54] Stirrer speed set to 0
[50:04] Datapoint id 40 collected
[50:04] Charge balance equation is out by 32.9%
[50:04] Stirrer speed set to 55
[50:09] pH 3.93 -> 4.13
[50:09] Using cautious pH adjust
[50:09] Dispensed 0.000541 mL of Base (0.5 M KOH)
[50:14] Stepping pH = 4.19
[50:30] Stirrer speed set to 0
[50:42] Datapoint id 41 collected
[50:42] Charge balance equation is out by 50.0%
[50:42] Stirrer speed set to 55
[50:47] pH 4.16 -> 4.36
[50:47] Using cautious pH adjust
[50:47] Dispensed 0.000353 mL of Base (0.5 M KOH)
[50:52] Stepping pH = 4.33
[50:52] Dispensed 0.000047 mL of Base (0.5 M KOH)
[50:57] Stepping pH = 4.33
[50:57] Dispensed 0.000235 mL of Base (0.5 M KOH)
[51:02] Stepping pH = 4.58

Sample name: **M12_octanol** Experiment start time: **3/3/2018 4:33:34 PM**
Assay name: **pH-metric high logP** Analyst: **Pion**
Assay ID: **18C-03013** Instrument ID: **T312060**
Filename: **C:\Sirius_T3\Mehtap\20180302_exp29_logP_T3-2\18C-03013_M12_octanol_pH-metric high logP.t3r**

### Experiment Log (continued)

[51:17] Stirrer speed set to 0
[52:03] Datapoint id 42 collected
[52:03] Charge balance equation is out by 10.7%
[52:03] Stirrer speed set to 55
[52:08] pH 4.50 -> 4.70
[52:08] Using charge balance adjust
[52:09] Dispensed 0.000353 mL of Base (0.5 M KOH)
[52:29] Stirrer speed set to 0
[53:21] Datapoint id 43 collected
[53:21] Charge balance equation is out by 15.6%
[53:21] Stirrer speed set to 55
[53:26] pH 4.76 -> 4.96
[53:26] Using cautious pH adjust
[53:26] Dispensed 0.000094 mL of Base (0.5 M KOH)
[53:31] Stepping pH = 4.80
[53:31] Dispensed 0.000188 mL of Base (0.5 M KOH)
[53:36] Stepping pH = 5.29
[53:51] Stirrer speed set to 0
[54:40] Datapoint id 44 collected
[54:40] Charge balance equation is out by -45.8%
[54:40] Stirrer speed set to 55
[54:45] pH 5.11 -> 5.31
[54:45] Using cautious pH adjust
[54:45] Dispensed 0.000047 mL of Base (0.5 M KOH)
[54:50] Stepping pH = 5.10
[54:50] Dispensed 0.000259 mL of Base (0.5 M KOH)
[54:55] Stepping pH = 6.40
[55:11] Stirrer speed set to 0
[56:11] Datapoint id 45 collected
[56:11] Charge balance equation is out by -213.5%
[56:11] Stirrer speed set to 55
[56:16] pH 6.18 -> 6.38
[56:16] Using cautious pH adjust
[56:16] Dispensed 0.000024 mL of Base (0.5 M KOH)
[56:21] Stepping pH = 6.15
[56:21] Dispensed 0.000141 mL of Base (0.5 M KOH)
[56:26] Stepping pH = 6.88
[56:41] Stirrer speed set to 0
[57:41] Datapoint id 46 collected
[57:41] Charge balance equation is out by -239.0%
[57:41] Stirrer speed set to 55
[57:46] pH 6.75 -> 6.95
[57:46] Using cautious pH adjust
[57:46] Dispensed 0.000024 mL of Base (0.5 M KOH)
[57:51] Stepping pH = 6.70
[57:51] Dispensed 0.000094 mL of Base (0.5 M KOH)
[57:56] Stepping pH = 7.05
[58:12] Stirrer speed set to 0
[59:12] Datapoint id 47 collected
[59:12] Charge balance equation is out by -295.8%
[59:12] Stirrer speed set to 55
[59:17] pH 6.96 -> 7.16
[59:17] Using cautious pH adjust
[59:17] Dispensed 0.000024 mL of Base (0.5 M KOH)
[59:22] Stepping pH = 6.90
[59:22] Dispensed 0.000071 mL of Base (0.5 M KOH)
[59:27] Stepping pH = 7.18
[59:42] Stirrer speed set to 0
[1:00:42] Datapoint id 48 collected

### Experiment Log (continued)

[1:00:42] Charge balance equation is out by -327.5%
[1:00:42] Stirrer speed set to 55
[1:00:47] pH 7.10 -> 7.30
[1:00:47] Using cautious pH adjust
[1:00:47] Dispensed 0.000024 mL of Base (0.5 M KOH)
[1:00:52] Stepping pH = 7.09
[1:00:52] Dispensed 0.000047 mL of Base (0.5 M KOH)
[1:00:57] Stepping pH = 7.27
[1:00:57] Dispensed 0.000024 mL of Base (0.5 M KOH)
[1:01:03] Stepping pH = 7.45
[1:01:18] Stirrer speed set to 0
[1:02:18] Datapoint id 49 collected
[1:02:18] Charge balance equation is out by -423.0%
[1:02:18] Stirrer speed set to 55
[1:02:23] pH 7.49 -> 7.69
[1:02:23] Using cautious pH adjust
[1:02:23] Dispensed 0.000024 mL of Base (0.5 M KOH)
[1:02:28] Stepping pH = 7.52
[1:02:28] Dispensed 0.000024 mL of Base (0.5 M KOH)
[1:02:33] Stepping pH = 7.64
[1:02:33] Dispensed 0.000024 mL of Base (0.5 M KOH)
[1:02:38] Stepping pH = 7.87
[1:02:53] Stirrer speed set to 0
[1:03:53] Datapoint id 50 collected
[1:03:53] Charge balance equation is out by -597.1%
[1:03:53] Stirrer speed set to 55
[1:03:58] pH 7.64 -> 7.84
[1:03:58] Using cautious pH adjust
[1:03:58] Dispensed 0.000024 mL of Base (0.5 M KOH)
[1:04:04] Stepping pH = 7.62
[1:04:04] Dispensed 0.000024 mL of Base (0.5 M KOH)
[1:04:09] Stepping pH = 7.72
[1:04:09] Dispensed 0.000024 mL of Base (0.5 M KOH)
[1:04:14] Stepping pH = 7.86
[1:04:29] Stirrer speed set to 0
[1:05:29] Datapoint id 51 collected
[1:05:29] Charge balance equation is out by -723.3%
[1:05:29] Stirrer speed set to 55
[1:05:34] pH 7.76 -> 7.96
[1:05:34] Using cautious pH adjust
[1:05:34] Dispensed 0.000024 mL of Base (0.5 M KOH)
[1:05:39] Stepping pH = 7.75
[1:05:39] Dispensed 0.000024 mL of Base (0.5 M KOH)
[1:05:44] Stepping pH = 7.91
[1:05:44] Dispensed 0.000024 mL of Base (0.5 M KOH)
[1:05:49] Stepping pH = 8.15
[1:06:05] Stirrer speed set to 0
[1:07:05] Datapoint id 52 collected
[1:07:05] Charge balance equation is out by -787.5%
[1:07:05] Stirrer speed set to 55
[1:07:10] pH 8.14 -> 8.34
[1:07:10] Using cautious pH adjust
[1:07:10] Dispensed 0.000024 mL of Base (0.5 M KOH)
[1:07:15] Stepping pH = 8.11
[1:07:15] Dispensed 0.000024 mL of Base (0.5 M KOH)
[1:07:20] Stepping pH = 8.17
[1:07:20] Dispensed 0.000024 mL of Base (0.5 M KOH)
[1:07:25] Stepping pH = 8.30
[1:07:25] Dispensed 0.000024 mL of Base (0.5 M KOH)

| | | | |
|---|---|---|---|
| Sample name: | **M12_octanol** | Experiment start time: | **3/3/2018 4:33:34 PM** |
| Assay name: | **pH-metric high logP** | Analyst: | **Pion** |
| Assay ID: | **18C-03013** | Instrument ID: | **T312060** |
| Filename: | **C:\Sirius_T3\Mehtap\20180302_exp29_logP_T3-2\18C-03013_M12_octanol_pH-metric high logP.t3r** | | |

### Experiment Log (continued)

[1:07:30] Stepping pH = 8.46
[1:07:45] Stirrer speed set to 0
[1:08:45] Datapoint id 53 collected
[1:08:45] Charge balance equation is out by -1,072.2%
[1:08:45] Stirrer speed set to 55
[1:08:50] pH 8.40 -> 8.60
[1:08:50] Using cautious pH adjust
[1:08:50] Dispensed 0.000024 mL of Base (0.5 M KOH)
[1:08:56] Stepping pH = 8.40
[1:08:56] Dispensed 0.000024 mL of Base (0.5 M KOH)
[1:09:01] Stepping pH = 8.48
[1:09:01] Dispensed 0.000024 mL of Base (0.5 M KOH)
[1:09:06] Stepping pH = 8.59
[1:09:06] Dispensed 0.000024 mL of Base (0.5 M KOH)
[1:09:11] Stepping pH = 8.69
[1:09:26] Stirrer speed set to 0
[1:10:06] Datapoint id 54 collected
[1:10:06] Charge balance equation is out by -722.4%
[1:10:06] Stirrer speed set to 55
[1:10:11] pH 8.66 -> 8.86
[1:10:11] Using cautious pH adjust
[1:10:11] Dispensed 0.000024 mL of Base (0.5 M KOH)
[1:10:16] Stepping pH = 8.66
[1:10:16] Dispensed 0.000024 mL of Base (0.5 M KOH)
[1:10:21] Stepping pH = 8.72
[1:10:21] Dispensed 0.000047 mL of Base (0.5 M KOH)
[1:10:26] Stepping pH = 8.90
[1:10:41] Stirrer speed set to 0
[1:11:35] Datapoint id 55 collected
[1:11:35] Charge balance equation is out by -320.7%
[1:11:35] Stirrer speed set to 55
[1:11:40] pH 8.88 -> 9.05
[1:11:40] Using cautious pH adjust
[1:11:40] Dispensed 0.000024 mL of Base (0.5 M KOH)
[1:11:45] Stepping pH = 8.87
[1:11:45] Dispensed 0.000071 mL of Base (0.5 M KOH)
[1:11:50] Stepping pH = 9.03
[1:12:05] Stirrer speed set to 0
[1:12:42] Datapoint id 56 collected
[1:12:42] Charge balance equation is out by -231.7%
[1:12:42] Titration 3 of 3
[1:12:42] Adding initial titrants
[1:12:42] Automatically add 0.20000 mL of Octanol
[1:12:47] Dispensed 0.200000 mL of Octanol
[1:12:47] Stirrer speed set to 10
[1:12:48] Stirrer speed set to 60
[1:12:48] Iterative adjust 9.05 -> 2.00
[1:12:48] pH 9.05 -> 2.00
[1:12:50] Dispensed 0.059784 mL of Acid (0.5 M HCl)
[1:12:55] pH 2.02 -> 2.00
[1:12:55] Dispensed 0.002728 mL of Acid (0.5 M HCl)
[1:13:45] Stirrer speed set to 0
[1:13:55] Datapoint id 57 collected
[1:13:55] Stirrer speed set to 60
[1:14:00] pH 1.99 -> 2.19
[1:14:00] Using cautious pH adjust
[1:14:01] Dispensed 0.009619 mL of Base (0.5 M KOH)
[1:14:06] Stepping pH = 2.07
[1:14:06] Dispensed 0.008843 mL of Base (0.5 M KOH)

| | | | |
|---|---|---|---|
| Sample name: | **M12_octanol** | Experiment start time: | **3/3/2018 4:33:34 PM** |
| Assay name: | **pH-metric high logP** | Analyst: | **Pion** |
| Assay ID: | **18C-03013** | Instrument ID: | **T312060** |
| Filename: | **C:\Sirius_T3\Mehtap\20180302_exp29_logP_T3-2\18C-03013_M12_octanol_pH-metric high logP.t3r** | | |

### Experiment Log (continued)

[1:14:11] Stepping pH = 2.16
[1:14:11] Dispensed 0.002375 mL of Base (0.5 M KOH)
[1:14:16] Stepping pH = 2.19
[1:14:32] Stirrer speed set to 0
[1:14:42] Datapoint id 58 collected
[1:14:42] Charge balance equation is out by -8.3%
[1:14:42] Stirrer speed set to 60
[1:14:47] pH 2.19 -> 2.39
[1:14:47] Using charge balance adjust
[1:14:47] Dispensed 0.012653 mL of Base (0.5 M KOH)
[1:15:07] Stirrer speed set to 0
[1:15:17] Datapoint id 59 collected
[1:15:17] Charge balance equation is out by -9.4%
[1:15:17] Stirrer speed set to 60
[1:15:22] pH 2.37 -> 2.57
[1:15:22] Using charge balance adjust
[1:15:23] Dispensed 0.008937 mL of Base (0.5 M KOH)
[1:15:43] Stirrer speed set to 0
[1:15:53] Datapoint id 60 collected
[1:15:53] Charge balance equation is out by -3.8%
[1:15:53] Stirrer speed set to 60
[1:15:59] pH 2.57 -> 2.77
[1:15:59] Using charge balance adjust
[1:15:59] Dispensed 0.006373 mL of Base (0.5 M KOH)
[1:16:19] Stirrer speed set to 0
[1:16:29] Datapoint id 61 collected
[1:16:29] Charge balance equation is out by -0.8%
[1:16:29] Stirrer speed set to 60
[1:16:34] pH 2.78 -> 2.98
[1:16:34] Using charge balance adjust
[1:16:34] Dispensed 0.004680 mL of Base (0.5 M KOH)
[1:16:55] Stirrer speed set to 0
[1:17:05] Datapoint id 62 collected
[1:17:05] Charge balance equation is out by 5.8%
[1:17:05] Stirrer speed set to 60
[1:17:10] pH 3.00 -> 3.20
[1:17:10] Using charge balance adjust
[1:17:10] Dispensed 0.003410 mL of Base (0.5 M KOH)
[1:17:30] Stirrer speed set to 0
[1:17:40] Datapoint id 63 collected
[1:17:40] Charge balance equation is out by 22.0%
[1:17:40] Stirrer speed set to 60
[1:17:45] pH 3.24 -> 3.44
[1:17:45] Using cautious pH adjust
[1:17:45] Dispensed 0.001152 mL of Base (0.5 M KOH)
[1:17:50] Stepping pH = 3.35
[1:17:51] Dispensed 0.000706 mL of Base (0.5 M KOH)
[1:17:56] Stepping pH = 3.43
[1:17:56] Dispensed 0.000141 mL of Base (0.5 M KOH)
[1:18:01] Stepping pH = 3.43
[1:18:01] Dispensed 0.000165 mL of Base (0.5 M KOH)
[1:18:06] Stepping pH = 3.44
[1:18:21] Stirrer speed set to 0
[1:18:31] Datapoint id 64 collected
[1:18:31] Charge balance equation is out by 6.4%
[1:18:31] Stirrer speed set to 60
[1:18:36] pH 3.45 -> 3.65
[1:18:36] Using charge balance adjust
[1:18:36] Dispensed 0.001623 mL of Base (0.5 M KOH)

| | | | |
|---|---|---|---|
| Sample name: | **M12_octanol** | Experiment start time: | **3/3/2018 4:33:34 PM** |
| Assay name: | **pH-metric high logP** | Analyst: | **Pion** |
| Assay ID: | **18C-03013** | Instrument ID: | **T312060** |
| Filename: | **C:\Sirius_T3\Mehtap\20180302_exp29_logP_T3-2\18C-03013_M12_octanol_pH-metric high logP.t3r** | | |

### Experiment Log (continued)

[1:18:57] Stirrer speed set to 0
[1:19:07] Datapoint id 65 collected
[1:19:07] Charge balance equation is out by 64.7%
[1:19:07] Stirrer speed set to 60
[1:19:12] pH 3.79 -> 3.99
[1:19:12] Using cautious pH adjust
[1:19:12] Dispensed 0.000423 mL of Base (0.5 M KOH)
[1:19:17] Stepping pH = 3.90
[1:19:18] Dispensed 0.000212 mL of Base (0.5 M KOH)
[1:19:23] Stepping pH = 3.99
[1:19:38] Stirrer speed set to 0
[1:19:48] Datapoint id 66 collected
[1:19:48] Charge balance equation is out by 25.8%
[1:19:48] Stirrer speed set to 60
[1:19:53] pH 4.00 -> 4.20
[1:19:53] Using cautious pH adjust
[1:19:53] Dispensed 0.000282 mL of Base (0.5 M KOH)
[1:19:58] Stepping pH = 4.15
[1:19:58] Dispensed 0.000071 mL of Base (0.5 M KOH)
[1:20:03] Stepping pH = 4.17
[1:20:03] Dispensed 0.000047 mL of Base (0.5 M KOH)
[1:20:08] Stepping pH = 4.18
[1:20:08] Dispensed 0.000024 mL of Base (0.5 M KOH)
[1:20:14] Stepping pH = 4.18
[1:20:14] Dispensed 0.000212 mL of Base (0.5 M KOH)
[1:20:19] Stepping pH = 4.45
[1:20:34] Stirrer speed set to 0
[1:20:45] Datapoint id 67 collected
[1:20:45] Charge balance equation is out by -17.7%
[1:20:45] Stirrer speed set to 60
[1:20:50] pH 4.46 -> 4.66
[1:20:50] Using cautious pH adjust
[1:20:50] Dispensed 0.000094 mL of Base (0.5 M KOH)
[1:20:55] Stepping pH = 4.56
[1:20:55] Dispensed 0.000071 mL of Base (0.5 M KOH)
[1:21:00] Stepping pH = 4.66
[1:21:15] Stirrer speed set to 0
[1:21:32] Datapoint id 68 collected
[1:21:32] Charge balance equation is out by 13.8%
[1:21:32] Stirrer speed set to 60
[1:21:37] pH 4.64 -> 4.84
[1:21:37] Using charge balance adjust
[1:21:38] Dispensed 0.000141 mL of Base (0.5 M KOH)
[1:21:58] Stirrer speed set to 0
[1:22:11] Datapoint id 69 collected
[1:22:11] Charge balance equation is out by 29.9%
[1:22:11] Stirrer speed set to 60
[1:22:16] pH 4.94 -> 5.14
[1:22:16] Using cautious pH adjust
[1:22:16] Dispensed 0.000047 mL of Base (0.5 M KOH)
[1:22:22] Stepping pH = 4.95
[1:22:22] Dispensed 0.000094 mL of Base (0.5 M KOH)
[1:22:27] Stepping pH = 5.40
[1:22:42] Stirrer speed set to 0
[1:22:57] Datapoint id 70 collected
[1:22:57] Charge balance equation is out by -92.0%
[1:22:57] Stirrer speed set to 60
[1:23:02] pH 5.25 -> 5.45
[1:23:02] Using cautious pH adjust

| | | | |
|---|---|---|---|
| Sample name: | **M12_octanol** | Experiment start time: | **3/3/2018 4:33:34 PM** |
| Assay name: | **pH-metric high logP** | Analyst: | **Pion** |
| Assay ID: | **18C-03013** | Instrument ID: | **T312060** |
| Filename: | **C:\Sirius_T3\Mehtap\20180302_exp29_logP_T3-2\18C-03013_M12_octanol_pH-metric high logP.t3r** | | |

### Experiment Log (continued)

[1:23:03] Dispensed 0.000024 mL of Base (0.5 M KOH)
[1:23:08] Stepping pH = 5.23
[1:23:08] Dispensed 0.000118 mL of Base (0.5 M KOH)
[1:23:13] Stepping pH = 6.55
[1:23:28] Stirrer speed set to 0
[1:24:28] Datapoint id 71 collected
[1:24:28] Charge balance equation is out by -224.6%
[1:24:28] Stirrer speed set to 60
[1:24:33] pH 6.43 -> 6.63
[1:24:33] Using cautious pH adjust
[1:24:33] Dispensed 0.000024 mL of Base (0.5 M KOH)
[1:24:38] Stepping pH = 6.42
[1:24:38] Dispensed 0.000118 mL of Base (0.5 M KOH)
[1:24:43] Stepping pH = 7.17
[1:24:58] Stirrer speed set to 0
[1:25:59] Datapoint id 72 collected
[1:25:59] Charge balance equation is out by -216.2%
[1:25:59] Stirrer speed set to 60
[1:26:04] pH 7.15 -> 7.35
[1:26:04] Using cautious pH adjust
[1:26:04] Dispensed 0.000024 mL of Base (0.5 M KOH)
[1:26:09] Stepping pH = 7.17
[1:26:09] Dispensed 0.000024 mL of Base (0.5 M KOH)
[1:26:14] Stepping pH = 7.23
[1:26:14] Dispensed 0.000024 mL of Base (0.5 M KOH)
[1:26:19] Stepping pH = 7.25
[1:26:19] Dispensed 0.000047 mL of Base (0.5 M KOH)
[1:26:24] Stepping pH = 7.50
[1:26:39] Stirrer speed set to 0
[1:27:39] Datapoint id 73 collected
[1:27:39] Charge balance equation is out by -583.7%
[1:27:39] Stirrer speed set to 60
[1:27:44] pH 7.42 -> 7.62
[1:27:44] Using cautious pH adjust
[1:27:44] Dispensed 0.000024 mL of Base (0.5 M KOH)
[1:27:49] Stepping pH = 7.42
[1:27:49] Dispensed 0.000024 mL of Base (0.5 M KOH)
[1:27:55] Stepping pH = 7.46
[1:27:55] Dispensed 0.000024 mL of Base (0.5 M KOH)
[1:28:00] Stepping pH = 7.58
[1:28:00] Dispensed 0.000024 mL of Base (0.5 M KOH)
[1:28:05] Stepping pH = 7.75
[1:28:20] Stirrer speed set to 0
[1:29:20] Datapoint id 74 collected
[1:29:20] Charge balance equation is out by -640.0%
[1:29:20] Stirrer speed set to 60
[1:29:25] pH 7.76 -> 7.96
[1:29:25] Using cautious pH adjust
[1:29:25] Dispensed 0.000024 mL of Base (0.5 M KOH)
[1:29:30] Stepping pH = 7.85
[1:29:30] Dispensed 0.000024 mL of Base (0.5 M KOH)
[1:29:35] Stepping pH = 7.93
[1:29:35] Dispensed 0.000024 mL of Base (0.5 M KOH)
[1:29:40] Stepping pH = 8.03
[1:29:56] Stirrer speed set to 0
[1:30:56] Datapoint id 75 collected
[1:30:56] Charge balance equation is out by -626.4%
[1:30:56] Stirrer speed set to 60
[1:31:01] pH 7.91 -> 8.11

| | | | |
|---|---|---|---|
| Sample name: | **M12_octanol** | Experiment start time: | **3/3/2018 4:33:34 PM** |
| Assay name: | **pH-metric high logP** | Analyst: | **Pion** |
| Assay ID: | **18C-03013** | Instrument ID: | **T312060** |
| Filename: | **C:\Sirius_T3\Mehtap\20180302_exp29_logP_T3-2\18C-03013_M12_octanol_pH-metric high logP.t3r** | | |

### Experiment Log (continued)

[1:31:01] Using cautious pH adjust
[1:31:01] Dispensed 0.000024 mL of Base (0.5 M KOH)
[1:31:06] Stepping pH = 7.96
[1:31:06] Dispensed 0.000024 mL of Base (0.5 M KOH)
[1:31:11] Stepping pH = 8.09
[1:31:11] Dispensed 0.000024 mL of Base (0.5 M KOH)
[1:31:16] Stepping pH = 8.32
[1:31:31] Stirrer speed set to 0
[1:32:31] Datapoint id 76 collected
[1:32:31] Charge balance equation is out by -645.7%
[1:32:31] Stirrer speed set to 60
[1:32:36] pH 8.27 -> 8.47
[1:32:36] Using cautious pH adjust
[1:32:36] Dispensed 0.000024 mL of Base (0.5 M KOH)
[1:32:41] Stepping pH = 8.32
[1:32:41] Dispensed 0.000024 mL of Base (0.5 M KOH)
[1:32:47] Stepping pH = 8.46
[1:32:47] Dispensed 0.000024 mL of Base (0.5 M KOH)
[1:32:52] Stepping pH = 8.58
[1:33:07] Stirrer speed set to 0
[1:33:44] Datapoint id 77 collected
[1:33:44] Charge balance equation is out by -426.1%
[1:33:44] Stirrer speed set to 60
[1:33:49] pH 8.51 -> 8.71
[1:33:49] Using cautious pH adjust
[1:33:49] Dispensed 0.000024 mL of Base (0.5 M KOH)
[1:33:54] Stepping pH = 8.53
[1:33:54] Dispensed 0.000024 mL of Base (0.5 M KOH)
[1:33:59] Stepping pH = 8.59
[1:33:59] Dispensed 0.000024 mL of Base (0.5 M KOH)
[1:34:04] Stepping pH = 8.72
[1:34:19] Stirrer speed set to 0
[1:35:17] Datapoint id 78 collected
[1:35:17] Charge balance equation is out by -316.1%
[1:35:17] Stirrer speed set to 60
[1:35:22] pH 8.65 -> 8.85
[1:35:22] Using cautious pH adjust
[1:35:22] Dispensed 0.000024 mL of Base (0.5 M KOH)
[1:35:27] Stepping pH = 8.63
[1:35:27] Dispensed 0.000071 mL of Base (0.5 M KOH)
[1:35:32] Stepping pH = 8.91
[1:35:48] Stirrer speed set to 0
[1:36:05] Datapoint id 79 collected
[1:36:05] Charge balance equation is out by -257.2%
[1:36:05] Stirrer speed set to 60
[1:36:10] pH 8.90 -> 9.05
[1:36:10] Using cautious pH adjust
[1:36:10] Dispensed 0.000024 mL of Base (0.5 M KOH)
[1:36:15] Stepping pH = 8.92
[1:36:15] Dispensed 0.000024 mL of Base (0.5 M KOH)
[1:36:20] Stepping pH = 8.94
[1:36:21] Dispensed 0.000094 mL of Base (0.5 M KOH)
[1:36:26] Stepping pH = 9.14
[1:36:41] Stirrer speed set to 0
[1:36:54] Datapoint id 80 collected
[1:36:54] Charge balance equation is out by -421.1%
[1:36:54] Argon flow rate set to 0
[1:36:58] Titrator arm moved over Titration position
