## Supplementary material for "Octanol-water partition coefficient measurements for the SAMPL6 Blind Prediction Challenge": SM12_18C-03014_M12_octanol_pH-metric high logP_report.pdf

| | | | |
|---|---|---|---|
| Sample name: | **M12_octanol** | Experiment start time: | **3/3/2018 6:11:21 PM** |
| Assay name: | **pH-metric high logP** | Analyst: | **Pion** |
| Assay ID: | **18C-03014** | Instrument ID: | **T312060** |
| Filename: | **C:\Sirius_T3\Mehtap\20180302_exp29_logP_T3-2\18C-03014_M12_octanol_pH-metric high logP.t3r** | | |

#### pH-metric Result

| | | |
|---|---|---|
| logP (XH +) | 0.62 | ±0.05 (n=50) |
| logP (neutral X) | 3.91 | ±0.01 (n=50) |

#### 18C-03014 Points 1 to 25

| | |
|---|---|
| M12_octanol concentration factor | 0.960 |
| Carbonate | 0.5327 mM |
| Acidity error | -1.77424 mM |

#### 18C-03014 Points 26 to 51

| | |
|---|---|
| M12_octanol concentration factor | 1.007 |
| Carbonate | 0.2455 mM |
| Acidity error | -1.88168 mM |

#### 18C-03014 Points 52 to 76

| | |
|---|---|
| M12_octanol concentration factor | 1.171 |
| Carbonate | 0.3091 mM |
| Acidity error | -3.49881 mM |

#### Warnings and errors

| | |
|---|---|
| Errors | None |
| Warnings | Excessive carbonate concentration present |

#### Sample logD and percent species

| pH | M12_octanol logD | M12_octanol M12_octanolH | M12_octanol M12_octanol | M12_octanol M12_octanolH* | M12_octanol M12_octanol* | Comment |
|---|---|---|---|---|---|---|
| 1.000 | 0.66 | 17.81 % | 0.00 % | 74.63 % | 7.56 % | |
| 1.200 | 0.69 | 17.06 % | 0.00 % | 71.47 % | 11.47 % | Stomach pH |
| 2.000 | 0.93 | 10.60 % | 0.01 % | 44.41 % | 44.98 % | |
| 3.000 | 1.67 | 2.10 % | 0.01 % | 8.80 % | 89.09 % | |
| 4.000 | 2.61 | 0.23 % | 0.01 % | 0.98 % | 98.78 % | |
| 5.000 | 3.45 | 0.02 % | 0.01 % | 0.10 % | 99.87 % | |
| 6.000 | 3.83 | 0.00 % | 0.01 % | 0.01 % | 99.98 % | |
| 6.500 | 3.88 | 0.00 % | 0.01 % | 0.00 % | 99.98 % | |
| 7.000 | 3.90 | 0.00 % | 0.01 % | 0.00 % | 99.99 % | |
| 7.400 | 3.90 | 0.00 % | 0.01 % | 0.00 % | 99.99 % | Blood pH |
| 8.000 | 3.91 | 0.00 % | 0.01 % | 0.00 % | 99.99 % | |
| 9.000 | 3.91 | 0.00 % | 0.01 % | 0.00 % | 99.99 % | |
| 10.000 | 3.91 | 0.00 % | 0.01 % | 0.00 % | 99.99 % | |
| 11.000 | 3.91 | 0.00 % | 0.01 % | 0.00 % | 99.99 % | |
| 12.000 | 3.91 | 0.00 % | 0.01 % | 0.00 % | 99.99 % | |

| | | | |
|---|---|---|---|
| Sample name: | **M12_octanol** | Experiment start time: | **3/3/2018 6:11:21 PM** |
| Assay name: | **pH-metric high logP** | Analyst: | **Pion** |
| Assay ID: | **18C-03014** | Instrument ID: | **T312060** |
| Filename: | **C:\Sirius_T3\Mehtap\20180302_exp29_logP_T3-2\18C-03014_M12_octanol_pH-metric high logP.t3r** | | |

#### Graphs

Charge balance residuals

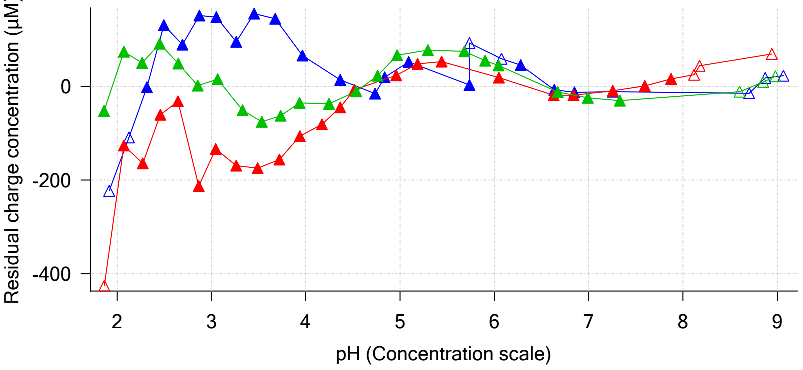


Ionic strength

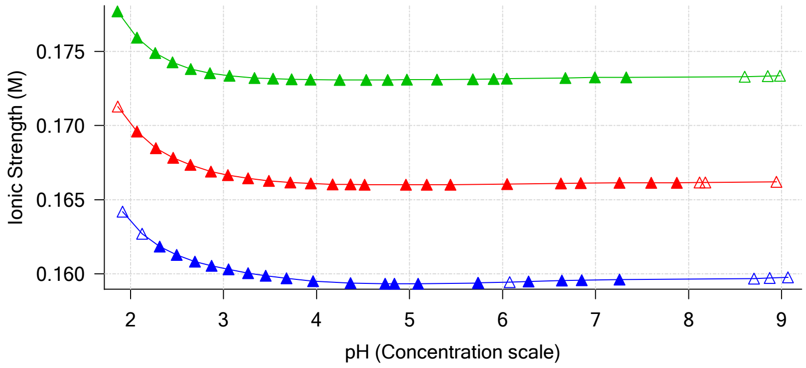


Buffer index profile

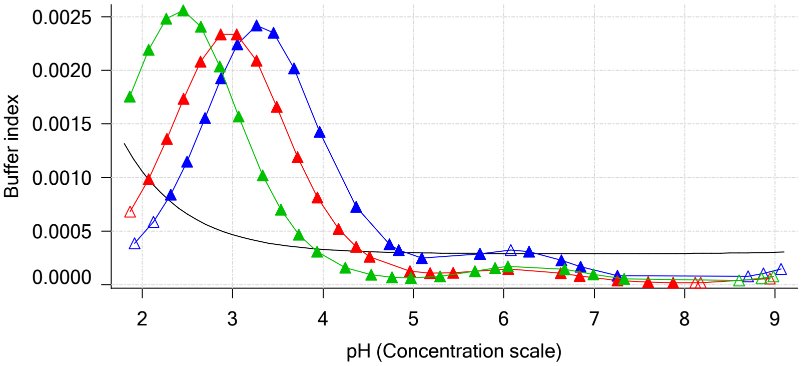


Sample M12_octanol Ionisation Graph

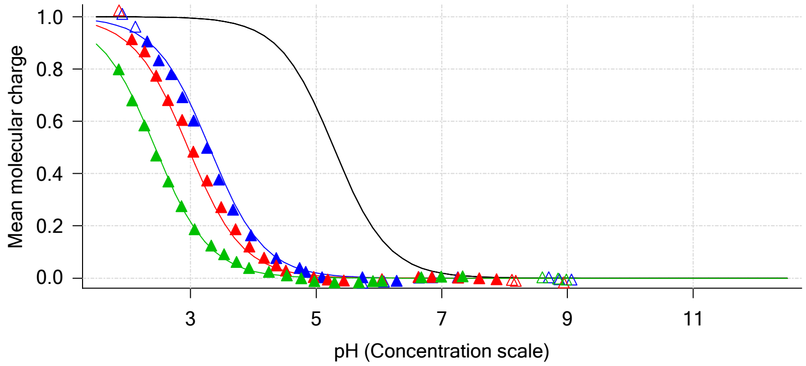


Distribution of species for sample M12_octanol

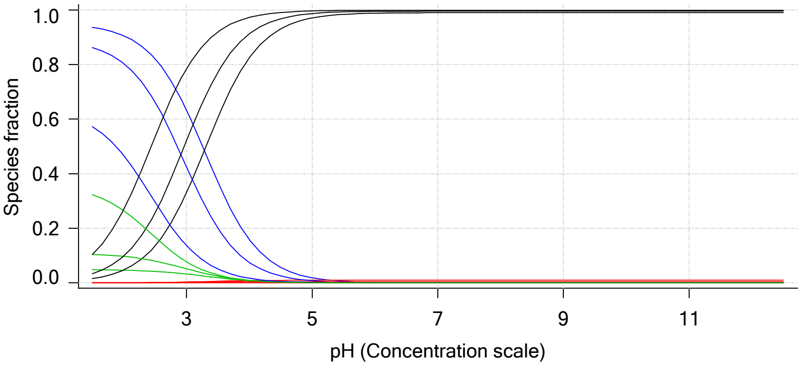


Lipophilicity Profile

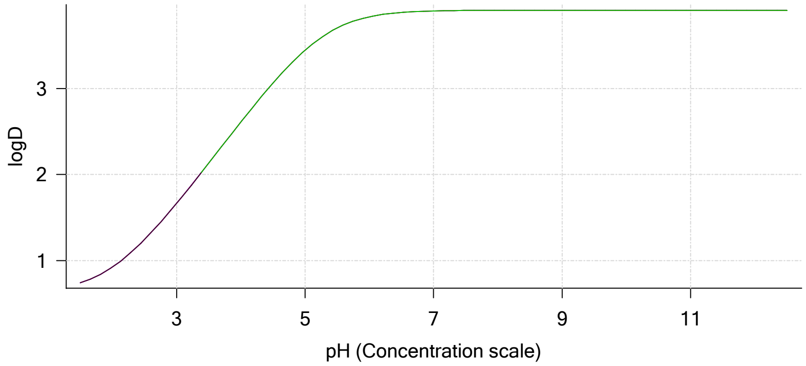


Log R Profile

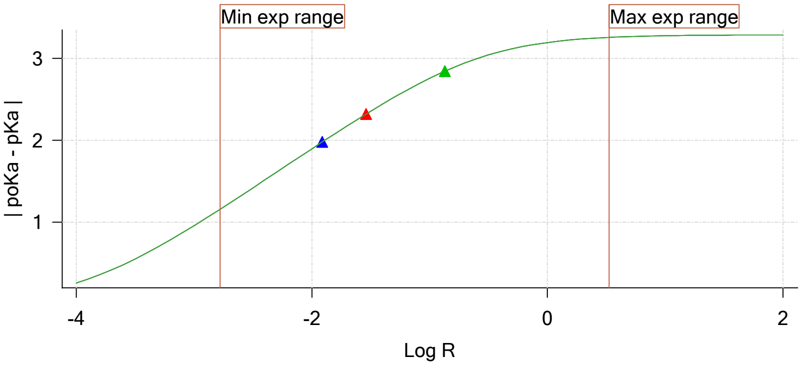


Sample Carbonate Ionisation Graph

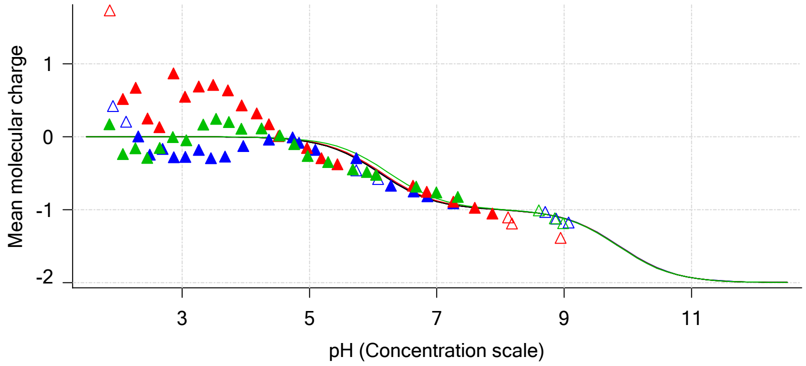

| | | | |
|---|---|---|---|
| Sample name: | **M12_octanol** | Experiment start time: | **3/3/2018 6:11:21 PM** |
| Assay name: | **pH-metric high logP** | Analyst: | **Pion** |
| Assay ID: | **18C-03014** | Instrument ID: | **T312060** |
| Filename: | **C:\Sirius_T3\Mehtap\20180302_exp29_logP_T3-2\18C-03014_M12_octanol_pH-metric high logP.t3r** | | |

#### Graphs (continued)

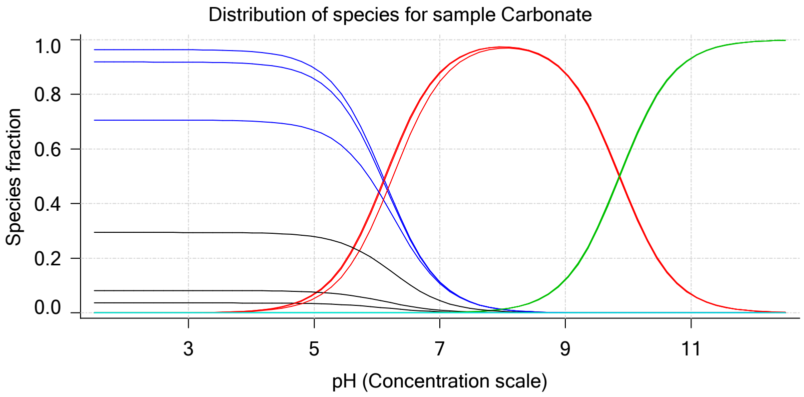

| | | | |
|---|---|---|---|
| Sample name: | **M12_octanol** | Experiment start time: | **3/3/2018 6:11:21 PM** |
| Assay name: | **pH-metric high logP** | Analyst: | **Pion** |
| Assay ID: | **18C-03014** | Instrument ID: | **T312060** |
| Filename: | **C:\Sirius_T3\Mehtap\20180302_exp29_logP_T3-2\18C-03014_M12_octanol_pH-metric high logP.t3r** | | |

### pH-metric high logP Titration 1 of 3 18C-03014 Points 1 to 25

#### Overall results

| | |
|---|---|
| RMSD | 0.070 |
| Average ionic strength | 0.160 M |
| Average temperature | 24.9°C |
| Partition ratio | 0.0123 : 1 |
| Analyte concentration range | 4301.0 μM to 4446.8 μM |
| Total points considered | 18 of 25 |

#### Warnings and errors

| | |
|---|---|
| Errors | None |
| Warnings | Excessive carbonate concentration present |
| | Excessive acidity error present |

#### Four-Plus parameters

| | | | |
|---|---|---|---|
| Alpha | 0.111 | 3/3/2018 6:11:21 PM | C:\Sirius_T3\HCl18C02.t3r |
| S | 0.9988 | 3/3/2018 6:11:21 PM | C:\Sirius_T3\HCl18C02.t3r |
| jH | 1.0 | 3/3/2018 6:11:21 PM | C:\Sirius_T3\HCl18C02.t3r |
| jOH | -0.8 | 3/3/2018 6:11:21 PM | C:\Sirius_T3\HCl18C02.t3r |

#### Titrants

| | | | |
|---|---|---|---|
| 0.50 M HCl | 0.999058 | 3/3/2018 6:11:21 PM | C:\Sirius_T3\HCl18C02.t3r |
| 0.50 M KOH | 0.999845 | 3/3/2018 6:11:21 PM | C:\Sirius_T3\KOH18B27.t3r |

#### Sample

| | |
|---|---|
| M12_octanol concentration factor | 0.960 |
| M12_octanol stoichiometry | 1.000 |
| Chloride stoichiometry | 1.000 |
| Base pKa 1 | 5.28 |
| logP (XH +) | 0.88 |
| logP (neutral X) | 3.99 |

#### Sample graphs

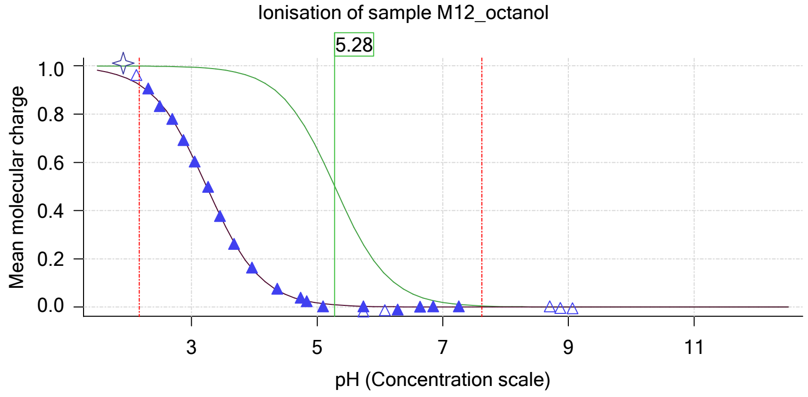


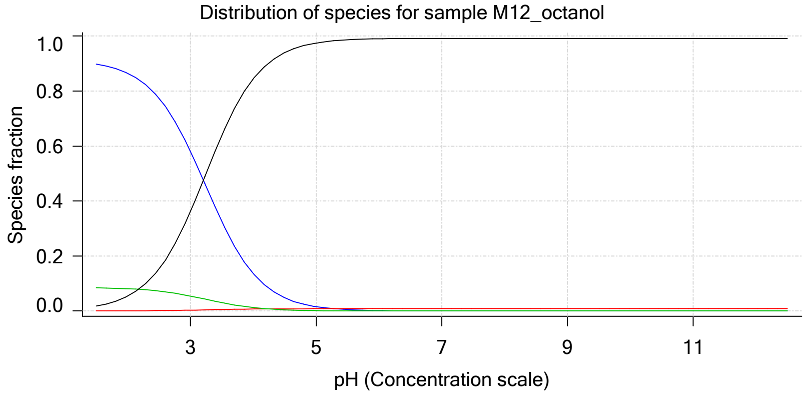

Sample name: **M12_octanol**
Assay name: **pH-metric high logP**
Assay ID: **18C-03014**
Filename: **C:\Sirius_T3\Mehtap\20180302_exp29_logP_T3-2\18C-03014_M12_octanol_pH-metric high logP.t3r**
Experiment start time: **3/3/2018 6:11:21 PM**
Analyst: **Pion**
Instrument ID: **T312060**

#### Sample graphs (continued)

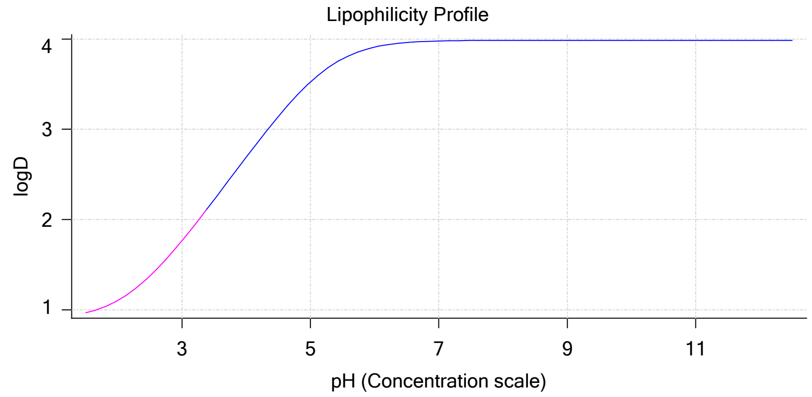


#### Sample logD and percent species

| pH | M12_octanol logD | M12_octanol M12_octanolH | M12_octanol M12_octanol | M12_octanol M12_octanolH* | M12_octanol M12_octanol* | Comment |
|---|---|---|---|---|---|---|
| 1.000 | 0.91 | 90.92 % | 0.00 % | 8.51 % | 0.57 % | |
| 1.200 | 0.93 | 90.62 % | 0.01 % | 8.48 % | 0.90 % | Stomach pH |
| 2.000 | 1.10 | 86.47 % | 0.05 % | 8.09 % | 5.39 % | |
| 3.000 | 1.76 | 58.05 % | 0.30 % | 5.43 % | 36.21 % | |
| 4.000 | 2.69 | 13.54 % | 0.71 % | 1.27 % | 84.48 % | |
| 5.000 | 3.52 | 1.56 % | 0.82 % | 0.15 % | 97.47 % | |
| 6.000 | 3.91 | 0.16 % | 0.83 % | 0.01 % | 98.99 % | |
| 6.500 | 3.96 | 0.05 % | 0.83 % | 0.00 % | 99.11 % | |
| 7.000 | 3.98 | 0.02 % | 0.83 % | 0.00 % | 99.15 % | |
| 7.400 | 3.98 | 0.01 % | 0.83 % | 0.00 % | 99.16 % | Blood pH |
| 8.000 | 3.99 | 0.00 % | 0.83 % | 0.00 % | 99.16 % | |
| 9.000 | 3.99 | 0.00 % | 0.83 % | 0.00 % | 99.17 % | |
| 10.000 | 3.99 | 0.00 % | 0.83 % | 0.00 % | 99.17 % | |
| 11.000 | 3.99 | 0.00 % | 0.83 % | 0.00 % | 99.17 % | |
| 12.000 | 3.99 | 0.00 % | 0.83 % | 0.00 % | 99.17 % | |

#### Carbonate and acidity

| | |
|---|---|
| Carbonate | 0.533 mM |
| Acidity error | -1.774 mM |

#### Other graphs

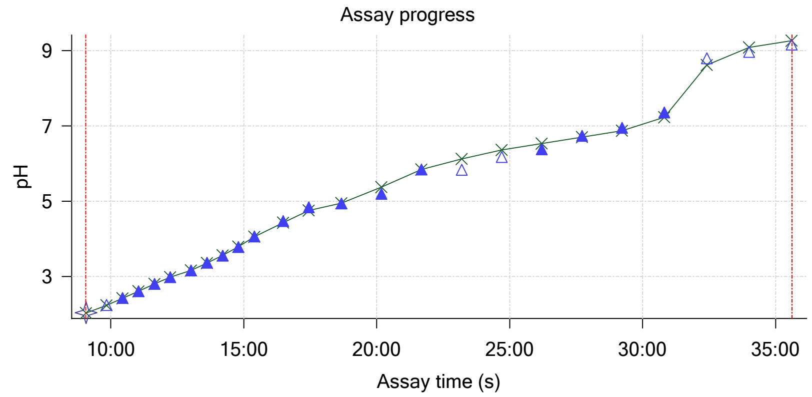


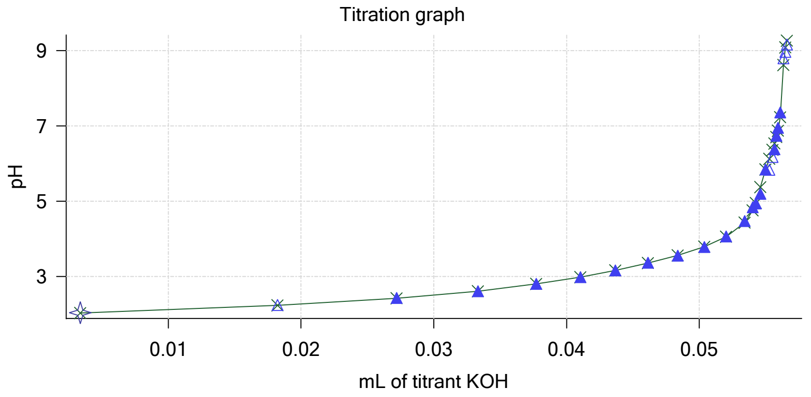

Sample name: **M12_octanol**
Assay name: **pH-metric high logP**
Assay ID: **18C-03014**
Filename: **C:\Sirius_T3\Mehtap\20180302_exp29_logP_T3-2\18C-03014_M12_octanol_pH-metric high logP.t3r**

Experiment start time: **3/3/2018 6:11:21 PM**
Analyst: **Pion**
Instrument ID: **T312060**

#### Other graphs (continued)

Charge balance residuals

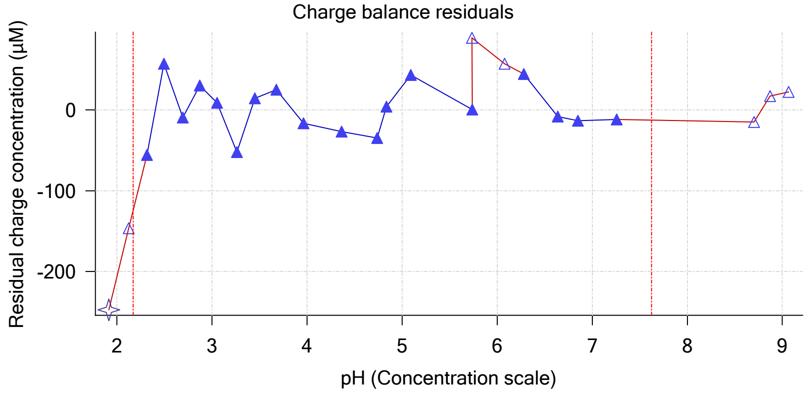


Buffer index profile

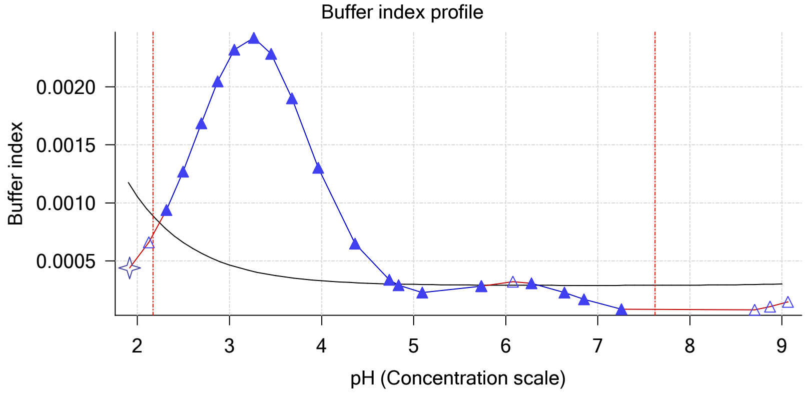


Ionic strength

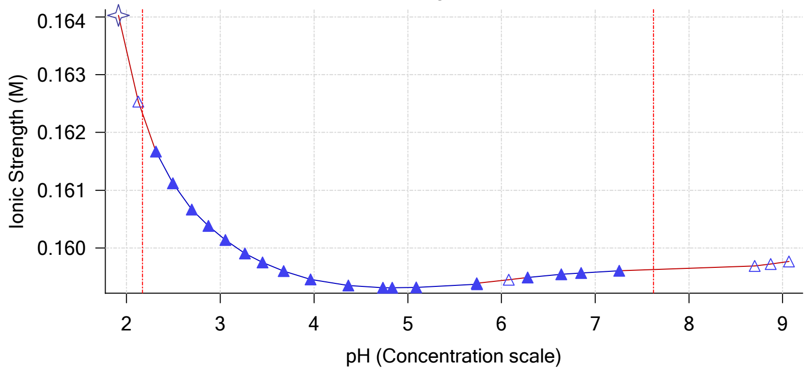


Log R Profile

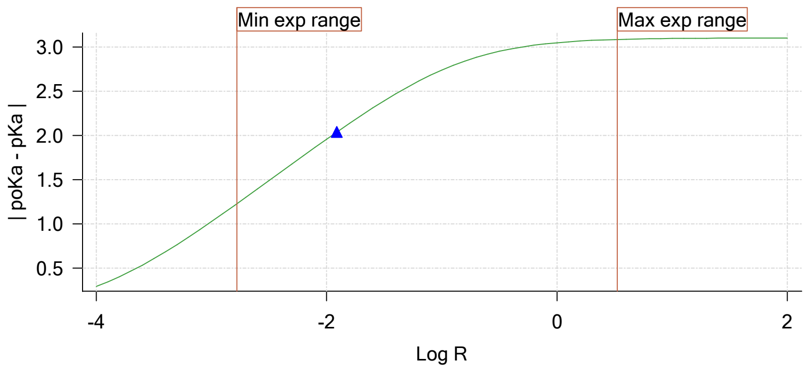


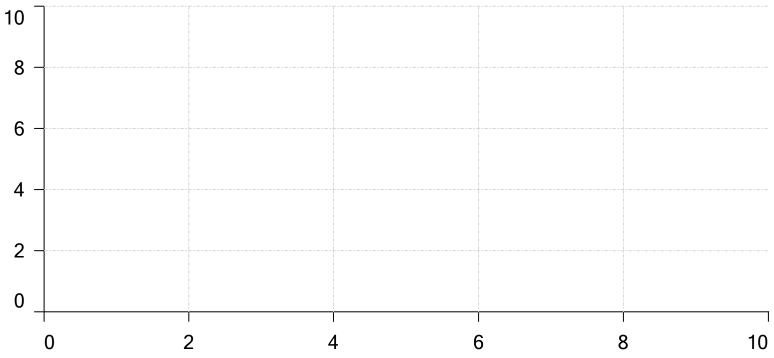


Ionisation of sample Carbonate

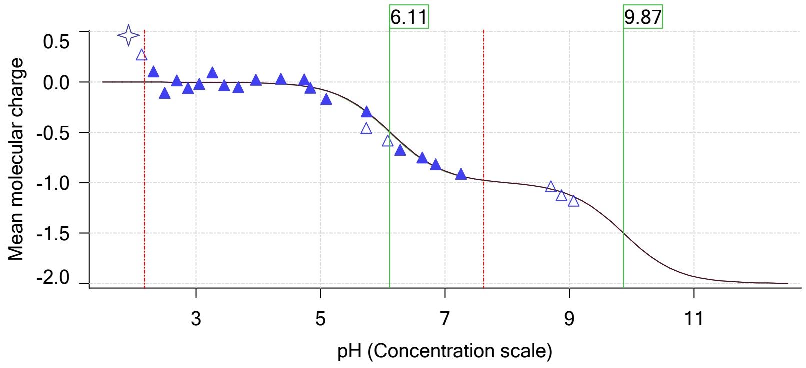

| | | | |
|---|---|---|---|
| Sample name: | **M12_octanol** | Experiment start time: | **3/3/2018 6:11:21 PM** |
| Assay name: | **pH-metric high logP** | Analyst: | **Pion** |
| Assay ID: | **18C-03014** | Instrument ID: | **T312060** |
| Filename: | **C:\Sirius_T3\Mehtap\20180302_exp29_logP_T3-2\18C-03014_M12_octanol_pH-metric high logP.t3r** | | |

# pH-metric high logP   Titration 2 of 3   18C-03014 Points 26 to 51

#### Overall results

| | |
|---|---|
| RMSD | 0.236 |
| Average ionic strength | 0.166 M |
| Average temperature | 25.0°C |
| Partition ratio | 0.0288 : 1 |
| Analyte concentration range | 3930.3 µM to 4070.1 µM |
| Total points considered | 22 of 26 |

#### Warnings and errors

| | |
|---|---|
| Errors | None |
| Warnings | Excessive acidity error present |

#### Four-Plus parameters

| | | | |
|---|---|---|---|
| Alpha | 0.111 | 3/3/2018 6:11:21 PM | C:\Sirius_T3\HCl18C02.t3r |
| S | 0.9988 | 3/3/2018 6:11:21 PM | C:\Sirius_T3\HCl18C02.t3r |
| jH | 1.0 | 3/3/2018 6:11:21 PM | C:\Sirius_T3\HCl18C02.t3r |
| jOH | -0.8 | 3/3/2018 6:11:21 PM | C:\Sirius_T3\HCl18C02.t3r |

#### Titrants

| | | | |
|---|---|---|---|
| 0.50 M HCl | 0.999058 | 3/3/2018 6:11:21 PM | C:\Sirius_T3\HCl18C02.t3r |
| 0.50 M KOH | 0.999845 | 3/3/2018 6:11:21 PM | C:\Sirius_T3\KOH18B27.t3r |

#### Sample

| | |
|---|---|
| M12_octanol concentration factor | 1.007 |
| M12_octanol stoichiometry | 1.000 |
| Chloride stoichiometry | 1.000 |
| Base pKa 1 | 5.28 |
| logP (XH +) | 1.02 |
| logP (neutral X) | 3.90 |

#### Sample graphs

Ionisation of sample M12_octanol

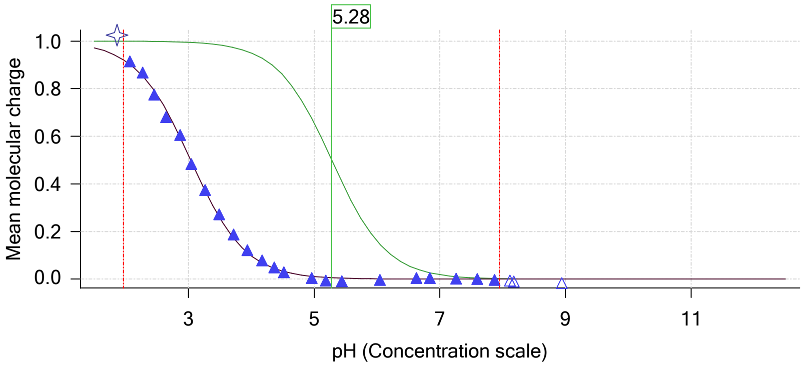


Distribution of species for sample M12_octanol

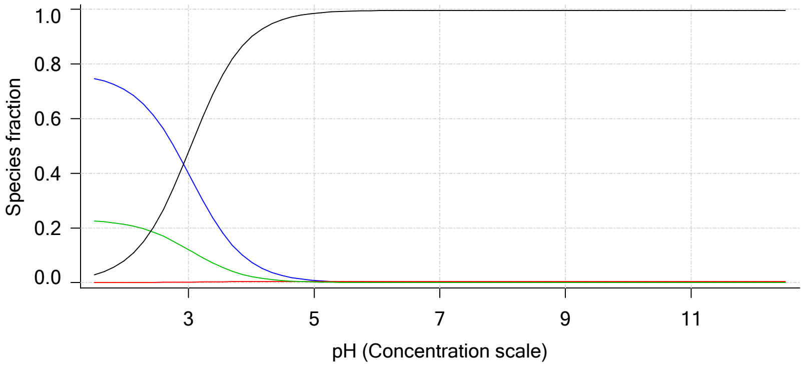

| | | | |
|---|---|---|---|
| Sample name: | **M12_octanol** | Experiment start time: | **3/3/2018 6:11:21 PM** |
| Assay name: | **pH-metric high logP** | Analyst: | **Pion** |
| Assay ID: | **18C-03014** | Instrument ID: | **T312060** |
| Filename: | **C:\Sirius_T3\Mehtap\20180302_exp29_logP_T3-2\18C-03014_M12_octanol_pH-metric high logP.t3r** | | |

#### Sample graphs (continued)

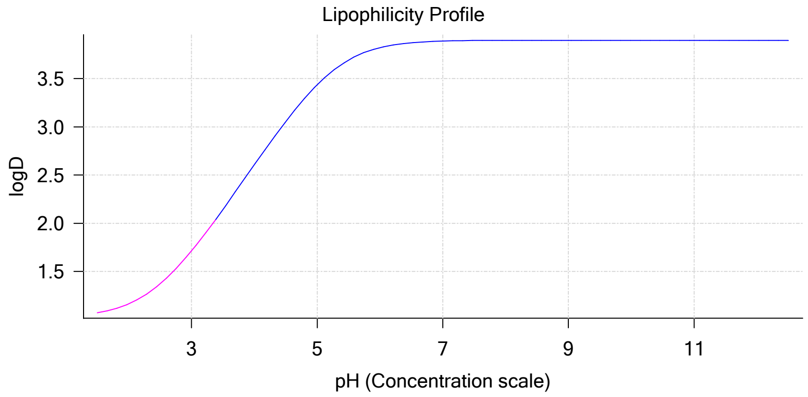


#### Sample logD and percent species

| pH | M12_octanol logD | M12_octanol M12_octanolH | M12_octanol M12_octanol | M12_octanol M12_octanolH* | M12_octanol M12_octanol* | Comment |
|---|---|---|---|---|---|---|
| 1.000 | 1.04 | 76.11 % | 0.00 % | 22.98 % | 0.91 % | |
| 1.200 | 1.05 | 75.71 % | 0.01 % | 22.85 % | 1.43 % | Stomach pH |
| 2.000 | 1.16 | 70.33 % | 0.04 % | 21.23 % | 8.40 % | |
| 3.000 | 1.71 | 39.98 % | 0.21 % | 12.07 % | 47.74 % | |
| 4.000 | 2.61 | 7.52 % | 0.39 % | 2.27 % | 89.81 % | |
| 5.000 | 3.44 | 0.82 % | 0.43 % | 0.25 % | 98.49 % | |
| 6.000 | 3.82 | 0.08 % | 0.44 % | 0.03 % | 99.45 % | |
| 6.500 | 3.87 | 0.03 % | 0.44 % | 0.01 % | 99.53 % | |
| 7.000 | 3.89 | 0.01 % | 0.44 % | 0.00 % | 99.55 % | |
| 7.400 | 3.89 | 0.00 % | 0.44 % | 0.00 % | 99.56 % | Blood pH |
| 8.000 | 3.90 | 0.00 % | 0.44 % | 0.00 % | 99.56 % | |
| 9.000 | 3.90 | 0.00 % | 0.44 % | 0.00 % | 99.56 % | |
| 10.000 | 3.90 | 0.00 % | 0.44 % | 0.00 % | 99.56 % | |
| 11.000 | 3.90 | 0.00 % | 0.44 % | 0.00 % | 99.56 % | |
| 12.000 | 3.90 | 0.00 % | 0.44 % | 0.00 % | 99.56 % | |

#### Carbonate and acidity

| | |
|---|---|
| Carbonate | 0.246 mM |
| Acidity error | -1.882 mM |

#### Other graphs

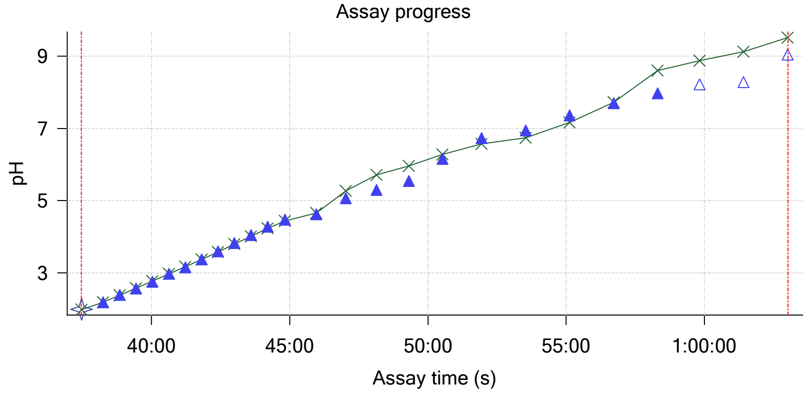


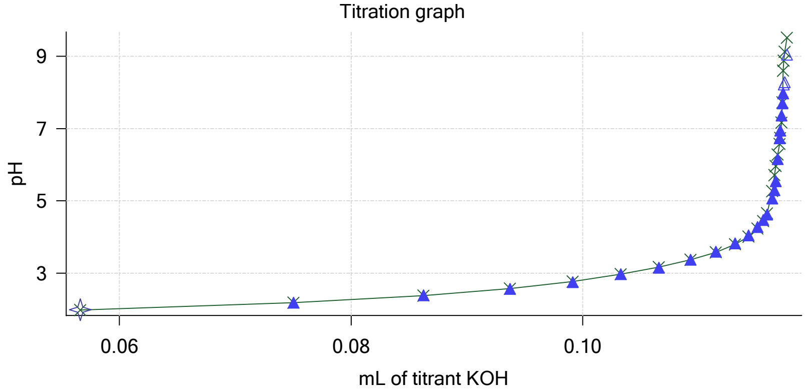

Sample name: **M12_octanol**
Assay name: **pH-metric high logP**
Assay ID: **18C-03014**
Filename: **C:\Sirius_T3\Mehtap\20180302_exp29_logP_T3-2\18C-03014_M12_octanol_pH-metric high logP.t3r**

Experiment start time: **3/3/2018 6:11:21 PM**
Analyst: **Pion**
Instrument ID: **T312060**

#### Other graphs (continued)

Charge balance residuals

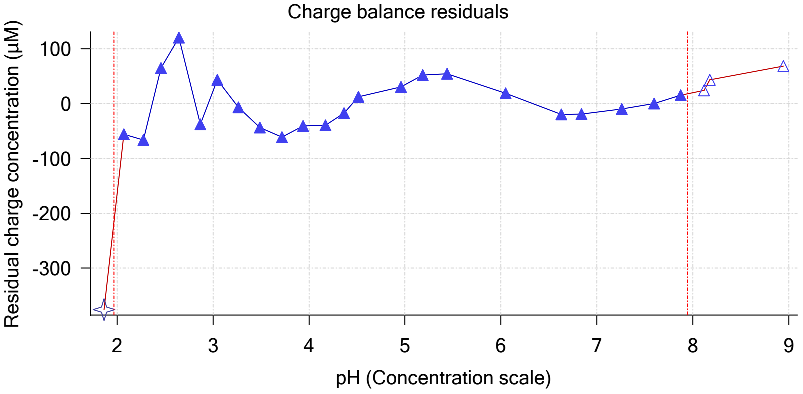


Buffer index profile

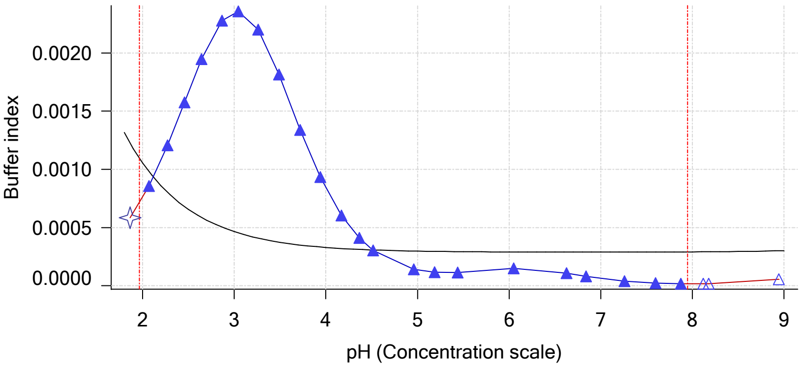


Ionic strength

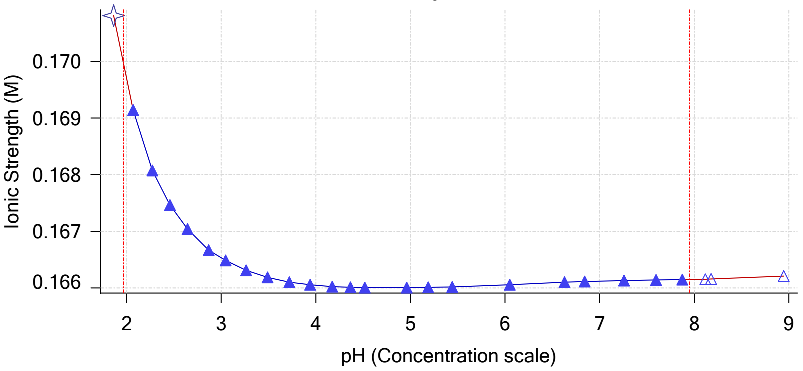


Log R Profile

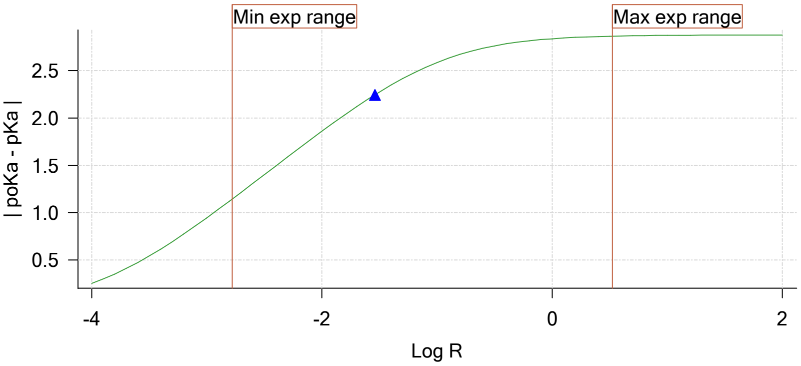


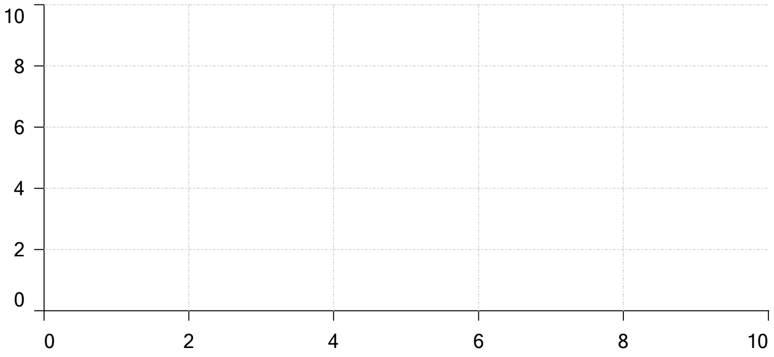


Ionisation of sample Carbonate

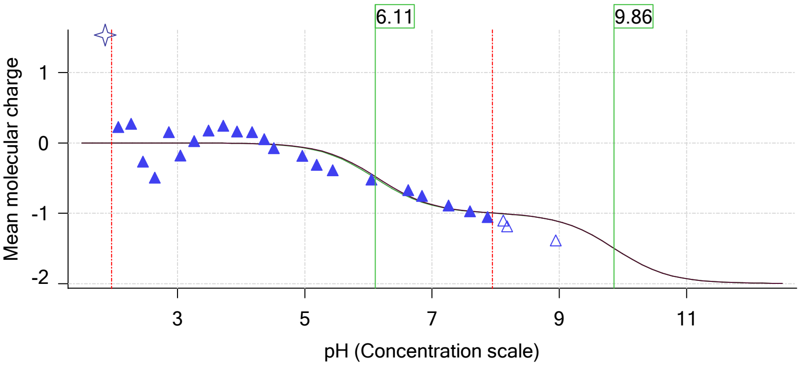

| | | | |
|---|---|---|---|
| Sample name: | **M12_octanol** | Experiment start time: | **3/3/2018 6:11:21 PM** |
| Assay name: | **pH-metric high logP** | Analyst: | **Pion** |
| Assay ID: | **18C-03014** | Instrument ID: | **T312060** |
| Filename: | **C:\Sirius_T3\Mehtap\20180302_exp29_logP_T3-2\18C-03014_M12_octanol_pH-metric high logP.t3r** | | |

### pH-metric high logP Titration 3 of 3 18C-03014 Points 52 to 76

#### Overall results

| | |
|---|---|
| RMSD | 0.325 |
| Average ionic strength | 0.174 M |
| Average temperature | 25.0°C |
| Partition ratio | 0.1350 : 1 |
| Analyte concentration range | 3312.8 µM to 3415.8 µM |
| Total points considered | 22 of 25 |

#### Warnings and errors

| | |
|---|---|
| Errors | None |
| Warnings | Excessive acidity error present |

#### Four-Plus parameters

| | | | |
|---|---|---|---|
| Alpha | 0.111 | 3/3/2018 6:11:21 PM | C:\Sirius_T3\HCl18C02.t3r |
| S | 0.9988 | 3/3/2018 6:11:21 PM | C:\Sirius_T3\HCl18C02.t3r |
| jH | 1.0 | 3/3/2018 6:11:21 PM | C:\Sirius_T3\HCl18C02.t3r |
| jOH | -0.8 | 3/3/2018 6:11:21 PM | C:\Sirius_T3\HCl18C02.t3r |

#### Titrants

| | | | |
|---|---|---|---|
| 0.50 M HCl | 0.999058 | 3/3/2018 6:11:21 PM | C:\Sirius_T3\HCl18C02.t3r |
| 0.50 M KOH | 0.999845 | 3/3/2018 6:11:21 PM | C:\Sirius_T3\KOH18B27.t3r |

#### Sample

| | |
|---|---|
| M12_octanol concentration factor | 1.171 |
| M12_octanol stoichiometry | 1.000 |
| Chloride stoichiometry | 1.000 |
| Base pKa 1 | 5.28 |
| logP (XH +) | 1.02 |
| logP (neutral X) | 4.11 |

#### Sample graphs

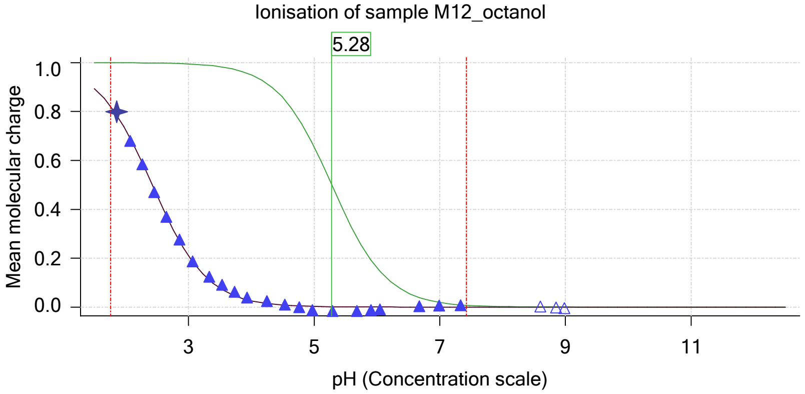


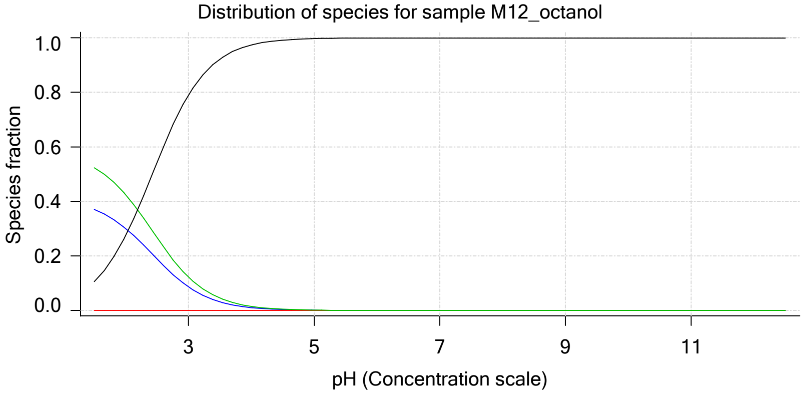

| | | | |
|---|---|---|---|
| Sample name: | **M12_octanol** | Experiment start time: | **3/3/2018 6:11:21 PM** |
| Assay name: | **pH-metric high logP** | Analyst: | **Pion** |
| Assay ID: | **18C-03014** | Instrument ID: | **T312060** |
| Filename: | **C:\Sirius_T3\Mehtap\20180302_exp29_logP_T3-2\18C-03014_M12_octanol_pH-metric high logP.t3r** | | |

#### Sample graphs (continued)

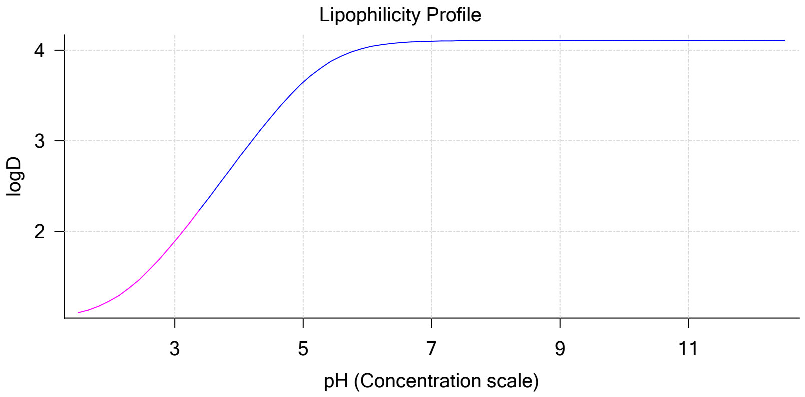


#### Sample logD and percent species

| pH | M12_octanol logD | M12_octanol M12_octanolH | M12_octanol M12_octanol | M12_octanol M12_octanolH* | M12_octanol M12_octanol* | Comment |
|---|---|---|---|---|---|---|
| 1.000 | 1.05 | 39.92 % | 0.00 % | 56.44 % | 3.63 % | |
| 1.200 | 1.06 | 39.09 % | 0.00 % | 55.27 % | 5.64 % | Stomach pH |
| 2.000 | 1.24 | 30.08 % | 0.02 % | 42.52 % | 27.38 % | |
| 3.000 | 1.89 | 8.68 % | 0.05 % | 12.27 % | 79.01 % | |
| 4.000 | 2.81 | 1.07 % | 0.06 % | 1.51 % | 97.36 % | |
| 5.000 | 3.65 | 0.11 % | 0.06 % | 0.15 % | 99.68 % | |
| 6.000 | 4.03 | 0.01 % | 0.06 % | 0.02 % | 99.92 % | |
| 6.500 | 4.08 | 0.00 % | 0.06 % | 0.00 % | 99.93 % | |
| 7.000 | 4.10 | 0.00 % | 0.06 % | 0.00 % | 99.94 % | |
| 7.400 | 4.11 | 0.00 % | 0.06 % | 0.00 % | 99.94 % | Blood pH |
| 8.000 | 4.11 | 0.00 % | 0.06 % | 0.00 % | 99.94 % | |
| 9.000 | 4.11 | 0.00 % | 0.06 % | 0.00 % | 99.94 % | |
| 10.000 | 4.11 | 0.00 % | 0.06 % | 0.00 % | 99.94 % | |
| 11.000 | 4.11 | 0.00 % | 0.06 % | 0.00 % | 99.94 % | |
| 12.000 | 4.11 | 0.00 % | 0.06 % | 0.00 % | 99.94 % | |

#### Carbonate and acidity

| | |
|---|---|
| Carbonate | 0.309 mM |
| Acidity error | -3.499 mM |

#### Other graphs

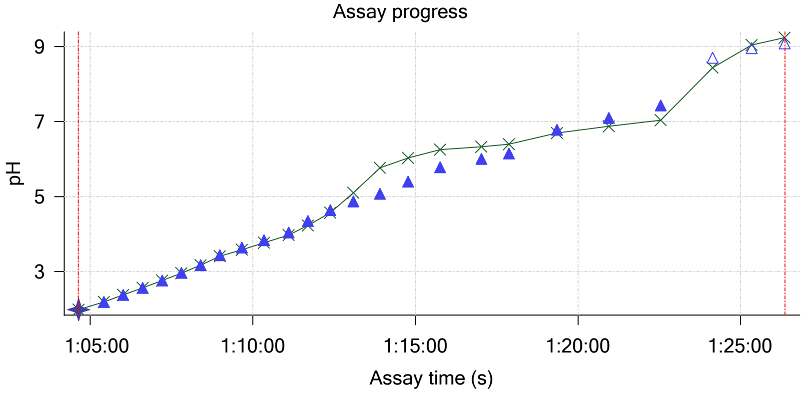


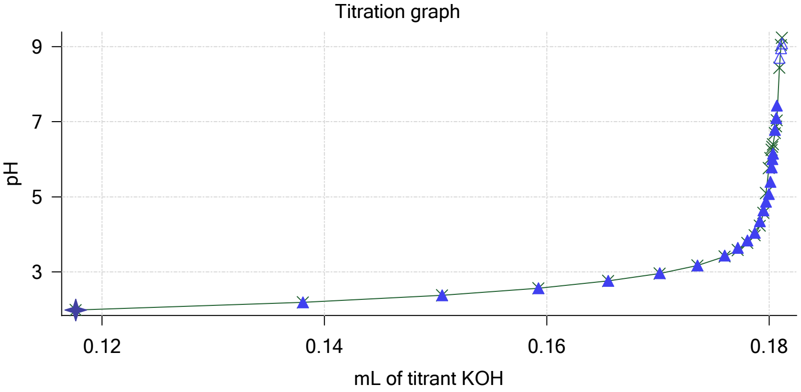

Sample name: **M12_octanol**
Assay name: **pH-metric high logP**
Assay ID: **18C-03014**
Filename: **C:\Sirius_T3\Mehtap\20180302_exp29_logP_T3-2\18C-03014_M12_octanol_pH-metric high logP.t3r**

Experiment start time: **3/3/2018 6:11:21 PM**
Analyst: **Pion**
Instrument ID: **T312060**

#### Other graphs (continued)

Charge balance residuals

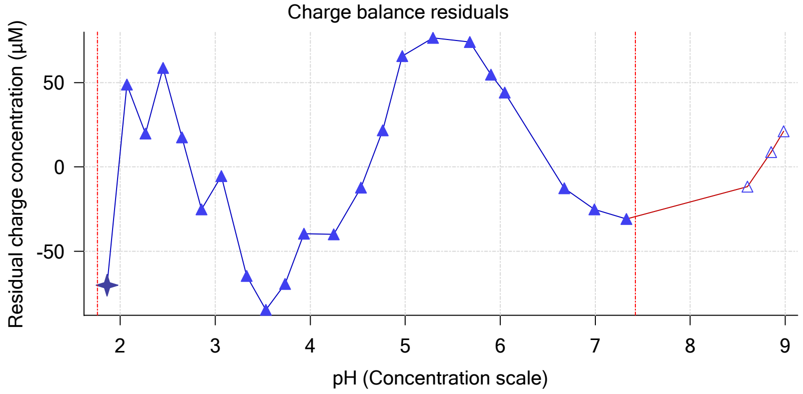


Buffer index profile

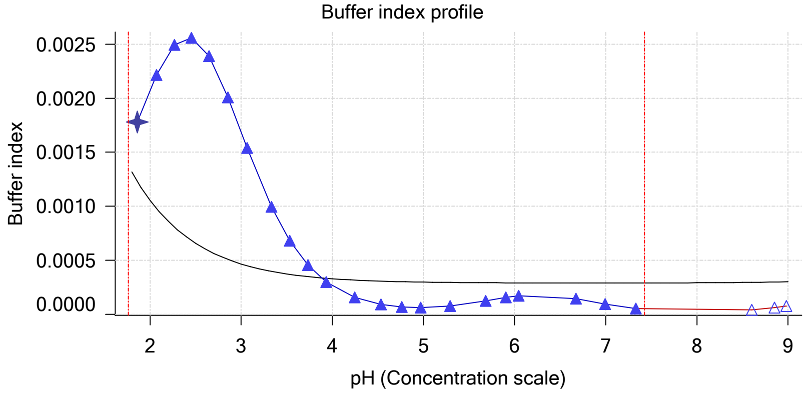


Ionic strength

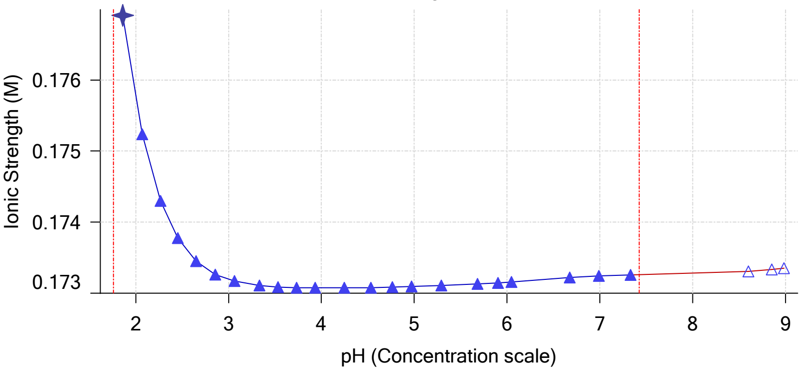


Log R Profile

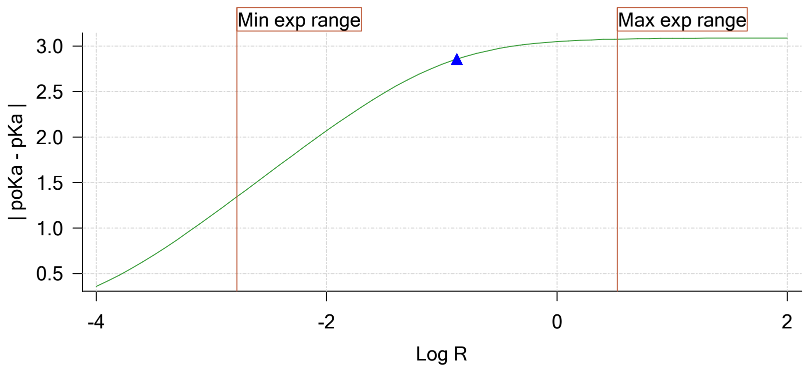


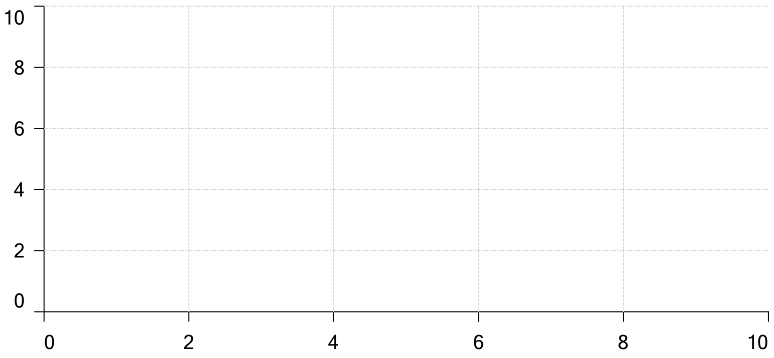


Ionisation of sample Carbonate

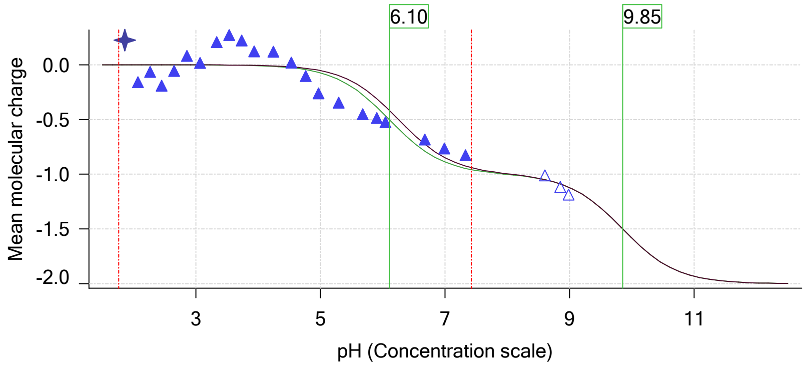

| | | | |
|---|---|---|---|
| Sample name: | **M12_octanol** | Experiment start time: | **3/3/2018 6:11:21 PM** |
| Assay name: | **pH-metric high logP** | Analyst: | **Pion** |
| Assay ID: | **18C-03014** | Instrument ID: | **T312060** |
| Filename: | **C:\Sirius_T3\Mehtap\20180302_exp29_logP_T3-2\18C-03014_M12_octanol_pH-metric high logP.t3r** | | |

#### Assay Model

| Settings | Value | Date/Time changed | Imported from |
|---|---|---|---|
| Sample name | M12_octanol | 2/28/2018 2:58:36 PM | User entered value |
| Sample by | Weight | | Default value |
| Sample weight | 0.002040 g | 3/2/2018 5:10:25 PM | User entered value |
| Formula weight | 292.16 g/mol | 2/28/2018 2:58:36 PM | User entered value |
| Solubility | Unknown | | Default value |
| Molecular weight | 255.70 | 2/28/2018 2:58:36 PM | User entered value |
| Individual pKa ionic environments | No | | Default value |
| Number of pKas | 1 | 2/28/2018 2:58:36 PM | User entered value |
| Sample is a | Base | 2/28/2018 2:58:36 PM | User entered value |
| pKa 1 | 5.28 | 2/28/2018 2:58:36 PM | User entered value |
| logp (XH +) | 1.02 | 3/2/2018 3:44:28 PM | User entered value |
| logP (neutral X) | 3.79 | 3/2/2018 3:44:35 PM | User entered value |
| Stoichiometry | 1.00000 | | Default value |
| Aprotic counterion name | Chloride | | From standards.xml file |
| Stoichiometry | 1.00 | | From standards.xml file |
| Charge per counterion | -1 | | From standards.xml file |

#### Events

| Time | Event | Water | Acid | Base | Octanol | pH | dpH/dt | pH R-squared | pH SD | dpH/dt time |
|---|---|---|---|---|---|---|---|---|---|---|
| 5:59.7 | Initial pH = 3.54 | | | | | | | | | |
| 9:04.2 | Data point 1 | 1.50000 mL | 0.04685 mL | 0.00336 mL | 0.01999 mL | 2.035 | -0.01024 | 0.61219 | 0.00065 | 10.0 s |
| 9:50.4 | Data point 2 | 1.50000 mL | 0.04685 mL | 0.01823 mL | 0.01999 mL | 2.240 | -0.00195 | 0.36606 | 0.00016 | 10.5 s |
| 10:26.5 | Data point 3 | 1.50000 mL | 0.04685 mL | 0.02721 mL | 0.01999 mL | 2.427 | -0.00750 | 0.72499 | 0.00043 | 10.5 s |
| 11:02.6 | Data point 4 | 1.50000 mL | 0.04685 mL | 0.03333 mL | 0.01999 mL | 2.606 | -0.00567 | 0.86217 | 0.00030 | 10.5 s |
| 11:38.7 | Data point 5 | 1.50000 mL | 0.04685 mL | 0.03772 mL | 0.01999 mL | 2.803 | -0.01366 | 0.90570 | 0.00071 | 10.0 s |
| 12:14.2 | Data point 6 | 1.50000 mL | 0.04685 mL | 0.04104 mL | 0.01999 mL | 2.980 | -0.01014 | 0.26097 | 0.00098 | 11.0 s |
| 13:01.1 | Data point 7 | 1.50000 mL | 0.04685 mL | 0.04367 mL | 0.01999 mL | 3.161 | -0.00882 | 0.80998 | 0.00048 | 10.5 s |
| 13:37.0 | Data point 8 | 1.50000 mL | 0.04685 mL | 0.04614 mL | 0.01999 mL | 3.370 | -0.00523 | 0.34459 | 0.00044 | 10.0 s |
| 14:12.5 | Data point 9 | 1.50000 mL | 0.04685 mL | 0.04838 mL | 0.01999 mL | 3.558 | -0.00999 | 0.44733 | 0.00074 | 10.0 s |
| 14:48.0 | Data point 10 | 1.50000 mL | 0.04685 mL | 0.05038 mL | 0.01999 mL | 3.784 | -0.01707 | 0.81320 | 0.00094 | 10.5 s |
| 15:24.0 | Data point 11 | 1.50000 mL | 0.04685 mL | 0.05202 mL | 0.01999 mL | 4.069 | -0.01895 | 0.92370 | 0.00097 | 29.5 s |
| 16:29.4 | Data point 12 | 1.50000 mL | 0.04685 mL | 0.05341 mL | 0.01999 mL | 4.470 | -0.01716 | 0.79592 | 0.00095 | 22.0 s |
| 17:27.1 | Data point 13 | 1.50000 mL | 0.04685 mL | 0.05402 mL | 0.01999 mL | 4.841 | -0.01868 | 0.91616 | 0.00096 | 47.5 s |
| 18:40.1 | Data point 14 | 1.50000 mL | 0.04685 mL | 0.05426 mL | 0.01999 mL | 4.941 | -0.02276 | 0.93191 | 0.00116 | Timed out at 59.5 s |
| 20:10.6 | Data point 15 | 1.50000 mL | 0.04685 mL | 0.05461 mL | 0.01999 mL | 5.196 | -0.04657 | 0.99269 | 0.00231 | Timed out at 59.5 s |
| 21:41.2 | Data point 16 | 1.50000 mL | 0.04685 mL | 0.05499 mL | 0.01999 mL | 5.842 | -0.13692 | 0.98959 | 0.00680 | Timed out at 59.5 s |
| 23:11.7 | Data point 17 | 1.50000 mL | 0.04685 mL | 0.05527 mL | 0.01999 mL | 5.839 | -0.10548 | 0.98431 | 0.00525 | Timed out at 59.5 s |
| 24:42.2 | Data point 18 | 1.50000 mL | 0.04685 mL | 0.05550 mL | 0.01999 mL | 6.181 | -0.07455 | 0.99489 | 0.00369 | Timed out at 59.5 s |
| 26:12.7 | Data point 19 | 1.50000 mL | 0.04685 mL | 0.05567 mL | 0.01999 mL | 6.383 | -0.02576 | 0.98099 | 0.00128 | Timed out at 59.5 s |
| 27:43.2 | Data point 20 | 1.50000 mL | 0.04685 mL | 0.05581 mL | 0.01999 mL | 6.739 | -0.03694 | 0.97684 | 0.00185 | Timed out at 59.5 s |
| 29:13.6 | Data point 21 | 1.50000 mL | 0.04685 mL | 0.05593 mL | 0.01999 mL | 6.952 | -0.03900 | 0.97414 | 0.00195 | Timed out at 59.5 s |
| 30:49.3 | Data point 22 | 1.50000 mL | 0.04685 mL | 0.05609 mL | 0.01999 mL | 7.359 | -0.06446 | 0.99560 | 0.00319 | Timed out at 59.5 s |
| 32:25.0 | Data point 23 | 1.50000 mL | 0.04685 mL | 0.05633 mL | 0.01999 mL | 8.802 | -0.02839 | 0.97337 | 0.00142 | Timed out at 59.5 s |
| 34:00.7 | Data point 24 | 1.50000 mL | 0.04685 mL | 0.05649 mL | 0.01999 mL | 8.971 | -0.03345 | 0.98026 | 0.00167 | Timed out at 59.5 s |
| 35:36.4 | Data point 25 | 1.50000 mL | 0.04685 mL | 0.05661 mL | 0.01999 mL | 9.166 | -0.01865 | 0.93346 | 0.00095 | 48.0 s |

Sample name: **M12_octanol**
Assay name: **pH-metric high logP**
Assay ID: **18C-03014**
Filename: **C:\Sirius_T3\Mehtap\20180302_exp29_logP_T3-2\18C-03014_M12_octanol_pH-metric high logP.t3r**

Experiment start time: **3/3/2018 6:11:21 PM**
Analyst: **Pion**
Instrument ID: **T312060**

#### Events (continued)

| Time | Event | Water | Acid | Base | Octanol | pH | dpH/dt | pH R-squared | pH SD | dpH/dt time |
|---|---|---|---|---|---|---|---|---|---|---|
| 37:28.6 | Data point 26 | 1.50000 mL | 0.10894 mL | 0.05661 mL | 0.05000 mL | 1.986 | -0.01183 | 0.86781 | 0.00063 | 10.0 s |
| 38:14.8 | Data point 27 | 1.50000 mL | 0.10894 mL | 0.07502 mL | 0.05000 mL | 2.187 | -0.01196 | 0.55172 | 0.00080 | 10.5 s |
| 38:51.0 | Data point 28 | 1.50000 mL | 0.10894 mL | 0.08624 mL | 0.05000 mL | 2.386 | -0.00645 | 0.60653 | 0.00041 | 10.0 s |
| 39:26.5 | Data point 29 | 1.50000 mL | 0.10894 mL | 0.09372 mL | 0.05000 mL | 2.568 | -0.00151 | 0.03213 | 0.00042 | 10.0 s |
| 40:02.1 | Data point 30 | 1.50000 mL | 0.10894 mL | 0.09913 mL | 0.05000 mL | 2.753 | -0.00324 | 0.58318 | 0.00021 | 10.5 s |
| 40:38.2 | Data point 31 | 1.50000 mL | 0.10894 mL | 0.10329 mL | 0.05000 mL | 2.974 | -0.00350 | 0.62117 | 0.00022 | 10.0 s |
| 41:13.7 | Data point 32 | 1.50000 mL | 0.10894 mL | 0.10656 mL | 0.05000 mL | 3.153 | -0.00315 | 0.45340 | 0.00023 | 10.0 s |
| 41:49.2 | Data point 33 | 1.50000 mL | 0.10894 mL | 0.10931 mL | 0.05000 mL | 3.370 | -0.00431 | 0.37822 | 0.00035 | 10.0 s |
| 42:24.7 | Data point 34 | 1.50000 mL | 0.10894 mL | 0.11150 mL | 0.05000 mL | 3.595 | -0.01024 | 0.34800 | 0.00086 | 10.0 s |
| 43:00.1 | Data point 35 | 1.50000 mL | 0.10894 mL | 0.11315 mL | 0.05000 mL | 3.826 | -0.01287 | 0.75168 | 0.00073 | 10.5 s |
| 43:36.1 | Data point 36 | 1.50000 mL | 0.10894 mL | 0.11432 mL | 0.05000 mL | 4.044 | -0.01264 | 0.72037 | 0.00074 | 11.0 s |
| 44:12.5 | Data point 37 | 1.50000 mL | 0.10894 mL | 0.11510 mL | 0.05000 mL | 4.277 | -0.00961 | 0.25227 | 0.00094 | 12.0 s |
| 44:49.9 | Data point 38 | 1.50000 mL | 0.10894 mL | 0.11559 mL | 0.05000 mL | 4.471 | -0.01870 | 0.92262 | 0.00096 | 42.5 s |
| 45:57.9 | Data point 39 | 1.50000 mL | 0.10894 mL | 0.11592 mL | 0.05000 mL | 4.621 | -0.01913 | 0.96775 | 0.00096 | 33.0 s |
| 47:01.5 | Data point 40 | 1.50000 mL | 0.10894 mL | 0.11637 mL | 0.05000 mL | 5.064 | -0.01981 | 0.97178 | 0.00099 | 36.5 s |
| 48:08.6 | Data point 41 | 1.50000 mL | 0.10894 mL | 0.11656 mL | 0.05000 mL | 5.290 | -0.01849 | 0.90940 | 0.00096 | 39.5 s |
| 49:18.8 | Data point 42 | 1.50000 mL | 0.10894 mL | 0.11668 mL | 0.05000 mL | 5.544 | -0.01813 | 0.90749 | 0.00094 | 42.5 s |
| 50:31.9 | Data point 43 | 1.50000 mL | 0.10894 mL | 0.11684 mL | 0.05000 mL | 6.152 | -0.01812 | 0.90297 | 0.00094 | 54.0 s |
| 51:56.5 | Data point 44 | 1.50000 mL | 0.10894 mL | 0.11698 mL | 0.05000 mL | 6.731 | -0.03629 | 0.94871 | 0.00184 | Timed out at 59.5 s |
| 53:32.0 | Data point 45 | 1.50000 mL | 0.10894 mL | 0.11705 mL | 0.05000 mL | 6.942 | -0.03907 | 0.98089 | 0.00195 | Timed out at 59.5 s |
| 55:07.6 | Data point 46 | 1.50000 mL | 0.10894 mL | 0.11717 mL | 0.05000 mL | 7.360 | -0.05485 | 0.98317 | 0.00273 | Timed out at 59.5 s |
| 56:43.2 | Data point 47 | 1.50000 mL | 0.10894 mL | 0.11724 mL | 0.05000 mL | 7.698 | -0.06448 | 0.99495 | 0.00319 | Timed out at 59.5 s |
| 58:18.8 | Data point 48 | 1.50000 mL | 0.10894 mL | 0.11731 mL | 0.05000 mL | 7.975 | -0.06889 | 0.99504 | 0.00341 | Timed out at 59.5 s |
| 59:49.3 | Data point 49 | 1.50000 mL | 0.10894 mL | 0.11736 mL | 0.05000 mL | 8.218 | -0.05194 | 0.98871 | 0.00258 | Timed out at 59.5 s |
| 1:01:24.9 | Data point 50 | 1.50000 mL | 0.10894 mL | 0.11743 mL | 0.05000 mL | 8.279 | -0.03256 | 0.93039 | 0.00167 | Timed out at 59.5 s |
| 1:03:00.6 | Data point 51 | 1.50000 mL | 0.10894 mL | 0.11764 mL | 0.05000 mL | 9.044 | -0.01889 | 0.94983 | 0.00096 | 35.0 s |
| 1:04:38.7 | Data point 52 | 1.50000 mL | 0.17653 mL | 0.11764 mL | 0.25000 mL | 1.983 | -0.01328 | 0.50845 | 0.00092 | 10.0 s |
| 1:05:25.0 | Data point 53 | 1.50000 mL | 0.17653 mL | 0.13808 mL | 0.25000 mL | 2.188 | -0.00421 | 0.56558 | 0.00028 | 10.0 s |
| 1:06:00.8 | Data point 54 | 1.50000 mL | 0.17653 mL | 0.15059 mL | 0.25000 mL | 2.379 | -0.01117 | 0.72575 | 0.00065 | 10.0 s |
| 1:06:36.4 | Data point 55 | 1.50000 mL | 0.17653 mL | 0.15924 mL | 0.25000 mL | 2.563 | 0.00955 | 0.39409 | 0.00075 | 10.5 s |
| 1:07:12.5 | Data point 56 | 1.50000 mL | 0.17653 mL | 0.16552 mL | 0.25000 mL | 2.758 | -0.00090 | 0.01671 | 0.00034 | 10.0 s |
| 1:07:48.0 | Data point 57 | 1.50000 mL | 0.17653 mL | 0.17016 mL | 0.25000 mL | 2.964 | -0.00646 | 0.68297 | 0.00039 | 10.0 s |
| 1:08:23.5 | Data point 58 | 1.50000 mL | 0.17653 mL | 0.17354 mL | 0.25000 mL | 3.172 | -0.01240 | 0.57154 | 0.00081 | 10.0 s |
| 1:08:59.1 | Data point 59 | 1.50000 mL | 0.17653 mL | 0.17601 mL | 0.25000 mL | 3.439 | -0.00851 | 0.81815 | 0.00046 | 10.0 s |
| 1:09:39.7 | Data point 60 | 1.50000 mL | 0.17653 mL | 0.17719 mL | 0.25000 mL | 3.639 | -0.00768 | 0.92817 | 0.00039 | 10.0 s |
| 1:10:20.4 | Data point 61 | 1.50000 mL | 0.17653 mL | 0.17806 mL | 0.25000 mL | 3.840 | -0.01302 | 0.73901 | 0.00075 | 10.0 s |
| 1:11:06.2 | Data point 62 | 1.50000 mL | 0.17653 mL | 0.17869 mL | 0.25000 mL | 4.040 | -0.01483 | 0.68740 | 0.00088 | 10.0 s |
| 1:11:41.6 | Data point 63 | 1.50000 mL | 0.17653 mL | 0.17916 mL | 0.25000 mL | 4.352 | -0.01359 | 0.79722 | 0.00075 | 11.0 s |
| 1:12:23.2 | Data point 64 | 1.50000 mL | 0.17653 mL | 0.17949 mL | 0.25000 mL | 4.639 | -0.01394 | 0.58554 | 0.00090 | 12.0 s |
| 1:13:05.6 | Data point 65 | 1.50000 mL | 0.17653 mL | 0.17973 mL | 0.25000 mL | 4.868 | -0.01439 | 0.70067 | 0.00085 | 13.0 s |
| 1:13:54.3 | Data point 66 | 1.50000 mL | 0.17653 mL | 0.17996 mL | 0.25000 mL | 5.075 | -0.01857 | 0.88631 | 0.00097 | 16.5 s |
| 1:14:46.4 | Data point 67 | 1.50000 mL | 0.17653 mL | 0.18010 mL | 0.25000 mL | 5.398 | -0.01592 | 0.73444 | 0.00092 | 28.5 s |
| 1:15:45.5 | Data point 68 | 1.50000 mL | 0.17653 mL | 0.18024 mL | 0.25000 mL | 5.785 | -0.01894 | 0.91475 | 0.00098 | 46.0 s |
| 1:17:02.1 | Data point 69 | 1.50000 mL | 0.17653 mL | 0.18029 mL | 0.25000 mL | 6.007 | -0.01848 | 0.84150 | 0.00100 | 25.0 s |
| 1:17:52.5 | Data point 70 | 1.50000 mL | 0.17653 mL | 0.18034 mL | 0.25000 mL | 6.149 | -0.01829 | 0.90450 | 0.00095 | 53.0 s |
| 1:19:21.4 | Data point 71 | 1.50000 mL | 0.17653 mL | 0.18053 mL | 0.25000 mL | 6.777 | -0.04095 | 0.98006 | 0.00204 | Timed out at 59.5 s |
| 1:20:57.0 | Data point 72 | 1.50000 mL | 0.17653 mL | 0.18062 mL | 0.25000 mL | 7.093 | -0.04542 | 0.98750 | 0.00225 | Timed out at 59.5 s |

| | | | |
|---|---|---|---|
| Sample name: | **M12_octanol** | Experiment start time: | **3/3/2018 6:11:21 PM** |
| Assay name: | **pH-metric high logP** | Analyst: | **Pion** |
| Assay ID: | **18C-03014** | Instrument ID: | **T312060** |
| Filename: | **C:\Sirius_T3\Mehtap\20180302_exp29_logP_T3-2\18C-03014_M12_octanol_pH-metric high logP.t3r** | | |

#### Events (continued)

| Time | Event | Water | Acid | Base | Octanol | pH | dpH/dt | pH R-squared | pH SD | dpH/dt time |
|---|---|---|---|---|---|---|---|---|---|---|
| 1:22:32.6 | Data point 73 | 1.50000 mL | 0.17653 mL | 0.18069 mL | 0.25000 mL | 7.432 | -0.06081 | 0.97336 | 0.00305 | Timed out at 59.5 s |
| 1:24:08.3 | Data point 74 | 1.50000 mL | 0.17653 mL | 0.18093 mL | 0.25000 mL | 8.702 | -0.01670 | 0.95705 | 0.00084 | 36.5 s |
| 1:25:20.5 | Data point 75 | 1.50000 mL | 0.17653 mL | 0.18107 mL | 0.25000 mL | 8.952 | -0.01969 | 0.97272 | 0.00099 | 25.0 s |
| 1:26:21.1 | Data point 76 | 1.50000 mL | 0.17653 mL | 0.18116 mL | 0.25000 mL | 9.082 | -0.01391 | 0.60511 | 0.00088 | 15.0 s |
| 1:26:45.2 | Assay volumes | 1.50000 mL | 0.17653 mL | 0.18116 mL | 0.25000 mL | | | | | |

| | | | |
|---|---|---|---|
| Sample name: | **M12_octanol** | Experiment start time: | **3/3/2018 6:11:21 PM** |
| Assay name: | **pH-metric high logP** | Analyst: | **Pion** |
| Assay ID: | **18C-03014** | Instrument ID: | **T312060** |
| Filename: | **C:\Sirius_T3\Mehtap\20180302_exp29_logP_T3-2\18C-03014_M12_octanol_pH-metric high logP.t3r** | | |

| | | | |
|---|---|---|---|
| Sample name: | **M12_octanol** | Experiment start time: | **3/3/2018 6:11:21 PM** |
| Assay name: | **pH-metric high logP** | Analyst: | **Pion** |
| Assay ID: | **18C-03014** | Instrument ID: | **T312060** |
| Filename: | **C:\Sirius_T3\Mehtap\20180302_exp29_logP_T3-2\18C-03014_M12_octanol_pH-metric high logP.t3r** | | |

#### Assay Settings (continued)

| Setting | Value | Original Value | Date/Time changed | Imported from |
|---|---|---|---|---|
| ***Titration 3*** | | | | |
| Titrate from | Low to high pH | | | |
| Add additional water | 0.00 mL | | | |
| Additional partition solvent volume | 0.200 mL | | | |
| Additional partition solvent added | Automatic | | | |
| After pH adjust stir for | 30 seconds | | | |
| Stir to allow partitioning for | 15 seconds | | | |
| Stirrer speed for partitioning | 60% | | | |
| ***Data Point Stability*** | | | | |
| Stir during data point collection | No | | | |
| Delay before data point collection | 0 seconds | | | |
| Number of points to average | 20 points | | | |
| Time interval between points | 0.50 seconds | | | |
| Required maximum standard deviation | 0.00100 dpH/dt | | | |
| Stability timeout after | 60 seconds | | | |

#### Calibration Settings

| Setting | Value | Date/Time changed | Imported from |
|---|---|---|---|
| Four-Plus alpha | 0.111 | 3/3/2018 6:11:21 PM | C:\Sirius_T3\HCl18C02.t3r |
| Four-Plus S | 0.9988 | 3/3/2018 6:11:21 PM | C:\Sirius_T3\HCl18C02.t3r |
| Four-Plus jH | 1.0 | 3/3/2018 6:11:21 PM | C:\Sirius_T3\HCl18C02.t3r |
| Four-Plus jOH | -0.8 | 3/3/2018 6:11:21 PM | C:\Sirius_T3\HCl18C02.t3r |
| Base concentration factor | 1.000 | 3/3/2018 6:11:21 PM | C:\Sirius_T3\KOH18B27.t3r |
| Acid concentration factor | 0.999 | 3/3/2018 6:11:21 PM | C:\Sirius_T3\HCl18C02.t3r |

#### Instrument Settings

| Setting | Value | Batch Id | Install date |
|---|---|---|---|
| Instrument owner | Merck | | |
| Instrument ID | T312060 | | |
| Instrument type | T3 Simulator | | |
| Software version | 1.1.3.0 | | |
| Dispenser module | | T3DM1200361 | 3/31/2009 5:24:52 AM |
| Dispenser 0 | Water | | 3/31/2009 5:25:05 AM |
| Syringe volume | 2.5 mL | | |
| Firmware version | 1.2.1(r2) | | |
| Titrant | Water (0.15 M KCl) | 02-06-2018 | 2/27/2018 10:05:59 AM |
| Dispenser 2 | Acid | | 3/31/2009 5:25:11 AM |
| Syringe volume | 0.5 mL | | |
| Firmware version | 1.2.1(r2) | | |
| Titrant | Acid (0.5 M HCl) | 02-27-2018 | 2/27/2018 10:27:22 AM |
| Dispenser 1 | Base | | 3/31/2009 5:25:21 AM |
| Syringe volume | 0.5 mL | | |
| Firmware version | 1.2.1(r2) | | |
| Titrant | Base (0.5 M KOH) | 9/22/2017 | 2/27/2018 10:21:22 AM |
| Dispenser 5 | Cosolvent | | 3/31/2009 5:26:24 AM |
| Syringe volume | 2.5 mL | | |
| Firmware version | 1.2.1(r2) | | |
| Distribution valve 5 | Distribution Valve | | 3/31/2009 5:28:19 AM |
| Firmware version | 1.1.3 | | |
| Port A | Methanol (80%, 0.15 M KCl) | 09-26-17 | 2/7/2018 9:42:01 AM |
| Port B | Cyclohexane | 11-01-17 | 2/27/2018 10:37:57 AM |
| Dispenser 3 | Buffer | | 8/3/2010 5:05:16 AM |
| Syringe volume | 0.5 mL | | |
| Firmware version | 1.2.1(r2) | | |
| Titrant | Dodecane | 2018/01/31 | 2/28/2018 10:18:04 AM |
| Dispenser 6 | Octanol | | 10/22/2010 10:52:43 AM |

| | | | |
|---|---|---|---|
| Sample name: | **M12_octanol** | Experiment start time: | **3/3/2018 6:11:21 PM** |
| Assay name: | **pH-metric high logP** | Analyst: | **Pion** |
| Assay ID: | **18C-03014** | Instrument ID: | **T312060** |
| Filename: | **C:\Sirius_T3\Mehtap\20180302_exp29_logP_T3-2\18C-03014_M12_octanol_pH-metric high logP.t3r** | | |

| | | | |
|---|---|---|---|
| Sample name: | **M12_octanol** | Experiment start time: | **3/3/2018 6:11:21 PM** |
| Assay name: | **pH-metric high logP** | Analyst: | **Pion** |
| Assay ID: | **18C-03014** | Instrument ID: | **T312060** |
| Filename: | **C:\Sirius_T3\Mehtap\20180302_exp29_logP_T3-2\18C-03014_M12_octanol_pH-metric high logP.t3r** | | |

#### Instrument Settings (continued)

| Setting | Value | Batch Id | Install date |
|---|---|---|---|
| Spectrometer calibration stir duration | 5 s | | |
| Spectrometer calibration stir speed | 30% | | |
| Spectrometer calibration wash pump volume | 20.0 mL | | |
| Spectrometer calibration wash stir duration | 5 s | | |
| Spectrometer calibration wash stir speed | 30% | | |
| Overhead dispense height | 10000 | | |

#### Refinement Settings

| Setting | Value | Default value |
|---|---|---|
| Turbidity detection method | None | None |
| Turbidity wavelength to assess | 500.0 nm | 500.0 nm |
| Turbidity maximum absorbance | 0.100 | 0.100 |
| Turbidity probe threshold | 50.00 | 50.00 |

#### Experiment Log

[2:37] Air gap created for Water (0.15 M KCl)
[2:38] Air gap created for Acid (0.5 M HCl)
[2:38] Air gap created for Base (0.5 M KOH)
[2:38] Air gap released for Water (0.15 M KCl)
[2:42] Titrator arm moved over Titration position
[2:42] Titration 1 of 3
[2:42] Adding initial titrants
[2:42] Automatically add 1.50000 mL of water
[3:07] Dispensed 1.500000 mL of Water (0.15 M KCl)
[3:11] Titrator arm moved over Drain
[5:53] Titrator arm moved to Titration position
[5:53] Argon flow rate set to 100
[5:53] Stirrer speed set to 10
[5:58] Automatically add 0.02000 mL of Octanol
[5:58] Dispensed 0.019991 mL of Octanol
[5:59] Initial pH = 3.54
[5:59] Iterative adjust 3.54 -> 2.00
[5:59] pH 3.54 -> 2.00
[6:01] Air gap released for Acid (0.5 M HCl)
[6:02] Dispensed 0.045296 mL of Acid (0.5 M HCl)
[6:07] pH 2.02 -> 2.00
[6:07] Dispensed 0.001552 mL of Acid (0.5 M HCl)
[6:12] Holding pH 2.00
[8:12] Stirrer speed set to 0
[8:12] Stirrer speed set to 50
[8:12] Iterative adjust 1.97 -> 2.00
[8:12] pH 1.97 -> 2.00
[8:13] Air gap released for Base (0.5 M KOH)
[8:14] Dispensed 0.003363 mL of Base (0.5 M KOH)
[9:04] Stirrer speed set to 0
[9:14] Datapoint id 1 collected
[9:14] Stirrer speed set to 50
[9:19] pH 2.04 -> 2.24
[9:19] Using cautious pH adjust
[9:19] Dispensed 0.007079 mL of Base (0.5 M KOH)
[9:24] Stepping pH = 2.12
[9:25] Dispensed 0.006444 mL of Base (0.5 M KOH)
[9:30] Stepping pH = 2.22
[9:30] Dispensed 0.001341 mL of Base (0.5 M KOH)
[9:35] Stepping pH = 2.24
[9:50] Stirrer speed set to 0
[10:01] Datapoint id 2 collected

Sample name: **M12_octanol**
Assay name: **pH-metric high logP**
Assay ID: **18C-03014**
Filename: **C:\Sirius_T3\Mehtap\20180302_exp29_logP_T3-2\18C-03014_M12_octanol_pH-metric high logP.t3r**
Experiment start time: **3/3/2018 6:11:21 PM**
Analyst: **Pion**
Instrument ID: **T312060**

#### Experiment Log (continued)

[10:01] Charge balance equation is out by -4.8%
[10:01] Stirrer speed set to 50
[10:06] pH 2.25 -> 2.45
[10:06] Using charge balance adjust
[10:06] Dispensed 0.008984 mL of Base (0.5 M KOH)
[10:26] Stirrer speed set to 0
[10:37] Datapoint id 3 collected
[10:37] Charge balance equation is out by -10.3%
[10:37] Stirrer speed set to 50
[10:42] pH 2.43 -> 2.63
[10:42] Using charge balance adjust
[10:42] Dispensed 0.006115 mL of Base (0.5 M KOH)
[11:02] Stirrer speed set to 0
[11:13] Datapoint id 4 collected
[11:13] Charge balance equation is out by -13.9%
[11:13] Stirrer speed set to 50
[11:18] pH 2.61 -> 2.81
[11:18] Using charge balance adjust
[11:18] Dispensed 0.004398 mL of Base (0.5 M KOH)
[11:38] Stirrer speed set to 0
[11:48] Datapoint id 5 collected
[11:48] Charge balance equation is out by -4.8%
[11:48] Stirrer speed set to 50
[11:54] pH 2.81 -> 3.01
[11:54] Using charge balance adjust
[11:54] Dispensed 0.003316 mL of Base (0.5 M KOH)
[12:14] Stirrer speed set to 0
[12:25] Datapoint id 6 collected
[12:25] Charge balance equation is out by -15.2%
[12:25] Stirrer speed set to 50
[12:30] pH 2.99 -> 3.19
[12:30] Using cautious pH adjust
[12:30] Dispensed 0.001388 mL of Base (0.5 M KOH)
[12:35] Stepping pH = 3.08
[12:35] Dispensed 0.000988 mL of Base (0.5 M KOH)
[12:41] Stepping pH = 3.16
[12:41] Dispensed 0.000259 mL of Base (0.5 M KOH)
[12:46] Stepping pH = 3.18
[13:01] Stirrer speed set to 0
[13:11] Datapoint id 7 collected
[13:11] Charge balance equation is out by 4.7%
[13:11] Stirrer speed set to 50
[13:16] pH 3.16 -> 3.36
[13:16] Using charge balance adjust
[13:17] Dispensed 0.002469 mL of Base (0.5 M KOH)
[13:37] Stirrer speed set to 0
[13:47] Datapoint id 8 collected
[13:47] Charge balance equation is out by 4.1%
[13:47] Stirrer speed set to 50
[13:52] pH 3.37 -> 3.57
[13:52] Using charge balance adjust
[13:52] Dispensed 0.002234 mL of Base (0.5 M KOH)
[14:12] Stirrer speed set to 0
[14:22] Datapoint id 9 collected
[14:22] Charge balance equation is out by -7.1%
[14:22] Stirrer speed set to 50
[14:27] pH 3.56 -> 3.76
[14:27] Using charge balance adjust
[14:28] Dispensed 0.001999 mL of Base (0.5 M KOH)

Sample name: **M12_octanol**
Assay name: **pH-metric high logP**
Assay ID: **18C-03014**
Filename: **C:\Sirius_T3\Mehtap\20180302_exp29_logP_T3-2\18C-03014_M12_octanol_pH-metric high logP.t3r**
Experiment start time: **3/3/2018 6:11:21 PM**
Analyst: **Pion**
Instrument ID: **T312060**

#### Experiment Log (continued)

[14:48] Stirrer speed set to 0
[14:58] Datapoint id 10 collected
[14:58] Charge balance equation is out by 13.2%
[14:58] Stirrer speed set to 50
[15:03] pH 3.78 -> 3.98
[15:03] Using charge balance adjust
[15:04] Dispensed 0.001646 mL of Base (0.5 M KOH)
[15:24] Stirrer speed set to 0
[15:53] Datapoint id 11 collected
[15:53] Charge balance equation is out by 42.3%
[15:53] Stirrer speed set to 50
[15:58] pH 4.06 -> 4.26
[15:58] Using cautious pH adjust
[15:59] Dispensed 0.000588 mL of Base (0.5 M KOH)
[16:04] Stepping pH = 4.24
[16:04] Dispensed 0.000047 mL of Base (0.5 M KOH)
[16:09] Stepping pH = 4.21
[16:09] Dispensed 0.000753 mL of Base (0.5 M KOH)
[16:14] Stepping pH = 4.64
[16:29] Stirrer speed set to 0
[16:51] Datapoint id 12 collected
[16:51] Charge balance equation is out by -18.3%
[16:51] Stirrer speed set to 50
[16:56] pH 4.49 -> 4.69
[16:56] Using cautious pH adjust
[16:56] Dispensed 0.000282 mL of Base (0.5 M KOH)
[17:01] Stepping pH = 4.67
[17:01] Dispensed 0.000024 mL of Base (0.5 M KOH)
[17:07] Stepping pH = 4.64
[17:07] Dispensed 0.000306 mL of Base (0.5 M KOH)
[17:12] Stepping pH = 4.98
[17:27] Stirrer speed set to 0
[18:14] Datapoint id 13 collected
[18:14] Charge balance equation is out by -11.2%
[18:14] Stirrer speed set to 50
[18:20] pH 4.89 -> 5.09
[18:20] Using charge balance adjust
[18:20] Dispensed 0.000235 mL of Base (0.5 M KOH)
[18:40] Stirrer speed set to 0
[19:40] Datapoint id 14 collected
[19:40] Charge balance equation is out by -77.2%
[19:40] Stirrer speed set to 50
[19:45] pH 5.03 -> 5.23
[19:45] Using cautious pH adjust
[19:45] Dispensed 0.000094 mL of Base (0.5 M KOH)
[19:50] Stepping pH = 5.04
[19:50] Dispensed 0.000259 mL of Base (0.5 M KOH)
[19:55] Stepping pH = 5.67
[20:10] Stirrer speed set to 0
[21:10] Datapoint id 15 collected
[21:10] Charge balance equation is out by -86.8%
[21:10] Stirrer speed set to 50
[21:16] pH 5.31 -> 5.51
[21:16] Using cautious pH adjust
[21:16] Dispensed 0.000047 mL of Base (0.5 M KOH)
[21:21] Stepping pH = 5.28
[21:21] Dispensed 0.000329 mL of Base (0.5 M KOH)
[21:26] Stepping pH = 6.63
[21:41] Stirrer speed set to 0

| | | | |
|---|---|---|---|
| Sample name: | **M12_octanol** | Experiment start time: | **3/3/2018 6:11:21 PM** |
| Assay name: | **pH-metric high logP** | Analyst: | **Pion** |
| Assay ID: | **18C-03014** | Instrument ID: | **T312060** |
| Filename: | **C:\Sirius_T3\Mehtap\20180302_exp29_logP_T3-2\18C-03014_M12_octanol_pH-metric high logP.t3r** | | |

#### Experiment Log (continued)

[22:41] Datapoint id 16 collected
[22:41] Charge balance equation is out by -249.2%
[22:41] Stirrer speed set to 50
[22:46] pH 5.92 -> 6.12
[22:46] Using cautious pH adjust
[22:46] Dispensed 0.000024 mL of Base (0.5 M KOH)
[22:51] Stepping pH = 5.77
[22:51] Dispensed 0.000259 mL of Base (0.5 M KOH)
[22:56] Stepping pH = 6.69
[23:11] Stirrer speed set to 0
[24:11] Datapoint id 17 collected
[24:11] Charge balance equation is out by -408.9%
[24:11] Stirrer speed set to 50
[24:17] pH 5.97 -> 6.17
[24:17] Using cautious pH adjust
[24:17] Dispensed 0.000024 mL of Base (0.5 M KOH)
[24:22] Stepping pH = 5.89
[24:22] Dispensed 0.000212 mL of Base (0.5 M KOH)
[24:27] Stepping pH = 6.77
[24:42] Stirrer speed set to 0
[25:42] Datapoint id 18 collected
[25:42] Charge balance equation is out by -301.9%
[25:42] Stirrer speed set to 50
[25:47] pH 6.27 -> 6.47
[25:47] Using cautious pH adjust
[25:47] Dispensed 0.000024 mL of Base (0.5 M KOH)
[25:52] Stepping pH = 6.23
[25:52] Dispensed 0.000141 mL of Base (0.5 M KOH)
[25:57] Stepping pH = 6.60
[26:12] Stirrer speed set to 0
[27:12] Datapoint id 19 collected
[27:12] Charge balance equation is out by -248.4%
[27:13] Stirrer speed set to 50
[27:18] pH 6.39 -> 6.59
[27:18] Using cautious pH adjust
[27:18] Dispensed 0.000024 mL of Base (0.5 M KOH)
[27:23] Stepping pH = 6.37
[27:23] Dispensed 0.000118 mL of Base (0.5 M KOH)
[27:28] Stepping pH = 6.70
[27:43] Stirrer speed set to 0
[28:43] Datapoint id 20 collected
[28:43] Charge balance equation is out by -226.3%
[28:43] Stirrer speed set to 50
[28:48] pH 6.69 -> 6.89
[28:48] Using cautious pH adjust
[28:48] Dispensed 0.000024 mL of Base (0.5 M KOH)
[28:53] Stepping pH = 6.65
[28:53] Dispensed 0.000094 mL of Base (0.5 M KOH)
[28:58] Stepping pH = 6.93
[29:13] Stirrer speed set to 0
[30:13] Datapoint id 21 collected
[30:13] Charge balance equation is out by -266.6%
[30:13] Stirrer speed set to 50
[30:19] pH 6.92 -> 7.12
[30:19] Using cautious pH adjust
[30:19] Dispensed 0.000024 mL of Base (0.5 M KOH)
[30:24] Stepping pH = 6.85
[30:24] Dispensed 0.000071 mL of Base (0.5 M KOH)
[30:29] Stepping pH = 6.95

Sample name: **M12_octanol** Experiment start time: **3/3/2018 6:11:21 PM**
Assay name: **pH-metric high logP** Analyst: **Pion**
Assay ID: **18C-03014** Instrument ID: **T312060**
Filename: **C:\Sirius_T3\Mehtap\20180302_exp29_logP_T3-2\18C-03014_M12_octanol_pH-metric high logP.t3r**

#### Experiment Log (continued)

[30:29] Dispensed 0.000071 mL of Base (0.5 M KOH)
[30:34] Stepping pH = 7.36
[30:49] Stirrer speed set to 0
[31:49] Datapoint id 22 collected
[31:49] Charge balance equation is out by -694.1%
[31:49] Stirrer speed set to 50
[31:54] pH 7.29 -> 7.49
[31:54] Using cautious pH adjust
[31:54] Dispensed 0.000024 mL of Base (0.5 M KOH)
[31:59] Stepping pH = 7.25
[31:59] Dispensed 0.000024 mL of Base (0.5 M KOH)
[32:05] Stepping pH = 7.22
[32:05] Dispensed 0.000188 mL of Base (0.5 M KOH)
[32:10] Stepping pH = 8.57
[32:25] Stirrer speed set to 0
[33:25] Datapoint id 23 collected
[33:25] Charge balance equation is out by -2,178.7%
[33:25] Stirrer speed set to 50
[33:30] pH 8.80 -> 9.00
[33:30] Using cautious pH adjust
[33:30] Dispensed 0.000024 mL of Base (0.5 M KOH)
[33:35] Stepping pH = 8.79
[33:35] Dispensed 0.000071 mL of Base (0.5 M KOH)
[33:40] Stepping pH = 8.89
[33:40] Dispensed 0.000071 mL of Base (0.5 M KOH)
[33:45] Stepping pH = 9.02
[34:00] Stirrer speed set to 0
[35:00] Datapoint id 24 collected
[35:01] Charge balance equation is out by -456.4%
[35:01] Stirrer speed set to 50
[35:06] pH 8.94 -> 9.05
[35:06] Using cautious pH adjust
[35:06] Dispensed 0.000024 mL of Base (0.5 M KOH)
[35:11] Stepping pH = 8.91
[35:11] Dispensed 0.000047 mL of Base (0.5 M KOH)
[35:16] Stepping pH = 8.96
[35:16] Dispensed 0.000047 mL of Base (0.5 M KOH)
[35:21] Stepping pH = 9.16
[35:36] Stirrer speed set to 0
[36:24] Datapoint id 25 collected
[36:24] Charge balance equation is out by -631.2%
[36:24] Titration 2 of 3
[36:24] Adding initial titrants
[36:24] Automatically add 0.03000 mL of Octanol
[36:25] Dispensed 0.030009 mL of Octanol
[36:25] Stirrer speed set to 10
[36:26] Stirrer speed set to 55
[36:26] Iterative adjust 9.17 -> 2.00
[36:26] pH 9.17 -> 2.00
[36:28] Dispensed 0.056138 mL of Acid (0.5 M HCl)
[36:33] pH 2.05 -> 2.00
[36:33] Dispensed 0.004633 mL of Acid (0.5 M HCl)
[36:38] pH 2.01 -> 2.00
[36:38] Dispensed 0.001317 mL of Acid (0.5 M HCl)
[37:28] Stirrer speed set to 0
[37:38] Datapoint id 26 collected
[37:38] Stirrer speed set to 55
[37:43] pH 1.99 -> 2.19
[37:43] Using cautious pH adjust

Sample name: **M12_octanol**
Assay name: **pH-metric high logP**
Assay ID: **18C-03014**
Filename: **C:\Sirius_T3\Mehtap\20180302_exp29_logP_T3-2\18C-03014_M12_octanol_pH-metric high logP.t3r**
Experiment start time: **3/3/2018 6:11:21 PM**
Analyst: **Pion**
Instrument ID: **T312060**

#### Experiment Log (continued)

[37:44] Dispensed 0.008725 mL of Base (0.5 M KOH)
[37:49] Stepping pH = 2.07
[37:49] Dispensed 0.007079 mL of Base (0.5 M KOH)
[37:54] Stepping pH = 2.15
[37:54] Dispensed 0.002611 mL of Base (0.5 M KOH)
[38:00] Stepping pH = 2.19
[38:15] Stirrer speed set to 0
[38:25] Datapoint id 27 collected
[38:25] Charge balance equation is out by -5.5%
[38:25] Stirrer speed set to 55
[38:30] pH 2.19 -> 2.39
[38:30] Using charge balance adjust
[38:31] Dispensed 0.011218 mL of Base (0.5 M KOH)
[38:51] Stirrer speed set to 0
[39:01] Datapoint id 28 collected
[39:01] Charge balance equation is out by -2.0%
[39:01] Stirrer speed set to 55
[39:06] pH 2.39 -> 2.59
[39:06] Using charge balance adjust
[39:06] Dispensed 0.007479 mL of Base (0.5 M KOH)
[39:26] Stirrer speed set to 0
[39:36] Datapoint id 29 collected
[39:36] Charge balance equation is out by -11.7%
[39:36] Stirrer speed set to 55
[39:41] pH 2.57 -> 2.77
[39:41] Using charge balance adjust
[39:42] Dispensed 0.005409 mL of Base (0.5 M KOH)
[40:02] Stirrer speed set to 0
[40:12] Datapoint id 30 collected
[40:12] Charge balance equation is out by -10.8%
[40:12] Stirrer speed set to 55
[40:17] pH 2.76 -> 2.96
[40:17] Using charge balance adjust
[40:18] Dispensed 0.004163 mL of Base (0.5 M KOH)
[40:38] Stirrer speed set to 0
[40:48] Datapoint id 31 collected
[40:48] Charge balance equation is out by 7.0%
[40:48] Stirrer speed set to 55
[40:53] pH 2.97 -> 3.17
[40:53] Using charge balance adjust
[40:53] Dispensed 0.003269 mL of Base (0.5 M KOH)
[41:13] Stirrer speed set to 0
[41:24] Datapoint id 32 collected
[41:24] Charge balance equation is out by -9.7%
[41:24] Stirrer speed set to 55
[41:29] pH 3.15 -> 3.35
[41:29] Using charge balance adjust
[41:29] Dispensed 0.002752 mL of Base (0.5 M KOH)
[41:49] Stirrer speed set to 0
[41:59] Datapoint id 33 collected
[41:59] Charge balance equation is out by 8.4%
[41:59] Stirrer speed set to 55
[42:04] pH 3.37 -> 3.57
[42:04] Using charge balance adjust
[42:04] Dispensed 0.002187 mL of Base (0.5 M KOH)
[42:24] Stirrer speed set to 0
[42:34] Datapoint id 34 collected
[42:34] Charge balance equation is out by 10.6%
[42:34] Stirrer speed set to 55

Sample name: **M12_octanol** Experiment start time: **3/3/2018 6:11:21 PM**
Assay name: **pH-metric high logP** Analyst: **Pion**
Assay ID: **18C-03014** Instrument ID: **T312060**
Filename: **C:\Sirius_T3\Mehtap\20180302_exp29_logP_T3-2\18C-03014_M12_octanol_pH-metric high logP.t3r**

#### Experiment Log (continued)

[42:40] pH 3.60 -> 3.80
[42:40] Using charge balance adjust
[42:40] Dispensed 0.001646 mL of Base (0.5 M KOH)
[43:00] Stirrer speed set to 0
[43:10] Datapoint id 35 collected
[43:10] Charge balance equation is out by 11.7%
[43:10] Stirrer speed set to 55
[43:16] pH 3.83 -> 4.03
[43:16] Using charge balance adjust
[43:16] Dispensed 0.001176 mL of Base (0.5 M KOH)
[43:36] Stirrer speed set to 0
[43:47] Datapoint id 36 collected
[43:47] Charge balance equation is out by 6.6%
[43:47] Stirrer speed set to 55
[43:52] pH 4.06 -> 4.26
[43:52] Using charge balance adjust
[43:52] Dispensed 0.000776 mL of Base (0.5 M KOH)
[44:12] Stirrer speed set to 0
[44:24] Datapoint id 37 collected
[44:24] Charge balance equation is out by 10.2%
[44:24] Stirrer speed set to 55
[44:29] pH 4.29 -> 4.49
[44:29] Using charge balance adjust
[44:29] Dispensed 0.000494 mL of Base (0.5 M KOH)
[44:50] Stirrer speed set to 0
[45:32] Datapoint id 38 collected
[45:32] Charge balance equation is out by -11.9%
[45:32] Stirrer speed set to 55
[45:37] pH 4.50 -> 4.70
[45:37] Using charge balance adjust
[45:37] Dispensed 0.000329 mL of Base (0.5 M KOH)
[45:58] Stirrer speed set to 0
[46:31] Datapoint id 39 collected
[46:31] Charge balance equation is out by -38.3%
[46:31] Stirrer speed set to 55
[46:36] pH 4.65 -> 4.85
[46:36] Using cautious pH adjust
[46:36] Dispensed 0.000118 mL of Base (0.5 M KOH)
[46:41] Stepping pH = 4.66
[46:41] Dispensed 0.000329 mL of Base (0.5 M KOH)
[46:46] Stepping pH = 5.18
[47:01] Stirrer speed set to 0
[47:38] Datapoint id 40 collected
[47:38] Charge balance equation is out by -91.6%
[47:38] Stirrer speed set to 55
[47:43] pH 5.10 -> 5.30
[47:43] Using cautious pH adjust
[47:43] Dispensed 0.000047 mL of Base (0.5 M KOH)
[47:48] Stepping pH = 5.11
[47:48] Dispensed 0.000141 mL of Base (0.5 M KOH)
[47:53] Stepping pH = 5.38
[48:08] Stirrer speed set to 0
[48:48] Datapoint id 41 collected
[48:48] Charge balance equation is out by -92.4%
[48:48] Stirrer speed set to 55
[48:53] pH 5.32 -> 5.52
[48:53] Using cautious pH adjust
[48:53] Dispensed 0.000024 mL of Base (0.5 M KOH)
[48:58] Stepping pH = 5.34

| | | | |
|---|---|---|---|
| Sample name: | **M12_octanol** | Experiment start time: | **3/3/2018 6:11:21 PM** |
| Assay name: | **pH-metric high logP** | Analyst: | **Pion** |
| Assay ID: | **18C-03014** | Instrument ID: | **T312060** |
| Filename: | **C:\Sirius_T3\Mehtap\20180302_exp29_logP_T3-2\18C-03014_M12_octanol_pH-metric high logP.t3r** | | |

#### Experiment Log (continued)

[48:58] Dispensed 0.000094 mL of Base (0.5 M KOH)
[49:03] Stepping pH = 5.61
[49:19] Stirrer speed set to 0
[50:01] Datapoint id 42 collected
[50:01] Charge balance equation is out by -86.2%
[50:01] Stirrer speed set to 55
[50:06] pH 5.58 -> 5.78
[50:06] Using cautious pH adjust
[50:06] Dispensed 0.000024 mL of Base (0.5 M KOH)
[50:11] Stepping pH = 5.58
[50:11] Dispensed 0.000141 mL of Base (0.5 M KOH)
[50:17] Stepping pH = 6.15
[50:32] Stirrer speed set to 0
[51:26] Datapoint id 43 collected
[51:26] Charge balance equation is out by -209.7%
[51:26] Stirrer speed set to 55
[51:31] pH 6.15 -> 6.35
[51:31] Using cautious pH adjust
[51:31] Dispensed 0.000024 mL of Base (0.5 M KOH)
[51:36] Stepping pH = 6.15
[51:36] Dispensed 0.000118 mL of Base (0.5 M KOH)
[51:41] Stepping pH = 6.66
[51:56] Stirrer speed set to 0
[52:56] Datapoint id 44 collected
[52:56] Charge balance equation is out by -201.5%
[52:56] Stirrer speed set to 55
[53:01] pH 6.73 -> 6.93
[53:01] Using cautious pH adjust
[53:01] Dispensed 0.000024 mL of Base (0.5 M KOH)
[53:06] Stepping pH = 6.77
[53:06] Dispensed 0.000024 mL of Base (0.5 M KOH)
[53:12] Stepping pH = 6.86
[53:12] Dispensed 0.000024 mL of Base (0.5 M KOH)
[53:17] Stepping pH = 6.98
[53:32] Stirrer speed set to 0
[54:32] Datapoint id 45 collected
[54:32] Charge balance equation is out by -131.1%
[54:32] Stirrer speed set to 55
[54:37] pH 6.91 -> 7.11
[54:37] Using cautious pH adjust
[54:37] Dispensed 0.000024 mL of Base (0.5 M KOH)
[54:42] Stepping pH = 6.88
[54:42] Dispensed 0.000071 mL of Base (0.5 M KOH)
[54:47] Stepping pH = 7.09
[54:47] Dispensed 0.000024 mL of Base (0.5 M KOH)
[54:52] Stepping pH = 7.32
[55:07] Stirrer speed set to 0
[56:07] Datapoint id 46 collected
[56:07] Charge balance equation is out by -381.0%
[56:07] Stirrer speed set to 55
[56:13] pH 7.34 -> 7.54
[56:13] Using cautious pH adjust
[56:13] Dispensed 0.000024 mL of Base (0.5 M KOH)
[56:18] Stepping pH = 7.34
[56:18] Dispensed 0.000024 mL of Base (0.5 M KOH)
[56:23] Stepping pH = 7.41
[56:23] Dispensed 0.000024 mL of Base (0.5 M KOH)
[56:28] Stepping pH = 7.58
[56:43] Stirrer speed set to 0

#### Experiment Log (continued)

[57:43] Datapoint id 47 collected
[57:43] Charge balance equation is out by -592.2%
[57:43] Stirrer speed set to 55
[57:48] pH 7.62 -> 7.82
[57:48] Using cautious pH adjust
[57:48] Dispensed 0.000024 mL of Base (0.5 M KOH)
[57:53] Stepping pH = 7.57
[57:53] Dispensed 0.000024 mL of Base (0.5 M KOH)
[57:58] Stepping pH = 7.67
[57:58] Dispensed 0.000024 mL of Base (0.5 M KOH)
[58:04] Stepping pH = 7.88
[58:19] Stirrer speed set to 0
[59:19] Datapoint id 48 collected
[59:19] Charge balance equation is out by -780.6%
[59:19] Stirrer speed set to 55
[59:24] pH 7.95 -> 8.15
[59:24] Using cautious pH adjust
[59:24] Dispensed 0.000024 mL of Base (0.5 M KOH)
[59:29] Stepping pH = 8.00
[59:29] Dispensed 0.000024 mL of Base (0.5 M KOH)
[59:34] Stepping pH = 8.15
[59:49] Stirrer speed set to 0
[1:00:49] Datapoint id 49 collected
[1:00:49] Charge balance equation is out by -477.9%
[1:00:49] Stirrer speed set to 55
[1:00:54] pH 8.16 -> 8.36
[1:00:54] Using cautious pH adjust
[1:00:54] Dispensed 0.000024 mL of Base (0.5 M KOH)
[1:00:59] Stepping pH = 8.15
[1:00:59] Dispensed 0.000024 mL of Base (0.5 M KOH)
[1:01:04] Stepping pH = 8.21
[1:01:05] Dispensed 0.000024 mL of Base (0.5 M KOH)
[1:01:10] Stepping pH = 8.35
[1:01:25] Stirrer speed set to 0
[1:02:25] Datapoint id 50 collected
[1:02:25] Charge balance equation is out by -765.8%
[1:02:25] Stirrer speed set to 55
[1:02:30] pH 8.21 -> 8.41
[1:02:30] Using cautious pH adjust
[1:02:30] Dispensed 0.000024 mL of Base (0.5 M KOH)
[1:02:35] Stepping pH = 8.16
[1:02:35] Dispensed 0.000024 mL of Base (0.5 M KOH)
[1:02:40] Stepping pH = 8.12
[1:02:40] Dispensed 0.000165 mL of Base (0.5 M KOH)
[1:02:45] Stepping pH = 8.99
[1:03:00] Stirrer speed set to 0
[1:03:35] Datapoint id 51 collected
[1:03:35] Charge balance equation is out by -2,119.2%
[1:03:35] Titration 3 of 3
[1:03:35] Adding initial titrants
[1:03:35] Automatically add 0.20000 mL of Octanol
[1:03:40] Dispensed 0.200000 mL of Octanol
[1:03:40] Stirrer speed set to 10
[1:03:41] Stirrer speed set to 60
[1:03:41] Iterative adjust 9.05 -> 2.00
[1:03:41] pH 9.05 -> 2.00
[1:03:43] Dispensed 0.058467 mL of Acid (0.5 M HCl)
[1:03:48] pH 2.09 -> 2.00
[1:03:48] Dispensed 0.009125 mL of Acid (0.5 M HCl)

Sample name: **M12_octanol**
Assay name: **pH-metric high logP**
Assay ID: **18C-03014**
Filename: **C:\Sirius_T3\Mehtap\20180302_exp29_logP_T3-2\18C-03014_M12_octanol_pH-metric high logP.t3r**

Experiment start time: **3/3/2018 6:11:21 PM**
Analyst: **Pion**
Instrument ID: **T312060**

#### Experiment Log (continued)

[1:04:38] Stirrer speed set to 0
[1:04:48] Datapoint id 52 collected
[1:04:48] Stirrer speed set to 60
[1:04:54] pH 1.99 -> 2.19
[1:04:54] Using cautious pH adjust
[1:04:54] Dispensed 0.009548 mL of Base (0.5 M KOH)
[1:04:59] Stepping pH = 2.07
[1:04:59] Dispensed 0.007973 mL of Base (0.5 M KOH)
[1:05:04] Stepping pH = 2.15
[1:05:05] Dispensed 0.002916 mL of Base (0.5 M KOH)
[1:05:10] Stepping pH = 2.19
[1:05:25] Stirrer speed set to 0
[1:05:35] Datapoint id 53 collected
[1:05:35] Charge balance equation is out by -7.1%
[1:05:35] Stirrer speed set to 60
[1:05:40] pH 2.19 -> 2.39
[1:05:40] Using charge balance adjust
[1:05:40] Dispensed 0.012512 mL of Base (0.5 M KOH)
[1:06:01] Stirrer speed set to 0
[1:06:11] Datapoint id 54 collected
[1:06:11] Charge balance equation is out by -6.0%
[1:06:11] Stirrer speed set to 60
[1:06:16] pH 2.38 -> 2.58
[1:06:16] Using charge balance adjust
[1:06:16] Dispensed 0.008655 mL of Base (0.5 M KOH)
[1:06:36] Stirrer speed set to 0
[1:06:47] Datapoint id 55 collected
[1:06:47] Charge balance equation is out by -9.8%
[1:06:47] Stirrer speed set to 60
[1:06:52] pH 2.57 -> 2.77
[1:06:52] Using charge balance adjust
[1:06:52] Dispensed 0.006279 mL of Base (0.5 M KOH)
[1:07:12] Stirrer speed set to 0
[1:07:22] Datapoint id 56 collected
[1:07:22] Charge balance equation is out by -5.8%
[1:07:22] Stirrer speed set to 60
[1:07:27] pH 2.76 -> 2.96
[1:07:27] Using charge balance adjust
[1:07:28] Dispensed 0.004633 mL of Base (0.5 M KOH)
[1:07:48] Stirrer speed set to 0
[1:07:58] Datapoint id 57 collected
[1:07:58] Charge balance equation is out by 0.1%
[1:07:58] Stirrer speed set to 60
[1:08:03] pH 2.97 -> 3.17
[1:08:03] Using charge balance adjust
[1:08:03] Dispensed 0.003387 mL of Base (0.5 M KOH)
[1:08:23] Stirrer speed set to 0
[1:08:33] Datapoint id 58 collected
[1:08:33] Charge balance equation is out by 0.6%
[1:08:33] Stirrer speed set to 60
[1:08:38] pH 3.18 -> 3.38
[1:08:38] Using charge balance adjust
[1:08:39] Dispensed 0.002469 mL of Base (0.5 M KOH)
[1:08:59] Stirrer speed set to 0
[1:09:09] Datapoint id 59 collected
[1:09:09] Charge balance equation is out by 31.4%
[1:09:09] Stirrer speed set to 60
[1:09:14] pH 3.45 -> 3.65
[1:09:14] Using cautious pH adjust

| | | | |
|---|---|---|---|
| Sample name: | **M12_octanol** | Experiment start time: | **3/3/2018 6:11:21 PM** |
| Assay name: | **pH-metric high logP** | Analyst: | **Pion** |
| Assay ID: | **18C-03014** | Instrument ID: | **T312060** |
| Filename: | **C:\Sirius_T3\Mehtap\20180302_exp29_logP_T3-2\18C-03014_M12_octanol_pH-metric high logP.t3r** | | |

#### Experiment Log (continued)

[1:09:14] Dispensed 0.000776 mL of Base (0.5 M KOH)
[1:09:19] Stepping pH = 3.56
[1:09:19] Dispensed 0.000400 mL of Base (0.5 M KOH)
[1:09:24] Stepping pH = 3.64
[1:09:39] Stirrer speed set to 0
[1:09:49] Datapoint id 60 collected
[1:09:49] Charge balance equation is out by 23.3%
[1:09:49] Stirrer speed set to 60
[1:09:55] pH 3.65 -> 3.85
[1:09:55] Using cautious pH adjust
[1:09:55] Dispensed 0.000517 mL of Base (0.5 M KOH)
[1:10:00] Stepping pH = 3.74
[1:10:00] Dispensed 0.000353 mL of Base (0.5 M KOH)
[1:10:05] Stepping pH = 3.84
[1:10:20] Stirrer speed set to 0
[1:10:30] Datapoint id 61 collected
[1:10:30] Charge balance equation is out by 16.0%
[1:10:30] Stirrer speed set to 60
[1:10:35] pH 3.85 -> 4.05
[1:10:35] Using cautious pH adjust
[1:10:35] Dispensed 0.000353 mL of Base (0.5 M KOH)
[1:10:41] Stepping pH = 3.95
[1:10:41] Dispensed 0.000235 mL of Base (0.5 M KOH)
[1:10:46] Stepping pH = 4.03
[1:10:46] Dispensed 0.000047 mL of Base (0.5 M KOH)
[1:10:51] Stepping pH = 4.04
[1:11:06] Stirrer speed set to 0
[1:11:16] Datapoint id 62 collected
[1:11:16] Charge balance equation is out by 12.2%
[1:11:16] Stirrer speed set to 60
[1:11:21] pH 4.05 -> 4.25
[1:11:21] Using charge balance adjust
[1:11:21] Dispensed 0.000470 mL of Base (0.5 M KOH)
[1:11:41] Stirrer speed set to 0
[1:11:52] Datapoint id 63 collected
[1:11:52] Charge balance equation is out by 53.3%
[1:11:52] Stirrer speed set to 60
[1:11:57] pH 4.37 -> 4.57
[1:11:57] Using cautious pH adjust
[1:11:58] Dispensed 0.000118 mL of Base (0.5 M KOH)
[1:12:03] Stepping pH = 4.40
[1:12:03] Dispensed 0.000212 mL of Base (0.5 M KOH)
[1:12:08] Stepping pH = 4.63
[1:12:23] Stirrer speed set to 0
[1:12:35] Datapoint id 64 collected
[1:12:35] Charge balance equation is out by -42.4%
[1:12:35] Stirrer speed set to 60
[1:12:40] pH 4.66 -> 4.86
[1:12:40] Using cautious pH adjust
[1:12:40] Dispensed 0.000071 mL of Base (0.5 M KOH)
[1:12:45] Stepping pH = 4.68
[1:12:45] Dispensed 0.000165 mL of Base (0.5 M KOH)
[1:12:50] Stepping pH = 4.86
[1:13:05] Stirrer speed set to 0
[1:13:18] Datapoint id 65 collected
[1:13:18] Charge balance equation is out by -88.6%
[1:13:18] Stirrer speed set to 60
[1:13:23] pH 4.89 -> 5.09
[1:13:24] Using cautious pH adjust

Sample name: **M12_octanol** Experiment start time: **3/3/2018 6:11:21 PM**
Assay name: **pH-metric high logP** Analyst: **Pion**
Assay ID: **18C-03014** Instrument ID: **T312060**
Filename: **C:\Sirius_T3\Mehtap\20180302_exp29_logP_T3-2\18C-03014_M12_octanol_pH-metric high logP.t3r**

#### Experiment Log (continued)

[1:13:24] Dispensed 0.000047 mL of Base (0.5 M KOH)
[1:13:29] Stepping pH = 4.90
[1:13:29] Dispensed 0.000118 mL of Base (0.5 M KOH)
[1:13:34] Stepping pH = 4.99
[1:13:34] Dispensed 0.000071 mL of Base (0.5 M KOH)
[1:13:39] Stepping pH = 5.09
[1:13:54] Stirrer speed set to 0
[1:14:11] Datapoint id 66 collected
[1:14:11] Charge balance equation is out by -190.1%
[1:14:11] Stirrer speed set to 60
[1:14:16] pH 5.10 -> 5.30
[1:14:16] Using cautious pH adjust
[1:14:16] Dispensed 0.000024 mL of Base (0.5 M KOH)
[1:14:21] Stepping pH = 5.11
[1:14:21] Dispensed 0.000071 mL of Base (0.5 M KOH)
[1:14:26] Stepping pH = 5.22
[1:14:26] Dispensed 0.000047 mL of Base (0.5 M KOH)
[1:14:31] Stepping pH = 5.38
[1:14:46] Stirrer speed set to 0
[1:15:15] Datapoint id 67 collected
[1:15:15] Charge balance equation is out by -164.6%
[1:15:15] Stirrer speed set to 60
[1:15:20] pH 5.41 -> 5.61
[1:15:20] Using cautious pH adjust
[1:15:20] Dispensed 0.000024 mL of Base (0.5 M KOH)
[1:15:25] Stepping pH = 5.39
[1:15:25] Dispensed 0.000118 mL of Base (0.5 M KOH)
[1:15:30] Stepping pH = 5.71
[1:15:45] Stirrer speed set to 0
[1:16:31] Datapoint id 68 collected
[1:16:31] Charge balance equation is out by -225.5%
[1:16:31] Stirrer speed set to 60
[1:16:36] pH 5.83 -> 6.03
[1:16:36] Using cautious pH adjust
[1:16:36] Dispensed 0.000024 mL of Base (0.5 M KOH)
[1:16:42] Stepping pH = 5.89
[1:16:42] Dispensed 0.000024 mL of Base (0.5 M KOH)
[1:16:47] Stepping pH = 6.03
[1:17:02] Stirrer speed set to 0
[1:17:27] Datapoint id 69 collected
[1:17:27] Charge balance equation is out by -8.4%
[1:17:27] Stirrer speed set to 60
[1:17:32] pH 5.99 -> 6.19
[1:17:32] Using charge balance adjust
[1:17:32] Dispensed 0.000047 mL of Base (0.5 M KOH)
[1:17:52] Stirrer speed set to 0
[1:18:45] Datapoint id 70 collected
[1:18:45] Charge balance equation is out by -19.7%
[1:18:45] Stirrer speed set to 60
[1:18:50] pH 6.13 -> 6.33
[1:18:50] Using cautious pH adjust
[1:18:50] Dispensed 0.000024 mL of Base (0.5 M KOH)
[1:18:56] Stepping pH = 6.13
[1:18:56] Dispensed 0.000071 mL of Base (0.5 M KOH)
[1:19:01] Stepping pH = 6.19
[1:19:01] Dispensed 0.000094 mL of Base (0.5 M KOH)
[1:19:06] Stepping pH = 6.80
[1:19:21] Stirrer speed set to 0
[1:20:21] Datapoint id 71 collected

Sample name: **M12_octanol**
Assay name: **pH-metric high logP**
Assay ID: **18C-03014**
Filename: **C:\Sirius_T3\Mehtap\20180302_exp29_logP_T3-2\18C-03014_M12_octanol_pH-metric high logP.t3r**

Experiment start time: **3/3/2018 6:11:21 PM**
Analyst: **Pion**
Instrument ID: **T312060**

#### Experiment Log (continued)

[1:20:21] Charge balance equation is out by -288.0%
[1:20:21] Stirrer speed set to 60
[1:20:26] pH 6.74 -> 6.94
[1:20:26] Using cautious pH adjust
[1:20:26] Dispensed 0.000024 mL of Base (0.5 M KOH)
[1:20:31] Stepping pH = 6.75
[1:20:31] Dispensed 0.000047 mL of Base (0.5 M KOH)
[1:20:37] Stepping pH = 6.90
[1:20:37] Dispensed 0.000024 mL of Base (0.5 M KOH)
[1:20:42] Stepping pH = 7.06
[1:20:57] Stirrer speed set to 0
[1:21:57] Datapoint id 72 collected
[1:21:57] Charge balance equation is out by -166.3%
[1:21:57] Stirrer speed set to 60
[1:22:02] pH 7.10 -> 7.30
[1:22:02] Using cautious pH adjust
[1:22:02] Dispensed 0.000024 mL of Base (0.5 M KOH)
[1:22:07] Stepping pH = 7.14
[1:22:07] Dispensed 0.000024 mL of Base (0.5 M KOH)
[1:22:12] Stepping pH = 7.26
[1:22:12] Dispensed 0.000024 mL of Base (0.5 M KOH)
[1:22:17] Stepping pH = 7.42
[1:22:32] Stirrer speed set to 0
[1:23:32] Datapoint id 73 collected
[1:23:32] Charge balance equation is out by -199.6%
[1:23:32] Stirrer speed set to 60
[1:23:37] pH 7.39 -> 7.59
[1:23:37] Using cautious pH adjust
[1:23:38] Dispensed 0.000024 mL of Base (0.5 M KOH)
[1:23:43] Stepping pH = 7.37
[1:23:43] Dispensed 0.000024 mL of Base (0.5 M KOH)
[1:23:48] Stepping pH = 7.35
[1:23:48] Dispensed 0.000188 mL of Base (0.5 M KOH)
[1:23:53] Stepping pH = 8.65
[1:24:08] Stirrer speed set to 0
[1:24:45] Datapoint id 74 collected
[1:24:45] Charge balance equation is out by -1,860.7%
[1:24:45] Stirrer speed set to 60
[1:24:50] pH 8.76 -> 8.96
[1:24:50] Using cautious pH adjust
[1:24:50] Dispensed 0.000024 mL of Base (0.5 M KOH)
[1:24:55] Stepping pH = 8.76
[1:24:55] Dispensed 0.000047 mL of Base (0.5 M KOH)
[1:25:00] Stepping pH = 8.82
[1:25:00] Dispensed 0.000071 mL of Base (0.5 M KOH)
[1:25:05] Stepping pH = 8.95
[1:25:20] Stirrer speed set to 0
[1:25:45] Datapoint id 75 collected
[1:25:45] Charge balance equation is out by -332.5%
[1:25:45] Stirrer speed set to 60
[1:25:50] pH 8.97 -> 9.05
[1:25:50] Using cautious pH adjust
[1:25:50] Dispensed 0.000024 mL of Base (0.5 M KOH)
[1:25:56] Stepping pH = 8.99
[1:25:56] Dispensed 0.000024 mL of Base (0.5 M KOH)
[1:26:01] Stepping pH = 8.99
[1:26:01] Dispensed 0.000047 mL of Base (0.5 M KOH)
[1:26:06] Stepping pH = 9.08
[1:26:21] Stirrer speed set to 0

| | | | |
|---|---|---|---|
| Sample name: | **M12_octanol** | Experiment start time: | **3/3/2018 6:11:21 PM** |
| Assay name: | **pH-metric high logP** | Analyst: | **Pion** |
| Assay ID: | **18C-03014** | Instrument ID: | **T312060** |
| Filename: | **C:\Sirius_T3\Mehtap\20180302_exp29_logP_T3-2\18C-03014_M12_octanol_pH-metric high logP.t3r** | | |

#### Experiment Log (continued)

[1:26:36] Datapoint id 76 collected
[1:26:36] Charge balance equation is out by -438.5%
[1:26:36] Argon flow rate set to 0
[1:26:40] Titrator arm moved over Titration position
