## Supplementary material for "Octanol-water partition coefficient measurements for the SAMPL6 Blind Prediction Challenge": SM12_18C-03015_M12_octanol_pH-metric high logP_report.pdf

| | | | |
|---|---|---|---|
| Sample name: | **M12_octanol** | Experiment start time: | **3/3/2018 7:38:51 PM** |
| Assay name: | **pH-metric high logP** | Analyst: | **Pion** |
| Assay ID: | **18C-03015** | Instrument ID: | **T312060** |
| Filename: | **C:\Sirius_T3\Mehtap\20180302_exp29_logP_T3-2\18C-03015_M12_octanol_pH-metric high logP.t3r** | | |

#### pH-metric Result

| | | |
|---|---|---|
| logP (XH +) | 0.40 | ±0.06 (n=50) |
| logP (neutral X) | 3.80 | ±0.01 (n=50) |

##### 18C-03015 Points 1 to 18

| | |
|---|---|
| M12_octanol concentration factor | 1.084 |
| Carbonate | 0.0827 mM |
| Acidity error | -3.00593 mM |

##### 18C-03015 Points 19 to 29

| | |
|---|---|
| M12_octanol concentration factor | 1.664 |
| Carbonate | 0.1338 mM |
| Acidity error | -2.74094 mM |

##### 18C-03015 Points 30 to 56

| | |
|---|---|
| M12_octanol concentration factor | 0.918 |
| Carbonate | 0.0569 mM |
| Acidity error | 8.57201 mM |

#### Warnings and errors

| | |
|---|---|
| Errors | None |
| Warnings | None |

#### Sample logD and percent species

| pH | M12_octanol logD | M12_octanol M12_octanolH | M12_octanol M12_octanol | M12_octanol M12_octanolH* | M12_octanol M12_octanol* | Comment |
|---|---|---|---|---|---|---|
| 1.000 | 0.45 | 26.11 % | 0.00 % | 65.29 % | 8.60 % | |
| 1.200 | 0.48 | 24.86 % | 0.00 % | 62.16 % | 12.97 % | Stomach pH |
| 2.000 | 0.76 | 14.72 % | 0.01 % | 36.81 % | 48.46 % | |
| 3.000 | 1.55 | 2.75 % | 0.01 % | 6.86 % | 90.38 % | |
| 4.000 | 2.50 | 0.30 % | 0.02 % | 0.75 % | 98.93 % | |
| 5.000 | 3.33 | 0.03 % | 0.02 % | 0.08 % | 99.88 % | |
| 6.000 | 3.72 | 0.00 % | 0.02 % | 0.01 % | 99.97 % | |
| 6.500 | 3.77 | 0.00 % | 0.02 % | 0.00 % | 99.98 % | |
| 7.000 | 3.79 | 0.00 % | 0.02 % | 0.00 % | 99.98 % | |
| 7.400 | 3.79 | 0.00 % | 0.02 % | 0.00 % | 99.98 % | Blood pH |
| 8.000 | 3.80 | 0.00 % | 0.02 % | 0.00 % | 99.98 % | |
| 9.000 | 3.80 | 0.00 % | 0.02 % | 0.00 % | 99.98 % | |
| 10.000 | 3.80 | 0.00 % | 0.02 % | 0.00 % | 99.98 % | |
| 11.000 | 3.80 | 0.00 % | 0.02 % | 0.00 % | 99.98 % | |
| 12.000 | 3.80 | 0.00 % | 0.02 % | 0.00 % | 99.98 % | |

| | | | |
|---|---|---|---|
| Sample name: | **M12_octanol** | Experiment start time: | **3/3/2018 7:38:51 PM** |
| Assay name: | **pH-metric high logP** | Analyst: | **Pion** |
| Assay ID: | **18C-03015** | Instrument ID: | **T312060** |
| Filename: | **C:\Sirius_T3\Mehtap\20180302_exp29_logP_T3-2\18C-03015_M12_octanol_pH-metric high logP.t3r** | | |

#### Graphs

Charge balance residuals

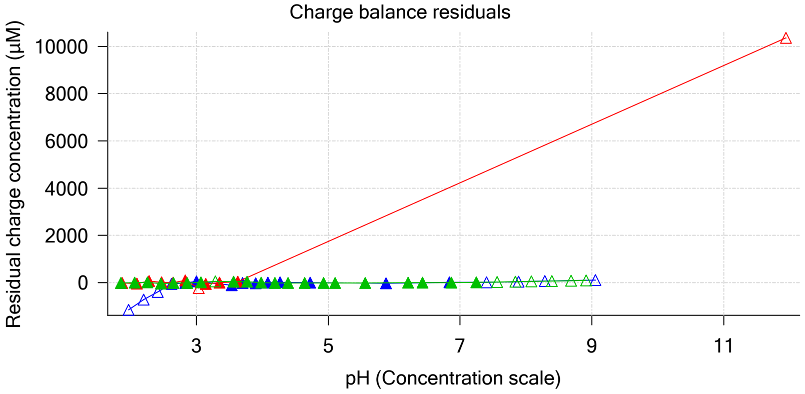


Ionic strength

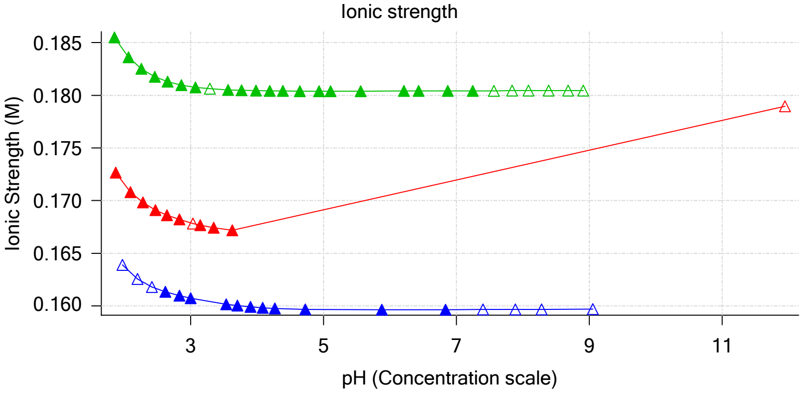


Buffer index profile

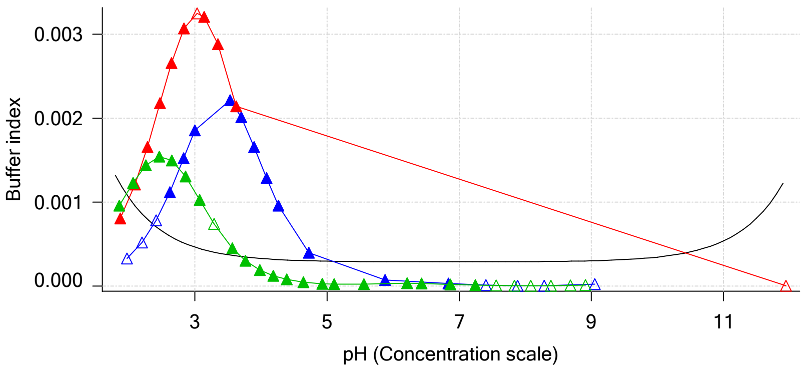


Sample M12_octanol Ionisation Graph

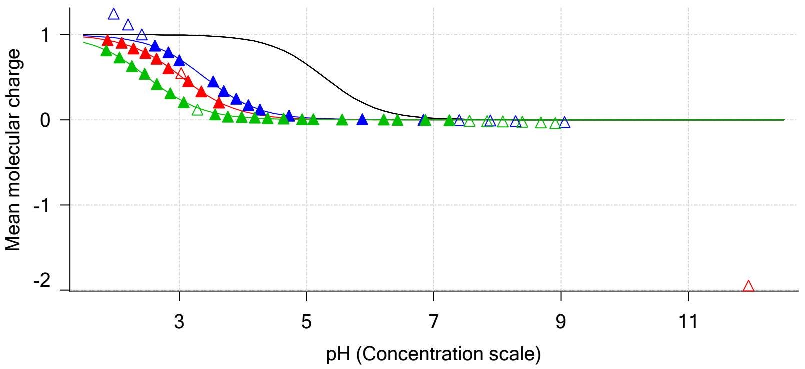


Distribution of species for sample M12_octanol

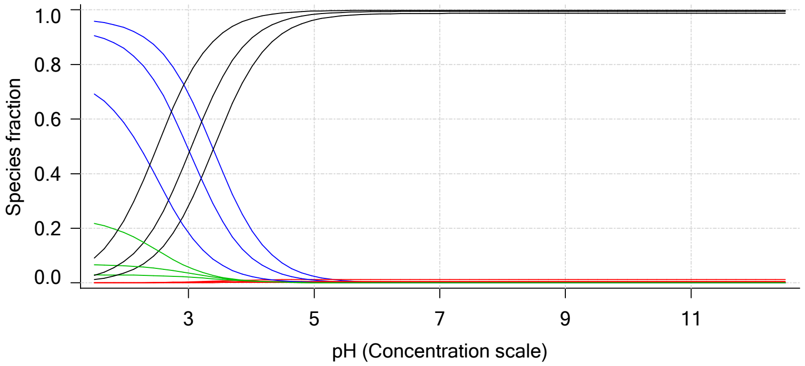


Lipophilicity Profile

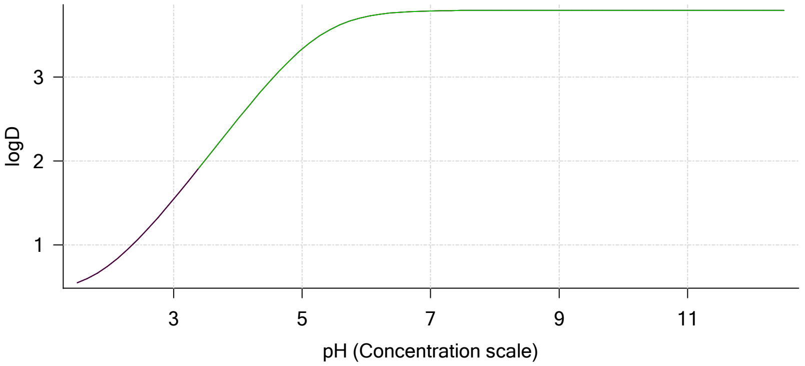


Log R Profile

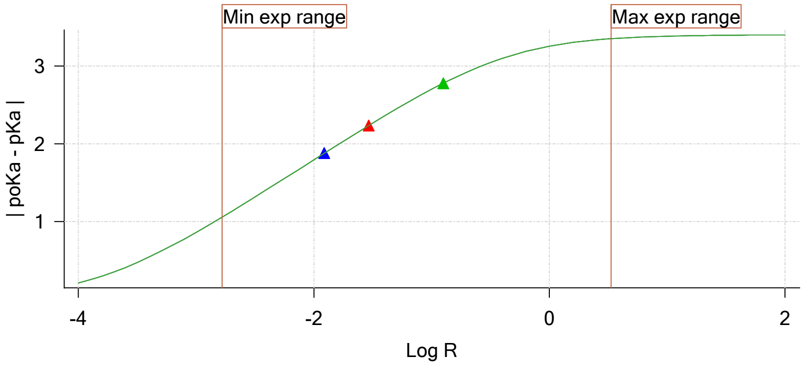


Sample Carbonate Ionisation Graph

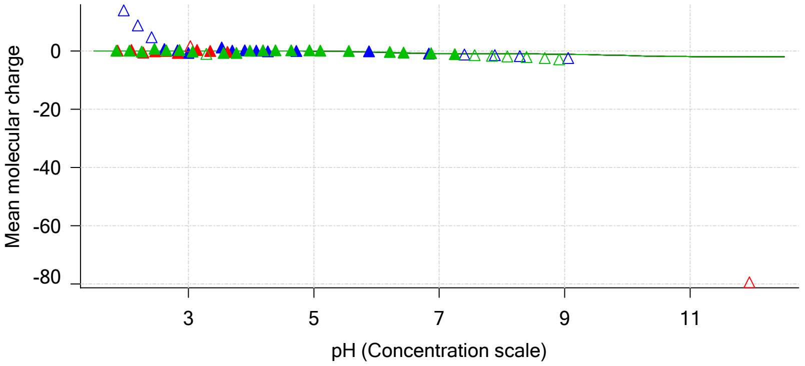

| | | | |
|---|---|---|---|
| Sample name: | **M12_octanol** | Experiment start time: | **3/3/2018 7:38:51 PM** |
| Assay name: | **pH-metric high logP** | Analyst: | **Pion** |
| Assay ID: | **18C-03015** | Instrument ID: | **T312060** |
| Filename: | **C:\Sirius_T3\Mehtap\20180302_exp29_logP_T3-2\18C-03015_M12_octanol_pH-metric high logP.t3r** | | |

#### Graphs (continued)

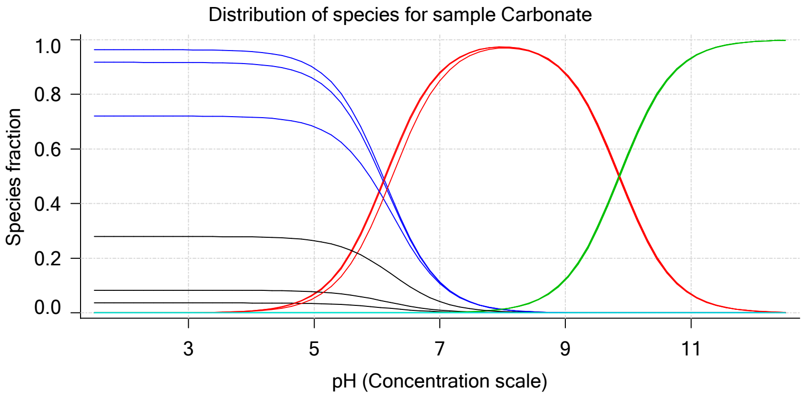

| | | | |
|---|---|---|---|
| Sample name: | **M12_octanol** | Experiment start time: | **3/3/2018 7:38:51 PM** |
| Assay name: | **pH-metric high logP** | Analyst: | **Pion** |
| Assay ID: | **18C-03015** | Instrument ID: | **T312060** |
| Filename: | **C:\Sirius_T3\Mehtap\20180302_exp29_logP_T3-2\18C-03015_M12_octanol_pH-metric high logP.t3r** | | |

# pH-metric high logP   Titration 1 of 3   18C-03015 Points 1 to 18

#### Overall results

| | |
|---|---|
| RMSD | 0.115 |
| Average ionic strength | 0.160 M |
| Average temperature | 24.9°C |
| Partition ratio | 0.0123 : 1 |
| Analyte concentration range | 3571.6 µM to 3683.7 µM |
| Total points considered | 11 of 18 |

#### Warnings and errors

| | |
|---|---|
| Errors | None |
| Warnings | Excessive acidity error present |

#### Four-Plus parameters

| | | | |
|---|---|---|---|
| Alpha | 0.111 | 3/3/2018 7:38:51 PM | C:\Sirius_T3\HCl18C02.t3r |
| S | 0.9988 | 3/3/2018 7:38:51 PM | C:\Sirius_T3\HCl18C02.t3r |
| jH | 1.0 | 3/3/2018 7:38:51 PM | C:\Sirius_T3\HCl18C02.t3r |
| jOH | -0.8 | 3/3/2018 7:38:51 PM | C:\Sirius_T3\HCl18C02.t3r |

#### Titrants

| | | | |
|---|---|---|---|
| 0.50 M HCl | 0.999058 | 3/3/2018 7:38:51 PM | C:\Sirius_T3\HCl18C02.t3r |
| 0.50 M KOH | 0.999845 | 3/3/2018 7:38:51 PM | C:\Sirius_T3\KOH18B27.t3r |

#### Sample

| | |
|---|---|
| M12_octanol concentration factor | 1.084 |
| M12_octanol stoichiometry | 1.000 |
| Chloride stoichiometry | 1.000 |
| Base pKa 1 | 5.28 |
| logP (XH +) | 1.02 |
| logP (neutral X) | 3.82 |

#### Sample graphs

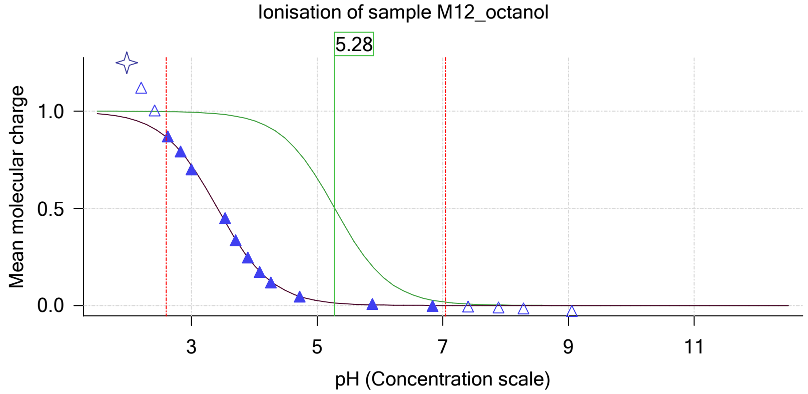


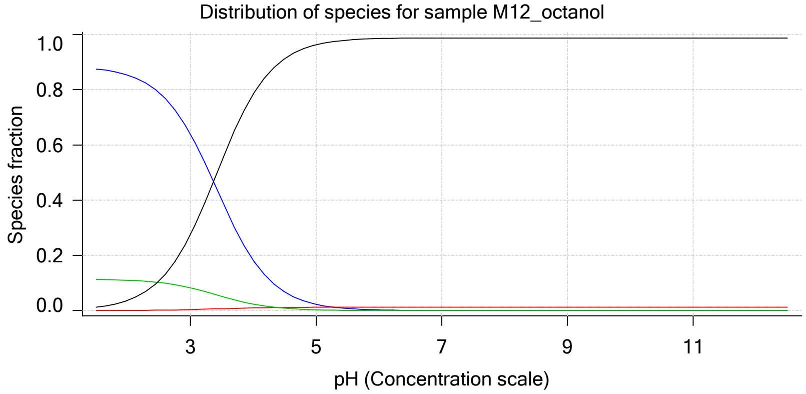

Sample name: **M12_octanol**
Assay name: **pH-metric high logP**
Assay ID: **18C-03015**
Filename: **C:\Sirius_T3\Mehtap\20180302_exp29_logP_T3-2\18C-03015_M12_octanol_pH-metric high logP.t3r**

Experiment start time: **3/3/2018 7:38:51 PM**
Analyst: **Pion**
Instrument ID: **T312060**

#### Sample graphs (continued)

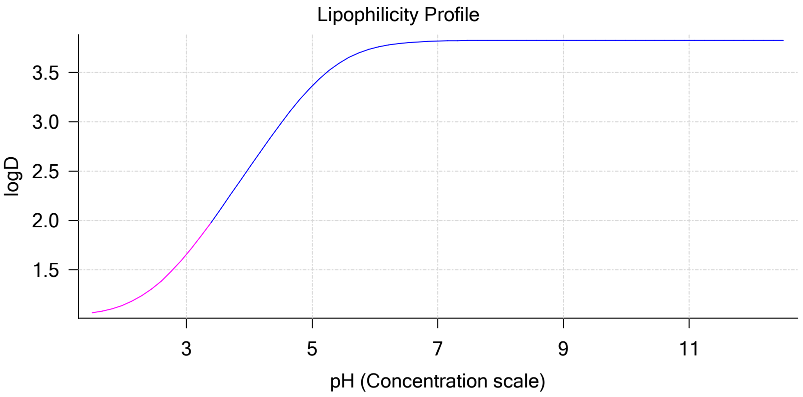


#### Sample logD and percent species

| pH | M12_octanol logD | M12_octanol M12_octanolH | M12_octanol M12_octanol | M12_octanol M12_octanolH* | M12_octanol M12_octanol* | Comment |
|---|---|---|---|---|---|---|
| 1.000 | 1.03 | 88.26 % | 0.00 % | 11.36 % | 0.38 % | |
| 1.200 | 1.04 | 88.06 % | 0.01 % | 11.33 % | 0.60 % | Stomach pH |
| 2.000 | 1.15 | 85.31 % | 0.04 % | 10.98 % | 3.67 % | |
| 3.000 | 1.66 | 63.92 % | 0.34 % | 8.22 % | 27.52 % | |
| 4.000 | 2.54 | 18.23 % | 0.96 % | 2.34 % | 78.47 % | |
| 5.000 | 3.36 | 2.24 % | 1.17 % | 0.29 % | 96.30 % | |
| 6.000 | 3.75 | 0.23 % | 1.20 % | 0.03 % | 98.54 % | |
| 6.500 | 3.80 | 0.07 % | 1.20 % | 0.01 % | 98.71 % | |
| 7.000 | 3.82 | 0.02 % | 1.20 % | 0.00 % | 98.77 % | |
| 7.400 | 3.82 | 0.01 % | 1.20 % | 0.00 % | 98.79 % | Blood pH |
| 8.000 | 3.82 | 0.00 % | 1.20 % | 0.00 % | 98.79 % | |
| 9.000 | 3.82 | 0.00 % | 1.20 % | 0.00 % | 98.80 % | |
| 10.000 | 3.82 | 0.00 % | 1.20 % | 0.00 % | 98.80 % | |
| 11.000 | 3.82 | 0.00 % | 1.20 % | 0.00 % | 98.80 % | |
| 12.000 | 3.82 | 0.00 % | 1.20 % | 0.00 % | 98.80 % | |

#### Carbonate and acidity

Carbonate 0.083 mM
Acidity error -3.006 mM

#### Other graphs

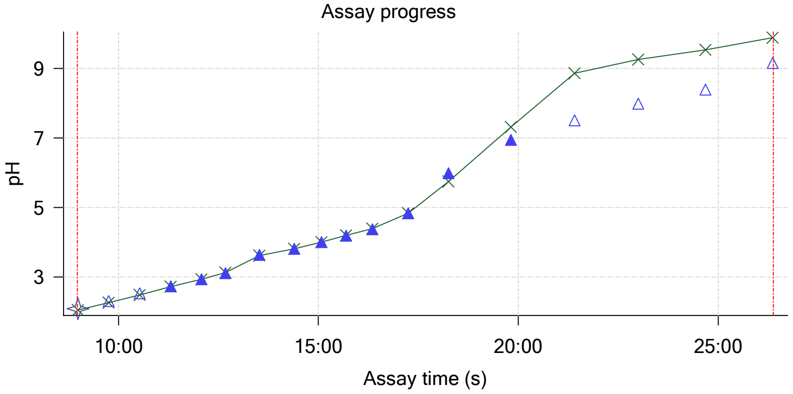


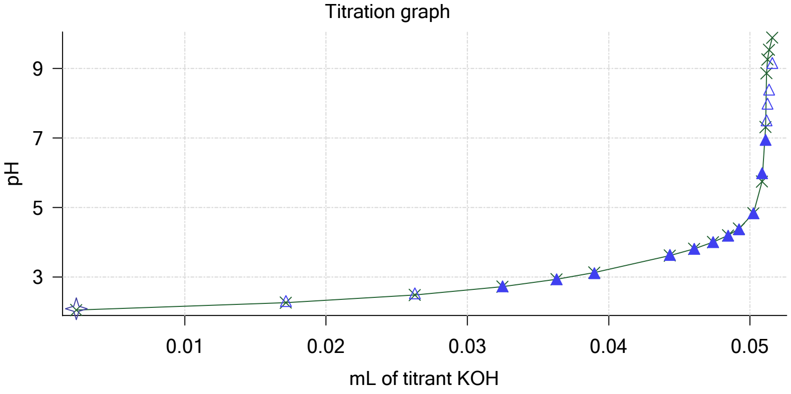

Sample name: **M12_octanol**
Assay name: **pH-metric high logP**
Assay ID: **18C-03015**
Filename: **C:\Sirius_T3\Mehtap\20180302_exp29_logP_T3-2\18C-03015_M12_octanol_pH-metric high logP.t3r**

Experiment start time: **3/3/2018 7:38:51 PM**
Analyst: **Pion**
Instrument ID: **T312060**

#### Other graphs (continued)

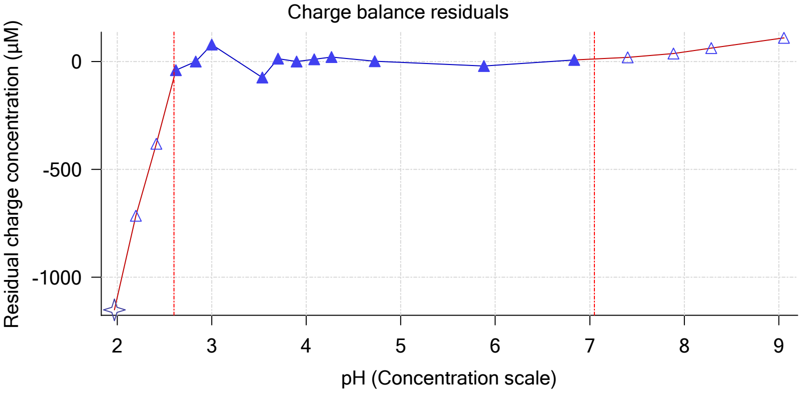


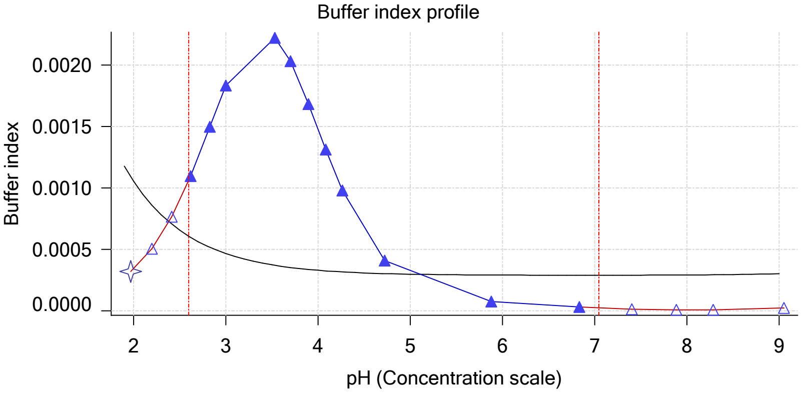


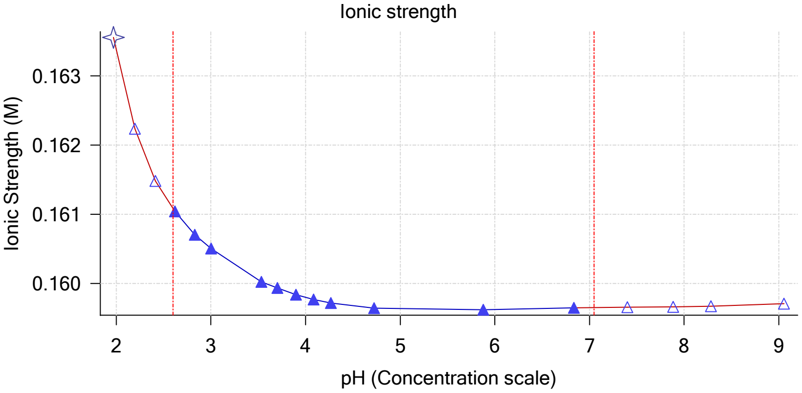


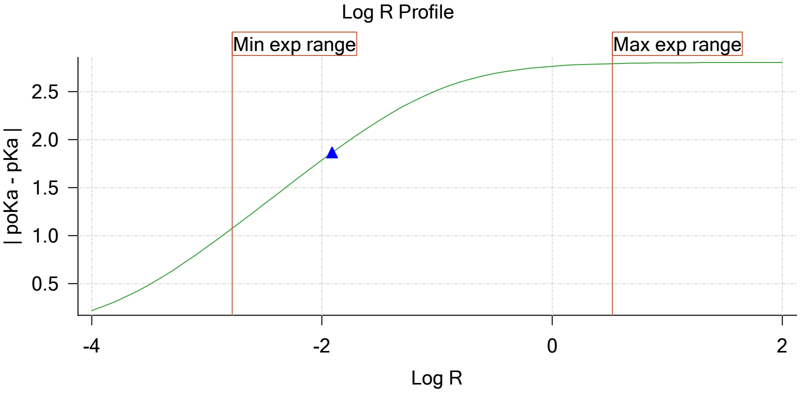


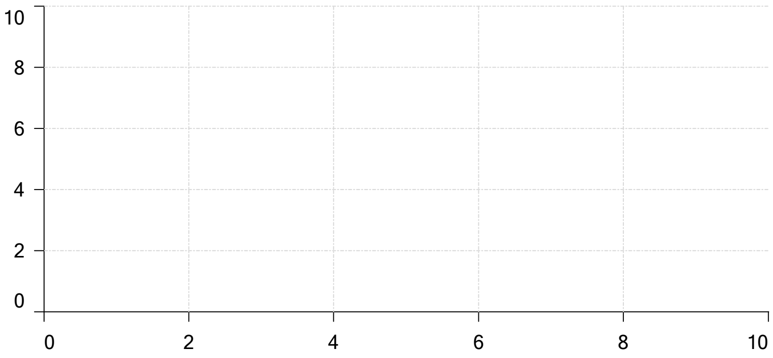


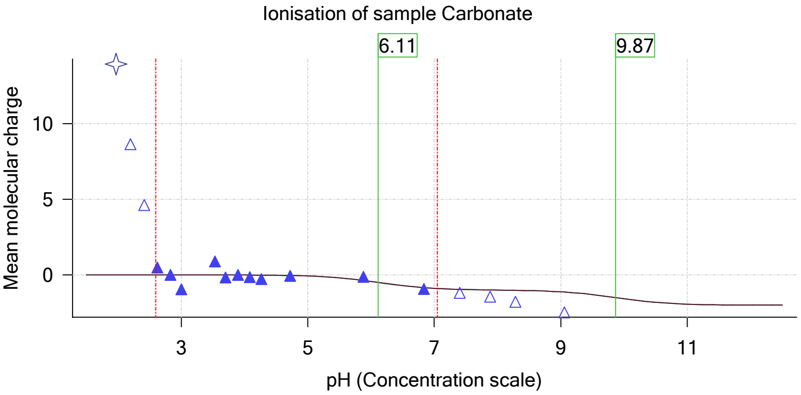

| | | | |
|---|---|---|---|
| Sample name: | **M12_octanol** | Experiment start time: | **3/3/2018 7:38:51 PM** |
| Assay name: | **pH-metric high logP** | Analyst: | **Pion** |
| Assay ID: | **18C-03015** | Instrument ID: | **T312060** |
| Filename: | **C:\Sirius_T3\Mehtap\20180302_exp29_logP_T3-2\18C-03015_M12_octanol_pH-metric high logP.t3r** | | |

### pH-metric high logP Titration 2 of 3 18C-03015 Points 19 to 29

#### Overall results

| | |
|---|---|
| RMSD | 0.008 |
| Average ionic strength | 0.168 M |
| Average temperature | 25.0°C |
| Partition ratio | 0.0293 : 1 |
| Analyte concentration range | 3109.3 µM to 3391.7 µM |
| Total points considered | 9 of 11 |

#### Warnings and errors

| | |
|---|---|
| Errors | None |
| Warnings | Sample concentration factor out of range<br>Excessive acidity error present |

#### Four-Plus parameters

| | | | |
|---|---|---|---|
| Alpha | 0.111 | 3/3/2018 7:38:51 PM | C:\Sirius_T3\HCl18C02.t3r |
| S | 0.9988 | 3/3/2018 7:38:51 PM | C:\Sirius_T3\HCl18C02.t3r |
| jH | 1.0 | 3/3/2018 7:38:51 PM | C:\Sirius_T3\HCl18C02.t3r |
| jOH | -0.8 | 3/3/2018 7:38:51 PM | C:\Sirius_T3\HCl18C02.t3r |

#### Titrants

| | | | |
|---|---|---|---|
| 0.50 M HCl | 0.999058 | 3/3/2018 7:38:51 PM | C:\Sirius_T3\HCl18C02.t3r |
| 0.50 M KOH | 0.999845 | 3/3/2018 7:38:51 PM | C:\Sirius_T3\KOH18B27.t3r |

#### Sample

| | |
|---|---|
| M12_octanol concentration factor | 1.664 |
| M12_octanol stoichiometry | 1.000 |
| Chloride stoichiometry | 1.000 |
| Base pKa 1 | 5.28 |
| logP (XH +) | 1.02 |
| logP (neutral X) | 3.89 |

#### Sample graphs

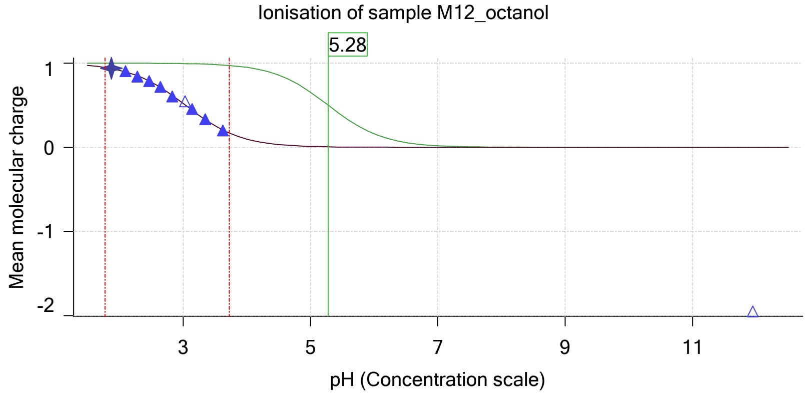


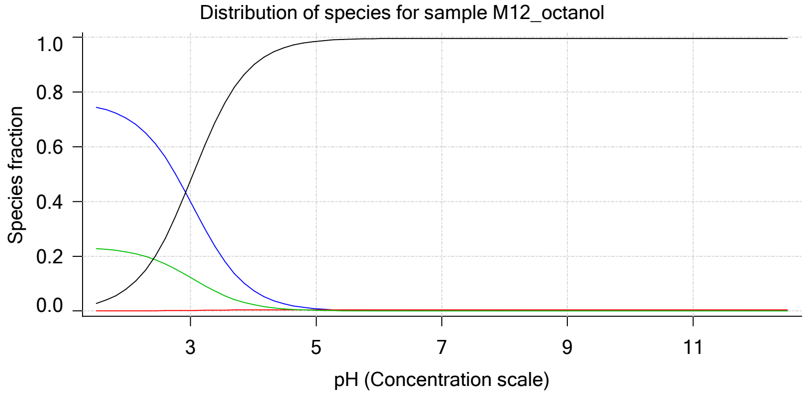

| | | | |
|---|---|---|---|
| Sample name: | **M12_octanol** | Experiment start time: | **3/3/2018 7:38:51 PM** |
| Assay name: | **pH-metric high logP** | Analyst: | **Pion** |
| Assay ID: | **18C-03015** | Instrument ID: | **T312060** |
| Filename: | **C:\Sirius_T3\Mehtap\20180302_exp29_logP_T3-2\18C-03015_M12_octanol_pH-metric high logP.t3r** | | |

#### Sample graphs (continued)

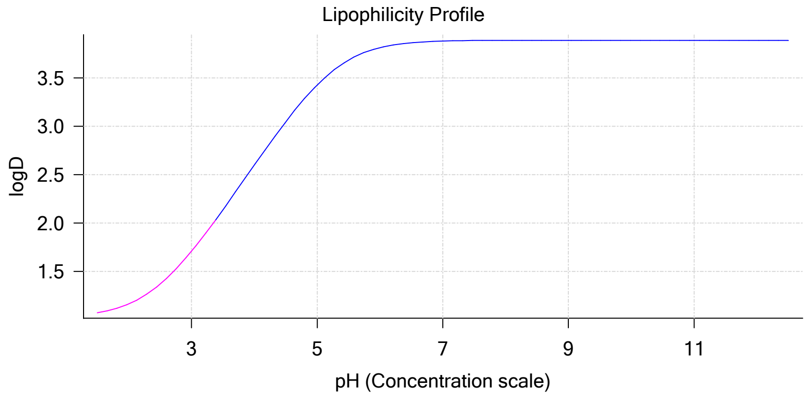


#### Sample logD and percent species

| pH | M12_octanol logD | M12_octanol M12_octanolH | M12_octanol M12_octanol | M12_octanol M12_octanolH* | M12_octanol M12_octanol* | Comment |
|---|---|---|---|---|---|---|
| 1.000 | 1.04 | 75.84 % | 0.00 % | 23.25 % | 0.90 % | |
| 1.200 | 1.05 | 75.44 % | 0.01 % | 23.13 % | 1.43 % | Stomach pH |
| 2.000 | 1.16 | 70.11 % | 0.04 % | 21.50 % | 8.36 % | |
| 3.000 | 1.71 | 39.94 % | 0.21 % | 12.25 % | 47.60 % | |
| 4.000 | 2.60 | 7.53 % | 0.40 % | 2.31 % | 89.76 % | |
| 5.000 | 3.43 | 0.83 % | 0.43 % | 0.25 % | 98.49 % | |
| 6.000 | 3.81 | 0.08 % | 0.44 % | 0.03 % | 99.45 % | |
| 6.500 | 3.86 | 0.03 % | 0.44 % | 0.01 % | 99.53 % | |
| 7.000 | 3.88 | 0.01 % | 0.44 % | 0.00 % | 99.55 % | |
| 7.400 | 3.89 | 0.00 % | 0.44 % | 0.00 % | 99.56 % | Blood pH |
| 8.000 | 3.89 | 0.00 % | 0.44 % | 0.00 % | 99.56 % | |
| 9.000 | 3.89 | 0.00 % | 0.44 % | 0.00 % | 99.56 % | |
| 10.000 | 3.89 | 0.00 % | 0.44 % | 0.00 % | 99.56 % | |
| 11.000 | 3.89 | 0.00 % | 0.44 % | 0.00 % | 99.56 % | |
| 12.000 | 3.89 | 0.00 % | 0.44 % | 0.00 % | 99.56 % | |

#### Carbonate and acidity

| | |
|---|---|
| Carbonate | 0.134 mM |
| Acidity error | -2.741 mM |

#### Other graphs

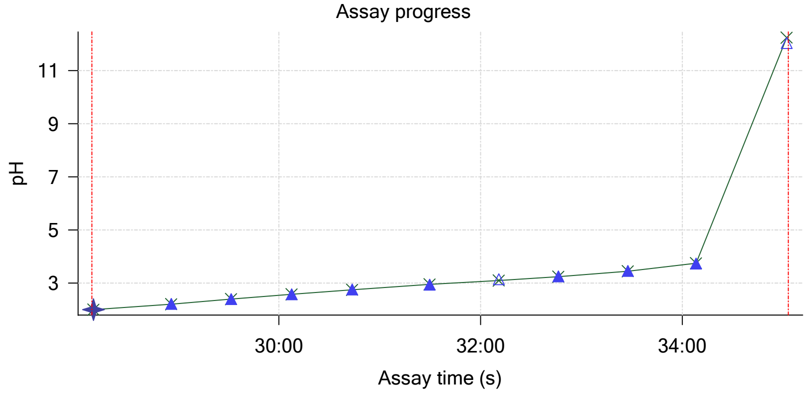


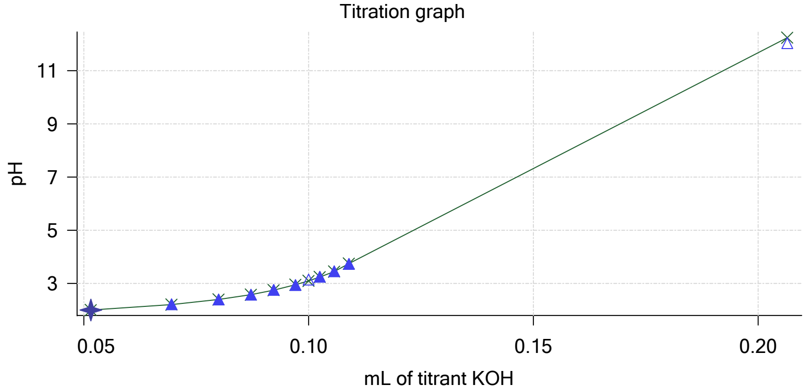

Sample name: **M12_octanol**
Assay name: **pH-metric high logP**
Assay ID: **18C-03015**
Filename: **C:\Sirius_T3\Mehtap\20180302_exp29_logP_T3-2\18C-03015_M12_octanol_pH-metric high logP.t3r**

Experiment start time: **3/3/2018 7:38:51 PM**
Analyst: **Pion**
Instrument ID: **T312060**

#### Other graphs (continued)

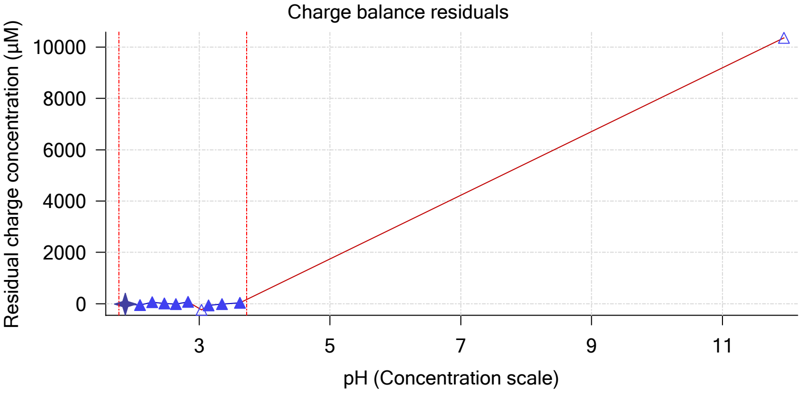


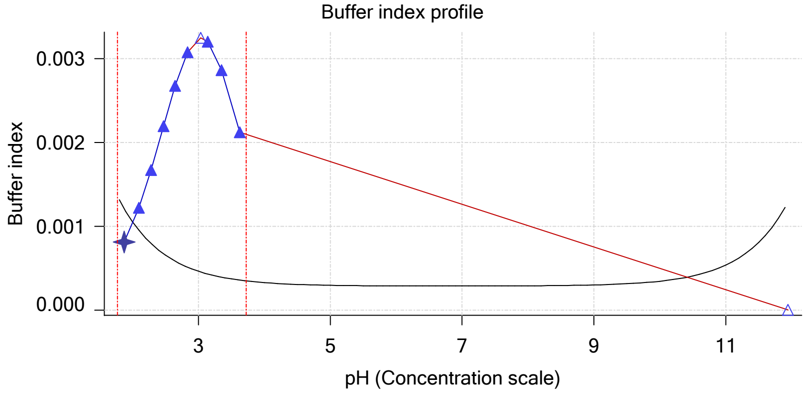


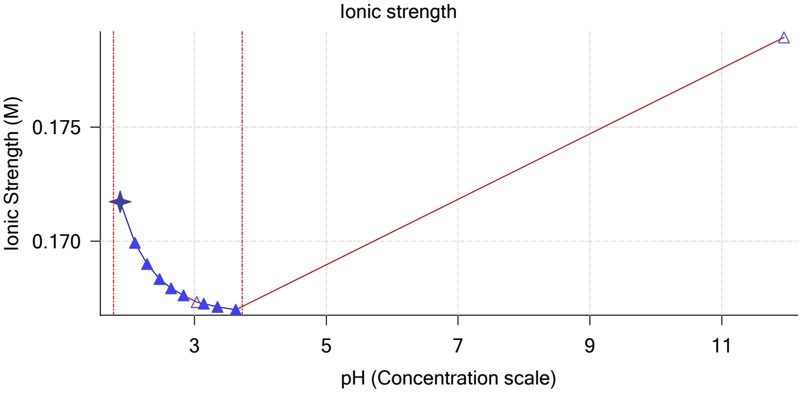


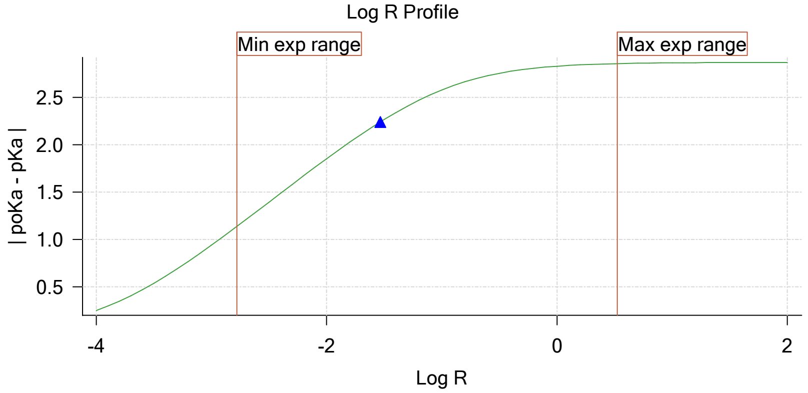


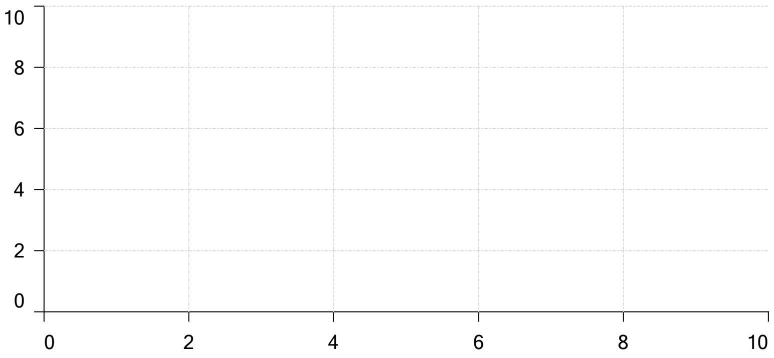


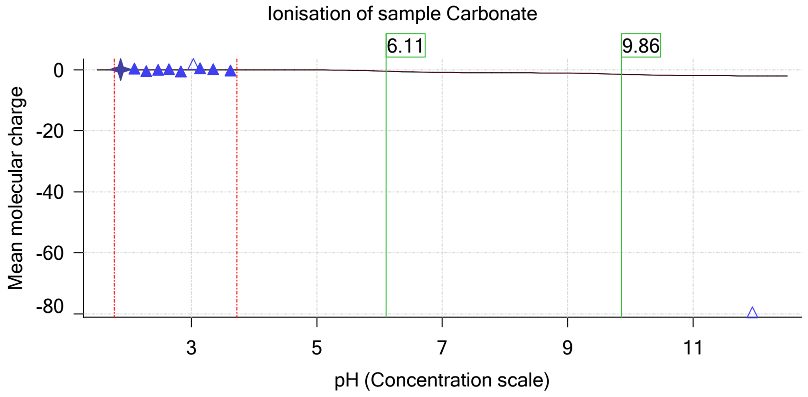

| | | | |
|---|---|---|---|
| Sample name: | **M12_octanol** | Experiment start time: | **3/3/2018 7:38:51 PM** |
| Assay name: | **pH-metric high logP** | Analyst: | **Pion** |
| Assay ID: | **18C-03015** | Instrument ID: | **T312060** |
| Filename: | **C:\Sirius_T3\Mehtap\20180302_exp29_logP_T3-2\18C-03015_M12_octanol_pH-metric high logP.t3r** | | |

# pH-metric high logP   Titration 3 of 3   18C-03015 Points 30 to 56

#### Overall results

| | |
|---|---|
| RMSD | 0.199 |
| Average ionic strength | 0.181 M |
| Average temperature | 25.0°C |
| Partition ratio | 0.1256 : 1 |
| Analyte concentration range | 2574.7 μM to 2650.8 μM |
| Total points considered | 20 of 27 |

#### Warnings and errors

| | |
|---|---|
| Errors | None |
| Warnings | Excessive acidity error present |

#### Four-Plus parameters

| | | | |
|---|---|---|---|
| Alpha | 0.111 | 3/3/2018 7:38:51 PM | C:\Sirius_T3\HCl18C02.t3r |
| S | 0.9988 | 3/3/2018 7:38:51 PM | C:\Sirius_T3\HCl18C02.t3r |
| jH | 1.0 | 3/3/2018 7:38:51 PM | C:\Sirius_T3\HCl18C02.t3r |
| jOH | -0.8 | 3/3/2018 7:38:51 PM | C:\Sirius_T3\HCl18C02.t3r |

#### Titrants

| | | | |
|---|---|---|---|
| 0.50 M HCl | 0.999058 | 3/3/2018 7:38:51 PM | C:\Sirius_T3\HCl18C02.t3r |
| 0.50 M KOH | 0.999845 | 3/3/2018 7:38:51 PM | C:\Sirius_T3\KOH18B27.t3r |

#### Sample

| | |
|---|---|
| M12_octanol concentration factor | 0.918 |
| M12_octanol stoichiometry | 1.000 |
| Chloride stoichiometry | 1.000 |
| Base pKa 1 | 5.28 |
| logP (XH +) | 1.02 |
| logP (neutral X) | 4.04 |

#### Sample graphs

Ionisation of sample M12_octanol

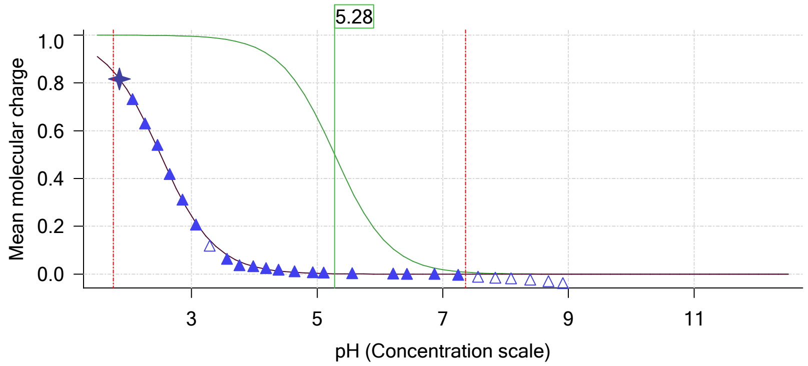


Distribution of species for sample M12_octanol

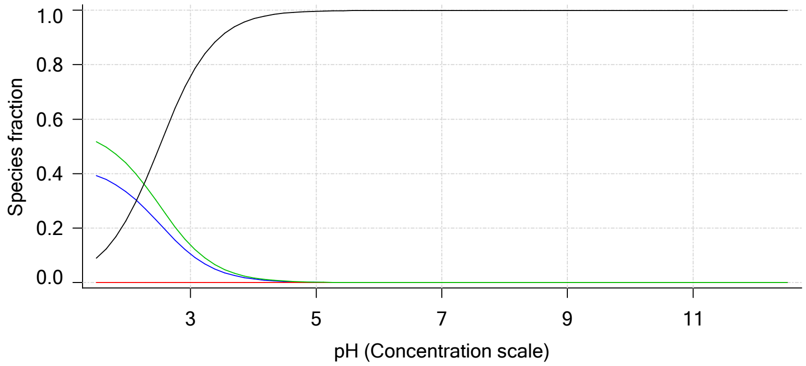

| | | | |
|---|---|---|---|
| Sample name: | **M12_octanol** | Experiment start time: | **3/3/2018 7:38:51 PM** |
| Assay name: | **pH-metric high logP** | Analyst: | **Pion** |
| Assay ID: | **18C-03015** | Instrument ID: | **T312060** |
| Filename: | **C:\Sirius_T3\Mehtap\20180302_exp29_logP_T3-2\18C-03015_M12_octanol_pH-metric high logP.t3r** | | |

#### Sample graphs (continued)

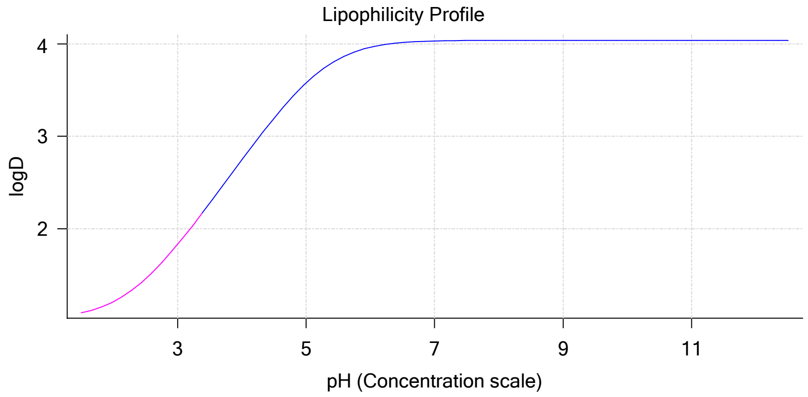


#### Sample logD and percent species

| pH | M12_octanol logD | M12_octanol M12_octanolH | M12_octanol M12_octanol | M12_octanol M12_octanolH* | M12_octanol M12_octanol* | Comment |
|---|---|---|---|---|---|---|
| 1.000 | 1.04 | 41.89 % | 0.00 % | 55.08 % | 3.02 % | |
| 1.200 | 1.06 | 41.16 % | 0.00 % | 54.12 % | 4.71 % | Stomach pH |
| 2.000 | 1.21 | 32.92 % | 0.02 % | 43.29 % | 23.77 % | |
| 3.000 | 1.83 | 10.48 % | 0.06 % | 13.78 % | 75.68 % | |
| 4.000 | 2.75 | 1.34 % | 0.07 % | 1.76 % | 96.83 % | |
| 5.000 | 3.58 | 0.14 % | 0.07 % | 0.18 % | 99.61 % | |
| 6.000 | 3.96 | 0.01 % | 0.07 % | 0.02 % | 99.90 % | |
| 6.500 | 4.01 | 0.00 % | 0.07 % | 0.01 % | 99.92 % | |
| 7.000 | 4.03 | 0.00 % | 0.07 % | 0.00 % | 99.92 % | |
| 7.400 | 4.04 | 0.00 % | 0.07 % | 0.00 % | 99.93 % | Blood pH |
| 8.000 | 4.04 | 0.00 % | 0.07 % | 0.00 % | 99.93 % | |
| 9.000 | 4.04 | 0.00 % | 0.07 % | 0.00 % | 99.93 % | |
| 10.000 | 4.04 | 0.00 % | 0.07 % | 0.00 % | 99.93 % | |
| 11.000 | 4.04 | 0.00 % | 0.07 % | 0.00 % | 99.93 % | |
| 12.000 | 4.04 | 0.00 % | 0.07 % | 0.00 % | 99.93 % | |

#### Carbonate and acidity

| | | |
|---|---|---|
| 💎 | Carbonate | 0.057 mM |
| 💎 | Acidity error | 8.572 mM |

#### Other graphs

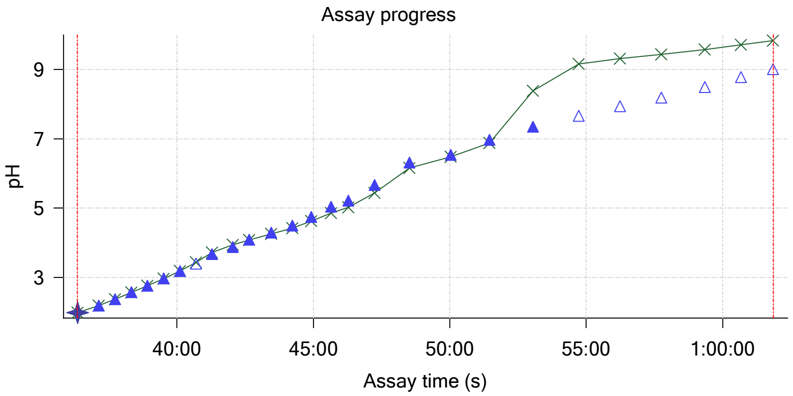


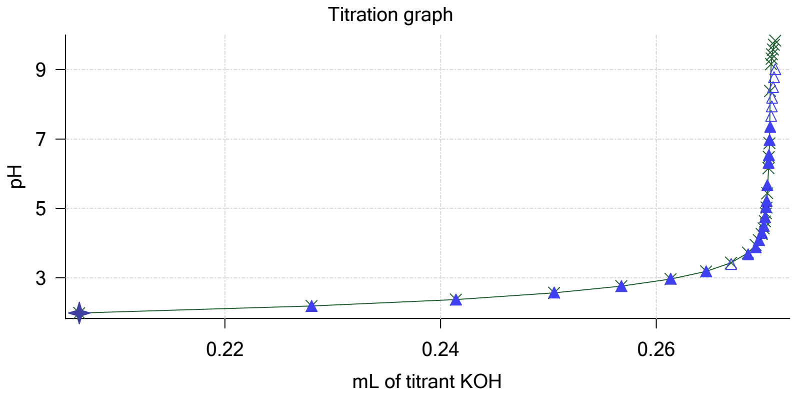

Sample name: **M12_octanol**
Assay name: **pH-metric high logP**
Assay ID: **18C-03015**
Filename: **C:\Sirius_T3\Mehtap\20180302_exp29_logP_T3-2\18C-03015_M12_octanol_pH-metric high logP.t3r**

Experiment start time: **3/3/2018 7:38:51 PM**
Analyst: **Pion**
Instrument ID: **T312060**

#### Other graphs (continued)

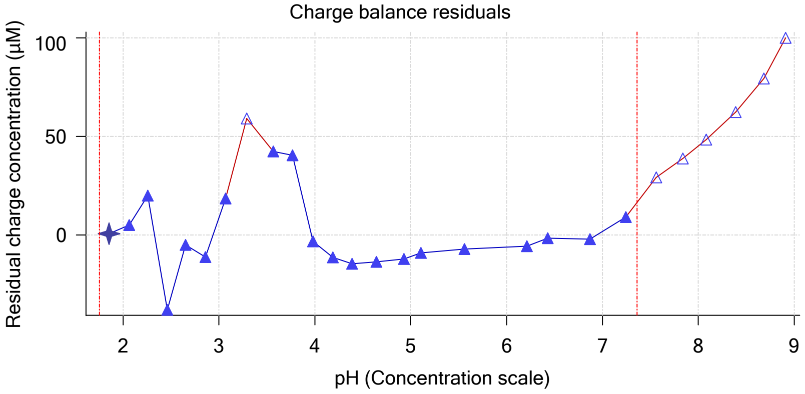


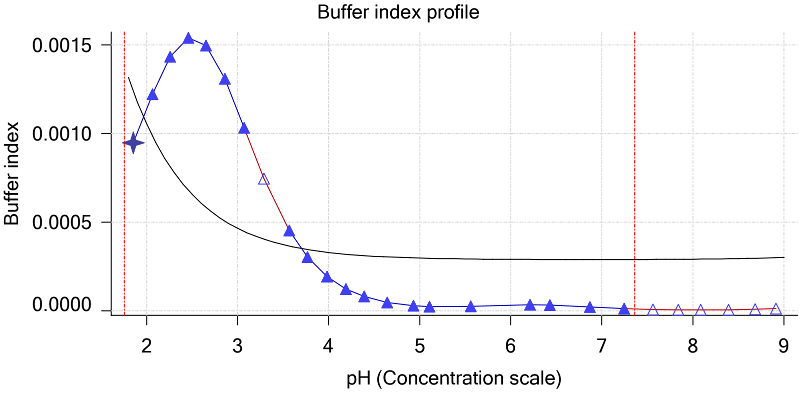


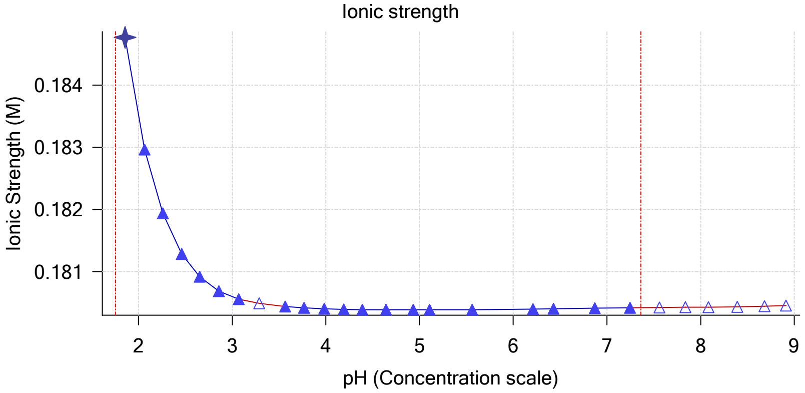


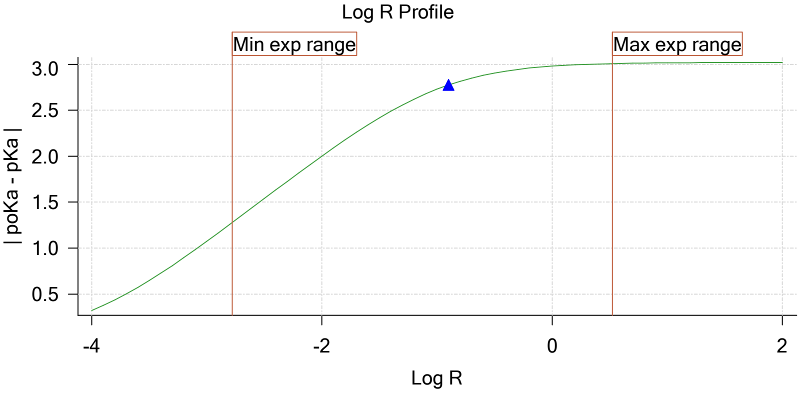


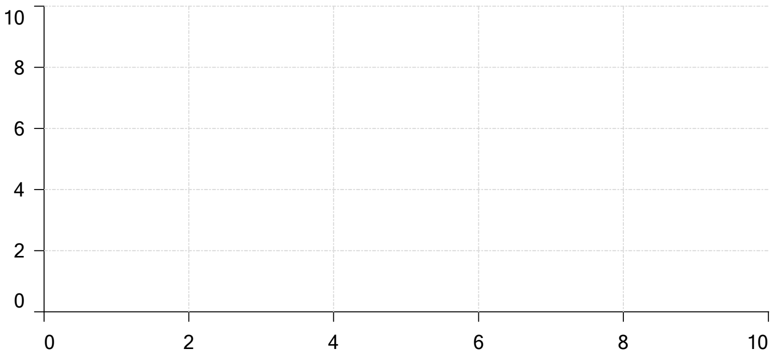


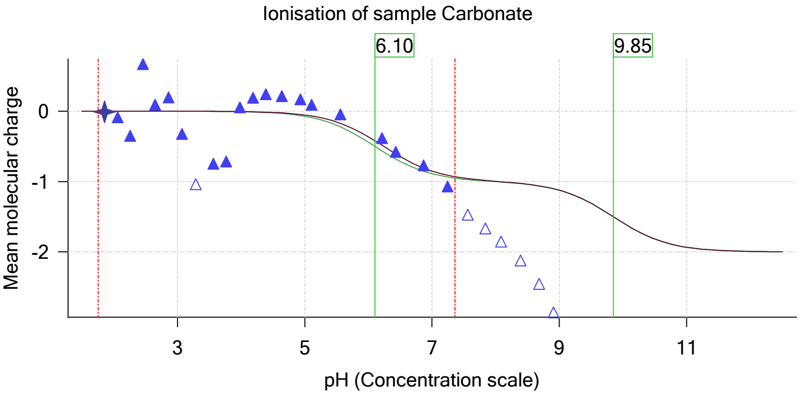

| | | | |
|---|---|---|---|
| Sample name: | **M12_octanol** | Experiment start time: | **3/3/2018 7:38:51 PM** |
| Assay name: | **pH-metric high logP** | Analyst: | **Pion** |
| Assay ID: | **18C-03015** | Instrument ID: | **T312060** |
| Filename: | **C:\Sirius_T3\Mehtap\20180302_exp29_logP_T3-2\18C-03015_M12_octanol_pH-metric high logP.t3r** | | |

#### Assay Model

| Settings | Value | Date/Time changed | Imported from |
|---|---|---|---|
| Sample name | M12_octanol | 2/28/2018 2:58:36 PM | User entered value |
| Sample by | Weight | | Default value |
| Sample weight | 0.001690 g | 3/2/2018 5:10:33 PM | User entered value |
| Formula weight | 292.16 g/mol | 2/28/2018 2:58:36 PM | User entered value |
| Solubility | Unknown | | Default value |
| Molecular weight | 255.70 | 2/28/2018 2:58:36 PM | User entered value |
| Individual pKa ionic environments | No | | Default value |
| Number of pKas | 1 | 2/28/2018 2:58:36 PM | User entered value |
| Sample is a | Base | 2/28/2018 2:58:36 PM | User entered value |
| pKa 1 | 5.28 | 2/28/2018 2:58:36 PM | User entered value |
| logp (XH +) | 1.02 | 3/2/2018 3:44:28 PM | User entered value |
| logP (neutral X) | 3.79 | 3/2/2018 3:44:35 PM | User entered value |
| Stoichiometry | 1.00000 | | Default value |
| Aprotic counterion name | Chloride | | From standards.xml file |
| Stoichiometry | 1.00 | | From standards.xml file |
| Charge per counterion | -1 | | From standards.xml file |

#### Events

| Time | Event | Water | Acid | Base | Octanol | pH | dpH/dt | pH R-squared | pH SD | dpH/dt time |
|---|---|---|---|---|---|---|---|---|---|---|
| 5:59.8 | Initial pH = 4.05 | | | | | | | | | |
| 8:59.2 | Data point 1 | 1.50000 mL | 0.04798 mL | 0.00233 mL | 0.01999 mL | 2.090 | -0.00070 | 0.00914 | 0.00036 | 10.0 s |
| 9:45.4 | Data point 2 | 1.50000 mL | 0.04798 mL | 0.01717 mL | 0.01999 mL | 2.312 | -0.00312 | 0.68871 | 0.00019 | 10.5 s |
| 10:31.9 | Data point 3 | 1.50000 mL | 0.04798 mL | 0.02629 mL | 0.01999 mL | 2.526 | 0.00243 | 0.06275 | 0.00048 | 10.5 s |
| 11:18.4 | Data point 4 | 1.50000 mL | 0.04798 mL | 0.03248 mL | 0.01999 mL | 2.730 | -0.00626 | 0.82301 | 0.00034 | 10.0 s |
| 12:04.3 | Data point 5 | 1.50000 mL | 0.04798 mL | 0.03631 mL | 0.01999 mL | 2.938 | -0.00734 | 0.94317 | 0.00037 | 10.5 s |
| 12:40.3 | Data point 6 | 1.50000 mL | 0.04798 mL | 0.03899 mL | 0.01999 mL | 3.110 | -0.01187 | 0.90433 | 0.00062 | 10.0 s |
| 13:31.5 | Data point 7 | 1.50000 mL | 0.04798 mL | 0.04433 mL | 0.01999 mL | 3.640 | -0.01726 | 0.73640 | 0.00099 | 11.5 s |
| 14:23.9 | Data point 8 | 1.50000 mL | 0.04798 mL | 0.04605 mL | 0.01999 mL | 3.809 | -0.01938 | 0.96322 | 0.00098 | 15.0 s |
| 15:04.4 | Data point 9 | 1.50000 mL | 0.04798 mL | 0.04739 mL | 0.01999 mL | 4.005 | -0.01944 | 0.95579 | 0.00098 | 12.0 s |
| 15:41.8 | Data point 10 | 1.50000 mL | 0.04798 mL | 0.04845 mL | 0.01999 mL | 4.191 | -0.01847 | 0.88746 | 0.00097 | 13.5 s |
| 16:20.8 | Data point 11 | 1.50000 mL | 0.04798 mL | 0.04922 mL | 0.01999 mL | 4.372 | -0.01878 | 0.94552 | 0.00095 | 23.5 s |
| 17:14.9 | Data point 12 | 1.50000 mL | 0.04798 mL | 0.05024 mL | 0.01999 mL | 4.829 | -0.01888 | 0.96108 | 0.00095 | 30.0 s |
| 18:15.5 | Data point 13 | 1.50000 mL | 0.04798 mL | 0.05085 mL | 0.01999 mL | 5.982 | -0.01949 | 0.93892 | 0.00099 | 52.5 s |
| 19:49.0 | Data point 14 | 1.50000 mL | 0.04798 mL | 0.05111 mL | 0.01999 mL | 6.938 | -0.03568 | 0.99301 | 0.00177 | Timed out at 59.5 s |
| 21:24.6 | Data point 15 | 1.50000 mL | 0.04798 mL | 0.05118 mL | 0.01999 mL | 7.503 | -0.03856 | 0.98751 | 0.00192 | Timed out at 59.5 s |
| 23:00.2 | Data point 16 | 1.50000 mL | 0.04798 mL | 0.05125 mL | 0.01999 mL | 7.986 | -0.04249 | 0.99373 | 0.00210 | Timed out at 59.5 s |
| 24:41.0 | Data point 17 | 1.50000 mL | 0.04798 mL | 0.05134 mL | 0.01999 mL | 8.384 | -0.03068 | 0.99526 | 0.00152 | Timed out at 59.5 s |
| 26:21.7 | Data point 18 | 1.50000 mL | 0.04798 mL | 0.05158 mL | 0.01999 mL | 9.153 | -0.01970 | 0.96174 | 0.00099 | 49.0 s |
| 28:09.7 | Data point 19 | 1.50000 mL | 0.10388 mL | 0.05158 mL | 0.05000 mL | 1.995 | -0.00675 | 0.60729 | 0.00043 | 10.0 s |
| 28:55.9 | Data point 20 | 1.50000 mL | 0.10388 mL | 0.06954 mL | 0.05000 mL | 2.211 | -0.00301 | 0.08351 | 0.00051 | 10.0 s |
| 29:31.6 | Data point 21 | 1.50000 mL | 0.10388 mL | 0.08001 mL | 0.05000 mL | 2.394 | 0.00217 | 0.25971 | 0.00021 | 10.5 s |
| 30:07.6 | Data point 22 | 1.50000 mL | 0.10388 mL | 0.08718 mL | 0.05000 mL | 2.582 | 0.01006 | 0.35818 | 0.00083 | 10.5 s |
| 30:43.7 | Data point 23 | 1.50000 mL | 0.10388 mL | 0.09226 mL | 0.05000 mL | 2.754 | -0.00412 | 0.68848 | 0.00025 | 10.0 s |
| 31:29.7 | Data point 24 | 1.50000 mL | 0.10388 mL | 0.09711 mL | 0.05000 mL | 2.942 | -0.00701 | 0.24786 | 0.00069 | 10.5 s |
| 32:10.9 | Data point 25 | 1.50000 mL | 0.10388 mL | 0.10002 mL | 0.05000 mL | 3.142 | -0.00388 | 0.12360 | 0.00054 | 10.0 s |
| 32:46.3 | Data point 26 | 1.50000 mL | 0.10388 mL | 0.10249 mL | 0.05000 mL | 3.250 | -0.00583 | 0.78197 | 0.00033 | 10.5 s |
| 33:27.5 | Data point 27 | 1.50000 mL | 0.10388 mL | 0.10574 mL | 0.05000 mL | 3.456 | -0.01149 | 0.59025 | 0.00074 | 10.0 s |
| 34:08.3 | Data point 28 | 1.50000 mL | 0.10388 mL | 0.10898 mL | 0.05000 mL | 3.732 | 0.00666 | 0.19651 | 0.00074 | 10.5 s |
| 35:02.1 | Data point 29 | 1.50000 mL | 0.10388 mL | 0.20652 mL | 0.05000 mL | 12.026 | 0.00788 | 0.91250 | 0.00041 | 10.5 s |
| 36:22.1 | Data point 30 | 1.50000 mL | 0.22564 mL | 0.20652 mL | 0.25000 mL | 1.977 | -0.01444 | 0.92481 | 0.00074 | 10.0 s |
| 37:08.5 | Data point 31 | 1.50000 mL | 0.22564 mL | 0.22803 mL | 0.25000 mL | 2.180 | -0.00826 | 0.64596 | 0.00051 | 10.0 s |
| 37:44.3 | Data point 32 | 1.50000 mL | 0.22564 mL | 0.24142 mL | 0.25000 mL | 2.372 | -0.00705 | 0.53601 | 0.00048 | 10.0 s |

Sample name: **M12_octanol**
Assay name: **pH-metric high logP**
Assay ID: **18C-03015**
Filename: **C:\Sirius_T3\Mehtap\20180302_exp29_logP_T3-2\18C-03015_M12_octanol_pH-metric high logP.t3r**

Experiment start time: **3/3/2018 7:38:51 PM**
Analyst: **Pion**
Instrument ID: **T312060**

#### Events (continued)

| Time | Event | Water | Acid | Base | Octanol | pH | dpH/dt | pH R-squared | pH SD | dpH/dt time |
|---|---|---|---|---|---|---|---|---|---|---|
| 38:19.9 | Data point 33 | 1.50000 mL | 0.22564 mL | 0.25052 mL | 0.25000 mL | 2.571 | -0.00201 | 0.05339 | 0.00043 | 10.0 s |
| 38:55.5 | Data point 34 | 1.50000 mL | 0.22564 mL | 0.25677 mL | 0.25000 mL | 2.761 | -0.00400 | 0.13797 | 0.00053 | 10.0 s |
| 39:31.1 | Data point 35 | 1.50000 mL | 0.22564 mL | 0.26134 mL | 0.25000 mL | 2.967 | -0.00538 | 0.70571 | 0.00032 | 10.0 s |
| 40:06.6 | Data point 36 | 1.50000 mL | 0.22564 mL | 0.26463 mL | 0.25000 mL | 3.179 | -0.00342 | 0.11319 | 0.00050 | 10.0 s |
| 40:42.1 | Data point 37 | 1.50000 mL | 0.22564 mL | 0.26693 mL | 0.25000 mL | 3.396 | -0.00748 | 0.56380 | 0.00049 | 10.0 s |
| 41:17.6 | Data point 38 | 1.50000 mL | 0.22564 mL | 0.26851 mL | 0.25000 mL | 3.672 | -0.00209 | 0.04091 | 0.00051 | 10.0 s |
| 42:03.3 | Data point 39 | 1.50000 mL | 0.22564 mL | 0.26921 mL | 0.25000 mL | 3.873 | -0.00392 | 0.08581 | 0.00066 | 10.0 s |
| 42:38.7 | Data point 40 | 1.50000 mL | 0.22564 mL | 0.26952 mL | 0.25000 mL | 4.087 | -0.00077 | 0.00175 | 0.00091 | 18.0 s |
| 43:27.2 | Data point 41 | 1.50000 mL | 0.22564 mL | 0.26978 mL | 0.25000 mL | 4.294 | -0.00113 | 0.00401 | 0.00088 | 10.5 s |
| 44:13.6 | Data point 42 | 1.50000 mL | 0.22564 mL | 0.26994 mL | 0.25000 mL | 4.494 | -0.00826 | 0.23363 | 0.00084 | 11.0 s |
| 44:55.1 | Data point 43 | 1.50000 mL | 0.22564 mL | 0.27009 mL | 0.25000 mL | 4.746 | -0.01490 | 0.56284 | 0.00098 | 18.5 s |
| 45:39.0 | Data point 44 | 1.50000 mL | 0.22564 mL | 0.27018 mL | 0.25000 mL | 5.035 | -0.01117 | 0.33968 | 0.00095 | 12.5 s |
| 46:16.9 | Data point 45 | 1.50000 mL | 0.22564 mL | 0.27023 mL | 0.25000 mL | 5.210 | -0.01900 | 0.90187 | 0.00099 | 27.5 s |
| 47:14.9 | Data point 46 | 1.50000 mL | 0.22564 mL | 0.27030 mL | 0.25000 mL | 5.664 | -0.01760 | 0.93799 | 0.00090 | 46.0 s |
| 48:31.5 | Data point 47 | 1.50000 mL | 0.22564 mL | 0.27039 mL | 0.25000 mL | 6.314 | -0.03485 | 0.93285 | 0.00178 | Timed out at 59.5 s |
| 50:02.1 | Data point 48 | 1.50000 mL | 0.22564 mL | 0.27044 mL | 0.25000 mL | 6.532 | -0.03321 | 0.95029 | 0.00168 | Timed out at 59.5 s |
| 51:27.4 | Data point 49 | 1.50000 mL | 0.22564 mL | 0.27048 mL | 0.25000 mL | 6.971 | -0.05095 | 0.98242 | 0.00254 | Timed out at 59.5 s |
| 53:03.0 | Data point 50 | 1.50000 mL | 0.22564 mL | 0.27055 mL | 0.25000 mL | 7.347 | -0.06715 | 0.99206 | 0.00333 | Timed out at 59.5 s |
| 54:43.9 | Data point 51 | 1.50000 mL | 0.22564 mL | 0.27065 mL | 0.25000 mL | 7.663 | -0.06618 | 0.98872 | 0.00329 | Timed out at 59.5 s |
| 56:14.4 | Data point 52 | 1.50000 mL | 0.22564 mL | 0.27070 mL | 0.25000 mL | 7.938 | -0.05473 | 0.99204 | 0.00271 | Timed out at 59.5 s |
| 57:44.9 | Data point 53 | 1.50000 mL | 0.22564 mL | 0.27074 mL | 0.25000 mL | 8.184 | -0.04284 | 0.94491 | 0.00218 | Timed out at 59.5 s |
| 59:20.6 | Data point 54 | 1.50000 mL | 0.22564 mL | 0.27081 mL | 0.25000 mL | 8.490 | -0.00848 | 0.37648 | 0.00068 | 38.5 s |
| 1:00:40.1 | Data point 55 | 1.50000 mL | 0.22564 mL | 0.27091 mL | 0.25000 mL | 8.782 | -0.00488 | 0.07901 | 0.00086 | 29.5 s |
| 1:01:50.4 | Data point 56 | 1.50000 mL | 0.22564 mL | 0.27103 mL | 0.25000 mL | 9.009 | -0.01631 | 0.80065 | 0.00090 | 14.5 s |
| 1:02:13.9 | Assay volumes | 1.50000 mL | 0.22564 mL | 0.27103 mL | 0.25000 mL | | | | | |

| | | | |
|---|---|---|---|
| Sample name: | **M12_octanol** | Experiment start time: | **3/3/2018 7:38:51 PM** |
| Assay name: | **pH-metric high logP** | Analyst: | **Pion** |
| Assay ID: | **18C-03015** | Instrument ID: | **T312060** |
| Filename: | **C:\Sirius_T3\Mehtap\20180302_exp29_logP_T3-2\18C-03015_M12_octanol_pH-metric high logP.t3r** | | |

| | | | |
|---|---|---|---|
| Sample name: | **M12_octanol** | Experiment start time: | **3/3/2018 7:38:51 PM** |
| Assay name: | **pH-metric high logP** | Analyst: | **Pion** |
| Assay ID: | **18C-03015** | Instrument ID: | **T312060** |
| Filename: | **C:\Sirius_T3\Mehtap\20180302_exp29_logP_T3-2\18C-03015_M12_octanol_pH-metric high logP.t3r** | | |

#### Assay Settings (continued)

| Setting | Value | Original Value | Date/Time changed | Imported from |
|---|---|---|---|---|
| ***Titration 3*** | | | | |
| Titrate from | Low to high pH | | | |
| Add additional water | 0.00 mL | | | |
| Additional partition solvent volume | 0.200 mL | | | |
| Additional partition solvent added | Automatic | | | |
| After pH adjust stir for | 30 seconds | | | |
| Stir to allow partitioning for | 15 seconds | | | |
| Stirrer speed for partitioning | 60% | | | |
| ***Data Point Stability*** | | | | |
| Stir during data point collection | No | | | |
| Delay before data point collection | 0 seconds | | | |
| Number of points to average | 20 points | | | |
| Time interval between points | 0.50 seconds | | | |
| Required maximum standard deviation | 0.00100 dpH/dt | | | |
| Stability timeout after | 60 seconds | | | |

#### Calibration Settings

| Setting | Value | Date/Time changed | Imported from |
|---|---|---|---|
| Four-Plus alpha | 0.111 | 3/3/2018 7:38:51 PM | C:\Sirius_T3\HCl18C02.t3r |
| Four-Plus S | 0.9988 | 3/3/2018 7:38:51 PM | C:\Sirius_T3\HCl18C02.t3r |
| Four-Plus jH | 1.0 | 3/3/2018 7:38:51 PM | C:\Sirius_T3\HCl18C02.t3r |
| Four-Plus jOH | -0.8 | 3/3/2018 7:38:51 PM | C:\Sirius_T3\HCl18C02.t3r |
| Base concentration factor | 1.000 | 3/3/2018 7:38:51 PM | C:\Sirius_T3\KOH18B27.t3r |
| Acid concentration factor | 0.999 | 3/3/2018 7:38:51 PM | C:\Sirius_T3\HCl18C02.t3r |

#### Instrument Settings

| Setting | Value | Batch Id | Install date |
|---|---|---|---|
| Instrument owner | Merck | | |
| Instrument ID | T312060 | | |
| Instrument type | T3 Simulator | | |
| Software version | 1.1.3.0 | | |
| Dispenser module | | T3DM1200361 | 3/31/2009 5:24:52 AM |
| Dispenser 0 | Water | | 3/31/2009 5:25:05 AM |
| Syringe volume | 2.5 mL | | |
| Firmware version | 1.2.1(r2) | | |
| Titrant | Water (0.15 M KCl) | 02-06-2018 | 2/27/2018 10:05:59 AM |
| Dispenser 2 | Acid | | 3/31/2009 5:25:11 AM |
| Syringe volume | 0.5 mL | | |
| Firmware version | 1.2.1(r2) | | |
| Titrant | Acid (0.5 M HCl) | 02-27-2018 | 2/27/2018 10:27:22 AM |
| Dispenser 1 | Base | | 3/31/2009 5:25:21 AM |
| Syringe volume | 0.5 mL | | |
| Firmware version | 1.2.1(r2) | | |
| Titrant | Base (0.5 M KOH) | 9/22/2017 | 2/27/2018 10:21:22 AM |
| Dispenser 5 | Cosolvent | | 3/31/2009 5:26:24 AM |
| Syringe volume | 2.5 mL | | |
| Firmware version | 1.2.1(r2) | | |
| Distribution valve 5 | Distribution Valve | | 3/31/2009 5:28:19 AM |
| Firmware version | 1.1.3 | | |
| Port A | Methanol (80%, 0.15 M KCl) | 09-26-17 | 2/7/2018 9:42:01 AM |
| Port B | Cyclohexane | 11-01-17 | 2/27/2018 10:37:57 AM |
| Dispenser 3 | Buffer | | 8/3/2010 5:05:16 AM |
| Syringe volume | 0.5 mL | | |
| Firmware version | 1.2.1(r2) | | |
| Titrant | Dodecane | 2018/01/31 | 2/28/2018 10:18:04 AM |
| Dispenser 6 | Octanol | | 10/22/2010 10:52:43 AM |

| | | | |
|---|---|---|---|
| Sample name: | **M12_octanol** | Experiment start time: | **3/3/2018 7:38:51 PM** |
| Assay name: | **pH-metric high logP** | Analyst: | **Pion** |
| Assay ID: | **18C-03015** | Instrument ID: | **T312060** |
| Filename: | **C:\Sirius_T3\Mehtap\20180302_exp29_logP_T3-2\18C-03015_M12_octanol_pH-metric high logP.t3r** | | |

| | | | |
|---|---|---|---|
| Sample name: | **M12_octanol** | Experiment start time: | **3/3/2018 7:38:51 PM** |
| Assay name: | **pH-metric high logP** | Analyst: | **Pion** |
| Assay ID: | **18C-03015** | Instrument ID: | **T312060** |
| Filename: | **C:\Sirius_T3\Mehtap\20180302_exp29_logP_T3-2\18C-03015_M12_octanol_pH-metric high logP.t3r** | | |

#### Instrument Settings (continued)

| Setting | Value | Batch Id | Install date |
|---|---|---|---|
| Spectrometer calibration stir duration | 5 s | | |
| Spectrometer calibration stir speed | 30% | | |
| Spectrometer calibration wash pump volume | 20.0 mL | | |
| Spectrometer calibration wash stir duration | 5 s | | |
| Spectrometer calibration wash stir speed | 30% | | |
| Overhead dispense height | 10000 | | |

#### Refinement Settings

| Setting | Value | Default value |
|---|---|---|
| Turbidity detection method | None | None |
| Turbidity wavelength to assess | 500.0 nm | 500.0 nm |
| Turbidity maximum absorbance | 0.100 | 0.100 |
| Turbidity probe threshold | 50.00 | 50.00 |

#### Experiment Log

[2:37] Air gap created for Water (0.15 M KCl)
[2:37] Air gap created for Acid (0.5 M HCl)
[2:38] Air gap created for Base (0.5 M KOH)
[2:38] Air gap released for Water (0.15 M KCl)
[2:42] Titrator arm moved over Titration position
[2:42] Titration 1 of 3
[2:42] Adding initial titrants
[2:42] Automatically add 1.50000 mL of water
[3:07] Dispensed 1.500000 mL of Water (0.15 M KCl)
[3:11] Titrator arm moved over Drain
[5:53] Titrator arm moved to Titration position
[5:53] Argon flow rate set to 100
[5:53] Stirrer speed set to 10
[5:58] Automatically add 0.02000 mL of Octanol
[5:58] Dispensed 0.019991 mL of Octanol
[6:00] Initial pH = 4.05
[6:00] Iterative adjust 4.05 -> 2.00
[6:00] pH 4.05 -> 2.00
[6:01] Air gap released for Acid (0.5 M HCl)
[6:02] Dispensed 0.047977 mL of Acid (0.5 M HCl)
[6:07] Holding pH 2.00
[8:07] Stirrer speed set to 0
[8:07] Stirrer speed set to 50
[8:07] Iterative adjust 1.98 -> 2.00
[8:07] pH 1.98 -> 2.00
[8:08] Air gap released for Base (0.5 M KOH)
[8:09] Dispensed 0.002328 mL of Base (0.5 M KOH)
[8:59] Stirrer speed set to 0
[9:09] Datapoint id 1 collected
[9:09] Stirrer speed set to 50
[9:14] pH 2.10 -> 2.30
[9:14] Using cautious pH adjust
[9:14] Dispensed 0.006232 mL of Base (0.5 M KOH)
[9:19] Stepping pH = 2.16
[9:20] Dispensed 0.006726 mL of Base (0.5 M KOH)
[9:25] Stepping pH = 2.26
[9:25] Dispensed 0.001881 mL of Base (0.5 M KOH)
[9:30] Stepping pH = 2.31
[9:45] Stirrer speed set to 0
[9:56] Datapoint id 2 collected
[9:56] Charge balance equation is out by -18.9%
[9:56] Stirrer speed set to 50

Sample name: **M12_octanol** Experiment start time: **3/3/2018 7:38:51 PM**
Assay name: **pH-metric high logP** Analyst: **Pion**
Assay ID: **18C-03015** Instrument ID: **T312060**
Filename: **C:\Sirius_T3\Mehtap\20180302_exp29_logP_T3-2\18C-03015_M12_octanol_pH-metric high logP.t3r**

#### Experiment Log (continued)

[10:01] pH 2.32 -> 2.52
[10:01] Using cautious pH adjust
[10:01] Dispensed 0.003810 mL of Base (0.5 M KOH)
[10:06] Stepping pH = 2.39
[10:06] Dispensed 0.004186 mL of Base (0.5 M KOH)
[10:11] Stepping pH = 2.49
[10:12] Dispensed 0.001129 mL of Base (0.5 M KOH)
[10:17] Stepping pH = 2.53
[10:32] Stirrer speed set to 0
[10:42] Datapoint id 3 collected
[10:42] Charge balance equation is out by -19.8%
[10:42] Stirrer speed set to 50
[10:47] pH 2.54 -> 2.74
[10:47] Using cautious pH adjust
[10:48] Dispensed 0.002469 mL of Base (0.5 M KOH)
[10:53] Stepping pH = 2.60
[10:53] Dispensed 0.002893 mL of Base (0.5 M KOH)
[10:58] Stepping pH = 2.70
[10:58] Dispensed 0.000823 mL of Base (0.5 M KOH)
[11:03] Stepping pH = 2.73
[11:18] Stirrer speed set to 0
[11:28] Datapoint id 4 collected
[11:28] Charge balance equation is out by -25.7%
[11:28] Stirrer speed set to 50
[11:33] pH 2.74 -> 2.94
[11:33] Using cautious pH adjust
[11:34] Dispensed 0.001740 mL of Base (0.5 M KOH)
[11:39] Stepping pH = 2.82
[11:39] Dispensed 0.001599 mL of Base (0.5 M KOH)
[11:44] Stepping pH = 2.90
[11:44] Dispensed 0.000494 mL of Base (0.5 M KOH)
[11:49] Stepping pH = 2.94
[12:04] Stirrer speed set to 0
[12:15] Datapoint id 5 collected
[12:15] Charge balance equation is out by -9.6%
[12:15] Stirrer speed set to 50
[12:20] pH 2.94 -> 3.14
[12:20] Using charge balance adjust
[12:20] Dispensed 0.002681 mL of Base (0.5 M KOH)
[12:40] Stirrer speed set to 0
[12:50] Datapoint id 6 collected
[12:50] Charge balance equation is out by -17.5%
[12:50] Stirrer speed set to 50
[12:55] pH 3.11 -> 3.31
[12:55] Using cautious pH adjust
[12:55] Dispensed 0.001152 mL of Base (0.5 M KOH)
[13:01] Stepping pH = 3.19
[13:01] Dispensed 0.001129 mL of Base (0.5 M KOH)
[13:06] Stepping pH = 3.27
[13:06] Dispensed 0.000447 mL of Base (0.5 M KOH)
[13:11] Stepping pH = 3.27
[13:11] Dispensed 0.002611 mL of Base (0.5 M KOH)
[13:16] Stepping pH = 3.68
[13:31] Stirrer speed set to 0
[13:43] Datapoint id 7 collected
[13:43] Charge balance equation is out by -131.1%
[13:43] Stirrer speed set to 50
[13:48] pH 3.65 -> 3.85
[13:48] Using cautious pH adjust

| | | | |
|---|---|---|---|
| Sample name: | **M12_octanol** | Experiment start time: | **3/3/2018 7:38:51 PM** |
| Assay name: | **pH-metric high logP** | Analyst: | **Pion** |
| Assay ID: | **18C-03015** | Instrument ID: | **T312060** |
| Filename: | **C:\Sirius_T3\Mehtap\20180302_exp29_logP_T3-2\18C-03015_M12_octanol_pH-metric high logP.t3r** | | |

#### Experiment Log (continued)

[13:48] Dispensed 0.000800 mL of Base (0.5 M KOH)
[13:53] Stepping pH = 3.73
[13:53] Dispensed 0.000682 mL of Base (0.5 M KOH)
[13:58] Stepping pH = 3.83
[13:58] Dispensed 0.000094 mL of Base (0.5 M KOH)
[14:03] Stepping pH = 3.83
[14:04] Dispensed 0.000141 mL of Base (0.5 M KOH)
[14:09] Stepping pH = 3.85
[14:24] Stirrer speed set to 0
[14:39] Datapoint id 8 collected
[14:39] Charge balance equation is out by -7.4%
[14:39] Stirrer speed set to 50
[14:44] pH 3.82 -> 4.02
[14:44] Using charge balance adjust
[14:44] Dispensed 0.001341 mL of Base (0.5 M KOH)
[15:04] Stirrer speed set to 0
[15:16] Datapoint id 9 collected
[15:16] Charge balance equation is out by -8.5%
[15:16] Stirrer speed set to 50
[15:21] pH 4.01 -> 4.21
[15:21] Using charge balance adjust
[15:21] Dispensed 0.001058 mL of Base (0.5 M KOH)
[15:42] Stirrer speed set to 0
[15:55] Datapoint id 10 collected
[15:55] Charge balance equation is out by -11.1%
[15:55] Stirrer speed set to 50
[16:00] pH 4.21 -> 4.41
[16:00] Using charge balance adjust
[16:00] Dispensed 0.000776 mL of Base (0.5 M KOH)
[16:21] Stirrer speed set to 0
[16:44] Datapoint id 11 collected
[16:44] Charge balance equation is out by -19.6%
[16:44] Stirrer speed set to 50
[16:49] pH 4.41 -> 4.61
[16:49] Using cautious pH adjust
[16:49] Dispensed 0.000282 mL of Base (0.5 M KOH)
[16:54] Stepping pH = 4.43
[16:55] Dispensed 0.000729 mL of Base (0.5 M KOH)
[17:00] Stepping pH = 4.88
[17:15] Stirrer speed set to 0
[17:45] Datapoint id 12 collected
[17:45] Charge balance equation is out by -84.8%
[17:45] Stirrer speed set to 50
[17:50] pH 4.90 -> 5.10
[17:50] Using cautious pH adjust
[17:50] Dispensed 0.000094 mL of Base (0.5 M KOH)
[17:55] Stepping pH = 4.89
[17:55] Dispensed 0.000517 mL of Base (0.5 M KOH)
[18:00] Stepping pH = 5.56
[18:15] Stirrer speed set to 0
[19:08] Datapoint id 13 collected
[19:08] Charge balance equation is out by -203.9%
[19:08] Stirrer speed set to 50
[19:13] pH 6.15 -> 6.35
[19:13] Using cautious pH adjust
[19:13] Dispensed 0.000024 mL of Base (0.5 M KOH)
[19:18] Stepping pH = 6.19
[19:18] Dispensed 0.000047 mL of Base (0.5 M KOH)
[19:23] Stepping pH = 6.22

| | | | |
|---|---|---|---|
| Sample name: | **M12_octanol** | Experiment start time: | **3/3/2018 7:38:51 PM** |
| Assay name: | **pH-metric high logP** | Analyst: | **Pion** |
| Assay ID: | **18C-03015** | Instrument ID: | **T312060** |
| Filename: | **C:\Sirius_T3\Mehtap\20180302_exp29_logP_T3-2\18C-03015_M12_octanol_pH-metric high logP.t3r** | | |

#### Experiment Log (continued)

[19:23] Dispensed 0.000094 mL of Base (0.5 M KOH)
[19:29] Stepping pH = 6.26
[19:29] Dispensed 0.000094 mL of Base (0.5 M KOH)
[19:34] Stepping pH = 6.43
[19:49] Stirrer speed set to 0
[20:49] Datapoint id 14 collected
[20:49] Charge balance equation is out by -426.3%
[20:49] Stirrer speed set to 50
[20:54] pH 7.06 -> 7.26
[20:54] Using cautious pH adjust
[20:54] Dispensed 0.000024 mL of Base (0.5 M KOH)
[20:59] Stepping pH = 7.13
[20:59] Dispensed 0.000024 mL of Base (0.5 M KOH)
[21:04] Stepping pH = 7.20
[21:04] Dispensed 0.000024 mL of Base (0.5 M KOH)
[21:09] Stepping pH = 7.31
[21:24] Stirrer speed set to 0
[22:24] Datapoint id 15 collected
[22:24] Charge balance equation is out by -279.1%
[22:24] Stirrer speed set to 50
[22:29] pH 7.62 -> 7.82
[22:29] Using cautious pH adjust
[22:30] Dispensed 0.000024 mL of Base (0.5 M KOH)
[22:35] Stepping pH = 7.71
[22:35] Dispensed 0.000024 mL of Base (0.5 M KOH)
[22:40] Stepping pH = 7.77
[22:40] Dispensed 0.000024 mL of Base (0.5 M KOH)
[22:45] Stepping pH = 7.83
[23:00] Stirrer speed set to 0
[24:00] Datapoint id 16 collected
[24:00] Charge balance equation is out by -785.0%
[24:00] Stirrer speed set to 50
[24:05] pH 8.16 -> 8.36
[24:05] Using cautious pH adjust
[24:05] Dispensed 0.000024 mL of Base (0.5 M KOH)
[24:10] Stepping pH = 8.25
[24:10] Dispensed 0.000024 mL of Base (0.5 M KOH)
[24:15] Stepping pH = 8.30
[24:15] Dispensed 0.000024 mL of Base (0.5 M KOH)
[24:21] Stepping pH = 8.32
[24:21] Dispensed 0.000024 mL of Base (0.5 M KOH)
[24:26] Stepping pH = 8.35
[24:41] Stirrer speed set to 0
[25:41] Datapoint id 17 collected
[25:41] Charge balance equation is out by -917.7%
[25:41] Stirrer speed set to 50
[25:46] pH 8.43 -> 8.63
[25:46] Using cautious pH adjust
[25:46] Dispensed 0.000024 mL of Base (0.5 M KOH)
[25:51] Stepping pH = 8.44
[25:51] Dispensed 0.000024 mL of Base (0.5 M KOH)
[25:56] Stepping pH = 8.44
[25:56] Dispensed 0.000047 mL of Base (0.5 M KOH)
[26:01] Stepping pH = 8.46
[26:01] Dispensed 0.000141 mL of Base (0.5 M KOH)
[26:06] Stepping pH = 8.83
[26:22] Stirrer speed set to 0
[27:11] Datapoint id 18 collected
[27:11] Charge balance equation is out by -1,849.5%

| | | | |
|---|---|---|---|
| Sample name: | **M12_octanol** | Experiment start time: | **3/3/2018 7:38:51 PM** |
| Assay name: | **pH-metric high logP** | Analyst: | **Pion** |
| Assay ID: | **18C-03015** | Instrument ID: | **T312060** |
| Filename: | **C:\Sirius_T3\Mehtap\20180302_exp29_logP_T3-2\18C-03015_M12_octanol_pH-metric high logP.t3r** | | |

#### Experiment Log (continued)

[27:11] Titration 2 of 3
[27:11] Adding initial titrants
[27:11] Automatically add 0.03000 mL of Octanol
[27:11] Dispensed 0.030009 mL of Octanol
[27:11] Stirrer speed set to 10
[27:13] Stirrer speed set to 55
[27:13] Iterative adjust 9.17 -> 2.00
[27:13] pH 9.17 -> 2.00
[27:14] Dispensed 0.053763 mL of Acid (0.5 M HCl)
[27:19] pH 2.02 -> 2.00
[27:19] Dispensed 0.002140 mL of Acid (0.5 M HCl)
[28:09] Stirrer speed set to 0
[28:19] Datapoint id 19 collected
[28:19] Stirrer speed set to 55
[28:25] pH 2.01 -> 2.21
[28:25] Using cautious pH adjust
[28:25] Dispensed 0.008208 mL of Base (0.5 M KOH)
[28:30] Stepping pH = 2.09
[28:30] Dispensed 0.007785 mL of Base (0.5 M KOH)
[28:35] Stepping pH = 2.18
[28:36] Dispensed 0.001976 mL of Base (0.5 M KOH)
[28:41] Stepping pH = 2.21
[28:56] Stirrer speed set to 0
[29:06] Datapoint id 20 collected
[29:06] Charge balance equation is out by -9.5%
[29:06] Stirrer speed set to 55
[29:11] pH 2.22 -> 2.42
[29:11] Using charge balance adjust
[29:11] Dispensed 0.010466 mL of Base (0.5 M KOH)
[29:31] Stirrer speed set to 0
[29:42] Datapoint id 21 collected
[29:42] Charge balance equation is out by -10.6%
[29:42] Stirrer speed set to 55
[29:47] pH 2.40 -> 2.60
[29:47] Using charge balance adjust
[29:47] Dispensed 0.007173 mL of Base (0.5 M KOH)
[30:07] Stirrer speed set to 0
[30:18] Datapoint id 22 collected
[30:18] Charge balance equation is out by -9.0%
[30:18] Stirrer speed set to 55
[30:23] pH 2.59 -> 2.79
[30:23] Using charge balance adjust
[30:23] Dispensed 0.005080 mL of Base (0.5 M KOH)
[30:43] Stirrer speed set to 0
[30:53] Datapoint id 23 collected
[30:53] Charge balance equation is out by -17.4%
[30:53] Stirrer speed set to 55
[30:59] pH 2.76 -> 2.96
[30:59] Using cautious pH adjust
[30:59] Dispensed 0.001952 mL of Base (0.5 M KOH)
[31:04] Stepping pH = 2.83
[31:04] Dispensed 0.002070 mL of Base (0.5 M KOH)
[31:09] Stepping pH = 2.91
[31:09] Dispensed 0.000823 mL of Base (0.5 M KOH)
[31:14] Stepping pH = 2.95
[31:29] Stirrer speed set to 0
[31:40] Datapoint id 24 collected
[31:40] Charge balance equation is out by -23.5%
[31:40] Stirrer speed set to 55

| | | | |
|---|---|---|---|
| Sample name: | **M12_octanol** | Experiment start time: | **3/3/2018 7:38:51 PM** |
| Assay name: | **pH-metric high logP** | Analyst: | **Pion** |
| Assay ID: | **18C-03015** | Instrument ID: | **T312060** |
| Filename: | **C:\Sirius_T3\Mehtap\20180302_exp29_logP_T3-2\18C-03015_M12_octanol_pH-metric high logP.t3r** | | |

#### Experiment Log (continued)

[31:45] pH 2.95 -> 3.15
[31:45] Using cautious pH adjust
[31:45] Dispensed 0.001552 mL of Base (0.5 M KOH)
[31:50] Stepping pH = 3.03
[31:50] Dispensed 0.001364 mL of Base (0.5 M KOH)
[31:56] Stepping pH = 3.15
[32:11] Stirrer speed set to 0
[32:21] Datapoint id 25 collected
[32:21] Charge balance equation is out by 5.6%
[32:21] Stirrer speed set to 55
[32:26] pH 3.15 -> 3.35
[32:26] Using charge balance adjust
[32:26] Dispensed 0.002469 mL of Base (0.5 M KOH)
[32:46] Stirrer speed set to 0
[32:57] Datapoint id 26 collected
[32:57] Charge balance equation is out by -50.0%
[32:57] Stirrer speed set to 55
[33:02] pH 3.25 -> 3.45
[33:02] Using cautious pH adjust
[33:02] Dispensed 0.001105 mL of Base (0.5 M KOH)
[33:07] Stepping pH = 3.29
[33:07] Dispensed 0.002140 mL of Base (0.5 M KOH)
[33:12] Stepping pH = 3.45
[33:27] Stirrer speed set to 0
[33:37] Datapoint id 27 collected
[33:37] Charge balance equation is out by -46.4%
[33:37] Stirrer speed set to 55
[33:42] pH 3.46 -> 3.66
[33:42] Using cautious pH adjust
[33:43] Dispensed 0.000870 mL of Base (0.5 M KOH)
[33:48] Stepping pH = 3.48
[33:48] Dispensed 0.002375 mL of Base (0.5 M KOH)
[33:53] Stepping pH = 3.75
[34:08] Stirrer speed set to 0
[34:19] Datapoint id 28 collected
[34:19] Charge balance equation is out by -87.2%
[34:19] Stirrer speed set to 55
[34:24] pH 3.74 -> 3.94
[34:24] Using cautious pH adjust
[34:24] Dispensed 0.000588 mL of Base (0.5 M KOH)
[34:29] Stepping pH = 3.73
[34:29] Dispensed 0.003057 mL of Base (0.5 M KOH)
[34:34] Stepping pH = 3.73
[34:35] Dispensed 0.015522 mL of Base (0.5 M KOH)
[34:40] Stepping pH = 3.73
[34:42] Dispensed 0.078363 mL of Base (0.5 M KOH)
[34:47] Stepping pH = 12.02
[35:02] Stirrer speed set to 0
[35:12] Datapoint id 29 collected
[35:12] Charge balance equation is out by -8,214.2%
[35:12] Titration 3 of 3
[35:12] Adding initial titrants
[35:12] Automatically add 0.20000 mL of Octanol
[35:17] Dispensed 0.200000 mL of Octanol
[35:17] Stirrer speed set to 10
[35:18] Stirrer speed set to 60
[35:18] Iterative adjust 12.03 -> 2.00
[35:18] pH 12.03 -> 2.00
[35:21] Dispensed 0.100000 mL of Acid (0.5 M HCl)

Sample name: **M12_octanol**
Assay name: **pH-metric high logP**
Assay ID: **18C-03015**
Filename: **C:\Sirius_T3\Mehtap\20180302_exp29_logP_T3-2\18C-03015_M12_octanol_pH-metric high logP.t3r**
Experiment start time: **3/3/2018 7:38:51 PM**
Analyst: **Pion**
Instrument ID: **T312060**

#### Experiment Log (continued)

[35:26] pH 2.19 -> 2.00
[35:26] Dispensed 0.018650 mL of Acid (0.5 M HCl)
[35:31] pH 2.03 -> 2.00
[35:32] Dispensed 0.003104 mL of Acid (0.5 M HCl)
[36:22] Stirrer speed set to 0
[36:32] Datapoint id 30 collected
[36:32] Stirrer speed set to 60
[36:37] pH 1.98 -> 2.18
[36:37] Using cautious pH adjust
[36:37] Dispensed 0.010348 mL of Base (0.5 M KOH)
[36:42] Stepping pH = 2.07
[36:43] Dispensed 0.008420 mL of Base (0.5 M KOH)
[36:48] Stepping pH = 2.15
[36:48] Dispensed 0.002752 mL of Base (0.5 M KOH)
[36:53] Stepping pH = 2.18
[37:08] Stirrer speed set to 0
[37:18] Datapoint id 31 collected
[37:18] Charge balance equation is out by -4.0%
[37:18] Stirrer speed set to 60
[37:23] pH 2.18 -> 2.38
[37:23] Using charge balance adjust
[37:24] Dispensed 0.013382 mL of Base (0.5 M KOH)
[37:44] Stirrer speed set to 0
[37:54] Datapoint id 32 collected
[37:54] Charge balance equation is out by -6.2%
[37:54] Stirrer speed set to 60
[37:59] pH 2.38 -> 2.58
[37:59] Using charge balance adjust
[38:00] Dispensed 0.009102 mL of Base (0.5 M KOH)
[38:20] Stirrer speed set to 0
[38:30] Datapoint id 33 collected
[38:30] Charge balance equation is out by -2.7%
[38:30] Stirrer speed set to 60
[38:35] pH 2.58 -> 2.78
[38:35] Using charge balance adjust
[38:35] Dispensed 0.006256 mL of Base (0.5 M KOH)
[38:55] Stirrer speed set to 0
[39:05] Datapoint id 34 collected
[39:05] Charge balance equation is out by -9.2%
[39:05] Stirrer speed set to 60
[39:10] pH 2.77 -> 2.97
[39:10] Using charge balance adjust
[39:11] Dispensed 0.004563 mL of Base (0.5 M KOH)
[39:31] Stirrer speed set to 0
[39:41] Datapoint id 35 collected
[39:41] Charge balance equation is out by -0.9%
[39:41] Stirrer speed set to 60
[39:46] pH 2.97 -> 3.17
[39:46] Using charge balance adjust
[39:46] Dispensed 0.003293 mL of Base (0.5 M KOH)
[40:06] Stirrer speed set to 0
[40:16] Datapoint id 36 collected
[40:16] Charge balance equation is out by 3.4%
[40:16] Stirrer speed set to 60
[40:22] pH 3.18 -> 3.38
[40:22] Using charge balance adjust
[40:22] Dispensed 0.002305 mL of Base (0.5 M KOH)
[40:42] Stirrer speed set to 0
[40:52] Datapoint id 37 collected

| | | | |
|---|---|---|---|
| Sample name: | **M12_octanol** | Experiment start time: | **3/3/2018 7:38:51 PM** |
| Assay name: | **pH-metric high logP** | Analyst: | **Pion** |
| Assay ID: | **18C-03015** | Instrument ID: | **T312060** |
| Filename: | **C:\Sirius_T3\Mehtap\20180302_exp29_logP_T3-2\18C-03015_M12_octanol_pH-metric high logP.t3r** | | |

#### Experiment Log (continued)

[40:52] Charge balance equation is out by 6.0%
[40:52] Stirrer speed set to 60
[40:57] pH 3.40 -> 3.60
[40:57] Using charge balance adjust
[40:57] Dispensed 0.001576 mL of Base (0.5 M KOH)
[41:17] Stirrer speed set to 0
[41:27] Datapoint id 38 collected
[41:27] Charge balance equation is out by 34.7%
[41:27] Stirrer speed set to 60
[41:32] pH 3.68 -> 3.88
[41:33] Using cautious pH adjust
[41:33] Dispensed 0.000447 mL of Base (0.5 M KOH)
[41:38] Stepping pH = 3.80
[41:38] Dispensed 0.000212 mL of Base (0.5 M KOH)
[41:43] Stepping pH = 3.87
[41:43] Dispensed 0.000047 mL of Base (0.5 M KOH)
[41:48] Stepping pH = 3.88
[42:03] Stirrer speed set to 0
[42:13] Datapoint id 39 collected
[42:13] Charge balance equation is out by 21.2%
[42:13] Stirrer speed set to 60
[42:18] pH 3.88 -> 4.08
[42:18] Using cautious pH adjust
[42:18] Dispensed 0.000306 mL of Base (0.5 M KOH)
[42:23] Stepping pH = 4.09
[42:38] Stirrer speed set to 0
[42:57] Datapoint id 40 collected
[42:57] Charge balance equation is out by 50.0%
[42:57] Stirrer speed set to 60
[43:02] pH 4.10 -> 4.30
[43:02] Using cautious pH adjust
[43:02] Dispensed 0.000188 mL of Base (0.5 M KOH)
[43:07] Stepping pH = 4.24
[43:07] Dispensed 0.000071 mL of Base (0.5 M KOH)
[43:12] Stepping pH = 4.29
[43:27] Stirrer speed set to 0
[43:38] Datapoint id 41 collected
[43:38] Charge balance equation is out by 32.9%
[43:38] Stirrer speed set to 60
[43:43] pH 4.32 -> 4.52
[43:43] Using cautious pH adjust
[43:43] Dispensed 0.000118 mL of Base (0.5 M KOH)
[43:48] Stepping pH = 4.48
[43:48] Dispensed 0.000024 mL of Base (0.5 M KOH)
[43:53] Stepping pH = 4.50
[43:53] Dispensed 0.000024 mL of Base (0.5 M KOH)
[43:58] Stepping pH = 4.51
[44:13] Stirrer speed set to 0
[44:24] Datapoint id 42 collected
[44:24] Charge balance equation is out by 33.2%
[44:24] Stirrer speed set to 60
[44:30] pH 4.50 -> 4.70
[44:30] Using cautious pH adjust
[44:30] Dispensed 0.000071 mL of Base (0.5 M KOH)
[44:35] Stepping pH = 4.58
[44:35] Dispensed 0.000071 mL of Base (0.5 M KOH)
[44:40] Stepping pH = 4.76
[44:55] Stirrer speed set to 0
[45:13] Datapoint id 43 collected

| | | | |
|---|---|---|---|
| Sample name: | **M12_octanol** | Experiment start time: | **3/3/2018 7:38:51 PM** |
| Assay name: | **pH-metric high logP** | Analyst: | **Pion** |
| Assay ID: | **18C-03015** | Instrument ID: | **T312060** |
| Filename: | **C:\Sirius_T3\Mehtap\20180302_exp29_logP_T3-2\18C-03015_M12_octanol_pH-metric high logP.t3r** | | |

#### Experiment Log (continued)

[45:13] Charge balance equation is out by 6.7%
[45:13] Stirrer speed set to 60
[45:19] pH 4.82 -> 5.02
[45:19] Using charge balance adjust
[45:19] Dispensed 0.000094 mL of Base (0.5 M KOH)
[45:39] Stirrer speed set to 0
[45:51] Datapoint id 44 collected
[45:51] Charge balance equation is out by 9.7%
[45:51] Stirrer speed set to 60
[45:56] pH 5.11 -> 5.31
[45:56] Using charge balance adjust
[45:56] Dispensed 0.000047 mL of Base (0.5 M KOH)
[46:17] Stirrer speed set to 0
[46:44] Datapoint id 45 collected
[46:44] Charge balance equation is out by -48.8%
[46:44] Stirrer speed set to 60
[46:49] pH 5.25 -> 5.45
[46:49] Using cautious pH adjust
[46:49] Dispensed 0.000024 mL of Base (0.5 M KOH)
[46:54] Stepping pH = 5.29
[46:55] Dispensed 0.000047 mL of Base (0.5 M KOH)
[47:00] Stepping pH = 5.71
[47:15] Stirrer speed set to 0
[48:01] Datapoint id 46 collected
[48:01] Charge balance equation is out by -35.1%
[48:01] Stirrer speed set to 60
[48:06] pH 5.76 -> 5.96
[48:06] Using cautious pH adjust
[48:06] Dispensed 0.000024 mL of Base (0.5 M KOH)
[48:11] Stepping pH = 5.77
[48:11] Dispensed 0.000071 mL of Base (0.5 M KOH)
[48:16] Stepping pH = 6.49
[48:31] Stirrer speed set to 0
[49:31] Datapoint id 47 collected
[49:31] Charge balance equation is out by -96.3%
[49:31] Stirrer speed set to 60
[49:36] pH 6.31 -> 6.51
[49:36] Using cautious pH adjust
[49:37] Dispensed 0.000024 mL of Base (0.5 M KOH)
[49:42] Stepping pH = 6.43
[49:42] Dispensed 0.000024 mL of Base (0.5 M KOH)
[49:47] Stepping pH = 6.61
[50:02] Stirrer speed set to 0
[51:02] Datapoint id 48 collected
[51:02] Charge balance equation is out by 8.4%
[51:02] Stirrer speed set to 60
[51:07] pH 6.70 -> 6.90
[51:07] Using charge balance adjust
[51:07] Dispensed 0.000047 mL of Base (0.5 M KOH)
[51:27] Stirrer speed set to 0
[52:27] Datapoint id 49 collected
[52:27] Charge balance equation is out by 34.8%
[52:27] Stirrer speed set to 60
[52:32] pH 7.04 -> 7.24
[52:32] Using cautious pH adjust
[52:32] Dispensed 0.000024 mL of Base (0.5 M KOH)
[52:37] Stepping pH = 7.10
[52:38] Dispensed 0.000024 mL of Base (0.5 M KOH)
[52:43] Stepping pH = 7.22

Sample name: **M12_octanol**
Assay name: **pH-metric high logP**
Assay ID: **18C-03015**
Filename: **C:\Sirius_T3\Mehtap\20180302_exp29_logP_T3-2\18C-03015_M12_octanol_pH-metric high logP.t3r**
Experiment start time: **3/3/2018 7:38:51 PM**
Analyst: **Pion**
Instrument ID: **T312060**

#### Experiment Log (continued)

[52:43] Dispensed 0.000024 mL of Base (0.5 M KOH)
[52:48] Stepping pH = 7.37
[53:03] Stirrer speed set to 0
[54:03] Datapoint id 50 collected
[54:03] Charge balance equation is out by -157.9%
[54:03] Stirrer speed set to 60
[54:08] pH 7.43 -> 7.63
[54:08] Using cautious pH adjust
[54:08] Dispensed 0.000024 mL of Base (0.5 M KOH)
[54:13] Stepping pH = 7.56
[54:13] Dispensed 0.000024 mL of Base (0.5 M KOH)
[54:18] Stepping pH = 7.60
[54:18] Dispensed 0.000024 mL of Base (0.5 M KOH)
[54:23] Stepping pH = 7.61
[54:24] Dispensed 0.000024 mL of Base (0.5 M KOH)
[54:29] Stepping pH = 7.67
[54:44] Stirrer speed set to 0
[55:44] Datapoint id 51 collected
[55:44] Charge balance equation is out by -542.8%
[55:44] Stirrer speed set to 60
[55:49] pH 7.65 -> 7.85
[55:49] Using cautious pH adjust
[55:49] Dispensed 0.000024 mL of Base (0.5 M KOH)
[55:54] Stepping pH = 7.78
[55:54] Dispensed 0.000024 mL of Base (0.5 M KOH)
[55:59] Stepping pH = 7.97
[56:14] Stirrer speed set to 0
[57:14] Datapoint id 52 collected
[57:14] Charge balance equation is out by -323.4%
[57:14] Stirrer speed set to 60
[57:19] pH 8.02 -> 8.22
[57:19] Using cautious pH adjust
[57:19] Dispensed 0.000024 mL of Base (0.5 M KOH)
[57:24] Stepping pH = 8.15
[57:25] Dispensed 0.000024 mL of Base (0.5 M KOH)
[57:30] Stepping pH = 8.27
[57:45] Stirrer speed set to 0
[58:45] Datapoint id 53 collected
[58:45] Charge balance equation is out by -340.7%
[58:45] Stirrer speed set to 60
[58:50] pH 8.22 -> 8.42
[58:50] Using cautious pH adjust
[58:50] Dispensed 0.000024 mL of Base (0.5 M KOH)
[58:55] Stepping pH = 8.29
[58:55] Dispensed 0.000024 mL of Base (0.5 M KOH)
[59:00] Stepping pH = 8.35
[59:00] Dispensed 0.000024 mL of Base (0.5 M KOH)
[59:05] Stepping pH = 8.51
[59:20] Stirrer speed set to 0
[59:59] Datapoint id 54 collected
[59:59] Charge balance equation is out by -425.0%
[59:59] Stirrer speed set to 60
[1:00:04] pH 8.57 -> 8.77
[1:00:04] Using cautious pH adjust
[1:00:04] Dispensed 0.000024 mL of Base (0.5 M KOH)
[1:00:09] Stepping pH = 8.60
[1:00:09] Dispensed 0.000024 mL of Base (0.5 M KOH)
[1:00:15] Stepping pH = 8.67
[1:00:15] Dispensed 0.000024 mL of Base (0.5 M KOH)

Sample name: **M12_octanol** Experiment start time: **3/3/2018 7:38:51 PM**
Assay name: **pH-metric high logP** Analyst: **Pion**
Assay ID: **18C-03015** Instrument ID: **T312060**
Filename: **C:\Sirius_T3\Mehtap\20180302_exp29_logP_T3-2\18C-03015_M12_octanol_pH-metric high logP.t3r**

#### Experiment Log (continued)

[1:00:20] Stepping pH = 8.72
[1:00:20] Dispensed 0.000024 mL of Base (0.5 M KOH)
[1:00:25] Stepping pH = 8.80
[1:00:40] Stirrer speed set to 0
[1:01:09] Datapoint id 55 collected
[1:01:09] Charge balance equation is out by -284.9%
[1:01:09] Stirrer speed set to 60
[1:01:15] pH 8.80 -> 9.00
[1:01:15] Using cautious pH adjust
[1:01:15] Dispensed 0.000024 mL of Base (0.5 M KOH)
[1:01:20] Stepping pH = 8.81
[1:01:20] Dispensed 0.000047 mL of Base (0.5 M KOH)
[1:01:25] Stepping pH = 8.92
[1:01:25] Dispensed 0.000024 mL of Base (0.5 M KOH)
[1:01:30] Stepping pH = 8.97
[1:01:30] Dispensed 0.000024 mL of Base (0.5 M KOH)
[1:01:35] Stepping pH = 9.01
[1:01:50] Stirrer speed set to 0
[1:02:05] Datapoint id 56 collected
[1:02:05] Charge balance equation is out by -258.0%
[1:02:05] Argon flow rate set to 0
[1:02:09] Titrator arm moved over Titration position
