## Supplementary material for "Octanol-water partition coefficient measurements for the SAMPL6 Blind Prediction Challenge": SM13_18C-09011_M13_octanol_pH-metric high logP_report.pdf

| | | | |
|---|---|---|---|
| Sample name: | **M13_octanol** | Experiment start time: | **3/9/2018 5:12:57 PM** |
| Assay name: | **pH-metric high logP** | Analyst: | **Pion** |
| Assay ID: | **18C-09011** | Instrument ID: | **T312060** |
| Filename: | **C:\Sirius_T3\Mehtap\20180309_exp31_logP_T3-2\18C-09011_M13_octanol_pH-metric high logP.t3r** | | |

#### pH-metric Result

| | | |
|---|---|---|
| logP (XH +) | -4.95 | ±1.76 (n=49) |
| logP (neutral X) | 2.87 | ±0.01 (n=49) |
| RMSD | 0.236 | |

#### 18C-09011 Points 2 to 18

| | |
|---|---|
| M13_octanol concentration factor | 0.835 |
| Carbonate | 0.0000 mM |
| Acidity error | 3.14531 mM |

#### 18C-09011 Points 19 to 34

| | |
|---|---|
| M13_octanol concentration factor | 0.903 |
| Carbonate | 0.0000 mM |
| Acidity error | 3.32532 mM |

#### 18C-09011 Points 35 to 65

| | |
|---|---|
| M13_octanol concentration factor | 0.690 |
| Carbonate | 0.2248 mM |
| Acidity error | 3.00531 mM |

#### Warnings and errors

| | |
|---|---|
| Errors | None |
| Warnings | One or more logP values out of range |

#### Sample logD and percent species

| pH | M13_octanol logD | M13_octanol M13_octanolH | M13_octanol M13_octanol | M13_octanol M13_octanolH* | M13_octanol M13_octanol* | Comment |
|---|---|---|---|---|---|---|
| 1.000 | -1.90 | 98.76 % | 0.00 % | 0.00 % | 1.24 % | |
| 1.200 | -1.70 | 98.05 % | 0.00 % | 0.00 % | 1.95 % | Stomach pH |
| 2.000 | -0.90 | 88.86 % | 0.02 % | 0.00 % | 11.13 % | |
| 3.000 | 0.10 | 44.37 % | 0.08 % | 0.00 % | 55.56 % | |
| 4.000 | 1.09 | 7.39 % | 0.13 % | 0.00 % | 92.49 % | |
| 5.000 | 2.03 | 0.79 % | 0.13 % | 0.00 % | 99.07 % | |
| 6.000 | 2.67 | 0.08 % | 0.14 % | 0.00 % | 99.78 % | |
| 6.500 | 2.79 | 0.03 % | 0.14 % | 0.00 % | 99.84 % | |
| 7.000 | 2.84 | 0.01 % | 0.14 % | 0.00 % | 99.86 % | |
| 7.400 | 2.86 | 0.00 % | 0.14 % | 0.00 % | 99.86 % | Blood pH |
| 8.000 | 2.87 | 0.00 % | 0.14 % | 0.00 % | 99.86 % | |
| 9.000 | 2.87 | 0.00 % | 0.14 % | 0.00 % | 99.86 % | |
| 10.000 | 2.87 | 0.00 % | 0.14 % | 0.00 % | 99.86 % | |
| 11.000 | 2.87 | 0.00 % | 0.14 % | 0.00 % | 99.86 % | |
| 12.000 | 2.87 | 0.00 % | 0.14 % | 0.00 % | 99.86 % | |

### pH-metric Result

| | | | |
|---|---|---|---|
| Sample name: | **M13_octanol** | Experiment start time: | **3/9/2018 5:12:57 PM** |
| Assay name: | **pH-metric high logP** | Analyst: | **Pion** |
| Assay ID: | **18C-09011** | Instrument ID: | **T312060** |
| Filename: | **C:\Sirius_T3\Mehtap\20180309_exp31_logP_T3-2\18C-09011_M13_octanol_pH-metric high logP.t3r** | | |

#### Graphs

Charge balance residuals

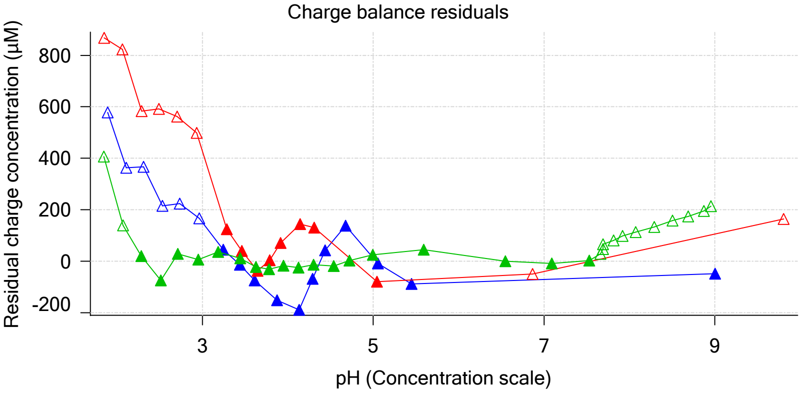


Ionic strength

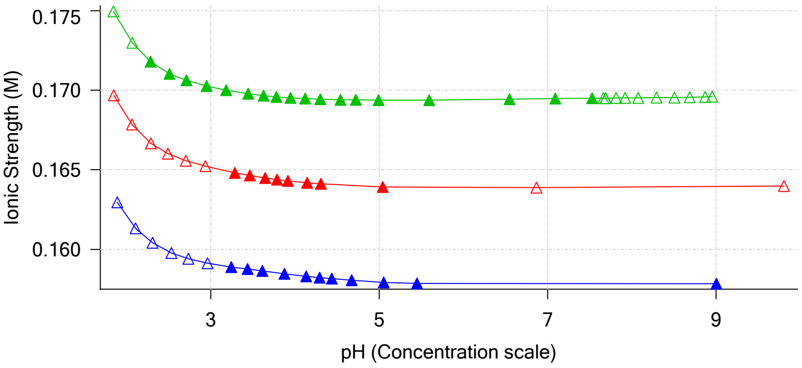


Buffer index profile

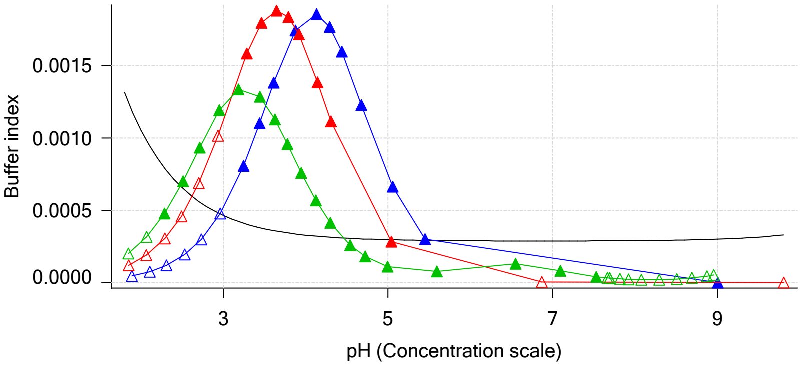


Sample M13_octanol Ionisation Graph

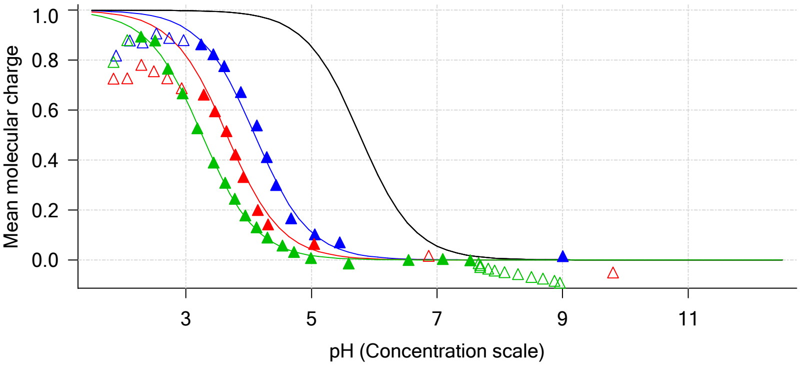


Distribution of species for sample M13_octanol

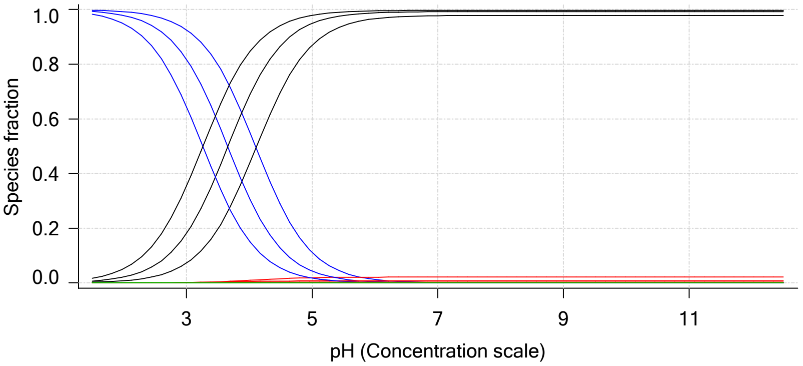


Lipophilicity Profile

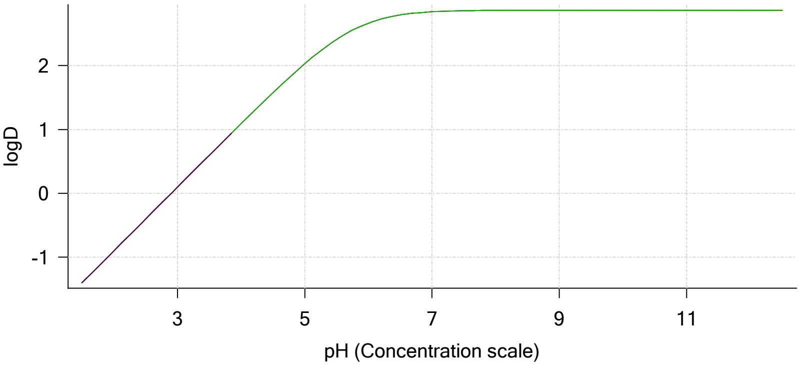


Log R Profile

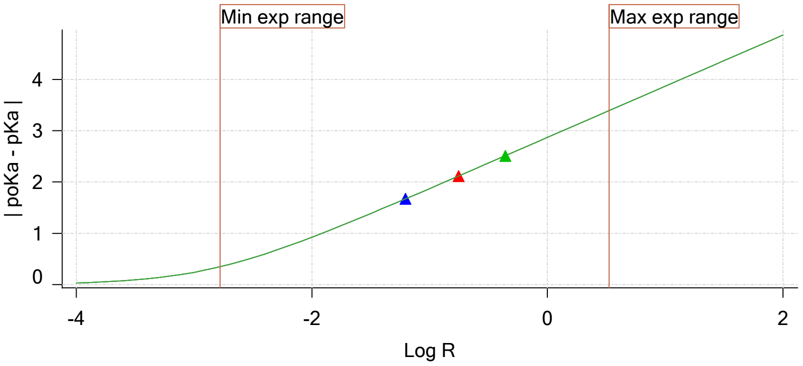


Sample Carbonate Ionisation Graph

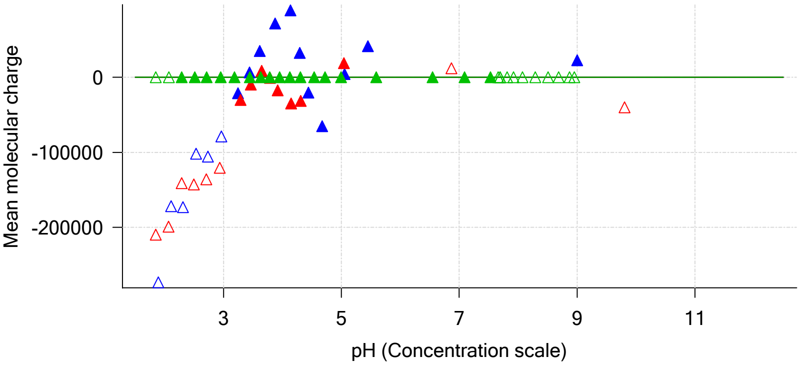

Sample name: **M13_octanol**
Assay name: **pH-metric high logP**
Assay ID: **18C-09011**
Filename: **C:\Sirius_T3\Mehtap\20180309_exp31_logP_T3-2\18C-09011_M13_octanol_pH-metric high logP.t3r**

Experiment start time: **3/9/2018 5:12:57 PM**
Analyst: **Pion**
Instrument ID: **T312060**

#### Graphs (continued)

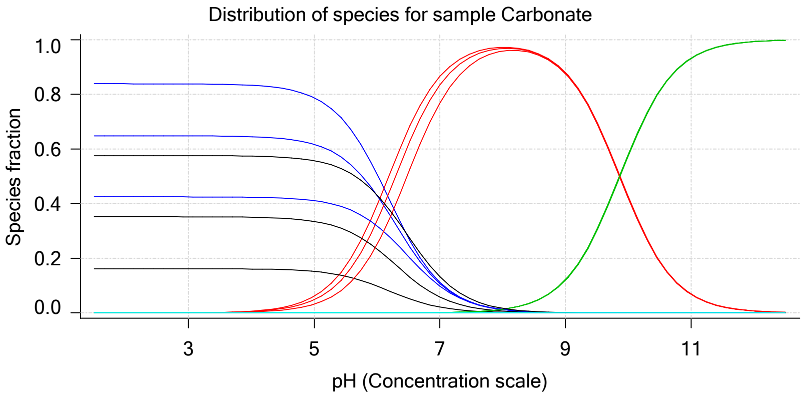

| | | | |
|---|---|---|---|
| Sample name: | **M13_octanol** | Experiment start time: | **3/9/2018 5:12:57 PM** |
| Assay name: | **pH-metric high logP** | Analyst: | **Pion** |
| Assay ID: | **18C-09011** | Instrument ID: | **T312060** |
| Filename: | **C:\Sirius_T3\Mehtap\20180309_exp31_logP_T3-2\18C-09011_M13_octanol_pH-metric high logP.t3r** | | |

### pH-metric high logP Titration 1 of 3 18C-09011 Points 2 to 18

#### Overall results

| | |
|---|---|
| RMSD | 0.383 |
| Average ionic strength | 0.158 M |
| Average temperature | 24.9°C |
| Partition ratio | 0.0624 : 1 |
| Analyte concentration range | 3623.6 μM to 3730.0 μM |
| Total points considered | 11 of 17 |

#### Warnings and errors

| | |
|---|---|
| Errors | None |
| Warnings | One or more logP values out of range<br>Excessive acidity error present |

#### Four-Plus parameters

| | | | |
|---|---|---|---|
| Alpha | 0.102 | 3/9/2018 5:12:57 PM | C:\Sirius_T3\HCl18C09.t3r |
| S | 0.9967 | 3/9/2018 5:12:57 PM | C:\Sirius_T3\HCl18C09.t3r |
| jH | 1.2 | 3/9/2018 5:12:57 PM | C:\Sirius_T3\HCl18C09.t3r |
| jOH | 0.0 | 3/9/2018 5:12:57 PM | C:\Sirius_T3\HCl18C09.t3r |

#### Titrants

| | | | |
|---|---|---|---|
| 0.50 M HCl | 0.999843 | 3/9/2018 5:12:57 PM | C:\Sirius_T3\HCl18C09.t3r |
| 0.50 M KOH | 0.999845 | 3/9/2018 5:12:57 PM | C:\Sirius_T3\KOH18B27.t3r |

#### Sample

| | |
|---|---|
| M13_octanol concentration factor | 0.835 |
| Base pKa 1 | 5.77 |
| logP (XH +) | -4.56 |
| logP (neutral X) | 2.83 |

#### Sample graphs

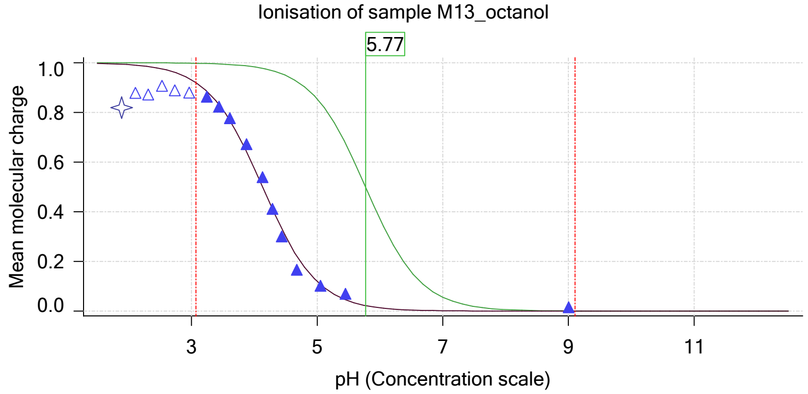


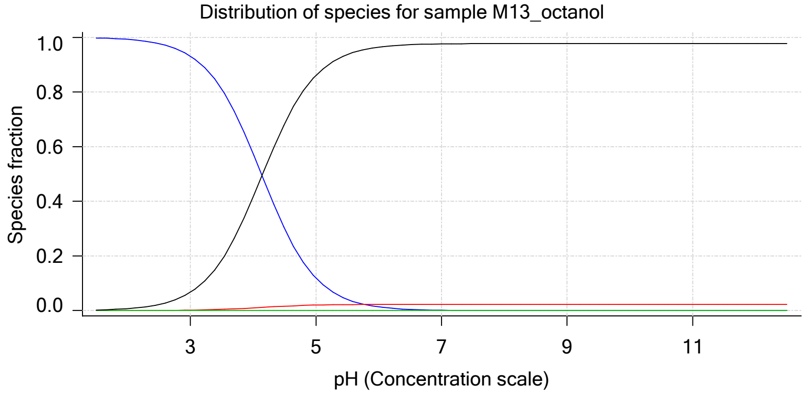

| | | | |
|---|---|---|---|
| Sample name: | **M13_octanol** | Experiment start time: | **3/9/2018 5:12:57 PM** |
| Assay name: | **pH-metric high logP** | Analyst: | **Pion** |
| Assay ID: | **18C-09011** | Instrument ID: | **T312060** |
| Filename: | **C:\Sirius_T3\Mehtap\20180309_exp31_logP_T3-2\18C-09011_M13_octanol_pH-metric high logP.t3r** | | |

#### Sample graphs (continued)

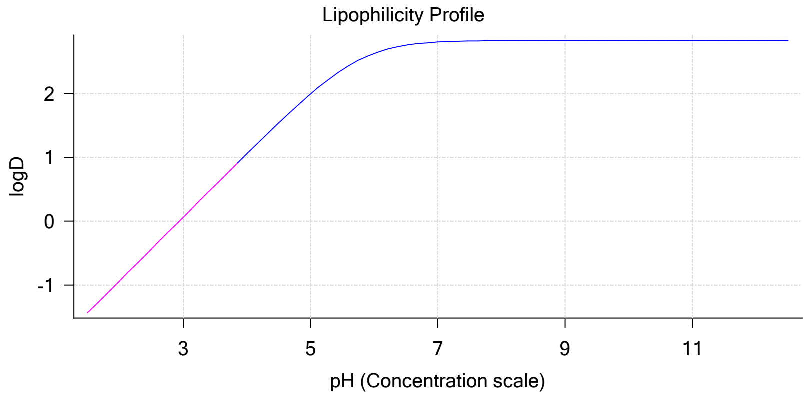


#### Sample logD and percent species

| pH | M13_octanol logD | M13_octanol M13_octanolH | M13_octanol M13_octanol | M13_octanol M13_octanolH* | M13_octanol M13_octanol* | Comment |
|---|---|---|---|---|---|---|
| 1.000 | -1.94 | 99.93 % | 0.00 % | 0.00 % | 0.07 % | |
| 1.200 | -1.74 | 99.88 % | 0.00 % | 0.00 % | 0.11 % | Stomach pH |
| 2.000 | -0.94 | 99.27 % | 0.02 % | 0.00 % | 0.72 % | |
| 3.000 | 0.06 | 93.11 % | 0.16 % | 0.00 % | 6.73 % | |
| 4.000 | 1.06 | 57.49 % | 0.98 % | 0.00 % | 41.53 % | |
| 5.000 | 2.00 | 11.91 % | 2.02 % | 0.00 % | 86.06 % | |
| 6.000 | 2.63 | 1.33 % | 2.27 % | 0.00 % | 96.40 % | |
| 6.500 | 2.76 | 0.43 % | 2.29 % | 0.00 % | 97.29 % | |
| 7.000 | 2.81 | 0.14 % | 2.29 % | 0.00 % | 97.57 % | |
| 7.400 | 2.82 | 0.05 % | 2.30 % | 0.00 % | 97.65 % | Blood pH |
| 8.000 | 2.83 | 0.01 % | 2.30 % | 0.00 % | 97.69 % | |
| 9.000 | 2.83 | 0.00 % | 2.30 % | 0.00 % | 97.70 % | |
| 10.000 | 2.83 | 0.00 % | 2.30 % | 0.00 % | 97.70 % | |
| 11.000 | 2.83 | 0.00 % | 2.30 % | 0.00 % | 97.70 % | |
| 12.000 | 2.83 | 0.00 % | 2.30 % | 0.00 % | 97.70 % | |

#### Carbonate and acidity

| | |
|---|---|
| Carbonate | 0.000 mM |
| Acidity error | 3.145 mM |

#### Other graphs

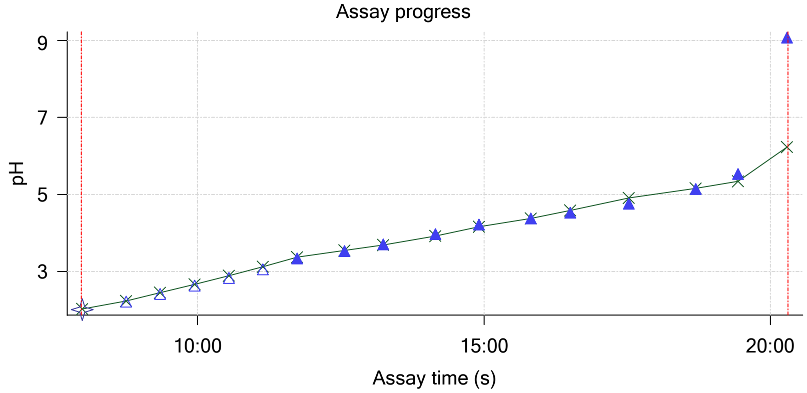


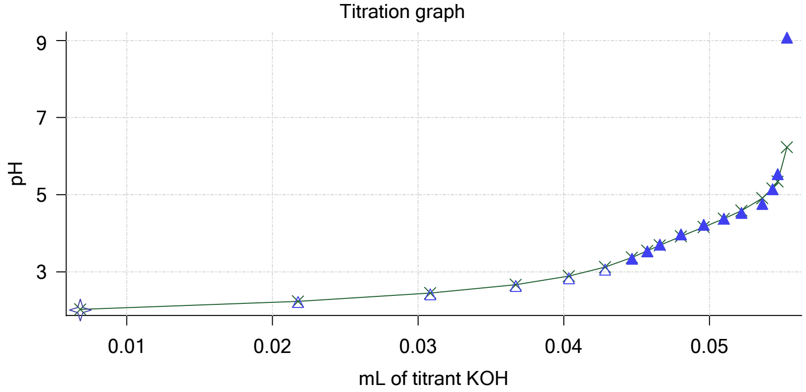

| | | | |
|---|---|---|---|
| Sample name: | **M13_octanol** | Experiment start time: | **3/9/2018 5:12:57 PM** |
| Assay name: | **pH-metric high logP** | Analyst: | **Pion** |
| Assay ID: | **18C-09011** | Instrument ID: | **T312060** |
| Filename: | **C:\Sirius_T3\Mehtap\20180309_exp31_logP_T3-2\18C-09011_M13_octanol_pH-metric high logP.t3r** | | |

#### Other graphs (continued)

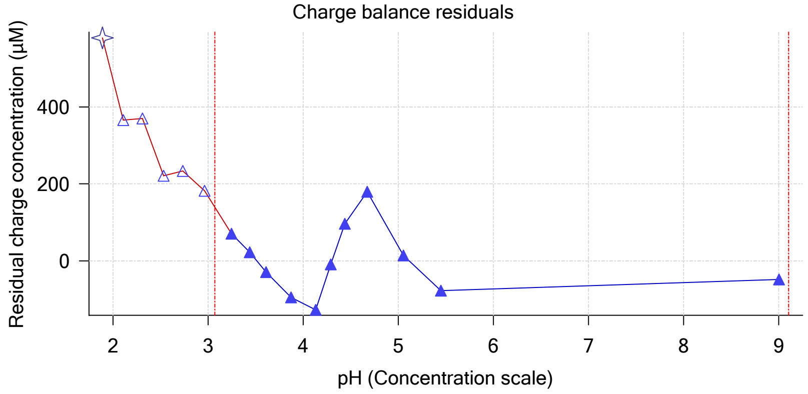


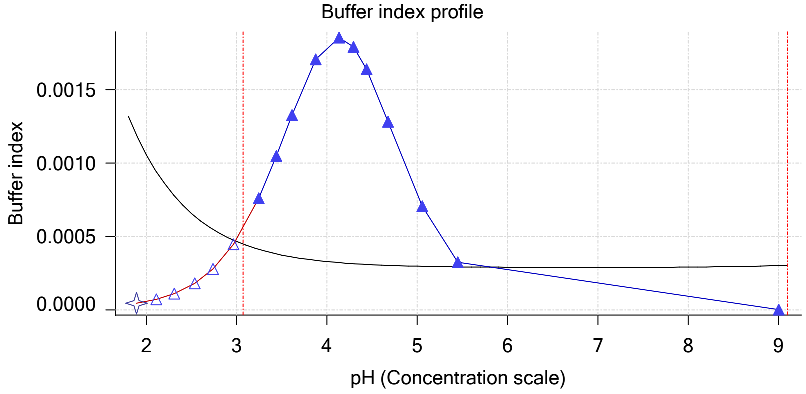


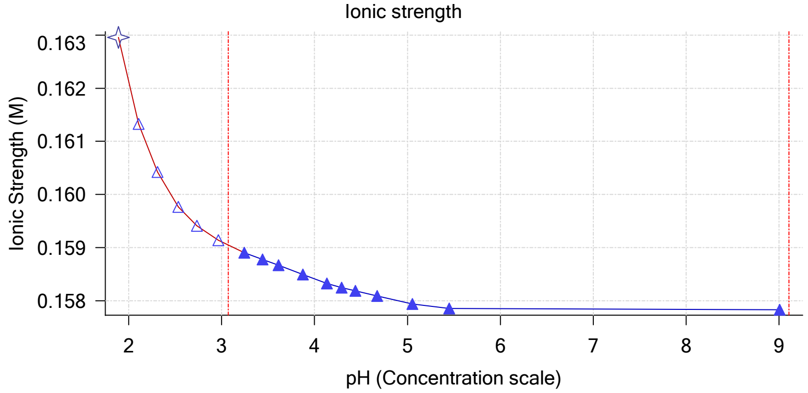


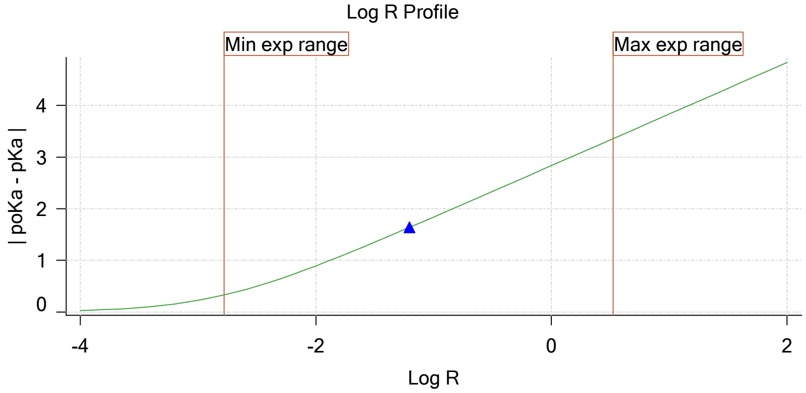


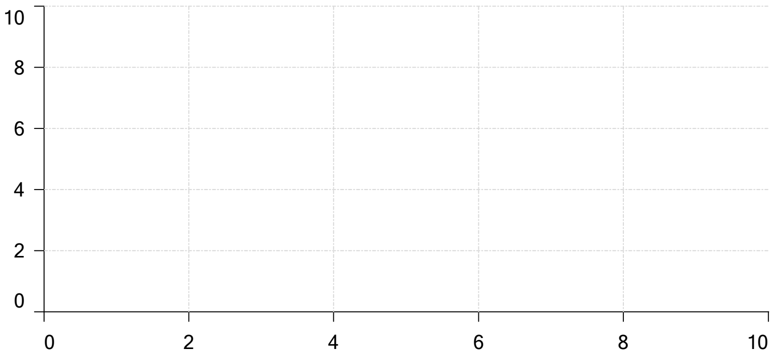

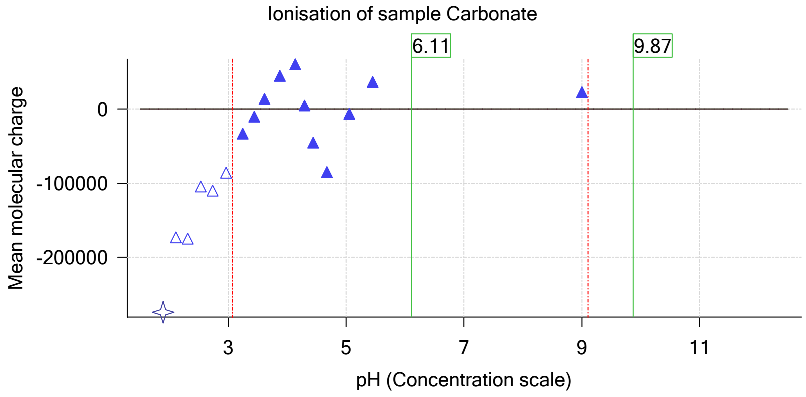

| | | | |
|---|---|---|---|
| Sample name: | **M13_octanol** | Experiment start time: | **3/9/2018 5:12:57 PM** |
| Assay name: | **pH-metric high logP** | Analyst: | **Pion** |
| Assay ID: | **18C-09011** | Instrument ID: | **T312060** |
| Filename: | **C:\Sirius_T3\Mehtap\20180309_exp31_logP_T3-2\18C-09011_M13_octanol_pH-metric high logP.t3r** | | |

# pH-metric high logP   Titration 2 of 3   18C-09011 Points 19 to 34

#### Overall results

| | |
|---|---|
| RMSD | 0.118 |
| Average ionic strength | 0.164 M |
| Average temperature | 24.9°C |
| Partition ratio | 0.1757 : 1 |
| Analyte concentration range | 3064.9 µM to 3153.0 µM |
| Total points considered | 8 of 16 |

#### Warnings and errors

| | |
|---|---|
| Errors | None |
| Warnings | One or more logP values out of range |
| | Excessive acidity error present |

#### Four-Plus parameters

| | | | |
|---|---|---|---|
| Alpha | 0.102 | 3/9/2018 5:12:57 PM | C:\Sirius_T3\HCl18C09.t3r |
| S | 0.9967 | 3/9/2018 5:12:57 PM | C:\Sirius_T3\HCl18C09.t3r |
| jH | 1.2 | 3/9/2018 5:12:57 PM | C:\Sirius_T3\HCl18C09.t3r |
| jOH | 0.0 | 3/9/2018 5:12:57 PM | C:\Sirius_T3\HCl18C09.t3r |

#### Titrants

| | | | |
|---|---|---|---|
| 0.50 M HCl | 0.999843 | 3/9/2018 5:12:57 PM | C:\Sirius_T3\HCl18C09.t3r |
| 0.50 M KOH | 0.999845 | 3/9/2018 5:12:57 PM | C:\Sirius_T3\KOH18B27.t3r |

#### Sample

| | |
|---|---|
| M13_octanol concentration factor | 0.903 |
| Base pKa 1 | 5.77 |
| logP (XH +) | -4.56 |
| logP (neutral X) | 2.90 |

#### Sample graphs

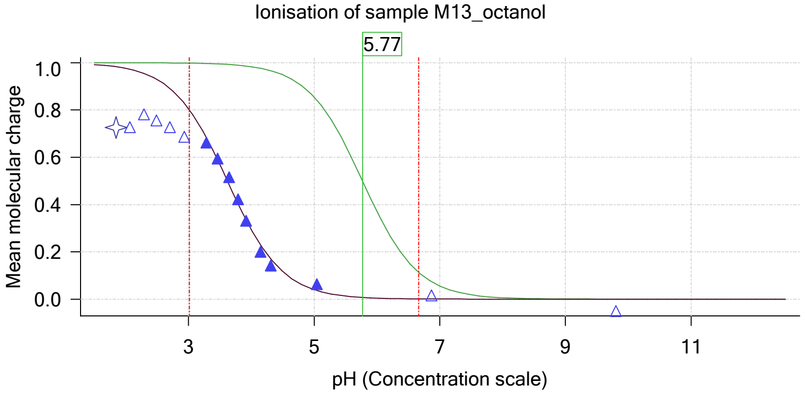


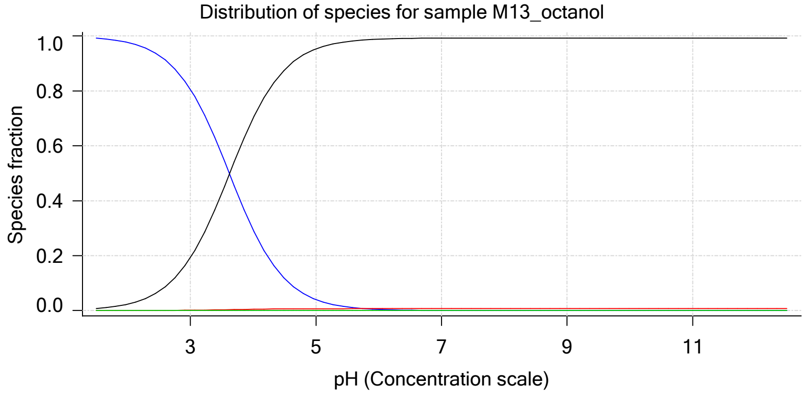

| | | | |
|---|---|---|---|
| Sample name: | **M13_octanol** | Experiment start time: | **3/9/2018 5:12:57 PM** |
| Assay name: | **pH-metric high logP** | Analyst: | **Pion** |
| Assay ID: | **18C-09011** | Instrument ID: | **T312060** |
| Filename: | **C:\Sirius_T3\Mehtap\20180309_exp31_logP_T3-2\18C-09011_M13_octanol_pH-metric high logP.t3r** | | |

#### Sample graphs (continued)

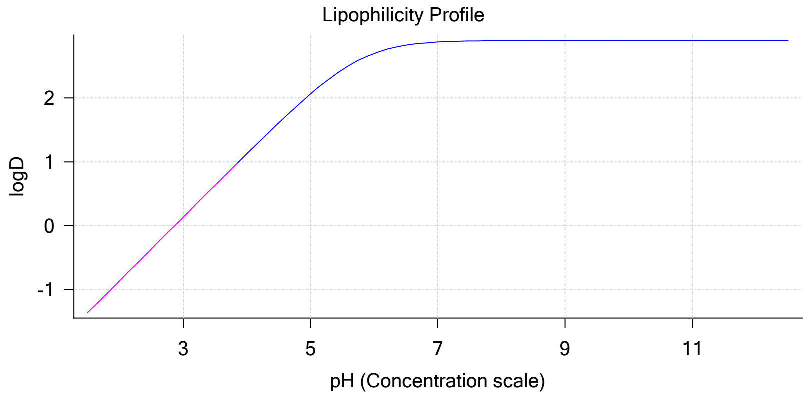


#### Sample logD and percent species

| pH | M13_octanol logD | M13_octanol M13_octanolH | M13_octanol M13_octanol | M13_octanol M13_octanolH* | M13_octanol M13_octanol* | Comment |
|---|---|---|---|---|---|---|
| 1.000 | -1.87 | 99.76 % | 0.00 % | 0.00 % | 0.24 % | |
| 1.200 | -1.67 | 99.62 % | 0.00 % | 0.00 % | 0.38 % | Stomach pH |
| 2.000 | -0.87 | 97.66 % | 0.02 % | 0.00 % | 2.33 % | |
| 3.000 | 0.13 | 80.65 % | 0.14 % | 0.00 % | 19.21 % | |
| 4.000 | 1.13 | 29.42 % | 0.50 % | 0.00 % | 70.08 % | |
| 5.000 | 2.06 | 4.00 % | 0.68 % | 0.00 % | 95.32 % | |
| 6.000 | 2.70 | 0.42 % | 0.70 % | 0.00 % | 98.88 % | |
| 6.500 | 2.83 | 0.13 % | 0.71 % | 0.00 % | 99.16 % | |
| 7.000 | 2.88 | 0.04 % | 0.71 % | 0.00 % | 99.25 % | |
| 7.400 | 2.89 | 0.02 % | 0.71 % | 0.00 % | 99.28 % | Blood pH |
| 8.000 | 2.90 | 0.00 % | 0.71 % | 0.00 % | 99.29 % | |
| 9.000 | 2.90 | 0.00 % | 0.71 % | 0.00 % | 99.29 % | |
| 10.000 | 2.90 | 0.00 % | 0.71 % | 0.00 % | 99.29 % | |
| 11.000 | 2.90 | 0.00 % | 0.71 % | 0.00 % | 99.29 % | |
| 12.000 | 2.90 | 0.00 % | 0.71 % | 0.00 % | 99.29 % | |

#### Carbonate and acidity

| | |
|---|---|
| Carbonate | 0.000 mM |
| Acidity error | 3.325 mM |

#### Other graphs

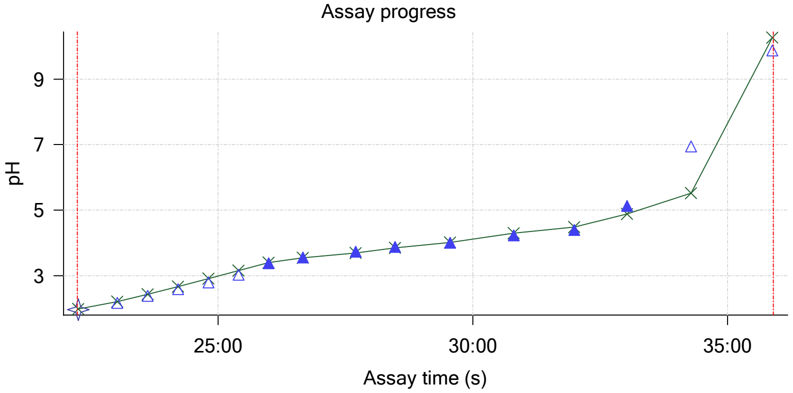


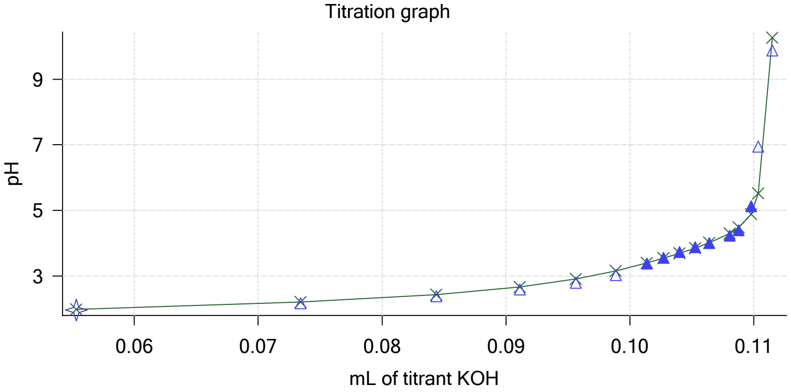

Sample name: **M13_octanol**
Assay name: **pH-metric high logP**
Assay ID: **18C-09011**
Filename: **C:\Sirius_T3\Mehtap\20180309_exp31_logP_T3-2\18C-09011_M13_octanol_pH-metric high logP.t3r**

Experiment start time: **3/9/2018 5:12:57 PM**
Analyst: **Pion**
Instrument ID: **T312060**

#### Other graphs (continued)

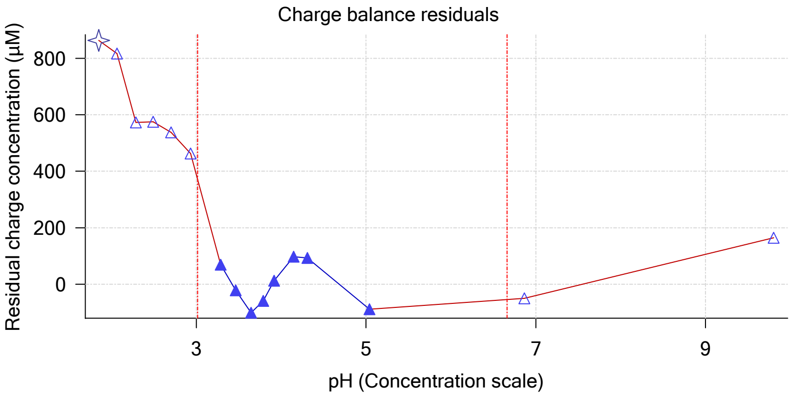


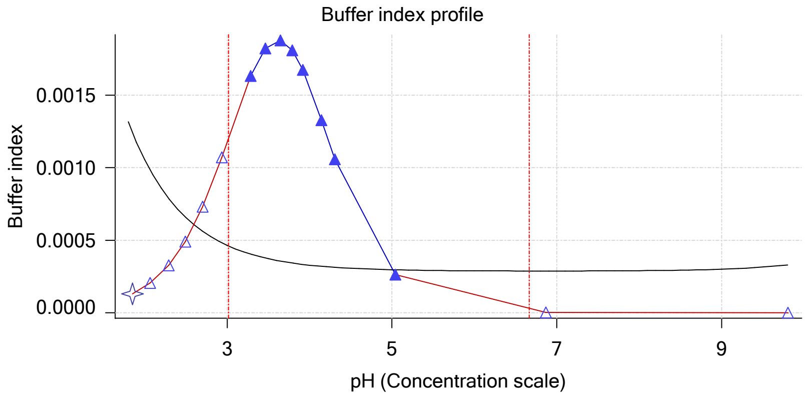


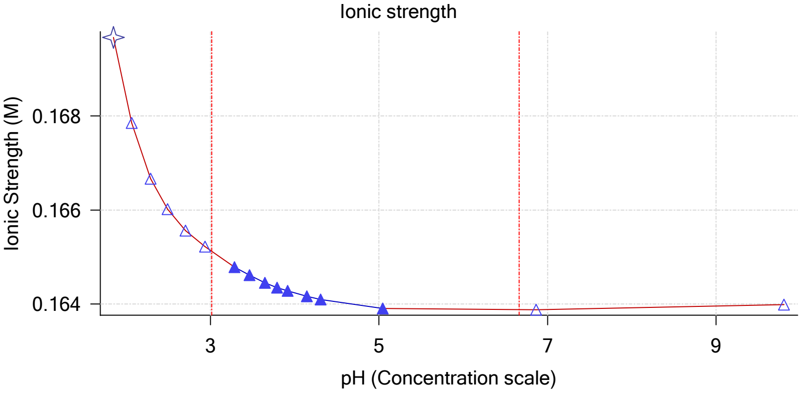


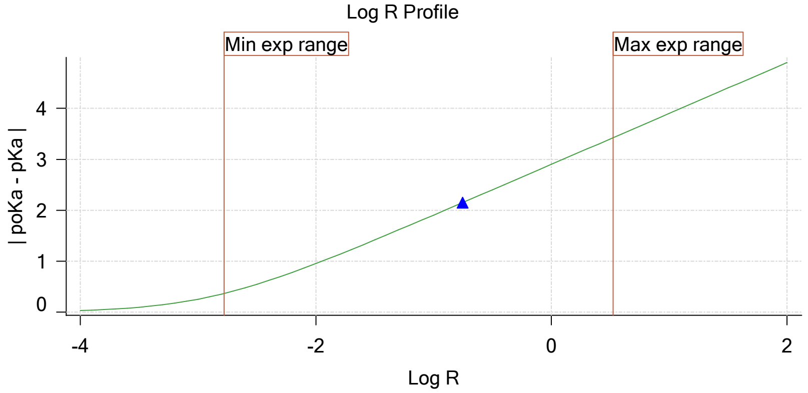


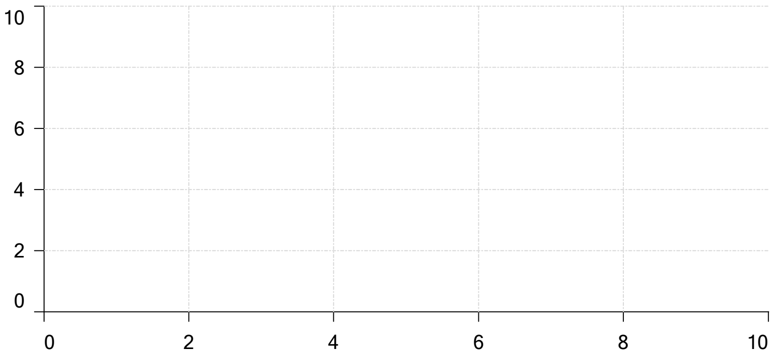


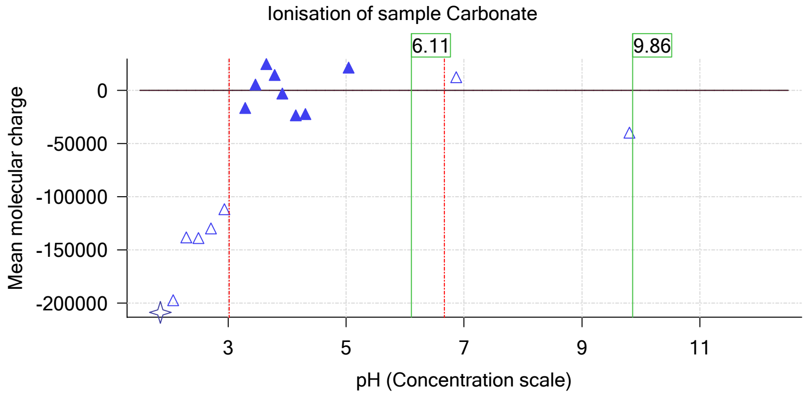

| | | | |
|---|---|---|---|
| Sample name: | **M13_octanol** | Experiment start time: | **3/9/2018 5:12:57 PM** |
| Assay name: | **pH-metric high logP** | Analyst: | **Pion** |
| Assay ID: | **18C-09011** | Instrument ID: | **T312060** |
| Filename: | **C:\Sirius_T3\Mehtap\20180309_exp31_logP_T3-2\18C-09011_M13_octanol_pH-metric high logP.t3r** | | |

### pH-metric high logP Titration 3 of 3 18C-09011 Points 35 to 65

#### Overall results

| | |
|---|---|
| RMSD | 0.137 |
| Average ionic strength | 0.170 M |
| Average temperature | 25.0°C |
| Partition ratio | 0.4387 : 1 |
| Analyte concentration range | 2344.1 μM to 2398.3 μM |
| Total points considered | 18 of 31 |

#### Warnings and errors

| | |
|---|---|
| Errors | None |
| Warnings | Sample concentration factor out of range |
| | One or more logP values out of range |
| | Excessive acidity error present |

#### Four-Plus parameters

| | | | |
|---|---|---|---|
| Alpha | 0.102 | 3/9/2018 5:12:57 PM | C:\Sirius_T3\HCl18C09.t3r |
| S | 0.9967 | 3/9/2018 5:12:57 PM | C:\Sirius_T3\HCl18C09.t3r |
| jH | 1.2 | 3/9/2018 5:12:57 PM | C:\Sirius_T3\HCl18C09.t3r |
| jOH | 0.0 | 3/9/2018 5:12:57 PM | C:\Sirius_T3\HCl18C09.t3r |

#### Titrants

| | | | |
|---|---|---|---|
| 0.50 M HCl | 0.999843 | 3/9/2018 5:12:57 PM | C:\Sirius_T3\HCl18C09.t3r |
| 0.50 M KOH | 0.999845 | 3/9/2018 5:12:57 PM | C:\Sirius_T3\KOH18B27.t3r |

#### Sample

| | |
|---|---|
| M13_octanol concentration factor | 0.690 |
| Base pKa 1 | 5.77 |
| logP (XH +) | -4.56 |
| logP (neutral X) | 2.86 |

#### Sample graphs

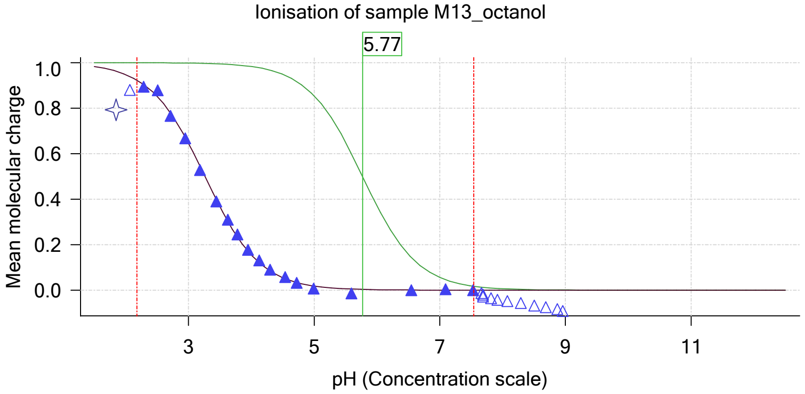


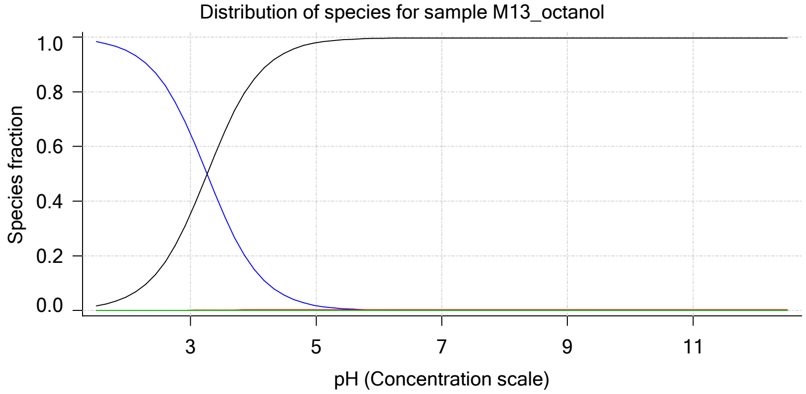

| | | | |
|---|---|---|---|
| Sample name: | **M13_octanol** | Experiment start time: | **3/9/2018 5:12:57 PM** |
| Assay name: | **pH-metric high logP** | Analyst: | **Pion** |
| Assay ID: | **18C-09011** | Instrument ID: | **T312060** |
| Filename: | **C:\Sirius_T3\Mehtap\20180309_exp31_logP_T3-2\18C-09011_M13_octanol_pH-metric high logP.t3r** | | |

#### Sample graphs (continued)

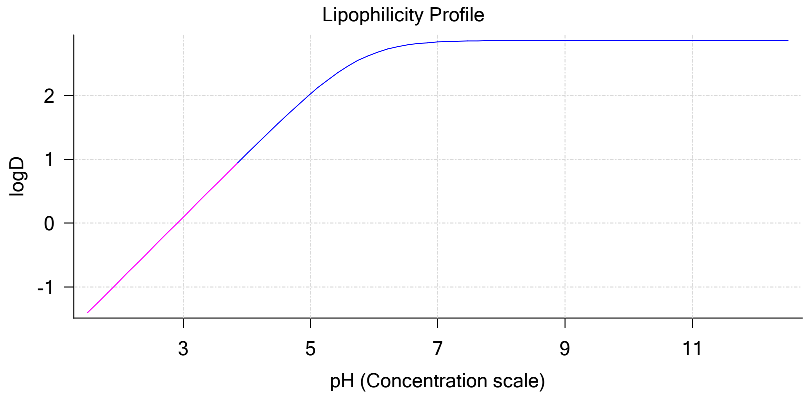


#### Sample logD and percent species

| pH | M13_octanol logD | M13_octanol M13_octanolH | M13_octanol M13_octanol | M13_octanol M13_octanolH* | M13_octanol M13_octanol* | Comment |
|---|---|---|---|---|---|---|
| 1.000 | -1.90 | 99.46 % | 0.00 % | 0.00 % | 0.54 % | |
| 1.200 | -1.71 | 99.14 % | 0.00 % | 0.00 % | 0.86 % | Stomach pH |
| 2.000 | -0.91 | 94.82 % | 0.02 % | 0.00 % | 5.17 % | |
| 3.000 | 0.09 | 64.66 % | 0.11 % | 0.00 % | 35.23 % | |
| 4.000 | 1.09 | 15.47 % | 0.26 % | 0.00 % | 84.27 % | |
| 5.000 | 2.03 | 1.80 % | 0.31 % | 0.00 % | 97.90 % | |
| 6.000 | 2.66 | 0.18 % | 0.31 % | 0.00 % | 99.51 % | |
| 6.500 | 2.79 | 0.06 % | 0.31 % | 0.00 % | 99.63 % | |
| 7.000 | 2.84 | 0.02 % | 0.31 % | 0.00 % | 99.67 % | |
| 7.400 | 2.85 | 0.01 % | 0.31 % | 0.00 % | 99.68 % | Blood pH |
| 8.000 | 2.86 | 0.00 % | 0.31 % | 0.00 % | 99.69 % | |
| 9.000 | 2.86 | 0.00 % | 0.31 % | 0.00 % | 99.69 % | |
| 10.000 | 2.86 | 0.00 % | 0.31 % | 0.00 % | 99.69 % | |
| 11.000 | 2.86 | 0.00 % | 0.31 % | 0.00 % | 99.69 % | |
| 12.000 | 2.86 | 0.00 % | 0.31 % | 0.00 % | 99.69 % | |

#### Carbonate and acidity

| | |
|---|---|
| Carbonate | 0.225 mM |
| Acidity error | 3.005 mM |

#### Other graphs

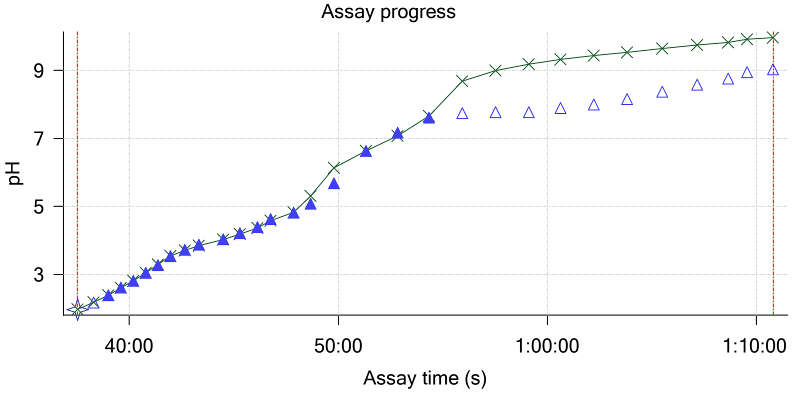


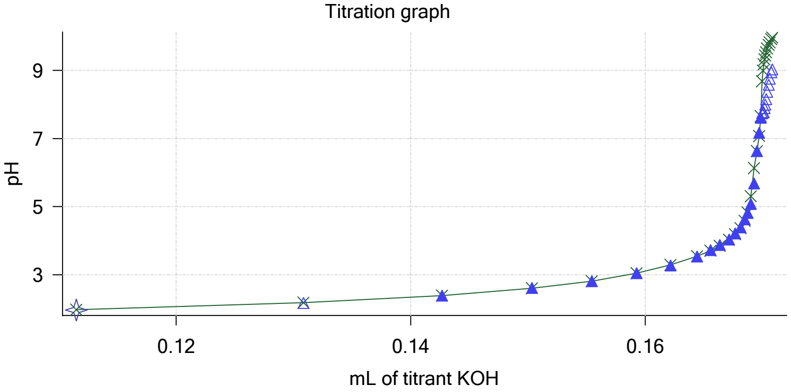

| | | | |
|---|---|---|---|
| Sample name: | **M13_octanol** | Experiment start time: | **3/9/2018 5:12:57 PM** |
| Assay name: | **pH-metric high logP** | Analyst: | **Pion** |
| Assay ID: | **18C-09011** | Instrument ID: | **T312060** |
| Filename: | **C:\Sirius_T3\Mehtap\20180309_exp31_logP_T3-2\18C-09011_M13_octanol_pH-metric high logP.t3r** | | |

#### Other graphs (continued)

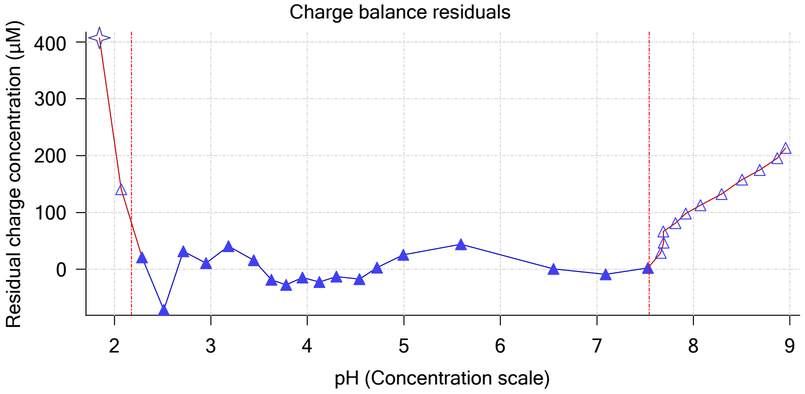


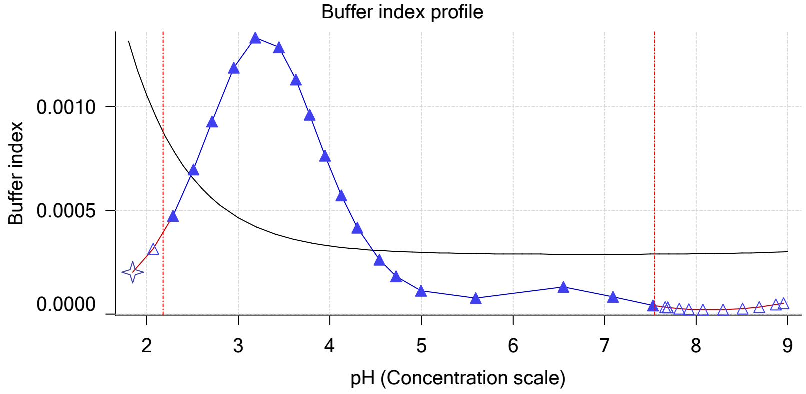


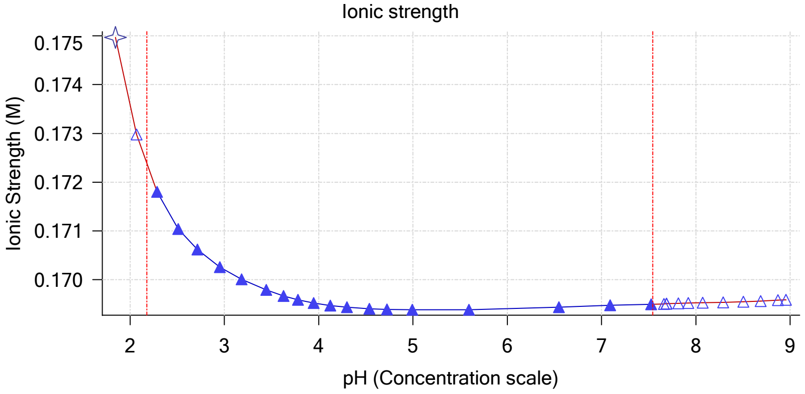


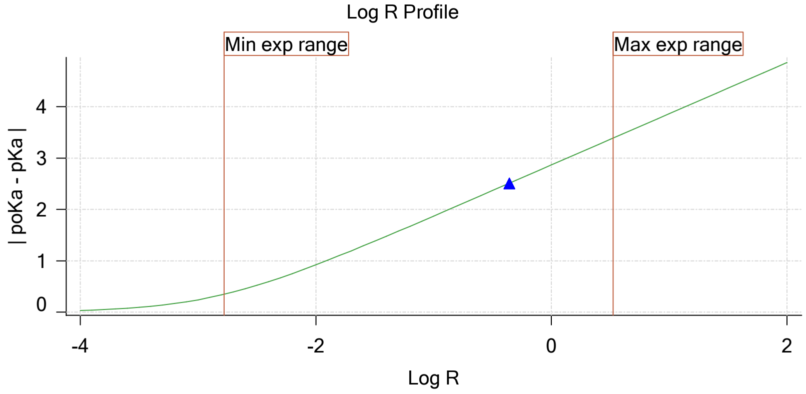


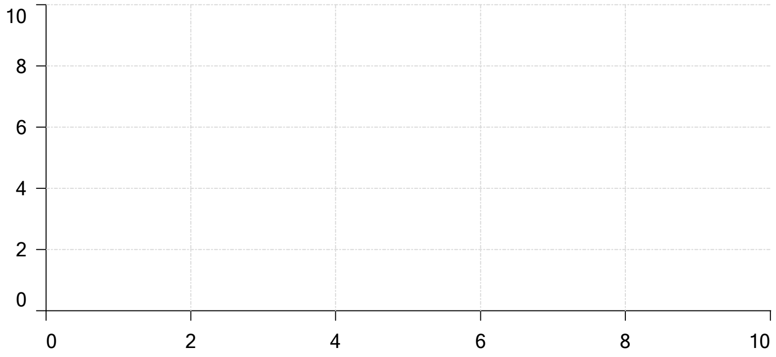


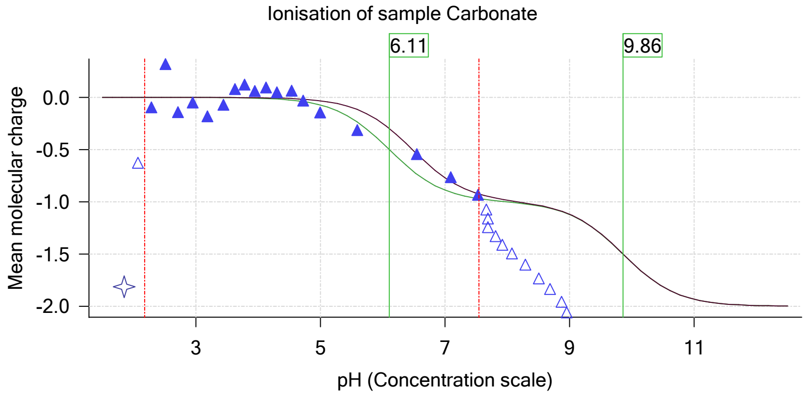

| | | | |
|---|---|---|---|
| Sample name: | **M13_octanol** | Experiment start time: | **3/9/2018 5:12:57 PM** |
| Assay name: | **pH-metric high logP** | Analyst: | **Pion** |
| Assay ID: | **18C-09011** | Instrument ID: | **T312060** |
| Filename: | **C:\Sirius_T3\Mehtap\20180309_exp31_logP_T3-2\18C-09011_M13_octanol_pH-metric high logP.t3r** | | |

#### Assay Model

| Settings | Value | Date/Time changed | Imported from |
|---|---|---|---|
| Sample name | M13_octanol | 2/27/2018 5:57:49 PM | User entered value |
| Sample by | Weight | | Default value |
| Sample weight | 0.001820 g | 3/9/2018 2:22:18 PM | User entered value |
| Formula weight | 295.34 g/mol | 2/27/2018 5:57:49 PM | User entered value |
| Solubility | Unknown | | Default value |
| Molecular weight | 295.34 | 2/27/2018 5:57:49 PM | User entered value |
| Individual pKa ionic environments | No | | Default value |
| Number of pKas | 1 | 2/27/2018 5:57:49 PM | User entered value |
| Sample is a | Base | 2/27/2018 5:57:49 PM | User entered value |
| pKa 1 | 5.77 | 2/27/2018 5:57:49 PM | User entered value |
| logp (XH +) | -4.56 | 3/2/2018 4:30:48 PM | User entered value |
| logP (neutral X) | 2.99 | 3/2/2018 4:30:43 PM | User entered value |

#### Events

| Time | Event | Water | Acid | Base | Octanol | pH | dpH/dt | pH R-squared | pH SD | dpH/[illegible] time |
|---|---|---|---|---|---|---|---|---|---|---|
| 4:58.3 | Manual volume addition | | | | 0.10000 mL | | | | | |
| 4:59.4 | Initial pH = 4.15 | | | | | | | | | |
| 7:58.9 | Data point 2 | 1.50000 mL | 0.04534 mL | 0.00680 mL | 0.10000 mL | 2.003 | -0.00733 | 0.78688 | 0.00041 | 10.0 s |
| 8:45.1 | Data point 3 | 1.50000 mL | 0.04534 mL | 0.02175 mL | 0.10000 mL | 2.213 | -0.00482 | 0.44185 | 0.00036 | 10.0 s |
| 9:20.7 | Data point 4 | 1.50000 mL | 0.04534 mL | 0.03083 mL | 0.10000 mL | 2.410 | -0.00055 | 0.04460 | 0.00013 | 10.5 s |
| 9:56.8 | Data point 5 | 1.50000 mL | 0.04534 mL | 0.03669 mL | 0.10000 mL | 2.631 | -0.00364 | 0.36884 | 0.00030 | 10.5 s |
| 10:32.8 | Data point 6 | 1.50000 mL | 0.04534 mL | 0.04033 mL | 0.10000 mL | 2.831 | -0.01005 | 0.38801 | 0.00080 | 10.0 s |
| 11:08.3 | Data point 7 | 1.50000 mL | 0.04534 mL | 0.04283 mL | 0.10000 mL | 3.058 | -0.01425 | 0.84558 | 0.00077 | 10.5 s |
| 11:44.3 | Data point 8 | 1.50000 mL | 0.04534 mL | 0.04466 mL | 0.10000 mL | 3.338 | -0.01848 | 0.91945 | 0.00095 | 19.0 s |
| 12:33.9 | Data point 9 | 1.50000 mL | 0.04534 mL | 0.04572 mL | 0.10000 mL | 3.531 | -0.01812 | 0.81824 | 0.00099 | 10.0 s |
| 13:14.4 | Data point 10 | 1.50000 mL | 0.04534 mL | 0.04659 mL | 0.10000 mL | 3.703 | -0.01908 | 0.89474 | 0.00100 | 19.0 s |
| 14:09.3 | Data point 11 | 1.50000 mL | 0.04534 mL | 0.04802 mL | 0.10000 mL | 3.965 | -0.01908 | 0.89676 | 0.00099 | 20.0 s |
| 14:54.7 | Data point 12 | 1.50000 mL | 0.04534 mL | 0.04960 mL | 0.10000 mL | 4.221 | -0.01943 | 0.92861 | 0.00100 | 19.0 s |
| 15:49.3 | Data point 13 | 1.50000 mL | 0.04534 mL | 0.05099 mL | 0.10000 mL | 4.380 | -0.01820 | 0.88224 | 0.00096 | 16.0 s |
| 16:30.5 | Data point 14 | 1.50000 mL | 0.04534 mL | 0.05219 mL | 0.10000 mL | 4.526 | -0.01937 | 0.97139 | 0.00097 | 25.5 s |
| 17:31.9 | Data point 15 | 1.50000 mL | 0.04534 mL | 0.05362 mL | 0.10000 mL | 4.762 | -0.01918 | 0.96005 | 0.00097 | 34.5 s |
| 18:42.1 | Data point 16 | 1.50000 mL | 0.04534 mL | 0.05433 mL | 0.10000 mL | 5.141 | -0.01371 | 0.48473 | 0.00097 | 19.0 s |
| 19:26.3 | Data point 17 | 1.50000 mL | 0.04534 mL | 0.05468 mL | 0.10000 mL | 5.535 | -0.00613 | 0.11557 | 0.00089 | 21.0 s |
| 20:17.7 | Data point 18 | 1.50000 mL | 0.04534 mL | 0.05532 mL | 0.10000 mL | 9.077 | -0.02136 | 0.97236 | 0.00107 | Time[illegible] out at |
| 22:15.2 | Data point 19 | 1.50000 mL | 0.09918 mL | 0.05532 mL | 0.30000 mL | 1.963 | -0.01099 | 0.97572 | 0.00055 | 10.0 s |
| 23:01.4 | Data point 20 | 1.50000 mL | 0.09918 mL | 0.07342 mL | 0.30000 mL | 2.171 | -0.01453 | 0.61507 | 0.00092 | 10.0 s |

| | | | |
|---|---|---|---|
| Sample name: | **M13_octanol** | Experiment start time: | **3/9/2018 5:12:57 PM** |
| Assay name: | **pH-metric high logP** | Analyst: | **Pion** |
| Assay ID: | **18C-09011** | Instrument ID: | **T312060** |
| Filename: | **C:\Sirius_T3\Mehtap\20180309_exp31_logP_T3-2\18C-09011_M13_octanol_pH-metric high logP.t3r** | | |

#### Events (continued)

| Time | Event | Water | Acid | Base | Octanol | pH | dpH/dt | pH R-squared | pH SD | dpH/dt time |
|---|---|---|---|---|---|---|---|---|---|---|
| 23:37.0 | Data point 21 | 1.50000 mL | 0.09918 mL | 0.08438 mL | 0.30000 mL | 2.390 | 0.00209 | 0.42447 | 0.00016 | 10.5 s |
| 24:13.0 | Data point 22 | 1.50000 mL | 0.09918 mL | 0.09111 mL | 0.30000 mL | 2.590 | -0.00255 | 0.12562 | 0.00035 | 10.0 s |
| 24:48.6 | Data point 23 | 1.50000 mL | 0.09918 mL | 0.09565 mL | 0.30000 mL | 2.800 | 0.00274 | 0.51472 | 0.00019 | 10.0 s |
| 25:24.0 | Data point 24 | 1.50000 mL | 0.09918 mL | 0.09887 mL | 0.30000 mL | 3.028 | -0.00247 | 0.23539 | 0.00025 | 10.0 s |
| 25:59.5 | Data point 25 | 1.50000 mL | 0.09918 mL | 0.10136 mL | 0.30000 mL | 3.378 | -0.01613 | 0.84514 | 0.00087 | 10.0 s |
| 26:40.1 | Data point 26 | 1.50000 mL | 0.09918 mL | 0.10271 mL | 0.30000 mL | 3.555 | -0.01471 | 0.68891 | 0.00088 | 26.5 s |
| 27:42.3 | Data point 27 | 1.50000 mL | 0.09918 mL | 0.10400 mL | 0.30000 mL | 3.737 | -0.00307 | 0.05380 | 0.00065 | 10.5 s |
| 28:28.5 | Data point 28 | 1.50000 mL | 0.09918 mL | 0.10527 mL | 0.30000 mL | 3.879 | -0.01698 | 0.74340 | 0.00098 | 29.0 s |
| 29:33.3 | Data point 29 | 1.50000 mL | 0.09918 mL | 0.10642 mL | 0.30000 mL | 4.007 | -0.01912 | 0.90936 | 0.00099 | 34.0 s |
| 30:48.3 | Data point 30 | 1.50000 mL | 0.09918 mL | 0.10807 mL | 0.30000 mL | 4.234 | -0.01945 | 0.95903 | 0.00098 | 40.5 s |
| 31:59.4 | Data point 31 | 1.50000 mL | 0.09918 mL | 0.10880 mL | 0.30000 mL | 4.397 | -0.01934 | 0.97445 | 0.00097 | 31.5 s |
| 33:01.6 | Data point 32 | 1.50000 mL | 0.09918 mL | 0.10981 mL | 0.30000 mL | 5.129 | -0.01995 | 0.97002 | 0.00100 | 45.0 s |
| 34:17.2 | Data point 33 | 1.50000 mL | 0.09918 mL | 0.11037 mL | 0.30000 mL | 6.946 | -0.12491 | 0.99710 | 0.00618 | Timed out at 59.5 s |
| 35:52.9 | Data point 34 | 1.50000 mL | 0.09918 mL | 0.11150 mL | 0.30000 mL | 9.875 | -0.00416 | 0.04424 | 0.00098 | 13.5 s |
| 37:31.9 | Data point 35 | 1.50000 mL | 0.15807 mL | 0.11150 mL | 0.80000 mL | 1.959 | -0.00284 | 0.16111 | 0.00035 | 10.0 s |
| 38:18.1 | Data point 36 | 1.50000 mL | 0.15807 mL | 0.13086 mL | 0.80000 mL | 2.175 | -0.00636 | 0.10985 | 0.00095 | 16.0 s |
| 38:59.8 | Data point 37 | 1.50000 mL | 0.15807 mL | 0.14266 mL | 0.80000 mL | 2.388 | -0.01190 | 0.63324 | 0.00074 | 10.0 s |
| 39:35.4 | Data point 38 | 1.50000 mL | 0.15807 mL | 0.15033 mL | 0.80000 mL | 2.610 | -0.00893 | 0.81503 | 0.00049 | 10.5 s |
| 40:11.5 | Data point 39 | 1.50000 mL | 0.15807 mL | 0.15543 mL | 0.80000 mL | 2.809 | -0.01069 | 0.89777 | 0.00056 | 10.0 s |
| 40:47.0 | Data point 40 | 1.50000 mL | 0.15807 mL | 0.15924 mL | 0.80000 mL | 3.044 | -0.00343 | 0.27090 | 0.00033 | 10.0 s |
| 41:22.5 | Data point 41 | 1.50000 mL | 0.15807 mL | 0.16214 mL | 0.80000 mL | 3.276 | -0.01814 | 0.88874 | 0.00095 | 10.5 s |
| 41:58.5 | Data point 42 | 1.50000 mL | 0.15807 mL | 0.16439 mL | 0.80000 mL | 3.535 | -0.01802 | 0.93392 | 0.00092 | 10.0 s |
| 42:39.1 | Data point 43 | 1.50000 mL | 0.15807 mL | 0.16555 mL | 0.80000 mL | 3.719 | -0.01345 | 0.67807 | 0.00081 | 10.0 s |
| 43:19.7 | Data point 44 | 1.50000 mL | 0.15807 mL | 0.16634 mL | 0.80000 mL | 3.870 | -0.01661 | 0.94479 | 0.00084 | 29.0 s |
| 44:29.7 | Data point 45 | 1.50000 mL | 0.15807 mL | 0.16712 mL | 0.80000 mL | 4.037 | -0.00694 | 0.13256 | 0.00094 | 11.5 s |
| 45:17.0 | Data point 46 | 1.50000 mL | 0.15807 mL | 0.16766 mL | 0.80000 mL | 4.213 | 0.00141 | 0.00562 | 0.00093 | 10.5 s |
| 46:08.4 | Data point 47 | 1.50000 mL | 0.15807 mL | 0.16811 mL | 0.80000 mL | 4.389 | -0.01790 | 0.90016 | 0.00093 | 11.5 s |
| 46:45.3 | Data point 48 | 1.50000 mL | 0.15807 mL | 0.16846 mL | 0.80000 mL | 4.627 | -0.01929 | 0.94369 | 0.00098 | 35.5 s |
| 47:51.4 | Data point 49 | 1.50000 mL | 0.15807 mL | 0.16872 mL | 0.80000 mL | 4.810 | -0.01458 | 0.57061 | 0.00095 | 19.0 s |
| 48:40.8 | Data point 50 | 1.50000 mL | 0.15807 mL | 0.16898 mL | 0.80000 mL | 5.080 | -0.01848 | 0.89699 | 0.00096 | 36.0 s |
| 49:47.4 | Data point 51 | 1.50000 mL | 0.15807 mL | 0.16926 mL | 0.80000 mL | 5.677 | -0.01986 | 0.97546 | 0.00099 | 56.0 s |
| 51:19.2 | Data point 52 | 1.50000 mL | 0.15807 mL | 0.16950 mL | 0.80000 mL | 6.630 | -0.10160 | 0.99674 | 0.00502 | Timed out at 59.5 s |
| 52:49.7 | Data point 53 | 1.50000 mL | 0.15807 mL | 0.16969 mL | 0.80000 mL | 7.171 | -0.11670 | 0.99695 | 0.00577 | Timed out at 59.5 s |
| 54:20.2 | Data point 54 | 1.50000 mL | 0.15807 mL | 0.16983 mL | 0.80000 mL | 7.606 | -0.10666 | 0.99532 | 0.00527 | Timed out at 59.5 s |
| 55:55.8 | Data point 55 | 1.50000 mL | 0.15807 mL | 0.16994 mL | 0.80000 mL | 7.740 | -0.06542 | 0.99344 | 0.00324 | Timed out at 59.5 s |
| 57:31.5 | Data point 56 | 1.50000 mL | 0.15807 mL | 0.17001 mL | 0.80000 mL | 7.767 | -0.04725 | 0.99135 | 0.00234 | Timed out at 59.5 s |
| 59:07.1 | Data point 57 | 1.50000 mL | 0.15807 mL | 0.17008 mL | 0.80000 mL | 7.767 | -0.03770 | 0.96936 | 0.00189 | Timed out at 59.5 s |
| 1:00:37.5 | Data point 58 | 1.50000 mL | 0.15807 mL | 0.17016 mL | 0.80000 mL | 7.892 | -0.03830 | 0.97684 | 0.00191 | Timed out at 59.5 s |
| 1:02:13.0 | Data point 59 | 1.50000 mL | 0.15807 mL | 0.17023 mL | 0.80000 mL | 7.996 | -0.03517 | 0.97225 | 0.00176 | Timed out at 59.5 s |
| 1:03:48.7 | Data point 60 | 1.50000 mL | 0.15807 mL | 0.17030 mL | 0.80000 mL | 8.151 | -0.02938 | 0.97141 | 0.00147 | Timed out at 59.5 s |
| 1:05:29.4 | Data point 61 | 1.50000 mL | 0.15807 mL | 0.17039 mL | 0.80000 mL | 8.367 | -0.02206 | 0.95552 | 0.00111 | Timed out at 59.5 s |
| 1:07:10.1 | Data point 62 | 1.50000 mL | 0.15807 mL | 0.17051 mL | 0.80000 mL | 8.580 | -0.01905 | 0.91928 | 0.00098 | 48.0 s |
| 1:08:39.0 | Data point 63 | 1.50000 mL | 0.15807 mL | 0.17060 mL | 0.80000 mL | 8.761 | -0.01127 | 0.33329 | 0.00096 | 18.5 s |
| 1:09:33.2 | Data point 64 | 1.50000 mL | 0.15807 mL | 0.17072 mL | 0.80000 mL | 8.944 | -0.01844 | 0.86480 | 0.00098 | 43.5 s |
| 1:10:47.3 | Data point 65 | 1.50000 mL | 0.15807 mL | 0.17081 mL | 0.80000 mL | 9.028 | -0.01594 | 0.74914 | 0.00091 | 17.0 s |
| 1:11:13.3 | Assay volumes | 1.50000 mL | 0.15807 mL | 0.17081 mL | 0.80000 mL | | | | | |

| | | | |
|---|---|---|---|
| Sample name: | **M13_octanol** | Experiment start time: | **3/9/2018 5:12:57 PM** |
| Assay name: | **pH-metric high logP** | Analyst: | **Pion** |
| Assay ID: | **18C-09011** | Instrument ID: | **T312060** |
| Filename: | **C:\Sirius_T3\Mehtap\20180309_exp31_logP_T3-2\18C-09011_M13_octanol_pH-metric high logP.t3r** | | |

| | | | |
|---|---|---|---|
| Sample name: | **M13_octanol** | Experiment start time: | **3/9/2018 5:12:57 PM** |
| Assay name: | **pH-metric high logP** | Analyst: | **Pion** |
| Assay ID: | **18C-09011** | Instrument ID: | **T312060** |
| Filename: | **C:\Sirius_T3\Mehtap\20180309_exp31_logP_T3-2\18C-09011_M13_octanol_pH-metric high logP.t3r** | | |

#### Assay Settings (continued)

| Setting | Value | Original Value | Date/Time changed | Imported from |
|---|---|---|---|---|
| ***Titration 3*** | | | | |
| Titrate from | Low to high pH | | | |
| Add additional water | 0.00 mL | | | |
| Additional partition solvent volume | 0.500 mL | | | |
| Additional partition solvent added | Automatic | | | |
| After pH adjust stir for | 30 seconds | | | |
| Stir to allow partitioning for | 15 seconds | | | |
| Stirrer speed for partitioning | 60% | | | |
| ***Data Point Stability*** | | | | |
| Stir during data point collection | No | | | |
| Delay before data point collection | 0 seconds | | | |
| Number of points to average | 20 points | | | |
| Time interval between points | 0.50 seconds | | | |
| Required maximum standard deviation | 0.00100 dpH/dt | | | |
| Stability timeout after | 60 seconds | | | |

#### Calibration Settings

| Setting | Value | Date/Time changed | Imported from |
|---|---|---|---|
| Four-Plus alpha | 0.102 | 3/9/2018 5:12:57 PM | C:\Sirius_T3\HCl18C09.t3r |
| Four-Plus S | 0.9967 | 3/9/2018 5:12:57 PM | C:\Sirius_T3\HCl18C09.t3r |
| Four-Plus jH | 1.2 | 3/9/2018 5:12:57 PM | C:\Sirius_T3\HCl18C09.t3r |
| Four-Plus jOH | 0.0 | 3/9/2018 5:12:57 PM | C:\Sirius_T3\HCl18C09.t3r |
| Base concentration factor | 1.000 | 3/9/2018 5:12:57 PM | C:\Sirius_T3\KOH18B27.t3r |
| Acid concentration factor | 1.000 | 3/9/2018 5:12:57 PM | C:\Sirius_T3\HCl18C09.t3r |

#### Instrument Settings

| Setting | Value | Batch Id | Install date |
|---|---|---|---|
| Instrument owner | Merck | | |
| Instrument ID | T312060 | | |
| Instrument type | T3 Simulator | | |
| Software version | 1.1.3.0 | | |
| Dispenser module | | T3DM1200361 | 3/31/2009 6:24:52 AM |
| Dispenser 0 | Water | | 3/31/2009 6:25:05 AM |
| Syringe volume | 2.5 mL | | |
| Firmware version | 1.2.1(r2) | | |
| Titrant | Water (0.15 M KCl) | 02-06-2018 | 2/27/2018 11:05:59 AM |
| Dispenser 2 | Acid | | 3/31/2009 6:25:11 AM |
| Syringe volume | 0.5 mL | | |
| Firmware version | 1.2.1(r2) | | |
| Titrant | Acid (0.5 M HCl) | 02-27-2018 | 2/27/2018 11:27:22 AM |
| Dispenser 1 | Base | | 3/31/2009 6:25:21 AM |
| Syringe volume | 0.5 mL | | |
| Firmware version | 1.2.1(r2) | | |
| Titrant | Base (0.5 M KOH) | 9/22/2017 | 2/27/2018 11:21:22 AM |
| Dispenser 5 | Cosolvent | | 3/31/2009 6:26:24 AM |
| Syringe volume | 2.5 mL | | |
| Firmware version | 1.2.1(r2) | | |
| Distribution valve 5 | Distribution Valve | | 3/31/2009 6:28:19 AM |
| Firmware version | 1.1.3 | | |
| Port A | Methanol (80%, 0.15 M KCl) | 02-08-2018 | 3/6/2018 10:28:59 AM |
| Port B | Cyclohexane | 11-01-17 | 2/27/2018 11:37:57 AM |
| Dispenser 3 | Buffer | | 8/3/2010 6:05:16 AM |
| Syringe volume | 0.5 mL | | |
| Firmware version | 1.2.1(r2) | | |
| Titrant | Dodecane | 2018/01/31 | 2/28/2018 11:18:04 AM |
| Dispenser 6 | Octanol | | 10/22/2010 11:52:43 AM |

| | | | |
|---|---|---|---|
| Sample name: | **M13_octanol** | Experiment start time: | **3/9/2018 5:12:57 PM** |
| Assay name: | **pH-metric high logP** | Analyst: | **Pion** |
| Assay ID: | **18C-09011** | Instrument ID: | **T312060** |
| Filename: | **C:\Sirius_T3\Mehtap\20180309_exp31_logP_T3-2\18C-09011_M13_octanol_pH-metric high logP.t3r** | | |

| | | | |
|---|---|---|---|
| Sample name: | **M13_octanol** | Experiment start time: | **3/9/2018 5:12:57 PM** |
| Assay name: | **pH-metric high logP** | Analyst: | **Pion** |
| Assay ID: | **18C-09011** | Instrument ID: | **T312060** |
| Filename: | **C:\Sirius_T3\Mehtap\20180309_exp31_logP_T3-2\18C-09011_M13_octanol_pH-metric high logP.t3r** | | |

#### Instrument Settings (continued)

| Setting | Value | Batch Id | Install date |
|---|---|---|---|
| Spectrometer calibration stir duration | 5 s | | |
| Spectrometer calibration stir speed | 30% | | |
| Spectrometer calibration wash pump volume | 20.0 mL | | |
| Spectrometer calibration wash stir duration | 5 s | | |
| Spectrometer calibration wash stir speed | 30% | | |
| Overhead dispense height | 10000 | | |

#### Refinement Settings

| Setting | Value | Default value |
|---|---|---|
| Turbidity detection method | None | None |
| Turbidity wavelength to assess | 500.0 nm | 500.0 nm |
| Turbidity maximum absorbance | 0.100 | 0.100 |
| Turbidity probe threshold | 50.00 | 50.00 |
