## Supplementary material for "Octanol-water partition coefficient measurements for the SAMPL6 Blind Prediction Challenge": SM13_18C-16016_M13_octanol_pH-metric high logP_report.pdf

| | | | |
|---|---|---|---|
| Sample name: | **M13_octanol** | Experiment start time: | **3/16/2018 8:34:17 PM** |
| Assay name: | **pH-metric high logP** | Analyst: | **Pion** |
| Assay ID: | **18C-16016** | Instrument ID: | **T311053** |
| Filename: | **C:\Sirius_T3\Mehtap\20180316_exp32_logP_T3-1\18C-16016_M13_octanol_pH-metric high logP.t3r** | | |

#### pH-metric Result

| | | |
|---|---|---|
| logP (XH +) | -7.57 | ±1.49 (n=50) |
| logP (neutral X) | 2.89 | ±0.02 (n=50) |
| RMSD | 3.854 | |

#### 18C-16016 Points 2 to 15

| | |
|---|---|
| M13_octanol concentration factor | 0.878 |
| Carbonate | 0.0003 mM |
| Acidity error | 2.29946 mM |

#### 18C-16016 Points 16 to 39

| | |
|---|---|
| M13_octanol concentration factor | 0.780 |
| Carbonate | 0.0000 mM |
| Acidity error | 2.26129 mM |

#### 18C-16016 Points 40 to 63

| | |
|---|---|
| M13_octanol concentration factor | 0.706 |
| Carbonate | 0.1381 mM |
| Acidity error | 2.26901 mM |

#### Warnings and errors

| | |
|---|---|
| Errors | None |
| Warnings | One or more logP values out of range |

#### Sample logD and percent species

| pH | M13_octanol logD | M13_octanol M13_octanolH | M13_octanol M13_octanol | M13_octanol M13_octanolH* | M13_octanol M13_octanol* | Comment |
|---|---|---|---|---|---|---|
| 1.000 | -1.88 | 98.68 % | 0.00 % | 0.00 % | 1.31 % | |
| 1.200 | -1.68 | 97.93 % | 0.00 % | 0.00 % | 2.07 % | Stomach pH |
| 2.000 | -0.88 | 88.23 % | 0.01 % | 0.00 % | 11.76 % | |
| 3.000 | 0.12 | 42.84 % | 0.07 % | 0.00 % | 57.08 % | |
| 4.000 | 1.12 | 6.97 % | 0.12 % | 0.00 % | 92.91 % | |
| 5.000 | 2.06 | 0.74 % | 0.13 % | 0.00 % | 99.13 % | |
| 6.000 | 2.69 | 0.07 % | 0.13 % | 0.00 % | 99.80 % | |
| 6.500 | 2.82 | 0.02 % | 0.13 % | 0.00 % | 99.85 % | |
| 7.000 | 2.87 | 0.01 % | 0.13 % | 0.00 % | 99.87 % | |
| 7.400 | 2.88 | 0.00 % | 0.13 % | 0.00 % | 99.87 % | Blood pH |
| 8.000 | 2.89 | 0.00 % | 0.13 % | 0.00 % | 99.87 % | |
| 9.000 | 2.89 | 0.00 % | 0.13 % | 0.00 % | 99.87 % | |
| 10.000 | 2.89 | 0.00 % | 0.13 % | 0.00 % | 99.87 % | |
| 11.000 | 2.89 | 0.00 % | 0.13 % | 0.00 % | 99.87 % | |
| 12.000 | 2.89 | 0.00 % | 0.13 % | 0.00 % | 99.87 % | |

Sample name: **M13_octanol**
Assay name: **pH-metric high logP**
Assay ID: **18C-16016**
Filename: **C:\Sirius_T3\Mehtap\20180316_exp32_logP_T3-1\18C-16016_M13_octanol_pH-metric high logP.t3r**

Experiment start time: **3/16/2018 8:34:17 PM**
Analyst: **Pion**
Instrument ID: **T311053**

#### Graphs

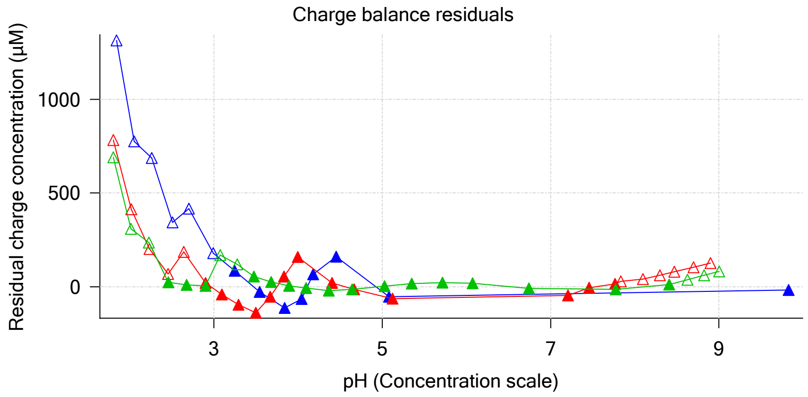


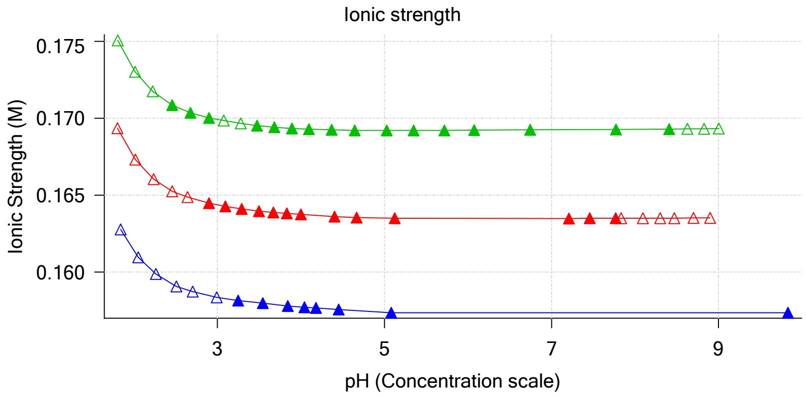


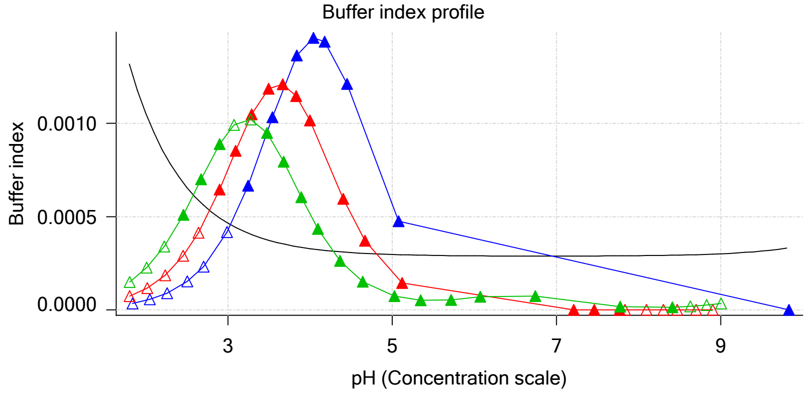


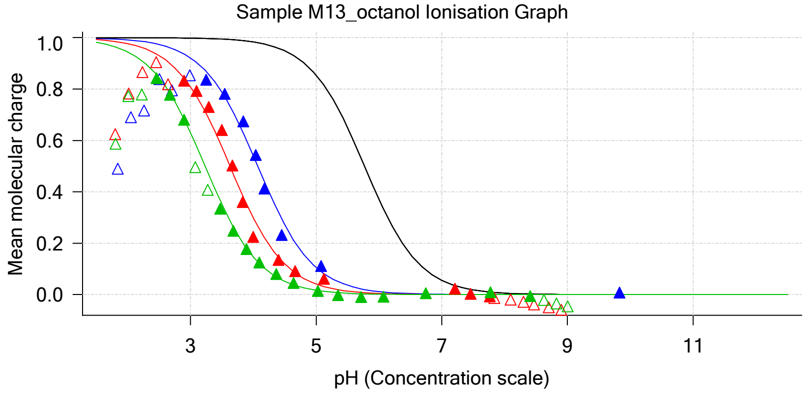


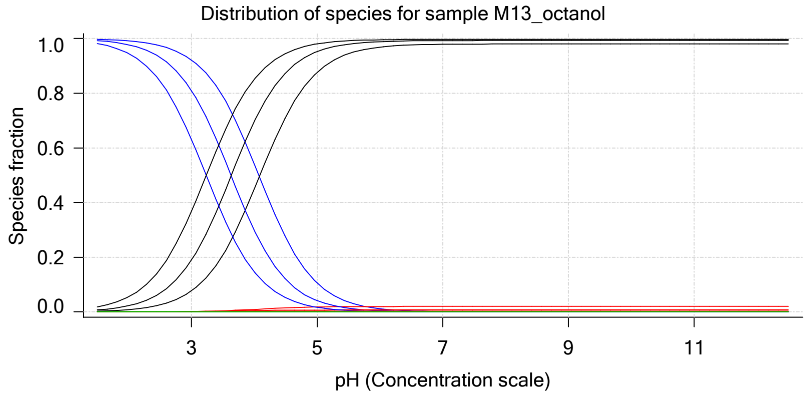


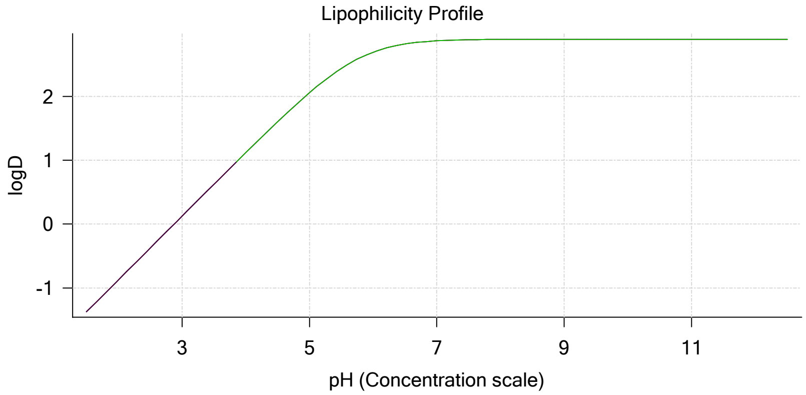


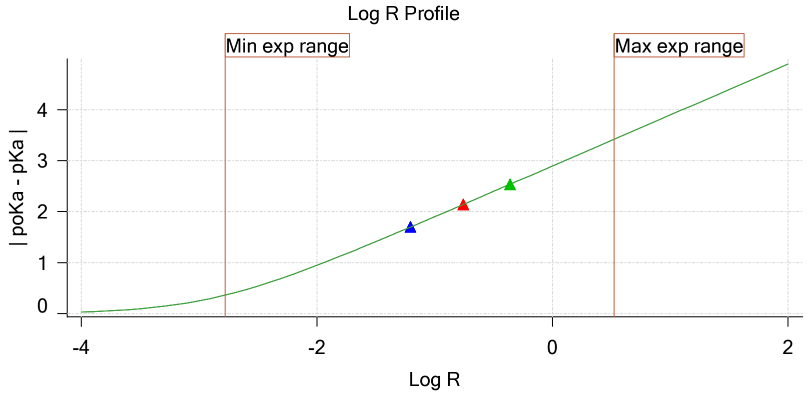


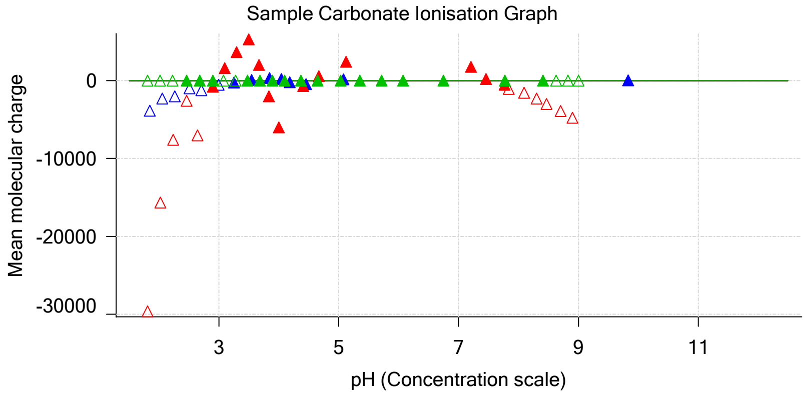

| | | | |
|---|---|---|---|
| Sample name: | **M13_octanol** | Experiment start time: | **3/16/2018 8:34:17 PM** |
| Assay name: | **pH-metric high logP** | Analyst: | **Pion** |
| Assay ID: | **18C-16016** | Instrument ID: | **T311053** |
| Filename: | **C:\Sirius_T3\Mehtap\20180316_exp32_logP_T3-1\18C-16016_M13_octanol_pH-metric high logP.t3r** | | |

#### Graphs (continued)

Distribution of species for sample Carbonate

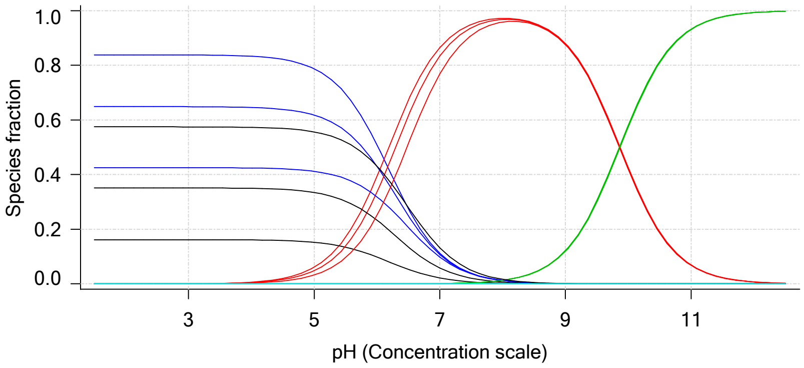

| | | | |
|---|---|---|---|
| Sample name: | **M13_octanol** | Experiment start time: | **3/16/2018 8:34:17 PM** |
| Assay name: | **pH-metric high logP** | Analyst: | **Pion** |
| Assay ID: | **18C-16016** | Instrument ID: | **T311053** |
| Filename: | **C:\Sirius_T3\Mehtap\20180316_exp32_logP_T3-1\18C-16016_M13_octanol_pH-metric high logP.t3r** | | |

### pH-metric high logP Titration 1 of 3 18C-16016 Points 2 to 15

#### Overall results

| | |
|---|---|
| RMSD | 0.073 |
| Average ionic strength | 0.158 M |
| Average temperature | 24.8°C |
| Partition ratio | 0.0624 : 1 |
| Analyte concentration range | 2710.7 μM to 2790.7 μM |
| Total points considered | 8 of 14 |

#### Warnings and errors

| | |
|---|---|
| Errors | None |
| Warnings | One or more logP values out of range<br>Excessive acidity error present |

#### Four-Plus parameters

| | | | |
|---|---|---|---|
| Alpha | 0.167 | 3/16/2018 8:34:16 PM | C:\Sirius_T3\HCl18C16.t3r |
| S | 0.9932 | 3/16/2018 8:34:16 PM | C:\Sirius_T3\HCl18C16.t3r |
| jH | 0.7 | 3/16/2018 8:34:16 PM | C:\Sirius_T3\HCl18C16.t3r |
| jOH | -0.9 | 3/16/2018 8:34:16 PM | C:\Sirius_T3\HCl18C16.t3r |

#### Titrants

| | | | |
|---|---|---|---|
| 0.50 M HCl | 0.990198 | 3/16/2018 8:34:16 PM | C:\Sirius_T3\HCl18C16.t3r |
| 0.50 M KOH | 1.009620 | 3/16/2018 8:34:17 PM | C:\Sirius_T3\KOH18C16.t3r |

#### Sample

| | |
|---|---|
| M13_octanol concentration factor | 0.878 |
| Base pKa 1 | 5.77 |
| logP (XH +) | -4.56 |
| logP (neutral X) | 2.90 |

#### Sample graphs

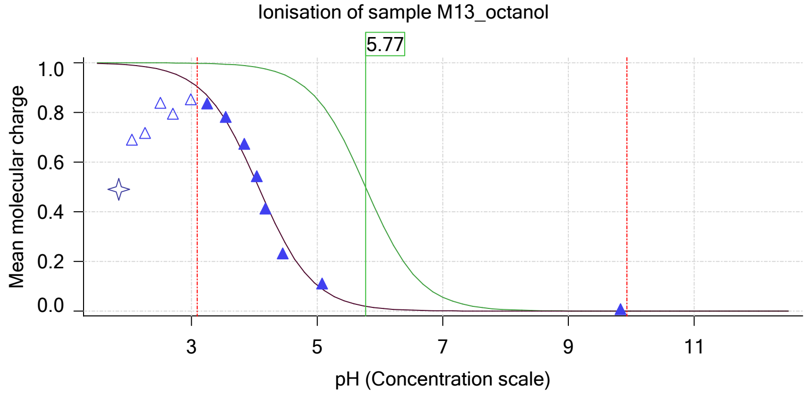


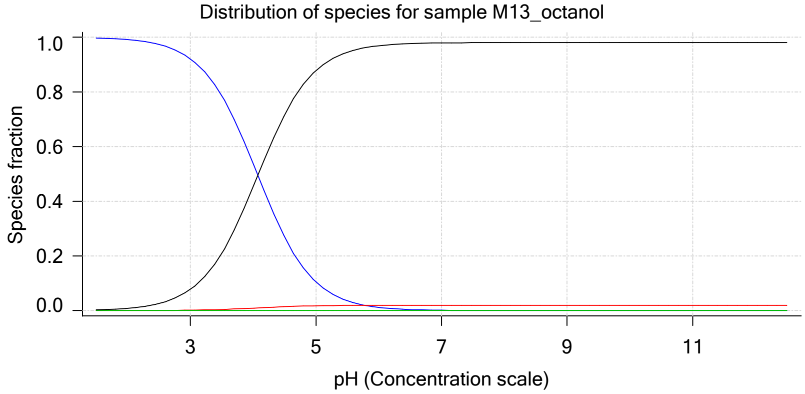

| | | | |
|---|---|---|---|
| Sample name: | **M13_octanol** | Experiment start time: | **3/16/2018 8:34:17 PM** |
| Assay name: | **pH-metric high logP** | Analyst: | **Pion** |
| Assay ID: | **18C-16016** | Instrument ID: | **T311053** |
| Filename: | **C:\Sirius_T3\Mehtap\20180316_exp32_logP_T3-1\18C-16016_M13_octanol_pH-metric high logP.t3r** | | |

#### Sample graphs (continued)

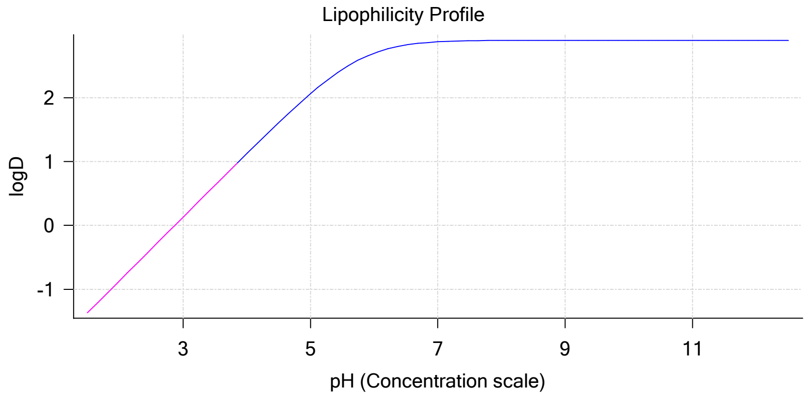


#### Sample logD and percent species

| pH | M13_octanol logD | M13_octanol M13_octanolH | M13_octanol M13_octanol | M13_octanol M13_octanolH* | M13_octanol M13_octanol* | Comment |
|---|---|---|---|---|---|---|
| 1.000 | -1.87 | 99.91 % | 0.00 % | 0.00 % | 0.08 % | |
| 1.200 | -1.67 | 99.86 % | 0.00 % | 0.00 % | 0.13 % | Stomach pH |
| 2.000 | -0.87 | 99.15 % | 0.02 % | 0.00 % | 0.83 % | |
| 3.000 | 0.13 | 92.10 % | 0.16 % | 0.00 % | 7.75 % | |
| 4.000 | 1.12 | 53.81 % | 0.91 % | 0.00 % | 45.27 % | |
| 5.000 | 2.06 | 10.43 % | 1.77 % | 0.00 % | 87.79 % | |
| 6.000 | 2.70 | 1.15 % | 1.96 % | 0.00 % | 96.89 % | |
| 6.500 | 2.83 | 0.37 % | 1.97 % | 0.00 % | 97.66 % | |
| 7.000 | 2.87 | 0.12 % | 1.98 % | 0.00 % | 97.91 % | |
| 7.400 | 2.89 | 0.05 % | 1.98 % | 0.00 % | 97.98 % | Blood pH |
| 8.000 | 2.90 | 0.01 % | 1.98 % | 0.00 % | 98.01 % | |
| 9.000 | 2.90 | 0.00 % | 1.98 % | 0.00 % | 98.02 % | |
| 10.000 | 2.90 | 0.00 % | 1.98 % | 0.00 % | 98.02 % | |
| 11.000 | 2.90 | 0.00 % | 1.98 % | 0.00 % | 98.02 % | |
| 12.000 | 2.90 | 0.00 % | 1.98 % | 0.00 % | 98.02 % | |

#### Carbonate and acidity

| | |
|---|---|
| Carbonate | 0.000 mM |
| Acidity error | 2.299 mM |

#### Other graphs

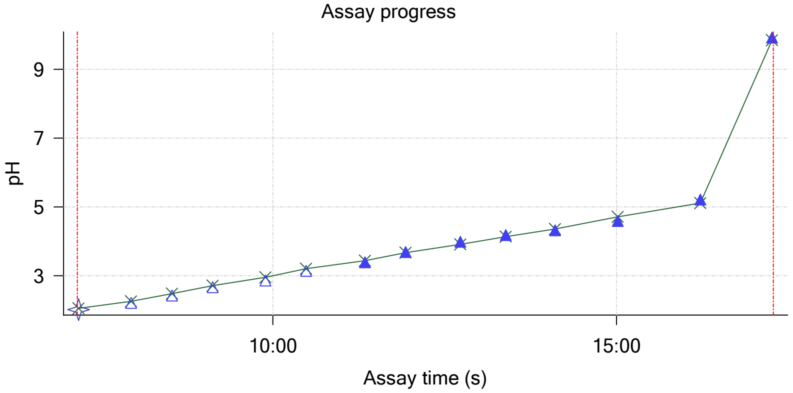


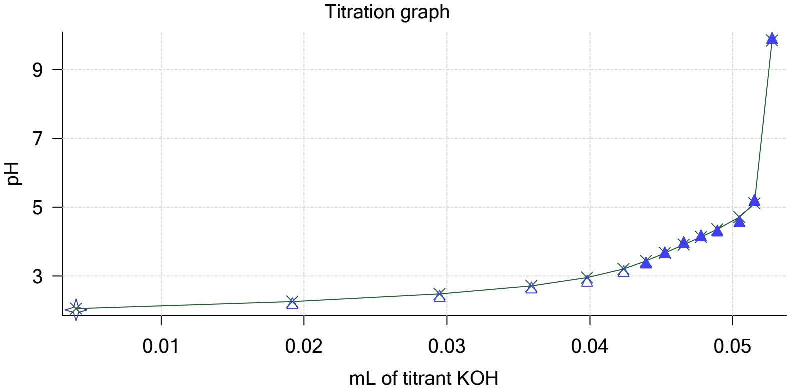

Sample name: **M13_octanol**
Assay name: **pH-metric high logP**
Assay ID: **18C-16016**
Filename: **C:\Sirius_T3\Mehtap\20180316_exp32_logP_T3-1\18C-16016_M13_octanol_pH-metric high logP.t3r**

Experiment start time: **3/16/2018 8:34:17 PM**
Analyst: **Pion**
Instrument ID: **T311053**

#### Other graphs (continued)

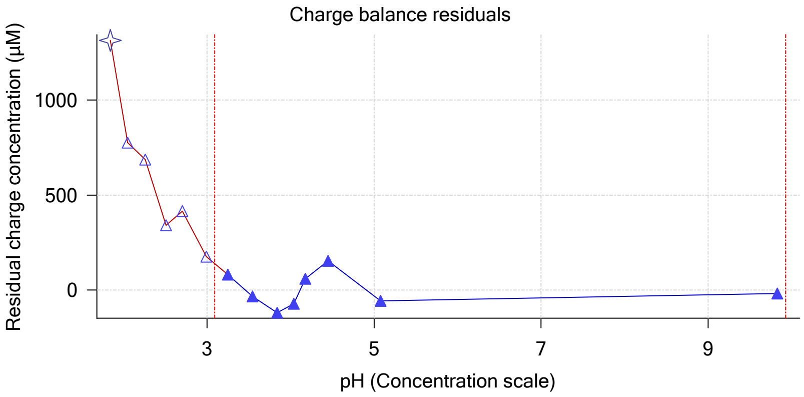


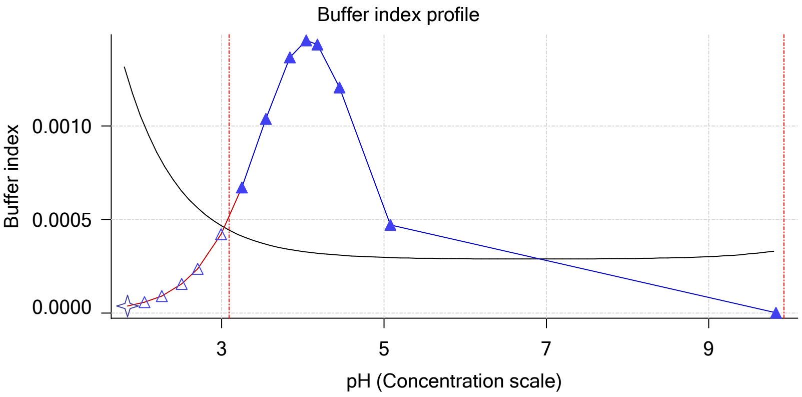


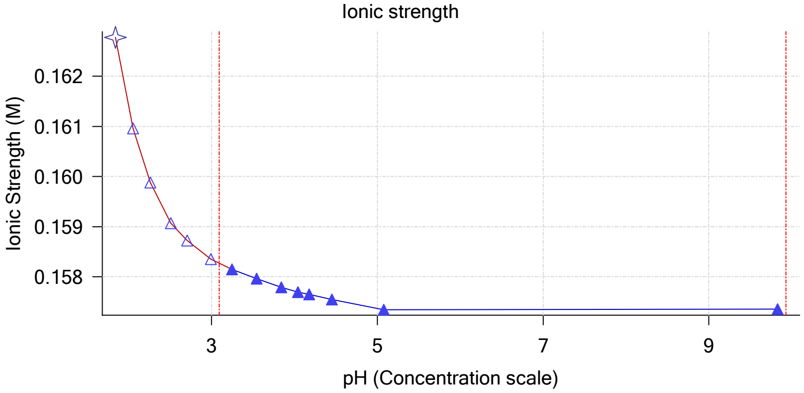


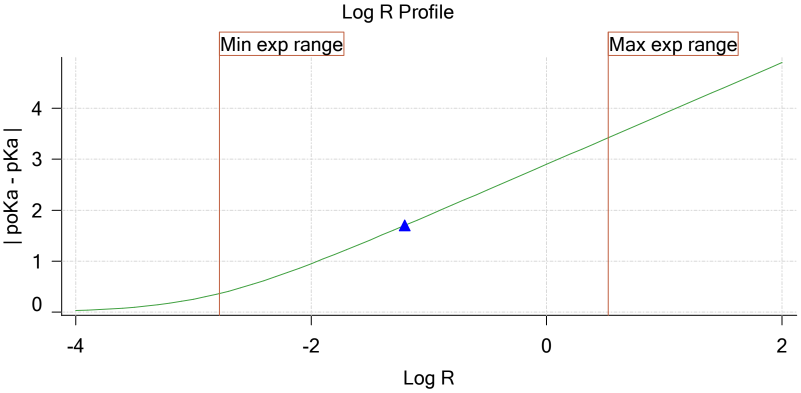


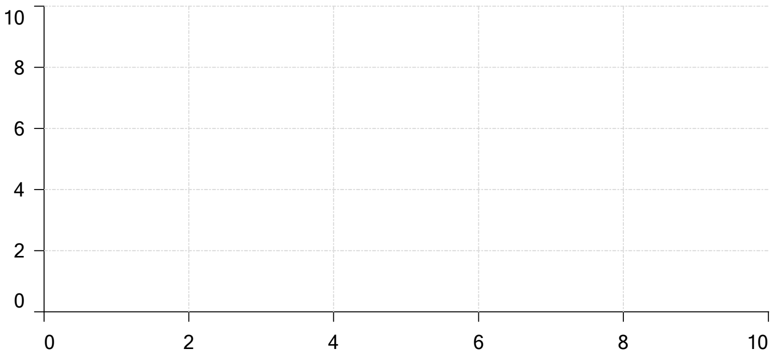


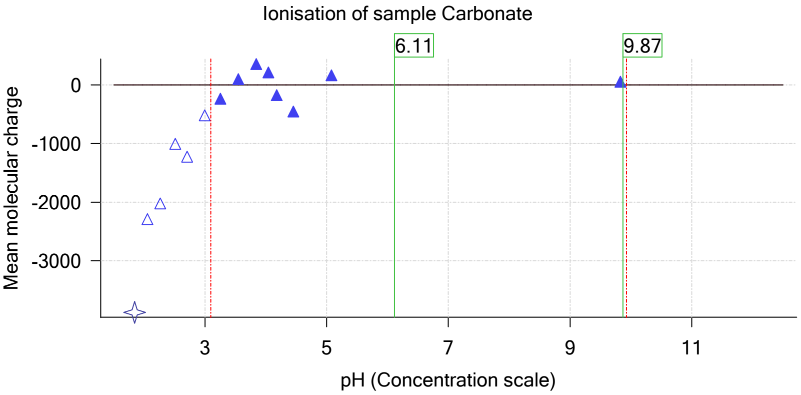

| | | | |
|---|---|---|---|
| Sample name: | **M13_octanol** | Experiment start time: | **3/16/2018 8:34:17 PM** |
| Assay name: | **pH-metric high logP** | Analyst: | **Pion** |
| Assay ID: | **18C-16016** | Instrument ID: | **T311053** |
| Filename: | **C:\Sirius_T3\Mehtap\20180316_exp32_logP_T3-1\18C-16016_M13_octanol_pH-metric high logP.t3r** | | |

### pH-metric high logP Titration 2 of 3 18C-16016 Points 16 to 39

#### Overall results

| | |
|---|---|
| RMSD | 6.380 |
| Average ionic strength | 0.164 M |
| Average temperature | 24.9°C |
| Partition ratio | 0.1752 : 1 |
| Analyte concentration range | 2287.9 µM to 2354.2 µM |
| Total points considered | 13 of 24 |

#### Warnings and errors

| | |
|---|---|
| Errors | None |
| Warnings | One or more logP values out of range |
| | Excessive acidity error present |

#### Four-Plus parameters

| | | | |
|---|---|---|---|
| Alpha | 0.167 | 3/16/2018 8:34:16 PM | C:\Sirius_T3\HCl18C16.t3r |
| S | 0.9932 | 3/16/2018 8:34:16 PM | C:\Sirius_T3\HCl18C16.t3r |
| jH | 0.7 | 3/16/2018 8:34:16 PM | C:\Sirius_T3\HCl18C16.t3r |
| jOH | -0.9 | 3/16/2018 8:34:16 PM | C:\Sirius_T3\HCl18C16.t3r |

#### Titrants

| | | | |
|---|---|---|---|
| 0.50 M HCl | 0.990198 | 3/16/2018 8:34:16 PM | C:\Sirius_T3\HCl18C16.t3r |
| 0.50 M KOH | 1.009620 | 3/16/2018 8:34:17 PM | C:\Sirius_T3\KOH18C16.t3r |

#### Sample

| | |
|---|---|
| M13_octanol concentration factor | 0.780 |
| Base pKa 1 | 5.77 |
| logP (XH +) | -4.56 |
| logP (neutral X) | 2.88 |

#### Sample graphs

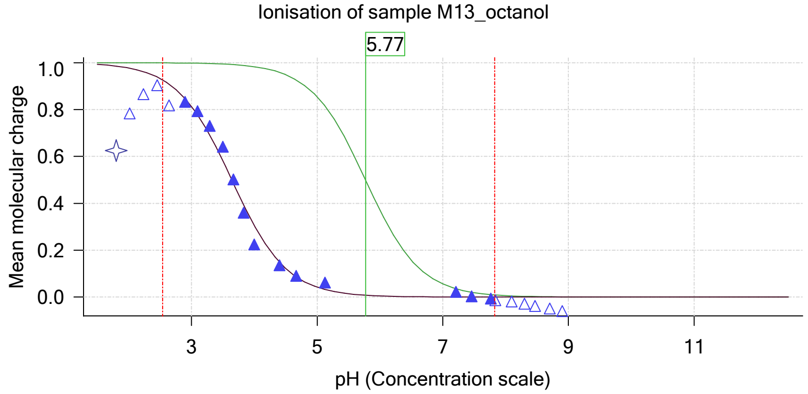


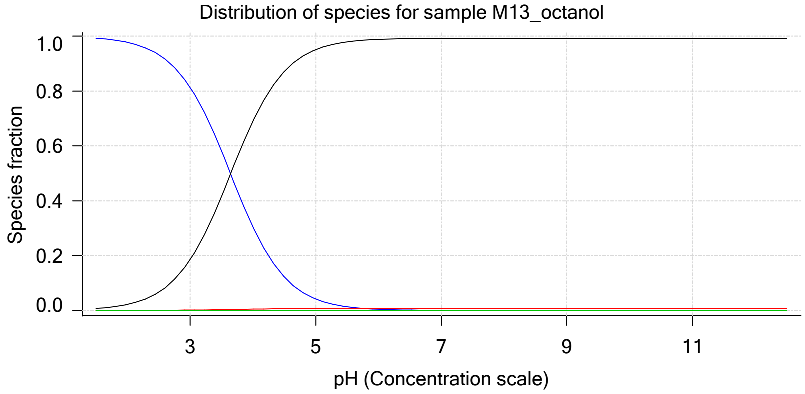

| | | | |
|---|---|---|---|
| Sample name: | **M13_octanol** | Experiment start time: | **3/16/2018 8:34:17 PM** |
| Assay name: | **pH-metric high logP** | Analyst: | **Pion** |
| Assay ID: | **18C-16016** | Instrument ID: | **T311053** |
| Filename: | **C:\Sirius_T3\Mehtap\20180316_exp32_logP_T3-1\18C-16016_M13_octanol_pH-metric high logP.t3r** | | |

#### Sample graphs (continued)

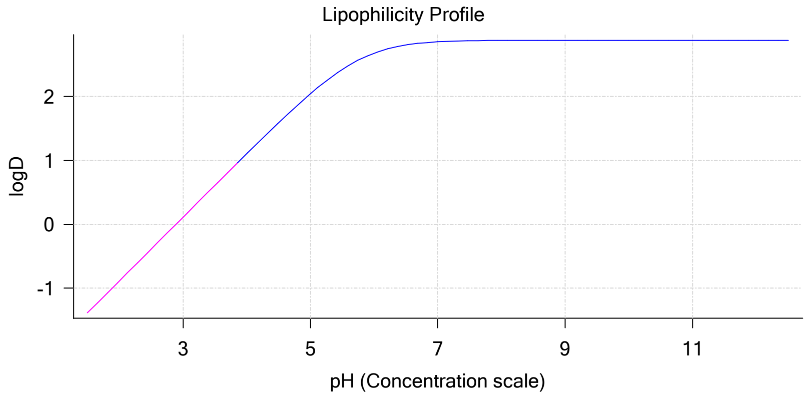


#### Sample logD and percent species

| pH | M13_octanol logD | M13_octanol M13_octanolH | M13_octanol M13_octanol | M13_octanol M13_octanolH* | M13_octanol M13_octanol* | Comment |
|---|---|---|---|---|---|---|
| 1.000 | -1.89 | 99.77 % | 0.00 % | 0.00 % | 0.23 % | |
| 1.200 | -1.69 | 99.64 % | 0.00 % | 0.00 % | 0.36 % | Stomach pH |
| 2.000 | -0.89 | 97.77 % | 0.02 % | 0.00 % | 2.22 % | |
| 3.000 | 0.11 | 81.41 % | 0.14 % | 0.00 % | 18.45 % | |
| 4.000 | 1.10 | 30.46 % | 0.52 % | 0.00 % | 69.02 % | |
| 5.000 | 2.04 | 4.20 % | 0.71 % | 0.00 % | 95.09 % | |
| 6.000 | 2.68 | 0.44 % | 0.74 % | 0.00 % | 98.82 % | |
| 6.500 | 2.81 | 0.14 % | 0.74 % | 0.00 % | 99.12 % | |
| 7.000 | 2.86 | 0.04 % | 0.74 % | 0.00 % | 99.21 % | |
| 7.400 | 2.87 | 0.02 % | 0.74 % | 0.00 % | 99.24 % | Blood pH |
| 8.000 | 2.88 | 0.00 % | 0.74 % | 0.00 % | 99.25 % | |
| 9.000 | 2.88 | 0.00 % | 0.74 % | 0.00 % | 99.26 % | |
| 10.000 | 2.88 | 0.00 % | 0.74 % | 0.00 % | 99.26 % | |
| 11.000 | 2.88 | 0.00 % | 0.74 % | 0.00 % | 99.26 % | |
| 12.000 | 2.88 | 0.00 % | 0.74 % | 0.00 % | 99.26 % | |

#### Carbonate and acidity

| | |
|---|---|
| Carbonate | 0.000 mM |
| Acidity error | 2.261 mM |

#### Other graphs

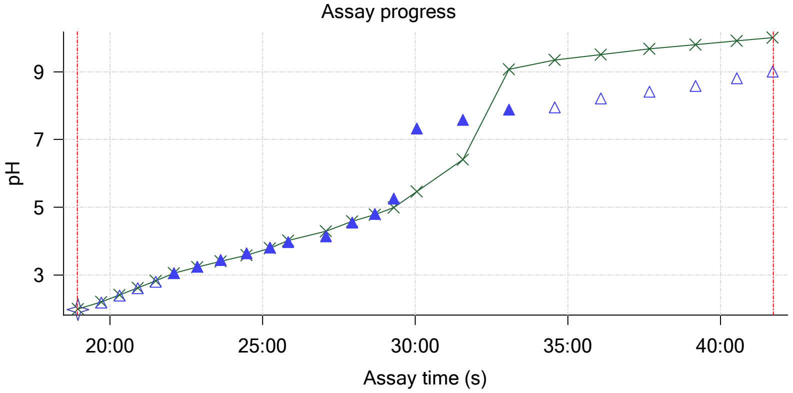


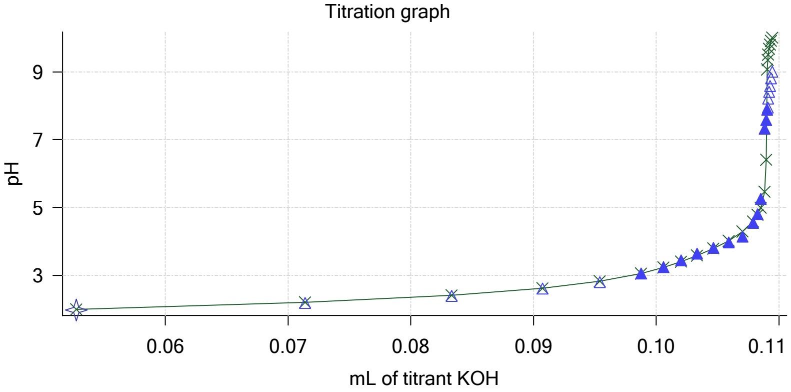

| | | | |
|---|---|---|---|
| Sample name: | **M13_octanol** | Experiment start time: | **3/16/2018 8:34:17 PM** |
| Assay name: | **pH-metric high logP** | Analyst: | **Pion** |
| Assay ID: | **18C-16016** | Instrument ID: | **T311053** |
| Filename: | **C:\Sirius_T3\Mehtap\20180316_exp32_logP_T3-1\18C-16016_M13_octanol_pH-metric high logP.t3r** | | |

#### Other graphs (continued)

Charge balance residuals

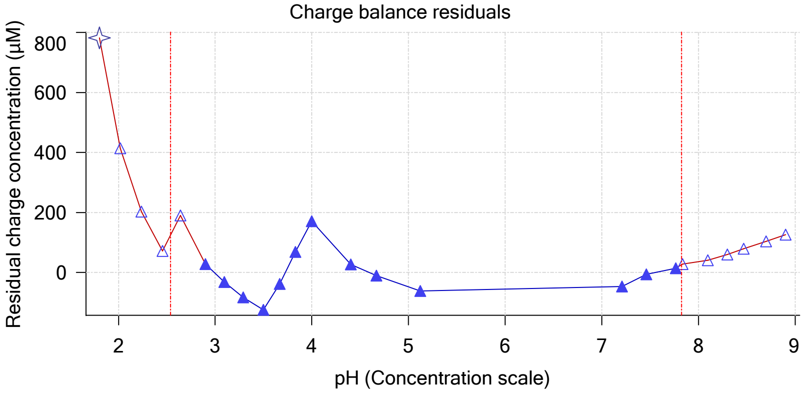


Buffer index profile

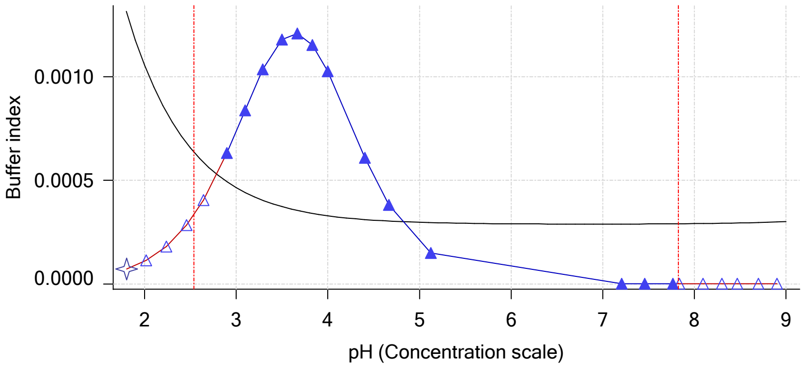


Ionic strength

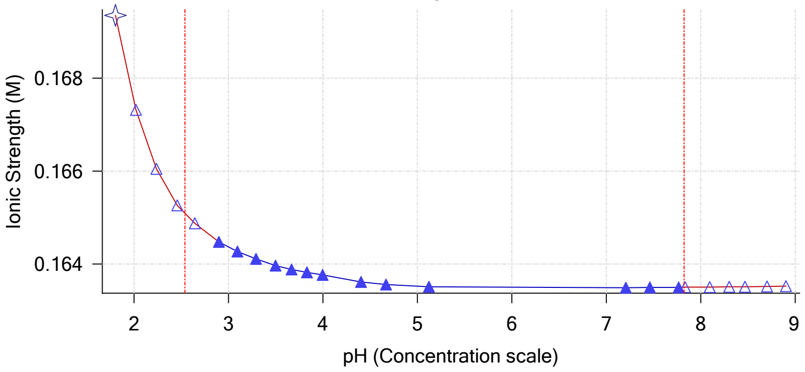


Log R Profile

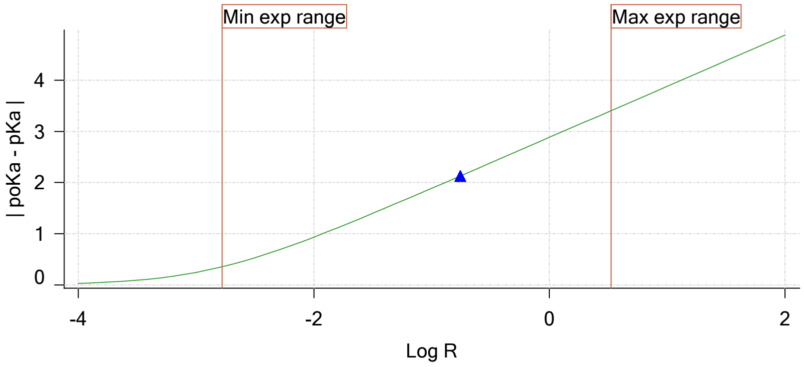


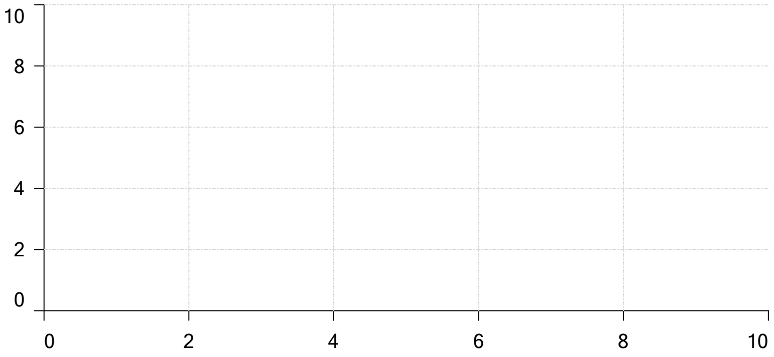


Ionisation of sample Carbonate

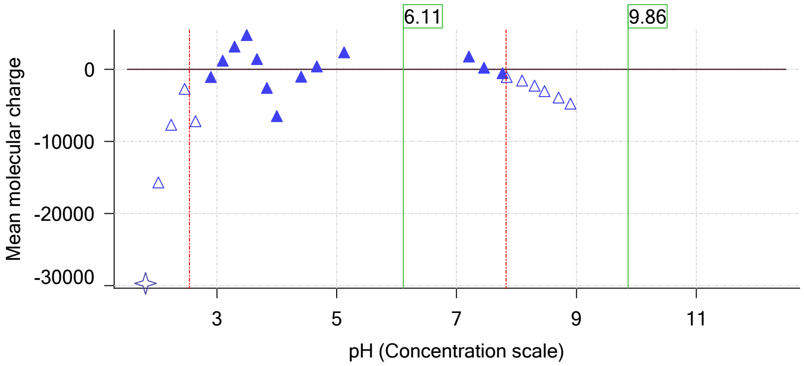

| | | | |
|---|---|---|---|
| Sample name: | **M13_octanol** | Experiment start time: | **3/16/2018 8:34:17 PM** |
| Assay name: | **pH-metric high logP** | Analyst: | **Pion** |
| Assay ID: | **18C-16016** | Instrument ID: | **T311053** |
| Filename: | **C:\Sirius_T3\Mehtap\20180316_exp32_logP_T3-1\18C-16016_M13_octanol_pH-metric high logP.t3r** | | |

### pH-metric high logP Titration 3 of 3 18C-16016 Points 40 to 63

#### Overall results

| | |
|---|---|
| RMSD | 0.265 |
| Average ionic strength | 0.169 M |
| Average temperature | 25.0°C |
| Partition ratio | 0.4371 : 1 |
| Analyte concentration range | 1749.3 µM to 1790.0 µM |
| Total points considered | 16 of 24 |

#### Warnings and errors

| | |
|---|---|
| Errors | None |
| Warnings | One or more logP values out of range<br>Excessive acidity error present |

#### Four-Plus parameters

| | | | |
|---|---|---|---|
| Alpha | 0.167 | 3/16/2018 8:34:16 PM | C:\Sirius_T3\HCl18C16.t3r |
| S | 0.9932 | 3/16/2018 8:34:16 PM | C:\Sirius_T3\HCl18C16.t3r |
| jH | 0.7 | 3/16/2018 8:34:16 PM | C:\Sirius_T3\HCl18C16.t3r |
| jOH | -0.9 | 3/16/2018 8:34:16 PM | C:\Sirius_T3\HCl18C16.t3r |

#### Titrants

| | | | |
|---|---|---|---|
| 0.50 M HCl | 0.990198 | 3/16/2018 8:34:16 PM | C:\Sirius_T3\HCl18C16.t3r |
| 0.50 M KOH | 1.009620 | 3/16/2018 8:34:17 PM | C:\Sirius_T3\KOH18C16.t3r |

#### Sample

| | |
|---|---|
| M13_octanol concentration factor | 0.706 |
| Base pKa 1 | 5.77 |
| logP (XH +) | -4.56 |
| logP (neutral X) | 2.92 |

#### Sample graphs

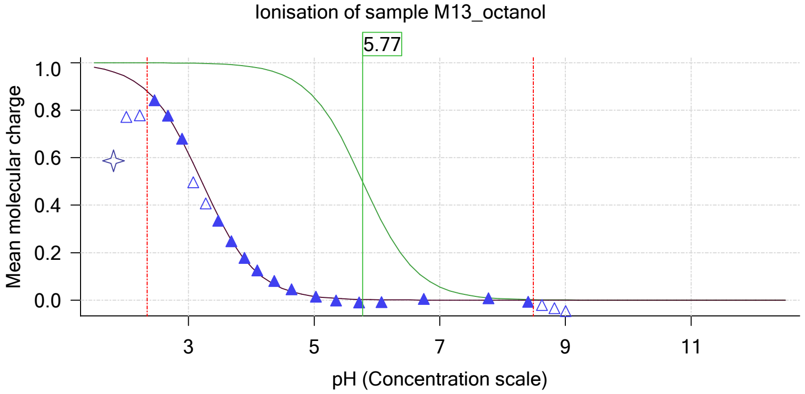


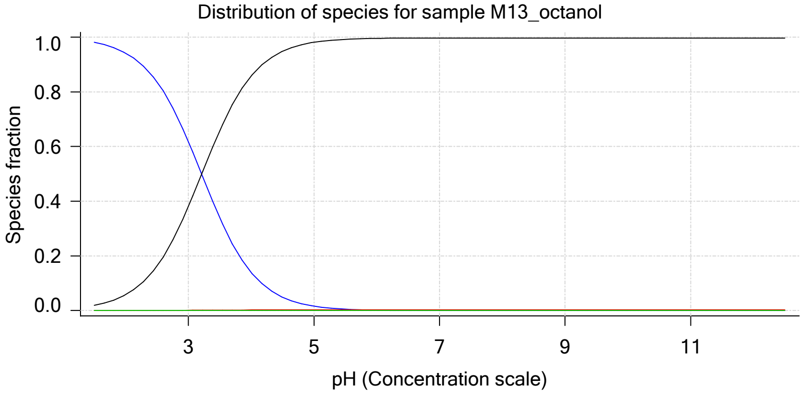

| | | | |
|---|---|---|---|
| Sample name: | **M13_octanol** | Experiment start time: | **3/16/2018 8:34:17 PM** |
| Assay name: | **pH-metric high logP** | Analyst: | **Pion** |
| Assay ID: | **18C-16016** | Instrument ID: | **T311053** |
| Filename: | **C:\Sirius_T3\Mehtap\20180316_exp32_logP_T3-1\18C-16016_M13_octanol_pH-metric high logP.t3r** | | |

#### Sample graphs (continued)

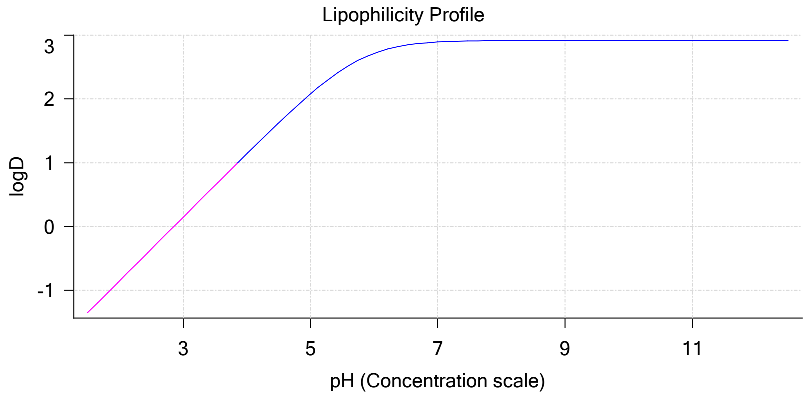


#### Sample logD and percent species

| pH | M13_octanol logD | M13_octanol M13_octanolH | M13_octanol M13_octanol | M13_octanol M13_octanolH* | M13_octanol M13_octanol* | Comment |
|---|---|---|---|---|---|---|
| 1.000 | -1.85 | 99.39 % | 0.00 % | 0.00 % | 0.61 % | |
| 1.200 | -1.65 | 99.03 % | 0.00 % | 0.00 % | 0.96 % | Stomach pH |
| 2.000 | -0.85 | 94.19 % | 0.02 % | 0.00 % | 5.79 % | |
| 3.000 | 0.15 | 61.87 % | 0.11 % | 0.00 % | 38.02 % | |
| 4.000 | 1.14 | 13.96 % | 0.24 % | 0.00 % | 85.80 % | |
| 5.000 | 2.08 | 1.60 % | 0.27 % | 0.00 % | 98.13 % | |
| 6.000 | 2.72 | 0.16 % | 0.28 % | 0.00 % | 99.56 % | |
| 6.500 | 2.84 | 0.05 % | 0.28 % | 0.00 % | 99.67 % | |
| 7.000 | 2.89 | 0.02 % | 0.28 % | 0.00 % | 99.71 % | |
| 7.400 | 2.91 | 0.01 % | 0.28 % | 0.00 % | 99.72 % | Blood pH |
| 8.000 | 2.92 | 0.00 % | 0.28 % | 0.00 % | 99.72 % | |
| 9.000 | 2.92 | 0.00 % | 0.28 % | 0.00 % | 99.72 % | |
| 10.000 | 2.92 | 0.00 % | 0.28 % | 0.00 % | 99.72 % | |
| 11.000 | 2.92 | 0.00 % | 0.28 % | 0.00 % | 99.72 % | |
| 12.000 | 2.92 | 0.00 % | 0.28 % | 0.00 % | 99.72 % | |

#### Carbonate and acidity

| | |
|---|---|
| Carbonate | 0.138 mM |
| Acidity error | 2.269 mM |

#### Other graphs

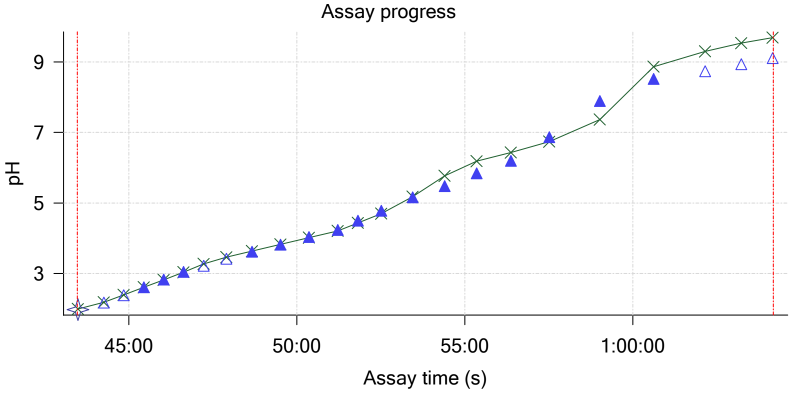


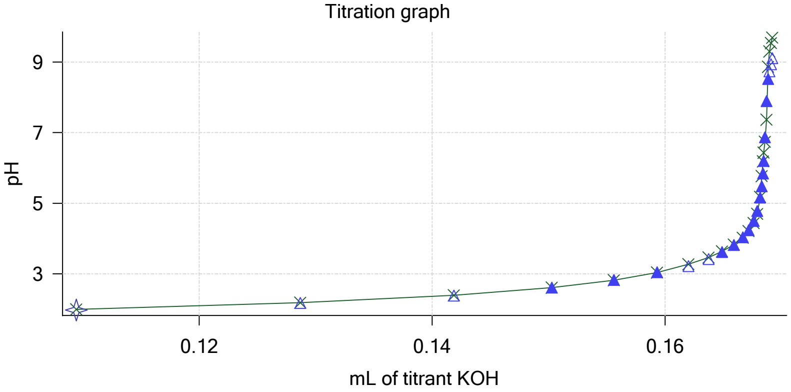

| | | | |
|---|---|---|---|
| Sample name: | **M13_octanol** | Experiment start time: | **3/16/2018 8:34:17 PM** |
| Assay name: | **pH-metric high logP** | Analyst: | **Pion** |
| Assay ID: | **18C-16016** | Instrument ID: | **T311053** |
| Filename: | **C:\Sirius_T3\Mehtap\20180316_exp32_logP_T3-1\18C-16016_M13_octanol_pH-metric high logP.t3r** | | |

#### Other graphs (continued)

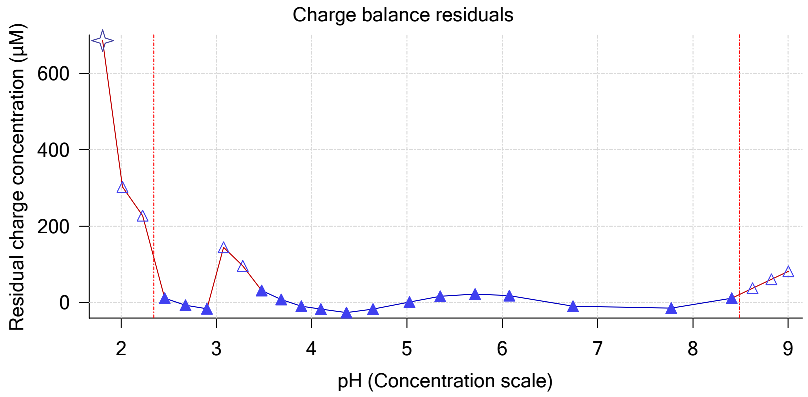


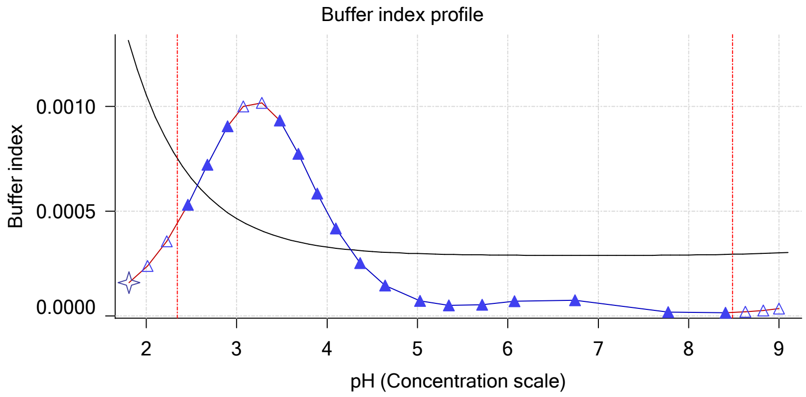


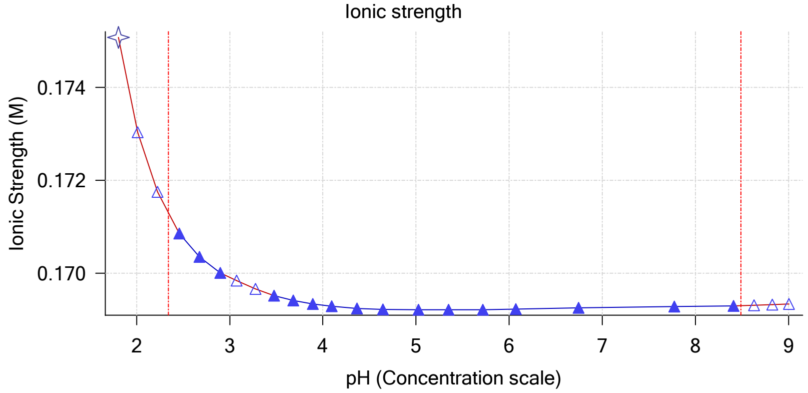


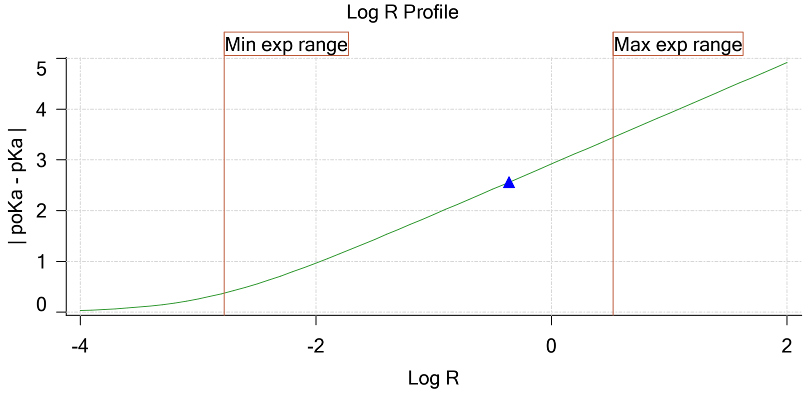


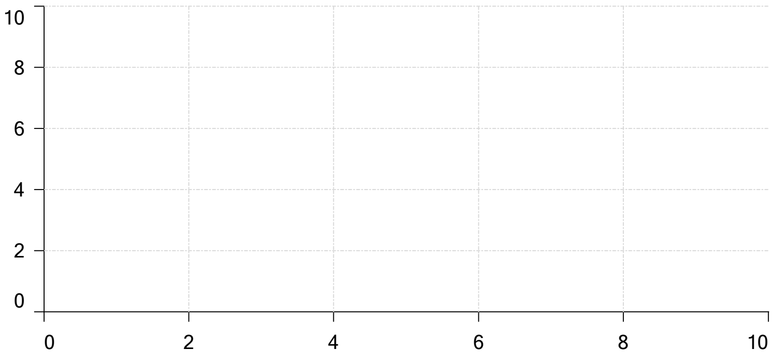


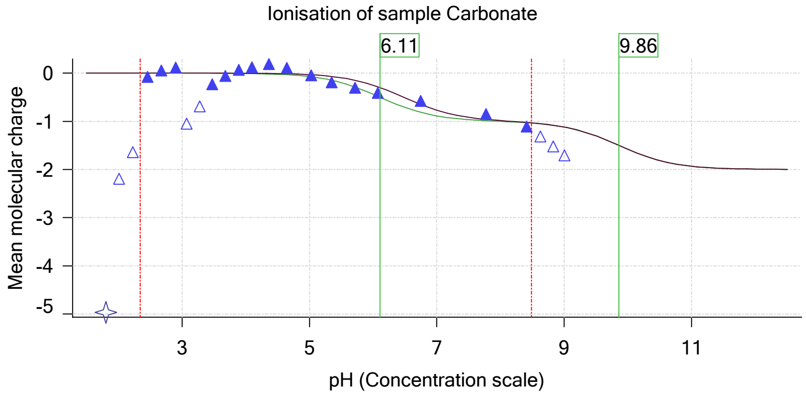

| | | | |
|---|---|---|---|
| Sample name: | **M13_octanol** | Experiment start time: | **3/16/2018 8:34:17 PM** |
| Assay name: | **pH-metric high logP** | Analyst: | **Pion** |
| Assay ID: | **18C-16016** | Instrument ID: | **T311053** |
| Filename: | **C:\Sirius_T3\Mehtap\20180316_exp32_logP_T3-1\18C-16016_M13_octanol_pH-metric high logP.t3r** | | |

#### Assay Model

| Settings | Value | Date/Time changed | Imported from |
|---|---|---|---|
| Sample name | M13_octanol | 2/27/2018 5:57:49 PM | User entered value |
| Sample by | Weight | | Default value |
| Sample weight | 0.001360 g | 3/16/2018 5:08:03 PM | User entered value |
| Formula weight | 295.34 g/mol | 2/27/2018 5:57:49 PM | User entered value |
| Solubility | Unknown | | Default value |
| Molecular weight | 295.34 | 2/27/2018 5:57:49 PM | User entered value |
| Individual pKa ionic environments | No | | Default value |
| Number of pKas | 1 | 2/27/2018 5:57:49 PM | User entered value |
| Sample is a | Base | 2/27/2018 5:57:49 PM | User entered value |
| pKa 1 | 5.77 | 2/27/2018 5:57:49 PM | User entered value |
| logp (XH +) | -4.56 | 3/2/2018 4:30:48 PM | User entered value |
| logP (neutral X) | 2.99 | 3/2/2018 4:30:43 PM | User entered value |

#### Events

| Time | Event | Water | Acid | Base | Octanol | pH | dpH/dt | pH R-squared | pH SD | dpH/[illegible] time |
|---|---|---|---|---|---|---|---|---|---|---|
| 4:09.8 | Manual volume addition | | | | 0.10000 mL | | | | | |
| 4:10.9 | Initial pH = 3.69 | | | | | | | | | |
| 7:10.4 | Data point 2 | 1.50000 mL | 0.04605 mL | 0.00405 mL | 0.10000 mL | 2.007 | 0.00659 | 0.78015 | 0.00037 | 10.0 s |
| 7:56.4 | Data point 3 | 1.50000 mL | 0.04605 mL | 0.01919 mL | 0.10000 mL | 2.211 | 0.00314 | 0.24211 | 0.00032 | 10.0 s |
| 8:32.1 | Data point 4 | 1.50000 mL | 0.04605 mL | 0.02949 mL | 0.10000 mL | 2.417 | -0.01053 | 0.38713 | 0.00084 | 10.0 s |
| 9:07.7 | Data point 5 | 1.50000 mL | 0.04605 mL | 0.03591 mL | 0.10000 mL | 2.660 | -0.00938 | 0.73483 | 0.00054 | 10.0 s |
| 9:53.5 | Data point 6 | 1.50000 mL | 0.04605 mL | 0.03982 mL | 0.10000 mL | 2.855 | -0.00211 | 0.30085 | 0.00019 | 10.0 s |
| 10:29.0 | Data point 7 | 1.50000 mL | 0.04605 mL | 0.04236 mL | 0.10000 mL | 3.139 | -0.00782 | 0.88102 | 0.00041 | 10.5 s |
| 11:20.4 | Data point 8 | 1.50000 mL | 0.04605 mL | 0.04393 mL | 0.10000 mL | 3.394 | -0.00736 | 0.41781 | 0.00056 | 10.0 s |
| 11:55.9 | Data point 9 | 1.50000 mL | 0.04605 mL | 0.04525 mL | 0.10000 mL | 3.686 | -0.01800 | 0.93937 | 0.00092 | 12.0 s |
| 12:43.6 | Data point 10 | 1.50000 mL | 0.04605 mL | 0.04657 mL | 0.10000 mL | 3.982 | -0.01971 | 0.97981 | 0.00098 | 14.5 s |
| 13:23.6 | Data point 11 | 1.50000 mL | 0.04605 mL | 0.04779 mL | 0.10000 mL | 4.181 | -0.01960 | 0.95605 | 0.00099 | 17.5 s |
| 14:06.5 | Data point 12 | 1.50000 mL | 0.04605 mL | 0.04892 mL | 0.10000 mL | 4.317 | -0.01919 | 0.90540 | 0.00100 | 19.0 s |
| 15:01.2 | Data point 13 | 1.50000 mL | 0.04605 mL | 0.05047 mL | 0.10000 mL | 4.588 | -0.01947 | 0.95881 | 0.00098 | 36.5 s |
| 16:13.4 | Data point 14 | 1.50000 mL | 0.04605 mL | 0.05153 mL | 0.10000 mL | 5.212 | 0.01166 | 0.44692 | 0.00086 | 32.0 s |
| 17:16.0 | Data point 15 | 1.50000 mL | 0.04605 mL | 0.05275 mL | 0.10000 mL | 9.929 | -0.01846 | 0.92278 | 0.00095 | 43.5 s |
| 18:57.2 | Data point 16 | 1.50000 mL | 0.10329 mL | 0.05275 mL | 0.30000 mL | 1.971 | -0.00110 | 0.01132 | 0.00051 | 10.0 s |
| 19:43.4 | Data point 17 | 1.50000 mL | 0.10329 mL | 0.07140 mL | 0.30000 mL | 2.179 | 0.00116 | 0.00555 | 0.00077 | 10.0 s |
| 20:19.1 | Data point 18 | 1.50000 mL | 0.10329 mL | 0.08333 mL | 0.30000 mL | 2.393 | -0.00552 | 0.08741 | 0.00092 | 10.0 s |
| 20:54.7 | Data point 19 | 1.50000 mL | 0.10329 mL | 0.09073 mL | 0.30000 mL | 2.610 | -0.00664 | 0.54896 | 0.00044 | 10.0 s |
| 21:30.3 | Data point 20 | 1.50000 mL | 0.10329 mL | 0.09541 mL | 0.30000 mL | 2.793 | -0.01670 | 0.91771 | 0.00086 | 10.5 s |

| | | | |
|---|---|---|---|
| Sample name: | **M13_octanol** | Experiment start time: | **3/16/2018 8:34:17 PM** |
| Assay name: | **pH-metric high logP** | Analyst: | **Pion** |
| Assay ID: | **18C-16016** | Instrument ID: | **T311053** |
| Filename: | **C:\Sirius_T3\Mehtap\20180316_exp32_logP_T3-1\18C-16016_M13_octanol_pH-metric high logP.t3r** | | |

#### Events (continued)

| Time | Event | Water | Acid | Base | Octanol | pH | dpH/dt | pH R-squared | pH SD | dpH/dt time |
|---|---|---|---|---|---|---|---|---|---|---|
| 22:06.3 | Data point 21 | 1.50000 mL | 0.10329 mL | 0.09875 mL | 0.30000 mL | 3.048 | -0.00688 | 0.24255 | 0.00069 | 10.0 s |
| 22:52.1 | Data point 22 | 1.50000 mL | 0.10329 mL | 0.10059 mL | 0.30000 mL | 3.242 | -0.00986 | 0.45925 | 0.00072 | 10.0 s |
| 23:37.8 | Data point 23 | 1.50000 mL | 0.10329 mL | 0.10202 mL | 0.30000 mL | 3.436 | -0.00502 | 0.22328 | 0.00052 | 10.0 s |
| 24:28.7 | Data point 24 | 1.50000 mL | 0.10329 mL | 0.10332 mL | 0.30000 mL | 3.641 | -0.01457 | 0.55071 | 0.00097 | 10.0 s |
| 25:14.5 | Data point 25 | 1.50000 mL | 0.10329 mL | 0.10466 mL | 0.30000 mL | 3.811 | -0.01728 | 0.77590 | 0.00097 | 10.5 s |
| 25:50.4 | Data point 26 | 1.50000 mL | 0.10329 mL | 0.10590 mL | 0.30000 mL | 3.972 | -0.02006 | 0.98341 | 0.00100 | 33.5 s |
| 27:04.9 | Data point 27 | 1.50000 mL | 0.10329 mL | 0.10703 mL | 0.30000 mL | 4.138 | -0.01923 | 0.92979 | 0.00098 | 26.0 s |
| 27:56.3 | Data point 28 | 1.50000 mL | 0.10329 mL | 0.10788 mL | 0.30000 mL | 4.540 | -0.01305 | 0.56589 | 0.00086 | 14.0 s |
| 28:40.9 | Data point 29 | 1.50000 mL | 0.10329 mL | 0.10826 mL | 0.30000 mL | 4.803 | -0.00985 | 0.36149 | 0.00081 | 12.0 s |
| 29:18.3 | Data point 30 | 1.50000 mL | 0.10329 mL | 0.10851 mL | 0.30000 mL | 5.255 | -0.01714 | 0.86889 | 0.00091 | 14.5 s |
| 30:03.2 | Data point 31 | 1.50000 mL | 0.10329 mL | 0.10882 mL | 0.30000 mL | 7.326 | -0.14623 | 0.99588 | 0.00723 | Timed out at 59.5 s |
| 31:33.7 | Data point 32 | 1.50000 mL | 0.10329 mL | 0.10896 mL | 0.30000 mL | 7.576 | -0.08507 | 0.99600 | 0.00421 | Timed out at 59.5 s |
| 33:04.1 | Data point 33 | 1.50000 mL | 0.10329 mL | 0.10903 mL | 0.30000 mL | 7.881 | -0.07193 | 0.99419 | 0.00357 | Timed out at 59.5 s |
| 34:34.5 | Data point 34 | 1.50000 mL | 0.10329 mL | 0.10908 mL | 0.30000 mL | 7.949 | -0.04523 | 0.99327 | 0.00224 | Timed out at 59.5 s |
| 36:05.0 | Data point 35 | 1.50000 mL | 0.10329 mL | 0.10913 mL | 0.30000 mL | 8.208 | -0.03919 | 0.98798 | 0.00195 | Timed out at 59.5 s |
| 37:40.6 | Data point 36 | 1.50000 mL | 0.10329 mL | 0.10920 mL | 0.30000 mL | 8.410 | -0.02748 | 0.97538 | 0.00137 | Timed out at 59.5 s |
| 39:11.0 | Data point 37 | 1.50000 mL | 0.10329 mL | 0.10927 mL | 0.30000 mL | 8.578 | -0.01916 | 0.95293 | 0.00097 | 45.5 s |
| 40:32.1 | Data point 38 | 1.50000 mL | 0.10329 mL | 0.10936 mL | 0.30000 mL | 8.808 | -0.01860 | 0.93287 | 0.00095 | 40.0 s |
| 41:42.6 | Data point 39 | 1.50000 mL | 0.10329 mL | 0.10945 mL | 0.30000 mL | 9.007 | -0.01934 | 0.95051 | 0.00098 | 26.0 s |
| 43:28.9 | Data point 40 | 1.50000 mL | 0.16319 mL | 0.10945 mL | 0.80000 mL | 1.973 | -0.00097 | 0.00987 | 0.00048 | 10.0 s |
| 44:15.2 | Data point 41 | 1.50000 mL | 0.16319 mL | 0.12869 mL | 0.80000 mL | 2.174 | -0.01641 | 0.68236 | 0.00098 | 10.0 s |
| 44:50.9 | Data point 42 | 1.50000 mL | 0.16319 mL | 0.14189 mL | 0.80000 mL | 2.382 | -0.00586 | 0.20619 | 0.00064 | 10.0 s |
| 45:26.6 | Data point 43 | 1.50000 mL | 0.16319 mL | 0.15028 mL | 0.80000 mL | 2.612 | -0.00646 | 0.51998 | 0.00044 | 10.0 s |
| 46:02.1 | Data point 44 | 1.50000 mL | 0.16319 mL | 0.15562 mL | 0.80000 mL | 2.826 | -0.01012 | 0.47373 | 0.00073 | 10.0 s |
| 46:37.6 | Data point 45 | 1.50000 mL | 0.16319 mL | 0.15931 mL | 0.80000 mL | 3.047 | -0.00998 | 0.83294 | 0.00054 | 10.5 s |
| 47:13.6 | Data point 46 | 1.50000 mL | 0.16319 mL | 0.16202 mL | 0.80000 mL | 3.221 | -0.00942 | 0.29407 | 0.00086 | 10.0 s |
| 47:54.2 | Data point 47 | 1.50000 mL | 0.16319 mL | 0.16373 mL | 0.80000 mL | 3.423 | -0.00386 | 0.10317 | 0.00059 | 10.0 s |
| 48:39.8 | Data point 48 | 1.50000 mL | 0.16319 mL | 0.16491 mL | 0.80000 mL | 3.619 | -0.00709 | 0.28131 | 0.00066 | 10.0 s |
| 49:30.7 | Data point 49 | 1.50000 mL | 0.16319 mL | 0.16592 mL | 0.80000 mL | 3.823 | -0.01281 | 0.70214 | 0.00075 | 10.0 s |
| 50:21.6 | Data point 50 | 1.50000 mL | 0.16319 mL | 0.16667 mL | 0.80000 mL | 4.033 | -0.00051 | 0.00324 | 0.00044 | 10.5 s |
| 51:12.9 | Data point 51 | 1.50000 mL | 0.16319 mL | 0.16719 mL | 0.80000 mL | 4.236 | -0.00921 | 0.79034 | 0.00051 | 10.5 s |
| 51:48.8 | Data point 52 | 1.50000 mL | 0.16319 mL | 0.16761 mL | 0.80000 mL | 4.504 | -0.01403 | 0.68052 | 0.00084 | 11.5 s |
| 52:30.8 | Data point 53 | 1.50000 mL | 0.16319 mL | 0.16792 mL | 0.80000 mL | 4.779 | -0.01789 | 0.81440 | 0.00098 | 25.5 s |
| 53:26.7 | Data point 54 | 1.50000 mL | 0.16319 mL | 0.16818 mL | 0.80000 mL | 5.161 | -0.01713 | 0.81703 | 0.00093 | 21.5 s |
| 54:23.9 | Data point 55 | 1.50000 mL | 0.16319 mL | 0.16832 mL | 0.80000 mL | 5.480 | -0.01521 | 0.57942 | 0.00099 | 21.5 s |
| 55:21.0 | Data point 56 | 1.50000 mL | 0.16319 mL | 0.16842 mL | 0.80000 mL | 5.844 | -0.01837 | 0.87683 | 0.00097 | 30.5 s |
| 56:22.0 | Data point 57 | 1.50000 mL | 0.16319 mL | 0.16849 mL | 0.80000 mL | 6.198 | -0.01775 | 0.86830 | 0.00094 | 38.0 s |
| 57:30.5 | Data point 58 | 1.50000 mL | 0.16319 mL | 0.16858 mL | 0.80000 mL | 6.865 | -0.07156 | 0.99469 | 0.00354 | Timed out at 59.5 s |
| 59:00.9 | Data point 59 | 1.50000 mL | 0.16319 mL | 0.16872 mL | 0.80000 mL | 7.887 | -0.09593 | 0.99713 | 0.00475 | Timed out at 59.5 s |
| 1:00:36.6 | Data point 60 | 1.50000 mL | 0.16319 mL | 0.16886 mL | 0.80000 mL | 8.519 | -0.01911 | 0.89364 | 0.00100 | 56.5 s |
| 1:02:08.7 | Data point 61 | 1.50000 mL | 0.16319 mL | 0.16898 mL | 0.80000 mL | 8.736 | -0.01754 | 0.87325 | 0.00093 | 29.0 s |
| 1:03:13.4 | Data point 62 | 1.50000 mL | 0.16319 mL | 0.16910 mL | 0.80000 mL | 8.932 | -0.01508 | 0.70146 | 0.00089 | 20.5 s |
| 1:04:09.5 | Data point 63 | 1.50000 mL | 0.16319 mL | 0.16921 mL | 0.80000 mL | 9.107 | -0.01562 | 0.77923 | 0.00087 | 14.5 s |
| 1:04:33.0 | Assay volumes | 1.50000 mL | 0.16319 mL | 0.16921 mL | 0.80000 mL | | | | | |

| | | | |
|---|---|---|---|
| Sample name: | **M13_octanol** | Experiment start time: | **3/16/2018 8:34:17 PM** |
| Assay name: | **pH-metric high logP** | Analyst: | **Pion** |
| Assay ID: | **18C-16016** | Instrument ID: | **T311053** |
| Filename: | **C:\Sirius_T3\Mehtap\20180316_exp32_logP_T3-1\18C-16016_M13_octanol_pH-metric high logP.t3r** | | |

| | | | |
|---|---|---|---|
| Sample name: | **M13_octanol** | Experiment start time: | **3/16/2018 8:34:17 PM** |
| Assay name: | **pH-metric high logP** | Analyst: | **Pion** |
| Assay ID: | **18C-16016** | Instrument ID: | **T311053** |
| Filename: | **C:\Sirius_T3\Mehtap\20180316_exp32_logP_T3-1\18C-16016_M13_octanol_pH-metric high logP.t3r** | | |

#### Assay Settings (continued)

| Setting | Value | Original Value | Date/Time changed | Imported from |
|---|---|---|---|---|
| ***Titration 3*** | | | | |
| Titrate from | Low to high pH | | | |
| Add additional water | 0.00 mL | | | |
| Additional partition solvent volume | 0.500 mL | | | |
| Additional partition solvent added | Automatic | | | |
| After pH adjust stir for | 30 seconds | | | |
| Stir to allow partitioning for | 15 seconds | | | |
| Stirrer speed for partitioning | 60% | | | |
| ***Data Point Stability*** | | | | |
| Stir during data point collection | No | | | |
| Delay before data point collection | 0 seconds | | | |
| Number of points to average | 20 points | | | |
| Time interval between points | 0.50 seconds | | | |
| Required maximum standard deviation | 0.00100 dpH/dt | | | |
| Stability timeout after | 60 seconds | | | |

#### Calibration Settings

| Setting | Value | Date/Time changed | Imported from |
|---|---|---|---|
| Four-Plus alpha | 0.167 | 3/16/2018 8:34:16 PM | C:\Sirius_T3\HCl18C16.t3r |
| Four-Plus S | 0.9932 | 3/16/2018 8:34:16 PM | C:\Sirius_T3\HCl18C16.t3r |
| Four-Plus jH | 0.7 | 3/16/2018 8:34:16 PM | C:\Sirius_T3\HCl18C16.t3r |
| Four-Plus jOH | -0.9 | 3/16/2018 8:34:16 PM | C:\Sirius_T3\HCl18C16.t3r |
| Base concentration factor | 1.010 | 3/16/2018 8:34:17 PM | C:\Sirius_T3\KOH18C16.t3r |
| Acid concentration factor | 0.990 | 3/16/2018 8:34:16 PM | C:\Sirius_T3\HCl18C16.t3r |

#### Instrument Settings

| Setting | Value | Batch Id | Install date |
|---|---|---|---|
| Instrument owner | Merck | | |
| Instrument ID | T311053 | | |
| Instrument type | T3 Simulator | | |
| Software version | 1.1.3.0 | | |
| Dispenser module | | T3DM1100253 | 3/31/2009 6:24:52 AM |
| Dispenser 0 | Water | | 3/31/2009 6:25:05 AM |
| Syringe volume | 2.5 mL | | |
| Firmware version | 1.2.1(r2) | | |
| Titrant | Water (0.15 M KCl) | 2-6-18 | 3/5/2018 11:53:26 AM |
| Dispenser 2 | Acid | | 3/31/2009 6:25:11 AM |
| Syringe volume | 0.5 mL | | |
| Firmware version | 1.2.1(r2) | | |
| Titrant | Acid (0.5 M HCl) | 03-16-2018 | 3/16/2018 11:26:11 AM |
| Dispenser 1 | Base | | 3/31/2009 6:25:21 AM |
| Syringe volume | 0.5 mL | | |
| Firmware version | 1.2.1(r2) | | |
| Titrant | Base (0.5 M KOH) | 09-22-17 | 3/16/2018 11:19:32 AM |
| Dispenser 5 | Cosolvent | | 3/31/2009 6:26:24 AM |
| Syringe volume | 2.5 mL | | |
| Firmware version | 1.2.1(r2) | | |
| Distribution valve 5 | Distribution Valve | | 3/31/2009 6:28:19 AM |
| Firmware version | 1.1.3 | | |
| Port A | Methanol (80%, 0.15 M KCl) | 03-16-2018 | 3/16/2018 11:40:37 AM |
| Port B | Cyclohexane | | 1/16/2018 1:42:16 PM |
| Port C | MeCN (50%, 0.15 M KCl) | 10-30-17 | 3/16/2018 11:52:45 AM |
| Dispenser 3 | Buffer | | 8/3/2010 6:05:16 AM |
| Syringe volume | 0.5 mL | | |
| Firmware version | 1.2.1(r2) | | |
| Titrant | Dodecane | 1-31-2018 | 2/28/2018 10:07:18 AM |

| | | | |
|---|---|---|---|
| Sample name: | **M13_octanol** | Experiment start time: | **3/16/2018 8:34:17 PM** |
| Assay name: | **pH-metric high logP** | Analyst: | **Pion** |
| Assay ID: | **18C-16016** | Instrument ID: | **T311053** |
| Filename: | **C:\Sirius_T3\Mehtap\20180316_exp32_logP_T3-1\18C-16016_M13_octanol_pH-metric high logP.t3r** | | |

#### Instrument Settings (continued)

| Setting | Value | Batch Id | Install date |
|---|---|---|---|
| Dispenser 6 | Octanol | | 10/22/2010 11:52:43 AM |
| Syringe volume | 0.5 mL | | |
| Firmware version | 1.2.1(r2) | | |
| Titrant | Octanol | 1-31-2018 | 1/31/2018 5:24:59 PM |
| Titrator | | T3TM1100153 | 3/31/2009 6:24:17 AM |
| Horizontal axis firmware version | 1.17 AI1DI2DO2 Stepper 2 | | |
| Vertical axis firmware version | 1.17 AI1DI2DO2 Stepper 2 | | |
| Chassis I/O firmware version | 1.11 AI1DI0DO4 Norgren I/O | | |
| Probe I/O firmware version | 1.1.1 | | |
| Electrode | T3 Electrode | T3E0769 | 8/15/2017 10:21:54 AM |
| E0 calibration | -10.42 mV | | 3/16/2018 8:34:44 PM |
| Filling solution | 3M KCl | KCL095 | 3/16/2018 11:15:26 AM |
| Liquids | | | |
| Wash 1 | 50% IPA:50% Water | | 3/16/2018 11:15:44 AM |
| Wash 2 | 0.5% Trition X-100 in H20 | | 3/16/2018 11:15:47 AM |
| Buffer position 1 | pH7 Wash | | 3/16/2018 11:15:49 AM |
| Buffer position 2 | pH 7 | | 3/16/2018 11:15:51 AM |
| Storage position | | | 3/16/2018 11:16:23 AM |
| Wash water | 4.3e+003 mL | 2-27-18 | 2/27/2018 10:29:26 AM |
| Waste | 5.9e+003 mL | | 2/27/2018 10:29:30 AM |
| Temperature controller | | | 8/5/2010 7:35:13 AM |
| Turbidity detector | | | 3/31/2009 6:24:45 AM |
| Spectrometer | | 072390 | 11/23/2010 12:22:28 PM |
| Dip probe | | 11086 | |
| Wavelength coefficient A0 | 185.563 | | |
| Wavelength coefficient A1 | 2.17439 | | |
| Wavelength coefficient A2 | -0.000285622 | | |
| Total lamp lit time | 804:59:49 | | 11/23/2010 12:22:28 PM |
| Calibrated on | 2/22/2018 5:36:24 PM | | |
| Integration time | 17 | | |
| Scans averaged | 10 | | |
| Autoloader | | T3AL1100237 | 11/10/2015 10:34:13 AM |
| Left-right axis firmware version | 1.17 AI1DI2DO2 Stepper 2 | | |
| Front-back axis firmware version | 1.17 AI1DI2DO2 Stepper 2 | | |
| Vertical axis firmware version | 1.17 AI1DI2DO2 Stepper 2 | | |
| Chassis I/O firmware version | 1.11 AI1DI0DO4 Norgren I/O | | |
| Configuration | | | |
| Alternate titration position | Titration position | | |
| Alternate reference position | Reference position | | |
| Maximum standard vial volume | 3.50 mL | | |
| Maximum alternate vial volume | 25.00 mL | | |
| Automatic action idle period | 5 minute(s) | | |
| Titrant tube volume | 1.3 mL | | |
| Syringe flush count | 3.50 | | |
| Flowing wash pump volume | 20.0 mL | | |
| Flowing wash stir duration | 5 s | | |
| Flowing wash stir speed | 30% | | |
| Solvent wash stir duration | 5 s | | |
| Solvent wash stir speed | 30% | | |
| Surfactant wash stir duration | 5 s | | |
| Surfactant wash stir speed | 30% | | |
| E0 calibration minimum number of points | 10 | | |
| E0 calibration maximum standard deviation | 0.01500 | | |
| E0 calibration timeout period | 60 s | | |
| E0 calibration stir duration | 5 s | | |
| E0 calibration preparation stir speed | 30% | | |
| E0 calibration buffer wash stir duration | 5 s | | |
| E0 calibration buffer wash stir speed | 30% | | |

| | | | |
|---|---|---|---|
| Sample name: | **M13_octanol** | Experiment start time: | **3/16/2018 8:34:17 PM** |
| Assay name: | **pH-metric high logP** | Analyst: | **Pion** |
| Assay ID: | **18C-16016** | Instrument ID: | **T311053** |
| Filename: | **C:\Sirius_T3\Mehtap\20180316_exp32_logP_T3-1\18C-16016_M13_octanol_pH-metric high logP.t3r** | | |

#### Instrument Settings (continued)

| Setting | Value | Batch Id | Install date |
|---|---|---|---|
| E0 calibration reading stir speed | 0% | | |
| Spectrometer calibration stir duration | 5 s | | |
| Spectrometer calibration stir speed | 30% | | |
| Spectrometer calibration wash pump volume | 20.0 mL | | |
| Spectrometer calibration wash stir duration | 5 s | | |
| Spectrometer calibration wash stir speed | 30% | | |
| Overhead dispense height | 10000 | | |

#### Refinement Settings

| Setting | Value | Default value |
|---|---|---|
| Turbidity detection method | None | None |
| Turbidity wavelength to assess | 500.0 nm | 500.0 nm |
| Turbidity maximum absorbance | 0.100 | 0.100 |
| Turbidity probe threshold | 50.00 | 50.00 |
