## Supplementary material for "Octanol-water partition coefficient measurements for the SAMPL6 Blind Prediction Challenge": SM13_18C-26010_M13_octanol_pH-metric high logP_report.pdf

| | | | |
|---|---|---|---|
| Sample name: | **M13_octanol** | Experiment start time: | **3/26/2018 4:16:35 PM** |
| Assay name: | **pH-metric high logP** | Analyst: | **Pion** |
| Assay ID: | **18C-26010** | Instrument ID: | **T312060** |
| Filename: | **C:\Sirius_T3\Mehtap\20180327_exp34_logP_T3-2\18C-26010_M13_octanol_pH-metric high logP.t3r** | | |

### pH-metric Result

| | | |
|---|---|---|
| logP (XH +) | 0.09 | ±0.08 (n=50) |
| logP (neutral X) | 2.99 | ±0.03 (n=50) |

### 18C-26010 Points 2 to 16

| | |
|---|---|
| M13_octanol concentration factor | 1.006 |
| Carbonate | 0.0000 mM |
| Acidity error | 2.38959 mM |

### 18C-26010 Points 17 to 40

| | |
|---|---|
| M13_octanol concentration factor | 0.639 |
| Carbonate | 0.1054 mM |
| Acidity error | 2.15628 mM |

### 18C-26010 Points 41 to 68

| | |
|---|---|
| M13_octanol concentration factor | 0.745 |
| Carbonate | 0.1125 mM |
| Acidity error | 2.15554 mM |

### Warnings and errors

| | |
|---|---|
| Errors | None |
| Warnings | None |

### Sample logD and percent species

| pH | M13_octanol logD | M13_octanol M13_octanolH | M13_octanol M13_octanol | M13_octanol M13_octanolH* | M13_octanol M13_octanol* | Comment |
|---|---|---|---|---|---|---|
| 1.000 | 0.09 | 44.67 % | 0.00 % | 54.60 % | 0.74 % | |
| 1.200 | 0.10 | 44.47 % | 0.00 % | 54.36 % | 1.17 % | Stomach pH |
| 2.000 | 0.14 | 41.88 % | 0.01 % | 51.19 % | 6.93 % | |
| 3.000 | 0.46 | 25.78 % | 0.04 % | 31.52 % | 42.65 % | |
| 4.000 | 1.24 | 5.32 % | 0.09 % | 6.51 % | 88.08 % | |
| 5.000 | 2.15 | 0.60 % | 0.10 % | 0.73 % | 98.57 % | |
| 6.000 | 2.79 | 0.06 % | 0.10 % | 0.07 % | 99.76 % | |
| 6.500 | 2.91 | 0.02 % | 0.10 % | 0.02 % | 99.86 % | |
| 7.000 | 2.96 | 0.01 % | 0.10 % | 0.01 % | 99.88 % | |
| 7.400 | 2.98 | 0.00 % | 0.10 % | 0.00 % | 99.89 % | Blood pH |
| 8.000 | 2.99 | 0.00 % | 0.10 % | 0.00 % | 99.90 % | |
| 9.000 | 2.99 | 0.00 % | 0.10 % | 0.00 % | 99.90 % | |
| 10.000 | 2.99 | 0.00 % | 0.10 % | 0.00 % | 99.90 % | |
| 11.000 | 2.99 | 0.00 % | 0.10 % | 0.00 % | 99.90 % | |
| 12.000 | 2.99 | 0.00 % | 0.10 % | 0.00 % | 99.90 % | |

Sample name: **M13_octanol**
Assay name: **pH-metric high logP**
Assay ID: **18C-26010**
Filename: **C:\Sirius_T3\Mehtap\20180327_exp34_logP_T3-2\18C-26010_M13_octanol_pH-metric high logP.t3r**

Experiment start time: **3/26/2018 4:16:35 PM**
Analyst: **Pion**
Instrument ID: **T312060**

### Graphs

Charge balance residuals

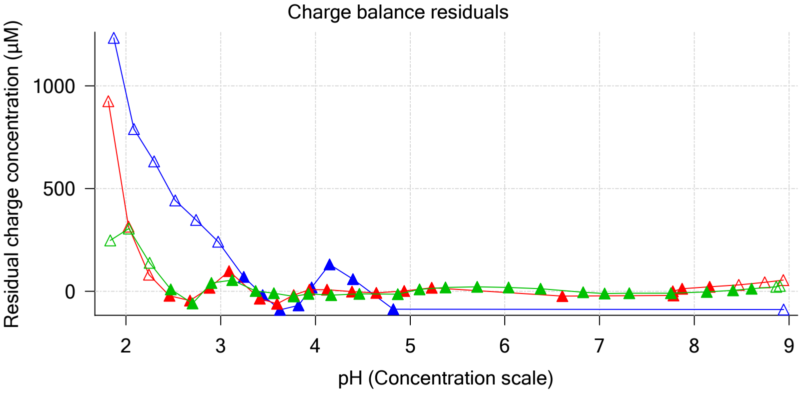


Ionic strength

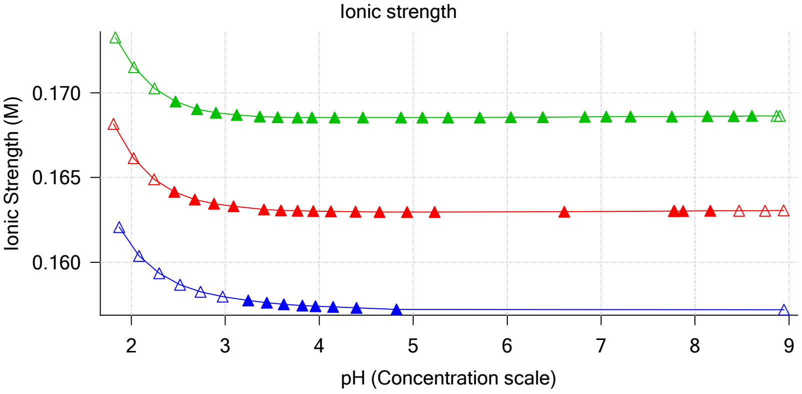


Buffer index profile

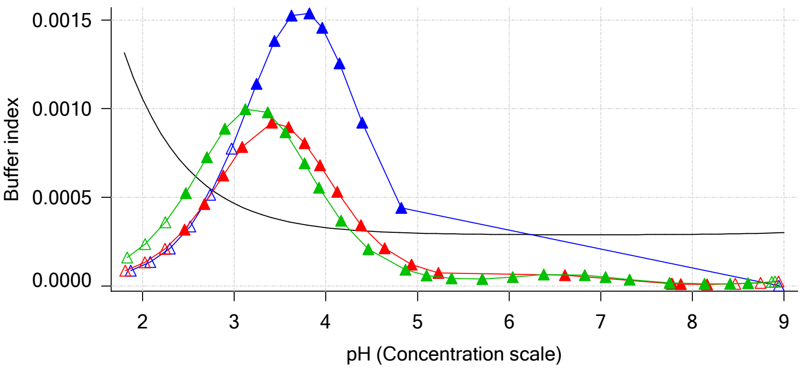


Sample M13_octanol Ionisation Graph

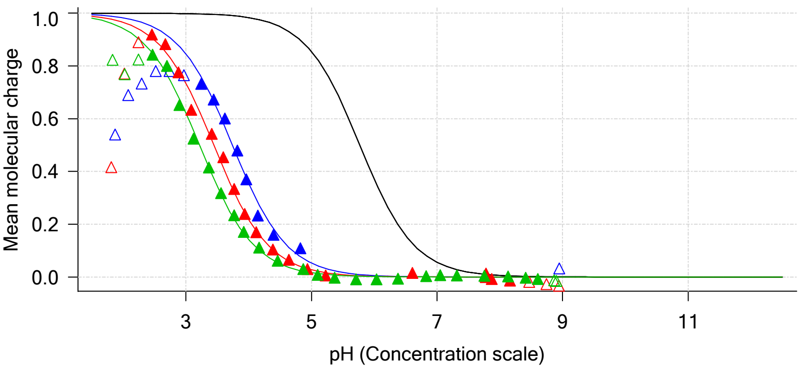


Distribution of species for sample M13_octanol

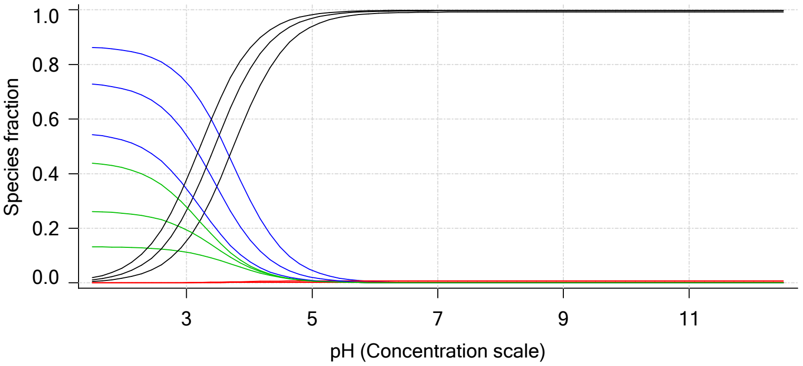


Lipophilicity Profile

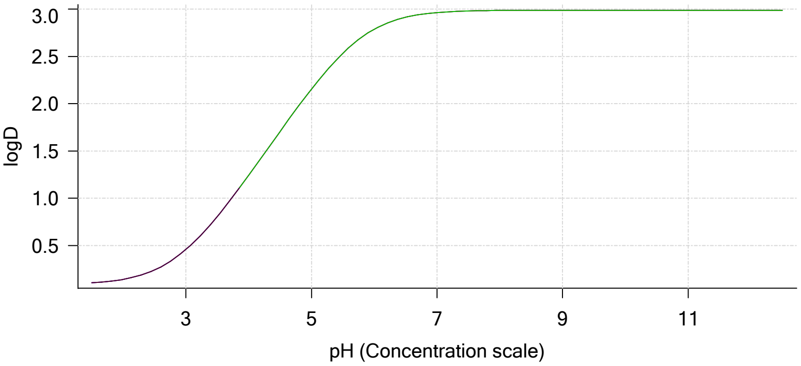


Log R Profile

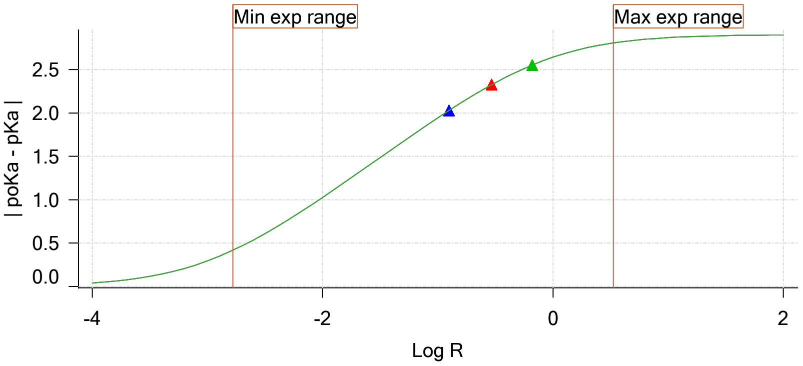


Sample Carbonate Ionisation Graph

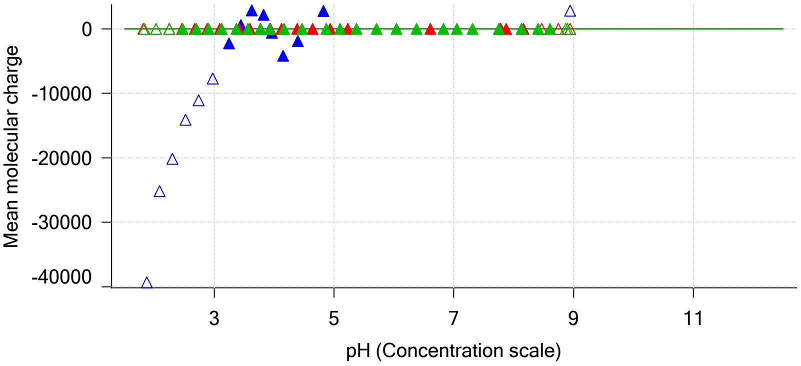

| | | | |
|---|---|---|---|
| Sample name: | **M13_octanol** | Experiment start time: | **3/26/2018 4:16:35 PM** |
| Assay name: | **pH-metric high logP** | Analyst: | **Pion** |
| Assay ID: | **18C-26010** | Instrument ID: | **T312060** |
| Filename: | **C:\Sirius_T3\Mehtap\20180327_exp34_logP_T3-2\18C-26010_M13_octanol_pH-metric high logP.t3r** | | |

### Graphs (continued)

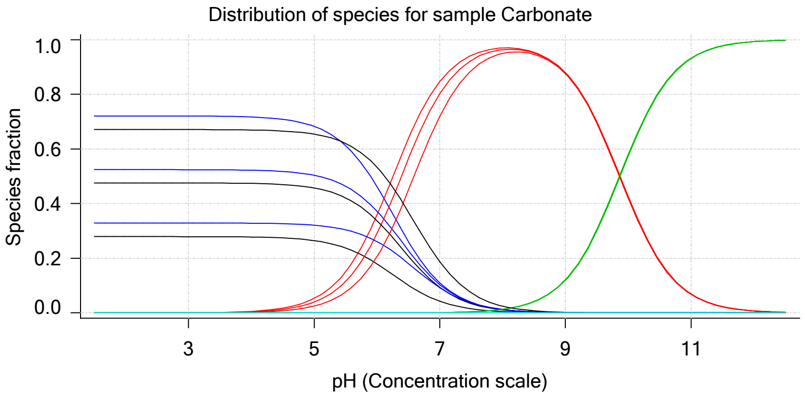

| | | | |
|---|---|---|---|
| Sample name: | **M13_octanol** | Experiment start time: | **3/26/2018 4:16:35 PM** |
| Assay name: | **pH-metric high logP** | Analyst: | **Pion** |
| Assay ID: | **18C-26010** | Instrument ID: | **T312060** |
| Filename: | **C:\Sirius_T3\Mehtap\20180327_exp34_logP_T3-2\18C-26010_M13_octanol_pH-metric high logP.t3r** | | |

# pH-metric high logP   Titration 1 of 3   18C-26010 Points 2 to 16

### Overall results

| | |
|---|---|
| RMSD | 0.079 |
| Average ionic strength | 0.158 M |
| Average temperature | 24.9°C |
| Partition ratio | 0.1253 : 1 |
| Analyte concentration range | 2374.9 μM to 2437.5 μM |
| Total points considered | 8 of 15 |

### Warnings and errors

| | |
|---|---|
| Errors | None |
| Warnings | One or more logP values out of range |
| | Excessive acidity error present |

### Four-Plus parameters

| | | | |
|---|---|---|---|
| Alpha | 0.122 | 3/26/2018 4:16:34 PM | C:\Sirius_T3\18C-26006_Blank standardisation.t3r |
| S | 0.9974 | 3/26/2018 4:16:34 PM | C:\Sirius_T3\18C-26006_Blank standardisation.t3r |
| jH | 1.1 | 3/26/2018 4:16:34 PM | C:\Sirius_T3\18C-26006_Blank standardisation.t3r |
| jOH | -0.5 | 3/26/2018 4:16:34 PM | C:\Sirius_T3\18C-26006_Blank standardisation.t3r |

### Titrants

| | | | |
|---|---|---|---|
| 0.50 M HCl | 0.994478 | 3/26/2018 4:16:34 PM | C:\Sirius_T3\18C-26006_Blank standardisation.t3r |
| 0.50 M KOH | 1.003190 | 3/26/2018 4:16:35 PM | C:\Sirius_T3\KOH18C23.t3r |

### Sample

| | |
|---|---|
| M13_octanol concentration factor | 1.006 |
| Base pKa 1 | 5.77 |
| logP (XH +) | -4.56 |
| logP (neutral X) | 2.93 |

### Sample graphs

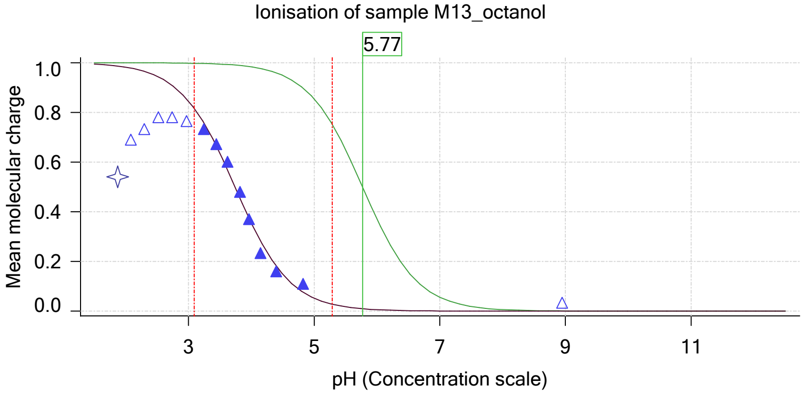


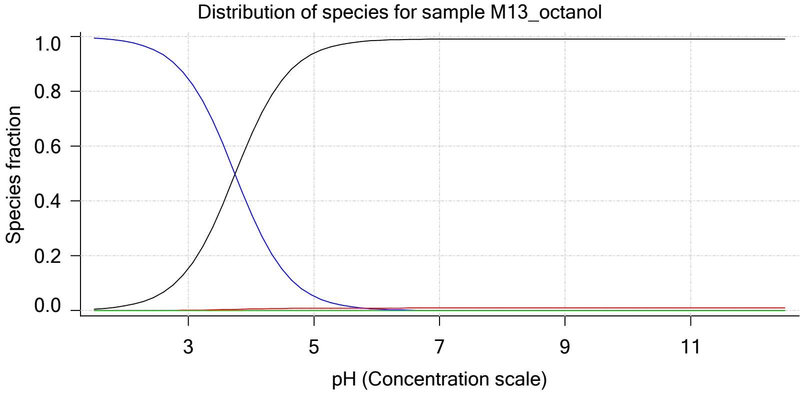

| | | | |
|---|---|---|---|
| Sample name: | **M13_octanol** | Experiment start time: | **3/26/2018 4:16:35 PM** |
| Assay name: | **pH-metric high logP** | Analyst: | **Pion** |
| Assay ID: | **18C-26010** | Instrument ID: | **T312060** |
| Filename: | **C:\Sirius_T3\Mehtap\20180327_exp34_logP_T3-2\18C-26010_M13_octanol_pH-metric high logP.t3r** | | |

### Sample graphs (continued)

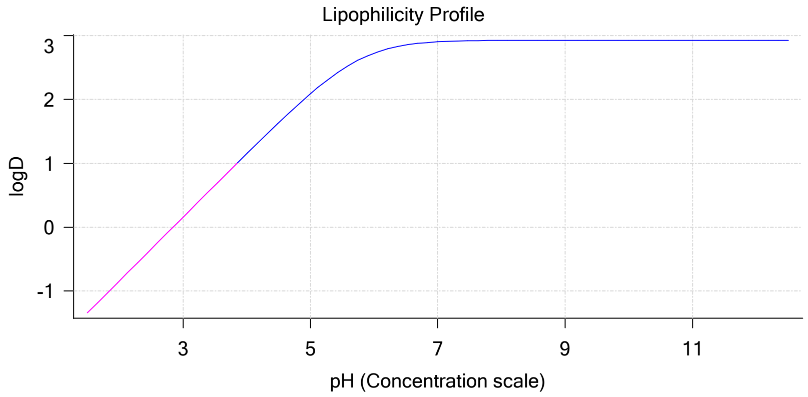


### Sample logD and percent species

| pH | M13_octanol logD | M13_octanol M13_octanolH | M13_octanol M13_octanol | M13_octanol M13_octanolH* | M13_octanol M13_octanol* | Comment |
|---|---|---|---|---|---|---|
| 1.000 | -1.84 | 99.82 % | 0.00 % | 0.00 % | 0.18 % | |
| 1.200 | -1.64 | 99.71 % | 0.00 % | 0.00 % | 0.28 % | Stomach pH |
| 2.000 | -0.84 | 98.21 % | 0.02 % | 0.00 % | 1.77 % | |
| 3.000 | 0.16 | 84.62 % | 0.14 % | 0.00 % | 15.23 % | |
| 4.000 | 1.15 | 35.50 % | 0.60 % | 0.00 % | 63.90 % | |
| 5.000 | 2.09 | 5.22 % | 0.89 % | 0.00 % | 93.90 % | |
| 6.000 | 2.73 | 0.55 % | 0.93 % | 0.00 % | 98.52 % | |
| 6.500 | 2.85 | 0.17 % | 0.93 % | 0.00 % | 98.89 % | |
| 7.000 | 2.90 | 0.05 % | 0.93 % | 0.00 % | 99.01 % | |
| 7.400 | 2.92 | 0.02 % | 0.93 % | 0.00 % | 99.04 % | Blood pH |
| 8.000 | 2.92 | 0.01 % | 0.93 % | 0.00 % | 99.06 % | |
| 9.000 | 2.93 | 0.00 % | 0.93 % | 0.00 % | 99.06 % | |
| 10.000 | 2.93 | 0.00 % | 0.93 % | 0.00 % | 99.07 % | |
| 11.000 | 2.93 | 0.00 % | 0.93 % | 0.00 % | 99.07 % | |
| 12.000 | 2.93 | 0.00 % | 0.93 % | 0.00 % | 99.07 % | |

### Carbonate and acidity

| | |
|---|---|
| Carbonate | 0.000 mM |
| Acidity error | 2.390 mM |

### Other graphs

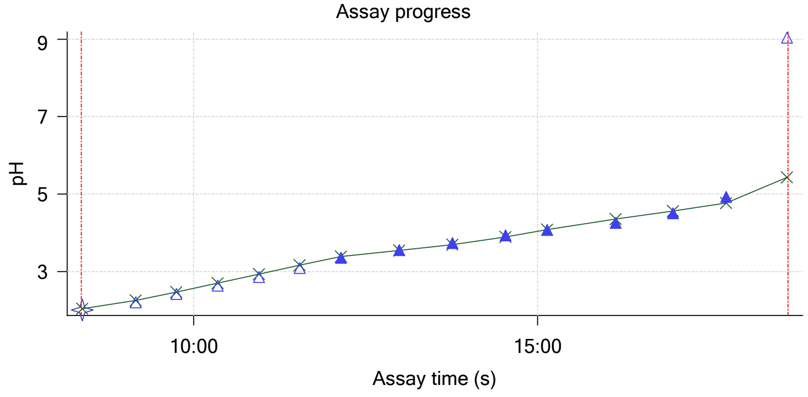


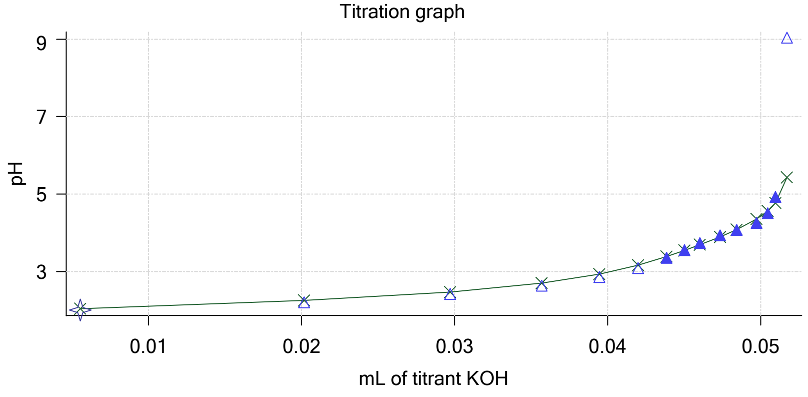

Sample name: **M13_octanol**
Assay name: **pH-metric high logP**
Assay ID: **18C-26010**
Filename: **C:\Sirius_T3\Mehtap\20180327_exp34_logP_T3-2\18C-26010_M13_octanol_pH-metric high logP.t3r**

Experiment start time: **3/26/2018 4:16:35 PM**
Analyst: **Pion**
Instrument ID: **T312060**

### Other graphs (continued)

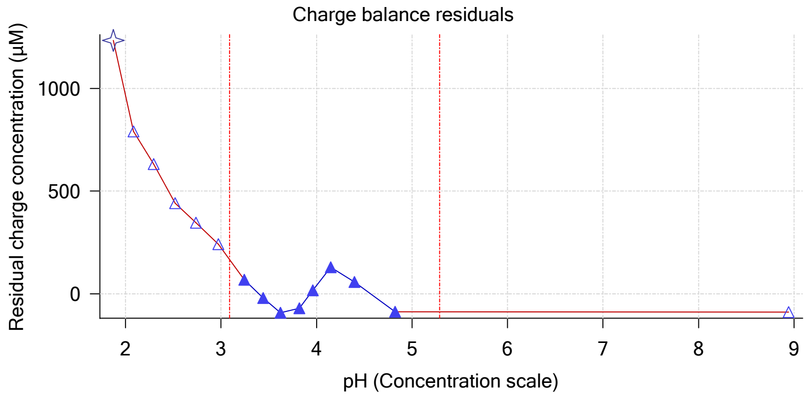


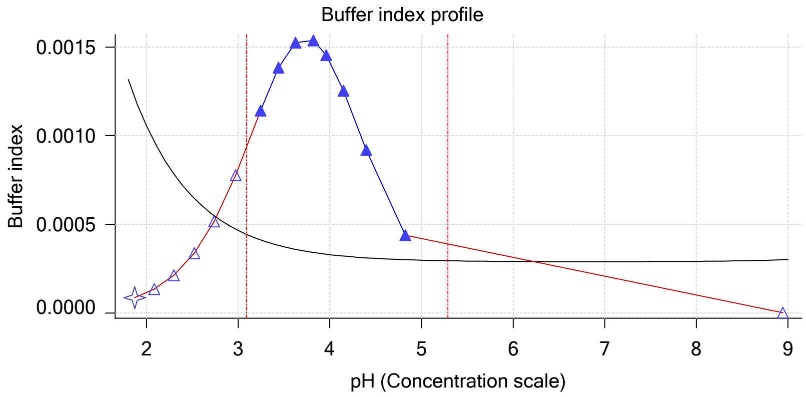


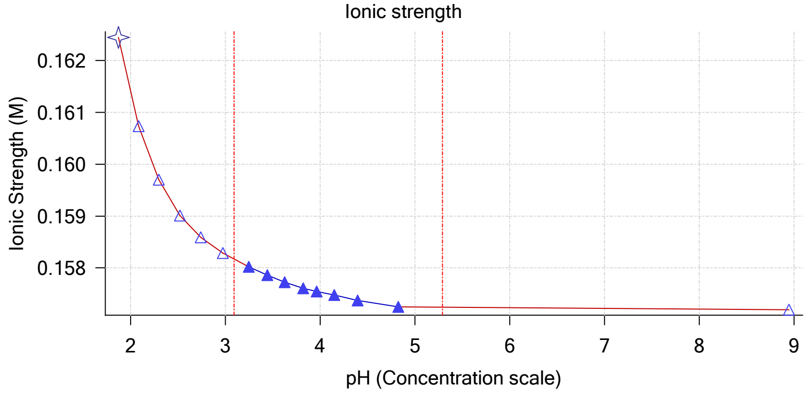


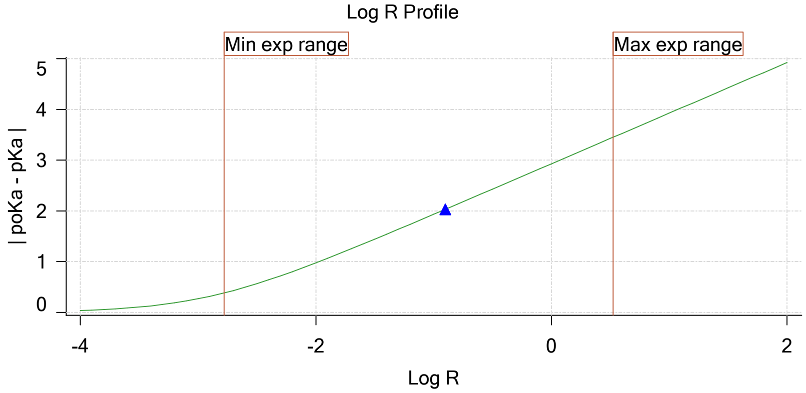


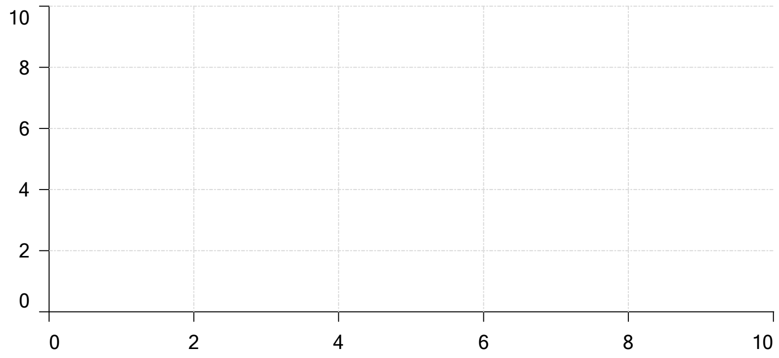


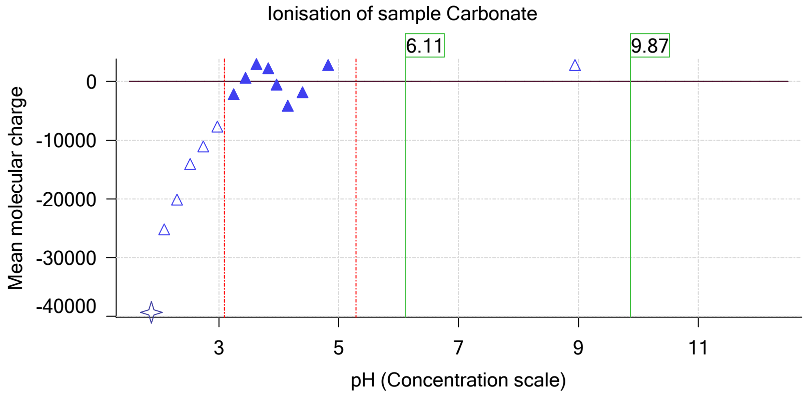

| | | | |
|---|---|---|---|
| Sample name: | **M13_octanol** | Experiment start time: | **3/26/2018 4:16:35 PM** |
| Assay name: | **pH-metric high logP** | Analyst: | **Pion** |
| Assay ID: | **18C-26010** | Instrument ID: | **T312060** |
| Filename: | **C:\Sirius_T3\Mehtap\20180327_exp34_logP_T3-2\18C-26010_M13_octanol_pH-metric high logP.t3r** | | |

# pH-metric high logP   Titration 2 of 3   18C-26010 Points 17 to 40

### Overall results

| | |
|---|---|
| RMSD | 0.567 |
| Average ionic strength | 0.163 M |
| Average temperature | 25.0°C |
| Partition ratio | 0.2937 : 1 |
| Analyte concentration range | 1935.6 μM to 1984.1 μM |
| Total points considered | 18 of 24 |

### Warnings and errors

| | |
|---|---|
| Errors | None |
| Warnings | Sample concentration factor out of range |
| | One or more logP values out of range |
| | Excessive acidity error present |

### Four-Plus parameters

| | | | |
|---|---|---|---|
| Alpha | 0.122 | 3/26/2018 4:16:34 PM | C:\Sirius_T3\18C-26006_Blank standardisation.t3r |
| S | 0.9974 | 3/26/2018 4:16:34 PM | C:\Sirius_T3\18C-26006_Blank standardisation.t3r |
| jH | 1.1 | 3/26/2018 4:16:34 PM | C:\Sirius_T3\18C-26006_Blank standardisation.t3r |
| jOH | -0.5 | 3/26/2018 4:16:34 PM | C:\Sirius_T3\18C-26006_Blank standardisation.t3r |

### Titrants

| | | | |
|---|---|---|---|
| 0.50 M HCl | 0.994478 | 3/26/2018 4:16:34 PM | C:\Sirius_T3\18C-26006_Blank standardisation.t3r |
| 0.50 M KOH | 1.003190 | 3/26/2018 4:16:35 PM | C:\Sirius_T3\KOH18C23.t3r |

### Sample

| | |
|---|---|
| M13_octanol concentration factor | 0.639 |
| Base pKa 1 | 5.77 |
| logP (XH +) | -4.56 |
| logP (neutral X) | 2.85 |

### Sample graphs

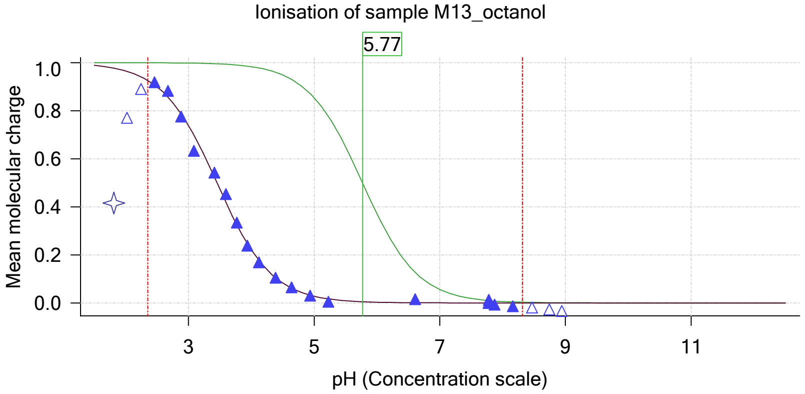


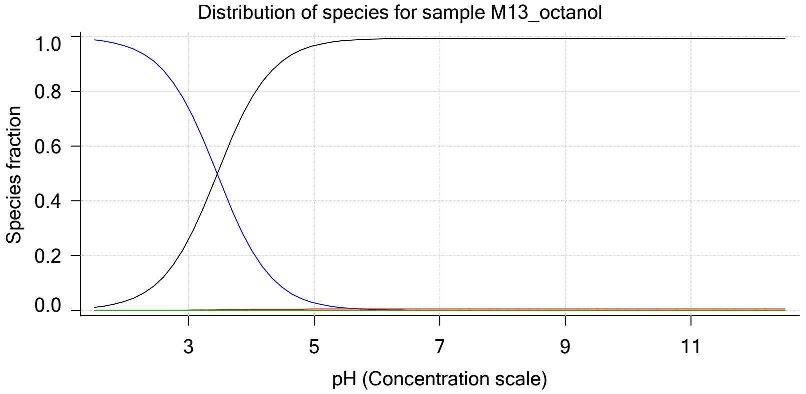

Sample name: **M13_octanol**
Assay name: **pH-metric high logP**
Assay ID: **18C-26010**
Filename: **C:\Sirius_T3\Mehtap\20180327_exp34_logP_T3-2\18C-26010_M13_octanol_pH-metric high logP.t3r**

Experiment start time: **3/26/2018 4:16:35 PM**
Analyst: **Pion**
Instrument ID: **T312060**

### Sample graphs (continued)

Lipophilicity Profile

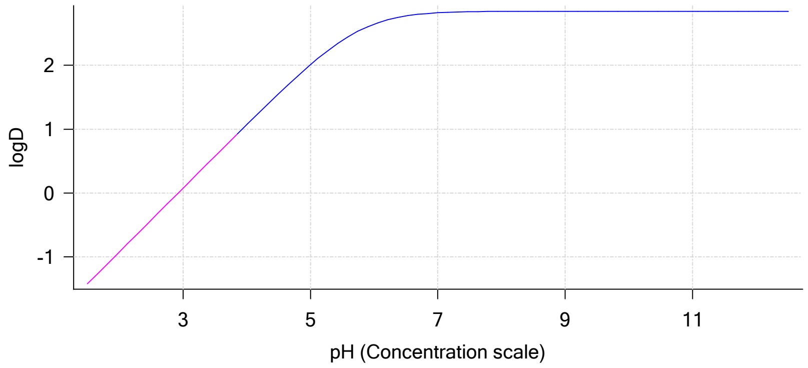


### Sample logD and percent species

| pH | M13_octanol logD | M13_octanol M13_octanolH | M13_octanol M13_octanol | M13_octanol M13_octanolH* | M13_octanol M13_octanol* | Comment |
|---|---|---|---|---|---|---|
| 1.000 | -1.92 | 99.65 % | 0.00 % | 0.00 % | 0.35 % | |
| 1.200 | -1.72 | 99.44 % | 0.00 % | 0.00 % | 0.55 % | Stomach pH |
| 2.000 | -0.92 | 96.60 % | 0.02 % | 0.00 % | 3.39 % | |
| 3.000 | 0.08 | 73.95 % | 0.13 % | 0.00 % | 25.93 % | |
| 4.000 | 1.07 | 22.11 % | 0.38 % | 0.00 % | 77.52 % | |
| 5.000 | 2.01 | 2.76 % | 0.47 % | 0.00 % | 96.77 % | |
| 6.000 | 2.65 | 0.28 % | 0.48 % | 0.00 % | 99.24 % | |
| 6.500 | 2.77 | 0.09 % | 0.48 % | 0.00 % | 99.43 % | |
| 7.000 | 2.82 | 0.03 % | 0.48 % | 0.00 % | 99.49 % | |
| 7.400 | 2.84 | 0.01 % | 0.48 % | 0.00 % | 99.51 % | Blood pH |
| 8.000 | 2.84 | 0.00 % | 0.48 % | 0.00 % | 99.52 % | |
| 9.000 | 2.85 | 0.00 % | 0.48 % | 0.00 % | 99.52 % | |
| 10.000 | 2.85 | 0.00 % | 0.48 % | 0.00 % | 99.52 % | |
| 11.000 | 2.85 | 0.00 % | 0.48 % | 0.00 % | 99.52 % | |
| 12.000 | 2.85 | 0.00 % | 0.48 % | 0.00 % | 99.52 % | |

### Carbonate and acidity

Carbonate 0.105 mM
Acidity error 2.156 mM

### Other graphs

Assay progress

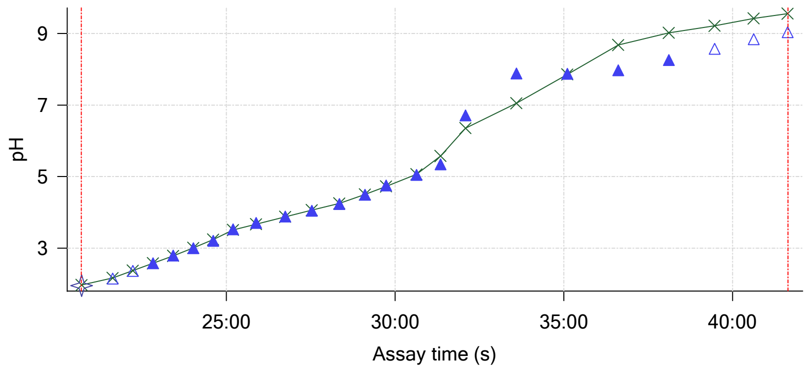


Titration graph

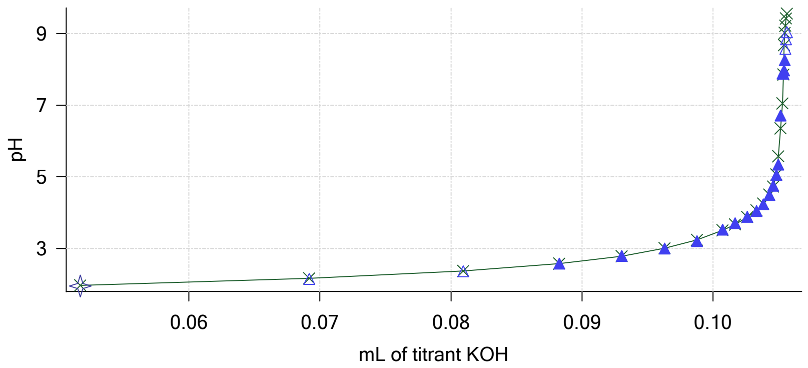

| | | | |
|---|---|---|---|
| Sample name: | **M13_octanol** | Experiment start time: | **3/26/2018 4:16:35 PM** |
| Assay name: | **pH-metric high logP** | Analyst: | **Pion** |
| Assay ID: | **18C-26010** | Instrument ID: | **T312060** |
| Filename: | **C:\Sirius_T3\Mehtap\20180327_exp34_logP_T3-2\18C-26010_M13_octanol_pH-metric high logP.t3r** | | |

### Other graphs (continued)

Charge balance residuals

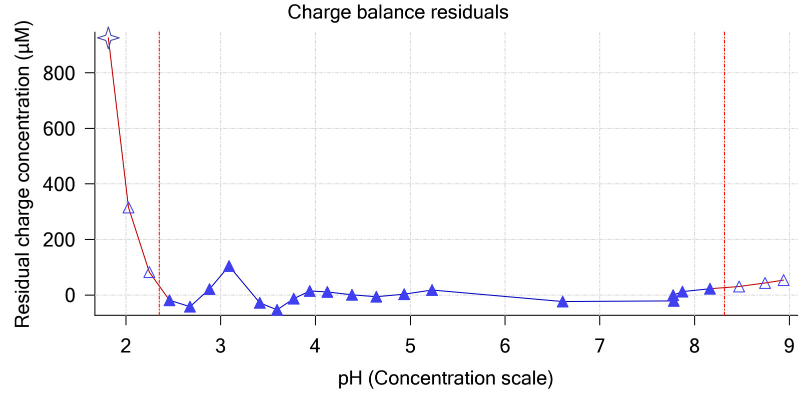


Buffer index profile

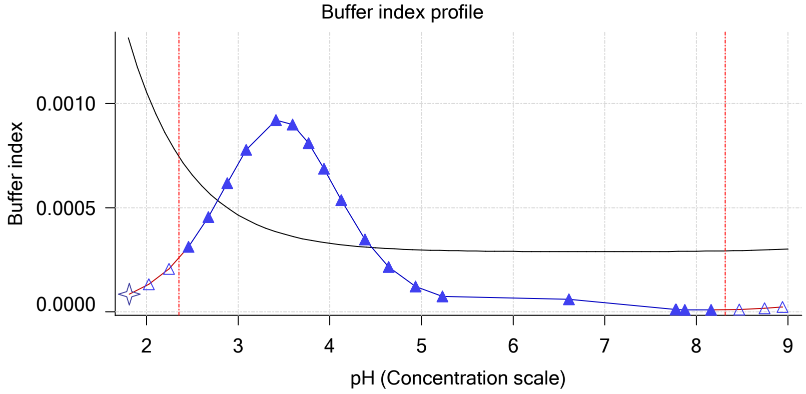


Ionic strength

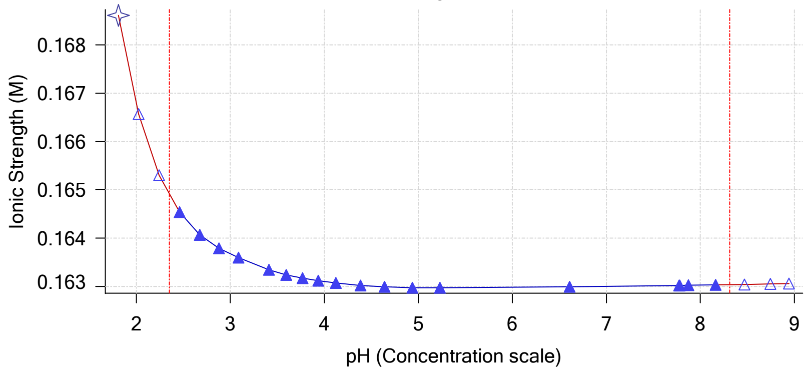


Log R Profile

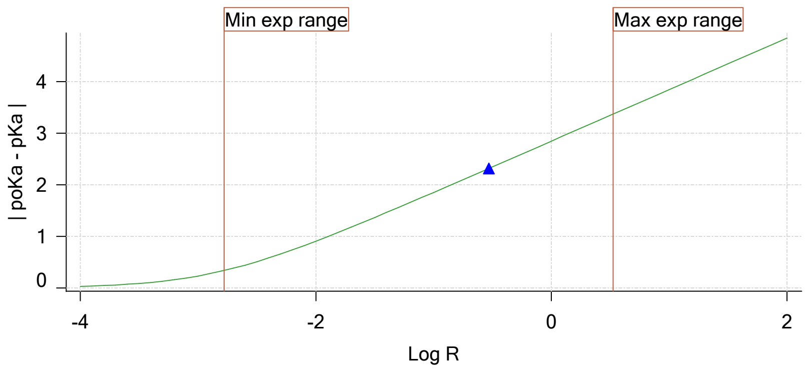


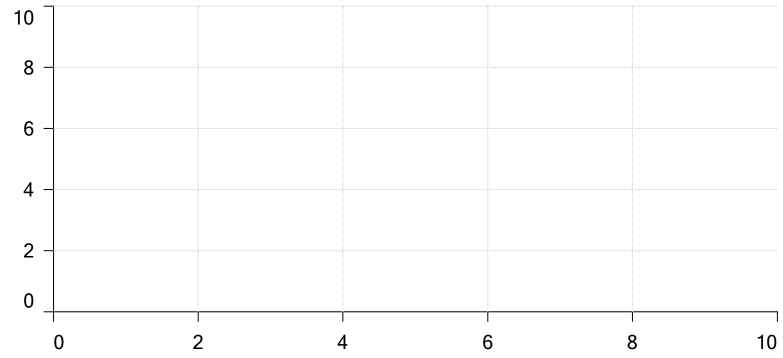


Ionisation of sample Carbonate

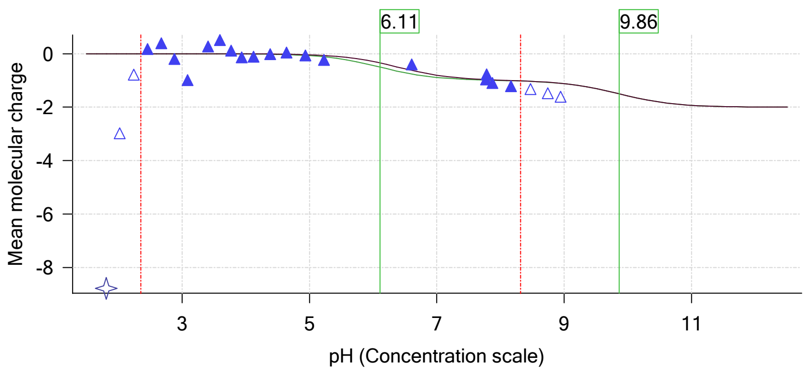

| | | | |
|---|---|---|---|
| Sample name: | **M13_octanol** | Experiment start time: | **3/26/2018 4:16:35 PM** |
| Assay name: | **pH-metric high logP** | Analyst: | **Pion** |
| Assay ID: | **18C-26010** | Instrument ID: | **T312060** |
| Filename: | **C:\Sirius_T3\Mehtap\20180327_exp34_logP_T3-2\18C-26010_M13_octanol_pH-metric high logP.t3r** | | |

# pH-metric high logP   Titration 3 of 3   18C-26010 Points 41 to 68

### Overall results

| | |
|---|---|
| RMSD | 0.220 |
| Average ionic strength | 0.169 M |
| Average temperature | 25.0°C |
| Partition ratio | 0.6600 : 1 |
| Analyte concentration range | 1412.7 μM to 1440.3 μM |
| Total points considered | 23 of 28 |

### Warnings and errors

| | |
|---|---|
| Errors | None |
| Warnings | One or more logP values out of range |
| | Excessive acidity error present |

### Four-Plus parameters

| | | | |
|---|---|---|---|
| Alpha | 0.122 | 3/26/2018 4:16:34 PM | C:\Sirius_T3\18C-26006_Blank standardisation.t3r |
| S | 0.9974 | 3/26/2018 4:16:34 PM | C:\Sirius_T3\18C-26006_Blank standardisation.t3r |
| jH | 1.1 | 3/26/2018 4:16:34 PM | C:\Sirius_T3\18C-26006_Blank standardisation.t3r |
| jOH | -0.5 | 3/26/2018 4:16:34 PM | C:\Sirius_T3\18C-26006_Blank standardisation.t3r |

### Titrants

| | | | |
|---|---|---|---|
| 0.50 M HCl | 0.994478 | 3/26/2018 4:16:34 PM | C:\Sirius_T3\18C-26006_Blank standardisation.t3r |
| 0.50 M KOH | 1.003190 | 3/26/2018 4:16:35 PM | C:\Sirius_T3\KOH18C23.t3r |

### Sample

| | |
|---|---|
| M13_octanol concentration factor | 0.745 |
| Base pKa 1 | 5.77 |
| logP (XH +) | -4.56 |
| logP (neutral X) | 2.73 |

### Sample graphs

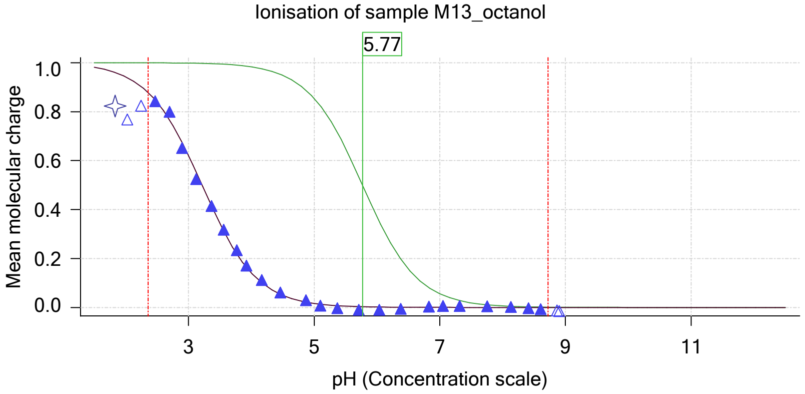


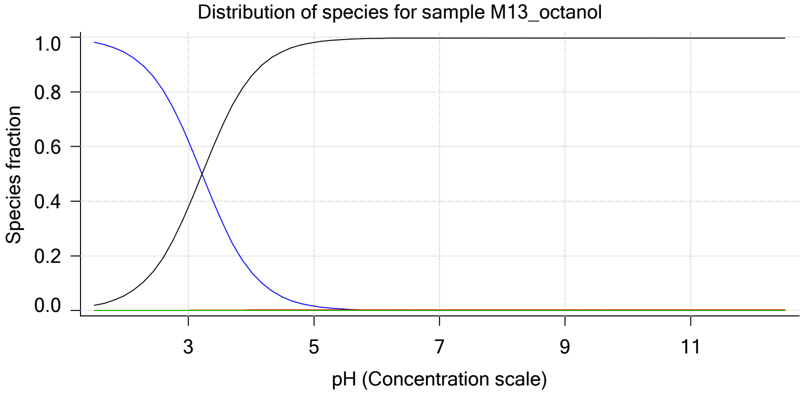

| | | | |
|---|---|---|---|
| Sample name: | **M13_octanol** | Experiment start time: | **3/26/2018 4:16:35 PM** |
| Assay name: | **pH-metric high logP** | Analyst: | **Pion** |
| Assay ID: | **18C-26010** | Instrument ID: | **T312060** |
| Filename: | **C:\Sirius_T3\Mehtap\20180327_exp34_logP_T3-2\18C-26010_M13_octanol_pH-metric high logP.t3r** | | |

### Sample graphs (continued)

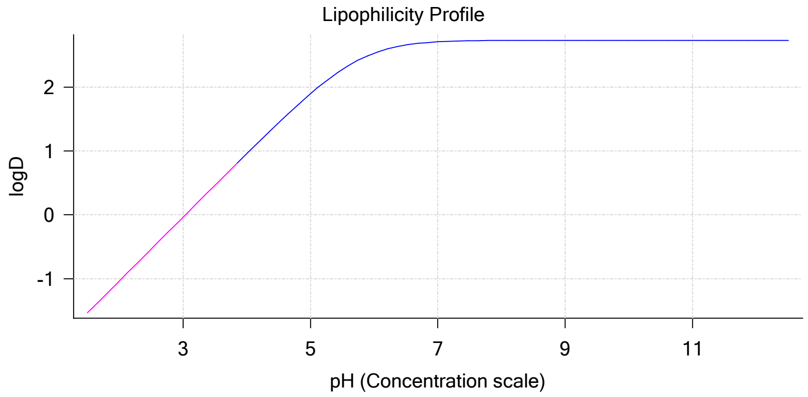


### Sample logD and percent species

| pH | M13_octanol logD | M13_octanol M13_octanolH | M13_octanol M13_octanol | M13_octanol M13_octanolH* | M13_octanol M13_octanol* | Comment |
|---|---|---|---|---|---|---|
| 1.000 | -2.04 | 99.39 % | 0.00 % | 0.00 % | 0.60 % | |
| 1.200 | -1.84 | 99.04 % | 0.00 % | 0.00 % | 0.95 % | Stomach pH |
| 2.000 | -1.04 | 94.26 % | 0.02 % | 0.00 % | 5.72 % | |
| 3.000 | -0.04 | 62.18 % | 0.11 % | 0.00 % | 37.72 % | |
| 4.000 | 0.96 | 14.12 % | 0.24 % | 0.00 % | 85.64 % | |
| 5.000 | 1.90 | 1.62 % | 0.27 % | 0.00 % | 98.11 % | |
| 6.000 | 2.53 | 0.16 % | 0.28 % | 0.00 % | 99.56 % | |
| 6.500 | 2.66 | 0.05 % | 0.28 % | 0.00 % | 99.67 % | |
| 7.000 | 2.71 | 0.02 % | 0.28 % | 0.00 % | 99.70 % | |
| 7.400 | 2.72 | 0.01 % | 0.28 % | 0.00 % | 99.71 % | Blood pH |
| 8.000 | 2.73 | 0.00 % | 0.28 % | 0.00 % | 99.72 % | |
| 9.000 | 2.73 | 0.00 % | 0.28 % | 0.00 % | 99.72 % | |
| 10.000 | 2.73 | 0.00 % | 0.28 % | 0.00 % | 99.72 % | |
| 11.000 | 2.73 | 0.00 % | 0.28 % | 0.00 % | 99.72 % | |
| 12.000 | 2.73 | 0.00 % | 0.28 % | 0.00 % | 99.72 % | |

### Carbonate and acidity

| | |
|---|---|
| Carbonate | 0.113 mM |
| Acidity error | 2.156 mM |

### Other graphs

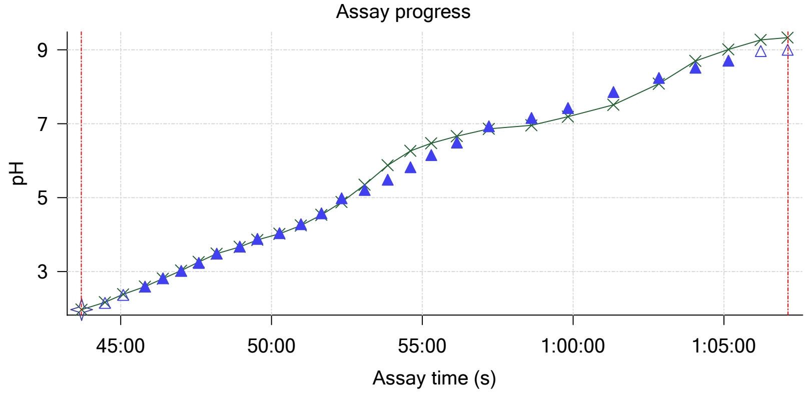


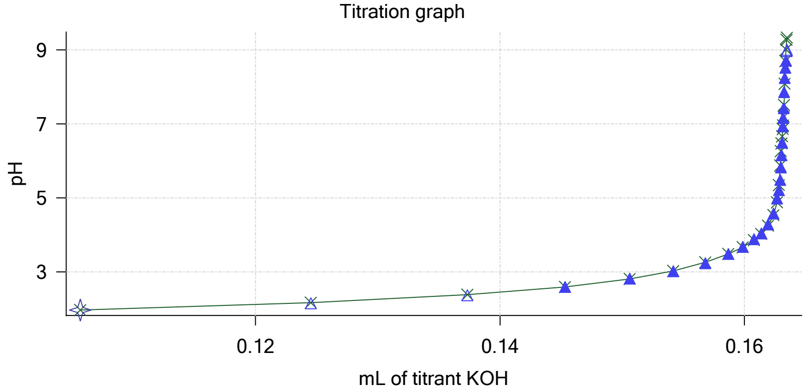

Sample name: **M13_octanol**
Assay name: **pH-metric high logP**
Assay ID: **18C-26010**
Filename: **C:\Sirius_T3\Mehtap\20180327_exp34_logP_T3-2\18C-26010_M13_octanol_pH-metric high logP.t3r**

Experiment start time: **3/26/2018 4:16:35 PM**
Analyst: **Pion**
Instrument ID: **T312060**

### Other graphs (continued)

Charge balance residuals

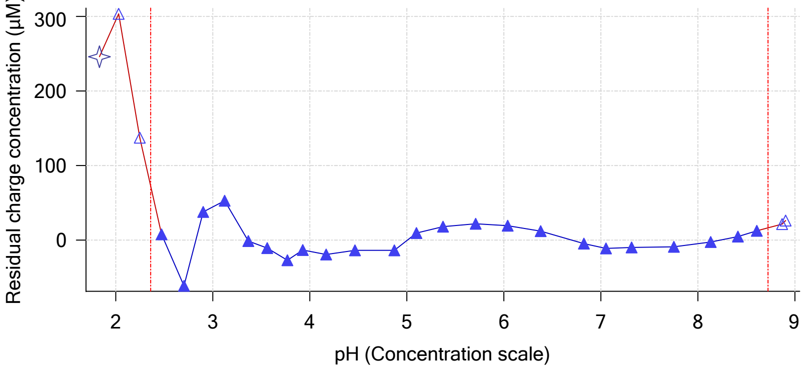


Buffer index profile

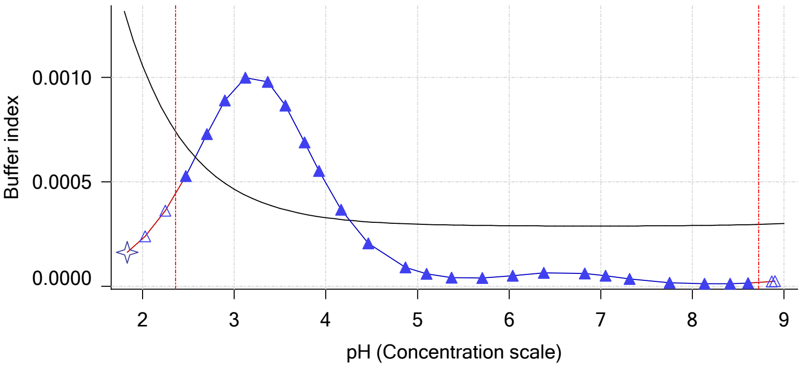


Ionic strength

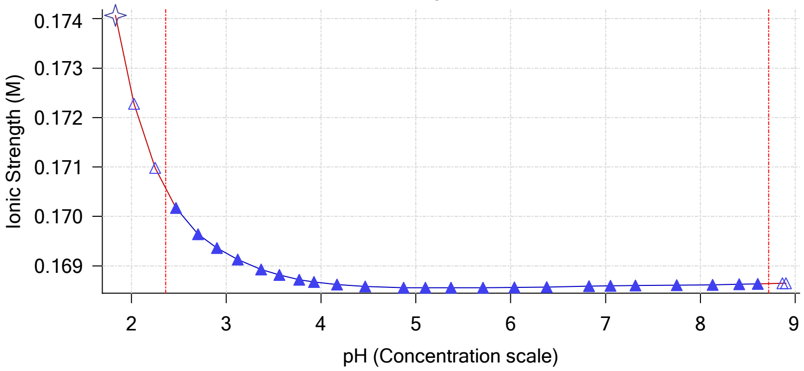


Log R Profile

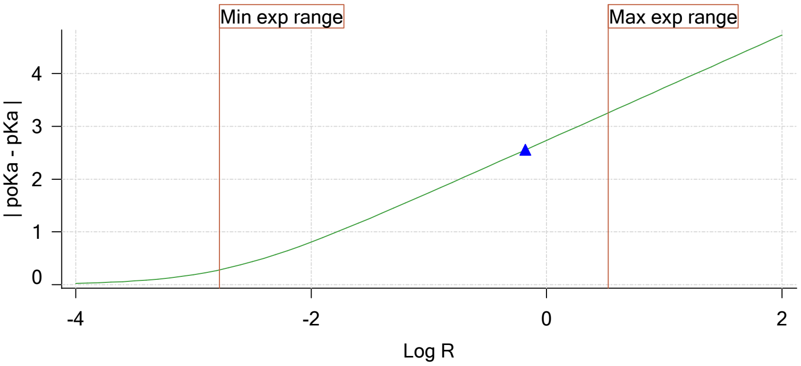


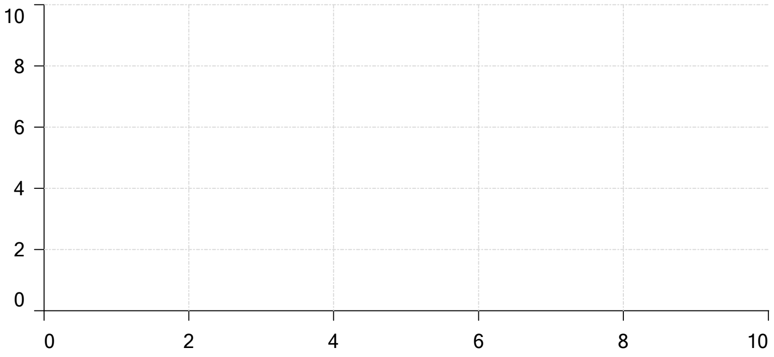


Ionisation of sample Carbonate

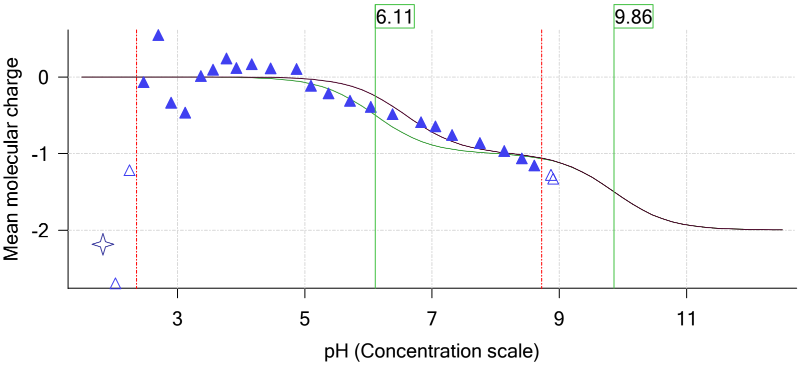

| | | | |
|---|---|---|---|
| Sample name: | **M13_octanol** | Experiment start time: | **3/26/2018 4:16:35 PM** |
| Assay name: | **pH-metric high logP** | Analyst: | **Pion** |
| Assay ID: | **18C-26010** | Instrument ID: | **T312060** |
| Filename: | **C:\Sirius_T3\Mehtap\20180327_exp34_logP_T3-2\18C-26010_M13_octanol_pH-metric high logP.t3r** | | |

### Assay Model

| Settings | Value | Date/Time changed | Imported from |
|---|---|---|---|
| Sample name | M13_octanol | 2/27/2018 5:57:49 PM | User entered value |
| Sample by | Weight | | Default value |
| Sample weight | 0.001260 g | 3/26/2018 4:15:03 PM | User entered value |
| Formula weight | 295.34 g/mol | 2/27/2018 5:57:49 PM | User entered value |
| Solubility | Unknown | | Default value |
| Molecular weight | 295.34 | 2/27/2018 5:57:49 PM | User entered value |
| Individual pKa ionic environments | No | | Default value |
| Number of pKas | 1 | 2/27/2018 5:57:49 PM | User entered value |
| Sample is a | Base | 2/27/2018 5:57:49 PM | User entered value |
| pKa 1 | 5.77 | 2/27/2018 5:57:49 PM | User entered value |
| logp (XH +) | -4.56 | 3/2/2018 4:30:48 PM | User entered value |
| logP (neutral X) | 2.99 | 3/2/2018 4:30:43 PM | User entered value |

### Events

| Time | Event | Water | Acid | Base | Octanol | pH | dpH/dt | pH R-squared | pH SD | dpH/d time |
|---|---|---|---|---|---|---|---|---|---|---|
| 5:17.0 | Manual volume addition | | | | 0.20000 mL | | | | | |
| 5:18.1 | Initial pH = 3.49 | | | | | | | | | |
| 8:22.9 | Data point 2 | 1.50000 mL | 0.04473 mL | 0.00553 mL | 0.20000 mL | 2.003 | -0.00959 | 0.73625 | 0.00055 | 10.5 s |
| 9:09.4 | Data point 3 | 1.50000 mL | 0.04473 mL | 0.02016 mL | 0.20000 mL | 2.209 | 0.00564 | 0.12526 | 0.00079 | 10.0 s |
| 9:45.0 | Data point 4 | 1.50000 mL | 0.04473 mL | 0.02970 mL | 0.20000 mL | 2.418 | 0.01231 | 0.87847 | 0.00065 | 10.5 s |
| 10:21.0 | Data point 5 | 1.50000 mL | 0.04473 mL | 0.03568 mL | 0.20000 mL | 2.638 | 0.00365 | 0.22847 | 0.00038 | 10.5 s |
| 10:57.1 | Data point 6 | 1.50000 mL | 0.04473 mL | 0.03944 mL | 0.20000 mL | 2.855 | -0.00035 | 0.00242 | 0.00035 | 10.0 s |
| 11:32.6 | Data point 7 | 1.50000 mL | 0.04473 mL | 0.04198 mL | 0.20000 mL | 3.087 | 0.00584 | 0.40770 | 0.00045 | 10.5 s |
| 12:08.5 | Data point 8 | 1.50000 mL | 0.04473 mL | 0.04384 mL | 0.20000 mL | 3.358 | -0.00076 | 0.00844 | 0.00041 | 10.0 s |
| 12:59.5 | Data point 9 | 1.50000 mL | 0.04473 mL | 0.04501 mL | 0.20000 mL | 3.555 | -0.00079 | 0.00582 | 0.00051 | 10.5 s |
| 13:45.7 | Data point 10 | 1.50000 mL | 0.04473 mL | 0.04603 mL | 0.20000 mL | 3.736 | -0.01231 | 0.48567 | 0.00087 | 10.5 s |
| 14:32.0 | Data point 11 | 1.50000 mL | 0.04473 mL | 0.04734 mL | 0.20000 mL | 3.933 | -0.00938 | 0.30311 | 0.00084 | 11.0 s |
| 15:08.5 | Data point 12 | 1.50000 mL | 0.04473 mL | 0.04842 mL | 0.20000 mL | 4.073 | -0.01778 | 0.77327 | 0.00100 | 13.5 s |
| 16:08.2 | Data point 13 | 1.50000 mL | 0.04473 mL | 0.04972 mL | 0.20000 mL | 4.259 | -0.01176 | 0.47760 | 0.00084 | 14.5 s |
| 16:58.3 | Data point 14 | 1.50000 mL | 0.04473 mL | 0.05045 mL | 0.20000 mL | 4.506 | 0.01843 | 0.89339 | 0.00096 | 21.0 s |
| 17:44.7 | Data point 15 | 1.50000 mL | 0.04473 mL | 0.05096 mL | 0.20000 mL | 4.931 | 0.01593 | 0.72074 | 0.00093 | 22.5 s |
| 18:37.6 | Data point 16 | 1.50000 mL | 0.04473 mL | 0.05172 mL | 0.20000 mL | 9.043 | -0.08412 | 0.99442 | 0.00416 | Time out at |
| 20:42.7 | Data point 17 | 1.50000 mL | 0.09854 mL | 0.05172 mL | 0.50000 mL | 1.945 | 0.00587 | 0.10684 | 0.00089 | 19.0 s |
| 21:38.0 | Data point 18 | 1.50000 mL | 0.09854 mL | 0.06919 mL | 0.50000 mL | 2.151 | -0.00296 | 0.10195 | 0.00046 | 10.0 s |
| 22:13.6 | Data point 19 | 1.50000 mL | 0.09854 mL | 0.08095 mL | 0.50000 mL | 2.366 | 0.00068 | 0.00245 | 0.00068 | 10.5 s |
| 22:49.6 | Data point 20 | 1.50000 mL | 0.09854 mL | 0.08826 mL | 0.50000 mL | 2.578 | -0.01356 | 0.80667 | 0.00075 | 10.0 s |

| | | | |
|---|---|---|---|
| Sample name: | **M13_octanol** | Experiment start time: | **3/26/2018 4:16:35 PM** |
| Assay name: | **pH-metric high logP** | Analyst: | **Pion** |
| Assay ID: | **18C-26010** | Instrument ID: | **T312060** |
| Filename: | **C:\Sirius_T3\Mehtap\20180327_exp34_logP_T3-2\18C-26010_M13_octanol_pH-metric high logP.t3r** | | |

### Events (continued)

| Time | Event | Water | Acid | Base | Octanol | pH | dpH/dt | pH R-squared | pH SD | dpH/dt time |
|---|---|---|---|---|---|---|---|---|---|---|
| 23:25.1 | Data point 21 | 1.50000 mL | 0.09854 mL | 0.09304 mL | 0.50000 mL | 2.792 | 0.00291 | 0.03033 | 0.00083 | 10.5 s |
| 24:01.1 | Data point 22 | 1.50000 mL | 0.09854 mL | 0.09631 mL | 0.50000 mL | 2.996 | -0.00625 | 0.16494 | 0.00076 | 10.0 s |
| 24:36.5 | Data point 23 | 1.50000 mL | 0.09854 mL | 0.09878 mL | 0.50000 mL | 3.202 | -0.00833 | 0.32865 | 0.00072 | 10.0 s |
| 25:11.7 | Data point 24 | 1.50000 mL | 0.09854 mL | 0.10073 mL | 0.50000 mL | 3.524 | -0.01622 | 0.91829 | 0.00084 | 10.5 s |
| 25:52.8 | Data point 25 | 1.50000 mL | 0.09854 mL | 0.10167 mL | 0.50000 mL | 3.706 | -0.01693 | 0.76864 | 0.00095 | 11.0 s |
| 26:44.5 | Data point 26 | 1.50000 mL | 0.09854 mL | 0.10261 mL | 0.50000 mL | 3.881 | -0.01256 | 0.44869 | 0.00093 | 12.0 s |
| 27:32.0 | Data point 27 | 1.50000 mL | 0.09854 mL | 0.10332 mL | 0.50000 mL | 4.048 | -0.01675 | 0.86339 | 0.00089 | 13.0 s |
| 28:20.7 | Data point 28 | 1.50000 mL | 0.09854 mL | 0.10383 mL | 0.50000 mL | 4.234 | -0.01972 | 0.95699 | 0.00100 | 10.0 s |
| 29:06.4 | Data point 29 | 1.50000 mL | 0.09854 mL | 0.10430 mL | 0.50000 mL | 4.495 | -0.01298 | 0.53508 | 0.00088 | 12.0 s |
| 29:43.8 | Data point 30 | 1.50000 mL | 0.09854 mL | 0.10459 mL | 0.50000 mL | 4.751 | 0.01337 | 0.62718 | 0.00083 | 23.5 s |
| 30:37.9 | Data point 31 | 1.50000 mL | 0.09854 mL | 0.10482 mL | 0.50000 mL | 5.045 | -0.01188 | 0.50324 | 0.00083 | 12.5 s |
| 31:21.0 | Data point 32 | 1.50000 mL | 0.09854 mL | 0.10499 mL | 0.50000 mL | 5.337 | -0.01679 | 0.75428 | 0.00095 | 13.5 s |
| 32:05.0 | Data point 33 | 1.50000 mL | 0.09854 mL | 0.10515 mL | 0.50000 mL | 6.712 | -0.05259 | 0.99413 | 0.00261 | Timed out at 59.5 s |
| 33:35.6 | Data point 34 | 1.50000 mL | 0.09854 mL | 0.10529 mL | 0.50000 mL | 7.882 | -0.07582 | 0.99362 | 0.00376 | Timed out at 59.5 s |
| 35:06.1 | Data point 35 | 1.50000 mL | 0.09854 mL | 0.10536 mL | 0.50000 mL | 7.874 | -0.03303 | 0.99176 | 0.00164 | Timed out at 59.5 s |
| 36:36.6 | Data point 36 | 1.50000 mL | 0.09854 mL | 0.10541 mL | 0.50000 mL | 7.975 | -0.02481 | 0.97940 | 0.00124 | Timed out at 59.5 s |
| 38:07.1 | Data point 37 | 1.50000 mL | 0.09854 mL | 0.10546 mL | 0.50000 mL | 8.262 | -0.01947 | 0.94137 | 0.00099 | 50.5 s |
| 39:28.1 | Data point 38 | 1.50000 mL | 0.09854 mL | 0.10550 mL | 0.50000 mL | 8.569 | -0.01705 | 0.77622 | 0.00096 | 34.0 s |
| 40:37.7 | Data point 39 | 1.50000 mL | 0.09854 mL | 0.10557 mL | 0.50000 mL | 8.843 | -0.01363 | 0.52717 | 0.00093 | 24.5 s |
| 41:37.8 | Data point 40 | 1.50000 mL | 0.09854 mL | 0.10564 mL | 0.50000 mL | 9.039 | -0.01553 | 0.70869 | 0.00091 | 19.0 s |
| 43:42.7 | Data point 41 | 1.50000 mL | 0.15644 mL | 0.10564 mL | 1.20000 mL | 1.964 | -0.01340 | 0.57402 | 0.00087 | 10.0 s |
| 44:28.9 | Data point 42 | 1.50000 mL | 0.15644 mL | 0.12453 mL | 1.20000 mL | 2.154 | 0.00096 | 0.01426 | 0.00040 | 10.5 s |
| 45:05.1 | Data point 43 | 1.50000 mL | 0.15644 mL | 0.13735 mL | 1.20000 mL | 2.370 | 0.01091 | 0.34363 | 0.00092 | 18.0 s |
| 45:48.7 | Data point 44 | 1.50000 mL | 0.15644 mL | 0.14532 mL | 1.20000 mL | 2.589 | -0.00436 | 0.09759 | 0.00069 | 10.0 s |
| 46:24.3 | Data point 45 | 1.50000 mL | 0.15644 mL | 0.15061 mL | 1.20000 mL | 2.818 | -0.00140 | 0.02683 | 0.00042 | 10.5 s |
| 47:00.3 | Data point 46 | 1.50000 mL | 0.15644 mL | 0.15419 mL | 1.20000 mL | 3.015 | -0.01292 | 0.43338 | 0.00097 | 10.0 s |
| 47:35.8 | Data point 47 | 1.50000 mL | 0.15644 mL | 0.15682 mL | 1.20000 mL | 3.235 | -0.00551 | 0.33990 | 0.00047 | 10.0 s |
| 48:11.2 | Data point 48 | 1.50000 mL | 0.15644 mL | 0.15870 mL | 1.20000 mL | 3.480 | -0.00486 | 0.44316 | 0.00036 | 10.0 s |
| 48:57.0 | Data point 49 | 1.50000 mL | 0.15644 mL | 0.15988 mL | 1.20000 mL | 3.672 | -0.01560 | 0.96977 | 0.00078 | 10.0 s |
| 49:32.4 | Data point 50 | 1.50000 mL | 0.15644 mL | 0.16079 mL | 1.20000 mL | 3.881 | 0.00315 | 0.02822 | 0.00093 | 18.5 s |
| 50:16.3 | Data point 51 | 1.50000 mL | 0.15644 mL | 0.16138 mL | 1.20000 mL | 4.036 | -0.01090 | 0.30494 | 0.00098 | 12.0 s |
| 50:58.8 | Data point 52 | 1.50000 mL | 0.15644 mL | 0.16195 mL | 1.20000 mL | 4.277 | -0.01737 | 0.75358 | 0.00099 | 10.0 s |
| 51:39.4 | Data point 53 | 1.50000 mL | 0.15644 mL | 0.16239 mL | 1.20000 mL | 4.574 | -0.01819 | 0.84303 | 0.00098 | 10.0 s |
| 52:19.8 | Data point 54 | 1.50000 mL | 0.15644 mL | 0.16268 mL | 1.20000 mL | 4.979 | -0.01369 | 0.66561 | 0.00083 | 10.0 s |
| 53:05.6 | Data point 55 | 1.50000 mL | 0.15644 mL | 0.16284 mL | 1.20000 mL | 5.207 | -0.00278 | 0.09353 | 0.00045 | 10.0 s |
| 53:51.2 | Data point 56 | 1.50000 mL | 0.15644 mL | 0.16294 mL | 1.20000 mL | 5.479 | -0.00914 | 0.49731 | 0.00064 | 10.0 s |
| 54:36.8 | Data point 57 | 1.50000 mL | 0.15644 mL | 0.16301 mL | 1.20000 mL | 5.816 | -0.01695 | 0.93689 | 0.00086 | 10.5 s |
| 55:17.8 | Data point 58 | 1.50000 mL | 0.15644 mL | 0.16305 mL | 1.20000 mL | 6.145 | -0.00680 | 0.14161 | 0.00089 | 20.5 s |
| 56:08.7 | Data point 59 | 1.50000 mL | 0.15644 mL | 0.16310 mL | 1.20000 mL | 6.484 | -0.01501 | 0.62688 | 0.00094 | 33.5 s |
| 57:12.7 | Data point 60 | 1.50000 mL | 0.15644 mL | 0.16315 mL | 1.20000 mL | 6.929 | -0.01950 | 0.93593 | 0.00100 | 59.0 s |
| 58:37.2 | Data point 61 | 1.50000 mL | 0.15644 mL | 0.16317 mL | 1.20000 mL | 7.157 | -0.01773 | 0.81355 | 0.00097 | 42.0 s |
| 59:49.7 | Data point 62 | 1.50000 mL | 0.15644 mL | 0.16322 mL | 1.20000 mL | 7.418 | -0.02771 | 0.99293 | 0.00137 | Timed out at 59.5 s |
| 1:01:20.3 | Data point 63 | 1.50000 mL | 0.15644 mL | 0.16326 mL | 1.20000 mL | 7.854 | -0.03775 | 0.96659 | 0.00190 | Timed out at 59.5 s |
| 1:02:50.9 | Data point 64 | 1.50000 mL | 0.15644 mL | 0.16331 mL | 1.20000 mL | 8.232 | -0.01636 | 0.69169 | 0.00097 | 42.0 s |
| 1:04:03.3 | Data point 65 | 1.50000 mL | 0.15644 mL | 0.16336 mL | 1.20000 mL | 8.510 | -0.01437 | 0.53636 | 0.00097 | 35.0 s |
| 1:05:08.9 | Data point 66 | 1.50000 mL | 0.15644 mL | 0.16340 mL | 1.20000 mL | 8.705 | -0.01715 | 0.74002 | 0.00098 | 28.5 s |
| 1:06:13.2 | Data point 67 | 1.50000 mL | 0.15644 mL | 0.16348 mL | 1.20000 mL | 8.966 | -0.00089 | 0.00198 | 0.00099 | 28.0 s |
| 1:07:06.6 | Data point 68 | 1.50000 mL | 0.15644 mL | 0.16350 mL | 1.20000 mL | 9.001 | 0.00342 | 0.03245 | 0.00094 | 26.5 s |
| 1:07:42.2 | Assay volumes | 1.50000 mL | 0.15644 mL | 0.16350 mL | 1.20000 mL | | | | | |

| | | | |
|---|---|---|---|
| Sample name: | **M13_octanol** | Experiment start time: | **3/26/2018 4:16:35 PM** |
| Assay name: | **pH-metric high logP** | Analyst: | **Pion** |
| Assay ID: | **18C-26010** | Instrument ID: | **T312060** |
| Filename: | **C:\Sirius_T3\Mehtap\20180327_exp34_logP_T3-2\18C-26010_M13_octanol_pH-metric high logP.t3r** | | |

| | | | |
|---|---|---|---|
| Sample name: | **M13_octanol** | Experiment start time: | **3/26/2018 4:16:35 PM** |
| Assay name: | **pH-metric high logP** | Analyst: | **Pion** |
| Assay ID: | **18C-26010** | Instrument ID: | **T312060** |
| Filename: | **C:\Sirius_T3\Mehtap\20180327_exp34_logP_T3-2\18C-26010_M13_octanol_pH-metric high logP.t3r** | | |

### Assay Settings (continued)

| Setting | Value | Original Value | Date/Time changed | Imported from |
|---|---|---|---|---|
| ***Titration 3*** | | | | |
| Titrate from | Low to high pH | | | |
| Add additional water | 0.00 mL | | | |
| Additional partition solvent volume | 0.700 mL | | | |
| Additional partition solvent added | Automatic | | | |
| After pH adjust stir for | 30 seconds | | | |
| Stir to allow partitioning for | 15 seconds | | | |
| Stirrer speed for partitioning | 60% | | | |
| ***Data Point Stability*** | | | | |
| Stir during data point collection | No | | | |
| Delay before data point collection | 0 seconds | | | |
| Number of points to average | 20 points | | | |
| Time interval between points | 0.50 seconds | | | |
| Required maximum standard deviation | 0.00100 dpH/dt | | | |
| Stability timeout after | 60 seconds | | | |

### Calibration Settings

| Setting | Value | Date/Time changed | Imported from |
|---|---|---|---|
| Four-Plus alpha | 0.122 | 3/26/2018 4:16:34 PM | C:\Sirius_T3\18C-26006_Blank standardisation.t3r |
| Four-Plus S | 0.9974 | 3/26/2018 4:16:34 PM | C:\Sirius_T3\18C-26006_Blank standardisation.t3r |
| Four-Plus jH | 1.1 | 3/26/2018 4:16:34 PM | C:\Sirius_T3\18C-26006_Blank standardisation.t3r |
| Four-Plus jOH | -0.5 | 3/26/2018 4:16:34 PM | C:\Sirius_T3\18C-26006_Blank standardisation.t3r |
| Base concentration factor | 1.003 | 3/26/2018 4:16:35 PM | C:\Sirius_T3\KOH18C23.t3r |
| Acid concentration factor | 0.994 | 3/26/2018 4:16:34 PM | C:\Sirius_T3\18C-26006_Blank standardisation.t3r |

### Instrument Settings

| Setting | Value | Batch Id | Install date |
|---|---|---|---|
| Instrument owner | Merck | | |
| Instrument ID | T312060 | | |
| Instrument type | T3 Simulator | | |
| Software version | 1.1.3.0 | | |
| Dispenser module | | T3DM1200361 | 3/31/2009 6:24:52 AM |
| Dispenser 0 | Water | | 3/31/2009 6:25:05 AM |
| Syringe volume | 2.5 mL | | |
| Firmware version | 1.2.1(r2) | | |
| Titrant | Water (0.15 M KCl) | 02-06-2018 | 3/16/2018 11:09:18 AM |
| Dispenser 2 | Acid | | 3/31/2009 6:25:11 AM |
| Syringe volume | 0.5 mL | | |
| Firmware version | 1.2.1(r2) | | |
| Titrant | Acid (0.5 M HCl) | 03-16-2018 | 3/16/2018 10:56:23 AM |
| Dispenser 1 | Base | | 3/31/2009 6:25:21 AM |
| Syringe volume | 0.5 mL | | |
| Firmware version | 1.2.1(r2) | | |
| Titrant | Base (0.5 M KOH) | 3/22/2018 | 3/23/2018 9:34:17 AM |
| Dispenser 5 | Cosolvent | | 3/31/2009 6:26:24 AM |
| Syringe volume | 2.5 mL | | |
| Firmware version | 1.2.1(r2) | | |
| Distribution valve 5 | Distribution Valve | | 3/31/2009 6:28:19 AM |
| Firmware version | 1.1.3 | | |
| Port A | Methanol (80%, 0.15 M KCl) | 02-08-2018 | 3/6/2018 10:28:59 AM |
| Port B | Cyclohexane | 11-01-17 | 2/27/2018 11:37:57 AM |
| Dispenser 3 | Buffer | | 8/3/2010 6:05:16 AM |
| Syringe volume | 0.5 mL | | |
| Firmware version | 1.2.1(r2) | | |
| Titrant | Dodecane | 2018/01/31 | 2/28/2018 11:18:04 AM |
| Dispenser 6 | Octanol | | 10/22/2010 11:52:43 AM |

| | | | |
|---|---|---|---|
| Sample name: | **M13_octanol** | Experiment start time: | **3/26/2018 4:16:35 PM** |
| Assay name: | **pH-metric high logP** | Analyst: | **Pion** |
| Assay ID: | **18C-26010** | Instrument ID: | **T312060** |
| Filename: | **C:\Sirius_T3\Mehtap\20180327_exp34_logP_T3-2\18C-26010_M13_octanol_pH-metric high logP.t3r** | | |

| | | | |
|---|---|---|---|
| Sample name: | **M13_octanol** | Experiment start time: | **3/26/2018 4:16:35 PM** |
| Assay name: | **pH-metric high logP** | Analyst: | **Pion** |
| Assay ID: | **18C-26010** | Instrument ID: | **T312060** |
| Filename: | **C:\Sirius_T3\Mehtap\20180327_exp34_logP_T3-2\18C-26010_M13_octanol_pH-metric high logP.t3r** | | |

### Instrument Settings (continued)

| Setting | Value | Batch Id | Install date |
|---|---|---|---|
| Spectrometer calibration stir duration | 5 s | | |
| Spectrometer calibration stir speed | 30% | | |
| Spectrometer calibration wash pump volume | 20.0 mL | | |
| Spectrometer calibration wash stir duration | 5 s | | |
| Spectrometer calibration wash stir speed | 30% | | |
| Overhead dispense height | 10000 | | |

### Refinement Settings

| Setting | Value | Default value |
|---|---|---|
| Turbidity detection method | None | None |
| Turbidity wavelength to assess | 500.0 nm | 500.0 nm |
| Turbidity maximum absorbance | 0.100 | 0.100 |
| Turbidity probe threshold | 50.00 | 50.00 |

### Experiment Log

[2:41] Air gap created for Water (0.15 M KCl)
[2:41] Air gap created for Acid (0.5 M HCl)
[2:41] Air gap created for Base (0.5 M KOH)
[2:42] Air gap released for Water (0.15 M KCl)
[2:46] Titrator arm moved over Titration position
[2:46] Titration 1 of 3
[2:46] Adding initial titrants
[2:46] Automatically add 1.50000 mL of water
[3:11] Dispensed 1.500000 mL of Water (0.15 M KCl)
[3:15] Titrator arm moved over Drain
[4:56] Titrator arm moved to Titration position
[4:56] Argon flow rate set to 100
[4:56] Stirrer speed set to 10
[5:18] Initial pH = 3.49
[5:18] Iterative adjust 3.49 -> 2.00
[5:18] pH 3.49 -> 2.00
[5:19] Air gap released for Acid (0.5 M HCl)
[5:20] Dispensed 0.042404 mL of Acid (0.5 M HCl)
[5:25] pH 2.02 -> 2.00
[5:25] Dispensed 0.002328 mL of Acid (0.5 M HCl)
[5:30] Holding pH 2.00
[7:30] Stirrer speed set to 0
[7:30] Stirrer speed set to 50
[7:30] Iterative adjust 1.95 -> 2.00
[7:30] pH 1.95 -> 2.00
[7:31] Air gap released for Base (0.5 M KOH)
[7:32] Dispensed 0.005527 mL of Base (0.5 M KOH)
[8:22] Stirrer speed set to 0
[8:33] Datapoint id 2 collected
[8:33] Stirrer speed set to 50
[8:38] pH 2.01 -> 2.21
[8:38] Using cautious pH adjust
[8:38] Dispensed 0.007808 mL of Base (0.5 M KOH)
[8:43] Stepping pH = 2.09
[8:44] Dispensed 0.006068 mL of Base (0.5 M KOH)
[8:49] Stepping pH = 2.19
[8:49] Dispensed 0.000753 mL of Base (0.5 M KOH)
[8:54] Stepping pH = 2.21
[9:09] Stirrer speed set to 0
[9:19] Datapoint id 3 collected
[9:19] Charge balance equation is out by 6.4%
[9:19] Stirrer speed set to 50

| | | | |
|---|---|---|---|
| Sample name: | **M13_octanol** | Experiment start time: | **3/26/2018 4:16:35 PM** |
| Assay name: | **pH-metric high logP** | Analyst: | **Pion** |
| Assay ID: | **18C-26010** | Instrument ID: | **T312060** |
| Filename: | **C:\Sirius_T3\Mehtap\20180327_exp34_logP_T3-2\18C-26010_M13_octanol_pH-metric high logP.t3r** | | |

### Experiment Log (continued)

[9:24] pH 2.22 -> 2.42
[9:24] Using charge balance adjust
[9:24] Dispensed 0.009548 mL of Base (0.5 M KOH)
[9:45] Stirrer speed set to 0
[9:55] Datapoint id 4 collected
[9:55] Charge balance equation is out by -0.4%
[9:55] Stirrer speed set to 50
[10:00] pH 2.43 -> 2.63
[10:00] Using charge balance adjust
[10:00] Dispensed 0.005974 mL of Base (0.5 M KOH)
[10:21] Stirrer speed set to 0
[10:31] Datapoint id 5 collected
[10:31] Charge balance equation is out by 5.5%
[10:31] Stirrer speed set to 50
[10:36] pH 2.65 -> 2.85
[10:36] Using charge balance adjust
[10:36] Dispensed 0.003763 mL of Base (0.5 M KOH)
[10:57] Stirrer speed set to 0
[11:07] Datapoint id 6 collected
[11:07] Charge balance equation is out by 4.1%
[11:07] Stirrer speed set to 50
[11:12] pH 2.87 -> 3.07
[11:12] Using charge balance adjust
[11:12] Dispensed 0.002540 mL of Base (0.5 M KOH)
[11:32] Stirrer speed set to 0
[11:43] Datapoint id 7 collected
[11:43] Charge balance equation is out by 10.5%
[11:43] Stirrer speed set to 50
[11:48] pH 3.10 -> 3.30
[11:48] Using charge balance adjust
[11:48] Dispensed 0.001858 mL of Base (0.5 M KOH)
[12:08] Stirrer speed set to 0
[12:18] Datapoint id 8 collected
[12:18] Charge balance equation is out by 30.9%
[12:18] Stirrer speed set to 50
[12:23] pH 3.36 -> 3.56
[12:23] Using cautious pH adjust
[12:23] Dispensed 0.000753 mL of Base (0.5 M KOH)
[12:28] Stepping pH = 3.52
[12:29] Dispensed 0.000212 mL of Base (0.5 M KOH)
[12:34] Stepping pH = 3.55
[12:34] Dispensed 0.000094 mL of Base (0.5 M KOH)
[12:39] Stepping pH = 3.55
[12:39] Dispensed 0.000118 mL of Base (0.5 M KOH)
[12:44] Stepping pH = 3.57
[12:59] Stirrer speed set to 0
[13:10] Datapoint id 9 collected
[13:10] Charge balance equation is out by 23.3%
[13:10] Stirrer speed set to 50
[13:15] pH 3.56 -> 3.76
[13:15] Using cautious pH adjust
[13:15] Dispensed 0.000682 mL of Base (0.5 M KOH)
[13:20] Stepping pH = 3.73
[13:20] Dispensed 0.000141 mL of Base (0.5 M KOH)
[13:25] Stepping pH = 3.74
[13:25] Dispensed 0.000188 mL of Base (0.5 M KOH)
[13:30] Stepping pH = 3.77
[13:45] Stirrer speed set to 0
[13:56] Datapoint id 10 collected

Sample name: **M13_octanol**
Assay name: **pH-metric high logP**
Assay ID: **18C-26010**
Filename: **C:\Sirius_T3\Mehtap\20180327_exp34_logP_T3-2\18C-26010_M13_octanol_pH-metric high logP.t3r**
Experiment start time: **3/26/2018 4:16:35 PM**
Analyst: **Pion**
Instrument ID: **T312060**

### Experiment Log (continued)

[13:56] Charge balance equation is out by 25.9%
[13:56] Stirrer speed set to 50
[14:01] pH 3.74 -> 3.94
[14:01] Using cautious pH adjust
[14:01] Dispensed 0.000635 mL of Base (0.5 M KOH)
[14:06] Stepping pH = 3.89
[14:06] Dispensed 0.000188 mL of Base (0.5 M KOH)
[14:11] Stepping pH = 3.90
[14:11] Dispensed 0.000494 mL of Base (0.5 M KOH)
[14:16] Stepping pH = 4.00
[14:32] Stirrer speed set to 0
[14:43] Datapoint id 11 collected
[14:43] Charge balance equation is out by -4.2%
[14:43] Stirrer speed set to 50
[14:48] pH 3.94 -> 4.14
[14:48] Using charge balance adjust
[14:48] Dispensed 0.001082 mL of Base (0.5 M KOH)
[15:08] Stirrer speed set to 0
[15:22] Datapoint id 12 collected
[15:22] Charge balance equation is out by -32.8%
[15:22] Stirrer speed set to 50
[15:27] pH 4.08 -> 4.28
[15:27] Using cautious pH adjust
[15:27] Dispensed 0.000470 mL of Base (0.5 M KOH)
[15:32] Stepping pH = 4.19
[15:32] Dispensed 0.000259 mL of Base (0.5 M KOH)
[15:37] Stepping pH = 4.23
[15:37] Dispensed 0.000212 mL of Base (0.5 M KOH)
[15:42] Stepping pH = 4.26
[15:42] Dispensed 0.000094 mL of Base (0.5 M KOH)
[15:47] Stepping pH = 4.26
[15:48] Dispensed 0.000259 mL of Base (0.5 M KOH)
[15:53] Stepping pH = 4.34
[16:08] Stirrer speed set to 0
[16:22] Datapoint id 13 collected
[16:22] Charge balance equation is out by -36.9%
[16:22] Stirrer speed set to 50
[16:27] pH 4.26 -> 4.46
[16:27] Using cautious pH adjust
[16:27] Dispensed 0.000376 mL of Base (0.5 M KOH)
[16:33] Stepping pH = 4.41
[16:33] Dispensed 0.000094 mL of Base (0.5 M KOH)
[16:38] Stepping pH = 4.41
[16:38] Dispensed 0.000259 mL of Base (0.5 M KOH)
[16:43] Stepping pH = 4.58
[16:58] Stirrer speed set to 0
[17:19] Datapoint id 14 collected
[17:19] Charge balance equation is out by -0.5%
[17:19] Stirrer speed set to 50
[17:24] pH 4.50 -> 4.70
[17:24] Using charge balance adjust
[17:24] Dispensed 0.000517 mL of Base (0.5 M KOH)
[17:44] Stirrer speed set to 0
[18:07] Datapoint id 15 collected
[18:07] Charge balance equation is out by 114.9%
[18:07] Stirrer speed set to 50
[18:12] pH 4.88 -> 5.08
[18:12] Using cautious pH adjust
[18:12] Dispensed 0.000118 mL of Base (0.5 M KOH)

| | | | |
|---|---|---|---|
| Sample name: | **M13_octanol** | Experiment start time: | **3/26/2018 4:16:35 PM** |
| Assay name: | **pH-metric high logP** | Analyst: | **Pion** |
| Assay ID: | **18C-26010** | Instrument ID: | **T312060** |
| Filename: | **C:\Sirius_T3\Mehtap\20180327_exp34_logP_T3-2\18C-26010_M13_octanol_pH-metric high logP.t3r** | | |

### Experiment Log (continued)

[18:17] Stepping pH = 4.87
[18:17] Dispensed 0.000635 mL of Base (0.5 M KOH)
[18:22] Stepping pH = 9.43
[18:37] Stirrer speed set to 0
[19:37] Datapoint id 16 collected
[19:37] Charge balance equation is out by -207.2%
[19:37] Titration 2 of 3
[19:37] Adding initial titrants
[19:37] Automatically add 0.30000 mL of Octanol
[19:44] Dispensed 0.300000 mL of Octanol
[19:44] Stirrer speed set to 10
[19:45] Stirrer speed set to 55
[19:45] Iterative adjust 9.03 -> 2.00
[19:45] pH 9.03 -> 2.00
[19:47] Dispensed 0.052140 mL of Acid (0.5 M HCl)
[19:52] pH 2.02 -> 2.00
[19:52] Dispensed 0.001670 mL of Acid (0.5 M HCl)
[20:42] Stirrer speed set to 0
[21:01] Datapoint id 17 collected
[21:01] Stirrer speed set to 55
[21:06] pH 1.95 -> 2.15
[21:06] Using cautious pH adjust
[21:07] Dispensed 0.009525 mL of Base (0.5 M KOH)
[21:12] Stepping pH = 2.05
[21:12] Dispensed 0.006279 mL of Base (0.5 M KOH)
[21:17] Stepping pH = 2.13
[21:17] Dispensed 0.001670 mL of Base (0.5 M KOH)
[21:22] Stepping pH = 2.15
[21:38] Stirrer speed set to 0
[21:48] Datapoint id 18 collected
[21:48] Charge balance equation is out by 8.3%
[21:48] Stirrer speed set to 55
[21:53] pH 2.16 -> 2.36
[21:53] Using charge balance adjust
[21:53] Dispensed 0.011759 mL of Base (0.5 M KOH)
[22:13] Stirrer speed set to 0
[22:24] Datapoint id 19 collected
[22:24] Charge balance equation is out by 2.3%
[22:24] Stirrer speed set to 55
[22:29] pH 2.38 -> 2.58
[22:29] Using charge balance adjust
[22:29] Dispensed 0.007314 mL of Base (0.5 M KOH)
[22:49] Stirrer speed set to 0
[22:59] Datapoint id 20 collected
[22:59] Charge balance equation is out by 0.9%
[22:59] Stirrer speed set to 55
[23:04] pH 2.59 -> 2.79
[23:04] Using charge balance adjust
[23:05] Dispensed 0.004774 mL of Base (0.5 M KOH)
[23:25] Stirrer speed set to 0
[23:35] Datapoint id 21 collected
[23:35] Charge balance equation is out by 2.4%
[23:35] Stirrer speed set to 55
[23:40] pH 2.81 -> 3.01
[23:40] Using charge balance adjust
[23:40] Dispensed 0.003269 mL of Base (0.5 M KOH)
[24:01] Stirrer speed set to 0
[24:11] Datapoint id 22 collected
[24:11] Charge balance equation is out by -4.9%

| | | | |
|---|---|---|---|
| Sample name: | **M13_octanol** | Experiment start time: | **3/26/2018 4:16:35 PM** |
| Assay name: | **pH-metric high logP** | Analyst: | **Pion** |
| Assay ID: | **18C-26010** | Instrument ID: | **T312060** |
| Filename: | **C:\Sirius_T3\Mehtap\20180327_exp34_logP_T3-2\18C-26010_M13_octanol_pH-metric high logP.t3r** | | |

### Experiment Log (continued)

[24:11] Stirrer speed set to 55
[24:16] pH 3.01 -> 3.21
[24:16] Using charge balance adjust
[24:16] Dispensed 0.002469 mL of Base (0.5 M KOH)
[24:36] Stirrer speed set to 0
[24:46] Datapoint id 23 collected
[24:46] Charge balance equation is out by -3.0%
[24:46] Stirrer speed set to 55
[24:51] pH 3.21 -> 3.41
[24:51] Using charge balance adjust
[24:51] Dispensed 0.001952 mL of Base (0.5 M KOH)
[25:11] Stirrer speed set to 0
[25:22] Datapoint id 24 collected
[25:22] Charge balance equation is out by 54.8%
[25:22] Stirrer speed set to 55
[25:27] pH 3.53 -> 3.73
[25:27] Using cautious pH adjust
[25:27] Dispensed 0.000706 mL of Base (0.5 M KOH)
[25:32] Stepping pH = 3.68
[25:32] Dispensed 0.000235 mL of Base (0.5 M KOH)
[25:37] Stepping pH = 3.73
[25:52] Stirrer speed set to 0
[26:03] Datapoint id 25 collected
[26:03] Charge balance equation is out by 34.0%
[26:03] Stirrer speed set to 55
[26:08] pH 3.72 -> 3.92
[26:08] Using cautious pH adjust
[26:09] Dispensed 0.000564 mL of Base (0.5 M KOH)
[26:14] Stepping pH = 3.85
[26:14] Dispensed 0.000188 mL of Base (0.5 M KOH)
[26:19] Stepping pH = 3.89
[26:19] Dispensed 0.000094 mL of Base (0.5 M KOH)
[26:24] Stepping pH = 3.90
[26:24] Dispensed 0.000094 mL of Base (0.5 M KOH)
[26:29] Stepping pH = 3.92
[26:44] Stirrer speed set to 0
[26:56] Datapoint id 26 collected
[26:56] Charge balance equation is out by 15.5%
[26:56] Stirrer speed set to 55
[27:01] pH 3.89 -> 4.09
[27:01] Using cautious pH adjust
[27:01] Dispensed 0.000447 mL of Base (0.5 M KOH)
[27:06] Stepping pH = 4.03
[27:06] Dispensed 0.000141 mL of Base (0.5 M KOH)
[27:11] Stepping pH = 4.06
[27:11] Dispensed 0.000118 mL of Base (0.5 M KOH)
[27:16] Stepping pH = 4.09
[27:32] Stirrer speed set to 0
[27:45] Datapoint id 27 collected
[27:45] Charge balance equation is out by 19.9%
[27:45] Stirrer speed set to 55
[27:50] pH 4.06 -> 4.26
[27:50] Using cautious pH adjust
[27:50] Dispensed 0.000329 mL of Base (0.5 M KOH)
[27:55] Stepping pH = 4.19
[27:55] Dispensed 0.000118 mL of Base (0.5 M KOH)
[28:00] Stepping pH = 4.23
[28:00] Dispensed 0.000071 mL of Base (0.5 M KOH)
[28:05] Stepping pH = 4.25

Sample name: **M13_octanol**
Assay name: **pH-metric high logP**
Assay ID: **18C-26010**
Filename: **C:\Sirius_T3\Mehtap\20180327_exp34_logP_T3-2\18C-26010_M13_octanol_pH-metric high logP.t3r**
Experiment start time: **3/26/2018 4:16:35 PM**
Analyst: **Pion**
Instrument ID: **T312060**

### Experiment Log (continued)

[28:20] Stirrer speed set to 0
[28:30] Datapoint id 28 collected
[28:30] Charge balance equation is out by 21.2%
[28:30] Stirrer speed set to 55
[28:35] pH 4.24 -> 4.44
[28:35] Using cautious pH adjust
[28:35] Dispensed 0.000235 mL of Base (0.5 M KOH)
[28:41] Stepping pH = 4.39
[28:41] Dispensed 0.000071 mL of Base (0.5 M KOH)
[28:46] Stepping pH = 4.39
[28:46] Dispensed 0.000165 mL of Base (0.5 M KOH)
[28:51] Stepping pH = 4.52
[29:06] Stirrer speed set to 0
[29:18] Datapoint id 29 collected
[29:18] Charge balance equation is out by -0.7%
[29:18] Stirrer speed set to 55
[29:23] pH 4.50 -> 4.70
[29:23] Using charge balance adjust
[29:23] Dispensed 0.000282 mL of Base (0.5 M KOH)
[29:43] Stirrer speed set to 0
[30:07] Datapoint id 30 collected
[30:07] Charge balance equation is out by 24.6%
[30:07] Stirrer speed set to 55
[30:12] pH 4.75 -> 4.95
[30:12] Using cautious pH adjust
[30:12] Dispensed 0.000094 mL of Base (0.5 M KOH)
[30:17] Stepping pH = 4.80
[30:17] Dispensed 0.000141 mL of Base (0.5 M KOH)
[30:22] Stepping pH = 5.09
[30:37] Stirrer speed set to 0
[30:50] Datapoint id 31 collected
[30:50] Charge balance equation is out by -26.9%
[30:50] Stirrer speed set to 55
[30:55] pH 5.05 -> 5.25
[30:55] Using cautious pH adjust
[30:55] Dispensed 0.000047 mL of Base (0.5 M KOH)
[31:00] Stepping pH = 5.07
[31:00] Dispensed 0.000118 mL of Base (0.5 M KOH)
[31:05] Stepping pH = 5.37
[31:21] Stirrer speed set to 0
[31:34] Datapoint id 32 collected
[31:34] Charge balance equation is out by -86.8%
[31:34] Stirrer speed set to 55
[31:39] pH 5.41 -> 5.61
[31:39] Using cautious pH adjust
[31:39] Dispensed 0.000024 mL of Base (0.5 M KOH)
[31:44] Stepping pH = 5.40
[31:44] Dispensed 0.000141 mL of Base (0.5 M KOH)
[31:49] Stepping pH = 6.97
[32:05] Stirrer speed set to 0
[33:05] Datapoint id 33 collected
[33:05] Charge balance equation is out by -201.9%
[33:05] Stirrer speed set to 55
[33:10] pH 6.45 -> 6.65
[33:10] Using cautious pH adjust
[33:10] Dispensed 0.000024 mL of Base (0.5 M KOH)
[33:15] Stepping pH = 6.42
[33:15] Dispensed 0.000118 mL of Base (0.5 M KOH)
[33:20] Stepping pH = 8.04

### Experiment Log (continued)

[33:35] Stirrer speed set to 0
[34:35] Datapoint id 34 collected
[34:35] Charge balance equation is out by -241.6%
[34:35] Stirrer speed set to 55
[34:40] pH 7.57 -> 7.77
[34:40] Using cautious pH adjust
[34:40] Dispensed 0.000024 mL of Base (0.5 M KOH)
[34:45] Stepping pH = 7.53
[34:45] Dispensed 0.000047 mL of Base (0.5 M KOH)
[34:51] Stepping pH = 7.84
[35:06] Stirrer speed set to 0
[36:06] Datapoint id 35 collected
[36:06] Charge balance equation is out by -393.8%
[36:06] Stirrer speed set to 55
[36:11] pH 7.75 -> 7.95
[36:11] Using cautious pH adjust
[36:11] Dispensed 0.000024 mL of Base (0.5 M KOH)
[36:16] Stepping pH = 7.80
[36:16] Dispensed 0.000024 mL of Base (0.5 M KOH)
[36:21] Stepping pH = 8.02
[36:36] Stirrer speed set to 0
[37:36] Datapoint id 36 collected
[37:36] Charge balance equation is out by -332.8%
[37:36] Stirrer speed set to 55
[37:41] pH 8.03 -> 8.23
[37:41] Using cautious pH adjust
[37:41] Dispensed 0.000024 mL of Base (0.5 M KOH)
[37:46] Stepping pH = 8.20
[37:46] Dispensed 0.000024 mL of Base (0.5 M KOH)
[37:52] Stepping pH = 8.36
[38:07] Stirrer speed set to 0
[38:57] Datapoint id 37 collected
[38:57] Charge balance equation is out by -361.1%
[38:57] Stirrer speed set to 55
[39:02] pH 8.24 -> 8.44
[39:02] Using cautious pH adjust
[39:02] Dispensed 0.000024 mL of Base (0.5 M KOH)
[39:07] Stepping pH = 8.35
[39:07] Dispensed 0.000024 mL of Base (0.5 M KOH)
[39:13] Stepping pH = 8.60
[39:28] Stirrer speed set to 0
[40:02] Datapoint id 38 collected
[40:02] Charge balance equation is out by -280.6%
[40:02] Stirrer speed set to 55
[40:07] pH 8.66 -> 8.86
[40:07] Using cautious pH adjust
[40:07] Dispensed 0.000024 mL of Base (0.5 M KOH)
[40:12] Stepping pH = 8.70
[40:12] Dispensed 0.000024 mL of Base (0.5 M KOH)
[40:17] Stepping pH = 8.80
[40:17] Dispensed 0.000024 mL of Base (0.5 M KOH)
[40:22] Stepping pH = 8.89
[40:37] Stirrer speed set to 0
[41:02] Datapoint id 39 collected
[41:02] Charge balance equation is out by -172.2%
[41:02] Stirrer speed set to 55
[41:07] pH 8.88 -> 9.05
[41:07] Using cautious pH adjust
[41:07] Dispensed 0.000024 mL of Base (0.5 M KOH)

Sample name: **M13_octanol** Experiment start time: **3/26/2018 4:16:35 PM**
Assay name: **pH-metric high logP** Analyst: **Pion**
Assay ID: **18C-26010** Instrument ID: **T312060**
Filename: **C:\Sirius_T3\Mehtap\20180327_exp34_logP_T3-2\18C-26010_M13_octanol_pH-metric high logP.t3r**

### Experiment Log (continued)

[41:12] Stepping pH = 8.90
[41:12] Dispensed 0.000024 mL of Base (0.5 M KOH)
[41:17] Stepping pH = 8.98
[41:17] Dispensed 0.000024 mL of Base (0.5 M KOH)
[41:22] Stepping pH = 9.07
[41:37] Stirrer speed set to 0
[41:56] Datapoint id 40 collected
[41:56] Charge balance equation is out by -170.5%
[41:56] Titration 3 of 3
[41:56] Adding initial titrants
[41:56] Automatically add 0.70000 mL of Octanol
[42:44] Dispensed 0.700000 mL of Octanol
[42:44] Stirrer speed set to 10
[42:45] Stirrer speed set to 60
[42:45] Iterative adjust 9.02 -> 2.00
[42:45] pH 9.02 -> 2.00
[42:47] Dispensed 0.054727 mL of Acid (0.5 M HCl)
[42:52] pH 2.03 -> 2.00
[42:52] Dispensed 0.003175 mL of Acid (0.5 M HCl)
[43:42] Stirrer speed set to 0
[43:52] Datapoint id 41 collected
[43:52] Stirrer speed set to 60
[43:57] pH 1.96 -> 2.16
[43:57] Using cautious pH adjust
[43:58] Dispensed 0.009972 mL of Base (0.5 M KOH)
[44:03] Stepping pH = 2.05
[44:03] Dispensed 0.007596 mL of Base (0.5 M KOH)
[44:08] Stepping pH = 2.14
[44:08] Dispensed 0.001317 mL of Base (0.5 M KOH)
[44:13] Stepping pH = 2.16
[44:28] Stirrer speed set to 0
[44:39] Datapoint id 42 collected
[44:39] Charge balance equation is out by 5.2%
[44:39] Stirrer speed set to 60
[44:44] pH 2.16 -> 2.36
[44:44] Using charge balance adjust
[44:44] Dispensed 0.012817 mL of Base (0.5 M KOH)
[45:05] Stirrer speed set to 0
[45:23] Datapoint id 43 collected
[45:23] Charge balance equation is out by 5.4%
[45:23] Stirrer speed set to 60
[45:28] pH 2.38 -> 2.58
[45:28] Using charge balance adjust
[45:28] Dispensed 0.007973 mL of Base (0.5 M KOH)
[45:48] Stirrer speed set to 0
[45:58] Datapoint id 44 collected
[45:58] Charge balance equation is out by 3.7%
[45:58] Stirrer speed set to 60
[46:03] pH 2.59 -> 2.79
[46:03] Using charge balance adjust
[46:04] Dispensed 0.005292 mL of Base (0.5 M KOH)
[46:24] Stirrer speed set to 0
[46:34] Datapoint id 45 collected
[46:34] Charge balance equation is out by 12.0%
[46:34] Stirrer speed set to 60
[46:39] pH 2.82 -> 3.02
[46:39] Using charge balance adjust
[46:40] Dispensed 0.003575 mL of Base (0.5 M KOH)
[47:00] Stirrer speed set to 0

| | | | |
|---|---|---|---|
| Sample name: | **M13_octanol** | Experiment start time: | **3/26/2018 4:16:35 PM** |
| Assay name: | **pH-metric high logP** | Analyst: | **Pion** |
| Assay ID: | **18C-26010** | Instrument ID: | **T312060** |
| Filename: | **C:\Sirius_T3\Mehtap\20180327_exp34_logP_T3-2\18C-26010_M13_octanol_pH-metric high logP.t3r** | | |

### Experiment Log (continued)

[47:10] Datapoint id 46 collected
[47:10] Charge balance equation is out by -4.8%
[47:10] Stirrer speed set to 60
[47:15] pH 3.02 -> 3.22
[47:15] Using charge balance adjust
[47:15] Dispensed 0.002634 mL of Base (0.5 M KOH)
[47:35] Stirrer speed set to 0
[47:45] Datapoint id 47 collected
[47:45] Charge balance equation is out by 7.7%
[47:45] Stirrer speed set to 60
[47:50] pH 3.25 -> 3.45
[47:50] Using charge balance adjust
[47:51] Dispensed 0.001881 mL of Base (0.5 M KOH)
[48:11] Stirrer speed set to 0
[48:21] Datapoint id 48 collected
[48:21] Charge balance equation is out by 17.5%
[48:21] Stirrer speed set to 60
[48:26] pH 3.49 -> 3.69
[48:26] Using cautious pH adjust
[48:26] Dispensed 0.000635 mL of Base (0.5 M KOH)
[48:31] Stepping pH = 3.59
[48:31] Dispensed 0.000400 mL of Base (0.5 M KOH)
[48:36] Stepping pH = 3.66
[48:36] Dispensed 0.000141 mL of Base (0.5 M KOH)
[48:41] Stepping pH = 3.68
[48:57] Stirrer speed set to 0
[49:07] Datapoint id 49 collected
[49:07] Charge balance equation is out by 7.9%
[49:07] Stirrer speed set to 60
[49:12] pH 3.68 -> 3.88
[49:12] Using charge balance adjust
[49:12] Dispensed 0.000917 mL of Base (0.5 M KOH)
[49:32] Stirrer speed set to 0
[49:51] Datapoint id 50 collected
[49:51] Charge balance equation is out by 2.2%
[49:51] Stirrer speed set to 60
[49:56] pH 3.90 -> 4.10
[49:56] Using charge balance adjust
[49:56] Dispensed 0.000588 mL of Base (0.5 M KOH)
[50:16] Stirrer speed set to 0
[50:28] Datapoint id 51 collected
[50:28] Charge balance equation is out by -30.8%
[50:28] Stirrer speed set to 60
[50:33] pH 4.05 -> 4.25
[50:33] Using cautious pH adjust
[50:33] Dispensed 0.000212 mL of Base (0.5 M KOH)
[50:38] Stepping pH = 4.09
[50:38] Dispensed 0.000353 mL of Base (0.5 M KOH)
[50:43] Stepping pH = 4.29
[50:58] Stirrer speed set to 0
[51:08] Datapoint id 52 collected
[51:08] Charge balance equation is out by -28.1%
[51:08] Stirrer speed set to 60
[51:14] pH 4.27 -> 4.47
[51:14] Using cautious pH adjust
[51:14] Dispensed 0.000141 mL of Base (0.5 M KOH)
[51:19] Stepping pH = 4.30
[51:19] Dispensed 0.000306 mL of Base (0.5 M KOH)
[51:24] Stepping pH = 4.59

### Experiment Log (continued)

[51:39] Stirrer speed set to 0
[51:49] Datapoint id 53 collected
[51:49] Charge balance equation is out by -65.4%
[51:49] Stirrer speed set to 60
[51:54] pH 4.58 -> 4.78
[51:54] Using cautious pH adjust
[51:54] Dispensed 0.000071 mL of Base (0.5 M KOH)
[51:59] Stepping pH = 4.59
[51:59] Dispensed 0.000212 mL of Base (0.5 M KOH)
[52:04] Stepping pH = 4.99
[52:19] Stirrer speed set to 0
[52:29] Datapoint id 54 collected
[52:29] Charge balance equation is out by -96.2%
[52:29] Stirrer speed set to 60
[52:35] pH 4.97 -> 5.17
[52:35] Using cautious pH adjust
[52:35] Dispensed 0.000024 mL of Base (0.5 M KOH)
[52:40] Stepping pH = 4.97
[52:40] Dispensed 0.000094 mL of Base (0.5 M KOH)
[52:45] Stepping pH = 5.07
[52:45] Dispensed 0.000047 mL of Base (0.5 M KOH)
[52:50] Stepping pH = 5.19
[53:05] Stirrer speed set to 0
[53:15] Datapoint id 55 collected
[53:15] Charge balance equation is out by -183.3%
[53:15] Stirrer speed set to 60
[53:20] pH 5.19 -> 5.39
[53:20] Using cautious pH adjust
[53:20] Dispensed 0.000024 mL of Base (0.5 M KOH)
[53:25] Stepping pH = 5.20
[53:25] Dispensed 0.000047 mL of Base (0.5 M KOH)
[53:31] Stepping pH = 5.35
[53:31] Dispensed 0.000024 mL of Base (0.5 M KOH)
[53:36] Stepping pH = 5.46
[53:51] Stirrer speed set to 0
[54:01] Datapoint id 56 collected
[54:01] Charge balance equation is out by -136.9%
[54:01] Stirrer speed set to 60
[54:06] pH 5.47 -> 5.67
[54:06] Using cautious pH adjust
[54:06] Dispensed 0.000024 mL of Base (0.5 M KOH)
[54:11] Stepping pH = 5.50
[54:11] Dispensed 0.000024 mL of Base (0.5 M KOH)
[54:16] Stepping pH = 5.63
[54:16] Dispensed 0.000024 mL of Base (0.5 M KOH)
[54:21] Stepping pH = 5.79
[54:36] Stirrer speed set to 0
[54:47] Datapoint id 57 collected
[54:47] Charge balance equation is out by -142.7%
[54:47] Stirrer speed set to 60
[54:52] pH 5.85 -> 6.05
[54:52] Using cautious pH adjust
[54:52] Dispensed 0.000024 mL of Base (0.5 M KOH)
[54:57] Stepping pH = 5.91
[54:57] Dispensed 0.000024 mL of Base (0.5 M KOH)
[55:02] Stepping pH = 6.14
[55:17] Stirrer speed set to 0
[55:38] Datapoint id 58 collected
[55:38] Charge balance equation is out by -37.8%

Sample name: **M13_octanol**
Assay name: **pH-metric high logP**
Assay ID: **18C-26010**
Filename: **C:\Sirius_T3\Mehtap\20180327_exp34_logP_T3-2\18C-26010_M13_octanol_pH-metric high logP.t3r**

Experiment start time: **3/26/2018 4:16:35 PM**
Analyst: **Pion**
Instrument ID: **T312060**

### Experiment Log (continued)

[55:38] Stirrer speed set to 60
[55:43] pH 6.09 -> 6.29
[55:43] Using cautious pH adjust
[55:43] Dispensed 0.000024 mL of Base (0.5 M KOH)
[55:48] Stepping pH = 6.20
[55:48] Dispensed 0.000024 mL of Base (0.5 M KOH)
[55:53] Stepping pH = 6.54
[56:08] Stirrer speed set to 0
[56:42] Datapoint id 59 collected
[56:42] Charge balance equation is out by -15.1%
[56:42] Stirrer speed set to 60
[56:47] pH 6.56 -> 6.76
[56:47] Using cautious pH adjust
[56:47] Dispensed 0.000024 mL of Base (0.5 M KOH)
[56:52] Stepping pH = 6.65
[56:52] Dispensed 0.000024 mL of Base (0.5 M KOH)
[56:57] Stepping pH = 6.97
[57:12] Stirrer speed set to 0
[58:11] Datapoint id 60 collected
[58:11] Charge balance equation is out by 3.8%
[58:11] Stirrer speed set to 60
[58:16] pH 6.99 -> 7.19
[58:16] Using charge balance adjust
[58:17] Dispensed 0.000024 mL of Base (0.5 M KOH)
[58:37] Stirrer speed set to 0
[59:19] Datapoint id 61 collected
[59:19] Charge balance equation is out by -18.2%
[59:19] Stirrer speed set to 60
[59:24] pH 6.94 -> 7.14
[59:24] Using cautious pH adjust
[59:24] Dispensed 0.000024 mL of Base (0.5 M KOH)
[59:29] Stepping pH = 6.99
[59:29] Dispensed 0.000024 mL of Base (0.5 M KOH)
[59:34] Stepping pH = 7.34
[59:49] Stirrer speed set to 0
[1:00:49] Datapoint id 62 collected
[1:00:49] Charge balance equation is out by -30.8%
[1:00:49] Stirrer speed set to 60
[1:00:54] pH 7.34 -> 7.54
[1:00:54] Using cautious pH adjust
[1:00:55] Dispensed 0.000024 mL of Base (0.5 M KOH)
[1:01:00] Stepping pH = 7.38
[1:01:00] Dispensed 0.000024 mL of Base (0.5 M KOH)
[1:01:05] Stepping pH = 7.84
[1:01:20] Stirrer speed set to 0
[1:02:20] Datapoint id 63 collected
[1:02:20] Charge balance equation is out by -73.9%
[1:02:20] Stirrer speed set to 60
[1:02:25] pH 7.88 -> 8.08
[1:02:25] Using cautious pH adjust
[1:02:25] Dispensed 0.000024 mL of Base (0.5 M KOH)
[1:02:30] Stepping pH = 7.97
[1:02:30] Dispensed 0.000024 mL of Base (0.5 M KOH)
[1:02:35] Stepping pH = 8.26
[1:02:50] Stirrer speed set to 0
[1:03:33] Datapoint id 64 collected
[1:03:33] Charge balance equation is out by -237.4%
[1:03:33] Stirrer speed set to 60
[1:03:38] pH 8.20 -> 8.40

| | | | |
|---|---|---|---|
| Sample name: | **M13_octanol** | Experiment start time: | **3/26/2018 4:16:35 PM** |
| Assay name: | **pH-metric high logP** | Analyst: | **Pion** |
| Assay ID: | **18C-26010** | Instrument ID: | **T312060** |
| Filename: | **C:\Sirius_T3\Mehtap\20180327_exp34_logP_T3-2\18C-26010_M13_octanol_pH-metric high logP.t3r** | | |

### Experiment Log (continued)

[1:03:38] Using cautious pH adjust
[1:03:38] Dispensed 0.000024 mL of Base (0.5 M KOH)
[1:03:43] Stepping pH = 8.28
[1:03:43] Dispensed 0.000024 mL of Base (0.5 M KOH)
[1:03:48] Stepping pH = 8.53
[1:04:03] Stirrer speed set to 0
[1:04:38] Datapoint id 65 collected
[1:04:38] Charge balance equation is out by -230.0%
[1:04:38] Stirrer speed set to 60
[1:04:43] pH 8.49 -> 8.69
[1:04:43] Using cautious pH adjust
[1:04:43] Dispensed 0.000024 mL of Base (0.5 M KOH)
[1:04:48] Stepping pH = 8.55
[1:04:48] Dispensed 0.000024 mL of Base (0.5 M KOH)
[1:04:53] Stepping pH = 8.71
[1:05:08] Stirrer speed set to 0
[1:05:37] Datapoint id 66 collected
[1:05:37] Charge balance equation is out by -127.2%
[1:05:37] Stirrer speed set to 60
[1:05:42] pH 8.71 -> 8.91
[1:05:42] Using cautious pH adjust
[1:05:42] Dispensed 0.000024 mL of Base (0.5 M KOH)
[1:05:47] Stepping pH = 8.74
[1:05:47] Dispensed 0.000024 mL of Base (0.5 M KOH)
[1:05:53] Stepping pH = 8.87
[1:05:53] Dispensed 0.000024 mL of Base (0.5 M KOH)
[1:05:58] Stepping pH = 8.98
[1:06:13] Stirrer speed set to 0
[1:06:41] Datapoint id 67 collected
[1:06:41] Charge balance equation is out by -152.4%
[1:06:41] Stirrer speed set to 60
[1:06:46] pH 8.97 -> 9.05
[1:06:46] Using cautious pH adjust
[1:06:46] Dispensed 0.000024 mL of Base (0.5 M KOH)
[1:06:51] Stepping pH = 9.00
[1:07:06] Stirrer speed set to 0
[1:07:33] Datapoint id 68 collected
[1:07:33] Charge balance equation is out by -27.6%
[1:07:33] Argon flow rate set to 0
[1:07:37] Titrator arm moved over Titration position
[1:07:58] The autoloader failed to pick at location "Sample position"
