## Supplementary material for "Octanol-water partition coefficient measurements for the SAMPL6 Blind Prediction Challenge": SM14_18B-28002_M15_octanol_pH-metric high logP_report.pdf

| | | | |
|---|---|---|---|
| Sample name: | **M15_octanol** | Experiment start time: | **2/28/2018 1:10:47 AM** |
| Assay name: | **pH-metric high logP** | Analyst: | **Pion** |
| Assay ID: | **18B-28002** | Instrument ID: | **T312060** |
| Filename: | **C:\Sirius_T3\Mehtap\20180227_exp27_logP_T3-2\18B-28002_M15_octanol_pH-metric high logP.t3r** | | |

#### pH-metric Result

| | | |
|---|---|---|
| logP (XH2 2+) | -2.38 | |
| logP (XH +) | -4.96 | ±2.15 (n=47) |
| logP (neutral X) | 1.92 | ±0.01 (n=47) |

#### 18B-28002 Points 1 to 35

| | |
|---|---|
| M15_octanol concentration factor | 0.761 |
| Carbonate | 0.1910 mM |
| Acidity error | 0.00862 mM |

#### 18B-28002 Points 36 to 72

| | |
|---|---|
| M15_octanol concentration factor | 0.839 |
| Carbonate | 0.2848 mM |
| Acidity error | -0.02801 mM |

#### 18B-28002 Points 73 to 106

| | |
|---|---|
| M15_octanol concentration factor | 0.903 |
| Carbonate | 0.2174 mM |
| Acidity error | 0.16739 mM |

#### Warnings and errors

| | |
|---|---|
| Errors | None |
| Warnings | One or more logP values out of range |

#### Sample logD and percent species

| pH | M15_octanol logD | M15_octanol M15_octanolH2 | M15_octanol M15_octanolH | M15_octanol M15_octanol | M15_octanol M15_octanolH2* | M15_octanol M15_octanolH* | M15_octanol M15_octanol* | Comment |
|---|---|---|---|---|---|---|---|---|
| 1.000 | -2.38 | 97.03 % | 2.55 % | 0.00 % | 0.40 % | 0.00 % | 0.01 % | |
| 1.200 | -2.37 | 95.59 % | 3.98 % | 0.00 % | 0.40 % | 0.00 % | 0.03 % | Stomach pH |
| 2.000 | -1.92 | 78.23 % | 20.58 % | 0.01 % | 0.33 % | 0.00 % | 0.86 % | |
| 3.000 | -0.52 | 21.05 % | 55.37 % | 0.28 % | 0.09 % | 0.00 % | 23.22 % | |
| 4.000 | 0.59 | 0.72 % | 18.93 % | 0.95 % | 0.00 % | 0.00 % | 79.40 % | |
| 5.000 | 1.45 | 0.01 % | 2.30 % | 1.15 % | 0.00 % | 0.00 % | 96.54 % | |
| 6.000 | 1.84 | 0.00 % | 0.24 % | 1.18 % | 0.00 % | 0.00 % | 98.59 % | |
| 6.500 | 1.90 | 0.00 % | 0.07 % | 1.18 % | 0.00 % | 0.00 % | 98.75 % | |
| 7.000 | 1.91 | 0.00 % | 0.02 % | 1.18 % | 0.00 % | 0.00 % | 98.80 % | |
| 7.400 | 1.92 | 0.00 % | 0.01 % | 1.18 % | 0.00 % | 0.00 % | 98.81 % | Blood pH |
| 8.000 | 1.92 | 0.00 % | 0.00 % | 1.18 % | 0.00 % | 0.00 % | 98.82 % | |
| 9.000 | 1.92 | 0.00 % | 0.00 % | 1.18 % | 0.00 % | 0.00 % | 98.82 % | |
| 10.000 | 1.92 | 0.00 % | 0.00 % | 1.18 % | 0.00 % | 0.00 % | 98.82 % | |
| 11.000 | 1.92 | 0.00 % | 0.00 % | 1.18 % | 0.00 % | 0.00 % | 98.82 % | |
| 12.000 | 1.92 | 0.00 % | 0.00 % | 1.18 % | 0.00 % | 0.00 % | 98.82 % | |

Sample name: **M15_octanol**
Assay name: **pH-metric high logP**
Assay ID: **18B-28002**
Filename: **C:\Sirius_T3\Mehtap\20180227_exp27_logP_T3-2\18B-28002_M15_octanol_pH-metric high logP.t3r**

Experiment start time: **2/28/2018 1:10:47 AM**
Analyst: **Pion**
Instrument ID: **T312060**

#### Graphs

Charge balance residuals

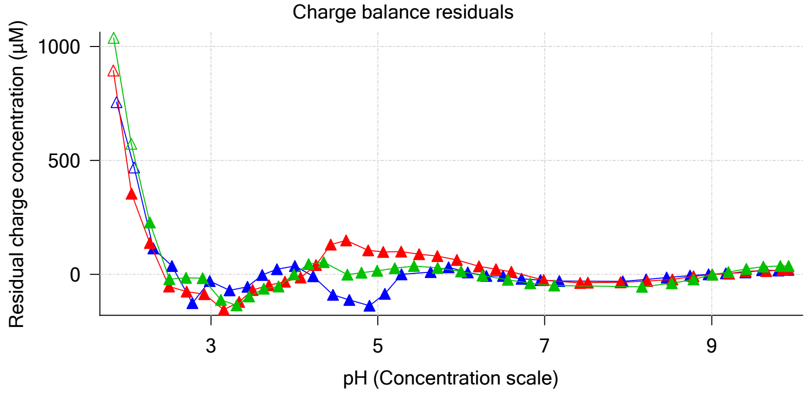


Ionic strength

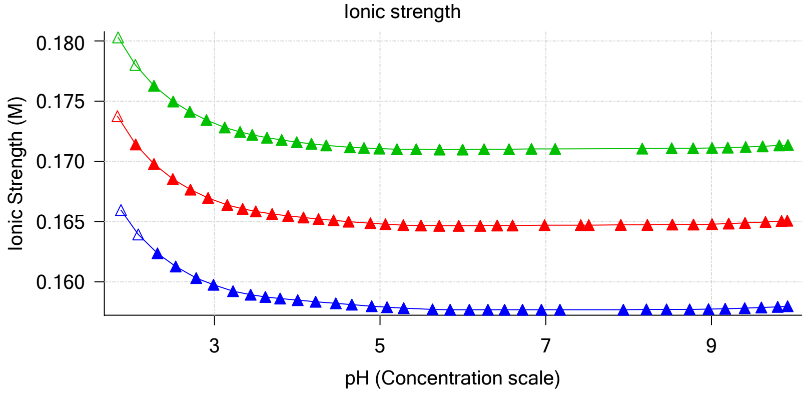


Buffer index profile

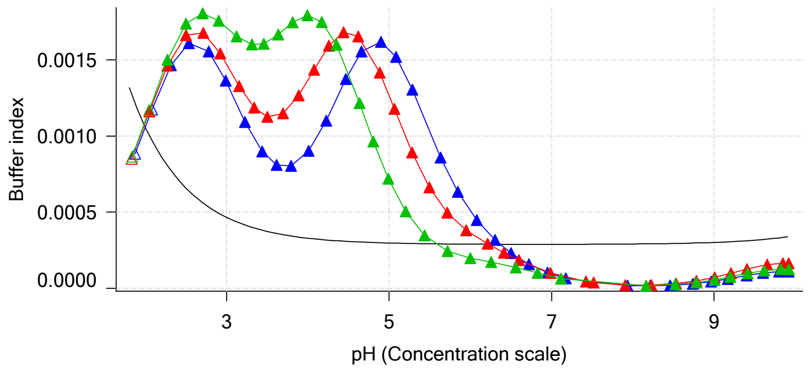


Sample M15_octanol Ionisation Graph

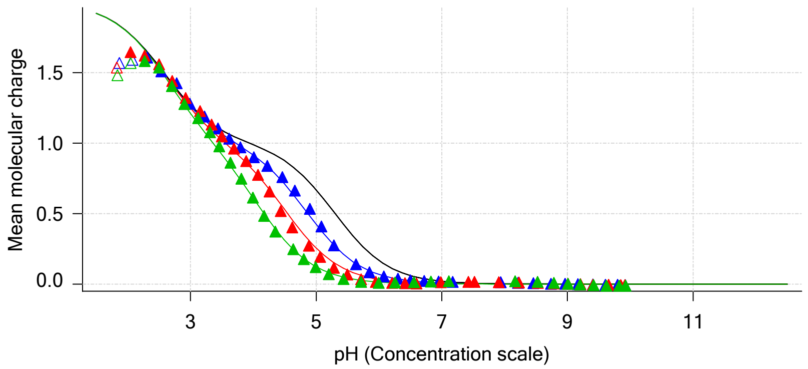


Distribution of species for sample M15_octanol

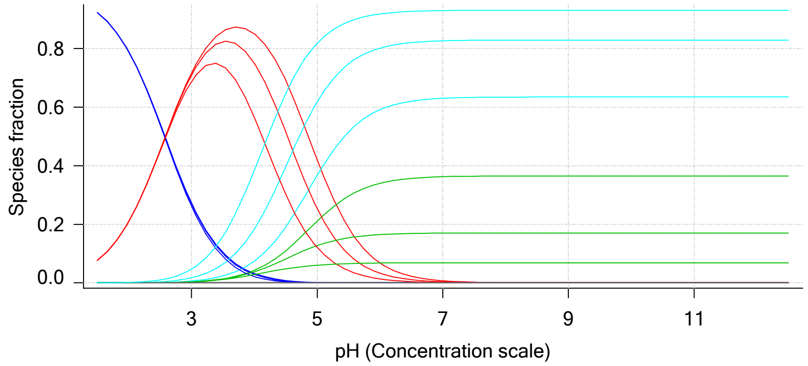


Lipophilicity Profile

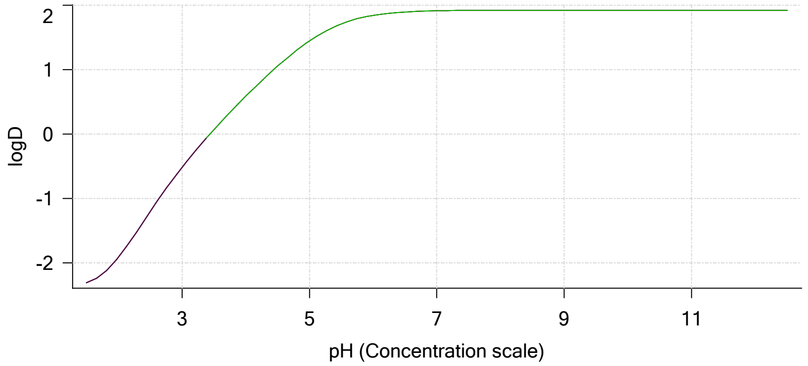


Log R Profile

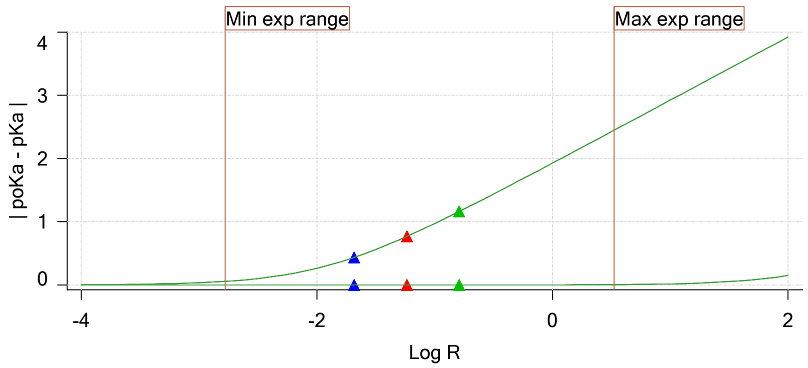


Sample Carbonate Ionisation Graph

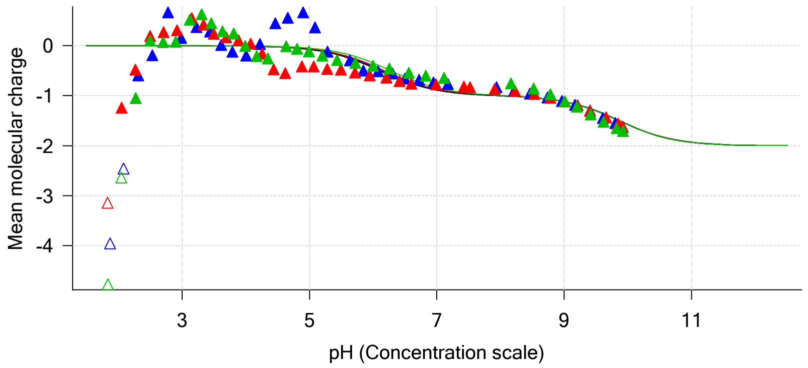

Sample name: **M15_octanol**
Assay name: **pH-metric high logP**
Assay ID: **18B-28002**
Filename: **C:\Sirius_T3\Mehtap\20180227_exp27_logP_T3-2\18B-28002_M15_octanol_pH-metric high logP.t3r**

Experiment start time: **2/28/2018 1:10:47 AM**
Analyst: **Pion**
Instrument ID: **T312060**

#### Graphs (continued)

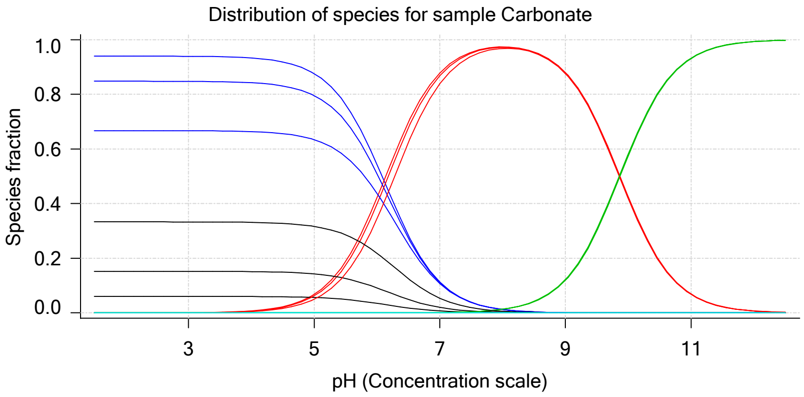

| | | | |
|---|---|---|---|
| Sample name: | **M15_octanol** | Experiment start time: | **2/28/2018 1:10:47 AM** |
| Assay name: | **pH-metric high logP** | Analyst: | **Pion** |
| Assay ID: | **18B-28002** | Instrument ID: | **T312060** |
| Filename: | **C:\Sirius_T3\Mehtap\20180227_exp27_logP_T3-2\18B-28002_M15_octanol_pH-metric high logP.t3r** | | |

### pH-metric high logP Titration 1 of 3 18B-28002 Points 1 to 35

#### Overall results

| | |
|---|---|
| RMSD | 0.336 |
| Average ionic strength | 0.158 M |
| Average temperature | 25.0°C |
| Partition ratio | 0.0208 : 1 |
| Analyte concentration range | 3493.3 μM to 3618.9 μM |
| Total points considered | 33 of 35 |

#### Warnings and errors

| | |
|---|---|
| Errors | None |
| Warnings | One or more logP values out of range |

#### Four-Plus parameters

| | | | |
|---|---|---|---|
| Alpha | 0.130 | 2/28/2018 1:10:47 AM | C:\Sirius_T3\HCl18B27.t3r |
| S | 0.9970 | 2/28/2018 1:10:47 AM | C:\Sirius_T3\HCl18B27.t3r |
| jH | 0.8 | 2/28/2018 1:10:47 AM | C:\Sirius_T3\HCl18B27.t3r |
| jOH | -0.4 | 2/28/2018 1:10:47 AM | C:\Sirius_T3\HCl18B27.t3r |

#### Titrants

| | | | |
|---|---|---|---|
| 0.50 M HCl | 0.993513 | 2/28/2018 1:10:47 AM | C:\Sirius_T3\HCl18B27.t3r |
| 0.50 M KOH | 0.999845 | 2/28/2018 1:10:47 AM | C:\Sirius_T3\KOH18B27.t3r |

#### Sample

| | |
|---|---|
| M15_octanol concentration factor | 0.761 |
| Base pKa 1 | 2.58 |
| Base pKa 2 | 5.30 |
| logP (XH2 2+) | -2.38 |
| logP (XH +) | -9.58 |
| logP (neutral X) | 1.85 |

#### Sample graphs

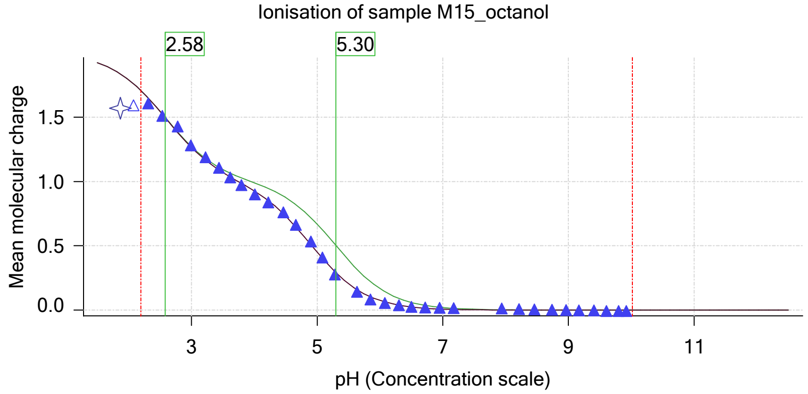


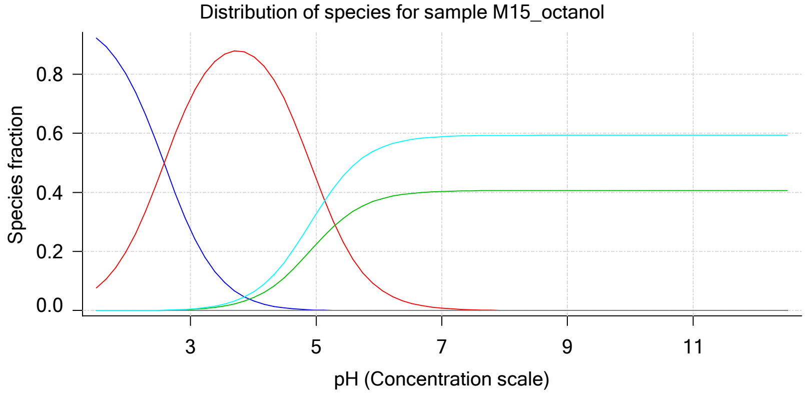

Sample name: **M15_octanol**
Assay name: **pH-metric high logP**
Assay ID: **18B-28002**
Filename: **C:\Sirius_T3\Mehtap\20180227_exp27_logP_T3-2\18B-28002_M15_octanol_pH-metric high logP.t3r**
Experiment start time: **2/28/2018 1:10:47 AM**
Analyst: **Pion**
Instrument ID: **T312060**

#### Sample graphs (continued)

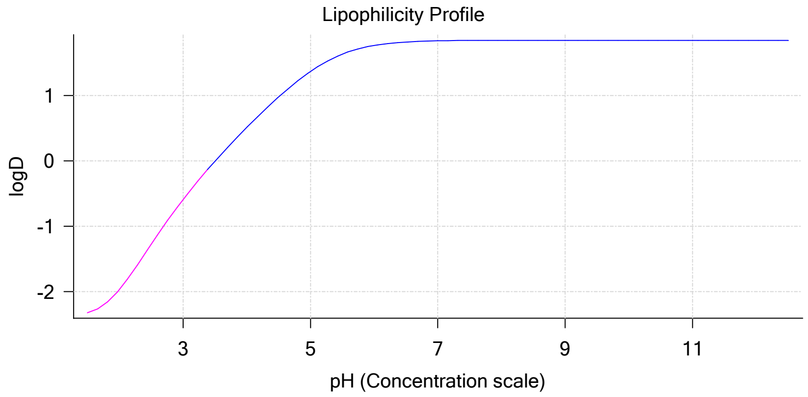


#### Sample logD and percent species

| pH | M15_octanol logD | M15_octanol M15_octanolH2 | M15_octanol M15_octanolH | M15_octanol M15_octanol | M15_octanol M15_octanolH2* | M15_octanol M15_octanolH* | M15_octanol M15_octanol* | Comment |
|---|---|---|---|---|---|---|---|---|
| 1.000 | -2.38 | 97.43 % | 2.56 % | 0.00 % | 0.01 % | 0.00 % | 0.00 % | |
| 1.200 | -2.37 | 95.99 % | 4.00 % | 0.00 % | 0.01 % | 0.00 % | 0.00 % | Stomach pH |
| 2.000 | -1.97 | 79.15 % | 20.82 % | 0.01 % | 0.01 % | 0.00 % | 0.02 % | |
| 3.000 | -0.59 | 27.30 % | 71.81 % | 0.36 % | 0.00 % | 0.00 % | 0.53 % | |
| 4.000 | 0.51 | 3.27 % | 86.11 % | 4.32 % | 0.00 % | 0.00 % | 6.30 % | |
| 5.000 | 1.37 | 0.17 % | 44.71 % | 22.41 % | 0.00 % | 0.00 % | 32.72 % | |
| 6.000 | 1.77 | 0.00 % | 7.50 % | 37.60 % | 0.00 % | 0.00 % | 54.90 % | |
| 6.500 | 1.82 | 0.00 % | 2.50 % | 39.63 % | 0.00 % | 0.00 % | 57.87 % | |
| 7.000 | 1.84 | 0.00 % | 0.80 % | 40.32 % | 0.00 % | 0.00 % | 58.87 % | |
| 7.400 | 1.84 | 0.00 % | 0.32 % | 40.52 % | 0.00 % | 0.00 % | 59.16 % | Blood pH |
| 8.000 | 1.85 | 0.00 % | 0.08 % | 40.62 % | 0.00 % | 0.00 % | 59.30 % | |
| 9.000 | 1.85 | 0.00 % | 0.01 % | 40.64 % | 0.00 % | 0.00 % | 59.35 % | |
| 10.000 | 1.85 | 0.00 % | 0.00 % | 40.65 % | 0.00 % | 0.00 % | 59.35 % | |
| 11.000 | 1.85 | 0.00 % | 0.00 % | 40.65 % | 0.00 % | 0.00 % | 59.35 % | |
| 12.000 | 1.85 | 0.00 % | 0.00 % | 40.65 % | 0.00 % | 0.00 % | 59.35 % | |

#### Carbonate and acidity

| | |
|---|---|
| Carbonate | 0.191 mM |
| Acidity error | 0.009 mM |

#### Other graphs

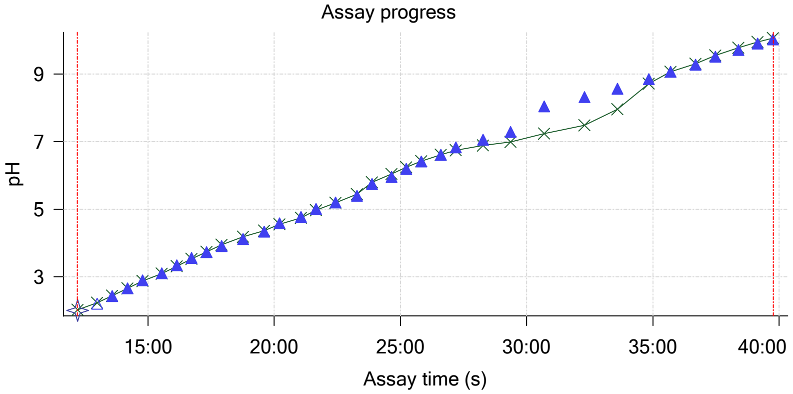


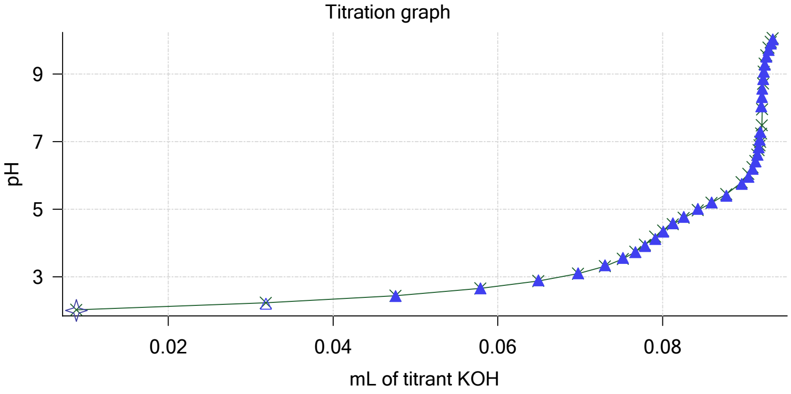

| | | | |
|---|---|---|---|
| Sample name: | **M15_octanol** | Experiment start time: | **2/28/2018 1:10:47 AM** |
| Assay name: | **pH-metric high logP** | Analyst: | **Pion** |
| Assay ID: | **18B-28002** | Instrument ID: | **T312060** |
| Filename: | **C:\Sirius_T3\Mehtap\20180227_exp27_logP_T3-2\18B-28002_M15_octanol_pH-metric high logP.t3r** | | |

#### Other graphs (continued)

Charge balance residuals

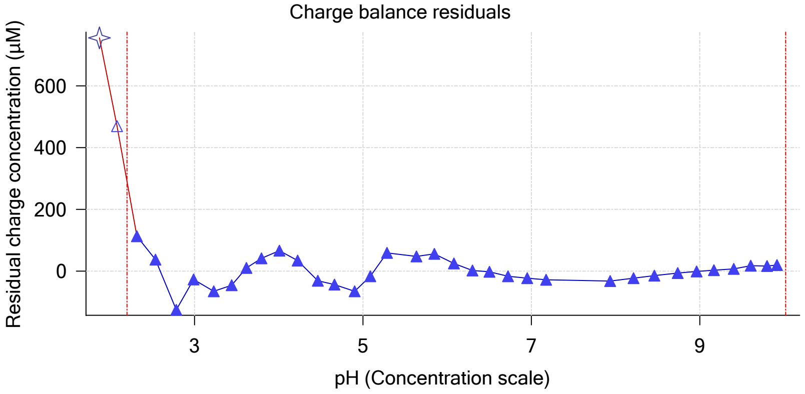


Buffer index profile

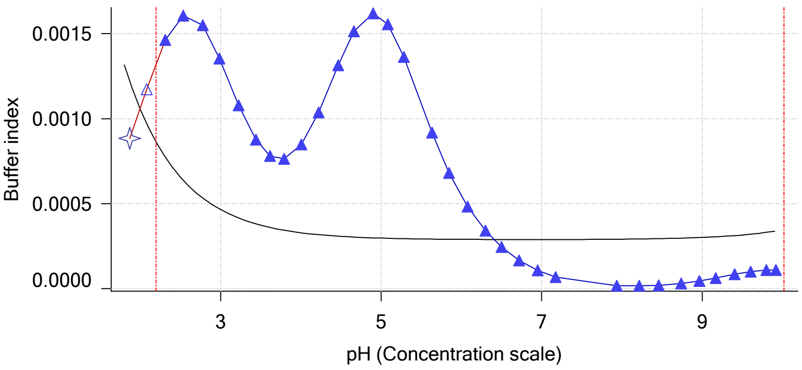


Ionic strength

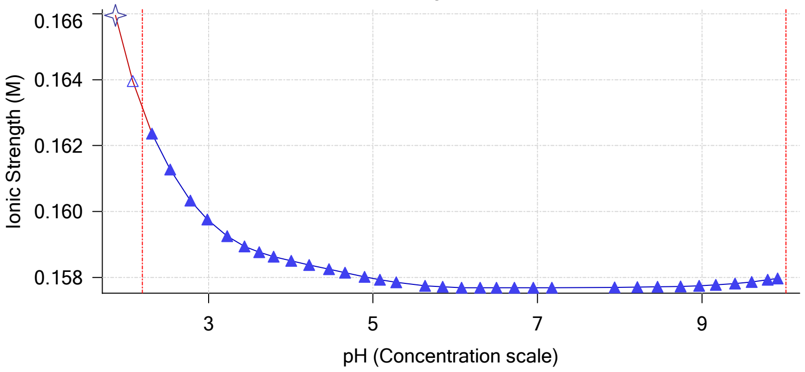


Log R Profile

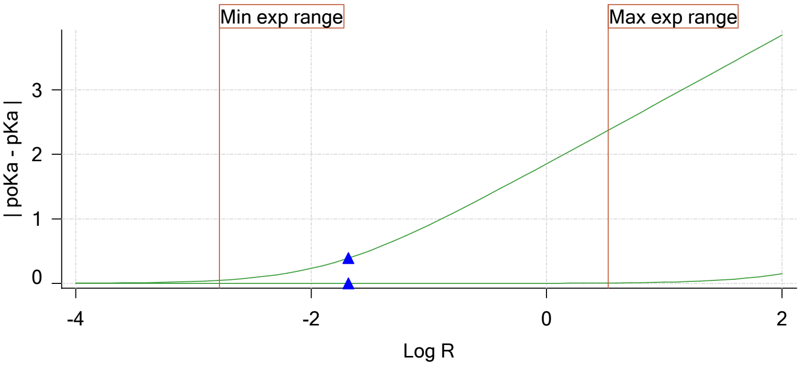


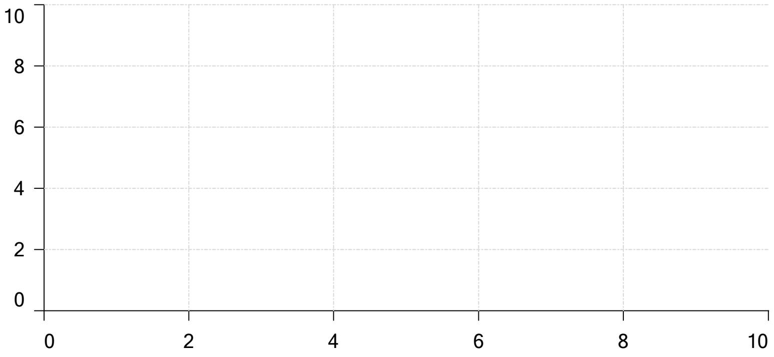


Ionisation of sample Carbonate

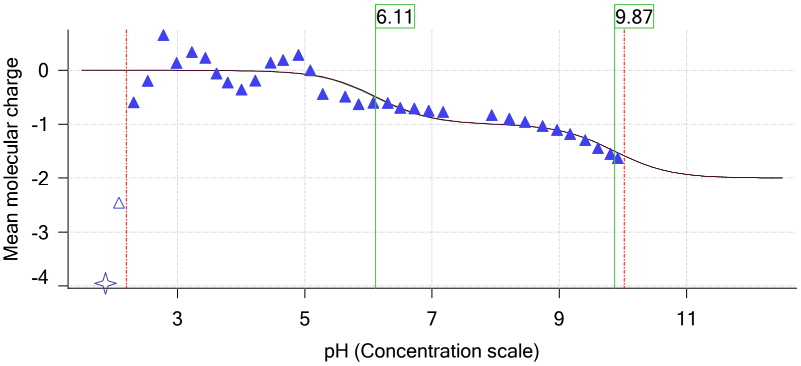

| | | | |
|---|---|---|---|
| Sample name: | **M15_octanol** | Experiment start time: | **2/28/2018 1:10:47 AM** |
| Assay name: | **pH-metric high logP** | Analyst: | **Pion** |
| Assay ID: | **18B-28002** | Instrument ID: | **T312060** |
| Filename: | **C:\Sirius_T3\Mehtap\20180227_exp27_logP_T3-2\18B-28002_M15_octanol_pH-metric high logP.t3r** | | |

# pH-metric high logP   Titration 2 of 3   18B-28002 Points 36 to 72

#### Overall results

| | |
|---|---|
| RMSD | 0.387 |
| Average ionic strength | 0.166 M |
| Average temperature | 25.0°C |
| Partition ratio | 0.0581 : 1 |
| Analyte concentration range | 3113.6 μM to 3230.1 μM |
| Total points considered | 36 of 37 |

#### Warnings and errors

| | |
|---|---|
| Errors | None |
| Warnings | One or more logP values out of range |

#### Four-Plus parameters

| | | | |
|---|---|---|---|
| Alpha | 0.130 | 2/28/2018 1:10:47 AM | C:\Sirius_T3\HCl18B27.t3r |
| S | 0.9970 | 2/28/2018 1:10:47 AM | C:\Sirius_T3\HCl18B27.t3r |
| jH | 0.8 | 2/28/2018 1:10:47 AM | C:\Sirius_T3\HCl18B27.t3r |
| jOH | -0.4 | 2/28/2018 1:10:47 AM | C:\Sirius_T3\HCl18B27.t3r |

#### Titrants

| | | | |
|---|---|---|---|
| 0.50 M HCl | 0.993513 | 2/28/2018 1:10:47 AM | C:\Sirius_T3\HCl18B27.t3r |
| 0.50 M KOH | 0.999845 | 2/28/2018 1:10:47 AM | C:\Sirius_T3\KOH18B27.t3r |

#### Sample

| | |
|---|---|
| M15_octanol concentration factor | 0.839 |
| Base pKa 1 | 2.58 |
| Base pKa 2 | 5.30 |
| logP (XH2 2+) | -2.38 |
| logP (XH +) | -3.43 |
| logP (neutral X) | 1.98 |

#### Sample graphs

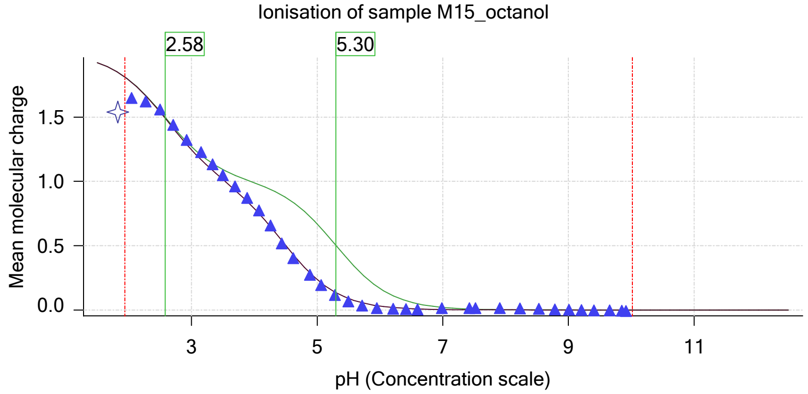


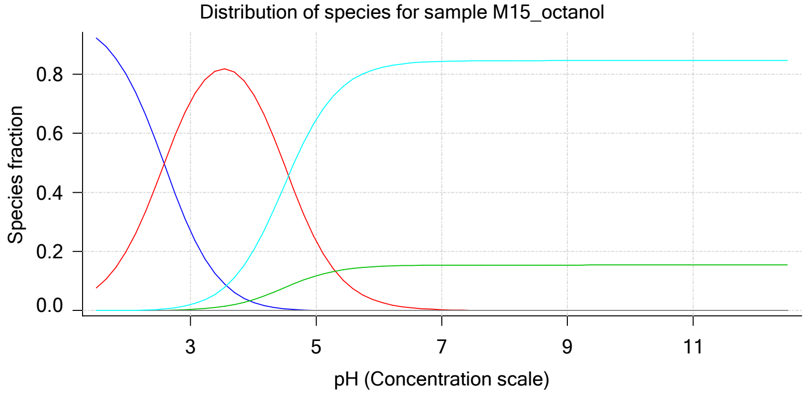

| | | | |
|---|---|---|---|
| Sample name: | **M15_octanol** | Experiment start time: | **2/28/2018 1:10:47 AM** |
| Assay name: | **pH-metric high logP** | Analyst: | **Pion** |
| Assay ID: | **18B-28002** | Instrument ID: | **T312060** |
| Filename: | **C:\Sirius_T3\Mehtap\20180227_exp27_logP_T3-2\18B-28002_M15_octanol_pH-metric high logP.t3r** | | |

#### Sample graphs (continued)

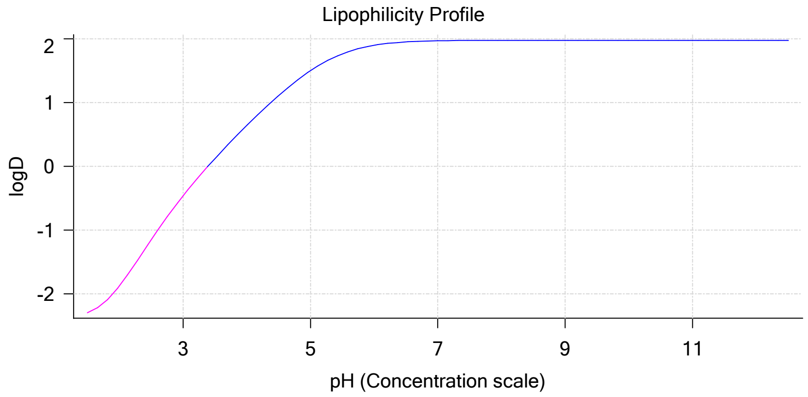


#### Sample logD and percent species

| pH | M15_octanol logD | M15_octanol M15_octanolH2 | M15_octanol M15_octanolH | M15_octanol M15_octanol | M15_octanol M15_octanolH2* | M15_octanol M15_octanolH* | M15_octanol M15_octanol* | Comment |
|---|---|---|---|---|---|---|---|---|
| 1.000 | -2.38 | 97.41 % | 2.56 % | 0.00 % | 0.02 % | 0.00 % | 0.00 % | |
| 1.200 | -2.36 | 95.97 % | 4.00 % | 0.00 % | 0.02 % | 0.00 % | 0.00 % | Stomach pH |
| 2.000 | -1.88 | 79.11 % | 20.81 % | 0.01 % | 0.02 % | 0.00 % | 0.06 % | |
| 3.000 | -0.46 | 26.91 % | 70.78 % | 0.35 % | 0.01 % | 0.00 % | 1.95 % | |
| 4.000 | 0.64 | 2.79 % | 73.34 % | 3.68 % | 0.00 % | 0.00 % | 20.19 % | |
| 5.000 | 1.50 | 0.09 % | 23.48 % | 11.77 % | 0.00 % | 0.00 % | 64.66 % | |
| 6.000 | 1.90 | 0.00 % | 2.98 % | 14.94 % | 0.00 % | 0.00 % | 82.08 % | |
| 6.500 | 1.95 | 0.00 % | 0.96 % | 15.25 % | 0.00 % | 0.00 % | 83.79 % | |
| 7.000 | 1.97 | 0.00 % | 0.31 % | 15.35 % | 0.00 % | 0.00 % | 84.34 % | |
| 7.400 | 1.97 | 0.00 % | 0.12 % | 15.38 % | 0.00 % | 0.00 % | 84.50 % | Blood pH |
| 8.000 | 1.97 | 0.00 % | 0.03 % | 15.39 % | 0.00 % | 0.00 % | 84.57 % | |
| 9.000 | 1.98 | 0.00 % | 0.00 % | 15.40 % | 0.00 % | 0.00 % | 84.60 % | |
| 10.000 | 1.98 | 0.00 % | 0.00 % | 15.40 % | 0.00 % | 0.00 % | 84.60 % | |
| 11.000 | 1.98 | 0.00 % | 0.00 % | 15.40 % | 0.00 % | 0.00 % | 84.60 % | |
| 12.000 | 1.98 | 0.00 % | 0.00 % | 15.40 % | 0.00 % | 0.00 % | 84.60 % | |

#### Carbonate and acidity

| | |
|---|---|
| Carbonate | 0.285 mM |
| Acidity error | -0.028 mM |

#### Other graphs

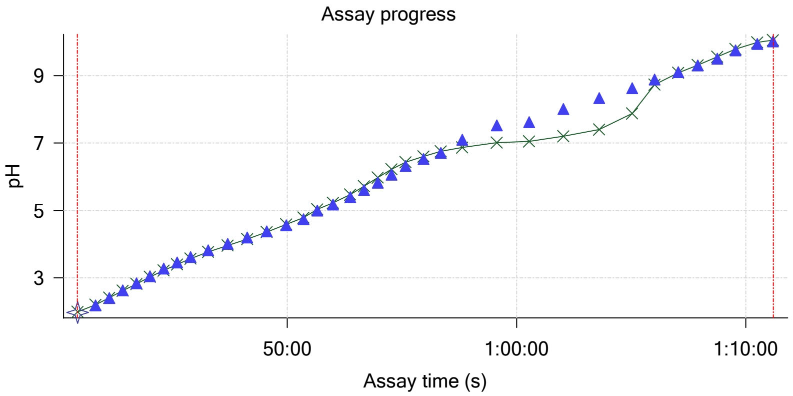


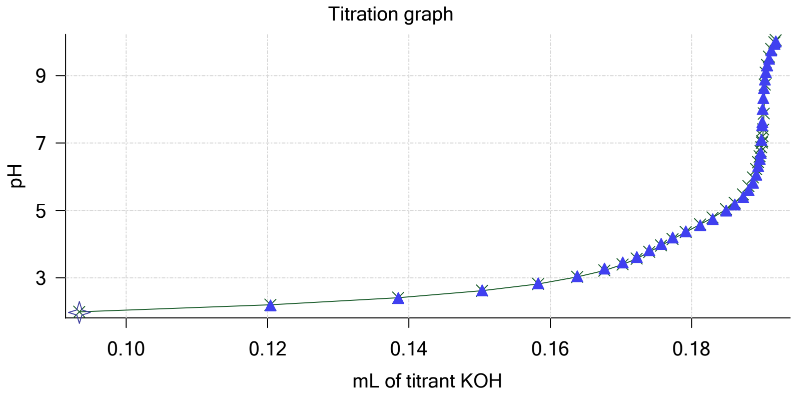

| | | | |
|---|---|---|---|
| Sample name: | **M15_octanol** | Experiment start time: | **2/28/2018 1:10:47 AM** |
| Assay name: | **pH-metric high logP** | Analyst: | **Pion** |
| Assay ID: | **18B-28002** | Instrument ID: | **T312060** |
| Filename: | **C:\Sirius_T3\Mehtap\20180227_exp27_logP_T3-2\18B-28002_M15_octanol_pH-metric high logP.t3r** | | |

#### Other graphs (continued)

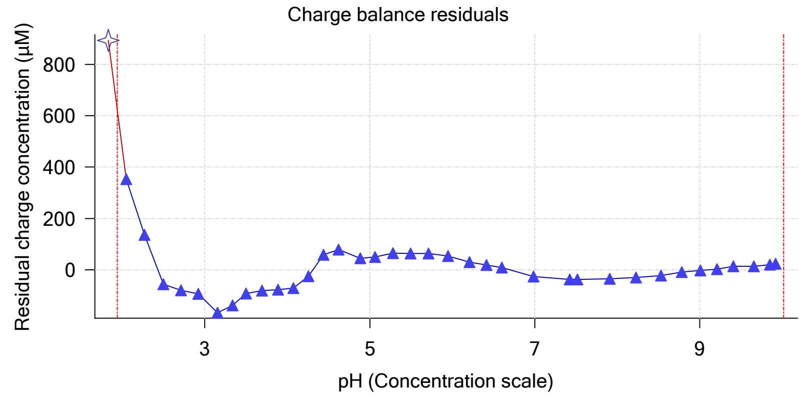


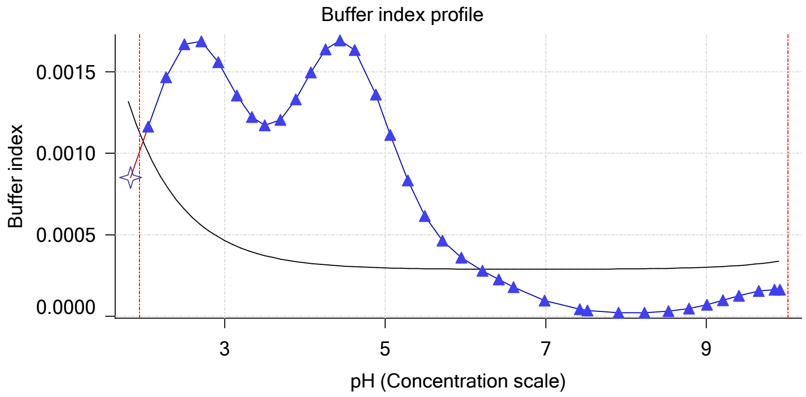


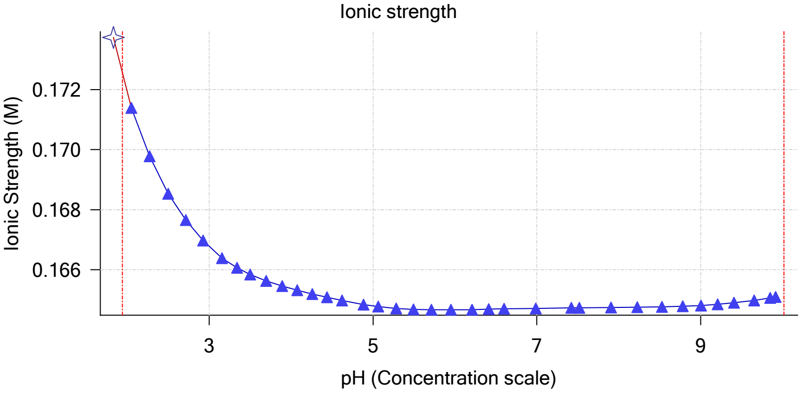


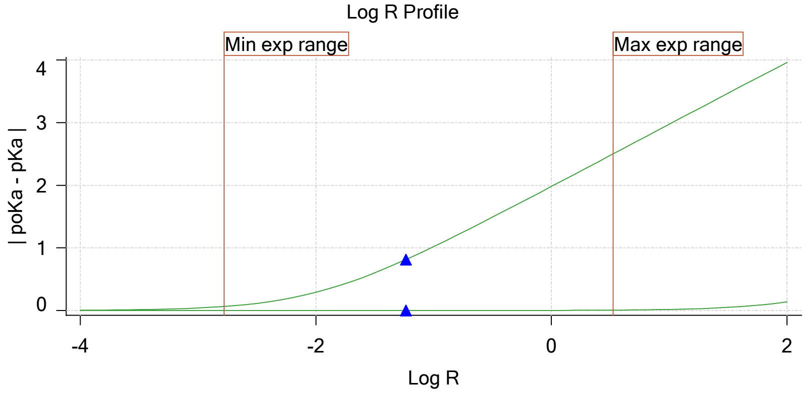


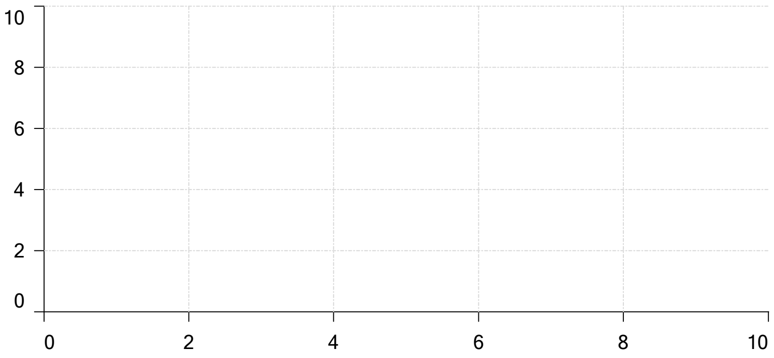


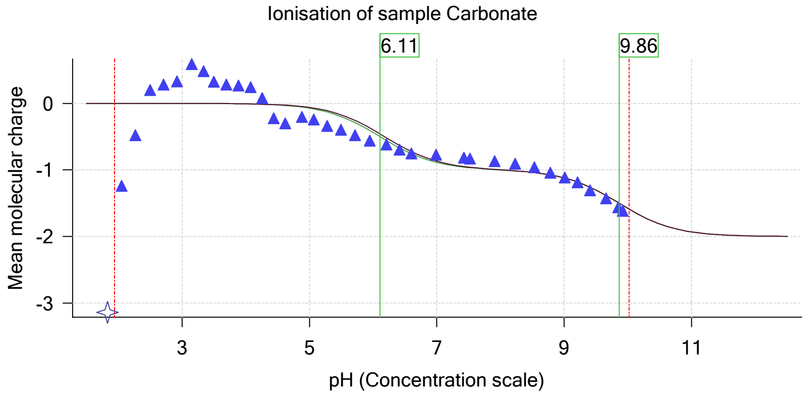

| | | | |
|---|---|---|---|
| Sample name: | **M15_octanol** | Experiment start time: | **2/28/2018 1:10:47 AM** |
| Assay name: | **pH-metric high logP** | Analyst: | **Pion** |
| Assay ID: | **18B-28002** | Instrument ID: | **T312060** |
| Filename: | **C:\Sirius_T3\Mehtap\20180227_exp27_logP_T3-2\18B-28002_M15_octanol_pH-metric high logP.t3r** | | |

### pH-metric high logP Titration 3 of 3 18B-28002 Points 73 to 106

#### Overall results

| | |
|---|---|
| RMSD | 0.501 |
| Average ionic strength | 0.172 M |
| Average temperature | 25.0°C |
| Partition ratio | 0.1616 : 1 |
| Analyte concentration range | 2625.5 μM to 2712.6 μM |
| Total points considered | 32 of 34 |

#### Warnings and errors

| | |
|---|---|
| Errors | None |
| Warnings | One or more logP values out of range |

#### Four-Plus parameters

| | | | |
|---|---|---|---|
| Alpha | 0.130 | 2/28/2018 1:10:47 AM | C:\Sirius_T3\HCl18B27.t3r |
| S | 0.9970 | 2/28/2018 1:10:47 AM | C:\Sirius_T3\HCl18B27.t3r |
| jH | 0.8 | 2/28/2018 1:10:47 AM | C:\Sirius_T3\HCl18B27.t3r |
| jOH | -0.4 | 2/28/2018 1:10:47 AM | C:\Sirius_T3\HCl18B27.t3r |

#### Titrants

| | | | |
|---|---|---|---|
| 0.50 M HCl | 0.993513 | 2/28/2018 1:10:47 AM | C:\Sirius_T3\HCl18B27.t3r |
| 0.50 M KOH | 0.999845 | 2/28/2018 1:10:47 AM | C:\Sirius_T3\KOH18B27.t3r |

#### Sample

| | |
|---|---|
| M15_octanol concentration factor | 0.903 |
| Base pKa 1 | 2.58 |
| Base pKa 2 | 5.30 |
| logP (XH2 2+) | -2.38 |
| logP (XH +) | -3.95 |
| logP (neutral X) | 1.91 |

#### Sample graphs

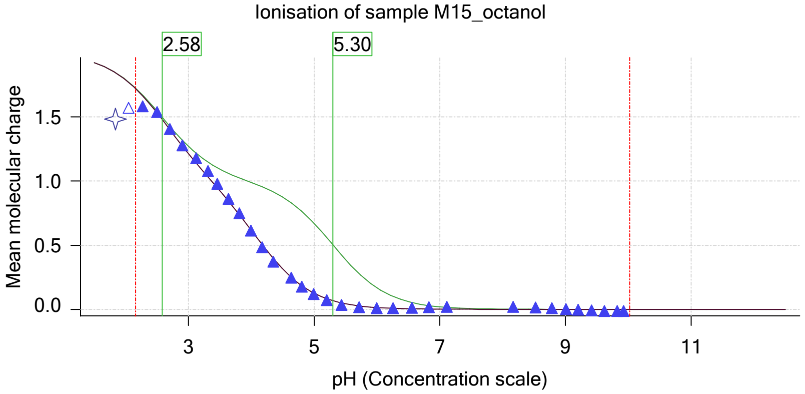


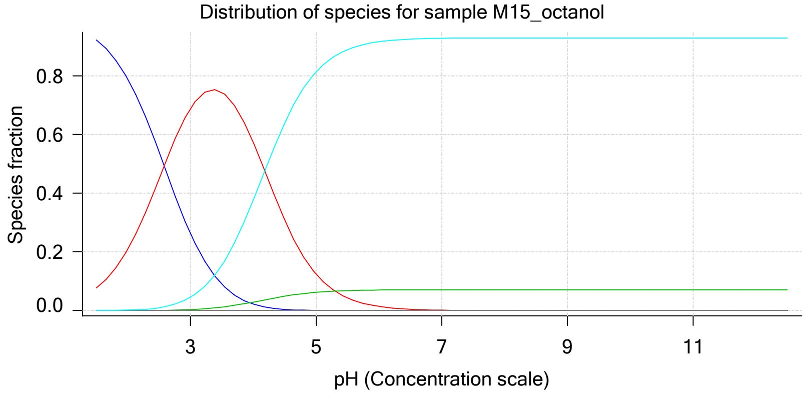

| | | | |
|---|---|---|---|
| Sample name: | **M15_octanol** | Experiment start time: | **2/28/2018 1:10:47 AM** |
| Assay name: | **pH-metric high logP** | Analyst: | **Pion** |
| Assay ID: | **18B-28002** | Instrument ID: | **T312060** |
| Filename: | **C:\Sirius_T3\Mehtap\20180227_exp27_logP_T3-2\18B-28002_M15_octanol_pH-metric high logP.t3r** | | |

#### Sample graphs (continued)

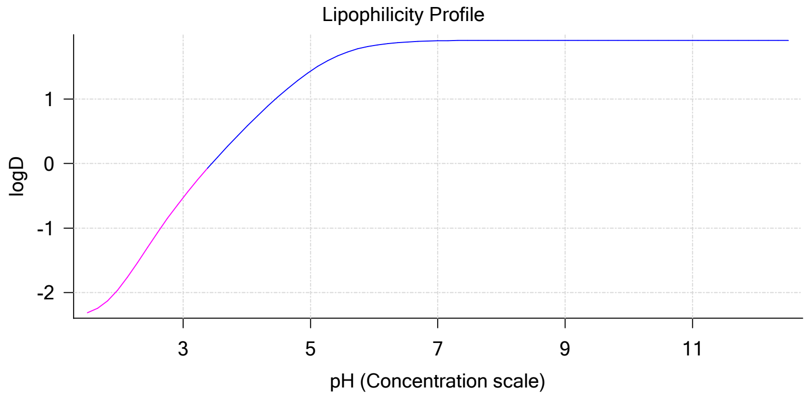


#### Sample logD and percent species

| pH | M15_octanol logD | M15_octanol M15_octanolH2 | M15_octanol M15_octanolH | M15_octanol M15_octanol | M15_octanol M15_octanolH2* | M15_octanol M15_octanolH* | M15_octanol M15_octanol* | Comment |
|---|---|---|---|---|---|---|---|---|
| 1.000 | -2.38 | 97.37 % | 2.56 % | 0.00 % | 0.07 % | 0.00 % | 0.00 % | |
| 1.200 | -2.37 | 95.93 % | 4.00 % | 0.00 % | 0.06 % | 0.00 % | 0.00 % | Stomach pH |
| 2.000 | -1.93 | 79.02 % | 20.78 % | 0.01 % | 0.05 % | 0.00 % | 0.14 % | |
| 3.000 | -0.53 | 26.20 % | 68.91 % | 0.35 % | 0.02 % | 0.00 % | 4.53 % | |
| 4.000 | 0.57 | 2.18 % | 57.29 % | 2.87 % | 0.00 % | 0.00 % | 37.65 % | |
| 5.000 | 1.43 | 0.05 % | 12.38 % | 6.21 % | 0.00 % | 0.00 % | 81.37 % | |
| 6.000 | 1.83 | 0.00 % | 1.39 % | 6.99 % | 0.00 % | 0.00 % | 91.62 % | |
| 6.500 | 1.88 | 0.00 % | 0.45 % | 7.05 % | 0.00 % | 0.00 % | 92.50 % | |
| 7.000 | 1.90 | 0.00 % | 0.14 % | 7.08 % | 0.00 % | 0.00 % | 92.78 % | |
| 7.400 | 1.91 | 0.00 % | 0.06 % | 7.08 % | 0.00 % | 0.00 % | 92.86 % | Blood pH |
| 8.000 | 1.91 | 0.00 % | 0.01 % | 7.08 % | 0.00 % | 0.00 % | 92.90 % | |
| 9.000 | 1.91 | 0.00 % | 0.00 % | 7.09 % | 0.00 % | 0.00 % | 92.91 % | |
| 10.000 | 1.91 | 0.00 % | 0.00 % | 7.09 % | 0.00 % | 0.00 % | 92.91 % | |
| 11.000 | 1.91 | 0.00 % | 0.00 % | 7.09 % | 0.00 % | 0.00 % | 92.91 % | |
| 12.000 | 1.91 | 0.00 % | 0.00 % | 7.09 % | 0.00 % | 0.00 % | 92.91 % | |

#### Carbonate and acidity

| | |
|---|---|
| Carbonate | 0.217 mM |
| Acidity error | 0.167 mM |

#### Other graphs

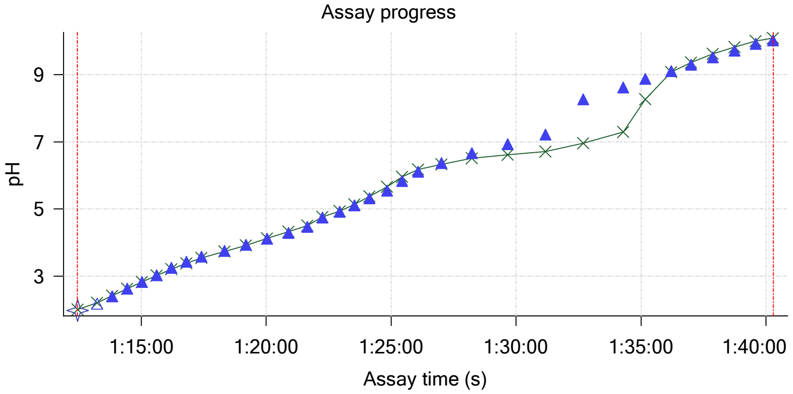


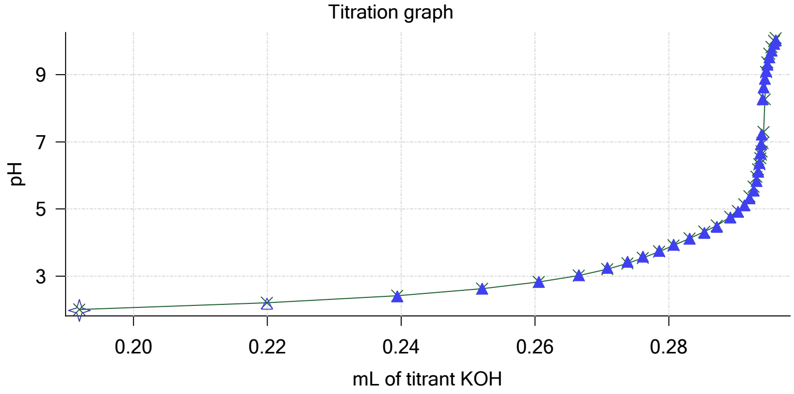

Sample name: **M15_octanol**
Assay name: **pH-metric high logP**
Assay ID: **18B-28002**
Filename: **C:\Sirius_T3\Mehtap\20180227_exp27_logP_T3-2\18B-28002_M15_octanol_pH-metric high logP.t3r**

Experiment start time: **2/28/2018 1:10:47 AM**
Analyst: **Pion**
Instrument ID: **T312060**

#### Other graphs (continued)

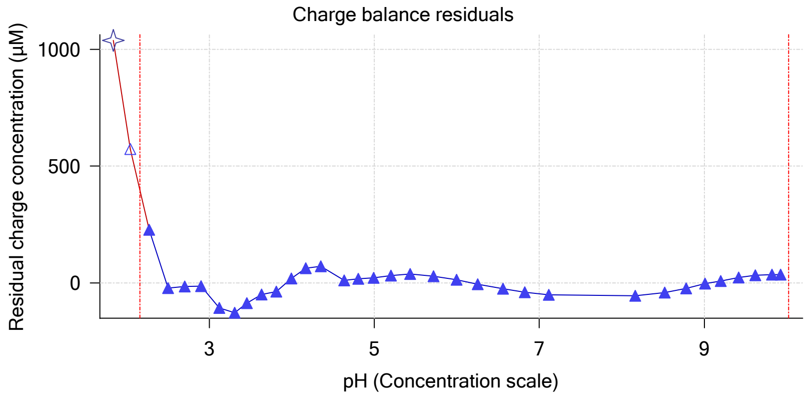


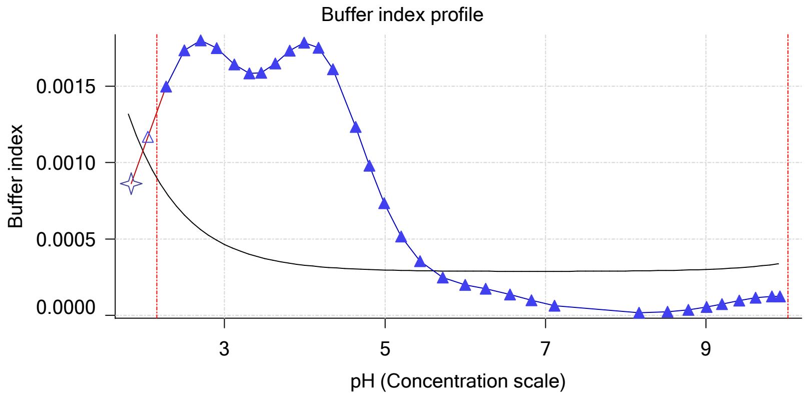


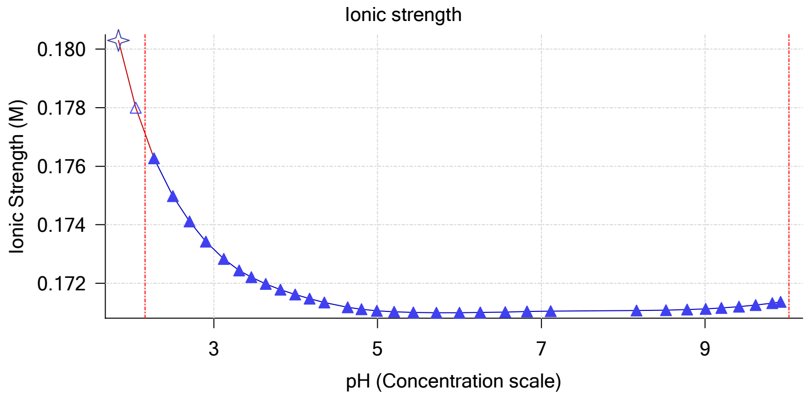


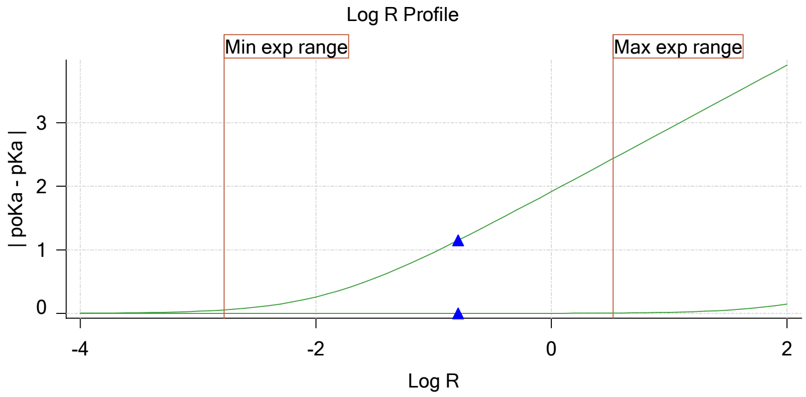


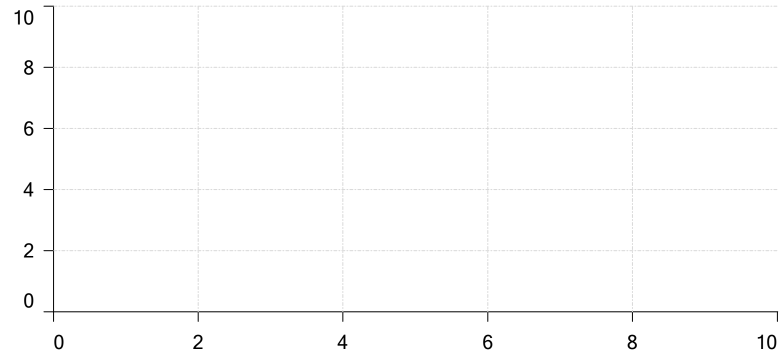


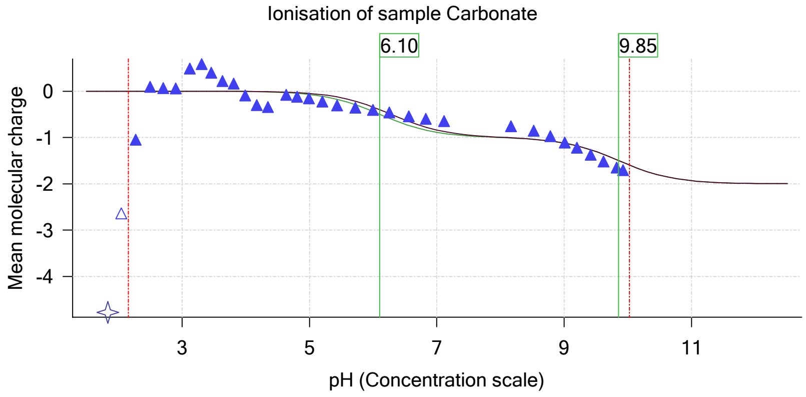

| | | | |
|---|---|---|---|
| Sample name: | **M15_octanol** | Experiment start time: | **2/28/2018 1:10:47 AM** |
| Assay name: | **pH-metric high logP** | Analyst: | **Pion** |
| Assay ID: | **18B-28002** | Instrument ID: | **T312060** |
| Filename: | **C:\Sirius_T3\Mehtap\20180227_exp27_logP_T3-2\18B-28002_M15_octanol_pH-metric high logP.t3r** | | |

#### Assay Model

| Settings | Value | Date/Time changed | Imported from |
|---|---|---|---|
| Sample name | M15_octanol | 2/27/2018 5:03:03 PM | User entered value |
| Sample by | Weight | | Default value |
| Sample weight | 0.001780 g | 2/27/2018 6:41:09 PM | User entered value |
| Formula weight | 209.25 g/mol | 2/27/2018 5:03:03 PM | User entered value |
| Solubility | Unknown | | Default value |
| Molecular weight | 209.25 | 2/27/2018 5:03:03 PM | User entered value |
| Individual pKa ionic environments | No | | Default value |
| Number of pKas | 2 | 2/27/2018 5:03:03 PM | User entered value |
| Sample is a | Base | 2/27/2018 5:03:03 PM | User entered value |
| pKa 1 | 2.58 | 2/27/2018 5:03:03 PM | User entered value |
| pKa 2 | 5.30 | 2/27/2018 5:03:03 PM | User entered value |
| logP (XH2 2+) | -2.38 | 2/27/2018 5:03:37 PM | User entered value |
| logp (XH +) | -1.38 | 2/27/2018 5:03:29 PM | User entered value |
| logP (neutral X) | 1.33 | 2/27/2018 5:03:21 PM | User entered value |

#### Events

| Time | Event | Water | Acid | Base | Octanol | pH | dpH/dt | pH R-squared | pH SD | dpH/dt time |
|---|---|---|---|---|---|---|---|---|---|---|
| 9:12.2 | Initial pH = 6.77 | | | | | | | | | |
| 12:12.8 | Data point 1 | 2.20002 mL | 0.09175 mL | 0.00884 mL | 0.05000 mL | 2.005 | -0.01157 | 0.40408 | 0.00090 | 10.0 s |
| 12:59.2 | Data point 2 | 2.20002 mL | 0.09175 mL | 0.03187 mL | 0.05000 mL | 2.209 | -0.00700 | 0.51322 | 0.00048 | 10.0 s |
| 13:35.0 | Data point 3 | 2.20002 mL | 0.09175 mL | 0.04760 mL | 0.05000 mL | 2.439 | -0.01430 | 0.68245 | 0.00086 | 10.5 s |
| 14:11.1 | Data point 4 | 2.20002 mL | 0.09175 mL | 0.05788 mL | 0.05000 mL | 2.658 | -0.01104 | 0.75037 | 0.00063 | 10.0 s |
| 14:46.7 | Data point 5 | 2.20002 mL | 0.09175 mL | 0.06491 mL | 0.05000 mL | 2.903 | -0.00268 | 0.03532 | 0.00070 | 10.0 s |
| 15:32.6 | Data point 6 | 2.20002 mL | 0.09175 mL | 0.06976 mL | 0.05000 mL | 3.109 | 0.00003 | 0.00001 | 0.00060 | 10.0 s |
| 16:08.1 | Data point 7 | 2.20002 mL | 0.09175 mL | 0.07300 mL | 0.05000 mL | 3.347 | -0.00484 | 0.65284 | 0.00030 | 10.0 s |
| 16:43.6 | Data point 8 | 2.20002 mL | 0.09175 mL | 0.07516 mL | 0.05000 mL | 3.559 | -0.01288 | 0.80904 | 0.00071 | 10.0 s |
| 17:19.1 | Data point 9 | 2.20002 mL | 0.09175 mL | 0.07669 mL | 0.05000 mL | 3.735 | -0.00362 | 0.61081 | 0.00023 | 10.0 s |
| 17:54.5 | Data point 10 | 2.20002 mL | 0.09175 mL | 0.07787 mL | 0.05000 mL | 3.912 | -0.00298 | 0.56183 | 0.00020 | 10.0 s |
| 18:45.5 | Data point 11 | 2.20002 mL | 0.09175 mL | 0.07909 mL | 0.05000 mL | 4.125 | 0.00402 | 0.19132 | 0.00045 | 10.0 s |
| 19:36.3 | Data point 12 | 2.20002 mL | 0.09175 mL | 0.08008 mL | 0.05000 mL | 4.341 | -0.00621 | 0.78463 | 0.00035 | 10.5 s |
| 20:12.2 | Data point 13 | 2.20002 mL | 0.09175 mL | 0.08123 mL | 0.05000 mL | 4.581 | -0.01069 | 0.76499 | 0.00060 | 10.0 s |
| 21:03.2 | Data point 14 | 2.20002 mL | 0.09175 mL | 0.08255 mL | 0.05000 mL | 4.776 | -0.00334 | 0.62508 | 0.00021 | 10.5 s |
| 21:39.2 | Data point 15 | 2.20002 mL | 0.09175 mL | 0.08429 mL | 0.05000 mL | 5.014 | 0.00455 | 0.39258 | 0.00036 | 10.5 s |
| 22:25.5 | Data point 16 | 2.20002 mL | 0.09175 mL | 0.08596 mL | 0.05000 mL | 5.200 | -0.00496 | 0.25675 | 0.00048 | 10.0 s |
| 23:16.5 | Data point 17 | 2.20002 mL | 0.09175 mL | 0.08772 mL | 0.05000 mL | 5.399 | -0.01087 | 0.54074 | 0.00073 | 10.5 s |
| 23:52.5 | Data point 18 | 2.20002 mL | 0.09175 mL | 0.08958 mL | 0.05000 mL | 5.748 | -0.00304 | 0.04364 | 0.00072 | 10.5 s |
| 24:38.7 | Data point 19 | 2.20002 mL | 0.09175 mL | 0.09043 mL | 0.05000 mL | 5.958 | -0.00868 | 0.75112 | 0.00049 | 10.0 s |
| 25:14.1 | Data point 20 | 2.20002 mL | 0.09175 mL | 0.09092 mL | 0.05000 mL | 6.190 | -0.01451 | 0.83075 | 0.00079 | 10.0 s |
| 25:49.5 | Data point 21 | 2.20002 mL | 0.09175 mL | 0.09125 mL | 0.05000 mL | 6.413 | -0.01920 | 0.95433 | 0.00097 | 10.5 s |
| 26:35.7 | Data point 22 | 2.20002 mL | 0.09175 mL | 0.09151 mL | 0.05000 mL | 6.612 | -0.01748 | 0.96637 | 0.00088 | 10.5 s |
| 27:11.6 | Data point 23 | 2.20002 mL | 0.09175 mL | 0.09165 mL | 0.05000 mL | 6.831 | -0.01570 | 0.64440 | 0.00097 | 34.0 s |
| 28:16.3 | Data point 24 | 2.20002 mL | 0.09175 mL | 0.09177 mL | 0.05000 mL | 7.058 | -0.01716 | 0.88880 | 0.00090 | 35.0 s |
| 29:21.8 | Data point 25 | 2.20002 mL | 0.09175 mL | 0.09184 mL | 0.05000 mL | 7.282 | -0.01947 | 0.92580 | 0.00100 | 49.5 s |
| 30:41.9 | Data point 26 | 2.20002 mL | 0.09175 mL | 0.09196 mL | 0.05000 mL | 8.043 | -0.03436 | 0.98140 | 0.00171 | Timed out at 59.5 s |
| 32:17.6 | Data point 27 | 2.20002 mL | 0.09175 mL | 0.09203 mL | 0.05000 mL | 8.319 | -0.01934 | 0.98179 | 0.00096 | 42.5 s |
| 33:35.8 | Data point 28 | 2.20002 mL | 0.09175 mL | 0.09210 mL | 0.05000 mL | 8.564 | -0.01948 | 0.95892 | 0.00098 | 39.0 s |
| 34:50.5 | Data point 29 | 2.20002 mL | 0.09175 mL | 0.09219 mL | 0.05000 mL | 8.842 | -0.01977 | 0.97108 | 0.00099 | 21.0 s |
| 35:42.1 | Data point 30 | 2.20002 mL | 0.09175 mL | 0.09229 mL | 0.05000 mL | 9.067 | -0.01861 | 0.91683 | 0.00096 | 29.0 s |
| 36:41.7 | Data point 31 | 2.20002 mL | 0.09175 mL | 0.09240 mL | 0.05000 mL | 9.270 | -0.01864 | 0.96902 | 0.00094 | 16.0 s |
| 37:28.2 | Data point 32 | 2.20002 mL | 0.09175 mL | 0.09259 mL | 0.05000 mL | 9.504 | -0.01908 | 0.97520 | 0.00095 | 14.0 s |
| 38:23.1 | Data point 33 | 2.20002 mL | 0.09175 mL | 0.09285 mL | 0.05000 mL | 9.705 | -0.01982 | 0.95760 | 0.00100 | 10.5 s |
| 39:09.4 | Data point 34 | 2.20002 mL | 0.09175 mL | 0.09313 mL | 0.05000 mL | 9.900 | -0.01284 | 0.77555 | 0.00072 | 10.5 s |
| 39:45.3 | Data point 35 | 2.20002 mL | 0.09175 mL | 0.09337 mL | 0.05000 mL | 10.019 | -0.01383 | 0.93336 | 0.00071 | 10.0 s |
| 40:51.5 | Data point 36 | 2.20002 mL | 0.19015 mL | 0.09337 mL | 0.15000 mL | 1.966 | -0.01118 | 0.80408 | 0.00062 | 10.0 s |
| 41:38.0 | Data point 37 | 2.20002 mL | 0.19015 mL | 0.12041 mL | 0.15000 mL | 2.180 | 0.01132 | 0.58782 | 0.00073 | 10.0 s |

| | | | |
|---|---|---|---|
| Sample name: | **M15_octanol** | Experiment start time: | **2/28/2018 1:10:47 AM** |
| Assay name: | **pH-metric high logP** | Analyst: | **Pion** |
| Assay ID: | **18B-28002** | Instrument ID: | **T312060** |
| Filename: | **C:\Sirius_T3\Mehtap\20180227_exp27_logP_T3-2\18B-28002_M15_octanol_pH-metric high logP.t3r** | | |

#### Events (continued)

| Time | Event | Water | Acid | Base | Octanol | pH | dpH/dt | pH R-squared | pH SD | dpH/dt time |
|---|---|---|---|---|---|---|---|---|---|---|
| 42:13.9 | Data point 38 | 2.20002 mL | 0.19015 mL | 0.13848 mL | 0.15000 mL | 2.397 | -0.00623 | 0.18166 | 0.00072 | 10.0 s |
| 42:49.6 | Data point 39 | 2.20002 mL | 0.19015 mL | 0.15038 mL | 0.15000 mL | 2.623 | -0.00443 | 0.19218 | 0.00050 | 10.0 s |
| 43:25.2 | Data point 40 | 2.20002 mL | 0.19015 mL | 0.15830 mL | 0.15000 mL | 2.834 | -0.00869 | 0.46386 | 0.00063 | 10.0 s |
| 44:00.8 | Data point 41 | 2.20002 mL | 0.19015 mL | 0.16381 mL | 0.15000 mL | 3.045 | 0.00047 | 0.00116 | 0.00068 | 10.0 s |
| 44:36.3 | Data point 42 | 2.20002 mL | 0.19015 mL | 0.16766 mL | 0.15000 mL | 3.275 | 0.00012 | 0.00005 | 0.00083 | 10.0 s |
| 45:11.8 | Data point 43 | 2.20002 mL | 0.19015 mL | 0.17027 mL | 0.15000 mL | 3.459 | -0.00226 | 0.43409 | 0.00017 | 10.0 s |
| 45:47.3 | Data point 44 | 2.20002 mL | 0.19015 mL | 0.17222 mL | 0.15000 mL | 3.620 | -0.01083 | 0.78989 | 0.00060 | 10.0 s |
| 46:33.1 | Data point 45 | 2.20002 mL | 0.19015 mL | 0.17404 mL | 0.15000 mL | 3.813 | 0.01106 | 0.77170 | 0.00062 | 10.0 s |
| 47:24.1 | Data point 46 | 2.20002 mL | 0.19015 mL | 0.17568 mL | 0.15000 mL | 4.007 | -0.00395 | 0.53793 | 0.00027 | 10.0 s |
| 48:15.0 | Data point 47 | 2.20002 mL | 0.19015 mL | 0.17730 mL | 0.15000 mL | 4.193 | -0.00655 | 0.22705 | 0.00068 | 10.0 s |
| 49:06.0 | Data point 48 | 2.20002 mL | 0.19015 mL | 0.17916 mL | 0.15000 mL | 4.376 | 0.00748 | 0.59754 | 0.00048 | 10.0 s |
| 49:57.1 | Data point 49 | 2.20002 mL | 0.19015 mL | 0.18123 mL | 0.15000 mL | 4.555 | 0.00753 | 0.36418 | 0.00062 | 10.0 s |
| 50:42.9 | Data point 50 | 2.20002 mL | 0.19015 mL | 0.18297 mL | 0.15000 mL | 4.737 | -0.00461 | 0.27683 | 0.00043 | 10.0 s |
| 51:18.3 | Data point 51 | 2.20002 mL | 0.19015 mL | 0.18490 mL | 0.15000 mL | 4.999 | -0.00646 | 0.54035 | 0.00043 | 10.5 s |
| 51:59.5 | Data point 52 | 2.20002 mL | 0.19015 mL | 0.18610 mL | 0.15000 mL | 5.180 | -0.00205 | 0.01692 | 0.00078 | 10.0 s |
| 52:45.2 | Data point 53 | 2.20002 mL | 0.19015 mL | 0.18728 mL | 0.15000 mL | 5.395 | 0.00304 | 0.03920 | 0.00076 | 10.0 s |
| 53:20.6 | Data point 54 | 2.20002 mL | 0.19015 mL | 0.18808 mL | 0.15000 mL | 5.608 | -0.01570 | 0.82199 | 0.00086 | 11.0 s |
| 53:57.1 | Data point 55 | 2.20002 mL | 0.19015 mL | 0.18869 mL | 0.15000 mL | 5.827 | -0.01657 | 0.86910 | 0.00088 | 10.5 s |
| 54:33.0 | Data point 56 | 2.20002 mL | 0.19015 mL | 0.18914 mL | 0.15000 mL | 6.061 | -0.01364 | 0.47158 | 0.00098 | 11.5 s |
| 55:09.9 | Data point 57 | 2.20002 mL | 0.19015 mL | 0.18944 mL | 0.15000 mL | 6.322 | -0.01837 | 0.84621 | 0.00099 | 16.0 s |
| 55:56.4 | Data point 58 | 2.20002 mL | 0.19015 mL | 0.18965 mL | 0.15000 mL | 6.525 | -0.01620 | 0.78360 | 0.00090 | 15.0 s |
| 56:42.0 | Data point 59 | 2.20002 mL | 0.19015 mL | 0.18979 mL | 0.15000 mL | 6.710 | -0.01910 | 0.93669 | 0.00097 | 31.0 s |
| 57:38.4 | Data point 60 | 2.20002 mL | 0.19015 mL | 0.18989 mL | 0.15000 mL | 7.095 | -0.01967 | 0.99319 | 0.00097 | 59.0 s |
| 59:08.1 | Data point 61 | 2.20002 mL | 0.19015 mL | 0.18998 mL | 0.15000 mL | 7.531 | -0.03741 | 0.97375 | 0.00187 | Timed out at 59.5 s |
| 1:00:33.4 | Data point 62 | 2.20002 mL | 0.19015 mL | 0.19001 mL | 0.15000 mL | 7.625 | -0.01967 | 0.98081 | 0.00098 | 53.5 s |
| 1:02:02.7 | Data point 63 | 2.20002 mL | 0.19015 mL | 0.19007 mL | 0.15000 mL | 8.014 | -0.01881 | 0.97922 | 0.00094 | 58.0 s |
| 1:03:36.4 | Data point 64 | 2.20002 mL | 0.19015 mL | 0.19015 mL | 0.15000 mL | 8.332 | -0.01820 | 0.91738 | 0.00094 | 45.0 s |
| 1:05:02.3 | Data point 65 | 2.20002 mL | 0.19015 mL | 0.19024 mL | 0.15000 mL | 8.633 | -0.01854 | 0.84927 | 0.00099 | 18.0 s |
| 1:06:01.1 | Data point 66 | 2.20002 mL | 0.19015 mL | 0.19038 mL | 0.15000 mL | 8.886 | -0.01878 | 0.92878 | 0.00096 | 21.5 s |
| 1:07:03.5 | Data point 67 | 2.20002 mL | 0.19015 mL | 0.19052 mL | 0.15000 mL | 9.109 | -0.01850 | 0.95816 | 0.00093 | 15.0 s |
| 1:07:54.1 | Data point 68 | 2.20002 mL | 0.19015 mL | 0.19069 mL | 0.15000 mL | 9.310 | -0.01772 | 0.81325 | 0.00097 | 10.5 s |
| 1:08:45.4 | Data point 69 | 2.20002 mL | 0.19015 mL | 0.19095 mL | 0.15000 mL | 9.509 | -0.01583 | 0.65574 | 0.00097 | 17.0 s |
| 1:09:33.1 | Data point 70 | 2.20002 mL | 0.19015 mL | 0.19130 mL | 0.15000 mL | 9.756 | -0.01460 | 0.55168 | 0.00097 | 16.5 s |
| 1:10:30.5 | Data point 71 | 2.20002 mL | 0.19015 mL | 0.19172 mL | 0.15000 mL | 9.948 | -0.00681 | 0.21656 | 0.00072 | 10.0 s |
| 1:11:11.1 | Data point 72 | 2.20002 mL | 0.19015 mL | 0.19191 mL | 0.15000 mL | 10.017 | 0.00087 | 0.01530 | 0.00035 | 10.0 s |
| 1:12:27.4 | Data point 73 | 2.20002 mL | 0.29403 mL | 0.19191 mL | 0.45000 mL | 1.972 | -0.01003 | 0.70683 | 0.00059 | 10.0 s |
| 1:13:13.9 | Data point 74 | 2.20002 mL | 0.29403 mL | 0.21997 mL | 0.45000 mL | 2.175 | 0.00579 | 0.71483 | 0.00034 | 10.5 s |
| 1:13:50.3 | Data point 75 | 2.20002 mL | 0.29403 mL | 0.23942 mL | 0.45000 mL | 2.398 | -0.00050 | 0.00121 | 0.00071 | 10.0 s |
| 1:14:26.0 | Data point 76 | 2.20002 mL | 0.29403 mL | 0.25212 mL | 0.45000 mL | 2.626 | -0.01002 | 0.42775 | 0.00076 | 10.0 s |
| 1:15:01.6 | Data point 77 | 2.20002 mL | 0.29403 mL | 0.26056 mL | 0.45000 mL | 2.826 | -0.00389 | 0.08255 | 0.00067 | 10.0 s |
| 1:15:37.2 | Data point 78 | 2.20002 mL | 0.29403 mL | 0.26653 mL | 0.45000 mL | 3.025 | 0.01417 | 0.79174 | 0.00079 | 10.0 s |
| 1:16:12.6 | Data point 79 | 2.20002 mL | 0.29403 mL | 0.27081 mL | 0.45000 mL | 3.243 | 0.00428 | 0.28164 | 0.00040 | 10.0 s |
| 1:16:48.2 | Data point 80 | 2.20002 mL | 0.29403 mL | 0.27382 mL | 0.45000 mL | 3.430 | 0.00870 | 0.54657 | 0.00058 | 10.5 s |
| 1:17:24.2 | Data point 81 | 2.20002 mL | 0.29403 mL | 0.27613 mL | 0.45000 mL | 3.577 | -0.01656 | 0.72331 | 0.00096 | 15.0 s |
| 1:18:20.2 | Data point 82 | 2.20002 mL | 0.29403 mL | 0.27858 mL | 0.45000 mL | 3.754 | 0.00239 | 0.04126 | 0.00058 | 10.0 s |
| 1:19:11.2 | Data point 83 | 2.20002 mL | 0.29403 mL | 0.28074 mL | 0.45000 mL | 3.931 | -0.00851 | 0.21217 | 0.00091 | 10.0 s |
| 1:20:02.2 | Data point 84 | 2.20002 mL | 0.29403 mL | 0.28311 mL | 0.45000 mL | 4.112 | 0.00633 | 0.50236 | 0.00044 | 10.0 s |
| 1:20:53.2 | Data point 85 | 2.20002 mL | 0.29403 mL | 0.28530 mL | 0.45000 mL | 4.288 | 0.00616 | 0.42147 | 0.00047 | 10.0 s |
| 1:21:39.1 | Data point 86 | 2.20002 mL | 0.29403 mL | 0.28716 mL | 0.45000 mL | 4.468 | 0.00653 | 0.41686 | 0.00050 | 10.5 s |
| 1:22:15.1 | Data point 87 | 2.20002 mL | 0.29403 mL | 0.28920 mL | 0.45000 mL | 4.750 | -0.01024 | 0.56608 | 0.00067 | 10.5 s |
| 1:22:56.2 | Data point 88 | 2.20002 mL | 0.29403 mL | 0.29033 mL | 0.45000 mL | 4.921 | -0.00504 | 0.30494 | 0.00045 | 10.5 s |
| 1:23:32.1 | Data point 89 | 2.20002 mL | 0.29403 mL | 0.29128 mL | 0.45000 mL | 5.107 | -0.00661 | 0.13487 | 0.00089 | 10.0 s |
| 1:24:07.6 | Data point 90 | 2.20002 mL | 0.29403 mL | 0.29207 mL | 0.45000 mL | 5.317 | -0.00338 | 0.02804 | 0.00100 | 17.0 s |
| 1:24:50.0 | Data point 91 | 2.20002 mL | 0.29403 mL | 0.29269 mL | 0.45000 mL | 5.549 | -0.00970 | 0.38066 | 0.00078 | 11.0 s |
| 1:25:26.4 | Data point 92 | 2.20002 mL | 0.29403 mL | 0.29311 mL | 0.45000 mL | 5.831 | -0.01048 | 0.30330 | 0.00094 | 12.5 s |
| 1:26:04.3 | Data point 93 | 2.20002 mL | 0.29403 mL | 0.29337 mL | 0.45000 mL | 6.110 | -0.01816 | 0.86837 | 0.00096 | 31.0 s |

| | | | |
|---|---|---|---|
| Sample name: | **M15_octanol** | Experiment start time: | **2/28/2018 1:10:47 AM** |
| Assay name: | **pH-metric high logP** | Analyst: | **Pion** |
| Assay ID: | **18B-28002** | Instrument ID: | **T312060** |
| Filename: | **C:\Sirius_T3\Mehtap\20180227_exp27_logP_T3-2\18B-28002_M15_octanol_pH-metric high logP.t3r** | | |

#### Events (continued)

| Time | Event | Water | Acid | Base | Octanol | pH | dpH/dt | pH R-squared | pH SD | dpH/dt time |
|---|---|---|---|---|---|---|---|---|---|---|
| 1:27:00.8 | Data point 94 | 2.20002 mL | 0.29403 mL | 0.29353 mL | 0.45000 mL | 6.366 | -0.01843 | 0.94543 | 0.00094 | 42.0 s |
| 1:28:13.4 | Data point 95 | 2.20002 mL | 0.29403 mL | 0.29370 mL | 0.45000 mL | 6.669 | -0.01870 | 0.98301 | 0.00093 | 56.0 s |
| 1:29:40.0 | Data point 96 | 2.20002 mL | 0.29403 mL | 0.29379 mL | 0.45000 mL | 6.935 | -0.02391 | 0.96793 | 0.00120 | Timed out at 59.5 s |
| 1:31:10.7 | Data point 97 | 2.20002 mL | 0.29403 mL | 0.29386 mL | 0.45000 mL | 7.222 | -0.03766 | 0.98462 | 0.00187 | Timed out at 59.5 s |
| 1:32:41.3 | Data point 98 | 2.20002 mL | 0.29403 mL | 0.29403 mL | 0.45000 mL | 8.270 | -0.03076 | 0.99007 | 0.00153 | Timed out at 59.5 s |
| 1:34:16.9 | Data point 99 | 2.20002 mL | 0.29403 mL | 0.29417 mL | 0.45000 mL | 8.626 | -0.01455 | 0.55806 | 0.00096 | 17.5 s |
| 1:35:10.1 | Data point 100 | 2.20002 mL | 0.29403 mL | 0.29433 mL | 0.45000 mL | 8.885 | 0.00677 | 0.11530 | 0.00099 | 21.5 s |
| 1:36:12.4 | Data point 101 | 2.20002 mL | 0.29403 mL | 0.29454 mL | 0.45000 mL | 9.111 | -0.00911 | 0.28833 | 0.00084 | 11.5 s |
| 1:36:59.8 | Data point 102 | 2.20002 mL | 0.29403 mL | 0.29473 mL | 0.45000 mL | 9.302 | -0.01144 | 0.38459 | 0.00091 | 12.0 s |
| 1:37:52.6 | Data point 103 | 2.20002 mL | 0.29403 mL | 0.29501 mL | 0.45000 mL | 9.516 | -0.01706 | 0.74556 | 0.00098 | 11.0 s |
| 1:38:44.4 | Data point 104 | 2.20002 mL | 0.29403 mL | 0.29534 mL | 0.45000 mL | 9.718 | -0.01826 | 0.87768 | 0.00096 | 10.0 s |
| 1:39:35.4 | Data point 105 | 2.20002 mL | 0.29403 mL | 0.29574 mL | 0.45000 mL | 9.922 | -0.00385 | 0.15864 | 0.00048 | 10.5 s |
| 1:40:16.4 | Data point 106 | 2.20002 mL | 0.29403 mL | 0.29598 mL | 0.45000 mL | 10.023 | -0.00776 | 0.63643 | 0.00048 | 10.5 s |
| 1:40:36.1 | Assay volumes | 2.20002 mL | 0.29403 mL | 0.29598 mL | 0.45000 mL | | | | | |

| | | | |
|---|---|---|---|
| Sample name: | **M15_octanol** | Experiment start time: | **2/28/2018 1:10:47 AM** |
| Assay name: | **pH-metric high logP** | Analyst: | **Pion** |
| Assay ID: | **18B-28002** | Instrument ID: | **T312060** |
| Filename: | **C:\Sirius_T3\Mehtap\20180227_exp27_logP_T3-2\18B-28002_M15_octanol_pH-metric high logP.t3r** | | |

#### Assay Settings

| Setting | Value | Original Value | Date/Time changed | Imported from |
|---|---|---|---|---|
| ***General Settings*** | | | | |
| Analyst name | Pion | | | |
| ***Standard Experiment Settings*** | | | | |
| Number of titrations | 3 | | | |
| Minimum pH | 2.000 | | | |
| Maximum pH | 10.000 | | | |
| pH step between points of | 0.200 | | | |
| Minimum titrant addition | 0.00002 mL | | | |
| Maximum titrant addition | 0.10000 mL | | | |
| Argon flow rate | 100% | | | |
| Start titration using | Cautious pH adjust | | | |
| ***Advanced General Settings*** | | | | |
| Detect turbidity using | None | | | |
| Collect turbidity sensor data | No | | | |
| Collect UV spectra | No | | | |
| Stir after titrant addition for | 5 seconds | | | |
| For titrant addition, stir at | 10% | | | |
| ***Titrant Pre-Dose*** | | | | |
| Titrant pre-dose | None | | | |
| ***Assay Medium*** | | | | |
| ISA water volume | 2.20 mL | | | |
| Water added | Automatic | | | |
| Partition solvent type | Octanol | | | |
| Partition volume | 0.050 mL | | | |
| Partition solvent added | Automatic | | | |
| After partition addition, stir for | 1 seconds | | | |
| ***Sample Sonication*** | | | | |
| Sonicate | Yes | | | |
| Adjust pH for sonication | No | | | |
| Sonicate for | 300 seconds | | | |
| After sonication stir for | 5 seconds | | | |
| ***Sample Dissolution*** | | | | |
| Perform a dissolution stage | Yes | | | |
| Adjust and hold pH for dissolution | To start pH | | | |
| Stir to dissolve for | 120 seconds | | | |
| For dissolution, stir at | 10% | | | |
| ***Carbonate purge*** | | | | |
| Perform a carbonate purge | No | | | |
| ***Temperature Control*** | | | | |
| Wait for temperature | Yes | | | |
| Required start temperature | 25.0°C | | | |
| Acceptable deviation | 0.5°C | | | |
| Time to wait | 60 seconds | | | |
| Stir speed of | 50% | | | |
| ***Titration 1*** | | | | |
| Titrate from | Low to high pH | | | |
| Adjust to start pH | Yes | | | |
| After pH adjust stir for | 30 seconds | | | |
| Stir to allow partitioning for | 15 seconds | | | |
| Stirrer speed for partitioning | 50% | | | |
| ***Titration 2*** | | | | |
| Titrate from | Low to high pH | | | |
| Add additional water | 0.00 mL | | | |
| Additional partition solvent volume | 0.100 mL | | | |
| Additional partition solvent added | Automatic | | | |
| After pH adjust stir for | 30 seconds | | | |
| Stir to allow partitioning for | 15 seconds | | | |
| Stirrer speed for partitioning | 55% | | | |

| | | | |
|---|---|---|---|
| Sample name: | **M15_octanol** | Experiment start time: | **2/28/2018 1:10:47 AM** |
| Assay name: | **pH-metric high logP** | Analyst: | **Pion** |
| Assay ID: | **18B-28002** | Instrument ID: | **T312060** |
| Filename: | **C:\Sirius_T3\Mehtap\20180227_exp27_logP_T3-2\18B-28002_M15_octanol_pH-metric high logP.t3r** | | |

#### Assay Settings (continued)

| Setting | Value | Original Value | Date/Time changed | Imported from |
|---|---|---|---|---|
| ***Titration 3*** | | | | |
| Titrate from | Low to high pH | | | |
| Add additional water | 0.00 mL | | | |
| Additional partition solvent volume | 0.300 mL | | | |
| Additional partition solvent added | Automatic | | | |
| After pH adjust stir for | 30 seconds | | | |
| Stir to allow partitioning for | 15 seconds | | | |
| Stirrer speed for partitioning | 60% | | | |
| ***Data Point Stability*** | | | | |
| Stir during data point collection | No | | | |
| Delay before data point collection | 0 seconds | | | |
| Number of points to average | 20 points | | | |
| Time interval between points | 0.50 seconds | | | |
| Required maximum standard deviation | 0.00100 dpH/dt | | | |
| Stability timeout after | 60 seconds | | | |

#### Calibration Settings

| Setting | Value | Date/Time changed | Imported from |
|---|---|---|---|
| Four-Plus alpha | 0.130 | 2/28/2018 1:10:47 AM | C:\Sirius_T3\HCl18B27.t3r |
| Four-Plus S | 0.9970 | 2/28/2018 1:10:47 AM | C:\Sirius_T3\HCl18B27.t3r |
| Four-Plus jH | 0.8 | 2/28/2018 1:10:47 AM | C:\Sirius_T3\HCl18B27.t3r |
| Four-Plus jOH | -0.4 | 2/28/2018 1:10:47 AM | C:\Sirius_T3\HCl18B27.t3r |
| Base concentration factor | 1.000 | 2/28/2018 1:10:47 AM | C:\Sirius_T3\KOH18B27.t3r |
| Acid concentration factor | 0.994 | 2/28/2018 1:10:47 AM | C:\Sirius_T3\HCl18B27.t3r |

#### Instrument Settings

| Setting | Value | Batch Id | Install date |
|---|---|---|---|
| Instrument owner | Merck | | |
| Instrument ID | T312060 | | |
| Instrument type | T3 Simulator | | |
| Software version | 1.1.3.0 | | |
| Dispenser module | | T3DM1200361 | 3/31/2009 5:24:52 AM |
| Dispenser 0 | Water | | 3/31/2009 5:25:05 AM |
| Syringe volume | 2.5 mL | | |
| Firmware version | 1.2.1(r2) | | |
| Titrant | Water (0.15 M KCl) | 02-06-2018 | 2/27/2018 10:05:59 AM |
| Dispenser 2 | Acid | | 3/31/2009 5:25:11 AM |
| Syringe volume | 0.5 mL | | |
| Firmware version | 1.2.1(r2) | | |
| Titrant | Acid (0.5 M HCl) | 02-27-2018 | 2/27/2018 10:27:22 AM |
| Dispenser 1 | Base | | 3/31/2009 5:25:21 AM |
| Syringe volume | 0.5 mL | | |
| Firmware version | 1.2.1(r2) | | |
| Titrant | Base (0.5 M KOH) | 9/22/2017 | 2/27/2018 10:21:22 AM |
| Dispenser 5 | Cosolvent | | 3/31/2009 5:26:24 AM |
| Syringe volume | 2.5 mL | | |
| Firmware version | 1.2.1(r2) | | |
| Distribution valve 5 | Distribution Valve | | 3/31/2009 5:28:19 AM |
| Firmware version | 1.1.3 | | |
| Port A | Methanol (80%, 0.15 M KCl) | 09-26-17 | 2/7/2018 9:42:01 AM |
| Port B | Cyclohexane | 11-01-17 | 2/27/2018 10:37:57 AM |
| Dispenser 3 | Buffer | | 8/3/2010 5:05:16 AM |
| Syringe volume | 0.5 mL | | |
| Firmware version | 1.2.1(r2) | | |
| Titrant | Dodecane | 2018/01/31 | 1/31/2018 12:26:26 PM |
| Dispenser 6 | Octanol | | 10/22/2010 10:52:43 AM |

| | | | |
|---|---|---|---|
| Sample name: | **M15_octanol** | Experiment start time: | **2/28/2018 1:10:47 AM** |
| Assay name: | **pH-metric high logP** | Analyst: | **Pion** |
| Assay ID: | **18B-28002** | Instrument ID: | **T312060** |
| Filename: | **C:\Sirius_T3\Mehtap\20180227_exp27_logP_T3-2\18B-28002_M15_octanol_pH-metric high logP.t3r** | | |

#### Instrument Settings (continued)

| Setting | Value | Batch Id | Install date |
|---|---|---|---|
| Syringe volume | 0.5 mL | | |
| Firmware version | 1.2.1(r2) | | |
| Titrant | Octanol | 01-31-2018 | 2/27/2018 9:59:35 AM |
| Titrator | | T3TM1200161 | 3/31/2009 5:24:17 AM |
| Horizontal axis firmware version | 1.17 AI1DI2DO2 Stepper 2 | | |
| Vertical axis firmware version | 1.17 AI1DI2DO2 Stepper 2 | | |
| Chassis I/O firmware version | 1.11 AI1DI0DO4 Norgren I/O | | |
| Probe I/O firmware version | 1.1.1 | | |
| Electrode | T3 Electrode | T3E0923 | 1/23/2018 2:01:00 PM |
| E0 calibration | +2.60 mV | | 2/28/2018 1:11:15 AM |
| Filling solution | 3M KCl | KCL097 | 2/27/2018 9:49:43 AM |
| Liquids | | | |
| Wash 1 | 50% IPA:50% Water | | 2/27/2018 9:49:58 AM |
| Wash 2 | 0.5% Trition X-100 in H20 | | 2/27/2018 9:50:01 AM |
| Buffer position 1 | pH7 Wash | | 2/27/2018 9:50:04 AM |
| Buffer position 2 | pH 7 | | 2/27/2018 9:50:06 AM |
| Storage position | | | 2/27/2018 9:55:12 AM |
| Wash water | 9.1e+003 mL | 02-27-2018 | 2/27/2018 9:54:39 AM |
| Waste | 6.3e+003 mL | | 11/28/2017 10:36:29 AM |
| Temperature controller | | | 8/5/2010 6:35:13 AM |
| Turbidity detector | | | 3/31/2009 5:24:45 AM |
| Spectrometer | | 074811 | 11/23/2010 11:22:28 AM |
| Dip probe | | 10196 | |
| Wavelength coefficient A0 | 183.333 | | |
| Wavelength coefficient A1 | 2.21568 | | |
| Wavelength coefficient A2 | -0.000289308 | | |
| Total lamp lit time | 110:52:56 | | 11/23/2010 11:22:28 AM |
| Calibrated on | 2/27/2018 10:40:38 AM | | |
| Integration time | 40 | | |
| Scans averaged | 10 | | |
| Autoloader | | T3AL1200345 | 11/10/2015 9:34:13 AM |
| Left-right axis firmware version | 1.17 AI1DI2DO2 Stepper 2 | | |
| Front-back axis firmware version | 1.17 AI1DI2DO2 Stepper 2 | | |
| Vertical axis firmware version | 1.17 AI1DI2DO2 Stepper 2 | | |
| Chassis I/O firmware version | 1.11 AI1DI0DO4 Norgren I/O | | |
| Configuration | | | |
| Alternate titration position | Titration position | | |
| Alternate reference position | Reference position | | |
| Maximum standard vial volume | 3.50 mL | | |
| Maximum alternate vial volume | 25.00 mL | | |
| Automatic action idle period | 5 minute(s) | | |
| Titrant tube volume | 1.3 mL | | |
| Syringe flush count | 3.50 | | |
| Flowing wash pump volume | 20.0 mL | | |
| Flowing wash stir duration | 5 s | | |
| Flowing wash stir speed | 30% | | |
| Solvent wash stir duration | 5 s | | |
| Solvent wash stir speed | 30% | | |
| Surfactant wash stir duration | 5 s | | |
| Surfactant wash stir speed | 30% | | |
| E0 calibration minimum number of points | 10 | | |
| E0 calibration maximum standard deviation | 0.01500 | | |
| E0 calibration timeout period | 60 s | | |
| E0 calibration stir duration | 5 s | | |
| E0 calibration preparation stir speed | 30% | | |
| E0 calibration buffer wash stir duration | 5 s | | |
| E0 calibration buffer wash stir speed | 30% | | |
| E0 calibration reading stir speed | 0% | | |

| | | | |
|---|---|---|---|
| Sample name: | **M15_octanol** | Experiment start time: | **2/28/2018 1:10:47 AM** |
| Assay name: | **pH-metric high logP** | Analyst: | **Pion** |
| Assay ID: | **18B-28002** | Instrument ID: | **T312060** |
| Filename: | **C:\Sirius_T3\Mehtap\20180227_exp27_logP_T3-2\18B-28002_M15_octanol_pH-metric high logP.t3r** | | |

#### Instrument Settings (continued)

| Setting | Value | Batch Id | Install date |
|---|---|---|---|
| Spectrometer calibration stir duration | 5 s | | |
| Spectrometer calibration stir speed | 30% | | |
| Spectrometer calibration wash pump volume | 20.0 mL | | |
| Spectrometer calibration wash stir duration | 5 s | | |
| Spectrometer calibration wash stir speed | 30% | | |
| Overhead dispense height | 10000 | | |

#### Refinement Settings

| Setting | Value | Default value |
|---|---|---|
| Turbidity detection method | None | None |
| Turbidity wavelength to assess | 500.0 nm | 500.0 nm |
| Turbidity maximum absorbance | 0.100 | 0.100 |
| Turbidity probe threshold | 50.00 | 50.00 |

#### Experiment Log

[2:37] Air gap created for Water (0.15 M KCl)
[2:37] Air gap created for Acid (0.5 M HCl)
[2:38] Air gap created for Base (0.5 M KOH)
[2:38] Air gap released for Water (0.15 M KCl)
[2:42] Titrator arm moved over Titration position
[2:42] Titration 1 of 3
[2:42] Adding initial titrants
[2:42] Automatically add 2.20000 mL of water
[3:19] Dispensed 2.200024 mL of Water (0.15 M KCl)
[3:23] Titrator arm moved over Drain
[9:05] Titrator arm moved to Titration position
[9:05] Argon flow rate set to 100
[9:05] Stirrer speed set to 10
[9:10] Automatically add 0.05000 mL of Octanol
[9:11] Dispensed 0.050000 mL of Octanol
[9:12] Initial pH = 6.77
[9:12] Iterative adjust 6.77 -> 2.00
[9:12] pH 6.77 -> 2.00
[9:15] Air gap released for Acid (0.5 M HCl)
[9:15] Dispensed 0.091745 mL of Acid (0.5 M HCl)
[9:21] Holding pH 2.00
[11:21] Stirrer speed set to 0
[11:21] Stirrer speed set to 50
[11:21] Iterative adjust 1.95 -> 2.00
[11:21] pH 1.95 -> 2.00
[11:21] Air gap released for Base (0.5 M KOH)
[11:22] Dispensed 0.008843 mL of Base (0.5 M KOH)
[12:13] Stirrer speed set to 0
[12:23] Datapoint id 1 collected
[12:23] Stirrer speed set to 50
[12:28] pH 2.01 -> 2.21
[12:28] Using cautious pH adjust
[12:28] Dispensed 0.012112 mL of Base (0.5 M KOH)
[12:33] Stepping pH = 2.10
[12:34] Dispensed 0.008631 mL of Base (0.5 M KOH)
[12:39] Stepping pH = 2.18
[12:39] Dispensed 0.002281 mL of Base (0.5 M KOH)
[12:44] Stepping pH = 2.21
[12:59] Stirrer speed set to 0
[13:09] Datapoint id 2 collected
[13:09] Charge balance equation is out by 4.9%
[13:09] Stirrer speed set to 50

| | | | |
|---|---|---|---|
| Sample name: | **M15_octanol** | Experiment start time: | **2/28/2018 1:10:47 AM** |
| Assay name: | **pH-metric high logP** | Analyst: | **Pion** |
| Assay ID: | **18B-28002** | Instrument ID: | **T312060** |
| Filename: | **C:\Sirius_T3\Mehtap\20180227_exp27_logP_T3-2\18B-28002_M15_octanol_pH-metric high logP.t3r** | | |

#### Experiment Log (continued)

[13:14] pH 2.22 -> 2.42
[13:14] Using charge balance adjust
[13:15] Dispensed 0.015734 mL of Base (0.5 M KOH)
[13:35] Stirrer speed set to 0
[13:45] Datapoint id 3 collected
[13:45] Charge balance equation is out by 9.6%
[13:45] Stirrer speed set to 50
[13:50] pH 2.44 -> 2.64
[13:50] Using charge balance adjust
[13:51] Dispensed 0.010278 mL of Base (0.5 M KOH)
[14:11] Stirrer speed set to 0
[14:21] Datapoint id 4 collected
[14:21] Charge balance equation is out by 6.7%
[14:21] Stirrer speed set to 50
[14:26] pH 2.66 -> 2.86
[14:26] Using charge balance adjust
[14:26] Dispensed 0.007032 mL of Base (0.5 M KOH)
[14:47] Stirrer speed set to 0
[14:57] Datapoint id 5 collected
[14:57] Charge balance equation is out by 19.7%
[14:57] Stirrer speed set to 50
[15:02] pH 2.91 -> 3.11
[15:02] Using cautious pH adjust
[15:02] Dispensed 0.002305 mL of Base (0.5 M KOH)
[15:07] Stepping pH = 2.99
[15:07] Dispensed 0.002023 mL of Base (0.5 M KOH)
[15:12] Stepping pH = 3.08
[15:12] Dispensed 0.000517 mL of Base (0.5 M KOH)
[15:17] Stepping pH = 3.11
[15:32] Stirrer speed set to 0
[15:42] Datapoint id 6 collected
[15:42] Charge balance equation is out by -4.8%
[15:42] Stirrer speed set to 50
[15:48] pH 3.12 -> 3.32
[15:48] Using charge balance adjust
[15:48] Dispensed 0.003246 mL of Base (0.5 M KOH)
[16:08] Stirrer speed set to 0
[16:18] Datapoint id 7 collected
[16:18] Charge balance equation is out by 13.6%
[16:18] Stirrer speed set to 50
[16:23] pH 3.35 -> 3.55
[16:23] Using charge balance adjust
[16:23] Dispensed 0.002164 mL of Base (0.5 M KOH)
[16:43] Stirrer speed set to 0
[16:53] Datapoint id 8 collected
[16:53] Charge balance equation is out by 3.6%
[16:53] Stirrer speed set to 50
[16:59] pH 3.56 -> 3.76
[16:59] Using charge balance adjust
[16:59] Dispensed 0.001529 mL of Base (0.5 M KOH)
[17:19] Stirrer speed set to 0
[17:29] Datapoint id 9 collected
[17:29] Charge balance equation is out by -14.3%
[17:29] Stirrer speed set to 50
[17:34] pH 3.75 -> 3.95
[17:34] Using charge balance adjust
[17:34] Dispensed 0.001176 mL of Base (0.5 M KOH)
[17:54] Stirrer speed set to 0
[18:04] Datapoint id 10 collected

Sample name: **M15_octanol** Experiment start time: **2/28/2018 1:10:47 AM**
Assay name: **pH-metric high logP** Analyst: **Pion**
Assay ID: **18B-28002** Instrument ID: **T312060**
Filename: **C:\Sirius_T3\Mehtap\20180227_exp27_logP_T3-2\18B-28002_M15_octanol_pH-metric high logP.t3r**

#### Experiment Log (continued)

[18:04] Charge balance equation is out by -20.3%
[18:04] Stirrer speed set to 50
[18:09] pH 3.92 -> 4.12
[18:09] Using cautious pH adjust
[18:10] Dispensed 0.000517 mL of Base (0.5 M KOH)
[18:15] Stepping pH = 4.00
[18:15] Dispensed 0.000470 mL of Base (0.5 M KOH)
[18:20] Stepping pH = 4.09
[18:20] Dispensed 0.000118 mL of Base (0.5 M KOH)
[18:25] Stepping pH = 4.10
[18:25] Dispensed 0.000118 mL of Base (0.5 M KOH)
[18:30] Stepping pH = 4.12
[18:45] Stirrer speed set to 0
[18:55] Datapoint id 11 collected
[18:55] Charge balance equation is out by -18.7%
[18:55] Stirrer speed set to 50
[19:00] pH 4.13 -> 4.33
[19:00] Using cautious pH adjust
[19:00] Dispensed 0.000517 mL of Base (0.5 M KOH)
[19:05] Stepping pH = 4.26
[19:06] Dispensed 0.000259 mL of Base (0.5 M KOH)
[19:11] Stepping pH = 4.31
[19:11] Dispensed 0.000071 mL of Base (0.5 M KOH)
[19:16] Stepping pH = 4.32
[19:16] Dispensed 0.000141 mL of Base (0.5 M KOH)
[19:21] Stepping pH = 4.35
[19:36] Stirrer speed set to 0
[19:47] Datapoint id 12 collected
[19:47] Charge balance equation is out by 3.7%
[19:47] Stirrer speed set to 50
[19:52] pH 4.35 -> 4.55
[19:52] Using charge balance adjust
[19:52] Dispensed 0.001152 mL of Base (0.5 M KOH)
[20:12] Stirrer speed set to 0
[20:22] Datapoint id 13 collected
[20:22] Charge balance equation is out by 17.7%
[20:22] Stirrer speed set to 50
[20:27] pH 4.59 -> 4.79
[20:27] Using cautious pH adjust
[20:27] Dispensed 0.000729 mL of Base (0.5 M KOH)
[20:32] Stepping pH = 4.71
[20:33] Dispensed 0.000400 mL of Base (0.5 M KOH)
[20:38] Stepping pH = 4.77
[20:38] Dispensed 0.000094 mL of Base (0.5 M KOH)
[20:43] Stepping pH = 4.77
[20:43] Dispensed 0.000094 mL of Base (0.5 M KOH)
[20:48] Stepping pH = 4.79
[21:03] Stirrer speed set to 0
[21:14] Datapoint id 14 collected
[21:14] Charge balance equation is out by 8.9%
[21:14] Stirrer speed set to 50
[21:19] pH 4.78 -> 4.98
[21:19] Using charge balance adjust
[21:19] Dispensed 0.001740 mL of Base (0.5 M KOH)
[21:39] Stirrer speed set to 0
[21:50] Datapoint id 15 collected
[21:50] Charge balance equation is out by 16.6%
[21:50] Stirrer speed set to 50
[21:55] pH 5.02 -> 5.22

#### Experiment Log (continued)

[21:55] Using cautious pH adjust
[21:55] Dispensed 0.000988 mL of Base (0.5 M KOH)
[22:00] Stepping pH = 5.14
[22:00] Dispensed 0.000494 mL of Base (0.5 M KOH)
[22:05] Stepping pH = 5.20
[22:05] Dispensed 0.000188 mL of Base (0.5 M KOH)
[22:10] Stepping pH = 5.21
[22:25] Stirrer speed set to 0
[22:35] Datapoint id 16 collected
[22:35] Charge balance equation is out by 15.1%
[22:35] Stirrer speed set to 50
[22:40] pH 5.20 -> 5.40
[22:40] Using cautious pH adjust
[22:41] Dispensed 0.001011 mL of Base (0.5 M KOH)
[22:46] Stepping pH = 5.32
[22:46] Dispensed 0.000517 mL of Base (0.5 M KOH)
[22:51] Stepping pH = 5.39
[22:51] Dispensed 0.000118 mL of Base (0.5 M KOH)
[22:56] Stepping pH = 5.39
[22:56] Dispensed 0.000118 mL of Base (0.5 M KOH)
[23:01] Stepping pH = 5.41
[23:16] Stirrer speed set to 0
[23:27] Datapoint id 17 collected
[23:27] Charge balance equation is out by 12.5%
[23:27] Stirrer speed set to 50
[23:32] pH 5.40 -> 5.60
[23:32] Using charge balance adjust
[23:32] Dispensed 0.001858 mL of Base (0.5 M KOH)
[23:52] Stirrer speed set to 0
[24:03] Datapoint id 18 collected
[24:03] Charge balance equation is out by 72.8%
[24:03] Stirrer speed set to 50
[24:08] pH 5.75 -> 5.95
[24:08] Using cautious pH adjust
[24:08] Dispensed 0.000659 mL of Base (0.5 M KOH)
[24:13] Stepping pH = 5.93
[24:13] Dispensed 0.000071 mL of Base (0.5 M KOH)
[24:18] Stepping pH = 5.93
[24:18] Dispensed 0.000118 mL of Base (0.5 M KOH)
[24:23] Stepping pH = 5.97
[24:38] Stirrer speed set to 0
[24:49] Datapoint id 19 collected
[24:49] Charge balance equation is out by 34.9%
[24:49] Stirrer speed set to 50
[24:54] pH 5.97 -> 6.17
[24:54] Using cautious pH adjust
[24:54] Dispensed 0.000494 mL of Base (0.5 M KOH)
[24:59] Stepping pH = 6.20
[25:14] Stirrer speed set to 0
[25:24] Datapoint id 20 collected
[25:24] Charge balance equation is out by 50.0%
[25:24] Stirrer speed set to 50
[25:29] pH 6.20 -> 6.40
[25:29] Using cautious pH adjust
[25:29] Dispensed 0.000329 mL of Base (0.5 M KOH)
[25:34] Stepping pH = 6.42
[25:49] Stirrer speed set to 0
[26:00] Datapoint id 21 collected
[26:00] Charge balance equation is out by 50.0%

Sample name: **M15_octanol** Experiment start time: **2/28/2018 1:10:47 AM**
Assay name: **pH-metric high logP** Analyst: **Pion**
Assay ID: **18B-28002** Instrument ID: **T312060**
Filename: **C:\Sirius_T3\Mehtap\20180227_exp27_logP_T3-2\18B-28002_M15_octanol_pH-metric high logP.t3r**

#### Experiment Log (continued)

[26:00] Stirrer speed set to 50
[26:05] pH 6.42 -> 6.62
[26:05] Using cautious pH adjust
[26:05] Dispensed 0.000212 mL of Base (0.5 M KOH)
[26:10] Stepping pH = 6.60
[26:10] Dispensed 0.000024 mL of Base (0.5 M KOH)
[26:15] Stepping pH = 6.61
[26:15] Dispensed 0.000024 mL of Base (0.5 M KOH)
[26:20] Stepping pH = 6.62
[26:36] Stirrer speed set to 0
[26:46] Datapoint id 22 collected
[26:46] Charge balance equation is out by 37.9%
[26:46] Stirrer speed set to 50
[26:51] pH 6.62 -> 6.82
[26:51] Using cautious pH adjust
[26:51] Dispensed 0.000141 mL of Base (0.5 M KOH)
[26:56] Stepping pH = 6.84
[27:11] Stirrer speed set to 0
[27:45] Datapoint id 23 collected
[27:45] Charge balance equation is out by 50.0%
[27:45] Stirrer speed set to 50
[27:51] pH 6.85 -> 7.05
[27:51] Using cautious pH adjust
[27:51] Dispensed 0.000094 mL of Base (0.5 M KOH)
[27:56] Stepping pH = 7.02
[27:56] Dispensed 0.000024 mL of Base (0.5 M KOH)
[28:01] Stepping pH = 7.06
[28:16] Stirrer speed set to 0
[28:51] Datapoint id 24 collected
[28:51] Charge balance equation is out by 39.1%
[28:51] Stirrer speed set to 50
[28:56] pH 7.08 -> 7.28
[28:56] Using cautious pH adjust
[28:56] Dispensed 0.000047 mL of Base (0.5 M KOH)
[29:01] Stepping pH = 7.19
[29:01] Dispensed 0.000024 mL of Base (0.5 M KOH)
[29:07] Stepping pH = 7.28
[29:22] Stirrer speed set to 0
[30:11] Datapoint id 25 collected
[30:11] Charge balance equation is out by 23.6%
[30:11] Stirrer speed set to 50
[30:16] pH 7.30 -> 7.50
[30:16] Using cautious pH adjust
[30:16] Dispensed 0.000024 mL of Base (0.5 M KOH)
[30:21] Stepping pH = 7.31
[30:22] Dispensed 0.000094 mL of Base (0.5 M KOH)
[30:27] Stepping pH = 8.01
[30:42] Stirrer speed set to 0
[31:42] Datapoint id 26 collected
[31:42] Charge balance equation is out by -92.6%
[31:42] Stirrer speed set to 50
[31:47] pH 8.09 -> 8.29
[31:47] Using cautious pH adjust
[31:47] Dispensed 0.000024 mL of Base (0.5 M KOH)
[31:52] Stepping pH = 8.11
[31:52] Dispensed 0.000024 mL of Base (0.5 M KOH)
[31:57] Stepping pH = 8.18
[31:57] Dispensed 0.000024 mL of Base (0.5 M KOH)
[32:02] Stepping pH = 8.30

| | | | |
|---|---|---|---|
| Sample name: | **M15_octanol** | Experiment start time: | **2/28/2018 1:10:47 AM** |
| Assay name: | **pH-metric high logP** | Analyst: | **Pion** |
| Assay ID: | **18B-28002** | Instrument ID: | **T312060** |
| Filename: | **C:\Sirius_T3\Mehtap\20180227_exp27_logP_T3-2\18B-28002_M15_octanol_pH-metric high logP.t3r** | | |

#### Experiment Log (continued)

[32:17] Stirrer speed set to 0
[33:00] Datapoint id 27 collected
[33:00] Charge balance equation is out by -279.3%
[33:00] Stirrer speed set to 50
[33:05] pH 8.36 -> 8.56
[33:05] Using cautious pH adjust
[33:05] Dispensed 0.000024 mL of Base (0.5 M KOH)
[33:10] Stepping pH = 8.38
[33:10] Dispensed 0.000024 mL of Base (0.5 M KOH)
[33:15] Stepping pH = 8.46
[33:15] Dispensed 0.000024 mL of Base (0.5 M KOH)
[33:21] Stepping pH = 8.56
[33:36] Stirrer speed set to 0
[34:15] Datapoint id 28 collected
[34:15] Charge balance equation is out by -251.0%
[34:15] Stirrer speed set to 50
[34:20] pH 8.60 -> 8.80
[34:20] Using cautious pH adjust
[34:20] Dispensed 0.000024 mL of Base (0.5 M KOH)
[34:25] Stepping pH = 8.62
[34:25] Dispensed 0.000047 mL of Base (0.5 M KOH)
[34:30] Stepping pH = 8.79
[34:30] Dispensed 0.000024 mL of Base (0.5 M KOH)
[34:35] Stepping pH = 8.85
[34:50] Stirrer speed set to 0
[35:11] Datapoint id 29 collected
[35:11] Charge balance equation is out by -177.5%
[35:11] Stirrer speed set to 50
[35:16] pH 8.86 -> 9.06
[35:16] Using cautious pH adjust
[35:17] Dispensed 0.000024 mL of Base (0.5 M KOH)
[35:22] Stepping pH = 8.87
[35:22] Dispensed 0.000071 mL of Base (0.5 M KOH)
[35:27] Stepping pH = 9.07
[35:42] Stirrer speed set to 0
[36:11] Datapoint id 30 collected
[36:11] Charge balance equation is out by -92.2%
[36:11] Stirrer speed set to 50
[36:16] pH 9.08 -> 9.28
[36:16] Using cautious pH adjust
[36:16] Dispensed 0.000047 mL of Base (0.5 M KOH)
[36:21] Stepping pH = 9.12
[36:21] Dispensed 0.000071 mL of Base (0.5 M KOH)
[36:26] Stepping pH = 9.27
[36:41] Stirrer speed set to 0
[36:58] Datapoint id 31 collected
[36:58] Charge balance equation is out by -49.9%
[36:58] Stirrer speed set to 50
[37:03] pH 9.28 -> 9.48
[37:03] Using cautious pH adjust
[37:03] Dispensed 0.000047 mL of Base (0.5 M KOH)
[37:08] Stepping pH = 9.30
[37:08] Dispensed 0.000141 mL of Base (0.5 M KOH)
[37:13] Stepping pH = 9.51
[37:28] Stirrer speed set to 0
[37:42] Datapoint id 32 collected
[37:42] Charge balance equation is out by -87.7%
[37:42] Stirrer speed set to 50
[37:47] pH 9.52 -> 9.72

Sample name: **M15_octanol** Experiment start time: **2/28/2018 1:10:47 AM**
Assay name: **pH-metric high logP** Analyst: **Pion**
Assay ID: **18B-28002** Instrument ID: **T312060**
Filename: **C:\Sirius_T3\Mehtap\20180227_exp27_logP_T3-2\18B-28002_M15_octanol_pH-metric high logP.t3r**

#### Experiment Log (continued)

[37:47] Using cautious pH adjust
[37:47] Dispensed 0.000094 mL of Base (0.5 M KOH)
[37:52] Stepping pH = 9.58
[37:52] Dispensed 0.000118 mL of Base (0.5 M KOH)
[37:57] Stepping pH = 9.69
[37:58] Dispensed 0.000024 mL of Base (0.5 M KOH)
[38:03] Stepping pH = 9.70
[38:03] Dispensed 0.000024 mL of Base (0.5 M KOH)
[38:08] Stepping pH = 9.71
[38:23] Stirrer speed set to 0
[38:33] Datapoint id 33 collected
[38:33] Charge balance equation is out by -52.5%
[38:33] Stirrer speed set to 50
[38:39] pH 9.71 -> 9.91
[38:39] Using cautious pH adjust
[38:39] Dispensed 0.000118 mL of Base (0.5 M KOH)
[38:44] Stepping pH = 9.80
[38:44] Dispensed 0.000118 mL of Base (0.5 M KOH)
[38:49] Stepping pH = 9.88
[38:49] Dispensed 0.000047 mL of Base (0.5 M KOH)
[38:54] Stepping pH = 9.91
[39:09] Stirrer speed set to 0
[39:20] Datapoint id 34 collected
[39:20] Charge balance equation is out by -10.0%
[39:20] Stirrer speed set to 50
[39:25] pH 9.91 -> 10.05
[39:25] Using charge balance adjust
[39:25] Dispensed 0.000235 mL of Base (0.5 M KOH)
[39:45] Stirrer speed set to 0
[39:55] Datapoint id 35 collected
[39:55] Charge balance equation is out by -45.0%
[39:55] Titration 2 of 3
[39:55] Adding initial titrants
[39:55] Automatically add 0.10000 mL of Octanol
[39:58] Dispensed 0.100000 mL of Octanol
[39:58] Stirrer speed set to 10
[39:59] Stirrer speed set to 55
[39:59] Iterative adjust 10.03 -> 2.00
[39:59] pH 10.03 -> 2.00
[40:01] Dispensed 0.098401 mL of Acid (0.5 M HCl)
[40:51] Stirrer speed set to 0
[41:01] Datapoint id 36 collected
[41:01] Stirrer speed set to 55
[41:06] pH 1.97 -> 2.17
[41:06] Using cautious pH adjust
[41:07] Dispensed 0.014111 mL of Base (0.5 M KOH)
[41:12] Stepping pH = 2.07
[41:12] Dispensed 0.010278 mL of Base (0.5 M KOH)
[41:17] Stepping pH = 2.15
[41:18] Dispensed 0.002658 mL of Base (0.5 M KOH)
[41:23] Stepping pH = 2.18
[41:38] Stirrer speed set to 0
[41:48] Datapoint id 37 collected
[41:48] Charge balance equation is out by 4.2%
[41:48] Stirrer speed set to 55
[41:53] pH 2.19 -> 2.39
[41:53] Using charge balance adjust
[41:53] Dispensed 0.018062 mL of Base (0.5 M KOH)
[42:14] Stirrer speed set to 0

Sample name: **M15_octanol** Experiment start time: **2/28/2018 1:10:47 AM**
Assay name: **pH-metric high logP** Analyst: **Pion**
Assay ID: **18B-28002** Instrument ID: **T312060**
Filename: **C:\Sirius_T3\Mehtap\20180227_exp27_logP_T3-2\18B-28002_M15_octanol_pH-metric high logP.t3r**

#### Experiment Log (continued)

[42:24] Datapoint id 38 collected
[42:24] Charge balance equation is out by 5.1%
[42:24] Stirrer speed set to 55
[42:29] pH 2.40 -> 2.60
[42:29] Using charge balance adjust
[42:29] Dispensed 0.011900 mL of Base (0.5 M KOH)
[42:49] Stirrer speed set to 0
[42:59] Datapoint id 39 collected
[42:59] Charge balance equation is out by 11.2%
[42:59] Stirrer speed set to 55
[43:05] pH 2.63 -> 2.83
[43:05] Using charge balance adjust
[43:05] Dispensed 0.007926 mL of Base (0.5 M KOH)
[43:25] Stirrer speed set to 0
[43:35] Datapoint id 40 collected
[43:35] Charge balance equation is out by 3.5%
[43:35] Stirrer speed set to 55
[43:40] pH 2.84 -> 3.04
[43:40] Using charge balance adjust
[43:40] Dispensed 0.005503 mL of Base (0.5 M KOH)
[44:01] Stirrer speed set to 0
[44:11] Datapoint id 41 collected
[44:11] Charge balance equation is out by 1.6%
[44:11] Stirrer speed set to 55
[44:16] pH 3.05 -> 3.25
[44:16] Using charge balance adjust
[44:16] Dispensed 0.003857 mL of Base (0.5 M KOH)
[44:36] Stirrer speed set to 0
[44:46] Datapoint id 42 collected
[44:46] Charge balance equation is out by 10.8%
[44:46] Stirrer speed set to 55
[44:51] pH 3.28 -> 3.48
[44:51] Using charge balance adjust
[44:51] Dispensed 0.002611 mL of Base (0.5 M KOH)
[45:12] Stirrer speed set to 0
[45:22] Datapoint id 43 collected
[45:22] Charge balance equation is out by -10.7%
[45:22] Stirrer speed set to 55
[45:27] pH 3.46 -> 3.66
[45:27] Using charge balance adjust
[45:27] Dispensed 0.001952 mL of Base (0.5 M KOH)
[45:47] Stirrer speed set to 0
[45:57] Datapoint id 44 collected
[45:57] Charge balance equation is out by -21.3%
[45:57] Stirrer speed set to 55
[46:02] pH 3.63 -> 3.83
[46:02] Using cautious pH adjust
[46:02] Dispensed 0.000776 mL of Base (0.5 M KOH)
[46:07] Stepping pH = 3.70
[46:07] Dispensed 0.000729 mL of Base (0.5 M KOH)
[46:13] Stepping pH = 3.78
[46:13] Dispensed 0.000306 mL of Base (0.5 M KOH)
[46:18] Stepping pH = 3.82
[46:33] Stirrer speed set to 0
[46:43] Datapoint id 45 collected
[46:43] Charge balance equation is out by -17.3%
[46:43] Stirrer speed set to 55
[46:48] pH 3.82 -> 4.02
[46:48] Using cautious pH adjust

Sample name: **M15_octanol** Experiment start time: **2/28/2018 1:10:47 AM**
Assay name: **pH-metric high logP** Analyst: **Pion**
Assay ID: **18B-28002** Instrument ID: **T312060**
Filename: **C:\Sirius_T3\Mehtap\20180227_exp27_logP_T3-2\18B-28002_M15_octanol_pH-metric high logP.t3r**

#### Experiment Log (continued)

[46:48] Dispensed 0.000659 mL of Base (0.5 M KOH)
[46:53] Stepping pH = 3.90
[46:53] Dispensed 0.000588 mL of Base (0.5 M KOH)
[46:58] Stepping pH = 3.98
[46:58] Dispensed 0.000282 mL of Base (0.5 M KOH)
[47:04] Stepping pH = 4.01
[47:04] Dispensed 0.000118 mL of Base (0.5 M KOH)
[47:09] Stepping pH = 4.02
[47:24] Stirrer speed set to 0
[47:34] Datapoint id 46 collected
[47:34] Charge balance equation is out by -25.1%
[47:34] Stirrer speed set to 55
[47:39] pH 4.01 -> 4.21
[47:39] Using cautious pH adjust
[47:39] Dispensed 0.000611 mL of Base (0.5 M KOH)
[47:44] Stepping pH = 4.09
[47:44] Dispensed 0.000588 mL of Base (0.5 M KOH)
[47:49] Stepping pH = 4.16
[47:49] Dispensed 0.000306 mL of Base (0.5 M KOH)
[47:55] Stepping pH = 4.20
[47:55] Dispensed 0.000118 mL of Base (0.5 M KOH)
[48:00] Stepping pH = 4.20
[48:15] Stirrer speed set to 0
[48:25] Datapoint id 47 collected
[48:25] Charge balance equation is out by -30.8%
[48:25] Stirrer speed set to 55
[48:30] pH 4.20 -> 4.40
[48:30] Using cautious pH adjust
[48:30] Dispensed 0.000659 mL of Base (0.5 M KOH)
[48:35] Stepping pH = 4.27
[48:35] Dispensed 0.000682 mL of Base (0.5 M KOH)
[48:40] Stepping pH = 4.34
[48:40] Dispensed 0.000376 mL of Base (0.5 M KOH)
[48:46] Stepping pH = 4.38
[48:46] Dispensed 0.000141 mL of Base (0.5 M KOH)
[48:51] Stepping pH = 4.39
[49:06] Stirrer speed set to 0
[49:16] Datapoint id 48 collected
[49:16] Charge balance equation is out by -41.1%
[49:16] Stirrer speed set to 55
[49:21] pH 4.38 -> 4.58
[49:21] Using cautious pH adjust
[49:21] Dispensed 0.000753 mL of Base (0.5 M KOH)
[49:26] Stepping pH = 4.45
[49:26] Dispensed 0.000800 mL of Base (0.5 M KOH)
[49:31] Stepping pH = 4.54
[49:32] Dispensed 0.000353 mL of Base (0.5 M KOH)
[49:37] Stepping pH = 4.56
[49:37] Dispensed 0.000165 mL of Base (0.5 M KOH)
[49:42] Stepping pH = 4.57
[49:57] Stirrer speed set to 0
[50:07] Datapoint id 49 collected
[50:07] Charge balance equation is out by -36.8%
[50:07] Stirrer speed set to 55
[50:12] pH 4.56 -> 4.76
[50:12] Using cautious pH adjust
[50:12] Dispensed 0.000870 mL of Base (0.5 M KOH)
[50:17] Stepping pH = 4.66
[50:17] Dispensed 0.000659 mL of Base (0.5 M KOH)

Sample name: **M15_octanol** Experiment start time: **2/28/2018 1:10:47 AM**
Assay name: **pH-metric high logP** Analyst: **Pion**
Assay ID: **18B-28002** Instrument ID: **T312060**
Filename: **C:\Sirius_T3\Mehtap\20180227_exp27_logP_T3-2\18B-28002_M15_octanol_pH-metric high logP.t3r**

#### Experiment Log (continued)

[50:22] Stepping pH = 4.74
[50:22] Dispensed 0.000212 mL of Base (0.5 M KOH)
[50:28] Stepping pH = 4.75
[50:43] Stirrer speed set to 0
[50:53] Datapoint id 50 collected
[50:53] Charge balance equation is out by 0.1%
[50:53] Stirrer speed set to 55
[50:58] pH 4.74 -> 4.94
[50:58] Using charge balance adjust
[50:58] Dispensed 0.001929 mL of Base (0.5 M KOH)
[51:18] Stirrer speed set to 0
[51:29] Datapoint id 51 collected
[51:29] Charge balance equation is out by 29.5%
[51:29] Stirrer speed set to 55
[51:34] pH 5.00 -> 5.20
[51:34] Using cautious pH adjust
[51:34] Dispensed 0.001011 mL of Base (0.5 M KOH)
[51:39] Stepping pH = 5.17
[51:39] Dispensed 0.000188 mL of Base (0.5 M KOH)
[51:44] Stepping pH = 5.20
[51:59] Stirrer speed set to 0
[52:09] Datapoint id 52 collected
[52:09] Charge balance equation is out by 41.1%
[52:09] Stirrer speed set to 55
[52:14] pH 5.19 -> 5.39
[52:14] Using cautious pH adjust
[52:14] Dispensed 0.000941 mL of Base (0.5 M KOH)
[52:20] Stepping pH = 5.37
[52:20] Dispensed 0.000071 mL of Base (0.5 M KOH)
[52:25] Stepping pH = 5.37
[52:25] Dispensed 0.000165 mL of Base (0.5 M KOH)
[52:30] Stepping pH = 5.41
[52:45] Stirrer speed set to 0
[52:55] Datapoint id 53 collected
[52:55] Charge balance equation is out by 37.1%
[52:55] Stirrer speed set to 55
[53:00] pH 5.40 -> 5.60
[53:00] Using cautious pH adjust
[53:00] Dispensed 0.000800 mL of Base (0.5 M KOH)
[53:05] Stepping pH = 5.62
[53:20] Stirrer speed set to 0
[53:31] Datapoint id 54 collected
[53:31] Charge balance equation is out by 50.0%
[53:31] Stirrer speed set to 55
[53:37] pH 5.62 -> 5.82
[53:37] Using cautious pH adjust
[53:37] Dispensed 0.000611 mL of Base (0.5 M KOH)
[53:42] Stepping pH = 5.83
[53:57] Stirrer speed set to 0
[54:07] Datapoint id 55 collected
[54:07] Charge balance equation is out by 50.0%
[54:07] Stirrer speed set to 55
[54:13] pH 5.84 -> 6.04
[54:13] Using cautious pH adjust
[54:13] Dispensed 0.000447 mL of Base (0.5 M KOH)
[54:18] Stepping pH = 6.07
[54:33] Stirrer speed set to 0
[54:44] Datapoint id 56 collected
[54:44] Charge balance equation is out by 50.0%

| | | | |
|---|---|---|---|
| Sample name: | **M15_octanol** | Experiment start time: | **2/28/2018 1:10:47 AM** |
| Assay name: | **pH-metric high logP** | Analyst: | **Pion** |
| Assay ID: | **18B-28002** | Instrument ID: | **T312060** |
| Filename: | **C:\Sirius_T3\Mehtap\20180227_exp27_logP_T3-2\18B-28002_M15_octanol_pH-metric high logP.t3r** | | |

#### Experiment Log (continued)

[54:44] Stirrer speed set to 55
[54:49] pH 6.07 -> 6.27
[54:49] Using cautious pH adjust
[54:50] Dispensed 0.000306 mL of Base (0.5 M KOH)
[54:55] Stepping pH = 6.34
[55:10] Stirrer speed set to 0
[55:26] Datapoint id 57 collected
[55:26] Charge balance equation is out by 50.0%
[55:26] Stirrer speed set to 55
[55:31] pH 6.34 -> 6.54
[55:31] Using cautious pH adjust
[55:31] Dispensed 0.000188 mL of Base (0.5 M KOH)
[55:36] Stepping pH = 6.51
[55:36] Dispensed 0.000024 mL of Base (0.5 M KOH)
[55:41] Stepping pH = 6.54
[55:56] Stirrer speed set to 0
[56:11] Datapoint id 58 collected
[56:11] Charge balance equation is out by 43.1%
[56:11] Stirrer speed set to 55
[56:16] pH 6.54 -> 6.74
[56:16] Using cautious pH adjust
[56:16] Dispensed 0.000118 mL of Base (0.5 M KOH)
[56:22] Stepping pH = 6.69
[56:22] Dispensed 0.000024 mL of Base (0.5 M KOH)
[56:27] Stepping pH = 6.73
[56:42] Stirrer speed set to 0
[57:13] Datapoint id 59 collected
[57:13] Charge balance equation is out by 37.4%
[57:13] Stirrer speed set to 55
[57:18] pH 6.75 -> 6.95
[57:18] Using cautious pH adjust
[57:18] Dispensed 0.000094 mL of Base (0.5 M KOH)
[57:23] Stepping pH = 7.12
[57:38] Stirrer speed set to 0
[58:37] Datapoint id 60 collected
[58:37] Charge balance equation is out by 50.0%
[58:37] Stirrer speed set to 55
[58:42] pH 7.14 -> 7.34
[58:42] Using cautious pH adjust
[58:42] Dispensed 0.000047 mL of Base (0.5 M KOH)
[58:47] Stepping pH = 7.20
[58:48] Dispensed 0.000047 mL of Base (0.5 M KOH)
[58:53] Stepping pH = 7.55
[59:08] Stirrer speed set to 0
[1:00:08] Datapoint id 61 collected
[1:00:08] Charge balance equation is out by -6.7%
[1:00:08] Stirrer speed set to 55
[1:00:13] pH 7.60 -> 7.80
[1:00:13] Using charge balance adjust
[1:00:13] Dispensed 0.000024 mL of Base (0.5 M KOH)
[1:00:33] Stirrer speed set to 0
[1:01:27] Datapoint id 62 collected
[1:01:27] Charge balance equation is out by -85.7%
[1:01:27] Stirrer speed set to 55
[1:01:32] pH 7.69 -> 7.89
[1:01:32] Using cautious pH adjust
[1:01:32] Dispensed 0.000024 mL of Base (0.5 M KOH)
[1:01:37] Stepping pH = 7.73
[1:01:37] Dispensed 0.000024 mL of Base (0.5 M KOH)

Sample name: **M15_octanol** Experiment start time: **2/28/2018 1:10:47 AM**
Assay name: **pH-metric high logP** Analyst: **Pion**
Assay ID: **18B-28002** Instrument ID: **T312060**
Filename: **C:\Sirius_T3\Mehtap\20180227_exp27_logP_T3-2\18B-28002_M15_octanol_pH-metric high logP.t3r**

#### Experiment Log (continued)

[1:01:42] Stepping pH = 7.88
[1:01:42] Dispensed 0.000024 mL of Base (0.5 M KOH)
[1:01:47] Stepping pH = 8.02
[1:02:02] Stirrer speed set to 0
[1:03:01] Datapoint id 63 collected
[1:03:01] Charge balance equation is out by -137.8%
[1:03:01] Stirrer speed set to 55
[1:03:06] pH 8.09 -> 8.29
[1:03:06] Using cautious pH adjust
[1:03:06] Dispensed 0.000024 mL of Base (0.5 M KOH)
[1:03:11] Stepping pH = 8.15
[1:03:11] Dispensed 0.000024 mL of Base (0.5 M KOH)
[1:03:16] Stepping pH = 8.22
[1:03:16] Dispensed 0.000024 mL of Base (0.5 M KOH)
[1:03:21] Stepping pH = 8.32
[1:03:36] Stirrer speed set to 0
[1:04:21] Datapoint id 64 collected
[1:04:21] Charge balance equation is out by -239.8%
[1:04:21] Stirrer speed set to 55
[1:04:26] pH 8.40 -> 8.60
[1:04:26] Using cautious pH adjust
[1:04:27] Dispensed 0.000024 mL of Base (0.5 M KOH)
[1:04:32] Stepping pH = 8.43
[1:04:32] Dispensed 0.000024 mL of Base (0.5 M KOH)
[1:04:37] Stepping pH = 8.48
[1:04:37] Dispensed 0.000024 mL of Base (0.5 M KOH)
[1:04:42] Stepping pH = 8.55
[1:04:42] Dispensed 0.000024 mL of Base (0.5 M KOH)
[1:04:47] Stepping pH = 8.62
[1:05:02] Stirrer speed set to 0
[1:05:20] Datapoint id 65 collected
[1:05:20] Charge balance equation is out by -340.7%
[1:05:20] Stirrer speed set to 55
[1:05:25] pH 8.67 -> 8.87
[1:05:25] Using cautious pH adjust
[1:05:25] Dispensed 0.000024 mL of Base (0.5 M KOH)
[1:05:30] Stepping pH = 8.68
[1:05:30] Dispensed 0.000047 mL of Base (0.5 M KOH)
[1:05:36] Stepping pH = 8.75
[1:05:36] Dispensed 0.000047 mL of Base (0.5 M KOH)
[1:05:41] Stepping pH = 8.85
[1:05:41] Dispensed 0.000024 mL of Base (0.5 M KOH)
[1:05:46] Stepping pH = 8.89
[1:06:01] Stirrer speed set to 0
[1:06:22] Datapoint id 66 collected
[1:06:22] Charge balance equation is out by -302.4%
[1:06:22] Stirrer speed set to 55
[1:06:28] pH 8.90 -> 9.10
[1:06:28] Using cautious pH adjust
[1:06:28] Dispensed 0.000024 mL of Base (0.5 M KOH)
[1:06:33] Stepping pH = 8.91
[1:06:33] Dispensed 0.000071 mL of Base (0.5 M KOH)
[1:06:38] Stepping pH = 9.05
[1:06:38] Dispensed 0.000024 mL of Base (0.5 M KOH)
[1:06:43] Stepping pH = 9.08
[1:06:43] Dispensed 0.000024 mL of Base (0.5 M KOH)
[1:06:48] Stepping pH = 9.11
[1:07:03] Stirrer speed set to 0
[1:07:18] Datapoint id 67 collected

| | | | |
|---|---|---|---|
| Sample name: | **M15_octanol** | Experiment start time: | **2/28/2018 1:10:47 AM** |
| Assay name: | **pH-metric high logP** | Analyst: | **Pion** |
| Assay ID: | **18B-28002** | Instrument ID: | **T312060** |
| Filename: | **C:\Sirius_T3\Mehtap\20180227_exp27_logP_T3-2\18B-28002_M15_octanol_pH-metric high logP.t3r** | | |

#### Experiment Log (continued)

[1:07:18] Charge balance equation is out by -180.0%
[1:07:18] Stirrer speed set to 55
[1:07:23] pH 9.12 -> 9.32
[1:07:23] Using cautious pH adjust
[1:07:23] Dispensed 0.000047 mL of Base (0.5 M KOH)
[1:07:28] Stepping pH = 9.16
[1:07:29] Dispensed 0.000094 mL of Base (0.5 M KOH)
[1:07:34] Stepping pH = 9.30
[1:07:34] Dispensed 0.000024 mL of Base (0.5 M KOH)
[1:07:39] Stepping pH = 9.31
[1:07:54] Stirrer speed set to 0
[1:08:04] Datapoint id 68 collected
[1:08:04] Charge balance equation is out by -73.1%
[1:08:04] Stirrer speed set to 55
[1:08:10] pH 9.32 -> 9.52
[1:08:10] Using cautious pH adjust
[1:08:10] Dispensed 0.000071 mL of Base (0.5 M KOH)
[1:08:15] Stepping pH = 9.36
[1:08:15] Dispensed 0.000118 mL of Base (0.5 M KOH)
[1:08:20] Stepping pH = 9.47
[1:08:20] Dispensed 0.000047 mL of Base (0.5 M KOH)
[1:08:25] Stepping pH = 9.50
[1:08:25] Dispensed 0.000024 mL of Base (0.5 M KOH)
[1:08:30] Stepping pH = 9.51
[1:08:45] Stirrer speed set to 0
[1:09:02] Datapoint id 69 collected
[1:09:02] Charge balance equation is out by -116.1%
[1:09:02] Stirrer speed set to 55
[1:09:07] pH 9.52 -> 9.72
[1:09:07] Using cautious pH adjust
[1:09:07] Dispensed 0.000094 mL of Base (0.5 M KOH)
[1:09:13] Stepping pH = 9.53
[1:09:13] Dispensed 0.000259 mL of Base (0.5 M KOH)
[1:09:18] Stepping pH = 9.76
[1:09:33] Stirrer speed set to 0
[1:09:49] Datapoint id 70 collected
[1:09:49] Charge balance equation is out by -89.8%
[1:09:49] Stirrer speed set to 55
[1:09:55] pH 9.76 -> 9.96
[1:09:55] Using cautious pH adjust
[1:09:55] Dispensed 0.000141 mL of Base (0.5 M KOH)
[1:10:00] Stepping pH = 9.82
[1:10:00] Dispensed 0.000212 mL of Base (0.5 M KOH)
[1:10:05] Stepping pH = 9.94
[1:10:05] Dispensed 0.000047 mL of Base (0.5 M KOH)
[1:10:10] Stepping pH = 9.95
[1:10:10] Dispensed 0.000024 mL of Base (0.5 M KOH)
[1:10:15] Stepping pH = 9.96
[1:10:30] Stirrer speed set to 0
[1:10:40] Datapoint id 71 collected
[1:10:40] Charge balance equation is out by -51.5%
[1:10:40] Stirrer speed set to 55
[1:10:45] pH 9.96 -> 10.05
[1:10:45] Using cautious pH adjust
[1:10:46] Dispensed 0.000094 mL of Base (0.5 M KOH)
[1:10:51] Stepping pH = 9.99
[1:10:51] Dispensed 0.000094 mL of Base (0.5 M KOH)
[1:10:56] Stepping pH = 10.02
[1:11:11] Stirrer speed set to 0

| | | | |
|---|---|---|---|
| Sample name: | **M15_octanol** | Experiment start time: | **2/28/2018 1:10:47 AM** |
| Assay name: | **pH-metric high logP** | Analyst: | **Pion** |
| Assay ID: | **18B-28002** | Instrument ID: | **T312060** |
| Filename: | **C:\Sirius_T3\Mehtap\20180227_exp27_logP_T3-2\18B-28002_M15_octanol_pH-metric high logP.t3r** | | |

#### Experiment Log (continued)

[1:11:21] Datapoint id 72 collected
[1:11:21] Charge balance equation is out by -8.6%
[1:11:21] Titration 3 of 3
[1:11:21] Adding initial titrants
[1:11:21] Automatically add 0.30000 mL of Octanol
[1:11:28] Dispensed 0.300000 mL of Octanol
[1:11:28] Stirrer speed set to 10
[1:11:29] Stirrer speed set to 60
[1:11:29] Iterative adjust 10.02 -> 2.00
[1:11:29] pH 10.02 -> 2.00
[1:11:32] Dispensed 0.100000 mL of Acid (0.5 M HCl)
[1:11:37] pH 2.02 -> 2.00
[1:11:37] Dispensed 0.003881 mL of Acid (0.5 M HCl)
[1:12:27] Stirrer speed set to 0
[1:12:37] Datapoint id 73 collected
[1:12:37] Stirrer speed set to 60
[1:12:42] pH 1.98 -> 2.18
[1:12:42] Using cautious pH adjust
[1:12:43] Dispensed 0.015122 mL of Base (0.5 M KOH)
[1:12:48] Stepping pH = 2.07
[1:12:48] Dispensed 0.010160 mL of Base (0.5 M KOH)
[1:12:53] Stepping pH = 2.15
[1:12:54] Dispensed 0.002775 mL of Base (0.5 M KOH)
[1:12:59] Stepping pH = 2.18
[1:13:14] Stirrer speed set to 0
[1:13:24] Datapoint id 74 collected
[1:13:24] Charge balance equation is out by 7.3%
[1:13:24] Stirrer speed set to 60
[1:13:29] pH 2.19 -> 2.39
[1:13:29] Using charge balance adjust
[1:13:30] Dispensed 0.019450 mL of Base (0.5 M KOH)
[1:13:50] Stirrer speed set to 0
[1:14:00] Datapoint id 75 collected
[1:14:00] Charge balance equation is out by 6.0%
[1:14:00] Stirrer speed set to 60
[1:14:05] pH 2.40 -> 2.60
[1:14:05] Using charge balance adjust
[1:14:06] Dispensed 0.012700 mL of Base (0.5 M KOH)
[1:14:26] Stirrer speed set to 0
[1:14:36] Datapoint id 76 collected
[1:14:36] Charge balance equation is out by 11.5%
[1:14:36] Stirrer speed set to 60
[1:14:41] pH 2.63 -> 2.83
[1:14:41] Using charge balance adjust
[1:14:41] Dispensed 0.008443 mL of Base (0.5 M KOH)
[1:15:01] Stirrer speed set to 0
[1:15:11] Datapoint id 77 collected
[1:15:11] Charge balance equation is out by -0.8%
[1:15:11] Stirrer speed set to 60
[1:15:17] pH 2.83 -> 3.03
[1:15:17] Using charge balance adjust
[1:15:17] Dispensed 0.005974 mL of Base (0.5 M KOH)
[1:15:37] Stirrer speed set to 0
[1:15:47] Datapoint id 78 collected
[1:15:47] Charge balance equation is out by -2.6%
[1:15:47] Stirrer speed set to 60
[1:15:52] pH 3.03 -> 3.23
[1:15:52] Using charge balance adjust
[1:15:52] Dispensed 0.004280 mL of Base (0.5 M KOH)

#### Experiment Log (continued)

[1:16:12] Stirrer speed set to 0
[1:16:22] Datapoint id 79 collected
[1:16:22] Charge balance equation is out by 6.7%
[1:16:22] Stirrer speed set to 60
[1:16:28] pH 3.25 -> 3.45
[1:16:28] Using charge balance adjust
[1:16:28] Dispensed 0.003010 mL of Base (0.5 M KOH)
[1:16:48] Stirrer speed set to 0
[1:16:59] Datapoint id 80 collected
[1:16:59] Charge balance equation is out by -9.4%
[1:16:59] Stirrer speed set to 60
[1:17:04] pH 3.43 -> 3.63
[1:17:04] Using charge balance adjust
[1:17:04] Dispensed 0.002305 mL of Base (0.5 M KOH)
[1:17:24] Stirrer speed set to 0
[1:17:39] Datapoint id 81 collected
[1:17:39] Charge balance equation is out by -27.6%
[1:17:39] Stirrer speed set to 60
[1:17:44] pH 3.57 -> 3.77
[1:17:44] Using cautious pH adjust
[1:17:44] Dispensed 0.000988 mL of Base (0.5 M KOH)
[1:17:49] Stepping pH = 3.65
[1:17:49] Dispensed 0.000917 mL of Base (0.5 M KOH)
[1:17:55] Stepping pH = 3.73
[1:17:55] Dispensed 0.000423 mL of Base (0.5 M KOH)
[1:18:00] Stepping pH = 3.76
[1:18:00] Dispensed 0.000118 mL of Base (0.5 M KOH)
[1:18:05] Stepping pH = 3.77
[1:18:20] Stirrer speed set to 0
[1:18:30] Datapoint id 82 collected
[1:18:30] Charge balance equation is out by -24.4%
[1:18:30] Stirrer speed set to 60
[1:18:35] pH 3.76 -> 3.96
[1:18:35] Using cautious pH adjust
[1:18:35] Dispensed 0.000870 mL of Base (0.5 M KOH)
[1:18:40] Stepping pH = 3.85
[1:18:40] Dispensed 0.000706 mL of Base (0.5 M KOH)
[1:18:46] Stepping pH = 3.91
[1:18:46] Dispensed 0.000400 mL of Base (0.5 M KOH)
[1:18:51] Stepping pH = 3.94
[1:18:51] Dispensed 0.000188 mL of Base (0.5 M KOH)
[1:18:56] Stepping pH = 3.95
[1:19:11] Stirrer speed set to 0
[1:19:21] Datapoint id 83 collected
[1:19:21] Charge balance equation is out by -24.7%
[1:19:21] Stirrer speed set to 60
[1:19:26] pH 3.94 -> 4.14
[1:19:26] Using cautious pH adjust
[1:19:26] Dispensed 0.000847 mL of Base (0.5 M KOH)
[1:19:31] Stepping pH = 4.01
[1:19:31] Dispensed 0.000917 mL of Base (0.5 M KOH)
[1:19:37] Stepping pH = 4.09
[1:19:37] Dispensed 0.000447 mL of Base (0.5 M KOH)
[1:19:42] Stepping pH = 4.12
[1:19:42] Dispensed 0.000165 mL of Base (0.5 M KOH)
[1:19:47] Stepping pH = 4.13
[1:20:02] Stirrer speed set to 0
[1:20:12] Datapoint id 84 collected
[1:20:12] Charge balance equation is out by -41.6%

Sample name: **M15_octanol** Experiment start time: **2/28/2018 1:10:47 AM**
Assay name: **pH-metric high logP** Analyst: **Pion**
Assay ID: **18B-28002** Instrument ID: **T312060**
Filename: **C:\Sirius_T3\Mehtap\20180227_exp27_logP_T3-2\18B-28002_M15_octanol_pH-metric high logP.t3r**

#### Experiment Log (continued)

[1:20:12] Stirrer speed set to 60
[1:20:17] pH 4.11 -> 4.31
[1:20:17] Using cautious pH adjust
[1:20:17] Dispensed 0.000894 mL of Base (0.5 M KOH)
[1:20:22] Stepping pH = 4.19
[1:20:22] Dispensed 0.000823 mL of Base (0.5 M KOH)
[1:20:28] Stepping pH = 4.27
[1:20:28] Dispensed 0.000329 mL of Base (0.5 M KOH)
[1:20:33] Stepping pH = 4.30
[1:20:33] Dispensed 0.000141 mL of Base (0.5 M KOH)
[1:20:38] Stepping pH = 4.30
[1:20:53] Stirrer speed set to 0
[1:21:03] Datapoint id 85 collected
[1:21:03] Charge balance equation is out by -23.0%
[1:21:03] Stirrer speed set to 60
[1:21:08] pH 4.29 -> 4.49
[1:21:08] Using cautious pH adjust
[1:21:08] Dispensed 0.000964 mL of Base (0.5 M KOH)
[1:21:13] Stepping pH = 4.39
[1:21:14] Dispensed 0.000706 mL of Base (0.5 M KOH)
[1:21:19] Stepping pH = 4.47
[1:21:19] Dispensed 0.000188 mL of Base (0.5 M KOH)
[1:21:24] Stepping pH = 4.49
[1:21:39] Stirrer speed set to 0
[1:21:49] Datapoint id 86 collected
[1:21:49] Charge balance equation is out by 3.5%
[1:21:49] Stirrer speed set to 60
[1:21:55] pH 4.47 -> 4.67
[1:21:55] Using charge balance adjust
[1:21:55] Dispensed 0.002046 mL of Base (0.5 M KOH)
[1:22:15] Stirrer speed set to 0
[1:22:25] Datapoint id 87 collected
[1:22:25] Charge balance equation is out by 40.8%
[1:22:25] Stirrer speed set to 60
[1:22:31] pH 4.75 -> 4.95
[1:22:31] Using cautious pH adjust
[1:22:31] Dispensed 0.001035 mL of Base (0.5 M KOH)
[1:22:36] Stepping pH = 4.93
[1:22:36] Dispensed 0.000094 mL of Base (0.5 M KOH)
[1:22:41] Stepping pH = 4.94
[1:22:56] Stirrer speed set to 0
[1:23:07] Datapoint id 88 collected
[1:23:07] Charge balance equation is out by 45.1%
[1:23:07] Stirrer speed set to 60
[1:23:12] pH 4.93 -> 5.13
[1:23:12] Using cautious pH adjust
[1:23:12] Dispensed 0.000941 mL of Base (0.5 M KOH)
[1:23:17] Stepping pH = 5.13
[1:23:32] Stirrer speed set to 0
[1:23:42] Datapoint id 89 collected
[1:23:42] Charge balance equation is out by 50.0%
[1:23:42] Stirrer speed set to 60
[1:23:47] pH 5.11 -> 5.31
[1:23:47] Using cautious pH adjust
[1:23:47] Dispensed 0.000800 mL of Base (0.5 M KOH)
[1:23:52] Stepping pH = 5.34
[1:24:07] Stirrer speed set to 0
[1:24:24] Datapoint id 90 collected
[1:24:24] Charge balance equation is out by 50.0%

| | | | |
|---|---|---|---|
| Sample name: | **M15_octanol** | Experiment start time: | **2/28/2018 1:10:47 AM** |
| Assay name: | **pH-metric high logP** | Analyst: | **Pion** |
| Assay ID: | **18B-28002** | Instrument ID: | **T312060** |
| Filename: | **C:\Sirius_T3\Mehtap\20180227_exp27_logP_T3-2\18B-28002_M15_octanol_pH-metric high logP.t3r** | | |

#### Experiment Log (continued)

[1:24:24] Stirrer speed set to 60
[1:24:30] pH 5.32 -> 5.52
[1:24:30] Using cautious pH adjust
[1:24:30] Dispensed 0.000611 mL of Base (0.5 M KOH)
[1:24:35] Stepping pH = 5.57
[1:24:50] Stirrer speed set to 0
[1:25:01] Datapoint id 91 collected
[1:25:01] Charge balance equation is out by 50.0%
[1:25:01] Stirrer speed set to 60
[1:25:06] pH 5.56 -> 5.76
[1:25:06] Using cautious pH adjust
[1:25:06] Dispensed 0.000423 mL of Base (0.5 M KOH)
[1:25:11] Stepping pH = 5.85
[1:25:26] Stirrer speed set to 0
[1:25:39] Datapoint id 92 collected
[1:25:39] Charge balance equation is out by 50.0%
[1:25:39] Stirrer speed set to 60
[1:25:44] pH 5.85 -> 6.05
[1:25:44] Using cautious pH adjust
[1:25:44] Dispensed 0.000259 mL of Base (0.5 M KOH)
[1:25:49] Stepping pH = 6.16
[1:26:04] Stirrer speed set to 0
[1:26:35] Datapoint id 93 collected
[1:26:35] Charge balance equation is out by 50.0%
[1:26:35] Stirrer speed set to 60
[1:26:40] pH 6.11 -> 6.31
[1:26:40] Using cautious pH adjust
[1:26:40] Dispensed 0.000165 mL of Base (0.5 M KOH)
[1:26:46] Stepping pH = 6.42
[1:27:01] Stirrer speed set to 0
[1:27:43] Datapoint id 94 collected
[1:27:43] Charge balance equation is out by 50.0%
[1:27:43] Stirrer speed set to 60
[1:27:48] pH 6.39 -> 6.59
[1:27:48] Using cautious pH adjust
[1:27:48] Dispensed 0.000094 mL of Base (0.5 M KOH)
[1:27:53] Stepping pH = 6.50
[1:27:53] Dispensed 0.000071 mL of Base (0.5 M KOH)
[1:27:58] Stepping pH = 6.74
[1:28:13] Stirrer speed set to 0
[1:29:09] Datapoint id 95 collected
[1:29:09] Charge balance equation is out by 21.9%
[1:29:09] Stirrer speed set to 60
[1:29:14] pH 6.69 -> 6.89
[1:29:14] Using cautious pH adjust
[1:29:14] Dispensed 0.000071 mL of Base (0.5 M KOH)
[1:29:20] Stepping pH = 6.88
[1:29:20] Dispensed 0.000024 mL of Base (0.5 M KOH)
[1:29:25] Stepping pH = 6.99
[1:29:40] Stirrer speed set to 0
[1:30:40] Datapoint id 96 collected
[1:30:40] Charge balance equation is out by 34.4%
[1:30:40] Stirrer speed set to 60
[1:30:45] pH 7.00 -> 7.20
[1:30:45] Using cautious pH adjust
[1:30:45] Dispensed 0.000047 mL of Base (0.5 M KOH)
[1:30:50] Stepping pH = 7.14
[1:30:50] Dispensed 0.000024 mL of Base (0.5 M KOH)
[1:30:55] Stepping pH = 7.32

| | | | |
|---|---|---|---|
| Sample name: | **M15_octanol** | Experiment start time: | **2/28/2018 1:10:47 AM** |
| Assay name: | **pH-metric high logP** | Analyst: | **Pion** |
| Assay ID: | **18B-28002** | Instrument ID: | **T312060** |
| Filename: | **C:\Sirius_T3\Mehtap\20180227_exp27_logP_T3-2\18B-28002_M15_octanol_pH-metric high logP.t3r** | | |

#### Experiment Log (continued)

[1:31:11] Stirrer speed set to 0
[1:32:11] Datapoint id 97 collected
[1:32:11] Charge balance equation is out by 22.4%
[1:32:11] Stirrer speed set to 60
[1:32:16] pH 7.20 -> 7.40
[1:32:16] Using cautious pH adjust
[1:32:16] Dispensed 0.000024 mL of Base (0.5 M KOH)
[1:32:21] Stepping pH = 7.16
[1:32:21] Dispensed 0.000141 mL of Base (0.5 M KOH)
[1:32:26] Stepping pH = 8.36
[1:32:41] Stirrer speed set to 0
[1:33:41] Datapoint id 98 collected
[1:33:41] Charge balance equation is out by -248.0%
[1:33:41] Stirrer speed set to 60
[1:33:46] pH 8.32 -> 8.52
[1:33:46] Using cautious pH adjust
[1:33:46] Dispensed 0.000024 mL of Base (0.5 M KOH)
[1:33:51] Stepping pH = 8.32
[1:33:51] Dispensed 0.000024 mL of Base (0.5 M KOH)
[1:33:56] Stepping pH = 8.34
[1:33:57] Dispensed 0.000094 mL of Base (0.5 M KOH)
[1:34:02] Stepping pH = 8.63
[1:34:17] Stirrer speed set to 0
[1:34:34] Datapoint id 99 collected
[1:34:34] Charge balance equation is out by -554.6%
[1:34:34] Stirrer speed set to 60
[1:34:39] pH 8.67 -> 8.87
[1:34:39] Using cautious pH adjust
[1:34:39] Dispensed 0.000024 mL of Base (0.5 M KOH)
[1:34:45] Stepping pH = 8.67
[1:34:45] Dispensed 0.000047 mL of Base (0.5 M KOH)
[1:34:50] Stepping pH = 8.72
[1:34:50] Dispensed 0.000094 mL of Base (0.5 M KOH)
[1:34:55] Stepping pH = 8.88
[1:35:10] Stirrer speed set to 0
[1:35:31] Datapoint id 100 collected
[1:35:31] Charge balance equation is out by -321.4%
[1:35:31] Stirrer speed set to 60
[1:35:37] pH 8.92 -> 9.12
[1:35:37] Using cautious pH adjust
[1:35:37] Dispensed 0.000024 mL of Base (0.5 M KOH)
[1:35:42] Stepping pH = 8.92
[1:35:42] Dispensed 0.000094 mL of Base (0.5 M KOH)
[1:35:47] Stepping pH = 9.04
[1:35:47] Dispensed 0.000047 mL of Base (0.5 M KOH)
[1:35:52] Stepping pH = 9.08
[1:35:52] Dispensed 0.000047 mL of Base (0.5 M KOH)
[1:35:57] Stepping pH = 9.13
[1:36:12] Stirrer speed set to 0
[1:36:24] Datapoint id 101 collected
[1:36:24] Charge balance equation is out by -240.5%
[1:36:24] Stirrer speed set to 60
[1:36:29] pH 9.12 -> 9.32
[1:36:29] Using cautious pH adjust
[1:36:29] Dispensed 0.000047 mL of Base (0.5 M KOH)
[1:36:34] Stepping pH = 9.14
[1:36:34] Dispensed 0.000118 mL of Base (0.5 M KOH)
[1:36:39] Stepping pH = 9.30
[1:36:39] Dispensed 0.000024 mL of Base (0.5 M KOH)

Sample name: **M15_octanol**
Assay name: **pH-metric high logP**
Assay ID: **18B-28002**
Filename: **C:\Sirius_T3\Mehtap\20180227_exp27_logP_T3-2\18B-28002_M15_octanol_pH-metric high logP.t3r**

Experiment start time: **2/28/2018 1:10:47 AM**
Analyst: **Pion**
Instrument ID: **T312060**

#### Experiment Log (continued)

[1:36:44] Stepping pH = 9.31
[1:37:00] Stirrer speed set to 0
[1:37:12] Datapoint id 102 collected
[1:37:12] Charge balance equation is out by -99.8%
[1:37:12] Stirrer speed set to 60
[1:37:17] pH 9.31 -> 9.51
[1:37:17] Using cautious pH adjust
[1:37:17] Dispensed 0.000071 mL of Base (0.5 M KOH)
[1:37:22] Stepping pH = 9.35
[1:37:22] Dispensed 0.000141 mL of Base (0.5 M KOH)
[1:37:27] Stepping pH = 9.48
[1:37:27] Dispensed 0.000047 mL of Base (0.5 M KOH)
[1:37:32] Stepping pH = 9.50
[1:37:32] Dispensed 0.000024 mL of Base (0.5 M KOH)
[1:37:37] Stepping pH = 9.52
[1:37:52] Stirrer speed set to 0
[1:38:03] Datapoint id 103 collected
[1:38:03] Charge balance equation is out by -92.7%
[1:38:03] Stirrer speed set to 60
[1:38:09] pH 9.52 -> 9.72
[1:38:09] Using cautious pH adjust
[1:38:09] Dispensed 0.000094 mL of Base (0.5 M KOH)
[1:38:14] Stepping pH = 9.59
[1:38:14] Dispensed 0.000141 mL of Base (0.5 M KOH)
[1:38:19] Stepping pH = 9.68
[1:38:19] Dispensed 0.000047 mL of Base (0.5 M KOH)
[1:38:24] Stepping pH = 9.70
[1:38:24] Dispensed 0.000047 mL of Base (0.5 M KOH)
[1:38:29] Stepping pH = 9.72
[1:38:44] Stirrer speed set to 0
[1:38:54] Datapoint id 104 collected
[1:38:54] Charge balance equation is out by -63.5%
[1:38:54] Stirrer speed set to 60
[1:38:59] pH 9.73 -> 9.93
[1:38:59] Using cautious pH adjust
[1:38:59] Dispensed 0.000141 mL of Base (0.5 M KOH)
[1:39:05] Stepping pH = 9.81
[1:39:05] Dispensed 0.000165 mL of Base (0.5 M KOH)
[1:39:10] Stepping pH = 9.90
[1:39:10] Dispensed 0.000071 mL of Base (0.5 M KOH)
[1:39:15] Stepping pH = 9.92
[1:39:15] Dispensed 0.000024 mL of Base (0.5 M KOH)
[1:39:20] Stepping pH = 9.93
[1:39:35] Stirrer speed set to 0
[1:39:46] Datapoint id 105 collected
[1:39:46] Charge balance equation is out by -32.3%
[1:39:46] Stirrer speed set to 60
[1:39:51] pH 9.93 -> 10.05
[1:39:51] Using cautious pH adjust
[1:39:51] Dispensed 0.000118 mL of Base (0.5 M KOH)
[1:39:56] Stepping pH = 9.98
[1:39:56] Dispensed 0.000118 mL of Base (0.5 M KOH)
[1:40:01] Stepping pH = 10.03
[1:40:16] Stirrer speed set to 0
[1:40:27] Datapoint id 106 collected
[1:40:27] Charge balance equation is out by 0.6%
[1:40:27] Argon flow rate set to 0
[1:40:31] Titrator arm moved over Titration position
