## Supplementary material for "Octanol-water partition coefficient measurements for the SAMPL6 Blind Prediction Challenge": SM14_18C-01004_M15_octanol_pH-metric high logP_report.pdf

| | | | |
|---|---|---|---|
| Sample name: | **M15_octanol** | Experiment start time: | **3/1/2018 4:33:50 AM** |
| Assay name: | **pH-metric high logP** | Analyst: | **Pion** |
| Assay ID: | **18C-01004** | Instrument ID: | **T312060** |
| Filename: | **C:\Sirius_T3\Mehtap\20180228_exp28_logP_T3-2\18C-01004_M15_octanol_pH-metric high logP.t3r** | | |

#### pH-metric Result

| | | |
|---|---|---|
| logP (XH2 2+) | -5.50 | |
| logP (XH +) | -0.11 | ±0.20 (n=50) |
| logP (neutral X) | 1.92 | ±0.01 (n=50) |

#### 18C-01004 Points 1 to 36

| | |
|---|---|
| M15_octanol concentration factor | 0.910 |
| Carbonate | 0.2391 mM |
| Acidity error | -0.31186 mM |

#### 18C-01004 Points 37 to 69

| | |
|---|---|
| M15_octanol concentration factor | 0.903 |
| Carbonate | 0.1847 mM |
| Acidity error | -0.25845 mM |

#### 18C-01004 Points 70 to 106

| | |
|---|---|
| M15_octanol concentration factor | 0.913 |
| Carbonate | 0.0779 mM |
| Acidity error | -0.09188 mM |

#### Warnings and errors

| | |
|---|---|
| Errors | None |
| Warnings | One or more logP values out of range |

#### Sample logD and percent species

| pH | M15_octanol logD | M15_octanol M15_octanolH2 | M15_octanol M15_octanolH | M15_octanol M15_octanol | M15_octanol M15_octanolH2* | M15_octanol M15_octanolH* | M15_octanol M15_octanol* | Comment |
|---|---|---|---|---|---|---|---|---|
| 1.000 | -1.70 | 95.54 % | 2.51 % | 0.00 % | 0.00 % | 1.94 % | 0.01 % | |
| 1.200 | -1.51 | 93.10 % | 3.88 % | 0.00 % | 0.00 % | 3.00 % | 0.03 % | Stomach pH |
| 2.000 | -0.77 | 67.69 % | 17.81 % | 0.01 % | 0.00 % | 13.75 % | 0.74 % | |
| 3.000 | -0.07 | 14.76 % | 38.83 % | 0.19 % | 0.00 % | 29.98 % | 16.23 % | |
| 4.000 | 0.66 | 0.63 % | 16.56 % | 0.83 % | 0.00 % | 12.79 % | 69.20 % | |
| 5.000 | 1.45 | 0.01 % | 2.27 % | 1.14 % | 0.00 % | 1.75 % | 94.83 % | |
| 6.000 | 1.84 | 0.00 % | 0.24 % | 1.18 % | 0.00 % | 0.18 % | 98.40 % | |
| 6.500 | 1.89 | 0.00 % | 0.07 % | 1.18 % | 0.00 % | 0.06 % | 98.68 % | |
| 7.000 | 1.91 | 0.00 % | 0.02 % | 1.18 % | 0.00 % | 0.02 % | 98.77 % | |
| 7.400 | 1.92 | 0.00 % | 0.01 % | 1.18 % | 0.00 % | 0.01 % | 98.80 % | Blood pH |
| 8.000 | 1.92 | 0.00 % | 0.00 % | 1.19 % | 0.00 % | 0.00 % | 98.81 % | |
| 9.000 | 1.92 | 0.00 % | 0.00 % | 1.19 % | 0.00 % | 0.00 % | 98.81 % | |
| 10.000 | 1.92 | 0.00 % | 0.00 % | 1.19 % | 0.00 % | 0.00 % | 98.81 % | |
| 11.000 | 1.92 | 0.00 % | 0.00 % | 1.19 % | 0.00 % | 0.00 % | 98.81 % | |
| 12.000 | 1.92 | 0.00 % | 0.00 % | 1.19 % | 0.00 % | 0.00 % | 98.81 % | |

### pH-metric Result

Sample name: **M15_octanol**
Assay name: **pH-metric high logP**
Assay ID: **18C-01004**
Filename: **C:\Sirius_T3\Mehtap\20180228_exp28_logP_T3-2\18C-01004_M15_octanol_pH-metric high logP.t3r**

Experiment start time: **3/1/2018 4:33:50 AM**
Analyst: **Pion**
Instrument ID: **T312060**

#### Graphs

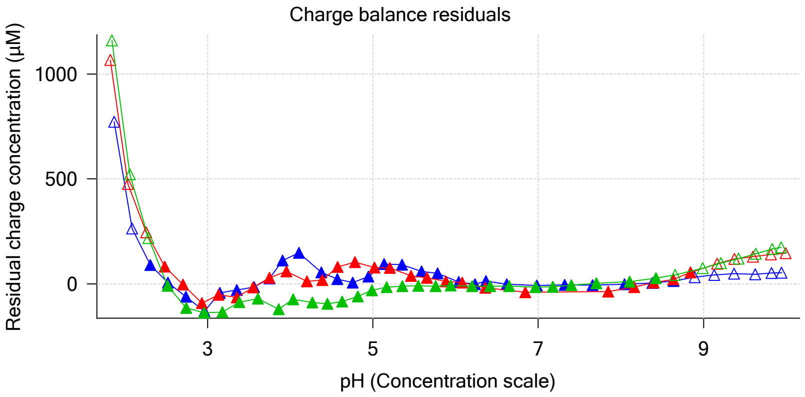


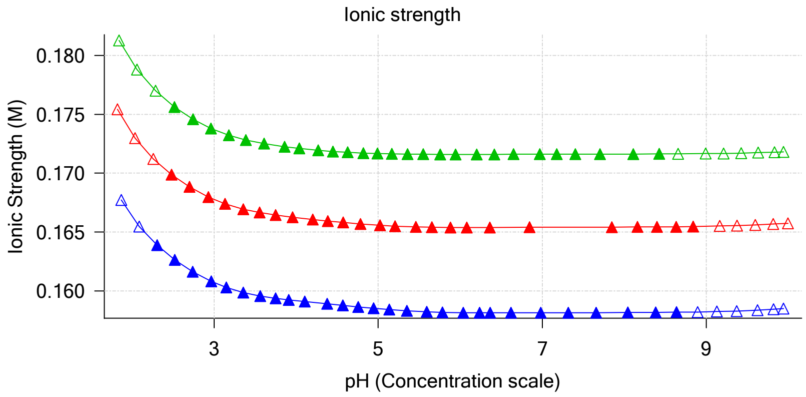


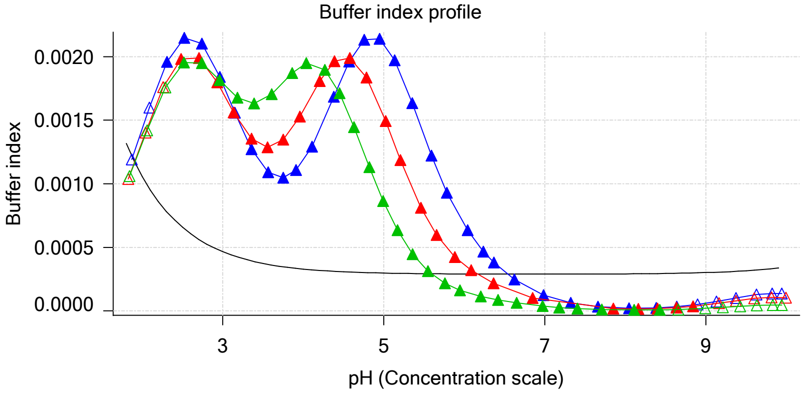


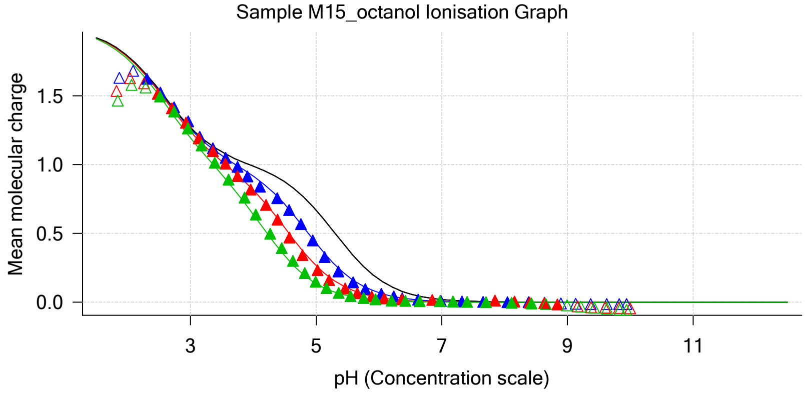


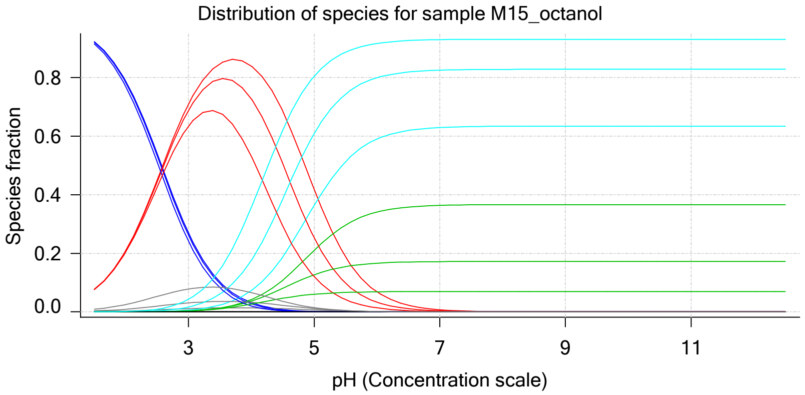


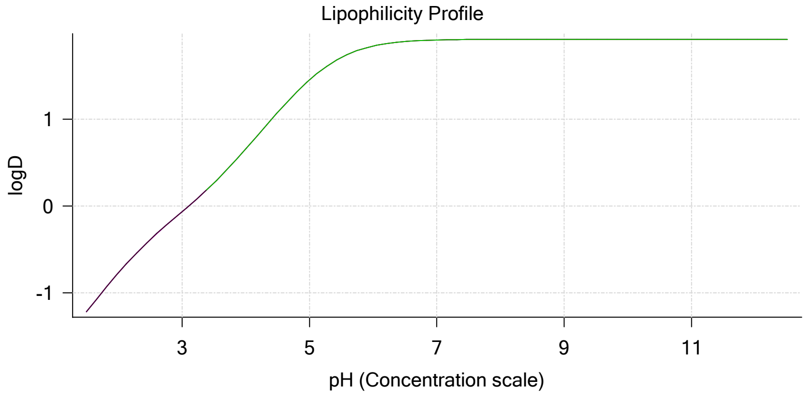


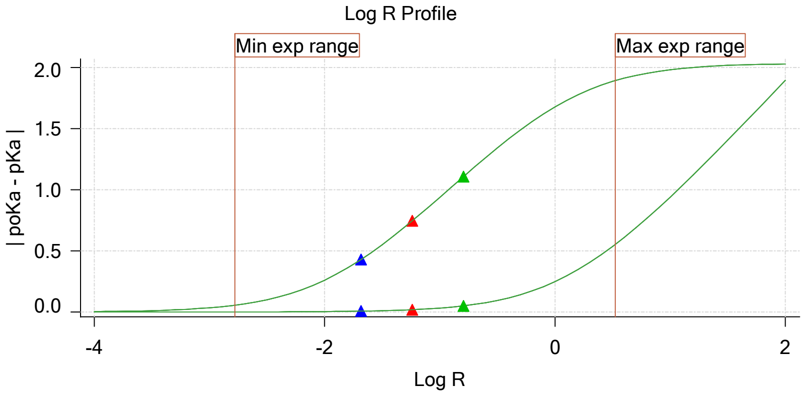


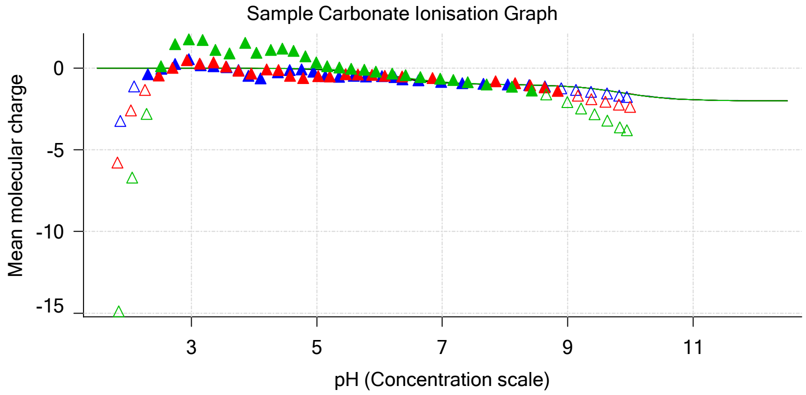

| | | | |
|---|---|---|---|
| Sample name: | **M15_octanol** | Experiment start time: | **3/1/2018 4:33:50 AM** |
| Assay name: | **pH-metric high logP** | Analyst: | **Pion** |
| Assay ID: | **18C-01004** | Instrument ID: | **T312060** |
| Filename: | **C:\Sirius_T3\Mehtap\20180228_exp28_logP_T3-2\18C-01004_M15_octanol_pH-metric high logP.t3r** | | |

#### Graphs (continued)

Distribution of species for sample Carbonate

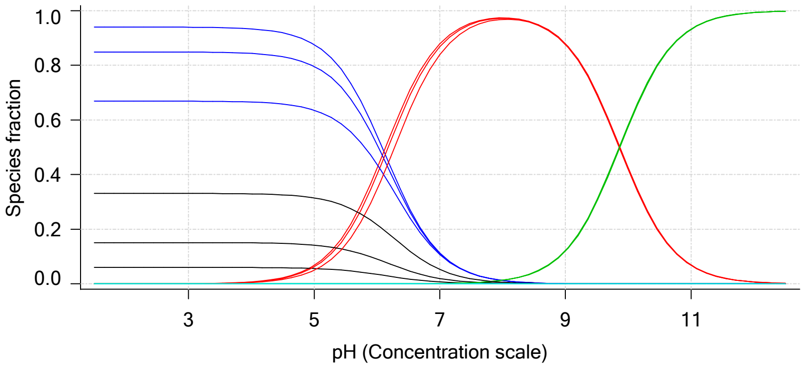

| | | | |
|---|---|---|---|
| Sample name: | **M15_octanol** | Experiment start time: | **3/1/2018 4:33:50 AM** |
| Assay name: | **pH-metric high logP** | Analyst: | **Pion** |
| Assay ID: | **18C-01004** | Instrument ID: | **T312060** |
| Filename: | **C:\Sirius_T3\Mehtap\20180228_exp28_logP_T3-2\18C-01004_M15_octanol_pH-metric high logP.t3r** | | |

### pH-metric high logP Titration 1 of 3 18C-01004 Points 1 to 36

#### Overall results

| | |
|---|---|
| RMSD | 0.075 |
| Average ionic strength | 0.159 M |
| Average temperature | 25.0°C |
| Partition ratio | 0.0208 : 1 |
| Analyte concentration range | 3893.3 μM to 4047.6 μM |
| Total points considered | 28 of 36 |

#### Warnings and errors

| | |
|---|---|
| Errors | None |
| Warnings | One or more logP values out of range |

#### Four-Plus parameters

| | | | |
|---|---|---|---|
| Alpha | 0.130 | 3/1/2018 4:33:50 AM | C:\Sirius_T3\HCl18B27.t3r |
| S | 0.9970 | 3/1/2018 4:33:50 AM | C:\Sirius_T3\HCl18B27.t3r |
| jH | 0.8 | 3/1/2018 4:33:50 AM | C:\Sirius_T3\HCl18B27.t3r |
| jOH | -0.4 | 3/1/2018 4:33:50 AM | C:\Sirius_T3\HCl18B27.t3r |

#### Titrants

| | | | |
|---|---|---|---|
| 0.50 M HCl | 0.993513 | 3/1/2018 4:33:50 AM | C:\Sirius_T3\HCl18B27.t3r |
| 0.50 M KOH | 0.999845 | 3/1/2018 4:33:50 AM | C:\Sirius_T3\KOH18B27.t3r |

#### Sample

| | |
|---|---|
| M15_octanol concentration factor | 0.910 |
| Base pKa 1 | 2.58 |
| Base pKa 2 | 5.30 |
| logP (XH2 2+) | -5.50 |
| logP (XH +) | -4.16 |
| logP (neutral X) | 1.96 |

#### Sample graphs

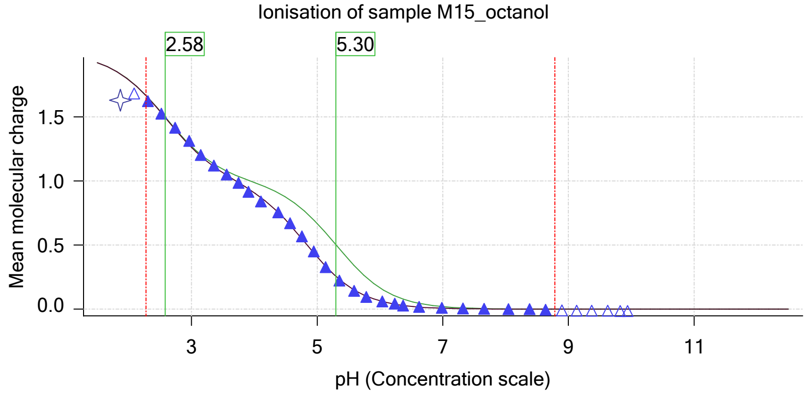


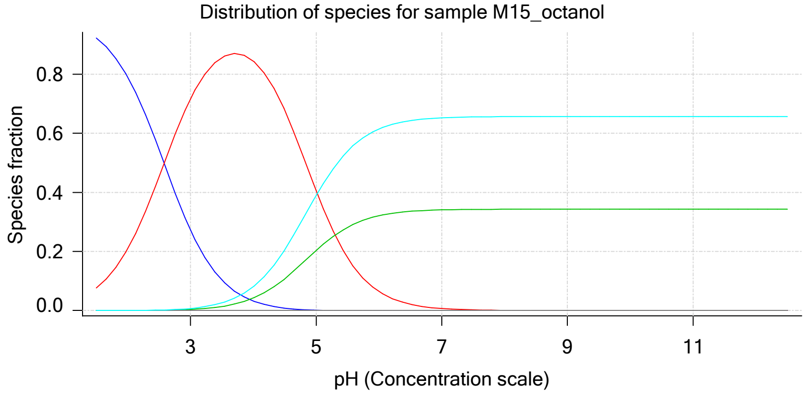

| | | | |
|---|---|---|---|
| Sample name: | **M15_octanol** | Experiment start time: | **3/1/2018 4:33:50 AM** |
| Assay name: | **pH-metric high logP** | Analyst: | **Pion** |
| Assay ID: | **18C-01004** | Instrument ID: | **T312060** |
| Filename: | **C:\Sirius_T3\Mehtap\20180228_exp28_logP_T3-2\18C-01004_M15_octanol_pH-metric high logP.t3r** | | |

#### Sample graphs (continued)

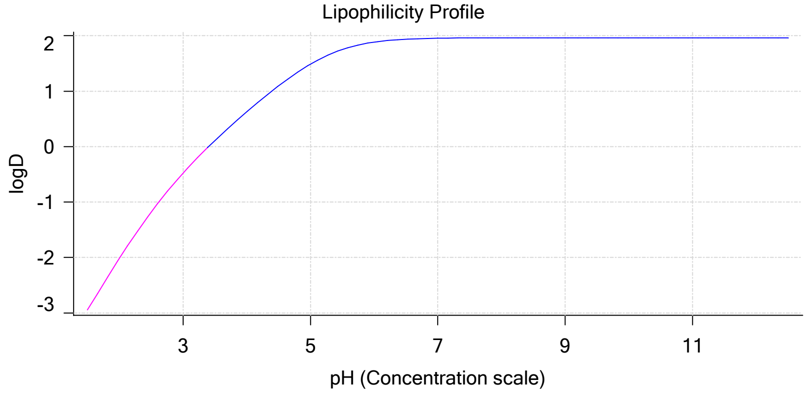


#### Sample logD and percent species

| pH | M15_octanol logD | M15_octanol M15_octanolH2 | M15_octanol M15_octanolH | M15_octanol M15_octanol | M15_octanol M15_octanolH2* | M15_octanol M15_octanolH* | M15_octanol M15_octanol* | Comment |
|---|---|---|---|---|---|---|---|---|
| 1.000 | -3.91 | 97.44 % | 2.56 % | 0.00 % | 0.00 % | 0.00 % | 0.00 % | |
| 1.200 | -3.52 | 96.00 % | 4.00 % | 0.00 % | 0.00 % | 0.00 % | 0.00 % | Stomach pH |
| 2.000 | -2.02 | 79.15 % | 20.82 % | 0.01 % | 0.00 % | 0.00 % | 0.02 % | |
| 3.000 | -0.48 | 27.26 % | 71.69 % | 0.36 % | 0.00 % | 0.00 % | 0.69 % | |
| 4.000 | 0.63 | 3.21 % | 84.45 % | 4.23 % | 0.00 % | 0.00 % | 8.10 % | |
| 5.000 | 1.49 | 0.15 % | 40.58 % | 20.34 % | 0.00 % | 0.00 % | 38.93 % | |
| 6.000 | 1.89 | 0.00 % | 6.41 % | 32.11 % | 0.00 % | 0.00 % | 61.48 % | |
| 6.500 | 1.94 | 0.00 % | 2.12 % | 33.58 % | 0.00 % | 0.00 % | 64.30 % | |
| 7.000 | 1.96 | 0.00 % | 0.68 % | 34.08 % | 0.00 % | 0.00 % | 65.24 % | |
| 7.400 | 1.96 | 0.00 % | 0.27 % | 34.22 % | 0.00 % | 0.00 % | 65.51 % | Blood pH |
| 8.000 | 1.96 | 0.00 % | 0.07 % | 34.29 % | 0.00 % | 0.00 % | 65.64 % | |
| 9.000 | 1.96 | 0.00 % | 0.01 % | 34.31 % | 0.00 % | 0.00 % | 65.68 % | |
| 10.000 | 1.96 | 0.00 % | 0.00 % | 34.31 % | 0.00 % | 0.00 % | 65.69 % | |
| 11.000 | 1.96 | 0.00 % | 0.00 % | 34.31 % | 0.00 % | 0.00 % | 65.69 % | |
| 12.000 | 1.96 | 0.00 % | 0.00 % | 34.31 % | 0.00 % | 0.00 % | 65.69 % | |

#### Carbonate and acidity

| | |
|---|---|
| Carbonate | 0.239 mM |
| Acidity error | -0.312 mM |

#### Other graphs

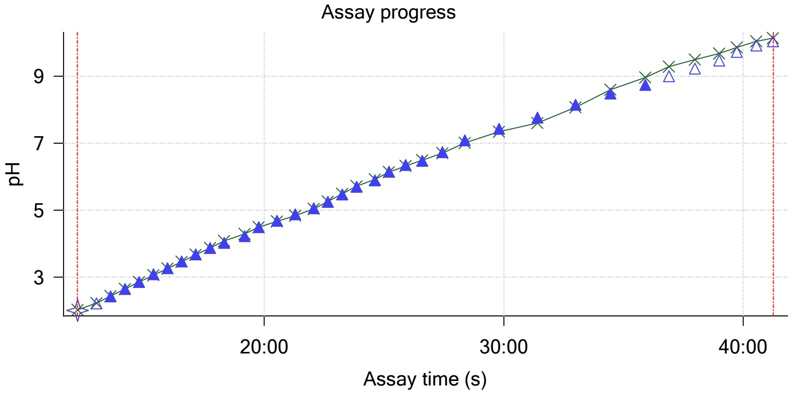


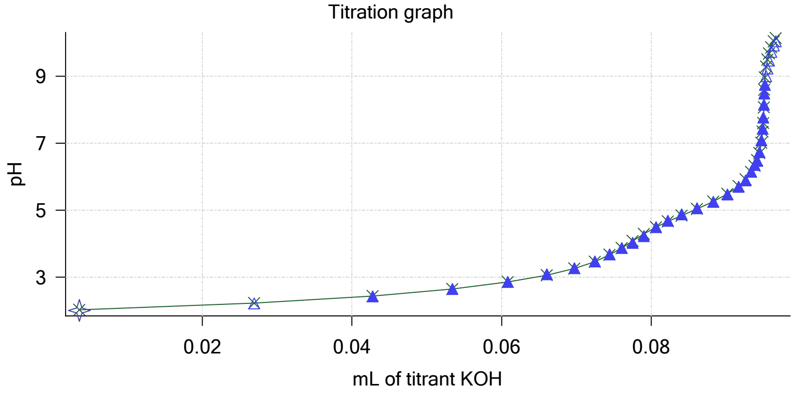

Sample name: **M15_octanol**
Assay name: **pH-metric high logP**
Assay ID: **18C-01004**
Filename: **C:\Sirius_T3\Mehtap\20180228_exp28_logP_T3-2\18C-01004_M15_octanol_pH-metric high logP.t3r**

Experiment start time: **3/1/2018 4:33:50 AM**
Analyst: **Pion**
Instrument ID: **T312060**

#### Other graphs (continued)

Charge balance residuals

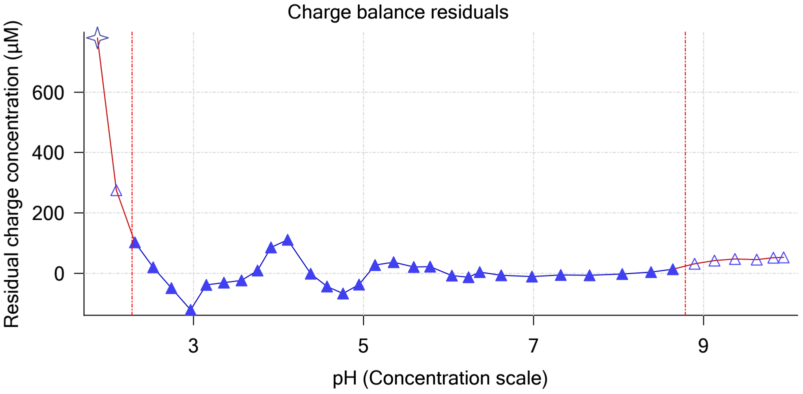


Buffer index profile

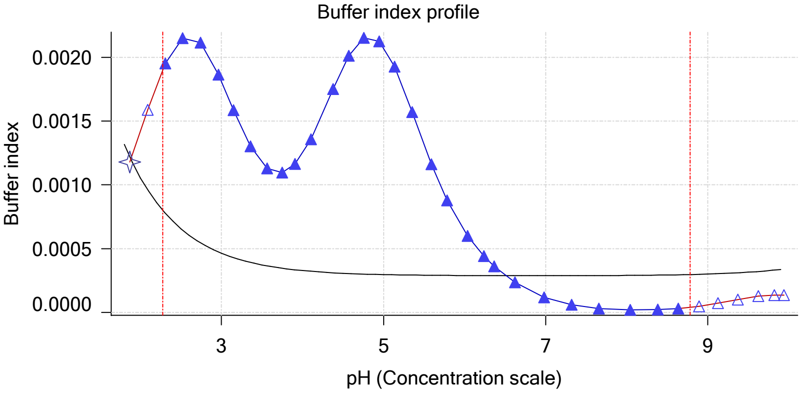


Ionic strength

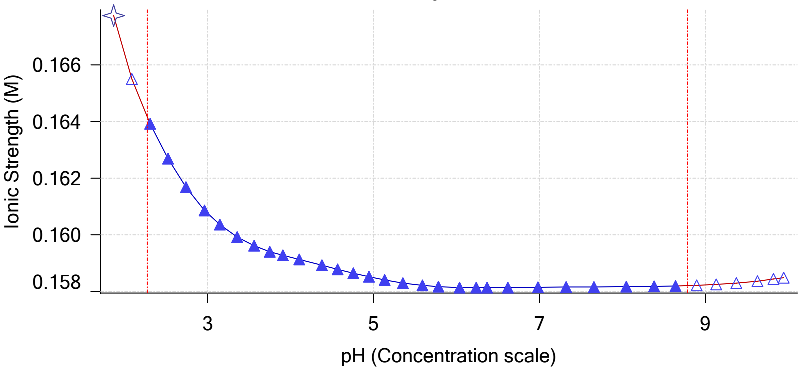


Log R Profile

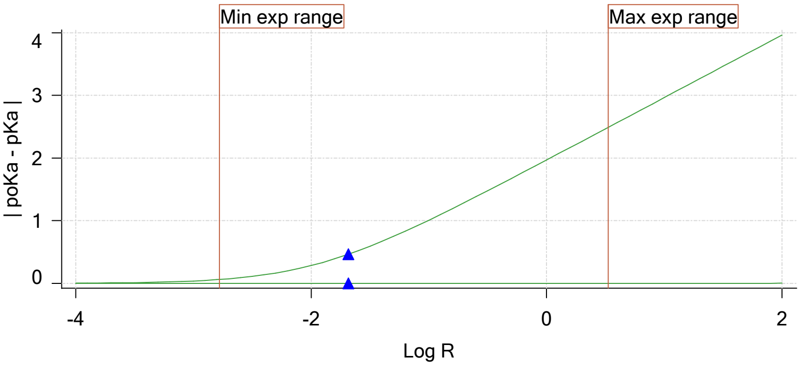


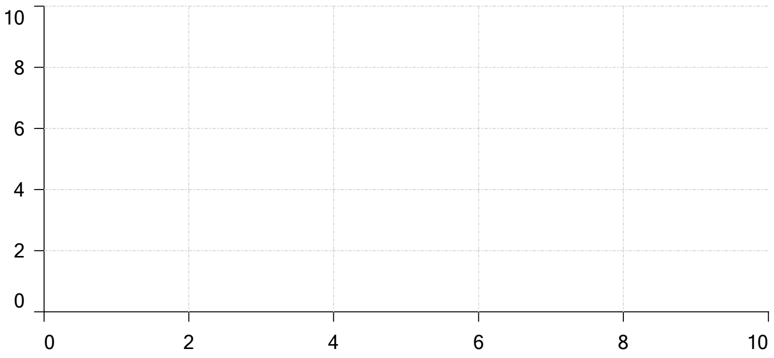


Ionisation of sample Carbonate

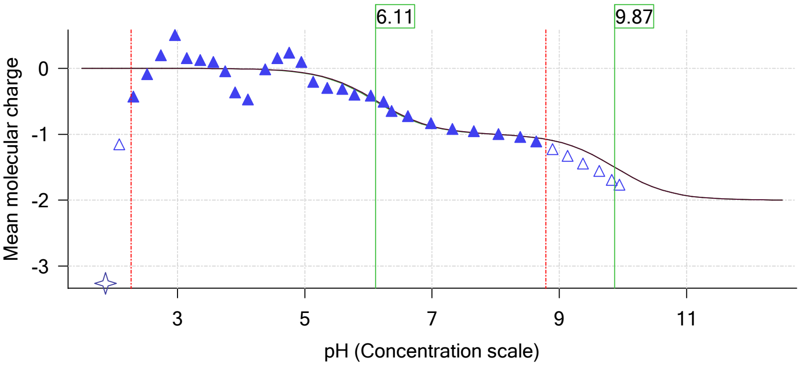

| | | | |
|---|---|---|---|
| Sample name: | **M15_octanol** | Experiment start time: | **3/1/2018 4:33:50 AM** |
| Assay name: | **pH-metric high logP** | Analyst: | **Pion** |
| Assay ID: | **18C-01004** | Instrument ID: | **T312060** |
| Filename: | **C:\Sirius_T3\Mehtap\20180228_exp28_logP_T3-2\18C-01004_M15_octanol_pH-metric high logP.t3r** | | |

# pH-metric high logP   Titration 2 of 3   18C-01004 Points 37 to 69

#### Overall results

| | |
|---|---|
| RMSD | 0.470 |
| Average ionic strength | 0.166 M |
| Average temperature | 25.0°C |
| Partition ratio | 0.0577 : 1 |
| Analyte concentration range | 3455.8 μM to 3592.3 μM |
| Total points considered | 25 of 33 |

#### Warnings and errors

| | |
|---|---|
| Errors | None |
| Warnings | One or more logP values out of range |

#### Four-Plus parameters

| | | | |
|---|---|---|---|
| Alpha | 0.130 | 3/1/2018 4:33:50 AM | C:\Sirius_T3\HCl18B27.t3r |
| S | 0.9970 | 3/1/2018 4:33:50 AM | C:\Sirius_T3\HCl18B27.t3r |
| jH | 0.8 | 3/1/2018 4:33:50 AM | C:\Sirius_T3\HCl18B27.t3r |
| jOH | -0.4 | 3/1/2018 4:33:50 AM | C:\Sirius_T3\HCl18B27.t3r |

#### Titrants

| | | | |
|---|---|---|---|
| 0.50 M HCl | 0.993513 | 3/1/2018 4:33:50 AM | C:\Sirius_T3\HCl18B27.t3r |
| 0.50 M KOH | 0.999845 | 3/1/2018 4:33:50 AM | C:\Sirius_T3\KOH18B27.t3r |

#### Sample

| | |
|---|---|
| M15_octanol concentration factor | 0.903 |
| Base pKa 1 | 2.58 |
| Base pKa 2 | 5.30 |
| logP (XH2 2+) | -5.50 |
| logP (XH +) | -3.58 |
| logP (neutral X) | 1.94 |

#### Sample graphs

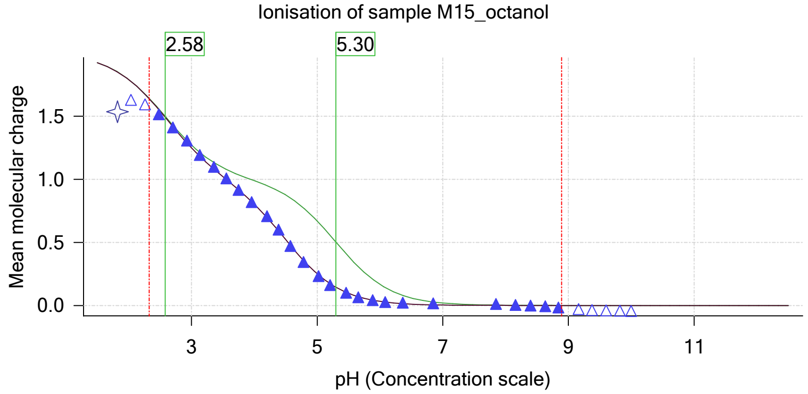


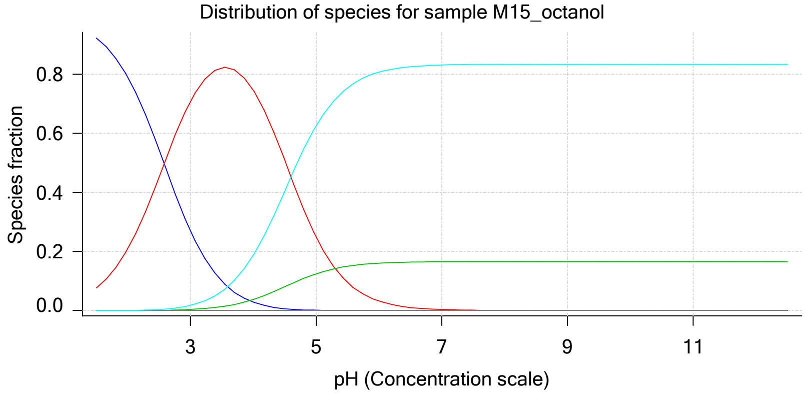

| | | | |
|---|---|---|---|
| Sample name: | **M15_octanol** | Experiment start time: | **3/1/2018 4:33:50 AM** |
| Assay name: | **pH-metric high logP** | Analyst: | **Pion** |
| Assay ID: | **18C-01004** | Instrument ID: | **T312060** |
| Filename: | **C:\Sirius_T3\Mehtap\20180228_exp28_logP_T3-2\18C-01004_M15_octanol_pH-metric high logP.t3r** | | |

#### Sample graphs (continued)

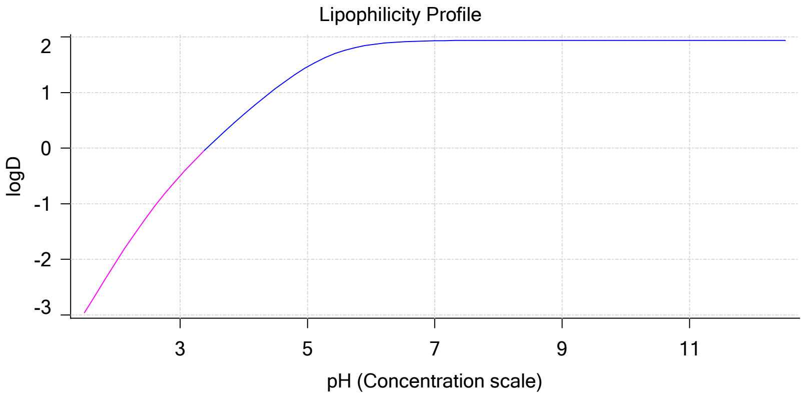


#### Sample logD and percent species

| pH | M15_octanol logD | M15_octanol M15_octanolH2 | M15_octanol M15_octanolH | M15_octanol M15_octanol | M15_octanol M15_octanolH2* | M15_octanol M15_octanolH* | M15_octanol M15_octanol* | Comment |
|---|---|---|---|---|---|---|---|---|
| 1.000 | -3.91 | 97.44 % | 2.56 % | 0.00 % | 0.00 % | 0.00 % | 0.00 % | |
| 1.200 | -3.54 | 96.00 % | 4.00 % | 0.00 % | 0.00 % | 0.00 % | 0.00 % | Stomach pH |
| 2.000 | -2.04 | 79.12 % | 20.81 % | 0.01 % | 0.00 % | 0.00 % | 0.05 % | |
| 3.000 | -0.50 | 26.96 % | 70.90 % | 0.36 % | 0.00 % | 0.00 % | 1.79 % | |
| 4.000 | 0.60 | 2.84 % | 74.62 % | 3.74 % | 0.00 % | 0.00 % | 18.80 % | |
| 5.000 | 1.46 | 0.09 % | 24.85 % | 12.46 % | 0.00 % | 0.00 % | 62.60 % | |
| 6.000 | 1.86 | 0.00 % | 3.20 % | 16.06 % | 0.00 % | 0.00 % | 80.73 % | |
| 6.500 | 1.91 | 0.00 % | 1.04 % | 16.42 % | 0.00 % | 0.00 % | 82.54 % | |
| 7.000 | 1.93 | 0.00 % | 0.33 % | 16.54 % | 0.00 % | 0.00 % | 83.13 % | |
| 7.400 | 1.94 | 0.00 % | 0.13 % | 16.57 % | 0.00 % | 0.00 % | 83.30 % | Blood pH |
| 8.000 | 1.94 | 0.00 % | 0.03 % | 16.59 % | 0.00 % | 0.00 % | 83.38 % | |
| 9.000 | 1.94 | 0.00 % | 0.00 % | 16.59 % | 0.00 % | 0.00 % | 83.40 % | |
| 10.000 | 1.94 | 0.00 % | 0.00 % | 16.60 % | 0.00 % | 0.00 % | 83.40 % | |
| 11.000 | 1.94 | 0.00 % | 0.00 % | 16.60 % | 0.00 % | 0.00 % | 83.40 % | |
| 12.000 | 1.94 | 0.00 % | 0.00 % | 16.60 % | 0.00 % | 0.00 % | 83.40 % | |

#### Carbonate and acidity

| | |
|---|---|
| Carbonate | 0.185 mM |
| Acidity error | -0.258 mM |

#### Other graphs

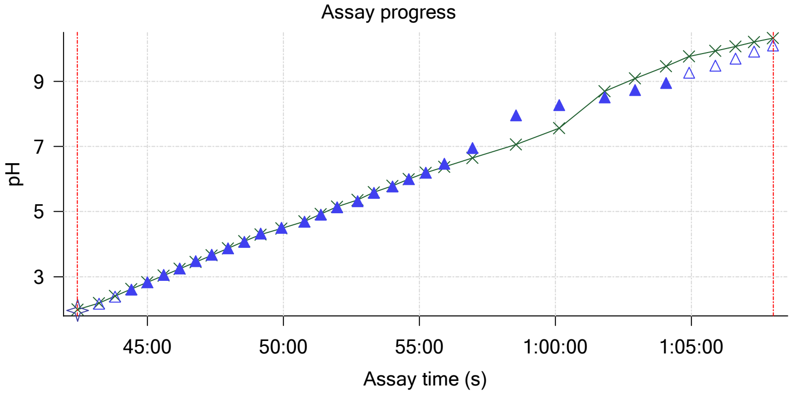


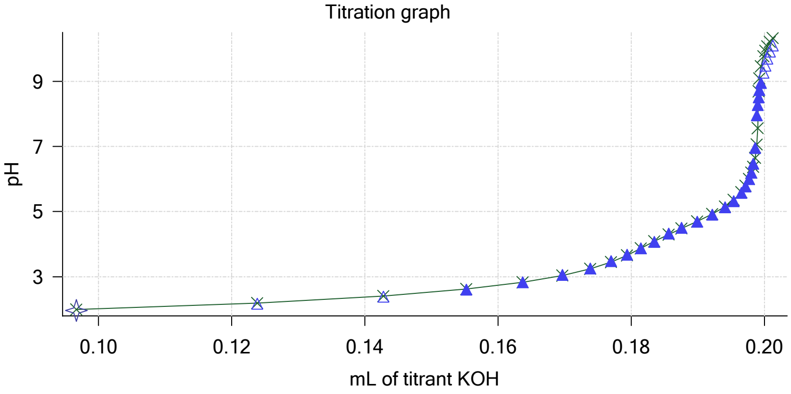

Sample name: **M15_octanol**
Assay name: **pH-metric high logP**
Assay ID: **18C-01004**
Filename: **C:\Sirius_T3\Mehtap\20180228_exp28_logP_T3-2\18C-01004_M15_octanol_pH-metric high logP.t3r**

Experiment start time: **3/1/2018 4:33:50 AM**
Analyst: **Pion**
Instrument ID: **T312060**

#### Other graphs (continued)

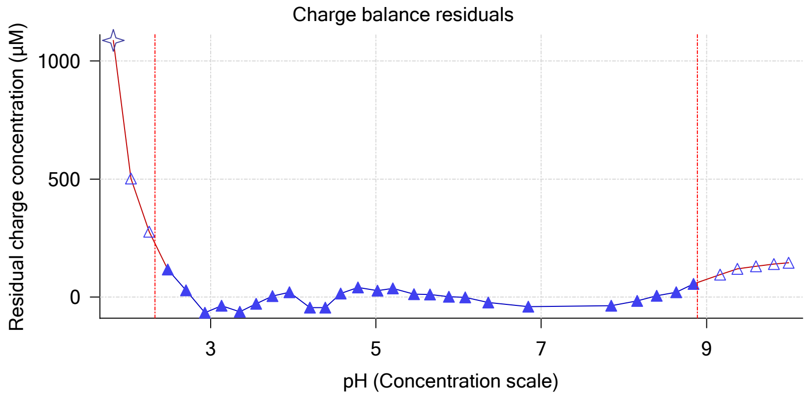


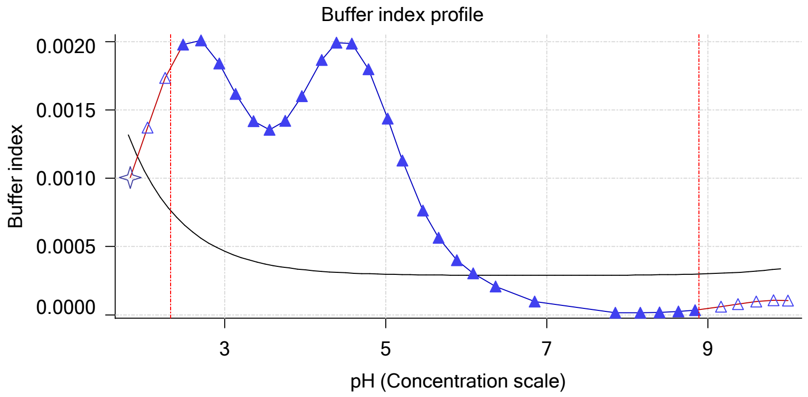


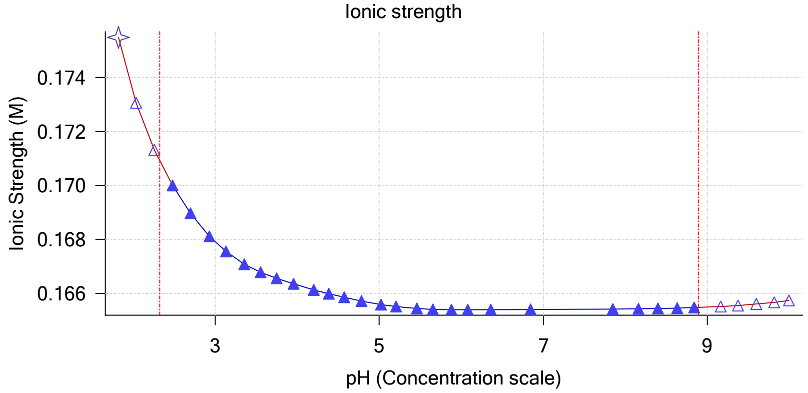


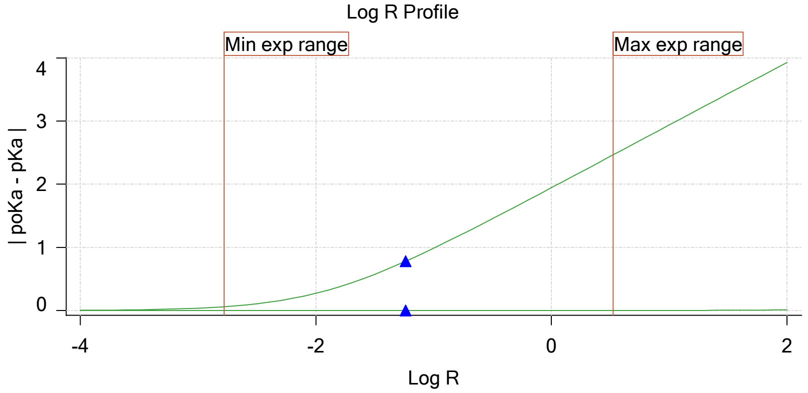


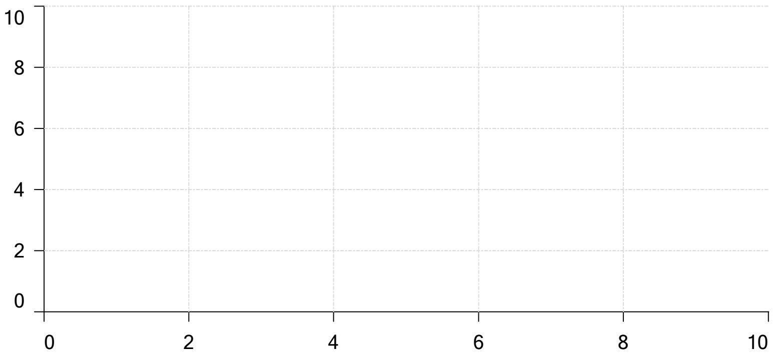


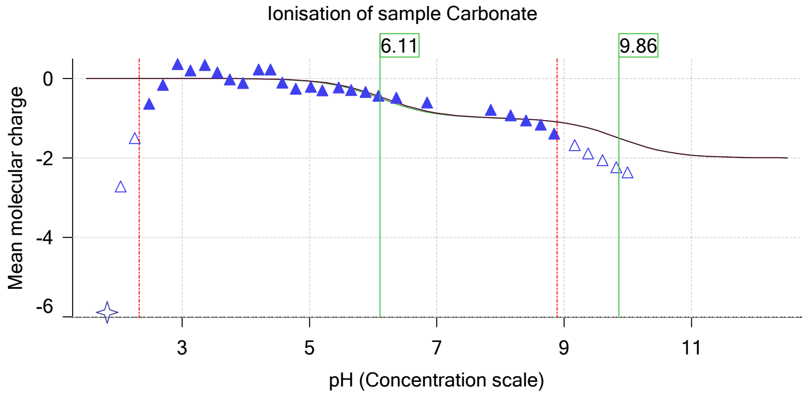

| | | | |
|---|---|---|---|
| Sample name: | **M15_octanol** | Experiment start time: | **3/1/2018 4:33:50 AM** |
| Assay name: | **pH-metric high logP** | Analyst: | **Pion** |
| Assay ID: | **18C-01004** | Instrument ID: | **T312060** |
| Filename: | **C:\Sirius_T3\Mehtap\20180228_exp28_logP_T3-2\18C-01004_M15_octanol_pH-metric high logP.t3r** | | |

# pH-metric high logP   Titration 3 of 3   18C-01004 Points 70 to 106

#### Overall results

| | |
|---|---|
| RMSD | 0.356 |
| Average ionic strength | 0.172 M |
| Average temperature | 25.0°C |
| Partition ratio | 0.1602 : 1 |
| Analyte concentration range | 2913.6 μM to 3011.2 μM |
| Total points considered | 27 of 37 |

#### Warnings and errors

| | |
|---|---|
| Errors | None |
| Warnings | One or more logP values out of range |

#### Four-Plus parameters

| | | | |
|---|---|---|---|
| Alpha | 0.130 | 3/1/2018 4:33:50 AM | C:\Sirius_T3\HCl18B27.t3r |
| S | 0.9970 | 3/1/2018 4:33:50 AM | C:\Sirius_T3\HCl18B27.t3r |
| jH | 0.8 | 3/1/2018 4:33:50 AM | C:\Sirius_T3\HCl18B27.t3r |
| jOH | -0.4 | 3/1/2018 4:33:50 AM | C:\Sirius_T3\HCl18B27.t3r |

#### Titrants

| | | | |
|---|---|---|---|
| 0.50 M HCl | 0.993513 | 3/1/2018 4:33:50 AM | C:\Sirius_T3\HCl18B27.t3r |
| 0.50 M KOH | 0.999845 | 3/1/2018 4:33:50 AM | C:\Sirius_T3\KOH18B27.t3r |

#### Sample

| | |
|---|---|
| M15_octanol concentration factor | 0.913 |
| Base pKa 1 | 2.58 |
| Base pKa 2 | 5.30 |
| logP (XH2 2+) | -5.50 |
| logP (XH +) | -1.60 |
| logP (neutral X) | 1.81 |

#### Sample graphs

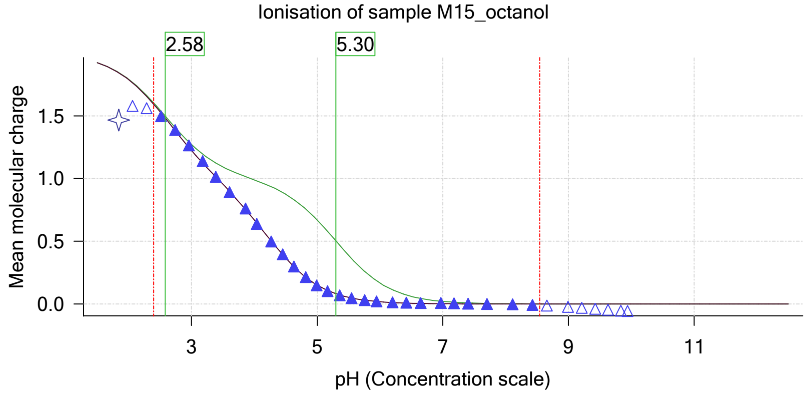


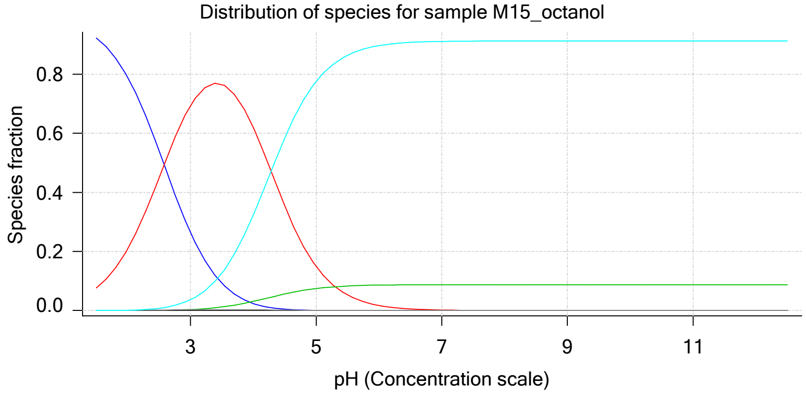

| | | | |
|---|---|---|---|
| Sample name: | **M15_octanol** | Experiment start time: | **3/1/2018 4:33:50 AM** |
| Assay name: | **pH-metric high logP** | Analyst: | **Pion** |
| Assay ID: | **18C-01004** | Instrument ID: | **T312060** |
| Filename: | **C:\Sirius_T3\Mehtap\20180228_exp28_logP_T3-2\18C-01004_M15_octanol_pH-metric high logP.t3r** | | |

#### Sample graphs (continued)

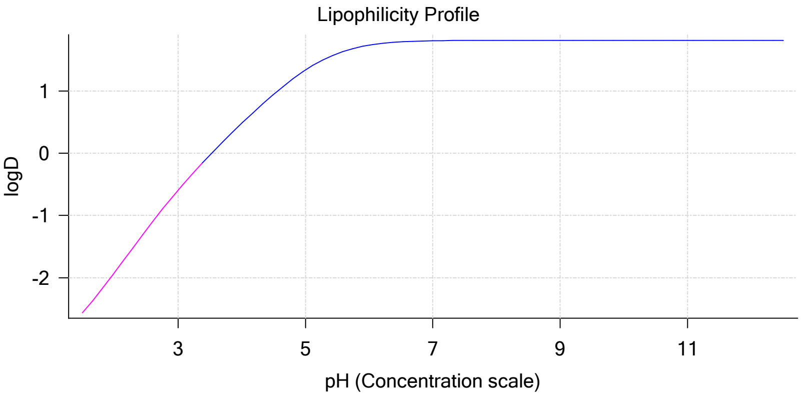


#### Sample logD and percent species

| pH | M15_octanol logD | M15_octanol M15_octanolH2 | M15_octanol M15_octanolH | M15_octanol M15_octanol | M15_octanol M15_octanolH2* | M15_octanol M15_octanolH* | M15_octanol M15_octanol* | Comment |
|---|---|---|---|---|---|---|---|---|
| 1.000 | -3.13 | 97.43 % | 2.56 % | 0.00 % | 0.00 % | 0.01 % | 0.00 % | |
| 1.200 | -2.91 | 95.98 % | 4.00 % | 0.00 % | 0.00 % | 0.02 % | 0.00 % | Stomach pH |
| 2.000 | -1.92 | 79.01 % | 20.78 % | 0.01 % | 0.00 % | 0.08 % | 0.11 % | |
| 3.000 | -0.60 | 26.38 % | 69.38 % | 0.35 % | 0.00 % | 0.28 % | 3.62 % | |
| 4.000 | 0.48 | 2.36 % | 61.96 % | 3.11 % | 0.00 % | 0.25 % | 32.33 % | |
| 5.000 | 1.34 | 0.06 % | 14.87 % | 7.45 % | 0.00 % | 0.06 % | 77.56 % | |
| 6.000 | 1.73 | 0.00 % | 1.72 % | 8.61 % | 0.00 % | 0.01 % | 89.66 % | |
| 6.500 | 1.79 | 0.00 % | 0.55 % | 8.72 % | 0.00 % | 0.00 % | 90.73 % | |
| 7.000 | 1.80 | 0.00 % | 0.17 % | 8.75 % | 0.00 % | 0.00 % | 91.08 % | |
| 7.400 | 1.81 | 0.00 % | 0.07 % | 8.76 % | 0.00 % | 0.00 % | 91.17 % | Blood pH |
| 8.000 | 1.81 | 0.00 % | 0.02 % | 8.76 % | 0.00 % | 0.00 % | 91.22 % | |
| 9.000 | 1.81 | 0.00 % | 0.00 % | 8.76 % | 0.00 % | 0.00 % | 91.23 % | |
| 10.000 | 1.81 | 0.00 % | 0.00 % | 8.76 % | 0.00 % | 0.00 % | 91.24 % | |
| 11.000 | 1.81 | 0.00 % | 0.00 % | 8.76 % | 0.00 % | 0.00 % | 91.24 % | |
| 12.000 | 1.81 | 0.00 % | 0.00 % | 8.76 % | 0.00 % | 0.00 % | 91.24 % | |

#### Carbonate and acidity

| | |
|---|---|
| Carbonate | 0.078 mM |
| Acidity error | -0.092 mM |

#### Other graphs

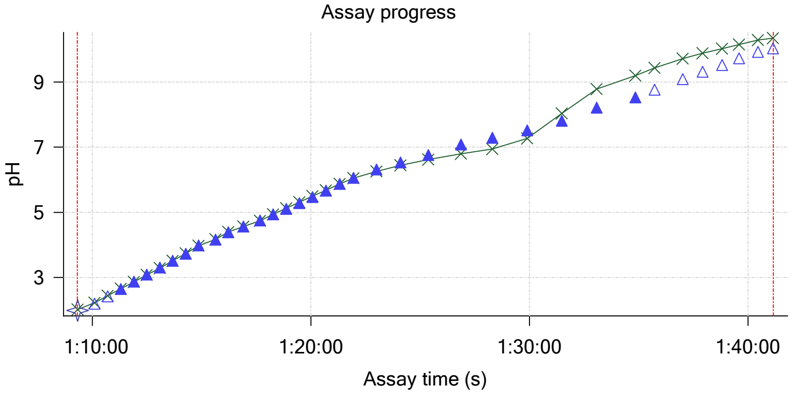


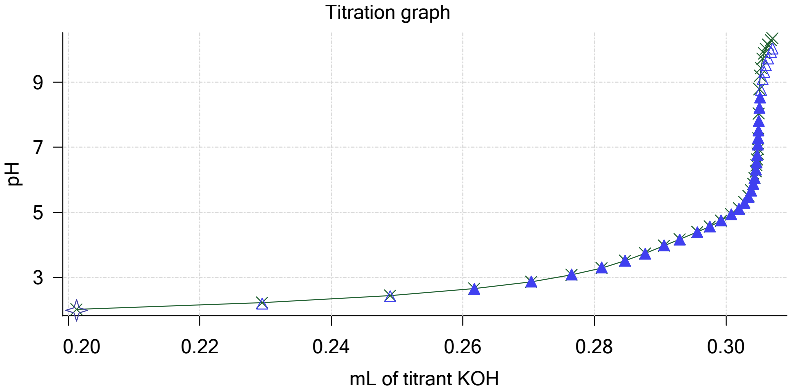

| | | | |
|---|---|---|---|
| Sample name: | **M15_octanol** | Experiment start time: | **3/1/2018 4:33:50 AM** |
| Assay name: | **pH-metric high logP** | Analyst: | **Pion** |
| Assay ID: | **18C-01004** | Instrument ID: | **T312060** |
| Filename: | **C:\Sirius_T3\Mehtap\20180228_exp28_logP_T3-2\18C-01004_M15_octanol_pH-metric high logP.t3r** | | |

#### Other graphs (continued)

Charge balance residuals

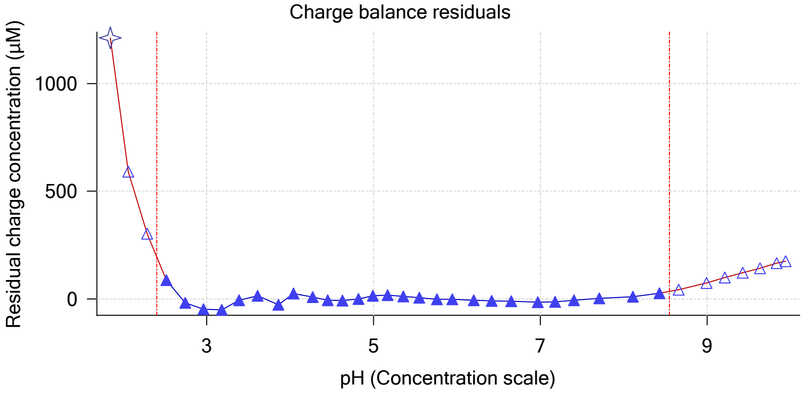


Buffer index profile

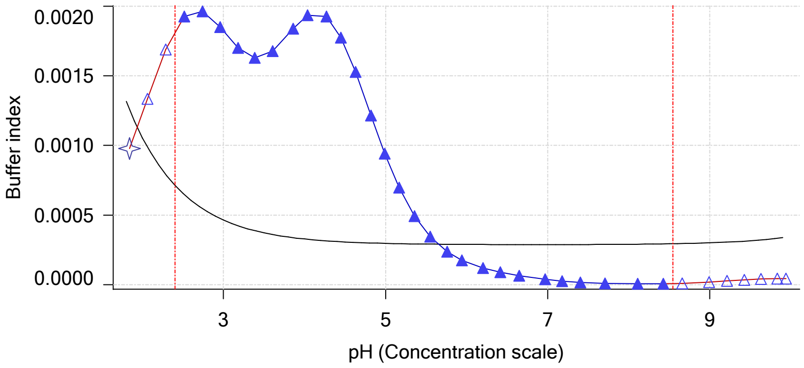


Ionic strength

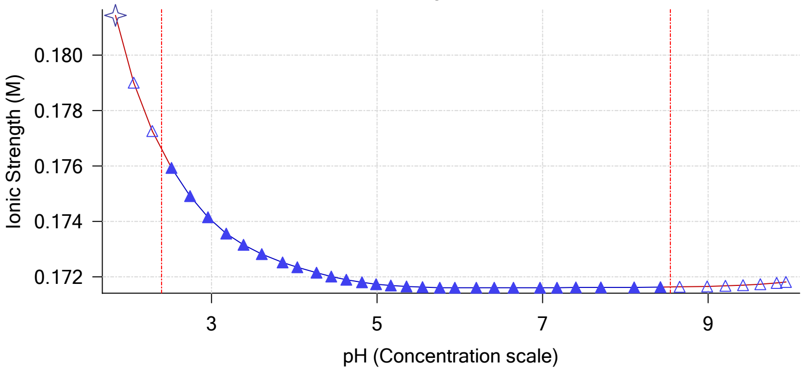


Log R Profile

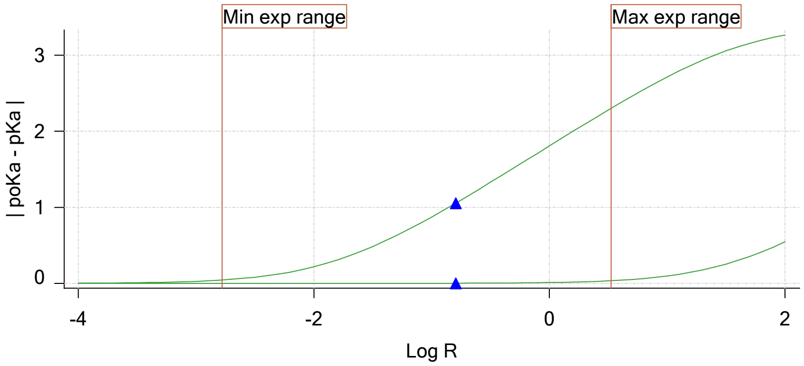


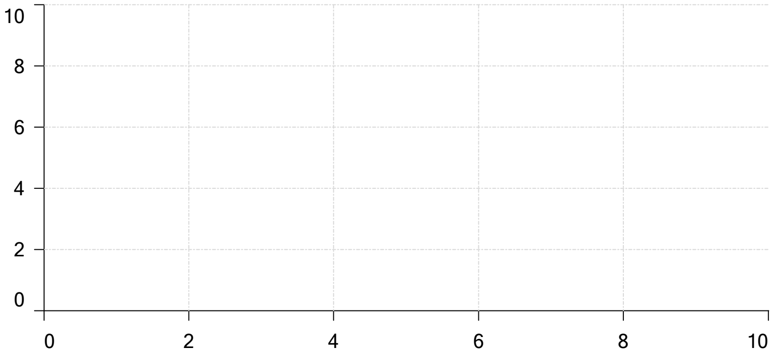


Ionisation of sample Carbonate

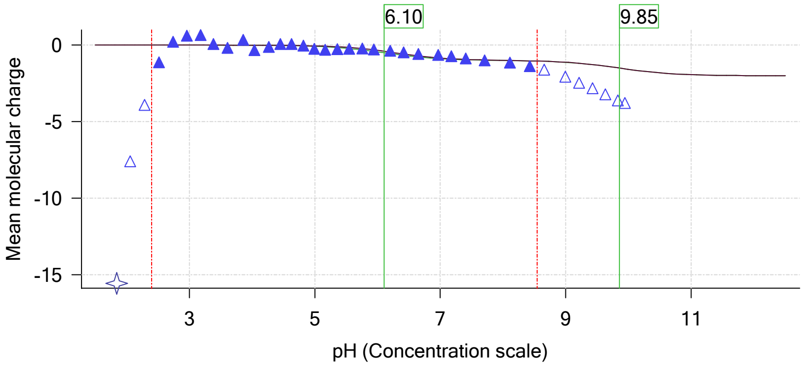

| | | | |
|---|---|---|---|
| Sample name: | **M15_octanol** | Experiment start time: | **3/1/2018 4:33:50 AM** |
| Assay name: | **pH-metric high logP** | Analyst: | **Pion** |
| Assay ID: | **18C-01004** | Instrument ID: | **T312060** |
| Filename: | **C:\Sirius_T3\Mehtap\20180228_exp28_logP_T3-2\18C-01004_M15_octanol_pH-metric high logP.t3r** | | |

#### Assay Model

| Settings | Value | Date/Time changed | Imported from |
|---|---|---|---|
| Sample name | M15_octanol | 2/27/2018 5:03:03 PM | User entered value |
| Sample by | Weight | | Default value |
| Sample weight | 0.001990 g | 2/28/2018 4:25:17 PM | User entered value |
| Formula weight | 209.25 g/mol | 2/27/2018 5:03:03 PM | User entered value |
| Solubility | Unknown | | Default value |
| Molecular weight | 209.25 | 2/27/2018 5:03:03 PM | User entered value |
| Individual pKa ionic environments | No | | Default value |
| Number of pKas | 2 | 2/27/2018 5:03:03 PM | User entered value |
| Sample is a | Base | 2/27/2018 5:03:03 PM | User entered value |
| pKa 1 | 2.58 | 2/27/2018 5:03:03 PM | User entered value |
| pKa 2 | 5.30 | 2/27/2018 5:03:03 PM | User entered value |
| logP (XH2 2+) | -5.50 | 2/28/2018 2:10:35 PM | User entered value |
| logp (XH +) | -4.96 | 2/28/2018 2:10:28 PM | User entered value |
| logP (neutral X) | 1.92 | 2/28/2018 2:10:15 PM | User entered value |

#### Events

| Time | Event | Water | Acid | Base | Octanol | pH | dpH/dt | pH R-squared | pH SD | dpH/dt time |
|---|---|---|---|---|---|---|---|---|---|---|
| 9:11.8 | Initial pH = 7.75 | | | | | | | | | |
| 12:12.6 | Data point 1 | 2.20002 mL | 0.09603 mL | 0.00355 mL | 0.05000 mL | 2.004 | -0.00004 | 0.00005 | 0.00026 | 10.5 s |
| 12:59.6 | Data point 2 | 2.20002 mL | 0.09603 mL | 0.02693 mL | 0.05000 mL | 2.218 | -0.00410 | 0.09174 | 0.00067 | 10.0 s |
| 13:35.4 | Data point 3 | 2.20002 mL | 0.09603 mL | 0.04276 mL | 0.05000 mL | 2.434 | -0.00622 | 0.41443 | 0.00048 | 10.0 s |
| 14:11.0 | Data point 4 | 2.20002 mL | 0.09603 mL | 0.05341 mL | 0.05000 mL | 2.647 | -0.00112 | 0.17193 | 0.00013 | 10.0 s |
| 14:46.6 | Data point 5 | 2.20002 mL | 0.09603 mL | 0.06084 mL | 0.05000 mL | 2.863 | -0.00762 | 0.16820 | 0.00092 | 10.0 s |
| 15:22.1 | Data point 6 | 2.20002 mL | 0.09603 mL | 0.06606 mL | 0.05000 mL | 3.086 | -0.00070 | 0.05715 | 0.00014 | 10.0 s |
| 15:57.6 | Data point 7 | 2.20002 mL | 0.09603 mL | 0.06973 mL | 0.05000 mL | 3.269 | -0.00182 | 0.13794 | 0.00024 | 10.0 s |
| 16:33.1 | Data point 8 | 2.20002 mL | 0.09603 mL | 0.07246 mL | 0.05000 mL | 3.476 | -0.00683 | 0.20917 | 0.00074 | 10.0 s |
| 17:08.6 | Data point 9 | 2.20002 mL | 0.09603 mL | 0.07446 mL | 0.05000 mL | 3.682 | -0.00883 | 0.24236 | 0.00089 | 10.0 s |
| 17:44.1 | Data point 10 | 2.20002 mL | 0.09603 mL | 0.07606 mL | 0.05000 mL | 3.870 | -0.00207 | 0.48492 | 0.00015 | 10.5 s |
| 18:20.0 | Data point 11 | 2.20002 mL | 0.09603 mL | 0.07752 mL | 0.05000 mL | 4.027 | -0.00297 | 0.52144 | 0.00020 | 10.0 s |
| 19:11.0 | Data point 12 | 2.20002 mL | 0.09603 mL | 0.07902 mL | 0.05000 mL | 4.224 | 0.00693 | 0.14332 | 0.00090 | 10.0 s |
| 19:46.5 | Data point 13 | 2.20002 mL | 0.09603 mL | 0.08067 mL | 0.05000 mL | 4.495 | -0.00297 | 0.63948 | 0.00018 | 10.0 s |
| 20:32.2 | Data point 14 | 2.20002 mL | 0.09603 mL | 0.08224 mL | 0.05000 mL | 4.685 | -0.00829 | 0.27907 | 0.00078 | 10.0 s |
| 21:18.0 | Data point 15 | 2.20002 mL | 0.09603 mL | 0.08405 mL | 0.05000 mL | 4.873 | -0.00361 | 0.81281 | 0.00020 | 10.5 s |
| 22:04.3 | Data point 16 | 2.20002 mL | 0.09603 mL | 0.08615 mL | 0.05000 mL | 5.061 | -0.00994 | 0.26388 | 0.00096 | 10.0 s |
| 22:39.8 | Data point 17 | 2.20002 mL | 0.09603 mL | 0.08831 mL | 0.05000 mL | 5.249 | -0.01621 | 0.90472 | 0.00084 | 10.0 s |
| 23:15.2 | Data point 18 | 2.20002 mL | 0.09603 mL | 0.09019 mL | 0.05000 mL | 5.466 | -0.00885 | 0.83876 | 0.00048 | 10.5 s |
| 23:51.2 | Data point 19 | 2.20002 mL | 0.09603 mL | 0.09167 mL | 0.05000 mL | 5.704 | -0.01077 | 0.55398 | 0.00071 | 10.0 s |
| 24:36.9 | Data point 20 | 2.20002 mL | 0.09603 mL | 0.09262 mL | 0.05000 mL | 5.896 | -0.01015 | 0.64626 | 0.00062 | 10.0 s |
| 25:12.4 | Data point 21 | 2.20002 mL | 0.09603 mL | 0.09337 mL | 0.05000 mL | 6.150 | -0.01917 | 0.94736 | 0.00097 | 11.5 s |
| 25:54.3 | Data point 22 | 2.20002 mL | 0.09603 mL | 0.09384 mL | 0.05000 mL | 6.348 | -0.01956 | 0.94838 | 0.00099 | 16.0 s |
| 26:35.7 | Data point 23 | 2.20002 mL | 0.09603 mL | 0.09417 mL | 0.05000 mL | 6.477 | -0.01852 | 0.92877 | 0.00095 | 20.5 s |
| 27:26.8 | Data point 24 | 2.20002 mL | 0.09603 mL | 0.09447 mL | 0.05000 mL | 6.731 | -0.01978 | 0.97668 | 0.00099 | 25.0 s |
| 28:22.5 | Data point 25 | 2.20002 mL | 0.09603 mL | 0.09476 mL | 0.05000 mL | 7.091 | -0.01939 | 0.96778 | 0.00097 | 50.5 s |
| 29:48.7 | Data point 26 | 2.20002 mL | 0.09603 mL | 0.09492 mL | 0.05000 mL | 7.427 | -0.02618 | 0.93832 | 0.00134 | Timed out at 59.5 s |
| 31:24.4 | Data point 27 | 2.20002 mL | 0.09603 mL | 0.09499 mL | 0.05000 mL | 7.765 | -0.02782 | 0.96820 | 0.00140 | Timed out at 59.5 s |
| 32:60.0 | Data point 28 | 2.20002 mL | 0.09603 mL | 0.09506 mL | 0.05000 mL | 8.150 | -0.01942 | 0.97507 | 0.00097 | 51.5 s |
| 34:27.2 | Data point 29 | 2.20002 mL | 0.09603 mL | 0.09513 mL | 0.05000 mL | 8.488 | -0.01822 | 0.88544 | 0.00096 | 47.0 s |
| 35:55.0 | Data point 30 | 2.20002 mL | 0.09603 mL | 0.09523 mL | 0.05000 mL | 8.742 | -0.01803 | 0.81814 | 0.00098 | 18.0 s |
| 36:53.9 | Data point 31 | 2.20002 mL | 0.09603 mL | 0.09539 mL | 0.05000 mL | 8.998 | -0.01906 | 0.91802 | 0.00098 | 29.5 s |
| 37:59.3 | Data point 32 | 2.20002 mL | 0.09603 mL | 0.09556 mL | 0.05000 mL | 9.231 | -0.01960 | 0.94492 | 0.00100 | 24.5 s |
| 38:59.8 | Data point 33 | 2.20002 mL | 0.09603 mL | 0.09577 mL | 0.05000 mL | 9.473 | -0.01978 | 0.98072 | 0.00099 | 14.0 s |
| 39:44.2 | Data point 34 | 2.20002 mL | 0.09603 mL | 0.09605 mL | 0.05000 mL | 9.727 | -0.01838 | 0.95975 | 0.00093 | 13.0 s |
| 40:32.9 | Data point 35 | 2.20002 mL | 0.09603 mL | 0.09640 mL | 0.05000 mL | 9.921 | -0.01891 | 0.96239 | 0.00095 | 11.0 s |
| 41:14.5 | Data point 36 | 2.20002 mL | 0.09603 mL | 0.09666 mL | 0.05000 mL | 10.042 | -0.01797 | 0.91702 | 0.00093 | 10.0 s |

| | | | |
|---|---|---|---|
| Sample name: | **M15_octanol** | Experiment start time: | **3/1/2018 4:33:50 AM** |
| Assay name: | **pH-metric high logP** | Analyst: | **Pion** |
| Assay ID: | **18C-01004** | Instrument ID: | **T312060** |
| Filename: | **C:\Sirius_T3\Mehtap\20180228_exp28_logP_T3-2\18C-01004_M15_octanol_pH-metric high logP.t3r** | | |

#### Events (continued)

| Time | Event | Water | Acid | Base | Octanol | pH | dpH/dt | pH R-squared | pH SD | dpH/dt time |
|---|---|---|---|---|---|---|---|---|---|---|
| 42:26.2 | Data point 37 | 2.20002 mL | 0.20071 mL | 0.09666 mL | 0.15000 mL | 1.959 | -0.00249 | 0.02483 | 0.00078 | 10.0 s |
| 43:12.8 | Data point 38 | 2.20002 mL | 0.20071 mL | 0.12385 mL | 0.15000 mL | 2.169 | 0.00115 | 0.01005 | 0.00057 | 10.0 s |
| 43:48.7 | Data point 39 | 2.20002 mL | 0.20071 mL | 0.14278 mL | 0.15000 mL | 2.387 | 0.00292 | 0.23411 | 0.00030 | 10.0 s |
| 44:24.4 | Data point 40 | 2.20002 mL | 0.20071 mL | 0.15522 mL | 0.15000 mL | 2.607 | 0.00278 | 0.64171 | 0.00017 | 10.0 s |
| 45:00.1 | Data point 41 | 2.20002 mL | 0.20071 mL | 0.16369 mL | 0.15000 mL | 2.825 | 0.00765 | 0.45122 | 0.00056 | 10.0 s |
| 45:35.6 | Data point 42 | 2.20002 mL | 0.20071 mL | 0.16964 mL | 0.15000 mL | 3.053 | 0.00261 | 0.02150 | 0.00088 | 10.0 s |
| 46:11.0 | Data point 43 | 2.20002 mL | 0.20071 mL | 0.17382 mL | 0.15000 mL | 3.254 | -0.00337 | 0.07118 | 0.00062 | 10.0 s |
| 46:46.5 | Data point 44 | 2.20002 mL | 0.20071 mL | 0.17695 mL | 0.15000 mL | 3.476 | -0.00111 | 0.17901 | 0.00013 | 10.0 s |
| 47:22.0 | Data point 45 | 2.20002 mL | 0.20071 mL | 0.17935 mL | 0.15000 mL | 3.673 | -0.00484 | 0.08087 | 0.00084 | 10.5 s |
| 47:58.0 | Data point 46 | 2.20002 mL | 0.20071 mL | 0.18144 mL | 0.15000 mL | 3.869 | -0.00085 | 0.00291 | 0.00078 | 10.5 s |
| 48:34.0 | Data point 47 | 2.20002 mL | 0.20071 mL | 0.18347 mL | 0.15000 mL | 4.074 | -0.00208 | 0.18891 | 0.00024 | 10.0 s |
| 49:09.5 | Data point 48 | 2.20002 mL | 0.20071 mL | 0.18561 mL | 0.15000 mL | 4.321 | 0.00313 | 0.03406 | 0.00084 | 10.5 s |
| 49:55.8 | Data point 49 | 2.20002 mL | 0.20071 mL | 0.18756 mL | 0.15000 mL | 4.505 | -0.00239 | 0.41560 | 0.00018 | 10.0 s |
| 50:46.8 | Data point 50 | 2.20002 mL | 0.20071 mL | 0.18989 mL | 0.15000 mL | 4.692 | 0.00658 | 0.10707 | 0.00099 | 10.0 s |
| 51:22.3 | Data point 51 | 2.20002 mL | 0.20071 mL | 0.19215 mL | 0.15000 mL | 4.901 | -0.00276 | 0.35548 | 0.00023 | 10.5 s |
| 51:58.2 | Data point 52 | 2.20002 mL | 0.20071 mL | 0.19410 mL | 0.15000 mL | 5.136 | -0.00855 | 0.19756 | 0.00095 | 10.0 s |
| 52:44.0 | Data point 53 | 2.20002 mL | 0.20071 mL | 0.19539 mL | 0.15000 mL | 5.320 | 0.01104 | 0.48343 | 0.00078 | 10.5 s |
| 53:19.9 | Data point 54 | 2.20002 mL | 0.20071 mL | 0.19652 mL | 0.15000 mL | 5.575 | -0.01538 | 0.61586 | 0.00097 | 10.0 s |
| 54:00.5 | Data point 55 | 2.20002 mL | 0.20071 mL | 0.19718 mL | 0.15000 mL | 5.767 | -0.01599 | 0.72745 | 0.00093 | 10.5 s |
| 54:36.4 | Data point 56 | 2.20002 mL | 0.20071 mL | 0.19770 mL | 0.15000 mL | 5.995 | -0.01720 | 0.89603 | 0.00090 | 12.5 s |
| 55:14.3 | Data point 57 | 2.20002 mL | 0.20071 mL | 0.19805 mL | 0.15000 mL | 6.194 | -0.01773 | 0.93943 | 0.00090 | 15.0 s |
| 55:54.8 | Data point 58 | 2.20002 mL | 0.20071 mL | 0.19831 mL | 0.15000 mL | 6.473 | -0.01702 | 0.93578 | 0.00087 | 32.0 s |
| 56:57.3 | Data point 59 | 2.20002 mL | 0.20071 mL | 0.19859 mL | 0.15000 mL | 6.955 | -0.02431 | 0.97693 | 0.00121 | Timed out at 59.5 s |
| 58:33.1 | Data point 60 | 2.20002 mL | 0.20071 mL | 0.19885 mL | 0.15000 mL | 7.955 | -0.04787 | 0.97420 | 0.00240 | Timed out at 59.5 s |
| 1:00:08.8 | Data point 61 | 2.20002 mL | 0.20071 mL | 0.19899 mL | 0.15000 mL | 8.265 | -0.01855 | 0.84395 | 0.00100 | 59.0 s |
| 1:01:48.7 | Data point 62 | 2.20002 mL | 0.20071 mL | 0.19913 mL | 0.15000 mL | 8.503 | -0.01414 | 0.67047 | 0.00085 | 31.5 s |
| 1:02:55.9 | Data point 63 | 2.20002 mL | 0.20071 mL | 0.19925 mL | 0.15000 mL | 8.736 | -0.01917 | 0.92795 | 0.00098 | 28.0 s |
| 1:04:04.8 | Data point 64 | 2.20002 mL | 0.20071 mL | 0.19948 mL | 0.15000 mL | 8.945 | -0.01399 | 0.56454 | 0.00092 | 14.5 s |
| 1:04:54.9 | Data point 65 | 2.20002 mL | 0.20071 mL | 0.19984 mL | 0.15000 mL | 9.265 | -0.01776 | 0.81338 | 0.00097 | 22.5 s |
| 1:05:53.3 | Data point 66 | 2.20002 mL | 0.20071 mL | 0.20012 mL | 0.15000 mL | 9.476 | -0.01942 | 0.96420 | 0.00098 | 13.5 s |
| 1:06:37.4 | Data point 67 | 2.20002 mL | 0.20071 mL | 0.20042 mL | 0.15000 mL | 9.700 | -0.01799 | 0.94238 | 0.00092 | 11.0 s |
| 1:07:18.9 | Data point 68 | 2.20002 mL | 0.20071 mL | 0.20082 mL | 0.15000 mL | 9.916 | -0.01678 | 0.93188 | 0.00086 | 10.5 s |
| 1:08:00.0 | Data point 69 | 2.20002 mL | 0.20071 mL | 0.20125 mL | 0.15000 mL | 10.094 | -0.01862 | 0.92012 | 0.00096 | 13.5 s |
| 1:09:19.8 | Data point 70 | 2.20002 mL | 0.30703 mL | 0.20125 mL | 0.45000 mL | 1.978 | -0.00661 | 0.17199 | 0.00079 | 10.0 s |
| 1:10:06.4 | Data point 71 | 2.20002 mL | 0.30703 mL | 0.22949 mL | 0.45000 mL | 2.191 | 0.00020 | 0.00450 | 0.00015 | 10.0 s |
| 1:10:42.3 | Data point 72 | 2.20002 mL | 0.30703 mL | 0.24894 mL | 0.45000 mL | 2.413 | -0.00624 | 0.12108 | 0.00089 | 10.5 s |
| 1:11:18.6 | Data point 73 | 2.20002 mL | 0.30703 mL | 0.26174 mL | 0.45000 mL | 2.641 | -0.00127 | 0.02629 | 0.00039 | 10.0 s |
| 1:11:54.2 | Data point 74 | 2.20002 mL | 0.30703 mL | 0.27039 mL | 0.45000 mL | 2.864 | 0.01123 | 0.56243 | 0.00074 | 10.0 s |
| 1:12:29.8 | Data point 75 | 2.20002 mL | 0.30703 mL | 0.27653 mL | 0.45000 mL | 3.082 | 0.00260 | 0.02083 | 0.00089 | 10.0 s |
| 1:13:05.3 | Data point 76 | 2.20002 mL | 0.30703 mL | 0.28109 mL | 0.45000 mL | 3.300 | 0.00115 | 0.17309 | 0.00014 | 10.0 s |
| 1:13:40.9 | Data point 77 | 2.20002 mL | 0.30703 mL | 0.28467 mL | 0.45000 mL | 3.507 | -0.00125 | 0.06719 | 0.00024 | 10.0 s |
| 1:14:16.4 | Data point 78 | 2.20002 mL | 0.30703 mL | 0.28772 mL | 0.45000 mL | 3.728 | -0.00003 | 0.00007 | 0.00016 | 10.0 s |
| 1:14:51.9 | Data point 79 | 2.20002 mL | 0.30703 mL | 0.29057 mL | 0.45000 mL | 3.979 | 0.00062 | 0.00143 | 0.00080 | 10.5 s |
| 1:15:38.3 | Data point 80 | 2.20002 mL | 0.30703 mL | 0.29299 mL | 0.45000 mL | 4.156 | 0.00694 | 0.12696 | 0.00096 | 10.0 s |
| 1:16:13.8 | Data point 81 | 2.20002 mL | 0.30703 mL | 0.29563 mL | 0.45000 mL | 4.386 | -0.00283 | 0.55635 | 0.00019 | 10.0 s |
| 1:16:54.4 | Data point 82 | 2.20002 mL | 0.30703 mL | 0.29753 mL | 0.45000 mL | 4.567 | 0.00299 | 0.02388 | 0.00096 | 10.5 s |
| 1:17:40.7 | Data point 83 | 2.20002 mL | 0.30703 mL | 0.29925 mL | 0.45000 mL | 4.745 | 0.01034 | 0.46849 | 0.00075 | 10.5 s |
| 1:18:16.7 | Data point 84 | 2.20002 mL | 0.30703 mL | 0.30080 mL | 0.45000 mL | 4.934 | -0.00687 | 0.30375 | 0.00062 | 10.5 s |
| 1:18:52.6 | Data point 85 | 2.20002 mL | 0.30703 mL | 0.30195 mL | 0.45000 mL | 5.106 | -0.00848 | 0.37580 | 0.00068 | 10.5 s |
| 1:19:28.6 | Data point 86 | 2.20002 mL | 0.30703 mL | 0.30280 mL | 0.45000 mL | 5.282 | -0.00416 | 0.04532 | 0.00097 | 10.5 s |
| 1:20:04.5 | Data point 87 | 2.20002 mL | 0.30703 mL | 0.30341 mL | 0.45000 mL | 5.472 | 0.00416 | 0.04632 | 0.00095 | 11.5 s |
| 1:20:41.4 | Data point 88 | 2.20002 mL | 0.30703 mL | 0.30383 mL | 0.45000 mL | 5.662 | 0.00750 | 0.22112 | 0.00079 | 12.5 s |
| 1:21:19.3 | Data point 89 | 2.20002 mL | 0.30703 mL | 0.30414 mL | 0.45000 mL | 5.871 | -0.01602 | 0.74413 | 0.00092 | 12.5 s |
| 1:21:57.2 | Data point 90 | 2.20002 mL | 0.30703 mL | 0.30435 mL | 0.45000 mL | 6.055 | -0.01729 | 0.85517 | 0.00092 | 32.5 s |
| 1:23:00.4 | Data point 91 | 2.20002 mL | 0.30703 mL | 0.30454 mL | 0.45000 mL | 6.313 | -0.01803 | 0.92861 | 0.00092 | 40.0 s |

| Sample name: | **M15_octanol** | Experiment start time: | **3/1/2018 4:33:50 AM** |
|---|---|---|---|
| Assay name: | **pH-metric high logP** | Analyst: | **Pion** |
| Assay ID: | **18C-01004** | Instrument ID: | **T312060** |
| Filename: | **C:\Sirius_T3\Mehtap\20180228_exp28_logP_T3-2\18C-01004_M15_octanol_pH-metric high logP.t3r** | | |

#### Events (continued)

| Time | Event | Water | Acid | Base | Octanol | pH | dpH/dt | pH R-squared | pH SD | dpH/dt time |
|---|---|---|---|---|---|---|---|---|---|---|
| 1:24:05.7 | Data point 92 | 2.20002 mL | 0.30703 mL | 0.30466 mL | 0.45000 mL | 6.527 | -0.01857 | 0.95760 | 0.00094 | 51.0 s |
| 1:25:22.1 | Data point 93 | 2.20002 mL | 0.30703 mL | 0.30475 mL | 0.45000 mL | 6.759 | -0.01815 | 0.93102 | 0.00093 | 59.5 s |
| 1:26:52.2 | Data point 94 | 2.20002 mL | 0.30703 mL | 0.30482 mL | 0.45000 mL | 7.078 | -0.04861 | 0.94386 | 0.00247 | Timed out at 59.5 s |
| 1:28:17.6 | Data point 95 | 2.20002 mL | 0.30703 mL | 0.30487 mL | 0.45000 mL | 7.285 | -0.04933 | 0.98898 | 0.00245 | Timed out at 59.5 s |
| 1:29:53.3 | Data point 96 | 2.20002 mL | 0.30703 mL | 0.30494 mL | 0.45000 mL | 7.511 | -0.04973 | 0.95602 | 0.00251 | Timed out at 59.5 s |
| 1:31:28.9 | Data point 97 | 2.20002 mL | 0.30703 mL | 0.30501 mL | 0.45000 mL | 7.812 | -0.04080 | 0.98347 | 0.00203 | Timed out at 59.5 s |
| 1:33:04.5 | Data point 98 | 2.20002 mL | 0.30703 mL | 0.30508 mL | 0.45000 mL | 8.214 | -0.02455 | 0.95406 | 0.00124 | Timed out at 59.5 s |
| 1:34:50.4 | Data point 99 | 2.20002 mL | 0.30703 mL | 0.30520 mL | 0.45000 mL | 8.531 | -0.01310 | 0.49687 | 0.00092 | 17.0 s |
| 1:35:43.0 | Data point 100 | 2.20002 mL | 0.30703 mL | 0.30532 mL | 0.45000 mL | 8.762 | -0.01738 | 0.82617 | 0.00094 | 31.5 s |
| 1:37:00.5 | Data point 101 | 2.20002 mL | 0.30703 mL | 0.30557 mL | 0.45000 mL | 9.094 | -0.00556 | 0.09210 | 0.00091 | 13.5 s |
| 1:37:54.9 | Data point 102 | 2.20002 mL | 0.30703 mL | 0.30581 mL | 0.45000 mL | 9.313 | -0.01317 | 0.45469 | 0.00096 | 18.0 s |
| 1:38:48.6 | Data point 103 | 2.20002 mL | 0.30703 mL | 0.30607 mL | 0.45000 mL | 9.526 | -0.01066 | 0.32164 | 0.00093 | 11.0 s |
| 1:39:35.3 | Data point 104 | 2.20002 mL | 0.30703 mL | 0.30640 mL | 0.45000 mL | 9.732 | -0.01018 | 0.39405 | 0.00080 | 11.0 s |
| 1:40:27.1 | Data point 105 | 2.20002 mL | 0.30703 mL | 0.30682 mL | 0.45000 mL | 9.932 | -0.01216 | 0.85368 | 0.00065 | 10.5 s |
| 1:41:08.2 | Data point 106 | 2.20002 mL | 0.30703 mL | 0.30708 mL | 0.45000 mL | 10.040 | -0.01217 | 0.84433 | 0.00065 | 10.0 s |
| 1:41:27.3 | Assay volumes | 2.20002 mL | 0.30703 mL | 0.30708 mL | 0.45000 mL | | | | | |

| | | | |
|---|---|---|---|
| Sample name: | **M15_octanol** | Experiment start time: | **3/1/2018 4:33:50 AM** |
| Assay name: | **pH-metric high logP** | Analyst: | **Pion** |
| Assay ID: | **18C-01004** | Instrument ID: | **T312060** |
| Filename: | **C:\Sirius_T3\Mehtap\20180228_exp28_logP_T3-2\18C-01004_M15_octanol_pH-metric high logP.t3r** | | |

| | | | |
|---|---|---|---|
| Sample name: | **M15_octanol** | Experiment start time: | **3/1/2018 4:33:50 AM** |
| Assay name: | **pH-metric high logP** | Analyst: | **Pion** |
| Assay ID: | **18C-01004** | Instrument ID: | **T312060** |
| Filename: | **C:\Sirius_T3\Mehtap\20180228_exp28_logP_T3-2\18C-01004_M15_octanol_pH-metric high logP.t3r** | | |

#### Assay Settings (continued)

| Setting | Value | Original Value | Date/Time changed | Imported from |
|---|---|---|---|---|
| ***Titration 3*** | | | | |
| Titrate from | Low to high pH | | | |
| Add additional water | 0.00 mL | | | |
| Additional partition solvent volume | 0.300 mL | | | |
| Additional partition solvent added | Automatic | | | |
| After pH adjust stir for | 30 seconds | | | |
| Stir to allow partitioning for | 15 seconds | | | |
| Stirrer speed for partitioning | 60% | | | |
| ***Data Point Stability*** | | | | |
| Stir during data point collection | No | | | |
| Delay before data point collection | 0 seconds | | | |
| Number of points to average | 20 points | | | |
| Time interval between points | 0.50 seconds | | | |
| Required maximum standard deviation | 0.00100 dpH/dt | | | |
| Stability timeout after | 60 seconds | | | |

#### Calibration Settings

| Setting | Value | Date/Time changed | Imported from |
|---|---|---|---|
| Four-Plus alpha | 0.130 | 3/1/2018 4:33:50 AM | C:\Sirius_T3\HCl18B27.t3r |
| Four-Plus S | 0.9970 | 3/1/2018 4:33:50 AM | C:\Sirius_T3\HCl18B27.t3r |
| Four-Plus jH | 0.8 | 3/1/2018 4:33:50 AM | C:\Sirius_T3\HCl18B27.t3r |
| Four-Plus jOH | -0.4 | 3/1/2018 4:33:50 AM | C:\Sirius_T3\HCl18B27.t3r |
| Base concentration factor | 1.000 | 3/1/2018 4:33:50 AM | C:\Sirius_T3\KOH18B27.t3r |
| Acid concentration factor | 0.994 | 3/1/2018 4:33:50 AM | C:\Sirius_T3\HCl18B27.t3r |

#### Instrument Settings

| Setting | Value | Batch Id | Install date |
|---|---|---|---|
| Instrument owner | Merck | | |
| Instrument ID | T312060 | | |
| Instrument type | T3 Simulator | | |
| Software version | 1.1.3.0 | | |
| Dispenser module | | T3DM1200361 | 3/31/2009 5:24:52 AM |
| Dispenser 0 | Water | | 3/31/2009 5:25:05 AM |
| Syringe volume | 2.5 mL | | |
| Firmware version | 1.2.1(r2) | | |
| Titrant | Water (0.15 M KCl) | 02-06-2018 | 2/27/2018 10:05:59 AM |
| Dispenser 2 | Acid | | 3/31/2009 5:25:11 AM |
| Syringe volume | 0.5 mL | | |
| Firmware version | 1.2.1(r2) | | |
| Titrant | Acid (0.5 M HCl) | 02-27-2018 | 2/27/2018 10:27:22 AM |
| Dispenser 1 | Base | | 3/31/2009 5:25:21 AM |
| Syringe volume | 0.5 mL | | |
| Firmware version | 1.2.1(r2) | | |
| Titrant | Base (0.5 M KOH) | 9/22/2017 | 2/27/2018 10:21:22 AM |
| Dispenser 5 | Cosolvent | | 3/31/2009 5:26:24 AM |
| Syringe volume | 2.5 mL | | |
| Firmware version | 1.2.1(r2) | | |
| Distribution valve 5 | Distribution Valve | | 3/31/2009 5:28:19 AM |
| Firmware version | 1.1.3 | | |
| Port A | Methanol (80%, 0.15 M KCl) | 09-26-17 | 2/7/2018 9:42:01 AM |
| Port B | Cyclohexane | 11-01-17 | 2/27/2018 10:37:57 AM |
| Dispenser 3 | Buffer | | 8/3/2010 5:05:16 AM |
| Syringe volume | 0.5 mL | | |
| Firmware version | 1.2.1(r2) | | |
| Titrant | Dodecane | 2018/01/31 | 2/28/2018 10:18:04 AM |
| Dispenser 6 | Octanol | | 10/22/2010 10:52:43 AM |

| | | | |
|---|---|---|---|
| Sample name: | **M15_octanol** | Experiment start time: | **3/1/2018 4:33:50 AM** |
| Assay name: | **pH-metric high logP** | Analyst: | **Pion** |
| Assay ID: | **18C-01004** | Instrument ID: | **T312060** |
| Filename: | **C:\Sirius_T3\Mehtap\20180228_exp28_logP_T3-2\18C-01004_M15_octanol_pH-metric high logP.t3r** | | |

Sample name: **M15_octanol** Experiment start time: **3/1/2018 4:33:50 AM**
Assay name: **pH-metric high logP** Analyst: **Pion**
Assay ID: **18C-01004** Instrument ID: **T312060**
Filename: **C:\Sirius_T3\Mehtap\20180228_exp28_logP_T3-2\18C-01004_M15_octanol_pH-metric high logP.t3r**

#### Instrument Settings (continued)

| Setting | Value | Batch Id | Install date |
|---|---|---|---|
| Spectrometer calibration stir duration | 5 s | | |
| Spectrometer calibration stir speed | 30% | | |
| Spectrometer calibration wash pump volume | 20.0 mL | | |
| Spectrometer calibration wash stir duration | 5 s | | |
| Spectrometer calibration wash stir speed | 30% | | |
| Overhead dispense height | 10000 | | |

#### Refinement Settings

| Setting | Value | Default value |
|---|---|---|
| Turbidity detection method | None | None |
| Turbidity wavelength to assess | 500.0 nm | 500.0 nm |
| Turbidity maximum absorbance | 0.100 | 0.100 |
| Turbidity probe threshold | 50.00 | 50.00 |

#### Experiment Log

[2:38] Air gap created for Water (0.15 M KCl)
[2:38] Air gap created for Acid (0.5 M HCl)
[2:38] Air gap created for Base (0.5 M KOH)
[2:39] Air gap released for Water (0.15 M KCl)
[2:42] Titrator arm moved over Titration position
[2:42] Titration 1 of 3
[2:42] Adding initial titrants
[2:42] Automatically add 2.20000 mL of water
[3:19] Dispensed 2.200024 mL of Water (0.15 M KCl)
[3:24] Titrator arm moved over Drain
[9:05] Titrator arm moved to Titration position
[9:05] Argon flow rate set to 100
[9:05] Stirrer speed set to 10
[9:10] Automatically add 0.05000 mL of Octanol
[9:11] Dispensed 0.050000 mL of Octanol
[9:12] Initial pH = 7.75
[9:12] Iterative adjust 7.75 -> 2.00
[9:12] pH 7.75 -> 2.00
[9:15] Air gap released for Acid (0.5 M HCl)
[9:16] Dispensed 0.096025 mL of Acid (0.5 M HCl)
[9:21] Holding pH 2.00
[11:21] Stirrer speed set to 0
[11:21] Stirrer speed set to 50
[11:21] Iterative adjust 1.98 -> 2.00
[11:21] pH 1.98 -> 2.00
[11:22] Air gap released for Base (0.5 M KOH)
[11:22] Dispensed 0.003551 mL of Base (0.5 M KOH)
[12:13] Stirrer speed set to 0
[12:23] Datapoint id 1 collected
[12:23] Stirrer speed set to 50
[12:29] pH 2.01 -> 2.21
[12:29] Using cautious pH adjust
[12:29] Dispensed 0.012159 mL of Base (0.5 M KOH)
[12:34] Stepping pH = 2.10
[12:34] Dispensed 0.009290 mL of Base (0.5 M KOH)
[12:40] Stepping pH = 2.19
[12:40] Dispensed 0.001929 mL of Base (0.5 M KOH)
[12:45] Stepping pH = 2.22
[13:00] Stirrer speed set to 0
[13:10] Datapoint id 2 collected
[13:10] Charge balance equation is out by 3.9%
[13:10] Stirrer speed set to 50

| | | | |
|---|---|---|---|
| Sample name: | **M15_octanol** | Experiment start time: | **3/1/2018 4:33:50 AM** |
| Assay name: | **pH-metric high logP** | Analyst: | **Pion** |
| Assay ID: | **18C-01004** | Instrument ID: | **T312060** |
| Filename: | **C:\Sirius_T3\Mehtap\20180228_exp28_logP_T3-2\18C-01004_M15_octanol_pH-metric high logP.t3r** | | |

#### Experiment Log (continued)

[13:15] pH 2.22 -> 2.42
[13:15] Using charge balance adjust
[13:15] Dispensed 0.015828 mL of Base (0.5 M KOH)
[13:36] Stirrer speed set to 0
[13:46] Datapoint id 3 collected
[13:46] Charge balance equation is out by 5.0%
[13:46] Stirrer speed set to 50
[13:51] pH 2.44 -> 2.64
[13:51] Using charge balance adjust
[13:51] Dispensed 0.010654 mL of Base (0.5 M KOH)
[14:11] Stirrer speed set to 0
[14:21] Datapoint id 4 collected
[14:21] Charge balance equation is out by 4.3%
[14:21] Stirrer speed set to 50
[14:26] pH 2.65 -> 2.85
[14:26] Using charge balance adjust
[14:27] Dispensed 0.007432 mL of Base (0.5 M KOH)
[14:47] Stirrer speed set to 0
[14:57] Datapoint id 5 collected
[14:57] Charge balance equation is out by 5.6%
[14:57] Stirrer speed set to 50
[15:02] pH 2.87 -> 3.07
[15:02] Using charge balance adjust
[15:02] Dispensed 0.005221 mL of Base (0.5 M KOH)
[15:22] Stirrer speed set to 0
[15:32] Datapoint id 6 collected
[15:32] Charge balance equation is out by 7.8%
[15:32] Stirrer speed set to 50
[15:38] pH 3.09 -> 3.29
[15:38] Using charge balance adjust
[15:38] Dispensed 0.003669 mL of Base (0.5 M KOH)
[15:58] Stirrer speed set to 0
[16:08] Datapoint id 7 collected
[16:08] Charge balance equation is out by -10.7%
[16:08] Stirrer speed set to 50
[16:13] pH 3.27 -> 3.47
[16:13] Using charge balance adjust
[16:13] Dispensed 0.002728 mL of Base (0.5 M KOH)
[16:33] Stirrer speed set to 0
[16:43] Datapoint id 8 collected
[16:43] Charge balance equation is out by 1.1%
[16:43] Stirrer speed set to 50
[16:49] pH 3.48 -> 3.68
[16:49] Using charge balance adjust
[16:49] Dispensed 0.001999 mL of Base (0.5 M KOH)
[17:09] Stirrer speed set to 0
[17:19] Datapoint id 9 collected
[17:19] Charge balance equation is out by 0.9%
[17:19] Stirrer speed set to 50
[17:24] pH 3.69 -> 3.89
[17:24] Using charge balance adjust
[17:24] Dispensed 0.001599 mL of Base (0.5 M KOH)
[17:44] Stirrer speed set to 0
[17:55] Datapoint id 10 collected
[17:55] Charge balance equation is out by -8.5%
[17:55] Stirrer speed set to 50
[18:00] pH 3.87 -> 4.07
[18:00] Using charge balance adjust
[18:00] Dispensed 0.001458 mL of Base (0.5 M KOH)

| | | | |
|---|---|---|---|
| Sample name: | **M15_octanol** | Experiment start time: | **3/1/2018 4:33:50 AM** |
| Assay name: | **pH-metric high logP** | Analyst: | **Pion** |
| Assay ID: | **18C-01004** | Instrument ID: | **T312060** |
| Filename: | **C:\Sirius_T3\Mehtap\20180228_exp28_logP_T3-2\18C-01004_M15_octanol_pH-metric high logP.t3r** | | |

#### Experiment Log (continued)

[18:20] Stirrer speed set to 0
[18:30] Datapoint id 11 collected
[18:30] Charge balance equation is out by -23.7%
[18:30] Stirrer speed set to 50
[18:35] pH 4.03 -> 4.23
[18:35] Using cautious pH adjust
[18:35] Dispensed 0.000729 mL of Base (0.5 M KOH)
[18:41] Stepping pH = 4.13
[18:41] Dispensed 0.000564 mL of Base (0.5 M KOH)
[18:46] Stepping pH = 4.21
[18:46] Dispensed 0.000118 mL of Base (0.5 M KOH)
[18:51] Stepping pH = 4.22
[18:51] Dispensed 0.000094 mL of Base (0.5 M KOH)
[18:56] Stepping pH = 4.23
[19:11] Stirrer speed set to 0
[19:21] Datapoint id 12 collected
[19:21] Charge balance equation is out by -3.6%
[19:21] Stirrer speed set to 50
[19:26] pH 4.23 -> 4.43
[19:26] Using charge balance adjust
[19:27] Dispensed 0.001646 mL of Base (0.5 M KOH)
[19:47] Stirrer speed set to 0
[19:57] Datapoint id 13 collected
[19:57] Charge balance equation is out by 33.3%
[19:57] Stirrer speed set to 50
[20:02] pH 4.50 -> 4.70
[20:02] Using cautious pH adjust
[20:02] Dispensed 0.001011 mL of Base (0.5 M KOH)
[20:07] Stepping pH = 4.63
[20:07] Dispensed 0.000447 mL of Base (0.5 M KOH)
[20:12] Stepping pH = 4.68
[20:12] Dispensed 0.000118 mL of Base (0.5 M KOH)
[20:17] Stepping pH = 4.69
[20:32] Stirrer speed set to 0
[20:42] Datapoint id 14 collected
[20:42] Charge balance equation is out by 22.0%
[20:42] Stirrer speed set to 50
[20:48] pH 4.69 -> 4.89
[20:48] Using cautious pH adjust
[20:48] Dispensed 0.001105 mL of Base (0.5 M KOH)
[20:53] Stepping pH = 4.82
[20:53] Dispensed 0.000517 mL of Base (0.5 M KOH)
[20:58] Stepping pH = 4.87
[20:58] Dispensed 0.000188 mL of Base (0.5 M KOH)
[21:03] Stepping pH = 4.88
[21:18] Stirrer speed set to 0
[21:29] Datapoint id 15 collected
[21:29] Charge balance equation is out by 17.8%
[21:29] Stirrer speed set to 50
[21:34] pH 4.88 -> 5.08
[21:34] Using cautious pH adjust
[21:34] Dispensed 0.001129 mL of Base (0.5 M KOH)
[21:39] Stepping pH = 4.99
[21:39] Dispensed 0.000706 mL of Base (0.5 M KOH)
[21:44] Stepping pH = 5.05
[21:44] Dispensed 0.000259 mL of Base (0.5 M KOH)
[21:49] Stepping pH = 5.07
[22:05] Stirrer speed set to 0
[22:15] Datapoint id 16 collected

Sample name: **M15_octanol**
Assay name: **pH-metric high logP**
Assay ID: **18C-01004**
Filename: **C:\Sirius_T3\Mehtap\20180228_exp28_logP_T3-2\18C-01004_M15_octanol_pH-metric high logP.t3r**
Experiment start time: **3/1/2018 4:33:50 AM**
Analyst: **Pion**
Instrument ID: **T312060**

#### Experiment Log (continued)

[22:15] Charge balance equation is out by 7.7%
[22:15] Stirrer speed set to 50
[22:20] pH 5.07 -> 5.27
[22:20] Using charge balance adjust
[22:20] Dispensed 0.002164 mL of Base (0.5 M KOH)
[22:40] Stirrer speed set to 0
[22:50] Datapoint id 17 collected
[22:50] Charge balance equation is out by -8.1%
[22:50] Stirrer speed set to 50
[22:55] pH 5.25 -> 5.45
[22:55] Using charge balance adjust
[22:55] Dispensed 0.001881 mL of Base (0.5 M KOH)
[23:15] Stirrer speed set to 0
[23:26] Datapoint id 18 collected
[23:26] Charge balance equation is out by 7.2%
[23:26] Stirrer speed set to 50
[23:31] pH 5.47 -> 5.67
[23:31] Using charge balance adjust
[23:31] Dispensed 0.001482 mL of Base (0.5 M KOH)
[23:51] Stirrer speed set to 0
[24:02] Datapoint id 19 collected
[24:02] Charge balance equation is out by 17.3%
[24:02] Stirrer speed set to 50
[24:07] pH 5.71 -> 5.91
[24:07] Using cautious pH adjust
[24:07] Dispensed 0.000517 mL of Base (0.5 M KOH)
[24:12] Stepping pH = 5.80
[24:12] Dispensed 0.000376 mL of Base (0.5 M KOH)
[24:17] Stepping pH = 5.90
[24:17] Dispensed 0.000047 mL of Base (0.5 M KOH)
[24:22] Stepping pH = 5.90
[24:37] Stirrer speed set to 0
[24:47] Datapoint id 20 collected
[24:47] Charge balance equation is out by 10.0%
[24:47] Stirrer speed set to 50
[24:52] pH 5.90 -> 6.10
[24:52] Using charge balance adjust
[24:52] Dispensed 0.000753 mL of Base (0.5 M KOH)
[25:13] Stirrer speed set to 0
[25:24] Datapoint id 21 collected
[25:24] Charge balance equation is out by 24.3%
[25:24] Stirrer speed set to 50
[25:29] pH 6.15 -> 6.35
[25:29] Using cautious pH adjust
[25:29] Dispensed 0.000235 mL of Base (0.5 M KOH)
[25:34] Stepping pH = 6.23
[25:34] Dispensed 0.000235 mL of Base (0.5 M KOH)
[25:39] Stepping pH = 6.35
[25:55] Stirrer speed set to 0
[26:11] Datapoint id 22 collected
[26:11] Charge balance equation is out by 3.5%
[26:11] Stirrer speed set to 50
[26:16] pH 6.36 -> 6.56
[26:16] Using charge balance adjust
[26:16] Dispensed 0.000329 mL of Base (0.5 M KOH)
[26:36] Stirrer speed set to 0
[26:57] Datapoint id 23 collected
[26:57] Charge balance equation is out by -39.1%
[26:57] Stirrer speed set to 50

| | | | |
|---|---|---|---|
| Sample name: | **M15_octanol** | Experiment start time: | **3/1/2018 4:33:50 AM** |
| Assay name: | **pH-metric high logP** | Analyst: | **Pion** |
| Assay ID: | **18C-01004** | Instrument ID: | **T312060** |
| Filename: | **C:\Sirius_T3\Mehtap\20180228_exp28_logP_T3-2\18C-01004_M15_octanol_pH-metric high logP.t3r** | | |

#### Experiment Log (continued)

[27:02] pH 6.48 -> 6.68
[27:02] Using cautious pH adjust
[27:02] Dispensed 0.000118 mL of Base (0.5 M KOH)
[27:07] Stepping pH = 6.53
[27:07] Dispensed 0.000188 mL of Base (0.5 M KOH)
[27:12] Stepping pH = 6.72
[27:27] Stirrer speed set to 0
[27:52] Datapoint id 24 collected
[27:52] Charge balance equation is out by -27.4%
[27:52] Stirrer speed set to 50
[27:57] pH 6.75 -> 6.95
[27:57] Using cautious pH adjust
[27:57] Dispensed 0.000071 mL of Base (0.5 M KOH)
[28:02] Stepping pH = 6.77
[28:03] Dispensed 0.000212 mL of Base (0.5 M KOH)
[28:08] Stepping pH = 7.07
[28:23] Stirrer speed set to 0
[29:13] Datapoint id 25 collected
[29:13] Charge balance equation is out by -83.1%
[29:13] Stirrer speed set to 50
[29:18] pH 7.12 -> 7.32
[29:18] Using cautious pH adjust
[29:18] Dispensed 0.000024 mL of Base (0.5 M KOH)
[29:24] Stepping pH = 7.13
[29:24] Dispensed 0.000094 mL of Base (0.5 M KOH)
[29:29] Stepping pH = 7.23
[29:29] Dispensed 0.000047 mL of Base (0.5 M KOH)
[29:34] Stepping pH = 7.42
[29:49] Stirrer speed set to 0
[30:49] Datapoint id 26 collected
[30:49] Charge balance equation is out by -166.1%
[30:49] Stirrer speed set to 50
[30:54] pH 7.49 -> 7.69
[30:54] Using cautious pH adjust
[30:54] Dispensed 0.000024 mL of Base (0.5 M KOH)
[30:59] Stepping pH = 7.55
[30:59] Dispensed 0.000024 mL of Base (0.5 M KOH)
[31:04] Stepping pH = 7.61
[31:04] Dispensed 0.000024 mL of Base (0.5 M KOH)
[31:10] Stepping pH = 7.74
[31:25] Stirrer speed set to 0
[32:25] Datapoint id 27 collected
[32:25] Charge balance equation is out by -89.7%
[32:25] Stirrer speed set to 50
[32:30] pH 7.81 -> 8.01
[32:30] Using cautious pH adjust
[32:30] Dispensed 0.000024 mL of Base (0.5 M KOH)
[32:35] Stepping pH = 7.86
[32:35] Dispensed 0.000024 mL of Base (0.5 M KOH)
[32:40] Stepping pH = 7.97
[32:40] Dispensed 0.000024 mL of Base (0.5 M KOH)
[32:45] Stepping pH = 8.12
[33:00] Stirrer speed set to 0
[33:52] Datapoint id 28 collected
[33:52] Charge balance equation is out by -214.6%
[33:52] Stirrer speed set to 50
[33:57] pH 8.22 -> 8.42
[33:57] Using cautious pH adjust
[33:57] Dispensed 0.000024 mL of Base (0.5 M KOH)

Sample name: **M15_octanol** Experiment start time: **3/1/2018 4:33:50 AM**
Assay name: **pH-metric high logP** Analyst: **Pion**
Assay ID: **18C-01004** Instrument ID: **T312060**
Filename: **C:\Sirius_T3\Mehtap\20180228_exp28_logP_T3-2\18C-01004_M15_octanol_pH-metric high logP.t3r**

#### Experiment Log (continued)

[34:02] Stepping pH = 8.27
[34:02] Dispensed 0.000024 mL of Base (0.5 M KOH)
[34:07] Stepping pH = 8.37
[34:07] Dispensed 0.000024 mL of Base (0.5 M KOH)
[34:12] Stepping pH = 8.48
[34:27] Stirrer speed set to 0
[35:15] Datapoint id 29 collected
[35:15] Charge balance equation is out by -257.9%
[35:15] Stirrer speed set to 50
[35:20] pH 8.53 -> 8.73
[35:20] Using cautious pH adjust
[35:20] Dispensed 0.000024 mL of Base (0.5 M KOH)
[35:25] Stepping pH = 8.56
[35:25] Dispensed 0.000024 mL of Base (0.5 M KOH)
[35:30] Stepping pH = 8.62
[35:30] Dispensed 0.000024 mL of Base (0.5 M KOH)
[35:35] Stepping pH = 8.68
[35:35] Dispensed 0.000024 mL of Base (0.5 M KOH)
[35:40] Stepping pH = 8.74
[35:55] Stirrer speed set to 0
[36:13] Datapoint id 30 collected
[36:13] Charge balance equation is out by -326.8%
[36:13] Stirrer speed set to 50
[36:18] pH 8.76 -> 8.96
[36:18] Using cautious pH adjust
[36:18] Dispensed 0.000024 mL of Base (0.5 M KOH)
[36:24] Stepping pH = 8.78
[36:24] Dispensed 0.000047 mL of Base (0.5 M KOH)
[36:29] Stepping pH = 8.83
[36:29] Dispensed 0.000071 mL of Base (0.5 M KOH)
[36:34] Stepping pH = 8.95
[36:34] Dispensed 0.000024 mL of Base (0.5 M KOH)
[36:39] Stepping pH = 9.00
[36:54] Stirrer speed set to 0
[37:24] Datapoint id 31 collected
[37:24] Charge balance equation is out by -326.2%
[37:24] Stirrer speed set to 50
[37:29] pH 9.02 -> 9.22
[37:29] Using cautious pH adjust
[37:29] Dispensed 0.000024 mL of Base (0.5 M KOH)
[37:34] Stepping pH = 9.03
[37:34] Dispensed 0.000094 mL of Base (0.5 M KOH)
[37:39] Stepping pH = 9.14
[37:39] Dispensed 0.000047 mL of Base (0.5 M KOH)
[37:44] Stepping pH = 9.23
[38:00] Stirrer speed set to 0
[38:24] Datapoint id 32 collected
[38:24] Charge balance equation is out by -182.2%
[38:24] Stirrer speed set to 50
[38:29] pH 9.24 -> 9.44
[38:29] Using cautious pH adjust
[38:29] Dispensed 0.000047 mL of Base (0.5 M KOH)
[38:34] Stepping pH = 9.26
[38:34] Dispensed 0.000141 mL of Base (0.5 M KOH)
[38:40] Stepping pH = 9.43
[38:40] Dispensed 0.000024 mL of Base (0.5 M KOH)
[38:45] Stepping pH = 9.47
[39:00] Stirrer speed set to 0
[39:14] Datapoint id 33 collected

| | | | |
|---|---|---|---|
| Sample name: | **M15_octanol** | Experiment start time: | **3/1/2018 4:33:50 AM** |
| Assay name: | **pH-metric high logP** | Analyst: | **Pion** |
| Assay ID: | **18C-01004** | Instrument ID: | **T312060** |
| Filename: | **C:\Sirius_T3\Mehtap\20180228_exp28_logP_T3-2\18C-01004_M15_octanol_pH-metric high logP.t3r** | | |

#### Experiment Log (continued)

[39:14] Charge balance equation is out by -108.6%
[39:14] Stirrer speed set to 50
[39:19] pH 9.49 -> 9.69
[39:19] Using cautious pH adjust
[39:19] Dispensed 0.000071 mL of Base (0.5 M KOH)
[39:24] Stepping pH = 9.51
[39:24] Dispensed 0.000212 mL of Base (0.5 M KOH)
[39:29] Stepping pH = 9.72
[39:45] Stirrer speed set to 0
[39:58] Datapoint id 34 collected
[39:58] Charge balance equation is out by -84.8%
[39:58] Stirrer speed set to 50
[40:03] pH 9.73 -> 9.93
[40:03] Using cautious pH adjust
[40:03] Dispensed 0.000141 mL of Base (0.5 M KOH)
[40:08] Stepping pH = 9.80
[40:08] Dispensed 0.000165 mL of Base (0.5 M KOH)
[40:13] Stepping pH = 9.90
[40:13] Dispensed 0.000047 mL of Base (0.5 M KOH)
[40:18] Stepping pH = 9.93
[40:33] Stirrer speed set to 0
[40:44] Datapoint id 35 collected
[40:44] Charge balance equation is out by -29.7%
[40:44] Stirrer speed set to 50
[40:49] pH 9.93 -> 10.05
[40:49] Using cautious pH adjust
[40:49] Dispensed 0.000118 mL of Base (0.5 M KOH)
[40:55] Stepping pH = 9.97
[40:55] Dispensed 0.000141 mL of Base (0.5 M KOH)
[41:00] Stepping pH = 10.04
[41:15] Stirrer speed set to 0
[41:25] Datapoint id 36 collected
[41:25] Charge balance equation is out by -12.1%
[41:25] Titration 2 of 3
[41:25] Adding initial titrants
[41:25] Automatically add 0.10000 mL of Octanol
[41:27] Dispensed 0.100000 mL of Octanol
[41:27] Stirrer speed set to 10
[41:28] Stirrer speed set to 55
[41:28] Iterative adjust 10.05 -> 2.00
[41:28] pH 10.05 -> 2.00
[41:31] Dispensed 0.100000 mL of Acid (0.5 M HCl)
[41:36] pH 2.03 -> 2.00
[41:36] Dispensed 0.004680 mL of Acid (0.5 M HCl)
[42:26] Stirrer speed set to 0
[42:36] Datapoint id 37 collected
[42:36] Stirrer speed set to 55
[42:42] pH 1.96 -> 2.16
[42:42] Using cautious pH adjust
[42:42] Dispensed 0.014675 mL of Base (0.5 M KOH)
[42:47] Stepping pH = 2.05
[42:48] Dispensed 0.010466 mL of Base (0.5 M KOH)
[42:53] Stepping pH = 2.14
[42:53] Dispensed 0.002046 mL of Base (0.5 M KOH)
[42:58] Stepping pH = 2.17
[43:13] Stirrer speed set to 0
[43:23] Datapoint id 38 collected
[43:23] Charge balance equation is out by 7.4%
[43:23] Stirrer speed set to 55

| | | | |
|---|---|---|---|
| Sample name: | **M15_octanol** | Experiment start time: | **3/1/2018 4:33:50 AM** |
| Assay name: | **pH-metric high logP** | Analyst: | **Pion** |
| Assay ID: | **18C-01004** | Instrument ID: | **T312060** |
| Filename: | **C:\Sirius_T3\Mehtap\20180228_exp28_logP_T3-2\18C-01004_M15_octanol_pH-metric high logP.t3r** | | |

#### Experiment Log (continued)

[43:28] pH 2.17 -> 2.37
[43:28] Using charge balance adjust
[43:29] Dispensed 0.018932 mL of Base (0.5 M KOH)
[43:49] Stirrer speed set to 0
[43:59] Datapoint id 39 collected
[43:59] Charge balance equation is out by 7.7%
[43:59] Stirrer speed set to 55
[44:04] pH 2.39 -> 2.59
[44:04] Using charge balance adjust
[44:05] Dispensed 0.012441 mL of Base (0.5 M KOH)
[44:25] Stirrer speed set to 0
[44:35] Datapoint id 40 collected
[44:35] Charge balance equation is out by 8.0%
[44:35] Stirrer speed set to 55
[44:40] pH 2.61 -> 2.81
[44:40] Using charge balance adjust
[44:40] Dispensed 0.008467 mL of Base (0.5 M KOH)
[45:00] Stirrer speed set to 0
[45:10] Datapoint id 41 collected
[45:10] Charge balance equation is out by 6.4%
[45:10] Stirrer speed set to 55
[45:15] pH 2.83 -> 3.03
[45:15] Using charge balance adjust
[45:16] Dispensed 0.005950 mL of Base (0.5 M KOH)
[45:36] Stirrer speed set to 0
[45:46] Datapoint id 42 collected
[45:46] Charge balance equation is out by 10.4%
[45:46] Stirrer speed set to 55
[45:51] pH 3.06 -> 3.26
[45:51] Using charge balance adjust
[45:51] Dispensed 0.004186 mL of Base (0.5 M KOH)
[46:11] Stirrer speed set to 0
[46:21] Datapoint id 43 collected
[46:21] Charge balance equation is out by -2.9%
[46:21] Stirrer speed set to 55
[46:26] pH 3.26 -> 3.46
[46:26] Using charge balance adjust
[46:27] Dispensed 0.003128 mL of Base (0.5 M KOH)
[46:47] Stirrer speed set to 0
[46:57] Datapoint id 44 collected
[46:57] Charge balance equation is out by 9.8%
[46:57] Stirrer speed set to 55
[47:02] pH 3.48 -> 3.68
[47:02] Using charge balance adjust
[47:02] Dispensed 0.002399 mL of Base (0.5 M KOH)
[47:22] Stirrer speed set to 0
[47:33] Datapoint id 45 collected
[47:33] Charge balance equation is out by -4.0%
[47:33] Stirrer speed set to 55
[47:38] pH 3.68 -> 3.88
[47:38] Using charge balance adjust
[47:38] Dispensed 0.002093 mL of Base (0.5 M KOH)
[47:58] Stirrer speed set to 0
[48:09] Datapoint id 46 collected
[48:09] Charge balance equation is out by -4.6%
[48:09] Stirrer speed set to 55
[48:14] pH 3.87 -> 4.07
[48:14] Using charge balance adjust
[48:14] Dispensed 0.002023 mL of Base (0.5 M KOH)

Sample name: **M15_octanol**
Assay name: **pH-metric high logP**
Assay ID: **18C-01004**
Filename: **C:\Sirius_T3\Mehtap\20180228_exp28_logP_T3-2\18C-01004_M15_octanol_pH-metric high logP.t3r**
Experiment start time: **3/1/2018 4:33:50 AM**
Analyst: **Pion**
Instrument ID: **T312060**

#### Experiment Log (continued)

[48:34] Stirrer speed set to 0
[48:44] Datapoint id 47 collected
[48:44] Charge balance equation is out by 0.6%
[48:44] Stirrer speed set to 55
[48:49] pH 4.07 -> 4.27
[48:49] Using charge balance adjust
[48:50] Dispensed 0.002140 mL of Base (0.5 M KOH)
[49:10] Stirrer speed set to 0
[49:20] Datapoint id 48 collected
[49:20] Charge balance equation is out by 22.9%
[49:20] Stirrer speed set to 55
[49:25] pH 4.32 -> 4.52
[49:25] Using cautious pH adjust
[49:25] Dispensed 0.001176 mL of Base (0.5 M KOH)
[49:31] Stepping pH = 4.44
[49:31] Dispensed 0.000635 mL of Base (0.5 M KOH)
[49:36] Stepping pH = 4.51
[49:36] Dispensed 0.000141 mL of Base (0.5 M KOH)
[49:41] Stepping pH = 4.52
[49:56] Stirrer speed set to 0
[50:06] Datapoint id 49 collected
[50:06] Charge balance equation is out by 16.8%
[50:06] Stirrer speed set to 55
[50:11] pH 4.51 -> 4.71
[50:11] Using cautious pH adjust
[50:11] Dispensed 0.001176 mL of Base (0.5 M KOH)
[50:16] Stepping pH = 4.61
[50:16] Dispensed 0.000800 mL of Base (0.5 M KOH)
[50:22] Stepping pH = 4.68
[50:22] Dispensed 0.000212 mL of Base (0.5 M KOH)
[50:27] Stepping pH = 4.70
[50:27] Dispensed 0.000141 mL of Base (0.5 M KOH)
[50:32] Stepping pH = 4.70
[50:47] Stirrer speed set to 0
[50:57] Datapoint id 50 collected
[50:57] Charge balance equation is out by 1.4%
[50:57] Stirrer speed set to 55
[51:02] pH 4.69 -> 4.89
[51:02] Using charge balance adjust
[51:02] Dispensed 0.002258 mL of Base (0.5 M KOH)
[51:23] Stirrer speed set to 0
[51:33] Datapoint id 51 collected
[51:33] Charge balance equation is out by 4.7%
[51:33] Stirrer speed set to 55
[51:38] pH 4.90 -> 5.10
[51:38] Using charge balance adjust
[51:38] Dispensed 0.001952 mL of Base (0.5 M KOH)
[51:58] Stirrer speed set to 0
[52:09] Datapoint id 52 collected
[52:09] Charge balance equation is out by 16.1%
[52:09] Stirrer speed set to 55
[52:14] pH 5.14 -> 5.34
[52:14] Using cautious pH adjust
[52:14] Dispensed 0.000753 mL of Base (0.5 M KOH)
[52:19] Stepping pH = 5.25
[52:19] Dispensed 0.000447 mL of Base (0.5 M KOH)
[52:24] Stepping pH = 5.32
[52:24] Dispensed 0.000094 mL of Base (0.5 M KOH)
[52:29] Stepping pH = 5.33

| | | | |
|---|---|---|---|
| Sample name: | **M15_octanol** | Experiment start time: | **3/1/2018 4:33:50 AM** |
| Assay name: | **pH-metric high logP** | Analyst: | **Pion** |
| Assay ID: | **18C-01004** | Instrument ID: | **T312060** |
| Filename: | **C:\Sirius_T3\Mehtap\20180228_exp28_logP_T3-2\18C-01004_M15_octanol_pH-metric high logP.t3r** | | |

#### Experiment Log (continued)

[52:44] Stirrer speed set to 0
[52:55] Datapoint id 53 collected
[52:55] Charge balance equation is out by 13.9%
[52:55] Stirrer speed set to 55
[53:00] pH 5.33 -> 5.53
[53:00] Using charge balance adjust
[53:00] Dispensed 0.001129 mL of Base (0.5 M KOH)
[53:20] Stirrer speed set to 0
[53:30] Datapoint id 54 collected
[53:30] Charge balance equation is out by 23.2%
[53:30] Stirrer speed set to 55
[53:35] pH 5.58 -> 5.78
[53:35] Using cautious pH adjust
[53:35] Dispensed 0.000376 mL of Base (0.5 M KOH)
[53:41] Stepping pH = 5.67
[53:41] Dispensed 0.000282 mL of Base (0.5 M KOH)
[53:46] Stepping pH = 5.77
[54:01] Stirrer speed set to 0
[54:11] Datapoint id 55 collected
[54:11] Charge balance equation is out by 11.5%
[54:11] Stirrer speed set to 55
[54:16] pH 5.77 -> 5.97
[54:16] Using charge balance adjust
[54:16] Dispensed 0.000517 mL of Base (0.5 M KOH)
[54:37] Stirrer speed set to 0
[54:49] Datapoint id 56 collected
[54:49] Charge balance equation is out by 10.2%
[54:49] Stirrer speed set to 55
[54:54] pH 6.00 -> 6.20
[54:54] Using charge balance adjust
[54:54] Dispensed 0.000353 mL of Base (0.5 M KOH)
[55:15] Stirrer speed set to 0
[55:30] Datapoint id 57 collected
[55:30] Charge balance equation is out by -5.4%
[55:30] Stirrer speed set to 55
[55:35] pH 6.21 -> 6.41
[55:35] Using charge balance adjust
[55:35] Dispensed 0.000259 mL of Base (0.5 M KOH)
[55:55] Stirrer speed set to 0
[56:27] Datapoint id 58 collected
[56:27] Charge balance equation is out by 31.8%
[56:27] Stirrer speed set to 55
[56:32] pH 6.50 -> 6.70
[56:32] Using cautious pH adjust
[56:32] Dispensed 0.000071 mL of Base (0.5 M KOH)
[56:37] Stepping pH = 6.51
[56:37] Dispensed 0.000212 mL of Base (0.5 M KOH)
[56:42] Stepping pH = 6.96
[56:58] Stirrer speed set to 0
[57:58] Datapoint id 59 collected
[57:58] Charge balance equation is out by -84.9%
[57:58] Stirrer speed set to 55
[58:03] pH 6.98 -> 7.18
[58:03] Using cautious pH adjust
[58:03] Dispensed 0.000024 mL of Base (0.5 M KOH)
[58:08] Stepping pH = 7.01
[58:08] Dispensed 0.000071 mL of Base (0.5 M KOH)
[58:13] Stepping pH = 7.02
[58:13] Dispensed 0.000165 mL of Base (0.5 M KOH)

| | | | |
|---|---|---|---|
| Sample name: | **M15_octanol** | Experiment start time: | **3/1/2018 4:33:50 AM** |
| Assay name: | **pH-metric high logP** | Analyst: | **Pion** |
| Assay ID: | **18C-01004** | Instrument ID: | **T312060** |
| Filename: | **C:\Sirius_T3\Mehtap\20180228_exp28_logP_T3-2\18C-01004_M15_octanol_pH-metric high logP.t3r** | | |

#### Experiment Log (continued)

[58:18] Stepping pH = 7.94
[58:33] Stirrer speed set to 0
[59:33] Datapoint id 60 collected
[59:33] Charge balance equation is out by -331.1%
[59:33] Stirrer speed set to 55
[59:39] pH 8.01 -> 8.21
[59:39] Using cautious pH adjust
[59:39] Dispensed 0.000024 mL of Base (0.5 M KOH)
[59:44] Stepping pH = 8.02
[59:44] Dispensed 0.000024 mL of Base (0.5 M KOH)
[59:49] Stepping pH = 8.02
[59:49] Dispensed 0.000094 mL of Base (0.5 M KOH)
[59:54] Stepping pH = 8.25
[1:00:09] Stirrer speed set to 0
[1:01:08] Datapoint id 61 collected
[1:01:08] Charge balance equation is out by -895.9%
[1:01:08] Stirrer speed set to 55
[1:01:13] pH 8.29 -> 8.49
[1:01:13] Using cautious pH adjust
[1:01:13] Dispensed 0.000024 mL of Base (0.5 M KOH)
[1:01:18] Stepping pH = 8.30
[1:01:19] Dispensed 0.000024 mL of Base (0.5 M KOH)
[1:01:24] Stepping pH = 8.31
[1:01:24] Dispensed 0.000071 mL of Base (0.5 M KOH)
[1:01:29] Stepping pH = 8.46
[1:01:29] Dispensed 0.000024 mL of Base (0.5 M KOH)
[1:01:34] Stepping pH = 8.51
[1:01:49] Stirrer speed set to 0
[1:02:21] Datapoint id 62 collected
[1:02:21] Charge balance equation is out by -675.6%
[1:02:21] Stirrer speed set to 55
[1:02:26] pH 8.53 -> 8.73
[1:02:26] Using cautious pH adjust
[1:02:26] Dispensed 0.000024 mL of Base (0.5 M KOH)
[1:02:31] Stepping pH = 8.55
[1:02:31] Dispensed 0.000024 mL of Base (0.5 M KOH)
[1:02:36] Stepping pH = 8.59
[1:02:36] Dispensed 0.000071 mL of Base (0.5 M KOH)
[1:02:41] Stepping pH = 8.73
[1:02:56] Stirrer speed set to 0
[1:03:24] Datapoint id 63 collected
[1:03:24] Charge balance equation is out by -363.9%
[1:03:24] Stirrer speed set to 55
[1:03:29] pH 8.75 -> 8.95
[1:03:29] Using cautious pH adjust
[1:03:29] Dispensed 0.000024 mL of Base (0.5 M KOH)
[1:03:34] Stepping pH = 8.77
[1:03:35] Dispensed 0.000047 mL of Base (0.5 M KOH)
[1:03:40] Stepping pH = 8.81
[1:03:40] Dispensed 0.000094 mL of Base (0.5 M KOH)
[1:03:45] Stepping pH = 8.88
[1:03:45] Dispensed 0.000071 mL of Base (0.5 M KOH)
[1:03:50] Stepping pH = 8.94
[1:04:05] Stirrer speed set to 0
[1:04:20] Datapoint id 64 collected
[1:04:20] Charge balance equation is out by -534.8%
[1:04:20] Stirrer speed set to 55
[1:04:25] pH 8.96 -> 9.16
[1:04:25] Using cautious pH adjust

| | | | |
|---|---|---|---|
| Sample name: | **M15_octanol** | Experiment start time: | **3/1/2018 4:33:50 AM** |
| Assay name: | **pH-metric high logP** | Analyst: | **Pion** |
| Assay ID: | **18C-01004** | Instrument ID: | **T312060** |
| Filename: | **C:\Sirius_T3\Mehtap\20180228_exp28_logP_T3-2\18C-01004_M15_octanol_pH-metric high logP.t3r** | | |

#### Experiment Log (continued)

[1:04:25] Dispensed 0.000024 mL of Base (0.5 M KOH)
[1:04:30] Stepping pH = 8.97
[1:04:30] Dispensed 0.000094 mL of Base (0.5 M KOH)
[1:04:35] Stepping pH = 8.99
[1:04:35] Dispensed 0.000235 mL of Base (0.5 M KOH)
[1:04:40] Stepping pH = 9.25
[1:04:55] Stirrer speed set to 0
[1:05:18] Datapoint id 65 collected
[1:05:18] Charge balance equation is out by -466.4%
[1:05:18] Stirrer speed set to 55
[1:05:23] pH 9.28 -> 9.48
[1:05:23] Using cautious pH adjust
[1:05:23] Dispensed 0.000047 mL of Base (0.5 M KOH)
[1:05:28] Stepping pH = 9.28
[1:05:28] Dispensed 0.000165 mL of Base (0.5 M KOH)
[1:05:33] Stepping pH = 9.42
[1:05:33] Dispensed 0.000071 mL of Base (0.5 M KOH)
[1:05:38] Stepping pH = 9.47
[1:05:54] Stirrer speed set to 0
[1:06:07] Datapoint id 66 collected
[1:06:07] Charge balance equation is out by -157.2%
[1:06:07] Stirrer speed set to 55
[1:06:12] pH 9.48 -> 9.68
[1:06:12] Using cautious pH adjust
[1:06:12] Dispensed 0.000094 mL of Base (0.5 M KOH)
[1:06:17] Stepping pH = 9.51
[1:06:17] Dispensed 0.000212 mL of Base (0.5 M KOH)
[1:06:23] Stepping pH = 9.69
[1:06:38] Stirrer speed set to 0
[1:06:49] Datapoint id 67 collected
[1:06:49] Charge balance equation is out by -70.6%
[1:06:49] Stirrer speed set to 55
[1:06:54] pH 9.71 -> 9.91
[1:06:54] Using cautious pH adjust
[1:06:54] Dispensed 0.000141 mL of Base (0.5 M KOH)
[1:06:59] Stepping pH = 9.75
[1:06:59] Dispensed 0.000259 mL of Base (0.5 M KOH)
[1:07:04] Stepping pH = 9.91
[1:07:19] Stirrer speed set to 0
[1:07:30] Datapoint id 68 collected
[1:07:30] Charge balance equation is out by -45.5%
[1:07:30] Stirrer speed set to 55
[1:07:35] pH 9.92 -> 10.05
[1:07:35] Using cautious pH adjust
[1:07:35] Dispensed 0.000118 mL of Base (0.5 M KOH)
[1:07:40] Stepping pH = 9.94
[1:07:40] Dispensed 0.000306 mL of Base (0.5 M KOH)
[1:07:45] Stepping pH = 10.09
[1:08:00] Stirrer speed set to 0
[1:08:14] Datapoint id 69 collected
[1:08:14] Charge balance equation is out by -72.3%
[1:08:14] Titration 3 of 3
[1:08:14] Adding initial titrants
[1:08:14] Automatically add 0.30000 mL of Octanol
[1:08:21] Dispensed 0.300000 mL of Octanol
[1:08:21] Stirrer speed set to 10
[1:08:22] Stirrer speed set to 60
[1:08:22] Iterative adjust 10.10 -> 2.00
[1:08:22] pH 10.10 -> 2.00

Sample name: **M15_octanol** Experiment start time: **3/1/2018 4:33:50 AM**
Assay name: **pH-metric high logP** Analyst: **Pion**
Assay ID: **18C-01004** Instrument ID: **T312060**
Filename: **C:\Sirius_T3\Mehtap\20180228_exp28_logP_T3-2\18C-01004_M15_octanol_pH-metric high logP.t3r**

#### Experiment Log (continued)

[1:08:24] Dispensed 0.100000 mL of Acid (0.5 M HCl)
[1:08:30] pH 2.04 -> 2.00
[1:08:30] Dispensed 0.006326 mL of Acid (0.5 M HCl)
[1:09:20] Stirrer speed set to 0
[1:09:30] Datapoint id 70 collected
[1:09:30] Stirrer speed set to 60
[1:09:35] pH 1.99 -> 2.19
[1:09:35] Using cautious pH adjust
[1:09:36] Dispensed 0.015052 mL of Base (0.5 M KOH)
[1:09:41] Stepping pH = 2.08
[1:09:41] Dispensed 0.010818 mL of Base (0.5 M KOH)
[1:09:46] Stepping pH = 2.16
[1:09:46] Dispensed 0.002375 mL of Base (0.5 M KOH)
[1:09:52] Stepping pH = 2.19
[1:10:07] Stirrer speed set to 0
[1:10:17] Datapoint id 71 collected
[1:10:17] Charge balance equation is out by 6.2%
[1:10:17] Stirrer speed set to 60
[1:10:22] pH 2.19 -> 2.39
[1:10:22] Using charge balance adjust
[1:10:22] Dispensed 0.019450 mL of Base (0.5 M KOH)
[1:10:43] Stirrer speed set to 0
[1:10:53] Datapoint id 72 collected
[1:10:53] Charge balance equation is out by 9.2%
[1:10:53] Stirrer speed set to 60
[1:10:58] pH 2.41 -> 2.61
[1:10:58] Using charge balance adjust
[1:10:59] Dispensed 0.012794 mL of Base (0.5 M KOH)
[1:11:19] Stirrer speed set to 0
[1:11:29] Datapoint id 73 collected
[1:11:29] Charge balance equation is out by 13.1%
[1:11:29] Stirrer speed set to 60
[1:11:34] pH 2.64 -> 2.84
[1:11:34] Using charge balance adjust
[1:11:34] Dispensed 0.008655 mL of Base (0.5 M KOH)
[1:11:54] Stirrer speed set to 0
[1:12:05] Datapoint id 74 collected
[1:12:05] Charge balance equation is out by 9.7%
[1:12:05] Stirrer speed set to 60
[1:12:10] pH 2.87 -> 3.07
[1:12:10] Using charge balance adjust
[1:12:10] Dispensed 0.006138 mL of Base (0.5 M KOH)
[1:12:30] Stirrer speed set to 0
[1:12:40] Datapoint id 75 collected
[1:12:40] Charge balance equation is out by 5.4%
[1:12:40] Stirrer speed set to 60
[1:12:45] pH 3.09 -> 3.29
[1:12:45] Using charge balance adjust
[1:12:45] Dispensed 0.004563 mL of Base (0.5 M KOH)
[1:13:06] Stirrer speed set to 0
[1:13:16] Datapoint id 76 collected
[1:13:16] Charge balance equation is out by 6.3%
[1:13:16] Stirrer speed set to 60
[1:13:21] pH 3.30 -> 3.50
[1:13:21] Using charge balance adjust
[1:13:21] Dispensed 0.003575 mL of Base (0.5 M KOH)
[1:13:41] Stirrer speed set to 0
[1:13:51] Datapoint id 77 collected
[1:13:51] Charge balance equation is out by 2.0%

| | | | |
|---|---|---|---|
| Sample name: | **M15_octanol** | Experiment start time: | **3/1/2018 4:33:50 AM** |
| Assay name: | **pH-metric high logP** | Analyst: | **Pion** |
| Assay ID: | **18C-01004** | Instrument ID: | **T312060** |
| Filename: | **C:\Sirius_T3\Mehtap\20180228_exp28_logP_T3-2\18C-01004_M15_octanol_pH-metric high logP.t3r** | | |

#### Experiment Log (continued)

[1:13:51] Stirrer speed set to 60
[1:13:56] pH 3.51 -> 3.71
[1:13:56] Using charge balance adjust
[1:13:57] Dispensed 0.003057 mL of Base (0.5 M KOH)
[1:14:17] Stirrer speed set to 0
[1:14:27] Datapoint id 78 collected
[1:14:27] Charge balance equation is out by 10.2%
[1:14:27] Stirrer speed set to 60
[1:14:32] pH 3.73 -> 3.93
[1:14:32] Using charge balance adjust
[1:14:32] Dispensed 0.002846 mL of Base (0.5 M KOH)
[1:14:52] Stirrer speed set to 0
[1:15:03] Datapoint id 79 collected
[1:15:03] Charge balance equation is out by 26.2%
[1:15:03] Stirrer speed set to 60
[1:15:08] pH 3.98 -> 4.18
[1:15:08] Using cautious pH adjust
[1:15:08] Dispensed 0.001364 mL of Base (0.5 M KOH)
[1:15:13] Stepping pH = 4.08
[1:15:13] Dispensed 0.000894 mL of Base (0.5 M KOH)
[1:15:18] Stepping pH = 4.16
[1:15:18] Dispensed 0.000165 mL of Base (0.5 M KOH)
[1:15:23] Stepping pH = 4.17
[1:15:39] Stirrer speed set to 0
[1:15:49] Datapoint id 80 collected
[1:15:49] Charge balance equation is out by 10.9%
[1:15:49] Stirrer speed set to 60
[1:15:54] pH 4.16 -> 4.36
[1:15:54] Using charge balance adjust
[1:15:54] Dispensed 0.002634 mL of Base (0.5 M KOH)
[1:16:14] Stirrer speed set to 0
[1:16:24] Datapoint id 81 collected
[1:16:24] Charge balance equation is out by 15.1%
[1:16:24] Stirrer speed set to 60
[1:16:29] pH 4.38 -> 4.58
[1:16:29] Using cautious pH adjust
[1:16:29] Dispensed 0.001152 mL of Base (0.5 M KOH)
[1:16:34] Stepping pH = 4.48
[1:16:34] Dispensed 0.000753 mL of Base (0.5 M KOH)
[1:16:40] Stepping pH = 4.57
[1:16:55] Stirrer speed set to 0
[1:17:05] Datapoint id 82 collected
[1:17:05] Charge balance equation is out by 17.4%
[1:17:05] Stirrer speed set to 60
[1:17:10] pH 4.56 -> 4.76
[1:17:10] Using cautious pH adjust
[1:17:10] Dispensed 0.000964 mL of Base (0.5 M KOH)
[1:17:16] Stepping pH = 4.67
[1:17:16] Dispensed 0.000611 mL of Base (0.5 M KOH)
[1:17:21] Stepping pH = 4.74
[1:17:21] Dispensed 0.000141 mL of Base (0.5 M KOH)
[1:17:26] Stepping pH = 4.76
[1:17:41] Stirrer speed set to 0
[1:17:52] Datapoint id 83 collected
[1:17:52] Charge balance equation is out by 12.3%
[1:17:52] Stirrer speed set to 60
[1:17:57] pH 4.75 -> 4.95
[1:17:57] Using charge balance adjust
[1:17:57] Dispensed 0.001552 mL of Base (0.5 M KOH)

Sample name: **M15_octanol**
Assay name: **pH-metric high logP**
Assay ID: **18C-01004**
Filename: **C:\Sirius_T3\Mehtap\20180228_exp28_logP_T3-2\18C-01004_M15_octanol_pH-metric high logP.t3r**

Experiment start time: **3/1/2018 4:33:50 AM**
Analyst: **Pion**
Instrument ID: **T312060**

#### Experiment Log (continued)

[1:18:17] Stirrer speed set to 0
[1:18:27] Datapoint id 84 collected
[1:18:27] Charge balance equation is out by -6.1%
[1:18:27] Stirrer speed set to 60
[1:18:33] pH 4.93 -> 5.13
[1:18:33] Using charge balance adjust
[1:18:33] Dispensed 0.001152 mL of Base (0.5 M KOH)
[1:18:53] Stirrer speed set to 0
[1:19:03] Datapoint id 85 collected
[1:19:03] Charge balance equation is out by -13.9%
[1:19:03] Stirrer speed set to 60
[1:19:09] pH 5.11 -> 5.31
[1:19:09] Using charge balance adjust
[1:19:09] Dispensed 0.000847 mL of Base (0.5 M KOH)
[1:19:29] Stirrer speed set to 0
[1:19:39] Datapoint id 86 collected
[1:19:39] Charge balance equation is out by -13.8%
[1:19:39] Stirrer speed set to 60
[1:19:44] pH 5.29 -> 5.49
[1:19:44] Using charge balance adjust
[1:19:45] Dispensed 0.000611 mL of Base (0.5 M KOH)
[1:20:05] Stirrer speed set to 0
[1:20:16] Datapoint id 87 collected
[1:20:16] Charge balance equation is out by -8.9%
[1:20:16] Stirrer speed set to 60
[1:20:21] pH 5.48 -> 5.68
[1:20:21] Using charge balance adjust
[1:20:21] Dispensed 0.000423 mL of Base (0.5 M KOH)
[1:20:42] Stirrer speed set to 0
[1:20:54] Datapoint id 88 collected
[1:20:54] Charge balance equation is out by -11.2%
[1:20:54] Stirrer speed set to 60
[1:20:59] pH 5.68 -> 5.88
[1:20:59] Using charge balance adjust
[1:20:59] Dispensed 0.000306 mL of Base (0.5 M KOH)
[1:21:20] Stirrer speed set to 0
[1:21:32] Datapoint id 89 collected
[1:21:32] Charge balance equation is out by -4.3%
[1:21:32] Stirrer speed set to 60
[1:21:37] pH 5.89 -> 6.09
[1:21:37] Using charge balance adjust
[1:21:37] Dispensed 0.000212 mL of Base (0.5 M KOH)
[1:21:58] Stirrer speed set to 0
[1:22:30] Datapoint id 90 collected
[1:22:30] Charge balance equation is out by -17.7%
[1:22:30] Stirrer speed set to 60
[1:22:35] pH 6.08 -> 6.28
[1:22:35] Using cautious pH adjust
[1:22:35] Dispensed 0.000094 mL of Base (0.5 M KOH)
[1:22:40] Stepping pH = 6.15
[1:22:40] Dispensed 0.000094 mL of Base (0.5 M KOH)
[1:22:46] Stepping pH = 6.33
[1:23:01] Stirrer speed set to 0
[1:23:41] Datapoint id 91 collected
[1:23:41] Charge balance equation is out by -5.1%
[1:23:41] Stirrer speed set to 60
[1:23:46] pH 6.35 -> 6.55
[1:23:46] Using charge balance adjust
[1:23:46] Dispensed 0.000118 mL of Base (0.5 M KOH)

| | | | |
|---|---|---|---|
| Sample name: | **M15_octanol** | Experiment start time: | **3/1/2018 4:33:50 AM** |
| Assay name: | **pH-metric high logP** | Analyst: | **Pion** |
| Assay ID: | **18C-01004** | Instrument ID: | **T312060** |
| Filename: | **C:\Sirius_T3\Mehtap\20180228_exp28_logP_T3-2\18C-01004_M15_octanol_pH-metric high logP.t3r** | | |

#### Experiment Log (continued)

[1:24:06] Stirrer speed set to 0
[1:24:57] Datapoint id 92 collected
[1:24:57] Charge balance equation is out by -13.5%
[1:24:57] Stirrer speed set to 60
[1:25:02] pH 6.61 -> 6.81
[1:25:02] Using charge balance adjust
[1:25:02] Dispensed 0.000094 mL of Base (0.5 M KOH)
[1:25:22] Stirrer speed set to 0
[1:26:22] Datapoint id 93 collected
[1:26:22] Charge balance equation is out by -26.4%
[1:26:22] Stirrer speed set to 60
[1:26:27] pH 6.83 -> 7.03
[1:26:27] Using cautious pH adjust
[1:26:27] Dispensed 0.000024 mL of Base (0.5 M KOH)
[1:26:32] Stepping pH = 6.90
[1:26:32] Dispensed 0.000047 mL of Base (0.5 M KOH)
[1:26:37] Stepping pH = 7.10
[1:26:53] Stirrer speed set to 0
[1:27:53] Datapoint id 94 collected
[1:27:53] Charge balance equation is out by -5.1%
[1:27:53] Stirrer speed set to 60
[1:27:58] pH 7.15 -> 7.35
[1:27:58] Using charge balance adjust
[1:27:58] Dispensed 0.000047 mL of Base (0.5 M KOH)
[1:28:18] Stirrer speed set to 0
[1:29:18] Datapoint id 95 collected
[1:29:18] Charge balance equation is out by -32.1%
[1:29:18] Stirrer speed set to 60
[1:29:23] pH 7.27 -> 7.47
[1:29:23] Using cautious pH adjust
[1:29:23] Dispensed 0.000024 mL of Base (0.5 M KOH)
[1:29:28] Stepping pH = 7.34
[1:29:28] Dispensed 0.000024 mL of Base (0.5 M KOH)
[1:29:33] Stepping pH = 7.40
[1:29:33] Dispensed 0.000024 mL of Base (0.5 M KOH)
[1:29:38] Stepping pH = 7.52
[1:29:54] Stirrer speed set to 0
[1:30:54] Datapoint id 96 collected
[1:30:54] Charge balance equation is out by -94.3%
[1:30:54] Stirrer speed set to 60
[1:30:59] pH 7.53 -> 7.73
[1:30:59] Using cautious pH adjust
[1:30:59] Dispensed 0.000024 mL of Base (0.5 M KOH)
[1:31:04] Stepping pH = 7.55
[1:31:04] Dispensed 0.000024 mL of Base (0.5 M KOH)
[1:31:09] Stepping pH = 7.63
[1:31:09] Dispensed 0.000024 mL of Base (0.5 M KOH)
[1:31:14] Stepping pH = 7.75
[1:31:29] Stirrer speed set to 0
[1:32:29] Datapoint id 97 collected
[1:32:29] Charge balance equation is out by -237.6%
[1:32:29] Stirrer speed set to 60
[1:32:34] pH 7.96 -> 8.16
[1:32:34] Using cautious pH adjust
[1:32:34] Dispensed 0.000024 mL of Base (0.5 M KOH)
[1:32:39] Stepping pH = 8.09
[1:32:39] Dispensed 0.000024 mL of Base (0.5 M KOH)
[1:32:45] Stepping pH = 8.13
[1:32:45] Dispensed 0.000024 mL of Base (0.5 M KOH)

| | | | |
|---|---|---|---|
| Sample name: | **M15_octanol** | Experiment start time: | **3/1/2018 4:33:50 AM** |
| Assay name: | **pH-metric high logP** | Analyst: | **Pion** |
| Assay ID: | **18C-01004** | Instrument ID: | **T312060** |
| Filename: | **C:\Sirius_T3\Mehtap\20180228_exp28_logP_T3-2\18C-01004_M15_octanol_pH-metric high logP.t3r** | | |

#### Experiment Log (continued)

[1:32:50] Stepping pH = 8.22
[1:33:05] Stirrer speed set to 0
[1:34:05] Datapoint id 98 collected
[1:34:05] Charge balance equation is out by -329.5%
[1:34:05] Stirrer speed set to 60
[1:34:10] pH 8.28 -> 8.48
[1:34:10] Using cautious pH adjust
[1:34:10] Dispensed 0.000024 mL of Base (0.5 M KOH)
[1:34:15] Stepping pH = 8.35
[1:34:15] Dispensed 0.000024 mL of Base (0.5 M KOH)
[1:34:20] Stepping pH = 8.41
[1:34:20] Dispensed 0.000024 mL of Base (0.5 M KOH)
[1:34:25] Stepping pH = 8.43
[1:34:25] Dispensed 0.000024 mL of Base (0.5 M KOH)
[1:34:30] Stepping pH = 8.44
[1:34:30] Dispensed 0.000024 mL of Base (0.5 M KOH)
[1:34:36] Stepping pH = 8.50
[1:34:51] Stirrer speed set to 0
[1:35:08] Datapoint id 99 collected
[1:35:08] Charge balance equation is out by -520.9%
[1:35:08] Stirrer speed set to 60
[1:35:13] pH 8.57 -> 8.77
[1:35:13] Using cautious pH adjust
[1:35:13] Dispensed 0.000024 mL of Base (0.5 M KOH)
[1:35:18] Stepping pH = 8.59
[1:35:18] Dispensed 0.000047 mL of Base (0.5 M KOH)
[1:35:23] Stepping pH = 8.67
[1:35:23] Dispensed 0.000047 mL of Base (0.5 M KOH)
[1:35:28] Stepping pH = 8.76
[1:35:43] Stirrer speed set to 0
[1:36:15] Datapoint id 100 collected
[1:36:15] Charge balance equation is out by -224.9%
[1:36:15] Stirrer speed set to 60
[1:36:20] pH 8.79 -> 8.99
[1:36:20] Using cautious pH adjust
[1:36:20] Dispensed 0.000024 mL of Base (0.5 M KOH)
[1:36:25] Stepping pH = 8.82
[1:36:25] Dispensed 0.000071 mL of Base (0.5 M KOH)
[1:36:30] Stepping pH = 8.90
[1:36:30] Dispensed 0.000047 mL of Base (0.5 M KOH)
[1:36:35] Stepping pH = 8.97
[1:36:35] Dispensed 0.000024 mL of Base (0.5 M KOH)
[1:36:41] Stepping pH = 8.96
[1:36:41] Dispensed 0.000094 mL of Base (0.5 M KOH)
[1:36:46] Stepping pH = 9.08
[1:37:01] Stirrer speed set to 0
[1:37:14] Datapoint id 101 collected
[1:37:14] Charge balance equation is out by -394.5%
[1:37:14] Stirrer speed set to 60
[1:37:19] pH 9.11 -> 9.31
[1:37:19] Using cautious pH adjust
[1:37:20] Dispensed 0.000047 mL of Base (0.5 M KOH)
[1:37:25] Stepping pH = 9.14
[1:37:25] Dispensed 0.000118 mL of Base (0.5 M KOH)
[1:37:30] Stepping pH = 9.26
[1:37:30] Dispensed 0.000047 mL of Base (0.5 M KOH)
[1:37:35] Stepping pH = 9.30
[1:37:35] Dispensed 0.000024 mL of Base (0.5 M KOH)
[1:37:40] Stepping pH = 9.31

| | | | |
|---|---|---|---|
| Sample name: | **M15_octanol** | Experiment start time: | **3/1/2018 4:33:50 AM** |
| Assay name: | **pH-metric high logP** | Analyst: | **Pion** |
| Assay ID: | **18C-01004** | Instrument ID: | **T312060** |
| Filename: | **C:\Sirius_T3\Mehtap\20180228_exp28_logP_T3-2\18C-01004_M15_octanol_pH-metric high logP.t3r** | | |

#### Experiment Log (continued)

[1:37:55] Stirrer speed set to 0
[1:38:13] Datapoint id 102 collected
[1:38:13] Charge balance equation is out by -152.8%
[1:38:13] Stirrer speed set to 60
[1:38:18] pH 9.33 -> 9.53
[1:38:18] Using cautious pH adjust
[1:38:18] Dispensed 0.000071 mL of Base (0.5 M KOH)
[1:38:23] Stepping pH = 9.37
[1:38:24] Dispensed 0.000141 mL of Base (0.5 M KOH)
[1:38:29] Stepping pH = 9.50
[1:38:29] Dispensed 0.000047 mL of Base (0.5 M KOH)
[1:38:34] Stepping pH = 9.52
[1:38:49] Stirrer speed set to 0
[1:39:00] Datapoint id 103 collected
[1:39:00] Charge balance equation is out by -71.5%
[1:39:00] Stirrer speed set to 60
[1:39:05] pH 9.54 -> 9.74
[1:39:05] Using cautious pH adjust
[1:39:05] Dispensed 0.000094 mL of Base (0.5 M KOH)
[1:39:10] Stepping pH = 9.58
[1:39:10] Dispensed 0.000188 mL of Base (0.5 M KOH)
[1:39:15] Stepping pH = 9.71
[1:39:15] Dispensed 0.000047 mL of Base (0.5 M KOH)
[1:39:20] Stepping pH = 9.73
[1:39:36] Stirrer speed set to 0
[1:39:47] Datapoint id 104 collected
[1:39:47] Charge balance equation is out by -52.8%
[1:39:47] Stirrer speed set to 60
[1:39:52] pH 9.74 -> 9.94
[1:39:52] Using cautious pH adjust
[1:39:52] Dispensed 0.000165 mL of Base (0.5 M KOH)
[1:39:57] Stepping pH = 9.82
[1:39:57] Dispensed 0.000165 mL of Base (0.5 M KOH)
[1:40:02] Stepping pH = 9.91
[1:40:02] Dispensed 0.000071 mL of Base (0.5 M KOH)
[1:40:07] Stepping pH = 9.93
[1:40:07] Dispensed 0.000024 mL of Base (0.5 M KOH)
[1:40:12] Stepping pH = 9.94
[1:40:27] Stirrer speed set to 0
[1:40:38] Datapoint id 105 collected
[1:40:38] Charge balance equation is out by -33.7%
[1:40:38] Stirrer speed set to 60
[1:40:43] pH 9.94 -> 10.05
[1:40:43] Using cautious pH adjust
[1:40:43] Dispensed 0.000118 mL of Base (0.5 M KOH)
[1:40:48] Stepping pH = 9.97
[1:40:48] Dispensed 0.000141 mL of Base (0.5 M KOH)
[1:40:53] Stepping pH = 10.04
[1:41:08] Stirrer speed set to 0
[1:41:18] Datapoint id 106 collected
[1:41:18] Charge balance equation is out by -10.7%
[1:41:18] Argon flow rate set to 0
[1:41:22] Titrator arm moved over Titration position
