## Supplementary material for "Octanol-water partition coefficient measurements for the SAMPL6 Blind Prediction Challenge": SM14_18C-01005_M15_octanol_pH-metric high logP_report.pdf

| | | | |
|---|---|---|---|
| Sample name: | **M15_octanol** | Experiment start time: | **3/1/2018 6:16:08 AM** |
| Assay name: | **pH-metric high logP** | Analyst: | **Pion** |
| Assay ID: | **18C-01005** | Instrument ID: | **T312060** |
| Filename: | **C:\Sirius_T3\Mehtap\20180228_exp28_logP_T3-2\18C-01005_M15_octanol_pH-metric high logP.t3r** | | |

#### pH-metric Result

| | | |
|---|---|---|
| logP (XH2 2+) | -5.50 | |
| logP (XH +) | -0.05 | ±0.07 (n=50) |
| logP (neutral X) | 1.93 | ±0.01 (n=50) |

#### 18C-01005 Points 1 to 35

| | |
|---|---|
| M15_octanol concentration factor | 0.875 |
| Carbonate | 0.2232 mM |
| Acidity error | -0.23667 mM |

#### 18C-01005 Points 36 to 71

| | |
|---|---|
| M15_octanol concentration factor | 0.909 |
| Carbonate | 0.1837 mM |
| Acidity error | -0.20987 mM |

#### 18C-01005 Points 72 to 105

| | |
|---|---|
| M15_octanol concentration factor | 0.947 |
| Carbonate | 0.0524 mM |
| Acidity error | -0.05926 mM |

#### Warnings and errors

| | |
|---|---|
| Errors | None |
| Warnings | One or more logP values out of range |

#### Sample logD and percent species

| pH | M15_octanol logD | M15_octanol M15_octanolH2 | M15_octanol M15_octanolH | M15_octanol M15_octanol | M15_octanol M15_octanolH2* | M15_octanol M15_octanolH* | M15_octanol M15_octanol* | Comment |
|---|---|---|---|---|---|---|---|---|
| 1.000 | -1.64 | 95.26 % | 2.51 % | 0.00 % | 0.00 % | 2.22 % | 0.01 % | |
| 1.200 | -1.45 | 92.68 % | 3.86 % | 0.00 % | 0.00 % | 3.43 % | 0.03 % | Stomach pH |
| 2.000 | -0.71 | 66.33 % | 17.45 % | 0.01 % | 0.00 % | 15.47 % | 0.75 % | |
| 3.000 | -0.02 | 14.08 % | 37.04 % | 0.19 % | 0.00 % | 32.84 % | 15.86 % | |
| 4.000 | 0.68 | 0.61 % | 15.98 % | 0.80 % | 0.00 % | 14.17 % | 68.44 % | |
| 5.000 | 1.46 | 0.01 % | 2.21 % | 1.11 % | 0.00 % | 1.96 % | 94.71 % | |
| 6.000 | 1.85 | 0.00 % | 0.23 % | 1.15 % | 0.00 % | 0.20 % | 98.41 % | |
| 6.500 | 1.91 | 0.00 % | 0.07 % | 1.16 % | 0.00 % | 0.06 % | 98.71 % | |
| 7.000 | 1.92 | 0.00 % | 0.02 % | 1.16 % | 0.00 % | 0.02 % | 98.80 % | |
| 7.400 | 1.93 | 0.00 % | 0.01 % | 1.16 % | 0.00 % | 0.01 % | 98.83 % | Blood pH |
| 8.000 | 1.93 | 0.00 % | 0.00 % | 1.16 % | 0.00 % | 0.00 % | 98.84 % | |
| 9.000 | 1.93 | 0.00 % | 0.00 % | 1.16 % | 0.00 % | 0.00 % | 98.84 % | |
| 10.000 | 1.93 | 0.00 % | 0.00 % | 1.16 % | 0.00 % | 0.00 % | 98.84 % | |
| 11.000 | 1.93 | 0.00 % | 0.00 % | 1.16 % | 0.00 % | 0.00 % | 98.84 % | |
| 12.000 | 1.93 | 0.00 % | 0.00 % | 1.16 % | 0.00 % | 0.00 % | 98.84 % | |

| | | | |
|---|---|---|---|
| Sample name: | **M15_octanol** | Experiment start time: | **3/1/2018 6:16:08 AM** |
| Assay name: | **pH-metric high logP** | Analyst: | **Pion** |
| Assay ID: | **18C-01005** | Instrument ID: | **T312060** |
| Filename: | **C:\Sirius_T3\Mehtap\20180228_exp28_logP_T3-2\18C-01005_M15_octanol_pH-metric high logP.t3r** | | |

#### Graphs

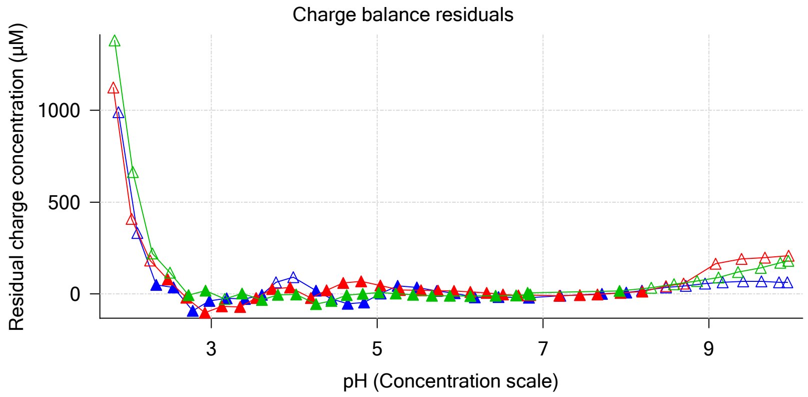


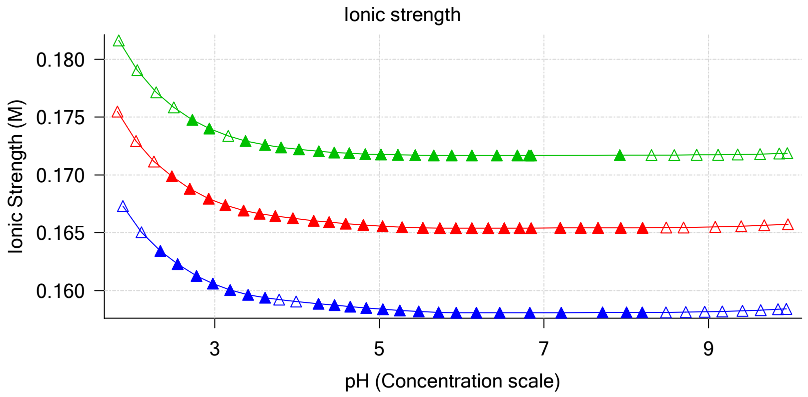


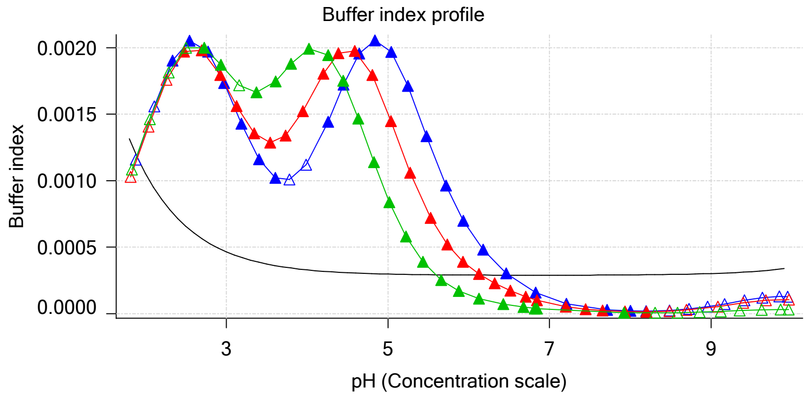


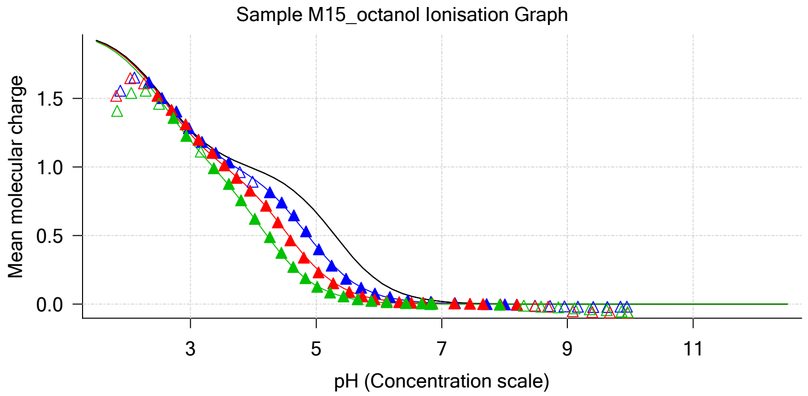


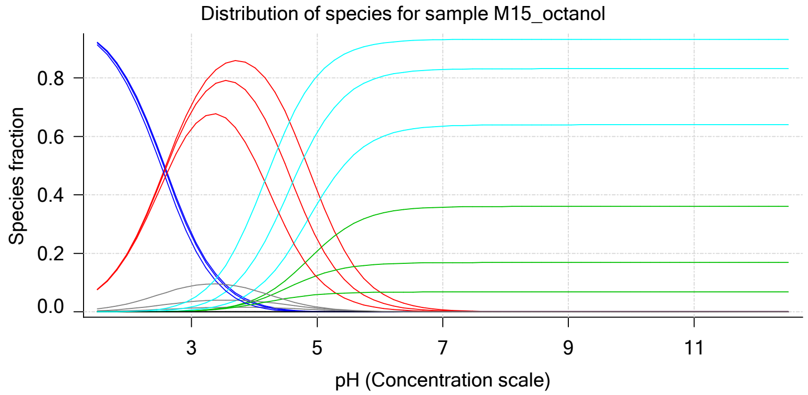


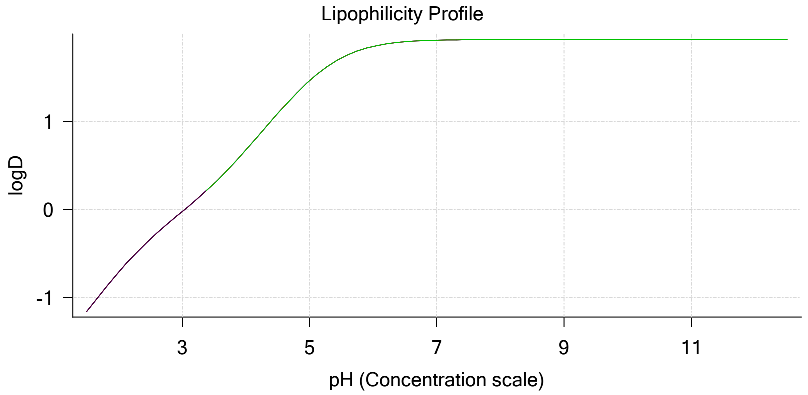


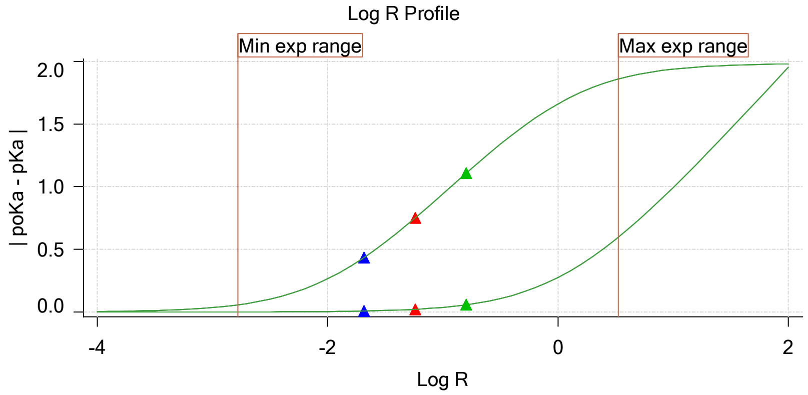


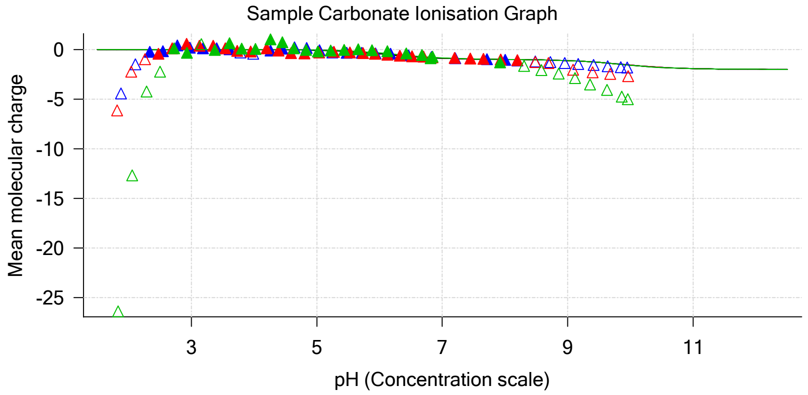

| | | | |
|---|---|---|---|
| Sample name: | **M15_octanol** | Experiment start time: | **3/1/2018 6:16:08 AM** |
| Assay name: | **pH-metric high logP** | Analyst: | **Pion** |
| Assay ID: | **18C-01005** | Instrument ID: | **T312060** |
| Filename: | **C:\Sirius_T3\Mehtap\20180228_exp28_logP_T3-2\18C-01005_M15_octanol_pH-metric high logP.t3r** | | |

#### Graphs (continued)

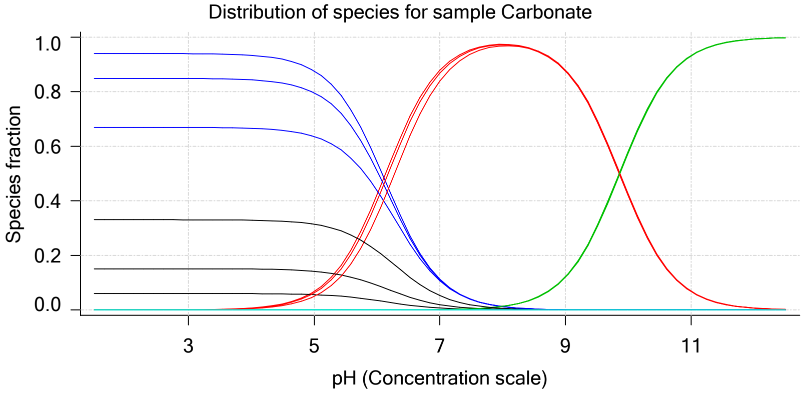

| | | | |
|---|---|---|---|
| Sample name: | **M15_octanol** | Experiment start time: | **3/1/2018 6:16:08 AM** |
| Assay name: | **pH-metric high logP** | Analyst: | **Pion** |
| Assay ID: | **18C-01005** | Instrument ID: | **T312060** |
| Filename: | **C:\Sirius_T3\Mehtap\20180228_exp28_logP_T3-2\18C-01005_M15_octanol_pH-metric high logP.t3r** | | |

### pH-metric high logP Titration 1 of 3 18C-01005 Points 1 to 35

#### Overall results

| | |
|---|---|
| RMSD | 0.176 |
| Average ionic strength | 0.159 M |
| Average temperature | 25.0°C |
| Partition ratio | 0.0208 : 1 |
| Analyte concentration range | 3855.0 μM to 4000.4 μM |
| Total points considered | 23 of 35 |

#### Warnings and errors

| | |
|---|---|
| Errors | None |
| Warnings | One or more logP values out of range |

#### Four-Plus parameters

| | | | |
|---|---|---|---|
| Alpha | 0.130 | 3/1/2018 6:16:08 AM | C:\Sirius_T3\HCl18B27.t3r |
| S | 0.9970 | 3/1/2018 6:16:08 AM | C:\Sirius_T3\HCl18B27.t3r |
| jH | 0.8 | 3/1/2018 6:16:08 AM | C:\Sirius_T3\HCl18B27.t3r |
| jOH | -0.4 | 3/1/2018 6:16:08 AM | C:\Sirius_T3\HCl18B27.t3r |

#### Titrants

| | | | |
|---|---|---|---|
| 0.50 M HCl | 0.993513 | 3/1/2018 6:16:08 AM | C:\Sirius_T3\HCl18B27.t3r |
| 0.50 M KOH | 0.999845 | 3/1/2018 6:16:08 AM | C:\Sirius_T3\KOH18B27.t3r |

#### Sample

| | |
|---|---|
| M15_octanol concentration factor | 0.875 |
| Base pKa 1 | 2.58 |
| Base pKa 2 | 5.30 |
| logP (XH2 2+) | -5.50 |
| logP (XH +) | -3.48 |
| logP (neutral X) | 1.91 |

#### Sample graphs

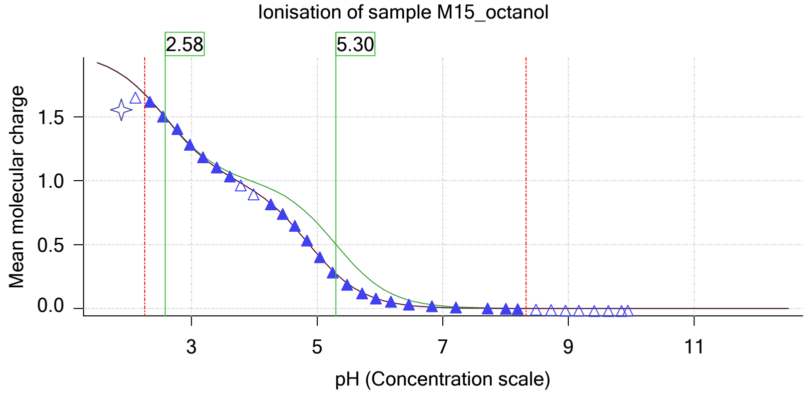


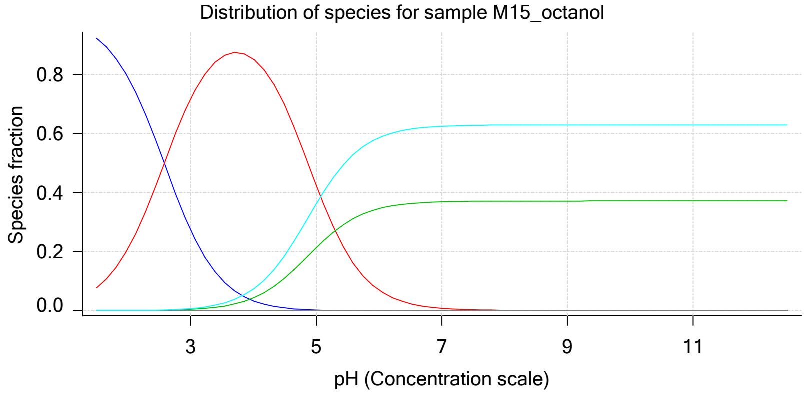

| | | | |
|---|---|---|---|
| Sample name: | **M15_octanol** | Experiment start time: | **3/1/2018 6:16:08 AM** |
| Assay name: | **pH-metric high logP** | Analyst: | **Pion** |
| Assay ID: | **18C-01005** | Instrument ID: | **T312060** |
| Filename: | **C:\Sirius_T3\Mehtap\20180228_exp28_logP_T3-2\18C-01005_M15_octanol_pH-metric high logP.t3r** | | |

#### Sample graphs (continued)

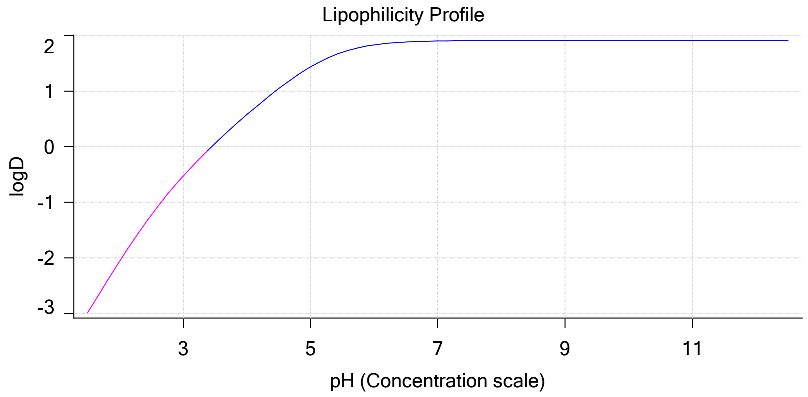


#### Sample logD and percent species

| pH | M15_octanol logD | M15_octanol M15_octanolH2 | M15_octanol M15_octanolH | M15_octanol M15_octanol | M15_octanol M15_octanolH2* | M15_octanol M15_octanolH* | M15_octanol M15_octanol* | Comment |
|---|---|---|---|---|---|---|---|---|
| 1.000 | -3.93 | 97.44 % | 2.56 % | 0.00 % | 0.00 % | 0.00 % | 0.00 % | |
| 1.200 | -3.56 | 96.00 % | 4.00 % | 0.00 % | 0.00 % | 0.00 % | 0.00 % | Stomach pH |
| 2.000 | -2.07 | 79.15 % | 20.82 % | 0.01 % | 0.00 % | 0.00 % | 0.02 % | |
| 3.000 | -0.53 | 27.28 % | 71.75 % | 0.36 % | 0.00 % | 0.00 % | 0.61 % | |
| 4.000 | 0.58 | 3.24 % | 85.24 % | 4.27 % | 0.00 % | 0.00 % | 7.24 % | |
| 5.000 | 1.43 | 0.16 % | 42.47 % | 21.28 % | 0.00 % | 0.00 % | 36.08 % | |
| 6.000 | 1.83 | 0.00 % | 6.89 % | 34.54 % | 0.00 % | 0.00 % | 58.56 % | |
| 6.500 | 1.89 | 0.00 % | 2.29 % | 36.25 % | 0.00 % | 0.00 % | 61.46 % | |
| 7.000 | 1.90 | 0.00 % | 0.73 % | 36.83 % | 0.00 % | 0.00 % | 62.44 % | |
| 7.400 | 1.91 | 0.00 % | 0.29 % | 36.99 % | 0.00 % | 0.00 % | 62.71 % | Blood pH |
| 8.000 | 1.91 | 0.00 % | 0.07 % | 37.07 % | 0.00 % | 0.00 % | 62.85 % | |
| 9.000 | 1.91 | 0.00 % | 0.01 % | 37.10 % | 0.00 % | 0.00 % | 62.89 % | |
| 10.000 | 1.91 | 0.00 % | 0.00 % | 37.10 % | 0.00 % | 0.00 % | 62.90 % | |
| 11.000 | 1.91 | 0.00 % | 0.00 % | 37.10 % | 0.00 % | 0.00 % | 62.90 % | |
| 12.000 | 1.91 | 0.00 % | 0.00 % | 37.10 % | 0.00 % | 0.00 % | 62.90 % | |

#### Carbonate and acidity

| | |
|---|---|
| Carbonate | 0.223 mM |
| Acidity error | -0.237 mM |

#### Other graphs

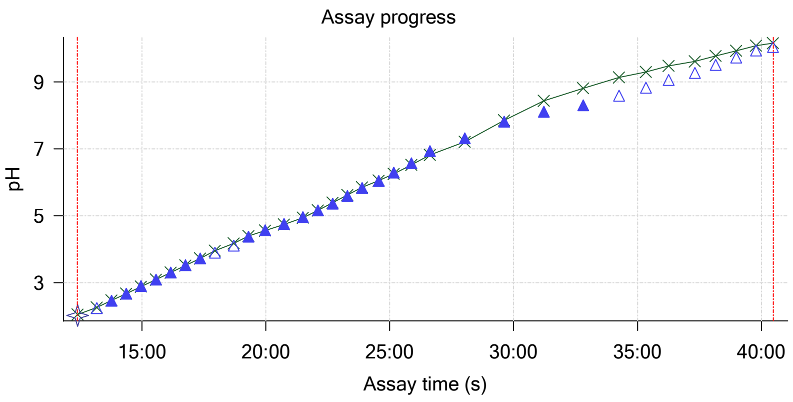


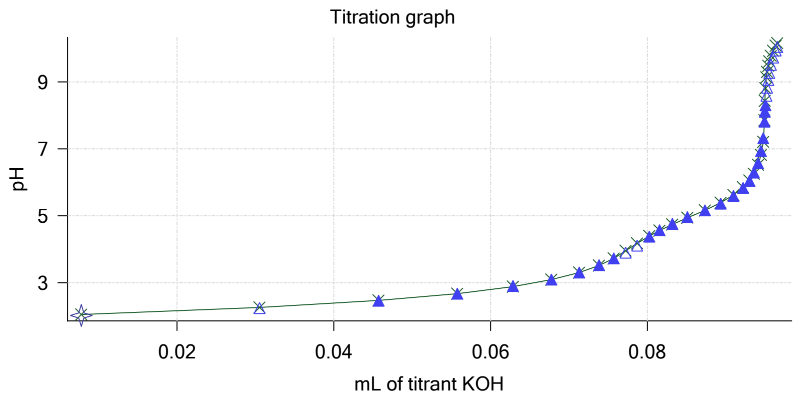

| | | | |
|---|---|---|---|
| Sample name: | **M15_octanol** | Experiment start time: | **3/1/2018 6:16:08 AM** |
| Assay name: | **pH-metric high logP** | Analyst: | **Pion** |
| Assay ID: | **18C-01005** | Instrument ID: | **T312060** |
| Filename: | **C:\Sirius_T3\Mehtap\20180228_exp28_logP_T3-2\18C-01005_M15_octanol_pH-metric high logP.t3r** | | |

#### Other graphs (continued)

Charge balance residuals

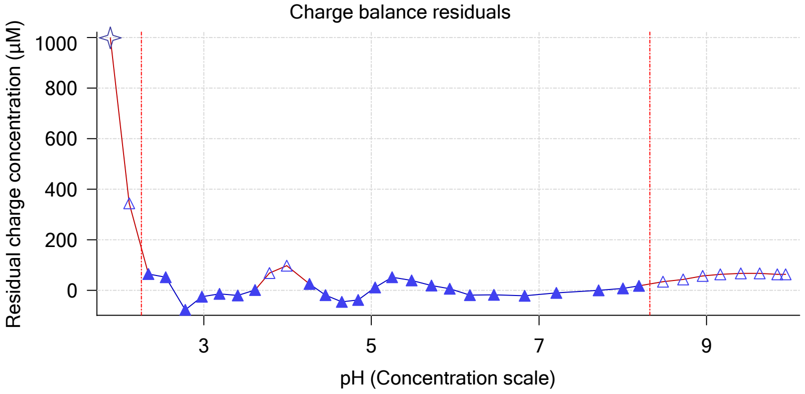


Buffer index profile

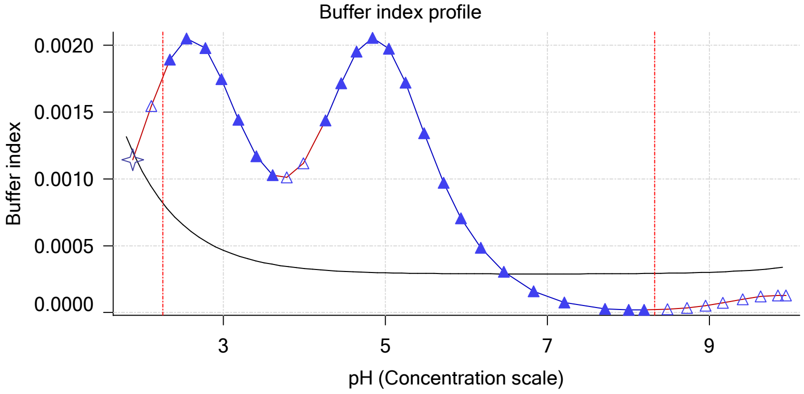


Ionic strength

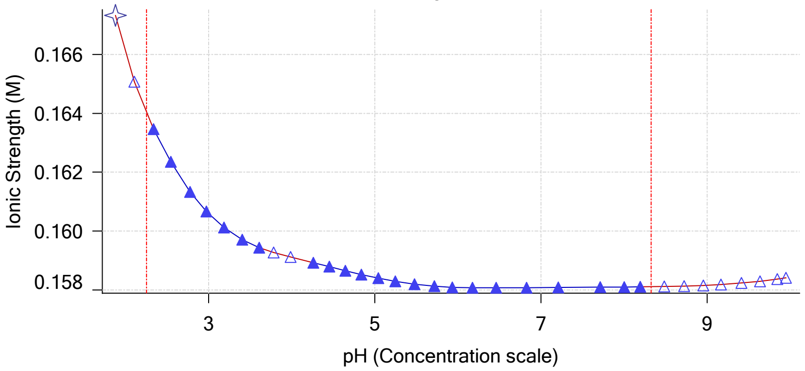


Log R Profile

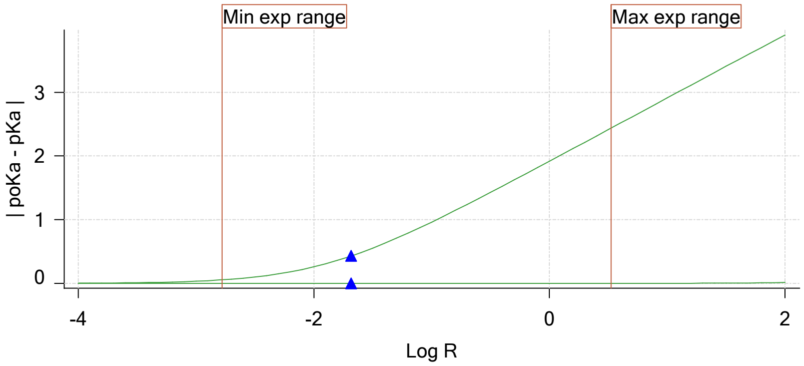


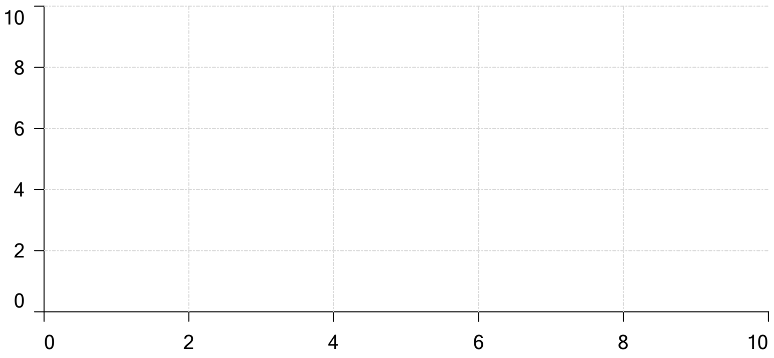


Ionisation of sample Carbonate

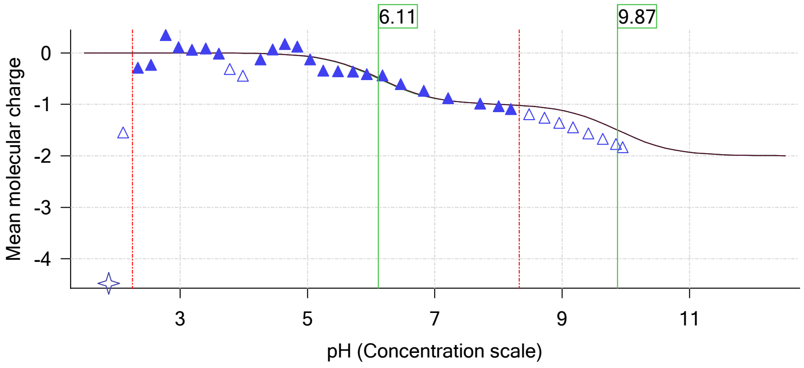

| | | | |
|---|---|---|---|
| Sample name: | **M15_octanol** | Experiment start time: | **3/1/2018 6:16:08 AM** |
| Assay name: | **pH-metric high logP** | Analyst: | **Pion** |
| Assay ID: | **18C-01005** | Instrument ID: | **T312060** |
| Filename: | **C:\Sirius_T3\Mehtap\20180228_exp28_logP_T3-2\18C-01005_M15_octanol_pH-metric high logP.t3r** | | |

### pH-metric high logP Titration 2 of 3 18C-01005 Points 36 to 71

#### Overall results

| | |
|---|---|
| RMSD | 0.148 |
| Average ionic strength | 0.166 M |
| Average temperature | 25.0°C |
| Partition ratio | 0.0576 : 1 |
| Analyte concentration range | 3420.0 µM to 3556.1 µM |
| Total points considered | 27 of 36 |

#### Warnings and errors

| | |
|---|---|
| Errors | None |
| Warnings | One or more logP values out of range |

#### Four-Plus parameters

| | | | |
|---|---|---|---|
| Alpha | 0.130 | 3/1/2018 6:16:08 AM | C:\Sirius_T3\HCl18B27.t3r |
| S | 0.9970 | 3/1/2018 6:16:08 AM | C:\Sirius_T3\HCl18B27.t3r |
| jH | 0.8 | 3/1/2018 6:16:08 AM | C:\Sirius_T3\HCl18B27.t3r |
| jOH | -0.4 | 3/1/2018 6:16:08 AM | C:\Sirius_T3\HCl18B27.t3r |

#### Titrants

| | | | |
|---|---|---|---|
| 0.50 M HCl | 0.993513 | 3/1/2018 6:16:08 AM | C:\Sirius_T3\HCl18B27.t3r |
| 0.50 M KOH | 0.999845 | 3/1/2018 6:16:08 AM | C:\Sirius_T3\KOH18B27.t3r |

#### Sample

| | |
|---|---|
| M15_octanol concentration factor | 0.909 |
| Base pKa 1 | 2.58 |
| Base pKa 2 | 5.30 |
| logP (XH2 2+) | -5.50 |
| logP (XH +) | -4.43 |
| logP (neutral X) | 1.93 |

#### Sample graphs

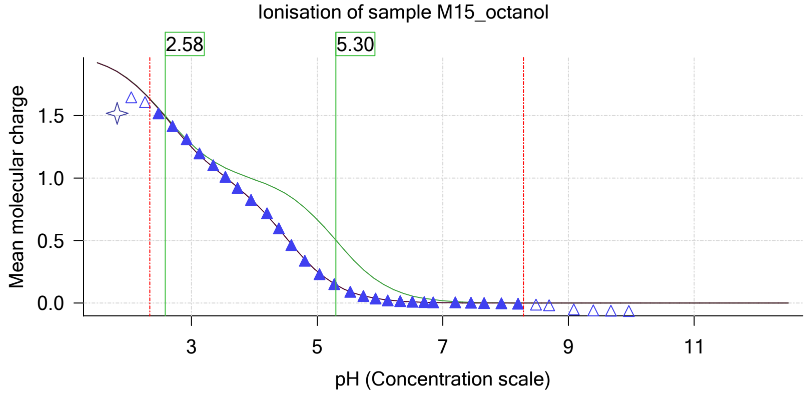


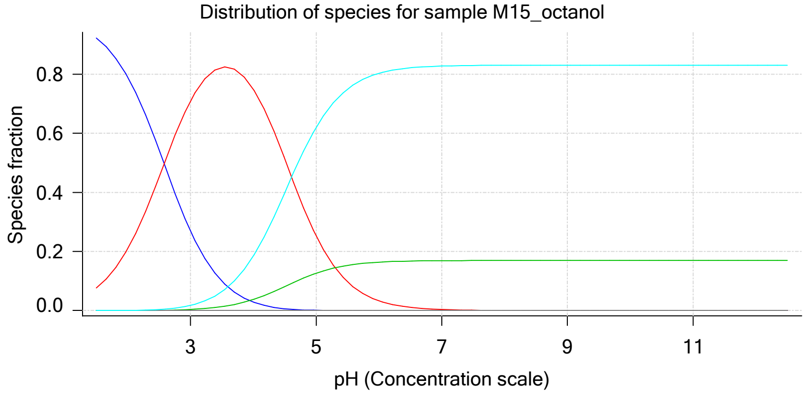

| | | | |
|---|---|---|---|
| Sample name: | **M15_octanol** | Experiment start time: | **3/1/2018 6:16:08 AM** |
| Assay name: | **pH-metric high logP** | Analyst: | **Pion** |
| Assay ID: | **18C-01005** | Instrument ID: | **T312060** |
| Filename: | **C:\Sirius_T3\Mehtap\20180228_exp28_logP_T3-2\18C-01005_M15_octanol_pH-metric high logP.t3r** | | |

#### Sample graphs (continued)

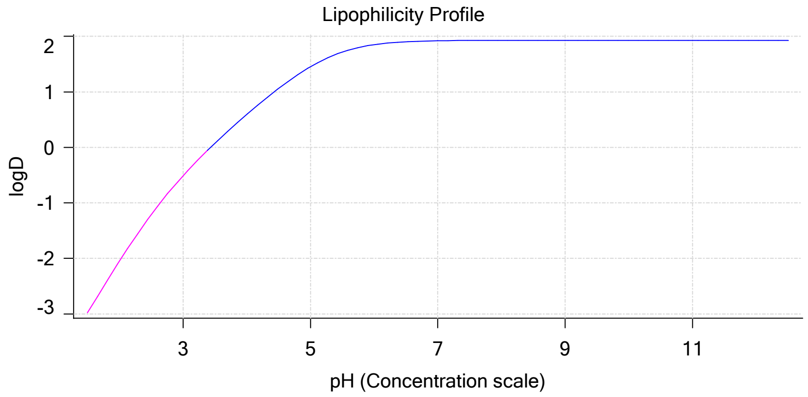


#### Sample logD and percent species

| pH | M15_octanol logD | M15_octanol M15_octanolH2 | M15_octanol M15_octanolH | M15_octanol M15_octanol | M15_octanol M15_octanolH2* | M15_octanol M15_octanolH* | M15_octanol M15_octanol* | Comment |
|---|---|---|---|---|---|---|---|---|
| 1.000 | -3.95 | 97.44 % | 2.56 % | 0.00 % | 0.00 % | 0.00 % | 0.00 % | |
| 1.200 | -3.56 | 96.00 % | 4.00 % | 0.00 % | 0.00 % | 0.00 % | 0.00 % | Stomach pH |
| 2.000 | -2.05 | 79.13 % | 20.81 % | 0.01 % | 0.00 % | 0.00 % | 0.05 % | |
| 3.000 | -0.51 | 26.97 % | 70.93 % | 0.36 % | 0.00 % | 0.00 % | 1.74 % | |
| 4.000 | 0.59 | 2.85 % | 74.97 % | 3.76 % | 0.00 % | 0.00 % | 18.42 % | |
| 5.000 | 1.45 | 0.10 % | 25.24 % | 12.65 % | 0.00 % | 0.00 % | 62.01 % | |
| 6.000 | 1.85 | 0.00 % | 3.27 % | 16.39 % | 0.00 % | 0.00 % | 80.34 % | |
| 6.500 | 1.90 | 0.00 % | 1.06 % | 16.76 % | 0.00 % | 0.00 % | 82.18 % | |
| 7.000 | 1.92 | 0.00 % | 0.34 % | 16.89 % | 0.00 % | 0.00 % | 82.78 % | |
| 7.400 | 1.93 | 0.00 % | 0.13 % | 16.92 % | 0.00 % | 0.00 % | 82.94 % | Blood pH |
| 8.000 | 1.93 | 0.00 % | 0.03 % | 16.94 % | 0.00 % | 0.00 % | 83.03 % | |
| 9.000 | 1.93 | 0.00 % | 0.00 % | 16.94 % | 0.00 % | 0.00 % | 83.05 % | |
| 10.000 | 1.93 | 0.00 % | 0.00 % | 16.94 % | 0.00 % | 0.00 % | 83.06 % | |
| 11.000 | 1.93 | 0.00 % | 0.00 % | 16.94 % | 0.00 % | 0.00 % | 83.06 % | |
| 12.000 | 1.93 | 0.00 % | 0.00 % | 16.94 % | 0.00 % | 0.00 % | 83.06 % | |

#### Carbonate and acidity

| | |
|---|---|
| Carbonate | 0.184 mM |
| Acidity error | -0.210 mM |

#### Other graphs

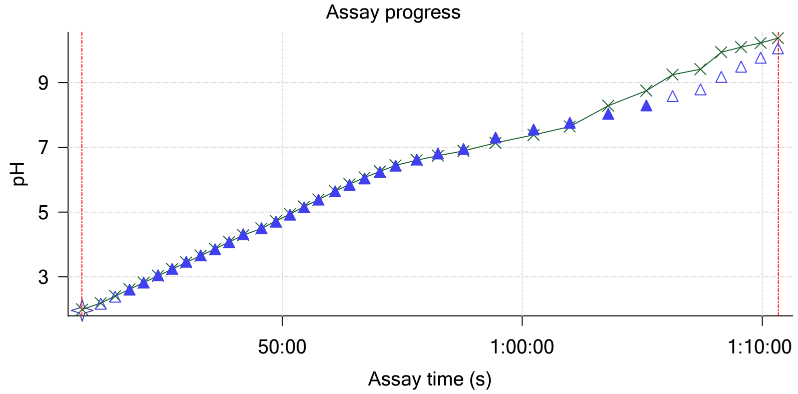


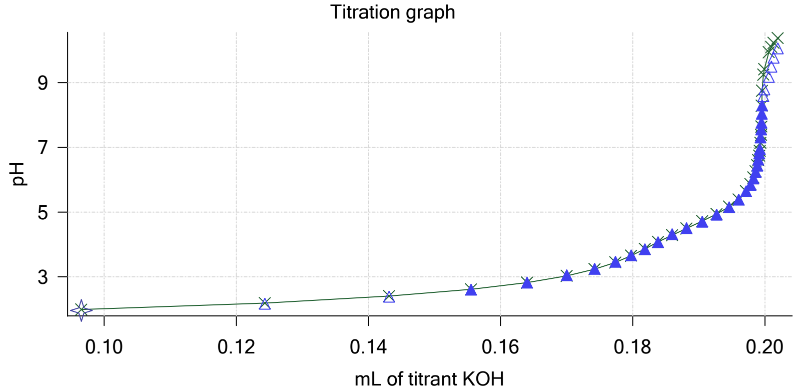

Sample name: **M15_octanol** Experiment start time: **3/1/2018 6:16:08 AM**
Assay name: **pH-metric high logP** Analyst: **Pion**
Assay ID: **18C-01005** Instrument ID: **T312060**
Filename: **C:\Sirius_T3\Mehtap\20180228_exp28_logP_T3-2\18C-01005_M15_octanol_pH-metric high logP.t3r**

#### Other graphs (continued)

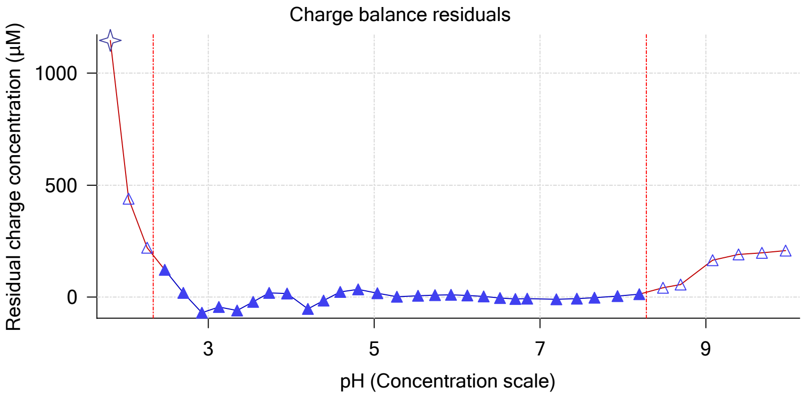


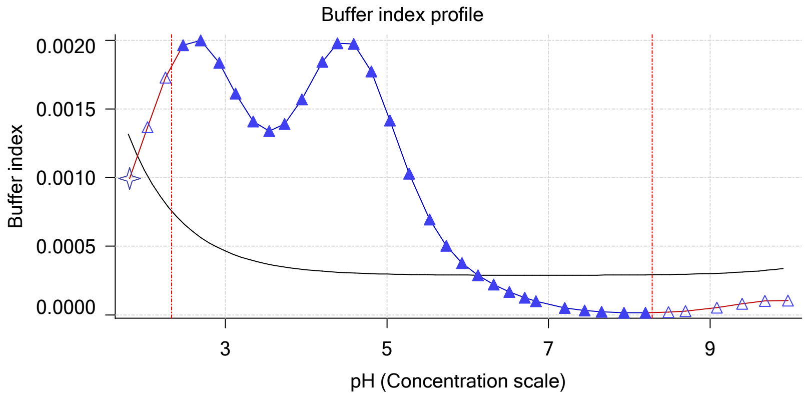


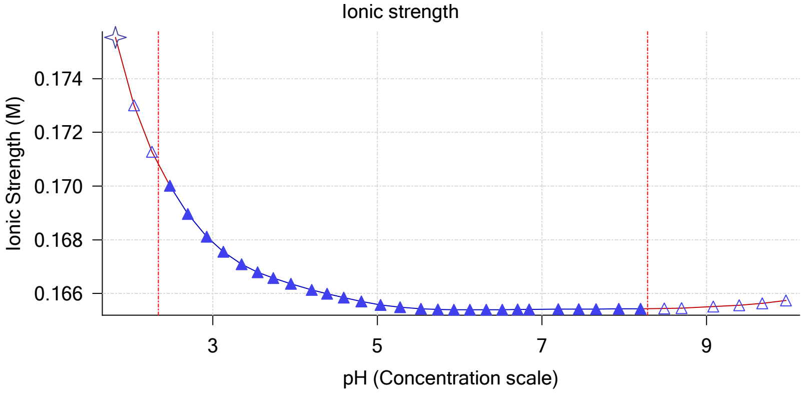


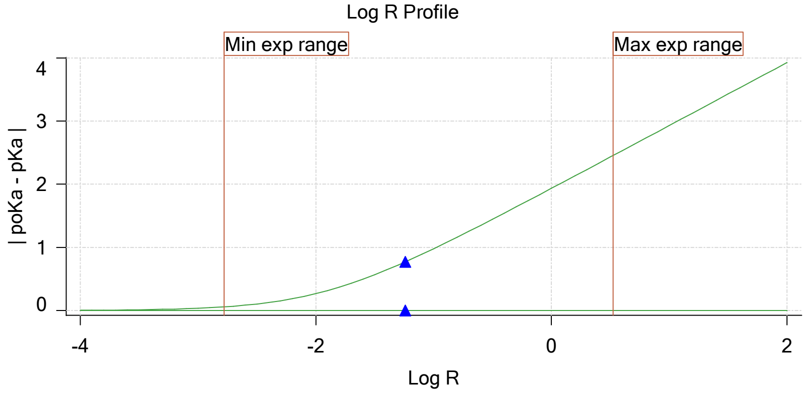


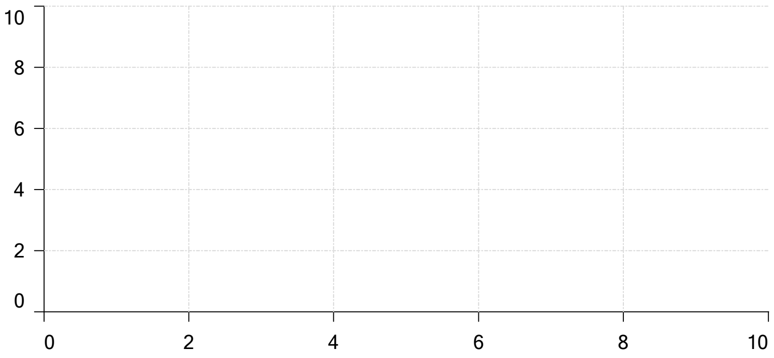


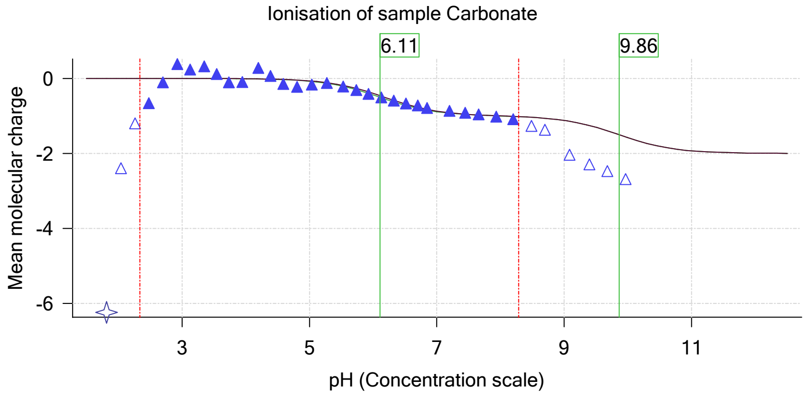

| | | | |
|---|---|---|---|
| Sample name: | **M15_octanol** | Experiment start time: | **3/1/2018 6:16:08 AM** |
| Assay name: | **pH-metric high logP** | Analyst: | **Pion** |
| Assay ID: | **18C-01005** | Instrument ID: | **T312060** |
| Filename: | **C:\Sirius_T3\Mehtap\20180228_exp28_logP_T3-2\18C-01005_M15_octanol_pH-metric high logP.t3r** | | |

### pH-metric high logP Titration 3 of 3 18C-01005 Points 72 to 105

#### Overall results

| | |
|---|---|
| RMSD | 0.401 |
| Average ionic strength | 0.172 M |
| Average temperature | 25.0°C |
| Partition ratio | 0.1600 : 1 |
| Analyte concentration range | 2881.0 µM to 2978.5 µM |
| Total points considered | 21 of 34 |

#### Warnings and errors

| | |
|---|---|
| Errors | None |
| Warnings | One or more logP values out of range |

#### Four-Plus parameters

| | | | |
|---|---|---|---|
| Alpha | 0.130 | 3/1/2018 6:16:08 AM | C:\Sirius_T3\HCl18B27.t3r |
| S | 0.9970 | 3/1/2018 6:16:08 AM | C:\Sirius_T3\HCl18B27.t3r |
| jH | 0.8 | 3/1/2018 6:16:08 AM | C:\Sirius_T3\HCl18B27.t3r |
| jOH | -0.4 | 3/1/2018 6:16:08 AM | C:\Sirius_T3\HCl18B27.t3r |

#### Titrants

| | | | |
|---|---|---|---|
| 0.50 M HCl | 0.993513 | 3/1/2018 6:16:08 AM | C:\Sirius_T3\HCl18B27.t3r |
| 0.50 M KOH | 0.999845 | 3/1/2018 6:16:08 AM | C:\Sirius_T3\KOH18B27.t3r |

#### Sample

| | |
|---|---|
| M15_octanol concentration factor | 0.947 |
| Base pKa 1 | 2.58 |
| Base pKa 2 | 5.30 |
| logP (XH2 2+) | -5.50 |
| logP (XH +) | -0.01 |
| logP (neutral X) | 1.92 |

#### Sample graphs

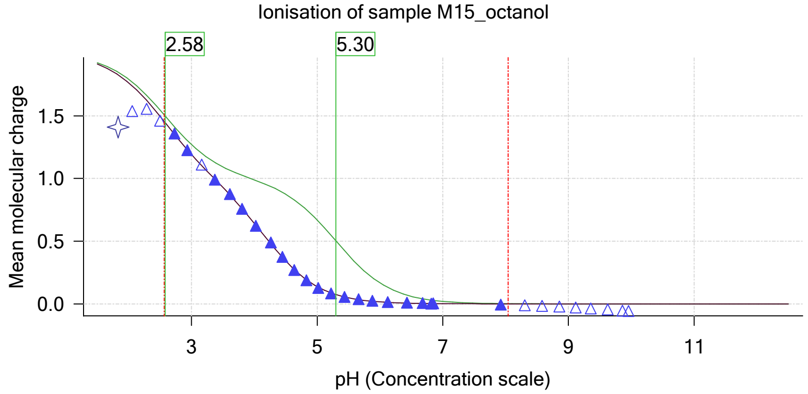


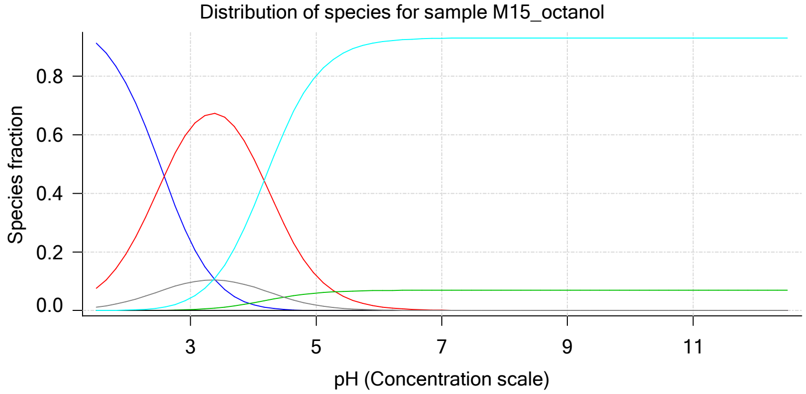

| | | | |
|---|---|---|---|
| Sample name: | **M15_octanol** | Experiment start time: | **3/1/2018 6:16:08 AM** |
| Assay name: | **pH-metric high logP** | Analyst: | **Pion** |
| Assay ID: | **18C-01005** | Instrument ID: | **T312060** |
| Filename: | **C:\Sirius_T3\Mehtap\20180228_exp28_logP_T3-2\18C-01005_M15_octanol_pH-metric high logP.t3r** | | |

#### Sample graphs (continued)

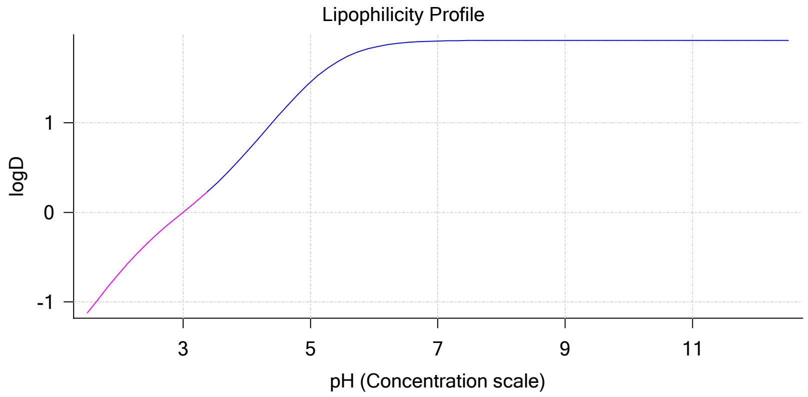


#### Sample logD and percent species

| pH | M15_octanol logD | M15_octanol M15_octanolH2 | M15_octanol M15_octanolH | M15_octanol M15_octanol | M15_octanol M15_octanolH2* | M15_octanol M15_octanolH* | M15_octanol M15_octanol* | Comment |
|---|---|---|---|---|---|---|---|---|
| 1.000 | -1.60 | 97.05 % | 2.55 % | 0.00 % | 0.00 % | 0.40 % | 0.00 % | |
| 1.200 | -1.41 | 95.40 % | 3.98 % | 0.00 % | 0.00 % | 0.62 % | 0.00 % | Stomach pH |
| 2.000 | -0.68 | 76.59 % | 20.15 % | 0.01 % | 0.00 % | 3.12 % | 0.14 % | |
| 3.000 | 0.00 | 23.66 % | 62.22 % | 0.31 % | 0.00 % | 9.63 % | 4.18 % | |
| 4.000 | 0.68 | 1.99 % | 52.22 % | 2.62 % | 0.00 % | 8.08 % | 35.10 % | |
| 5.000 | 1.46 | 0.05 % | 11.93 % | 5.98 % | 0.00 % | 1.85 % | 80.20 % | |
| 6.000 | 1.85 | 0.00 % | 1.36 % | 6.83 % | 0.00 % | 0.21 % | 91.60 % | |
| 6.500 | 1.90 | 0.00 % | 0.44 % | 6.90 % | 0.00 % | 0.07 % | 92.59 % | |
| 7.000 | 1.91 | 0.00 % | 0.14 % | 6.93 % | 0.00 % | 0.02 % | 92.91 % | |
| 7.400 | 1.92 | 0.00 % | 0.06 % | 6.93 % | 0.00 % | 0.01 % | 93.00 % | Blood pH |
| 8.000 | 1.92 | 0.00 % | 0.01 % | 6.94 % | 0.00 % | 0.00 % | 93.05 % | |
| 9.000 | 1.92 | 0.00 % | 0.00 % | 6.94 % | 0.00 % | 0.00 % | 93.06 % | |
| 10.000 | 1.92 | 0.00 % | 0.00 % | 6.94 % | 0.00 % | 0.00 % | 93.06 % | |
| 11.000 | 1.92 | 0.00 % | 0.00 % | 6.94 % | 0.00 % | 0.00 % | 93.06 % | |
| 12.000 | 1.92 | 0.00 % | 0.00 % | 6.94 % | 0.00 % | 0.00 % | 93.06 % | |

#### Carbonate and acidity

| | |
|---|---|
| Carbonate | 0.052 mM |
| Acidity error | -0.059 mM |

#### Other graphs

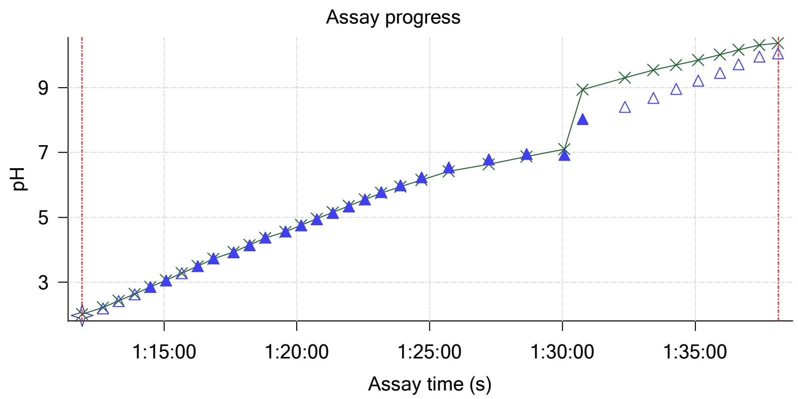


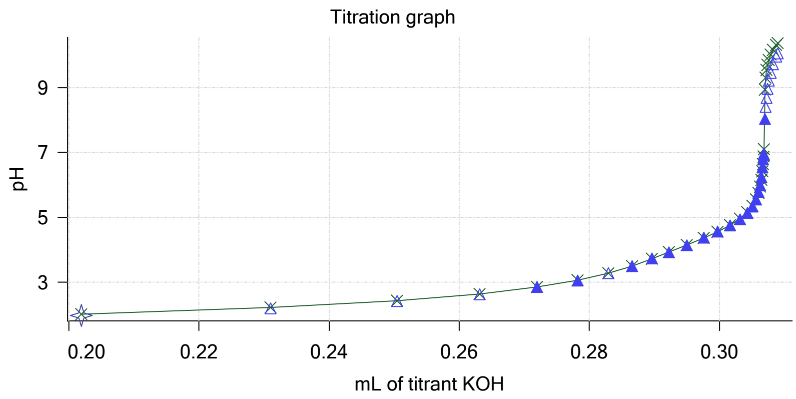

| | | | |
|---|---|---|---|
| Sample name: | **M15_octanol** | Experiment start time: | **3/1/2018 6:16:08 AM** |
| Assay name: | **pH-metric high logP** | Analyst: | **Pion** |
| Assay ID: | **18C-01005** | Instrument ID: | **T312060** |
| Filename: | **C:\Sirius_T3\Mehtap\20180228_exp28_logP_T3-2\18C-01005_M15_octanol_pH-metric high logP.t3r** | | |

#### Other graphs (continued)

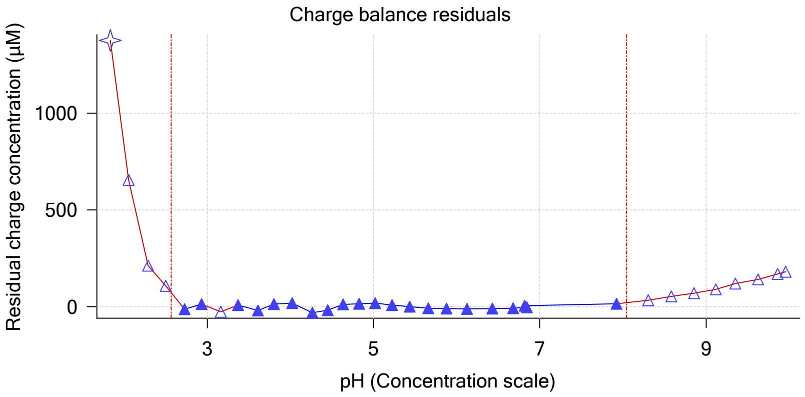


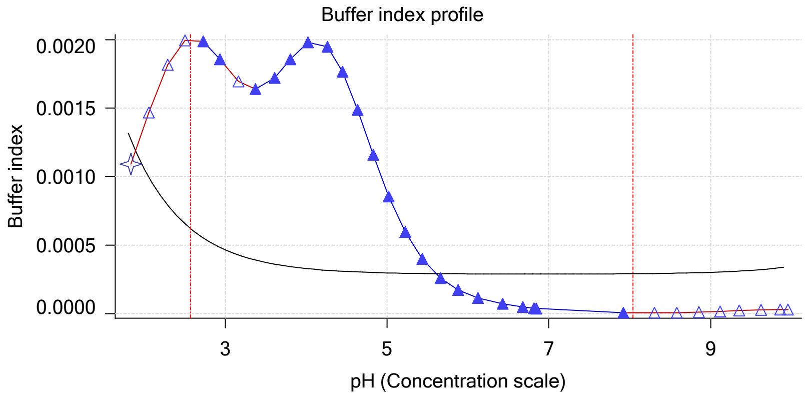


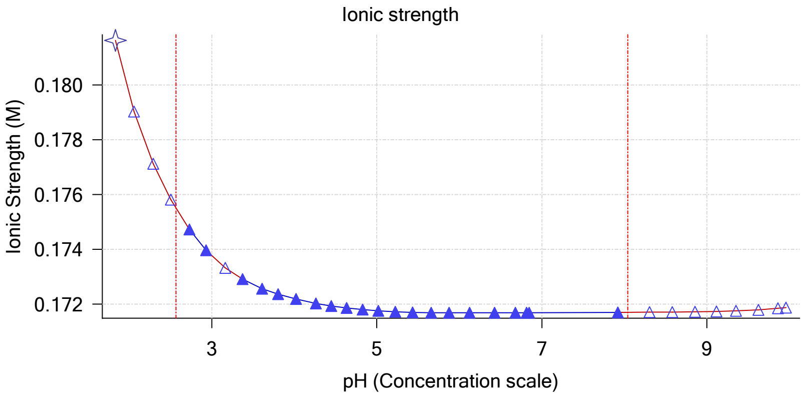


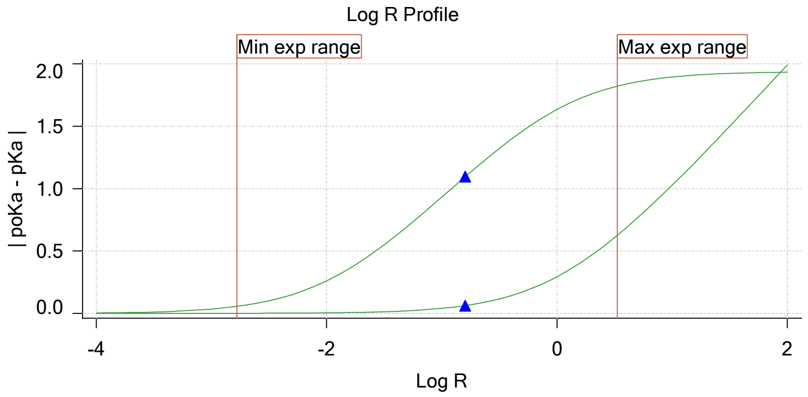


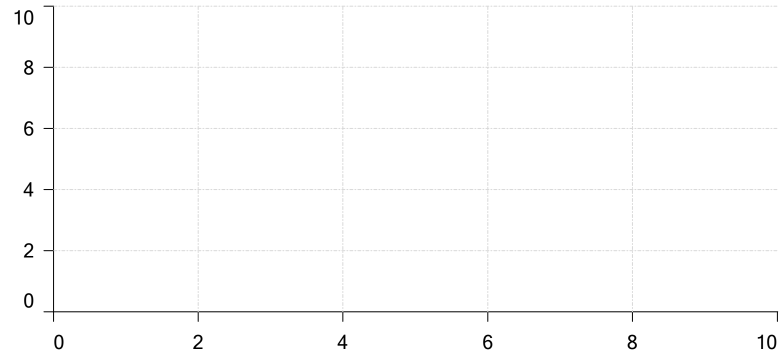


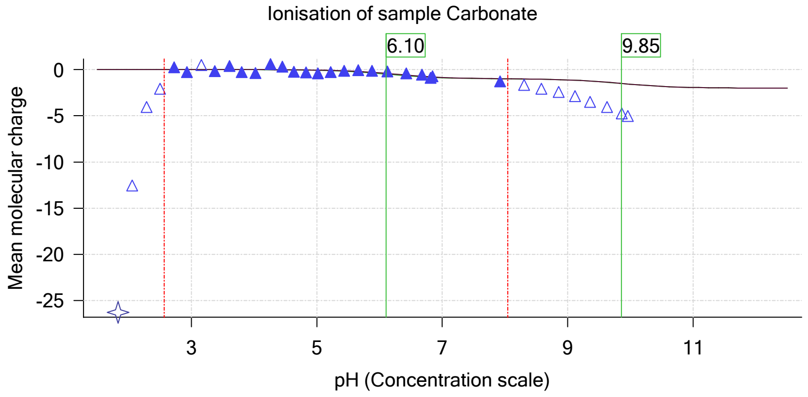

Sample name: **M15_octanol**
Assay name: **pH-metric high logP**
Assay ID: **18C-01005**
Filename: **C:\Sirius_T3\Mehtap\20180228_exp28_logP_T3-2\18C-01005_M15_octanol_pH-metric high logP.t3r**

Experiment start time: **3/1/2018 6:16:08 AM**
Analyst: **Pion**
Instrument ID: **T312060**

#### Assay Model

| Settings | Value | Date/Time changed | Imported from |
|---|---|---|---|
| Sample name | M15_octanol | 2/27/2018 5:03:03 PM | User entered value |
| Sample by | Weight | | Default value |
| Sample weight | 0.001970 g | 2/28/2018 4:25:24 PM | User entered value |
| Formula weight | 209.25 g/mol | 2/27/2018 5:03:03 PM | User entered value |
| Solubility | Unknown | | Default value |
| Molecular weight | 209.25 | 2/27/2018 5:03:03 PM | User entered value |
| Individual pKa ionic environments | No | | Default value |
| Number of pKas | 2 | 2/27/2018 5:03:03 PM | User entered value |
| Sample is a | Base | 2/27/2018 5:03:03 PM | User entered value |
| pKa 1 | 2.58 | 2/27/2018 5:03:03 PM | User entered value |
| pKa 2 | 5.30 | 2/27/2018 5:03:03 PM | User entered value |
| logP (XH2 2+) | -5.50 | 2/28/2018 2:10:35 PM | User entered value |
| logp (XH +) | -4.96 | 2/28/2018 2:10:28 PM | User entered value |
| logP (neutral X) | 1.92 | 2/28/2018 2:10:15 PM | User entered value |

#### Events

| Time | Event | Water | Acid | Base | Octanol | pH | dpH/dt | pH R-squared | pH SD | dpH/dt time |
|---|---|---|---|---|---|---|---|---|---|---|
| 9:23.6 | Initial pH = 7.47 | | | | | | | | | |
| 12:24.5 | Data point 1 | 2.20002 mL | 0.09563 mL | 0.00778 mL | 0.05000 mL | 2.016 | 0.00623 | 0.50576 | 0.00043 | 10.0 s |
| 13:10.8 | Data point 2 | 2.20002 mL | 0.09563 mL | 0.03050 mL | 0.05000 mL | 2.239 | -0.00953 | 0.35232 | 0.00079 | 10.0 s |
| 13:46.7 | Data point 3 | 2.20002 mL | 0.09563 mL | 0.04570 mL | 0.05000 mL | 2.464 | -0.00246 | 0.19027 | 0.00028 | 10.0 s |
| 14:22.3 | Data point 4 | 2.20002 mL | 0.09563 mL | 0.05576 mL | 0.05000 mL | 2.670 | 0.01234 | 0.53634 | 0.00083 | 10.0 s |
| 14:58.0 | Data point 5 | 2.20002 mL | 0.09563 mL | 0.06286 mL | 0.05000 mL | 2.900 | 0.00043 | 0.01947 | 0.00015 | 10.5 s |
| 15:34.0 | Data point 6 | 2.20002 mL | 0.09563 mL | 0.06776 mL | 0.05000 mL | 3.097 | -0.00497 | 0.18772 | 0.00057 | 10.0 s |
| 16:09.5 | Data point 7 | 2.20002 mL | 0.09563 mL | 0.07131 mL | 0.05000 mL | 3.307 | -0.00870 | 0.18868 | 0.00099 | 10.0 s |
| 16:45.1 | Data point 8 | 2.20002 mL | 0.09563 mL | 0.07385 mL | 0.05000 mL | 3.525 | -0.00071 | 0.08489 | 0.00012 | 10.5 s |
| 17:21.0 | Data point 9 | 2.20002 mL | 0.09563 mL | 0.07571 mL | 0.05000 mL | 3.728 | -0.00085 | 0.13611 | 0.00011 | 10.0 s |
| 17:56.5 | Data point 10 | 2.20002 mL | 0.09563 mL | 0.07723 mL | 0.05000 mL | 3.901 | -0.00101 | 0.13288 | 0.00014 | 10.0 s |
| 18:42.4 | Data point 11 | 2.20002 mL | 0.09563 mL | 0.07869 mL | 0.05000 mL | 4.105 | -0.00256 | 0.29892 | 0.00023 | 10.0 s |
| 19:17.9 | Data point 12 | 2.20002 mL | 0.09563 mL | 0.08022 mL | 0.05000 mL | 4.378 | -0.00592 | 0.25762 | 0.00058 | 10.0 s |
| 19:58.5 | Data point 13 | 2.20002 mL | 0.09563 mL | 0.08154 mL | 0.05000 mL | 4.569 | -0.00172 | 0.03384 | 0.00046 | 10.0 s |
| 20:44.3 | Data point 14 | 2.20002 mL | 0.09563 mL | 0.08316 mL | 0.05000 mL | 4.761 | -0.00067 | 0.00810 | 0.00037 | 10.0 s |
| 21:30.1 | Data point 15 | 2.20002 mL | 0.09563 mL | 0.08511 mL | 0.05000 mL | 4.956 | -0.00175 | 0.25727 | 0.00017 | 10.5 s |
| 22:06.1 | Data point 16 | 2.20002 mL | 0.09563 mL | 0.08732 mL | 0.05000 mL | 5.158 | -0.00440 | 0.41157 | 0.00034 | 10.0 s |
| 22:41.6 | Data point 17 | 2.20002 mL | 0.09563 mL | 0.08935 mL | 0.05000 mL | 5.362 | 0.00350 | 0.06911 | 0.00066 | 10.5 s |
| 23:17.6 | Data point 18 | 2.20002 mL | 0.09563 mL | 0.09099 mL | 0.05000 mL | 5.592 | -0.01026 | 0.39926 | 0.00080 | 10.0 s |
| 23:53.0 | Data point 19 | 2.20002 mL | 0.09563 mL | 0.09222 mL | 0.05000 mL | 5.831 | -0.00232 | 0.01329 | 0.00099 | 10.0 s |
| 24:33.6 | Data point 20 | 2.20002 mL | 0.09563 mL | 0.09302 mL | 0.05000 mL | 6.046 | -0.01229 | 0.83805 | 0.00066 | 10.5 s |
| 25:09.5 | Data point 21 | 2.20002 mL | 0.09563 mL | 0.09358 mL | 0.05000 mL | 6.288 | -0.01579 | 0.76334 | 0.00089 | 12.5 s |
| 25:52.6 | Data point 22 | 2.20002 mL | 0.09563 mL | 0.09412 mL | 0.05000 mL | 6.573 | -0.01546 | 0.70398 | 0.00091 | 14.5 s |
| 26:37.6 | Data point 23 | 2.20002 mL | 0.09563 mL | 0.09450 mL | 0.05000 mL | 6.938 | -0.01891 | 0.87947 | 0.00100 | 48.5 s |
| 28:01.9 | Data point 24 | 2.20002 mL | 0.09563 mL | 0.09476 mL | 0.05000 mL | 7.316 | -0.02494 | 0.93500 | 0.00127 | Timed out at 59.5 s |
| 29:37.6 | Data point 25 | 2.20002 mL | 0.09563 mL | 0.09492 mL | 0.05000 mL | 7.818 | -0.04146 | 0.96788 | 0.00208 | Timed out at 59.5 s |
| 31:13.2 | Data point 26 | 2.20002 mL | 0.09563 mL | 0.09499 mL | 0.05000 mL | 8.108 | -0.02610 | 0.95271 | 0.00132 | Timed out at 59.5 s |
| 32:48.8 | Data point 27 | 2.20002 mL | 0.09563 mL | 0.09506 mL | 0.05000 mL | 8.299 | -0.01949 | 0.95727 | 0.00098 | 45.5 s |
| 34:15.2 | Data point 28 | 2.20002 mL | 0.09563 mL | 0.09518 mL | 0.05000 mL | 8.587 | -0.01701 | 0.74040 | 0.00098 | 24.5 s |
| 35:20.4 | Data point 29 | 2.20002 mL | 0.09563 mL | 0.09527 mL | 0.05000 mL | 8.827 | -0.01958 | 0.96654 | 0.00098 | 19.5 s |
| 36:15.6 | Data point 30 | 2.20002 mL | 0.09563 mL | 0.09541 mL | 0.05000 mL | 9.056 | -0.01834 | 0.92738 | 0.00094 | 28.0 s |
| 37:19.3 | Data point 31 | 2.20002 mL | 0.09563 mL | 0.09556 mL | 0.05000 mL | 9.269 | -0.01816 | 0.91609 | 0.00094 | 14.5 s |
| 38:09.5 | Data point 32 | 2.20002 mL | 0.09563 mL | 0.09577 mL | 0.05000 mL | 9.513 | -0.01968 | 0.96171 | 0.00099 | 14.0 s |
| 38:59.2 | Data point 33 | 2.20002 mL | 0.09563 mL | 0.09603 mL | 0.05000 mL | 9.735 | -0.01921 | 0.97472 | 0.00096 | 12.5 s |
| 39:47.4 | Data point 34 | 2.20002 mL | 0.09563 mL | 0.09635 mL | 0.05000 mL | 9.945 | -0.00595 | 0.27714 | 0.00056 | 10.0 s |
| 40:27.9 | Data point 35 | 2.20002 mL | 0.09563 mL | 0.09657 mL | 0.05000 mL | 10.046 | -0.01774 | 0.92128 | 0.00091 | 10.0 s |

| | | | |
|---|---|---|---|
| Sample name: | **M15_octanol** | Experiment start time: | **3/1/2018 6:16:08 AM** |
| Assay name: | **pH-metric high logP** | Analyst: | **Pion** |
| Assay ID: | **18C-01005** | Instrument ID: | **T312060** |
| Filename: | **C:\Sirius_T3\Mehtap\20180228_exp28_logP_T3-2\18C-01005_M15_octanol_pH-metric high logP.t3r** | | |

#### Events (continued)

| Time | Event | Water | Acid | Base | Octanol | pH | dpH/dt | pH R-squared | pH SD | dpH/dt time |
|---|---|---|---|---|---|---|---|---|---|---|
| 41:39.6 | Data point 36 | 2.20002 mL | 0.20089 mL | 0.09657 mL | 0.15000 mL | 1.954 | -0.00850 | 0.92951 | 0.00043 | 10.0 s |
| 42:26.1 | Data point 37 | 2.20002 mL | 0.20089 mL | 0.12434 mL | 0.15000 mL | 2.171 | 0.00527 | 0.50918 | 0.00036 | 10.0 s |
| 43:02.0 | Data point 38 | 2.20002 mL | 0.20089 mL | 0.14313 mL | 0.15000 mL | 2.389 | -0.01330 | 0.58147 | 0.00086 | 10.0 s |
| 43:37.7 | Data point 39 | 2.20002 mL | 0.20089 mL | 0.15550 mL | 0.15000 mL | 2.602 | -0.00134 | 0.03743 | 0.00034 | 10.0 s |
| 44:13.3 | Data point 40 | 2.20002 mL | 0.20089 mL | 0.16402 mL | 0.15000 mL | 2.821 | -0.00056 | 0.01526 | 0.00022 | 10.0 s |
| 44:48.9 | Data point 41 | 2.20002 mL | 0.20089 mL | 0.16999 mL | 0.15000 mL | 3.047 | 0.00129 | 0.22583 | 0.00013 | 10.0 s |
| 45:24.4 | Data point 42 | 2.20002 mL | 0.20089 mL | 0.17420 mL | 0.15000 mL | 3.248 | 0.00052 | 0.05566 | 0.00011 | 10.0 s |
| 45:59.9 | Data point 43 | 2.20002 mL | 0.20089 mL | 0.17735 mL | 0.15000 mL | 3.468 | -0.00387 | 0.16203 | 0.00047 | 10.0 s |
| 46:35.4 | Data point 44 | 2.20002 mL | 0.20089 mL | 0.17975 mL | 0.15000 mL | 3.662 | -0.00150 | 0.13762 | 0.00020 | 10.5 s |
| 47:11.4 | Data point 45 | 2.20002 mL | 0.20089 mL | 0.18182 mL | 0.15000 mL | 3.852 | 0.00050 | 0.03983 | 0.00012 | 10.0 s |
| 47:46.8 | Data point 46 | 2.20002 mL | 0.20089 mL | 0.18382 mL | 0.15000 mL | 4.067 | -0.00502 | 0.37193 | 0.00041 | 10.0 s |
| 48:22.3 | Data point 47 | 2.20002 mL | 0.20089 mL | 0.18594 mL | 0.15000 mL | 4.318 | -0.00521 | 0.16247 | 0.00064 | 10.0 s |
| 49:08.2 | Data point 48 | 2.20002 mL | 0.20089 mL | 0.18812 mL | 0.15000 mL | 4.506 | -0.00615 | 0.38511 | 0.00049 | 10.0 s |
| 49:43.7 | Data point 49 | 2.20002 mL | 0.20089 mL | 0.19048 mL | 0.15000 mL | 4.707 | -0.00269 | 0.27582 | 0.00025 | 10.0 s |
| 50:19.1 | Data point 50 | 2.20002 mL | 0.20089 mL | 0.19269 mL | 0.15000 mL | 4.921 | -0.01206 | 0.44085 | 0.00090 | 10.0 s |
| 50:54.6 | Data point 51 | 2.20002 mL | 0.20089 mL | 0.19457 mL | 0.15000 mL | 5.153 | 0.00999 | 0.53425 | 0.00068 | 10.5 s |
| 51:30.6 | Data point 52 | 2.20002 mL | 0.20089 mL | 0.19600 mL | 0.15000 mL | 5.389 | -0.01513 | 0.66733 | 0.00091 | 10.5 s |
| 52:11.7 | Data point 53 | 2.20002 mL | 0.20089 mL | 0.19715 mL | 0.15000 mL | 5.641 | -0.01166 | 0.35160 | 0.00097 | 11.0 s |
| 52:48.2 | Data point 54 | 2.20002 mL | 0.20089 mL | 0.19781 mL | 0.15000 mL | 5.848 | -0.00111 | 0.00552 | 0.00073 | 12.0 s |
| 53:25.6 | Data point 55 | 2.20002 mL | 0.20089 mL | 0.19826 mL | 0.15000 mL | 6.041 | -0.01512 | 0.65316 | 0.00092 | 13.0 s |
| 54:04.0 | Data point 56 | 2.20002 mL | 0.20089 mL | 0.19859 mL | 0.15000 mL | 6.237 | -0.01714 | 0.87476 | 0.00091 | 14.0 s |
| 54:43.4 | Data point 57 | 2.20002 mL | 0.20089 mL | 0.19882 mL | 0.15000 mL | 6.433 | -0.01867 | 0.91161 | 0.00097 | 27.5 s |
| 55:36.4 | Data point 58 | 2.20002 mL | 0.20089 mL | 0.19899 mL | 0.15000 mL | 6.627 | -0.01996 | 0.98353 | 0.00099 | 27.0 s |
| 56:28.8 | Data point 59 | 2.20002 mL | 0.20089 mL | 0.19911 mL | 0.15000 mL | 6.813 | -0.01534 | 0.97399 | 0.00077 | 39.0 s |
| 57:33.2 | Data point 60 | 2.20002 mL | 0.20089 mL | 0.19920 mL | 0.15000 mL | 6.955 | -0.01953 | 0.94508 | 0.00099 | 49.0 s |
| 58:52.9 | Data point 61 | 2.20002 mL | 0.20089 mL | 0.19932 mL | 0.15000 mL | 7.307 | -0.03681 | 0.97518 | 0.00184 | Timed out at 59.5 s |
| 1:00:28.6 | Data point 62 | 2.20002 mL | 0.20089 mL | 0.19939 mL | 0.15000 mL | 7.553 | -0.03803 | 0.97454 | 0.00190 | Timed out at 59.5 s |
| 1:01:59.1 | Data point 63 | 2.20002 mL | 0.20089 mL | 0.19944 mL | 0.15000 mL | 7.763 | -0.03211 | 0.96281 | 0.00162 | Timed out at 59.5 s |
| 1:03:34.7 | Data point 64 | 2.20002 mL | 0.20089 mL | 0.19951 mL | 0.15000 mL | 8.039 | -0.02677 | 0.95360 | 0.00135 | Timed out at 59.5 s |
| 1:05:10.3 | Data point 65 | 2.20002 mL | 0.20089 mL | 0.19958 mL | 0.15000 mL | 8.302 | -0.01665 | 0.74155 | 0.00095 | 24.5 s |
| 1:06:15.7 | Data point 66 | 2.20002 mL | 0.20089 mL | 0.19977 mL | 0.15000 mL | 8.588 | -0.01458 | 0.68624 | 0.00087 | 29.0 s |
| 1:07:25.6 | Data point 67 | 2.20002 mL | 0.20089 mL | 0.19988 mL | 0.15000 mL | 8.798 | -0.01709 | 0.86919 | 0.00091 | 16.5 s |
| 1:08:17.8 | Data point 68 | 2.20002 mL | 0.20089 mL | 0.20059 mL | 0.15000 mL | 9.185 | -0.01489 | 0.69562 | 0.00088 | 14.0 s |
| 1:09:07.5 | Data point 69 | 2.20002 mL | 0.20089 mL | 0.20094 mL | 0.15000 mL | 9.496 | -0.01158 | 0.36700 | 0.00094 | 18.0 s |
| 1:09:56.1 | Data point 70 | 2.20002 mL | 0.20089 mL | 0.20132 mL | 0.15000 mL | 9.775 | -0.01160 | 0.42092 | 0.00088 | 12.5 s |
| 1:10:39.2 | Data point 71 | 2.20002 mL | 0.20089 mL | 0.20193 mL | 0.15000 mL | 10.062 | -0.01596 | 0.67334 | 0.00096 | 10.0 s |
| 1:11:55.5 | Data point 72 | 2.20002 mL | 0.30889 mL | 0.20193 mL | 0.45000 mL | 1.969 | 0.00067 | 0.00237 | 0.00068 | 10.0 s |
| 1:12:42.0 | Data point 73 | 2.20002 mL | 0.30889 mL | 0.23100 mL | 0.45000 mL | 2.187 | 0.01750 | 0.85670 | 0.00093 | 10.0 s |
| 1:13:17.9 | Data point 74 | 2.20002 mL | 0.30889 mL | 0.25045 mL | 0.45000 mL | 2.416 | 0.01285 | 0.80261 | 0.00071 | 10.0 s |
| 1:13:53.6 | Data point 75 | 2.20002 mL | 0.30889 mL | 0.26312 mL | 0.45000 mL | 2.628 | 0.00730 | 0.64955 | 0.00045 | 10.0 s |
| 1:14:29.3 | Data point 76 | 2.20002 mL | 0.30889 mL | 0.27197 mL | 0.45000 mL | 2.851 | 0.00165 | 0.38392 | 0.00013 | 10.0 s |
| 1:15:04.8 | Data point 77 | 2.20002 mL | 0.30889 mL | 0.27822 mL | 0.45000 mL | 3.054 | 0.00066 | 0.05640 | 0.00014 | 10.0 s |
| 1:15:40.4 | Data point 78 | 2.20002 mL | 0.30889 mL | 0.28293 mL | 0.45000 mL | 3.283 | -0.00318 | 0.06439 | 0.00062 | 10.0 s |
| 1:16:15.9 | Data point 79 | 2.20002 mL | 0.30889 mL | 0.28655 mL | 0.45000 mL | 3.492 | 0.00558 | 0.58627 | 0.00036 | 10.0 s |
| 1:16:51.4 | Data point 80 | 2.20002 mL | 0.30889 mL | 0.28961 mL | 0.45000 mL | 3.729 | -0.00357 | 0.40444 | 0.00028 | 10.0 s |
| 1:17:37.3 | Data point 81 | 2.20002 mL | 0.30889 mL | 0.29222 mL | 0.45000 mL | 3.922 | 0.00000 | 0.00000 | 0.00050 | 10.5 s |
| 1:18:13.3 | Data point 82 | 2.20002 mL | 0.30889 mL | 0.29497 mL | 0.45000 mL | 4.141 | -0.00271 | 0.20156 | 0.00030 | 10.0 s |
| 1:18:48.8 | Data point 83 | 2.20002 mL | 0.30889 mL | 0.29758 mL | 0.45000 mL | 4.380 | -0.00694 | 0.22006 | 0.00073 | 10.0 s |
| 1:19:34.6 | Data point 84 | 2.20002 mL | 0.30889 mL | 0.29972 mL | 0.45000 mL | 4.565 | 0.00192 | 0.03274 | 0.00052 | 10.0 s |
| 1:20:10.1 | Data point 85 | 2.20002 mL | 0.30889 mL | 0.30165 mL | 0.45000 mL | 4.750 | -0.00572 | 0.22132 | 0.00060 | 10.0 s |
| 1:20:45.6 | Data point 86 | 2.20002 mL | 0.30889 mL | 0.30315 mL | 0.45000 mL | 4.943 | -0.00051 | 0.00238 | 0.00052 | 10.5 s |
| 1:21:21.6 | Data point 87 | 2.20002 mL | 0.30889 mL | 0.30428 mL | 0.45000 mL | 5.134 | -0.00660 | 0.13315 | 0.00089 | 10.5 s |
| 1:21:57.5 | Data point 88 | 2.20002 mL | 0.30889 mL | 0.30508 mL | 0.45000 mL | 5.338 | -0.00192 | 0.01187 | 0.00087 | 11.0 s |

| | | | |
|---|---|---|---|
| Sample name: | **M15_octanol** | Experiment start time: | **3/1/2018 6:16:08 AM** |
| Assay name: | **pH-metric high logP** | Analyst: | **Pion** |
| Assay ID: | **18C-01005** | Instrument ID: | **T312060** |
| Filename: | **C:\Sirius_T3\Mehtap\20180228_exp28_logP_T3-2\18C-01005_M15_octanol_pH-metric high logP.t3r** | | |

#### Events (continued)

| Time | Event | Water | Acid | Base | Octanol | pH | dpH/dt | pH R-squared | pH SD | dpH/dt time |
|---|---|---|---|---|---|---|---|---|---|---|
| 1:22:33.9 | Data point 89 | 2.20002 mL | 0.30889 mL | 0.30562 mL | 0.45000 mL | 5.547 | -0.00385 | 0.05953 | 0.00078 | 11.5 s |
| 1:23:10.9 | Data point 90 | 2.20002 mL | 0.30889 mL | 0.30600 mL | 0.45000 mL | 5.773 | -0.01234 | 0.38543 | 0.00098 | 18.0 s |
| 1:23:54.3 | Data point 91 | 2.20002 mL | 0.30889 mL | 0.30626 mL | 0.45000 mL | 5.990 | -0.01294 | 0.62761 | 0.00081 | 22.0 s |
| 1:24:41.7 | Data point 92 | 2.20002 mL | 0.30889 mL | 0.30644 mL | 0.45000 mL | 6.237 | -0.01660 | 0.92533 | 0.00085 | 30.5 s |
| 1:25:42.9 | Data point 93 | 2.20002 mL | 0.30889 mL | 0.30661 mL | 0.45000 mL | 6.538 | -0.02270 | 0.93887 | 0.00116 | Timed out at 59.5 s |
| 1:27:13.4 | Data point 94 | 2.20002 mL | 0.30889 mL | 0.30670 mL | 0.45000 mL | 6.787 | -0.03247 | 0.95665 | 0.00164 | Timed out at 59.5 s |
| 1:28:38.8 | Data point 95 | 2.20002 mL | 0.30889 mL | 0.30677 mL | 0.45000 mL | 6.955 | -0.02853 | 0.95075 | 0.00145 | Timed out at 59.5 s |
| 1:30:04.2 | Data point 96 | 2.20002 mL | 0.30889 mL | 0.30682 mL | 0.45000 mL | 6.926 | -0.00738 | 0.28183 | 0.00069 | 10.5 s |
| 1:30:45.2 | Data point 97 | 2.20002 mL | 0.30889 mL | 0.30699 mL | 0.45000 mL | 8.028 | -0.06462 | 0.99131 | 0.00321 | Timed out at 59.5 s |
| 1:32:20.9 | Data point 98 | 2.20002 mL | 0.30889 mL | 0.30710 mL | 0.45000 mL | 8.410 | -0.01600 | 0.62423 | 0.00100 | 23.5 s |
| 1:33:25.2 | Data point 99 | 2.20002 mL | 0.30889 mL | 0.30724 mL | 0.45000 mL | 8.685 | -0.01186 | 0.41042 | 0.00092 | 15.5 s |
| 1:34:16.4 | Data point 100 | 2.20002 mL | 0.30889 mL | 0.30739 mL | 0.45000 mL | 8.958 | -0.00728 | 0.18983 | 0.00082 | 14.5 s |
| 1:35:06.5 | Data point 101 | 2.20002 mL | 0.30889 mL | 0.30757 mL | 0.45000 mL | 9.218 | -0.01378 | 0.67757 | 0.00083 | 13.0 s |
| 1:35:55.3 | Data point 102 | 2.20002 mL | 0.30889 mL | 0.30785 mL | 0.45000 mL | 9.455 | -0.01183 | 0.35300 | 0.00098 | 11.5 s |
| 1:36:37.4 | Data point 103 | 2.20002 mL | 0.30889 mL | 0.30821 mL | 0.45000 mL | 9.725 | -0.00923 | 0.21248 | 0.00099 | 11.5 s |
| 1:37:24.7 | Data point 104 | 2.20002 mL | 0.30889 mL | 0.30870 mL | 0.45000 mL | 9.960 | -0.01830 | 0.91815 | 0.00094 | 11.0 s |
| 1:38:06.3 | Data point 105 | 2.20002 mL | 0.30889 mL | 0.30896 mL | 0.45000 mL | 10.057 | -0.01398 | 0.91248 | 0.00072 | 10.5 s |
| 1:38:25.8 | Assay volumes | 2.20002 mL | 0.30889 mL | 0.30896 mL | 0.45000 mL | | | | | |

| | | | |
|---|---|---|---|
| Sample name: | **M15_octanol** | Experiment start time: | **3/1/2018 6:16:08 AM** |
| Assay name: | **pH-metric high logP** | Analyst: | **Pion** |
| Assay ID: | **18C-01005** | Instrument ID: | **T312060** |
| Filename: | **C:\Sirius_T3\Mehtap\20180228_exp28_logP_T3-2\18C-01005_M15_octanol_pH-metric high logP.t3r** | | |

| | | | |
|---|---|---|---|
| Sample name: | **M15_octanol** | Experiment start time: | **3/1/2018 6:16:08 AM** |
| Assay name: | **pH-metric high logP** | Analyst: | **Pion** |
| Assay ID: | **18C-01005** | Instrument ID: | **T312060** |
| Filename: | **C:\Sirius_T3\Mehtap\20180228_exp28_logP_T3-2\18C-01005_M15_octanol_pH-metric high logP.t3r** | | |

#### Assay Settings (continued)

| Setting | Value | Original Value | Date/Time changed | Imported from |
|---|---|---|---|---|
| ***Titration 3*** | | | | |
| Titrate from | Low to high pH | | | |
| Add additional water | 0.00 mL | | | |
| Additional partition solvent volume | 0.300 mL | | | |
| Additional partition solvent added | Automatic | | | |
| After pH adjust stir for | 30 seconds | | | |
| Stir to allow partitioning for | 15 seconds | | | |
| Stirrer speed for partitioning | 60% | | | |
| ***Data Point Stability*** | | | | |
| Stir during data point collection | No | | | |
| Delay before data point collection | 0 seconds | | | |
| Number of points to average | 20 points | | | |
| Time interval between points | 0.50 seconds | | | |
| Required maximum standard deviation | 0.00100 dpH/dt | | | |
| Stability timeout after | 60 seconds | | | |

#### Calibration Settings

| Setting | Value | Date/Time changed | Imported from |
|---|---|---|---|
| Four-Plus alpha | 0.130 | 3/1/2018 6:16:08 AM | C:\Sirius_T3\HCl18B27.t3r |
| Four-Plus S | 0.9970 | 3/1/2018 6:16:08 AM | C:\Sirius_T3\HCl18B27.t3r |
| Four-Plus jH | 0.8 | 3/1/2018 6:16:08 AM | C:\Sirius_T3\HCl18B27.t3r |
| Four-Plus jOH | -0.4 | 3/1/2018 6:16:08 AM | C:\Sirius_T3\HCl18B27.t3r |
| Base concentration factor | 1.000 | 3/1/2018 6:16:08 AM | C:\Sirius_T3\KOH18B27.t3r |
| Acid concentration factor | 0.994 | 3/1/2018 6:16:08 AM | C:\Sirius_T3\HCl18B27.t3r |

#### Instrument Settings

| Setting | Value | Batch Id | Install date |
|---|---|---|---|
| Instrument owner | Merck | | |
| Instrument ID | T312060 | | |
| Instrument type | T3 Simulator | | |
| Software version | 1.1.3.0 | | |
| Dispenser module | | T3DM1200361 | 3/31/2009 5:24:52 AM |
| Dispenser 0 | Water | | 3/31/2009 5:25:05 AM |
| Syringe volume | 2.5 mL | | |
| Firmware version | 1.2.1(r2) | | |
| Titrant | Water (0.15 M KCl) | 02-06-2018 | 2/27/2018 10:05:59 AM |
| Dispenser 2 | Acid | | 3/31/2009 5:25:11 AM |
| Syringe volume | 0.5 mL | | |
| Firmware version | 1.2.1(r2) | | |
| Titrant | Acid (0.5 M HCl) | 02-27-2018 | 2/27/2018 10:27:22 AM |
| Dispenser 1 | Base | | 3/31/2009 5:25:21 AM |
| Syringe volume | 0.5 mL | | |
| Firmware version | 1.2.1(r2) | | |
| Titrant | Base (0.5 M KOH) | 9/22/2017 | 2/27/2018 10:21:22 AM |
| Dispenser 5 | Cosolvent | | 3/31/2009 5:26:24 AM |
| Syringe volume | 2.5 mL | | |
| Firmware version | 1.2.1(r2) | | |
| Distribution valve 5 | Distribution Valve | | 3/31/2009 5:28:19 AM |
| Firmware version | 1.1.3 | | |
| Port A | Methanol (80%, 0.15 M KCl) | 09-26-17 | 2/7/2018 9:42:01 AM |
| Port B | Cyclohexane | 11-01-17 | 2/27/2018 10:37:57 AM |
| Dispenser 3 | Buffer | | 8/3/2010 5:05:16 AM |
| Syringe volume | 0.5 mL | | |
| Firmware version | 1.2.1(r2) | | |
| Titrant | Dodecane | 2018/01/31 | 2/28/2018 10:18:04 AM |
| Dispenser 6 | Octanol | | 10/22/2010 10:52:43 AM |

| | | | |
|---|---|---|---|
| Sample name: | **M15_octanol** | Experiment start time: | **3/1/2018 6:16:08 AM** |
| Assay name: | **pH-metric high logP** | Analyst: | **Pion** |
| Assay ID: | **18C-01005** | Instrument ID: | **T312060** |
| Filename: | **C:\Sirius_T3\Mehtap\20180228_exp28_logP_T3-2\18C-01005_M15_octanol_pH-metric high logP.t3r** | | |

| | | | |
|---|---|---|---|
| Sample name: | **M15_octanol** | Experiment start time: | **3/1/2018 6:16:08 AM** |
| Assay name: | **pH-metric high logP** | Analyst: | **Pion** |
| Assay ID: | **18C-01005** | Instrument ID: | **T312060** |
| Filename: | **C:\Sirius_T3\Mehtap\20180228_exp28_logP_T3-2\18C-01005_M15_octanol_pH-metric high logP.t3r** | | |

#### Instrument Settings (continued)

| Setting | Value | Batch Id | Install date |
|---|---|---|---|
| Spectrometer calibration stir duration | 5 s | | |
| Spectrometer calibration stir speed | 30% | | |
| Spectrometer calibration wash pump volume | 20.0 mL | | |
| Spectrometer calibration wash stir duration | 5 s | | |
| Spectrometer calibration wash stir speed | 30% | | |
| Overhead dispense height | 10000 | | |

#### Refinement Settings

| Setting | Value | Default value |
|---|---|---|
| Turbidity detection method | None | None |
| Turbidity wavelength to assess | 500.0 nm | 500.0 nm |
| Turbidity maximum absorbance | 0.100 | 0.100 |
| Turbidity probe threshold | 50.00 | 50.00 |

#### Experiment Log

[2:49] Air gap created for Water (0.15 M KCl)
[2:49] Air gap created for Acid (0.5 M HCl)
[2:50] Air gap created for Base (0.5 M KOH)
[2:50] Air gap released for Water (0.15 M KCl)
[2:54] Titrator arm moved over Titration position
[2:54] Titration 1 of 3
[2:54] Adding initial titrants
[2:54] Automatically add 2.20000 mL of water
[3:31] Dispensed 2.200024 mL of Water (0.15 M KCl)
[3:35] Titrator arm moved over Drain
[9:16] Titrator arm moved to Titration position
[9:16] Argon flow rate set to 100
[9:16] Stirrer speed set to 10
[9:21] Automatically add 0.05000 mL of Octanol
[9:23] Dispensed 0.050000 mL of Octanol
[9:24] Initial pH = 7.47
[9:24] Iterative adjust 7.47 -> 2.00
[9:24] pH 7.47 -> 2.00
[9:27] Air gap released for Acid (0.5 M HCl)
[9:27] Dispensed 0.095626 mL of Acid (0.5 M HCl)
[9:32] Holding pH 2.00
[11:33] Stirrer speed set to 0
[11:33] Stirrer speed set to 50
[11:33] Iterative adjust 1.95 -> 2.00
[11:33] pH 1.95 -> 2.00
[11:33] Air gap released for Base (0.5 M KOH)
[11:34] Dispensed 0.007785 mL of Base (0.5 M KOH)
[12:25] Stirrer speed set to 0
[12:35] Datapoint id 1 collected
[12:35] Stirrer speed set to 50
[12:40] pH 2.03 -> 2.23
[12:40] Using cautious pH adjust
[12:40] Dispensed 0.011783 mL of Base (0.5 M KOH)
[12:45] Stepping pH = 2.11
[12:46] Dispensed 0.009196 mL of Base (0.5 M KOH)
[12:51] Stepping pH = 2.21
[12:51] Dispensed 0.001740 mL of Base (0.5 M KOH)
[12:56] Stepping pH = 2.23
[13:11] Stirrer speed set to 0
[13:21] Datapoint id 2 collected
[13:21] Charge balance equation is out by 3.7%
[13:21] Stirrer speed set to 50

Sample name: **M15_octanol** Experiment start time: **3/1/2018 6:16:08 AM**
Assay name: **pH-metric high logP** Analyst: **Pion**
Assay ID: **18C-01005** Instrument ID: **T312060**
Filename: **C:\Sirius_T3\Mehtap\20180228_exp28_logP_T3-2\18C-01005_M15_octanol_pH-metric high logP.t3r**

#### Experiment Log (continued)

[13:26] pH 2.24 -> 2.44
[13:26] Using charge balance adjust
[13:27] Dispensed 0.015193 mL of Base (0.5 M KOH)
[13:47] Stirrer speed set to 0
[13:57] Datapoint id 3 collected
[13:57] Charge balance equation is out by 9.8%
[13:57] Stirrer speed set to 50
[14:02] pH 2.47 -> 2.67
[14:02] Using charge balance adjust
[14:02] Dispensed 0.010066 mL of Base (0.5 M KOH)
[14:22] Stirrer speed set to 0
[14:32] Datapoint id 4 collected
[14:32] Charge balance equation is out by 0.9%
[14:32] Stirrer speed set to 50
[14:38] pH 2.68 -> 2.88
[14:38] Using charge balance adjust
[14:38] Dispensed 0.007103 mL of Base (0.5 M KOH)
[14:58] Stirrer speed set to 0
[15:09] Datapoint id 5 collected
[15:09] Charge balance equation is out by 10.7%
[15:09] Stirrer speed set to 50
[15:14] pH 2.91 -> 3.11
[15:14] Using charge balance adjust
[15:14] Dispensed 0.004892 mL of Base (0.5 M KOH)
[15:34] Stirrer speed set to 0
[15:44] Datapoint id 6 collected
[15:44] Charge balance equation is out by -6.0%
[15:44] Stirrer speed set to 50
[15:49] pH 3.11 -> 3.31
[15:49] Using charge balance adjust
[15:49] Dispensed 0.003551 mL of Base (0.5 M KOH)
[16:10] Stirrer speed set to 0
[16:20] Datapoint id 7 collected
[16:20] Charge balance equation is out by 0.0%
[16:20] Stirrer speed set to 50
[16:25] pH 3.31 -> 3.51
[16:25] Using charge balance adjust
[16:25] Dispensed 0.002540 mL of Base (0.5 M KOH)
[16:45] Stirrer speed set to 0
[16:56] Datapoint id 8 collected
[16:56] Charge balance equation is out by 4.9%
[16:56] Stirrer speed set to 50
[17:01] pH 3.53 -> 3.73
[17:01] Using charge balance adjust
[17:01] Dispensed 0.001858 mL of Base (0.5 M KOH)
[17:21] Stirrer speed set to 0
[17:31] Datapoint id 9 collected
[17:31] Charge balance equation is out by -2.1%
[17:31] Stirrer speed set to 50
[17:36] pH 3.74 -> 3.94
[17:36] Using charge balance adjust
[17:36] Dispensed 0.001529 mL of Base (0.5 M KOH)
[17:57] Stirrer speed set to 0
[18:07] Datapoint id 10 collected
[18:07] Charge balance equation is out by -17.7%
[18:07] Stirrer speed set to 50
[18:12] pH 3.91 -> 4.11
[18:12] Using cautious pH adjust
[18:12] Dispensed 0.000729 mL of Base (0.5 M KOH)

Sample name: **M15_octanol** Experiment start time: **3/1/2018 6:16:08 AM**
Assay name: **pH-metric high logP** Analyst: **Pion**
Assay ID: **18C-01005** Instrument ID: **T312060**
Filename: **C:\Sirius_T3\Mehtap\20180228_exp28_logP_T3-2\18C-01005_M15_octanol_pH-metric high logP.t3r**

#### Experiment Log (continued)

[18:17] Stepping pH = 3.99
[18:17] Dispensed 0.000635 mL of Base (0.5 M KOH)
[18:22] Stepping pH = 4.09
[18:22] Dispensed 0.000094 mL of Base (0.5 M KOH)
[18:27] Stepping pH = 4.11
[18:43] Stirrer speed set to 0
[18:53] Datapoint id 11 collected
[18:53] Charge balance equation is out by -0.4%
[18:53] Stirrer speed set to 50
[18:58] pH 4.11 -> 4.31
[18:58] Using charge balance adjust
[18:58] Dispensed 0.001529 mL of Base (0.5 M KOH)
[19:18] Stirrer speed set to 0
[19:28] Datapoint id 12 collected
[19:28] Charge balance equation is out by 33.2%
[19:28] Stirrer speed set to 50
[19:33] pH 4.38 -> 4.58
[19:33] Using cautious pH adjust
[19:33] Dispensed 0.000917 mL of Base (0.5 M KOH)
[19:38] Stepping pH = 4.51
[19:38] Dispensed 0.000400 mL of Base (0.5 M KOH)
[19:44] Stepping pH = 4.57
[19:59] Stirrer speed set to 0
[20:09] Datapoint id 13 collected
[20:09] Charge balance equation is out by 28.2%
[20:09] Stirrer speed set to 50
[20:14] pH 4.57 -> 4.77
[20:14] Using cautious pH adjust
[20:14] Dispensed 0.001035 mL of Base (0.5 M KOH)
[20:19] Stepping pH = 4.70
[20:19] Dispensed 0.000494 mL of Base (0.5 M KOH)
[20:24] Stepping pH = 4.76
[20:24] Dispensed 0.000094 mL of Base (0.5 M KOH)
[20:29] Stepping pH = 4.77
[20:44] Stirrer speed set to 0
[20:54] Datapoint id 14 collected
[20:54] Charge balance equation is out by 22.3%
[20:54] Stirrer speed set to 50
[21:00] pH 4.77 -> 4.97
[21:00] Using cautious pH adjust
[21:00] Dispensed 0.001129 mL of Base (0.5 M KOH)
[21:05] Stepping pH = 4.88
[21:05] Dispensed 0.000635 mL of Base (0.5 M KOH)
[21:10] Stepping pH = 4.95
[21:10] Dispensed 0.000188 mL of Base (0.5 M KOH)
[21:15] Stepping pH = 4.96
[21:30] Stirrer speed set to 0
[21:41] Datapoint id 15 collected
[21:41] Charge balance equation is out by 12.8%
[21:41] Stirrer speed set to 50
[21:46] pH 4.96 -> 5.16
[21:46] Using charge balance adjust
[21:46] Dispensed 0.002211 mL of Base (0.5 M KOH)
[22:06] Stirrer speed set to 0
[22:16] Datapoint id 16 collected
[22:16] Charge balance equation is out by -2.8%
[22:16] Stirrer speed set to 50
[22:21] pH 5.16 -> 5.36
[22:21] Using charge balance adjust

Sample name: **M15_octanol** Experiment start time: **3/1/2018 6:16:08 AM**
Assay name: **pH-metric high logP** Analyst: **Pion**
Assay ID: **18C-01005** Instrument ID: **T312060**
Filename: **C:\Sirius_T3\Mehtap\20180228_exp28_logP_T3-2\18C-01005_M15_octanol_pH-metric high logP.t3r**

#### Experiment Log (continued)

[22:22] Dispensed 0.002023 mL of Base (0.5 M KOH)
[22:42] Stirrer speed set to 0
[22:52] Datapoint id 17 collected
[22:52] Charge balance equation is out by -0.1%
[22:52] Stirrer speed set to 50
[22:57] pH 5.37 -> 5.57
[22:57] Using charge balance adjust
[22:57] Dispensed 0.001646 mL of Base (0.5 M KOH)
[23:18] Stirrer speed set to 0
[23:28] Datapoint id 18 collected
[23:28] Charge balance equation is out by 12.1%
[23:28] Stirrer speed set to 50
[23:33] pH 5.60 -> 5.80
[23:33] Using charge balance adjust
[23:33] Dispensed 0.001223 mL of Base (0.5 M KOH)
[23:53] Stirrer speed set to 0
[24:03] Datapoint id 19 collected
[24:03] Charge balance equation is out by 16.5%
[24:03] Stirrer speed set to 50
[24:08] pH 5.84 -> 6.04
[24:08] Using cautious pH adjust
[24:08] Dispensed 0.000423 mL of Base (0.5 M KOH)
[24:13] Stepping pH = 5.92
[24:14] Dispensed 0.000376 mL of Base (0.5 M KOH)
[24:19] Stepping pH = 6.03
[24:34] Stirrer speed set to 0
[24:44] Datapoint id 20 collected
[24:44] Charge balance equation is out by 3.8%
[24:44] Stirrer speed set to 50
[24:49] pH 6.05 -> 6.25
[24:49] Using charge balance adjust
[24:49] Dispensed 0.000564 mL of Base (0.5 M KOH)
[25:10] Stirrer speed set to 0
[25:22] Datapoint id 21 collected
[25:22] Charge balance equation is out by 16.9%
[25:22] Stirrer speed set to 50
[25:27] pH 6.29 -> 6.49
[25:27] Using cautious pH adjust
[25:27] Dispensed 0.000188 mL of Base (0.5 M KOH)
[25:32] Stepping pH = 6.33
[25:33] Dispensed 0.000353 mL of Base (0.5 M KOH)
[25:38] Stepping pH = 6.52
[25:53] Stirrer speed set to 0
[26:07] Datapoint id 22 collected
[26:07] Charge balance equation is out by -49.2%
[26:07] Stirrer speed set to 50
[26:12] pH 6.58 -> 6.78
[26:12] Using cautious pH adjust
[26:12] Dispensed 0.000094 mL of Base (0.5 M KOH)
[26:17] Stepping pH = 6.60
[26:18] Dispensed 0.000282 mL of Base (0.5 M KOH)
[26:23] Stepping pH = 6.84
[26:38] Stirrer speed set to 0
[27:26] Datapoint id 23 collected
[27:26] Charge balance equation is out by -83.4%
[27:26] Stirrer speed set to 50
[27:31] pH 6.94 -> 7.14
[27:31] Using cautious pH adjust
[27:32] Dispensed 0.000047 mL of Base (0.5 M KOH)

Sample name: **M15_octanol** Experiment start time: **3/1/2018 6:16:08 AM**
Assay name: **pH-metric high logP** Analyst: **Pion**
Assay ID: **18C-01005** Instrument ID: **T312060**
Filename: **C:\Sirius_T3\Mehtap\20180228_exp28_logP_T3-2\18C-01005_M15_octanol_pH-metric high logP.t3r**

#### Experiment Log (continued)

[27:37] Stepping pH = 6.96
[27:37] Dispensed 0.000118 mL of Base (0.5 M KOH)
[27:42] Stepping pH = 7.04
[27:42] Dispensed 0.000094 mL of Base (0.5 M KOH)
[27:47] Stepping pH = 7.24
[28:02] Stirrer speed set to 0
[29:02] Datapoint id 24 collected
[29:02] Charge balance equation is out by -171.1%
[29:02] Stirrer speed set to 50
[29:07] pH 7.30 -> 7.50
[29:07] Using cautious pH adjust
[29:07] Dispensed 0.000024 mL of Base (0.5 M KOH)
[29:12] Stepping pH = 7.32
[29:12] Dispensed 0.000047 mL of Base (0.5 M KOH)
[29:17] Stepping pH = 7.36
[29:17] Dispensed 0.000094 mL of Base (0.5 M KOH)
[29:23] Stepping pH = 7.64
[29:38] Stirrer speed set to 0
[30:38] Datapoint id 25 collected
[30:38] Charge balance equation is out by -247.2%
[30:38] Stirrer speed set to 50
[30:43] pH 7.80 -> 8.00
[30:43] Using cautious pH adjust
[30:43] Dispensed 0.000024 mL of Base (0.5 M KOH)
[30:48] Stepping pH = 7.88
[30:48] Dispensed 0.000024 mL of Base (0.5 M KOH)
[30:53] Stepping pH = 7.97
[30:53] Dispensed 0.000024 mL of Base (0.5 M KOH)
[30:58] Stepping pH = 8.09
[31:13] Stirrer speed set to 0
[32:13] Datapoint id 26 collected
[32:13] Charge balance equation is out by -210.8%
[32:13] Stirrer speed set to 50
[32:18] pH 8.06 -> 8.26
[32:18] Using cautious pH adjust
[32:19] Dispensed 0.000024 mL of Base (0.5 M KOH)
[32:24] Stepping pH = 8.11
[32:24] Dispensed 0.000024 mL of Base (0.5 M KOH)
[32:29] Stepping pH = 8.16
[32:29] Dispensed 0.000024 mL of Base (0.5 M KOH)
[32:34] Stepping pH = 8.26
[32:49] Stirrer speed set to 0
[33:35] Datapoint id 27 collected
[33:35] Charge balance equation is out by -276.2%
[33:35] Stirrer speed set to 50
[33:40] pH 8.28 -> 8.48
[33:40] Using cautious pH adjust
[33:40] Dispensed 0.000024 mL of Base (0.5 M KOH)
[33:45] Stepping pH = 8.29
[33:45] Dispensed 0.000024 mL of Base (0.5 M KOH)
[33:50] Stepping pH = 8.33
[33:50] Dispensed 0.000047 mL of Base (0.5 M KOH)
[33:55] Stepping pH = 8.43
[33:55] Dispensed 0.000024 mL of Base (0.5 M KOH)
[34:00] Stepping pH = 8.55
[34:15] Stirrer speed set to 0
[34:40] Datapoint id 28 collected
[34:40] Charge balance equation is out by -522.0%
[34:40] Stirrer speed set to 50

Sample name: **M15_octanol** Experiment start time: **3/1/2018 6:16:08 AM**
Assay name: **pH-metric high logP** Analyst: **Pion**
Assay ID: **18C-01005** Instrument ID: **T312060**
Filename: **C:\Sirius_T3\Mehtap\20180228_exp28_logP_T3-2\18C-01005_M15_octanol_pH-metric high logP.t3r**

#### Experiment Log (continued)

[34:45] pH 8.59 -> 8.79
[34:45] Using cautious pH adjust
[34:45] Dispensed 0.000024 mL of Base (0.5 M KOH)
[34:50] Stepping pH = 8.63
[34:50] Dispensed 0.000024 mL of Base (0.5 M KOH)
[34:55] Stepping pH = 8.69
[34:55] Dispensed 0.000024 mL of Base (0.5 M KOH)
[35:00] Stepping pH = 8.75
[35:00] Dispensed 0.000024 mL of Base (0.5 M KOH)
[35:05] Stepping pH = 8.81
[35:21] Stirrer speed set to 0
[35:40] Datapoint id 29 collected
[35:40] Charge balance equation is out by -241.0%
[35:40] Stirrer speed set to 50
[35:45] pH 8.84 -> 9.04
[35:45] Using cautious pH adjust
[35:45] Dispensed 0.000024 mL of Base (0.5 M KOH)
[35:50] Stepping pH = 8.86
[35:50] Dispensed 0.000071 mL of Base (0.5 M KOH)
[35:55] Stepping pH = 8.93
[35:56] Dispensed 0.000047 mL of Base (0.5 M KOH)
[36:01] Stepping pH = 9.03
[36:16] Stirrer speed set to 0
[36:44] Datapoint id 30 collected
[36:44] Charge balance equation is out by -213.1%
[36:44] Stirrer speed set to 50
[36:49] pH 9.07 -> 9.27
[36:49] Using cautious pH adjust
[36:49] Dispensed 0.000047 mL of Base (0.5 M KOH)
[36:54] Stepping pH = 9.11
[36:54] Dispensed 0.000071 mL of Base (0.5 M KOH)
[36:59] Stepping pH = 9.21
[36:59] Dispensed 0.000024 mL of Base (0.5 M KOH)
[37:04] Stepping pH = 9.27
[37:19] Stirrer speed set to 0
[37:34] Datapoint id 31 collected
[37:34] Charge balance equation is out by -90.2%
[37:34] Stirrer speed set to 50
[37:39] pH 9.28 -> 9.48
[37:39] Using cautious pH adjust
[37:39] Dispensed 0.000047 mL of Base (0.5 M KOH)
[37:44] Stepping pH = 9.31
[37:44] Dispensed 0.000141 mL of Base (0.5 M KOH)
[37:49] Stepping pH = 9.46
[37:49] Dispensed 0.000024 mL of Base (0.5 M KOH)
[37:55] Stepping pH = 9.51
[38:10] Stirrer speed set to 0
[38:24] Datapoint id 32 collected
[38:24] Charge balance equation is out by -88.1%
[38:24] Stirrer speed set to 50
[38:29] pH 9.52 -> 9.72
[38:29] Using cautious pH adjust
[38:29] Dispensed 0.000094 mL of Base (0.5 M KOH)
[38:34] Stepping pH = 9.57
[38:34] Dispensed 0.000141 mL of Base (0.5 M KOH)
[38:39] Stepping pH = 9.70
[38:39] Dispensed 0.000024 mL of Base (0.5 M KOH)
[38:44] Stepping pH = 9.73
[38:59] Stirrer speed set to 0

Sample name: **M15_octanol**
Assay name: **pH-metric high logP**
Assay ID: **18C-01005**
Filename: **C:\Sirius_T3\Mehtap\20180228_exp28_logP_T3-2\18C-01005_M15_octanol_pH-metric high logP.t3r**
Experiment start time: **3/1/2018 6:16:08 AM**
Analyst: **Pion**
Instrument ID: **T312060**

#### Experiment Log (continued)

[39:12] Datapoint id 33 collected
[39:12] Charge balance equation is out by -50.1%
[39:12] Stirrer speed set to 50
[39:17] pH 9.74 -> 9.94
[39:17] Using cautious pH adjust
[39:17] Dispensed 0.000141 mL of Base (0.5 M KOH)
[39:22] Stepping pH = 9.82
[39:22] Dispensed 0.000141 mL of Base (0.5 M KOH)
[39:27] Stepping pH = 9.91
[39:27] Dispensed 0.000047 mL of Base (0.5 M KOH)
[39:32] Stepping pH = 9.94
[39:48] Stirrer speed set to 0
[39:58] Datapoint id 34 collected
[39:58] Charge balance equation is out by -18.1%
[39:58] Stirrer speed set to 50
[40:03] pH 9.95 -> 10.05
[40:03] Using cautious pH adjust
[40:03] Dispensed 0.000094 mL of Base (0.5 M KOH)
[40:08] Stepping pH = 9.98
[40:08] Dispensed 0.000118 mL of Base (0.5 M KOH)
[40:13] Stepping pH = 10.04
[40:28] Stirrer speed set to 0
[40:38] Datapoint id 35 collected
[40:38] Charge balance equation is out by -23.6%
[40:38] Titration 2 of 3
[40:38] Adding initial titrants
[40:38] Automatically add 0.10000 mL of Octanol
[40:41] Dispensed 0.100000 mL of Octanol
[40:41] Stirrer speed set to 10
[40:42] Stirrer speed set to 55
[40:42] Iterative adjust 10.05 -> 2.00
[40:42] pH 10.05 -> 2.00
[40:44] Dispensed 0.100000 mL of Acid (0.5 M HCl)
[40:49] pH 2.03 -> 2.00
[40:49] Dispensed 0.005268 mL of Acid (0.5 M HCl)
[41:40] Stirrer speed set to 0
[41:50] Datapoint id 36 collected
[41:50] Stirrer speed set to 55
[41:55] pH 1.96 -> 2.16
[41:55] Using cautious pH adjust
[41:55] Dispensed 0.014722 mL of Base (0.5 M KOH)
[42:00] Stepping pH = 2.05
[42:01] Dispensed 0.011054 mL of Base (0.5 M KOH)
[42:06] Stepping pH = 2.14
[42:06] Dispensed 0.001999 mL of Base (0.5 M KOH)
[42:11] Stepping pH = 2.17
[42:26] Stirrer speed set to 0
[42:36] Datapoint id 37 collected
[42:36] Charge balance equation is out by 5.8%
[42:36] Stirrer speed set to 55
[42:41] pH 2.17 -> 2.37
[42:41] Using charge balance adjust
[42:42] Dispensed 0.018791 mL of Base (0.5 M KOH)
[43:02] Stirrer speed set to 0
[43:12] Datapoint id 38 collected
[43:12] Charge balance equation is out by 6.9%
[43:12] Stirrer speed set to 55
[43:17] pH 2.39 -> 2.59
[43:17] Using charge balance adjust

Sample name: **M15_octanol**
Assay name: **pH-metric high logP**
Assay ID: **18C-01005**
Filename: **C:\Sirius_T3\Mehtap\20180228_exp28_logP_T3-2\18C-01005_M15_octanol_pH-metric high logP.t3r**

Experiment start time: **3/1/2018 6:16:08 AM**
Analyst: **Pion**
Instrument ID: **T312060**

#### Experiment Log (continued)

[43:18] Dispensed 0.012371 mL of Base (0.5 M KOH)
[43:38] Stirrer speed set to 0
[43:48] Datapoint id 39 collected
[43:48] Charge balance equation is out by 4.3%
[43:48] Stirrer speed set to 55
[43:53] pH 2.61 -> 2.81
[43:53] Using charge balance adjust
[43:53] Dispensed 0.008514 mL of Base (0.5 M KOH)
[44:13] Stirrer speed set to 0
[44:23] Datapoint id 40 collected
[44:23] Charge balance equation is out by 6.2%
[44:23] Stirrer speed set to 55
[44:29] pH 2.83 -> 3.03
[44:29] Using charge balance adjust
[44:29] Dispensed 0.005974 mL of Base (0.5 M KOH)
[44:49] Stirrer speed set to 0
[44:59] Datapoint id 41 collected
[44:59] Charge balance equation is out by 9.4%
[44:59] Stirrer speed set to 55
[45:04] pH 3.05 -> 3.25
[45:04] Using charge balance adjust
[45:04] Dispensed 0.004210 mL of Base (0.5 M KOH)
[45:25] Stirrer speed set to 0
[45:35] Datapoint id 42 collected
[45:35] Charge balance equation is out by -1.8%
[45:35] Stirrer speed set to 55
[45:40] pH 3.25 -> 3.45
[45:40] Using charge balance adjust
[45:40] Dispensed 0.003151 mL of Base (0.5 M KOH)
[46:00] Stirrer speed set to 0
[46:10] Datapoint id 43 collected
[46:10] Charge balance equation is out by 8.8%
[46:10] Stirrer speed set to 55
[46:15] pH 3.47 -> 3.67
[46:15] Using charge balance adjust
[46:15] Dispensed 0.002399 mL of Base (0.5 M KOH)
[46:36] Stirrer speed set to 0
[46:46] Datapoint id 44 collected
[46:46] Charge balance equation is out by -5.2%
[46:46] Stirrer speed set to 55
[46:51] pH 3.67 -> 3.87
[46:51] Using charge balance adjust
[46:51] Dispensed 0.002070 mL of Base (0.5 M KOH)
[47:11] Stirrer speed set to 0
[47:22] Datapoint id 45 collected
[47:22] Charge balance equation is out by -7.7%
[47:22] Stirrer speed set to 55
[47:27] pH 3.86 -> 4.06
[47:27] Using charge balance adjust
[47:27] Dispensed 0.001999 mL of Base (0.5 M KOH)
[47:47] Stirrer speed set to 0
[47:57] Datapoint id 46 collected
[47:57] Charge balance equation is out by 5.7%
[47:57] Stirrer speed set to 55
[48:02] pH 4.07 -> 4.27
[48:02] Using charge balance adjust
[48:02] Dispensed 0.002117 mL of Base (0.5 M KOH)
[48:22] Stirrer speed set to 0
[48:32] Datapoint id 47 collected

| | | | |
|---|---|---|---|
| Sample name: | **M15_octanol** | Experiment start time: | **3/1/2018 6:16:08 AM** |
| Assay name: | **pH-metric high logP** | Analyst: | **Pion** |
| Assay ID: | **18C-01005** | Instrument ID: | **T312060** |
| Filename: | **C:\Sirius_T3\Mehtap\20180228_exp28_logP_T3-2\18C-01005_M15_octanol_pH-metric high logP.t3r** | | |

#### Experiment Log (continued)

[48:32] Charge balance equation is out by 24.4%
[48:32] Stirrer speed set to 55
[48:38] pH 4.31 -> 4.51
[48:38] Using cautious pH adjust
[48:38] Dispensed 0.001152 mL of Base (0.5 M KOH)
[48:43] Stepping pH = 4.41
[48:43] Dispensed 0.000870 mL of Base (0.5 M KOH)
[48:48] Stepping pH = 4.50
[48:48] Dispensed 0.000165 mL of Base (0.5 M KOH)
[48:53] Stepping pH = 4.51
[49:08] Stirrer speed set to 0
[49:18] Datapoint id 48 collected
[49:18] Charge balance equation is out by 5.4%
[49:18] Stirrer speed set to 55
[49:23] pH 4.51 -> 4.71
[49:23] Using charge balance adjust
[49:24] Dispensed 0.002352 mL of Base (0.5 M KOH)
[49:44] Stirrer speed set to 0
[49:54] Datapoint id 49 collected
[49:54] Charge balance equation is out by -0.6%
[49:54] Stirrer speed set to 55
[49:59] pH 4.71 -> 4.91
[49:59] Using charge balance adjust
[49:59] Dispensed 0.002211 mL of Base (0.5 M KOH)
[50:19] Stirrer speed set to 0
[50:29] Datapoint id 50 collected
[50:29] Charge balance equation is out by 5.6%
[50:29] Stirrer speed set to 55
[50:34] pH 4.93 -> 5.13
[50:34] Using charge balance adjust
[50:35] Dispensed 0.001881 mL of Base (0.5 M KOH)
[50:55] Stirrer speed set to 0
[51:05] Datapoint id 51 collected
[51:05] Charge balance equation is out by 13.1%
[51:05] Stirrer speed set to 55
[51:10] pH 5.16 -> 5.36
[51:10] Using charge balance adjust
[51:11] Dispensed 0.001435 mL of Base (0.5 M KOH)
[51:31] Stirrer speed set to 0
[51:41] Datapoint id 52 collected
[51:41] Charge balance equation is out by 15.8%
[51:41] Stirrer speed set to 55
[51:46] pH 5.39 -> 5.59
[51:46] Using cautious pH adjust
[51:46] Dispensed 0.000517 mL of Base (0.5 M KOH)
[51:52] Stepping pH = 5.45
[51:52] Dispensed 0.000635 mL of Base (0.5 M KOH)
[51:57] Stepping pH = 5.60
[52:12] Stirrer speed set to 0
[52:23] Datapoint id 53 collected
[52:23] Charge balance equation is out by -12.6%
[52:23] Stirrer speed set to 55
[52:28] pH 5.65 -> 5.85
[52:28] Using charge balance adjust
[52:28] Dispensed 0.000659 mL of Base (0.5 M KOH)
[52:48] Stirrer speed set to 0
[53:00] Datapoint id 54 collected
[53:00] Charge balance equation is out by 0.5%
[53:00] Stirrer speed set to 55

Sample name: **M15_octanol**
Assay name: **pH-metric high logP**
Assay ID: **18C-01005**
Filename: **C:\Sirius_T3\Mehtap\20180228_exp28_logP_T3-2\18C-01005_M15_octanol_pH-metric high logP.t3r**

Experiment start time: **3/1/2018 6:16:08 AM**
Analyst: **Pion**
Instrument ID: **T312060**

#### Experiment Log (continued)

[53:05] pH 5.86 -> 6.06
[53:05] Using charge balance adjust
[53:06] Dispensed 0.000447 mL of Base (0.5 M KOH)
[53:26] Stirrer speed set to 0
[53:39] Datapoint id 55 collected
[53:39] Charge balance equation is out by -8.3%
[53:39] Stirrer speed set to 55
[53:44] pH 6.05 -> 6.25
[53:44] Using charge balance adjust
[53:44] Dispensed 0.000329 mL of Base (0.5 M KOH)
[54:04] Stirrer speed set to 0
[54:18] Datapoint id 56 collected
[54:18] Charge balance equation is out by -4.8%
[54:18] Stirrer speed set to 55
[54:23] pH 6.24 -> 6.44
[54:23] Using charge balance adjust
[54:23] Dispensed 0.000235 mL of Base (0.5 M KOH)
[54:44] Stirrer speed set to 0
[55:11] Datapoint id 57 collected
[55:11] Charge balance equation is out by -4.5%
[55:11] Stirrer speed set to 55
[55:16] pH 6.43 -> 6.63
[55:16] Using charge balance adjust
[55:16] Dispensed 0.000165 mL of Base (0.5 M KOH)
[55:37] Stirrer speed set to 0
[56:04] Datapoint id 58 collected
[56:04] Charge balance equation is out by -2.3%
[56:04] Stirrer speed set to 55
[56:09] pH 6.63 -> 6.83
[56:09] Using charge balance adjust
[56:09] Dispensed 0.000118 mL of Base (0.5 M KOH)
[56:29] Stirrer speed set to 0
[57:08] Datapoint id 59 collected
[57:08] Charge balance equation is out by -7.5%
[57:08] Stirrer speed set to 55
[57:13] pH 6.81 -> 7.01
[57:13] Using charge balance adjust
[57:13] Dispensed 0.000094 mL of Base (0.5 M KOH)
[57:33] Stirrer speed set to 0
[58:22] Datapoint id 60 collected
[58:22] Charge balance equation is out by -25.4%
[58:22] Stirrer speed set to 55
[58:28] pH 6.94 -> 7.14
[58:28] Using cautious pH adjust
[58:28] Dispensed 0.000024 mL of Base (0.5 M KOH)
[58:33] Stepping pH = 6.96
[58:33] Dispensed 0.000094 mL of Base (0.5 M KOH)
[58:38] Stepping pH = 7.14
[58:53] Stirrer speed set to 0
[59:53] Datapoint id 61 collected
[59:53] Charge balance equation is out by -82.5%
[59:53] Stirrer speed set to 55
[59:58] pH 7.28 -> 7.48
[59:58] Using cautious pH adjust
[59:58] Dispensed 0.000024 mL of Base (0.5 M KOH)
[1:00:03] Stepping pH = 7.34
[1:00:03] Dispensed 0.000024 mL of Base (0.5 M KOH)
[1:00:08] Stepping pH = 7.41
[1:00:09] Dispensed 0.000024 mL of Base (0.5 M KOH)

Sample name: **M15_octanol**
Assay name: **pH-metric high logP**
Assay ID: **18C-01005**
Filename: **C:\Sirius_T3\Mehtap\20180228_exp28_logP_T3-2\18C-01005_M15_octanol_pH-metric high logP.t3r**
Experiment start time: **3/1/2018 6:16:08 AM**
Analyst: **Pion**
Instrument ID: **T312060**

#### Experiment Log (continued)

[1:00:14] Stepping pH = 7.51
[1:00:29] Stirrer speed set to 0
[1:01:29] Datapoint id 62 collected
[1:01:29] Charge balance equation is out by -78.6%
[1:01:29] Stirrer speed set to 55
[1:01:34] pH 7.53 -> 7.73
[1:01:34] Using cautious pH adjust
[1:01:34] Dispensed 0.000024 mL of Base (0.5 M KOH)
[1:01:39] Stepping pH = 7.61
[1:01:39] Dispensed 0.000024 mL of Base (0.5 M KOH)
[1:01:44] Stepping pH = 7.72
[1:01:59] Stirrer speed set to 0
[1:02:59] Datapoint id 63 collected
[1:02:59] Charge balance equation is out by -84.1%
[1:02:59] Stirrer speed set to 55
[1:03:04] pH 7.73 -> 7.93
[1:03:04] Using cautious pH adjust
[1:03:04] Dispensed 0.000024 mL of Base (0.5 M KOH)
[1:03:09] Stepping pH = 7.79
[1:03:10] Dispensed 0.000024 mL of Base (0.5 M KOH)
[1:03:15] Stepping pH = 7.89
[1:03:15] Dispensed 0.000024 mL of Base (0.5 M KOH)
[1:03:20] Stepping pH = 7.99
[1:03:35] Stirrer speed set to 0
[1:04:35] Datapoint id 64 collected
[1:04:35] Charge balance equation is out by -269.2%
[1:04:35] Stirrer speed set to 55
[1:04:40] pH 8.03 -> 8.23
[1:04:40] Using cautious pH adjust
[1:04:40] Dispensed 0.000024 mL of Base (0.5 M KOH)
[1:04:45] Stepping pH = 8.10
[1:04:45] Dispensed 0.000024 mL of Base (0.5 M KOH)
[1:04:50] Stepping pH = 8.18
[1:04:50] Dispensed 0.000024 mL of Base (0.5 M KOH)
[1:04:55] Stepping pH = 8.27
[1:05:10] Stirrer speed set to 0
[1:05:35] Datapoint id 65 collected
[1:05:35] Charge balance equation is out by -326.7%
[1:05:35] Stirrer speed set to 55
[1:05:40] pH 8.29 -> 8.49
[1:05:40] Using cautious pH adjust
[1:05:40] Dispensed 0.000024 mL of Base (0.5 M KOH)
[1:05:45] Stepping pH = 8.32
[1:05:45] Dispensed 0.000024 mL of Base (0.5 M KOH)
[1:05:50] Stepping pH = 8.36
[1:05:51] Dispensed 0.000047 mL of Base (0.5 M KOH)
[1:05:56] Stepping pH = 8.37
[1:05:56] Dispensed 0.000094 mL of Base (0.5 M KOH)
[1:06:01] Stepping pH = 8.53
[1:06:16] Stirrer speed set to 0
[1:06:45] Datapoint id 66 collected
[1:06:45] Charge balance equation is out by -901.8%
[1:06:45] Stirrer speed set to 55
[1:06:50] pH 8.59 -> 8.79
[1:06:50] Using cautious pH adjust
[1:06:50] Dispensed 0.000024 mL of Base (0.5 M KOH)
[1:06:55] Stepping pH = 8.62
[1:06:55] Dispensed 0.000024 mL of Base (0.5 M KOH)
[1:07:00] Stepping pH = 8.66

| | | | |
|---|---|---|---|
| Sample name: | **M15_octanol** | Experiment start time: | **3/1/2018 6:16:08 AM** |
| Assay name: | **pH-metric high logP** | Analyst: | **Pion** |
| Assay ID: | **18C-01005** | Instrument ID: | **T312060** |
| Filename: | **C:\Sirius_T3\Mehtap\20180228_exp28_logP_T3-2\18C-01005_M15_octanol_pH-metric high logP.t3r** | | |

#### Experiment Log (continued)

[1:07:00] Dispensed 0.000047 mL of Base (0.5 M KOH)
[1:07:05] Stepping pH = 8.73
[1:07:05] Dispensed 0.000024 mL of Base (0.5 M KOH)
[1:07:11] Stepping pH = 8.78
[1:07:26] Stirrer speed set to 0
[1:07:42] Datapoint id 67 collected
[1:07:42] Charge balance equation is out by -378.7%
[1:07:42] Stirrer speed set to 55
[1:07:47] pH 8.81 -> 9.01
[1:07:47] Using cautious pH adjust
[1:07:47] Dispensed 0.000024 mL of Base (0.5 M KOH)
[1:07:52] Stepping pH = 8.81
[1:07:53] Dispensed 0.000118 mL of Base (0.5 M KOH)
[1:07:58] Stepping pH = 8.81
[1:07:58] Dispensed 0.000564 mL of Base (0.5 M KOH)
[1:08:03] Stepping pH = 9.06
[1:08:18] Stirrer speed set to 0
[1:08:32] Datapoint id 68 collected
[1:08:32] Charge balance equation is out by -1,458.4%
[1:08:32] Stirrer speed set to 55
[1:08:37] pH 9.20 -> 9.40
[1:08:37] Using cautious pH adjust
[1:08:37] Dispensed 0.000047 mL of Base (0.5 M KOH)
[1:08:42] Stepping pH = 9.21
[1:08:42] Dispensed 0.000141 mL of Base (0.5 M KOH)
[1:08:47] Stepping pH = 9.28
[1:08:47] Dispensed 0.000165 mL of Base (0.5 M KOH)
[1:08:53] Stepping pH = 9.44
[1:09:08] Stirrer speed set to 0
[1:09:26] Datapoint id 69 collected
[1:09:26] Charge balance equation is out by -266.5%
[1:09:26] Stirrer speed set to 55
[1:09:31] pH 9.51 -> 9.71
[1:09:31] Using cautious pH adjust
[1:09:31] Dispensed 0.000094 mL of Base (0.5 M KOH)
[1:09:36] Stepping pH = 9.51
[1:09:36] Dispensed 0.000282 mL of Base (0.5 M KOH)
[1:09:41] Stepping pH = 9.70
[1:09:56] Stirrer speed set to 0
[1:10:09] Datapoint id 70 collected
[1:10:09] Charge balance equation is out by -98.1%
[1:10:09] Stirrer speed set to 55
[1:10:14] pH 9.79 -> 9.99
[1:10:14] Using cautious pH adjust
[1:10:14] Dispensed 0.000165 mL of Base (0.5 M KOH)
[1:10:19] Stepping pH = 9.80
[1:10:19] Dispensed 0.000447 mL of Base (0.5 M KOH)
[1:10:24] Stepping pH = 10.04
[1:10:39] Stirrer speed set to 0
[1:10:49] Datapoint id 71 collected
[1:10:49] Charge balance equation is out by -90.6%
[1:10:49] Titration 3 of 3
[1:10:49] Adding initial titrants
[1:10:49] Automatically add 0.30000 mL of Octanol
[1:10:56] Dispensed 0.300000 mL of Octanol
[1:10:56] Stirrer speed set to 10
[1:10:58] Stirrer speed set to 60
[1:10:58] Iterative adjust 10.07 -> 2.00
[1:10:58] pH 10.07 -> 2.00

Sample name: **M15_octanol**
Assay name: **pH-metric high logP**
Assay ID: **18C-01005**
Filename: **C:\Sirius_T3\Mehtap\20180228_exp28_logP_T3-2\18C-01005_M15_octanol_pH-metric high logP.t3r**

Experiment start time: **3/1/2018 6:16:08 AM**
Analyst: **Pion**
Instrument ID: **T312060**

#### Experiment Log (continued)

[1:11:00] Dispensed 0.100000 mL of Acid (0.5 M HCl)
[1:11:05] pH 2.05 -> 2.00
[1:11:05] Dispensed 0.007996 mL of Acid (0.5 M HCl)
[1:11:56] Stirrer speed set to 0
[1:12:06] Datapoint id 72 collected
[1:12:06] Stirrer speed set to 60
[1:12:11] pH 1.98 -> 2.18
[1:12:11] Using cautious pH adjust
[1:12:11] Dispensed 0.015334 mL of Base (0.5 M KOH)
[1:12:16] Stepping pH = 2.06
[1:12:17] Dispensed 0.011900 mL of Base (0.5 M KOH)
[1:12:22] Stepping pH = 2.16
[1:12:22] Dispensed 0.001834 mL of Base (0.5 M KOH)
[1:12:27] Stepping pH = 2.18
[1:12:42] Stirrer speed set to 0
[1:12:52] Datapoint id 73 collected
[1:12:52] Charge balance equation is out by 5.2%
[1:12:52] Stirrer speed set to 60
[1:12:57] pH 2.19 -> 2.39
[1:12:57] Using charge balance adjust
[1:12:58] Dispensed 0.019450 mL of Base (0.5 M KOH)
[1:13:18] Stirrer speed set to 0
[1:13:28] Datapoint id 74 collected
[1:13:28] Charge balance equation is out by 10.4%
[1:13:28] Stirrer speed set to 60
[1:13:33] pH 2.42 -> 2.62
[1:13:33] Using charge balance adjust
[1:13:34] Dispensed 0.012676 mL of Base (0.5 M KOH)
[1:13:54] Stirrer speed set to 0
[1:14:04] Datapoint id 75 collected
[1:14:04] Charge balance equation is out by 3.9%
[1:14:04] Stirrer speed set to 60
[1:14:09] pH 2.63 -> 2.83
[1:14:09] Using charge balance adjust
[1:14:09] Dispensed 0.008843 mL of Base (0.5 M KOH)
[1:14:29] Stirrer speed set to 0
[1:14:39] Datapoint id 76 collected
[1:14:39] Charge balance equation is out by 9.8%
[1:14:39] Stirrer speed set to 60
[1:14:45] pH 2.86 -> 3.06
[1:14:45] Using charge balance adjust
[1:14:45] Dispensed 0.006256 mL of Base (0.5 M KOH)
[1:15:05] Stirrer speed set to 0
[1:15:15] Datapoint id 77 collected
[1:15:15] Charge balance equation is out by -0.7%
[1:15:15] Stirrer speed set to 60
[1:15:20] pH 3.06 -> 3.26
[1:15:20] Using charge balance adjust
[1:15:20] Dispensed 0.004704 mL of Base (0.5 M KOH)
[1:15:41] Stirrer speed set to 0
[1:15:51] Datapoint id 78 collected
[1:15:51] Charge balance equation is out by 11.5%
[1:15:51] Stirrer speed set to 60
[1:15:56] pH 3.28 -> 3.48
[1:15:56] Using charge balance adjust
[1:15:56] Dispensed 0.003622 mL of Base (0.5 M KOH)
[1:16:16] Stirrer speed set to 0
[1:16:26] Datapoint id 79 collected
[1:16:26] Charge balance equation is out by 4.3%

| | | | |
|---|---|---|---|
| Sample name: | **M15_octanol** | Experiment start time: | **3/1/2018 6:16:08 AM** |
| Assay name: | **pH-metric high logP** | Analyst: | **Pion** |
| Assay ID: | **18C-01005** | Instrument ID: | **T312060** |
| Filename: | **C:\Sirius_T3\Mehtap\20180228_exp28_logP_T3-2\18C-01005_M15_octanol_pH-metric high logP.t3r** | | |

#### Experiment Log (continued)

[1:16:26] Stirrer speed set to 60
[1:16:31] pH 3.50 -> 3.70
[1:16:31] Using charge balance adjust
[1:16:31] Dispensed 0.003057 mL of Base (0.5 M KOH)
[1:16:52] Stirrer speed set to 0
[1:17:02] Datapoint id 80 collected
[1:17:02] Charge balance equation is out by 17.0%
[1:17:02] Stirrer speed set to 60
[1:17:07] pH 3.73 -> 3.93
[1:17:07] Using cautious pH adjust
[1:17:07] Dispensed 0.001411 mL of Base (0.5 M KOH)
[1:17:12] Stepping pH = 3.83
[1:17:12] Dispensed 0.001011 mL of Base (0.5 M KOH)
[1:17:17] Stepping pH = 3.91
[1:17:17] Dispensed 0.000188 mL of Base (0.5 M KOH)
[1:17:22] Stepping pH = 3.93
[1:17:37] Stirrer speed set to 0
[1:17:48] Datapoint id 81 collected
[1:17:48] Charge balance equation is out by 7.0%
[1:17:48] Stirrer speed set to 60
[1:17:53] pH 3.92 -> 4.12
[1:17:53] Using charge balance adjust
[1:17:53] Dispensed 0.002752 mL of Base (0.5 M KOH)
[1:18:13] Stirrer speed set to 0
[1:18:23] Datapoint id 82 collected
[1:18:23] Charge balance equation is out by 8.4%
[1:18:23] Stirrer speed set to 60
[1:18:29] pH 4.14 -> 4.34
[1:18:29] Using charge balance adjust
[1:18:29] Dispensed 0.002611 mL of Base (0.5 M KOH)
[1:18:49] Stirrer speed set to 0
[1:18:59] Datapoint id 83 collected
[1:18:59] Charge balance equation is out by 20.8%
[1:18:59] Stirrer speed set to 60
[1:19:04] pH 4.38 -> 4.58
[1:19:04] Using cautious pH adjust
[1:19:04] Dispensed 0.001152 mL of Base (0.5 M KOH)
[1:19:09] Stepping pH = 4.47
[1:19:09] Dispensed 0.000847 mL of Base (0.5 M KOH)
[1:19:15] Stepping pH = 4.56
[1:19:15] Dispensed 0.000141 mL of Base (0.5 M KOH)
[1:19:20] Stepping pH = 4.58
[1:19:35] Stirrer speed set to 0
[1:19:45] Datapoint id 84 collected
[1:19:45] Charge balance equation is out by 7.2%
[1:19:45] Stirrer speed set to 60
[1:19:50] pH 4.57 -> 4.77
[1:19:50] Using charge balance adjust
[1:19:50] Dispensed 0.001929 mL of Base (0.5 M KOH)
[1:20:10] Stirrer speed set to 0
[1:20:20] Datapoint id 85 collected
[1:20:20] Charge balance equation is out by -8.5%
[1:20:20] Stirrer speed set to 60
[1:20:25] pH 4.75 -> 4.95
[1:20:25] Using charge balance adjust
[1:20:26] Dispensed 0.001505 mL of Base (0.5 M KOH)
[1:20:46] Stirrer speed set to 0
[1:20:56] Datapoint id 86 collected
[1:20:56] Charge balance equation is out by -4.5%

Sample name: **M15_octanol**
Assay name: **pH-metric high logP**
Assay ID: **18C-01005**
Filename: **C:\Sirius_T3\Mehtap\20180228_exp28_logP_T3-2\18C-01005_M15_octanol_pH-metric high logP.t3r**
Experiment start time: **3/1/2018 6:16:08 AM**
Analyst: **Pion**
Instrument ID: **T312060**

#### Experiment Log (continued)

[1:20:56] Stirrer speed set to 60
[1:21:01] pH 4.95 -> 5.15
[1:21:01] Using charge balance adjust
[1:21:01] Dispensed 0.001129 mL of Base (0.5 M KOH)
[1:21:22] Stirrer speed set to 0
[1:21:32] Datapoint id 87 collected
[1:21:32] Charge balance equation is out by -5.7%
[1:21:32] Stirrer speed set to 60
[1:21:37] pH 5.14 -> 5.34
[1:21:37] Using charge balance adjust
[1:21:37] Dispensed 0.000800 mL of Base (0.5 M KOH)
[1:21:58] Stirrer speed set to 0
[1:22:09] Datapoint id 88 collected
[1:22:09] Charge balance equation is out by -0.8%
[1:22:09] Stirrer speed set to 60
[1:22:14] pH 5.34 -> 5.54
[1:22:14] Using charge balance adjust
[1:22:14] Dispensed 0.000541 mL of Base (0.5 M KOH)
[1:22:34] Stirrer speed set to 0
[1:22:46] Datapoint id 89 collected
[1:22:46] Charge balance equation is out by 1.2%
[1:22:46] Stirrer speed set to 60
[1:22:51] pH 5.55 -> 5.75
[1:22:51] Using charge balance adjust
[1:22:51] Dispensed 0.000376 mL of Base (0.5 M KOH)
[1:23:11] Stirrer speed set to 0
[1:23:29] Datapoint id 90 collected
[1:23:29] Charge balance equation is out by 9.5%
[1:23:29] Stirrer speed set to 60
[1:23:34] pH 5.77 -> 5.97
[1:23:34] Using charge balance adjust
[1:23:34] Dispensed 0.000259 mL of Base (0.5 M KOH)
[1:23:54] Stirrer speed set to 0
[1:24:16] Datapoint id 91 collected
[1:24:16] Charge balance equation is out by 7.6%
[1:24:16] Stirrer speed set to 60
[1:24:22] pH 6.00 -> 6.20
[1:24:22] Using charge balance adjust
[1:24:22] Dispensed 0.000188 mL of Base (0.5 M KOH)
[1:24:42] Stirrer speed set to 0
[1:25:12] Datapoint id 92 collected
[1:25:12] Charge balance equation is out by 18.0%
[1:25:12] Stirrer speed set to 60
[1:25:18] pH 6.23 -> 6.43
[1:25:18] Using cautious pH adjust
[1:25:18] Dispensed 0.000071 mL of Base (0.5 M KOH)
[1:25:23] Stepping pH = 6.29
[1:25:23] Dispensed 0.000094 mL of Base (0.5 M KOH)
[1:25:28] Stepping pH = 6.50
[1:25:43] Stirrer speed set to 0
[1:26:43] Datapoint id 93 collected
[1:26:43] Charge balance equation is out by -15.4%
[1:26:43] Stirrer speed set to 60
[1:26:48] pH 6.54 -> 6.74
[1:26:48] Using cautious pH adjust
[1:26:48] Dispensed 0.000047 mL of Base (0.5 M KOH)
[1:26:53] Stepping pH = 6.63
[1:26:53] Dispensed 0.000047 mL of Base (0.5 M KOH)
[1:26:58] Stepping pH = 6.80

Sample name: **M15_octanol**
Assay name: **pH-metric high logP**
Assay ID: **18C-01005**
Filename: **C:\Sirius_T3\Mehtap\20180228_exp28_logP_T3-2\18C-01005_M15_octanol_pH-metric high logP.t3r**
Experiment start time: **3/1/2018 6:16:08 AM**
Analyst: **Pion**
Instrument ID: **T312060**

#### Experiment Log (continued)

[1:27:14] Stirrer speed set to 0
[1:28:14] Datapoint id 94 collected
[1:28:14] Charge balance equation is out by 12.0%
[1:28:14] Stirrer speed set to 60
[1:28:19] pH 6.78 -> 6.98
[1:28:19] Using charge balance adjust
[1:28:19] Dispensed 0.000071 mL of Base (0.5 M KOH)
[1:28:39] Stirrer speed set to 0
[1:29:39] Datapoint id 95 collected
[1:29:39] Charge balance equation is out by -11.1%
[1:29:39] Stirrer speed set to 60
[1:29:44] pH 6.97 -> 7.17
[1:29:44] Using charge balance adjust
[1:29:44] Dispensed 0.000047 mL of Base (0.5 M KOH)
[1:30:04] Stirrer speed set to 0
[1:30:15] Datapoint id 96 collected
[1:30:15] Charge balance equation is out by -76.1%
[1:30:15] Stirrer speed set to 60
[1:30:20] pH 6.91 -> 7.11
[1:30:20] Using cautious pH adjust
[1:30:20] Dispensed 0.000024 mL of Base (0.5 M KOH)
[1:30:25] Stepping pH = 6.90
[1:30:25] Dispensed 0.000141 mL of Base (0.5 M KOH)
[1:30:30] Stepping pH = 7.91
[1:30:45] Stirrer speed set to 0
[1:31:45] Datapoint id 97 collected
[1:31:45] Charge balance equation is out by -214.0%
[1:31:45] Stirrer speed set to 60
[1:31:50] pH 8.09 -> 8.29
[1:31:50] Using cautious pH adjust
[1:31:51] Dispensed 0.000024 mL of Base (0.5 M KOH)
[1:31:56] Stepping pH = 8.14
[1:31:56] Dispensed 0.000024 mL of Base (0.5 M KOH)
[1:32:01] Stepping pH = 8.14
[1:32:01] Dispensed 0.000071 mL of Base (0.5 M KOH)
[1:32:06] Stepping pH = 8.36
[1:32:21] Stirrer speed set to 0
[1:32:45] Datapoint id 98 collected
[1:32:45] Charge balance equation is out by -580.1%
[1:32:45] Stirrer speed set to 60
[1:32:50] pH 8.43 -> 8.63
[1:32:50] Using cautious pH adjust
[1:32:50] Dispensed 0.000024 mL of Base (0.5 M KOH)
[1:32:55] Stepping pH = 8.45
[1:32:55] Dispensed 0.000024 mL of Base (0.5 M KOH)
[1:33:00] Stepping pH = 8.48
[1:33:00] Dispensed 0.000071 mL of Base (0.5 M KOH)
[1:33:05] Stepping pH = 8.61
[1:33:05] Dispensed 0.000024 mL of Base (0.5 M KOH)
[1:33:10] Stepping pH = 8.68
[1:33:25] Stirrer speed set to 0
[1:33:41] Datapoint id 99 collected
[1:33:41] Charge balance equation is out by -464.7%
[1:33:41] Stirrer speed set to 60
[1:33:46] pH 8.69 -> 8.89
[1:33:46] Using cautious pH adjust
[1:33:46] Dispensed 0.000024 mL of Base (0.5 M KOH)
[1:33:51] Stepping pH = 8.71
[1:33:51] Dispensed 0.000047 mL of Base (0.5 M KOH)

Sample name: **M15_octanol**
Assay name: **pH-metric high logP**
Assay ID: **18C-01005**
Filename: **C:\Sirius_T3\Mehtap\20180228_exp28_logP_T3-2\18C-01005_M15_octanol_pH-metric high logP.t3r**
Experiment start time: **3/1/2018 6:16:08 AM**
Analyst: **Pion**
Instrument ID: **T312060**

#### Experiment Log (continued)

[1:33:56] Stepping pH = 8.77
[1:33:56] Dispensed 0.000071 mL of Base (0.5 M KOH)
[1:34:01] Stepping pH = 8.90
[1:34:17] Stirrer speed set to 0
[1:34:31] Datapoint id 100 collected
[1:34:31] Charge balance equation is out by -252.1%
[1:34:31] Stirrer speed set to 60
[1:34:36] pH 8.96 -> 9.16
[1:34:36] Using cautious pH adjust
[1:34:36] Dispensed 0.000024 mL of Base (0.5 M KOH)
[1:34:41] Stepping pH = 8.98
[1:34:41] Dispensed 0.000094 mL of Base (0.5 M KOH)
[1:34:46] Stepping pH = 9.09
[1:34:46] Dispensed 0.000071 mL of Base (0.5 M KOH)
[1:34:52] Stepping pH = 9.18
[1:35:07] Stirrer speed set to 0
[1:35:20] Datapoint id 101 collected
[1:35:20] Charge balance equation is out by -179.3%
[1:35:20] Stirrer speed set to 60
[1:35:25] pH 9.23 -> 9.43
[1:35:25] Using cautious pH adjust
[1:35:25] Dispensed 0.000047 mL of Base (0.5 M KOH)
[1:35:30] Stepping pH = 9.24
[1:35:30] Dispensed 0.000165 mL of Base (0.5 M KOH)
[1:35:35] Stepping pH = 9.36
[1:35:35] Dispensed 0.000071 mL of Base (0.5 M KOH)
[1:35:40] Stepping pH = 9.43
[1:35:55] Stirrer speed set to 0
[1:36:07] Datapoint id 102 collected
[1:36:07] Charge balance equation is out by -161.3%
[1:36:07] Stirrer speed set to 60
[1:36:12] pH 9.46 -> 9.66
[1:36:12] Using cautious pH adjust
[1:36:12] Dispensed 0.000094 mL of Base (0.5 M KOH)
[1:36:17] Stepping pH = 9.47
[1:36:17] Dispensed 0.000259 mL of Base (0.5 M KOH)
[1:36:22] Stepping pH = 9.67
[1:36:38] Stirrer speed set to 0
[1:36:49] Datapoint id 103 collected
[1:36:49] Charge balance equation is out by -94.3%
[1:36:49] Stirrer speed set to 60
[1:36:54] pH 9.73 -> 9.93
[1:36:54] Using cautious pH adjust
[1:36:54] Dispensed 0.000141 mL of Base (0.5 M KOH)
[1:36:59] Stepping pH = 9.76
[1:36:59] Dispensed 0.000306 mL of Base (0.5 M KOH)
[1:37:05] Stepping pH = 9.91
[1:37:05] Dispensed 0.000047 mL of Base (0.5 M KOH)
[1:37:10] Stepping pH = 9.95
[1:37:25] Stirrer speed set to 0
[1:37:36] Datapoint id 104 collected
[1:37:36] Charge balance equation is out by -63.7%
[1:37:36] Stirrer speed set to 60
[1:37:41] pH 9.97 -> 10.05
[1:37:41] Using cautious pH adjust
[1:37:41] Dispensed 0.000094 mL of Base (0.5 M KOH)
[1:37:46] Stepping pH = 9.98
[1:37:46] Dispensed 0.000165 mL of Base (0.5 M KOH)
[1:37:51] Stepping pH = 10.04

| | | | |
|---|---|---|---|
| Sample name: | **M15_octanol** | Experiment start time: | **3/1/2018 6:16:08 AM** |
| Assay name: | **pH-metric high logP** | Analyst: | **Pion** |
| Assay ID: | **18C-01005** | Instrument ID: | **T312060** |
| Filename: | **C:\Sirius_T3\Mehtap\20180228_exp28_logP_T3-2\18C-01005_M15_octanol_pH-metric high logP.t3r** | | |

#### Experiment Log (continued)

[1:38:06] Stirrer speed set to 0
[1:38:17] Datapoint id 105 collected
[1:38:17] Charge balance equation is out by -43.3%
[1:38:17] Argon flow rate set to 0
[1:38:21] Titrator arm moved over Titration position
