## Supplementary material for "Octanol-water partition coefficient measurements for the SAMPL6 Blind Prediction Challenge": SM14_18C-03006_M15_octanol_pH-metric high logP_report.pdf

| | | | |
|---|---|---|---|
| Sample name: | **M15_octanol** | Experiment start time: | **3/3/2018 8:26:22 AM** |
| Assay name: | **pH-metric high logP** | Analyst: | **Pion** |
| Assay ID: | **18C-03006** | Instrument ID: | **T312060** |
| Filename: | **C:\Sirius_T3\Mehtap\20180302_exp29_logP_T3-2\18C-03006_M15_octanol_pH-metric high logP.t3r** | | |

#### pH-metric Result

| | | |
|---|---|---|
| logP (XH2 2+) | -5.50 | |
| logP (XH +) | 0.28 | ±0.16 (n=50) |
| logP (neutral X) | 2.04 | ±0.02 (n=50) |

##### 18C-03006 Points 1 to 32

| | |
|---|---|
| M15_octanol concentration factor | 0.894 |
| Carbonate | 0.2191 mM |
| Acidity error | -0.25037 mM |

##### 18C-03006 Points 33 to 65

| | |
|---|---|
| M15_octanol concentration factor | 0.950 |
| Carbonate | 0.2177 mM |
| Acidity error | -0.29629 mM |

##### 18C-03006 Points 66 to 99

| | |
|---|---|
| M15_octanol concentration factor | 1.015 |
| Carbonate | 0.2265 mM |
| Acidity error | -0.47876 mM |

#### Warnings and errors

| | |
|---|---|
| Errors | None |
| Warnings | One or more logP values out of range |

#### Sample logD and percent species

| pH | M15_octanol logD | M15_octanol M15_octanolH2 | M15_octanol M15_octanolH | M15_octanol M15_octanol | M15_octanol M15_octanolH2* | M15_octanol M15_octanolH* | M15_octanol M15_octanol* | Comment |
|---|---|---|---|---|---|---|---|---|
| 1.000 | -1.31 | 92.86 % | 2.44 % | 0.00 % | 0.00 % | 4.68 % | 0.01 % | |
| 1.200 | -1.11 | 89.13 % | 3.72 % | 0.00 % | 0.00 % | 7.12 % | 0.03 % | Stomach pH |
| 2.000 | -0.39 | 56.12 % | 14.76 % | 0.01 % | 0.00 % | 28.29 % | 0.82 % | |
| 3.000 | 0.25 | 9.86 % | 25.93 % | 0.13 % | 0.00 % | 49.68 % | 14.41 % | |
| 4.000 | 0.84 | 0.44 % | 11.68 % | 0.59 % | 0.00 % | 22.38 % | 64.91 % | |
| 5.000 | 1.58 | 0.01 % | 1.70 % | 0.85 % | 0.00 % | 3.25 % | 94.20 % | |
| 6.000 | 1.97 | 0.00 % | 0.18 % | 0.89 % | 0.00 % | 0.34 % | 98.59 % | |
| 6.500 | 2.02 | 0.00 % | 0.06 % | 0.89 % | 0.00 % | 0.11 % | 98.94 % | |
| 7.000 | 2.04 | 0.00 % | 0.02 % | 0.89 % | 0.00 % | 0.03 % | 99.05 % | |
| 7.400 | 2.04 | 0.00 % | 0.01 % | 0.89 % | 0.00 % | 0.01 % | 99.09 % | Blood pH |
| 8.000 | 2.04 | 0.00 % | 0.00 % | 0.89 % | 0.00 % | 0.00 % | 99.10 % | |
| 9.000 | 2.04 | 0.00 % | 0.00 % | 0.89 % | 0.00 % | 0.00 % | 99.11 % | |
| 10.000 | 2.04 | 0.00 % | 0.00 % | 0.89 % | 0.00 % | 0.00 % | 99.11 % | |
| 11.000 | 2.04 | 0.00 % | 0.00 % | 0.89 % | 0.00 % | 0.00 % | 99.11 % | |
| 12.000 | 2.04 | 0.00 % | 0.00 % | 0.89 % | 0.00 % | 0.00 % | 99.11 % | |

| | | | |
|---|---|---|---|
| Sample name: | **M15_octanol** | Experiment start time: | **3/3/2018 8:26:22 AM** |
| Assay name: | **pH-metric high logP** | Analyst: | **Pion** |
| Assay ID: | **18C-03006** | Instrument ID: | **T312060** |
| Filename: | **C:\Sirius_T3\Mehtap\20180302_exp29_logP_T3-2\18C-03006_M15_octanol_pH-metric high logP.t3r** | | |

#### Graphs

Charge balance residuals

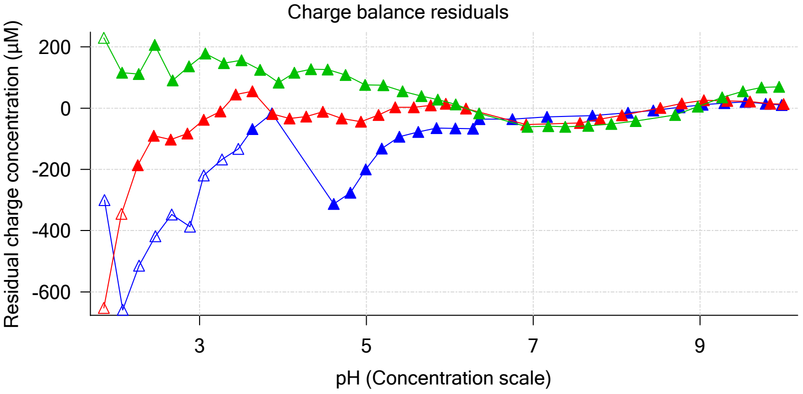


Ionic strength

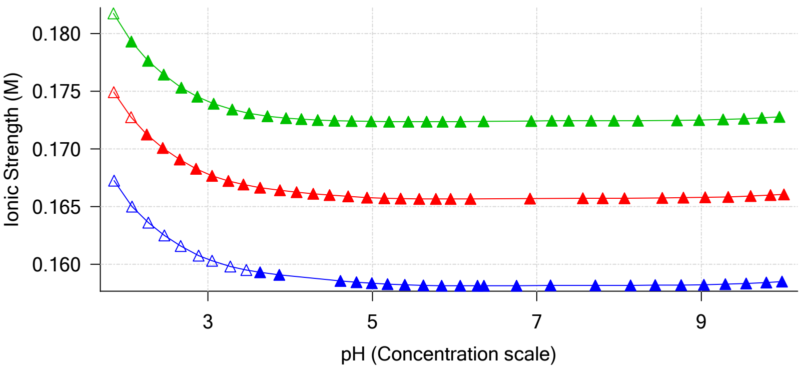


Buffer index profile

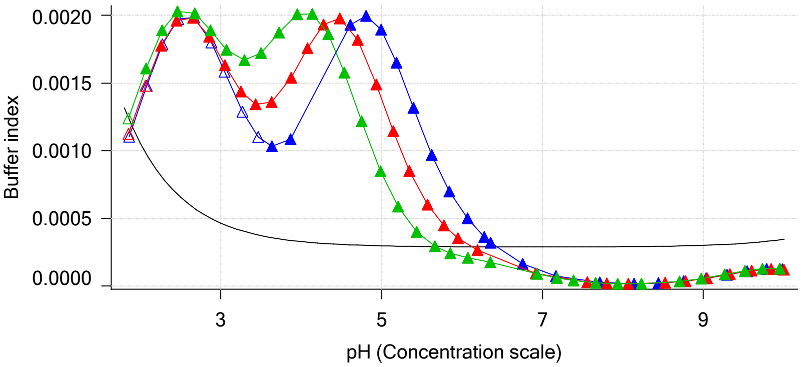


Sample M15_octanol Ionisation Graph

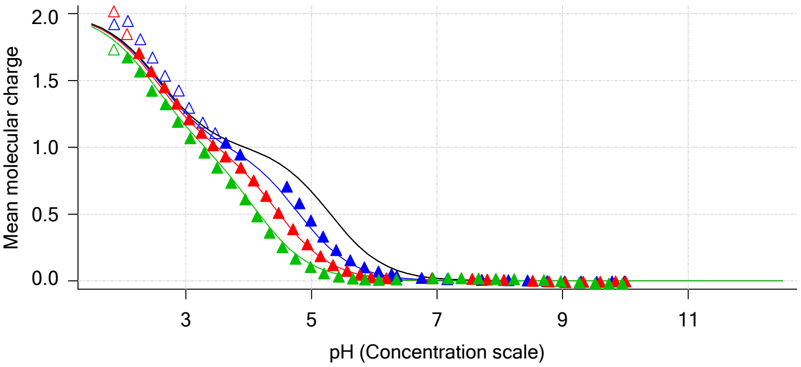


Distribution of species for sample M15_octanol

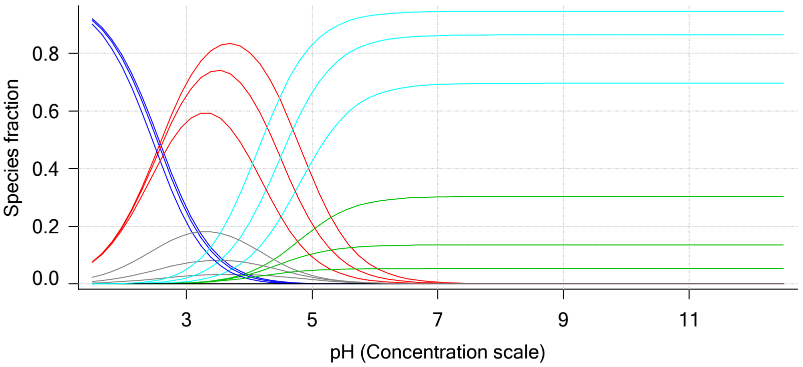


Lipophilicity Profile

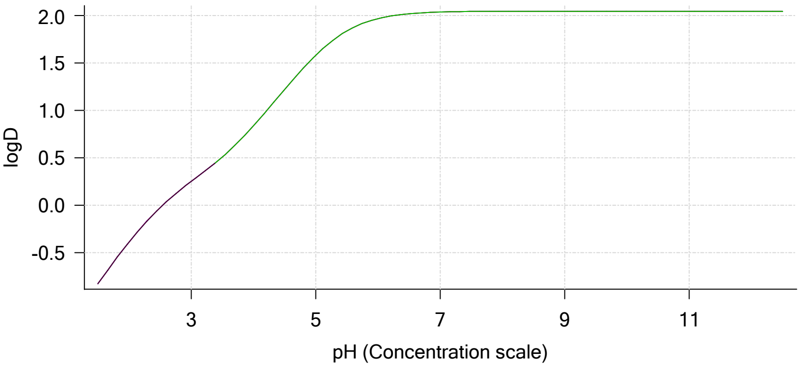


Log R Profile

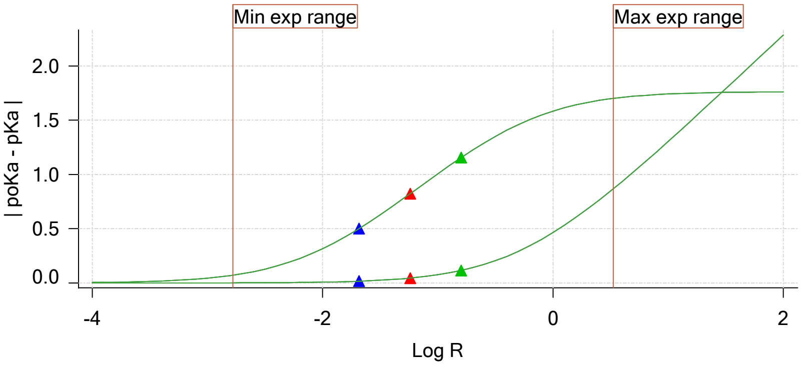


Sample Carbonate Ionisation Graph

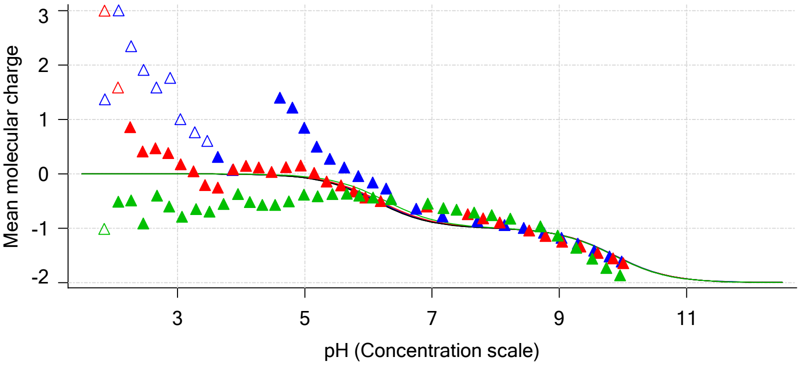

| | | | |
|---|---|---|---|
| Sample name: | **M15_octanol** | Experiment start time: | **3/3/2018 8:26:22 AM** |
| Assay name: | **pH-metric high logP** | Analyst: | **Pion** |
| Assay ID: | **18C-03006** | Instrument ID: | **T312060** |
| Filename: | **C:\Sirius_T3\Mehtap\20180302_exp29_logP_T3-2\18C-03006_M15_octanol_pH-metric high logP.t3r** | | |

#### Graphs (continued)

Distribution of species for sample Carbonate

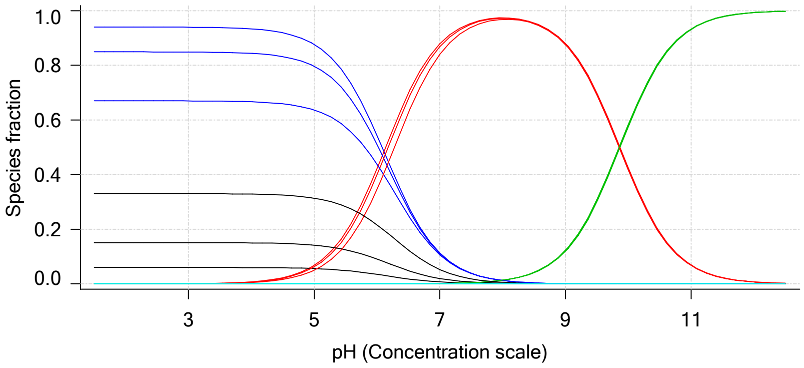

| | | | |
|---|---|---|---|
| Sample name: | **M15_octanol** | Experiment start time: | **3/3/2018 8:26:22 AM** |
| Assay name: | **pH-metric high logP** | Analyst: | **Pion** |
| Assay ID: | **18C-03006** | Instrument ID: | **T312060** |
| Filename: | **C:\Sirius_T3\Mehtap\20180302_exp29_logP_T3-2\18C-03006_M15_octanol_pH-metric high logP.t3r** | | |

# pH-metric high logP   Titration 1 of 3   18C-03006 Points 1 to 32

#### Overall results

| | |
|---|---|
| RMSD | 0.221 |
| Average ionic strength | 0.158 M |
| Average temperature | 25.0°C |
| Partition ratio | 0.0207 : 1 |
| Analyte concentration range | 3659.4 µM to 3810.2 µM |
| Total points considered | 23 of 32 |

#### Warnings and errors

| | |
|---|---|
| Errors | None |
| Warnings | One or more logP values out of range |

#### Four-Plus parameters

| | | | |
|---|---|---|---|
| Alpha | 0.111 | 3/3/2018 8:26:22 AM | C:\Sirius_T3\HCl18C02.t3r |
| S | 0.9988 | 3/3/2018 8:26:22 AM | C:\Sirius_T3\HCl18C02.t3r |
| jH | 1.0 | 3/3/2018 8:26:22 AM | C:\Sirius_T3\HCl18C02.t3r |
| jOH | -0.8 | 3/3/2018 8:26:22 AM | C:\Sirius_T3\HCl18C02.t3r |

#### Titrants

| | | | |
|---|---|---|---|
| 0.50 M HCl | 0.999058 | 3/3/2018 8:26:22 AM | C:\Sirius_T3\HCl18C02.t3r |
| 0.50 M KOH | 0.999845 | 3/3/2018 8:26:22 AM | C:\Sirius_T3\KOH18B27.t3r |

#### Sample

| | |
|---|---|
| M15_octanol concentration factor | 0.894 |
| Base pKa 1 | 2.58 |
| Base pKa 2 | 5.30 |
| logP (XH2 2+) | -5.50 |
| logP (XH +) | -2.63 |
| logP (neutral X) | 1.84 |

#### Sample graphs

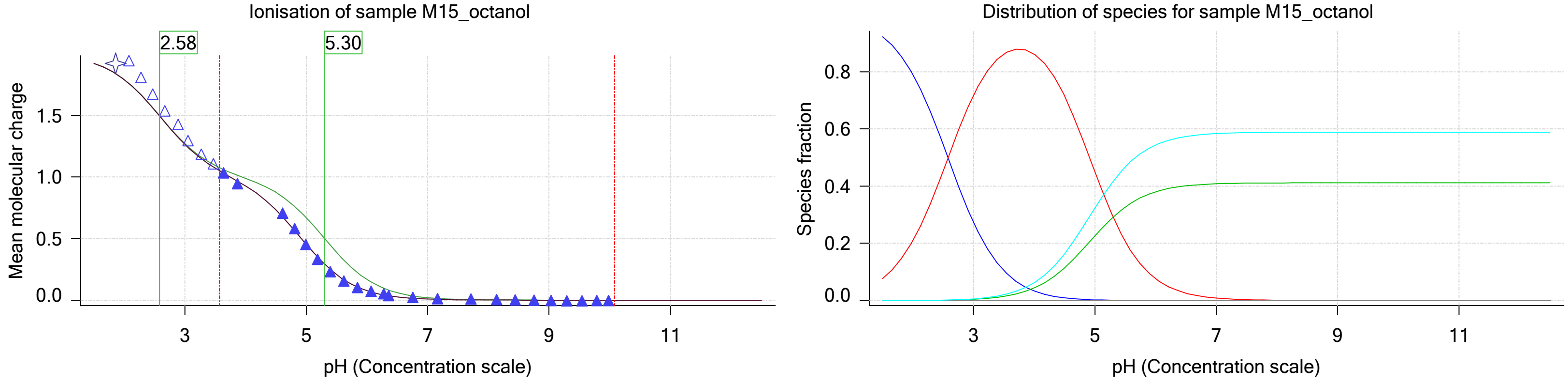

| | | | |
|---|---|---|---|
| Sample name: | **M15_octanol** | Experiment start time: | **3/3/2018 8:26:22 AM** |
| Assay name: | **pH-metric high logP** | Analyst: | **Pion** |
| Assay ID: | **18C-03006** | Instrument ID: | **T312060** |
| Filename: | **C:\Sirius_T3\Mehtap\20180302_exp29_logP_T3-2\18C-03006_M15_octanol_pH-metric high logP.t3r** | | |

#### Sample graphs (continued)

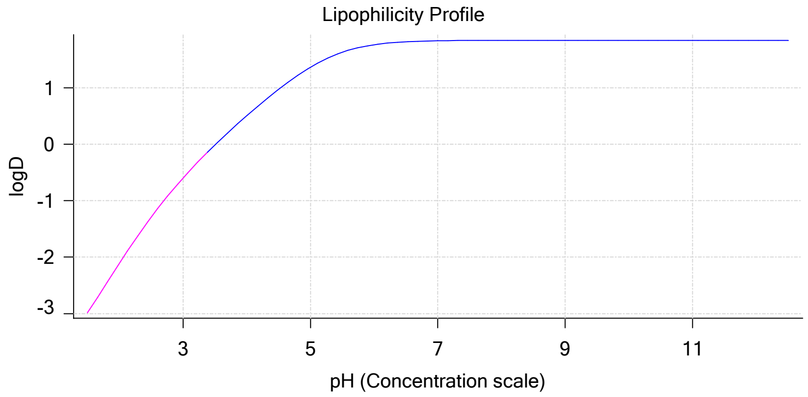


#### Sample logD and percent species

| pH | M15_octanol logD | M15_octanol M15_octanolH2 | M15_octanol M15_octanolH | M15_octanol M15_octanol | M15_octanol M15_octanolH2* | M15_octanol M15_octanolH* | M15_octanol M15_octanol* | Comment |
|---|---|---|---|---|---|---|---|---|
| 1.000 | -3.82 | 97.44 % | 2.56 % | 0.00 % | 0.00 % | 0.00 % | 0.00 % | |
| 1.200 | -3.50 | 96.00 % | 4.00 % | 0.00 % | 0.00 % | 0.00 % | 0.00 % | Stomach pH |
| 2.000 | -2.11 | 79.15 % | 20.82 % | 0.01 % | 0.00 % | 0.00 % | 0.01 % | |
| 3.000 | -0.60 | 27.30 % | 71.82 % | 0.36 % | 0.00 % | 0.00 % | 0.52 % | |
| 4.000 | 0.50 | 3.28 % | 86.21 % | 4.32 % | 0.00 % | 0.00 % | 6.18 % | |
| 5.000 | 1.36 | 0.17 % | 45.00 % | 22.55 % | 0.00 % | 0.00 % | 32.28 % | |
| 6.000 | 1.76 | 0.00 % | 7.58 % | 38.01 % | 0.00 % | 0.00 % | 54.40 % | |
| 6.500 | 1.81 | 0.00 % | 2.53 % | 40.09 % | 0.00 % | 0.00 % | 57.38 % | |
| 7.000 | 1.83 | 0.00 % | 0.81 % | 40.80 % | 0.00 % | 0.00 % | 58.39 % | |
| 7.400 | 1.84 | 0.00 % | 0.33 % | 41.00 % | 0.00 % | 0.00 % | 58.68 % | Blood pH |
| 8.000 | 1.84 | 0.00 % | 0.08 % | 41.10 % | 0.00 % | 0.00 % | 58.82 % | |
| 9.000 | 1.84 | 0.00 % | 0.01 % | 41.13 % | 0.00 % | 0.00 % | 58.86 % | |
| 10.000 | 1.84 | 0.00 % | 0.00 % | 41.13 % | 0.00 % | 0.00 % | 58.87 % | |
| 11.000 | 1.84 | 0.00 % | 0.00 % | 41.13 % | 0.00 % | 0.00 % | 58.87 % | |
| 12.000 | 1.84 | 0.00 % | 0.00 % | 41.13 % | 0.00 % | 0.00 % | 58.87 % | |

#### Carbonate and acidity

| | |
|---|---|
| Carbonate | 0.219 mM |
| Acidity error | -0.250 mM |

#### Other graphs

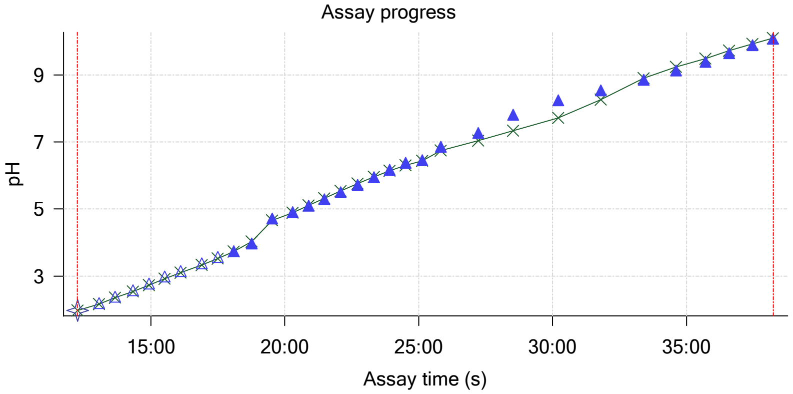


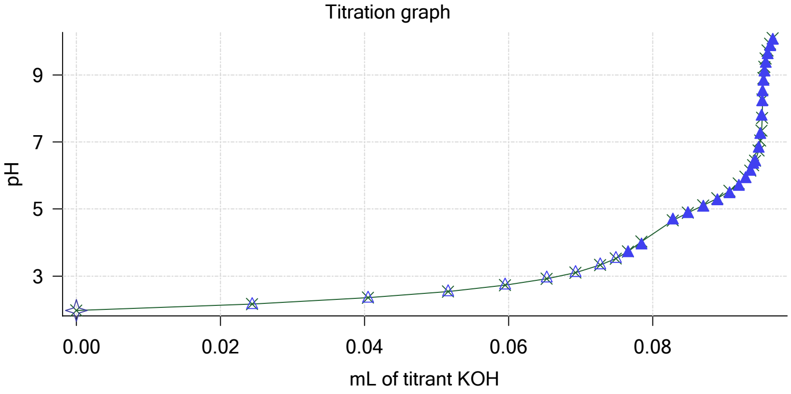

| | | | |
|---|---|---|---|
| Sample name: | **M15_octanol** | Experiment start time: | **3/3/2018 8:26:22 AM** |
| Assay name: | **pH-metric high logP** | Analyst: | **Pion** |
| Assay ID: | **18C-03006** | Instrument ID: | **T312060** |
| Filename: | **C:\Sirius_T3\Mehtap\20180302_exp29_logP_T3-2\18C-03006_M15_octanol_pH-metric high logP.t3r** | | |

#### Other graphs (continued)

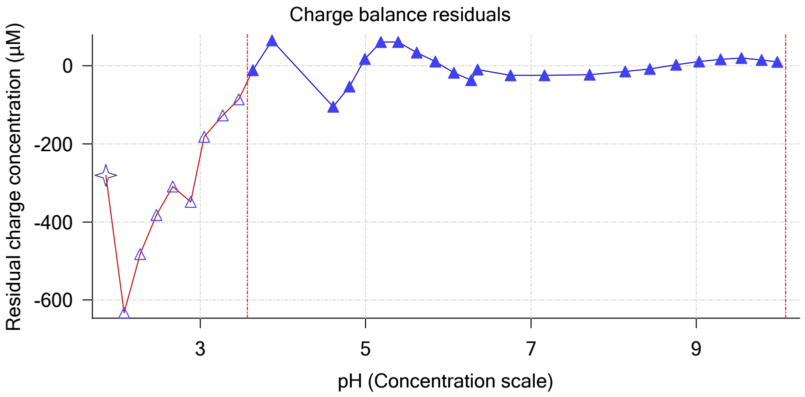


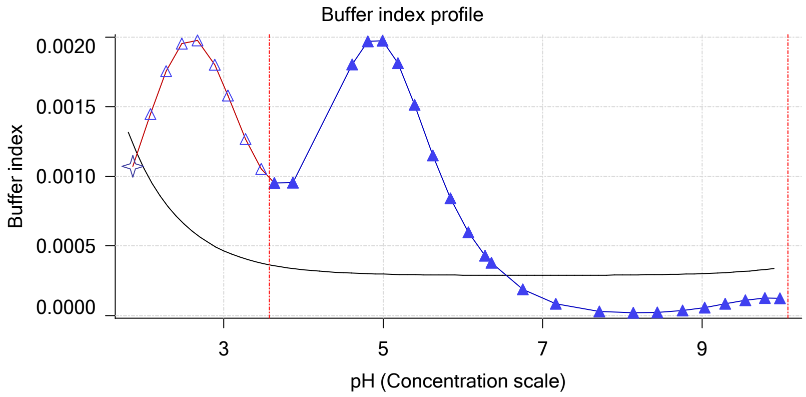


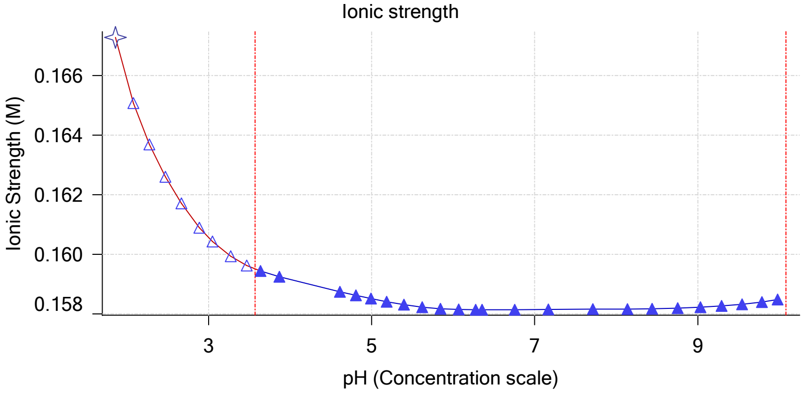


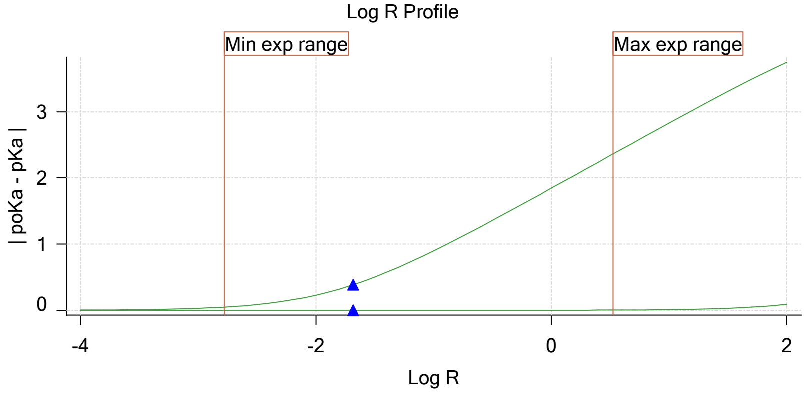


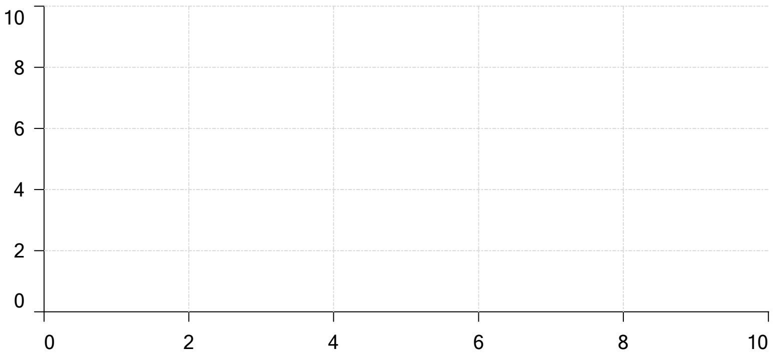


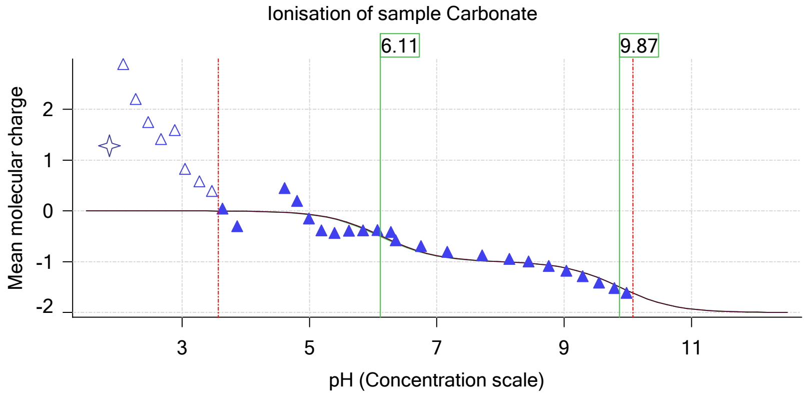

| | | | |
|---|---|---|---|
| Sample name: | **M15_octanol** | Experiment start time: | **3/3/2018 8:26:22 AM** |
| Assay name: | **pH-metric high logP** | Analyst: | **Pion** |
| Assay ID: | **18C-03006** | Instrument ID: | **T312060** |
| Filename: | **C:\Sirius_T3\Mehtap\20180302_exp29_logP_T3-2\18C-03006_M15_octanol_pH-metric high logP.t3r** | | |

### pH-metric high logP Titration 2 of 3 18C-03006 Points 33 to 65

#### Overall results

| | |
|---|---|
| RMSD | 0.481 |
| Average ionic strength | 0.167 M |
| Average temperature | 25.0°C |
| Partition ratio | 0.0576 : 1 |
| Analyte concentration range | 3242.7 µM to 3373.5 µM |
| Total points considered | 31 of 33 |

#### Warnings and errors

| | |
|---|---|
| Errors | None |
| Warnings | One or more logP values out of range |

#### Four-Plus parameters

| | | | |
|---|---|---|---|
| Alpha | 0.111 | 3/3/2018 8:26:22 AM | C:\Sirius_T3\HCl18C02.t3r |
| S | 0.9988 | 3/3/2018 8:26:22 AM | C:\Sirius_T3\HCl18C02.t3r |
| jH | 1.0 | 3/3/2018 8:26:22 AM | C:\Sirius_T3\HCl18C02.t3r |
| jOH | -0.8 | 3/3/2018 8:26:22 AM | C:\Sirius_T3\HCl18C02.t3r |

#### Titrants

| | | | |
|---|---|---|---|
| 0.50 M HCl | 0.999058 | 3/3/2018 8:26:22 AM | C:\Sirius_T3\HCl18C02.t3r |
| 0.50 M KOH | 0.999845 | 3/3/2018 8:26:22 AM | C:\Sirius_T3\KOH18B27.t3r |

#### Sample

| | |
|---|---|
| M15_octanol concentration factor | 0.950 |
| Base pKa 1 | 2.58 |
| Base pKa 2 | 5.30 |
| logP (XH2 2+) | -5.50 |
| logP (XH +) | -2.46 |
| logP (neutral X) | 1.99 |

#### Sample graphs

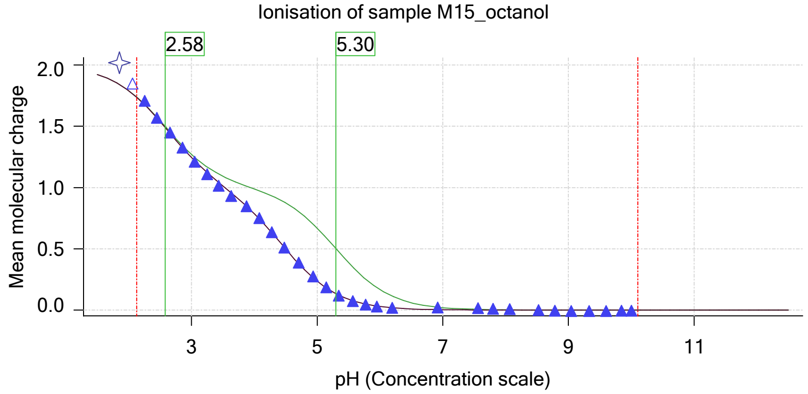


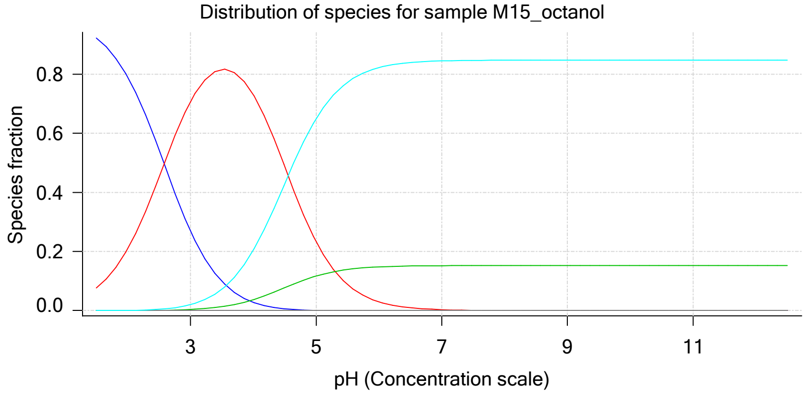

| | | | |
|---|---|---|---|
| Sample name: | **M15_octanol** | Experiment start time: | **3/3/2018 8:26:22 AM** |
| Assay name: | **pH-metric high logP** | Analyst: | **Pion** |
| Assay ID: | **18C-03006** | Instrument ID: | **T312060** |
| Filename: | **C:\Sirius_T3\Mehtap\20180302_exp29_logP_T3-2\18C-03006_M15_octanol_pH-metric high logP.t3r** | | |

#### Sample graphs (continued)

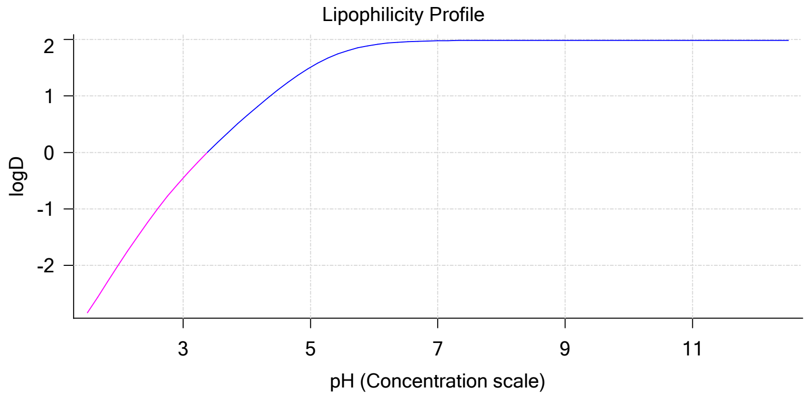


#### Sample logD and percent species

| pH | M15_octanol logD | M15_octanol M15_octanolH2 | M15_octanol M15_octanolH | M15_octanol M15_octanol | M15_octanol M15_octanolH2* | M15_octanol M15_octanolH* | M15_octanol M15_octanol* | Comment |
|---|---|---|---|---|---|---|---|---|
| 1.000 | -3.66 | 97.44 % | 2.56 % | 0.00 % | 0.00 % | 0.00 % | 0.00 % | |
| 1.200 | -3.35 | 96.00 % | 4.00 % | 0.00 % | 0.00 % | 0.00 % | 0.00 % | Stomach pH |
| 2.000 | -1.97 | 79.12 % | 20.81 % | 0.01 % | 0.00 % | 0.00 % | 0.06 % | |
| 3.000 | -0.45 | 26.90 % | 70.76 % | 0.35 % | 0.00 % | 0.01 % | 1.97 % | |
| 4.000 | 0.65 | 2.78 % | 73.13 % | 3.67 % | 0.00 % | 0.01 % | 20.41 % | |
| 5.000 | 1.51 | 0.09 % | 23.28 % | 11.67 % | 0.00 % | 0.00 % | 64.96 % | |
| 6.000 | 1.91 | 0.00 % | 2.95 % | 14.77 % | 0.00 % | 0.00 % | 82.28 % | |
| 6.500 | 1.96 | 0.00 % | 0.95 % | 15.08 % | 0.00 % | 0.00 % | 83.97 % | |
| 7.000 | 1.98 | 0.00 % | 0.30 % | 15.18 % | 0.00 % | 0.00 % | 84.52 % | |
| 7.400 | 1.98 | 0.00 % | 0.12 % | 15.21 % | 0.00 % | 0.00 % | 84.67 % | Blood pH |
| 8.000 | 1.98 | 0.00 % | 0.03 % | 15.22 % | 0.00 % | 0.00 % | 84.75 % | |
| 9.000 | 1.99 | 0.00 % | 0.00 % | 15.22 % | 0.00 % | 0.00 % | 84.77 % | |
| 10.000 | 1.99 | 0.00 % | 0.00 % | 15.22 % | 0.00 % | 0.00 % | 84.78 % | |
| 11.000 | 1.99 | 0.00 % | 0.00 % | 15.22 % | 0.00 % | 0.00 % | 84.78 % | |
| 12.000 | 1.99 | 0.00 % | 0.00 % | 15.22 % | 0.00 % | 0.00 % | 84.78 % | |

#### Carbonate and acidity

| | |
|---|---|
| Carbonate | 0.218 mM |
| Acidity error | -0.296 mM |

#### Other graphs

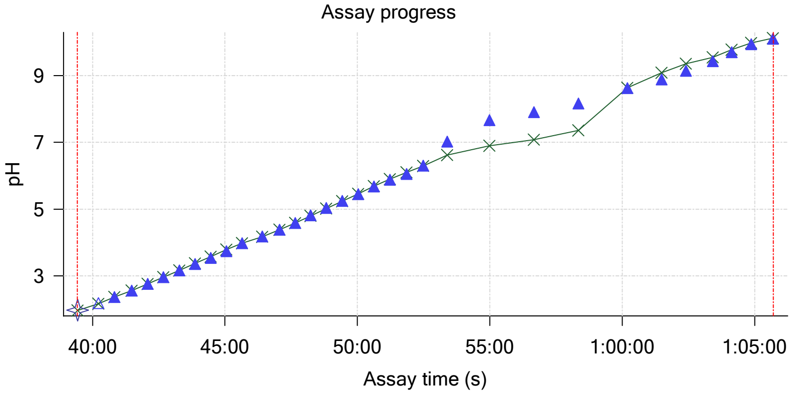


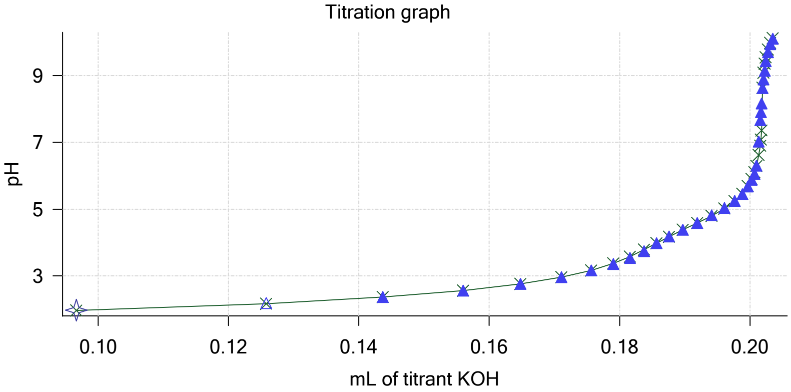

| | | | |
|---|---|---|---|
| Sample name: | **M15_octanol** | Experiment start time: | **3/3/2018 8:26:22 AM** |
| Assay name: | **pH-metric high logP** | Analyst: | **Pion** |
| Assay ID: | **18C-03006** | Instrument ID: | **T312060** |
| Filename: | **C:\Sirius_T3\Mehtap\20180302_exp29_logP_T3-2\18C-03006_M15_octanol_pH-metric high logP.t3r** | | |

#### Other graphs (continued)

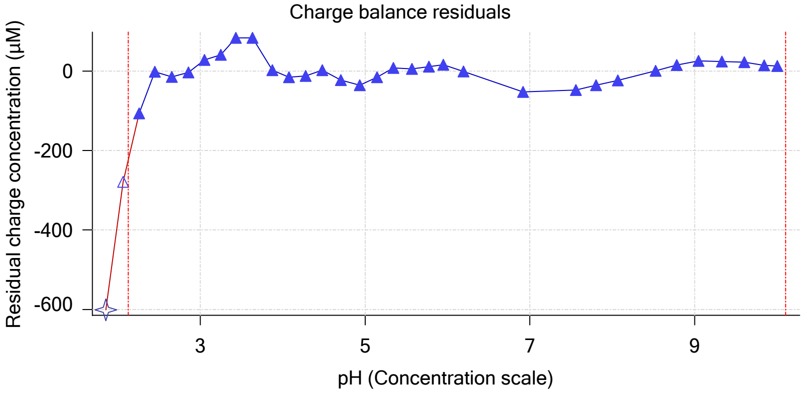


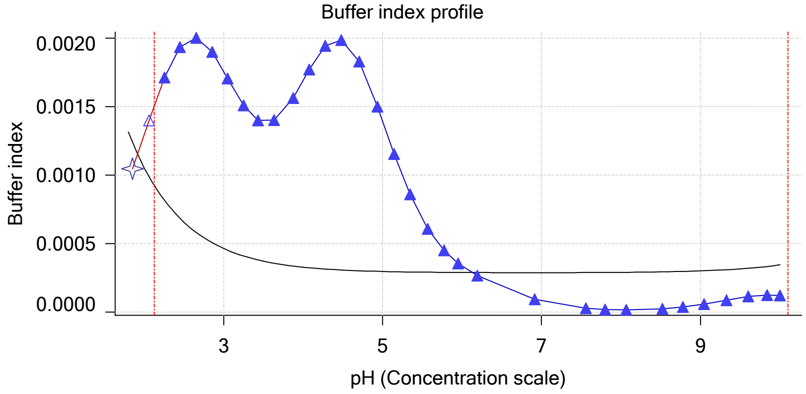


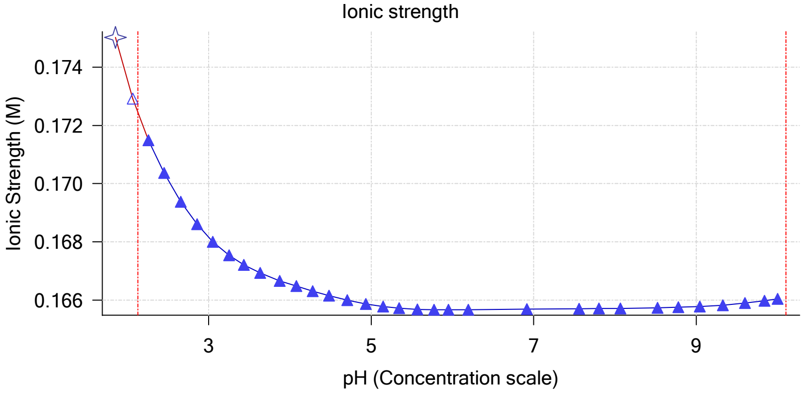


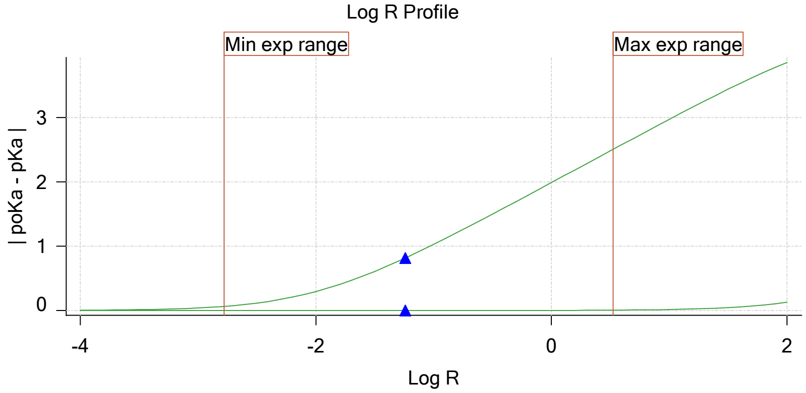


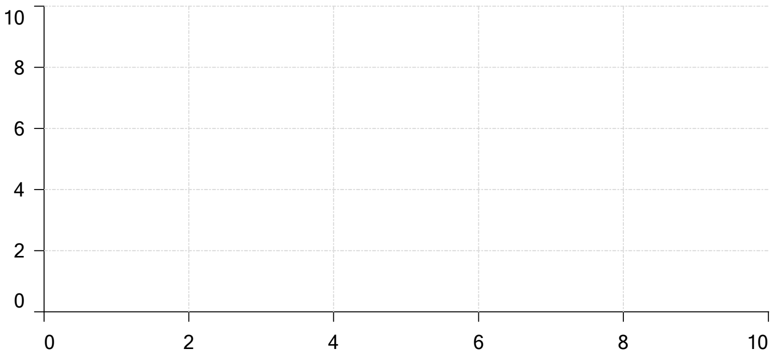


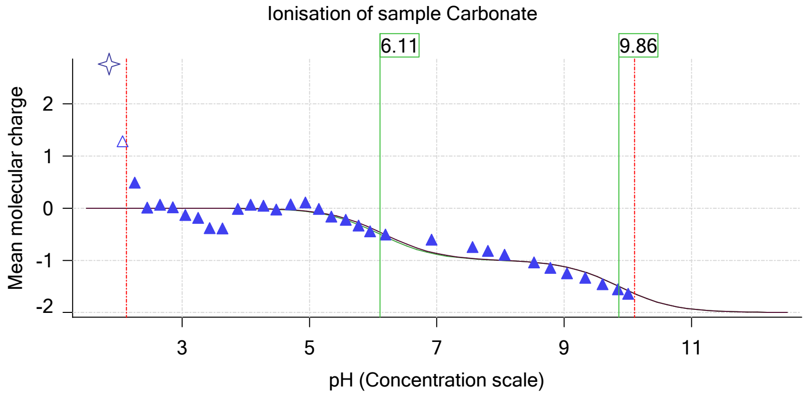

| | | | |
|---|---|---|---|
| Sample name: | **M15_octanol** | Experiment start time: | **3/3/2018 8:26:22 AM** |
| Assay name: | **pH-metric high logP** | Analyst: | **Pion** |
| Assay ID: | **18C-03006** | Instrument ID: | **T312060** |
| Filename: | **C:\Sirius_T3\Mehtap\20180302_exp29_logP_T3-2\18C-03006_M15_octanol_pH-metric high logP.t3r** | | |

### pH-metric high logP Titration 3 of 3 18C-03006 Points 66 to 99

#### Overall results

| | |
|---|---|
| RMSD | 0.728 |
| Average ionic strength | 0.173 M |
| Average temperature | 25.0°C |
| Partition ratio | 0.1596 : 1 |
| Analyte concentration range | 2725.9 μM to 2821.3 μM |
| Total points considered | 33 of 34 |

#### Warnings and errors

| | |
|---|---|
| Errors | None |
| Warnings | One or more logP values out of range |

#### Four-Plus parameters

| | | | |
|---|---|---|---|
| Alpha | 0.111 | 3/3/2018 8:26:22 AM | C:\Sirius_T3\HCl18C02.t3r |
| S | 0.9988 | 3/3/2018 8:26:22 AM | C:\Sirius_T3\HCl18C02.t3r |
| jH | 1.0 | 3/3/2018 8:26:22 AM | C:\Sirius_T3\HCl18C02.t3r |
| jOH | -0.8 | 3/3/2018 8:26:22 AM | C:\Sirius_T3\HCl18C02.t3r |

#### Titrants

| | | | |
|---|---|---|---|
| 0.50 M HCl | 0.999058 | 3/3/2018 8:26:22 AM | C:\Sirius_T3\HCl18C02.t3r |
| 0.50 M KOH | 0.999845 | 3/3/2018 8:26:22 AM | C:\Sirius_T3\KOH18B27.t3r |

#### Sample

| | |
|---|---|
| M15_octanol concentration factor | 1.015 |
| Base pKa 1 | 2.58 |
| Base pKa 2 | 5.30 |
| logP (XH2 2+) | -5.50 |
| logP (XH +) | 0.54 |
| logP (neutral X) | 2.20 |

#### Sample graphs

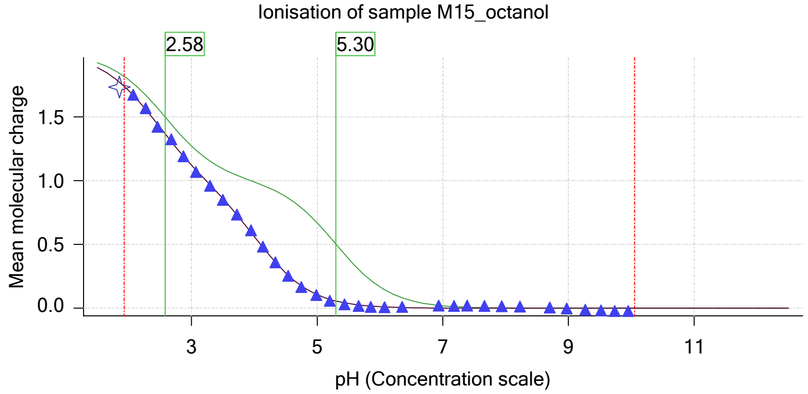


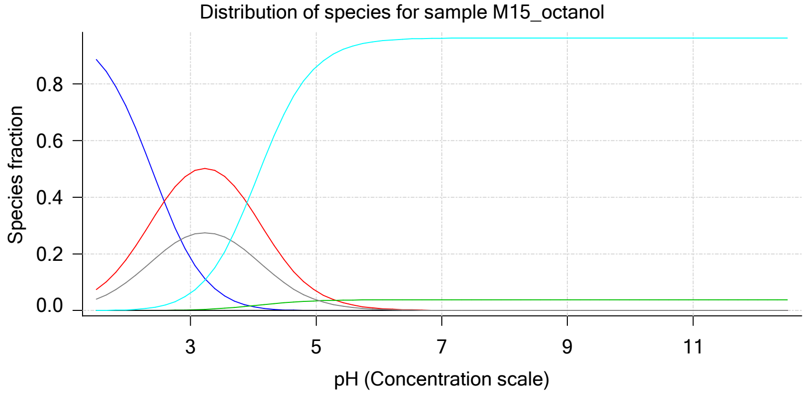

| | | | |
|---|---|---|---|
| Sample name: | **M15_octanol** | Experiment start time: | **3/3/2018 8:26:22 AM** |
| Assay name: | **pH-metric high logP** | Analyst: | **Pion** |
| Assay ID: | **18C-03006** | Instrument ID: | **T312060** |
| Filename: | **C:\Sirius_T3\Mehtap\20180302_exp29_logP_T3-2\18C-03006_M15_octanol_pH-metric high logP.t3r** | | |

#### Sample graphs (continued)

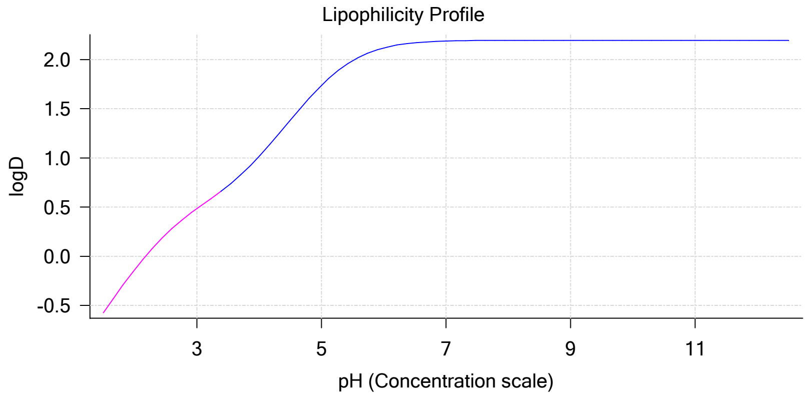


#### Sample logD and percent species

| pH | M15_octanol logD | M15_octanol M15_octanolH2 | M15_octanol M15_octanolH | M15_octanol M15_octanol | M15_octanol M15_octanolH2* | M15_octanol M15_octanolH* | M15_octanol M15_octanol* | Comment |
|---|---|---|---|---|---|---|---|---|
| 1.000 | -1.05 | 96.09 % | 2.53 % | 0.00 % | 0.00 % | 1.38 % | 0.00 % | |
| 1.200 | -0.86 | 93.93 % | 3.92 % | 0.00 % | 0.00 % | 2.14 % | 0.01 % | Stomach pH |
| 2.000 | -0.14 | 70.90 % | 18.65 % | 0.01 % | 0.00 % | 10.21 % | 0.23 % | |
| 3.000 | 0.48 | 18.47 % | 48.58 % | 0.24 % | 0.00 % | 26.59 % | 6.11 % | |
| 4.000 | 1.02 | 1.31 % | 34.56 % | 1.73 % | 0.00 % | 18.91 % | 43.49 % | |
| 5.000 | 1.74 | 0.03 % | 6.83 % | 3.42 % | 0.00 % | 3.74 % | 85.98 % | |
| 6.000 | 2.12 | 0.00 % | 0.76 % | 3.79 % | 0.00 % | 0.41 % | 95.05 % | |
| 6.500 | 2.17 | 0.00 % | 0.24 % | 3.82 % | 0.00 % | 0.13 % | 95.81 % | |
| 7.000 | 2.19 | 0.00 % | 0.08 % | 3.83 % | 0.00 % | 0.04 % | 96.06 % | |
| 7.400 | 2.19 | 0.00 % | 0.03 % | 3.83 % | 0.00 % | 0.02 % | 96.12 % | Blood pH |
| 8.000 | 2.20 | 0.00 % | 0.01 % | 3.83 % | 0.00 % | 0.00 % | 96.16 % | |
| 9.000 | 2.20 | 0.00 % | 0.00 % | 3.83 % | 0.00 % | 0.00 % | 96.17 % | |
| 10.000 | 2.20 | 0.00 % | 0.00 % | 3.83 % | 0.00 % | 0.00 % | 96.17 % | |
| 11.000 | 2.20 | 0.00 % | 0.00 % | 3.83 % | 0.00 % | 0.00 % | 96.17 % | |
| 12.000 | 2.20 | 0.00 % | 0.00 % | 3.83 % | 0.00 % | 0.00 % | 96.17 % | |

#### Carbonate and acidity

| | |
|---|---|
| Carbonate | 0.227 mM |
| Acidity error | -0.479 mM |

#### Other graphs

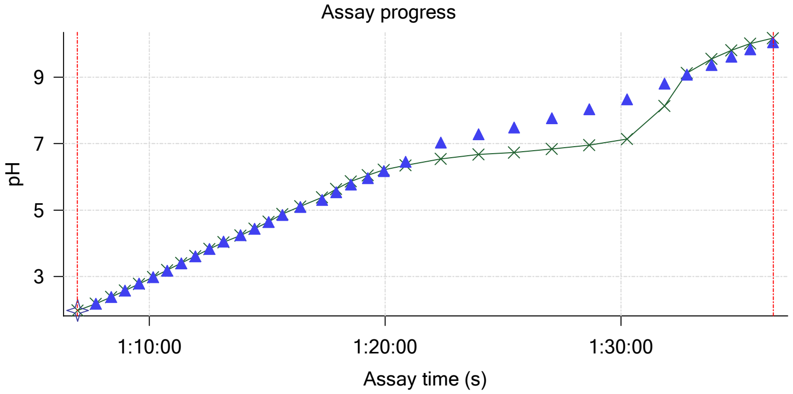


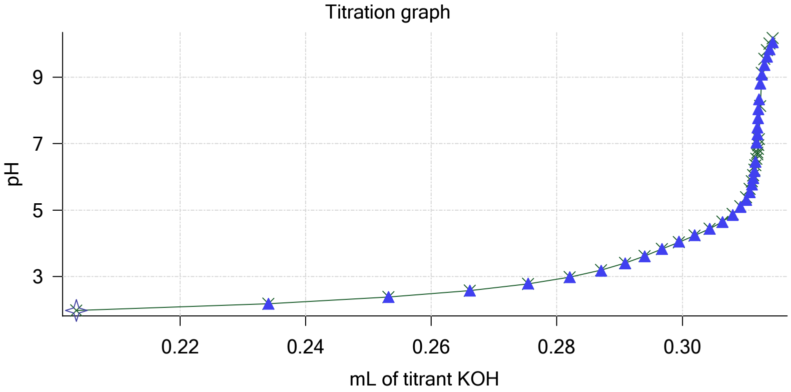

| | | | |
|---|---|---|---|
| Sample name: | **M15_octanol** | Experiment start time: | **3/3/2018 8:26:22 AM** |
| Assay name: | **pH-metric high logP** | Analyst: | **Pion** |
| Assay ID: | **18C-03006** | Instrument ID: | **T312060** |
| Filename: | **C:\Sirius_T3\Mehtap\20180302_exp29_logP_T3-2\18C-03006_M15_octanol_pH-metric high logP.t3r** | | |

#### Other graphs (continued)

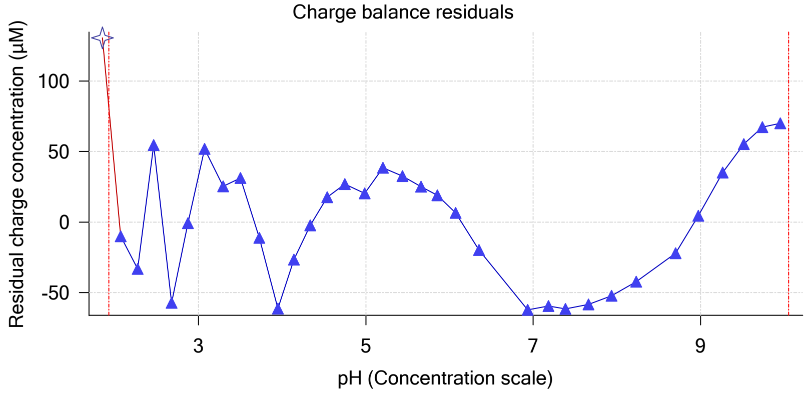


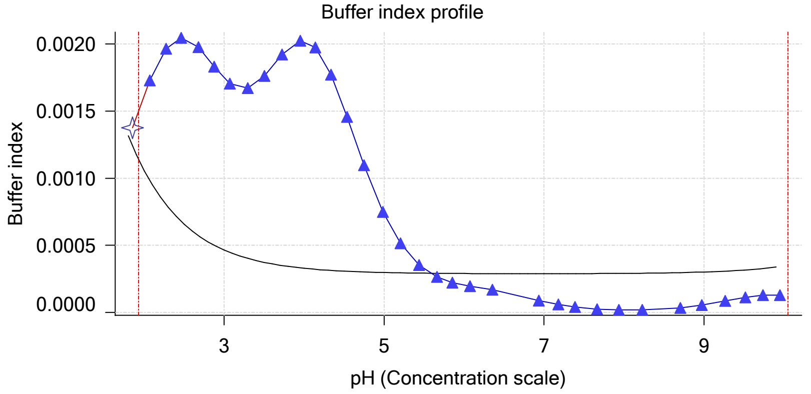


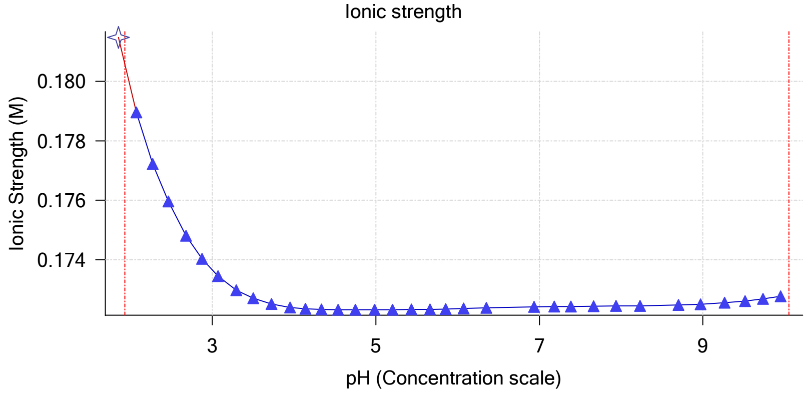


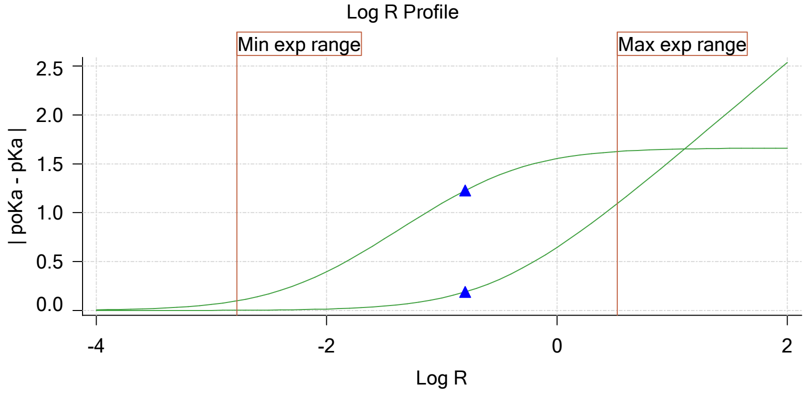


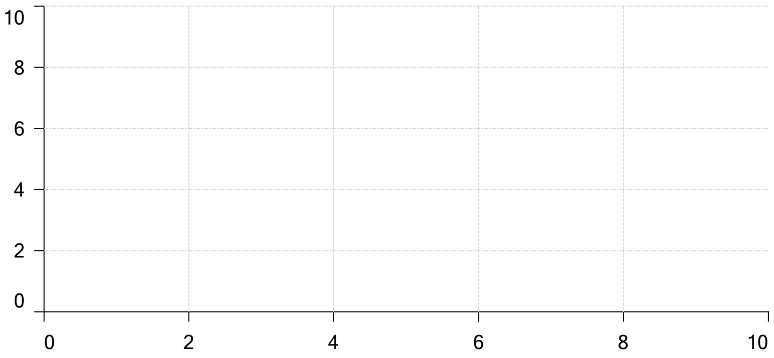

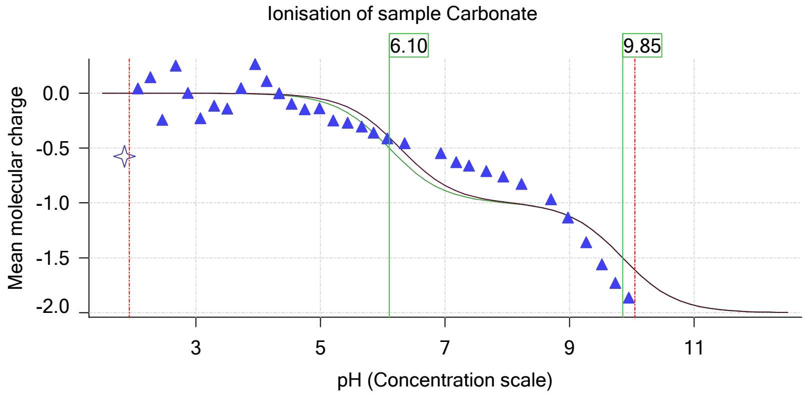

| | | | |
|---|---|---|---|
| Sample name: | **M15_octanol** | Experiment start time: | **3/3/2018 8:26:22 AM** |
| Assay name: | **pH-metric high logP** | Analyst: | **Pion** |
| Assay ID: | **18C-03006** | Instrument ID: | **T312060** |
| Filename: | **C:\Sirius_T3\Mehtap\20180302_exp29_logP_T3-2\18C-03006_M15_octanol_pH-metric high logP.t3r** | | |

#### Assay Model

| Settings | Value | Date/Time changed | Imported from |
|---|---|---|---|
| Sample name | M15_octanol | 2/27/2018 5:03:03 PM | User entered value |
| Sample by | Weight | | Default value |
| Sample weight | 0.001870 g | 3/2/2018 5:09:20 PM | User entered value |
| Formula weight | 209.25 g/mol | 2/27/2018 5:03:03 PM | User entered value |
| Solubility | Unknown | | Default value |
| Molecular weight | 209.25 | 2/27/2018 5:03:03 PM | User entered value |
| Individual pKa ionic environments | No | | Default value |
| Number of pKas | 2 | 2/27/2018 5:03:03 PM | User entered value |
| Sample is a | Base | 2/27/2018 5:03:03 PM | User entered value |
| pKa 1 | 2.58 | 2/27/2018 5:03:03 PM | User entered value |
| pKa 2 | 5.30 | 2/27/2018 5:03:03 PM | User entered value |
| logP (XH2 2+) | -5.50 | 2/28/2018 2:10:35 PM | User entered value |
| logp (XH +) | -4.96 | 2/28/2018 2:10:28 PM | User entered value |
| logP (neutral X) | 1.92 | 2/28/2018 2:10:15 PM | User entered value |

#### Events

| Time | Event | Water | Acid | Base | Octanol | pH | dpH/dt | pH R-squared | pH SD | dpH/dt time |
|---|---|---|---|---|---|---|---|---|---|---|
| 9:11.9 | Initial pH = 6.94 | | | | | | | | | |
| 12:16.5 | Data point 1 | 2.20002 mL | 0.09546 mL | 0.00000 mL | 0.05000 mL | 1.982 | 0.00078 | 0.01682 | 0.00030 | 10.0 s |
| 13:04.2 | Data point 2 | 2.20002 mL | 0.09546 mL | 0.02439 mL | 0.05000 mL | 2.195 | 0.00638 | 0.59345 | 0.00041 | 10.0 s |
| 13:40.0 | Data point 3 | 2.20002 mL | 0.09546 mL | 0.04048 mL | 0.05000 mL | 2.388 | 0.01407 | 0.48508 | 0.00100 | 14.5 s |
| 14:20.2 | Data point 4 | 2.20002 mL | 0.09546 mL | 0.05160 mL | 0.05000 mL | 2.582 | -0.00048 | 0.03777 | 0.00012 | 10.0 s |
| 14:55.8 | Data point 5 | 2.20002 mL | 0.09546 mL | 0.05952 mL | 0.05000 mL | 2.777 | -0.00803 | 0.16330 | 0.00098 | 10.0 s |
| 15:31.3 | Data point 6 | 2.20002 mL | 0.09546 mL | 0.06526 mL | 0.05000 mL | 2.994 | -0.00078 | 0.13803 | 0.00010 | 10.0 s |
| 16:06.8 | Data point 7 | 2.20002 mL | 0.09546 mL | 0.06928 mL | 0.05000 mL | 3.157 | -0.01081 | 0.35970 | 0.00089 | 11.5 s |
| 16:54.2 | Data point 8 | 2.20002 mL | 0.09546 mL | 0.07272 mL | 0.05000 mL | 3.378 | -0.00291 | 0.60595 | 0.00018 | 10.5 s |
| 17:30.2 | Data point 9 | 2.20002 mL | 0.09546 mL | 0.07491 mL | 0.05000 mL | 3.574 | 0.00106 | 0.01066 | 0.00051 | 10.0 s |
| 18:05.7 | Data point 10 | 2.20002 mL | 0.09546 mL | 0.07658 mL | 0.05000 mL | 3.742 | -0.00515 | 0.52674 | 0.00035 | 10.0 s |
| 18:46.3 | Data point 11 | 2.20002 mL | 0.09546 mL | 0.07843 mL | 0.05000 mL | 3.974 | -0.00048 | 0.00212 | 0.00052 | 10.0 s |
| 19:32.3 | Data point 12 | 2.20002 mL | 0.09546 mL | 0.08281 mL | 0.05000 mL | 4.715 | 0.01154 | 0.48986 | 0.00081 | 10.0 s |
| 20:18.1 | Data point 13 | 2.20002 mL | 0.09546 mL | 0.08488 mL | 0.05000 mL | 4.912 | -0.00828 | 0.71346 | 0.00048 | 10.0 s |
| 20:53.6 | Data point 14 | 2.20002 mL | 0.09546 mL | 0.08699 mL | 0.05000 mL | 5.098 | -0.00493 | 0.05980 | 0.00099 | 10.0 s |
| 21:29.0 | Data point 15 | 2.20002 mL | 0.09546 mL | 0.08897 mL | 0.05000 mL | 5.290 | -0.01048 | 0.79475 | 0.00058 | 10.5 s |
| 22:05.1 | Data point 16 | 2.20002 mL | 0.09546 mL | 0.09064 mL | 0.05000 mL | 5.499 | 0.01607 | 0.65427 | 0.00098 | 13.0 s |
| 22:43.5 | Data point 17 | 2.20002 mL | 0.09546 mL | 0.09196 mL | 0.05000 mL | 5.725 | -0.00818 | 0.73332 | 0.00047 | 10.5 s |
| 23:19.5 | Data point 18 | 2.20002 mL | 0.09546 mL | 0.09290 mL | 0.05000 mL | 5.946 | -0.01316 | 0.44366 | 0.00098 | 10.0 s |
| 23:55.0 | Data point 19 | 2.20002 mL | 0.09546 mL | 0.09353 mL | 0.05000 mL | 6.171 | -0.01198 | 0.37219 | 0.00097 | 10.5 s |
| 24:30.9 | Data point 20 | 2.20002 mL | 0.09546 mL | 0.09396 mL | 0.05000 mL | 6.380 | -0.00989 | 0.27105 | 0.00094 | 11.5 s |
| 25:07.8 | Data point 21 | 2.20002 mL | 0.09546 mL | 0.09424 mL | 0.05000 mL | 6.459 | -0.00347 | 0.03282 | 0.00095 | 11.5 s |
| 25:49.9 | Data point 22 | 2.20002 mL | 0.09546 mL | 0.09468 mL | 0.05000 mL | 6.856 | -0.01929 | 0.91505 | 0.00100 | 47.5 s |
| 27:13.3 | Data point 23 | 2.20002 mL | 0.09546 mL | 0.09494 mL | 0.05000 mL | 7.267 | -0.01675 | 0.68623 | 0.00100 | 42.5 s |
| 28:31.5 | Data point 24 | 2.20002 mL | 0.09546 mL | 0.09508 mL | 0.05000 mL | 7.813 | -0.04110 | 0.97438 | 0.00206 | Timed out at 59.5 s |
| 30:12.4 | Data point 25 | 2.20002 mL | 0.09546 mL | 0.09518 mL | 0.05000 mL | 8.240 | -0.03638 | 0.97370 | 0.00182 | Timed out at 59.5 s |
| 31:48.0 | Data point 26 | 2.20002 mL | 0.09546 mL | 0.09525 mL | 0.05000 mL | 8.540 | -0.01949 | 0.94537 | 0.00099 | 55.0 s |
| 33:23.8 | Data point 27 | 2.20002 mL | 0.09546 mL | 0.09537 mL | 0.05000 mL | 8.856 | -0.01251 | 0.96457 | 0.00063 | 36.5 s |
| 34:36.2 | Data point 28 | 2.20002 mL | 0.09546 mL | 0.09551 mL | 0.05000 mL | 9.132 | -0.01942 | 0.95772 | 0.00098 | 30.0 s |
| 35:42.0 | Data point 29 | 2.20002 mL | 0.09546 mL | 0.09570 mL | 0.05000 mL | 9.391 | -0.01949 | 0.95986 | 0.00098 | 17.5 s |
| 36:35.3 | Data point 30 | 2.20002 mL | 0.09546 mL | 0.09595 mL | 0.05000 mL | 9.643 | -0.01556 | 0.97273 | 0.00078 | 17.0 s |
| 37:28.0 | Data point 31 | 2.20002 mL | 0.09546 mL | 0.09628 mL | 0.05000 mL | 9.887 | -0.01964 | 0.96938 | 0.00098 | 14.5 s |
| 38:13.3 | Data point 32 | 2.20002 mL | 0.09546 mL | 0.09666 mL | 0.05000 mL | 10.078 | -0.01833 | 0.96056 | 0.00092 | 11.5 s |
| 39:26.5 | Data point 33 | 2.20002 mL | 0.20245 mL | 0.09666 mL | 0.15000 mL | 1.977 | -0.01121 | 0.68100 | 0.00067 | 10.0 s |
| 40:13.0 | Data point 34 | 2.20002 mL | 0.20245 mL | 0.12578 mL | 0.15000 mL | 2.181 | -0.00937 | 0.91245 | 0.00048 | 10.5 s |
| 40:49.3 | Data point 35 | 2.20002 mL | 0.20245 mL | 0.14367 mL | 0.15000 mL | 2.372 | 0.01563 | 0.60347 | 0.00099 | 14.0 s |
| 41:29.0 | Data point 36 | 2.20002 mL | 0.20245 mL | 0.15602 mL | 0.15000 mL | 2.562 | -0.00403 | 0.15580 | 0.00050 | 10.0 s |

Sample name: **M15_octanol**
Assay name: **pH-metric high logP**
Assay ID: **18C-03006**
Filename: **C:\Sirius_T3\Mehtap\20180302_exp29_logP_T3-2\18C-03006_M15_octanol_pH-metric high logP.t3r**
Experiment start time: **3/3/2018 8:26:22 AM**
Analyst: **Pion**
Instrument ID: **T312060**

#### Events (continued)

| Time | Event | Water | Acid | Base | Octanol | pH | dpH/dt | pH R-squared | pH SD | dpH/dt time |
|---|---|---|---|---|---|---|---|---|---|---|
| 42:04.6 | Data point 37 | 2.20002 mL | 0.20245 mL | 0.16482 mL | 0.15000 mL | 2.766 | -0.00114 | 0.29314 | 0.00010 | 10.5 s |
| 42:40.6 | Data point 38 | 2.20002 mL | 0.20245 mL | 0.17107 mL | 0.15000 mL | 2.965 | 0.00091 | 0.07557 | 0.00016 | 10.5 s |
| 43:16.5 | Data point 39 | 2.20002 mL | 0.20245 mL | 0.17564 mL | 0.15000 mL | 3.160 | -0.00686 | 0.19692 | 0.00076 | 10.0 s |
| 43:52.1 | Data point 40 | 2.20002 mL | 0.20245 mL | 0.17902 mL | 0.15000 mL | 3.358 | -0.00225 | 0.26531 | 0.00022 | 10.0 s |
| 44:27.6 | Data point 41 | 2.20002 mL | 0.20245 mL | 0.18161 mL | 0.15000 mL | 3.541 | -0.00311 | 0.56195 | 0.00020 | 10.0 s |
| 45:03.0 | Data point 42 | 2.20002 mL | 0.20245 mL | 0.18375 mL | 0.15000 mL | 3.740 | -0.00336 | 0.76299 | 0.00019 | 10.0 s |
| 45:38.5 | Data point 43 | 2.20002 mL | 0.20245 mL | 0.18568 mL | 0.15000 mL | 3.982 | -0.00227 | 0.41703 | 0.00017 | 10.0 s |
| 46:24.3 | Data point 44 | 2.20002 mL | 0.20245 mL | 0.18758 mL | 0.15000 mL | 4.184 | -0.00909 | 0.29388 | 0.00083 | 13.5 s |
| 47:03.3 | Data point 45 | 2.20002 mL | 0.20245 mL | 0.18970 mL | 0.15000 mL | 4.384 | -0.00440 | 0.08998 | 0.00072 | 10.0 s |
| 47:38.8 | Data point 46 | 2.20002 mL | 0.20245 mL | 0.19193 mL | 0.15000 mL | 4.585 | 0.00070 | 0.00237 | 0.00071 | 10.0 s |
| 48:14.2 | Data point 47 | 2.20002 mL | 0.20245 mL | 0.19412 mL | 0.15000 mL | 4.810 | -0.00945 | 0.23883 | 0.00096 | 10.0 s |
| 48:49.7 | Data point 48 | 2.20002 mL | 0.20245 mL | 0.19610 mL | 0.15000 mL | 5.040 | -0.00439 | 0.29196 | 0.00040 | 10.5 s |
| 49:25.6 | Data point 49 | 2.20002 mL | 0.20245 mL | 0.19767 mL | 0.15000 mL | 5.250 | -0.00627 | 0.48785 | 0.00044 | 10.5 s |
| 50:01.6 | Data point 50 | 2.20002 mL | 0.20245 mL | 0.19885 mL | 0.15000 mL | 5.450 | -0.01045 | 0.65521 | 0.00064 | 10.5 s |
| 50:37.5 | Data point 51 | 2.20002 mL | 0.20245 mL | 0.19969 mL | 0.15000 mL | 5.674 | -0.01202 | 0.60887 | 0.00076 | 11.0 s |
| 51:13.9 | Data point 52 | 2.20002 mL | 0.20245 mL | 0.20028 mL | 0.15000 mL | 5.877 | -0.01959 | 0.97062 | 0.00098 | 12.0 s |
| 51:51.4 | Data point 53 | 2.20002 mL | 0.20245 mL | 0.20068 mL | 0.15000 mL | 6.055 | -0.01017 | 0.29811 | 0.00092 | 12.5 s |
| 52:29.4 | Data point 54 | 2.20002 mL | 0.20245 mL | 0.20099 mL | 0.15000 mL | 6.297 | -0.01877 | 0.95054 | 0.00095 | 23.5 s |
| 53:23.6 | Data point 55 | 2.20002 mL | 0.20245 mL | 0.20136 mL | 0.15000 mL | 7.018 | -0.05573 | 0.98191 | 0.00278 | Timed out at 59.5 s |
| 54:59.2 | Data point 56 | 2.20002 mL | 0.20245 mL | 0.20158 mL | 0.15000 mL | 7.661 | -0.08712 | 0.98891 | 0.00433 | Timed out at 59.5 s |
| 56:40.0 | Data point 57 | 2.20002 mL | 0.20245 mL | 0.20167 mL | 0.15000 mL | 7.903 | -0.06287 | 0.97801 | 0.00314 | Timed out at 59.5 s |
| 58:20.7 | Data point 58 | 2.20002 mL | 0.20245 mL | 0.20176 mL | 0.15000 mL | 8.166 | -0.04207 | 0.98021 | 0.00210 | Timed out at 59.5 s |
| 1:00:11.8 | Data point 59 | 2.20002 mL | 0.20245 mL | 0.20195 mL | 0.15000 mL | 8.624 | -0.01762 | 0.98142 | 0.00088 | 36.5 s |
| 1:01:29.3 | Data point 60 | 2.20002 mL | 0.20245 mL | 0.20209 mL | 0.15000 mL | 8.881 | -0.01366 | 0.48711 | 0.00097 | 19.5 s |
| 1:02:24.3 | Data point 61 | 2.20002 mL | 0.20245 mL | 0.20226 mL | 0.15000 mL | 9.145 | -0.00856 | 0.31850 | 0.00075 | 30.0 s |
| 1:03:24.9 | Data point 62 | 2.20002 mL | 0.20245 mL | 0.20245 mL | 0.15000 mL | 9.427 | -0.01587 | 0.72413 | 0.00092 | 12.5 s |
| 1:04:08.0 | Data point 63 | 2.20002 mL | 0.20245 mL | 0.20275 mL | 0.15000 mL | 9.699 | -0.01697 | 0.88469 | 0.00089 | 13.5 s |
| 1:04:52.1 | Data point 64 | 2.20002 mL | 0.20245 mL | 0.20313 mL | 0.15000 mL | 9.940 | -0.01307 | 0.46035 | 0.00095 | 18.5 s |
| 1:05:41.2 | Data point 65 | 2.20002 mL | 0.20245 mL | 0.20350 mL | 0.15000 mL | 10.101 | -0.01920 | 0.91102 | 0.00099 | 10.5 s |
| 1:06:58.1 | Data point 66 | 2.20002 mL | 0.31409 mL | 0.20350 mL | 0.45000 mL | 1.975 | -0.01558 | 0.84604 | 0.00084 | 10.0 s |
| 1:07:44.6 | Data point 67 | 2.20002 mL | 0.31409 mL | 0.23408 mL | 0.45000 mL | 2.187 | 0.01489 | 0.60774 | 0.00094 | 12.5 s |
| 1:08:23.0 | Data point 68 | 2.20002 mL | 0.31409 mL | 0.25320 mL | 0.45000 mL | 2.385 | 0.00088 | 0.09557 | 0.00014 | 10.0 s |
| 1:08:58.7 | Data point 69 | 2.20002 mL | 0.31409 mL | 0.26618 mL | 0.45000 mL | 2.573 | -0.00649 | 0.12500 | 0.00091 | 10.0 s |
| 1:09:34.4 | Data point 70 | 2.20002 mL | 0.31409 mL | 0.27545 mL | 0.45000 mL | 2.787 | -0.00191 | 0.06611 | 0.00037 | 10.0 s |
| 1:10:09.9 | Data point 71 | 2.20002 mL | 0.31409 mL | 0.28206 mL | 0.45000 mL | 2.981 | 0.00572 | 0.62627 | 0.00036 | 10.5 s |
| 1:10:46.0 | Data point 72 | 2.20002 mL | 0.31409 mL | 0.28704 mL | 0.45000 mL | 3.178 | -0.01361 | 0.49274 | 0.00096 | 10.5 s |
| 1:11:21.9 | Data point 73 | 2.20002 mL | 0.31409 mL | 0.29090 mL | 0.45000 mL | 3.401 | 0.00911 | 0.20975 | 0.00098 | 10.0 s |
| 1:11:57.5 | Data point 74 | 2.20002 mL | 0.31409 mL | 0.29400 mL | 0.45000 mL | 3.608 | -0.00440 | 0.13243 | 0.00060 | 10.0 s |
| 1:12:32.9 | Data point 75 | 2.20002 mL | 0.31409 mL | 0.29678 mL | 0.45000 mL | 3.831 | 0.00458 | 0.65602 | 0.00028 | 10.5 s |
| 1:13:09.0 | Data point 76 | 2.20002 mL | 0.31409 mL | 0.29944 mL | 0.45000 mL | 4.056 | -0.00707 | 0.18756 | 0.00081 | 18.0 s |
| 1:13:52.5 | Data point 77 | 2.20002 mL | 0.31409 mL | 0.30198 mL | 0.45000 mL | 4.244 | -0.00994 | 0.26743 | 0.00095 | 10.0 s |
| 1:14:28.0 | Data point 78 | 2.20002 mL | 0.31409 mL | 0.30435 mL | 0.45000 mL | 4.441 | -0.00154 | 0.01632 | 0.00059 | 10.0 s |
| 1:15:03.5 | Data point 79 | 2.20002 mL | 0.31409 mL | 0.30640 mL | 0.45000 mL | 4.642 | -0.00829 | 0.23500 | 0.00084 | 10.0 s |
| 1:15:38.9 | Data point 80 | 2.20002 mL | 0.31409 mL | 0.30804 mL | 0.45000 mL | 4.854 | 0.01636 | 0.66406 | 0.00099 | 20.5 s |
| 1:16:24.9 | Data point 81 | 2.20002 mL | 0.31409 mL | 0.30927 mL | 0.45000 mL | 5.090 | 0.00586 | 0.12896 | 0.00081 | 19.0 s |
| 1:17:19.6 | Data point 82 | 2.20002 mL | 0.31409 mL | 0.31016 mL | 0.45000 mL | 5.308 | -0.00746 | 0.23824 | 0.00075 | 11.0 s |
| 1:17:56.0 | Data point 83 | 2.20002 mL | 0.31409 mL | 0.31070 mL | 0.45000 mL | 5.539 | -0.01384 | 0.63849 | 0.00086 | 11.5 s |
| 1:18:32.9 | Data point 84 | 2.20002 mL | 0.31409 mL | 0.31105 mL | 0.45000 mL | 5.765 | 0.00783 | 0.17459 | 0.00093 | 18.0 s |
| 1:19:16.4 | Data point 85 | 2.20002 mL | 0.31409 mL | 0.31129 mL | 0.45000 mL | 5.959 | -0.01816 | 0.90344 | 0.00094 | 15.0 s |
| 1:19:56.8 | Data point 86 | 2.20002 mL | 0.31409 mL | 0.31148 mL | 0.45000 mL | 6.176 | -0.00051 | 0.00066 | 0.00098 | 30.0 s |
| 1:20:52.2 | Data point 87 | 2.20002 mL | 0.31409 mL | 0.31162 mL | 0.45000 mL | 6.455 | -0.01711 | 0.79155 | 0.00095 | 59.5 s |
| 1:22:22.2 | Data point 88 | 2.20002 mL | 0.31409 mL | 0.31181 mL | 0.45000 mL | 7.035 | -0.09484 | 0.98262 | 0.00473 | Timed out at 59.5 s |

| | | | |
|---|---|---|---|
| Sample name: | **M15_octanol** | Experiment start time: | **3/3/2018 8:26:22 AM** |
| Assay name: | **pH-metric high logP** | Analyst: | **Pion** |
| Assay ID: | **18C-03006** | Instrument ID: | **T312060** |
| Filename: | **C:\Sirius_T3\Mehtap\20180302_exp29_logP_T3-2\18C-03006_M15_octanol_pH-metric high logP.t3r** | | |

#### Events (continued)

| Time | Event | Water | Acid | Base | Octanol | pH | dpH/dt | pH R-squared | pH SD | dpH/dt time |
|---|---|---|---|---|---|---|---|---|---|---|
| 1:23:57.9 | Data point 89 | 2.20002 mL | 0.31409 mL | 0.31192 mL | 0.45000 mL | 7.282 | -0.07791 | 0.98221 | 0.00388 | Timed out at 59.5 s |
| 1:25:28.4 | Data point 90 | 2.20002 mL | 0.31409 mL | 0.31197 mL | 0.45000 mL | 7.488 | -0.07538 | 0.98886 | 0.00374 | Timed out at 59.5 s |
| 1:27:04.1 | Data point 91 | 2.20002 mL | 0.31409 mL | 0.31204 mL | 0.45000 mL | 7.764 | -0.08258 | 0.98709 | 0.00411 | Timed out at 59.5 s |
| 1:28:39.7 | Data point 92 | 2.20002 mL | 0.31409 mL | 0.31211 mL | 0.45000 mL | 8.037 | -0.06736 | 0.98943 | 0.00334 | Timed out at 59.5 s |
| 1:30:15.3 | Data point 93 | 2.20002 mL | 0.31409 mL | 0.31221 mL | 0.45000 mL | 8.330 | -0.04664 | 0.96334 | 0.00235 | Timed out at 59.5 s |
| 1:31:51.0 | Data point 94 | 2.20002 mL | 0.31409 mL | 0.31242 mL | 0.45000 mL | 8.802 | -0.01725 | 0.85754 | 0.00092 | 21.0 s |
| 1:32:47.7 | Data point 95 | 2.20002 mL | 0.31409 mL | 0.31268 mL | 0.45000 mL | 9.073 | -0.00613 | 0.09179 | 0.00100 | 27.5 s |
| 1:33:50.9 | Data point 96 | 2.20002 mL | 0.31409 mL | 0.31305 mL | 0.45000 mL | 9.365 | -0.01654 | 0.78160 | 0.00093 | 14.5 s |
| 1:34:41.2 | Data point 97 | 2.20002 mL | 0.31409 mL | 0.31345 mL | 0.45000 mL | 9.616 | -0.01302 | 0.60064 | 0.00083 | 12.5 s |
| 1:35:29.4 | Data point 98 | 2.20002 mL | 0.31409 mL | 0.31388 mL | 0.45000 mL | 9.835 | 0.00468 | 0.05381 | 0.00100 | 21.5 s |
| 1:36:26.7 | Data point 99 | 2.20002 mL | 0.31409 mL | 0.31439 mL | 0.45000 mL | 10.049 | 0.00989 | 0.27168 | 0.00094 | 11.5 s |
| 1:36:47.2 | Assay volumes | 2.20002 mL | 0.31409 mL | 0.31439 mL | 0.45000 mL | | | | | |

| | | | |
|---|---|---|---|
| Sample name: | **M15_octanol** | Experiment start time: | **3/3/2018 8:26:22 AM** |
| Assay name: | **pH-metric high logP** | Analyst: | **Pion** |
| Assay ID: | **18C-03006** | Instrument ID: | **T312060** |
| Filename: | **C:\Sirius_T3\Mehtap\20180302_exp29_logP_T3-2\18C-03006_M15_octanol_pH-metric high logP.t3r** | | |

| | | | |
|---|---|---|---|
| Sample name: | **M15_octanol** | Experiment start time: | **3/3/2018 8:26:22 AM** |
| Assay name: | **pH-metric high logP** | Analyst: | **Pion** |
| Assay ID: | **18C-03006** | Instrument ID: | **T312060** |
| Filename: | **C:\Sirius_T3\Mehtap\20180302_exp29_logP_T3-2\18C-03006_M15_octanol_pH-metric high logP.t3r** | | |

#### Assay Settings (continued)

| Setting | Value | Original Value | Date/Time changed | Imported from |
|---|---|---|---|---|
| ***Titration 3*** | | | | |
| Titrate from | Low to high pH | | | |
| Add additional water | 0.00 mL | | | |
| Additional partition solvent volume | 0.300 mL | | | |
| Additional partition solvent added | Automatic | | | |
| After pH adjust stir for | 30 seconds | | | |
| Stir to allow partitioning for | 15 seconds | | | |
| Stirrer speed for partitioning | 60% | | | |
| ***Data Point Stability*** | | | | |
| Stir during data point collection | No | | | |
| Delay before data point collection | 0 seconds | | | |
| Number of points to average | 20 points | | | |
| Time interval between points | 0.50 seconds | | | |
| Required maximum standard deviation | 0.00100 dpH/dt | | | |
| Stability timeout after | 60 seconds | | | |

#### Calibration Settings

| Setting | Value | Date/Time changed | Imported from |
|---|---|---|---|
| Four-Plus alpha | 0.111 | 3/3/2018 8:26:22 AM | C:\Sirius_T3\HCl18C02.t3r |
| Four-Plus S | 0.9988 | 3/3/2018 8:26:22 AM | C:\Sirius_T3\HCl18C02.t3r |
| Four-Plus jH | 1.0 | 3/3/2018 8:26:22 AM | C:\Sirius_T3\HCl18C02.t3r |
| Four-Plus jOH | -0.8 | 3/3/2018 8:26:22 AM | C:\Sirius_T3\HCl18C02.t3r |
| Base concentration factor | 1.000 | 3/3/2018 8:26:22 AM | C:\Sirius_T3\KOH18B27.t3r |
| Acid concentration factor | 0.999 | 3/3/2018 8:26:22 AM | C:\Sirius_T3\HCl18C02.t3r |

#### Instrument Settings

| Setting | Value | Batch Id | Install date |
|---|---|---|---|
| Instrument owner | Merck | | |
| Instrument ID | T312060 | | |
| Instrument type | T3 Simulator | | |
| Software version | 1.1.3.0 | | |
| Dispenser module | | T3DM1200361 | 3/31/2009 5:24:52 AM |
| Dispenser 0 | Water | | 3/31/2009 5:25:05 AM |
| Syringe volume | 2.5 mL | | |
| Firmware version | 1.2.1(r2) | | |
| Titrant | Water (0.15 M KCl) | 02-06-2018 | 2/27/2018 10:05:59 AM |
| Dispenser 2 | Acid | | 3/31/2009 5:25:11 AM |
| Syringe volume | 0.5 mL | | |
| Firmware version | 1.2.1(r2) | | |
| Titrant | Acid (0.5 M HCl) | 02-27-2018 | 2/27/2018 10:27:22 AM |
| Dispenser 1 | Base | | 3/31/2009 5:25:21 AM |
| Syringe volume | 0.5 mL | | |
| Firmware version | 1.2.1(r2) | | |
| Titrant | Base (0.5 M KOH) | 9/22/2017 | 2/27/2018 10:21:22 AM |
| Dispenser 5 | Cosolvent | | 3/31/2009 5:26:24 AM |
| Syringe volume | 2.5 mL | | |
| Firmware version | 1.2.1(r2) | | |
| Distribution valve 5 | Distribution Valve | | 3/31/2009 5:28:19 AM |
| Firmware version | 1.1.3 | | |
| Port A | Methanol (80%, 0.15 M KCl) | 09-26-17 | 2/7/2018 9:42:01 AM |
| Port B | Cyclohexane | 11-01-17 | 2/27/2018 10:37:57 AM |
| Dispenser 3 | Buffer | | 8/3/2010 5:05:16 AM |
| Syringe volume | 0.5 mL | | |
| Firmware version | 1.2.1(r2) | | |
| Titrant | Dodecane | 2018/01/31 | 2/28/2018 10:18:04 AM |
| Dispenser 6 | Octanol | | 10/22/2010 10:52:43 AM |

| | | | |
|---|---|---|---|
| Sample name: | **M15_octanol** | Experiment start time: | **3/3/2018 8:26:22 AM** |
| Assay name: | **pH-metric high logP** | Analyst: | **Pion** |
| Assay ID: | **18C-03006** | Instrument ID: | **T312060** |
| Filename: | **C:\Sirius_T3\Mehtap\20180302_exp29_logP_T3-2\18C-03006_M15_octanol_pH-metric high logP.t3r** | | |

| | | | |
|---|---|---|---|
| Sample name: | **M15_octanol** | Experiment start time: | **3/3/2018 8:26:22 AM** |
| Assay name: | **pH-metric high logP** | Analyst: | **Pion** |
| Assay ID: | **18C-03006** | Instrument ID: | **T312060** |
| Filename: | **C:\Sirius_T3\Mehtap\20180302_exp29_logP_T3-2\18C-03006_M15_octanol_pH-metric high logP.t3r** | | |

#### Instrument Settings (continued)

| Setting | Value | Batch Id | Install date |
|---|---|---|---|
| Spectrometer calibration stir duration | 5 s | | |
| Spectrometer calibration stir speed | 30% | | |
| Spectrometer calibration wash pump volume | 20.0 mL | | |
| Spectrometer calibration wash stir duration | 5 s | | |
| Spectrometer calibration wash stir speed | 30% | | |
| Overhead dispense height | 10000 | | |

#### Refinement Settings

| Setting | Value | Default value |
|---|---|---|
| Turbidity detection method | None | None |
| Turbidity wavelength to assess | 500.0 nm | 500.0 nm |
| Turbidity maximum absorbance | 0.100 | 0.100 |
| Turbidity probe threshold | 50.00 | 50.00 |

#### Experiment Log

[2:38] Air gap created for Water (0.15 M KCl)
[2:38] Air gap created for Acid (0.5 M HCl)
[2:38] Air gap created for Base (0.5 M KOH)
[2:39] Air gap released for Water (0.15 M KCl)
[2:43] Titrator arm moved over Titration position
[2:43] Titration 1 of 3
[2:43] Adding initial titrants
[2:43] Automatically add 2.20000 mL of water
[3:19] Dispensed 2.200024 mL of Water (0.15 M KCl)
[3:24] Titrator arm moved over Drain
[9:05] Titrator arm moved to Titration position
[9:05] Argon flow rate set to 100
[9:05] Stirrer speed set to 10
[9:10] Automatically add 0.05000 mL of Octanol
[9:11] Dispensed 0.050000 mL of Octanol
[9:12] Initial pH = 6.94
[9:12] Iterative adjust 6.94 -> 2.00
[9:12] pH 6.94 -> 2.00
[9:15] Air gap released for Acid (0.5 M HCl)
[9:16] Dispensed 0.091322 mL of Acid (0.5 M HCl)
[9:21] pH 2.02 -> 2.00
[9:21] Dispensed 0.002563 mL of Acid (0.5 M HCl)
[9:26] pH 2.01 -> 2.00
[9:26] Dispensed 0.001576 mL of Acid (0.5 M HCl)
[9:31] Holding pH 2.00
[11:32] Stirrer speed set to 0
[11:32] Stirrer speed set to 50
[11:32] Iterative adjust 1.99 -> 2.00
[12:17] Stirrer speed set to 0
[12:27] Datapoint id 1 collected
[12:27] Stirrer speed set to 50
[12:32] pH 1.99 -> 2.19
[12:32] Using cautious pH adjust
[12:33] Air gap released for Base (0.5 M KOH)
[12:34] Dispensed 0.012465 mL of Base (0.5 M KOH)
[12:39] Stepping pH = 2.08
[12:39] Dispensed 0.009243 mL of Base (0.5 M KOH)
[12:44] Stepping pH = 2.16
[12:44] Dispensed 0.002681 mL of Base (0.5 M KOH)
[12:49] Stepping pH = 2.19
[13:05] Stirrer speed set to 0
[13:15] Datapoint id 2 collected

Sample name: **M15_octanol** Experiment start time: **3/3/2018 8:26:22 AM**
Assay name: **pH-metric high logP** Analyst: **Pion**
Assay ID: **18C-03006** Instrument ID: **T312060**
Filename: **C:\Sirius_T3\Mehtap\20180302_exp29_logP_T3-2\18C-03006_M15_octanol_pH-metric high logP.t3r**

#### Experiment Log (continued)

[13:15] Charge balance equation is out by 2.1%
[13:15] Stirrer speed set to 50
[13:20] pH 2.20 -> 2.40
[13:20] Using charge balance adjust
[13:20] Dispensed 0.016087 mL of Base (0.5 M KOH)
[13:40] Stirrer speed set to 0
[13:55] Datapoint id 3 collected
[13:55] Charge balance equation is out by -5.3%
[13:55] Stirrer speed set to 50
[14:00] pH 2.39 -> 2.59
[14:00] Using charge balance adjust
[14:00] Dispensed 0.011124 mL of Base (0.5 M KOH)
[14:21] Stirrer speed set to 0
[14:31] Datapoint id 4 collected
[14:31] Charge balance equation is out by -5.5%
[14:31] Stirrer speed set to 50
[14:36] pH 2.59 -> 2.79
[14:36] Using charge balance adjust
[14:36] Dispensed 0.007926 mL of Base (0.5 M KOH)
[14:56] Stirrer speed set to 0
[15:06] Datapoint id 5 collected
[15:06] Charge balance equation is out by -5.2%
[15:06] Stirrer speed set to 50
[15:11] pH 2.78 -> 2.98
[15:11] Using charge balance adjust
[15:12] Dispensed 0.005738 mL of Base (0.5 M KOH)
[15:32] Stirrer speed set to 0
[15:42] Datapoint id 6 collected
[15:42] Charge balance equation is out by 5.1%
[15:42] Stirrer speed set to 50
[15:47] pH 3.00 -> 3.20
[15:47] Using charge balance adjust
[15:47] Dispensed 0.004022 mL of Base (0.5 M KOH)
[16:07] Stirrer speed set to 0
[16:19] Datapoint id 7 collected
[16:19] Charge balance equation is out by -21.3%
[16:19] Stirrer speed set to 50
[16:24] pH 3.16 -> 3.36
[16:24] Using cautious pH adjust
[16:24] Dispensed 0.001529 mL of Base (0.5 M KOH)
[16:29] Stepping pH = 3.23
[16:29] Dispensed 0.001693 mL of Base (0.5 M KOH)
[16:34] Stepping pH = 3.35
[16:34] Dispensed 0.000212 mL of Base (0.5 M KOH)
[16:39] Stepping pH = 3.38
[16:55] Stirrer speed set to 0
[17:05] Datapoint id 8 collected
[17:05] Charge balance equation is out by -11.9%
[17:05] Stirrer speed set to 50
[17:10] pH 3.38 -> 3.58
[17:10] Using charge balance adjust
[17:10] Dispensed 0.002187 mL of Base (0.5 M KOH)
[17:31] Stirrer speed set to 0
[17:41] Datapoint id 9 collected
[17:41] Charge balance equation is out by -4.7%
[17:41] Stirrer speed set to 50
[17:46] pH 3.58 -> 3.78
[17:46] Using charge balance adjust
[17:46] Dispensed 0.001670 mL of Base (0.5 M KOH)

| | | | |
|---|---|---|---|
| Sample name: | **M15_octanol** | Experiment start time: | **3/3/2018 8:26:22 AM** |
| Assay name: | **pH-metric high logP** | Analyst: | **Pion** |
| Assay ID: | **18C-03006** | Instrument ID: | **T312060** |
| Filename: | **C:\Sirius_T3\Mehtap\20180302_exp29_logP_T3-2\18C-03006_M15_octanol_pH-metric high logP.t3r** | | |

#### Experiment Log (continued)

[18:06] Stirrer speed set to 0
[18:16] Datapoint id 10 collected
[18:16] Charge balance equation is out by -19.9%
[18:16] Stirrer speed set to 50
[18:21] pH 3.75 -> 3.95
[18:21] Using cautious pH adjust
[18:21] Dispensed 0.000729 mL of Base (0.5 M KOH)
[18:26] Stepping pH = 3.79
[18:26] Dispensed 0.001129 mL of Base (0.5 M KOH)
[18:32] Stepping pH = 3.95
[18:47] Stirrer speed set to 0
[18:57] Datapoint id 11 collected
[18:57] Charge balance equation is out by -27.1%
[18:57] Stirrer speed set to 50
[19:02] pH 3.98 -> 4.18
[19:02] Using cautious pH adjust
[19:02] Dispensed 0.000706 mL of Base (0.5 M KOH)
[19:07] Stepping pH = 4.04
[19:07] Dispensed 0.000917 mL of Base (0.5 M KOH)
[19:12] Stepping pH = 4.04
[19:12] Dispensed 0.002752 mL of Base (0.5 M KOH)
[19:18] Stepping pH = 4.69
[19:33] Stirrer speed set to 0
[19:43] Datapoint id 12 collected
[19:43] Charge balance equation is out by -214.2%
[19:43] Stirrer speed set to 50
[19:48] pH 4.72 -> 4.92
[19:48] Using cautious pH adjust
[19:48] Dispensed 0.001058 mL of Base (0.5 M KOH)
[19:53] Stepping pH = 4.82
[19:53] Dispensed 0.000800 mL of Base (0.5 M KOH)
[19:58] Stepping pH = 4.90
[19:58] Dispensed 0.000212 mL of Base (0.5 M KOH)
[20:03] Stepping pH = 4.92
[20:18] Stirrer speed set to 0
[20:29] Datapoint id 13 collected
[20:29] Charge balance equation is out by 2.5%
[20:29] Stirrer speed set to 50
[20:34] pH 4.92 -> 5.12
[20:34] Using charge balance adjust
[20:34] Dispensed 0.002117 mL of Base (0.5 M KOH)
[20:54] Stirrer speed set to 0
[21:04] Datapoint id 14 collected
[21:04] Charge balance equation is out by -10.3%
[21:04] Stirrer speed set to 50
[21:09] pH 5.10 -> 5.30
[21:09] Using charge balance adjust
[21:09] Dispensed 0.001976 mL of Base (0.5 M KOH)
[21:29] Stirrer speed set to 0
[21:40] Datapoint id 15 collected
[21:40] Charge balance equation is out by -6.8%
[21:40] Stirrer speed set to 50
[21:45] pH 5.30 -> 5.50
[21:45] Using charge balance adjust
[21:45] Dispensed 0.001670 mL of Base (0.5 M KOH)
[22:05] Stirrer speed set to 0
[22:19] Datapoint id 16 collected
[22:19] Charge balance equation is out by 1.3%
[22:19] Stirrer speed set to 50

| | | | |
|---|---|---|---|
| Sample name: | **M15_octanol** | Experiment start time: | **3/3/2018 8:26:22 AM** |
| Assay name: | **pH-metric high logP** | Analyst: | **Pion** |
| Assay ID: | **18C-03006** | Instrument ID: | **T312060** |
| Filename: | **C:\Sirius_T3\Mehtap\20180302_exp29_logP_T3-2\18C-03006_M15_octanol_pH-metric high logP.t3r** | | |

#### Experiment Log (continued)

[22:24] pH 5.50 -> 5.70
[22:24] Using charge balance adjust
[22:24] Dispensed 0.001317 mL of Base (0.5 M KOH)
[22:44] Stirrer speed set to 0
[22:54] Datapoint id 17 collected
[22:54] Charge balance equation is out by 10.1%
[22:54] Stirrer speed set to 50
[23:00] pH 5.73 -> 5.93
[23:00] Using charge balance adjust
[23:00] Dispensed 0.000941 mL of Base (0.5 M KOH)
[23:20] Stirrer speed set to 0
[23:30] Datapoint id 18 collected
[23:30] Charge balance equation is out by 7.6%
[23:30] Stirrer speed set to 50
[23:35] pH 5.95 -> 6.15
[23:35] Using charge balance adjust
[23:35] Dispensed 0.000635 mL of Base (0.5 M KOH)
[23:55] Stirrer speed set to 0
[24:06] Datapoint id 19 collected
[24:06] Charge balance equation is out by 8.2%
[24:06] Stirrer speed set to 50
[24:11] pH 6.18 -> 6.38
[24:11] Using charge balance adjust
[24:11] Dispensed 0.000423 mL of Base (0.5 M KOH)
[24:31] Stirrer speed set to 0
[24:43] Datapoint id 20 collected
[24:43] Charge balance equation is out by 0.3%
[24:43] Stirrer speed set to 50
[24:48] pH 6.39 -> 6.59
[24:48] Using charge balance adjust
[24:48] Dispensed 0.000282 mL of Base (0.5 M KOH)
[25:08] Stirrer speed set to 0
[25:20] Datapoint id 21 collected
[25:20] Charge balance equation is out by -64.1%
[25:20] Stirrer speed set to 50
[25:25] pH 6.47 -> 6.67
[25:25] Using cautious pH adjust
[25:25] Dispensed 0.000118 mL of Base (0.5 M KOH)
[25:30] Stepping pH = 6.48
[25:30] Dispensed 0.000329 mL of Base (0.5 M KOH)
[25:35] Stepping pH = 6.72
[25:50] Stirrer speed set to 0
[26:38] Datapoint id 22 collected
[26:38] Charge balance equation is out by -84.6%
[26:38] Stirrer speed set to 50
[26:43] pH 6.88 -> 7.08
[26:43] Using cautious pH adjust
[26:43] Dispensed 0.000047 mL of Base (0.5 M KOH)
[26:48] Stepping pH = 6.89
[26:48] Dispensed 0.000141 mL of Base (0.5 M KOH)
[26:53] Stepping pH = 6.99
[26:53] Dispensed 0.000071 mL of Base (0.5 M KOH)
[26:59] Stepping pH = 7.16
[27:14] Stirrer speed set to 0
[27:56] Datapoint id 23 collected
[27:56] Charge balance equation is out by -162.3%
[27:56] Stirrer speed set to 50
[28:01] pH 7.29 -> 7.49
[28:01] Using cautious pH adjust

Sample name: **M15_octanol**
Assay name: **pH-metric high logP**
Assay ID: **18C-03006**
Filename: **C:\Sirius_T3\Mehtap\20180302_exp29_logP_T3-2\18C-03006_M15_octanol_pH-metric high logP.t3r**
Experiment start time: **3/3/2018 8:26:22 AM**
Analyst: **Pion**
Instrument ID: **T312060**

#### Experiment Log (continued)

[28:01] Dispensed 0.000024 mL of Base (0.5 M KOH)
[28:07] Stepping pH = 7.31
[28:07] Dispensed 0.000071 mL of Base (0.5 M KOH)
[28:12] Stepping pH = 7.38
[28:12] Dispensed 0.000047 mL of Base (0.5 M KOH)
[28:17] Stepping pH = 7.59
[28:32] Stirrer speed set to 0
[29:32] Datapoint id 24 collected
[29:32] Charge balance equation is out by -197.2%
[29:32] Stirrer speed set to 50
[29:37] pH 7.82 -> 8.02
[29:37] Using cautious pH adjust
[29:37] Dispensed 0.000024 mL of Base (0.5 M KOH)
[29:42] Stepping pH = 7.84
[29:42] Dispensed 0.000024 mL of Base (0.5 M KOH)
[29:47] Stepping pH = 7.88
[29:47] Dispensed 0.000024 mL of Base (0.5 M KOH)
[29:53] Stepping pH = 7.99
[29:53] Dispensed 0.000024 mL of Base (0.5 M KOH)
[29:58] Stepping pH = 8.15
[30:13] Stirrer speed set to 0
[31:13] Datapoint id 25 collected
[31:13] Charge balance equation is out by -439.5%
[31:13] Stirrer speed set to 50
[31:18] pH 8.28 -> 8.48
[31:18] Using cautious pH adjust
[31:18] Dispensed 0.000024 mL of Base (0.5 M KOH)
[31:23] Stepping pH = 8.34
[31:23] Dispensed 0.000024 mL of Base (0.5 M KOH)
[31:28] Stepping pH = 8.41
[31:28] Dispensed 0.000024 mL of Base (0.5 M KOH)
[31:33] Stepping pH = 8.51
[31:48] Stirrer speed set to 0
[32:44] Datapoint id 26 collected
[32:44] Charge balance equation is out by -243.3%
[32:44] Stirrer speed set to 50
[32:49] pH 8.54 -> 8.74
[32:49] Using cautious pH adjust
[32:49] Dispensed 0.000024 mL of Base (0.5 M KOH)
[32:54] Stepping pH = 8.57
[32:54] Dispensed 0.000024 mL of Base (0.5 M KOH)
[32:59] Stepping pH = 8.63
[32:59] Dispensed 0.000047 mL of Base (0.5 M KOH)
[33:04] Stepping pH = 8.72
[33:04] Dispensed 0.000024 mL of Base (0.5 M KOH)
[33:09] Stepping pH = 8.81
[33:24] Stirrer speed set to 0
[34:01] Datapoint id 27 collected
[34:01] Charge balance equation is out by -320.9%
[34:01] Stirrer speed set to 50
[34:06] pH 8.87 -> 9.07
[34:06] Using cautious pH adjust
[34:06] Dispensed 0.000024 mL of Base (0.5 M KOH)
[34:11] Stepping pH = 8.90
[34:11] Dispensed 0.000071 mL of Base (0.5 M KOH)
[34:16] Stepping pH = 8.98
[34:16] Dispensed 0.000047 mL of Base (0.5 M KOH)
[34:21] Stepping pH = 9.08
[34:37] Stirrer speed set to 0

#### Experiment Log (continued)

[35:07] Datapoint id 28 collected
[35:07] Charge balance equation is out by -173.9%
[35:07] Stirrer speed set to 50
[35:12] pH 9.15 -> 9.35
[35:12] Using cautious pH adjust
[35:12] Dispensed 0.000047 mL of Base (0.5 M KOH)
[35:17] Stepping pH = 9.18
[35:17] Dispensed 0.000094 mL of Base (0.5 M KOH)
[35:22] Stepping pH = 9.28
[35:22] Dispensed 0.000047 mL of Base (0.5 M KOH)
[35:27] Stepping pH = 9.36
[35:42] Stirrer speed set to 0
[36:00] Datapoint id 29 collected
[36:00] Charge balance equation is out by -130.4%
[36:00] Stirrer speed set to 50
[36:05] pH 9.40 -> 9.60
[36:05] Using cautious pH adjust
[36:05] Dispensed 0.000071 mL of Base (0.5 M KOH)
[36:10] Stepping pH = 9.42
[36:10] Dispensed 0.000165 mL of Base (0.5 M KOH)
[36:15] Stepping pH = 9.57
[36:15] Dispensed 0.000024 mL of Base (0.5 M KOH)
[36:21] Stepping pH = 9.63
[36:36] Stirrer speed set to 0
[36:53] Datapoint id 30 collected
[36:53] Charge balance equation is out by -89.0%
[36:53] Stirrer speed set to 50
[36:58] pH 9.65 -> 9.85
[36:58] Using cautious pH adjust
[36:58] Dispensed 0.000118 mL of Base (0.5 M KOH)
[37:03] Stepping pH = 9.69
[37:03] Dispensed 0.000188 mL of Base (0.5 M KOH)
[37:08] Stepping pH = 9.83
[37:08] Dispensed 0.000024 mL of Base (0.5 M KOH)
[37:13] Stepping pH = 9.88
[37:28] Stirrer speed set to 0
[37:43] Datapoint id 31 collected
[37:43] Charge balance equation is out by -50.9%
[37:43] Stirrer speed set to 50
[37:48] pH 9.89 -> 10.05
[37:48] Using cautious pH adjust
[37:48] Dispensed 0.000141 mL of Base (0.5 M KOH)
[37:53] Stepping pH = 9.93
[37:53] Dispensed 0.000235 mL of Base (0.5 M KOH)
[37:59] Stepping pH = 10.04
[38:14] Stirrer speed set to 0
[38:25] Datapoint id 32 collected
[38:25] Charge balance equation is out by -35.0%
[38:25] Titration 2 of 3
[38:25] Adding initial titrants
[38:25] Automatically add 0.10000 mL of Octanol
[38:28] Dispensed 0.100000 mL of Octanol
[38:28] Stirrer speed set to 10
[38:29] Stirrer speed set to 55
[38:29] Iterative adjust 10.08 -> 2.00
[38:29] pH 10.08 -> 2.00
[38:31] Dispensed 0.097789 mL of Acid (0.5 M HCl)
[38:36] pH 2.06 -> 2.00
[38:37] Dispensed 0.009196 mL of Acid (0.5 M HCl)

#### Experiment Log (continued)

[39:27] Stirrer speed set to 0
[39:37] Datapoint id 33 collected
[39:37] Stirrer speed set to 55
[39:42] pH 1.98 -> 2.18
[39:42] Using cautious pH adjust
[39:42] Dispensed 0.013711 mL of Base (0.5 M KOH)
[39:48] Stepping pH = 2.06
[39:48] Dispensed 0.011947 mL of Base (0.5 M KOH)
[39:53] Stepping pH = 2.15
[39:53] Dispensed 0.003457 mL of Base (0.5 M KOH)
[39:58] Stepping pH = 2.18
[40:13] Stirrer speed set to 0
[40:24] Datapoint id 34 collected
[40:24] Charge balance equation is out by -6.2%
[40:24] Stirrer speed set to 55
[40:29] pH 2.18 -> 2.38
[40:29] Using charge balance adjust
[40:30] Dispensed 0.017897 mL of Base (0.5 M KOH)
[40:50] Stirrer speed set to 0
[41:04] Datapoint id 35 collected
[41:04] Charge balance equation is out by -6.3%
[41:04] Stirrer speed set to 55
[41:09] pH 2.38 -> 2.58
[41:09] Using charge balance adjust
[41:09] Dispensed 0.012347 mL of Base (0.5 M KOH)
[41:29] Stirrer speed set to 0
[41:39] Datapoint id 36 collected
[41:39] Charge balance equation is out by -7.2%
[41:39] Stirrer speed set to 55
[41:44] pH 2.57 -> 2.77
[41:44] Using charge balance adjust
[41:45] Dispensed 0.008796 mL of Base (0.5 M KOH)
[42:05] Stirrer speed set to 0
[42:16] Datapoint id 37 collected
[42:16] Charge balance equation is out by -0.8%
[42:16] Stirrer speed set to 55
[42:21] pH 2.77 -> 2.97
[42:21] Using charge balance adjust
[42:21] Dispensed 0.006256 mL of Base (0.5 M KOH)
[42:41] Stirrer speed set to 0
[42:51] Datapoint id 38 collected
[42:51] Charge balance equation is out by -4.0%
[42:51] Stirrer speed set to 55
[42:56] pH 2.97 -> 3.17
[42:56] Using charge balance adjust
[42:57] Dispensed 0.004563 mL of Base (0.5 M KOH)
[43:17] Stirrer speed set to 0
[43:27] Datapoint id 39 collected
[43:27] Charge balance equation is out by -5.3%
[43:27] Stirrer speed set to 55
[43:32] pH 3.16 -> 3.36
[43:32] Using charge balance adjust
[43:32] Dispensed 0.003387 mL of Base (0.5 M KOH)
[43:52] Stirrer speed set to 0
[44:02] Datapoint id 40 collected
[44:02] Charge balance equation is out by -3.5%
[44:03] Stirrer speed set to 55
[44:08] pH 3.36 -> 3.56
[44:08] Using charge balance adjust

Sample name: **M15_octanol** Experiment start time: **3/3/2018 8:26:22 AM**
Assay name: **pH-metric high logP** Analyst: **Pion**
Assay ID: **18C-03006** Instrument ID: **T312060**
Filename: **C:\Sirius_T3\Mehtap\20180302_exp29_logP_T3-2\18C-03006_M15_octanol_pH-metric high logP.t3r**

#### Experiment Log (continued)

[44:08] Dispensed 0.002587 mL of Base (0.5 M KOH)
[44:28] Stirrer speed set to 0
[44:38] Datapoint id 41 collected
[44:38] Charge balance equation is out by -11.2%
[44:38] Stirrer speed set to 55
[44:43] pH 3.55 -> 3.75
[44:43] Using charge balance adjust
[44:43] Dispensed 0.002140 mL of Base (0.5 M KOH)
[45:03] Stirrer speed set to 0
[45:13] Datapoint id 42 collected
[45:13] Charge balance equation is out by -2.6%
[45:13] Stirrer speed set to 55
[45:19] pH 3.75 -> 3.95
[45:19] Using charge balance adjust
[45:19] Dispensed 0.001929 mL of Base (0.5 M KOH)
[45:39] Stirrer speed set to 0
[45:49] Datapoint id 43 collected
[45:49] Charge balance equation is out by 18.3%
[45:49] Stirrer speed set to 55
[45:54] pH 3.98 -> 4.18
[45:54] Using cautious pH adjust
[45:54] Dispensed 0.000988 mL of Base (0.5 M KOH)
[45:59] Stepping pH = 4.08
[45:59] Dispensed 0.000753 mL of Base (0.5 M KOH)
[46:04] Stepping pH = 4.16
[46:05] Dispensed 0.000165 mL of Base (0.5 M KOH)
[46:10] Stepping pH = 4.19
[46:25] Stirrer speed set to 0
[46:38] Datapoint id 44 collected
[46:38] Charge balance equation is out by 2.9%
[46:38] Stirrer speed set to 55
[46:43] pH 4.19 -> 4.39
[46:43] Using charge balance adjust
[46:44] Dispensed 0.002117 mL of Base (0.5 M KOH)
[47:04] Stirrer speed set to 0
[47:14] Datapoint id 45 collected
[47:14] Charge balance equation is out by -2.0%
[47:14] Stirrer speed set to 55
[47:19] pH 4.39 -> 4.59
[47:19] Using charge balance adjust
[47:19] Dispensed 0.002234 mL of Base (0.5 M KOH)
[47:39] Stirrer speed set to 0
[47:49] Datapoint id 46 collected
[47:49] Charge balance equation is out by -1.0%
[47:49] Stirrer speed set to 55
[47:54] pH 4.59 -> 4.79
[47:54] Using charge balance adjust
[47:54] Dispensed 0.002187 mL of Base (0.5 M KOH)
[48:15] Stirrer speed set to 0
[48:25] Datapoint id 47 collected
[48:25] Charge balance equation is out by 11.4%
[48:25] Stirrer speed set to 55
[48:30] pH 4.81 -> 5.01
[48:30] Using charge balance adjust
[48:30] Dispensed 0.001976 mL of Base (0.5 M KOH)
[48:50] Stirrer speed set to 0
[49:01] Datapoint id 48 collected
[49:01] Charge balance equation is out by 14.0%
[49:01] Stirrer speed set to 55

#### Experiment Log (continued)

[49:06] pH 5.04 -> 5.24
[49:06] Using charge balance adjust
[49:06] Dispensed 0.001576 mL of Base (0.5 M KOH)
[49:26] Stirrer speed set to 0
[49:37] Datapoint id 49 collected
[49:37] Charge balance equation is out by 3.3%
[49:37] Stirrer speed set to 55
[49:42] pH 5.26 -> 5.46
[49:42] Using charge balance adjust
[49:42] Dispensed 0.001176 mL of Base (0.5 M KOH)
[50:02] Stirrer speed set to 0
[50:13] Datapoint id 50 collected
[50:13] Charge balance equation is out by -3.5%
[50:13] Stirrer speed set to 55
[50:18] pH 5.46 -> 5.66
[50:18] Using charge balance adjust
[50:18] Dispensed 0.000847 mL of Base (0.5 M KOH)
[50:38] Stirrer speed set to 0
[50:49] Datapoint id 51 collected
[50:49] Charge balance equation is out by 7.1%
[50:49] Stirrer speed set to 55
[50:54] pH 5.68 -> 5.88
[50:54] Using charge balance adjust
[50:54] Dispensed 0.000588 mL of Base (0.5 M KOH)
[51:14] Stirrer speed set to 0
[51:26] Datapoint id 52 collected
[51:26] Charge balance equation is out by -2.4%
[51:26] Stirrer speed set to 55
[51:31] pH 5.88 -> 6.08
[51:31] Using charge balance adjust
[51:32] Dispensed 0.000400 mL of Base (0.5 M KOH)
[51:52] Stirrer speed set to 0
[52:04] Datapoint id 53 collected
[52:04] Charge balance equation is out by -12.9%
[52:04] Stirrer speed set to 55
[52:09] pH 6.06 -> 6.26
[52:09] Using charge balance adjust
[52:10] Dispensed 0.000306 mL of Base (0.5 M KOH)
[52:30] Stirrer speed set to 0
[52:53] Datapoint id 54 collected
[52:53] Charge balance equation is out by 17.6%
[52:53] Stirrer speed set to 55
[52:58] pH 6.30 -> 6.50
[52:58] Using cautious pH adjust
[52:59] Dispensed 0.000094 mL of Base (0.5 M KOH)
[53:04] Stepping pH = 6.32
[53:04] Dispensed 0.000282 mL of Base (0.5 M KOH)
[53:09] Stepping pH = 6.75
[53:24] Stirrer speed set to 0
[54:24] Datapoint id 55 collected
[54:24] Charge balance equation is out by -87.2%
[54:24] Stirrer speed set to 55
[54:29] pH 7.05 -> 7.25
[54:29] Using cautious pH adjust
[54:29] Dispensed 0.000024 mL of Base (0.5 M KOH)
[54:34] Stepping pH = 7.07
[54:34] Dispensed 0.000071 mL of Base (0.5 M KOH)
[54:39] Stepping pH = 7.11
[54:39] Dispensed 0.000118 mL of Base (0.5 M KOH)

| | | | |
|---|---|---|---|
| Sample name: | **M15_octanol** | Experiment start time: | **3/3/2018 8:26:22 AM** |
| Assay name: | **pH-metric high logP** | Analyst: | **Pion** |
| Assay ID: | **18C-03006** | Instrument ID: | **T312060** |
| Filename: | **C:\Sirius_T3\Mehtap\20180302_exp29_logP_T3-2\18C-03006_M15_octanol_pH-metric high logP.t3r** | | |

#### Experiment Log (continued)

[54:45] Stepping pH = 7.46
[55:00] Stirrer speed set to 0
[56:00] Datapoint id 56 collected
[56:00] Charge balance equation is out by -289.0%
[56:00] Stirrer speed set to 55
[56:05] pH 7.69 -> 7.89
[56:05] Using cautious pH adjust
[56:05] Dispensed 0.000024 mL of Base (0.5 M KOH)
[56:10] Stepping pH = 7.74
[56:10] Dispensed 0.000024 mL of Base (0.5 M KOH)
[56:15] Stepping pH = 7.79
[56:15] Dispensed 0.000024 mL of Base (0.5 M KOH)
[56:20] Stepping pH = 7.84
[56:20] Dispensed 0.000024 mL of Base (0.5 M KOH)
[56:25] Stepping pH = 7.93
[56:40] Stirrer speed set to 0
[57:40] Datapoint id 57 collected
[57:40] Charge balance equation is out by -381.5%
[57:40] Stirrer speed set to 55
[57:45] pH 7.90 -> 8.10
[57:45] Using cautious pH adjust
[57:46] Dispensed 0.000024 mL of Base (0.5 M KOH)
[57:51] Stepping pH = 7.94
[57:51] Dispensed 0.000024 mL of Base (0.5 M KOH)
[57:56] Stepping pH = 8.00
[57:56] Dispensed 0.000024 mL of Base (0.5 M KOH)
[58:01] Stepping pH = 8.08
[58:01] Dispensed 0.000024 mL of Base (0.5 M KOH)
[58:06] Stepping pH = 8.17
[58:21] Stirrer speed set to 0
[59:21] Datapoint id 58 collected
[59:21] Charge balance equation is out by -472.6%
[59:21] Stirrer speed set to 55
[59:26] pH 8.19 -> 8.39
[59:26] Using cautious pH adjust
[59:26] Dispensed 0.000024 mL of Base (0.5 M KOH)
[59:31] Stepping pH = 8.22
[59:31] Dispensed 0.000024 mL of Base (0.5 M KOH)
[59:37] Stepping pH = 8.27
[59:37] Dispensed 0.000024 mL of Base (0.5 M KOH)
[59:42] Stepping pH = 8.33
[59:42] Dispensed 0.000024 mL of Base (0.5 M KOH)
[59:47] Stepping pH = 8.37
[59:47] Dispensed 0.000024 mL of Base (0.5 M KOH)
[59:52] Stepping pH = 8.35
[59:52] Dispensed 0.000071 mL of Base (0.5 M KOH)
[59:57] Stepping pH = 8.58
[1:00:12] Stirrer speed set to 0
[1:00:49] Datapoint id 59 collected
[1:00:49] Charge balance equation is out by -1,013.4%
[1:00:49] Stirrer speed set to 55
[1:00:54] pH 8.64 -> 8.84
[1:00:54] Using cautious pH adjust
[1:00:54] Dispensed 0.000024 mL of Base (0.5 M KOH)
[1:00:59] Stepping pH = 8.66
[1:00:59] Dispensed 0.000047 mL of Base (0.5 M KOH)
[1:01:04] Stepping pH = 8.73
[1:01:04] Dispensed 0.000047 mL of Base (0.5 M KOH)
[1:01:09] Stepping pH = 8.83

| | | | |
|---|---|---|---|
| Sample name: | **M15_octanol** | Experiment start time: | **3/3/2018 8:26:22 AM** |
| Assay name: | **pH-metric high logP** | Analyst: | **Pion** |
| Assay ID: | **18C-03006** | Instrument ID: | **T312060** |
| Filename: | **C:\Sirius_T3\Mehtap\20180302_exp29_logP_T3-2\18C-03006_M15_octanol_pH-metric high logP.t3r** | | |

#### Experiment Log (continued)

[1:01:09] Dispensed 0.000024 mL of Base (0.5 M KOH)
[1:01:15] Stepping pH = 8.89
[1:01:30] Stirrer speed set to 0
[1:01:49] Datapoint id 60 collected
[1:01:49] Charge balance equation is out by -319.6%
[1:01:49] Stirrer speed set to 55
[1:01:54] pH 8.89 -> 9.09
[1:01:54] Using cautious pH adjust
[1:01:54] Dispensed 0.000024 mL of Base (0.5 M KOH)
[1:01:59] Stepping pH = 8.91
[1:01:59] Dispensed 0.000071 mL of Base (0.5 M KOH)
[1:02:04] Stepping pH = 8.99
[1:02:05] Dispensed 0.000071 mL of Base (0.5 M KOH)
[1:02:10] Stepping pH = 9.11
[1:02:25] Stirrer speed set to 0
[1:02:55] Datapoint id 61 collected
[1:02:55] Charge balance equation is out by -216.7%
[1:02:55] Stirrer speed set to 55
[1:03:00] pH 9.15 -> 9.35
[1:03:00] Using cautious pH adjust
[1:03:00] Dispensed 0.000047 mL of Base (0.5 M KOH)
[1:03:05] Stepping pH = 9.17
[1:03:05] Dispensed 0.000141 mL of Base (0.5 M KOH)
[1:03:10] Stepping pH = 9.37
[1:03:25] Stirrer speed set to 0
[1:03:38] Datapoint id 62 collected
[1:03:38] Charge balance equation is out by -91.0%
[1:03:38] Stirrer speed set to 55
[1:03:43] pH 9.43 -> 9.63
[1:03:43] Using cautious pH adjust
[1:03:43] Dispensed 0.000071 mL of Base (0.5 M KOH)
[1:03:48] Stepping pH = 9.45
[1:03:48] Dispensed 0.000235 mL of Base (0.5 M KOH)
[1:03:53] Stepping pH = 9.64
[1:04:08] Stirrer speed set to 0
[1:04:22] Datapoint id 63 collected
[1:04:22] Charge balance equation is out by -92.6%
[1:04:22] Stirrer speed set to 55
[1:04:27] pH 9.71 -> 9.91
[1:04:27] Using cautious pH adjust
[1:04:27] Dispensed 0.000141 mL of Base (0.5 M KOH)
[1:04:32] Stepping pH = 9.75
[1:04:32] Dispensed 0.000235 mL of Base (0.5 M KOH)
[1:04:37] Stepping pH = 9.91
[1:04:53] Stirrer speed set to 0
[1:05:11] Datapoint id 64 collected
[1:05:11] Charge balance equation is out by -37.0%
[1:05:11] Stirrer speed set to 55
[1:05:16] pH 9.95 -> 10.05
[1:05:16] Using cautious pH adjust
[1:05:16] Dispensed 0.000094 mL of Base (0.5 M KOH)
[1:05:21] Stepping pH = 9.95
[1:05:21] Dispensed 0.000282 mL of Base (0.5 M KOH)
[1:05:26] Stepping pH = 10.07
[1:05:42] Stirrer speed set to 0
[1:05:52] Datapoint id 65 collected
[1:05:52] Charge balance equation is out by -89.5%
[1:05:52] Titration 3 of 3
[1:05:52] Adding initial titrants

Sample name: **M15_octanol**
Assay name: **pH-metric high logP**
Assay ID: **18C-03006**
Filename: **C:\Sirius_T3\Mehtap\20180302_exp29_logP_T3-2\18C-03006_M15_octanol_pH-metric high logP.t3r**
Experiment start time: **3/3/2018 8:26:22 AM**
Analyst: **Pion**
Instrument ID: **T312060**

#### Experiment Log (continued)

[1:05:52] Automatically add 0.30000 mL of Octanol
[1:05:59] Dispensed 0.300000 mL of Octanol
[1:05:59] Stirrer speed set to 10
[1:06:00] Stirrer speed set to 60
[1:06:00] Iterative adjust 10.11 -> 2.00
[1:06:00] pH 10.11 -> 2.00
[1:06:03] Dispensed 0.100000 mL of Acid (0.5 M HCl)
[1:06:08] pH 2.07 -> 2.00
[1:06:08] Dispensed 0.011642 mL of Acid (0.5 M HCl)
[1:06:58] Stirrer speed set to 0
[1:07:08] Datapoint id 66 collected
[1:07:08] Stirrer speed set to 60
[1:07:14] pH 1.98 -> 2.18
[1:07:14] Using cautious pH adjust
[1:07:14] Dispensed 0.014887 mL of Base (0.5 M KOH)
[1:07:19] Stepping pH = 2.06
[1:07:20] Dispensed 0.012512 mL of Base (0.5 M KOH)
[1:07:25] Stepping pH = 2.15
[1:07:25] Dispensed 0.003175 mL of Base (0.5 M KOH)
[1:07:30] Stepping pH = 2.18
[1:07:45] Stirrer speed set to 0
[1:07:58] Datapoint id 67 collected
[1:07:58] Charge balance equation is out by -2.6%
[1:07:58] Stirrer speed set to 60
[1:08:03] pH 2.19 -> 2.39
[1:08:03] Using charge balance adjust
[1:08:03] Dispensed 0.019120 mL of Base (0.5 M KOH)
[1:08:23] Stirrer speed set to 0
[1:08:33] Datapoint id 68 collected
[1:08:33] Charge balance equation is out by -2.3%
[1:08:33] Stirrer speed set to 60
[1:08:39] pH 2.39 -> 2.59
[1:08:39] Using charge balance adjust
[1:08:39] Dispensed 0.012982 mL of Base (0.5 M KOH)
[1:08:59] Stirrer speed set to 0
[1:09:09] Datapoint id 69 collected
[1:09:09] Charge balance equation is out by -8.3%
[1:09:09] Stirrer speed set to 60
[1:09:14] pH 2.58 -> 2.78
[1:09:14] Using charge balance adjust
[1:09:15] Dispensed 0.009266 mL of Base (0.5 M KOH)
[1:09:35] Stirrer speed set to 0
[1:09:45] Datapoint id 70 collected
[1:09:45] Charge balance equation is out by 2.2%
[1:09:45] Stirrer speed set to 60
[1:09:50] pH 2.79 -> 2.99
[1:09:50] Using charge balance adjust
[1:09:50] Dispensed 0.006609 mL of Base (0.5 M KOH)
[1:10:10] Stirrer speed set to 0
[1:10:21] Datapoint id 71 collected
[1:10:21] Charge balance equation is out by -5.3%
[1:10:21] Stirrer speed set to 60
[1:10:26] pH 2.98 -> 3.18
[1:10:26] Using charge balance adjust
[1:10:26] Dispensed 0.004986 mL of Base (0.5 M KOH)
[1:10:46] Stirrer speed set to 0
[1:10:57] Datapoint id 72 collected
[1:10:57] Charge balance equation is out by -2.9%
[1:10:57] Stirrer speed set to 60

| | | | |
|---|---|---|---|
| Sample name: | **M15_octanol** | Experiment start time: | **3/3/2018 8:26:22 AM** |
| Assay name: | **pH-metric high logP** | Analyst: | **Pion** |
| Assay ID: | **18C-03006** | Instrument ID: | **T312060** |
| Filename: | **C:\Sirius_T3\Mehtap\20180302_exp29_logP_T3-2\18C-03006_M15_octanol_pH-metric high logP.t3r** | | |

#### Experiment Log (continued)

[1:11:02] pH 3.18 -> 3.38
[1:11:02] Using charge balance adjust
[1:11:02] Dispensed 0.003857 mL of Base (0.5 M KOH)
[1:11:22] Stirrer speed set to 0
[1:11:32] Datapoint id 73 collected
[1:11:32] Charge balance equation is out by 9.5%
[1:11:32] Stirrer speed set to 60
[1:11:37] pH 3.41 -> 3.61
[1:11:37] Using charge balance adjust
[1:11:38] Dispensed 0.003104 mL of Base (0.5 M KOH)
[1:11:58] Stirrer speed set to 0
[1:12:08] Datapoint id 74 collected
[1:12:08] Charge balance equation is out by 0.7%
[1:12:08] Stirrer speed set to 60
[1:12:13] pH 3.61 -> 3.81
[1:12:13] Using charge balance adjust
[1:12:13] Dispensed 0.002775 mL of Base (0.5 M KOH)
[1:12:33] Stirrer speed set to 0
[1:12:44] Datapoint id 75 collected
[1:12:44] Charge balance equation is out by 12.4%
[1:12:44] Stirrer speed set to 60
[1:12:49] pH 3.83 -> 4.03
[1:12:49] Using charge balance adjust
[1:12:49] Dispensed 0.002658 mL of Base (0.5 M KOH)
[1:13:09] Stirrer speed set to 0
[1:13:27] Datapoint id 76 collected
[1:13:27] Charge balance equation is out by 13.3%
[1:13:27] Stirrer speed set to 60
[1:13:33] pH 4.05 -> 4.25
[1:13:33] Using charge balance adjust
[1:13:33] Dispensed 0.002540 mL of Base (0.5 M KOH)
[1:13:53] Stirrer speed set to 0
[1:14:03] Datapoint id 77 collected
[1:14:03] Charge balance equation is out by -4.8%
[1:14:03] Stirrer speed set to 60
[1:14:08] pH 4.24 -> 4.44
[1:14:08] Using charge balance adjust
[1:14:08] Dispensed 0.002375 mL of Base (0.5 M KOH)
[1:14:28] Stirrer speed set to 0
[1:14:38] Datapoint id 78 collected
[1:14:38] Charge balance equation is out by 0.7%
[1:14:38] Stirrer speed set to 60
[1:14:44] pH 4.44 -> 4.64
[1:14:44] Using charge balance adjust
[1:14:44] Dispensed 0.002046 mL of Base (0.5 M KOH)
[1:15:04] Stirrer speed set to 0
[1:15:14] Datapoint id 79 collected
[1:15:14] Charge balance equation is out by 0.9%
[1:15:14] Stirrer speed set to 60
[1:15:19] pH 4.64 -> 4.84
[1:15:19] Using charge balance adjust
[1:15:19] Dispensed 0.001646 mL of Base (0.5 M KOH)
[1:15:39] Stirrer speed set to 0
[1:16:00] Datapoint id 80 collected
[1:16:00] Charge balance equation is out by 5.4%
[1:16:00] Stirrer speed set to 60
[1:16:05] pH 4.85 -> 5.05
[1:16:05] Using charge balance adjust
[1:16:05] Dispensed 0.001223 mL of Base (0.5 M KOH)

| | | | |
|---|---|---|---|
| Sample name: | **M15_octanol** | Experiment start time: | **3/3/2018 8:26:22 AM** |
| Assay name: | **pH-metric high logP** | Analyst: | **Pion** |
| Assay ID: | **18C-03006** | Instrument ID: | **T312060** |
| Filename: | **C:\Sirius_T3\Mehtap\20180302_exp29_logP_T3-2\18C-03006_M15_octanol_pH-metric high logP.t3r** | | |

#### Experiment Log (continued)

[1:16:25] Stirrer speed set to 0
[1:16:44] Datapoint id 81 collected
[1:16:44] Charge balance equation is out by 18.8%
[1:16:44] Stirrer speed set to 60
[1:16:49] pH 5.09 -> 5.29
[1:16:49] Using cautious pH adjust
[1:16:49] Dispensed 0.000400 mL of Base (0.5 M KOH)
[1:16:55] Stepping pH = 5.16
[1:16:55] Dispensed 0.000447 mL of Base (0.5 M KOH)
[1:17:00] Stepping pH = 5.28
[1:17:00] Dispensed 0.000047 mL of Base (0.5 M KOH)
[1:17:05] Stepping pH = 5.31
[1:17:20] Stirrer speed set to 0
[1:17:31] Datapoint id 82 collected
[1:17:31] Charge balance equation is out by -9.2%
[1:17:31] Stirrer speed set to 60
[1:17:36] pH 5.32 -> 5.52
[1:17:36] Using charge balance adjust
[1:17:36] Dispensed 0.000541 mL of Base (0.5 M KOH)
[1:17:56] Stirrer speed set to 0
[1:18:08] Datapoint id 83 collected
[1:18:08] Charge balance equation is out by 10.6%
[1:18:08] Stirrer speed set to 60
[1:18:13] pH 5.55 -> 5.75
[1:18:13] Using charge balance adjust
[1:18:13] Dispensed 0.000353 mL of Base (0.5 M KOH)
[1:18:33] Stirrer speed set to 0
[1:18:51] Datapoint id 84 collected
[1:18:51] Charge balance equation is out by 9.2%
[1:18:51] Stirrer speed set to 60
[1:18:56] pH 5.77 -> 5.97
[1:18:56] Using charge balance adjust
[1:18:57] Dispensed 0.000235 mL of Base (0.5 M KOH)
[1:19:17] Stirrer speed set to 0
[1:19:32] Datapoint id 85 collected
[1:19:32] Charge balance equation is out by -5.5%
[1:19:32] Stirrer speed set to 60
[1:19:37] pH 5.96 -> 6.16
[1:19:37] Using charge balance adjust
[1:19:37] Dispensed 0.000188 mL of Base (0.5 M KOH)
[1:19:57] Stirrer speed set to 0
[1:20:27] Datapoint id 86 collected
[1:20:27] Charge balance equation is out by 5.3%
[1:20:27] Stirrer speed set to 60
[1:20:32] pH 6.18 -> 6.38
[1:20:32] Using charge balance adjust
[1:20:32] Dispensed 0.000141 mL of Base (0.5 M KOH)
[1:20:53] Stirrer speed set to 0
[1:21:52] Datapoint id 87 collected
[1:21:52] Charge balance equation is out by 37.0%
[1:21:52] Stirrer speed set to 60
[1:21:57] pH 6.45 -> 6.65
[1:21:57] Using cautious pH adjust
[1:21:57] Dispensed 0.000047 mL of Base (0.5 M KOH)
[1:22:02] Stepping pH = 6.47
[1:22:02] Dispensed 0.000141 mL of Base (0.5 M KOH)
[1:22:08] Stepping pH = 6.91
[1:22:23] Stirrer speed set to 0
[1:23:23] Datapoint id 88 collected

| | | | |
|---|---|---|---|
| Sample name: | **M15_octanol** | Experiment start time: | **3/3/2018 8:26:22 AM** |
| Assay name: | **pH-metric high logP** | Analyst: | **Pion** |
| Assay ID: | **18C-03006** | Instrument ID: | **T312060** |
| Filename: | **C:\Sirius_T3\Mehtap\20180302_exp29_logP_T3-2\18C-03006_M15_octanol_pH-metric high logP.t3r** | | |

#### Experiment Log (continued)

[1:23:23] Charge balance equation is out by -88.9%
[1:23:23] Stirrer speed set to 60
[1:23:28] pH 7.01 -> 7.21
[1:23:28] Using cautious pH adjust
[1:23:28] Dispensed 0.000024 mL of Base (0.5 M KOH)
[1:23:33] Stepping pH = 7.02
[1:23:33] Dispensed 0.000071 mL of Base (0.5 M KOH)
[1:23:38] Stepping pH = 7.16
[1:23:38] Dispensed 0.000024 mL of Base (0.5 M KOH)
[1:23:43] Stepping pH = 7.31
[1:23:58] Stirrer speed set to 0
[1:24:58] Datapoint id 89 collected
[1:24:58] Charge balance equation is out by -137.7%
[1:24:58] Stirrer speed set to 60
[1:25:03] pH 7.32 -> 7.52
[1:25:03] Using cautious pH adjust
[1:25:03] Dispensed 0.000024 mL of Base (0.5 M KOH)
[1:25:09] Stepping pH = 7.42
[1:25:09] Dispensed 0.000024 mL of Base (0.5 M KOH)
[1:25:14] Stepping pH = 7.52
[1:25:29] Stirrer speed set to 0
[1:26:29] Datapoint id 90 collected
[1:26:29] Charge balance equation is out by -46.2%
[1:26:29] Stirrer speed set to 60
[1:26:34] pH 7.52 -> 7.72
[1:26:34] Using cautious pH adjust
[1:26:34] Dispensed 0.000024 mL of Base (0.5 M KOH)
[1:26:39] Stepping pH = 7.60
[1:26:39] Dispensed 0.000024 mL of Base (0.5 M KOH)
[1:26:44] Stepping pH = 7.66
[1:26:44] Dispensed 0.000024 mL of Base (0.5 M KOH)
[1:26:49] Stepping pH = 7.76
[1:27:04] Stirrer speed set to 0
[1:28:04] Datapoint id 91 collected
[1:28:04] Charge balance equation is out by -201.7%
[1:28:04] Stirrer speed set to 60
[1:28:10] pH 7.83 -> 8.03
[1:28:10] Using cautious pH adjust
[1:28:10] Dispensed 0.000024 mL of Base (0.5 M KOH)
[1:28:15] Stepping pH = 7.86
[1:28:15] Dispensed 0.000024 mL of Base (0.5 M KOH)
[1:28:20] Stepping pH = 7.99
[1:28:20] Dispensed 0.000024 mL of Base (0.5 M KOH)
[1:28:25] Stepping pH = 8.13
[1:28:40] Stirrer speed set to 0
[1:29:40] Datapoint id 92 collected
[1:29:40] Charge balance equation is out by -321.0%
[1:29:40] Stirrer speed set to 60
[1:29:45] pH 8.07 -> 8.27
[1:29:45] Using cautious pH adjust
[1:29:45] Dispensed 0.000024 mL of Base (0.5 M KOH)
[1:29:50] Stepping pH = 8.07
[1:29:50] Dispensed 0.000047 mL of Base (0.5 M KOH)
[1:29:55] Stepping pH = 8.24
[1:29:56] Dispensed 0.000024 mL of Base (0.5 M KOH)
[1:30:01] Stepping pH = 8.36
[1:30:16] Stirrer speed set to 0
[1:31:16] Datapoint id 93 collected
[1:31:16] Charge balance equation is out by -434.1%

| | | | |
|---|---|---|---|
| Sample name: | **M15_octanol** | Experiment start time: | **3/3/2018 8:26:22 AM** |
| Assay name: | **pH-metric high logP** | Analyst: | **Pion** |
| Assay ID: | **18C-03006** | Instrument ID: | **T312060** |
| Filename: | **C:\Sirius_T3\Mehtap\20180302_exp29_logP_T3-2\18C-03006_M15_octanol_pH-metric high logP.t3r** | | |

#### Experiment Log (continued)

[1:31:16] Stirrer speed set to 60
[1:31:21] pH 8.37 -> 8.57
[1:31:21] Using cautious pH adjust
[1:31:21] Dispensed 0.000024 mL of Base (0.5 M KOH)
[1:31:26] Stepping pH = 8.37
[1:31:26] Dispensed 0.000024 mL of Base (0.5 M KOH)
[1:31:31] Stepping pH = 8.37
[1:31:31] Dispensed 0.000165 mL of Base (0.5 M KOH)
[1:31:36] Stepping pH = 8.72
[1:31:51] Stirrer speed set to 0
[1:32:12] Datapoint id 94 collected
[1:32:12] Charge balance equation is out by -893.2%
[1:32:12] Stirrer speed set to 60
[1:32:18] pH 8.83 -> 9.03
[1:32:18] Using cautious pH adjust
[1:32:18] Dispensed 0.000024 mL of Base (0.5 M KOH)
[1:32:23] Stepping pH = 8.84
[1:32:23] Dispensed 0.000071 mL of Base (0.5 M KOH)
[1:32:28] Stepping pH = 8.87
[1:32:28] Dispensed 0.000165 mL of Base (0.5 M KOH)
[1:32:33] Stepping pH = 9.03
[1:32:48] Stirrer speed set to 0
[1:33:16] Datapoint id 95 collected
[1:33:16] Charge balance equation is out by -410.9%
[1:33:16] Stirrer speed set to 60
[1:33:21] pH 9.09 -> 9.29
[1:33:21] Using cautious pH adjust
[1:33:21] Dispensed 0.000047 mL of Base (0.5 M KOH)
[1:33:26] Stepping pH = 9.10
[1:33:26] Dispensed 0.000118 mL of Base (0.5 M KOH)
[1:33:31] Stepping pH = 9.14
[1:33:31] Dispensed 0.000212 mL of Base (0.5 M KOH)
[1:33:36] Stepping pH = 9.32
[1:33:51] Stirrer speed set to 0
[1:34:06] Datapoint id 96 collected
[1:34:06] Charge balance equation is out by -341.4%
[1:34:06] Stirrer speed set to 60
[1:34:11] pH 9.38 -> 9.58
[1:34:11] Using cautious pH adjust
[1:34:11] Dispensed 0.000071 mL of Base (0.5 M KOH)
[1:34:16] Stepping pH = 9.39
[1:34:16] Dispensed 0.000235 mL of Base (0.5 M KOH)
[1:34:21] Stepping pH = 9.52
[1:34:21] Dispensed 0.000094 mL of Base (0.5 M KOH)
[1:34:26] Stepping pH = 9.60
[1:34:42] Stirrer speed set to 0
[1:34:54] Datapoint id 97 collected
[1:34:54] Charge balance equation is out by -153.1%
[1:34:54] Stirrer speed set to 60
[1:34:59] pH 9.62 -> 9.82
[1:34:59] Using cautious pH adjust
[1:34:59] Dispensed 0.000118 mL of Base (0.5 M KOH)
[1:35:04] Stepping pH = 9.66
[1:35:04] Dispensed 0.000235 mL of Base (0.5 M KOH)
[1:35:10] Stepping pH = 9.78
[1:35:10] Dispensed 0.000071 mL of Base (0.5 M KOH)
[1:35:15] Stepping pH = 9.83
[1:35:30] Stirrer speed set to 0
[1:35:51] Datapoint id 98 collected

Sample name: **M15_octanol**
Assay name: **pH-metric high logP**
Assay ID: **18C-03006**
Filename: **C:\Sirius_T3\Mehtap\20180302_exp29_logP_T3-2\18C-03006_M15_octanol_pH-metric high logP.t3r**
Experiment start time: **3/3/2018 8:26:22 AM**
Analyst: **Pion**
Instrument ID: **T312060**

#### Experiment Log (continued)

[1:35:51] Charge balance equation is out by -75.4%
[1:35:51] Stirrer speed set to 60
[1:35:57] pH 9.84 -> 10.04
[1:35:57] Using cautious pH adjust
[1:35:57] Dispensed 0.000188 mL of Base (0.5 M KOH)
[1:36:02] Stepping pH = 9.90
[1:36:02] Dispensed 0.000235 mL of Base (0.5 M KOH)
[1:36:07] Stepping pH = 10.00
[1:36:07] Dispensed 0.000094 mL of Base (0.5 M KOH)
[1:36:12] Stepping pH = 10.04
[1:36:27] Stirrer speed set to 0
[1:36:39] Datapoint id 99 collected
[1:36:39] Charge balance equation is out by -35.9%
[1:36:39] Argon flow rate set to 0
[1:36:43] Titrator arm moved over Titration position
