## Supplementary material for "Octanol-water partition coefficient measurements for the SAMPL6 Blind Prediction Challenge": SM15_18C-01007_M16_octanol_pH-metric high logP_report.pdf

| | | | |
|---|---|---|---|
| Sample name: | **M16_octanol** | Experiment start time: | **3/1/2018 9:25:34 AM** |
| Assay name: | **pH-metric high logP** | Analyst: | **Pion** |
| Assay ID: | **18C-01007** | Instrument ID: | **T312060** |
| Filename: | **C:\Sirius_T3\Mehtap\20180228_exp28_logP_T3-2\18C-01007_M16_octanol_pH-metric high logP.t3r** | | |

### pH-metric Result

| | | |
|---|---|---|
| logP (XH2 +) | 1.38 | ±1.07 (n=50) |
| logP (neutral XH) | 3.14 | ±0.01 (n=50) |
| logP (X -) | -1.62 | |

### 18C-01007 Points 1 to 30

| | |
|---|---|
| M16_octanol concentration factor | 0.780 |
| Carbonate | 0.1052 mM |
| Acidity error | 3.38405 mM |

### 18C-01007 Points 31 to 58

| | |
|---|---|
| M16_octanol concentration factor | 0.751 |
| Carbonate | 0.8332 mM |
| Acidity error | 2.20126 mM |

### 18C-01007 Points 59 to 86

| | |
|---|---|
| M16_octanol concentration factor | 0.772 |
| Carbonate | 0.5481 mM |
| Acidity error | 2.29676 mM |

### Warnings and errors

| | |
|---|---|
| Errors | None |
| Warnings | Excessive carbonate concentration present |

### Sample logD and percent species

| pH | M16_octanol logD | M16_octanol M16_octanolH2 | M16_octanol M16_octanolH | M16_octanol M16_octanol | M16_octanol M16_octanolH2* | M16_octanol M16_octanolH* | M16_octanol M16_octanol* | Comment |
|---|---|---|---|---|---|---|---|---|
| 1.000 | 1.39 | 3.94 % | 0.00 % | 0.00 % | 94.96 % | 1.10 % | 0.00 % | |
| 1.200 | 1.39 | 3.92 % | 0.00 % | 0.00 % | 94.35 % | 1.73 % | 0.00 % | Stomach pH |
| 2.000 | 1.43 | 3.59 % | 0.01 % | 0.00 % | 86.42 % | 9.99 % | 0.00 % | |
| 3.000 | 1.71 | 1.89 % | 0.04 % | 0.00 % | 45.50 % | 52.58 % | 0.00 % | |
| 4.000 | 2.40 | 0.33 % | 0.07 % | 0.00 % | 7.93 % | 91.67 % | 0.00 % | |
| 5.000 | 2.97 | 0.04 % | 0.07 % | 0.00 % | 0.86 % | 99.04 % | 0.00 % | |
| 6.000 | 3.12 | 0.00 % | 0.07 % | 0.00 % | 0.09 % | 99.84 % | 0.00 % | |
| 6.500 | 3.14 | 0.00 % | 0.07 % | 0.00 % | 0.03 % | 99.90 % | 0.00 % | |
| 7.000 | 3.14 | 0.00 % | 0.07 % | 0.00 % | 0.01 % | 99.92 % | 0.00 % | |
| 7.400 | 3.13 | 0.00 % | 0.07 % | 0.00 % | 0.00 % | 99.92 % | 0.00 % | Blood pH |
| 8.000 | 3.10 | 0.00 % | 0.07 % | 0.01 % | 0.00 % | 99.92 % | 0.00 % | |
| 9.000 | 2.81 | 0.00 % | 0.07 % | 0.08 % | 0.00 % | 99.84 % | 0.00 % | |
| 10.000 | 2.05 | 0.00 % | 0.07 % | 0.82 % | 0.00 % | 99.09 % | 0.02 % | |
| 11.000 | 1.08 | 0.00 % | 0.07 % | 7.59 % | 0.00 % | 92.16 % | 0.18 % | |
| 12.000 | 0.09 | 0.00 % | 0.04 % | 44.66 % | 0.00 % | 54.23 % | 1.07 % | |

| | | | |
|---|---|---|---|
| Sample name: | **M16_octanol** | Experiment start time: | **3/1/2018 9:25:34 AM** |
| Assay name: | **pH-metric high logP** | Analyst: | **Pion** |
| Assay ID: | **18C-01007** | Instrument ID: | **T312060** |
| Filename: | **C:\Sirius_T3\Mehtap\20180228_exp28_logP_T3-2\18C-01007_M16_octanol_pH-metric high logP.t3r** | | |

### Graphs

Charge balance residuals

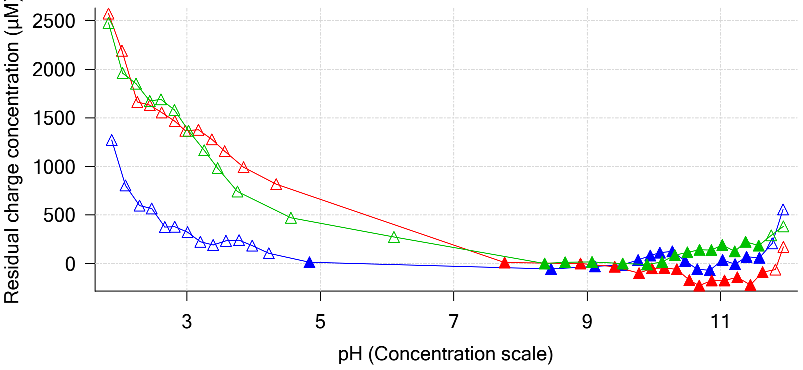


Ionic strength

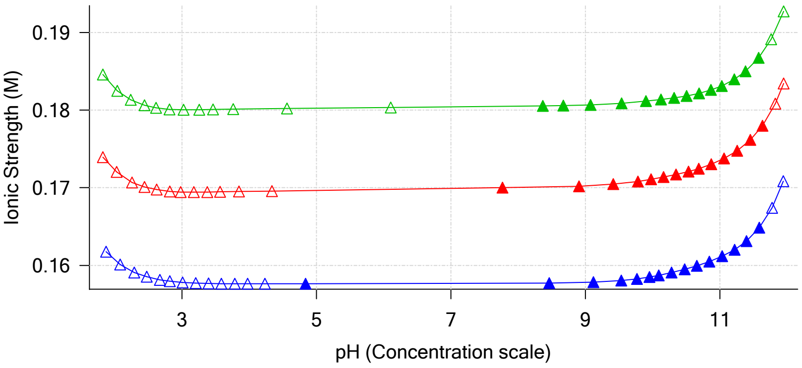


Buffer index profile

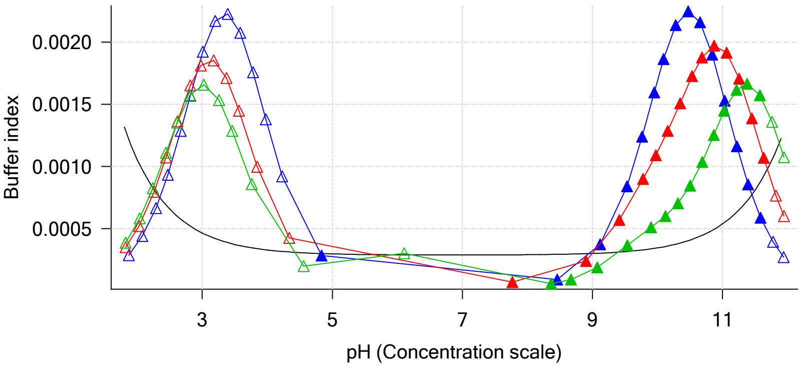


Sample M16_octanol Ionisation Graph

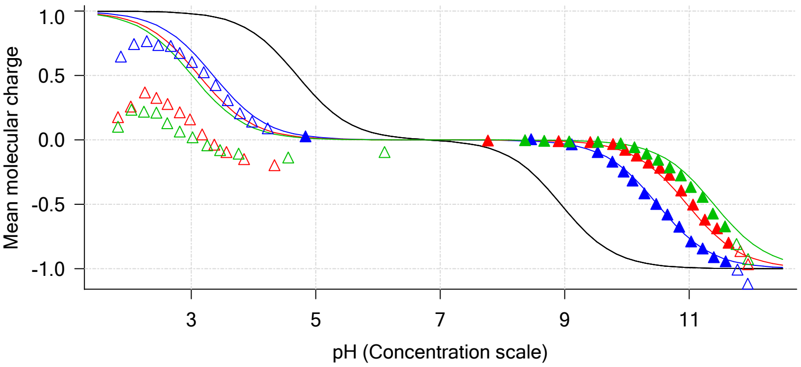


Distribution of species for sample M16_octanol

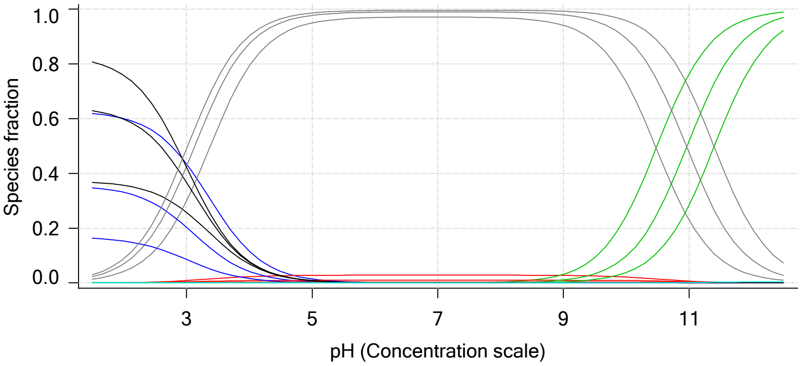


Lipophilicity Profile

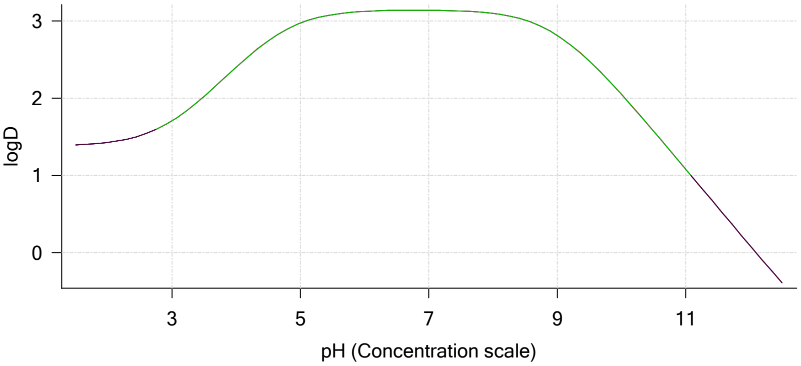


Log R Profile

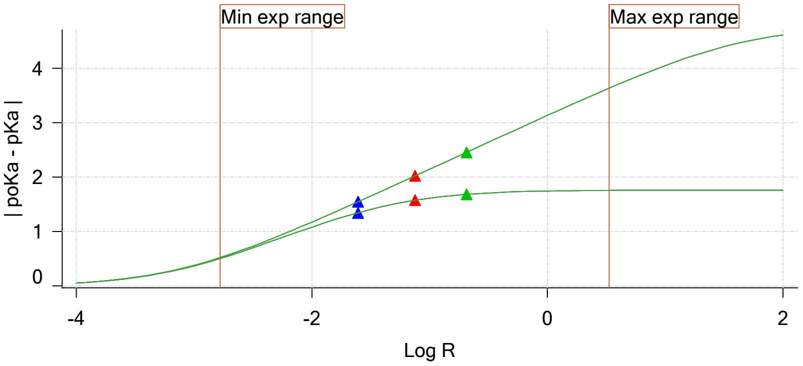


Sample Carbonate Ionisation Graph

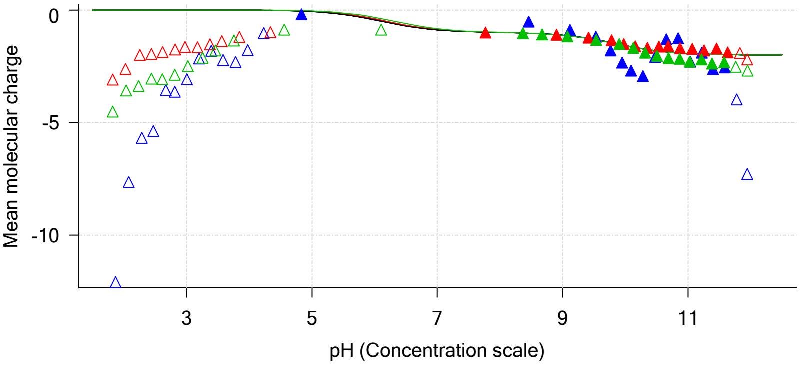

Sample name: **M16_octanol**
Assay name: **pH-metric high logP**
Assay ID: **18C-01007**
Filename: **C:\Sirius_T3\Mehtap\20180228_exp28_logP_T3-2\18C-01007_M16_octanol_pH-metric high logP.t3r**

Experiment start time: **3/1/2018 9:25:34 AM**
Analyst: **Pion**
Instrument ID: **T312060**

### Graphs (continued)

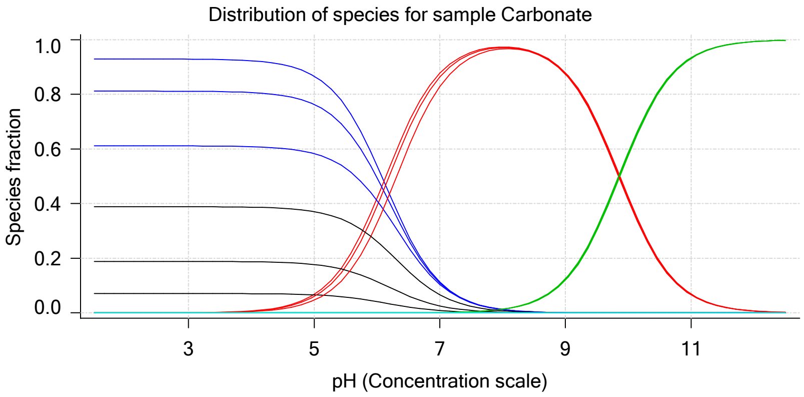

| | | | |
|---|---|---|---|
| Sample name: | **M16_octanol** | Experiment start time: | **3/1/2018 9:25:34 AM** |
| Assay name: | **pH-metric high logP** | Analyst: | **Pion** |
| Assay ID: | **18C-01007** | Instrument ID: | **T312060** |
| Filename: | **C:\Sirius_T3\Mehtap\20180228_exp28_logP_T3-2\18C-01007_M16_octanol_pH-metric high logP.t3r** | | |

# pH-metric high logP   Titration 1 of 3   18C-01007 Points 1 to 30

### Overall results

| | |
|---|---|
| RMSD | 0.139 |
| Average ionic strength | 0.160 M |
| Average temperature | 25.0°C |
| Partition ratio | 0.0246 : 1 |
| Analyte concentration range | 4642.2 μM to 4978.9 μM |
| Total points considered | 15 of 30 |

### Warnings and errors

| | |
|---|---|
| Errors | None |
| Warnings | Excessive acidity error present |

### Four-Plus parameters

| | | | |
|---|---|---|---|
| Alpha | 0.130 | 3/1/2018 9:25:33 AM | C:\Sirius_T3\HCl18B27.t3r |
| S | 0.9970 | 3/1/2018 9:25:33 AM | C:\Sirius_T3\HCl18B27.t3r |
| jH | 0.8 | 3/1/2018 9:25:33 AM | C:\Sirius_T3\HCl18B27.t3r |
| jOH | -0.4 | 3/1/2018 9:25:33 AM | C:\Sirius_T3\HCl18B27.t3r |

### Titrants

| | | | |
|---|---|---|---|
| 0.50 M HCl | 0.993513 | 3/1/2018 9:25:34 AM | C:\Sirius_T3\HCl18B27.t3r |
| 0.50 M KOH | 0.999845 | 3/1/2018 9:25:34 AM | C:\Sirius_T3\KOH18B27.t3r |

### Sample

| | |
|---|---|
| M16_octanol concentration factor | 0.780 |
| Base pKa 1 | 4.70 |
| Acid pKa 2 | 8.94 |
| logP (XH2 +) | 1.38 |
| logP (neutral XH) | 3.12 |
| logP (X -) | -1.62 |

### Sample graphs

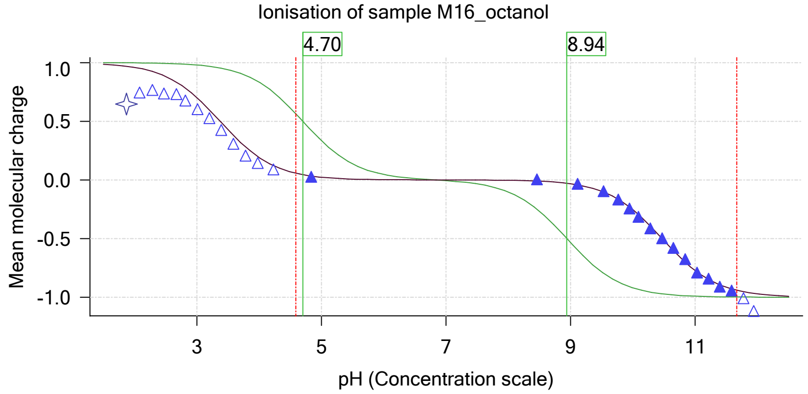


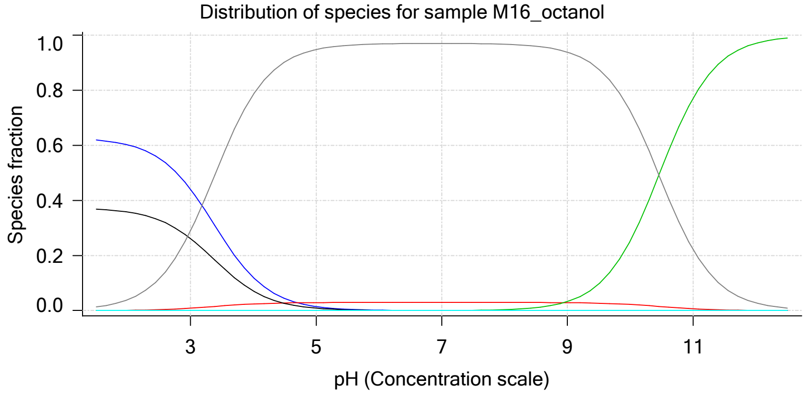

| | | | |
|---|---|---|---|
| Sample name: | **M16_octanol** | Experiment start time: | **3/1/2018 9:25:34 AM** |
| Assay name: | **pH-metric high logP** | Analyst: | **Pion** |
| Assay ID: | **18C-01007** | Instrument ID: | **T312060** |
| Filename: | **C:\Sirius_T3\Mehtap\20180228_exp28_logP_T3-2\18C-01007_M16_octanol_pH-metric high logP.t3r** | | |

### Sample graphs (continued)

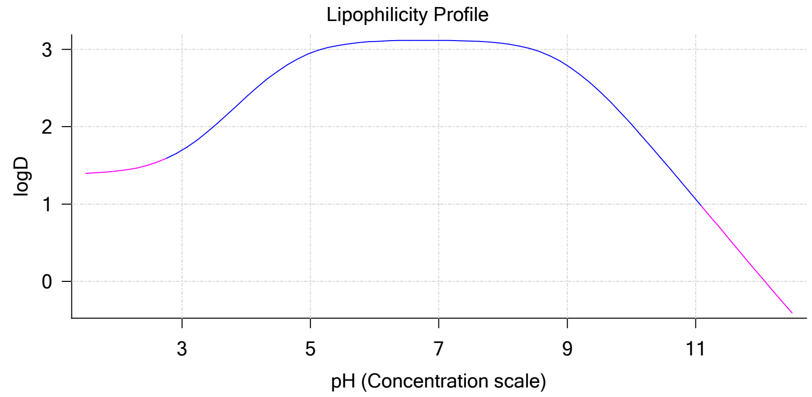


### Sample logD and percent species

| pH | M16_octanol logD | M16_octanol M16_octanolH2 | M16_octanol M16_octanolH | M16_octanol M16_octanol | M16_octanol M16_octanolH2* | M16_octanol M16_octanolH* | M16_octanol M16_octanol* | Comment |
|---|---|---|---|---|---|---|---|---|
| 1.000 | 1.39 | 62.47 % | 0.01 % | 0.00 % | 37.11 % | 0.41 % | 0.00 % | |
| 1.200 | 1.39 | 62.31 % | 0.02 % | 0.00 % | 37.02 % | 0.65 % | 0.00 % | Stomach pH |
| 2.000 | 1.43 | 60.18 % | 0.12 % | 0.00 % | 35.75 % | 3.94 % | 0.00 % | |
| 3.000 | 1.70 | 44.07 % | 0.88 % | 0.00 % | 26.18 % | 28.87 % | 0.00 % | |
| 4.000 | 2.38 | 11.99 % | 2.39 % | 0.00 % | 7.12 % | 78.50 % | 0.00 % | |
| 5.000 | 2.95 | 1.45 % | 2.89 % | 0.00 % | 0.86 % | 94.80 % | 0.00 % | |
| 6.000 | 3.10 | 0.15 % | 2.95 % | 0.00 % | 0.09 % | 96.81 % | 0.00 % | |
| 6.500 | 3.12 | 0.05 % | 2.95 % | 0.01 % | 0.03 % | 96.96 % | 0.00 % | |
| 7.000 | 3.12 | 0.01 % | 2.95 % | 0.03 % | 0.01 % | 96.99 % | 0.00 % | |
| 7.400 | 3.11 | 0.01 % | 2.95 % | 0.09 % | 0.00 % | 96.95 % | 0.00 % | Blood pH |
| 8.000 | 3.08 | 0.00 % | 2.95 % | 0.34 % | 0.00 % | 96.71 % | 0.00 % | |
| 9.000 | 2.79 | 0.00 % | 2.86 % | 3.28 % | 0.00 % | 93.86 % | 0.00 % | |
| 10.000 | 2.03 | 0.00 % | 2.21 % | 25.34 % | 0.00 % | 72.44 % | 0.01 % | |
| 11.000 | 1.06 | 0.00 % | 0.67 % | 77.21 % | 0.00 % | 22.08 % | 0.05 % | |
| 12.000 | 0.07 | 0.00 % | 0.08 % | 97.08 % | 0.00 % | 2.78 % | 0.06 % | |

### Carbonate and acidity

| | |
|---|---|
| Carbonate | 0.105 mM |
| Acidity error | 3.384 mM |

### Other graphs

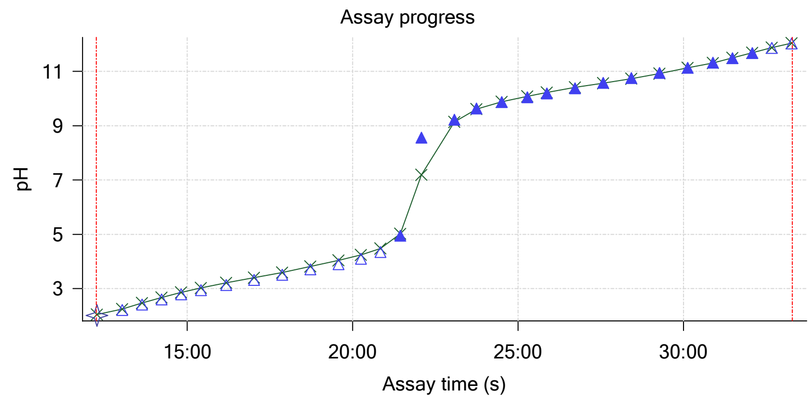


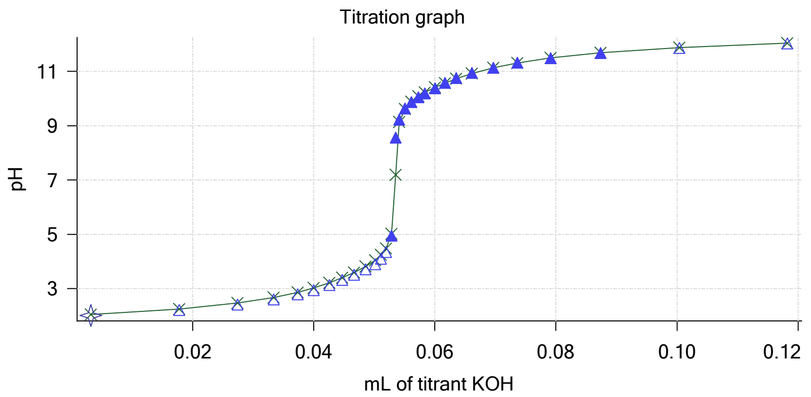

Sample name: **M16_octanol** Experiment start time: **3/1/2018 9:25:34 AM**
Assay name: **pH-metric high logP** Analyst: **Pion**
Assay ID: **18C-01007** Instrument ID: **T312060**
Filename: **C:\Sirius_T3\Mehtap\20180228_exp28_logP_T3-2\18C-01007_M16_octanol_pH-metric high logP.t3r**

### Other graphs (continued)

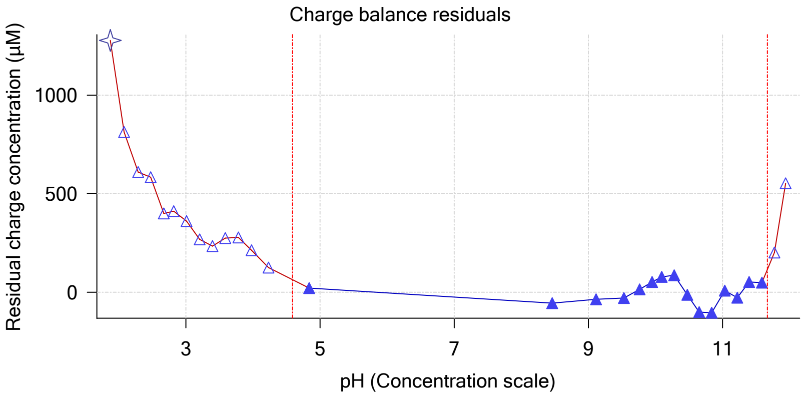


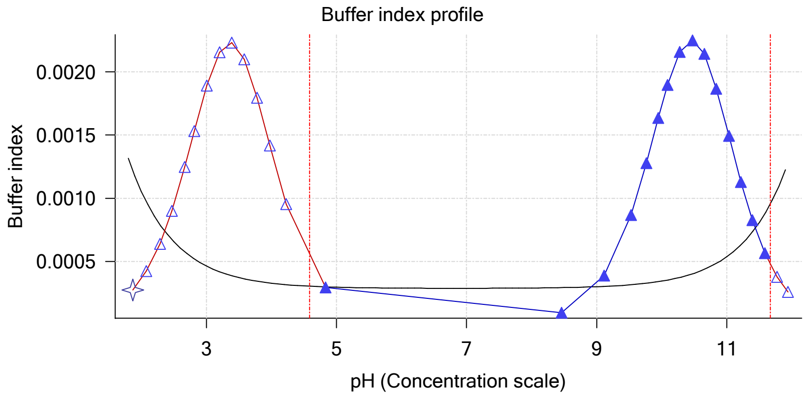


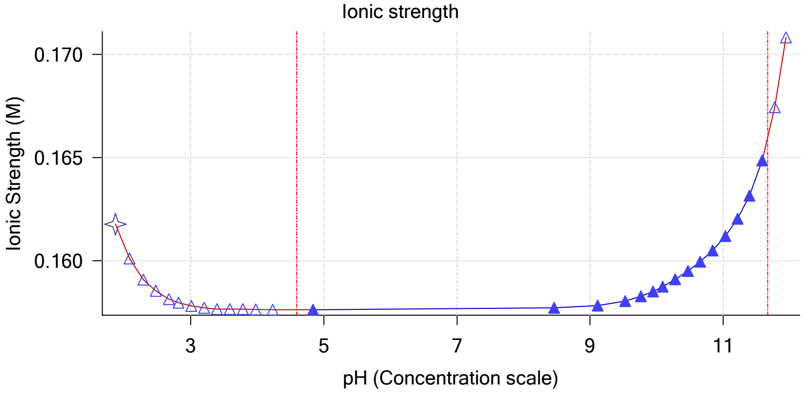


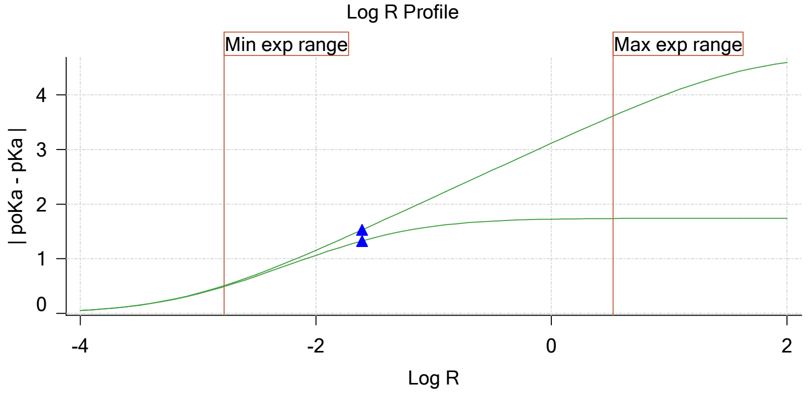


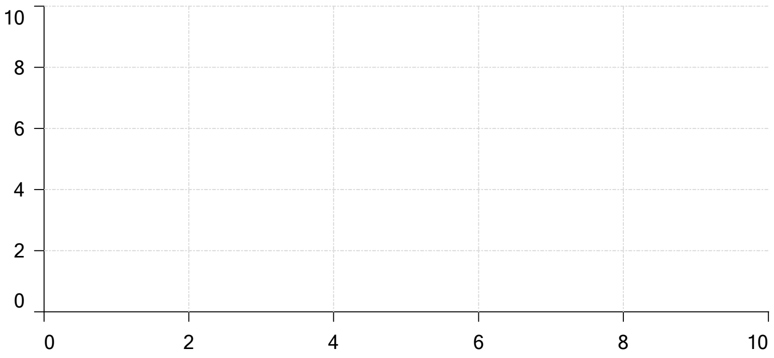


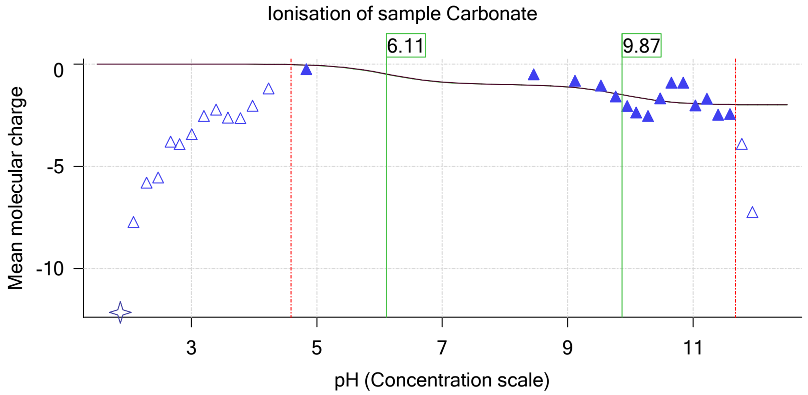

| | | | |
|---|---|---|---|
| Sample name: | **M16_octanol** | Experiment start time: | **3/1/2018 9:25:34 AM** |
| Assay name: | **pH-metric high logP** | Analyst: | **Pion** |
| Assay ID: | **18C-01007** | Instrument ID: | **T312060** |
| Filename: | **C:\Sirius_T3\Mehtap\20180228_exp28_logP_T3-2\18C-01007_M16_octanol_pH-metric high logP.t3r** | | |

# pH-metric high logP   Titration 2 of 3   18C-01007 Points 31 to 58

### Overall results

| | |
|---|---|
| RMSD | 0.048 |
| Average ionic strength | 0.173 M |
| Average temperature | 25.0°C |
| Partition ratio | 0.0752 : 1 |
| Analyte concentration range | 3835.8 μM to 4100.8 μM |
| Total points considered | 14 of 28 |

### Warnings and errors

| | |
|---|---|
| Errors | None |
| Warnings | One or more logP values out of range |
| | Excessive carbonate concentration present |
| | Excessive acidity error present |

### Four-Plus parameters

| | | | |
|---|---|---|---|
| Alpha | 0.130 | 3/1/2018 9:25:33 AM | C:\Sirius_T3\HCl18B27.t3r |
| S | 0.9970 | 3/1/2018 9:25:33 AM | C:\Sirius_T3\HCl18B27.t3r |
| jH | 0.8 | 3/1/2018 9:25:33 AM | C:\Sirius_T3\HCl18B27.t3r |
| jOH | -0.4 | 3/1/2018 9:25:33 AM | C:\Sirius_T3\HCl18B27.t3r |

### Titrants

| | | | |
|---|---|---|---|
| 0.50 M HCl | 0.993513 | 3/1/2018 9:25:34 AM | C:\Sirius_T3\HCl18B27.t3r |
| 0.50 M KOH | 0.999845 | 3/1/2018 9:25:34 AM | C:\Sirius_T3\KOH18B27.t3r |

### Sample

| | |
|---|---|
| M16_octanol concentration factor | 0.751 |
| Base pKa 1 | 4.70 |
| Acid pKa 2 | 8.94 |
| logP (XH2 +) | -5.98 |
| logP (neutral XH) | 3.25 |
| logP (X -) | -1.62 |

### Sample graphs

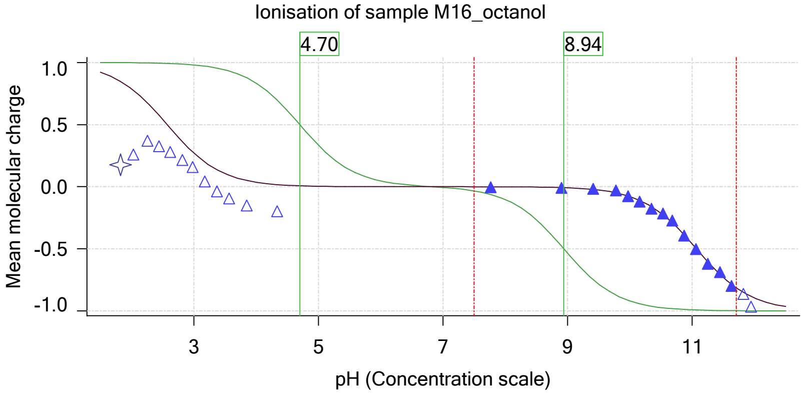


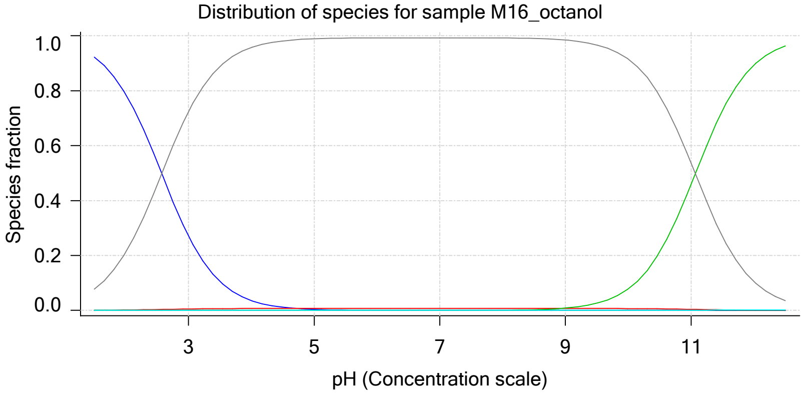

| | | | |
|---|---|---|---|
| Sample name: | **M16_octanol** | Experiment start time: | **3/1/2018 9:25:34 AM** |
| Assay name: | **pH-metric high logP** | Analyst: | **Pion** |
| Assay ID: | **18C-01007** | Instrument ID: | **T312060** |
| Filename: | **C:\Sirius_T3\Mehtap\20180228_exp28_logP_T3-2\18C-01007_M16_octanol_pH-metric high logP.t3r** | | |

### Sample graphs (continued)

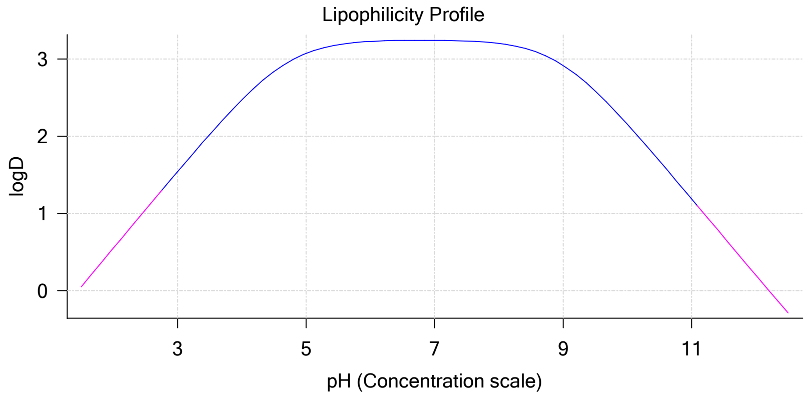


### Sample logD and percent species

| pH | M16_octanol logD | M16_octanol M16_octanolH2 | M16_octanol M16_octanolH | M16_octanol M16_octanol | M16_octanol M16_octanolH2* | M16_octanol M16_octanolH* | M16_octanol M16_octanol* | Comment |
|---|---|---|---|---|---|---|---|---|
| 1.000 | -0.45 | 97.38 % | 0.02 % | 0.00 % | 0.00 % | 2.60 % | 0.00 % | |
| 1.200 | -0.25 | 95.91 % | 0.03 % | 0.00 % | 0.00 % | 4.06 % | 0.00 % | Stomach pH |
| 2.000 | 0.55 | 78.80 % | 0.16 % | 0.00 % | 0.00 % | 21.04 % | 0.00 % | |
| 3.000 | 1.54 | 27.10 % | 0.54 % | 0.00 % | 0.00 % | 72.36 % | 0.00 % | |
| 4.000 | 2.47 | 3.58 % | 0.72 % | 0.00 % | 0.00 % | 95.70 % | 0.00 % | |
| 5.000 | 3.07 | 0.37 % | 0.74 % | 0.00 % | 0.00 % | 98.89 % | 0.00 % | |
| 6.000 | 3.23 | 0.04 % | 0.74 % | 0.00 % | 0.00 % | 99.22 % | 0.00 % | |
| 6.500 | 3.24 | 0.01 % | 0.74 % | 0.00 % | 0.00 % | 99.24 % | 0.00 % | |
| 7.000 | 3.24 | 0.00 % | 0.74 % | 0.01 % | 0.00 % | 99.25 % | 0.00 % | |
| 7.400 | 3.24 | 0.00 % | 0.74 % | 0.02 % | 0.00 % | 99.24 % | 0.00 % | Blood pH |
| 8.000 | 3.20 | 0.00 % | 0.74 % | 0.09 % | 0.00 % | 99.17 % | 0.00 % | |
| 9.000 | 2.92 | 0.00 % | 0.74 % | 0.84 % | 0.00 % | 98.42 % | 0.00 % | |
| 10.000 | 2.15 | 0.00 % | 0.68 % | 7.85 % | 0.00 % | 91.45 % | 0.01 % | |
| 11.000 | 1.19 | 0.00 % | 0.40 % | 45.96 % | 0.00 % | 53.56 % | 0.08 % | |
| 12.000 | 0.20 | 0.00 % | 0.08 % | 89.35 % | 0.00 % | 10.41 % | 0.16 % | |

### Carbonate and acidity

| | |
|---|---|
| Carbonate | 0.833 mM |
| Acidity error | 2.201 mM |

### Other graphs

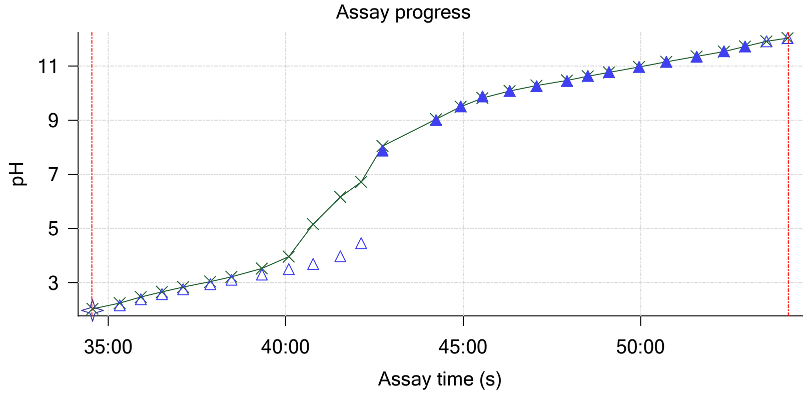


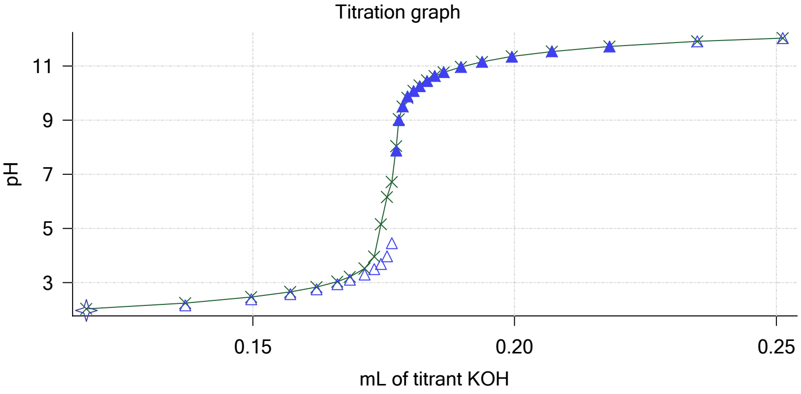

Sample name: **M16_octanol**
Assay name: **pH-metric high logP**
Assay ID: **18C-01007**
Filename: **C:\Sirius_T3\Mehtap\20180228_exp28_logP_T3-2\18C-01007_M16_octanol_pH-metric high logP.t3r**

Experiment start time: **3/1/2018 9:25:34 AM**
Analyst: **Pion**
Instrument ID: **T312060**

### Other graphs (continued)

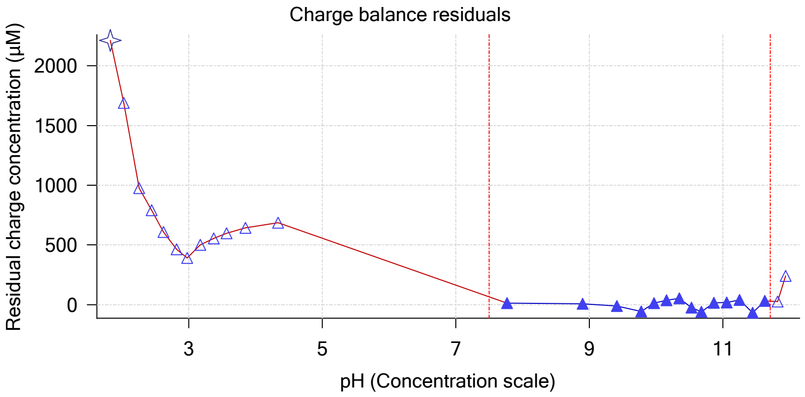


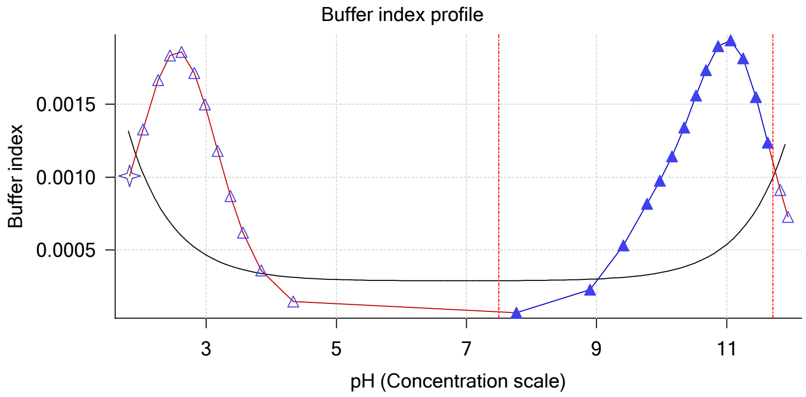


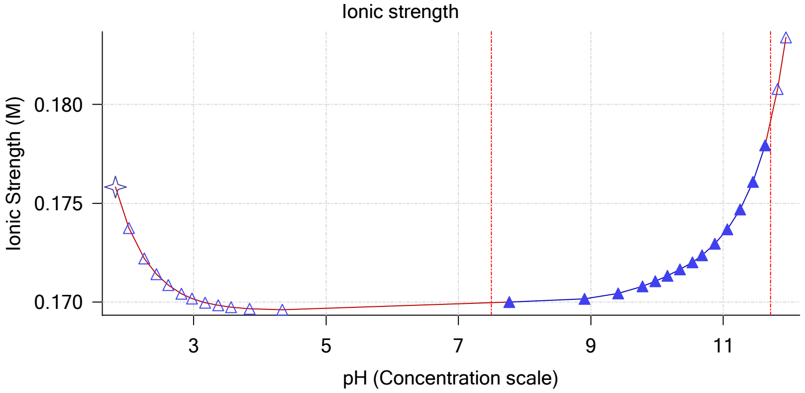


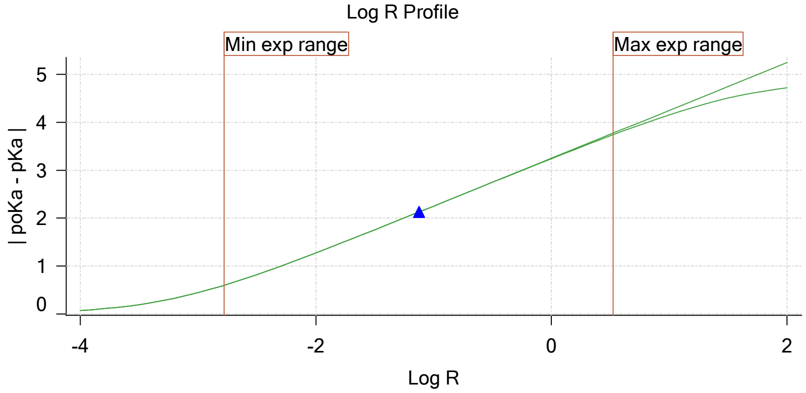


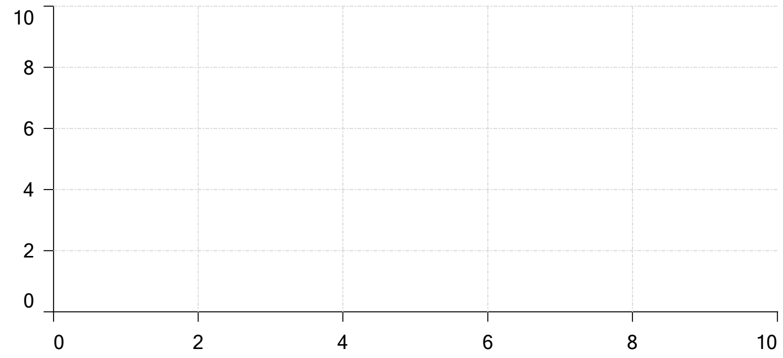


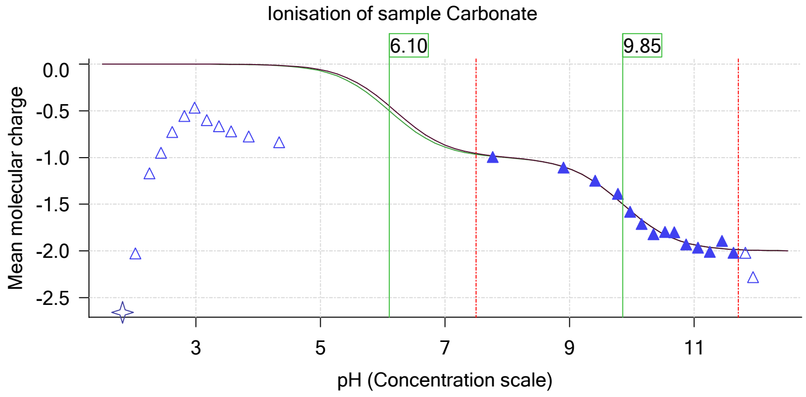

| | | | |
|---|---|---|---|
| Sample name: | **M16_octanol** | Experiment start time: | **3/1/2018 9:25:34 AM** |
| Assay name: | **pH-metric high logP** | Analyst: | **Pion** |
| Assay ID: | **18C-01007** | Instrument ID: | **T312060** |
| Filename: | **C:\Sirius_T3\Mehtap\20180228_exp28_logP_T3-2\18C-01007_M16_octanol_pH-metric high logP.t3r** | | |

# pH-metric high logP   Titration 3 of 3   18C-01007 Points 59 to 86

### Overall results

| | |
|---|---|
| RMSD | 0.033 |
| Average ionic strength | 0.182 M |
| Average temperature | 25.0°C |
| Partition ratio | 0.2059 : 1 |
| Analyte concentration range | 2984.0 μM to 3160.6 μM |
| Total points considered | 14 of 28 |

### Warnings and errors

| | |
|---|---|
| Errors | None |
| Warnings | Excessive carbonate concentration present |
| | Excessive acidity error present |

### Four-Plus parameters

| | | | |
|---|---|---|---|
| Alpha | 0.130 | 3/1/2018 9:25:33 AM | C:\Sirius_T3\HCl18B27.t3r |
| S | 0.9970 | 3/1/2018 9:25:33 AM | C:\Sirius_T3\HCl18B27.t3r |
| jH | 0.8 | 3/1/2018 9:25:33 AM | C:\Sirius_T3\HCl18B27.t3r |
| jOH | -0.4 | 3/1/2018 9:25:33 AM | C:\Sirius_T3\HCl18B27.t3r |

### Titrants

| | | | |
|---|---|---|---|
| 0.50 M HCl | 0.993513 | 3/1/2018 9:25:34 AM | C:\Sirius_T3\HCl18B27.t3r |
| 0.50 M KOH | 0.999845 | 3/1/2018 9:25:34 AM | C:\Sirius_T3\KOH18B27.t3r |

### Sample

| | |
|---|---|
| M16_octanol concentration factor | 0.772 |
| Base pKa 1 | 4.70 |
| Acid pKa 2 | 8.94 |
| logP (XH2 +) | -0.97 |
| logP (neutral XH) | 3.02 |
| logP (X -) | -1.62 |

### Sample graphs

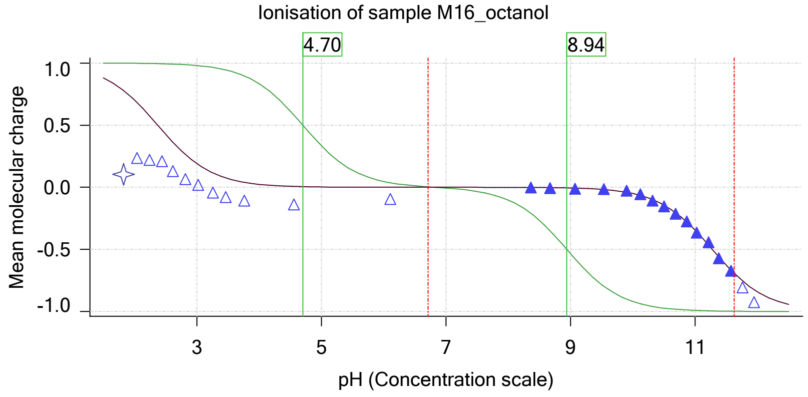


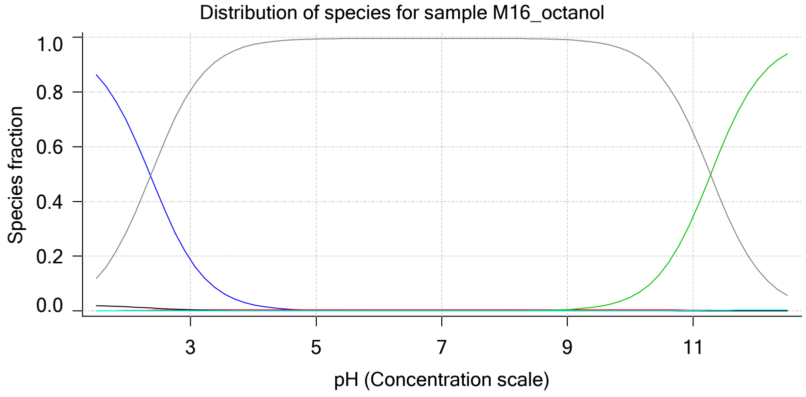

| | | | |
|---|---|---|---|
| Sample name: | **M16_octanol** | Experiment start time: | **3/1/2018 9:25:34 AM** |
| Assay name: | **pH-metric high logP** | Analyst: | **Pion** |
| Assay ID: | **18C-01007** | Instrument ID: | **T312060** |
| Filename: | **C:\Sirius_T3\Mehtap\20180228_exp28_logP_T3-2\18C-01007_M16_octanol_pH-metric high logP.t3r** | | |

### Sample graphs (continued)

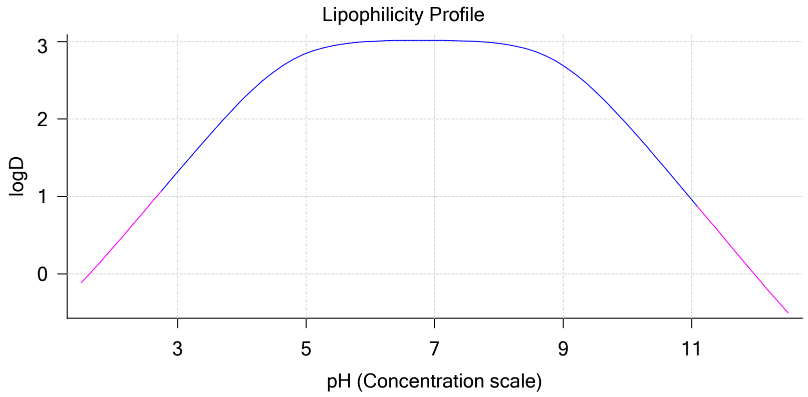


### Sample logD and percent species

| pH | M16_octanol logD | M16_octanol M16_octanolH2 | M16_octanol M16_octanolH | M16_octanol M16_octanol | M16_octanol M16_octanolH2* | M16_octanol M16_octanolH* | M16_octanol M16_octanol* | Comment |
|---|---|---|---|---|---|---|---|---|
| 1.000 | -0.50 | 93.84 % | 0.02 % | 0.00 % | 2.07 % | 4.07 % | 0.00 % | |
| 1.200 | -0.36 | 91.65 % | 0.03 % | 0.00 % | 2.02 % | 6.29 % | 0.00 % | Stomach pH |
| 2.000 | 0.34 | 68.62 % | 0.14 % | 0.00 % | 1.51 % | 29.73 % | 0.00 % | |
| 3.000 | 1.32 | 18.60 % | 0.37 % | 0.00 % | 0.41 % | 80.61 % | 0.00 % | |
| 4.000 | 2.24 | 2.24 % | 0.45 % | 0.00 % | 0.05 % | 97.26 % | 0.00 % | |
| 5.000 | 2.85 | 0.23 % | 0.46 % | 0.00 % | 0.01 % | 99.31 % | 0.00 % | |
| 6.000 | 3.00 | 0.02 % | 0.46 % | 0.00 % | 0.00 % | 99.52 % | 0.00 % | |
| 6.500 | 3.01 | 0.01 % | 0.46 % | 0.00 % | 0.00 % | 99.53 % | 0.00 % | |
| 7.000 | 3.02 | 0.00 % | 0.46 % | 0.01 % | 0.00 % | 99.53 % | 0.00 % | |
| 7.400 | 3.01 | 0.00 % | 0.46 % | 0.01 % | 0.00 % | 99.53 % | 0.00 % | Blood pH |
| 8.000 | 2.98 | 0.00 % | 0.46 % | 0.05 % | 0.00 % | 99.49 % | 0.00 % | |
| 9.000 | 2.69 | 0.00 % | 0.46 % | 0.52 % | 0.00 % | 99.02 % | 0.00 % | |
| 10.000 | 1.93 | 0.00 % | 0.44 % | 5.00 % | 0.00 % | 94.54 % | 0.02 % | |
| 11.000 | 0.96 | 0.00 % | 0.30 % | 34.42 % | 0.00 % | 65.11 % | 0.17 % | |
| 12.000 | -0.03 | 0.00 % | 0.07 % | 83.68 % | 0.00 % | 15.83 % | 0.41 % | |

### Carbonate and acidity

| | |
|---|---|
| Carbonate | 0.548 mM |
| Acidity error | 2.297 mM |

### Other graphs

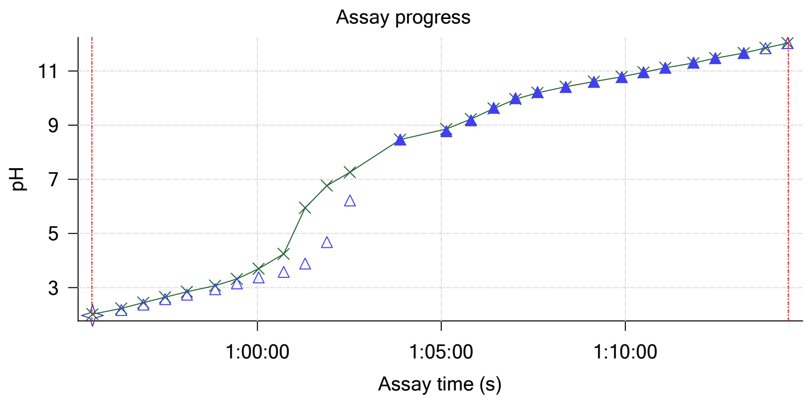


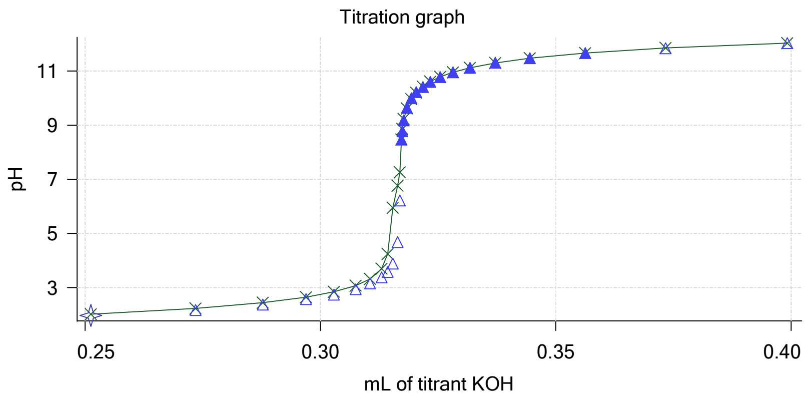

| | | | |
|---|---|---|---|
| Sample name: | **M16_octanol** | Experiment start time: | **3/1/2018 9:25:34 AM** |
| Assay name: | **pH-metric high logP** | Analyst: | **Pion** |
| Assay ID: | **18C-01007** | Instrument ID: | **T312060** |
| Filename: | **C:\Sirius_T3\Mehtap\20180228_exp28_logP_T3-2\18C-01007_M16_octanol_pH-metric high logP.t3r** | | |

### Other graphs (continued)

Charge balance residuals

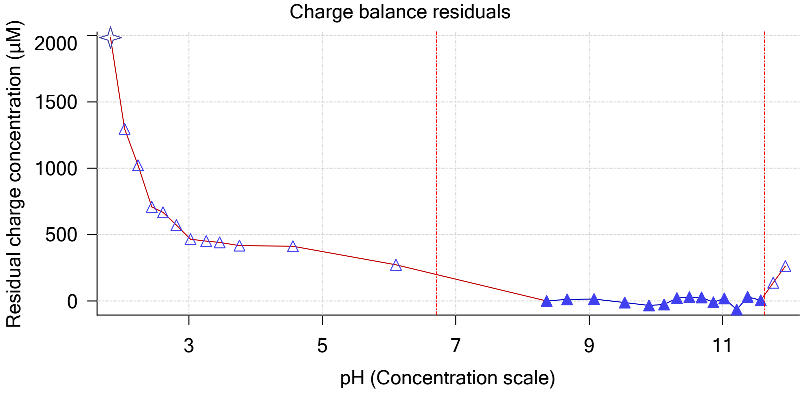


Buffer index profile

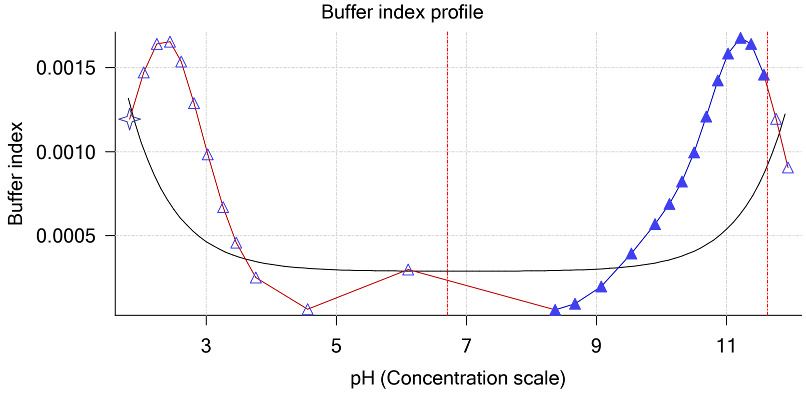


Ionic strength

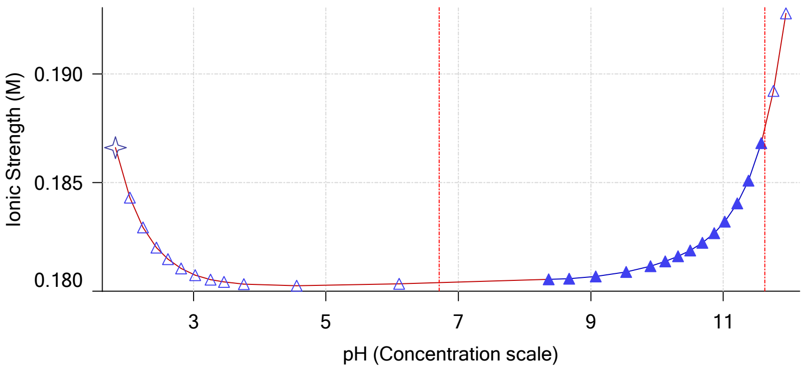


Log R Profile

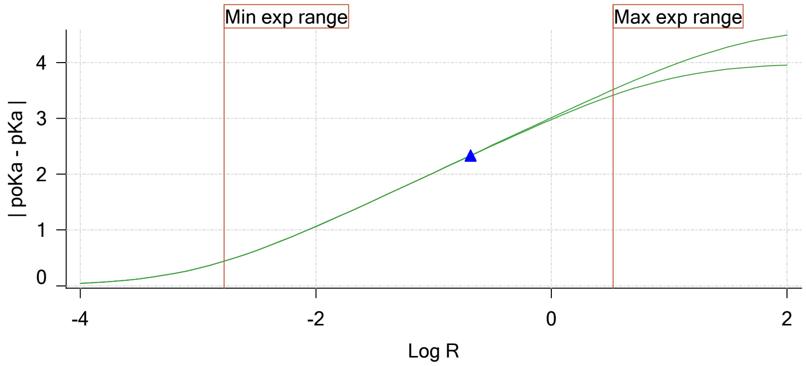


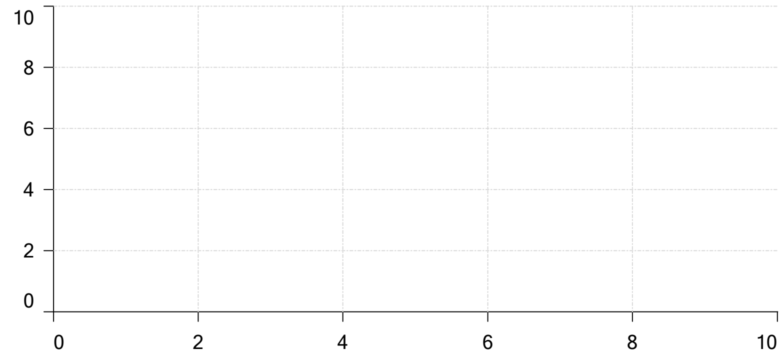

Ionisation of sample Carbonate

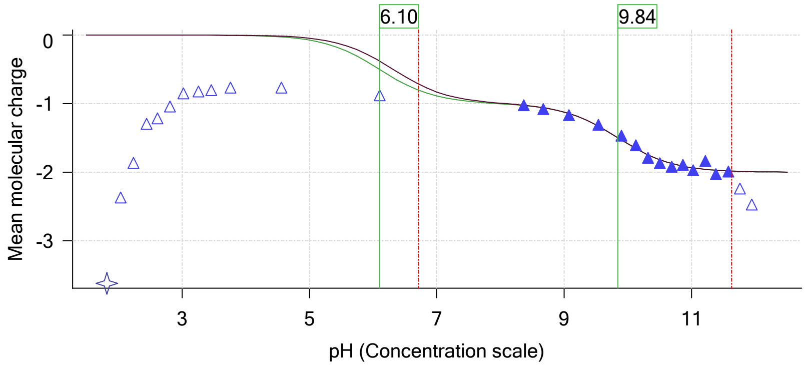

| | | | |
|---|---|---|---|
| Sample name: | **M16_octanol** | Experiment start time: | **3/1/2018 9:25:34 AM** |
| Assay name: | **pH-metric high logP** | Analyst: | **Pion** |
| Assay ID: | **18C-01007** | Instrument ID: | **T312060** |
| Filename: | **C:\Sirius_T3\Mehtap\20180228_exp28_logP_T3-2\18C-01007_M16_octanol_pH-metric high logP.t3r** | | |

### Assay Model

| Settings | Value | Date/Time changed | Imported from |
|---|---|---|---|
| Sample name | M16_octanol | 2/27/2018 6:14:13 PM | User entered value |
| Sample by | Weight | | Default value |
| Sample weight | 0.001660 g | 2/28/2018 4:25:40 PM | User entered value |
| Formula weight | 210.23 g/mol | 2/27/2018 5:08:55 PM | User entered value |
| Solubility | Unknown | | Default value |
| Molecular weight | 210.23 | 2/27/2018 5:08:55 PM | User entered value |
| Individual pKa ionic environments | No | | Default value |
| Number of pKas | 2 | 2/27/2018 5:08:55 PM | User entered value |
| Sample is a | Ampholyte | 2/27/2018 5:08:55 PM | User entered value |
| pKa 1 | 4.70 | 2/27/2018 5:08:55 PM | User entered value |
| Type | Base | 2/27/2018 5:08:55 PM | User entered value |
| pKa 2 | 8.94 | 2/27/2018 5:08:55 PM | User entered value |
| Type | Acid | 2/27/2018 5:08:55 PM | User entered value |
| logp (XH2 +) | -0.97 | 2/27/2018 5:09:30 PM | User entered value |
| logP (neutral XH) | 1.47 | 2/27/2018 5:09:44 PM | User entered value |
| logP (X -) | -1.62 | 2/27/2018 5:09:50 PM | User entered value |

### Events

| Time | Event | Water | Acid | Base | Octanol | pH | dpH/dt | pH R-squared | pH SD | dpH/dt time |
|---|---|---|---|---|---|---|---|---|---|---|
| 9:11.9 | Initial pH = 3.56 | | | | | | | | | |
| 12:16.4 | Data point 1 | 1.50000 mL | 0.04271 mL | 0.00320 mL | 0.04000 mL | 2.005 | -0.00710 | 0.21574 | 0.00076 | 10.0 s |
| 13:02.6 | Data point 2 | 1.50000 mL | 0.04271 mL | 0.01778 mL | 0.04000 mL | 2.209 | -0.00559 | 0.12657 | 0.00078 | 10.0 s |
| 13:38.3 | Data point 3 | 1.50000 mL | 0.04271 mL | 0.02740 mL | 0.04000 mL | 2.414 | 0.00086 | 0.04511 | 0.00020 | 10.0 s |
| 14:13.8 | Data point 4 | 1.50000 mL | 0.04271 mL | 0.03340 mL | 0.04000 mL | 2.598 | -0.00678 | 0.14397 | 0.00088 | 10.0 s |
| 14:49.3 | Data point 5 | 1.50000 mL | 0.04271 mL | 0.03737 mL | 0.04000 mL | 2.793 | -0.00172 | 0.13364 | 0.00023 | 10.0 s |
| 15:24.9 | Data point 6 | 1.50000 mL | 0.04271 mL | 0.03998 mL | 0.04000 mL | 2.938 | -0.00257 | 0.34140 | 0.00022 | 10.0 s |
| 16:10.7 | Data point 7 | 1.50000 mL | 0.04271 mL | 0.04262 mL | 0.04000 mL | 3.130 | -0.00330 | 0.56608 | 0.00022 | 10.0 s |
| 17:01.6 | Data point 8 | 1.50000 mL | 0.04271 mL | 0.04471 mL | 0.04000 mL | 3.323 | -0.00522 | 0.10478 | 0.00080 | 10.0 s |
| 17:52.6 | Data point 9 | 1.50000 mL | 0.04271 mL | 0.04666 mL | 0.04000 mL | 3.513 | -0.00408 | 0.55895 | 0.00027 | 10.0 s |
| 18:43.6 | Data point 10 | 1.50000 mL | 0.04271 mL | 0.04859 mL | 0.04000 mL | 3.704 | -0.00652 | 0.61139 | 0.00041 | 10.0 s |
| 19:34.5 | Data point 11 | 1.50000 mL | 0.04271 mL | 0.05014 mL | 0.04000 mL | 3.901 | -0.01015 | 0.66408 | 0.00061 | 10.0 s |
| 20:15.1 | Data point 12 | 1.50000 mL | 0.04271 mL | 0.05113 mL | 0.04000 mL | 4.094 | -0.01157 | 0.56816 | 0.00076 | 10.0 s |
| 20:50.6 | Data point 13 | 1.50000 mL | 0.04271 mL | 0.05195 mL | 0.04000 mL | 4.347 | -0.01522 | 0.64804 | 0.00093 | 10.5 s |
| 21:26.5 | Data point 14 | 1.50000 mL | 0.04271 mL | 0.05287 mL | 0.04000 mL | 4.951 | -0.01799 | 0.87526 | 0.00095 | 13.0 s |
| 22:05.0 | Data point 15 | 1.50000 mL | 0.04271 mL | 0.05355 mL | 0.04000 mL | 8.563 | -0.01946 | 0.98111 | 0.00097 | 34.0 s |
| 23:04.4 | Data point 16 | 1.50000 mL | 0.04271 mL | 0.05412 mL | 0.04000 mL | 9.218 | -0.01863 | 0.95964 | 0.00094 | 14.5 s |
| 23:44.4 | Data point 17 | 1.50000 mL | 0.04271 mL | 0.05506 mL | 0.04000 mL | 9.631 | -0.01929 | 0.94749 | 0.00098 | 10.5 s |
| 24:30.7 | Data point 18 | 1.50000 mL | 0.04271 mL | 0.05614 mL | 0.04000 mL | 9.865 | -0.00801 | 0.59789 | 0.00051 | 10.5 s |
| 25:17.1 | Data point 19 | 1.50000 mL | 0.04271 mL | 0.05729 mL | 0.04000 mL | 10.048 | -0.00508 | 0.13625 | 0.00068 | 10.0 s |
| 25:52.5 | Data point 20 | 1.50000 mL | 0.04271 mL | 0.05837 mL | 0.04000 mL | 10.190 | -0.01000 | 0.36786 | 0.00081 | 10.0 s |
| 26:43.4 | Data point 21 | 1.50000 mL | 0.04271 mL | 0.06002 mL | 0.04000 mL | 10.377 | -0.00716 | 0.23433 | 0.00073 | 10.0 s |
| 27:34.3 | Data point 22 | 1.50000 mL | 0.04271 mL | 0.06167 mL | 0.04000 mL | 10.570 | -0.00128 | 0.01797 | 0.00047 | 10.0 s |
| 28:25.3 | Data point 23 | 1.50000 mL | 0.04271 mL | 0.06355 mL | 0.04000 mL | 10.750 | -0.00897 | 0.69945 | 0.00053 | 10.0 s |
| 29:16.4 | Data point 24 | 1.50000 mL | 0.04271 mL | 0.06611 mL | 0.04000 mL | 10.936 | -0.00376 | 0.05852 | 0.00077 | 10.0 s |
| 30:07.5 | Data point 25 | 1.50000 mL | 0.04271 mL | 0.06966 mL | 0.04000 mL | 11.128 | -0.00187 | 0.33816 | 0.00016 | 10.0 s |
| 30:53.5 | Data point 26 | 1.50000 mL | 0.04271 mL | 0.07361 mL | 0.04000 mL | 11.313 | -0.00175 | 0.34200 | 0.00015 | 10.0 s |
| 31:29.0 | Data point 27 | 1.50000 mL | 0.04271 mL | 0.07912 mL | 0.04000 mL | 11.488 | 0.00172 | 0.30827 | 0.00015 | 10.0 s |
| 32:04.7 | Data point 28 | 1.50000 mL | 0.04271 mL | 0.08739 mL | 0.04000 mL | 11.678 | 0.00337 | 0.74144 | 0.00019 | 10.0 s |
| 32:40.4 | Data point 29 | 1.50000 mL | 0.04271 mL | 0.10040 mL | 0.04000 mL | 11.865 | -0.00296 | 0.02218 | 0.00098 | 10.0 s |
| 33:16.3 | Data point 30 | 1.50000 mL | 0.04271 mL | 0.11820 mL | 0.04000 mL | 12.028 | 0.01264 | 0.63373 | 0.00078 | 10.0 s |
| 34:33.6 | Data point 31 | 1.50000 mL | 0.16729 mL | 0.11820 mL | 0.14000 mL | 1.958 | -0.01998 | 0.98240 | 0.00100 | 10.0 s |
| 35:19.9 | Data point 32 | 1.50000 mL | 0.16729 mL | 0.13709 mL | 0.14000 mL | 2.157 | 0.00252 | 0.43182 | 0.00019 | 10.0 s |
| 35:55.6 | Data point 33 | 1.50000 mL | 0.16729 mL | 0.14972 mL | 0.14000 mL | 2.384 | 0.00020 | 0.00012 | 0.00092 | 10.0 s |
| 36:31.2 | Data point 34 | 1.50000 mL | 0.16729 mL | 0.15720 mL | 0.14000 mL | 2.567 | 0.00609 | 0.52065 | 0.00042 | 10.0 s |
| 37:06.8 | Data point 35 | 1.50000 mL | 0.16729 mL | 0.16218 mL | 0.14000 mL | 2.744 | 0.00186 | 0.31955 | 0.00016 | 10.0 s |
| 37:52.7 | Data point 36 | 1.50000 mL | 0.16729 mL | 0.16616 mL | 0.14000 mL | 2.940 | 0.00060 | 0.05168 | 0.00013 | 10.5 s |

| | | | |
|---|---|---|---|
| Sample name: | **M16_octanol** | Experiment start time: | **3/1/2018 9:25:34 AM** |
| Assay name: | **pH-metric high logP** | Analyst: | **Pion** |
| Assay ID: | **18C-01007** | Instrument ID: | **T312060** |
| Filename: | **C:\Sirius_T3\Mehtap\20180228_exp28_logP_T3-2\18C-01007_M16_octanol_pH-metric high logP.t3r** | | |

### Events (continued)

| Time | Event | Water | Acid | Base | Octanol | pH | dpH/dt | pH R-squared | pH SD | dpH/dt time |
|---|---|---|---|---|---|---|---|---|---|---|
| 38:28.7 | Data point 37 | 1.50000 mL | 0.16729 mL | 0.16856 mL | 0.14000 mL | 3.099 | -0.00077 | 0.04289 | 0.00018 | 10.0 s |
| 39:19.8 | Data point 38 | 1.50000 mL | 0.16729 mL | 0.17136 mL | 0.14000 mL | 3.295 | -0.00378 | 0.07695 | 0.00067 | 10.0 s |
| 40:05.6 | Data point 39 | 1.50000 mL | 0.16729 mL | 0.17321 mL | 0.14000 mL | 3.493 | -0.00681 | 0.31042 | 0.00060 | 10.5 s |
| 40:46.7 | Data point 40 | 1.50000 mL | 0.16729 mL | 0.17446 mL | 0.14000 mL | 3.684 | -0.00803 | 0.18101 | 0.00093 | 10.0 s |
| 41:32.4 | Data point 41 | 1.50000 mL | 0.16729 mL | 0.17561 mL | 0.14000 mL | 3.965 | 0.00188 | 0.07002 | 0.00035 | 10.0 s |
| 42:07.9 | Data point 42 | 1.50000 mL | 0.16729 mL | 0.17655 mL | 0.14000 mL | 4.455 | -0.01704 | 0.79788 | 0.00094 | 10.5 s |
| 42:43.8 | Data point 43 | 1.50000 mL | 0.16729 mL | 0.17742 mL | 0.14000 mL | 7.873 | -0.08043 | 0.99747 | 0.00398 | Timed out at 59.5 s |
| 44:14.3 | Data point 44 | 1.50000 mL | 0.16729 mL | 0.17789 mL | 0.14000 mL | 9.006 | -0.01224 | 0.48541 | 0.00087 | 16.0 s |
| 44:55.8 | Data point 45 | 1.50000 mL | 0.16729 mL | 0.17862 mL | 0.14000 mL | 9.515 | -0.01874 | 0.92592 | 0.00096 | 11.5 s |
| 45:32.7 | Data point 46 | 1.50000 mL | 0.16729 mL | 0.17959 mL | 0.14000 mL | 9.879 | -0.00549 | 0.23712 | 0.00056 | 10.0 s |
| 46:18.6 | Data point 47 | 1.50000 mL | 0.16729 mL | 0.18071 mL | 0.14000 mL | 10.073 | -0.01138 | 0.44990 | 0.00084 | 10.0 s |
| 47:04.3 | Data point 48 | 1.50000 mL | 0.16729 mL | 0.18187 mL | 0.14000 mL | 10.258 | -0.00900 | 0.33929 | 0.00076 | 10.0 s |
| 47:55.4 | Data point 49 | 1.50000 mL | 0.16729 mL | 0.18330 mL | 0.14000 mL | 10.446 | -0.00333 | 0.51756 | 0.00023 | 10.0 s |
| 48:30.9 | Data point 50 | 1.50000 mL | 0.16729 mL | 0.18476 mL | 0.14000 mL | 10.630 | -0.00872 | 0.81800 | 0.00048 | 10.0 s |
| 49:06.3 | Data point 51 | 1.50000 mL | 0.16729 mL | 0.18648 mL | 0.14000 mL | 10.779 | -0.00578 | 0.38967 | 0.00046 | 10.0 s |
| 49:57.4 | Data point 52 | 1.50000 mL | 0.16729 mL | 0.18977 mL | 0.14000 mL | 10.967 | 0.00098 | 0.03975 | 0.00024 | 10.0 s |
| 50:43.3 | Data point 53 | 1.50000 mL | 0.16729 mL | 0.19381 mL | 0.14000 mL | 11.158 | -0.00373 | 0.09690 | 0.00059 | 10.0 s |
| 51:34.5 | Data point 54 | 1.50000 mL | 0.16729 mL | 0.19948 mL | 0.14000 mL | 11.348 | -0.00214 | 0.18325 | 0.00025 | 10.0 s |
| 52:20.5 | Data point 55 | 1.50000 mL | 0.16729 mL | 0.20715 mL | 0.14000 mL | 11.543 | 0.00767 | 0.67130 | 0.00046 | 10.0 s |
| 52:56.2 | Data point 56 | 1.50000 mL | 0.16729 mL | 0.21816 mL | 0.14000 mL | 11.723 | 0.00230 | 0.01310 | 0.00099 | 10.5 s |
| 53:32.6 | Data point 57 | 1.50000 mL | 0.16729 mL | 0.23492 mL | 0.14000 mL | 11.912 | 0.00035 | 0.00040 | 0.00087 | 10.0 s |
| 54:08.4 | Data point 58 | 1.50000 mL | 0.16729 mL | 0.25125 mL | 0.14000 mL | 12.032 | -0.00066 | 0.00154 | 0.00082 | 10.0 s |
| 55:30.7 | Data point 59 | 1.50000 mL | 0.30701 mL | 0.25125 mL | 0.44000 mL | 1.958 | -0.01398 | 0.59830 | 0.00089 | 10.5 s |
| 56:17.5 | Data point 60 | 1.50000 mL | 0.30701 mL | 0.27349 mL | 0.44000 mL | 2.167 | 0.01516 | 0.86723 | 0.00080 | 10.5 s |
| 56:53.8 | Data point 61 | 1.50000 mL | 0.30701 mL | 0.28777 mL | 0.44000 mL | 2.365 | 0.00123 | 0.04714 | 0.00028 | 10.0 s |
| 57:29.4 | Data point 62 | 1.50000 mL | 0.30701 mL | 0.29692 mL | 0.44000 mL | 2.567 | 0.00332 | 0.71527 | 0.00019 | 10.0 s |
| 58:05.0 | Data point 63 | 1.50000 mL | 0.30701 mL | 0.30287 mL | 0.44000 mL | 2.737 | -0.00087 | 0.02536 | 0.00027 | 10.0 s |
| 58:50.9 | Data point 64 | 1.50000 mL | 0.30701 mL | 0.30748 mL | 0.44000 mL | 2.936 | 0.00264 | 0.18981 | 0.00030 | 10.0 s |
| 59:26.4 | Data point 65 | 1.50000 mL | 0.30701 mL | 0.31054 mL | 0.44000 mL | 3.144 | -0.00133 | 0.27873 | 0.00012 | 10.0 s |
| 1:00:01.9 | Data point 66 | 1.50000 mL | 0.30701 mL | 0.31298 mL | 0.44000 mL | 3.378 | -0.00152 | 0.05010 | 0.00033 | 10.0 s |
| 1:00:42.5 | Data point 67 | 1.50000 mL | 0.30701 mL | 0.31430 mL | 0.44000 mL | 3.580 | -0.00279 | 0.07835 | 0.00049 | 10.0 s |
| 1:01:17.9 | Data point 68 | 1.50000 mL | 0.30701 mL | 0.31538 mL | 0.44000 mL | 3.878 | -0.00524 | 0.64775 | 0.00032 | 10.0 s |
| 1:01:53.4 | Data point 69 | 1.50000 mL | 0.30701 mL | 0.31642 mL | 0.44000 mL | 4.674 | -0.00689 | 0.18723 | 0.00079 | 12.0 s |
| 1:02:30.8 | Data point 70 | 1.50000 mL | 0.30701 mL | 0.31686 mL | 0.44000 mL | 6.215 | -0.01858 | 0.91294 | 0.00096 | 51.5 s |
| 1:03:52.9 | Data point 71 | 1.50000 mL | 0.30701 mL | 0.31724 mL | 0.44000 mL | 8.470 | -0.01798 | 0.89260 | 0.00094 | 50.0 s |
| 1:05:08.4 | Data point 72 | 1.50000 mL | 0.30701 mL | 0.31740 mL | 0.44000 mL | 8.775 | -0.01405 | 0.63869 | 0.00087 | 14.5 s |
| 1:05:48.3 | Data point 73 | 1.50000 mL | 0.30701 mL | 0.31771 mL | 0.44000 mL | 9.175 | -0.01421 | 0.63693 | 0.00088 | 12.0 s |
| 1:06:25.7 | Data point 74 | 1.50000 mL | 0.30701 mL | 0.31834 mL | 0.44000 mL | 9.635 | -0.00550 | 0.11202 | 0.00081 | 10.0 s |
| 1:07:01.2 | Data point 75 | 1.50000 mL | 0.30701 mL | 0.31933 mL | 0.44000 mL | 10.000 | 0.00089 | 0.00892 | 0.00046 | 10.5 s |
| 1:07:37.1 | Data point 76 | 1.50000 mL | 0.30701 mL | 0.32037 mL | 0.44000 mL | 10.223 | -0.00159 | 0.02094 | 0.00054 | 10.0 s |
| 1:08:23.0 | Data point 77 | 1.50000 mL | 0.30701 mL | 0.32175 mL | 0.44000 mL | 10.415 | 0.00362 | 0.09900 | 0.00057 | 10.0 s |
| 1:09:08.8 | Data point 78 | 1.50000 mL | 0.30701 mL | 0.32333 mL | 0.44000 mL | 10.600 | 0.00093 | 0.00262 | 0.00090 | 10.0 s |
| 1:09:54.6 | Data point 79 | 1.50000 mL | 0.30701 mL | 0.32545 mL | 0.44000 mL | 10.784 | 0.01040 | 0.85611 | 0.00056 | 10.0 s |
| 1:10:30.1 | Data point 80 | 1.50000 mL | 0.30701 mL | 0.32815 mL | 0.44000 mL | 10.962 | 0.00448 | 0.80191 | 0.00025 | 10.0 s |
| 1:11:05.6 | Data point 81 | 1.50000 mL | 0.30701 mL | 0.33177 mL | 0.44000 mL | 11.121 | 0.00570 | 0.85507 | 0.00030 | 10.0 s |
| 1:11:51.6 | Data point 82 | 1.50000 mL | 0.30701 mL | 0.33711 mL | 0.44000 mL | 11.309 | 0.00647 | 0.21129 | 0.00070 | 10.0 s |
| 1:12:27.2 | Data point 83 | 1.50000 mL | 0.30701 mL | 0.34450 mL | 0.44000 mL | 11.473 | 0.00664 | 0.82184 | 0.00036 | 10.0 s |
| 1:13:13.4 | Data point 84 | 1.50000 mL | 0.30701 mL | 0.35626 mL | 0.44000 mL | 11.668 | 0.00657 | 0.91736 | 0.00034 | 10.0 s |
| 1:13:49.2 | Data point 85 | 1.50000 mL | 0.30701 mL | 0.37333 mL | 0.44000 mL | 11.850 | 0.00586 | 0.31317 | 0.00052 | 10.0 s |
| 1:14:25.2 | Data point 86 | 1.50000 mL | 0.30701 mL | 0.39915 mL | 0.44000 mL | 12.033 | 0.00565 | 0.29376 | 0.00051 | 10.0 s |
| 1:14:44.3 | Assay volumes | 1.50000 mL | 0.30701 mL | 0.39915 mL | 0.44000 mL | | | | | |

| | | | |
|---|---|---|---|
| Sample name: | **M16_octanol** | Experiment start time: | **3/1/2018 9:25:34 AM** |
| Assay name: | **pH-metric high logP** | Analyst: | **Pion** |
| Assay ID: | **18C-01007** | Instrument ID: | **T312060** |
| Filename: | **C:\Sirius_T3\Mehtap\20180228_exp28_logP_T3-2\18C-01007_M16_octanol_pH-metric high logP.t3r** | | |

| | | | |
|---|---|---|---|
| Sample name: | **M16_octanol** | Experiment start time: | **3/1/2018 9:25:34 AM** |
| Assay name: | **pH-metric high logP** | Analyst: | **Pion** |
| Assay ID: | **18C-01007** | Instrument ID: | **T312060** |
| Filename: | **C:\Sirius_T3\Mehtap\20180228_exp28_logP_T3-2\18C-01007_M16_octanol_pH-metric high logP.t3r** | | |

### Assay Settings (continued)

| Setting | Value | Original Value | Date/Time changed | Imported from |
|---|---|---|---|---|
| ***Titration 3*** | | | | |
| Titrate from | Low to high pH | | | |
| Add additional water | 0.00 mL | | | |
| Additional partition solvent volume | 0.300 mL | | | |
| Additional partition solvent added | Automatic | | | |
| After pH adjust stir for | 30 seconds | | | |
| Stir to allow partitioning for | 15 seconds | | | |
| Stirrer speed for partitioning | 60% | | | |
| ***Data Point Stability*** | | | | |
| Stir during data point collection | No | | | |
| Delay before data point collection | 0 seconds | | | |
| Number of points to average | 20 points | | | |
| Time interval between points | 0.50 seconds | | | |
| Required maximum standard deviation | 0.00100 dpH/dt | | | |
| Stability timeout after | 60 seconds | | | |

### Calibration Settings

| Setting | Value | Date/Time changed | Imported from |
|---|---|---|---|
| Four-Plus alpha | 0.130 | 3/1/2018 9:25:33 AM | C:\Sirius_T3\HCl18B27.t3r |
| Four-Plus S | 0.9970 | 3/1/2018 9:25:33 AM | C:\Sirius_T3\HCl18B27.t3r |
| Four-Plus jH | 0.8 | 3/1/2018 9:25:33 AM | C:\Sirius_T3\HCl18B27.t3r |
| Four-Plus jOH | -0.4 | 3/1/2018 9:25:33 AM | C:\Sirius_T3\HCl18B27.t3r |
| Base concentration factor | 1.000 | 3/1/2018 9:25:34 AM | C:\Sirius_T3\KOH18B27.t3r |
| Acid concentration factor | 0.994 | 3/1/2018 9:25:34 AM | C:\Sirius_T3\HCl18B27.t3r |

### Instrument Settings

| Setting | Value | Batch Id | Install date |
|---|---|---|---|
| Instrument owner | Merck | | |
| Instrument ID | T312060 | | |
| Instrument type | T3 Simulator | | |
| Software version | 1.1.3.0 | | |
| Dispenser module | | T3DM1200361 | 3/31/2009 5:24:52 AM |
| Dispenser 0 | Water | | 3/31/2009 5:25:05 AM |
| Syringe volume | 2.5 mL | | |
| Firmware version | 1.2.1(r2) | | |
| Titrant | Water (0.15 M KCl) | 02-06-2018 | 2/27/2018 10:05:59 AM |
| Dispenser 2 | Acid | | 3/31/2009 5:25:11 AM |
| Syringe volume | 0.5 mL | | |
| Firmware version | 1.2.1(r2) | | |
| Titrant | Acid (0.5 M HCl) | 02-27-2018 | 2/27/2018 10:27:22 AM |
| Dispenser 1 | Base | | 3/31/2009 5:25:21 AM |
| Syringe volume | 0.5 mL | | |
| Firmware version | 1.2.1(r2) | | |
| Titrant | Base (0.5 M KOH) | 9/22/2017 | 2/27/2018 10:21:22 AM |
| Dispenser 5 | Cosolvent | | 3/31/2009 5:26:24 AM |
| Syringe volume | 2.5 mL | | |
| Firmware version | 1.2.1(r2) | | |
| Distribution valve 5 | Distribution Valve | | 3/31/2009 5:28:19 AM |
| Firmware version | 1.1.3 | | |
| Port A | Methanol (80%, 0.15 M KCl) | 09-26-17 | 2/7/2018 9:42:01 AM |
| Port B | Cyclohexane | 11-01-17 | 2/27/2018 10:37:57 AM |
| Dispenser 3 | Buffer | | 8/3/2010 5:05:16 AM |
| Syringe volume | 0.5 mL | | |
| Firmware version | 1.2.1(r2) | | |
| Titrant | Dodecane | 2018/01/31 | 2/28/2018 10:18:04 AM |
| Dispenser 6 | Octanol | | 10/22/2010 10:52:43 AM |

| | | | |
|---|---|---|---|
| Sample name: | **M16_octanol** | Experiment start time: | **3/1/2018 9:25:34 AM** |
| Assay name: | **pH-metric high logP** | Analyst: | **Pion** |
| Assay ID: | **18C-01007** | Instrument ID: | **T312060** |
| Filename: | **C:\Sirius_T3\Mehtap\20180228_exp28_logP_T3-2\18C-01007_M16_octanol_pH-metric high logP.t3r** | | |

| | | | |
|---|---|---|---|
| Sample name: | **M16_octanol** | Experiment start time: | **3/1/2018 9:25:34 AM** |
| Assay name: | **pH-metric high logP** | Analyst: | **Pion** |
| Assay ID: | **18C-01007** | Instrument ID: | **T312060** |
| Filename: | **C:\Sirius_T3\Mehtap\20180228_exp28_logP_T3-2\18C-01007_M16_octanol_pH-metric high logP.t3r** | | |

### Instrument Settings (continued)

| Setting | Value | Batch Id | Install date |
|---|---|---|---|
| Spectrometer calibration stir duration | 5 s | | |
| Spectrometer calibration stir speed | 30% | | |
| Spectrometer calibration wash pump volume | 20.0 mL | | |
| Spectrometer calibration wash stir duration | 5 s | | |
| Spectrometer calibration wash stir speed | 30% | | |
| Overhead dispense height | 10000 | | |

### Refinement Settings

| Setting | Value | Default value |
|---|---|---|
| Turbidity detection method | None | None |
| Turbidity wavelength to assess | 500.0 nm | 500.0 nm |
| Turbidity maximum absorbance | 0.100 | 0.100 |
| Turbidity probe threshold | 50.00 | 50.00 |

### Experiment Log

[2:48] Air gap created for Water (0.15 M KCl)
[2:49] Air gap created for Acid (0.5 M HCl)
[2:49] Air gap created for Base (0.5 M KOH)
[2:49] Air gap released for Water (0.15 M KCl)
[2:53] Titrator arm moved over Titration position
[2:53] Titration 1 of 3
[2:53] Adding initial titrants
[2:53] Automatically add 1.50000 mL of water
[3:18] Dispensed 1.500000 mL of Water (0.15 M KCl)
[3:23] Titrator arm moved over Drain
[9:04] Titrator arm moved to Titration position
[9:04] Argon flow rate set to 100
[9:04] Stirrer speed set to 10
[9:09] Automatically add 0.04000 mL of Octanol
[9:10] Dispensed 0.040005 mL of Octanol
[9:11] Initial pH = 3.56
[9:11] Iterative adjust 3.56 -> 2.00
[9:11] pH 3.56 -> 2.00
[9:13] Air gap released for Acid (0.5 M HCl)
[9:14] Dispensed 0.041439 mL of Acid (0.5 M HCl)
[9:19] pH 2.01 -> 2.00
[9:19] Dispensed 0.001270 mL of Acid (0.5 M HCl)
[9:24] Holding pH 2.00
[11:24] Stirrer speed set to 0
[11:24] Stirrer speed set to 50
[11:24] Iterative adjust 1.97 -> 2.00
[11:24] pH 1.97 -> 2.00
[11:25] Air gap released for Base (0.5 M KOH)
[11:26] Dispensed 0.003198 mL of Base (0.5 M KOH)
[12:16] Stirrer speed set to 0
[12:26] Datapoint id 1 collected
[12:26] Stirrer speed set to 50
[12:31] pH 2.01 -> 2.21
[12:31] Using cautious pH adjust
[12:31] Dispensed 0.007714 mL of Base (0.5 M KOH)
[12:37] Stepping pH = 2.10
[12:37] Dispensed 0.005903 mL of Base (0.5 M KOH)
[12:42] Stepping pH = 2.19
[12:42] Dispensed 0.000964 mL of Base (0.5 M KOH)
[12:47] Stepping pH = 2.21
[13:02] Stirrer speed set to 0
[13:12] Datapoint id 2 collected

Sample name: **M16_octanol**
Assay name: **pH-metric high logP**
Assay ID: **18C-01007**
Filename: **C:\Sirius_T3\Mehtap\20180228_exp28_logP_T3-2\18C-01007_M16_octanol_pH-metric high logP.t3r**

Experiment start time: **3/1/2018 9:25:34 AM**
Analyst: **Pion**
Instrument ID: **T312060**

### Experiment Log (continued)

[13:12] Charge balance equation is out by 5.6%
[13:12] Stirrer speed set to 50
[13:17] pH 2.21 -> 2.41
[13:17] Using charge balance adjust
[13:18] Dispensed 0.009619 mL of Base (0.5 M KOH)
[13:38] Stirrer speed set to 0
[13:48] Datapoint id 3 collected
[13:48] Charge balance equation is out by -0.3%
[13:48] Stirrer speed set to 50
[13:53] pH 2.42 -> 2.62
[13:53] Using charge balance adjust
[13:53] Dispensed 0.005997 mL of Base (0.5 M KOH)
[14:13] Stirrer speed set to 0
[14:23] Datapoint id 4 collected
[14:23] Charge balance equation is out by -11.6%
[14:23] Stirrer speed set to 50
[14:29] pH 2.60 -> 2.80
[14:29] Using charge balance adjust
[14:29] Dispensed 0.003975 mL of Base (0.5 M KOH)
[14:49] Stirrer speed set to 0
[14:59] Datapoint id 5 collected
[14:59] Charge balance equation is out by -5.5%
[14:59] Stirrer speed set to 50
[15:04] pH 2.80 -> 3.00
[15:04] Using charge balance adjust
[15:04] Dispensed 0.002611 mL of Base (0.5 M KOH)
[15:24] Stirrer speed set to 0
[15:34] Datapoint id 6 collected
[15:34] Charge balance equation is out by -31.1%
[15:34] Stirrer speed set to 50
[15:40] pH 2.94 -> 3.14
[15:40] Using cautious pH adjust
[15:40] Dispensed 0.000988 mL of Base (0.5 M KOH)
[15:45] Stepping pH = 3.01
[15:45] Dispensed 0.001152 mL of Base (0.5 M KOH)
[15:50] Stepping pH = 3.09
[15:50] Dispensed 0.000494 mL of Base (0.5 M KOH)
[15:55] Stepping pH = 3.13
[16:10] Stirrer speed set to 0
[16:20] Datapoint id 7 collected
[16:20] Charge balance equation is out by -33.7%
[16:20] Stirrer speed set to 50
[16:25] pH 3.14 -> 3.34
[16:25] Using cautious pH adjust
[16:25] Dispensed 0.000729 mL of Base (0.5 M KOH)
[16:31] Stepping pH = 3.20
[16:31] Dispensed 0.000823 mL of Base (0.5 M KOH)
[16:36] Stepping pH = 3.28
[16:36] Dispensed 0.000400 mL of Base (0.5 M KOH)
[16:41] Stepping pH = 3.32
[16:41] Dispensed 0.000141 mL of Base (0.5 M KOH)
[16:46] Stepping pH = 3.33
[17:01] Stirrer speed set to 0
[17:11] Datapoint id 8 collected
[17:11] Charge balance equation is out by -44.8%
[17:11] Stirrer speed set to 50
[17:16] pH 3.33 -> 3.53
[17:16] Using cautious pH adjust
[17:16] Dispensed 0.000588 mL of Base (0.5 M KOH)

| | | | |
|---|---|---|---|
| Sample name: | **M16_octanol** | Experiment start time: | **3/1/2018 9:25:34 AM** |
| Assay name: | **pH-metric high logP** | Analyst: | **Pion** |
| Assay ID: | **18C-01007** | Instrument ID: | **T312060** |
| Filename: | **C:\Sirius_T3\Mehtap\20180228_exp28_logP_T3-2\18C-01007_M16_octanol_pH-metric high logP.t3r** | | |

### Experiment Log (continued)

[17:22] Stepping pH = 3.39
[17:22] Dispensed 0.000729 mL of Base (0.5 M KOH)
[17:27] Stepping pH = 3.46
[17:27] Dispensed 0.000447 mL of Base (0.5 M KOH)
[17:32] Stepping pH = 3.51
[17:32] Dispensed 0.000188 mL of Base (0.5 M KOH)
[17:37] Stepping pH = 3.52
[17:52] Stirrer speed set to 0
[18:02] Datapoint id 9 collected
[18:02] Charge balance equation is out by -64.9%
[18:02] Stirrer speed set to 50
[18:07] pH 3.52 -> 3.72
[18:07] Using cautious pH adjust
[18:07] Dispensed 0.000564 mL of Base (0.5 M KOH)
[18:13] Stepping pH = 3.58
[18:13] Dispensed 0.000706 mL of Base (0.5 M KOH)
[18:18] Stepping pH = 3.64
[18:18] Dispensed 0.000564 mL of Base (0.5 M KOH)
[18:23] Stepping pH = 3.71
[18:23] Dispensed 0.000094 mL of Base (0.5 M KOH)
[18:28] Stepping pH = 3.71
[18:43] Stirrer speed set to 0
[18:53] Datapoint id 10 collected
[18:53] Charge balance equation is out by -71.8%
[18:53] Stirrer speed set to 50
[18:58] pH 3.71 -> 3.91
[18:58] Using cautious pH adjust
[18:58] Dispensed 0.000611 mL of Base (0.5 M KOH)
[19:03] Stepping pH = 3.79
[19:04] Dispensed 0.000588 mL of Base (0.5 M KOH)
[19:09] Stepping pH = 3.87
[19:09] Dispensed 0.000259 mL of Base (0.5 M KOH)
[19:14] Stepping pH = 3.90
[19:14] Dispensed 0.000094 mL of Base (0.5 M KOH)
[19:19] Stepping pH = 3.91
[19:34] Stirrer speed set to 0
[19:44] Datapoint id 11 collected
[19:44] Charge balance equation is out by -28.6%
[19:44] Stirrer speed set to 50
[19:49] pH 3.91 -> 4.11
[19:49] Using cautious pH adjust
[19:49] Dispensed 0.000706 mL of Base (0.5 M KOH)
[19:54] Stepping pH = 4.05
[19:55] Dispensed 0.000282 mL of Base (0.5 M KOH)
[20:00] Stepping pH = 4.10
[20:15] Stirrer speed set to 0
[20:25] Datapoint id 12 collected
[20:25] Charge balance equation is out by 30.4%
[20:25] Stirrer speed set to 50
[20:30] pH 4.10 -> 4.30
[20:30] Using cautious pH adjust
[20:30] Dispensed 0.000823 mL of Base (0.5 M KOH)
[20:35] Stepping pH = 4.37
[20:50] Stirrer speed set to 0
[21:01] Datapoint id 13 collected
[21:01] Charge balance equation is out by 50.0%
[21:01] Stirrer speed set to 50
[21:06] pH 4.36 -> 4.56
[21:06] Using cautious pH adjust

| | | | |
|---|---|---|---|
| Sample name: | **M16_octanol** | Experiment start time: | **3/1/2018 9:25:34 AM** |
| Assay name: | **pH-metric high logP** | Analyst: | **Pion** |
| Assay ID: | **18C-01007** | Instrument ID: | **T312060** |
| Filename: | **C:\Sirius_T3\Mehtap\20180228_exp28_logP_T3-2\18C-01007_M16_octanol_pH-metric high logP.t3r** | | |

### Experiment Log (continued)

[21:06] Dispensed 0.000917 mL of Base (0.5 M KOH)
[21:11] Stepping pH = 4.99
[21:26] Stirrer speed set to 0
[21:39] Datapoint id 14 collected
[21:39] Charge balance equation is out by 50.0%
[21:39] Stirrer speed set to 50
[21:44] pH 4.97 -> 5.17
[21:44] Using cautious pH adjust
[21:44] Dispensed 0.000682 mL of Base (0.5 M KOH)
[21:49] Stepping pH = 8.17
[22:05] Stirrer speed set to 0
[22:39] Datapoint id 15 collected
[22:39] Charge balance equation is out by 50.0%
[22:39] Stirrer speed set to 50
[22:44] pH 8.55 -> 8.75
[22:44] Using cautious pH adjust
[22:44] Dispensed 0.000564 mL of Base (0.5 M KOH)
[22:49] Stepping pH = 9.23
[23:04] Stirrer speed set to 0
[23:19] Datapoint id 16 collected
[23:19] Charge balance equation is out by 50.0%
[23:19] Stirrer speed set to 50
[23:24] pH 9.22 -> 9.42
[23:24] Using cautious pH adjust
[23:24] Dispensed 0.000941 mL of Base (0.5 M KOH)
[23:29] Stepping pH = 9.65
[23:44] Stirrer speed set to 0
[23:54] Datapoint id 17 collected
[23:54] Charge balance equation is out by 50.0%
[23:54] Stirrer speed set to 50
[24:00] pH 9.63 -> 9.83
[24:00] Using cautious pH adjust
[24:00] Dispensed 0.000776 mL of Base (0.5 M KOH)
[24:05] Stepping pH = 9.82
[24:05] Dispensed 0.000047 mL of Base (0.5 M KOH)
[24:10] Stepping pH = 9.82
[24:10] Dispensed 0.000259 mL of Base (0.5 M KOH)
[24:15] Stepping pH = 9.88
[24:30] Stirrer speed set to 0
[24:41] Datapoint id 18 collected
[24:41] Charge balance equation is out by 30.2%
[24:41] Stirrer speed set to 50
[24:46] pH 9.87 -> 10.07
[24:46] Using cautious pH adjust
[24:46] Dispensed 0.000635 mL of Base (0.5 M KOH)
[24:51] Stepping pH = 9.99
[24:51] Dispensed 0.000282 mL of Base (0.5 M KOH)
[24:56] Stepping pH = 10.02
[24:56] Dispensed 0.000235 mL of Base (0.5 M KOH)
[25:02] Stepping pH = 10.06
[25:17] Stirrer speed set to 0
[25:27] Datapoint id 19 collected
[25:27] Charge balance equation is out by 7.6%
[25:27] Stirrer speed set to 50
[25:32] pH 10.05 -> 10.25
[25:32] Using charge balance adjust
[25:32] Dispensed 0.001082 mL of Base (0.5 M KOH)
[25:52] Stirrer speed set to 0
[26:02] Datapoint id 20 collected

| | | | |
|---|---|---|---|
| Sample name: | **M16_octanol** | Experiment start time: | **3/1/2018 9:25:34 AM** |
| Assay name: | **pH-metric high logP** | Analyst: | **Pion** |
| Assay ID: | **18C-01007** | Instrument ID: | **T312060** |
| Filename: | **C:\Sirius_T3\Mehtap\20180228_exp28_logP_T3-2\18C-01007_M16_octanol_pH-metric high logP.t3r** | | |

### Experiment Log (continued)

[26:02] Charge balance equation is out by -29.3%
[26:02] Stirrer speed set to 50
[26:07] pH 10.19 -> 10.39
[26:07] Using cautious pH adjust
[26:07] Dispensed 0.000517 mL of Base (0.5 M KOH)
[26:12] Stepping pH = 10.25
[26:12] Dispensed 0.000635 mL of Base (0.5 M KOH)
[26:18] Stepping pH = 10.34
[26:18] Dispensed 0.000329 mL of Base (0.5 M KOH)
[26:23] Stepping pH = 10.37
[26:23] Dispensed 0.000165 mL of Base (0.5 M KOH)
[26:28] Stepping pH = 10.39
[26:43] Stirrer speed set to 0
[26:53] Datapoint id 21 collected
[26:53] Charge balance equation is out by -61.1%
[26:53] Stirrer speed set to 50
[26:58] pH 10.38 -> 10.58
[26:58] Using cautious pH adjust
[26:58] Dispensed 0.000517 mL of Base (0.5 M KOH)
[27:03] Stepping pH = 10.45
[27:03] Dispensed 0.000611 mL of Base (0.5 M KOH)
[27:08] Stepping pH = 10.52
[27:08] Dispensed 0.000353 mL of Base (0.5 M KOH)
[27:14] Stepping pH = 10.56
[27:14] Dispensed 0.000165 mL of Base (0.5 M KOH)
[27:19] Stepping pH = 10.58
[27:34] Stirrer speed set to 0
[27:44] Datapoint id 22 collected
[27:44] Charge balance equation is out by -57.1%
[27:44] Stirrer speed set to 50
[27:49] pH 10.57 -> 10.77
[27:49] Using cautious pH adjust
[27:49] Dispensed 0.000611 mL of Base (0.5 M KOH)
[27:54] Stepping pH = 10.64
[27:54] Dispensed 0.000706 mL of Base (0.5 M KOH)
[27:59] Stepping pH = 10.72
[28:00] Dispensed 0.000400 mL of Base (0.5 M KOH)
[28:05] Stepping pH = 10.75
[28:05] Dispensed 0.000165 mL of Base (0.5 M KOH)
[28:10] Stepping pH = 10.76
[28:25] Stirrer speed set to 0
[28:35] Datapoint id 23 collected
[28:35] Charge balance equation is out by -50.4%
[28:35] Stirrer speed set to 50
[28:40] pH 10.75 -> 10.95
[28:40] Using cautious pH adjust
[28:40] Dispensed 0.000823 mL of Base (0.5 M KOH)
[28:45] Stepping pH = 10.83
[28:45] Dispensed 0.000941 mL of Base (0.5 M KOH)
[28:50] Stepping pH = 10.90
[28:51] Dispensed 0.000588 mL of Base (0.5 M KOH)
[28:56] Stepping pH = 10.94
[28:56] Dispensed 0.000212 mL of Base (0.5 M KOH)
[29:01] Stepping pH = 10.94
[29:16] Stirrer speed set to 0
[29:26] Datapoint id 24 collected
[29:26] Charge balance equation is out by -55.0%
[29:26] Stirrer speed set to 50
[29:31] pH 10.94 -> 11.14

Sample name: **M16_octanol** Experiment start time: **3/1/2018 9:25:34 AM**
Assay name: **pH-metric high logP** Analyst: **Pion**
Assay ID: **18C-01007** Instrument ID: **T312060**
Filename: **C:\Sirius_T3\Mehtap\20180228_exp28_logP_T3-2\18C-01007_M16_octanol_pH-metric high logP.t3r**

### Experiment Log (continued)

[29:31] Using cautious pH adjust
[29:31] Dispensed 0.001199 mL of Base (0.5 M KOH)
[29:36] Stepping pH = 11.01
[29:36] Dispensed 0.001458 mL of Base (0.5 M KOH)
[29:42] Stepping pH = 11.09
[29:42] Dispensed 0.000706 mL of Base (0.5 M KOH)
[29:47] Stepping pH = 11.13
[29:47] Dispensed 0.000188 mL of Base (0.5 M KOH)
[29:52] Stepping pH = 11.14
[30:07] Stirrer speed set to 0
[30:17] Datapoint id 25 collected
[30:17] Charge balance equation is out by -48.0%
[30:17] Stirrer speed set to 50
[30:22] pH 11.13 -> 11.33
[30:22] Using cautious pH adjust
[30:22] Dispensed 0.001787 mL of Base (0.5 M KOH)
[30:27] Stepping pH = 11.22
[30:28] Dispensed 0.001552 mL of Base (0.5 M KOH)
[30:33] Stepping pH = 11.30
[30:33] Dispensed 0.000611 mL of Base (0.5 M KOH)
[30:38] Stepping pH = 11.32
[30:53] Stirrer speed set to 0
[31:03] Datapoint id 26 collected
[31:03] Charge balance equation is out by -10.0%
[31:03] Stirrer speed set to 50
[31:08] pH 11.32 -> 11.52
[31:08] Using charge balance adjust
[31:08] Dispensed 0.005503 mL of Base (0.5 M KOH)
[31:29] Stirrer speed set to 0
[31:39] Datapoint id 27 collected
[31:39] Charge balance equation is out by -14.2%
[31:39] Stirrer speed set to 50
[31:44] pH 11.49 -> 11.69
[31:44] Using charge balance adjust
[31:44] Dispensed 0.008278 mL of Base (0.5 M KOH)
[32:04] Stirrer speed set to 0
[32:14] Datapoint id 28 collected
[32:14] Charge balance equation is out by -7.1%
[32:14] Stirrer speed set to 50
[32:19] pH 11.68 -> 11.88
[32:19] Using charge balance adjust
[32:20] Dispensed 0.013006 mL of Base (0.5 M KOH)
[32:40] Stirrer speed set to 0
[32:50] Datapoint id 29 collected
[32:50] Charge balance equation is out by -8.7%
[32:50] Stirrer speed set to 50
[32:55] pH 11.87 -> 12.05
[32:55] Using charge balance adjust
[32:56] Dispensed 0.017803 mL of Base (0.5 M KOH)
[33:16] Stirrer speed set to 0
[33:26] Datapoint id 30 collected
[33:26] Charge balance equation is out by -21.5%
[33:26] Titration 2 of 3
[33:26] Adding initial titrants
[33:26] Automatically add 0.10000 mL of Octanol
[33:28] Dispensed 0.100000 mL of Octanol
[33:28] Stirrer speed set to 10
[33:29] Stirrer speed set to 55
[33:29] Iterative adjust 12.04 -> 2.00

| | | | |
|---|---|---|---|
| Sample name: | **M16_octanol** | Experiment start time: | **3/1/2018 9:25:34 AM** |
| Assay name: | **pH-metric high logP** | Analyst: | **Pion** |
| Assay ID: | **18C-01007** | Instrument ID: | **T312060** |
| Filename: | **C:\Sirius_T3\Mehtap\20180228_exp28_logP_T3-2\18C-01007_M16_octanol_pH-metric high logP.t3r** | | |

### Experiment Log (continued)

[33:29] pH 12.04 -> 2.00
[33:32] Dispensed 0.100000 mL of Acid (0.5 M HCl)
[33:37] pH 2.26 -> 2.00
[33:38] Dispensed 0.022437 mL of Acid (0.5 M HCl)
[33:43] pH 2.02 -> 2.00
[33:43] Dispensed 0.002140 mL of Acid (0.5 M HCl)
[34:33] Stirrer speed set to 0
[34:43] Datapoint id 31 collected
[34:43] Stirrer speed set to 55
[34:48] pH 1.96 -> 2.16
[34:48] Using cautious pH adjust
[34:49] Dispensed 0.010113 mL of Base (0.5 M KOH)
[34:54] Stepping pH = 2.06
[34:54] Dispensed 0.006703 mL of Base (0.5 M KOH)
[34:59] Stepping pH = 2.13
[34:59] Dispensed 0.002070 mL of Base (0.5 M KOH)
[35:04] Stepping pH = 2.16
[35:19] Stirrer speed set to 0
[35:30] Datapoint id 32 collected
[35:30] Charge balance equation is out by 6.7%
[35:30] Stirrer speed set to 55
[35:35] pH 2.16 -> 2.36
[35:35] Using charge balance adjust
[35:35] Dispensed 0.012629 mL of Base (0.5 M KOH)
[35:55] Stirrer speed set to 0
[36:05] Datapoint id 33 collected
[36:05] Charge balance equation is out by 11.2%
[36:05] Stirrer speed set to 55
[36:10] pH 2.39 -> 2.59
[36:10] Using charge balance adjust
[36:11] Dispensed 0.007479 mL of Base (0.5 M KOH)
[36:31] Stirrer speed set to 0
[36:41] Datapoint id 34 collected
[36:41] Charge balance equation is out by -11.6%
[36:41] Stirrer speed set to 55
[36:46] pH 2.57 -> 2.77
[36:46] Using charge balance adjust
[36:46] Dispensed 0.004986 mL of Base (0.5 M KOH)
[37:06] Stirrer speed set to 0
[37:16] Datapoint id 35 collected
[37:16] Charge balance equation is out by -15.2%
[37:16] Stirrer speed set to 55
[37:21] pH 2.75 -> 2.95
[37:21] Using cautious pH adjust
[37:22] Dispensed 0.001740 mL of Base (0.5 M KOH)
[37:27] Stepping pH = 2.82
[37:27] Dispensed 0.001740 mL of Base (0.5 M KOH)
[37:32] Stepping pH = 2.91
[37:32] Dispensed 0.000494 mL of Base (0.5 M KOH)
[37:37] Stepping pH = 2.94
[37:52] Stirrer speed set to 0
[38:03] Datapoint id 36 collected
[38:03] Charge balance equation is out by -14.7%
[38:03] Stirrer speed set to 55
[38:08] pH 2.94 -> 3.14
[38:08] Using charge balance adjust
[38:08] Dispensed 0.002399 mL of Base (0.5 M KOH)
[38:28] Stirrer speed set to 0
[38:38] Datapoint id 37 collected

| | | | |
|---|---|---|---|
| Sample name: | **M16_octanol** | Experiment start time: | **3/1/2018 9:25:34 AM** |
| Assay name: | **pH-metric high logP** | Analyst: | **Pion** |
| Assay ID: | **18C-01007** | Instrument ID: | **T312060** |
| Filename: | **C:\Sirius_T3\Mehtap\20180228_exp28_logP_T3-2\18C-01007_M16_octanol_pH-metric high logP.t3r** | | |

### Experiment Log (continued)

[38:38] Charge balance equation is out by -22.9%
[38:38] Stirrer speed set to 55
[38:43] pH 3.10 -> 3.30
[38:43] Using cautious pH adjust
[38:44] Dispensed 0.000941 mL of Base (0.5 M KOH)
[38:49] Stepping pH = 3.17
[38:49] Dispensed 0.001105 mL of Base (0.5 M KOH)
[38:54] Stepping pH = 3.25
[38:54] Dispensed 0.000611 mL of Base (0.5 M KOH)
[38:59] Stepping pH = 3.29
[38:59] Dispensed 0.000141 mL of Base (0.5 M KOH)
[39:04] Stepping pH = 3.30
[39:19] Stirrer speed set to 0
[39:29] Datapoint id 38 collected
[39:29] Charge balance equation is out by -46.9%
[39:29] Stirrer speed set to 55
[39:34] pH 3.30 -> 3.50
[39:34] Using cautious pH adjust
[39:35] Dispensed 0.000800 mL of Base (0.5 M KOH)
[39:40] Stepping pH = 3.38
[39:40] Dispensed 0.000776 mL of Base (0.5 M KOH)
[39:45] Stepping pH = 3.46
[39:45] Dispensed 0.000282 mL of Base (0.5 M KOH)
[39:50] Stepping pH = 3.50
[40:05] Stirrer speed set to 0
[40:16] Datapoint id 39 collected
[40:16] Charge balance equation is out by -15.2%
[40:16] Stirrer speed set to 55
[40:21] pH 3.50 -> 3.70
[40:21] Using cautious pH adjust
[40:21] Dispensed 0.000776 mL of Base (0.5 M KOH)
[40:26] Stepping pH = 3.61
[40:26] Dispensed 0.000470 mL of Base (0.5 M KOH)
[40:31] Stepping pH = 3.69
[40:46] Stirrer speed set to 0
[40:56] Datapoint id 40 collected
[40:56] Charge balance equation is out by 19.5%
[40:56] Stirrer speed set to 55
[41:01] pH 3.69 -> 3.89
[41:01] Using cautious pH adjust
[41:02] Dispensed 0.000823 mL of Base (0.5 M KOH)
[41:07] Stepping pH = 3.88
[41:07] Dispensed 0.000047 mL of Base (0.5 M KOH)
[41:12] Stepping pH = 3.88
[41:12] Dispensed 0.000282 mL of Base (0.5 M KOH)
[41:17] Stepping pH = 3.96
[41:32] Stirrer speed set to 0
[41:42] Datapoint id 41 collected
[41:42] Charge balance equation is out by 29.0%
[41:42] Stirrer speed set to 55
[41:47] pH 3.97 -> 4.17
[41:47] Using cautious pH adjust
[41:47] Dispensed 0.000941 mL of Base (0.5 M KOH)
[41:52] Stepping pH = 4.46
[42:07] Stirrer speed set to 0
[42:18] Datapoint id 42 collected
[42:18] Charge balance equation is out by 50.0%
[42:18] Stirrer speed set to 55
[42:23] pH 4.46 -> 4.66

| | | | |
|---|---|---|---|
| Sample name: | **M16_octanol** | Experiment start time: | **3/1/2018 9:25:34 AM** |
| Assay name: | **pH-metric high logP** | Analyst: | **Pion** |
| Assay ID: | **18C-01007** | Instrument ID: | **T312060** |
| Filename: | **C:\Sirius_T3\Mehtap\20180228_exp28_logP_T3-2\18C-01007_M16_octanol_pH-metric high logP.t3r** | | |

### Experiment Log (continued)

[42:23] Using cautious pH adjust
[42:23] Dispensed 0.000870 mL of Base (0.5 M KOH)
[42:28] Stepping pH = 7.47
[42:43] Stirrer speed set to 0
[43:43] Datapoint id 43 collected
[43:43] Charge balance equation is out by 50.0%
[43:43] Stirrer speed set to 55
[43:48] pH 7.65 -> 7.85
[43:48] Using cautious pH adjust
[43:49] Dispensed 0.000071 mL of Base (0.5 M KOH)
[43:54] Stepping pH = 7.58
[43:54] Dispensed 0.000400 mL of Base (0.5 M KOH)
[43:59] Stepping pH = 9.04
[44:14] Stirrer speed set to 0
[44:30] Datapoint id 44 collected
[44:30] Charge balance equation is out by -270.4%
[44:30] Stirrer speed set to 55
[44:35] pH 9.00 -> 9.20
[44:35] Using cautious pH adjust
[44:35] Dispensed 0.000729 mL of Base (0.5 M KOH)
[44:40] Stepping pH = 9.52
[44:55] Stirrer speed set to 0
[45:07] Datapoint id 45 collected
[45:07] Charge balance equation is out by 50.0%
[45:07] Stirrer speed set to 55
[45:12] pH 9.51 -> 9.71
[45:12] Using cautious pH adjust
[45:12] Dispensed 0.000964 mL of Base (0.5 M KOH)
[45:17] Stepping pH = 9.88
[45:32] Stirrer speed set to 0
[45:42] Datapoint id 46 collected
[45:42] Charge balance equation is out by 50.0%
[45:42] Stirrer speed set to 55
[45:47] pH 9.88 -> 10.08
[45:47] Using cautious pH adjust
[45:48] Dispensed 0.000847 mL of Base (0.5 M KOH)
[45:53] Stepping pH = 10.05
[45:53] Dispensed 0.000118 mL of Base (0.5 M KOH)
[45:58] Stepping pH = 10.06
[45:58] Dispensed 0.000165 mL of Base (0.5 M KOH)
[46:03] Stepping pH = 10.08
[46:18] Stirrer speed set to 0
[46:28] Datapoint id 47 collected
[46:28] Charge balance equation is out by 32.6%
[46:28] Stirrer speed set to 55
[46:33] pH 10.08 -> 10.28
[46:33] Using cautious pH adjust
[46:33] Dispensed 0.000753 mL of Base (0.5 M KOH)
[46:38] Stepping pH = 10.22
[46:39] Dispensed 0.000259 mL of Base (0.5 M KOH)
[46:44] Stepping pH = 10.25
[46:44] Dispensed 0.000141 mL of Base (0.5 M KOH)
[46:49] Stepping pH = 10.27
[47:04] Stirrer speed set to 0
[47:14] Datapoint id 48 collected
[47:14] Charge balance equation is out by 22.5%
[47:14] Stirrer speed set to 55
[47:19] pH 10.26 -> 10.46
[47:19] Using cautious pH adjust

Sample name: **M16_octanol**
Assay name: **pH-metric high logP**
Assay ID: **18C-01007**
Filename: **C:\Sirius_T3\Mehtap\20180228_exp28_logP_T3-2\18C-01007_M16_octanol_pH-metric high logP.t3r**
Experiment start time: **3/1/2018 9:25:34 AM**
Analyst: **Pion**
Instrument ID: **T312060**

### Experiment Log (continued)

[47:19] Dispensed 0.000706 mL of Base (0.5 M KOH)
[47:24] Stepping pH = 10.37
[47:24] Dispensed 0.000423 mL of Base (0.5 M KOH)
[47:29] Stepping pH = 10.43
[47:30] Dispensed 0.000235 mL of Base (0.5 M KOH)
[47:35] Stepping pH = 10.45
[47:35] Dispensed 0.000071 mL of Base (0.5 M KOH)
[47:40] Stepping pH = 10.45
[47:55] Stirrer speed set to 0
[48:05] Datapoint id 49 collected
[48:05] Charge balance equation is out by -1.7%
[48:05] Stirrer speed set to 55
[48:10] pH 10.45 -> 10.65
[48:10] Using charge balance adjust
[48:10] Dispensed 0.001458 mL of Base (0.5 M KOH)
[48:30] Stirrer speed set to 0
[48:40] Datapoint id 50 collected
[48:40] Charge balance equation is out by -9.7%
[48:40] Stirrer speed set to 55
[48:46] pH 10.63 -> 10.83
[48:46] Using charge balance adjust
[48:46] Dispensed 0.001717 mL of Base (0.5 M KOH)
[49:06] Stirrer speed set to 0
[49:16] Datapoint id 51 collected
[49:16] Charge balance equation is out by -26.7%
[49:16] Stirrer speed set to 55
[49:21] pH 10.78 -> 10.98
[49:21] Using cautious pH adjust
[49:21] Dispensed 0.001058 mL of Base (0.5 M KOH)
[49:26] Stepping pH = 10.85
[49:26] Dispensed 0.001199 mL of Base (0.5 M KOH)
[49:31] Stepping pH = 10.92
[49:32] Dispensed 0.000753 mL of Base (0.5 M KOH)
[49:37] Stepping pH = 10.96
[49:37] Dispensed 0.000282 mL of Base (0.5 M KOH)
[49:42] Stepping pH = 10.97
[49:57] Stirrer speed set to 0
[50:07] Datapoint id 52 collected
[50:07] Charge balance equation is out by -54.7%
[50:07] Stirrer speed set to 55
[50:12] pH 10.97 -> 11.17
[50:12] Using cautious pH adjust
[50:12] Dispensed 0.001529 mL of Base (0.5 M KOH)
[50:17] Stepping pH = 11.05
[50:18] Dispensed 0.001646 mL of Base (0.5 M KOH)
[50:23] Stepping pH = 11.13
[50:23] Dispensed 0.000870 mL of Base (0.5 M KOH)
[50:28] Stepping pH = 11.16
[50:43] Stirrer speed set to 0
[50:53] Datapoint id 53 collected
[50:53] Charge balance equation is out by -31.7%
[50:53] Stirrer speed set to 55
[50:58] pH 11.16 -> 11.36
[50:58] Using cautious pH adjust
[50:58] Dispensed 0.002258 mL of Base (0.5 M KOH)
[51:03] Stepping pH = 11.25
[51:03] Dispensed 0.001929 mL of Base (0.5 M KOH)
[51:09] Stepping pH = 11.32
[51:09] Dispensed 0.001152 mL of Base (0.5 M KOH)

| | | | |
|---|---|---|---|
| Sample name: | **M16_octanol** | Experiment start time: | **3/1/2018 9:25:34 AM** |
| Assay name: | **pH-metric high logP** | Analyst: | **Pion** |
| Assay ID: | **18C-01007** | Instrument ID: | **T312060** |
| Filename: | **C:\Sirius_T3\Mehtap\20180228_exp28_logP_T3-2\18C-01007_M16_octanol_pH-metric high logP.t3r** | | |

### Experiment Log (continued)

[51:14] Stepping pH = 11.35
[51:14] Dispensed 0.000329 mL of Base (0.5 M KOH)
[51:19] Stepping pH = 11.36
[51:34] Stirrer speed set to 0
[51:44] Datapoint id 54 collected
[51:44] Charge balance equation is out by -25.3%
[51:44] Stirrer speed set to 55
[51:49] pH 11.35 -> 11.55
[51:49] Using cautious pH adjust
[51:49] Dispensed 0.003504 mL of Base (0.5 M KOH)
[51:54] Stepping pH = 11.45
[51:55] Dispensed 0.002987 mL of Base (0.5 M KOH)
[52:00] Stepping pH = 11.52
[52:00] Dispensed 0.001176 mL of Base (0.5 M KOH)
[52:05] Stepping pH = 11.55
[52:20] Stirrer speed set to 0
[52:30] Datapoint id 55 collected
[52:30] Charge balance equation is out by -9.2%
[52:30] Stirrer speed set to 55
[52:35] pH 11.55 -> 11.75
[52:35] Using charge balance adjust
[52:36] Dispensed 0.011007 mL of Base (0.5 M KOH)
[52:56] Stirrer speed set to 0
[53:06] Datapoint id 56 collected
[53:06] Charge balance equation is out by -12.5%
[53:06] Stirrer speed set to 55
[53:11] pH 11.73 -> 11.93
[53:11] Using charge balance adjust
[53:12] Dispensed 0.016769 mL of Base (0.5 M KOH)
[53:32] Stirrer speed set to 0
[53:42] Datapoint id 57 collected
[53:42] Charge balance equation is out by -8.0%
[53:42] Stirrer speed set to 55
[53:47] pH 11.92 -> 12.05
[53:47] Using charge balance adjust
[53:48] Dispensed 0.016322 mL of Base (0.5 M KOH)
[54:08] Stirrer speed set to 0
[54:18] Datapoint id 58 collected
[54:18] Charge balance equation is out by -42.2%
[54:18] Titration 3 of 3
[54:18] Adding initial titrants
[54:18] Automatically add 0.30000 mL of Octanol
[54:25] Dispensed 0.300000 mL of Octanol
[54:25] Stirrer speed set to 10
[54:26] Stirrer speed set to 60
[54:26] Iterative adjust 12.04 -> 2.00
[54:26] pH 12.04 -> 2.00
[54:29] Dispensed 0.100000 mL of Acid (0.5 M HCl)
[54:34] pH 2.44 -> 2.00
[54:35] Dispensed 0.035842 mL of Acid (0.5 M HCl)
[54:40] pH 2.03 -> 2.00
[54:40] Dispensed 0.003881 mL of Acid (0.5 M HCl)
[55:30] Stirrer speed set to 0
[55:41] Datapoint id 59 collected
[55:41] Stirrer speed set to 60
[55:46] pH 1.97 -> 2.17
[55:46] Using cautious pH adjust
[55:46] Dispensed 0.011477 mL of Base (0.5 M KOH)
[55:51] Stepping pH = 2.05

Sample name: **M16_octanol** Experiment start time: **3/1/2018 9:25:34 AM**
Assay name: **pH-metric high logP** Analyst: **Pion**
Assay ID: **18C-01007** Instrument ID: **T312060**
Filename: **C:\Sirius_T3\Mehtap\20180228_exp28_logP_T3-2\18C-01007_M16_octanol_pH-metric high logP.t3r**

### Experiment Log (continued)

[55:52] Dispensed 0.008937 mL of Base (0.5 M KOH)
[55:57] Stepping pH = 2.14
[55:57] Dispensed 0.001834 mL of Base (0.5 M KOH)
[56:02] Stepping pH = 2.17
[56:17] Stirrer speed set to 0
[56:28] Datapoint id 60 collected
[56:28] Charge balance equation is out by 3.1%
[56:28] Stirrer speed set to 60
[56:33] pH 2.17 -> 2.37
[56:33] Using charge balance adjust
[56:33] Dispensed 0.014276 mL of Base (0.5 M KOH)
[56:53] Stirrer speed set to 0
[57:03] Datapoint id 61 collected
[57:03] Charge balance equation is out by -3.3%
[57:03] Stirrer speed set to 60
[57:08] pH 2.37 -> 2.57
[57:08] Using charge balance adjust
[57:09] Dispensed 0.009149 mL of Base (0.5 M KOH)
[57:29] Stirrer speed set to 0
[57:39] Datapoint id 62 collected
[57:39] Charge balance equation is out by -1.3%
[57:39] Stirrer speed set to 60
[57:44] pH 2.57 -> 2.77
[57:44] Using charge balance adjust
[57:44] Dispensed 0.005950 mL of Base (0.5 M KOH)
[58:05] Stirrer speed set to 0
[58:15] Datapoint id 63 collected
[58:15] Charge balance equation is out by -16.5%
[58:15] Stirrer speed set to 60
[58:20] pH 2.74 -> 2.94
[58:20] Using cautious pH adjust
[58:20] Dispensed 0.002117 mL of Base (0.5 M KOH)
[58:25] Stepping pH = 2.82
[58:25] Dispensed 0.001952 mL of Base (0.5 M KOH)
[58:30] Stepping pH = 2.91
[58:30] Dispensed 0.000541 mL of Base (0.5 M KOH)
[58:35] Stepping pH = 2.94
[58:50] Stirrer speed set to 0
[59:00] Datapoint id 64 collected
[59:00] Charge balance equation is out by -8.5%
[59:00] Stirrer speed set to 60
[59:06] pH 2.94 -> 3.14
[59:06] Using charge balance adjust
[59:06] Dispensed 0.003057 mL of Base (0.5 M KOH)
[59:26] Stirrer speed set to 0
[59:36] Datapoint id 65 collected
[59:36] Charge balance equation is out by 2.6%
[59:36] Stirrer speed set to 60
[59:41] pH 3.15 -> 3.35
[59:41] Using charge balance adjust
[59:41] Dispensed 0.002446 mL of Base (0.5 M KOH)
[1:00:01] Stirrer speed set to 0
[1:00:11] Datapoint id 66 collected
[1:00:11] Charge balance equation is out by 15.7%
[1:00:11] Stirrer speed set to 60
[1:00:17] pH 3.38 -> 3.58
[1:00:17] Using cautious pH adjust
[1:00:17] Dispensed 0.001105 mL of Base (0.5 M KOH)
[1:00:22] Stepping pH = 3.54

Sample name: **M16_octanol** Experiment start time: **3/1/2018 9:25:34 AM**
Assay name: **pH-metric high logP** Analyst: **Pion**
Assay ID: **18C-01007** Instrument ID: **T312060**
Filename: **C:\Sirius_T3\Mehtap\20180228_exp28_logP_T3-2\18C-01007_M16_octanol_pH-metric high logP.t3r**

### Experiment Log (continued)

[1:00:22] Dispensed 0.000212 mL of Base (0.5 M KOH)
[1:00:27] Stepping pH = 3.58
[1:00:42] Stirrer speed set to 0
[1:00:52] Datapoint id 67 collected
[1:00:52] Charge balance equation is out by 40.2%
[1:00:52] Stirrer speed set to 60
[1:00:57] pH 3.58 -> 3.78
[1:00:57] Using cautious pH adjust
[1:00:57] Dispensed 0.001082 mL of Base (0.5 M KOH)
[1:01:02] Stepping pH = 3.88
[1:01:18] Stirrer speed set to 0
[1:01:28] Datapoint id 68 collected
[1:01:28] Charge balance equation is out by 50.0%
[1:01:28] Stirrer speed set to 60
[1:01:33] pH 3.88 -> 4.08
[1:01:33] Using cautious pH adjust
[1:01:33] Dispensed 0.001035 mL of Base (0.5 M KOH)
[1:01:38] Stepping pH = 4.71
[1:01:53] Stirrer speed set to 0
[1:02:05] Datapoint id 69 collected
[1:02:05] Charge balance equation is out by 50.0%
[1:02:05] Stirrer speed set to 60
[1:02:10] pH 4.69 -> 4.89
[1:02:10] Using cautious pH adjust
[1:02:10] Dispensed 0.000447 mL of Base (0.5 M KOH)
[1:02:15] Stepping pH = 6.55
[1:02:30] Stirrer speed set to 0
[1:03:22] Datapoint id 70 collected
[1:03:22] Charge balance equation is out by 50.0%
[1:03:22] Stirrer speed set to 60
[1:03:27] pH 6.17 -> 6.37
[1:03:27] Using cautious pH adjust
[1:03:27] Dispensed 0.000047 mL of Base (0.5 M KOH)
[1:03:32] Stepping pH = 6.09
[1:03:32] Dispensed 0.000329 mL of Base (0.5 M KOH)
[1:03:37] Stepping pH = 8.68
[1:03:52] Stirrer speed set to 0
[1:04:43] Datapoint id 71 collected
[1:04:43] Charge balance equation is out by -318.8%
[1:04:43] Stirrer speed set to 60
[1:04:48] pH 8.44 -> 8.64
[1:04:48] Using cautious pH adjust
[1:04:48] Dispensed 0.000165 mL of Base (0.5 M KOH)
[1:04:53] Stepping pH = 8.82
[1:05:08] Stirrer speed set to 0
[1:05:22] Datapoint id 72 collected
[1:05:22] Charge balance equation is out by 50.0%
[1:05:22] Stirrer speed set to 60
[1:05:28] pH 8.77 -> 8.97
[1:05:28] Using cautious pH adjust
[1:05:28] Dispensed 0.000306 mL of Base (0.5 M KOH)
[1:05:33] Stepping pH = 9.20
[1:05:48] Stirrer speed set to 0
[1:06:00] Datapoint id 73 collected
[1:06:00] Charge balance equation is out by 50.0%
[1:06:00] Stirrer speed set to 60
[1:06:05] pH 9.18 -> 9.38
[1:06:05] Using cautious pH adjust
[1:06:05] Dispensed 0.000635 mL of Base (0.5 M KOH)

Sample name: **M16_octanol** Experiment start time: **3/1/2018 9:25:34 AM**
Assay name: **pH-metric high logP** Analyst: **Pion**
Assay ID: **18C-01007** Instrument ID: **T312060**
Filename: **C:\Sirius_T3\Mehtap\20180228_exp28_logP_T3-2\18C-01007_M16_octanol_pH-metric high logP.t3r**

### Experiment Log (continued)

[1:06:10] Stepping pH = 9.65
[1:06:25] Stirrer speed set to 0
[1:06:35] Datapoint id 74 collected
[1:06:35] Charge balance equation is out by 50.0%
[1:06:35] Stirrer speed set to 60
[1:06:40] pH 9.64 -> 9.84
[1:06:40] Using cautious pH adjust
[1:06:41] Dispensed 0.000988 mL of Base (0.5 M KOH)
[1:06:46] Stepping pH = 10.01
[1:07:01] Stirrer speed set to 0
[1:07:11] Datapoint id 75 collected
[1:07:11] Charge balance equation is out by 50.0%
[1:07:11] Stirrer speed set to 60
[1:07:16] pH 10.00 -> 10.20
[1:07:16] Using cautious pH adjust
[1:07:16] Dispensed 0.001035 mL of Base (0.5 M KOH)
[1:07:22] Stepping pH = 10.23
[1:07:37] Stirrer speed set to 0
[1:07:47] Datapoint id 76 collected
[1:07:47] Charge balance equation is out by 50.0%
[1:07:47] Stirrer speed set to 60
[1:07:52] pH 10.22 -> 10.42
[1:07:52] Using cautious pH adjust
[1:07:52] Dispensed 0.001011 mL of Base (0.5 M KOH)
[1:07:57] Stepping pH = 10.39
[1:07:57] Dispensed 0.000212 mL of Base (0.5 M KOH)
[1:08:02] Stepping pH = 10.40
[1:08:02] Dispensed 0.000165 mL of Base (0.5 M KOH)
[1:08:07] Stepping pH = 10.42
[1:08:23] Stirrer speed set to 0
[1:08:33] Datapoint id 77 collected
[1:08:33] Charge balance equation is out by 31.5%
[1:08:33] Stirrer speed set to 60
[1:08:38] pH 10.42 -> 10.62
[1:08:38] Using cautious pH adjust
[1:08:38] Dispensed 0.001011 mL of Base (0.5 M KOH)
[1:08:43] Stepping pH = 10.55
[1:08:43] Dispensed 0.000376 mL of Base (0.5 M KOH)
[1:08:48] Stepping pH = 10.59
[1:08:48] Dispensed 0.000188 mL of Base (0.5 M KOH)
[1:08:53] Stepping pH = 10.61
[1:09:08] Stirrer speed set to 0
[1:09:18] Datapoint id 78 collected
[1:09:18] Charge balance equation is out by 21.6%
[1:09:18] Stirrer speed set to 60
[1:09:23] pH 10.60 -> 10.80
[1:09:23] Using cautious pH adjust
[1:09:24] Dispensed 0.001105 mL of Base (0.5 M KOH)
[1:09:29] Stepping pH = 10.71
[1:09:29] Dispensed 0.000729 mL of Base (0.5 M KOH)
[1:09:34] Stepping pH = 10.77
[1:09:34] Dispensed 0.000282 mL of Base (0.5 M KOH)
[1:09:39] Stepping pH = 10.79
[1:09:54] Stirrer speed set to 0
[1:10:04] Datapoint id 79 collected
[1:10:04] Charge balance equation is out by 5.1%
[1:10:04] Stirrer speed set to 60
[1:10:09] pH 10.78 -> 10.98
[1:10:09] Using charge balance adjust

| | | | |
|---|---|---|---|
| Sample name: | **M16_octanol** | Experiment start time: | **3/1/2018 9:25:34 AM** |
| Assay name: | **pH-metric high logP** | Analyst: | **Pion** |
| Assay ID: | **18C-01007** | Instrument ID: | **T312060** |
| Filename: | **C:\Sirius_T3\Mehtap\20180228_exp28_logP_T3-2\18C-01007_M16_octanol_pH-metric high logP.t3r** | | |

### Experiment Log (continued)

[1:10:09] Dispensed 0.002705 mL of Base (0.5 M KOH)
[1:10:30] Stirrer speed set to 0
[1:10:40] Datapoint id 80 collected
[1:10:40] Charge balance equation is out by -11.6%
[1:10:40] Stirrer speed set to 60
[1:10:45] pH 10.96 -> 11.16
[1:10:45] Using charge balance adjust
[1:10:45] Dispensed 0.003622 mL of Base (0.5 M KOH)
[1:11:05] Stirrer speed set to 0
[1:11:15] Datapoint id 81 collected
[1:11:15] Charge balance equation is out by -20.9%
[1:11:15] Stirrer speed set to 60
[1:11:20] pH 11.12 -> 11.32
[1:11:20] Using cautious pH adjust
[1:11:21] Dispensed 0.002469 mL of Base (0.5 M KOH)
[1:11:26] Stepping pH = 11.21
[1:11:26] Dispensed 0.002140 mL of Base (0.5 M KOH)
[1:11:31] Stepping pH = 11.29
[1:11:31] Dispensed 0.000729 mL of Base (0.5 M KOH)
[1:11:36] Stepping pH = 11.31
[1:11:51] Stirrer speed set to 0
[1:12:01] Datapoint id 82 collected
[1:12:01] Charge balance equation is out by -7.8%
[1:12:01] Stirrer speed set to 60
[1:12:06] pH 11.31 -> 11.51
[1:12:06] Using charge balance adjust
[1:12:07] Dispensed 0.007385 mL of Base (0.5 M KOH)
[1:12:27] Stirrer speed set to 0
[1:12:37] Datapoint id 83 collected
[1:12:37] Charge balance equation is out by -18.4%
[1:12:37] Stirrer speed set to 60
[1:12:42] pH 11.48 -> 11.68
[1:12:42] Using cautious pH adjust
[1:12:42] Dispensed 0.005386 mL of Base (0.5 M KOH)
[1:12:47] Stepping pH = 11.57
[1:12:47] Dispensed 0.004586 mL of Base (0.5 M KOH)
[1:12:53] Stepping pH = 11.65
[1:12:53] Dispensed 0.001787 mL of Base (0.5 M KOH)
[1:12:58] Stepping pH = 11.67
[1:13:13] Stirrer speed set to 0
[1:13:23] Datapoint id 84 collected
[1:13:23] Charge balance equation is out by -8.5%
[1:13:23] Stirrer speed set to 60
[1:13:28] pH 11.67 -> 11.87
[1:13:28] Using charge balance adjust
[1:13:29] Dispensed 0.017074 mL of Base (0.5 M KOH)
[1:13:49] Stirrer speed set to 0
[1:13:59] Datapoint id 85 collected
[1:13:59] Charge balance equation is out by -11.5%
[1:13:59] Stirrer speed set to 60
[1:14:04] pH 11.85 -> 12.05
[1:14:04] Using charge balance adjust
[1:14:05] Dispensed 0.025823 mL of Base (0.5 M KOH)
[1:14:25] Stirrer speed set to 0
[1:14:35] Datapoint id 86 collected
[1:14:35] Charge balance equation is out by -9.9%
[1:14:35] Argon flow rate set to 0
[1:14:39] Titrator arm moved over Titration position
[1:15:00] The autoloader failed to pick at location "Sample position"
