## Supplementary material for "Octanol-water partition coefficient measurements for the SAMPL6 Blind Prediction Challenge": SM15_18C-01008_M16_octanol_pH-metric high logP_report.pdf

| | | | |
|---|---|---|---|
| Sample name: | **M16_octanol** | Experiment start time: | **3/1/2018 10:41:05 AM** |
| Assay name: | **pH-metric high logP** | Analyst: | **Pion** |
| Assay ID: | **18C-01008** | Instrument ID: | **T312060** |
| Filename: | **C:\Sirius_T3\Mehtap\20180228_exp28_logP_T3-2\18C-01008_M16_octanol_pH-metric high logP.t3r** | | |

#### pH-metric Result

| | |
|---|---|
| logP (XH2 +) | -1.34 |
| logP (neutral XH) | 3.04 |
| logP (X -) | -1.62 |

#### 18C-01008 Points 1 to 32

| | |
|---|---|
| M16_octanol concentration factor | 0.795 |
| Carbonate | 0.0308 mM |
| Acidity error | 3.10957 mM |

#### 18C-01008 Points 33 to 60

| | |
|---|---|
| M16_octanol concentration factor | 0.841 |
| Carbonate | 0.2347 mM |
| Acidity error | 2.72359 mM |

#### 18C-01008 Points 61 to 95

| | |
|---|---|
| M16_octanol concentration factor | 0.890 |
| Carbonate | 0.3111 mM |
| Acidity error | 2.55032 mM |

#### Warnings and errors

| | |
|---|---|
| Errors | None |
| Warnings | User stopped refinement before final result convergence occured |

#### Sample logD and percent species

| pH | M16_octanol logD | M16_octanol M16_octanolH2 | M16_octanol M16_octanolH | M16_octanol M16_octanol | M16_octanol M16_octanolH2* | M16_octanol M16_octanolH* | M16_octanol M16_octanol* | Comment |
|---|---|---|---|---|---|---|---|---|
| 1.000 | -0.58 | 79.04 % | 0.02 % | 0.00 % | 3.57 % | 17.37 % | 0.00 % | |
| 1.200 | -0.41 | 71.75 % | 0.02 % | 0.00 % | 3.24 % | 24.99 % | 0.00 % | Stomach pH |
| 2.000 | 0.35 | 30.82 % | 0.06 % | 0.00 % | 1.39 % | 67.73 % | 0.00 % | |
| 3.000 | 1.33 | 4.34 % | 0.09 % | 0.00 % | 0.20 % | 95.38 % | 0.00 % | |
| 4.000 | 2.26 | 0.45 % | 0.09 % | 0.00 % | 0.02 % | 99.44 % | 0.00 % | |
| 5.000 | 2.87 | 0.05 % | 0.09 % | 0.00 % | 0.00 % | 99.86 % | 0.00 % | |
| 6.000 | 3.02 | 0.00 % | 0.09 % | 0.00 % | 0.00 % | 99.90 % | 0.00 % | |
| 6.500 | 3.03 | 0.00 % | 0.09 % | 0.00 % | 0.00 % | 99.91 % | 0.00 % | |
| 7.000 | 3.03 | 0.00 % | 0.09 % | 0.00 % | 0.00 % | 99.91 % | 0.00 % | |
| 7.400 | 3.03 | 0.00 % | 0.09 % | 0.00 % | 0.00 % | 99.91 % | 0.00 % | Blood pH |
| 8.000 | 2.99 | 0.00 % | 0.09 % | 0.01 % | 0.00 % | 99.90 % | 0.00 % | |
| 9.000 | 2.71 | 0.00 % | 0.09 % | 0.10 % | 0.00 % | 99.80 % | 0.00 % | |
| 10.000 | 1.95 | 0.00 % | 0.09 % | 1.03 % | 0.00 % | 98.85 % | 0.02 % | |
| 11.000 | 0.98 | 0.00 % | 0.08 % | 9.41 % | 0.00 % | 90.28 % | 0.23 % | |
| 12.000 | -0.01 | 0.00 % | 0.04 % | 50.40 % | 0.00 % | 48.35 % | 1.21 % | |

Sample name: **M16_octanol**
Assay name: **pH-metric high logP**
Assay ID: **18C-01008**
Filename: **C:\Sirius_T3\Mehtap\20180228_exp28_logP_T3-2\18C-01008_M16_octanol_pH-metric high logP.t3r**

Experiment start time: **3/1/2018 10:41:05 AM**
Analyst: **Pion**
Instrument ID: **T312060**

#### Graphs

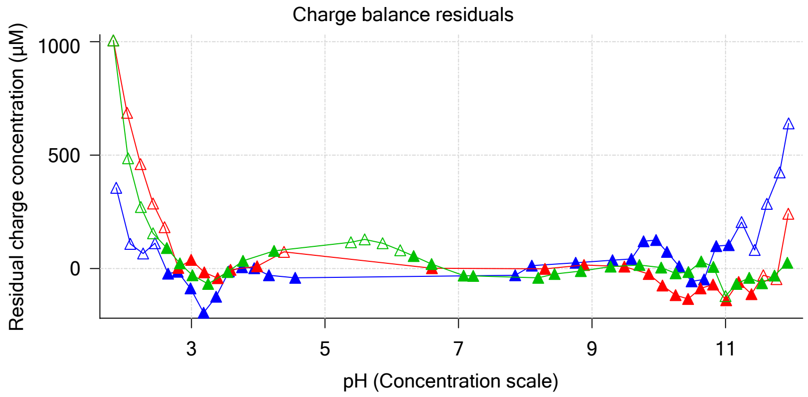


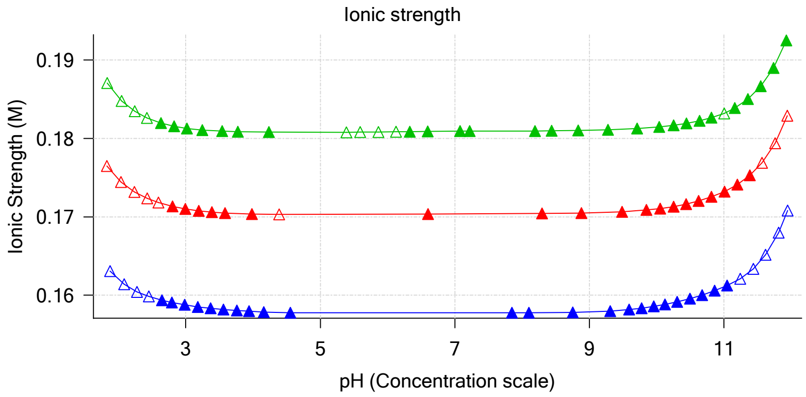


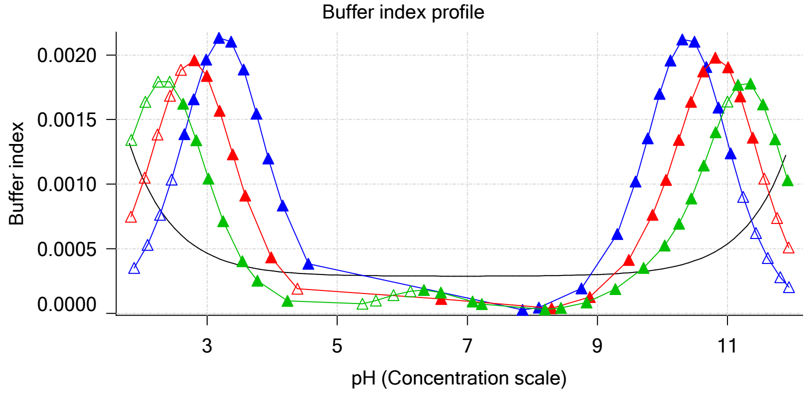


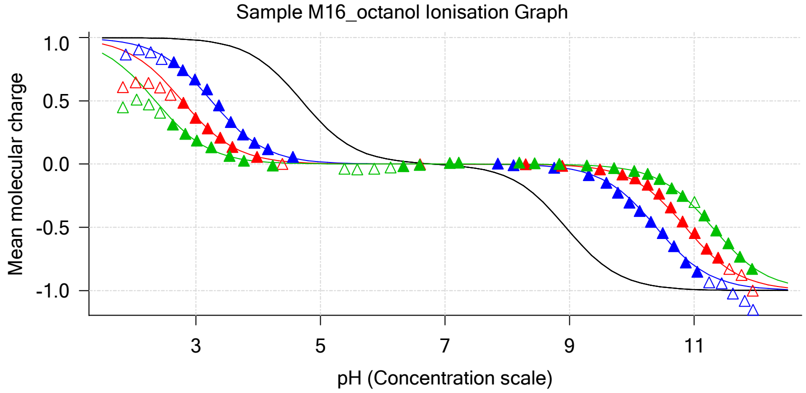


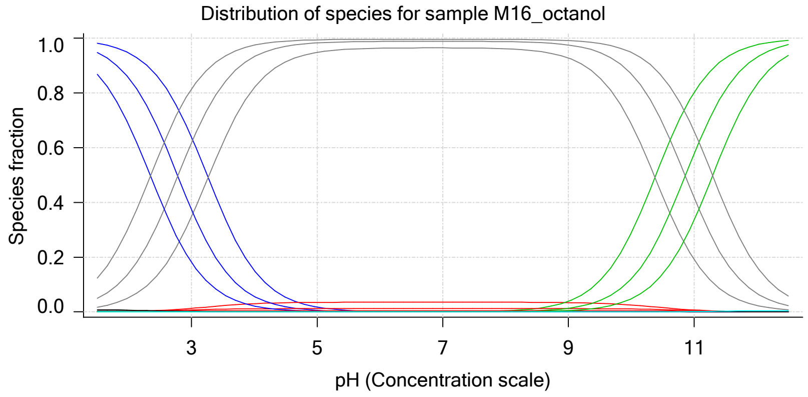


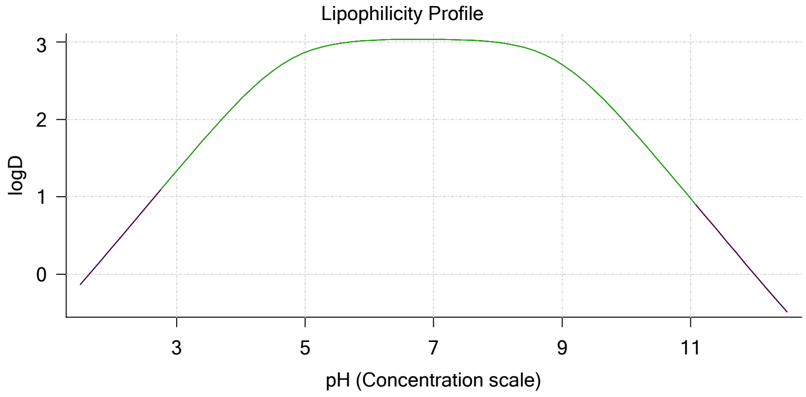


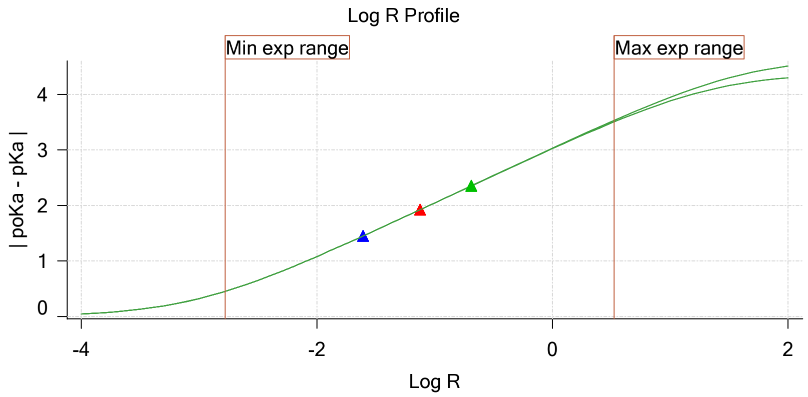


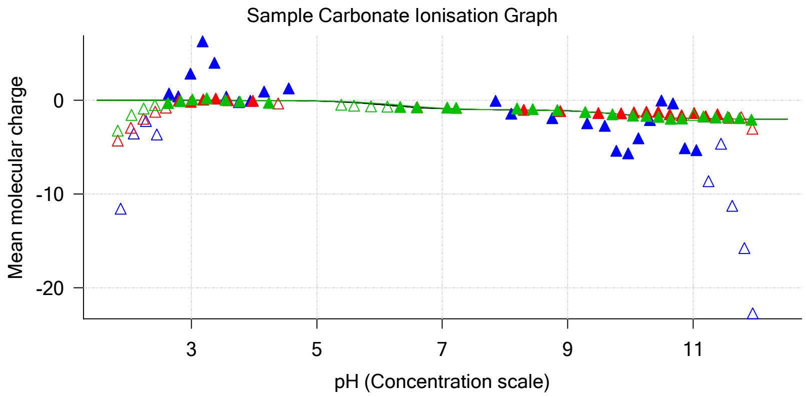

| | | | |
|---|---|---|---|
| Sample name: | **M16_octanol** | Experiment start time: | **3/1/2018 10:41:05 AM** |
| Assay name: | **pH-metric high logP** | Analyst: | **Pion** |
| Assay ID: | **18C-01008** | Instrument ID: | **T312060** |
| Filename: | **C:\Sirius_T3\Mehtap\20180228_exp28_logP_T3-2\18C-01008_M16_octanol_pH-metric high logP.t3r** | | |

#### Graphs (continued)

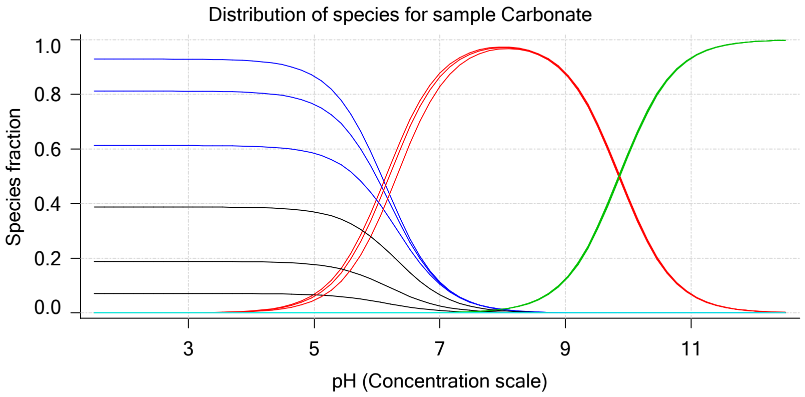

| | | | |
|---|---|---|---|
| Sample name: | **M16_octanol** | Experiment start time: | **3/1/2018 10:41:05 AM** |
| Assay name: | **pH-metric high logP** | Analyst: | **Pion** |
| Assay ID: | **18C-01008** | Instrument ID: | **T312060** |
| Filename: | **C:\Sirius_T3\Mehtap\20180228_exp28_logP_T3-2\18C-01008_M16_octanol_pH-metric high logP.t3r** | | |

### pH-metric high logP Titration 1 of 3 18C-01008 Points 1 to 32

#### Overall results

| | |
|---|---|
| RMSD | 0.203 |
| Average ionic strength | 0.159 M |
| Average temperature | 25.0°C |
| Partition ratio | 0.0247 : 1 |
| Analyte concentration range | 4379.7 µM to 4702.9 µM |
| Total points considered | 23 of 32 |

#### Warnings and errors

| | |
|---|---|
| Errors | None |
| Warnings | Excessive acidity error present |

#### Four-Plus parameters

| | | | |
|---|---|---|---|
| Alpha | 0.130 | 3/1/2018 10:41:04 AM | C:\Sirius_T3\HCl18B27.t3r |
| S | 0.9970 | 3/1/2018 10:41:04 AM | C:\Sirius_T3\HCl18B27.t3r |
| jH | 0.8 | 3/1/2018 10:41:04 AM | C:\Sirius_T3\HCl18B27.t3r |
| jOH | -0.4 | 3/1/2018 10:41:04 AM | C:\Sirius_T3\HCl18B27.t3r |

#### Titrants

| | | | |
|---|---|---|---|
| 0.50 M HCl | 0.993513 | 3/1/2018 10:41:05 AM | C:\Sirius_T3\HCl18B27.t3r |
| 0.50 M KOH | 0.999845 | 3/1/2018 10:41:05 AM | C:\Sirius_T3\KOH18B27.t3r |

#### Sample

| | |
|---|---|
| M16_octanol concentration factor | 0.795 |
| Base pKa 1 | 4.70 |
| Acid pKa 2 | 8.94 |
| logP (XH2 +) | -0.95 |
| logP (neutral XH) | 3.01 |
| logP (X -) | -1.62 |

#### Sample graphs

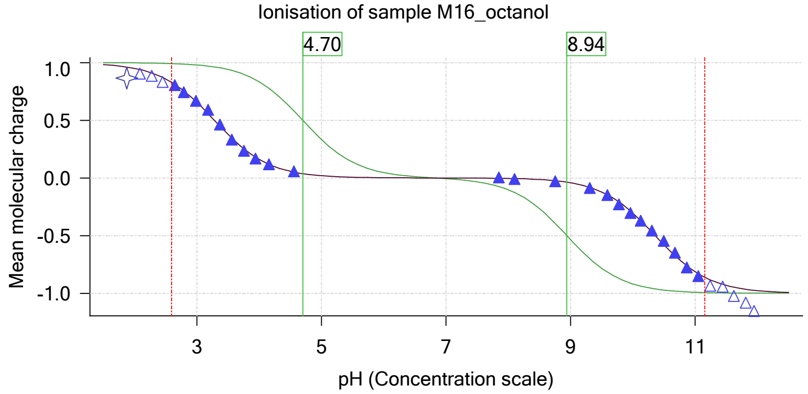


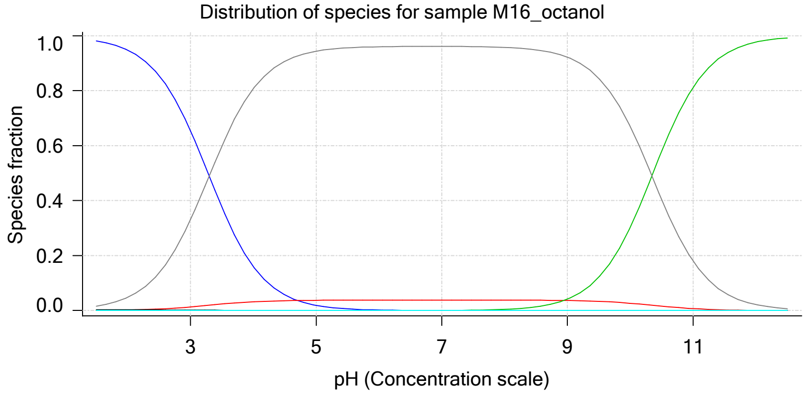

| | | | |
|---|---|---|---|
| Sample name: | **M16_octanol** | Experiment start time: | **3/1/2018 10:41:05 AM** |
| Assay name: | **pH-metric high logP** | Analyst: | **Pion** |
| Assay ID: | **18C-01008** | Instrument ID: | **T312060** |
| Filename: | **C:\Sirius_T3\Mehtap\20180228_exp28_logP_T3-2\18C-01008_M16_octanol_pH-metric high logP.t3r** | | |

#### Sample graphs (continued)

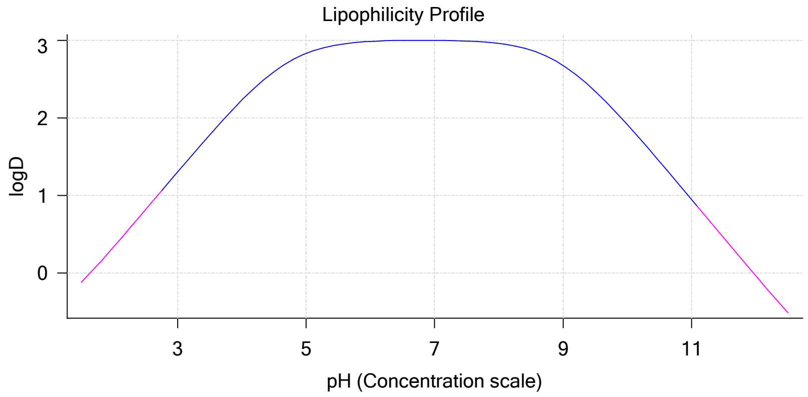


#### Sample logD and percent species

| pH | M16_octanol logD | M16_octanol M16_octanolH2 | M16_octanol M16_octanolH | M16_octanol M16_octanol | M16_octanol M16_octanolH2* | M16_octanol M16_octanolH* | M16_octanol M16_octanol* | Comment |
|---|---|---|---|---|---|---|---|---|
| 1.000 | -0.50 | 99.20 % | 0.02 % | 0.00 % | 0.28 % | 0.50 % | 0.00 % | |
| 1.200 | -0.36 | 98.90 % | 0.03 % | 0.00 % | 0.27 % | 0.79 % | 0.00 % | Stomach pH |
| 2.000 | 0.33 | 94.77 % | 0.19 % | 0.00 % | 0.26 % | 4.78 % | 0.00 % | |
| 3.000 | 1.30 | 65.49 % | 1.31 % | 0.00 % | 0.18 % | 33.02 % | 0.00 % | |
| 4.000 | 2.23 | 16.01 % | 3.19 % | 0.00 % | 0.04 % | 80.75 % | 0.00 % | |
| 5.000 | 2.83 | 1.87 % | 3.73 % | 0.00 % | 0.01 % | 94.39 % | 0.00 % | |
| 6.000 | 2.99 | 0.19 % | 3.80 % | 0.00 % | 0.00 % | 96.01 % | 0.00 % | |
| 6.500 | 3.00 | 0.06 % | 3.80 % | 0.01 % | 0.00 % | 96.12 % | 0.00 % | |
| 7.000 | 3.00 | 0.02 % | 3.80 % | 0.04 % | 0.00 % | 96.13 % | 0.00 % | |
| 7.400 | 3.00 | 0.01 % | 3.80 % | 0.11 % | 0.00 % | 96.08 % | 0.00 % | Blood pH |
| 8.000 | 2.96 | 0.00 % | 3.79 % | 0.44 % | 0.00 % | 95.77 % | 0.00 % | |
| 9.000 | 2.68 | 0.00 % | 3.65 % | 4.19 % | 0.00 % | 92.16 % | 0.00 % | |
| 10.000 | 1.91 | 0.00 % | 2.65 % | 30.40 % | 0.00 % | 66.93 % | 0.02 % | |
| 11.000 | 0.95 | 0.00 % | 0.71 % | 81.34 % | 0.00 % | 17.91 % | 0.05 % | |
| 12.000 | -0.04 | 0.00 % | 0.09 % | 97.71 % | 0.00 % | 2.15 % | 0.06 % | |

#### Carbonate and acidity

| | |
|---|---|
| Carbonate | 0.031 mM |
| Acidity error | 3.110 mM |

#### Other graphs

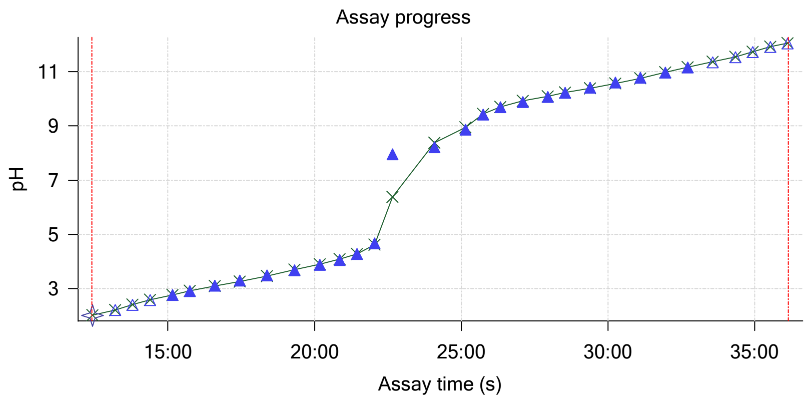


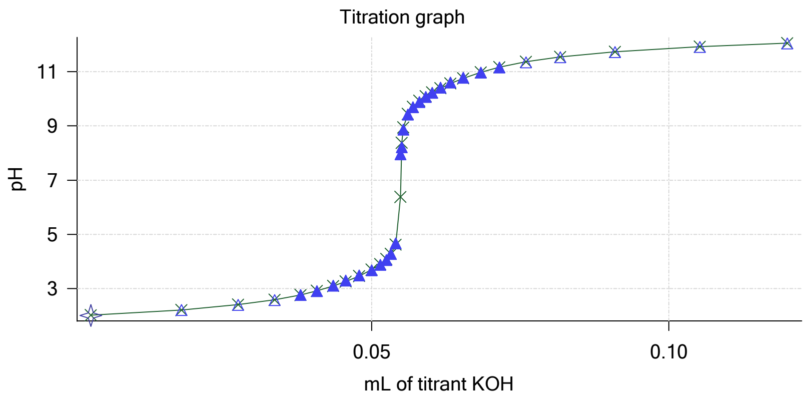

| | | | |
|---|---|---|---|
| Sample name: | **M16_octanol** | Experiment start time: | **3/1/2018 10:41:05 AM** |
| Assay name: | **pH-metric high logP** | Analyst: | **Pion** |
| Assay ID: | **18C-01008** | Instrument ID: | **T312060** |
| Filename: | **C:\Sirius_T3\Mehtap\20180228_exp28_logP_T3-2\18C-01008_M16_octanol_pH-metric high logP.t3r** | | |

#### Other graphs (continued)

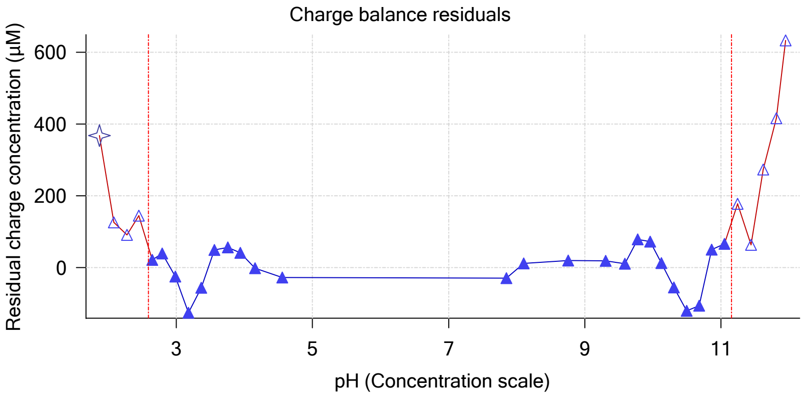


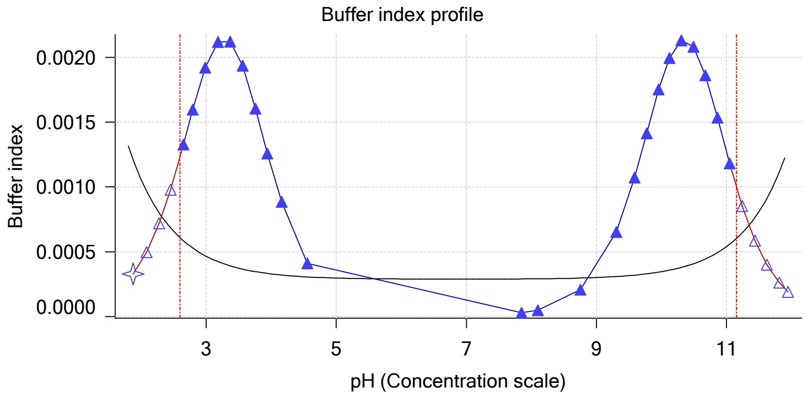


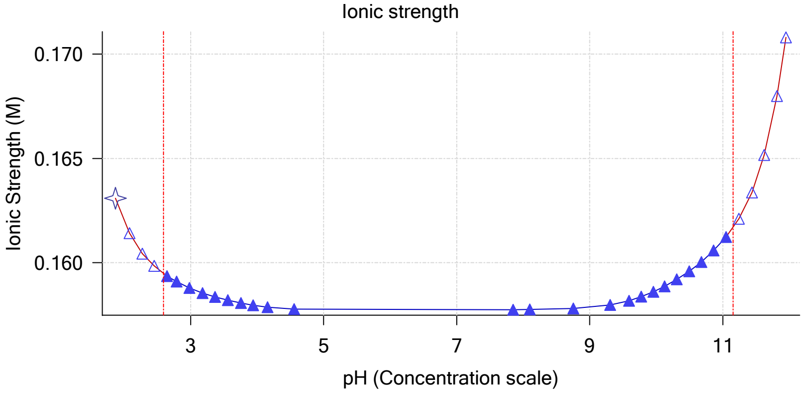


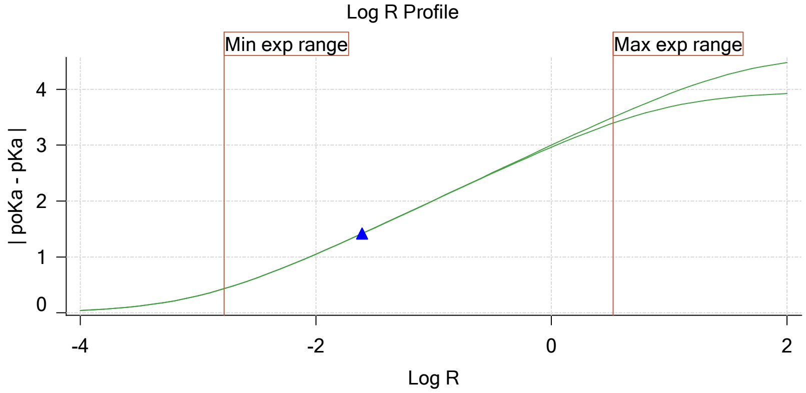


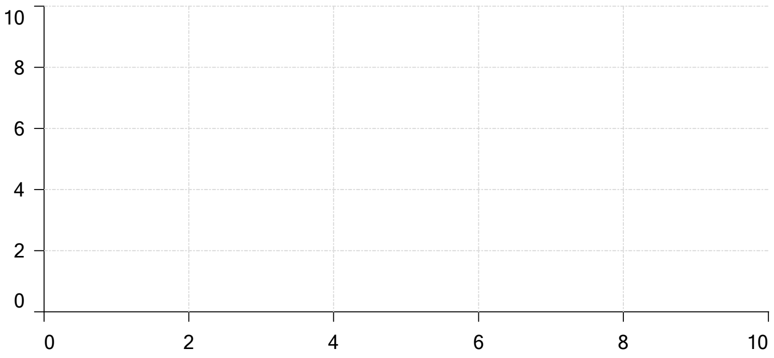

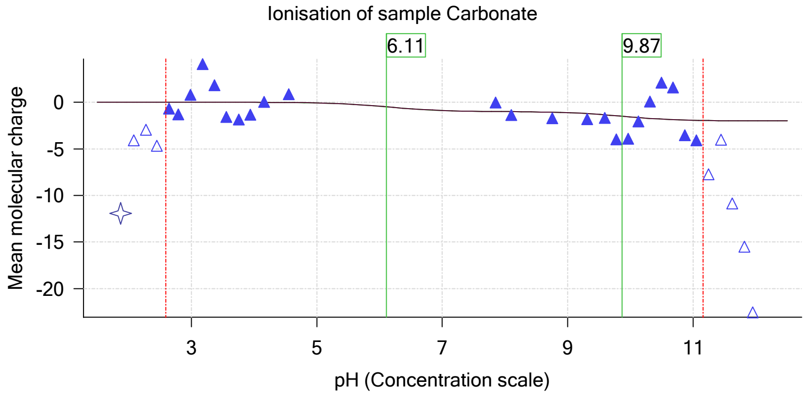

| | | | |
|---|---|---|---|
| Sample name: | **M16_octanol** | Experiment start time: | **3/1/2018 10:41:05 AM** |
| Assay name: | **pH-metric high logP** | Analyst: | **Pion** |
| Assay ID: | **18C-01008** | Instrument ID: | **T312060** |
| Filename: | **C:\Sirius_T3\Mehtap\20180228_exp28_logP_T3-2\18C-01008_M16_octanol_pH-metric high logP.t3r** | | |

# pH-metric high logP   Titration 2 of 3   18C-01008 Points 33 to 60

#### Overall results

| | |
|---|---|
| RMSD | 0.036 |
| Average ionic strength | 0.171 M |
| Average temperature | 25.0°C |
| Partition ratio | 0.0751 : 1 |
| Analyte concentration range | 3612.7 µM to 3864.7 µM |
| Total points considered | 19 of 28 |

#### Warnings and errors

| | |
|---|---|
| Errors | None |
| Warnings | Excessive acidity error present |

#### Four-Plus parameters

| | | | |
|---|---|---|---|
| Alpha | 0.130 | 3/1/2018 10:41:04 AM | C:\Sirius_T3\HCl18B27.t3r |
| S | 0.9970 | 3/1/2018 10:41:04 AM | C:\Sirius_T3\HCl18B27.t3r |
| jH | 0.8 | 3/1/2018 10:41:04 AM | C:\Sirius_T3\HCl18B27.t3r |
| jOH | -0.4 | 3/1/2018 10:41:04 AM | C:\Sirius_T3\HCl18B27.t3r |

#### Titrants

| | | | |
|---|---|---|---|
| 0.50 M HCl | 0.993513 | 3/1/2018 10:41:05 AM | C:\Sirius_T3\HCl18B27.t3r |
| 0.50 M KOH | 0.999845 | 3/1/2018 10:41:05 AM | C:\Sirius_T3\KOH18B27.t3r |

#### Sample

| | |
|---|---|
| M16_octanol concentration factor | 0.841 |
| Base pKa 1 | 4.70 |
| Acid pKa 2 | 8.94 |
| logP (XH2 +) | 0.31 |
| logP (neutral XH) | 3.10 |
| logP (X -) | -1.62 |

#### Sample graphs

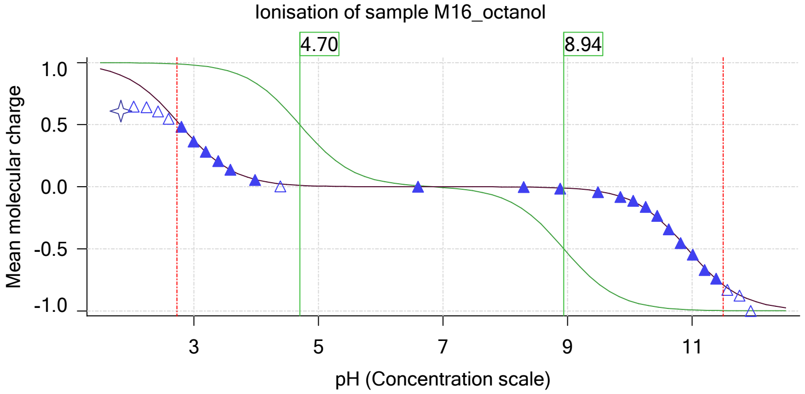


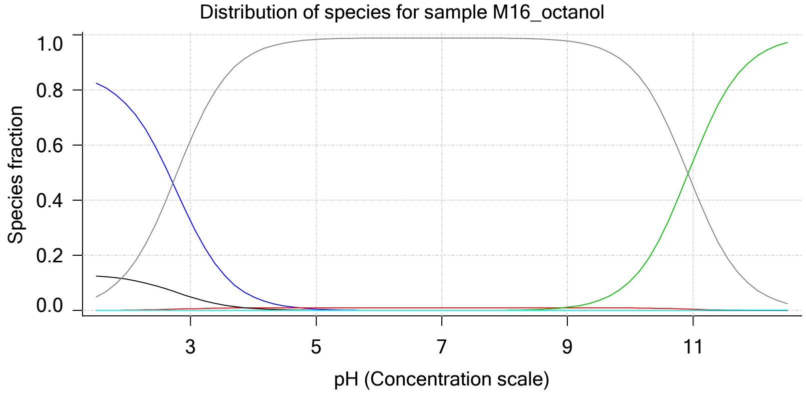

| | | | |
|---|---|---|---|
| Sample name: | **M16_octanol** | Experiment start time: | **3/1/2018 10:41:05 AM** |
| Assay name: | **pH-metric high logP** | Analyst: | **Pion** |
| Assay ID: | **18C-01008** | Instrument ID: | **T312060** |
| Filename: | **C:\Sirius_T3\Mehtap\20180228_exp28_logP_T3-2\18C-01008_M16_octanol_pH-metric high logP.t3r** | | |

#### Sample graphs (continued)

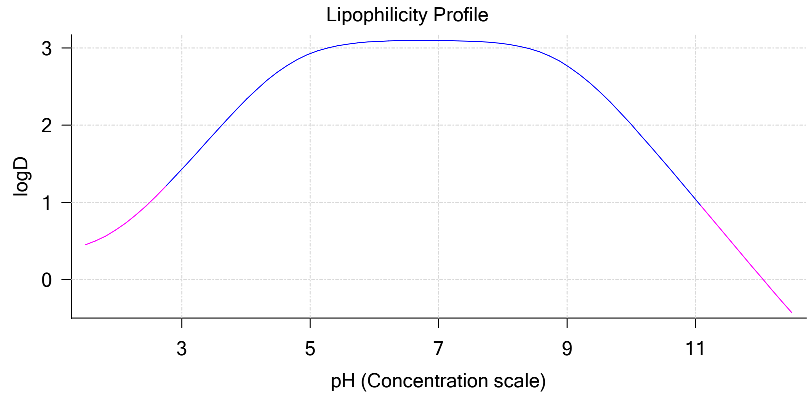


#### Sample logD and percent species

| pH | M16_octanol logD | M16_octanol M16_octanolH2 | M16_octanol M16_octanolH | M16_octanol M16_octanol | M16_octanol M16_octanolH2* | M16_octanol M16_octanolH* | M16_octanol M16_octanol* | Comment |
|---|---|---|---|---|---|---|---|---|
| 1.000 | 0.36 | 85.40 % | 0.02 % | 0.00 % | 12.96 % | 1.62 % | 0.00 % | |
| 1.200 | 0.38 | 84.59 % | 0.03 % | 0.00 % | 12.83 % | 2.55 % | 0.00 % | Stomach pH |
| 2.000 | 0.66 | 74.43 % | 0.15 % | 0.00 % | 11.29 % | 14.13 % | 0.00 % | |
| 3.000 | 1.43 | 32.57 % | 0.65 % | 0.00 % | 4.94 % | 61.84 % | 0.00 % | |
| 4.000 | 2.33 | 4.92 % | 0.98 % | 0.00 % | 0.75 % | 93.36 % | 0.00 % | |
| 5.000 | 2.93 | 0.52 % | 1.03 % | 0.00 % | 0.08 % | 98.37 % | 0.00 % | |
| 6.000 | 3.08 | 0.05 % | 1.04 % | 0.00 % | 0.01 % | 98.90 % | 0.00 % | |
| 6.500 | 3.09 | 0.02 % | 1.04 % | 0.00 % | 0.00 % | 98.94 % | 0.00 % | |
| 7.000 | 3.10 | 0.01 % | 1.04 % | 0.01 % | 0.00 % | 98.94 % | 0.00 % | |
| 7.400 | 3.09 | 0.00 % | 1.04 % | 0.03 % | 0.00 % | 98.93 % | 0.00 % | Blood pH |
| 8.000 | 3.06 | 0.00 % | 1.04 % | 0.12 % | 0.00 % | 98.84 % | 0.00 % | |
| 9.000 | 2.77 | 0.00 % | 1.03 % | 1.18 % | 0.00 % | 97.79 % | 0.00 % | |
| 10.000 | 2.01 | 0.00 % | 0.93 % | 10.66 % | 0.00 % | 88.39 % | 0.02 % | |
| 11.000 | 1.04 | 0.00 % | 0.47 % | 54.37 % | 0.00 % | 45.06 % | 0.10 % | |
| 12.000 | 0.05 | 0.00 % | 0.08 % | 92.12 % | 0.00 % | 7.64 % | 0.17 % | |

#### Carbonate and acidity

| | |
|---|---|
| Carbonate | 0.235 mM |
| Acidity error | 2.724 mM |

#### Other graphs

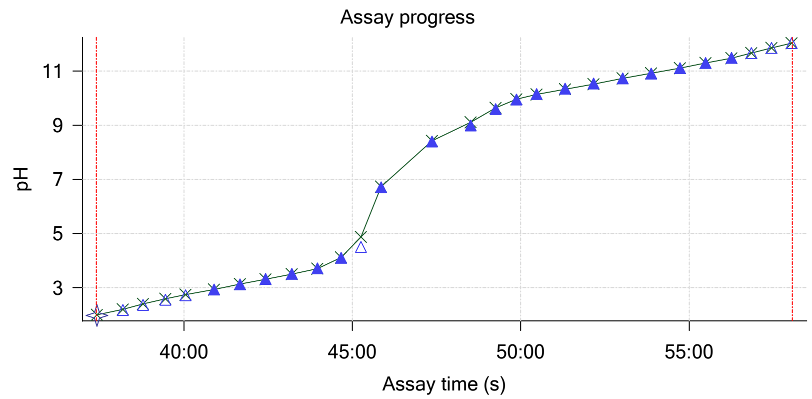


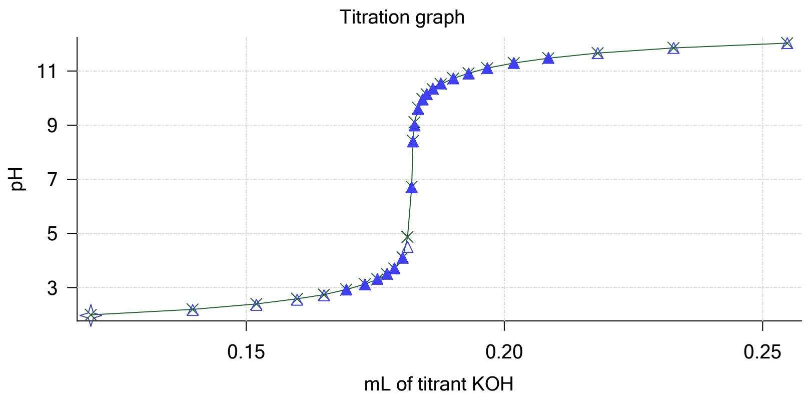

Sample name: **M16_octanol** Experiment start time: **3/1/2018 10:41:05 AM**
Assay name: **pH-metric high logP** Analyst: **Pion**
Assay ID: **18C-01008** Instrument ID: **T312060**
Filename: **C:\Sirius_T3\Mehtap\20180228_exp28_logP_T3-2\18C-01008_M16_octanol_pH-metric high logP.t3r**

#### Other graphs (continued)

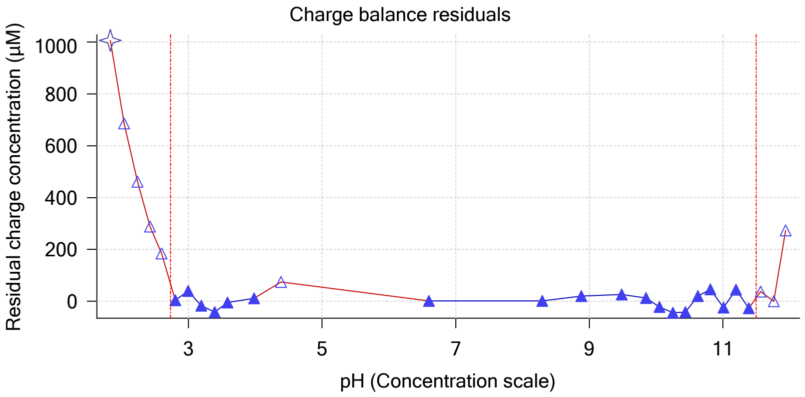


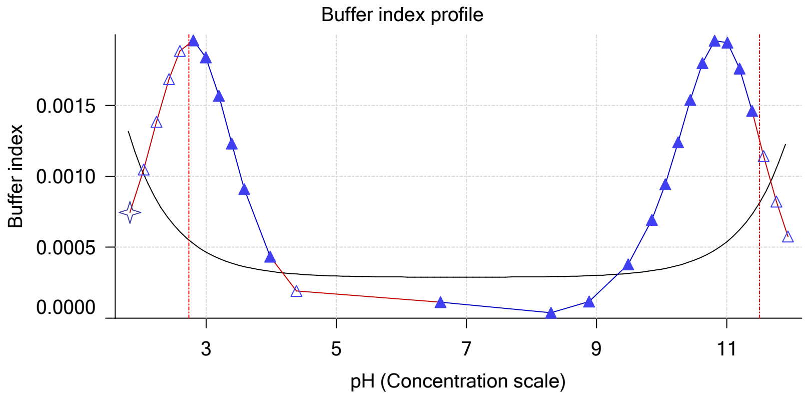


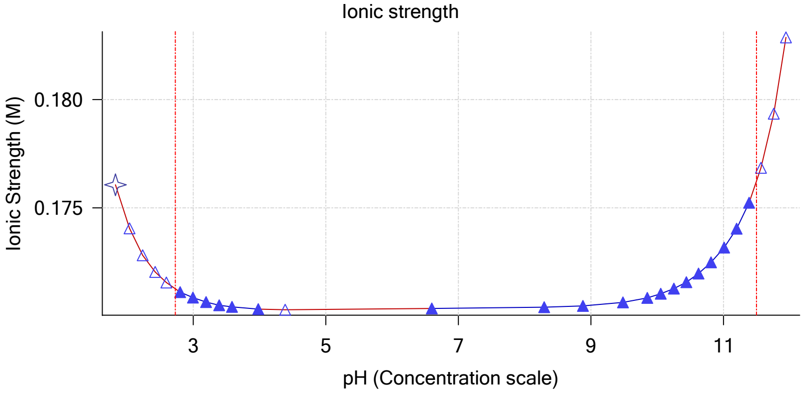


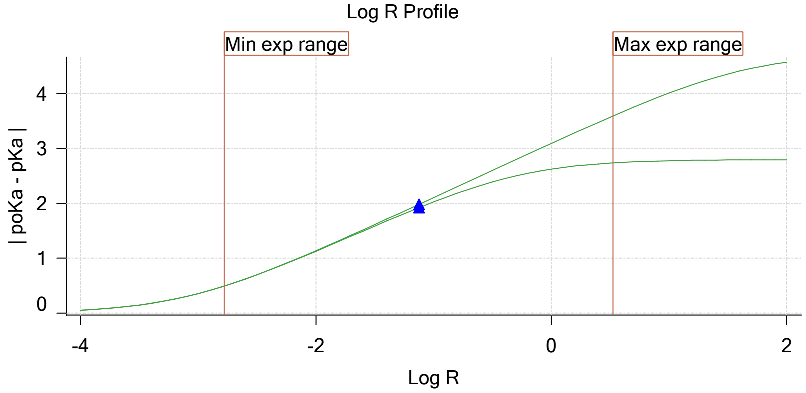


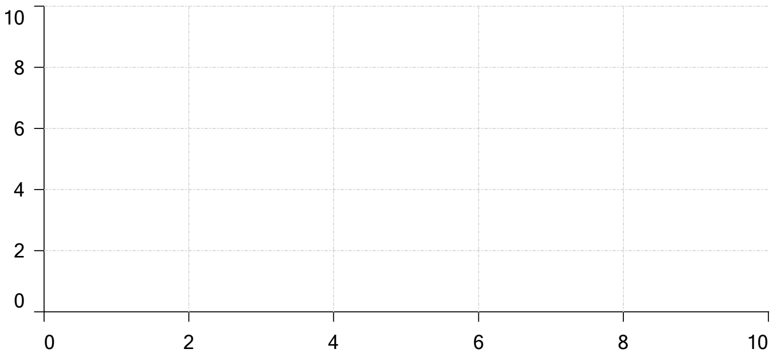


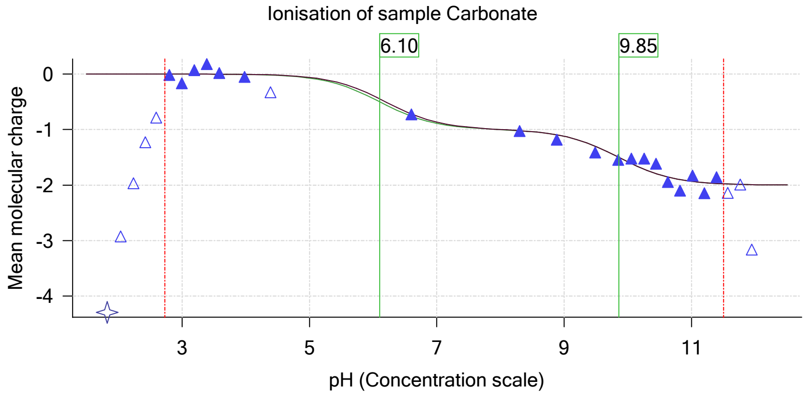

| | | | |
|---|---|---|---|
| Sample name: | **M16_octanol** | Experiment start time: | **3/1/2018 10:41:05 AM** |
| Assay name: | **pH-metric high logP** | Analyst: | **Pion** |
| Assay ID: | **18C-01008** | Instrument ID: | **T312060** |
| Filename: | **C:\Sirius_T3\Mehtap\20180228_exp28_logP_T3-2\18C-01008_M16_octanol_pH-metric high logP.t3r** | | |

# pH-metric high logP   Titration 3 of 3   18C-01008 Points 61 to 95

#### Overall results

| | |
|---|---|
| RMSD | 0.274 |
| Average ionic strength | 0.182 M |
| Average temperature | 25.0°C |
| Partition ratio | 0.2049 : 1 |
| Analyte concentration range | 2812.2 µM to 2977.4 µM |
| Total points considered | 26 of 35 |

#### Warnings and errors

| | |
|---|---|
| Errors | None |
| Warnings | One or more logP values out of range |
| | Excessive acidity error present |

#### Four-Plus parameters

| | | | |
|---|---|---|---|
| Alpha | 0.130 | 3/1/2018 10:41:04 AM | C:\Sirius_T3\HCl18B27.t3r |
| S | 0.9970 | 3/1/2018 10:41:04 AM | C:\Sirius_T3\HCl18B27.t3r |
| jH | 0.8 | 3/1/2018 10:41:04 AM | C:\Sirius_T3\HCl18B27.t3r |
| jOH | -0.4 | 3/1/2018 10:41:04 AM | C:\Sirius_T3\HCl18B27.t3r |

#### Titrants

| | | | |
|---|---|---|---|
| 0.50 M HCl | 0.993513 | 3/1/2018 10:41:05 AM | C:\Sirius_T3\HCl18B27.t3r |
| 0.50 M KOH | 0.999845 | 3/1/2018 10:41:05 AM | C:\Sirius_T3\KOH18B27.t3r |

#### Sample

| | |
|---|---|
| M16_octanol concentration factor | 0.890 |
| Base pKa 1 | 4.70 |
| Acid pKa 2 | 8.94 |
| logP (XH2 +) | -3.82 |
| logP (neutral XH) | 3.06 |
| logP (X -) | -1.62 |

#### Sample graphs

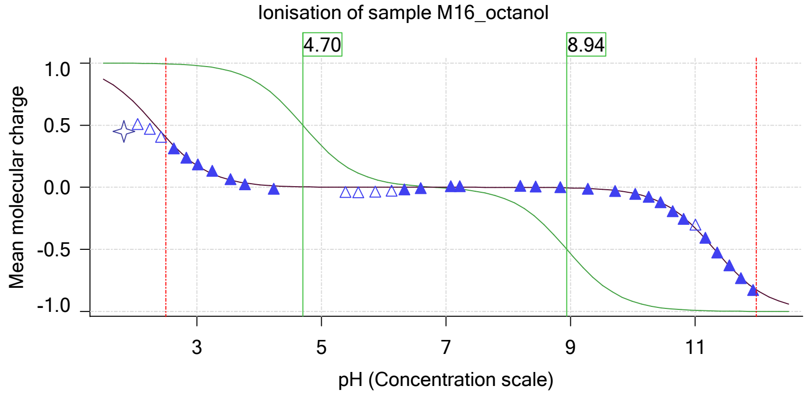


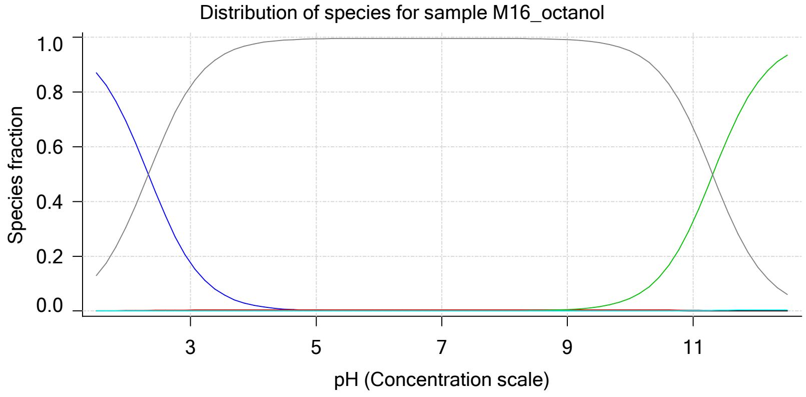

Sample name: **M16_octanol**
Assay name: **pH-metric high logP**
Assay ID: **18C-01008**
Filename: **C:\Sirius_T3\Mehtap\20180228_exp28_logP_T3-2\18C-01008_M16_octanol_pH-metric high logP.t3r**

Experiment start time: **3/1/2018 10:41:05 AM**
Analyst: **Pion**
Instrument ID: **T312060**

#### Sample graphs (continued)

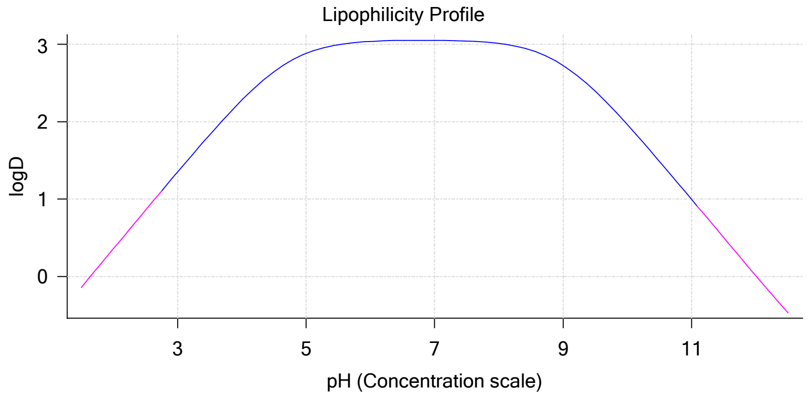


#### Sample logD and percent species

| pH | M16_octanol logD | M16_octanol M16_octanolH2 | M16_octanol M16_octanolH | M16_octanol M16_octanol | M16_octanol M16_octanolH2* | M16_octanol M16_octanolH* | M16_octanol M16_octanol* | Comment |
|---|---|---|---|---|---|---|---|---|
| 1.000 | -0.64 | 95.51 % | 0.02 % | 0.00 % | 0.00 % | 4.47 % | 0.00 % | |
| 1.200 | -0.44 | 93.06 % | 0.03 % | 0.00 % | 0.00 % | 6.91 % | 0.00 % | Stomach pH |
| 2.000 | 0.36 | 68.01 % | 0.14 % | 0.00 % | 0.00 % | 31.85 % | 0.00 % | |
| 3.000 | 1.35 | 17.53 % | 0.35 % | 0.00 % | 0.00 % | 82.12 % | 0.00 % | |
| 4.000 | 2.28 | 2.08 % | 0.42 % | 0.00 % | 0.00 % | 97.50 % | 0.00 % | |
| 5.000 | 2.88 | 0.21 % | 0.42 % | 0.00 % | 0.00 % | 99.36 % | 0.00 % | |
| 6.000 | 3.04 | 0.02 % | 0.42 % | 0.00 % | 0.00 % | 99.55 % | 0.00 % | |
| 6.500 | 3.05 | 0.01 % | 0.42 % | 0.00 % | 0.00 % | 99.57 % | 0.00 % | |
| 7.000 | 3.05 | 0.00 % | 0.42 % | 0.00 % | 0.00 % | 99.57 % | 0.00 % | |
| 7.400 | 3.05 | 0.00 % | 0.42 % | 0.01 % | 0.00 % | 99.56 % | 0.00 % | Blood pH |
| 8.000 | 3.01 | 0.00 % | 0.42 % | 0.05 % | 0.00 % | 99.53 % | 0.00 % | |
| 9.000 | 2.73 | 0.00 % | 0.42 % | 0.48 % | 0.00 % | 99.09 % | 0.00 % | |
| 10.000 | 1.96 | 0.00 % | 0.40 % | 4.64 % | 0.00 % | 94.93 % | 0.02 % | |
| 11.000 | 1.00 | 0.00 % | 0.28 % | 32.70 % | 0.00 % | 66.85 % | 0.16 % | |
| 12.000 | 0.01 | 0.00 % | 0.07 % | 82.63 % | 0.00 % | 16.89 % | 0.41 % | |

#### Carbonate and acidity

Carbonate 0.311 mM
Acidity error 2.550 mM

#### Other graphs

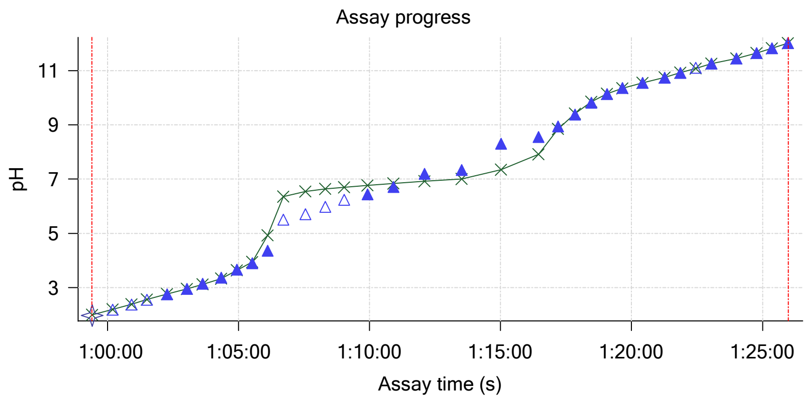


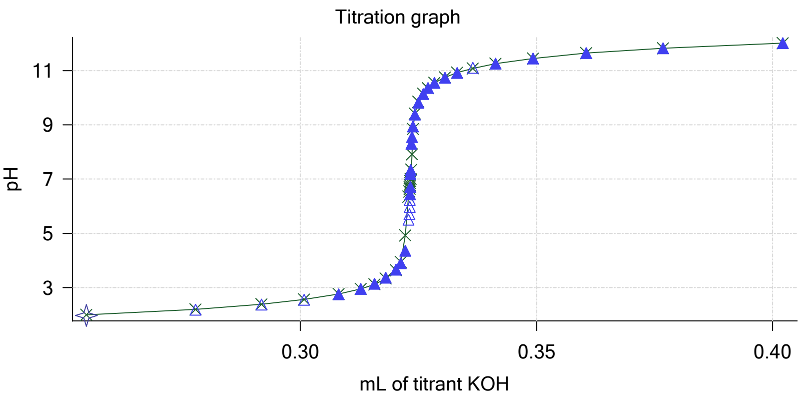

| | | | |
|---|---|---|---|
| Sample name: | **M16_octanol** | Experiment start time: | **3/1/2018 10:41:05 AM** |
| Assay name: | **pH-metric high logP** | Analyst: | **Pion** |
| Assay ID: | **18C-01008** | Instrument ID: | **T312060** |
| Filename: | **C:\Sirius_T3\Mehtap\20180228_exp28_logP_T3-2\18C-01008_M16_octanol_pH-metric high logP.t3r** | | |

#### Other graphs (continued)

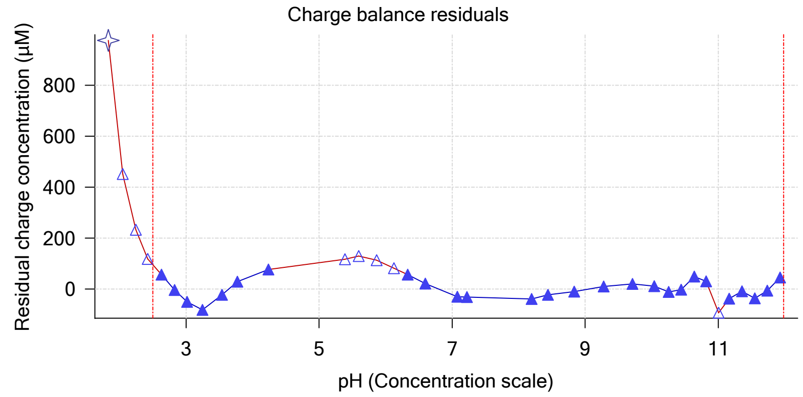


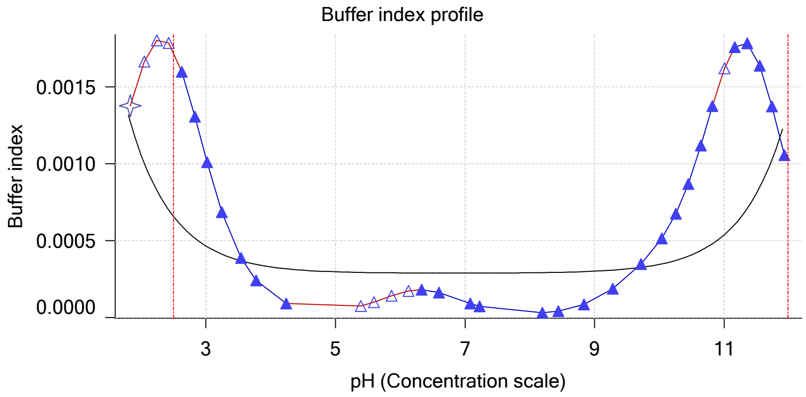


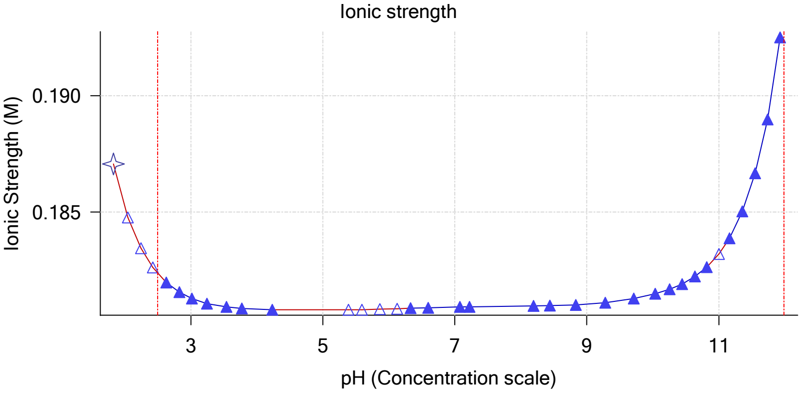


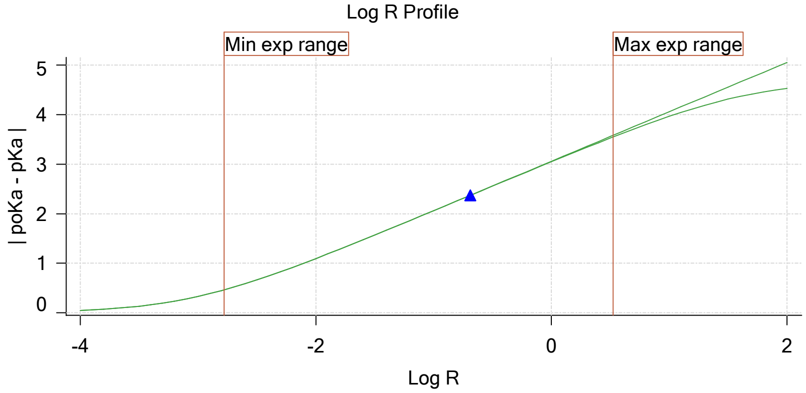


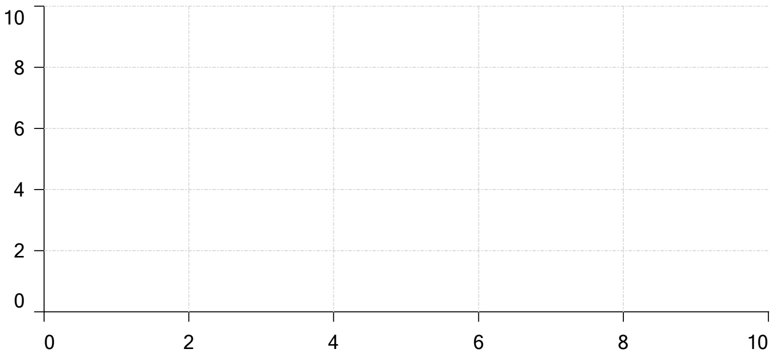


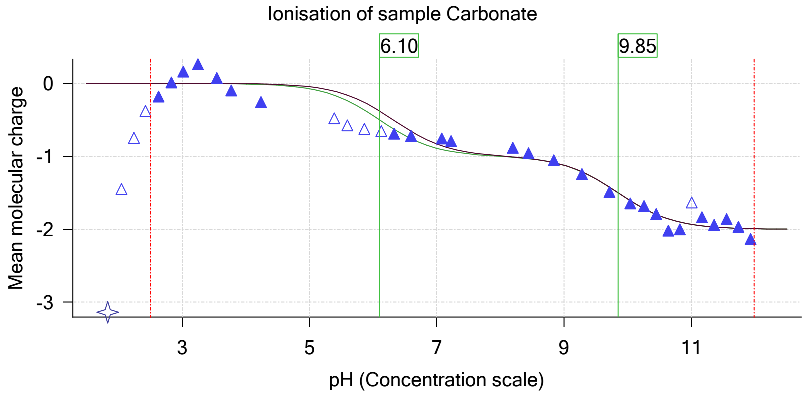

| | | | |
|---|---|---|---|
| Sample name: | **M16_octanol** | Experiment start time: | **3/1/2018 10:41:05 AM** |
| Assay name: | **pH-metric high logP** | Analyst: | **Pion** |
| Assay ID: | **18C-01008** | Instrument ID: | **T312060** |
| Filename: | **C:\Sirius_T3\Mehtap\20180228_exp28_logP_T3-2\18C-01008_M16_octanol_pH-metric high logP.t3r** | | |

#### Assay Model

| Settings | Value | Date/Time changed | Imported from |
|---|---|---|---|
| Sample name | M16_octanol | 2/27/2018 6:14:13 PM | User entered value |
| Sample by | Weight | | Default value |
| Sample weight | 0.001570 g | 2/28/2018 4:25:47 PM | User entered value |
| Formula weight | 210.23 g/mol | 2/27/2018 5:08:55 PM | User entered value |
| Solubility | Unknown | | Default value |
| Molecular weight | 210.23 | 2/27/2018 5:08:55 PM | User entered value |
| Individual pKa ionic environments | No | | Default value |
| Number of pKas | 2 | 2/27/2018 5:08:55 PM | User entered value |
| Sample is a | Ampholyte | 2/27/2018 5:08:55 PM | User entered value |
| pKa 1 | 4.70 | 2/27/2018 5:08:55 PM | User entered value |
| Type | Base | 2/27/2018 5:08:55 PM | User entered value |
| pKa 2 | 8.94 | 2/27/2018 5:08:55 PM | User entered value |
| Type | Acid | 2/27/2018 5:08:55 PM | User entered value |
| logp (XH2 +) | -0.97 | 2/27/2018 5:09:30 PM | User entered value |
| logP (neutral XH) | 1.47 | 2/27/2018 5:09:44 PM | User entered value |
| logP (X -) | -1.62 | 2/27/2018 5:09:50 PM | User entered value |

#### Events

| Time | Event | Water | Acid | Base | Octanol | pH | dpH/dt | pH R-squared | pH SD | dpH/dt time |
|---|---|---|---|---|---|---|---|---|---|---|
| 9:20.9 | Initial pH = 3.61 | | | | | | | | | |
| 12:25.5 | Data point 1 | 1.50000 mL | 0.04518 mL | 0.00275 mL | 0.04000 mL | 2.007 | -0.00290 | 0.67446 | 0.00017 | 10.5 s |
| 13:12.3 | Data point 2 | 1.50000 mL | 0.04518 mL | 0.01799 mL | 0.04000 mL | 2.212 | 0.00158 | 0.11489 | 0.00023 | 10.0 s |
| 13:47.9 | Data point 3 | 1.50000 mL | 0.04518 mL | 0.02752 mL | 0.04000 mL | 2.403 | 0.00139 | 0.25998 | 0.00013 | 10.0 s |
| 14:23.5 | Data point 4 | 1.50000 mL | 0.04518 mL | 0.03368 mL | 0.04000 mL | 2.577 | -0.00059 | 0.10502 | 0.00009 | 10.0 s |
| 15:09.4 | Data point 5 | 1.50000 mL | 0.04518 mL | 0.03805 mL | 0.04000 mL | 2.771 | -0.00565 | 0.10551 | 0.00086 | 10.0 s |
| 15:44.9 | Data point 6 | 1.50000 mL | 0.04518 mL | 0.04080 mL | 0.04000 mL | 2.914 | -0.00330 | 0.57968 | 0.00021 | 10.0 s |
| 16:35.9 | Data point 7 | 1.50000 mL | 0.04518 mL | 0.04353 mL | 0.04000 mL | 3.106 | -0.00387 | 0.67760 | 0.00023 | 10.0 s |
| 17:26.9 | Data point 8 | 1.50000 mL | 0.04518 mL | 0.04567 mL | 0.04000 mL | 3.301 | -0.00110 | 0.02661 | 0.00033 | 10.0 s |
| 18:23.0 | Data point 9 | 1.50000 mL | 0.04518 mL | 0.04793 mL | 0.04000 mL | 3.489 | -0.01147 | 0.32477 | 0.00099 | 10.0 s |
| 19:19.1 | Data point 10 | 1.50000 mL | 0.04518 mL | 0.05000 mL | 0.04000 mL | 3.679 | -0.00492 | 0.73539 | 0.00028 | 10.5 s |
| 20:10.5 | Data point 11 | 1.50000 mL | 0.04518 mL | 0.05148 mL | 0.04000 mL | 3.876 | -0.01175 | 0.51526 | 0.00081 | 10.0 s |
| 20:51.3 | Data point 12 | 1.50000 mL | 0.04518 mL | 0.05247 mL | 0.04000 mL | 4.059 | -0.01315 | 0.67954 | 0.00079 | 10.0 s |
| 21:26.7 | Data point 13 | 1.50000 mL | 0.04518 mL | 0.05322 mL | 0.04000 mL | 4.276 | -0.00889 | 0.68480 | 0.00053 | 10.5 s |
| 22:02.7 | Data point 14 | 1.50000 mL | 0.04518 mL | 0.05407 mL | 0.04000 mL | 4.672 | -0.01660 | 0.79604 | 0.00092 | 11.5 s |
| 22:39.6 | Data point 15 | 1.50000 mL | 0.04518 mL | 0.05489 mL | 0.04000 mL | 7.954 | -0.05119 | 0.98315 | 0.00255 | Timed out at 59.5 s |
| 24:04.9 | Data point 16 | 1.50000 mL | 0.04518 mL | 0.05506 mL | 0.04000 mL | 8.206 | -0.02013 | 0.98810 | 0.00100 | 38.0 s |
| 25:08.3 | Data point 17 | 1.50000 mL | 0.04518 mL | 0.05534 mL | 0.04000 mL | 8.859 | -0.01483 | 0.91186 | 0.00077 | 10.5 s |
| 25:44.3 | Data point 18 | 1.50000 mL | 0.04518 mL | 0.05611 mL | 0.04000 mL | 9.410 | -0.01450 | 0.98052 | 0.00072 | 10.5 s |
| 26:20.3 | Data point 19 | 1.50000 mL | 0.04518 mL | 0.05696 mL | 0.04000 mL | 9.691 | -0.01710 | 0.91406 | 0.00088 | 10.0 s |
| 27:06.1 | Data point 20 | 1.50000 mL | 0.04518 mL | 0.05804 mL | 0.04000 mL | 9.876 | -0.01009 | 0.92731 | 0.00052 | 10.0 s |
| 27:57.0 | Data point 21 | 1.50000 mL | 0.04518 mL | 0.05915 mL | 0.04000 mL | 10.060 | -0.00874 | 0.33867 | 0.00074 | 10.0 s |
| 28:32.5 | Data point 22 | 1.50000 mL | 0.04518 mL | 0.06018 mL | 0.04000 mL | 10.223 | -0.01285 | 0.40934 | 0.00099 | 10.0 s |
| 29:23.5 | Data point 23 | 1.50000 mL | 0.04518 mL | 0.06159 mL | 0.04000 mL | 10.406 | -0.00727 | 0.93231 | 0.00037 | 10.5 s |
| 30:14.9 | Data point 24 | 1.50000 mL | 0.04518 mL | 0.06329 mL | 0.04000 mL | 10.593 | -0.00688 | 0.41094 | 0.00053 | 10.0 s |
| 31:06.0 | Data point 25 | 1.50000 mL | 0.04518 mL | 0.06543 mL | 0.04000 mL | 10.776 | -0.01364 | 0.90456 | 0.00071 | 10.5 s |
| 31:57.5 | Data point 26 | 1.50000 mL | 0.04518 mL | 0.06839 mL | 0.04000 mL | 10.961 | -0.01272 | 0.56692 | 0.00083 | 10.0 s |
| 32:43.3 | Data point 27 | 1.50000 mL | 0.04518 mL | 0.07150 mL | 0.04000 mL | 11.147 | 0.00312 | 0.33680 | 0.00027 | 10.0 s |
| 33:34.4 | Data point 28 | 1.50000 mL | 0.04518 mL | 0.07596 mL | 0.04000 mL | 11.337 | -0.00151 | 0.26906 | 0.00014 | 10.0 s |
| 34:20.4 | Data point 29 | 1.50000 mL | 0.04518 mL | 0.08177 mL | 0.04000 mL | 11.533 | -0.00131 | 0.26464 | 0.00013 | 10.0 s |
| 34:56.1 | Data point 30 | 1.50000 mL | 0.04518 mL | 0.09099 mL | 0.04000 mL | 11.712 | 0.01582 | 0.69550 | 0.00094 | 10.0 s |
| 35:31.9 | Data point 31 | 1.50000 mL | 0.04518 mL | 0.10525 mL | 0.04000 mL | 11.903 | -0.01334 | 0.54570 | 0.00089 | 10.0 s |
| 36:07.6 | Data point 32 | 1.50000 mL | 0.04518 mL | 0.11997 mL | 0.04000 mL | 12.036 | -0.00001 | 0.00000 | 0.00060 | 10.0 s |
| 37:25.0 | Data point 33 | 1.50000 mL | 0.17239 mL | 0.11997 mL | 0.14000 mL | 1.962 | -0.01011 | 0.31324 | 0.00089 | 10.0 s |
| 38:11.4 | Data point 34 | 1.50000 mL | 0.17239 mL | 0.13967 mL | 0.14000 mL | 2.167 | -0.01017 | 0.51336 | 0.00070 | 10.0 s |
| 38:47.1 | Data point 35 | 1.50000 mL | 0.17239 mL | 0.15202 mL | 0.14000 mL | 2.364 | 0.01692 | 0.77326 | 0.00095 | 14.5 s |

| | | | |
|---|---|---|---|
| Sample name: | **M16_octanol** | Experiment start time: | **3/1/2018 10:41:05 AM** |
| Assay name: | **pH-metric high logP** | Analyst: | **Pion** |
| Assay ID: | **18C-01008** | Instrument ID: | **T312060** |
| Filename: | **C:\Sirius_T3\Mehtap\20180228_exp28_logP_T3-2\18C-01008_M16_octanol_pH-metric high logP.t3r** | | |

#### Events (continued)

| Time | Event | Water | Acid | Base | Octanol | pH | dpH/dt | pH R-squared | pH SD | dpH/dt time |
|---|---|---|---|---|---|---|---|---|---|---|
| 39:27.2 | Data point 36 | 1.50000 mL | 0.17239 mL | 0.15988 mL | 0.14000 mL | 2.550 | -0.00131 | 0.00551 | 0.00087 | 10.0 s |
| 40:02.8 | Data point 37 | 1.50000 mL | 0.17239 mL | 0.16508 mL | 0.14000 mL | 2.719 | 0.01355 | 0.83353 | 0.00073 | 10.0 s |
| 40:53.9 | Data point 38 | 1.50000 mL | 0.17239 mL | 0.16943 mL | 0.14000 mL | 2.923 | -0.00343 | 0.05295 | 0.00074 | 10.0 s |
| 41:39.9 | Data point 39 | 1.50000 mL | 0.17239 mL | 0.17302 mL | 0.14000 mL | 3.116 | 0.00068 | 0.02704 | 0.00020 | 10.0 s |
| 42:25.7 | Data point 40 | 1.50000 mL | 0.17239 mL | 0.17545 mL | 0.14000 mL | 3.313 | 0.01297 | 0.52106 | 0.00089 | 10.5 s |
| 43:12.1 | Data point 41 | 1.50000 mL | 0.17239 mL | 0.17726 mL | 0.14000 mL | 3.511 | 0.00068 | 0.03379 | 0.00018 | 10.0 s |
| 43:57.8 | Data point 42 | 1.50000 mL | 0.17239 mL | 0.17872 mL | 0.14000 mL | 3.704 | -0.00606 | 0.15775 | 0.00075 | 16.5 s |
| 44:39.8 | Data point 43 | 1.50000 mL | 0.17239 mL | 0.18032 mL | 0.14000 mL | 4.100 | -0.00521 | 0.09608 | 0.00083 | 10.0 s |
| 45:15.3 | Data point 44 | 1.50000 mL | 0.17239 mL | 0.18123 mL | 0.14000 mL | 4.503 | -0.00893 | 0.23890 | 0.00090 | 10.5 s |
| 45:51.2 | Data point 45 | 1.50000 mL | 0.17239 mL | 0.18203 mL | 0.14000 mL | 6.710 | -0.05359 | 0.98654 | 0.00266 | Timed out at 59.5 s |
| 47:21.8 | Data point 46 | 1.50000 mL | 0.17239 mL | 0.18234 mL | 0.14000 mL | 8.402 | -0.02008 | 0.98880 | 0.00100 | 43.5 s |
| 48:30.8 | Data point 47 | 1.50000 mL | 0.17239 mL | 0.18260 mL | 0.14000 mL | 8.986 | -0.01599 | 0.67601 | 0.00096 | 19.0 s |
| 49:15.2 | Data point 48 | 1.50000 mL | 0.17239 mL | 0.18328 mL | 0.14000 mL | 9.587 | -0.01391 | 0.57721 | 0.00090 | 11.5 s |
| 49:52.1 | Data point 49 | 1.50000 mL | 0.17239 mL | 0.18420 mL | 0.14000 mL | 9.950 | -0.01192 | 0.86726 | 0.00063 | 10.5 s |
| 50:28.1 | Data point 50 | 1.50000 mL | 0.17239 mL | 0.18497 mL | 0.14000 mL | 10.154 | -0.00728 | 0.19670 | 0.00081 | 10.0 s |
| 51:19.0 | Data point 51 | 1.50000 mL | 0.17239 mL | 0.18612 mL | 0.14000 mL | 10.353 | -0.01567 | 0.63815 | 0.00097 | 10.0 s |
| 52:10.0 | Data point 52 | 1.50000 mL | 0.17239 mL | 0.18772 mL | 0.14000 mL | 10.539 | -0.00172 | 0.22733 | 0.00018 | 10.5 s |
| 53:01.4 | Data point 53 | 1.50000 mL | 0.17239 mL | 0.19010 mL | 0.14000 mL | 10.724 | -0.00528 | 0.43270 | 0.00040 | 10.0 s |
| 53:52.4 | Data point 54 | 1.50000 mL | 0.17239 mL | 0.19306 mL | 0.14000 mL | 10.912 | 0.00664 | 0.52199 | 0.00045 | 10.0 s |
| 54:43.5 | Data point 55 | 1.50000 mL | 0.17239 mL | 0.19668 mL | 0.14000 mL | 11.108 | -0.00005 | 0.00018 | 0.00019 | 10.0 s |
| 55:29.4 | Data point 56 | 1.50000 mL | 0.17239 mL | 0.20186 mL | 0.14000 mL | 11.294 | 0.00132 | 0.14476 | 0.00017 | 10.0 s |
| 56:15.3 | Data point 57 | 1.50000 mL | 0.17239 mL | 0.20851 mL | 0.14000 mL | 11.482 | -0.00055 | 0.02091 | 0.00019 | 10.0 s |
| 56:51.0 | Data point 58 | 1.50000 mL | 0.17239 mL | 0.21809 mL | 0.14000 mL | 11.660 | 0.01284 | 0.83806 | 0.00069 | 10.0 s |
| 57:26.7 | Data point 59 | 1.50000 mL | 0.17239 mL | 0.23274 mL | 0.14000 mL | 11.852 | 0.00826 | 0.93309 | 0.00042 | 10.0 s |
| 58:02.7 | Data point 60 | 1.50000 mL | 0.17239 mL | 0.25475 mL | 0.14000 mL | 12.031 | 0.00778 | 0.92215 | 0.00040 | 10.0 s |
| 59:24.9 | Data point 61 | 1.50000 mL | 0.31343 mL | 0.25475 mL | 0.44000 mL | 1.965 | -0.00463 | 0.40371 | 0.00036 | 10.0 s |
| 1:00:11.3 | Data point 62 | 1.50000 mL | 0.31343 mL | 0.27785 mL | 0.44000 mL | 2.178 | 0.01276 | 0.89016 | 0.00067 | 17.5 s |
| 1:00:54.6 | Data point 63 | 1.50000 mL | 0.31343 mL | 0.29184 mL | 0.44000 mL | 2.370 | 0.01126 | 0.59387 | 0.00072 | 10.0 s |
| 1:01:30.1 | Data point 64 | 1.50000 mL | 0.31343 mL | 0.30085 mL | 0.44000 mL | 2.548 | 0.01630 | 0.71883 | 0.00095 | 10.0 s |
| 1:02:16.1 | Data point 65 | 1.50000 mL | 0.31343 mL | 0.30814 mL | 0.44000 mL | 2.753 | -0.00161 | 0.01660 | 0.00062 | 10.0 s |
| 1:03:02.1 | Data point 66 | 1.50000 mL | 0.31343 mL | 0.31279 mL | 0.44000 mL | 2.951 | 0.01037 | 0.37883 | 0.00083 | 10.0 s |
| 1:03:37.6 | Data point 67 | 1.50000 mL | 0.31343 mL | 0.31573 mL | 0.44000 mL | 3.137 | 0.00768 | 0.16641 | 0.00093 | 17.5 s |
| 1:04:20.5 | Data point 68 | 1.50000 mL | 0.31343 mL | 0.31813 mL | 0.44000 mL | 3.367 | 0.00085 | 0.00280 | 0.00079 | 10.0 s |
| 1:04:56.0 | Data point 69 | 1.50000 mL | 0.31343 mL | 0.32025 mL | 0.44000 mL | 3.661 | 0.00260 | 0.02567 | 0.00080 | 10.0 s |
| 1:05:31.4 | Data point 70 | 1.50000 mL | 0.31343 mL | 0.32128 mL | 0.44000 mL | 3.891 | -0.00269 | 0.34212 | 0.00023 | 10.0 s |
| 1:06:06.9 | Data point 71 | 1.50000 mL | 0.31343 mL | 0.32227 mL | 0.44000 mL | 4.353 | -0.00164 | 0.02000 | 0.00057 | 10.0 s |
| 1:06:42.3 | Data point 72 | 1.50000 mL | 0.31343 mL | 0.32295 mL | 0.44000 mL | 5.501 | -0.01091 | 0.30965 | 0.00097 | 20.5 s |
| 1:07:33.3 | Data point 73 | 1.50000 mL | 0.31343 mL | 0.32310 mL | 0.44000 mL | 5.706 | 0.01056 | 0.34237 | 0.00089 | 19.5 s |
| 1:08:18.3 | Data point 74 | 1.50000 mL | 0.31343 mL | 0.32317 mL | 0.44000 mL | 5.975 | -0.01091 | 0.35546 | 0.00090 | 18.0 s |
| 1:09:01.7 | Data point 75 | 1.50000 mL | 0.31343 mL | 0.32321 mL | 0.44000 mL | 6.238 | -0.01377 | 0.50757 | 0.00095 | 28.0 s |
| 1:09:55.2 | Data point 76 | 1.50000 mL | 0.31343 mL | 0.32326 mL | 0.44000 mL | 6.441 | -0.01475 | 0.53518 | 0.00100 | 34.0 s |
| 1:10:54.6 | Data point 77 | 1.50000 mL | 0.31343 mL | 0.32331 mL | 0.44000 mL | 6.706 | -0.01933 | 0.97748 | 0.00097 | 40.5 s |
| 1:12:05.6 | Data point 78 | 1.50000 mL | 0.31343 mL | 0.32335 mL | 0.44000 mL | 7.187 | -0.05389 | 0.98362 | 0.00268 | Timed out at 59.5 s |
| 1:13:31.0 | Data point 79 | 1.50000 mL | 0.31343 mL | 0.32340 mL | 0.44000 mL | 7.330 | -0.03553 | 0.97995 | 0.00177 | Timed out at 59.5 s |
| 1:15:01.5 | Data point 80 | 1.50000 mL | 0.31343 mL | 0.32354 mL | 0.44000 mL | 8.298 | -0.03259 | 0.96414 | 0.00164 | Timed out at 59.5 s |
| 1:16:26.9 | Data point 81 | 1.50000 mL | 0.31343 mL | 0.32366 mL | 0.44000 mL | 8.546 | -0.01671 | 0.68295 | 0.00100 | 19.0 s |
| 1:17:11.3 | Data point 82 | 1.50000 mL | 0.31343 mL | 0.32385 mL | 0.44000 mL | 8.939 | -0.01490 | 0.60712 | 0.00094 | 14.0 s |
| 1:17:50.8 | Data point 83 | 1.50000 mL | 0.31343 mL | 0.32427 mL | 0.44000 mL | 9.381 | 0.00173 | 0.00806 | 0.00095 | 12.0 s |
| 1:18:28.1 | Data point 84 | 1.50000 mL | 0.31343 mL | 0.32505 mL | 0.44000 mL | 9.814 | 0.00015 | 0.00006 | 0.00099 | 10.5 s |
| 1:19:03.9 | Data point 85 | 1.50000 mL | 0.31343 mL | 0.32604 mL | 0.44000 mL | 10.137 | 0.00630 | 0.10167 | 0.00098 | 10.0 s |
| 1:19:39.4 | Data point 86 | 1.50000 mL | 0.31343 mL | 0.32702 mL | 0.44000 mL | 10.354 | 0.00292 | 0.02377 | 0.00094 | 10.0 s |
| 1:20:25.1 | Data point 87 | 1.50000 mL | 0.31343 mL | 0.32843 mL | 0.44000 mL | 10.543 | 0.00828 | 0.57942 | 0.00054 | 10.0 s |
| 1:21:16.2 | Data point 88 | 1.50000 mL | 0.31343 mL | 0.33067 mL | 0.44000 mL | 10.736 | 0.01106 | 0.70690 | 0.00065 | 10.0 s |

| | | | |
|---|---|---|---|
| Sample name: | **M16_octanol** | Experiment start time: | **3/1/2018 10:41:05 AM** |
| Assay name: | **pH-metric high logP** | Analyst: | **Pion** |
| Assay ID: | **18C-01008** | Instrument ID: | **T312060** |
| Filename: | **C:\Sirius_T3\Mehtap\20180228_exp28_logP_T3-2\18C-01008_M16_octanol_pH-metric high logP.t3r** | | |

#### Events (continued)

| Time | Event | Water | Acid | Base | Octanol | pH | dpH/dt | pH R-squared | pH SD | dpH/dt time |
|---|---|---|---|---|---|---|---|---|---|---|
| 1:21:51.7 | Data point 89 | 1.50000 mL | 0.31343 mL | 0.33321 mL | 0.44000 mL | 10.914 | 0.00908 | 0.75807 | 0.00051 | 10.0 s |
| 1:22:27.2 | Data point 90 | 1.50000 mL | 0.31343 mL | 0.33655 mL | 0.44000 mL | 11.100 | 0.00124 | 0.00385 | 0.00099 | 10.0 s |
| 1:23:02.7 | Data point 91 | 1.50000 mL | 0.31343 mL | 0.34132 mL | 0.44000 mL | 11.258 | 0.01768 | 0.78273 | 0.00099 | 16.0 s |
| 1:23:60.0 | Data point 92 | 1.50000 mL | 0.31343 mL | 0.34927 mL | 0.44000 mL | 11.450 | 0.01237 | 0.90868 | 0.00064 | 10.0 s |
| 1:24:46.2 | Data point 93 | 1.50000 mL | 0.31343 mL | 0.36051 mL | 0.44000 mL | 11.643 | 0.01058 | 0.60009 | 0.00067 | 10.0 s |
| 1:25:21.9 | Data point 94 | 1.50000 mL | 0.31343 mL | 0.37681 mL | 0.44000 mL | 11.828 | 0.01552 | 0.68214 | 0.00093 | 10.0 s |
| 1:25:58.0 | Data point 95 | 1.50000 mL | 0.31343 mL | 0.40214 mL | 0.44000 mL | 12.016 | 0.00685 | 0.34934 | 0.00057 | 10.0 s |
| 1:26:17.0 | Assay volumes | 1.50000 mL | 0.31343 mL | 0.40214 mL | 0.44000 mL | | | | | |

| | | | |
|---|---|---|---|
| Sample name: | **M16_octanol** | Experiment start time: | **3/1/2018 10:41:05 AM** |
| Assay name: | **pH-metric high logP** | Analyst: | **Pion** |
| Assay ID: | **18C-01008** | Instrument ID: | **T312060** |
| Filename: | **C:\Sirius_T3\Mehtap\20180228_exp28_logP_T3-2\18C-01008_M16_octanol_pH-metric high logP.t3r** | | |

| | | | |
|---|---|---|---|
| Sample name: | **M16_octanol** | Experiment start time: | **3/1/2018 10:41:05 AM** |
| Assay name: | **pH-metric high logP** | Analyst: | **Pion** |
| Assay ID: | **18C-01008** | Instrument ID: | **T312060** |
| Filename: | **C:\Sirius_T3\Mehtap\20180228_exp28_logP_T3-2\18C-01008_M16_octanol_pH-metric high logP.t3r** | | |

#### Assay Settings (continued)

| Setting | Value | Original Value | Date/Time changed | Imported from |
|---|---|---|---|---|
| ***Titration 3*** | | | | |
| Titrate from | Low to high pH | | | |
| Add additional water | 0.00 mL | | | |
| Additional partition solvent volume | 0.300 mL | | | |
| Additional partition solvent added | Automatic | | | |
| After pH adjust stir for | 30 seconds | | | |
| Stir to allow partitioning for | 15 seconds | | | |
| Stirrer speed for partitioning | 60% | | | |
| ***Data Point Stability*** | | | | |
| Stir during data point collection | No | | | |
| Delay before data point collection | 0 seconds | | | |
| Number of points to average | 20 points | | | |
| Time interval between points | 0.50 seconds | | | |
| Required maximum standard deviation | 0.00100 dpH/dt | | | |
| Stability timeout after | 60 seconds | | | |

#### Calibration Settings

| Setting | Value | Date/Time changed | Imported from |
|---|---|---|---|
| Four-Plus alpha | 0.130 | 3/1/2018 10:41:04 AM | C:\Sirius_T3\HCl18B27.t3r |
| Four-Plus S | 0.9970 | 3/1/2018 10:41:04 AM | C:\Sirius_T3\HCl18B27.t3r |
| Four-Plus jH | 0.8 | 3/1/2018 10:41:04 AM | C:\Sirius_T3\HCl18B27.t3r |
| Four-Plus jOH | -0.4 | 3/1/2018 10:41:04 AM | C:\Sirius_T3\HCl18B27.t3r |
| Base concentration factor | 1.000 | 3/1/2018 10:41:05 AM | C:\Sirius_T3\KOH18B27.t3r |
| Acid concentration factor | 0.994 | 3/1/2018 10:41:05 AM | C:\Sirius_T3\HCl18B27.t3r |

#### Instrument Settings

| Setting | Value | Batch Id | Install date |
|---|---|---|---|
| Instrument owner | Merck | | |
| Instrument ID | T312060 | | |
| Instrument type | T3 Simulator | | |
| Software version | 1.1.3.0 | | |
| Dispenser module | | T3DM1200361 | 3/31/2009 5:24:52 AM |
| Dispenser 0 | Water | | 3/31/2009 5:25:05 AM |
| Syringe volume | 2.5 mL | | |
| Firmware version | 1.2.1(r2) | | |
| Titrant | Water (0.15 M KCl) | 02-06-2018 | 2/27/2018 10:05:59 AM |
| Dispenser 2 | Acid | | 3/31/2009 5:25:11 AM |
| Syringe volume | 0.5 mL | | |
| Firmware version | 1.2.1(r2) | | |
| Titrant | Acid (0.5 M HCl) | 02-27-2018 | 2/27/2018 10:27:22 AM |
| Dispenser 1 | Base | | 3/31/2009 5:25:21 AM |
| Syringe volume | 0.5 mL | | |
| Firmware version | 1.2.1(r2) | | |
| Titrant | Base (0.5 M KOH) | 9/22/2017 | 2/27/2018 10:21:22 AM |
| Dispenser 5 | Cosolvent | | 3/31/2009 5:26:24 AM |
| Syringe volume | 2.5 mL | | |
| Firmware version | 1.2.1(r2) | | |
| Distribution valve 5 | Distribution Valve | | 3/31/2009 5:28:19 AM |
| Firmware version | 1.1.3 | | |
| Port A | Methanol (80%, 0.15 M KCl) | 09-26-17 | 2/7/2018 9:42:01 AM |
| Port B | Cyclohexane | 11-01-17 | 2/27/2018 10:37:57 AM |
| Dispenser 3 | Buffer | | 8/3/2010 5:05:16 AM |
| Syringe volume | 0.5 mL | | |
| Firmware version | 1.2.1(r2) | | |
| Titrant | Dodecane | 2018/01/31 | 2/28/2018 10:18:04 AM |
| Dispenser 6 | Octanol | | 10/22/2010 10:52:43 AM |

| | | | |
|---|---|---|---|
| Sample name: | **M16_octanol** | Experiment start time: | **3/1/2018 10:41:05 AM** |
| Assay name: | **pH-metric high logP** | Analyst: | **Pion** |
| Assay ID: | **18C-01008** | Instrument ID: | **T312060** |
| Filename: | **C:\Sirius_T3\Mehtap\20180228_exp28_logP_T3-2\18C-01008_M16_octanol_pH-metric high logP.t3r** | | |

| | | | |
|---|---|---|---|
| Sample name: | **M16_octanol** | Experiment start time: | **3/1/2018 10:41:05 AM** |
| Assay name: | **pH-metric high logP** | Analyst: | **Pion** |
| Assay ID: | **18C-01008** | Instrument ID: | **T312060** |
| Filename: | **C:\Sirius_T3\Mehtap\20180228_exp28_logP_T3-2\18C-01008_M16_octanol_pH-metric high logP.t3r** | | |

#### Instrument Settings (continued)

| Setting | Value | Batch Id | Install date |
|---|---|---|---|
| Spectrometer calibration stir duration | 5 s | | |
| Spectrometer calibration stir speed | 30% | | |
| Spectrometer calibration wash pump volume | 20.0 mL | | |
| Spectrometer calibration wash stir duration | 5 s | | |
| Spectrometer calibration wash stir speed | 30% | | |
| Overhead dispense height | 10000 | | |

#### Refinement Settings

| Setting | Value | Default value |
|---|---|---|
| Turbidity detection method | None | None |
| Turbidity wavelength to assess | 500.0 nm | 500.0 nm |
| Turbidity maximum absorbance | 0.100 | 0.100 |
| Turbidity probe threshold | 50.00 | 50.00 |

#### Experiment Log

[2:58] Air gap created for Water (0.15 M KCl)
[2:58] Air gap created for Acid (0.5 M HCl)
[2:58] Air gap created for Base (0.5 M KOH)
[2:59] Air gap released for Water (0.15 M KCl)
[3:03] Titrator arm moved over Titration position
[3:03] Titration 1 of 3
[3:03] Adding initial titrants
[3:03] Automatically add 1.50000 mL of water
[3:28] Dispensed 1.500000 mL of Water (0.15 M KCl)
[3:32] Titrator arm moved over Drain
[9:13] Titrator arm moved to Titration position
[9:13] Argon flow rate set to 100
[9:13] Stirrer speed set to 10
[9:18] Automatically add 0.04000 mL of Octanol
[9:19] Dispensed 0.040005 mL of Octanol
[9:21] Initial pH = 3.61
[9:21] Iterative adjust 3.61 -> 2.00
[9:21] pH 3.61 -> 2.00
[9:22] Air gap released for Acid (0.5 M HCl)
[9:23] Dispensed 0.041627 mL of Acid (0.5 M HCl)
[9:28] pH 2.04 -> 2.00
[9:28] Dispensed 0.003551 mL of Acid (0.5 M HCl)
[9:33] Holding pH 2.00
[11:33] Stirrer speed set to 0
[11:33] Stirrer speed set to 50
[11:33] Iterative adjust 1.97 -> 2.00
[11:33] pH 1.97 -> 2.00
[11:34] Air gap released for Base (0.5 M KOH)
[11:35] Dispensed 0.002752 mL of Base (0.5 M KOH)
[12:25] Stirrer speed set to 0
[12:36] Datapoint id 1 collected
[12:36] Stirrer speed set to 50
[12:41] pH 2.02 -> 2.22
[12:41] Using cautious pH adjust
[12:41] Dispensed 0.007667 mL of Base (0.5 M KOH)
[12:46] Stepping pH = 2.10
[12:46] Dispensed 0.005974 mL of Base (0.5 M KOH)
[12:52] Stepping pH = 2.19
[12:52] Dispensed 0.001599 mL of Base (0.5 M KOH)
[12:57] Stepping pH = 2.22
[13:12] Stirrer speed set to 0
[13:22] Datapoint id 2 collected

| | | | |
|---|---|---|---|
| Sample name: | **M16_octanol** | Experiment start time: | **3/1/2018 10:41:05 AM** |
| Assay name: | **pH-metric high logP** | Analyst: | **Pion** |
| Assay ID: | **18C-01008** | Instrument ID: | **T312060** |
| Filename: | **C:\Sirius_T3\Mehtap\20180228_exp28_logP_T3-2\18C-01008_M16_octanol_pH-metric high logP.t3r** | | |

#### Experiment Log (continued)

[13:22] Charge balance equation is out by 0.5%
[13:22] Stirrer speed set to 50
[13:27] pH 2.22 -> 2.42
[13:27] Using charge balance adjust
[13:27] Dispensed 0.009525 mL of Base (0.5 M KOH)
[13:48] Stirrer speed set to 0
[13:58] Datapoint id 3 collected
[13:58] Charge balance equation is out by -8.2%
[13:58] Stirrer speed set to 50
[14:03] pH 2.41 -> 2.61
[14:03] Using charge balance adjust
[14:03] Dispensed 0.006162 mL of Base (0.5 M KOH)
[14:23] Stirrer speed set to 0
[14:33] Datapoint id 4 collected
[14:33] Charge balance equation is out by -16.7%
[14:33] Stirrer speed set to 50
[14:38] pH 2.58 -> 2.78
[14:38] Using cautious pH adjust
[14:38] Dispensed 0.002093 mL of Base (0.5 M KOH)
[14:43] Stepping pH = 2.67
[14:44] Dispensed 0.001623 mL of Base (0.5 M KOH)
[14:49] Stepping pH = 2.74
[14:49] Dispensed 0.000659 mL of Base (0.5 M KOH)
[14:54] Stepping pH = 2.78
[15:09] Stirrer speed set to 0
[15:19] Datapoint id 5 collected
[15:19] Charge balance equation is out by -4.9%
[15:19] Stirrer speed set to 50
[15:24] pH 2.78 -> 2.98
[15:24] Using charge balance adjust
[15:24] Dispensed 0.002752 mL of Base (0.5 M KOH)
[15:45] Stirrer speed set to 0
[15:55] Datapoint id 6 collected
[15:55] Charge balance equation is out by -31.1%
[15:55] Stirrer speed set to 50
[16:00] pH 2.92 -> 3.12
[16:00] Using cautious pH adjust
[16:00] Dispensed 0.001035 mL of Base (0.5 M KOH)
[16:05] Stepping pH = 2.99
[16:05] Dispensed 0.001129 mL of Base (0.5 M KOH)
[16:10] Stepping pH = 3.07
[16:10] Dispensed 0.000447 mL of Base (0.5 M KOH)
[16:15] Stepping pH = 3.11
[16:15] Dispensed 0.000118 mL of Base (0.5 M KOH)
[16:20] Stepping pH = 3.12
[16:36] Stirrer speed set to 0
[16:46] Datapoint id 7 collected
[16:46] Charge balance equation is out by -32.9%
[16:46] Stirrer speed set to 50
[16:51] pH 3.11 -> 3.31
[16:51] Using cautious pH adjust
[16:51] Dispensed 0.000729 mL of Base (0.5 M KOH)
[16:56] Stepping pH = 3.18
[16:56] Dispensed 0.000776 mL of Base (0.5 M KOH)
[17:01] Stepping pH = 3.25
[17:01] Dispensed 0.000494 mL of Base (0.5 M KOH)
[17:06] Stepping pH = 3.30
[17:06] Dispensed 0.000141 mL of Base (0.5 M KOH)
[17:11] Stepping pH = 3.31

| | | | |
|---|---|---|---|
| Sample name: | **M16_octanol** | Experiment start time: | **3/1/2018 10:41:05 AM** |
| Assay name: | **pH-metric high logP** | Analyst: | **Pion** |
| Assay ID: | **18C-01008** | Instrument ID: | **T312060** |
| Filename: | **C:\Sirius_T3\Mehtap\20180228_exp28_logP_T3-2\18C-01008_M16_octanol_pH-metric high logP.t3r** | | |

#### Experiment Log (continued)

[17:26] Stirrer speed set to 0
[17:37] Datapoint id 8 collected
[17:37] Charge balance equation is out by -46.4%
[17:37] Stirrer speed set to 50
[17:42] pH 3.30 -> 3.50
[17:42] Using cautious pH adjust
[17:42] Dispensed 0.000588 mL of Base (0.5 M KOH)
[17:47] Stepping pH = 3.36
[17:47] Dispensed 0.000800 mL of Base (0.5 M KOH)
[17:52] Stepping pH = 3.43
[17:52] Dispensed 0.000541 mL of Base (0.5 M KOH)
[17:57] Stepping pH = 3.48
[17:57] Dispensed 0.000188 mL of Base (0.5 M KOH)
[18:02] Stepping pH = 3.49
[18:02] Dispensed 0.000141 mL of Base (0.5 M KOH)
[18:08] Stepping pH = 3.50
[18:23] Stirrer speed set to 0
[18:33] Datapoint id 9 collected
[18:33] Charge balance equation is out by -90.2%
[18:33] Stirrer speed set to 50
[18:38] pH 3.49 -> 3.69
[18:38] Using cautious pH adjust
[18:38] Dispensed 0.000541 mL of Base (0.5 M KOH)
[18:43] Stepping pH = 3.55
[18:43] Dispensed 0.000753 mL of Base (0.5 M KOH)
[18:48] Stepping pH = 3.63
[18:48] Dispensed 0.000447 mL of Base (0.5 M KOH)
[18:53] Stepping pH = 3.67
[18:53] Dispensed 0.000188 mL of Base (0.5 M KOH)
[18:58] Stepping pH = 3.68
[18:59] Dispensed 0.000141 mL of Base (0.5 M KOH)
[19:04] Stepping pH = 3.69
[19:19] Stirrer speed set to 0
[19:29] Datapoint id 10 collected
[19:29] Charge balance equation is out by -90.4%
[19:29] Stirrer speed set to 50
[19:34] pH 3.68 -> 3.88
[19:34] Using cautious pH adjust
[19:34] Dispensed 0.000564 mL of Base (0.5 M KOH)
[19:40] Stepping pH = 3.76
[19:40] Dispensed 0.000564 mL of Base (0.5 M KOH)
[19:45] Stepping pH = 3.85
[19:45] Dispensed 0.000212 mL of Base (0.5 M KOH)
[19:50] Stepping pH = 3.87
[19:50] Dispensed 0.000141 mL of Base (0.5 M KOH)
[19:55] Stepping pH = 3.89
[20:10] Stirrer speed set to 0
[20:20] Datapoint id 11 collected
[20:20] Charge balance equation is out by -31.4%
[20:20] Stirrer speed set to 50
[20:25] pH 3.88 -> 4.08
[20:25] Using cautious pH adjust
[20:26] Dispensed 0.000659 mL of Base (0.5 M KOH)
[20:31] Stepping pH = 4.01
[20:31] Dispensed 0.000329 mL of Base (0.5 M KOH)
[20:36] Stepping pH = 4.07
[20:51] Stirrer speed set to 0
[21:01] Datapoint id 12 collected
[21:01] Charge balance equation is out by 25.7%

| | | | |
|---|---|---|---|
| Sample name: | **M16_octanol** | Experiment start time: | **3/1/2018 10:41:05 AM** |
| Assay name: | **pH-metric high logP** | Analyst: | **Pion** |
| Assay ID: | **18C-01008** | Instrument ID: | **T312060** |
| Filename: | **C:\Sirius_T3\Mehtap\20180228_exp28_logP_T3-2\18C-01008_M16_octanol_pH-metric high logP.t3r** | | |

#### Experiment Log (continued)

[21:01] Stirrer speed set to 50
[21:06] pH 4.07 -> 4.27
[21:06] Using cautious pH adjust
[21:06] Dispensed 0.000753 mL of Base (0.5 M KOH)
[21:11] Stepping pH = 4.29
[21:26] Stirrer speed set to 0
[21:37] Datapoint id 13 collected
[21:37] Charge balance equation is out by 50.0%
[21:37] Stirrer speed set to 50
[21:42] pH 4.28 -> 4.48
[21:42] Using cautious pH adjust
[21:42] Dispensed 0.000847 mL of Base (0.5 M KOH)
[21:47] Stepping pH = 4.69
[22:02] Stirrer speed set to 0
[22:14] Datapoint id 14 collected
[22:14] Charge balance equation is out by 50.0%
[22:14] Stirrer speed set to 50
[22:19] pH 4.68 -> 4.88
[22:19] Using cautious pH adjust
[22:19] Dispensed 0.000823 mL of Base (0.5 M KOH)
[22:24] Stepping pH = 7.46
[22:39] Stirrer speed set to 0
[23:39] Datapoint id 15 collected
[23:39] Charge balance equation is out by 50.0%
[23:39] Stirrer speed set to 50
[23:44] pH 7.86 -> 8.06
[23:44] Using cautious pH adjust
[23:44] Dispensed 0.000165 mL of Base (0.5 M KOH)
[23:49] Stepping pH = 8.28
[24:04] Stirrer speed set to 0
[24:43] Datapoint id 16 collected
[24:43] Charge balance equation is out by 50.0%
[24:43] Stirrer speed set to 50
[24:48] pH 8.16 -> 8.36
[24:48] Using cautious pH adjust
[24:48] Dispensed 0.000282 mL of Base (0.5 M KOH)
[24:53] Stepping pH = 8.87
[25:08] Stirrer speed set to 0
[25:18] Datapoint id 17 collected
[25:18] Charge balance equation is out by 50.0%
[25:18] Stirrer speed set to 50
[25:24] pH 8.85 -> 9.05
[25:24] Using cautious pH adjust
[25:24] Dispensed 0.000776 mL of Base (0.5 M KOH)
[25:29] Stepping pH = 9.43
[25:44] Stirrer speed set to 0
[25:54] Datapoint id 18 collected
[25:54] Charge balance equation is out by 50.0%
[25:54] Stirrer speed set to 50
[26:00] pH 9.41 -> 9.61
[26:00] Using cautious pH adjust
[26:00] Dispensed 0.000847 mL of Base (0.5 M KOH)
[26:05] Stepping pH = 9.70
[26:20] Stirrer speed set to 0
[26:30] Datapoint id 19 collected
[26:30] Charge balance equation is out by 50.0%
[26:30] Stirrer speed set to 50
[26:35] pH 9.69 -> 9.89
[26:35] Using cautious pH adjust

| | | | |
|---|---|---|---|
| Sample name: | **M16_octanol** | Experiment start time: | **3/1/2018 10:41:05 AM** |
| Assay name: | **pH-metric high logP** | Analyst: | **Pion** |
| Assay ID: | **18C-01008** | Instrument ID: | **T312060** |
| Filename: | **C:\Sirius_T3\Mehtap\20180228_exp28_logP_T3-2\18C-01008_M16_octanol_pH-metric high logP.t3r** | | |

#### Experiment Log (continued)

[26:35] Dispensed 0.000706 mL of Base (0.5 M KOH)
[26:40] Stepping pH = 9.85
[26:40] Dispensed 0.000165 mL of Base (0.5 M KOH)
[26:45] Stepping pH = 9.86
[26:45] Dispensed 0.000212 mL of Base (0.5 M KOH)
[26:51] Stepping pH = 9.89
[27:06] Stirrer speed set to 0
[27:16] Datapoint id 20 collected
[27:16] Charge balance equation is out by 23.1%
[27:16] Stirrer speed set to 50
[27:21] pH 9.87 -> 10.07
[27:21] Using cautious pH adjust
[27:21] Dispensed 0.000588 mL of Base (0.5 M KOH)
[27:26] Stepping pH = 10.00
[27:26] Dispensed 0.000282 mL of Base (0.5 M KOH)
[27:31] Stepping pH = 10.04
[27:31] Dispensed 0.000165 mL of Base (0.5 M KOH)
[27:36] Stepping pH = 10.06
[27:36] Dispensed 0.000071 mL of Base (0.5 M KOH)
[27:42] Stepping pH = 10.07
[27:57] Stirrer speed set to 0
[28:07] Datapoint id 21 collected
[28:07] Charge balance equation is out by 5.9%
[28:07] Stirrer speed set to 50
[28:12] pH 10.06 -> 10.26
[28:12] Using charge balance adjust
[28:12] Dispensed 0.001035 mL of Base (0.5 M KOH)
[28:32] Stirrer speed set to 0
[28:42] Datapoint id 22 collected
[28:42] Charge balance equation is out by -19.8%
[28:42] Stirrer speed set to 50
[28:47] pH 10.23 -> 10.43
[28:47] Using cautious pH adjust
[28:47] Dispensed 0.000494 mL of Base (0.5 M KOH)
[28:52] Stepping pH = 10.30
[28:53] Dispensed 0.000517 mL of Base (0.5 M KOH)
[28:58] Stepping pH = 10.37
[28:58] Dispensed 0.000282 mL of Base (0.5 M KOH)
[29:03] Stepping pH = 10.41
[29:03] Dispensed 0.000118 mL of Base (0.5 M KOH)
[29:08] Stepping pH = 10.42
[29:23] Stirrer speed set to 0
[29:34] Datapoint id 23 collected
[29:34] Charge balance equation is out by -43.7%
[29:34] Stirrer speed set to 50
[29:39] pH 10.41 -> 10.61
[29:39] Using cautious pH adjust
[29:39] Dispensed 0.000517 mL of Base (0.5 M KOH)
[29:44] Stepping pH = 10.48
[29:44] Dispensed 0.000611 mL of Base (0.5 M KOH)
[29:49] Stepping pH = 10.55
[29:49] Dispensed 0.000376 mL of Base (0.5 M KOH)
[29:54] Stepping pH = 10.59
[29:54] Dispensed 0.000188 mL of Base (0.5 M KOH)
[29:59] Stepping pH = 10.60
[30:15] Stirrer speed set to 0
[30:25] Datapoint id 24 collected
[30:25] Charge balance equation is out by -63.3%
[30:25] Stirrer speed set to 50

Sample name: **M16_octanol** Experiment start time: **3/1/2018 10:41:05 AM**
Assay name: **pH-metric high logP** Analyst: **Pion**
Assay ID: **18C-01008** Instrument ID: **T312060**
Filename: **C:\Sirius_T3\Mehtap\20180228_exp28_logP_T3-2\18C-01008_M16_octanol_pH-metric high logP.t3r**

#### Experiment Log (continued)

[30:30] pH 10.60 -> 10.80
[30:30] Using cautious pH adjust
[30:30] Dispensed 0.000635 mL of Base (0.5 M KOH)
[30:35] Stepping pH = 10.66
[30:35] Dispensed 0.000776 mL of Base (0.5 M KOH)
[30:40] Stepping pH = 10.73
[30:40] Dispensed 0.000541 mL of Base (0.5 M KOH)
[30:45] Stepping pH = 10.78
[30:45] Dispensed 0.000188 mL of Base (0.5 M KOH)
[30:51] Stepping pH = 10.79
[31:06] Stirrer speed set to 0
[31:16] Datapoint id 25 collected
[31:16] Charge balance equation is out by -66.0%
[31:16] Stirrer speed set to 50
[31:21] pH 10.78 -> 10.98
[31:21] Using cautious pH adjust
[31:21] Dispensed 0.000870 mL of Base (0.5 M KOH)
[31:26] Stepping pH = 10.84
[31:27] Dispensed 0.001082 mL of Base (0.5 M KOH)
[31:32] Stepping pH = 10.91
[31:32] Dispensed 0.000800 mL of Base (0.5 M KOH)
[31:37] Stepping pH = 10.96
[31:37] Dispensed 0.000212 mL of Base (0.5 M KOH)
[31:42] Stepping pH = 10.97
[31:57] Stirrer speed set to 0
[32:07] Datapoint id 26 collected
[32:07] Charge balance equation is out by -71.2%
[32:07] Stirrer speed set to 50
[32:12] pH 10.96 -> 11.16
[32:12] Using cautious pH adjust
[32:12] Dispensed 0.001246 mL of Base (0.5 M KOH)
[32:17] Stepping pH = 11.04
[32:17] Dispensed 0.001270 mL of Base (0.5 M KOH)
[32:23] Stepping pH = 11.12
[32:23] Dispensed 0.000588 mL of Base (0.5 M KOH)
[32:28] Stepping pH = 11.15
[32:43] Stirrer speed set to 0
[32:53] Datapoint id 27 collected
[32:53] Charge balance equation is out by -24.0%
[32:53] Stirrer speed set to 50
[32:58] pH 11.15 -> 11.35
[32:58] Using cautious pH adjust
[32:58] Dispensed 0.001881 mL of Base (0.5 M KOH)
[33:03] Stepping pH = 11.25
[33:03] Dispensed 0.001505 mL of Base (0.5 M KOH)
[33:09] Stepping pH = 11.31
[33:09] Dispensed 0.000729 mL of Base (0.5 M KOH)
[33:14] Stepping pH = 11.34
[33:14] Dispensed 0.000353 mL of Base (0.5 M KOH)
[33:19] Stepping pH = 11.34
[33:34] Stirrer speed set to 0
[33:44] Datapoint id 28 collected
[33:44] Charge balance equation is out by -18.0%
[33:44] Stirrer speed set to 50
[33:49] pH 11.34 -> 11.54
[33:49] Using cautious pH adjust
[33:49] Dispensed 0.002916 mL of Base (0.5 M KOH)
[33:54] Stepping pH = 11.44
[33:55] Dispensed 0.002328 mL of Base (0.5 M KOH)

| | | | |
|---|---|---|---|
| Sample name: | **M16_octanol** | Experiment start time: | **3/1/2018 10:41:05 AM** |
| Assay name: | **pH-metric high logP** | Analyst: | **Pion** |
| Assay ID: | **18C-01008** | Instrument ID: | **T312060** |
| Filename: | **C:\Sirius_T3\Mehtap\20180228_exp28_logP_T3-2\18C-01008_M16_octanol_pH-metric high logP.t3r** | | |

#### Experiment Log (continued)

[34:00] Stepping pH = 11.52
[34:00] Dispensed 0.000564 mL of Base (0.5 M KOH)
[34:05] Stepping pH = 11.54
[34:20] Stirrer speed set to 0
[34:30] Datapoint id 29 collected
[34:30] Charge balance equation is out by 0.7%
[34:30] Stirrer speed set to 50
[34:35] pH 11.54 -> 11.74
[34:35] Using charge balance adjust
[34:36] Dispensed 0.009219 mL of Base (0.5 M KOH)
[34:56] Stirrer speed set to 0
[35:06] Datapoint id 30 collected
[35:06] Charge balance equation is out by -12.5%
[35:06] Stirrer speed set to 50
[35:11] pH 11.72 -> 11.92
[35:11] Using charge balance adjust
[35:11] Dispensed 0.014252 mL of Base (0.5 M KOH)
[35:31] Stirrer speed set to 0
[35:42] Datapoint id 31 collected
[35:42] Charge balance equation is out by -7.9%
[35:42] Stirrer speed set to 50
[35:47] pH 11.91 -> 12.05
[35:47] Using charge balance adjust
[35:47] Dispensed 0.014722 mL of Base (0.5 M KOH)
[36:07] Stirrer speed set to 0
[36:17] Datapoint id 32 collected
[36:17] Charge balance equation is out by -36.5%
[36:17] Titration 2 of 3
[36:17] Adding initial titrants
[36:17] Automatically add 0.10000 mL of Octanol
[36:20] Dispensed 0.100000 mL of Octanol
[36:20] Stirrer speed set to 10
[36:21] Stirrer speed set to 55
[36:21] Iterative adjust 12.04 -> 2.00
[36:21] pH 12.04 -> 2.00
[36:23] Dispensed 0.100000 mL of Acid (0.5 M HCl)
[36:28] pH 2.29 -> 2.00
[36:29] Dispensed 0.024106 mL of Acid (0.5 M HCl)
[36:34] pH 2.03 -> 2.00
[36:34] Dispensed 0.003104 mL of Acid (0.5 M HCl)
[37:25] Stirrer speed set to 0
[37:35] Datapoint id 33 collected
[37:35] Stirrer speed set to 55
[37:40] pH 1.97 -> 2.17
[37:40] Using cautious pH adjust
[37:40] Dispensed 0.009995 mL of Base (0.5 M KOH)
[37:45] Stepping pH = 2.06
[37:46] Dispensed 0.007291 mL of Base (0.5 M KOH)
[37:51] Stepping pH = 2.13
[37:51] Dispensed 0.002422 mL of Base (0.5 M KOH)
[37:56] Stepping pH = 2.17
[38:11] Stirrer speed set to 0
[38:21] Datapoint id 34 collected
[38:21] Charge balance equation is out by 1.4%
[38:21] Stirrer speed set to 55
[38:26] pH 2.17 -> 2.37
[38:26] Using charge balance adjust
[38:26] Dispensed 0.012347 mL of Base (0.5 M KOH)
[38:47] Stirrer speed set to 0

| | | | |
|---|---|---|---|
| Sample name: | **M16_octanol** | Experiment start time: | **3/1/2018 10:41:05 AM** |
| Assay name: | **pH-metric high logP** | Analyst: | **Pion** |
| Assay ID: | **18C-01008** | Instrument ID: | **T312060** |
| Filename: | **C:\Sirius_T3\Mehtap\20180228_exp28_logP_T3-2\18C-01008_M16_octanol_pH-metric high logP.t3r** | | |

#### Experiment Log (continued)

[39:01] Datapoint id 35 collected
[39:01] Charge balance equation is out by -4.4%
[39:01] Stirrer speed set to 55
[39:06] pH 2.37 -> 2.57
[39:06] Using charge balance adjust
[39:07] Dispensed 0.007855 mL of Base (0.5 M KOH)
[39:27] Stirrer speed set to 0
[39:37] Datapoint id 36 collected
[39:37] Charge balance equation is out by -10.3%
[39:37] Stirrer speed set to 55
[39:42] pH 2.56 -> 2.76
[39:42] Using charge balance adjust
[39:42] Dispensed 0.005198 mL of Base (0.5 M KOH)
[40:02] Stirrer speed set to 0
[40:12] Datapoint id 37 collected
[40:12] Charge balance equation is out by -18.8%
[40:12] Stirrer speed set to 55
[40:18] pH 2.73 -> 2.93
[40:18] Using cautious pH adjust
[40:18] Dispensed 0.001811 mL of Base (0.5 M KOH)
[40:23] Stepping pH = 2.81
[40:23] Dispensed 0.001552 mL of Base (0.5 M KOH)
[40:28] Stepping pH = 2.88
[40:28] Dispensed 0.000729 mL of Base (0.5 M KOH)
[40:33] Stepping pH = 2.91
[40:33] Dispensed 0.000259 mL of Base (0.5 M KOH)
[40:38] Stepping pH = 2.93
[40:54] Stirrer speed set to 0
[41:04] Datapoint id 38 collected
[41:04] Charge balance equation is out by -19.6%
[41:04] Stirrer speed set to 55
[41:09] pH 2.93 -> 3.13
[41:09] Using cautious pH adjust
[41:09] Dispensed 0.001223 mL of Base (0.5 M KOH)
[41:14] Stepping pH = 2.98
[41:14] Dispensed 0.001576 mL of Base (0.5 M KOH)
[41:19] Stepping pH = 3.07
[41:19] Dispensed 0.000800 mL of Base (0.5 M KOH)
[41:24] Stepping pH = 3.12
[41:40] Stirrer speed set to 0
[41:50] Datapoint id 39 collected
[41:50] Charge balance equation is out by -46.4%
[41:50] Stirrer speed set to 55
[41:55] pH 3.12 -> 3.32
[41:55] Using cautious pH adjust
[41:55] Dispensed 0.000917 mL of Base (0.5 M KOH)
[42:00] Stepping pH = 3.18
[42:00] Dispensed 0.001105 mL of Base (0.5 M KOH)
[42:05] Stepping pH = 3.28
[42:05] Dispensed 0.000400 mL of Base (0.5 M KOH)
[42:10] Stepping pH = 3.32
[42:25] Stirrer speed set to 0
[42:36] Datapoint id 40 collected
[42:36] Charge balance equation is out by -32.0%
[42:36] Stirrer speed set to 55
[42:41] pH 3.32 -> 3.52
[42:41] Using cautious pH adjust
[42:41] Dispensed 0.000776 mL of Base (0.5 M KOH)
[42:46] Stepping pH = 3.40

| | | | |
|---|---|---|---|
| Sample name: | **M16_octanol** | Experiment start time: | **3/1/2018 10:41:05 AM** |
| Assay name: | **pH-metric high logP** | Analyst: | **Pion** |
| Assay ID: | **18C-01008** | Instrument ID: | **T312060** |
| Filename: | **C:\Sirius_T3\Mehtap\20180228_exp28_logP_T3-2\18C-01008_M16_octanol_pH-metric high logP.t3r** | | |

#### Experiment Log (continued)

[42:46] Dispensed 0.000753 mL of Base (0.5 M KOH)
[42:51] Stepping pH = 3.48
[42:51] Dispensed 0.000282 mL of Base (0.5 M KOH)
[42:57] Stepping pH = 3.51
[43:12] Stirrer speed set to 0
[43:22] Datapoint id 41 collected
[43:22] Charge balance equation is out by -16.3%
[43:22] Stirrer speed set to 55
[43:27] pH 3.52 -> 3.72
[43:27] Using cautious pH adjust
[43:27] Dispensed 0.000753 mL of Base (0.5 M KOH)
[43:32] Stepping pH = 3.61
[43:32] Dispensed 0.000564 mL of Base (0.5 M KOH)
[43:37] Stepping pH = 3.69
[43:37] Dispensed 0.000141 mL of Base (0.5 M KOH)
[43:42] Stepping pH = 3.71
[43:57] Stirrer speed set to 0
[44:14] Datapoint id 42 collected
[44:14] Charge balance equation is out by 4.1%
[44:14] Stirrer speed set to 55
[44:19] pH 3.71 -> 3.91
[44:19] Using charge balance adjust
[44:19] Dispensed 0.001599 mL of Base (0.5 M KOH)
[44:39] Stirrer speed set to 0
[44:49] Datapoint id 43 collected
[44:49] Charge balance equation is out by 96.0%
[44:49] Stirrer speed set to 55
[44:55] pH 4.11 -> 4.31
[44:55] Using cautious pH adjust
[44:55] Dispensed 0.000917 mL of Base (0.5 M KOH)
[45:00] Stepping pH = 4.50
[45:15] Stirrer speed set to 0
[45:25] Datapoint id 44 collected
[45:25] Charge balance equation is out by 50.0%
[45:25] Stirrer speed set to 55
[45:31] pH 4.51 -> 4.71
[45:31] Using cautious pH adjust
[45:31] Dispensed 0.000800 mL of Base (0.5 M KOH)
[45:36] Stepping pH = 6.63
[45:51] Stirrer speed set to 0
[46:51] Datapoint id 45 collected
[46:51] Charge balance equation is out by 50.0%
[46:51] Stirrer speed set to 55
[46:56] pH 6.57 -> 6.77
[46:56] Using cautious pH adjust
[46:56] Dispensed 0.000047 mL of Base (0.5 M KOH)
[47:01] Stepping pH = 6.51
[47:01] Dispensed 0.000259 mL of Base (0.5 M KOH)
[47:06] Stepping pH = 8.44
[47:21] Stirrer speed set to 0
[48:05] Datapoint id 46 collected
[48:05] Charge balance equation is out by -289.9%
[48:05] Stirrer speed set to 55
[48:10] pH 8.37 -> 8.57
[48:10] Using cautious pH adjust
[48:10] Dispensed 0.000259 mL of Base (0.5 M KOH)
[48:15] Stepping pH = 9.01
[48:30] Stirrer speed set to 0
[48:49] Datapoint id 47 collected

| | | | |
|---|---|---|---|
| Sample name: | **M16_octanol** | Experiment start time: | **3/1/2018 10:41:05 AM** |
| Assay name: | **pH-metric high logP** | Analyst: | **Pion** |
| Assay ID: | **18C-01008** | Instrument ID: | **T312060** |
| Filename: | **C:\Sirius_T3\Mehtap\20180228_exp28_logP_T3-2\18C-01008_M16_octanol_pH-metric high logP.t3r** | | |

#### Experiment Log (continued)

[48:49] Charge balance equation is out by 50.0%
[48:49] Stirrer speed set to 55
[48:55] pH 8.98 -> 9.18
[48:55] Using cautious pH adjust
[48:55] Dispensed 0.000682 mL of Base (0.5 M KOH)
[49:00] Stepping pH = 9.60
[49:15] Stirrer speed set to 0
[49:26] Datapoint id 48 collected
[49:26] Charge balance equation is out by 50.0%
[49:26] Stirrer speed set to 55
[49:31] pH 9.59 -> 9.79
[49:31] Using cautious pH adjust
[49:32] Dispensed 0.000917 mL of Base (0.5 M KOH)
[49:37] Stepping pH = 9.96
[49:52] Stirrer speed set to 0
[50:02] Datapoint id 49 collected
[50:02] Charge balance equation is out by 50.0%
[50:02] Stirrer speed set to 55
[50:07] pH 9.95 -> 10.15
[50:07] Using cautious pH adjust
[50:07] Dispensed 0.000776 mL of Base (0.5 M KOH)
[50:13] Stepping pH = 10.16
[50:28] Stirrer speed set to 0
[50:38] Datapoint id 50 collected
[50:38] Charge balance equation is out by 50.0%
[50:38] Stirrer speed set to 55
[50:43] pH 10.16 -> 10.36
[50:43] Using cautious pH adjust
[50:43] Dispensed 0.000706 mL of Base (0.5 M KOH)
[50:48] Stepping pH = 10.30
[50:48] Dispensed 0.000259 mL of Base (0.5 M KOH)
[50:53] Stepping pH = 10.34
[50:53] Dispensed 0.000094 mL of Base (0.5 M KOH)
[50:58] Stepping pH = 10.35
[50:58] Dispensed 0.000094 mL of Base (0.5 M KOH)
[51:03] Stepping pH = 10.36
[51:19] Stirrer speed set to 0
[51:29] Datapoint id 51 collected
[51:29] Charge balance equation is out by 19.2%
[51:29] Stirrer speed set to 55
[51:34] pH 10.35 -> 10.55
[51:34] Using cautious pH adjust
[51:34] Dispensed 0.000682 mL of Base (0.5 M KOH)
[51:39] Stepping pH = 10.45
[51:39] Dispensed 0.000517 mL of Base (0.5 M KOH)
[51:44] Stepping pH = 10.51
[51:44] Dispensed 0.000282 mL of Base (0.5 M KOH)
[51:49] Stepping pH = 10.54
[51:49] Dispensed 0.000118 mL of Base (0.5 M KOH)
[51:55] Stepping pH = 10.55
[52:10] Stirrer speed set to 0
[52:20] Datapoint id 52 collected
[52:20] Charge balance equation is out by -16.9%
[52:20] Stirrer speed set to 55
[52:25] pH 10.54 -> 10.74
[52:25] Using cautious pH adjust
[52:25] Dispensed 0.000753 mL of Base (0.5 M KOH)
[52:30] Stepping pH = 10.61
[52:30] Dispensed 0.000823 mL of Base (0.5 M KOH)

#### Experiment Log (continued)

[52:36] Stepping pH = 10.68
[52:36] Dispensed 0.000588 mL of Base (0.5 M KOH)
[52:41] Stepping pH = 10.72
[52:41] Dispensed 0.000212 mL of Base (0.5 M KOH)
[52:46] Stepping pH = 10.73
[53:01] Stirrer speed set to 0
[53:11] Datapoint id 53 collected
[53:11] Charge balance equation is out by -57.5%
[53:11] Stirrer speed set to 55
[53:16] pH 10.73 -> 10.93
[53:16] Using cautious pH adjust
[53:16] Dispensed 0.000964 mL of Base (0.5 M KOH)
[53:21] Stepping pH = 10.79
[53:21] Dispensed 0.001176 mL of Base (0.5 M KOH)
[53:27] Stepping pH = 10.87
[53:27] Dispensed 0.000659 mL of Base (0.5 M KOH)
[53:32] Stepping pH = 10.92
[53:32] Dispensed 0.000165 mL of Base (0.5 M KOH)
[53:37] Stepping pH = 10.92
[53:52] Stirrer speed set to 0
[54:02] Datapoint id 54 collected
[54:02] Charge balance equation is out by -53.2%
[54:02] Stirrer speed set to 55
[54:07] pH 10.92 -> 11.12
[54:07] Using cautious pH adjust
[54:07] Dispensed 0.001364 mL of Base (0.5 M KOH)
[54:12] Stepping pH = 11.00
[54:13] Dispensed 0.001317 mL of Base (0.5 M KOH)
[54:18] Stepping pH = 11.07
[54:18] Dispensed 0.000706 mL of Base (0.5 M KOH)
[54:23] Stepping pH = 11.11
[54:23] Dispensed 0.000235 mL of Base (0.5 M KOH)
[54:28] Stepping pH = 11.12
[54:43] Stirrer speed set to 0
[54:53] Datapoint id 55 collected
[54:53] Charge balance equation is out by -31.5%
[54:53] Stirrer speed set to 55
[54:58] pH 11.11 -> 11.31
[54:58] Using cautious pH adjust
[54:58] Dispensed 0.002023 mL of Base (0.5 M KOH)
[55:03] Stepping pH = 11.19
[55:04] Dispensed 0.002093 mL of Base (0.5 M KOH)
[55:09] Stepping pH = 11.27
[55:09] Dispensed 0.001058 mL of Base (0.5 M KOH)
[55:14] Stepping pH = 11.30
[55:29] Stirrer speed set to 0
[55:39] Datapoint id 56 collected
[55:39] Charge balance equation is out by -27.4%
[55:39] Stirrer speed set to 55
[55:44] pH 11.30 -> 11.50
[55:44] Using cautious pH adjust
[55:44] Dispensed 0.003081 mL of Base (0.5 M KOH)
[55:49] Stepping pH = 11.40
[55:50] Dispensed 0.002446 mL of Base (0.5 M KOH)
[55:55] Stepping pH = 11.46
[55:55] Dispensed 0.001129 mL of Base (0.5 M KOH)
[56:00] Stepping pH = 11.49
[56:15] Stirrer speed set to 0
[56:25] Datapoint id 57 collected

Sample name: **M16_octanol** Experiment start time: **3/1/2018 10:41:05 AM**
Assay name: **pH-metric high logP** Analyst: **Pion**
Assay ID: **18C-01008** Instrument ID: **T312060**
Filename: **C:\Sirius_T3\Mehtap\20180228_exp28_logP_T3-2\18C-01008_M16_octanol_pH-metric high logP.t3r**

#### Experiment Log (continued)

[56:25] Charge balance equation is out by -7.6%
[56:25] Stirrer speed set to 55
[56:30] pH 11.49 -> 11.69
[56:30] Using charge balance adjust
[56:30] Dispensed 0.009572 mL of Base (0.5 M KOH)
[56:51] Stirrer speed set to 0
[57:01] Datapoint id 58 collected
[57:01] Charge balance equation is out by -13.0%
[57:01] Stirrer speed set to 55
[57:06] pH 11.67 -> 11.87
[57:06] Using charge balance adjust
[57:06] Dispensed 0.014652 mL of Base (0.5 M KOH)
[57:26] Stirrer speed set to 0
[57:36] Datapoint id 59 collected
[57:36] Charge balance equation is out by -6.6%
[57:36] Stirrer speed set to 55
[57:41] pH 11.86 -> 12.05
[57:41] Using charge balance adjust
[57:42] Dispensed 0.022013 mL of Base (0.5 M KOH)
[58:02] Stirrer speed set to 0
[58:12] Datapoint id 60 collected
[58:12] Charge balance equation is out by -13.7%
[58:12] Titration 3 of 3
[58:12] Adding initial titrants
[58:12] Automatically add 0.30000 mL of Octanol
[58:19] Dispensed 0.300000 mL of Octanol
[58:19] Stirrer speed set to 10
[58:20] Stirrer speed set to 60
[58:20] Iterative adjust 12.04 -> 2.00
[58:20] pH 12.04 -> 2.00
[58:23] Dispensed 0.100000 mL of Acid (0.5 M HCl)
[58:28] pH 2.44 -> 2.00
[58:29] Dispensed 0.036007 mL of Acid (0.5 M HCl)
[58:34] pH 2.04 -> 2.00
[58:34] Dispensed 0.005033 mL of Acid (0.5 M HCl)
[59:25] Stirrer speed set to 0
[59:35] Datapoint id 61 collected
[59:35] Stirrer speed set to 60
[59:40] pH 1.97 -> 2.17
[59:40] Using cautious pH adjust
[59:40] Dispensed 0.011359 mL of Base (0.5 M KOH)
[59:45] Stepping pH = 2.05
[59:45] Dispensed 0.009595 mL of Base (0.5 M KOH)
[59:51] Stepping pH = 2.15
[59:51] Dispensed 0.002140 mL of Base (0.5 M KOH)
[59:56] Stepping pH = 2.17
[1:00:11] Stirrer speed set to 0
[1:00:28] Datapoint id 62 collected
[1:00:28] Charge balance equation is out by -1.7%
[1:00:28] Stirrer speed set to 60
[1:00:34] pH 2.18 -> 2.38
[1:00:34] Using charge balance adjust
[1:00:34] Dispensed 0.013993 mL of Base (0.5 M KOH)
[1:00:54] Stirrer speed set to 0
[1:01:04] Datapoint id 63 collected
[1:01:04] Charge balance equation is out by -6.3%
[1:01:04] Stirrer speed set to 60
[1:01:09] pH 2.38 -> 2.58
[1:01:09] Using charge balance adjust

| | | | |
|---|---|---|---|
| Sample name: | **M16_octanol** | Experiment start time: | **3/1/2018 10:41:05 AM** |
| Assay name: | **pH-metric high logP** | Analyst: | **Pion** |
| Assay ID: | **18C-01008** | Instrument ID: | **T312060** |
| Filename: | **C:\Sirius_T3\Mehtap\20180228_exp28_logP_T3-2\18C-01008_M16_octanol_pH-metric high logP.t3r** | | |

#### Experiment Log (continued)

[1:01:10] Dispensed 0.009008 mL of Base (0.5 M KOH)
[1:01:30] Stirrer speed set to 0
[1:01:40] Datapoint id 64 collected
[1:01:40] Charge balance equation is out by -15.0%
[1:01:40] Stirrer speed set to 60
[1:01:45] pH 2.56 -> 2.76
[1:01:45] Using cautious pH adjust
[1:01:45] Dispensed 0.003057 mL of Base (0.5 M KOH)
[1:01:50] Stepping pH = 2.62
[1:01:50] Dispensed 0.003198 mL of Base (0.5 M KOH)
[1:01:55] Stepping pH = 2.72
[1:01:56] Dispensed 0.001035 mL of Base (0.5 M KOH)
[1:02:01] Stepping pH = 2.75
[1:02:16] Stirrer speed set to 0
[1:02:26] Datapoint id 65 collected
[1:02:26] Charge balance equation is out by -19.0%
[1:02:26] Stirrer speed set to 60
[1:02:31] pH 2.76 -> 2.96
[1:02:31] Using cautious pH adjust
[1:02:31] Dispensed 0.002046 mL of Base (0.5 M KOH)
[1:02:36] Stepping pH = 2.84
[1:02:36] Dispensed 0.001834 mL of Base (0.5 M KOH)
[1:02:41] Stepping pH = 2.92
[1:02:41] Dispensed 0.000776 mL of Base (0.5 M KOH)
[1:02:47] Stepping pH = 2.95
[1:03:02] Stirrer speed set to 0
[1:03:12] Datapoint id 66 collected
[1:03:12] Charge balance equation is out by -13.9%
[1:03:12] Stirrer speed set to 60
[1:03:17] pH 2.96 -> 3.16
[1:03:17] Using charge balance adjust
[1:03:17] Dispensed 0.002940 mL of Base (0.5 M KOH)
[1:03:37] Stirrer speed set to 0
[1:03:55] Datapoint id 67 collected
[1:03:55] Charge balance equation is out by -11.3%
[1:03:55] Stirrer speed set to 60
[1:04:00] pH 3.14 -> 3.34
[1:04:00] Using charge balance adjust
[1:04:00] Dispensed 0.002399 mL of Base (0.5 M KOH)
[1:04:20] Stirrer speed set to 0
[1:04:30] Datapoint id 68 collected
[1:04:30] Charge balance equation is out by 13.3%
[1:04:30] Stirrer speed set to 60
[1:04:35] pH 3.37 -> 3.57
[1:04:35] Using charge balance adjust
[1:04:35] Dispensed 0.002117 mL of Base (0.5 M KOH)
[1:04:56] Stirrer speed set to 0
[1:05:06] Datapoint id 69 collected
[1:05:06] Charge balance equation is out by 44.0%
[1:05:06] Stirrer speed set to 60
[1:05:11] pH 3.67 -> 3.87
[1:05:11] Using cautious pH adjust
[1:05:11] Dispensed 0.001035 mL of Base (0.5 M KOH)
[1:05:16] Stepping pH = 3.90
[1:05:31] Stirrer speed set to 0
[1:05:41] Datapoint id 70 collected
[1:05:41] Charge balance equation is out by 50.0%
[1:05:41] Stirrer speed set to 60
[1:05:46] pH 3.90 -> 4.10

| | | | |
|---|---|---|---|
| Sample name: | **M16_octanol** | Experiment start time: | **3/1/2018 10:41:05 AM** |
| Assay name: | **pH-metric high logP** | Analyst: | **Pion** |
| Assay ID: | **18C-01008** | Instrument ID: | **T312060** |
| Filename: | **C:\Sirius_T3\Mehtap\20180228_exp28_logP_T3-2\18C-01008_M16_octanol_pH-metric high logP.t3r** | | |

#### Experiment Log (continued)

[1:05:46] Using cautious pH adjust
[1:05:46] Dispensed 0.000988 mL of Base (0.5 M KOH)
[1:05:51] Stepping pH = 4.36
[1:06:06] Stirrer speed set to 0
[1:06:17] Datapoint id 71 collected
[1:06:17] Charge balance equation is out by 50.0%
[1:06:17] Stirrer speed set to 60
[1:06:22] pH 4.36 -> 4.56
[1:06:22] Using cautious pH adjust
[1:06:22] Dispensed 0.000682 mL of Base (0.5 M KOH)
[1:06:27] Stepping pH = 5.66
[1:06:42] Stirrer speed set to 0
[1:07:02] Datapoint id 72 collected
[1:07:02] Charge balance equation is out by 50.0%
[1:07:02] Stirrer speed set to 60
[1:07:08] pH 5.46 -> 5.66
[1:07:08] Using cautious pH adjust
[1:07:08] Dispensed 0.000094 mL of Base (0.5 M KOH)
[1:07:13] Stepping pH = 5.60
[1:07:13] Dispensed 0.000047 mL of Base (0.5 M KOH)
[1:07:18] Stepping pH = 5.79
[1:07:33] Stirrer speed set to 0
[1:07:53] Datapoint id 73 collected
[1:07:53] Charge balance equation is out by 32.1%
[1:07:53] Stirrer speed set to 60
[1:07:58] pH 5.73 -> 5.93
[1:07:58] Using cautious pH adjust
[1:07:58] Dispensed 0.000071 mL of Base (0.5 M KOH)
[1:08:03] Stepping pH = 6.04
[1:08:18] Stirrer speed set to 0
[1:08:36] Datapoint id 74 collected
[1:08:36] Charge balance equation is out by 50.0%
[1:08:36] Stirrer speed set to 60
[1:08:41] pH 5.99 -> 6.19
[1:08:41] Using cautious pH adjust
[1:08:41] Dispensed 0.000047 mL of Base (0.5 M KOH)
[1:08:46] Stepping pH = 6.27
[1:09:01] Stirrer speed set to 0
[1:09:29] Datapoint id 75 collected
[1:09:29] Charge balance equation is out by 50.0%
[1:09:29] Stirrer speed set to 60
[1:09:34] pH 6.09 -> 6.29
[1:09:34] Using cautious pH adjust
[1:09:35] Dispensed 0.000047 mL of Base (0.5 M KOH)
[1:09:40] Stepping pH = 6.48
[1:09:55] Stirrer speed set to 0
[1:10:29] Datapoint id 76 collected
[1:10:29] Charge balance equation is out by 50.0%
[1:10:29] Stirrer speed set to 60
[1:10:34] pH 6.42 -> 6.62
[1:10:34] Using cautious pH adjust
[1:10:34] Dispensed 0.000047 mL of Base (0.5 M KOH)
[1:10:39] Stepping pH = 6.76
[1:10:54] Stirrer speed set to 0
[1:11:35] Datapoint id 77 collected
[1:11:35] Charge balance equation is out by 50.0%
[1:11:35] Stirrer speed set to 60
[1:11:40] pH 6.79 -> 6.99
[1:11:40] Using cautious pH adjust

| | | | |
|---|---|---|---|
| Sample name: | **M16_octanol** | Experiment start time: | **3/1/2018 10:41:05 AM** |
| Assay name: | **pH-metric high logP** | Analyst: | **Pion** |
| Assay ID: | **18C-01008** | Instrument ID: | **T312060** |
| Filename: | **C:\Sirius_T3\Mehtap\20180228_exp28_logP_T3-2\18C-01008_M16_octanol_pH-metric high logP.t3r** | | |

#### Experiment Log (continued)

[1:11:40] Dispensed 0.000024 mL of Base (0.5 M KOH)
[1:11:45] Stepping pH = 6.86
[1:11:45] Dispensed 0.000024 mL of Base (0.5 M KOH)
[1:11:50] Stepping pH = 7.15
[1:12:05] Stirrer speed set to 0
[1:13:05] Datapoint id 78 collected
[1:13:05] Charge balance equation is out by -4.3%
[1:13:05] Stirrer speed set to 60
[1:13:10] pH 6.87 -> 7.07
[1:13:10] Using charge balance adjust
[1:13:10] Dispensed 0.000047 mL of Base (0.5 M KOH)
[1:13:31] Stirrer speed set to 0
[1:14:31] Datapoint id 79 collected
[1:14:31] Charge balance equation is out by 129.5%
[1:14:31] Stirrer speed set to 60
[1:14:36] pH 7.26 -> 7.46
[1:14:36] Using cautious pH adjust
[1:14:36] Dispensed 0.000024 mL of Base (0.5 M KOH)
[1:14:41] Stepping pH = 7.23
[1:14:41] Dispensed 0.000118 mL of Base (0.5 M KOH)
[1:14:46] Stepping pH = 8.40
[1:15:01] Stirrer speed set to 0
[1:16:01] Datapoint id 80 collected
[1:16:01] Charge balance equation is out by -238.7%
[1:16:01] Stirrer speed set to 60
[1:16:06] pH 8.30 -> 8.50
[1:16:06] Using cautious pH adjust
[1:16:06] Dispensed 0.000118 mL of Base (0.5 M KOH)
[1:16:11] Stepping pH = 8.60
[1:16:27] Stirrer speed set to 0
[1:16:46] Datapoint id 81 collected
[1:16:46] Charge balance equation is out by 50.0%
[1:16:46] Stirrer speed set to 60
[1:16:51] pH 8.54 -> 8.74
[1:16:51] Using cautious pH adjust
[1:16:51] Dispensed 0.000188 mL of Base (0.5 M KOH)
[1:16:56] Stepping pH = 8.96
[1:17:11] Stirrer speed set to 0
[1:17:25] Datapoint id 82 collected
[1:17:25] Charge balance equation is out by 50.0%
[1:17:25] Stirrer speed set to 60
[1:17:30] pH 8.94 -> 9.14
[1:17:30] Using cautious pH adjust
[1:17:30] Dispensed 0.000423 mL of Base (0.5 M KOH)
[1:17:35] Stepping pH = 9.41
[1:17:50] Stirrer speed set to 0
[1:18:02] Datapoint id 83 collected
[1:18:02] Charge balance equation is out by 50.0%
[1:18:02] Stirrer speed set to 60
[1:18:07] pH 9.39 -> 9.59
[1:18:07] Using cautious pH adjust
[1:18:08] Dispensed 0.000776 mL of Base (0.5 M KOH)
[1:18:13] Stepping pH = 9.83
[1:18:28] Stirrer speed set to 0
[1:18:38] Datapoint id 84 collected
[1:18:38] Charge balance equation is out by 50.0%
[1:18:38] Stirrer speed set to 60
[1:18:43] pH 9.82 -> 10.02
[1:18:43] Using cautious pH adjust

Sample name: **M16_octanol** Experiment start time: **3/1/2018 10:41:05 AM**
Assay name: **pH-metric high logP** Analyst: **Pion**
Assay ID: **18C-01008** Instrument ID: **T312060**
Filename: **C:\Sirius_T3\Mehtap\20180228_exp28_logP_T3-2\18C-01008_M16_octanol_pH-metric high logP.t3r**

#### Experiment Log (continued)

[1:18:43] Dispensed 0.000988 mL of Base (0.5 M KOH)
[1:18:48] Stepping pH = 10.15
[1:19:04] Stirrer speed set to 0
[1:19:14] Datapoint id 85 collected
[1:19:14] Charge balance equation is out by 50.0%
[1:19:14] Stirrer speed set to 60
[1:19:19] pH 10.14 -> 10.34
[1:19:19] Using cautious pH adjust
[1:19:19] Dispensed 0.000988 mL of Base (0.5 M KOH)
[1:19:24] Stepping pH = 10.37
[1:19:39] Stirrer speed set to 0
[1:19:49] Datapoint id 86 collected
[1:19:49] Charge balance equation is out by 50.1%
[1:19:49] Stirrer speed set to 60
[1:19:54] pH 10.36 -> 10.56
[1:19:54] Using cautious pH adjust
[1:19:54] Dispensed 0.000964 mL of Base (0.5 M KOH)
[1:19:59] Stepping pH = 10.51
[1:19:59] Dispensed 0.000306 mL of Base (0.5 M KOH)
[1:20:05] Stepping pH = 10.54
[1:20:05] Dispensed 0.000141 mL of Base (0.5 M KOH)
[1:20:10] Stepping pH = 10.55
[1:20:25] Stirrer speed set to 0
[1:20:35] Datapoint id 87 collected
[1:20:35] Charge balance equation is out by 27.0%
[1:20:35] Stirrer speed set to 60
[1:20:40] pH 10.55 -> 10.75
[1:20:40] Using cautious pH adjust
[1:20:40] Dispensed 0.001058 mL of Base (0.5 M KOH)
[1:20:45] Stepping pH = 10.66
[1:20:45] Dispensed 0.000659 mL of Base (0.5 M KOH)
[1:20:50] Stepping pH = 10.71
[1:20:50] Dispensed 0.000329 mL of Base (0.5 M KOH)
[1:20:56] Stepping pH = 10.73
[1:20:56] Dispensed 0.000188 mL of Base (0.5 M KOH)
[1:21:01] Stepping pH = 10.74
[1:21:16] Stirrer speed set to 0
[1:21:26] Datapoint id 88 collected
[1:21:26] Charge balance equation is out by -5.7%
[1:21:26] Stirrer speed set to 60
[1:21:31] pH 10.74 -> 10.94
[1:21:31] Using charge balance adjust
[1:21:31] Dispensed 0.002540 mL of Base (0.5 M KOH)
[1:21:51] Stirrer speed set to 0
[1:22:01] Datapoint id 89 collected
[1:22:01] Charge balance equation is out by -13.9%
[1:22:01] Stirrer speed set to 60
[1:22:06] pH 10.92 -> 11.12
[1:22:06] Using charge balance adjust
[1:22:07] Dispensed 0.003340 mL of Base (0.5 M KOH)
[1:22:27] Stirrer speed set to 0
[1:22:37] Datapoint id 90 collected
[1:22:37] Charge balance equation is out by -9.0%
[1:22:37] Stirrer speed set to 60
[1:22:42] pH 11.10 -> 11.30
[1:22:42] Using charge balance adjust
[1:22:42] Dispensed 0.004774 mL of Base (0.5 M KOH)
[1:23:02] Stirrer speed set to 0
[1:23:18] Datapoint id 91 collected

| | | | |
|---|---|---|---|
| Sample name: | **M16_octanol** | Experiment start time: | **3/1/2018 10:41:05 AM** |
| Assay name: | **pH-metric high logP** | Analyst: | **Pion** |
| Assay ID: | **18C-01008** | Instrument ID: | **T312060** |
| Filename: | **C:\Sirius_T3\Mehtap\20180228_exp28_logP_T3-2\18C-01008_M16_octanol_pH-metric high logP.t3r** | | |

#### Experiment Log (continued)

[1:23:18] Charge balance equation is out by -23.1%
[1:23:18] Stirrer speed set to 60
[1:23:23] pH 11.26 -> 11.46
[1:23:23] Using cautious pH adjust
[1:23:24] Dispensed 0.003340 mL of Base (0.5 M KOH)
[1:23:29] Stepping pH = 11.35
[1:23:29] Dispensed 0.002893 mL of Base (0.5 M KOH)
[1:23:34] Stepping pH = 11.43
[1:23:34] Dispensed 0.001270 mL of Base (0.5 M KOH)
[1:23:39] Stepping pH = 11.45
[1:23:39] Dispensed 0.000447 mL of Base (0.5 M KOH)
[1:23:45] Stepping pH = 11.46
[1:24:00] Stirrer speed set to 0
[1:24:10] Datapoint id 92 collected
[1:24:10] Charge balance equation is out by -19.0%
[1:24:10] Stirrer speed set to 60
[1:24:15] pH 11.46 -> 11.66
[1:24:15] Using cautious pH adjust
[1:24:15] Dispensed 0.005174 mL of Base (0.5 M KOH)
[1:24:20] Stepping pH = 11.56
[1:24:20] Dispensed 0.004092 mL of Base (0.5 M KOH)
[1:24:25] Stepping pH = 11.62
[1:24:26] Dispensed 0.001976 mL of Base (0.5 M KOH)
[1:24:31] Stepping pH = 11.65
[1:24:46] Stirrer speed set to 0
[1:24:56] Datapoint id 93 collected
[1:24:56] Charge balance equation is out by -7.9%
[1:24:56] Stirrer speed set to 60
[1:25:01] pH 11.65 -> 11.85
[1:25:01] Using charge balance adjust
[1:25:01] Dispensed 0.016298 mL of Base (0.5 M KOH)
[1:25:22] Stirrer speed set to 0
[1:25:32] Datapoint id 94 collected
[1:25:32] Charge balance equation is out by -11.2%
[1:25:32] Stirrer speed set to 60
[1:25:37] pH 11.84 -> 12.04
[1:25:37] Using charge balance adjust
[1:25:37] Dispensed 0.025329 mL of Base (0.5 M KOH)
[1:25:58] Stirrer speed set to 0
[1:26:08] Datapoint id 95 collected
[1:26:08] Charge balance equation is out by -9.5%
[1:26:08] Argon flow rate set to 0
[1:26:12] Titrator arm moved over Titration position
