## Supplementary material for "Octanol-water partition coefficient measurements for the SAMPL6 Blind Prediction Challenge": SM15_18C-01009_M16_octanol_pH-metric high logP_report.pdf

| | | | |
|---|---|---|---|
| Sample name: | **M16_octanol** | Experiment start time: | **3/1/2018 12:08:09 PM** |
| Assay name: | **pH-metric high logP** | Analyst: | **Pion** |
| Assay ID: | **18C-01009** | Instrument ID: | **T312060** |
| Filename: | **C:\Sirius_T3\Mehtap\20180228_exp28_logP_T3-2\18C-01009_M16_octanol_pH-metric high logP.t3r** | | |

#### pH-metric Result

| | | |
|---|---|---|
| logP (XH2 +) | -0.17 | ±0.20 (n=50) |
| logP (neutral XH) | 3.04 | ±0.01 (n=50) |
| logP (X -) | -1.62 | |

#### 18C-01009 Points 1 to 31

| | |
|---|---|
| M16_octanol concentration factor | 0.767 |
| Carbonate | 0.0840 mM |
| Acidity error | 3.05344 mM |

#### 18C-01009 Points 32 to 62

| | |
|---|---|
| M16_octanol concentration factor | 0.794 |
| Carbonate | 0.1938 mM |
| Acidity error | 2.68082 mM |

#### 18C-01009 Points 63 to 92

| | |
|---|---|
| M16_octanol concentration factor | 0.836 |
| Carbonate | 0.2810 mM |
| Acidity error | 2.47693 mM |

#### Warnings and errors

| | |
|---|---|
| Errors | None |
| Warnings | None |

#### Sample logD and percent species

| pH | M16_octanol logD | M16_octanol M16_octanolH2 | M16_octanol M16_octanolH | M16_octanol M16_octanol | M16_octanol M16_octanolH2* | M16_octanol M16_octanolH* | M16_octanol M16_octanol* | Comment |
|---|---|---|---|---|---|---|---|---|
| 1.000 | -0.05 | 52.62 % | 0.01 % | 0.00 % | 35.72 % | 11.64 % | 0.00 % | |
| 1.200 | 0.01 | 49.26 % | 0.02 % | 0.00 % | 33.44 % | 17.28 % | 0.00 % | Stomach pH |
| 2.000 | 0.46 | 25.68 % | 0.05 % | 0.00 % | 17.44 % | 56.83 % | 0.00 % | |
| 3.000 | 1.35 | 4.20 % | 0.08 % | 0.00 % | 2.85 % | 92.87 % | 0.00 % | |
| 4.000 | 2.27 | 0.45 % | 0.09 % | 0.00 % | 0.30 % | 99.16 % | 0.00 % | |
| 5.000 | 2.87 | 0.05 % | 0.09 % | 0.00 % | 0.03 % | 99.83 % | 0.00 % | |
| 6.000 | 3.02 | 0.00 % | 0.09 % | 0.00 % | 0.00 % | 99.90 % | 0.00 % | |
| 6.500 | 3.04 | 0.00 % | 0.09 % | 0.00 % | 0.00 % | 99.91 % | 0.00 % | |
| 7.000 | 3.04 | 0.00 % | 0.09 % | 0.00 % | 0.00 % | 99.91 % | 0.00 % | |
| 7.400 | 3.03 | 0.00 % | 0.09 % | 0.00 % | 0.00 % | 99.91 % | 0.00 % | Blood pH |
| 8.000 | 3.00 | 0.00 % | 0.09 % | 0.01 % | 0.00 % | 99.90 % | 0.00 % | |
| 9.000 | 2.71 | 0.00 % | 0.09 % | 0.10 % | 0.00 % | 99.80 % | 0.00 % | |
| 10.000 | 1.95 | 0.00 % | 0.09 % | 1.02 % | 0.00 % | 98.86 % | 0.02 % | |
| 11.000 | 0.98 | 0.00 % | 0.08 % | 9.35 % | 0.00 % | 90.34 % | 0.22 % | |
| 12.000 | -0.00 | 0.00 % | 0.04 % | 50.23 % | 0.00 % | 48.52 % | 1.20 % | |

| | | | |
|---|---|---|---|
| Sample name: | **M16_octanol** | Experiment start time: | **3/1/2018 12:08:09 PM** |
| Assay name: | **pH-metric high logP** | Analyst: | **Pion** |
| Assay ID: | **18C-01009** | Instrument ID: | **T312060** |
| Filename: | **C:\Sirius_T3\Mehtap\20180228_exp28_logP_T3-2\18C-01009_M16_octanol_pH-metric high logP.t3r** | | |

#### Graphs

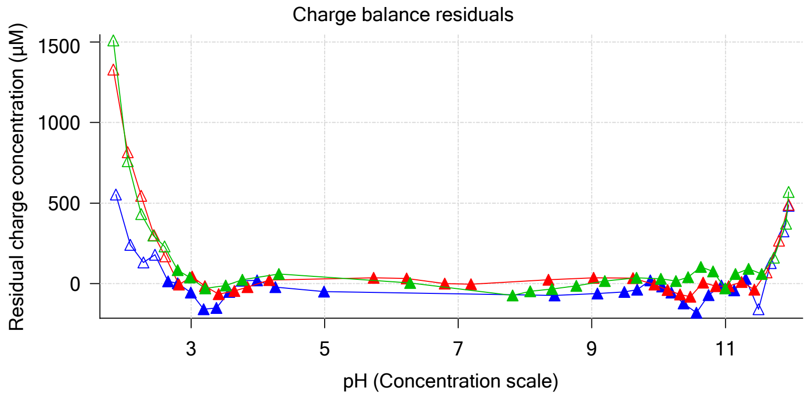


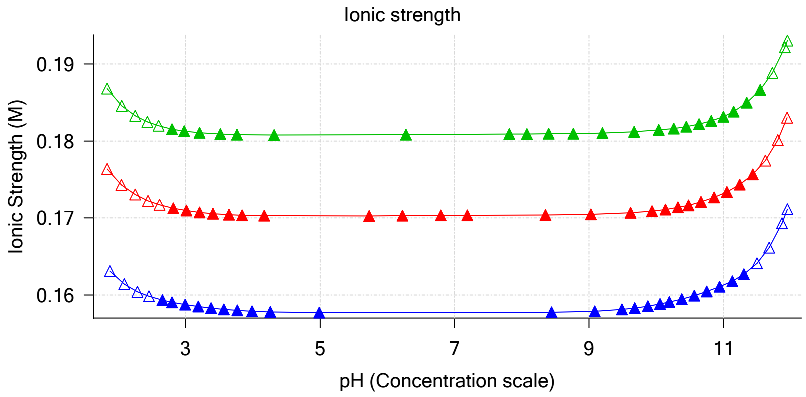


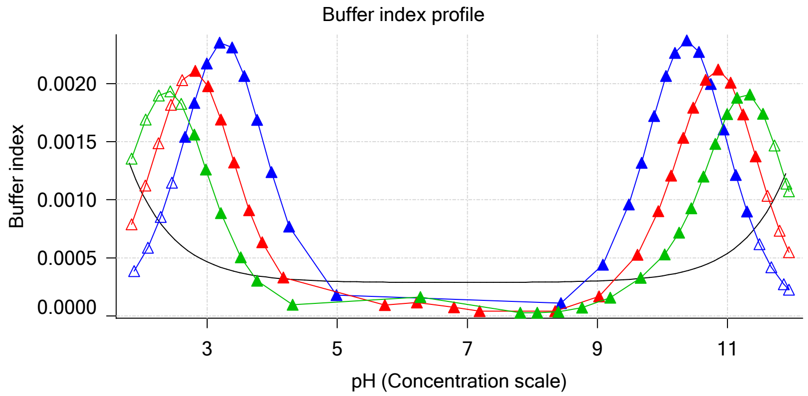


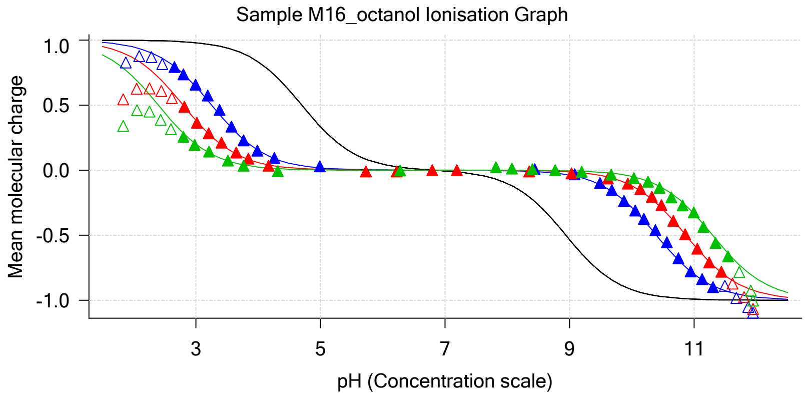


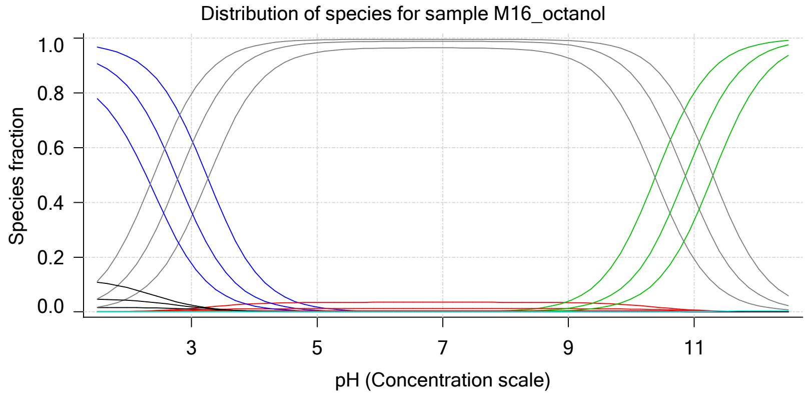


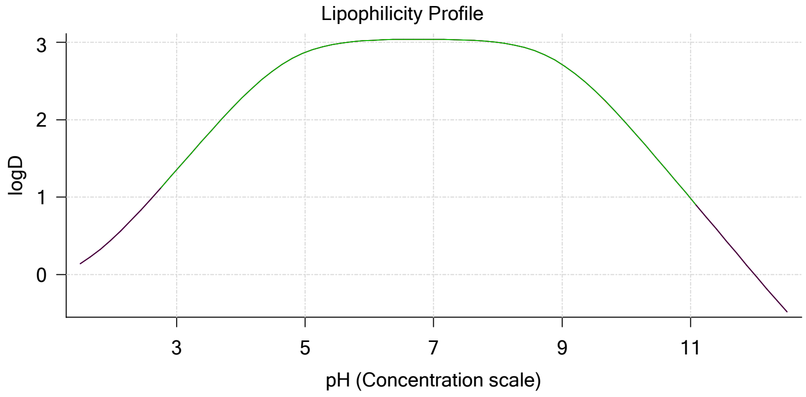


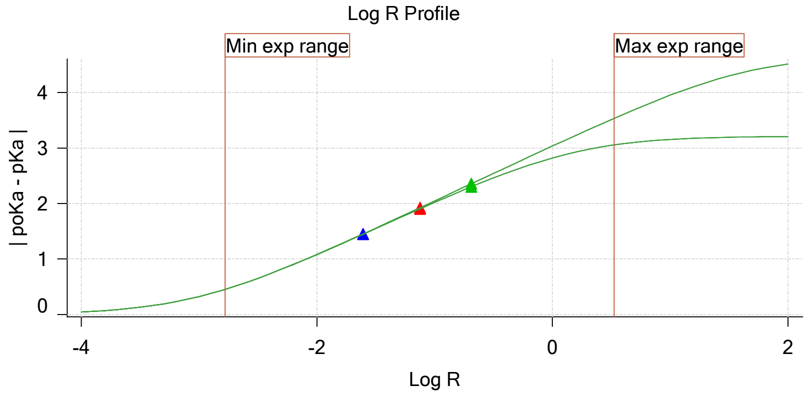


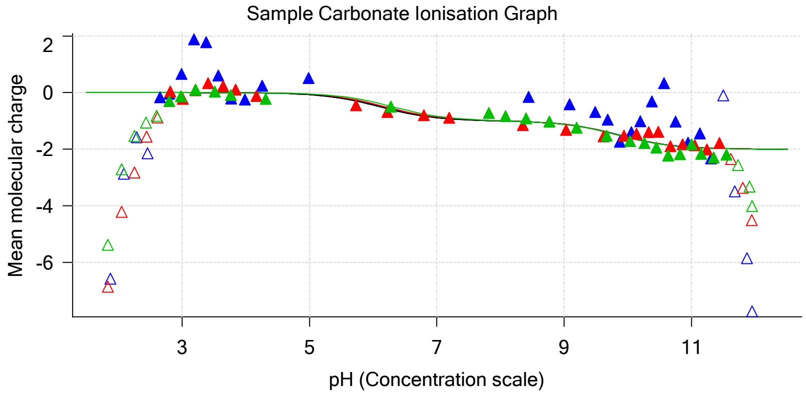

| | | | |
|---|---|---|---|
| Sample name: | **M16_octanol** | Experiment start time: | **3/1/2018 12:08:09 PM** |
| Assay name: | **pH-metric high logP** | Analyst: | **Pion** |
| Assay ID: | **18C-01009** | Instrument ID: | **T312060** |
| Filename: | **C:\Sirius_T3\Mehtap\20180228_exp28_logP_T3-2\18C-01009_M16_octanol_pH-metric high logP.t3r** | | |

#### Graphs (continued)

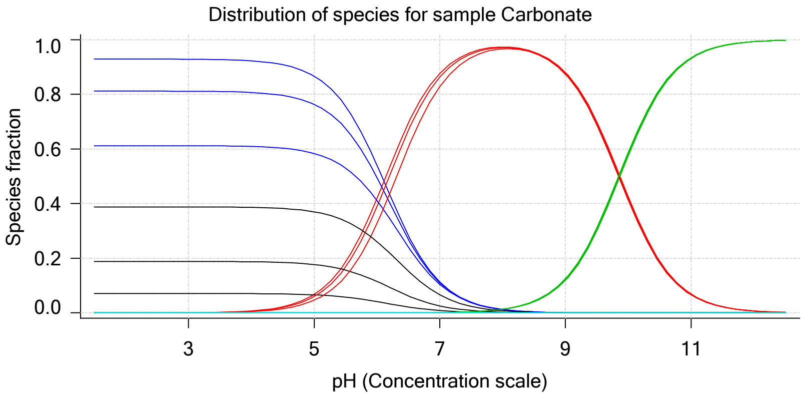

| | | | |
|---|---|---|---|
| Sample name: | **M16_octanol** | Experiment start time: | **3/1/2018 12:08:09 PM** |
| Assay name: | **pH-metric high logP** | Analyst: | **Pion** |
| Assay ID: | **18C-01009** | Instrument ID: | **T312060** |
| Filename: | **C:\Sirius_T3\Mehtap\20180228_exp28_logP_T3-2\18C-01009_M16_octanol_pH-metric high logP.t3r** | | |

# pH-metric high logP   Titration 1 of 3   18C-01009 Points 1 to 31

#### Overall results

| | |
|---|---|
| RMSD | 0.138 |
| Average ionic strength | 0.159 M |
| Average temperature | 25.0°C |
| Partition ratio | 0.0247 : 1 |
| Analyte concentration range | 4991.2 µM to 5365.1 µM |
| Total points considered | 23 of 31 |

#### Warnings and errors

| | |
|---|---|
| Errors | None |
| Warnings | Excessive acidity error present |

#### Four-Plus parameters

| | | | |
|---|---|---|---|
| Alpha | 0.130 | 3/1/2018 12:08:08 PM | C:\Sirius_T3\HCl18B27.t3r |
| S | 0.9970 | 3/1/2018 12:08:08 PM | C:\Sirius_T3\HCl18B27.t3r |
| jH | 0.8 | 3/1/2018 12:08:08 PM | C:\Sirius_T3\HCl18B27.t3r |
| jOH | -0.4 | 3/1/2018 12:08:08 PM | C:\Sirius_T3\HCl18B27.t3r |

#### Titrants

| | | | |
|---|---|---|---|
| 0.50 M HCl | 0.993513 | 3/1/2018 12:08:08 PM | C:\Sirius_T3\HCl18B27.t3r |
| 0.50 M KOH | 0.999845 | 3/1/2018 12:08:09 PM | C:\Sirius_T3\KOH18B27.t3r |

#### Sample

| | |
|---|---|
| M16_octanol concentration factor | 0.767 |
| Base pKa 1 | 4.70 |
| Acid pKa 2 | 8.94 |
| logP (XH2 +) | 0.85 |
| logP (neutral XH) | 3.08 |
| logP (X -) | -1.62 |

#### Sample graphs

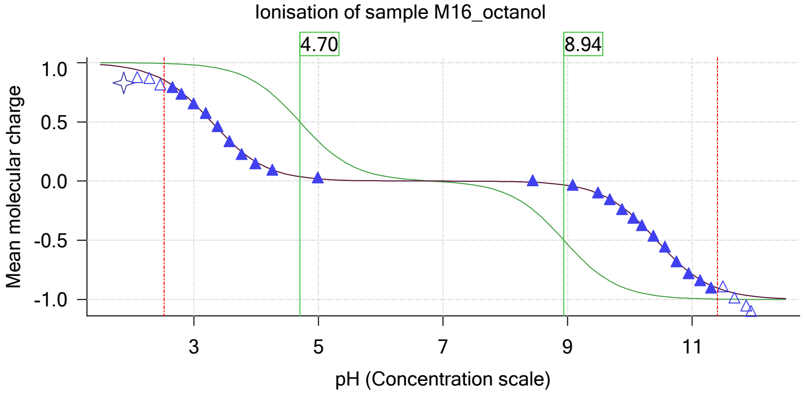


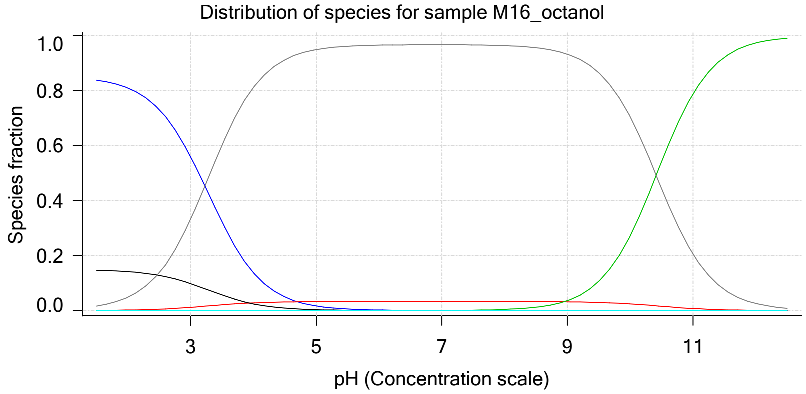

| | | | |
|---|---|---|---|
| Sample name: | **M16_octanol** | Experiment start time: | **3/1/2018 12:08:09 PM** |
| Assay name: | **pH-metric high logP** | Analyst: | **Pion** |
| Assay ID: | **18C-01009** | Instrument ID: | **T312060** |
| Filename: | **C:\Sirius_T3\Mehtap\20180228_exp28_logP_T3-2\18C-01009_M16_octanol_pH-metric high logP.t3r** | | |

#### Sample graphs (continued)

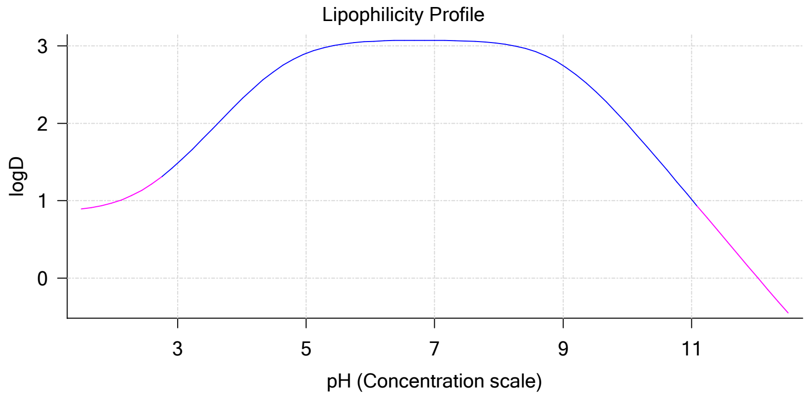


#### Sample logD and percent species

| pH | M16_octanol logD | M16_octanol M16_octanolH2 | M16_octanol M16_octanolH | M16_octanol M16_octanol | M16_octanol M16_octanolH2* | M16_octanol M16_octanolH* | M16_octanol M16_octanol* | Comment |
|---|---|---|---|---|---|---|---|---|
| 1.000 | 0.86 | 84.71 % | 0.02 % | 0.00 % | 14.77 % | 0.50 % | 0.00 % | |
| 1.200 | 0.87 | 84.45 % | 0.03 % | 0.00 % | 14.73 % | 0.79 % | 0.00 % | Stomach pH |
| 2.000 | 0.97 | 80.93 % | 0.16 % | 0.00 % | 14.12 % | 4.80 % | 0.00 % | |
| 3.000 | 1.48 | 55.95 % | 1.12 % | 0.00 % | 9.76 % | 33.17 % | 0.00 % | |
| 4.000 | 2.31 | 13.69 % | 2.73 % | 0.00 % | 2.39 % | 81.18 % | 0.00 % | |
| 5.000 | 2.90 | 1.60 % | 3.19 % | 0.00 % | 0.28 % | 94.92 % | 0.00 % | |
| 6.000 | 3.06 | 0.16 % | 3.25 % | 0.00 % | 0.03 % | 96.56 % | 0.00 % | |
| 6.500 | 3.07 | 0.05 % | 3.25 % | 0.01 % | 0.01 % | 96.67 % | 0.00 % | |
| 7.000 | 3.07 | 0.02 % | 3.25 % | 0.04 % | 0.00 % | 96.69 % | 0.00 % | |
| 7.400 | 3.07 | 0.01 % | 3.25 % | 0.09 % | 0.00 % | 96.65 % | 0.00 % | Blood pH |
| 8.000 | 3.03 | 0.00 % | 3.24 % | 0.37 % | 0.00 % | 96.38 % | 0.00 % | |
| 9.000 | 2.75 | 0.00 % | 3.14 % | 3.60 % | 0.00 % | 93.26 % | 0.00 % | |
| 10.000 | 1.98 | 0.00 % | 2.37 % | 27.21 % | 0.00 % | 70.41 % | 0.02 % | |
| 11.000 | 1.02 | 0.00 % | 0.69 % | 78.86 % | 0.00 % | 20.41 % | 0.05 % | |
| 12.000 | 0.03 | 0.00 % | 0.08 % | 97.34 % | 0.00 % | 2.52 % | 0.06 % | |

#### Carbonate and acidity

| | |
|---|---|
| Carbonate | 0.084 mM |
| Acidity error | 3.053 mM |

#### Other graphs

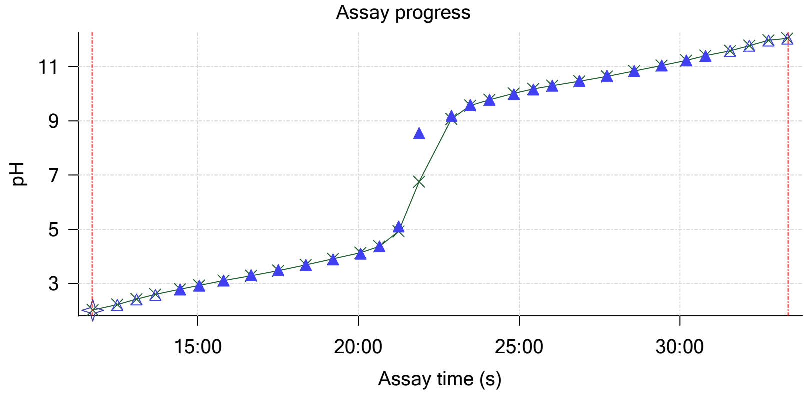


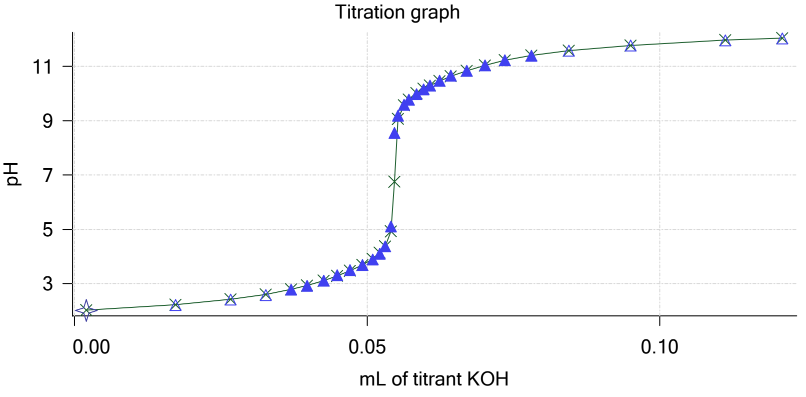

| | | | |
|---|---|---|---|
| Sample name: | **M16_octanol** | Experiment start time: | **3/1/2018 12:08:09 PM** |
| Assay name: | **pH-metric high logP** | Analyst: | **Pion** |
| Assay ID: | **18C-01009** | Instrument ID: | **T312060** |
| Filename: | **C:\Sirius_T3\Mehtap\20180228_exp28_logP_T3-2\18C-01009_M16_octanol_pH-metric high logP.t3r** | | |

#### Other graphs (continued)

Charge balance residuals

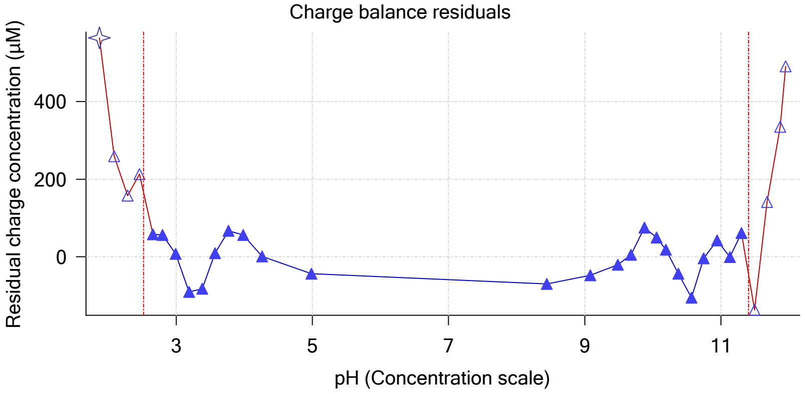


Buffer index profile

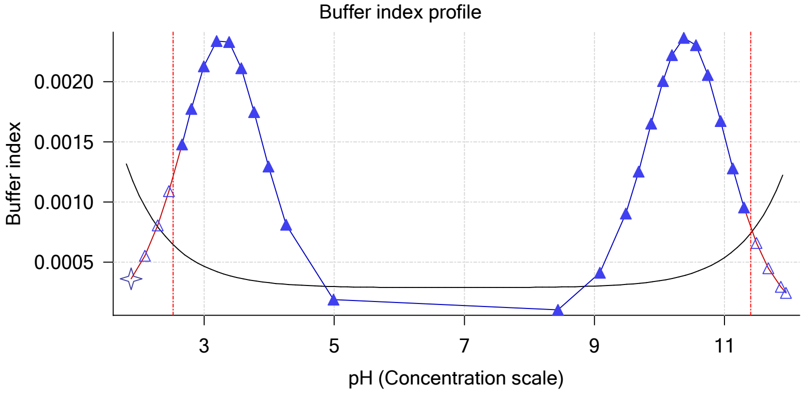


Ionic strength

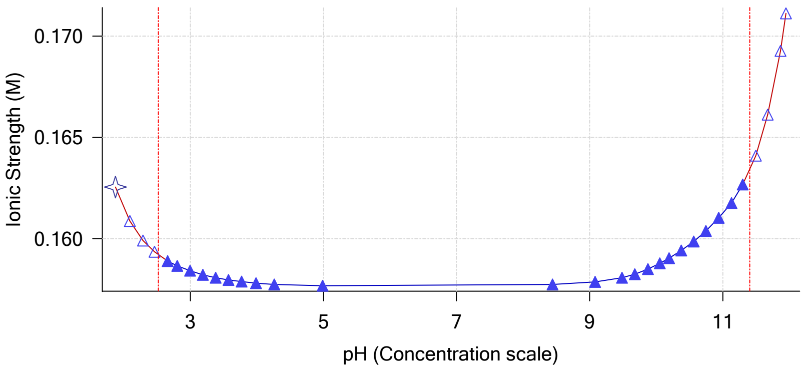


Log R Profile

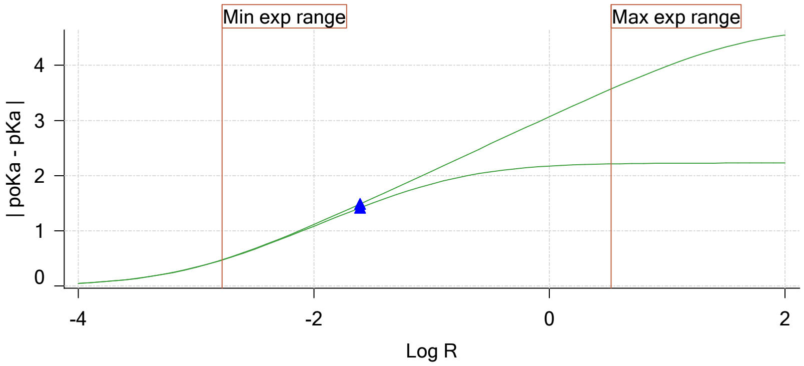


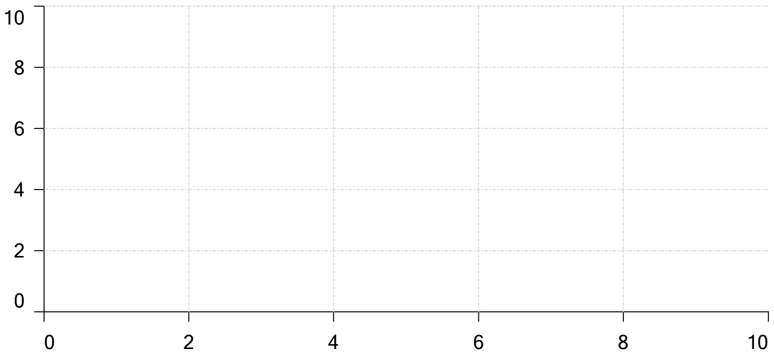


Ionisation of sample Carbonate

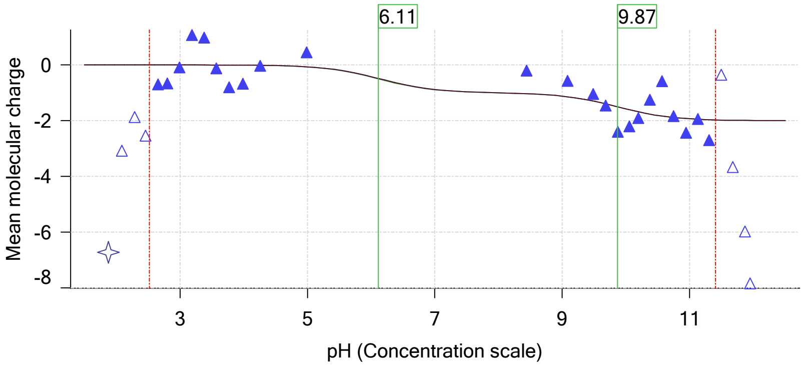

| | | | |
|---|---|---|---|
| Sample name: | **M16_octanol** | Experiment start time: | **3/1/2018 12:08:09 PM** |
| Assay name: | **pH-metric high logP** | Analyst: | **Pion** |
| Assay ID: | **18C-01009** | Instrument ID: | **T312060** |
| Filename: | **C:\Sirius_T3\Mehtap\20180228_exp28_logP_T3-2\18C-01009_M16_octanol_pH-metric high logP.t3r** | | |

### pH-metric high logP Titration 2 of 3 18C-01009 Points 32 to 62

#### Overall results

| | |
|---|---|
| RMSD | 0.147 |
| Average ionic strength | 0.171 M |
| Average temperature | 25.0°C |
| Partition ratio | 0.0751 : 1 |
| Analyte concentration range | 4115.3 µM to 4404.0 µM |
| Total points considered | 23 of 31 |

#### Warnings and errors

| | |
|---|---|
| Errors | None |
| Warnings | Excessive acidity error present |

#### Four-Plus parameters

| | | | |
|---|---|---|---|
| Alpha | 0.130 | 3/1/2018 12:08:08 PM | C:\Sirius_T3\HCl18B27.t3r |
| S | 0.9970 | 3/1/2018 12:08:08 PM | C:\Sirius_T3\HCl18B27.t3r |
| jH | 0.8 | 3/1/2018 12:08:08 PM | C:\Sirius_T3\HCl18B27.t3r |
| jOH | -0.4 | 3/1/2018 12:08:08 PM | C:\Sirius_T3\HCl18B27.t3r |

#### Titrants

| | | | |
|---|---|---|---|
| 0.50 M HCl | 0.993513 | 3/1/2018 12:08:08 PM | C:\Sirius_T3\HCl18B27.t3r |
| 0.50 M KOH | 0.999845 | 3/1/2018 12:08:09 PM | C:\Sirius_T3\KOH18B27.t3r |

#### Sample

| | |
|---|---|
| M16_octanol concentration factor | 0.794 |
| Base pKa 1 | 4.70 |
| Acid pKa 2 | 8.94 |
| logP (XH2 +) | 0.03 |
| logP (neutral XH) | 3.06 |
| logP (X -) | -1.62 |

#### Sample graphs

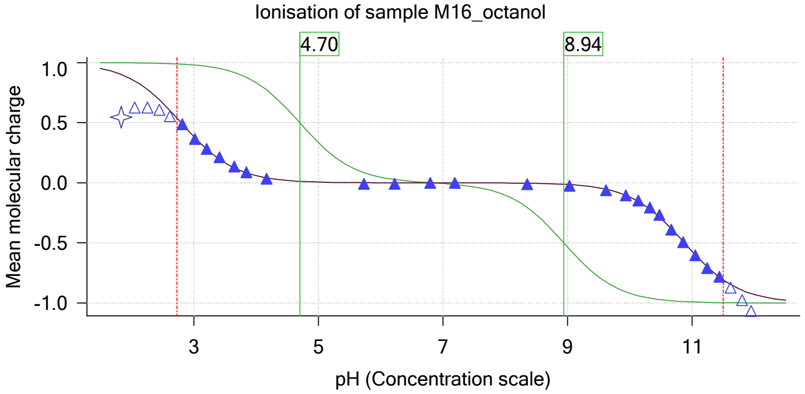


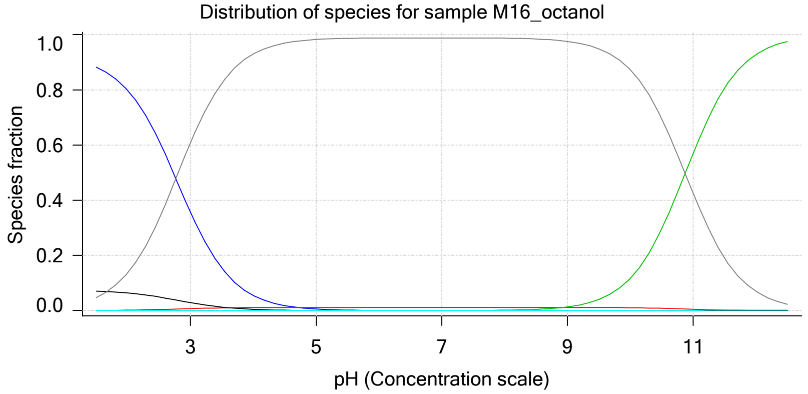

| | | | |
|---|---|---|---|
| Sample name: | **M16_octanol** | Experiment start time: | **3/1/2018 12:08:09 PM** |
| Assay name: | **pH-metric high logP** | Analyst: | **Pion** |
| Assay ID: | **18C-01009** | Instrument ID: | **T312060** |
| Filename: | **C:\Sirius_T3\Mehtap\20180228_exp28_logP_T3-2\18C-01009_M16_octanol_pH-metric high logP.t3r** | | |

#### Sample graphs (continued)

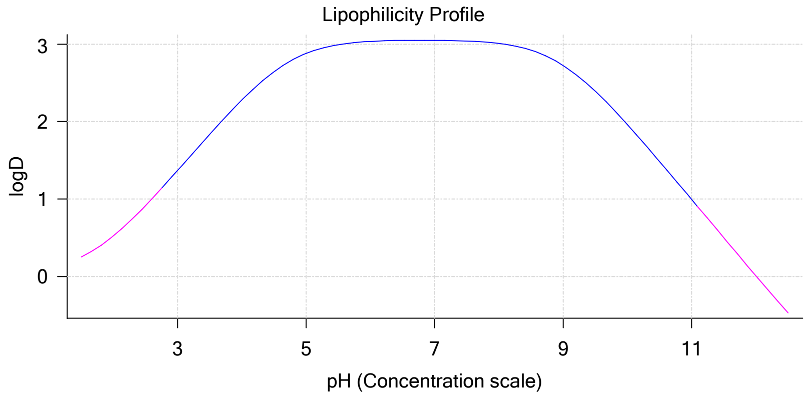


#### Sample logD and percent species

| pH | M16_octanol logD | M16_octanol M16_octanolH2 | M16_octanol M16_octanolH | M16_octanol M16_octanol | M16_octanol M16_octanolH2* | M16_octanol M16_octanolH* | M16_octanol M16_octanol* | Comment |
|---|---|---|---|---|---|---|---|---|
| 1.000 | 0.11 | 91.17 % | 0.02 % | 0.00 % | 7.26 % | 1.55 % | 0.00 % | |
| 1.200 | 0.15 | 90.33 % | 0.03 % | 0.00 % | 7.20 % | 2.44 % | 0.00 % | Stomach pH |
| 2.000 | 0.52 | 79.86 % | 0.16 % | 0.00 % | 6.36 % | 13.62 % | 0.00 % | |
| 3.000 | 1.37 | 35.65 % | 0.71 % | 0.00 % | 2.84 % | 60.80 % | 0.00 % | |
| 4.000 | 2.28 | 5.45 % | 1.09 % | 0.00 % | 0.43 % | 93.02 % | 0.00 % | |
| 5.000 | 2.88 | 0.58 % | 1.15 % | 0.00 % | 0.05 % | 98.23 % | 0.00 % | |
| 6.000 | 3.03 | 0.06 % | 1.16 % | 0.00 % | 0.00 % | 98.78 % | 0.00 % | |
| 6.500 | 3.05 | 0.02 % | 1.16 % | 0.00 % | 0.00 % | 98.82 % | 0.00 % | |
| 7.000 | 3.05 | 0.01 % | 1.16 % | 0.01 % | 0.00 % | 98.82 % | 0.00 % | |
| 7.400 | 3.04 | 0.00 % | 1.16 % | 0.03 % | 0.00 % | 98.81 % | 0.00 % | Blood pH |
| 8.000 | 3.01 | 0.00 % | 1.15 % | 0.13 % | 0.00 % | 98.71 % | 0.00 % | |
| 9.000 | 2.72 | 0.00 % | 1.14 % | 1.31 % | 0.00 % | 97.55 % | 0.00 % | |
| 10.000 | 1.96 | 0.00 % | 1.02 % | 11.72 % | 0.00 % | 87.24 % | 0.02 % | |
| 11.000 | 0.99 | 0.00 % | 0.50 % | 56.98 % | 0.00 % | 42.42 % | 0.10 % | |
| 12.000 | 0.01 | 0.00 % | 0.08 % | 92.84 % | 0.00 % | 6.91 % | 0.17 % | |

#### Carbonate and acidity

| | |
|---|---|
| Carbonate | 0.194 mM |
| Acidity error | 2.681 mM |

#### Other graphs

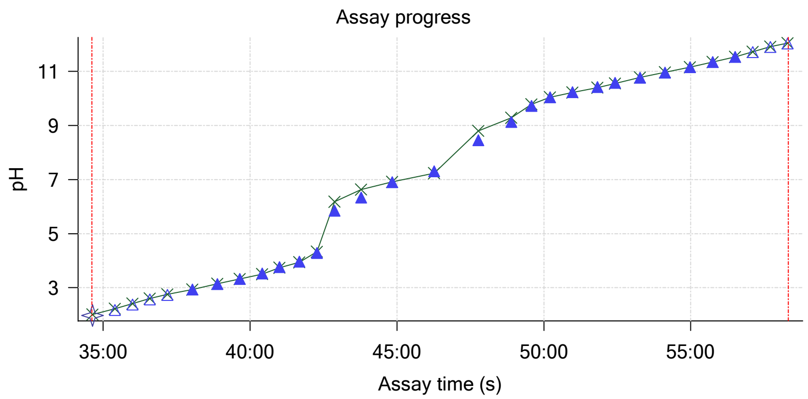


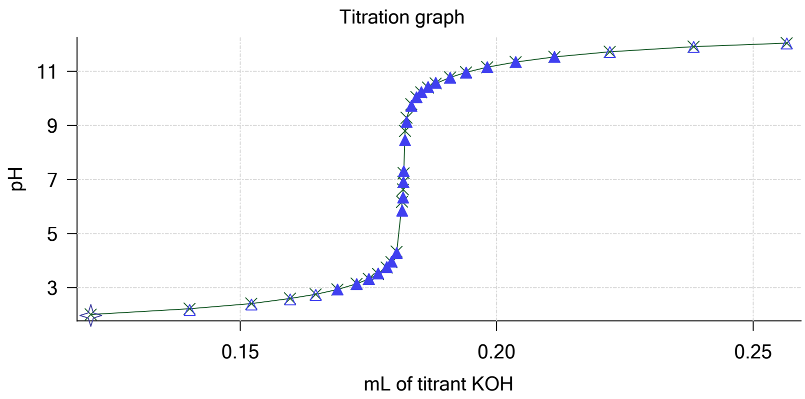

Sample name: **M16_octanol**
Assay name: **pH-metric high logP**
Assay ID: **18C-01009**
Filename: **C:\Sirius_T3\Mehtap\20180228_exp28_logP_T3-2\18C-01009_M16_octanol_pH-metric high logP.t3r**

Experiment start time: **3/1/2018 12:08:09 PM**
Analyst: **Pion**
Instrument ID: **T312060**

#### Other graphs (continued)

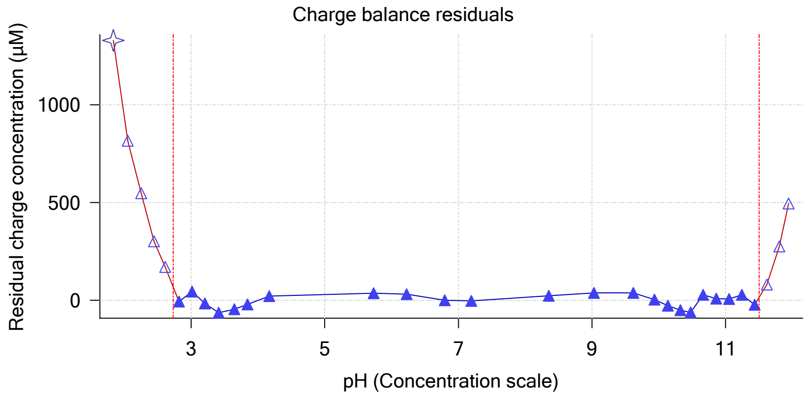


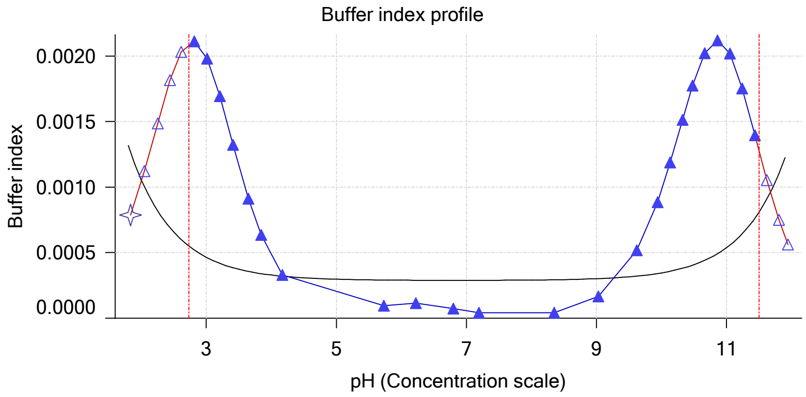


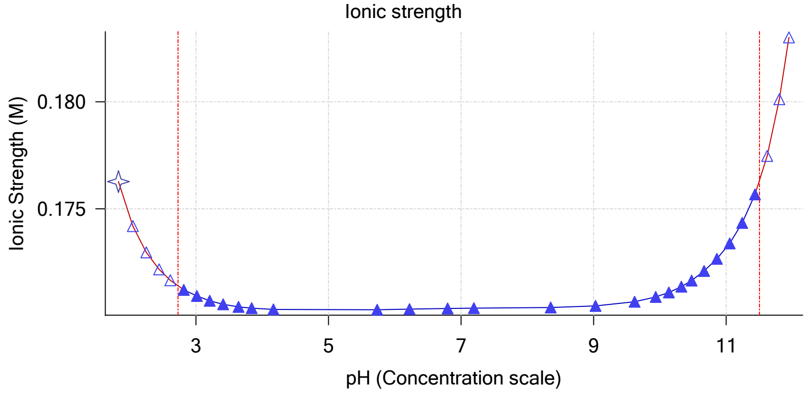


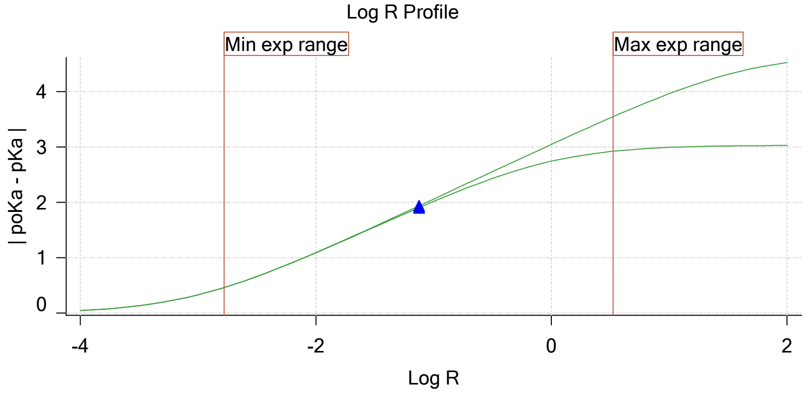


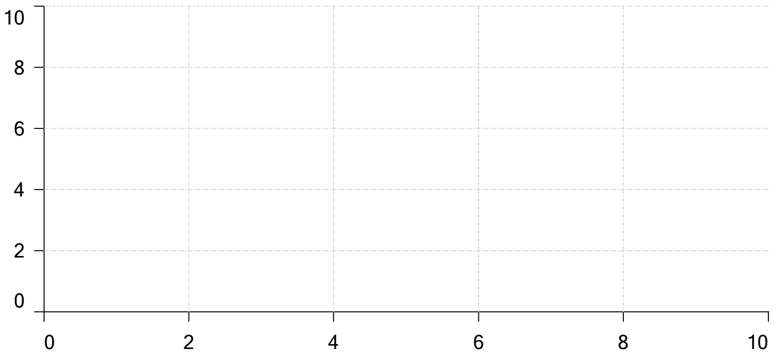


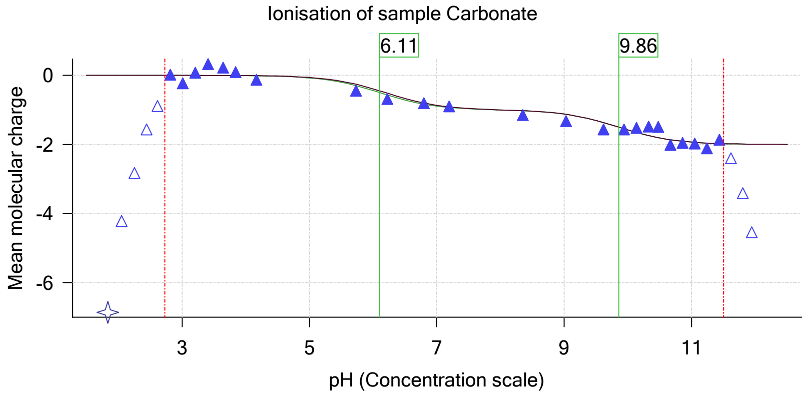

Sample name: **M16_octanol**
Assay name: **pH-metric high logP**
Assay ID: **18C-01009**
Filename: **C:\Sirius_T3\Mehtap\20180228_exp28_logP_T3-2\18C-01009_M16_octanol_pH-metric high logP.t3r**

Experiment start time: **3/1/2018 12:08:09 PM**
Analyst: **Pion**
Instrument ID: **T312060**

### pH-metric high logP Titration 3 of 3 18C-01009 Points 63 to 92

#### Overall results

| | |
|---|---|
| RMSD | 0.627 |
| Average ionic strength | 0.182 M |
| Average temperature | 25.0°C |
| Partition ratio | 0.2052 : 1 |
| Analyte concentration range | 3198.3 μM to 3391.8 μM |
| Total points considered | 22 of 30 |

#### Warnings and errors

| | |
|---|---|
| Errors | None |
| Warnings | One or more logP values out of range<br>Excessive acidity error present |

#### Four-Plus parameters

| | | | |
|---|---|---|---|
| Alpha | 0.130 | 3/1/2018 12:08:08 PM | C:\Sirius_T3\HCl18B27.t3r |
| S | 0.9970 | 3/1/2018 12:08:08 PM | C:\Sirius_T3\HCl18B27.t3r |
| jH | 0.8 | 3/1/2018 12:08:08 PM | C:\Sirius_T3\HCl18B27.t3r |
| jOH | -0.4 | 3/1/2018 12:08:08 PM | C:\Sirius_T3\HCl18B27.t3r |

#### Titrants

| | | | |
|---|---|---|---|
| 0.50 M HCl | 0.993513 | 3/1/2018 12:08:08 PM | C:\Sirius_T3\HCl18B27.t3r |
| 0.50 M KOH | 0.999845 | 3/1/2018 12:08:09 PM | C:\Sirius_T3\KOH18B27.t3r |

#### Sample

| | |
|---|---|
| M16_octanol concentration factor | 0.836 |
| Base pKa 1 | 4.70 |
| Acid pKa 2 | 8.94 |
| logP (XH2 +) | -5.10 |
| logP (neutral XH) | 3.01 |
| logP (X -) | -1.62 |

#### Sample graphs

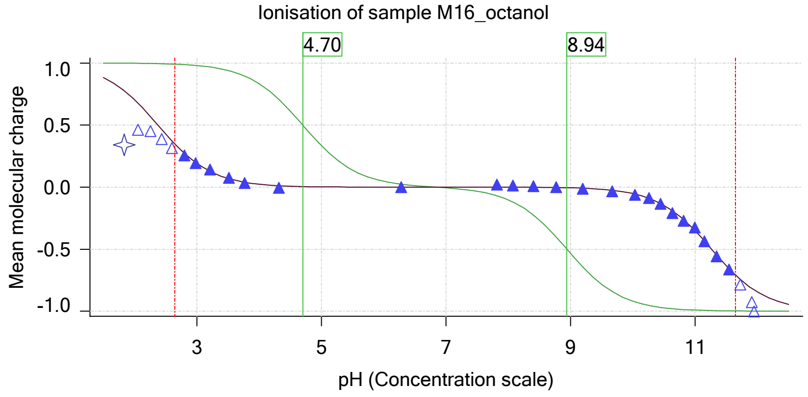


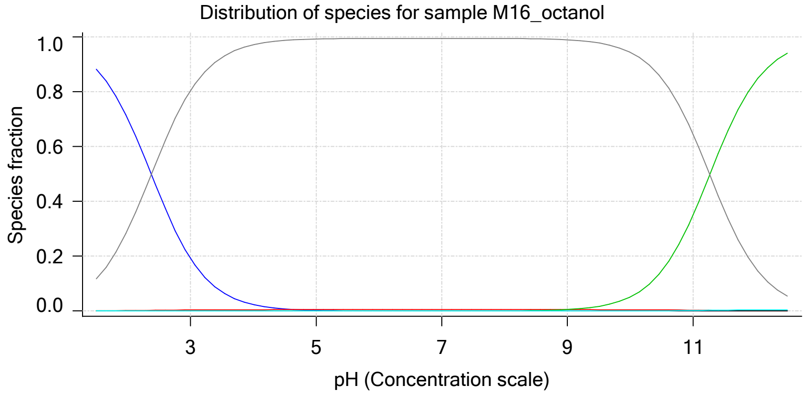

| | | | |
|---|---|---|---|
| Sample name: | **M16_octanol** | Experiment start time: | **3/1/2018 12:08:09 PM** |
| Assay name: | **pH-metric high logP** | Analyst: | **Pion** |
| Assay ID: | **18C-01009** | Instrument ID: | **T312060** |
| Filename: | **C:\Sirius_T3\Mehtap\20180228_exp28_logP_T3-2\18C-01009_M16_octanol_pH-metric high logP.t3r** | | |

#### Sample graphs (continued)

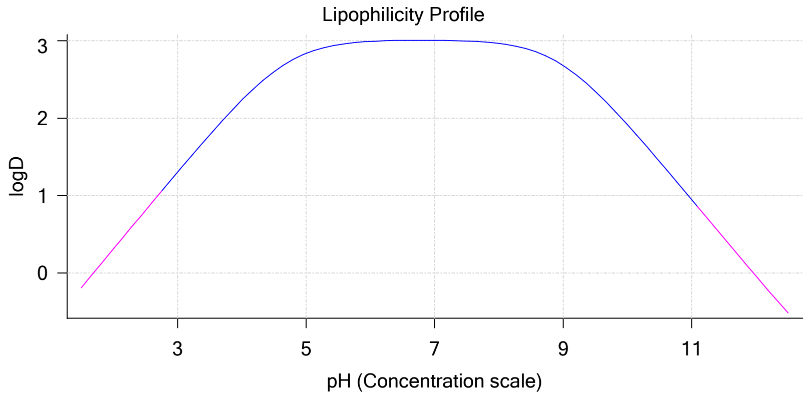


#### Sample logD and percent species

| pH | M16_octanol logD | M16_octanol M16_octanolH2 | M16_octanol M16_octanolH | M16_octanol M16_octanol | M16_octanol M16_octanolH2* | M16_octanol M16_octanolH* | M16_octanol M16_octanol* | Comment |
|---|---|---|---|---|---|---|---|---|
| 1.000 | -0.69 | 95.95 % | 0.02 % | 0.00 % | 0.00 % | 4.03 % | 0.00 % | |
| 1.200 | -0.49 | 93.73 % | 0.03 % | 0.00 % | 0.00 % | 6.24 % | 0.00 % | Stomach pH |
| 2.000 | 0.31 | 70.32 % | 0.14 % | 0.00 % | 0.00 % | 29.54 % | 0.00 % | |
| 3.000 | 1.30 | 19.16 % | 0.38 % | 0.00 % | 0.00 % | 80.46 % | 0.00 % | |
| 4.000 | 2.23 | 2.31 % | 0.46 % | 0.00 % | 0.00 % | 97.22 % | 0.00 % | |
| 5.000 | 2.83 | 0.24 % | 0.47 % | 0.00 % | 0.00 % | 99.29 % | 0.00 % | |
| 6.000 | 2.99 | 0.02 % | 0.47 % | 0.00 % | 0.00 % | 99.50 % | 0.00 % | |
| 6.500 | 3.00 | 0.01 % | 0.47 % | 0.00 % | 0.00 % | 99.52 % | 0.00 % | |
| 7.000 | 3.00 | 0.00 % | 0.47 % | 0.01 % | 0.00 % | 99.52 % | 0.00 % | |
| 7.400 | 3.00 | 0.00 % | 0.47 % | 0.01 % | 0.00 % | 99.51 % | 0.00 % | Blood pH |
| 8.000 | 2.96 | 0.00 % | 0.47 % | 0.05 % | 0.00 % | 99.47 % | 0.00 % | |
| 9.000 | 2.68 | 0.00 % | 0.47 % | 0.54 % | 0.00 % | 98.99 % | 0.00 % | |
| 10.000 | 1.92 | 0.00 % | 0.45 % | 5.15 % | 0.00 % | 94.38 % | 0.03 % | |
| 11.000 | 0.95 | 0.00 % | 0.31 % | 35.12 % | 0.00 % | 64.40 % | 0.17 % | |
| 12.000 | -0.04 | 0.00 % | 0.07 % | 84.09 % | 0.00 % | 15.42 % | 0.41 % | |

#### Carbonate and acidity

| | |
|---|---|
| Carbonate | 0.281 mM |
| Acidity error | 2.477 mM |

#### Other graphs

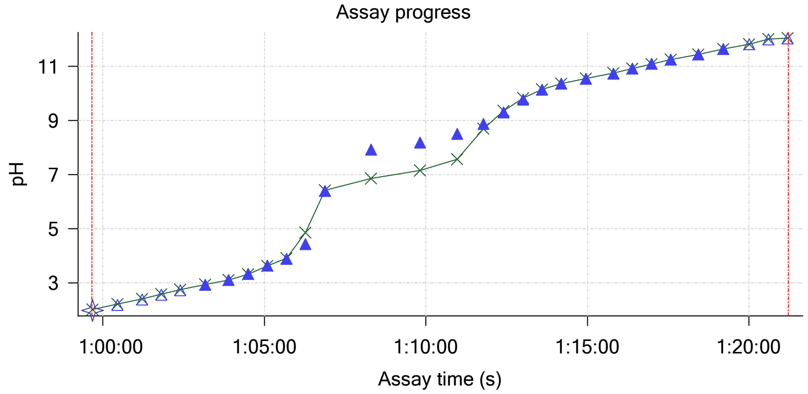


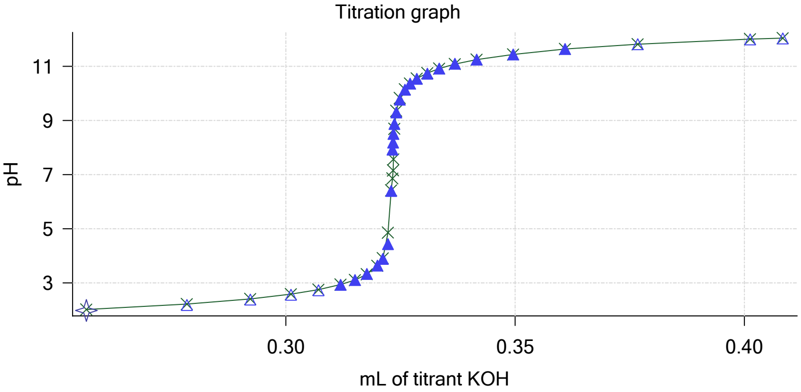

| | | | |
|---|---|---|---|
| Sample name: | **M16_octanol** | Experiment start time: | **3/1/2018 12:08:09 PM** |
| Assay name: | **pH-metric high logP** | Analyst: | **Pion** |
| Assay ID: | **18C-01009** | Instrument ID: | **T312060** |
| Filename: | **C:\Sirius_T3\Mehtap\20180228_exp28_logP_T3-2\18C-01009_M16_octanol_pH-metric high logP.t3r** | | |

#### Other graphs (continued)

Charge balance residuals

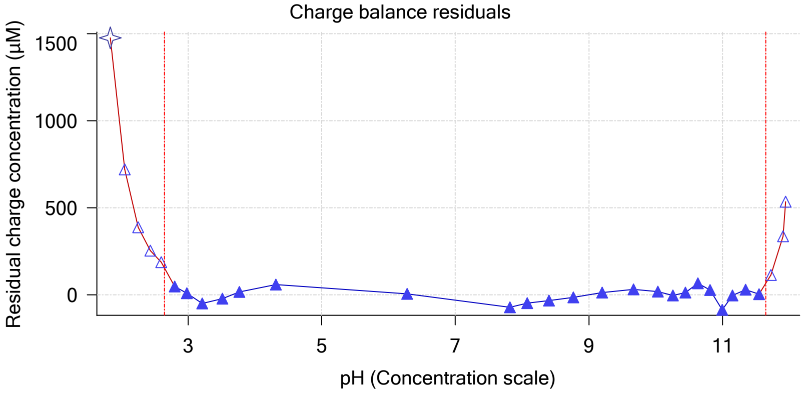


Buffer index profile

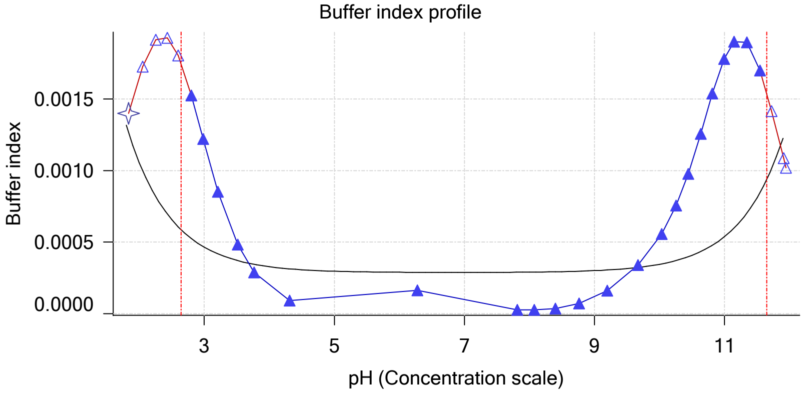


Ionic strength

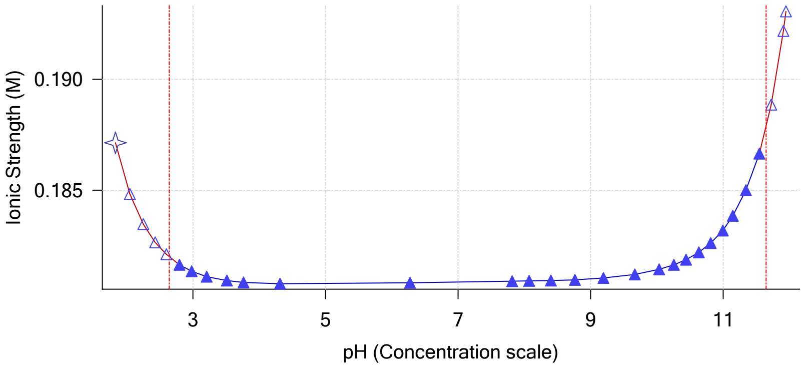


Log R Profile

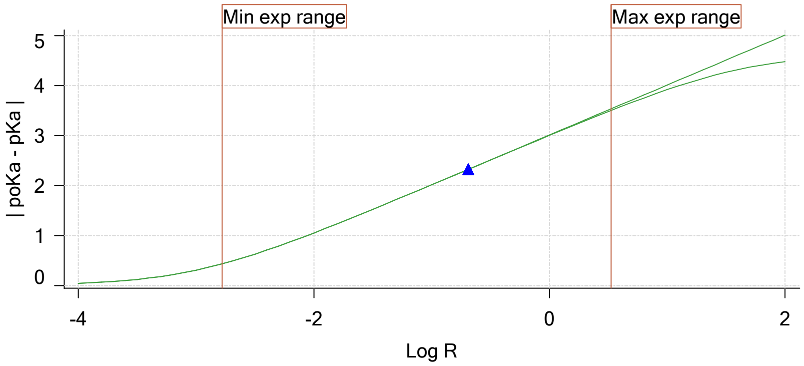


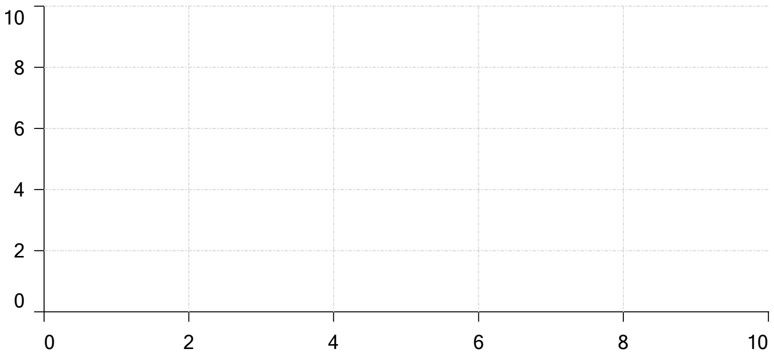


Ionisation of sample Carbonate

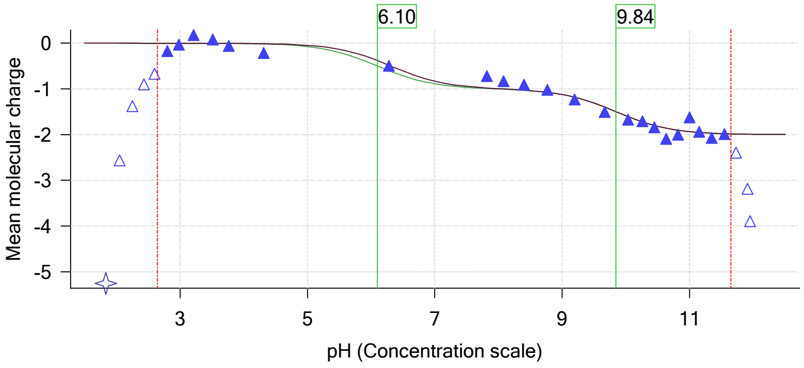

| | |
|---|---|
| Sample name: | **M16_octanol** |
| Assay name: | **pH-metric high logP** |
| Assay ID: | **18C-01009** |
| Filename: | **C:\Sirius_T3\Mehtap\20180228_exp28_logP_T3-2\18C-01009_M16_octanol_pH-metric high logP.t3r** |
| Experiment start time: | **3/1/2018 12:08:09 PM** |
| Analyst: | **Pion** |
| Instrument ID: | **T312060** |

#### Assay Model

| Settings | Value | Date/Time changed | Imported from |
|---|---|---|---|
| Sample name | M16_octanol | 2/27/2018 6:14:13 PM | User entered value |
| Sample by | Weight | | Default value |
| Sample weight | 0.001790 g | 2/28/2018 4:25:54 PM | User entered value |
| Formula weight | 210.23 g/mol | 2/27/2018 5:08:55 PM | User entered value |
| Solubility | Unknown | | Default value |
| Molecular weight | 210.23 | 2/27/2018 5:08:55 PM | User entered value |
| Individual pKa ionic environments | No | | Default value |
| Number of pKas | 2 | 2/27/2018 5:08:55 PM | User entered value |
| Sample is a | Ampholyte | 2/27/2018 5:08:55 PM | User entered value |
| pKa 1 | 4.70 | 2/27/2018 5:08:55 PM | User entered value |
| Type | Base | 2/27/2018 5:08:55 PM | User entered value |
| pKa 2 | 8.94 | 2/27/2018 5:08:55 PM | User entered value |
| Type | Acid | 2/27/2018 5:08:55 PM | User entered value |
| logp (XH2 +) | -0.97 | 2/27/2018 5:09:30 PM | User entered value |
| logP (neutral XH) | 1.47 | 2/27/2018 5:09:44 PM | User entered value |
| logP (X -) | -1.62 | 2/27/2018 5:09:50 PM | User entered value |

#### Events

| Time | Event | Water | Acid | Base | Octanol | pH | dpH/dt | pH R-squared | pH SD | dpH/dt time |
|---|---|---|---|---|---|---|---|---|---|---|
| 8:39.5 | Initial pH = 3.74 | | | | | | | | | |
| 11:44.1 | Data point 1 | 1.50000 mL | 0.04499 mL | 0.00202 mL | 0.04000 mL | 2.009 | -0.00786 | 0.25633 | 0.00077 | 10.0 s |
| 12:30.3 | Data point 2 | 1.50000 mL | 0.04499 mL | 0.01726 mL | 0.04000 mL | 2.218 | -0.00029 | 0.00763 | 0.00016 | 10.0 s |
| 13:06.0 | Data point 3 | 1.50000 mL | 0.04499 mL | 0.02667 mL | 0.04000 mL | 2.413 | -0.00594 | 0.13958 | 0.00079 | 10.0 s |
| 13:41.5 | Data point 4 | 1.50000 mL | 0.04499 mL | 0.03269 mL | 0.04000 mL | 2.584 | -0.00084 | 0.06954 | 0.00016 | 10.0 s |
| 14:27.4 | Data point 5 | 1.50000 mL | 0.04499 mL | 0.03702 mL | 0.04000 mL | 2.780 | -0.00378 | 0.06388 | 0.00074 | 10.0 s |
| 15:02.9 | Data point 6 | 1.50000 mL | 0.04499 mL | 0.03972 mL | 0.04000 mL | 2.922 | -0.00866 | 0.21191 | 0.00093 | 10.0 s |
| 15:48.8 | Data point 7 | 1.50000 mL | 0.04499 mL | 0.04259 mL | 0.04000 mL | 3.115 | -0.00483 | 0.80380 | 0.00027 | 10.0 s |
| 16:39.8 | Data point 8 | 1.50000 mL | 0.04499 mL | 0.04485 mL | 0.04000 mL | 3.309 | -0.00755 | 0.21309 | 0.00081 | 10.0 s |
| 17:30.8 | Data point 9 | 1.50000 mL | 0.04499 mL | 0.04706 mL | 0.04000 mL | 3.501 | -0.00612 | 0.78470 | 0.00034 | 10.0 s |
| 18:21.9 | Data point 10 | 1.50000 mL | 0.04499 mL | 0.04920 mL | 0.04000 mL | 3.691 | -0.00337 | 0.03338 | 0.00091 | 10.0 s |
| 19:12.8 | Data point 11 | 1.50000 mL | 0.04499 mL | 0.05092 mL | 0.04000 mL | 3.887 | -0.00649 | 0.12208 | 0.00092 | 10.0 s |
| 20:03.7 | Data point 12 | 1.50000 mL | 0.04499 mL | 0.05216 mL | 0.04000 mL | 4.105 | -0.01178 | 0.65123 | 0.00072 | 10.0 s |
| 20:39.2 | Data point 13 | 1.50000 mL | 0.04499 mL | 0.05306 mL | 0.04000 mL | 4.377 | -0.01852 | 0.90826 | 0.00096 | 10.5 s |
| 21:15.1 | Data point 14 | 1.50000 mL | 0.04499 mL | 0.05407 mL | 0.04000 mL | 5.103 | -0.01578 | 0.79081 | 0.00088 | 13.0 s |
| 21:53.5 | Data point 15 | 1.50000 mL | 0.04499 mL | 0.05468 mL | 0.04000 mL | 8.545 | -0.01967 | 0.99135 | 0.00098 | 34.5 s |
| 22:53.5 | Data point 16 | 1.50000 mL | 0.04499 mL | 0.05527 mL | 0.04000 mL | 9.186 | -0.01692 | 0.94412 | 0.00086 | 10.0 s |
| 23:28.9 | Data point 17 | 1.50000 mL | 0.04499 mL | 0.05628 mL | 0.04000 mL | 9.588 | -0.01207 | 0.94821 | 0.00061 | 10.0 s |
| 24:04.3 | Data point 18 | 1.50000 mL | 0.04499 mL | 0.05713 mL | 0.04000 mL | 9.780 | -0.01206 | 0.54950 | 0.00080 | 10.0 s |
| 24:50.2 | Data point 19 | 1.50000 mL | 0.04499 mL | 0.05842 mL | 0.04000 mL | 9.975 | -0.00092 | 0.02620 | 0.00028 | 10.5 s |
| 25:26.2 | Data point 20 | 1.50000 mL | 0.04499 mL | 0.05962 mL | 0.04000 mL | 10.154 | -0.00816 | 0.82899 | 0.00044 | 10.0 s |
| 26:01.6 | Data point 21 | 1.50000 mL | 0.04499 mL | 0.06070 mL | 0.04000 mL | 10.292 | -0.01383 | 0.62163 | 0.00087 | 10.0 s |
| 26:52.5 | Data point 22 | 1.50000 mL | 0.04499 mL | 0.06235 mL | 0.04000 mL | 10.475 | -0.00623 | 0.21568 | 0.00066 | 10.0 s |
| 27:43.4 | Data point 23 | 1.50000 mL | 0.04499 mL | 0.06430 mL | 0.04000 mL | 10.663 | -0.01087 | 0.60836 | 0.00069 | 10.0 s |
| 28:34.5 | Data point 24 | 1.50000 mL | 0.04499 mL | 0.06700 mL | 0.04000 mL | 10.845 | 0.00564 | 0.16265 | 0.00069 | 10.5 s |
| 29:26.1 | Data point 25 | 1.50000 mL | 0.04499 mL | 0.07013 mL | 0.04000 mL | 11.038 | -0.00670 | 0.84026 | 0.00036 | 10.0 s |
| 30:12.0 | Data point 26 | 1.50000 mL | 0.04499 mL | 0.07354 mL | 0.04000 mL | 11.226 | -0.00717 | 0.22192 | 0.00075 | 10.0 s |
| 30:47.5 | Data point 27 | 1.50000 mL | 0.04499 mL | 0.07806 mL | 0.04000 mL | 11.397 | 0.00185 | 0.09507 | 0.00030 | 10.0 s |
| 31:33.5 | Data point 28 | 1.50000 mL | 0.04499 mL | 0.08443 mL | 0.04000 mL | 11.589 | -0.00034 | 0.00072 | 0.00063 | 10.0 s |
| 32:09.1 | Data point 29 | 1.50000 mL | 0.04499 mL | 0.09501 mL | 0.04000 mL | 11.769 | -0.00357 | 0.09765 | 0.00056 | 10.0 s |
| 32:45.0 | Data point 30 | 1.50000 mL | 0.04499 mL | 0.11117 mL | 0.04000 mL | 11.958 | 0.00119 | 0.00998 | 0.00059 | 10.0 s |
| 33:20.6 | Data point 31 | 1.50000 mL | 0.04499 mL | 0.12091 mL | 0.04000 mL | 12.036 | 0.00213 | 0.35467 | 0.00018 | 10.0 s |
| 34:38.0 | Data point 32 | 1.50000 mL | 0.17241 mL | 0.12091 mL | 0.14000 mL | 1.969 | -0.00814 | 0.79043 | 0.00045 | 10.0 s |
| 35:24.3 | Data point 33 | 1.50000 mL | 0.17241 mL | 0.14019 mL | 0.14000 mL | 2.180 | -0.00761 | 0.40848 | 0.00059 | 10.0 s |
| 35:59.9 | Data point 34 | 1.50000 mL | 0.17241 mL | 0.15219 mL | 0.14000 mL | 2.379 | 0.01002 | 0.94066 | 0.00051 | 10.0 s |
| 36:35.6 | Data point 35 | 1.50000 mL | 0.17241 mL | 0.15974 mL | 0.14000 mL | 2.569 | 0.00480 | 0.68312 | 0.00029 | 10.0 s |
| 37:11.1 | Data point 36 | 1.50000 mL | 0.17241 mL | 0.16472 mL | 0.14000 mL | 2.737 | 0.00482 | 0.88092 | 0.00025 | 10.0 s |

| | | | |
|---|---|---|---|
| Sample name: | **M16_octanol** | Experiment start time: | **3/1/2018 12:08:09 PM** |
| Assay name: | **pH-metric high logP** | Analyst: | **Pion** |
| Assay ID: | **18C-01009** | Instrument ID: | **T312060** |
| Filename: | **C:\Sirius_T3\Mehtap\20180228_exp28_logP_T3-2\18C-01009_M16_octanol_pH-metric high logP.t3r** | | |

#### Events (continued)

| Time | Event | Water | Acid | Base | Octanol | pH | dpH/dt | pH R-squared | pH SD | dpH/dt time |
|---|---|---|---|---|---|---|---|---|---|---|
| 38:02.2 | Data point 37 | 1.50000 mL | 0.17241 mL | 0.16898 mL | 0.14000 mL | 2.939 | -0.00481 | 0.19222 | 0.00054 | 10.0 s |
| 38:53.3 | Data point 38 | 1.50000 mL | 0.17241 mL | 0.17272 mL | 0.14000 mL | 3.135 | 0.00012 | 0.00403 | 0.00010 | 10.0 s |
| 39:39.2 | Data point 39 | 1.50000 mL | 0.17241 mL | 0.17512 mL | 0.14000 mL | 3.328 | 0.00518 | 0.27140 | 0.00049 | 10.0 s |
| 40:25.0 | Data point 40 | 1.50000 mL | 0.17241 mL | 0.17693 mL | 0.14000 mL | 3.530 | -0.00192 | 0.46039 | 0.00014 | 10.0 s |
| 41:00.4 | Data point 41 | 1.50000 mL | 0.17241 mL | 0.17858 mL | 0.14000 mL | 3.764 | -0.00045 | 0.04173 | 0.00011 | 10.0 s |
| 41:41.0 | Data point 42 | 1.50000 mL | 0.17241 mL | 0.17954 mL | 0.14000 mL | 3.961 | 0.01048 | 0.42558 | 0.00079 | 10.0 s |
| 42:16.4 | Data point 43 | 1.50000 mL | 0.17241 mL | 0.18055 mL | 0.14000 mL | 4.287 | 0.00274 | 0.05649 | 0.00057 | 10.5 s |
| 42:52.4 | Data point 44 | 1.50000 mL | 0.17241 mL | 0.18156 mL | 0.14000 mL | 5.845 | -0.01422 | 0.55499 | 0.00094 | 24.0 s |
| 43:47.0 | Data point 45 | 1.50000 mL | 0.17241 mL | 0.18175 mL | 0.14000 mL | 6.336 | -0.01876 | 0.92228 | 0.00096 | 33.0 s |
| 44:50.5 | Data point 46 | 1.50000 mL | 0.17241 mL | 0.18184 mL | 0.14000 mL | 6.906 | -0.02960 | 0.95500 | 0.00150 | Timed out at 59.5 s |
| 46:15.9 | Data point 47 | 1.50000 mL | 0.17241 mL | 0.18191 mL | 0.14000 mL | 7.300 | -0.02968 | 0.97977 | 0.00148 | Timed out at 59.5 s |
| 47:46.5 | Data point 48 | 1.50000 mL | 0.17241 mL | 0.18215 mL | 0.14000 mL | 8.457 | -0.01552 | 0.97452 | 0.00078 | 41.5 s |
| 48:53.6 | Data point 49 | 1.50000 mL | 0.17241 mL | 0.18248 mL | 0.14000 mL | 9.131 | -0.01766 | 0.85050 | 0.00095 | 15.5 s |
| 49:34.5 | Data point 50 | 1.50000 mL | 0.17241 mL | 0.18337 mL | 0.14000 mL | 9.719 | -0.00134 | 0.00466 | 0.00097 | 13.0 s |
| 50:13.0 | Data point 51 | 1.50000 mL | 0.17241 mL | 0.18436 mL | 0.14000 mL | 10.037 | -0.00738 | 0.26859 | 0.00070 | 10.0 s |
| 50:58.7 | Data point 52 | 1.50000 mL | 0.17241 mL | 0.18532 mL | 0.14000 mL | 10.233 | 0.00095 | 0.02065 | 0.00033 | 10.0 s |
| 51:49.7 | Data point 53 | 1.50000 mL | 0.17241 mL | 0.18667 mL | 0.14000 mL | 10.421 | -0.00379 | 0.74501 | 0.00022 | 10.5 s |
| 52:25.6 | Data point 54 | 1.50000 mL | 0.17241 mL | 0.18815 mL | 0.14000 mL | 10.575 | -0.00945 | 0.32573 | 0.00082 | 10.0 s |
| 53:16.7 | Data point 55 | 1.50000 mL | 0.17241 mL | 0.19092 mL | 0.14000 mL | 10.762 | -0.00222 | 0.36351 | 0.00018 | 10.0 s |
| 54:07.8 | Data point 56 | 1.50000 mL | 0.17241 mL | 0.19405 mL | 0.14000 mL | 10.955 | -0.00583 | 0.12472 | 0.00082 | 10.0 s |
| 54:58.8 | Data point 57 | 1.50000 mL | 0.17241 mL | 0.19817 mL | 0.14000 mL | 11.147 | -0.00232 | 0.32508 | 0.00020 | 10.0 s |
| 55:44.8 | Data point 58 | 1.50000 mL | 0.17241 mL | 0.20372 mL | 0.14000 mL | 11.336 | -0.00054 | 0.03917 | 0.00013 | 10.0 s |
| 56:30.8 | Data point 59 | 1.50000 mL | 0.17241 mL | 0.21129 mL | 0.14000 mL | 11.529 | 0.00672 | 0.53103 | 0.00046 | 10.0 s |
| 57:06.5 | Data point 60 | 1.50000 mL | 0.17241 mL | 0.22204 mL | 0.14000 mL | 11.710 | 0.00008 | 0.00005 | 0.00057 | 10.0 s |
| 57:42.4 | Data point 61 | 1.50000 mL | 0.17241 mL | 0.23838 mL | 0.14000 mL | 11.894 | 0.00523 | 0.08361 | 0.00089 | 10.0 s |
| 58:18.3 | Data point 62 | 1.50000 mL | 0.17241 mL | 0.25654 mL | 0.14000 mL | 12.032 | 0.00844 | 0.20969 | 0.00091 | 10.0 s |
| 59:40.6 | Data point 63 | 1.50000 mL | 0.31378 mL | 0.25654 mL | 0.44000 mL | 1.970 | -0.00419 | 0.48669 | 0.00030 | 10.0 s |
| 1:00:27.0 | Data point 64 | 1.50000 mL | 0.31378 mL | 0.27848 mL | 0.44000 mL | 2.183 | -0.01035 | 0.62763 | 0.00065 | 20.0 s |
| 1:01:12.7 | Data point 65 | 1.50000 mL | 0.31378 mL | 0.29229 mL | 0.44000 mL | 2.381 | -0.00423 | 0.05683 | 0.00088 | 10.0 s |
| 1:01:48.4 | Data point 66 | 1.50000 mL | 0.31378 mL | 0.30115 mL | 0.44000 mL | 2.559 | 0.00526 | 0.10194 | 0.00081 | 10.0 s |
| 1:02:24.0 | Data point 67 | 1.50000 mL | 0.31378 mL | 0.30717 mL | 0.44000 mL | 2.724 | 0.00672 | 0.26884 | 0.00064 | 10.0 s |
| 1:03:09.9 | Data point 68 | 1.50000 mL | 0.31378 mL | 0.31195 mL | 0.44000 mL | 2.924 | 0.01126 | 0.42408 | 0.00085 | 18.0 s |
| 1:03:53.4 | Data point 69 | 1.50000 mL | 0.31378 mL | 0.31512 mL | 0.44000 mL | 3.103 | 0.00666 | 0.70178 | 0.00039 | 10.5 s |
| 1:04:29.5 | Data point 70 | 1.50000 mL | 0.31378 mL | 0.31771 mL | 0.44000 mL | 3.331 | 0.00922 | 0.44168 | 0.00069 | 10.5 s |
| 1:05:05.5 | Data point 71 | 1.50000 mL | 0.31378 mL | 0.32001 mL | 0.44000 mL | 3.635 | 0.00860 | 0.24150 | 0.00086 | 10.0 s |
| 1:05:41.0 | Data point 72 | 1.50000 mL | 0.31378 mL | 0.32117 mL | 0.44000 mL | 3.886 | 0.00769 | 0.25705 | 0.00075 | 10.0 s |
| 1:06:16.4 | Data point 73 | 1.50000 mL | 0.31378 mL | 0.32227 mL | 0.44000 mL | 4.432 | 0.00226 | 0.01769 | 0.00084 | 10.5 s |
| 1:06:52.3 | Data point 74 | 1.50000 mL | 0.31378 mL | 0.32298 mL | 0.44000 mL | 6.389 | -0.01886 | 0.93280 | 0.00097 | 55.5 s |
| 1:08:18.4 | Data point 75 | 1.50000 mL | 0.31378 mL | 0.32326 mL | 0.44000 mL | 7.922 | -0.05162 | 0.99435 | 0.00256 | Timed out at 59.5 s |
| 1:09:48.9 | Data point 76 | 1.50000 mL | 0.31378 mL | 0.32340 mL | 0.44000 mL | 8.182 | -0.01649 | 0.68209 | 0.00099 | 44.0 s |
| 1:10:58.4 | Data point 77 | 1.50000 mL | 0.31378 mL | 0.32352 mL | 0.44000 mL | 8.506 | -0.01726 | 0.75067 | 0.00098 | 23.0 s |
| 1:11:46.7 | Data point 78 | 1.50000 mL | 0.31378 mL | 0.32371 mL | 0.44000 mL | 8.870 | -0.01698 | 0.74469 | 0.00097 | 12.0 s |
| 1:12:24.1 | Data point 79 | 1.50000 mL | 0.31378 mL | 0.32411 mL | 0.44000 mL | 9.299 | -0.01727 | 0.74481 | 0.00099 | 11.0 s |
| 1:13:00.6 | Data point 80 | 1.50000 mL | 0.31378 mL | 0.32491 mL | 0.44000 mL | 9.770 | -0.01163 | 0.54195 | 0.00078 | 10.0 s |
| 1:13:36.0 | Data point 81 | 1.50000 mL | 0.31378 mL | 0.32601 mL | 0.44000 mL | 10.135 | -0.00037 | 0.00108 | 0.00056 | 10.0 s |
| 1:14:11.5 | Data point 82 | 1.50000 mL | 0.31378 mL | 0.32709 mL | 0.44000 mL | 10.357 | -0.00019 | 0.00290 | 0.00018 | 10.0 s |
| 1:14:57.2 | Data point 83 | 1.50000 mL | 0.31378 mL | 0.32858 mL | 0.44000 mL | 10.542 | 0.00649 | 0.22886 | 0.00067 | 10.0 s |
| 1:15:48.3 | Data point 84 | 1.50000 mL | 0.31378 mL | 0.33088 mL | 0.44000 mL | 10.732 | 0.00792 | 0.45250 | 0.00058 | 10.0 s |
| 1:16:23.8 | Data point 85 | 1.50000 mL | 0.31378 mL | 0.33349 mL | 0.44000 mL | 10.912 | 0.00857 | 0.47694 | 0.00061 | 10.0 s |
| 1:16:59.3 | Data point 86 | 1.50000 mL | 0.31378 mL | 0.33688 mL | 0.44000 mL | 11.093 | 0.00726 | 0.16909 | 0.00087 | 10.0 s |
| 1:17:34.9 | Data point 87 | 1.50000 mL | 0.31378 mL | 0.34163 mL | 0.44000 mL | 11.245 | 0.00646 | 0.13436 | 0.00087 | 10.0 s |
| 1:18:26.0 | Data point 88 | 1.50000 mL | 0.31378 mL | 0.34962 mL | 0.44000 mL | 11.440 | 0.00609 | 0.16576 | 0.00074 | 10.0 s |
| 1:19:12.1 | Data point 89 | 1.50000 mL | 0.31378 mL | 0.36091 mL | 0.44000 mL | 11.639 | 0.01231 | 0.61053 | 0.00078 | 22.5 s |
| 1:20:00.5 | Data point 90 | 1.50000 mL | 0.31378 mL | 0.37676 mL | 0.44000 mL | 11.816 | 0.00711 | 0.92311 | 0.00037 | 10.0 s |

| | | | |
|---|---|---|---|
| Sample name: | **M16_octanol** | Experiment start time: | **3/1/2018 12:08:09 PM** |
| Assay name: | **pH-metric high logP** | Analyst: | **Pion** |
| Assay ID: | **18C-01009** | Instrument ID: | **T312060** |
| Filename: | **C:\Sirius_T3\Mehtap\20180228_exp28_logP_T3-2\18C-01009_M16_octanol_pH-metric high logP.t3r** | | |

#### Events (continued)

| Time | Event | Water | Acid | Base | Octanol | pH | dpH/dt | pH R-squared | pH SD | dpH/dt time |
|---|---|---|---|---|---|---|---|---|---|---|
| 1:20:36.5 | Data point 91 | 1.50000 mL | 0.31378 mL | 0.40127 mL | 0.44000 mL | 11.997 | 0.01097 | 0.61510 | 0.00069 | 10.0 s |
| 1:21:12.1 | Data point 92 | 1.50000 mL | 0.31378 mL | 0.40837 mL | 0.44000 mL | 12.034 | 0.01274 | 0.67396 | 0.00077 | 10.0 s |
| 1:21:31.3 | Assay volumes | 1.50000 mL | 0.31378 mL | 0.40837 mL | 0.44000 mL | | | | | |

| | | | |
|---|---|---|---|
| Sample name: | **M16_octanol** | Experiment start time: | **3/1/2018 12:08:09 PM** |
| Assay name: | **pH-metric high logP** | Analyst: | **Pion** |
| Assay ID: | **18C-01009** | Instrument ID: | **T312060** |
| Filename: | **C:\Sirius_T3\Mehtap\20180228_exp28_logP_T3-2\18C-01009_M16_octanol_pH-metric high logP.t3r** | | |

| | | | |
|---|---|---|---|
| Sample name: | **M16_octanol** | Experiment start time: | **3/1/2018 12:08:09 PM** |
| Assay name: | **pH-metric high logP** | Analyst: | **Pion** |
| Assay ID: | **18C-01009** | Instrument ID: | **T312060** |
| Filename: | **C:\Sirius_T3\Mehtap\20180228_exp28_logP_T3-2\18C-01009_M16_octanol_pH-metric high logP.t3r** | | |

#### Assay Settings (continued)

| Setting | Value | Original Value | Date/Time changed | Imported from |
|---|---|---|---|---|
| ***Titration 3*** | | | | |
| Titrate from | Low to high pH | | | |
| Add additional water | 0.00 mL | | | |
| Additional partition solvent volume | 0.300 mL | | | |
| Additional partition solvent added | Automatic | | | |
| After pH adjust stir for | 30 seconds | | | |
| Stir to allow partitioning for | 15 seconds | | | |
| Stirrer speed for partitioning | 60% | | | |
| ***Data Point Stability*** | | | | |
| Stir during data point collection | No | | | |
| Delay before data point collection | 0 seconds | | | |
| Number of points to average | 20 points | | | |
| Time interval between points | 0.50 seconds | | | |
| Required maximum standard deviation | 0.00100 dpH/dt | | | |
| Stability timeout after | 60 seconds | | | |

#### Calibration Settings

| Setting | Value | Date/Time changed | Imported from |
|---|---|---|---|
| Four-Plus alpha | 0.130 | 3/1/2018 12:08:08 PM | C:\Sirius_T3\HCl18B27.t3r |
| Four-Plus S | 0.9970 | 3/1/2018 12:08:08 PM | C:\Sirius_T3\HCl18B27.t3r |
| Four-Plus jH | 0.8 | 3/1/2018 12:08:08 PM | C:\Sirius_T3\HCl18B27.t3r |
| Four-Plus jOH | -0.4 | 3/1/2018 12:08:08 PM | C:\Sirius_T3\HCl18B27.t3r |
| Base concentration factor | 1.000 | 3/1/2018 12:08:09 PM | C:\Sirius_T3\KOH18B27.t3r |
| Acid concentration factor | 0.994 | 3/1/2018 12:08:08 PM | C:\Sirius_T3\HCl18B27.t3r |

#### Instrument Settings

| Setting | Value | Batch Id | Install date |
|---|---|---|---|
| Instrument owner | Merck | | |
| Instrument ID | T312060 | | |
| Instrument type | T3 Simulator | | |
| Software version | 1.1.3.0 | | |
| Dispenser module | | T3DM1200361 | 3/31/2009 5:24:52 AM |
| Dispenser 0 | Water | | 3/31/2009 5:25:05 AM |
| Syringe volume | 2.5 mL | | |
| Firmware version | 1.2.1(r2) | | |
| Titrant | Water (0.15 M KCl) | 02-06-2018 | 2/27/2018 10:05:59 AM |
| Dispenser 2 | Acid | | 3/31/2009 5:25:11 AM |
| Syringe volume | 0.5 mL | | |
| Firmware version | 1.2.1(r2) | | |
| Titrant | Acid (0.5 M HCl) | 02-27-2018 | 2/27/2018 10:27:22 AM |
| Dispenser 1 | Base | | 3/31/2009 5:25:21 AM |
| Syringe volume | 0.5 mL | | |
| Firmware version | 1.2.1(r2) | | |
| Titrant | Base (0.5 M KOH) | 9/22/2017 | 2/27/2018 10:21:22 AM |
| Dispenser 5 | Cosolvent | | 3/31/2009 5:26:24 AM |
| Syringe volume | 2.5 mL | | |
| Firmware version | 1.2.1(r2) | | |
| Distribution valve 5 | Distribution Valve | | 3/31/2009 5:28:19 AM |
| Firmware version | 1.1.3 | | |
| Port A | Methanol (80%, 0.15 M KCl) | 09-26-17 | 2/7/2018 9:42:01 AM |
| Port B | Cyclohexane | 11-01-17 | 2/27/2018 10:37:57 AM |
| Dispenser 3 | Buffer | | 8/3/2010 5:05:16 AM |
| Syringe volume | 0.5 mL | | |
| Firmware version | 1.2.1(r2) | | |
| Titrant | Dodecane | 2018/01/31 | 2/28/2018 10:18:04 AM |
| Dispenser 6 | Octanol | | 10/22/2010 10:52:43 AM |

| | | | |
|---|---|---|---|
| Sample name: | **M16_octanol** | Experiment start time: | **3/1/2018 12:08:09 PM** |
| Assay name: | **pH-metric high logP** | Analyst: | **Pion** |
| Assay ID: | **18C-01009** | Instrument ID: | **T312060** |
| Filename: | **C:\Sirius_T3\Mehtap\20180228_exp28_logP_T3-2\18C-01009_M16_octanol_pH-metric high logP.t3r** | | |

| | | | |
|---|---|---|---|
| Sample name: | **M16_octanol** | Experiment start time: | **3/1/2018 12:08:09 PM** |
| Assay name: | **pH-metric high logP** | Analyst: | **Pion** |
| Assay ID: | **18C-01009** | Instrument ID: | **T312060** |
| Filename: | **C:\Sirius_T3\Mehtap\20180228_exp28_logP_T3-2\18C-01009_M16_octanol_pH-metric high logP.t3r** | | |

#### Instrument Settings (continued)

| Setting | Value | Batch Id | Install date |
|---|---|---|---|
| Spectrometer calibration stir duration | 5 s | | |
| Spectrometer calibration stir speed | 30% | | |
| Spectrometer calibration wash pump volume | 20.0 mL | | |
| Spectrometer calibration wash stir duration | 5 s | | |
| Spectrometer calibration wash stir speed | 30% | | |
| Overhead dispense height | 10000 | | |

#### Refinement Settings

| Setting | Value | Default value |
|---|---|---|
| Turbidity detection method | None | None |
| Turbidity wavelength to assess | 500.0 nm | 500.0 nm |
| Turbidity maximum absorbance | 0.100 | 0.100 |
| Turbidity probe threshold | 50.00 | 50.00 |

#### Experiment Log

[2:16] Air gap created for Water (0.15 M KCl)
[2:17] Air gap created for Acid (0.5 M HCl)
[2:17] Air gap created for Base (0.5 M KOH)
[2:17] Air gap released for Water (0.15 M KCl)
[2:21] Titrator arm moved over Titration position
[2:21] Titration 1 of 3
[2:21] Adding initial titrants
[2:21] Automatically add 1.50000 mL of water
[2:46] Dispensed 1.500000 mL of Water (0.15 M KCl)
[2:51] Titrator arm moved over Drain
[8:32] Titrator arm moved to Titration position
[8:32] Argon flow rate set to 100
[8:32] Stirrer speed set to 10
[8:37] Automatically add 0.04000 mL of Octanol
[8:38] Dispensed 0.040005 mL of Octanol
[8:39] Initial pH = 3.74
[8:39] Iterative adjust 3.74 -> 2.00
[8:39] pH 3.74 -> 2.00
[8:41] Air gap released for Acid (0.5 M HCl)
[8:41] Dispensed 0.042615 mL of Acid (0.5 M HCl)
[8:46] pH 2.02 -> 2.00
[8:47] Dispensed 0.002375 mL of Acid (0.5 M HCl)
[8:52] Holding pH 2.00
[10:52] Stirrer speed set to 0
[10:52] Stirrer speed set to 50
[10:52] Iterative adjust 1.98 -> 2.00
[10:52] pH 1.98 -> 2.00
[10:53] Air gap released for Base (0.5 M KOH)
[10:53] Dispensed 0.002023 mL of Base (0.5 M KOH)
[11:44] Stirrer speed set to 0
[11:54] Datapoint id 1 collected
[11:54] Stirrer speed set to 50
[11:59] pH 2.02 -> 2.22
[11:59] Using cautious pH adjust
[11:59] Dispensed 0.007596 mL of Base (0.5 M KOH)
[12:04] Stepping pH = 2.10
[12:04] Dispensed 0.005997 mL of Base (0.5 M KOH)
[12:10] Stepping pH = 2.19
[12:10] Dispensed 0.001646 mL of Base (0.5 M KOH)
[12:15] Stepping pH = 2.22
[12:30] Stirrer speed set to 0
[12:40] Datapoint id 2 collected

Sample name: **M16_octanol**
Assay name: **pH-metric high logP**
Assay ID: **18C-01009**
Filename: **C:\Sirius_T3\Mehtap\20180228_exp28_logP_T3-2\18C-01009_M16_octanol_pH-metric high logP.t3r**

Experiment start time: **3/1/2018 12:08:09 PM**
Analyst: **Pion**
Instrument ID: **T312060**

#### Experiment Log (continued)

[12:40] Charge balance equation is out by -0.2%
[12:40] Stirrer speed set to 50
[12:45] pH 2.23 -> 2.43
[12:45] Using charge balance adjust
[12:45] Dispensed 0.009407 mL of Base (0.5 M KOH)
[13:06] Stirrer speed set to 0
[13:16] Datapoint id 3 collected
[13:16] Charge balance equation is out by -6.0%
[13:16] Stirrer speed set to 50
[13:21] pH 2.42 -> 2.62
[13:21] Using charge balance adjust
[13:21] Dispensed 0.006021 mL of Base (0.5 M KOH)
[13:41] Stirrer speed set to 0
[13:51] Datapoint id 4 collected
[13:51] Charge balance equation is out by -18.1%
[13:51] Stirrer speed set to 50
[13:56] pH 2.59 -> 2.79
[13:56] Using cautious pH adjust
[13:56] Dispensed 0.002046 mL of Base (0.5 M KOH)
[14:01] Stepping pH = 2.67
[14:02] Dispensed 0.001787 mL of Base (0.5 M KOH)
[14:07] Stepping pH = 2.76
[14:07] Dispensed 0.000494 mL of Base (0.5 M KOH)
[14:12] Stepping pH = 2.78
[14:27] Stirrer speed set to 0
[14:37] Datapoint id 5 collected
[14:37] Charge balance equation is out by -5.4%
[14:37] Stirrer speed set to 50
[14:42] pH 2.79 -> 2.99
[14:42] Using charge balance adjust
[14:42] Dispensed 0.002705 mL of Base (0.5 M KOH)
[15:02] Stirrer speed set to 0
[15:12] Datapoint id 6 collected
[15:12] Charge balance equation is out by -31.5%
[15:12] Stirrer speed set to 50
[15:18] pH 2.93 -> 3.13
[15:18] Using cautious pH adjust
[15:18] Dispensed 0.001035 mL of Base (0.5 M KOH)
[15:23] Stepping pH = 2.99
[15:23] Dispensed 0.001294 mL of Base (0.5 M KOH)
[15:28] Stepping pH = 3.08
[15:28] Dispensed 0.000541 mL of Base (0.5 M KOH)
[15:33] Stepping pH = 3.12
[15:48] Stirrer speed set to 0
[15:58] Datapoint id 7 collected
[15:58] Charge balance equation is out by -39.0%
[15:58] Stirrer speed set to 50
[16:03] pH 3.12 -> 3.32
[16:03] Using cautious pH adjust
[16:04] Dispensed 0.000753 mL of Base (0.5 M KOH)
[16:09] Stepping pH = 3.18
[16:09] Dispensed 0.000917 mL of Base (0.5 M KOH)
[16:14] Stepping pH = 3.27
[16:14] Dispensed 0.000423 mL of Base (0.5 M KOH)
[16:19] Stepping pH = 3.30
[16:19] Dispensed 0.000165 mL of Base (0.5 M KOH)
[16:24] Stepping pH = 3.32
[16:39] Stirrer speed set to 0
[16:49] Datapoint id 8 collected

| | | | |
|---|---|---|---|
| Sample name: | **M16_octanol** | Experiment start time: | **3/1/2018 12:08:09 PM** |
| Assay name: | **pH-metric high logP** | Analyst: | **Pion** |
| Assay ID: | **18C-01009** | Instrument ID: | **T312060** |
| Filename: | **C:\Sirius_T3\Mehtap\20180228_exp28_logP_T3-2\18C-01009_M16_octanol_pH-metric high logP.t3r** | | |

#### Experiment Log (continued)

[16:49] Charge balance equation is out by -50.3%
[16:49] Stirrer speed set to 50
[16:55] pH 3.32 -> 3.52
[16:55] Using cautious pH adjust
[16:55] Dispensed 0.000611 mL of Base (0.5 M KOH)
[17:00] Stepping pH = 3.37
[17:00] Dispensed 0.000823 mL of Base (0.5 M KOH)
[17:05] Stepping pH = 3.45
[17:05] Dispensed 0.000541 mL of Base (0.5 M KOH)
[17:10] Stepping pH = 3.49
[17:10] Dispensed 0.000235 mL of Base (0.5 M KOH)
[17:15] Stepping pH = 3.51
[17:30] Stirrer speed set to 0
[17:40] Datapoint id 9 collected
[17:40] Charge balance equation is out by -79.1%
[17:40] Stirrer speed set to 50
[17:45] pH 3.51 -> 3.71
[17:45] Using cautious pH adjust
[17:46] Dispensed 0.000588 mL of Base (0.5 M KOH)
[17:51] Stepping pH = 3.56
[17:51] Dispensed 0.000800 mL of Base (0.5 M KOH)
[17:56] Stepping pH = 3.63
[17:56] Dispensed 0.000635 mL of Base (0.5 M KOH)
[18:01] Stepping pH = 3.69
[18:01] Dispensed 0.000118 mL of Base (0.5 M KOH)
[18:06] Stepping pH = 3.70
[18:21] Stirrer speed set to 0
[18:31] Datapoint id 10 collected
[18:31] Charge balance equation is out by -81.1%
[18:31] Stirrer speed set to 50
[18:37] pH 3.70 -> 3.90
[18:37] Using cautious pH adjust
[18:37] Dispensed 0.000635 mL of Base (0.5 M KOH)
[18:42] Stepping pH = 3.77
[18:42] Dispensed 0.000729 mL of Base (0.5 M KOH)
[18:47] Stepping pH = 3.86
[18:47] Dispensed 0.000259 mL of Base (0.5 M KOH)
[18:52] Stepping pH = 3.89
[18:52] Dispensed 0.000094 mL of Base (0.5 M KOH)
[18:57] Stepping pH = 3.89
[19:12] Stirrer speed set to 0
[19:22] Datapoint id 11 collected
[19:22] Charge balance equation is out by -36.1%
[19:22] Stirrer speed set to 50
[19:27] pH 3.89 -> 4.09
[19:27] Using cautious pH adjust
[19:28] Dispensed 0.000753 mL of Base (0.5 M KOH)
[19:33] Stepping pH = 4.04
[19:33] Dispensed 0.000282 mL of Base (0.5 M KOH)
[19:38] Stepping pH = 4.08
[19:38] Dispensed 0.000071 mL of Base (0.5 M KOH)
[19:43] Stepping pH = 4.08
[19:43] Dispensed 0.000141 mL of Base (0.5 M KOH)
[19:48] Stepping pH = 4.11
[20:03] Stirrer speed set to 0
[20:13] Datapoint id 12 collected
[20:13] Charge balance equation is out by 17.6%
[20:13] Stirrer speed set to 50
[20:18] pH 4.11 -> 4.31

| | | | |
|---|---|---|---|
| Sample name: | **M16_octanol** | Experiment start time: | **3/1/2018 12:08:09 PM** |
| Assay name: | **pH-metric high logP** | Analyst: | **Pion** |
| Assay ID: | **18C-01009** | Instrument ID: | **T312060** |
| Filename: | **C:\Sirius_T3\Mehtap\20180228_exp28_logP_T3-2\18C-01009_M16_octanol_pH-metric high logP.t3r** | | |

#### Experiment Log (continued)

[20:18] Using cautious pH adjust
[20:18] Dispensed 0.000894 mL of Base (0.5 M KOH)
[20:24] Stepping pH = 4.40
[20:39] Stirrer speed set to 0
[20:49] Datapoint id 13 collected
[20:49] Charge balance equation is out by 50.0%
[20:49] Stirrer speed set to 50
[20:54] pH 4.39 -> 4.59
[20:54] Using cautious pH adjust
[20:54] Dispensed 0.001011 mL of Base (0.5 M KOH)
[21:00] Stepping pH = 5.11
[21:15] Stirrer speed set to 0
[21:28] Datapoint id 14 collected
[21:28] Charge balance equation is out by 50.0%
[21:28] Stirrer speed set to 50
[21:33] pH 5.12 -> 5.32
[21:33] Using cautious pH adjust
[21:33] Dispensed 0.000611 mL of Base (0.5 M KOH)
[21:38] Stepping pH = 8.25
[21:53] Stirrer speed set to 0
[22:28] Datapoint id 15 collected
[22:28] Charge balance equation is out by 50.0%
[22:28] Stirrer speed set to 50
[22:33] pH 8.53 -> 8.73
[22:33] Using cautious pH adjust
[22:33] Dispensed 0.000588 mL of Base (0.5 M KOH)
[22:38] Stepping pH = 9.19
[22:53] Stirrer speed set to 0
[23:03] Datapoint id 16 collected
[23:03] Charge balance equation is out by 50.0%
[23:03] Stirrer speed set to 50
[23:08] pH 9.18 -> 9.38
[23:08] Using cautious pH adjust
[23:08] Dispensed 0.001011 mL of Base (0.5 M KOH)
[23:13] Stepping pH = 9.60
[23:28] Stirrer speed set to 0
[23:38] Datapoint id 17 collected
[23:38] Charge balance equation is out by 50.0%
[23:38] Stirrer speed set to 50
[23:44] pH 9.59 -> 9.79
[23:44] Using cautious pH adjust
[23:44] Dispensed 0.000847 mL of Base (0.5 M KOH)
[23:49] Stepping pH = 9.79
[24:04] Stirrer speed set to 0
[24:14] Datapoint id 18 collected
[24:14] Charge balance equation is out by 50.0%
[24:14] Stirrer speed set to 50
[24:19] pH 9.78 -> 9.98
[24:19] Using cautious pH adjust
[24:19] Dispensed 0.000729 mL of Base (0.5 M KOH)
[24:24] Stepping pH = 9.91
[24:24] Dispensed 0.000282 mL of Base (0.5 M KOH)
[24:29] Stepping pH = 9.94
[24:30] Dispensed 0.000282 mL of Base (0.5 M KOH)
[24:35] Stepping pH = 9.98
[24:50] Stirrer speed set to 0
[25:00] Datapoint id 19 collected
[25:00] Charge balance equation is out by 10.9%
[25:00] Stirrer speed set to 50

#### Experiment Log (continued)

[25:05] pH 9.98 -> 10.18
[25:05] Using charge balance adjust
[25:05] Dispensed 0.001199 mL of Base (0.5 M KOH)
[25:26] Stirrer speed set to 0
[25:36] Datapoint id 20 collected
[25:36] Charge balance equation is out by -11.3%
[25:36] Stirrer speed set to 50
[25:41] pH 10.16 -> 10.36
[25:41] Using charge balance adjust
[25:41] Dispensed 0.001082 mL of Base (0.5 M KOH)
[26:01] Stirrer speed set to 0
[26:11] Datapoint id 21 collected
[26:11] Charge balance equation is out by -32.3%
[26:11] Stirrer speed set to 50
[26:16] pH 10.29 -> 10.49
[26:16] Using cautious pH adjust
[26:16] Dispensed 0.000517 mL of Base (0.5 M KOH)
[26:21] Stepping pH = 10.36
[26:21] Dispensed 0.000611 mL of Base (0.5 M KOH)
[26:27] Stepping pH = 10.43
[26:27] Dispensed 0.000353 mL of Base (0.5 M KOH)
[26:32] Stepping pH = 10.47
[26:32] Dispensed 0.000165 mL of Base (0.5 M KOH)
[26:37] Stepping pH = 10.49
[26:52] Stirrer speed set to 0
[27:02] Datapoint id 22 collected
[27:02] Charge balance equation is out by -58.4%
[27:02] Stirrer speed set to 50
[27:07] pH 10.48 -> 10.68
[27:07] Using cautious pH adjust
[27:07] Dispensed 0.000588 mL of Base (0.5 M KOH)
[27:12] Stepping pH = 10.55
[27:12] Dispensed 0.000682 mL of Base (0.5 M KOH)
[27:18] Stepping pH = 10.61
[27:18] Dispensed 0.000494 mL of Base (0.5 M KOH)
[27:23] Stepping pH = 10.66
[27:23] Dispensed 0.000188 mL of Base (0.5 M KOH)
[27:28] Stepping pH = 10.68
[27:43] Stirrer speed set to 0
[27:53] Datapoint id 23 collected
[27:53] Charge balance equation is out by -67.1%
[27:53] Stirrer speed set to 50
[27:58] pH 10.67 -> 10.87
[27:58] Using cautious pH adjust
[27:58] Dispensed 0.000729 mL of Base (0.5 M KOH)
[28:03] Stepping pH = 10.73
[28:03] Dispensed 0.000964 mL of Base (0.5 M KOH)
[28:08] Stepping pH = 10.80
[28:09] Dispensed 0.000706 mL of Base (0.5 M KOH)
[28:14] Stepping pH = 10.84
[28:14] Dispensed 0.000306 mL of Base (0.5 M KOH)
[28:19] Stepping pH = 10.86
[28:34] Stirrer speed set to 0
[28:45] Datapoint id 24 collected
[28:45] Charge balance equation is out by -85.3%
[28:45] Stirrer speed set to 50
[28:50] pH 10.85 -> 11.05
[28:50] Using cautious pH adjust
[28:50] Dispensed 0.001011 mL of Base (0.5 M KOH)

| | | | |
|---|---|---|---|
| Sample name: | **M16_octanol** | Experiment start time: | **3/1/2018 12:08:09 PM** |
| Assay name: | **pH-metric high logP** | Analyst: | **Pion** |
| Assay ID: | **18C-01009** | Instrument ID: | **T312060** |
| Filename: | **C:\Sirius_T3\Mehtap\20180228_exp28_logP_T3-2\18C-01009_M16_octanol_pH-metric high logP.t3r** | | |

#### Experiment Log (continued)

[28:55] Stepping pH = 10.91
[28:55] Dispensed 0.001270 mL of Base (0.5 M KOH)
[29:00] Stepping pH = 11.00
[29:00] Dispensed 0.000682 mL of Base (0.5 M KOH)
[29:05] Stepping pH = 11.04
[29:05] Dispensed 0.000165 mL of Base (0.5 M KOH)
[29:11] Stepping pH = 11.05
[29:26] Stirrer speed set to 0
[29:36] Datapoint id 25 collected
[29:36] Charge balance equation is out by -55.8%
[29:36] Stirrer speed set to 50
[29:41] pH 11.04 -> 11.24
[29:41] Using cautious pH adjust
[29:41] Dispensed 0.001505 mL of Base (0.5 M KOH)
[29:46] Stepping pH = 11.13
[29:46] Dispensed 0.001364 mL of Base (0.5 M KOH)
[29:51] Stepping pH = 11.21
[29:51] Dispensed 0.000541 mL of Base (0.5 M KOH)
[29:56] Stepping pH = 11.23
[30:11] Stirrer speed set to 0
[30:22] Datapoint id 26 collected
[30:22] Charge balance equation is out by -13.7%
[30:22] Stirrer speed set to 50
[30:27] pH 11.23 -> 11.43
[30:27] Using charge balance adjust
[30:27] Dispensed 0.004516 mL of Base (0.5 M KOH)
[30:47] Stirrer speed set to 0
[30:57] Datapoint id 27 collected
[30:57] Charge balance equation is out by -16.5%
[30:57] Stirrer speed set to 50
[31:02] pH 11.40 -> 11.60
[31:02] Using cautious pH adjust
[31:02] Dispensed 0.003340 mL of Base (0.5 M KOH)
[31:07] Stepping pH = 11.51
[31:08] Dispensed 0.002187 mL of Base (0.5 M KOH)
[31:13] Stepping pH = 11.57
[31:13] Dispensed 0.000847 mL of Base (0.5 M KOH)
[31:18] Stepping pH = 11.59
[31:33] Stirrer speed set to 0
[31:43] Datapoint id 28 collected
[31:43] Charge balance equation is out by 4.5%
[31:43] Stirrer speed set to 50
[31:48] pH 11.59 -> 11.79
[31:48] Using charge balance adjust
[31:48] Dispensed 0.010583 mL of Base (0.5 M KOH)
[32:09] Stirrer speed set to 0
[32:19] Datapoint id 29 collected
[32:19] Charge balance equation is out by -12.6%
[32:19] Stirrer speed set to 50
[32:24] pH 11.77 -> 11.97
[32:24] Using charge balance adjust
[32:24] Dispensed 0.016157 mL of Base (0.5 M KOH)
[32:44] Stirrer speed set to 0
[32:55] Datapoint id 30 collected
[32:55] Charge balance equation is out by -7.8%
[32:55] Stirrer speed set to 50
[33:00] pH 11.96 -> 12.05
[33:00] Using charge balance adjust
[33:00] Dispensed 0.009737 mL of Base (0.5 M KOH)

| | | | |
|---|---|---|---|
| Sample name: | **M16_octanol** | Experiment start time: | **3/1/2018 12:08:09 PM** |
| Assay name: | **pH-metric high logP** | Analyst: | **Pion** |
| Assay ID: | **18C-01009** | Instrument ID: | **T312060** |
| Filename: | **C:\Sirius_T3\Mehtap\20180228_exp28_logP_T3-2\18C-01009_M16_octanol_pH-metric high logP.t3r** | | |

#### Experiment Log (continued)

[33:20] Stirrer speed set to 0
[33:30] Datapoint id 31 collected
[33:30] Charge balance equation is out by -63.2%
[33:30] Titration 2 of 3
[33:30] Adding initial titrants
[33:30] Automatically add 0.10000 mL of Octanol
[33:33] Dispensed 0.100000 mL of Octanol
[33:33] Stirrer speed set to 10
[33:34] Stirrer speed set to 55
[33:34] Iterative adjust 12.04 -> 2.00
[33:34] pH 12.04 -> 2.00
[33:36] Dispensed 0.100000 mL of Acid (0.5 M HCl)
[33:41] pH 2.31 -> 2.00
[33:42] Dispensed 0.025141 mL of Acid (0.5 M HCl)
[33:47] pH 2.02 -> 2.00
[33:47] Dispensed 0.002281 mL of Acid (0.5 M HCl)
[34:38] Stirrer speed set to 0
[34:48] Datapoint id 32 collected
[34:48] Stirrer speed set to 55
[34:53] pH 1.98 -> 2.18
[34:53] Using cautious pH adjust
[34:53] Dispensed 0.009784 mL of Base (0.5 M KOH)
[34:58] Stepping pH = 2.06
[34:58] Dispensed 0.007502 mL of Base (0.5 M KOH)
[35:03] Stepping pH = 2.15
[35:04] Dispensed 0.001999 mL of Base (0.5 M KOH)
[35:09] Stepping pH = 2.18
[35:24] Stirrer speed set to 0
[35:34] Datapoint id 33 collected
[35:34] Charge balance equation is out by 1.4%
[35:34] Stirrer speed set to 55
[35:39] pH 2.19 -> 2.39
[35:39] Using charge balance adjust
[35:39] Dispensed 0.011994 mL of Base (0.5 M KOH)
[35:59] Stirrer speed set to 0
[36:10] Datapoint id 34 collected
[36:10] Charge balance equation is out by -3.2%
[36:10] Stirrer speed set to 55
[36:15] pH 2.39 -> 2.59
[36:15] Using charge balance adjust
[36:15] Dispensed 0.007549 mL of Base (0.5 M KOH)
[36:35] Stirrer speed set to 0
[36:45] Datapoint id 35 collected
[36:45] Charge balance equation is out by -9.6%
[36:45] Stirrer speed set to 55
[36:50] pH 2.58 -> 2.78
[36:50] Using charge balance adjust
[36:50] Dispensed 0.004986 mL of Base (0.5 M KOH)
[37:11] Stirrer speed set to 0
[37:21] Datapoint id 36 collected
[37:21] Charge balance equation is out by -20.0%
[37:21] Stirrer speed set to 55
[37:26] pH 2.74 -> 2.94
[37:26] Using cautious pH adjust
[37:26] Dispensed 0.001764 mL of Base (0.5 M KOH)
[37:31] Stepping pH = 2.82
[37:31] Dispensed 0.001623 mL of Base (0.5 M KOH)
[37:36] Stepping pH = 2.90
[37:36] Dispensed 0.000659 mL of Base (0.5 M KOH)

Sample name: **M16_octanol** Experiment start time: **3/1/2018 12:08:09 PM**
Assay name: **pH-metric high logP** Analyst: **Pion**
Assay ID: **18C-01009** Instrument ID: **T312060**
Filename: **C:\Sirius_T3\Mehtap\20180228_exp28_logP_T3-2\18C-01009_M16_octanol_pH-metric high logP.t3r**

#### Experiment Log (continued)

[37:41] Stepping pH = 2.93
[37:42] Dispensed 0.000212 mL of Base (0.5 M KOH)
[37:47] Stepping pH = 2.94
[38:02] Stirrer speed set to 0
[38:12] Datapoint id 37 collected
[38:12] Charge balance equation is out by -20.5%
[38:12] Stirrer speed set to 55
[38:17] pH 2.94 -> 3.14
[38:17] Using cautious pH adjust
[38:17] Dispensed 0.001223 mL of Base (0.5 M KOH)
[38:22] Stepping pH = 3.00
[38:22] Dispensed 0.001529 mL of Base (0.5 M KOH)
[38:27] Stepping pH = 3.08
[38:27] Dispensed 0.000823 mL of Base (0.5 M KOH)
[38:33] Stepping pH = 3.13
[38:33] Dispensed 0.000165 mL of Base (0.5 M KOH)
[38:38] Stepping pH = 3.14
[38:53] Stirrer speed set to 0
[39:03] Datapoint id 38 collected
[39:03] Charge balance equation is out by -53.0%
[39:03] Stirrer speed set to 55
[39:08] pH 3.14 -> 3.34
[39:08] Using cautious pH adjust
[39:08] Dispensed 0.000941 mL of Base (0.5 M KOH)
[39:13] Stepping pH = 3.21
[39:13] Dispensed 0.001035 mL of Base (0.5 M KOH)
[39:18] Stepping pH = 3.30
[39:18] Dispensed 0.000423 mL of Base (0.5 M KOH)
[39:24] Stepping pH = 3.33
[39:39] Stirrer speed set to 0
[39:49] Datapoint id 39 collected
[39:49] Charge balance equation is out by -27.4%
[39:49] Stirrer speed set to 55
[39:54] pH 3.34 -> 3.54
[39:54] Using cautious pH adjust
[39:54] Dispensed 0.000823 mL of Base (0.5 M KOH)
[39:59] Stepping pH = 3.42
[39:59] Dispensed 0.000753 mL of Base (0.5 M KOH)
[40:04] Stepping pH = 3.51
[40:04] Dispensed 0.000235 mL of Base (0.5 M KOH)
[40:09] Stepping pH = 3.53
[40:25] Stirrer speed set to 0
[40:35] Datapoint id 40 collected
[40:35] Charge balance equation is out by -8.9%
[40:35] Stirrer speed set to 55
[40:40] pH 3.53 -> 3.73
[40:40] Using charge balance adjust
[40:40] Dispensed 0.001646 mL of Base (0.5 M KOH)
[41:00] Stirrer speed set to 0
[41:10] Datapoint id 41 collected
[41:10] Charge balance equation is out by 16.2%
[41:10] Stirrer speed set to 55
[41:15] pH 3.77 -> 3.97
[41:15] Using cautious pH adjust
[41:15] Dispensed 0.000917 mL of Base (0.5 M KOH)
[41:20] Stepping pH = 3.96
[41:20] Dispensed 0.000047 mL of Base (0.5 M KOH)
[41:25] Stepping pH = 3.96
[41:41] Stirrer speed set to 0

| | | | |
|---|---|---|---|
| Sample name: | **M16_octanol** | Experiment start time: | **3/1/2018 12:08:09 PM** |
| Assay name: | **pH-metric high logP** | Analyst: | **Pion** |
| Assay ID: | **18C-01009** | Instrument ID: | **T312060** |
| Filename: | **C:\Sirius_T3\Mehtap\20180228_exp28_logP_T3-2\18C-01009_M16_octanol_pH-metric high logP.t3r** | | |

#### Experiment Log (continued)

[41:51] Datapoint id 42 collected
[41:51] Charge balance equation is out by 47.0%
[41:51] Stirrer speed set to 55
[41:56] pH 3.97 -> 4.17
[41:56] Using cautious pH adjust
[41:56] Dispensed 0.001011 mL of Base (0.5 M KOH)
[42:01] Stepping pH = 4.29
[42:16] Stirrer speed set to 0
[42:27] Datapoint id 43 collected
[42:27] Charge balance equation is out by 50.0%
[42:27] Stirrer speed set to 55
[42:32] pH 4.29 -> 4.49
[42:32] Using cautious pH adjust
[42:32] Dispensed 0.001011 mL of Base (0.5 M KOH)
[42:37] Stepping pH = 5.79
[42:52] Stirrer speed set to 0
[43:16] Datapoint id 44 collected
[43:16] Charge balance equation is out by 50.0%
[43:16] Stirrer speed set to 55
[43:21] pH 5.84 -> 6.04
[43:21] Using cautious pH adjust
[43:21] Dispensed 0.000118 mL of Base (0.5 M KOH)
[43:26] Stepping pH = 5.94
[43:26] Dispensed 0.000071 mL of Base (0.5 M KOH)
[43:31] Stepping pH = 6.18
[43:46] Stirrer speed set to 0
[44:20] Datapoint id 45 collected
[44:20] Charge balance equation is out by 18.2%
[44:20] Stirrer speed set to 55
[44:25] pH 6.29 -> 6.49
[44:25] Using cautious pH adjust
[44:25] Dispensed 0.000047 mL of Base (0.5 M KOH)
[44:30] Stepping pH = 6.38
[44:30] Dispensed 0.000047 mL of Base (0.5 M KOH)
[44:35] Stepping pH = 6.89
[44:50] Stirrer speed set to 0
[45:50] Datapoint id 46 collected
[45:50] Charge balance equation is out by 7.4%
[45:50] Stirrer speed set to 55
[45:55] pH 6.83 -> 7.03
[45:55] Using charge balance adjust
[45:55] Dispensed 0.000071 mL of Base (0.5 M KOH)
[46:15] Stirrer speed set to 0
[47:15] Datapoint id 47 collected
[47:15] Charge balance equation is out by 136.7%
[47:15] Stirrer speed set to 55
[47:21] pH 7.23 -> 7.43
[47:21] Using cautious pH adjust
[47:21] Dispensed 0.000024 mL of Base (0.5 M KOH)
[47:26] Stepping pH = 7.19
[47:26] Dispensed 0.000212 mL of Base (0.5 M KOH)
[47:31] Stepping pH = 8.55
[47:46] Stirrer speed set to 0
[48:28] Datapoint id 48 collected
[48:28] Charge balance equation is out by -241.0%
[48:28] Stirrer speed set to 55
[48:33] pH 8.44 -> 8.64
[48:33] Using cautious pH adjust
[48:33] Dispensed 0.000329 mL of Base (0.5 M KOH)

| | | | |
|---|---|---|---|
| Sample name: | **M16_octanol** | Experiment start time: | **3/1/2018 12:08:09 PM** |
| Assay name: | **pH-metric high logP** | Analyst: | **Pion** |
| Assay ID: | **18C-01009** | Instrument ID: | **T312060** |
| Filename: | **C:\Sirius_T3\Mehtap\20180228_exp28_logP_T3-2\18C-01009_M16_octanol_pH-metric high logP.t3r** | | |

#### Experiment Log (continued)

[48:38] Stepping pH = 9.15
[48:53] Stirrer speed set to 0
[49:09] Datapoint id 49 collected
[49:09] Charge balance equation is out by 50.0%
[49:09] Stirrer speed set to 55
[49:14] pH 9.13 -> 9.33
[49:14] Using cautious pH adjust
[49:14] Dispensed 0.000894 mL of Base (0.5 M KOH)
[49:19] Stepping pH = 9.72
[49:34] Stirrer speed set to 0
[49:47] Datapoint id 50 collected
[49:47] Charge balance equation is out by 50.0%
[49:47] Stirrer speed set to 55
[49:52] pH 9.72 -> 9.92
[49:52] Using cautious pH adjust
[49:52] Dispensed 0.000988 mL of Base (0.5 M KOH)
[49:57] Stepping pH = 10.04
[50:12] Stirrer speed set to 0
[50:23] Datapoint id 51 collected
[50:23] Charge balance equation is out by 50.0%
[50:23] Stirrer speed set to 55
[50:28] pH 10.04 -> 10.24
[50:28] Using cautious pH adjust
[50:28] Dispensed 0.000800 mL of Base (0.5 M KOH)
[50:33] Stepping pH = 10.22
[50:33] Dispensed 0.000071 mL of Base (0.5 M KOH)
[50:38] Stepping pH = 10.23
[50:38] Dispensed 0.000094 mL of Base (0.5 M KOH)
[50:43] Stepping pH = 10.24
[50:58] Stirrer speed set to 0
[51:08] Datapoint id 52 collected
[51:08] Charge balance equation is out by 39.5%
[51:08] Stirrer speed set to 55
[51:13] pH 10.24 -> 10.44
[51:13] Using cautious pH adjust
[51:13] Dispensed 0.000729 mL of Base (0.5 M KOH)
[51:19] Stepping pH = 10.36
[51:19] Dispensed 0.000376 mL of Base (0.5 M KOH)
[51:24] Stepping pH = 10.41
[51:24] Dispensed 0.000165 mL of Base (0.5 M KOH)
[51:29] Stepping pH = 10.43
[51:29] Dispensed 0.000071 mL of Base (0.5 M KOH)
[51:34] Stepping pH = 10.43
[51:49] Stirrer speed set to 0
[52:00] Datapoint id 53 collected
[52:00] Charge balance equation is out by 9.5%
[52:00] Stirrer speed set to 55
[52:05] pH 10.42 -> 10.62
[52:05] Using charge balance adjust
[52:05] Dispensed 0.001482 mL of Base (0.5 M KOH)
[52:25] Stirrer speed set to 0
[52:35] Datapoint id 54 collected
[52:35] Charge balance equation is out by -24.8%
[52:35] Stirrer speed set to 55
[52:40] pH 10.58 -> 10.78
[52:40] Using cautious pH adjust
[52:40] Dispensed 0.000823 mL of Base (0.5 M KOH)
[52:45] Stepping pH = 10.64
[52:46] Dispensed 0.001011 mL of Base (0.5 M KOH)

#### Experiment Log (continued)

[52:51] Stepping pH = 10.71
[52:51] Dispensed 0.000682 mL of Base (0.5 M KOH)
[52:56] Stepping pH = 10.76
[52:56] Dispensed 0.000259 mL of Base (0.5 M KOH)
[53:01] Stepping pH = 10.77
[53:16] Stirrer speed set to 0
[53:26] Datapoint id 55 collected
[53:26] Charge balance equation is out by -67.6%
[53:26] Stirrer speed set to 55
[53:31] pH 10.77 -> 10.97
[53:31] Using cautious pH adjust
[53:32] Dispensed 0.001058 mL of Base (0.5 M KOH)
[53:37] Stepping pH = 10.84
[53:37] Dispensed 0.001223 mL of Base (0.5 M KOH)
[53:42] Stepping pH = 10.91
[53:42] Dispensed 0.000682 mL of Base (0.5 M KOH)
[53:47] Stepping pH = 10.96
[53:47] Dispensed 0.000165 mL of Base (0.5 M KOH)
[53:52] Stepping pH = 10.96
[54:07] Stirrer speed set to 0
[54:17] Datapoint id 56 collected
[54:17] Charge balance equation is out by -47.4%
[54:17] Stirrer speed set to 55
[54:22] pH 10.96 -> 11.16
[54:22] Using cautious pH adjust
[54:23] Dispensed 0.001482 mL of Base (0.5 M KOH)
[54:28] Stepping pH = 11.04
[54:28] Dispensed 0.001458 mL of Base (0.5 M KOH)
[54:33] Stepping pH = 11.11
[54:33] Dispensed 0.000847 mL of Base (0.5 M KOH)
[54:38] Stepping pH = 11.14
[54:38] Dispensed 0.000329 mL of Base (0.5 M KOH)
[54:43] Stepping pH = 11.15
[54:58] Stirrer speed set to 0
[55:08] Datapoint id 57 collected
[55:08] Charge balance equation is out by -38.3%
[55:08] Stirrer speed set to 55
[55:14] pH 11.15 -> 11.35
[55:14] Using cautious pH adjust
[55:14] Dispensed 0.002234 mL of Base (0.5 M KOH)
[55:19] Stepping pH = 11.23
[55:19] Dispensed 0.002258 mL of Base (0.5 M KOH)
[55:24] Stepping pH = 11.31
[55:24] Dispensed 0.001058 mL of Base (0.5 M KOH)
[55:29] Stepping pH = 11.34
[55:44] Stirrer speed set to 0
[55:54] Datapoint id 58 collected
[55:54] Charge balance equation is out by -24.3%
[55:54] Stirrer speed set to 55
[55:59] pH 11.34 -> 11.54
[55:59] Using cautious pH adjust
[56:00] Dispensed 0.003434 mL of Base (0.5 M KOH)
[56:05] Stepping pH = 11.44
[56:05] Dispensed 0.002775 mL of Base (0.5 M KOH)
[56:10] Stepping pH = 11.50
[56:10] Dispensed 0.001364 mL of Base (0.5 M KOH)
[56:15] Stepping pH = 11.53
[56:30] Stirrer speed set to 0
[56:40] Datapoint id 59 collected

| | | | |
|---|---|---|---|
| Sample name: | **M16_octanol** | Experiment start time: | **3/1/2018 12:08:09 PM** |
| Assay name: | **pH-metric high logP** | Analyst: | **Pion** |
| Assay ID: | **18C-01009** | Instrument ID: | **T312060** |
| Filename: | **C:\Sirius_T3\Mehtap\20180228_exp28_logP_T3-2\18C-01009_M16_octanol_pH-metric high logP.t3r** | | |

#### Experiment Log (continued)

[56:40] Charge balance equation is out by -10.4%
[56:40] Stirrer speed set to 55
[56:45] pH 11.54 -> 11.74
[56:45] Using charge balance adjust
[56:46] Dispensed 0.010748 mL of Base (0.5 M KOH)
[57:06] Stirrer speed set to 0
[57:16] Datapoint id 60 collected
[57:16] Charge balance equation is out by -12.6%
[57:16] Stirrer speed set to 55
[57:21] pH 11.71 -> 11.91
[57:21] Using charge balance adjust
[57:22] Dispensed 0.016345 mL of Base (0.5 M KOH)
[57:42] Stirrer speed set to 0
[57:52] Datapoint id 61 collected
[57:52] Charge balance equation is out by -9.7%
[57:52] Stirrer speed set to 55
[57:57] pH 11.90 -> 12.05
[57:57] Using charge balance adjust
[57:58] Dispensed 0.018156 mL of Base (0.5 M KOH)
[58:18] Stirrer speed set to 0
[58:28] Datapoint id 62 collected
[58:28] Charge balance equation is out by -33.1%
[58:28] Titration 3 of 3
[58:28] Adding initial titrants
[58:28] Automatically add 0.30000 mL of Octanol
[58:35] Dispensed 0.300000 mL of Octanol
[58:35] Stirrer speed set to 10
[58:36] Stirrer speed set to 60
[58:36] Iterative adjust 12.04 -> 2.00
[58:36] pH 12.04 -> 2.00
[58:38] Dispensed 0.100000 mL of Acid (0.5 M HCl)
[58:44] pH 2.47 -> 2.00
[58:45] Dispensed 0.037465 mL of Acid (0.5 M HCl)
[58:50] pH 2.03 -> 2.00
[58:50] Dispensed 0.003904 mL of Acid (0.5 M HCl)
[59:40] Stirrer speed set to 0
[59:50] Datapoint id 63 collected
[59:50] Stirrer speed set to 60
[59:55] pH 1.98 -> 2.18
[59:55] Using cautious pH adjust
[59:56] Dispensed 0.011265 mL of Base (0.5 M KOH)
[1:00:01] Stepping pH = 2.06
[1:00:01] Dispensed 0.008796 mL of Base (0.5 M KOH)
[1:00:06] Stepping pH = 2.15
[1:00:06] Dispensed 0.001881 mL of Base (0.5 M KOH)
[1:00:11] Stepping pH = 2.18
[1:00:26] Stirrer speed set to 0
[1:00:47] Datapoint id 64 collected
[1:00:47] Charge balance equation is out by 2.6%
[1:00:47] Stirrer speed set to 60
[1:00:52] pH 2.19 -> 2.39
[1:00:52] Using charge balance adjust
[1:00:52] Dispensed 0.013805 mL of Base (0.5 M KOH)
[1:01:12] Stirrer speed set to 0
[1:01:22] Datapoint id 65 collected
[1:01:22] Charge balance equation is out by -4.3%
[1:01:22] Stirrer speed set to 60
[1:01:27] pH 2.39 -> 2.59
[1:01:27] Using charge balance adjust

| | | | |
|---|---|---|---|
| Sample name: | **M16_octanol** | Experiment start time: | **3/1/2018 12:08:09 PM** |
| Assay name: | **pH-metric high logP** | Analyst: | **Pion** |
| Assay ID: | **18C-01009** | Instrument ID: | **T312060** |
| Filename: | **C:\Sirius_T3\Mehtap\20180228_exp28_logP_T3-2\18C-01009_M16_octanol_pH-metric high logP.t3r** | | |

#### Experiment Log (continued)

[1:01:28] Dispensed 0.008866 mL of Base (0.5 M KOH)
[1:01:48] Stirrer speed set to 0
[1:01:58] Datapoint id 66 collected
[1:01:58] Charge balance equation is out by -13.7%
[1:01:58] Stirrer speed set to 60
[1:02:03] pH 2.57 -> 2.77
[1:02:03] Using charge balance adjust
[1:02:03] Dispensed 0.006021 mL of Base (0.5 M KOH)
[1:02:23] Stirrer speed set to 0
[1:02:34] Datapoint id 67 collected
[1:02:34] Charge balance equation is out by -21.7%
[1:02:34] Stirrer speed set to 60
[1:02:39] pH 2.73 -> 2.93
[1:02:39] Using cautious pH adjust
[1:02:39] Dispensed 0.002187 mL of Base (0.5 M KOH)
[1:02:44] Stepping pH = 2.82
[1:02:44] Dispensed 0.001834 mL of Base (0.5 M KOH)
[1:02:49] Stepping pH = 2.89
[1:02:49] Dispensed 0.000753 mL of Base (0.5 M KOH)
[1:02:54] Stepping pH = 2.92
[1:03:09] Stirrer speed set to 0
[1:03:27] Datapoint id 68 collected
[1:03:27] Charge balance equation is out by -9.5%
[1:03:27] Stirrer speed set to 60
[1:03:33] pH 2.93 -> 3.13
[1:03:33] Using charge balance adjust
[1:03:33] Dispensed 0.003175 mL of Base (0.5 M KOH)
[1:03:53] Stirrer speed set to 0
[1:04:03] Datapoint id 69 collected
[1:04:03] Charge balance equation is out by -12.8%
[1:04:03] Stirrer speed set to 60
[1:04:09] pH 3.11 -> 3.31
[1:04:09] Using charge balance adjust
[1:04:09] Dispensed 0.002587 mL of Base (0.5 M KOH)
[1:04:29] Stirrer speed set to 0
[1:04:40] Datapoint id 70 collected
[1:04:40] Charge balance equation is out by 10.9%
[1:04:40] Stirrer speed set to 60
[1:04:45] pH 3.34 -> 3.54
[1:04:45] Using charge balance adjust
[1:04:45] Dispensed 0.002305 mL of Base (0.5 M KOH)
[1:05:05] Stirrer speed set to 0
[1:05:15] Datapoint id 71 collected
[1:05:15] Charge balance equation is out by 48.3%
[1:05:15] Stirrer speed set to 60
[1:05:20] pH 3.64 -> 3.84
[1:05:20] Using cautious pH adjust
[1:05:20] Dispensed 0.001152 mL of Base (0.5 M KOH)
[1:05:25] Stepping pH = 3.89
[1:05:40] Stirrer speed set to 0
[1:05:51] Datapoint id 72 collected
[1:05:51] Charge balance equation is out by 50.0%
[1:05:51] Stirrer speed set to 60
[1:05:56] pH 3.89 -> 4.09
[1:05:56] Using cautious pH adjust
[1:05:56] Dispensed 0.001105 mL of Base (0.5 M KOH)
[1:06:01] Stepping pH = 4.44
[1:06:16] Stirrer speed set to 0
[1:06:26] Datapoint id 73 collected

| | | | |
|---|---|---|---|
| Sample name: | **M16_octanol** | Experiment start time: | **3/1/2018 12:08:09 PM** |
| Assay name: | **pH-metric high logP** | Analyst: | **Pion** |
| Assay ID: | **18C-01009** | Instrument ID: | **T312060** |
| Filename: | **C:\Sirius_T3\Mehtap\20180228_exp28_logP_T3-2\18C-01009_M16_octanol_pH-metric high logP.t3r** | | |

#### Experiment Log (continued)

[1:06:26] Charge balance equation is out by 50.0%
[1:06:26] Stirrer speed set to 60
[1:06:32] pH 4.44 -> 4.64
[1:06:32] Using cautious pH adjust
[1:06:32] Dispensed 0.000706 mL of Base (0.5 M KOH)
[1:06:37] Stepping pH = 6.57
[1:06:52] Stirrer speed set to 0
[1:07:47] Datapoint id 74 collected
[1:07:47] Charge balance equation is out by 50.0%
[1:07:47] Stirrer speed set to 60
[1:07:53] pH 6.31 -> 6.51
[1:07:53] Using cautious pH adjust
[1:07:53] Dispensed 0.000047 mL of Base (0.5 M KOH)
[1:07:58] Stepping pH = 6.28
[1:07:58] Dispensed 0.000235 mL of Base (0.5 M KOH)
[1:08:03] Stepping pH = 8.21
[1:08:18] Stirrer speed set to 0
[1:09:18] Datapoint id 75 collected
[1:09:18] Charge balance equation is out by -228.6%
[1:09:18] Stirrer speed set to 60
[1:09:23] pH 7.81 -> 8.01
[1:09:23] Using cautious pH adjust
[1:09:23] Dispensed 0.000047 mL of Base (0.5 M KOH)
[1:09:28] Stepping pH = 7.85
[1:09:28] Dispensed 0.000094 mL of Base (0.5 M KOH)
[1:09:33] Stepping pH = 8.33
[1:09:48] Stirrer speed set to 0
[1:10:33] Datapoint id 76 collected
[1:10:33] Charge balance equation is out by -40.8%
[1:10:33] Stirrer speed set to 60
[1:10:38] pH 8.20 -> 8.40
[1:10:38] Using cautious pH adjust
[1:10:38] Dispensed 0.000118 mL of Base (0.5 M KOH)
[1:10:43] Stepping pH = 8.56
[1:10:58] Stirrer speed set to 0
[1:11:21] Datapoint id 77 collected
[1:11:21] Charge balance equation is out by 50.0%
[1:11:21] Stirrer speed set to 60
[1:11:26] pH 8.49 -> 8.69
[1:11:26] Using cautious pH adjust
[1:11:26] Dispensed 0.000188 mL of Base (0.5 M KOH)
[1:11:31] Stepping pH = 8.91
[1:11:46] Stirrer speed set to 0
[1:11:58] Datapoint id 78 collected
[1:11:58] Charge balance equation is out by 50.0%
[1:11:58] Stirrer speed set to 60
[1:12:03] pH 8.87 -> 9.07
[1:12:03] Using cautious pH adjust
[1:12:03] Dispensed 0.000400 mL of Base (0.5 M KOH)
[1:12:09] Stepping pH = 9.33
[1:12:24] Stirrer speed set to 0
[1:12:35] Datapoint id 79 collected
[1:12:35] Charge balance equation is out by 50.0%
[1:12:35] Stirrer speed set to 60
[1:12:40] pH 9.30 -> 9.50
[1:12:40] Using cautious pH adjust
[1:12:40] Dispensed 0.000800 mL of Base (0.5 M KOH)
[1:12:45] Stepping pH = 9.79
[1:13:00] Stirrer speed set to 0

| | | | |
|---|---|---|---|
| Sample name: | **M16_octanol** | Experiment start time: | **3/1/2018 12:08:09 PM** |
| Assay name: | **pH-metric high logP** | Analyst: | **Pion** |
| Assay ID: | **18C-01009** | Instrument ID: | **T312060** |
| Filename: | **C:\Sirius_T3\Mehtap\20180228_exp28_logP_T3-2\18C-01009_M16_octanol_pH-metric high logP.t3r** | | |

#### Experiment Log (continued)

[1:13:10] Datapoint id 80 collected
[1:13:10] Charge balance equation is out by 50.0%
[1:13:10] Stirrer speed set to 60
[1:13:15] pH 9.77 -> 9.97
[1:13:15] Using cautious pH adjust
[1:13:15] Dispensed 0.001105 mL of Base (0.5 M KOH)
[1:13:20] Stepping pH = 10.15
[1:13:36] Stirrer speed set to 0
[1:13:46] Datapoint id 81 collected
[1:13:46] Charge balance equation is out by 50.0%
[1:13:46] Stirrer speed set to 60
[1:13:51] pH 10.14 -> 10.34
[1:13:51] Using cautious pH adjust
[1:13:51] Dispensed 0.001082 mL of Base (0.5 M KOH)
[1:13:56] Stepping pH = 10.37
[1:14:11] Stirrer speed set to 0
[1:14:21] Datapoint id 82 collected
[1:14:21] Charge balance equation is out by 50.0%
[1:14:21] Stirrer speed set to 60
[1:14:26] pH 10.36 -> 10.56
[1:14:26] Using cautious pH adjust
[1:14:26] Dispensed 0.001058 mL of Base (0.5 M KOH)
[1:14:31] Stepping pH = 10.51
[1:14:31] Dispensed 0.000306 mL of Base (0.5 M KOH)
[1:14:37] Stepping pH = 10.55
[1:14:37] Dispensed 0.000118 mL of Base (0.5 M KOH)
[1:14:42] Stepping pH = 10.55
[1:14:57] Stirrer speed set to 0
[1:15:07] Datapoint id 83 collected
[1:15:07] Charge balance equation is out by 29.7%
[1:15:07] Stirrer speed set to 60
[1:15:12] pH 10.55 -> 10.75
[1:15:12] Using cautious pH adjust
[1:15:12] Dispensed 0.001105 mL of Base (0.5 M KOH)
[1:15:17] Stepping pH = 10.66
[1:15:17] Dispensed 0.000682 mL of Base (0.5 M KOH)
[1:15:22] Stepping pH = 10.71
[1:15:22] Dispensed 0.000329 mL of Base (0.5 M KOH)
[1:15:27] Stepping pH = 10.73
[1:15:28] Dispensed 0.000188 mL of Base (0.5 M KOH)
[1:15:33] Stepping pH = 10.74
[1:15:48] Stirrer speed set to 0
[1:15:58] Datapoint id 84 collected
[1:15:58] Charge balance equation is out by -4.8%
[1:15:58] Stirrer speed set to 60
[1:16:03] pH 10.74 -> 10.94
[1:16:03] Using charge balance adjust
[1:16:03] Dispensed 0.002611 mL of Base (0.5 M KOH)
[1:16:23] Stirrer speed set to 0
[1:16:33] Datapoint id 85 collected
[1:16:33] Charge balance equation is out by -12.8%
[1:16:33] Stirrer speed set to 60
[1:16:38] pH 10.92 -> 11.12
[1:16:38] Using charge balance adjust
[1:16:39] Dispensed 0.003387 mL of Base (0.5 M KOH)
[1:16:59] Stirrer speed set to 0
[1:17:09] Datapoint id 86 collected
[1:17:09] Charge balance equation is out by -11.7%
[1:17:09] Stirrer speed set to 60

| | | | |
|---|---|---|---|
| Sample name: | **M16_octanol** | Experiment start time: | **3/1/2018 12:08:09 PM** |
| Assay name: | **pH-metric high logP** | Analyst: | **Pion** |
| Assay ID: | **18C-01009** | Instrument ID: | **T312060** |
| Filename: | **C:\Sirius_T3\Mehtap\20180228_exp28_logP_T3-2\18C-01009_M16_octanol_pH-metric high logP.t3r** | | |

#### Experiment Log (continued)

[1:17:14] pH 11.10 -> 11.30
[1:17:14] Using charge balance adjust
[1:17:14] Dispensed 0.004751 mL of Base (0.5 M KOH)
[1:17:34] Stirrer speed set to 0
[1:17:44] Datapoint id 87 collected
[1:17:44] Charge balance equation is out by -27.1%
[1:17:44] Stirrer speed set to 60
[1:17:50] pH 11.25 -> 11.45
[1:17:50] Using cautious pH adjust
[1:17:50] Dispensed 0.003269 mL of Base (0.5 M KOH)
[1:17:55] Stepping pH = 11.34
[1:17:55] Dispensed 0.002893 mL of Base (0.5 M KOH)
[1:18:00] Stepping pH = 11.41
[1:18:00] Dispensed 0.001364 mL of Base (0.5 M KOH)
[1:18:05] Stepping pH = 11.44
[1:18:05] Dispensed 0.000470 mL of Base (0.5 M KOH)
[1:18:10] Stepping pH = 11.45
[1:18:26] Stirrer speed set to 0
[1:18:36] Datapoint id 88 collected
[1:18:36] Charge balance equation is out by -22.0%
[1:18:36] Stirrer speed set to 60
[1:18:41] pH 11.45 -> 11.65
[1:18:41] Using cautious pH adjust
[1:18:41] Dispensed 0.005103 mL of Base (0.5 M KOH)
[1:18:46] Stepping pH = 11.54
[1:18:46] Dispensed 0.004163 mL of Base (0.5 M KOH)
[1:18:51] Stepping pH = 11.61
[1:18:51] Dispensed 0.002023 mL of Base (0.5 M KOH)
[1:18:57] Stepping pH = 11.64
[1:19:12] Stirrer speed set to 0
[1:19:34] Datapoint id 89 collected
[1:19:34] Charge balance equation is out by -10.3%
[1:19:34] Stirrer speed set to 60
[1:19:39] pH 11.64 -> 11.84
[1:19:39] Using charge balance adjust
[1:19:40] Dispensed 0.015851 mL of Base (0.5 M KOH)
[1:20:00] Stirrer speed set to 0
[1:20:10] Datapoint id 90 collected
[1:20:10] Charge balance equation is out by -12.5%
[1:20:10] Stirrer speed set to 60
[1:20:15] pH 11.82 -> 12.02
[1:20:15] Using charge balance adjust
[1:20:16] Dispensed 0.024506 mL of Base (0.5 M KOH)
[1:20:36] Stirrer speed set to 0
[1:20:46] Datapoint id 91 collected
[1:20:46] Charge balance equation is out by -12.2%
[1:20:46] Stirrer speed set to 60
[1:20:51] pH 12.01 -> 12.05
[1:20:51] Using charge balance adjust
[1:20:51] Dispensed 0.007103 mL of Base (0.5 M KOH)
[1:21:12] Stirrer speed set to 0
[1:21:22] Datapoint id 92 collected
[1:21:22] Charge balance equation is out by -85.4%
[1:21:22] Argon flow rate set to 0
[1:21:26] Titrator arm moved over Titration position
