## Supplementary material for "Octanol-water partition coefficient measurements for the SAMPL6 Blind Prediction Challenge": SM16_18C-09014_M18_octanol_pH-metric high logP_report.pdf

| | | | |
|---|---|---|---|
| Sample name: | **M18_octanol** | Experiment start time: | **3/9/2018 8:37:21 PM** |
| Assay name: | **pH-metric high logP** | Analyst: | **Pion** |
| Assay ID: | **18C-09014** | Instrument ID: | **T312060** |
| Filename: | **C:\Sirius_T3\Mehtap\20180309_exp31_logP_T3-2\18C-09014_M18_octanol_pH-metric high logP.t3r** | | |

#### pH-metric Result

| | | |
|---|---|---|
| logP (XH2 +) | -0.24 | ±0.09 (n=50) |
| logP (neutral XH) | 2.63 | ±0.01 (n=50) |
| logP (X -) | 0.46 | |
| RMSD | 0.266 | |

#### 18C-09014 Points 2 to 24

| | |
|---|---|
| M18_octanol concentration factor | 0.988 |
| Carbonate | 0.1577 mM |
| Acidity error | -0.03698 mM |

#### 18C-09014 Points 25 to 48

| | |
|---|---|
| M18_octanol concentration factor | 0.944 |
| Carbonate | 0.1161 mM |
| Acidity error | -0.54687 mM |

#### 18C-09014 Points 49 to 73

| | |
|---|---|
| M18_octanol concentration factor | 0.993 |
| Carbonate | 0.1503 mM |
| Acidity error | -1.10219 mM |

#### Warnings and errors

| | |
|---|---|
| Errors | None |
| Warnings | None |

#### Sample logD and percent species

| pH | M18_octanol logD | M18_octanol M18_octanolH2 | M18_octanol M18_octanolH | M18_octanol M18_octanol | M18_octanol M18_octanolH2* | M18_octanol M18_octanolH* | M18_octanol M18_octanol* | Comment |
|---|---|---|---|---|---|---|---|---|
| 1.000 | -0.23 | 63.00 % | 0.00 % | 0.00 % | 35.85 % | 1.14 % | 0.00 % | |
| 1.200 | -0.22 | 62.58 % | 0.00 % | 0.00 % | 35.61 % | 1.80 % | 0.00 % | Stomach pH |
| 2.000 | -0.12 | 57.11 % | 0.02 % | 0.00 % | 32.50 % | 10.37 % | 0.00 % | |
| 3.000 | 0.38 | 29.51 % | 0.13 % | 0.00 % | 16.79 % | 53.57 % | 0.00 % | |
| 4.000 | 1.25 | 5.06 % | 0.22 % | 0.00 % | 2.88 % | 91.85 % | 0.00 % | |
| 5.000 | 2.11 | 0.54 % | 0.23 % | 0.00 % | 0.31 % | 98.91 % | 0.00 % | |
| 6.000 | 2.54 | 0.05 % | 0.23 % | 0.00 % | 0.03 % | 99.68 % | 0.00 % | |
| 6.500 | 2.60 | 0.02 % | 0.23 % | 0.00 % | 0.01 % | 99.74 % | 0.00 % | |
| 7.000 | 2.62 | 0.01 % | 0.23 % | 0.00 % | 0.00 % | 99.76 % | 0.00 % | |
| 7.400 | 2.62 | 0.00 % | 0.23 % | 0.00 % | 0.00 % | 99.76 % | 0.00 % | Blood pH |
| 8.000 | 2.63 | 0.00 % | 0.23 % | 0.00 % | 0.00 % | 99.76 % | 0.00 % | |
| 9.000 | 2.62 | 0.00 % | 0.23 % | 0.01 % | 0.00 % | 99.75 % | 0.02 % | |
| 10.000 | 2.54 | 0.00 % | 0.23 % | 0.05 % | 0.00 % | 99.56 % | 0.15 % | |
| 11.000 | 2.13 | 0.00 % | 0.23 % | 0.51 % | 0.00 % | 97.77 % | 1.48 % | |
| 12.000 | 1.32 | 0.00 % | 0.19 % | 4.36 % | 0.00 % | 82.87 % | 12.57 % | |

Sample name: **M18_octanol**
Assay name: **pH-metric high logP**
Assay ID: **18C-09014**
Filename: **C:\Sirius_T3\Mehtap\20180309_exp31_logP_T3-2\18C-09014_M18_octanol_pH-metric high logP.t3r**

Experiment start time: **3/9/2018 8:37:21 PM**
Analyst: **Pion**
Instrument ID: **T312060**

#### Graphs

Charge balance residuals

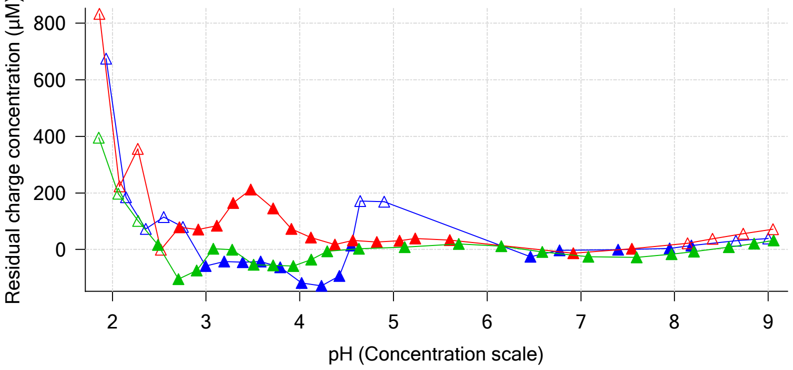


Ionic strength

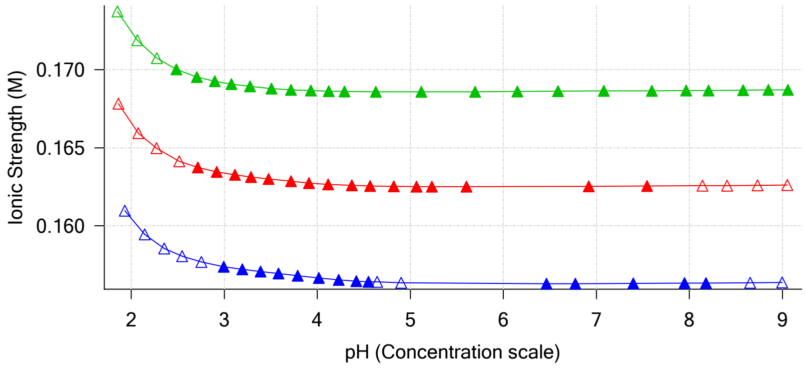


Buffer index profile

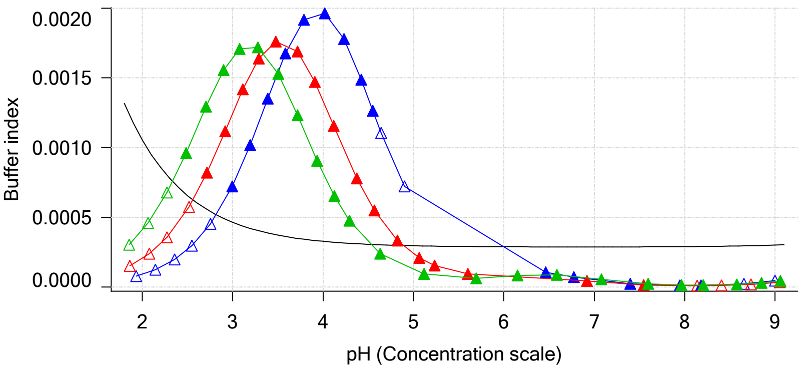


Sample M18_octanol Ionisation Graph

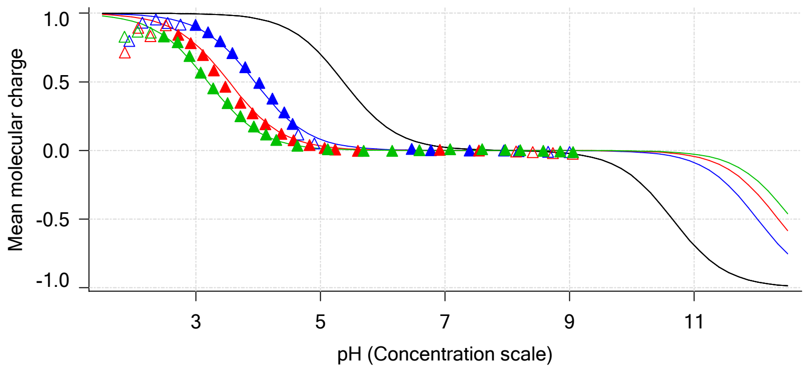


Distribution of species for sample M18_octanol

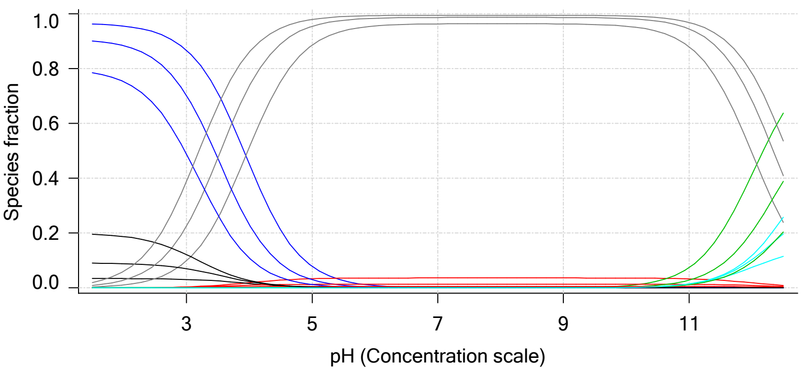


Lipophilicity Profile

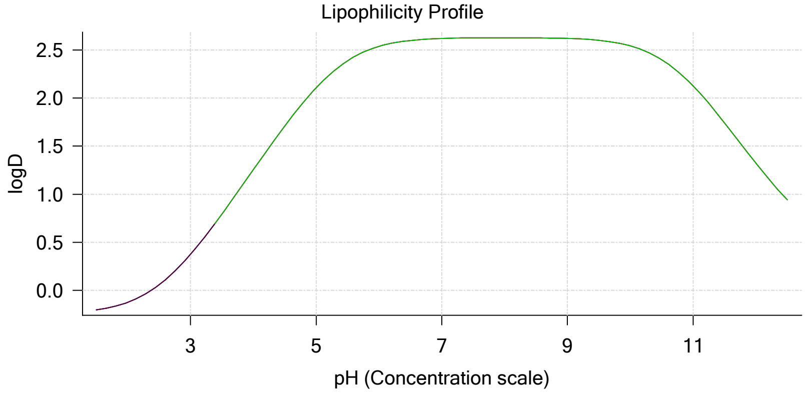


Log R Profile

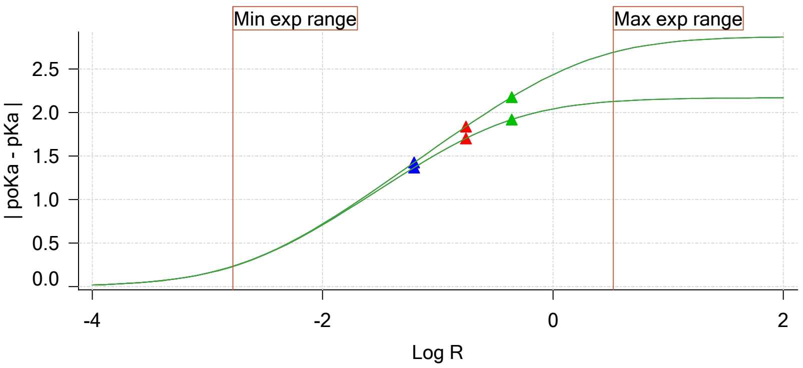


Sample Carbonate Ionisation Graph

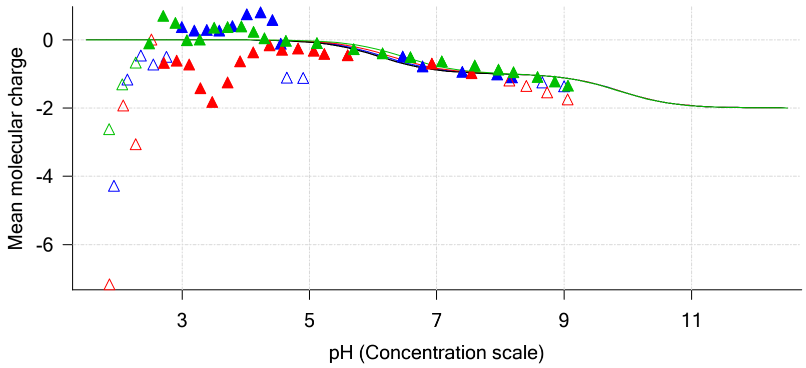

| | | | |
|---|---|---|---|
| Sample name: | **M18_octanol** | Experiment start time: | **3/9/2018 8:37:21 PM** |
| Assay name: | **pH-metric high logP** | Analyst: | **Pion** |
| Assay ID: | **18C-09014** | Instrument ID: | **T312060** |
| Filename: | **C:\Sirius_T3\Mehtap\20180309_exp31_logP_T3-2\18C-09014_M18_octanol_pH-metric high logP.t3r** | | |

#### Graphs (continued)

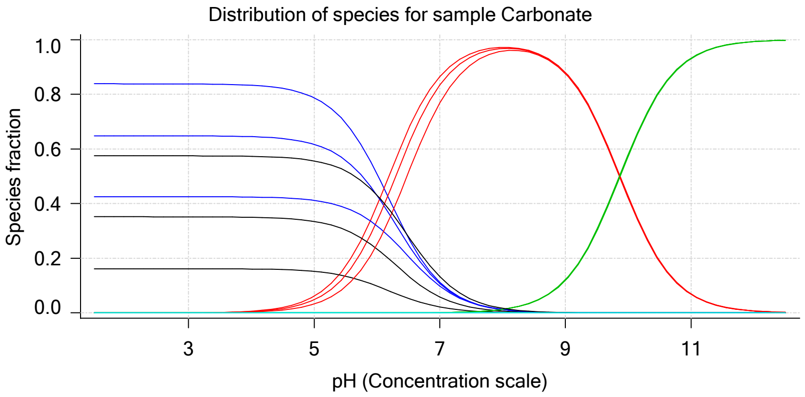

| | | | |
|---|---|---|---|
| Sample name: | **M18_octanol** | Experiment start time: | **3/9/2018 8:37:21 PM** |
| Assay name: | **pH-metric high logP** | Analyst: | **Pion** |
| Assay ID: | **18C-09014** | Instrument ID: | **T312060** |
| Filename: | **C:\Sirius_T3\Mehtap\20180309_exp31_logP_T3-2\18C-09014_M18_octanol_pH-metric high logP.t3r** | | |

### pH-metric high logP Titration 1 of 3 18C-09014 Points 2 to 24

#### Overall results

| | |
|---|---|
| RMSD | 0.227 |
| Average ionic strength | 0.157 M |
| Average temperature | 24.9°C |
| Partition ratio | 0.0624 : 1 |
| Analyte concentration range | 3258.0 μM to 3348.4 μM |
| Total points considered | 14 of 23 |

#### Warnings and errors

| | |
|---|---|
| Errors | None |
| Warnings | None |

#### Four-Plus parameters

| | | | |
|---|---|---|---|
| Alpha | 0.102 | 3/9/2018 8:37:21 PM | C:\Sirius_T3\HCl18C09.t3r |
| S | 0.9967 | 3/9/2018 8:37:21 PM | C:\Sirius_T3\HCl18C09.t3r |
| jH | 1.2 | 3/9/2018 8:37:21 PM | C:\Sirius_T3\HCl18C09.t3r |
| jOH | 0.0 | 3/9/2018 8:37:21 PM | C:\Sirius_T3\HCl18C09.t3r |

#### Titrants

| | | | |
|---|---|---|---|
| 0.50 M HCl | 0.999843 | 3/9/2018 8:37:21 PM | C:\Sirius_T3\HCl18C09.t3r |
| 0.50 M KOH | 0.999845 | 3/9/2018 8:37:21 PM | C:\Sirius_T3\KOH18B27.t3r |

#### Sample

| | |
|---|---|
| M18_octanol concentration factor | 0.988 |
| Base pKa 1 | 5.37 |
| Acid pKa 2 | 10.65 |
| logP (XH2 +) | -1.08 |
| logP (neutral XH) | 2.57 |
| logP (X -) | 0.46 |

#### Sample graphs

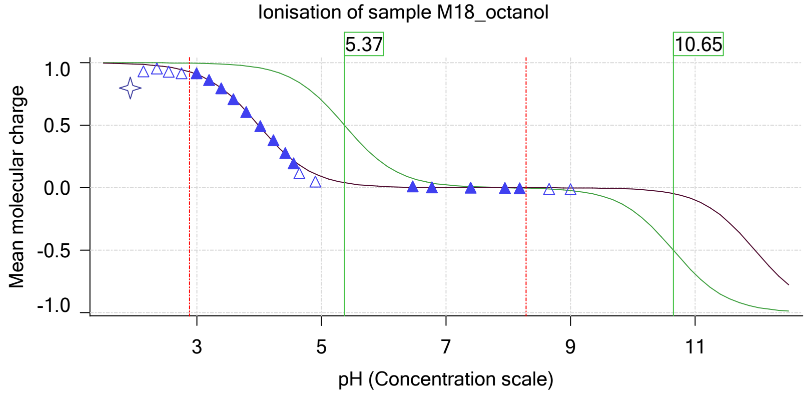


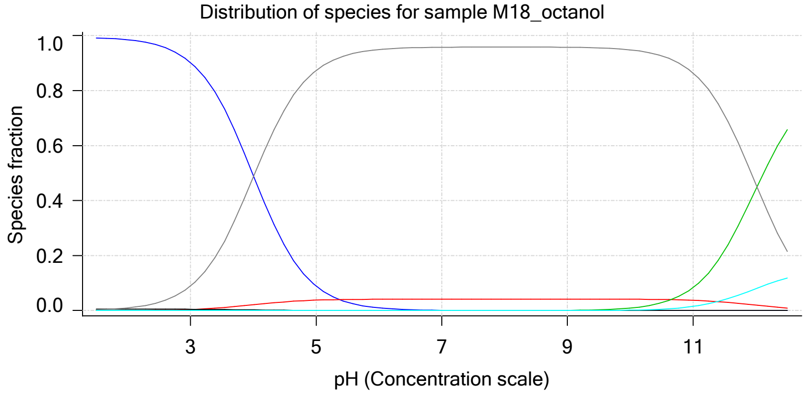

| | | | |
|---|---|---|---|
| Sample name: | **M18_octanol** | Experiment start time: | **3/9/2018 8:37:21 PM** |
| Assay name: | **pH-metric high logP** | Analyst: | **Pion** |
| Assay ID: | **18C-09014** | Instrument ID: | **T312060** |
| Filename: | **C:\Sirius_T3\Mehtap\20180309_exp31_logP_T3-2\18C-09014_M18_octanol_pH-metric high logP.t3r** | | |

#### Sample graphs (continued)

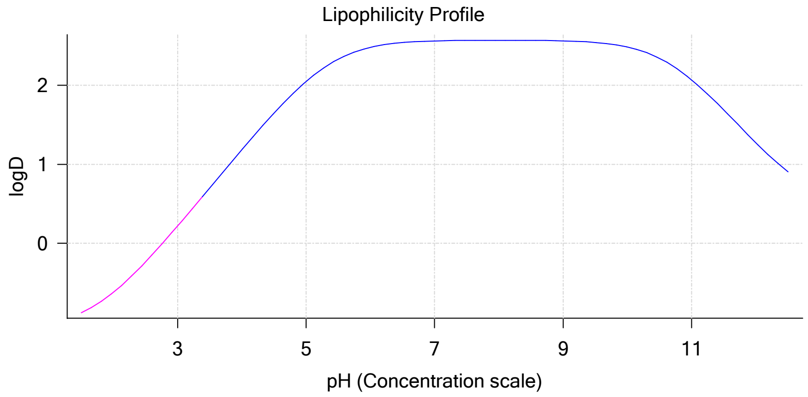


#### Sample logD and percent species

| pH | M18_octanol logD | M18_octanol M18_octanolH2 | M18_octanol M18_octanolH | M18_octanol M18_octanol | M18_octanol M18_octanolH2* | M18_octanol M18_octanolH* | M18_octanol M18_octanol* | Comment |
|---|---|---|---|---|---|---|---|---|
| 1.000 | -1.01 | 99.39 % | 0.00 % | 0.00 % | 0.51 % | 0.10 % | 0.00 % | |
| 1.200 | -0.97 | 99.33 % | 0.01 % | 0.00 % | 0.51 % | 0.16 % | 0.00 % | Stomach pH |
| 2.000 | -0.62 | 98.48 % | 0.04 % | 0.00 % | 0.51 % | 0.97 % | 0.00 % | |
| 3.000 | 0.22 | 90.24 % | 0.38 % | 0.00 % | 0.46 % | 8.91 % | 0.00 % | |
| 4.000 | 1.18 | 49.14 % | 2.10 % | 0.00 % | 0.25 % | 48.51 % | 0.00 % | |
| 5.000 | 2.05 | 8.85 % | 3.77 % | 0.00 % | 0.05 % | 87.33 % | 0.00 % | |
| 6.000 | 2.48 | 0.96 % | 4.10 % | 0.00 % | 0.00 % | 94.93 % | 0.00 % | |
| 6.500 | 2.54 | 0.31 % | 4.13 % | 0.00 % | 0.00 % | 95.56 % | 0.00 % | |
| 7.000 | 2.56 | 0.10 % | 4.14 % | 0.00 % | 0.00 % | 95.76 % | 0.00 % | |
| 7.400 | 2.56 | 0.04 % | 4.14 % | 0.00 % | 0.00 % | 95.82 % | 0.00 % | Blood pH |
| 8.000 | 2.57 | 0.01 % | 4.14 % | 0.01 % | 0.00 % | 95.84 % | 0.00 % | |
| 9.000 | 2.56 | 0.00 % | 4.14 % | 0.09 % | 0.00 % | 95.75 % | 0.02 % | |
| 10.000 | 2.48 | 0.00 % | 4.10 % | 0.92 % | 0.00 % | 94.82 % | 0.17 % | |
| 11.000 | 2.07 | 0.00 % | 3.73 % | 8.36 % | 0.00 % | 86.40 % | 1.50 % | |
| 12.000 | 1.27 | 0.00 % | 1.98 % | 44.28 % | 0.00 % | 45.77 % | 7.97 % | |

#### Carbonate and acidity

| | |
|---|---|
| Carbonate | 0.158 mM |
| Acidity error | -0.037 mM |

#### Other graphs

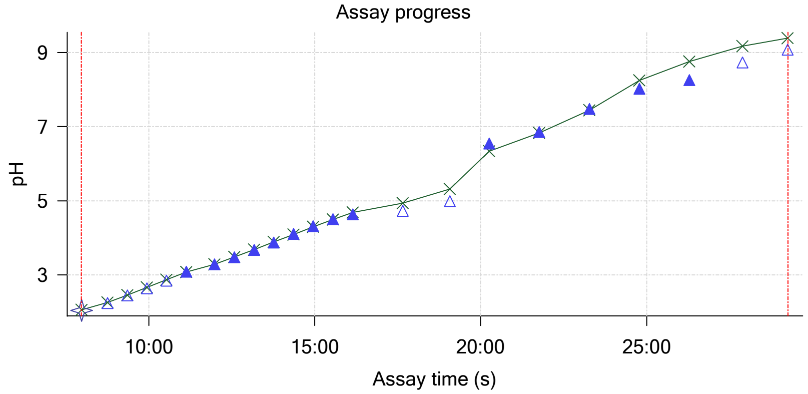


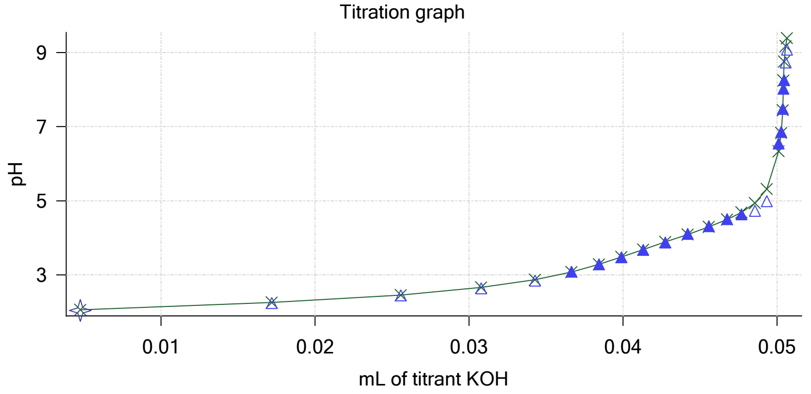

| | | | |
|---|---|---|---|
| Sample name: | **M18_octanol** | Experiment start time: | **3/9/2018 8:37:21 PM** |
| Assay name: | **pH-metric high logP** | Analyst: | **Pion** |
| Assay ID: | **18C-09014** | Instrument ID: | **T312060** |
| Filename: | **C:\Sirius_T3\Mehtap\20180309_exp31_logP_T3-2\18C-09014_M18_octanol_pH-metric high logP.t3r** | | |

#### Other graphs (continued)

Charge balance residuals

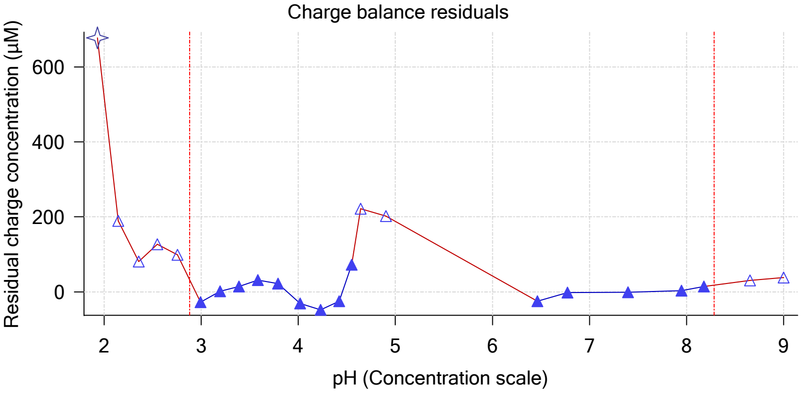


Buffer index profile

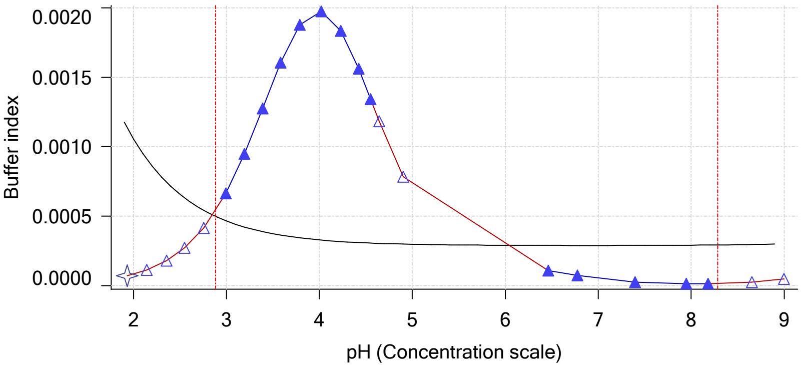


Ionic strength

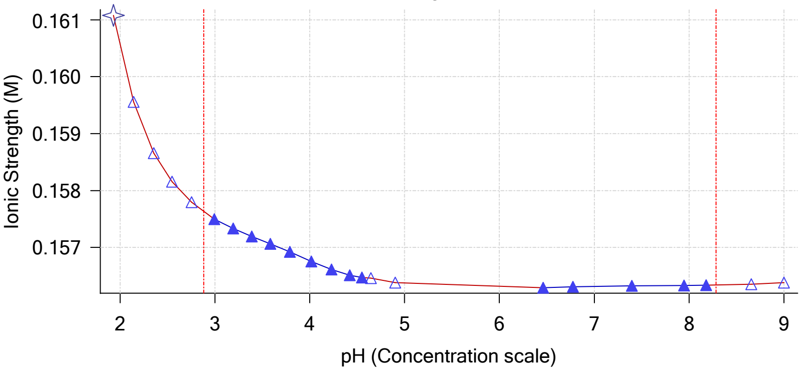


Log R Profile

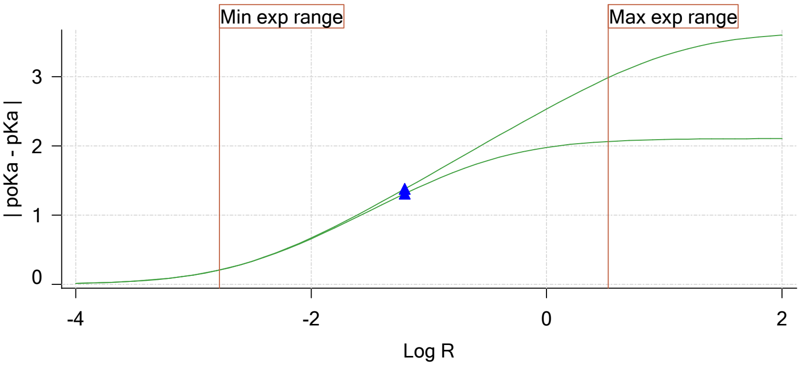


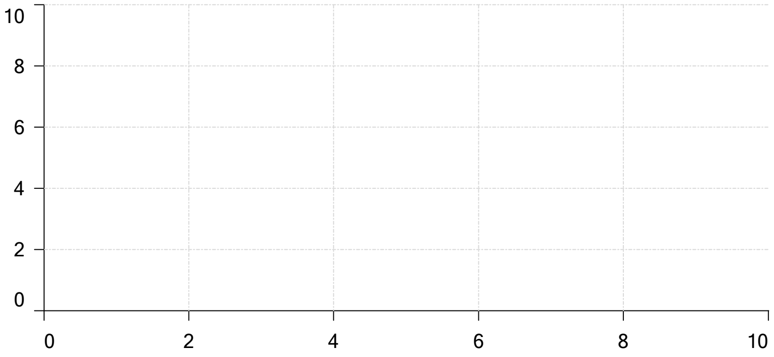

Ionisation of sample Carbonate

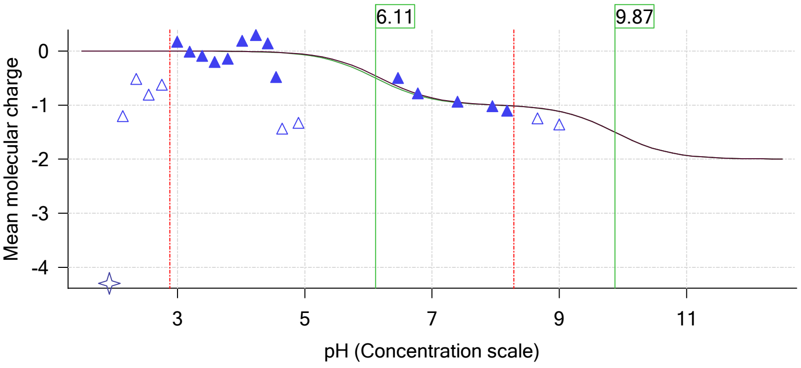

| | | | |
|---|---|---|---|
| Sample name: | **M18_octanol** | Experiment start time: | **3/9/2018 8:37:21 PM** |
| Assay name: | **pH-metric high logP** | Analyst: | **Pion** |
| Assay ID: | **18C-09014** | Instrument ID: | **T312060** |
| Filename: | **C:\Sirius_T3\Mehtap\20180309_exp31_logP_T3-2\18C-09014_M18_octanol_pH-metric high logP.t3r** | | |

# pH-metric high logP   Titration 2 of 3   18C-09014 Points 25 to 48

#### Overall results

| | |
|---|---|
| RMSD | 0.131 |
| Average ionic strength | 0.163 M |
| Average temperature | 25.0°C |
| Partition ratio | 0.1756 : 1 |
| Analyte concentration range | 2756.3 µM to 2832.3 µM |
| Total points considered | 16 of 24 |

#### Warnings and errors

| | |
|---|---|
| Errors | None |
| Warnings | None |

#### Four-Plus parameters

| | | | |
|---|---|---|---|
| Alpha | 0.102 | 3/9/2018 8:37:21 PM | C:\Sirius_T3\HCl18C09.t3r |
| S | 0.9967 | 3/9/2018 8:37:21 PM | C:\Sirius_T3\HCl18C09.t3r |
| jH | 1.2 | 3/9/2018 8:37:21 PM | C:\Sirius_T3\HCl18C09.t3r |
| jOH | 0.0 | 3/9/2018 8:37:21 PM | C:\Sirius_T3\HCl18C09.t3r |

#### Titrants

| | | | |
|---|---|---|---|
| 0.50 M HCl | 0.999843 | 3/9/2018 8:37:21 PM | C:\Sirius_T3\HCl18C09.t3r |
| 0.50 M KOH | 0.999845 | 3/9/2018 8:37:21 PM | C:\Sirius_T3\KOH18B27.t3r |

#### Sample

| | |
|---|---|
| M18_octanol concentration factor | 0.944 |
| Base pKa 1 | 5.37 |
| Acid pKa 2 | 10.65 |
| logP (XH2 +) | -1.92 |
| logP (neutral XH) | 2.67 |
| logP (X -) | 0.46 |

#### Sample graphs

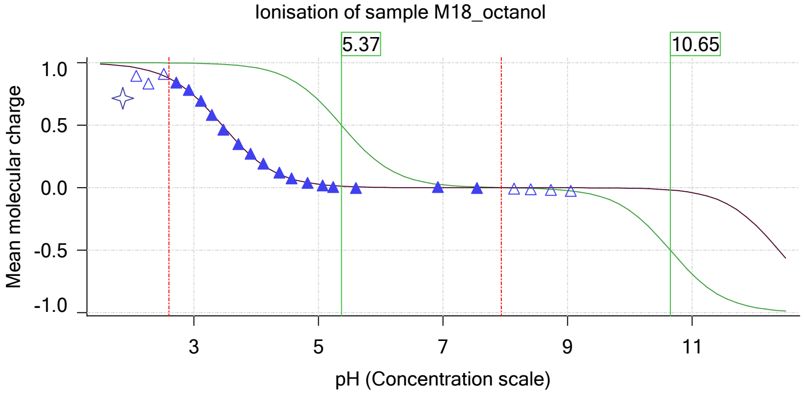


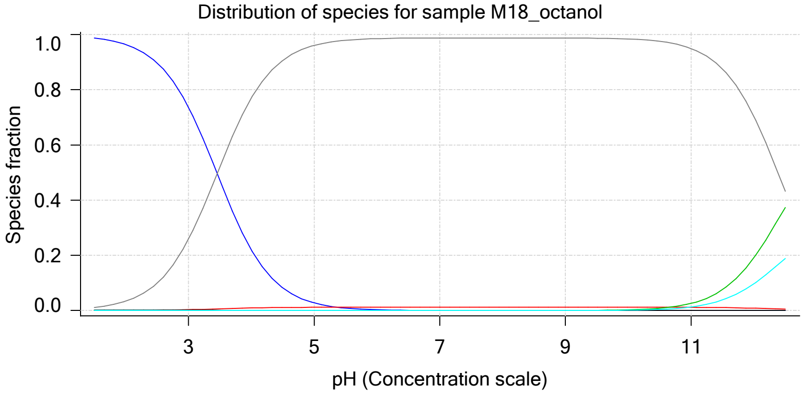

| | | | |
|---|---|---|---|
| Sample name: | **M18_octanol** | Experiment start time: | **3/9/2018 8:37:21 PM** |
| Assay name: | **pH-metric high logP** | Analyst: | **Pion** |
| Assay ID: | **18C-09014** | Instrument ID: | **T312060** |
| Filename: | **C:\Sirius_T3\Mehtap\20180309_exp31_logP_T3-2\18C-09014_M18_octanol_pH-metric high logP.t3r** | | |

#### Sample graphs (continued)

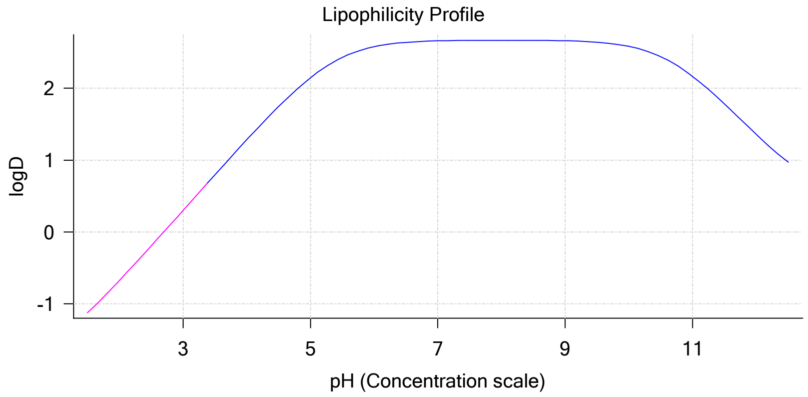


#### Sample logD and percent species

| pH | M18_octanol logD | M18_octanol M18_octanolH2 | M18_octanol M18_octanolH | M18_octanol M18_octanol | M18_octanol M18_octanolH2* | M18_octanol M18_octanolH* | M18_octanol M18_octanol* | Comment |
|---|---|---|---|---|---|---|---|---|
| 1.000 | -1.50 | 99.44 % | 0.00 % | 0.00 % | 0.21 % | 0.35 % | 0.00 % | |
| 1.200 | -1.36 | 99.23 % | 0.01 % | 0.00 % | 0.21 % | 0.55 % | 0.00 % | Stomach pH |
| 2.000 | -0.68 | 96.38 % | 0.04 % | 0.00 % | 0.20 % | 3.37 % | 0.00 % | |
| 3.000 | 0.30 | 73.74 % | 0.31 % | 0.00 % | 0.16 % | 25.79 % | 0.00 % | |
| 4.000 | 1.28 | 22.02 % | 0.94 % | 0.00 % | 0.05 % | 76.99 % | 0.00 % | |
| 5.000 | 2.14 | 2.75 % | 1.17 % | 0.00 % | 0.01 % | 96.07 % | 0.00 % | |
| 6.000 | 2.58 | 0.28 % | 1.20 % | 0.00 % | 0.00 % | 98.52 % | 0.00 % | |
| 6.500 | 2.64 | 0.09 % | 1.20 % | 0.00 % | 0.00 % | 98.71 % | 0.00 % | |
| 7.000 | 2.66 | 0.03 % | 1.20 % | 0.00 % | 0.00 % | 98.77 % | 0.00 % | |
| 7.400 | 2.66 | 0.01 % | 1.21 % | 0.00 % | 0.00 % | 98.78 % | 0.00 % | Blood pH |
| 8.000 | 2.67 | 0.00 % | 1.21 % | 0.00 % | 0.00 % | 98.79 % | 0.00 % | |
| 9.000 | 2.66 | 0.00 % | 1.20 % | 0.03 % | 0.00 % | 98.75 % | 0.01 % | |
| 10.000 | 2.58 | 0.00 % | 1.20 % | 0.27 % | 0.00 % | 98.39 % | 0.14 % | |
| 11.000 | 2.16 | 0.00 % | 1.16 % | 2.59 % | 0.00 % | 94.94 % | 1.31 % | |
| 12.000 | 1.36 | 0.00 % | 0.86 % | 19.18 % | 0.00 % | 70.24 % | 9.72 % | |

#### Carbonate and acidity

| | |
|---|---|
| Carbonate | 0.116 mM |
| Acidity error | -0.547 mM |

#### Other graphs

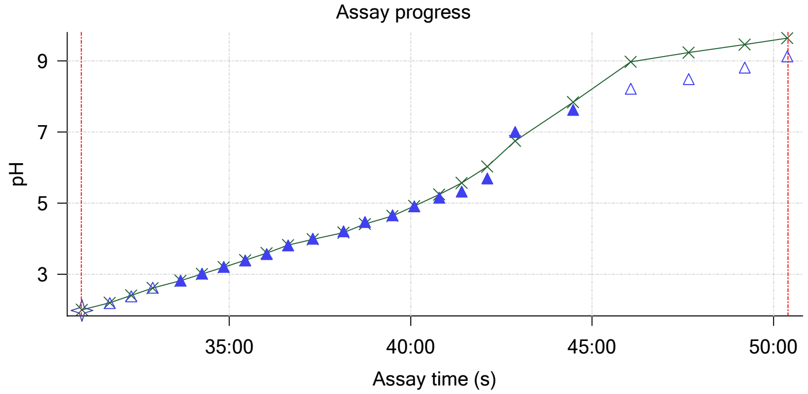


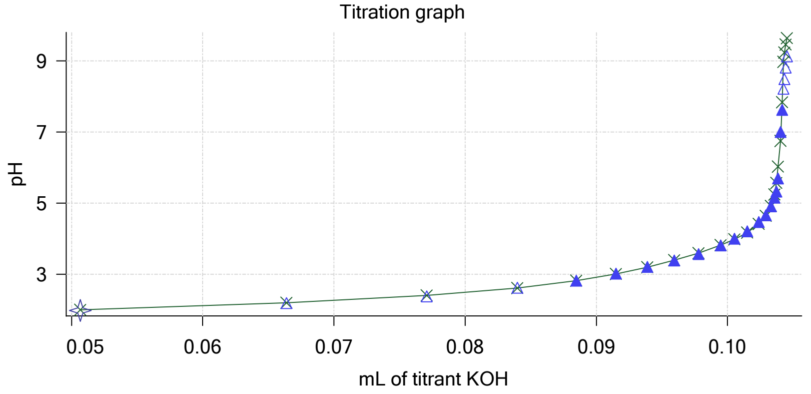

Sample name: **M18_octanol** Experiment start time: **3/9/2018 8:37:21 PM**
Assay name: **pH-metric high logP** Analyst: **Pion**
Assay ID: **18C-09014** Instrument ID: **T312060**
Filename: **C:\Sirius_T3\Mehtap\20180309_exp31_logP_T3-2\18C-09014_M18_octanol_pH-metric high logP.t3r**

#### Other graphs (continued)

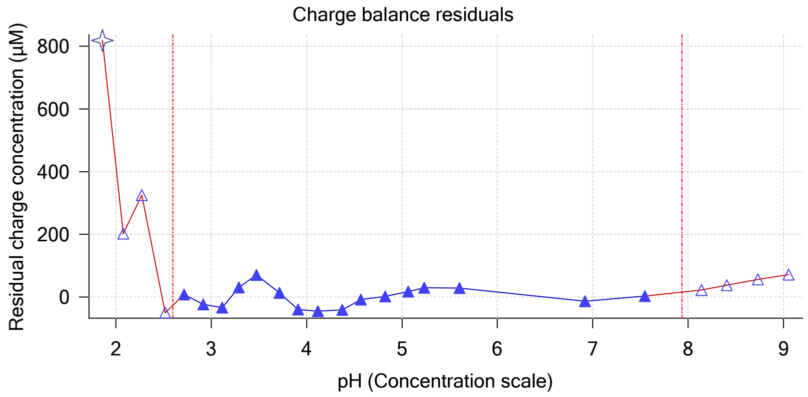


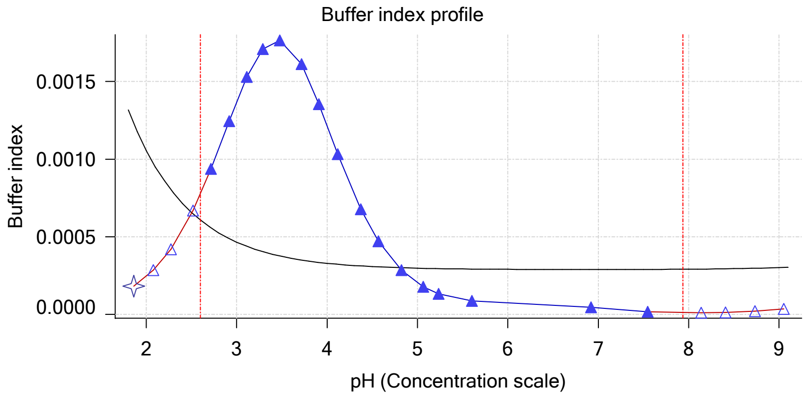


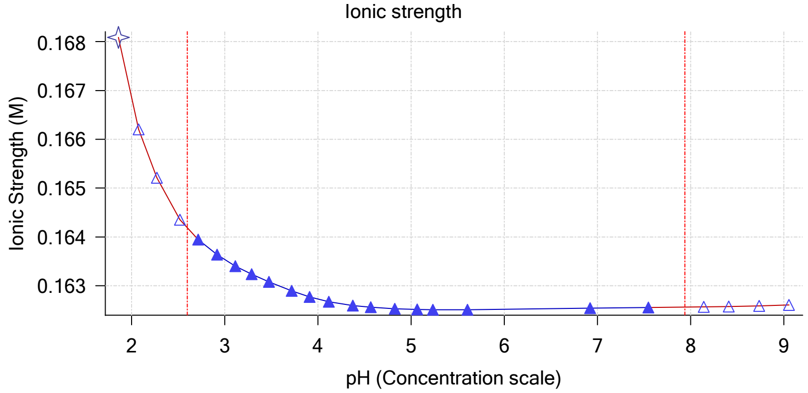


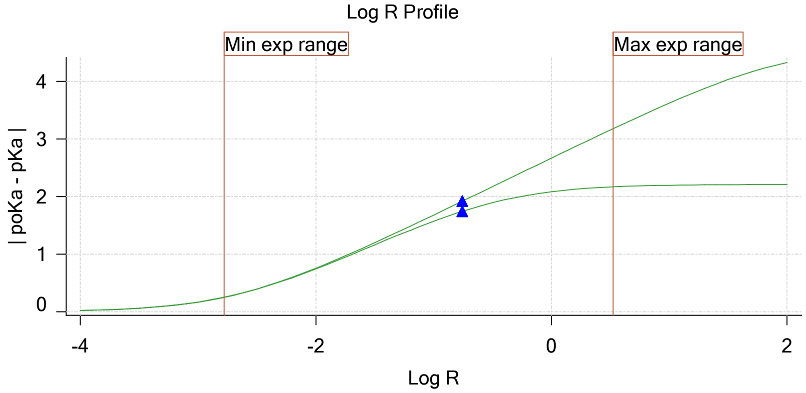


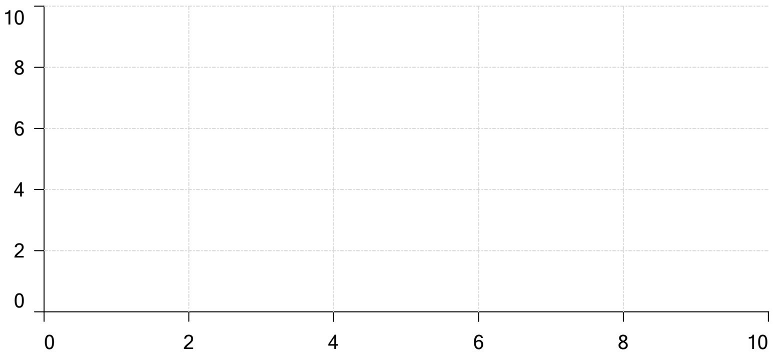


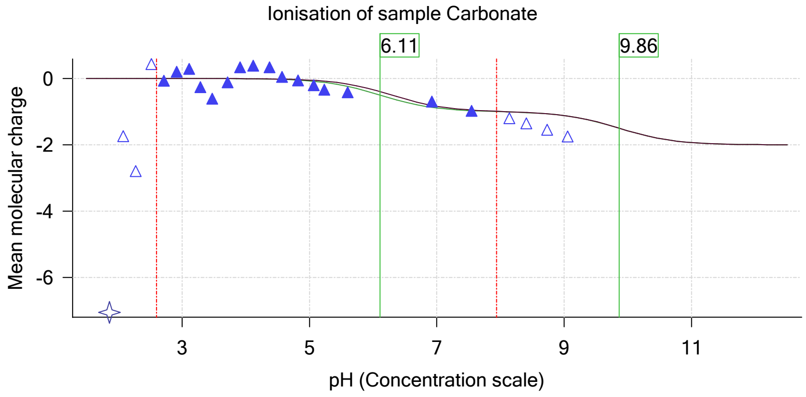

| | | | |
|---|---|---|---|
| Sample name: | **M18_octanol** | Experiment start time: | **3/9/2018 8:37:21 PM** |
| Assay name: | **pH-metric high logP** | Analyst: | **Pion** |
| Assay ID: | **18C-09014** | Instrument ID: | **T312060** |
| Filename: | **C:\Sirius_T3\Mehtap\20180309_exp31_logP_T3-2\18C-09014_M18_octanol_pH-metric high logP.t3r** | | |

### pH-metric high logP Titration 3 of 3 18C-09014 Points 49 to 73

#### Overall results

| | |
|---|---|
| RMSD | 0.342 |
| Average ionic strength | 0.169 M |
| Average temperature | 25.0°C |
| Partition ratio | 0.4375 : 1 |
| Analyte concentration range | 2105.7 µM to 2154.4 µM |
| Total points considered | 22 of 25 |

#### Warnings and errors

| | |
|---|---|
| Errors | None |
| Warnings | One or more logP values out of range<br>Excessive acidity error present |

#### Four-Plus parameters

| | | | |
|---|---|---|---|
| Alpha | 0.102 | 3/9/2018 8:37:21 PM | C:\Sirius_T3\HCl18C09.t3r |
| S | 0.9967 | 3/9/2018 8:37:21 PM | C:\Sirius_T3\HCl18C09.t3r |
| jH | 1.2 | 3/9/2018 8:37:21 PM | C:\Sirius_T3\HCl18C09.t3r |
| jOH | 0.0 | 3/9/2018 8:37:21 PM | C:\Sirius_T3\HCl18C09.t3r |

#### Titrants

| | | | |
|---|---|---|---|
| 0.50 M HCl | 0.999843 | 3/9/2018 8:37:21 PM | C:\Sirius_T3\HCl18C09.t3r |
| 0.50 M KOH | 0.999845 | 3/9/2018 8:37:21 PM | C:\Sirius_T3\KOH18B27.t3r |

#### Sample

| | |
|---|---|
| M18_octanol concentration factor | 0.993 |
| Base pKa 1 | 5.37 |
| Acid pKa 2 | 10.65 |
| logP (XH2 +) | -2.03 |
| logP (neutral XH) | 2.50 |
| logP (X -) | 0.46 |

#### Sample graphs

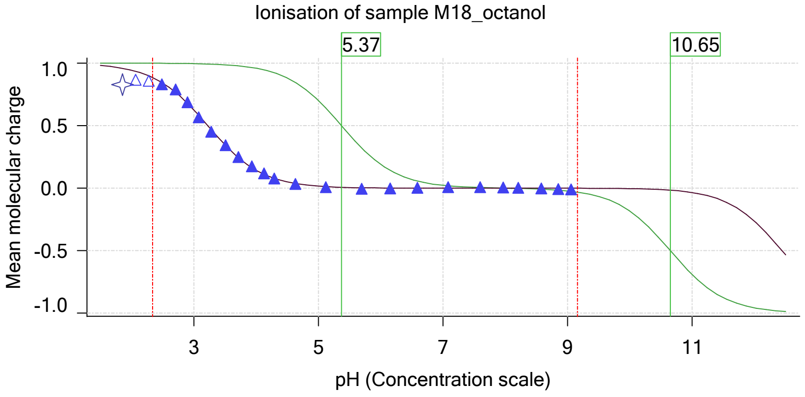


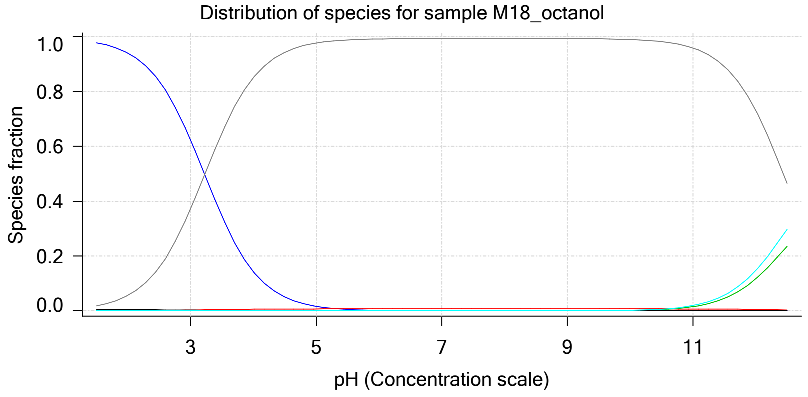

| | | | |
|---|---|---|---|
| Sample name: | **M18_octanol** | Experiment start time: | **3/9/2018 8:37:21 PM** |
| Assay name: | **pH-metric high logP** | Analyst: | **Pion** |
| Assay ID: | **18C-09014** | Instrument ID: | **T312060** |
| Filename: | **C:\Sirius_T3\Mehtap\20180309_exp31_logP_T3-2\18C-09014_M18_octanol_pH-metric high logP.t3r** | | |

#### Sample graphs (continued)

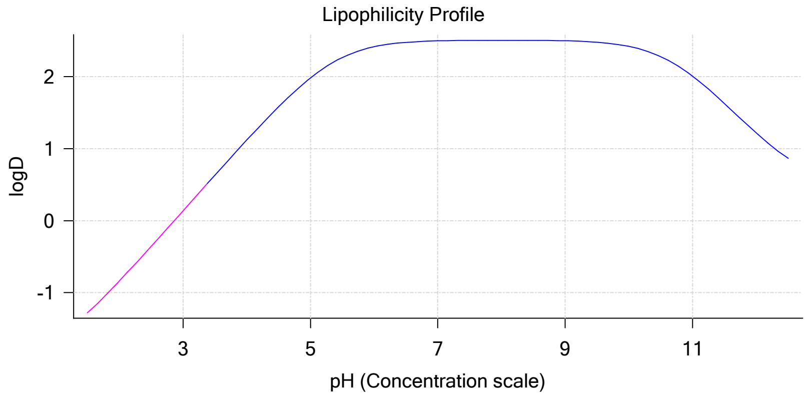


#### Sample logD and percent species

| pH | M18_octanol logD | M18_octanol M18_octanolH2 | M18_octanol M18_octanolH | M18_octanol M18_octanol | M18_octanol M18_octanolH2* | M18_octanol M18_octanolH* | M18_octanol M18_octanol* | Comment |
|---|---|---|---|---|---|---|---|---|
| 1.000 | -1.64 | 99.00 % | 0.00 % | 0.00 % | 0.40 % | 0.59 % | 0.00 % | |
| 1.200 | -1.51 | 98.66 % | 0.01 % | 0.00 % | 0.40 % | 0.93 % | 0.00 % | Stomach pH |
| 2.000 | -0.84 | 93.97 % | 0.04 % | 0.00 % | 0.38 % | 5.61 % | 0.00 % | |
| 3.000 | 0.14 | 62.30 % | 0.27 % | 0.00 % | 0.25 % | 37.18 % | 0.00 % | |
| 4.000 | 1.12 | 14.25 % | 0.61 % | 0.00 % | 0.06 % | 85.08 % | 0.00 % | |
| 5.000 | 1.98 | 1.64 % | 0.70 % | 0.00 % | 0.01 % | 97.66 % | 0.00 % | |
| 6.000 | 2.41 | 0.17 % | 0.71 % | 0.00 % | 0.00 % | 99.12 % | 0.00 % | |
| 6.500 | 2.47 | 0.05 % | 0.71 % | 0.00 % | 0.00 % | 99.24 % | 0.00 % | |
| 7.000 | 2.49 | 0.02 % | 0.71 % | 0.00 % | 0.00 % | 99.27 % | 0.00 % | |
| 7.400 | 2.50 | 0.01 % | 0.71 % | 0.00 % | 0.00 % | 99.28 % | 0.00 % | Blood pH |
| 8.000 | 2.50 | 0.00 % | 0.71 % | 0.00 % | 0.00 % | 99.29 % | 0.00 % | |
| 9.000 | 2.50 | 0.00 % | 0.71 % | 0.02 % | 0.00 % | 99.25 % | 0.02 % | |
| 10.000 | 2.42 | 0.00 % | 0.71 % | 0.16 % | 0.00 % | 98.93 % | 0.20 % | |
| 11.000 | 2.00 | 0.00 % | 0.69 % | 1.53 % | 0.00 % | 95.85 % | 1.93 % | |
| 12.000 | 1.22 | 0.00 % | 0.52 % | 11.69 % | 0.00 % | 73.05 % | 14.75 % | |

#### Carbonate and acidity

| | |
|---|---|
| Carbonate | 0.150 mM |
| Acidity error | -1.102 mM |

#### Other graphs

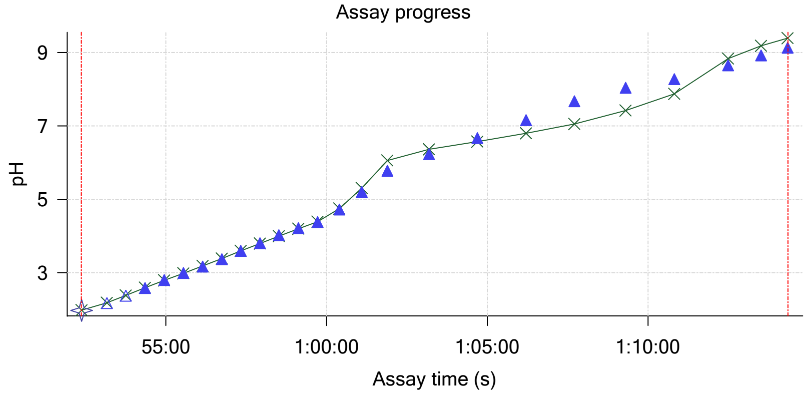


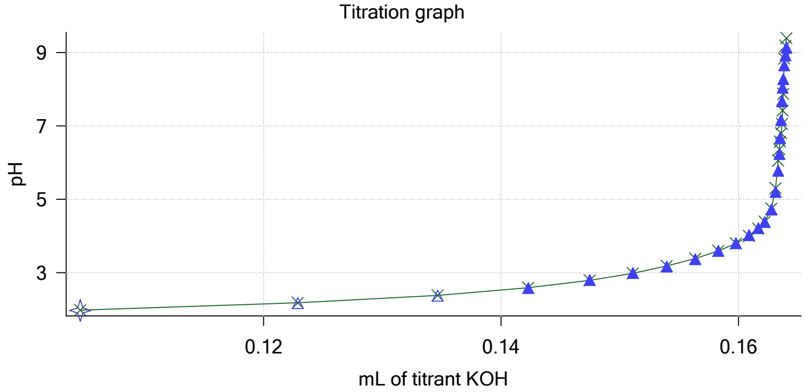

| | | | |
|---|---|---|---|
| Sample name: | **M18_octanol** | Experiment start time: | **3/9/2018 8:37:21 PM** |
| Assay name: | **pH-metric high logP** | Analyst: | **Pion** |
| Assay ID: | **18C-09014** | Instrument ID: | **T312060** |
| Filename: | **C:\Sirius_T3\Mehtap\20180309_exp31_logP_T3-2\18C-09014_M18_octanol_pH-metric high logP.t3r** | | |

#### Other graphs (continued)

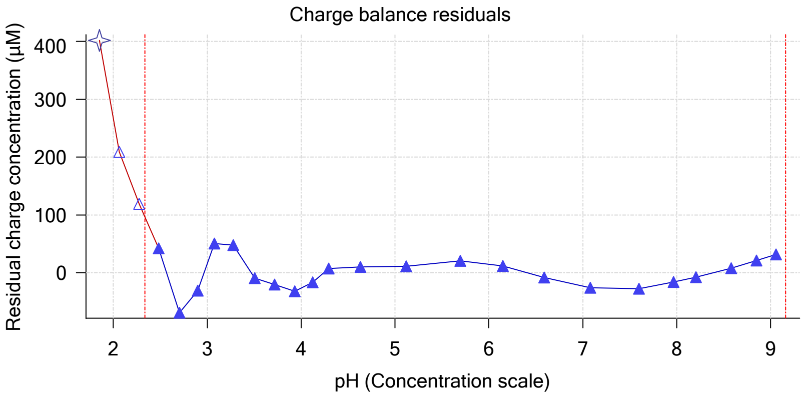


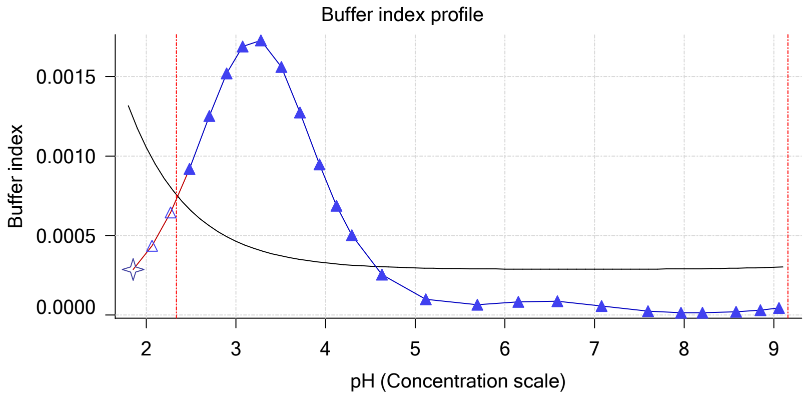


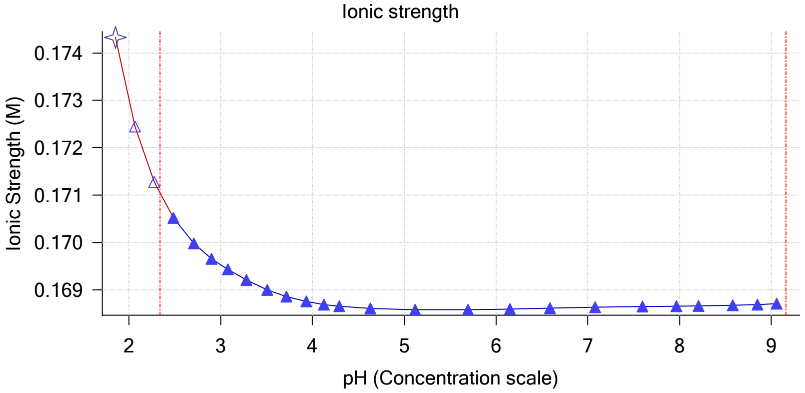


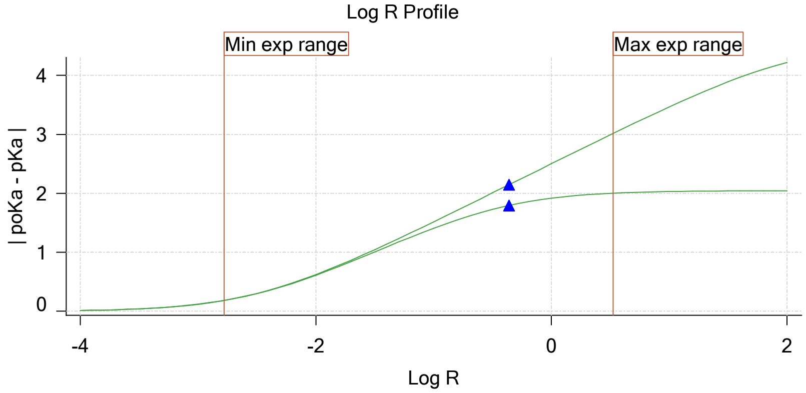


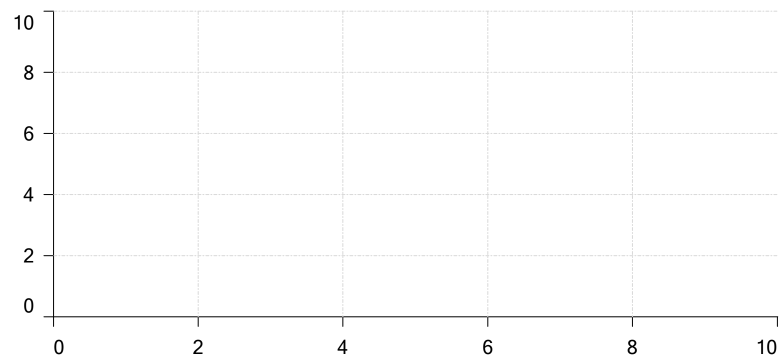

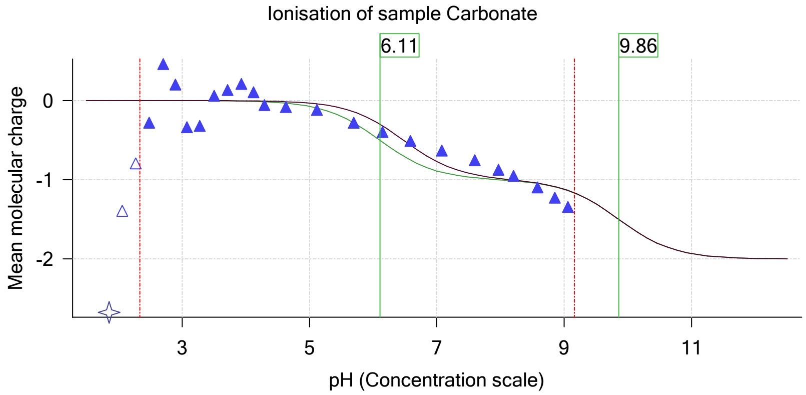

| | | | |
|---|---|---|---|
| Sample name: | **M18_octanol** | Experiment start time: | **3/9/2018 8:37:21 PM** |
| Assay name: | **pH-metric high logP** | Analyst: | **Pion** |
| Assay ID: | **18C-09014** | Instrument ID: | **T312060** |
| Filename: | **C:\Sirius_T3\Mehtap\20180309_exp31_logP_T3-2\18C-09014_M18_octanol_pH-metric high logP.t3r** | | |

#### Assay Model

| Settings | Value | Date/Time changed | Imported from |
|---|---|---|---|
| Sample name | M18_octanol | 2/27/2018 7:08:39 PM | User entered value |
| Sample by | Weight | | Default value |
| Sample weight | 0.001480 g | 3/9/2018 2:22:36 PM | User entered value |
| Formula weight | 267.11 g/mol | 2/27/2018 7:08:39 PM | User entered value |
| Solubility | Unknown | | Default value |
| Molecular weight | 267.11 | 2/27/2018 7:08:39 PM | User entered value |
| Individual pKa ionic environments | No | | Default value |
| Number of pKas | 2 | 2/27/2018 7:08:39 PM | User entered value |
| Sample is a | Ampholyte | 2/27/2018 7:08:39 PM | User entered value |
| pKa 1 | 5.37 | 2/27/2018 7:08:39 PM | User entered value |
| Type | Base | 2/27/2018 7:08:39 PM | User entered value |
| pKa 2 | 10.65 | 2/27/2018 7:08:39 PM | User entered value |
| Type | Acid | 2/27/2018 7:08:39 PM | User entered value |
| logp (XH2 +) | -0.35 | 2/28/2018 3:20:28 PM | User entered value |
| logP (neutral XH) | 2.57 | 3/2/2018 4:34:50 PM | User entered value |
| logP (X -) | 0.46 | 2/27/2018 7:09:34 PM | User entered value |

#### Events

| Time | Event | Water | Acid | Base | Octanol | pH | dpH/dt | pH R-squared | pH SD | dpH/ time |
|---|---|---|---|---|---|---|---|---|---|---|
| 4:57.9 | Manual volume addition | | | | 0.10000 mL | | | | | |
| 4:59.0 | Initial pH = 7.17 | | | | | | | | | |
| 7:58.6 | Data point 2 | 1.50000 mL | 0.05002 mL | 0.00475 mL | 0.10000 mL | 2.041 | -0.00537 | 0.75188 | 0.00031 | 10.5 s |
| 8:45.2 | Data point 3 | 1.50000 mL | 0.05002 mL | 0.01719 mL | 0.10000 mL | 2.248 | -0.00630 | 0.13591 | 0.00084 | 10.0 s |
| 9:20.9 | Data point 4 | 1.50000 mL | 0.05002 mL | 0.02556 mL | 0.10000 mL | 2.455 | -0.00491 | 0.10647 | 0.00074 | 10.0 s |
| 9:56.5 | Data point 5 | 1.50000 mL | 0.05002 mL | 0.03079 mL | 0.10000 mL | 2.645 | -0.00008 | 0.00076 | 0.00014 | 10.0 s |
| 10:31.9 | Data point 6 | 1.50000 mL | 0.05002 mL | 0.03429 mL | 0.10000 mL | 2.849 | -0.00867 | 0.76331 | 0.00049 | 10.5 s |
| 11:07.9 | Data point 7 | 1.50000 mL | 0.05002 mL | 0.03667 mL | 0.10000 mL | 3.088 | -0.00680 | 0.26943 | 0.00065 | 10.0 s |
| 11:58.8 | Data point 8 | 1.50000 mL | 0.05002 mL | 0.03845 mL | 0.10000 mL | 3.287 | -0.00858 | 0.64011 | 0.00053 | 10.5 s |
| 12:34.8 | Data point 9 | 1.50000 mL | 0.05002 mL | 0.03991 mL | 0.10000 mL | 3.481 | -0.00517 | 0.71303 | 0.00030 | 10.0 s |
| 13:10.2 | Data point 10 | 1.50000 mL | 0.05002 mL | 0.04132 mL | 0.10000 mL | 3.675 | -0.01100 | 0.68415 | 0.00066 | 10.0 s |
| 13:45.7 | Data point 11 | 1.50000 mL | 0.05002 mL | 0.04276 mL | 0.10000 mL | 3.880 | -0.01228 | 0.70505 | 0.00072 | 10.5 s |
| 14:21.7 | Data point 12 | 1.50000 mL | 0.05002 mL | 0.04421 mL | 0.10000 mL | 4.108 | -0.01324 | 0.78765 | 0.00074 | 10.0 s |
| 14:57.1 | Data point 13 | 1.50000 mL | 0.05002 mL | 0.04558 mL | 0.10000 mL | 4.320 | -0.00760 | 0.32722 | 0.00066 | 10.0 s |
| 15:32.6 | Data point 14 | 1.50000 mL | 0.05002 mL | 0.04675 mL | 0.10000 mL | 4.509 | -0.01875 | 0.88887 | 0.00098 | 10.5 s |
| 16:08.5 | Data point 15 | 1.50000 mL | 0.05002 mL | 0.04770 mL | 0.10000 mL | 4.638 | -0.09531 | 0.99824 | 0.00471 | Time out at |
| 17:39.0 | Data point 16 | 1.50000 mL | 0.05002 mL | 0.04857 mL | 0.10000 mL | 4.731 | -0.03485 | 0.99545 | 0.00173 | Time out at |
| 19:04.3 | Data point 17 | 1.50000 mL | 0.05002 mL | 0.04934 mL | 0.10000 mL | 4.989 | -0.01963 | 0.98594 | 0.00098 | 40.5 s |

| | | | |
|---|---|---|---|
| Sample name: | **M18_octanol** | Experiment start time: | **3/9/2018 8:37:21 PM** |
| Assay name: | **pH-metric high logP** | Analyst: | **Pion** |
| Assay ID: | **18C-09014** | Instrument ID: | **T312060** |
| Filename: | **C:\Sirius_T3\Mehtap\20180309_exp31_logP_T3-2\18C-09014_M18_octanol_pH-metric high logP.t3r** | | |

#### Events (continued)

| Time | Event | Water | Acid | Base | Octanol | pH | dpH/dt | pH R-squared | pH SD | dpH/dt time |
|---|---|---|---|---|---|---|---|---|---|---|
| 20:15.5 | Data point 18 | 1.50000 mL | 0.05002 mL | 0.05012 mL | 0.10000 mL | 6.544 | -0.03976 | 0.99389 | 0.00197 | Timed out at 59.5 s |
| 21:46.0 | Data point 19 | 1.50000 mL | 0.05002 mL | 0.05028 mL | 0.10000 mL | 6.855 | -0.03344 | 0.99646 | 0.00165 | Timed out at 59.5 s |
| 23:16.5 | Data point 20 | 1.50000 mL | 0.05002 mL | 0.05038 mL | 0.10000 mL | 7.476 | -0.05450 | 0.99713 | 0.00270 | Timed out at 59.5 s |
| 24:47.0 | Data point 21 | 1.50000 mL | 0.05002 mL | 0.05042 mL | 0.10000 mL | 8.023 | -0.06285 | 0.99743 | 0.00311 | Timed out at 59.5 s |
| 26:17.5 | Data point 22 | 1.50000 mL | 0.05002 mL | 0.05047 mL | 0.10000 mL | 8.256 | -0.04162 | 0.99520 | 0.00206 | Timed out at 59.5 s |
| 27:53.2 | Data point 23 | 1.50000 mL | 0.05002 mL | 0.05056 mL | 0.10000 mL | 8.730 | -0.01987 | 0.99215 | 0.00099 | 51.0 s |
| 29:14.8 | Data point 24 | 1.50000 mL | 0.05002 mL | 0.05066 mL | 0.10000 mL | 9.072 | -0.01966 | 0.96316 | 0.00099 | 38.5 s |
| 30:56.2 | Data point 25 | 1.50000 mL | 0.10564 mL | 0.05066 mL | 0.30000 mL | 1.973 | -0.00826 | 0.81865 | 0.00045 | 10.0 s |
| 31:42.4 | Data point 26 | 1.50000 mL | 0.10564 mL | 0.06639 mL | 0.30000 mL | 2.182 | -0.00740 | 0.49850 | 0.00052 | 10.0 s |
| 32:18.0 | Data point 27 | 1.50000 mL | 0.10564 mL | 0.07707 mL | 0.30000 mL | 2.373 | -0.00564 | 0.17659 | 0.00066 | 10.0 s |
| 32:53.6 | Data point 28 | 1.50000 mL | 0.10564 mL | 0.08398 mL | 0.30000 mL | 2.615 | 0.00002 | 0.00001 | 0.00050 | 10.0 s |
| 33:39.6 | Data point 29 | 1.50000 mL | 0.10564 mL | 0.08848 mL | 0.30000 mL | 2.812 | -0.00441 | 0.74067 | 0.00025 | 10.0 s |
| 34:15.1 | Data point 30 | 1.50000 mL | 0.10564 mL | 0.09151 mL | 0.30000 mL | 3.012 | -0.01157 | 0.77811 | 0.00065 | 10.5 s |
| 34:51.1 | Data point 31 | 1.50000 mL | 0.10564 mL | 0.09389 mL | 0.30000 mL | 3.207 | -0.00365 | 0.80988 | 0.00020 | 10.0 s |
| 35:26.5 | Data point 32 | 1.50000 mL | 0.10564 mL | 0.09593 mL | 0.30000 mL | 3.381 | -0.00664 | 0.44901 | 0.00049 | 10.0 s |
| 36:01.9 | Data point 33 | 1.50000 mL | 0.10564 mL | 0.09779 mL | 0.30000 mL | 3.567 | -0.01484 | 0.65999 | 0.00090 | 10.0 s |
| 36:37.4 | Data point 34 | 1.50000 mL | 0.10564 mL | 0.09948 mL | 0.30000 mL | 3.807 | -0.00593 | 0.14880 | 0.00076 | 10.0 s |
| 37:18.0 | Data point 35 | 1.50000 mL | 0.10564 mL | 0.10052 mL | 0.30000 mL | 3.999 | -0.00035 | 0.00113 | 0.00051 | 10.0 s |
| 38:08.9 | Data point 36 | 1.50000 mL | 0.10564 mL | 0.10151 mL | 0.30000 mL | 4.207 | -0.00527 | 0.79747 | 0.00029 | 10.0 s |
| 38:44.4 | Data point 37 | 1.50000 mL | 0.10564 mL | 0.10237 mL | 0.30000 mL | 4.462 | -0.00775 | 0.68238 | 0.00046 | 10.0 s |
| 39:30.1 | Data point 38 | 1.50000 mL | 0.10564 mL | 0.10292 mL | 0.30000 mL | 4.655 | -0.00577 | 0.49479 | 0.00040 | 10.5 s |
| 40:06.0 | Data point 39 | 1.50000 mL | 0.10564 mL | 0.10332 mL | 0.30000 mL | 4.910 | -0.01228 | 0.58176 | 0.00079 | 10.5 s |
| 40:47.1 | Data point 40 | 1.50000 mL | 0.10564 mL | 0.10358 mL | 0.30000 mL | 5.151 | -0.01553 | 0.76936 | 0.00087 | 12.0 s |
| 41:24.5 | Data point 41 | 1.50000 mL | 0.10564 mL | 0.10372 mL | 0.30000 mL | 5.319 | -0.01184 | 0.46775 | 0.00086 | 11.5 s |
| 42:06.6 | Data point 42 | 1.50000 mL | 0.10564 mL | 0.10386 mL | 0.30000 mL | 5.687 | -0.01865 | 0.92517 | 0.00096 | 15.5 s |
| 42:52.8 | Data point 43 | 1.50000 mL | 0.10564 mL | 0.10405 mL | 0.30000 mL | 7.000 | -0.08496 | 0.99448 | 0.00421 | Timed out at 59.5 s |
| 44:28.4 | Data point 44 | 1.50000 mL | 0.10564 mL | 0.10416 mL | 0.30000 mL | 7.624 | -0.06603 | 0.99772 | 0.00326 | Timed out at 59.5 s |
| 46:04.1 | Data point 45 | 1.50000 mL | 0.10564 mL | 0.10426 mL | 0.30000 mL | 8.216 | -0.04320 | 0.99574 | 0.00214 | Timed out at 59.5 s |
| 47:39.9 | Data point 46 | 1.50000 mL | 0.10564 mL | 0.10433 mL | 0.30000 mL | 8.482 | -0.01966 | 0.98330 | 0.00098 | 57.0 s |
| 49:12.6 | Data point 47 | 1.50000 mL | 0.10564 mL | 0.10442 mL | 0.30000 mL | 8.807 | -0.02001 | 0.98580 | 0.00100 | 40.0 s |
| 50:23.1 | Data point 48 | 1.50000 mL | 0.10564 mL | 0.10454 mL | 0.30000 mL | 9.125 | -0.01779 | 0.90618 | 0.00092 | 34.5 s |
| 52:23.2 | Data point 49 | 1.50000 mL | 0.16729 mL | 0.10454 mL | 0.80000 mL | 1.966 | -0.01371 | 0.95560 | 0.00069 | 10.5 s |
| 53:10.0 | Data point 50 | 1.50000 mL | 0.16729 mL | 0.12288 mL | 0.80000 mL | 2.170 | 0.00380 | 0.50246 | 0.00026 | 10.0 s |
| 53:45.8 | Data point 51 | 1.50000 mL | 0.16729 mL | 0.13467 mL | 0.80000 mL | 2.375 | 0.01474 | 0.76262 | 0.00083 | 10.0 s |
| 54:21.4 | Data point 52 | 1.50000 mL | 0.16729 mL | 0.14229 mL | 0.80000 mL | 2.582 | -0.00433 | 0.22604 | 0.00045 | 10.0 s |
| 54:56.9 | Data point 53 | 1.50000 mL | 0.16729 mL | 0.14746 mL | 0.80000 mL | 2.802 | -0.01110 | 0.57112 | 0.00072 | 10.5 s |
| 55:32.9 | Data point 54 | 1.50000 mL | 0.16729 mL | 0.15110 mL | 0.80000 mL | 2.992 | -0.00395 | 0.04126 | 0.00096 | 10.5 s |
| 56:09.0 | Data point 55 | 1.50000 mL | 0.16729 mL | 0.15398 mL | 0.80000 mL | 3.169 | 0.00169 | 0.00837 | 0.00091 | 10.0 s |
| 56:44.4 | Data point 56 | 1.50000 mL | 0.16729 mL | 0.15637 mL | 0.80000 mL | 3.371 | -0.00442 | 0.51991 | 0.00030 | 10.0 s |
| 57:20.0 | Data point 57 | 1.50000 mL | 0.16729 mL | 0.15832 mL | 0.80000 mL | 3.598 | -0.00476 | 0.13870 | 0.00063 | 10.5 s |
| 57:55.8 | Data point 58 | 1.50000 mL | 0.16729 mL | 0.15981 mL | 0.80000 mL | 3.807 | -0.00790 | 0.80262 | 0.00044 | 10.0 s |
| 58:31.3 | Data point 59 | 1.50000 mL | 0.16729 mL | 0.16091 mL | 0.80000 mL | 4.022 | 0.00289 | 0.02519 | 0.00090 | 10.5 s |
| 59:07.2 | Data point 60 | 1.50000 mL | 0.16729 mL | 0.16169 mL | 0.80000 mL | 4.213 | 0.00323 | 0.17283 | 0.00038 | 10.5 s |
| 59:43.0 | Data point 61 | 1.50000 mL | 0.16729 mL | 0.16223 mL | 0.80000 mL | 4.381 | -0.00427 | 0.08186 | 0.00074 | 10.5 s |
| 1:00:24.2 | Data point 62 | 1.50000 mL | 0.16729 mL | 0.16279 mL | 0.80000 mL | 4.718 | -0.01425 | 0.87866 | 0.00075 | 11.0 s |
| 1:01:05.7 | Data point 63 | 1.50000 mL | 0.16729 mL | 0.16315 mL | 0.80000 mL | 5.204 | -0.00635 | 0.10706 | 0.00096 | 12.0 s |
| 1:01:53.3 | Data point 64 | 1.50000 mL | 0.16729 mL | 0.16336 mL | 0.80000 mL | 5.779 | -0.01830 | 0.86406 | 0.00097 | 47.0 s |
| 1:03:10.9 | Data point 65 | 1.50000 mL | 0.16729 mL | 0.16345 mL | 0.80000 mL | 6.233 | -0.02014 | 0.98026 | 0.00100 | Timed out at 59.5 s |

Sample name: **M18_octanol**
Assay name: **pH-metric high logP**
Assay ID: **18C-09014**
Filename: **C:\Sirius_T3\Mehtap\20180309_exp31_logP_T3-2\18C-09014_M18_octanol_pH-metric high logP.t3r**

Experiment start time: **3/9/2018 8:37:21 PM**
Analyst: **Pion**
Instrument ID: **T312060**

#### Events (continued)

| Time | Event | Water | Acid | Base | Octanol | pH | dpH/dt | pH R-squared | pH SD | dpH/dt time |
|---|---|---|---|---|---|---|---|---|---|---|
| 1:04:41.4 | Data point 66 | 1.50000 mL | 0.16729 mL | 0.16352 mL | 0.80000 mL | 6.668 | -0.04896 | 0.99508 | 0.00242 | Timed out at 59.5 s |
| 1:06:12.0 | Data point 67 | 1.50000 mL | 0.16729 mL | 0.16359 mL | 0.80000 mL | 7.159 | -0.06899 | 0.99765 | 0.00341 | Timed out at 59.5 s |
| 1:07:42.4 | Data point 68 | 1.50000 mL | 0.16729 mL | 0.16366 mL | 0.80000 mL | 7.675 | -0.06553 | 0.99682 | 0.00324 | Timed out at 59.5 s |
| 1:09:18.0 | Data point 69 | 1.50000 mL | 0.16729 mL | 0.16373 mL | 0.80000 mL | 8.043 | -0.04078 | 0.98943 | 0.00202 | Timed out at 59.5 s |
| 1:10:48.5 | Data point 70 | 1.50000 mL | 0.16729 mL | 0.16378 mL | 0.80000 mL | 8.282 | -0.02444 | 0.97943 | 0.00122 | Timed out at 59.5 s |
| 1:12:29.2 | Data point 71 | 1.50000 mL | 0.16729 mL | 0.16388 mL | 0.80000 mL | 8.654 | -0.01571 | 0.67890 | 0.00094 | 25.5 s |
| 1:13:30.4 | Data point 72 | 1.50000 mL | 0.16729 mL | 0.16397 mL | 0.80000 mL | 8.923 | -0.00987 | 0.25217 | 0.00097 | 14.0 s |
| 1:14:20.0 | Data point 73 | 1.50000 mL | 0.16729 mL | 0.16406 mL | 0.80000 mL | 9.132 | -0.01934 | 0.95656 | 0.00098 | 25.0 s |
| 1:14:54.1 | Assay volumes | 1.50000 mL | 0.16729 mL | 0.16406 mL | 0.80000 mL | | | | | |

| | | | |
|---|---|---|---|
| Sample name: | **M18_octanol** | Experiment start time: | **3/9/2018 8:37:21 PM** |
| Assay name: | **pH-metric high logP** | Analyst: | **Pion** |
| Assay ID: | **18C-09014** | Instrument ID: | **T312060** |
| Filename: | **C:\Sirius_T3\Mehtap\20180309_exp31_logP_T3-2\18C-09014_M18_octanol_pH-metric high logP.t3r** | | |

| | | | |
|---|---|---|---|
| Sample name: | **M18_octanol** | Experiment start time: | **3/9/2018 8:37:21 PM** |
| Assay name: | **pH-metric high logP** | Analyst: | **Pion** |
| Assay ID: | **18C-09014** | Instrument ID: | **T312060** |
| Filename: | **C:\Sirius_T3\Mehtap\20180309_exp31_logP_T3-2\18C-09014_M18_octanol_pH-metric high logP.t3r** | | |

#### Assay Settings (continued)

| Setting | Value | Original Value | Date/Time changed | Imported from |
|---|---|---|---|---|
| ***Titration 3*** | | | | |
| Titrate from | Low to high pH | | | |
| Add additional water | 0.00 mL | | | |
| Additional partition solvent volume | 0.500 mL | | | |
| Additional partition solvent added | Automatic | | | |
| After pH adjust stir for | 30 seconds | | | |
| Stir to allow partitioning for | 15 seconds | | | |
| Stirrer speed for partitioning | 60% | | | |
| ***Data Point Stability*** | | | | |
| Stir during data point collection | No | | | |
| Delay before data point collection | 0 seconds | | | |
| Number of points to average | 20 points | | | |
| Time interval between points | 0.50 seconds | | | |
| Required maximum standard deviation | 0.00100 dpH/dt | | | |
| Stability timeout after | 60 seconds | | | |

#### Calibration Settings

| Setting | Value | Date/Time changed | Imported from |
|---|---|---|---|
| Four-Plus alpha | 0.102 | 3/9/2018 8:37:21 PM | C:\Sirius_T3\HCl18C09.t3r |
| Four-Plus S | 0.9967 | 3/9/2018 8:37:21 PM | C:\Sirius_T3\HCl18C09.t3r |
| Four-Plus jH | 1.2 | 3/9/2018 8:37:21 PM | C:\Sirius_T3\HCl18C09.t3r |
| Four-Plus jOH | 0.0 | 3/9/2018 8:37:21 PM | C:\Sirius_T3\HCl18C09.t3r |
| Base concentration factor | 1.000 | 3/9/2018 8:37:21 PM | C:\Sirius_T3\KOH18B27.t3r |
| Acid concentration factor | 1.000 | 3/9/2018 8:37:21 PM | C:\Sirius_T3\HCl18C09.t3r |

#### Instrument Settings

| Setting | Value | Batch Id | Install date |
|---|---|---|---|
| Instrument owner | Merck | | |
| Instrument ID | T312060 | | |
| Instrument type | T3 Simulator | | |
| Software version | 1.1.3.0 | | |
| Dispenser module | | T3DM1200361 | 3/31/2009 6:24:52 AM |
| Dispenser 0 | Water | | 3/31/2009 6:25:05 AM |
| Syringe volume | 2.5 mL | | |
| Firmware version | 1.2.1(r2) | | |
| Titrant | Water (0.15 M KCl) | 02-06-2018 | 2/27/2018 11:05:59 AM |
| Dispenser 2 | Acid | | 3/31/2009 6:25:11 AM |
| Syringe volume | 0.5 mL | | |
| Firmware version | 1.2.1(r2) | | |
| Titrant | Acid (0.5 M HCl) | 02-27-2018 | 2/27/2018 11:27:22 AM |
| Dispenser 1 | Base | | 3/31/2009 6:25:21 AM |
| Syringe volume | 0.5 mL | | |
| Firmware version | 1.2.1(r2) | | |
| Titrant | Base (0.5 M KOH) | 9/22/2017 | 2/27/2018 11:21:22 AM |
| Dispenser 5 | Cosolvent | | 3/31/2009 6:26:24 AM |
| Syringe volume | 2.5 mL | | |
| Firmware version | 1.2.1(r2) | | |
| Distribution valve 5 | Distribution Valve | | 3/31/2009 6:28:19 AM |
| Firmware version | 1.1.3 | | |
| Port A | Methanol (80%, 0.15 M KCl) | 02-08-2018 | 3/6/2018 10:28:59 AM |
| Port B | Cyclohexane | 11-01-17 | 2/27/2018 11:37:57 AM |
| Dispenser 3 | Buffer | | 8/3/2010 6:05:16 AM |
| Syringe volume | 0.5 mL | | |
| Firmware version | 1.2.1(r2) | | |
| Titrant | Dodecane | 2018/01/31 | 2/28/2018 11:18:04 AM |
| Dispenser 6 | Octanol | | 10/22/2010 11:52:43 AM |

| | | | |
|---|---|---|---|
| Sample name: | **M18_octanol** | Experiment start time: | **3/9/2018 8:37:21 PM** |
| Assay name: | **pH-metric high logP** | Analyst: | **Pion** |
| Assay ID: | **18C-09014** | Instrument ID: | **T312060** |
| Filename: | **C:\Sirius_T3\Mehtap\20180309_exp31_logP_T3-2\18C-09014_M18_octanol_pH-metric high logP.t3r** | | |

| | | | |
|---|---|---|---|
| Sample name: | **M18_octanol** | Experiment start time: | **3/9/2018 8:37:21 PM** |
| Assay name: | **pH-metric high logP** | Analyst: | **Pion** |
| Assay ID: | **18C-09014** | Instrument ID: | **T312060** |
| Filename: | **C:\Sirius_T3\Mehtap\20180309_exp31_logP_T3-2\18C-09014_M18_octanol_pH-metric high logP.t3r** | | |

#### Instrument Settings (continued)

| Setting | Value | Batch Id | Install date |
|---|---|---|---|
| Spectrometer calibration stir duration | 5 s | | |
| Spectrometer calibration stir speed | 30% | | |
| Spectrometer calibration wash pump volume | 20.0 mL | | |
| Spectrometer calibration wash stir duration | 5 s | | |
| Spectrometer calibration wash stir speed | 30% | | |
| Overhead dispense height | 10000 | | |

#### Refinement Settings

| Setting | Value | Default value |
|---|---|---|
| Turbidity detection method | None | None |
| Turbidity wavelength to assess | 500.0 nm | 500.0 nm |
| Turbidity maximum absorbance | 0.100 | 0.100 |
| Turbidity probe threshold | 50.00 | 50.00 |
