## Supplementary material for "Octanol-water partition coefficient measurements for the SAMPL6 Blind Prediction Challenge": SM16_18C-09015_M18_octanol_pH-metric high logP_report.pdf

| | | | |
|---|---|---|---|
| Sample name: | **M18_octanol** | Experiment start time: | **3/9/2018 9:53:01 PM** |
| Assay name: | **pH-metric high logP** | Analyst: | **Pion** |
| Assay ID: | **18C-09015** | Instrument ID: | **T312060** |
| Filename: | **C:\Sirius_T3\Mehtap\20180309_exp31_logP_T3-2\18C-09015_M18_octanol_pH-metric high logP.t3r** | | |

### pH-metric Result

| | | |
|---|---|---|
| logP (XH2 +) | -0.73 | ±0.56 (n=50) |
| logP (neutral XH) | 2.59 | ±0.01 (n=50) |
| logP (X -) | 0.46 | |
| RMSD | 1.511 | |

#### 18C-09015 Points 2 to 20

| | |
|---|---|
| M18_octanol concentration factor | 1.192 |
| Carbonate | 0.0000 mM |
| Acidity error | 0.28168 mM |

#### 18C-09015 Points 21 to 41

| | |
|---|---|
| M18_octanol concentration factor | 1.079 |
| Carbonate | 0.1008 mM |
| Acidity error | -0.00618 mM |

#### 18C-09015 Points 42 to 69

| | |
|---|---|
| M18_octanol concentration factor | 1.146 |
| Carbonate | 0.1707 mM |
| Acidity error | 0.14218 mM |

### Warnings and errors

| | |
|---|---|
| Errors | None |
| Warnings | None |

### Sample logD and percent species

| pH | M18_octanol logD | M18_octanol M18_octanolH2 | M18_octanol M18_octanolH | M18_octanol M18_octanol | M18_octanol M18_octanolH2* | M18_octanol M18_octanolH* | M18_octanol M18_octanol* | Comment |
|---|---|---|---|---|---|---|---|---|
| 1.000 | -0.69 | 83.14 % | 0.00 % | 0.00 % | 15.49 % | 1.37 % | 0.00 % | |
| 1.200 | -0.67 | 82.48 % | 0.01 % | 0.00 % | 15.36 % | 2.15 % | 0.00 % | Stomach pH |
| 2.000 | -0.46 | 74.02 % | 0.03 % | 0.00 % | 13.79 % | 12.16 % | 0.00 % | |
| 3.000 | 0.26 | 35.30 % | 0.15 % | 0.00 % | 6.57 % | 57.98 % | 0.00 % | |
| 4.000 | 1.20 | 5.66 % | 0.24 % | 0.00 % | 1.05 % | 93.04 % | 0.00 % | |
| 5.000 | 2.06 | 0.60 % | 0.26 % | 0.00 % | 0.11 % | 99.03 % | 0.00 % | |
| 6.000 | 2.49 | 0.06 % | 0.26 % | 0.00 % | 0.01 % | 99.67 % | 0.00 % | |
| 6.500 | 2.55 | 0.02 % | 0.26 % | 0.00 % | 0.00 % | 99.72 % | 0.00 % | |
| 7.000 | 2.58 | 0.01 % | 0.26 % | 0.00 % | 0.00 % | 99.73 % | 0.00 % | |
| 7.400 | 2.58 | 0.00 % | 0.26 % | 0.00 % | 0.00 % | 99.74 % | 0.00 % | Blood pH |
| 8.000 | 2.58 | 0.00 % | 0.26 % | 0.00 % | 0.00 % | 99.74 % | 0.00 % | |
| 9.000 | 2.58 | 0.00 % | 0.26 % | 0.01 % | 0.00 % | 99.72 % | 0.02 % | |
| 10.000 | 2.50 | 0.00 % | 0.26 % | 0.06 % | 0.00 % | 99.52 % | 0.17 % | |
| 11.000 | 2.08 | 0.00 % | 0.25 % | 0.57 % | 0.00 % | 97.54 % | 1.64 % | |
| 12.000 | 1.28 | 0.00 % | 0.21 % | 4.73 % | 0.00 % | 81.41 % | 13.65 % | |

| | | | |
|---|---|---|---|
| Sample name: | **M18_octanol** | Experiment start time: | **3/9/2018 9:53:01 PM** |
| Assay name: | **pH-metric high logP** | Analyst: | **Pion** |
| Assay ID: | **18C-09015** | Instrument ID: | **T312060** |
| Filename: | **C:\Sirius_T3\Mehtap\20180309_exp31_logP_T3-2\18C-09015_M18_octanol_pH-metric high logP.t3r** | | |

### Graphs

Charge balance residuals

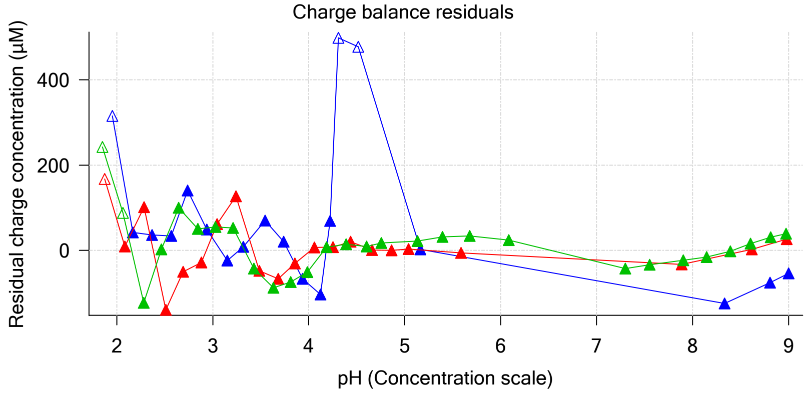


Ionic strength

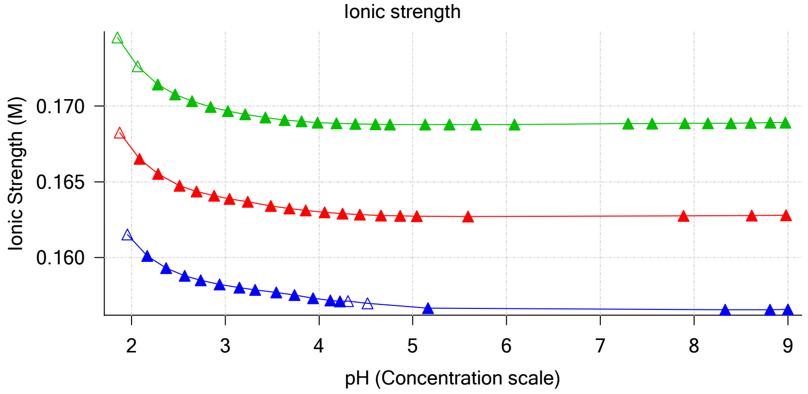


Buffer index profile

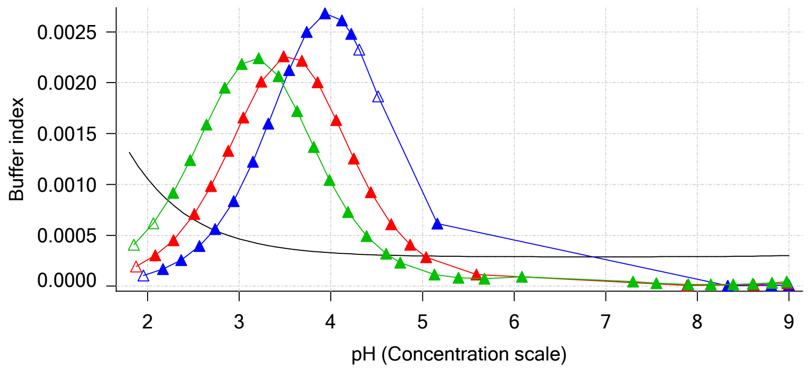


Sample M18_octanol Ionisation Graph

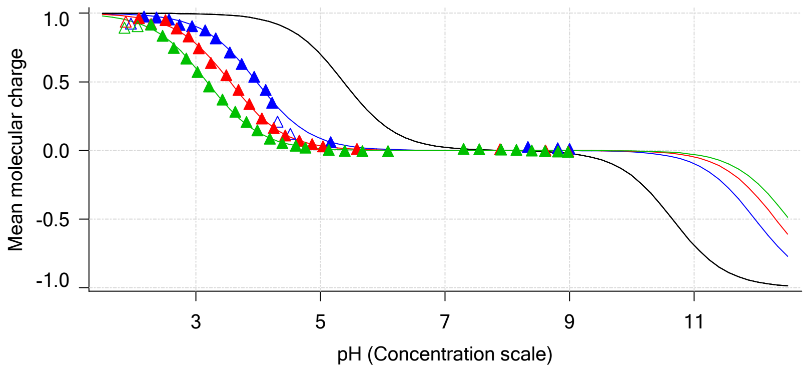


Distribution of species for sample M18_octanol

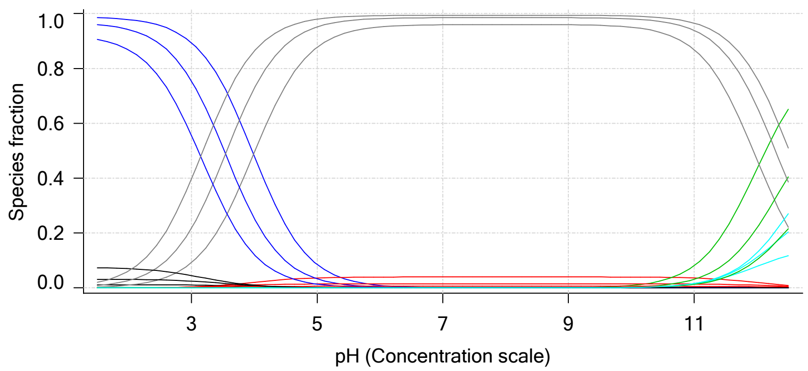


Lipophilicity Profile

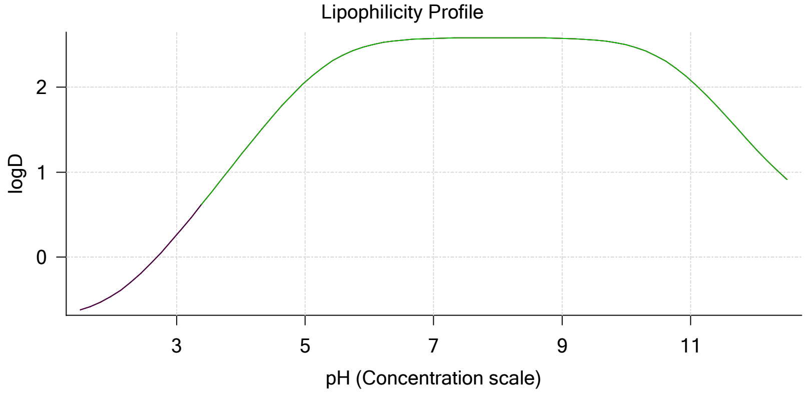


Log R Profile

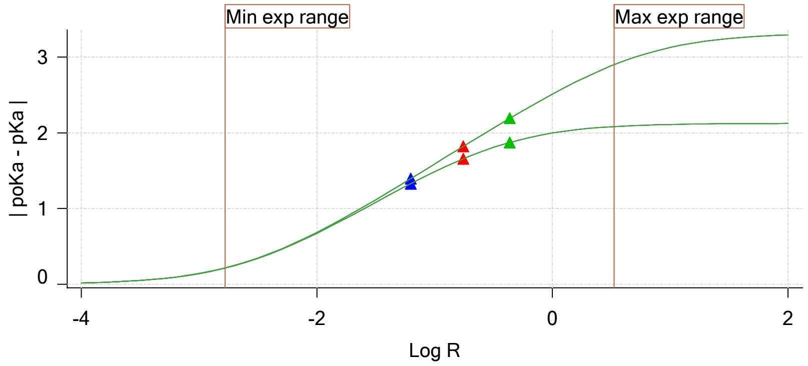


Sample Carbonate Ionisation Graph

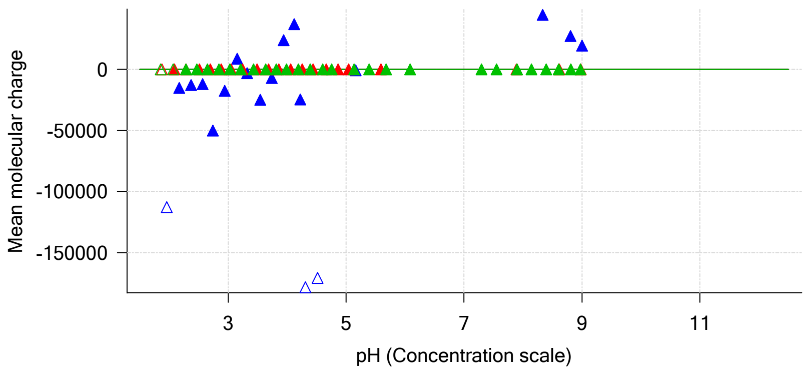

| | | | |
|---|---|---|---|
| Sample name: | **M18_octanol** | Experiment start time: | **3/9/2018 9:53:01 PM** |
| Assay name: | **pH-metric high logP** | Analyst: | **Pion** |
| Assay ID: | **18C-09015** | Instrument ID: | **T312060** |
| Filename: | **C:\Sirius_T3\Mehtap\20180309_exp31_logP_T3-2\18C-09015_M18_octanol_pH-metric high logP.t3r** | | |

### Graphs (continued)

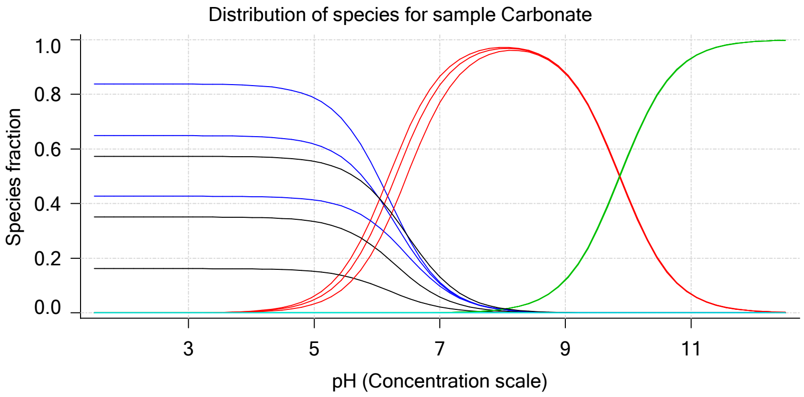

| | | | |
|---|---|---|---|
| Sample name: | **M18_octanol** | Experiment start time: | **3/9/2018 9:53:01 PM** |
| Assay name: | **pH-metric high logP** | Analyst: | **Pion** |
| Assay ID: | **18C-09015** | Instrument ID: | **T312060** |
| Filename: | **C:\Sirius_T3\Mehtap\20180309_exp31_logP_T3-2\18C-09015_M18_octanol_pH-metric high logP.t3r** | | |

# pH-metric high logP   Titration 1 of 3   18C-09015 Points 2 to 20

### Overall results

| | |
|---|---|
| RMSD | 2.905 |
| Average ionic strength | 0.158 M |
| Average temperature | 24.9°C |
| Partition ratio | 0.0626 : 1 |
| Analyte concentration range | 3669.6 µM to 3776.5 µM |
| Total points considered | 16 of 19 |

### Warnings and errors

| | |
|---|---|
| Errors | None |
| Warnings | One or more logP values out of range |

### Four-Plus parameters

| | | | |
|---|---|---|---|
| Alpha | 0.102 | 3/9/2018 9:53:01 PM | C:\Sirius_T3\HCl18C09.t3r |
| S | 0.9967 | 3/9/2018 9:53:01 PM | C:\Sirius_T3\HCl18C09.t3r |
| jH | 1.2 | 3/9/2018 9:53:01 PM | C:\Sirius_T3\HCl18C09.t3r |
| jOH | 0.0 | 3/9/2018 9:53:01 PM | C:\Sirius_T3\HCl18C09.t3r |

### Titrants

| | | | |
|---|---|---|---|
| 0.50 M HCl | 0.999843 | 3/9/2018 9:53:01 PM | C:\Sirius_T3\HCl18C09.t3r |
| 0.50 M KOH | 0.999845 | 3/9/2018 9:53:01 PM | C:\Sirius_T3\KOH18B27.t3r |

### Sample

| | |
|---|---|
| M18_octanol concentration factor | 1.192 |
| Base pKa 1 | 5.37 |
| Acid pKa 2 | 10.65 |
| logP (XH2 +) | -7.94 |
| logP (neutral XH) | 2.63 |
| logP (X -) | 0.46 |

### Sample graphs

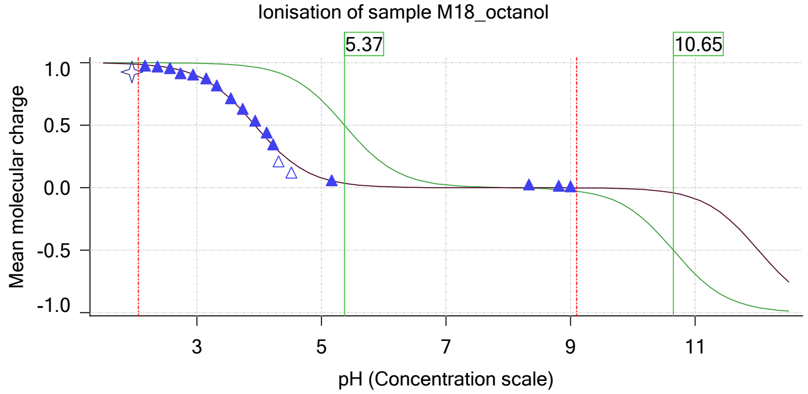


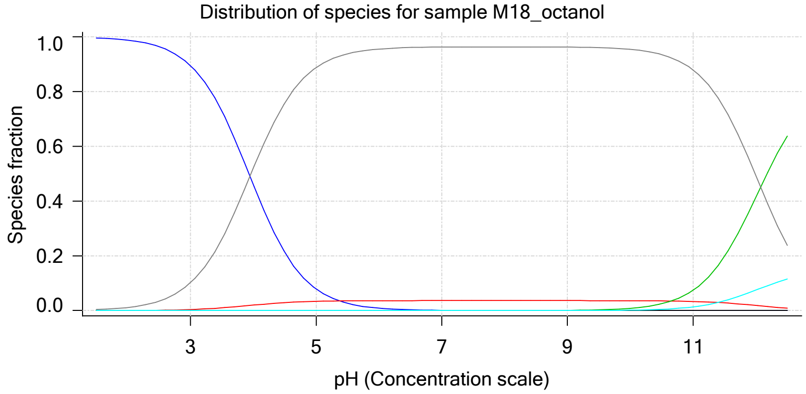

| | | | |
|---|---|---|---|
| Sample name: | **M18_octanol** | Experiment start time: | **3/9/2018 9:53:01 PM** |
| Assay name: | **pH-metric high logP** | Analyst: | **Pion** |
| Assay ID: | **18C-09015** | Instrument ID: | **T312060** |
| Filename: | **C:\Sirius_T3\Mehtap\20180309_exp31_logP_T3-2\18C-09015_M18_octanol_pH-metric high logP.t3r** | | |

### Sample graphs (continued)

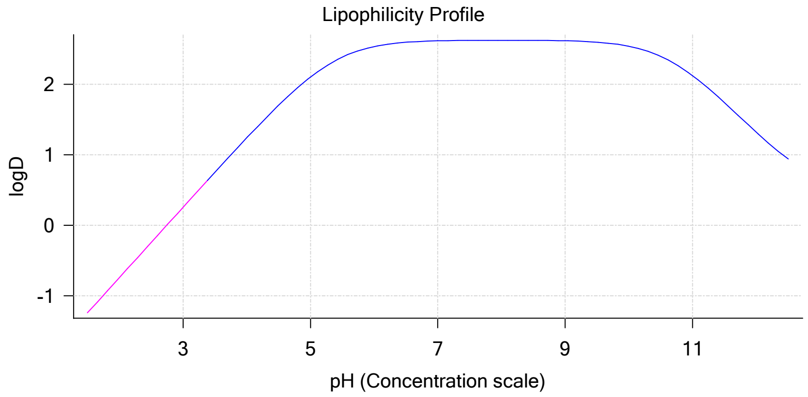


### Sample logD and percent species

| pH | M18_octanol logD | M18_octanol M18_octanolH2 | M18_octanol M18_octanolH | M18_octanol M18_octanol | M18_octanol M18_octanolH2* | M18_octanol M18_octanolH* | M18_octanol M18_octanol* | Comment |
|---|---|---|---|---|---|---|---|---|
| 1.000 | -1.74 | 99.88 % | 0.00 % | 0.00 % | 0.00 % | 0.11 % | 0.00 % | |
| 1.200 | -1.54 | 99.81 % | 0.01 % | 0.00 % | 0.00 % | 0.18 % | 0.00 % | Stomach pH |
| 2.000 | -0.74 | 98.84 % | 0.04 % | 0.00 % | 0.00 % | 1.12 % | 0.00 % | |
| 3.000 | 0.26 | 89.50 % | 0.38 % | 0.00 % | 0.00 % | 10.12 % | 0.00 % | |
| 4.000 | 1.24 | 46.01 % | 1.96 % | 0.00 % | 0.00 % | 52.03 % | 0.00 % | |
| 5.000 | 2.10 | 7.85 % | 3.35 % | 0.00 % | 0.00 % | 88.80 % | 0.00 % | |
| 6.000 | 2.54 | 0.84 % | 3.60 % | 0.00 % | 0.00 % | 95.55 % | 0.00 % | |
| 6.500 | 2.60 | 0.27 % | 3.63 % | 0.00 % | 0.00 % | 96.11 % | 0.00 % | |
| 7.000 | 2.62 | 0.09 % | 3.63 % | 0.00 % | 0.00 % | 96.28 % | 0.00 % | |
| 7.400 | 2.62 | 0.03 % | 3.63 % | 0.00 % | 0.00 % | 96.33 % | 0.00 % | Blood pH |
| 8.000 | 2.63 | 0.01 % | 3.63 % | 0.01 % | 0.00 % | 96.35 % | 0.00 % | |
| 9.000 | 2.62 | 0.00 % | 3.63 % | 0.08 % | 0.00 % | 96.27 % | 0.01 % | |
| 10.000 | 2.54 | 0.00 % | 3.60 % | 0.81 % | 0.00 % | 95.45 % | 0.15 % | |
| 11.000 | 2.12 | 0.00 % | 3.32 % | 7.42 % | 0.00 % | 87.92 % | 1.34 % | |
| 12.000 | 1.32 | 0.00 % | 1.85 % | 41.51 % | 0.00 % | 49.15 % | 7.49 % | |

### Carbonate and acidity

Carbonate 0.000 mM
Acidity error 0.282 mM

### Other graphs

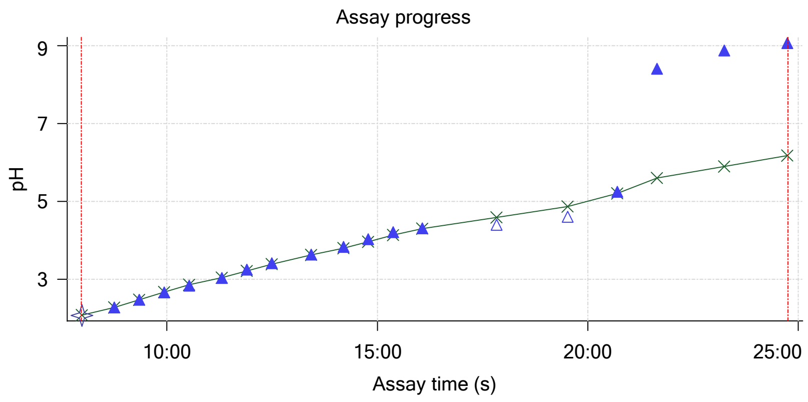


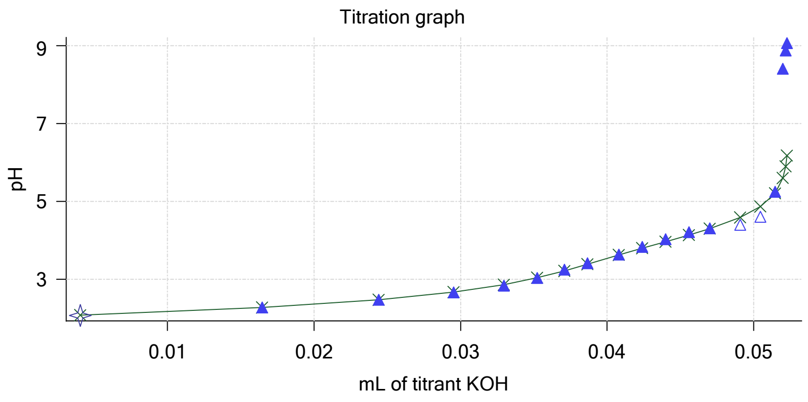

Sample name: **M18_octanol**
Assay name: **pH-metric high logP**
Assay ID: **18C-09015**
Filename: **C:\Sirius_T3\Mehtap\20180309_exp31_logP_T3-2\18C-09015_M18_octanol_pH-metric high logP.t3r**

Experiment start time: **3/9/2018 9:53:01 PM**
Analyst: **Pion**
Instrument ID: **T312060**

### Other graphs (continued)

Charge balance residuals

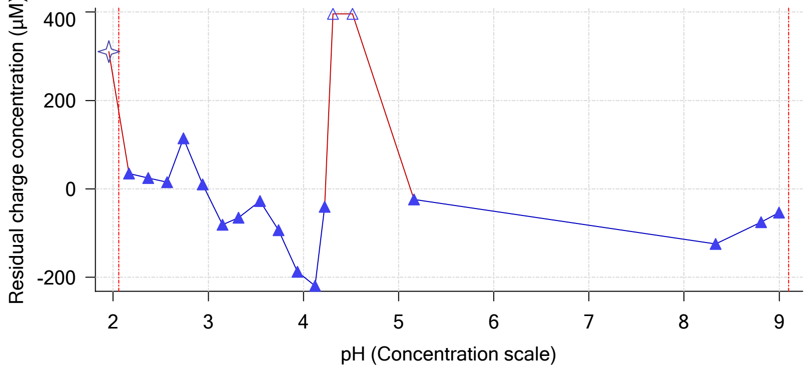


Buffer index profile

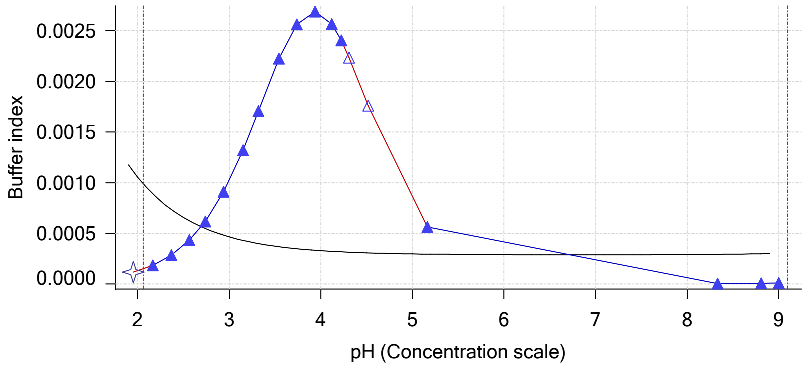


Ionic strength

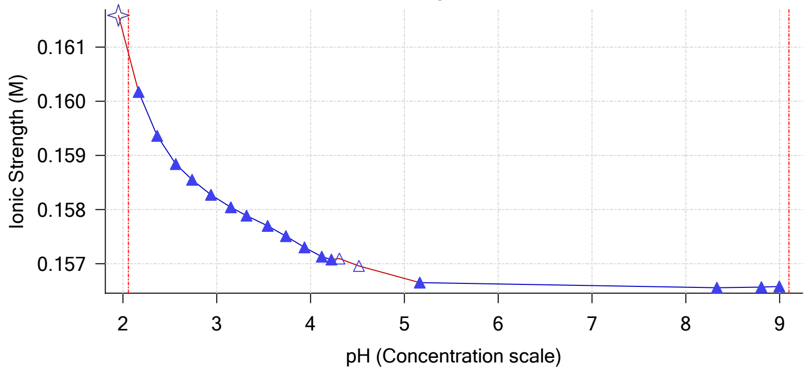


Log R Profile

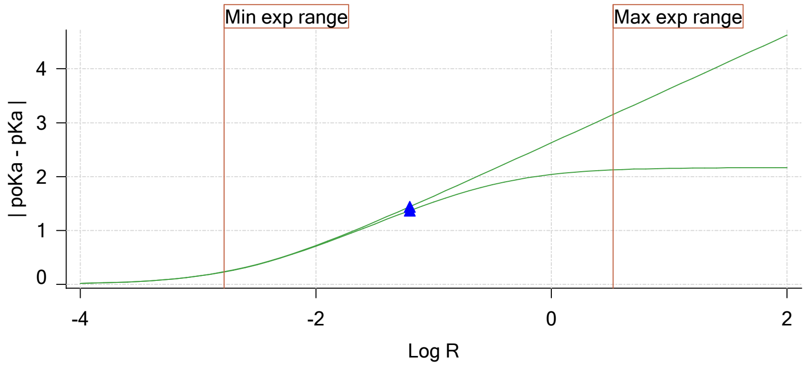


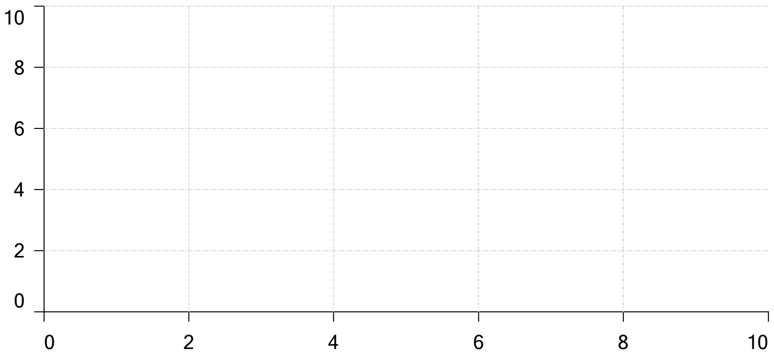


Ionisation of sample Carbonate

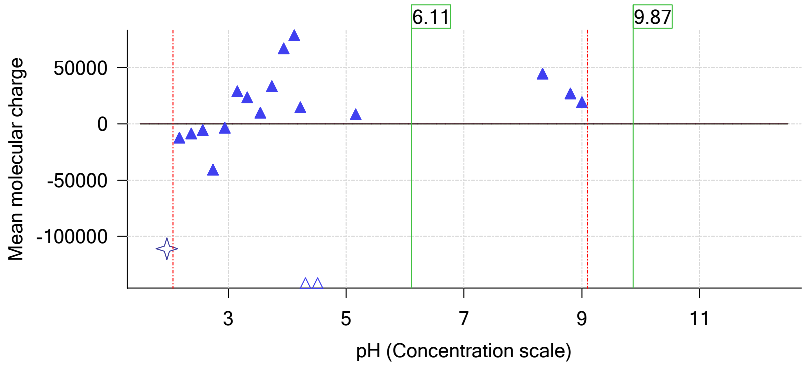

| | | | |
|---|---|---|---|
| Sample name: | **M18_octanol** | Experiment start time: | **3/9/2018 9:53:01 PM** |
| Assay name: | **pH-metric high logP** | Analyst: | **Pion** |
| Assay ID: | **18C-09015** | Instrument ID: | **T312060** |
| Filename: | **C:\Sirius_T3\Mehtap\20180309_exp31_logP_T3-2\18C-09015_M18_octanol_pH-metric high logP.t3r** | | |

# pH-metric high logP   Titration 2 of 3   18C-09015 Points 21 to 41

### Overall results

| | |
|---|---|
| RMSD | 0.624 |
| Average ionic strength | 0.164 M |
| Average temperature | 25.0°C |
| Partition ratio | 0.1752 : 1 |
| Analyte concentration range | 3095.9 µM to 3187.0 µM |
| Total points considered | 20 of 21 |

### Warnings and errors

| | |
|---|---|
| Errors | None |
| Warnings | One or more logP values out of range |

### Four-Plus parameters

| | | | |
|---|---|---|---|
| Alpha | 0.102 | 3/9/2018 9:53:01 PM | C:\Sirius_T3\HCl18C09.t3r |
| S | 0.9967 | 3/9/2018 9:53:01 PM | C:\Sirius_T3\HCl18C09.t3r |
| jH | 1.2 | 3/9/2018 9:53:01 PM | C:\Sirius_T3\HCl18C09.t3r |
| jOH | 0.0 | 3/9/2018 9:53:01 PM | C:\Sirius_T3\HCl18C09.t3r |

### Titrants

| | | | |
|---|---|---|---|
| 0.50 M HCl | 0.999843 | 3/9/2018 9:53:01 PM | C:\Sirius_T3\HCl18C09.t3r |
| 0.50 M KOH | 0.999845 | 3/9/2018 9:53:01 PM | C:\Sirius_T3\KOH18B27.t3r |

### Sample

| | |
|---|---|
| M18_octanol concentration factor | 1.079 |
| Base pKa 1 | 5.37 |
| Acid pKa 2 | 10.65 |
| logP (XH2 +) | -4.21 |
| logP (neutral XH) | 2.57 |
| logP (X -) | 0.46 |

### Sample graphs

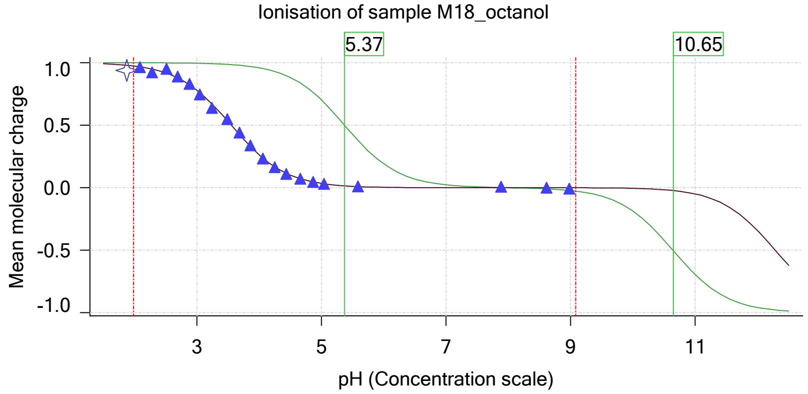


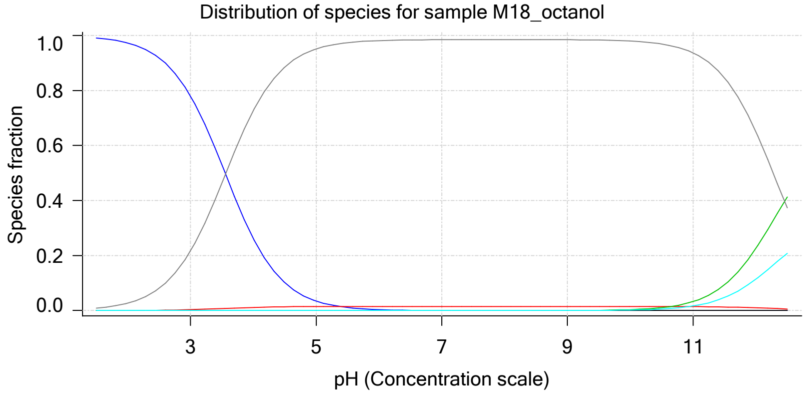

| | | | |
|---|---|---|---|
| Sample name: | **M18_octanol** | Experiment start time: | **3/9/2018 9:53:01 PM** |
| Assay name: | **pH-metric high logP** | Analyst: | **Pion** |
| Assay ID: | **18C-09015** | Instrument ID: | **T312060** |
| Filename: | **C:\Sirius_T3\Mehtap\20180309_exp31_logP_T3-2\18C-09015_M18_octanol_pH-metric high logP.t3r** | | |

### Sample graphs (continued)

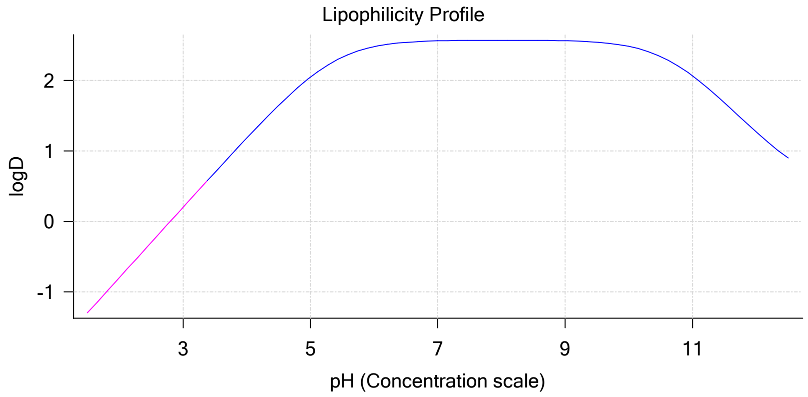


### Sample logD and percent species

| pH | M18_octanol logD | M18_octanol M18_octanolH2 | M18_octanol M18_octanolH | M18_octanol M18_octanol | M18_octanol M18_octanolH2* | M18_octanol M18_octanolH* | M18_octanol M18_octanol* | Comment |
|---|---|---|---|---|---|---|---|---|
| 1.000 | -1.80 | 99.72 % | 0.00 % | 0.00 % | 0.00 % | 0.28 % | 0.00 % | |
| 1.200 | -1.60 | 99.55 % | 0.01 % | 0.00 % | 0.00 % | 0.44 % | 0.00 % | Stomach pH |
| 2.000 | -0.80 | 97.25 % | 0.04 % | 0.00 % | 0.00 % | 2.71 % | 0.00 % | |
| 3.000 | 0.20 | 77.96 % | 0.33 % | 0.00 % | 0.00 % | 21.70 % | 0.00 % | |
| 4.000 | 1.18 | 26.13 % | 1.11 % | 0.00 % | 0.00 % | 72.75 % | 0.00 % | |
| 5.000 | 2.05 | 3.42 % | 1.46 % | 0.00 % | 0.00 % | 95.12 % | 0.00 % | |
| 6.000 | 2.48 | 0.35 % | 1.50 % | 0.00 % | 0.00 % | 98.14 % | 0.00 % | |
| 6.500 | 2.54 | 0.11 % | 1.51 % | 0.00 % | 0.00 % | 98.38 % | 0.00 % | |
| 7.000 | 2.56 | 0.04 % | 1.51 % | 0.00 % | 0.00 % | 98.46 % | 0.00 % | |
| 7.400 | 2.57 | 0.01 % | 1.51 % | 0.00 % | 0.00 % | 98.48 % | 0.00 % | Blood pH |
| 8.000 | 2.57 | 0.00 % | 1.51 % | 0.00 % | 0.00 % | 98.48 % | 0.00 % | |
| 9.000 | 2.56 | 0.00 % | 1.51 % | 0.03 % | 0.00 % | 98.44 % | 0.02 % | |
| 10.000 | 2.48 | 0.00 % | 1.50 % | 0.34 % | 0.00 % | 97.99 % | 0.17 % | |
| 11.000 | 2.07 | 0.00 % | 1.44 % | 3.22 % | 0.00 % | 93.72 % | 1.62 % | |
| 12.000 | 1.27 | 0.00 % | 1.00 % | 22.40 % | 0.00 % | 65.29 % | 11.31 % | |

### Carbonate and acidity

| | |
|---|---|
| Carbonate | 0.101 mM |
| Acidity error | -0.006 mM |

### Other graphs

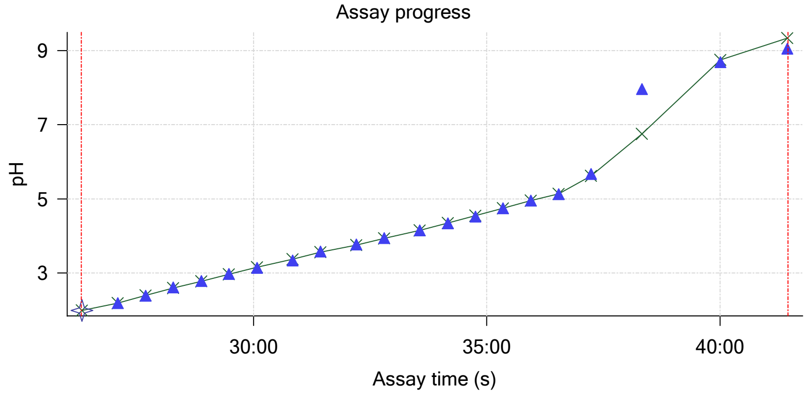


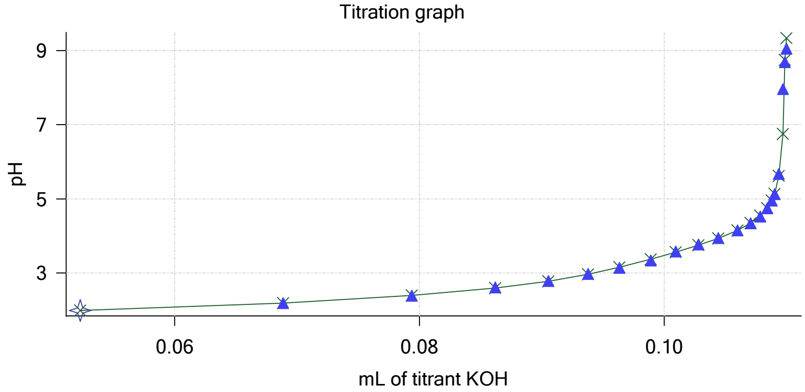

Sample name: **M18_octanol**
Assay name: **pH-metric high logP**
Assay ID: **18C-09015**
Filename: **C:\Sirius_T3\Mehtap\20180309_exp31_logP_T3-2\18C-09015_M18_octanol_pH-metric high logP.t3r**

Experiment start time: **3/9/2018 9:53:01 PM**
Analyst: **Pion**
Instrument ID: **T312060**

### Other graphs (continued)

Charge balance residuals

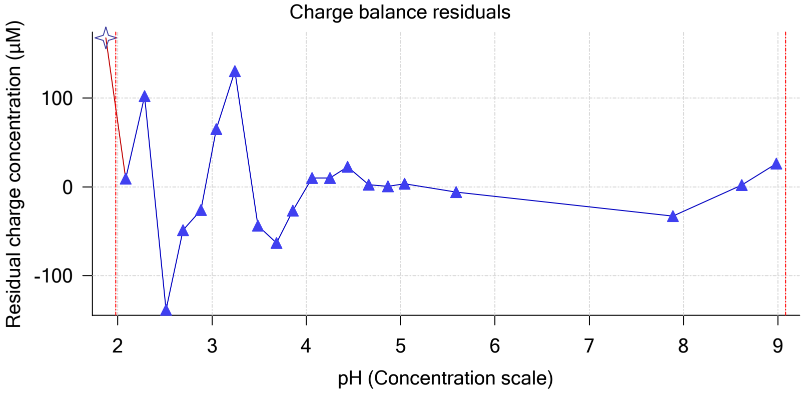


Buffer index profile

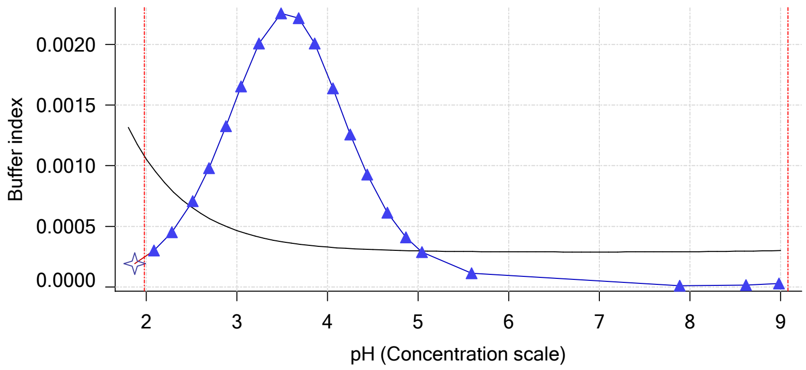


Ionic strength

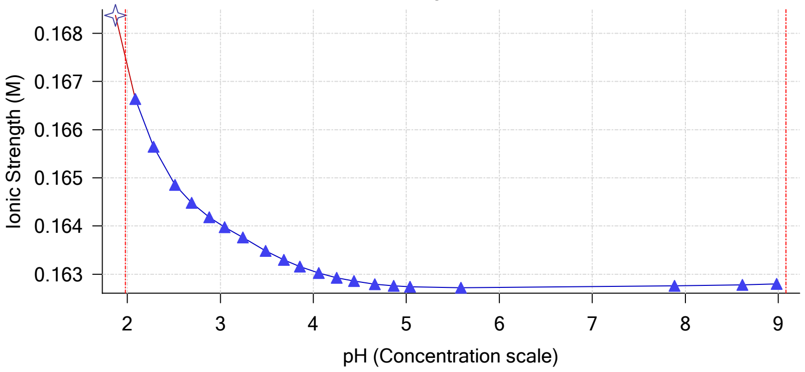


Log R Profile

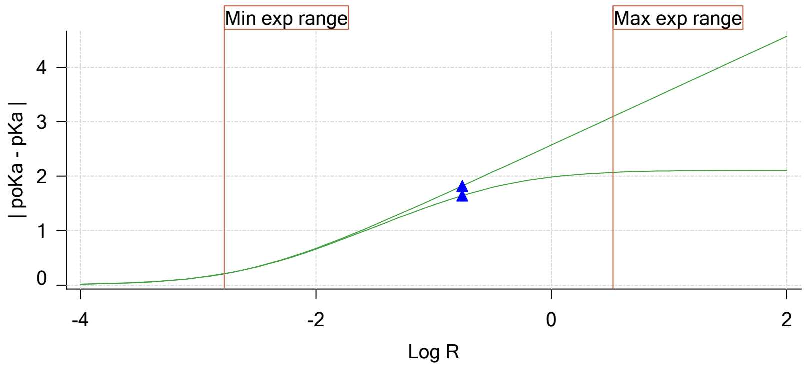


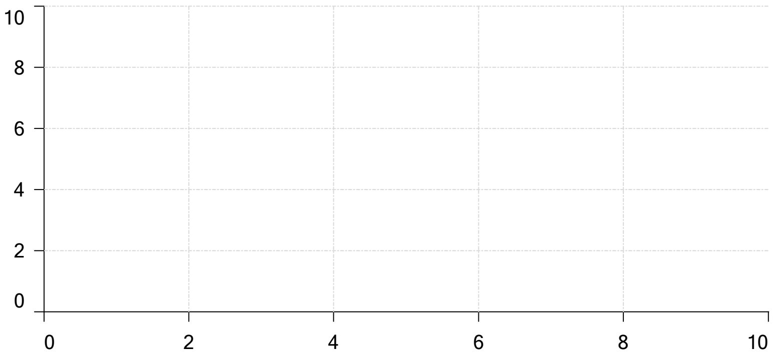


Ionisation of sample Carbonate

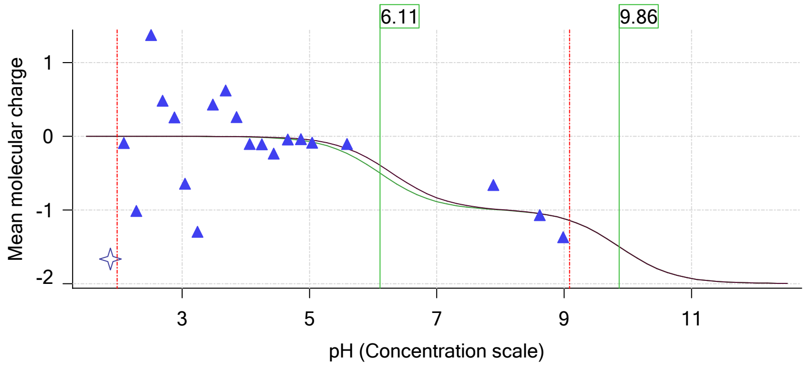

| | | | |
|---|---|---|---|
| Sample name: | **M18_octanol** | Experiment start time: | **3/9/2018 9:53:01 PM** |
| Assay name: | **pH-metric high logP** | Analyst: | **Pion** |
| Assay ID: | **18C-09015** | Instrument ID: | **T312060** |
| Filename: | **C:\Sirius_T3\Mehtap\20180309_exp31_logP_T3-2\18C-09015_M18_octanol_pH-metric high logP.t3r** | | |

# pH-metric high logP   Titration 3 of 3   18C-09015 Points 42 to 69

### Overall results

| | |
|---|---|
| RMSD | 0.425 |
| Average ionic strength | 0.169 M |
| Average temperature | 25.0°C |
| Partition ratio | 0.4343 : 1 |
| Analyte concentration range | 2362.4 µM to 2420.9 µM |
| Total points considered | 26 of 28 |

### Warnings and errors

| | |
|---|---|
| Errors | None |
| Warnings | One or more logP values out of range |

### Four-Plus parameters

| | | | |
|---|---|---|---|
| Alpha | 0.102 | 3/9/2018 9:53:01 PM | C:\Sirius_T3\HCl18C09.t3r |
| S | 0.9967 | 3/9/2018 9:53:01 PM | C:\Sirius_T3\HCl18C09.t3r |
| jH | 1.2 | 3/9/2018 9:53:01 PM | C:\Sirius_T3\HCl18C09.t3r |
| jOH | 0.0 | 3/9/2018 9:53:01 PM | C:\Sirius_T3\HCl18C09.t3r |

### Titrants

| | | | |
|---|---|---|---|
| 0.50 M HCl | 0.999843 | 3/9/2018 9:53:01 PM | C:\Sirius_T3\HCl18C09.t3r |
| 0.50 M KOH | 0.999845 | 3/9/2018 9:53:01 PM | C:\Sirius_T3\KOH18B27.t3r |

### Sample

| | |
|---|---|
| M18_octanol concentration factor | 1.146 |
| Base pKa 1 | 5.37 |
| Acid pKa 2 | 10.65 |
| logP (XH2 +) | -3.41 |
| logP (neutral XH) | 2.55 |
| logP (X -) | 0.46 |

### Sample graphs

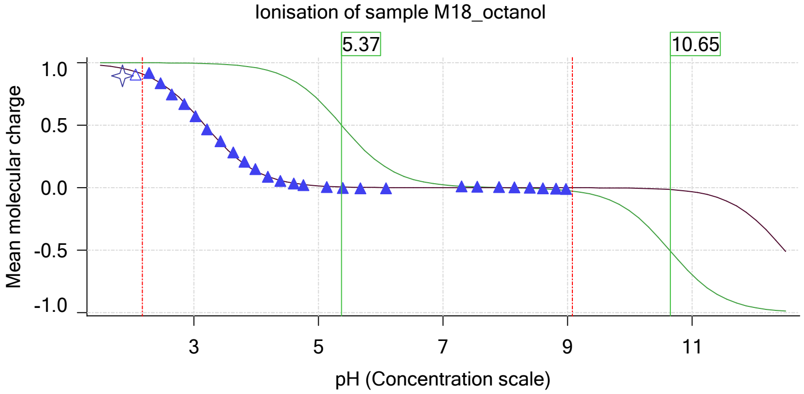


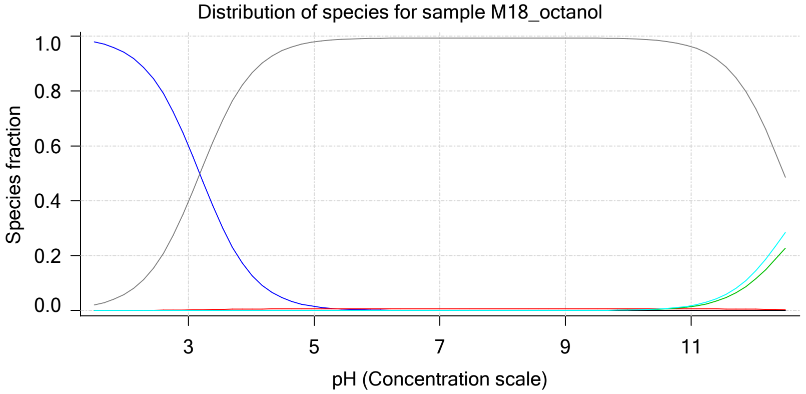

| | | | |
|---|---|---|---|
| Sample name: | **M18_octanol** | Experiment start time: | **3/9/2018 9:53:01 PM** |
| Assay name: | **pH-metric high logP** | Analyst: | **Pion** |
| Assay ID: | **18C-09015** | Instrument ID: | **T312060** |
| Filename: | **C:\Sirius_T3\Mehtap\20180309_exp31_logP_T3-2\18C-09015_M18_octanol_pH-metric high logP.t3r** | | |

### Sample graphs (continued)

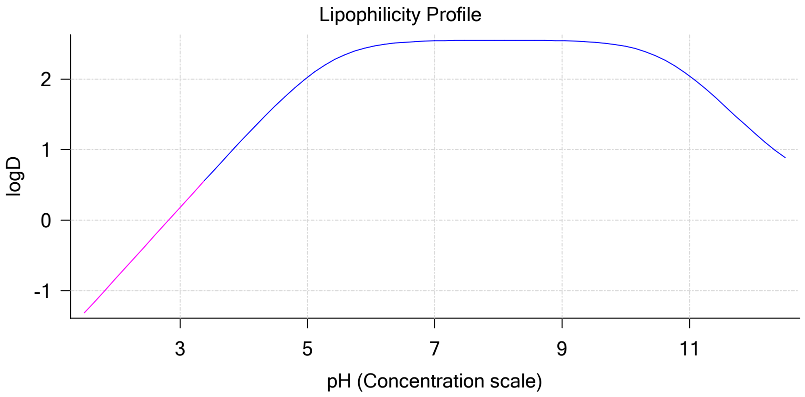


### Sample logD and percent species

| pH | M18_octanol logD | M18_octanol M18_octanolH2 | M18_octanol M18_octanolH | M18_octanol M18_octanol | M18_octanol M18_octanolH2* | M18_octanol M18_octanolH* | M18_octanol M18_octanol* | Comment |
|---|---|---|---|---|---|---|---|---|
| 1.000 | -1.80 | 99.32 % | 0.00 % | 0.00 % | 0.02 % | 0.66 % | 0.00 % | |
| 1.200 | -1.61 | 98.94 % | 0.01 % | 0.00 % | 0.02 % | 1.04 % | 0.00 % | Stomach pH |
| 2.000 | -0.81 | 93.72 % | 0.04 % | 0.00 % | 0.02 % | 6.22 % | 0.00 % | |
| 3.000 | 0.18 | 59.93 % | 0.26 % | 0.00 % | 0.01 % | 39.80 % | 0.00 % | |
| 4.000 | 1.17 | 13.02 % | 0.56 % | 0.00 % | 0.00 % | 86.43 % | 0.00 % | |
| 5.000 | 2.03 | 1.47 % | 0.63 % | 0.00 % | 0.00 % | 97.90 % | 0.00 % | |
| 6.000 | 2.46 | 0.15 % | 0.64 % | 0.00 % | 0.00 % | 99.21 % | 0.00 % | |
| 6.500 | 2.52 | 0.05 % | 0.64 % | 0.00 % | 0.00 % | 99.31 % | 0.00 % | |
| 7.000 | 2.54 | 0.01 % | 0.64 % | 0.00 % | 0.00 % | 99.35 % | 0.00 % | |
| 7.400 | 2.55 | 0.01 % | 0.64 % | 0.00 % | 0.00 % | 99.35 % | 0.00 % | Blood pH |
| 8.000 | 2.55 | 0.00 % | 0.64 % | 0.00 % | 0.00 % | 99.36 % | 0.00 % | |
| 9.000 | 2.54 | 0.00 % | 0.64 % | 0.01 % | 0.00 % | 99.33 % | 0.02 % | |
| 10.000 | 2.47 | 0.00 % | 0.64 % | 0.14 % | 0.00 % | 99.04 % | 0.18 % | |
| 11.000 | 2.05 | 0.00 % | 0.62 % | 1.38 % | 0.00 % | 96.26 % | 1.73 % | |
| 12.000 | 1.26 | 0.00 % | 0.48 % | 10.81 % | 0.00 % | 75.17 % | 13.54 % | |

### Carbonate and acidity

| | |
|---|---|
| Carbonate | 0.171 mM |
| Acidity error | 0.142 mM |

### Other graphs

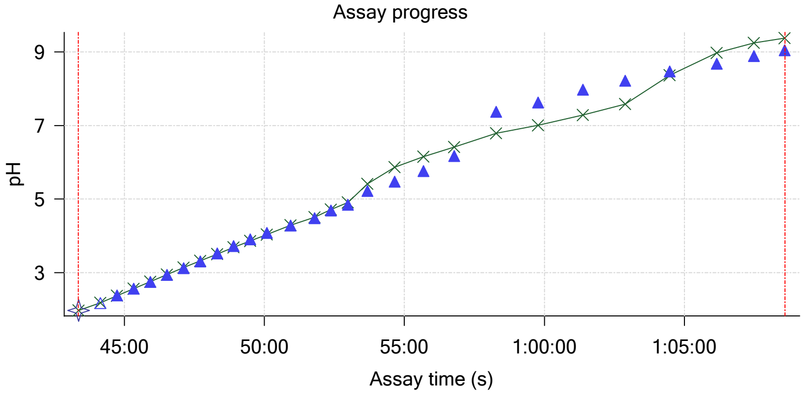


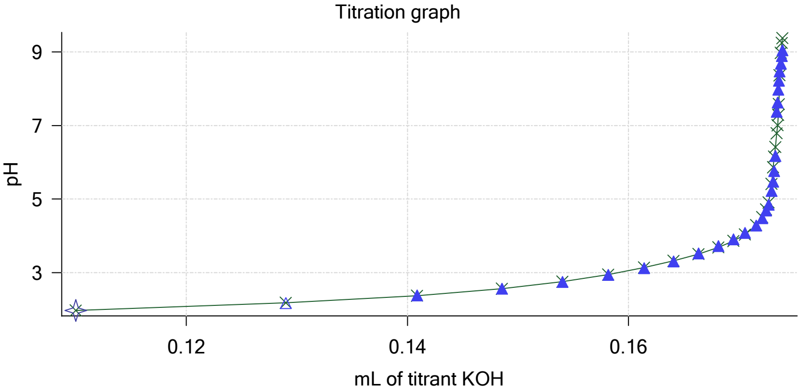

| | | | |
|---|---|---|---|
| Sample name: | **M18_octanol** | Experiment start time: | **3/9/2018 9:53:01 PM** |
| Assay name: | **pH-metric high logP** | Analyst: | **Pion** |
| Assay ID: | **18C-09015** | Instrument ID: | **T312060** |
| Filename: | **C:\Sirius_T3\Mehtap\20180309_exp31_logP_T3-2\18C-09015_M18_octanol_pH-metric high logP.t3r** | | |

### Other graphs (continued)

Charge balance residuals

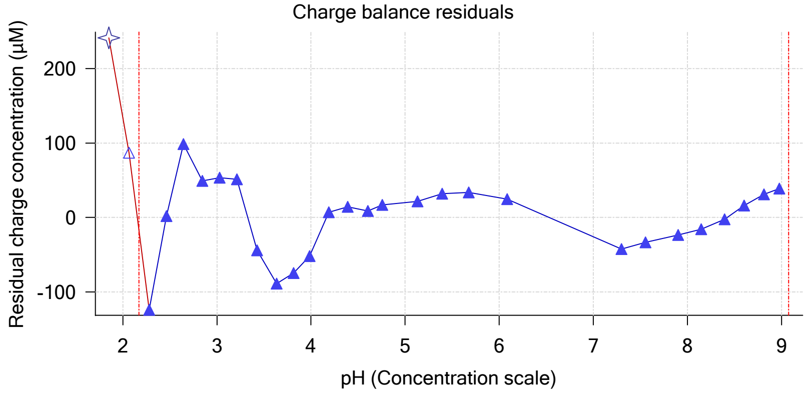


Buffer index profile

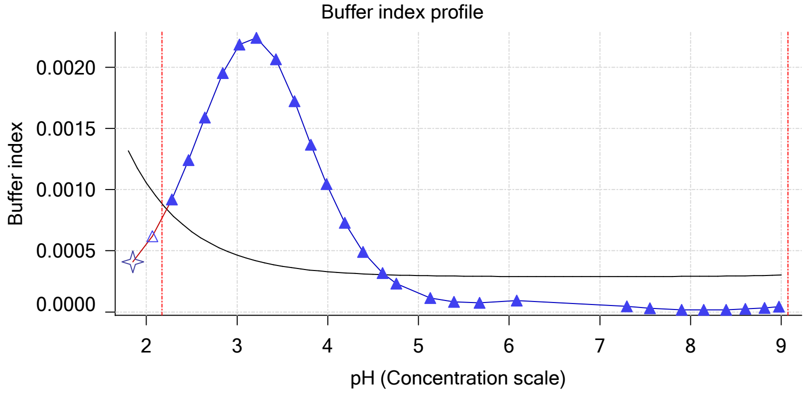


Ionic strength

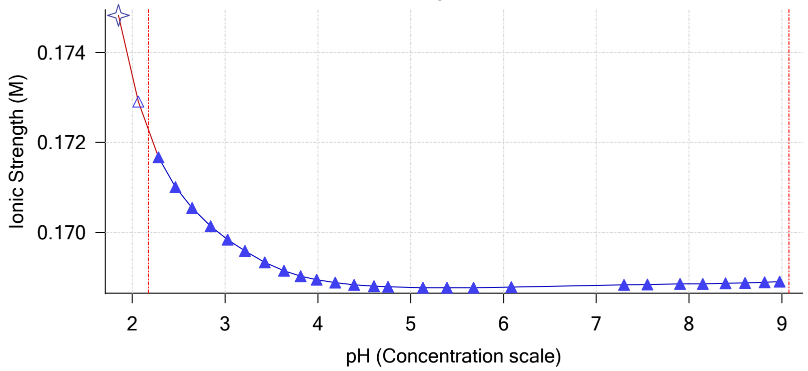


Log R Profile

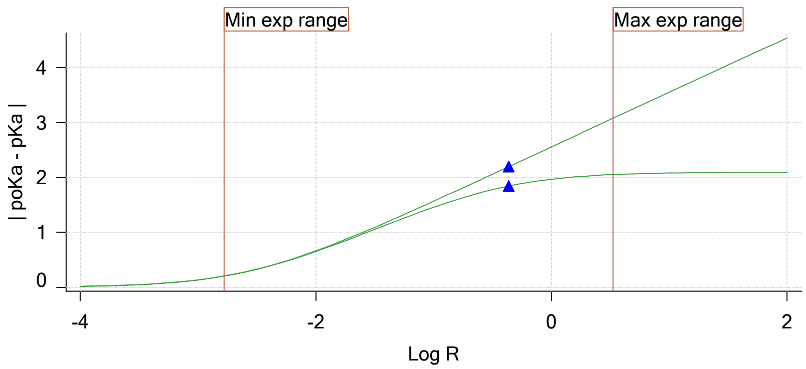


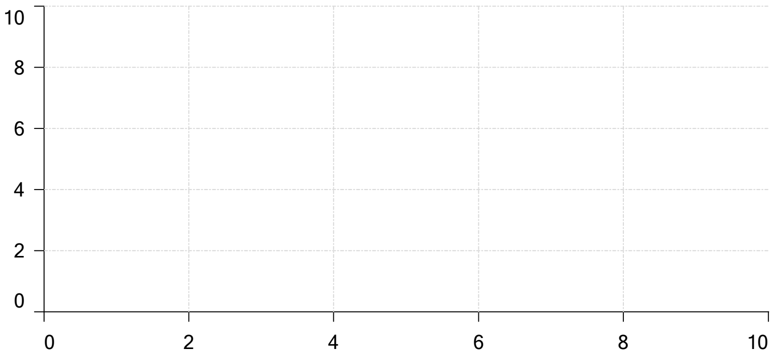


Ionisation of sample Carbonate

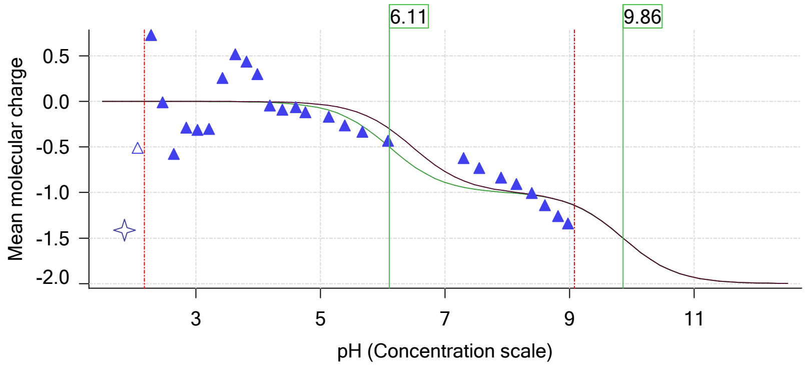

| | | | |
|---|---|---|---|
| Sample name: | **M18_octanol** | Experiment start time: | **3/9/2018 9:53:01 PM** |
| Assay name: | **pH-metric high logP** | Analyst: | **Pion** |
| Assay ID: | **18C-09015** | Instrument ID: | **T312060** |
| Filename: | **C:\Sirius_T3\Mehtap\20180309_exp31_logP_T3-2\18C-09015_M18_octanol_pH-metric high logP.t3r** | | |

### Assay Model

| Settings | Value | Date/Time changed | Imported from |
|---|---|---|---|
| Sample name | M18_octanol | 2/27/2018 7:08:39 PM | User entered value |
| Sample by | Weight | | Default value |
| Sample weight | 0.001670 g | 3/9/2018 2:22:44 PM | User entered value |
| Formula weight | 267.11 g/mol | 2/27/2018 7:08:39 PM | User entered value |
| Solubility | Unknown | | Default value |
| Molecular weight | 267.11 | 2/27/2018 7:08:39 PM | User entered value |
| Individual pKa ionic environments | No | | Default value |
| Number of pKas | 2 | 2/27/2018 7:08:39 PM | User entered value |
| Sample is a | Ampholyte | 2/27/2018 7:08:39 PM | User entered value |
| pKa 1 | 5.37 | 2/27/2018 7:08:39 PM | User entered value |
| Type | Base | 2/27/2018 7:08:39 PM | User entered value |
| pKa 2 | 10.65 | 2/27/2018 7:08:39 PM | User entered value |
| Type | Acid | 2/27/2018 7:08:39 PM | User entered value |
| logp (XH2 +) | -0.35 | 2/28/2018 3:20:28 PM | User entered value |
| logP (neutral XH) | 2.57 | 3/2/2018 4:34:50 PM | User entered value |
| logP (X -) | 0.46 | 2/27/2018 7:09:34 PM | User entered value |

### Events

| Time | Event | Water | Acid | Base | Octanol | pH | dpH/dt | pH R-squared | pH SD | dpH/[illegible] time |
|---|---|---|---|---|---|---|---|---|---|---|
| 4:58.2 | Manual volume addition | | | | 0.10000 mL | | | | | |
| 4:59.3 | Initial pH = 8.13 | | | | | | | | | |
| 7:58.9 | Data point 2 | 1.50000 mL | 0.05148 mL | 0.00405 mL | 0.10000 mL | 2.064 | -0.00398 | 0.61752 | 0.00025 | 10.0 s |
| 8:45.1 | Data point 3 | 1.50000 mL | 0.05148 mL | 0.01646 mL | 0.10000 mL | 2.270 | 0.00032 | 0.01048 | 0.00015 | 10.0 s |
| 9:20.7 | Data point 4 | 1.50000 mL | 0.05148 mL | 0.02441 mL | 0.10000 mL | 2.467 | 0.00127 | 0.33156 | 0.00011 | 10.0 s |
| 9:56.2 | Data point 5 | 1.50000 mL | 0.05148 mL | 0.02954 mL | 0.10000 mL | 2.663 | -0.00044 | 0.05449 | 0.00009 | 10.0 s |
| 10:31.9 | Data point 6 | 1.50000 mL | 0.05148 mL | 0.03295 mL | 0.10000 mL | 2.835 | -0.00106 | 0.08335 | 0.00018 | 10.5 s |
| 11:18.2 | Data point 7 | 1.50000 mL | 0.05148 mL | 0.03523 mL | 0.10000 mL | 3.034 | -0.00637 | 0.44601 | 0.00047 | 10.0 s |
| 11:53.8 | Data point 8 | 1.50000 mL | 0.05148 mL | 0.03709 mL | 0.10000 mL | 3.244 | -0.00918 | 0.56183 | 0.00060 | 10.5 s |
| 12:29.7 | Data point 9 | 1.50000 mL | 0.05148 mL | 0.03866 mL | 0.10000 mL | 3.411 | -0.01473 | 0.74716 | 0.00084 | 10.0 s |
| 13:25.9 | Data point 10 | 1.50000 mL | 0.05148 mL | 0.04080 mL | 0.10000 mL | 3.635 | -0.01009 | 0.71263 | 0.00059 | 10.0 s |
| 14:11.7 | Data point 11 | 1.50000 mL | 0.05148 mL | 0.04240 mL | 0.10000 mL | 3.828 | -0.01257 | 0.80770 | 0.00069 | 10.0 s |
| 14:47.2 | Data point 12 | 1.50000 mL | 0.05148 mL | 0.04400 mL | 0.10000 mL | 4.027 | -0.00532 | 0.89260 | 0.00028 | 10.0 s |
| 15:22.6 | Data point 13 | 1.50000 mL | 0.05148 mL | 0.04558 mL | 0.10000 mL | 4.210 | -0.01823 | 0.90318 | 0.00095 | 16.0 s |
| 16:04.1 | Data point 14 | 1.50000 mL | 0.05148 mL | 0.04701 mL | 0.10000 mL | 4.312 | -0.10209 | 0.99736 | 0.00505 | Time[illegible] out at |
| 17:50.2 | Data point 15 | 1.50000 mL | 0.05148 mL | 0.04908 mL | 0.10000 mL | 4.398 | -0.03232 | 0.99319 | 0.00160 | Time[illegible] out at |
| 19:31.1 | Data point 16 | 1.50000 mL | 0.05148 mL | 0.05047 mL | 0.10000 mL | 4.604 | -0.02003 | 0.98669 | 0.00100 | 45.5 s |
| 20:42.0 | Data point 17 | 1.50000 mL | 0.05148 mL | 0.05146 mL | 0.10000 mL | 5.249 | -0.01859 | 0.86523 | 0.00099 | 26.0 s |

| | | | |
|---|---|---|---|
| Sample name: | **M18_octanol** | Experiment start time: | **3/9/2018 9:53:01 PM** |
| Assay name: | **pH-metric high logP** | Analyst: | **Pion** |
| Assay ID: | **18C-09015** | Instrument ID: | **T312060** |
| Filename: | **C:\Sirius_T3\Mehtap\20180309_exp31_logP_T3-2\18C-09015_M18_octanol_pH-metric high logP.t3r** | | |

### Events (continued)

| Time | Event | Water | Acid | Base | Octanol | pH | dpH/dt | pH R-squared | pH SD | dpH/dt time |
|---|---|---|---|---|---|---|---|---|---|---|
| 21:38.6 | Data point 18 | 1.50000 mL | 0.05148 mL | 0.05200 mL | 0.10000 mL | 8.408 | -0.02877 | 0.97980 | 0.00144 | Timed out at 59.5 s |
| 23:14.2 | Data point 19 | 1.50000 mL | 0.05148 mL | 0.05219 mL | 0.10000 mL | 8.882 | -0.01997 | 0.97577 | 0.00100 | 54.5 s |
| 24:44.4 | Data point 20 | 1.50000 mL | 0.05148 mL | 0.05228 mL | 0.10000 mL | 9.072 | -0.01991 | 0.96826 | 0.00100 | 32.0 s |
| 26:19.4 | Data point 21 | 1.50000 mL | 0.10948 mL | 0.05228 mL | 0.30000 mL | 1.985 | -0.00807 | 0.73649 | 0.00046 | 10.0 s |
| 27:05.6 | Data point 22 | 1.50000 mL | 0.10948 mL | 0.06886 mL | 0.30000 mL | 2.191 | -0.00302 | 0.06536 | 0.00058 | 10.0 s |
| 27:41.3 | Data point 23 | 1.50000 mL | 0.10948 mL | 0.07937 mL | 0.30000 mL | 2.384 | 0.00289 | 0.58885 | 0.00019 | 10.0 s |
| 28:16.9 | Data point 24 | 1.50000 mL | 0.10948 mL | 0.08619 mL | 0.30000 mL | 2.610 | -0.00234 | 0.02222 | 0.00078 | 10.5 s |
| 28:52.9 | Data point 25 | 1.50000 mL | 0.10948 mL | 0.09055 mL | 0.30000 mL | 2.789 | -0.00887 | 0.28002 | 0.00083 | 10.0 s |
| 29:28.4 | Data point 26 | 1.50000 mL | 0.10948 mL | 0.09377 mL | 0.30000 mL | 2.975 | -0.00368 | 0.74326 | 0.00021 | 10.5 s |
| 30:04.5 | Data point 27 | 1.50000 mL | 0.10948 mL | 0.09633 mL | 0.30000 mL | 3.139 | -0.00884 | 0.23194 | 0.00091 | 10.0 s |
| 30:50.3 | Data point 28 | 1.50000 mL | 0.10948 mL | 0.09889 mL | 0.30000 mL | 3.334 | -0.01064 | 0.80030 | 0.00059 | 10.5 s |
| 31:26.3 | Data point 29 | 1.50000 mL | 0.10948 mL | 0.10094 mL | 0.30000 mL | 3.577 | -0.01028 | 0.41136 | 0.00079 | 10.0 s |
| 32:12.0 | Data point 30 | 1.50000 mL | 0.10948 mL | 0.10280 mL | 0.30000 mL | 3.773 | -0.00666 | 0.87855 | 0.00035 | 10.5 s |
| 32:48.0 | Data point 31 | 1.50000 mL | 0.10948 mL | 0.10442 mL | 0.30000 mL | 3.947 | -0.01424 | 0.74500 | 0.00081 | 10.0 s |
| 33:33.8 | Data point 32 | 1.50000 mL | 0.10948 mL | 0.10600 mL | 0.30000 mL | 4.149 | -0.00724 | 0.88195 | 0.00038 | 10.5 s |
| 34:09.8 | Data point 33 | 1.50000 mL | 0.10948 mL | 0.10706 mL | 0.30000 mL | 4.340 | -0.00713 | 0.59165 | 0.00046 | 10.0 s |
| 34:45.3 | Data point 34 | 1.50000 mL | 0.10948 mL | 0.10786 mL | 0.30000 mL | 4.525 | -0.01579 | 0.72847 | 0.00091 | 10.0 s |
| 35:20.7 | Data point 35 | 1.50000 mL | 0.10948 mL | 0.10842 mL | 0.30000 mL | 4.747 | -0.01335 | 0.78441 | 0.00074 | 10.5 s |
| 35:56.6 | Data point 36 | 1.50000 mL | 0.10948 mL | 0.10880 mL | 0.30000 mL | 4.951 | -0.00800 | 0.24231 | 0.00080 | 10.5 s |
| 36:32.5 | Data point 37 | 1.50000 mL | 0.10948 mL | 0.10903 mL | 0.30000 mL | 5.127 | -0.01493 | 0.70300 | 0.00088 | 11.0 s |
| 37:14.0 | Data point 38 | 1.50000 mL | 0.10948 mL | 0.10936 mL | 0.30000 mL | 5.672 | -0.01880 | 0.88256 | 0.00099 | 29.5 s |
| 38:19.4 | Data point 39 | 1.50000 mL | 0.10948 mL | 0.10969 mL | 0.30000 mL | 7.963 | -0.08034 | 0.99469 | 0.00398 | Timed out at 59.5 s |
| 40:00.2 | Data point 40 | 1.50000 mL | 0.10948 mL | 0.10985 mL | 0.30000 mL | 8.691 | -0.01737 | 0.90449 | 0.00090 | 50.5 s |
| 41:26.4 | Data point 41 | 1.50000 mL | 0.10948 mL | 0.11000 mL | 0.30000 mL | 9.055 | -0.02009 | 0.98442 | 0.00100 | 30.0 s |
| 43:22.0 | Data point 42 | 1.50000 mL | 0.17251 mL | 0.11000 mL | 0.80000 mL | 1.964 | -0.00989 | 0.81980 | 0.00054 | 10.0 s |
| 44:08.3 | Data point 43 | 1.50000 mL | 0.17251 mL | 0.12900 mL | 0.80000 mL | 2.170 | 0.00443 | 0.37114 | 0.00036 | 10.0 s |
| 44:44.0 | Data point 44 | 1.50000 mL | 0.17251 mL | 0.14088 mL | 0.80000 mL | 2.382 | 0.00263 | 0.03724 | 0.00067 | 10.5 s |
| 45:20.1 | Data point 45 | 1.50000 mL | 0.17251 mL | 0.14857 mL | 0.80000 mL | 2.564 | -0.00827 | 0.25892 | 0.00080 | 10.0 s |
| 45:55.7 | Data point 46 | 1.50000 mL | 0.17251 mL | 0.15404 mL | 0.80000 mL | 2.741 | -0.00871 | 0.52375 | 0.00059 | 10.0 s |
| 46:31.2 | Data point 47 | 1.50000 mL | 0.17251 mL | 0.15818 mL | 0.80000 mL | 2.939 | 0.00202 | 0.06322 | 0.00040 | 10.5 s |
| 47:07.2 | Data point 48 | 1.50000 mL | 0.17251 mL | 0.16141 mL | 0.80000 mL | 3.121 | -0.00814 | 0.90284 | 0.00042 | 10.0 s |
| 47:42.7 | Data point 49 | 1.50000 mL | 0.17251 mL | 0.16409 mL | 0.80000 mL | 3.304 | 0.00201 | 0.05752 | 0.00041 | 10.5 s |
| 48:18.7 | Data point 50 | 1.50000 mL | 0.17251 mL | 0.16634 mL | 0.80000 mL | 3.520 | -0.01362 | 0.46718 | 0.00098 | 10.0 s |
| 48:54.2 | Data point 51 | 1.50000 mL | 0.17251 mL | 0.16813 mL | 0.80000 mL | 3.724 | -0.00424 | 0.07060 | 0.00079 | 10.0 s |
| 49:29.6 | Data point 52 | 1.50000 mL | 0.17251 mL | 0.16950 mL | 0.80000 mL | 3.903 | -0.00679 | 0.94080 | 0.00035 | 10.5 s |
| 50:05.6 | Data point 53 | 1.50000 mL | 0.17251 mL | 0.17053 mL | 0.80000 mL | 4.076 | 0.00436 | 0.17804 | 0.00051 | 10.0 s |
| 50:56.5 | Data point 54 | 1.50000 mL | 0.17251 mL | 0.17154 mL | 0.80000 mL | 4.276 | -0.00388 | 0.13097 | 0.00053 | 10.0 s |
| 51:47.4 | Data point 55 | 1.50000 mL | 0.17251 mL | 0.17211 mL | 0.80000 mL | 4.477 | -0.01051 | 0.37200 | 0.00085 | 10.0 s |
| 52:22.7 | Data point 56 | 1.50000 mL | 0.17251 mL | 0.17246 mL | 0.80000 mL | 4.690 | -0.01333 | 0.59091 | 0.00086 | 11.0 s |
| 52:59.1 | Data point 57 | 1.50000 mL | 0.17251 mL | 0.17267 mL | 0.80000 mL | 4.843 | -0.01303 | 0.56430 | 0.00086 | 11.0 s |
| 53:40.6 | Data point 58 | 1.50000 mL | 0.17251 mL | 0.17295 mL | 0.80000 mL | 5.218 | -0.01867 | 0.86298 | 0.00099 | 28.0 s |
| 54:39.2 | Data point 59 | 1.50000 mL | 0.17251 mL | 0.17310 mL | 0.80000 mL | 5.477 | -0.01831 | 0.92742 | 0.00094 | 31.0 s |
| 55:40.7 | Data point 60 | 1.50000 mL | 0.17251 mL | 0.17319 mL | 0.80000 mL | 5.760 | -0.01927 | 0.92709 | 0.00099 | 35.5 s |
| 56:46.8 | Data point 61 | 1.50000 mL | 0.17251 mL | 0.17328 mL | 0.80000 mL | 6.166 | -0.01962 | 0.94684 | 0.00100 | 59.0 s |
| 58:16.4 | Data point 62 | 1.50000 mL | 0.17251 mL | 0.17342 mL | 0.80000 mL | 7.377 | -0.12941 | 0.99771 | 0.00639 | Timed out at 59.5 s |
| 59:46.9 | Data point 63 | 1.50000 mL | 0.17251 mL | 0.17350 mL | 0.80000 mL | 7.630 | -0.07477 | 0.98222 | 0.00373 | Timed out at 59.5 s |
| 1:01:22.5 | Data point 64 | 1.50000 mL | 0.17251 mL | 0.17357 mL | 0.80000 mL | 7.978 | -0.05777 | 0.98086 | 0.00288 | Timed out at 59.5 s |
| 1:02:53.1 | Data point 65 | 1.50000 mL | 0.17251 mL | 0.17361 mL | 0.80000 mL | 8.222 | -0.03185 | 0.98956 | 0.00158 | Timed out at 59.5 s |
| 1:04:28.8 | Data point 66 | 1.50000 mL | 0.17251 mL | 0.17368 mL | 0.80000 mL | 8.468 | -0.02381 | 0.91426 | 0.00123 | Timed out at 59.5 s |
| 1:06:09.5 | Data point 67 | 1.50000 mL | 0.17251 mL | 0.17378 mL | 0.80000 mL | 8.677 | -0.01737 | 0.85617 | 0.00093 | 43.5 s |

| | | | |
|---|---|---|---|
| Sample name: | **M18_octanol** | Experiment start time: | **3/9/2018 9:53:01 PM** |
| Assay name: | **pH-metric high logP** | Analyst: | **Pion** |
| Assay ID: | **18C-09015** | Instrument ID: | **T312060** |
| Filename: | **C:\Sirius_T3\Mehtap\20180309_exp31_logP_T3-2\18C-09015_M18_octanol_pH-metric high logP.t3r** | | |

### Events (continued)

| Time | Event | Water | Acid | Base | Octanol | pH | dpH/dt | pH R-squared | pH SD | dpH/dt time |
|---|---|---|---|---|---|---|---|---|---|---|
| 1:07:28.7 | Data point 68 | 1.50000 mL | 0.17251 mL | 0.17387 mL | 0.80000 mL | 8.888 | -0.01877 | 0.89665 | 0.00098 | 35.5 s |
| 1:08:34.8 | Data point 69 | 1.50000 mL | 0.17251 mL | 0.17394 mL | 0.80000 mL | 9.048 | -0.01938 | 0.95152 | 0.00098 | 14.0 s |
| 1:08:57.9 | Assay volumes | 1.50000 mL | 0.17251 mL | 0.17394 mL | 0.80000 mL | | | | | |

| | | | |
|---|---|---|---|
| Sample name: | **M18_octanol** | Experiment start time: | **3/9/2018 9:53:01 PM** |
| Assay name: | **pH-metric high logP** | Analyst: | **Pion** |
| Assay ID: | **18C-09015** | Instrument ID: | **T312060** |
| Filename: | **C:\Sirius_T3\Mehtap\20180309_exp31_logP_T3-2\18C-09015_M18_octanol_pH-metric high logP.t3r** | | |

| | | | |
|---|---|---|---|
| Sample name: | **M18_octanol** | Experiment start time: | **3/9/2018 9:53:01 PM** |
| Assay name: | **pH-metric high logP** | Analyst: | **Pion** |
| Assay ID: | **18C-09015** | Instrument ID: | **T312060** |
| Filename: | **C:\Sirius_T3\Mehtap\20180309_exp31_logP_T3-2\18C-09015_M18_octanol_pH-metric high logP.t3r** | | |

### Assay Settings (continued)

| Setting | Value | Original Value | Date/Time changed | Imported from |
|---|---|---|---|---|
| ***Titration 3*** | | | | |
| Titrate from | Low to high pH | | | |
| Add additional water | 0.00 mL | | | |
| Additional partition solvent volume | 0.500 mL | | | |
| Additional partition solvent added | Automatic | | | |
| After pH adjust stir for | 30 seconds | | | |
| Stir to allow partitioning for | 15 seconds | | | |
| Stirrer speed for partitioning | 60% | | | |
| ***Data Point Stability*** | | | | |
| Stir during data point collection | No | | | |
| Delay before data point collection | 0 seconds | | | |
| Number of points to average | 20 points | | | |
| Time interval between points | 0.50 seconds | | | |
| Required maximum standard deviation | 0.00100 dpH/dt | | | |
| Stability timeout after | 60 seconds | | | |

### Calibration Settings

| Setting | Value | Date/Time changed | Imported from |
|---|---|---|---|
| Four-Plus alpha | 0.102 | 3/9/2018 9:53:01 PM | C:\Sirius_T3\HCl18C09.t3r |
| Four-Plus S | 0.9967 | 3/9/2018 9:53:01 PM | C:\Sirius_T3\HCl18C09.t3r |
| Four-Plus jH | 1.2 | 3/9/2018 9:53:01 PM | C:\Sirius_T3\HCl18C09.t3r |
| Four-Plus jOH | 0.0 | 3/9/2018 9:53:01 PM | C:\Sirius_T3\HCl18C09.t3r |
| Base concentration factor | 1.000 | 3/9/2018 9:53:01 PM | C:\Sirius_T3\KOH18B27.t3r |
| Acid concentration factor | 1.000 | 3/9/2018 9:53:01 PM | C:\Sirius_T3\HCl18C09.t3r |

### Instrument Settings

| Setting | Value | Batch Id | Install date |
|---|---|---|---|
| Instrument owner | Merck | | |
| Instrument ID | T312060 | | |
| Instrument type | T3 Simulator | | |
| Software version | 1.1.3.0 | | |
| Dispenser module | | T3DM1200361 | 3/31/2009 6:24:52 AM |
| Dispenser 0 | Water | | 3/31/2009 6:25:05 AM |
| Syringe volume | 2.5 mL | | |
| Firmware version | 1.2.1(r2) | | |
| Titrant | Water (0.15 M KCl) | 02-06-2018 | 2/27/2018 11:05:59 AM |
| Dispenser 2 | Acid | | 3/31/2009 6:25:11 AM |
| Syringe volume | 0.5 mL | | |
| Firmware version | 1.2.1(r2) | | |
| Titrant | Acid (0.5 M HCl) | 02-27-2018 | 2/27/2018 11:27:22 AM |
| Dispenser 1 | Base | | 3/31/2009 6:25:21 AM |
| Syringe volume | 0.5 mL | | |
| Firmware version | 1.2.1(r2) | | |
| Titrant | Base (0.5 M KOH) | 9/22/2017 | 2/27/2018 11:21:22 AM |
| Dispenser 5 | Cosolvent | | 3/31/2009 6:26:24 AM |
| Syringe volume | 2.5 mL | | |
| Firmware version | 1.2.1(r2) | | |
| Distribution valve 5 | Distribution Valve | | 3/31/2009 6:28:19 AM |
| Firmware version | 1.1.3 | | |
| Port A | Methanol (80%, 0.15 M KCl) | 02-08-2018 | 3/6/2018 10:28:59 AM |
| Port B | Cyclohexane | 11-01-17 | 2/27/2018 11:37:57 AM |
| Dispenser 3 | Buffer | | 8/3/2010 6:05:16 AM |
| Syringe volume | 0.5 mL | | |
| Firmware version | 1.2.1(r2) | | |
| Titrant | Dodecane | 2018/01/31 | 2/28/2018 11:18:04 AM |
| Dispenser 6 | Octanol | | 10/22/2010 11:52:43 AM |

| | | | |
|---|---|---|---|
| Sample name: | **M18_octanol** | Experiment start time: | **3/9/2018 9:53:01 PM** |
| Assay name: | **pH-metric high logP** | Analyst: | **Pion** |
| Assay ID: | **18C-09015** | Instrument ID: | **T312060** |
| Filename: | **C:\Sirius_T3\Mehtap\20180309_exp31_logP_T3-2\18C-09015_M18_octanol_pH-metric high logP.t3r** | | |

| | | | |
|---|---|---|---|
| Sample name: | **M18_octanol** | Experiment start time: | **3/9/2018 9:53:01 PM** |
| Assay name: | **pH-metric high logP** | Analyst: | **Pion** |
| Assay ID: | **18C-09015** | Instrument ID: | **T312060** |
| Filename: | **C:\Sirius_T3\Mehtap\20180309_exp31_logP_T3-2\18C-09015_M18_octanol_pH-metric high logP.t3r** | | |

### Instrument Settings (continued)

| Setting | Value | Batch Id | Install date |
|---|---|---|---|
| Spectrometer calibration stir duration | 5 s | | |
| Spectrometer calibration stir speed | 30% | | |
| Spectrometer calibration wash pump volume | 20.0 mL | | |
| Spectrometer calibration wash stir duration | 5 s | | |
| Spectrometer calibration wash stir speed | 30% | | |
| Overhead dispense height | 10000 | | |

### Refinement Settings

| Setting | Value | Default value |
|---|---|---|
| Turbidity detection method | None | None |
| Turbidity wavelength to assess | 500.0 nm | 500.0 nm |
| Turbidity maximum absorbance | 0.100 | 0.100 |
| Turbidity probe threshold | 50.00 | 50.00 |
