## Supplementary material for "Octanol-water partition coefficient measurements for the SAMPL6 Blind Prediction Challenge": SM16_18C-09016_M18_octanol_pH-metric high logP_report.pdf

| | | | |
|---|---|---|---|
| Sample name: | **M18_octanol** | Experiment start time: | **3/9/2018 11:02:44 PM** |
| Assay name: | **pH-metric high logP** | Analyst: | **Pion** |
| Assay ID: | **18C-09016** | Instrument ID: | **T312060** |
| Filename: | **C:\Sirius_T3\Mehtap\20180309_exp31_logP_T3-2\18C-09016_M18_octanol_pH-metric high logP.t3r** | | |

#### pH-metric Result

| | | |
|---|---|---|
| logP (XH2 +) | -0.21 | ±0.12 (n=50) |
| logP (neutral XH) | 2.63 | ±0.02 (n=50) |
| logP (X -) | 0.46 | |
| RMSD | 0.583 | |

#### 18C-09016 Points 2 to 28

| | |
|---|---|
| M18_octanol concentration factor | 1.091 |
| Carbonate | 0.1016 mM |
| Acidity error | -0.00215 mM |

#### 18C-09016 Points 29 to 56

| | |
|---|---|
| M18_octanol concentration factor | 1.051 |
| Carbonate | 0.0884 mM |
| Acidity error | -0.10411 mM |

#### 18C-09016 Points 57 to 86

| | |
|---|---|
| M18_octanol concentration factor | 1.063 |
| Carbonate | 0.1449 mM |
| Acidity error | 0.00187 mM |

#### Warnings and errors

| | |
|---|---|
| Errors | None |
| Warnings | None |

#### Sample logD and percent species

| pH | M18_octanol logD | M18_octanol M18_octanolH2 | M18_octanol M18_octanolH | M18_octanol M18_octanol | M18_octanol M18_octanolH2* | M18_octanol M18_octanolH* | M18_octanol M18_octanol* | Comment |
|---|---|---|---|---|---|---|---|---|
| 1.000 | -0.19 | 61.03 % | 0.00 % | 0.00 % | 37.85 % | 1.12 % | 0.00 % | |
| 1.200 | -0.19 | 60.63 % | 0.00 % | 0.00 % | 37.60 % | 1.76 % | 0.00 % | Stomach pH |
| 2.000 | -0.10 | 55.44 % | 0.02 % | 0.00 % | 34.39 % | 10.15 % | 0.00 % | |
| 3.000 | 0.39 | 28.95 % | 0.12 % | 0.00 % | 17.95 % | 52.98 % | 0.00 % | |
| 4.000 | 1.26 | 5.01 % | 0.21 % | 0.00 % | 3.11 % | 91.67 % | 0.00 % | |
| 5.000 | 2.11 | 0.54 % | 0.23 % | 0.00 % | 0.34 % | 98.89 % | 0.00 % | |
| 6.000 | 2.54 | 0.05 % | 0.23 % | 0.00 % | 0.03 % | 99.68 % | 0.00 % | |
| 6.500 | 2.60 | 0.02 % | 0.23 % | 0.00 % | 0.01 % | 99.74 % | 0.00 % | |
| 7.000 | 2.62 | 0.01 % | 0.23 % | 0.00 % | 0.00 % | 99.76 % | 0.00 % | |
| 7.400 | 2.63 | 0.00 % | 0.23 % | 0.00 % | 0.00 % | 99.76 % | 0.00 % | Blood pH |
| 8.000 | 2.63 | 0.00 % | 0.23 % | 0.00 % | 0.00 % | 99.76 % | 0.00 % | |
| 9.000 | 2.62 | 0.00 % | 0.23 % | 0.01 % | 0.00 % | 99.75 % | 0.02 % | |
| 10.000 | 2.55 | 0.00 % | 0.23 % | 0.05 % | 0.00 % | 99.57 % | 0.15 % | |
| 11.000 | 2.13 | 0.00 % | 0.23 % | 0.51 % | 0.00 % | 97.79 % | 1.47 % | |
| 12.000 | 1.32 | 0.00 % | 0.19 % | 4.33 % | 0.00 % | 82.99 % | 12.49 % | |

| | | | |
|---|---|---|---|
| Sample name: | **M18_octanol** | Experiment start time: | **3/9/2018 11:02:44 PM** |
| Assay name: | **pH-metric high logP** | Analyst: | **Pion** |
| Assay ID: | **18C-09016** | Instrument ID: | **T312060** |
| Filename: | **C:\Sirius_T3\Mehtap\20180309_exp31_logP_T3-2\18C-09016_M18_octanol_pH-metric high logP.t3r** | | |

#### Graphs

Charge balance residuals

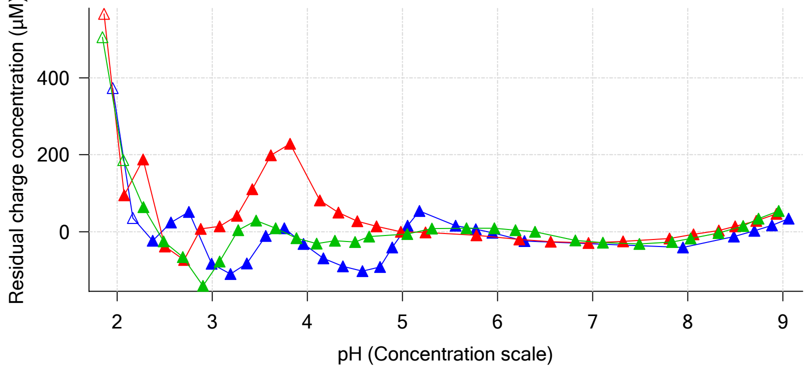


Ionic strength

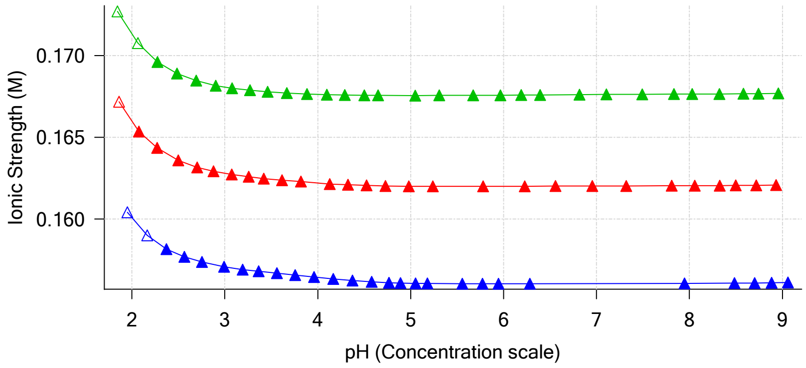


Buffer index profile

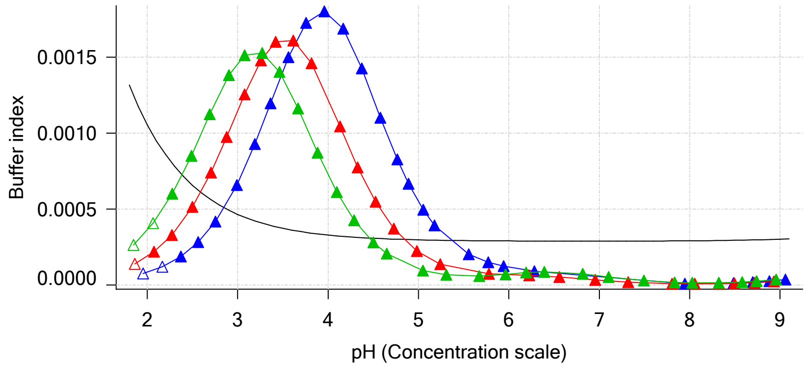


Sample M18_octanol Ionisation Graph

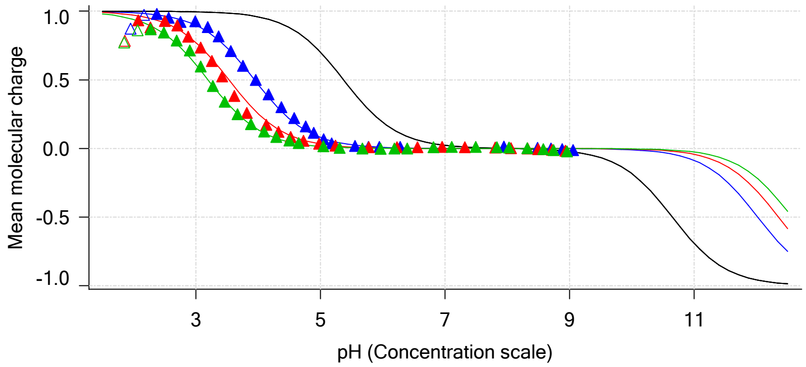


Distribution of species for sample M18_octanol

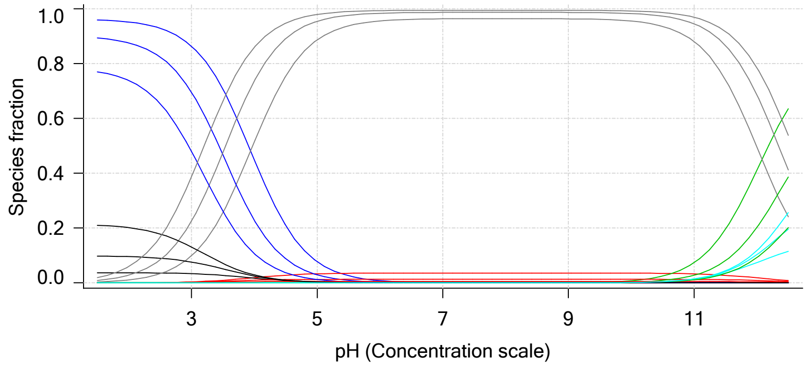


Lipophilicity Profile

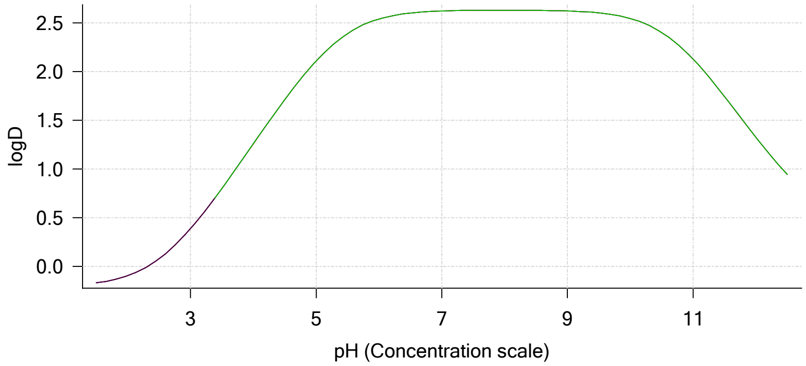


Log R Profile

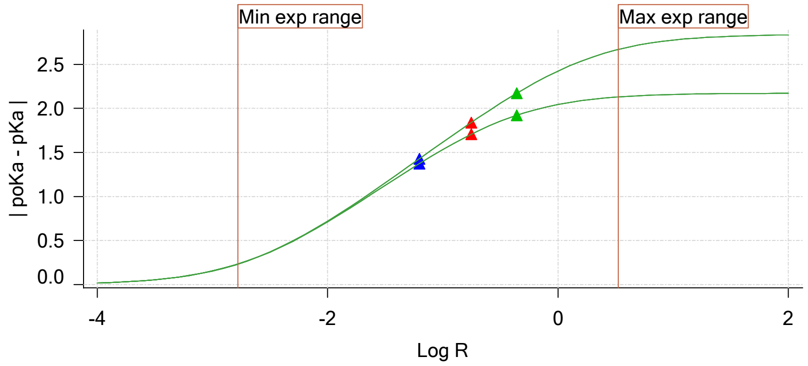


Sample Carbonate Ionisation Graph

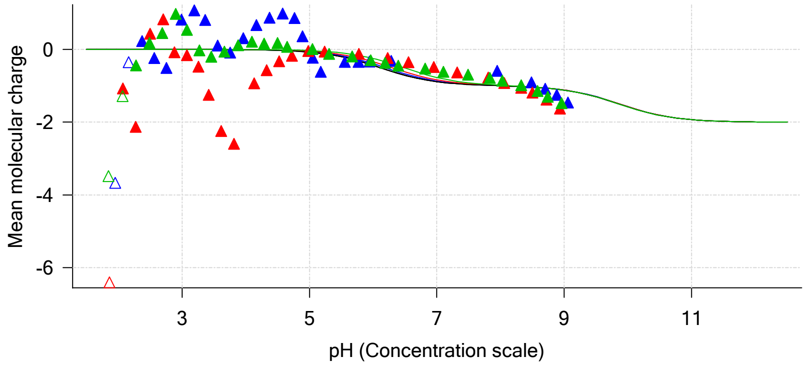

Sample name: **M18_octanol**
Assay name: **pH-metric high logP**
Assay ID: **18C-09016**
Filename: **C:\Sirius_T3\Mehtap\20180309_exp31_logP_T3-2\18C-09016_M18_octanol_pH-metric high logP.t3r**

Experiment start time: **3/9/2018 11:02:44 PM**
Analyst: **Pion**
Instrument ID: **T312060**

#### Graphs (continued)

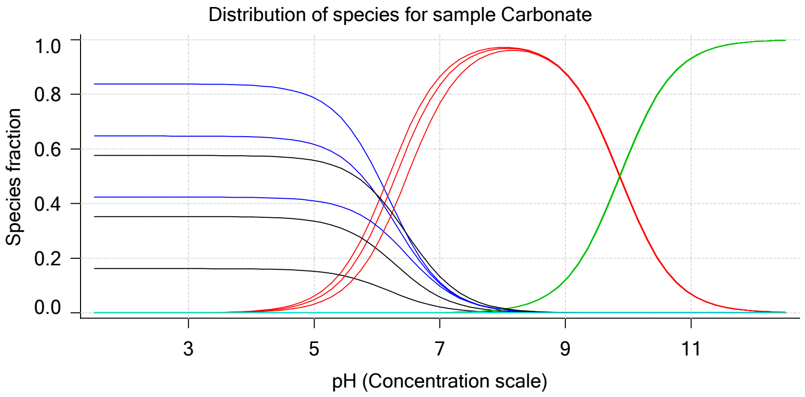

| | | | |
|---|---|---|---|
| Sample name: | **M18_octanol** | Experiment start time: | **3/9/2018 11:02:44 PM** |
| Assay name: | **pH-metric high logP** | Analyst: | **Pion** |
| Assay ID: | **18C-09016** | Instrument ID: | **T312060** |
| Filename: | **C:\Sirius_T3\Mehtap\20180309_exp31_logP_T3-2\18C-09016_M18_octanol_pH-metric high logP.t3r** | | |

### pH-metric high logP Titration 1 of 3 18C-09016 Points 2 to 28

#### Overall results

| | |
|---|---|
| RMSD | 0.731 |
| Average ionic strength | 0.156 M |
| Average temperature | 24.9°C |
| Partition ratio | 0.0626 : 1 |
| Analyte concentration range | 2691.8 µM to 2763.3 µM |
| Total points considered | 25 of 27 |

#### Warnings and errors

| | |
|---|---|
| Errors | None |
| Warnings | None |

#### Four-Plus parameters

| | | | |
|---|---|---|---|
| Alpha | 0.102 | 3/9/2018 11:02:43 PM | C:\Sirius_T3\HCl18C09.t3r |
| S | 0.9967 | 3/9/2018 11:02:43 PM | C:\Sirius_T3\HCl18C09.t3r |
| jH | 1.2 | 3/9/2018 11:02:43 PM | C:\Sirius_T3\HCl18C09.t3r |
| jOH | 0.0 | 3/9/2018 11:02:43 PM | C:\Sirius_T3\HCl18C09.t3r |

#### Titrants

| | | | |
|---|---|---|---|
| 0.50 M HCl | 0.999843 | 3/9/2018 11:02:44 PM | C:\Sirius_T3\HCl18C09.t3r |
| 0.50 M KOH | 0.999845 | 3/9/2018 11:02:44 PM | C:\Sirius_T3\KOH18B27.t3r |

#### Sample

| | |
|---|---|
| M18_octanol concentration factor | 1.091 |
| Base pKa 1 | 5.37 |
| Acid pKa 2 | 10.65 |
| logP (XH2 +) | -1.02 |
| logP (neutral XH) | 2.58 |
| logP (X -) | 0.46 |

#### Sample graphs

Ionisation of sample M18_octanol

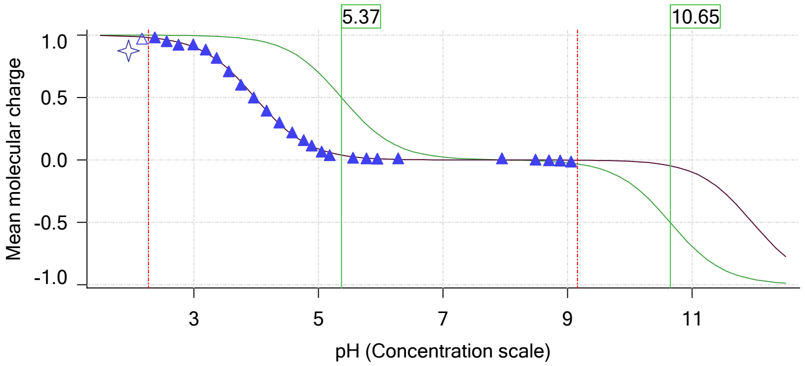


Distribution of species for sample M18_octanol

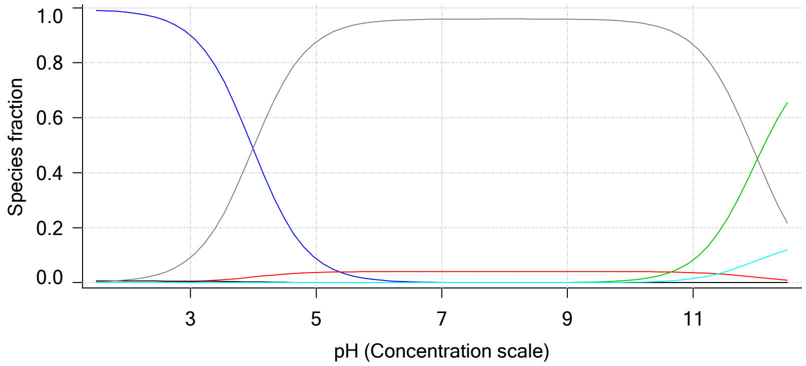

| | | | |
|---|---|---|---|
| Sample name: | **M18_octanol** | Experiment start time: | **3/9/2018 11:02:44 PM** |
| Assay name: | **pH-metric high logP** | Analyst: | **Pion** |
| Assay ID: | **18C-09016** | Instrument ID: | **T312060** |
| Filename: | **C:\Sirius_T3\Mehtap\20180309_exp31_logP_T3-2\18C-09016_M18_octanol_pH-metric high logP.t3r** | | |

#### Sample graphs (continued)

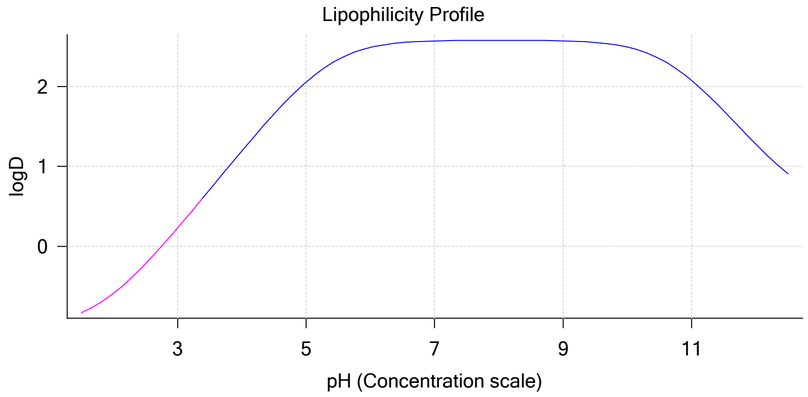


#### Sample logD and percent species

| pH | M18_octanol logD | M18_octanol M18_octanolH2 | M18_octanol M18_octanolH | M18_octanol M18_octanol | M18_octanol M18_octanolH2* | M18_octanol M18_octanolH* | M18_octanol M18_octanol* | Comment |
|---|---|---|---|---|---|---|---|---|
| 1.000 | -0.95 | 99.30 % | 0.00 % | 0.00 % | 0.60 % | 0.10 % | 0.00 % | |
| 1.200 | -0.91 | 99.24 % | 0.01 % | 0.00 % | 0.60 % | 0.16 % | 0.00 % | Stomach pH |
| 2.000 | -0.59 | 98.37 % | 0.04 % | 0.00 % | 0.59 % | 1.00 % | 0.00 % | |
| 3.000 | 0.23 | 89.97 % | 0.38 % | 0.00 % | 0.54 % | 9.11 % | 0.00 % | |
| 4.000 | 1.19 | 48.52 % | 2.07 % | 0.00 % | 0.29 % | 49.12 % | 0.00 % | |
| 5.000 | 2.06 | 8.65 % | 3.69 % | 0.00 % | 0.05 % | 87.60 % | 0.00 % | |
| 6.000 | 2.49 | 0.94 % | 4.00 % | 0.00 % | 0.01 % | 95.05 % | 0.00 % | |
| 6.500 | 2.55 | 0.30 % | 4.03 % | 0.00 % | 0.00 % | 95.67 % | 0.00 % | |
| 7.000 | 2.57 | 0.09 % | 4.04 % | 0.00 % | 0.00 % | 95.86 % | 0.00 % | |
| 7.400 | 2.57 | 0.04 % | 4.04 % | 0.00 % | 0.00 % | 95.92 % | 0.00 % | Blood pH |
| 8.000 | 2.58 | 0.01 % | 4.04 % | 0.01 % | 0.00 % | 95.94 % | 0.00 % | |
| 9.000 | 2.57 | 0.00 % | 4.04 % | 0.09 % | 0.00 % | 95.85 % | 0.02 % | |
| 10.000 | 2.49 | 0.00 % | 4.00 % | 0.90 % | 0.00 % | 94.94 % | 0.16 % | |
| 11.000 | 2.08 | 0.00 % | 3.65 % | 8.18 % | 0.00 % | 86.70 % | 1.48 % | |
| 12.000 | 1.28 | 0.00 % | 1.95 % | 43.76 % | 0.00 % | 46.39 % | 7.90 % | |

#### Carbonate and acidity

| | |
|---|---|
| Carbonate | 0.102 mM |
| Acidity error | -0.002 mM |

#### Other graphs

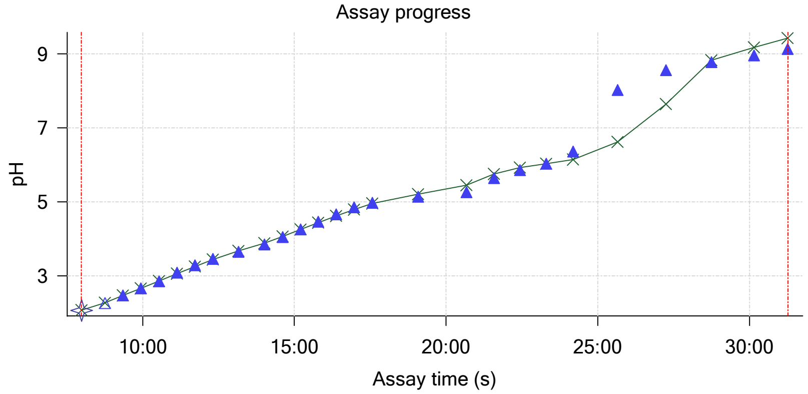


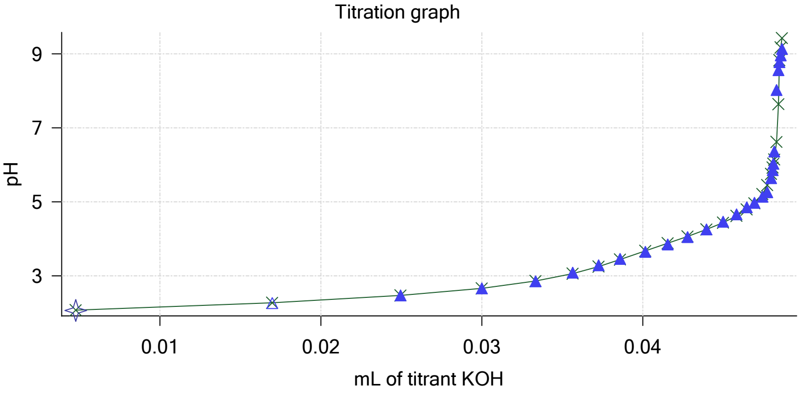

| | | | |
|---|---|---|---|
| Sample name: | **M18_octanol** | Experiment start time: | **3/9/2018 11:02:44 PM** |
| Assay name: | **pH-metric high logP** | Analyst: | **Pion** |
| Assay ID: | **18C-09016** | Instrument ID: | **T312060** |
| Filename: | **C:\Sirius_T3\Mehtap\20180309_exp31_logP_T3-2\18C-09016_M18_octanol_pH-metric high logP.t3r** | | |

#### Other graphs (continued)

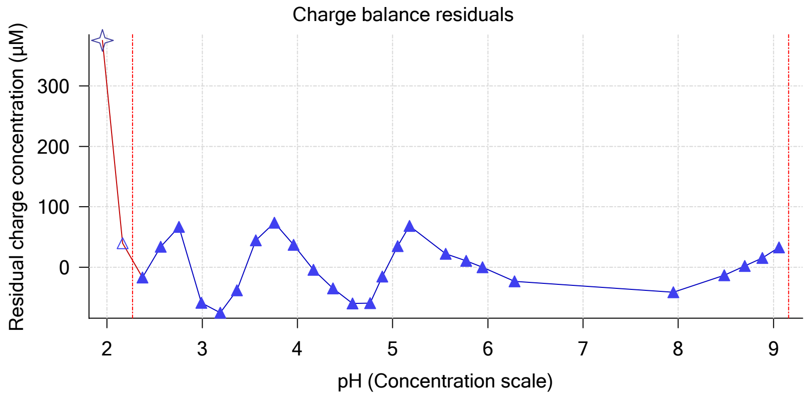


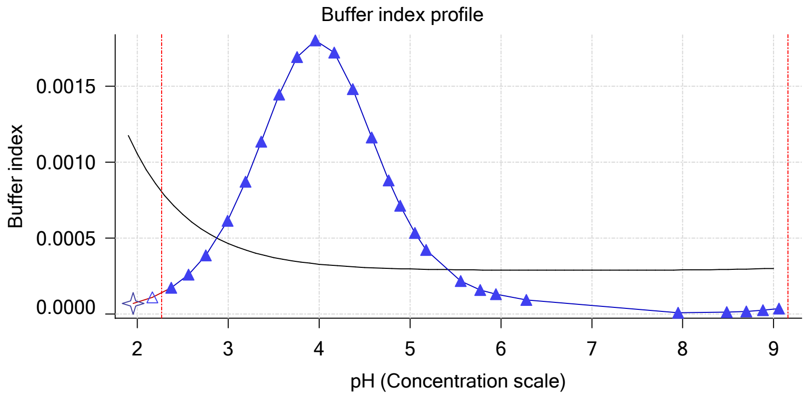


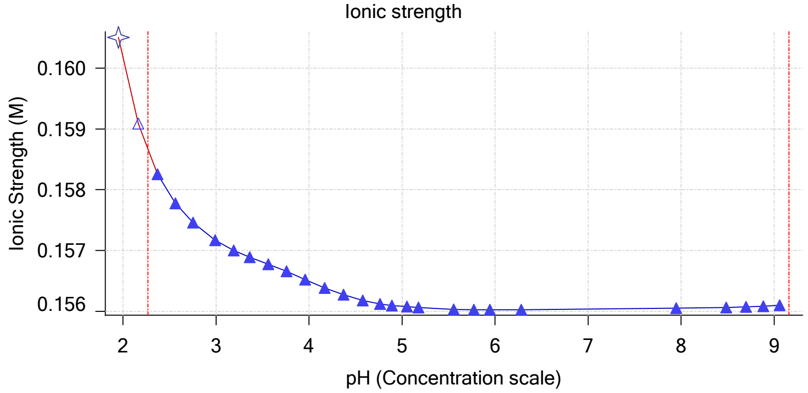


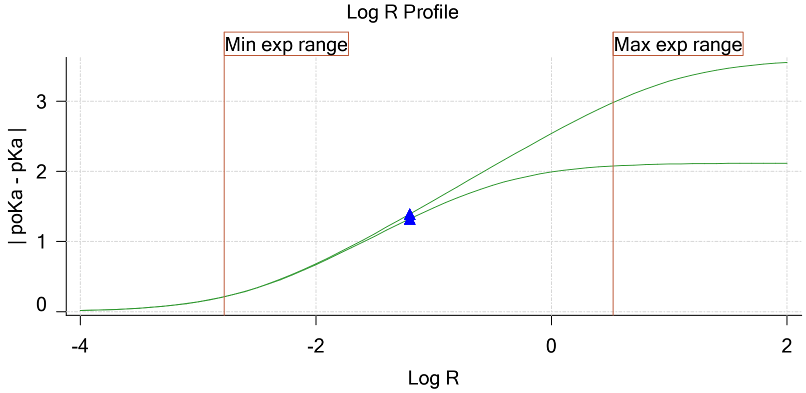


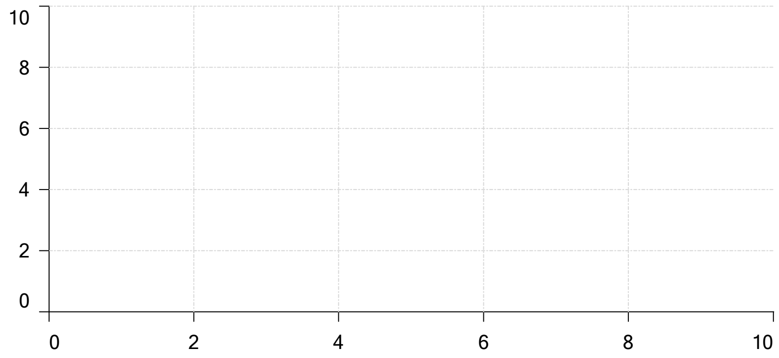

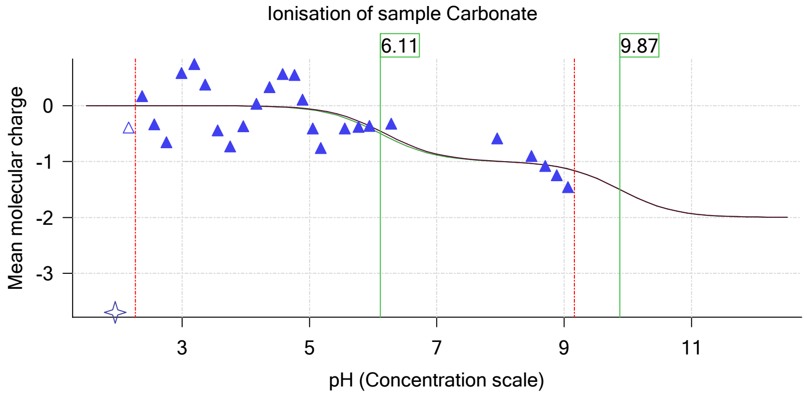

| | | | |
|---|---|---|---|
| Sample name: | **M18_octanol** | Experiment start time: | **3/9/2018 11:02:44 PM** |
| Assay name: | **pH-metric high logP** | Analyst: | **Pion** |
| Assay ID: | **18C-09016** | Instrument ID: | **T312060** |
| Filename: | **C:\Sirius_T3\Mehtap\20180309_exp31_logP_T3-2\18C-09016_M18_octanol_pH-metric high logP.t3r** | | |

# pH-metric high logP   Titration 2 of 3   18C-09016 Points 29 to 56

#### Overall results

| | |
|---|---|
| RMSD | 0.550 |
| Average ionic strength | 0.162 M |
| Average temperature | 25.0°C |
| Partition ratio | 0.1763 : 1 |
| Analyte concentration range | 2278.7 µM to 2341.3 µM |
| Total points considered | 27 of 28 |

#### Warnings and errors

| | |
|---|---|
| Errors | None |
| Warnings | One or more logP values out of range |

#### Four-Plus parameters

| | | | |
|---|---|---|---|
| Alpha | 0.102 | 3/9/2018 11:02:43 PM | C:\Sirius_T3\HCl18C09.t3r |
| S | 0.9967 | 3/9/2018 11:02:43 PM | C:\Sirius_T3\HCl18C09.t3r |
| jH | 1.2 | 3/9/2018 11:02:43 PM | C:\Sirius_T3\HCl18C09.t3r |
| jOH | 0.0 | 3/9/2018 11:02:43 PM | C:\Sirius_T3\HCl18C09.t3r |

#### Titrants

| | | | |
|---|---|---|---|
| 0.50 M HCl | 0.999843 | 3/9/2018 11:02:44 PM | C:\Sirius_T3\HCl18C09.t3r |
| 0.50 M KOH | 0.999845 | 3/9/2018 11:02:44 PM | C:\Sirius_T3\KOH18B27.t3r |

#### Sample

| | |
|---|---|
| M18_octanol concentration factor | 1.051 |
| Base pKa 1 | 5.37 |
| Acid pKa 2 | 10.65 |
| logP (XH2 +) | 7.44 |
| logP (neutral XH) | 9.35 |
| logP (X -) | 0.46 |

#### Sample graphs

Ionisation of sample M18_octanol

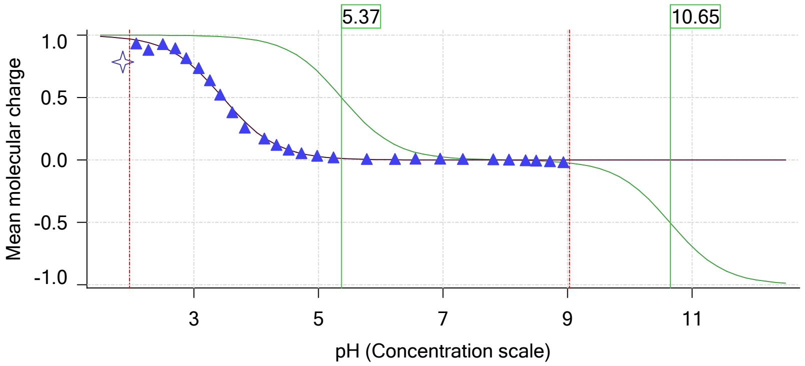


Distribution of species for sample M18_octanol

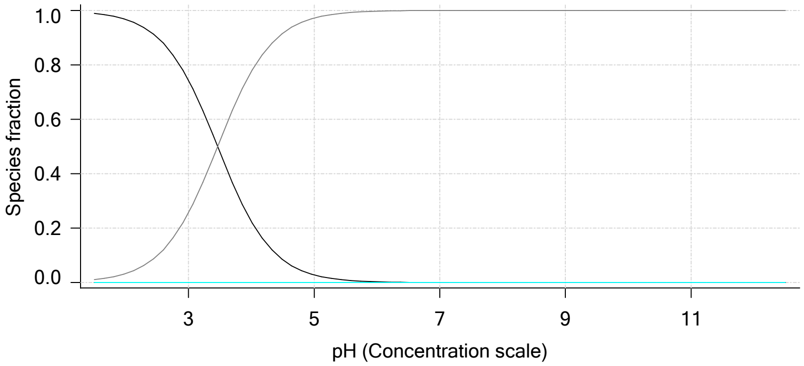

| | | | |
|---|---|---|---|
| Sample name: | **M18_octanol** | Experiment start time: | **3/9/2018 11:02:44 PM** |
| Assay name: | **pH-metric high logP** | Analyst: | **Pion** |
| Assay ID: | **18C-09016** | Instrument ID: | **T312060** |
| Filename: | **C:\Sirius_T3\Mehtap\20180309_exp31_logP_T3-2\18C-09016_M18_octanol_pH-metric high logP.t3r** | | |

#### Sample graphs (continued)

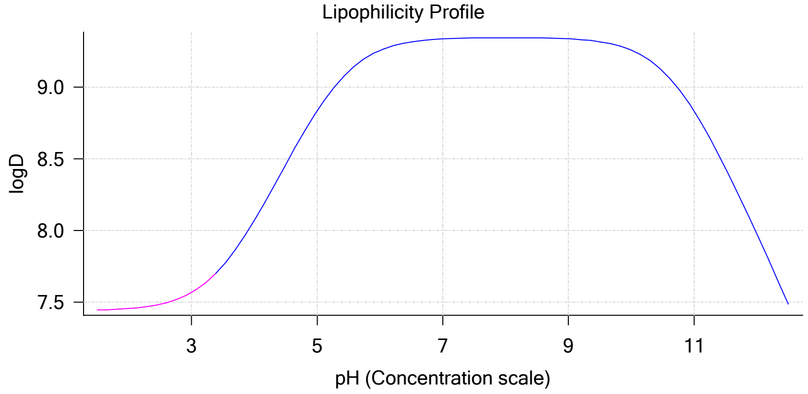


#### Sample logD and percent species

| pH | M18_octanol logD | M18_octanol M18_octanolH2 | M18_octanol M18_octanolH | M18_octanol M18_octanol | M18_octanol M18_octanolH2* | M18_octanol M18_octanolH* | M18_octanol M18_octanol* | Comment |
|---|---|---|---|---|---|---|---|---|
| 1.000 | 7.44 | 0.00 % | 0.00 % | 0.00 % | 99.66 % | 0.34 % | 0.00 % | |
| 1.200 | 7.44 | 0.00 % | 0.00 % | 0.00 % | 99.46 % | 0.54 % | 0.00 % | Stomach pH |
| 2.000 | 7.45 | 0.00 % | 0.00 % | 0.00 % | 96.66 % | 3.34 % | 0.00 % | |
| 3.000 | 7.57 | 0.00 % | 0.00 % | 0.00 % | 74.32 % | 25.68 % | 0.00 % | |
| 4.000 | 8.07 | 0.00 % | 0.00 % | 0.00 % | 22.45 % | 77.55 % | 0.00 % | |
| 5.000 | 8.84 | 0.00 % | 0.00 % | 0.00 % | 2.81 % | 97.19 % | 0.00 % | |
| 6.000 | 9.26 | 0.00 % | 0.00 % | 0.00 % | 0.29 % | 99.71 % | 0.00 % | |
| 6.500 | 9.32 | 0.00 % | 0.00 % | 0.00 % | 0.09 % | 99.91 % | 0.00 % | |
| 7.000 | 9.34 | 0.00 % | 0.00 % | 0.00 % | 0.03 % | 99.97 % | 0.00 % | |
| 7.400 | 9.34 | 0.00 % | 0.00 % | 0.00 % | 0.01 % | 99.99 % | 0.00 % | Blood pH |
| 8.000 | 9.35 | 0.00 % | 0.00 % | 0.00 % | 0.00 % | 100.00 % | 0.00 % | |
| 9.000 | 9.34 | 0.00 % | 0.00 % | 0.00 % | 0.00 % | 100.00 % | 0.00 % | |
| 10.000 | 9.26 | 0.00 % | 0.00 % | 0.00 % | 0.00 % | 100.00 % | 0.00 % | |
| 11.000 | 8.84 | 0.00 % | 0.00 % | 0.00 % | 0.00 % | 100.00 % | 0.00 % | |
| 12.000 | 7.98 | 0.00 % | 0.00 % | 0.00 % | 0.00 % | 100.00 % | 0.00 % | |

#### Carbonate and acidity

| | |
|---|---|
| Carbonate | 0.088 mM |
| Acidity error | -0.104 mM |

#### Other graphs

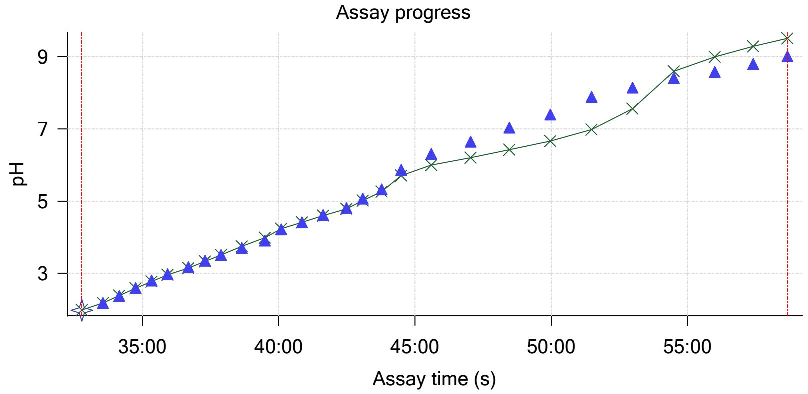


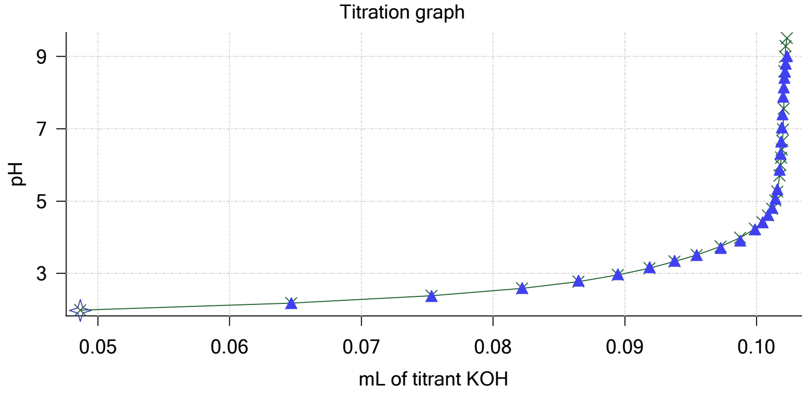

Sample name: **M18_octanol**
Assay name: **pH-metric high logP**
Assay ID: **18C-09016**
Filename: **C:\Sirius_T3\Mehtap\20180309_exp31_logP_T3-2\18C-09016_M18_octanol_pH-metric high logP.t3r**

Experiment start time: **3/9/2018 11:02:44 PM**
Analyst: **Pion**
Instrument ID: **T312060**

#### Other graphs (continued)

Charge balance residuals

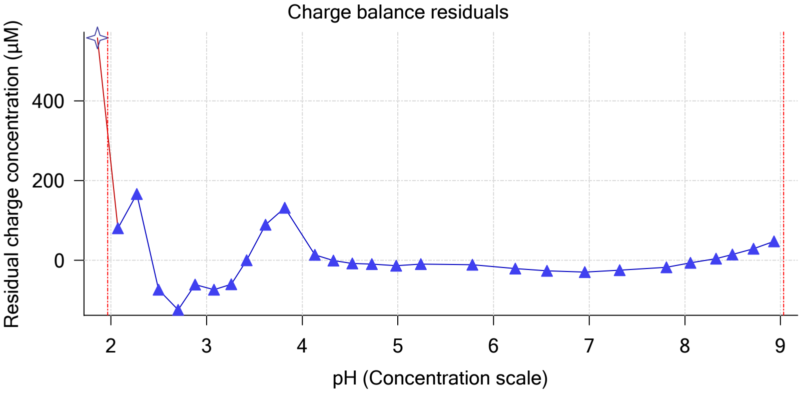


Buffer index profile

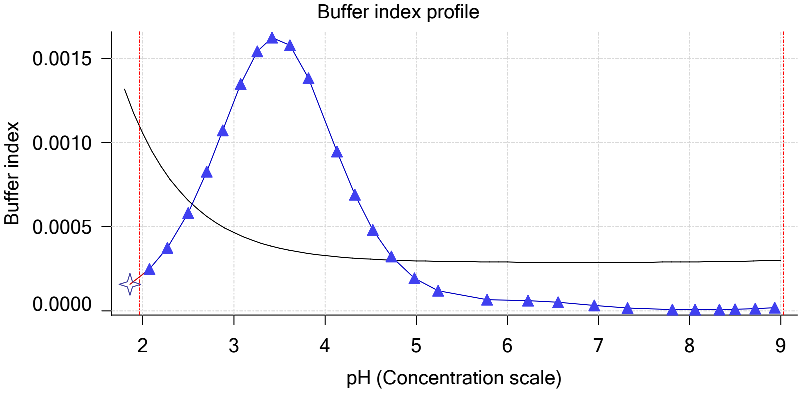


Ionic strength

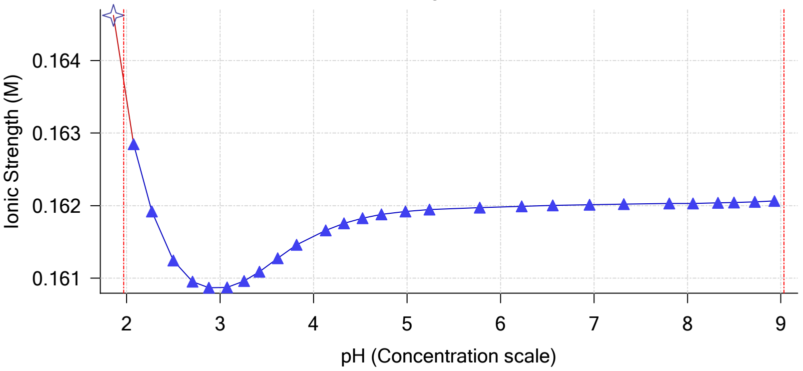


Log R Profile

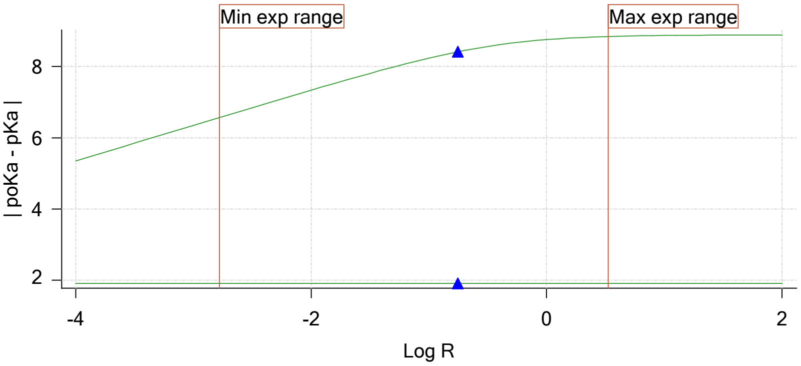


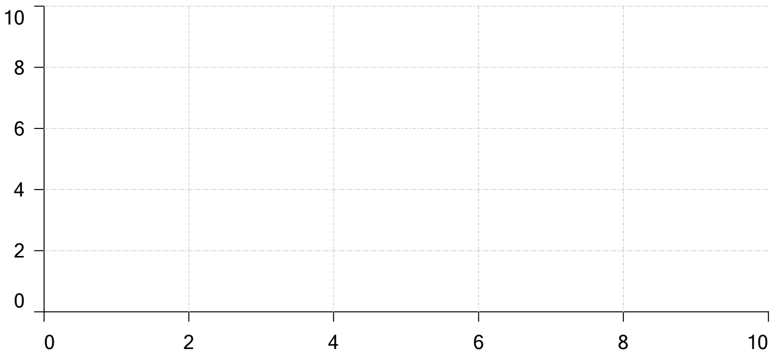

Ionisation of sample Carbonate

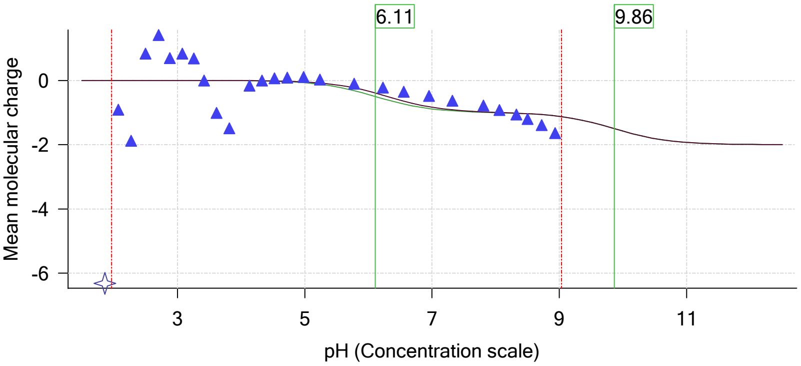

| | | | |
|---|---|---|---|
| Sample name: | **M18_octanol** | Experiment start time: | **3/9/2018 11:02:44 PM** |
| Assay name: | **pH-metric high logP** | Analyst: | **Pion** |
| Assay ID: | **18C-09016** | Instrument ID: | **T312060** |
| Filename: | **C:\Sirius_T3\Mehtap\20180309_exp31_logP_T3-2\18C-09016_M18_octanol_pH-metric high logP.t3r** | | |

# pH-metric high logP   Titration 3 of 3   18C-09016 Points 57 to 86

#### Overall results

| | |
|---|---|
| RMSD | 0.447 |
| Average ionic strength | 0.168 M |
| Average temperature | 25.0°C |
| Partition ratio | 0.4400 : 1 |
| Analyte concentration range | 1742.6 µM to 1782.5 µM |
| Total points considered | 28 of 30 |

#### Warnings and errors

| | |
|---|---|
| Errors | None |
| Warnings | One or more logP values out of range |

#### Four-Plus parameters

| | | | |
|---|---|---|---|
| Alpha | 0.102 | 3/9/2018 11:02:43 PM | C:\Sirius_T3\HCl18C09.t3r |
| S | 0.9967 | 3/9/2018 11:02:43 PM | C:\Sirius_T3\HCl18C09.t3r |
| jH | 1.2 | 3/9/2018 11:02:43 PM | C:\Sirius_T3\HCl18C09.t3r |
| jOH | 0.0 | 3/9/2018 11:02:43 PM | C:\Sirius_T3\HCl18C09.t3r |

#### Titrants

| | | | |
|---|---|---|---|
| 0.50 M HCl | 0.999843 | 3/9/2018 11:02:44 PM | C:\Sirius_T3\HCl18C09.t3r |
| 0.50 M KOH | 0.999845 | 3/9/2018 11:02:44 PM | C:\Sirius_T3\KOH18B27.t3r |

#### Sample

| | |
|---|---|
| M18_octanol concentration factor | 1.063 |
| Base pKa 1 | 5.37 |
| Acid pKa 2 | 10.65 |
| logP (XH2 +) | -3.10 |
| logP (neutral XH) | 2.50 |
| logP (X -) | 0.46 |

#### Sample graphs

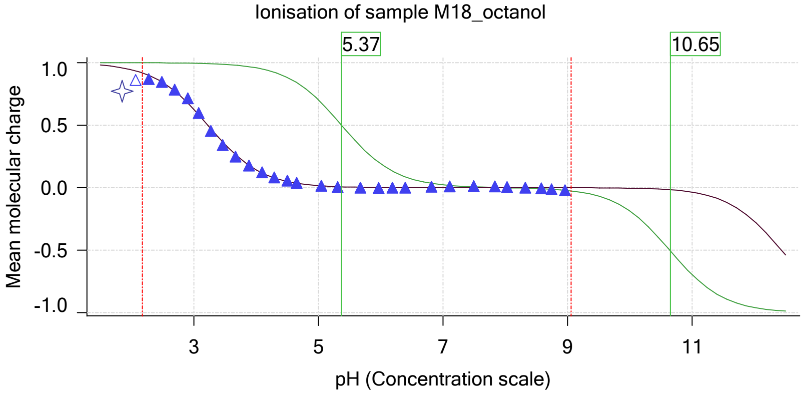


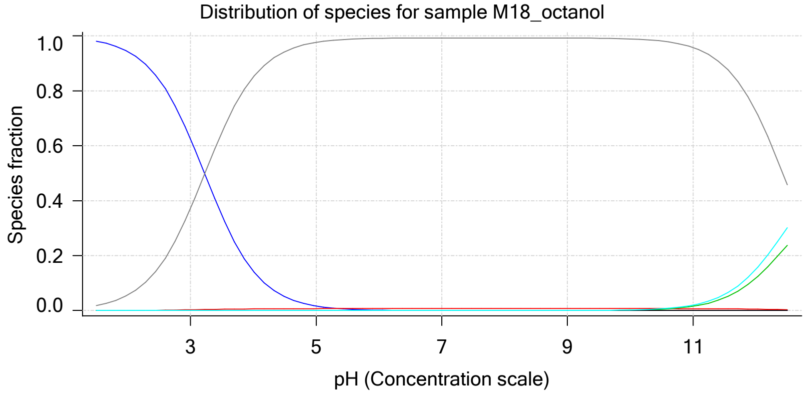

| | | | |
|---|---|---|---|
| Sample name: | **M18_octanol** | Experiment start time: | **3/9/2018 11:02:44 PM** |
| Assay name: | **pH-metric high logP** | Analyst: | **Pion** |
| Assay ID: | **18C-09016** | Instrument ID: | **T312060** |
| Filename: | **C:\Sirius_T3\Mehtap\20180309_exp31_logP_T3-2\18C-09016_M18_octanol_pH-metric high logP.t3r** | | |

#### Sample graphs (continued)

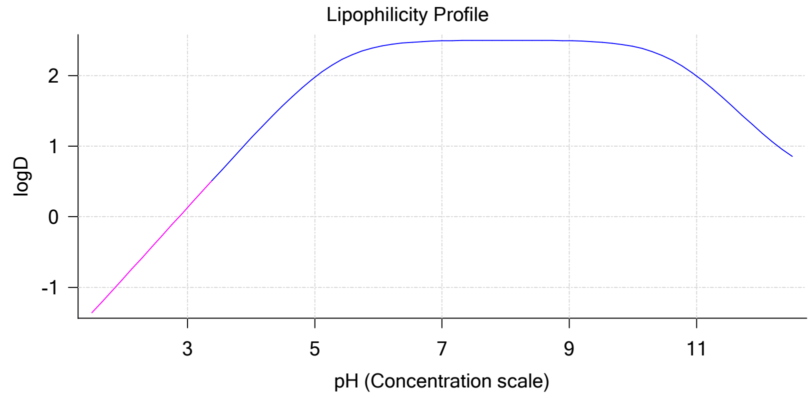


#### Sample logD and percent species

| pH | M18_octanol logD | M18_octanol M18_octanolH2 | M18_octanol M18_octanolH | M18_octanol M18_octanol | M18_octanol M18_octanolH2* | M18_octanol M18_octanolH* | M18_octanol M18_octanol* | Comment |
|---|---|---|---|---|---|---|---|---|
| 1.000 | -1.84 | 99.37 % | 0.00 % | 0.00 % | 0.04 % | 0.59 % | 0.00 % | |
| 1.200 | -1.65 | 99.02 % | 0.01 % | 0.00 % | 0.03 % | 0.94 % | 0.00 % | Stomach pH |
| 2.000 | -0.87 | 94.30 % | 0.04 % | 0.00 % | 0.03 % | 5.63 % | 0.00 % | |
| 3.000 | 0.13 | 62.45 % | 0.27 % | 0.00 % | 0.02 % | 37.26 % | 0.00 % | |
| 4.000 | 1.11 | 14.26 % | 0.61 % | 0.00 % | 0.01 % | 85.12 % | 0.00 % | |
| 5.000 | 1.98 | 1.64 % | 0.70 % | 0.00 % | 0.00 % | 97.66 % | 0.00 % | |
| 6.000 | 2.41 | 0.17 % | 0.71 % | 0.00 % | 0.00 % | 99.13 % | 0.00 % | |
| 6.500 | 2.47 | 0.05 % | 0.71 % | 0.00 % | 0.00 % | 99.24 % | 0.00 % | |
| 7.000 | 2.49 | 0.02 % | 0.71 % | 0.00 % | 0.00 % | 99.27 % | 0.00 % | |
| 7.400 | 2.50 | 0.01 % | 0.71 % | 0.00 % | 0.00 % | 99.28 % | 0.00 % | Blood pH |
| 8.000 | 2.50 | 0.00 % | 0.71 % | 0.00 % | 0.00 % | 99.28 % | 0.00 % | |
| 9.000 | 2.49 | 0.00 % | 0.71 % | 0.02 % | 0.00 % | 99.25 % | 0.02 % | |
| 10.000 | 2.42 | 0.00 % | 0.71 % | 0.16 % | 0.00 % | 98.93 % | 0.20 % | |
| 11.000 | 2.00 | 0.00 % | 0.69 % | 1.53 % | 0.00 % | 95.84 % | 1.95 % | |
| 12.000 | 1.21 | 0.00 % | 0.52 % | 11.68 % | 0.00 % | 72.98 % | 14.82 % | |

#### Carbonate and acidity

| | |
|---|---|
| Carbonate | 0.145 mM |
| Acidity error | 0.002 mM |

#### Other graphs

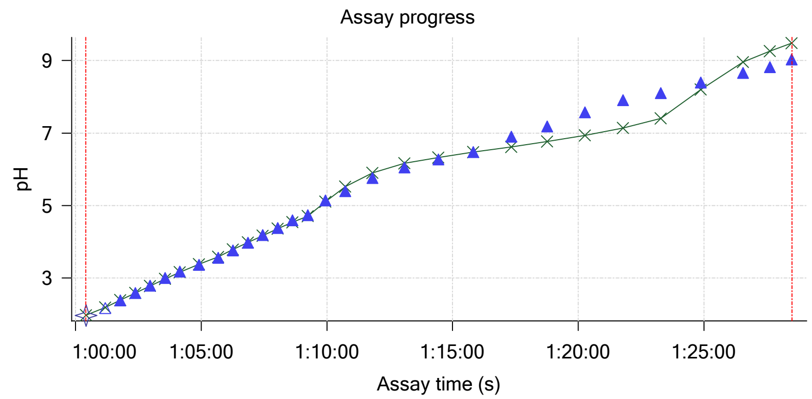


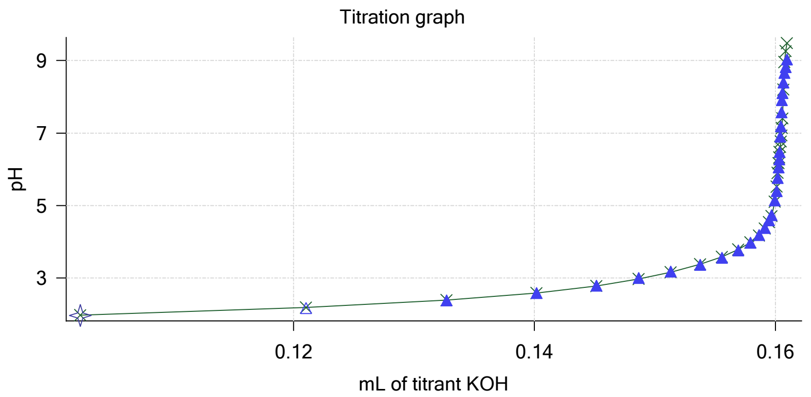

| | | | |
|---|---|---|---|
| Sample name: | **M18_octanol** | Experiment start time: | **3/9/2018 11:02:44 PM** |
| Assay name: | **pH-metric high logP** | Analyst: | **Pion** |
| Assay ID: | **18C-09016** | Instrument ID: | **T312060** |
| Filename: | **C:\Sirius_T3\Mehtap\20180309_exp31_logP_T3-2\18C-09016_M18_octanol_pH-metric high logP.t3r** | | |

#### Other graphs (continued)

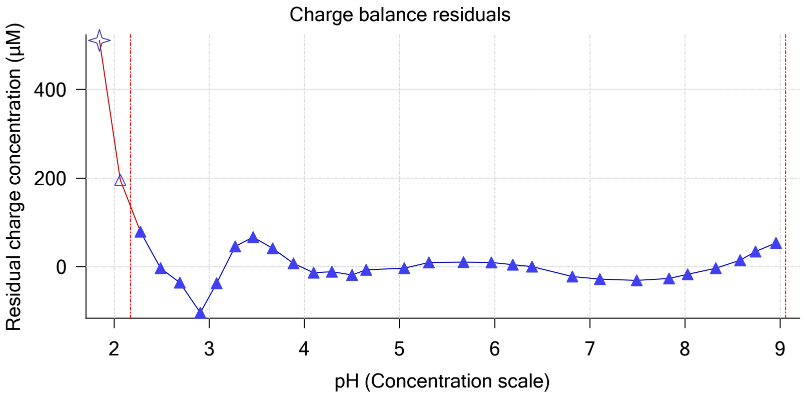


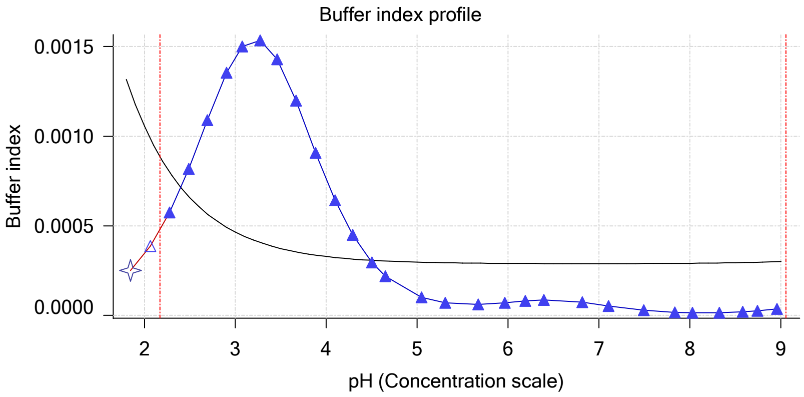


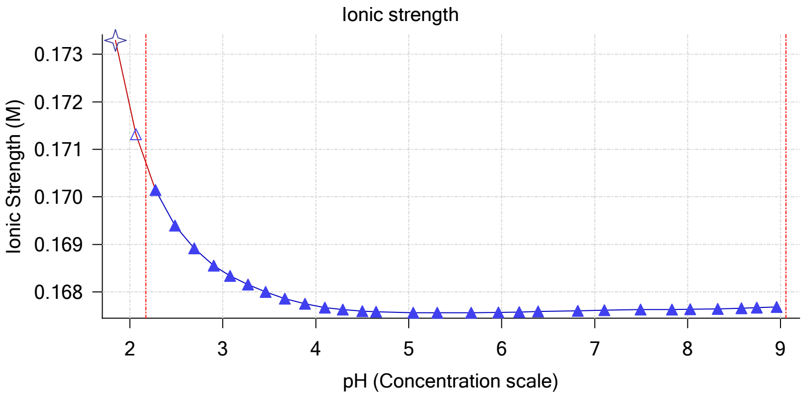


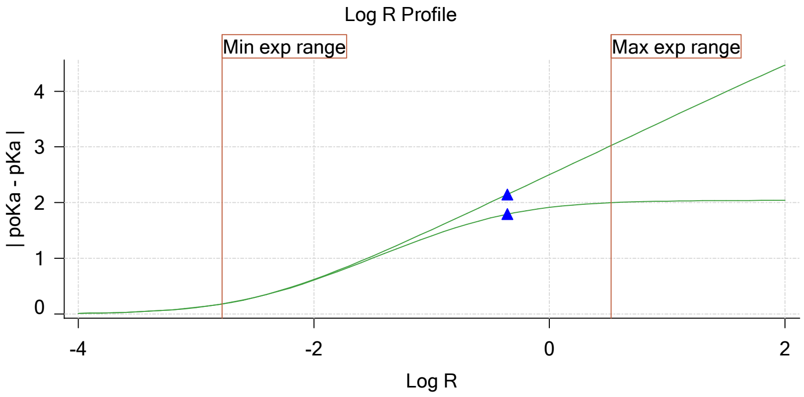


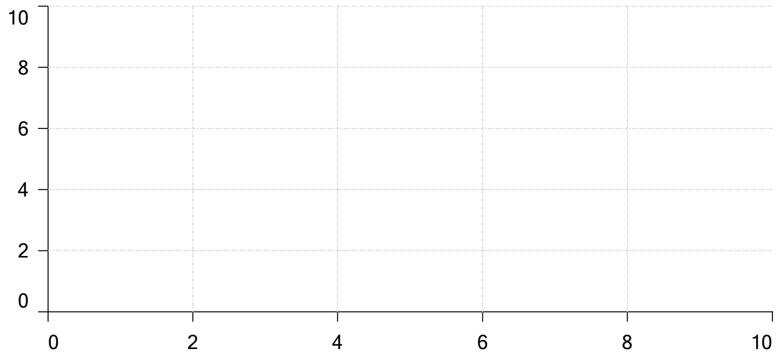

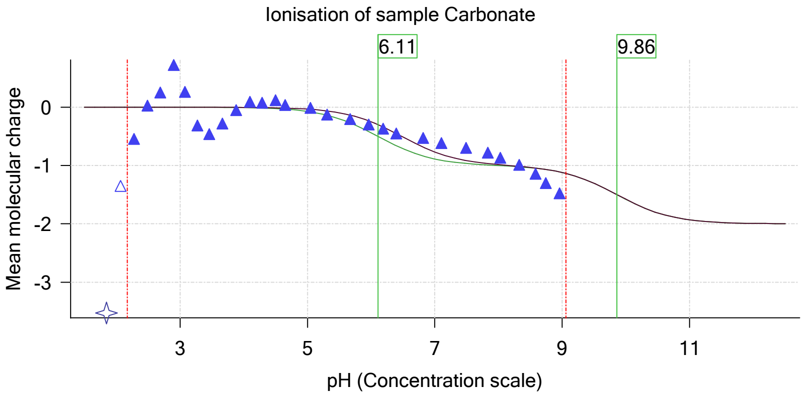

| | | | |
|---|---|---|---|
| Sample name: | **M18_octanol** | Experiment start time: | **3/9/2018 11:02:44 PM** |
| Assay name: | **pH-metric high logP** | Analyst: | **Pion** |
| Assay ID: | **18C-09016** | Instrument ID: | **T312060** |
| Filename: | **C:\Sirius_T3\Mehtap\20180309_exp31_logP_T3-2\18C-09016_M18_octanol_pH-metric high logP.t3r** | | |

#### Assay Model

| Settings | Value | Date/Time changed | Imported from |
|---|---|---|---|
| Sample name | M18_octanol | 2/27/2018 7:08:39 PM | User entered value |
| Sample by | Weight | | Default value |
| Sample weight | 0.001220 g | 3/9/2018 2:22:52 PM | User entered value |
| Formula weight | 267.11 g/mol | 2/27/2018 7:08:39 PM | User entered value |
| Solubility | Unknown | | Default value |
| Molecular weight | 267.11 | 2/27/2018 7:08:39 PM | User entered value |
| Individual pKa ionic environments | No | | Default value |
| Number of pKas | 2 | 2/27/2018 7:08:39 PM | User entered value |
| Sample is a | Ampholyte | 2/27/2018 7:08:39 PM | User entered value |
| pKa 1 | 5.37 | 2/27/2018 7:08:39 PM | User entered value |
| Type | Base | 2/27/2018 7:08:39 PM | User entered value |
| pKa 2 | 10.65 | 2/27/2018 7:08:39 PM | User entered value |
| Type | Acid | 2/27/2018 7:08:39 PM | User entered value |
| logp (XH2 +) | -0.35 | 2/28/2018 3:20:28 PM | User entered value |
| logP (neutral XH) | 2.57 | 3/2/2018 4:34:50 PM | User entered value |
| logP (X -) | 0.46 | 2/27/2018 7:09:34 PM | User entered value |

#### Events

| Time | Event | Water | Acid | Base | Octanol | pH | dpH/dt | pH R-squared | pH SD | dpH/d time |
|---|---|---|---|---|---|---|---|---|---|---|
| 4:58.7 | Manual volume addition | | | | 0.10000 mL | | | | | |
| 4:59.8 | Initial pH = 7.99 | | | | | | | | | |
| 7:59.3 | Data point 2 | 1.50000 mL | 0.04812 mL | 0.00477 mL | 0.10000 mL | 2.062 | -0.00681 | 0.27502 | 0.00064 | 10.0 s |
| 8:45.5 | Data point 3 | 1.50000 mL | 0.04812 mL | 0.01698 mL | 0.10000 mL | 2.268 | 0.00821 | 0.44883 | 0.00061 | 10.0 s |
| 9:21.1 | Data point 4 | 1.50000 mL | 0.04812 mL | 0.02495 mL | 0.10000 mL | 2.471 | -0.00977 | 0.70424 | 0.00058 | 10.0 s |
| 9:56.6 | Data point 5 | 1.50000 mL | 0.04812 mL | 0.03001 mL | 0.10000 mL | 2.662 | -0.00069 | 0.07464 | 0.00012 | 10.0 s |
| 10:32.1 | Data point 6 | 1.50000 mL | 0.04812 mL | 0.03335 mL | 0.10000 mL | 2.850 | -0.00288 | 0.33859 | 0.00024 | 10.0 s |
| 11:07.6 | Data point 7 | 1.50000 mL | 0.04812 mL | 0.03565 mL | 0.10000 mL | 3.085 | -0.01383 | 0.83922 | 0.00074 | 10.0 s |
| 11:43.3 | Data point 8 | 1.50000 mL | 0.04812 mL | 0.03725 mL | 0.10000 mL | 3.283 | -0.00418 | 0.74326 | 0.00024 | 10.0 s |
| 12:18.8 | Data point 9 | 1.50000 mL | 0.04812 mL | 0.03859 mL | 0.10000 mL | 3.456 | -0.01450 | 0.85812 | 0.00077 | 10.0 s |
| 13:09.7 | Data point 10 | 1.50000 mL | 0.04812 mL | 0.04017 mL | 0.10000 mL | 3.653 | -0.01030 | 0.40559 | 0.00080 | 10.0 s |
| 14:00.8 | Data point 11 | 1.50000 mL | 0.04812 mL | 0.04156 mL | 0.10000 mL | 3.848 | -0.00945 | 0.70466 | 0.00056 | 10.5 s |
| 14:36.7 | Data point 12 | 1.50000 mL | 0.04812 mL | 0.04278 mL | 0.10000 mL | 4.049 | -0.00976 | 0.86426 | 0.00052 | 10.0 s |
| 15:12.1 | Data point 13 | 1.50000 mL | 0.04812 mL | 0.04396 mL | 0.10000 mL | 4.256 | -0.00996 | 0.79638 | 0.00055 | 10.0 s |
| 15:47.4 | Data point 14 | 1.50000 mL | 0.04812 mL | 0.04499 mL | 0.10000 mL | 4.460 | -0.01429 | 0.84980 | 0.00077 | 10.0 s |
| 16:22.8 | Data point 15 | 1.50000 mL | 0.04812 mL | 0.04584 mL | 0.10000 mL | 4.667 | -0.01676 | 0.72668 | 0.00097 | 10.0 s |
| 16:58.2 | Data point 16 | 1.50000 mL | 0.04812 mL | 0.04647 mL | 0.10000 mL | 4.852 | -0.01624 | 0.82288 | 0.00088 | 10.5 s |
| 17:34.1 | Data point 17 | 1.50000 mL | 0.04812 mL | 0.04694 mL | 0.10000 mL | 4.977 | -0.06194 | 0.99731 | 0.00306 | Timed out at |

| | | | |
|---|---|---|---|
| Sample name: | **M18_octanol** | Experiment start time: | **3/9/2018 11:02:44 PM** |
| Assay name: | **pH-metric high logP** | Analyst: | **Pion** |
| Assay ID: | **18C-09016** | Instrument ID: | **T312060** |
| Filename: | **C:\Sirius_T3\Mehtap\20180309_exp31_logP_T3-2\18C-09016_M18_octanol_pH-metric high logP.t3r** | | |

#### Events (continued)

| Time | Event | Water | Acid | Base | Octanol | pH | dpH/dt | pH R-squared | pH SD | dpH/dt time |
|---|---|---|---|---|---|---|---|---|---|---|
| 19:04.6 | Data point 18 | 1.50000 mL | 0.04812 mL | 0.04744 mL | 0.10000 mL | 5.138 | -0.02805 | 0.99638 | 0.00139 | Timed out at 59.5 s |
| 20:40.3 | Data point 19 | 1.50000 mL | 0.04812 mL | 0.04774 mL | 0.10000 mL | 5.264 | -0.01845 | 0.96248 | 0.00093 | 29.0 s |
| 21:34.7 | Data point 20 | 1.50000 mL | 0.04812 mL | 0.04798 mL | 0.10000 mL | 5.641 | -0.01992 | 0.97402 | 0.00100 | 21.0 s |
| 22:26.3 | Data point 21 | 1.50000 mL | 0.04812 mL | 0.04807 mL | 0.10000 mL | 5.855 | -0.01912 | 0.93098 | 0.00098 | 26.5 s |
| 23:18.2 | Data point 22 | 1.50000 mL | 0.04812 mL | 0.04812 mL | 0.10000 mL | 6.027 | -0.01851 | 0.97469 | 0.00093 | 27.5 s |
| 24:11.1 | Data point 23 | 1.50000 mL | 0.04812 mL | 0.04817 mL | 0.10000 mL | 6.363 | -0.01958 | 0.96477 | 0.00099 | 57.5 s |
| 25:39.1 | Data point 24 | 1.50000 mL | 0.04812 mL | 0.04831 mL | 0.10000 mL | 8.024 | -0.07733 | 0.99662 | 0.00383 | Timed out at 59.5 s |
| 27:14.7 | Data point 25 | 1.50000 mL | 0.04812 mL | 0.04842 mL | 0.10000 mL | 8.560 | -0.02632 | 0.98996 | 0.00131 | Timed out at 59.5 s |
| 28:45.1 | Data point 26 | 1.50000 mL | 0.04812 mL | 0.04849 mL | 0.10000 mL | 8.774 | -0.02007 | 0.98634 | 0.00100 | 48.0 s |
| 30:08.9 | Data point 27 | 1.50000 mL | 0.04812 mL | 0.04857 mL | 0.10000 mL | 8.957 | -0.01923 | 0.96603 | 0.00097 | 36.0 s |
| 31:15.5 | Data point 28 | 1.50000 mL | 0.04812 mL | 0.04866 mL | 0.10000 mL | 9.132 | -0.01929 | 0.95230 | 0.00098 | 29.0 s |
| 32:47.3 | Data point 29 | 1.50000 mL | 0.10212 mL | 0.04866 mL | 0.30000 mL | 1.972 | 0.00050 | 0.00545 | 0.00033 | 10.5 s |
| 33:34.0 | Data point 30 | 1.50000 mL | 0.10212 mL | 0.06468 mL | 0.30000 mL | 2.179 | -0.01398 | 0.77392 | 0.00078 | 10.0 s |
| 34:09.7 | Data point 31 | 1.50000 mL | 0.10212 mL | 0.07533 mL | 0.30000 mL | 2.373 | 0.00146 | 0.41139 | 0.00011 | 10.0 s |
| 34:45.3 | Data point 32 | 1.50000 mL | 0.10212 mL | 0.08220 mL | 0.30000 mL | 2.598 | -0.00003 | 0.00017 | 0.00012 | 10.0 s |
| 35:20.8 | Data point 33 | 1.50000 mL | 0.10212 mL | 0.08648 mL | 0.30000 mL | 2.799 | -0.00882 | 0.36125 | 0.00072 | 10.0 s |
| 35:56.2 | Data point 34 | 1.50000 mL | 0.10212 mL | 0.08946 mL | 0.30000 mL | 2.973 | -0.00669 | 0.54661 | 0.00045 | 10.0 s |
| 36:42.0 | Data point 35 | 1.50000 mL | 0.10212 mL | 0.09186 mL | 0.30000 mL | 3.170 | -0.01175 | 0.75450 | 0.00067 | 10.5 s |
| 37:18.0 | Data point 36 | 1.50000 mL | 0.10212 mL | 0.09377 mL | 0.30000 mL | 3.350 | -0.00456 | 0.45623 | 0.00033 | 10.0 s |
| 37:53.4 | Data point 37 | 1.50000 mL | 0.10212 mL | 0.09544 mL | 0.30000 mL | 3.512 | -0.00080 | 0.01535 | 0.00032 | 10.0 s |
| 38:39.1 | Data point 38 | 1.50000 mL | 0.10212 mL | 0.09727 mL | 0.30000 mL | 3.706 | -0.00719 | 0.81084 | 0.00039 | 10.0 s |
| 39:30.1 | Data point 39 | 1.50000 mL | 0.10212 mL | 0.09875 mL | 0.30000 mL | 3.908 | -0.01381 | 0.88552 | 0.00072 | 10.5 s |
| 40:06.1 | Data point 40 | 1.50000 mL | 0.10212 mL | 0.09986 mL | 0.30000 mL | 4.220 | -0.00595 | 0.67659 | 0.00036 | 10.0 s |
| 40:51.7 | Data point 41 | 1.50000 mL | 0.10212 mL | 0.10045 mL | 0.30000 mL | 4.415 | -0.00682 | 0.73657 | 0.00039 | 10.5 s |
| 41:38.0 | Data point 42 | 1.50000 mL | 0.10212 mL | 0.10087 mL | 0.30000 mL | 4.612 | -0.01104 | 0.72091 | 0.00064 | 10.5 s |
| 42:29.5 | Data point 43 | 1.50000 mL | 0.10212 mL | 0.10118 mL | 0.30000 mL | 4.814 | -0.01440 | 0.75675 | 0.00082 | 10.5 s |
| 43:05.4 | Data point 44 | 1.50000 mL | 0.10212 mL | 0.10141 mL | 0.30000 mL | 5.068 | -0.01600 | 0.81524 | 0.00088 | 11.5 s |
| 43:47.3 | Data point 45 | 1.50000 mL | 0.10212 mL | 0.10158 mL | 0.30000 mL | 5.325 | -0.01213 | 0.64755 | 0.00074 | 12.0 s |
| 44:30.0 | Data point 46 | 1.50000 mL | 0.10212 mL | 0.10174 mL | 0.30000 mL | 5.861 | -0.01983 | 0.98013 | 0.00099 | 35.5 s |
| 45:36.0 | Data point 47 | 1.50000 mL | 0.10212 mL | 0.10181 mL | 0.30000 mL | 6.309 | -0.01961 | 0.97472 | 0.00098 | 56.0 s |
| 47:02.6 | Data point 48 | 1.50000 mL | 0.10212 mL | 0.10186 mL | 0.30000 mL | 6.639 | -0.03066 | 0.99363 | 0.00152 | Timed out at 59.5 s |
| 48:28.0 | Data point 49 | 1.50000 mL | 0.10212 mL | 0.10191 mL | 0.30000 mL | 7.034 | -0.05072 | 0.99593 | 0.00251 | Timed out at 59.5 s |
| 49:58.5 | Data point 50 | 1.50000 mL | 0.10212 mL | 0.10195 mL | 0.30000 mL | 7.398 | -0.06220 | 0.99680 | 0.00308 | Timed out at 59.5 s |
| 51:29.0 | Data point 51 | 1.50000 mL | 0.10212 mL | 0.10200 mL | 0.30000 mL | 7.885 | -0.06202 | 0.99692 | 0.00307 | Timed out at 59.5 s |
| 52:59.4 | Data point 52 | 1.50000 mL | 0.10212 mL | 0.10205 mL | 0.30000 mL | 8.135 | -0.04192 | 0.98906 | 0.00208 | Timed out at 59.5 s |
| 54:30.0 | Data point 53 | 1.50000 mL | 0.10212 mL | 0.10209 mL | 0.30000 mL | 8.402 | -0.02237 | 0.99033 | 0.00111 | Timed out at 59.5 s |
| 56:00.4 | Data point 54 | 1.50000 mL | 0.10212 mL | 0.10214 mL | 0.30000 mL | 8.573 | -0.01945 | 0.95229 | 0.00098 | 53.5 s |
| 57:24.5 | Data point 55 | 1.50000 mL | 0.10212 mL | 0.10221 mL | 0.30000 mL | 8.793 | -0.01809 | 0.98417 | 0.00090 | 34.5 s |
| 58:39.9 | Data point 56 | 1.50000 mL | 0.10212 mL | 0.10230 mL | 0.30000 mL | 9.005 | -0.01939 | 0.93292 | 0.00099 | 19.5 s |
| 1:00:24.9 | Data point 57 | 1.50000 mL | 0.16011 mL | 0.10230 mL | 0.80000 mL | 1.959 | -0.01114 | 0.87304 | 0.00059 | 10.0 s |
| 1:01:11.2 | Data point 58 | 1.50000 mL | 0.16011 mL | 0.12105 mL | 0.80000 mL | 2.170 | 0.00081 | 0.04813 | 0.00018 | 10.0 s |
| 1:01:46.9 | Data point 59 | 1.50000 mL | 0.16011 mL | 0.13271 mL | 0.80000 mL | 2.378 | 0.00136 | 0.09971 | 0.00021 | 10.0 s |
| 1:02:22.5 | Data point 60 | 1.50000 mL | 0.16011 mL | 0.14017 mL | 0.80000 mL | 2.585 | 0.01209 | 0.68746 | 0.00072 | 10.0 s |
| 1:02:58.1 | Data point 61 | 1.50000 mL | 0.16011 mL | 0.14513 mL | 0.80000 mL | 2.787 | 0.00716 | 0.56892 | 0.00047 | 10.0 s |
| 1:03:33.6 | Data point 62 | 1.50000 mL | 0.16011 mL | 0.14866 mL | 0.80000 mL | 2.998 | -0.00487 | 0.59781 | 0.00031 | 10.0 s |
| 1:04:09.0 | Data point 63 | 1.50000 mL | 0.16011 mL | 0.15129 mL | 0.80000 mL | 3.171 | 0.00223 | 0.08135 | 0.00039 | 10.0 s |
| 1:04:54.7 | Data point 64 | 1.50000 mL | 0.16011 mL | 0.15376 mL | 0.80000 mL | 3.364 | -0.00945 | 0.67088 | 0.00057 | 10.0 s |
| 1:05:40.5 | Data point 65 | 1.50000 mL | 0.16011 mL | 0.15555 mL | 0.80000 mL | 3.553 | -0.00773 | 0.66839 | 0.00047 | 10.0 s |

| | | | |
|---|---|---|---|
| Sample name: | **M18_octanol** | Experiment start time: | **3/9/2018 11:02:44 PM** |
| Assay name: | **pH-metric high logP** | Analyst: | **Pion** |
| Assay ID: | **18C-09016** | Instrument ID: | **T312060** |
| Filename: | **C:\Sirius_T3\Mehtap\20180309_exp31_logP_T3-2\18C-09016_M18_octanol_pH-metric high logP.t3r** | | |

#### Events (continued)

| Time | Event | Water | Acid | Base | Octanol | pH | dpH/dt | pH R-squared | pH SD | dpH/dt time |
|---|---|---|---|---|---|---|---|---|---|---|
| 1:06:16.0 | Data point 66 | 1.50000 mL | 0.16011 mL | 0.15691 mL | 0.80000 mL | 3.758 | -0.00478 | 0.09544 | 0.00076 | 10.0 s |
| 1:06:51.4 | Data point 67 | 1.50000 mL | 0.16011 mL | 0.15793 mL | 0.80000 mL | 3.973 | 0.00088 | 0.00414 | 0.00068 | 10.0 s |
| 1:07:26.8 | Data point 68 | 1.50000 mL | 0.16011 mL | 0.15863 mL | 0.80000 mL | 4.186 | -0.01639 | 0.67438 | 0.00099 | 10.0 s |
| 1:08:02.3 | Data point 69 | 1.50000 mL | 0.16011 mL | 0.15912 mL | 0.80000 mL | 4.378 | -0.01336 | 0.45806 | 0.00098 | 10.0 s |
| 1:08:37.7 | Data point 70 | 1.50000 mL | 0.16011 mL | 0.15945 mL | 0.80000 mL | 4.589 | -0.00985 | 0.60220 | 0.00063 | 11.0 s |
| 1:09:14.1 | Data point 71 | 1.50000 mL | 0.16011 mL | 0.15967 mL | 0.80000 mL | 4.736 | -0.00475 | 0.10626 | 0.00072 | 11.5 s |
| 1:09:56.1 | Data point 72 | 1.50000 mL | 0.16011 mL | 0.15995 mL | 0.80000 mL | 5.134 | -0.01189 | 0.37022 | 0.00096 | 12.0 s |
| 1:10:43.8 | Data point 73 | 1.50000 mL | 0.16011 mL | 0.16009 mL | 0.80000 mL | 5.393 | -0.01905 | 0.93142 | 0.00098 | 34.5 s |
| 1:11:48.9 | Data point 74 | 1.50000 mL | 0.16011 mL | 0.16018 mL | 0.80000 mL | 5.755 | -0.01516 | 0.77806 | 0.00085 | 45.5 s |
| 1:13:05.0 | Data point 75 | 1.50000 mL | 0.16011 mL | 0.16025 mL | 0.80000 mL | 6.047 | -0.01855 | 0.94284 | 0.00094 | 50.5 s |
| 1:14:26.2 | Data point 76 | 1.50000 mL | 0.16011 mL | 0.16030 mL | 0.80000 mL | 6.272 | -0.01969 | 0.96730 | 0.00099 | 58.0 s |
| 1:15:49.6 | Data point 77 | 1.50000 mL | 0.16011 mL | 0.16035 mL | 0.80000 mL | 6.474 | -0.02760 | 0.99228 | 0.00137 | Timed out at 59.5 s |
| 1:17:20.1 | Data point 78 | 1.50000 mL | 0.16011 mL | 0.16040 mL | 0.80000 mL | 6.897 | -0.06395 | 0.99250 | 0.00317 | Timed out at 59.5 s |
| 1:18:45.5 | Data point 79 | 1.50000 mL | 0.16011 mL | 0.16044 mL | 0.80000 mL | 7.185 | -0.06453 | 0.99687 | 0.00319 | Timed out at 59.5 s |
| 1:20:16.1 | Data point 80 | 1.50000 mL | 0.16011 mL | 0.16049 mL | 0.80000 mL | 7.570 | -0.08020 | 0.98798 | 0.00398 | Timed out at 59.5 s |
| 1:21:46.6 | Data point 81 | 1.50000 mL | 0.16011 mL | 0.16054 mL | 0.80000 mL | 7.910 | -0.05115 | 0.99413 | 0.00253 | Timed out at 59.5 s |
| 1:23:17.1 | Data point 82 | 1.50000 mL | 0.16011 mL | 0.16058 mL | 0.80000 mL | 8.105 | -0.04177 | 0.93249 | 0.00214 | Timed out at 59.5 s |
| 1:24:52.7 | Data point 83 | 1.50000 mL | 0.16011 mL | 0.16065 mL | 0.80000 mL | 8.398 | -0.02637 | 0.89962 | 0.00137 | Timed out at 59.5 s |
| 1:26:33.5 | Data point 84 | 1.50000 mL | 0.16011 mL | 0.16075 mL | 0.80000 mL | 8.655 | -0.01834 | 0.88375 | 0.00096 | 28.0 s |
| 1:27:37.2 | Data point 85 | 1.50000 mL | 0.16011 mL | 0.16084 mL | 0.80000 mL | 8.817 | -0.01433 | 0.54002 | 0.00096 | 16.5 s |
| 1:28:29.4 | Data point 86 | 1.50000 mL | 0.16011 mL | 0.16096 mL | 0.80000 mL | 9.030 | -0.01224 | 0.37247 | 0.00099 | 14.0 s |
| 1:28:52.4 | Assay volumes | 1.50000 mL | 0.16011 mL | 0.16096 mL | 0.80000 mL | | | | | |

| | | | |
|---|---|---|---|
| Sample name: | **M18_octanol** | Experiment start time: | **3/9/2018 11:02:44 PM** |
| Assay name: | **pH-metric high logP** | Analyst: | **Pion** |
| Assay ID: | **18C-09016** | Instrument ID: | **T312060** |
| Filename: | **C:\Sirius_T3\Mehtap\20180309_exp31_logP_T3-2\18C-09016_M18_octanol_pH-metric high logP.t3r** | | |

| | | | |
|---|---|---|---|
| Sample name: | **M18_octanol** | Experiment start time: | **3/9/2018 11:02:44 PM** |
| Assay name: | **pH-metric high logP** | Analyst: | **Pion** |
| Assay ID: | **18C-09016** | Instrument ID: | **T312060** |
| Filename: | **C:\Sirius_T3\Mehtap\20180309_exp31_logP_T3-2\18C-09016_M18_octanol_pH-metric high logP.t3r** | | |

#### Assay Settings (continued)

| Setting | Value | Original Value | Date/Time changed | Imported from |
|---|---|---|---|---|
| ***Titration 3*** | | | | |
| Titrate from | Low to high pH | | | |
| Add additional water | 0.00 mL | | | |
| Additional partition solvent volume | 0.500 mL | | | |
| Additional partition solvent added | Automatic | | | |
| After pH adjust stir for | 30 seconds | | | |
| Stir to allow partitioning for | 15 seconds | | | |
| Stirrer speed for partitioning | 60% | | | |
| ***Data Point Stability*** | | | | |
| Stir during data point collection | No | | | |
| Delay before data point collection | 0 seconds | | | |
| Number of points to average | 20 points | | | |
| Time interval between points | 0.50 seconds | | | |
| Required maximum standard deviation | 0.00100 dpH/dt | | | |
| Stability timeout after | 60 seconds | | | |

#### Calibration Settings

| Setting | Value | Date/Time changed | Imported from |
|---|---|---|---|
| Four-Plus alpha | 0.102 | 3/9/2018 11:02:43 PM | C:\Sirius_T3\HCl18C09.t3r |
| Four-Plus S | 0.9967 | 3/9/2018 11:02:43 PM | C:\Sirius_T3\HCl18C09.t3r |
| Four-Plus jH | 1.2 | 3/9/2018 11:02:43 PM | C:\Sirius_T3\HCl18C09.t3r |
| Four-Plus jOH | 0.0 | 3/9/2018 11:02:43 PM | C:\Sirius_T3\HCl18C09.t3r |
| Base concentration factor | 1.000 | 3/9/2018 11:02:44 PM | C:\Sirius_T3\KOH18B27.t3r |
| Acid concentration factor | 1.000 | 3/9/2018 11:02:44 PM | C:\Sirius_T3\HCl18C09.t3r |

#### Instrument Settings

| Setting | Value | Batch Id | Install date |
|---|---|---|---|
| Instrument owner | Merck | | |
| Instrument ID | T312060 | | |
| Instrument type | T3 Simulator | | |
| Software version | 1.1.3.0 | | |
| Dispenser module | | T3DM1200361 | 3/31/2009 6:24:52 AM |
| Dispenser 0 | Water | | 3/31/2009 6:25:05 AM |
| Syringe volume | 2.5 mL | | |
| Firmware version | 1.2.1(r2) | | |
| Titrant | Water (0.15 M KCl) | 02-06-2018 | 2/27/2018 11:05:59 AM |
| Dispenser 2 | Acid | | 3/31/2009 6:25:11 AM |
| Syringe volume | 0.5 mL | | |
| Firmware version | 1.2.1(r2) | | |
| Titrant | Acid (0.5 M HCl) | 02-27-2018 | 2/27/2018 11:27:22 AM |
| Dispenser 1 | Base | | 3/31/2009 6:25:21 AM |
| Syringe volume | 0.5 mL | | |
| Firmware version | 1.2.1(r2) | | |
| Titrant | Base (0.5 M KOH) | 9/22/2017 | 2/27/2018 11:21:22 AM |
| Dispenser 5 | Cosolvent | | 3/31/2009 6:26:24 AM |
| Syringe volume | 2.5 mL | | |
| Firmware version | 1.2.1(r2) | | |
| Distribution valve 5 | Distribution Valve | | 3/31/2009 6:28:19 AM |
| Firmware version | 1.1.3 | | |
| Port A | Methanol (80%, 0.15 M KCl) | 02-08-2018 | 3/6/2018 10:28:59 AM |
| Port B | Cyclohexane | 11-01-17 | 2/27/2018 11:37:57 AM |
| Dispenser 3 | Buffer | | 8/3/2010 6:05:16 AM |
| Syringe volume | 0.5 mL | | |
| Firmware version | 1.2.1(r2) | | |
| Titrant | Dodecane | 2018/01/31 | 2/28/2018 11:18:04 AM |
| Dispenser 6 | Octanol | | 10/22/2010 11:52:43 AM |

| | | | |
|---|---|---|---|
| Sample name: | **M18_octanol** | Experiment start time: | **3/9/2018 11:02:44 PM** |
| Assay name: | **pH-metric high logP** | Analyst: | **Pion** |
| Assay ID: | **18C-09016** | Instrument ID: | **T312060** |
| Filename: | **C:\Sirius_T3\Mehtap\20180309_exp31_logP_T3-2\18C-09016_M18_octanol_pH-metric high logP.t3r** | | |

| | | | |
|---|---|---|---|
| Sample name: | **M18_octanol** | Experiment start time: | **3/9/2018 11:02:44 PM** |
| Assay name: | **pH-metric high logP** | Analyst: | **Pion** |
| Assay ID: | **18C-09016** | Instrument ID: | **T312060** |
| Filename: | **C:\Sirius_T3\Mehtap\20180309_exp31_logP_T3-2\18C-09016_M18_octanol_pH-metric high logP.t3r** | | |

#### Instrument Settings (continued)

| Setting | Value | Batch Id | Install date |
|---|---|---|---|
| Spectrometer calibration stir duration | 5 s | | |
| Spectrometer calibration stir speed | 30% | | |
| Spectrometer calibration wash pump volume | 20.0 mL | | |
| Spectrometer calibration wash stir duration | 5 s | | |
| Spectrometer calibration wash stir speed | 30% | | |
| Overhead dispense height | 10000 | | |

#### Refinement Settings

| Setting | Value | Default value |
|---|---|---|
| Turbidity detection method | None | None |
| Turbidity wavelength to assess | 500.0 nm | 500.0 nm |
| Turbidity maximum absorbance | 0.100 | 0.100 |
| Turbidity probe threshold | 50.00 | 50.00 |
