## Supplementary material for "Octanol-water partition coefficient measurements for the SAMPL6 Blind Prediction Challenge": SM02_17L-14020_M02_octanol_pH-metric high logP.pdf

| | | | |
|---|---|---|---|
| Sample name: | **M02_octanol** | Experiment start time: | **12/14/2017 9:40:49 PM** |
| Assay name: | **pH-metric high logP** | Analyst: | **Dorothy Levorse** |
| Assay ID: | **17L-14020** | Instrument ID: | **T311053** |
| Filename: | **G:\OpenLab_PFA_Sirius-T3-1\Sirius-T3-1\Data2017\17L-14020_M02_octanol_pH-metric high logP.t3r** | | |

### pH-metric Result

| | |
|---|---|
| logP (XH +) | 1.38 |
| logP (neutral X) | 4.32 |
| RMSD | 0.298 |

### 17L-14020 Points 1 to 14

| | |
|---|---|
| M02_octanol concentration factor | 0.941 |
| Carbonate | 0.0792 mM |
| Acidity error | 0.00723 mM |

### 17L-14020 Points 15 to 31

| | |
|---|---|
| M02_octanol concentration factor | 0.825 |
| Carbonate | 0.1960 mM |
| Acidity error | 0.05972 mM |

### 17L-14020 Points 32 to 50

| | |
|---|---|
| M02_octanol concentration factor | 0.307 |
| Carbonate | 0.2201 mM |
| Acidity error | 0.61969 mM |

### Warnings and errors

| | |
|---|---|
| Errors | None |
| Warnings | None |

### Sample logD and percent species

| pH | M02_octanol logD | M02_octanol M02_octanolH | M02_octanol M02_octanol | M02_octanol M02_octanolH* | M02_octanol M02_octanol* | Comment |
|---|---|---|---|---|---|---|
| 1.000 | 1.41 | 3.71 % | 0.00 % | 88.98 % | 7.31 % | |
| 1.200 | 1.43 | 3.56 % | 0.00 % | 85.33 % | 11.11 % | Stomach pH |
| 2.000 | 1.64 | 2.24 % | 0.00 % | 53.67 % | 44.09 % | |
| 3.000 | 2.34 | 0.45 % | 0.00 % | 10.80 % | 88.74 % | |
| 4.000 | 3.26 | 0.05 % | 0.00 % | 1.20 % | 98.74 % | |
| 5.000 | 4.01 | 0.01 % | 0.00 % | 0.12 % | 99.87 % | |
| 6.000 | 4.28 | 0.00 % | 0.00 % | 0.01 % | 99.98 % | |
| 6.500 | 4.31 | 0.00 % | 0.00 % | 0.00 % | 99.99 % | |
| 7.000 | 4.32 | 0.00 % | 0.00 % | 0.00 % | 99.99 % | |
| 7.400 | 4.32 | 0.00 % | 0.00 % | 0.00 % | 99.99 % | Blood pH |
| 8.000 | 4.32 | 0.00 % | 0.00 % | 0.00 % | 100.00 % | |
| 9.000 | 4.32 | 0.00 % | 0.00 % | 0.00 % | 100.00 % | |
| 10.000 | 4.32 | 0.00 % | 0.00 % | 0.00 % | 100.00 % | |
| 11.000 | 4.32 | 0.00 % | 0.00 % | 0.00 % | 100.00 % | |
| 12.000 | 4.32 | 0.00 % | 0.00 % | 0.00 % | 100.00 % | |

| | | | |
|---|---|---|---|
| Sample name: | **M02_octanol** | Experiment start time: | **12/14/2017 9:40:49 PM** |
| Assay name: | **pH-metric high logP** | Analyst: | **Dorothy Levorse** |
| Assay ID: | **17L-14020** | Instrument ID: | **T311053** |
| Filename: | **G:\OpenLab_PFA_Sirius-T3-1\Sirius-T3-1\Data2017\17L-14020_M02_octanol_pH-metric high logP.t3r** | | |

### Graphs

Charge balance residuals

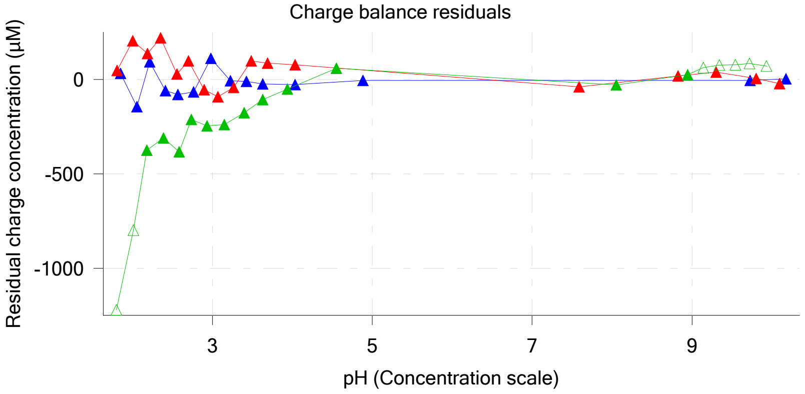


Ionic strength

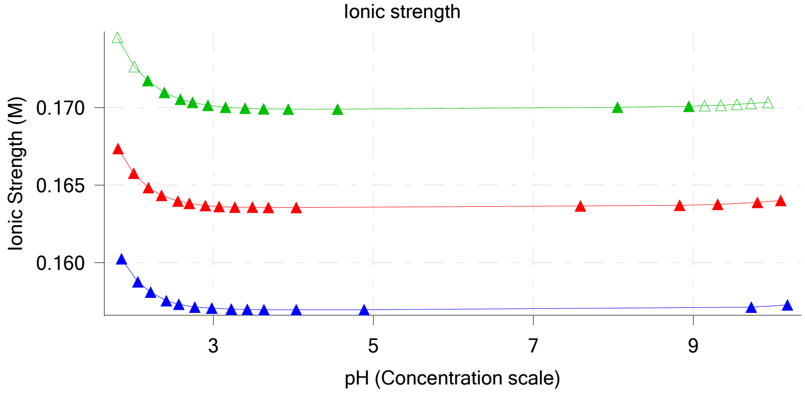


Buffer index profile

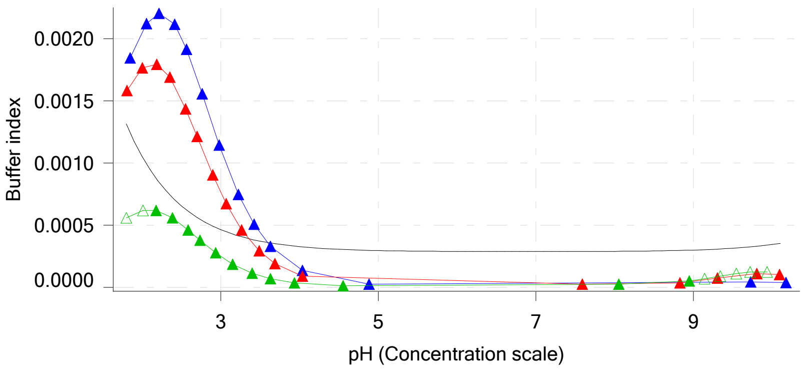


Sample M02_octanol Ionisation Graph

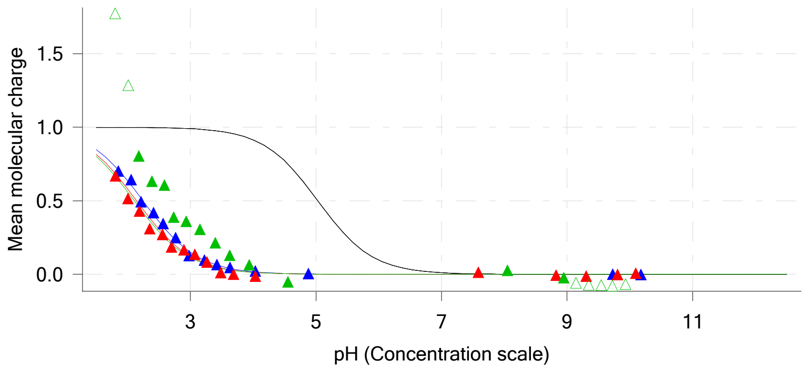


Distribution of species for sample M02_octanol

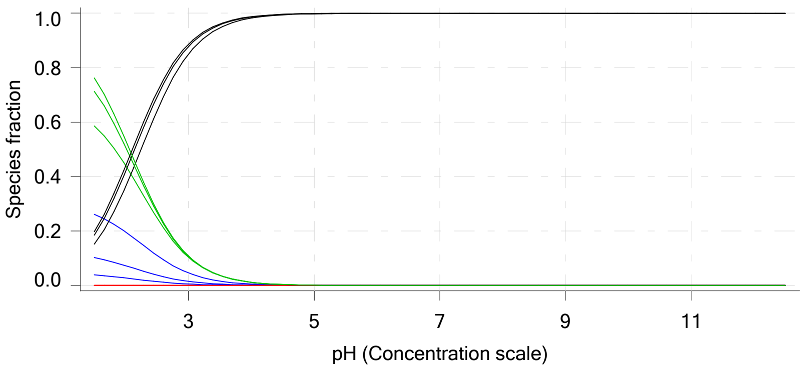


Lipophilicity Profile

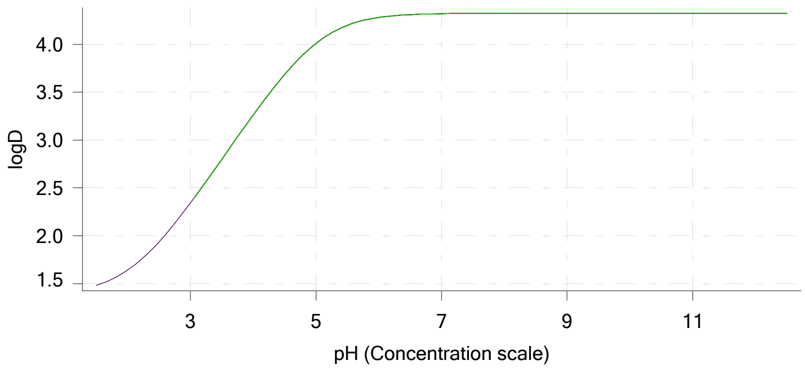


Log R Profile

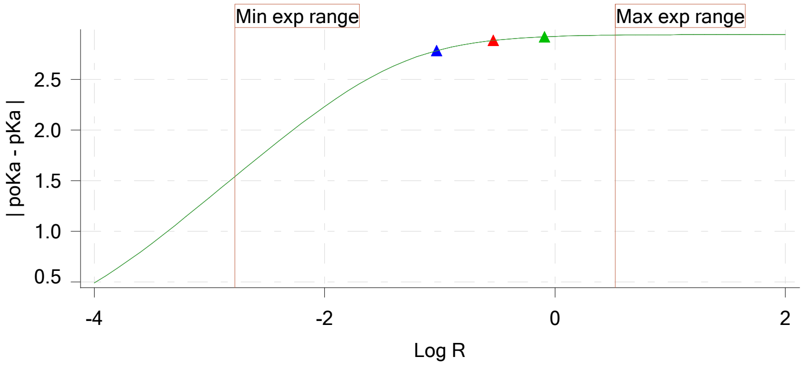


Sample Carbonate Ionisation Graph

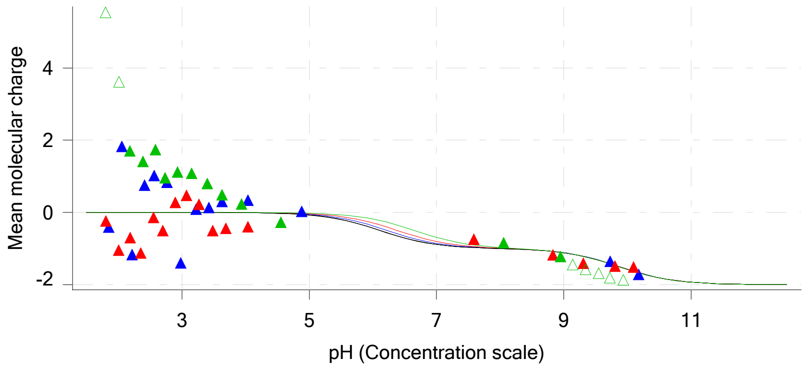

Sample name: **M02_octanol**
Assay name: **pH-metric high logP**
Assay ID: **17L-14020**
Filename: **G:\OpenLab_PFA_Sirius-T3-1\Sirius-T3-1\Data2017\17L-14020_M02_octanol_pH-metric high logP.t3r**

Experiment start time: **12/14/2017 9:40:49 PM**
Analyst: **Dorothy Levorse**
Instrument ID: **T311053**

### Graphs (continued)

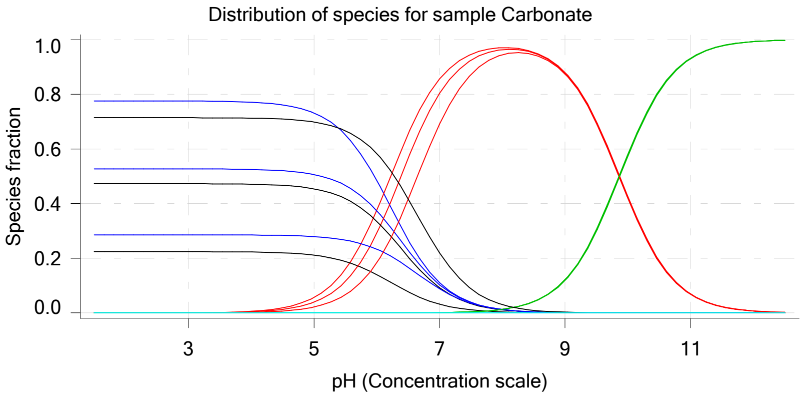

| | | | |
|---|---|---|---|
| Sample name: | **M02_octanol** | Experiment start time: | **12/14/2017 9:40:49 PM** |
| Assay name: | **pH-metric high logP** | Analyst: | **Dorothy Levorse** |
| Assay ID: | **17L-14020** | Instrument ID: | **T311053** |
| Filename: | **G:\OpenLab_PFA_Sirius-T3-1\Sirius-T3-1\Data2017\17L-14020_M02_octanol_pH-metric high logP.t3r** | | |

# pH-metric high logP   Titration 1 of 3   17L-14020 Points 1 to 14

### Overall results

| | |
|---|---|
| RMSD | 0.035 |
| Average ionic strength | 0.158 M |
| Average temperature | 24.8°C |
| Partition ratio | 0.0935 : 1 |
| Analyte concentration range | 3669.1 µM to 3783.1 µM |
| Total points considered | 14 of 14 |

### Warnings and errors

| | |
|---|---|
| Errors | None |
| Warnings | None |

### Four-Plus parameters

| | | | |
|---|---|---|---|
| Alpha | 0.156 | 12/14/2017 9:40:49 PM | C:\Sirius_T3\17L-14015_Blank standardisation.t3r |
| S | 0.9945 | 12/14/2017 9:40:49 PM | C:\Sirius_T3\17L-14015_Blank standardisation.t3r |
| jH | 0.6 | 12/14/2017 9:40:49 PM | C:\Sirius_T3\17L-14015_Blank standardisation.t3r |
| jOH | -0.7 | 12/14/2017 9:40:49 PM | C:\Sirius_T3\17L-14015_Blank standardisation.t3r |

### Titrants

| | | | |
|---|---|---|---|
| 0.50 M HCl | 1.001830 | 12/14/2017 9:40:49 PM | C:\Sirius_T3\17L-14015_Blank standardisation.t3r |
| 0.50 M KOH | 1.010240 | 12/14/2017 9:40:49 PM | C:\Sirius_T3\17L-11009_KHP_Base standardisation using KHP.t3r |

### Sample

| | |
|---|---|
| M02_octanol concentration factor | 0.941 |
| Base pKa 1 | 5.03 |
| logP (XH +) | -0.57 |
| logP (neutral X) | 3.81 |

### Sample graphs

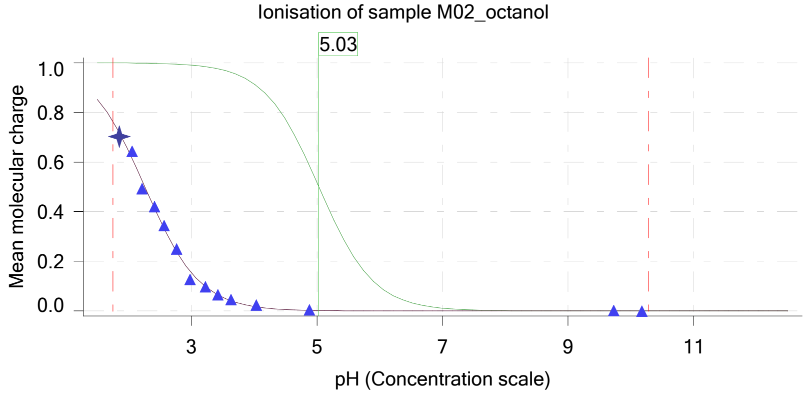


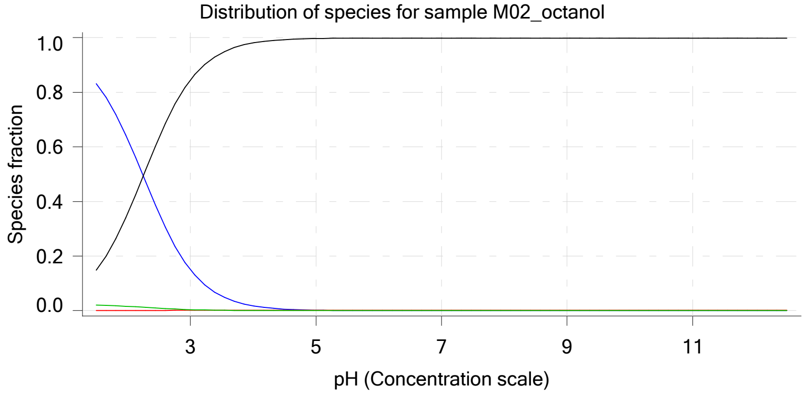

Sample name: **M02_octanol** Experiment start time: **12/14/2017 9:40:49 PM**
Assay name: **pH-metric high logP** Analyst: **Dorothy Levorse**
Assay ID: **17L-14020** Instrument ID: **T311053**
Filename: **G:\OpenLab_PFA_Sirius-T3-1\Sirius-T3-1\Data2017\17L-14020_M02_octanol_pH-metric high logP.t3r**

### Sample graphs (continued)

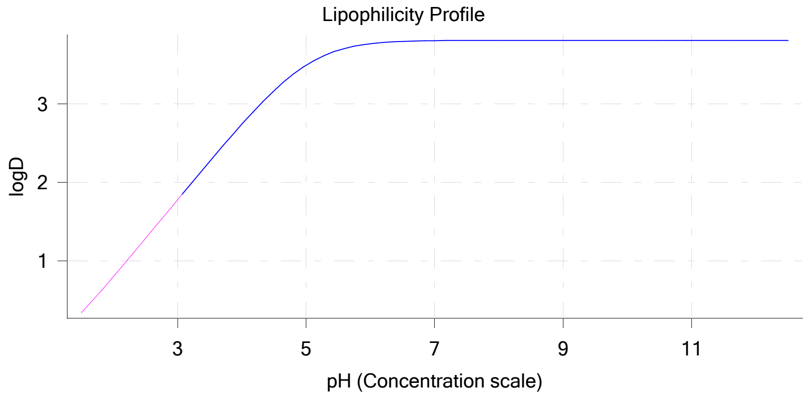


### Sample logD and percent species

| pH | M02_octanol logD | M02_octanol M02_octanolH | M02_octanol M02_octanol | M02_octanol M02_octanolH* | M02_octanol M02_octanol* | Comment |
|---|---|---|---|---|---|---|
| 1.000 | -0.06 | 92.45 % | 0.01 % | 2.33 % | 5.21 % | |
| 1.200 | 0.09 | 89.71 % | 0.01 % | 2.26 % | 8.02 % | Stomach pH |
| 2.000 | 0.80 | 62.89 % | 0.06 % | 1.58 % | 35.46 % | |
| 3.000 | 1.78 | 14.99 % | 0.14 % | 0.38 % | 84.50 % | |
| 4.000 | 2.74 | 1.74 % | 0.16 % | 0.04 % | 98.05 % | |
| 5.000 | 3.49 | 0.18 % | 0.16 % | 0.00 % | 99.65 % | |
| 6.000 | 3.77 | 0.02 % | 0.17 % | 0.00 % | 99.82 % | |
| 6.500 | 3.80 | 0.01 % | 0.17 % | 0.00 % | 99.83 % | |
| 7.000 | 3.81 | 0.00 % | 0.17 % | 0.00 % | 99.83 % | |
| 7.400 | 3.81 | 0.00 % | 0.17 % | 0.00 % | 99.83 % | Blood pH |
| 8.000 | 3.81 | 0.00 % | 0.17 % | 0.00 % | 99.83 % | |
| 9.000 | 3.81 | 0.00 % | 0.17 % | 0.00 % | 99.83 % | |
| 10.000 | 3.81 | 0.00 % | 0.17 % | 0.00 % | 99.83 % | |
| 11.000 | 3.81 | 0.00 % | 0.17 % | 0.00 % | 99.83 % | |
| 12.000 | 3.81 | 0.00 % | 0.17 % | 0.00 % | 99.83 % | |

### Carbonate and acidity

Carbonate 0.079 mM
Acidity error 0.007 mM

### Other graphs

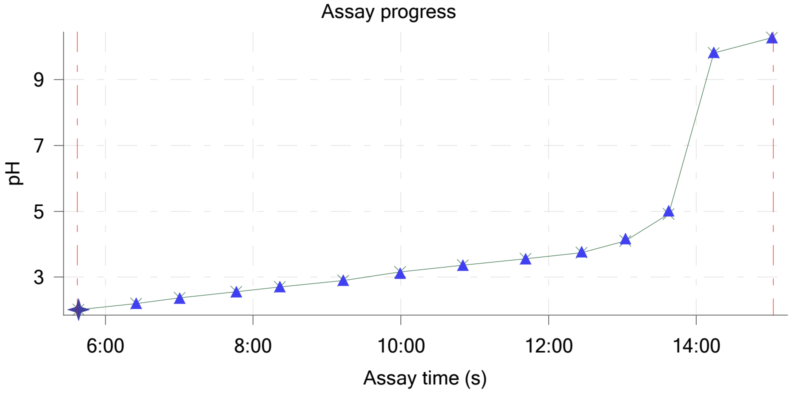


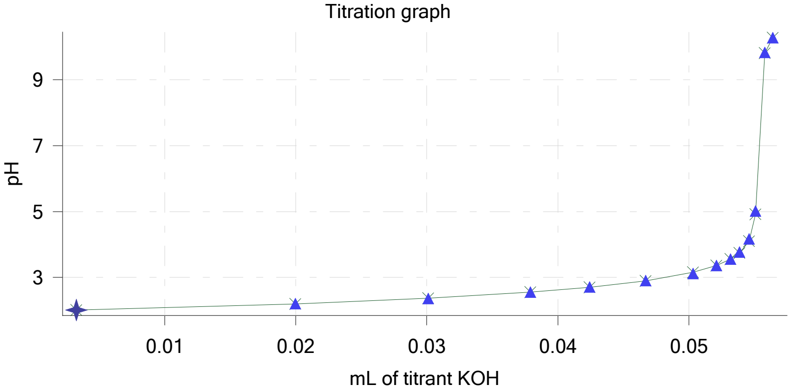

| | | | |
|---|---|---|---|
| Sample name: | **M02_octanol** | Experiment start time: | **12/14/2017 9:40:49 PM** |
| Assay name: | **pH-metric high logP** | Analyst: | **Dorothy Levorse** |
| Assay ID: | **17L-14020** | Instrument ID: | **T311053** |
| Filename: | **G:\OpenLab_PFA_Sirius-T3-1\Sirius-T3-1\Data2017\17L-14020_M02_octanol_pH-metric high logP.t3r** | | |

### Other graphs (continued)

Charge balance residuals

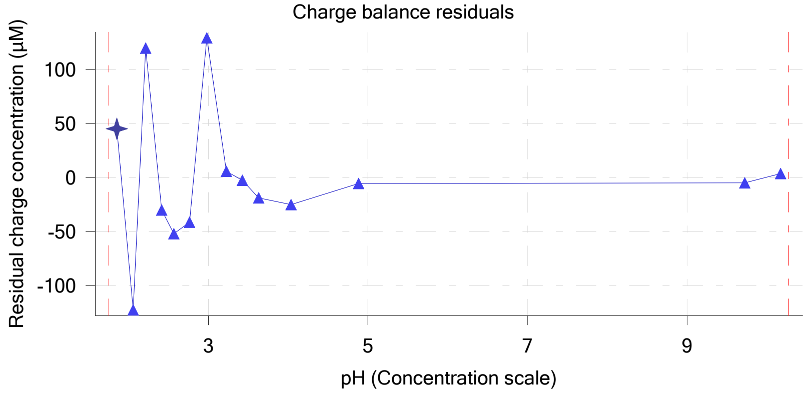


Buffer index profile

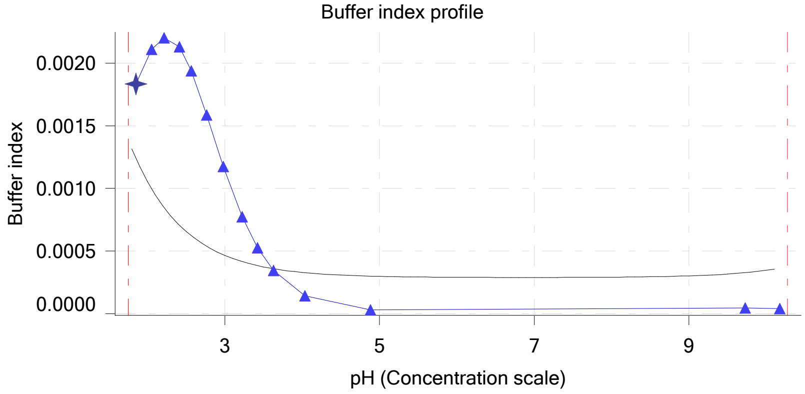


Ionic strength

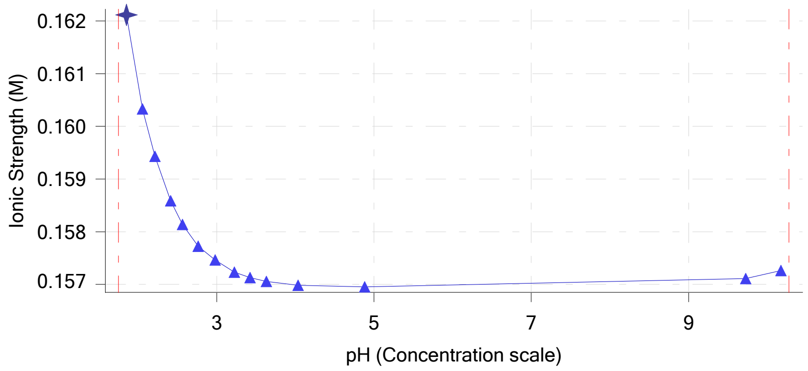


Log R Profile

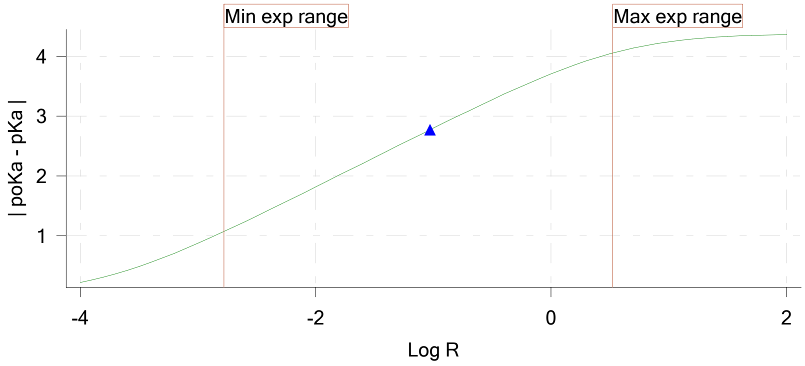


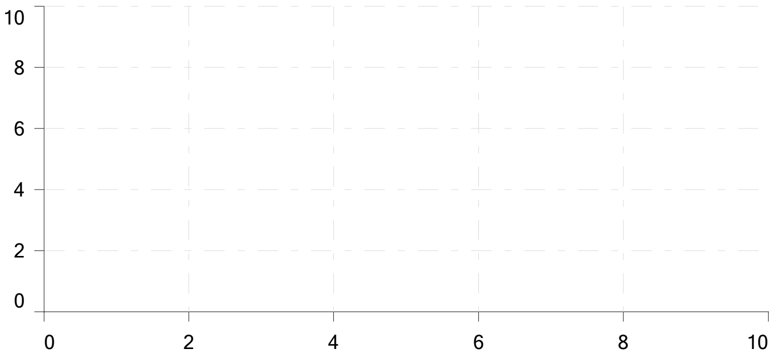


Ionisation of sample Carbonate

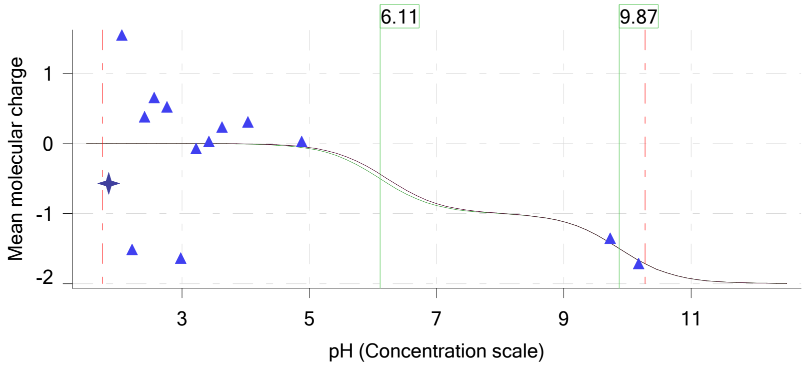

| | | | |
|---|---|---|---|
| Sample name: | **M02_octanol** | Experiment start time: | **12/14/2017 9:40:49 PM** |
| Assay name: | **pH-metric high logP** | Analyst: | **Dorothy Levorse** |
| Assay ID: | **17L-14020** | Instrument ID: | **T311053** |
| Filename: | **G:\OpenLab_PFA_Sirius-T3-1\Sirius-T3-1\Data2017\17L-14020_M02_octanol_pH-metric high logP.t3r** | | |

# pH-metric high logP   Titration 2 of 3   17L-14020 Points 15 to 31

### Overall results

| | |
|---|---|
| RMSD | 0.353 |
| Average ionic strength | 0.165 M |
| Average temperature | 24.9°C |
| Partition ratio | 0.2903 : 1 |
| Analyte concentration range | 2895.7 µM to 2976.2 µM |
| Total points considered | 17 of 17 |

### Warnings and errors

| | |
|---|---|
| Errors | None |
| Warnings | None |

### Four-Plus parameters

| | | | |
|---|---|---|---|
| Alpha | 0.156 | 12/14/2017 9:40:49 PM | C:\Sirius_T3\17L-14015_Blank standardisation.t3r |
| S | 0.9945 | 12/14/2017 9:40:49 PM | C:\Sirius_T3\17L-14015_Blank standardisation.t3r |
| jH | 0.6 | 12/14/2017 9:40:49 PM | C:\Sirius_T3\17L-14015_Blank standardisation.t3r |
| jOH | -0.7 | 12/14/2017 9:40:49 PM | C:\Sirius_T3\17L-14015_Blank standardisation.t3r |

### Titrants

| | | | |
|---|---|---|---|
| 0.50 M HCl | 1.001830 | 12/14/2017 9:40:49 PM | C:\Sirius_T3\17L-14015_Blank standardisation.t3r |
| 0.50 M KOH | 1.010240 | 12/14/2017 9:40:49 PM | C:\Sirius_T3\17L-11009_KHP_Base standardisation using KHP.t3r |

### Sample

| | |
|---|---|
| M02_octanol concentration factor | 0.825 |
| Base pKa 1 | 5.03 |
| logP (XH +) | -0.57 |
| logP (neutral X) | 3.52 |

### Sample graphs

Ionisation of sample M02_octanol

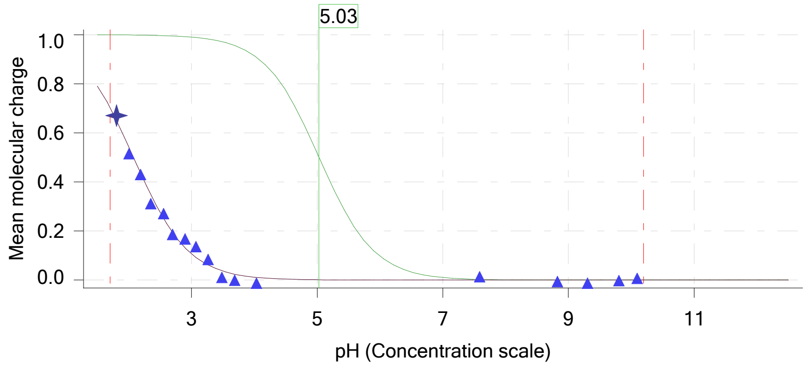


Distribution of species for sample M02_octanol

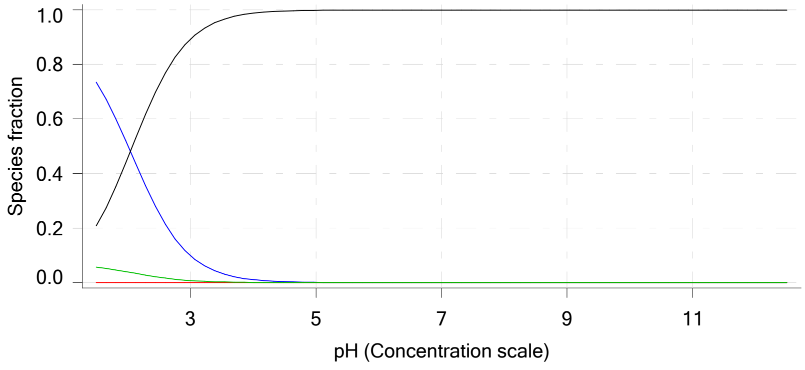

| | | | |
|---|---|---|---|
| Sample name: | **M02_octanol** | Experiment start time: | **12/14/2017 9:40:49 PM** |
| Assay name: | **pH-metric high logP** | Analyst: | **Dorothy Levorse** |
| Assay ID: | **17L-14020** | Instrument ID: | **T311053** |
| Filename: | **G:\OpenLab_PFA_Sirius-T3-1\Sirius-T3-1\Data2017\17L-14020_M02_octanol_pH-metric high logP.t3r** | | |

### Sample graphs (continued)

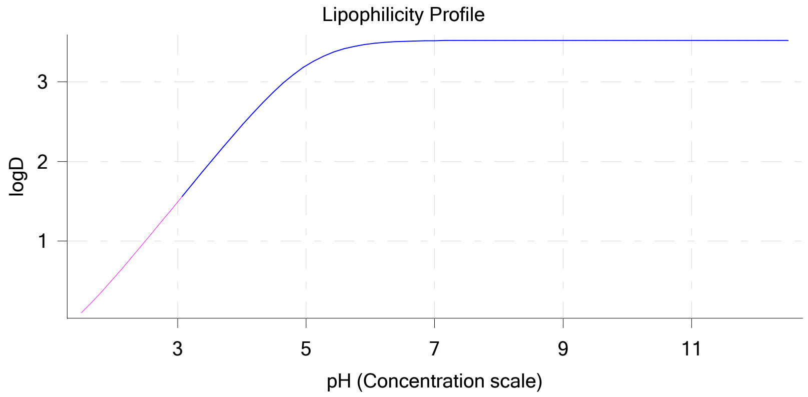


### Sample logD and percent species

| pH | M02_octanol logD | M02_octanol M02_octanolH | M02_octanol M02_octanol | M02_octanol M02_octanolH* | M02_octanol M02_octanol* | Comment |
|---|---|---|---|---|---|---|
| 1.000 | -0.24 | 85.58 % | 0.01 % | 6.69 % | 7.72 % | |
| 1.200 | -0.12 | 81.88 % | 0.01 % | 6.40 % | 11.71 % | Stomach pH |
| 2.000 | 0.53 | 50.46 % | 0.05 % | 3.94 % | 45.55 % | |
| 3.000 | 1.49 | 9.89 % | 0.09 % | 0.77 % | 89.25 % | |
| 4.000 | 2.45 | 1.09 % | 0.10 % | 0.09 % | 98.72 % | |
| 5.000 | 3.21 | 0.11 % | 0.10 % | 0.01 % | 99.78 % | |
| 6.000 | 3.48 | 0.01 % | 0.10 % | 0.00 % | 99.88 % | |
| 6.500 | 3.51 | 0.00 % | 0.10 % | 0.00 % | 99.89 % | |
| 7.000 | 3.52 | 0.00 % | 0.10 % | 0.00 % | 99.90 % | |
| 7.400 | 3.52 | 0.00 % | 0.10 % | 0.00 % | 99.90 % | Blood pH |
| 8.000 | 3.52 | 0.00 % | 0.10 % | 0.00 % | 99.90 % | |
| 9.000 | 3.52 | 0.00 % | 0.10 % | 0.00 % | 99.90 % | |
| 10.000 | 3.52 | 0.00 % | 0.10 % | 0.00 % | 99.90 % | |
| 11.000 | 3.52 | 0.00 % | 0.10 % | 0.00 % | 99.90 % | |
| 12.000 | 3.52 | 0.00 % | 0.10 % | 0.00 % | 99.90 % | |

### Carbonate and acidity

| | |
|---|---|
| Carbonate | 0.196 mM |
| Acidity error | 0.060 mM |

### Other graphs

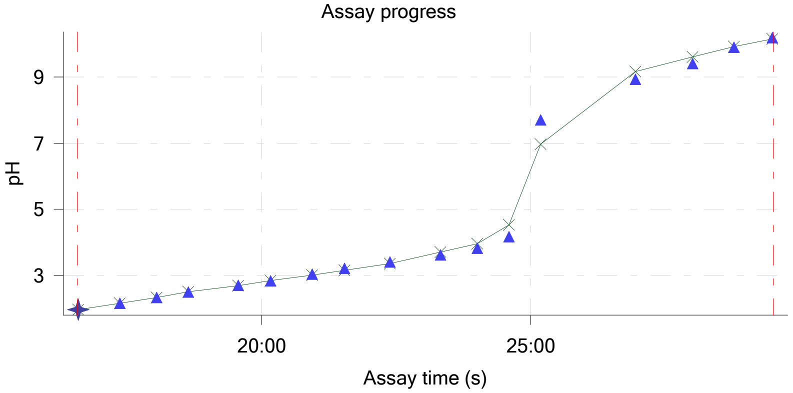


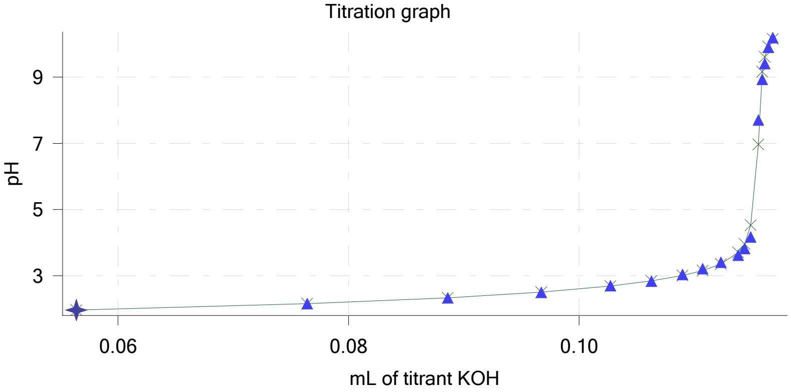

Sample name: **M02_octanol**
Assay name: **pH-metric high logP**
Assay ID: **17L-14020**
Filename: **G:\OpenLab_PFA_Sirius-T3-1\Sirius-T3-1\Data2017\17L-14020_M02_octanol_pH-metric high logP.t3r**

Experiment start time: **12/14/2017 9:40:49 PM**
Analyst: **Dorothy Levorse**
Instrument ID: **T311053**

### Other graphs (continued)

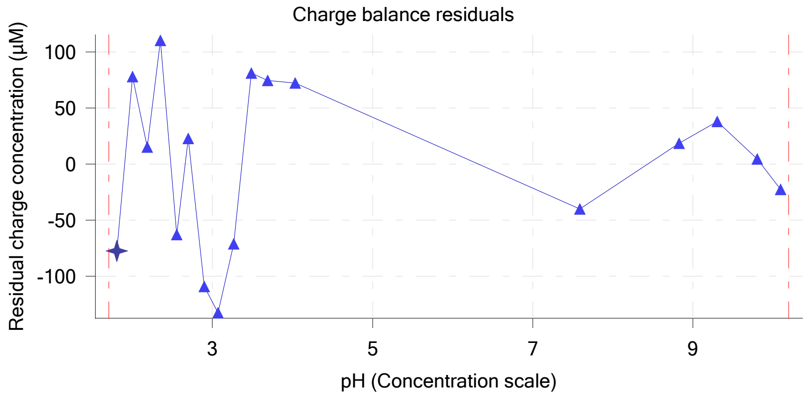


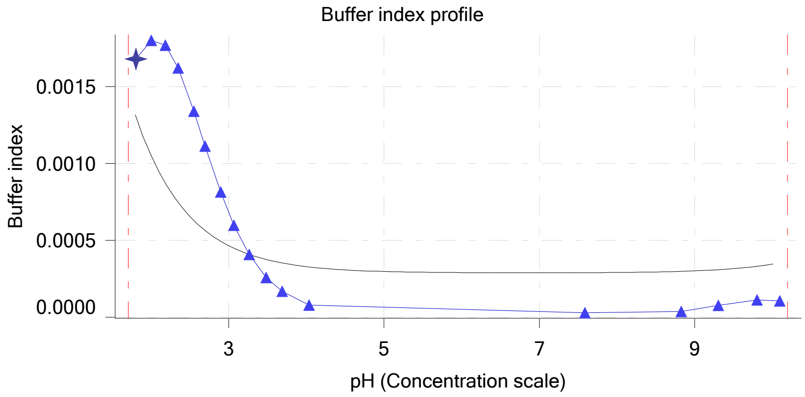


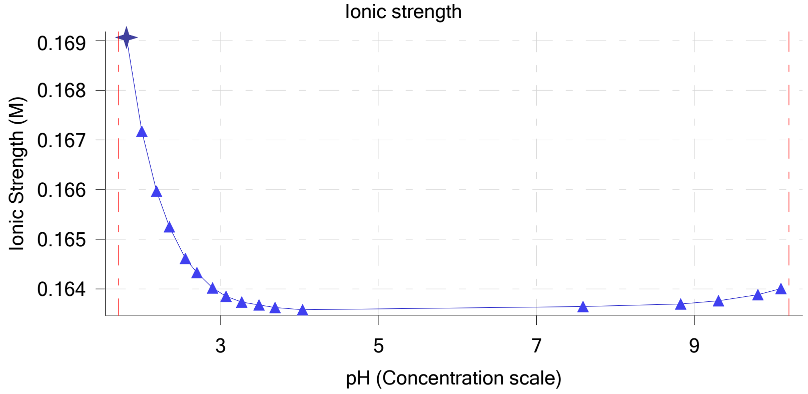


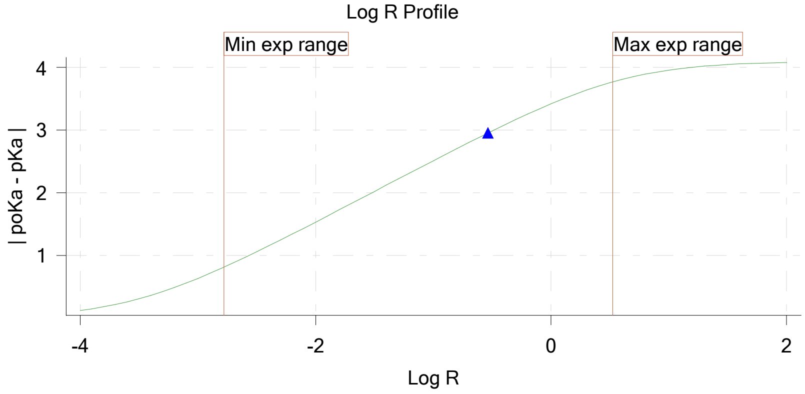


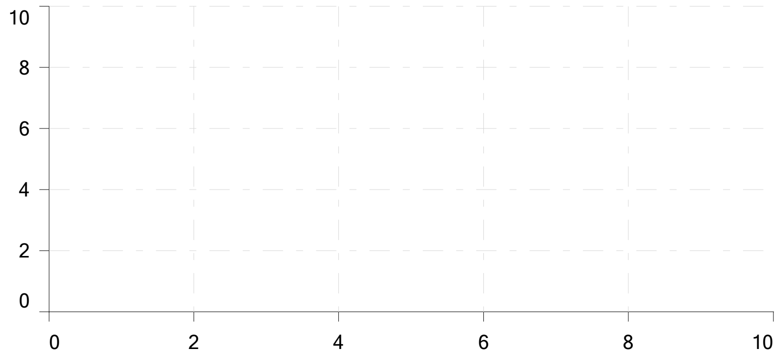


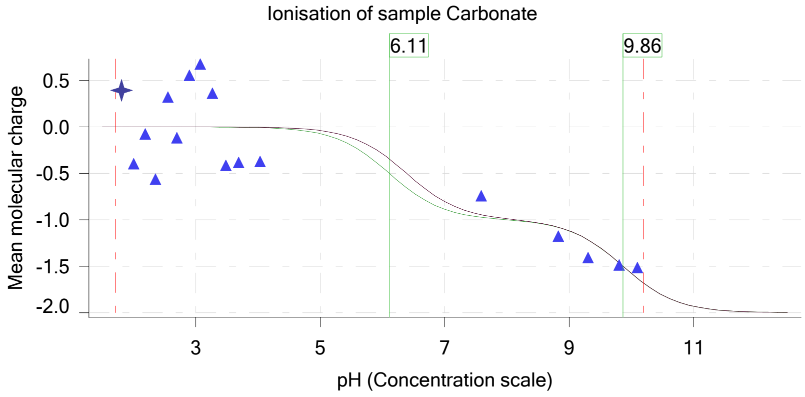

| | | | |
|---|---|---|---|
| Sample name: | **M02_octanol** | Experiment start time: | **12/14/2017 9:40:49 PM** |
| Assay name: | **pH-metric high logP** | Analyst: | **Dorothy Levorse** |
| Assay ID: | **17L-14020** | Instrument ID: | **T311053** |
| Filename: | **G:\OpenLab_PFA_Sirius-T3-1\Sirius-T3-1\Data2017\17L-14020_M02_octanol_pH-metric high logP.t3r** | | |

# pH-metric high logP   Titration 3 of 3   17L-14020 Points 32 to 50

### Overall results

| | |
|---|---|
| RMSD | 0.347 |
| Average ionic strength | 0.170 M |
| Average temperature | 25.0°C |
| Partition ratio | 0.8090 : 1 |
| Analyte concentration range | 1923.2 µM to 1961.3 µM |
| Total points considered | 12 of 19 |

### Warnings and errors

| | |
|---|---|
| Errors | None |
| Warnings | Sample concentration factor out of range |

### Four-Plus parameters

| | | | |
|---|---|---|---|
| Alpha | 0.156 | 12/14/2017 9:40:49 PM | C:\Sirius_T3\17L-14015_Blank standardisation.t3r |
| S | 0.9945 | 12/14/2017 9:40:49 PM | C:\Sirius_T3\17L-14015_Blank standardisation.t3r |
| jH | 0.6 | 12/14/2017 9:40:49 PM | C:\Sirius_T3\17L-14015_Blank standardisation.t3r |
| jOH | -0.7 | 12/14/2017 9:40:49 PM | C:\Sirius_T3\17L-14015_Blank standardisation.t3r |

### Titrants

| | | | |
|---|---|---|---|
| 0.50 M HCl | 1.001830 | 12/14/2017 9:40:49 PM | C:\Sirius_T3\17L-14015_Blank standardisation.t3r |
| 0.50 M KOH | 1.010240 | 12/14/2017 9:40:49 PM | C:\Sirius_T3\17L-11009_KHP_Base standardisation using KHP.t3r |

### Sample

| | |
|---|---|
| M02_octanol concentration factor | 0.307 |
| Base pKa 1 | 5.03 |
| logP (XH +) | -0.57 |
| logP (neutral X) | 2.51 |

### Sample graphs

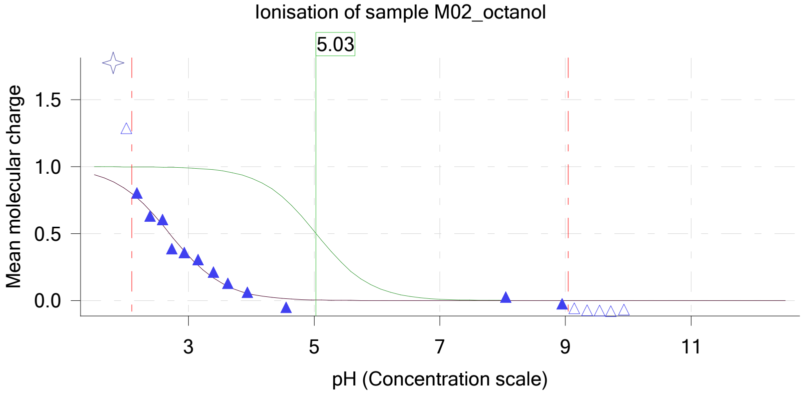


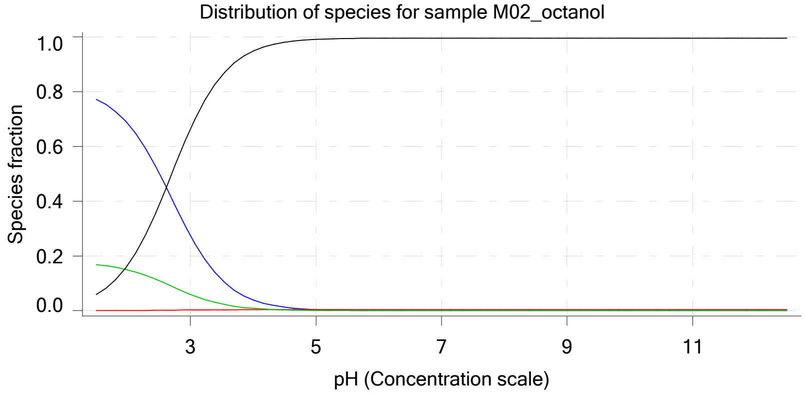

| | | | |
|---|---|---|---|
| Sample name: | **M02_octanol** | Experiment start time: | **12/14/2017 9:40:49 PM** |
| Assay name: | **pH-metric high logP** | Analyst: | **Dorothy Levorse** |
| Assay ID: | **17L-14020** | Instrument ID: | **T311053** |
| Filename: | **G:\OpenLab_PFA_Sirius-T3-1\Sirius-T3-1\Data2017\17L-14020_M02_octanol_pH-metric high logP.t3r** | | |

### Sample graphs (continued)

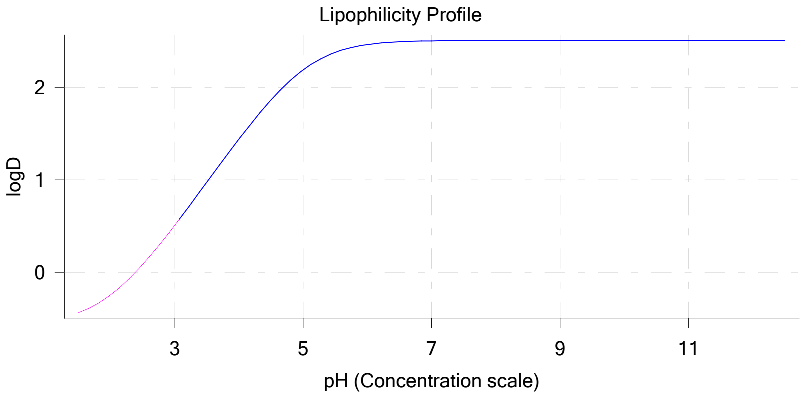


### Sample logD and percent species

| pH | M02_octanol logD | M02_octanol M02_octanolH | M02_octanol M02_octanol | M02_octanol M02_octanolH* | M02_octanol M02_octanol* | Comment |
|---|---|---|---|---|---|---|
| 1.000 | -0.52 | 80.51 % | 0.01 % | 17.53 % | 1.96 % | |
| 1.200 | -0.50 | 79.59 % | 0.01 % | 17.33 % | 3.07 % | Stomach pH |
| 2.000 | -0.24 | 68.41 % | 0.06 % | 14.90 % | 16.63 % | |
| 3.000 | 0.51 | 27.34 % | 0.26 % | 5.95 % | 66.45 % | |
| 4.000 | 1.44 | 3.90 % | 0.36 % | 0.85 % | 94.88 % | |
| 5.000 | 2.19 | 0.41 % | 0.38 % | 0.09 % | 99.12 % | |
| 6.000 | 2.46 | 0.04 % | 0.38 % | 0.01 % | 99.57 % | |
| 6.500 | 2.49 | 0.01 % | 0.38 % | 0.00 % | 99.60 % | |
| 7.000 | 2.50 | 0.00 % | 0.38 % | 0.00 % | 99.61 % | |
| 7.400 | 2.51 | 0.00 % | 0.38 % | 0.00 % | 99.62 % | Blood pH |
| 8.000 | 2.51 | 0.00 % | 0.38 % | 0.00 % | 99.62 % | |
| 9.000 | 2.51 | 0.00 % | 0.38 % | 0.00 % | 99.62 % | |
| 10.000 | 2.51 | 0.00 % | 0.38 % | 0.00 % | 99.62 % | |
| 11.000 | 2.51 | 0.00 % | 0.38 % | 0.00 % | 99.62 % | |
| 12.000 | 2.51 | 0.00 % | 0.38 % | 0.00 % | 99.62 % | |

### Carbonate and acidity

| | |
|---|---|
| Carbonate | 0.220 mM |
| Acidity error | 0.620 mM |

### Other graphs

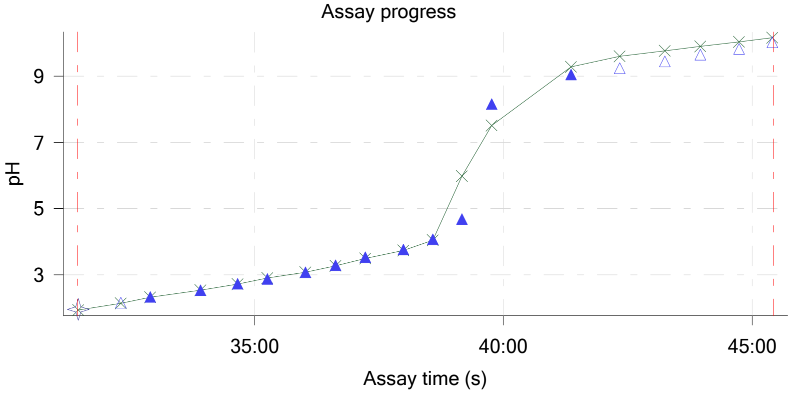


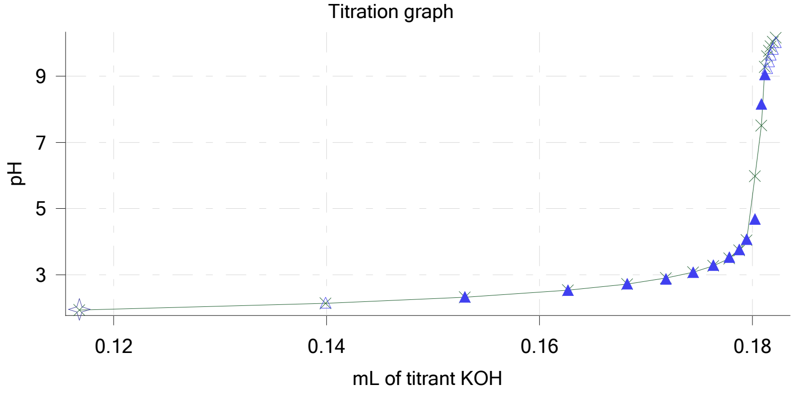

| | | | |
|---|---|---|---|
| Sample name: | **M02_octanol** | Experiment start time: | **12/14/2017 9:40:49 PM** |
| Assay name: | **pH-metric high logP** | Analyst: | **Dorothy Levorse** |
| Assay ID: | **17L-14020** | Instrument ID: | **T311053** |
| Filename: | **G:\OpenLab_PFA_Sirius-T3-1\Sirius-T3-1\Data2017\17L-14020_M02_octanol_pH-metric high logP.t3r** | | |

### Other graphs (continued)

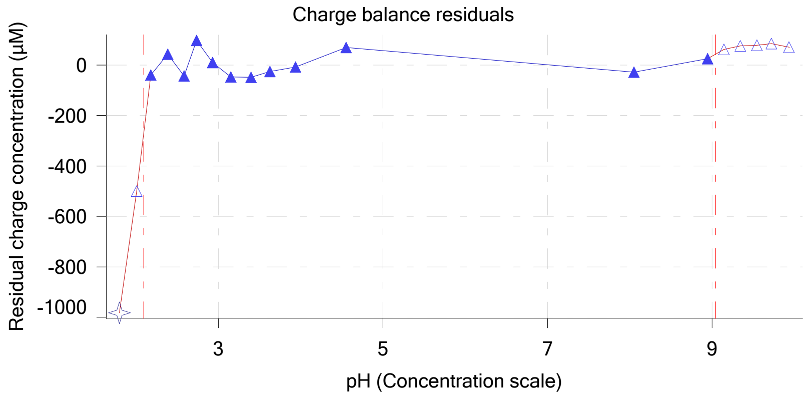


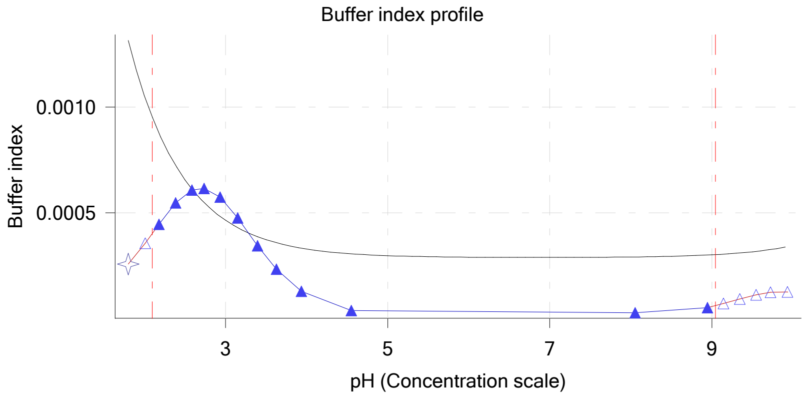


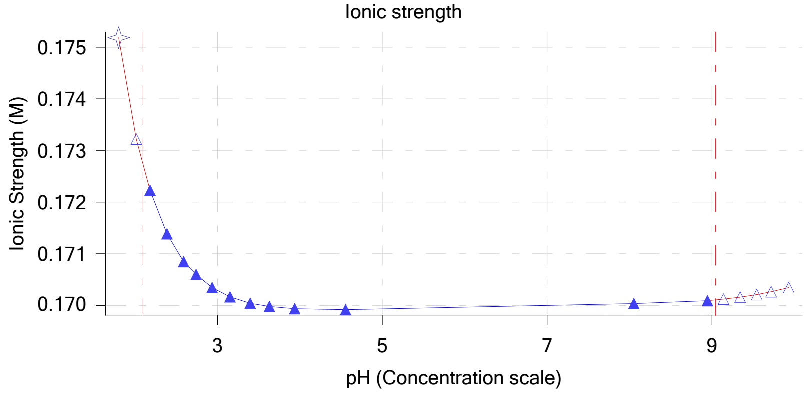


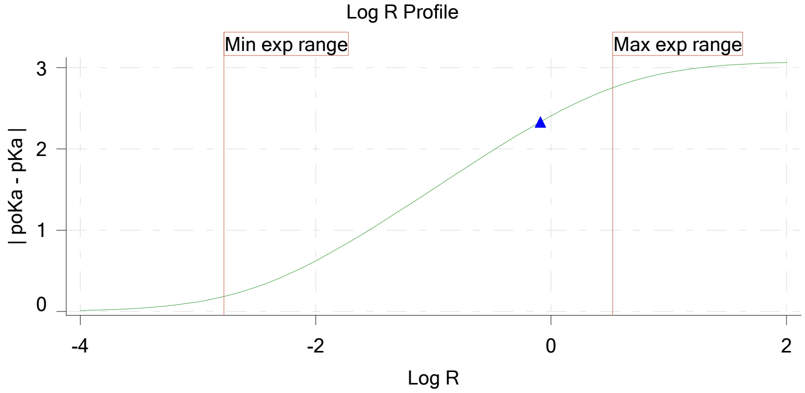


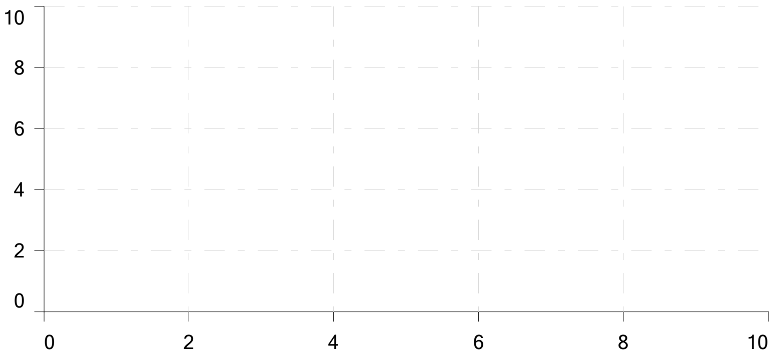

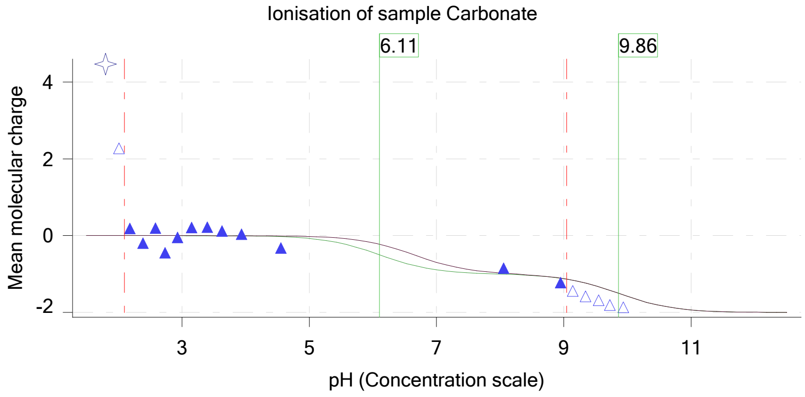

| | | | |
|---|---|---|---|
| Sample name: | **M02_octanol** | Experiment start time: | **12/14/2017 9:40:49 PM** |
| Assay name: | **pH-metric high logP** | Analyst: | **Dorothy Levorse** |
| Assay ID: | **17L-14020** | Instrument ID: | **T311053** |
| Filename: | **G:\OpenLab_PFA_Sirius-T3-1\Sirius-T3-1\Data2017\17L-14020_M02_octanol_pH-metric high logP.t3r** | | |

### Assay Model

| Settings | Value | Date/Time changed | Imported from |
|---|---|---|---|
| Sample name | M02_octanol | 12/6/2017 5:20:03 PM | User entered value |
| Sample by | Weight | | Default value |
| Sample weight | 0.001870 g | 12/14/2017 4:31:54 PM | User entered value |
| Formula weight | 289.26 g/mol | 12/6/2017 5:20:03 PM | User entered value |
| Solubility | Unknown | | Default value |
| Molecular weight | 289.26 | 12/6/2017 5:20:03 PM | User entered value |
| Individual pKa ionic environments | No | | Default value |
| Number of pKas | 1 | 12/6/2017 5:20:03 PM | User entered value |
| Sample is a | Base | 12/6/2017 5:20:03 PM | User entered value |
| pKa 1 | 5.03 | 12/6/2017 5:20:03 PM | User entered value |
| logp (XH +) | -0.57 | 12/14/2017 3:31:39 PM | User entered value |
| logP (neutral X) | 0.94 | 12/14/2017 3:31:32 PM | User entered value |

### Events

| Time | Event | Water | Acid | Base | Octanol | pH | dpH/dt | pH R-squared | pH SD | dpH/dt time |
|---|---|---|---|---|---|---|---|---|---|---|
| 2:38.7 | Initial pH = 6.57 | | | | | | | | | |
| 5:38.2 | Data point 1 | 1.50000 mL | 0.05560 mL | 0.00327 mL | 0.15000 mL | 2.007 | 0.00139 | 0.03368 | 0.00037 | 10.5 s |
| 6:24.9 | Data point 2 | 1.50000 mL | 0.05560 mL | 0.01997 mL | 0.15000 mL | 2.206 | -0.00747 | 0.39684 | 0.00059 | 10.0 s |
| 7:00.4 | Data point 3 | 1.50000 mL | 0.05560 mL | 0.03010 mL | 0.15000 mL | 2.363 | -0.00559 | 0.19734 | 0.00062 | 10.0 s |
| 7:46.3 | Data point 4 | 1.50000 mL | 0.05560 mL | 0.03791 mL | 0.15000 mL | 2.558 | -0.01487 | 0.86800 | 0.00079 | 10.0 s |
| 8:21.8 | Data point 5 | 1.50000 mL | 0.05560 mL | 0.04243 mL | 0.15000 mL | 2.710 | -0.00425 | 0.44243 | 0.00032 | 10.5 s |
| 9:13.3 | Data point 6 | 1.50000 mL | 0.05560 mL | 0.04671 mL | 0.15000 mL | 2.907 | -0.00837 | 0.81909 | 0.00046 | 10.5 s |
| 9:59.6 | Data point 7 | 1.50000 mL | 0.05560 mL | 0.05031 mL | 0.15000 mL | 3.120 | -0.01056 | 0.38594 | 0.00084 | 10.0 s |
| 10:50.5 | Data point 8 | 1.50000 mL | 0.05560 mL | 0.05209 mL | 0.15000 mL | 3.363 | -0.01094 | 0.53189 | 0.00074 | 10.0 s |
| 11:41.4 | Data point 9 | 1.50000 mL | 0.05560 mL | 0.05318 mL | 0.15000 mL | 3.562 | -0.00971 | 0.40417 | 0.00075 | 10.0 s |
| 12:27.0 | Data point 10 | 1.50000 mL | 0.05560 mL | 0.05386 mL | 0.15000 mL | 3.767 | -0.01049 | 0.64757 | 0.00064 | 10.0 s |
| 13:02.4 | Data point 11 | 1.50000 mL | 0.05560 mL | 0.05459 mL | 0.15000 mL | 4.170 | -0.00778 | 0.67581 | 0.00047 | 10.0 s |
| 13:37.7 | Data point 12 | 1.50000 mL | 0.05560 mL | 0.05508 mL | 0.15000 mL | 5.011 | 0.00369 | 0.07856 | 0.00065 | 11.0 s |
| 14:14.1 | Data point 13 | 1.50000 mL | 0.05560 mL | 0.05579 mL | 0.15000 mL | 9.827 | -0.01899 | 0.90855 | 0.00098 | 17.0 s |
| 15:01.6 | Data point 14 | 1.50000 mL | 0.05560 mL | 0.05640 mL | 0.15000 mL | 10.274 | -0.01908 | 0.91794 | 0.00098 | 17.0 s |
| 16:35.5 | Data point 15 | 1.50000 mL | 0.11580 mL | 0.05640 mL | 0.50000 mL | 1.964 | -0.00642 | 0.19348 | 0.00072 | 10.0 s |
| 17:21.8 | Data point 16 | 1.50000 mL | 0.11580 mL | 0.07641 mL | 0.50000 mL | 2.155 | -0.01440 | 0.57008 | 0.00094 | 15.5 s |
| 18:03.0 | Data point 17 | 1.50000 mL | 0.11580 mL | 0.08862 mL | 0.50000 mL | 2.336 | -0.01761 | 0.86346 | 0.00094 | 10.0 s |
| 18:38.5 | Data point 18 | 1.50000 mL | 0.11580 mL | 0.09671 mL | 0.50000 mL | 2.499 | -0.01747 | 0.85915 | 0.00093 | 14.5 s |
| 19:34.1 | Data point 19 | 1.50000 mL | 0.11580 mL | 0.10271 mL | 0.50000 mL | 2.699 | -0.01738 | 0.83571 | 0.00094 | 10.5 s |
| 20:10.1 | Data point 20 | 1.50000 mL | 0.11580 mL | 0.10626 mL | 0.50000 mL | 2.841 | -0.01842 | 0.94159 | 0.00094 | 10.5 s |
| 20:56.5 | Data point 21 | 1.50000 mL | 0.11580 mL | 0.10896 mL | 0.50000 mL | 3.039 | -0.00529 | 0.32126 | 0.00046 | 10.5 s |
| 21:32.5 | Data point 22 | 1.50000 mL | 0.11580 mL | 0.11070 mL | 0.50000 mL | 3.210 | -0.01298 | 0.72113 | 0.00075 | 10.0 s |
| 22:23.3 | Data point 23 | 1.50000 mL | 0.11580 mL | 0.11230 mL | 0.50000 mL | 3.404 | -0.01202 | 0.61736 | 0.00075 | 10.0 s |
| 23:19.3 | Data point 24 | 1.50000 mL | 0.11580 mL | 0.11381 mL | 0.50000 mL | 3.623 | -0.00105 | 0.02529 | 0.00033 | 10.5 s |
| 24:00.4 | Data point 25 | 1.50000 mL | 0.11580 mL | 0.11435 mL | 0.50000 mL | 3.825 | -0.01396 | 0.59332 | 0.00089 | 10.0 s |
| 24:35.7 | Data point 26 | 1.50000 mL | 0.11580 mL | 0.11486 mL | 0.50000 mL | 4.169 | -0.00892 | 0.71187 | 0.00052 | 10.0 s |
| 25:11.0 | Data point 27 | 1.50000 mL | 0.11580 mL | 0.11555 mL | 0.50000 mL | 7.702 | -0.10242 | 0.99687 | 0.00506 | Timed out at 59.5 s |
| 26:56.8 | Data point 28 | 1.50000 mL | 0.11580 mL | 0.11588 mL | 0.50000 mL | 8.934 | -0.01413 | 0.48746 | 0.00100 | 28.0 s |
| 28:00.6 | Data point 29 | 1.50000 mL | 0.11580 mL | 0.11611 mL | 0.50000 mL | 9.408 | -0.01497 | 0.88842 | 0.00078 | 15.5 s |
| 28:46.6 | Data point 30 | 1.50000 mL | 0.11580 mL | 0.11642 mL | 0.50000 mL | 9.903 | -0.01712 | 0.83019 | 0.00093 | 12.5 s |
| 29:29.6 | Data point 31 | 1.50000 mL | 0.11580 mL | 0.11679 mL | 0.50000 mL | 10.194 | -0.01474 | 0.64645 | 0.00091 | 10.0 s |
| 31:27.3 | Data point 32 | 1.50000 mL | 0.17935 mL | 0.11679 mL | 1.50000 mL | 1.957 | -0.01138 | 0.54149 | 0.00076 | 10.0 s |
| 32:18.7 | Data point 33 | 1.50000 mL | 0.17935 mL | 0.13991 mL | 1.50000 mL | 2.162 | -0.00826 | 0.34190 | 0.00070 | 10.0 s |
| 32:54.3 | Data point 34 | 1.50000 mL | 0.17935 mL | 0.15299 mL | 1.50000 mL | 2.329 | 0.00841 | 0.21086 | 0.00090 | 19.5 s |
| 33:54.9 | Data point 35 | 1.50000 mL | 0.17935 mL | 0.16268 mL | 1.50000 mL | 2.533 | -0.01444 | 0.56679 | 0.00095 | 19.0 s |
| 34:39.5 | Data point 36 | 1.50000 mL | 0.17935 mL | 0.16825 mL | 1.50000 mL | 2.729 | -0.01081 | 0.59250 | 0.00069 | 10.5 s |
| 35:15.5 | Data point 37 | 1.50000 mL | 0.17935 mL | 0.17190 mL | 1.50000 mL | 2.879 | 0.00523 | 0.07679 | 0.00093 | 10.0 s |
| 36:01.3 | Data point 38 | 1.50000 mL | 0.17935 mL | 0.17444 mL | 1.50000 mL | 3.074 | -0.01576 | 0.64444 | 0.00097 | 11.0 s |
| 36:37.8 | Data point 39 | 1.50000 mL | 0.17935 mL | 0.17634 mL | 1.50000 mL | 3.290 | -0.01372 | 0.74324 | 0.00079 | 10.5 s |

| | | | |
|---|---|---|---|
| Sample name: | **M02_octanol** | Experiment start time: | **12/14/2017 9:40:49 PM** |
| Assay name: | **pH-metric high logP** | Analyst: | **Dorothy Levorse** |
| Assay ID: | **17L-14020** | Instrument ID: | **T311053** |
| Filename: | **G:\OpenLab_PFA_Sirius-T3-1\Sirius-T3-1\Data2017\17L-14020_M02_octanol_pH-metric high logP.t3r** | | |

### Events (continued)

| Time | Event | Water | Acid | Base | Octanol | pH | dpH/dt | pH R-squared | pH SD | dpH/dt time |
|---|---|---|---|---|---|---|---|---|---|---|
| 37:13.7 | Data point 40 | 1.50000 mL | 0.17935 mL | 0.17782 mL | 1.50000 mL | 3.534 | -0.00886 | 0.57323 | 0.00058 | 10.0 s |
| 37:59.4 | Data point 41 | 1.50000 mL | 0.17935 mL | 0.17876 mL | 1.50000 mL | 3.765 | -0.00909 | 0.36165 | 0.00075 | 10.0 s |
| 38:34.8 | Data point 42 | 1.50000 mL | 0.17935 mL | 0.17947 mL | 1.50000 mL | 4.071 | -0.00570 | 0.25273 | 0.00056 | 10.0 s |
| 39:10.1 | Data point 43 | 1.50000 mL | 0.17935 mL | 0.18024 mL | 1.50000 mL | 4.684 | -0.00391 | 0.12958 | 0.00054 | 10.5 s |
| 39:46.0 | Data point 44 | 1.50000 mL | 0.17935 mL | 0.18083 mL | 1.50000 mL | 8.163 | -0.13728 | 0.99581 | 0.00680 | Timed out at 59.5 s |
| 41:21.6 | Data point 45 | 1.50000 mL | 0.17935 mL | 0.18119 mL | 1.50000 mL | 9.052 | -0.01846 | 0.83706 | 0.00100 | 23.0 s |
| 42:20.2 | Data point 46 | 1.50000 mL | 0.17935 mL | 0.18140 mL | 1.50000 mL | 9.247 | -0.00723 | 0.15620 | 0.00090 | 19.0 s |
| 43:14.9 | Data point 47 | 1.50000 mL | 0.17935 mL | 0.18156 mL | 1.50000 mL | 9.446 | -0.00873 | 0.24834 | 0.00086 | 12.0 s |
| 43:57.4 | Data point 48 | 1.50000 mL | 0.17935 mL | 0.18173 mL | 1.50000 mL | 9.647 | -0.00896 | 0.20472 | 0.00098 | 11.0 s |
| 44:44.1 | Data point 49 | 1.50000 mL | 0.17935 mL | 0.18194 mL | 1.50000 mL | 9.822 | 0.00182 | 0.01449 | 0.00075 | 10.0 s |
| 45:24.5 | Data point 50 | 1.50000 mL | 0.17935 mL | 0.18220 mL | 1.50000 mL | 10.033 | 0.01248 | 0.45958 | 0.00091 | 15.0 s |
| 45:48.4 | Assay volumes | 1.50000 mL | 0.17935 mL | 0.18220 mL | 1.50000 mL | | | | | |

| | | | |
|---|---|---|---|
| Sample name: | **M02_octanol** | Experiment start time: | **12/14/2017 9:40:49 PM** |
| Assay name: | **pH-metric high logP** | Analyst: | **Dorothy Levorse** |
| Assay ID: | **17L-14020** | Instrument ID: | **T311053** |
| Filename: | **G:\OpenLab_PFA_Sirius-T3-1\Sirius-T3-1\Data2017\17L-14020_M02_octanol_pH-metric high logP.t3r** | | |

| | | | |
|---|---|---|---|
| Sample name: | **M02_octanol** | Experiment start time: | **12/14/2017 9:40:49 PM** |
| Assay name: | **pH-metric high logP** | Analyst: | **Dorothy Levorse** |
| Assay ID: | **17L-14020** | Instrument ID: | **T311053** |
| Filename: | **G:\OpenLab_PFA_Sirius-T3-1\Sirius-T3-1\Data2017\17L-14020_M02_octanol_pH-metric high logP.t3r** | | |

### Assay Settings (continued)

| Setting | Value | Original Value | Date/Time changed | Imported from |
|---|---|---|---|---|
| Titrate from | Low to high pH | | | |
| Add additional water | 0.00 mL | | | |
| Additional partition solvent volume | 1.000 mL | | | |
| Additional partition solvent added | Automatic | | | |
| After pH adjust stir for | 30 seconds | | | |
| Stir to allow partitioning for | 15 seconds | | | |
| Stirrer speed for partitioning | 60% | | | |
| ***Data Point Stability*** | | | | |
| Stir during data point collection | No | | | |
| Delay before data point collection | 0 seconds | | | |
| Number of points to average | 20 points | | | |
| Time interval between points | 0.50 seconds | | | |
| Required maximum standard deviation | 0.00100 dpH/dt | | | |
| Stability timeout after | 60 seconds | | | |

### Calibration Settings

| Setting | Value | Date/Time changed | Imported from |
|---|---|---|---|
| Four-Plus alpha | 0.156 | 12/14/2017 9:40:49 PM | C:\Sirius_T3\17L-14015_Blank standardisation.t3r |
| Four-Plus S | 0.9945 | 12/14/2017 9:40:49 PM | C:\Sirius_T3\17L-14015_Blank standardisation.t3r |
| Four-Plus jH | 0.6 | 12/14/2017 9:40:49 PM | C:\Sirius_T3\17L-14015_Blank standardisation.t3r |
| Four-Plus jOH | -0.7 | 12/14/2017 9:40:49 PM | C:\Sirius_T3\17L-14015_Blank standardisation.t3r |
| Base concentration factor | 1.010 | 12/14/2017 9:40:49 PM | C:\Sirius_T3\17L-11009_KHP_Base standardisation using KHP.t3r |
| Acid concentration factor | 1.002 | 12/14/2017 9:40:49 PM | C:\Sirius_T3\17L-14015_Blank standardisation.t3r |

### Instrument Settings

| Setting | Value | Batch Id | Install date |
|---|---|---|---|
| Instrument owner | Merck | | |
| Instrument ID | T311053 | | |
| Instrument type | T3 Simulator | | |
| Software version | 1.1.3.0 | | |
| Dispenser module | | T3DM1100253 | 3/31/2009 6:24:52 AM |
| Dispenser 0 | Water | | 3/31/2009 6:25:05 AM |
| Syringe volume | 2.5 mL | | |
| Firmware version | 1.2.1(r2) | | |
| Titrant | Water (0.15 M KCl) | 10-10-2017 | 12/6/2017 10:48:42 AM |
| Dispenser 2 | Acid | | 3/31/2009 6:25:11 AM |
| Syringe volume | 0.5 mL | | |
| Firmware version | 1.2.1(r2) | | |
| Titrant | Acid (0.5 M HCl) | 12-6-2017 | 12/6/2017 10:50:31 AM |
| Dispenser 1 | Base | | 3/31/2009 6:25:21 AM |
| Syringe volume | 0.5 mL | | |
| Firmware version | 1.2.1(r2) | | |
| Titrant | Base (0.5 M KOH) | 09-22-17 | 11/27/2017 10:30:45 AM |
| Dispenser 5 | Cosolvent | | 3/31/2009 6:26:24 AM |
| Syringe volume | 2.5 mL | | |
| Firmware version | 1.2.1(r2) | | |
| Distribution valve 5 | Distribution Valve | | 3/31/2009 6:28:19 AM |
| Firmware version | 1.1.3 | | |
| Port A | Methanol (80%, 0.15 M KCl) | 9-26-17 | 12/11/2017 11:29:08 AM |
| Port B | Cyclohexane | | 10/19/2017 3:11:05 PM |
| Port C | MeCN (50%, 0.15 M KCl) | 10-30-17 | 11/27/2017 10:21:50 AM |
| Dispenser 3 | Buffer | | 8/3/2010 6:05:16 AM |
| Syringe volume | 0.5 mL | | |
| Firmware version | 1.2.1(r2) | | |
| Titrant | Dodecane | | 12/6/2017 10:50:59 AM |
| Dispenser 6 | Octanol | | 10/22/2010 11:52:43 AM |

| | | | |
|---|---|---|---|
| Sample name: | **M02_octanol** | Experiment start time: | **12/14/2017 9:40:49 PM** |
| Assay name: | **pH-metric high logP** | Analyst: | **Dorothy Levorse** |
| Assay ID: | **17L-14020** | Instrument ID: | **T311053** |
| Filename: | **G:\OpenLab_PFA_Sirius-T3-1\Sirius-T3-1\Data2017\17L-14020_M02_octanol_pH-metric high logP.t3r** | | |

### Instrument Settings (continued)

| Setting | Value | Batch Id | Install date |
|---|---|---|---|
| Syringe volume | 0.5 mL | | |
| Firmware version | 1.2.1(r2) | | |
| Titrant | Octanol | 9-14-17 | 12/11/2017 11:32:26 AM |
| Titrator | | T3TM1100153 | 3/31/2009 6:24:17 AM |
| Horizontal axis firmware version | 1.17 AI1DI2DO2 Stepper 2 | | |
| Vertical axis firmware version | 1.17 AI1DI2DO2 Stepper 2 | | |
| Chassis I/O firmware version | 1.11 AI1DI0DO4 Norgren I/O | | |
| Probe I/O firmware version | 1.1.1 | | |
| Electrode | T3 Electrode | T3E0769 | 8/15/2017 10:21:54 AM |
| E0 calibration | -5.35 mV | | 12/14/2017 9:41:17 PM |
| Filling solution | 3M KCl | KCL095 | 12/14/2017 3:53:24 PM |
| Liquids | | | |
| Wash 1 | 50% IPA:50% Water | | 12/14/2017 3:56:50 PM |
| Wash 2 | 0.5% Trition X-100 in H20 | | 12/14/2017 3:56:52 PM |
| Buffer position 1 | pH7 Wash | | 12/14/2017 3:56:54 PM |
| Buffer position 2 | pH 7 | | 12/14/2017 3:56:57 PM |
| Storage position | | | 12/14/2017 3:57:36 PM |
| Wash water | 6.6e+003 mL | 12-11-2017 | 12/11/2017 3:16:20 PM |
| Waste | 3.5e+003 mL | | 12/11/2017 3:16:25 PM |
| Temperature controller | | | 8/5/2010 7:35:13 AM |
| Turbidity detector | | | 3/31/2009 6:24:45 AM |
| Spectrometer | | 072390 | 11/23/2010 12:22:28 PM |
| Dip probe | | 11086 | |
| Wavelength coefficient A0 | 185.563 | | |
| Wavelength coefficient A1 | 2.17439 | | |
| Wavelength coefficient A2 | -0.000285622 | | |
| Total lamp lit time | 644:54:39 | | 11/23/2010 12:22:28 PM |
| Calibrated on | 12/4/2017 11:26:25 AM | | |
| Integration time | 11 | | |
| Scans averaged | 10 | | |
| Autoloader | | T3AL1100237 | 11/10/2015 10:34:13 AM |
| Left-right axis firmware version | 1.17 AI1DI2DO2 Stepper 2 | | |
| Front-back axis firmware version | 1.17 AI1DI2DO2 Stepper 2 | | |
| Vertical axis firmware version | 1.17 AI1DI2DO2 Stepper 2 | | |
| Chassis I/O firmware version | 1.11 AI1DI0DO4 Norgren I/O | | |
| Configuration | | | |
| Alternate titration position | Titration position | | |
| Alternate reference position | Reference position | | |
| Maximum standard vial volume | 3.50 mL | | |
| Maximum alternate vial volume | 25.00 mL | | |
| Automatic action idle period | 5 minute(s) | | |
| Titrant tube volume | 1.3 mL | | |
| Syringe flush count | 3.50 | | |
| Flowing wash pump volume | 20.0 mL | | |
| Flowing wash stir duration | 5 s | | |
| Flowing wash stir speed | 30% | | |
| Solvent wash stir duration | 5 s | | |
| Solvent wash stir speed | 30% | | |
| Surfactant wash stir duration | 5 s | | |
| Surfactant wash stir speed | 30% | | |
| E0 calibration minimum number of points | 10 | | |
| E0 calibration maximum standard deviation | 0.01500 | | |
| E0 calibration timeout period | 60 s | | |
| E0 calibration stir duration | 5 s | | |
| E0 calibration preparation stir speed | 30% | | |
| E0 calibration buffer wash stir duration | 5 s | | |
| E0 calibration buffer wash stir speed | 30% | | |
| E0 calibration reading stir speed | 0% | | |

| | | | |
|---|---|---|---|
| Sample name: | **M02_octanol** | Experiment start time: | **12/14/2017 9:40:49 PM** |
| Assay name: | **pH-metric high logP** | Analyst: | **Dorothy Levorse** |
| Assay ID: | **17L-14020** | Instrument ID: | **T311053** |
| Filename: | **G:\OpenLab_PFA_Sirius-T3-1\Sirius-T3-1\Data2017\17L-14020_M02_octanol_pH-metric high logP.t3r** | | |

### Instrument Settings (continued)

| Setting | Value | Batch Id | Install date |
|---|---|---|---|
| Spectrometer calibration stir duration | 5 s | | |
| Spectrometer calibration stir speed | 30% | | |
| Spectrometer calibration wash pump volume | 20.0 mL | | |
| Spectrometer calibration wash stir duration | 5 s | | |
| Spectrometer calibration wash stir speed | 30% | | |
| Overhead dispense height | 10000 | | |

### Refinement Settings

| Setting | Value | Default value |
|---|---|---|
| Turbidity detection method | None | None |
| Turbidity wavelength to assess | 500.0 nm | 500.0 nm |
| Turbidity maximum absorbance | 0.100 | 0.100 |
| Turbidity probe threshold | 50.00 | 50.00 |
