## Supplementary material for "Octanol-water partition coefficient measurements for the SAMPL6 Blind Prediction Challenge": SM08_18B-28014_M08_octanol_pH-metric high logP.pdf

| | | | |
|---|---|---|---|
| Sample name: | **M08_octanol** | Experiment start time: | **2/28/2018 9:57:03 PM** |
| Assay name: | **pH-metric high logP** | Analyst: | **Pion** |
| Assay ID: | **18B-28014** | Instrument ID: | **T312060** |
| Filename: | **G:\OpenLab_PFA_Sirius-T3-2\Data2018\18B-28014_M08_octanol_pH-metric high logP.t3r** | | |

#### pH-metric Result

| | |
|---|---|
| logP (neutral XH) | 3.97 |
| logP (X -) | 1.86 |
| RMSD | 0.921 |

#### 18B-28014 Points 2 to 29

| | |
|---|---|
| M08_octanol concentration factor | 0.885 |
| Carbonate | 0.0000 mM |
| Acidity error | -0.44323 mM |

#### 18B-28014 Points 30 to 68

| | |
|---|---|
| M08_octanol concentration factor | 1.148 |
| Carbonate | 0.0724 mM |
| Acidity error | -0.45335 mM |

#### 18B-28014 Points 69 to 110

| | |
|---|---|
| M08_octanol concentration factor | 0.843 |
| Carbonate | 0.4970 mM |
| Acidity error | -0.17938 mM |

#### Warnings and errors

| | |
|---|---|
| Errors | None |
| Warnings | None |

#### Sample logD and percent species

| pH | M08_octanol logD | M08_octanol M08_octanolH | M08_octanol M08_octanol | M08_octanol M08_octanolH* | M08_octanol M08_octanol* | Comment |
|---|---|---|---|---|---|---|
| 1.000 | 3.97 | 0.01 % | 0.00 % | 99.99 % | 0.00 % | |
| 1.200 | 3.97 | 0.01 % | 0.00 % | 99.99 % | 0.00 % | Stomach pH |
| 2.000 | 3.97 | 0.01 % | 0.00 % | 99.98 % | 0.00 % | |
| 3.000 | 3.95 | 0.01 % | 0.00 % | 99.94 % | 0.05 % | |
| 4.000 | 3.77 | 0.01 % | 0.01 % | 99.52 % | 0.46 % | |
| 5.000 | 3.15 | 0.01 % | 0.06 % | 95.49 % | 4.44 % | |
| 6.000 | 2.35 | 0.01 % | 0.43 % | 67.96 % | 31.59 % | |
| 6.500 | 2.08 | 0.00 % | 0.81 % | 40.15 % | 59.03 % | |
| 7.000 | 1.95 | 0.00 % | 1.12 % | 17.50 % | 81.37 % | |
| 7.400 | 1.90 | 0.00 % | 1.25 % | 7.79 % | 90.96 % | Blood pH |
| 8.000 | 1.87 | 0.00 % | 1.33 % | 2.08 % | 96.59 % | |
| 9.000 | 1.86 | 0.00 % | 1.35 % | 0.21 % | 98.43 % | |
| 10.000 | 1.86 | 0.00 % | 1.36 % | 0.02 % | 98.62 % | |
| 11.000 | 1.86 | 0.00 % | 1.36 % | 0.00 % | 98.64 % | |
| 12.000 | 1.86 | 0.00 % | 1.36 % | 0.00 % | 98.64 % | |

| | | | |
|---|---|---|---|
| Sample name: | **M08_octanol** | Experiment start time: | **2/28/2018 9:57:03 PM** |
| Assay name: | **pH-metric high logP** | Analyst: | **Pion** |
| Assay ID: | **18B-28014** | Instrument ID: | **T312060** |
| Filename: | **G:\OpenLab_PFA_Sirius-T3-2\Data2018\18B-28014_M08_octanol_pH-metric high logP.t3r** | | |

#### Graphs

Charge balance residuals

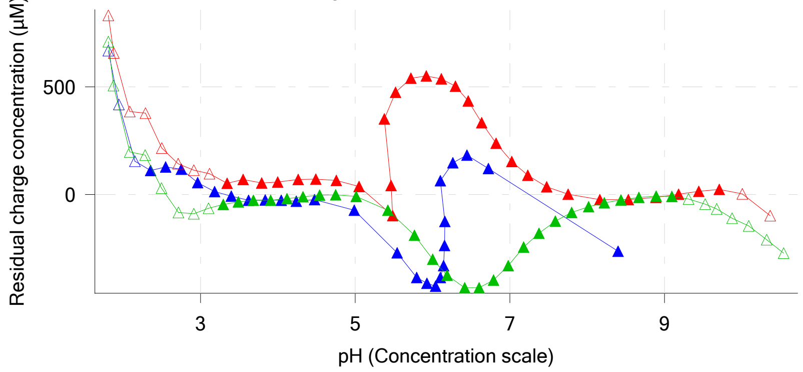


Ionic strength

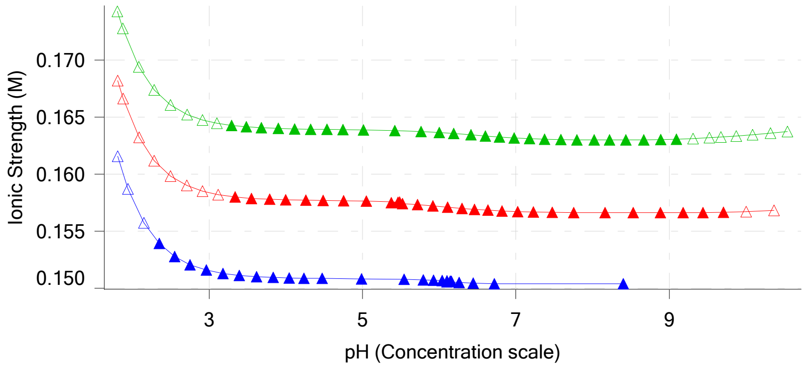


Buffer index profile

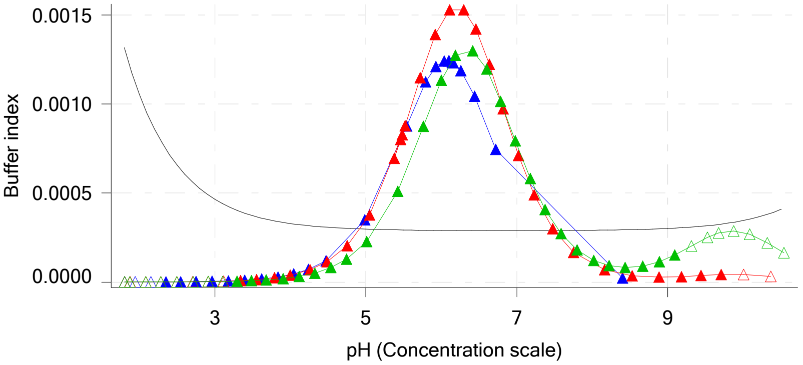


Sample M08_octanol Ionisation Graph

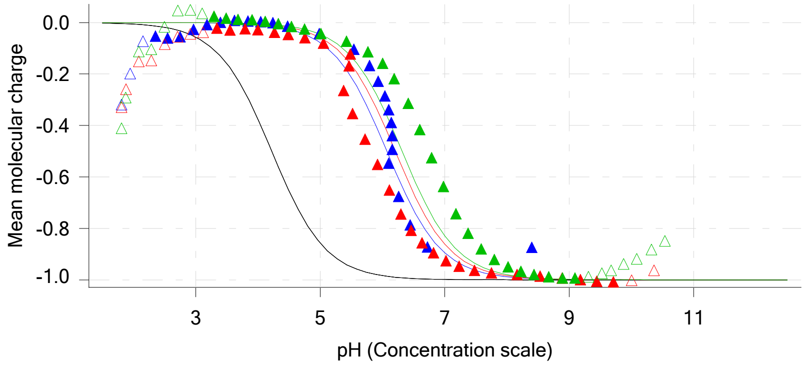


Distribution of species for sample M08_octanol

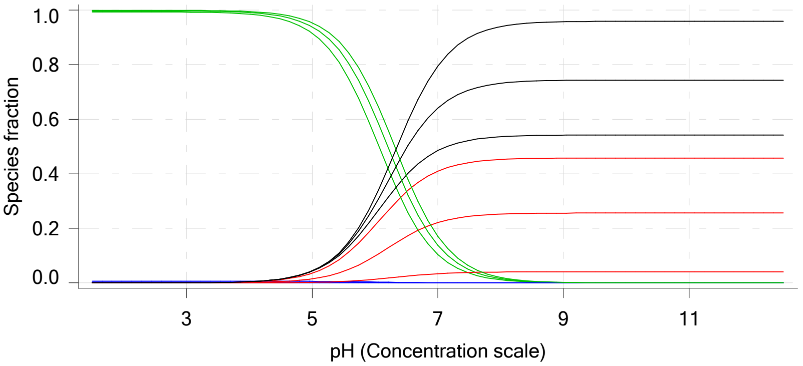


Lipophilicity Profile

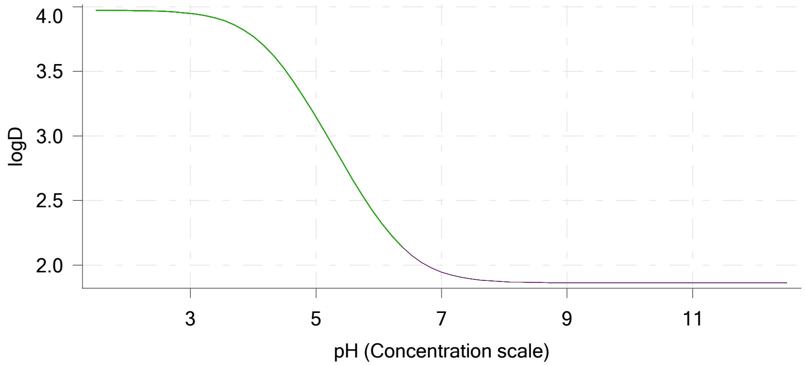


Log R Profile

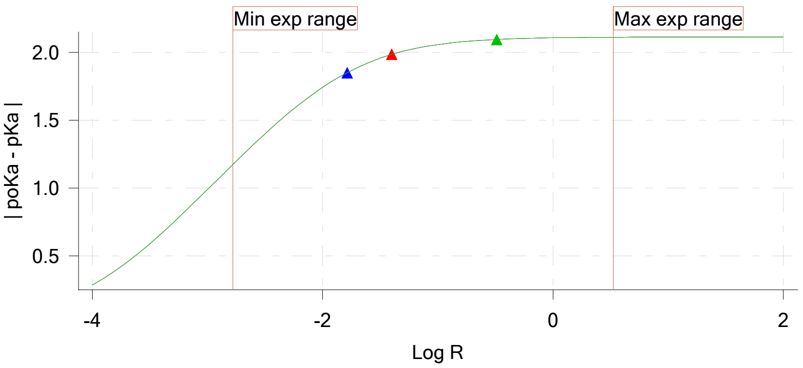


Sample Carbonate Ionisation Graph

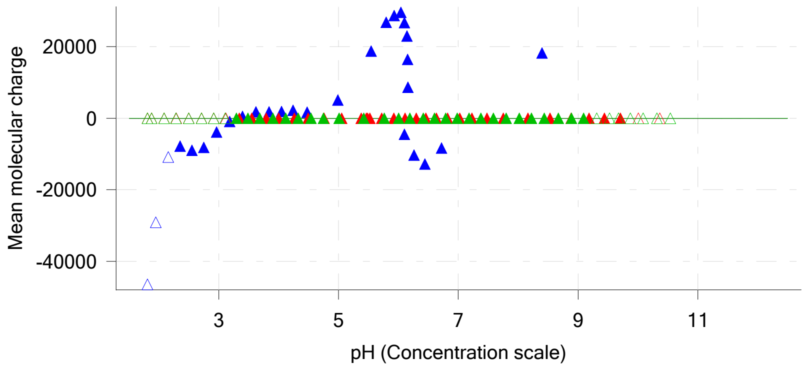

| | | | |
|---|---|---|---|
| Sample name: | **M08_octanol** | Experiment start time: | **2/28/2018 9:57:03 PM** |
| Assay name: | **pH-metric high logP** | Analyst: | **Pion** |
| Assay ID: | **18B-28014** | Instrument ID: | **T312060** |
| Filename: | **G:\OpenLab_PFA_Sirius-T3-2\Data2018\18B-28014_M08_octanol_pH-metric high logP.t3r** | | |

#### Graphs (continued)

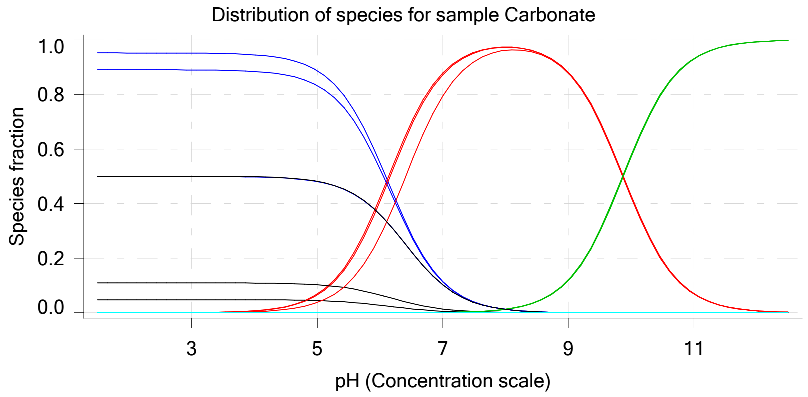

| | | | |
|---|---|---|---|
| Sample name: | **M08_octanol** | Experiment start time: | **2/28/2018 9:57:03 PM** |
| Assay name: | **pH-metric high logP** | Analyst: | **Pion** |
| Assay ID: | **18B-28014** | Instrument ID: | **T312060** |
| Filename: | **G:\OpenLab_PFA_Sirius-T3-2\Data2018\18B-28014_M08_octanol_pH-metric high logP.t3r** | | |

### pH-metric high logP Titration 1 of 3 18B-28014 Points 2 to 29

#### Overall results

| | |
|---|---|
| RMSD | 1.233 |
| Average ionic strength | 0.151 M |
| Average temperature | 24.9°C |
| Partition ratio | 0.0163 : 1 |
| Analyte concentration range | 2327.5 µM to 2407.5 µM |
| Total points considered | 25 of 28 |

#### Warnings and errors

| | |
|---|---|
| Errors | None |
| Warnings | None |

#### Four-Plus parameters

| | | | |
|---|---|---|---|
| Alpha | 0.130 | 2/28/2018 9:57:03 PM | C:\Sirius_T3\HCl18B27.t3r |
| S | 0.9970 | 2/28/2018 9:57:03 PM | C:\Sirius_T3\HCl18B27.t3r |
| jH | 0.8 | 2/28/2018 9:57:03 PM | C:\Sirius_T3\HCl18B27.t3r |
| jOH | -0.4 | 2/28/2018 9:57:03 PM | C:\Sirius_T3\HCl18B27.t3r |

#### Titrants

| | | | |
|---|---|---|---|
| 0.50 M HCl | 0.993513 | 2/28/2018 9:57:03 PM | C:\Sirius_T3\HCl18B27.t3r |
| 0.50 M KOH | 0.999845 | 2/28/2018 9:57:03 PM | C:\Sirius_T3\KOH18B27.t3r |

#### Sample

| | |
|---|---|
| M08_octanol concentration factor | 0.885 |
| Acid pKa 1 | 4.22 |
| logP (neutral XH) | 4.09 |
| logP (X -) | 1.80 |

#### Sample graphs

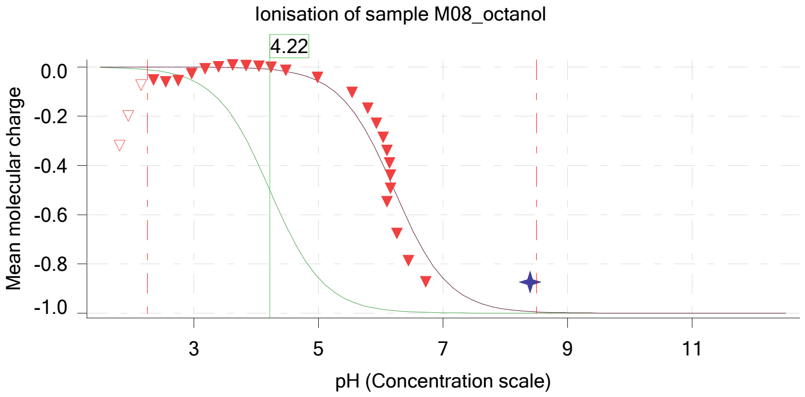


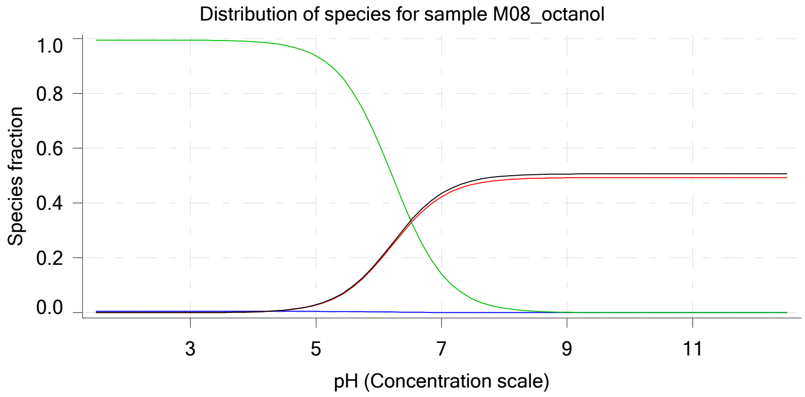

| | | | |
|---|---|---|---|
| Sample name: | **M08_octanol** | Experiment start time: | **2/28/2018 9:57:03 PM** |
| Assay name: | **pH-metric high logP** | Analyst: | **Pion** |
| Assay ID: | **18B-28014** | Instrument ID: | **T312060** |
| Filename: | **G:\OpenLab_PFA_Sirius-T3-2\Data2018\18B-28014_M08_octanol_pH-metric high logP.t3r** | | |

#### Sample graphs (continued)

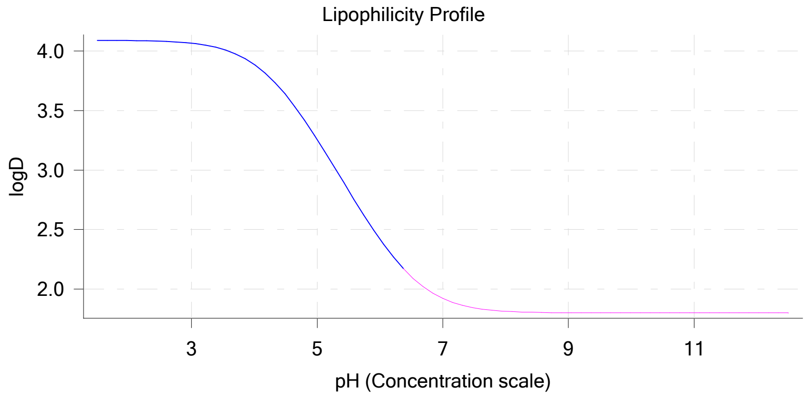


#### Sample logD and percent species

| pH | M08_octanol logD | M08_octanol M08_octanolH | M08_octanol M08_octanol | M08_octanol M08_octanolH* | M08_octanol M08_octanol* | Comment |
|---|---|---|---|---|---|---|
| 1.000 | 4.09 | 0.49 % | 0.00 % | 99.50 % | 0.00 % | |
| 1.200 | 4.09 | 0.49 % | 0.00 % | 99.50 % | 0.00 % | Stomach pH |
| 2.000 | 4.09 | 0.49 % | 0.00 % | 99.50 % | 0.00 % | |
| 3.000 | 4.07 | 0.49 % | 0.03 % | 99.44 % | 0.03 % | |
| 4.000 | 3.89 | 0.49 % | 0.30 % | 98.91 % | 0.30 % | |
| 5.000 | 3.26 | 0.47 % | 2.81 % | 93.83 % | 2.89 % | |
| 6.000 | 2.42 | 0.31 % | 18.58 % | 62.00 % | 19.11 % | |
| 6.500 | 2.10 | 0.17 % | 32.38 % | 34.16 % | 33.29 % | |
| 7.000 | 1.92 | 0.07 % | 42.31 % | 14.12 % | 43.50 % | |
| 7.400 | 1.85 | 0.03 % | 46.26 % | 6.14 % | 47.56 % | Blood pH |
| 8.000 | 1.81 | 0.01 % | 48.51 % | 1.62 % | 49.87 % | |
| 9.000 | 1.80 | 0.00 % | 49.23 % | 0.16 % | 50.61 % | |
| 10.000 | 1.80 | 0.00 % | 49.30 % | 0.02 % | 50.68 % | |
| 11.000 | 1.80 | 0.00 % | 49.31 % | 0.00 % | 50.69 % | |
| 12.000 | 1.80 | 0.00 % | 49.31 % | 0.00 % | 50.69 % | |

#### Carbonate and acidity

| | |
|---|---|
| Carbonate | 0.000 mM |
| Acidity error | -0.443 mM |

#### Other graphs

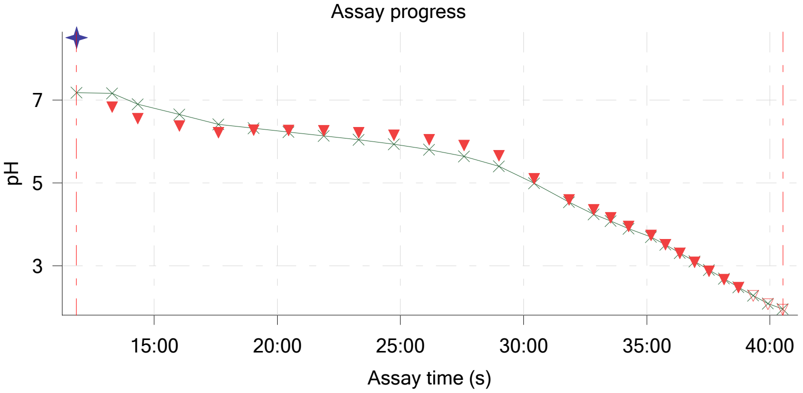


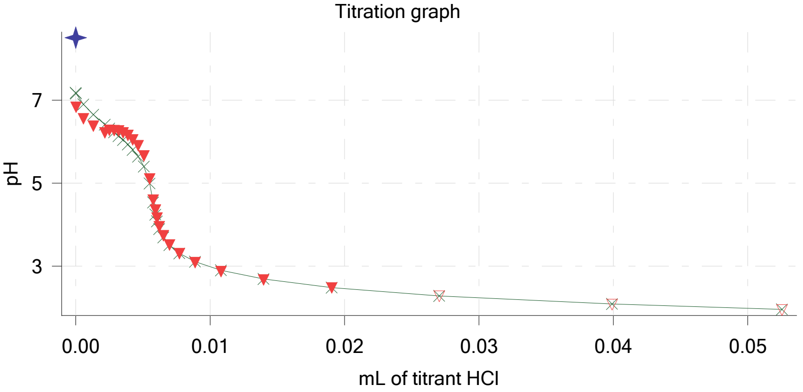

Sample name: **M08_octanol**
Assay name: **pH-metric high logP**
Assay ID: **18B-28014**
Filename: **G:\OpenLab_PFA_Sirius-T3-2\Data2018\18B-28014_M08_octanol_pH-metric high logP.t3r**

Experiment start time: **2/28/2018 9:57:03 PM**
Analyst: **Pion**
Instrument ID: **T312060**

#### Other graphs (continued)

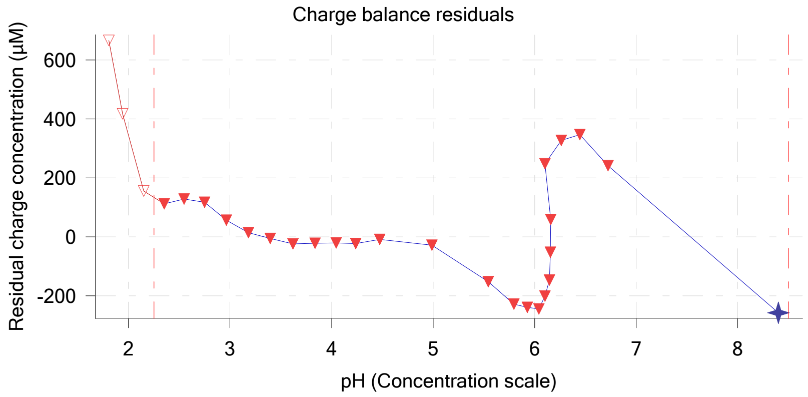


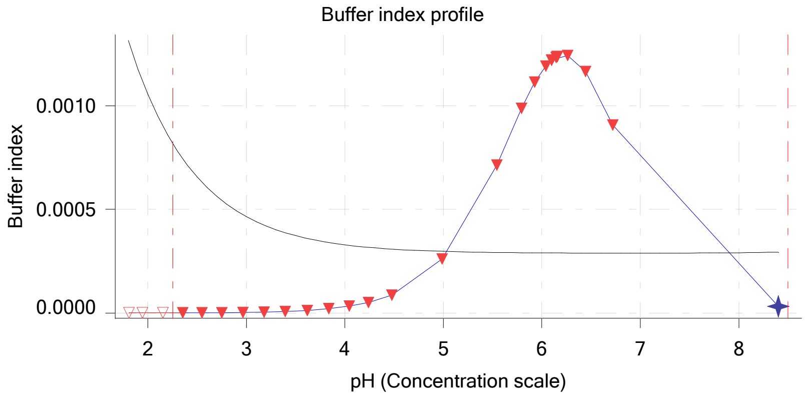


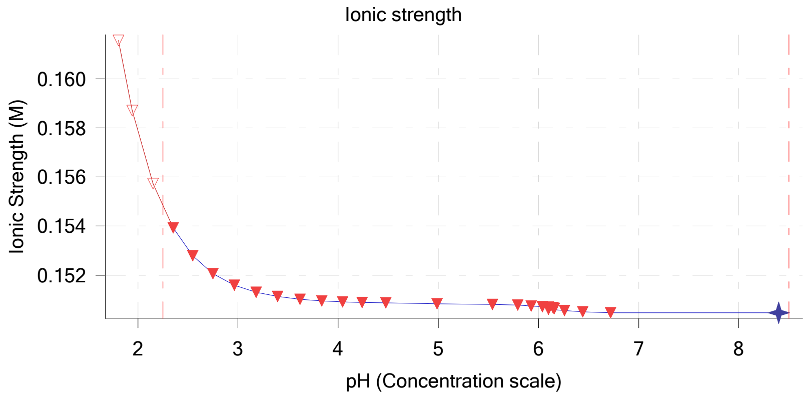


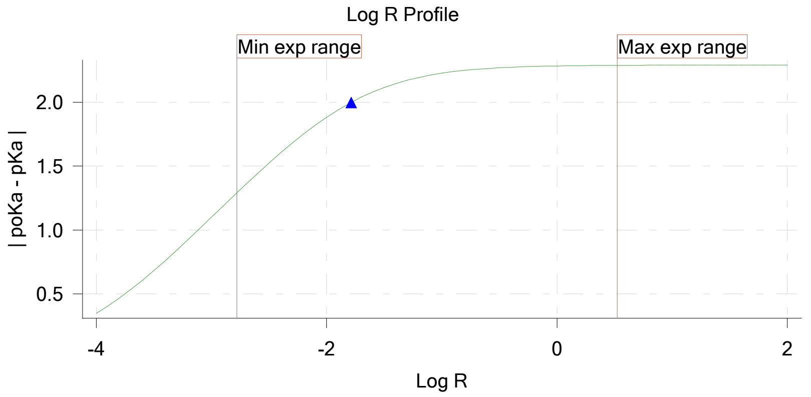


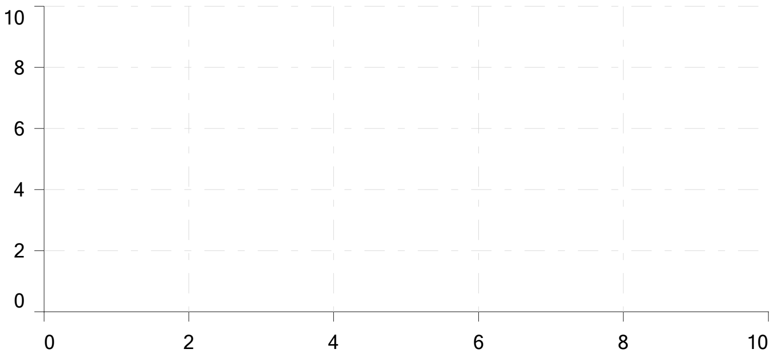


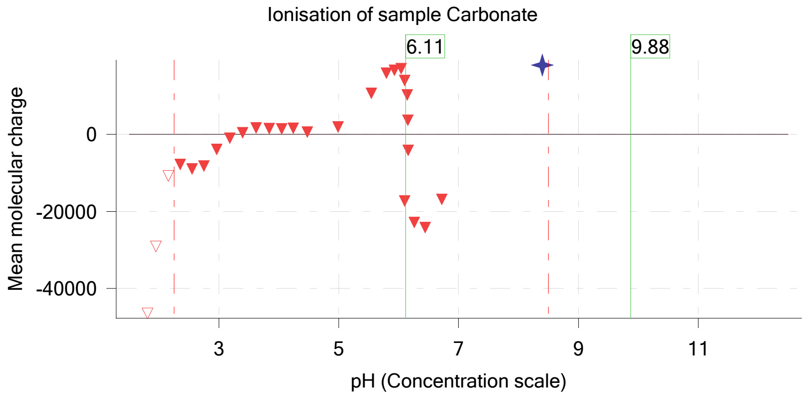

| | | | |
|---|---|---|---|
| Sample name: | **M08_octanol** | Experiment start time: | **2/28/2018 9:57:03 PM** |
| Assay name: | **pH-metric high logP** | Analyst: | **Pion** |
| Assay ID: | **18B-28014** | Instrument ID: | **T312060** |
| Filename: | **G:\OpenLab_PFA_Sirius-T3-2\Data2018\18B-28014_M08_octanol_pH-metric high logP.t3r** | | |

### pH-metric high logP Titration 2 of 3 18B-28014 Points 30 to 68

#### Overall results

| | |
|---|---|
| RMSD | 0.274 |
| Average ionic strength | 0.157 M |
| Average temperature | 25.0°C |
| Partition ratio | 0.0399 : 1 |
| Analyte concentration range | 2118.8 µM to 2194.1 µM |
| Total points considered | 29 of 39 |

#### Warnings and errors

| | |
|---|---|
| Errors | None |
| Warnings | None |

#### Four-Plus parameters

| | | | |
|---|---|---|---|
| Alpha | 0.130 | 2/28/2018 9:57:03 PM | C:\Sirius_T3\HCl18B27.t3r |
| S | 0.9970 | 2/28/2018 9:57:03 PM | C:\Sirius_T3\HCl18B27.t3r |
| jH | 0.8 | 2/28/2018 9:57:03 PM | C:\Sirius_T3\HCl18B27.t3r |
| jOH | -0.4 | 2/28/2018 9:57:03 PM | C:\Sirius_T3\HCl18B27.t3r |

#### Titrants

| | | | |
|---|---|---|---|
| 0.50 M HCl | 0.993513 | 2/28/2018 9:57:03 PM | C:\Sirius_T3\HCl18B27.t3r |
| 0.50 M KOH | 0.999845 | 2/28/2018 9:57:03 PM | C:\Sirius_T3\KOH18B27.t3r |

#### Sample

| | |
|---|---|
| M08_octanol concentration factor | 1.148 |
| Acid pKa 1 | 4.22 |
| logP (neutral XH) | 3.62 |
| logP (X -) | 1.80 |

#### Sample graphs

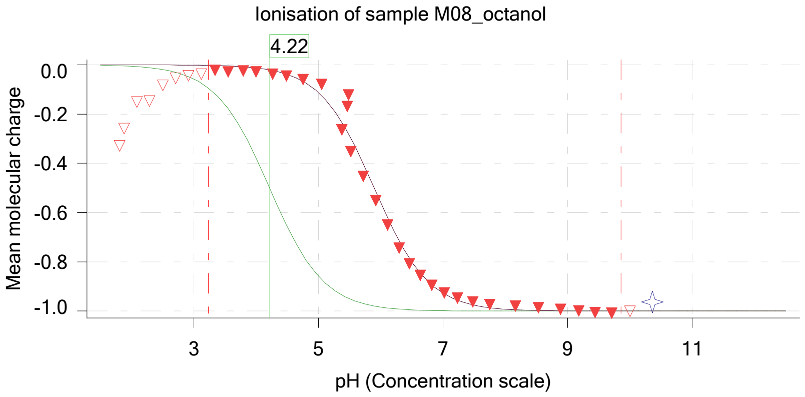


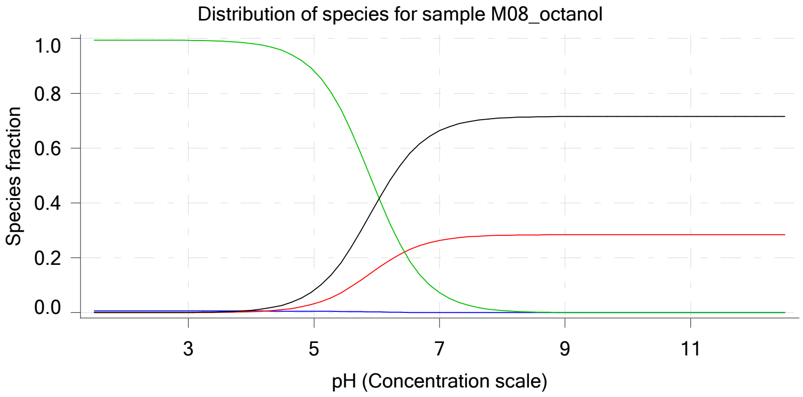

| | | | |
|---|---|---|---|
| Sample name: | **M08_octanol** | Experiment start time: | **2/28/2018 9:57:03 PM** |
| Assay name: | **pH-metric high logP** | Analyst: | **Pion** |
| Assay ID: | **18B-28014** | Instrument ID: | **T312060** |
| Filename: | **G:\OpenLab_PFA_Sirius-T3-2\Data2018\18B-28014_M08_octanol_pH-metric high logP.t3r** | | |

#### Sample graphs (continued)

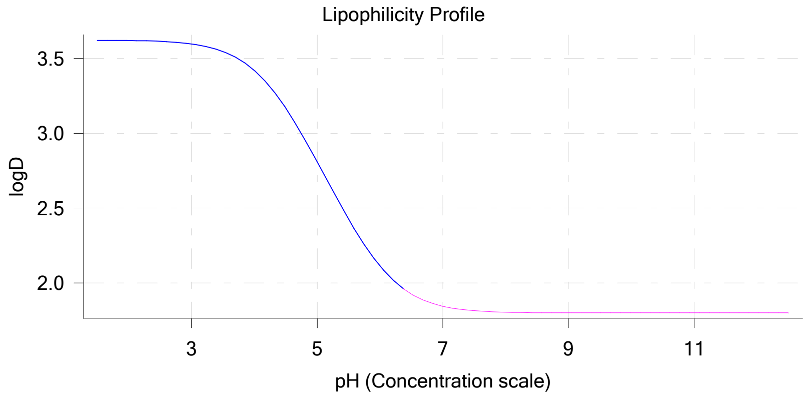


#### Sample logD and percent species

| pH | M08_octanol logD | M08_octanol M08_octanolH | M08_octanol M08_octanol | M08_octanol M08_octanolH* | M08_octanol M08_octanol* | Comment |
|---|---|---|---|---|---|---|
| 1.000 | 3.62 | 0.59 % | 0.00 % | 99.40 % | 0.00 % | |
| 1.200 | 3.62 | 0.59 % | 0.00 % | 99.40 % | 0.00 % | Stomach pH |
| 2.000 | 3.62 | 0.59 % | 0.00 % | 99.39 % | 0.01 % | |
| 3.000 | 3.60 | 0.59 % | 0.04 % | 99.28 % | 0.09 % | |
| 4.000 | 3.42 | 0.59 % | 0.35 % | 98.17 % | 0.89 % | |
| 5.000 | 2.81 | 0.53 % | 3.18 % | 88.28 % | 8.01 % | |
| 6.000 | 2.12 | 0.26 % | 15.85 % | 43.97 % | 39.91 % | |
| 6.500 | 1.93 | 0.12 % | 22.73 % | 19.93 % | 57.22 % | |
| 7.000 | 1.84 | 0.04 % | 26.34 % | 7.31 % | 66.31 % | |
| 7.400 | 1.82 | 0.02 % | 27.56 % | 3.04 % | 69.38 % | Blood pH |
| 8.000 | 1.80 | 0.00 % | 28.21 % | 0.78 % | 71.01 % | |
| 9.000 | 1.80 | 0.00 % | 28.41 % | 0.08 % | 71.51 % | |
| 10.000 | 1.80 | 0.00 % | 28.43 % | 0.01 % | 71.56 % | |
| 11.000 | 1.80 | 0.00 % | 28.43 % | 0.00 % | 71.57 % | |
| 12.000 | 1.80 | 0.00 % | 28.43 % | 0.00 % | 71.57 % | |

#### Carbonate and acidity

| | |
|---|---|
| Carbonate | 0.072 mM |
| Acidity error | -0.453 mM |

#### Other graphs

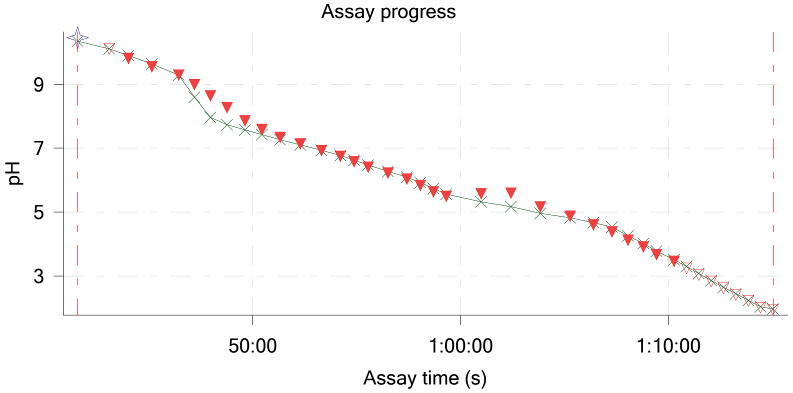


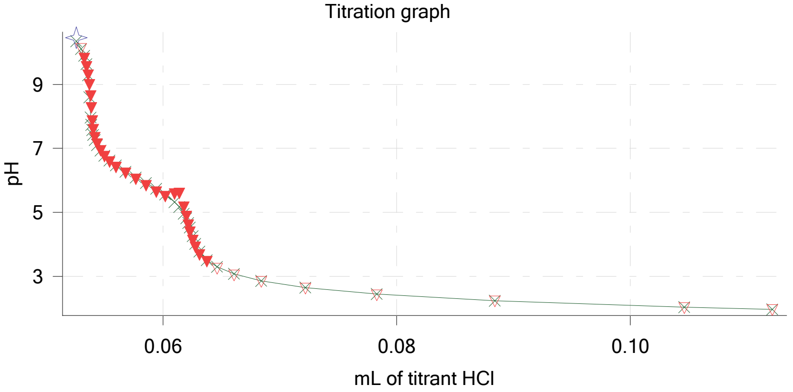

Sample name: **M08_octanol**
Assay name: **pH-metric high logP**
Assay ID: **18B-28014**
Filename: **G:\OpenLab_PFA_Sirius-T3-2\Data2018\18B-28014_M08_octanol_pH-metric high logP.t3r**

Experiment start time: **2/28/2018 9:57:03 PM**
Analyst: **Pion**
Instrument ID: **T312060**

#### Other graphs (continued)

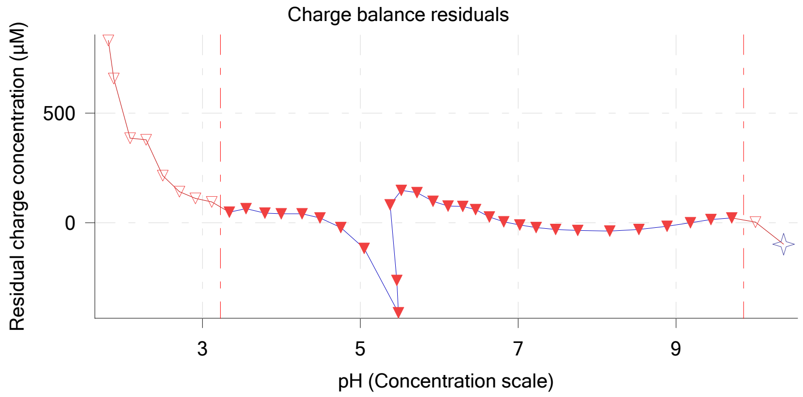


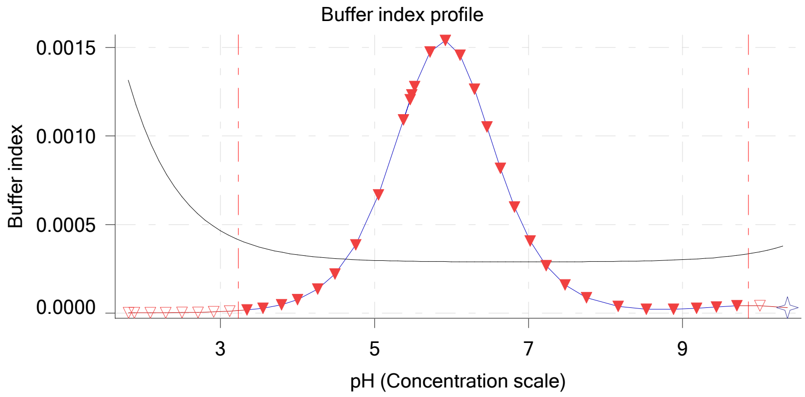


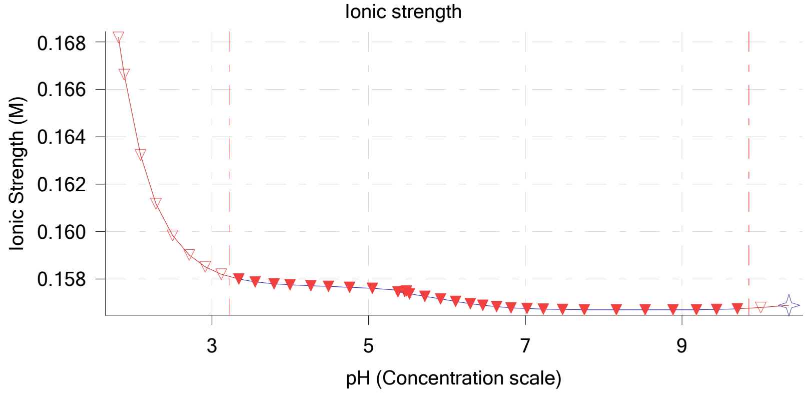


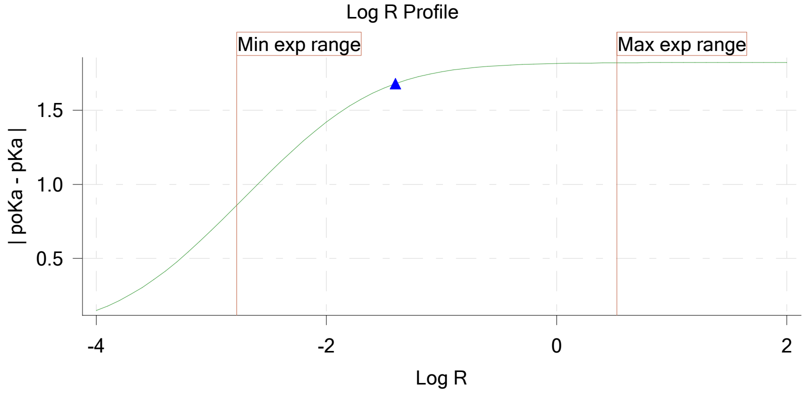


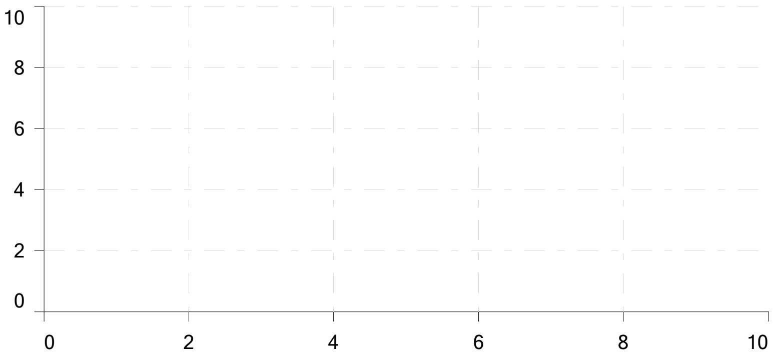


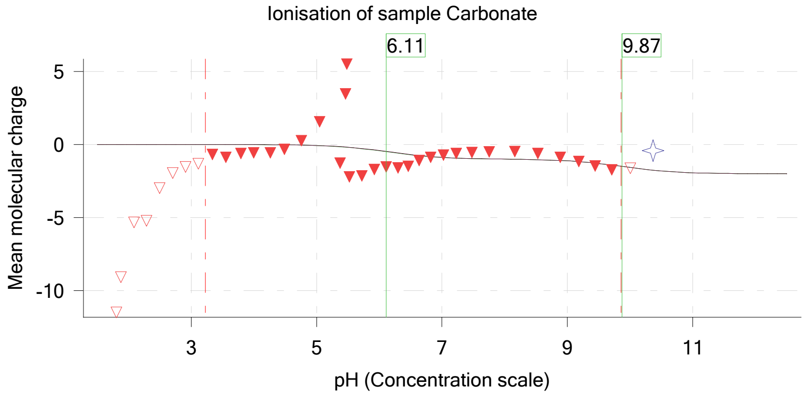

| | | | |
|---|---|---|---|
| Sample name: | **M08_octanol** | Experiment start time: | **2/28/2018 9:57:03 PM** |
| Assay name: | **pH-metric high logP** | Analyst: | **Pion** |
| Assay ID: | **18B-28014** | Instrument ID: | **T312060** |
| Filename: | **G:\OpenLab_PFA_Sirius-T3-2\Data2018\18B-28014_M08_octanol_pH-metric high logP.t3r** | | |

# pH-metric high logP   Titration 3 of 3   18B-28014 Points 69 to 110

#### Overall results

| | |
|---|---|
| RMSD | 0.091 |
| Average ionic strength | 0.164 M |
| Average temperature | 25.0°C |
| Partition ratio | 0.3243 : 1 |
| Analyte concentration range | 1558.1 µM to 1601.9 µM |
| Total points considered | 27 of 42 |

#### Warnings and errors

| | |
|---|---|
| Errors | None |
| Warnings | None |

#### Four-Plus parameters

| | | | |
|---|---|---|---|
| Alpha | 0.130 | 2/28/2018 9:57:03 PM | C:\Sirius_T3\HCl18B27.t3r |
| S | 0.9970 | 2/28/2018 9:57:03 PM | C:\Sirius_T3\HCl18B27.t3r |
| jH | 0.8 | 2/28/2018 9:57:03 PM | C:\Sirius_T3\HCl18B27.t3r |
| jOH | -0.4 | 2/28/2018 9:57:03 PM | C:\Sirius_T3\HCl18B27.t3r |

#### Titrants

| | | | |
|---|---|---|---|
| 0.50 M HCl | 0.993513 | 2/28/2018 9:57:03 PM | C:\Sirius_T3\HCl18B27.t3r |
| 0.50 M KOH | 0.999845 | 2/28/2018 9:57:03 PM | C:\Sirius_T3\KOH18B27.t3r |

#### Sample

| | |
|---|---|
| M08_octanol concentration factor | 0.843 |
| Acid pKa 1 | 4.22 |
| logP (neutral XH) | 4.34 |
| logP (X -) | 1.80 |

#### Sample graphs

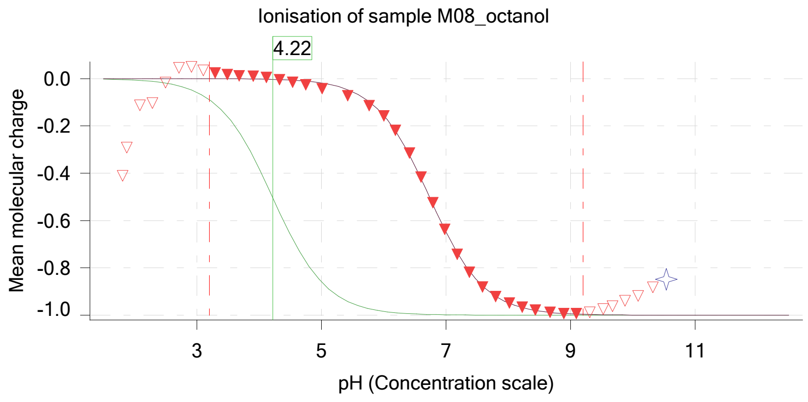


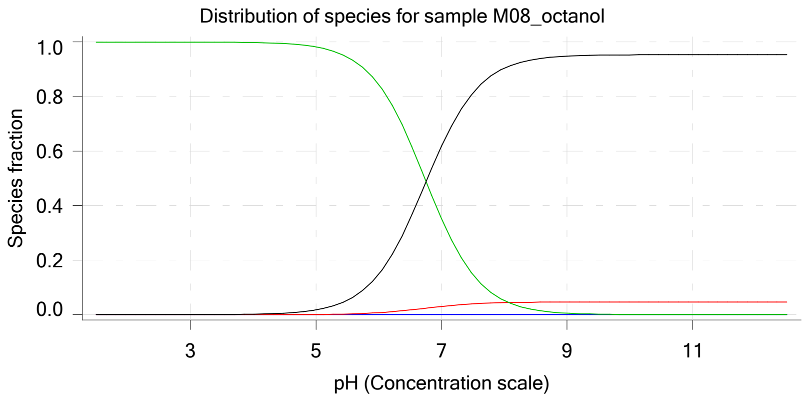

| | | | |
|---|---|---|---|
| Sample name: | **M08_octanol** | Experiment start time: | **2/28/2018 9:57:03 PM** |
| Assay name: | **pH-metric high logP** | Analyst: | **Pion** |
| Assay ID: | **18B-28014** | Instrument ID: | **T312060** |
| Filename: | **G:\OpenLab_PFA_Sirius-T3-2\Data2018\18B-28014_M08_octanol_pH-metric high logP.t3r** | | |

#### Sample graphs (continued)

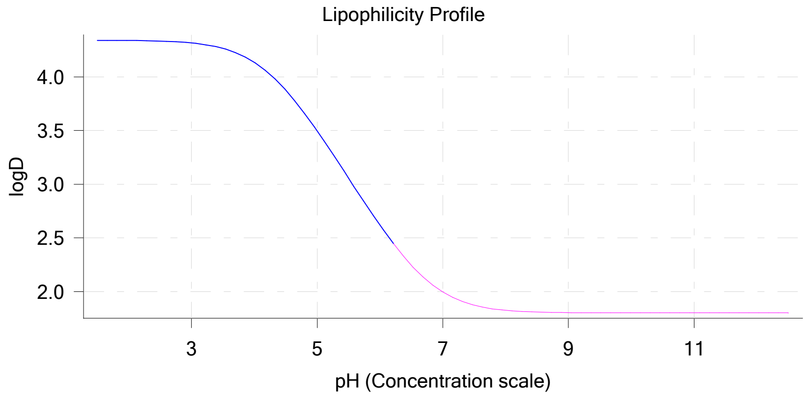


#### Sample logD and percent species

| pH | M08_octanol logD | M08_octanol M08_octanolH | M08_octanol M08_octanol | M08_octanol M08_octanolH* | M08_octanol M08_octanol* | Comment |
|---|---|---|---|---|---|---|
| 1.000 | 4.34 | 0.01 % | 0.00 % | 99.99 % | 0.00 % | |
| 1.200 | 4.34 | 0.01 % | 0.00 % | 99.99 % | 0.00 % | Stomach pH |
| 2.000 | 4.34 | 0.01 % | 0.00 % | 99.98 % | 0.00 % | |
| 3.000 | 4.32 | 0.01 % | 0.00 % | 99.97 % | 0.02 % | |
| 4.000 | 4.14 | 0.01 % | 0.01 % | 99.81 % | 0.17 % | |
| 5.000 | 3.50 | 0.01 % | 0.08 % | 98.21 % | 1.69 % | |
| 6.000 | 2.63 | 0.01 % | 0.71 % | 84.66 % | 14.61 % | |
| 6.500 | 2.25 | 0.01 % | 1.70 % | 63.59 % | 34.70 % | |
| 7.000 | 2.00 | 0.00 % | 3.00 % | 35.59 % | 61.41 % | |
| 7.400 | 1.89 | 0.00 % | 3.82 % | 18.03 % | 78.15 % | Blood pH |
| 8.000 | 1.82 | 0.00 % | 4.42 % | 5.24 % | 90.35 % | |
| 9.000 | 1.80 | 0.00 % | 4.63 % | 0.55 % | 94.82 % | |
| 10.000 | 1.80 | 0.00 % | 4.66 % | 0.06 % | 95.29 % | |
| 11.000 | 1.80 | 0.00 % | 4.66 % | 0.01 % | 95.33 % | |
| 12.000 | 1.80 | 0.00 % | 4.66 % | 0.00 % | 95.34 % | |

#### Carbonate and acidity

| | |
|---|---|
| Carbonate | 0.497 mM |
| Acidity error | -0.179 mM |

#### Other graphs

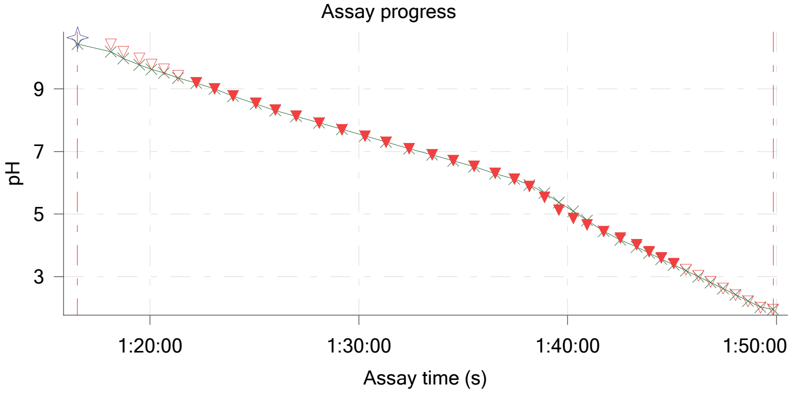


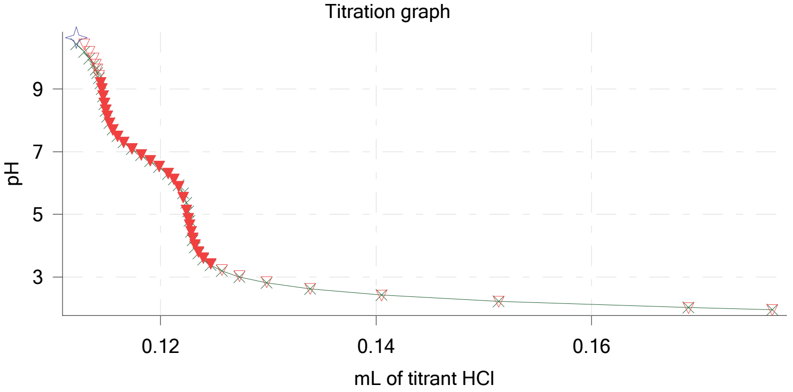

| | | | |
|---|---|---|---|
| Sample name: | **M08_octanol** | Experiment start time: | **2/28/2018 9:57:03 PM** |
| Assay name: | **pH-metric high logP** | Analyst: | **Pion** |
| Assay ID: | **18B-28014** | Instrument ID: | **T312060** |
| Filename: | **G:\OpenLab_PFA_Sirius-T3-2\Data2018\18B-28014_M08_octanol_pH-metric high logP.t3r** | | |

#### Other graphs (continued)

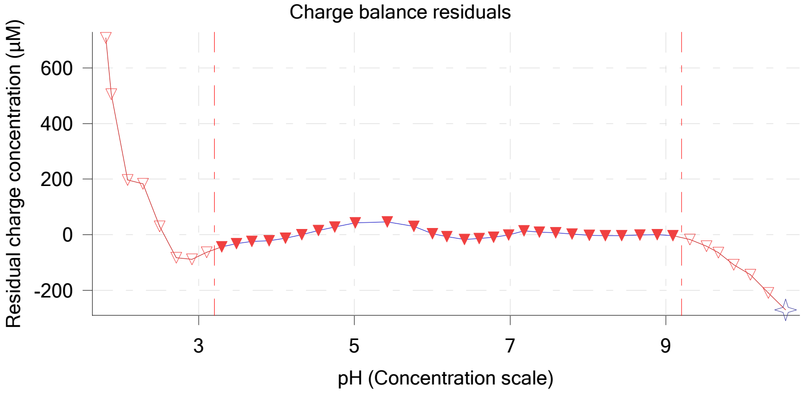


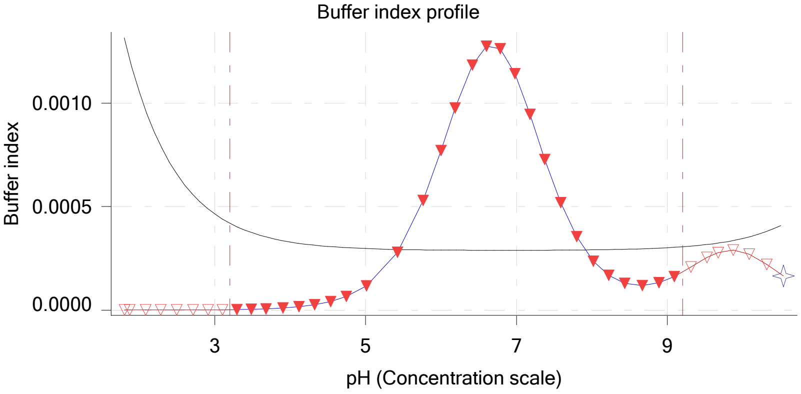


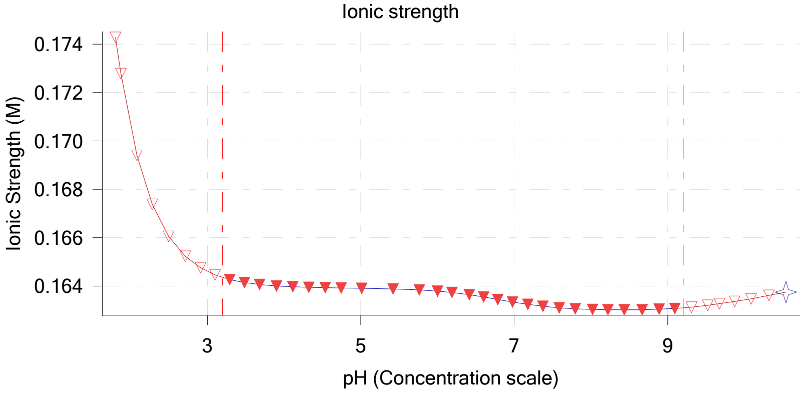


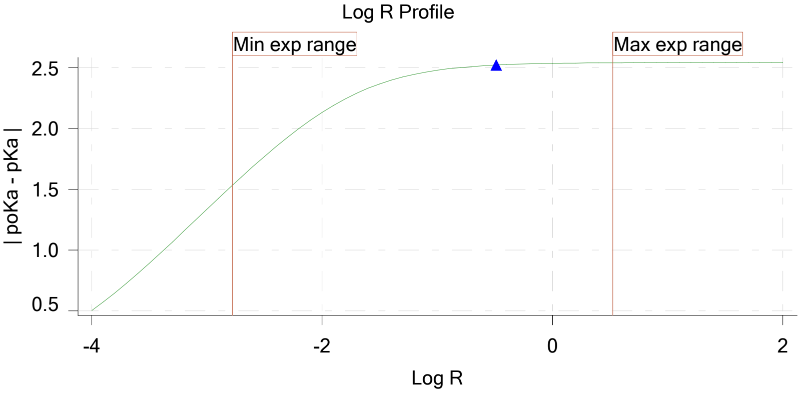


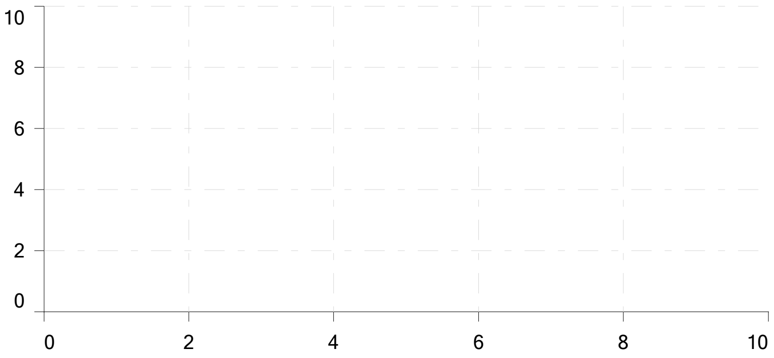


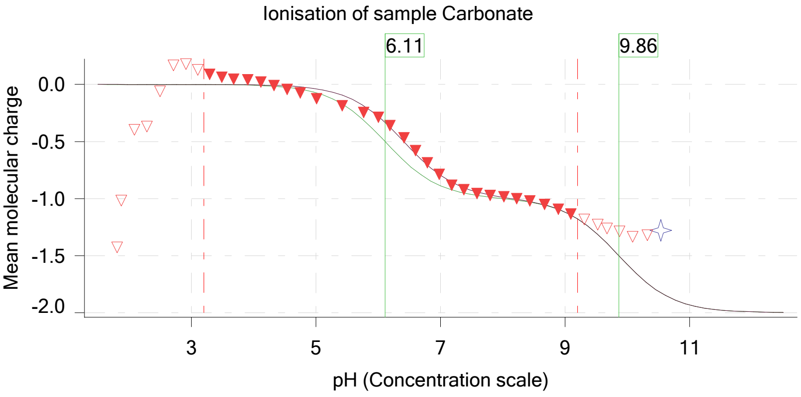

| | | | |
|---|---|---|---|
| Sample name: | **M08_octanol** | Experiment start time: | **2/28/2018 9:57:03 PM** |
| Assay name: | **pH-metric high logP** | Analyst: | **Pion** |
| Assay ID: | **18B-28014** | Instrument ID: | **T312060** |
| Filename: | **G:\OpenLab_PFA_Sirius-T3-2\Data2018\18B-28014_M08_octanol_pH-metric high logP.t3r** | | |

#### Assay Model

| Settings | Value | Date/Time changed | Imported from |
|---|---|---|---|
| Sample name | M08_octanol | 2/27/2018 5:33:51 PM | User entered value |
| Sample by | Weight | | Default value |
| Sample weight | 0.001080 g | 2/28/2018 5:24:25 PM | User entered value |
| Formula weight | 293.32 g/mol | 2/27/2018 5:33:51 PM | User entered value |
| Solubility | Unknown | | Default value |
| Molecular weight | 293.32 | 2/27/2018 5:33:51 PM | User entered value |
| Individual pKa ionic environments | No | | Default value |
| Number of pKas | 1 | 2/27/2018 5:33:51 PM | User entered value |
| Sample is a | Acid | 2/27/2018 5:33:51 PM | User entered value |
| pKa 1 | 4.22 | 2/27/2018 5:33:51 PM | User entered value |
| logP (neutral XH) | 3.45 | 2/28/2018 2:34:15 PM | User entered value |
| logP (X -) | 1.80 | 2/28/2018 2:34:50 PM | User entered value |

#### Events

| Time | Event | Water | Acid | Base | Octanol | pH | dpH/dt | pH R-squared | pH SD | dpH/[illegible] time |
|---|---|---|---|---|---|---|---|---|---|---|
| 8:58.1 | Manual volume addition | | | | 0.02500 mL | | | | | |
| 8:59.2 | Initial pH = 5.75 | | | | | | | | | |
| 11:51.1 | Data point 2 | 1.50000 mL | 0.00000 mL | 0.00437 mL | 0.02500 mL | 8.505 | -0.38369 | 0.99778 | 0.01897 | Timed out at |
| 13:17.8 | Data point 3 | 1.50000 mL | 0.00002 mL | 0.00437 mL | 0.02500 mL | 6.831 | 0.01922 | 0.90141 | 0.00100 | 32.0 s |
| 14:20.5 | Data point 4 | 1.50000 mL | 0.00059 mL | 0.00437 mL | 0.02500 mL | 6.554 | 0.05114 | 0.99769 | 0.00253 | Timed out at |
| 16:01.3 | Data point 5 | 1.50000 mL | 0.00132 mL | 0.00437 mL | 0.02500 mL | 6.372 | 0.07459 | 0.99483 | 0.00369 | Timed out at |
| 17:37.0 | Data point 6 | 1.50000 mL | 0.00216 mL | 0.00437 mL | 0.02500 mL | 6.212 | 0.09858 | 0.99770 | 0.00487 | Timed out at |
| 19:02.5 | Data point 7 | 1.50000 mL | 0.00252 mL | 0.00437 mL | 0.02500 mL | 6.270 | 0.08713 | 0.99504 | 0.00431 | Timed out at |
| 20:27.8 | Data point 8 | 1.50000 mL | 0.00287 mL | 0.00437 mL | 0.02500 mL | 6.267 | 0.08243 | 0.99748 | 0.00408 | Timed out at |
| 21:53.3 | Data point 9 | 1.50000 mL | 0.00320 mL | 0.00437 mL | 0.02500 mL | 6.255 | 0.08261 | 0.99774 | 0.00409 | Timed out at |
| 23:18.7 | Data point 10 | 1.50000 mL | 0.00353 mL | 0.00437 mL | 0.02500 mL | 6.212 | 0.07670 | 0.99623 | 0.00380 | Timed out at |
| 24:44.1 | Data point 11 | 1.50000 mL | 0.00388 mL | 0.00437 mL | 0.02500 mL | 6.152 | 0.08592 | 0.99776 | 0.00425 | Timed out at |
| 26:09.5 | Data point 12 | 1.50000 mL | 0.00426 mL | 0.00437 mL | 0.02500 mL | 6.040 | 0.08353 | 0.99742 | 0.00413 | Timed out at |
| 27:34.9 | Data point 13 | 1.50000 mL | 0.00466 mL | 0.00437 mL | 0.02500 mL | 5.906 | 0.08469 | 0.99736 | 0.00419 | Timed out at |
| 29:00.3 | Data point 14 | 1.50000 mL | 0.00508 mL | 0.00437 mL | 0.02500 mL | 5.655 | 0.08514 | 0.99629 | 0.00421 | Timed out at |

| | | | |
|---|---|---|---|
| Sample name: | **M08_octanol** | Experiment start time: | **2/28/2018 9:57:03 PM** |
| Assay name: | **pH-metric high logP** | Analyst: | **Pion** |
| Assay ID: | **18B-28014** | Instrument ID: | **T312060** |
| Filename: | **G:\OpenLab_PFA_Sirius-T3-2\Data2018\18B-28014_M08_octanol_pH-metric high logP.t3r** | | |

#### Events (continued)

| Time | Event | Water | Acid | Base | Octanol | pH | dpH/dt | pH R-squared | pH SD | dpH/dt time |
|---|---|---|---|---|---|---|---|---|---|---|
| 30:25.6 | Data point 15 | 1.50000 mL | 0.00550 mL | 0.00437 mL | 0.02500 mL | 5.102 | 0.04617 | 0.99374 | 0.00229 | Timed out at 59.5 s |
| 31:51.0 | Data point 16 | 1.50000 mL | 0.00576 mL | 0.00437 mL | 0.02500 mL | 4.592 | 0.01831 | 0.87650 | 0.00097 | 24.0 s |
| 32:50.6 | Data point 17 | 1.50000 mL | 0.00593 mL | 0.00437 mL | 0.02500 mL | 4.356 | 0.01879 | 0.87405 | 0.00099 | 16.0 s |
| 33:32.0 | Data point 18 | 1.50000 mL | 0.00604 mL | 0.00437 mL | 0.02500 mL | 4.163 | 0.01848 | 0.95772 | 0.00093 | 13.5 s |
| 34:16.1 | Data point 19 | 1.50000 mL | 0.00623 mL | 0.00437 mL | 0.02500 mL | 3.955 | 0.01959 | 0.95038 | 0.00099 | 13.0 s |
| 35:10.1 | Data point 20 | 1.50000 mL | 0.00654 mL | 0.00437 mL | 0.02500 mL | 3.739 | 0.01840 | 0.93630 | 0.00094 | 10.0 s |
| 35:45.5 | Data point 21 | 1.50000 mL | 0.00698 mL | 0.00437 mL | 0.02500 mL | 3.515 | 0.01107 | 0.91474 | 0.00057 | 10.0 s |
| 36:20.9 | Data point 22 | 1.50000 mL | 0.00771 mL | 0.00437 mL | 0.02500 mL | 3.301 | -0.00792 | 0.20574 | 0.00086 | 10.0 s |
| 36:56.3 | Data point 23 | 1.50000 mL | 0.00889 mL | 0.00437 mL | 0.02500 mL | 3.086 | -0.00002 | 0.00010 | 0.00009 | 10.0 s |
| 37:31.8 | Data point 24 | 1.50000 mL | 0.01082 mL | 0.00437 mL | 0.02500 mL | 2.873 | -0.00479 | 0.63485 | 0.00030 | 10.5 s |
| 38:07.8 | Data point 25 | 1.50000 mL | 0.01399 mL | 0.00437 mL | 0.02500 mL | 2.674 | -0.01143 | 0.54403 | 0.00077 | 10.0 s |
| 38:43.4 | Data point 26 | 1.50000 mL | 0.01905 mL | 0.00437 mL | 0.02500 mL | 2.478 | -0.01045 | 0.80609 | 0.00057 | 10.0 s |
| 39:19.0 | Data point 27 | 1.50000 mL | 0.02705 mL | 0.00437 mL | 0.02500 mL | 2.280 | -0.01095 | 0.68427 | 0.00065 | 10.0 s |
| 39:54.7 | Data point 28 | 1.50000 mL | 0.03991 mL | 0.00437 mL | 0.02500 mL | 2.076 | -0.01774 | 0.88391 | 0.00093 | 10.5 s |
| 40:31.0 | Data point 29 | 1.50000 mL | 0.05256 mL | 0.00437 mL | 0.02500 mL | 1.944 | -0.01689 | 0.91021 | 0.00087 | 10.0 s |
| 41:34.8 | Data point 30 | 1.50000 mL | 0.05256 mL | 0.06058 mL | 0.06500 mL | 10.463 | 0.01671 | 0.80475 | 0.00092 | 55.5 s |
| 43:06.2 | Data point 31 | 1.50000 mL | 0.05299 mL | 0.06058 mL | 0.06500 mL | 10.105 | -0.00088 | 0.00194 | 0.00098 | 30.0 s |
| 44:01.6 | Data point 32 | 1.50000 mL | 0.05325 mL | 0.06058 mL | 0.06500 mL | 9.808 | -0.01675 | 0.76998 | 0.00094 | 37.0 s |
| 45:09.2 | Data point 33 | 1.50000 mL | 0.05343 mL | 0.06058 mL | 0.06500 mL | 9.543 | -0.01717 | 0.74125 | 0.00098 | 41.5 s |
| 46:26.4 | Data point 34 | 1.50000 mL | 0.05357 mL | 0.06058 mL | 0.06500 mL | 9.285 | -0.00869 | 0.87535 | 0.00046 | 10.0 s |
| 47:12.0 | Data point 35 | 1.50000 mL | 0.05369 mL | 0.06058 mL | 0.06500 mL | 8.988 | 0.00397 | 0.06470 | 0.00077 | 10.0 s |
| 47:57.7 | Data point 36 | 1.50000 mL | 0.05379 mL | 0.06058 mL | 0.06500 mL | 8.634 | 0.01554 | 0.63385 | 0.00096 | 13.0 s |
| 48:46.3 | Data point 37 | 1.50000 mL | 0.05386 mL | 0.06058 mL | 0.06500 mL | 8.268 | 0.01699 | 0.72054 | 0.00099 | 16.0 s |
| 49:38.0 | Data point 38 | 1.50000 mL | 0.05393 mL | 0.06058 mL | 0.06500 mL | 7.860 | 0.01563 | 0.77436 | 0.00088 | 18.0 s |
| 50:26.5 | Data point 39 | 1.50000 mL | 0.05402 mL | 0.06058 mL | 0.06500 mL | 7.583 | 0.01366 | 0.60795 | 0.00087 | 17.5 s |
| 51:19.7 | Data point 40 | 1.50000 mL | 0.05416 mL | 0.06058 mL | 0.06500 mL | 7.337 | 0.01707 | 0.87652 | 0.00090 | 17.5 s |
| 52:18.0 | Data point 41 | 1.50000 mL | 0.05435 mL | 0.06058 mL | 0.06500 mL | 7.132 | 0.01352 | 0.65718 | 0.00082 | 15.0 s |
| 53:19.0 | Data point 42 | 1.50000 mL | 0.05463 mL | 0.06058 mL | 0.06500 mL | 6.929 | 0.01590 | 0.72033 | 0.00093 | 18.0 s |
| 54:12.7 | Data point 43 | 1.50000 mL | 0.05499 mL | 0.06058 mL | 0.06500 mL | 6.745 | 0.01686 | 0.80981 | 0.00093 | 15.0 s |
| 54:53.1 | Data point 44 | 1.50000 mL | 0.05541 mL | 0.06058 mL | 0.06500 mL | 6.570 | 0.01319 | 0.58565 | 0.00085 | 15.0 s |
| 55:33.5 | Data point 45 | 1.50000 mL | 0.05597 mL | 0.06058 mL | 0.06500 mL | 6.408 | 0.01755 | 0.91751 | 0.00090 | 16.0 s |
| 56:30.4 | Data point 46 | 1.50000 mL | 0.05680 mL | 0.06058 mL | 0.06500 mL | 6.222 | 0.01824 | 0.88012 | 0.00096 | 14.0 s |
| 57:25.4 | Data point 47 | 1.50000 mL | 0.05767 mL | 0.06058 mL | 0.06500 mL | 6.032 | 0.01601 | 0.70284 | 0.00094 | 13.0 s |
| 58:03.8 | Data point 48 | 1.50000 mL | 0.05854 mL | 0.06058 mL | 0.06500 mL | 5.832 | 0.01586 | 0.73469 | 0.00091 | 12.0 s |
| 58:41.4 | Data point 49 | 1.50000 mL | 0.05941 mL | 0.06058 mL | 0.06500 mL | 5.634 | 0.01736 | 0.78548 | 0.00097 | 12.0 s |
| 59:18.8 | Data point 50 | 1.50000 mL | 0.06018 mL | 0.06058 mL | 0.06500 mL | 5.490 | 0.10872 | 0.99860 | 0.00537 | Timed out at 59.5 s |
| 1:00:59.7 | Data point 51 | 1.50000 mL | 0.06098 mL | 0.06058 mL | 0.06500 mL | 5.576 | 0.16163 | 0.99800 | 0.00799 | Timed out at 59.5 s |
| 1:02:25.1 | Data point 52 | 1.50000 mL | 0.06138 mL | 0.06058 mL | 0.06500 mL | 5.597 | 0.09780 | 0.99796 | 0.00484 | Timed out at 59.5 s |
| 1:03:50.6 | Data point 53 | 1.50000 mL | 0.06178 mL | 0.06058 mL | 0.06500 mL | 5.164 | 0.03874 | 0.98187 | 0.00193 | Timed out at 59.5 s |
| 1:05:15.9 | Data point 54 | 1.50000 mL | 0.06199 mL | 0.06058 mL | 0.06500 mL | 4.871 | 0.01890 | 0.95104 | 0.00096 | 37.5 s |
| 1:06:24.0 | Data point 55 | 1.50000 mL | 0.06216 mL | 0.06058 mL | 0.06500 mL | 4.605 | 0.02009 | 0.98775 | 0.00100 | 22.5 s |
| 1:07:17.2 | Data point 56 | 1.50000 mL | 0.06230 mL | 0.06058 mL | 0.06500 mL | 4.379 | 0.01733 | 0.86039 | 0.00092 | 15.5 s |
| 1:08:03.2 | Data point 57 | 1.50000 mL | 0.06254 mL | 0.06058 mL | 0.06500 mL | 4.117 | 0.01748 | 0.84391 | 0.00094 | 14.0 s |
| 1:08:47.7 | Data point 58 | 1.50000 mL | 0.06277 mL | 0.06058 mL | 0.06500 mL | 3.910 | 0.01951 | 0.98119 | 0.00097 | 12.5 s |
| 1:09:25.6 | Data point 59 | 1.50000 mL | 0.06312 mL | 0.06058 mL | 0.06500 mL | 3.670 | 0.01603 | 0.74774 | 0.00092 | 10.0 s |
| 1:10:16.5 | Data point 60 | 1.50000 mL | 0.06376 mL | 0.06058 mL | 0.06500 mL | 3.462 | 0.00750 | 0.80522 | 0.00041 | 10.0 s |
| 1:10:52.0 | Data point 61 | 1.50000 mL | 0.06463 mL | 0.06058 mL | 0.06500 mL | 3.238 | -0.00145 | 0.00710 | 0.00085 | 10.0 s |
| 1:11:27.4 | Data point 62 | 1.50000 mL | 0.06609 mL | 0.06058 mL | 0.06500 mL | 3.035 | -0.00146 | 0.02277 | 0.00048 | 10.5 s |
| 1:12:03.4 | Data point 63 | 1.50000 mL | 0.06841 mL | 0.06058 mL | 0.06500 mL | 2.831 | -0.00808 | 0.27293 | 0.00076 | 10.0 s |
| 1:12:38.9 | Data point 64 | 1.50000 mL | 0.07218 mL | 0.06058 mL | 0.06500 mL | 2.624 | -0.00574 | 0.80349 | 0.00032 | 10.0 s |
| 1:13:14.6 | Data point 65 | 1.50000 mL | 0.07832 mL | 0.06058 mL | 0.06500 mL | 2.413 | -0.00805 | 0.73778 | 0.00046 | 10.0 s |
| 1:13:50.2 | Data point 66 | 1.50000 mL | 0.08841 mL | 0.06058 mL | 0.06500 mL | 2.216 | -0.00955 | 0.79291 | 0.00053 | 10.0 s |

sirius

| | | | |
|---|---|---|---|
| Sample name: | **M08_octanol** | Experiment start time: | **2/28/2018 9:57:03 PM** |
| Assay name: | **pH-metric high logP** | Analyst: | **Pion** |
| Assay ID: | **18B-28014** | Instrument ID: | **T312060** |
| Filename: | **G:\OpenLab_PFA_Sirius-T3-2\Data2018\18B-28014_M08_octanol_pH-metric high logP.t3r** | | |

#### Events (continued)

| Time | Event | Water | Acid | Base | Octanol | pH | dpH/dt | pH R-squared | pH SD | dpH/dt time |
|---|---|---|---|---|---|---|---|---|---|---|
| 1:14:26.0 | Data point 67 | 1.50000 mL | 0.10463 mL | 0.06058 mL | 0.06500 mL | 2.013 | -0.01476 | 0.90024 | 0.00077 | 10.0 s |
| 1:15:01.6 | Data point 68 | 1.50000 mL | 0.11218 mL | 0.06058 mL | 0.06500 mL | 1.944 | -0.00748 | 0.85449 | 0.00040 | 10.0 s |
| 1:16:32.0 | Data point 69 | 1.50000 mL | 0.11218 mL | 0.12131 mL | 0.56500 mL | 10.638 | -0.02692 | 0.97300 | 0.00135 | Timed out at 59.5 s |
| 1:18:07.8 | Data point 70 | 1.50000 mL | 0.11291 mL | 0.12131 mL | 0.56500 mL | 10.418 | -0.01629 | 0.90234 | 0.00085 | 10.5 s |
| 1:18:43.7 | Data point 71 | 1.50000 mL | 0.11341 mL | 0.12131 mL | 0.56500 mL | 10.185 | -0.00481 | 0.66052 | 0.00029 | 10.0 s |
| 1:19:29.4 | Data point 72 | 1.50000 mL | 0.11378 mL | 0.12131 mL | 0.56500 mL | 9.973 | 0.00415 | 0.69795 | 0.00025 | 10.0 s |
| 1:20:04.7 | Data point 73 | 1.50000 mL | 0.11399 mL | 0.12131 mL | 0.56500 mL | 9.775 | 0.00051 | 0.00415 | 0.00039 | 10.0 s |
| 1:20:40.0 | Data point 74 | 1.50000 mL | 0.11414 mL | 0.12131 mL | 0.56500 mL | 9.625 | 0.01049 | 0.80316 | 0.00058 | 10.0 s |
| 1:21:20.4 | Data point 75 | 1.50000 mL | 0.11430 mL | 0.12131 mL | 0.56500 mL | 9.413 | 0.01849 | 0.91674 | 0.00095 | 16.5 s |
| 1:22:12.7 | Data point 76 | 1.50000 mL | 0.11444 mL | 0.12131 mL | 0.56500 mL | 9.196 | 0.01358 | 0.49556 | 0.00095 | 17.5 s |
| 1:23:05.8 | Data point 77 | 1.50000 mL | 0.11456 mL | 0.12131 mL | 0.56500 mL | 8.994 | 0.01898 | 0.97137 | 0.00095 | 12.0 s |
| 1:23:58.6 | Data point 78 | 1.50000 mL | 0.11468 mL | 0.12131 mL | 0.56500 mL | 8.773 | 0.00423 | 0.05467 | 0.00089 | 19.5 s |
| 1:25:04.0 | Data point 79 | 1.50000 mL | 0.11479 mL | 0.12131 mL | 0.56500 mL | 8.540 | 0.01751 | 0.95118 | 0.00089 | 15.5 s |
| 1:26:00.5 | Data point 80 | 1.50000 mL | 0.11491 mL | 0.12131 mL | 0.56500 mL | 8.327 | 0.00999 | 0.26899 | 0.00095 | 18.5 s |
| 1:26:59.9 | Data point 81 | 1.50000 mL | 0.11505 mL | 0.12131 mL | 0.56500 mL | 8.125 | 0.00630 | 0.12857 | 0.00087 | 20.0 s |
| 1:28:05.9 | Data point 82 | 1.50000 mL | 0.11526 mL | 0.12131 mL | 0.56500 mL | 7.904 | 0.00961 | 0.25771 | 0.00093 | 19.5 s |
| 1:29:11.3 | Data point 83 | 1.50000 mL | 0.11557 mL | 0.12131 mL | 0.56500 mL | 7.692 | 0.01317 | 0.53307 | 0.00089 | 20.5 s |
| 1:30:17.8 | Data point 84 | 1.50000 mL | 0.11602 mL | 0.12131 mL | 0.56500 mL | 7.484 | 0.01066 | 0.33152 | 0.00091 | 19.5 s |
| 1:31:18.1 | Data point 85 | 1.50000 mL | 0.11658 mL | 0.12131 mL | 0.56500 mL | 7.286 | 0.01482 | 0.73005 | 0.00086 | 20.5 s |
| 1:32:24.8 | Data point 86 | 1.50000 mL | 0.11736 mL | 0.12131 mL | 0.56500 mL | 7.089 | 0.01351 | 0.56423 | 0.00089 | 19.5 s |
| 1:33:30.4 | Data point 87 | 1.50000 mL | 0.11820 mL | 0.12131 mL | 0.56500 mL | 6.896 | 0.01748 | 0.83538 | 0.00095 | 20.0 s |
| 1:34:31.3 | Data point 88 | 1.50000 mL | 0.11905 mL | 0.12131 mL | 0.56500 mL | 6.712 | 0.01401 | 0.48910 | 0.00099 | 19.0 s |
| 1:35:31.3 | Data point 89 | 1.50000 mL | 0.11987 mL | 0.12131 mL | 0.56500 mL | 6.525 | 0.00912 | 0.22976 | 0.00094 | 19.5 s |
| 1:36:31.7 | Data point 90 | 1.50000 mL | 0.12070 mL | 0.12131 mL | 0.56500 mL | 6.300 | 0.01131 | 0.37052 | 0.00092 | 19.5 s |
| 1:37:27.0 | Data point 91 | 1.50000 mL | 0.12124 mL | 0.12131 mL | 0.56500 mL | 6.113 | 0.01242 | 0.37784 | 0.00100 | 17.5 s |
| 1:38:09.9 | Data point 92 | 1.50000 mL | 0.12168 mL | 0.12131 mL | 0.56500 mL | 5.877 | 0.01326 | 0.59032 | 0.00085 | 18.0 s |
| 1:38:53.3 | Data point 93 | 1.50000 mL | 0.12211 mL | 0.12131 mL | 0.56500 mL | 5.536 | 0.01531 | 0.70704 | 0.00090 | 16.5 s |
| 1:39:35.3 | Data point 94 | 1.50000 mL | 0.12241 mL | 0.12131 mL | 0.56500 mL | 5.126 | 0.01004 | 0.33249 | 0.00086 | 15.5 s |
| 1:40:16.1 | Data point 95 | 1.50000 mL | 0.12258 mL | 0.12131 mL | 0.56500 mL | 4.862 | 0.01660 | 0.85540 | 0.00089 | 14.0 s |
| 1:40:55.6 | Data point 96 | 1.50000 mL | 0.12270 mL | 0.12131 mL | 0.56500 mL | 4.655 | 0.01286 | 0.68795 | 0.00077 | 12.5 s |
| 1:41:43.7 | Data point 97 | 1.50000 mL | 0.12284 mL | 0.12131 mL | 0.56500 mL | 4.442 | 0.01567 | 0.70157 | 0.00092 | 12.0 s |
| 1:42:31.3 | Data point 98 | 1.50000 mL | 0.12300 mL | 0.12131 mL | 0.56500 mL | 4.233 | 0.00457 | 0.07507 | 0.00082 | 11.0 s |
| 1:43:17.9 | Data point 99 | 1.50000 mL | 0.12321 mL | 0.12131 mL | 0.56500 mL | 4.023 | 0.01009 | 0.34406 | 0.00085 | 10.5 s |
| 1:43:53.8 | Data point 100 | 1.50000 mL | 0.12352 mL | 0.12131 mL | 0.56500 mL | 3.800 | 0.00859 | 0.33185 | 0.00074 | 10.0 s |
| 1:44:29.3 | Data point 101 | 1.50000 mL | 0.12397 mL | 0.12131 mL | 0.56500 mL | 3.606 | 0.01096 | 0.49543 | 0.00077 | 10.0 s |
| 1:45:04.7 | Data point 102 | 1.50000 mL | 0.12465 mL | 0.12131 mL | 0.56500 mL | 3.415 | -0.00112 | 0.00437 | 0.00083 | 10.0 s |
| 1:45:40.2 | Data point 103 | 1.50000 mL | 0.12571 mL | 0.12131 mL | 0.56500 mL | 3.225 | -0.00516 | 0.50756 | 0.00036 | 10.0 s |
| 1:46:15.7 | Data point 104 | 1.50000 mL | 0.12733 mL | 0.12131 mL | 0.56500 mL | 3.035 | 0.00000 | 0.00000 | 0.00038 | 10.0 s |
| 1:46:51.1 | Data point 105 | 1.50000 mL | 0.12984 mL | 0.12131 mL | 0.56500 mL | 2.835 | -0.01164 | 0.95088 | 0.00059 | 10.0 s |
| 1:47:26.7 | Data point 106 | 1.50000 mL | 0.13389 mL | 0.12131 mL | 0.56500 mL | 2.621 | -0.00308 | 0.11765 | 0.00044 | 10.0 s |
| 1:48:02.3 | Data point 107 | 1.50000 mL | 0.14052 mL | 0.12131 mL | 0.56500 mL | 2.411 | -0.01254 | 0.94329 | 0.00064 | 10.0 s |
| 1:48:38.0 | Data point 108 | 1.50000 mL | 0.15139 mL | 0.12131 mL | 0.56500 mL | 2.215 | -0.01773 | 0.98451 | 0.00088 | 10.0 s |
| 1:49:13.9 | Data point 109 | 1.50000 mL | 0.16900 mL | 0.12131 mL | 0.56500 mL | 2.009 | -0.01817 | 0.96311 | 0.00091 | 10.5 s |
| 1:49:50.1 | Data point 110 | 1.50000 mL | 0.17679 mL | 0.12131 mL | 0.56500 mL | 1.942 | -0.01341 | 0.97758 | 0.00067 | 10.5 s |
| 1:50:09.6 | Assay volumes | 1.50000 mL | 0.17679 mL | 0.12131 mL | 0.56500 mL | | | | | |

| | | | |
|---|---|---|---|
| Sample name: | **M08_octanol** | Experiment start time: | **2/28/2018 9:57:03 PM** |
| Assay name: | **pH-metric high logP** | Analyst: | **Pion** |
| Assay ID: | **18B-28014** | Instrument ID: | **T312060** |
| Filename: | **G:\OpenLab_PFA_Sirius-T3-2\Data2018\18B-28014_M08_octanol_pH-metric high logP.t3r** | | |

| | | | |
|---|---|---|---|
| Sample name: | **M08_octanol** | Experiment start time: | **2/28/2018 9:57:03 PM** |
| Assay name: | **pH-metric high logP** | Analyst: | **Pion** |
| Assay ID: | **18B-28014** | Instrument ID: | **T312060** |
| Filename: | **G:\OpenLab_PFA_Sirius-T3-2\Data2018\18B-28014_M08_octanol_pH-metric high logP.t3r** | | |

#### Assay Settings (continued)

| Setting | Value | Original Value | Date/Time changed | Imported from |
|---|---|---|---|---|
| ***Titration 3*** | | | | |
| Titrate from | High to low pH | | | |
| Add additional water | 0.00 mL | | | |
| Additional partition solvent volume | 0.500 mL | | | |
| Additional partition solvent added | Automatic | | | |
| After pH adjust stir for | 30 seconds | | | |
| Stir to allow partitioning for | 15 seconds | | | |
| Stirrer speed for partitioning | 60% | | | |
| ***Data Point Stability*** | | | | |
| Stir during data point collection | No | | | |
| Delay before data point collection | 0 seconds | | | |
| Number of points to average | 20 points | | | |
| Time interval between points | 0.50 seconds | | | |
| Required maximum standard deviation | 0.00100 dpH/dt | | | |
| Stability timeout after | 60 seconds | | | |

#### Calibration Settings

| Setting | Value | Date/Time changed | Imported from |
|---|---|---|---|
| Four-Plus alpha | 0.130 | 2/28/2018 9:57:03 PM | C:\Sirius_T3\HCl18B27.t3r |
| Four-Plus S | 0.9970 | 2/28/2018 9:57:03 PM | C:\Sirius_T3\HCl18B27.t3r |
| Four-Plus jH | 0.8 | 2/28/2018 9:57:03 PM | C:\Sirius_T3\HCl18B27.t3r |
| Four-Plus jOH | -0.4 | 2/28/2018 9:57:03 PM | C:\Sirius_T3\HCl18B27.t3r |
| Base concentration factor | 1.000 | 2/28/2018 9:57:03 PM | C:\Sirius_T3\KOH18B27.t3r |
| Acid concentration factor | 0.994 | 2/28/2018 9:57:03 PM | C:\Sirius_T3\HCl18B27.t3r |

#### Instrument Settings

| Setting | Value | Batch Id | Install date |
|---|---|---|---|
| Instrument owner | Merck | | |
| Instrument ID | T312060 | | |
| Instrument type | T3 Simulator | | |
| Software version | 1.1.3.0 | | |
| Dispenser module | | T3DM1200361 | 3/31/2009 6:24:52 AM |
| Dispenser 0 | Water | | 3/31/2009 6:25:05 AM |
| Syringe volume | 2.5 mL | | |
| Firmware version | 1.2.1(r2) | | |
| Titrant | Water (0.15 M KCl) | 02-06-2018 | 2/27/2018 11:05:59 AM |
| Dispenser 2 | Acid | | 3/31/2009 6:25:11 AM |
| Syringe volume | 0.5 mL | | |
| Firmware version | 1.2.1(r2) | | |
| Titrant | Acid (0.5 M HCl) | 02-27-2018 | 2/27/2018 11:27:22 AM |
| Dispenser 1 | Base | | 3/31/2009 6:25:21 AM |
| Syringe volume | 0.5 mL | | |
| Firmware version | 1.2.1(r2) | | |
| Titrant | Base (0.5 M KOH) | 9/22/2017 | 2/27/2018 11:21:22 AM |
| Dispenser 5 | Cosolvent | | 3/31/2009 6:26:24 AM |
| Syringe volume | 2.5 mL | | |
| Firmware version | 1.2.1(r2) | | |
| Distribution valve 5 | Distribution Valve | | 3/31/2009 6:28:19 AM |
| Firmware version | 1.1.3 | | |
| Port A | Methanol (80%, 0.15 M KCl) | 09-26-17 | 2/7/2018 10:42:01 AM |
| Port B | Cyclohexane | 11-01-17 | 2/27/2018 11:37:57 AM |
| Dispenser 3 | Buffer | | 8/3/2010 6:05:16 AM |
| Syringe volume | 0.5 mL | | |
| Firmware version | 1.2.1(r2) | | |
| Titrant | Dodecane | 2018/01/31 | 2/28/2018 11:18:04 AM |
| Dispenser 6 | Octanol | | 10/22/2010 11:52:43 AM |

| | | | |
|---|---|---|---|
| Sample name: | **M08_octanol** | Experiment start time: | **2/28/2018 9:57:03 PM** |
| Assay name: | **pH-metric high logP** | Analyst: | **Pion** |
| Assay ID: | **18B-28014** | Instrument ID: | **T312060** |
| Filename: | **G:\OpenLab_PFA_Sirius-T3-2\Data2018\18B-28014_M08_octanol_pH-metric high logP.t3r** | | |

| | | | |
|---|---|---|---|
| Sample name: | **M08_octanol** | Experiment start time: | **2/28/2018 9:57:03 PM** |
| Assay name: | **pH-metric high logP** | Analyst: | **Pion** |
| Assay ID: | **18B-28014** | Instrument ID: | **T312060** |
| Filename: | **G:\OpenLab_PFA_Sirius-T3-2\Data2018\18B-28014_M08_octanol_pH-metric high logP.t3r** | | |

#### Instrument Settings (continued)

| Setting | Value | Batch Id | Install date |
|---|---|---|---|
| Spectrometer calibration stir duration | 5 s | | |
| Spectrometer calibration stir speed | 30% | | |
| Spectrometer calibration wash pump volume | 20.0 mL | | |
| Spectrometer calibration wash stir duration | 5 s | | |
| Spectrometer calibration wash stir speed | 30% | | |
| Overhead dispense height | 10000 | | |

#### Refinement Settings

| Setting | Value | Default value |
|---|---|---|
| Turbidity detection method | None | None |
| Turbidity wavelength to assess | 500.0 nm | 500.0 nm |
| Turbidity maximum absorbance | 0.100 | 0.100 |
| Turbidity probe threshold | 50.00 | 50.00 |
