## Supplementary material for "Octanol-water partition coefficient measurements for the SAMPL6 Blind Prediction Challenge": SM14_17J-12008_M15_UV-metric psKa_report.pdf

| | | | |
|---|---|---|---|
| Sample name: | **M15** | Experiment start time: | **10/12/2017 9:40:50 AM** |
| Assay name: | **UV-metric psKa** | Analyst: | **Dorothy Levorse** |
| Assay ID: | **17J-12008** | Instrument ID: | **T311053** |
| Filename: | **C:\Sirius_T3\Mehtap\20171011_exp15_pKa\17J-12008_M15_UV-metric psKa.t3r** | | |

#### Yasuda-Shedlovsky result

| Extrapolation type | pKa 0% | SD | Intercept | Slope | R² | Ionic strength | Temperature |
|---|---|---|---|---|---|---|---|
| Yasuda-Shedlovsky | 2.56 | ±0.06 | 7.65 | -262.9812 | 0.9959 | 0.165 M | 24.9°C |
| Yasuda-Shedlovsky | 5.29 | ±0.06 | 9.46 | -190.6287 | 0.9935 | 0.165 M | 24.9°C |

#### Component assay results

| Titration | Methanol weight% | Direction | Result type | Dielectric constant | [H2O] | Ionic strength | Temperature | | psKa 1 | | psKa 2 |
|---|---|---|---|---|---|---|---|---|---|---|---|
| 17J-12008 Points 4 to 39 | 49.51 % | Up | UV-metric pKa | 56.6 | 24.7 M | 0.157 M | 24.9°C | ✓ | 1.62 | ✓ | 4.71 |
| 17J-12008 Points 41 to 77 | 40.06 % | Up | UV-metric pKa | 61.0 | 30.0 M | 0.166 M | 24.9°C | ✓ | 1.84 | ✓ | 4.83 |
| 17J-12008 Points 79 to 116 | 30.24 % | Up | UV-metric pKa | 65.4 | 35.7 M | 0.172 M | 24.9°C | ✓ | 2.09 | ✓ | 5.00 |

#### Graphs

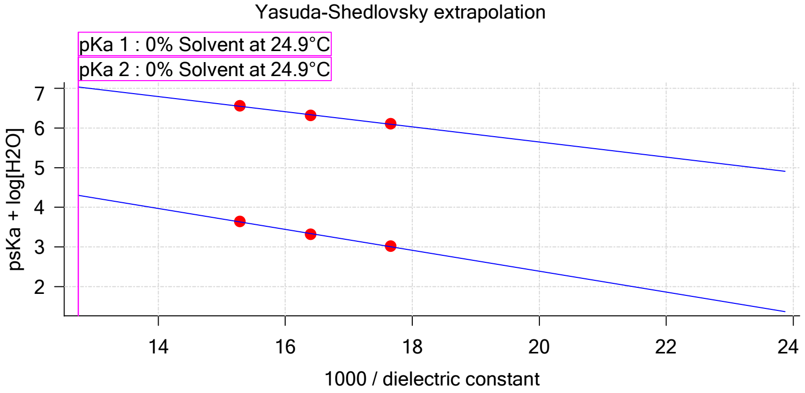


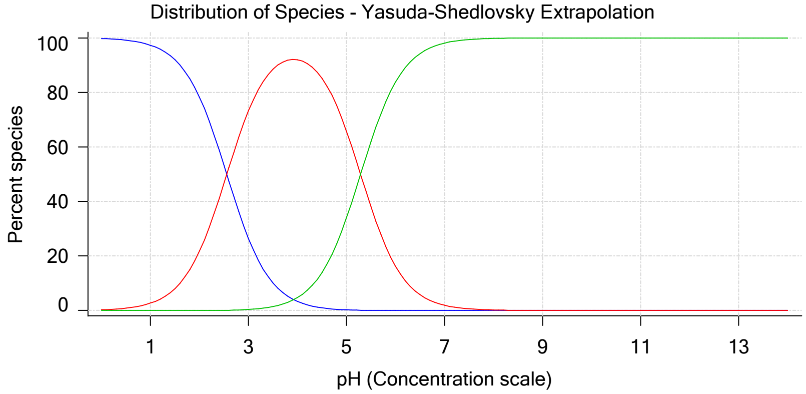


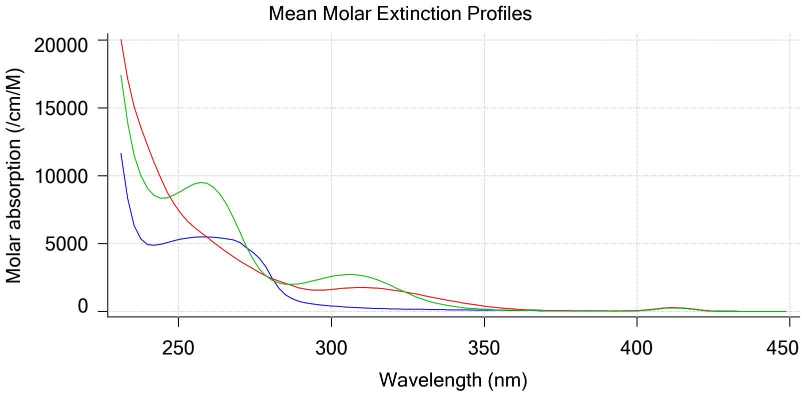


### UV-metric psKa Titration 1 of 3 17J-12008 Points 4 to 39

#### Results

| | |
|---|---|
| pKa 1 | **1.62** |
| pKa 2 | **4.71** |
| RMSD | **0.003 0.003 0.005** |
| Chi squared | **0.0250** |
| PCA calculated number of pKas | **3** |
| Average ionic strength | **0.157 M** |
| Average temperature | **24.9°C** |
| Analyte concentration range | **28.5 µM to 26.8 µM** |
| Methanol weight % | **49.5 %** |
| Dielectric constant | **56.6** |
| Water concentration | **24.7 M** |

| | | | |
|---|---|---|---|
| Sample name: | **M15** | Experiment start time: | **10/12/2017 9:40:50 AM** |
| Assay name: | **UV-metric psKa** | Analyst: | **Dorothy Levorse** |
| Assay ID: | **17J-12008** | Instrument ID: | **T311053** |
| Filename: | **C:\Sirius_T3\Mehtap\20171011_exp15_pKa\17J-12008_M15_UV-metric psKa.t3r** | | |

#### Results (continued)

| | |
|---|---|
| Number of pKas source | **Predicted** |
| Wavelength clipping | **230.0 nm to 450.0 nm** |
| pH clipping | **1.467 to 12.528** |

#### Warnings and errors

| | |
|---|---|
| Errors | None |
| Warnings | PCA calculation disagrees with predicted number of pKas |

#### Assay Settings

| Setting | Value | Original Value | Date/Time changed | Imported from |
|---|---|---|---|---|
| Buffer in use | Yes | | | |
| Buffer type | Phosphate Buffer | | | |
| ***Assay Medium*** | | | | |
| Volume of buffer introduced | 0.025000 mL | | | |
| Add buffer manually | Manual | | | |

#### Graphs

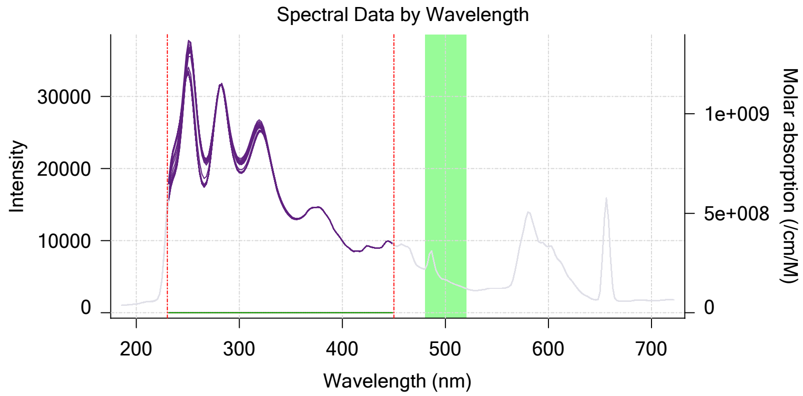


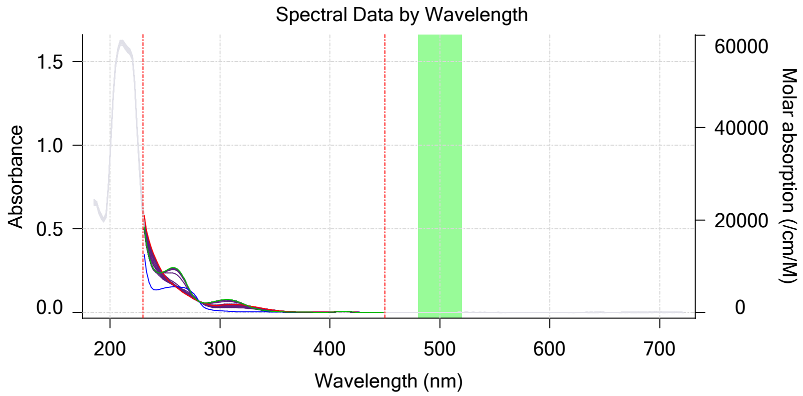


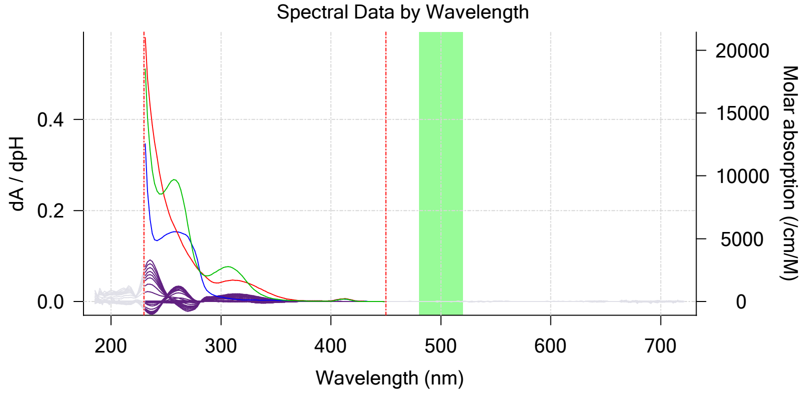


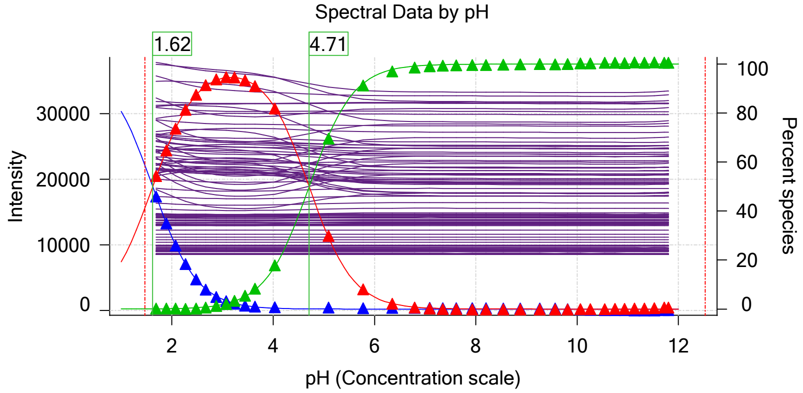

Sample name: **M15**
Assay name: **UV-metric psKa**
Assay ID: **17J-12008**
Filename: **C:\Sirius_T3\Mehtap\20171011_exp15_pKa\17J-12008_M15_UV-metric psKa.t3r**

Experiment start time: **10/12/2017 9:40:50 AM**
Analyst: **Dorothy Levorse**
Instrument ID: **T311053**

#### Graphs (continued)

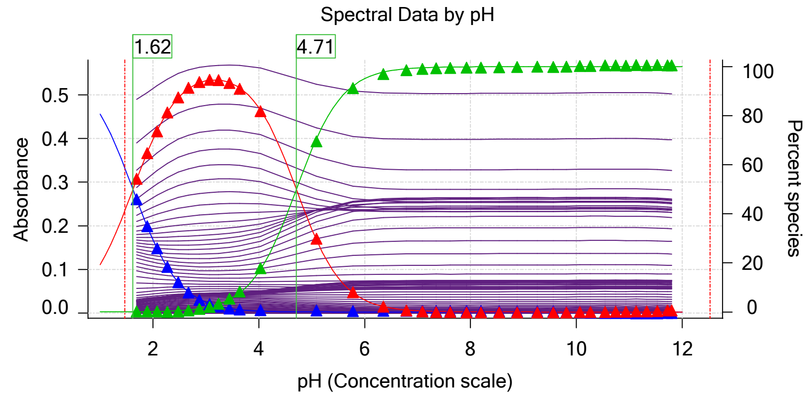


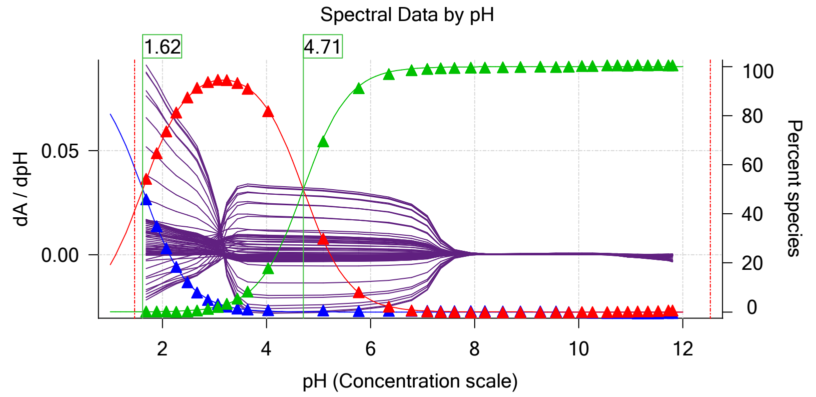


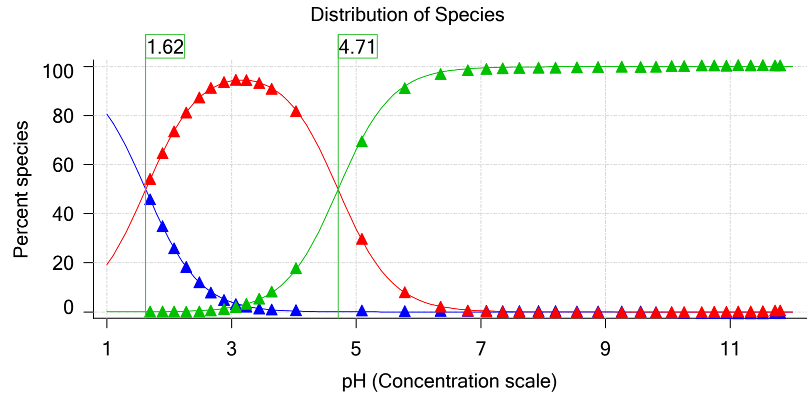


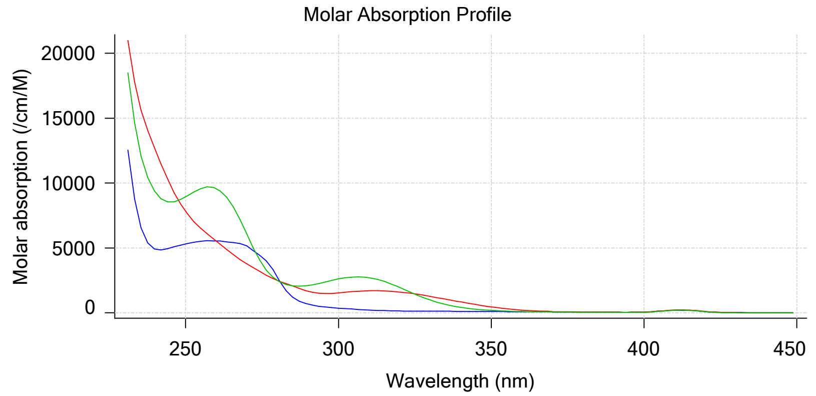


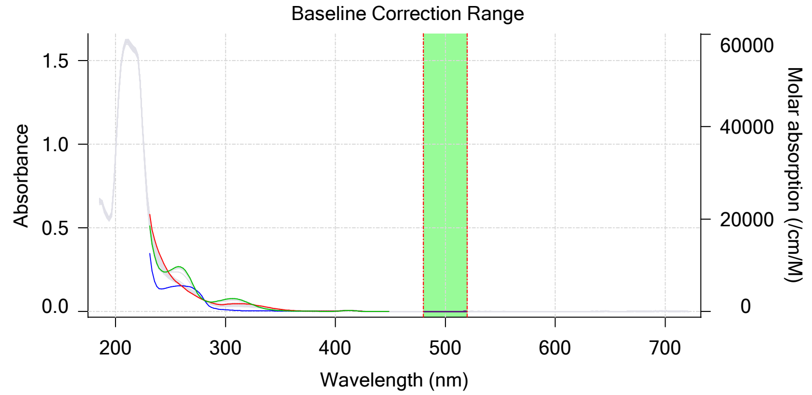


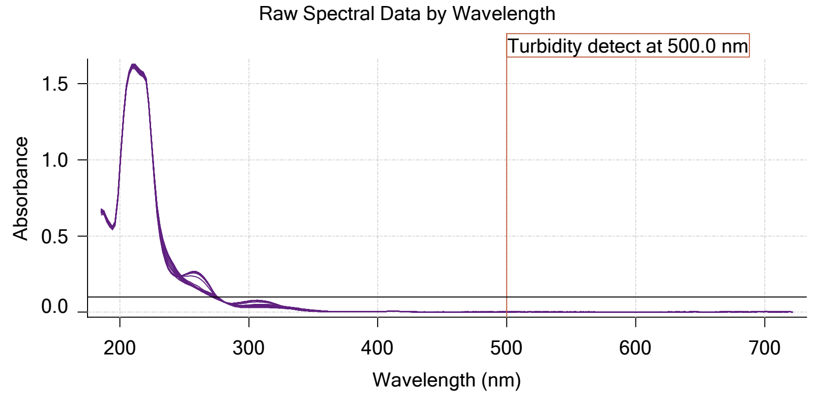


### UV-metric psKa Titration 2 of 3 17J-12008 Points 41 to 77

#### Results

| | |
|---|---|
| pKa 1 | **1.84** |
| pKa 2 | **4.83** |
| RMSD | **0.004 0.005 0.003** |
| Chi squared | **0.0275** |
| PCA calculated number of pKas | **3** |
| Average ionic strength | **0.166 M** |
| Average temperature | **24.9°C** |
| Analyte concentration range | **23.4 µM to 22.1 µM** |
| Methanol weight % | **40.1 %** |
| Dielectric constant | **61.0** |
| Water concentration | **30.0 M** |

| | | | |
|---|---|---|---|
| Sample name: | **M15** | Experiment start time: | **10/12/2017 9:40:50 AM** |
| Assay name: | **UV-metric psKa** | Analyst: | **Dorothy Levorse** |
| Assay ID: | **17J-12008** | Instrument ID: | **T311053** |
| Filename: | **C:\Sirius_T3\Mehtap\20171011_exp15_pKa\17J-12008_M15_UV-metric psKa.t3r** | | |

#### Results (continued)

| | |
|---|---|
| Number of pKas source | **Predicted** |
| Wavelength clipping | **230.0 nm to 450.0 nm** |
| pH clipping | **1.510 to 12.508** |

#### Warnings and errors

| | |
|---|---|
| Errors | None |
| Warnings | PCA calculation disagrees with predicted number of pKas |

#### Assay Settings

| Setting | Value | Original Value | Date/Time changed | Imported from |
|---|---|---|---|---|
| Buffer in use | Yes | | | |
| Buffer type | Phosphate Buffer | | | |
| ***Assay Medium*** | | | | |
| Volume of buffer introduced | 0.025000 mL | | | |
| Add buffer manually | Manual | | | |

#### Graphs

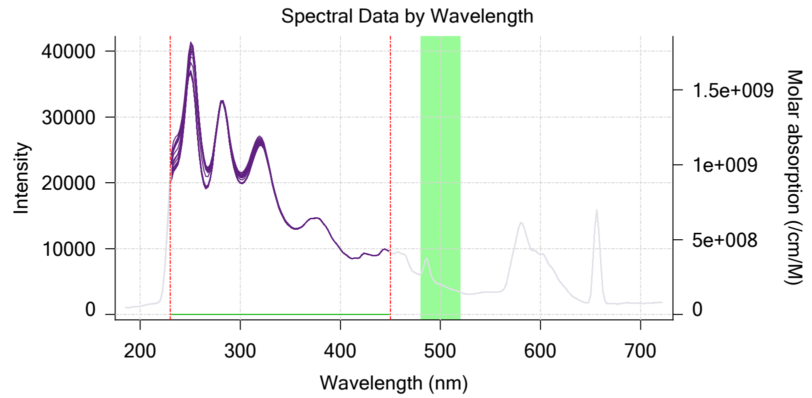


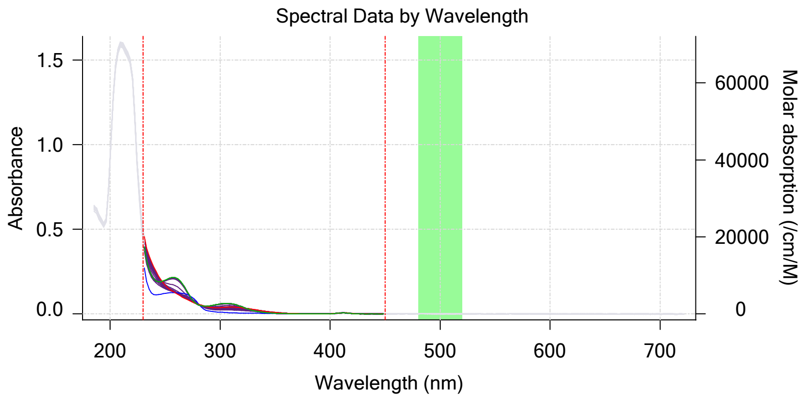


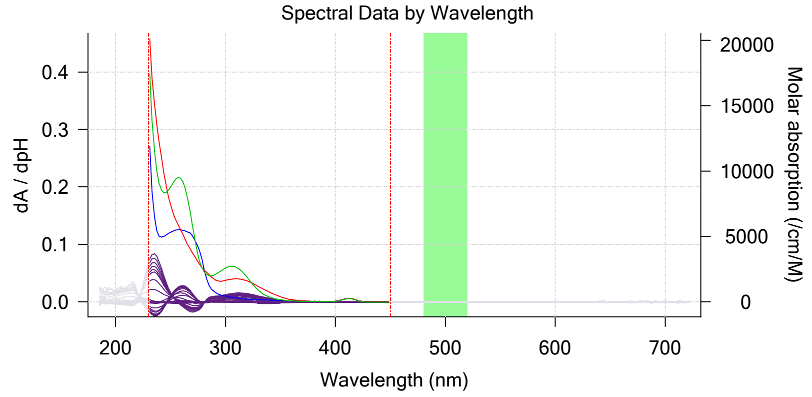


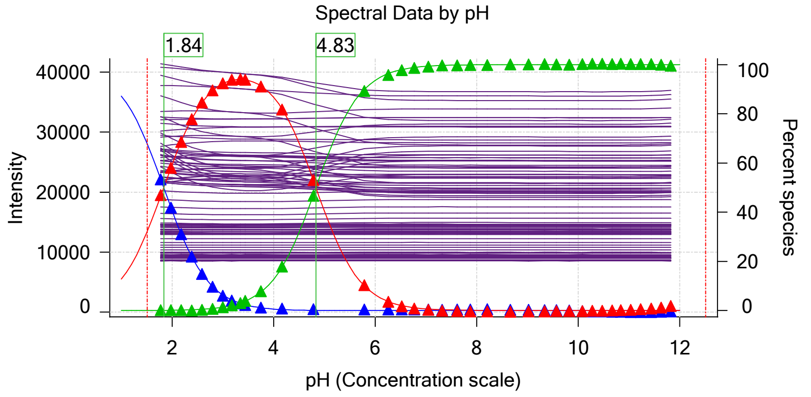

| | | | |
|---|---|---|---|
| Sample name: | **M15** | Experiment start time: | **10/12/2017 9:40:50 AM** |
| Assay name: | **UV-metric psKa** | Analyst: | **Dorothy Levorse** |
| Assay ID: | **17J-12008** | Instrument ID: | **T311053** |
| Filename: | **C:\Sirius_T3\Mehtap\20171011_exp15_pKa\17J-12008_M15_UV-metric psKa.t3r** | | |

#### Graphs (continued)

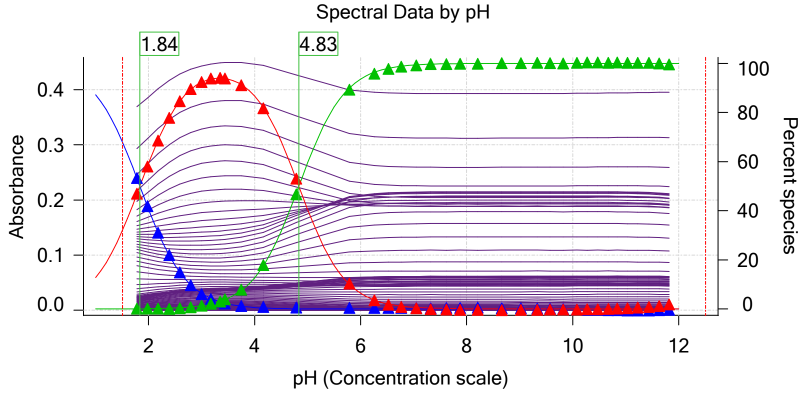


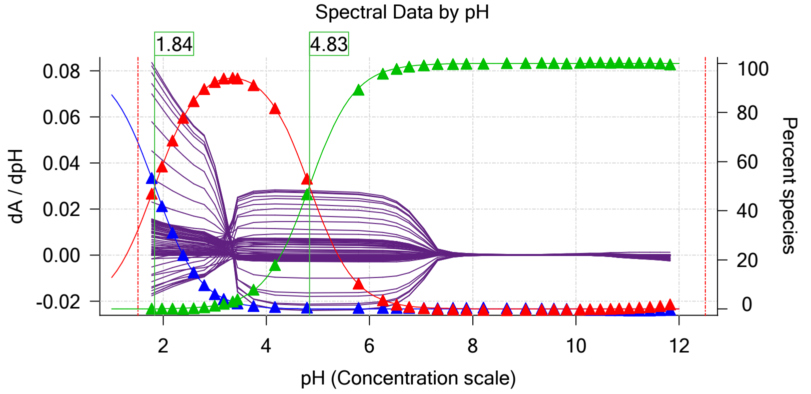


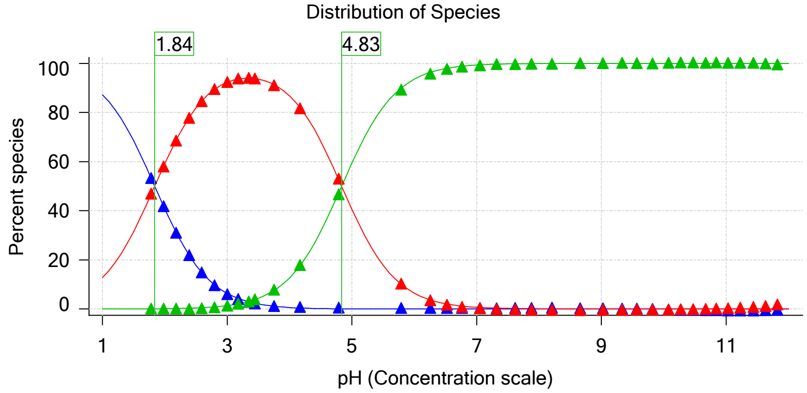


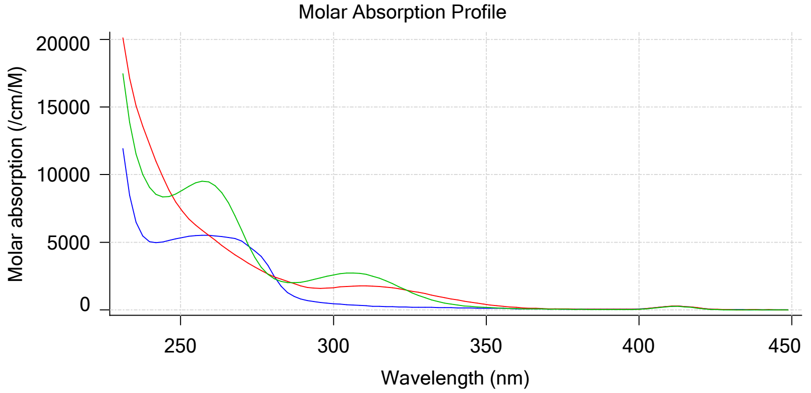


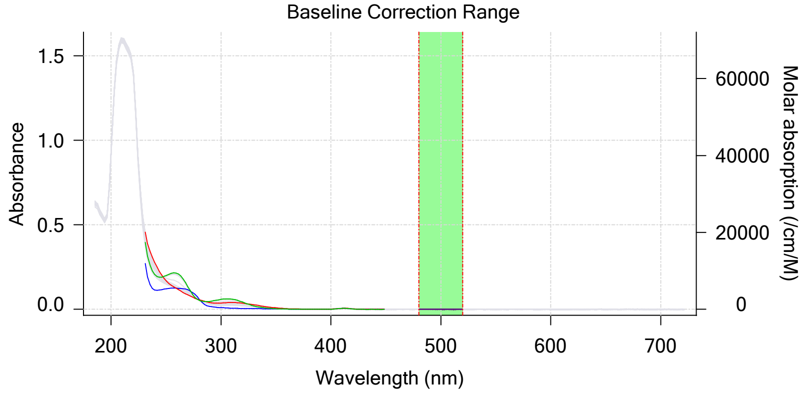


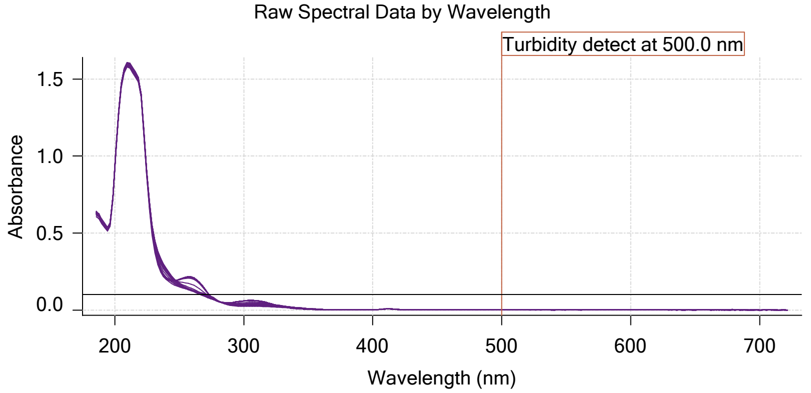


### UV-metric psKa Titration 3 of 3 17J-12008 Points 79 to 116

#### Results

| | |
|---|---|
| pKa 1 | **2.09** |
| pKa 2 | **5.00** |
| RMSD | **0.006 0.009 0.004** |
| Chi squared | **0.0538** |
| PCA calculated number of pKas | **3** |
| Average ionic strength | **0.172 M** |
| Average temperature | **24.9°C** |
| Analyte concentration range | **18.0 µM to 17.1 µM** |
| Methanol weight % | **30.2 %** |
| Dielectric constant | **65.4** |
| Water concentration | **35.7 M** |

Sample name: **M15**
Assay name: **UV-metric psKa**
Assay ID: **17J-12008**
Filename: **C:\Sirius_T3\Mehtap\20171011_exp15_pKa\17J-12008_M15_UV-metric psKa.t3r**

Experiment start time: **10/12/2017 9:40:50 AM**
Analyst: **Dorothy Levorse**
Instrument ID: **T311053**

#### Results (continued)

| | |
|---|---|
| Number of pKas source | **Predicted** |
| Wavelength clipping | **230.0 nm to 450.0 nm** |
| pH clipping | **1.516 to 12.527** |

#### Warnings and errors

| | |
|---|---|
| Errors | None |
| Warnings | PCA calculation disagrees with predicted number of pKas |

#### Assay Settings

| Setting | Value | Original Value | Date/Time changed | Imported from |
|---|---|---|---|---|
| Buffer in use | Yes | | | |
| Buffer type | Phosphate Buffer | | | |
| ***Assay Medium*** | | | | |
| Volume of buffer introduced | 0.025000 mL | | | |
| Add buffer manually | Manual | | | |

#### Graphs

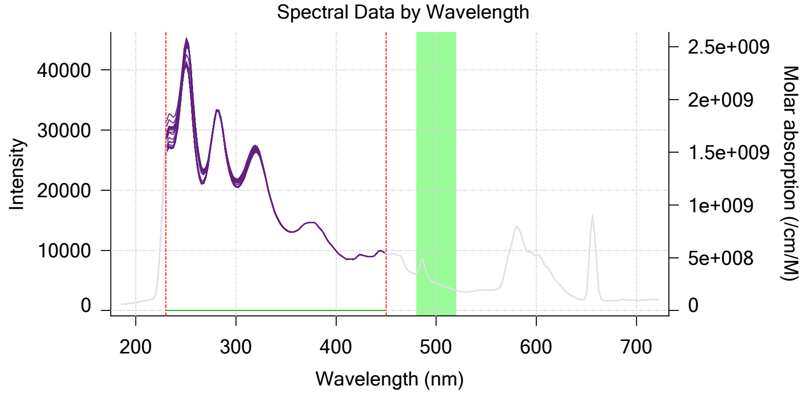


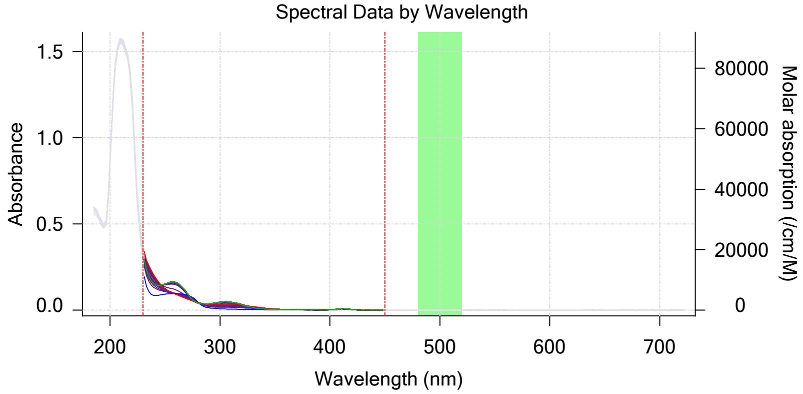


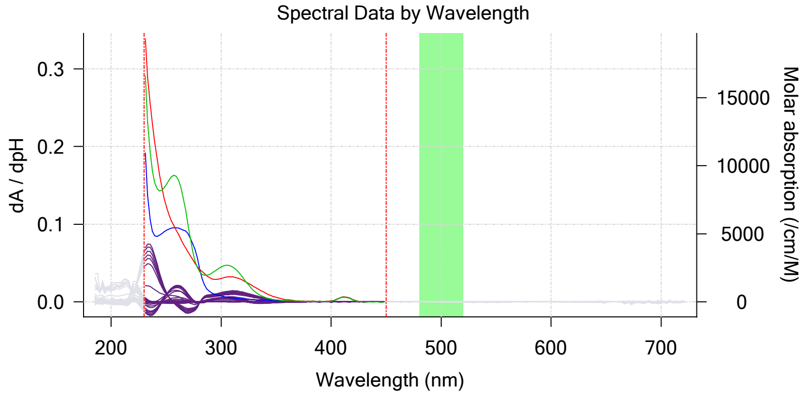


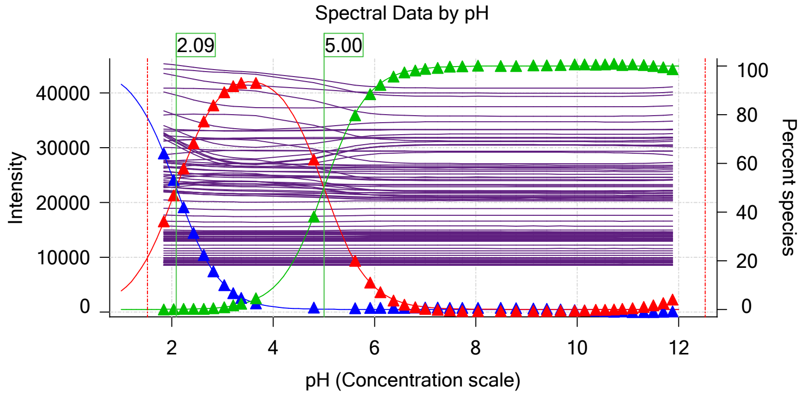

| | | | |
|---|---|---|---|
| Sample name: | **M15** | Experiment start time: | **10/12/2017 9:40:50 AM** |
| Assay name: | **UV-metric psKa** | Analyst: | **Dorothy Levorse** |
| Assay ID: | **17J-12008** | Instrument ID: | **T311053** |
| Filename: | **C:\Sirius_T3\Mehtap\20171011_exp15_pKa\17J-12008_M15_UV-metric psKa.t3r** | | |

#### Graphs (continued)

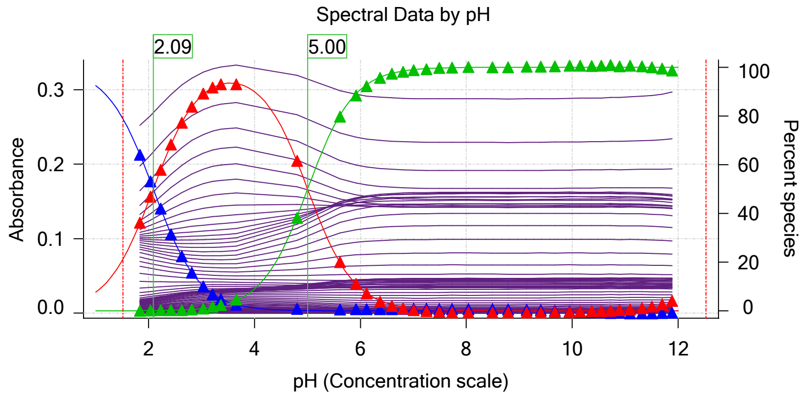


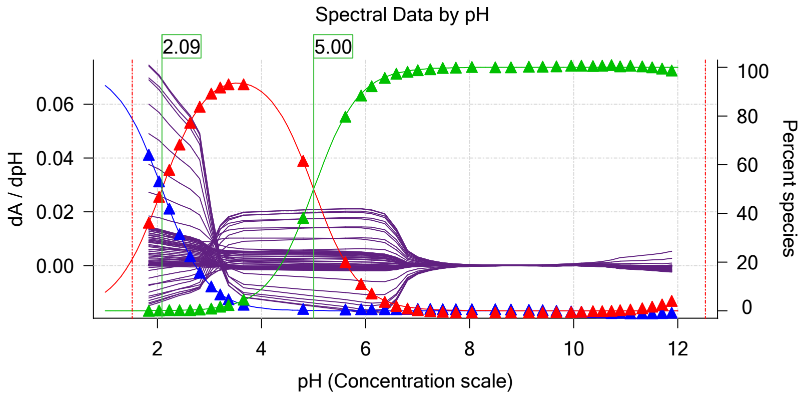


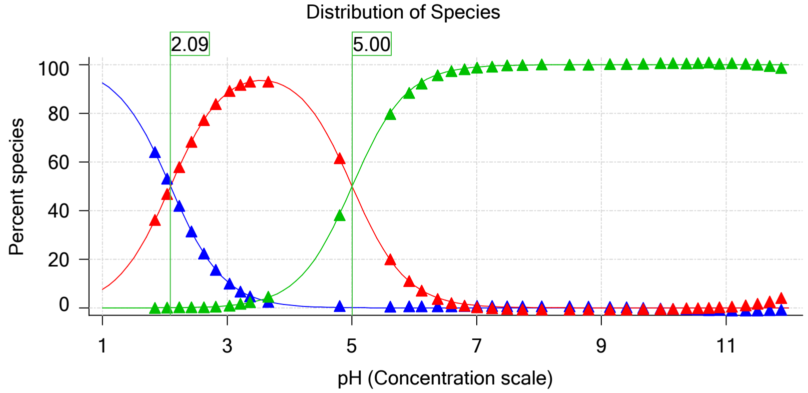


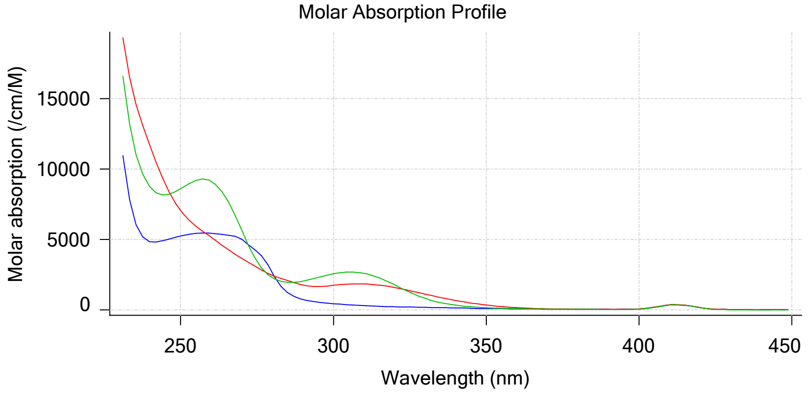


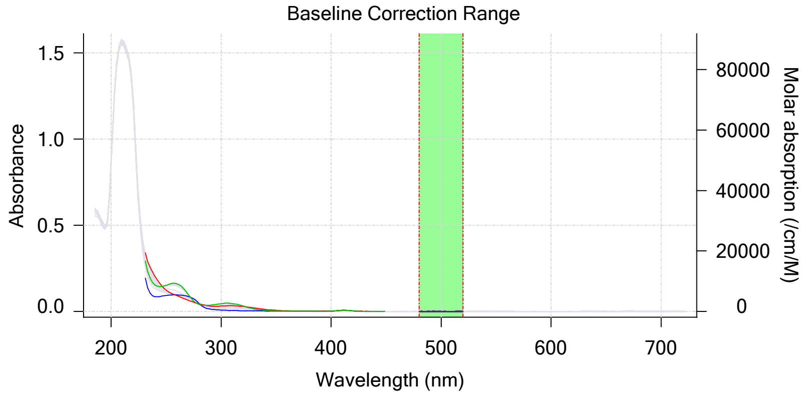


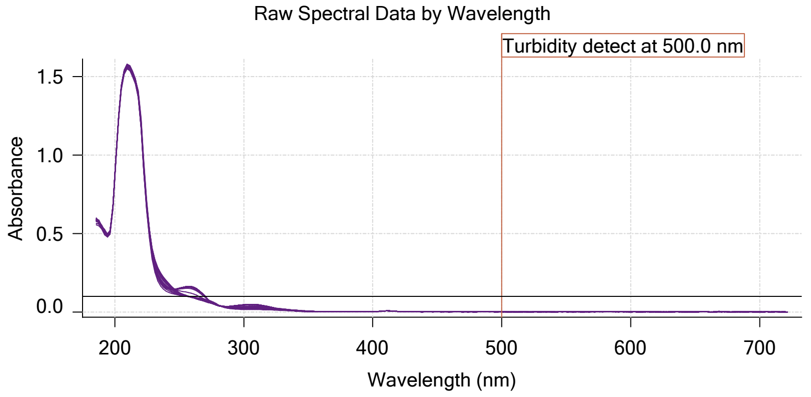


#### Assay Model

| Settings | Value | Date/Time changed | Imported from |
|---|---|---|---|
| Sample name | M15 | 10/11/2017 4:19:59 PM | User entered value |
| Sample by | Volume | | Default value |
| Sample volume | 0.0010 mL | 10/11/2017 4:19:59 PM | User entered value |
| Solvent | DMSO | | Default value |
| Sample concentration | 0.045400 M | 10/11/2017 4:19:59 PM | User entered value |
| Solubility | Unknown | | Default value |
| Molecular weight | 209.25 | 10/11/2017 4:20:11 PM | User entered value |
| Individual pKa ionic environments | No | | Default value |
| Number of pKas | 2 | 10/11/2017 4:19:59 PM | User entered value |
| Sample is a | Base | 10/11/2017 4:19:59 PM | User entered value |
| pKa 1 | 2.94 | 10/11/2017 4:19:59 PM | User entered value |
| pKa 2 | 5.25 | 10/11/2017 4:19:59 PM | User entered value |
| logP (XH2 2+) | -10.00 | | Default value |
| logp (XH +) | -10.00 | | Default value |

| | | | |
|---|---|---|---|
| Sample name: | **M15** | Experiment start time: | **10/12/2017 9:40:50 AM** |
| Assay name: | **UV-metric psKa** | Analyst: | **Dorothy Levorse** |
| Assay ID: | **17J-12008** | Instrument ID: | **T311053** |
| Filename: | **C:\Sirius_T3\Mehtap\20171011_exp15_pKa\17J-12008_M15_UV-metric psKa.t3r** | | |

#### Assay Model (continued)

| Settings | Value | Date/Time changed | Imported from |
|---|---|---|---|
| logP (neutral X) | -10.00 | 10/11/2017 4:19:59 PM | User entered value |

#### Events

| Time | Event | Water | Acid | Base | Methanol | Buffer | pH | dpH/dt | pH R-squar |
|---|---|---|---|---|---|---|---|---|---|
| 3:31.4 | Dark spectrum | | | | | | | | |
| 3:32.8 | Reference spectrum | | | | | | | | |
| 4:00.4 | Volume reset due to vial change | | | | | | | | |
| 4:44.6 | Initial pH = 8.34 | | | | | | | | |
| 5:50.1 | Data point 4 | 0.34995 mL | 0.06874 mL | 0.00000 mL | 1.15005 mL | 0.02500 mL | 1.967 | -0.00525 | 0.44790 |
| 6:19.0 | Data point 5 | 0.34995 mL | 0.06874 mL | 0.02491 mL | 1.15005 mL | 0.02500 mL | 2.169 | -0.00954 | 0.41836 |
| 6:35.8 | Data point 6 | 0.34995 mL | 0.06874 mL | 0.03996 mL | 1.15005 mL | 0.02500 mL | 2.352 | 0.01979 | 0.90627 |
| 6:52.7 | Data point 7 | 0.34995 mL | 0.06874 mL | 0.04976 mL | 1.15005 mL | 0.02500 mL | 2.545 | 0.02536 | 0.91550 |
| 7:09.4 | Data point 8 | 0.34995 mL | 0.06874 mL | 0.05614 mL | 1.15005 mL | 0.02500 mL | 2.755 | 0.01329 | 0.83619 |
| 7:26.1 | Data point 9 | 0.34995 mL | 0.06874 mL | 0.06011 mL | 1.15005 mL | 0.02500 mL | 2.944 | 0.01099 | 0.84718 |
| 7:42.7 | Data point 10 | 0.34995 mL | 0.06874 mL | 0.06270 mL | 1.15005 mL | 0.02500 mL | 3.150 | 0.00700 | 0.76597 |
| 7:59.4 | Data point 11 | 0.34995 mL | 0.06874 mL | 0.06432 mL | 1.15005 mL | 0.02500 mL | 3.342 | 0.00970 | 0.82302 |
| 8:15.9 | Data point 12 | 0.34995 mL | 0.06874 mL | 0.06536 mL | 1.15005 mL | 0.02500 mL | 3.510 | 0.01419 | 0.84907 |
| 8:47.9 | Data point 13 | 0.34995 mL | 0.06874 mL | 0.06637 mL | 1.15005 mL | 0.02500 mL | 3.713 | 0.02970 | 0.98099 |
| 9:25.1 | Data point 14 | 0.34995 mL | 0.06874 mL | 0.06729 mL | 1.15005 mL | 0.02500 mL | 3.910 | 0.03689 | 0.97433 |
| 9:46.8 | Data point 15 | 0.34995 mL | 0.06874 mL | 0.06780 mL | 1.15005 mL | 0.02500 mL | 4.300 | 0.07909 | 0.98985 |
| 10:13.5 | Data point 16 | 0.34995 mL | 0.06874 mL | 0.06823 mL | 1.15005 mL | 0.02500 mL | 5.353 | 0.09957 | 0.99392 |
| 11:21.8 | Data point 17 | 0.34995 mL | 0.06874 mL | 0.06841 mL | 1.15005 mL | 0.02500 mL | 6.034 | 0.13850 | 0.99424 |
| 12:38.5 | Data point 18 | 0.34995 mL | 0.06874 mL | 0.06851 mL | 1.15005 mL | 0.02500 mL | 6.608 | 0.09912 | 0.99017 |
| 13:50.4 | Data point 19 | 0.34995 mL | 0.06874 mL | 0.06863 mL | 1.15005 mL | 0.02500 mL | 7.042 | 0.10030 | 0.99113 |
| 14:43.6 | Data point 20 | 0.34995 mL | 0.06874 mL | 0.06874 mL | 1.15005 mL | 0.02500 mL | 7.340 | 0.09933 | 0.99340 |
| 15:31.6 | Data point 21 | 0.34995 mL | 0.06874 mL | 0.06886 mL | 1.15005 mL | 0.02500 mL | 7.598 | 0.09999 | 0.99204 |
| 16:15.0 | Data point 22 | 0.34995 mL | 0.06874 mL | 0.06900 mL | 1.15005 mL | 0.02500 mL | 7.863 | 0.09906 | 0.99051 |
| 17:05.4 | Data point 23 | 0.34995 mL | 0.06874 mL | 0.06914 mL | 1.15005 mL | 0.02500 mL | 8.174 | 0.09949 | 0.99270 |
| 17:55.3 | Data point 24 | 0.34995 mL | 0.06874 mL | 0.06926 mL | 1.15005 mL | 0.02500 mL | 8.445 | 0.09598 | 0.90762 |
| 18:31.8 | Data point 25 | 0.34995 mL | 0.06874 mL | 0.06936 mL | 1.15005 mL | 0.02500 mL | 8.783 | 0.09877 | 0.96284 |
| 19:21.0 | Data point 26 | 0.34995 mL | 0.06874 mL | 0.06945 mL | 1.15005 mL | 0.02500 mL | 9.120 | 0.09548 | 0.96493 |
| 20:12.5 | Data point 27 | 0.34995 mL | 0.06874 mL | 0.06957 mL | 1.15005 mL | 0.02500 mL | 9.500 | 0.09442 | 0.94739 |
| 20:53.2 | Data point 28 | 0.34995 mL | 0.06874 mL | 0.06968 mL | 1.15005 mL | 0.02500 mL | 9.802 | 0.09795 | 0.96406 |
| 21:26.0 | Data point 29 | 0.34995 mL | 0.06874 mL | 0.06983 mL | 1.15005 mL | 0.02500 mL | 10.047 | 0.09929 | 0.96654 |
| 21:48.5 | Data point 30 | 0.34995 mL | 0.06874 mL | 0.07001 mL | 1.15005 mL | 0.02500 mL | 10.298 | 0.02890 | 0.82114 |
| 22:20.3 | Data point 31 | 0.34995 mL | 0.06874 mL | 0.07030 mL | 1.15005 mL | 0.02500 mL | 10.497 | 0.01960 | 0.93055 |
| 22:37.0 | Data point 32 | 0.34995 mL | 0.06874 mL | 0.07072 mL | 1.15005 mL | 0.02500 mL | 10.777 | 0.00296 | 0.24851 |
| 23:08.8 | Data point 33 | 0.34995 mL | 0.06874 mL | 0.07166 mL | 1.15005 mL | 0.02500 mL | 10.976 | -0.00336 | 0.40552 |
| 23:41.0 | Data point 34 | 0.34995 mL | 0.06874 mL | 0.07314 mL | 1.15005 mL | 0.02500 mL | 11.169 | -0.01019 | 0.82632 |
| 24:13.2 | Data point 35 | 0.34995 mL | 0.06874 mL | 0.07563 mL | 1.15005 mL | 0.02500 mL | 11.359 | -0.00614 | 0.72743 |
| 24:40.3 | Data point 36 | 0.34995 mL | 0.06874 mL | 0.07949 mL | 1.15005 mL | 0.02500 mL | 11.558 | -0.01212 | 0.86864 |
| 25:12.5 | Data point 37 | 0.34995 mL | 0.06874 mL | 0.08544 mL | 1.15005 mL | 0.02500 mL | 11.754 | -0.00675 | 0.73695 |
| 25:45.0 | Data point 38 | 0.34995 mL | 0.06874 mL | 0.09544 mL | 1.15005 mL | 0.02500 mL | 11.945 | -0.00910 | 0.76255 |
| 26:06.9 | Data point 39 | 0.34995 mL | 0.06874 mL | 0.10085 mL | 1.15005 mL | 0.02500 mL | 12.028 | -0.00626 | 0.60483 |
| 27:48.4 | Reference spectrum | | | | | | | | |
| 28:52.4 | Data point 41 | 0.50000 mL | 0.16634 mL | 0.10087 mL | 1.15005 mL | 0.02500 mL | 2.010 | -0.05068 | 0.89643 |
| 29:19.9 | Data point 42 | 0.50000 mL | 0.16634 mL | 0.12429 mL | 1.15005 mL | 0.02500 mL | 2.208 | 0.00950 | 0.75308 |
| 29:36.9 | Data point 43 | 0.50000 mL | 0.16634 mL | 0.13930 mL | 1.15005 mL | 0.02500 mL | 2.410 | 0.01380 | 0.69492 |
| 29:53.8 | Data point 44 | 0.50000 mL | 0.16634 mL | 0.14880 mL | 1.15005 mL | 0.02500 mL | 2.617 | 0.00757 | 0.36152 |
| 30:10.6 | Data point 45 | 0.50000 mL | 0.16634 mL | 0.15475 mL | 1.15005 mL | 0.02500 mL | 2.816 | 0.02783 | 0.86319 |
| 30:27.4 | Data point 46 | 0.50000 mL | 0.16634 mL | 0.15849 mL | 1.15005 mL | 0.02500 mL | 3.023 | 0.00277 | 0.13850 |
| 30:44.1 | Data point 47 | 0.50000 mL | 0.16634 mL | 0.16082 mL | 1.15005 mL | 0.02500 mL | 3.227 | 0.01696 | 0.91503 |
| 31:00.8 | Data point 48 | 0.50000 mL | 0.16634 mL | 0.16228 mL | 1.15005 mL | 0.02500 mL | 3.402 | 0.01558 | 0.90902 |
| 31:17.4 | Data point 49 | 0.50000 mL | 0.16634 mL | 0.16326 mL | 1.15005 mL | 0.02500 mL | 3.572 | 0.02207 | 0.96804 |
| 31:33.9 | Data point 50 | 0.50000 mL | 0.16634 mL | 0.16392 mL | 1.15005 mL | 0.02500 mL | 3.664 | 0.02780 | 0.95210 |
| 31:55.6 | Data point 51 | 0.50000 mL | 0.16634 mL | 0.16477 mL | 1.15005 mL | 0.02500 mL | 3.975 | 0.03761 | 0.97378 |

sirius

| | | | |
|---|---|---|---|
| Sample name: | **M15** | Experiment start time: | **10/12/2017 9:40:50 AM** |
| Assay name: | **UV-metric psKa** | Analyst: | **Dorothy Levorse** |
| Assay ID: | **17J-12008** | Instrument ID: | **T311053** |
| Filename: | **C:\Sirius_T3\Mehtap\20171011_exp15_pKa\17J-12008_M15_UV-metric psKa.t3r** | | |

#### Events (continued)

| Time | Event | Water | Acid | Base | Methanol | Buffer | pH | dpH/dt | pH R-squared | [illegible] |
|---|---|---|---|---|---|---|---|---|---|---|
| 32:17.4 | Data point 52 | 0.50000 mL | 0.16634 mL | 0.16526 mL | 1.15005 mL | 0.02500 mL | 4.386 | 0.08947 | 0.99208 | 0. |
| 32:44.2 | Data point 53 | 0.50000 mL | 0.16634 mL | 0.16562 mL | 1.15005 mL | 0.02500 mL | 5.005 | 0.10041 | 0.99004 | 0. |
| 33:34.0 | Data point 54 | 0.50000 mL | 0.16634 mL | 0.16583 mL | 1.15005 mL | 0.02500 mL | 6.003 | 0.09781 | 0.99468 | 0. |
| 34:37.8 | Data point 55 | 0.50000 mL | 0.16634 mL | 0.16597 mL | 1.15005 mL | 0.02500 mL | 6.472 | 0.09955 | 0.99445 | 0. |
| 35:26.0 | Data point 56 | 0.50000 mL | 0.16634 mL | 0.16606 mL | 1.15005 mL | 0.02500 mL | 6.735 | 0.09893 | 0.98708 | 0. |
| 36:04.1 | Data point 57 | 0.50000 mL | 0.16634 mL | 0.16616 mL | 1.15005 mL | 0.02500 mL | 6.978 | 0.09618 | 0.98868 | 0. |
| 36:40.5 | Data point 58 | 0.50000 mL | 0.16634 mL | 0.16628 mL | 1.15005 mL | 0.02500 mL | 7.260 | 0.10005 | 0.98767 | 0. |
| 37:14.8 | Data point 59 | 0.50000 mL | 0.16634 mL | 0.16639 mL | 1.15005 mL | 0.02500 mL | 7.530 | 0.09972 | 0.98386 | 0. |
| 37:55.7 | Data point 60 | 0.50000 mL | 0.16634 mL | 0.16651 mL | 1.15005 mL | 0.02500 mL | 7.824 | 0.10067 | 0.99226 | 0. |
| 38:36.6 | Data point 61 | 0.50000 mL | 0.16634 mL | 0.16660 mL | 1.15005 mL | 0.02500 mL | 8.083 | 0.09648 | 0.97074 | 0. |
| 39:20.7 | Data point 62 | 0.50000 mL | 0.16634 mL | 0.16670 mL | 1.15005 mL | 0.02500 mL | 8.412 | 0.09956 | 0.98382 | 0. |
| 40:10.9 | Data point 63 | 0.50000 mL | 0.16634 mL | 0.16679 mL | 1.15005 mL | 0.02500 mL | 8.863 | 0.09764 | 0.98633 | 0. |
| 40:57.2 | Data point 64 | 0.50000 mL | 0.16634 mL | 0.16686 mL | 1.15005 mL | 0.02500 mL | 9.224 | 0.09288 | 0.94299 | 0. |
| 41:37.4 | Data point 65 | 0.50000 mL | 0.16634 mL | 0.16693 mL | 1.15005 mL | 0.02500 mL | 9.534 | 0.10038 | 0.98616 | 0. |
| 42:07.1 | Data point 66 | 0.50000 mL | 0.16634 mL | 0.16700 mL | 1.15005 mL | 0.02500 mL | 9.770 | 0.09351 | 0.96224 | 0. |
| 42:32.7 | Data point 67 | 0.50000 mL | 0.16634 mL | 0.16712 mL | 1.15005 mL | 0.02500 mL | 10.020 | 0.06284 | 0.97651 | 0. |
| 42:49.2 | Data point 68 | 0.50000 mL | 0.16634 mL | 0.16731 mL | 1.15005 mL | 0.02500 mL | 10.275 | 0.01695 | 0.81940 | 0. |
| 43:20.9 | Data point 69 | 0.50000 mL | 0.16634 mL | 0.16773 mL | 1.15005 mL | 0.02500 mL | 10.470 | 0.00881 | 0.64616 | 0. |
| 43:53.0 | Data point 70 | 0.50000 mL | 0.16634 mL | 0.16834 mL | 1.15005 mL | 0.02500 mL | 10.666 | 0.00308 | 0.32860 | 0. |
| 44:09.6 | Data point 71 | 0.50000 mL | 0.16634 mL | 0.16919 mL | 1.15005 mL | 0.02500 mL | 10.865 | -0.00524 | 0.65148 | 0. |
| 44:26.2 | Data point 72 | 0.50000 mL | 0.16634 mL | 0.17053 mL | 1.15005 mL | 0.02500 mL | 11.033 | -0.00879 | 0.79184 | 0. |
| 44:58.4 | Data point 73 | 0.50000 mL | 0.16634 mL | 0.17317 mL | 1.15005 mL | 0.02500 mL | 11.225 | -0.00979 | 0.78876 | 0. |
| 45:25.5 | Data point 74 | 0.50000 mL | 0.16634 mL | 0.17681 mL | 1.15005 mL | 0.02500 mL | 11.418 | -0.01504 | 0.90841 | 0. |
| 46:03.1 | Data point 75 | 0.50000 mL | 0.16634 mL | 0.18380 mL | 1.15005 mL | 0.02500 mL | 11.626 | -0.01108 | 0.62860 | 0. |
| 46:35.2 | Data point 76 | 0.50000 mL | 0.16634 mL | 0.19365 mL | 1.15005 mL | 0.02500 mL | 11.822 | -0.01119 | 0.87568 | 0. |
| 47:02.5 | Data point 77 | 0.50000 mL | 0.16634 mL | 0.20884 mL | 1.15005 mL | 0.02500 mL | 12.008 | -0.00839 | 0.82304 | 0. |
| 48:47.3 | Reference spectrum | | | | | | | | | |
| 50:10.6 | Data point 79 | 0.83996 mL | 0.29523 mL | 0.20887 mL | 1.15005 mL | 0.02500 mL | 2.016 | -0.01620 | 0.88855 | 0. |
| 50:38.2 | Data point 80 | 0.83996 mL | 0.29523 mL | 0.23544 mL | 1.15005 mL | 0.02500 mL | 2.210 | 0.01120 | 0.73558 | 0. |
| 50:55.3 | Data point 81 | 0.83996 mL | 0.29523 mL | 0.25249 mL | 1.15005 mL | 0.02500 mL | 2.406 | 0.00019 | 0.00022 | 0. |
| 51:12.1 | Data point 82 | 0.83996 mL | 0.29523 mL | 0.26352 mL | 1.15005 mL | 0.02500 mL | 2.600 | -0.01948 | 0.79039 | 0. |
| 51:28.9 | Data point 83 | 0.83996 mL | 0.29523 mL | 0.27067 mL | 1.15005 mL | 0.02500 mL | 2.802 | -0.00148 | 0.04503 | 0. |
| 51:45.6 | Data point 84 | 0.83996 mL | 0.29523 mL | 0.27514 mL | 1.15005 mL | 0.02500 mL | 2.990 | -0.01144 | 0.54674 | 0. |
| 52:02.2 | Data point 85 | 0.83996 mL | 0.29523 mL | 0.27803 mL | 1.15005 mL | 0.02500 mL | 3.209 | -0.01542 | 0.69490 | 0. |
| 52:18.8 | Data point 86 | 0.83996 mL | 0.29523 mL | 0.27980 mL | 1.15005 mL | 0.02500 mL | 3.383 | -0.00720 | 0.38977 | 0. |
| 52:35.4 | Data point 87 | 0.83996 mL | 0.29523 mL | 0.28097 mL | 1.15005 mL | 0.02500 mL | 3.538 | -0.00290 | 0.08907 | 0. |
| 53:07.5 | Data point 88 | 0.83996 mL | 0.29523 mL | 0.28140 mL | 1.15005 mL | 0.02500 mL | 3.830 | -0.01232 | 0.83931 | 0. |
| 53:34.5 | Data point 89 | 0.83996 mL | 0.29523 mL | 0.28429 mL | 1.15005 mL | 0.02500 mL | 4.972 | -0.08188 | 0.86107 | 0. |
| 54:03.1 | Data point 90 | 0.83996 mL | 0.29523 mL | 0.28457 mL | 1.15005 mL | 0.02500 mL | 5.781 | 0.01061 | 0.01674 | 0. |
| 54:25.4 | Data point 91 | 0.83996 mL | 0.29523 mL | 0.28471 mL | 1.15005 mL | 0.02500 mL | 6.080 | -0.05291 | 0.40901 | 0. |
| 54:47.5 | Data point 92 | 0.83996 mL | 0.29523 mL | 0.28481 mL | 1.15005 mL | 0.02500 mL | 6.283 | -0.06378 | 0.75328 | 0. |
| 55:09.2 | Data point 93 | 0.83996 mL | 0.29523 mL | 0.28492 mL | 1.15005 mL | 0.02500 mL | 6.535 | -0.07363 | 0.74563 | 0. |
| 55:35.7 | Data point 94 | 0.83996 mL | 0.29523 mL | 0.28504 mL | 1.15005 mL | 0.02500 mL | 6.755 | -0.09453 | 0.90047 | 0. |
| 56:06.4 | Data point 95 | 0.83996 mL | 0.29523 mL | 0.28516 mL | 1.15005 mL | 0.02500 mL | 6.966 | 0.01849 | 0.44080 | 0. |
| 56:33.3 | Data point 96 | 0.83996 mL | 0.29523 mL | 0.28528 mL | 1.15005 mL | 0.02500 mL | 7.165 | 0.05052 | 0.90661 | 0. |
| 57:05.2 | Data point 97 | 0.83996 mL | 0.29523 mL | 0.28542 mL | 1.15005 mL | 0.02500 mL | 7.406 | 0.09582 | 0.94132 | 0. |
| 57:37.1 | Data point 98 | 0.83996 mL | 0.29523 mL | 0.28554 mL | 1.15005 mL | 0.02500 mL | 7.646 | 0.09448 | 0.95830 | 0. |
| 58:11.7 | Data point 99 | 0.83996 mL | 0.29523 mL | 0.28563 mL | 1.15005 mL | 0.02500 mL | 7.893 | 0.09904 | 0.97936 | 0. |
| 58:51.1 | Data point 100 | 0.83996 mL | 0.29523 mL | 0.28572 mL | 1.15005 mL | 0.02500 mL | 8.200 | 0.09751 | 0.97855 | 0. |
| 59:35.9 | Data point 101 | 0.83996 mL | 0.29523 mL | 0.28582 mL | 1.15005 mL | 0.02500 mL | 8.648 | 0.09789 | 0.95408 | 0. |
| 1:00:19.7 | Data point 102 | 0.83996 mL | 0.29523 mL | 0.28589 mL | 1.15005 mL | 0.02500 mL | 8.948 | 0.09695 | 0.94482 | 0. |
| 1:01:00.0 | Data point 103 | 0.83996 mL | 0.29523 mL | 0.28598 mL | 1.15005 mL | 0.02500 mL | 9.292 | 0.09665 | 0.97234 | 0. |
| 1:01:30.1 | Data point 104 | 0.83996 mL | 0.29523 mL | 0.28608 mL | 1.15005 mL | 0.02500 mL | 9.557 | 0.09671 | 0.97597 | 0. |
| 1:01:55.9 | Data point 105 | 0.83996 mL | 0.29523 mL | 0.28619 mL | 1.15005 mL | 0.02500 mL | 9.823 | 0.05116 | 0.98408 | 0. |
| 1:02:12.4 | Data point 106 | 0.83996 mL | 0.29523 mL | 0.28638 mL | 1.15005 mL | 0.02500 mL | 10.102 | -0.00513 | 0.37093 | 0. |
| 1:02:44.3 | Data point 107 | 0.83996 mL | 0.29523 mL | 0.28676 mL | 1.15005 mL | 0.02500 mL | 10.299 | 0.00942 | 0.46416 | 0. |
| 1:03:00.9 | Data point 108 | 0.83996 mL | 0.29523 mL | 0.28735 mL | 1.15005 mL | 0.02500 mL | 10.517 | -0.02188 | 0.89061 | 0. |

| | | | |
|---|---|---|---|
| Sample name: | **M15** | Experiment start time: | **10/12/2017 9:40:50 AM** |
| Assay name: | **UV-metric psKa** | Analyst: | **Dorothy Levorse** |
| Assay ID: | **17J-12008** | Instrument ID: | **T311053** |
| Filename: | **C:\Sirius_T3\Mehtap\20171011_exp15_pKa\17J-12008_M15_UV-metric psKa.t3r** | | |

#### Events (continued)

| Time | Event | Water | Acid | Base | Methanol | Buffer | pH | dpH/dt | pH R-squared | pH SD |
|---|---|---|---|---|---|---|---|---|---|---|
| 1:03:17.3 | Data point 109 | 0.83996 mL | 0.29523 mL | 0.28829 mL | 1.15005 mL | 0.02500 mL | 10.700 | -0.02234 | 0.91518 | 0.00115 |
| 1:03:34.0 | Data point 110 | 0.83996 mL | 0.29523 mL | 0.28970 mL | 1.15005 mL | 0.02500 mL | 10.875 | -0.02240 | 0.88731 | 0.00117 |
| 1:03:50.7 | Data point 111 | 0.83996 mL | 0.29523 mL | 0.29177 mL | 1.15005 mL | 0.02500 mL | 11.040 | -0.02425 | 0.93339 | 0.00124 |
| 1:04:17.7 | Data point 112 | 0.83996 mL | 0.29523 mL | 0.29565 mL | 1.15005 mL | 0.02500 mL | 11.240 | -0.01758 | 0.89791 | 0.00092 |
| 1:04:55.2 | Data point 113 | 0.83996 mL | 0.29523 mL | 0.30287 mL | 1.15005 mL | 0.02500 mL | 11.461 | -0.01669 | 0.89086 | 0.00087 |
| 1:05:27.6 | Data point 114 | 0.83996 mL | 0.29523 mL | 0.31244 mL | 1.15005 mL | 0.02500 mL | 11.653 | -0.01628 | 0.88701 | 0.00083 |
| 1:05:54.9 | Data point 115 | 0.83996 mL | 0.29523 mL | 0.32733 mL | 1.15005 mL | 0.02500 mL | 11.844 | -0.01423 | 0.90910 | 0.00074 |
| 1:06:22.6 | Data point 116 | 0.83996 mL | 0.29523 mL | 0.35089 mL | 1.15005 mL | 0.02500 mL | 12.027 | -0.01514 | 0.88771 | 0.00079 |
| 1:08:22.0 | Assay volumes | 1.08996 mL | 0.43822 mL | 0.35089 mL | 1.15005 mL | 0.02500 mL | | | | |

#### Assay Settings

| Setting | Value | Original Value | Date/Time changed | Imported from |
|---|---|---|---|---|
| ***General Settings*** | | | | |
| Analyst name | Dorothy Levorse | | | |
| Separate reference vial | Yes | | | |
| ***Standard Experiment Settings*** | | | | |
| Number of titrations | 3 | | | |
| Minimum pH | 2.000 | | | |
| Maximum pH | 12.000 | | | |
| pH step between points of | 0.200 | | | |
| Minimum titrant addition | 0.00002 mL | | | |
| Maximum titrant addition | 0.10000 mL | | | |
| Argon flow rate | 100% | | | |
| Start titration using | Cautious pH adjust | | | |
| ***Advanced General Settings*** | | | | |
| Detect turbidity using | Spectrometer | | | |
| Monitor at a wavelength of | 500.0 nm | | | |
| Absorbance threshold of | 0.100 | | | |
| Collect turbidity sensor data | No | | | |
| Stir after titrant addition for | 5 seconds | | | |
| For titrant addition, stir at | 15% | | | |
| ***Titrant Pre-Dose*** | | | | |
| Titrant pre-dose | None | | | |
| ***Assay Medium*** | | | | |
| Cosolvent in use | Yes | | | |
| Cosolvent type | Methanol | | | |
| Cosolvent volume | 1.15 mL | | | |
| Cosolvent added | Automatic | | | |
| ISA water volume | 0.35 mL | | | |
| Water added | Automatic | | | |
| After water addition, stir for | 5 seconds | | | |
| At a speed of | 15% | | | |
| Buffer in use | Yes | | | |
| Buffer type | Phosphate Buffer | | | |
| Volume of buffer introduced | 0.025000 mL | | | |
| Add buffer manually | Manual | | | |
| After medium addition, stir for | 5 seconds | | | |
| ***Sample Sonication*** | | | | |
| Sonicate | No | | | |
| ***Sample Dissolution*** | | | | |
| Perform a dissolution stage | No | | | |
| ***Carbonate purge*** | | | | |
| Perform a carbonate purge | No | | | |
| ***Temperature Control*** | | | | |
| Wait for temperature | Yes | | | |
| Required start temperature | 25.0°C | | | |
| Acceptable deviation | 0.5°C | | | |

| | | | |
|---|---|---|---|
| Sample name: | **M15** | Experiment start time: | **10/12/2017 9:40:50 AM** |
| Assay name: | **UV-metric psKa** | Analyst: | **Dorothy Levorse** |
| Assay ID: | **17J-12008** | Instrument ID: | **T311053** |
| Filename: | **C:\Sirius_T3\Mehtap\20171011_exp15_pKa\17J-12008_M15_UV-metric psKa.t3r** | | |

#### Assay Settings (continued)

| Setting | Value | Original Value | Date/Time changed | Imported from |
|---|---|---|---|---|
| Time to wait | 60 seconds | | | |
| Stir speed of | 15% | | | |
| ***Titration 1*** | | | | |
| Titrate from | Low to high pH | | | |
| Adjust to start pH | Yes | | | |
| After pH adjust stir for | 10 seconds | | | |
| ***Titration 2*** | | | | |
| Titrate from | Low to high pH | | | |
| Additional cosolvent volume | 0.00 mL | | | |
| Add additional water | 0.15 mL | | | |
| Additional water added | Automatic | | | |
| After pH adjust stir for | 10 seconds | | | |
| ***Titration 3*** | | | | |
| Titrate from | Low to high pH | | | |
| Additional cosolvent volume | 0.00 mL | | | |
| Add additional water | 0.34 mL | | | |
| Additional water added | Automatic | | | |
| After pH adjust stir for | 10 seconds | | | |
| ***Data Point Stability*** | | | | |
| Stir during data point collection | Yes | | | |
| For point collection, stir at | 15% | | | |
| Delay before data point collection | 0 seconds | | | |
| Number of points to average | 20 points | | | |
| Time interval between points | 0.50 seconds | | | |
| Required maximum standard deviation | 0.00500 dpH/dt | | | |
| Stability timeout after | 60 seconds | | | |
| ***Experiment cleanup*** | | | | |
| Adjust pH to cleanup | To start pH | | | |
| And then stir for | 60 seconds | | | |
| For cleaning, stir at | 20% | | | |
| Then add water volume | 0.25 mL | | | |
| And then stir for | 30 seconds | | | |

#### Calibration Settings

| Setting | Value | Date/Time changed | Imported from |
|---|---|---|---|
| Four-Plus alpha | 0.109 | 10/12/2017 9:40:50 AM | C:\Sirius_T3\17J-11006_Blank standardisation.t3r |
| Four-Plus S | 1.0007 | 10/12/2017 9:40:50 AM | C:\Sirius_T3\17J-11006_Blank standardisation.t3r |
| Four-Plus jH | 0.3 | 10/12/2017 9:40:50 AM | C:\Sirius_T3\17J-11006_Blank standardisation.t3r |
| Four-Plus jOH | -0.2 | 10/12/2017 9:40:50 AM | C:\Sirius_T3\17J-11006_Blank standardisation.t3r |
| Base concentration factor | 1.011 | 10/12/2017 9:40:50 AM | C:\Sirius_T3\KOH17I22.t3r |
| Acid concentration factor | 0.995 | 10/12/2017 9:40:50 AM | C:\Sirius_T3\17J-11005_Blank standardisation.t3r |

#### Instrument Settings

| Setting | Value | Batch Id | Install date |
|---|---|---|---|
| Instrument owner | Merck | | |
| Instrument ID | T311053 | | |
| Instrument type | T3 Simulator | | |
| Software version | 1.1.3.0 | | |
| Dispenser module | | T3DM1100253 | 3/31/2009 6:24:52 AM |
| Dispenser 0 | Water | | 3/31/2009 6:25:05 AM |
| Syringe volume | 2.5 mL | | |
| Firmware version | 1.2.1(r2) | | |
| Titrant | Water (0.15 M KCl) | 10-10-2017 | 10/10/2017 10:48:53 AM |
| Dispenser 2 | Acid | | 3/31/2009 6:25:11 AM |
| Syringe volume | 0.5 mL | | |
| Firmware version | 1.2.1(r2) | | |

| | | | |
|---|---|---|---|
| Sample name: | **M15** | Experiment start time: | **10/12/2017 9:40:50 AM** |
| Assay name: | **UV-metric psKa** | Analyst: | **Dorothy Levorse** |
| Assay ID: | **17J-12008** | Instrument ID: | **T311053** |
| Filename: | **C:\Sirius_T3\Mehtap\20171011_exp15_pKa\17J-12008_M15_UV-metric psKa.t3r** | | |

#### Instrument Settings (continued)

| Setting | Value | Batch Id | Install date |
|---|---|---|---|
| Titrant | Acid (0.5 M HCl) | 166940 and 172875 | 10/6/2017 2:55:40 PM |
| Dispenser 1 | Base | | 3/31/2009 6:25:21 AM |
| Syringe volume | 0.5 mL | | |
| Firmware version | 1.2.1(r2) | | |
| Titrant | Base (0.5 M KOH) | 9-22-17 | 9/22/2017 4:02:42 PM |
| Dispenser 5 | Cosolvent | | 3/31/2009 6:26:24 AM |
| Syringe volume | 2.5 mL | | |
| Firmware version | 1.2.1(r2) | | |
| Distribution valve 5 | Distribution Valve | | 3/31/2009 6:28:19 AM |
| Firmware version | 1.1.3 | | |
| Port A | Methanol (80%, 0.15 M KCl) | 9-26-17 | 10/5/2017 5:02:03 PM |
| Port B | Cyclohexane | | 9/19/2017 2:15:02 PM |
| Port C | MeCN (50%, 0.15 M KCl) | 10-2-17 | 10/2/2017 11:28:55 AM |
| Dispenser 3 | Buffer | | 8/3/2010 6:05:16 AM |
| Syringe volume | 0.5 mL | | |
| Firmware version | 1.2.1(r2) | | |
| Titrant | Phosphate Buffer | | 10/10/2017 9:57:33 AM |
| Dispenser 6 | Octanol | | 10/22/2010 11:52:43 AM |
| Syringe volume | 0.5 mL | | |
| Firmware version | 1.2.1(r2) | | |
| Titrant | Octanol | 9-14-17 | 9/14/2017 10:30:38 AM |
| Titrator | | T3TM1100153 | 3/31/2009 6:24:17 AM |
| Horizontal axis firmware version | 1.17 AI1DI2DO2 Stepper 2 | | |
| Vertical axis firmware version | 1.17 AI1DI2DO2 Stepper 2 | | |
| Chassis I/O firmware version | 1.11 AI1DI0DO4 Norgren I/O | | |
| Probe I/O firmware version | 1.1.1 | | |
| Electrode | T3 Electrode | T3E0769 | 8/15/2017 10:21:54 AM |
| E0 calibration | -9.19 mV | | 10/12/2017 9:41:14 AM |
| Filling solution | 3M KCl | KCL095 | 10/10/2017 9:58:43 AM |
| Liquids | | | |
| Wash 1 | 50% IPA:50% Water | | 10/11/2017 8:31:15 AM |
| Wash 2 | 0.5% Trition X-100 in H20 | | 10/11/2017 8:31:17 AM |
| Buffer position 1 | pH7 Wash | | 10/11/2017 8:31:21 AM |
| Buffer position 2 | pH 7 | | 10/11/2017 8:31:23 AM |
| Storage position | | | 10/11/2017 8:31:26 AM |
| Wash water | 4.2e+003 mL | 10-6-17 | 10/6/2017 3:04:25 PM |
| Waste | 5.9e+003 mL | | 10/6/2017 3:04:33 PM |
| Temperature controller | | | 8/5/2010 7:35:13 AM |
| Turbidity detector | | | 3/31/2009 6:24:45 AM |
| Spectrometer | | 072390 | 11/23/2010 12:22:28 PM |
| Dip probe | | 11086 | |
| Wavelength coefficient A0 | 185.563 | | |
| Wavelength coefficient A1 | 2.17439 | | |
| Wavelength coefficient A2 | -0.000285622 | | |
| Total lamp lit time | 419:28:33 | | 11/23/2010 12:22:28 PM |
| Calibrated on | 10/11/2017 8:30:19 AM | | |
| Integration time | 10 | | |
| Scans averaged | 10 | | |
| Autoloader | | T3AL1100237 | 11/10/2015 10:34:13 AM |
| Left-right axis firmware version | 1.17 AI1DI2DO2 Stepper 2 | | |
| Front-back axis firmware version | 1.17 AI1DI2DO2 Stepper 2 | | |
| Vertical axis firmware version | 1.17 AI1DI2DO2 Stepper 2 | | |
| Chassis I/O firmware version | 1.11 AI1DI0DO4 Norgren I/O | | |
| Configuration | | | |
| Alternate titration position | Titration position | | |
| Alternate reference position | Reference position | | |
| Maximum standard vial volume | 3.50 mL | | |
| Maximum alternate vial volume | 25.00 mL | | |

| | | | |
|---|---|---|---|
| Sample name: | **M15** | Experiment start time: | **10/12/2017 9:40:50 AM** |
| Assay name: | **UV-metric psKa** | Analyst: | **Dorothy Levorse** |
| Assay ID: | **17J-12008** | Instrument ID: | **T311053** |
| Filename: | **C:\Sirius_T3\Mehtap\20171011_exp15_pKa\17J-12008_M15_UV-metric psKa.t3r** | | |

#### Instrument Settings (continued)

| Setting | Value | Batch Id | Install date |
|---|---|---|---|
| Automatic action idle period | 5 minute(s) | | |
| Titrant tube volume | 1.3 mL | | |
| Syringe flush count | 3.50 | | |
| Flowing wash pump volume | 20.0 mL | | |
| Flowing wash stir duration | 5 s | | |
| Flowing wash stir speed | 30% | | |
| Solvent wash stir duration | 5 s | | |
| Solvent wash stir speed | 30% | | |
| Surfactant wash stir duration | 5 s | | |
| Surfactant wash stir speed | 30% | | |
| E0 calibration minimum number of points | 10 | | |
| E0 calibration maximum standard deviation | 0.01500 | | |
| E0 calibration timeout period | 60 s | | |
| E0 calibration stir duration | 5 s | | |
| E0 calibration preparation stir speed | 30% | | |
| E0 calibration buffer wash stir duration | 5 s | | |
| E0 calibration buffer wash stir speed | 30% | | |
| E0 calibration reading stir speed | 0% | | |
| Spectrometer calibration stir duration | 5 s | | |
| Spectrometer calibration stir speed | 30% | | |
| Spectrometer calibration wash pump volume | 20.0 mL | | |
| Spectrometer calibration wash stir duration | 5 s | | |
| Spectrometer calibration wash stir speed | 30% | | |
| Overhead dispense height | 10000 | | |

#### Refinement Settings

| Setting | Value | Default value |
|---|---|---|
| Turbidity detection method | Spectrometer | Spectrometer |
| Turbidity wavelength to assess | 500.0 nm | 500.0 nm |
| Turbidity maximum absorbance | 0.100 | 0.100 |
| Turbidity probe threshold | 50.00 | 50.00 |
| Exclude turbid points | Yes | Yes |
| Low intensity warning threshold | 100 | 100 |
| Minimum absorbance change threshold | 0.100 | 0.100 |
| Eigenvector autocorrelation threshold | 0.80 | 0.80 |
| Maximum RMSD severe warning | 0.250 | 0.250 |
| Maximum RMSD warning | 0.050 | 0.050 |

#### Tray Information

Title

Location E1
