## Supplementary material for "Octanol-water partition coefficient measurements for the SAMPL6 Blind Prediction Challenge": SM15_17J-12009_M16_UV-metric psKa_report.pdf

| | | | |
|---|---|---|---|
| Sample name: | **M16** | Experiment start time: | **10/12/2017 10:50:14 AM** |
| Assay name: | **UV-metric psKa** | Analyst: | **Dorothy Levorse** |
| Assay ID: | **17J-12009** | Instrument ID: | **T311053** |
| Filename: | **C:\Sirius_T3\Mehtap\20171011_exp15_pKa\17J-12009_M16_UV-metric psKa.t3r** | | |

#### Yasuda-Shedlovsky result

| Extrapolation type | pKa 0% | SD | Intercept | Slope | R² | Ionic strength | Temperature |
|---|---|---|---|---|---|---|---|
| Yasuda-Shedlovsky | 4.67 | ±0.06 | 9.50 | -242.2362 | 0.9963 | 0.165 M | 24.9°C |
| Yasuda-Shedlovsky | 9.01 | ±0.02 | 10.19 | 44.3207 | 0.9912 | 0.165 M | 24.9°C |

#### Component assay results

| Titration | Methanol weight% | Direction | Result type | Dielectric constant | [H2O] | Ionic strength | Temperature | | psKa 1 | | psKa 2 |
|---|---|---|---|---|---|---|---|---|---|---|---|
| 17J-12009 Points 4 to 42 | 49.50 % | Up | UV-metric pKa | 56.6 | 24.7 M | 0.157 M | 24.9°C | ☑ | 3.84 | ☑ | 9.58 |
| 17J-12009 Points 44 to 84 | 39.91 % | Up | UV-metric pKa | 61.0 | 30.1 M | 0.166 M | 24.9°C | ☑ | 4.03 | ☑ | 9.45 |
| 17J-12009 Points 86 to 128 | 30.13 % | Up | UV-metric pKa | 65.5 | 35.8 M | 0.172 M | 24.9°C | ☑ | 4.26 | ☑ | 9.31 |

#### Graphs

Yasuda-Shedlovsky extrapolation

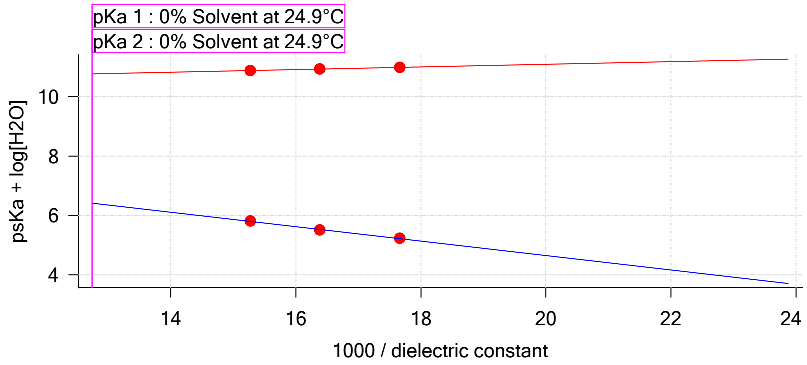


Distribution of Species - Yasuda-Shedlovsky Extrapolation

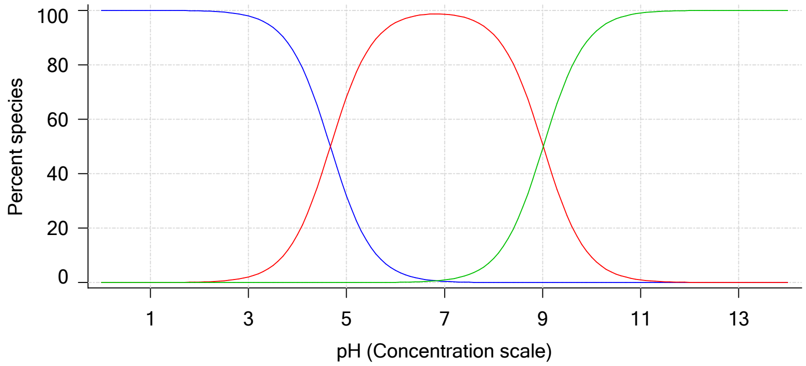


Mean Molar Extinction Profiles

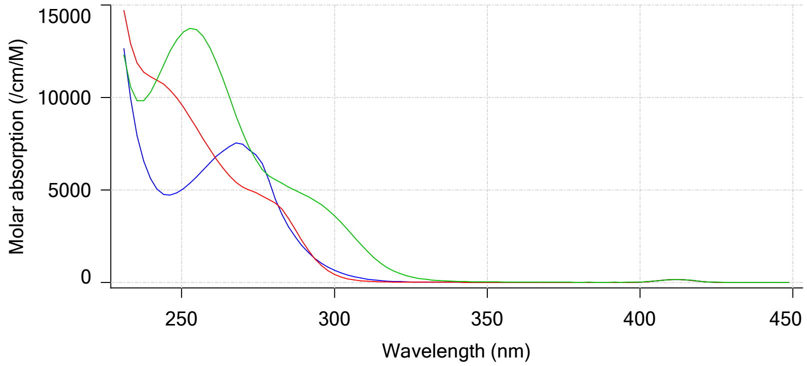


# UV-metric psKa   Titration 1 of 3   17J-12009 Points 4 to 42

#### Results

| | |
|---|---|
| pKa 1 | **3.84** |
| pKa 2 | **9.58** |
| RMSD | **0.001 0.001 0.001** |
| Chi squared | **0.0049** |
| PCA calculated number of pKas | **2** |
| Average ionic strength | **0.157 M** |
| Average temperature | **24.9°C** |
| Analyte concentration range | **68.6 µM to 64.5 µM** |
| Methanol weight % | **49.5 %** |
| Dielectric constant | **56.6** |
| Water concentration | **24.7 M** |

Sample name: **M16**
Assay name: **UV-metric psKa**
Assay ID: **17J-12009**
Filename: **C:\Sirius_T3\Mehtap\20171011_exp15_pKa\17J-12009_M16_UV-metric psKa.t3r**

Experiment start time: **10/12/2017 10:50:14 AM**
Analyst: **Dorothy Levorse**
Instrument ID: **T311053**

#### Results (continued)

| | |
|---|---|
| Number of pKas source | **Predicted** |
| Wavelength clipping | **230.0 nm to 450.0 nm** |
| pH clipping | **1.471 to 12.517** |

#### Warnings and errors

| | |
|---|---|
| Errors | None |
| Warnings | None |

#### Assay Settings

| Setting | Value | Original Value | Date/Time changed | Imported from |
|---|---|---|---|---|
| Buffer in use | Yes | | | |
| Buffer type | Phosphate Buffer | | | |
| ***Assay Medium*** | | | | |
| Volume of buffer introduced | 0.025000 mL | | | |
| Add buffer manually | Manual | | | |

#### Graphs

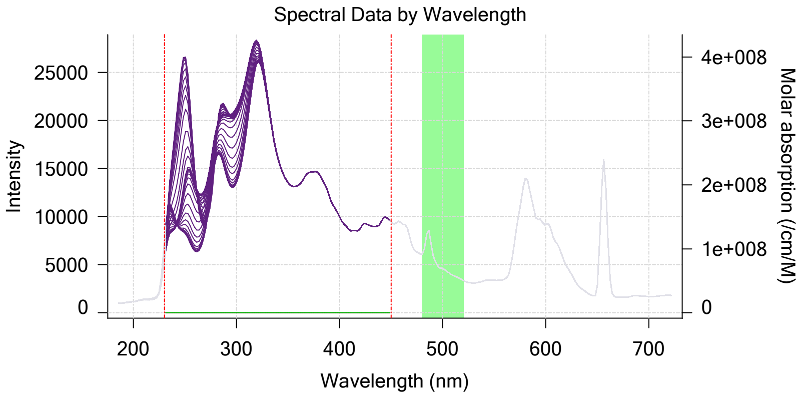


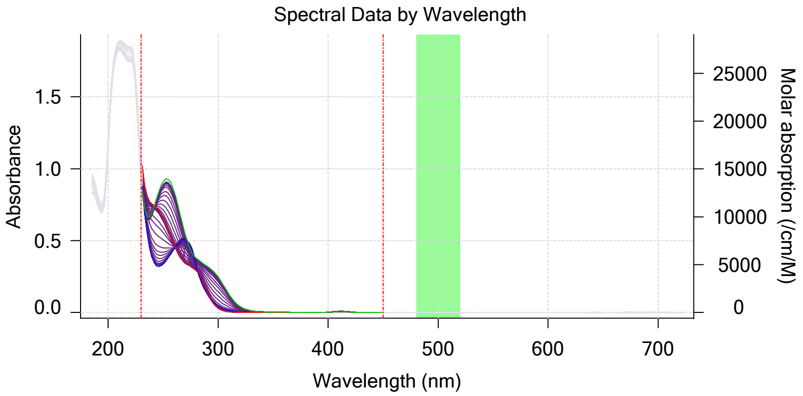


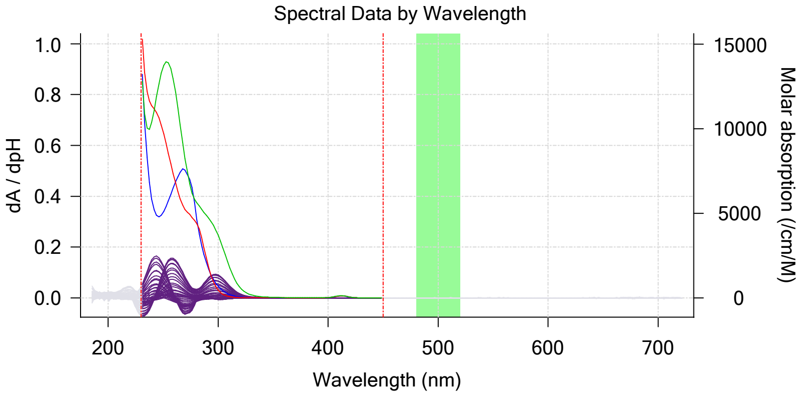


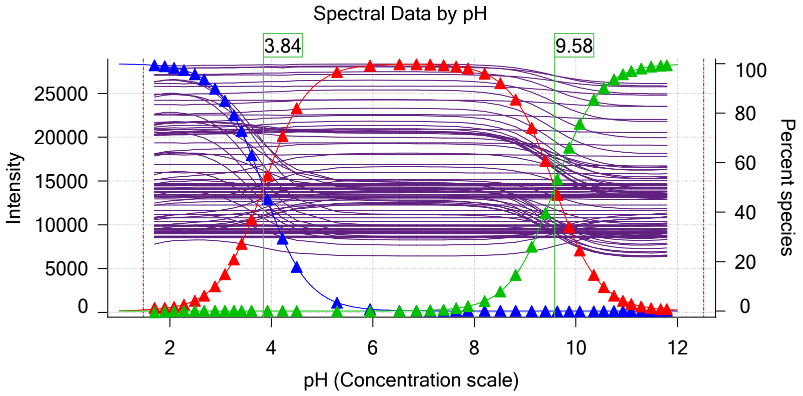

Sample name: **M16**
Assay name: **UV-metric psKa**
Assay ID: **17J-12009**
Filename: **C:\Sirius_T3\Mehtap\20171011_exp15_pKa\17J-12009_M16_UV-metric psKa.t3r**

Experiment start time: **10/12/2017 10:50:14 AM**
Analyst: **Dorothy Levorse**
Instrument ID: **T311053**

#### Graphs (continued)

Spectral Data by pH

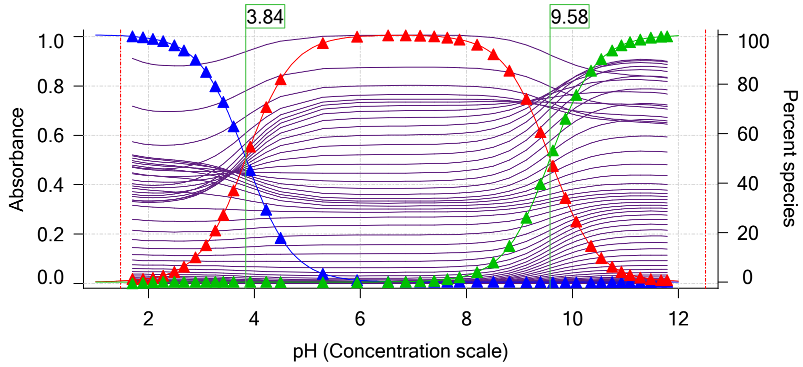


Spectral Data by pH

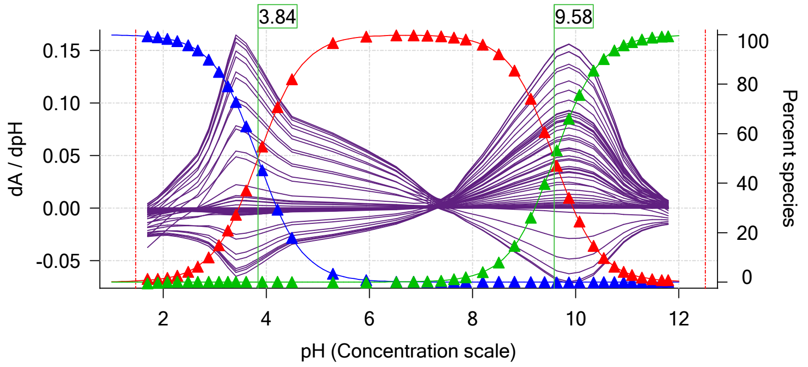


Distribution of Species

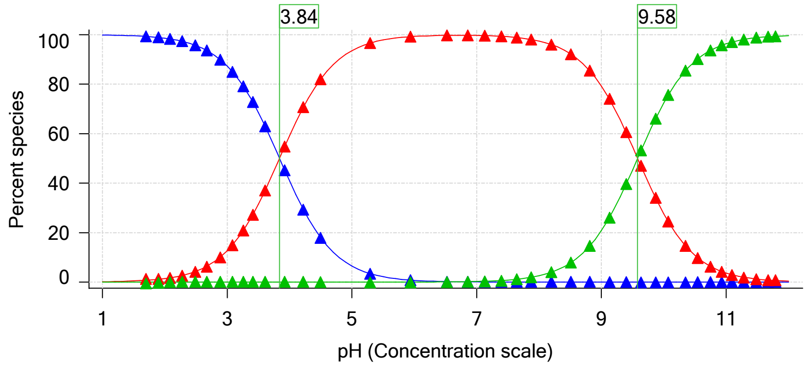


Molar Absorption Profile

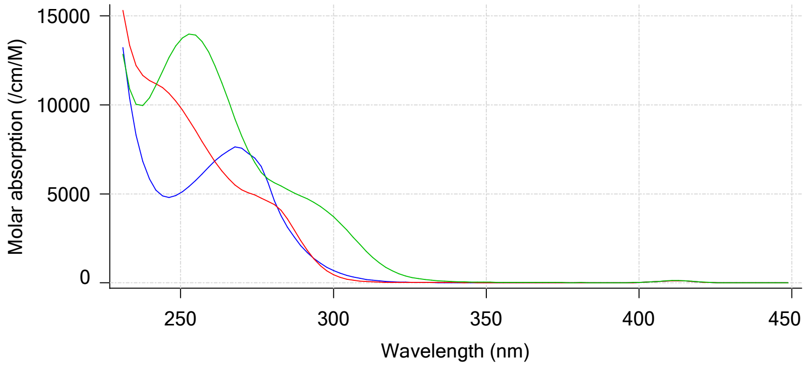


Baseline Correction Range

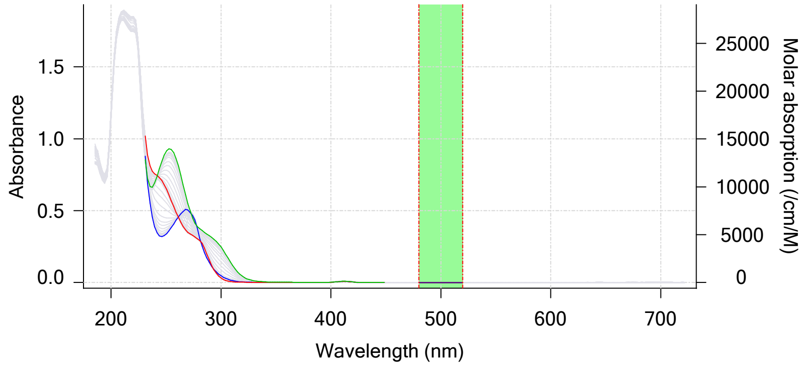


Raw Spectral Data by Wavelength

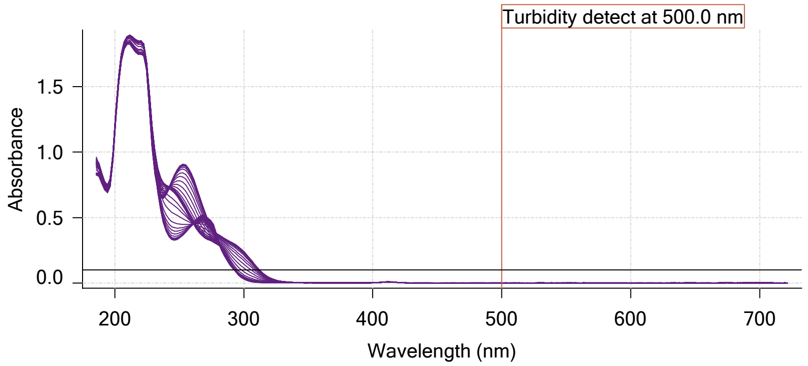


### UV-metric psKa Titration 2 of 3 17J-12009 Points 44 to 84

#### Results

| | |
|---|---|
| pKa 1 | **4.03** |
| pKa 2 | **9.45** |
| RMSD | **0.001 0.002 0.002** |
| Chi squared | **0.0062** |
| PCA calculated number of pKas | **2** |
| Average ionic strength | **0.166 M** |
| Average temperature | **24.9°C** |
| Analyte concentration range | **56.2 µM to 53.1 µM** |
| Methanol weight % | **39.9 %** |
| Dielectric constant | **61.0** |
| Water concentration | **30.1 M** |

| | | | |
|---|---|---|---|
| Sample name: | **M16** | Experiment start time: | **10/12/2017 10:50:14 AM** |
| Assay name: | **UV-metric psKa** | Analyst: | **Dorothy Levorse** |
| Assay ID: | **17J-12009** | Instrument ID: | **T311053** |
| Filename: | **C:\Sirius_T3\Mehtap\20171011_exp15_pKa\17J-12009_M16_UV-metric psKa.t3r** | | |

#### Results (continued)

| | |
|---|---|
| Number of pKas source | **Predicted** |
| Wavelength clipping | **230.0 nm to 450.0 nm** |
| pH clipping | **1.487 to 12.519** |

#### Warnings and errors

| | |
|---|---|
| Errors | None |
| Warnings | None |

#### Assay Settings

| **Setting** | **Value** | **Original Value** | **Date/Time changed** | **Imported from** |
|---|---|---|---|---|
| Buffer in use | Yes | | | |
| Buffer type | Phosphate Buffer | | | |
| ***Assay Medium*** | | | | |
| Volume of buffer introduced | 0.025000 mL | | | |
| Add buffer manually | Manual | | | |

#### Graphs

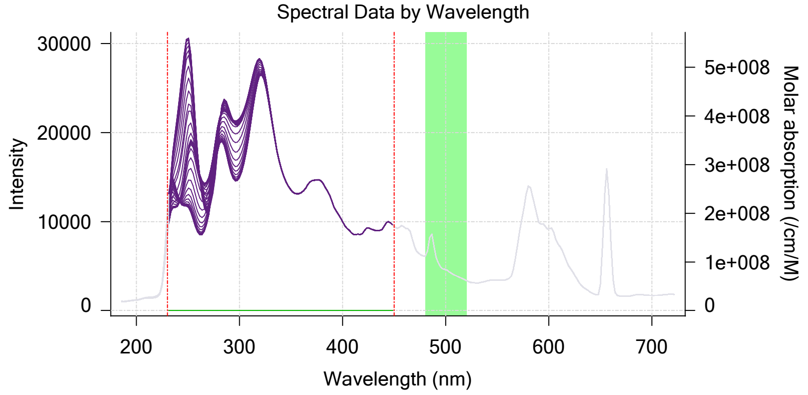


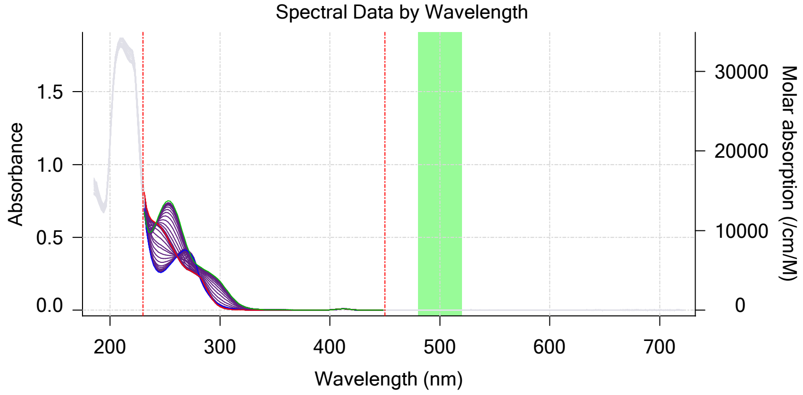


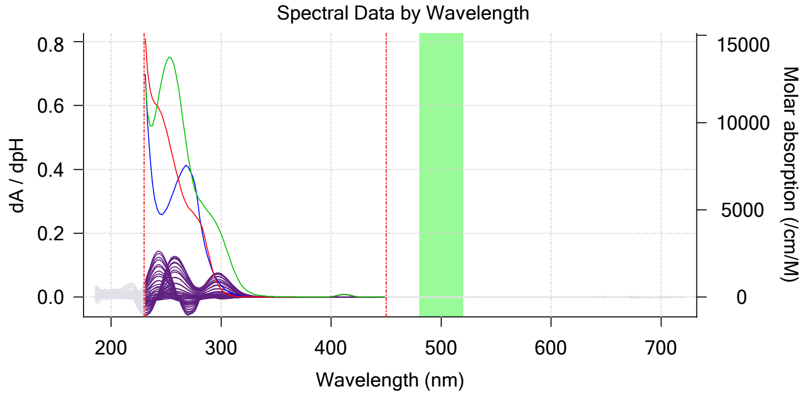


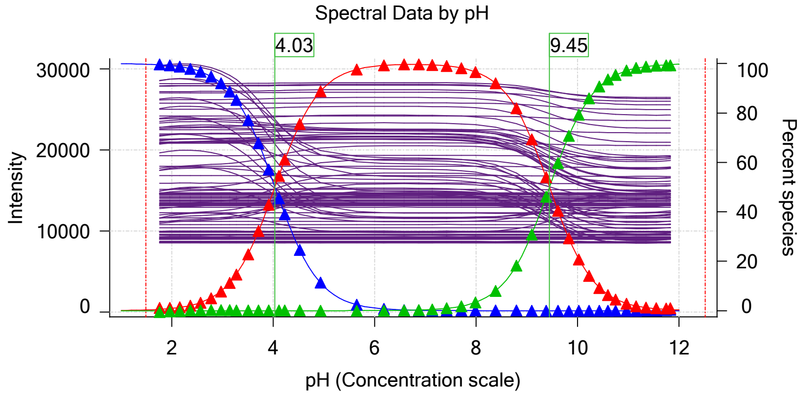

| | | | |
|---|---|---|---|
| Sample name: | **M16** | Experiment start time: | **10/12/2017 10:50:14 AM** |
| Assay name: | **UV-metric psKa** | Analyst: | **Dorothy Levorse** |
| Assay ID: | **17J-12009** | Instrument ID: | **T311053** |
| Filename: | **C:\Sirius_T3\Mehtap\20171011_exp15_pKa\17J-12009_M16_UV-metric psKa.t3r** | | |

#### Graphs (continued)

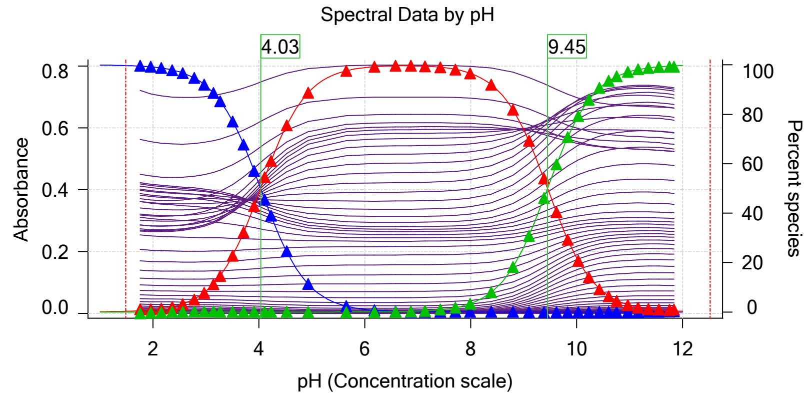


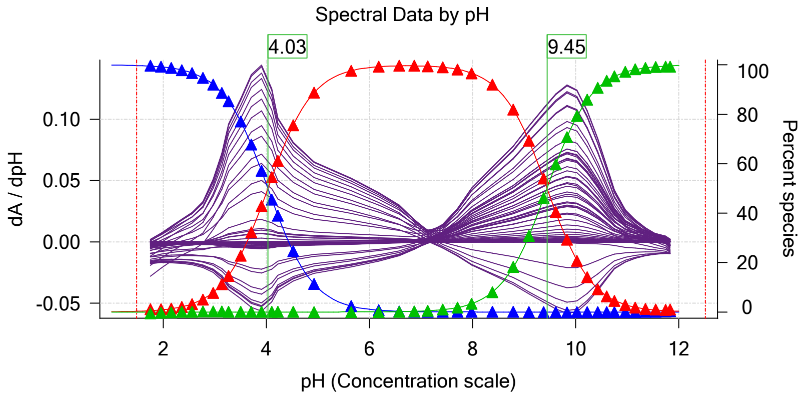


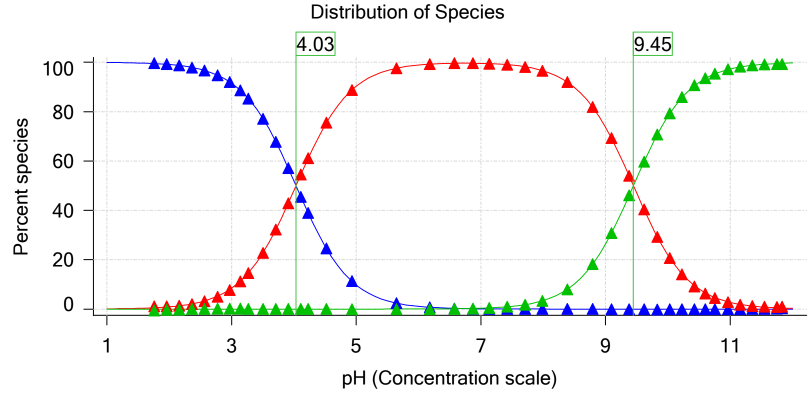


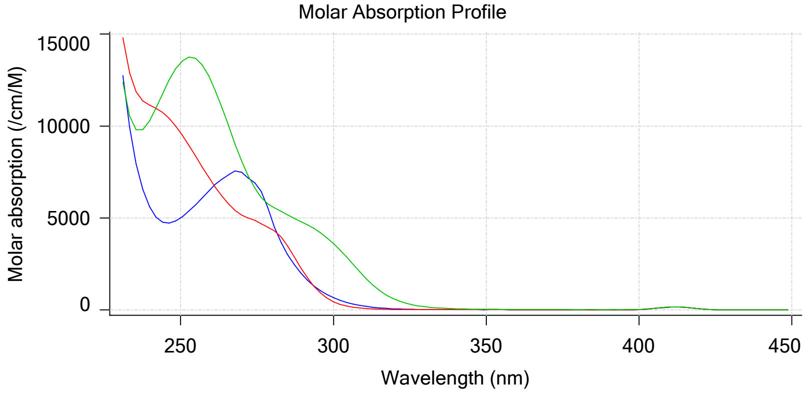


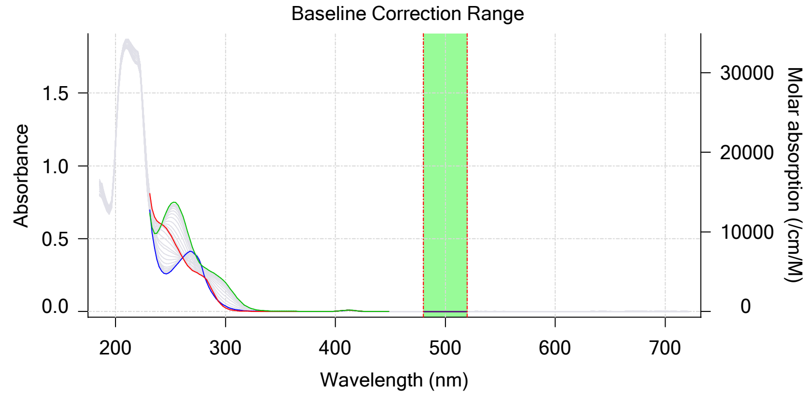


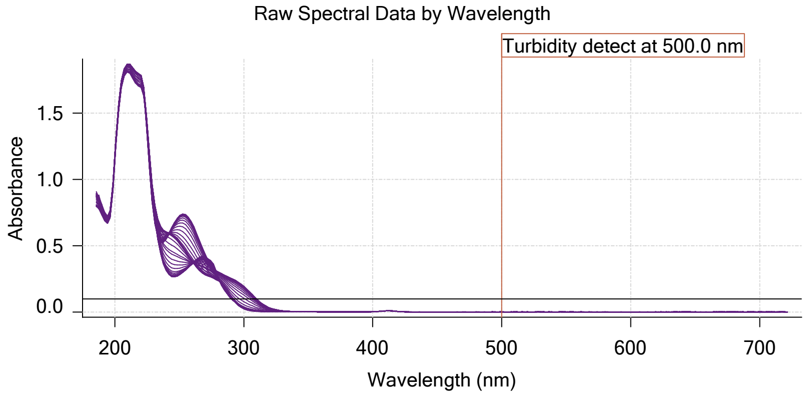


### UV-metric psKa Titration 3 of 3 17J-12009 Points 86 to 128

#### Results

| | |
|---|---|
| pKa 1 | **4.26** |
| pKa 2 | **9.31** |
| RMSD | **0.002 0.003 0.003** |
| Chi squared | **0.0098** |
| PCA calculated number of pKas | **3** |
| Average ionic strength | **0.172 M** |
| Average temperature | **24.9°C** |
| Analyte concentration range | **43.3 µM to 40.9 µM** |
| Methanol weight % | **30.1 %** |
| Dielectric constant | **65.5** |
| Water concentration | **35.8 M** |

| | | | |
|---|---|---|---|
| Sample name: | **M16** | Experiment start time: | **10/12/2017 10:50:14 AM** |
| Assay name: | **UV-metric psKa** | Analyst: | **Dorothy Levorse** |
| Assay ID: | **17J-12009** | Instrument ID: | **T311053** |
| Filename: | **C:\Sirius_T3\Mehtap\20171011_exp15_pKa\17J-12009_M16_UV-metric psKa.t3r** | | |

#### Results (continued)

| | |
|---|---|
| Number of pKas source | **Predicted** |
| Wavelength clipping | **230.0 nm to 450.0 nm** |
| pH clipping | **1.494 to 12.527** |

#### Warnings and errors

| | |
|---|---|
| Errors | None |
| Warnings | PCA calculation disagrees with predicted number of pKas |

#### Assay Settings

| Setting | Value | Original Value | Date/Time changed | Imported from |
|---|---|---|---|---|
| Buffer in use | Yes | | | |
| Buffer type | Phosphate Buffer | | | |
| ***Assay Medium*** | | | | |
| Volume of buffer introduced | 0.025000 mL | | | |
| Add buffer manually | Manual | | | |

#### Graphs

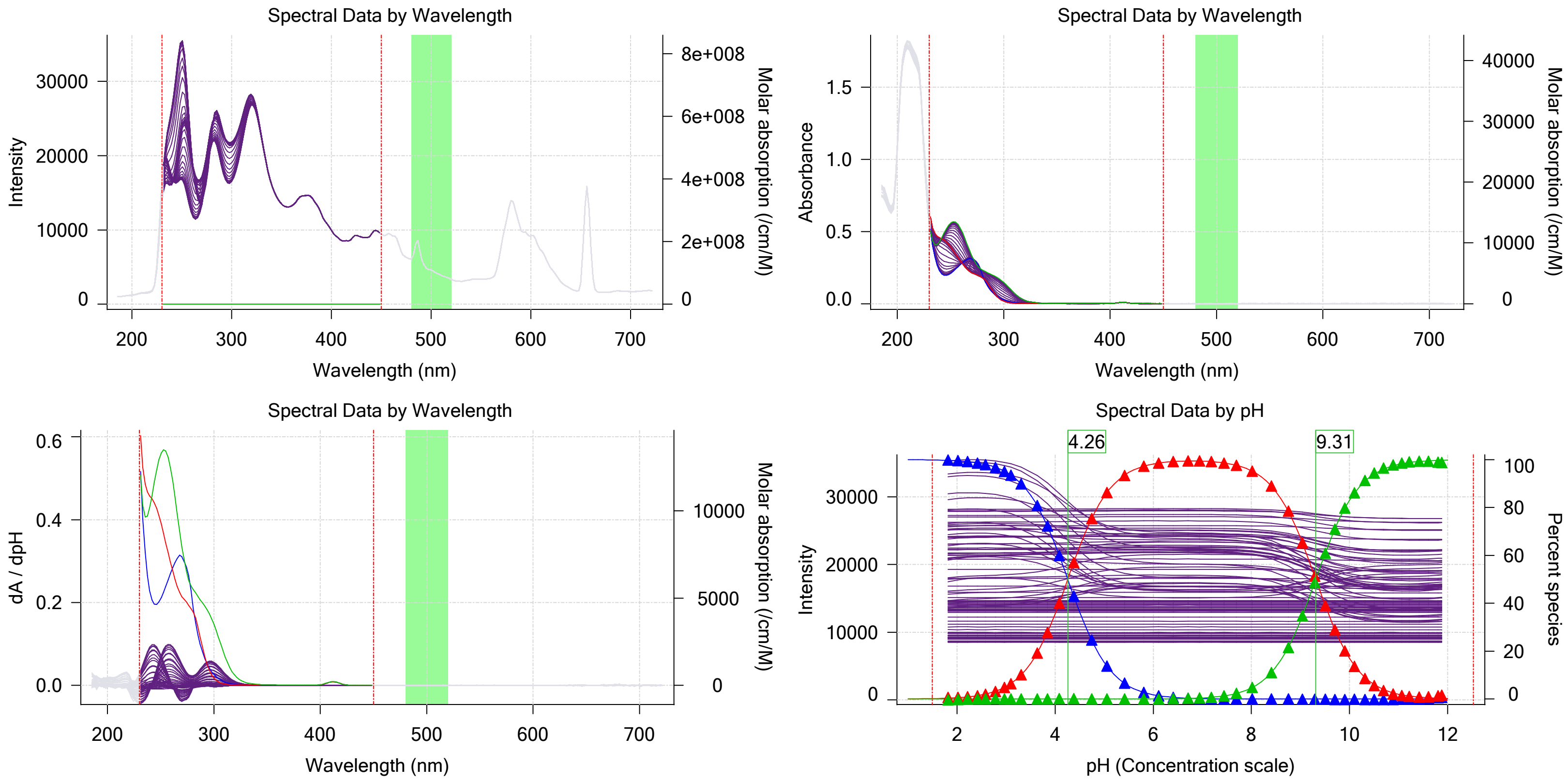

| | | | |
|---|---|---|---|
| Sample name: | **M16** | Experiment start time: | **10/12/2017 10:50:14 AM** |
| Assay name: | **UV-metric psKa** | Analyst: | **Dorothy Levorse** |
| Assay ID: | **17J-12009** | Instrument ID: | **T311053** |
| Filename: | **C:\Sirius_T3\Mehtap\20171011_exp15_pKa\17J-12009_M16_UV-metric psKa.t3r** | | |

#### Graphs (continued)

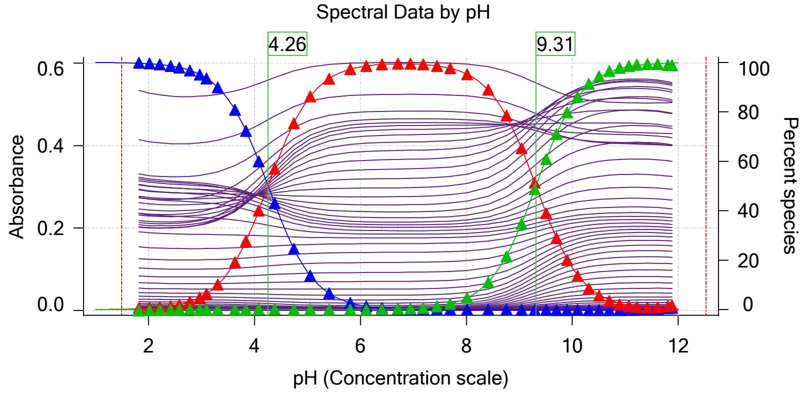


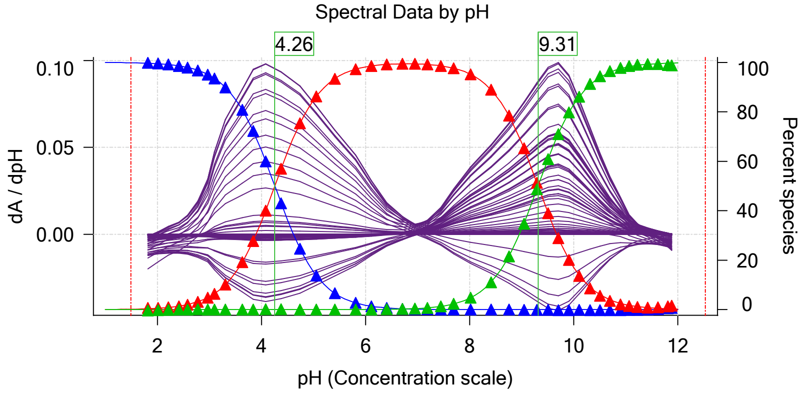


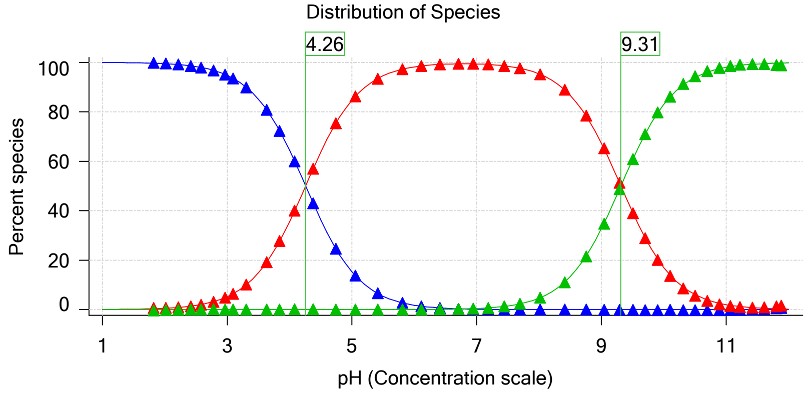


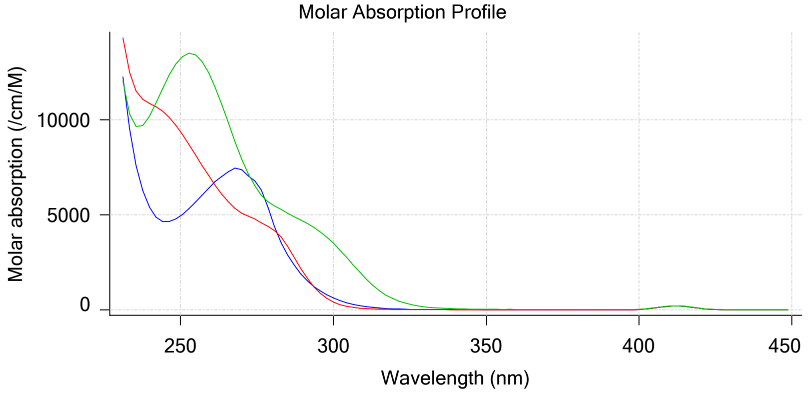


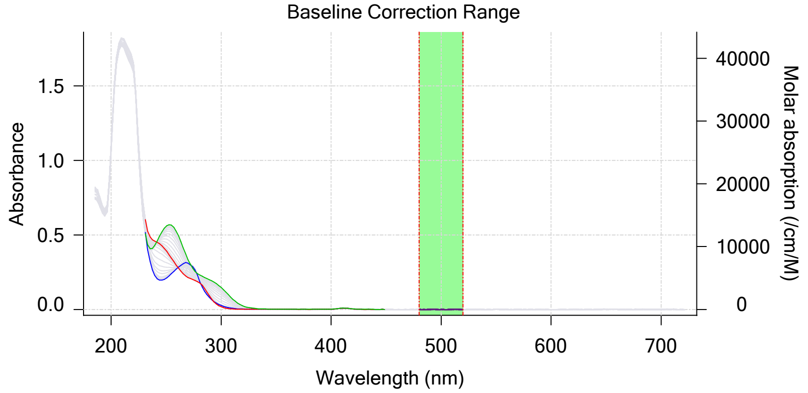


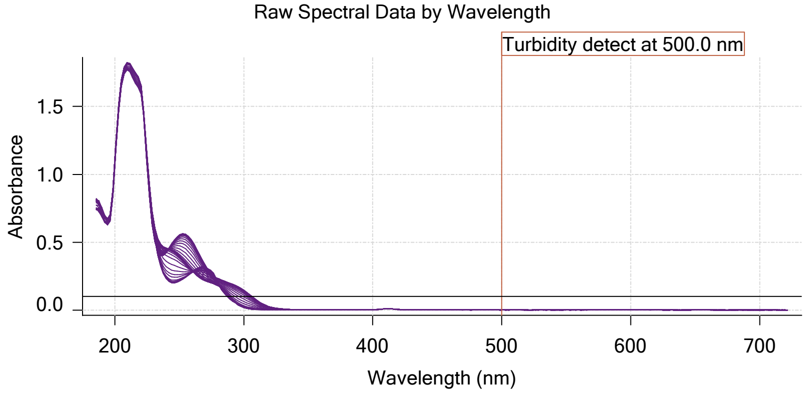


#### Assay Model

| Settings | Value | Date/Time changed | Imported from |
|---|---|---|---|
| Sample name | M16 | 10/11/2017 4:21:12 PM | User entered value |
| Sample by | Volume | | Default value |
| Sample volume | 0.0020 mL | 10/11/2017 4:21:12 PM | User entered value |
| Solvent | DMSO | | Default value |
| Sample concentration | 0.054700 M | 10/11/2017 4:21:12 PM | User entered value |
| Solubility | Unknown | | Default value |
| Molecular weight | 210.23 | 10/11/2017 4:21:22 PM | User entered value |
| Individual pKa ionic environments | No | | Default value |
| Number of pKas | 2 | 10/11/2017 4:21:12 PM | User entered value |
| Sample is a | Ampholyte | 10/11/2017 4:21:12 PM | User entered value |
| pKa 1 | 4.86 | 10/11/2017 4:21:12 PM | User entered value |
| Type | Base | 10/11/2017 4:21:12 PM | User entered value |
| pKa 2 | 9.07 | 10/11/2017 4:21:12 PM | User entered value |
| Type | Acid | 10/11/2017 4:21:12 PM | User entered value |

Sample name: **M16**
Assay name: **UV-metric psKa**
Assay ID: **17J-12009**
Filename: **C:\Sirius_T3\Mehtap\20171011_exp15_pKa\17J-12009_M16_UV-metric psKa.t3r**

Experiment start time: **10/12/2017 10:50:14 AM**
Analyst: **Dorothy Levorse**
Instrument ID: **T311053**

#### Assay Model (continued)

| Settings | Value | Date/Time changed | Imported from |
|---|---|---|---|
| logp (XH2 +) | -10.00 | | Default value |
| logP (neutral XH) | -10.00 | 10/11/2017 4:21:12 PM | User entered value |
| logP (X -) | -10.00 | | Default value |

#### Events

| Time | Event | Water | Acid | Base | Methanol | Buffer | pH | dpH/dt | pH R-squar |
|---|---|---|---|---|---|---|---|---|---|
| 3:40.5 | Dark spectrum | | | | | | | | |
| 3:42.0 | Reference spectrum | | | | | | | | |
| 4:09.7 | Volume reset due to vial change | | | | | | | | |
| 4:53.9 | Initial pH = 8.07 | | | | | | | | |
| 5:59.4 | Data point 4 | 0.34995 mL | 0.06860 mL | 0.00000 mL | 1.15005 mL | 0.02500 mL | 1.971 | -0.00967 | 0.69536 |
| 6:28.2 | Data point 5 | 0.34995 mL | 0.06860 mL | 0.02465 mL | 1.15005 mL | 0.02500 mL | 2.172 | -0.01785 | 0.72737 |
| 6:45.1 | Data point 6 | 0.34995 mL | 0.06860 mL | 0.03963 mL | 1.15005 mL | 0.02500 mL | 2.357 | 0.01480 | 0.92168 |
| 7:01.9 | Data point 7 | 0.34995 mL | 0.06860 mL | 0.04936 mL | 1.15005 mL | 0.02500 mL | 2.552 | 0.02578 | 0.84828 |
| 7:18.6 | Data point 8 | 0.34995 mL | 0.06860 mL | 0.05567 mL | 1.15005 mL | 0.02500 mL | 2.764 | 0.01968 | 0.80361 |
| 7:35.2 | Data point 9 | 0.34995 mL | 0.06860 mL | 0.05955 mL | 1.15005 mL | 0.02500 mL | 2.944 | 0.01316 | 0.85269 |
| 7:51.8 | Data point 10 | 0.34995 mL | 0.06860 mL | 0.06211 mL | 1.15005 mL | 0.02500 mL | 3.159 | 0.00948 | 0.75581 |
| 8:08.3 | Data point 11 | 0.34995 mL | 0.06860 mL | 0.06369 mL | 1.15005 mL | 0.02500 mL | 3.357 | 0.01152 | 0.78578 |
| 8:24.9 | Data point 12 | 0.34995 mL | 0.06860 mL | 0.06468 mL | 1.15005 mL | 0.02500 mL | 3.530 | 0.01310 | 0.90115 |
| 8:41.4 | Data point 13 | 0.34995 mL | 0.06860 mL | 0.06533 mL | 1.15005 mL | 0.02500 mL | 3.681 | 0.03201 | 0.96874 |
| 9:08.3 | Data point 14 | 0.34995 mL | 0.06860 mL | 0.06595 mL | 1.15005 mL | 0.02500 mL | 3.877 | 0.03400 | 0.97813 |
| 9:29.9 | Data point 15 | 0.34995 mL | 0.06860 mL | 0.06651 mL | 1.15005 mL | 0.02500 mL | 4.187 | 0.08154 | 0.99377 |
| 10:01.9 | Data point 16 | 0.34995 mL | 0.06860 mL | 0.06736 mL | 1.15005 mL | 0.02500 mL | 4.485 | 0.09919 | 0.99482 |
| 10:39.6 | Data point 17 | 0.34995 mL | 0.06860 mL | 0.06754 mL | 1.15005 mL | 0.02500 mL | 4.761 | 0.09769 | 0.98949 |
| 11:15.2 | Data point 18 | 0.34995 mL | 0.06860 mL | 0.06773 mL | 1.15005 mL | 0.02500 mL | 5.551 | 0.11230 | 0.99525 |
| 12:32.1 | Data point 19 | 0.34995 mL | 0.06860 mL | 0.06783 mL | 1.15005 mL | 0.02500 mL | 6.196 | 0.11157 | 0.99182 |
| 13:49.0 | Data point 20 | 0.34995 mL | 0.06860 mL | 0.06794 mL | 1.15005 mL | 0.02500 mL | 6.777 | 0.10014 | 0.99485 |
| 14:49.2 | Data point 21 | 0.34995 mL | 0.06860 mL | 0.06804 mL | 1.15005 mL | 0.02500 mL | 7.108 | 0.09753 | 0.98842 |
| 15:38.5 | Data point 22 | 0.34995 mL | 0.06860 mL | 0.06816 mL | 1.15005 mL | 0.02500 mL | 7.378 | 0.10099 | 0.99390 |
| 16:26.8 | Data point 23 | 0.34995 mL | 0.06860 mL | 0.06830 mL | 1.15005 mL | 0.02500 mL | 7.644 | 0.09933 | 0.99346 |
| 17:09.7 | Data point 24 | 0.34995 mL | 0.06860 mL | 0.06844 mL | 1.15005 mL | 0.02500 mL | 7.889 | 0.09867 | 0.98767 |
| 17:50.5 | Data point 25 | 0.34995 mL | 0.06860 mL | 0.06858 mL | 1.15005 mL | 0.02500 mL | 8.119 | 0.08217 | 0.72434 |
| 18:28.2 | Data point 26 | 0.34995 mL | 0.06860 mL | 0.06872 mL | 1.15005 mL | 0.02500 mL | 8.445 | 0.09452 | 0.96667 |
| 19:06.5 | Data point 27 | 0.34995 mL | 0.06860 mL | 0.06884 mL | 1.15005 mL | 0.02500 mL | 8.752 | 0.09935 | 0.99241 |
| 19:44.8 | Data point 28 | 0.34995 mL | 0.06860 mL | 0.06893 mL | 1.15005 mL | 0.02500 mL | 9.053 | 0.09736 | 0.98825 |
| 20:27.1 | Data point 29 | 0.34995 mL | 0.06860 mL | 0.06905 mL | 1.15005 mL | 0.02500 mL | 9.370 | 0.09941 | 0.98377 |
| 21:08.0 | Data point 30 | 0.34995 mL | 0.06860 mL | 0.06917 mL | 1.15005 mL | 0.02500 mL | 9.636 | 0.09723 | 0.97756 |
| 21:41.7 | Data point 31 | 0.34995 mL | 0.06860 mL | 0.06931 mL | 1.15005 mL | 0.02500 mL | 9.873 | 0.10040 | 0.98895 |
| 22:06.8 | Data point 32 | 0.34995 mL | 0.06860 mL | 0.06947 mL | 1.15005 mL | 0.02500 mL | 10.105 | 0.07554 | 0.97877 |
| 22:33.5 | Data point 33 | 0.34995 mL | 0.06860 mL | 0.06968 mL | 1.15005 mL | 0.02500 mL | 10.307 | 0.04184 | 0.93948 |
| 22:50.0 | Data point 34 | 0.34995 mL | 0.06860 mL | 0.06997 mL | 1.15005 mL | 0.02500 mL | 10.584 | 0.01590 | 0.90148 |
| 23:11.7 | Data point 35 | 0.34995 mL | 0.06860 mL | 0.07044 mL | 1.15005 mL | 0.02500 mL | 10.778 | 0.00406 | 0.39940 |
| 23:28.3 | Data point 36 | 0.34995 mL | 0.06860 mL | 0.07124 mL | 1.15005 mL | 0.02500 mL | 10.983 | -0.00419 | 0.36656 |
| 23:44.8 | Data point 37 | 0.34995 mL | 0.06860 mL | 0.07253 mL | 1.15005 mL | 0.02500 mL | 11.159 | -0.00606 | 0.49103 |
| 24:01.5 | Data point 38 | 0.34995 mL | 0.06860 mL | 0.07444 mL | 1.15005 mL | 0.02500 mL | 11.320 | -0.00736 | 0.63490 |
| 24:28.6 | Data point 39 | 0.34995 mL | 0.06860 mL | 0.07827 mL | 1.15005 mL | 0.02500 mL | 11.513 | -0.00938 | 0.89098 |
| 25:00.7 | Data point 40 | 0.34995 mL | 0.06860 mL | 0.08323 mL | 1.15005 mL | 0.02500 mL | 11.709 | -0.00915 | 0.76515 |
| 25:27.8 | Data point 41 | 0.34995 mL | 0.06860 mL | 0.09170 mL | 1.15005 mL | 0.02500 mL | 11.899 | -0.00817 | 0.77154 |
| 25:49.8 | Data point 42 | 0.34995 mL | 0.06860 mL | 0.09955 mL | 1.15005 mL | 0.02500 mL | 12.017 | -0.00561 | 0.60495 |
| 27:31.3 | Reference spectrum | | | | | | | | |
| 28:35.4 | Data point 44 | 0.50000 mL | 0.16928 mL | 0.09958 mL | 1.15005 mL | 0.02500 mL | 1.987 | -0.04742 | 0.92670 |
| 29:02.9 | Data point 45 | 0.50000 mL | 0.16928 mL | 0.12439 mL | 1.15005 mL | 0.02500 mL | 2.188 | 0.00964 | 0.81669 |
| 29:19.9 | Data point 46 | 0.50000 mL | 0.16928 mL | 0.14012 mL | 1.15005 mL | 0.02500 mL | 2.386 | 0.01430 | 0.79035 |
| 29:36.7 | Data point 47 | 0.50000 mL | 0.16928 mL | 0.15014 mL | 1.15005 mL | 0.02500 mL | 2.592 | 0.00124 | 0.02477 |
| 29:53.3 | Data point 48 | 0.50000 mL | 0.16928 mL | 0.15644 mL | 1.15005 mL | 0.02500 mL | 2.790 | 0.02782 | 0.86790 |
| 30:10.0 | Data point 49 | 0.50000 mL | 0.16928 mL | 0.16042 mL | 1.15005 mL | 0.02500 mL | 3.002 | 0.01190 | 0.82364 |

| | | | |
|---|---|---|---|
| Sample name: | **M16** | Experiment start time: | **10/12/2017 10:50:14 AM** |
| Assay name: | **UV-metric psKa** | Analyst: | **Dorothy Levorse** |
| Assay ID: | **17J-12009** | Instrument ID: | **T311053** |
| Filename: | **C:\Sirius_T3\Mehtap\20171011_exp15_pKa\17J-12009_M16_UV-metric psKa.t3r** | | |

#### Events (continued)

| Time | Event | Water | Acid | Base | Methanol | Buffer | pH | dpH/dt | pH R-squared | p[illegible] S[illegible] |
|---|---|---|---|---|---|---|---|---|---|---|
| 30:26.6 | Data point 50 | 0.50000 mL | 0.16928 mL | 0.16287 mL | 1.15005 mL | 0.02500 mL | 3.194 | 0.01465 | 0.92113 | 0. |
| 30:43.3 | Data point 51 | 0.50000 mL | 0.16928 mL | 0.16444 mL | 1.15005 mL | 0.02500 mL | 3.366 | 0.00927 | 0.80485 | 0. |
| 30:59.8 | Data point 52 | 0.50000 mL | 0.16928 mL | 0.16550 mL | 1.15005 mL | 0.02500 mL | 3.495 | 0.02401 | 0.92832 | 0. |
| 31:31.7 | Data point 53 | 0.50000 mL | 0.16928 mL | 0.16660 mL | 1.15005 mL | 0.02500 mL | 3.726 | 0.01485 | 0.78700 | 0. |
| 32:03.8 | Data point 54 | 0.50000 mL | 0.16928 mL | 0.16722 mL | 1.15005 mL | 0.02500 mL | 3.931 | 0.04364 | 0.98886 | 0. |
| 32:25.5 | Data point 55 | 0.50000 mL | 0.16928 mL | 0.16750 mL | 1.15005 mL | 0.02500 mL | 4.129 | 0.06512 | 0.98133 | 0. |
| 32:42.1 | Data point 56 | 0.50000 mL | 0.16928 mL | 0.16769 mL | 1.15005 mL | 0.02500 mL | 4.331 | 0.09337 | 0.99119 | 0. |
| 32:58.6 | Data point 57 | 0.50000 mL | 0.16928 mL | 0.16780 mL | 1.15005 mL | 0.02500 mL | 4.448 | 0.09931 | 0.98052 | 0. |
| 33:23.2 | Data point 58 | 0.50000 mL | 0.16928 mL | 0.16794 mL | 1.15005 mL | 0.02500 mL | 4.742 | 0.09999 | 0.99406 | 0. |
| 34:00.5 | Data point 59 | 0.50000 mL | 0.16928 mL | 0.16806 mL | 1.15005 mL | 0.02500 mL | 5.148 | 0.09881 | 0.97125 | 0. |
| 34:46.2 | Data point 60 | 0.50000 mL | 0.16928 mL | 0.16816 mL | 1.15005 mL | 0.02500 mL | 5.862 | 0.09761 | 0.98120 | 0. |
| 35:42.3 | Data point 61 | 0.50000 mL | 0.16928 mL | 0.16823 mL | 1.15005 mL | 0.02500 mL | 6.393 | 0.10013 | 0.99317 | 0. |
| 36:35.5 | Data point 62 | 0.50000 mL | 0.16928 mL | 0.16830 mL | 1.15005 mL | 0.02500 mL | 6.788 | 0.10099 | 0.99485 | 0. |
| 37:12.1 | Data point 63 | 0.50000 mL | 0.16928 mL | 0.16837 mL | 1.15005 mL | 0.02500 mL | 7.080 | 0.09927 | 0.98962 | 0. |
| 37:49.7 | Data point 64 | 0.50000 mL | 0.16928 mL | 0.16846 mL | 1.15005 mL | 0.02500 mL | 7.341 | 0.10010 | 0.98426 | 0. |
| 38:29.6 | Data point 65 | 0.50000 mL | 0.16928 mL | 0.16858 mL | 1.15005 mL | 0.02500 mL | 7.630 | 0.09718 | 0.98219 | 0. |
| 39:08.4 | Data point 66 | 0.50000 mL | 0.16928 mL | 0.16870 mL | 1.15005 mL | 0.02500 mL | 7.916 | 0.09773 | 0.97502 | 0. |
| 39:50.3 | Data point 67 | 0.50000 mL | 0.16928 mL | 0.16879 mL | 1.15005 mL | 0.02500 mL | 8.191 | 0.09950 | 0.98346 | 0. |
| 40:32.7 | Data point 68 | 0.50000 mL | 0.16928 mL | 0.16889 mL | 1.15005 mL | 0.02500 mL | 8.588 | 0.09613 | 0.96963 | 0. |
| 41:16.5 | Data point 69 | 0.50000 mL | 0.16928 mL | 0.16898 mL | 1.15005 mL | 0.02500 mL | 8.995 | 0.09558 | 0.97693 | 0. |
| 41:52.6 | Data point 70 | 0.50000 mL | 0.16928 mL | 0.16907 mL | 1.15005 mL | 0.02500 mL | 9.298 | 0.09442 | 0.96942 | 0. |
| 42:24.7 | Data point 71 | 0.50000 mL | 0.16928 mL | 0.16919 mL | 1.15005 mL | 0.02500 mL | 9.580 | 0.09533 | 0.93985 | 0. |
| 42:51.8 | Data point 72 | 0.50000 mL | 0.16928 mL | 0.16933 mL | 1.15005 mL | 0.02500 mL | 9.816 | 0.08831 | 0.97822 | 0. |
| 43:23.7 | Data point 73 | 0.50000 mL | 0.16928 mL | 0.16952 mL | 1.15005 mL | 0.02500 mL | 10.029 | 0.04734 | 0.94350 | 0. |
| 43:55.4 | Data point 74 | 0.50000 mL | 0.16928 mL | 0.16978 mL | 1.15005 mL | 0.02500 mL | 10.224 | 0.03020 | 0.94846 | 0. |
| 44:22.0 | Data point 75 | 0.50000 mL | 0.16928 mL | 0.17013 mL | 1.15005 mL | 0.02500 mL | 10.421 | 0.00656 | 0.39990 | 0. |
| 44:38.6 | Data point 76 | 0.50000 mL | 0.16928 mL | 0.17063 mL | 1.15005 mL | 0.02500 mL | 10.623 | 0.00409 | 0.37662 | 0. |
| 44:55.0 | Data point 77 | 0.50000 mL | 0.16928 mL | 0.17140 mL | 1.15005 mL | 0.02500 mL | 10.796 | -0.00702 | 0.60318 | 0. |
| 45:11.6 | Data point 78 | 0.50000 mL | 0.16928 mL | 0.17255 mL | 1.15005 mL | 0.02500 mL | 10.948 | -0.00557 | 0.68474 | 0. |
| 45:48.7 | Data point 79 | 0.50000 mL | 0.16928 mL | 0.17495 mL | 1.15005 mL | 0.02500 mL | 11.159 | -0.00838 | 0.90101 | 0. |
| 46:15.8 | Data point 80 | 0.50000 mL | 0.16928 mL | 0.17808 mL | 1.15005 mL | 0.02500 mL | 11.349 | -0.00924 | 0.85101 | 0. |
| 46:48.0 | Data point 81 | 0.50000 mL | 0.16928 mL | 0.18342 mL | 1.15005 mL | 0.02500 mL | 11.544 | -0.01208 | 0.87745 | 0. |
| 47:20.2 | Data point 82 | 0.50000 mL | 0.16928 mL | 0.19182 mL | 1.15005 mL | 0.02500 mL | 11.741 | -0.00938 | 0.82417 | 0. |
| 47:52.7 | Data point 83 | 0.50000 mL | 0.16928 mL | 0.20562 mL | 1.15005 mL | 0.02500 mL | 11.936 | -0.01072 | 0.80276 | 0. |
| 48:14.7 | Data point 84 | 0.50000 mL | 0.16928 mL | 0.21383 mL | 1.15005 mL | 0.02500 mL | 12.019 | -0.00860 | 0.76892 | 0. |
| 49:59.7 | Reference spectrum | | | | | | | | | |
| 51:23.3 | Data point 86 | 0.83996 mL | 0.29421 mL | 0.21385 mL | 1.15005 mL | 0.02500 mL | 1.994 | -0.02349 | 0.94569 | 0. |
| 51:50.9 | Data point 87 | 0.83996 mL | 0.29421 mL | 0.24224 mL | 1.15005 mL | 0.02500 mL | 2.191 | 0.01188 | 0.78011 | 0. |
| 52:07.8 | Data point 88 | 0.83996 mL | 0.29421 mL | 0.26011 mL | 1.15005 mL | 0.02500 mL | 2.388 | 0.00437 | 0.31795 | 0. |
| 52:24.7 | Data point 89 | 0.83996 mL | 0.29421 mL | 0.27161 mL | 1.15005 mL | 0.02500 mL | 2.589 | -0.02030 | 0.82416 | 0. |
| 52:57.1 | Data point 90 | 0.83996 mL | 0.29421 mL | 0.27895 mL | 1.15005 mL | 0.02500 mL | 2.756 | 0.00976 | 0.65367 | 0. |
| 53:24.2 | Data point 91 | 0.83996 mL | 0.29421 mL | 0.28335 mL | 1.15005 mL | 0.02500 mL | 2.951 | 0.00170 | 0.14583 | 0. |
| 53:40.9 | Data point 92 | 0.83996 mL | 0.29421 mL | 0.28652 mL | 1.15005 mL | 0.02500 mL | 3.140 | -0.00336 | 0.21309 | 0. |
| 53:57.5 | Data point 93 | 0.83996 mL | 0.29421 mL | 0.28857 mL | 1.15005 mL | 0.02500 mL | 3.268 | 0.00344 | 0.27724 | 0. |
| 54:29.4 | Data point 94 | 0.83996 mL | 0.29421 mL | 0.29022 mL | 1.15005 mL | 0.02500 mL | 3.477 | 0.01189 | 0.87382 | 0. |
| 54:45.9 | Data point 95 | 0.83996 mL | 0.29421 mL | 0.29116 mL | 1.15005 mL | 0.02500 mL | 3.802 | -0.00531 | 0.16094 | 0. |
| 55:17.7 | Data point 96 | 0.83996 mL | 0.29421 mL | 0.29168 mL | 1.15005 mL | 0.02500 mL | 4.009 | 0.03675 | 0.96951 | 0. |
| 55:34.1 | Data point 97 | 0.83996 mL | 0.29421 mL | 0.29196 mL | 1.15005 mL | 0.02500 mL | 4.250 | 0.04055 | 0.97110 | 0. |
| 55:55.7 | Data point 98 | 0.83996 mL | 0.29421 mL | 0.29224 mL | 1.15005 mL | 0.02500 mL | 4.547 | 0.07033 | 0.94381 | 0. |
| 56:17.3 | Data point 99 | 0.83996 mL | 0.29421 mL | 0.29243 mL | 1.15005 mL | 0.02500 mL | 4.910 | 0.09976 | 0.97446 | 0. |
| 56:50.0 | Data point 100 | 0.83996 mL | 0.29421 mL | 0.29255 mL | 1.15005 mL | 0.02500 mL | 5.222 | 0.09875 | 0.97662 | 0. |
| 57:25.7 | Data point 101 | 0.83996 mL | 0.29421 mL | 0.29264 mL | 1.15005 mL | 0.02500 mL | 5.577 | 0.09753 | 0.98708 | 0. |
| 58:07.8 | Data point 102 | 0.83996 mL | 0.29421 mL | 0.29273 mL | 1.15005 mL | 0.02500 mL | 5.971 | 0.10016 | 0.99125 | 0. |
| 58:46.4 | Data point 103 | 0.83996 mL | 0.29421 mL | 0.29283 mL | 1.15005 mL | 0.02500 mL | 6.275 | -0.03907 | 0.55717 | 0. |
| 59:08.1 | Data point 104 | 0.83996 mL | 0.29421 mL | 0.29292 mL | 1.15005 mL | 0.02500 mL | 6.571 | -0.08584 | 0.94201 | 0. |
| 59:29.6 | Data point 105 | 0.83996 mL | 0.29421 mL | 0.29304 mL | 1.15005 mL | 0.02500 mL | 6.871 | -0.09713 | 0.94088 | 0. |
| 1:00:05.2 | Data point 106 | 0.83996 mL | 0.29421 mL | 0.29318 mL | 1.15005 mL | 0.02500 mL | 7.107 | 0.09337 | 0.95943 | 0. |

| | | | |
|---|---|---|---|
| Sample name: | **M16** | Experiment start time: | **10/12/2017 10:50:14 AM** |
| Assay name: | **UV-metric psKa** | Analyst: | **Dorothy Levorse** |
| Assay ID: | **17J-12009** | Instrument ID: | **T311053** |
| Filename: | **C:\Sirius_T3\Mehtap\20171011_exp15_pKa\17J-12009_M16_UV-metric psKa.t3r** | | |

#### Events (continued)

| Time | Event | Water | Acid | Base | Methanol | Buffer | pH | dpH/dt | pH R-squared | pH SD |
|---|---|---|---|---|---|---|---|---|---|---|
| 1:00:37.2 | Data point 107 | 0.83996 mL | 0.29421 mL | 0.29332 mL | 1.15005 mL | 0.02500 mL | 7.346 | 0.08825 | 0.95666 | 0.0044[illegible] |
| 1:01:14.5 | Data point 108 | 0.83996 mL | 0.29421 mL | 0.29346 mL | 1.15005 mL | 0.02500 mL | 7.603 | 0.09465 | 0.96757 | 0.0047[illegible] |
| 1:01:49.7 | Data point 109 | 0.83996 mL | 0.29421 mL | 0.29358 mL | 1.15005 mL | 0.02500 mL | 7.851 | 0.09726 | 0.98075 | 0.0048[illegible] |
| 1:02:28.5 | Data point 110 | 0.83996 mL | 0.29421 mL | 0.29370 mL | 1.15005 mL | 0.02500 mL | 8.175 | 0.09974 | 0.98165 | 0.0049[illegible] |
| 1:03:12.6 | Data point 111 | 0.83996 mL | 0.29421 mL | 0.29381 mL | 1.15005 mL | 0.02500 mL | 8.568 | 0.09761 | 0.94615 | 0.0049[illegible] |
| 1:03:53.5 | Data point 112 | 0.83996 mL | 0.29421 mL | 0.29393 mL | 1.15005 mL | 0.02500 mL | 8.911 | 0.09881 | 0.98625 | 0.0049[illegible] |
| 1:04:30.8 | Data point 113 | 0.83996 mL | 0.29421 mL | 0.29405 mL | 1.15005 mL | 0.02500 mL | 9.197 | 0.09677 | 0.98197 | 0.0048[illegible] |
| 1:05:04.0 | Data point 114 | 0.83996 mL | 0.29421 mL | 0.29419 mL | 1.15005 mL | 0.02500 mL | 9.447 | 0.09347 | 0.95203 | 0.0047[illegible] |
| 1:05:26.6 | Data point 115 | 0.83996 mL | 0.29421 mL | 0.29436 mL | 1.15005 mL | 0.02500 mL | 9.661 | 0.05370 | 0.97656 | 0.0026[illegible] |
| 1:05:48.1 | Data point 116 | 0.83996 mL | 0.29421 mL | 0.29457 mL | 1.15005 mL | 0.02500 mL | 9.854 | 0.03229 | 0.87936 | 0.0017[illegible] |
| 1:06:20.0 | Data point 117 | 0.83996 mL | 0.29421 mL | 0.29490 mL | 1.15005 mL | 0.02500 mL | 10.055 | 0.02379 | 0.93949 | 0.0012[illegible] |
| 1:06:41.8 | Data point 118 | 0.83996 mL | 0.29421 mL | 0.29532 mL | 1.15005 mL | 0.02500 mL | 10.252 | -0.00051 | 0.01469 | 0.0002[illegible] |
| 1:07:18.8 | Data point 119 | 0.83996 mL | 0.29421 mL | 0.29654 mL | 1.15005 mL | 0.02500 mL | 10.464 | -0.00045 | 0.01218 | 0.0002[illegible] |
| 1:07:50.8 | Data point 120 | 0.83996 mL | 0.29421 mL | 0.29758 mL | 1.15005 mL | 0.02500 mL | 10.654 | -0.01243 | 0.71782 | 0.0007[illegible] |
| 1:08:23.0 | Data point 121 | 0.83996 mL | 0.29421 mL | 0.29908 mL | 1.15005 mL | 0.02500 mL | 10.846 | -0.00991 | 0.73895 | 0.0005[illegible] |
| 1:08:55.1 | Data point 122 | 0.83996 mL | 0.29421 mL | 0.30101 mL | 1.15005 mL | 0.02500 mL | 11.043 | -0.01277 | 0.71584 | 0.0007[illegible] |
| 1:09:11.8 | Data point 123 | 0.83996 mL | 0.29421 mL | 0.30407 mL | 1.15005 mL | 0.02500 mL | 11.214 | -0.01678 | 0.90057 | 0.0008[illegible] |
| 1:09:28.6 | Data point 124 | 0.83996 mL | 0.29421 mL | 0.30858 mL | 1.15005 mL | 0.02500 mL | 11.369 | -0.02172 | 0.91506 | 0.0011[illegible] |
| 1:10:00.9 | Data point 125 | 0.83996 mL | 0.29421 mL | 0.31733 mL | 1.15005 mL | 0.02500 mL | 11.568 | -0.01344 | 0.85187 | 0.0007[illegible] |
| 1:10:28.1 | Data point 126 | 0.83996 mL | 0.29421 mL | 0.32952 mL | 1.15005 mL | 0.02500 mL | 11.759 | -0.01385 | 0.84697 | 0.0007[illegible] |
| 1:10:55.7 | Data point 127 | 0.83996 mL | 0.29421 mL | 0.34993 mL | 1.15005 mL | 0.02500 mL | 11.949 | -0.01210 | 0.82759 | 0.0006[illegible] |
| 1:11:17.8 | Data point 128 | 0.83996 mL | 0.29421 mL | 0.36197 mL | 1.15005 mL | 0.02500 mL | 12.027 | -0.01064 | 0.72154 | 0.0006[illegible] |
| 1:13:17.2 | Assay volumes | 1.08996 mL | 0.43921 mL | 0.36197 mL | 1.15005 mL | 0.02500 mL | | | | |

#### Assay Settings

| Setting | Value | Original Value | Date/Time changed | Imported from |
|---|---|---|---|---|
| ***General Settings*** | | | | |
| Analyst name | Dorothy Levorse | | | |
| Separate reference vial | Yes | | | |
| ***Standard Experiment Settings*** | | | | |
| Number of titrations | 3 | | | |
| Minimum pH | 2.000 | | | |
| Maximum pH | 12.000 | | | |
| pH step between points of | 0.200 | | | |
| Minimum titrant addition | 0.00002 mL | | | |
| Maximum titrant addition | 0.10000 mL | | | |
| Argon flow rate | 100% | | | |
| Start titration using | Cautious pH adjust | | | |
| ***Advanced General Settings*** | | | | |
| Detect turbidity using | Spectrometer | | | |
| Monitor at a wavelength of | 500.0 nm | | | |
| Absorbance threshold of | 0.100 | | | |
| Collect turbidity sensor data | No | | | |
| Stir after titrant addition for | 5 seconds | | | |
| For titrant addition, stir at | 15% | | | |
| ***Titrant Pre-Dose*** | | | | |
| Titrant pre-dose | None | | | |
| ***Assay Medium*** | | | | |
| Cosolvent in use | Yes | | | |
| Cosolvent type | Methanol | | | |
| Cosolvent volume | 1.15 mL | | | |
| Cosolvent added | Automatic | | | |
| ISA water volume | 0.35 mL | | | |
| Water added | Automatic | | | |
| After water addition, stir for | 5 seconds | | | |
| At a speed of | 15% | | | |
| Buffer in use | Yes | | | |

| | | | |
|---|---|---|---|
| Sample name: | **M16** | Experiment start time: | **10/12/2017 10:50:14 AM** |
| Assay name: | **UV-metric psKa** | Analyst: | **Dorothy Levorse** |
| Assay ID: | **17J-12009** | Instrument ID: | **T311053** |
| Filename: | **C:\Sirius_T3\Mehtap\20171011_exp15_pKa\17J-12009_M16_UV-metric psKa.t3r** | | |

#### Calibration Settings

| Setting | Value | Date/Time changed | Imported from |
|---|---|---|---|
| Four-Plus alpha | 0.109 | 10/12/2017 10:50:14 AM | C:\Sirius_T3\17J-11006_Blank standardisation.t3r |
| Four-Plus S | 1.0007 | 10/12/2017 10:50:14 AM | C:\Sirius_T3\17J-11006_Blank standardisation.t3r |
| Four-Plus jH | 0.3 | 10/12/2017 10:50:14 AM | C:\Sirius_T3\17J-11006_Blank standardisation.t3r |
| Four-Plus jOH | -0.2 | 10/12/2017 10:50:14 AM | C:\Sirius_T3\17J-11006_Blank standardisation.t3r |
| Base concentration factor | 1.011 | 10/12/2017 10:50:14 AM | C:\Sirius_T3\KOH17I22.t3r |
| Acid concentration factor | 0.995 | 10/12/2017 10:50:14 AM | C:\Sirius_T3\17J-11005_Blank standardisation.t3r |

| | | | |
|---|---|---|---|
| Sample name: | **M16** | Experiment start time: | **10/12/2017 10:50:14 AM** |
| Assay name: | **UV-metric psKa** | Analyst: | **Dorothy Levorse** |
| Assay ID: | **17J-12009** | Instrument ID: | **T311053** |
| Filename: | **C:\Sirius_T3\Mehtap\20171011_exp15_pKa\17J-12009_M16_UV-metric psKa.t3r** | | |

| | | | |
|---|---|---|---|
| Sample name: | **M16** | Experiment start time: | **10/12/2017 10:50:14 AM** |
| Assay name: | **UV-metric psKa** | Analyst: | **Dorothy Levorse** |
| Assay ID: | **17J-12009** | Instrument ID: | **T311053** |
| Filename: | **C:\Sirius_T3\Mehtap\20171011_exp15_pKa\17J-12009_M16_UV-metric psKa.t3r** | | |

#### Tray Information

Title
Location F1
