## Supplementary material for "Octanol-water partition coefficient measurements for the SAMPL6 Blind Prediction Challenge": SM16_17J-12010_M18_UV-metric psKa_report.pdf

### Yasuda-Shedlovsky Result

| | | | |
|---|---|---|---|
| Sample name: | **M18** | Experiment start time: | **10/12/2017 12:04:33 PM** |
| Assay name: | **UV-metric psKa** | Analyst: | **Dorothy Levorse** |
| Assay ID: | **17J-12010** | Instrument ID: | **T311053** |
| Filename: | **C:\Sirius_T3\Mehtap\20171011_exp15_pKa\17J-12010_M18_UV-metric psKa.t3r** | | |

#### Yasuda-Shedlovsky result

| Extrapolation type | pKa 0% | SD | Intercept | Slope | R² | Ionic strength | Temperature |
|---|---|---|---|---|---|---|---|
| Yasuda-Shedlovsky | 5.35 | ±0.06 | 9.97 | -225.9587 | 0.9950 | 0.165 M | 24.9°C |
| Yasuda-Shedlovsky | 11.01 | ±0.05 | 11.71 | 81.7937 | 0.9745 | 0.165 M | 24.9°C |

#### Component assay results

| Titration | Methanol weight% | Direction | Result type | Dielectric constant | [H2O] | Ionic strength | Temperature | | psKa 1 | | psKa 2 |
|---|---|---|---|---|---|---|---|---|---|---|---|
| 17J-12010 Points 4 to 39 | 49.47 % | Up | UV-metric pKa | 56.6 | 24.7 M | 0.157 M | 24.9°C | ☑ | 4.60 | ☑ | 11.75 |
| 17J-12010 Points 41 to 78 | 40.05 % | Up | UV-metric pKa | 61.0 | 30.0 M | 0.166 M | 25.0°C | ☑ | 4.76 | ☑ | 11.59 |
| 17J-12010 Points 80 to 116 | 30.24 % | Up | UV-metric pKa | 65.4 | 35.7 M | 0.172 M | 24.9°C | ☑ | 4.97 | ☑ | 11.40 |

#### Graphs

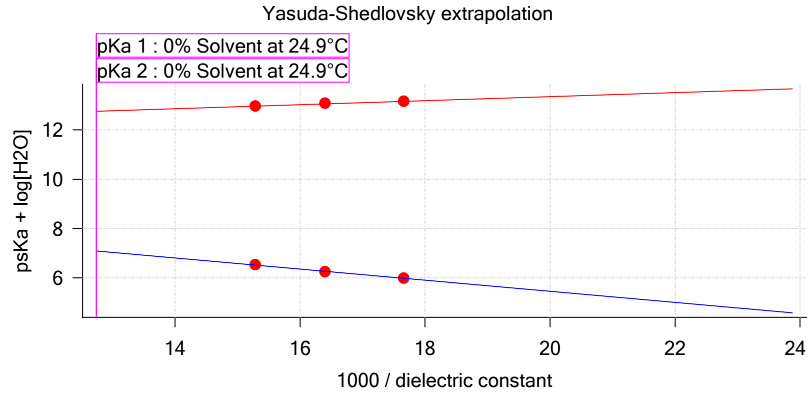


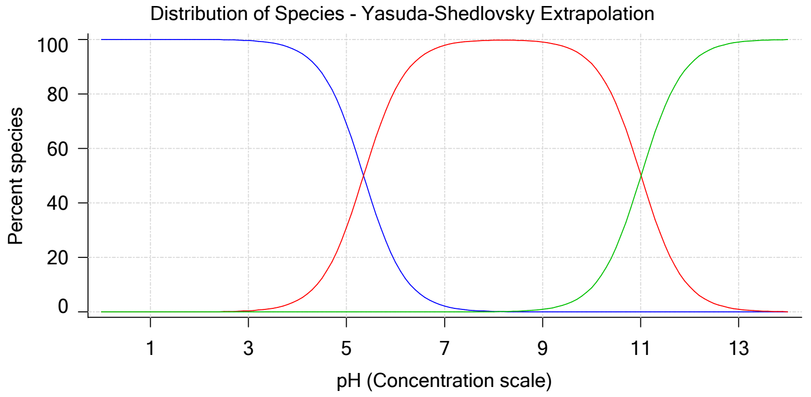


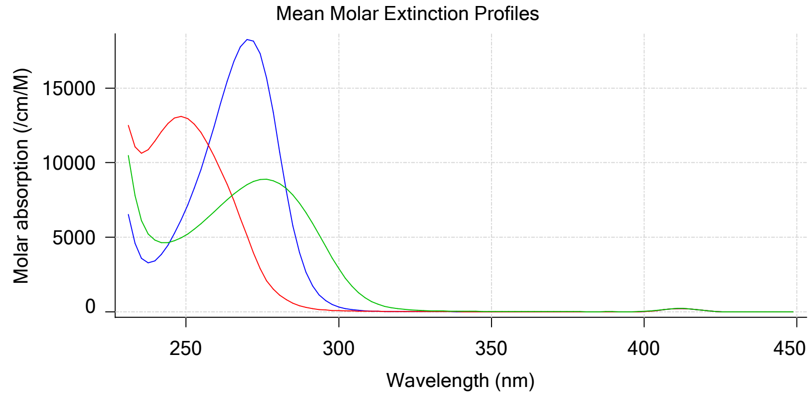


# UV-metric psKa   Titration 1 of 3   17J-12010 Points 4 to 39

#### Results

| | |
|---|---|
| pKa 1 | **4.60** |
| pKa 2 | **11.75** |
| RMSD | **0.003 0.002 0.001** |
| Chi squared | **0.0250** |
| PCA calculated number of pKas | **2** |
| Average ionic strength | **0.157 M** |
| Average temperature | **24.9°C** |
| Analyte concentration range | **48.1 µM to 45.2 µM** |
| Methanol weight % | **49.5 %** |
| Dielectric constant | **56.6** |
| Water concentration | **24.7 M** |

| | | | |
|---|---|---|---|
| Sample name: | **M18** | Experiment start time: | **10/12/2017 12:04:33 PM** |
| Assay name: | **UV-metric psKa** | Analyst: | **Dorothy Levorse** |
| Assay ID: | **17J-12010** | Instrument ID: | **T311053** |
| Filename: | **C:\Sirius_T3\Mehtap\20171011_exp15_pKa\17J-12010_M18_UV-metric psKa.t3r** | | |

#### Results (continued)

| | |
|---|---|
| Number of pKas source | **Predicted** |
| Wavelength clipping | **230.0 nm to 450.0 nm** |
| pH clipping | **1.468 to 12.516** |

#### Warnings and errors

| | |
|---|---|
| Errors | None |
| Warnings | None |

#### Assay Settings

| Setting | Value | Original Value | Date/Time changed | Imported from |
|---|---|---|---|---|
| Buffer in use | Yes | | | |
| Buffer type | Phosphate Buffer | | | |
| ***Assay Medium*** | | | | |
| Volume of buffer introduced | 0.025000 mL | | | |
| Add buffer manually | Manual | | | |

#### Graphs

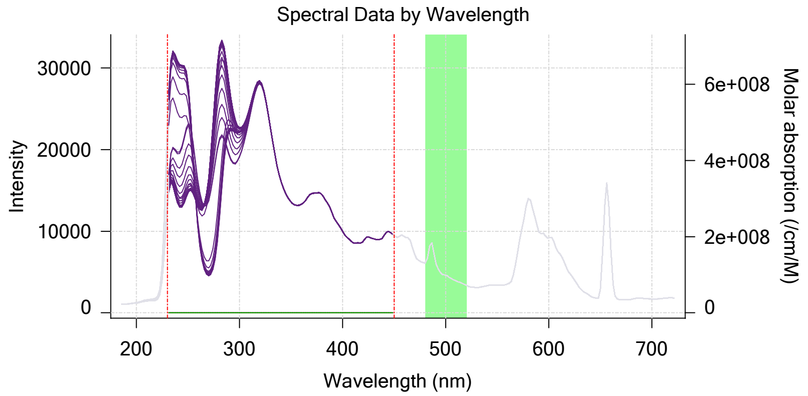


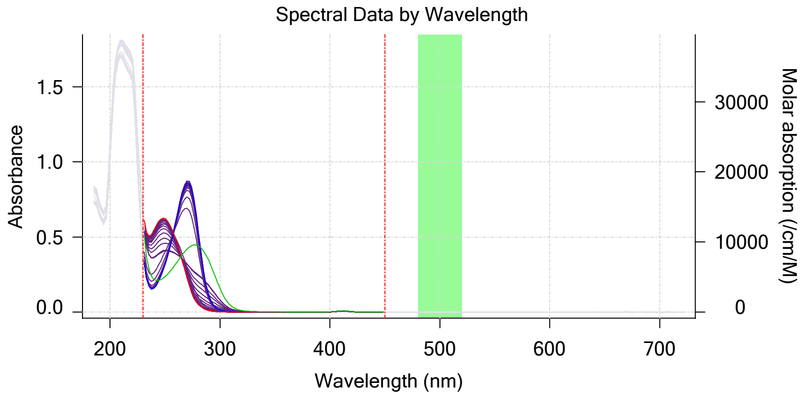


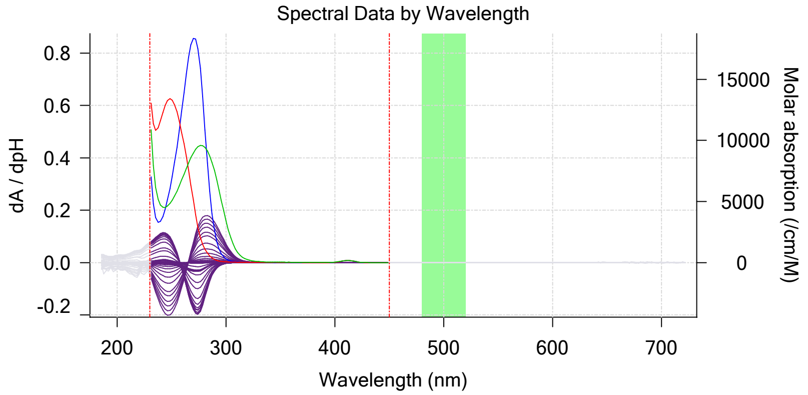


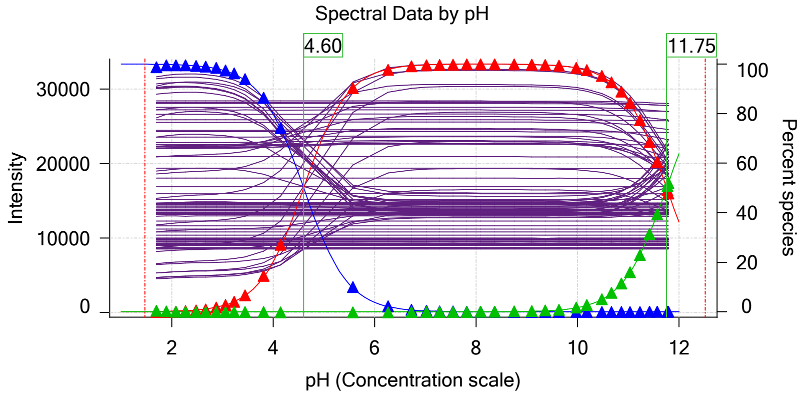

| | | | |
|---|---|---|---|
| Sample name: | **M18** | Experiment start time: | **10/12/2017 12:04:33 PM** |
| Assay name: | **UV-metric psKa** | Analyst: | **Dorothy Levorse** |
| Assay ID: | **17J-12010** | Instrument ID: | **T311053** |
| Filename: | **C:\Sirius_T3\Mehtap\20171011_exp15_pKa\17J-12010_M18_UV-metric psKa.t3r** | | |

#### Graphs (continued)

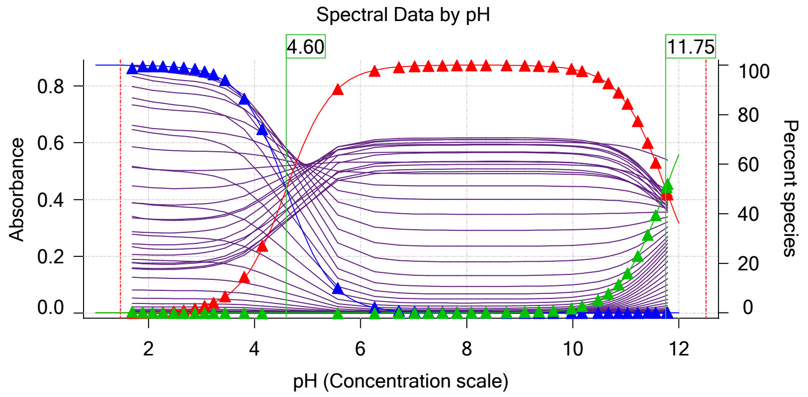


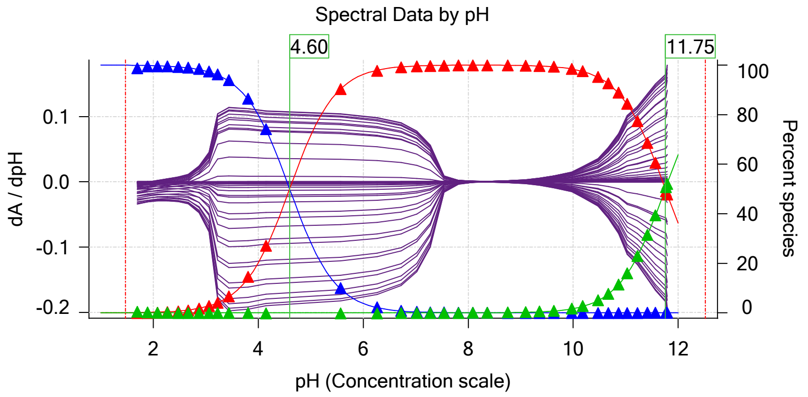


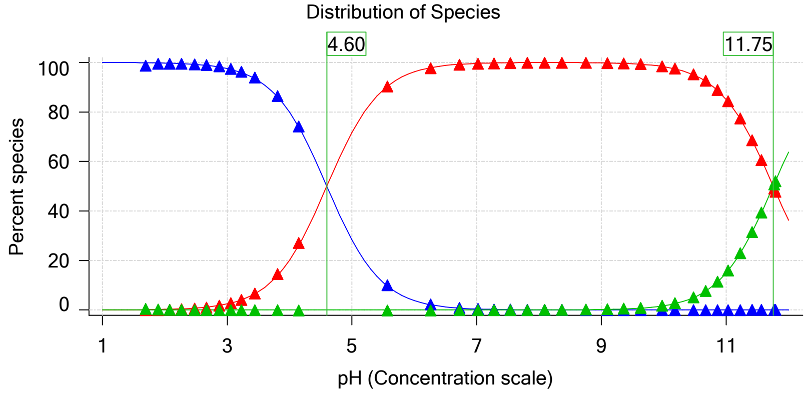


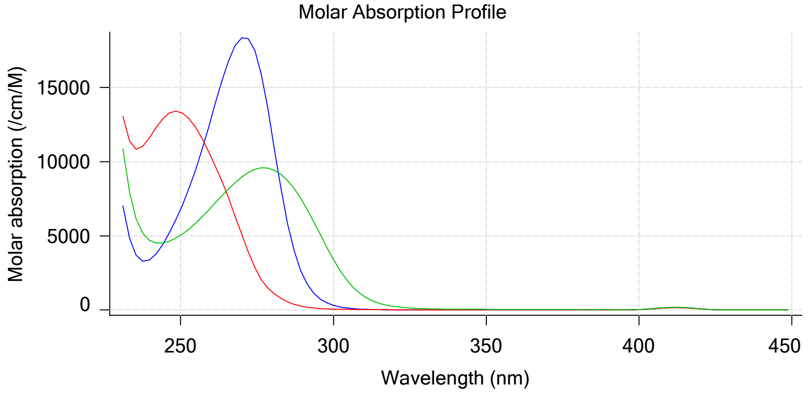


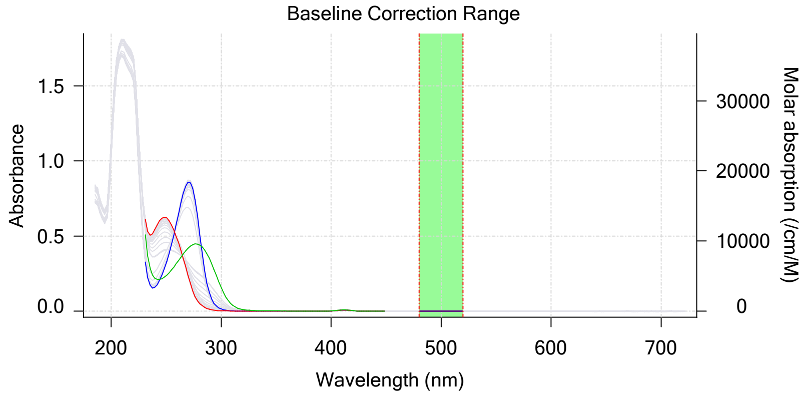


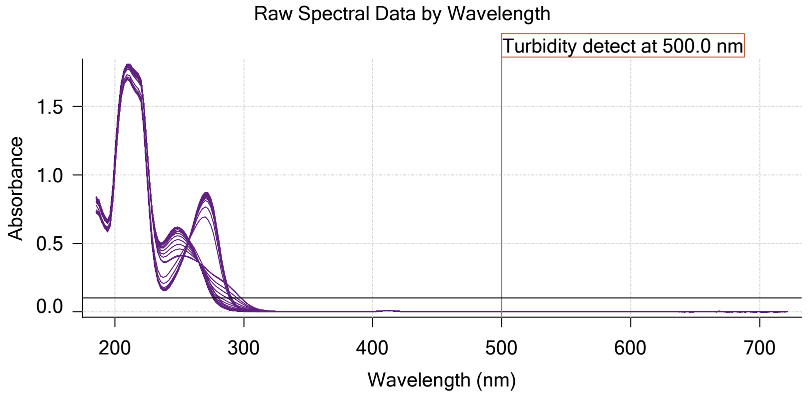


### UV-metric psKa Titration 2 of 3 17J-12010 Points 41 to 78

#### Results

| | |
|---|---|
| pKa 1 | **4.76** |
| pKa 2 | **11.59** |
| RMSD | **0.001 0.002 0.001** |
| Chi squared | **0.0136** |
| PCA calculated number of pKas | **2** |
| Average ionic strength | **0.166 M** |
| Average temperature | **25.0°C** |
| Analyte concentration range | **39.5 µM to 37.4 µM** |
| Methanol weight % | **40.0 %** |
| Dielectric constant | **61.0** |
| Water concentration | **30.0 M** |

| | | | |
|---|---|---|---|
| Sample name: | **M18** | Experiment start time: | **10/12/2017 12:04:33 PM** |
| Assay name: | **UV-metric psKa** | Analyst: | **Dorothy Levorse** |
| Assay ID: | **17J-12010** | Instrument ID: | **T311053** |
| Filename: | **C:\Sirius_T3\Mehtap\20171011_exp15_pKa\17J-12010_M18_UV-metric psKa.t3r** | | |

#### Results (continued)

| | |
|---|---|
| Number of pKas source | **Predicted** |
| Wavelength clipping | **230.0 nm to 450.0 nm** |
| pH clipping | **1.512 to 12.504** |

#### Warnings and errors

| | |
|---|---|
| Errors | None |
| Warnings | None |

#### Assay Settings

| Setting | Value | Original Value | Date/Time changed | Imported from |
|---|---|---|---|---|
| Buffer in use | Yes | | | |
| Buffer type | Phosphate Buffer | | | |
| ***Assay Medium*** | | | | |
| Volume of buffer introduced | 0.025000 mL | | | |
| Add buffer manually | Manual | | | |

#### Graphs

Spectral Data by Wavelength

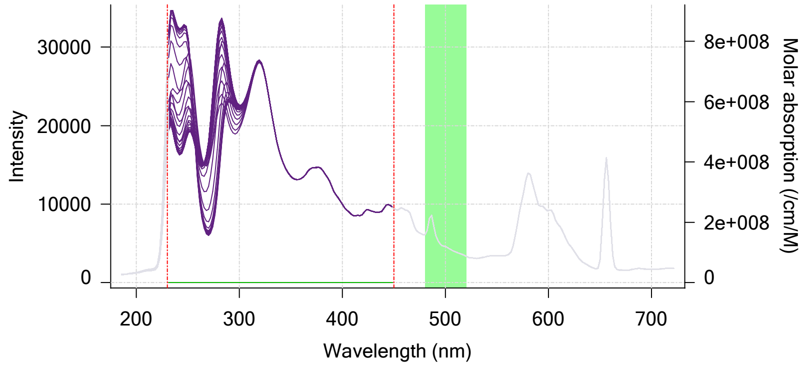


Spectral Data by Wavelength

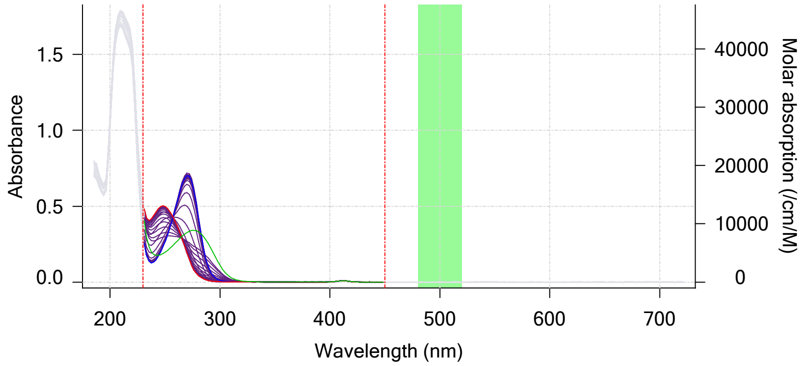


Spectral Data by Wavelength

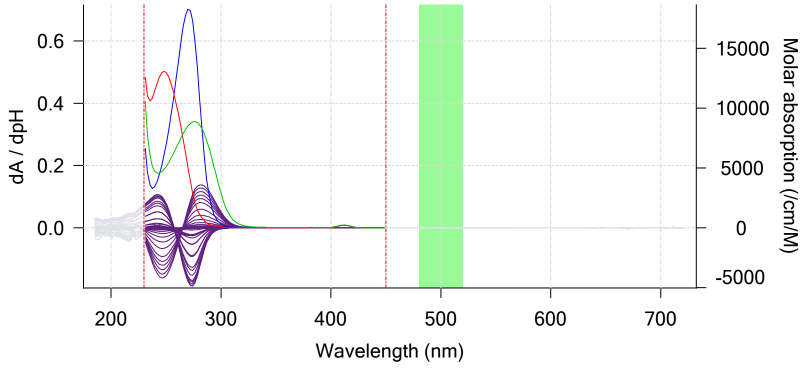


Spectral Data by pH

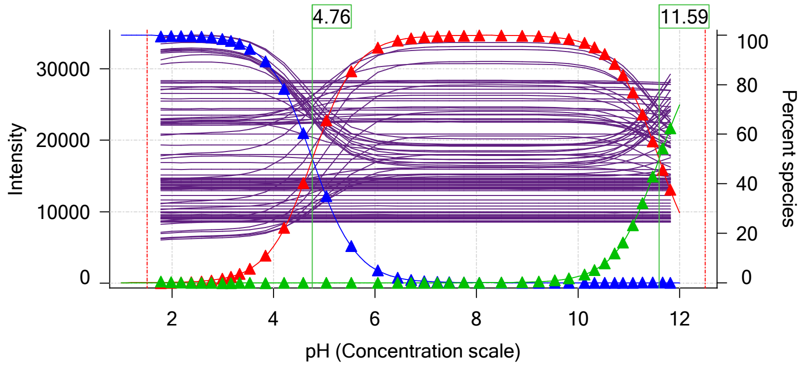

| | | | |
|---|---|---|---|
| Sample name: | **M18** | Experiment start time: | **10/12/2017 12:04:33 PM** |
| Assay name: | **UV-metric psKa** | Analyst: | **Dorothy Levorse** |
| Assay ID: | **17J-12010** | Instrument ID: | **T311053** |
| Filename: | **C:\Sirius_T3\Mehtap\20171011_exp15_pKa\17J-12010_M18_UV-metric psKa.t3r** | | |

#### Graphs (continued)

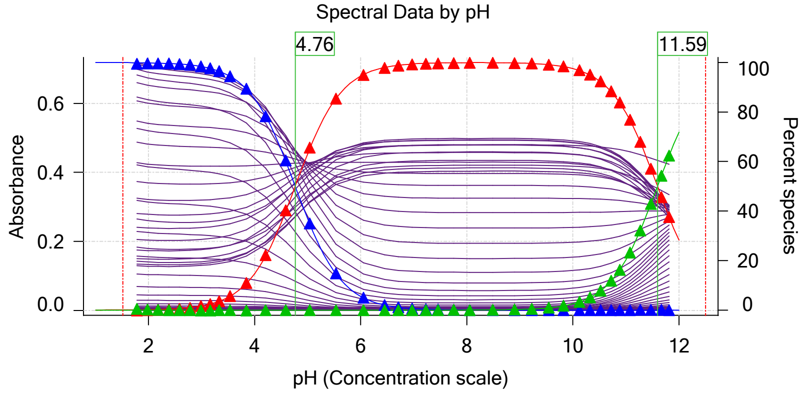


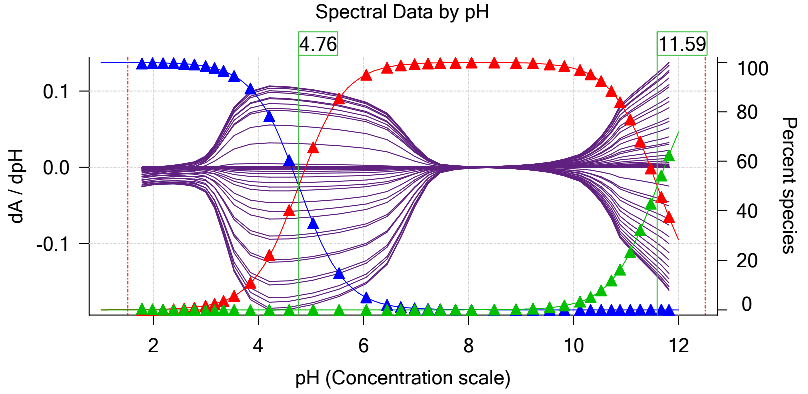


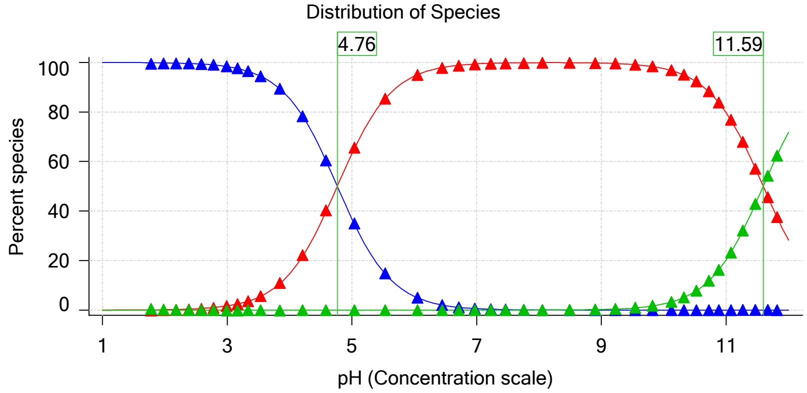


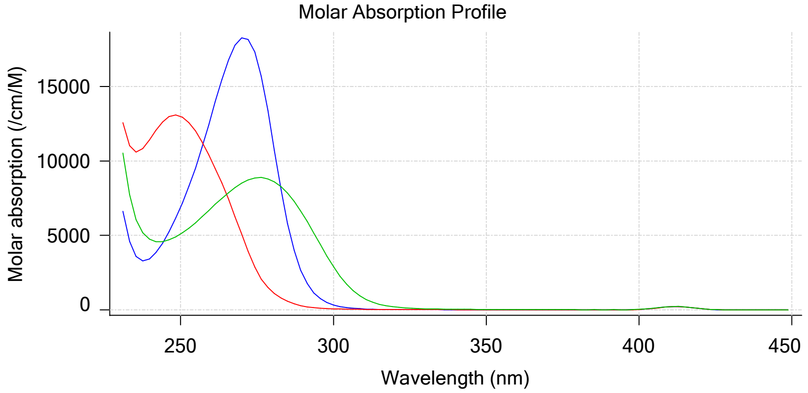


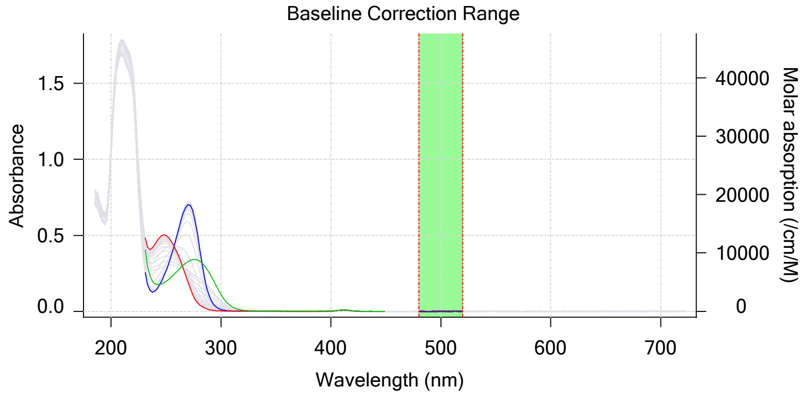


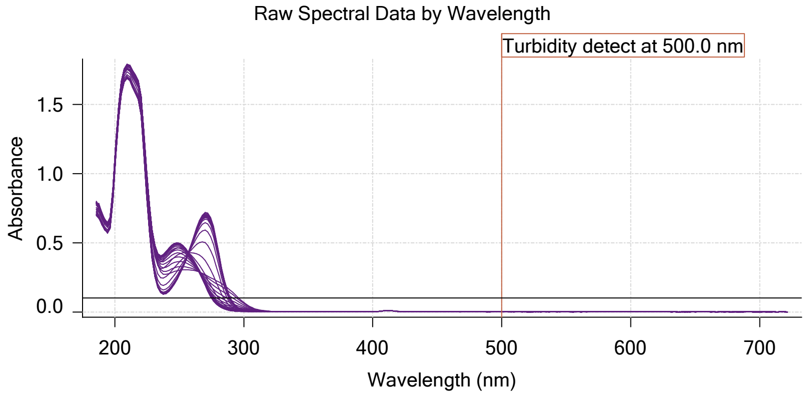


### UV-metric psKa Titration 3 of 3 17J-12010 Points 80 to 116

#### Results

| | |
|---|---|
| pKa 1 | **4.97** |
| pKa 2 | **11.40** |
| RMSD | **0.001 0.002 0.001** |
| Chi squared | **0.0077** |
| PCA calculated number of pKas | **2** |
| Average ionic strength | **0.172 M** |
| Average temperature | **24.9°C** |
| Analyte concentration range | **30.4 µM to 28.8 µM** |
| Methanol weight % | **30.2 %** |
| Dielectric constant | **65.4** |
| Water concentration | **35.7 M** |

| | | | |
|---|---|---|---|
| Sample name: | **M18** | Experiment start time: | **10/12/2017 12:04:33 PM** |
| Assay name: | **UV-metric psKa** | Analyst: | **Dorothy Levorse** |
| Assay ID: | **17J-12010** | Instrument ID: | **T311053** |
| Filename: | **C:\Sirius_T3\Mehtap\20171011_exp15_pKa\17J-12010_M18_UV-metric psKa.t3r** | | |

#### Results (continued)

| | |
|---|---|
| Number of pKas source | **Predicted** |
| Wavelength clipping | **230.0 nm to 450.0 nm** |
| pH clipping | **1.513 to 12.513** |

#### Warnings and errors

| | |
|---|---|
| Errors | None |
| Warnings | None |

#### Assay Settings

| Setting | Value | Original Value | Date/Time changed | Imported from |
|---|---|---|---|---|
| Buffer in use | Yes | | | |
| Buffer type | Phosphate Buffer | | | |
| ***Assay Medium*** | | | | |
| Volume of buffer introduced | 0.025000 mL | | | |
| Add buffer manually | Manual | | | |

#### Graphs

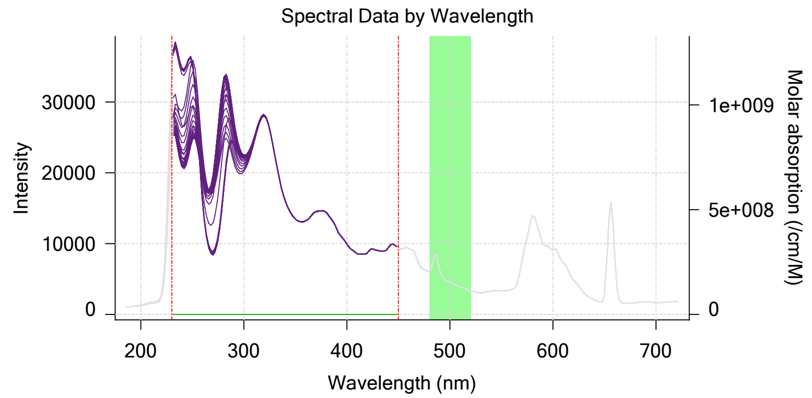


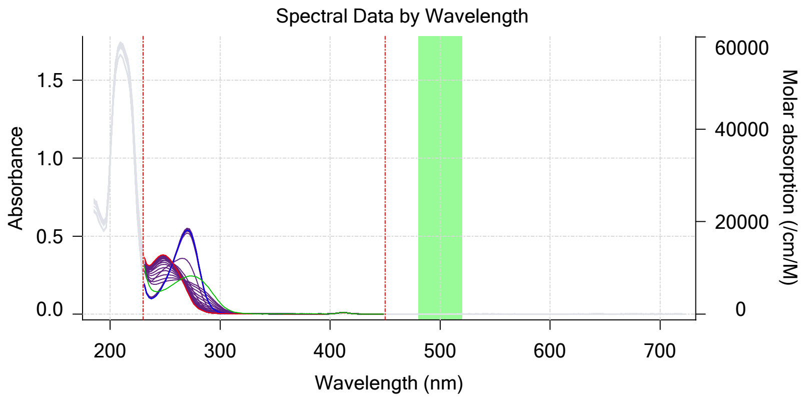


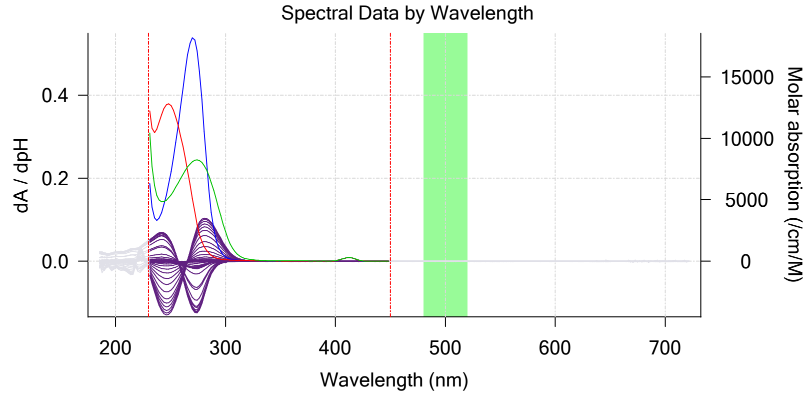


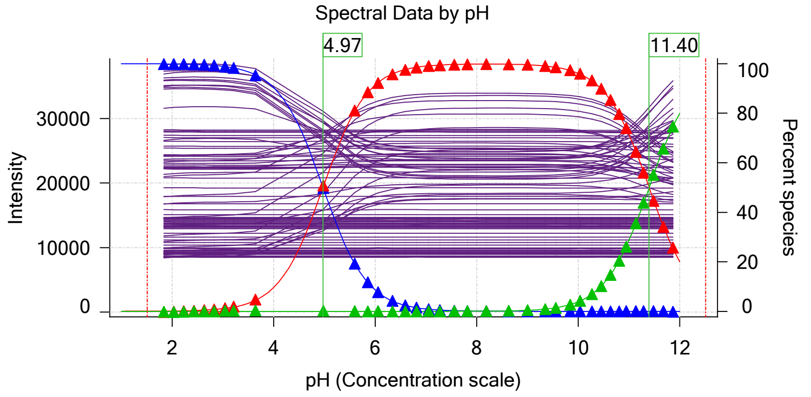

Sample name: **M18**
Assay name: **UV-metric psKa**
Assay ID: **17J-12010**
Filename: **C:\Sirius_T3\Mehtap\20171011_exp15_pKa\17J-12010_M18_UV-metric psKa.t3r**

Experiment start time: **10/12/2017 12:04:33 PM**
Analyst: **Dorothy Levorse**
Instrument ID: **T311053**

#### Graphs (continued)

Spectral Data by pH

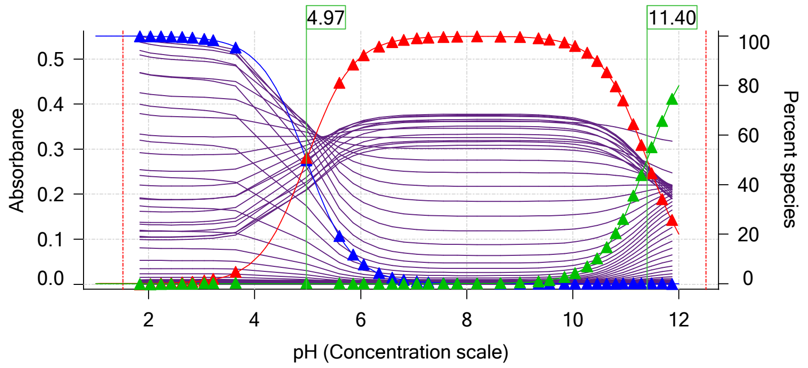


Spectral Data by pH

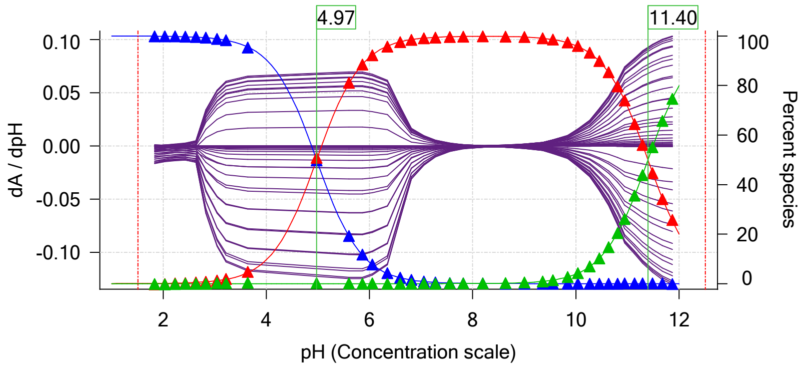


Distribution of Species

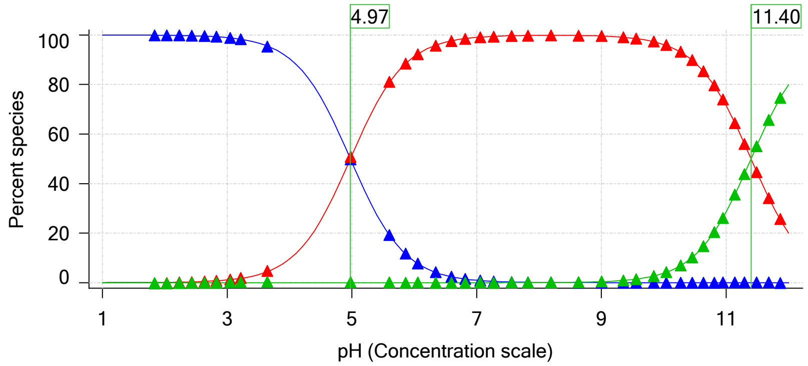


Molar Absorption Profile

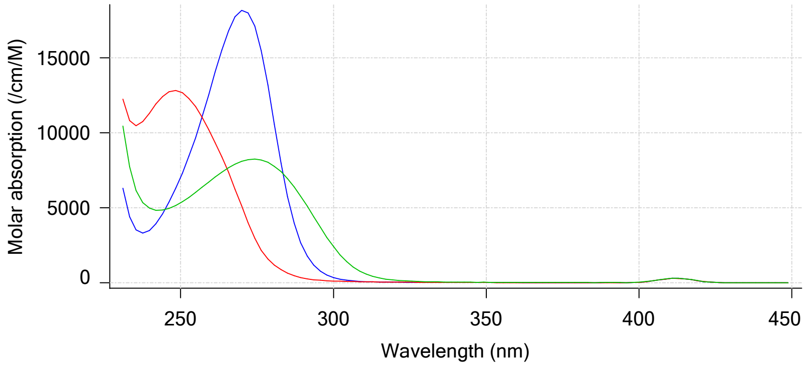


Baseline Correction Range

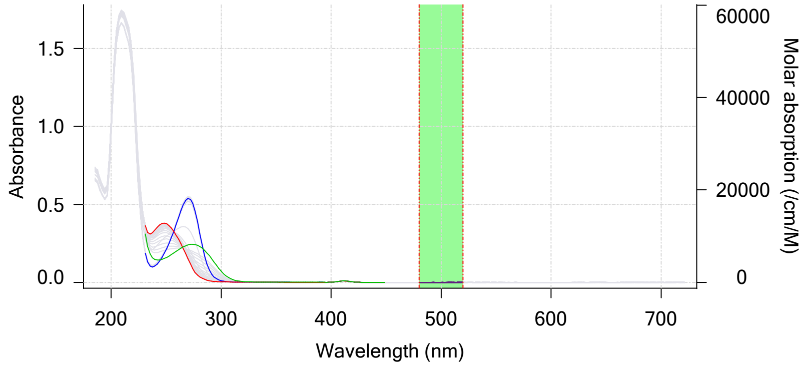


Raw Spectral Data by Wavelength

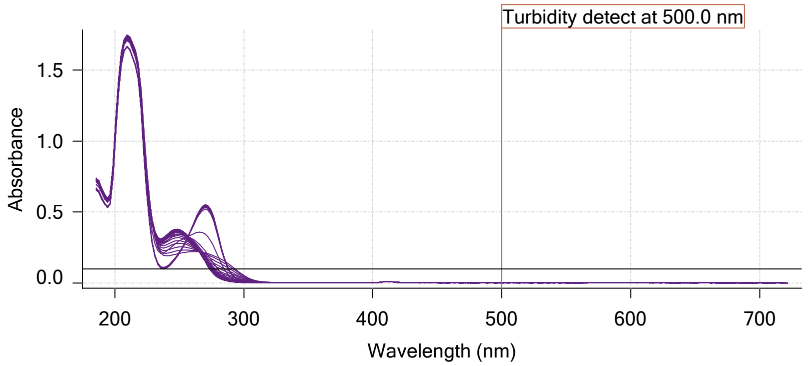


#### Assay Model

| Settings | Value | Date/Time changed | Imported from |
|---|---|---|---|
| Sample name | M18 | 10/11/2017 4:22:27 PM | User entered value |
| Sample by | Volume | | Default value |
| Sample volume | 0.0010 mL | 10/11/2017 4:22:27 PM | User entered value |
| Solvent | DMSO | | Default value |
| Sample concentration | 0.076700 M | 10/11/2017 4:22:27 PM | User entered value |
| Solubility | Unknown | | Default value |
| Molecular weight | 267.11 | 10/11/2017 4:22:36 PM | User entered value |
| Individual pKa ionic environments | No | | Default value |
| Number of pKas | 2 | 10/11/2017 4:22:27 PM | User entered value |
| Sample is a | Ampholyte | 10/11/2017 4:22:27 PM | User entered value |
| pKa 1 | 5.19 | 10/11/2017 4:22:27 PM | User entered value |
| Type | Base | 10/11/2017 4:22:27 PM | User entered value |
| pKa 2 | 10.85 | 10/11/2017 4:22:27 PM | User entered value |
| Type | Acid | 10/11/2017 4:22:27 PM | User entered value |

Sample name: **M18**
Assay name: **UV-metric psKa**
Assay ID: **17J-12010**
Filename: **C:\Sirius_T3\Mehtap\20171011_exp15_pKa\17J-12010_M18_UV-metric psKa.t3r**
Experiment start time: **10/12/2017 12:04:33 PM**
Analyst: **Dorothy Levorse**
Instrument ID: **T311053**

#### Assay Model (continued)

| Settings | Value | Date/Time changed | Imported from |
|---|---|---|---|
| logp (XH2 +) | -10.00 | | Default value |
| logP (neutral XH) | -10.00 | 10/11/2017 4:22:27 PM | User entered value |
| logP (X -) | -10.00 | | Default value |

#### Events

| Time | Event | Water | Acid | Base | Methanol | Buffer | pH | dpH/dt | pH R-squar |
|---|---|---|---|---|---|---|---|---|---|
| 3:31.4 | Dark spectrum | | | | | | | | |
| 3:32.7 | Reference spectrum | | | | | | | | |
| 4:00.4 | Volume reset due to vial change | | | | | | | | |
| 4:44.7 | Initial pH = 8.35 | | | | | | | | |
| 5:39.1 | Data point 4 | 0.34995 mL | 0.06919 mL | 0.00000 mL | 1.15005 mL | 0.02500 mL | 1.968 | -0.02216 | 0.93220 |
| 6:07.9 | Data point 5 | 0.34995 mL | 0.06919 mL | 0.02491 mL | 1.15005 mL | 0.02500 mL | 2.168 | -0.01140 | 0.51070 |
| 6:24.8 | Data point 6 | 0.34995 mL | 0.06919 mL | 0.04008 mL | 1.15005 mL | 0.02500 mL | 2.352 | 0.02310 | 0.86227 |
| 6:41.6 | Data point 7 | 0.34995 mL | 0.06919 mL | 0.04988 mL | 1.15005 mL | 0.02500 mL | 2.544 | 0.02591 | 0.86248 |
| 6:58.3 | Data point 8 | 0.34995 mL | 0.06919 mL | 0.05628 mL | 1.15005 mL | 0.02500 mL | 2.751 | 0.02119 | 0.81319 |
| 7:15.0 | Data point 9 | 0.34995 mL | 0.06919 mL | 0.06028 mL | 1.15005 mL | 0.02500 mL | 2.940 | 0.00531 | 0.69289 |
| 7:31.6 | Data point 10 | 0.34995 mL | 0.06919 mL | 0.06289 mL | 1.15005 mL | 0.02500 mL | 3.145 | 0.01086 | 0.71651 |
| 7:48.3 | Data point 11 | 0.34995 mL | 0.06919 mL | 0.06451 mL | 1.15005 mL | 0.02500 mL | 3.330 | 0.00949 | 0.76498 |
| 8:05.0 | Data point 12 | 0.34995 mL | 0.06919 mL | 0.06557 mL | 1.15005 mL | 0.02500 mL | 3.497 | 0.01228 | 0.83695 |
| 8:37.1 | Data point 13 | 0.34995 mL | 0.06919 mL | 0.06670 mL | 1.15005 mL | 0.02500 mL | 3.711 | 0.02264 | 0.95333 |
| 9:04.0 | Data point 14 | 0.34995 mL | 0.06919 mL | 0.06783 mL | 1.15005 mL | 0.02500 mL | 4.076 | 0.06944 | 0.98551 |
| 9:25.7 | Data point 15 | 0.34995 mL | 0.06919 mL | 0.06820 mL | 1.15005 mL | 0.02500 mL | 4.411 | 0.09971 | 0.99228 |
| 9:55.5 | Data point 16 | 0.34995 mL | 0.06919 mL | 0.06863 mL | 1.15005 mL | 0.02500 mL | 5.828 | 0.09719 | 0.97988 |
| 11:03.8 | Data point 17 | 0.34995 mL | 0.06919 mL | 0.06884 mL | 1.15005 mL | 0.02500 mL | 6.518 | 0.09753 | 0.98953 |
| 12:08.8 | Data point 18 | 0.34995 mL | 0.06919 mL | 0.06898 mL | 1.15005 mL | 0.02500 mL | 6.976 | 0.09815 | 0.98834 |
| 12:57.4 | Data point 19 | 0.34995 mL | 0.06919 mL | 0.06910 mL | 1.15005 mL | 0.02500 mL | 7.271 | 0.09986 | 0.98326 |
| 13:44.1 | Data point 20 | 0.34995 mL | 0.06919 mL | 0.06921 mL | 1.15005 mL | 0.02500 mL | 7.527 | 0.10032 | 0.99359 |
| 14:31.0 | Data point 21 | 0.34995 mL | 0.06919 mL | 0.06936 mL | 1.15005 mL | 0.02500 mL | 7.790 | 0.09965 | 0.99091 |
| 15:15.2 | Data point 22 | 0.34995 mL | 0.06919 mL | 0.06950 mL | 1.15005 mL | 0.02500 mL | 8.060 | 0.09971 | 0.98237 |
| 16:03.2 | Data point 23 | 0.34995 mL | 0.06919 mL | 0.06961 mL | 1.15005 mL | 0.02500 mL | 8.333 | 0.09994 | 0.97608 |
| 16:48.5 | Data point 24 | 0.34995 mL | 0.06919 mL | 0.06971 mL | 1.15005 mL | 0.02500 mL | 8.608 | 0.03826 | 0.16675 |
| 17:32.0 | Data point 25 | 0.34995 mL | 0.06919 mL | 0.06980 mL | 1.15005 mL | 0.02500 mL | 8.994 | 0.09896 | 0.98406 |
| 18:21.9 | Data point 26 | 0.34995 mL | 0.06919 mL | 0.06990 mL | 1.15005 mL | 0.02500 mL | 9.334 | 0.10061 | 0.98780 |
| 19:02.0 | Data point 27 | 0.34995 mL | 0.06919 mL | 0.06999 mL | 1.15005 mL | 0.02500 mL | 9.597 | 0.09781 | 0.98928 |
| 19:38.9 | Data point 28 | 0.34995 mL | 0.06919 mL | 0.07011 mL | 1.15005 mL | 0.02500 mL | 9.868 | 0.09614 | 0.98123 |
| 20:05.7 | Data point 29 | 0.34995 mL | 0.06919 mL | 0.07027 mL | 1.15005 mL | 0.02500 mL | 10.212 | 0.07694 | 0.96069 |
| 20:32.5 | Data point 30 | 0.34995 mL | 0.06919 mL | 0.07051 mL | 1.15005 mL | 0.02500 mL | 10.414 | 0.03263 | 0.92759 |
| 20:49.1 | Data point 31 | 0.34995 mL | 0.06919 mL | 0.07088 mL | 1.15005 mL | 0.02500 mL | 10.709 | 0.00864 | 0.78861 |
| 21:21.1 | Data point 32 | 0.34995 mL | 0.06919 mL | 0.07171 mL | 1.15005 mL | 0.02500 mL | 10.902 | -0.00050 | 0.01117 |
| 21:53.0 | Data point 33 | 0.34995 mL | 0.06919 mL | 0.07295 mL | 1.15005 mL | 0.02500 mL | 11.094 | -0.00659 | 0.71687 |
| 22:09.6 | Data point 34 | 0.34995 mL | 0.06919 mL | 0.07462 mL | 1.15005 mL | 0.02500 mL | 11.262 | -0.00654 | 0.74297 |
| 22:36.6 | Data point 35 | 0.34995 mL | 0.06919 mL | 0.07803 mL | 1.15005 mL | 0.02500 mL | 11.455 | -0.00627 | 0.67172 |
| 23:03.5 | Data point 36 | 0.34995 mL | 0.06919 mL | 0.08208 mL | 1.15005 mL | 0.02500 mL | 11.644 | -0.00712 | 0.70022 |
| 23:20.2 | Data point 37 | 0.34995 mL | 0.06919 mL | 0.08794 mL | 1.15005 mL | 0.02500 mL | 11.794 | -0.00717 | 0.74788 |
| 23:52.6 | Data point 38 | 0.34995 mL | 0.06919 mL | 0.09899 mL | 1.15005 mL | 0.02500 mL | 11.995 | 0.00175 | 0.11667 |
| 24:09.3 | Data point 39 | 0.34995 mL | 0.06919 mL | 0.10052 mL | 1.15005 mL | 0.02500 mL | 12.016 | -0.00742 | 0.79563 |
| 25:50.8 | Reference spectrum | | | | | | | | |
| 26:54.8 | Data point 41 | 0.50000 mL | 0.16660 mL | 0.10054 mL | 1.15005 mL | 0.02500 mL | 2.012 | -0.04740 | 0.93961 |
| 27:22.3 | Data point 42 | 0.50000 mL | 0.16660 mL | 0.12408 mL | 1.15005 mL | 0.02500 mL | 2.211 | 0.00909 | 0.74636 |
| 27:39.3 | Data point 43 | 0.50000 mL | 0.16660 mL | 0.13902 mL | 1.15005 mL | 0.02500 mL | 2.409 | 0.01364 | 0.85662 |
| 27:56.1 | Data point 44 | 0.50000 mL | 0.16660 mL | 0.14852 mL | 1.15005 mL | 0.02500 mL | 2.611 | 0.00818 | 0.78053 |
| 28:12.8 | Data point 45 | 0.50000 mL | 0.16660 mL | 0.15452 mL | 1.15005 mL | 0.02500 mL | 2.812 | 0.02851 | 0.91765 |
| 28:29.3 | Data point 46 | 0.50000 mL | 0.16660 mL | 0.15826 mL | 1.15005 mL | 0.02500 mL | 3.007 | 0.00158 | 0.05553 |
| 28:45.9 | Data point 47 | 0.50000 mL | 0.16660 mL | 0.16068 mL | 1.15005 mL | 0.02500 mL | 3.216 | 0.01208 | 0.86564 |
| 29:02.5 | Data point 48 | 0.50000 mL | 0.16660 mL | 0.16218 mL | 1.15005 mL | 0.02500 mL | 3.390 | 0.01997 | 0.93839 |
| 29:19.1 | Data point 49 | 0.50000 mL | 0.16660 mL | 0.16317 mL | 1.15005 mL | 0.02500 mL | 3.559 | 0.01585 | 0.88400 |

| | | | |
|---|---|---|---|
| Sample name: | **M18** | Experiment start time: | **10/12/2017 12:04:33 PM** |
| Assay name: | **UV-metric psKa** | Analyst: | **Dorothy Levorse** |
| Assay ID: | **17J-12010** | Instrument ID: | **T311053** |
| Filename: | **C:\Sirius_T3\Mehtap\20171011_exp15_pKa\17J-12010_M18_UV-metric psKa.t3r** | | |

#### Events (continued)

| Time | Event | Water | Acid | Base | Methanol | Buffer | pH | dpH/dt | pH R-squared | [illegible] |
|---|---|---|---|---|---|---|---|---|---|---|
| 29:46.0 | Data point 50 | 0.50000 mL | 0.16660 mL | 0.16432 mL | 1.15005 mL | 0.02500 mL | 3.760 | 0.02862 | 0.97119 | 0. |
| 30:07.8 | Data point 51 | 0.50000 mL | 0.16660 mL | 0.16498 mL | 1.15005 mL | 0.02500 mL | 4.068 | 0.04928 | 0.98585 | 0. |
| 30:29.6 | Data point 52 | 0.50000 mL | 0.16660 mL | 0.16538 mL | 1.15005 mL | 0.02500 mL | 4.427 | 0.09688 | 0.99530 | 0. |
| 30:56.4 | Data point 53 | 0.50000 mL | 0.16660 mL | 0.16571 mL | 1.15005 mL | 0.02500 mL | 4.805 | 0.09983 | 0.99043 | 0. |
| 31:36.3 | Data point 54 | 0.50000 mL | 0.16660 mL | 0.16590 mL | 1.15005 mL | 0.02500 mL | 5.259 | 0.09611 | 0.95932 | 0. |
| 32:34.2 | Data point 55 | 0.50000 mL | 0.16660 mL | 0.16602 mL | 1.15005 mL | 0.02500 mL | 5.750 | 0.09659 | 0.98588 | 0. |
| 33:30.9 | Data point 56 | 0.50000 mL | 0.16660 mL | 0.16611 mL | 1.15005 mL | 0.02500 mL | 6.267 | 0.09792 | 0.98701 | 0. |
| 34:25.6 | Data point 57 | 0.50000 mL | 0.16660 mL | 0.16620 mL | 1.15005 mL | 0.02500 mL | 6.660 | 0.09938 | 0.97707 | 0. |
| 35:03.3 | Data point 58 | 0.50000 mL | 0.16660 mL | 0.16630 mL | 1.15005 mL | 0.02500 mL | 6.923 | 0.10057 | 0.99505 | 0. |
| 35:42.0 | Data point 59 | 0.50000 mL | 0.16660 mL | 0.16639 mL | 1.15005 mL | 0.02500 mL | 7.179 | 0.09757 | 0.99359 | 0. |
| 36:22.5 | Data point 60 | 0.50000 mL | 0.16660 mL | 0.16651 mL | 1.15005 mL | 0.02500 mL | 7.435 | 0.09462 | 0.97663 | 0. |
| 37:01.3 | Data point 61 | 0.50000 mL | 0.16660 mL | 0.16663 mL | 1.15005 mL | 0.02500 mL | 7.668 | 0.09781 | 0.99247 | 0. |
| 37:45.8 | Data point 62 | 0.50000 mL | 0.16660 mL | 0.16675 mL | 1.15005 mL | 0.02500 mL | 7.960 | 0.09633 | 0.98309 | 0. |
| 38:28.7 | Data point 63 | 0.50000 mL | 0.16660 mL | 0.16684 mL | 1.15005 mL | 0.02500 mL | 8.260 | 0.09940 | 0.98649 | 0. |
| 39:14.2 | Data point 64 | 0.50000 mL | 0.16660 mL | 0.16693 mL | 1.15005 mL | 0.02500 mL | 8.692 | 0.09572 | 0.97048 | 0. |
| 40:04.2 | Data point 65 | 0.50000 mL | 0.16660 mL | 0.16703 mL | 1.15005 mL | 0.02500 mL | 9.104 | 0.09854 | 0.98929 | 0. |
| 40:47.4 | Data point 66 | 0.50000 mL | 0.16660 mL | 0.16712 mL | 1.15005 mL | 0.02500 mL | 9.427 | 0.09929 | 0.96392 | 0. |
| 41:20.1 | Data point 67 | 0.50000 mL | 0.16660 mL | 0.16722 mL | 1.15005 mL | 0.02500 mL | 9.744 | 0.09517 | 0.97511 | 0. |
| 41:44.8 | Data point 68 | 0.50000 mL | 0.16660 mL | 0.16733 mL | 1.15005 mL | 0.02500 mL | 10.020 | 0.06934 | 0.95893 | 0. |
| 42:01.3 | Data point 69 | 0.50000 mL | 0.16660 mL | 0.16755 mL | 1.15005 mL | 0.02500 mL | 10.322 | 0.01941 | 0.88056 | 0. |
| 42:33.2 | Data point 70 | 0.50000 mL | 0.16660 mL | 0.16804 mL | 1.15005 mL | 0.02500 mL | 10.519 | 0.00425 | 0.52856 | 0. |
| 43:05.3 | Data point 71 | 0.50000 mL | 0.16660 mL | 0.16874 mL | 1.15005 mL | 0.02500 mL | 10.718 | -0.00079 | 0.04953 | 0. |
| 43:22.0 | Data point 72 | 0.50000 mL | 0.16660 mL | 0.16973 mL | 1.15005 mL | 0.02500 mL | 10.915 | -0.00777 | 0.77415 | 0. |
| 43:38.6 | Data point 73 | 0.50000 mL | 0.16660 mL | 0.17126 mL | 1.15005 mL | 0.02500 mL | 11.078 | -0.00404 | 0.40114 | 0. |
| 44:10.7 | Data point 74 | 0.50000 mL | 0.16660 mL | 0.17415 mL | 1.15005 mL | 0.02500 mL | 11.270 | -0.01018 | 0.91389 | 0. |
| 44:37.7 | Data point 75 | 0.50000 mL | 0.16660 mL | 0.17806 mL | 1.15005 mL | 0.02500 mL | 11.461 | -0.01191 | 0.91645 | 0. |
| 45:10.1 | Data point 76 | 0.50000 mL | 0.16660 mL | 0.18535 mL | 1.15005 mL | 0.02500 mL | 11.659 | -0.00968 | 0.85015 | 0. |
| 45:42.6 | Data point 77 | 0.50000 mL | 0.16660 mL | 0.19617 mL | 1.15005 mL | 0.02500 mL | 11.857 | -0.00576 | 0.66939 | 0. |
| 46:04.7 | Data point 78 | 0.50000 mL | 0.16660 mL | 0.20889 mL | 1.15005 mL | 0.02500 mL | 12.004 | -0.00665 | 0.65235 | 0. |
| 47:49.5 | Reference spectrum | | | | | | | | | |
| 49:12.9 | Data point 80 | 0.83996 mL | 0.29548 mL | 0.20891 mL | 1.15005 mL | 0.02500 mL | 2.013 | -0.03025 | 0.95923 | 0. |
| 49:40.6 | Data point 81 | 0.83996 mL | 0.29548 mL | 0.23584 mL | 1.15005 mL | 0.02500 mL | 2.209 | 0.01165 | 0.71144 | 0. |
| 49:57.5 | Data point 82 | 0.83996 mL | 0.29548 mL | 0.25299 mL | 1.15005 mL | 0.02500 mL | 2.408 | 0.00774 | 0.43737 | 0. |
| 50:14.3 | Data point 83 | 0.83996 mL | 0.29548 mL | 0.26392 mL | 1.15005 mL | 0.02500 mL | 2.604 | -0.03359 | 0.83216 | 0. |
| 50:31.1 | Data point 84 | 0.83996 mL | 0.29548 mL | 0.27107 mL | 1.15005 mL | 0.02500 mL | 2.808 | 0.00980 | 0.53331 | 0. |
| 50:47.8 | Data point 85 | 0.83996 mL | 0.29548 mL | 0.27545 mL | 1.15005 mL | 0.02500 mL | 3.002 | -0.01498 | 0.82438 | 0. |
| 51:04.3 | Data point 86 | 0.83996 mL | 0.29548 mL | 0.27827 mL | 1.15005 mL | 0.02500 mL | 3.220 | -0.00773 | 0.40725 | 0. |
| 51:20.9 | Data point 87 | 0.83996 mL | 0.29548 mL | 0.27996 mL | 1.15005 mL | 0.02500 mL | 3.388 | 0.00254 | 0.11739 | 0. |
| 52:02.4 | Data point 88 | 0.83996 mL | 0.29548 mL | 0.28140 mL | 1.15005 mL | 0.02500 mL | 3.812 | -0.01429 | 0.83687 | 0. |
| 52:29.4 | Data point 89 | 0.83996 mL | 0.29548 mL | 0.28443 mL | 1.15005 mL | 0.02500 mL | 5.145 | -0.06232 | 0.60070 | 0. |
| 52:58.1 | Data point 90 | 0.83996 mL | 0.29548 mL | 0.28469 mL | 1.15005 mL | 0.02500 mL | 5.765 | 0.09540 | 0.95384 | 0. |
| 53:30.6 | Data point 91 | 0.83996 mL | 0.29548 mL | 0.28481 mL | 1.15005 mL | 0.02500 mL | 6.024 | 0.09838 | 0.96697 | 0. |
| 54:00.4 | Data point 92 | 0.83996 mL | 0.29548 mL | 0.28490 mL | 1.15005 mL | 0.02500 mL | 6.221 | -0.03478 | 0.34071 | 0. |
| 54:22.1 | Data point 93 | 0.83996 mL | 0.29548 mL | 0.28502 mL | 1.15005 mL | 0.02500 mL | 6.509 | -0.07557 | 0.70744 | 0. |
| 54:48.1 | Data point 94 | 0.83996 mL | 0.29548 mL | 0.28514 mL | 1.15005 mL | 0.02500 mL | 6.757 | -0.09151 | 0.92688 | 0. |
| 55:19.4 | Data point 95 | 0.83996 mL | 0.29548 mL | 0.28525 mL | 1.15005 mL | 0.02500 mL | 6.975 | 0.03002 | 0.62417 | 0. |
| 55:51.4 | Data point 96 | 0.83996 mL | 0.29548 mL | 0.28540 mL | 1.15005 mL | 0.02500 mL | 7.218 | 0.08354 | 0.85166 | 0. |
| 56:18.3 | Data point 97 | 0.83996 mL | 0.29548 mL | 0.28551 mL | 1.15005 mL | 0.02500 mL | 7.435 | 0.09524 | 0.94381 | 0. |
| 56:51.3 | Data point 98 | 0.83996 mL | 0.29548 mL | 0.28563 mL | 1.15005 mL | 0.02500 mL | 7.717 | 0.09909 | 0.98255 | 0. |
| 57:26.7 | Data point 99 | 0.83996 mL | 0.29548 mL | 0.28572 mL | 1.15005 mL | 0.02500 mL | 7.979 | 0.09940 | 0.97982 | 0. |
| 58:07.1 | Data point 100 | 0.83996 mL | 0.29548 mL | 0.28582 mL | 1.15005 mL | 0.02500 mL | 8.351 | 0.09745 | 0.98546 | 0. |
| 58:55.6 | Data point 101 | 0.83996 mL | 0.29548 mL | 0.28591 mL | 1.15005 mL | 0.02500 mL | 8.791 | 0.08853 | 0.90914 | 0. |
| 59:39.5 | Data point 102 | 0.83996 mL | 0.29548 mL | 0.28601 mL | 1.15005 mL | 0.02500 mL | 9.159 | 0.09262 | 0.96900 | 0. |
| 1:00:10.2 | Data point 103 | 0.83996 mL | 0.29548 mL | 0.28610 mL | 1.15005 mL | 0.02500 mL | 9.509 | 0.09447 | 0.94462 | 0. |
| 1:00:35.9 | Data point 104 | 0.83996 mL | 0.29548 mL | 0.28619 mL | 1.15005 mL | 0.02500 mL | 9.712 | 0.08737 | 0.95857 | 0. |
| 1:00:52.4 | Data point 105 | 0.83996 mL | 0.29548 mL | 0.28636 mL | 1.15005 mL | 0.02500 mL | 9.994 | 0.01804 | 0.91104 | 0. |
| 1:01:14.0 | Data point 106 | 0.83996 mL | 0.29548 mL | 0.28664 mL | 1.15005 mL | 0.02500 mL | 10.194 | 0.00435 | 0.41151 | 0. |

| | | | |
|---|---|---|---|
| Sample name: | **M18** | Experiment start time: | **10/12/2017 12:04:33 PM** |
| Assay name: | **UV-metric psKa** | Analyst: | **Dorothy Levorse** |
| Assay ID: | **17J-12010** | Instrument ID: | **T311053** |
| Filename: | **C:\Sirius_T3\Mehtap\20171011_exp15_pKa\17J-12010_M18_UV-metric psKa.t3r** | | |

#### Events (continued)

| Time | Event | Water | Acid | Base | Methanol | Buffer | pH | dpH/dt | pH R-squared | pH SD |
|---|---|---|---|---|---|---|---|---|---|---|
| 1:01:30.6 | Data point 107 | 0.83996 mL | 0.29548 mL | 0.28711 mL | 1.15005 mL | 0.02500 mL | 10.424 | -0.01587 | 0.92501 | 0.0008[illegible] |
| 1:01:47.3 | Data point 108 | 0.83996 mL | 0.29548 mL | 0.28789 mL | 1.15005 mL | 0.02500 mL | 10.607 | -0.01835 | 0.89412 | 0.0009[illegible] |
| 1:02:03.9 | Data point 109 | 0.83996 mL | 0.29548 mL | 0.28906 mL | 1.15005 mL | 0.02500 mL | 10.788 | -0.02060 | 0.94692 | 0.0010[illegible] |
| 1:02:20.6 | Data point 110 | 0.83996 mL | 0.29548 mL | 0.29080 mL | 1.15005 mL | 0.02500 mL | 10.960 | -0.02285 | 0.92858 | 0.0011[illegible] |
| 1:02:37.2 | Data point 111 | 0.83996 mL | 0.29548 mL | 0.29334 mL | 1.15005 mL | 0.02500 mL | 11.097 | -0.02119 | 0.92066 | 0.0010[illegible] |
| 1:03:04.2 | Data point 112 | 0.83996 mL | 0.29548 mL | 0.29673 mL | 1.15005 mL | 0.02500 mL | 11.287 | -0.01817 | 0.95686 | 0.0009[illegible] |
| 1:03:20.9 | Data point 113 | 0.83996 mL | 0.29548 mL | 0.30207 mL | 1.15005 mL | 0.02500 mL | 11.439 | -0.01973 | 0.92653 | 0.0010[illegible] |
| 1:03:53.3 | Data point 114 | 0.83996 mL | 0.29548 mL | 0.31101 mL | 1.15005 mL | 0.02500 mL | 11.634 | -0.01396 | 0.92329 | 0.0007[illegible] |
| 1:04:20.6 | Data point 115 | 0.83996 mL | 0.29548 mL | 0.32587 mL | 1.15005 mL | 0.02500 mL | 11.828 | -0.01381 | 0.93854 | 0.0007[illegible] |
| 1:04:48.3 | Data point 116 | 0.83996 mL | 0.29548 mL | 0.34875 mL | 1.15005 mL | 0.02500 mL | 12.013 | -0.00961 | 0.71039 | 0.0005[illegible] |
| 1:06:47.6 | Assay volumes | 1.08996 mL | 0.43631 mL | 0.34875 mL | 1.15005 mL | 0.02500 mL | | | | |

#### Assay Settings

| Setting | Value | Original Value | Date/Time changed | Imported from |
|---|---|---|---|---|
| ***General Settings*** | | | | |
| Analyst name | Dorothy Levorse | | | |
| Separate reference vial | Yes | | | |
| ***Standard Experiment Settings*** | | | | |
| Number of titrations | 3 | | | |
| Minimum pH | 2.000 | | | |
| Maximum pH | 12.000 | | | |
| pH step between points of | 0.200 | | | |
| Minimum titrant addition | 0.00002 mL | | | |
| Maximum titrant addition | 0.10000 mL | | | |
| Argon flow rate | 100% | | | |
| Start titration using | Cautious pH adjust | | | |
| ***Advanced General Settings*** | | | | |
| Detect turbidity using | Spectrometer | | | |
| Monitor at a wavelength of | 500.0 nm | | | |
| Absorbance threshold of | 0.100 | | | |
| Collect turbidity sensor data | No | | | |
| Stir after titrant addition for | 5 seconds | | | |
| For titrant addition, stir at | 15% | | | |
| ***Titrant Pre-Dose*** | | | | |
| Titrant pre-dose | None | | | |
| ***Assay Medium*** | | | | |
| Cosolvent in use | Yes | | | |
| Cosolvent type | Methanol | | | |
| Cosolvent volume | 1.15 mL | | | |
| Cosolvent added | Automatic | | | |
| ISA water volume | 0.35 mL | | | |
| Water added | Automatic | | | |
| After water addition, stir for | 5 seconds | | | |
| At a speed of | 15% | | | |
| Buffer in use | Yes | | | |
| Buffer type | Phosphate Buffer | | | |
| Volume of buffer introduced | 0.025000 mL | | | |
| Add buffer manually | Manual | | | |
| After medium addition, stir for | 5 seconds | | | |
| ***Sample Sonication*** | | | | |
| Sonicate | No | | | |
| ***Sample Dissolution*** | | | | |
| Perform a dissolution stage | No | | | |
| ***Carbonate purge*** | | | | |
| Perform a carbonate purge | No | | | |
| ***Temperature Control*** | | | | |
| Wait for temperature | Yes | | | |

| | | | |
|---|---|---|---|
| Sample name: | **M18** | Experiment start time: | **10/12/2017 12:04:33 PM** |
| Assay name: | **UV-metric psKa** | Analyst: | **Dorothy Levorse** |
| Assay ID: | **17J-12010** | Instrument ID: | **T311053** |
| Filename: | **C:\Sirius_T3\Mehtap\20171011_exp15_pKa\17J-12010_M18_UV-metric psKa.t3r** | | |

#### Assay Settings (continued)

| Setting | Value | Original Value | Date/Time changed | Imported from |
|---|---|---|---|---|
| Required start temperature | 25.0°C | | | |
| Acceptable deviation | 0.5°C | | | |
| Time to wait | 60 seconds | | | |
| Stir speed of | 15% | | | |
| ***Titration 1*** | | | | |
| Titrate from | Low to high pH | | | |
| Adjust to start pH | Yes | | | |
| After pH adjust stir for | 10 seconds | | | |
| ***Titration 2*** | | | | |
| Titrate from | Low to high pH | | | |
| Additional cosolvent volume | 0.00 mL | | | |
| Add additional water | 0.15 mL | | | |
| Additional water added | Automatic | | | |
| After pH adjust stir for | 10 seconds | | | |
| ***Titration 3*** | | | | |
| Titrate from | Low to high pH | | | |
| Additional cosolvent volume | 0.00 mL | | | |
| Add additional water | 0.34 mL | | | |
| Additional water added | Automatic | | | |
| After pH adjust stir for | 10 seconds | | | |
| ***Data Point Stability*** | | | | |
| Stir during data point collection | Yes | | | |
| For point collection, stir at | 15% | | | |
| Delay before data point collection | 0 seconds | | | |
| Number of points to average | 20 points | | | |
| Time interval between points | 0.50 seconds | | | |
| Required maximum standard deviation | 0.00500 dpH/dt | | | |
| Stability timeout after | 60 seconds | | | |
| ***Experiment cleanup*** | | | | |
| Adjust pH to cleanup | To start pH | | | |
| And then stir for | 60 seconds | | | |
| For cleaning, stir at | 20% | | | |
| Then add water volume | 0.25 mL | | | |
| And then stir for | 30 seconds | | | |

#### Calibration Settings

| Setting | Value | Date/Time changed | Imported from |
|---|---|---|---|
| Four-Plus alpha | 0.109 | 10/12/2017 12:04:33 PM | C:\Sirius_T3\17J-11006_Blank standardisation.t3r |
| Four-Plus S | 1.0007 | 10/12/2017 12:04:33 PM | C:\Sirius_T3\17J-11006_Blank standardisation.t3r |
| Four-Plus jH | 0.3 | 10/12/2017 12:04:33 PM | C:\Sirius_T3\17J-11006_Blank standardisation.t3r |
| Four-Plus jOH | -0.2 | 10/12/2017 12:04:33 PM | C:\Sirius_T3\17J-11006_Blank standardisation.t3r |
| Base concentration factor | 1.011 | 10/12/2017 12:04:33 PM | C:\Sirius_T3\KOH17I22.t3r |
| Acid concentration factor | 0.995 | 10/12/2017 12:04:33 PM | C:\Sirius_T3\17J-11005_Blank standardisation.t3r |

#### Instrument Settings

| Setting | Value | Batch Id | Install date |
|---|---|---|---|
| Instrument owner | Merck | | |
| Instrument ID | T311053 | | |
| Instrument type | T3 Simulator | | |
| Software version | 1.1.3.0 | | |
| Dispenser module | | T3DM1100253 | 3/31/2009 6:24:52 AM |
| Dispenser 0 | Water | | 3/31/2009 6:25:05 AM |
| Syringe volume | 2.5 mL | | |
| Firmware version | 1.2.1(r2) | | |
| Titrant | Water (0.15 M KCl) | 10-10-2017 | 10/10/2017 10:48:53 AM |
| Dispenser 2 | Acid | | 3/31/2009 6:25:11 AM |

| | | | |
|---|---|---|---|
| Sample name: | **M18** | Experiment start time: | **10/12/2017 12:04:33 PM** |
| Assay name: | **UV-metric psKa** | Analyst: | **Dorothy Levorse** |
| Assay ID: | **17J-12010** | Instrument ID: | **T311053** |
| Filename: | **C:\Sirius_T3\Mehtap\20171011_exp15_pKa\17J-12010_M18_UV-metric psKa.t3r** | | |

| | | | |
|---|---|---|---|
| Sample name: | **M18** | Experiment start time: | **10/12/2017 12:04:33 PM** |
| Assay name: | **UV-metric psKa** | Analyst: | **Dorothy Levorse** |
| Assay ID: | **17J-12010** | Instrument ID: | **T311053** |
| Filename: | **C:\Sirius_T3\Mehtap\20171011_exp15_pKa\17J-12010_M18_UV-metric psKa.t3r** | | |

#### Instrument Settings (continued)

| Setting | Value | Batch Id | Install date |
|---|---|---|---|
| Maximum standard vial volume | 3.50 mL | | |
| Maximum alternate vial volume | 25.00 mL | | |
| Automatic action idle period | 5 minute(s) | | |
| Titrant tube volume | 1.3 mL | | |
| Syringe flush count | 3.50 | | |
| Flowing wash pump volume | 20.0 mL | | |
| Flowing wash stir duration | 5 s | | |
| Flowing wash stir speed | 30% | | |
| Solvent wash stir duration | 5 s | | |
| Solvent wash stir speed | 30% | | |
| Surfactant wash stir duration | 5 s | | |
| Surfactant wash stir speed | 30% | | |
| E0 calibration minimum number of points | 10 | | |
| E0 calibration maximum standard deviation | 0.01500 | | |
| E0 calibration timeout period | 60 s | | |
| E0 calibration stir duration | 5 s | | |
| E0 calibration preparation stir speed | 30% | | |
| E0 calibration buffer wash stir duration | 5 s | | |
| E0 calibration buffer wash stir speed | 30% | | |
| E0 calibration reading stir speed | 0% | | |
| Spectrometer calibration stir duration | 5 s | | |
| Spectrometer calibration stir speed | 30% | | |
| Spectrometer calibration wash pump volume | 20.0 mL | | |
| Spectrometer calibration wash stir duration | 5 s | | |
| Spectrometer calibration wash stir speed | 30% | | |
| Overhead dispense height | 10000 | | |

#### Refinement Settings

| Setting | Value | Default value |
|---|---|---|
| Turbidity detection method | Spectrometer | Spectrometer |
| Turbidity wavelength to assess | 500.0 nm | 500.0 nm |
| Turbidity maximum absorbance | 0.100 | 0.100 |
| Turbidity probe threshold | 50.00 | 50.00 |
| Exclude turbid points | Yes | Yes |
| Low intensity warning threshold | 100 | 100 |
| Minimum absorbance change threshold | 0.100 | 0.100 |
| Eigenvector autocorrelation threshold | 0.80 | 0.80 |
| Maximum RMSD severe warning | 0.250 | 0.250 |
| Maximum RMSD warning | 0.050 | 0.050 |

#### Tray Information

Title

Location G1
